# Discovery of two structurally distinct classes of inhibitors targeting the nuclease MUS81 and enhancing efficacy of chemotherapy in cancer cells

**DOI:** 10.1101/2025.07.25.666773

**Authors:** Jana Prochazkova, Benoit Carbain, Victoria Marini, Fedor Nikulenkov, Stepan Havel, Naresh Akavaram, Prashant Khirsariya, Alexandra Sisakova, Jakub Cibulka, Michala Boudova, Magdalena Zacpalova, Magdalena Kalovska, Joana Rodrigues, Lukas Daniel, Jan Brezovsky, Claus Azzalin, Kamil Paruch, Lumir Krejci

## Abstract

Nucleases are emerging as promising pharmacological targets due to their essential role in maintaining genomic stability, which is crucial for cellular viability and can be exploited in the prevention and treatment of various diseases, including cancer. The conserved structure-specific endonuclease MUS81 is required for resolving branched DNA intermediates during replication, repair, and recombination. Aberrant activity of MUS81 leads to DNA damage, chromosomal abnormalities and genome instability, and contributes to oncogenesis. Pharmacological targeting of MUS81 thus represents an attractive underexplored therapeutic approach. Here we describe the discovery of two chemically distinct classes of small-molecule inhibitors of MUS81, exemplified by the compounds MU262 and MU876. Both compounds can effectively inhibit MUS81 *in vitro* and in the cell-based context and sensitize cancer cells to DNA-damaging agents through impairing their ability to repair DNA lesions. These compounds can be also used as chemical biology tools for further exploration of MUS81 function, and as leads in the process of drug discovery focused on development of new therapies that exploit DNA repair vulnerabilities in the treatment of cancer.

## INTRODUCTION

Genomic integrity is fundamental for cell survival and prevention of diseases associated with uncontrolled cellular proliferation, such as cancer. To maintain genomic stability, cells rely on various DNA repair pathways, including homologous recombination (HR), where nucleases play critical roles. Among them, the structure-specific endonuclease MUS81, belonging to the highly conserved XPF/MUS81 family, emerges as one of the key players in maintaining the integrity of the genome [1]. MUS81 forms a heterodimer with one of two catalytically inactive counterparts, EME1 or EME2 [2–4]. This complex performs multifaceted functions in DNA repair and replication fork processing. The importance of MUS81 for cellular homeostasis has been illustrated by the reports revealing that MUS81-knockout mice exhibit an increased tendency to form sporadic tumours, indicating the nuclease’s role as a tumour suppressor [5, 6]. In line with this, reduced MUS81 expression has been found to correlate with poor prognosis in various cancers, such as hepatocellular carcinoma and colorectal cancer [7, 8]. Conversely, elevated MUS81 levels have been associated with enhanced tumour cell migration and metastasis in gastric and ovarian cancer[9, 10], indicating its context-dependent dual role in cancer biology. Similarly, EME1 is upregulated in gastric cancer and other tumour types, and its depletion leads to cell cycle arrest and apoptosis [11], further confirming the reliance of some cancers on MUS81-EME1 activity. Collectively, these observations provide significant evidence that aberrant MUS81 activity leads to genomic instability and cancer development. This nuclease has thus emerged as an attractive therapeutic target [7, 12–17].

At the molecular level, MUS81 preferentially cleaves branched DNA structures, such as 3’flap, replication forks and nicked Holliday junctions – intermediates frequently formed during recombination or replication. By resolving these structures, MUS81 ensures accurate segregation of chromosomes during mitosis [18–21]. Its dysfunction leads to chromosomal abnormalities including micronuclei, anaphase and ultrafine bridges, and sister chromatin exchanges - hallmarks of genome instability [19, 22–24].

MUS81 also plays a critical role in processing stalled replication forks [25–27]. When replication forks stall due to obstacles or DNA lesions, MUS81-mediated cleavage enables their restart or resolution [24, 28–34], often through the generation of transient double-strand breaks (DSBs) that are subsequently repaired by break-induced replication repair pathway [32, 35]. Cells lacking MUS81-EME1 are hypersensitive to replication stress-inducing agents such as hydroxyurea (HU) or camptothecin (CPT) [23, 32], further linking the nuclease activity to the replication stress response. Dysregulation of replication fork processing, including replication-transcription collision, is increasingly recognised as a contributing factor to cancer development [36, 37], with MUS81 playing a key role in facilitating replication restart in these contexts [38, 39].

As described above, due to its essential role in DNA repair and replication fork processing, MUS81 has gained attention as a potential target for cancer therapy. Selective inhibition of MUS81 may induce synthetic lethality in cancer cells with compromised DNA repair pathways [40–43], and may enhance the efficacy of existing chemotherapies and radiotherapies [15, 44–46]. Despite the therapeutic potential of targeting MUS81, development of small-molecule MUS81 inhibitors remains rather underexplored. To date, only two reports describe small-molecule inhibitors capable of targeting MUS81 *in vitro*, but with limited evidence of their efficacy in cellular models [47, 48]. Our study identifies two classes of novel small-molecule inhibitors of MUS81 using two complementary high-throughput screening approaches. We describe structure-activity relationship (SAR) development in both series leading to the identification of the most potent compounds. In addition, we assessed the compounds’ impact on MUS81 nuclease activity *in vitro* and elucidated the mechanistic aspects of MUS81 inhibition, including the compounds’ binding to the target protein and a possible mode of action. We also show the impact of the newly discovered inhibitors on MUS81-dependent cellular processes, such as recombination-dependent repair pathways and formation of chromosomal aberrations. Importantly, the inhibitors reported herein potentiate the effects of the chemotherapeutic drug cisplatin and prevent cancer cells proliferation. In conclusion, the MUS81 inhibitors described in this study can be used as both molecular biology tools for the elucidation of the biological functions of MUS81 in genomic maintenance, and as lead compounds for developing targeted cancer therapies that exploit the inherent genomic instability of tumour cells.

## RESULTS

### Identification of small-molecule inhibitors of MUS81

To identify small-molecule inhibitors of MUS81, we used two complementary screening approaches. The first involved a structure-based virtual screening, where over 140.000 small molecules were docked into the active site of MUS81 [49] using AutoDock Vina software [50]. The predicted binding energies of these compounds ranged from -9.6 to -3.0 kcal.mol^-1^. Compounds with binding energy lower than - 8.0 kcal.mol^-1^ were selected for further analysis, yielding a set of 9074 candidates. These compounds were grouped into 27 clusters based on their predicted interactions with MUS81 using AuposSOM tool [51]. Each cluster represented a unique type of binding mode within the enzyme’s active site. Visual inspection of the top candidates using PyMol [52] confirmed their potential to efficiently block the catalytic site of MUS81. From this initial screening, 99 commercially available molecules, representing these clusters proportionally to the cluster sizes, were selected for further evaluation based on their predicted dissociation constants (K_D_) using neural network scoring function NNScore 2.0 [53] prioritising compounds with values below 1 μM. A final set of 20 compounds (Supplementary table 1) were purchased and evaluated in an *in vitro* nuclease assay using a 3’flap substrate, a well-established and relevant substrate for MUS81 [18]. The cleavage by purified MUS81-EME1 was monitored by gel electrophoresis, and compound activity was quantified by measuring the reduction in cleavage product intensity. Several compounds showed inhibitory activity with IC_50_ below 30 μM. Among them, the compound **1** (Scheme 1) exhibited the most potent inhibition, with IC_50_ ∼ 5 μM, and was selected for further optimisation and characterisation.

The second approach included a high-throughput *in vitro* screening of approximately 100.000 small molecules from the UCLA Molecular Screening Shared Resource compound libraries. For this, we designed a fluorogenic assay specific for MUS81-EME1, consisting of a 3’flap DNA substrate labelled with fluorescein (FAM) and a Black Hole Quencher (BHQ1; Fig. S1A). Upon endonucleolytic cleavage by recombinant MUS81-EME1, the fluorophore is separated from the quencher, resulting in a significant fluorescence increase, which provides a direct and quantitative readout of the enzymatic activity (Fig. S1A). Inhibitor activity is reflected by a corresponding reduction in fluorescent signal. From the initial screen, several hundred compounds showing reduced fluorescence were selected for validation in a dose-response assay. Of these, 72 compounds were confirmed to inhibit MUS81-EME1 nuclease activity. Based on the potency (IC_50_ values) and availability, 23 hits were purchased for further testing (Supplementary table 2). Among these, compound **2** (Scheme 2) was selected as the lead candidate for further development based on its favourable *in vitro* activity profile.

The candidate compounds **1** and **2** from both screens are both relatively small (MW = 373 and 428 g/mol, respectively), providing opportunities for further chemical modification and optimization of biological and physicochemical properties (e.g. activity, selectivity and aqueous solubility). These candidate compounds were thus subjected to medicinal chemistry optimization and SAR development, as described below.

### SAR optimization of the lead compounds from the *in silico* screening

The hit **1** was first resynthesized using the route depicted in Scheme 1. Specifically, condensation of properly substituted β-ketonitrile with hydrazine, followed by diazotization, reaction with cyanomethylbenzimidazole, and final cyclization provided the target compound (Scheme 1). Upon confirmation of the activity of the resynthesized compound **1**, the methodology was used for the preparation of additional analogues and development of the SAR in the series.

**Scheme 1.**
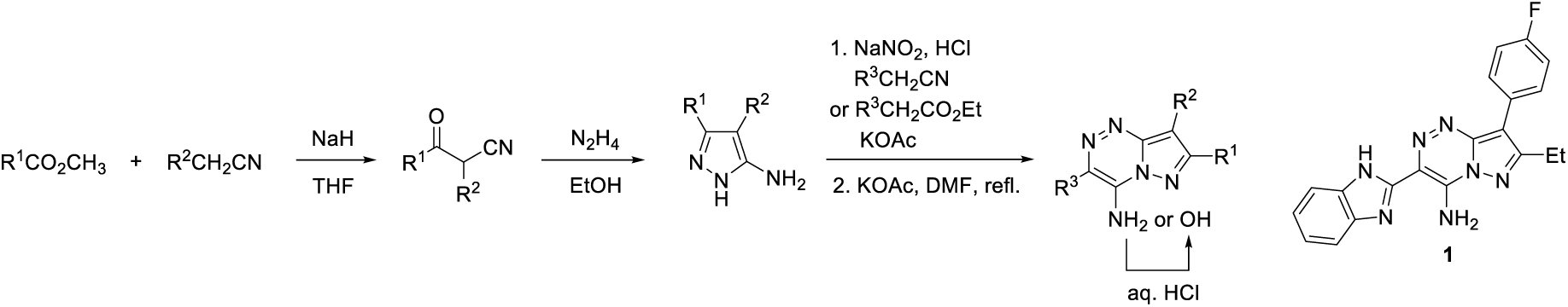
Synthetic route used for the preparation of the compound **1** and its analogues.

The SAR development in this series consisted of changing all four substituents around the pyrazolo[5,1-*c*][1,2,4]triazine scaffold (Table 1 and Table S1). As illustrated by the compounds **3**, **4** and **5** in Table 1, deletion of the substituents R^1^ (ethyl in **1**) and/or R^2^ (4-fluorophenyl in **1**) led to a significant decrease of MUS81 inhibitory activity. Less potent were also the analogues with alternatively substituted R^2^ phenyls (compounds **6**, **7**, **8** and **9**), cyclohexyl analogue **10**, and the compound **11** with R^1^ ethyl replaced with isopropyl (Table 1). Interestingly, the lack of 4-fluorophenyl motif could be compensated by larger substituents R^1^, exemplified by the compounds **12, 13**, **14**, and **15** (Table 1). However, these compounds were found to be only sparingly soluble in aqueous DMSO.

Deletion of the amino group in **1** led to the significantly less active compound **16** (Table 1). The methylated analogue **17** was also similarly inactive. In contrast, the replacement by hydroxy group led to the analogue **18** (**MU262**) (Table 1) that showed significantly improved activity in the primary biochemical nuclease assay.

Our attempts to replace the benzimidazole motif (i.e. substituent R^3^) were only partly successful, as illustrated by the compounds **19**, **20**, and **21** in Table 1. Similarly, numerous compounds with modified benzimidazole motif were significantly less potent - e.g. **22**, **23**, **24**, **25** and **26** (Table 1); except for the imidazole analogue **27**, which was however less soluble in aqueous DMSO. Finally, replacement of the pyrazolo[5,1-*c*][1,2,4]triazine core in **1** by the isosteric pyrazolo[1,5-*a*]pyridine scaffold resulted in the comparatively less potent compound **28** (Table 1).

**Table 1.**
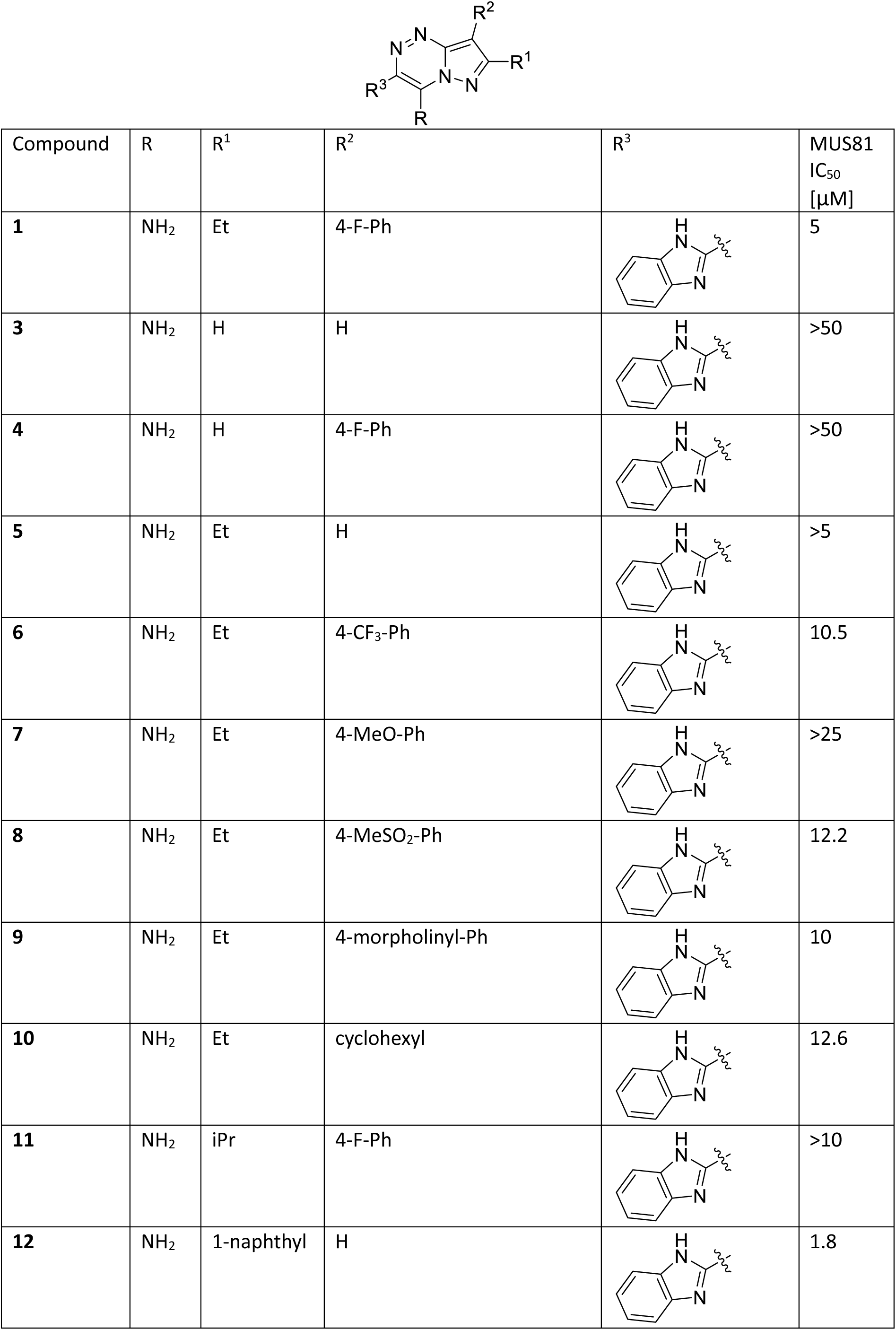
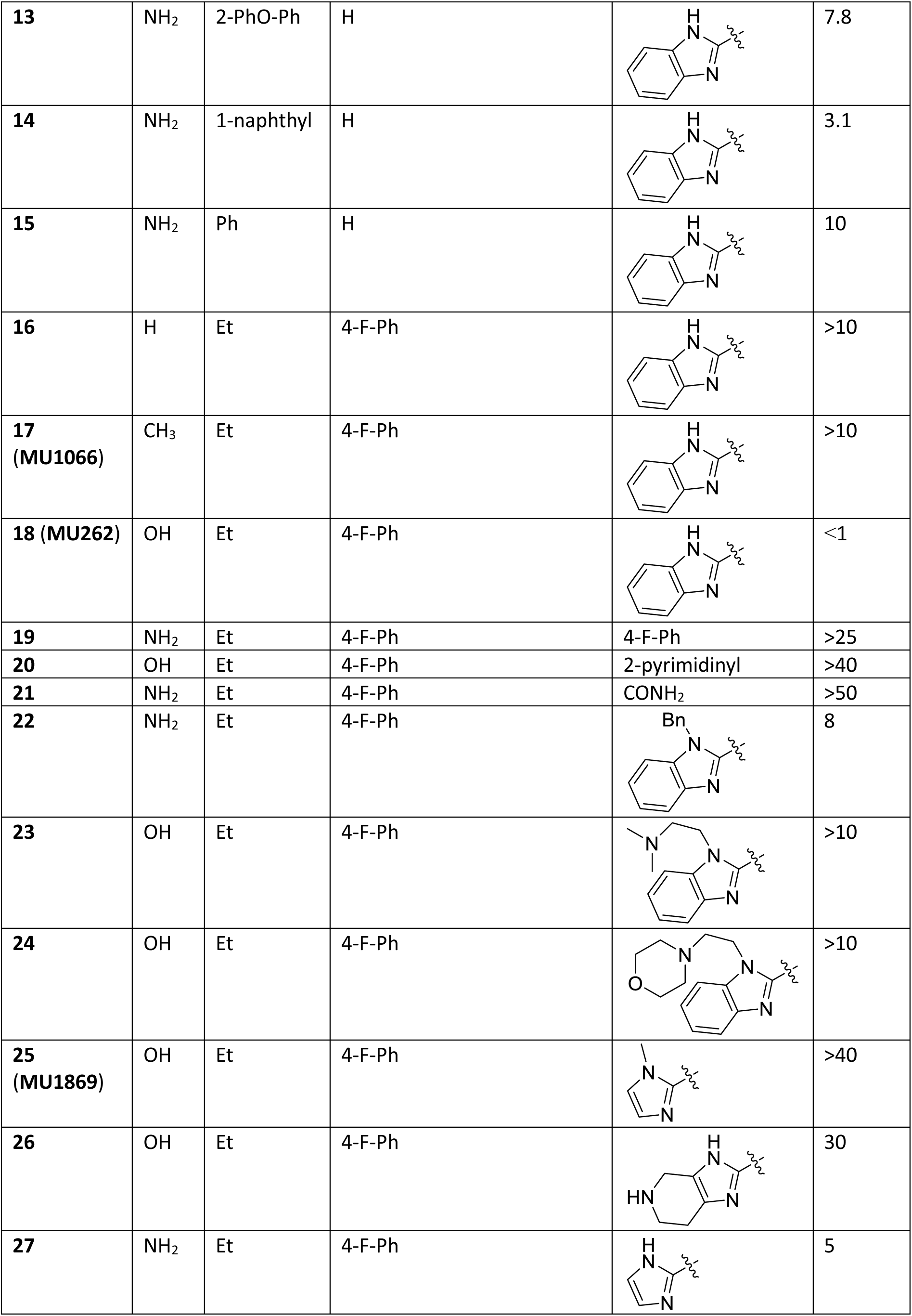
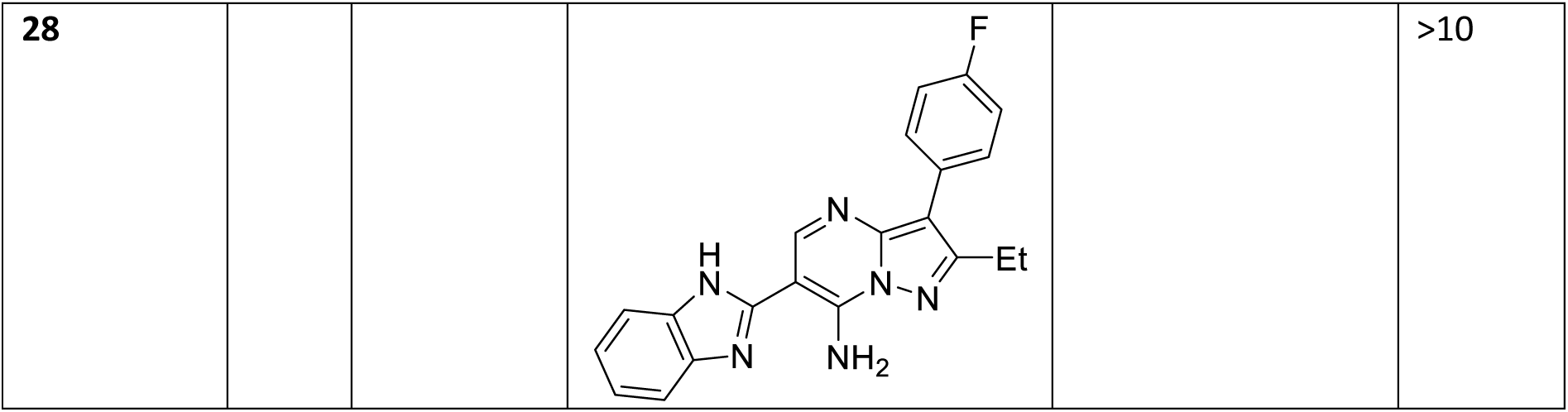
Structures and MUS81 *in vitro* IC50 values of the compound 1 and its selected analogues. Table including the complete set is included in Supporting Information (Supplementary table 3).

| Compound | R | R<sup>1</sup> | R<sup>2</sup> | R<sup>3</sup> | MUS81<br>IC<sub>50</sub><br>[μM] |
| --- | --- | --- | --- | --- | --- |
| **1** | NH<sub>2</sub> | Et | 4-F-Ph |  | 5 |
| **3** | NH<sub>2</sub> | H | H |  | >50 |
| **4** | NH<sub>2</sub> | H | 4-F-Ph |  | >50 |
| **5** | NH<sub>2</sub> | Et | H |  | >5 |
| **6** | NH<sub>2</sub> | Et | 4-CF<sub>3</sub>-Ph |  | 10.5 |
| **7** | NH<sub>2</sub> | Et | 4-MeO-Ph |  | >25 |
| **8** | NH<sub>2</sub> | Et | 4-MeSO<sub>2</sub>-Ph |  | 12.2 |
| **9** | NH<sub>2</sub> | Et | 4-morpholinyl-Ph |  | 10 |
| **10** | NH<sub>2</sub> | Et | cyclohexyl |  | 12.6 |
| **11** | NH<sub>2</sub> | iPr | 4-F-Ph |  | >10 |
| **12** | NH<sub>2</sub> | 1-naphthyl | H |  | 1.8 |
| **13** | NH<sub>2</sub> | 2-PhO-Ph | H |  | 7.8 |
| **14** | NH<sub>2</sub> | 1-naphthyl | H |  | 3.1 |
| **15** | NH<sub>2</sub> | Ph | H |  | 10 |
| **16** | H | Et | 4-F-Ph |  | >10 |
| **17**<br>(MU1066) | CH<sub>3</sub> | Et | 4-F-Ph |  | >10 |
| **18 (MU262)** | OH | Et | 4-F-Ph |  | <1 |
| **19** | NH<sub>2</sub> | Et | 4-F-Ph | 4-F-Ph | >25 |
| **20** | OH | Et | 4-F-Ph | 2-pyrimidinyl | >40 |
| **21** | NH<sub>2</sub> | Et | 4-F-Ph | CONH<sub>2</sub> | >50 |
| **22** | NH<sub>2</sub> | Et | 4-F-Ph |  | 8 |
| **23** | OH | Et | 4-F-Ph |  | >10 |
| **24** | OH | Et | 4-F-Ph |  | >10 |
| **25**<br>(MU1869) | OH | Et | 4-F-Ph |  | >40 |
| **26** | OH | Et | 4-F-Ph |  | 30 |
| **27** | NH<sub>2</sub> | Et | 4-F-Ph |  | 5 |
| 28 |  |  |  |  | >10 |

### SAR optimization of the lead compounds from *in vitro* screening

In the second series, the hit **2** served as the starting point in the SAR development. Along this line, we synthesized 53 analogues with the preserved central aminopyrazole scaffold (Table 2 and Table S2). Majority of the target compounds were prepared via condensation of properly substituted β-ketonitriles with heterocyclic hydrazines (typically possessing pyrimidine substituents), followed by acylation or alkylation of the NH_2_ group, as shown in Scheme 2.

**Scheme 2.**
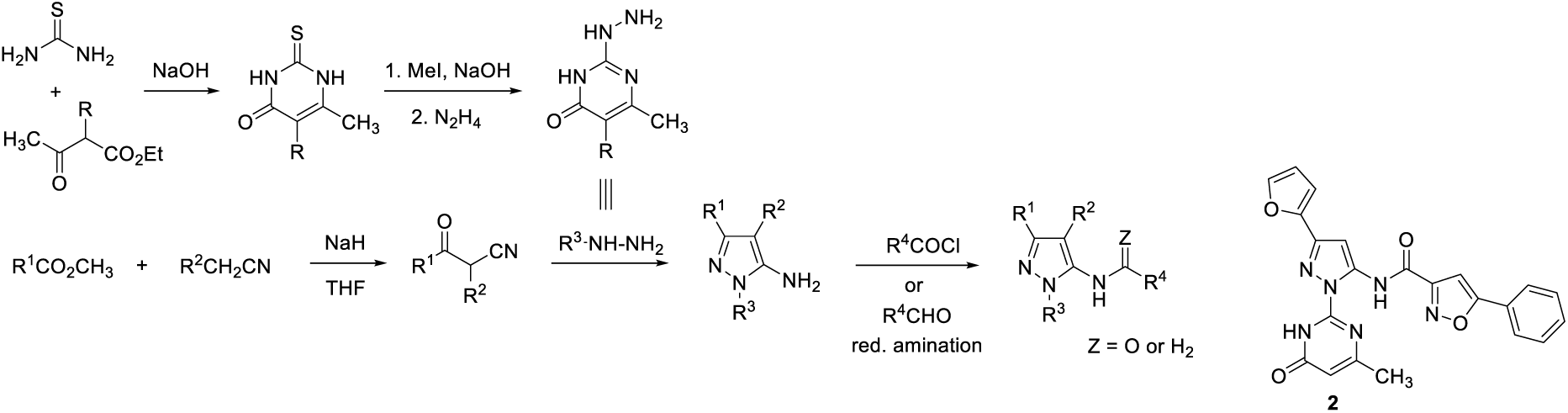
Synthesis of the compound **2** and its analogues.

Modification of R^1^ afforded several analogues with activity superior to that of **2**, namely compounds **29**, **30**, **31**, **32** (**MU876**) and **33** with the IC_50_ values 1.5, 4.5, 4.5, 0.5, and 4.5 μM, respectively (Table 2). In contrast, variation of the substituent R^4^ (typically substituted isoxazole) was less productive, with nearly all corresponding analogues exhibiting lower potency than the leads **2** and **32** (**MU876**) (Table 2). Less potent were also compounds with isosteres of the isoxazole motif, exemplified by the compounds **41**, **42**, **43** in Table 2, and the analogues **44** and **45** lacking the amidic carbonyl group and possessing methylated amidic nitrogen, respectively. Replacement/elaboration of the methylpyrimidinone motif R^3^ demonstrated that this moiety also needs to be preserved, and even minor modifications or substitutions resulted in significant decrease in the activity (**49**, **50**, **51**, **52, MU1003=53**). Of this subset, only the ethylated analogue **55** showed activity comparable to that of **32** (**MU876**) (Table 2).

**Table 2.**
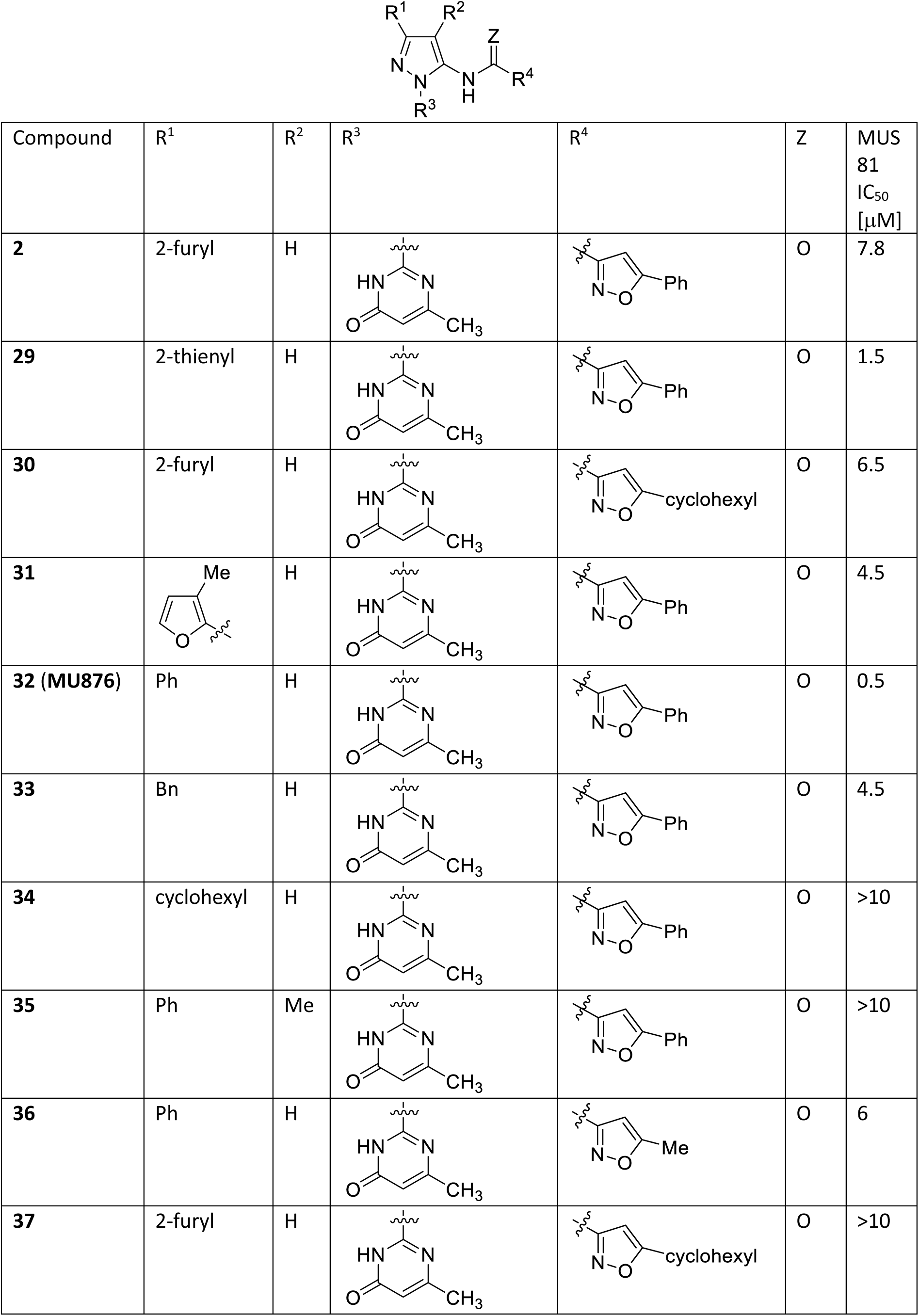
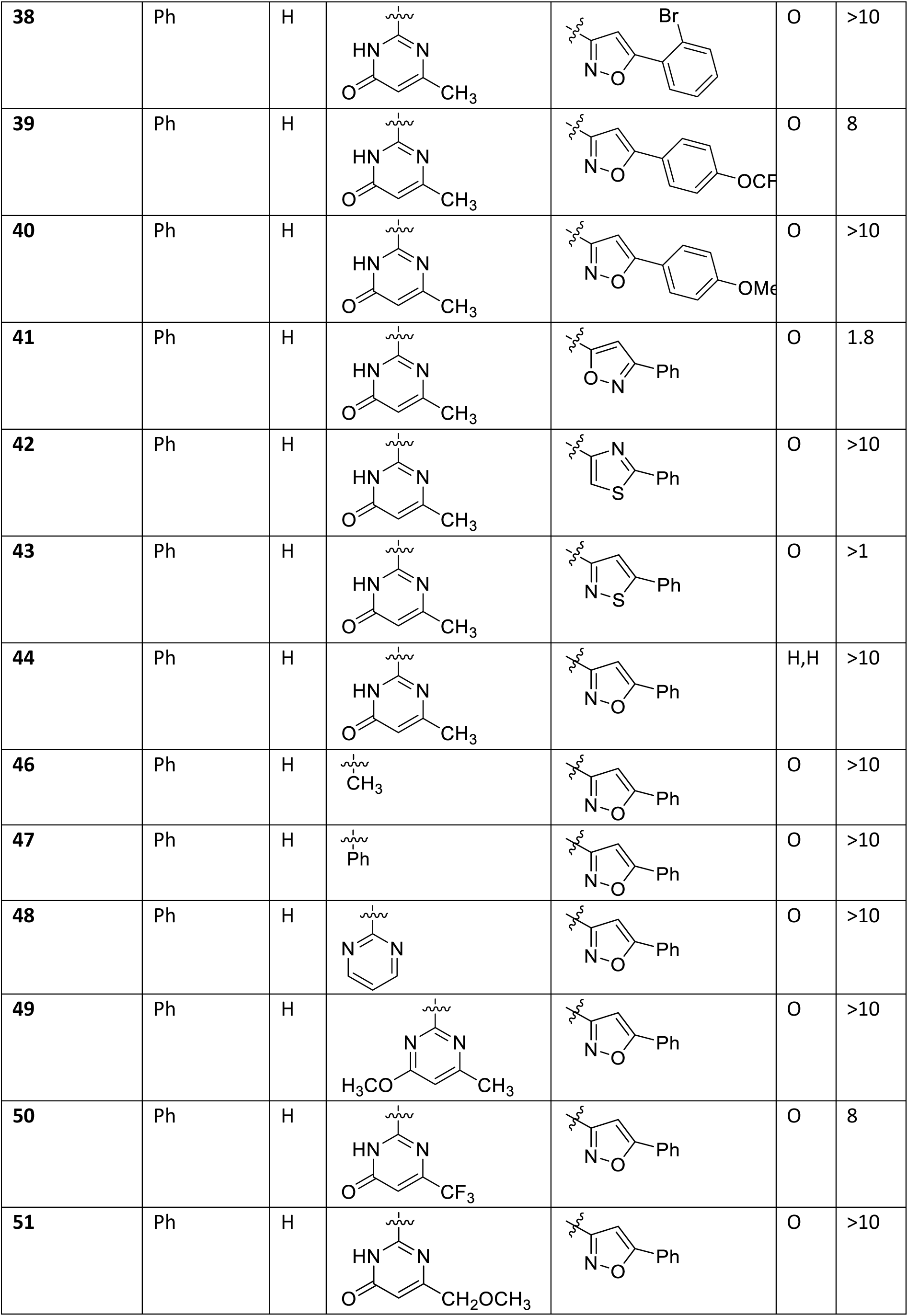
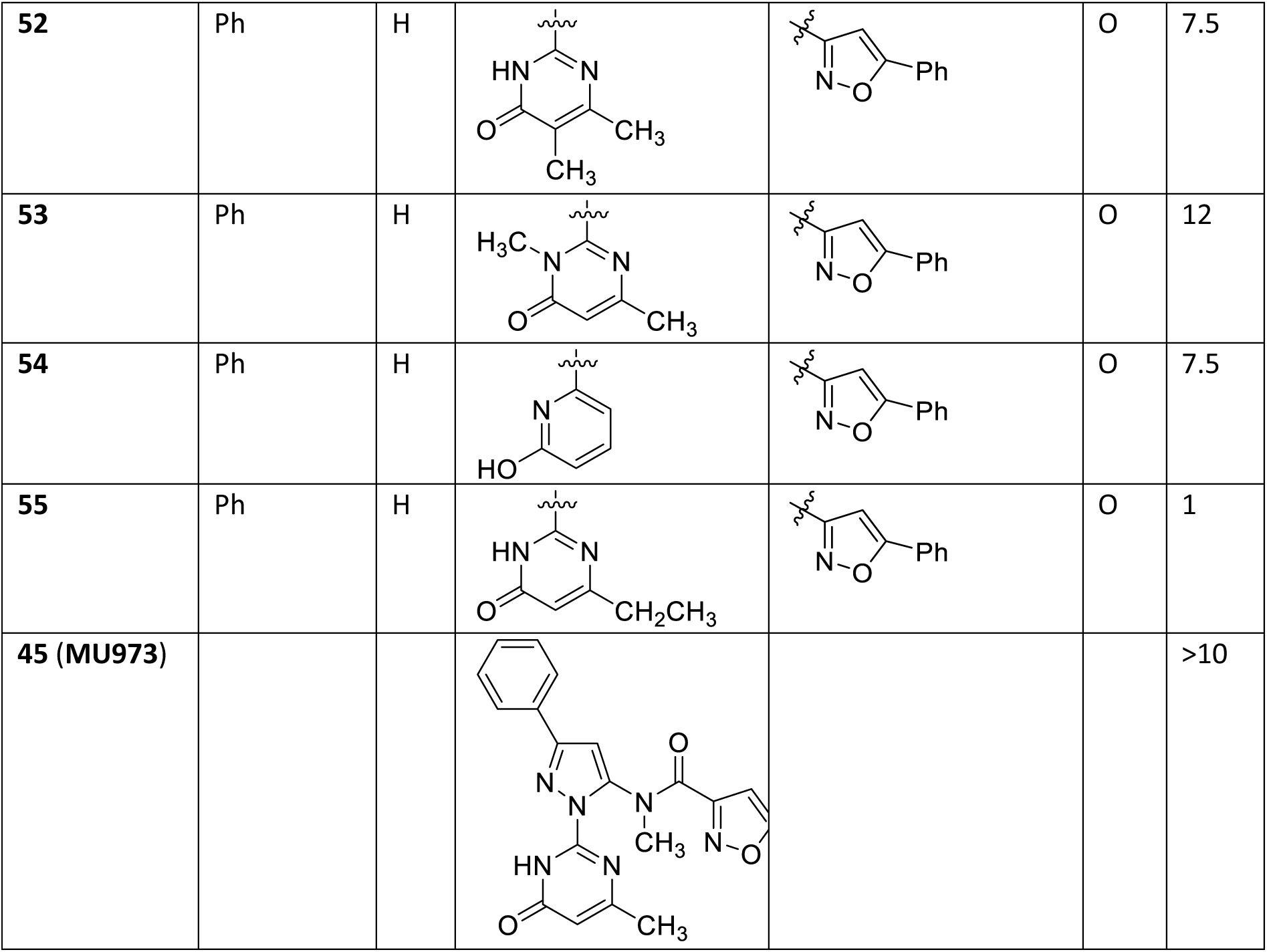
Structures and MUS81 IC50 values of the compound 2 and its selected analogues. Table including the complete set is included in Supporting Information (Supplementary table 4).

| Compound | R<sup>1</sup> | R<sup>2</sup> | R<sup>3</sup> | R<sup>4</sup> | Z | MUS<br>81<br>IC<sub>50</sub><br>[μM] |
| --- | --- | --- | --- | --- | --- | --- |
| **2** | 2-furyl | H |  |  | O | 7.8 |
| **29** | 2-thienyl | H |  |  | O | 1.5 |
| **30** | 2-furyl | H |  |  | O | 6.5 |
| **31** |  | H |  |  | O | 4.5 |
| **32 (MU876)** | Ph | H |  |  | O | 0.5 |
| **33** | Bn | H |  |  | O | 4.5 |
| **34** | cyclohexyl | H |  |  | O | >10 |
| **35** | Ph | Me |  |  | O | >10 |
| **36** | Ph | H |  |  | O | 6 |
| **37** | 2-furyl | H |  |  | O | >10 |
| 38 | Ph | H |  |  | O | >10 |
| 39 | Ph | H |  |  | O | 8 |
| 40 | Ph | H |  |  | O | >10 |
| 41 | Ph | H |  |  | O | 1.8 |
| 42 | Ph | H |  |  | O | >10 |
| 43 | Ph | H |  |  | O | >1 |
| 44 | Ph | H |  |  | H,H | >10 |
| 46 | Ph | H |  |  | O | >10 |
| 47 | Ph | H |  |  | O | >10 |
| 48 | Ph | H |  |  | O | >10 |
| 49 | Ph | H |  |  | O | >10 |
| 50 | Ph | H |  |  | O | 8 |
| 51 | Ph | H |  |  | O | >10 |
| 52 | Ph | H |  |  | O | 7.5 |
| 53 | Ph | H |  |  | O | 12 |
| 54 | Ph | H |  |  | O | 7.5 |
| 55 | Ph | H |  |  | O | 1 |
| 45 (MU973) |  |  |  |  |  | >10 |

### Two chemically diverse compounds, 18 (MU262) and 32 (MU876), inhibit MUS81-mediated DNA repair pathways in cells

To extend the SAR findings into a cellular context, selected derivatives from both chemical series that showed sufficient *in vitro* activity were tested in cell-based assays. Due to the current lack of assays that allow for direct and selective monitoring of MUS81 activity in cells, we used green fluorescent protein (GFP)-based reporter cell lines that detect DSB repair by homologous recombination (HR) or break-induced replication (BIR), two pathways in which MUS81 plays an important role [54–56] (Supplementary table 3 and 4).

These reporter assays enabled functional validation of MUS81 inhibition within a cellular setting and guided compound prioritisation based on biological efficacy. Of the tested compounds, **18** (**MU262**) and **32** (**MU876**) represented the most potent analogues within their respective chemical series in both *in vitro* as well as *in cellulo* settings (Table 1 and 2) and were selected for further profiling (Fig. 1A and D). Both compounds showed potent and dose-dependent inhibition of MUS81 with IC_50_ 0.87 μM for MU262 and 0.52 μM for MU876 in the *in vitro* assay (Fig. 1F-I). For each active compound a structurally similar negative control was also selected, **17** (**MU1066**) and **25** (**MU1869**) for **18**, and **45** (**MU973**) for **32** (Fig. 1B-C and 1E). These compounds show no activity in the *in vitro* assay (Fig. 1G and I, S2A-C).

**Figure 1.**
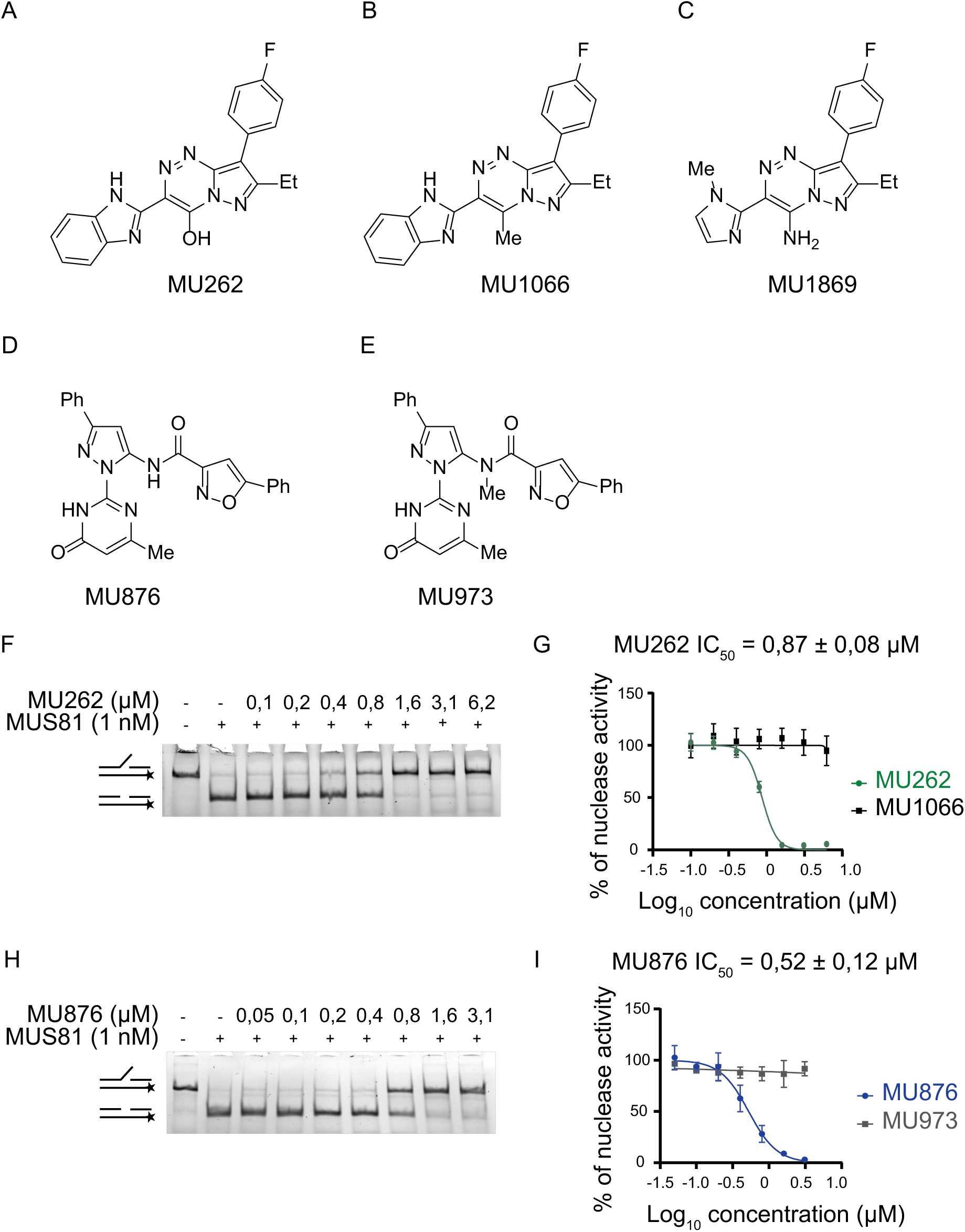
Two chemically diverse compounds inhibit the nuclease activity of MUS81-EME1 *in vitro*. (A) Chemical structure of MU262=18. (B) Chemical structure of MU1066=17. (C) Chemical structure of MU1869=25. (D) Chemical structure of MU876=32. (E) Chemical structure of MU973=45. (F) Purified MUS81-EME1 (hereafter labelled MUS81, 1 nM) was incubated with increasing concentrations of MU262, followed by the addition of 3 nM fluorescently labelled 3’ flap DNA. The reaction products were resolved on a native PAGE. Schematics on the side of the gel represent substrate and expected product of reaction. n = 3. (G) Quantification curve of MU262 and its negative control compound MU1066. 3ʹ flap DNA substrate processed by MUS81 was quantified using Multi Gauge software. Non-linear regression fitting model was used to calculate IC50 (mean ± s.e.), n = 3 or more. (H) Purified MUS81-EME1 (1 nM) was incubated with increasing concentrations of MU876, followed by the addition of 3 nM fluorescently labelled 3’ flap DNA. The reaction products were resolved on a native PAGE gel. Schematics on the side of the gel represent substrate and expected product of reaction. n = 3. (I) Quantification curve of MU876 and its negative control compound MU973. 3ʹ flap DNA substrate processed by MUS81 was quantified using Multi Gauge software. Non-linear regression fitting model was used to calculate IC50 (mean ± s.e.), n = 3 or more.

In cell-based assays assessing BIR we have first validated MUS81 dependency by using two different siRNAs to deplete MUS81 and observed a marked reduction in BIR efficiency (Fig. 2A). Notably, both compounds decrease BIR to the same extent as the siRNA-mediated depletion of MUS81, with an IC_50_ of 1.53 μM for **MU262** and 0.24 μM for **MU876**, respectively, while inactive control compounds showed no measurable effects (Fig. 2A-B). In addition to inhibiting BIR, both **MU262** and **MU876** significantly reduced the efficiency of an HR repair, with IC_50_ values in the range of 0.5-1 μM (Fig. 2C). This inhibition was again absent with their respective control analogues, further supporting the specificity of these inhibitors towards MUS81-dependent repair pathways.

To investigate their potential off-target effects, both compounds were also profiled in the MUS81-independent cell-based assay monitoring non-homologous end joining (NHEJ) pathway. In accordance with previous reports showing that inhibition of HR factors enhances NHEJ as a compensatory mechanism [57, 58], we observed that siRNA-depletion of BRCA2 or MUS81 modestly increased NHEJ activity (Fig. S3B). Correspondingly, the treatment with the MUS81 inhibitors **MU262** and **MU876** had no negative effect on NHEJ (Fig. 2D), indicating that they do not broadly suppress DNA repair but impair only HR/BIR pathways. Therefore, both compounds can be used as chemically orthogonal chemical probes for targeting the nuclease MUS81 in the cellular context.

**Figure 2.**
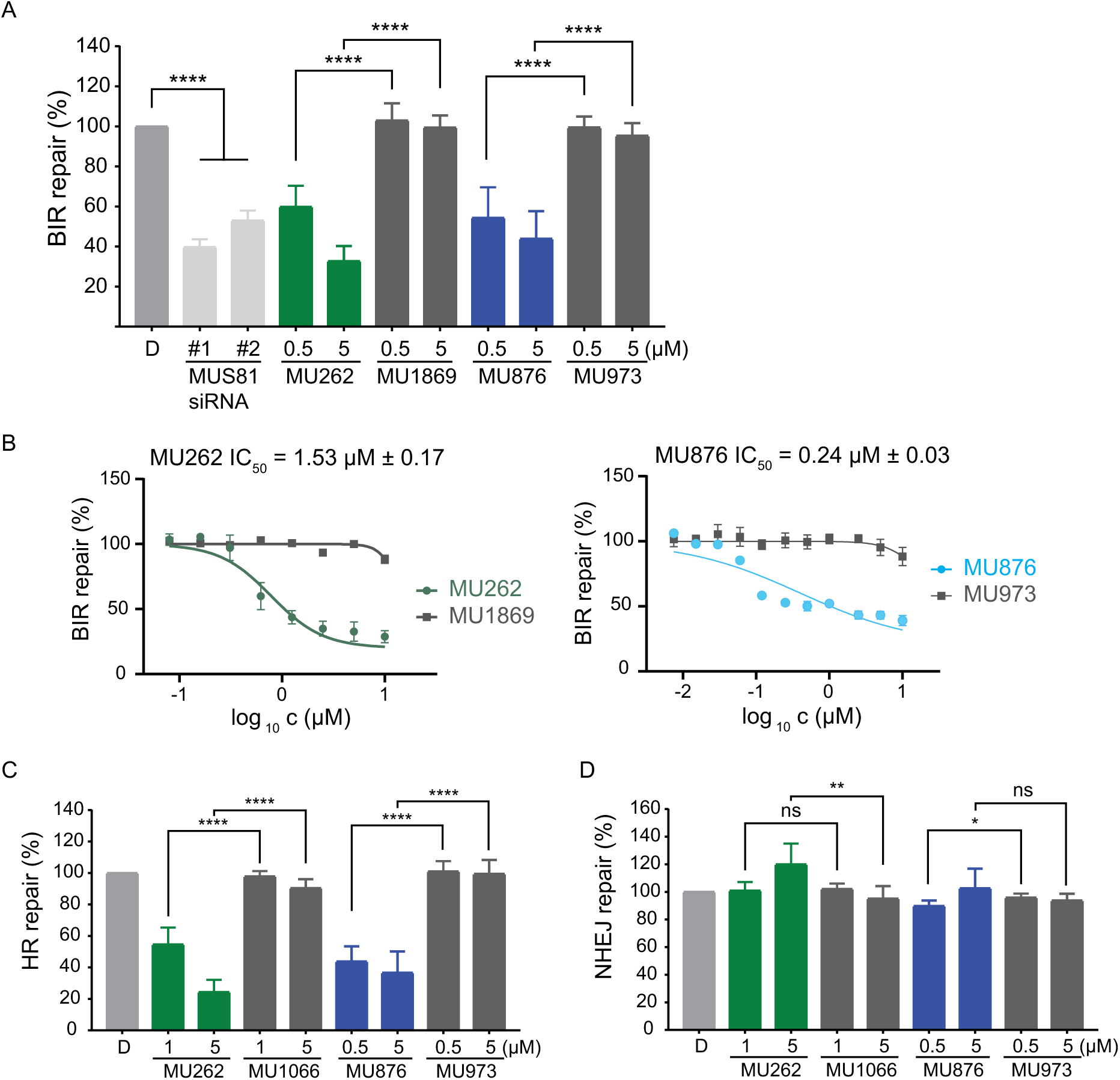
Small molecule inhibitors of MUS81 efficiently suppress the DNA repair pathways. (A) I-SceI-based BIR repair efficiency was measured in U2OS BIR-GFP cells treated with DMSO, two different siRNAs targeting MUS81, or the indicated concentrations of MU262 and MU876, and their control analogues for 72 hours. The percentage of repair was normalised to the DMSO-treated control (D). *n* = 3. error bars represent standard deviation (s.d.); **** *p* < 0.0001 (unpaired, two-tailed *t*-test). (B) Experiment was done as in (A), except a concentration range was wider to determine the IC50 value for MU262 and MU876 via a non-linear regression fitting model. *n* = 3. (C-D) I-SceI-based HR (C) and NHEJ (D) repair efficiency was measured using U2OS DR-GFP and EJ5-GFP cells, respectively, following treatment with DMSO, MU262, MU876, or their control analogues for 72 hours. The percentage of repair was normalised to the DMSO-treated control. *n* = 3. error bars represent s.d.; * *p* < 0.05, ** *p* < 0.01, **** *p* < 0.0001 (unpaired, two-tailed *t*-test).

### Inhibition of MUS81 does not induce cytotoxicity or disrupt cell cycle progression

Given these promising findings, we further investigated **MU262** and **MU876** specificity and potential cytotoxicity. Following a 3-day treatment, matching the duration of BIR and HR assays, Annexin V staining measured by flow cytometry showed no increase in apoptosis following treatment with **MU262** or **MU876**, nor with MUS81 depletion by siRNA, in contrast to the robust apoptosis induced by CPT (Fig. S4A, B). To evaluate possible effects on the cell cycle, we performed propidium iodide (PI) staining and 5-Ethynyl-2’-deoxyuridine (EdU) incorporation, as a marker of S phase. While CPT reduced the proportion of S phase cells, due to its anti-proliferative effect, MUS81 depletion only modestly increased the proportion of cells in the G1 phase, consistent with reduced proliferation (Fig. S4C). The cells treated with 1 μM **MU262** and **MU876** showed a mild increase in S phase (Fig. S4C), consistent with a defect in recombination-dependent repair during replication. The control compounds elicited no effect on cell cycle at this concentration. However, at 0.5 μM concentration of **MU262** and **MU876**, neither compound significantly affected the cell cycle distribution (Fig. S4C), in line with the previous studies reporting that depletion of MUS81 does not severely affect progression through S phase [42, 59, 60].

To evaluate the long-term effects of the inhibitors on the cellular viability, we treated cells twice a week for almost three weeks with low doses of **MU262** and **MU876**. **MU262** caused a mild reduction in viability at concentration of 1-3 μM while **MU876** more strongly suppressed proliferation at 0.25 and 0.5 μM (Fig. S5A-B). This mirrors the effect of the siRNA-mediated depletion of MUS81 (Fig. S5C-D).

### MU262 and MU876 inhibit MUS81 via a different mode of action

To elucidate the mechanism underlying the inhibitory activity of **MU262** and **MU876**, we performed several *in vitro* interaction assays. Using an electrophoretic mobility shift assay (EMSA), we examined how these compounds affect the binding of purified MUS81-EME1 complex to a 3’ flap DNA substrate. As expected, MUS81-EME1 efficiently bound to 3’flap DNA substrate (Fig. 3A). However, **MU876** disrupted this interaction in a concentration-dependent manner, effectively dissociating MUS81-EME1 from the DNA. In contrast, the control compound **MU973** had no significant effect (Fig. 3A and S6A). Interestingly, neither **MU262** nor its inactive analogue **MU1066** disrupted the MUS81-DNA interaction (Fig. 3B and S6), suggesting a different mechanism of action for **MU262** compared to **MU876**. This suggests that these two structurally different compounds interact with MUS81 through different binding modes. A previous study has shown that multiple MUS81 domains, including the nuclease active site, contribute to DNA substrate binding [61], offering the possibility that **MU876** may block the binding by targeting the protein-DNA interface.

To further confirm these findings, we employed bio-layer interferometry (BLI) to monitor the binding kinetics of MUS81 to immobilised DNA substrate in the presence of the inhibitors. Consistent with the results obtained by the EMSA assay, **MU876** impaired MUS81-DNA binding in a dose-dependent manner (Fig. 3C), whereas **MU262** had only negligible effect on this interaction (Fig. 3D).

In summary, while **MU876** appears to inhibit the nuclease MUS81 via interfering with DNA binding, **MU262** likely impairs the MUS81 enzymatic activity through an alternative, DNA binding-independent mechanism.

**Figure 3.**
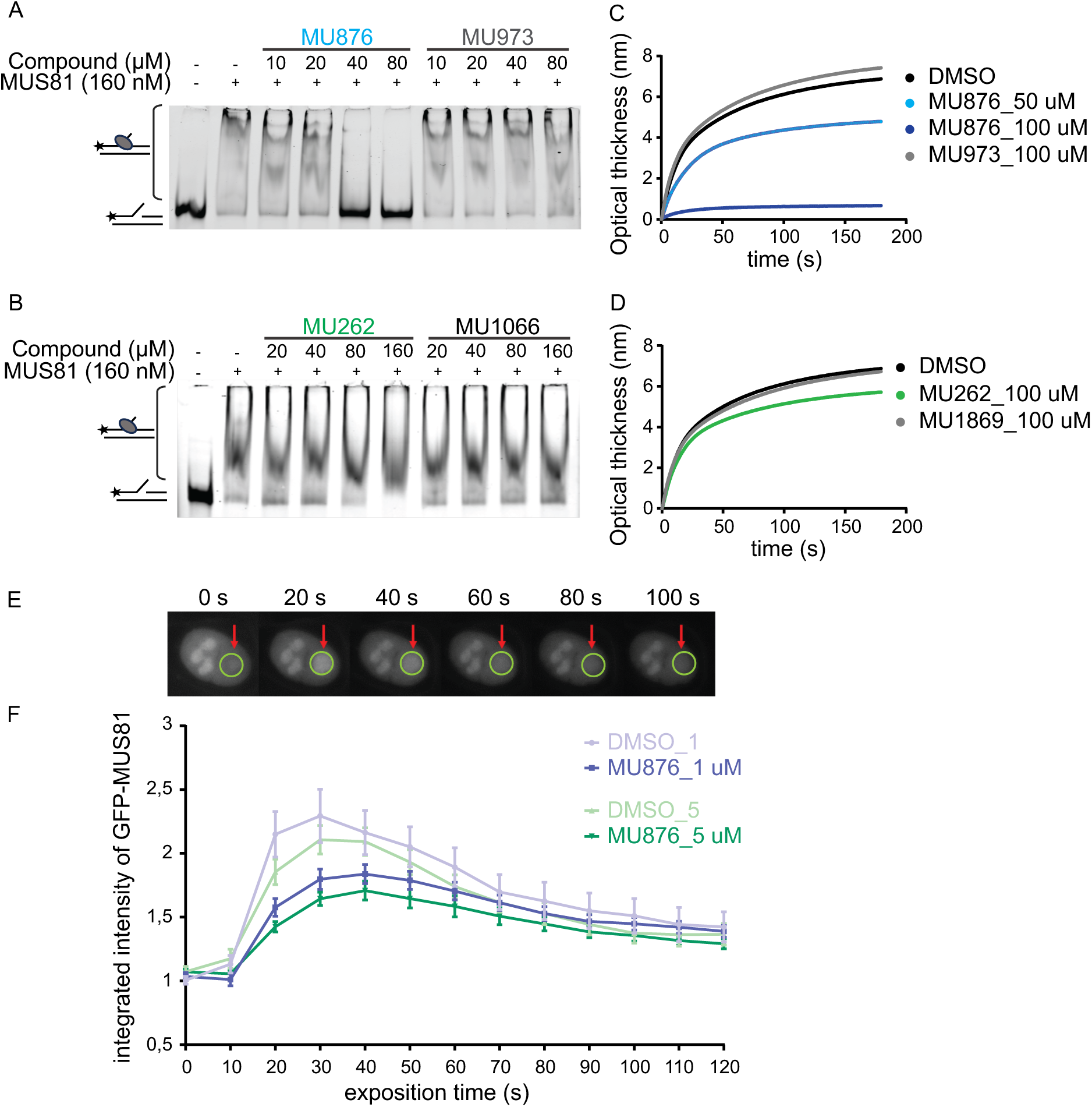
MU262 and MU876 have a different mode of action. (A-B) Purified MUS81-EME1 (160 nM) was incubated with increasing concentrations of MU262, MU876, or their respective control analogues, followed by the addition of 3 nM fluorescently labelled 3’ flap DNA substrate. MgCl2 was omitted from the reaction buffer to prevent substrate cleavage. The reaction products were resolved on a native PAGE gel. The lower band represents free DNA substrate, while the upper smear corresponds to the MUS81-DNA complex. n = 3. (C-D) A 3’ flap cleavage resistant DNA substrate was immobilised on a streptavidin-coated (SAX) sensor and incubated with truncated MUS81 (200 nM) alone, or with MUS81 pre-mixed with DMSO, MU876, MU262, or their respective control analogues. The real-time binding kinetics were measured as a change in optical thickness over time. Representative plots shown from two independent experiments. (E) Representative images of a DMSO-treated cell within 100 s after micro-irradiation. GFP-MUS81 U2OS cells were pre-incubated with DMSO or MU876 at 1 and 5 μM final concentration for 2 hours. Localised DNA damage was induced using a 355 nm laser micro-irradiation at specific sites in individual nuclei. The recruitment and retention of GFP-MUS81 at damage sites was traced by a sequential live-cell imaging for 120 seconds after micro-irradiation. (F) The integrated intensity of a GFP-MUS81 signal accumulation ± S.E. from (E). n = 3.

### Direct engagement of MU876 with MUS81 in cells

To confirm that **MU876** directly targets MUS81 in cellular setting, we generated HEK293 cells expressing GFP-tagged MUS81. As expected, GFP-MUS81 localised to the nucleus, with increased accumulation in nucleoli, in accordance with the previously observed nucleolar retention which is even enhanced upon UV treatment [64]. To assess DNA damage recruitment dynamics, we used laser micro-irradiation. GFP-MUS81 rapidly accumulated at damage sites, forming transient foci that disappeared shortly after recruitment (Fig. 3E). Treatment with **MU876** led to a concentration-dependent reduction of MUS81 recruitment to damage sites, however not affecting the nuclease retention once recruited (Fig. 3F). This corresponded to our *in vitro* experiments and supported the concept that **MU876** impairs the binding of MUS81 to damaged DNA in cells.

### MU262 and MU876 phenocopy siRNA-mediated MUS81 depletion

Next, we explored the phenotypic consequences of MUS81 inhibition by **MU262** and **MU876** in cells. One phenotype linked to MUS81 depletion is the formation of micronuclei, a hallmark of chromosomal aberrations [23]. Accordingly, we observed an increased number of micronuclei upon depletion of MUS81 by siRNA in U2OS cells (Fig. 4A). The treatment with **MU262** and **MU876** also significantly increased micronuclei formation, similarly to the effect of MUS81 depletion (Fig. 4A-B). The inactive control compounds had no effect on the formation of micronuclei (Fig. 4A), confirming the specificity of the observed phenotype.

In addition to its role in DNA repair, MUS81 has been implicated in telomere maintenance in cells that use alternative lengthening of telomeres (ALT) pathway, where depletion of MUS81 reduces telomere recombination and increases telomere loss [27, 65, 66]. Using fluorescence *in situ* hybridisation (FISH) on metaphase spreads, we demonstrated that both **MU262** and **MU876** significantly increased the presence of telomere-free ends (TFEs) (unlike the negative control compounds) in ALT-positive U2OS cells, but not in telomerase-positive and ALT-negative HeLa cells (Fig. 4C-D). This confirmed that MUS81 inhibition impairs ALT-dependent telomere maintenance and further verified the ability of the compounds to inhibit the MUS81 activity in the cell.

MUS81 plays a critical role in DNA repair, especially during replication and cell division [27]. Persistent DNA lesions are marked by phosphorylation of histone H2AX (γH2AX), a widely used marker of DNA damage that can be visualised by fluorescence microscopy [67]. To evaluate the impact of the MUS81 inhibitors **MU262** and **MU876** on DNA damage accumulation, we treated U2OS cells with them for 72h, and quantified γH2AX foci. Similarly to the cisplatin treatment, both inhibitors significantly increased the number of DNA damage foci in a concentration-dependent manner (Figs. 5A-B and S7A-B), indicating accumulation of unresolved DNA damage. In comparison, siRNA-mediated MUS81 depletion resulted in only a modest increase in γH2AX foci, likely due to the limited time of the protein absence upon the knock-down compared to a three-day treatment with the inhibitors (Fig. S7A-B).

To determine whether this persistent damage translates to reduced viability, we used colony formation assay (CFA). As expected, MUS81 depletion reduced the clonogenic survival of U2OS cells (Fig. 5C-D). Similarly, **MU262** and **MU876** significantly decreased cell survival, whereas the inactive control compounds had no effect (Fig. 5E-H). These results demonstrate that inhibition of MUS81 impairs the resolution of DNA lesions, and leads to increased DNA damage, chromosomal aberrations and reduced survival. These observations correlate with the previous report on impaired viability of MUS81-depleted cells under replication stress [42].

**Figure 4.**
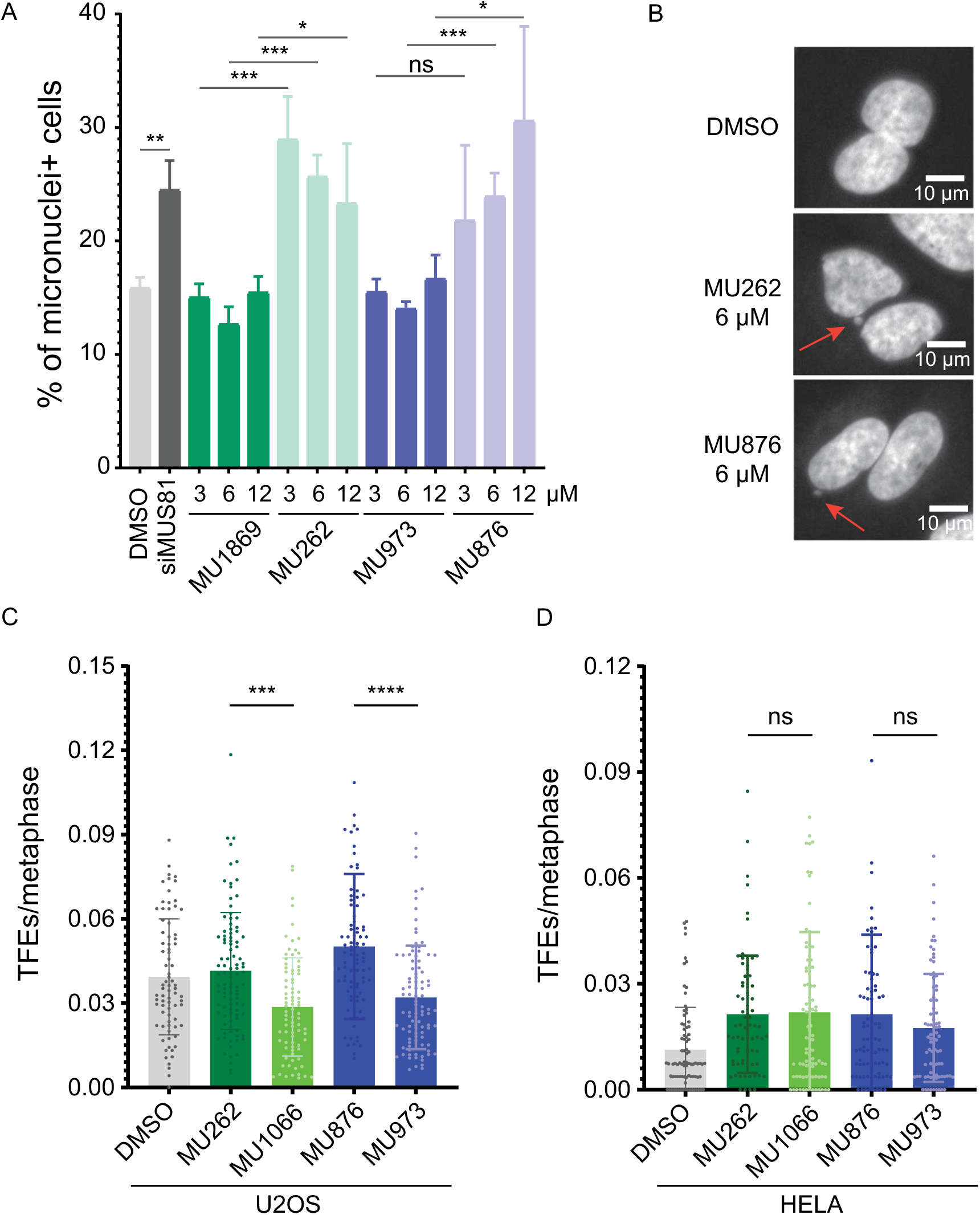
Small molecule inhibitors of MUS81 mimic the effect of siRNA-mediated MUS81 depletion in cells. (A) U2OS cells were treated with DMSO, MUS81-targeting siRNA, MU262, MU876, or their respective control analogues for 48 hours. Cytochalasin B (1,25 μg/mL) was added 16 hours before harvest to prevent cytokinesis. The graph shows the percentage of binucleated cells with micronuclei. n = 3; error bars represent s.d.; *, *p* < 0.05, **, *p* < 0.01 ***, *p* < 0.001 (unpaired, two-tailed *t*-test). (B) Representative images corresponding to the selected conditions from (A). (C) U2OS and (D) HeLa cells were incubated with DMSO, the indicated inhibitors, and their respective control analogues for 48 hours. Nocodazole (200 ng/mL) was added for 5 hours prior to the end of treatment to arrest cells in mitosis. Mitotic cells were collected by shake-off, and telomere fragility events (TFEs) were quantified manually for each metaphase spread. At least 70 metaphase spreads were quantified per condition. n = 3; error bars represent s.d.; ***, *p* < 0.001 ****, *p* < 0.0001 (ordinary One-way ANOVA).

**Figure 5.**
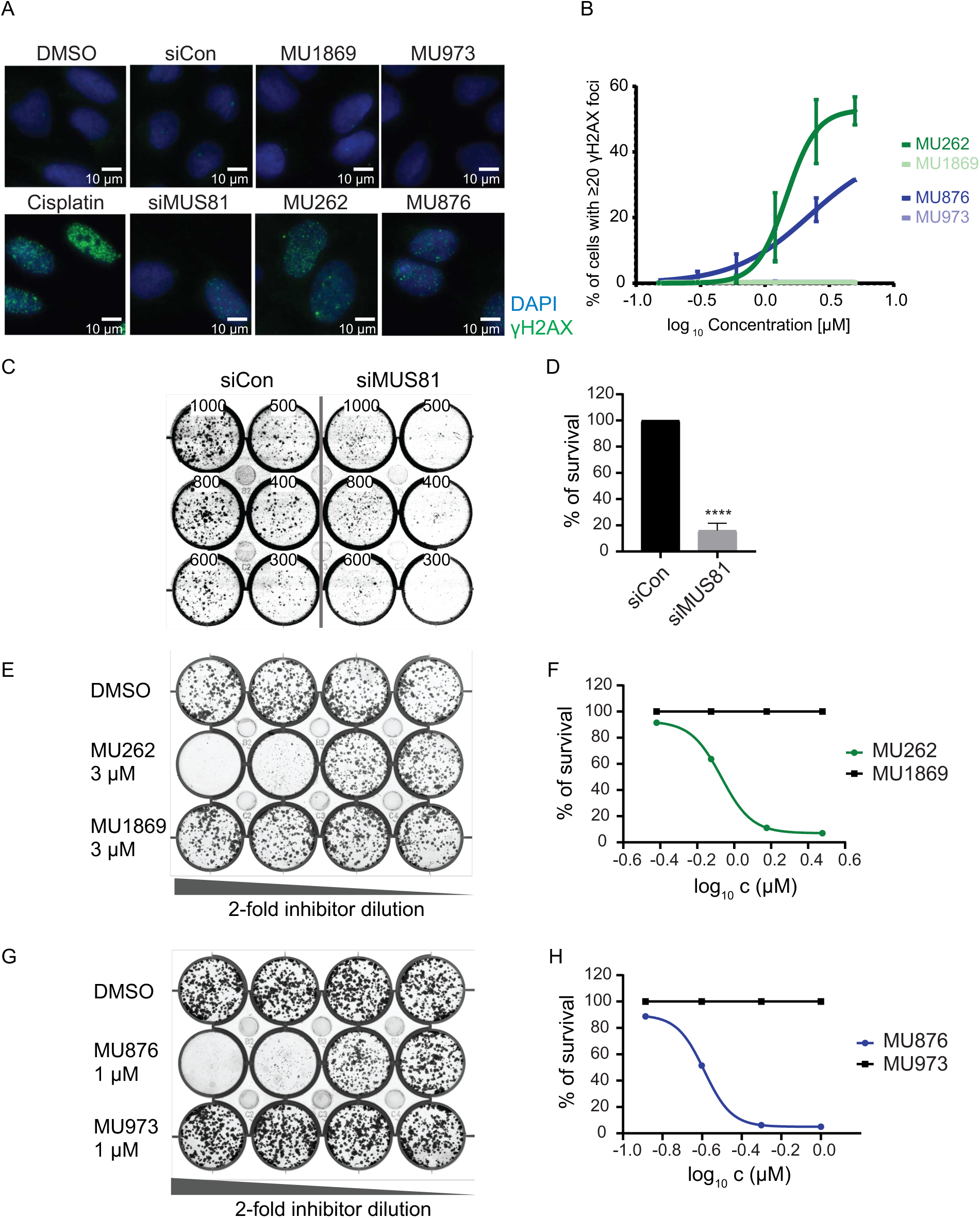
Endogenous DNA damage persists following treatment with MUS81 inhibitors. (A) U2OS cells were treated with DMSO, cisplatin as a positive control, control or MUS81-targeting siRNA, MU262, MU876 and their negative control counterparts at the indicated concentrations for 72 hours. Representative images show staining of a DNA damage marker γH2AX in green with counterstaining of DAPI in blue. (B) DNA damage was assessed by quantifying γH2AX foci per nucleus from (A) using CellProfiller software. The graph represents percentage (%) of cells with ≥20 γH2AX foci per nucleus across different concentrations of inhibitors. n = 3; error bars represent s.d. (C) Indicated number of U2OS cells was transfected with a non-targeting siRNA (siCon) or MUS81-targeting siRNA (siMUS81) seeded for a CFA. Cells were grown for 10-12 days, and colonies were visualised by crystal violet staining after harvest. n = 3, representative picture is shown. (D) Quantification of (C), error bars represent s.d.; ****, *p* < 0.0001 (unpaired, two-tailed *t*-test). (E) U2OS cells was treated with MU262 and its negative control MU1869 and seeded for a CFA. Cells were grown for 10-12 days, and colonies were visualised by crystal violet staining after harvest. n = 3, representative picture is shown. (F) Quantification of (E) using a non-linear regression fitting model. *n* = 3. (G) U2OS cells was treated with MU876 and its negative control MU973 and seeded for a CFA. Cells were grown for 10-12 days, and colonies were visualised by crystal violet staining after harvest. n = 3, representative picture is shown. (H) Quantification of (G) using a non-linear regression fitting model. *n* = 3.

### MUS81 inhibitors MU262 and MU876 potentiate the effect of chemotherapy and impair DNA repair

Despite the severe side effects associated with radio- and chemotherapy, cisplatin remains the primary option for numerous cancer patients worldwide [68]. Since MUS81-depleted cells are more sensitive to cisplatin [69], we explored whether MUS81 inhibitors could potentiate its cytotoxic effect. We treated various cancer cell lines, including U2OS, HEK293 and CAL51, with increasing concentration of **MU262** and **MU876** in combination with cisplatin and monitored cell survival 5 days after the treatment. As expected, cisplatin induced a dose-dependent decrease of viability, which was further enhanced by co-treatment with both MUS81 inhibitors in a dose-dependent manner (Fig. 6A-B, Fig. S8A-D). In contrast, no additional sensitisation was observed in CAL51 MUS81^-/-^ cells (Fig. S8E-F), supporting the on-target activity of the MUS81 inhibitors.

Mechanistically, the nuclease activity of MUS81 is critical for the repair of DNA damage induced by cross-linking agents such as mitomycin C (MMC) and cisplatin [5]. To assess whether our MUS81 inhibitors impact DNA repair induced by cisplatin treatment, we examined γH2AX foci formation following the cisplatin treatment in the presence or absence of **MU262** and **MU876**. Consistent with siRNA-mediated MUS81 depletion in U2OS (Fig. S9A) and genetic deletion in CAL51 cells (Fig. S9B), both compounds significantly reduced the number of γH2AX foci in a concentration-dependent manner compared to the DMSO controls (Fig. 6C and S9C). These findings suggest that MUS81 activity is required for cisplatin-induced DNA breaks and inhibition of MUS81 mimics genetic loss of function, effectively impairing cellular DNA damage response and enhancing the efficacy of cisplatin.

**Figure 6.**
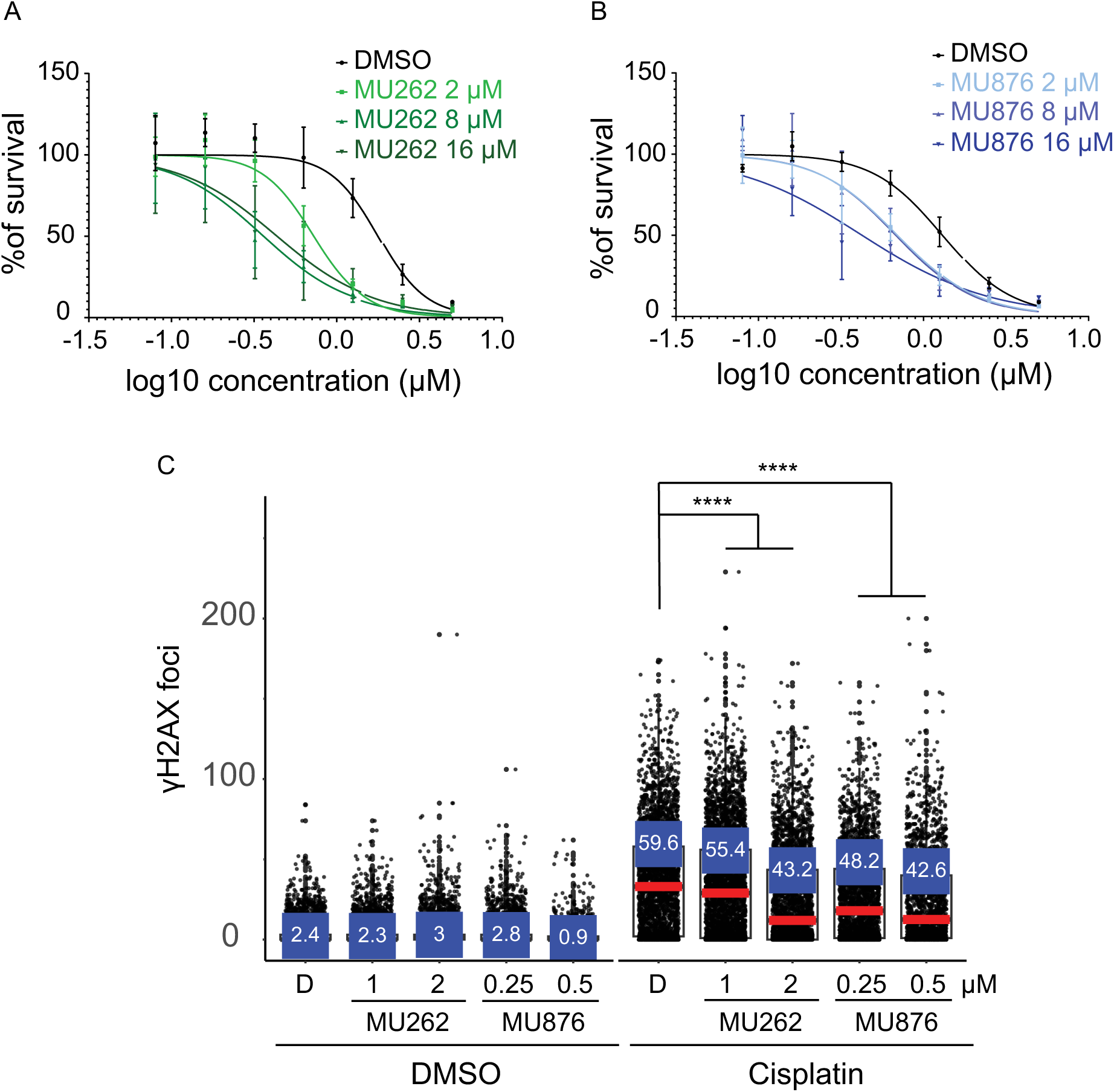
MUS81 inhibitors potentiate the effect of chemotherapy. (A-B) U2OS cells were treated with DMSO, MU262, or MU876 at the indicated concentrations in combination with range of cisplatin (0-5 μM) for 96 – 120 hours. Cell viability was assessed by the Cy-Quant assay. The survival was normalised to the DMSO-treated control. Graphs were generated in Graphpad Prism using the non-linear regression fitting model. n = 3, error bars represent s.d. (C) CAL51 WT cells were treated with DMSO, MU262, or MU876 at the indicated concentrations at each passage over two weeks. Cisplatin (12 μM) was added for the last 24 hours before harvest. DNA damage was assessed by quantifying γH2AX foci using fluorescence microscopy and CellProfiller software. Data were visualised plotted in R software. Red markers represent the median of γH2AX foci number per nucleus; blue markers represent the percentage of cells with ≥20 γH2AX foci per nucleus. n = 3., error bars represent s.d.; ****, *p* < 0.0001 (unpaired, two-tailed *t*-test).

## DISCUSSION AND CONCLUSIONS

Genomic instability is a hallmark of cancer and other diseases, often arising from defects in DNA repair pathways. While this vulnerability can be therapeutically exploited, as demonstrated by PARP inhibition in BRCA-mutated tumours [70], the number of clinically used agents that directly target DNA repair proteins remains limited [71]. In this respect, DNA nucleases, in particular those involved in the processing of stalled or collapsed replication forks, represent a promising yet underexplored class of targets. Among these, the endonuclease MUS81 plays a crucial role in safeguarding genome stability, especially during S phase and mitosis.

In our study, we have identified and characterised two distinct small-molecule inhibitors of MUS81 endonuclease, **MU262** and **MU876**. These compounds efficiently inhibit the *in vitro* nuclease activity of purified MUS81-EME1 complex at sub-micromolar concentrations. Importantly, both inhibitors phenocopy the effects of MUS81 depletion in cell-based assays, establishing their functional relevance and validating MUS81 as a druggable target. The comprehensive set of cell-based assays used in this study includes also detection of direct engagement of **MU876** with the nuclease MUS81 in living cells via recruitment assays.

Our characterisation of **MU262** and **MU876** represent a significant advancement over previously published studies on MUS81 inhibition, which lacked in-depth cellular validation [47, 48]. In this study, we used a comprehensive set of biochemical, biophysical, and cell-based analyses, demonstrating that both inhibitors **MU262** and **MU876** can effectively modulate MUS81-dependent processes, including BIR and HR pathways. Notably, treatment with either compound led to the accumulation of chromosomal aberrations such as formation of micronuclei and loss of telomeric DNA, i.e. phenotypes that closely mirror those observed in this study and previously also in MUS81-deficent mouse or human cells [22, 23, 72]. The structurally orthogonal MUS81 inhibitors reported herein also provide a unique opportunity to target the nuclease through different modes of inhibition: while **MU876** directly disrupts MUS81-DNA binding both *in vitro* and in cells, **MU262** appears to inhibit the nuclease via a distinct mechanism impairing downstream repair functions.

Despite their mild toxicity under short-term treatment, prolonged exposure to either **MU262** or **MU876** led to a marked reduction in cell proliferation and survival, closely mirroring the effect observed in this study but also previously upon MUS81 knockdown or knockout [29, 59, 73]. These effects are likely due to the accumulation of unresolved DNA structures and checkpoint activation, as observed previously in MUS81-deficient models, including transgenic mice [5]. However, we cannot exclude the possibility that part of the observed phenotypes may be caused by off-target effects. In fact, given the structural similarity and functional redundancy among human nucleases, dual inhibition can be both mechanistically plausible and therapeutically advantageous. For instance, synthetic lethality has been demonstrated in HEK 293 cells lacking both MUS81 and GEN1 [74]. Thus, future efforts focused on exploration of combined nuclease inhibition may lead to new therapeutic options with increased efficacy and lower risk of acquired resistance [75].

Importantly, we demonstrate that MUS81 inhibition sensitises cells to cisplatin, a routinely used chemotherapeutic agent whose efficacy is often limited by acquired resistance [76–78]. Both **MU262** and **MU876** enhance cisplatin-induced cytotoxicity across multiple cancer cell lines, underscoring the therapeutic potential of MUS81 inhibition in combination with chemotherapy. This effect was absent in MUS81-knockout cells and confirmed the specificity of the compounds. Mechanistically, we show that these inhibitors reduce the formation of γH2AX foci following cisplatin treatment, which is consistent with the model where MUS81 facilitates the generation of DSBs during interstrand crosslink (ICL) repair [69]. This further validates MUS81 as a critical component of the DNA damage response and suggests that its inhibition may potentiate the effects of genotoxic therapies. In contrast to single-agent therapies that are frequently associated with acquired resistance [79], combination therapies including MUS81 inhibitors may provide a more robust and durable clinical outcome.

Collectively, the data described in this report define the newly discovered compounds **MU262** and **MU876** as first-in-class inhibitors of the nuclease MUS81 with confirmed cellular activity that can be used as valuable tools in molecular biology studies focused on this nuclease. The findings reported herein also provide a foundation for further development of new single-agent or combination anti-cancer therapies that would utilize pharmacological targeting of MUS81.

## EXPERIMENTAL SECTION

### *In silico* ligand-based screening for MUS81-EME1 inhibitors

The three-dimensional structures of almost 150.000 molecules for virtual screening were downloaded from the clean drug-like subset of the ZINC database[80]. Only structures matching the selection criteria (xlogP ≤ 5, molecular weight ≤ 500 g/mol, number of H-bond donors ≤ 5 and number of H-bond acceptors ≤ 10) were selected for the screening. Furthermore, only molecules with the similarity lower than 0.8 (Tanimoto coefficient) were selected. Input files in Sybyl mol2 format were converted into AutoDock compliant format by MGLTools[81].

The crystal structure of human MUS81-EME1 was not described at the time of the initial screen. Therefore, the crystal structure of a chimerical complex of zebrafish MUS81/human EME1 was used as a template for homology modelling (PDB ID: 1J25; [61]). The chimeric complex was crystallised with truncated MUS81 (aa 303-612) and EME1 (aa 246-570) which preserves the nuclease activity of the complex. Since MUS81 nuclease activity is known to be dependent on the presence of bivalent metal ion, the Mn^2+^ was added into the active site of MUS81 protein based on superposition with structure of the Hef nuclease domain (PDB-ID:1J25; [82]). Corresponding amino acid sequences of MUS81 (residues 246-551) and EME1 (residues 246-570) was downloaded in FASTA format from the UniProtKB/Swiss-Prot database (MUS81 ID-Q96NY9, EME1 ID-Q96AY2). The model of MUS81-EME1 complex was built by using SWISS-MODEL [83] and was verified I-TASSER web server [84] . The active site of MUS81 included in this structure was used for molecular docking using the AutoDock Vina [50]. The region of the active site selected for molecular docking was set to 33 × 27 × 30 Å centred at the Mn^2+^ ion.

The docked conformations were scored by the AutoDock Vina software and rescored using NNScore 2.0 software [53]. A consensus score for each conformation was calculated by averaging the ranks obtained when AutoDock Vina score and the final NNScore 2.0 score. The common features in the binding modes of the docked conformations were used as a ground for molecule clustering. The clustering analysis was performed by AuposSOM [51] using default parameters where the map size was changed to 6 x 5 to increase the maximal number of clusters.

### *In vitro* high-throughput screening

The HTS was performed at the Molecular Screening Shared Resource (University of California Los Angeles) with approximately 100.000 compounds from their drug discovery libraries. All compounds were dissolved in DMSO. The compound transfer was done using BioMek FX (Beckman Coulter) equipped with a 0.5 µL pin tool, and other solutions were added by BioTek EL406 dispenser. The different components of the reaction were dispensed to black low-volume and flat-bottom 384-well plates (Polystyrene NBS; Corning 3820) in this order: 1) 4 µL master mix, 2) 50 nL compounds, 3) 1 µL recombinant MUS81-EME1. After a 30 min pre-incubation at room temperature, the reaction was initiated by adding 1 µL of 3’flap DNA substrate labelled with 6-carboxyfluorescein (FAM) and a Black Hole Quencher (BHQ1) (Figure S1). The plates were then incubated at room temperature for 1 hour before measuring fluorescence with Acquest 384-1536 plate reader (Molecular Devices). The final concentrations in the reaction were: 50 mM Tris-HCl pH 7.5, 1 mM DTT, 5 mM MgCl_2_, 0.1 mg/mL BSA, 100 mM KCl, 17 nM MUS81-EME1, 40 nM DNA, 8.3 µM compound (0.83% DMSO)

### Dose response assay (CZ OPENSCREEN IMG CAS)

The dose-response assay was performed in triplicates with 10 concentration points ranging from 5 nM to 10 µM. The reaction was done similarly to the primary HTS, with some modifications. The reaction volume was 5 µL and samples were prepared in 1536-plates. The compounds were transferred with a liquid handler Echo 550 (Labcyte, Beckman Coulter). Other reaction components were dispensed by Multidrop. Plates were scanned in an EnVision reader (PerkinElmer).

### Expression and purification of human MUS81

The MUS81-EME1 full-length expression plasmid was obtained from Stephen C. West (The Francis Crick Institute, US) was purified as previously described [85]. Briefly, 8 g of *E. coli* cell pellet was sonicated in 40 mL buffer C (50 mM Tris-HCl pH 7.5, 10% sucrose, 10 mM EDTA, 1 mM DTT, 0.01% NP40 and protease inhibitors) containing 150 mM KCl and clarified by centrifugation (100,000×g, 60 min). The cleared lysate was applied sequentially onto a 7-mL Q Sepharose and SP Sepharose columns (GE Healthcare Life Sciences). The SP Sepharose column was developed with a 70-mL gradient of 100-800 mM KCl in buffer K (20 mM K_2_HPO_4_ pH 7.5, 10% sucrose, 10 mM EDTA, 1 mM β-mercaptoethanol and 0.01% NP40). The peak fractions were pooled and mixed with 1.5-mL of His-Select Nickel Affinity Gel (Sigma Aldrich). The beads were washed with 10 column volumes of buffer K containing 150 mM KCl and 5 mM imidazole and bound proteins were eluted using 10 - 1000 mM imidazole in buffer K containing 150 mM KCl. The imidazole fractions were pooled and further fractionated using 0.5 mL heparin (GE Healthcare Life Sciences) with a 10 mL gradient of 250-900 mM KCl in buffer K. The fractions containing purified MUS81 were pooled, concentrated using a Vivaspin (Sartorius Stedim Biotech) concentrator and stored in 5 μL aliquots at -80 °C.

A truncated version of MUS81-EME1 complex, encompassing aminoacids 246-551 of MUS81 and a 178-570 of EME1 was generated. Plasmid co-expressing both proteins was transformed into bacteria and purified as described above.

### DNA substrates

All oligonucleotides used in this study were purchased from Eurofins Genomics and are listed in Supplementary table 5. The fluorescently labelled DNA substrates were prepared as previously described [86]. In short, equimolar amounts of individual oligonucleotides were annealed in hybridisation buffer H (50 mM Tris–HCl pH 7.5, 100 mM NaCl, 10 mM MgCl_2_). The mixture was heated to 70 °C for 3 min and cooled slowly to room temperature. The annealed DNA substrates were purified by fractionation on a 1-mL Mono Q column (GE Healthcare Life Sciences) with a 20-mL gradient of 50-1000 mM NaCl in 10 mM Tris–HCl pH 7.5. Fractions containing the DNA substrate were concentrated using a Vivaspin concentrator (Sartorius Stedim Biotech) with a 5 kDa cut-off and washed with lower salt buffer. The concentration of the DNA substrates was determined by absorbance measurement at 260 nm.

### Gel-based nuclease assay

Nuclease assays were performed in reaction buffer containing 50 mM Tris pH 7.5, 10 mM MgCl_2_, 1 mM DTT, 85 mM KCl and 20% glycerol. Purified MUS81-EME1 (1 nM) was pre-incubated in 9 μL reaction with indicated concentrations of inhibitors for 15 min at room temperature, followed by the addition of 3 nM of a 3’flap DNA substrate (listed in Supplementary table 5) and a further incubated for 15 min at 37 °C. The final concentration of DMSO in the reactions was 2%. The reactions were stopped by the addition of 0.1% SDS and 500 μg/mL of proteinase K, followed by incubation for 5 min at 37 °C. Each reaction was then mixed with 1/5 volume of loading buffer (60% glycerol, 10 mM Tris-HCl pH 7.5, 60 mM EDTA, and 0.1% Orange G) and loaded on a native PAGE gel (12% acrylamide gel in 1xTBE). The fluorescent DNA was visualised using a FLA-9000 (Fujifilm). Bands corresponding to cleaved substrates were selected for quantification using the MultiGauge 3.2 software. Final data represent integrated density of these bands with background subtraction. Fitting of the data with sigmoid curves were performed in OriginPro (OriginLab).

### Electrophoretic mobility shift assay

Purified MUS81-EME1 (160 nM) was incubated in a modified assay buffer without MgCl_2_ (50 mM Tris-HCl pH 7.4, 150 mM NaCl and 0.05 % Tween-20) with 15 nM of a 3’flap DNA substrate (listed in Supplementary table 5) in the absence or presence of selected inhibitors at RT for 10 minutes, followed by an incubation at 25 °C for 10 minutes. The reaction was stopped on ice and products were resolved using native PAGE (7.5% acrylamide gel in 0.5% TBE) at 4°C for 50 min (6.5 V/cm). Gels images were captured using a FLA-9000 (Fujifilm) and Typhoon (Amersham) scanners and quantified with Multi Gauge V3.2 software (Fujifilm).

### Biolayer Interferometry measurements (BLI)

Measurements were performed on a BLItz instrument (ForteBio) in a buffer containing 50 mM Tris-HCl pH7.5, 50 mM KCl, 10 mM MgCl_2_, and 0.05% Tween 20. A concentration of 15 nM biotinylated 3‘-flap (blocked) DNA substrate was immobilised on streptavidin-coated (SAX) sensors and incubated with 200 nM MUS81-EME1 fragment alone or premixed with indicated amount of inhibitors/DMSO. The real-time kinetics of protein association were measured as changes in optical thickness. The data were plotted in GraphPad Prism (GraphPad Software).

### Cell lines and culture conditions

U2OS WT cells were obtained from the European Collection of Authenticated Cell Cultures, U2OS-DR and U2OS-EJ5 cells were a kind gift from Dr. Jeremy Stark (City of Hope National Medical Center), while BIR-GFP cells were generously provided by Dr. Thanos D Halazonetis (University of Geneva). The U2OS GFP-MUS81 stable cell line was established using GFP-MUS81 WT or nuclease-dead (ND) DNA cloned into a pAIO plasmid [87] and co-transfected with a Flp-In recombinase plasmid. CAL51 cells were a kind gift from Dr. Martin Mistrik (Palacky University, Olomouc) and were used to generate the CAL51 MUS81^-/-^ cell line by CRISPR-Cas9 as described previously [88].

All cells were cultured in Dulbecco’s modified Eagle’s medium (DMEM, Gibco) supplemented with 10% (v/v) foetal bovine serum (FBS, Gibco), 100 U/mL penicillin and 100 μg/mL streptomycin (both from Biosera). All cells were grown at 37°C in humidified atmosphere in 5% CO2.

### RNAi

Cells were reverse-transfected with Lipofectamine RNAiMax (Life Technologies) and 30 nM of siRNA according to manufacturer’s instructions. Control siRNA used as a negative control (XWNeg9) and MUS81 siRNA (s37038) were obtained from ThermoFisher Scientific.

### Cell proliferation assay

CAL51 WT, CAL51 MUS81^-/-^, U2OS and HEK293 cells were seeded in a 96-well plate and treated with the respective concentration of selected inhibitors. After incubation for 4-5 days, cells were harvested while the untreated control was still sub-confluent. The plate was then incubated at -80 °C to achieve cell lysis. Cell proliferation was assessed using the CyQUANT™ Cell Proliferation Assay kit (Thermo Fisher Scientific) according to manufacturer’s instructions. Absorbance was measured using a plate reader (Infinite F500, Tecan Austria GmbH).

### Clonogenic survival assays

Cells were trypsinised, plated at the desired density in 6-well plates for colony forming assay, and then incubated for 10 days. Inhibitors at varying concentrations were added to the media after 24 hours. After incubation, colonies were fixed and stained with 0.5% crystal violet before scanning the plates. The colonies were then solubilised using 10% acetic acid solution and absorbance was measured with a plate reader. The percentage of survival of colonies was plotted with GraphPad Prism (GraphPad Software).

### Immunoblotting

To prepare whole cell extracts, cells were harvested by trypsinisation, washed with cold PBS and re-suspended in SDS-PAGE loading buffer. The samples were then sonicated and boiled at 70 °C for 10 min. Equal amounts of protein (50-100 μg) were separated on a 10% SDS-PAGE at 100 V, followed by transfer of proteins to nitrocellulose membrane using the semi-dry Trans-blot turbo Transfer system (1704150; Biorad). After transfer, membranes were blocked in 5% milk/PBST for 1 hour at room temperature and then incubated at 4 °C on a rocker overnight with the corresponding primary antibodies (Actin – ab184220, Abcam; MUS81 - ab14387, Abcam). The next day, the membranes were washed with PBST and incubated with the corresponding secondary antibodies (Anti-Rabbit IgG - A6154, Sigma-Aldrich; Anti-Mouse IgG, A0168, Sigma-Aldrich) for 1 hour at room temperature. Finally, the blots were developed by the Immobilon Western Chemiluminescent horseradish peroxidase (HRP) Substrate (WBKLS0500; MERCK Millipore), and images were acquired using the Luminescent Image Analyser (ImageQuant™ LAS 4000; Fujifilm).

### HR, BIR and NHEJ assays

Reporter DR-GFP and EJ2-GFP U2OS cells [89] along with BIR-GFP cells [90] were transfected with 2.5 μg of I-SceI-expressing pCAGGS vector and subsequently treated with MUS81 inhibitors at the indicated concentrations. After transfection (72 hours), cells were trypsinised and resuspended in 3% BSA in PBS. GFP fluorescence detection was carried out using a BD FACSVerse flow cytometer, and the data were analysed with FlowJo software (BD Life Sciences). Data presented in the graph were normalised to the DMSO-treated sample.

### EdU Cell cycle assay

U2OS WT cells (300.000) were seeded in a 6-well plate, a subset of them was treated with either control siRNA or siRNA targeting MUS81. After adhesion, cells were treated with the respective inhibitors for 72 hours during which CPT (100 μM, Sigma-Aldrich) was added for the last 16 h and 10 μM EdU (Sigma-Aldrich) was added for the last 1 hour. Cells were then harvested and stained using the Click-iT™ EdU Alexa Fluor™ 647 Flow Cytometry Assay Kit (Thermo Fisher Scientific). Finally, cells were resuspended in 1xPBS containing 5 μg/mL of propidium iodide (PI) and analysed using a BD FACSVerse flow cytometer (Becton Dickinson) and FlowJo software (BD Life Sciences). At least 20,000 cells were used for each measurement, and experiments were performed in duplicates.

### Annexin V apoptotic assay

U2OS WT cells (300.000) were seeded in a 6-well plate, a subset of them was treated with either control siRNA or siRNA targeting MUS81. After adhesion, cells were treated with the respective inhibitors for 72 hours during which CPT (100 μM, Sigma-Aldrich) was added for the last 2 h. Cells were harvested, washed in ice-cold PBS, and collected by centrifugation at 500×g for 5 min. Cells were simultaneously stained with FITC-labelled annexin V antibody (556419, Becton Dickinson), and PI (5 μL) at room temperature for 15 min, protected from light. Stained cells were analysed using a BD FACSVerse flow cytometer (Becton Dickinson) and FlowJo software (BD Life Sciences). At least 20.000 cells were used for each measurement, and experiments were performed in duplicates.

### Immunofluorescence

U2OS, CAL51 WT and CAL51 MUS81^-/-^ cells (10.000) were seeded in a 96-well plate 24 hours before the treatment. Cells were treated with the respective MUS81 inhibitor according to the assay scheme for either 2 weeks or 24 hours. In certain settings cells were treated with cisplatin (12 μM, Sigma-Aldrich) for the last 24 hours. After washing, cells were fixed with 3% PFA for 10 min at room temperature and permeabilised with 0,5% Triton X-100 (Sigma-Aldrich). Subsequently, cells were blocked with 5% bovine serum albumin (BSA, Sigma), incubated with γH2AX antibody (05-636, MERCK Millipore), and Alexa Fluor 488 anti-mouse secondary antibodies (Thermo Fischer, 1:1000). Images of the cells were taken using Nikon Eclipse microscope with nuclear staining by DAPI. The γH2AX foci were detected and quantified using CellProfiler (Broad Institute of MIT and Harvard) [91]. The analysed data were plotted in RStudio software using R programming language [92].

### Laser Microirradiation

HEK293 cells with GFP-MUS81 stable expression (cloned into a pAIO-based vector; a kind gift from Josef Jiricny) were grown in 35 mm ibidi’s μ-Dishes and pre-incubated with MU876=32 at 1 and 5 μM final concentration for two hours. Damage was induced using 355 nm laser line (UGA-42 Firefly, Rapp OptoElectronic, 20% output power, 3 iterations) connected to Delta Vision Elite Pro microscope (GE Healthcare). Recruitment and retention of GFP-MUS81 signal at damage site was traced with a 100x/1.4 (Olympus) objective at the indicated times after irradiation. GFP-MUS81 accumulation at the damage site was compared with an undamaged region within the same micro-irradiated cell. The average accumulation ± S.D. of GFP-MUS81 of three biological replicates is plotted.

### Solubility

Analysis of aqueous solubility was provided by Bienta Enamine Biology Services. Kinetic solubility assay was performed according to Enamine’s aqueous solubility SOP. Briefly, using a 20 mM stock solution of the compound in 100% DMSO, dilutions were prepared to a theoretical concentration of 400 μM in duplicates in PBS pH 7.4 (138 mM NaCl, 2.7 mM KCl, 10 mM K-phosphate) with 2% final DMSO. The experimental compound dilutions in PBS were further allowed to equilibrate at 25 °C in a thermostatic shaker for 2 hours and filtered through HTS filter plates using a vacuum manifold. The filtrates of test compounds were diluted 2-fold with acetonitrile with 2% DMSO. In parallel, compound dilutions in 50% acetonitrile/PBS were prepared to theoretical concentrations of 0.1, 1, 50, 100, and 200 μM with 2% final DMSO to generate calibration curves. Ondansetron was used as reference compound to control proper assay performance. All 22 samples were diluted 100-fold with 50% acetonitrile/water (v/v) mixes before LC-MS/MS measurement. The effective range of this assay is 0.2 – 400 μM and can be changed depending on the chemical nature of the compounds.

The thermodynamic solubility assay was performed according to Enamine’s aqueous solubility SOP. Briefly, the dry powder forms of the test compounds were mixed with phosphate-buffer (138 mM NaCl, 2.7 mM KCl, 10 mM K-phosphate, pH 7.4) to the theoretical concentration of 4 mM and further allowed to equilibrate at 25 °C in a thermostatic shaker. After 4 and 24 hours shaking, incubation mixtures were filtered through HTS filter plates using a vacuum manifold. The filtrates of test compounds were diluted 2-fold with acetonitrile. In parallel, using a 20 mM stock solution, compound dilutions in 50% acetonitrile/PBS mixes were prepared to the theoretical concentrations of 1, 25, 100, and 200 μM to generate calibration curves. Ondansetron was used as reference compound to control proper assay performance. All samples were diluted 100-fold with 50% methanol/water (v/v) mixes before LC-MS/MS measurement. The effective range of this assay is 2-400 μM and can be changed depending on the chemical nature of the compounds.

### Analysis of micronuclei

CAL51 WT and CAL51 MUS81^-/-^ cells were seeded in a 96-well plate 24 hours before the treatment. Alternatively, cells were transfected with 30 nM siRNA (control or MUS81) for 48 hours before treatment. Subsequently, cells were treated with respective inhibitors for 48 hours. During the last 16 hours cytochalasin B (Sigma-Aldrich) was added to the cells at a concentration of 1.25 μg/mL. At the end of the treatment, cells were washed in 1XPBS, fixed with 4% formaldehyde and washed again. DAPI (5 μg/mL, Panreac AppliChem) was added for nuclear staining. Images were acquired using Nikon Eclipse fluorescence microscope and analysed using ImageJ software [93].

### Cell survival assays

CAL51 WT cells were treated with DMSO, MU262=18, MU876=32, non-targeting siRNA or siRNA targeting MUS81. Cells were trypsinised, counted and re-plated twice a week and retreated with inhibitors or siRNA with every passage. The cell number was plotted for each time point. Data were plotted in Graphpad Prism (GraphPad Software) and a 2-way ANOVA test was used to assess significance.

### Detection of TFEs by DNA fluorescence *in situ* hybridisation (FISH)

U2OS and HeLa cells were seeded in 10 cm dishes. The next day, media was changed, and compounds were added at desired concentrations. This process was repeated after 43 hours except that compounds were added together with 200 ng/mL of nocodazole. Cells were collected 5 hours later by shake-off and incubated in 75 mM KCl at 37 °C for 10 min. Chromosomes were fixed in ice-cold methanol/acetic acid (3:1) and spread on glass slides. Slides were then treated with 20 μg/mL RNAse A (Sigma-Aldrich), in 1x PBS at 37 °C for 1 h, fixed in 4% formaldehyde (Sigma-Aldrich) in 1x PBS for 2 min, and treated with 70 μg/mL pepsin (Sigma-Aldrich) in 2 mM glycine, pH 2 (Sigma-Aldrich) at 37°C for 5 min. Slides were fixed again with 4% formaldehyde in 1x PBS for 2 min, incubated subsequently in 70%, 90% and 100% ethanol for 5 min each, and air-dried. A C-rich telomeric PNA probe (5’-AF568-OO *-p4*CCCTAACCCTAACCCTAA-3’; Panagene) diluted in hybridisation solution (10 mM Tris-HCl pH 7.2, 70% formamide, 0.5% blocking solution (Roche)) was applied onto the slides followed by incubation at 80°C for 5 min and at room temperature for 2 h. Slides were washed twice in 10 mM Tris-HCl pH 7.2, 70% formamide, 0.1% BSA and three times in 100 mM Tris-HCl pH 7.2, 150 mM NaCl, 0.08% Tween-20 at room temperature for 10 min each. DNA was counterstained with 100 ng/mL DAPI (Sigma-Aldrich) in 1x PBS and slides were mounted in Vectashield (Vectorlabs). Images were acquired with a Zeiss Cell Observer equipped with a cooled Axiocam 506 m camera and a 63X/1.4NA oil DIC M27 PlanApo N objective. Image analysis was performed using ImageJ [93] and Photoshop software (Adobe Inc.). TFEs were identified as chromatid ends lacking a detectable telomeric signal and the number of TFEs per metaphase is displayed.

## Supporting information

Supporting information_NMR spectra

Supporting information_Experimental procedures

## SUPPORTING INFORMATION

Additional experimental details, materials, and methods, and synthesis with NMR data for all compounds are provided in three separate files (Supplementary Figures and Tables, Experimental procedures, and NMR spectra).

## ACKNOWLEDGEMENTS

This work was supported by the following grants: SoMoPro (6SA17846) and the Czech Science Foundation (21-22593X). The authors also gratefully acknowledge the support provided by the following sources of funding: European Structural and Investment Funds, Operational Programme Research, Development and Education „Preclinical Progression of New Organic Compounds with Targeted Biological Activity” (Preclinprogress) CZ.02.1.01/0.0/0.0/16_025/0007381, the project CZ-OPENSCREEN: National Infrastructure for Chemical Biology (CZ-OPENSCREEN LM2023052), and Bader Philanthropies. Work in the Azzalin laboratory was supported by LaCaixa Foundation (project LCF/PR/HP21/52310016). We thank Vit Vsiansky, Veronika Weisova, Kristyna Slavikova and Barbora Barankova for help with some experiments. We also thank the company Artios Pharma for a productive collaboration.

## AUTHOR CONTRIBUTIONS

L.K., K.P., J.P., and B.C. designed the study and the experiments. L.K., K.P., J.P. wrote the paper.

J.P. performed most of the cell experiments. J.P., A.S. and M.Z. performed nuclease assays. L.D. and J.B. performed initial *in silico* screening. V.M. performed the *in vitro* HTS. B.C., S.H., N.A., and P.K. performed the synthesis of the most potent hits and derivatives. J.R. and C.A. performed detection of TFEs.

## ABBREVIATIONS USED

HR: homologous recombination
HU: hydroxyurea
CPT: camptothecin
SAR: structure-activity relationship
BIR: break-induced replication
NHEJ: non-homologous end joining
PI: propidium iodide
EdU: 5-Ethynyl-2’-deoxyuridine
EMSA: electrophoretic mobility shift assay
BLI: bio-layer interferometry
GFP: green fluorescent protein
ALT: alternative lengthening of telomeres
FISH: fluorescence *in-situ* hybridization
TFEs: telomere-free ends
CFA: colony formation assay
MMC: mitomycin C
ICL: interstrand crosslink
FAM: 6-carboxyfluorescein
BHQ1: black hole quencher
HRP: horseradish peroxidase

