## Supporting information_Figures for "Discovery of two structurally distinct classes of inhibitors targeting the nuclease MUS81 and enhancing efficacy of chemotherapy in cancer cells"

Republic

2 Department of Chemistry, Faculty of Science, Masaryk University, 62500 Brno, Czech

Republic

3 International Clinical Research Center, St. Anne’s University Hospital, Brno 656 91, Czech Republic

4 NCBR, Faculty of Science, Masaryk University, 62500 Brno, Czech Republic

5 GIMM - Gulbenkian Institute for Molecular Medicine, 1649-035 Lisbon, Portugal

6 Loschmidt Laboratories, Department of Experimental Biology and RECETOX, Faculty of

Science, Masaryk University, 62500 Brno, Czech Republic

7 Faculty of Medicine, University of Lisbon, 1649-028 Lisbon, Portugal

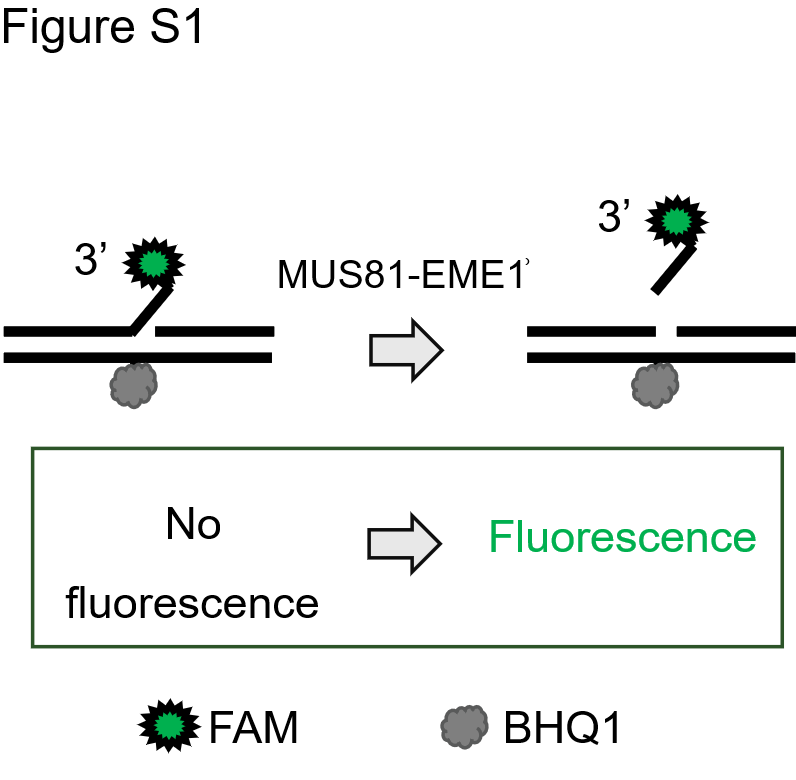

**Figure S1: A schematic representation of a fluorogenic assay specific for MUS81-EME1**

The assay consists of a 3’flap DNA substrate labelled with fluorescein (FAM) and a Black Hole Quencher. Upon endonucleolytic cleavage of the substrate by recombinant MUS81-EME1, FAM separates from BHQ1, resulting in a significant fluorescence increase, providing a direct readout of the enzymatic activity.

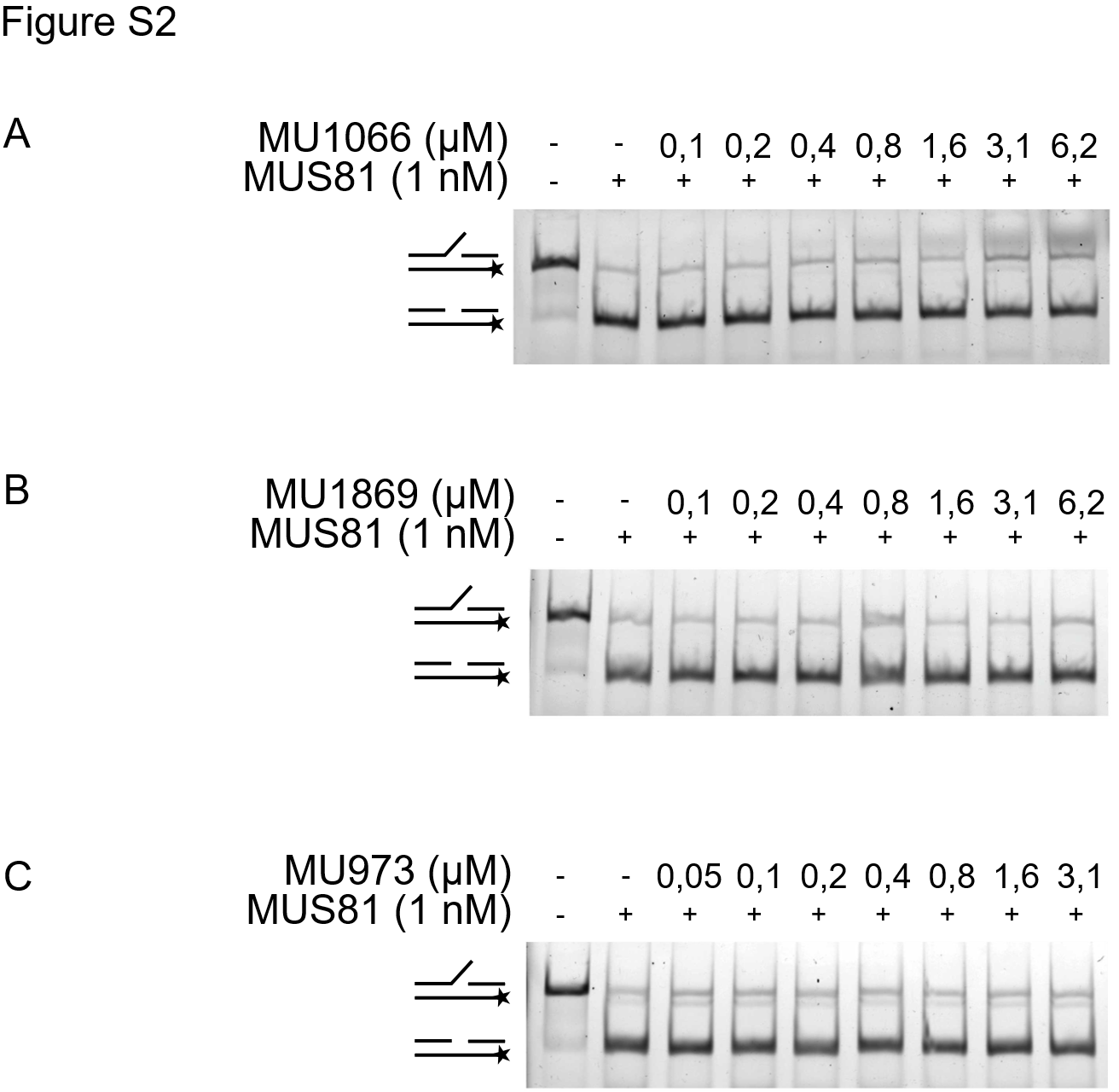

**Figure S2: In vitro nuclease assay of negative control compounds**

Purified human MUS81-EME1 (1 nM) was incubated with increasing concentrations of MU1066=17 (A), MU1869=25 (B) or MU973=45 (C), followed by the addition of 3 nm of fluorescently labelled 3’flap DNA. The reaction products were resolved on a native PAGE gel. The lower band corresponds to the cleaved DNA. Representative picture shown, n = 3.

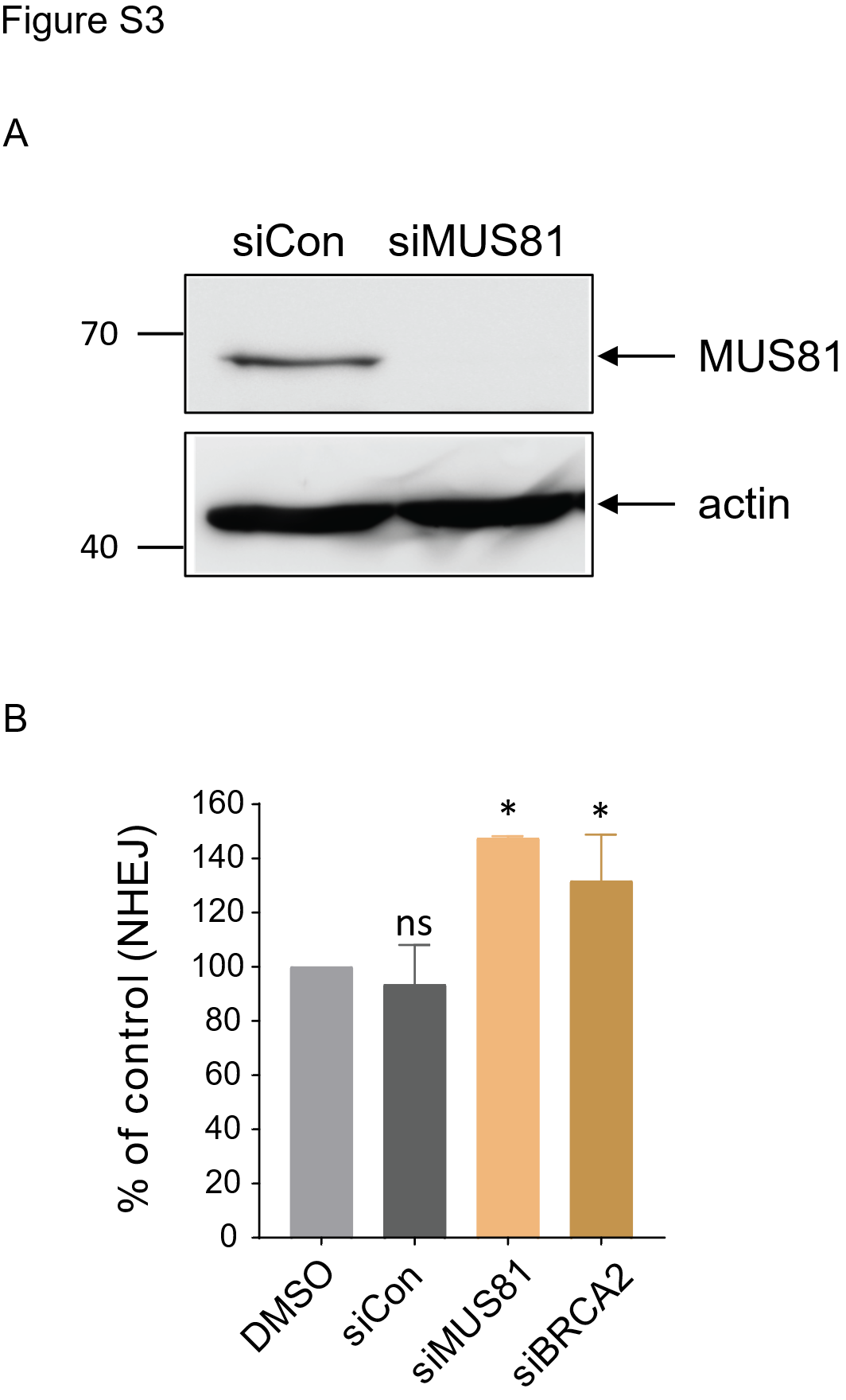

**Figure S3: MUS81 depletion does not affect NHEJ**

(A) U2OS BIR-GFP cells were treated with non-targeting siRNA (siCon) or siRNA against MUS81. Cells were harvested after three days, and WB was done using actin as a loading control.

(B) I-SceI-based NHEJ repair efficiency was measured using U2OS EJ5-GFP cells, treated with DMSO or siRNAs targeting MUS81 and BRCA2, respectively for 72 hours. The percentage of repair was normalised to the DMSO treated control. *n* = at least 2. error bars, s.d., **, *p* < 0.01 (unpaired, two-tailed *t*-test).

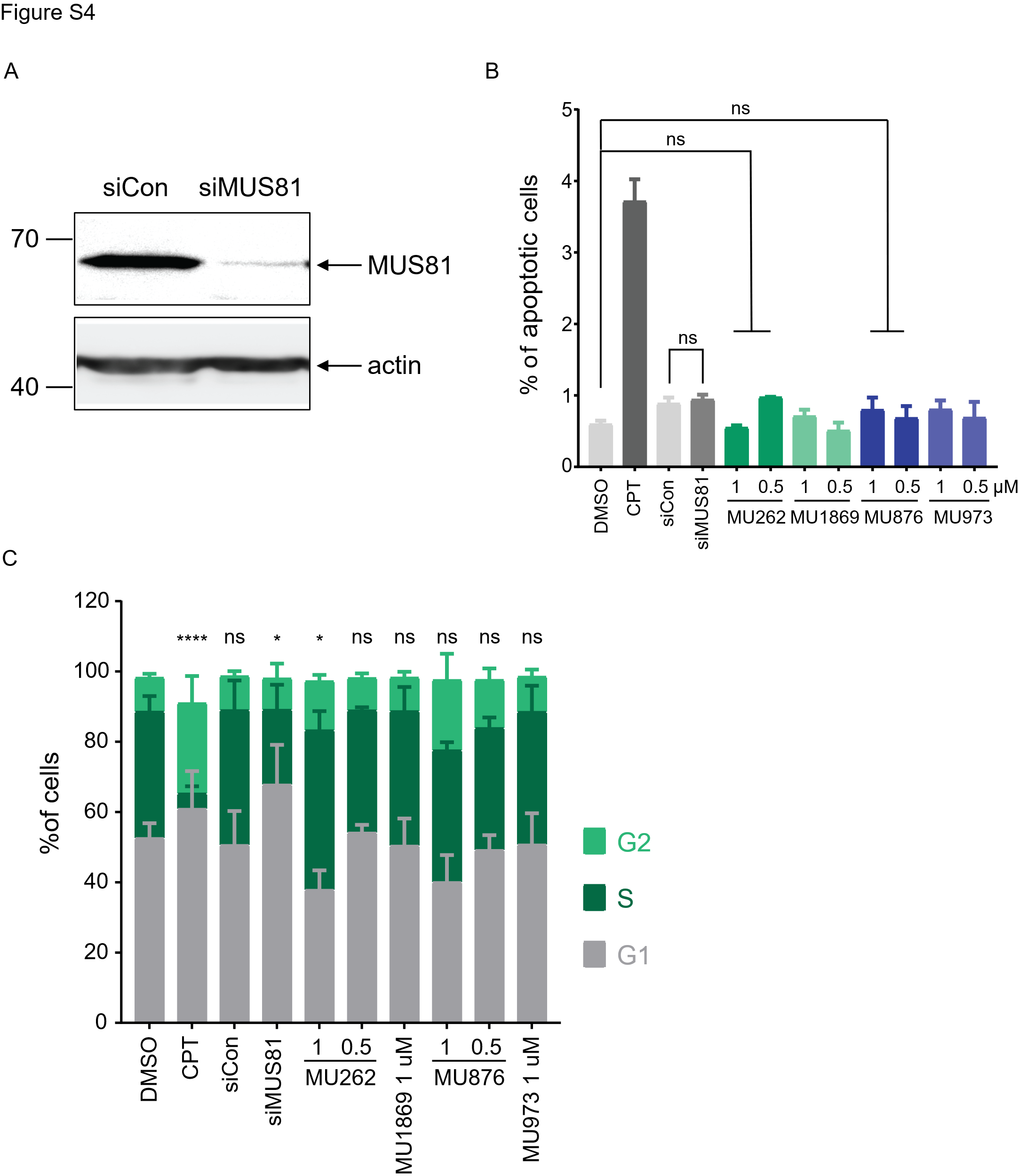

**Figure S4: Small molecule inhibitors of MUS81 do not have adverse side effects**

(A) U2OS WT cells were treated with non-targeting siRNA (siCon) or siRNA against MUS81. Cells were harvested after three, days and WB was done using actin as a loading control.

(B) U2OS WT cells were treated with DMSO, CPT, control or MUS81-targetting siRNA or the indicated concentrations of MU262=18, MU1869=25, MU876=32 and MU973=45 for 72 hours. Apoptotic cells were determined by Annexin V staining and measured by flow cytometry. n = at least 2. error bars, s.d. (unpaired, two-tailed *t*-test).

(C) U2OS WT cells were treated with DMSO, CPT, control or MUS81-targetting siRNA or the indicated concentrations of MU262=18, MU1869=25, MU876=32 and MU973=45 for 72 hours. Cells were labelled with EdU 30 minutes before harvest. Cell cycle was analysed using EdU and PI staining. Data were acquired by flow cytometry. n = 3. Error bars represent comparison to DMSO, s.d., *, *p* < 0.05 (unpaired, two-tailed *t*-test).

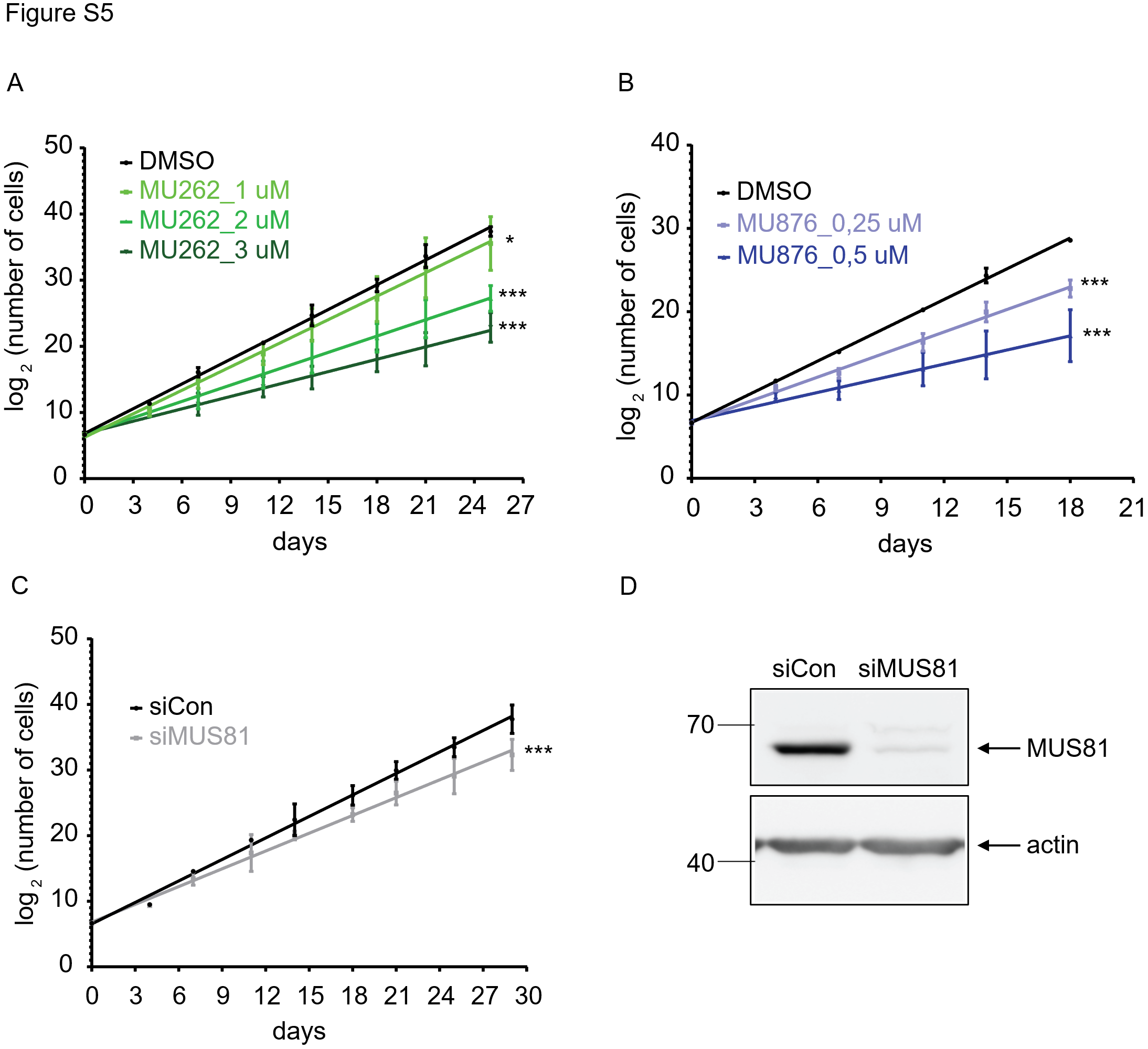

**Figure S5: Prolonged MUS81 depletion or inhibition slows cell growth**

(A) CAL51 WT cells were treated with DMSO and the indicated concentrations of MU262. Cells were left to grow for the indicated time and counted after each passage. The plots show non-linear fit of the exponential cell growth. n = at least 3. *, *p* < 0.05, ***, *p* < 0.001 (two-way ANOVA test).

(B) CAL51 WT cells were treated with DMSO and the indicated concentrations of MU876. Cells were left to grow for the indicated time and counted after each passage. The plots show non-linear fit of the exponential cell growth. n = 2. ***, *p* < 0.001 (two-way ANOVA test).

(C) CAL51 WT cells were treated with the non-targeting siRNA (siCon) and siRNA targeting MUS81. Cells were left to grow for the indicated time and counted after each passage. The plots show non-linear fit of the exponential cell growth. n = 2. ***, *p* < 0.001 (two-way ANOVA test).

(D) CAL51 WT cells were treated with control siRNA or siRNA against MUS81. Cells were harvested after three days, and WB was done using actin as a loading control.

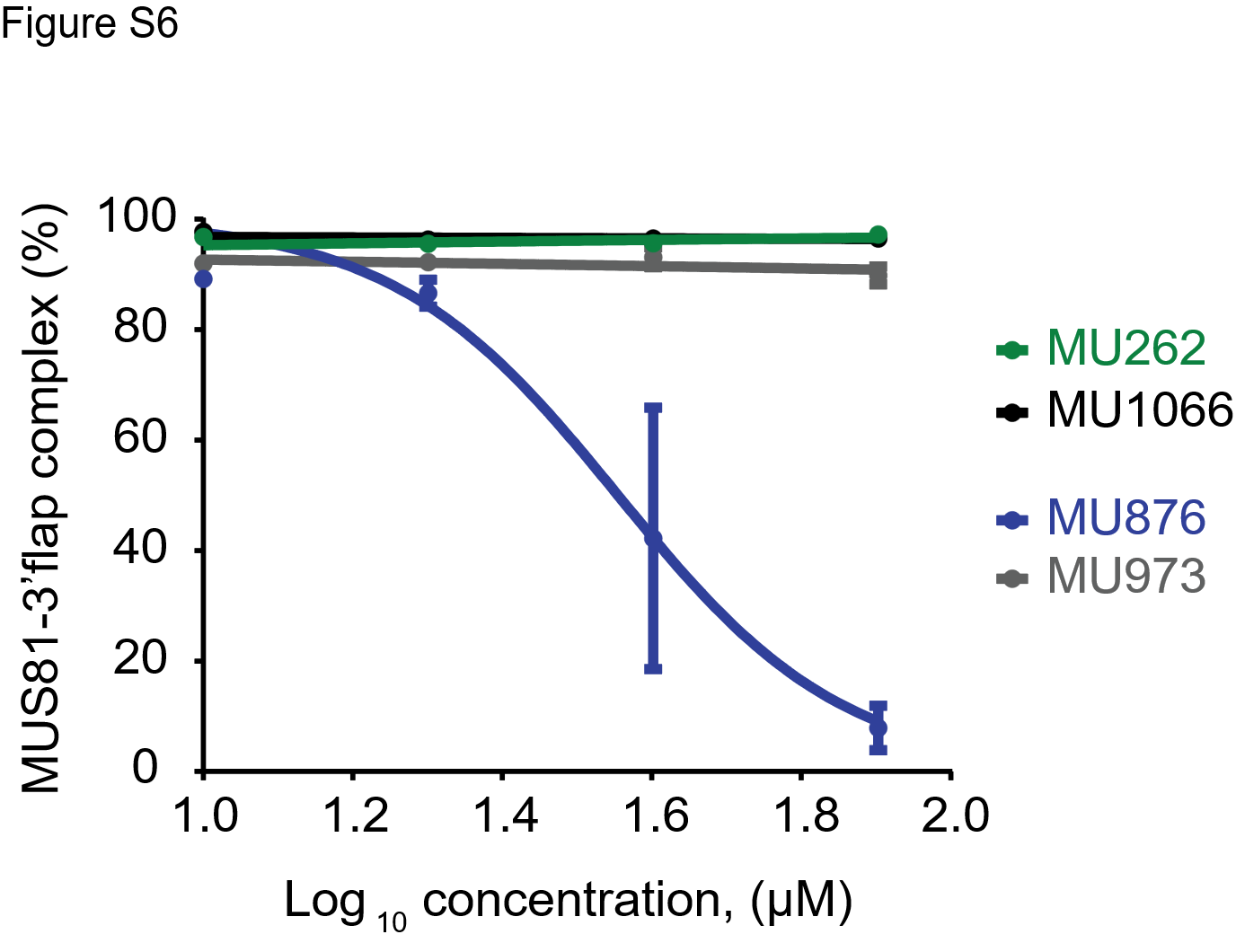

**Figure S6: Mechanism of function of the small molecules**

(A) Quantification of Fig. 4 A-B. Purified MUS81 (160 nM) was incubated with increasing concentrations of MU262, MU876, or their respective control analogues, followed by the addition of 3 nM fluorescently labelled 3’ flap DNA substrate. MgCl_2_ was omitted from the reaction buffer to prevent substrate cleavage. The reaction products were resolved on a native PAGE gel. The lower DNA band corresponding to the free DNA was used for quantification. Values were normalised to the negative control containing no enzyme. The graph represents % of 3’flap DNA substrate bound to MUS81 fitted to a sigmoid curve. n = 3.

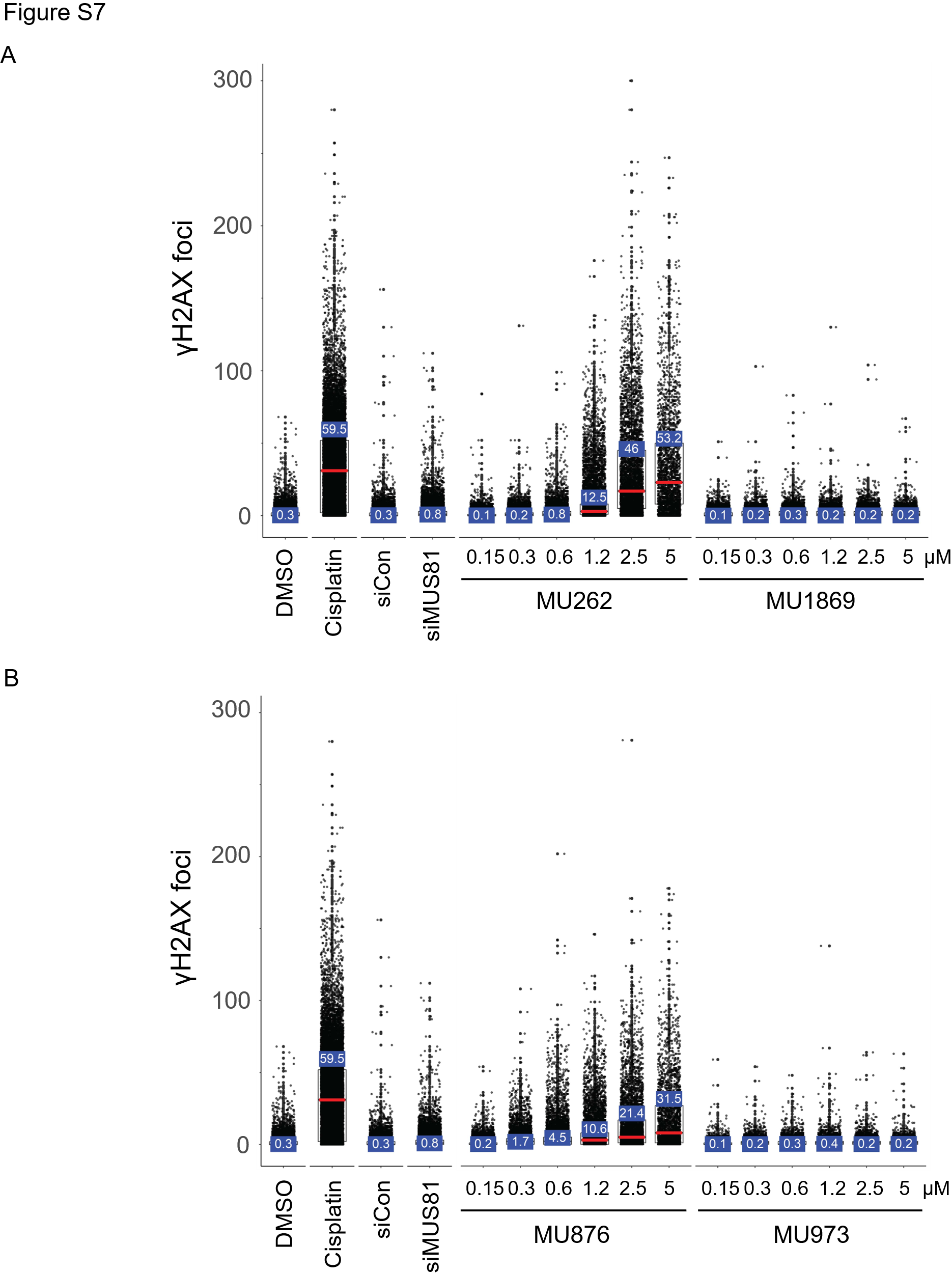

**Figure S7: Endogenous DNA damage persists following treatment with MUS81 inhibitors**

(A)U2OS cells were treated with DMSO, cisplatin as a positive control, non-targeting siRNA (siCon) or MUS81-targeting siRNA, MU262, and its negative control counterpart at the indicated concentrations for 72 hours. (B) U2OS cells were treated with DMSO, cisplatin as a positive control, non-targeting siRNA (siCon) or MUS81-targeting siRNA, MU876, and its negative control counterpart at the indicated concentrations for 72 hours. The number of γH2AX foci was assessed by fluorescence microscopy and quantified by a CellProfiller software. Data are plotted in R software. Red marker represents median of γH2AX foci number per nucleus. n = 3. (F) CAL51 MUS81^-/-^ cells and CAL51 cells treated with control siRNA. Cells were harvested after three days, and WB was done using actin as a loading control. (F) CAL51 WT and MUS81-/- cells were treated with DMSO, MU262 and MU876 at 16 μM concentration together with the indicated concentration of cisplatin for 96 – 120 hours. Cell viability was assessed by the Cy-Quant assay. The survival rate of cells was normalised to the DMSO control. Graphs are plotted in Graphpad Prism using the non-linear regression fitting model. n = 3.

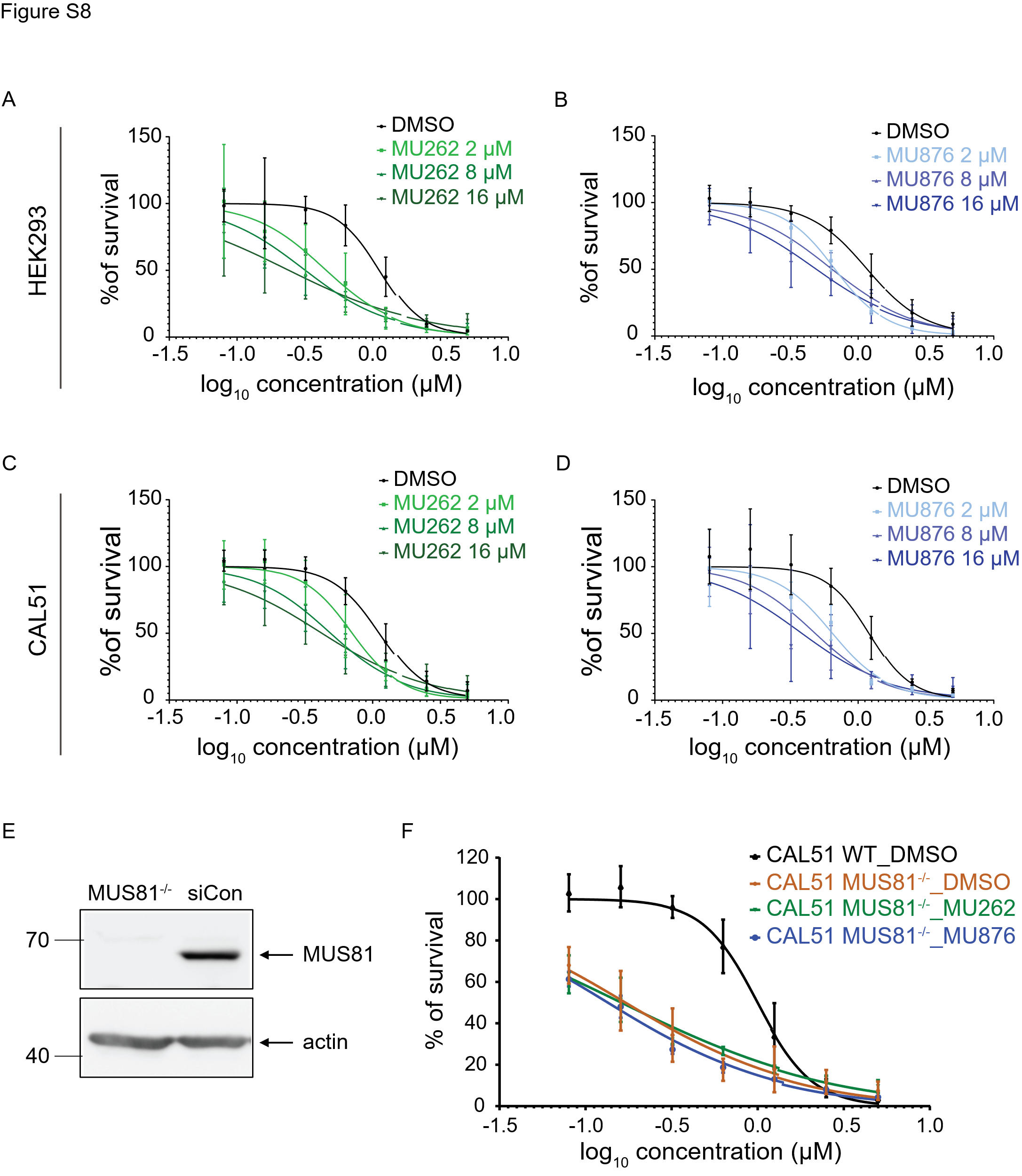

**Figure S8: MUS81 depletion sensitizes cells to cisplatin treatment**

HEK293 cells (A-B) and CAL51 cells (C-D) were treated with DMSO, MU262, or MU876 at the indicated concentrations in combination with range of cisplatin (0 – 5 μM) for 96 – 120 hours. Cell viability was assessed by the Cy-Quant assay. The survival was normalised to the DMSO-treated control. Graphs were generated in Graphpad Prism using the non-linear regression fitting model. n = 3, error bars represent s.d.

(E) CAL51 MUS81^-/-^ cells and CAL51 cells treated with control siRNA. Cells were harvested after three days and WB was done using actin as a loading control.

(F) CAL51 WT and MUS81-/- cells were treated with DMSO, MU262 and MU876 at 16 μM concentration together with the indicated concentration of cisplatin for 96 – 120 hours. Cell viability was assessed by the Cy-Quant assay. The survival rate of cells was normalised to the DMSO control. Graphs are plotted in Graphpad Prism using the non-linear regression fitting model. n = 3.

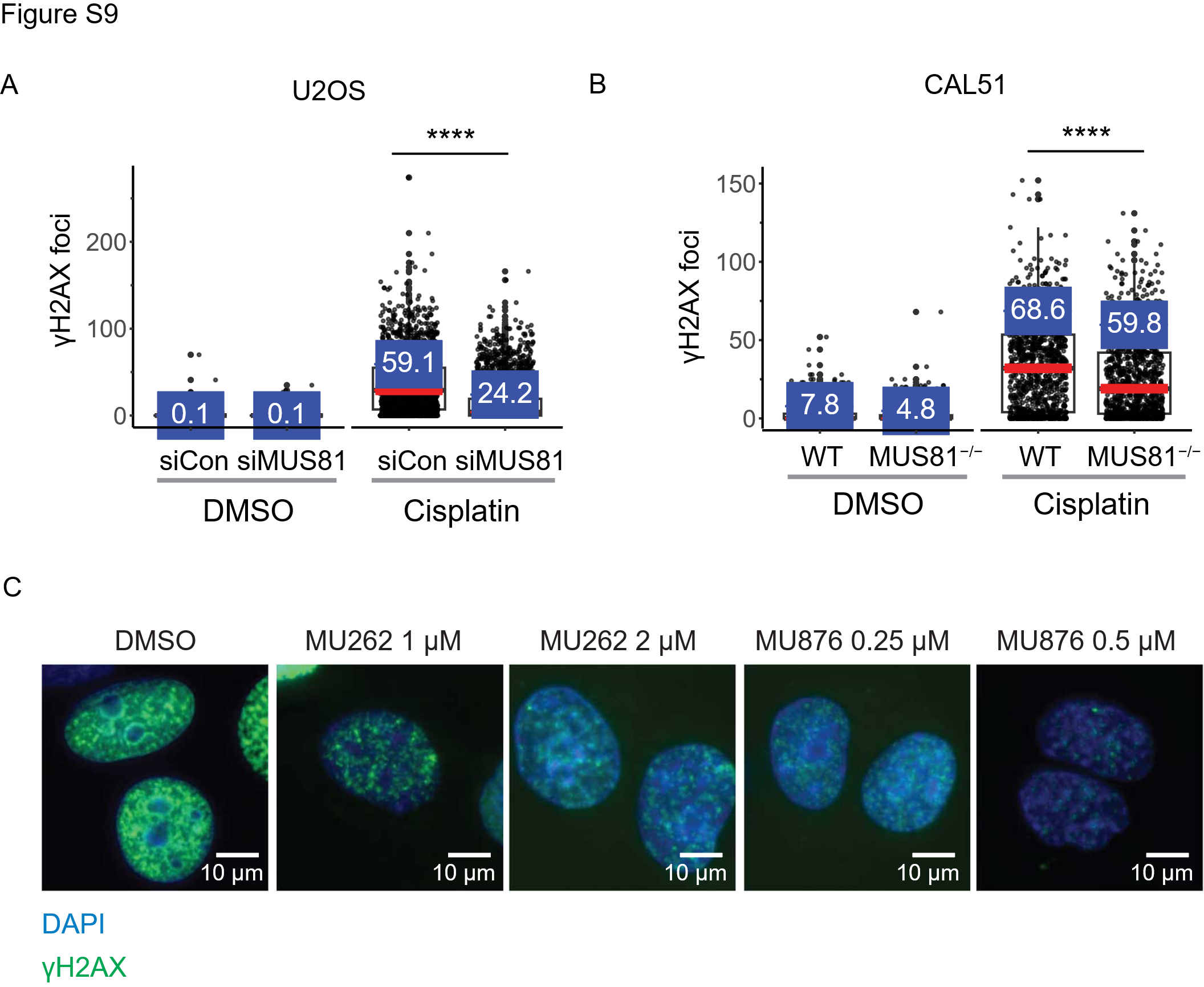

**Figure S9: MUS81 depletion potentiates the effect of chemotherapy**

(A) U2OS cells were treated with DMSO, non-targeting siRNA (siCon) or siRNA targeting MUS81 with every passage for a total length of two weeks. After that cisplatin (12 μM) was added for the last 24 hours before harvest. The number of foci of a DNA damage marker γH2AX was assessed by fluorescence microscopy and quantified by a CellProfiller software. Data are plotted in R software. Red marker represents median of γH2AX foci number per nucleus. n = 1., error bars, s.d., *, *p* < 0.05, **, *p* < 0.01 ***, *p* < 0.001 (unpaired, two-tailed *t*-test).

(B) CAL51 WT and MUS81^-/-^ cells were passaged for a total length of two weeks. After that cisplatin (12 μM) was added for the last 24 hours before harvest. The number of foci of a DNA damage marker γH2AX was assessed by fluorescence microscopy and quantified by a CellProfiller software. Data are plotted in R software. Red marker represents median of γH2AX foci number per nucleus. n = 1., error bars, s.d., *, *p* < 0.05, **, *p* < 0.01 ***, *p* < 0.001 (unpaired, two-tailed *t*-test).

(C) CAL51 WT cells were treated with DMSO, MU262, or MU876 at the indicated concentrations at each passage over two weeks. Cisplatin (12 μM) was added for the last 24 hours before harvest. DNA damage was assessed by visualising γH2AX foci using fluorescence microscopy, representative pictures are shown.

**Supplementary table 1: List of compounds identified and shortlisted in the *in silico* screening** ZINC code of the shortlisted compounds that were selected from in the silico screening. The values of the predicted kD and predicted binding energy are shown.

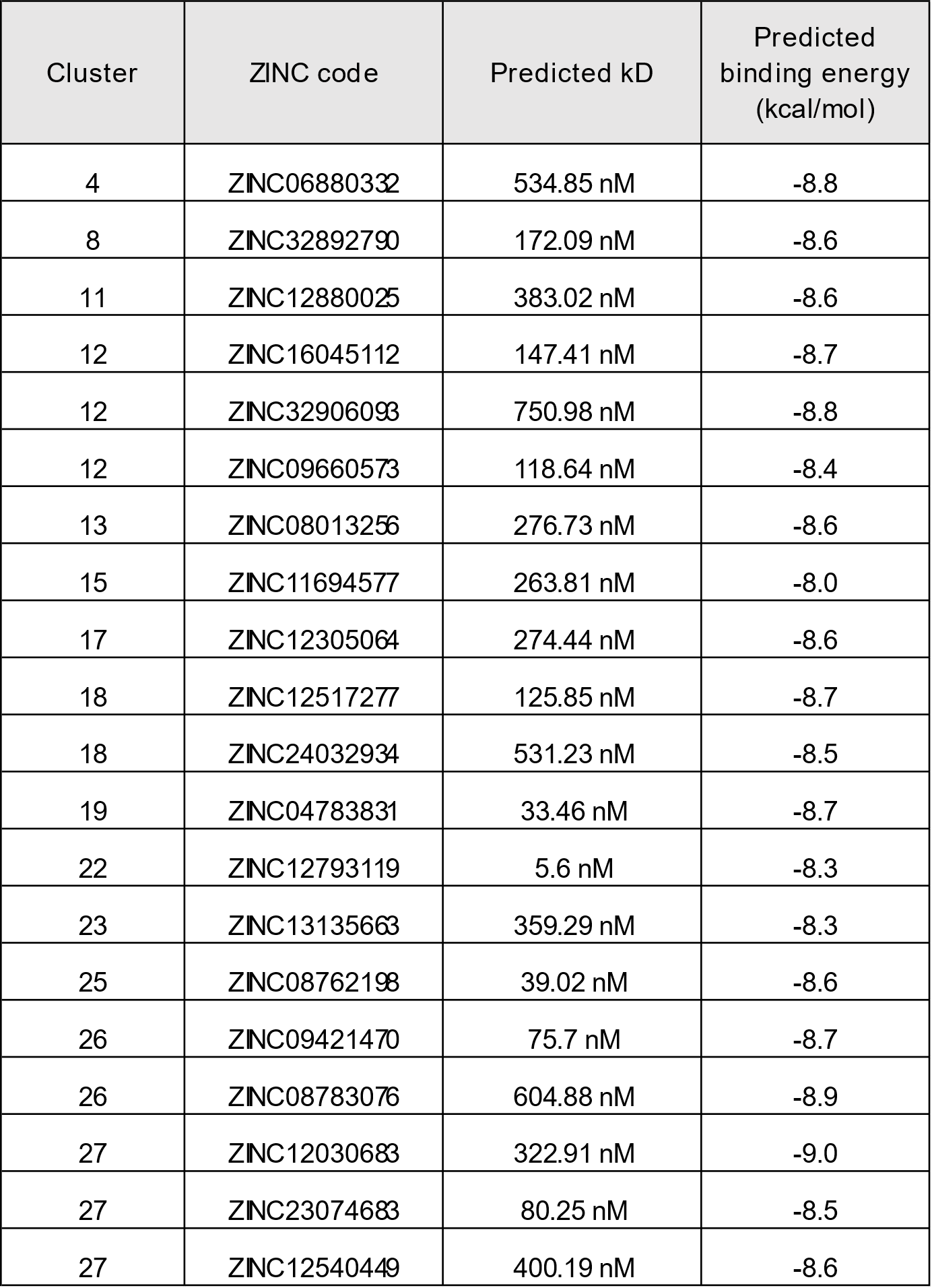

**Supplementary table 2: List of compounds identified and shortlisted in the i*n vitro* screening** Compound ID of the shortlisted compounds that were identified in the high-throughput screening. The compounds were purchased and a dose-response curve of the activity was assessed. The IC_50_ values are shown.

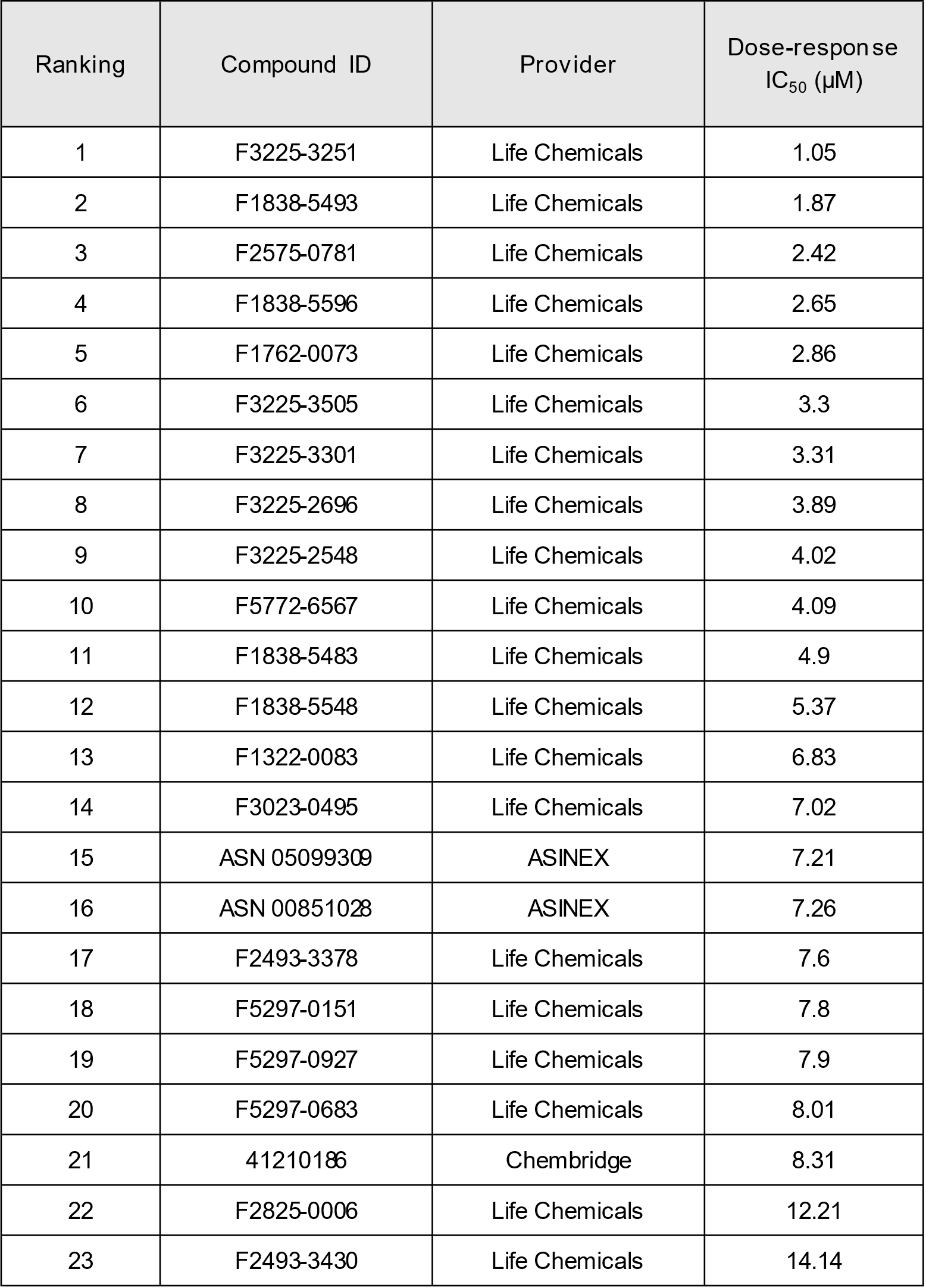

**Supplementary table 3: Structures and MUS81 *in vitro* and in cellulo IC_50_ values of the compound 1 and its analogues**

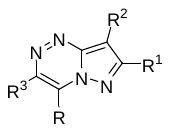

| **compound numbers** | **R** | **R^1^** | **R^2^** | **R^3^** | **In vitro IC_50_ interval**  **(μM)** | **Cell based IC_50_ interval (μM)** |
| --- | --- | --- | --- | --- | --- | --- |
| **1** | NH_2_ | Et | 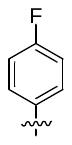 | 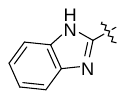 | 5-10 | >25 |
| **3** | NH_2_ | H | H | 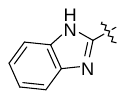 | >20 | N.A. |
| **4** | NH_2_ | H | 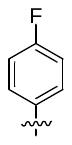 | 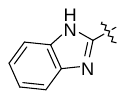 | >20 | N.A. |
| **5** | NH_2_ | Et | H | 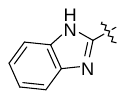 | 10-20 | N.A. |
| **6** | NH_2_ | Et | 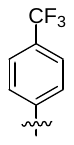 | 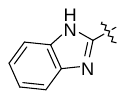 | 1-5 | >25 |
| **7** | NH_2_ | Et | 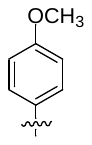 | 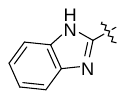 | >20 | >25 |
| **8** | NH_2_ | Et | 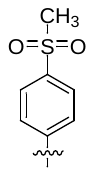 | 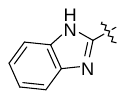 | >20 | 5-25 |
| **9** | NH_2_ | Et | 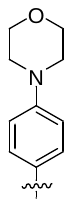 | 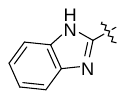 | >5 | >25 |
| **10** | NH_2_ | Et | 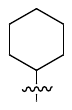 | 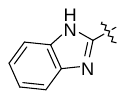 | 10-20 | >25 |
| **11** | NH_2_ | iPr | 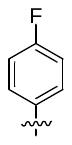 | 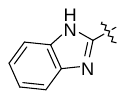 | 10-20 | N.A. |
| **12** | NH_2_ | 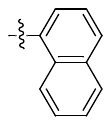 | H | 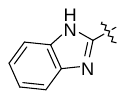 | 1-5 | >25 |
| **13** | NH_2_ | 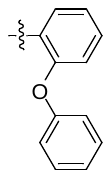 | H | 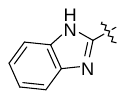 | 5-10 | >25 |
| **14** | NH_2_ | 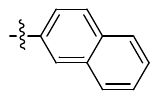 | H | 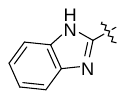 | 1-5 | >25 |
| **15** | NH_2_ | Ph | H | 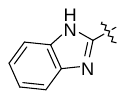 | 10-20 | >25 |
| **16** | H | Et | 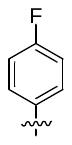 | 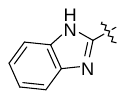 | >10 | 5-25 |
| **17** | CH_3_ | Et | 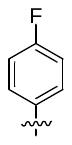 | 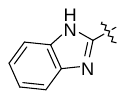 | >20 | >25 |
| **18** | OH | Et | 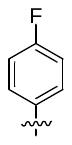 | 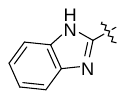 | <1 | <5 |
| **19** | NH_2_ | Et | 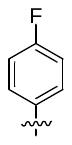 | CN | >20 | N.A. |
| **20** | OH | Et | 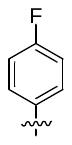 | 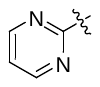 | >20 | 5-25 |
| **21** | NH_2_ | Et | 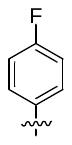 | CONH_2_ | >20 | N.A. |
| **22** | NH_2_ | Et | 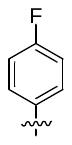 | 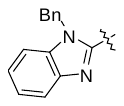 | 1-5 | >25 |
| **23** | OH | Et | 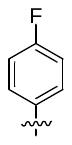 | 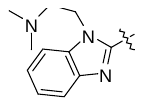 | >10 | 5-25 |
| **24** | OH | Et | 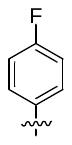 | 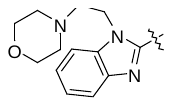 | >10 | 5-25 |
| **25** | OH | Et | 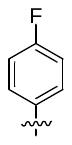 | 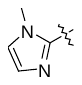 | >20 | >25 |
| **26** | OH | Et | 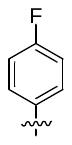 | 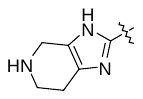 | >20 | N.A. |
| **27** | NH_2_ | Et | 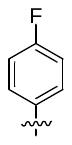 | 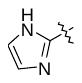 | 5-10 | >25 |
| **56** | NH_2_ | 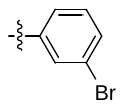 | H | 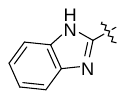 | 5-10 | 5-25 |
| **57** | NH_2_ | 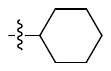 | H | 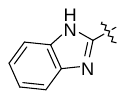 | >20 | N.A. |
| **58** | NH_2_ | iPr | H | 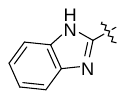 | >20 | N.A. |
| **59** | NH_2_ | 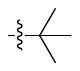 | H | 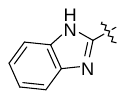 | >20 | N.A. |
| **60** | NH_2_ | 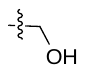 | H | 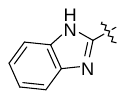 | >20 | N.A. |
| **61** | NH_2_ | 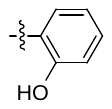 | H | 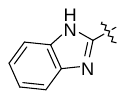 | 10-20 | >25 |
| **62** | NH_2_ | 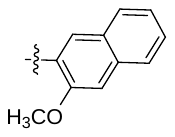 | H | 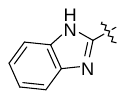 | 1-5 | >25 |
| **63** | NH_2_ | 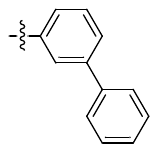 | H | 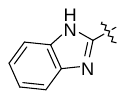 | >20 | N.A. |
| **64** | NH_2_ | 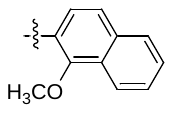 | H | 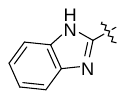 | >10 | N.A. |
| **65** | NH_2_ | 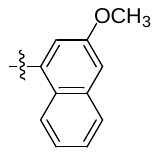 | H | 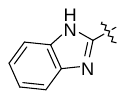 | 1-5 | >25 |
| **66** | NH_2_ | 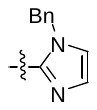 | H | 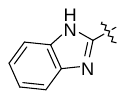 | 10-20 | >25 |
| **67** | NH_2_ | 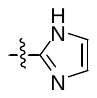 | H | 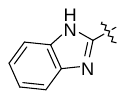 | >10 | N.A. |
| **68** | NH_2_ | 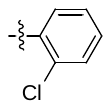 | H | 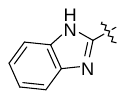 | 10-20 | N.A. |
| **69** | NH_2_ | 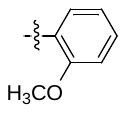 | H | 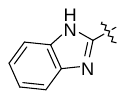 | 1-5 | >25 |
| **70** | NH_2_ | Et | H | 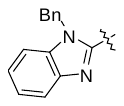 | >20 | N.A. |
| **71** | NH_2_ | Et | 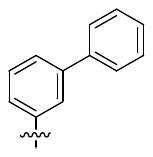 | 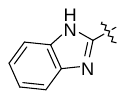 | 5-10 | >25 |
| **72** | NH_2_ | Et | 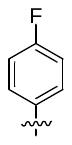 | 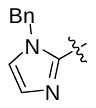 | >5 | >25 |
| **73** | NH_2_ | Et | 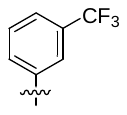 | 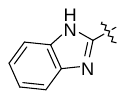 | 5-10 | 5-25 |
| **74** | NH_2_ | Et | 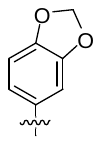 | 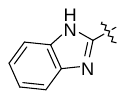 | >20 | >25 |
| **75** | NH_2_ | Et | 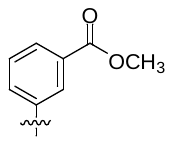 | 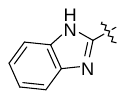 | 1-5 | 5-25 |
| **76** | NH_2_ | Et | 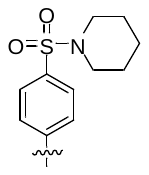 | 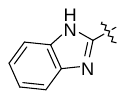 | >5 | 5-25 |
| **77** | NH_2_ | Et | 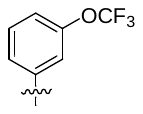 | 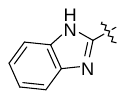 | 1-5 | >25 |
| **78** | NH_2_ | Et | 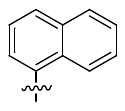 | 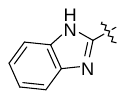 | 1-5 | >25 |
| **79** | OH | 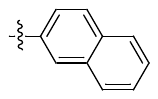 | H | 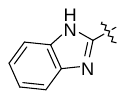 | 5-10 | >25 |
| **80** | OH | 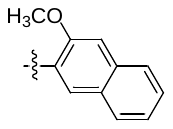 | H | 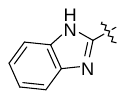 | 1-5 | >25 |
| **81** | OH | Et | H | 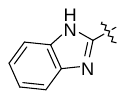 | >10 | N.A. |
| **82** | OH | 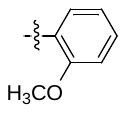 | H | 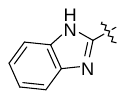 | 5-10 | 5-25 |
| **83** | OH | 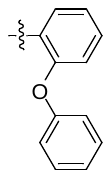 | H |  | 1-5 | <5 |
| **84** | OH |  | H |  | <1 | 5-25 |
| **85** | OH |  | H |  | 1-5 | >25 |
| **86** | OH |  | H |  | >40 | >25 |
| **87** | OH |  | H |  | 5-10 | 5-25 |
| **88** | OH |  | H |  | 1-5 | 5-25 |
| **89** | OH |  | H |  | 1-5 | 5-25 |
| **90** | OH | H |  |  | 1-5 | 5-25 |
| **91** | OH | H |  |  | 5-10 | >25 |
| **92** | OH | H |  |  | 1-5 | 5-25 |
| **93** | OH | Et |  |  | 10-20 | 5-25 |
| **94** | OH | Et |  |  | 1-5 | <5 |
| **95** | OH | Et |  |  | >10 | N.A. |
| **96** | OH | Et |  |  | 1-5 | 5-25 |
| **97** | OH | Et |  |  | 1-5 | <5 |
| **98** | NH_2_ | Et |  |  | 1-5 | N.A. |
| **99** | OH | Et |  |  | 1-5 | 5-25 |
| **100** | OH | Et |  |  | 1-5 | 5-25 |
| **101** | OH | Et |  |  | >20 | N.A. |
| **102** | OH | Et |  |  | 1-5 | <5 |
| **103** | OH | Et |  |  | >20 | N.A. |
| **104** | OH | Et |  |  | 1-5 | <5 |
| **105** | OH | Et |  |  | 10-20 | >25 |
| **106** | OH |  |  |  | 1-5 | 5-25 |
| **108** | OH | Et |  |  | >20 | >25 |
| **109** | OH | Et |  |  | <1 | <5 |
| **110** | OH | Et |  |  | <1 | <5 |
| **111** | OH | Et |  |  | <1 | 5-25 |
| **112** | OH | Et |  |  | >20 | N.A. |
| **113** | OH | Et |  |  | <1 | <5 |
| **114** | OH | Et |  |  | <1 | <5 |
| **115** | OH | Et |  |  | >20 | N.A. |
| **116** | OH | Et |  |  | >20 | N.A. |

**Miscellaneous targets**:

| **28** |  | >20 | >25 |
| --- | --- | --- | --- |
| **117** |  | >20 | N.A. |
| **107** |  | 10-20 | >25 |

*N.A. = not assessed

**Supplementary table 4: Structures and MUS81 *in vitro* and in cellulo IC_50_ values of the compound 2 and its analogues**

| **compound numbers** | **R^1^** | **R^2^** | **R^3^** | **R^4^** | **In vitro IC_50_ interval (μM)** | **Cell based IC_50_ interval (μM)** |
| --- | --- | --- | --- | --- | --- | --- |
| **36** |  | H |  |  | 5-10 | <5 |
| **32** |  | H |  |  | <1 | <1 |
| **35** |  | CH_3_ |  |  | >10 | <5 |
| **39** |  | H |  |  | 5-10 | 5-25 |
| **38** |  | H |  |  | >10 | <5 |
| **118** |  | H |  |  | 5-10 | <5 |
| **37** |  | H |  |  | >10 | 5-25 |
| **40** |  | H |  |  | >10 | 5-25 |
| **41** |  | H |  |  | 1-5 | <5 |
| **42** |  | H |  |  | 5-10 | <5 |
| **43** |  | H |  |  | <1 | <5 |
| **44** |  | H |  |  | >10 | 5-25 |
| **119** |  | H |  |  | >10 | <5 |
| **120** |  | H |  |  | >10 | N.A. |
| **121** |  | H |  |  | >10 | N.A. |
| **122** |  | H |  |  | >10 | <5 |
| **123** |  | H |  |  | >10 | N.A. |
| **140** |  | H |  |  | >10 | N.A. |
| **2** |  | H |  |  | <1 | 5-25 |
| **124** |  | H |  |  | >10 | 5-25 |
| **30** |  | H |  |  | 5-10 | <5 |
| **125** |  | H |  |  | >10 | N.A. |
| **126** |  | H |  |  | >10 | >25 |
| **127** |  | H |  |  | >10 | >25 |
| **128** |  | H |  |  | <1 | 5-25 |
| **29** |  | H |  |  | 1-5 | 5-25 |
| **129** |  | H |  |  | 1-5 | >25 |
| **34** |  | H |  |  | >10 | <5 |
| **130** |  | H |  |  | >10 | N.A. |
| **131** |  | H |  |  | 1-5 | <5 |
| **132** |  | H |  |  | >10 | <5 |
| **33** |  | H |  |  | 1-5 | <5 |
| **31** |  | H |  |  | 1-5 | 5-25 |
| **133** |  | H |  |  | 1-5 | 5-25 |
| **134** |  | H |  |  | 1-5 | <5 |
| **142** |  | H |  |  | 1-5 | <5 |
| **48** |  | H |  |  | >10 | 5-25 |
| **49** |  | H |  |  | >10 | 5-25 |
| **51** |  | H |  |  | >10 | N.A. |
| **46** |  | H | Methyl |  | >10 | >25 |
| **50** |  | H |  |  | 5-10 | <5 |
| **55** |  | H |  |  | <1 | <5 |
| **135** |  | H |  |  | >10 | <5 |
| **52** |  | H |  |  | 5-10 | <5 |
| **136** |  | H |  |  | >10 | >25 |
| **53** |  | H |  |  | 10-20 | 5-25 |
| **137** |  | H |  |  | >10 | >25 |
| **54** |  | H |  |  | 5-10 | 5-25 |
| **47** |  | H |  |  | >10 | N.A. |
| **139** |  | H |  |  | >10 | >25 |
| **138** |  | H |  |  | >10 | >25 |
| **143** |  | H |  |  | >10 | >25 |

**Miscellaneous targets**:

| **141** |  | >10 | <5 |
| --- | --- | --- | --- |
| **45** |  | >20 | >25 |

*N.A. = not assessed

**Supplementary table 5: List of DNA substrates**

DNA substate used for *in vitro* nuclease assay, EMSA and BLI.

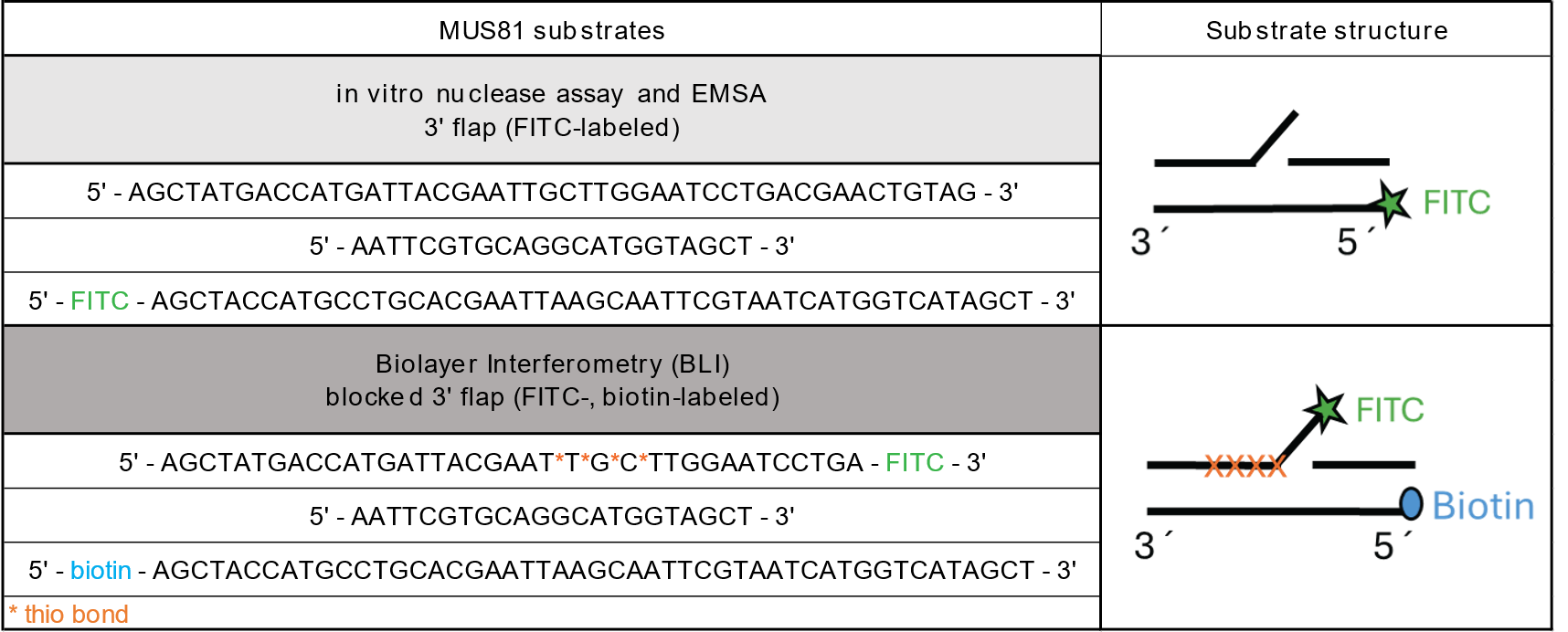
