## Supporting information_NMR spectra for "Discovery of two structurally distinct classes of inhibitors targeting the nuclease MUS81 and enhancing efficacy of chemotherapy in cancer cells"

**[1]H NMR, [13]C NMR and HRMS spectra of compound S1-S108**

[1]H (500 MHz) and [13]C NMR (126 MHz) spectra of **S1** in chloroform-*d*

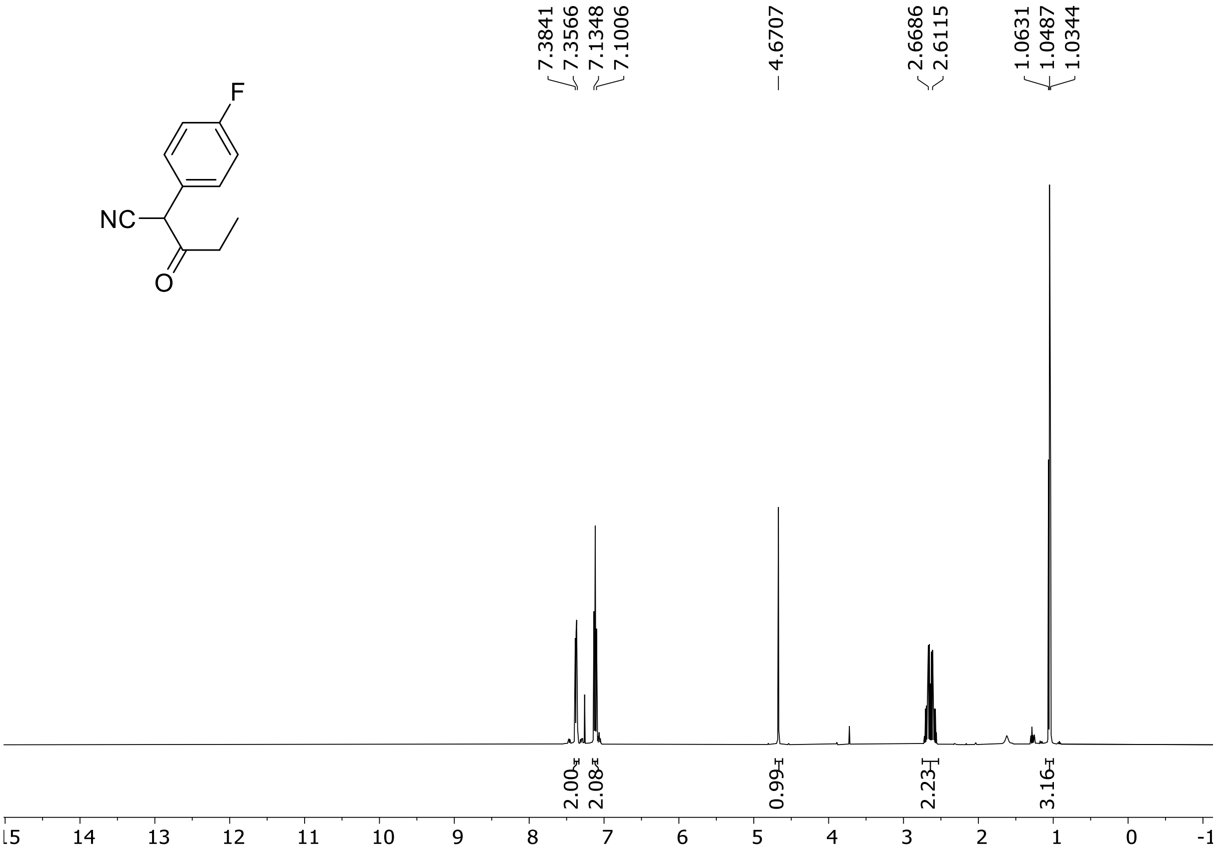

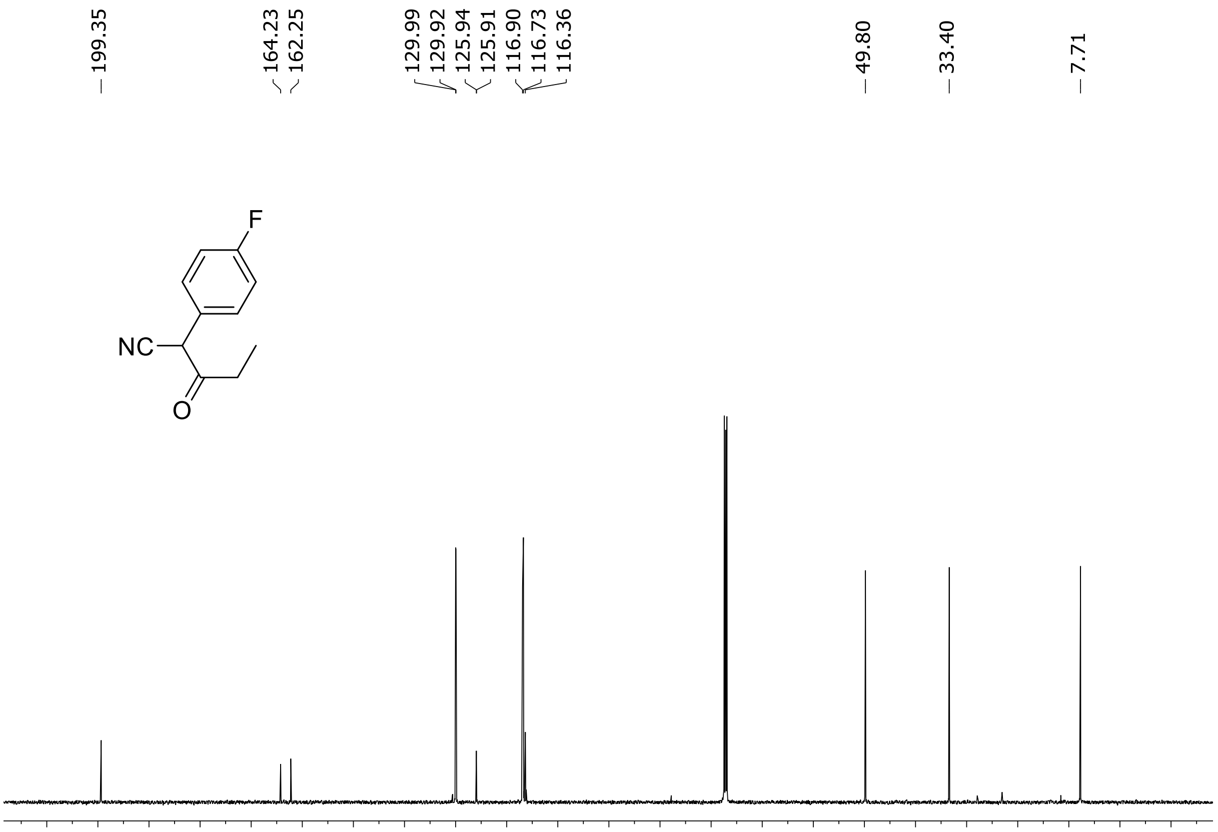

HRMS spectrum of **S1**

$C_{11}H_{10}FNO$ **mono m/z = 191,0746**

**ESI -** (MMI)
nitrogen flow 5 L/min, gas temperature 300°C, vaporizer 250°C,
nebulizer 45 psi, Vcap -2000 V, skimmer **65** V, fragmentor **40** V,
dissolved in MeOH

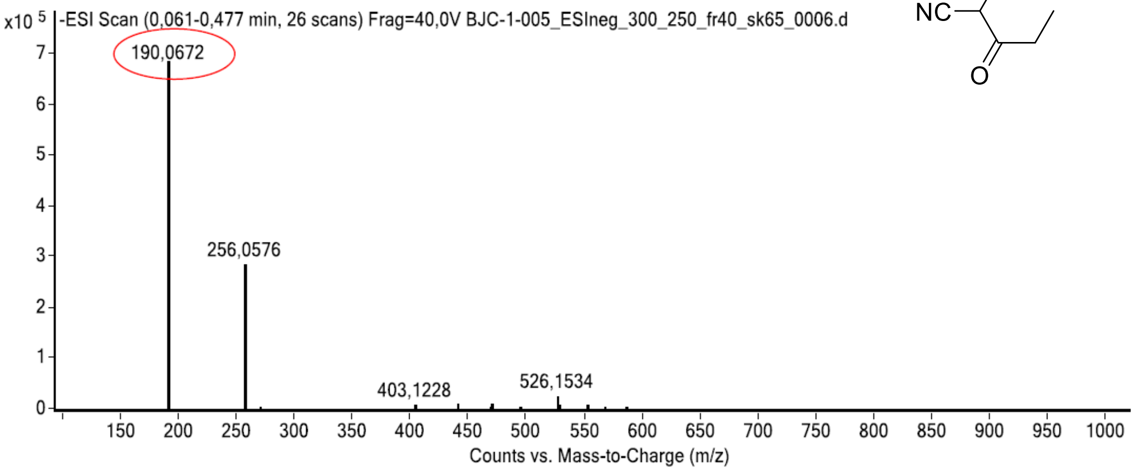

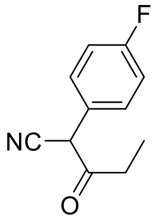

calculated mass:
$[M\text{-}H]^-$ = 190,0674 observed: $[M\text{-}H]^-$ = 190,0672 max. mass error = 1,1 ppm

[1]H (500 MHz) and [13]C NMR (126 MHz) spectra of **S2** in chloroform-*d*

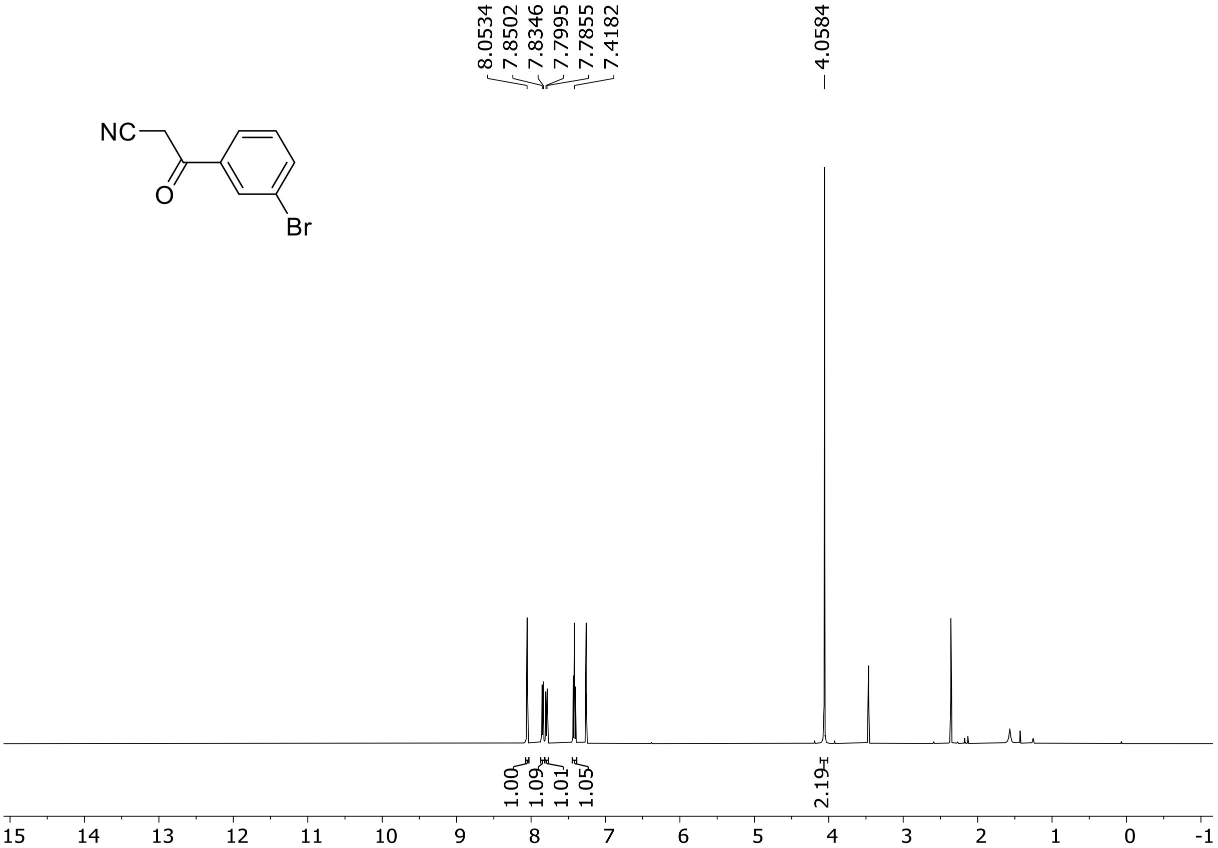

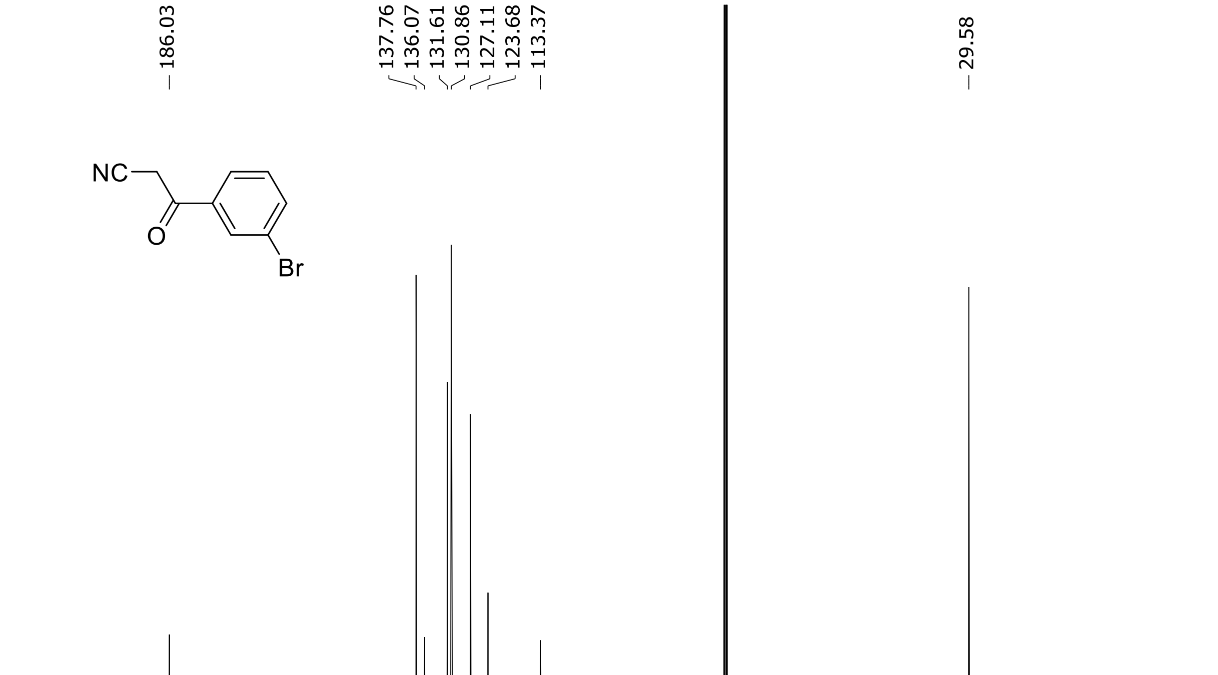

[1]H (500 MHz) and [13]C NMR (126 MHz) spectra of **S3** in chloroform-*d*

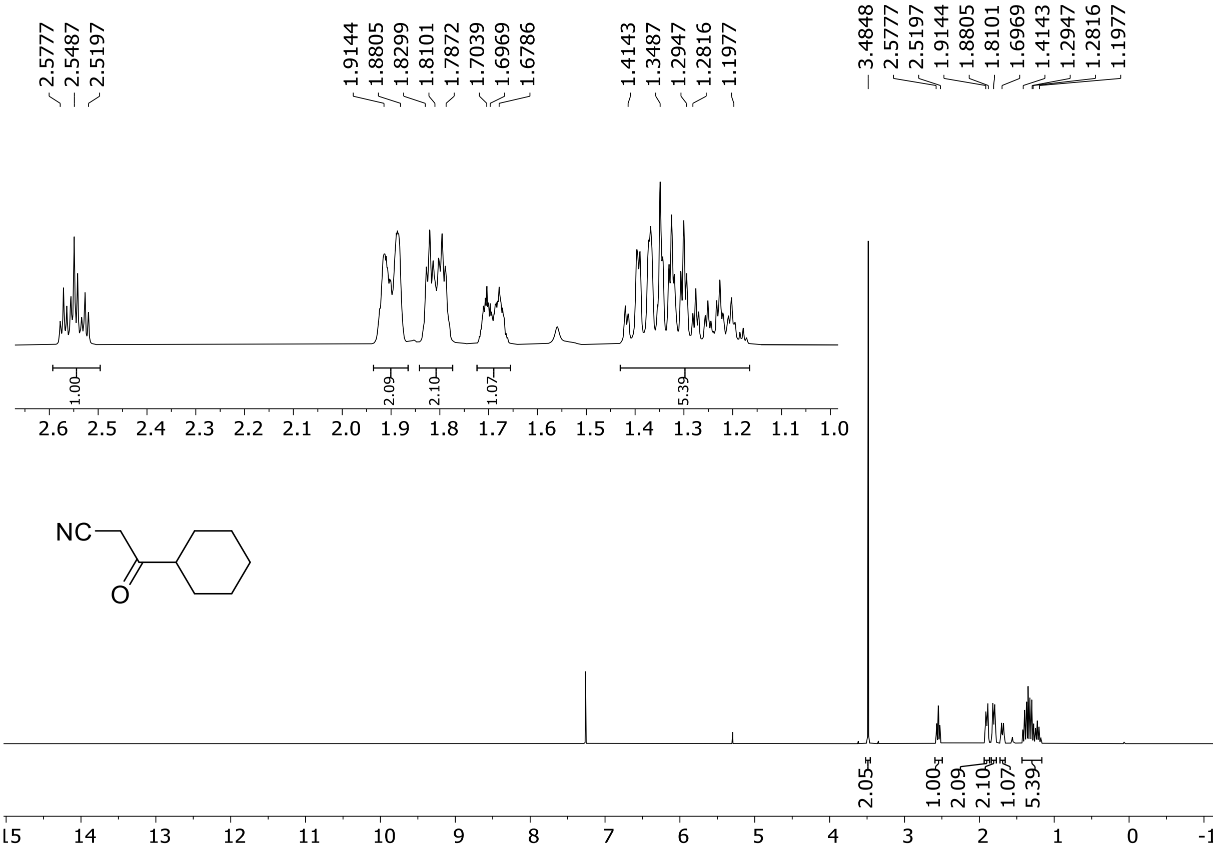

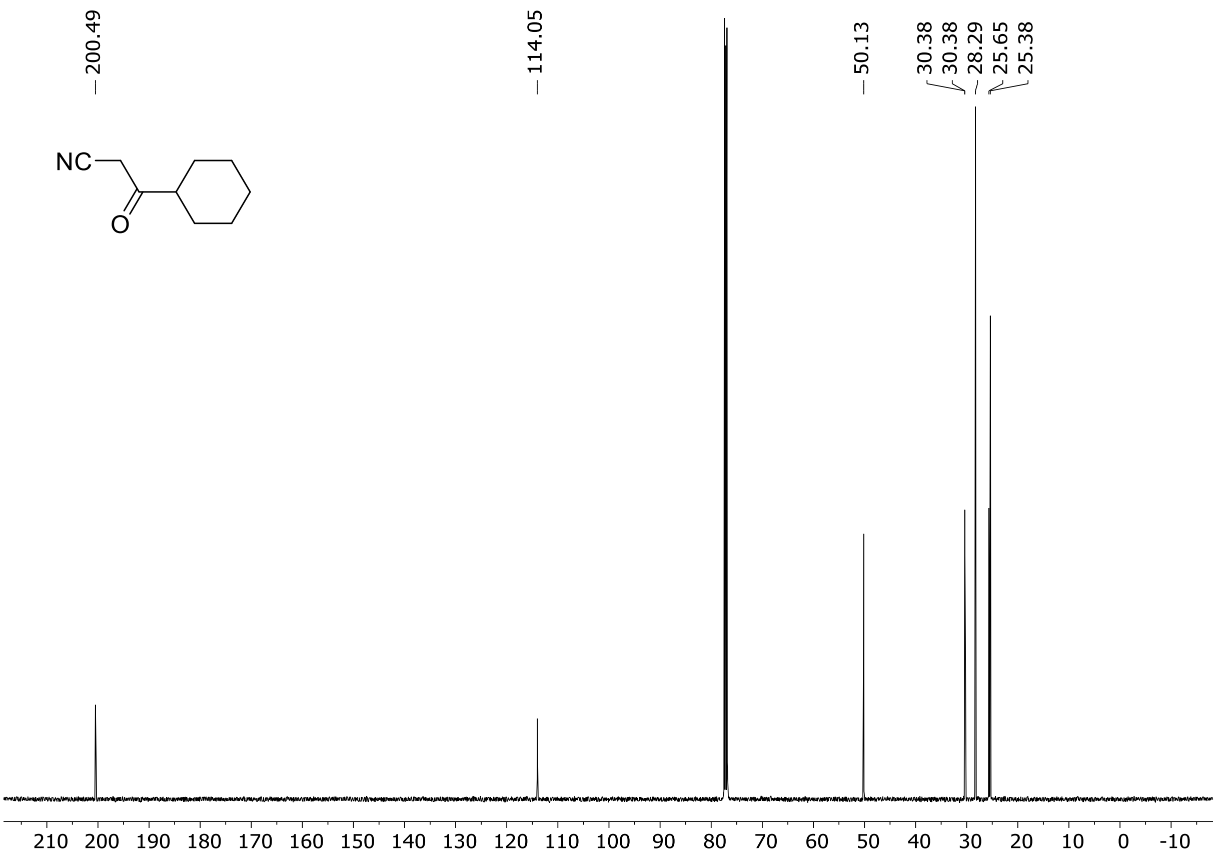

HRMS spectrum of **S3**

**$C_9H_{13}NO$** **mono m/z = 151.0997**

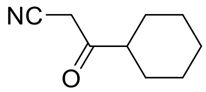

**APCI +** (MMI)
nitrogen flow 5 L/min, gas temperature 300°C, nebulizer 45 psi, skimmer 60 V, fragmentor 25 V, dissolved in methanol

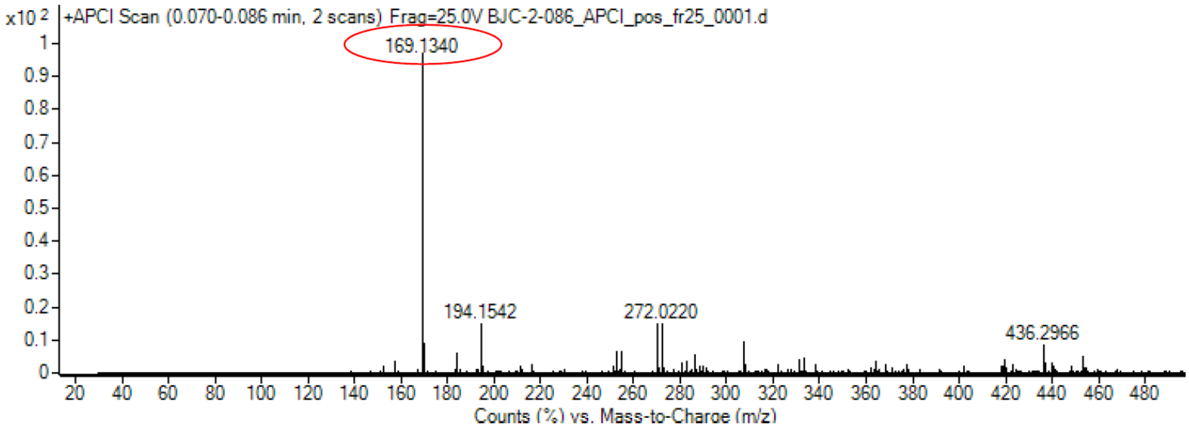

calculated mass: $[M+NH_4]^+$ = 169.1335 observed: $[M+H]^+$ = 169.1340 max. mass error = 2.9 ppm

FT-IR spectrum (neat) of **S3**

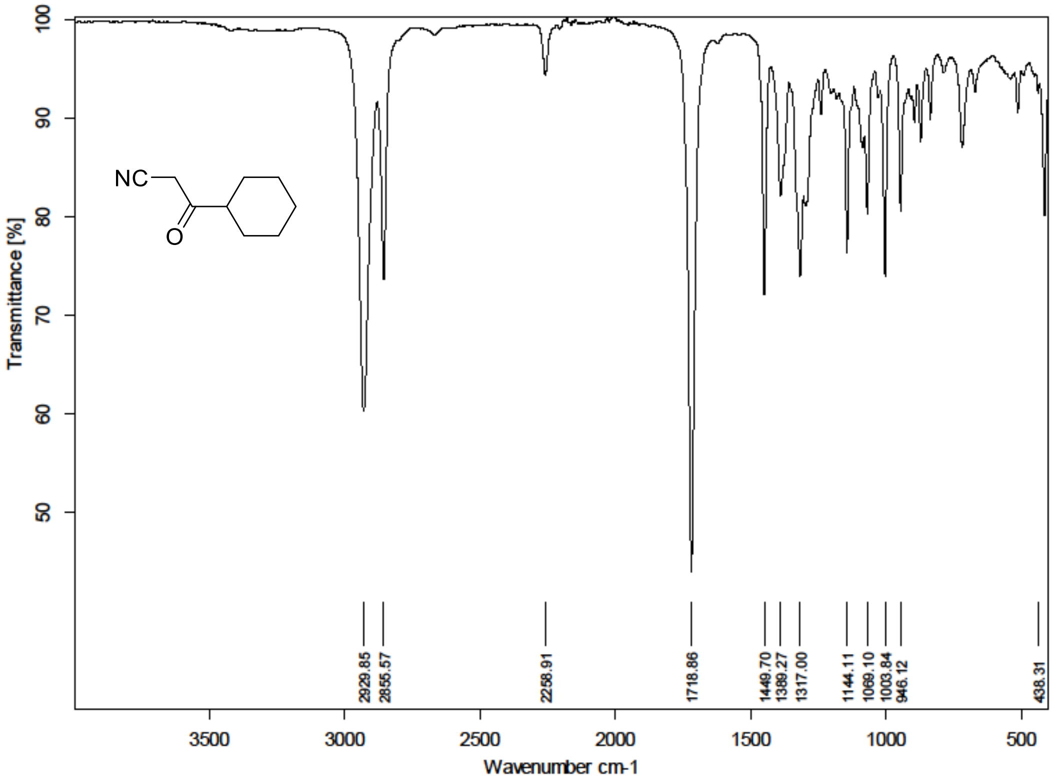

$^1$H (500 MHz) and $^{13}$C NMR (126 MHz) spectra of **S4** in chloroform-*d*

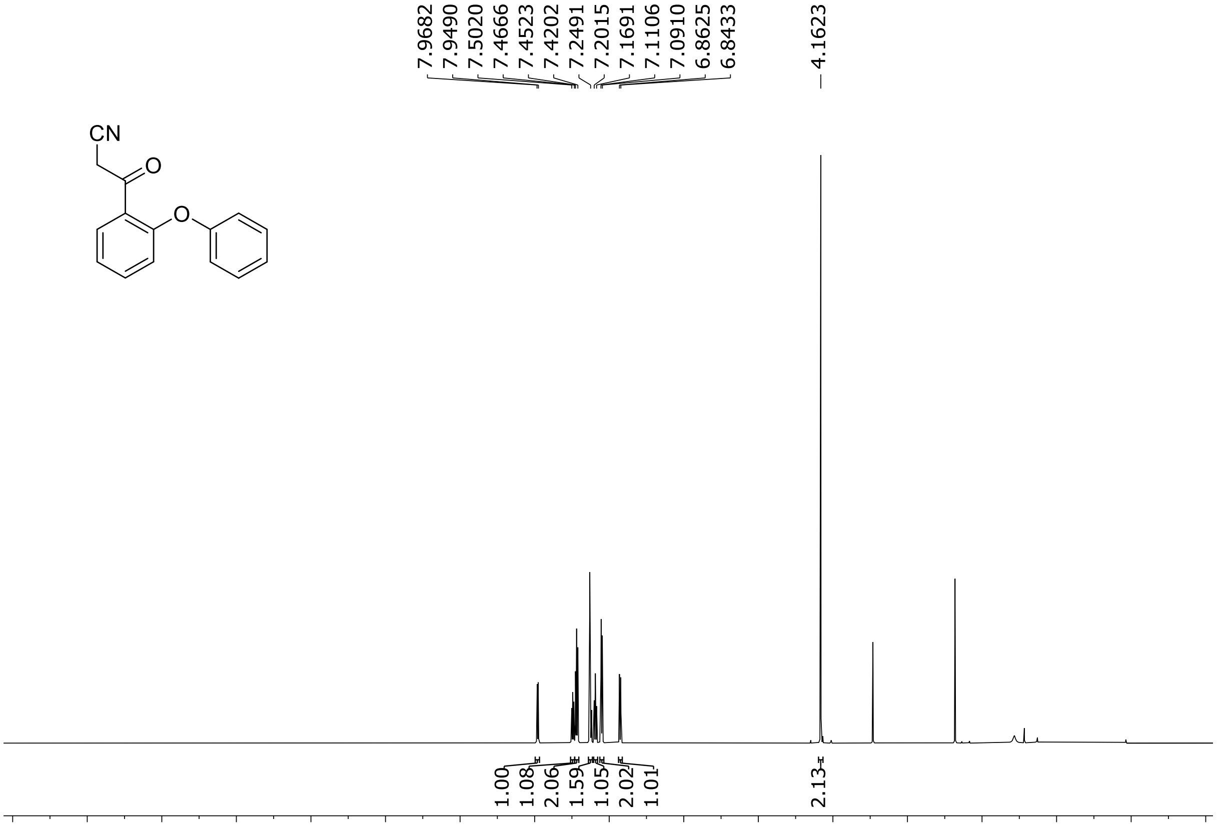

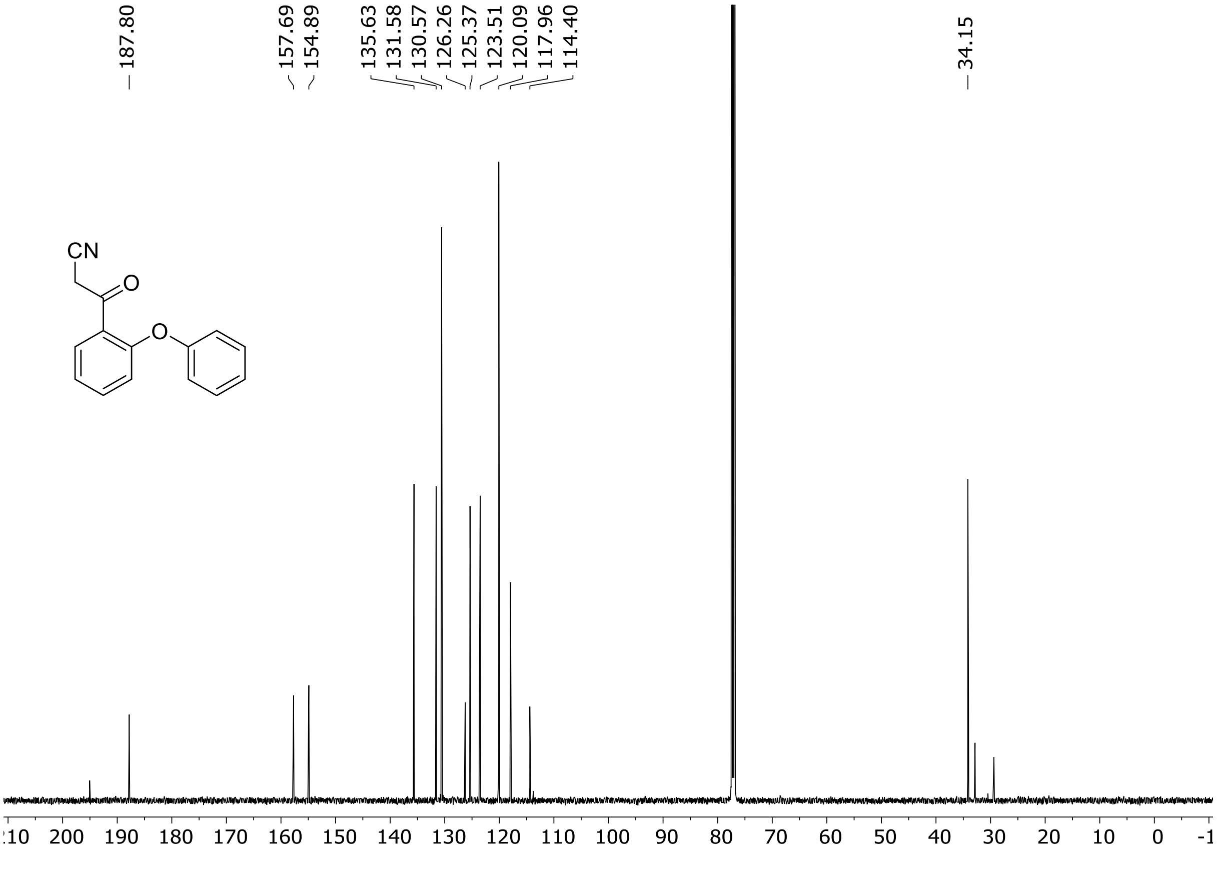

HRMS spectrum of **S4**

$C_{15}H_{11}NO_2$ **mono m/z = 237.079**

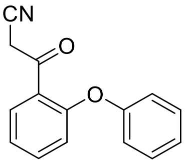

**APCI +** (MMI)
nitrogen flow 5 L/min, gas temperature 300°C, nebulizer 45 psi,
skimmer 60 V, fragmentor 23 V, dissolved in methanol

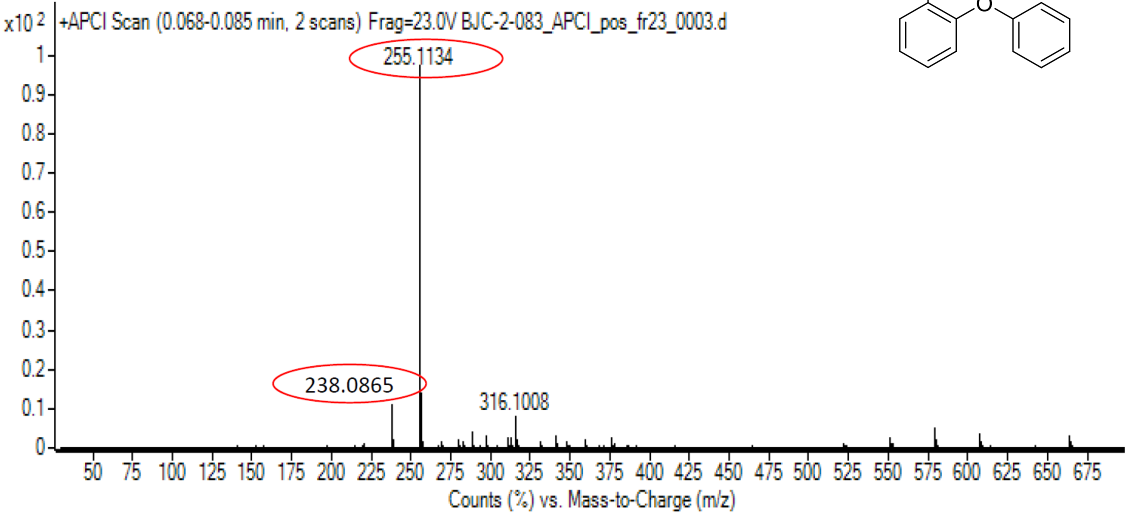

calculated mass: $[M+H]^+$ = 238.0863 observed: $[M+H]^+$ = 238.0865 max. mass error = 0.8 ppm

calculated mass: $[M+NH_4]^+$ = 255.1128 observed: $[M+NH_4]^+$ = 255.1134 max. mass error = 2.4 ppm

<sup></sup>[1]H (500 MHz) and [13]C NMR (126 MHz) spectra of **S5** in chloroform-*d*

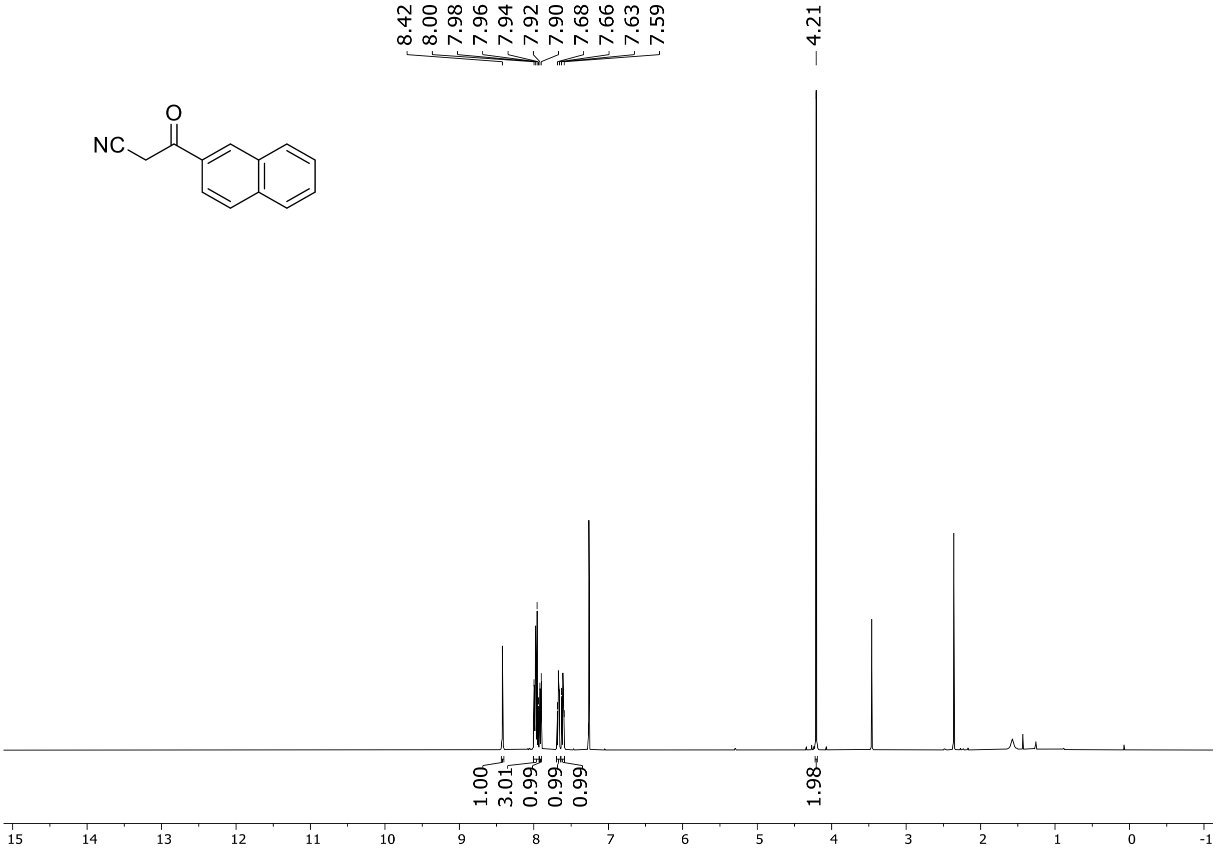

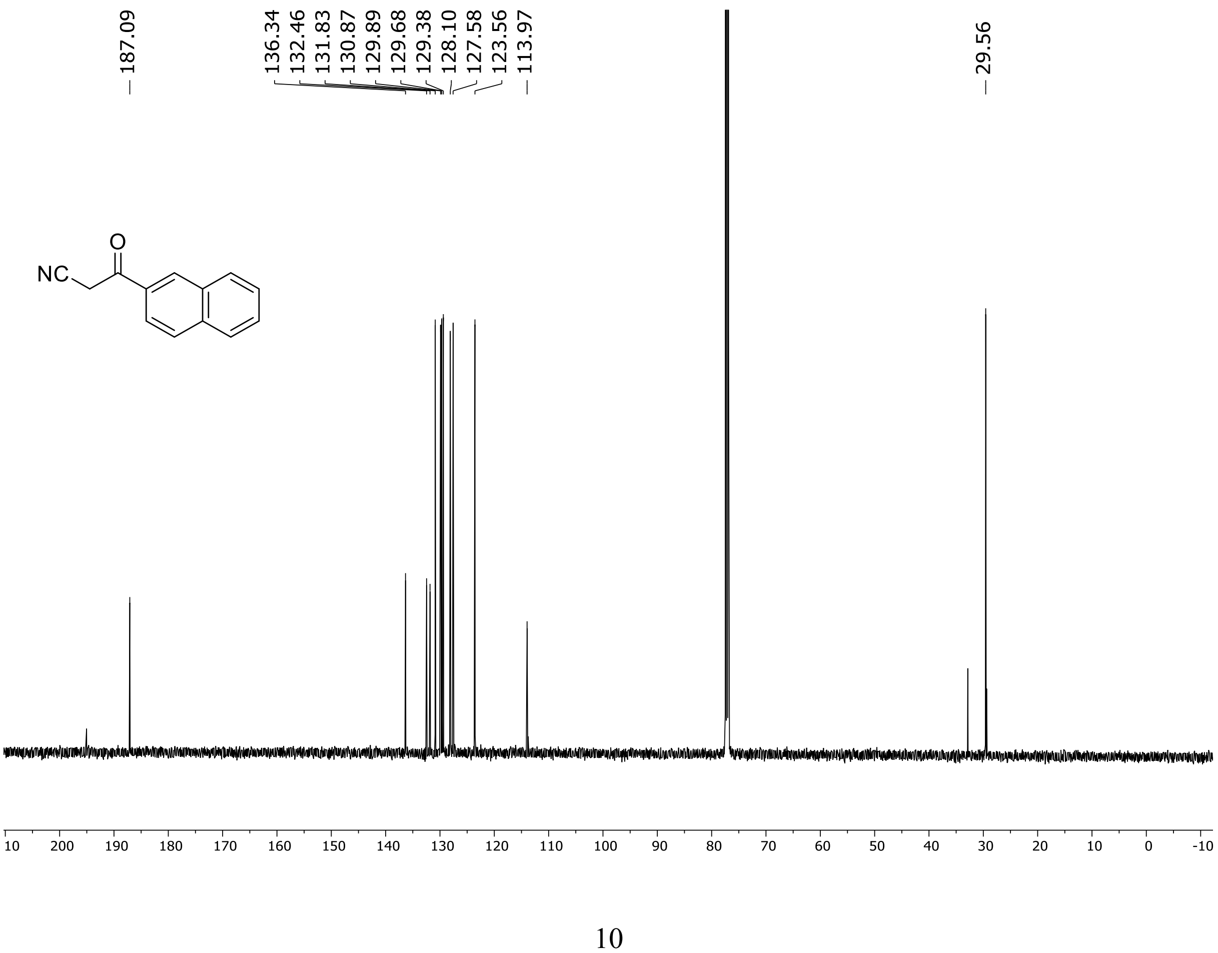

HRMS spectrum of **S5**

**$C_{13}H_9NO$ mono m/z = 195.0684**

**APCI -** (MMI)
nitrogen flow 5 L/min, gas temperature 300°C, nebulizer 45 psi,
skimmer 60 V, fragmentor 40 V, dissolved in methanol

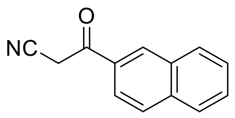

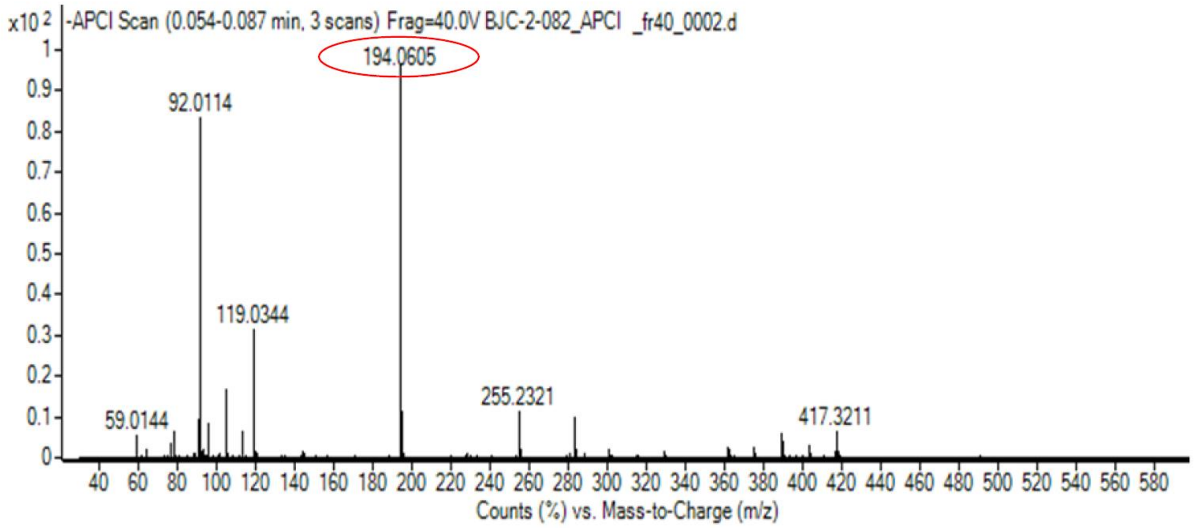

calculated mass: [M-H]$^-$ = 194.0611 observed: [M-H]$^-$ = 194.0605 max. mass error = 3.1 ppm

[1]H (500 MHz) and [13]C NMR (126 MHz) spectra of **S7** in chloroform-*d*

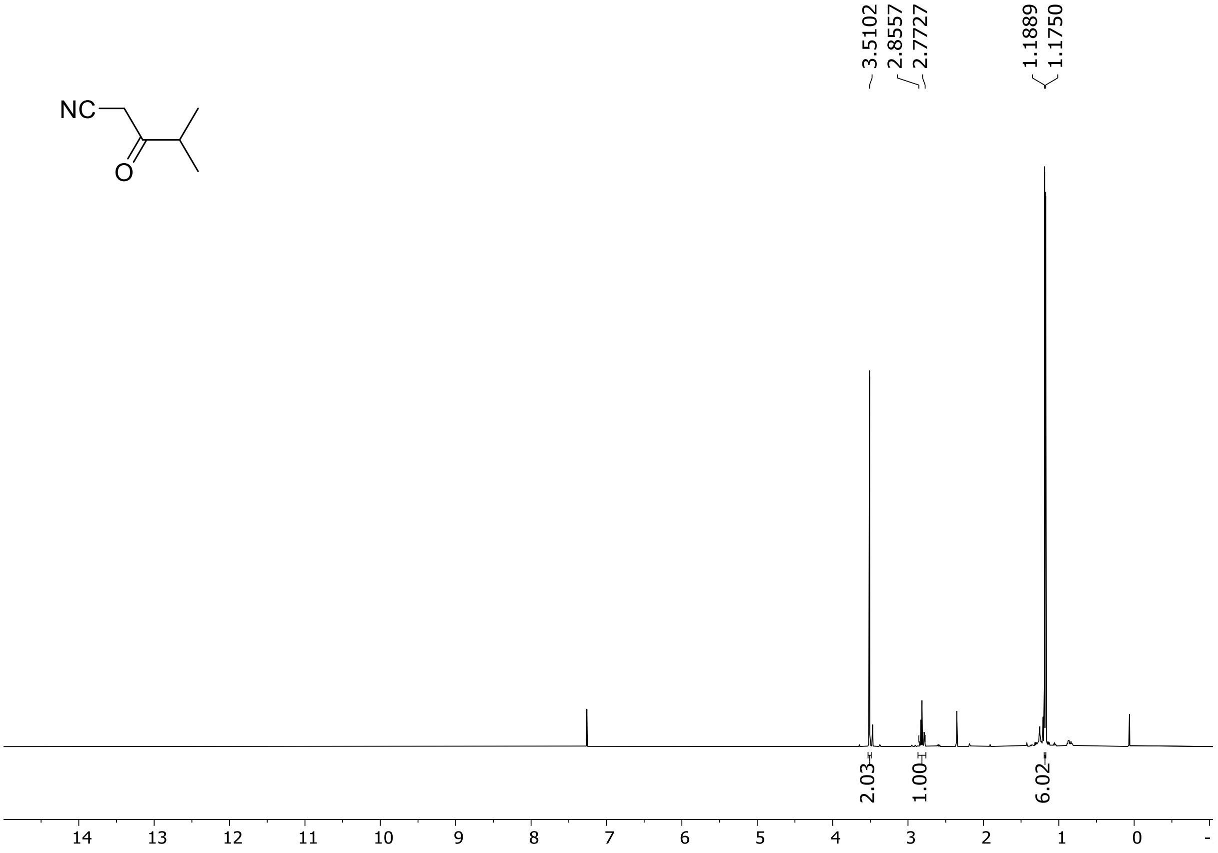

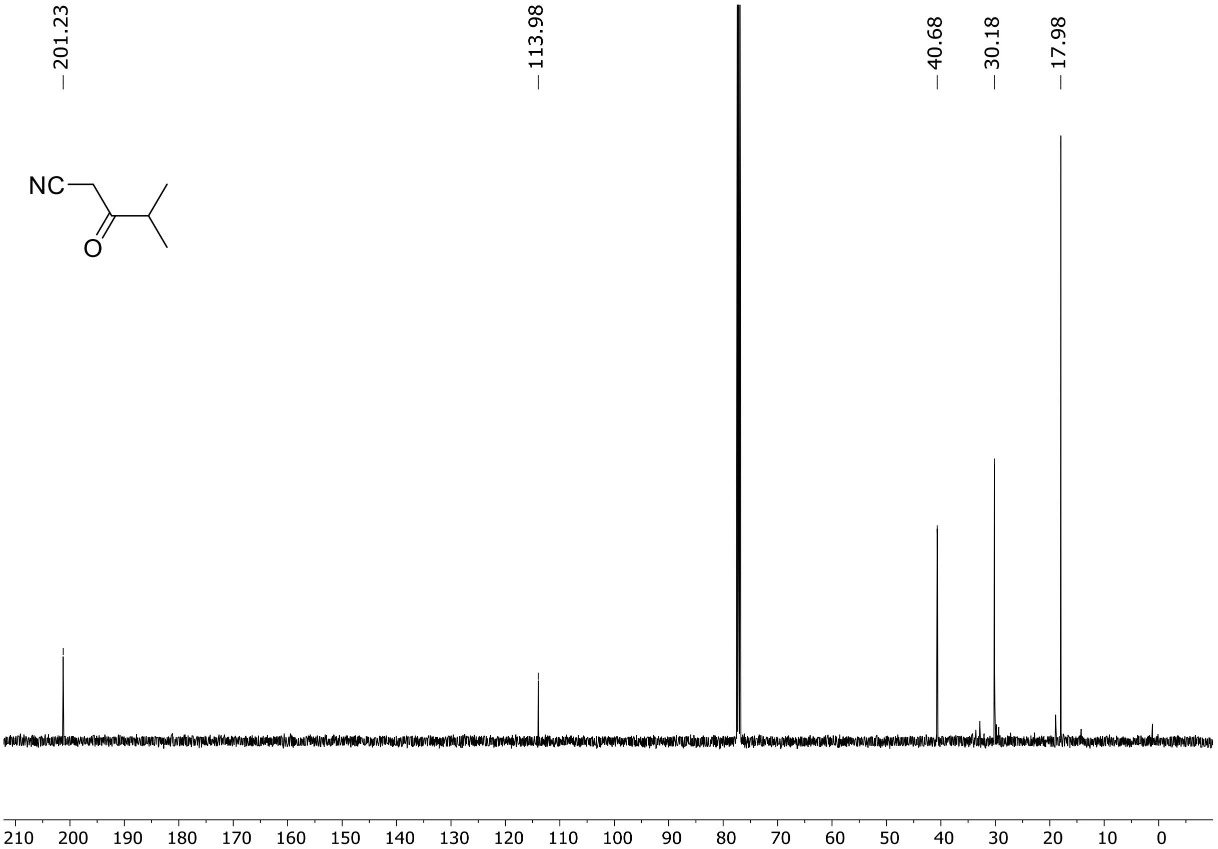

HRMS spectrum of **S7**

$C_6H_9NO$ **mono m/z = 111.0684**

**APCI -** (MMI)
nitrogen flow 5 L/min, gas temperature 300°C, nebulizer 45 psi, vaporizer 200°C
skimmer **65** V, fragmentor **40** V, dissolved in methanol

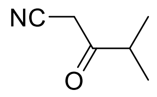

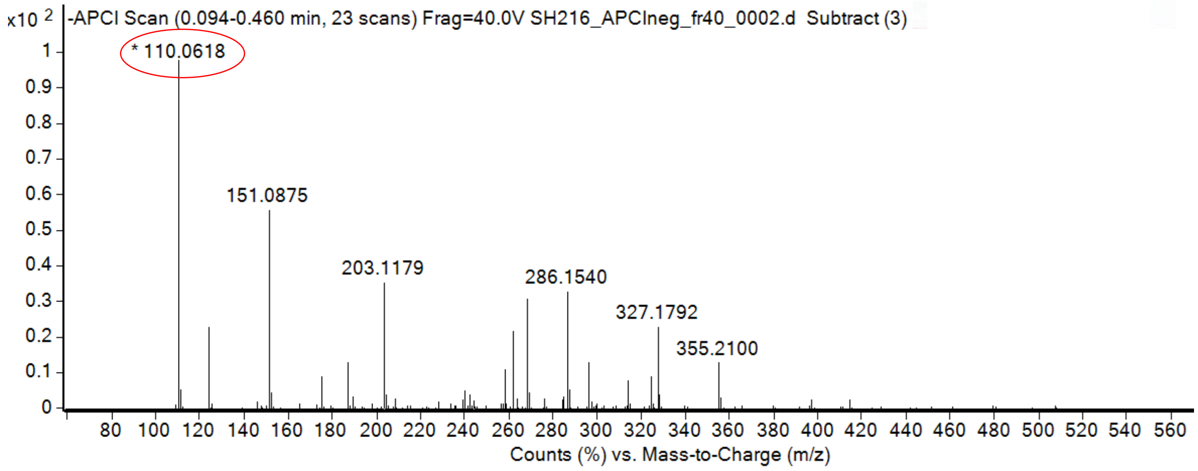

calculated mass: [M-H]$^-$ = 110.0611 observed: [M-H]$^-$ = 110.1618 max. mass error = 6.3 ppm

[1]H (500 MHz) and [13]C NMR (126 MHz) spectra of **S8** in chloroform-*d*

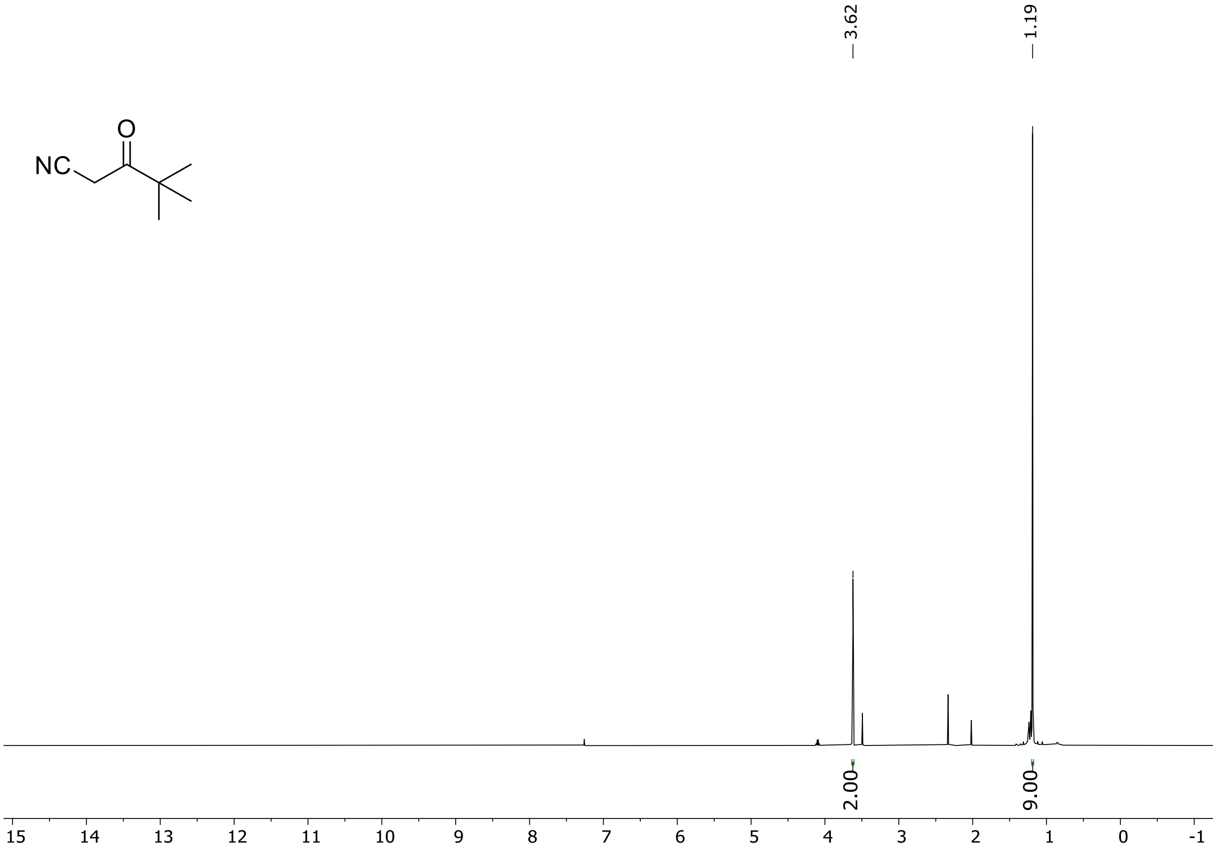

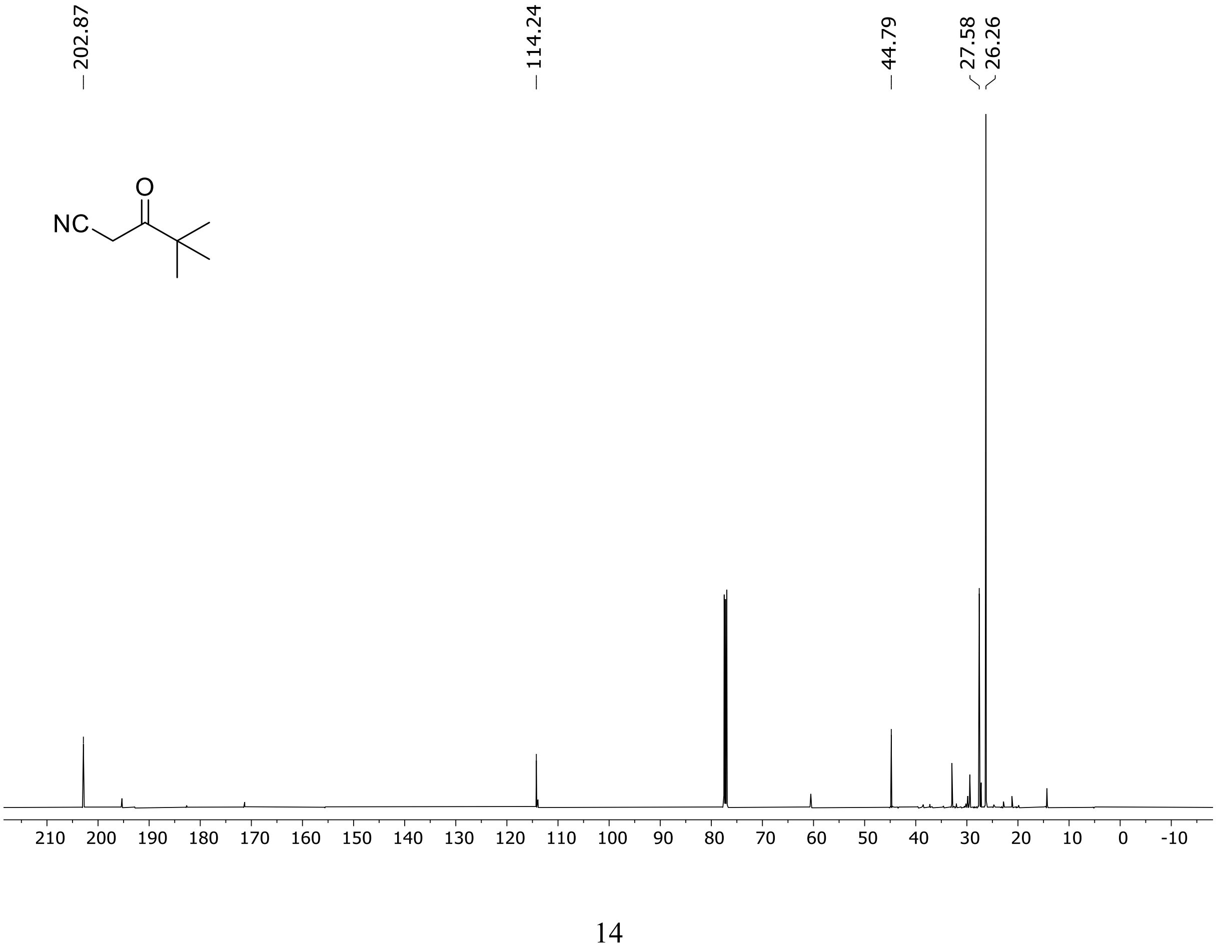

HRMS spectrum of **S8**

$C_7H_{11}NO$ **mono m/z = 125.0841**

**APCI -** (MMI)
nitrogen flow 5 L/min, gas temperature 300°C, nebulizer 45 psi, vaporizer 200°C
skimmer **65** V, fragmentor **27** V, dissolved in methanol

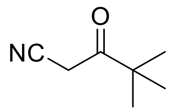

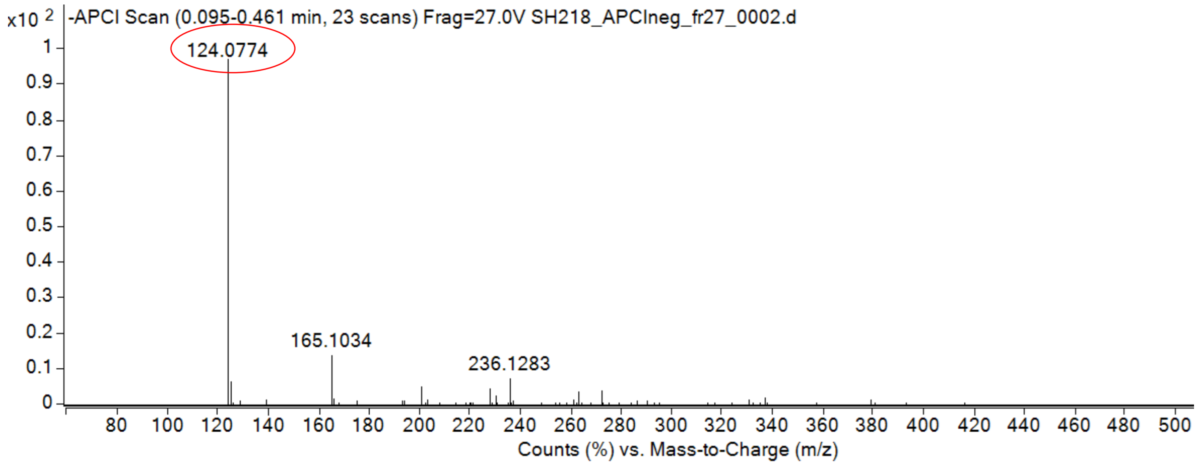

calculated mass: $[M-H]^-$ = 124.0768 observed: $[M-H]^-$ = 124.0774 max. mass error = 4.8 ppm

[1]H (500 MHz) and [13]C NMR (126 MHz) spectra of **S9** in chloroform-*d*

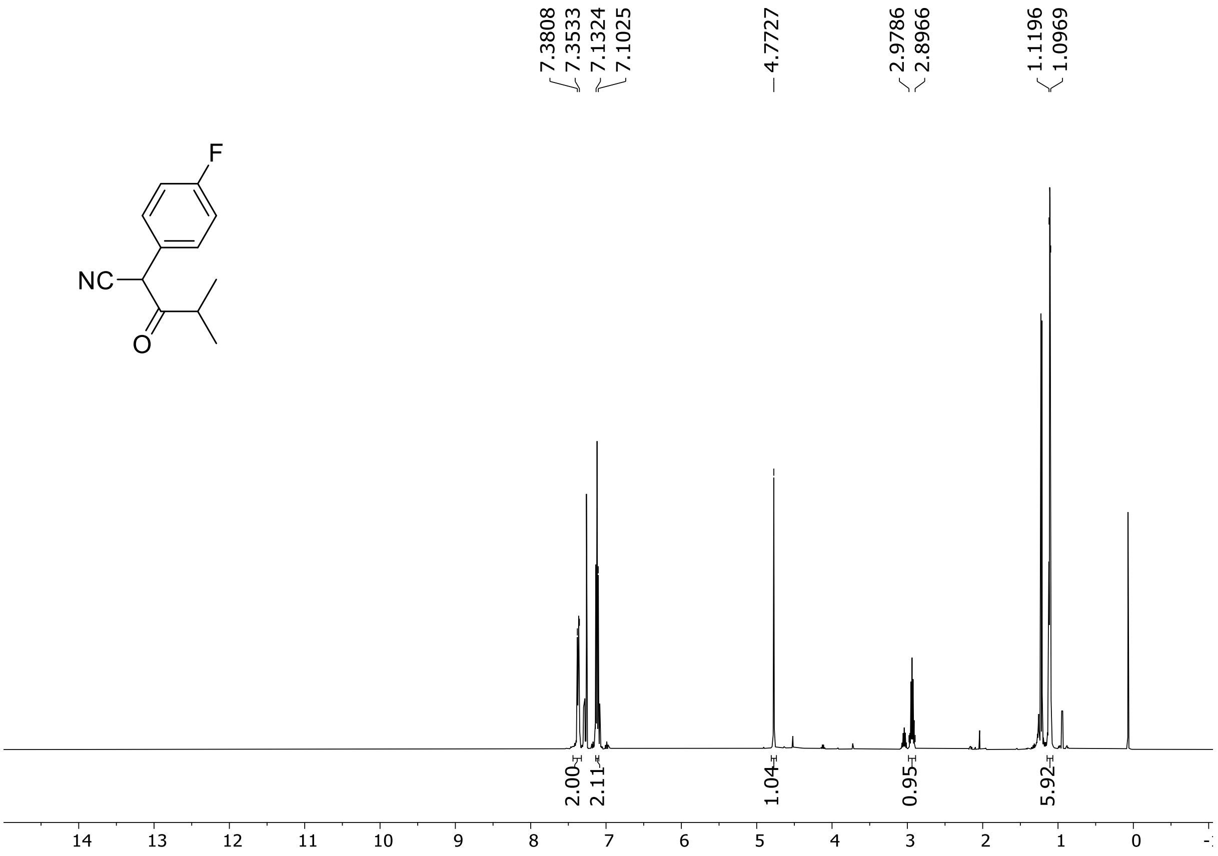

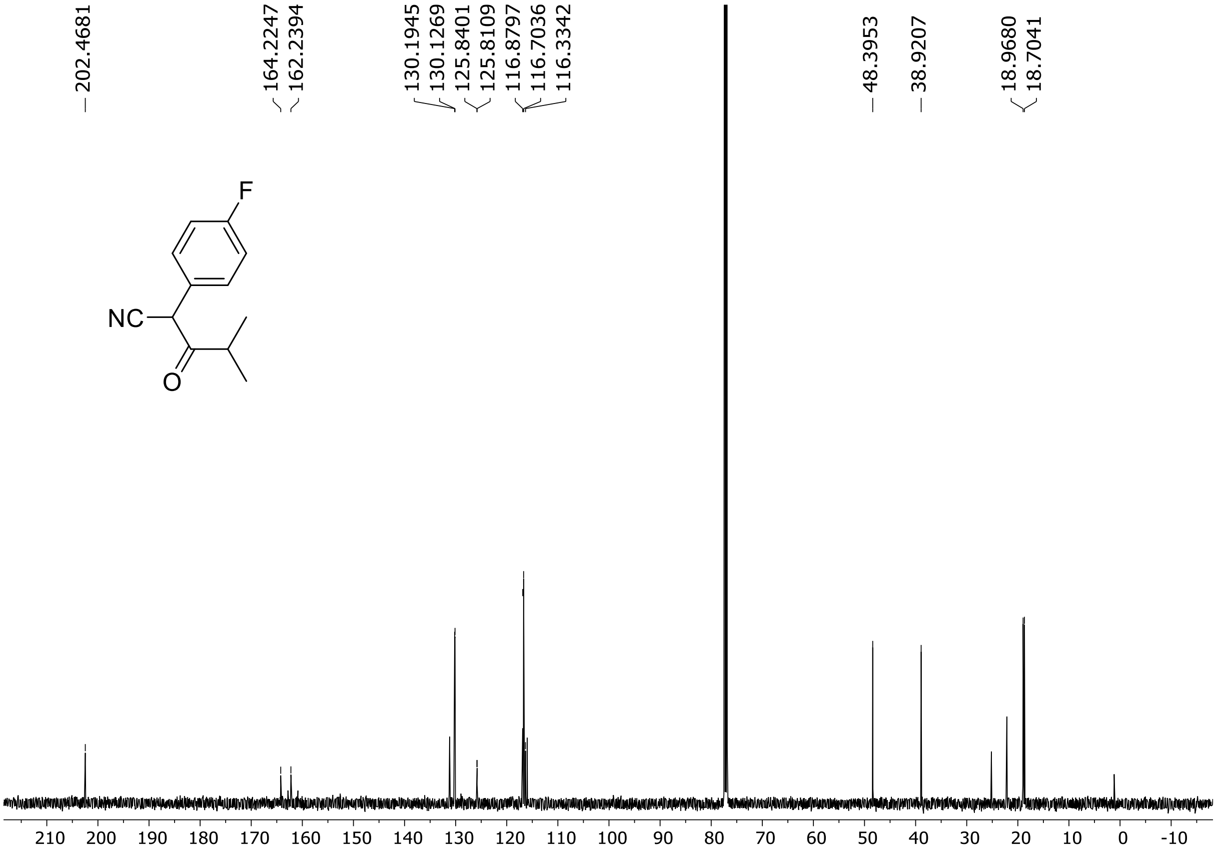

HRMS spectrum of **S9**

**$C_{12}H_{12}NOF$** **mono m/z = 205.0903**

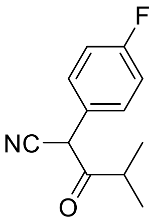

**APCI -** (MMI)
nitrogen flow 5 L/min, gas temperature 300°C, nebulizer 45 psi, vaporizer 200°C,
skimmer **65** V, fragmentor **36** V, dissolved in methanol

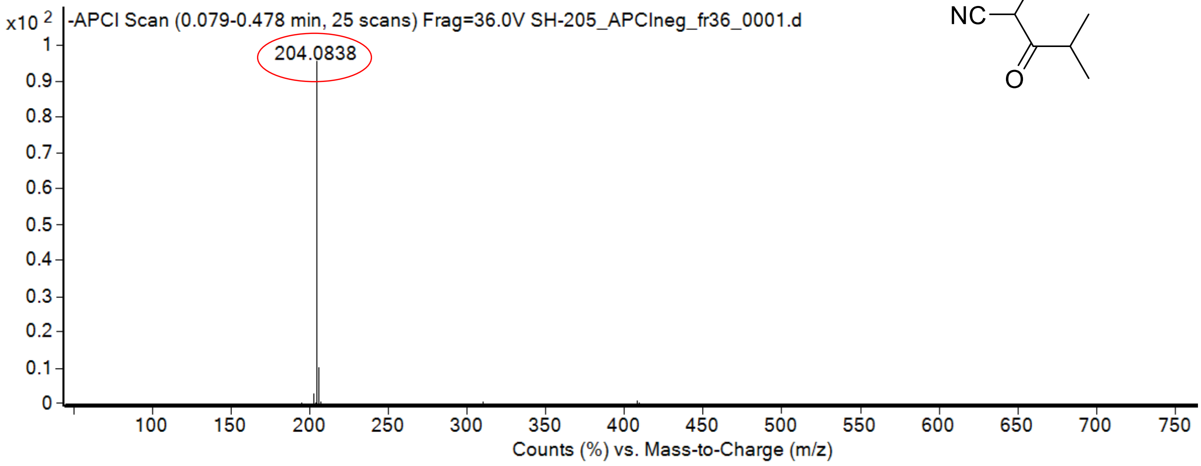

calculated mass: $[M\text{-}H]^-$ = 204.0830 observed: $[M+H]^+$ = 204.0838 max. mass error = 3.9 ppm

$^{1}H$ (500 MHz) and $^{13}C$ NMR (126 MHz) spectra of **S10** in chloroform-*d*

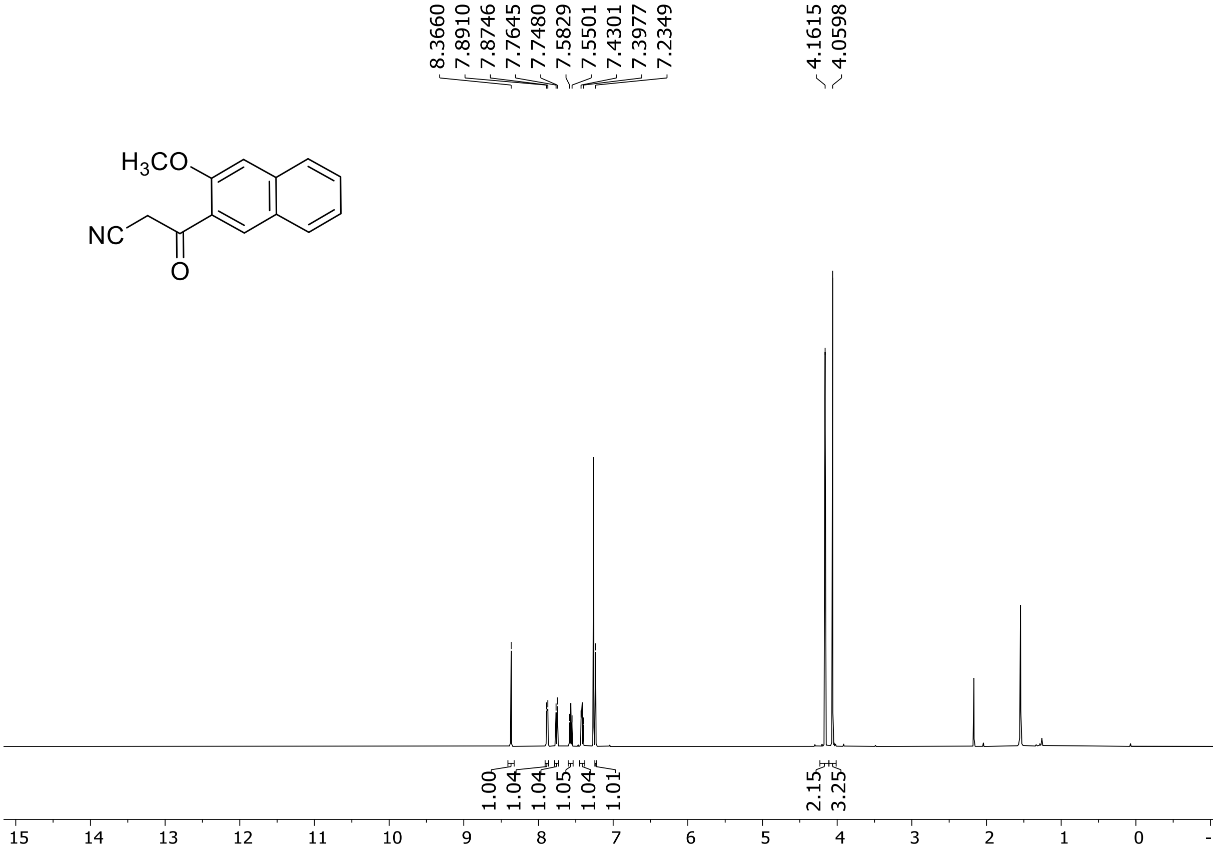

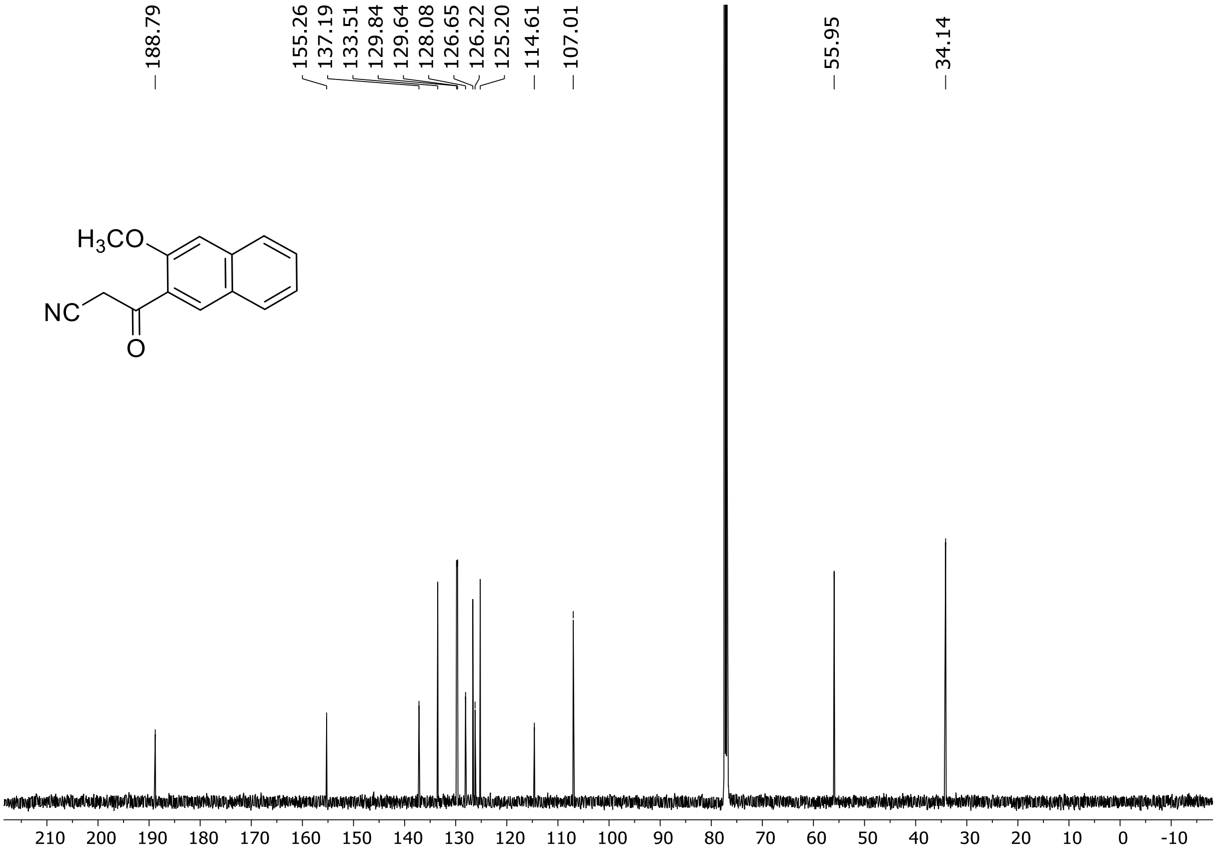

$^{1}H$ (500 MHz) and $^{13}C$ NMR (126 MHz) spectra of **S11** in chloroform-*d*

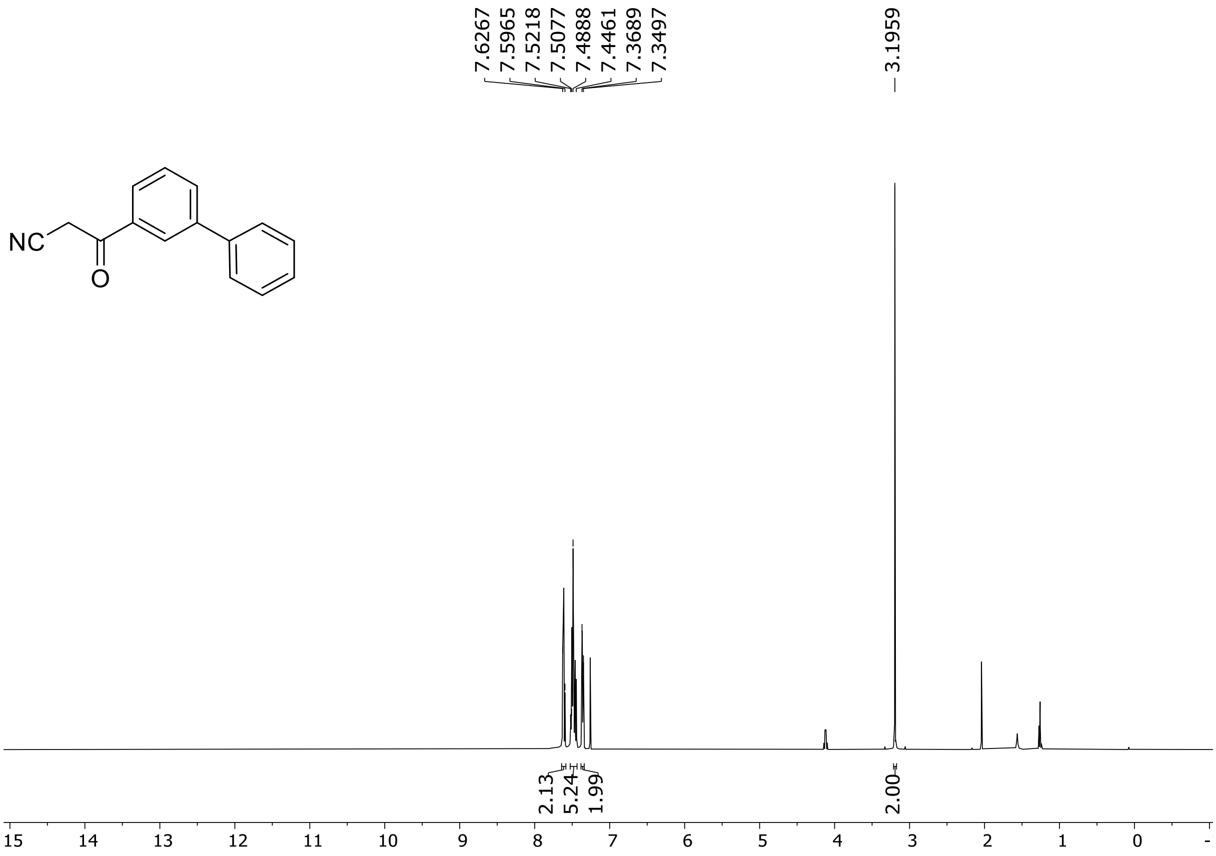

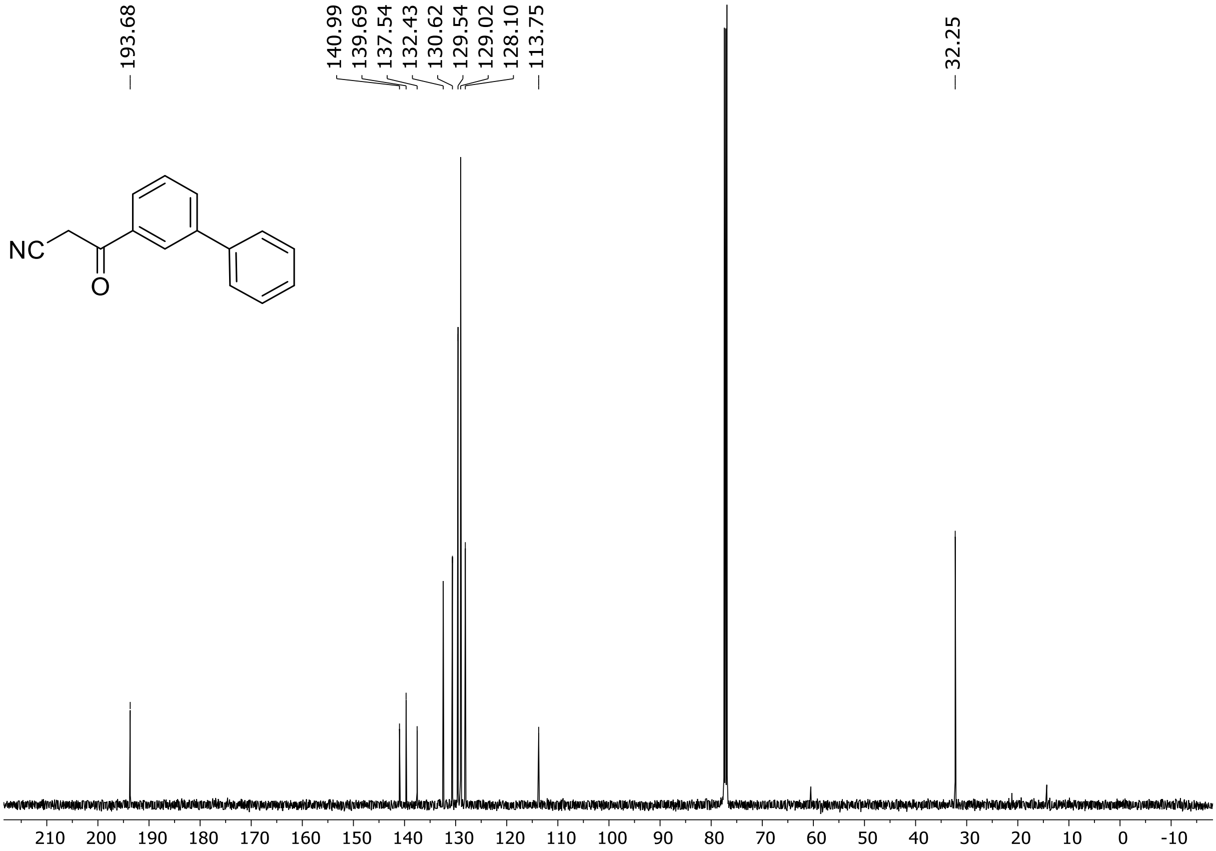

HRMS spectrum of **S11**

$C_{15}H_{11}NO$ **mono m/z = 221.0841**

**APCI -** (MMI)
nitrogen flow 5 L/min, gas temperature 300°C, nebulizer 45 psi, vaporizer 200°C
skimmer **65** V, fragmentor **15** V, dissolved in methanol

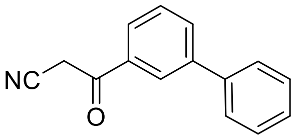

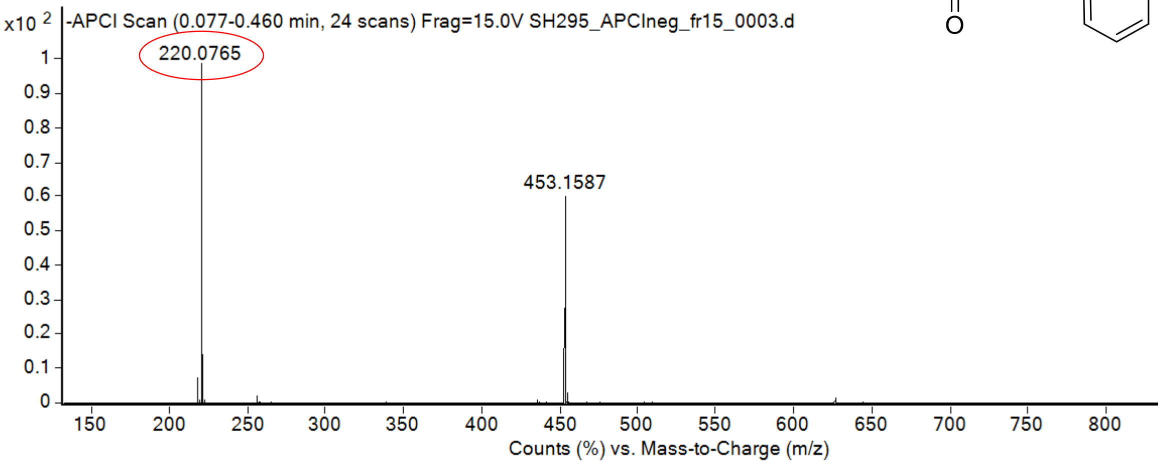

calculated mass: $[M-H]^-$ = 220.0768 observed: $[M-H]^-$ = 220.0765 max. mass error = 1.3 ppm

[1]H (500 MHz) and [13]C NMR (126 MHz) spectra of **S12** in chloroform-*d*

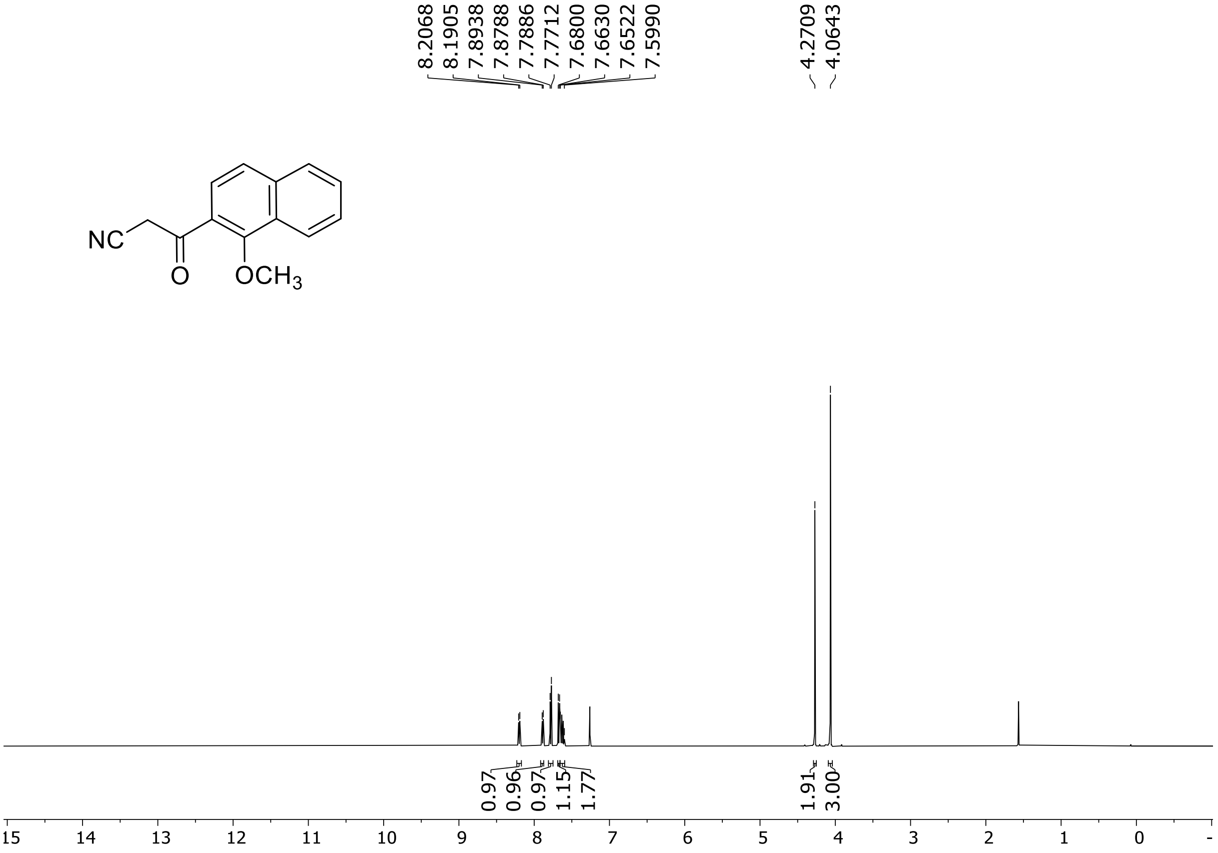

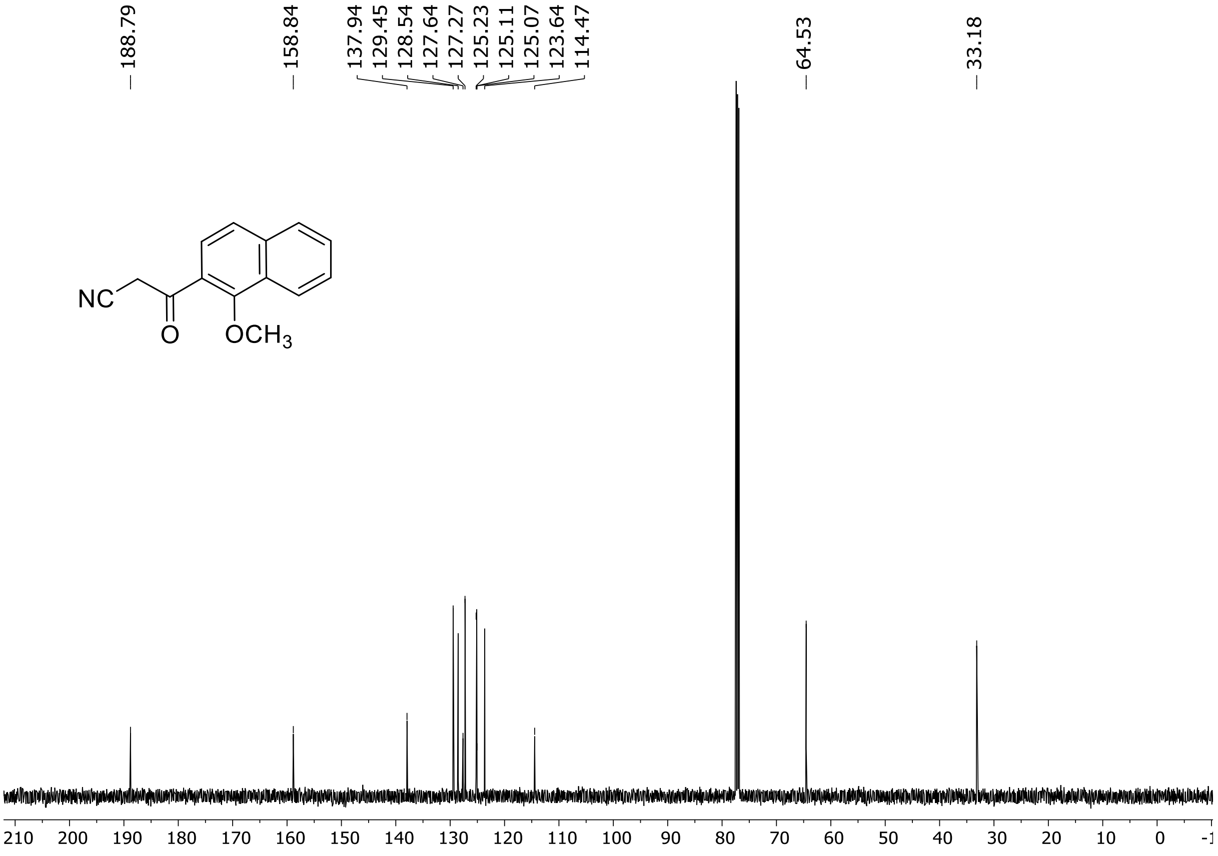

$^1H$ (500 MHz) and $^{13}C$ NMR (126 MHz) spectra of **S13** in chloroform-*d*

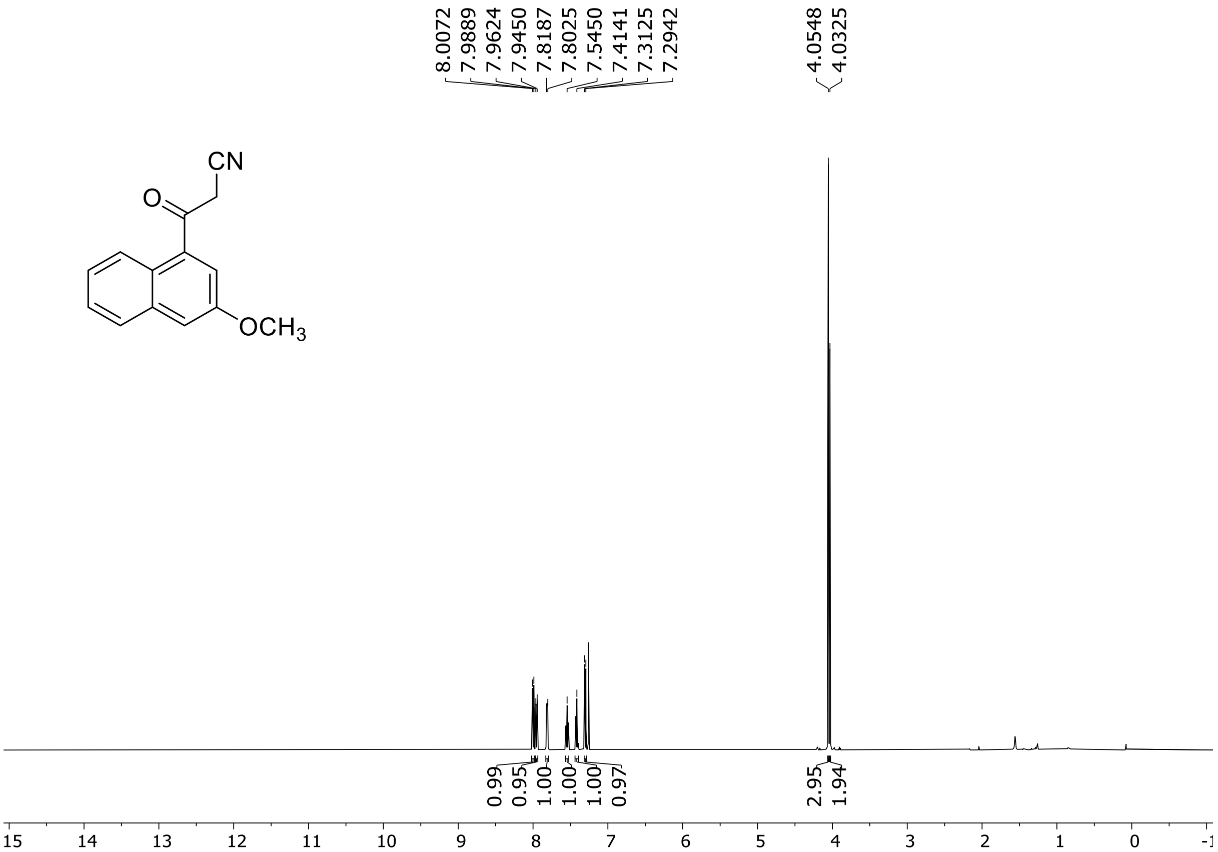

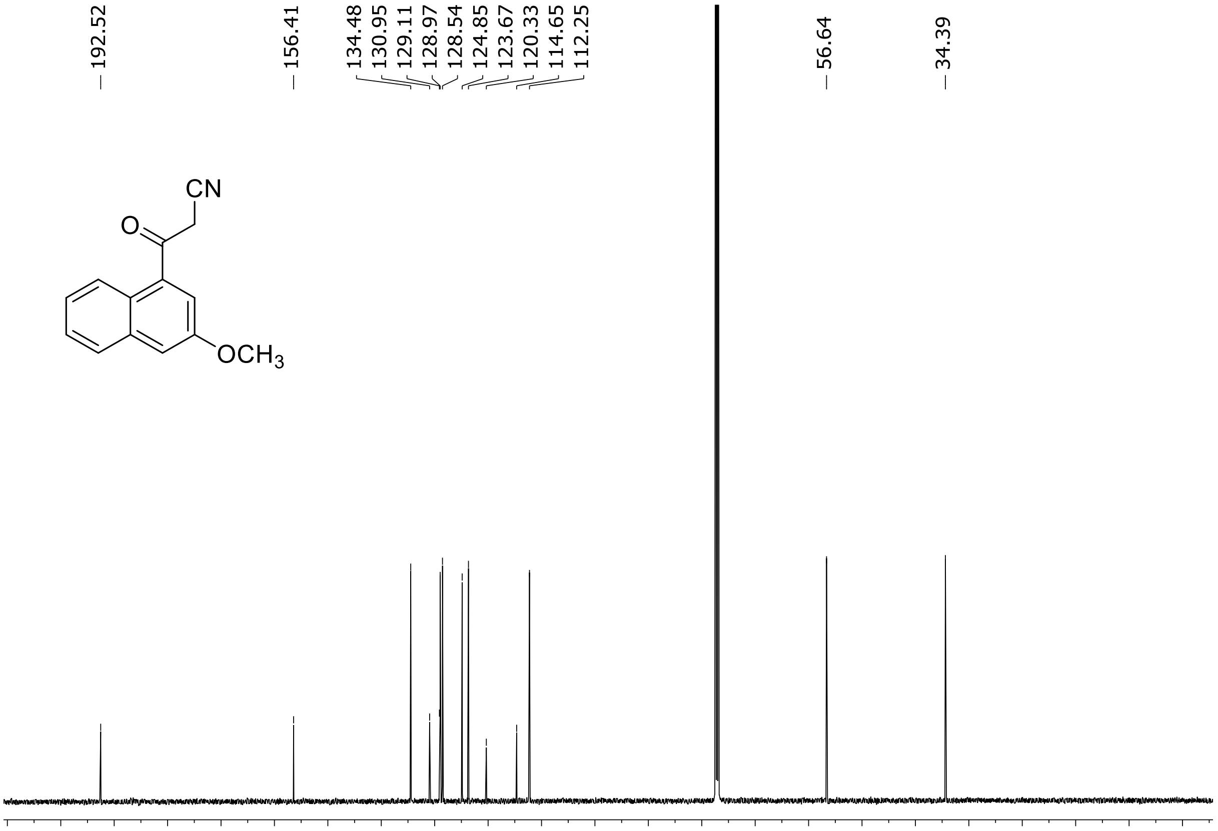

[1]H (500 MHz) and [13]C NMR (126 MHz) spectra of **S14** in chloroform-*d*

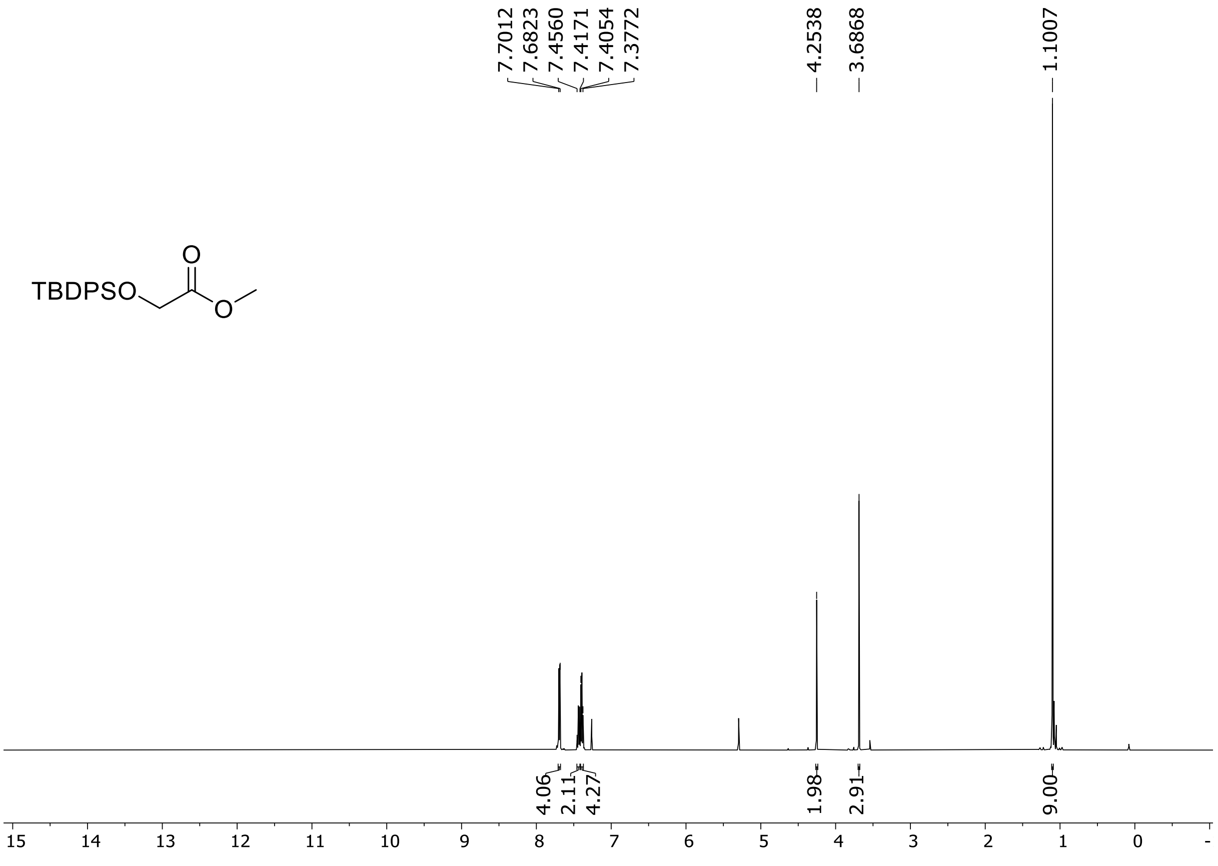

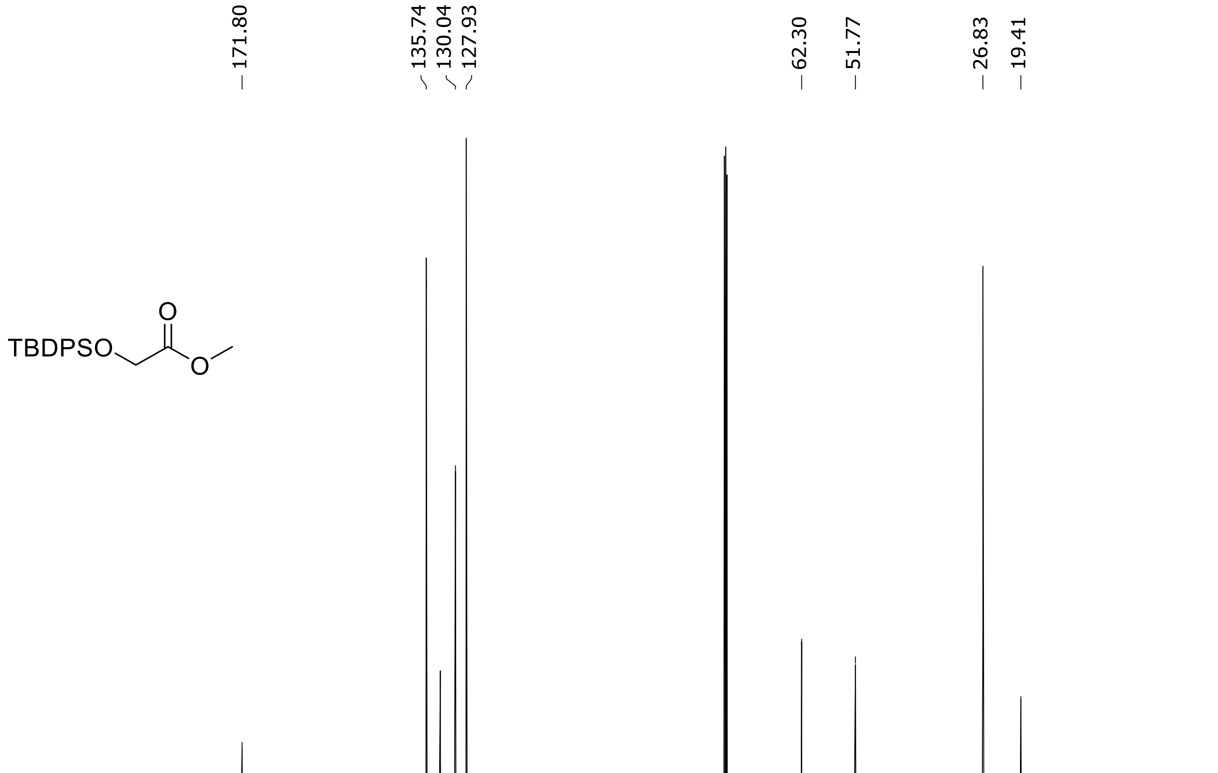

$^{1}$H (500 MHz) and $^{13}$C NMR (126 MHz) spectra of **S15** in chloroform-*d*

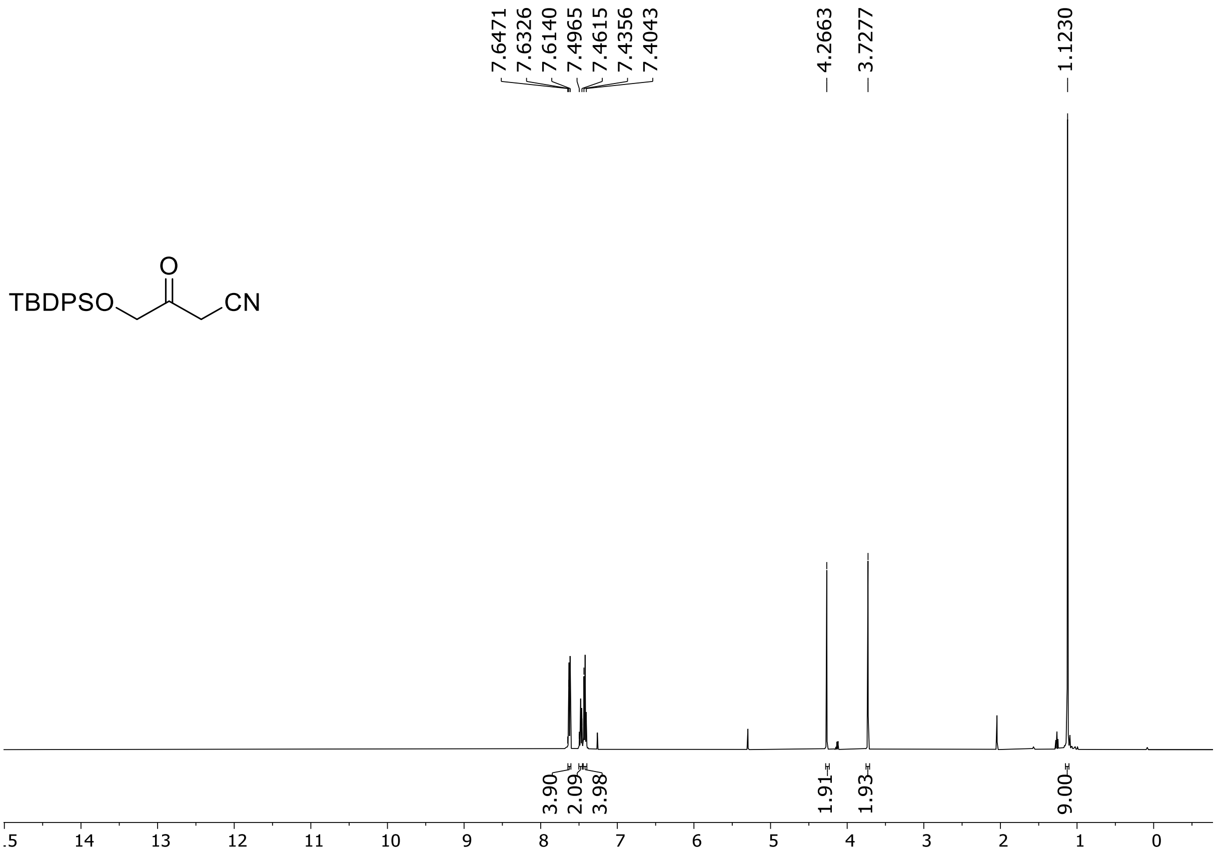

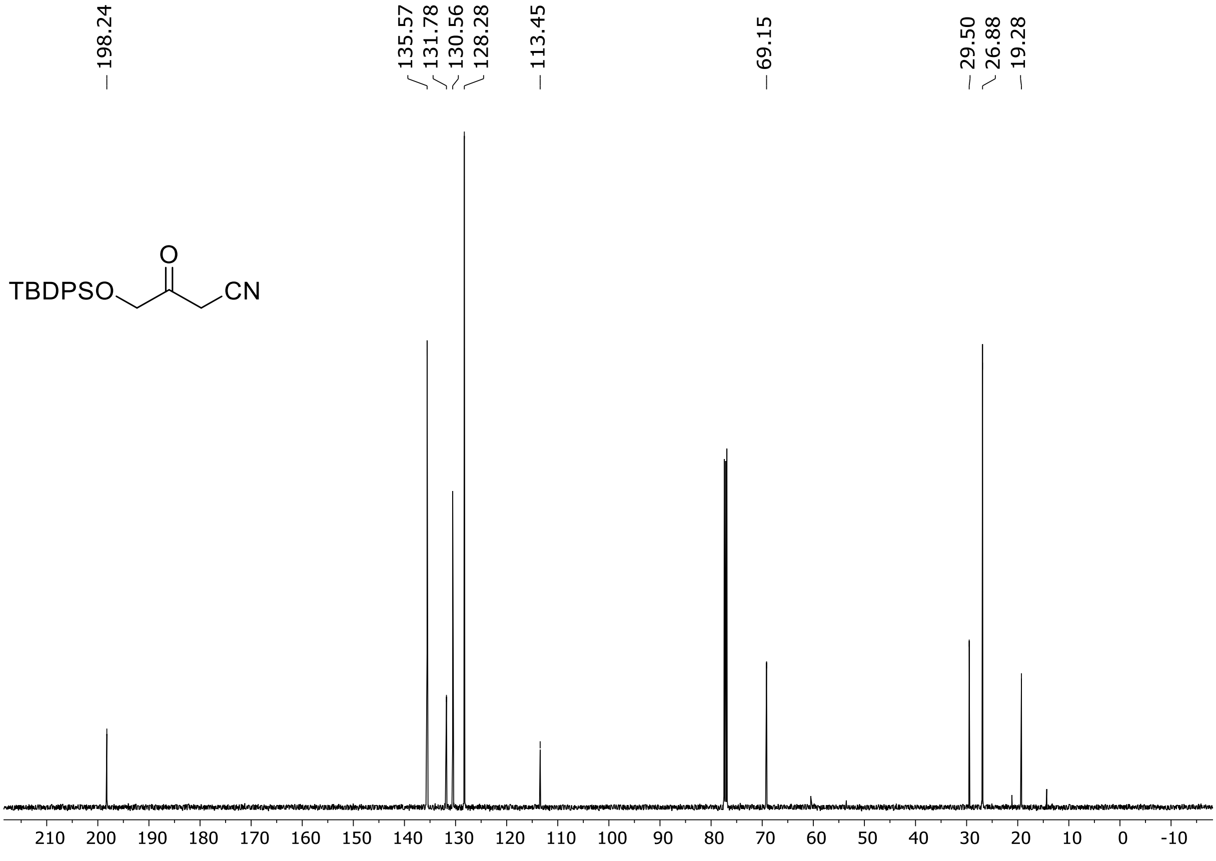

HRMS spectrum of **S15**

**$C_{20}H_{23}NO_2Si$** **mono m/z = 337.1498**

**APCI -** (MMI)
nitrogen flow 5 L/min, gas temperature 300°C, nebulizer 45 psi, vaporizer 200°C
skimmer **65** V, fragmentor **18** V, dissolved in methanol

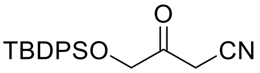

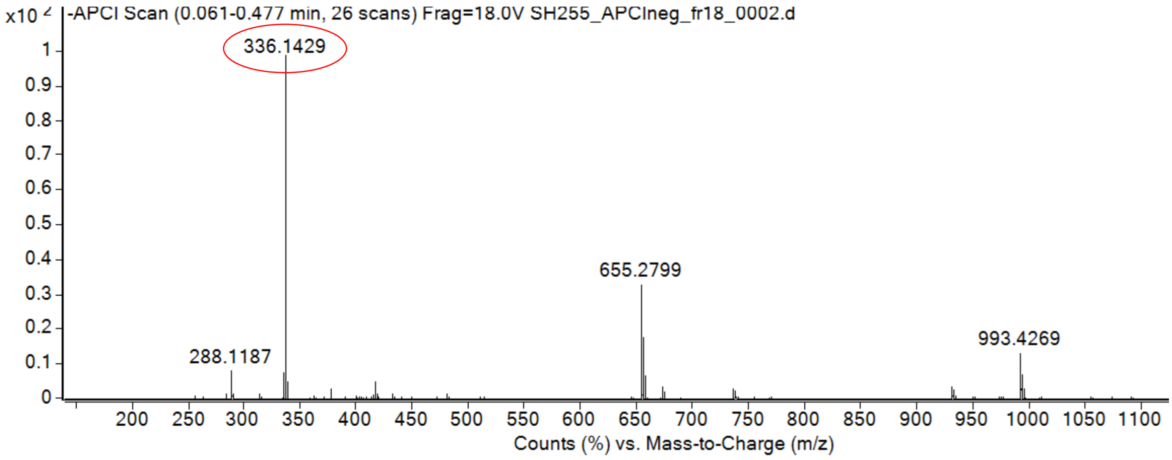

calculated mass: [M-H]$^-$ = 336.1425 observed: [M-H]$^-$ = 336.1429 max. mass error = 1.1 ppm

[1]H (500 MHz) spectrum of **S16** in chloroform-*d*

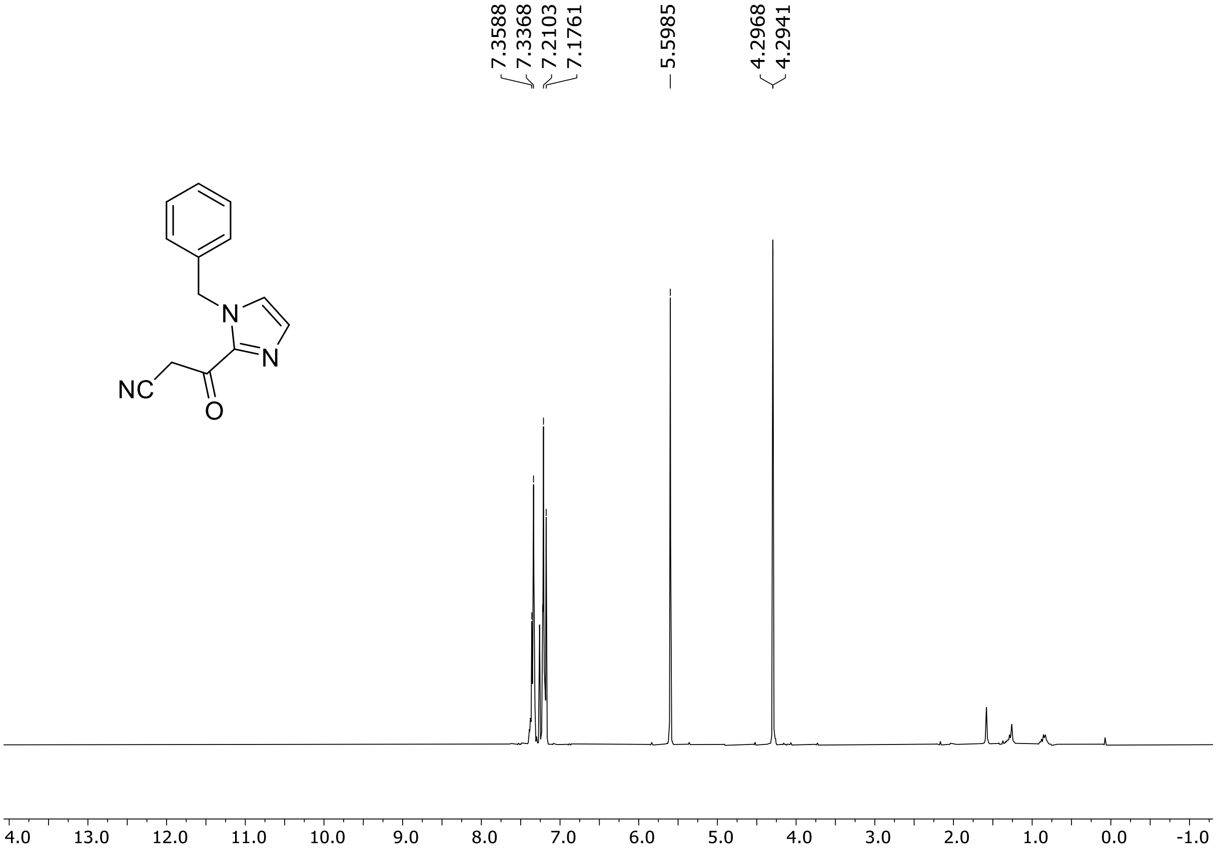

[1]H (500 MHz) and [13]C NMR (126 MHz) spectra of **S17** in chloroform-*d*

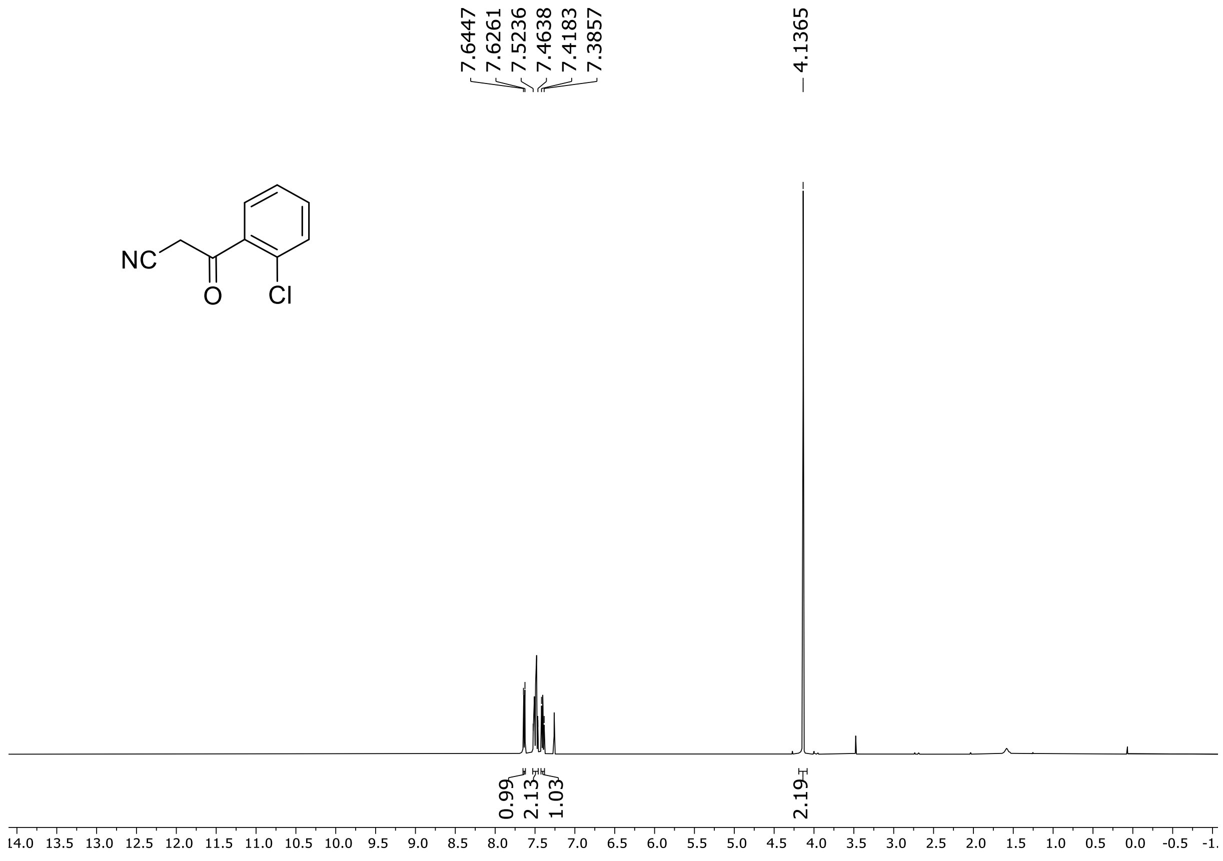

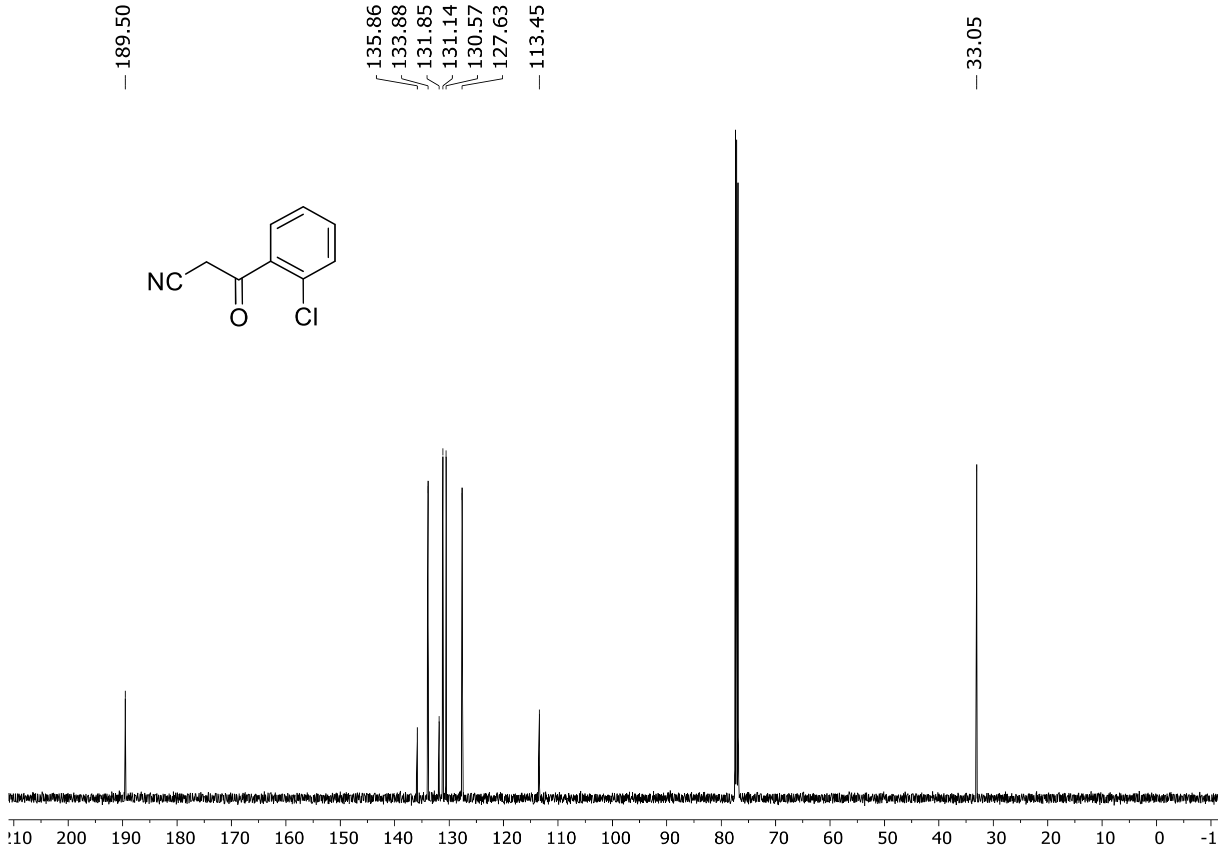

HRMS spectrum of **S17**

$C_9H_6ClNO$ **mono m/z = 179.0138**

**APCI -** (MMI)
nitrogen flow 5 L/min, gas temperature 300°C, nebulizer 45 psi, vaporizer 200°C
skimmer **65** V, fragmentor **25** V, dissolved in methanol

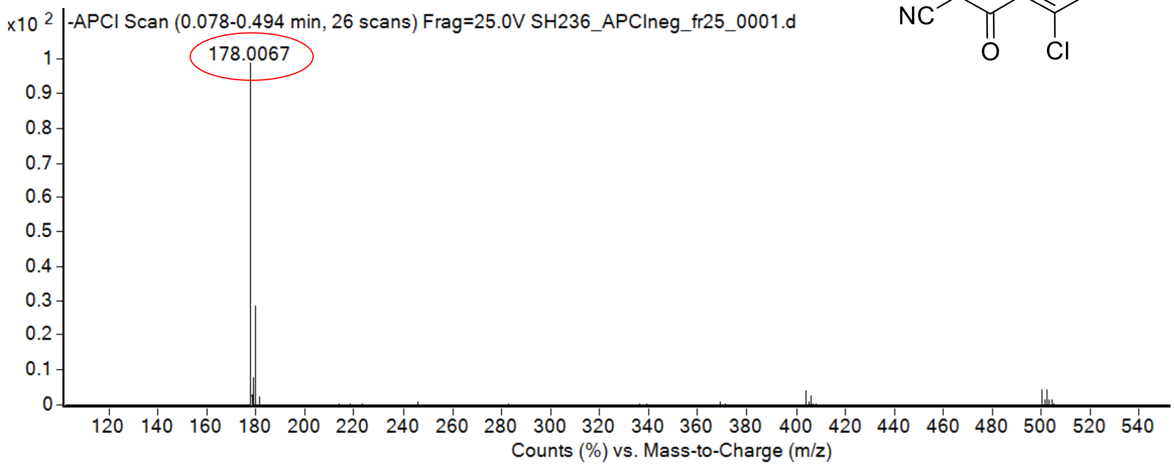

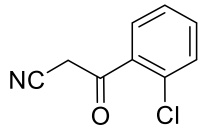

calculated mass: $[M\text{-}H]^-$ = 178.0065 observed: $[M\text{-}H]^-$ = 178.0067 max. mass error = 1.1 ppm

[1]H (500 MHz) and [13]C NMR (126 MHz) spectra of **S18** in chloroform-*d*

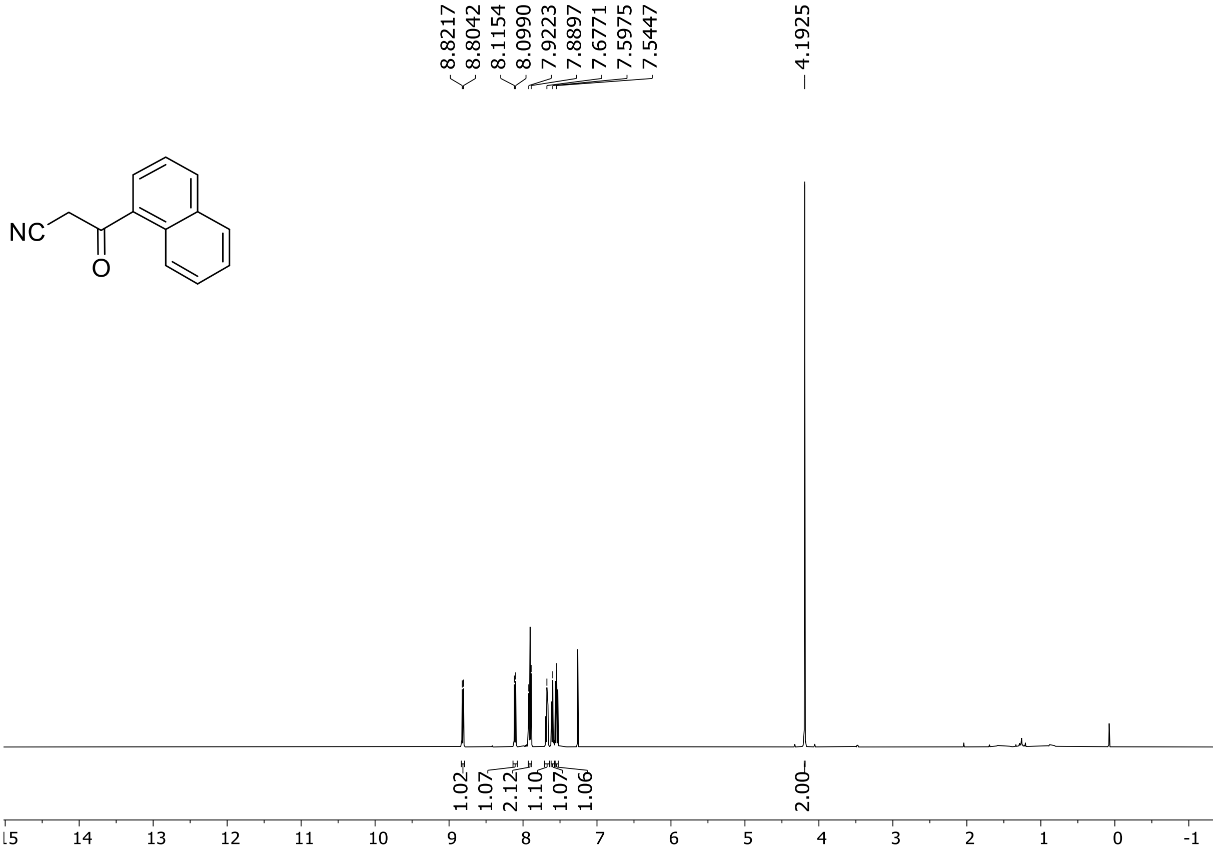

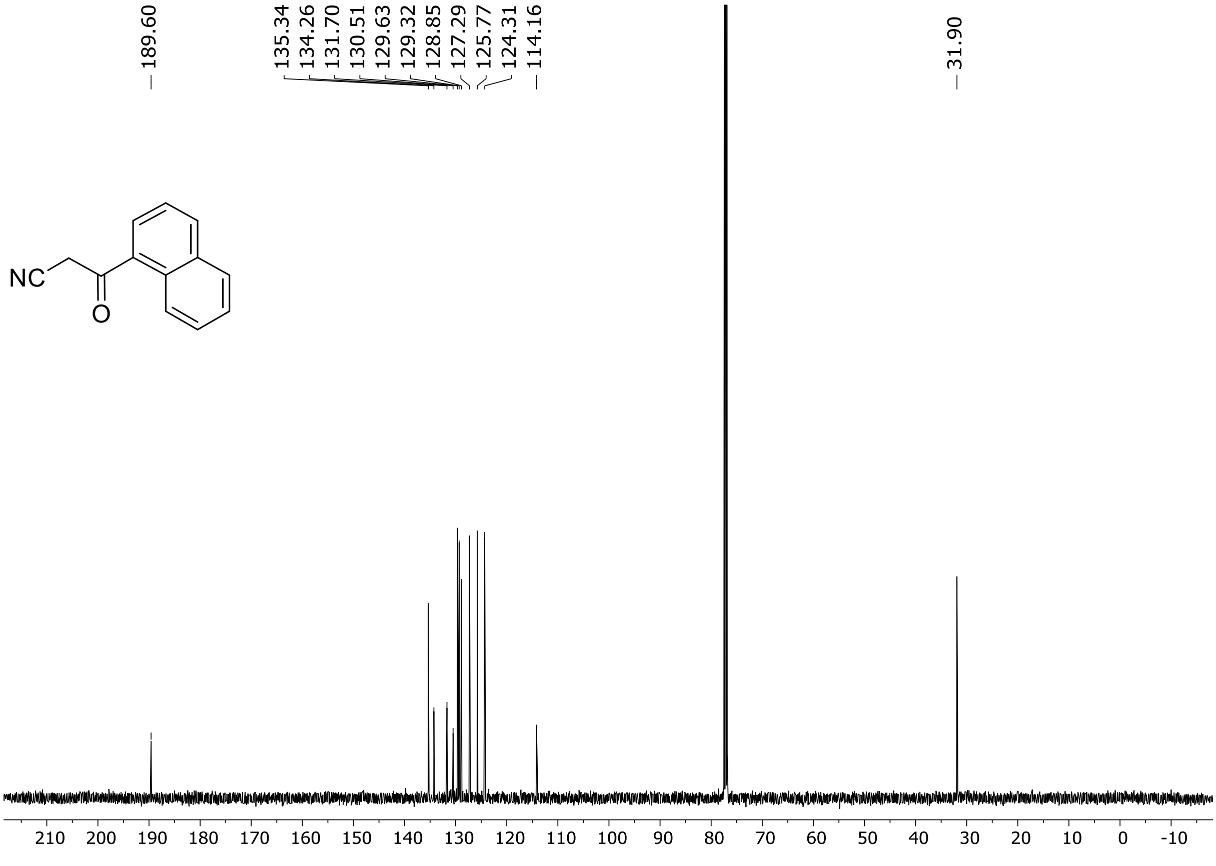

HRMS spectrum of **S18**

**$C_{13}H_9NO$** **mono m/z = 195.0684**

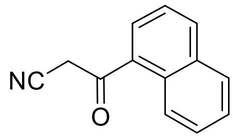

**APCI -** (MMI)
nitrogen flow 5 L/min, gas temperature 300°C, nebulizer 45 psi, vaporizer 200°C
skimmer **65** V, fragmentor **20** V, dissolved in methanol

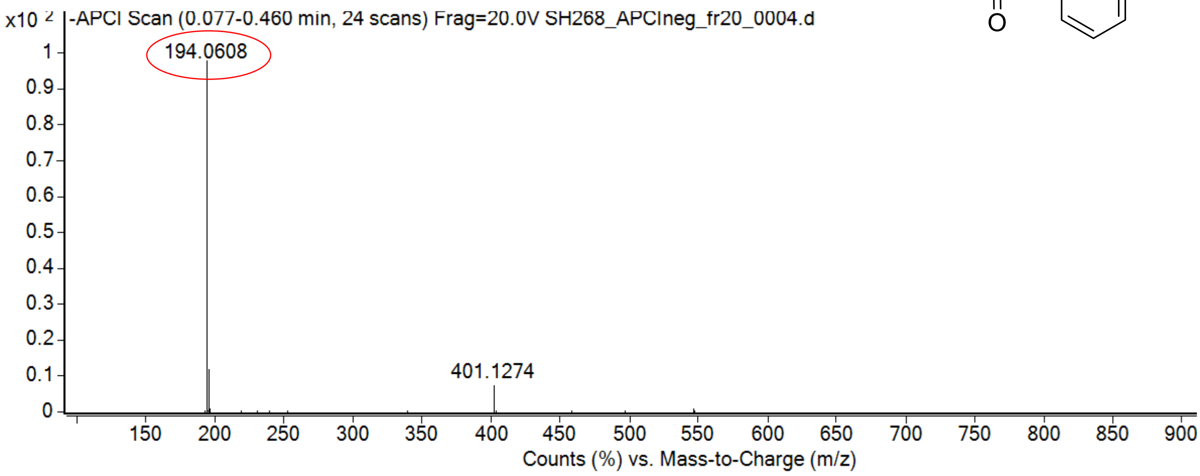

calculated mass: [M-H]$^-$ = 194.0611 observed: [M-H]$^-$ = 194.0608 max. mass error = 1.5 ppm

[1]H (300 MHz) NMR spectrum of **S19** in DMSO-$d_6$

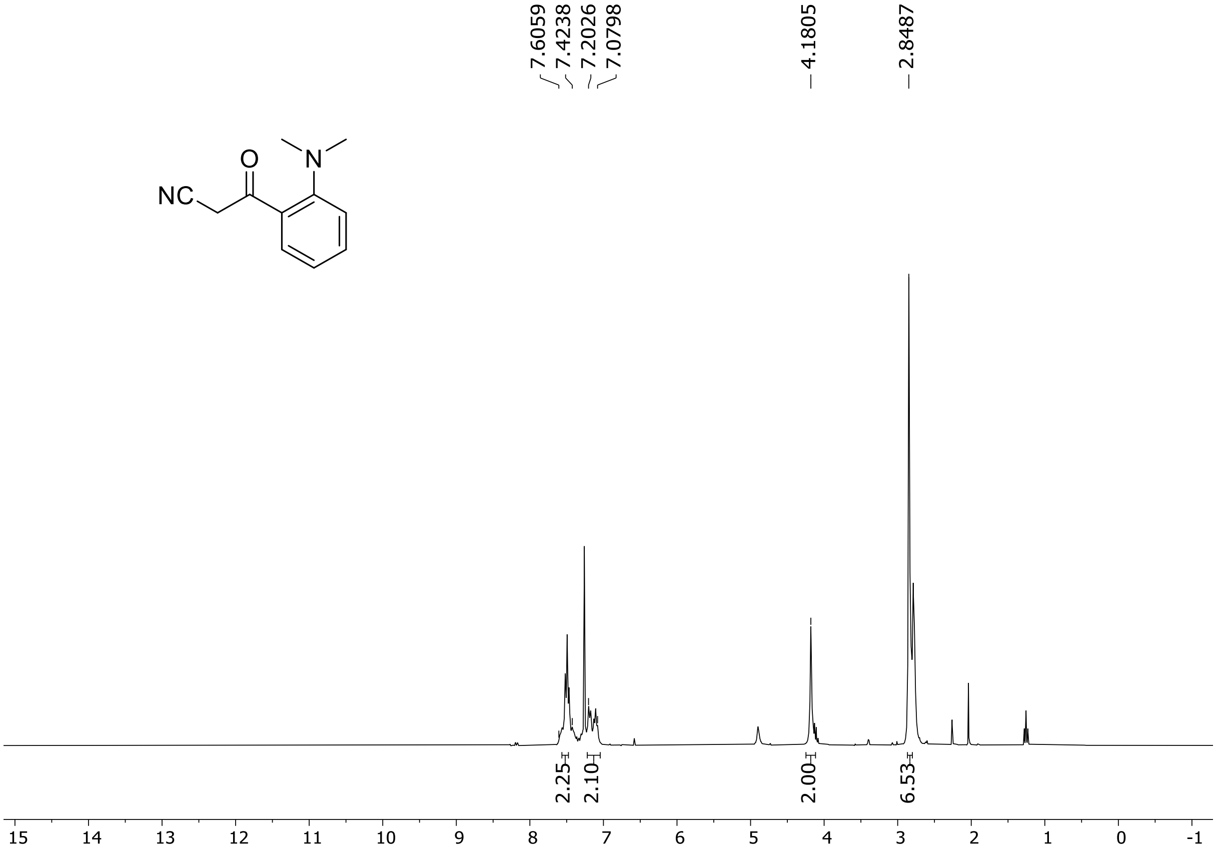

$^{1}H$ (500 MHz) and $^{13}C$ NMR (126 MHz) spectra of **S21** in chloroform-*d*

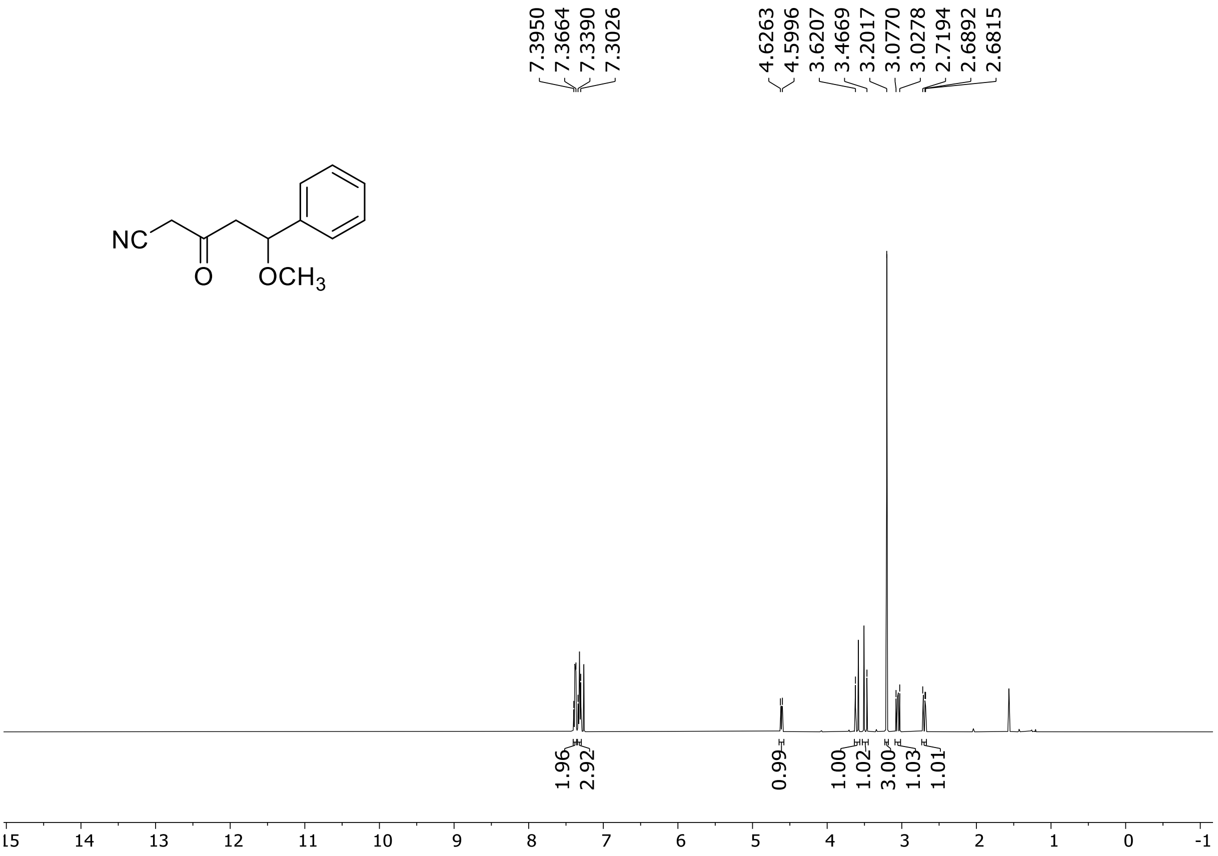

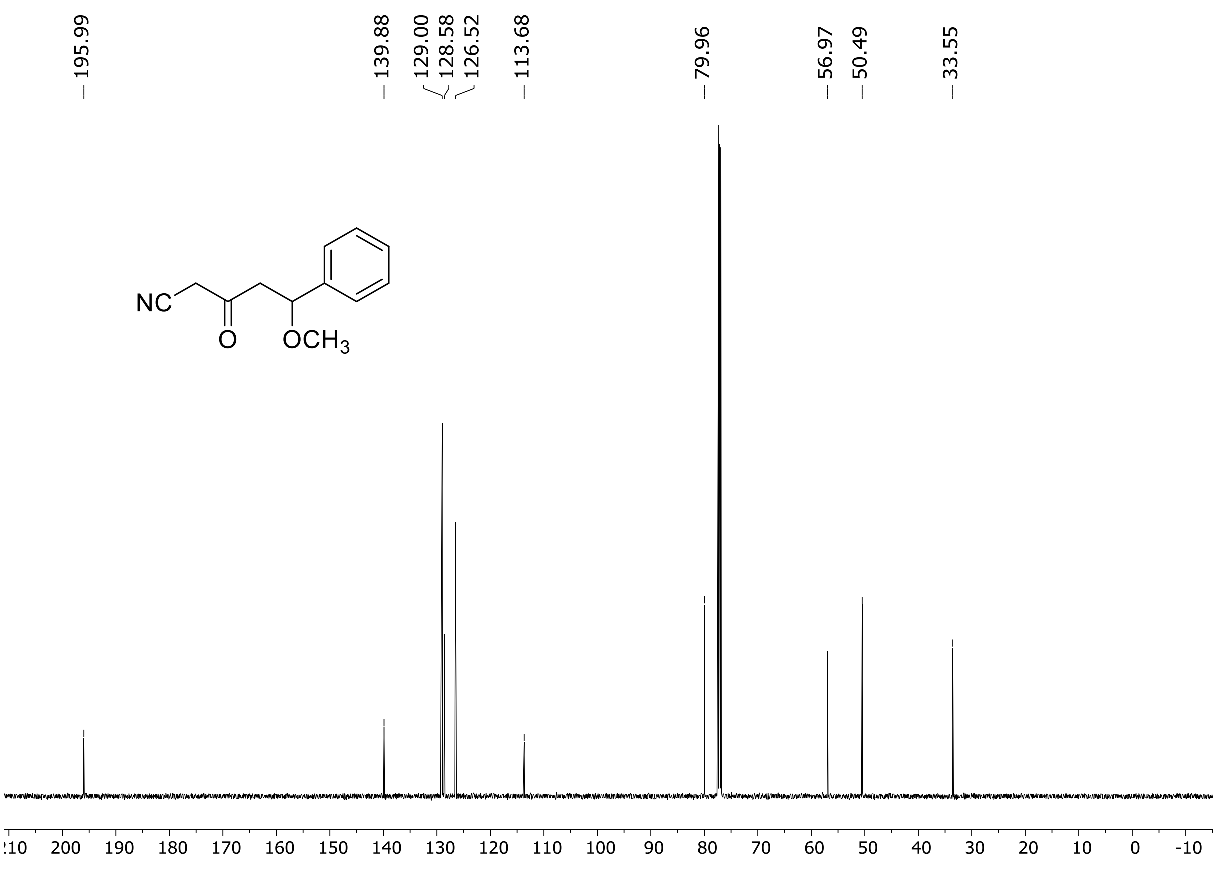

$^{1}H$ (500 MHz) and $^{13}C$ NMR (126 MHz) spectra of **S22** in chloroform-*d*

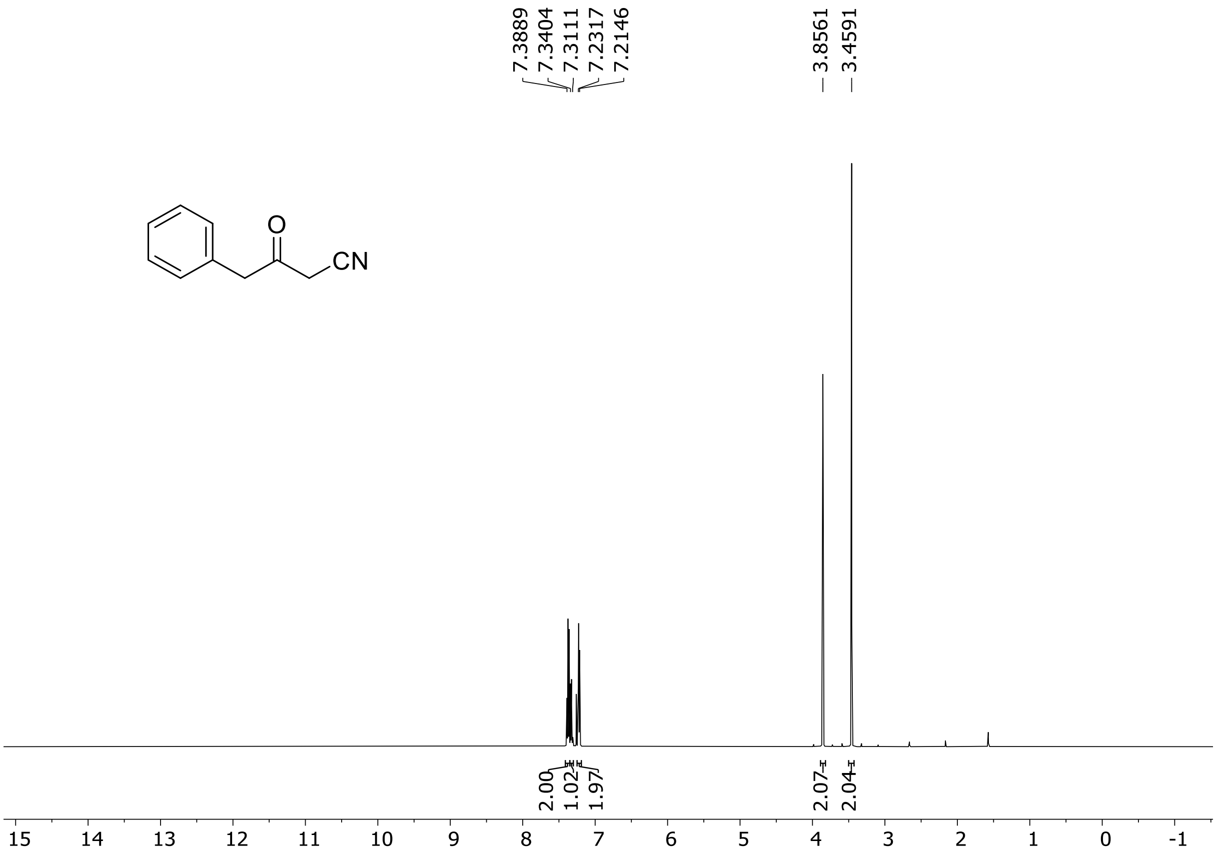

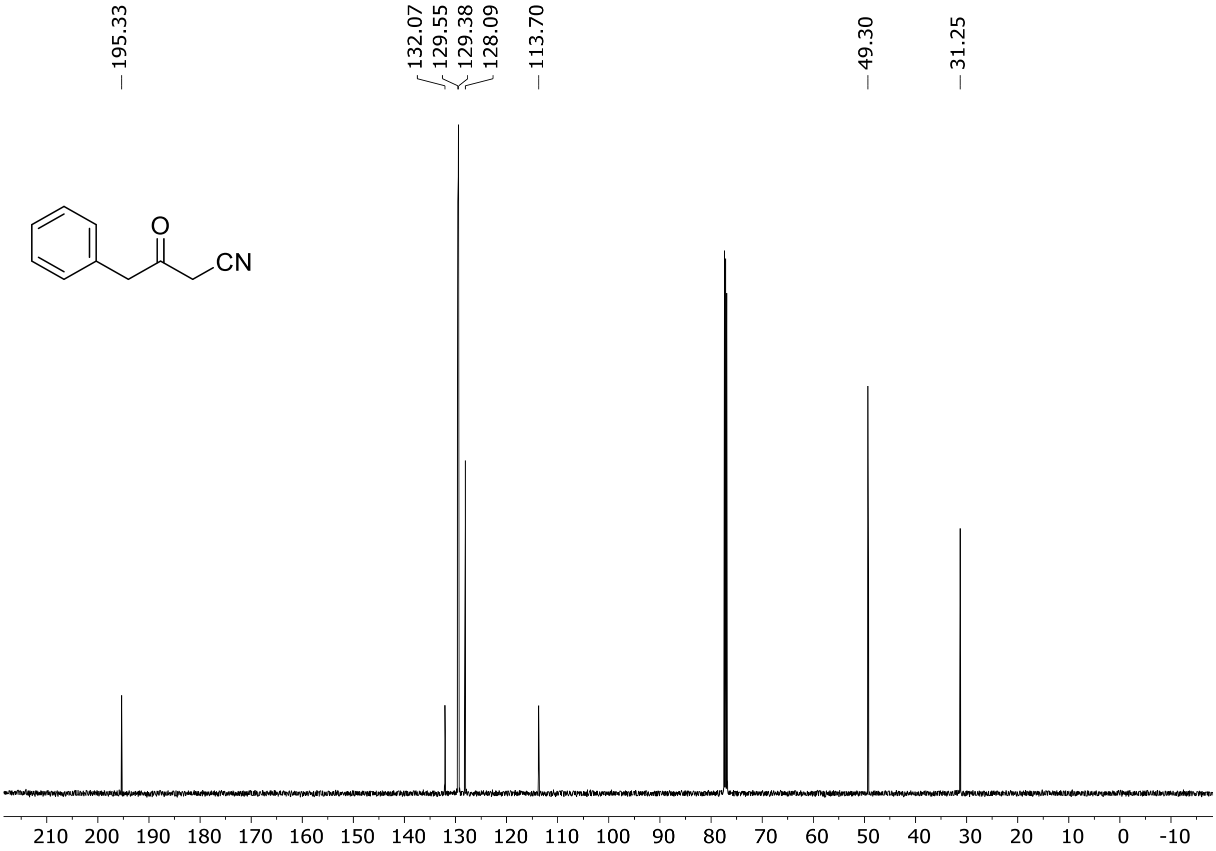

$^{1}$H (500 MHz) and $^{13}$C NMR (126 MHz) spectra of **S23** in chloroform-*d*

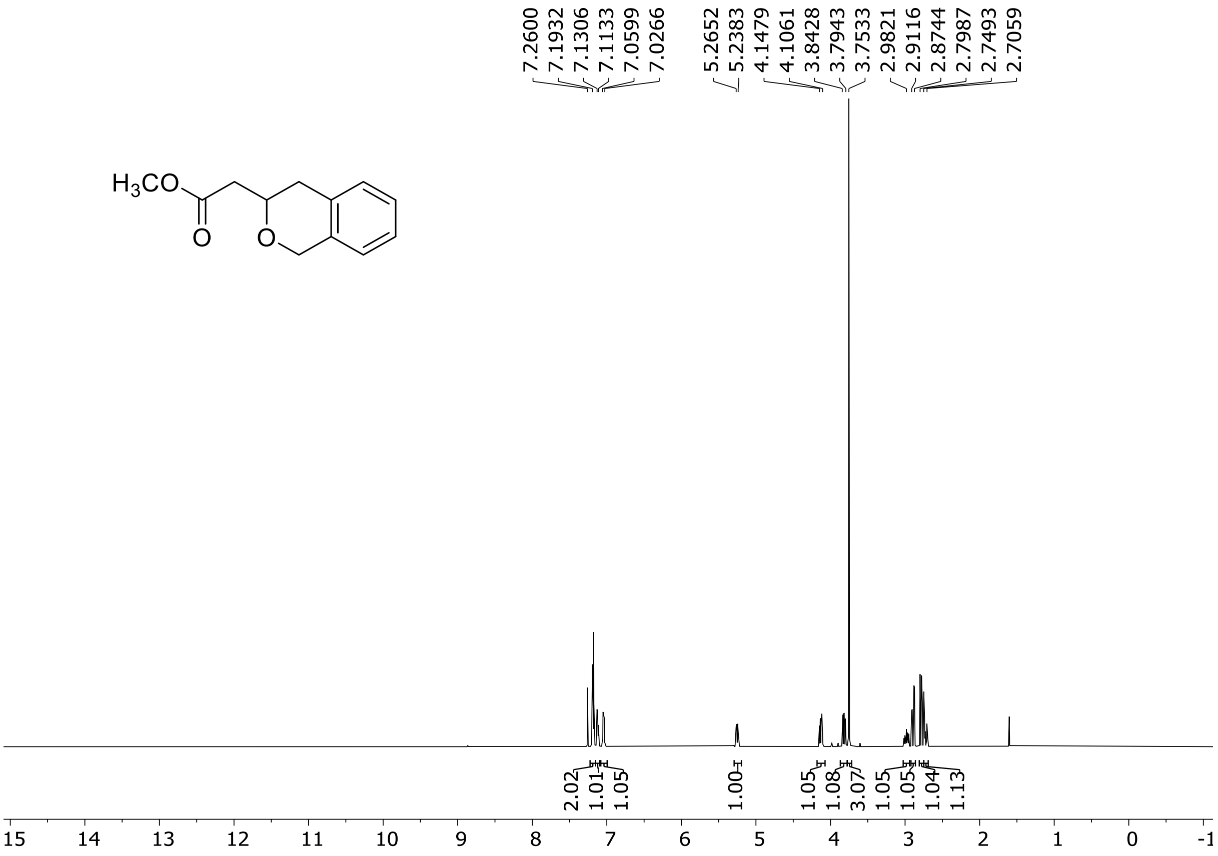

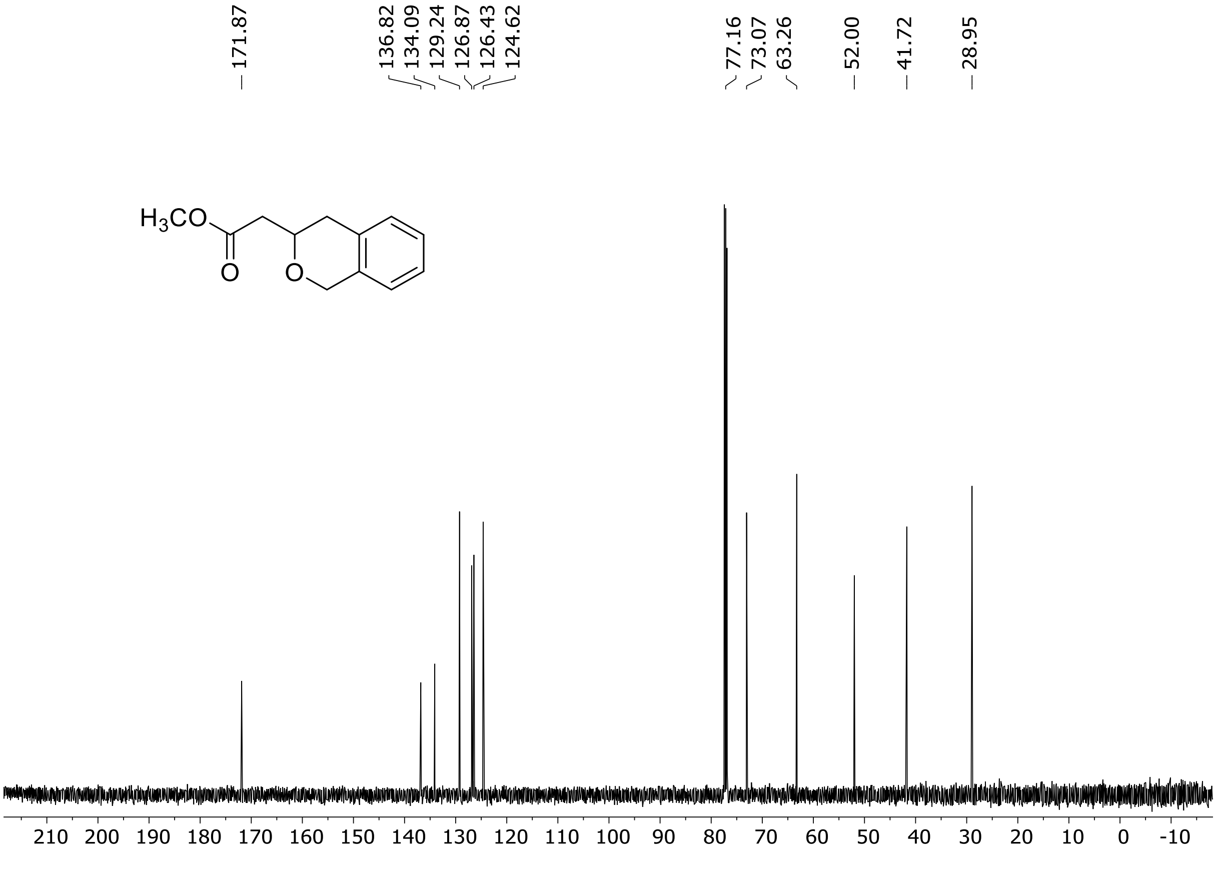

HRMS spectrum of **S23**

**$C_{12}H_{14}O_3$** **exact mass: 206.0943**

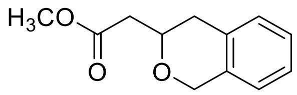

**APCI +** (MMI)
nitrogen flow 3 L/min, gas temperature 325°C, nebulizer 45 psig, skimmer 65 V, vaporizer 200°C, fragmentor 22 V, dissolved in methanol

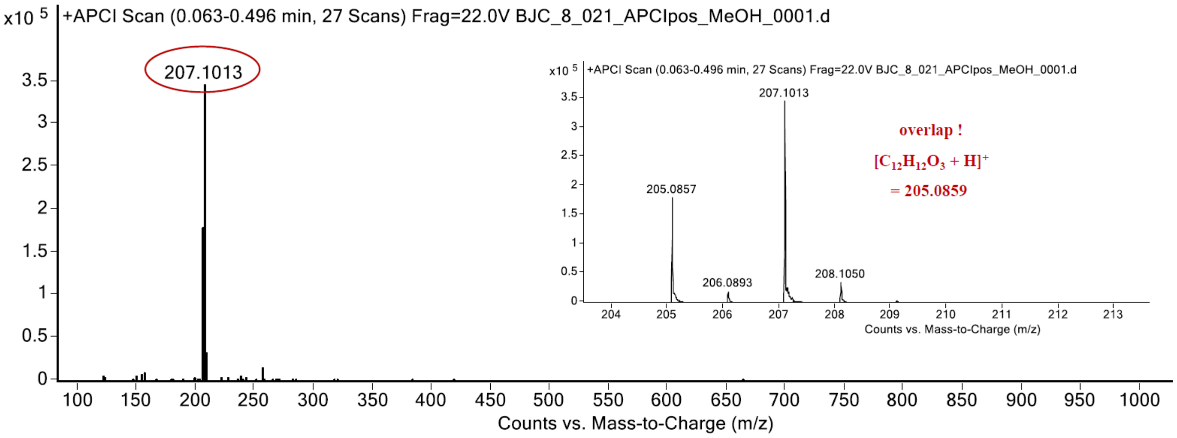

expected mass: $[M+H]^+ = 207.1016$ observed mass: $[M+H]^+ = 207.1013$ mass accuracy = - 1.4 ppm

[1]H (500 MHz) and [13]C NMR (126 MHz) spectra of **S24** in chloroform-*d*

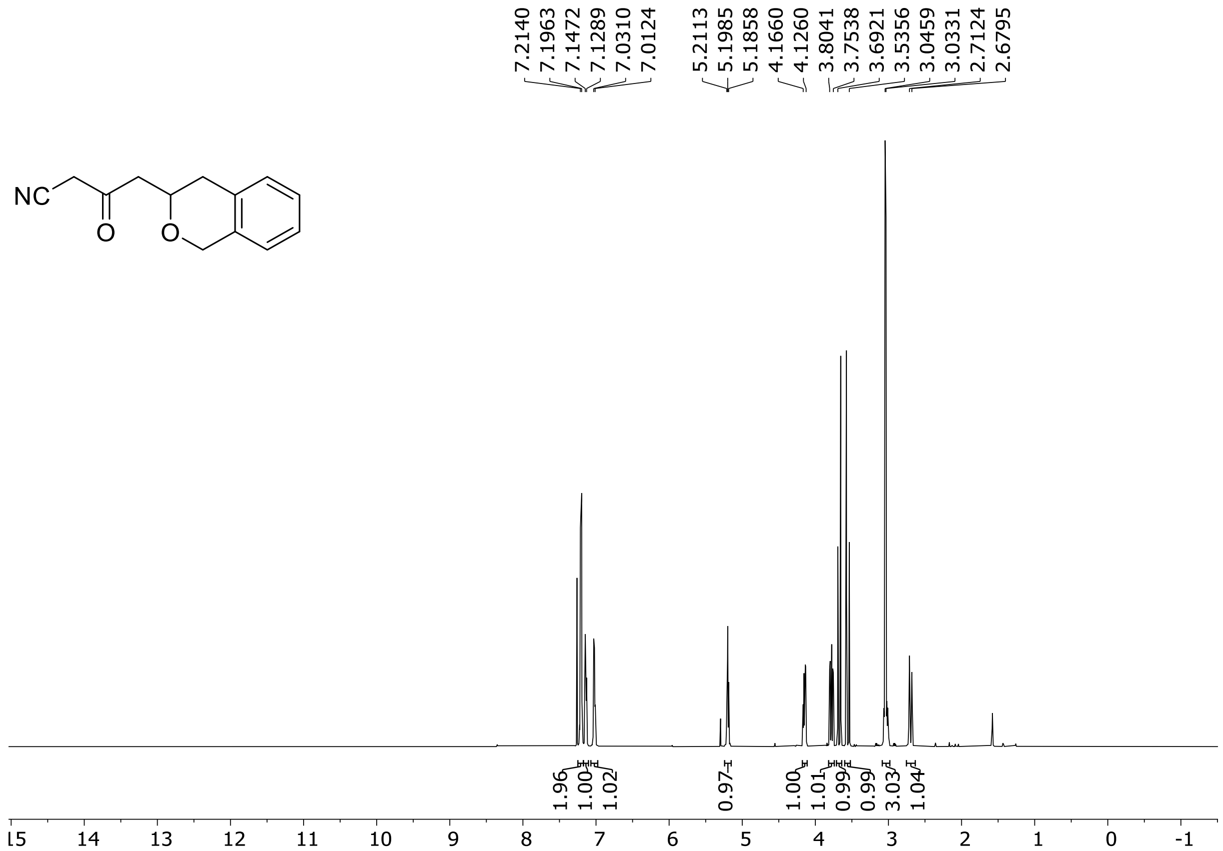

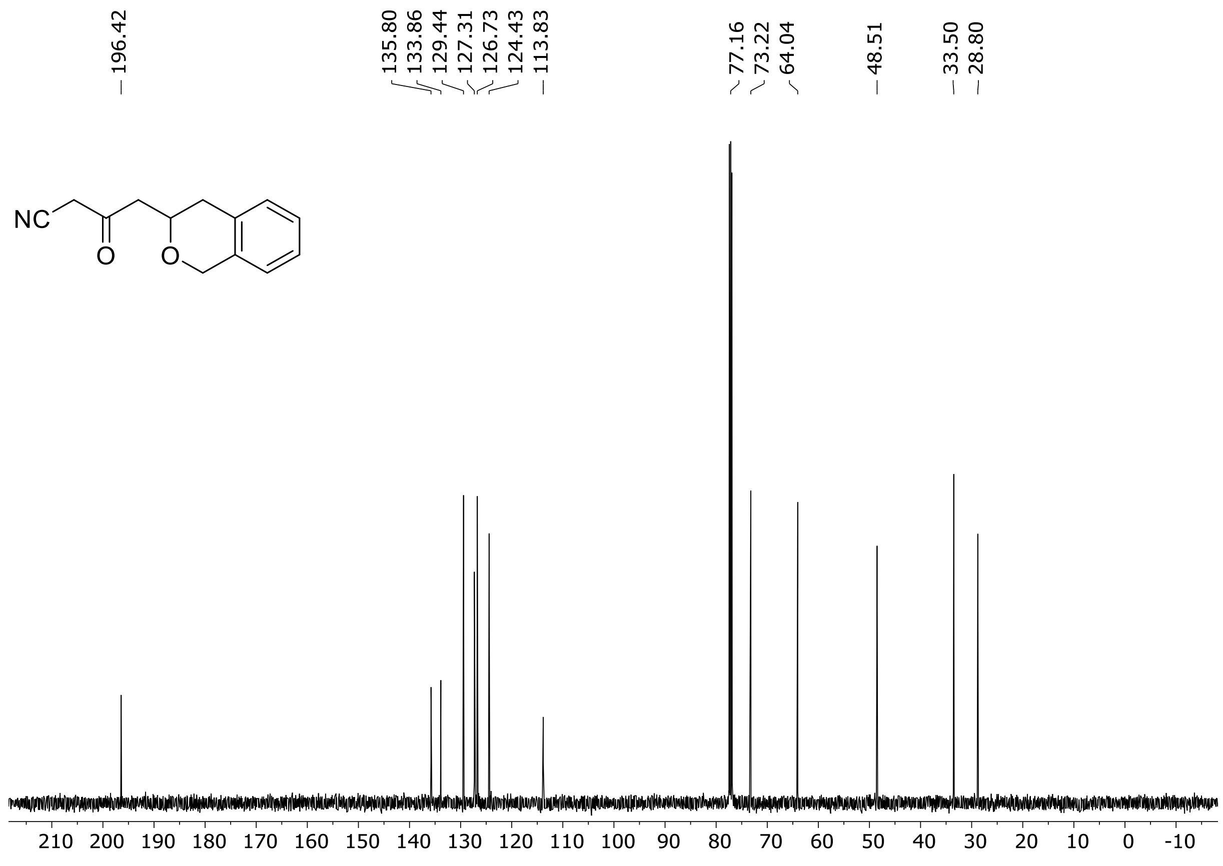

HRMS spectrum of **S24**

$C_{13}H_{13}NO_2$ **exact mass: 215.0946**

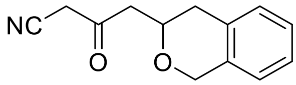

**APCI +** (MMI)
nitrogen flow 3 L/min, gas temperature 325°C, nebulizer 45 psig, skimmer 65 V, vaporizer 200°C, fragmentor 20 V, dissolved in methanol

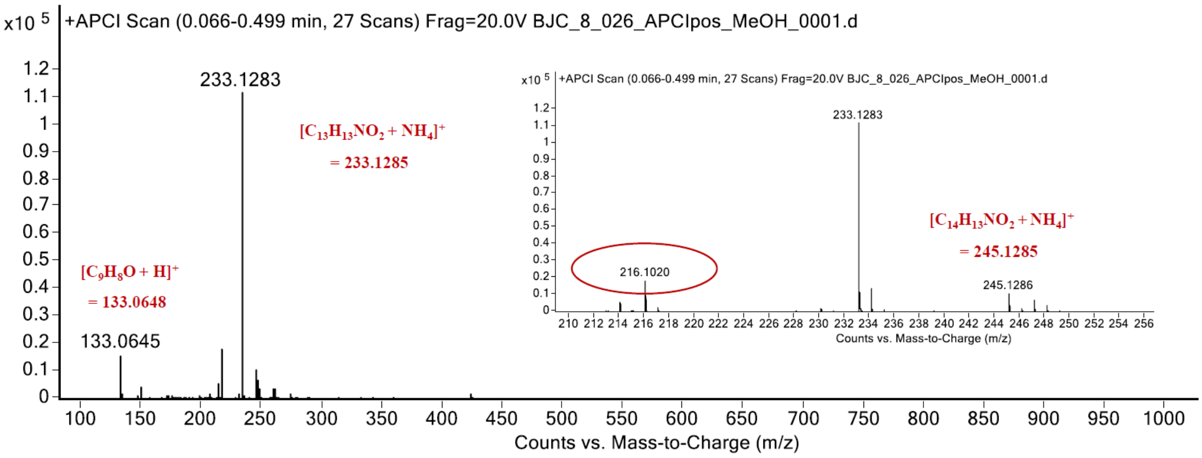

expected mass: $[M+H]^+$ = 216.1019 observed mass: $[M+H]^+$ = 216.1020 mass accuracy = 0.5 ppm

[1]H (500 MHz) NMR spectrum of **S26** in chloroform-*d*

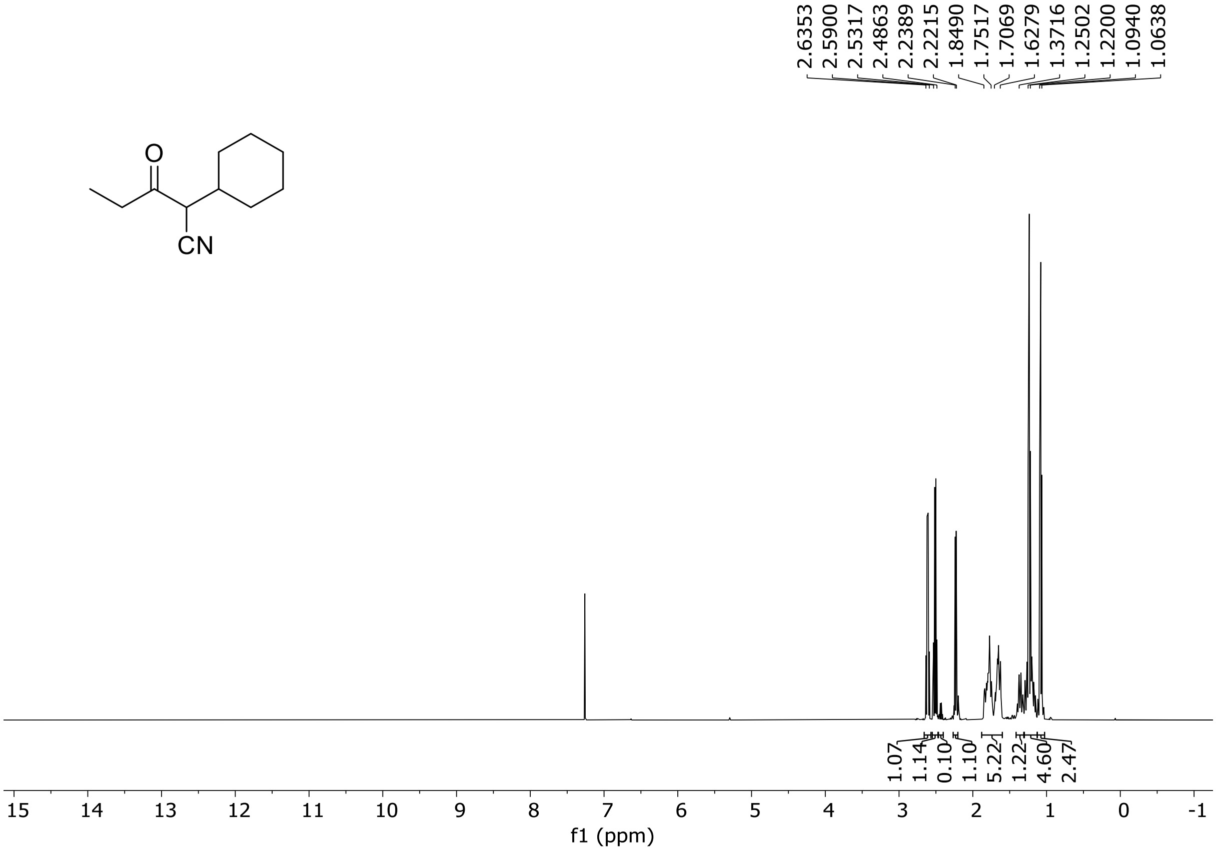

HRMS spectrum of **S26**

**$C_{11}H_{17}NO$** **exact mass: 179.1310**

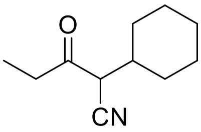

**APCI +** (MMI)
nitrogen flow 3 L/min, gas temperature 325°C, nebulizer 45 psig, skimmer 65 V, vaporizer 200°C, fragmentor 30 V, dissolved in methanol

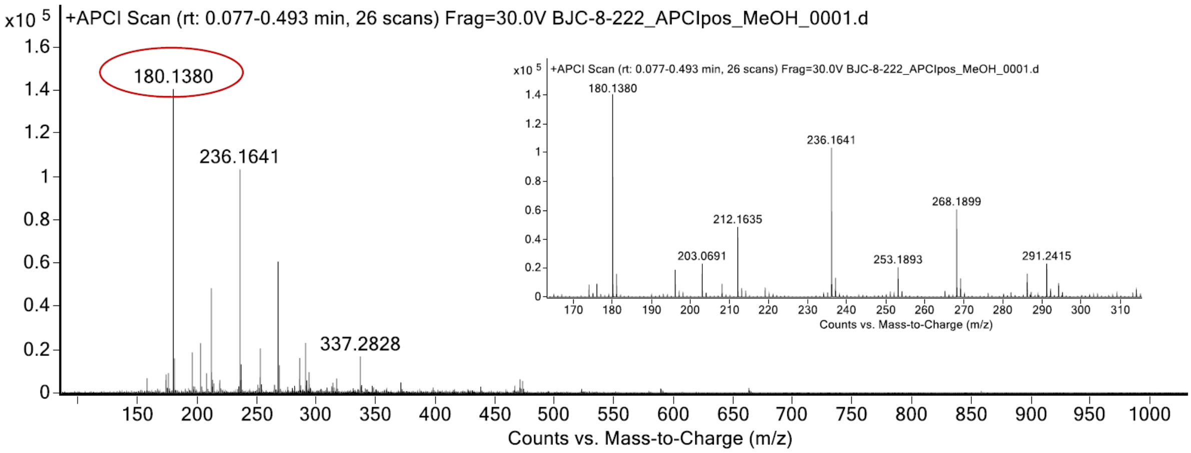

expected mass: $[M+H]^+$ = 180.1383 observed mass: $[M+H]^+$ = 180.1380 mass accuracy = - 1.7 ppm

[1]H (500 MHz) NMR spectrum of **S27** in chloroform-*d*

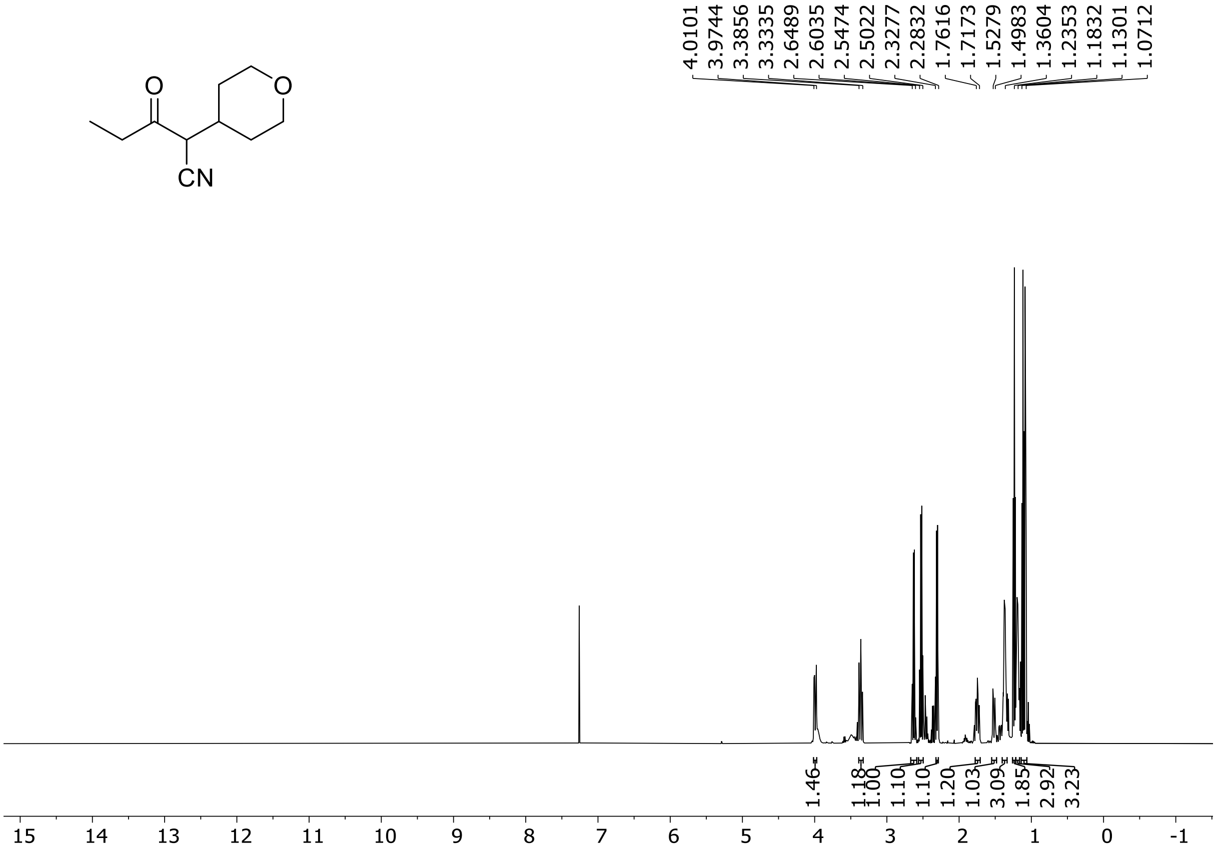

$^1$H (500 MHz) NMR spectrum of **S28** in chloroform-*d*

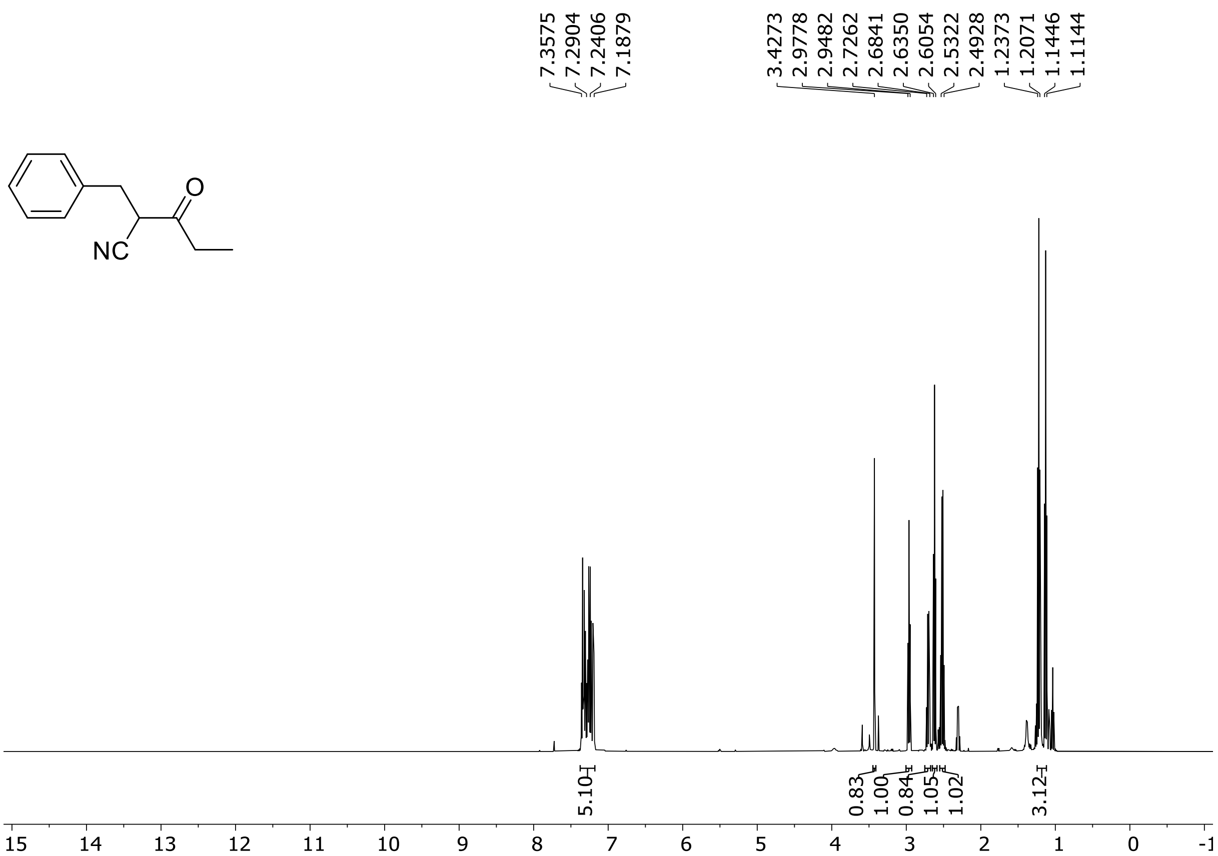

$^{1}H$ (500 MHz) and $^{13}C$ NMR (126 MHz) spectra of **S35** in chloroform-*d*

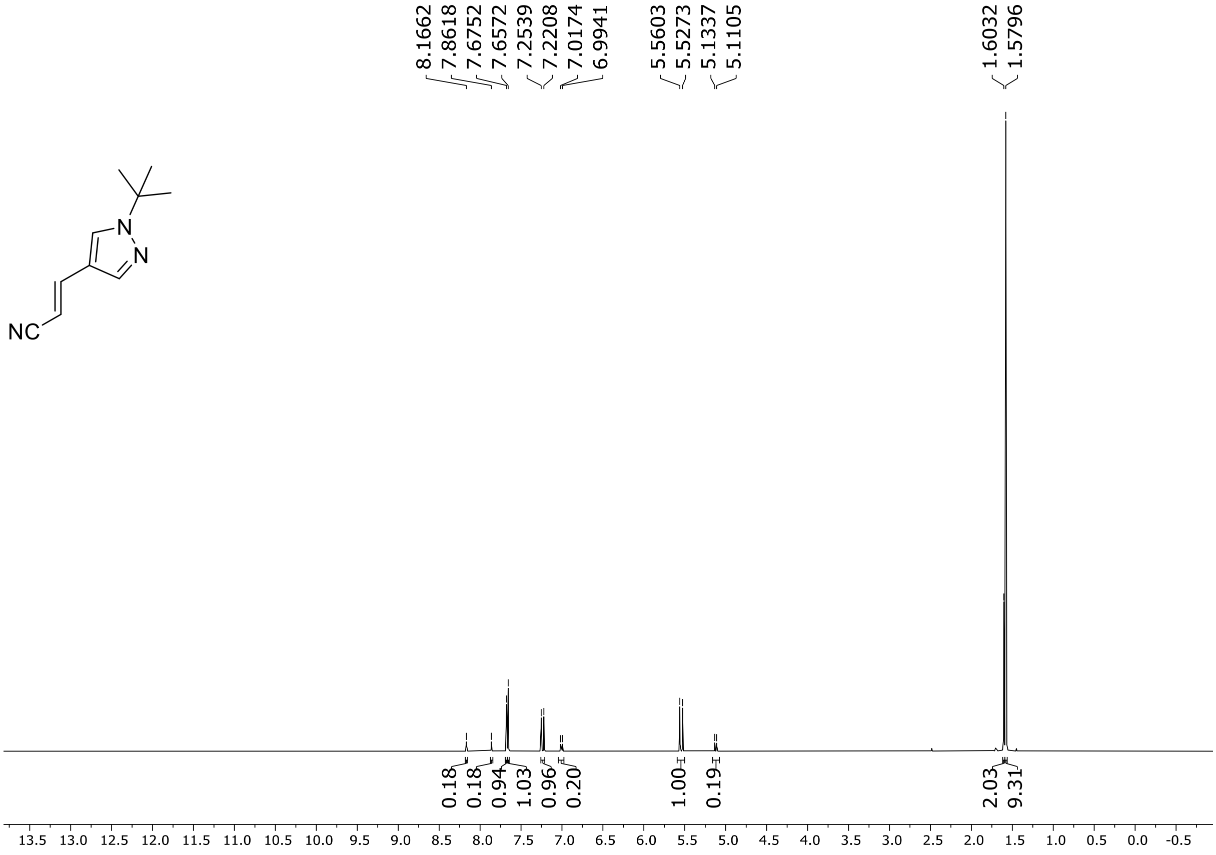

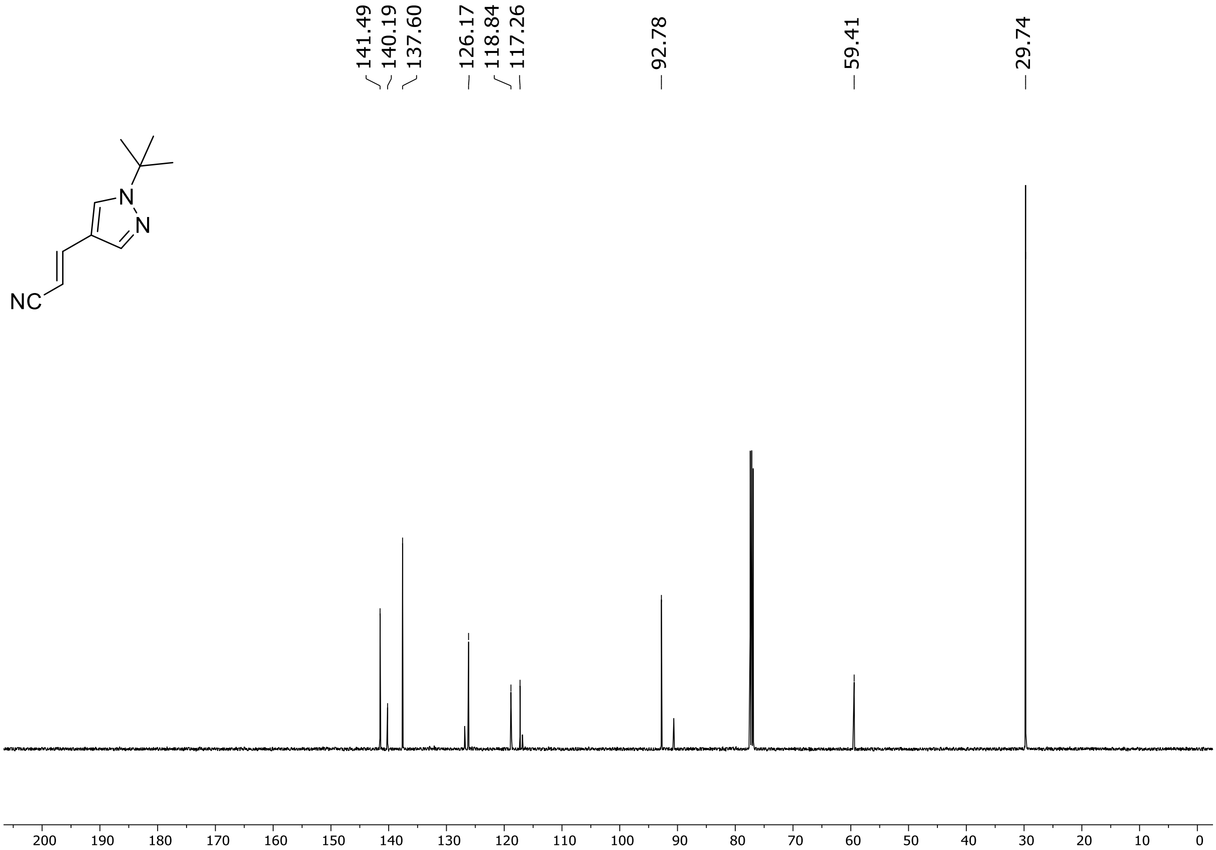

HRMS spectrum of **S35**

**$C_{10}H_{13}N_3$** **exact mass: 175.1109**

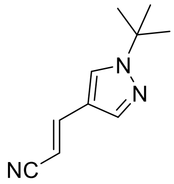

**APCI +** (MMI)
nitrogen flow 5 L/min, gas temperature 325°C, nebulizer 45 psig, skimmer 65 V, vaporizer 200°C, fragmentor 18 V, dissolved in methanol

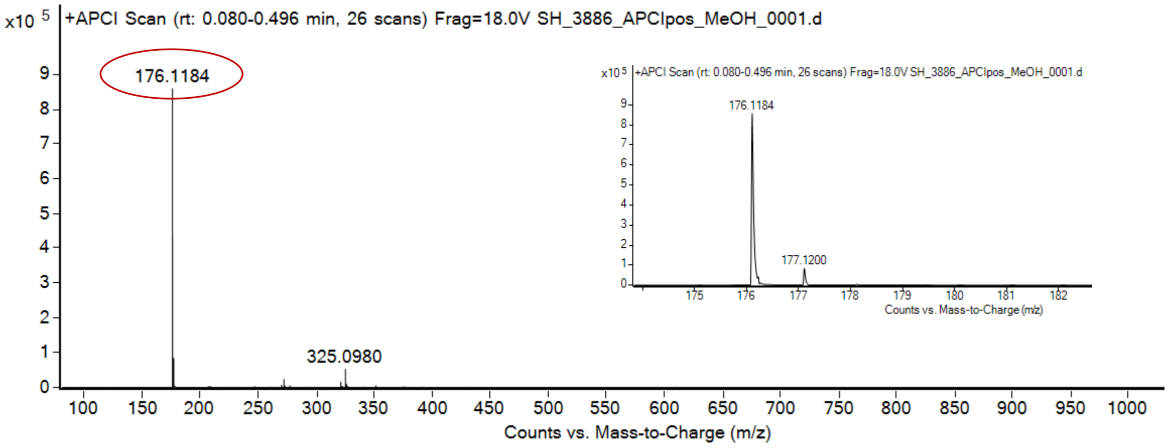

expected mass: $[M+H]^+$ = 176.1182 observed mass: $[M+H]^+$ = 176.1184 mass accuracy = 1.1 ppm

$^{1}H$ (500 MHz) and $^{13}C$ NMR (126 MHz) spectra of **S36** in chloroform-*d*

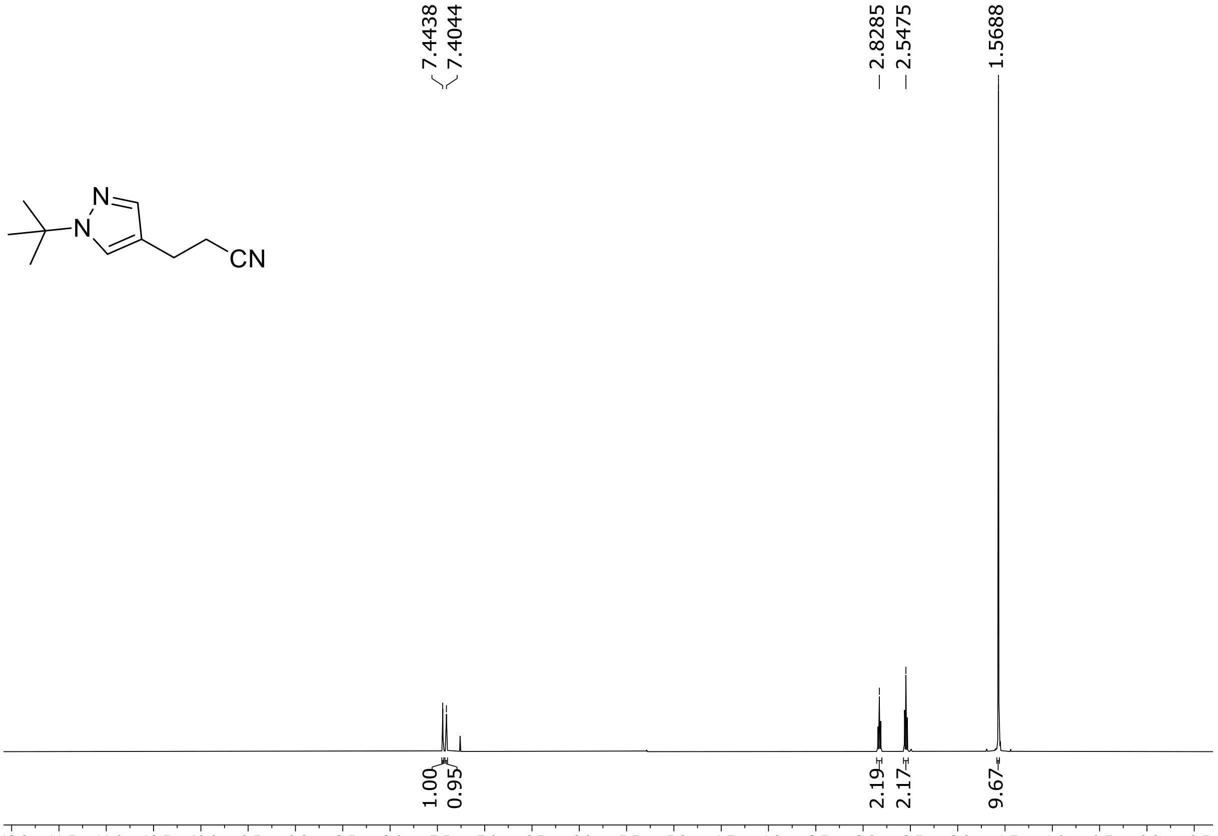

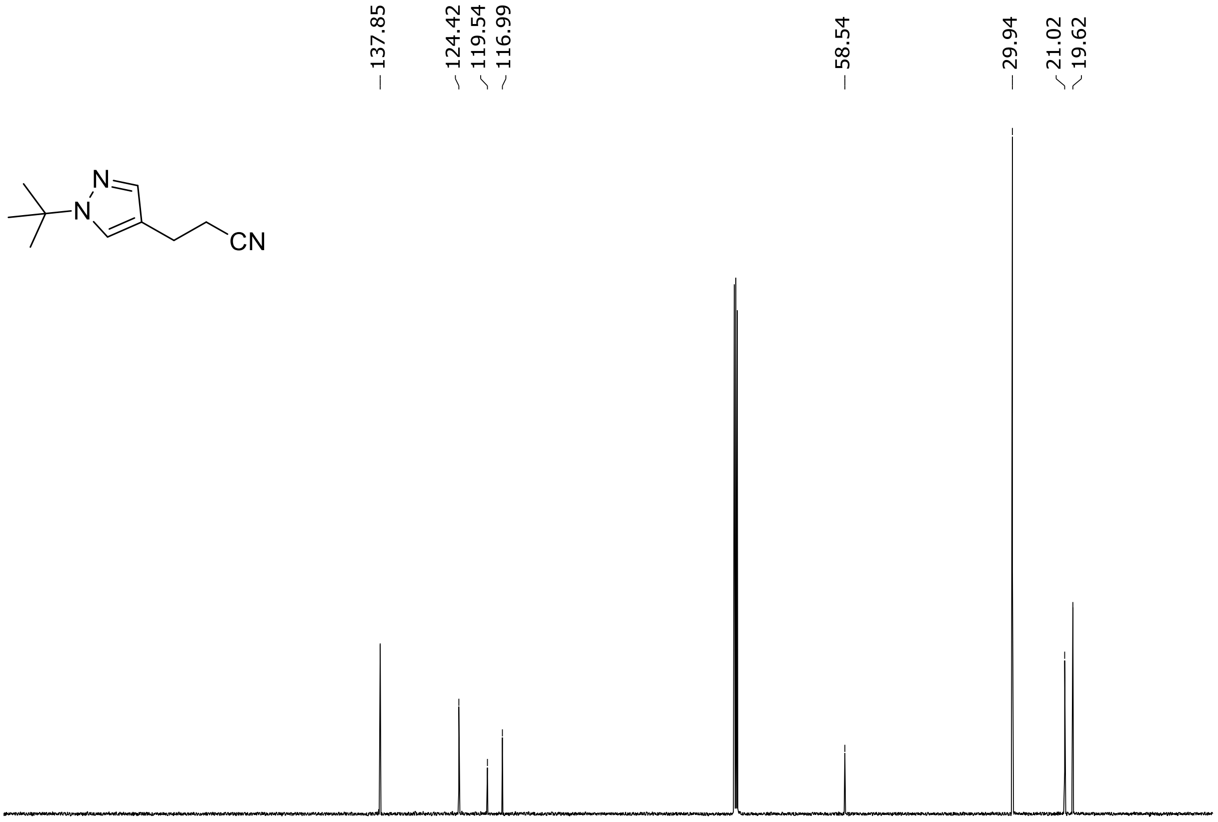

HRMS spectrum of **S36**

**$C_{10}H_{15}N_3$** **exact mass: 177.1266**

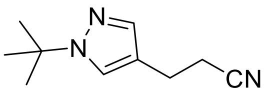

**APCI +** (MMI)
nitrogen flow 5 L/min, gas temperature 325°C, nebulizer 45 psig, skimmer 65 V, vaporizer 200°C, fragmentor 20 V, dissolved in methanol

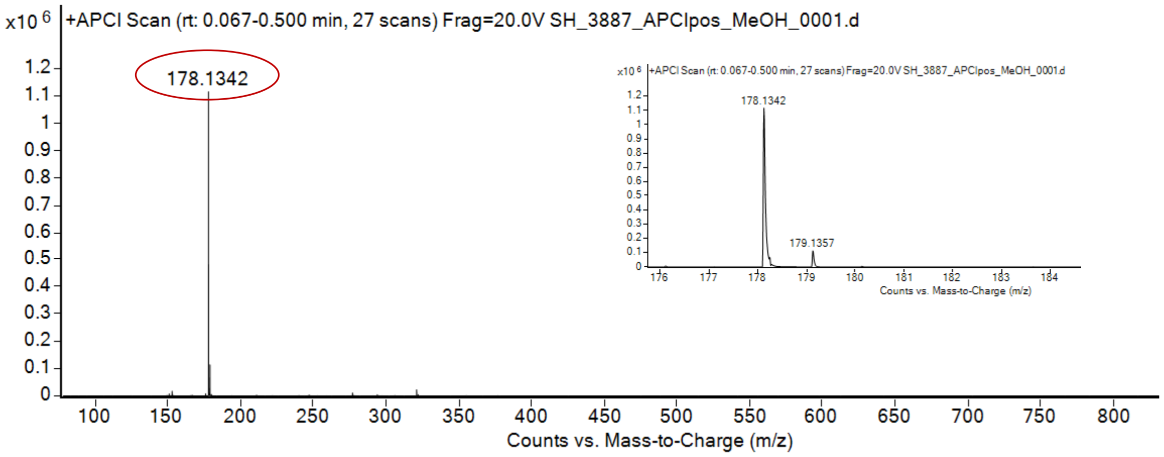

expected mass: [M+H]$^+$ = 178.1339 observed mass: [M+H]$^+$ = 178.1342 mass accuracy = - 1.7 ppm

[1]H (500 MHz) and [13]C NMR (126 MHz) spectra of **S38** in DMSO-$d_6$

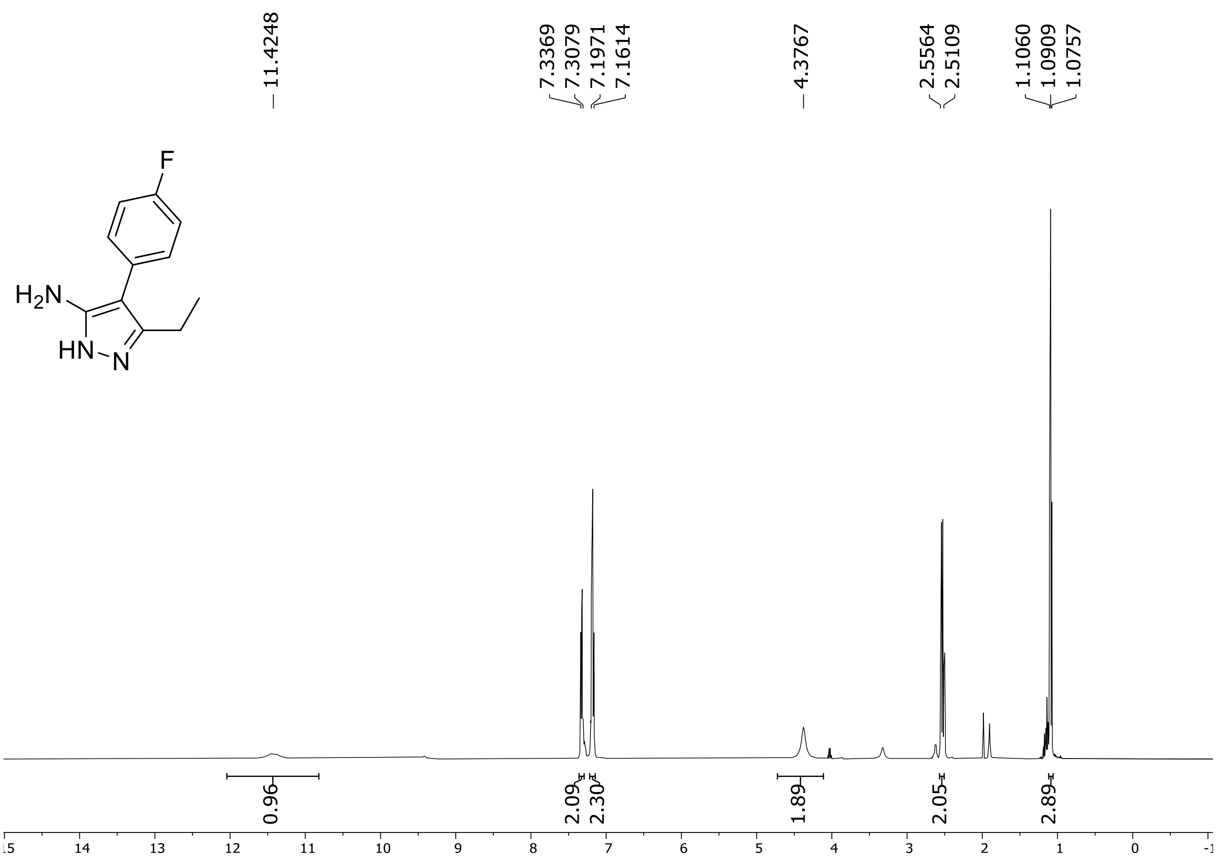

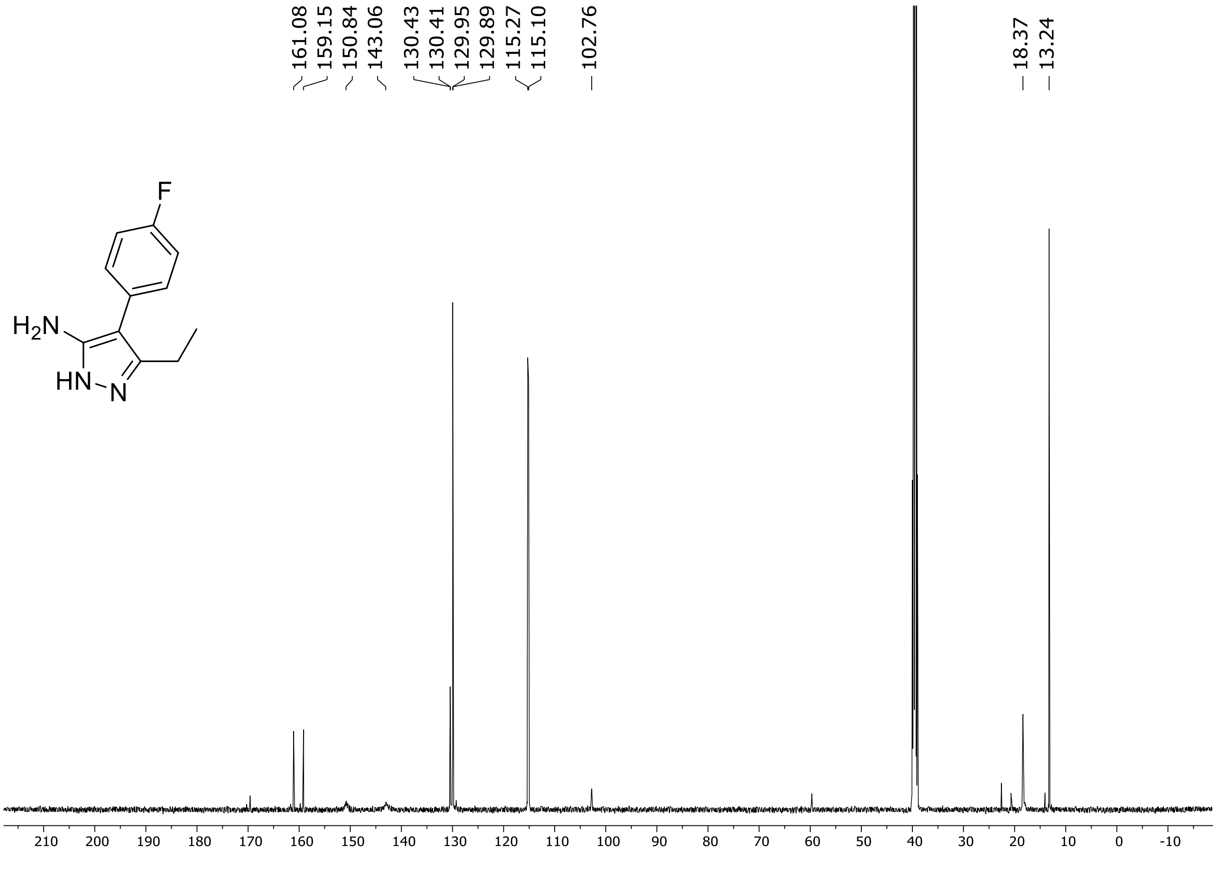

$^{19}$F (471 MHz) NMR spectrum of **S38** DMSO-$d_6$

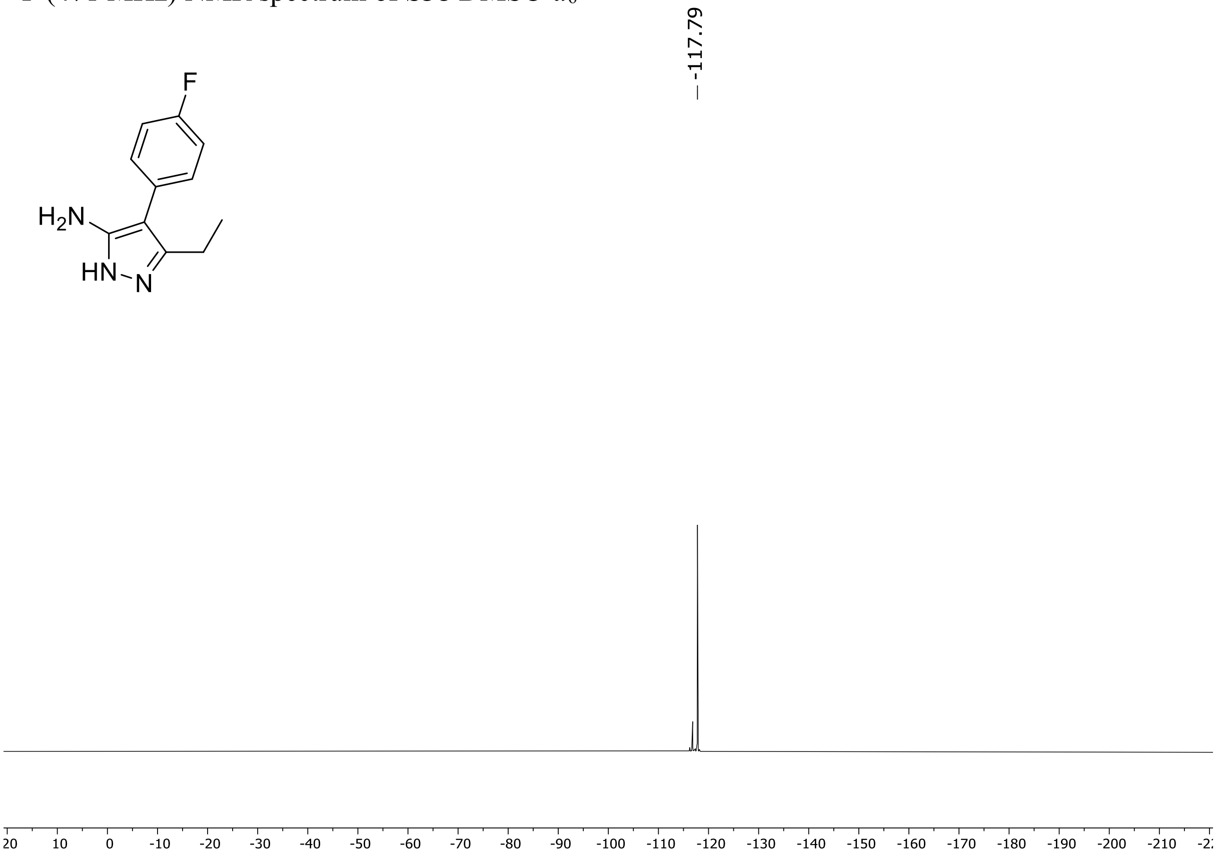

HRMS spectrum of **S38**

**$C_{11}H_{12}FN_3$** **mono m/z = 205,1001**

**ESI-** (MMI)
nitrogen flow 5 L/min, gas temperature 300°C, vaporizer 250°C,
nebulizer 45 psi, corona current 4 uA, Vcap -2000 V,
skimmer **65** V, fragmentor **150** V, dissolved in methanol (CH3OH)

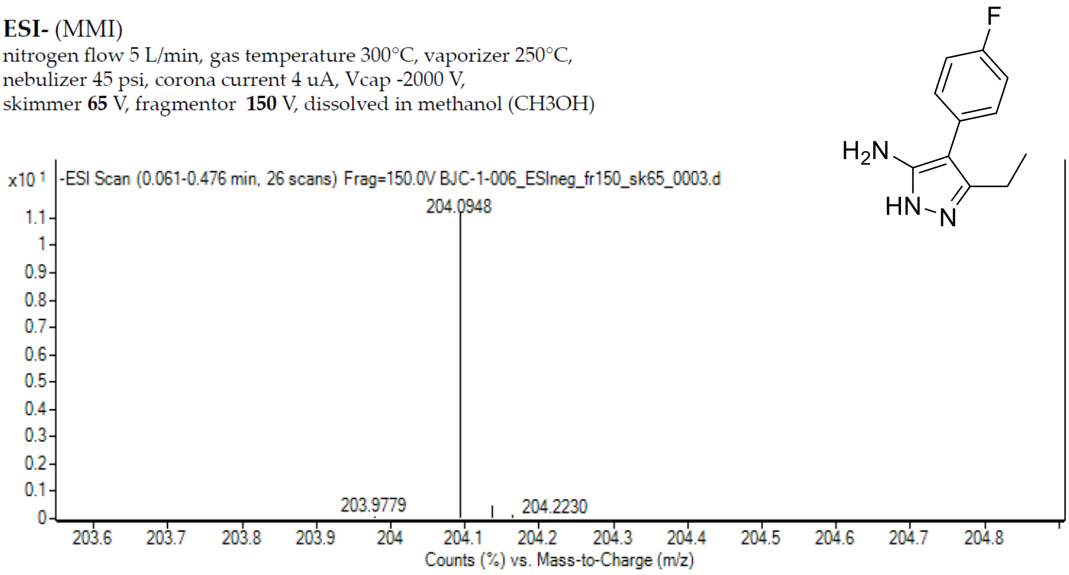

calculated mass: [M-H]$^-$ = 204,0942 observed: [M-H]$^-$ = 204,0948 max. mass error 2,9 ppm

$^{1}H$ (500 MHz) and $^{13}C$ NMR (126 MHz) spectra of **S39** in DMSO-$d_6$

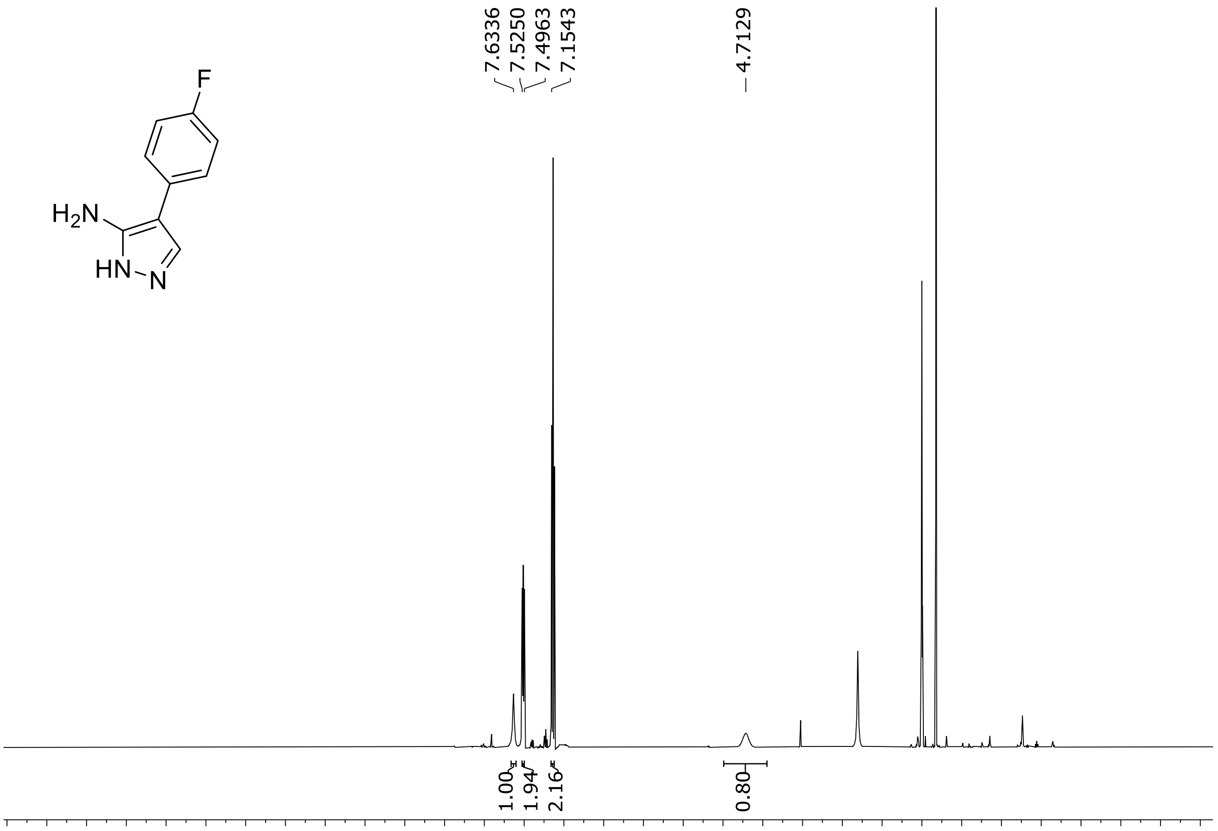

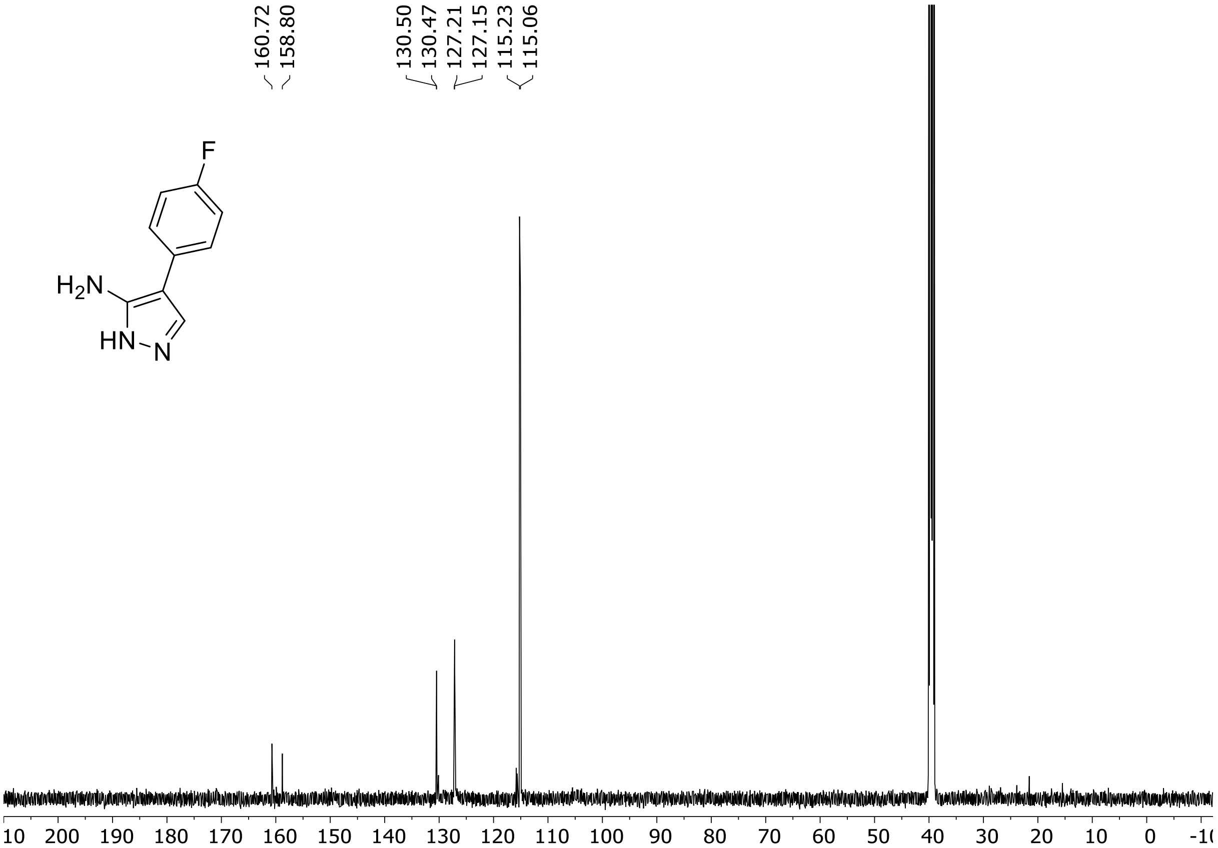

HRMS spectrum **S39**

**$C_9H_8FN_3$** **mono m/z = 177,0702**

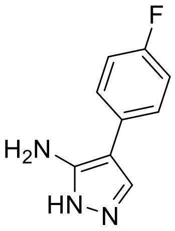

**ESI -** (MMI)
nitrogen flow 5 L/min, gas temperature 300°C, vaporizer 200°C,
nebulizer 45 psi, Vcap -2000 V, skimmer **65** V, fragmentor **40**V,
dissolved in DMSO

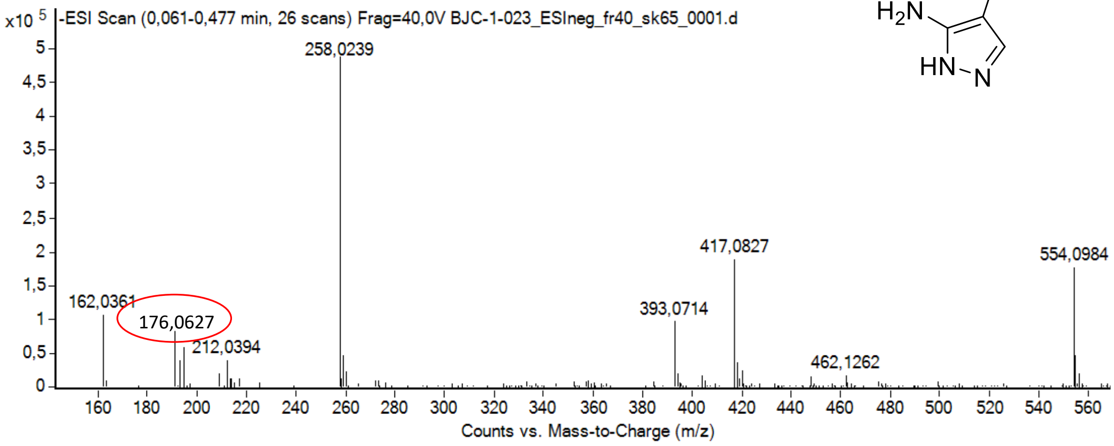

calculated mass:
$[M-H]^-$ = 176,0629 observed: $[M-H]^-$ = 176,0627 max. mass error = 1,1 ppm
$[M+Cl]^-$ = 212,0396 observed: $[M+Cl]^-$ = 212,0394 max. mass error = 0,9 ppm

$^{1}$H (500 MHz) and $^{13}$C NMR (126 MHz) spectra of **S40** in DMSO-$d_6$

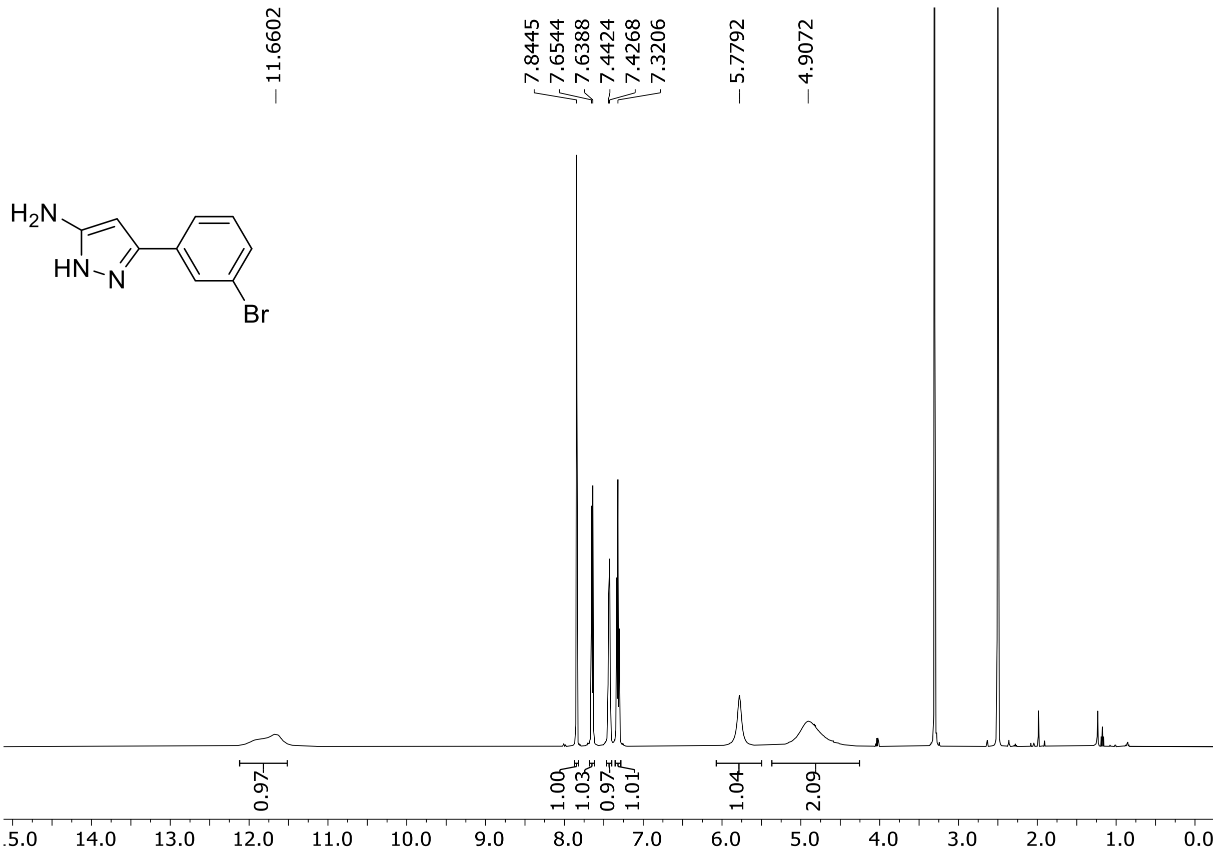

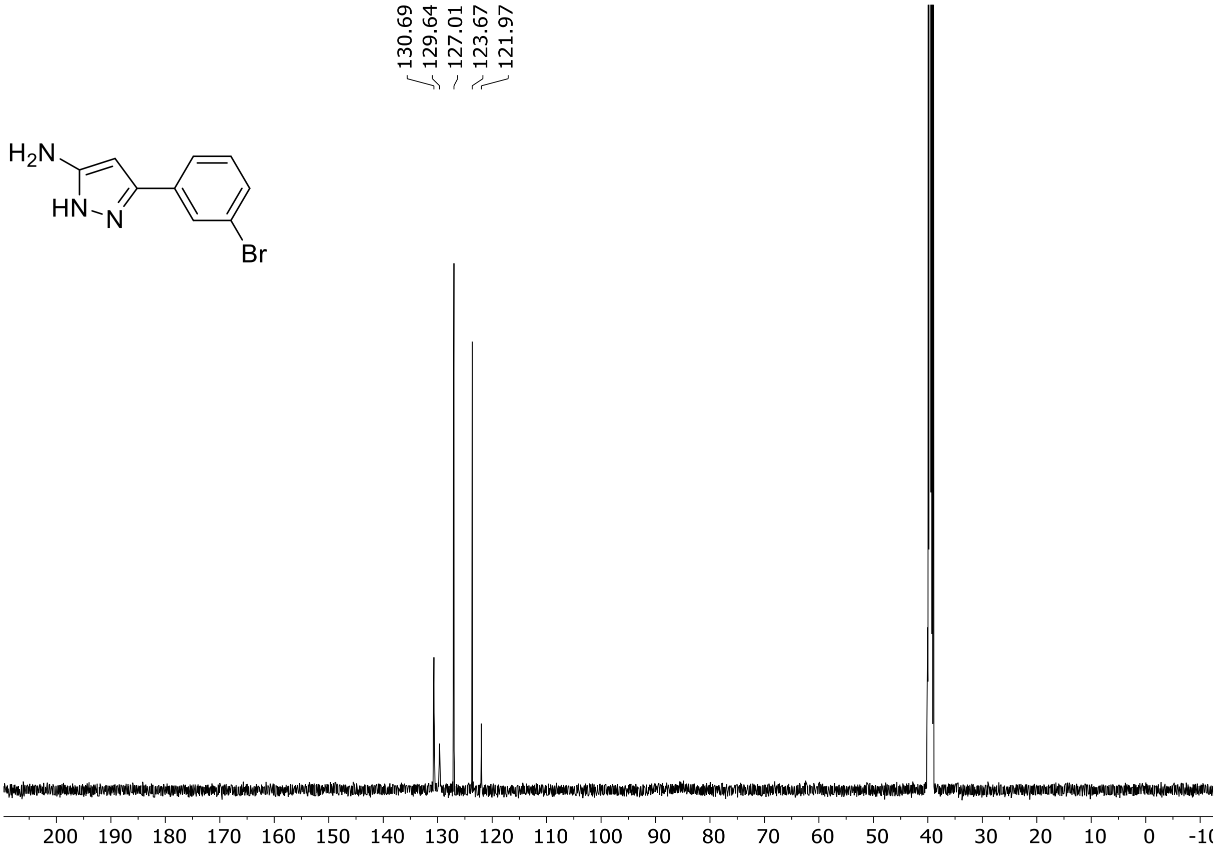

HRMS spectrum of **S40**

**$C_9H_8BrN_3$ mono m/z = 236.9902**

**APCI +** (MMI)
nitrogen flow 5 L/min, gas temperature 300°C, nebulizer 45 psi,
skimmer 60 V, fragmentor 32 V, dissolved in methanol

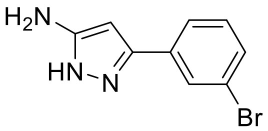

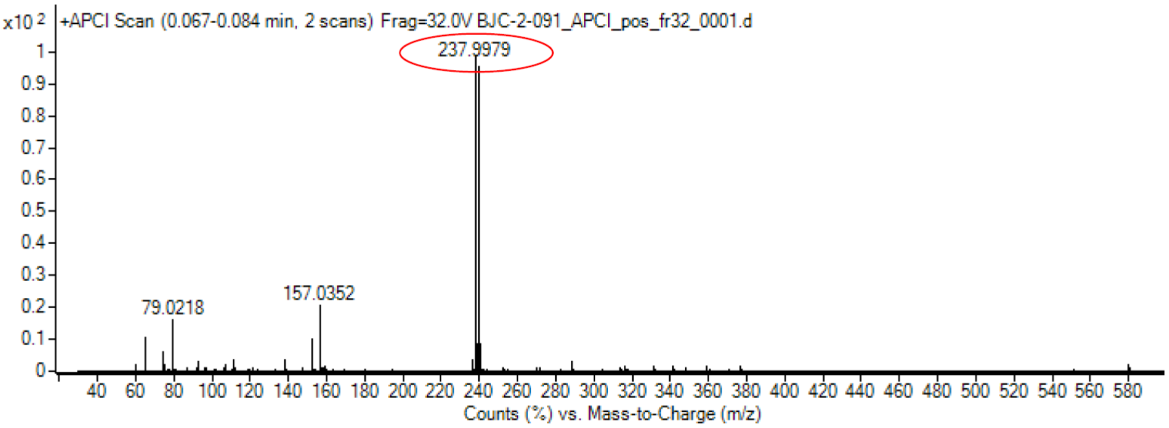

calculated mass: $[M+H]^+$ = 237.9974 observed: $[M+H]^+$ = 237.9979 max. mass error = 2.1 ppm

[1]H (500 MHz) and [13]C NMR (126 MHz) spectra of **S41** in chloroform-*d*

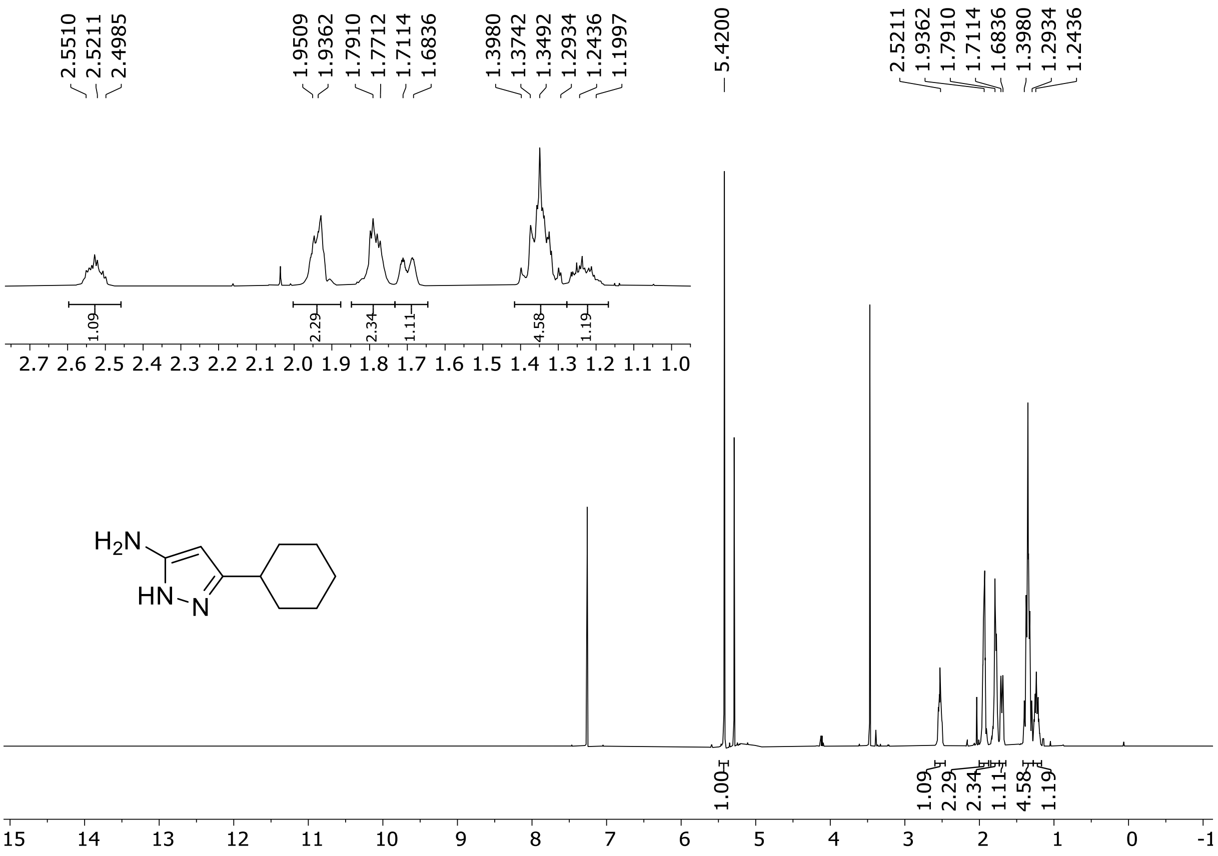

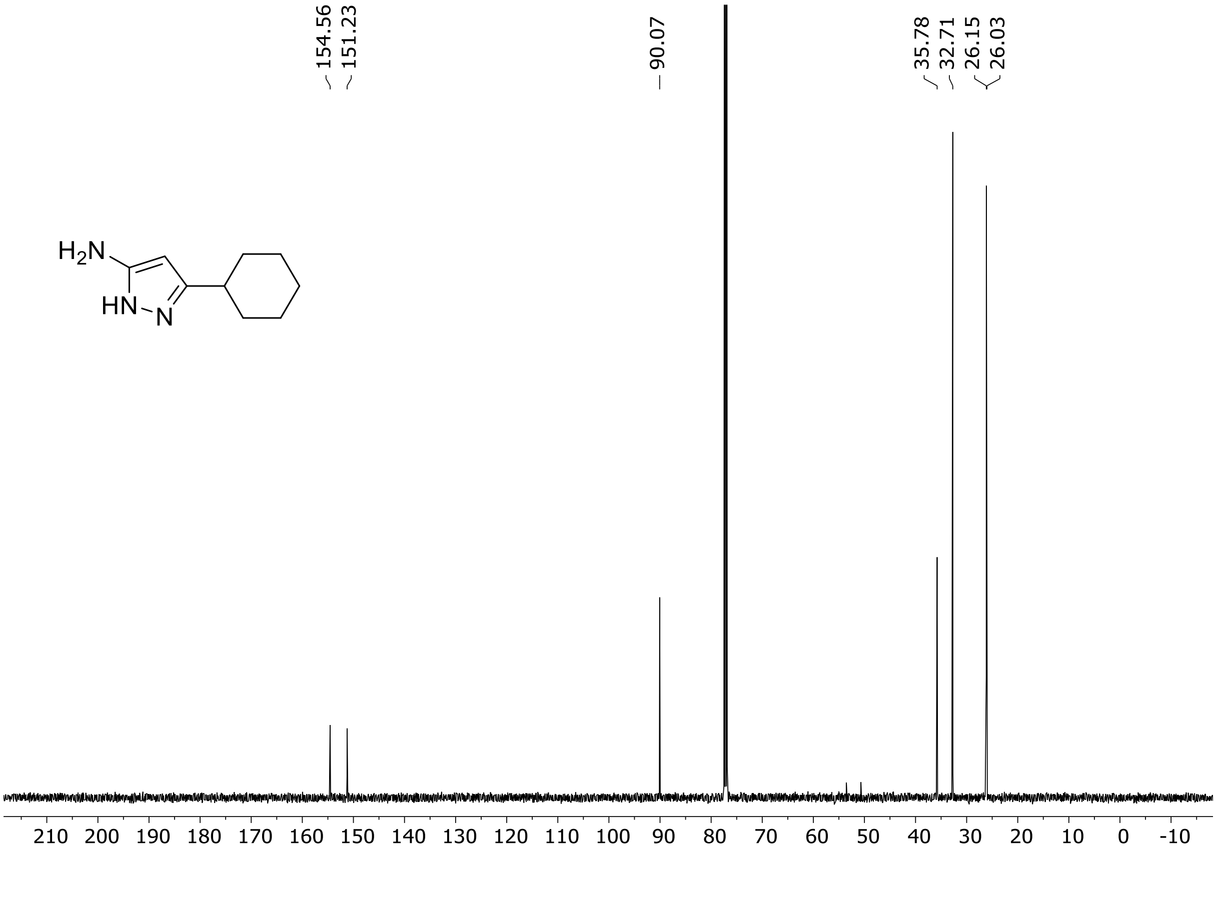

HRMS spectrum of **S41**

$C_9H_{15}N_3$ **mono m/z = 165.1266**

**APCI +** (MMI)
nitrogen flow 5 L/min, gas temperature 300°C, nebulizer 45 psi, skimmer 60 V, fragmentor 22 V, dissolved in methanol

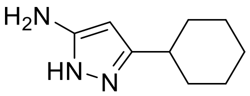

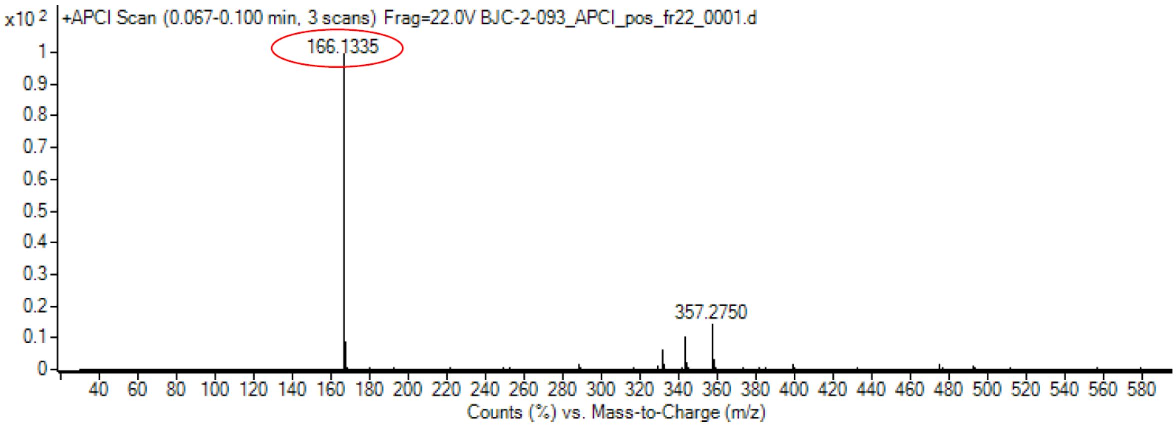

calculated mass: $[M+H]^+$= 166.1339 observed: $[M+H]^+$ = 166.1335 max. mass error = 2.4 ppm

[1]H (500 MHz) and [13]C NMR (126 MHz) spectra of **S42** in DMSO-$d_6$

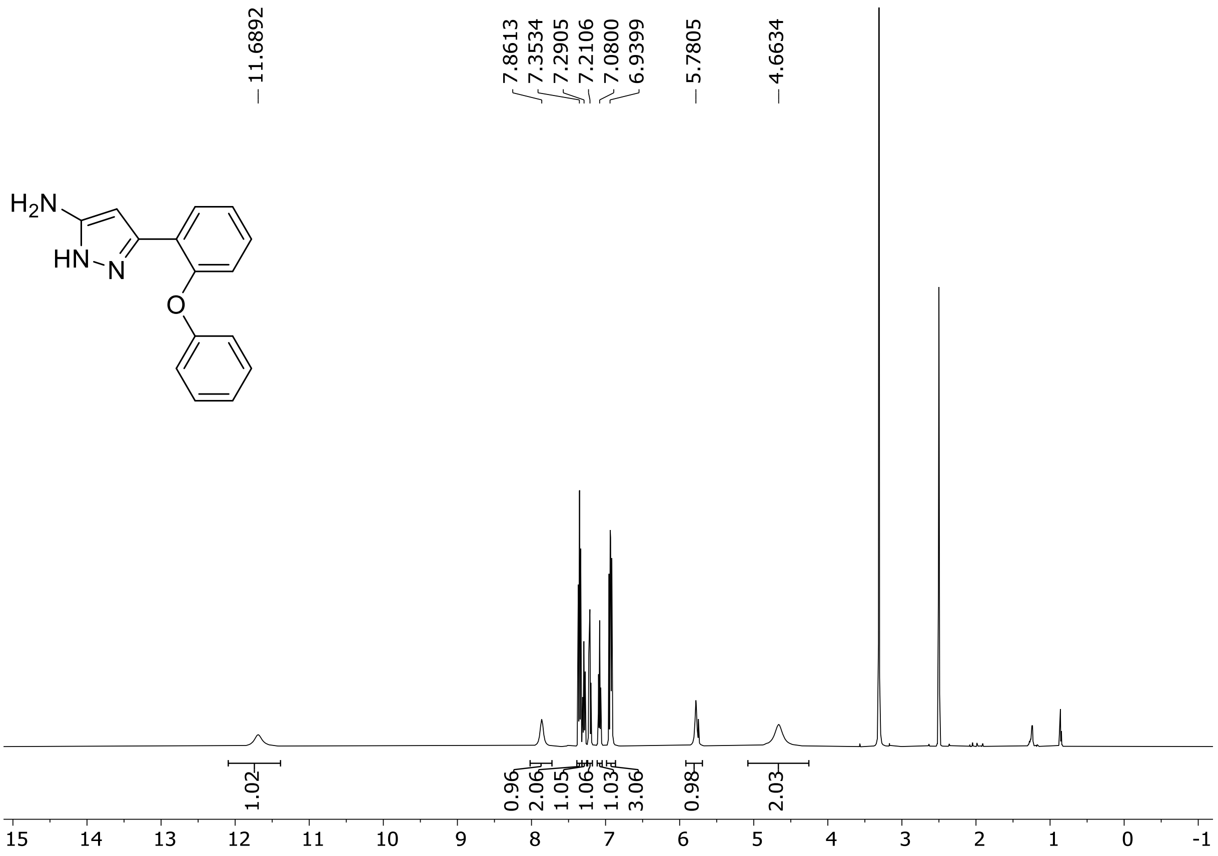

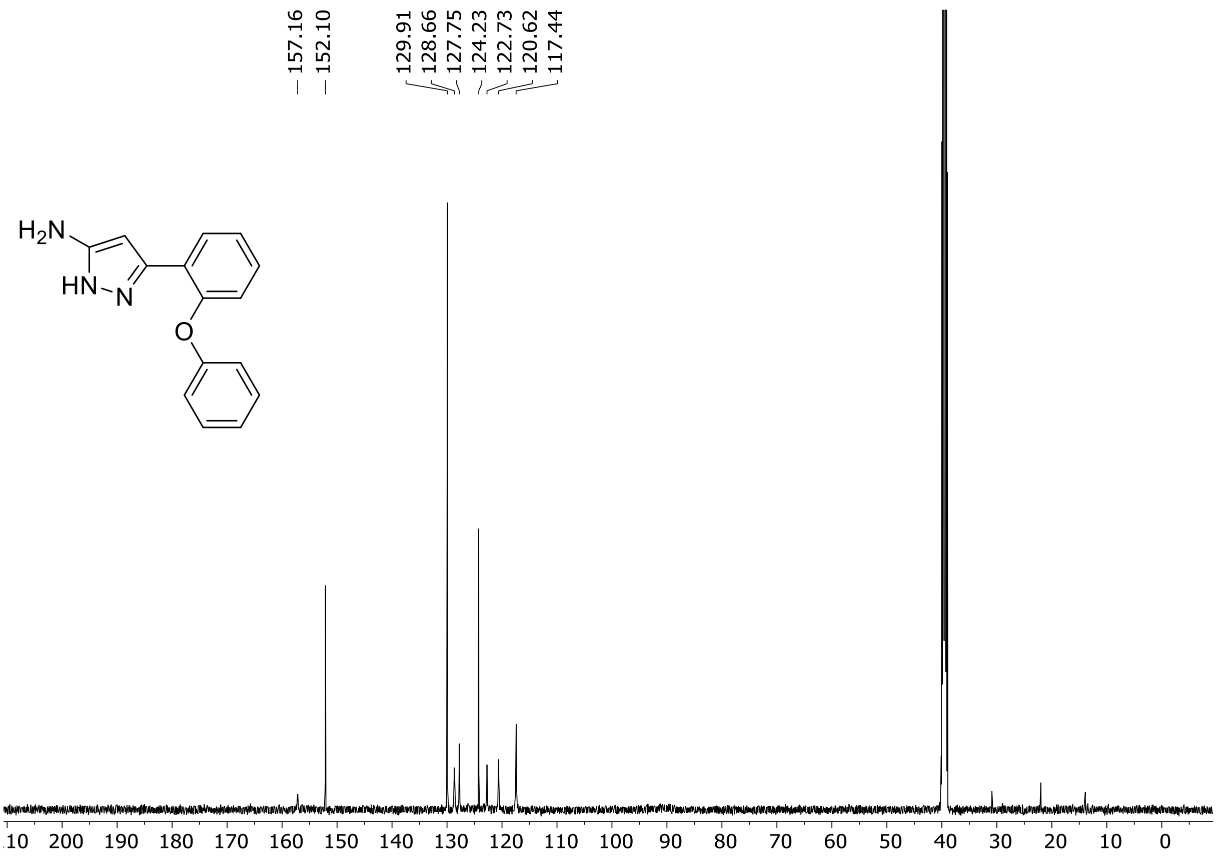

HRMS spectrum of **S42**

**$C_{15}H_{13}N_3O$ mono m/z = 251.1059**

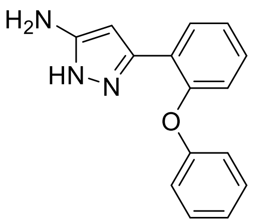

**APCI +** (MMI)
nitrogen flow 5 L/min, gas temperature 300°C, nebulizer 45 psi,
skimmer 60 V, fragmentor 25 V, dissolved in methanol

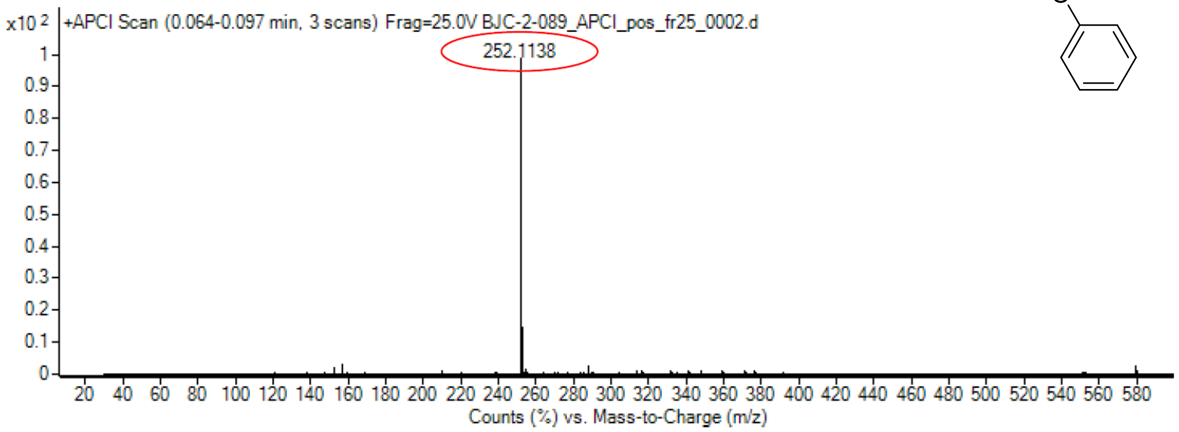

calculated mass: $[M+H]^+$ = 252.1131 observed: $[M+H]^+$ = 252.1138 max. mass error = 2.8 ppm

$^{1}H$ (500 MHz) and $^{13}C$ NMR (126 MHz) spectra of **S43** in DMSO-$d_6$

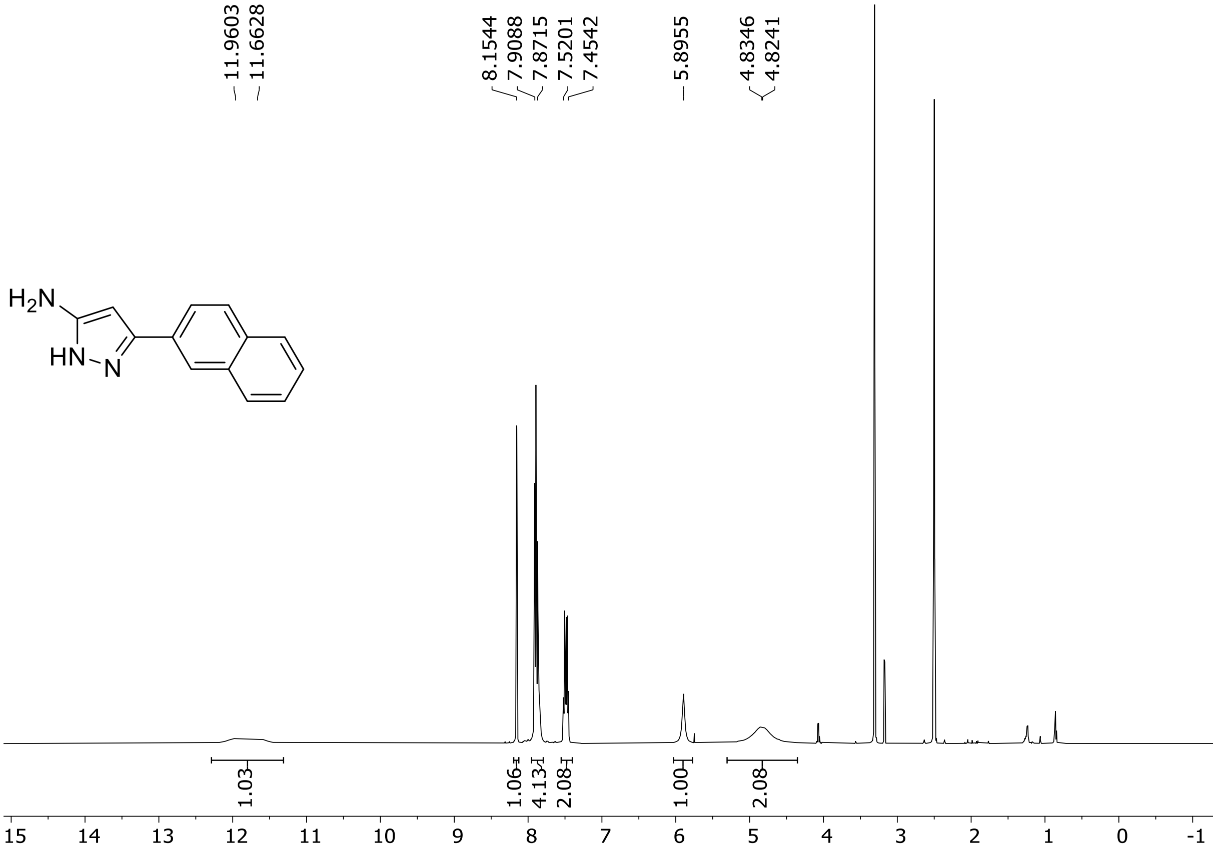

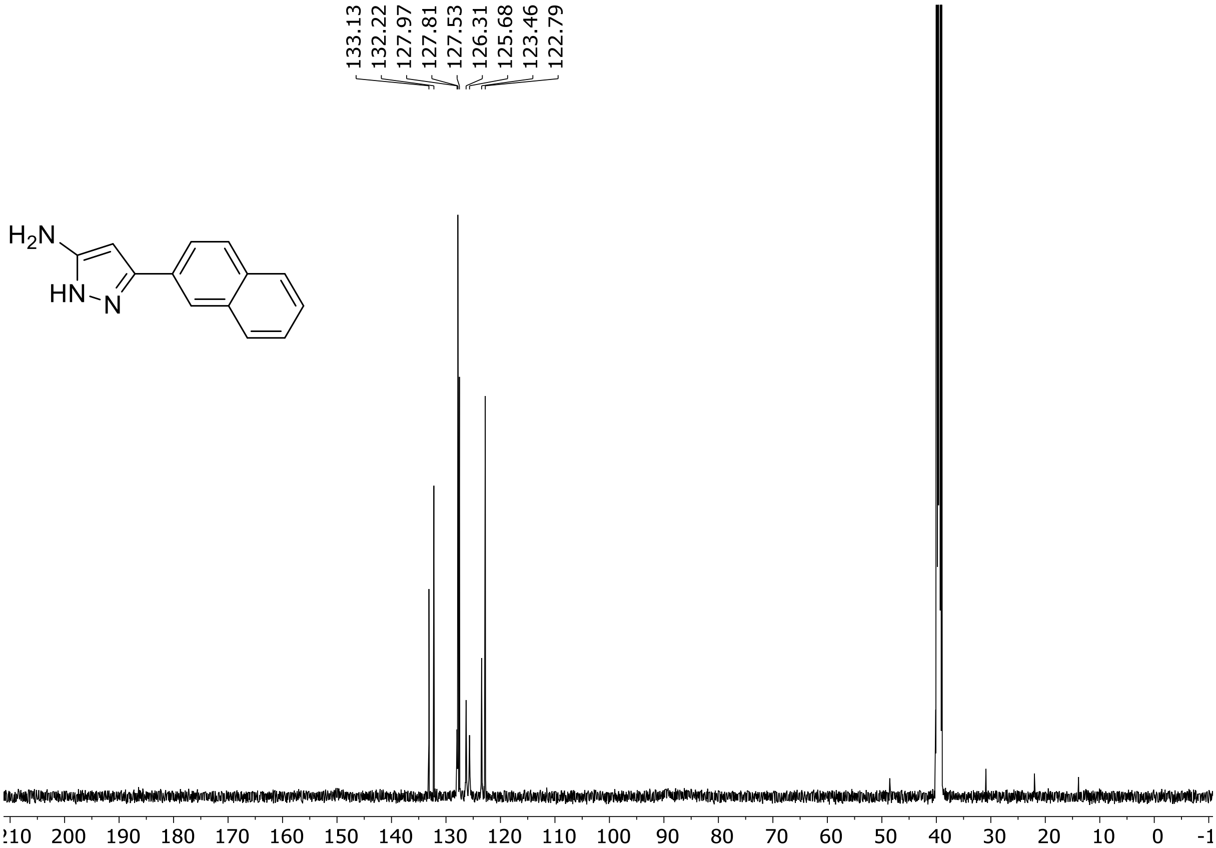

HRMS spectrum of **S43**

$C_{13}H_{11}N_3$ **mono m/z = 209.0953**

**ESI +** (MMI)
nitrogen flow 5 L/min, gas temperature 300°C, nebulizer 45 psi, skimmer 60 V, fragmentor 40 V, dissolved in methanol

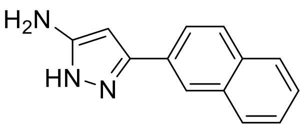

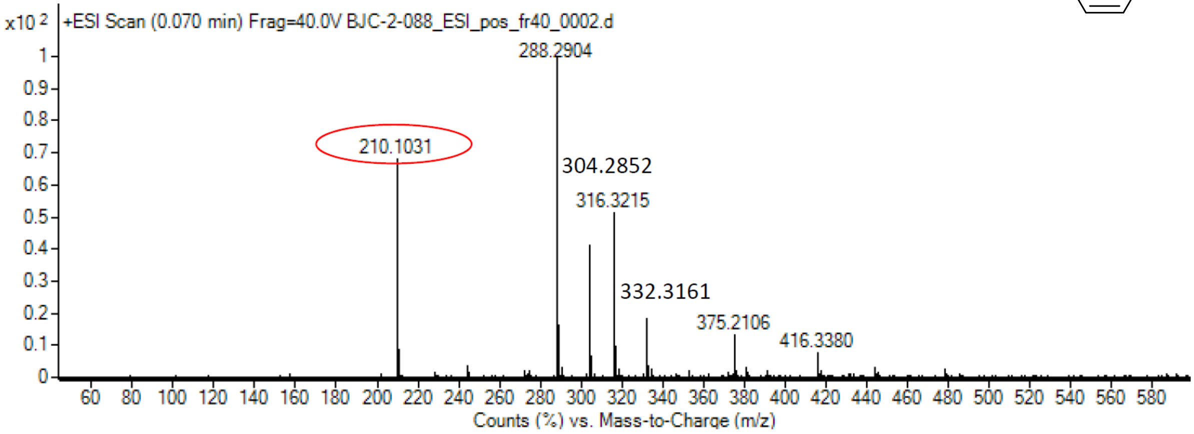

calculated mass: $[M+H]^+ = 210.1026$ observed: $[M+H]^+ = 210.1031$ max. mass error = 2.4 ppm

$^{1}H$ (500 MHz) and $^{13}C$ NMR (126 MHz) spectra of **S44** in chloroform-*d*

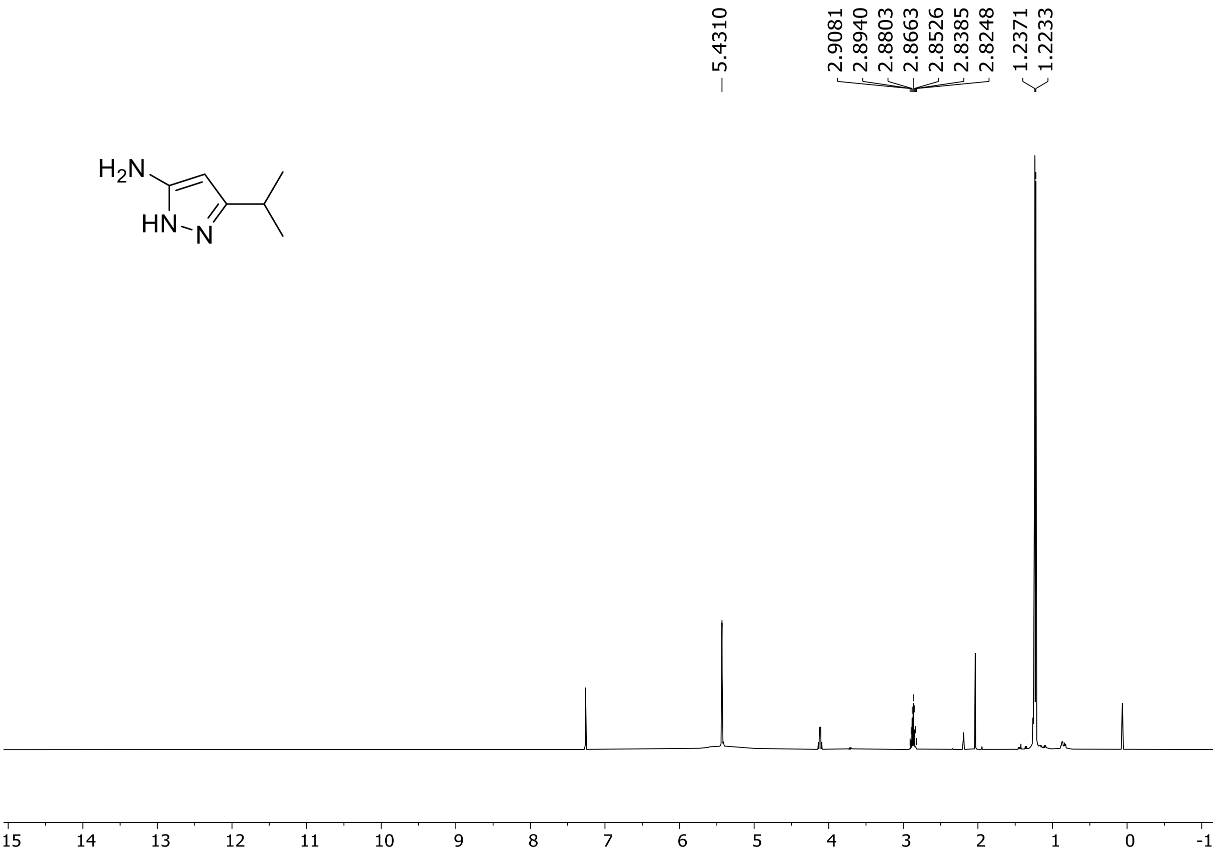

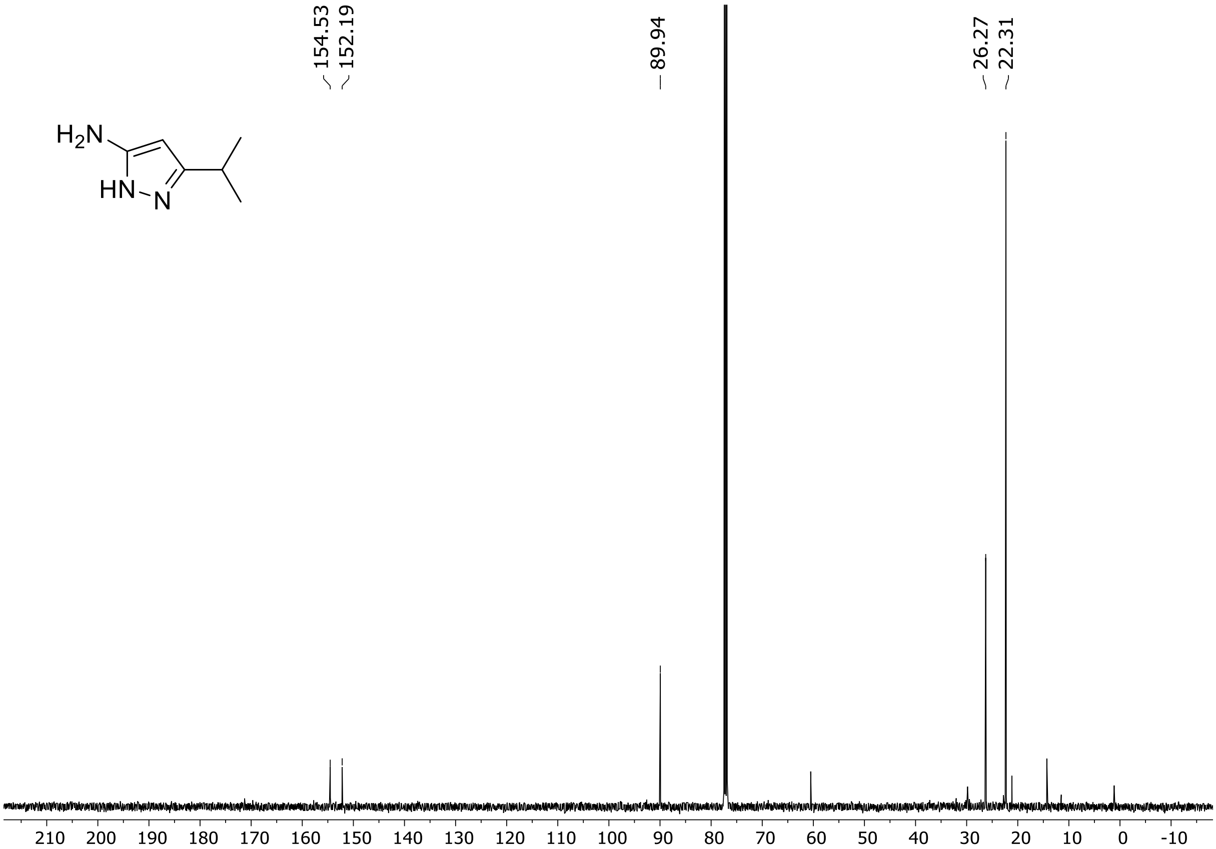

$^{1}H$ (500 MHz) and $^{13}C$ NMR (126 MHz) spectra of **S45** in DMSO-$d_6$

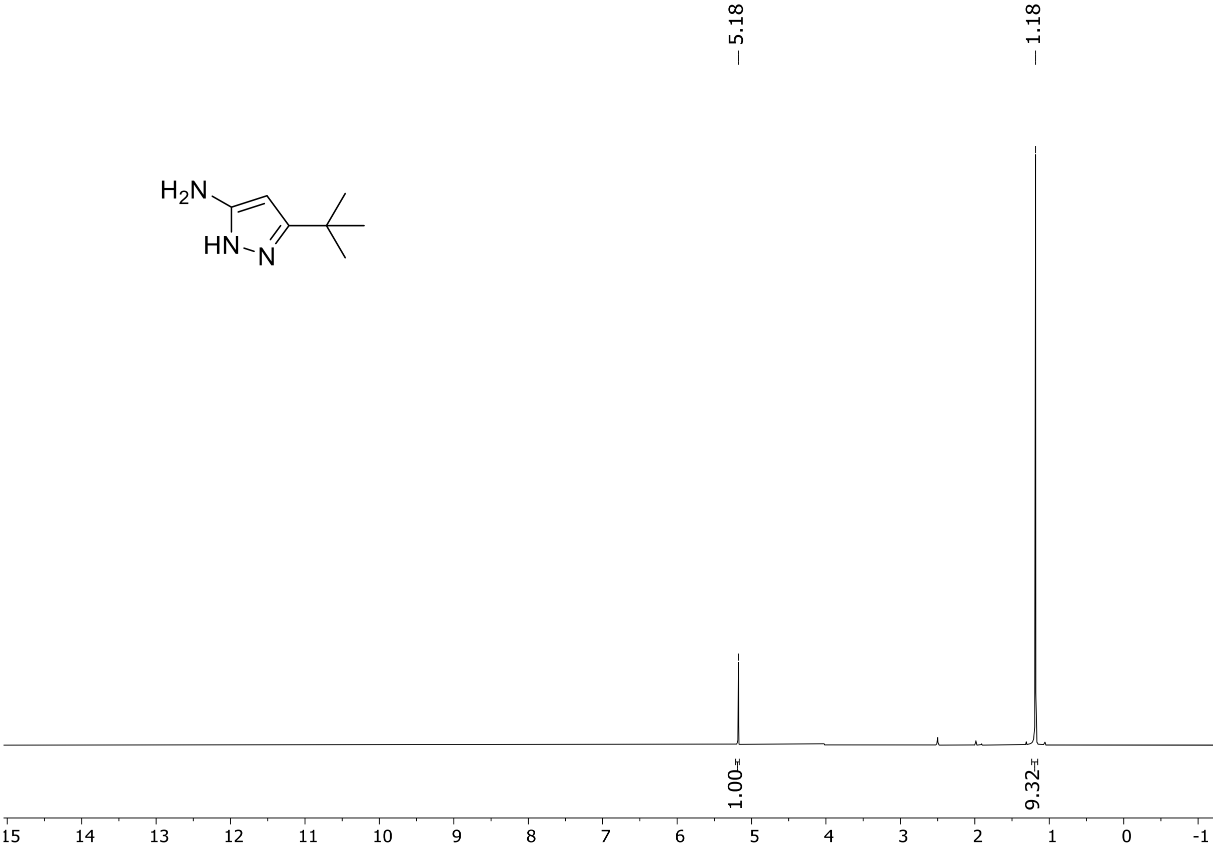

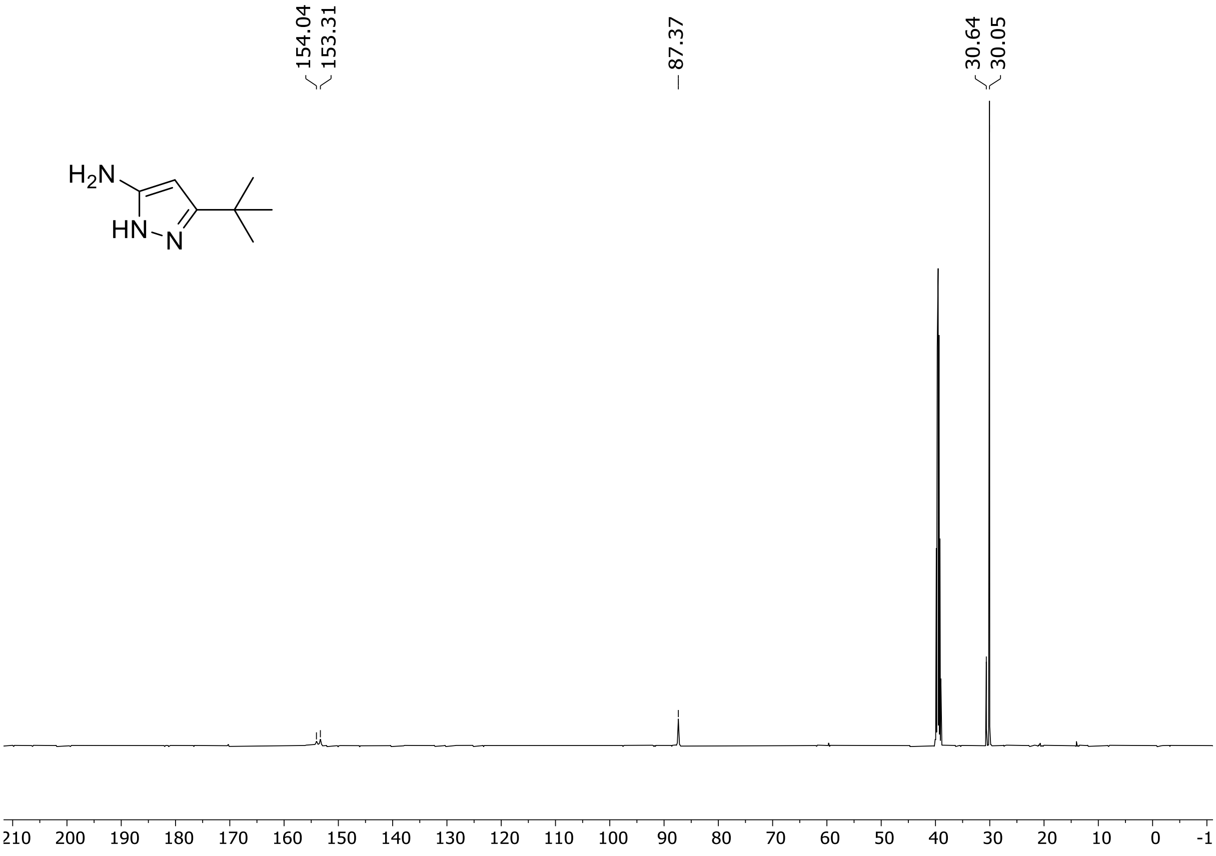

HRMS spectrum of **S45**

$C_7H_{13}N_3$ **mono m/z = 139.1109**

**APCI +** (MMI)
nitrogen flow 5 L/min, gas temperature 300°C, nebulizer 45 psi, vaporizer 200°C
skimmer **65** V, fragmentor **20** V, dissolved in methanol

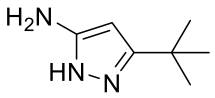

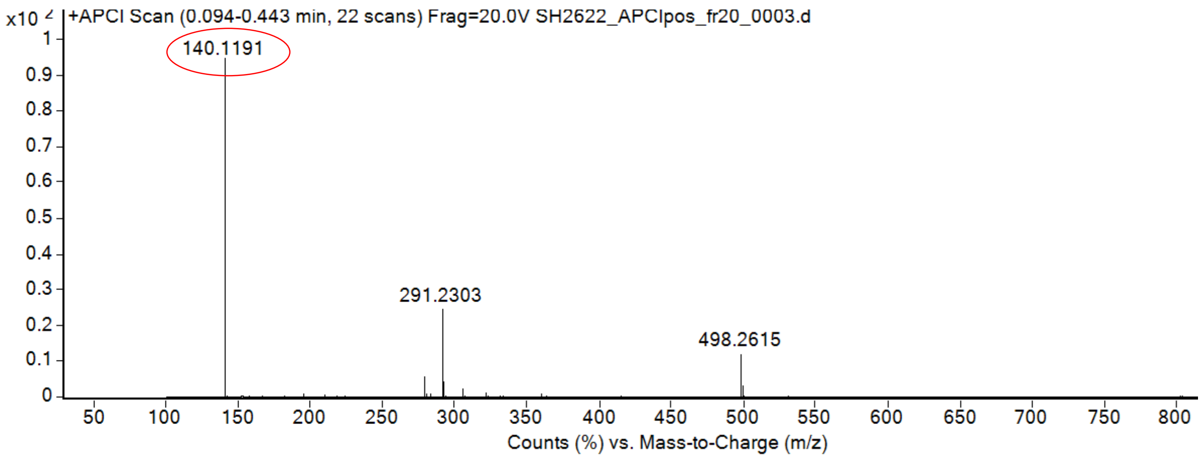

calculated mass: $[M+H]^+$ = 140.1182 observed: $[M+H]^+$ = 140.1191 max. mass error = 6.4 ppm

[1]H (500 MHz) and [13]C NMR (126 MHz) spectra of **S46** in chloroform-*d*

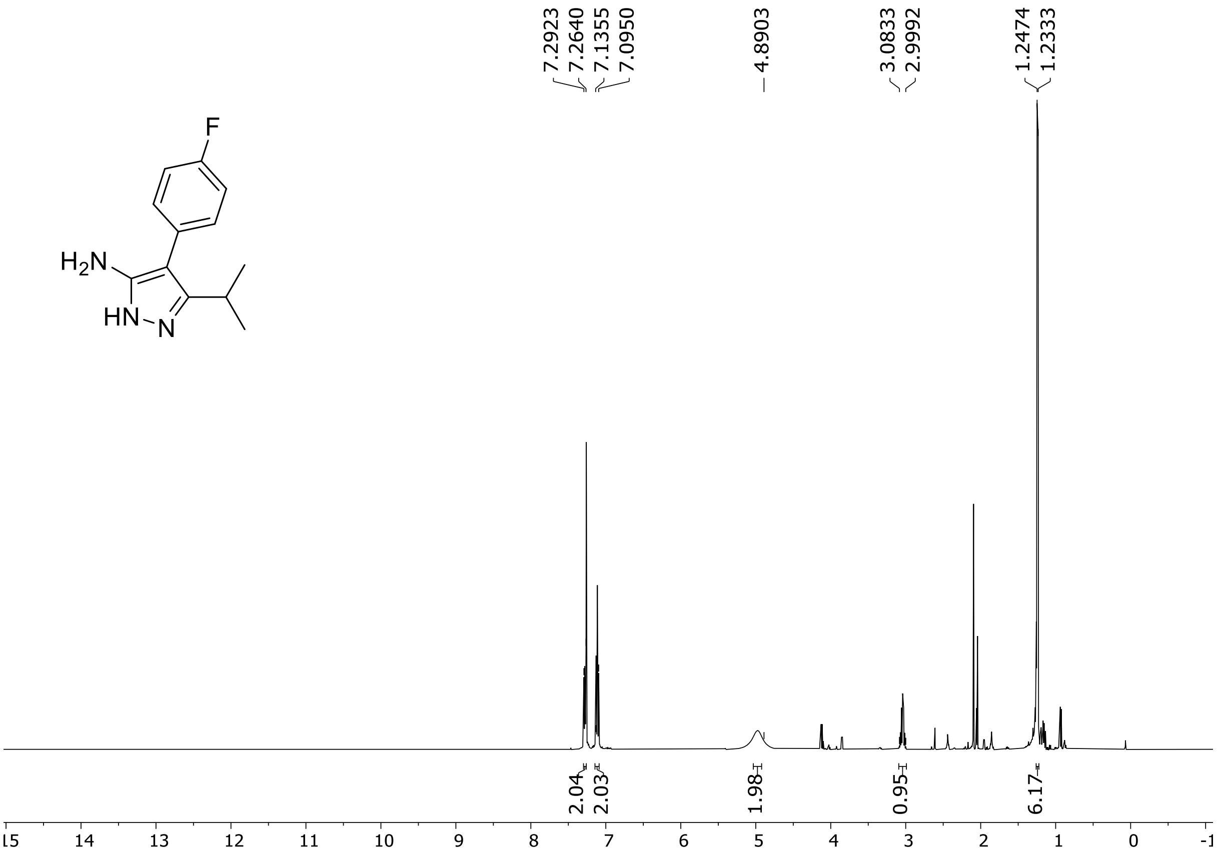

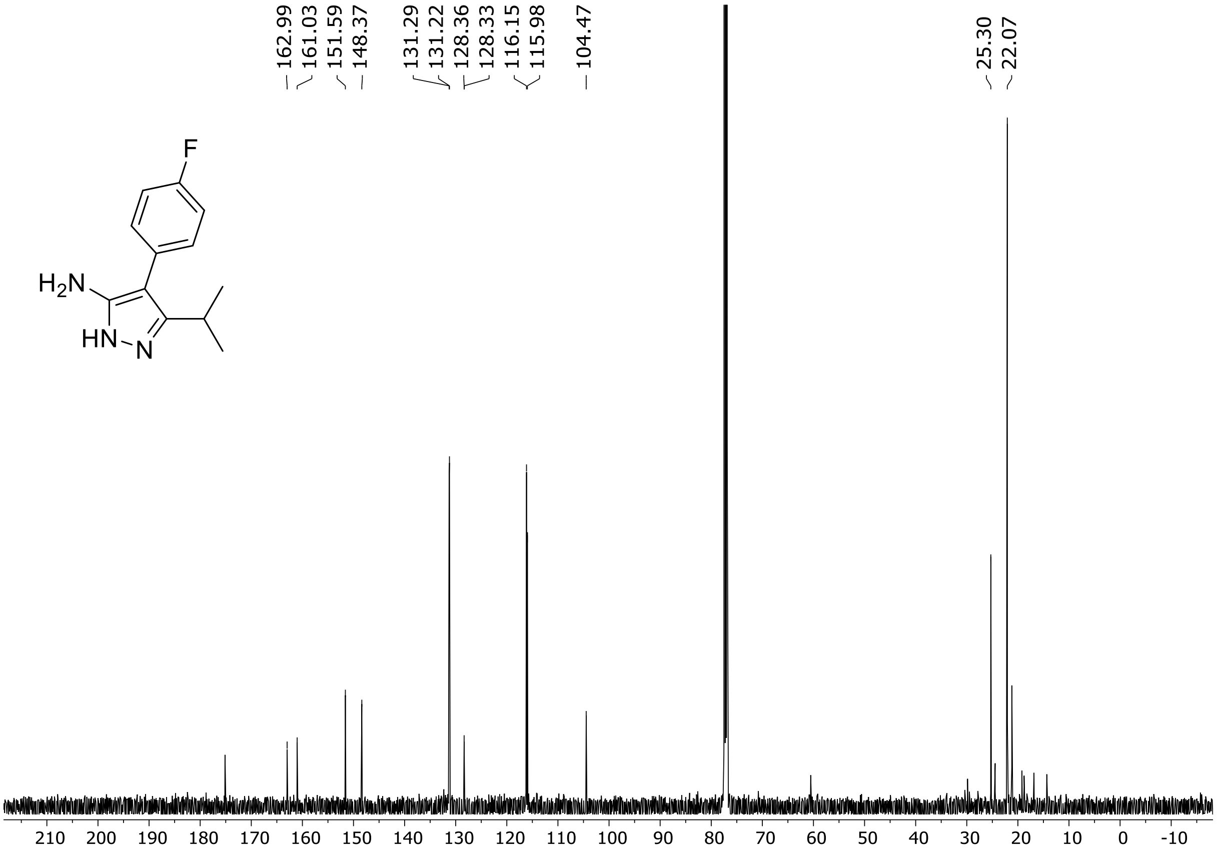

$^{1}$H (500 MHz) and $^{13}$C NMR (126 MHz) spectra of **S47** in chloroform-*d*

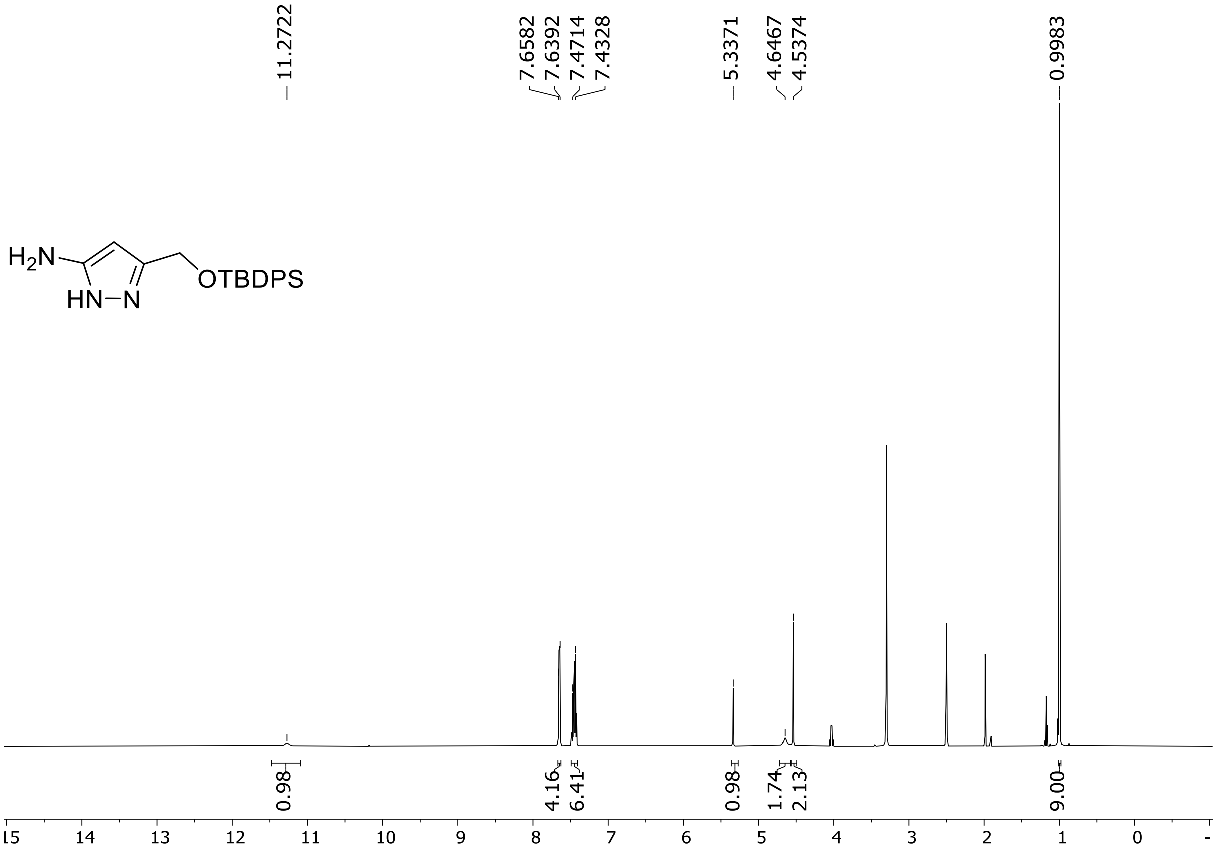

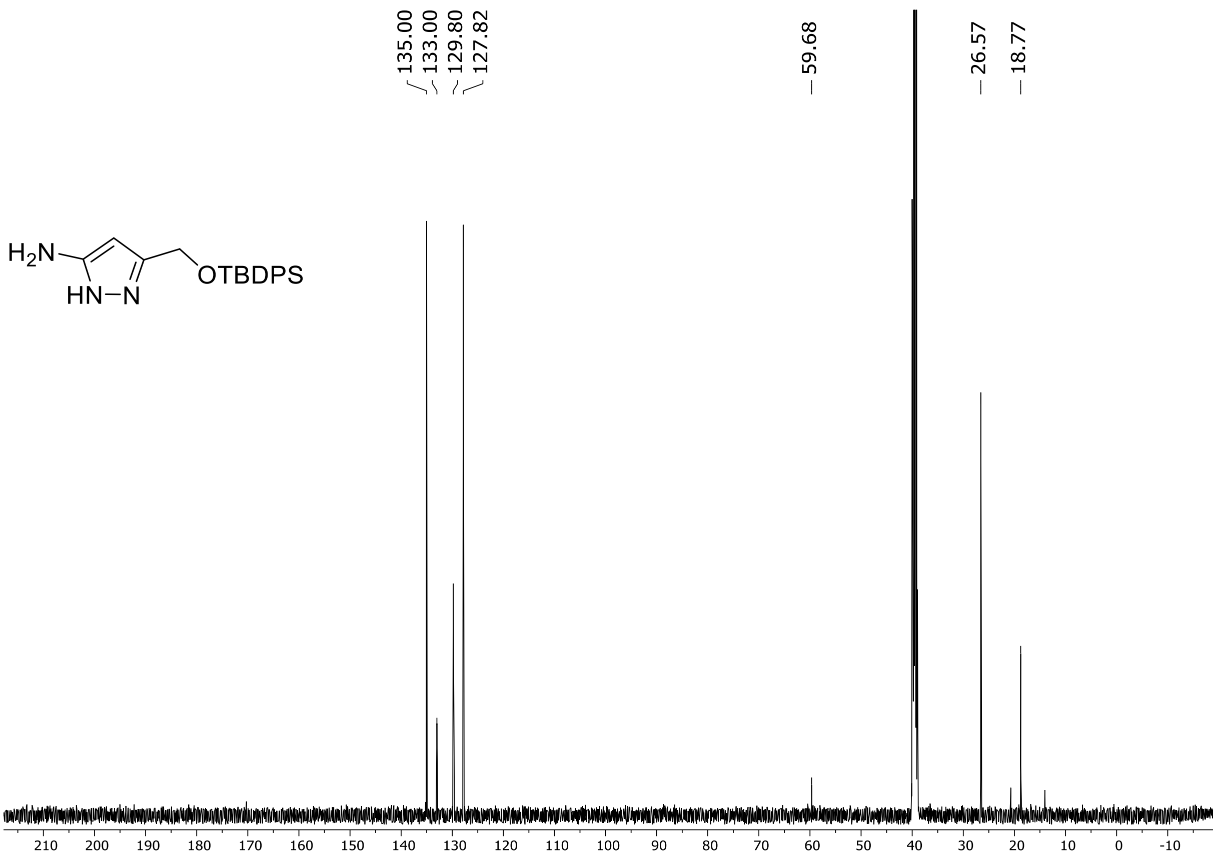

HRMS spectrum of **S47**

$C_{20}H_{25}N_{3}OSi$ **mono m/z = 351.1767**

**APCI +** (MMI)
nitrogen flow 5 L/min, gas temperature 300°C, nebulizer 45 psi, vaporizer 200°C
skimmer **65** V, fragmentor **5** V, dissolved in methanol

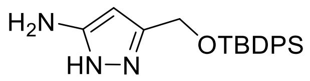

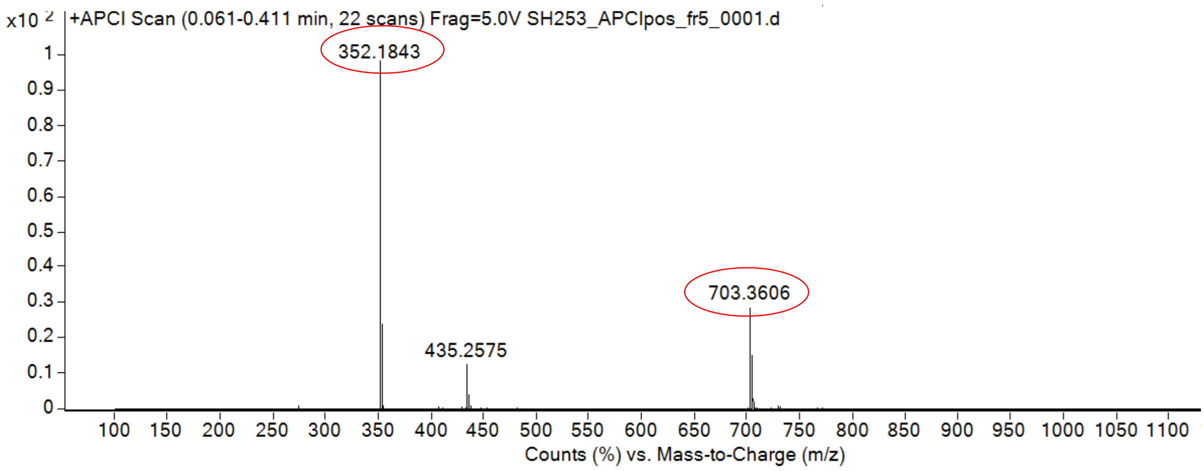

calculated mass: $[M+H]^+$ = 352.1840 observed: $[M+H]^+$ = 352.1843 max. mass error = 0.8 ppm

calculated mass: $[2xM+H]^+$ = 703.3607 observed: $[2xM+H]^+$ = 703.3606 max. mass error = 0.1 ppm

$^{1}H$ (500 MHz) and $^{13}C$ NMR (126 MHz) spectra of **S48** in DMSO-$d_6$

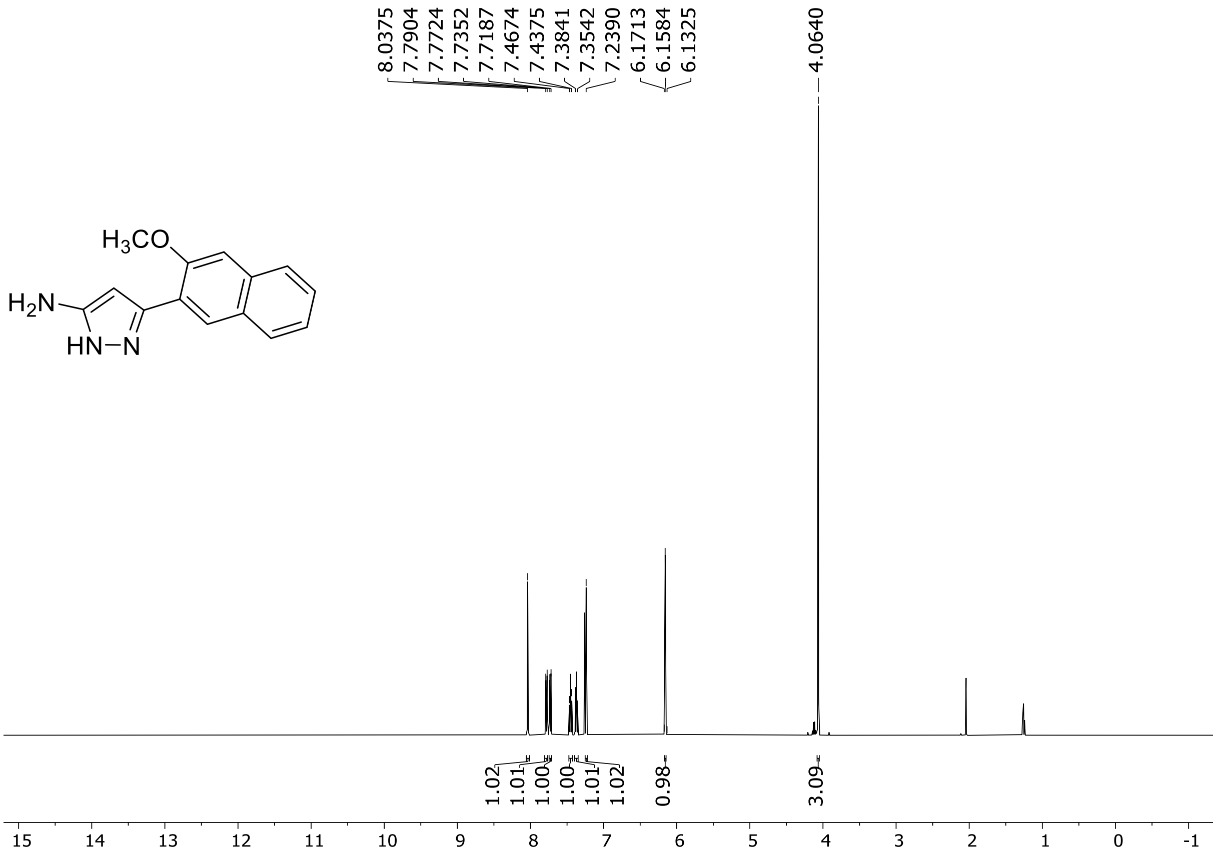

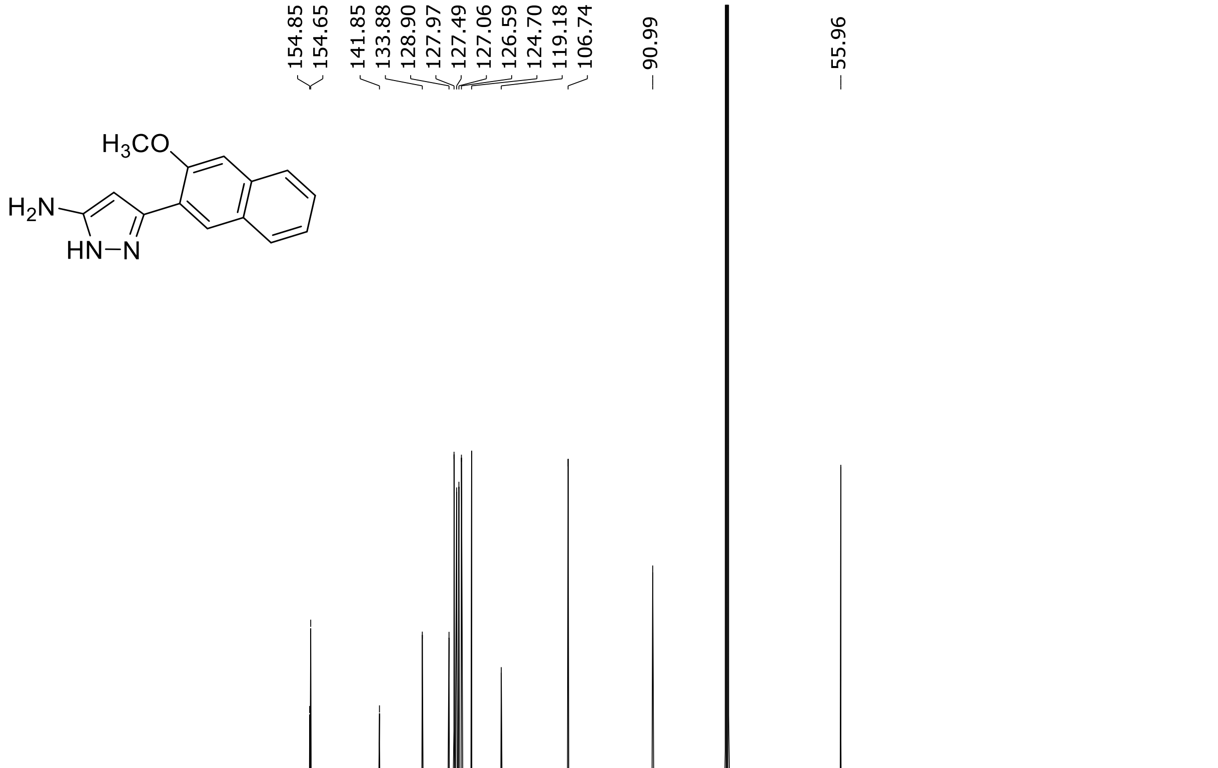

$^{1}H$ (500 MHz) and $^{13}C$ NMR (126 MHz) spectra of **S49** in DMSO-$d_6$

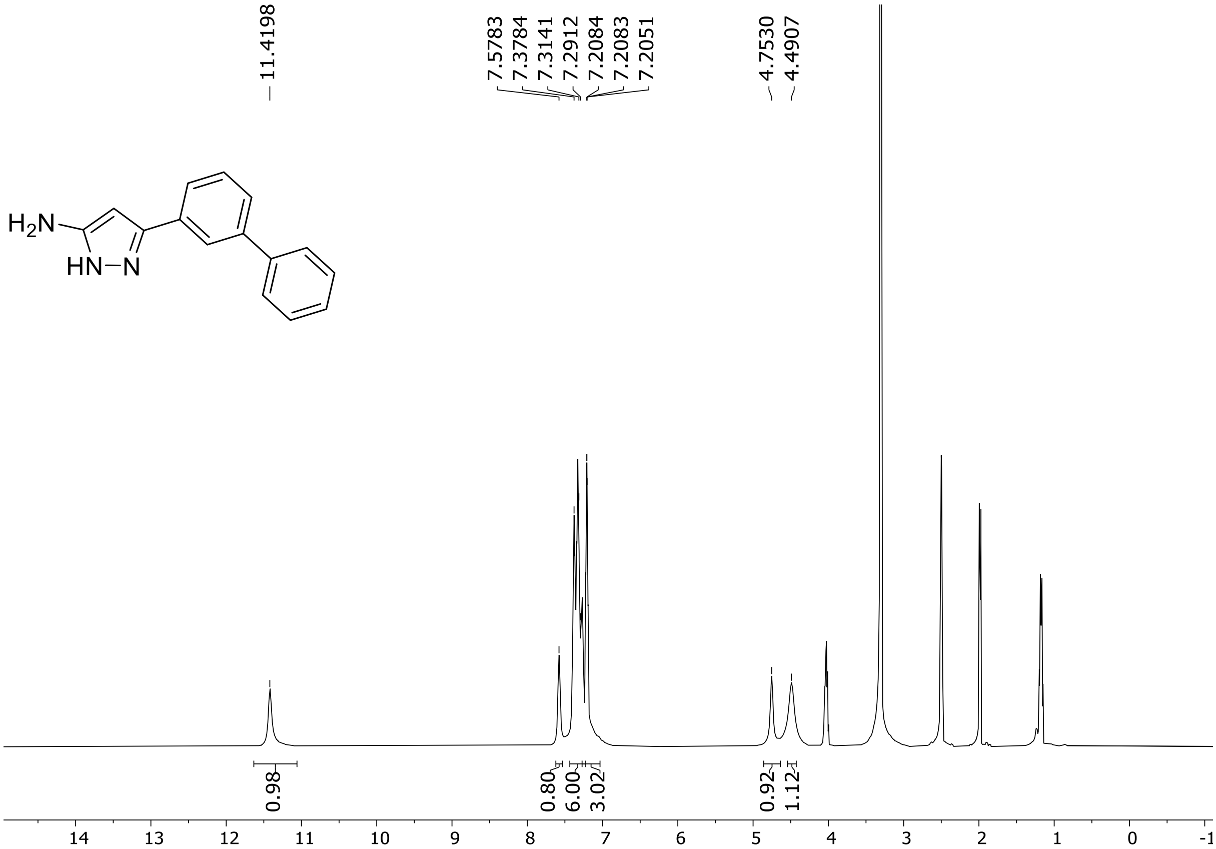

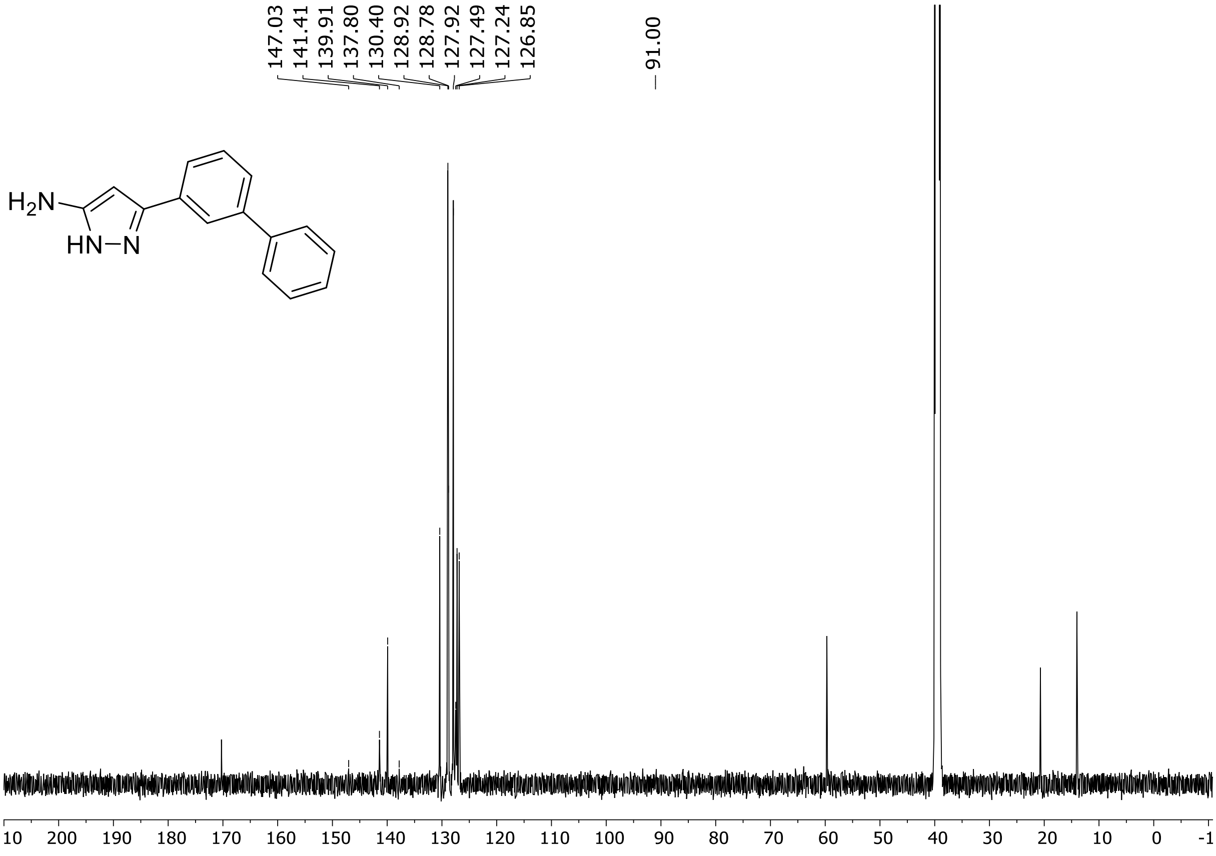

HRMS spectrum of **S49**

$C_{15}H_{13}N_3$ **mono m/z = 235.1109**

**APCI +** (MMI)
nitrogen flow 5 L/min, gas temperature 300°C, nebulizer 45 psi, vaporizer 200°C
skimmer **65** V, fragmentor **16** V, dissolved in methanol

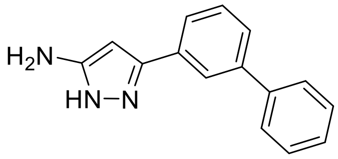

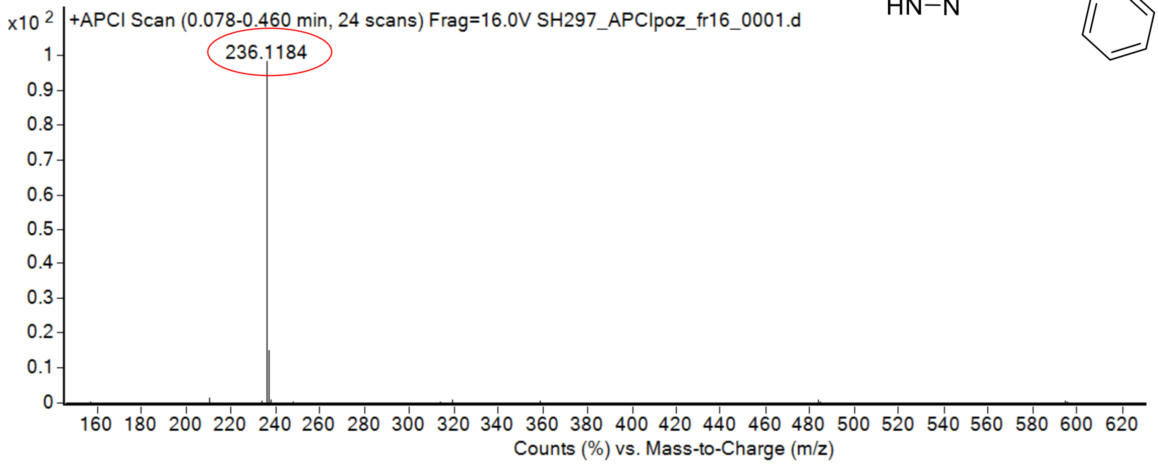

calculated mass: $[M+H]^+$ = 236.1182 observed: $[M+H]^+$ = 236.1184 max. mass error = 0.8 ppm

$^{1}H$ (500 MHz) and $^{13}C$ NMR (126 MHz) spectra of **S50** in DMSO-$d_6$

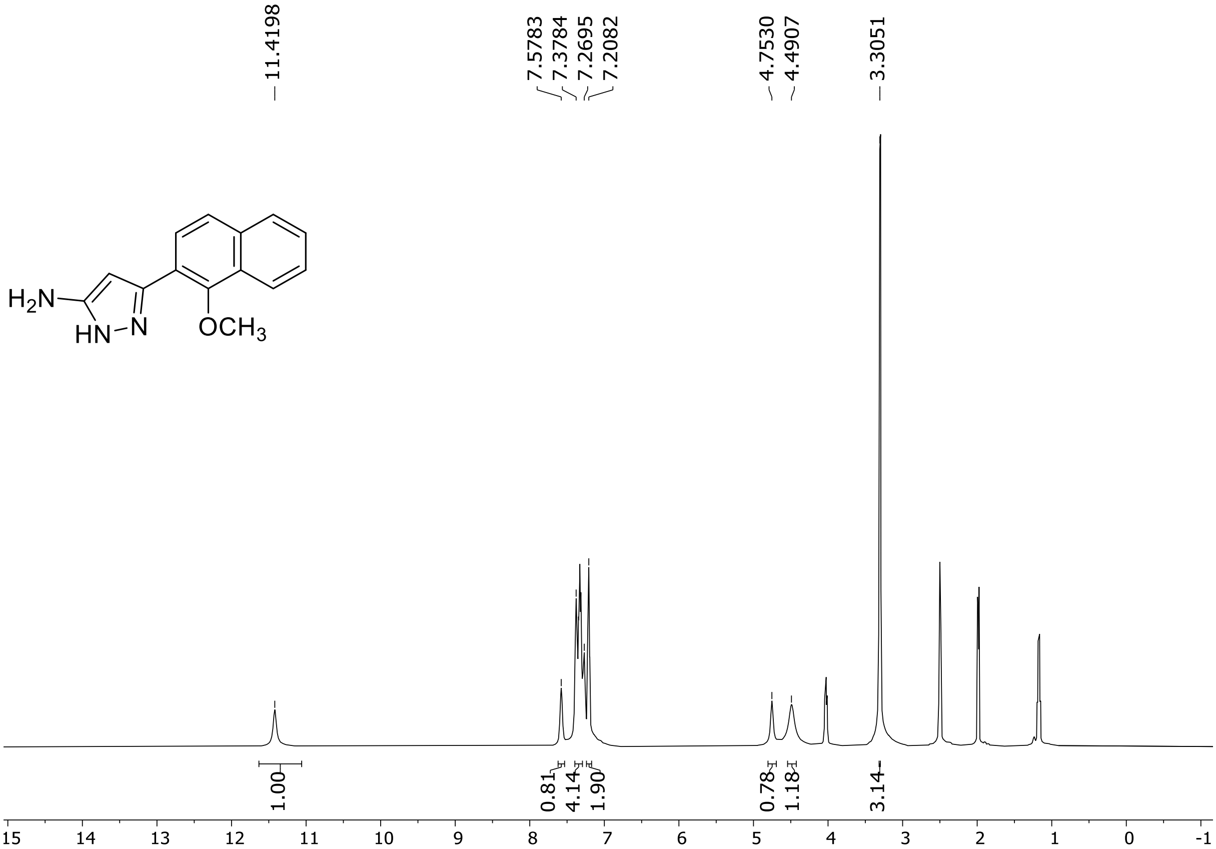

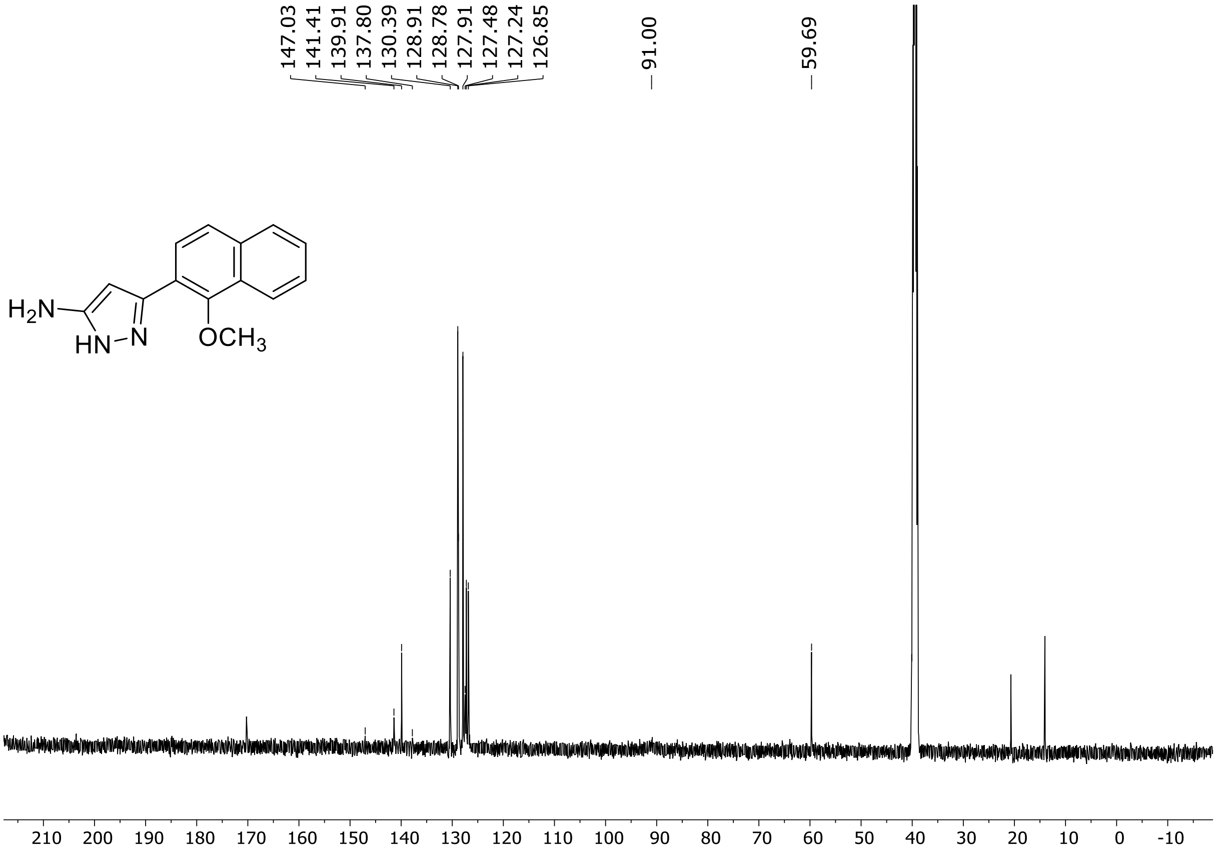

[1]H (500 MHz) and [13]C NMR (126 MHz) spectra of **S51** in DMSO-$d_6$

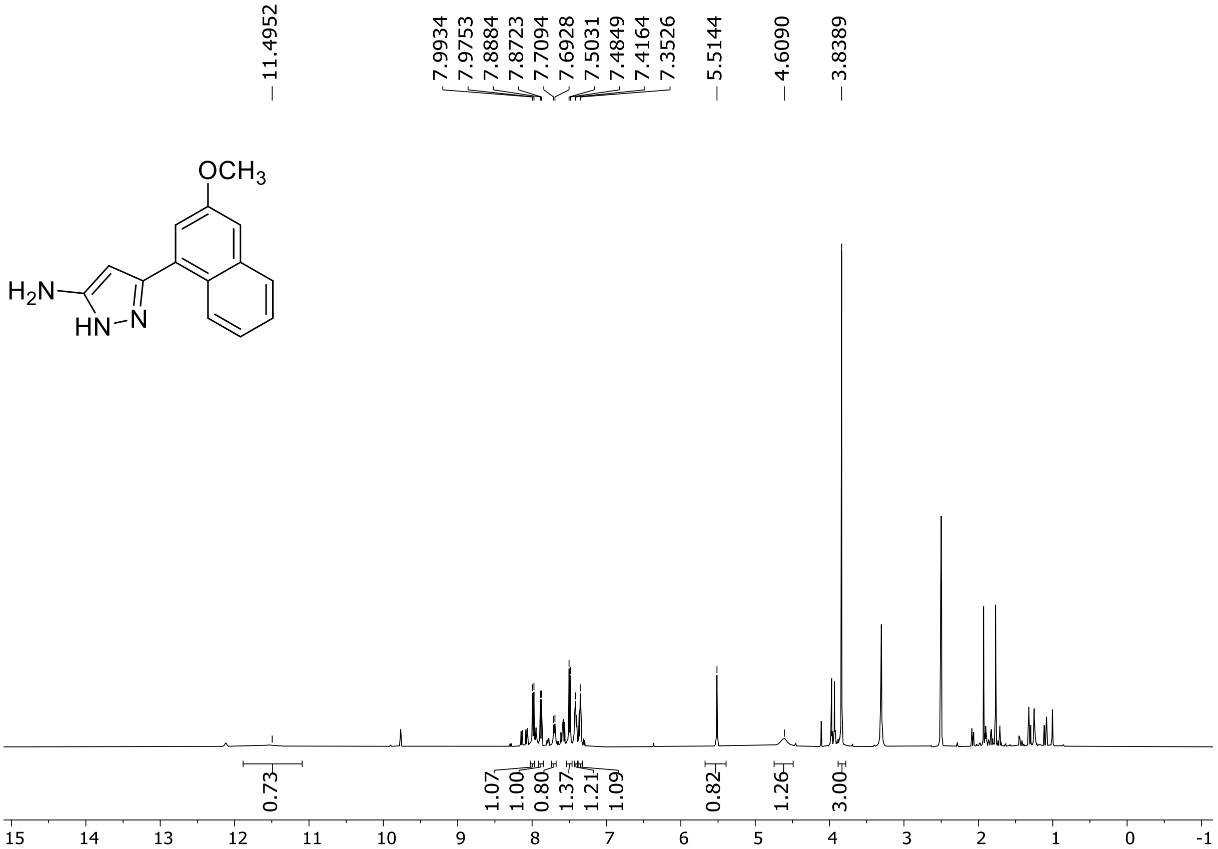

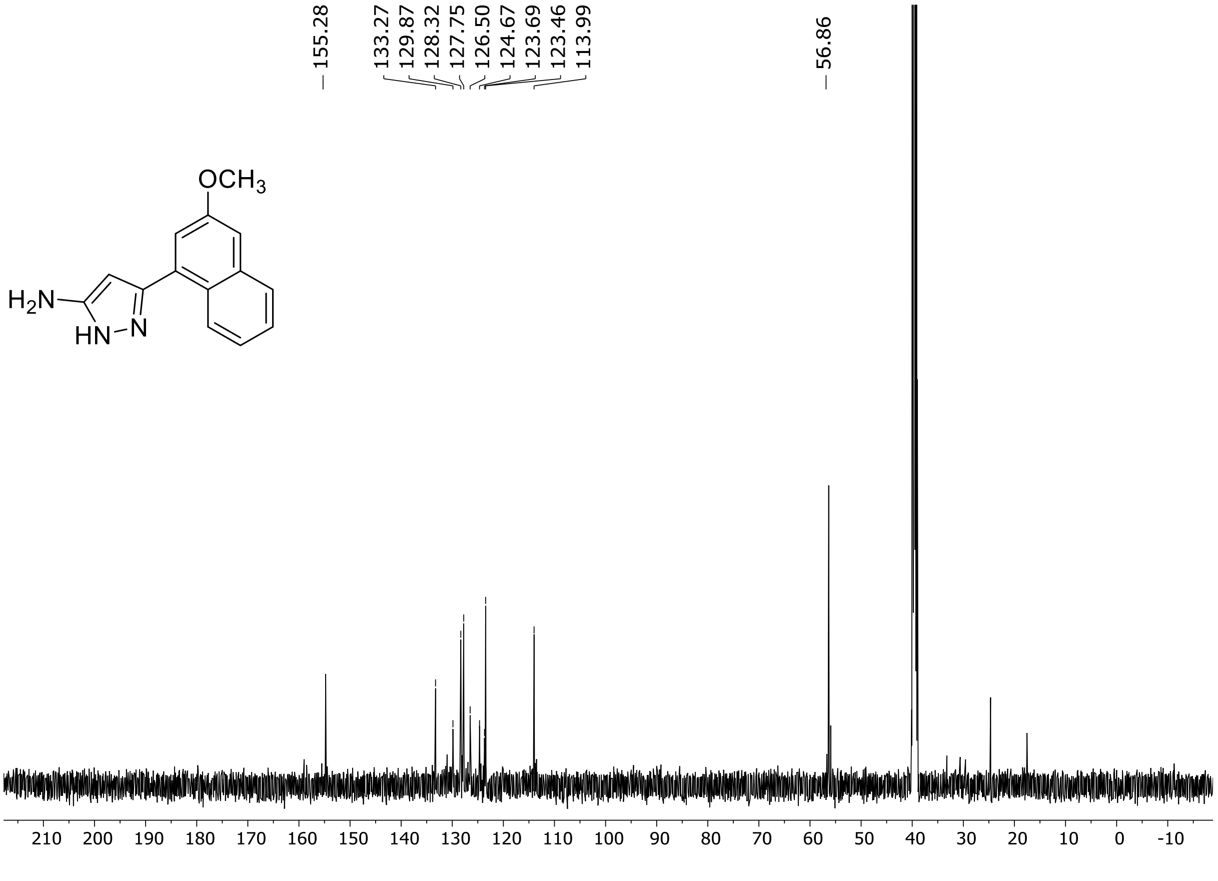

$^{1}H$ (500 MHz) and $^{13}C$ NMR (126 MHz) spectra of **S52** in DMSO-$d_6$

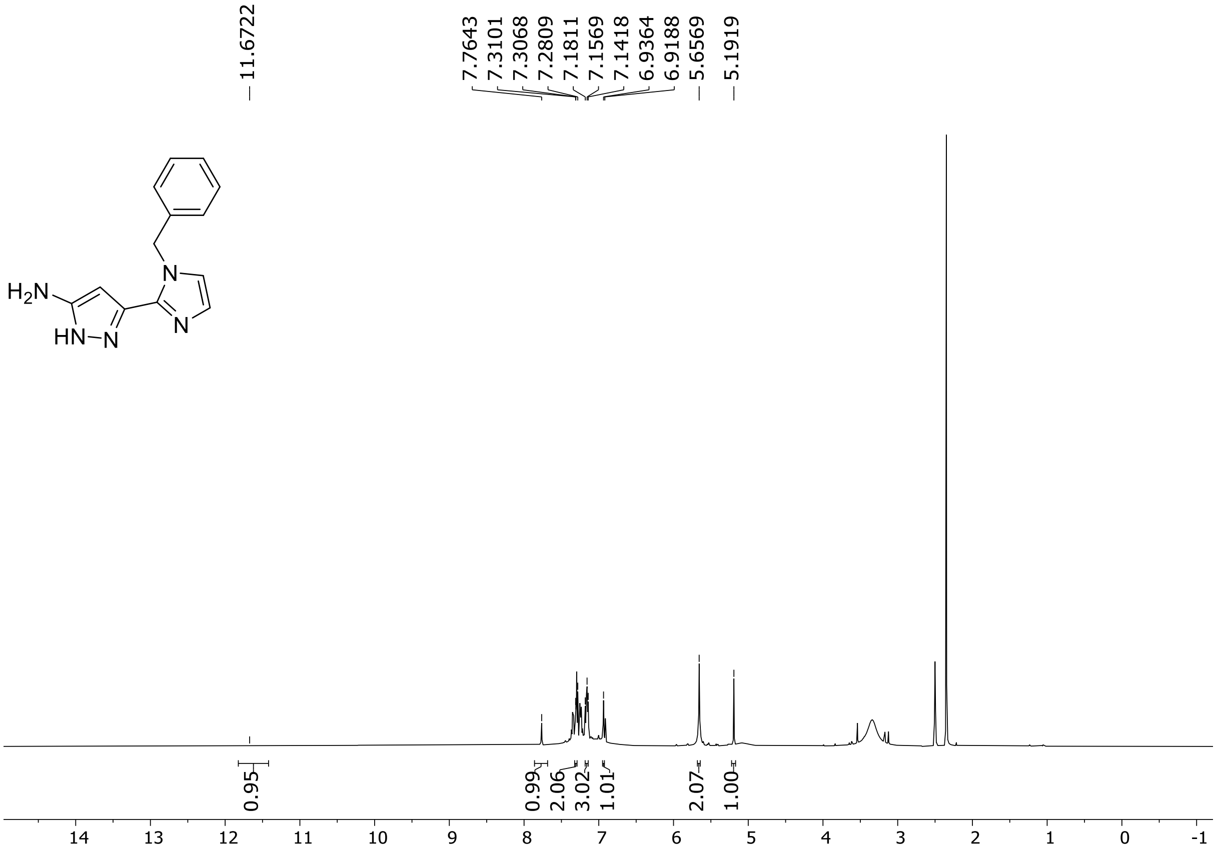

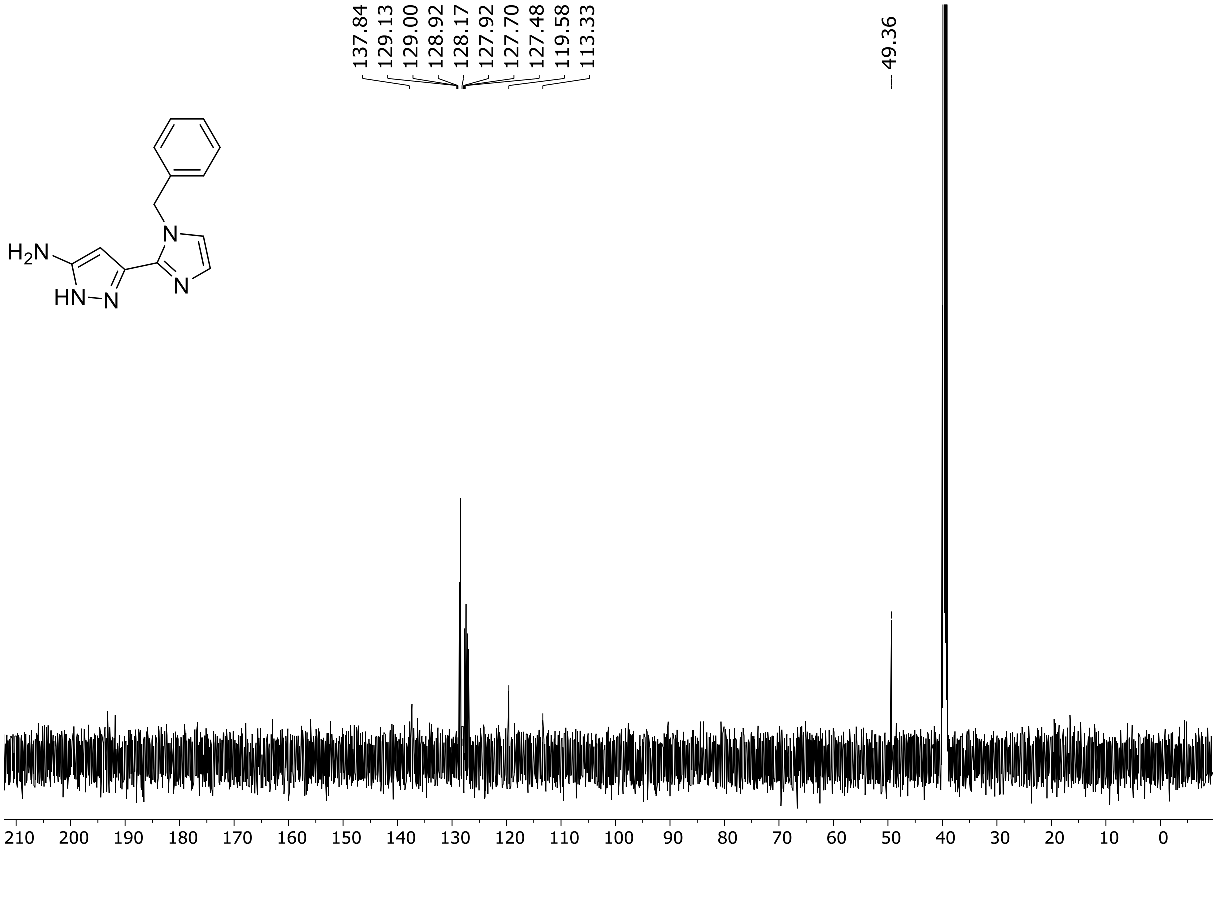

$^1H$ (500 MHz) and $^{13}C$ NMR (126 MHz) spectra of **S53** in DMSO-$d_6$

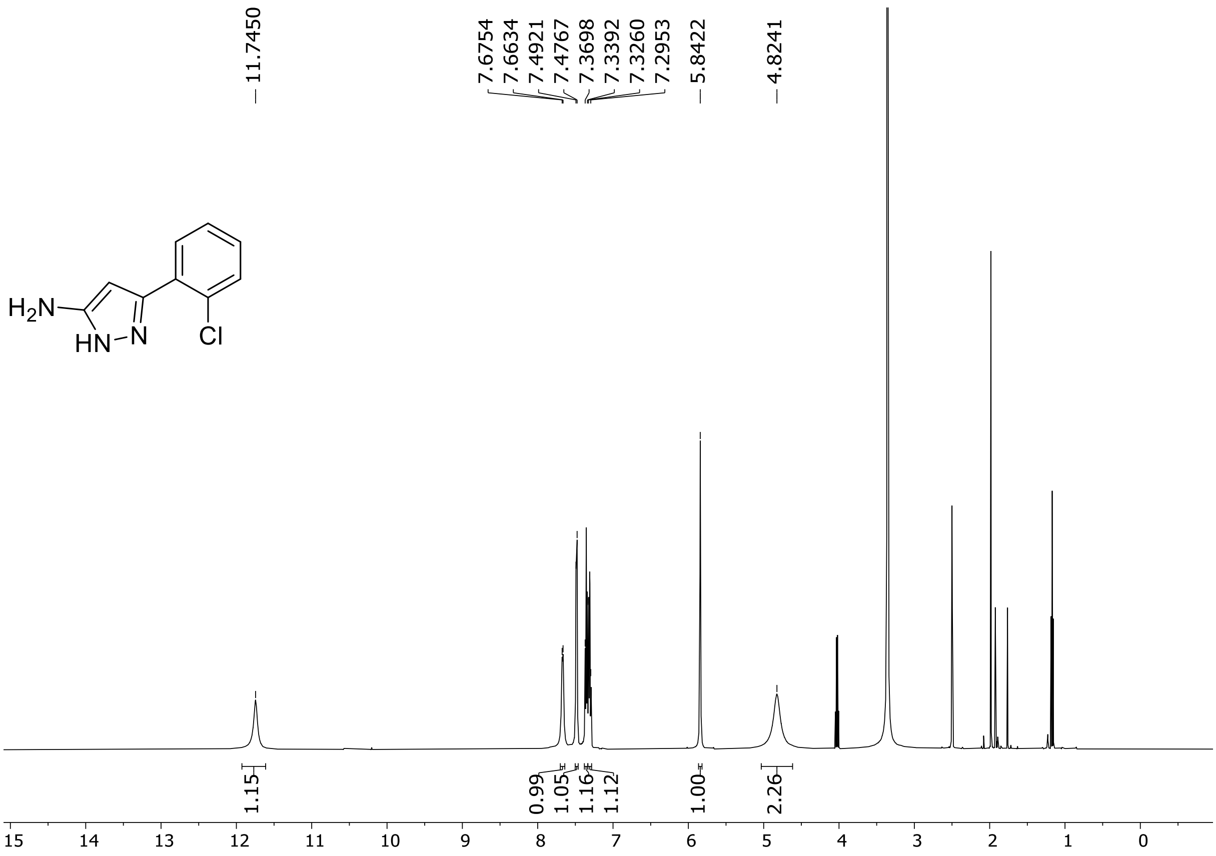

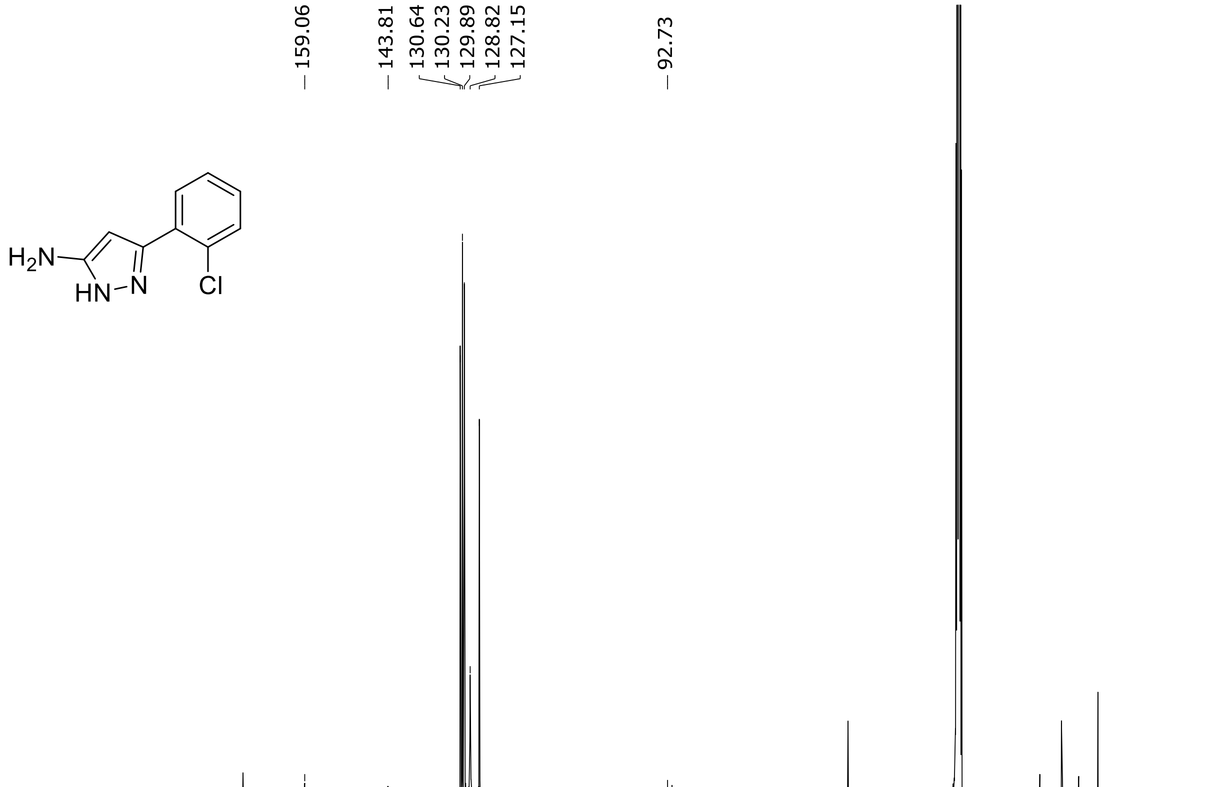

HRMS spectrum of **S53**

$C_9H_8ClN_3$ **mono m/z = 193.0407**

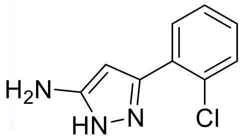

**APCI +** (MMI)
nitrogen flow 5 L/min, gas temperature 300°C, nebulizer 45 psi, vaporizer 200°C
skimmer **65** V, fragmentor **17** V, dissolved in methanol

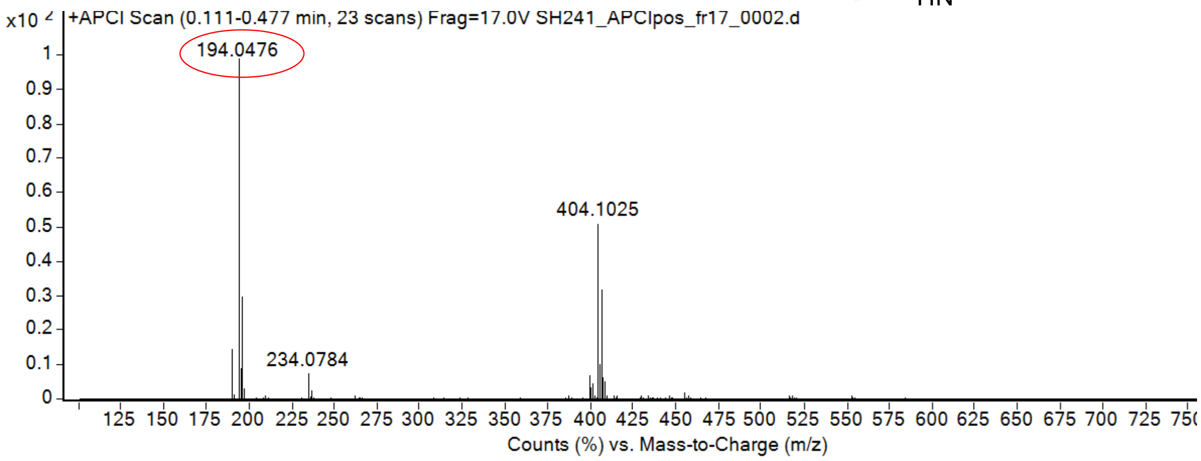

calculated mass: $[M+H]^+$ = 194.0480 observed: $[M+H]^+$ = 194.0476 max. mass error = 2 ppm

$^{1}H$ (500 MHz) and $^{13}C$ NMR (126 MHz) spectra of **S54** in DMSO-$d_6$

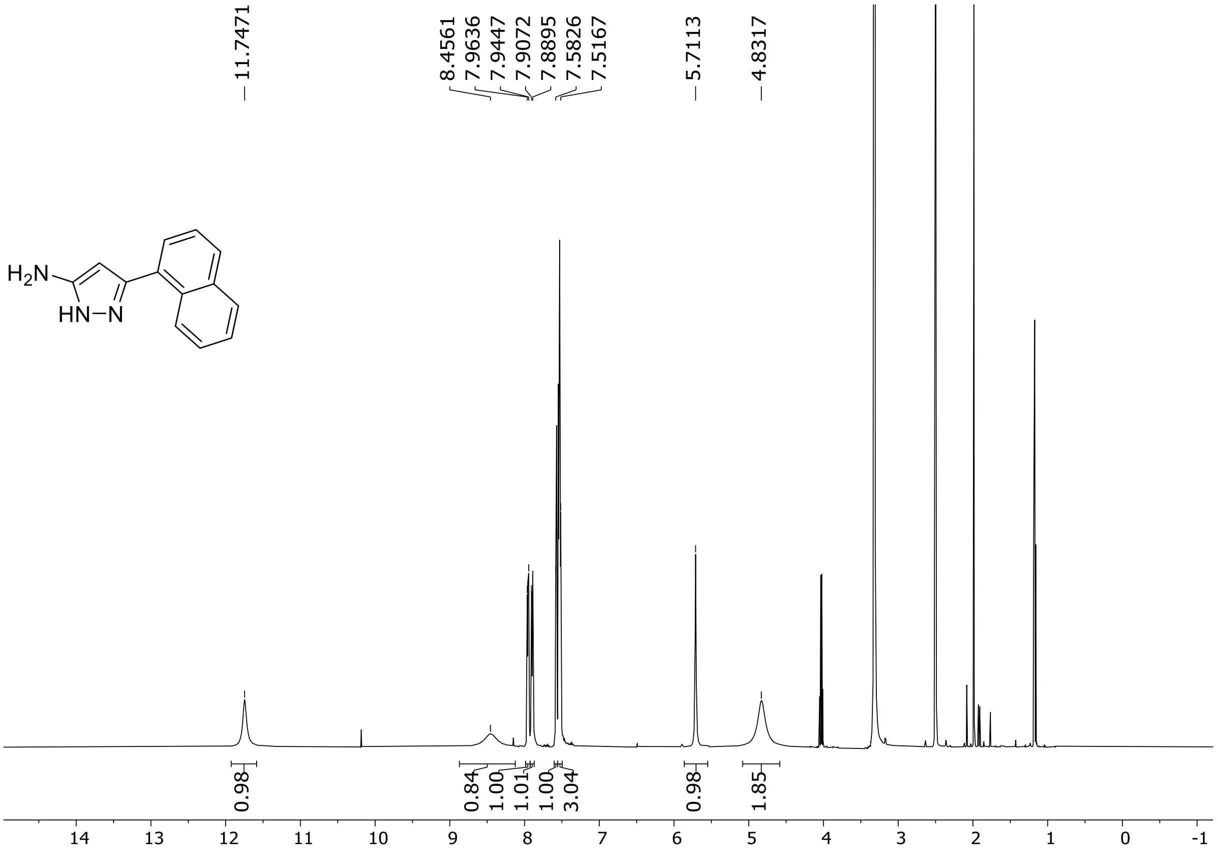

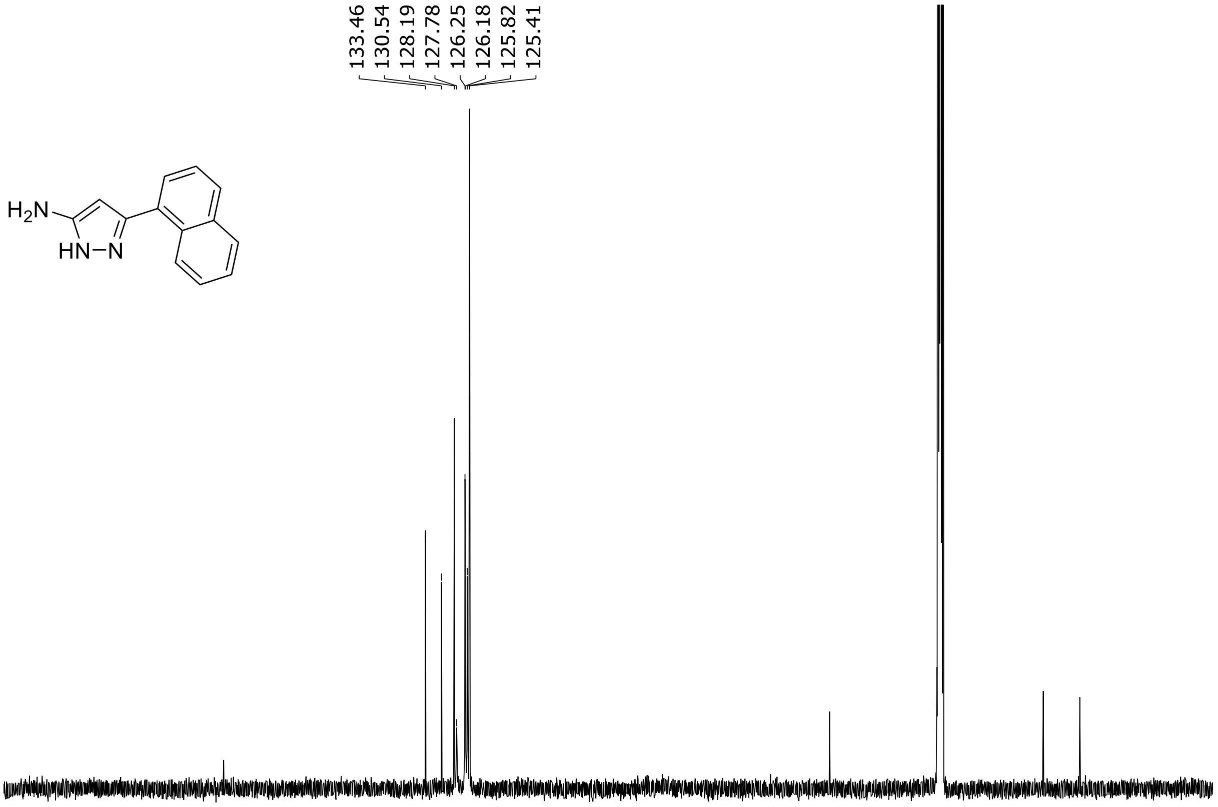

HRMS spectrum of **S54**

$C_{13}H_{11}N_3$ **mono m/z = 209.0953**

**APCI +** (MMI)
nitrogen flow 5 L/min, gas temperature 300°C, nebulizer 45 psi, vaporizer 200°C
skimmer **65** V, fragmentor **18** V, dissolved in methanol

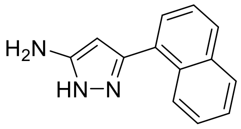

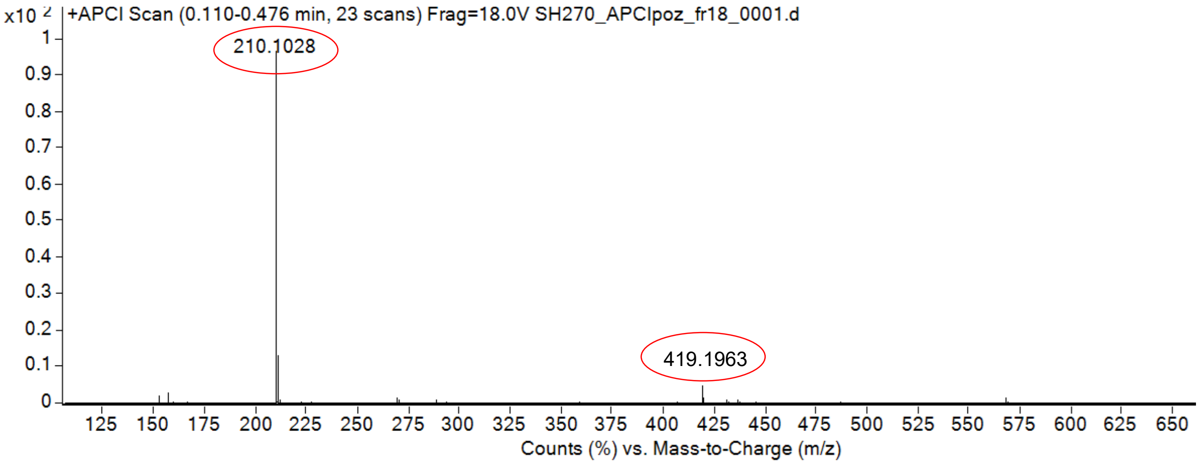

calculated mass: $[M+H]^+$ = 210.1026 observed: $[M+H]^+$ = 210.1028 max. mass error = 0.9 ppm

calculated mass: $[2xM+H]^+$ = 419.1979 observed: $[2xM+H]^+$ = 419.1963 max. mass error = 3.3 ppm

$^{1}H$ (500 MHz) and $^{13}C$ NMR (126 MHz) spectra of **S55** in DMSO-$d_6$

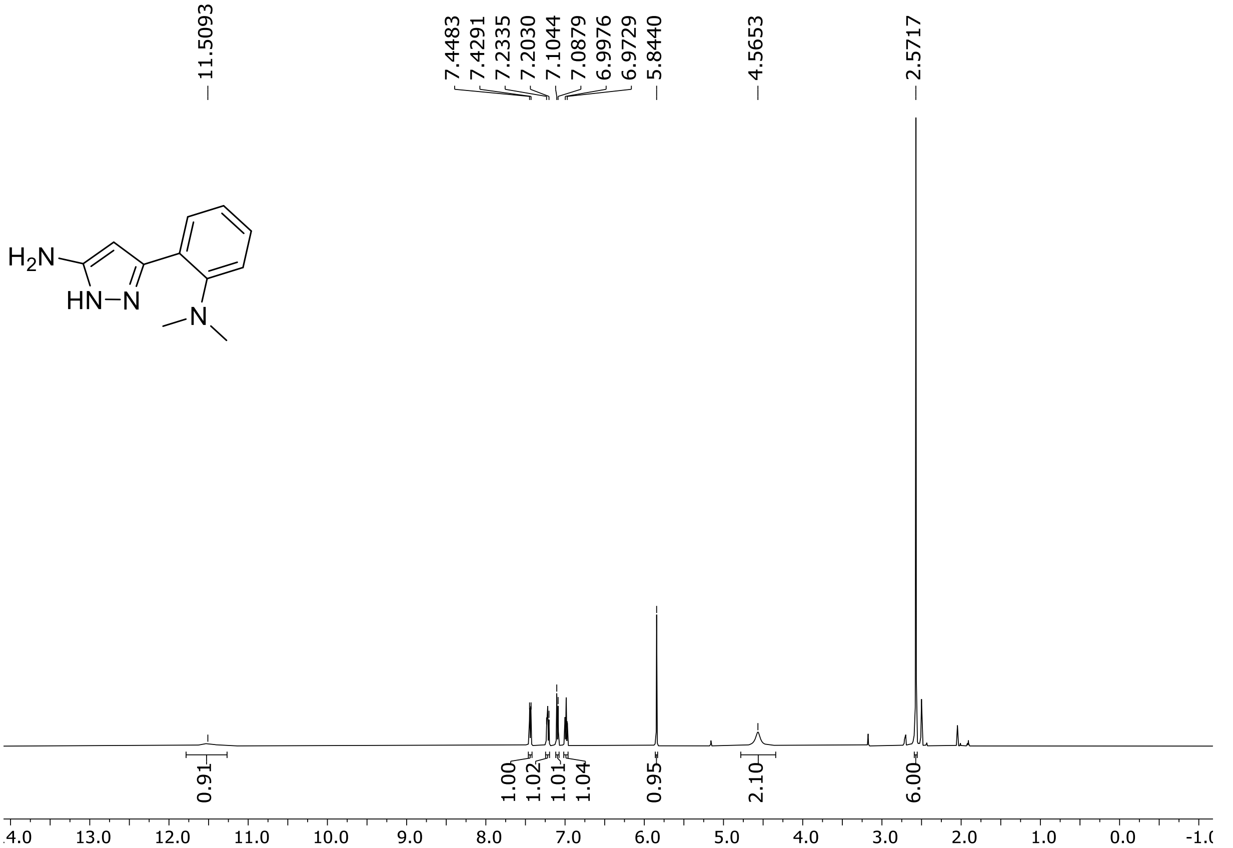

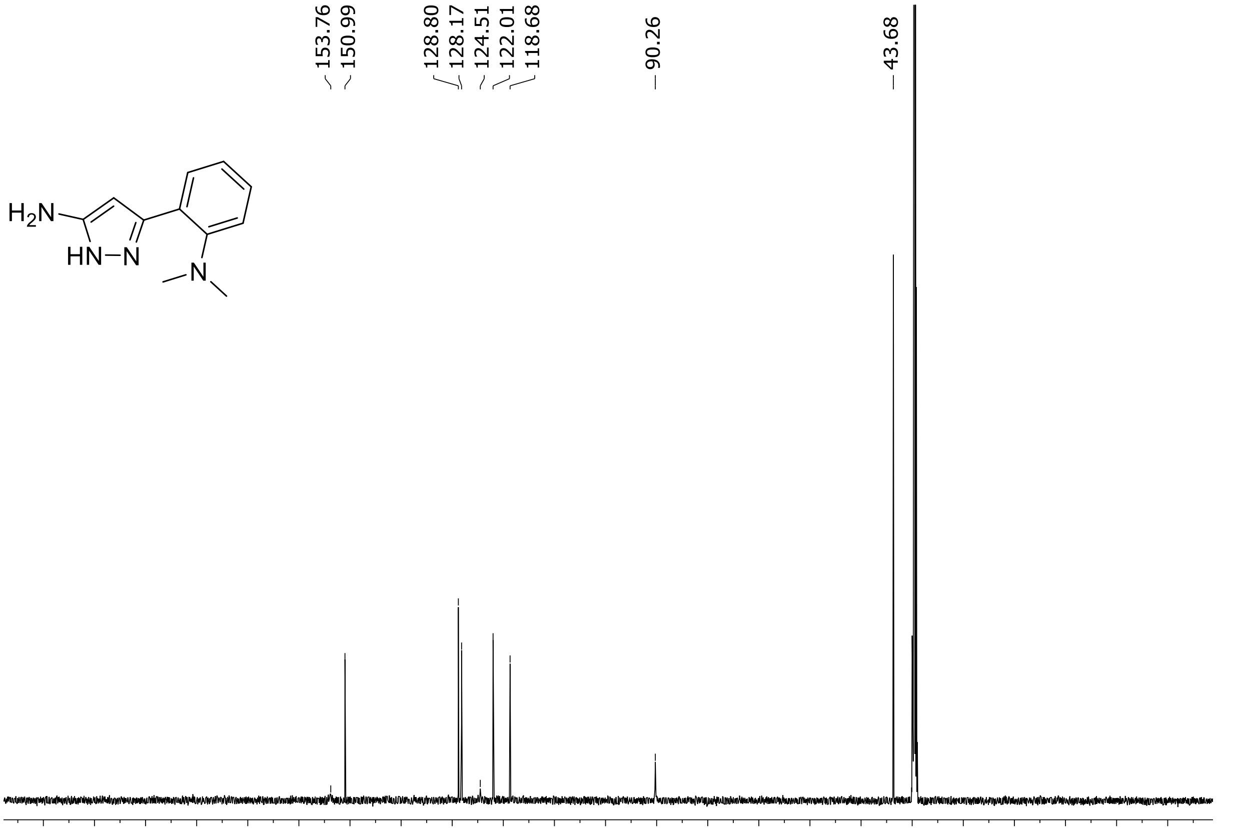

$^{1}H$ (500 MHz) and $^{13}C$ NMR (126 MHz) spectra of **S56** in chloroform-*d*

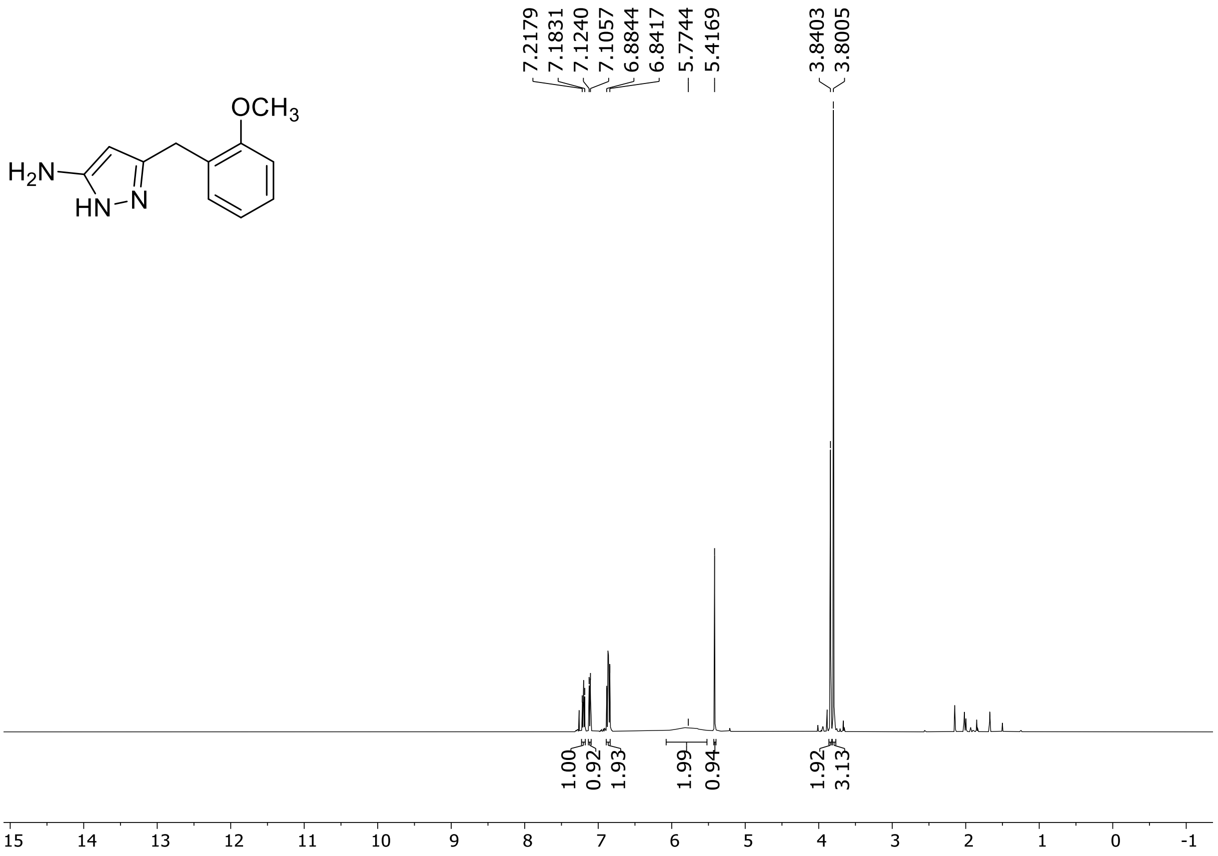

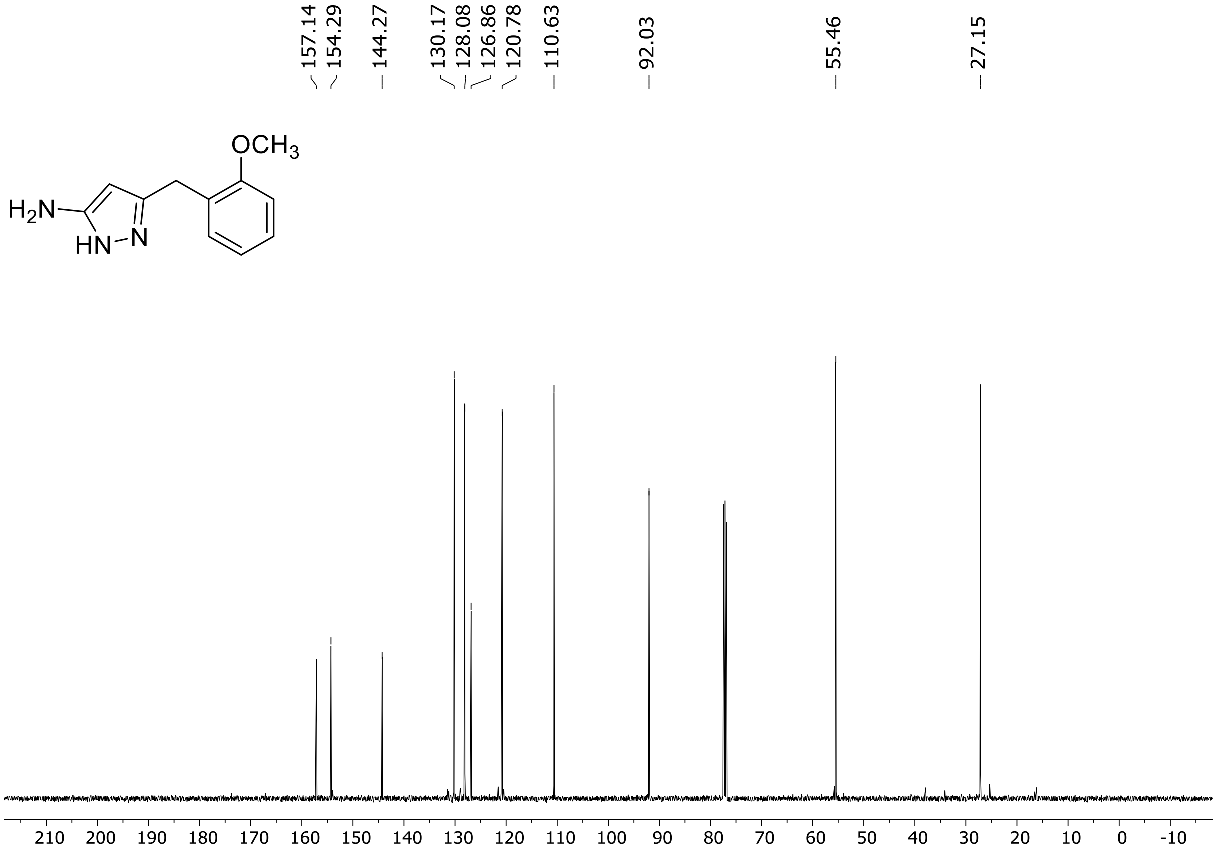

[1]H (500 MHz) and [13]C NMR (126 MHz) spectra of **S57** in chloroform-*d*

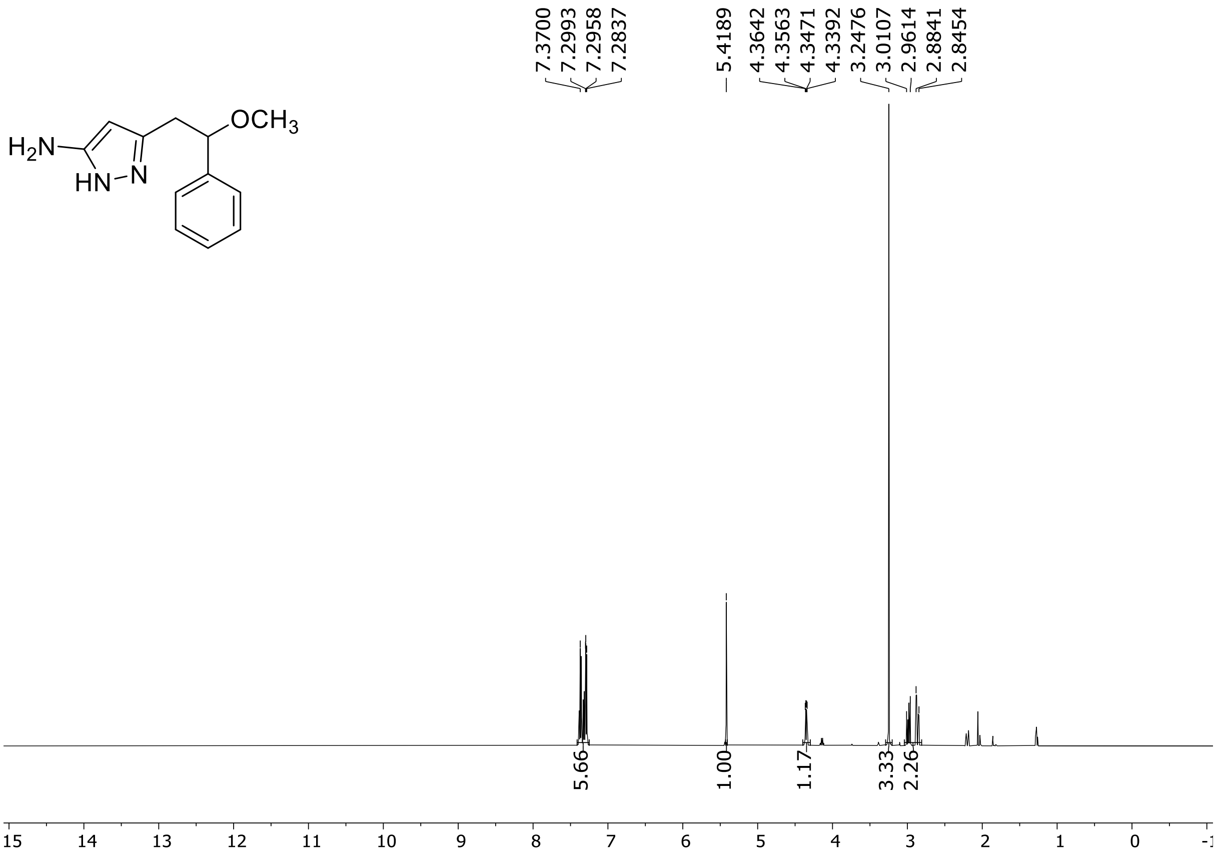

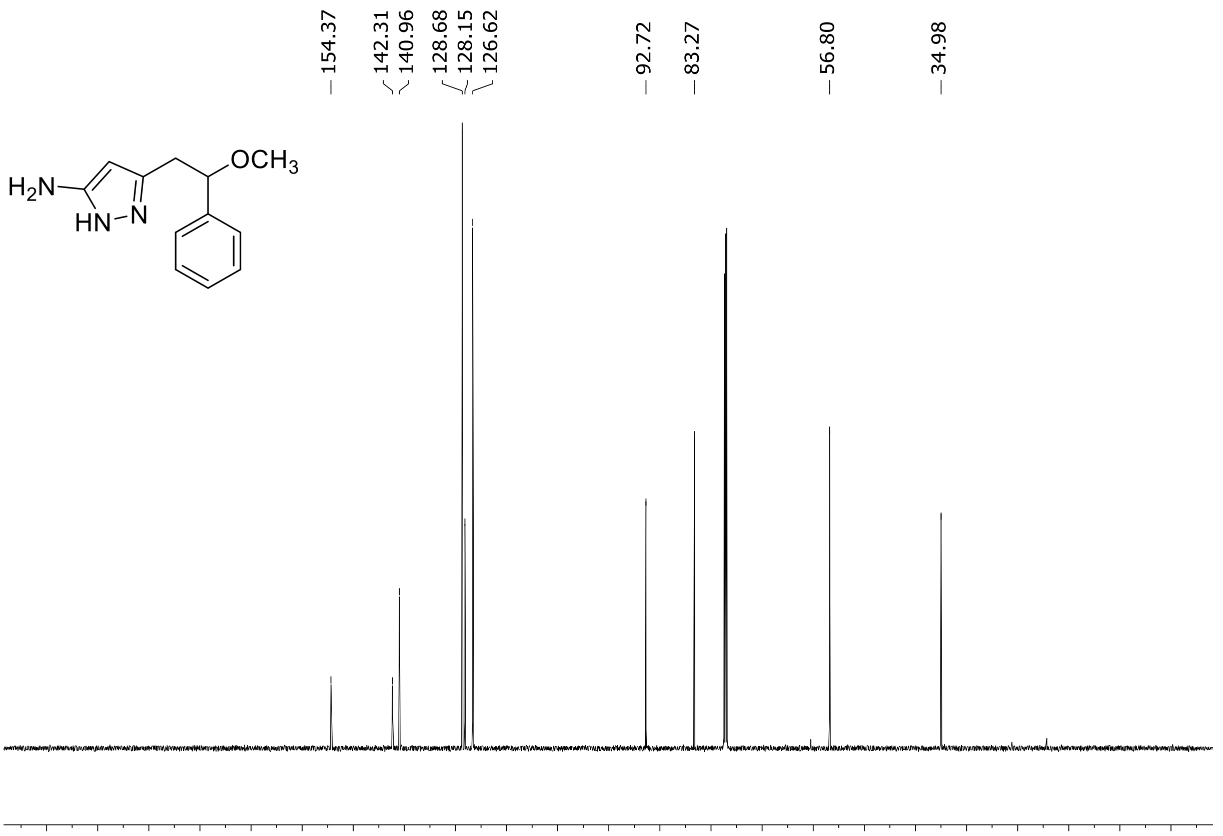

$^{1}H$ (500 MHz) and $^{13}C$ NMR (126 MHz) spectra of **S58** in chloroform-*d*

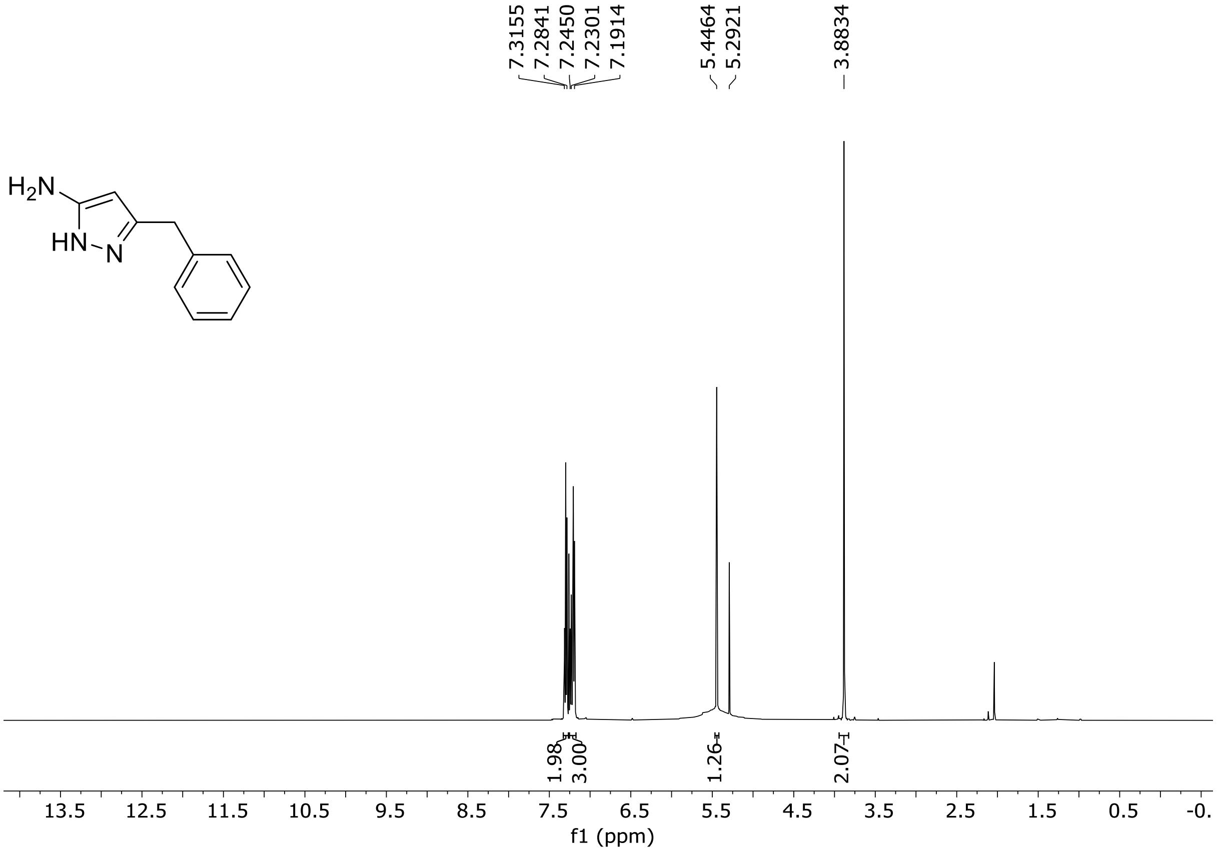

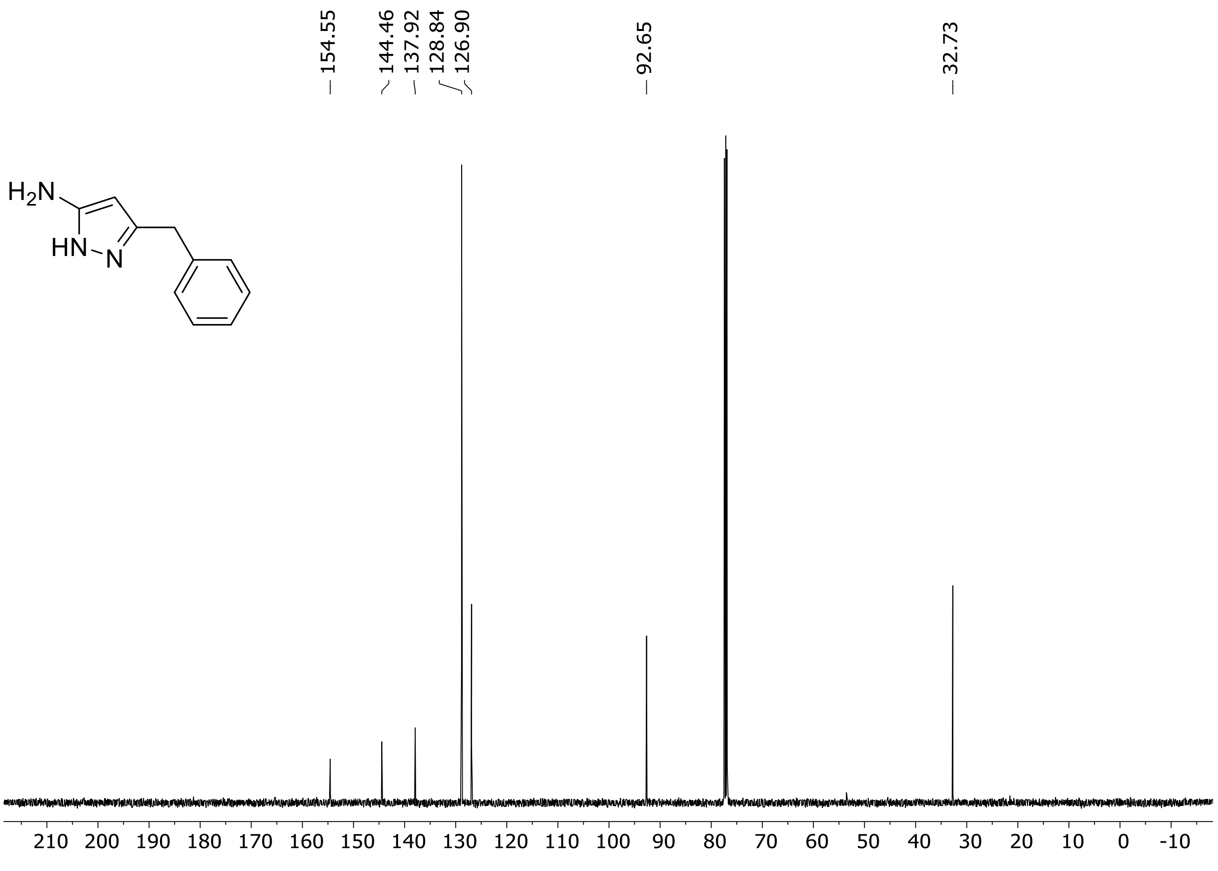

HRMS spectrum of **S58**

$C_{10}H_{11}N_3$ **exact mass: 173.0953**

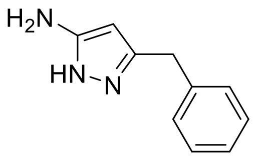

**APCI +** (MMI)
nitrogen flow 3 L/min, gas temperature 325°C, nebulizer 45 psig, skimmer 65 V, vaporizer 200°C, fragmentor 20 V, dissolved in methanol

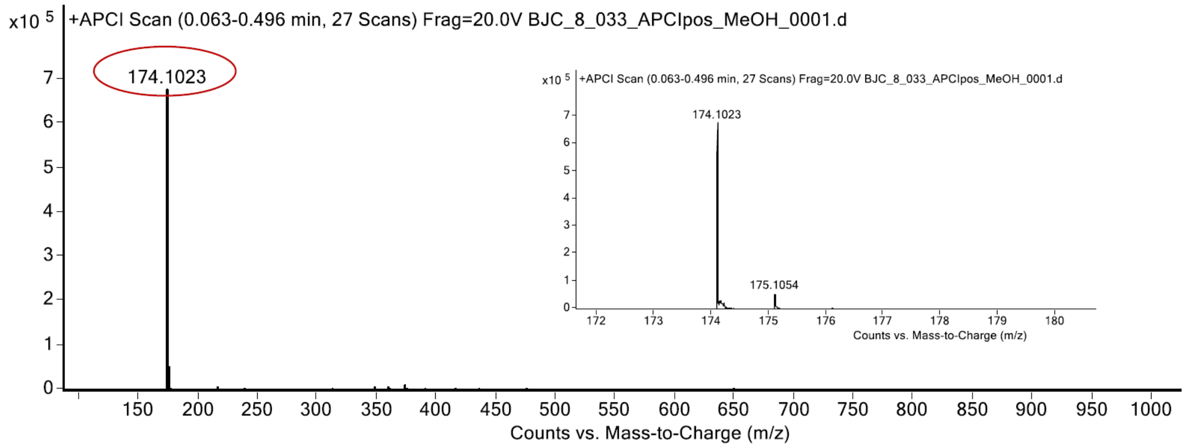

expected mass: $[M+H]^+$ = 174.1026 observed mass: $[M+H]^+$ = 174.1023 mass accuracy = - 1.7 ppm

$^{1}H$ (500 MHz) and $^{13}C$ NMR (126 MHz) spectra of **S59** in chloroform-*d*

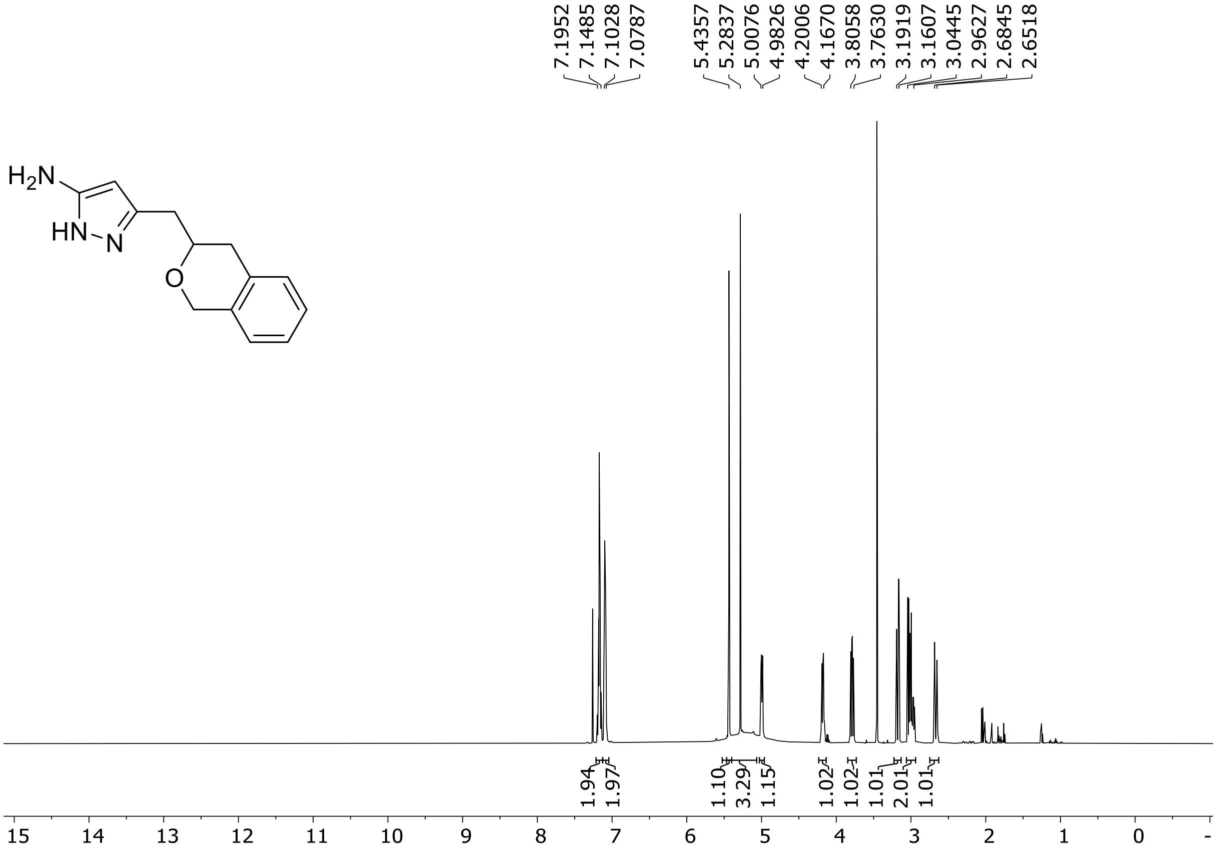

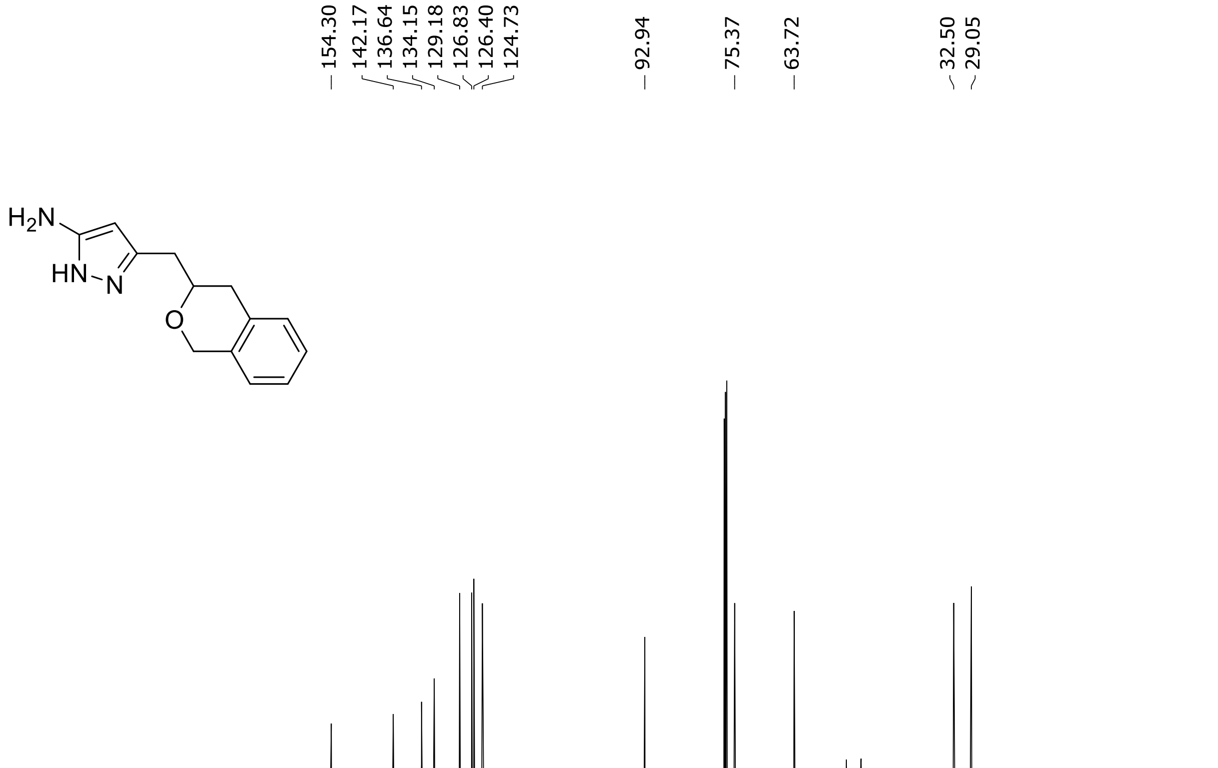

HRMS spectrum of **S59**

**$C_{13}H_{15}N_3O$** **exact mass: 229.1215**

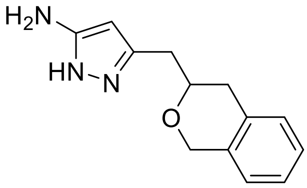

**APCI +** (MMI)
nitrogen flow 3 L/min, gas temperature 325°C, nebulizer 45 psig, skimmer 65 V, vaporizer 200°C, fragmentor 20 V, dissolved in methanol

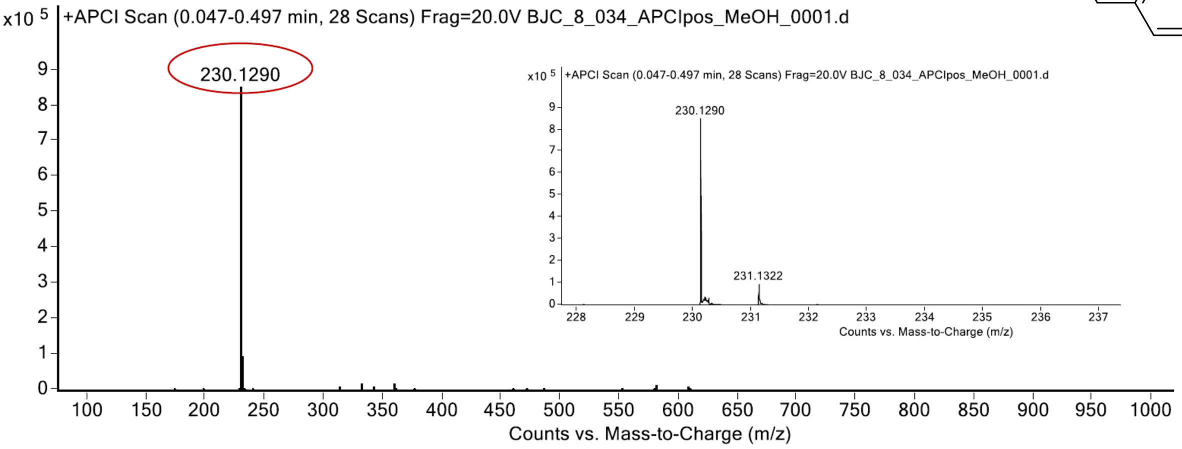

expected mass: $[M+H]^+$ = 230.1288 observed mass: $[M+H]^+$ = 230.1290 mass accuracy = 0.8 ppm

[1]H (500 MHz) and [13]C NMR (126 MHz) spectra of **S60** in chloroform-*d*

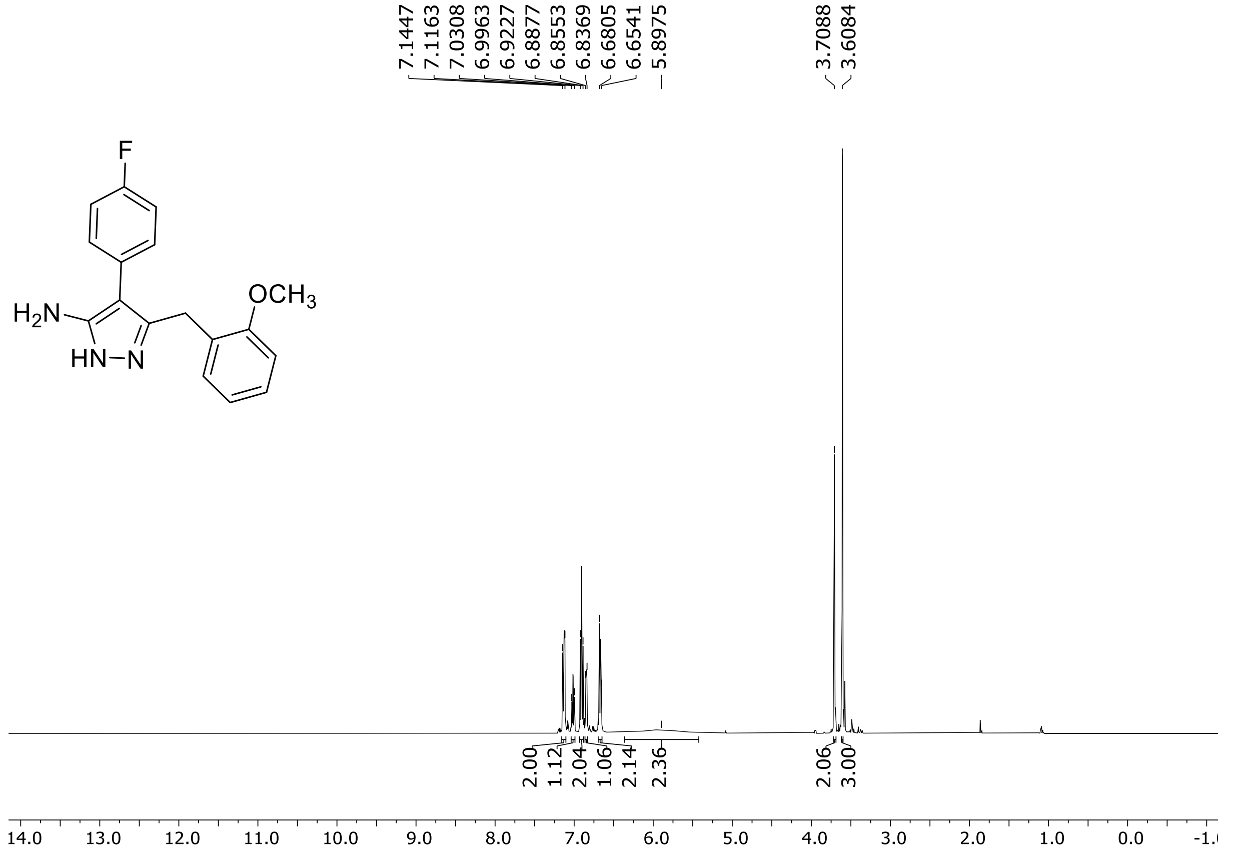

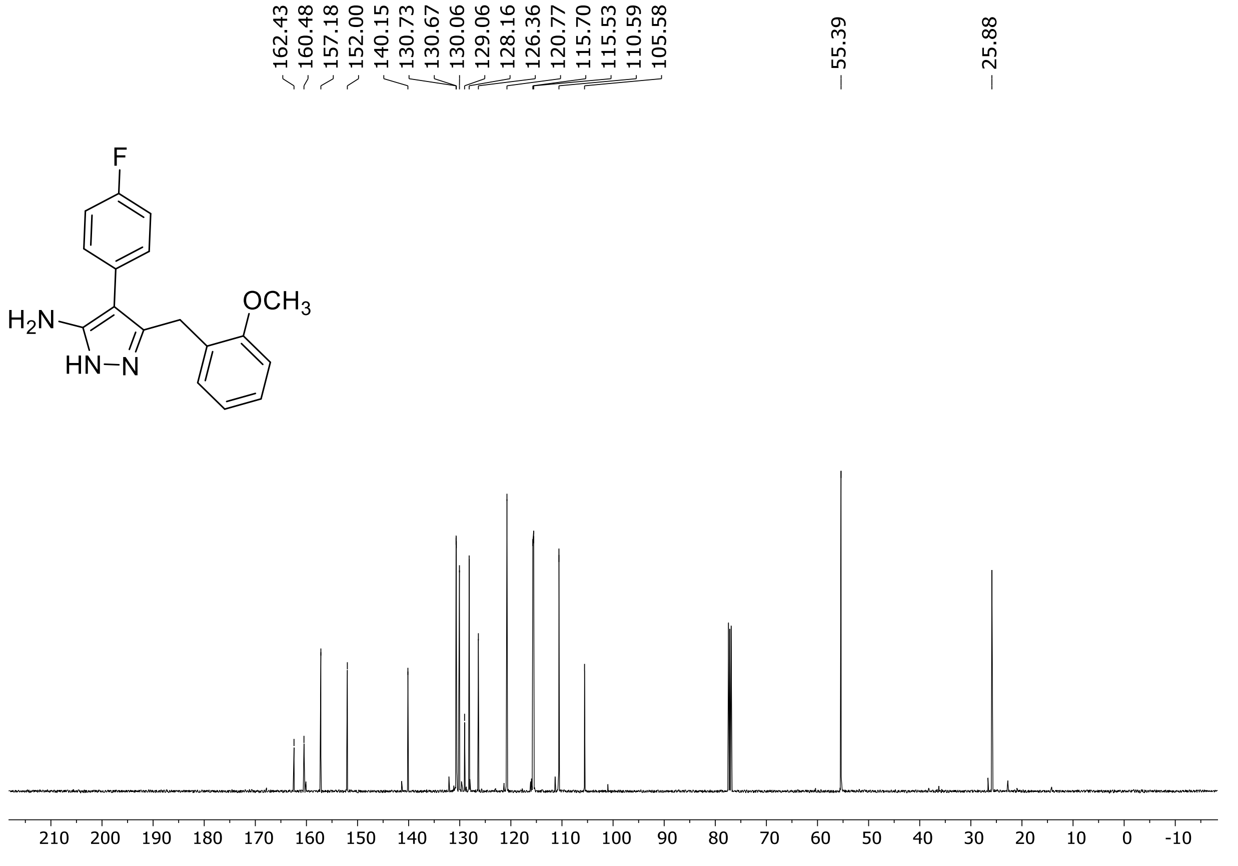

[19]F (471 MHz) NMR spectrum of **S60** in chloroform-*d*

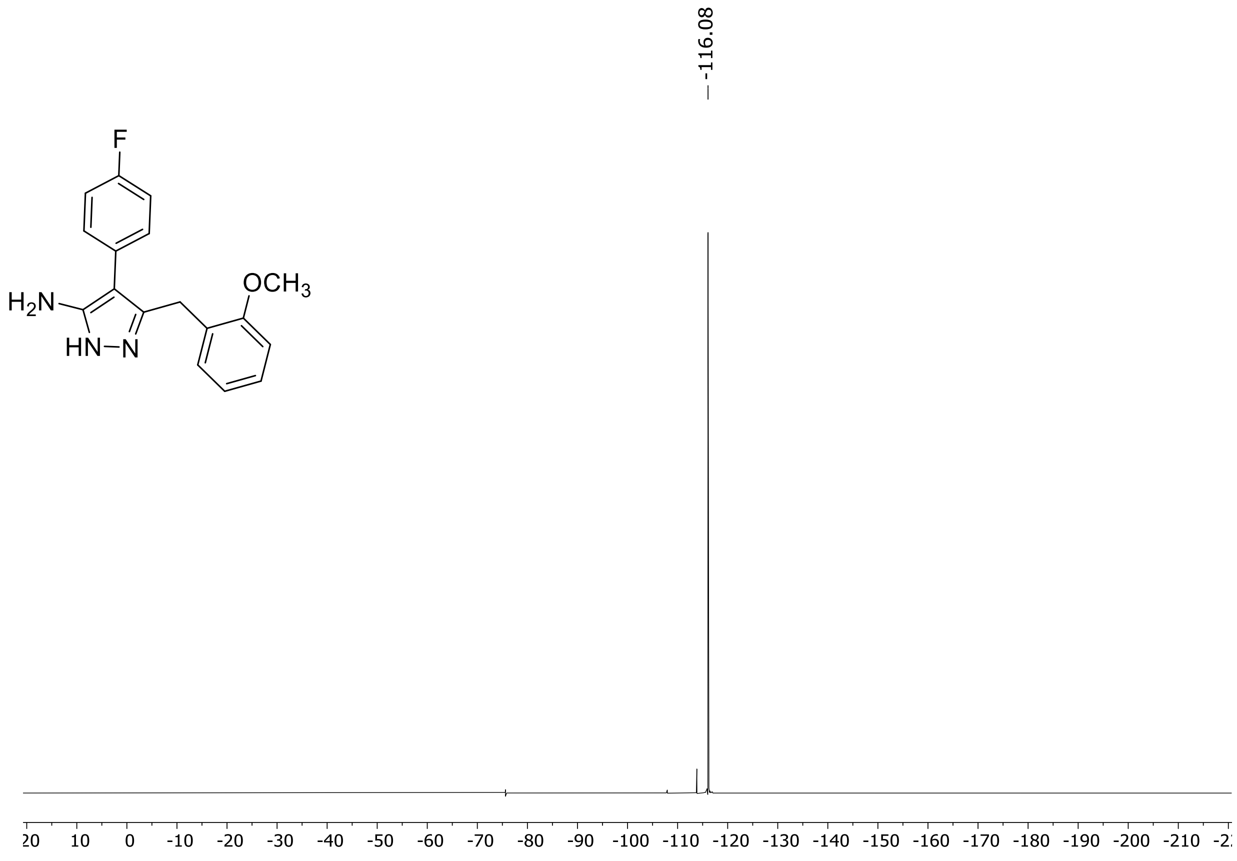

$^{1}H$ (500 MHz) and $^{13}C$ NMR (126 MHz) spectra of **S61** in chloroform-*d*

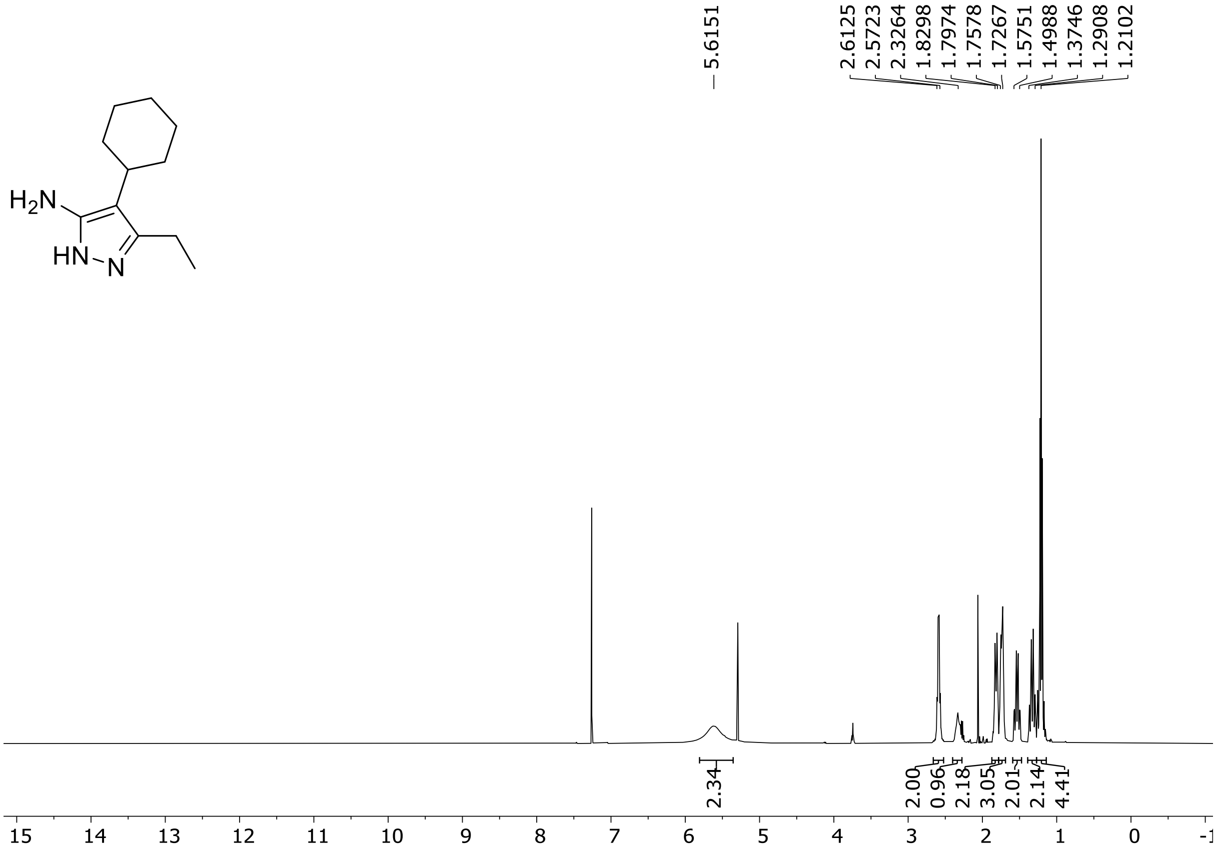

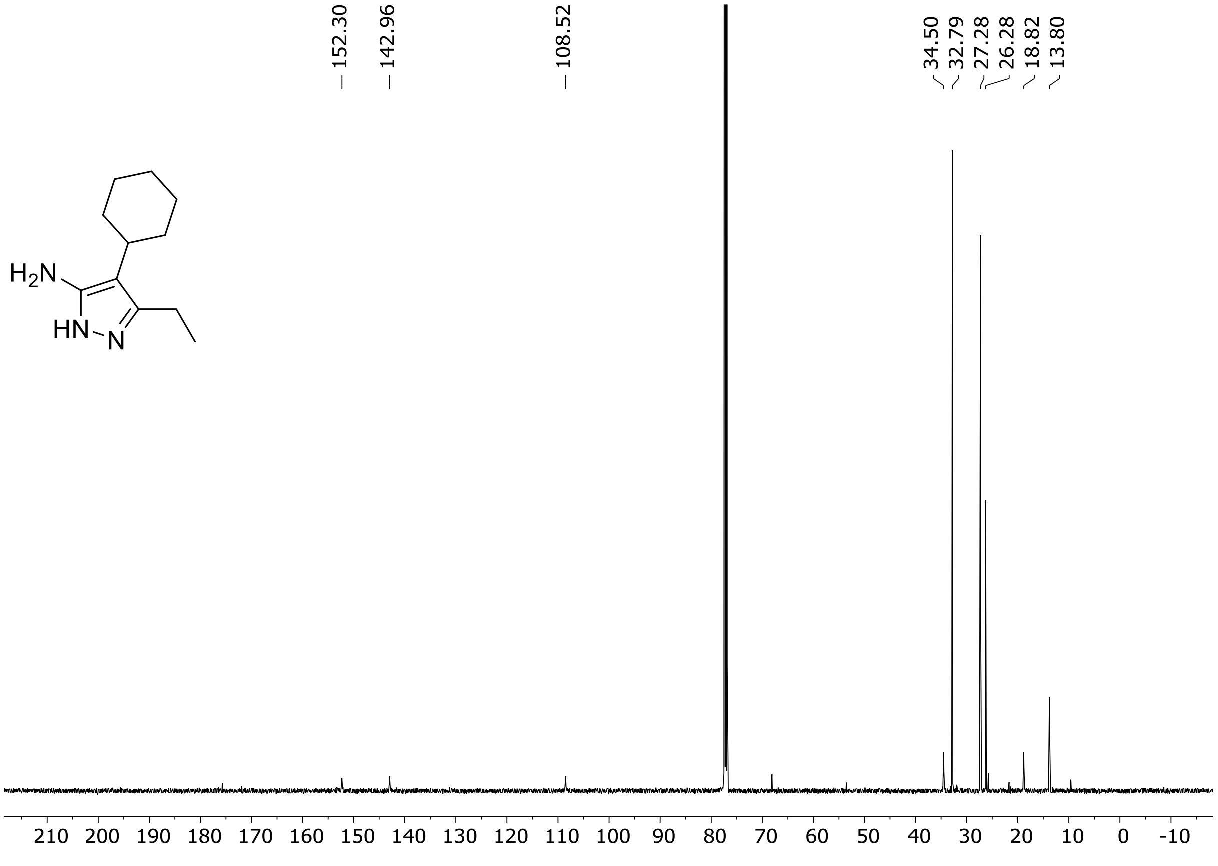

HRMS spectrum of **S61**

$C_{11}H_{19}N_3$ **exact mass: 193.1579**

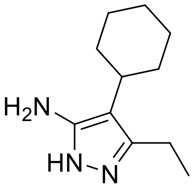

**APCI +** (MMI)
nitrogen flow 3 L/min, gas temperature 325°C, nebulizer 45 psig, skimmer 65 V, vaporizer 200°C, fragmentor 30 V, dissolved in methanol

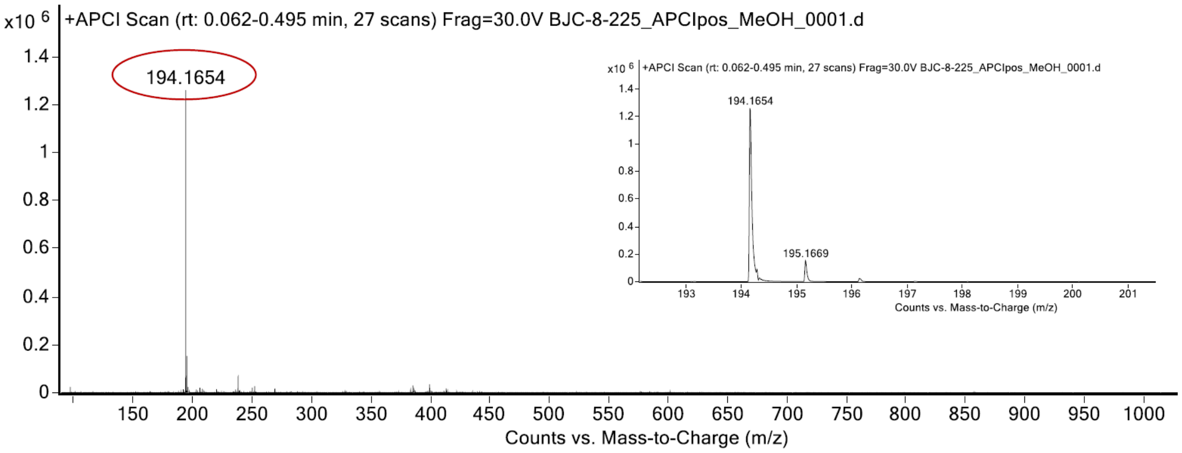

expected mass: $[M+H]^+$ = 194.1652 observed mass: $[M+H]^+$ = 194.1654 mass accuracy = 1.0 ppm

$^{1}H$ (500 MHz) and $^{13}C$ NMR (126 MHz) spectra of **S62** in chloroform-*d*

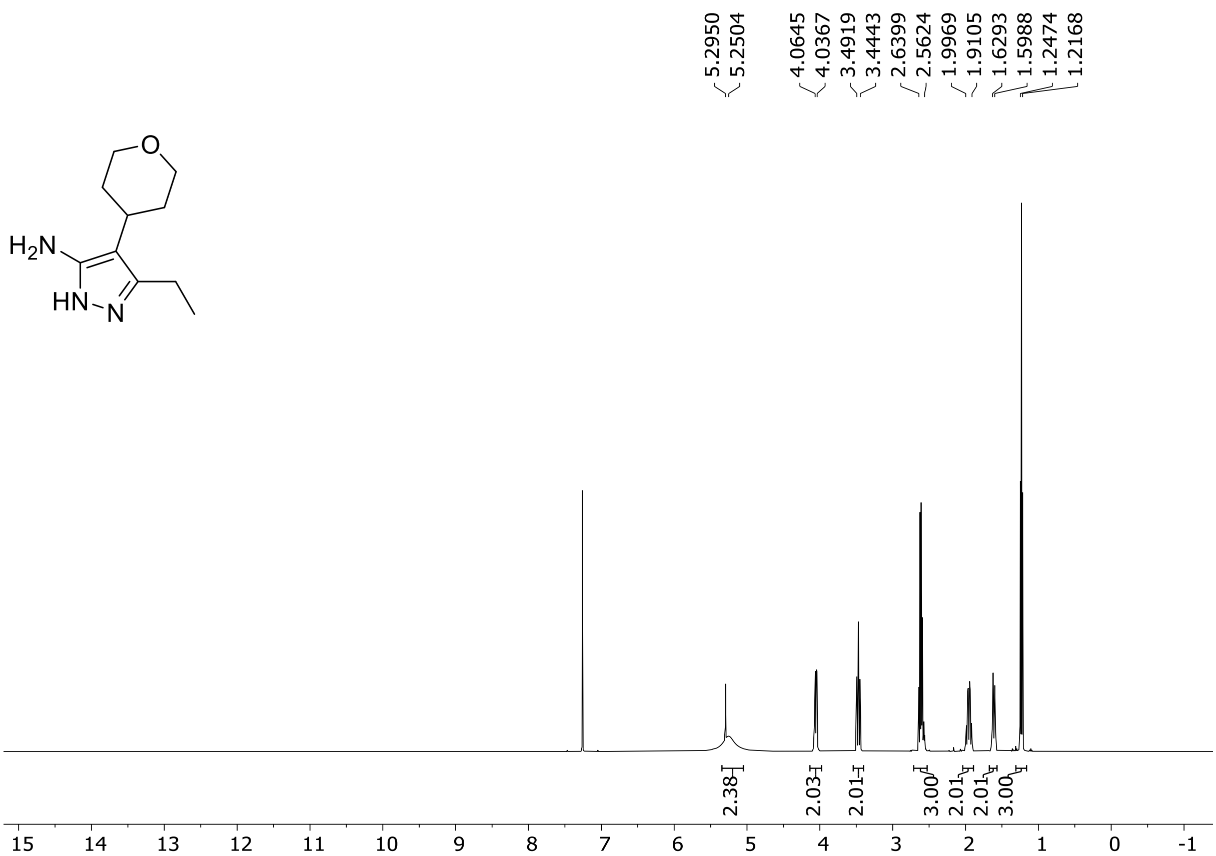

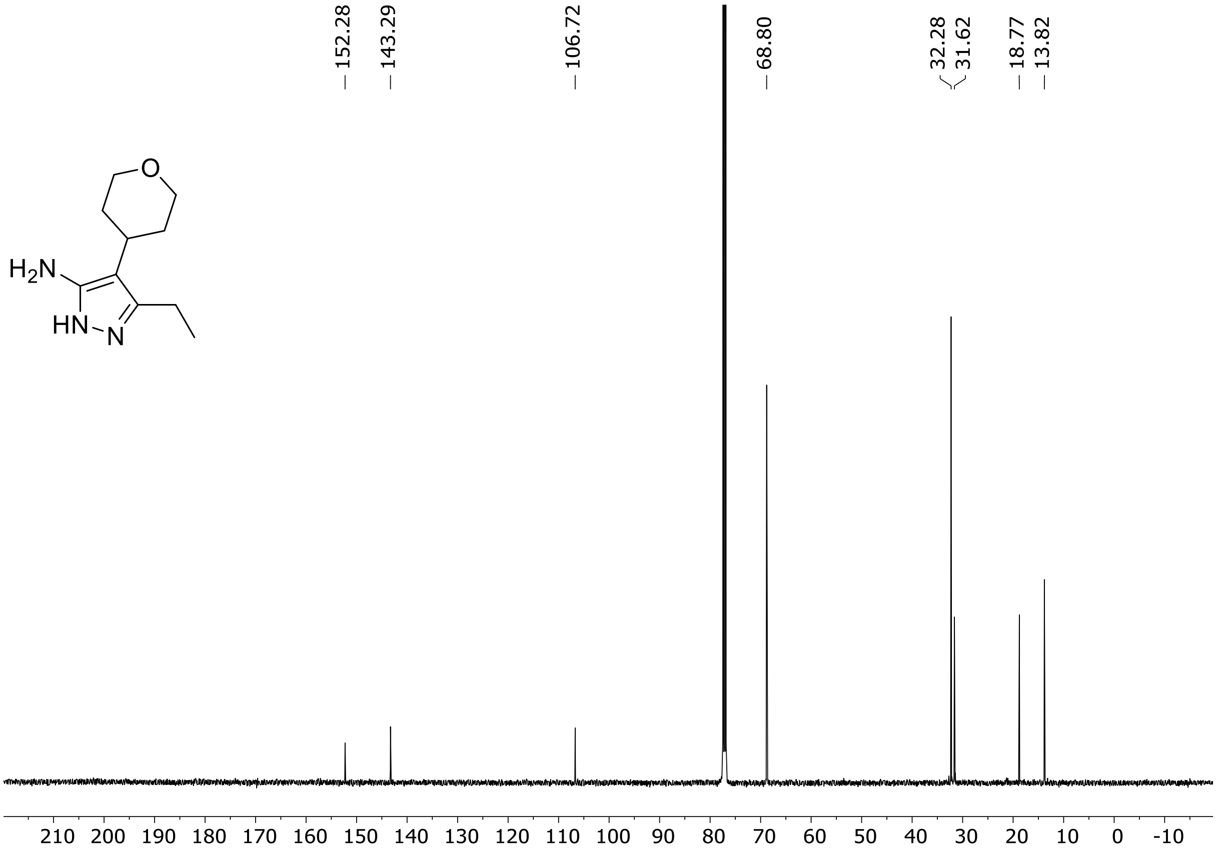

HRMS spectrum of **S62**

**$C_{10}H_{17}N_3O$** **exact mass: 193.1579**

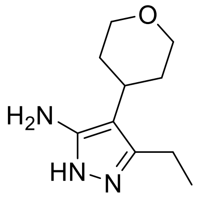

**APCI +** (MMI)
nitrogen flow 3 L/min, gas temperature 325°C, nebulizer 45 psig, skimmer 65 V, vaporizer 200°C, fragmentor 25 V, dissolved in methanol

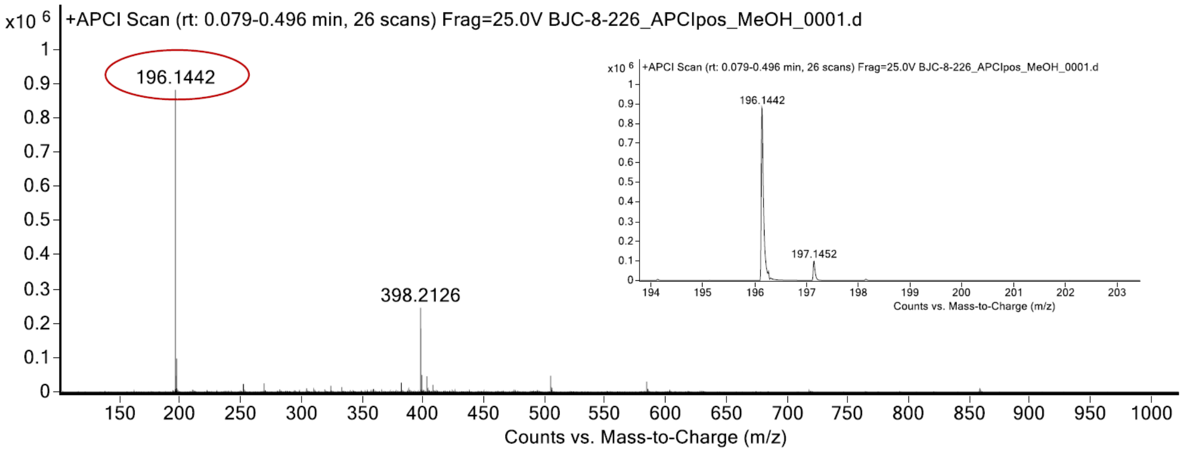

expected mass: $[M+H]^+$ = 196.1444 observed mass: $[M+H]^+$ = 196.1442 mass accuracy = - 1.0 ppm

[1]H (500 MHz) and [13]C NMR (126 MHz) spectra of **S63** in chloroform-*d*

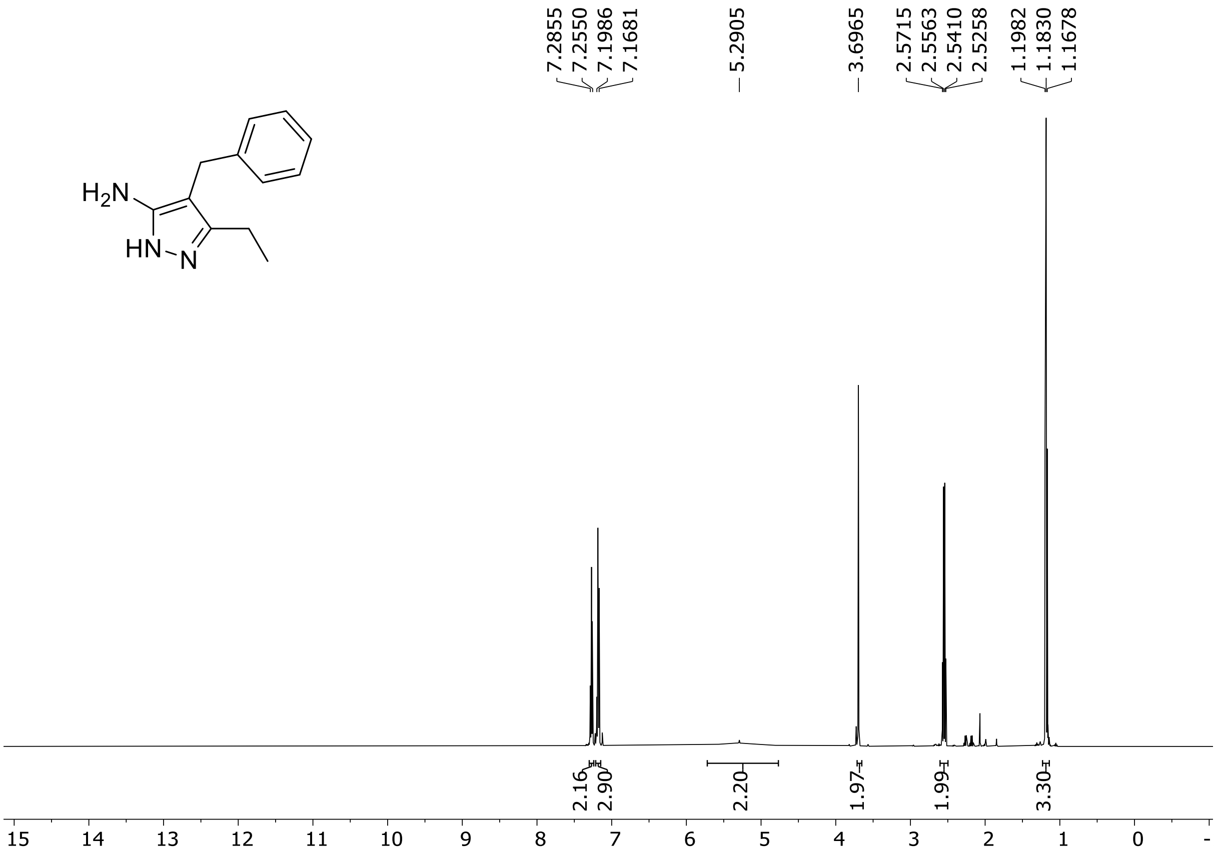

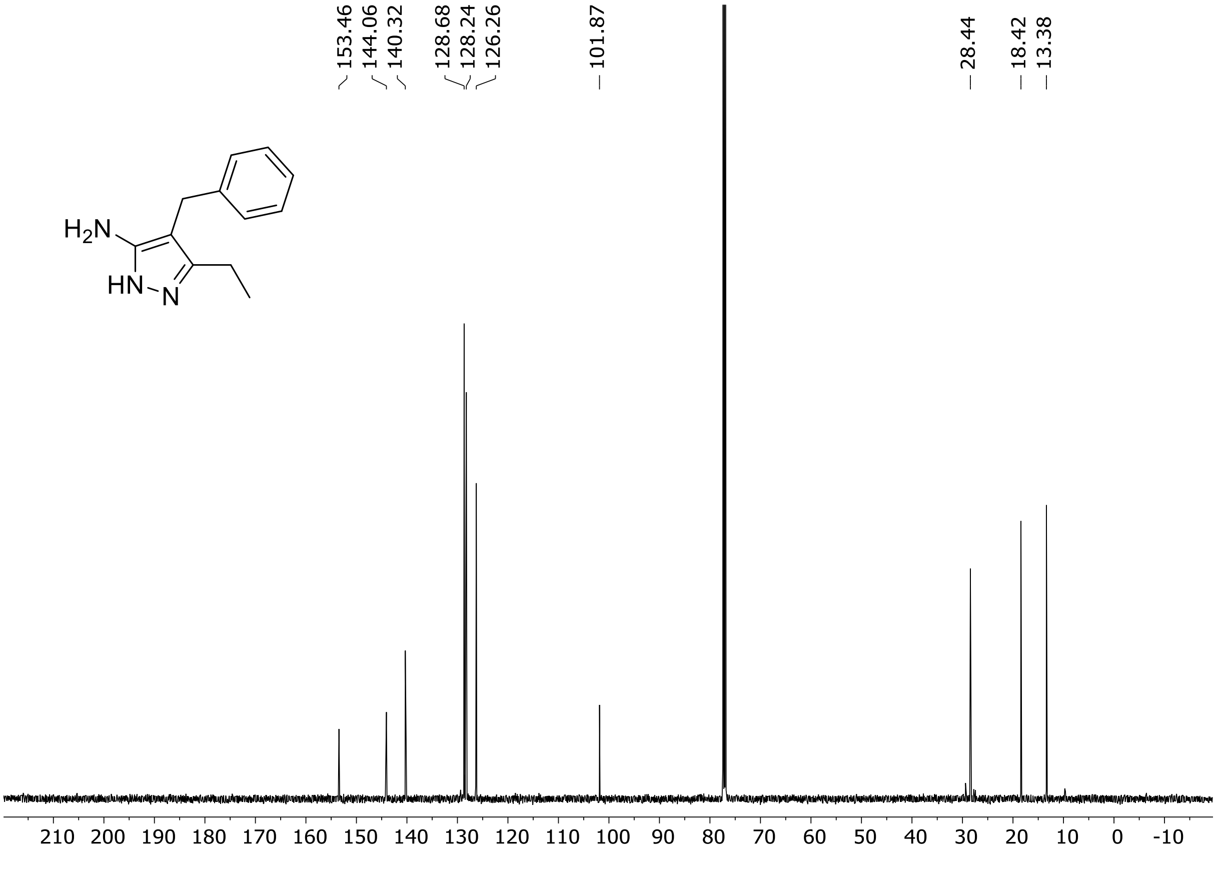

HRMS spectrum of **S63**

$C_{12}H_{15}N_3$ **exact mass: 201.1266**

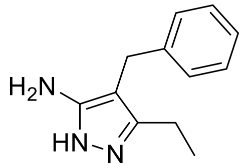

**APCI +** (MMI)
nitrogen flow 3 L/min, gas temperature 325°C, nebulizer 45 psig, skimmer 65 V, vaporizer 200°C, fragmentor 20 V, dissolved in methanol

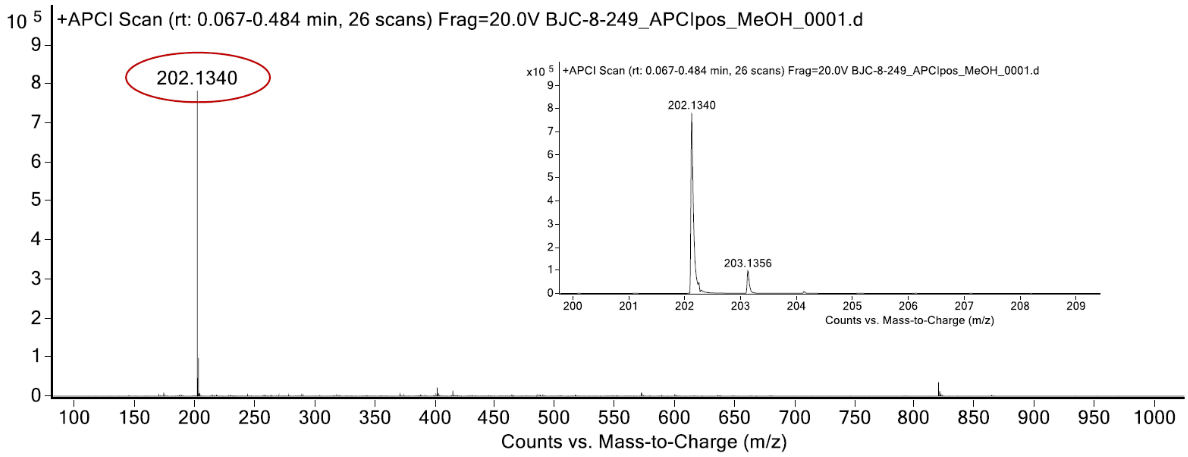

expected mass: $[M+H]^+ = 202.1339$ observed mass: $[M+H]^+ = 202.1340$ mass accuracy = 0.5 ppm

$^{1}H$ (500 MHz) and $^{13}C$ NMR (126 MHz) spectra of **S64** in chloroform-*d*

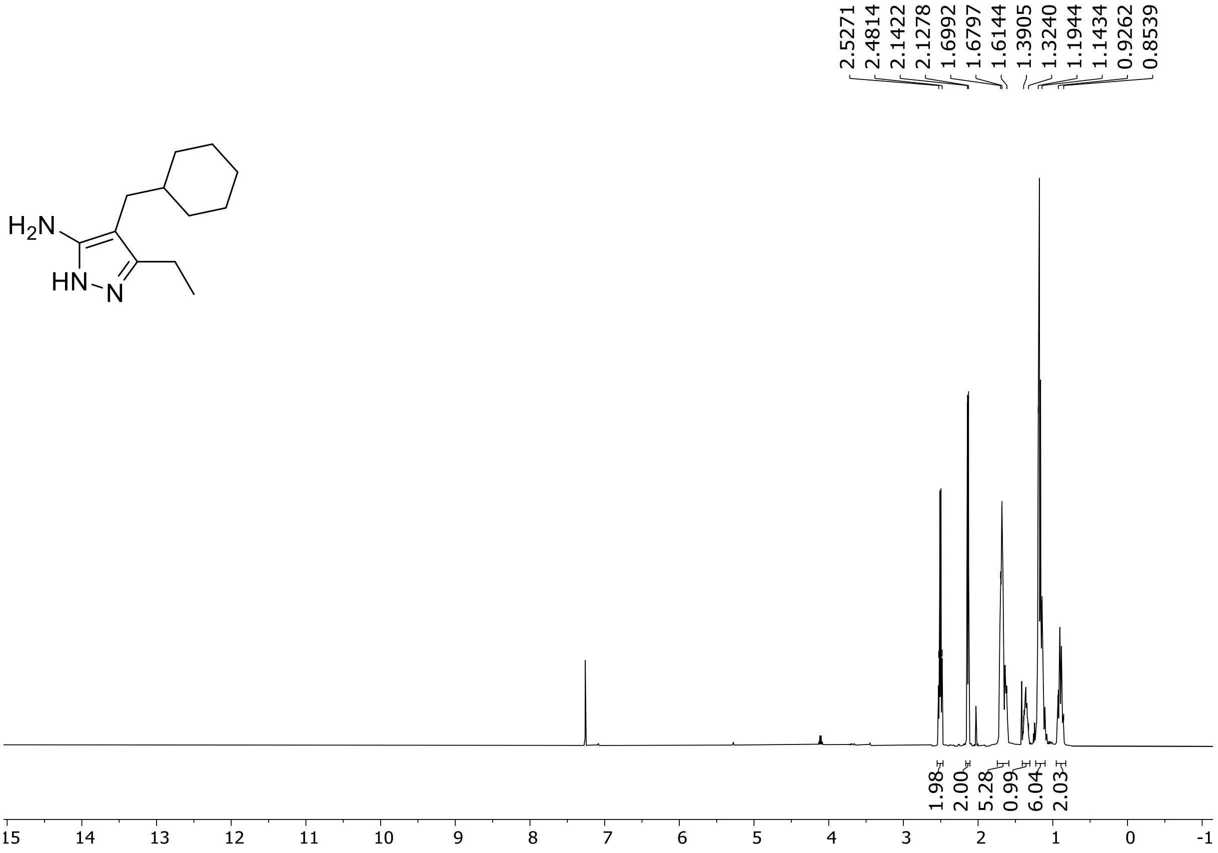

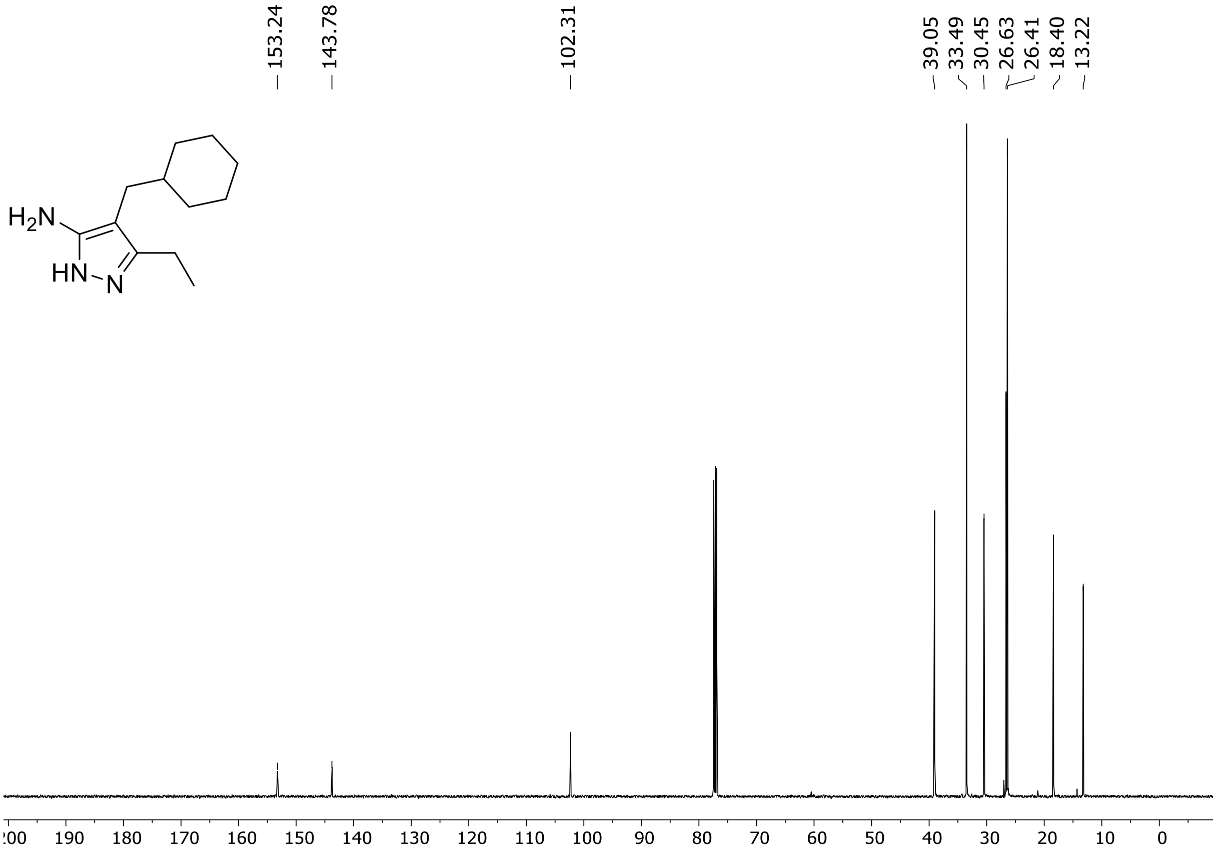

HRMS spectrum of **S64**

$C_{12}H_{21}N_3$ **exact mass: 207.1735**

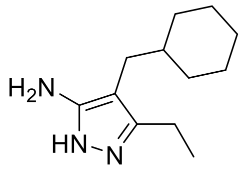

**APCI +** (MMI)
nitrogen flow 5 L/min, gas temperature 325°C, nebulizer 45 psig, skimmer 65 V,
vaporizer 200°C, fragmentor 20 V, dissolved in methanol

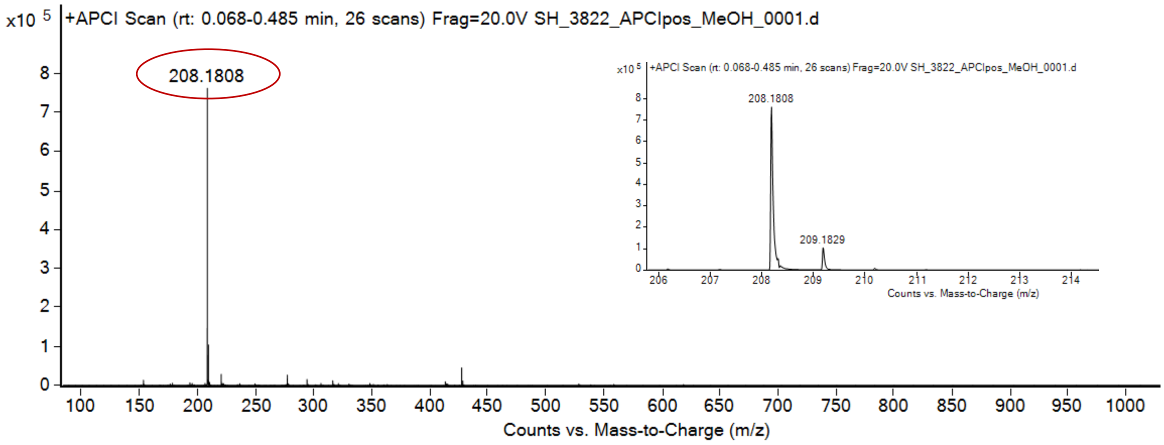

expected mass: $[M+H]^+ = 208.1808$ observed mass: $[M+H]^+ = 208.1808$ mass accuracy < 0.1 ppm

$^{1}H$ (500 MHz) and $^{13}C$ NMR (126 MHz) spectra of **S65** in chloroform-*d*

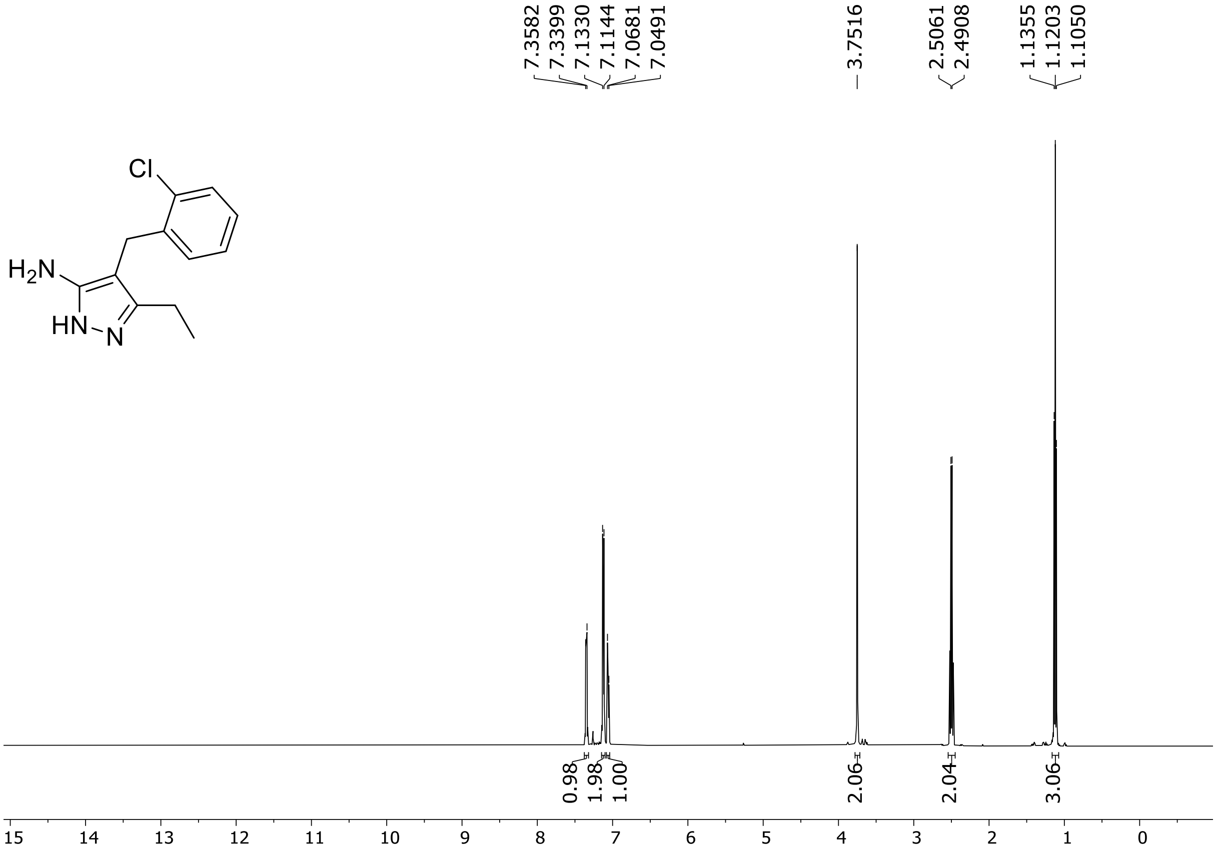

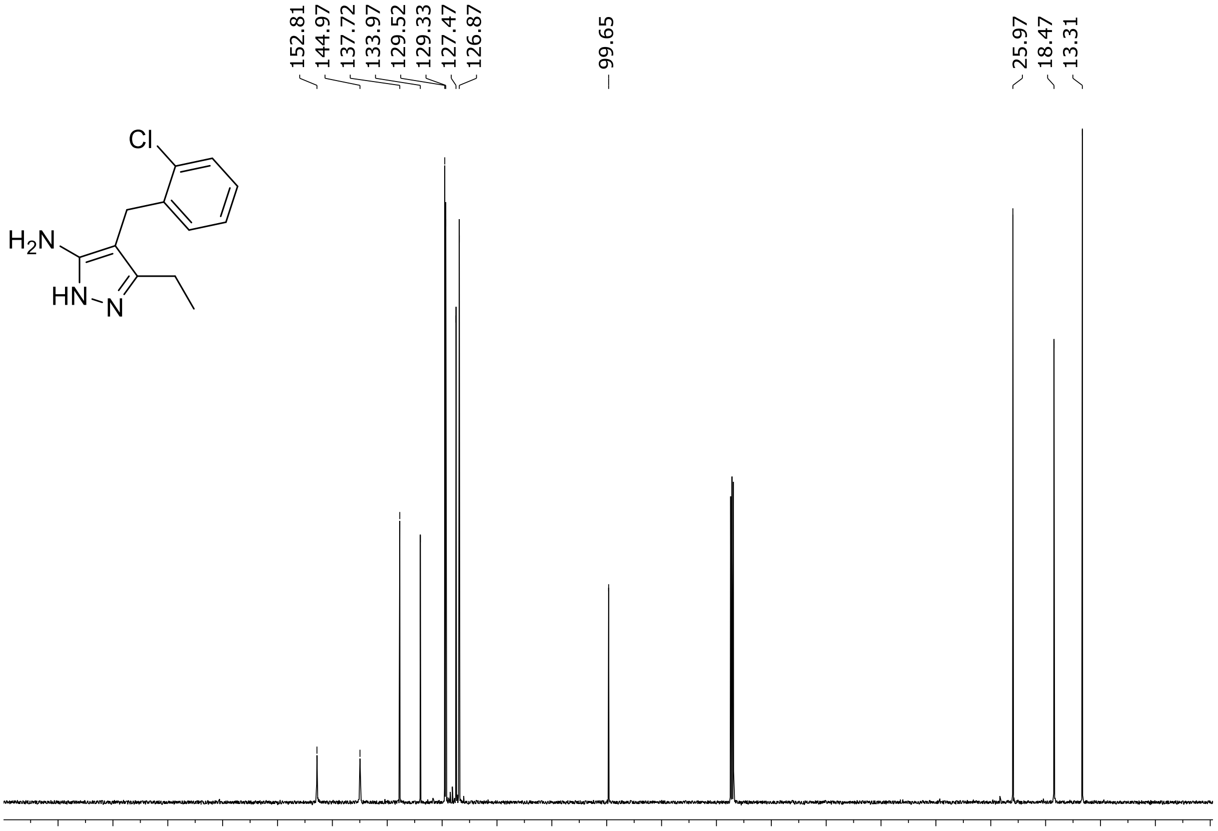

HRMS spectrum of **S65**

**$C_{12}H_{14}ClN_3$** **exact mass: 235.0876**

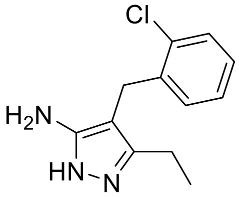

**APCI +** (MMI)
nitrogen flow 5 L/min, gas temperature 325°C, nebulizer 45 psig, skimmer 65 V, vaporizer 200°C, fragmentor 25 V, dissolved in methanol

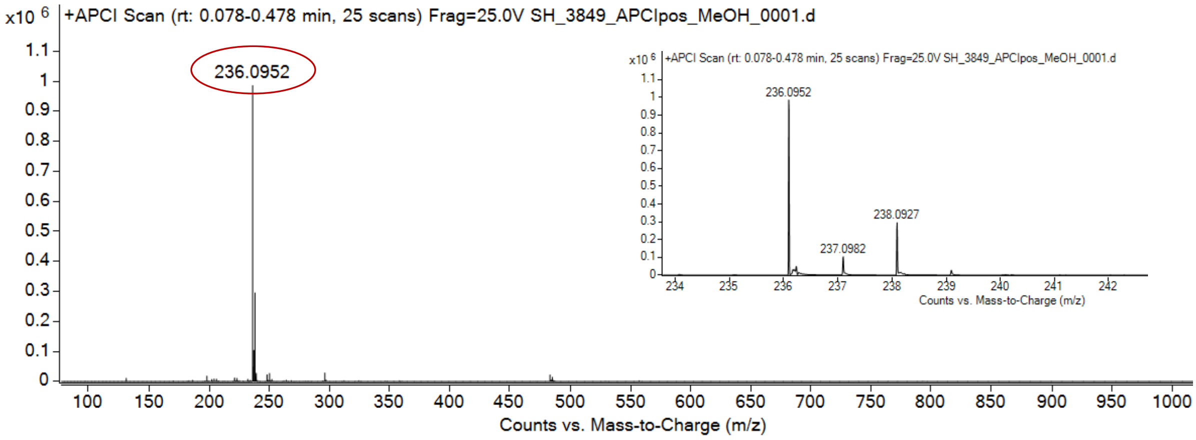

expected mass: $[M+H]^+$ = 236.0949 observed mass : $[M+H]^+$ = 236.0952 mass accuracy = 1.3 ppm

[1]H (500 MHz) and [13]C NMR (126 MHz) spectra of **S66** in chloroform-*d*

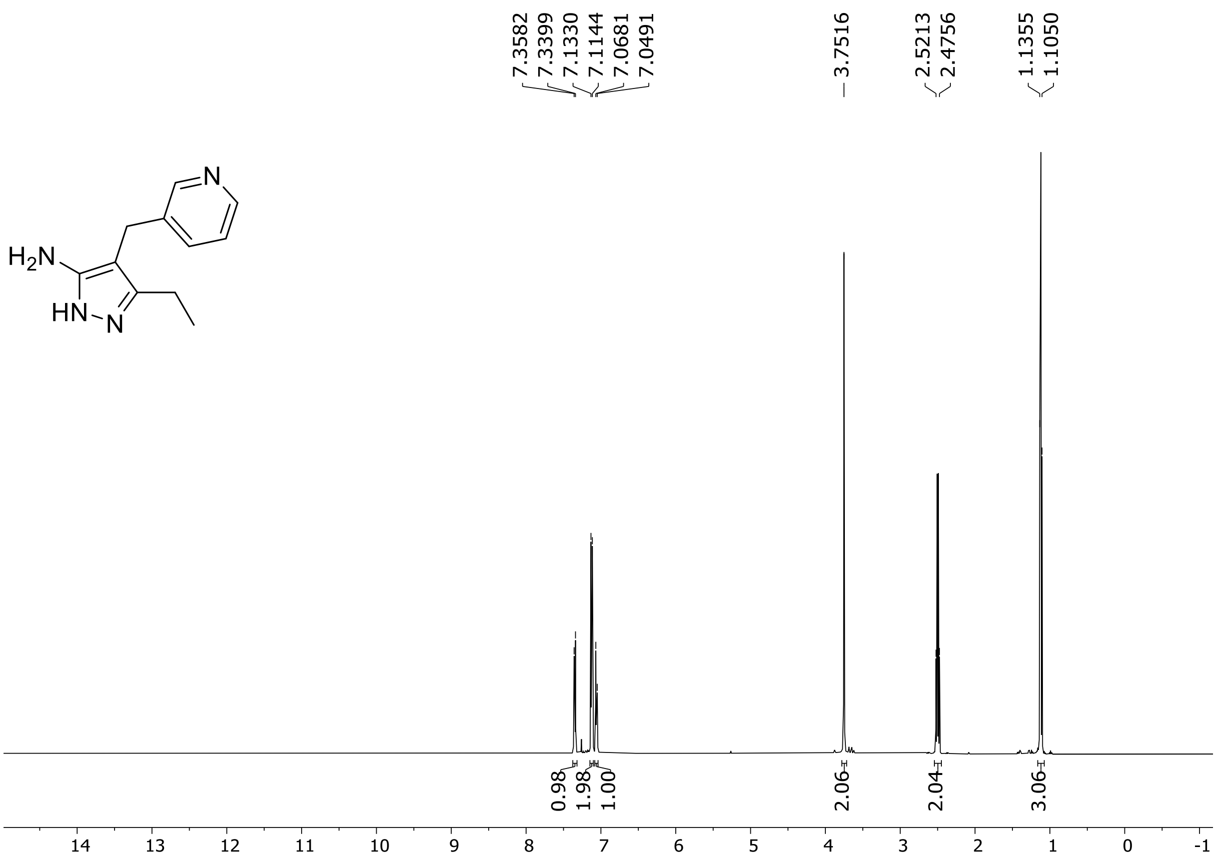

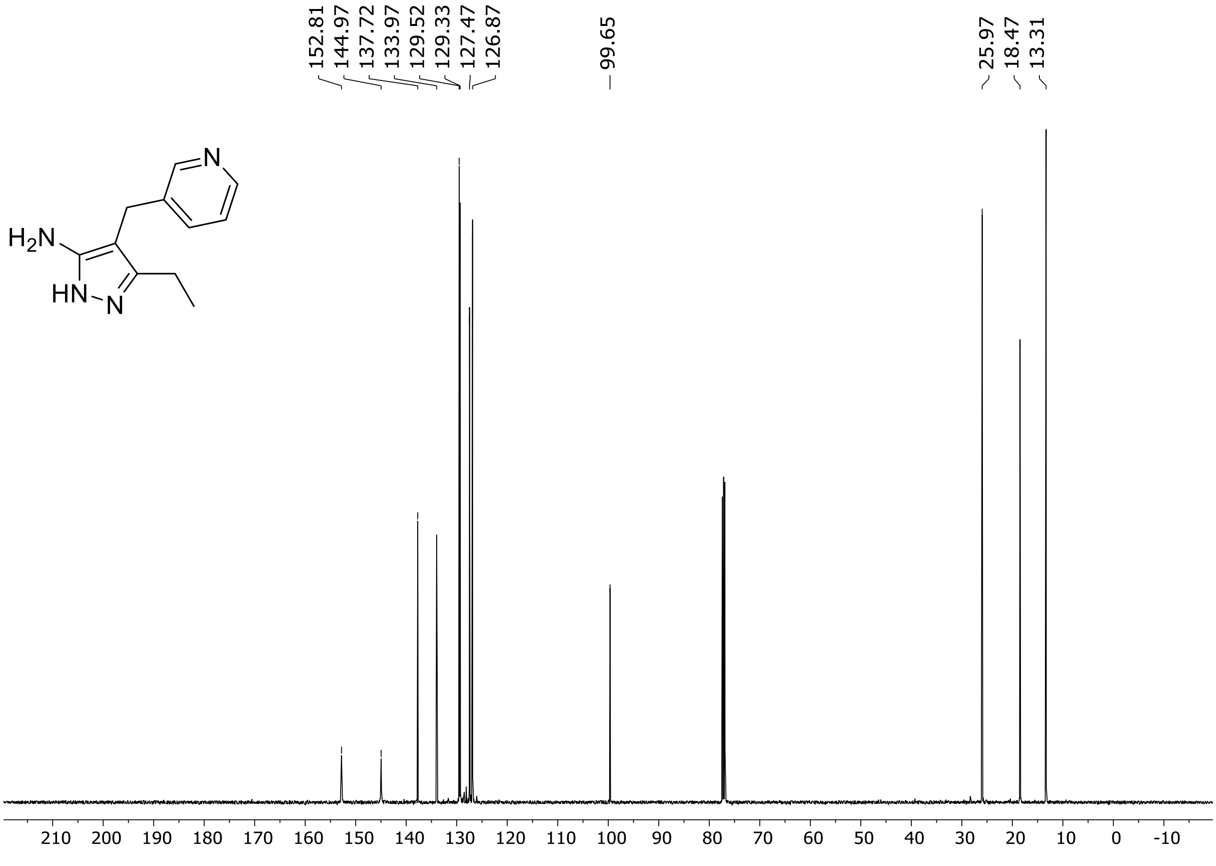

HRMS spectrum of **S66**

**$C_{11}H_{14}N_4$** **exact mass: 202.1218**

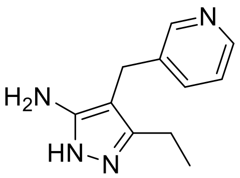

**APCI +** (MMI)
nitrogen flow 5 L/min, gas temperature 325°C, nebulizer 45 psig, skimmer 65 V, vaporizer 200°C, fragmentor 18 V, dissolved in methanol

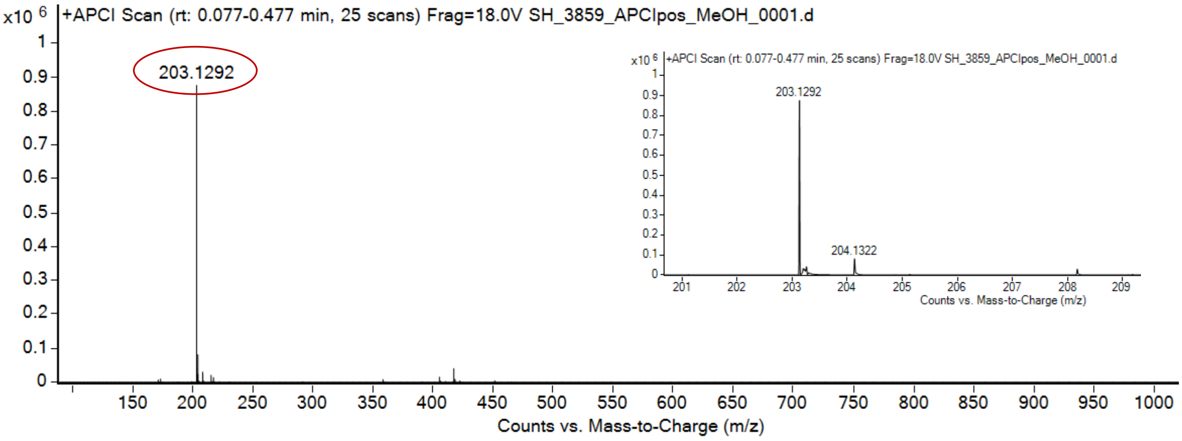

expected mass: $[M+H]^+$ = 203.1291 observed mass : $[M+H]^+$ = 203.1292 mass accuracy = 0.5 ppm

[1]H (500 MHz) and [13]C NMR (126 MHz) spectra of **S67** in chloroform-*d*

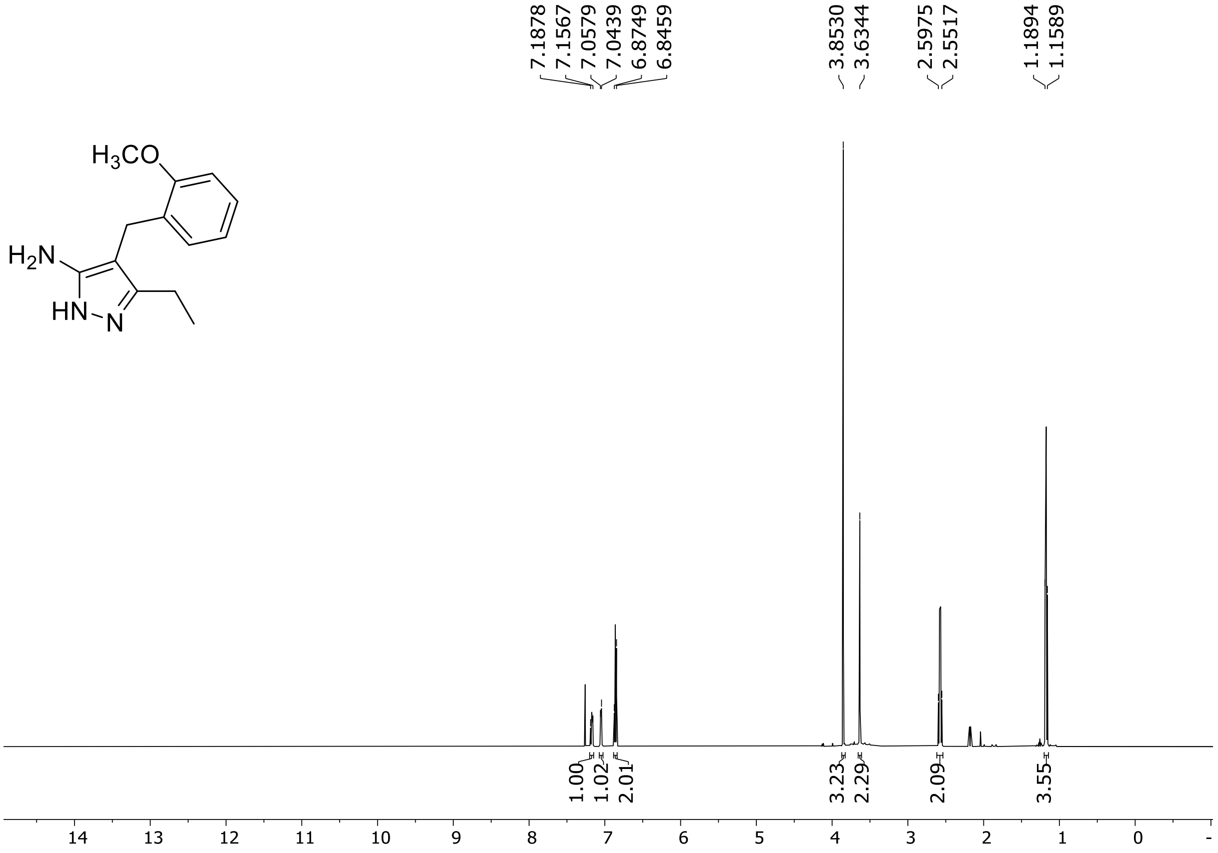

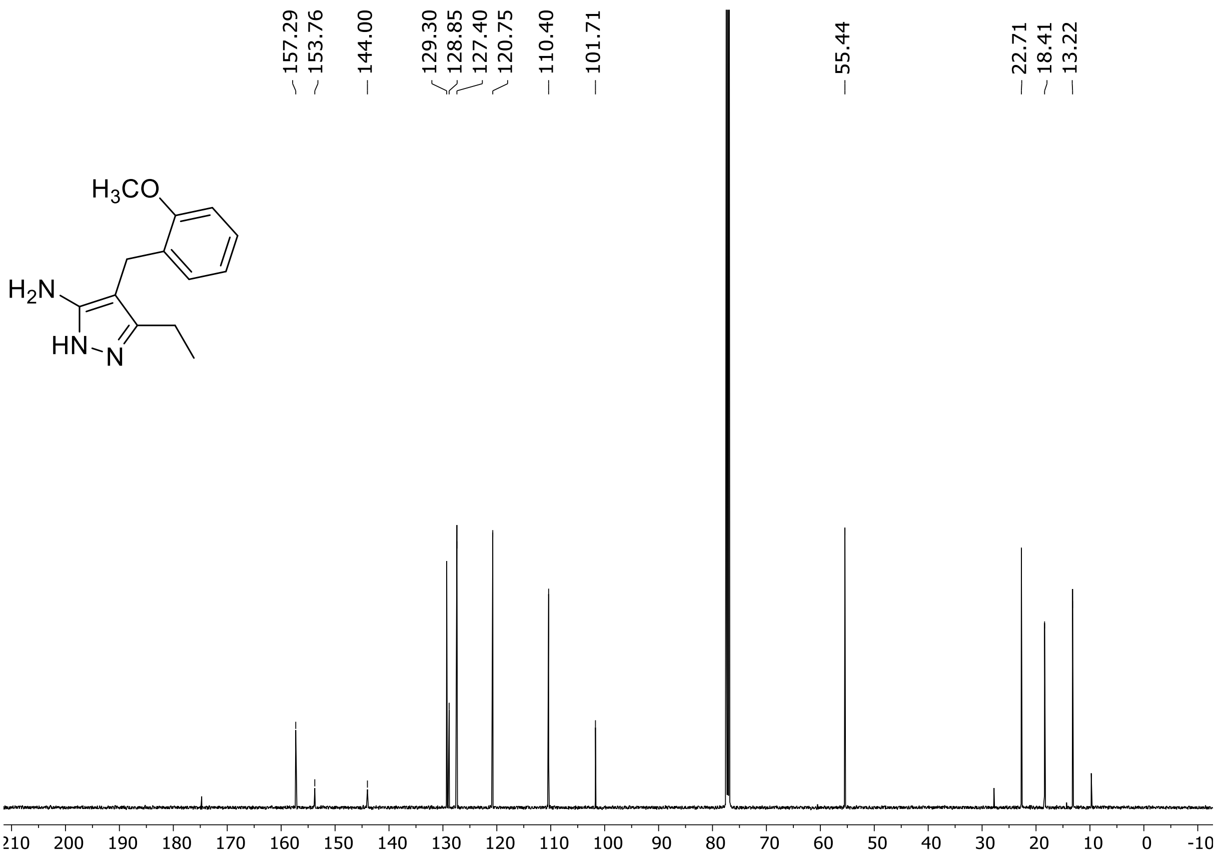

$^{1}H$ (500 MHz) and $^{13}C$ NMR (126 MHz) spectra of **S68** in chloroform-*d*

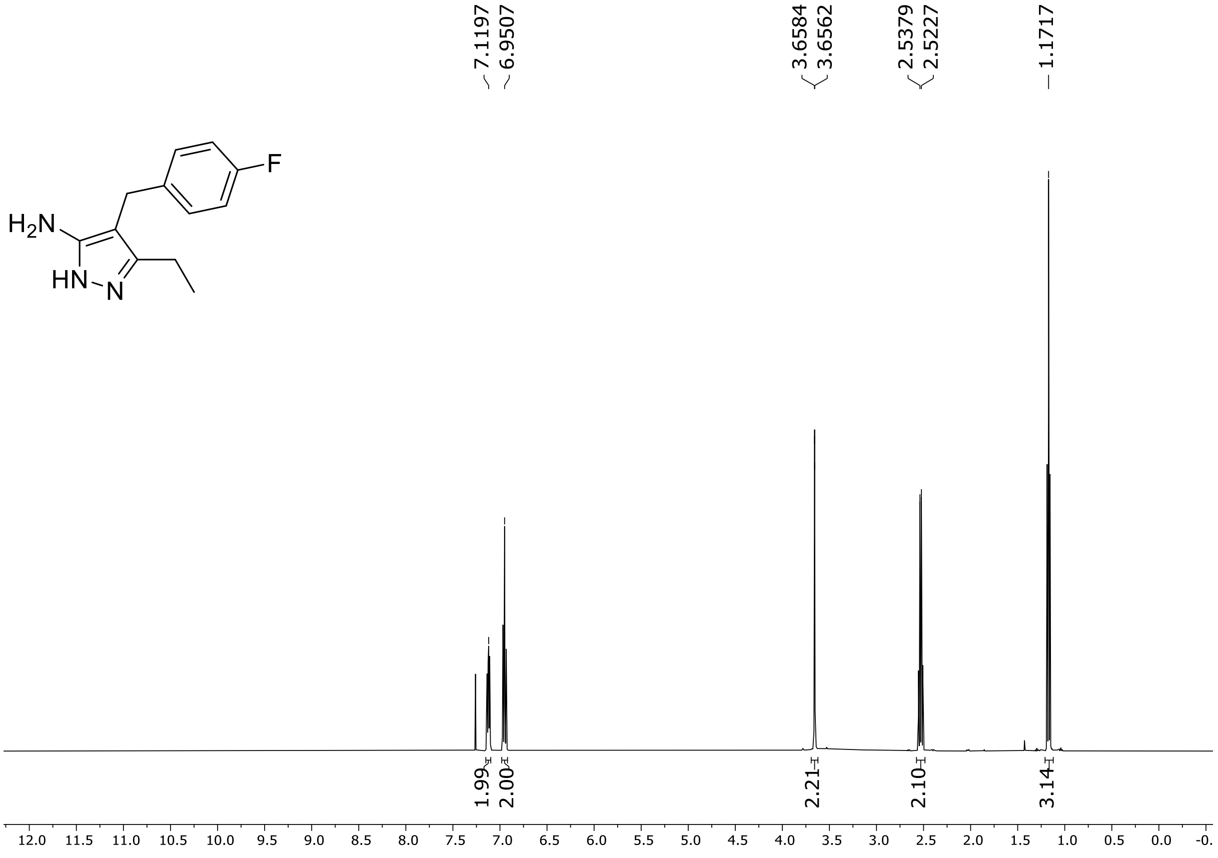

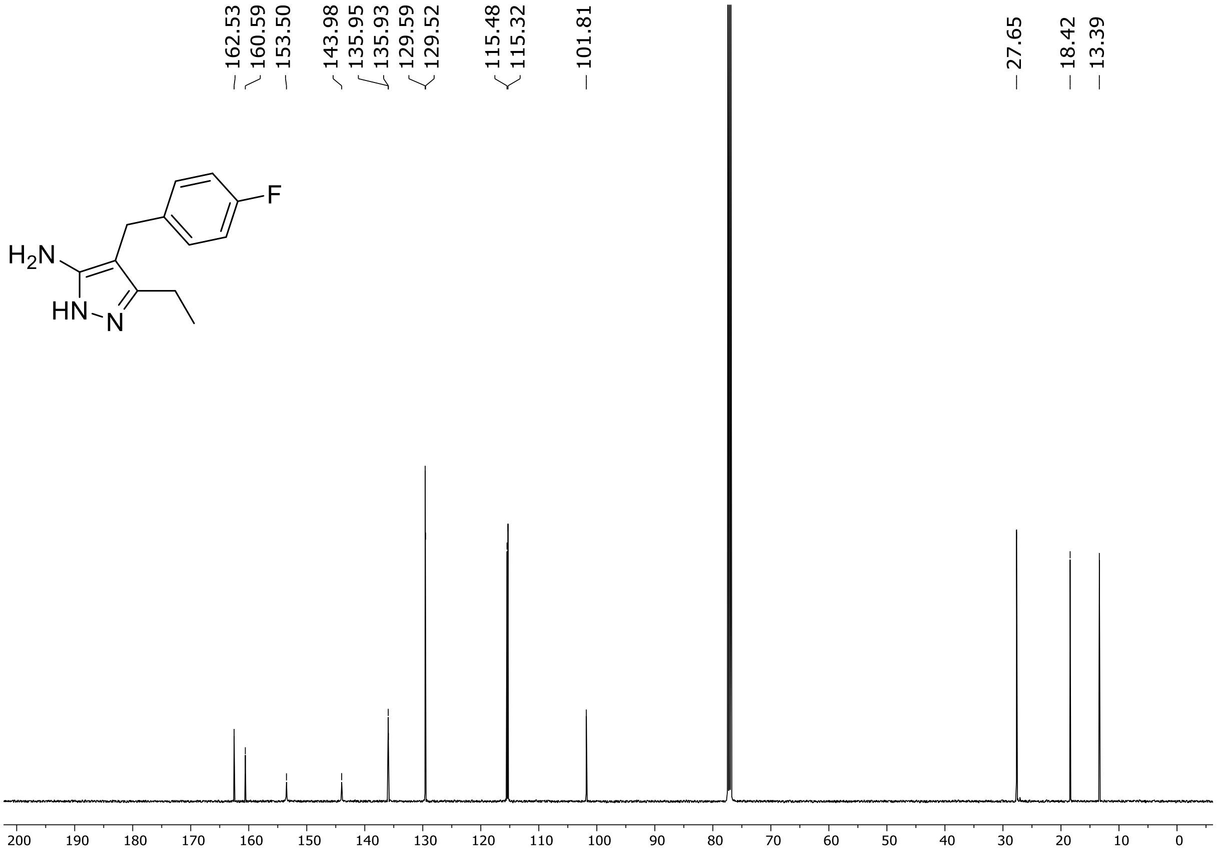

$^{19}$F (471 MHz) NMR spectrum of **S68** in chloroform-*d*

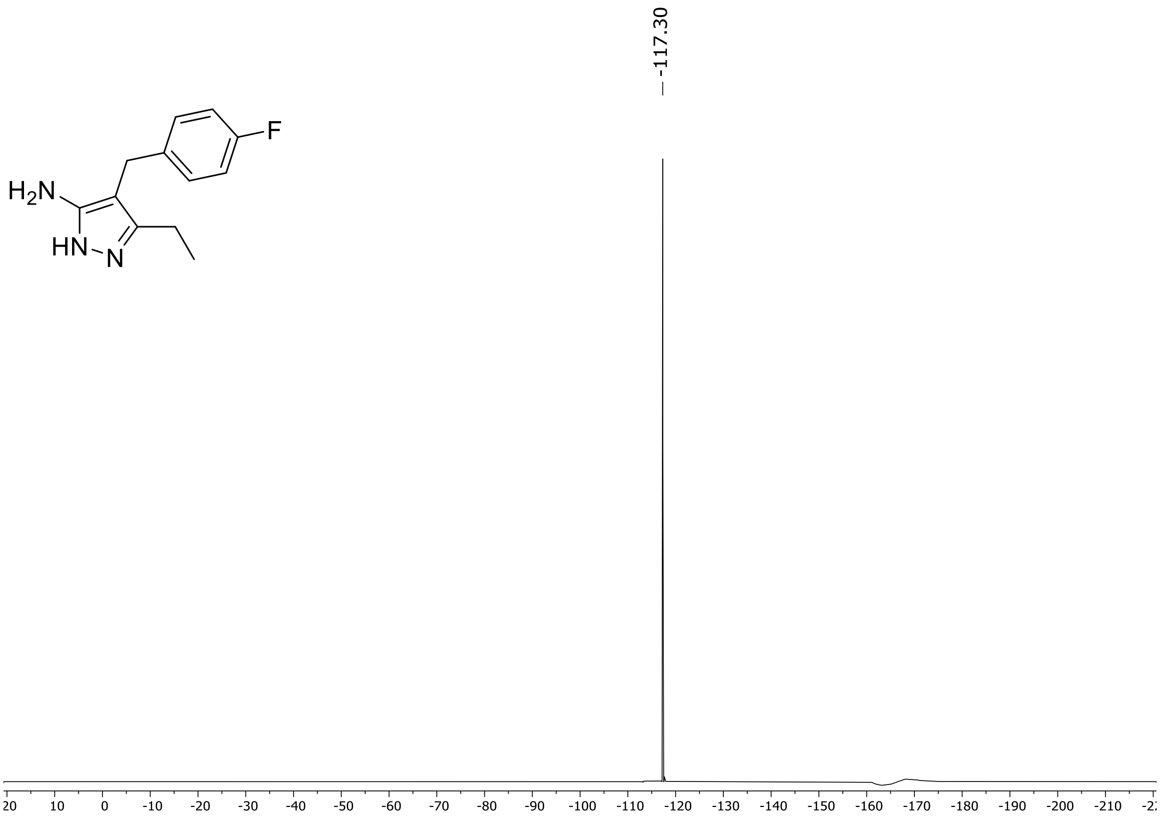

HRMS spectrum of **S68**

**$C_{12}H_{14}FN_3$** **exact mass: 219.1172**

**APCI +** (MMI)
nitrogen flow 5 L/min, gas temperature 325°C, nebulizer 45 psig, skimmer 65 V, vaporizer 200°C, fragmentor 18 V, dissolved in methanol

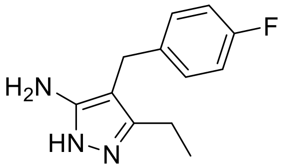

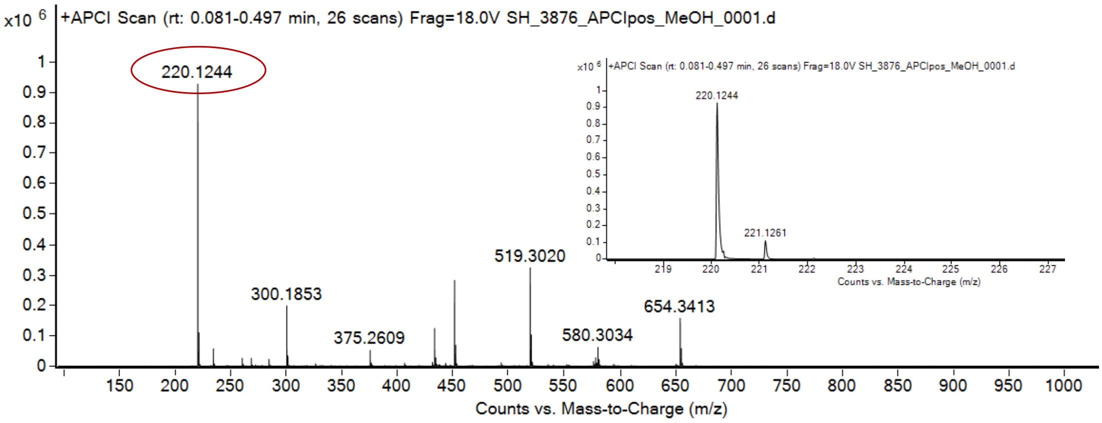

expected mass: [M+H]$^+$ = 220.1245 observed mass: [M+H]$^+$ = 220.1244 mass accuracy = - 0.5 ppm

$^{1}H$ (500 MHz) and $^{13}C$ NMR (126 MHz) spectra of **S69** in chloroform-*d*

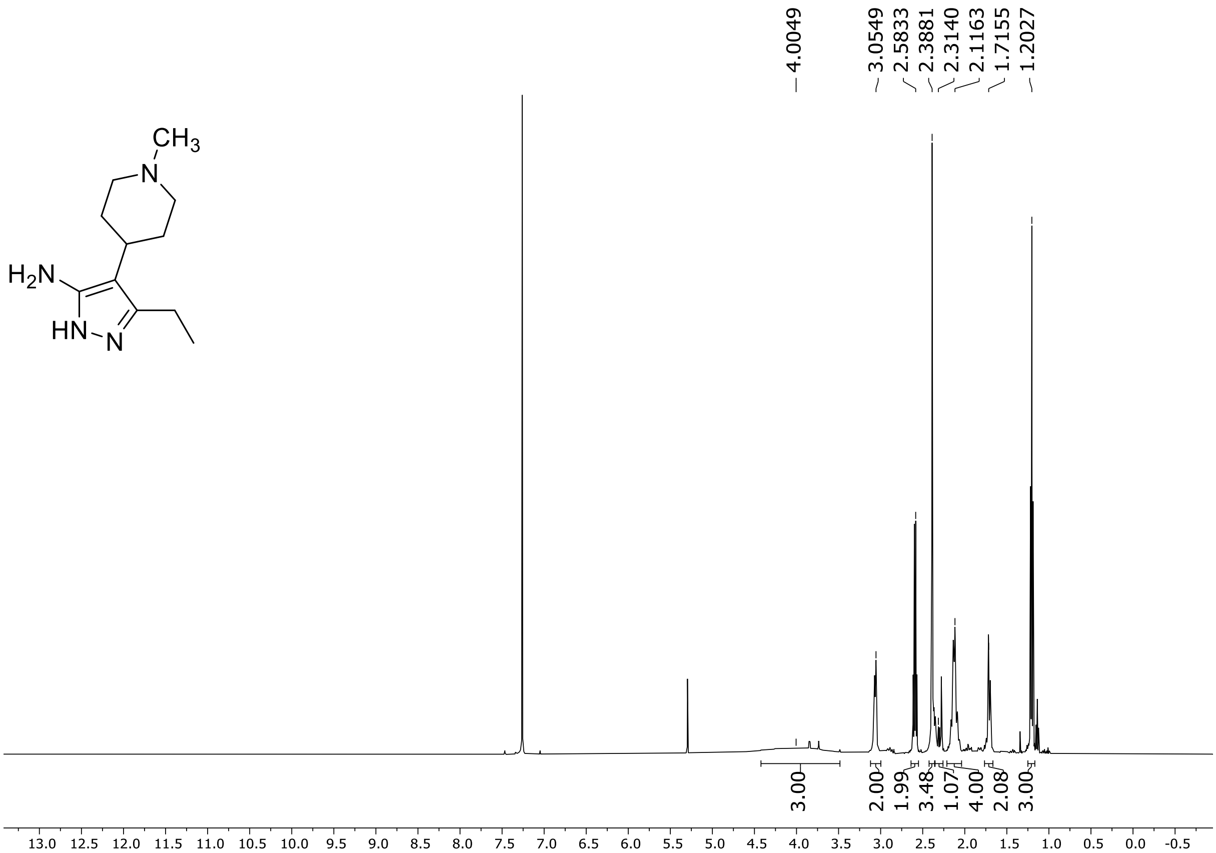

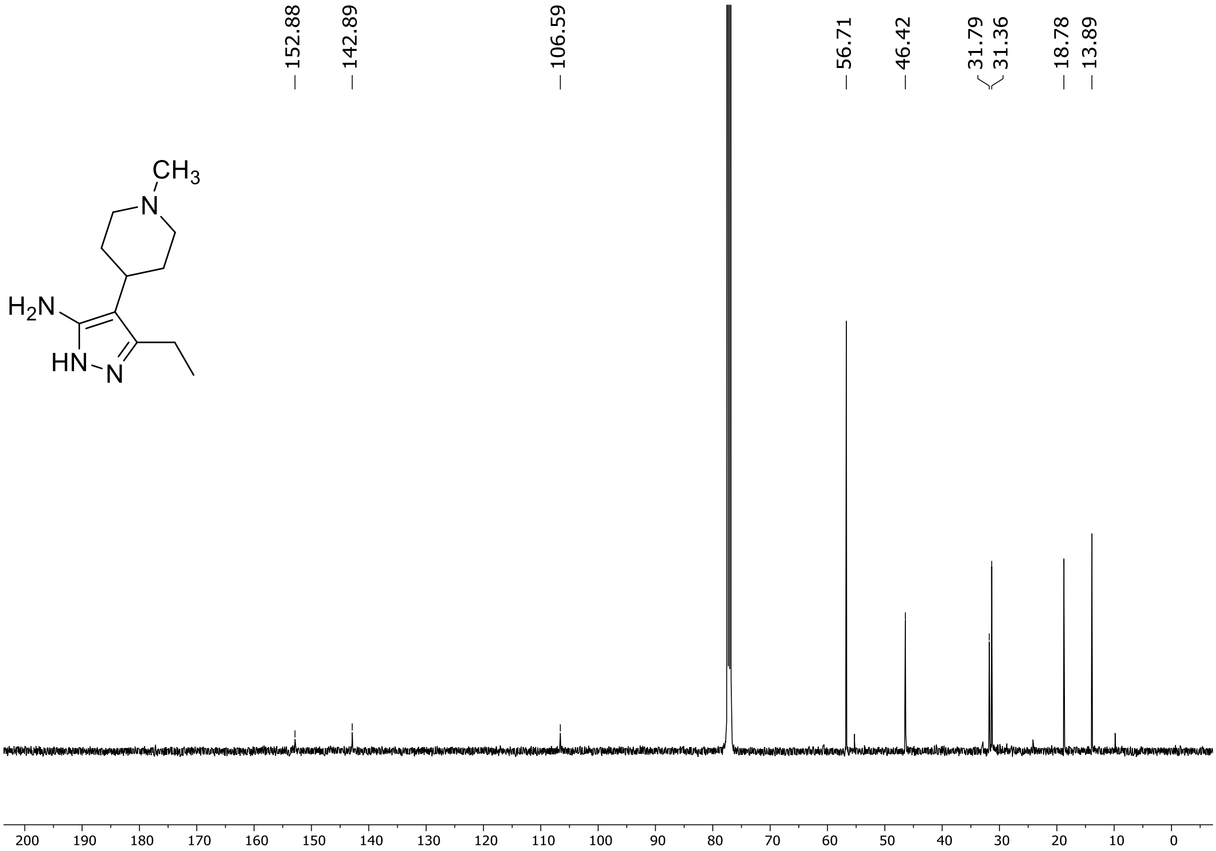

[1]H (500 MHz) and [13]C NMR (126 MHz) spectra of **S70** in chloroform-*d*

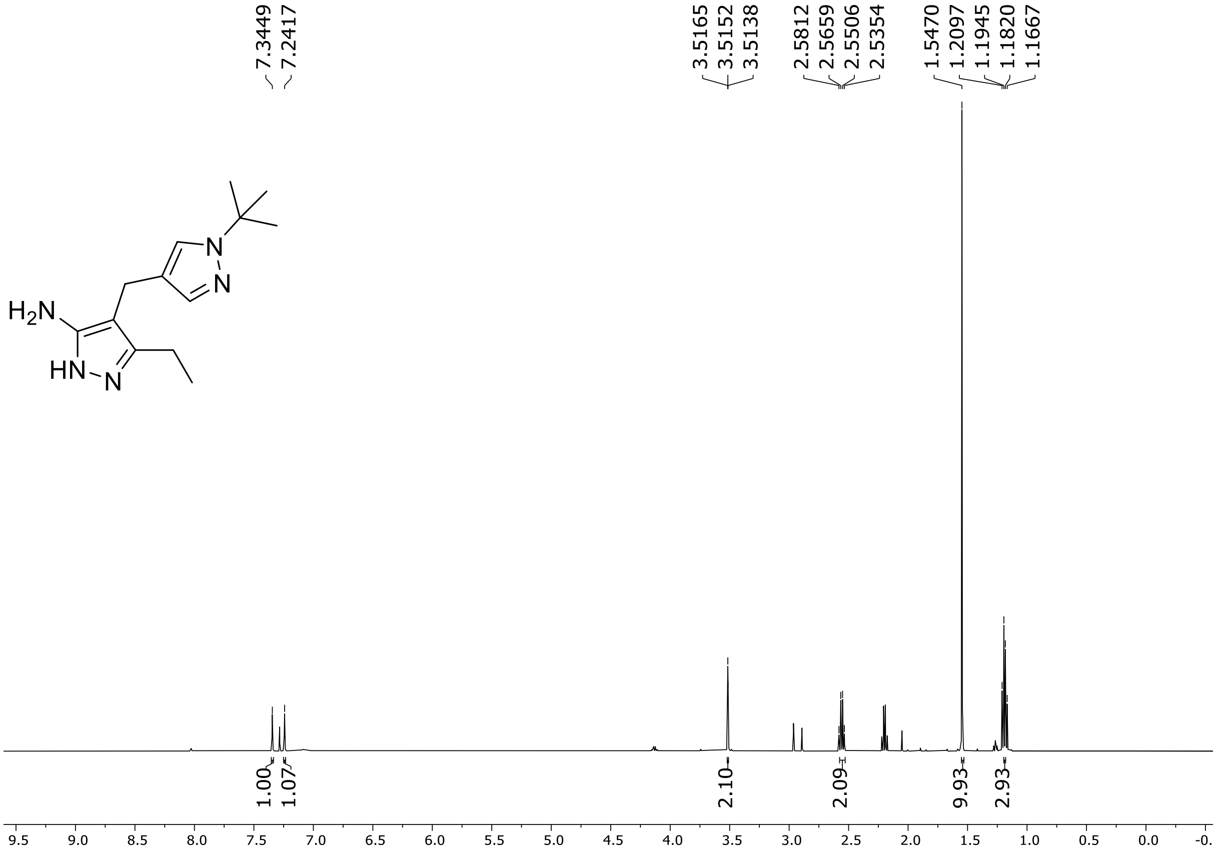

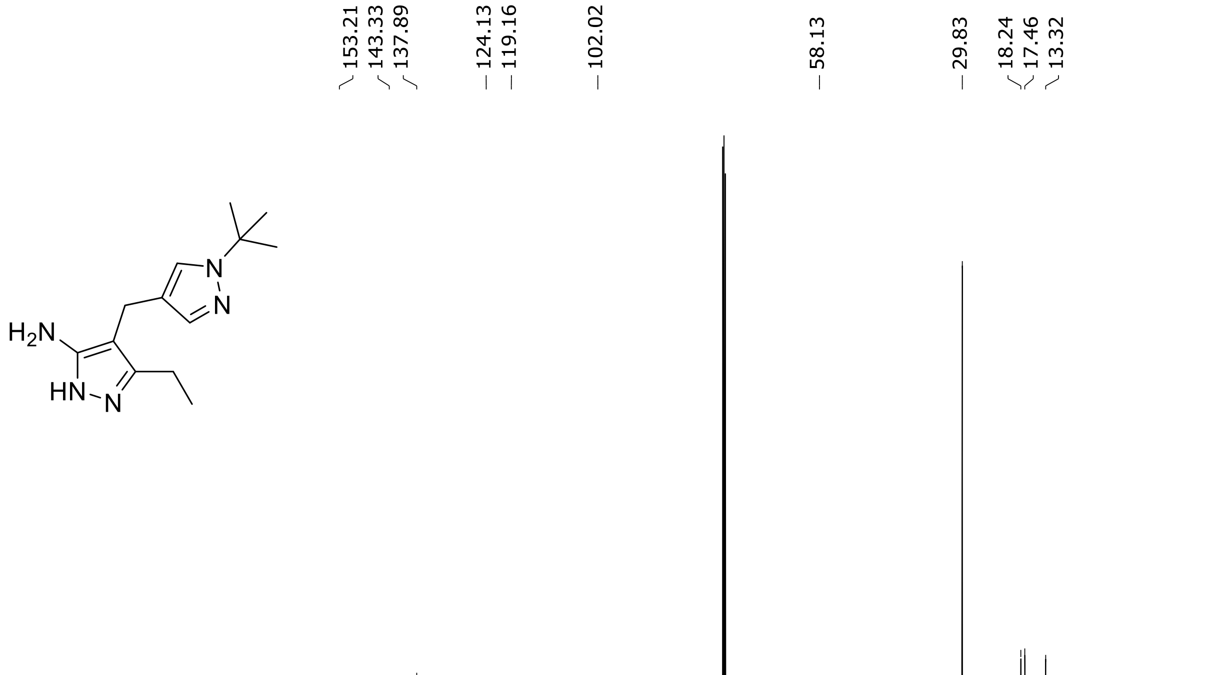

$^{1}H$ (500 MHz) and $^{13}C$ NMR (126 MHz) spectra of **S82** in DMSO-$d_6$

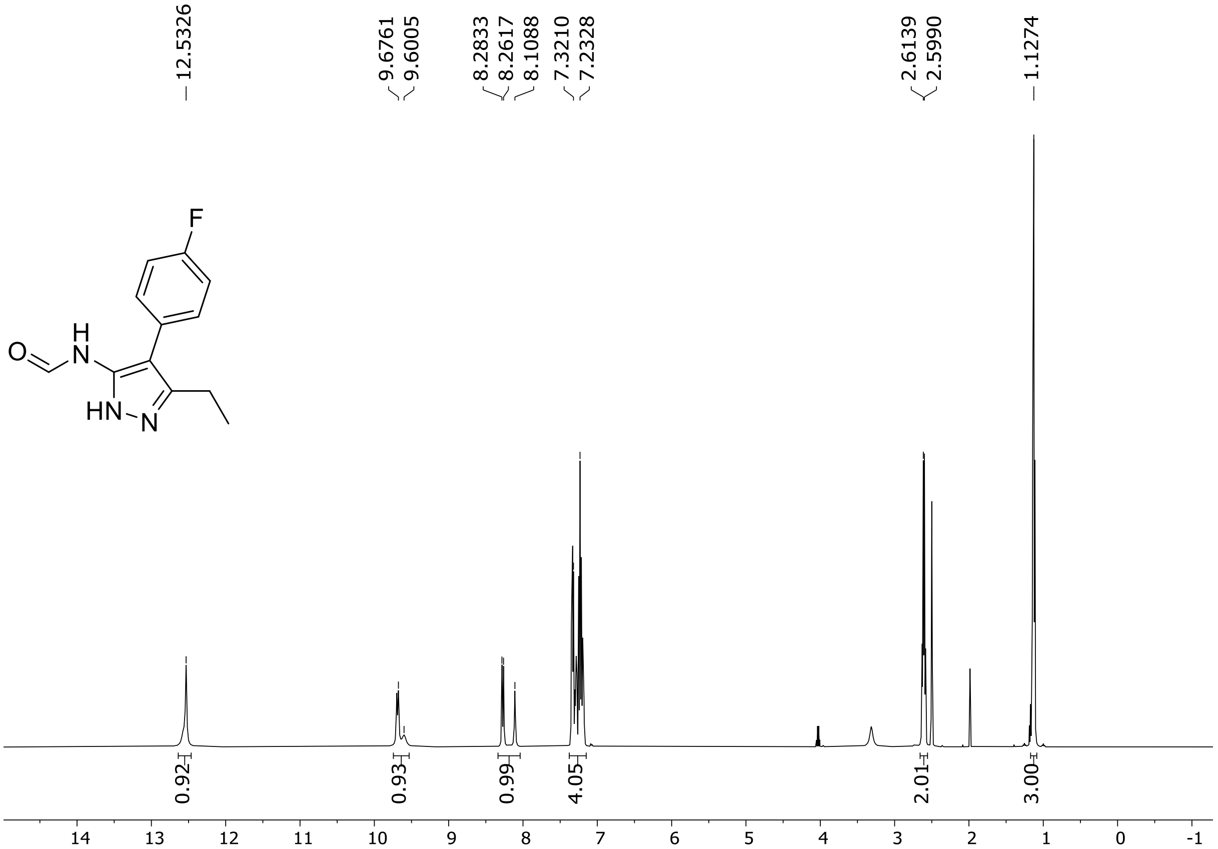

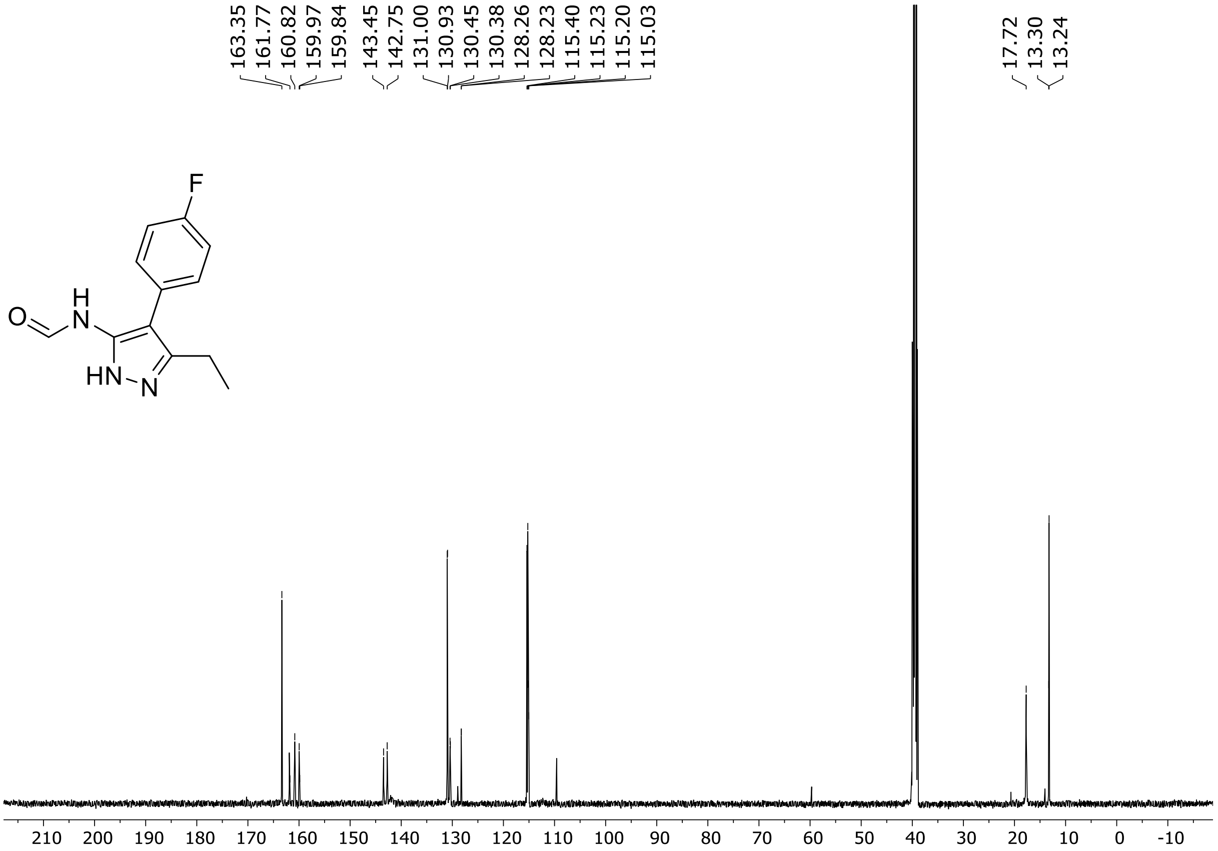

$^{19}F$ (471 MHz) NMR spectrum of **S82** in DMSO-$d_6$

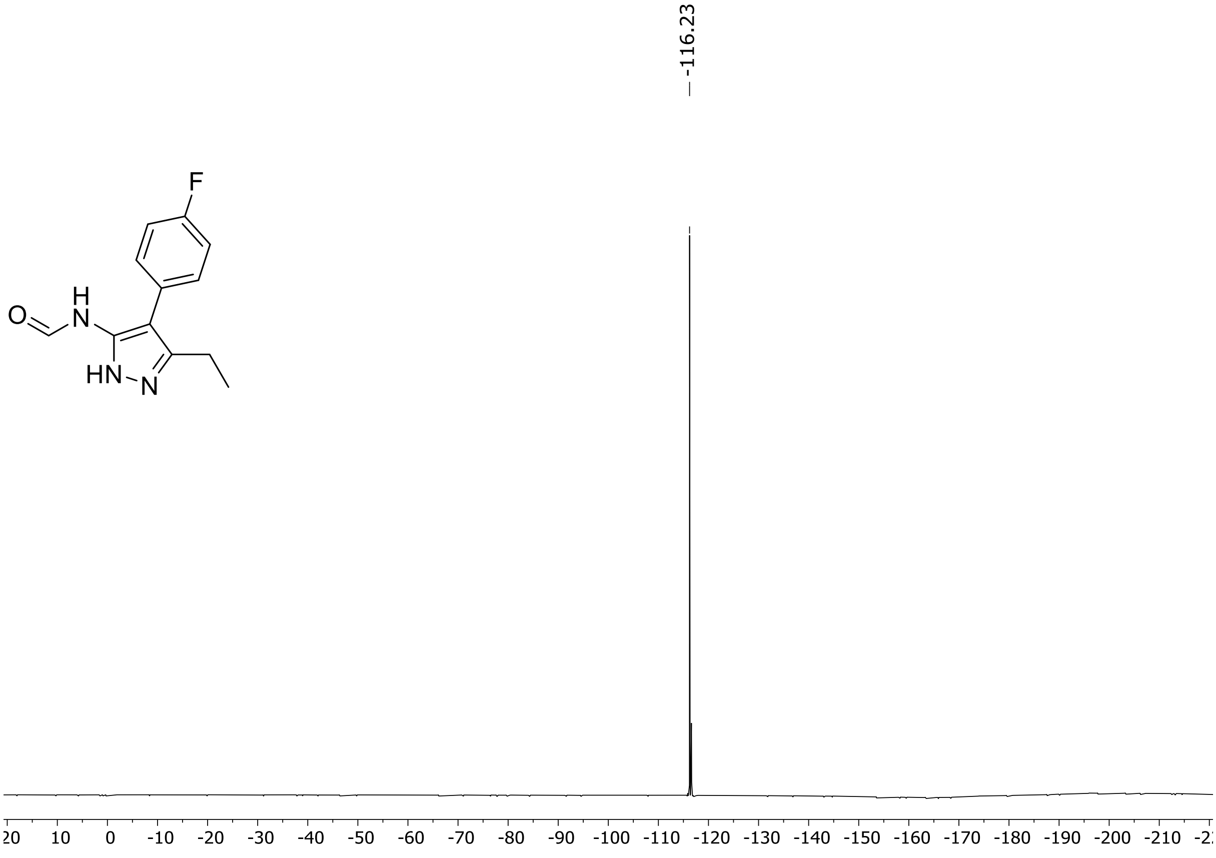

[1]H (500 MHz) and [13]C NMR (126 MHz) spectra of **S83** in DMSO-$d_6$

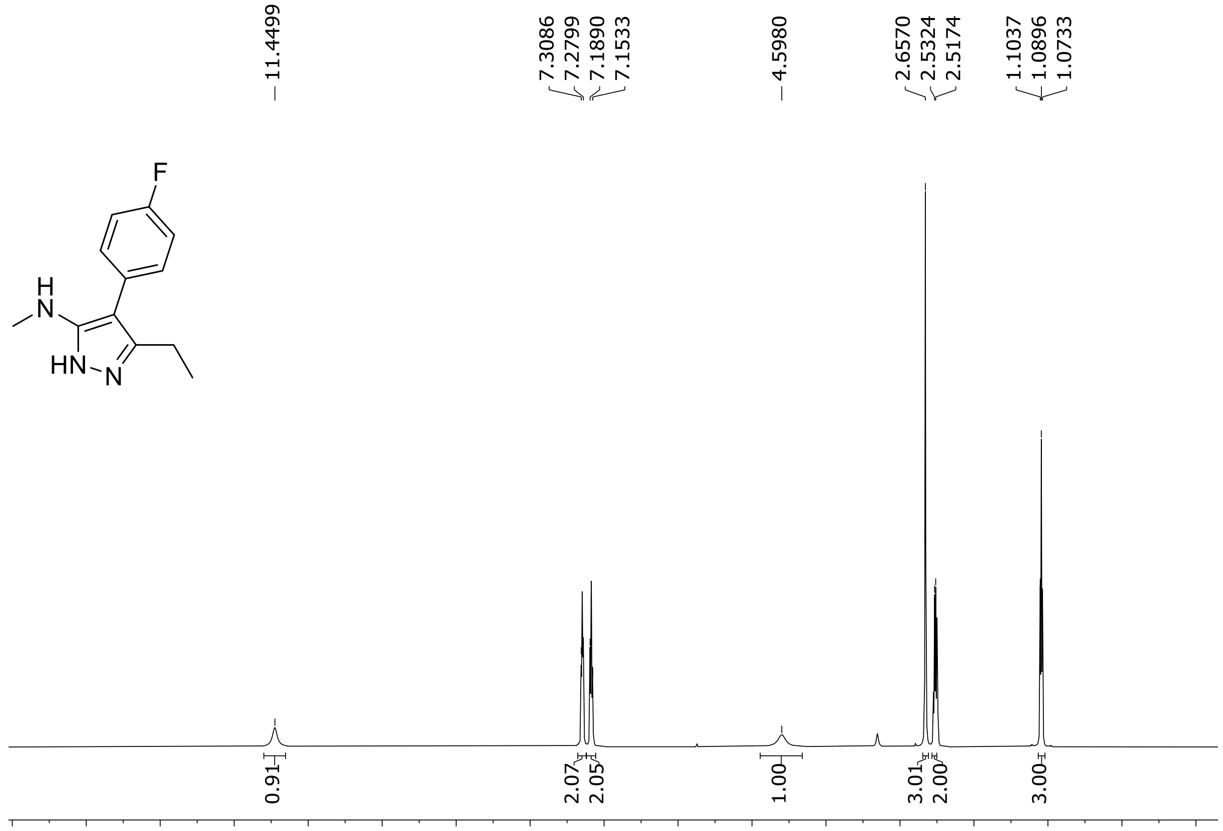

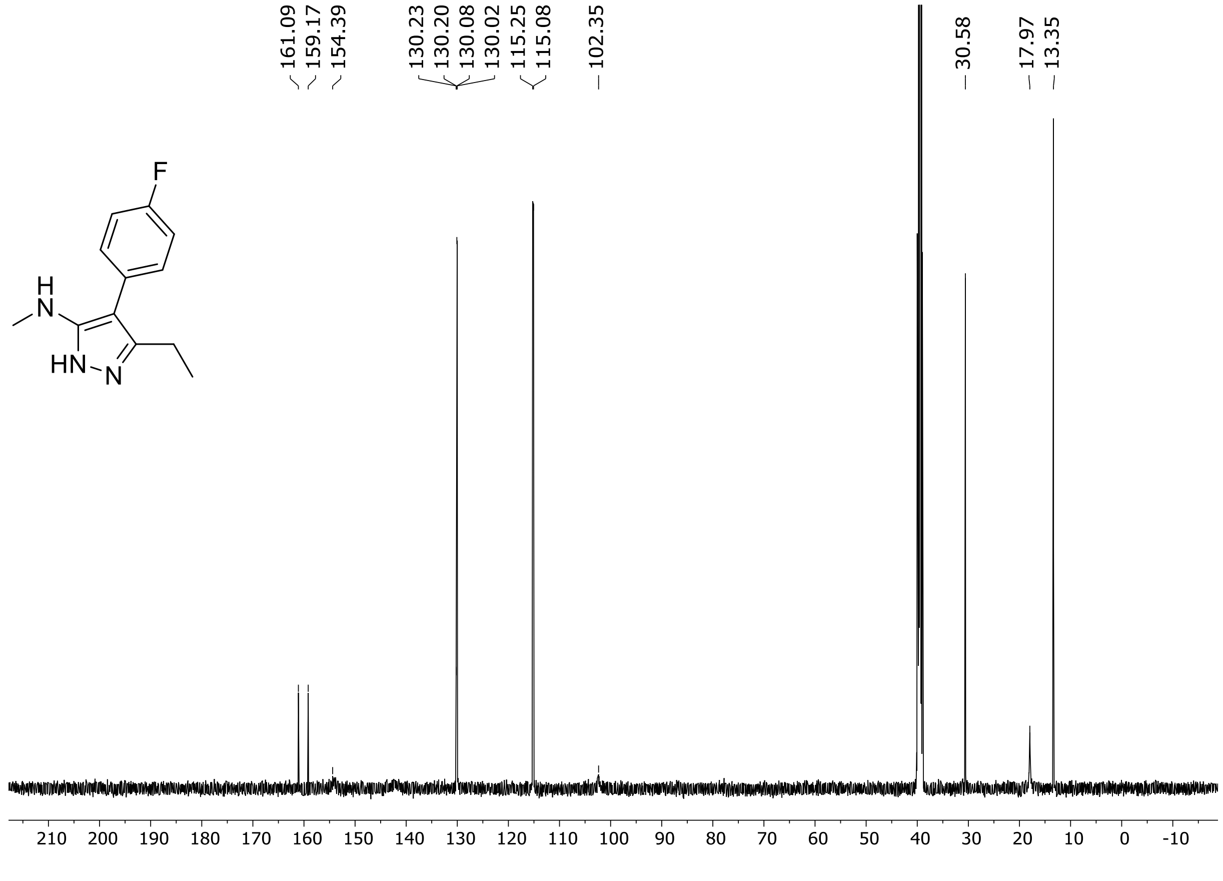

$^{19}F$ (471 MHz) NMR spectrum of **S83** in DMSO-$d_6$

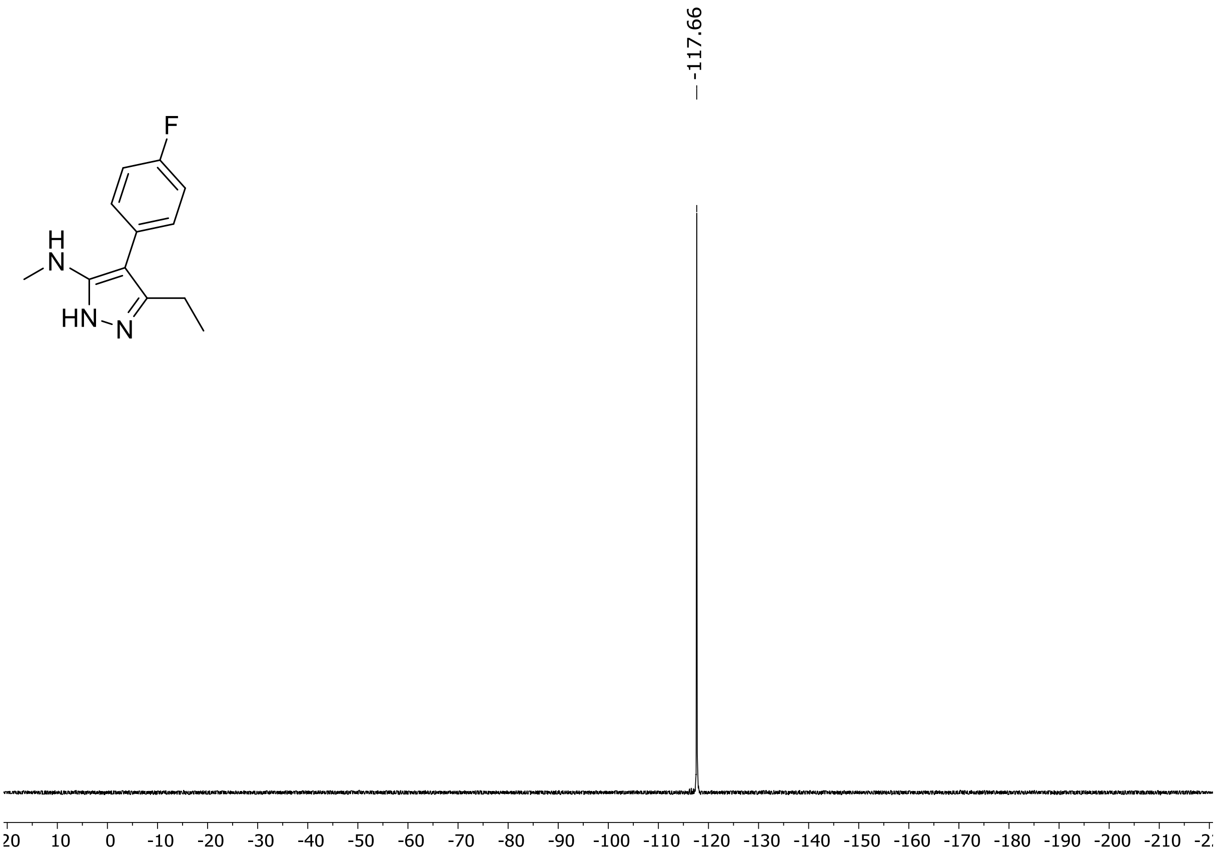

$^{1}H$ (500 MHz) and $^{13}C$ NMR (126 MHz) spectra of **S84** in DMSO-$d_6$

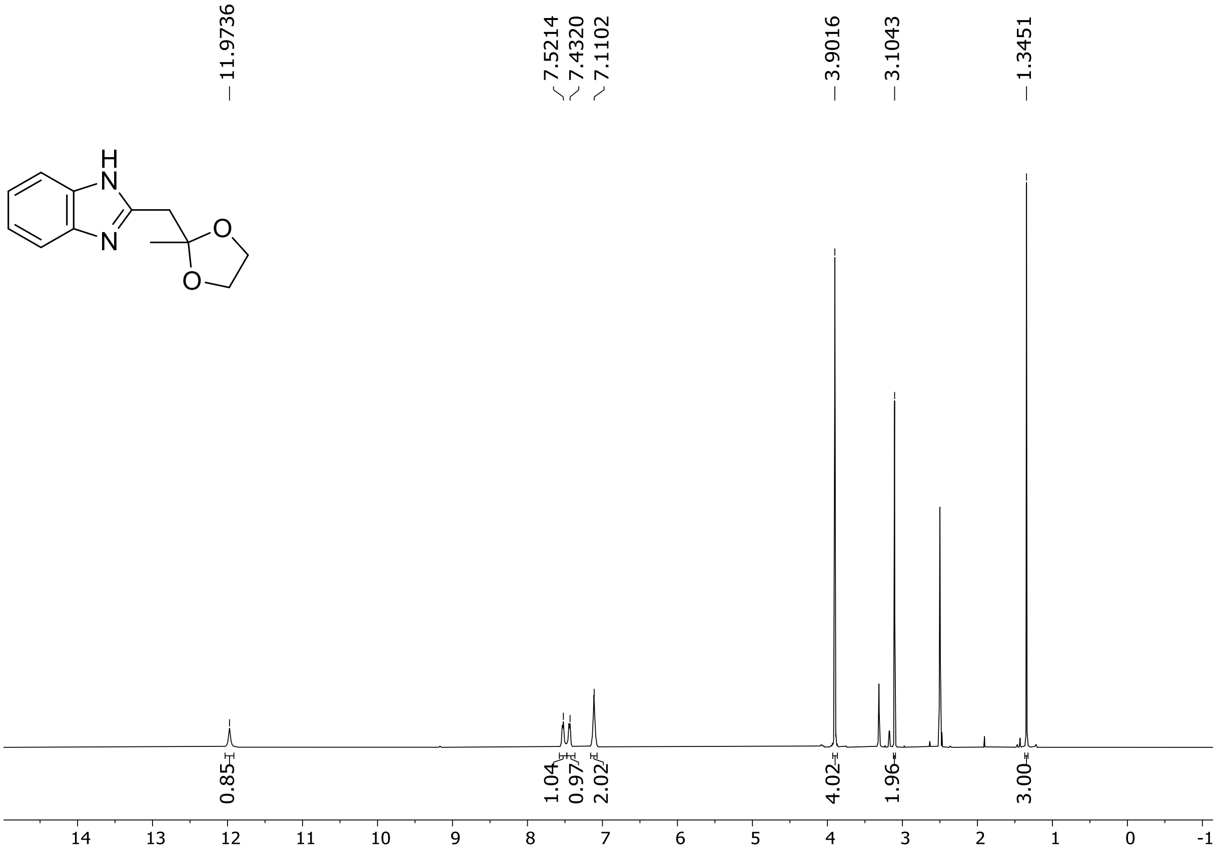

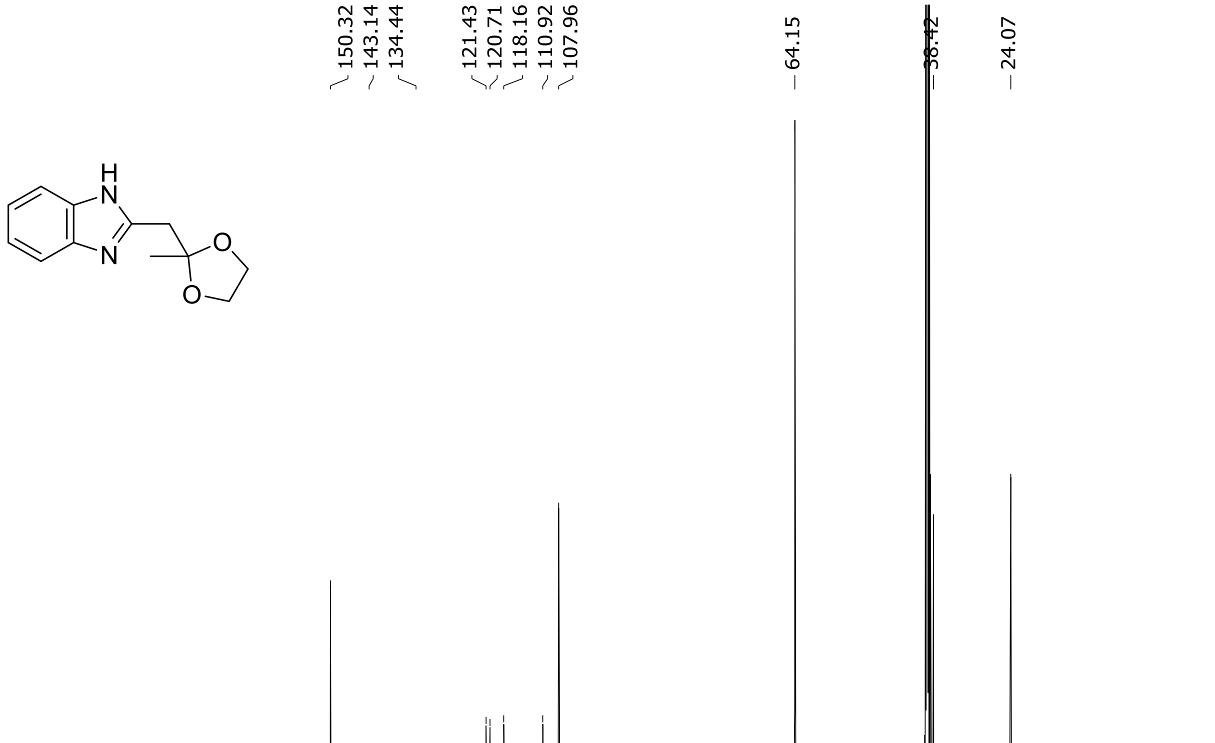

[1]H (500 MHz) and [13]C NMR (126 MHz) spectra of **S86** in DMSO-$d_6$

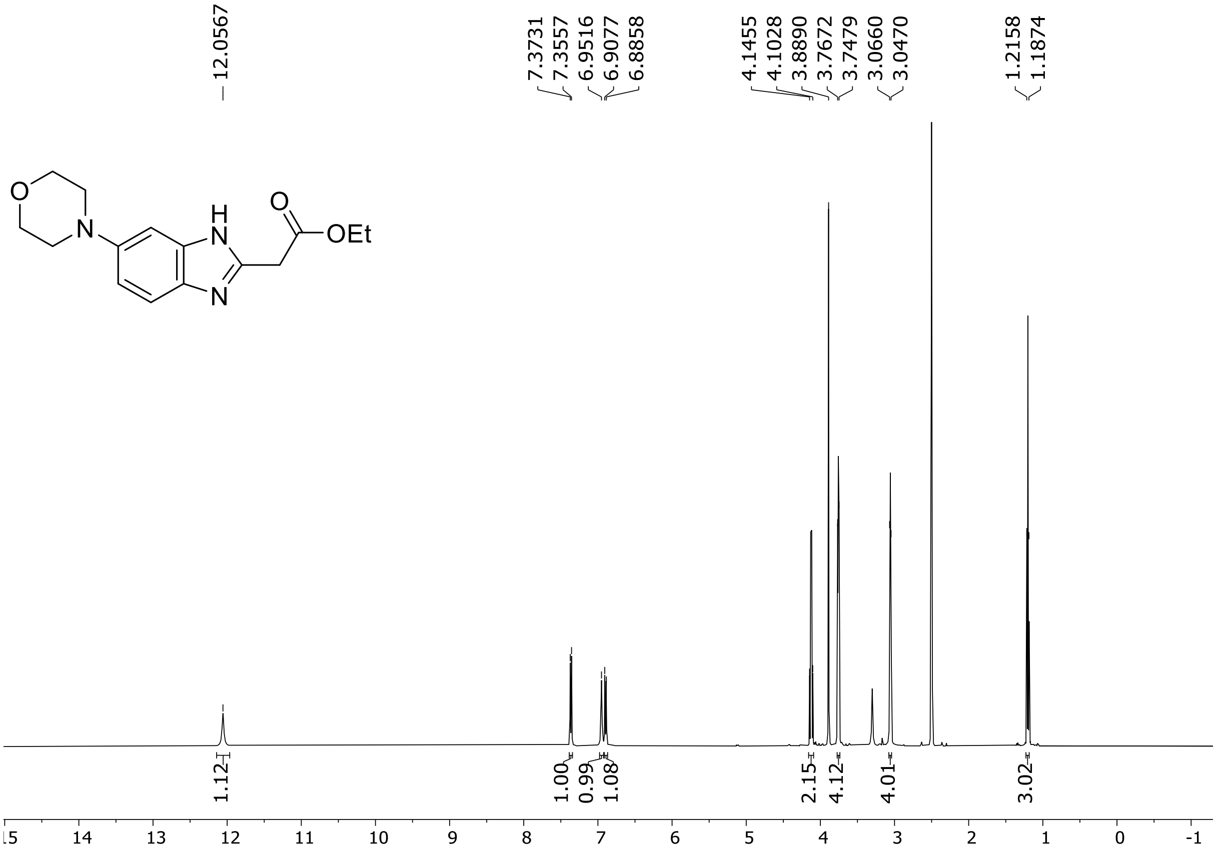

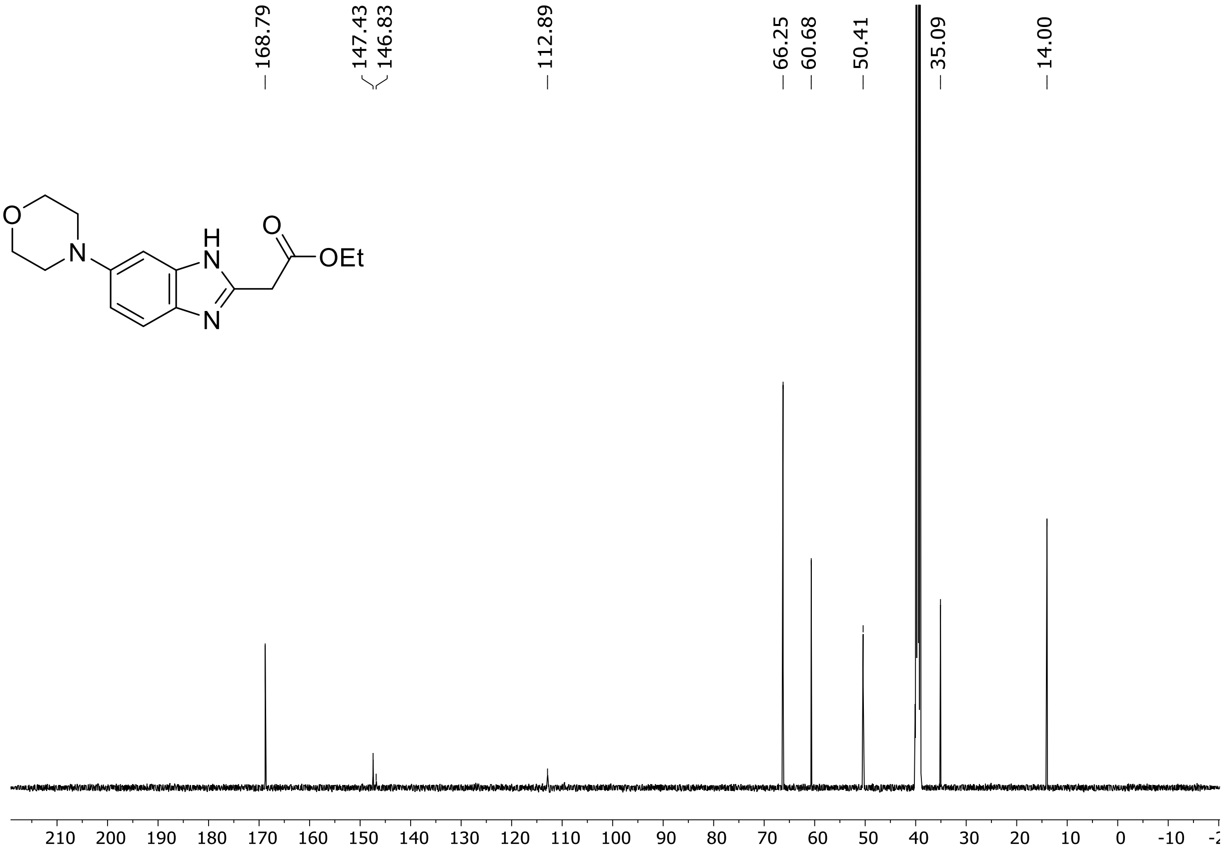

$^{1}H$ (500 MHz) and $^{13}C$ NMR (126 MHz) spectra of **S87** in DMSO-$d_6$

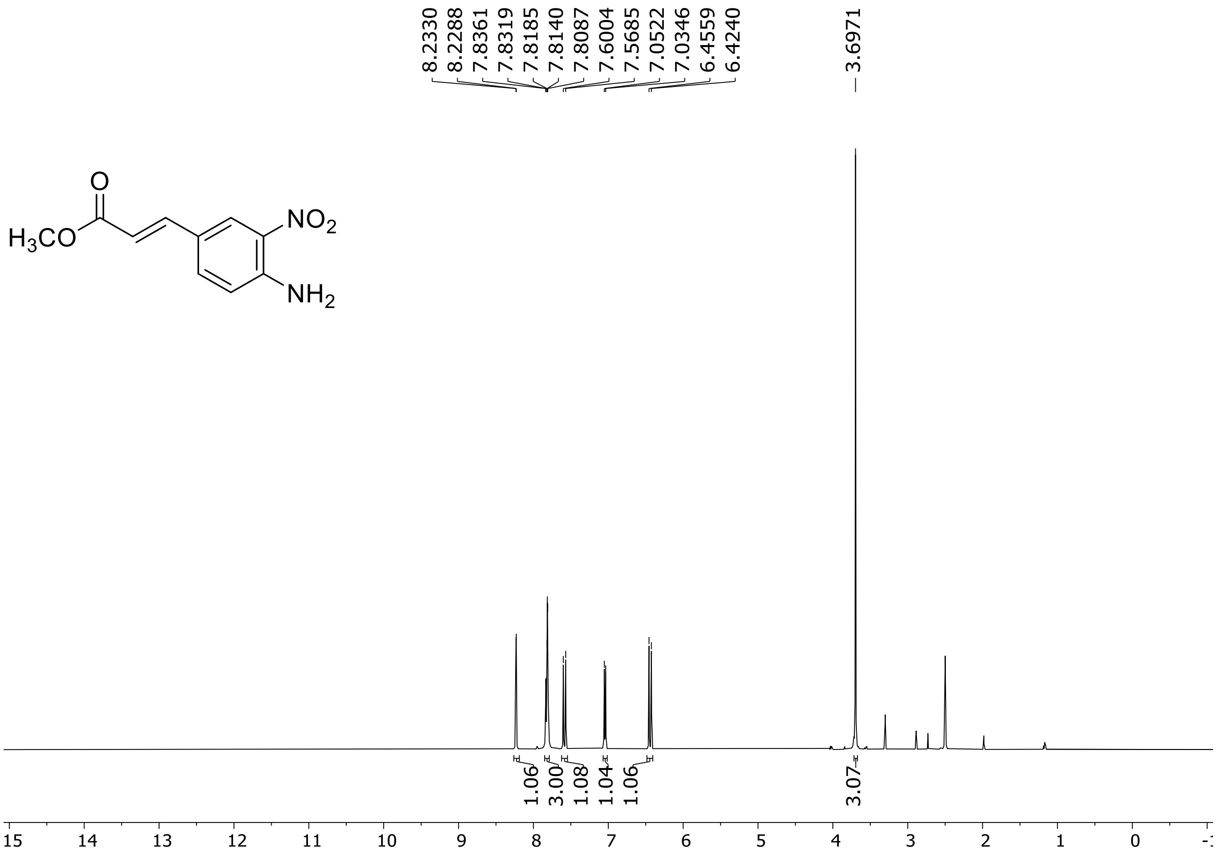

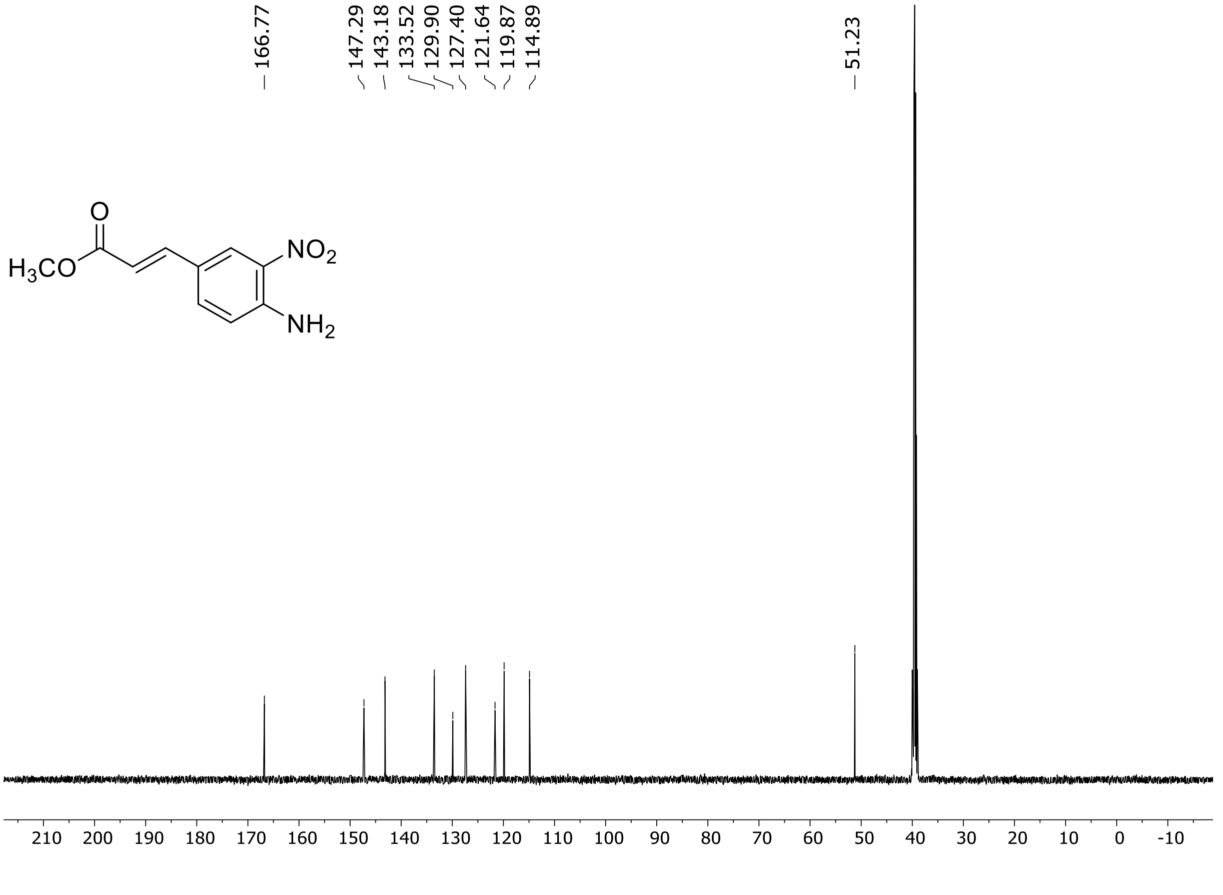

[1]H (500 MHz) NMR spectrum of **S88** in DMSO-$d_6$

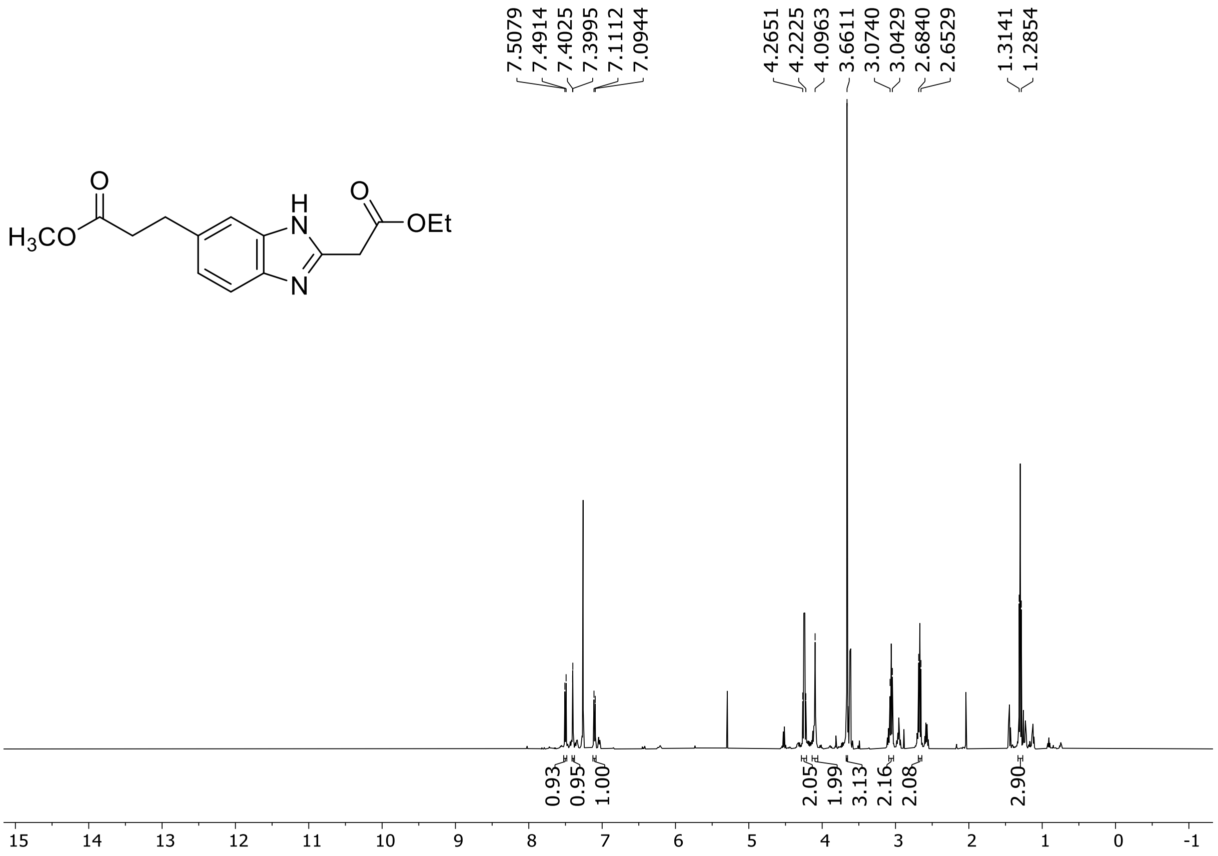

[1]H (500 MHz) and [13]C NMR (126 MHz) spectra of **S89** in chloroform-*d*

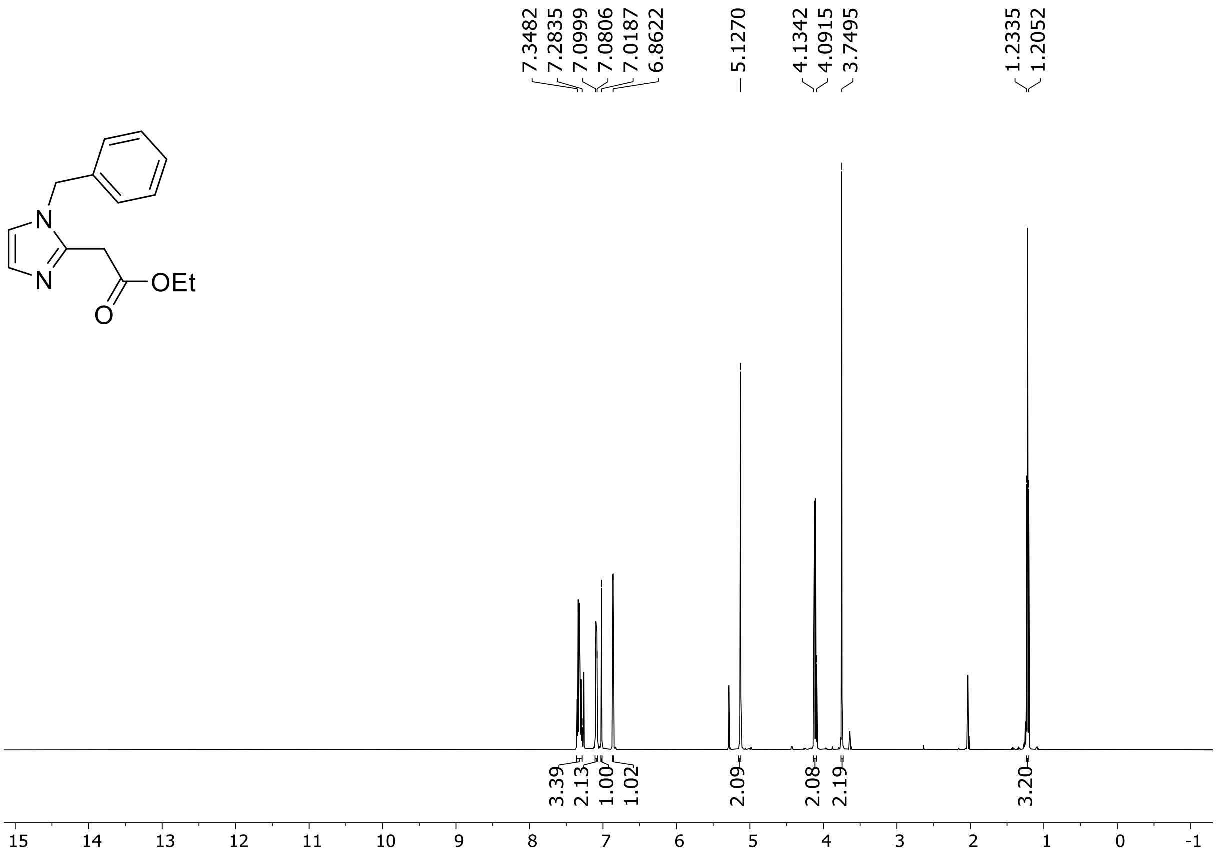

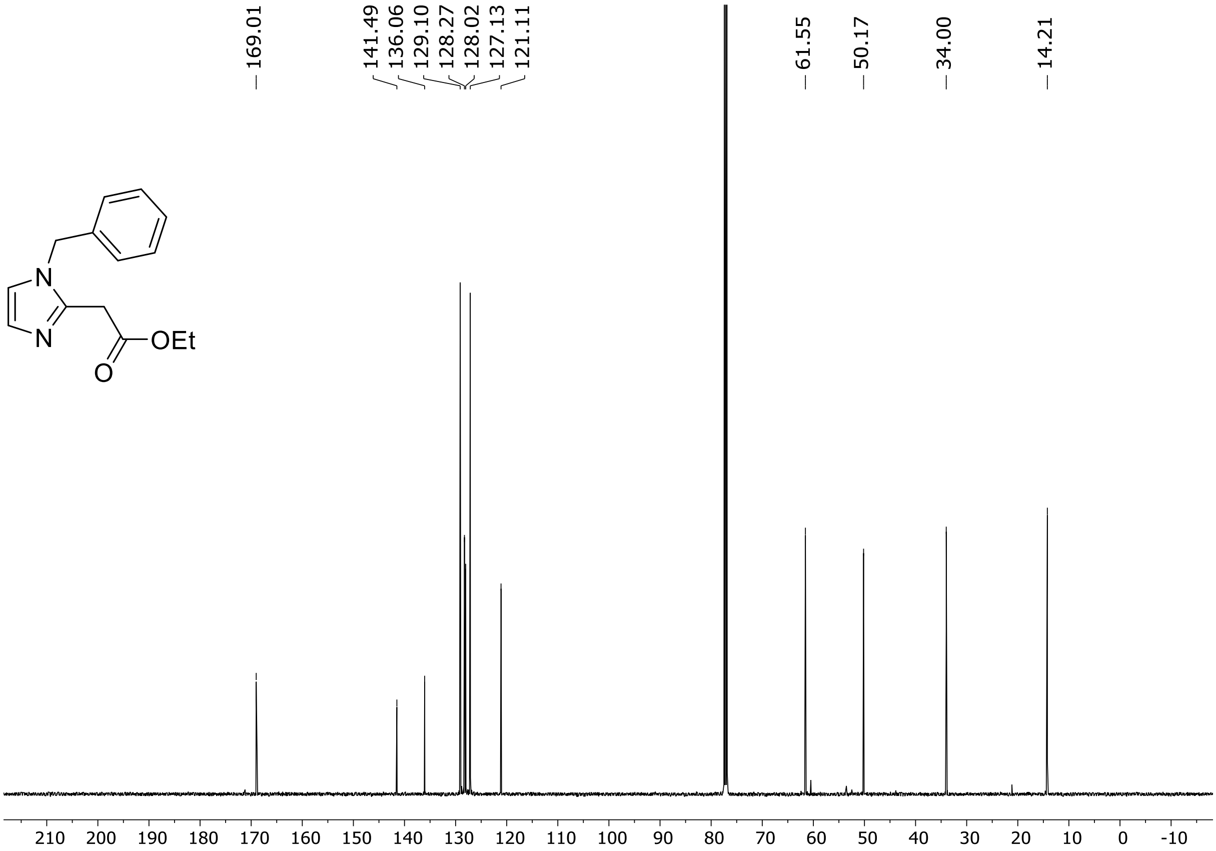

$^{1}$H (500 MHz) and $^{13}$C NMR (126 MHz) spectra of **S90** in chloroform-*d*

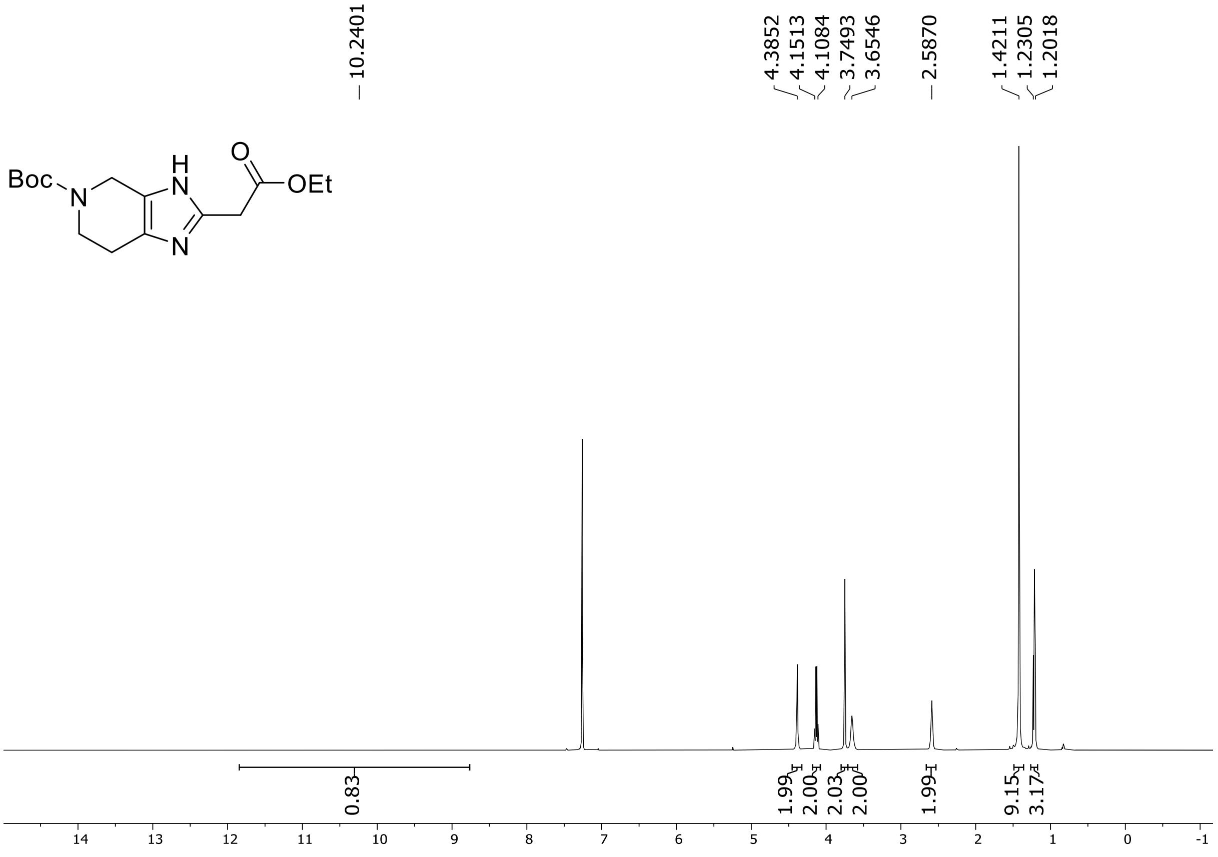

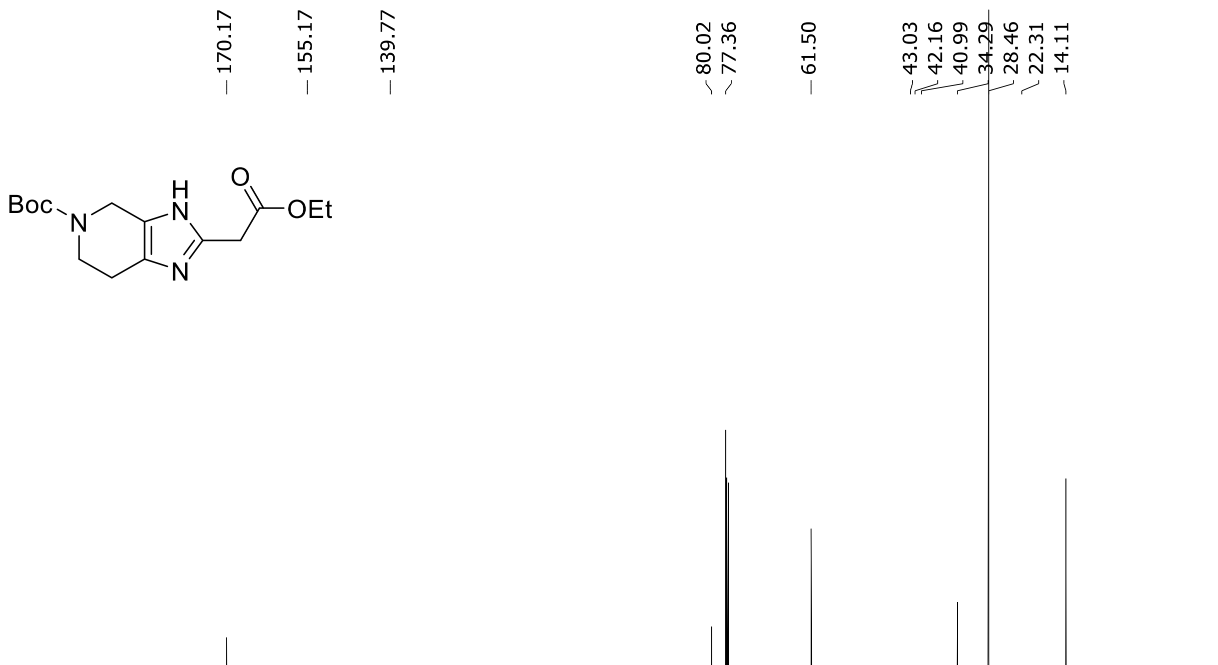

HRMS spectrum of **S90**

**$C_{15}H_{23}N_3O_4$** **exact mass: 309.1689**

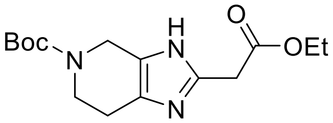

**APCI +** (MMI)
nitrogen flow 5 L/min, gas temperature 325°C, nebulizer 45 psig, skimmer 65 V, vaporizer 200°C, fragmentor 15 V, dissolved in methanol

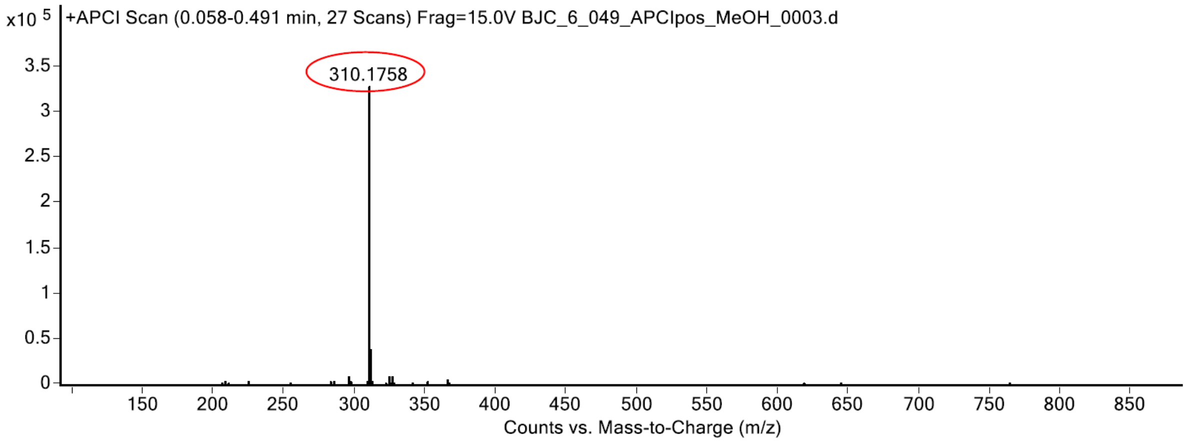

expected mass: $[M+H]^+$ = 310.1761 observed mass: $[M+H]^+$ = 310.1758 mass accuracy = - 1.0 ppm

[1]H (500 MHz) spectrum of **S91** in chloroform-*d*

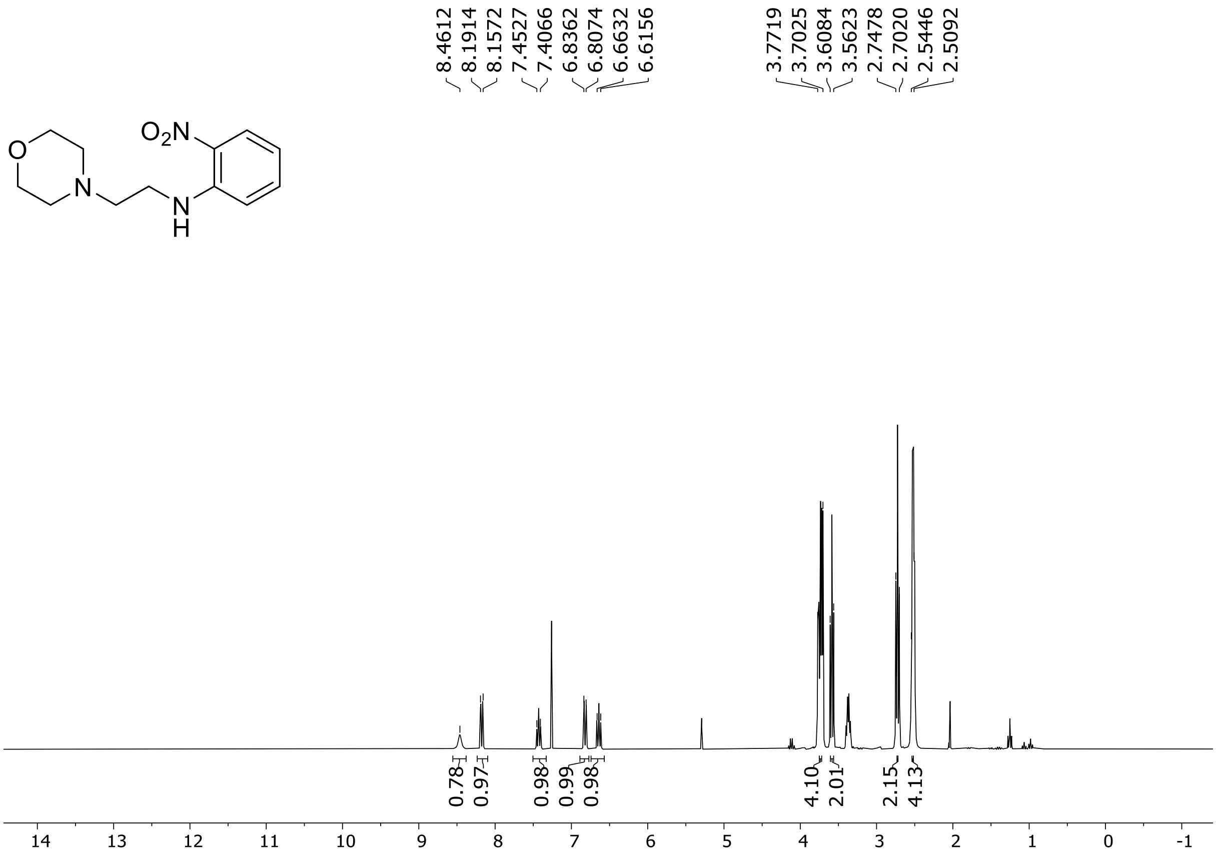

$^{1}H$ (500 MHz) and $^{13}C$ NMR (126 MHz) spectra of **S92** in chloroform-*d*

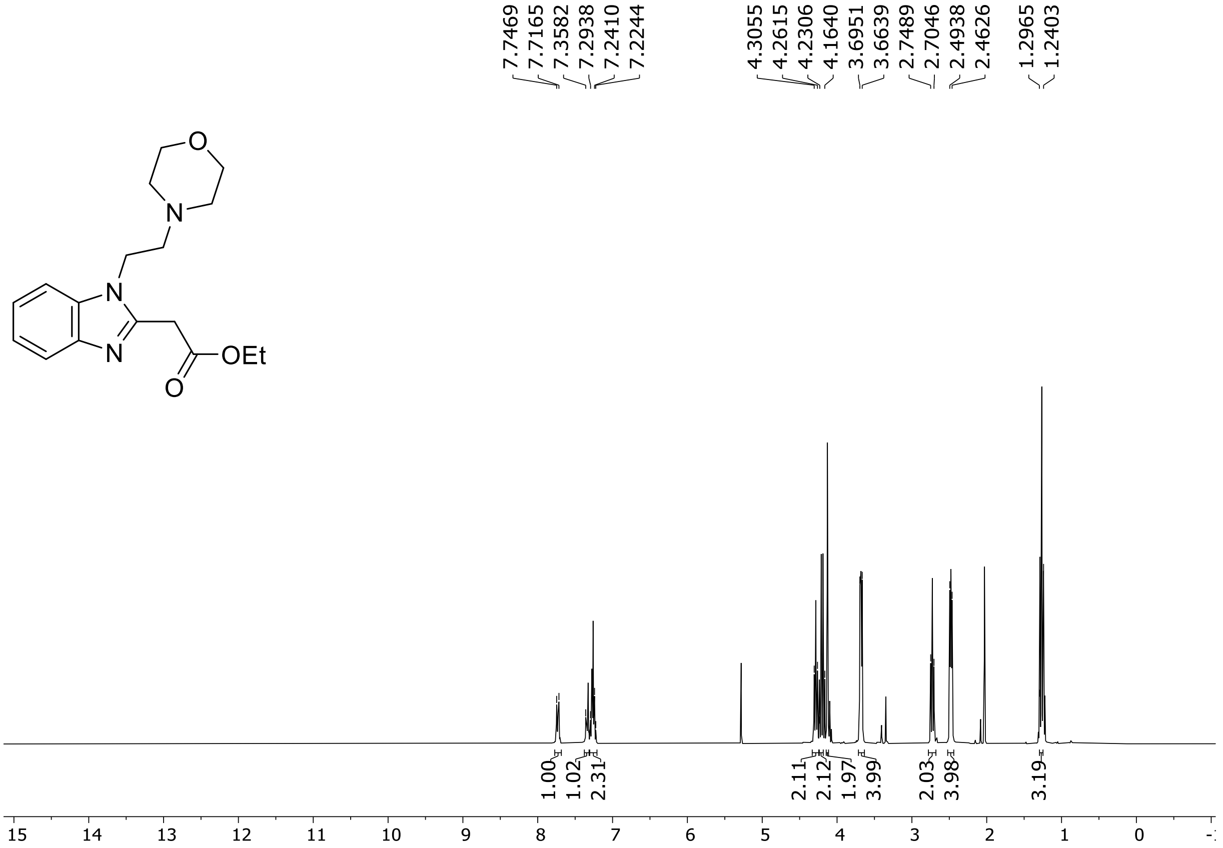

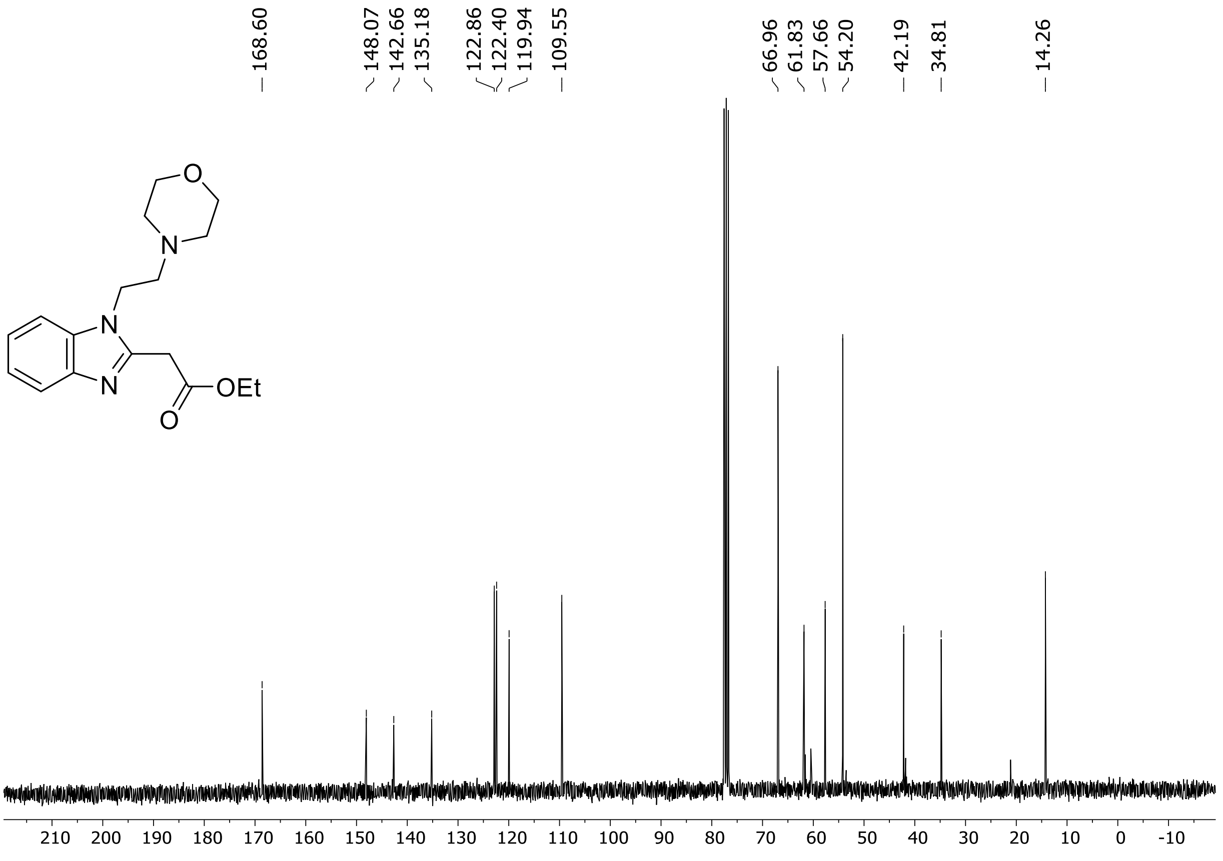

$^1$H (500 MHz) and $^{13}$C NMR (126 MHz) spectra of **S93** in chloroform-*d*

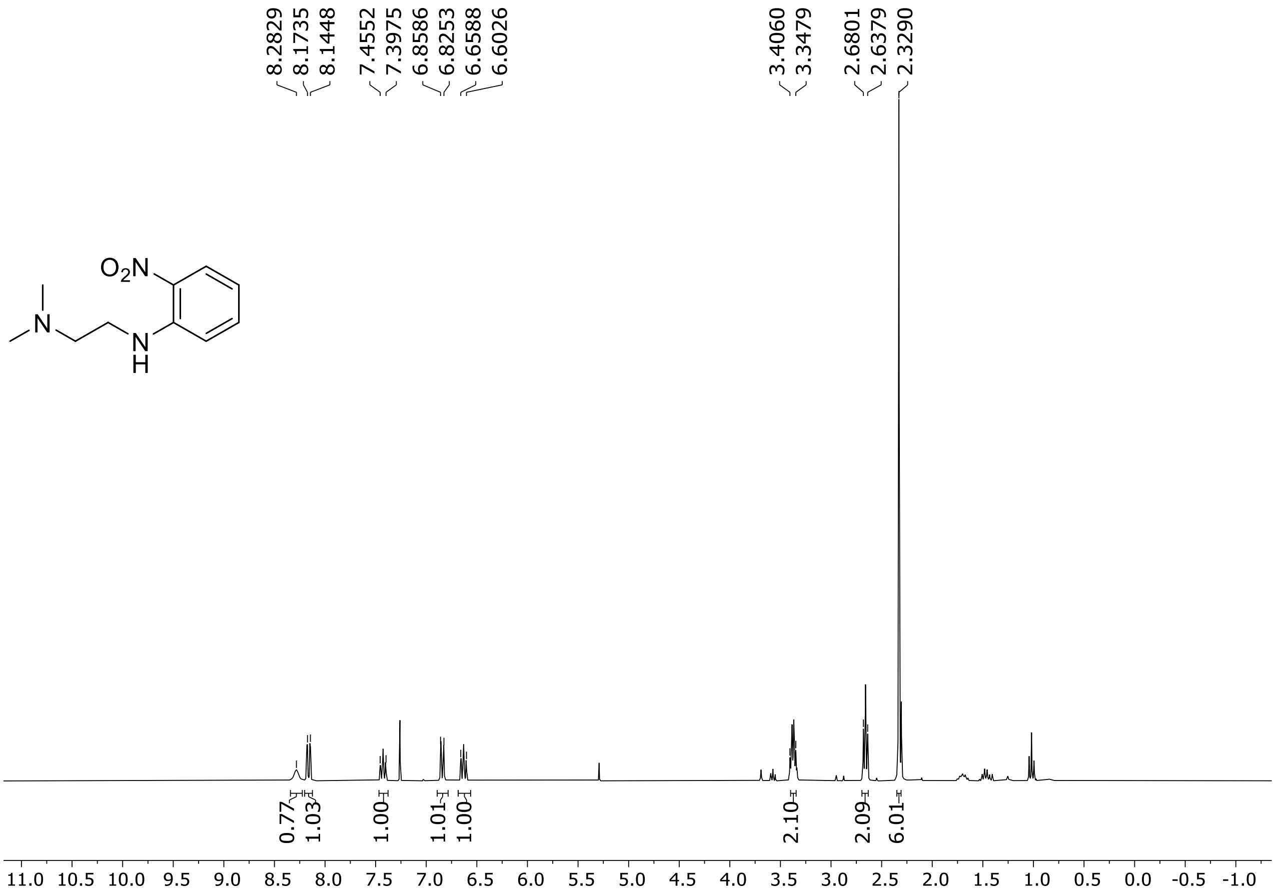

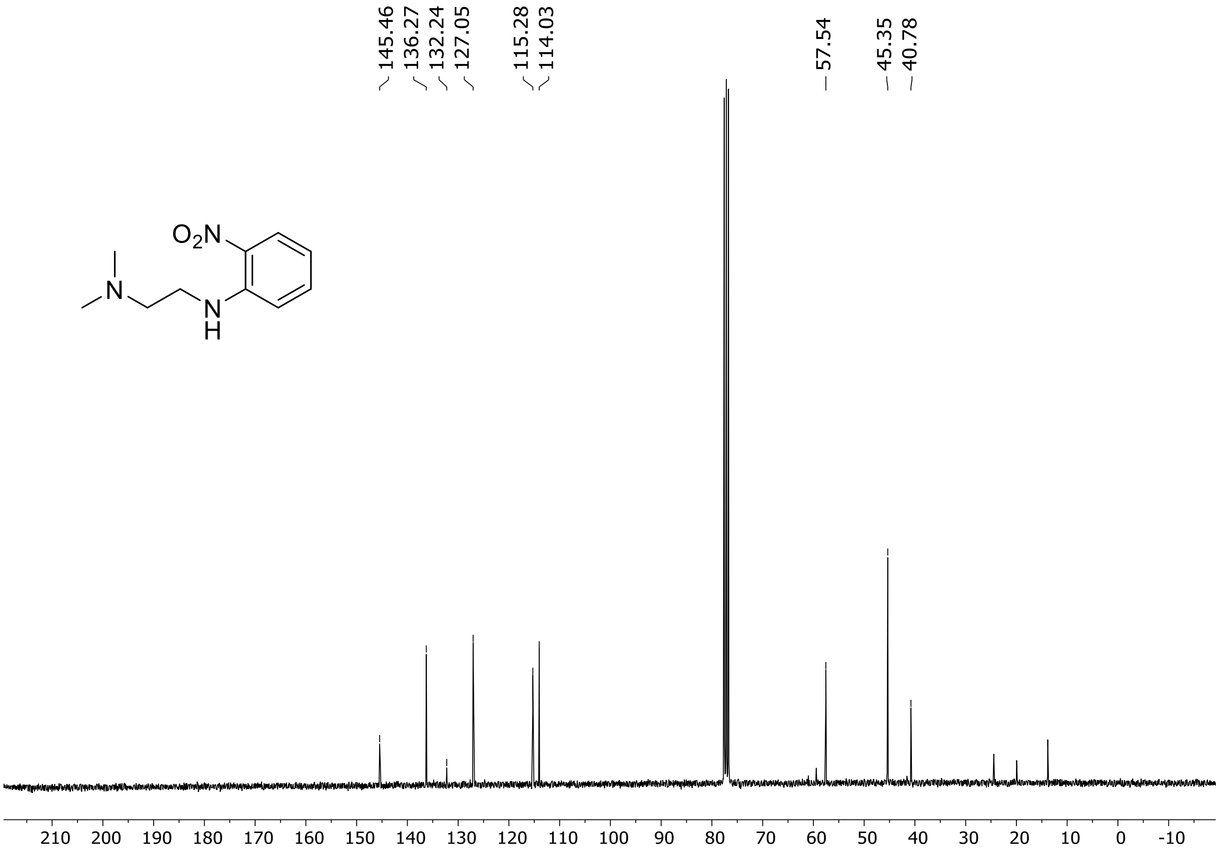

$^1$H (500 MHz) and $^{13}$C NMR (126 MHz) spectra of **S94** in chloroform-*d*

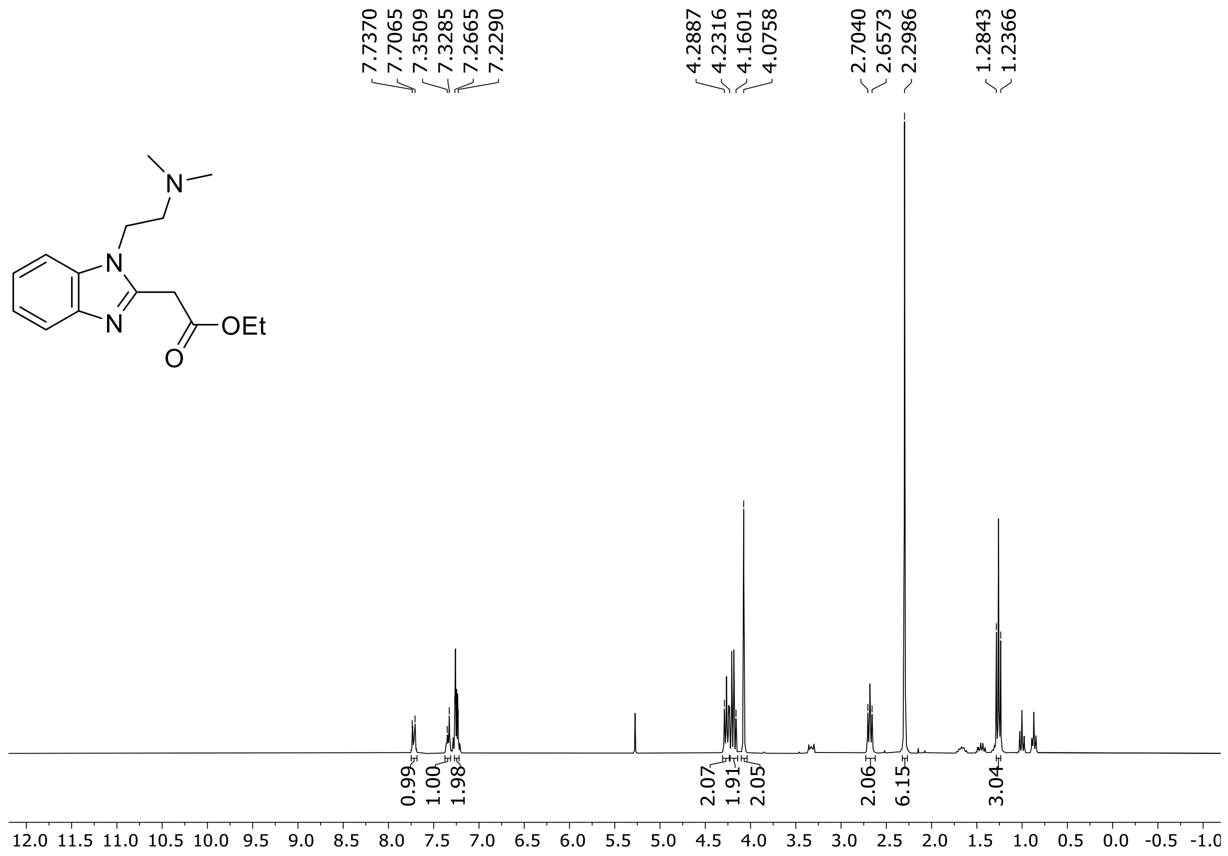

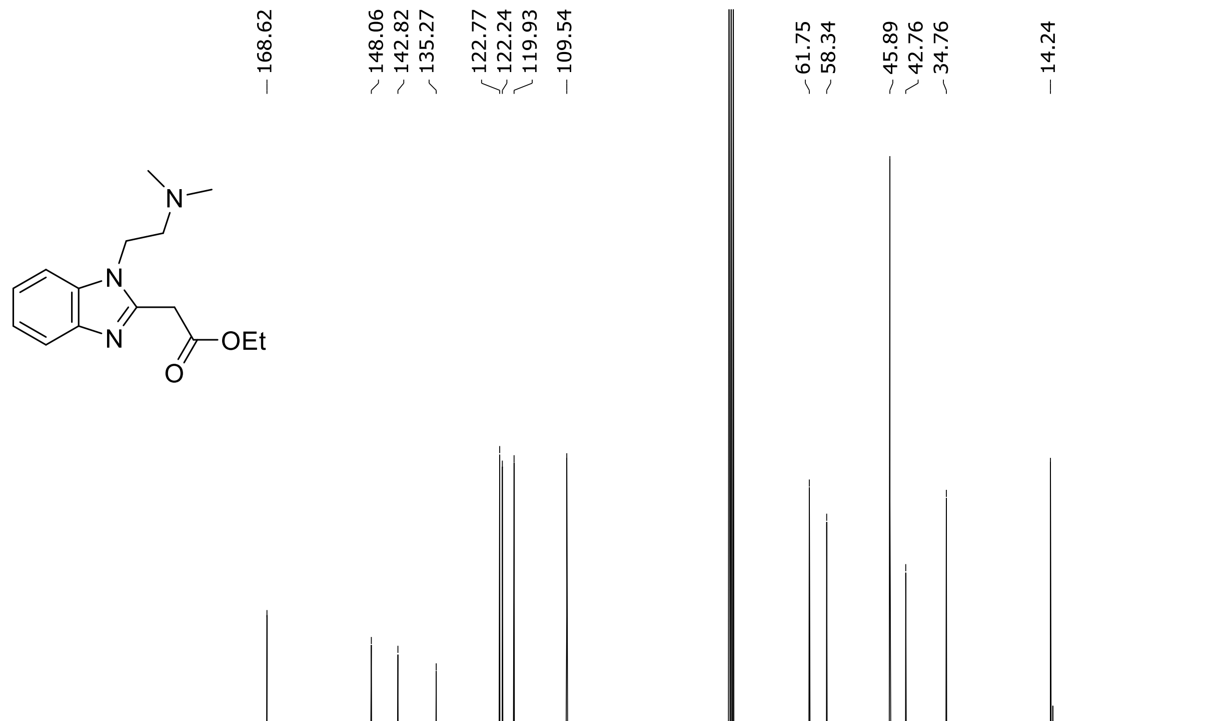

$^{1}$H (300 MHz) NMR spectrum of **S95** in chloroform-*d*

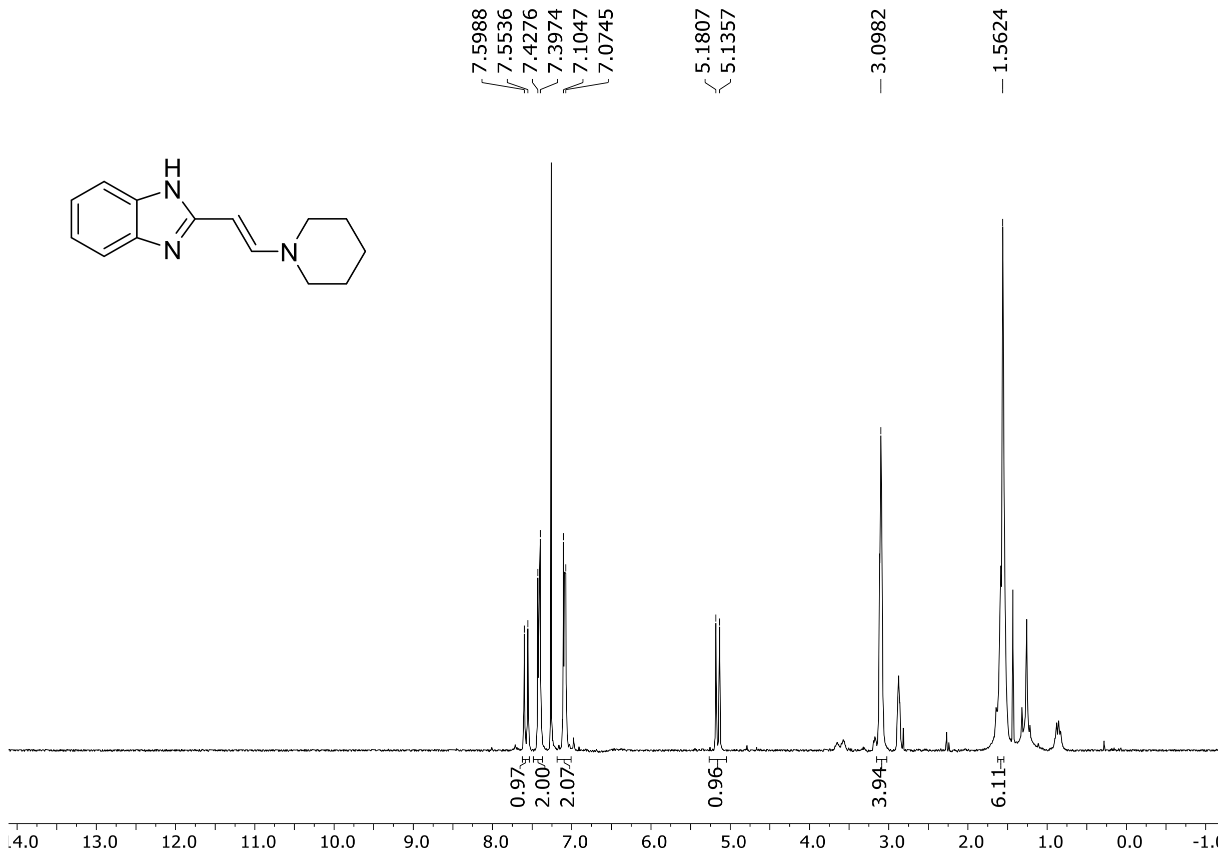

$^1$H (500 MHz) and $^{13}$C NMR (126 MHz) spectra of **S96** in chloroform-*d*

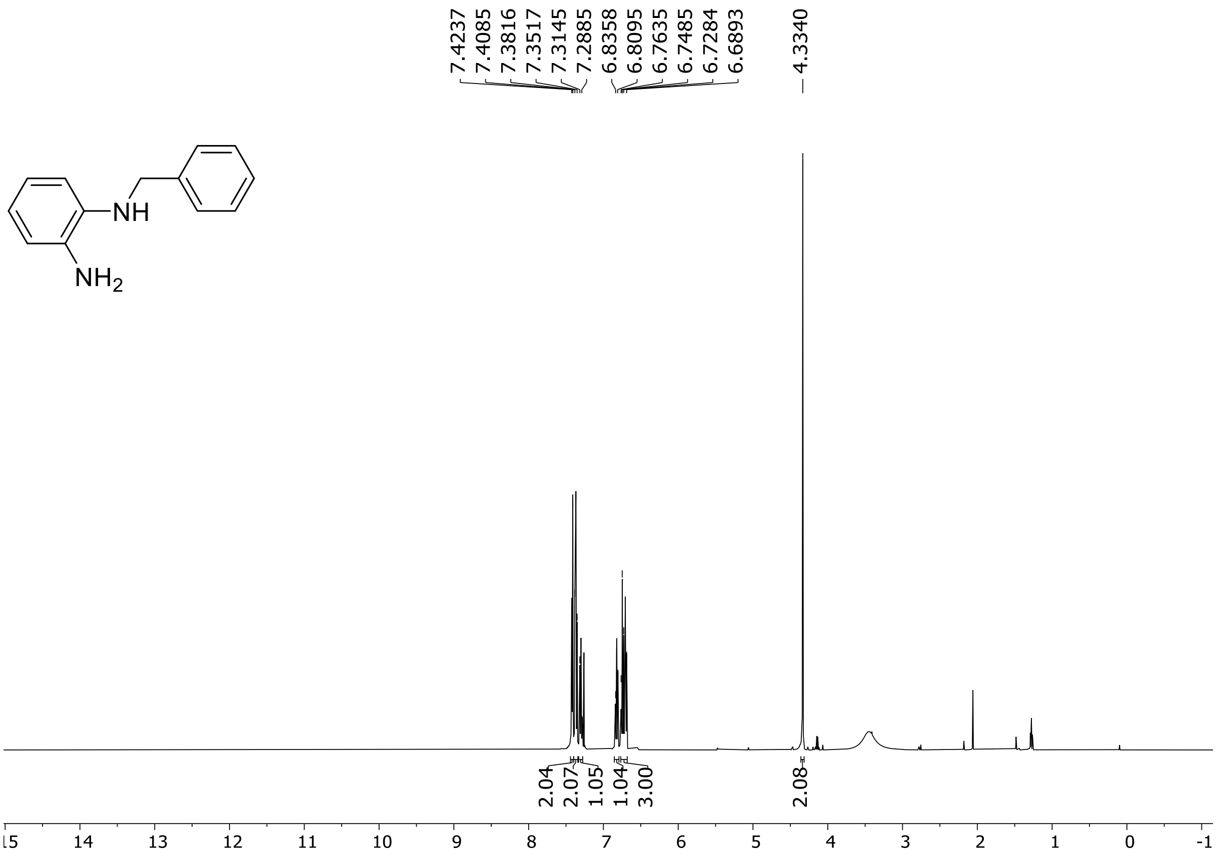

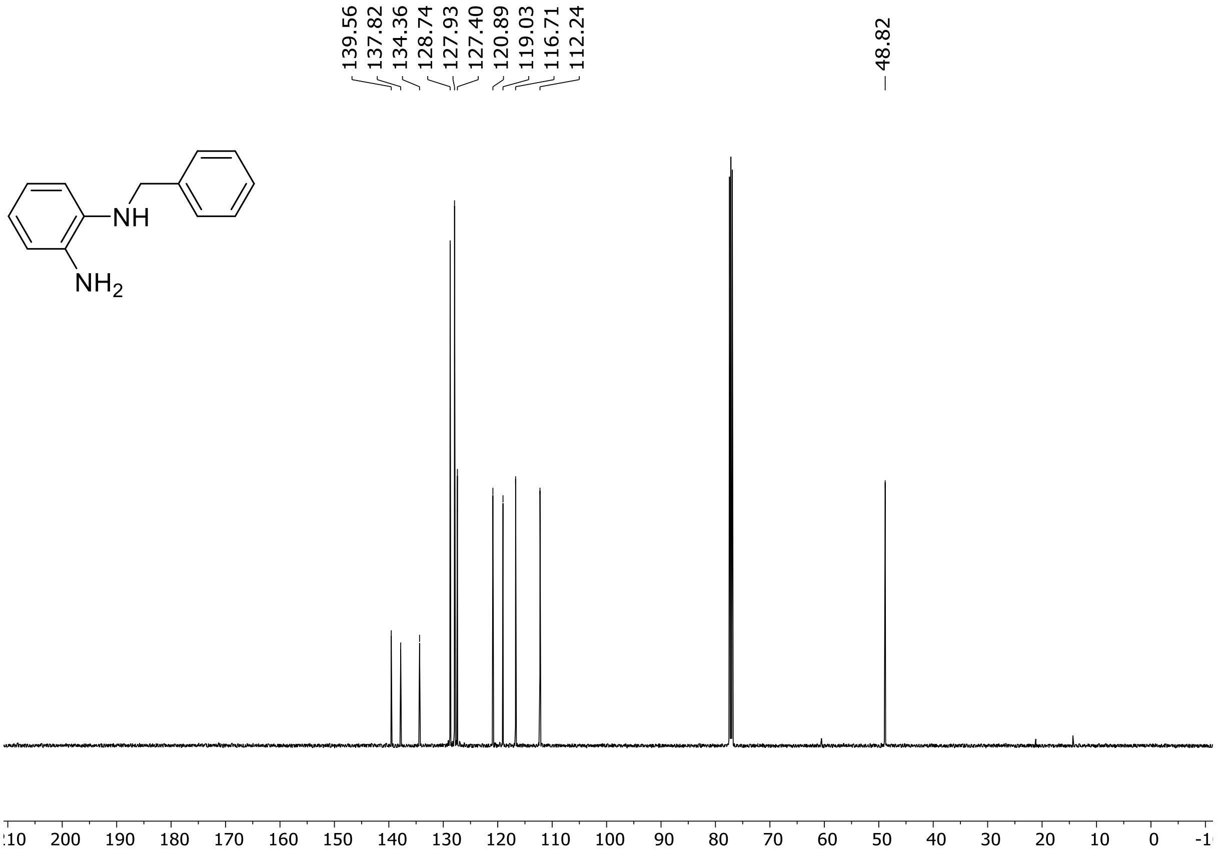

$^{1}$H (500 MHz) and $^{13}$C NMR (126 MHz) spectra of **S97** in chloroform-*d*

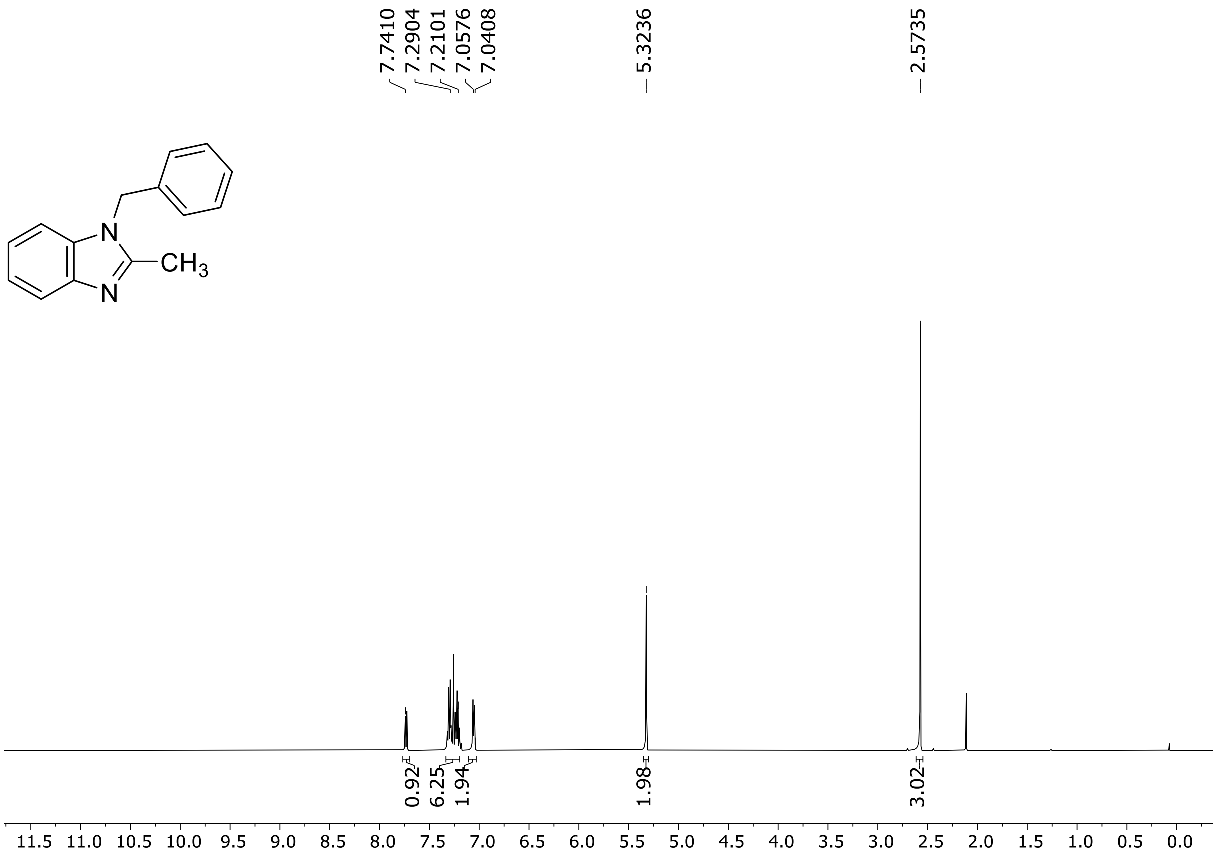

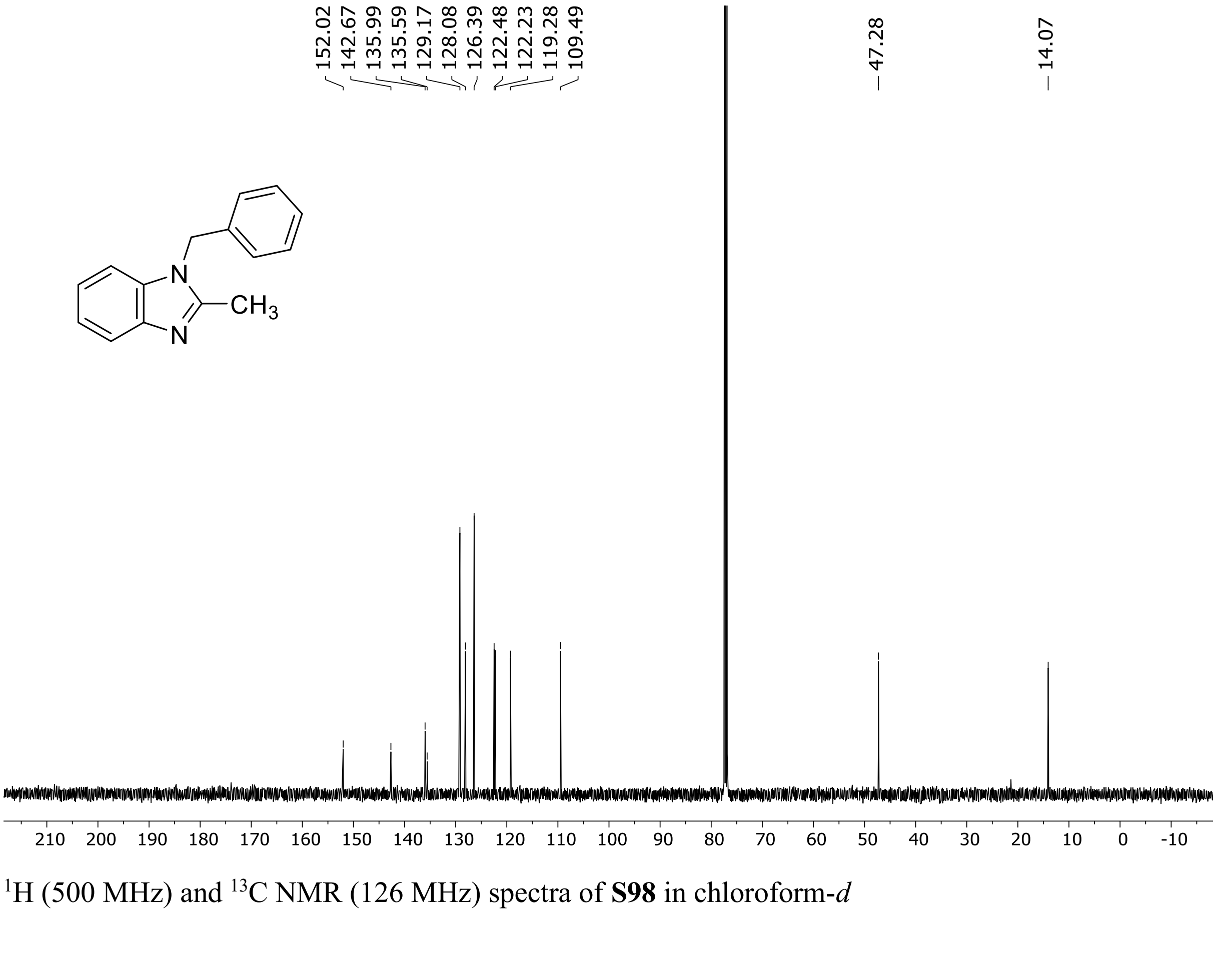

$^{1}H$ (500 MHz) and $^{13}C$ NMR (126 MHz) spectra of **S98** in chloroform-*d*

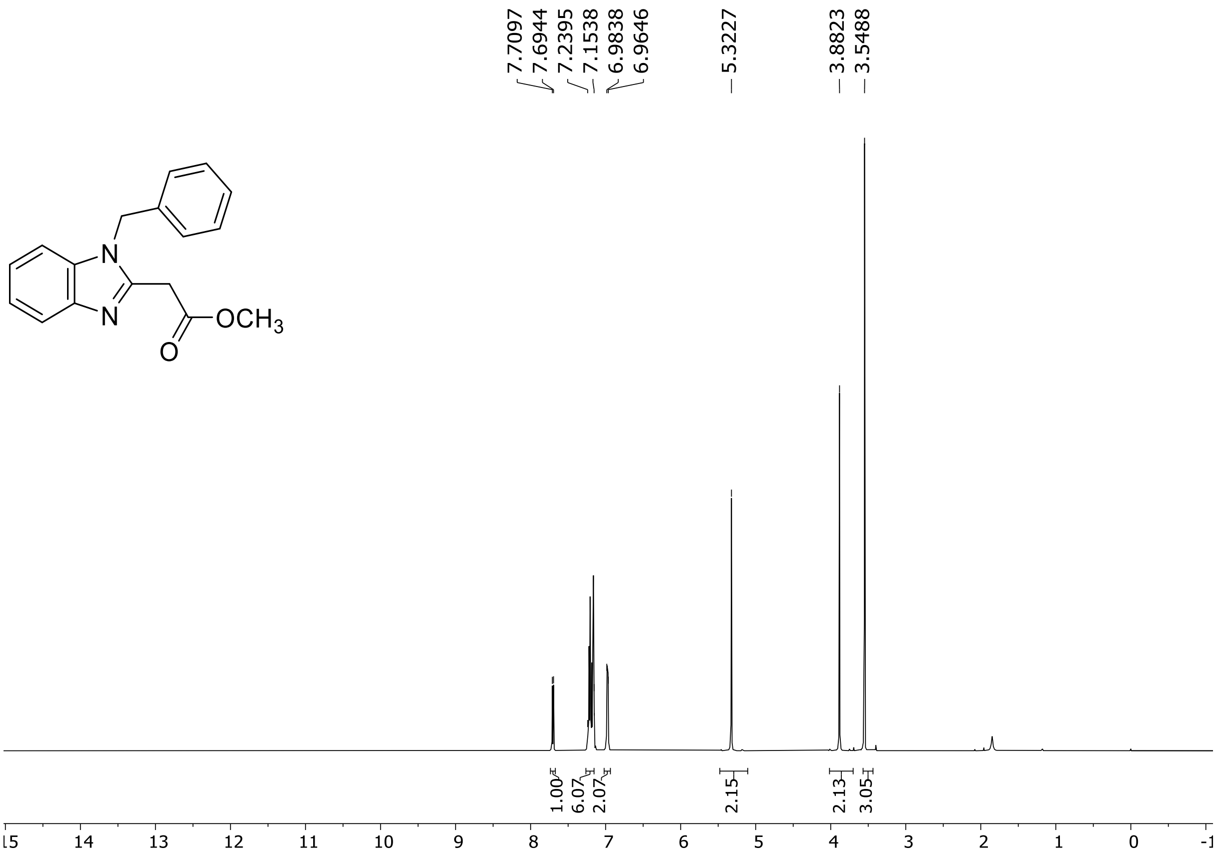

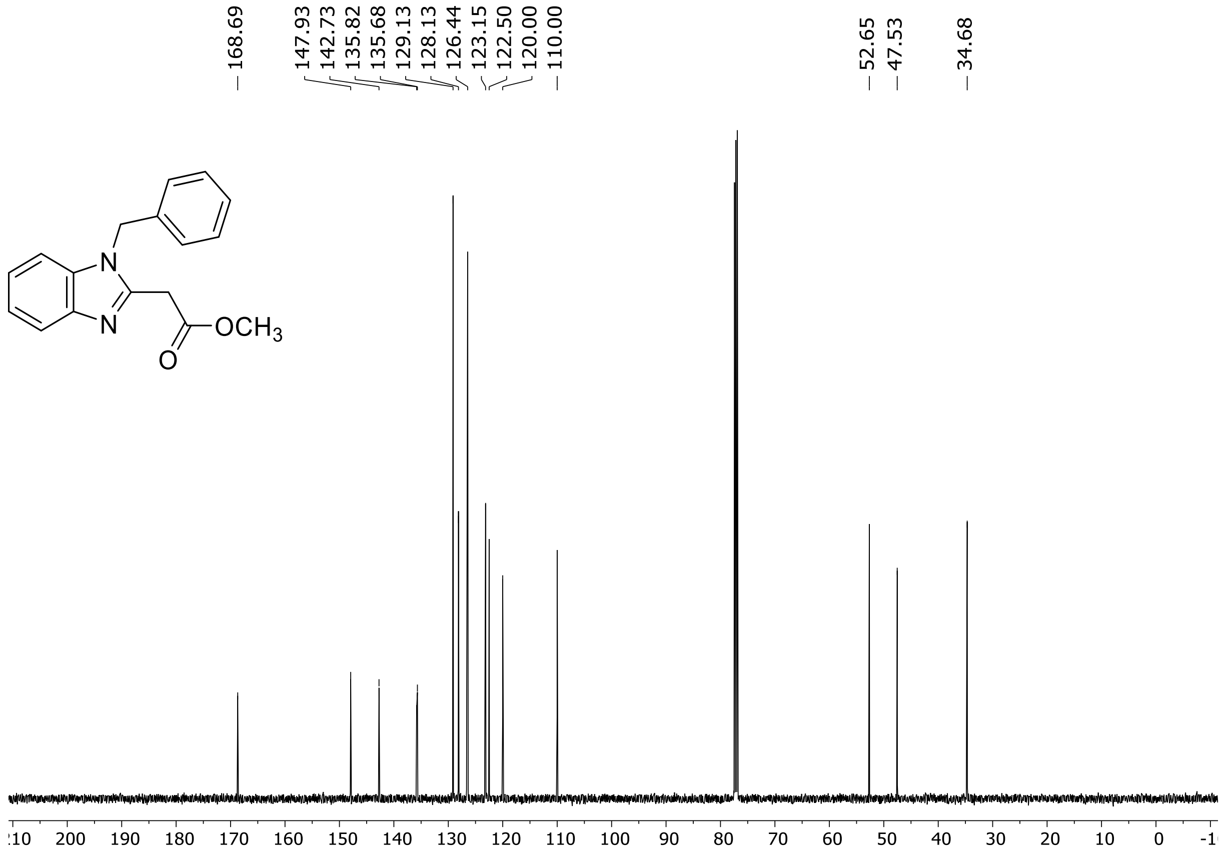

$^{1}$H (500 MHz) and $^{13}$C NMR (126 MHz) spectra of **S99** in chloroform-*d*

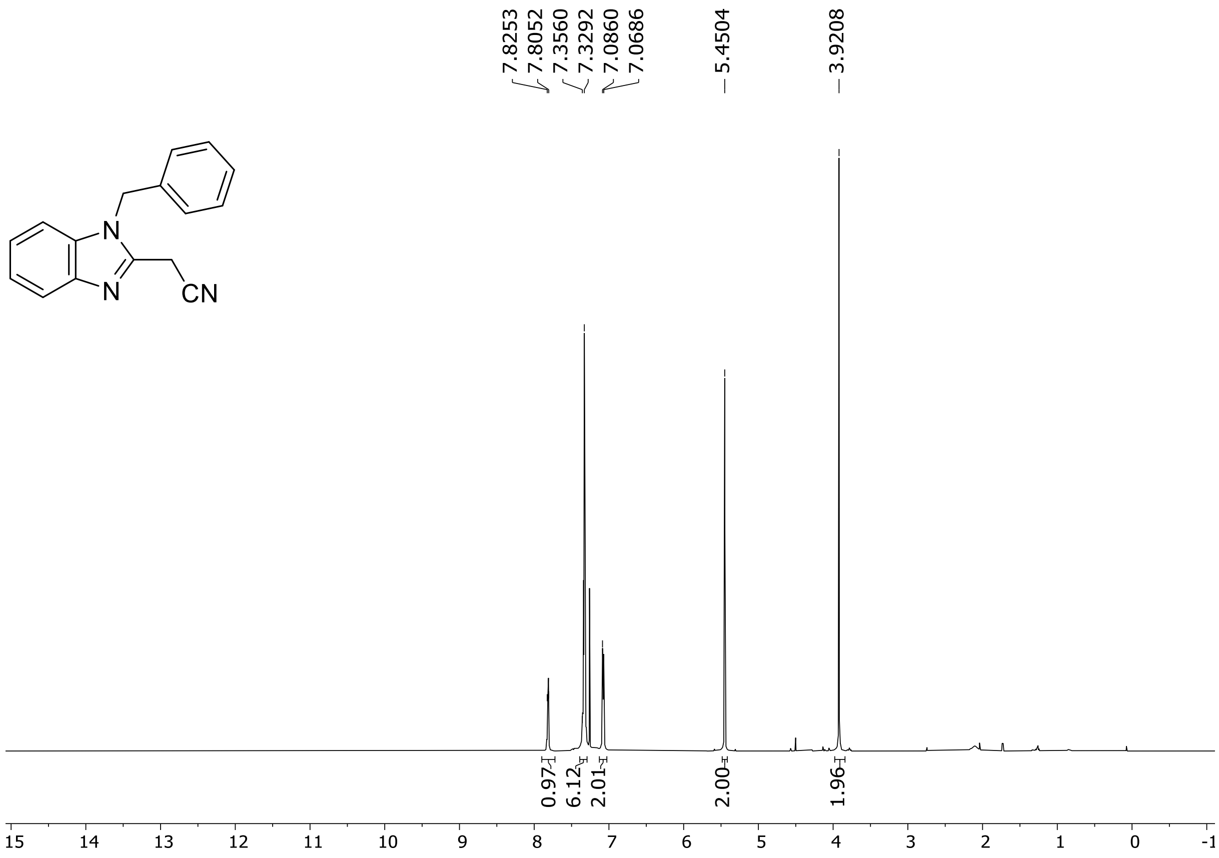

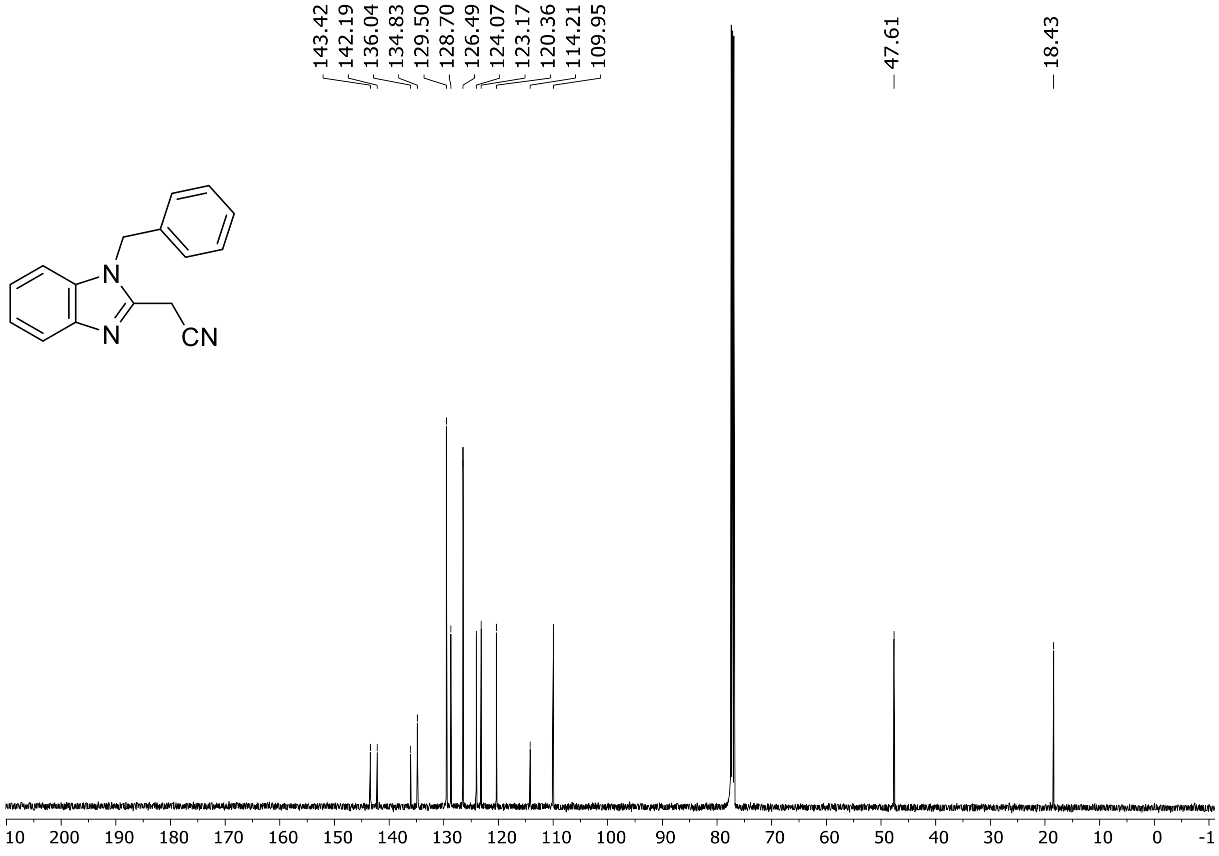

HRMS spectrum of **S99**

$C_{16}H_{13}N_3$ **mono m/z = 247.1109**

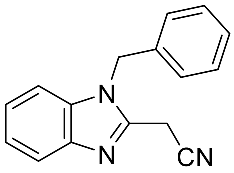

**APCI -** (MMI)
nitrogen flow 5 L/min, gas temperature 300°C, nebulizer 45 psi, vaporizer 200°C
skimmer **65** V, fragmentor **18** V, dissolved in methanol

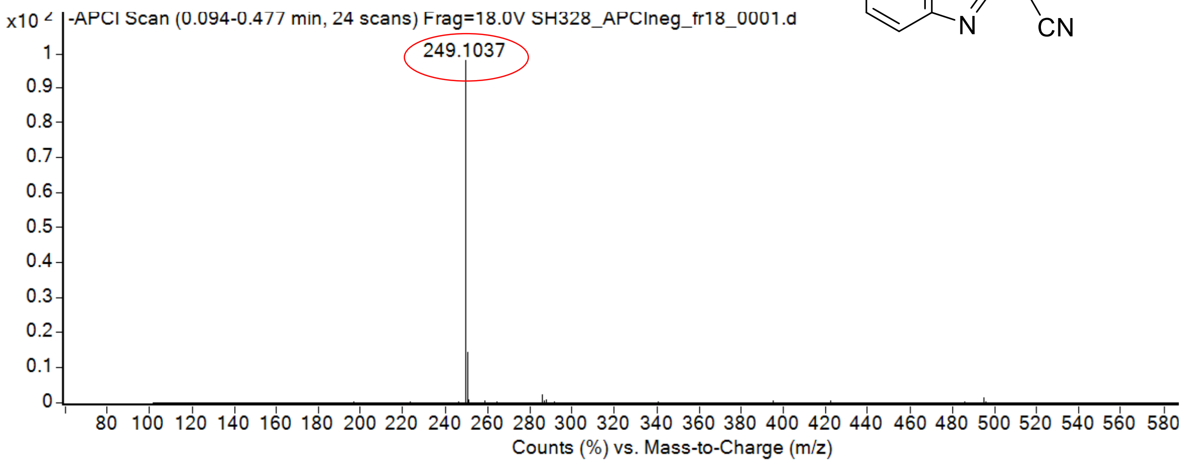

calculated mass: [M-H]$^-$ = 246.1037 observed: [M-H]$^-$ = 246.1037 max. mass error = < 0.1 ppm

$^1$H (500 MHz) and $^{13}$C NMR (126 MHz) spectra of **S100** in chloroform-*d*

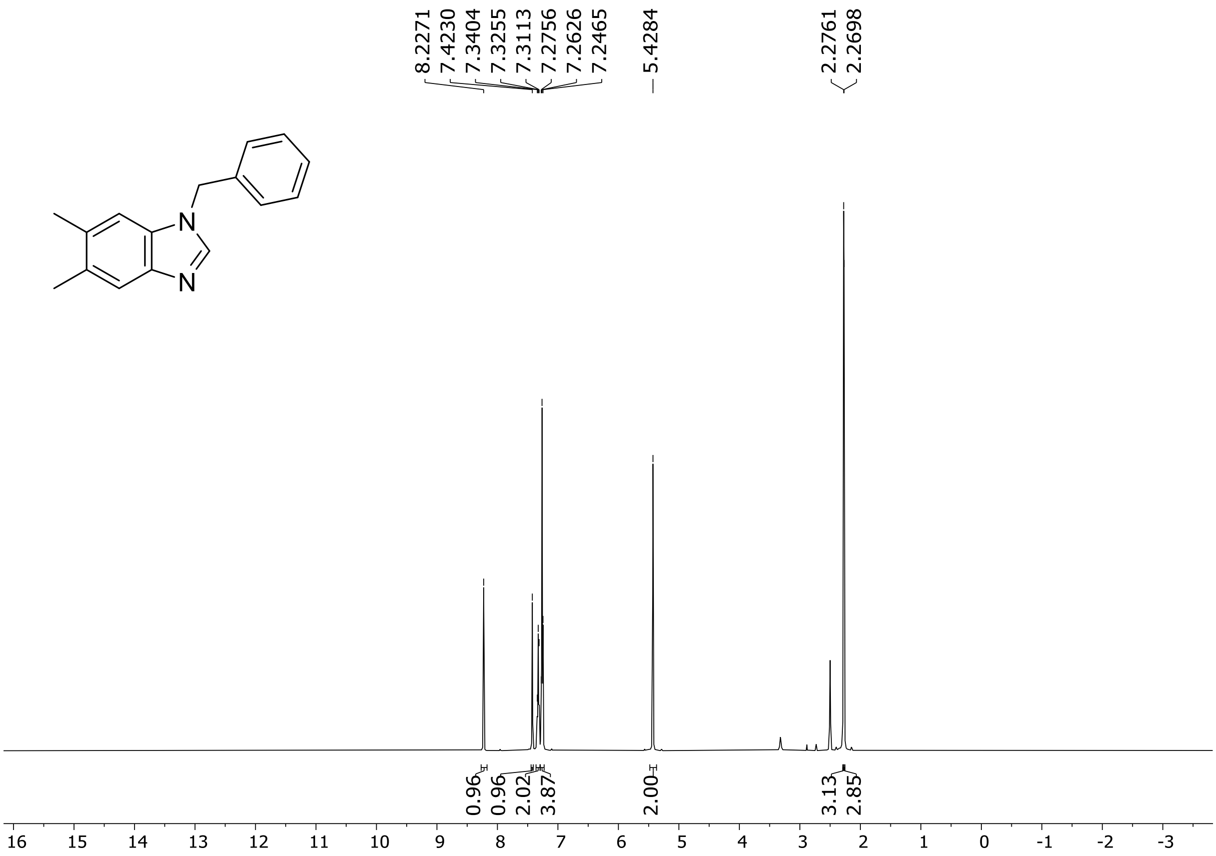
8.2271
7.4230
7.3404
7.3255
7.3113
7.2756
7.2626
7.2465
5.4284
2.2761
2.2698
N
N
0.96
0.96
2.02
3.87
2.00
3.13
2.85
16 15 14 13 12 11 10 9 8 7 6 5 4 3 2 1 0 -1 -2 -3

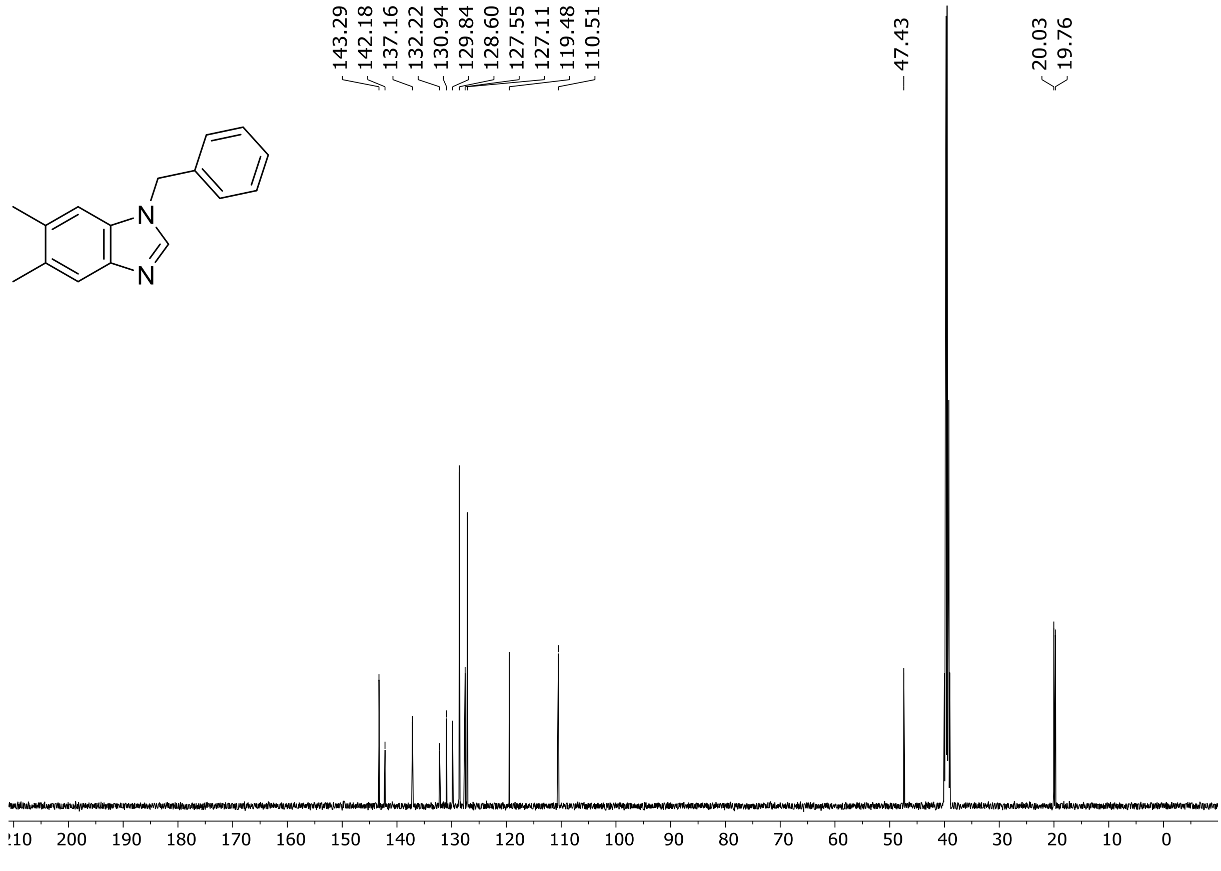
143.29
142.18
137.16
132.22
130.94
129.84
128.60
127.55
127.11
119.48
110.51
47.43
20.03
19.76
N
N
210 200 190 180 170 160 150 140 130 120 110 100 90 80 70 60 50 40 30 20 10 0

[1]H (500 MHz) and [13]C NMR (126 MHz) spectra of **S101** in chloroform-*d*

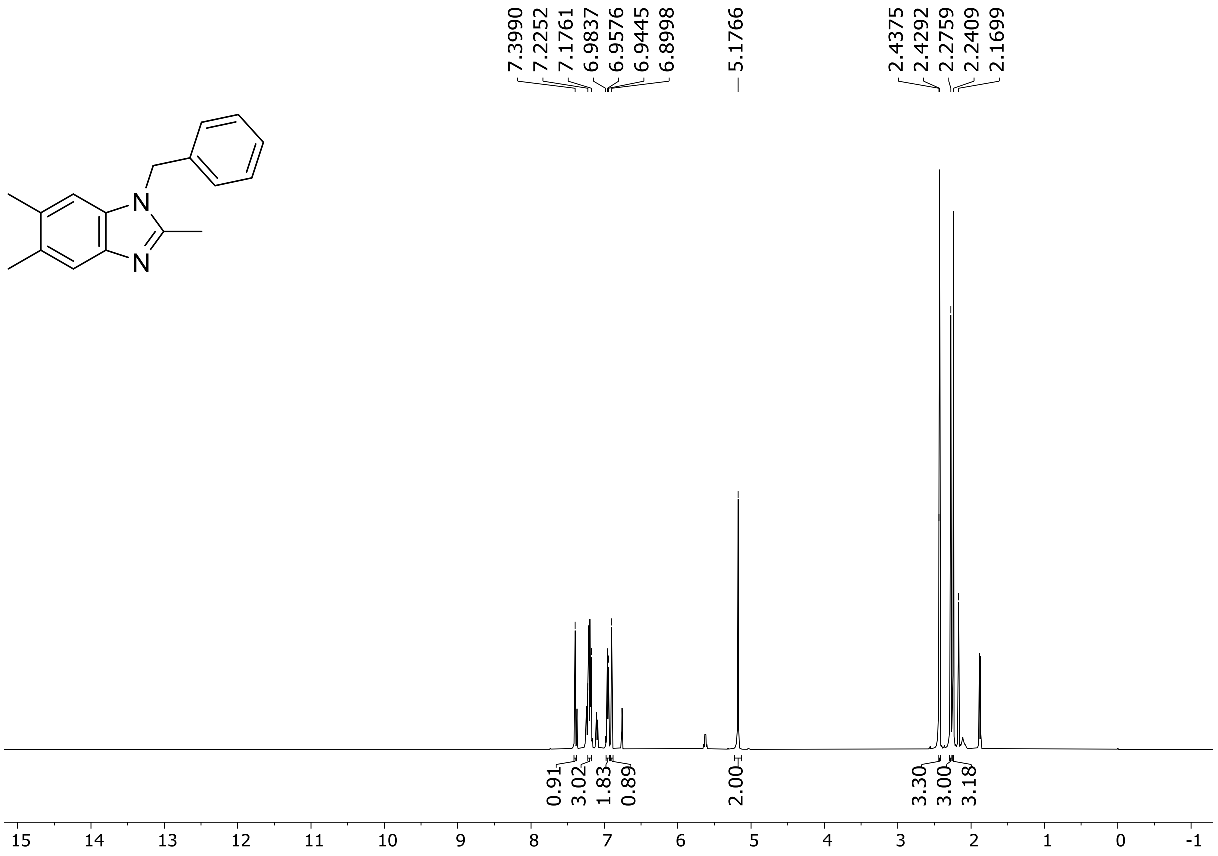

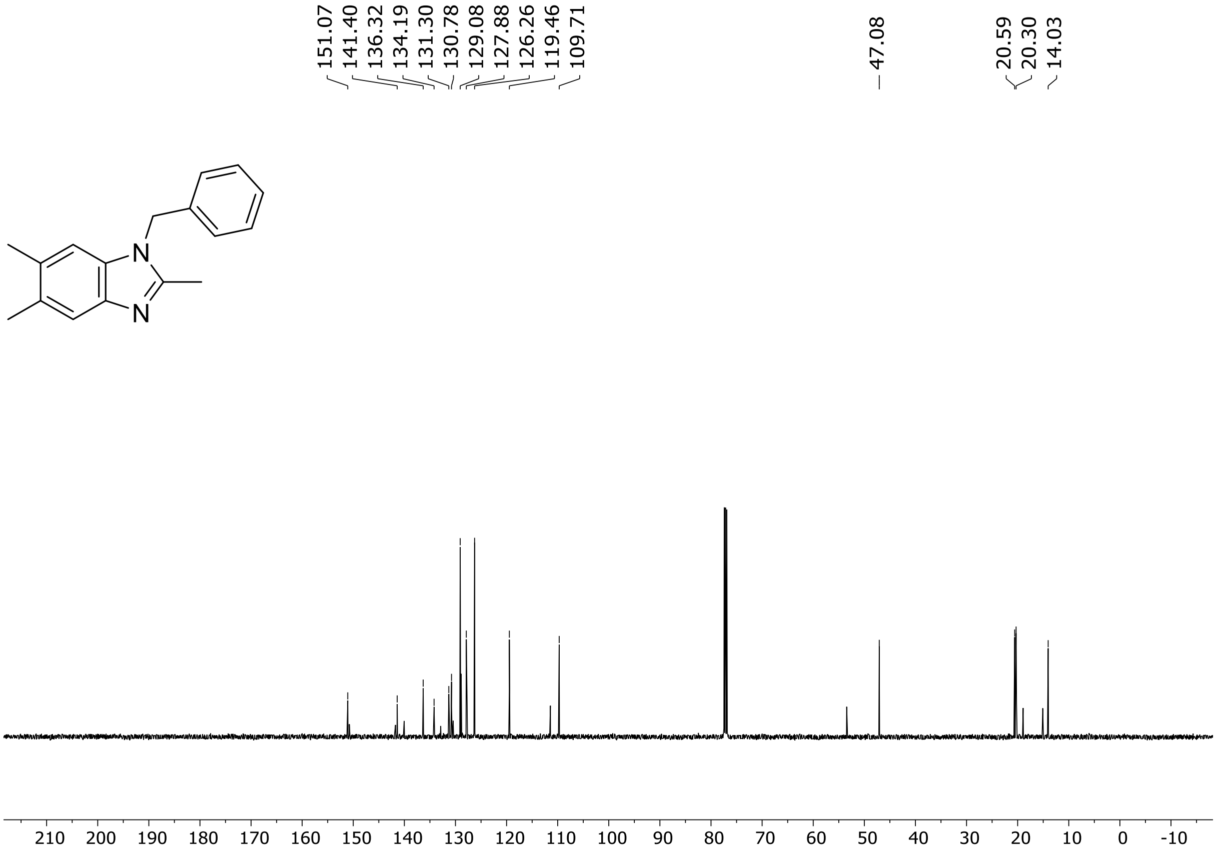

[1]H (500 MHz) and [13]C NMR (126 MHz) spectra of **S102** in chloroform-*d*

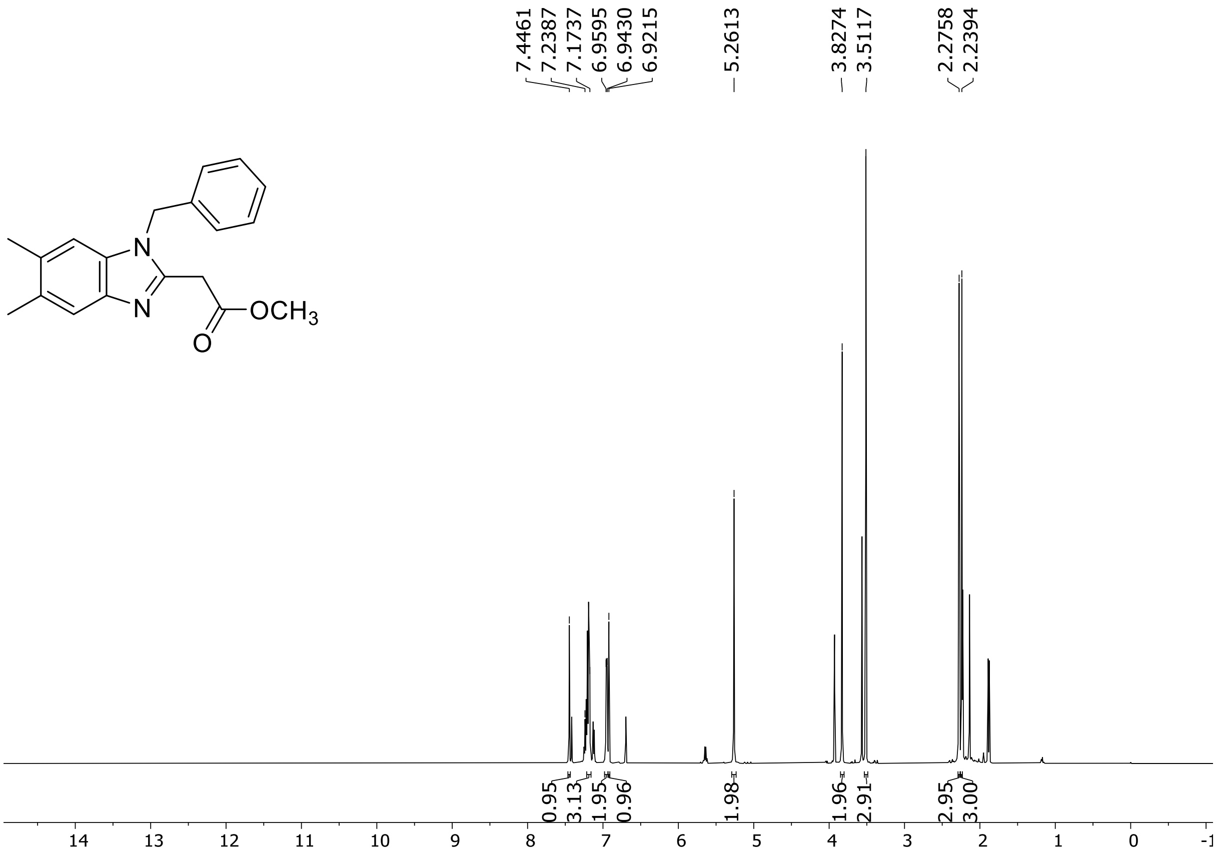

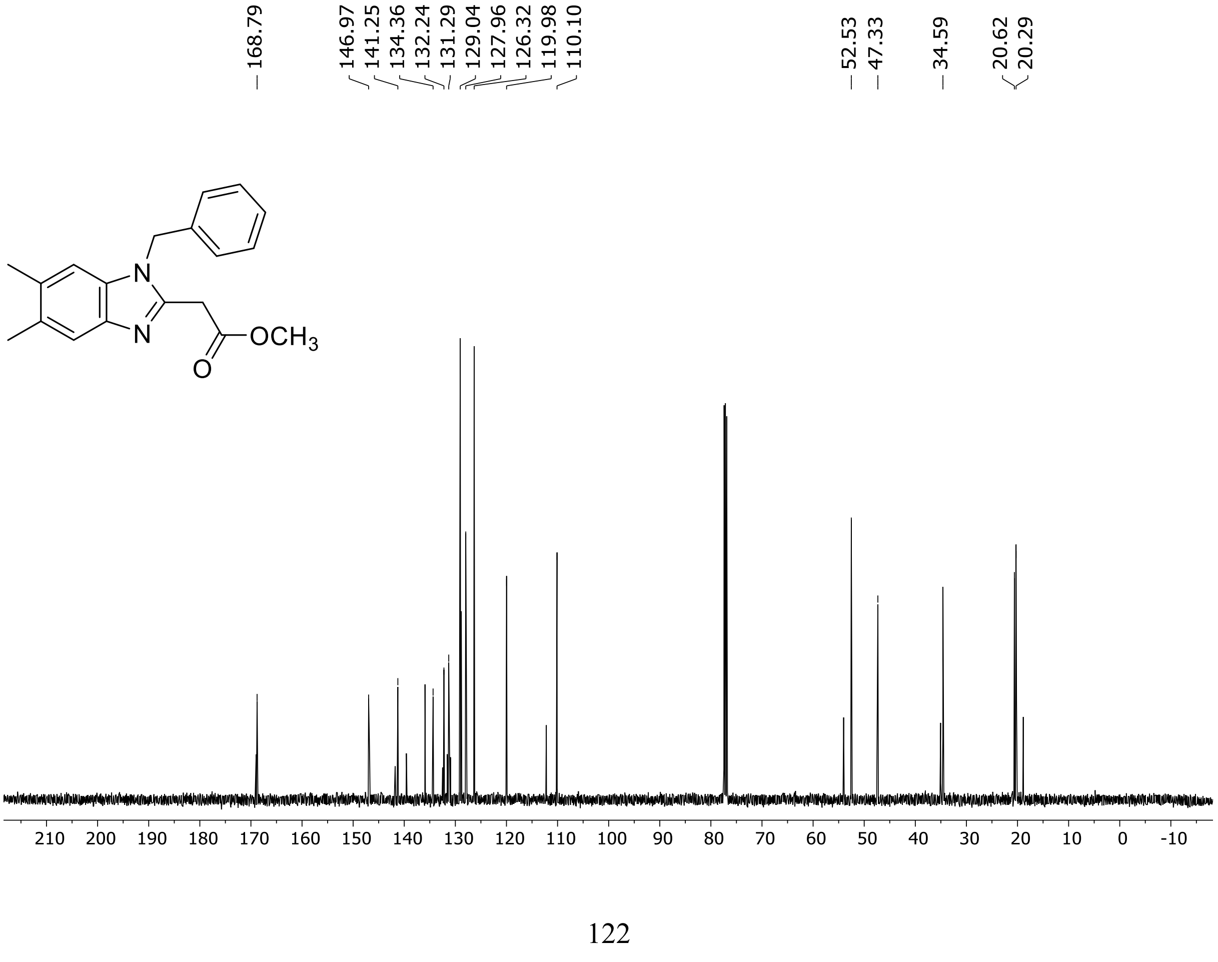

$^{1}H$ (500 MHz) and $^{13}C$ NMR (126 MHz) spectra of **S103** in DMSO-$d_6$

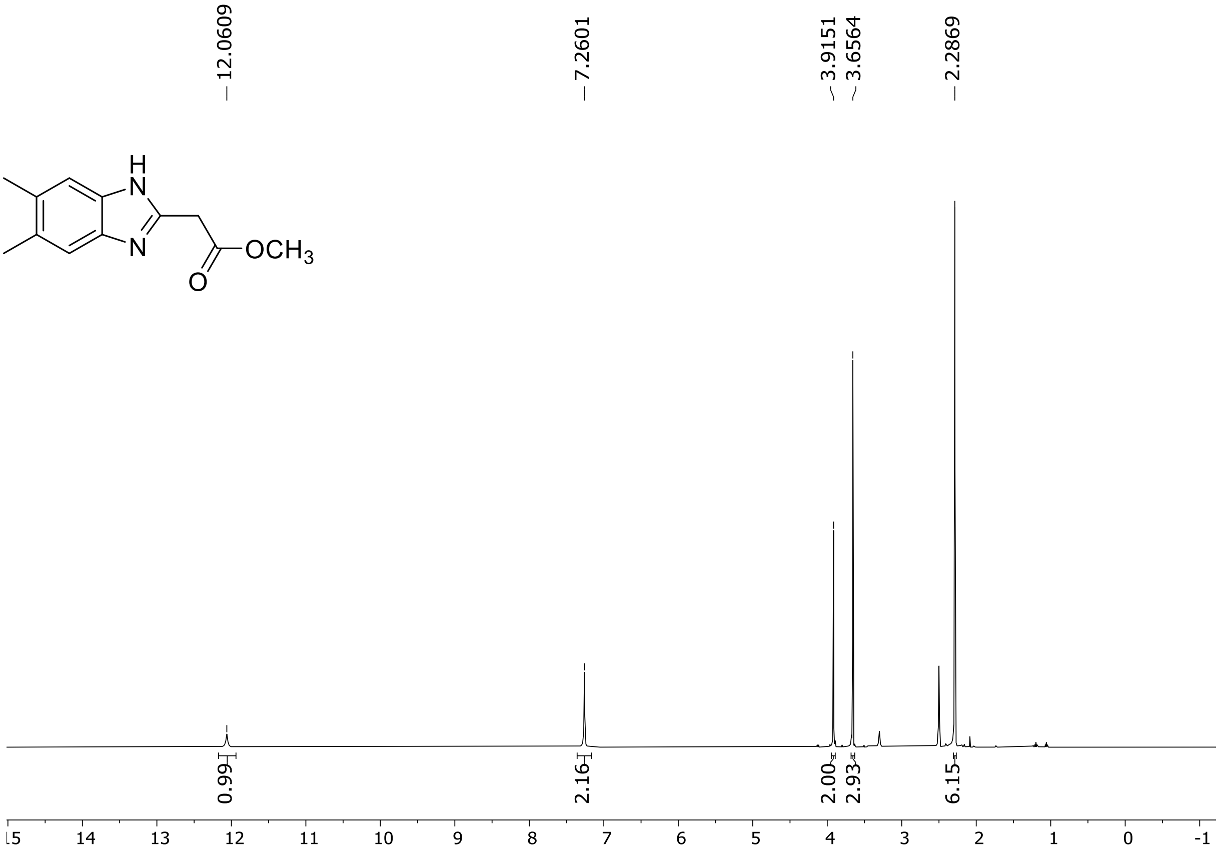

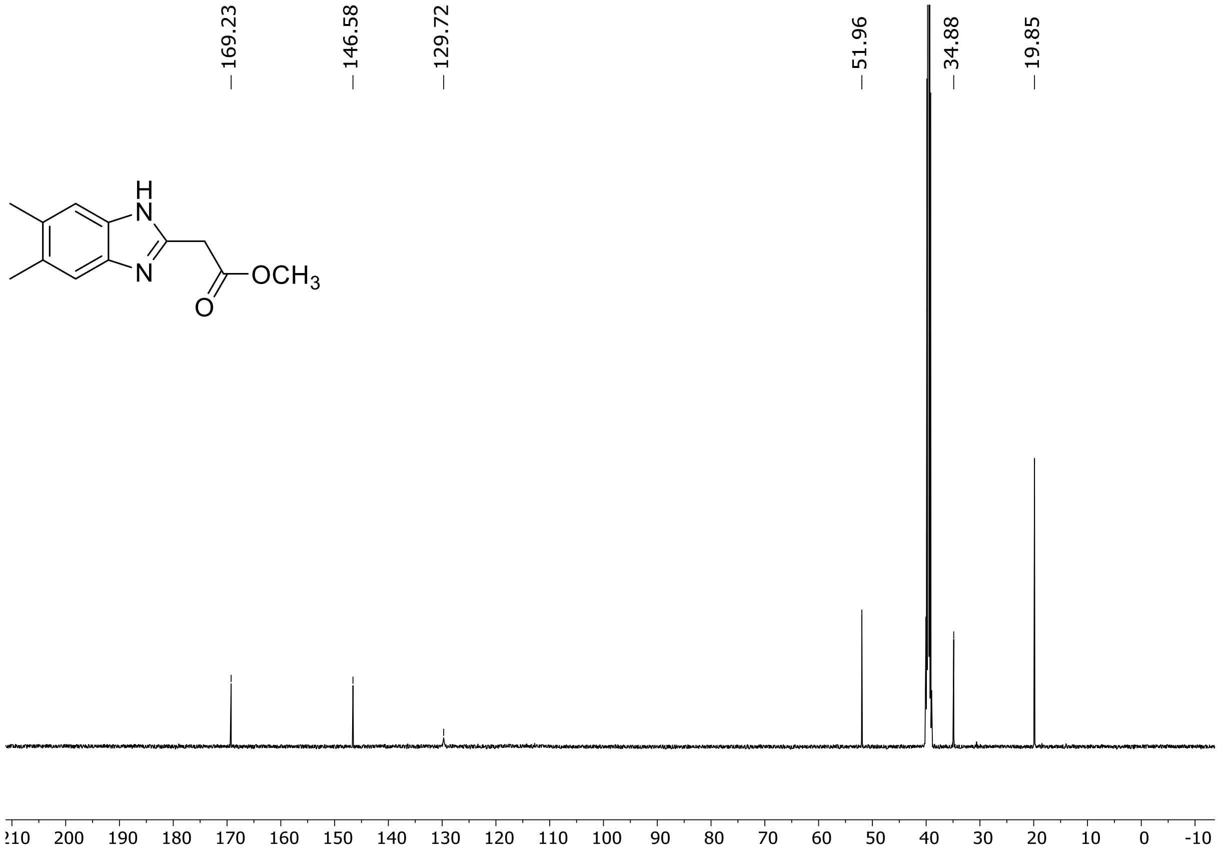

[1]H (500 MHz) and [13]C NMR (126 MHz) spectra of **S104** in DMSO-$d_6$

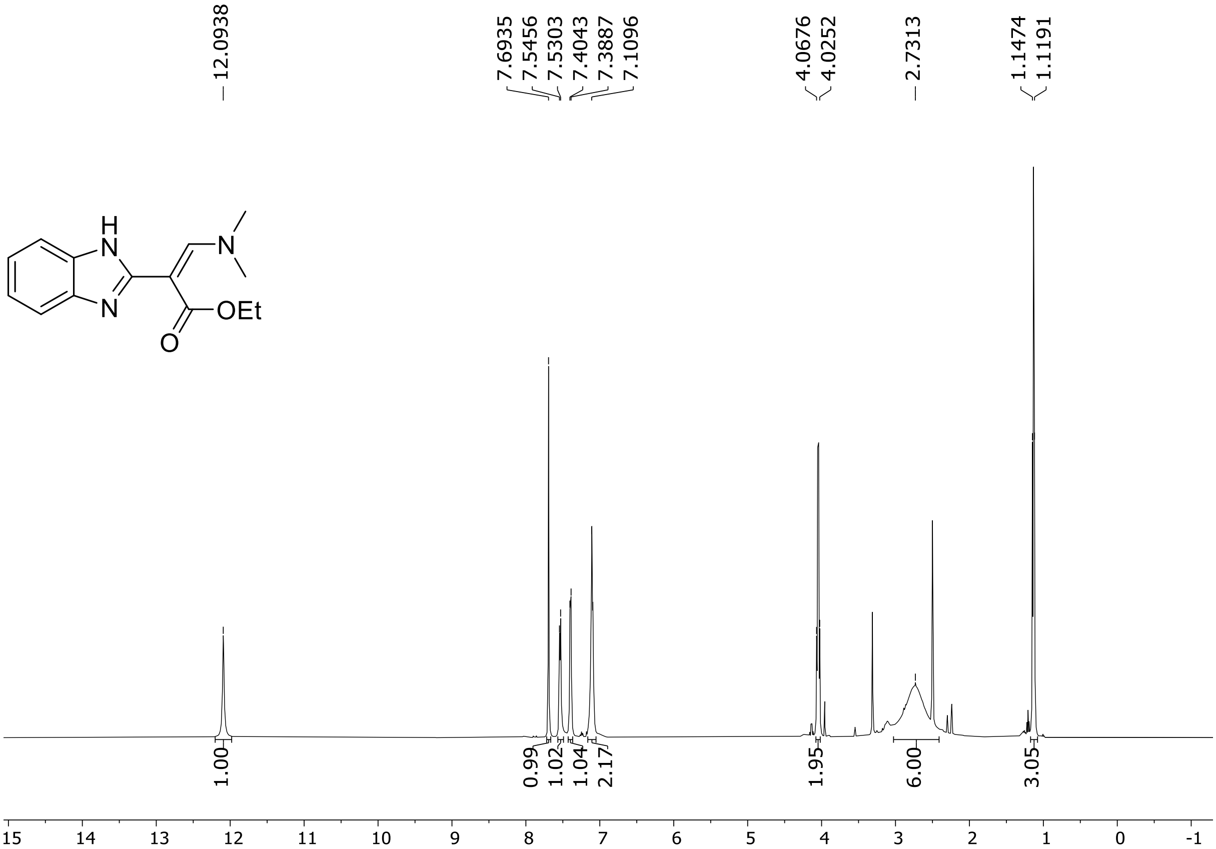

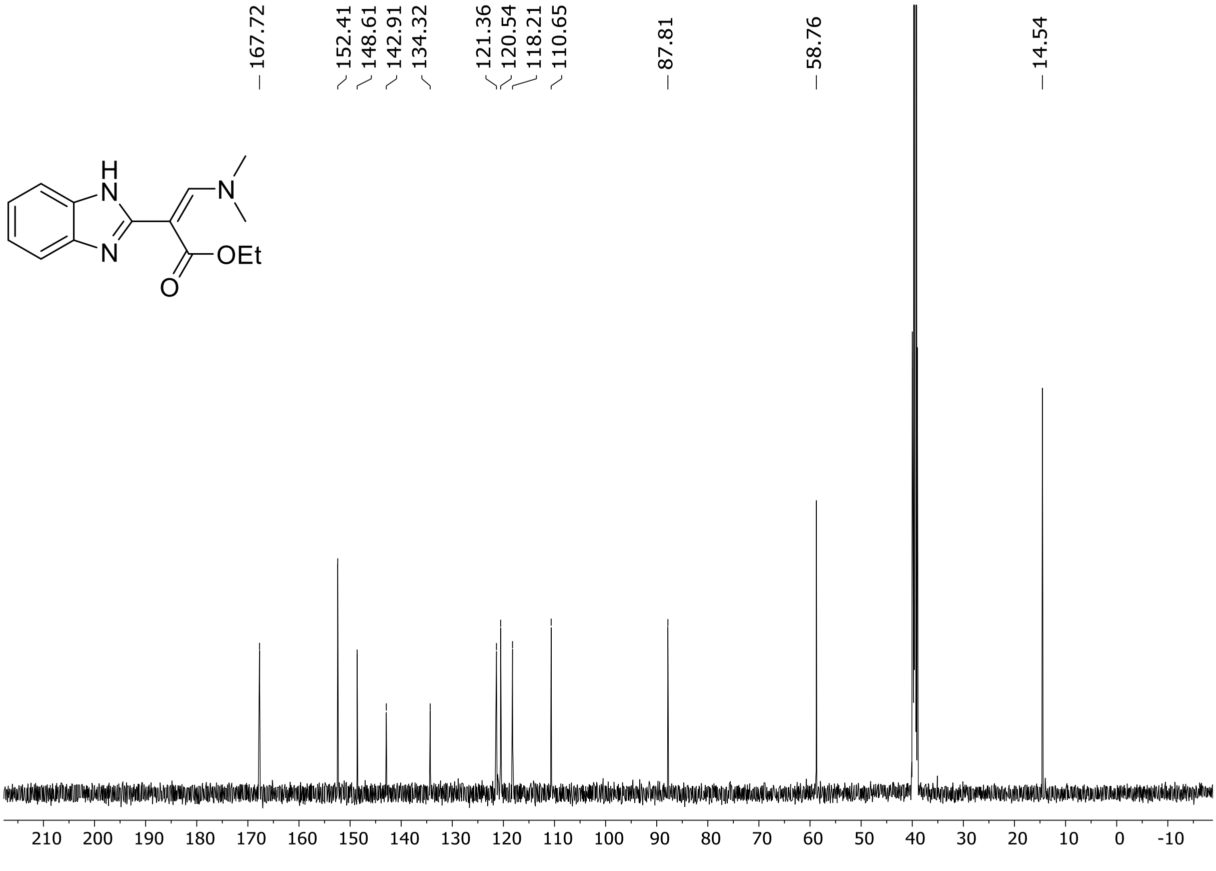

$^{1}H$ (500 MHz) and $^{13}C$ NMR (126 MHz) spectra of **S105** in DMSO-$d_6$

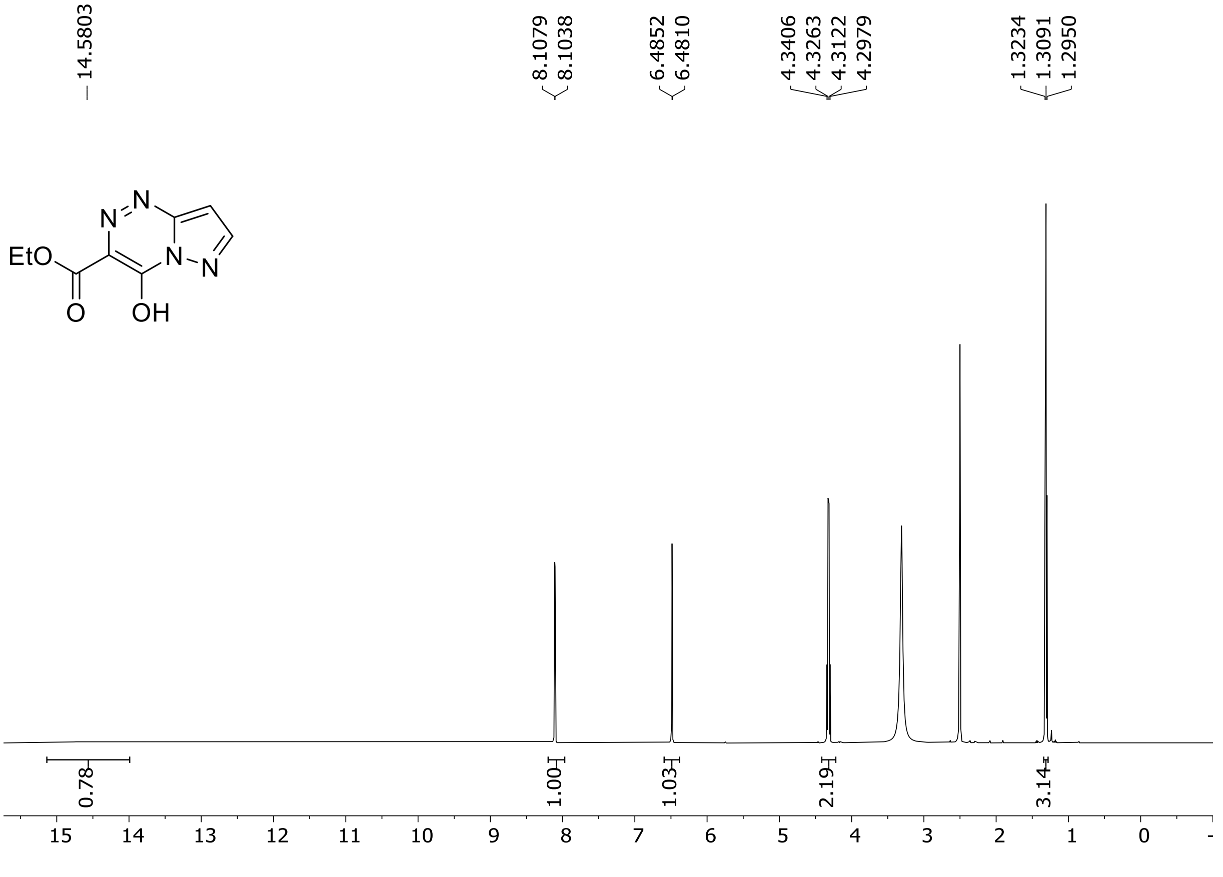

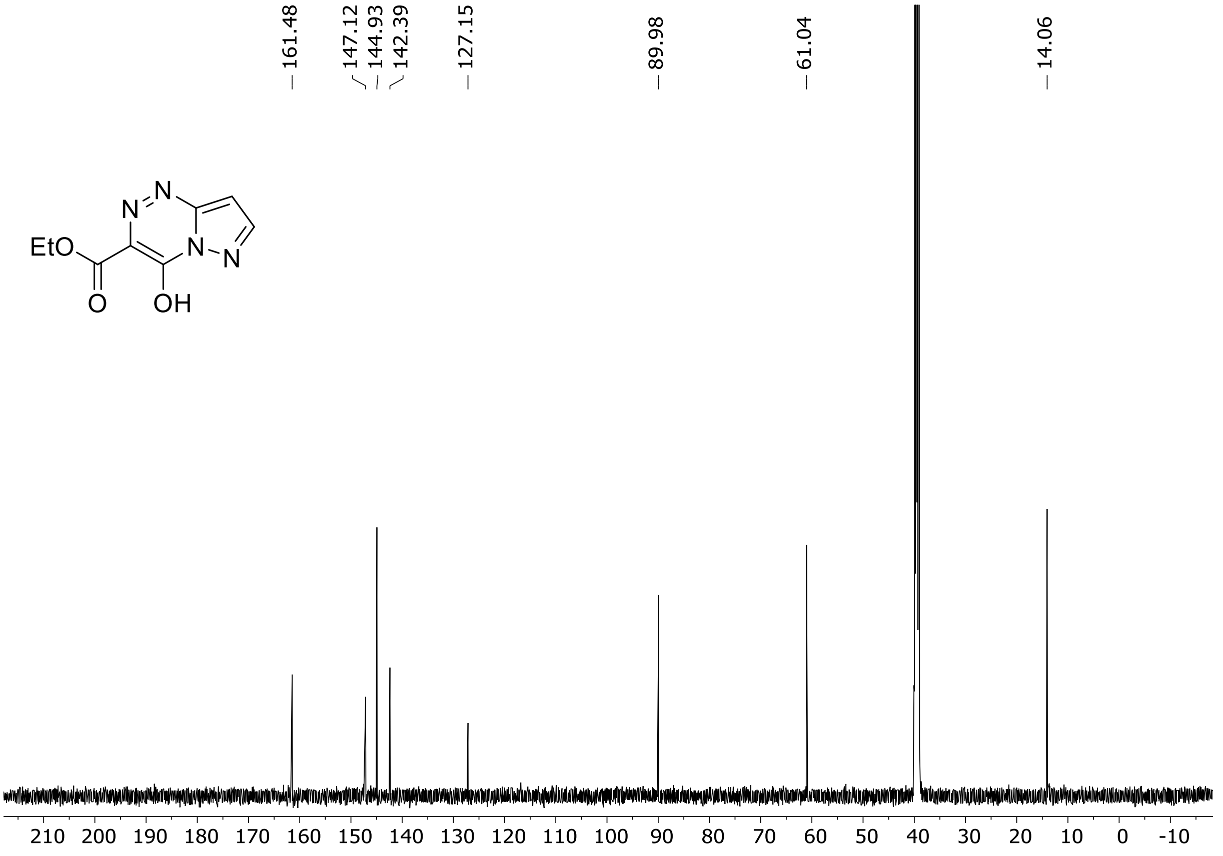

HRMS spectrum of **S105**

**$C_8H_8N_4O_3$ mono m/z = 208.17**

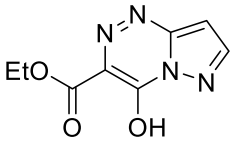

**APCI +** (MMI)

nitrogen flow 5 L/min, gas temperature 300°C, vaporizer 200°C, nebulizer 45 psi, skimmer **65** V, fragmentor 35 V, dissolved in MeOH

**ZOOM – range of interest**

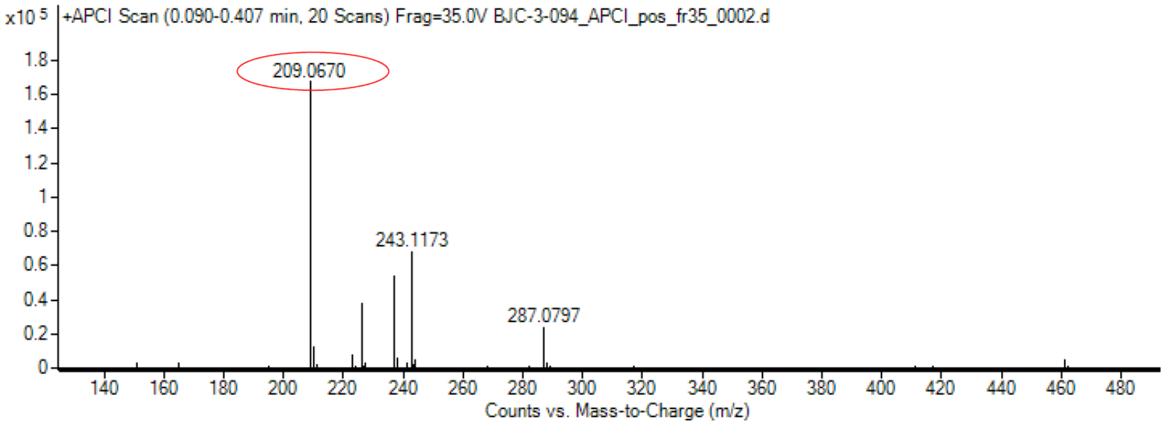

calculated mass: $[M+H]^+$ = 209.0669 observed: $[M+H]^+$ = 209.0670 max. mass error = 0.5 ppm

$^{1}H$ (500 MHz) and $^{13}C$ NMR (126 MHz) spectra of **S106** in DMSO-$d_6$

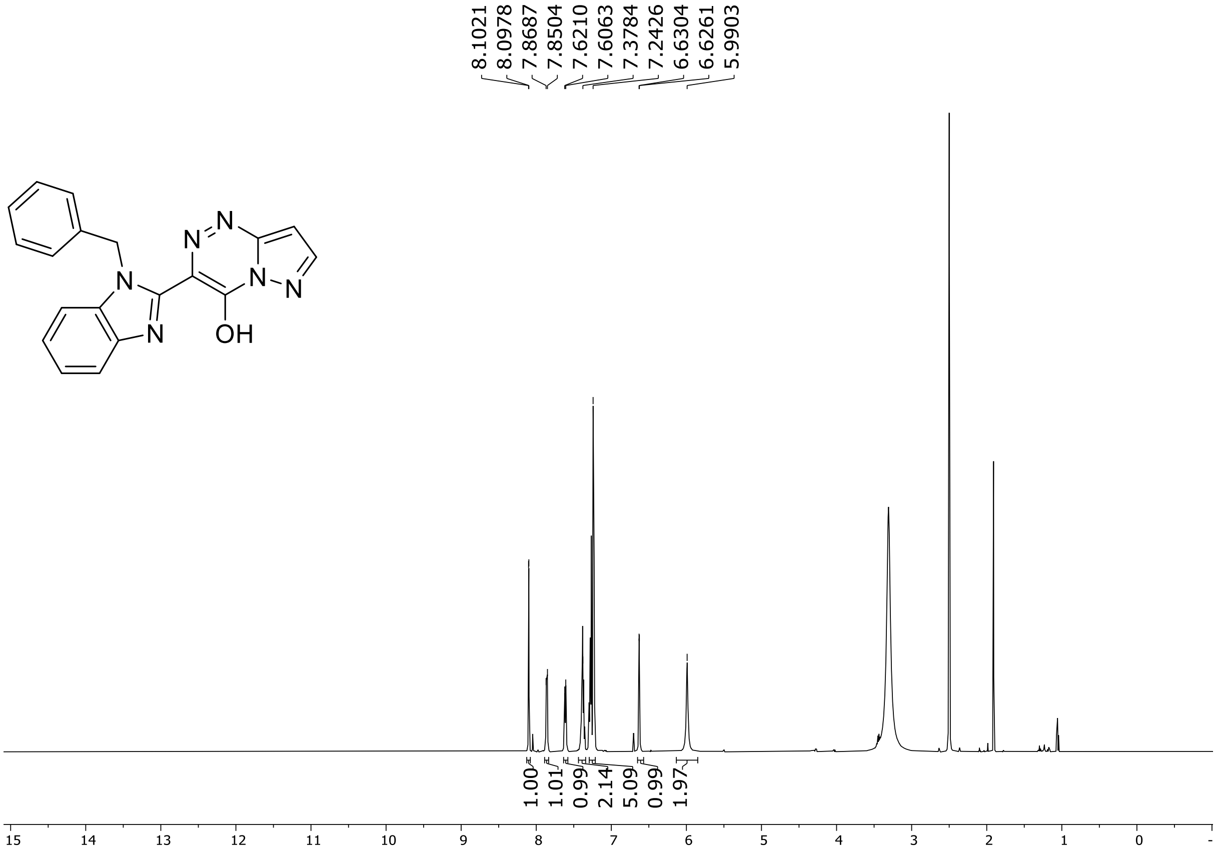

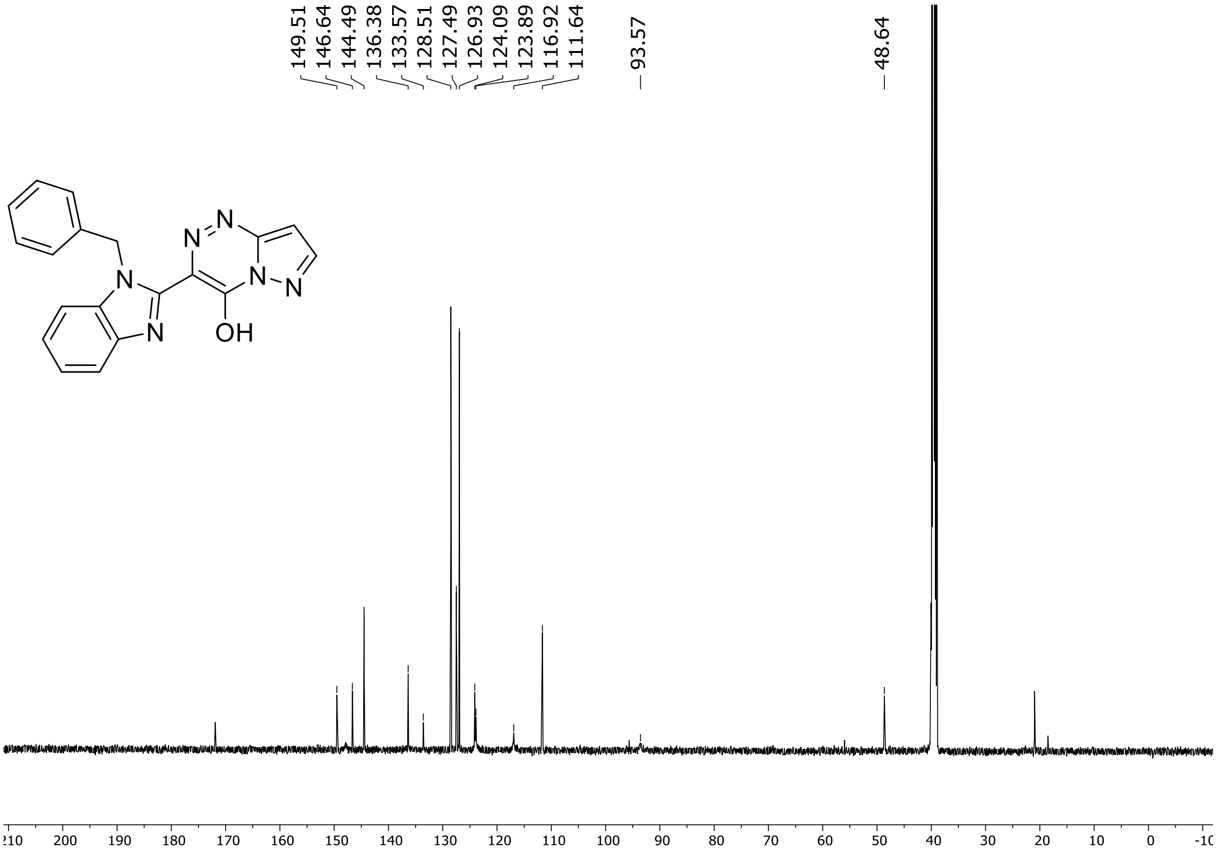

HRMS spectrum of **S106**

$C_{19}H_{14}N_6O$ mono m/z = 342.35

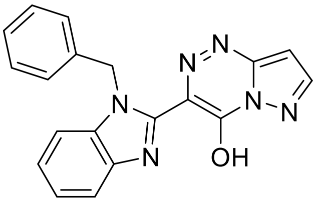

**APCI +** (MMI)

nitrogen flow 5 L/min, gas temperature 300°C, vaporizer 200°C, nebulizer 45 psi, skimmer **65** V, fragmentor 35 V, dissolved in MeOH

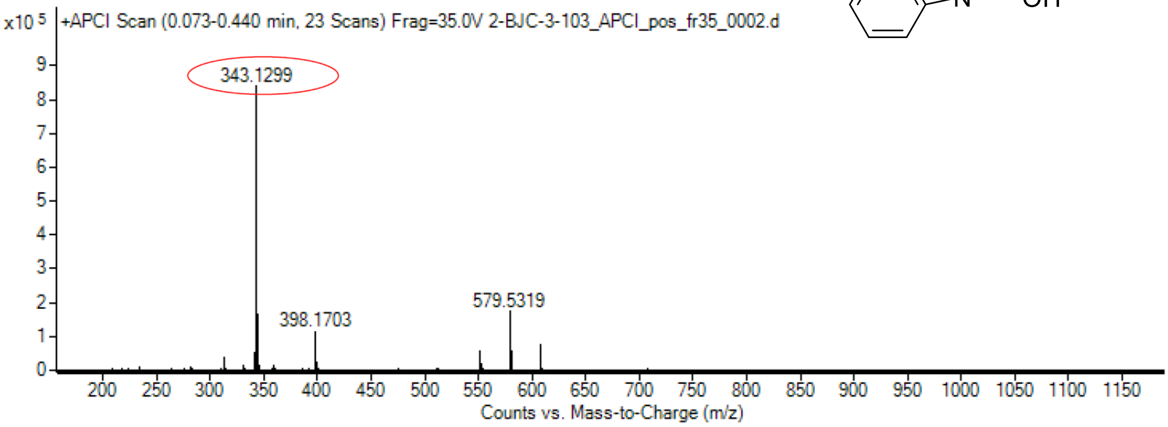

calculated mass: $[M+H]^+$ =343.1302 observed: $[M+H]^+$ = 343.1299 max. mass error = 0.8 ppm

$^{1}H$ (500 MHz) and $^{13}C$ NMR (126 MHz) spectra of **S107** in DMSO-$d_6$

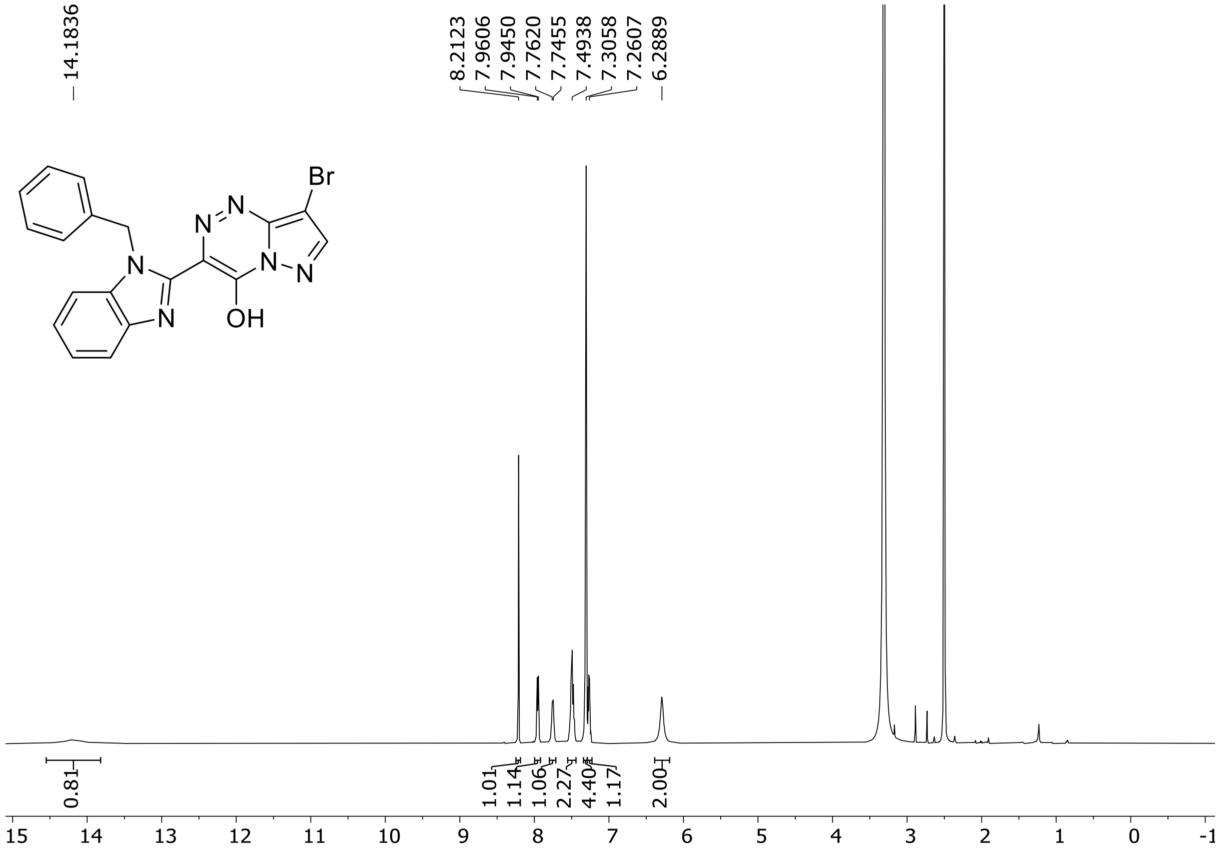

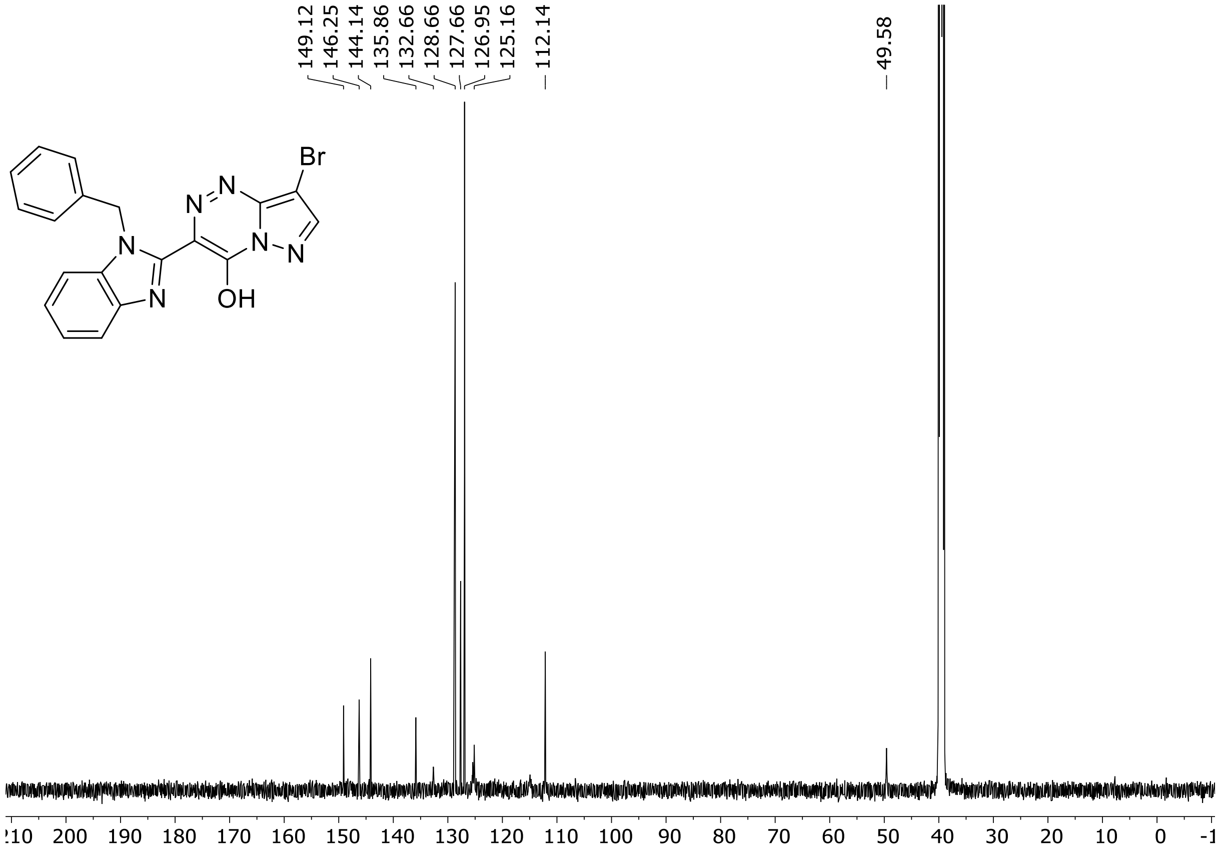

HRMS spectrum of **S107**

**$C_{19}H_{13}BrN_6O$ mono m/z = 420.03**

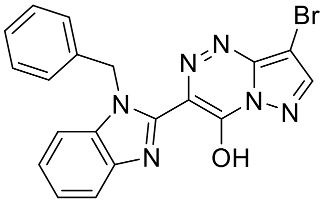

**ESI +** (MMI)

nitrogen flow 5 L/min, gas temperature 300°C, vaporizer 200°C, nebulizer 45 psi, skimmer **65** V, fragmentor 45 V, dissolved in MeOH

**ZOOM – range of interest**

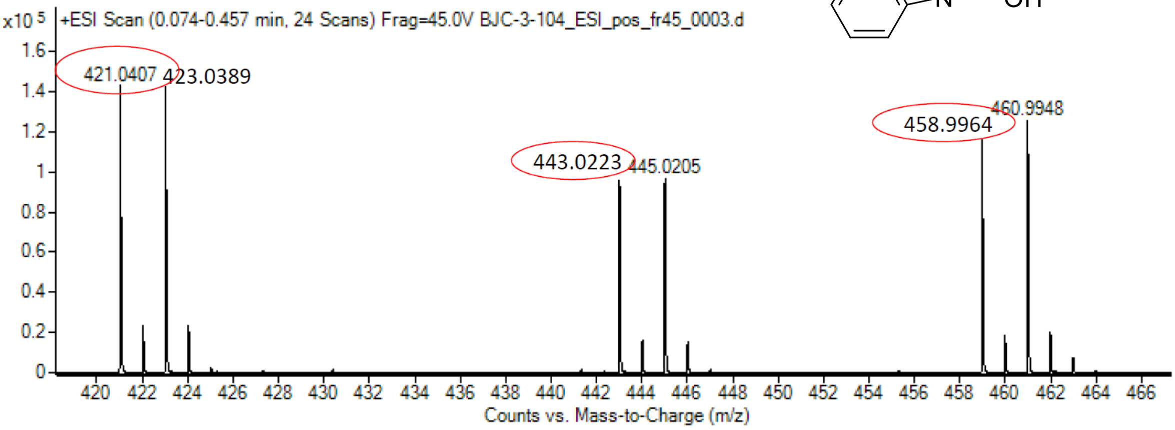

| | | | | |
|---|---|---|---|---|
| calculated mass: | $[M+H]^+$ = 421.0407 | observed: | $[M+H]^+$ = 421.0407 | max. mass error < 0.1 ppm |
| calculated mass: | $[M+Na]^+$ = 443.0226 | observed: | $[M+Na]^+$ = 443.0223 | max. mass error = 0.7 ppm |
| calculated mass: | $[M+K]^+$ = 458.9966 | observed: | $[M+K]^+$ = 458.9964 | max. mass error = 0.4 ppm |

[1]H (500 MHz) and [13]C NMR (126 MHz) spectra of **S108** in DMSO-$d_6$

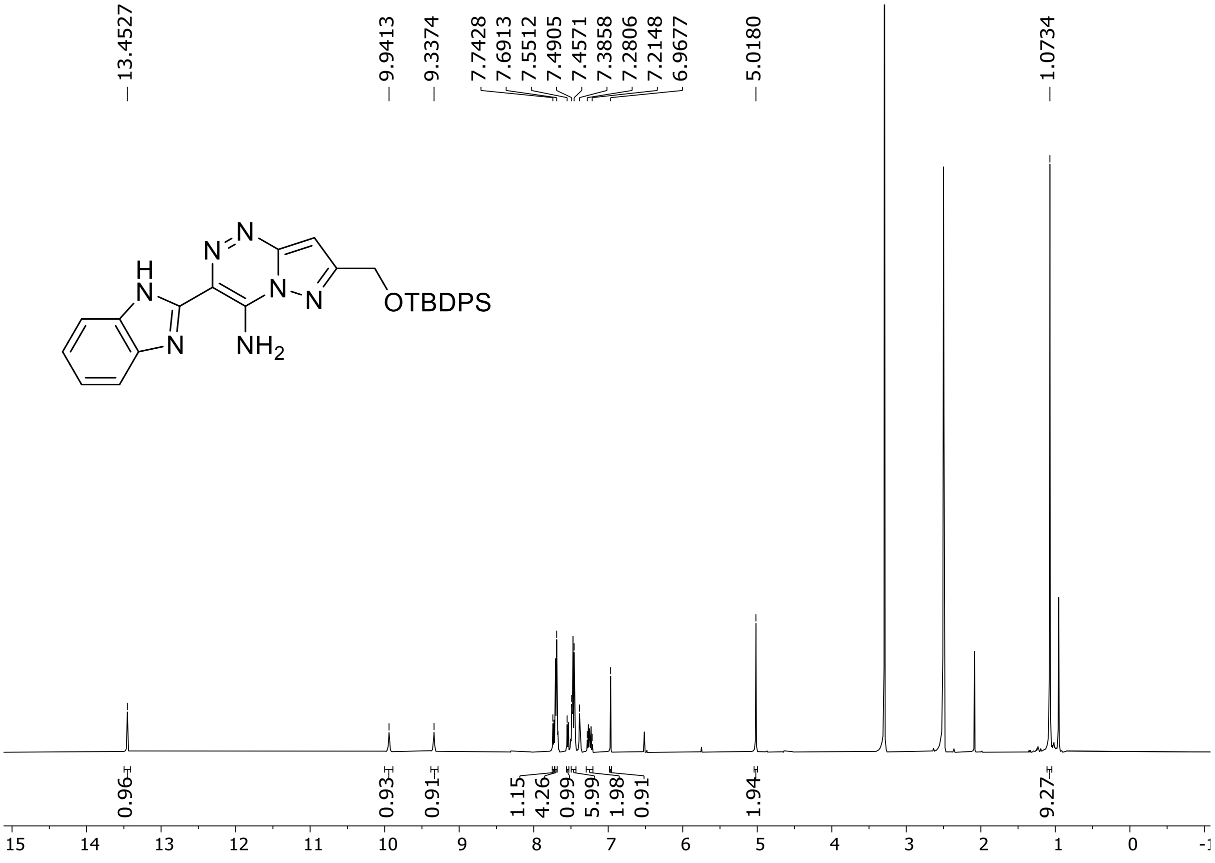

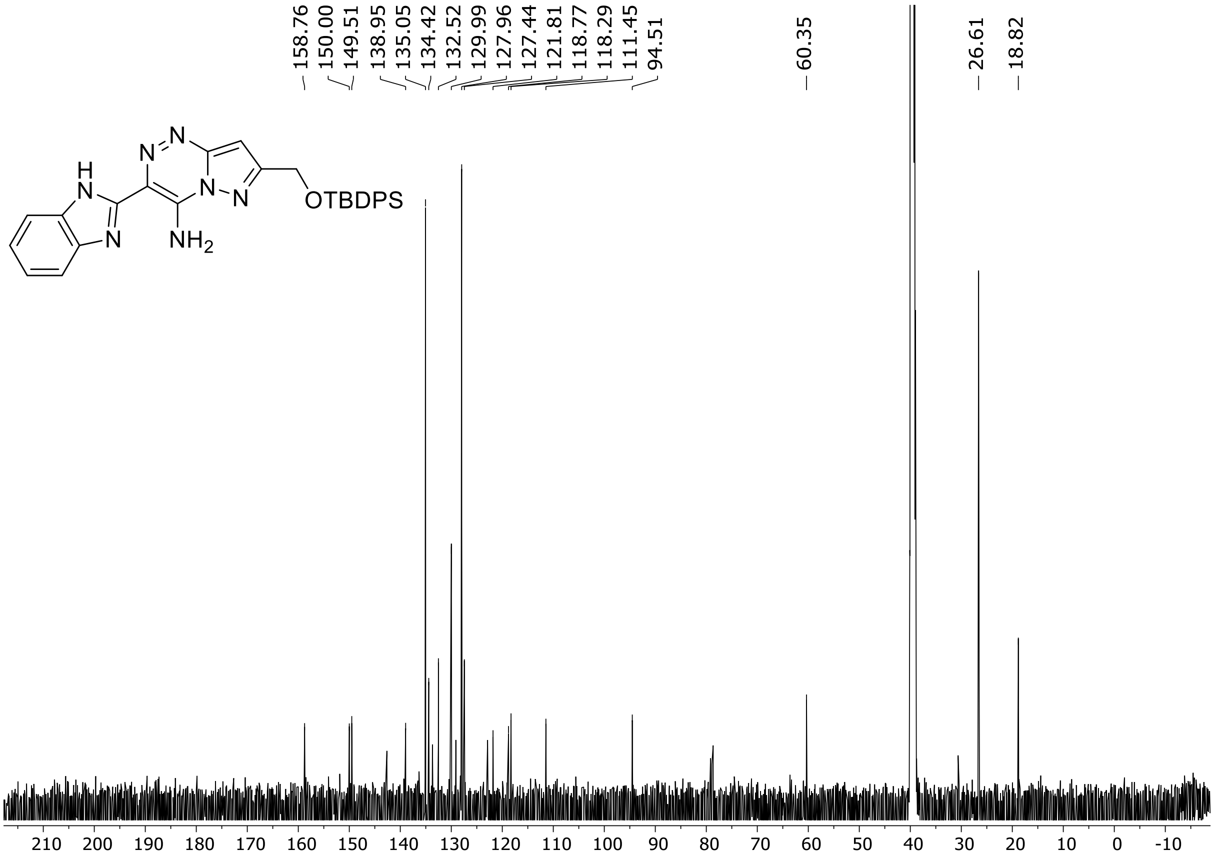

HRMS spectrum of **S108**

**$C_{29}H_{29}N_7OSi$** **mono m/z = 519.2203**

**APCI +** (MMI)
nitrogen flow 5 L/min, gas temperature 300°C, nebulizer 45 psi, vaporizer 200°C
skimmer **65** V, fragmentor **20** V, dissolved in methanol

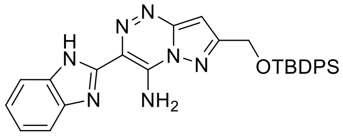

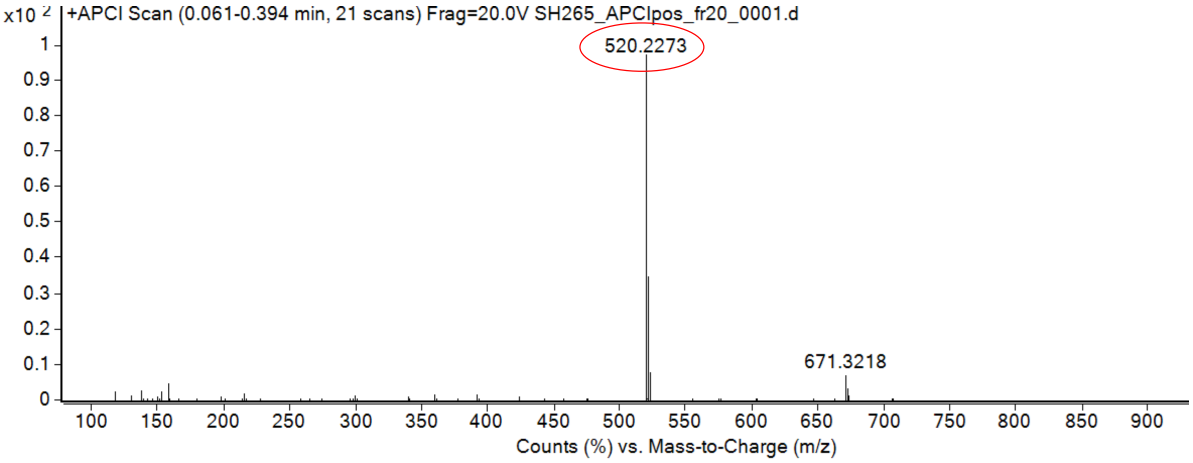

calculated mass: $[M+H]^+$ = 520.2276 observed: $[M+H]^+$ = 520.2273 max. mass error = 0.5 ppm

**[1]H NMR, [13]C NMR and HRMS spectra of compound 1 and 3-28**

[1]H (500 MHz) and [13]C NMR (126 MHz) spectra of **1** in DMSO-$d_6$

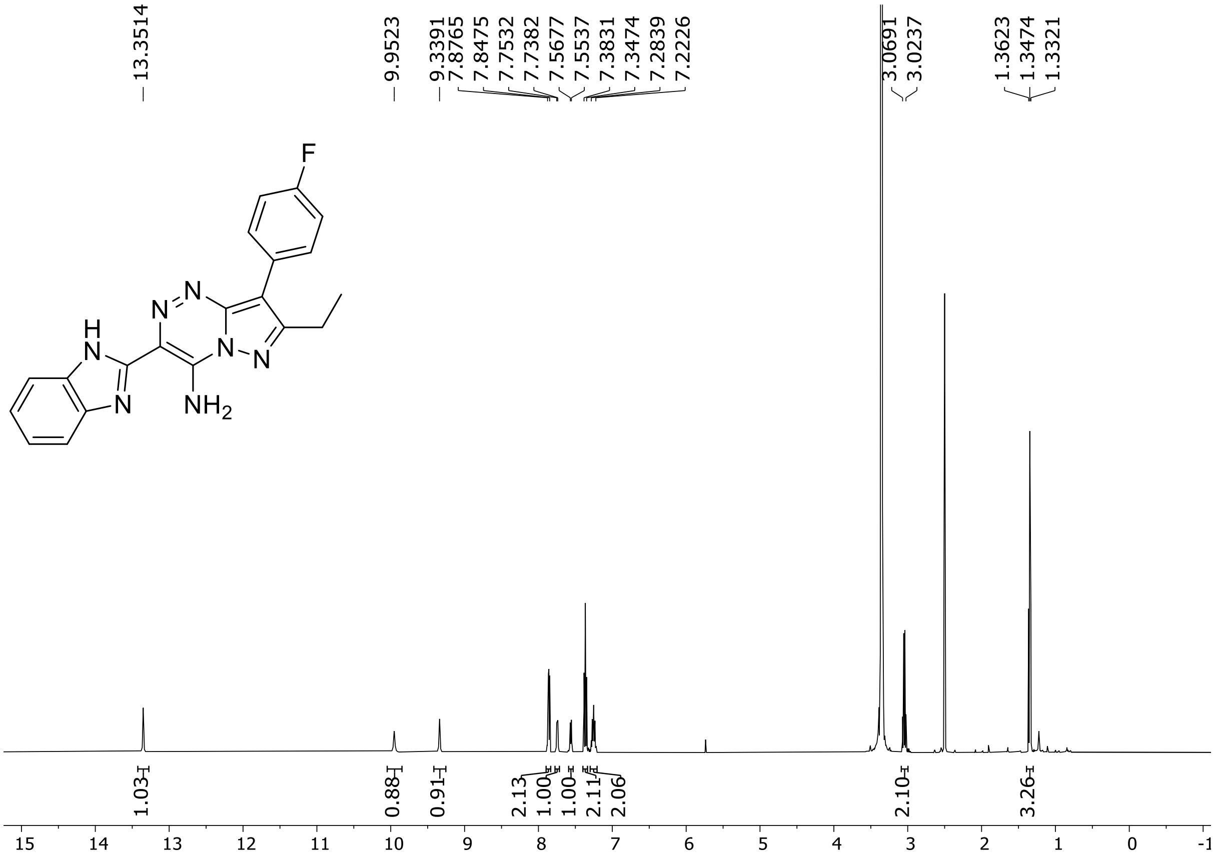

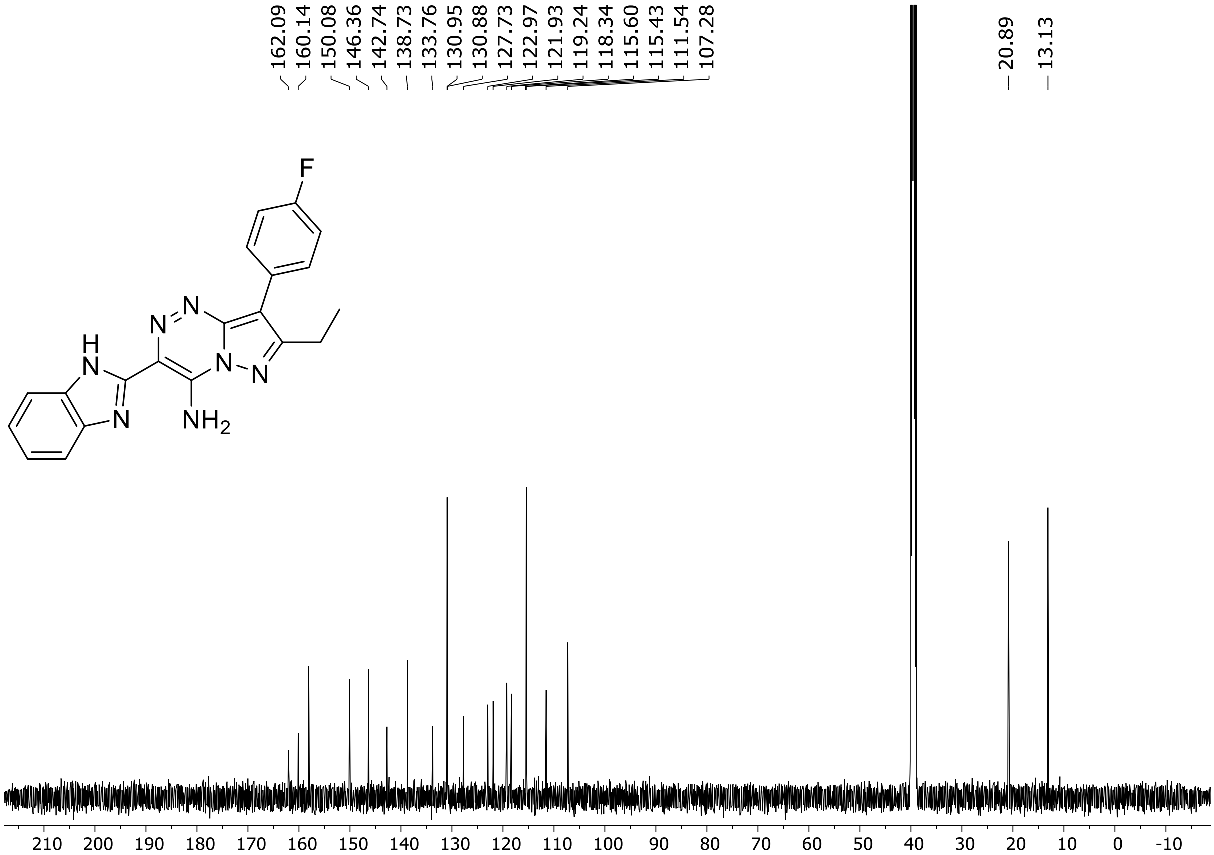

HRMS spectrum of **1**

$C_{20}H_{16}FN_7$ **mono m/z = 373,1446**

**APCI +** (MMI)
nitrogen flow 5 L/min, gas temperature 300°C, vaporizer 250°C,
nebulizer 45 psi, corona current 4 uA, Vcap -2000 V,
skimmer **65** V, fragmentor **35** V, dissolved in methanol (CH3OH)

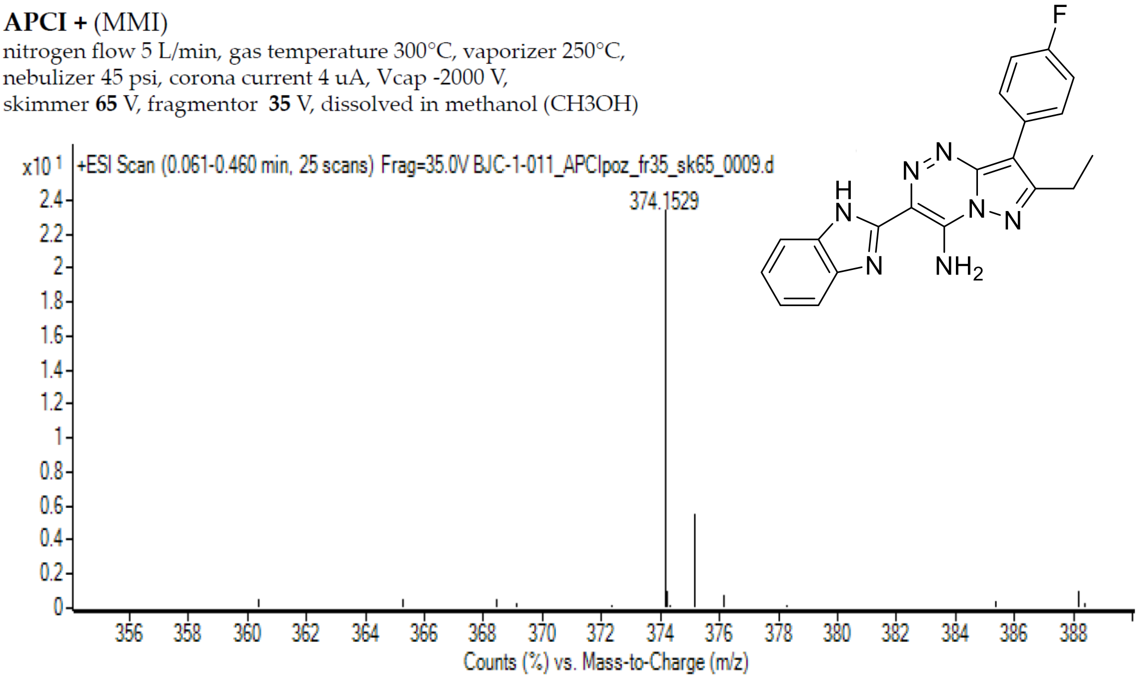

calculated mass: $[M+H]^+$ = 374,1524 observed: $[M+H]^+$ =374,1529 max. mass error 1,3 ppm

[1]H (500 MHz) and [13]C NMR (126 MHz) spectra of **3** in DMSO-$d_6$

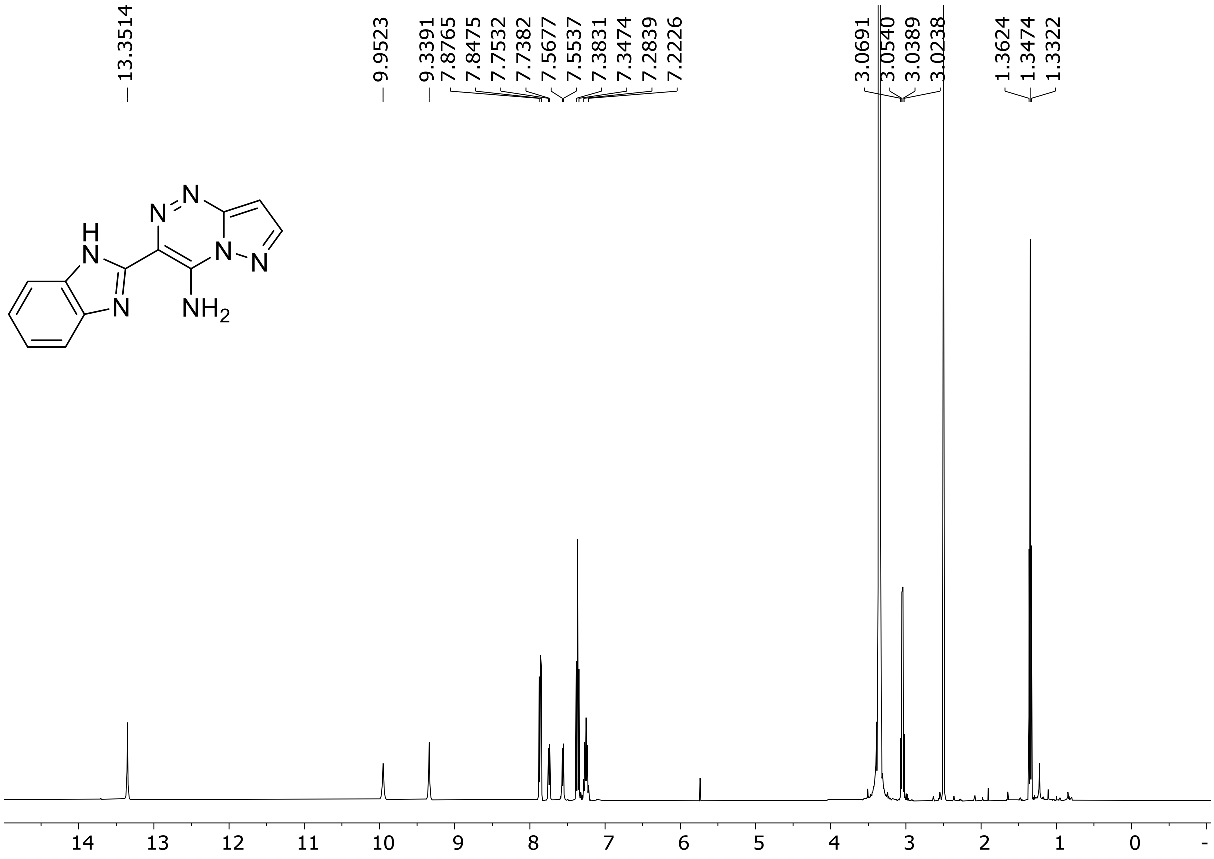

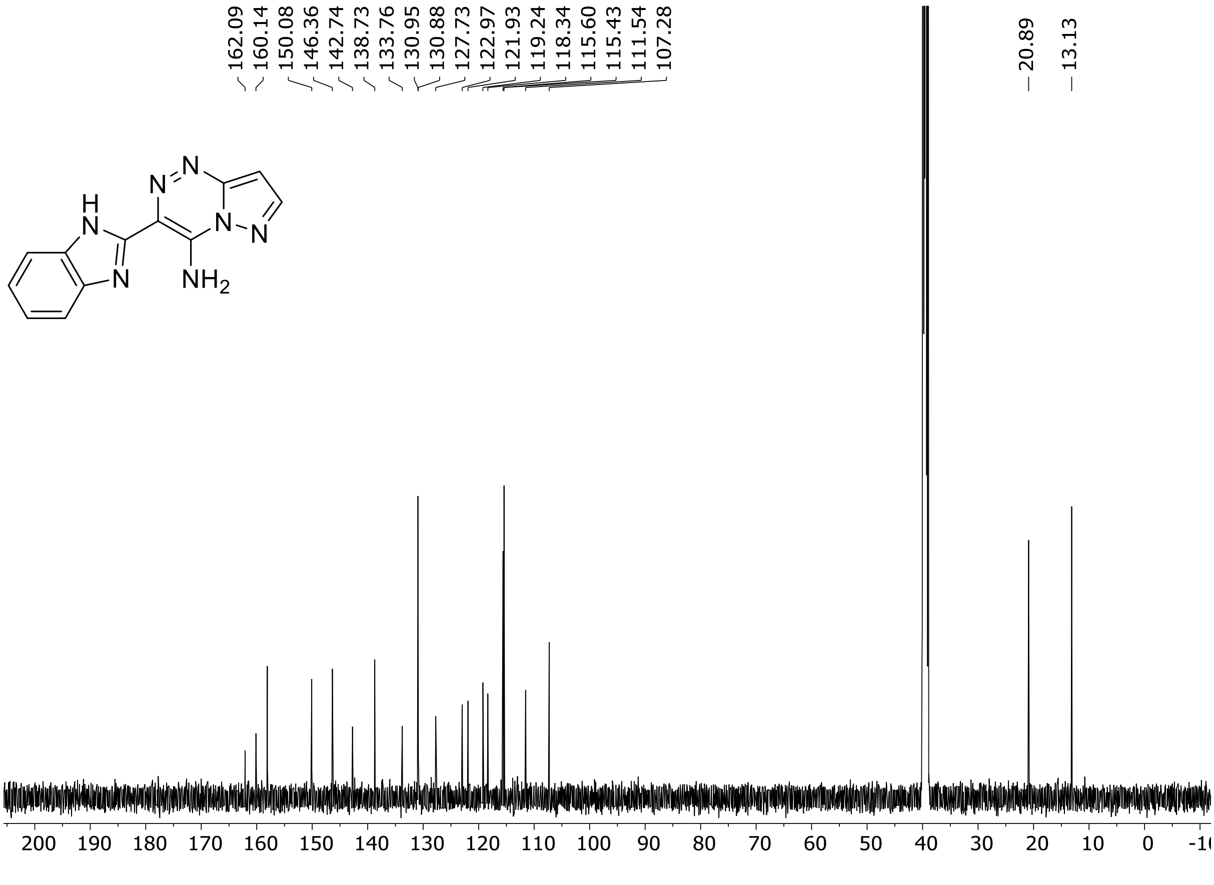

HRMS spectrum of **3**

$C_{12}H_9N_7$ **mono m/z = 251,2467**

**ESI -** (MMI)
nitrogen flow 5 L/min, gas temperature 300°C, vaporizer 200°C, nebulizer 45 psi, Vcap -2000 V, skimmer **65** V, fragmentor **35** V, dissolved in DMSO

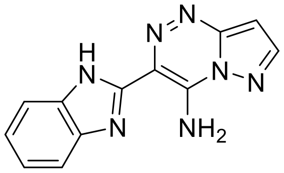

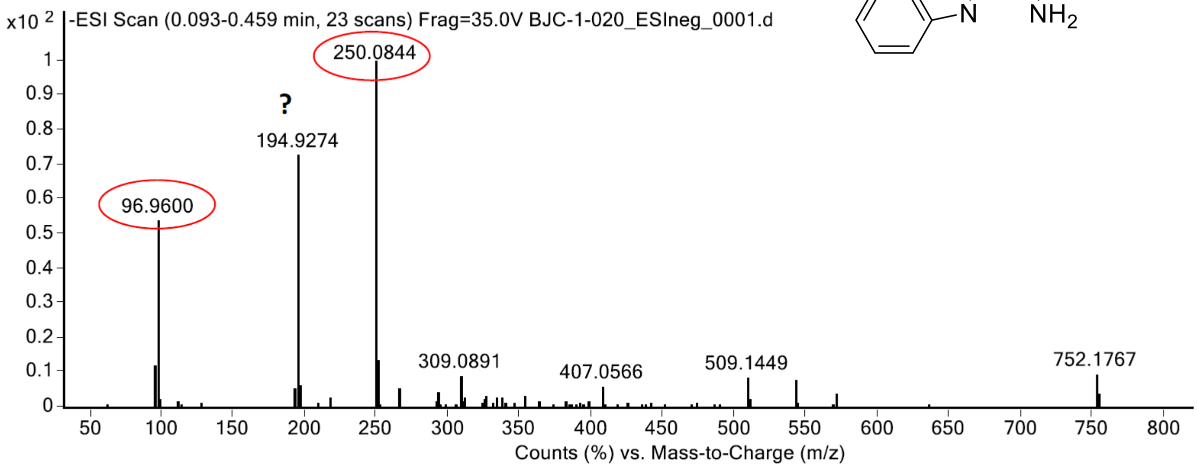

calculated mass: $[M-H]^-$ = 250,0846 observed: $[M-H]^-$ = 250,0844 max. mass error = 0,7 ppm

sulphuric acid
calculated mass: $[M-H]^-$ = 96,9601 observed: $[M-H]^-$ = 96,960 max. mass error = 1 ppm

[1]H (500 MHz) and [13]C NMR (126 MHz) spectra of **4** in DMSO-$d_6$

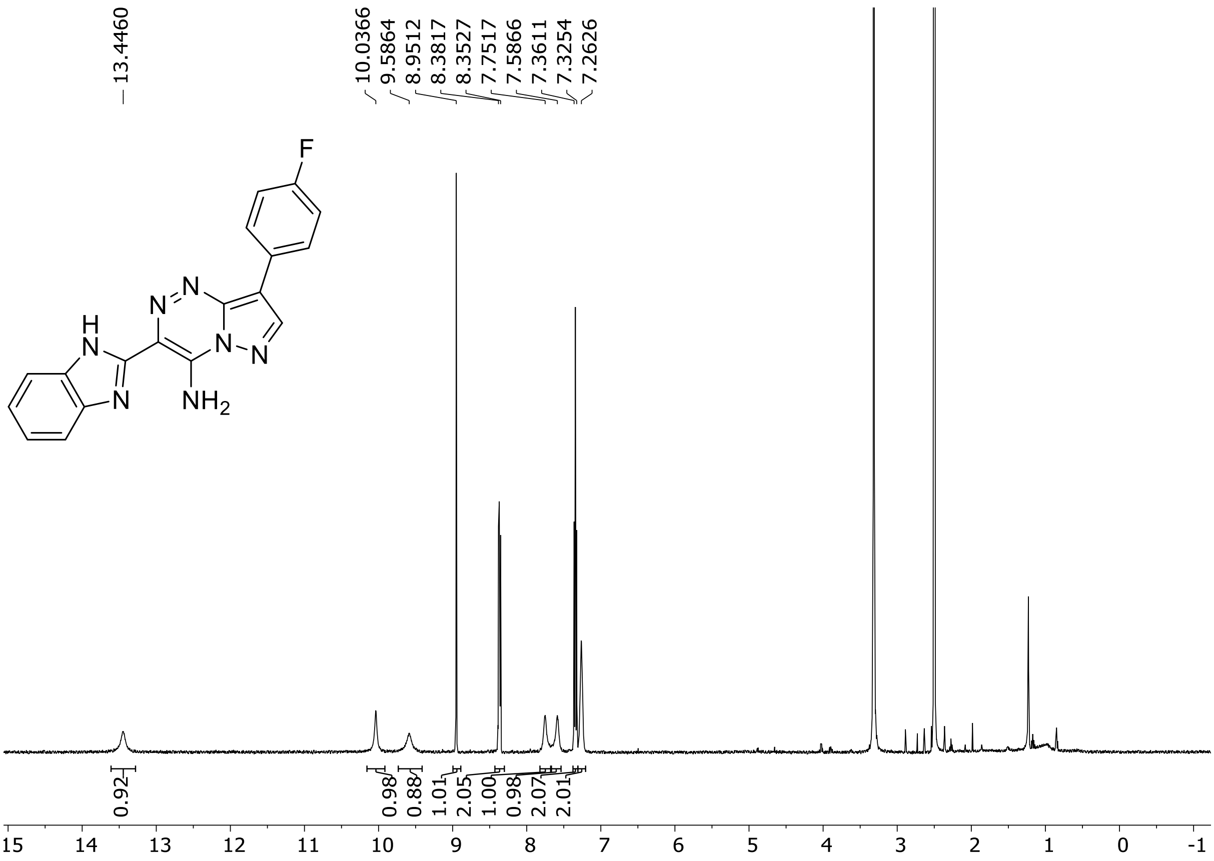

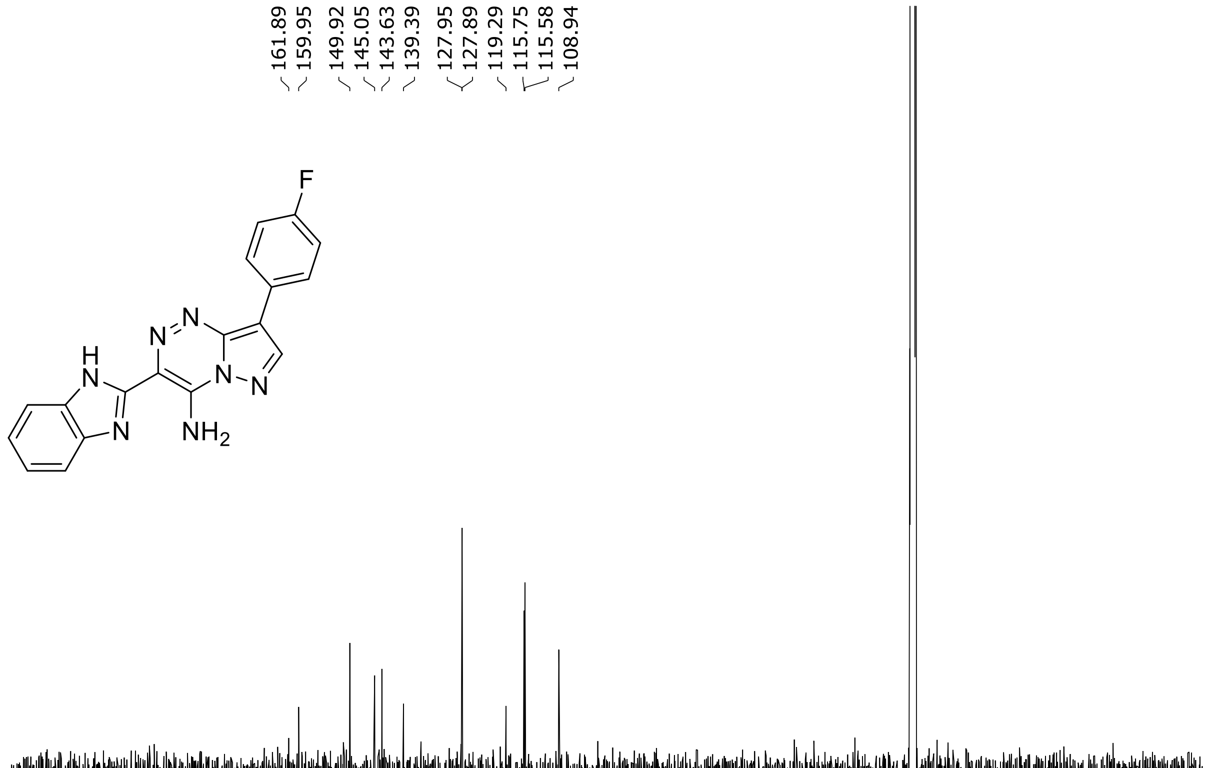

$^{19}F$ (282 MHz) NMR spectrum of **4** DMSO-$d_6$

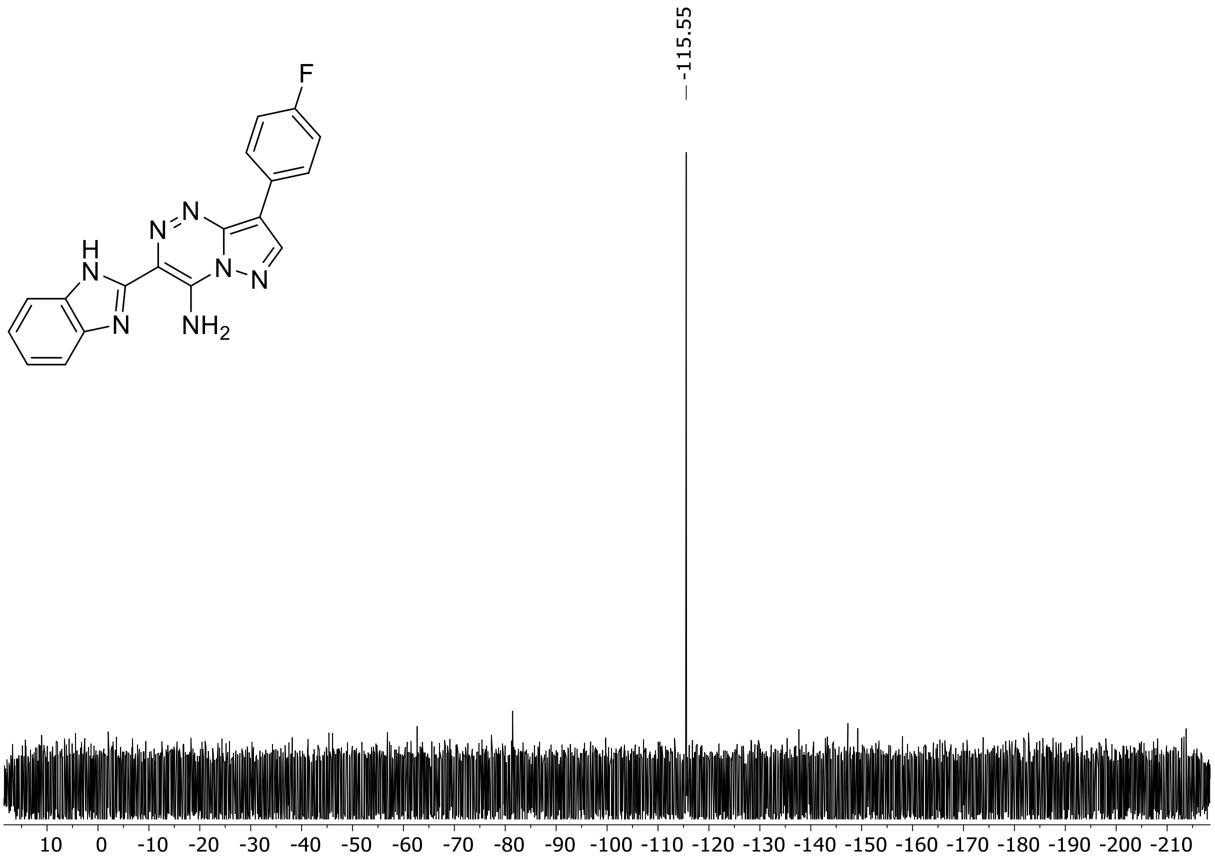

HRMS Spectra of **4**

**$C_{18}H_{12}FN_7$** **mono m/z = 345,1138**

**ESI -** (MMI)
nitrogen flow 5 L/min, gas temperature 300°C, vaporizer 200°C,
nebulizer 30 psi, skimmer **40** V, fragmentor **60** V, dissolved in DMSO

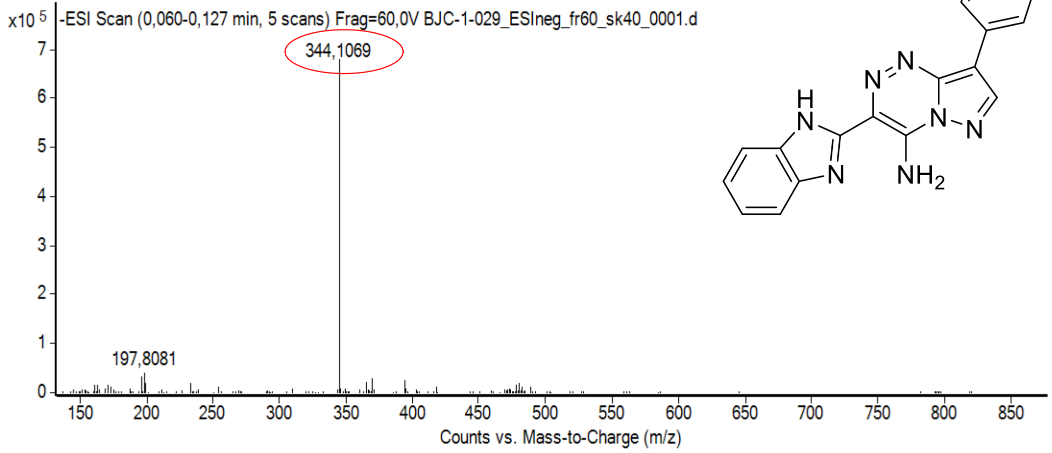

calculated mass: $[M-H]^-$ = 344,1065 observed: $[M-H]^-$ = 344,1069 max. mass error = 1,2 ppm

[1]H (500 MHz) and [13]C NMR (126 MHz) spectra of **5** in DMSO-$d_6$

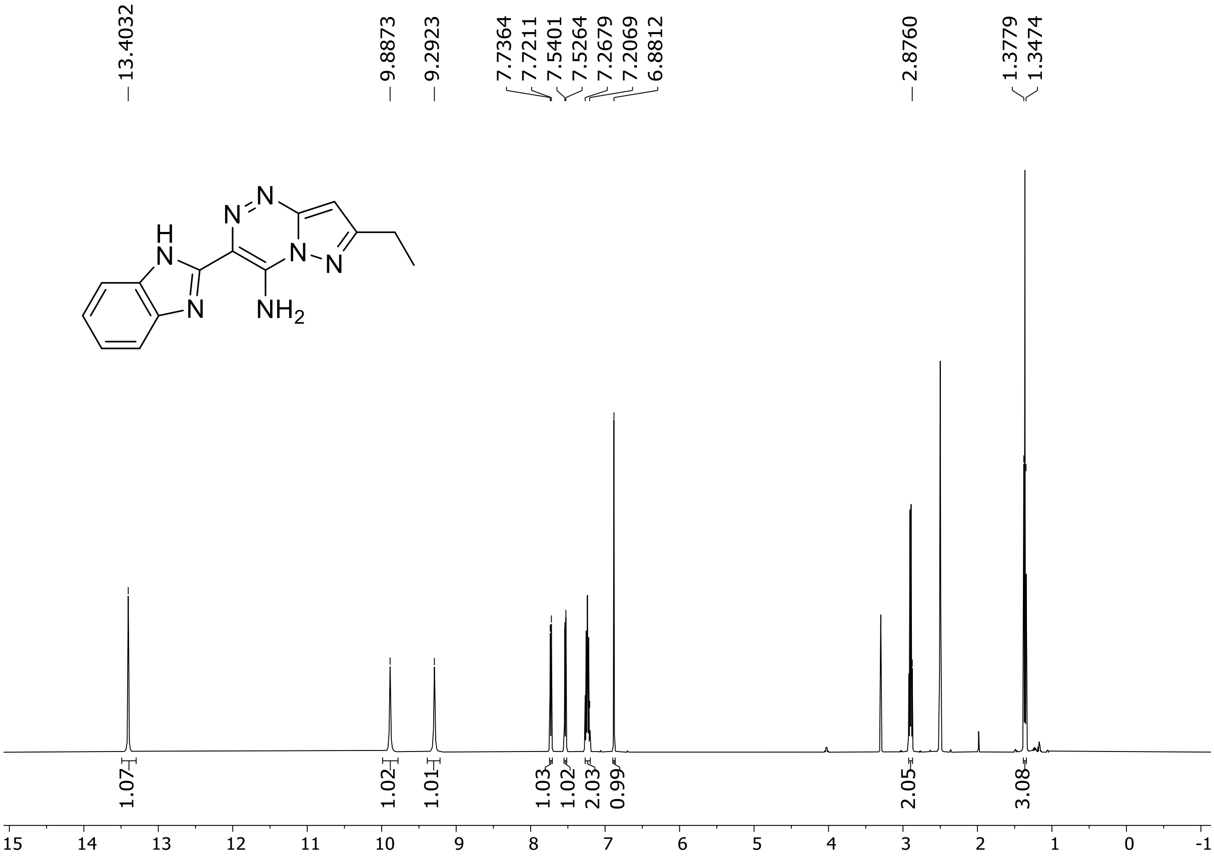

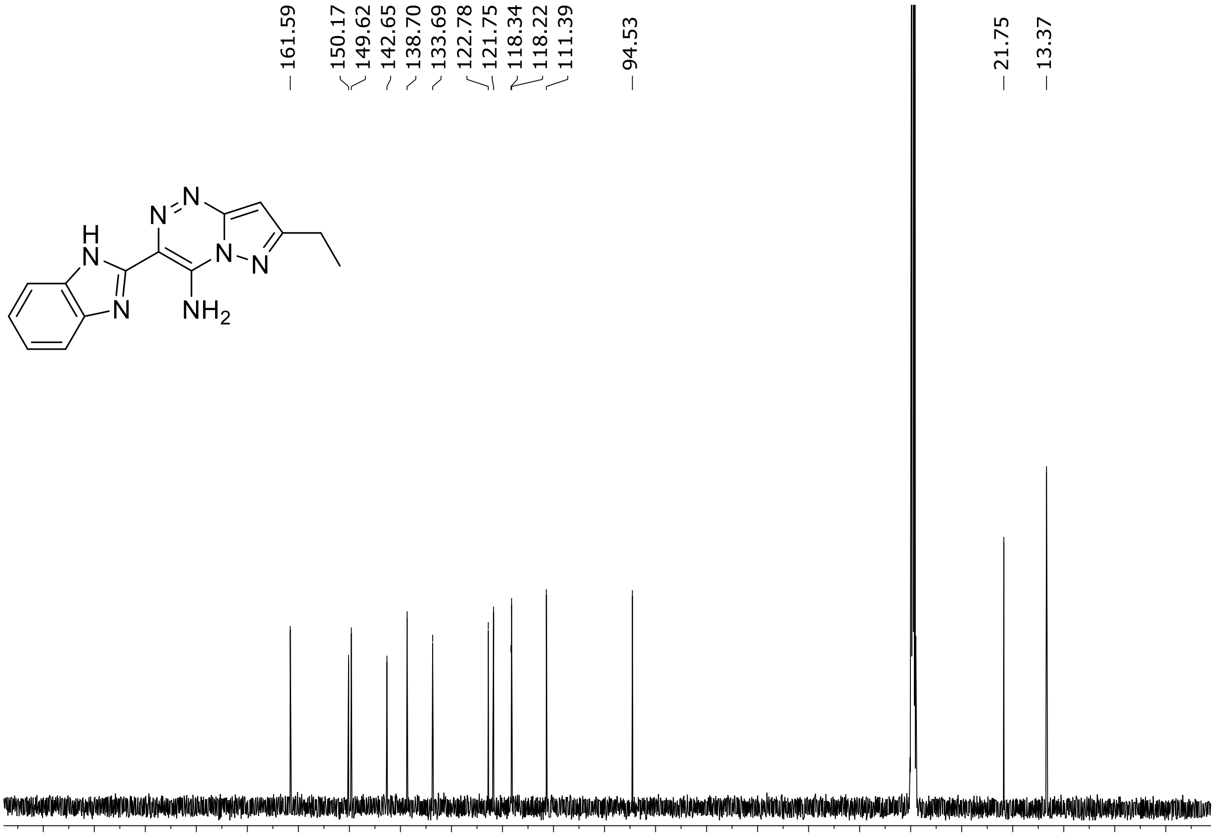

HRMS spectrum of **5**

**$C_{14}H_{13}N_7$** **mono m/z = 279,1232**

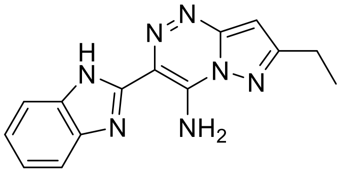

**APCI -** (MMI)
nitrogen flow 5 L/min, gas temperature 300°C, vaporizer 250°C,
nebulizer 45 psi, skimmer **65** V, fragmentor **30** V, dissolved in DMSO

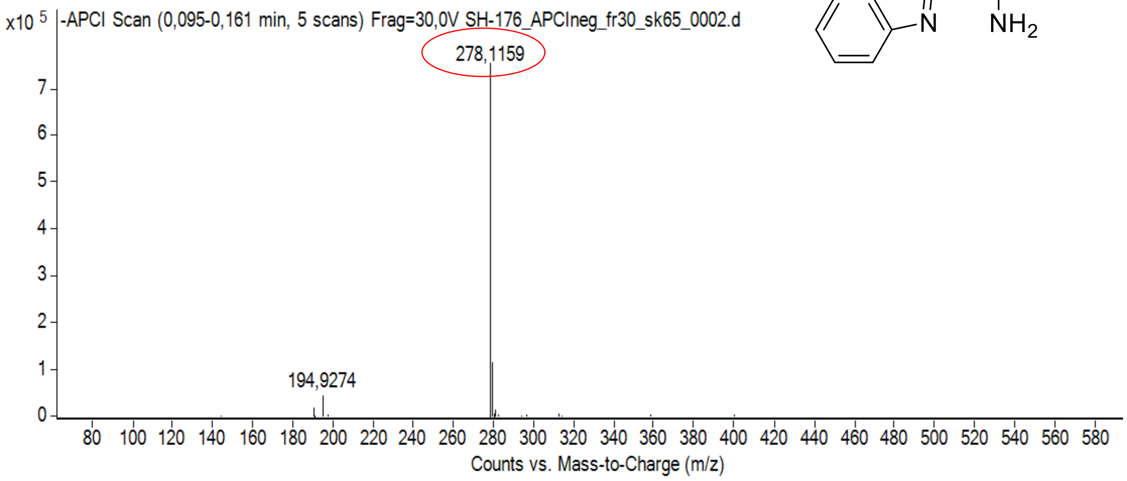

calculated mass: $[M\text{-}H]^-$ = 278,1160 observed: $[M\text{-}H]^-$ = 278,1159 max. mass error = 0,4 ppm

$^{1}H$ (500 MHz) and $^{13}C$ NMR (126 MHz) spectra of **6** in DMSO-$d_6$

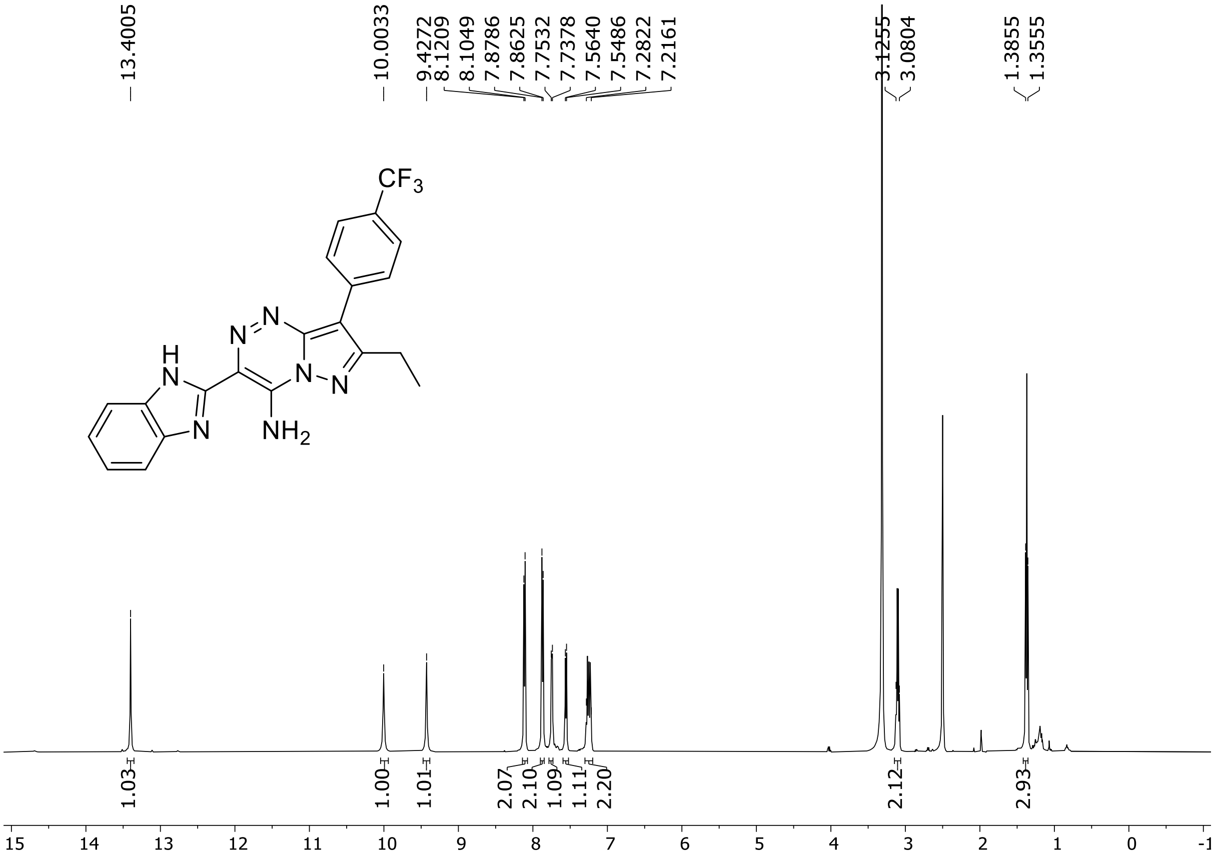

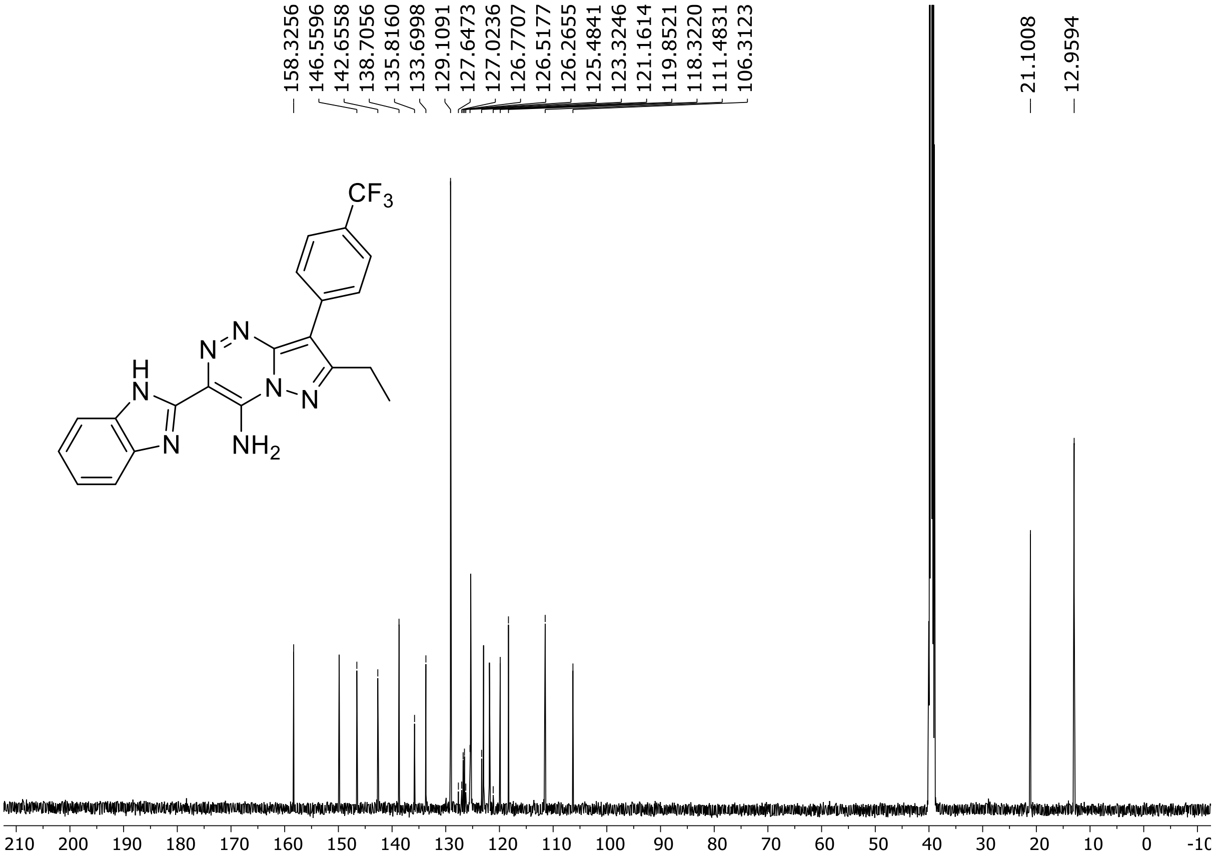

$^{19}$F (471 MHz) NMR spectrum of **6** DMSO-$d_6$

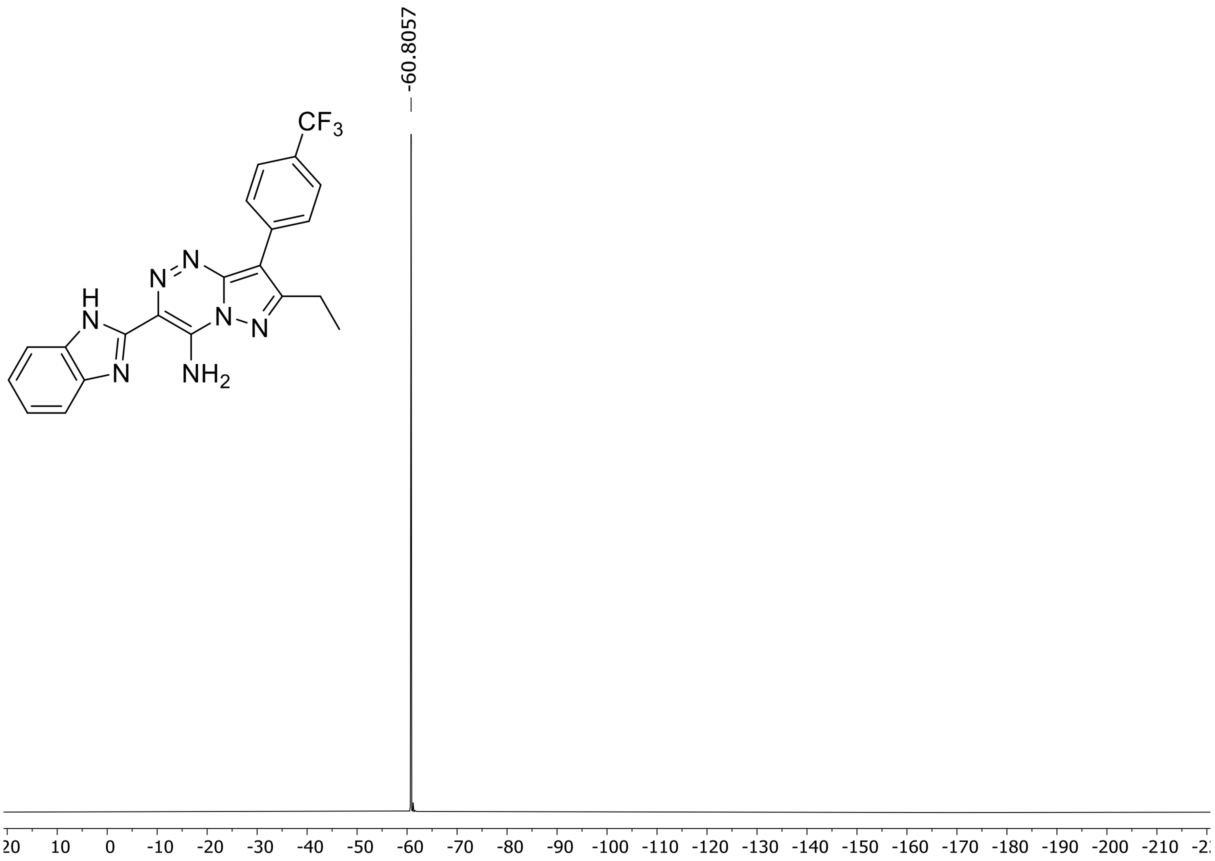

HRMS spectrum of **6**

$C_{21}H_{16}F_3N_7$ **mono *m/z* 423.1419**

**APCI +** (MMI)
nitrogen flow 5 L/min, gas temperature 325°C, nebulizer 45 psi, skimmer 65 V, vaporizer 250°C, fragmentor 35 V, dissolved in methanol

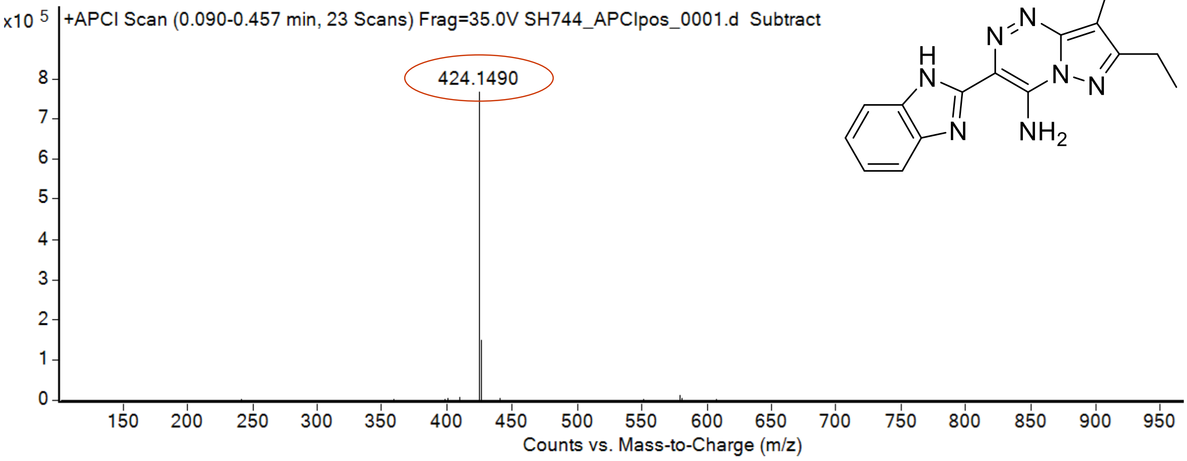

calculated mass: $[M+H]^+ = 424.1492$ observed: $[M+H]^+ = 424.1490$ mass accuracy = - 0.4 ppm

[1]H (500 MHz) and [13]C NMR (126 MHz) spectra of **7** in DMSO-$d_6$

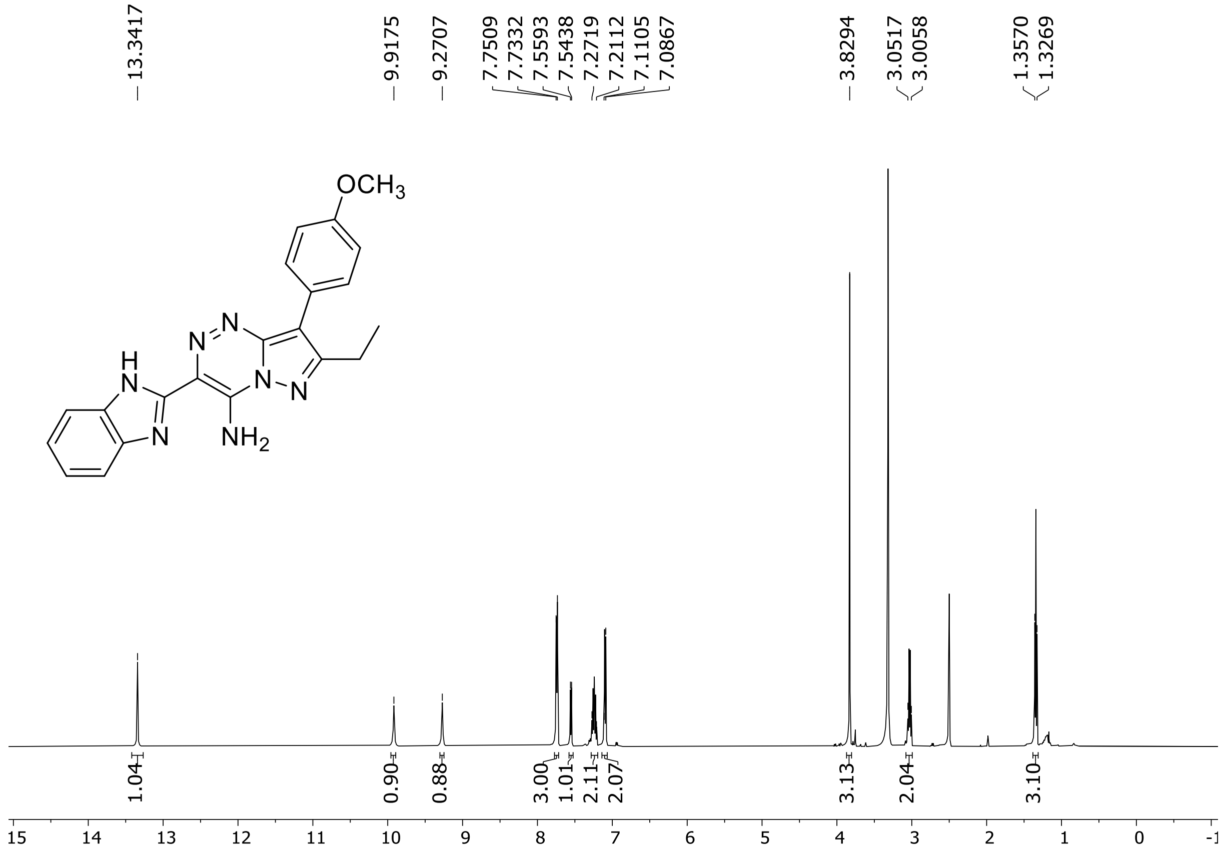

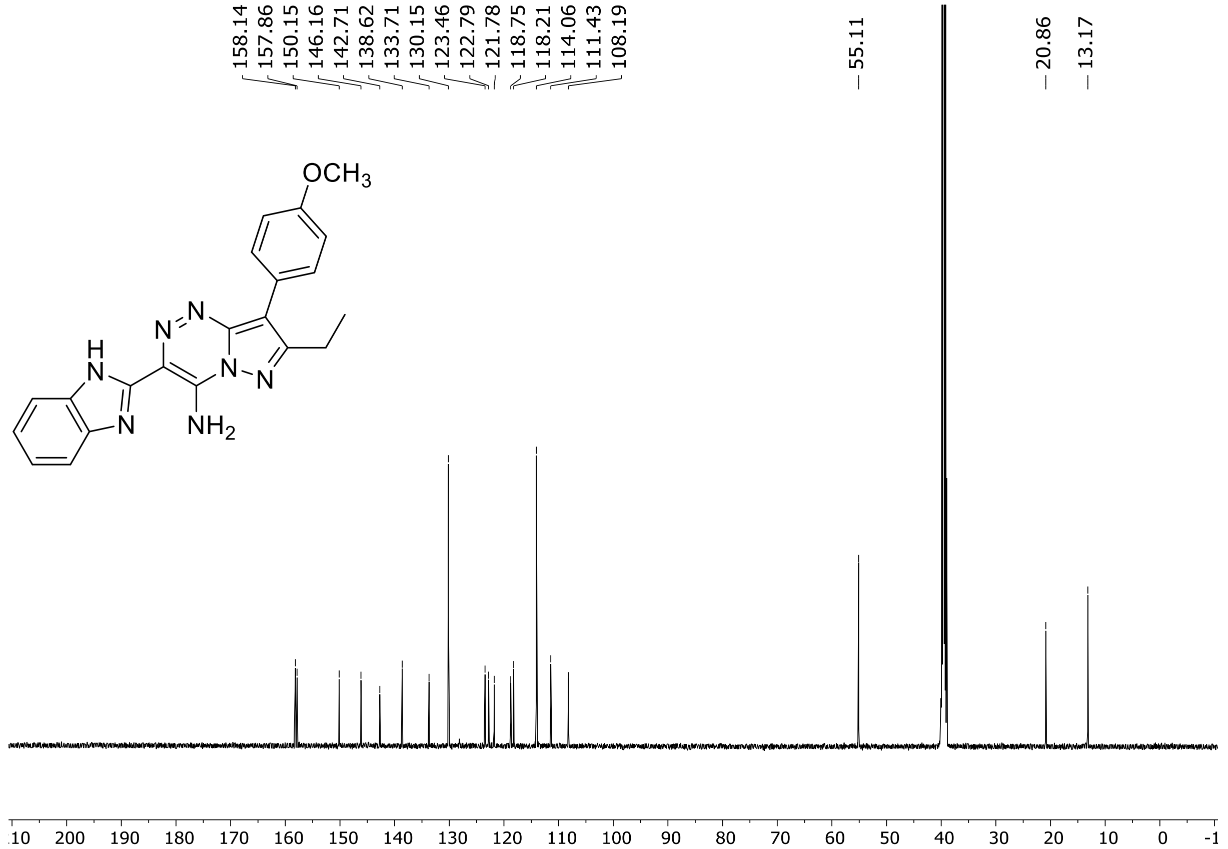

HRMS spectrum of **7**

**$C_{21}H_{19}N_7O$** **mono *m/z* 385.1651**

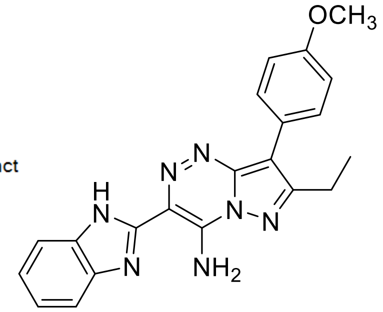

**APCI +** (MMI)
nitrogen flow 5 L/min, gas temperature 325°C, nebulizer 45 psi, skimmer 65 V, vaporizer 250°C, fragmentor 45 V, dissolved in methanol

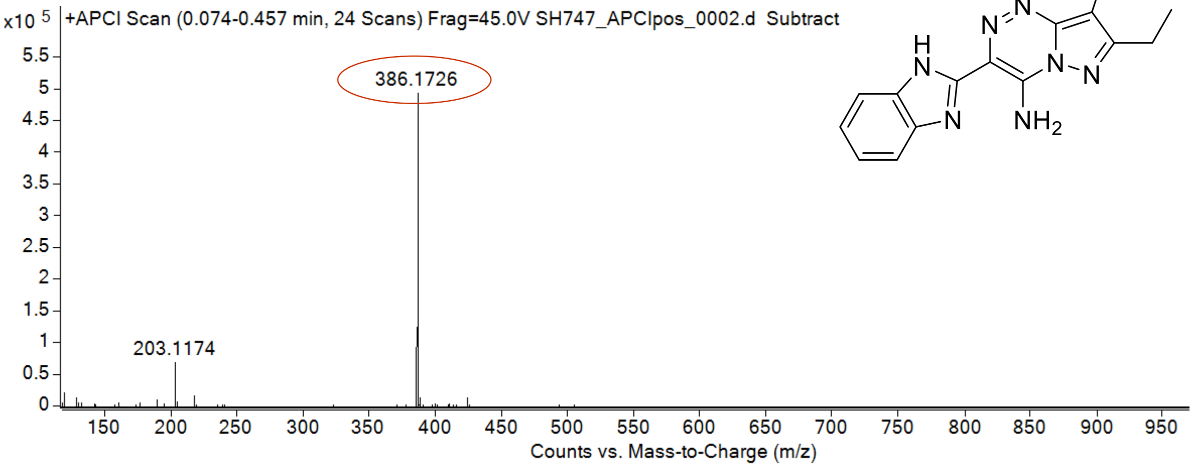

calculated mass: $[M+H]^+$ = 386.1724 observed: $[M+H]^+$ = 386.1726 mass accuracy = 0.5 ppm

$^{1}H$ (500 MHz) and $^{13}C$ NMR (126 MHz) spectra of **8** in DMSO-$d_6$

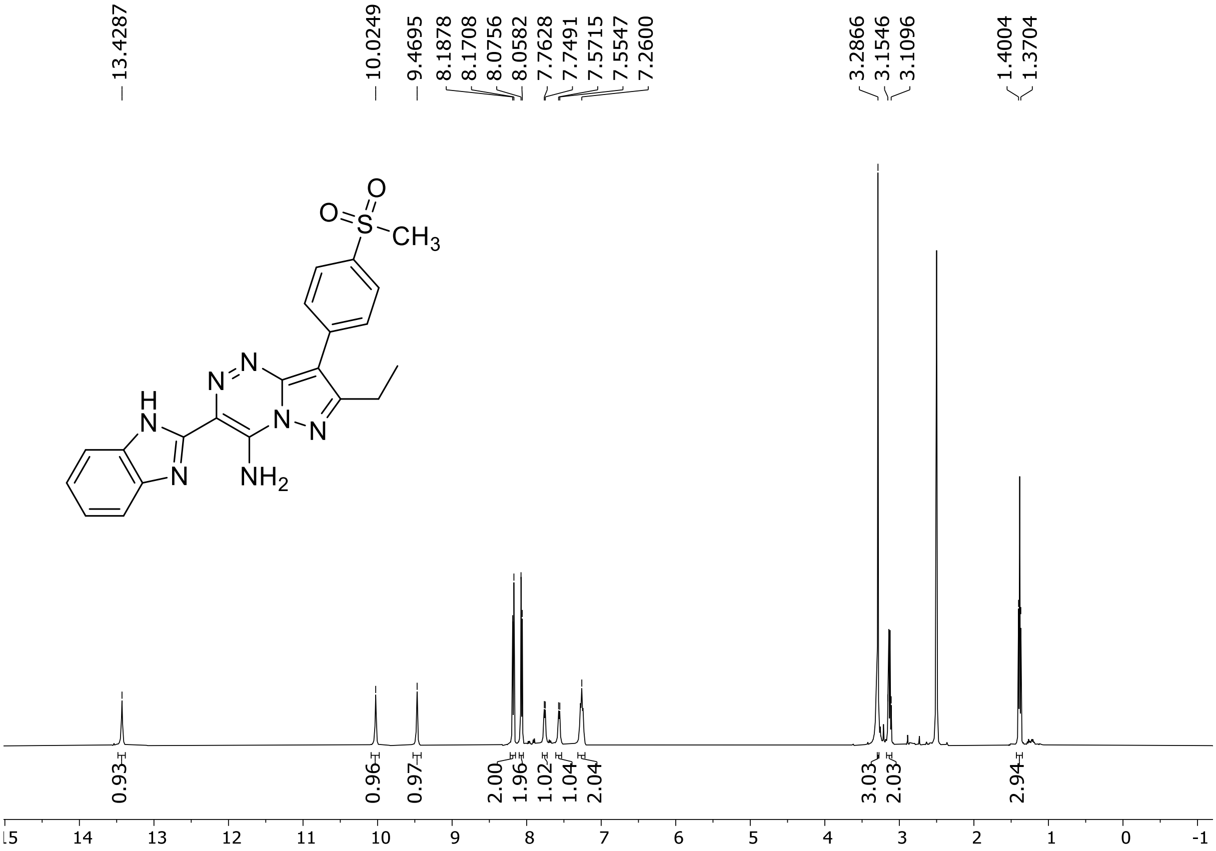

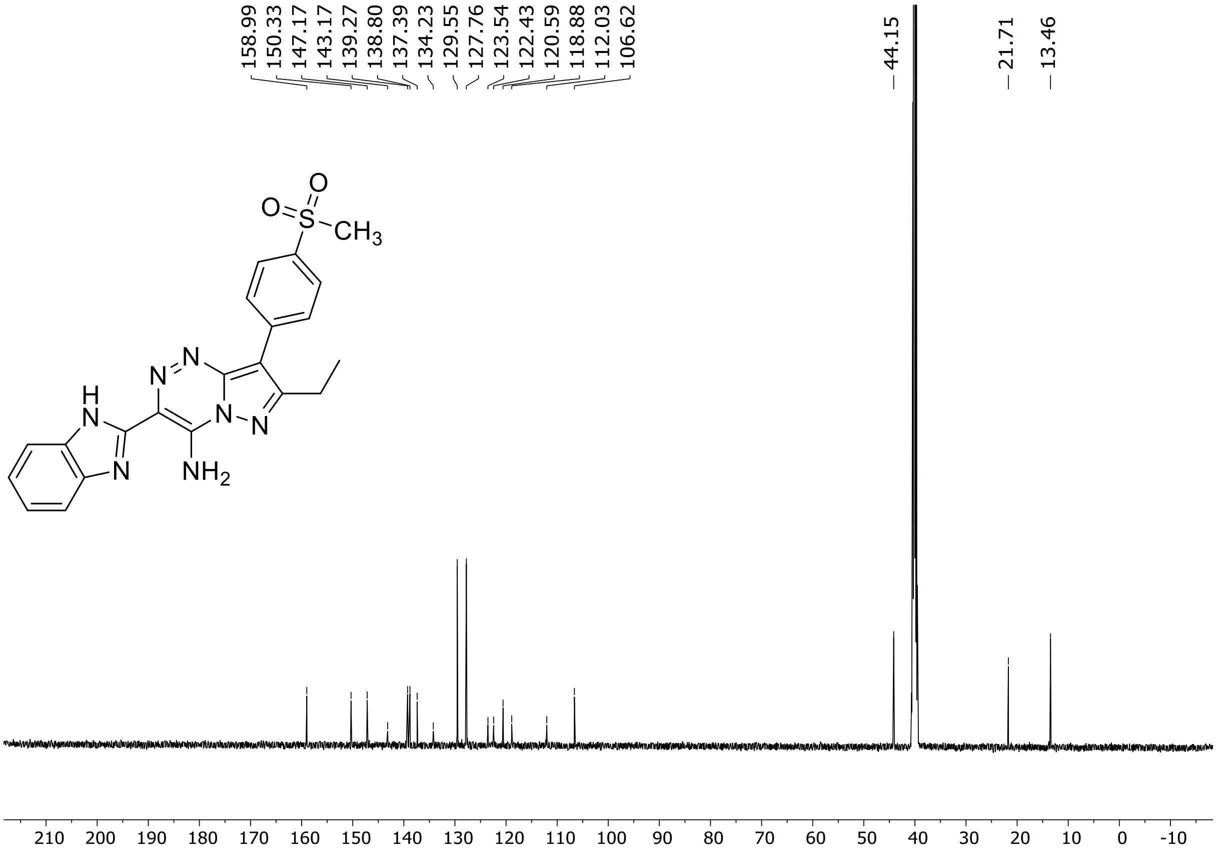

HRMS spectrum of **8**

**$C_{21}H_{19}N_7O_2S$** **mono *m*/*z* 433.1321**

**ESI -** (MMI)
nitrogen flow 5 L/min, gas temperature 325°C, nebulizer 45 psi, skimmer 65 V, capillary voltage 2500V, fragmentor 200 V, dissolved in methanol

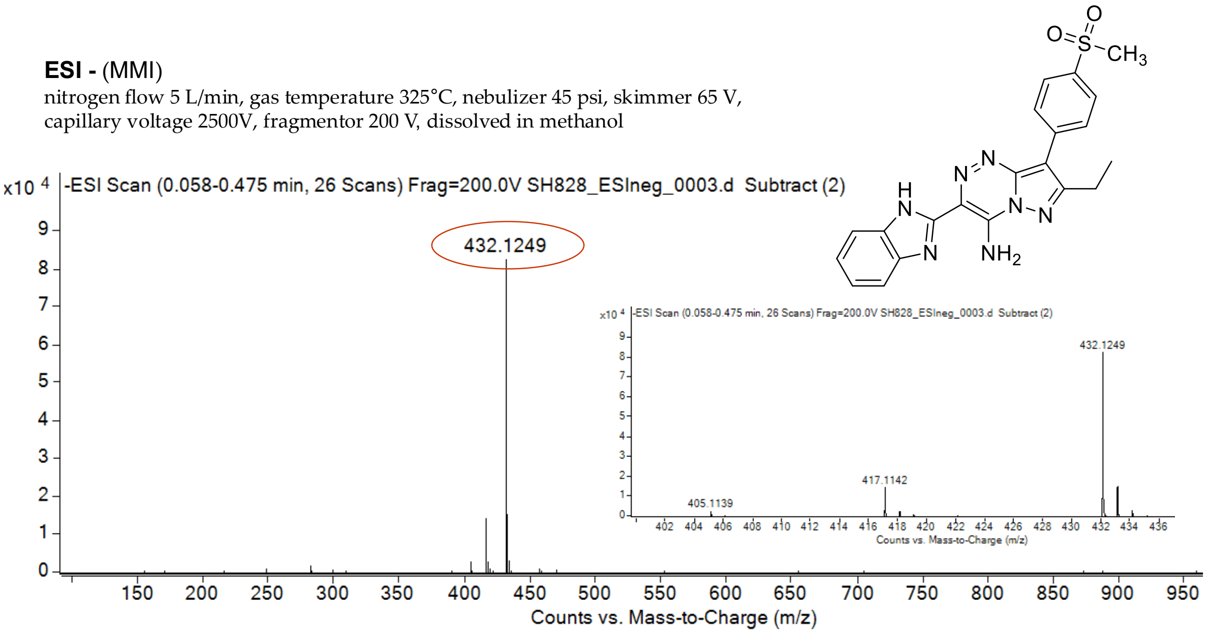

calculated mass: $[M-H]^-$ = 432.1248 observed: $[M-H]^-$ = 432.1249 mass accuracy = 0.2 ppm

$^{1}H$ (500 MHz) and $^{13}C$ NMR (126 MHz) spectra of **9** in DMSO-$d_6$

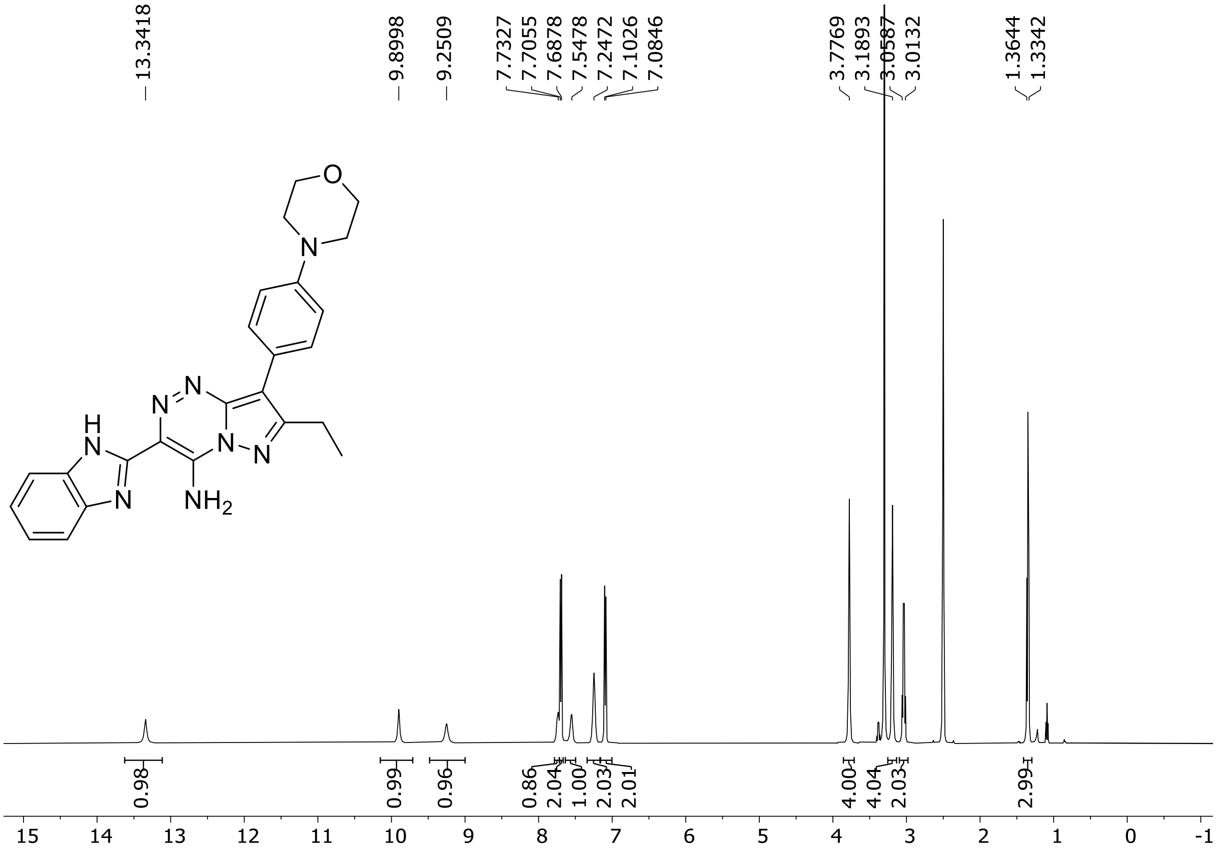

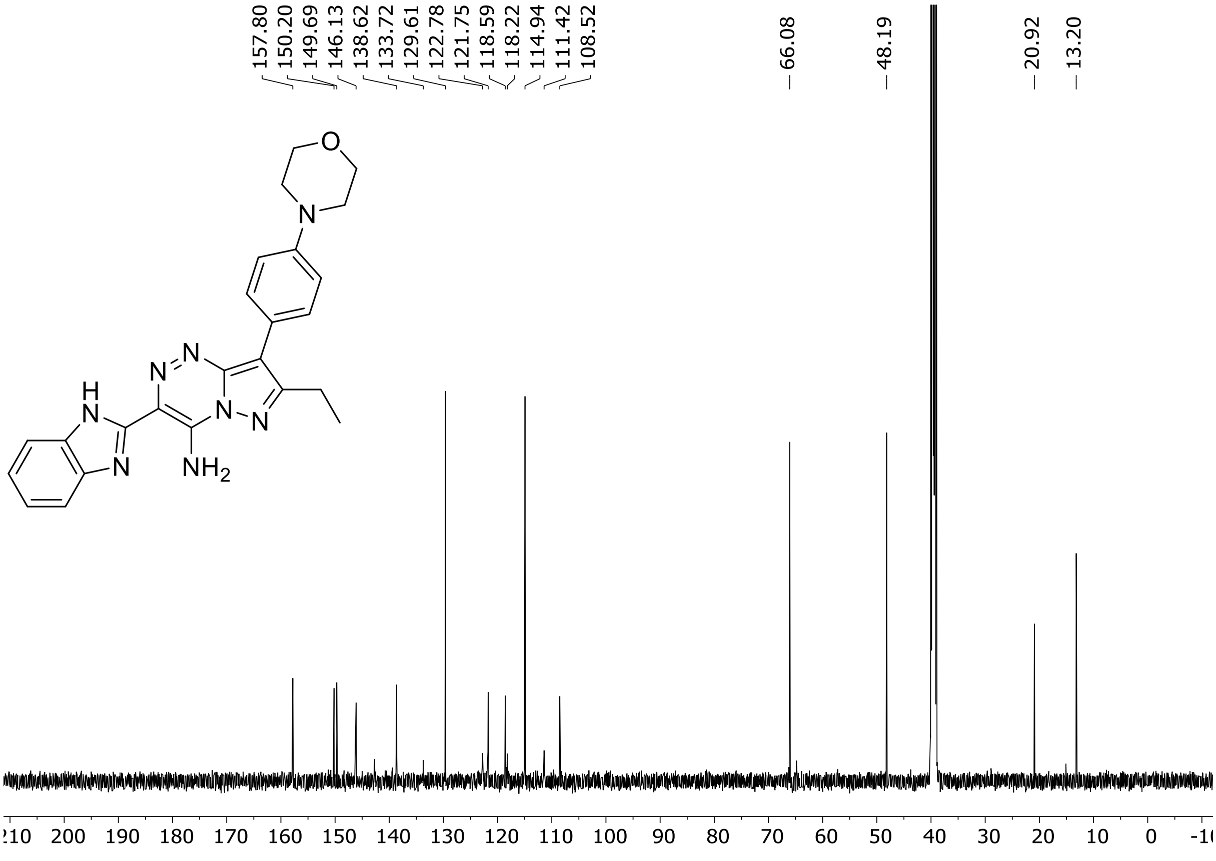

HRMS spectrum of **9**

$C_{24}H_{24}N_8O$ **mono *m/z* 440.2073**

**ESI -** (MMI)
nitrogen flow 5 L/min, gas temperature 325°C, nebulizer 45 psi, skimmer **65** V, capillary voltage 2500V, fragmentor 35 V, dissolved in methanol

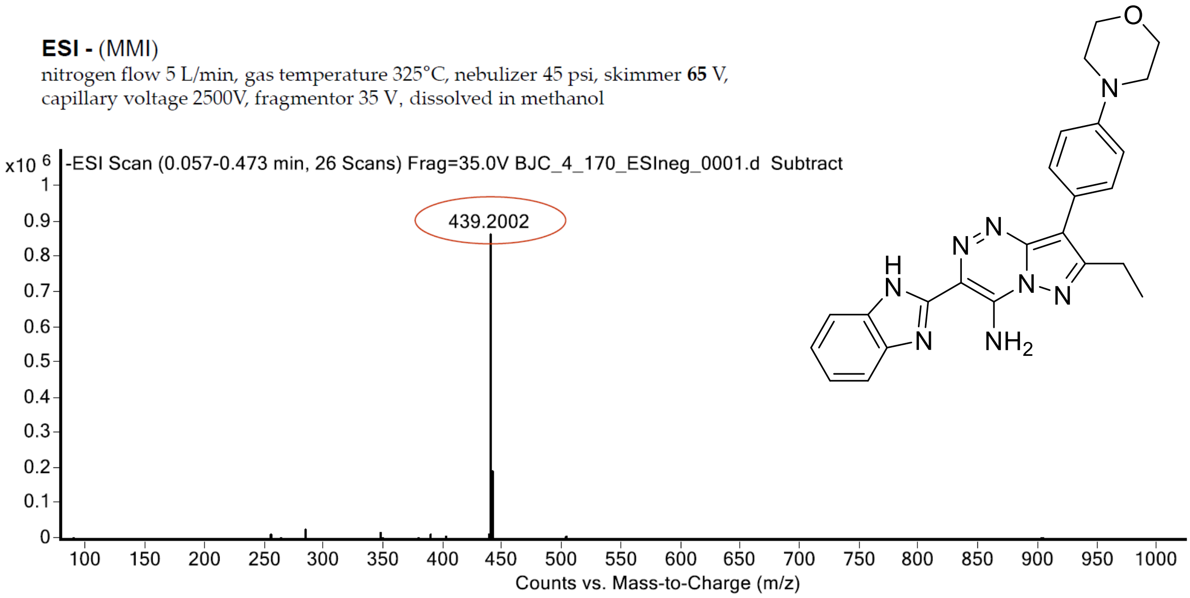

calculated mass: $[M\text{-}H]^-$ = 439.2000 observed: $[M\text{-}H]^-$ = 439.2002 mass accuracy = 0.4 ppm

$^{1}H$ (500 MHz) and $^{13}C$ NMR (126 MHz) spectra of **10** in DMSO-$d_6$

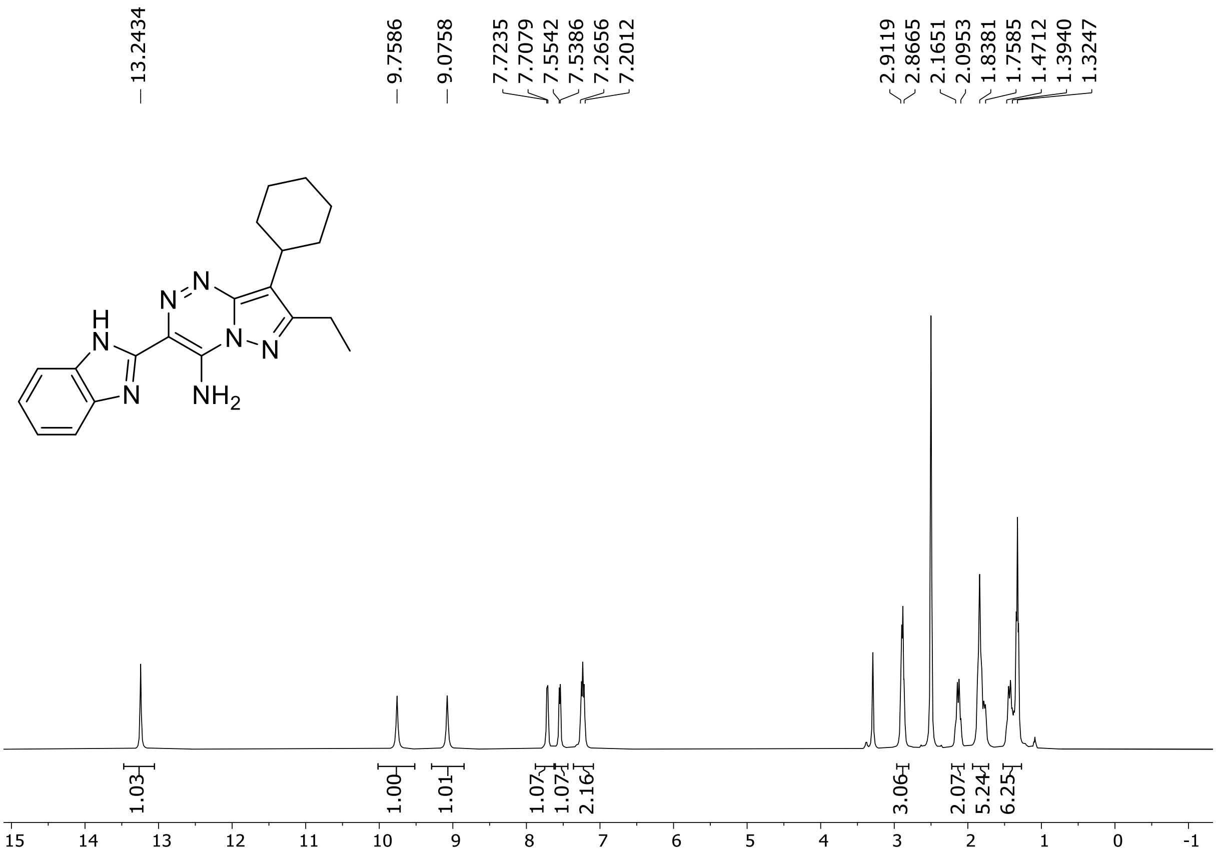

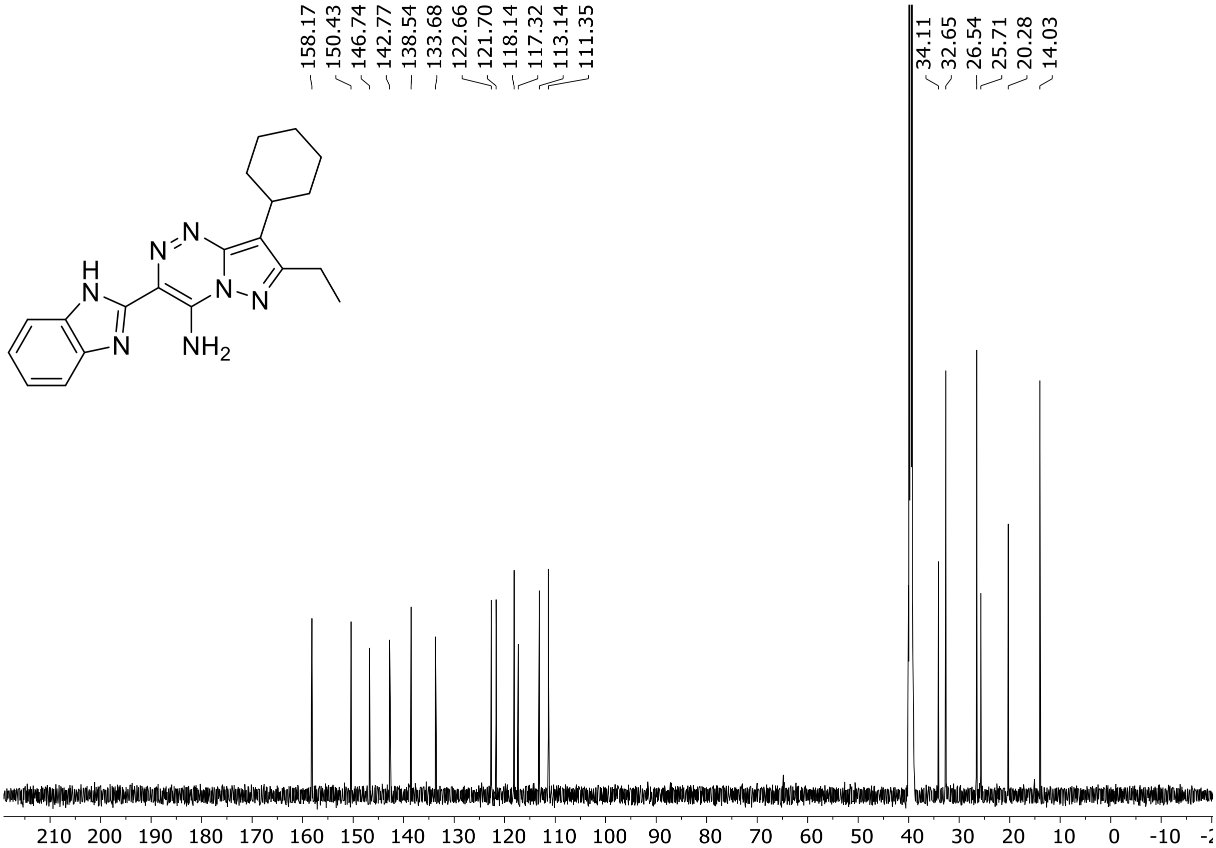

HRMS spectrum of **10**

$C_{20}H_{23}N_7$ **exact mass: 361.2015**

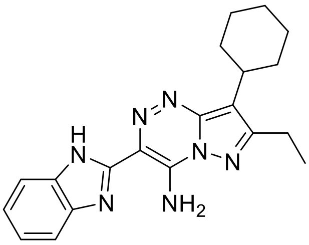

**APCI +** (MMI)
nitrogen flow 3 L/min, gas temperature 325°C, nebulizer 45 psig, skimmer 65 V, vaporizer 200°C, fragmentor 10 V, dissolved in methanol

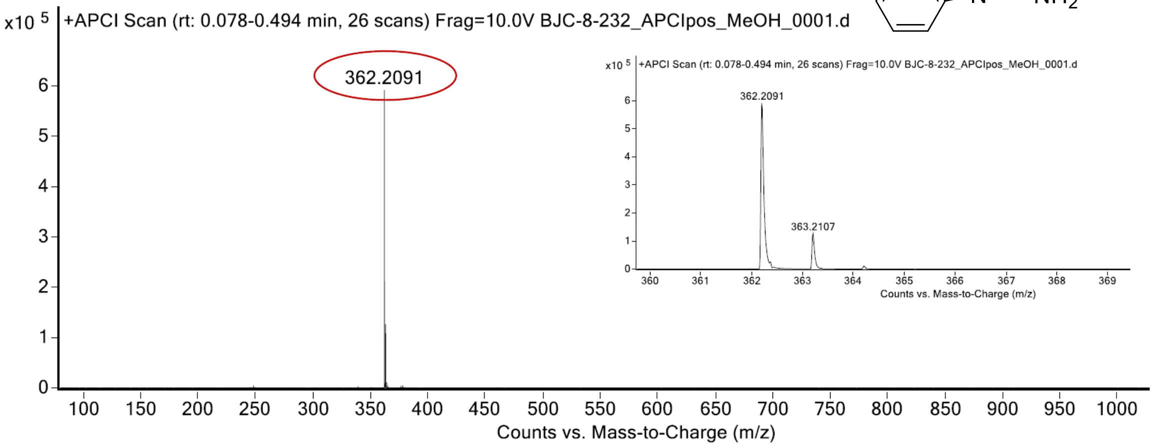

expected mass: $[M+H]^+ = 362.2088$ observed mass: $[M+H]^+ = 362.2091$ mass accuracy = 0.8 ppm

[1]H (500 MHz) and [13]C NMR (126 MHz) spectra of **11** in DMSO-$d_6$

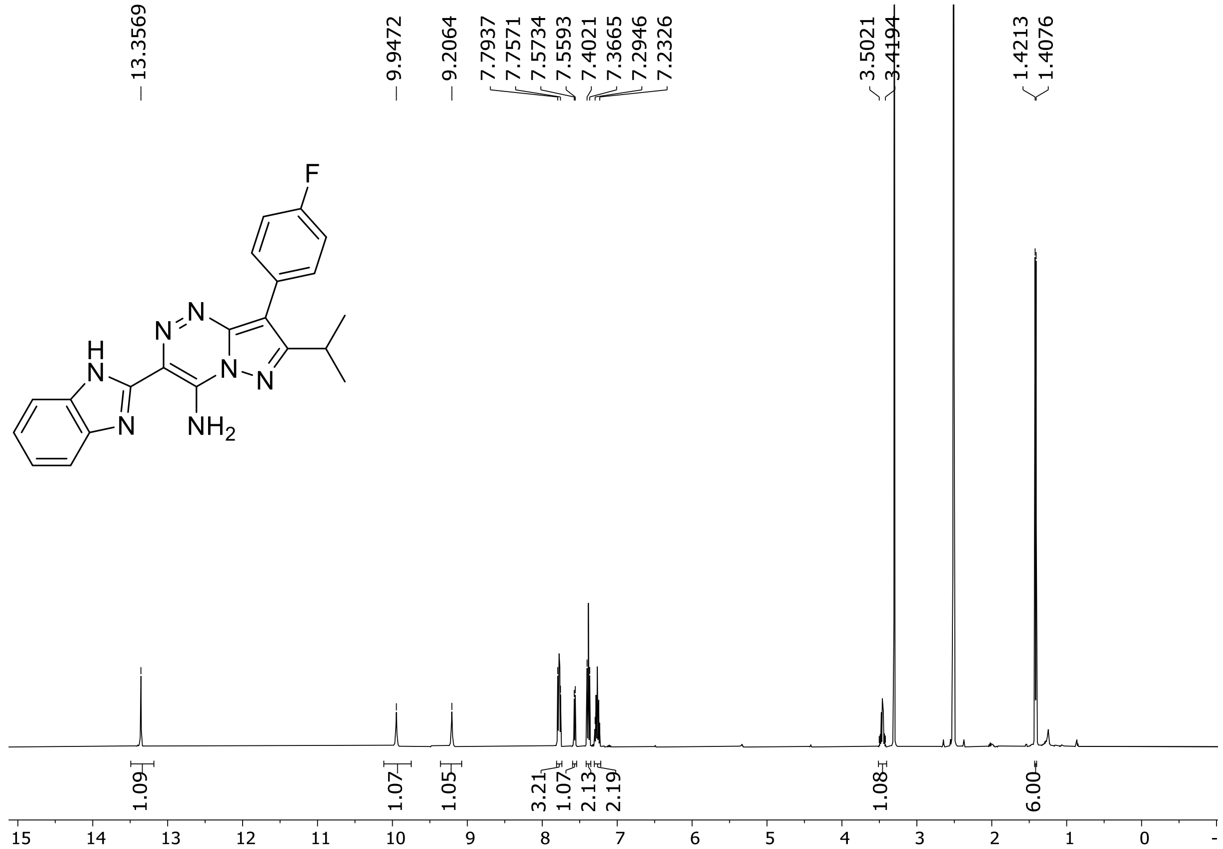

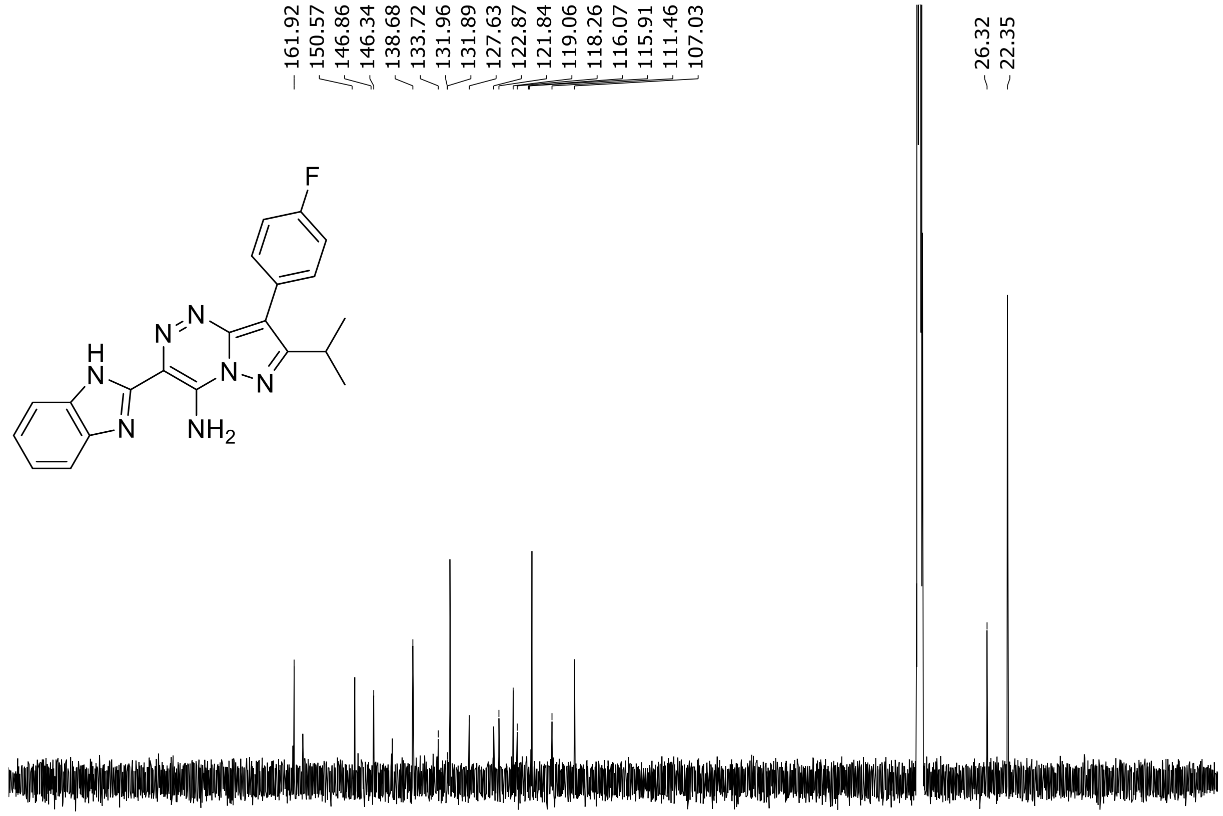

HRMS spectrum of **11**

**$C_{21}H_{18}FN_7$** **mono m/z = 387.1602**

**ESI -** (MMI)
nitrogen flow 5 L/min, gas temperature 300°C, vaporizer 250°C,
nebulizer 45 psi, skimmer **70** V, fragmentor 40 V, dissolved in DMSO

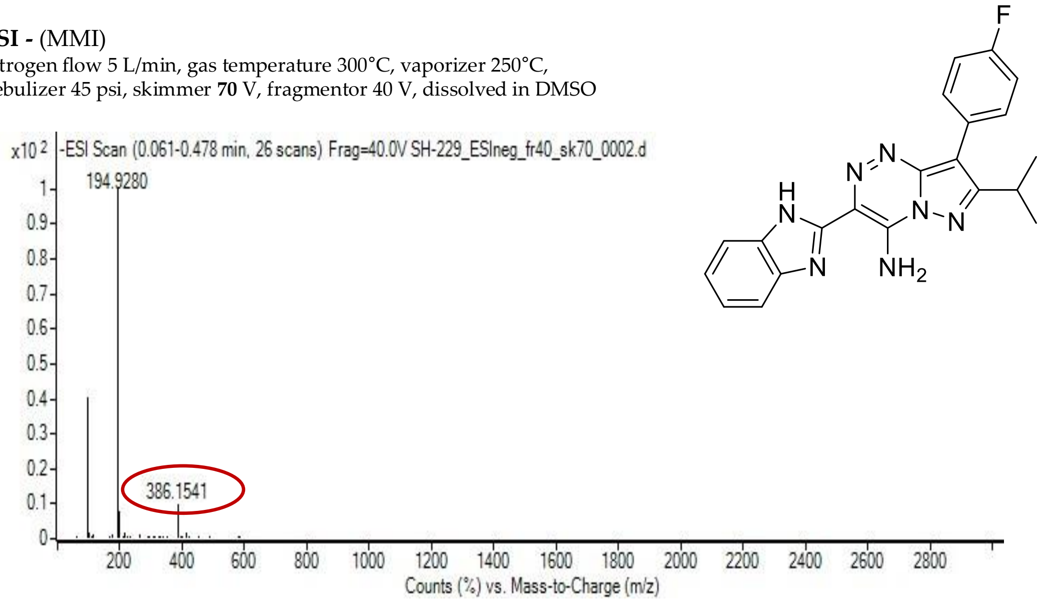

calculated mass: [M-H]$^-$ = 386.1535 observed: [M-H]$^-$ = 386.1541 m ax. mass error = 1.6 ppm

$^{1}H$ (500 MHz) and $^{13}C$ NMR (126 MHz) spectra of **12** in DMSO-$d_6$

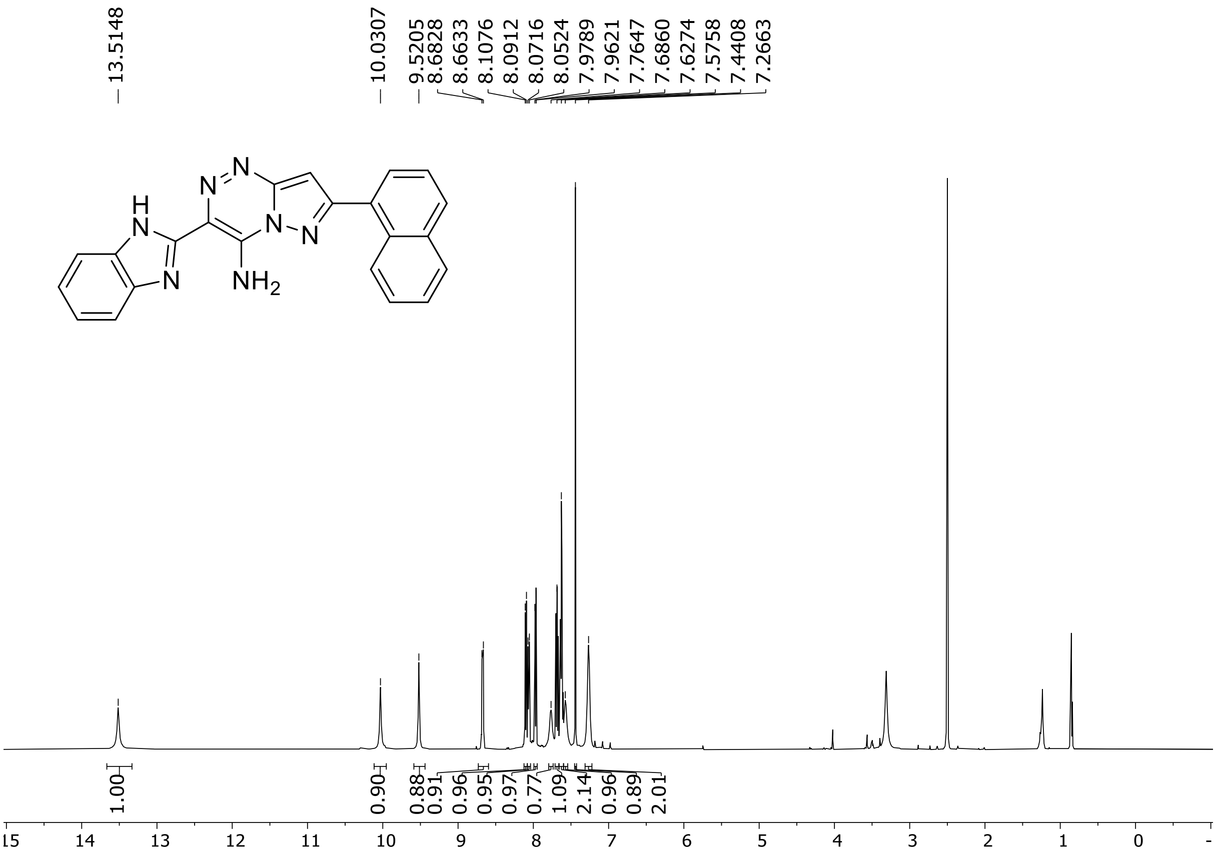

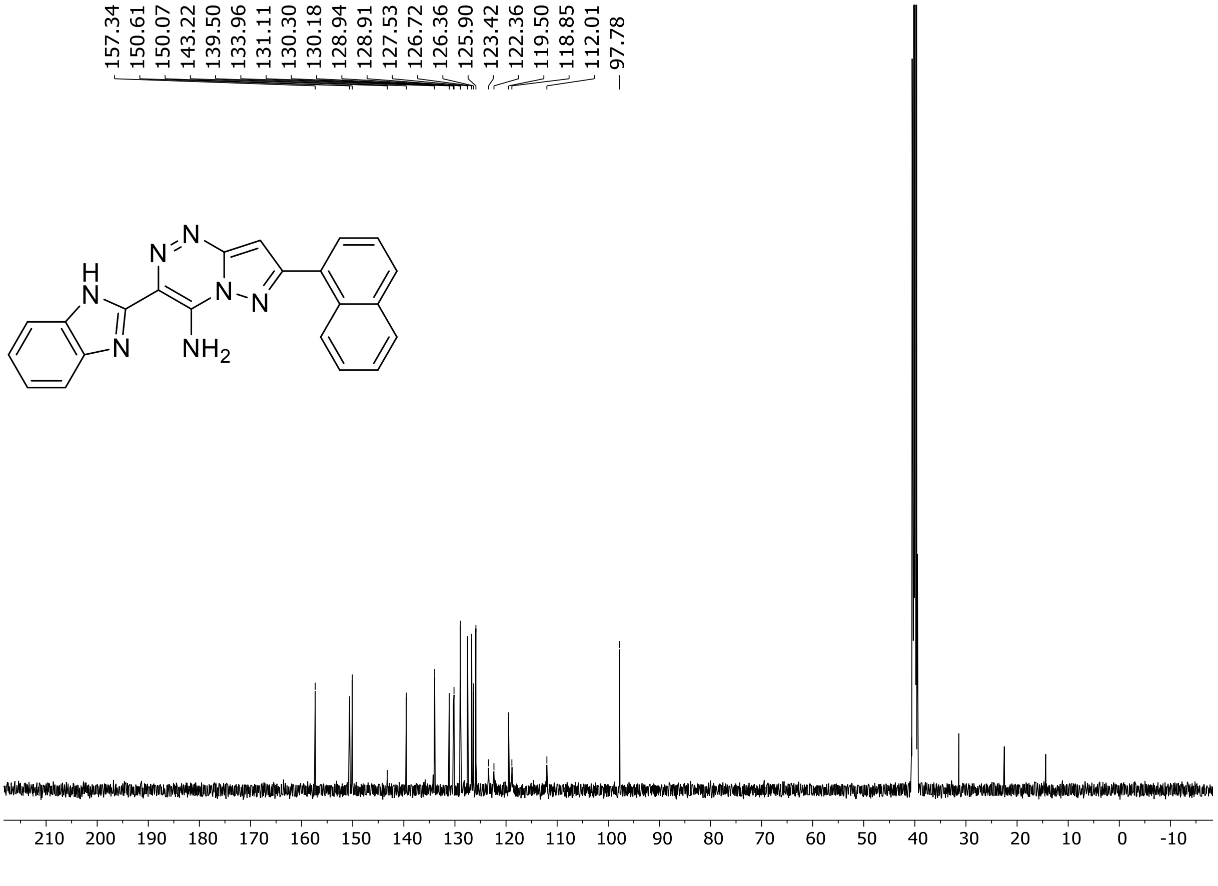

HRMS spectrum of **12**

$C_{22}H_{15}N_7$ **mono m/z = 377.1389**

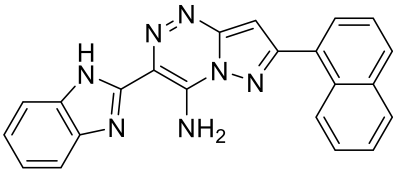

**APCI +** (MMI)
nitrogen flow 5 L/min, gas temperature 300°C, nebulizer 45 psi, vaporizer 200°C
skimmer **65** V, fragmentor **23** V, dissolved in methanol

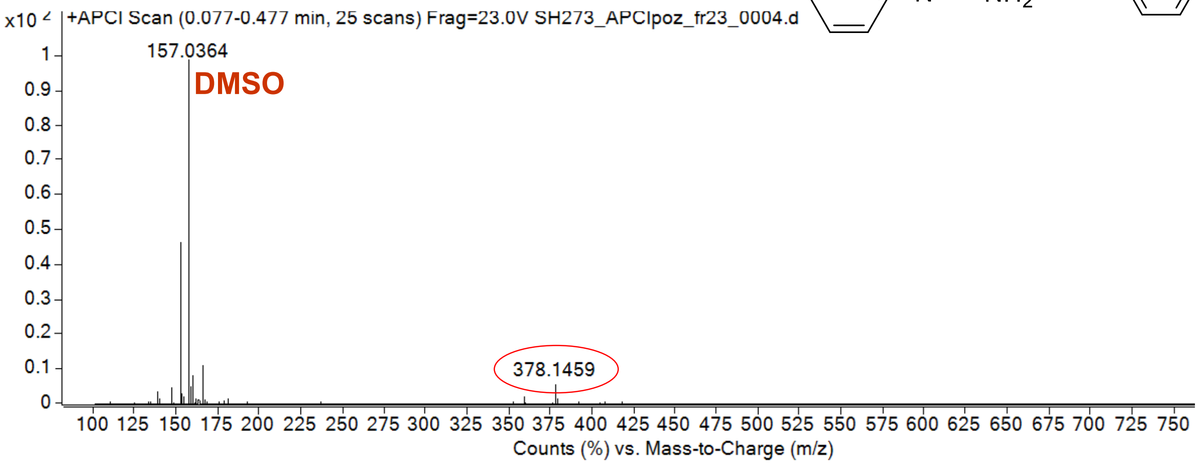

calculated mass: $[M+H]^+ = 378.1462$ observed: $[M+H]^+ = 378.1459$ max. mass error = 0.7 ppm

$^{1}H$ (500 MHz) and $^{13}C$ NMR (126 MHz) spectra of **13** in DMSO-$d_6$

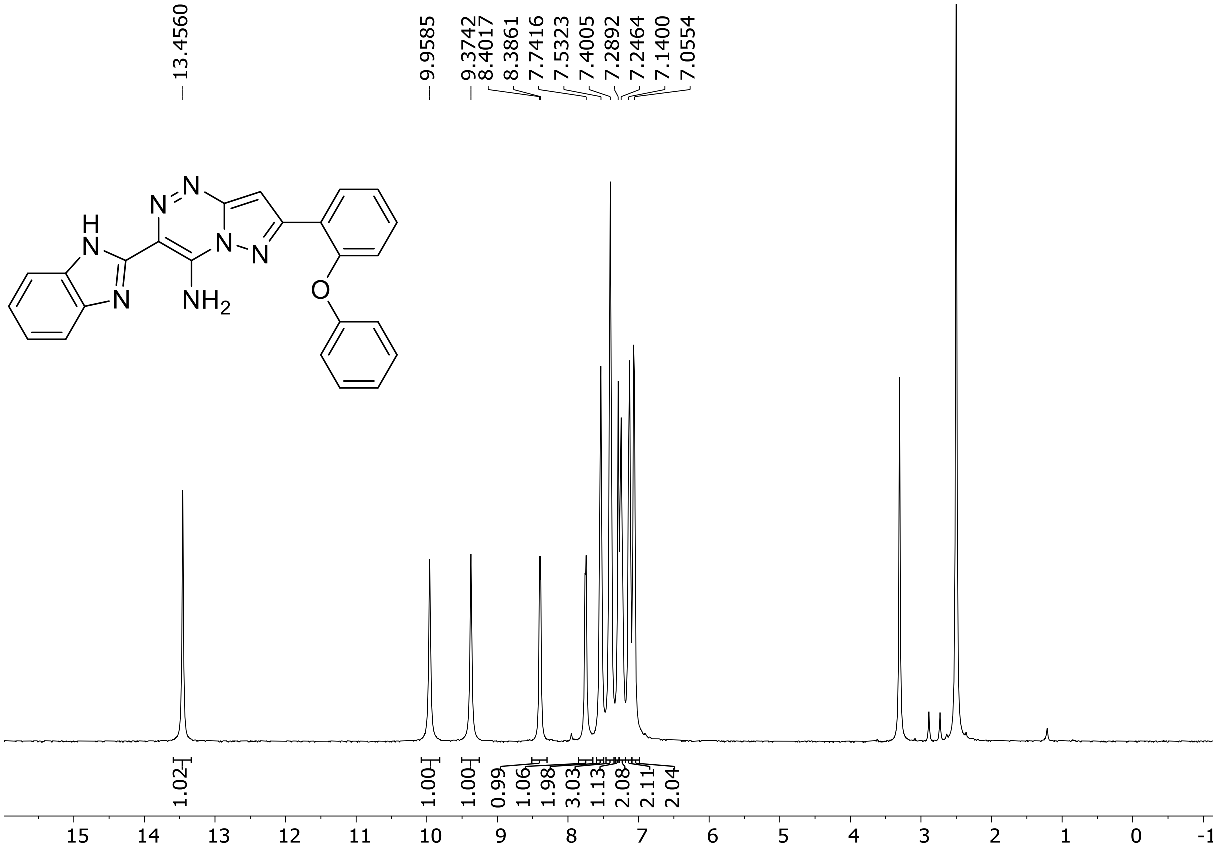

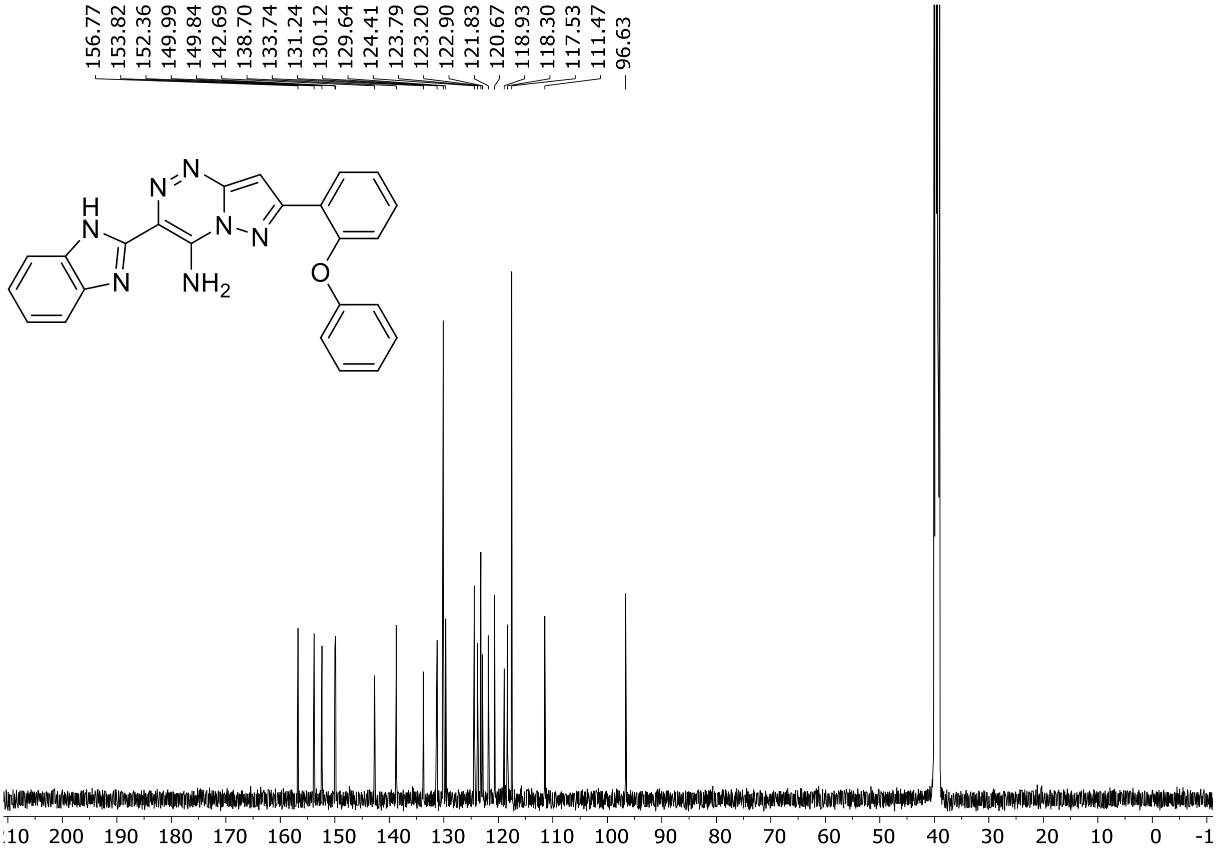

HRMS spectrum of **13**

$C_{24}H_{17}N_7O$ **mono m/z = 419.1495**

**ESI +** (MMI)
nitrogen flow 5 L/min, gas temperature 300°C, nebulizer 45 psi, skimmer 60 V, fragmentor 30 V, dissolved in methanol

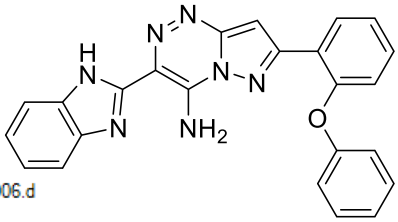

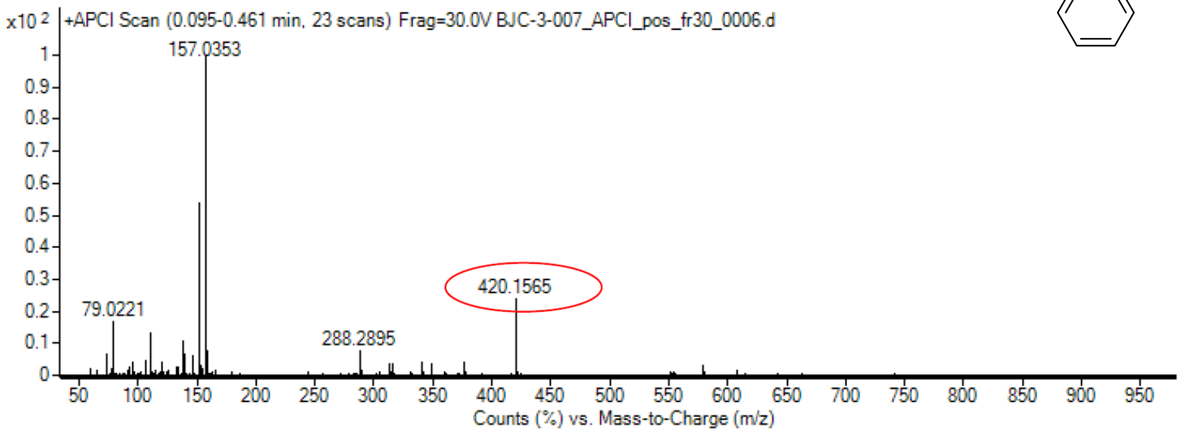

calculated mass: $[M+H]^+$ = 420.1567 observed: $[M+H]^+$ = 420.1565 max. mass error = 0.5 ppm

[1]H (500 MHz) and [13]C NMR (126 MHz) spectra of **14** in DMSO-$d_6$

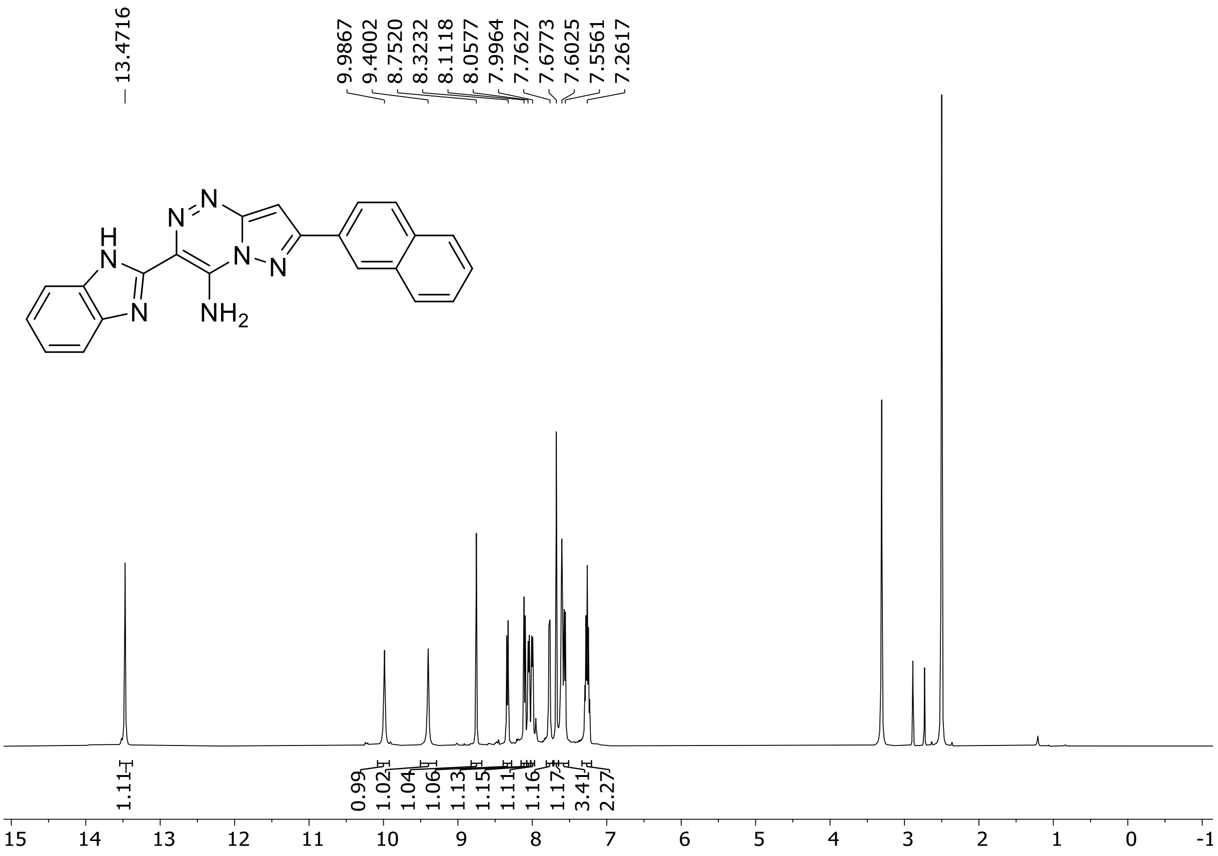

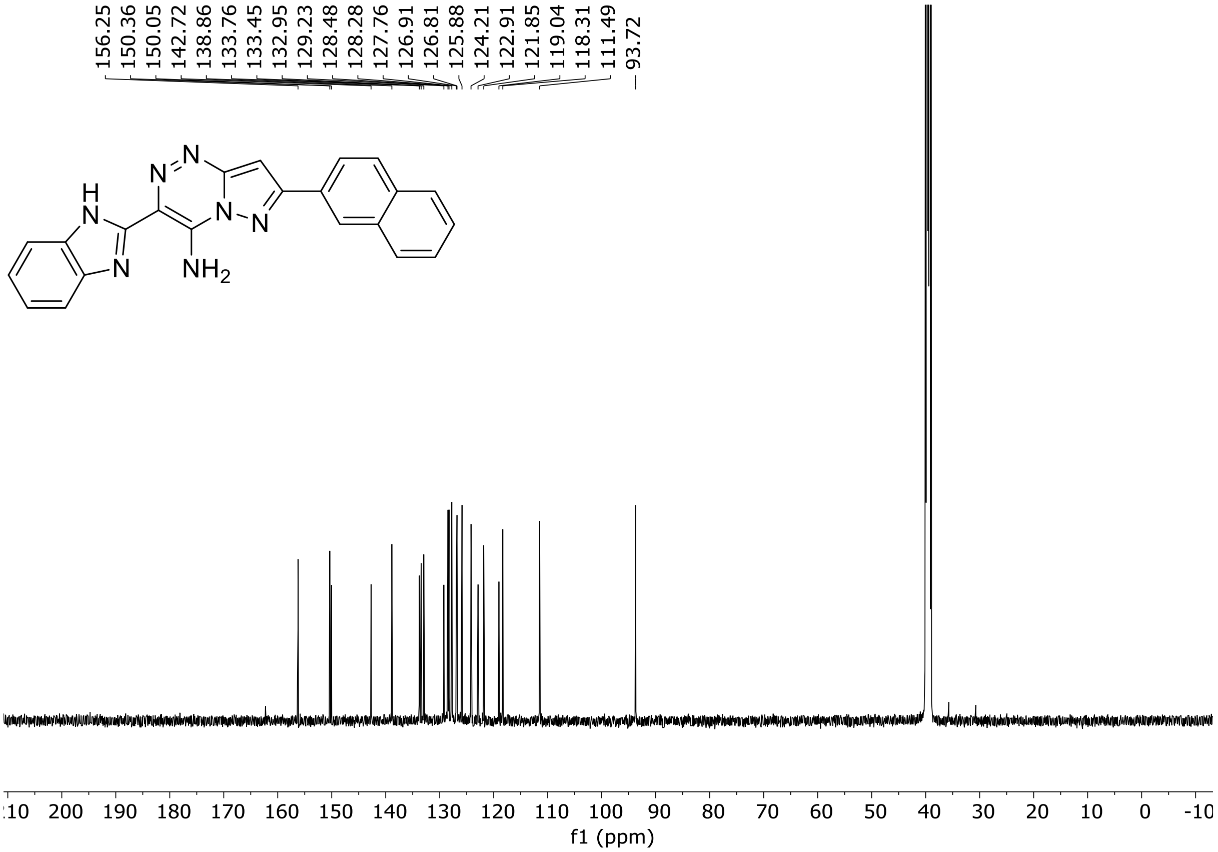

HRMS spectrum of **14**

$C_{22}H_{15}N_7$ **mono m/z = 377.1389**

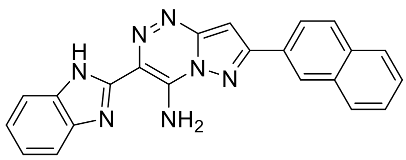

**APCI +** (MMI)
nitrogen flow 5 L/min, gas temperature 300°C, nebulizer 45 psi, skimmer 60 V, fragmentor 35 V, dissolved in methanol

ZOOM – range of interest

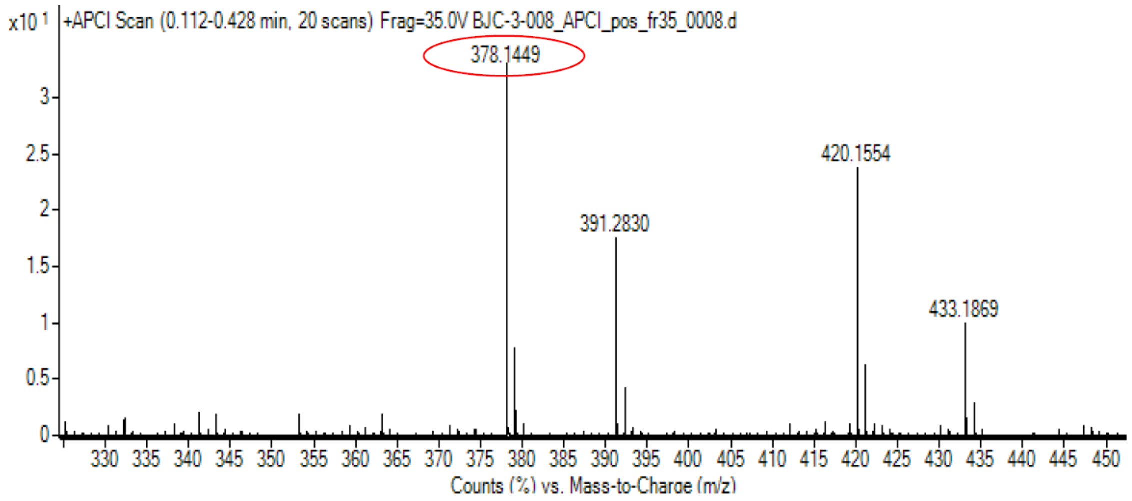

calculated mass: $[M+H]^+$= 378.1462 observed: $[M+H]^+$ = 378.1449 max. mass error = 3.4 ppm

[1]H (500 MHz) and [13]C NMR (126 MHz) spectra of **15** in DMSO-$d_6$

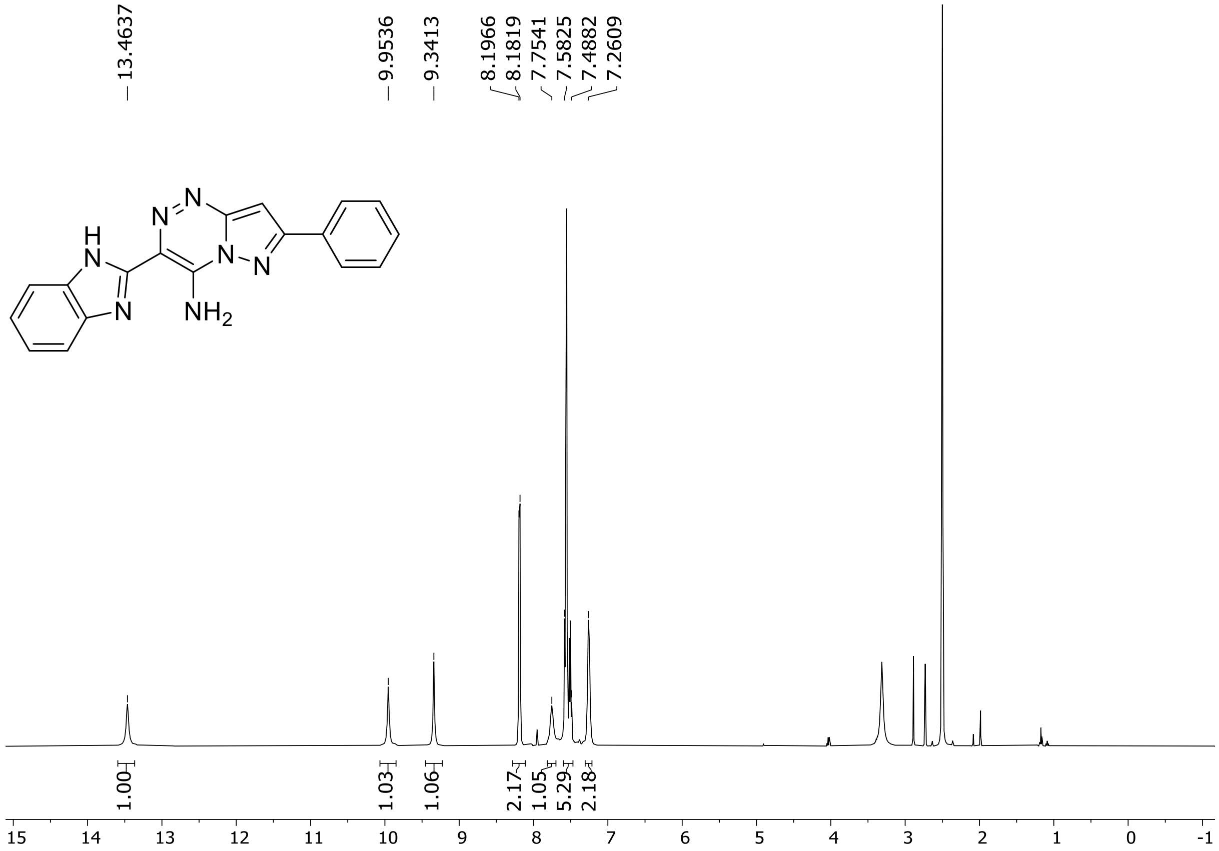

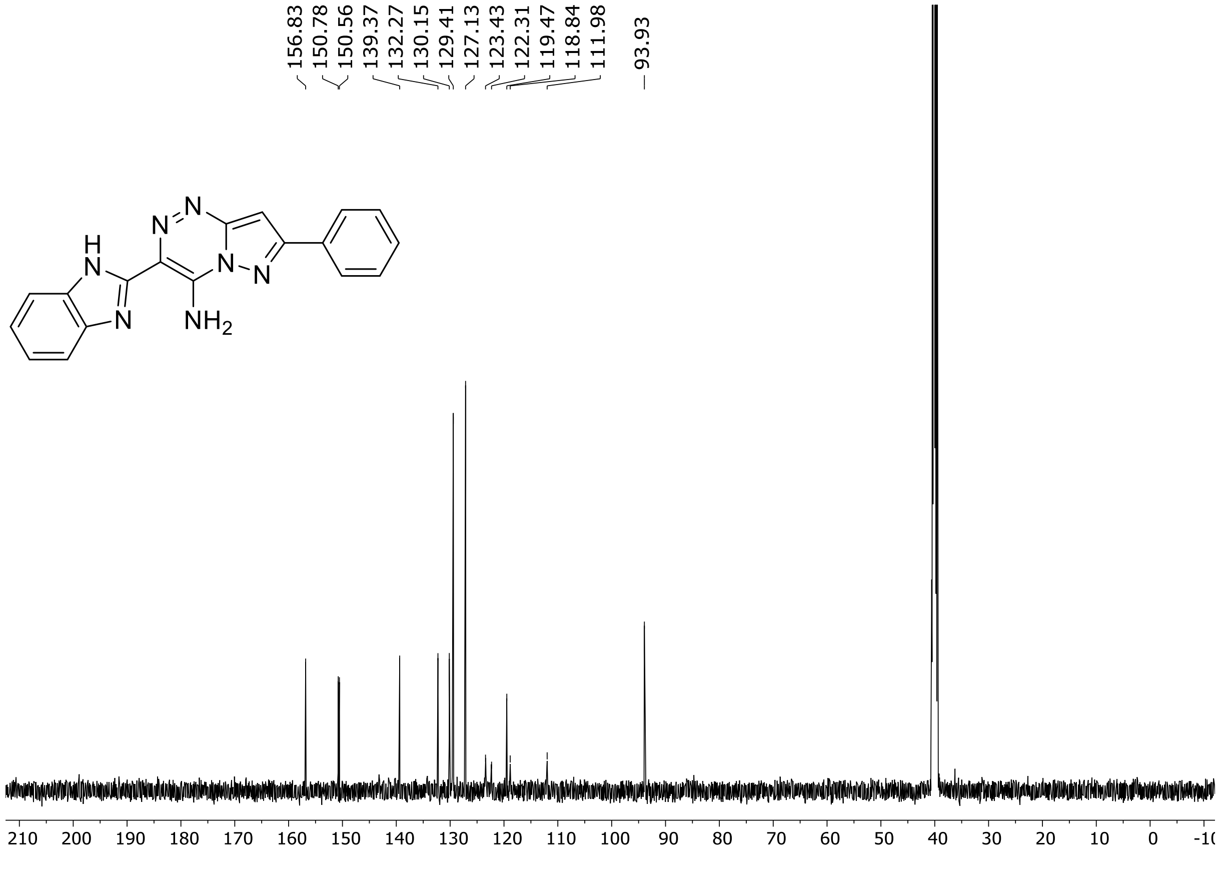

HRMS spectrum of **15**

**$C_{18}H_{13}N_7$** **exact mass: 327.1232**

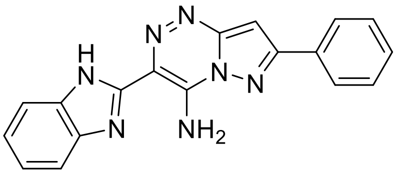

**APCI +** (MMI)
nitrogen flow 5 L/min, gas temperature 325°C, nebulizer 45 psig, skimmer 65 V, vaporizer 200°C, fragmentor 5 V, dissolved in methanol

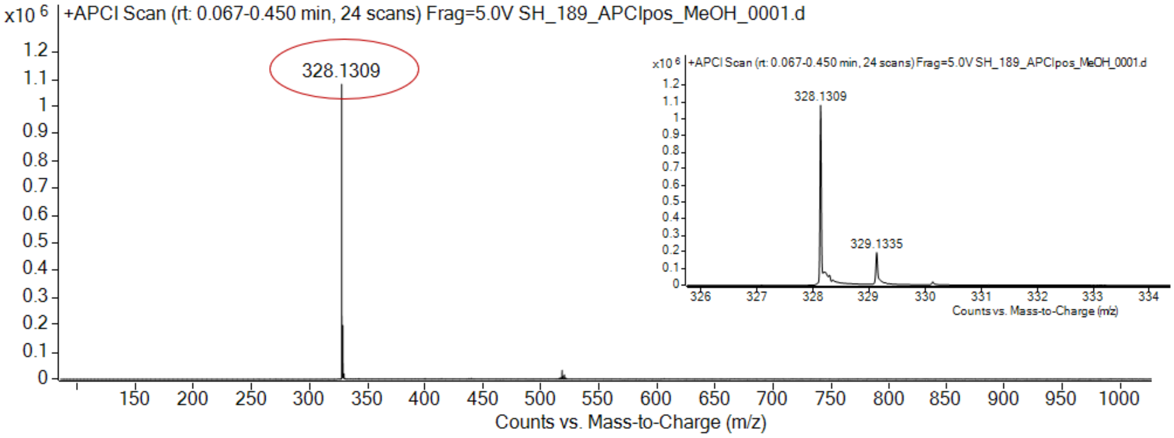

expected mass: $[M+H]^+$ = 328.1305 observed mass : $[M+H]^+$ = 328.1309 mass accuracy = 1.2 ppm

$^{1}H$ (300 MHz) and $^{13}C$ NMR (126 MHz) spectra of **16** in DMSO-$d_6$

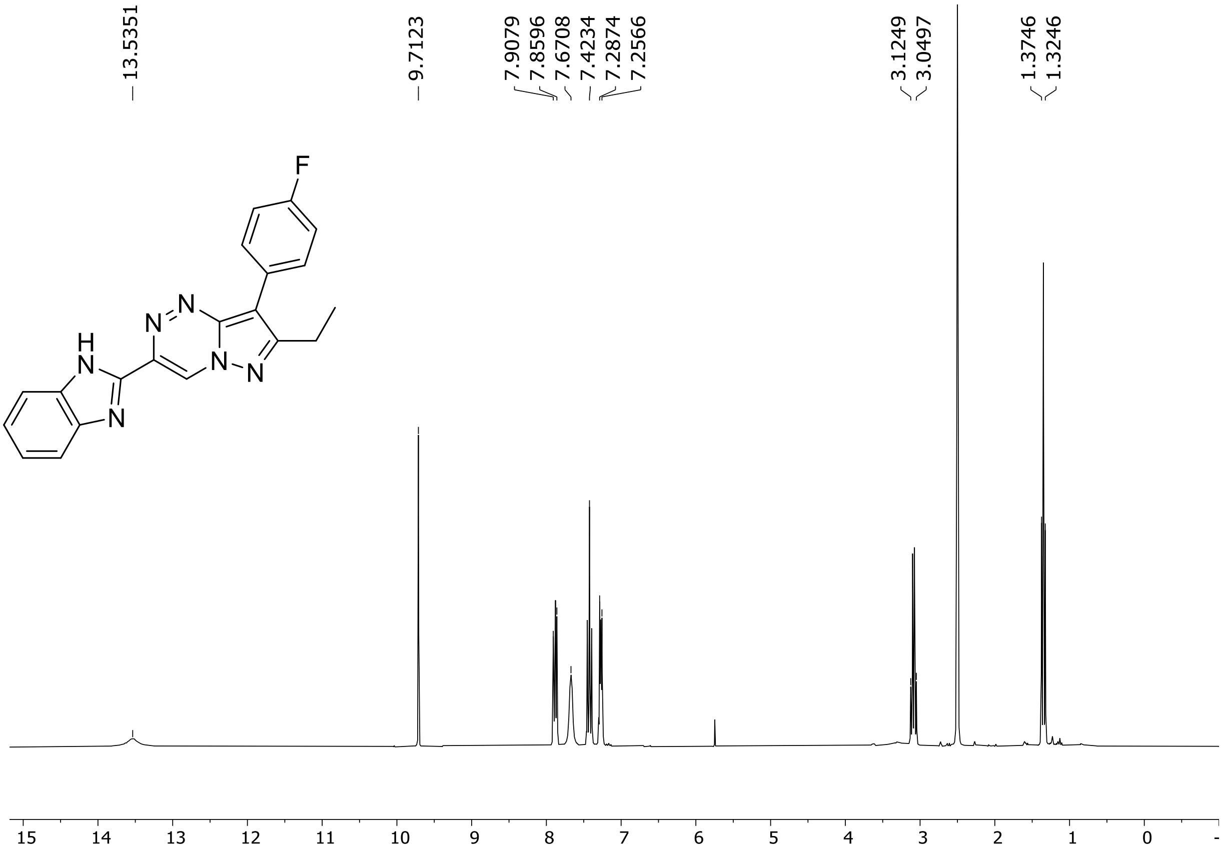

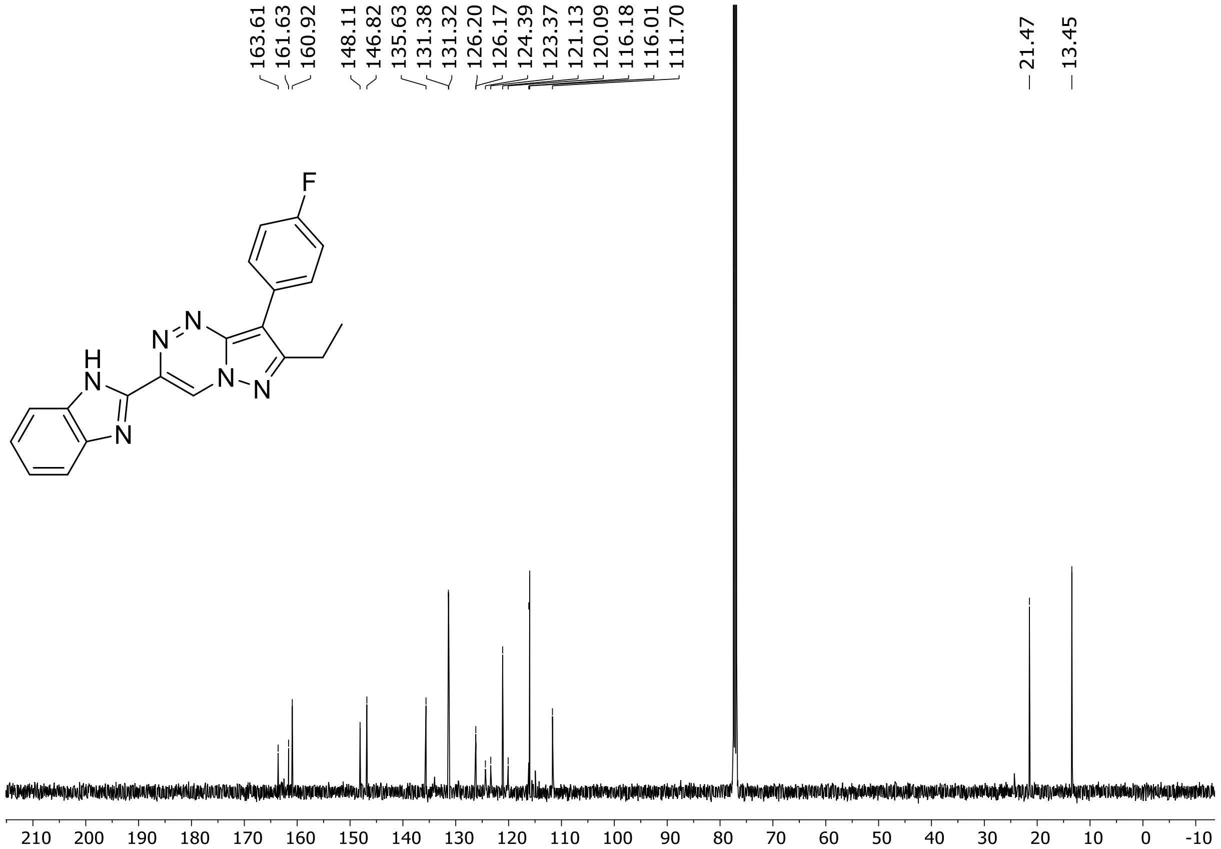

$^{19}F$ (282 MHz) NMR spectrum of **16** DMSO-$d_6$

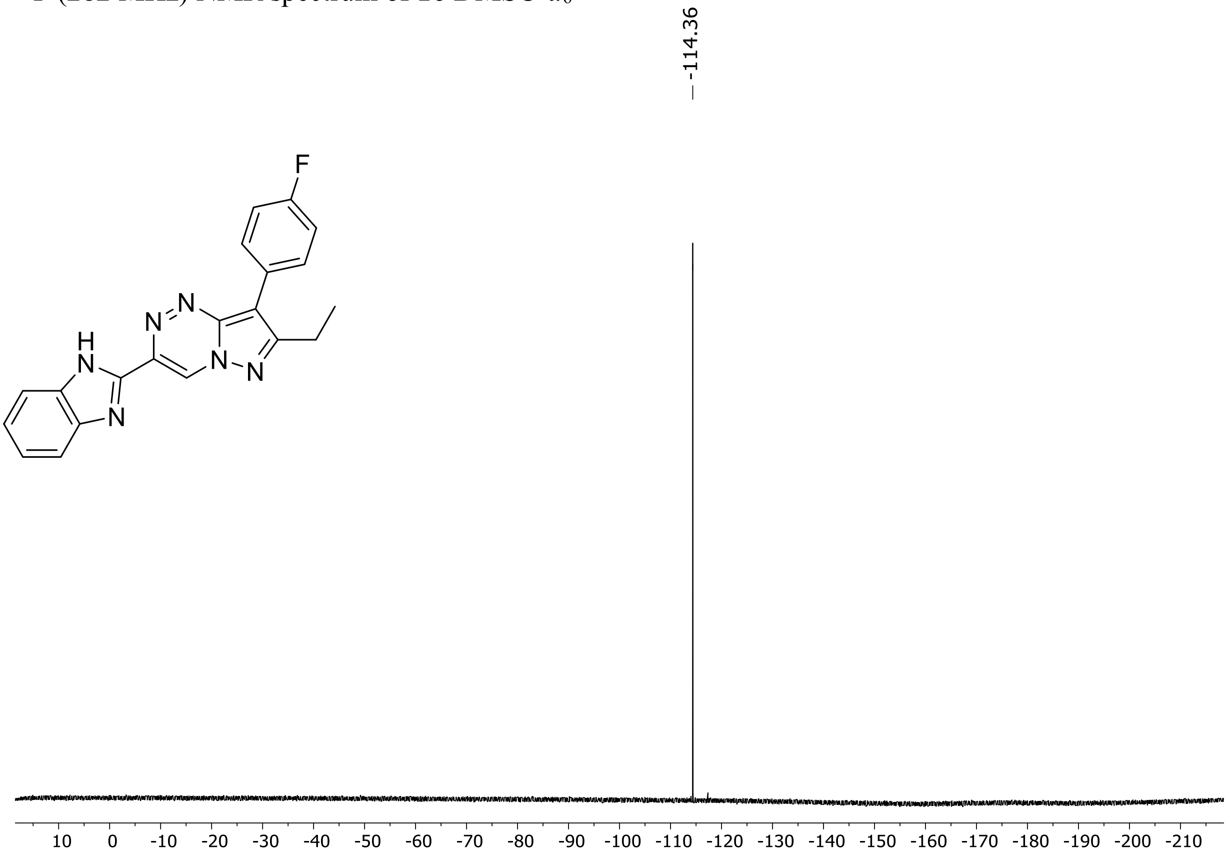

HRMS spectrum of **16**

**$C_{20}H_{15}FN_6$** **exact mass: 358.1342**

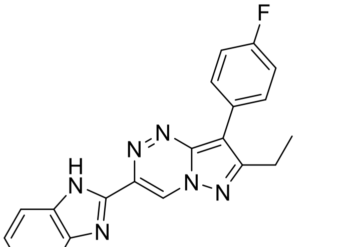

**APCI +** (MMI)
nitrogen flow 5 L/min, gas temperature 325°C, nebulizer 45 psig, skimmer 65 V, vaporizer 200°C, fragmentor 20 V, dissolved in methanol

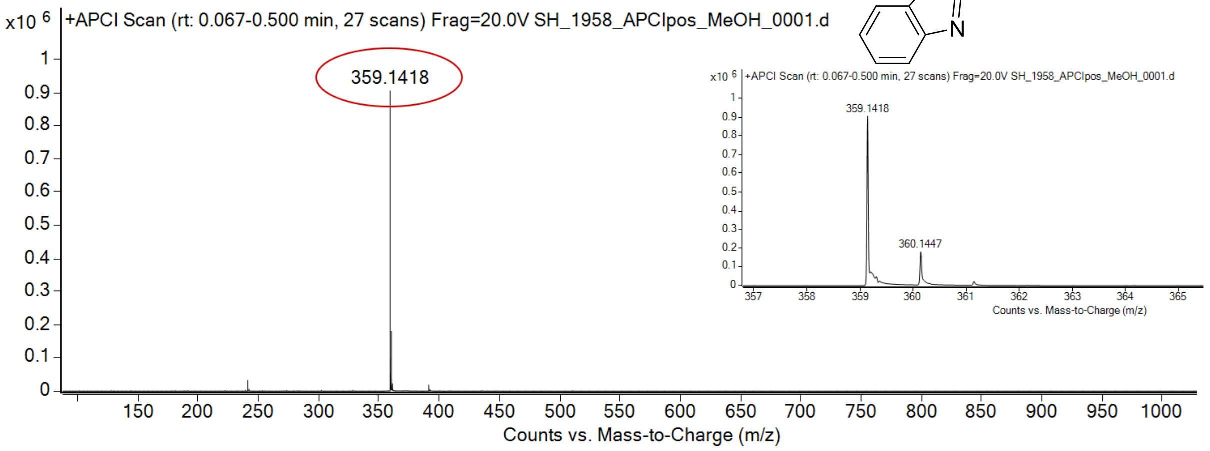

expected mass: [M+H]$^+$ = 359.1415 observed mass : [M+H]$^+$ = 359.1418 mass accuracy = 0.8 ppm

[1]H (500 MHz) and [13]C NMR (126 MHz) spectra of **17** in DMSO-$d_6$

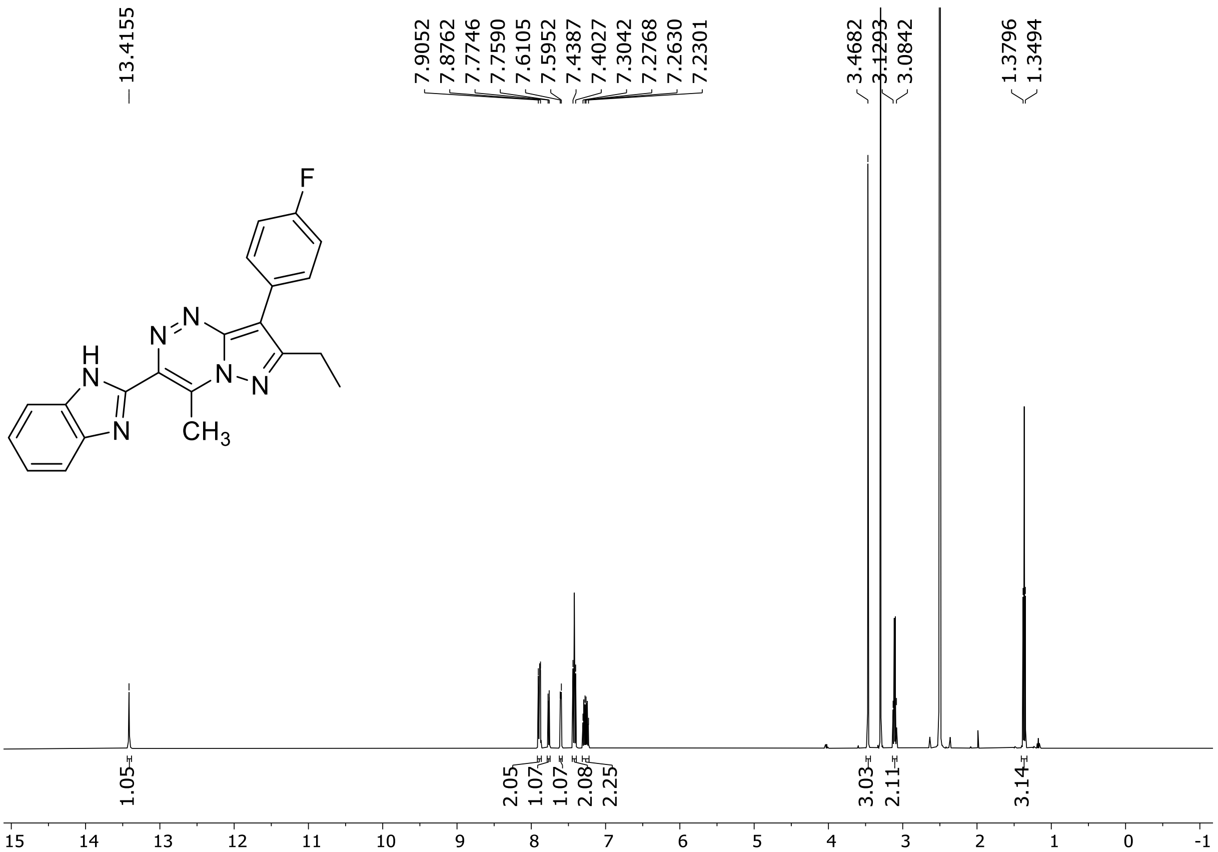

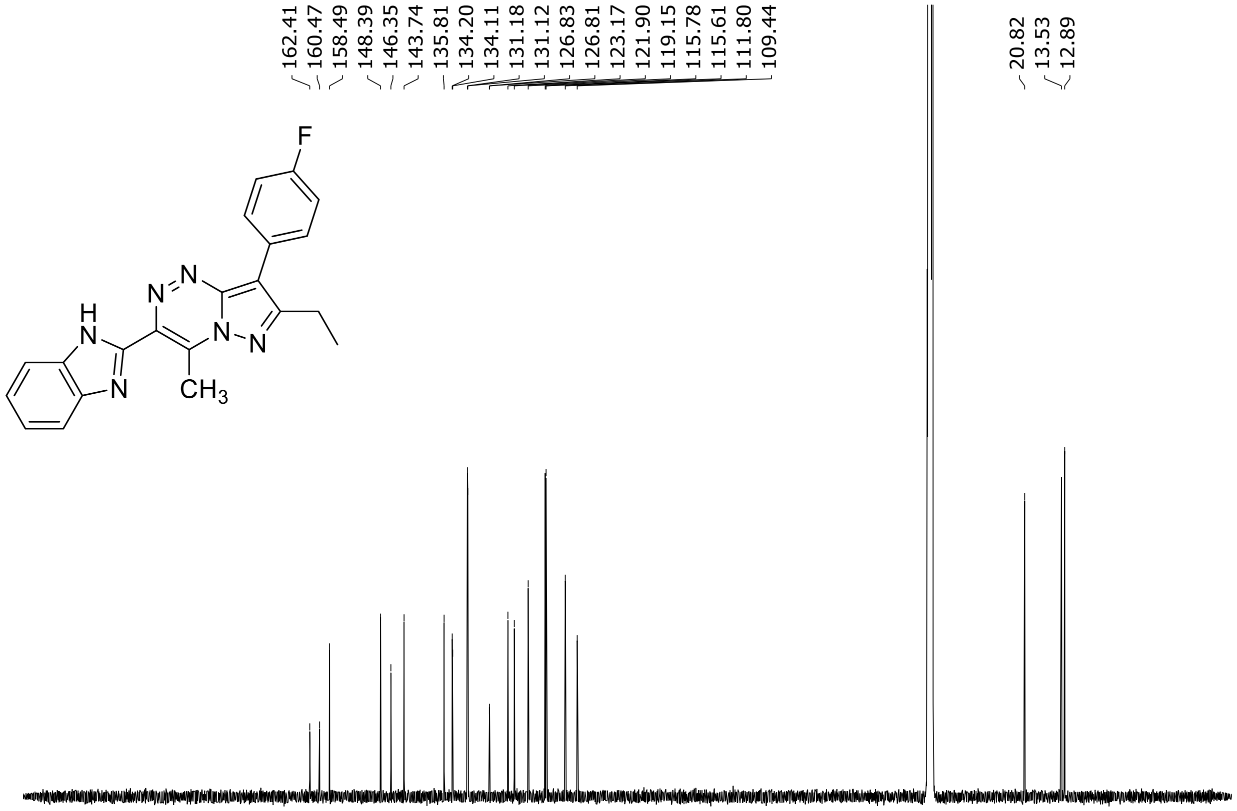

$^{19}F$ (282 MHz) NMR spectrum of **17** DMSO-$d_6$

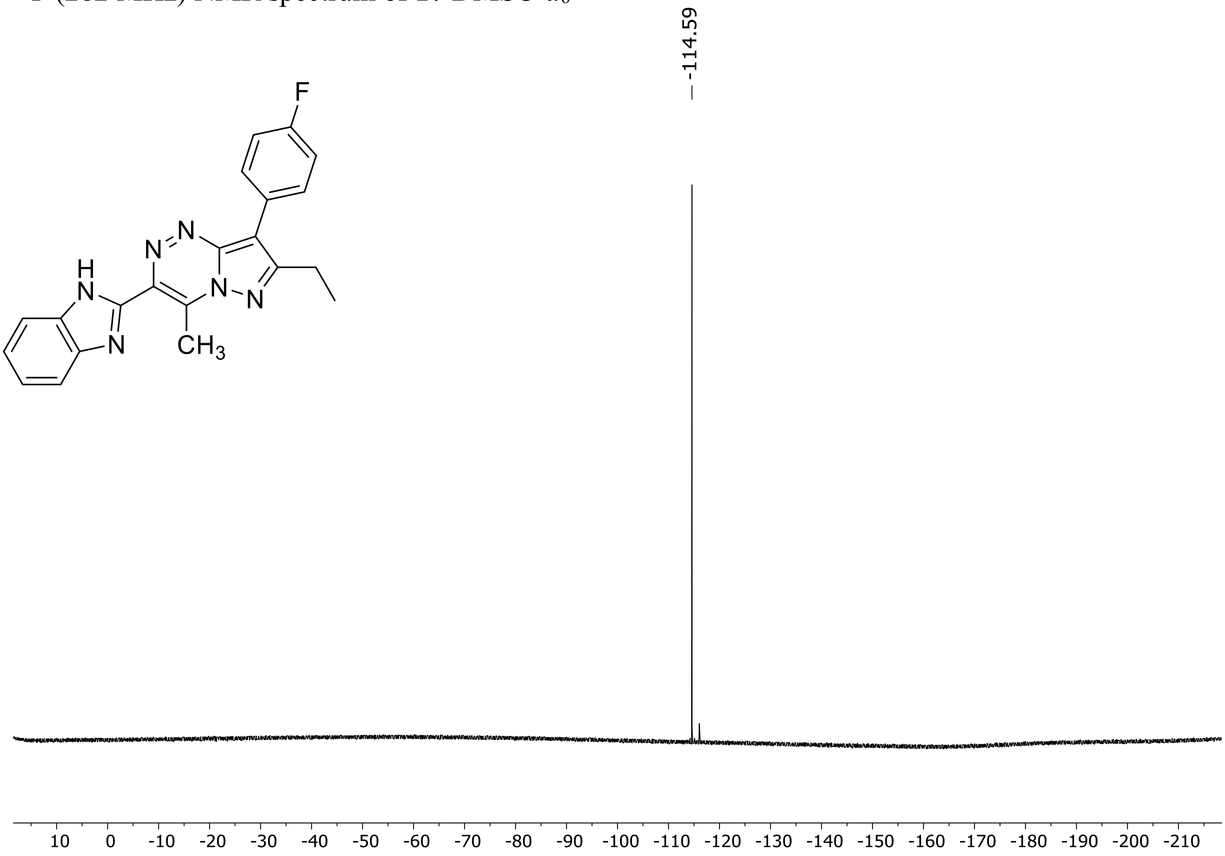

HRMS spectrum of **17**

**$C_{21}H_{17}FN_6$** **exact mass: 372.1499**

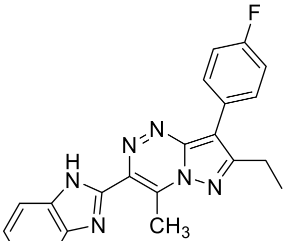

**APCI +** (MMI)
nitrogen flow 5 L/min, gas temperature 325°C, nebulizer 45 psig, skimmer 65 V, vaporizer 200°C, fragmentor 10 V, dissolved in methanol

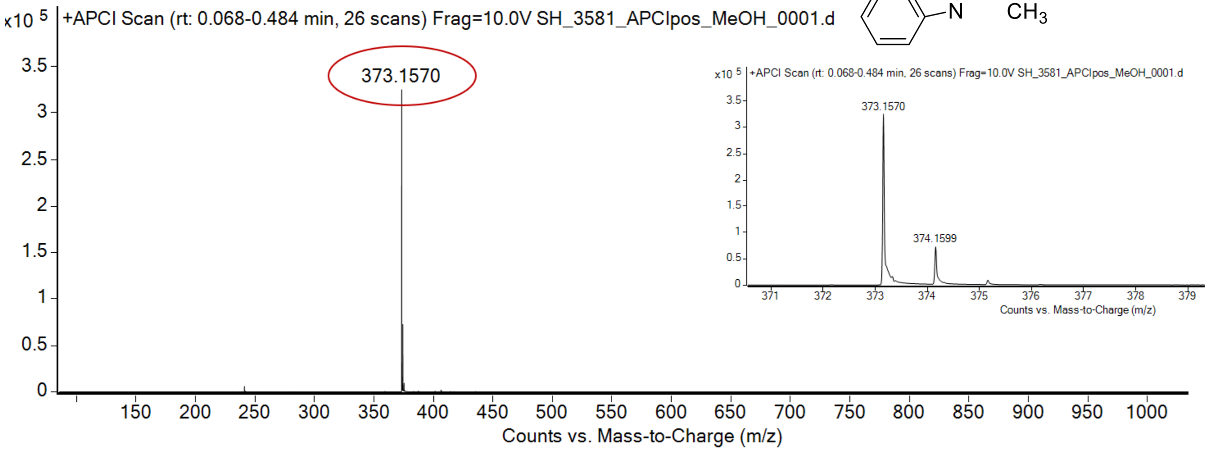

expected mass: $[M+H]^+$ = 373.1571 observed mass : $[M+H]^+$ = 373.1570 mass accuracy = - 0.3 ppm

$^{1}H$ (500 MHz) and $^{13}C$ NMR (126 MHz) spectra of **18** in DMSO-$d_6$

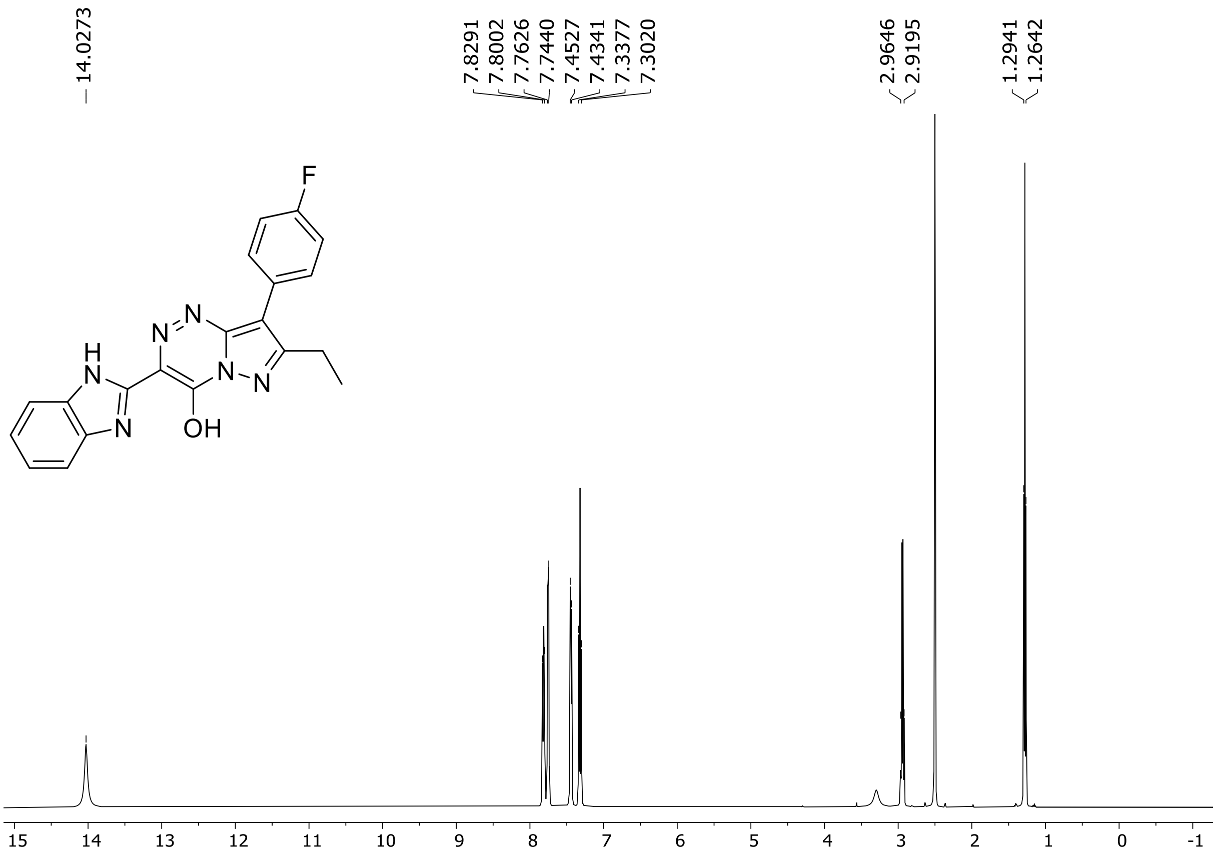

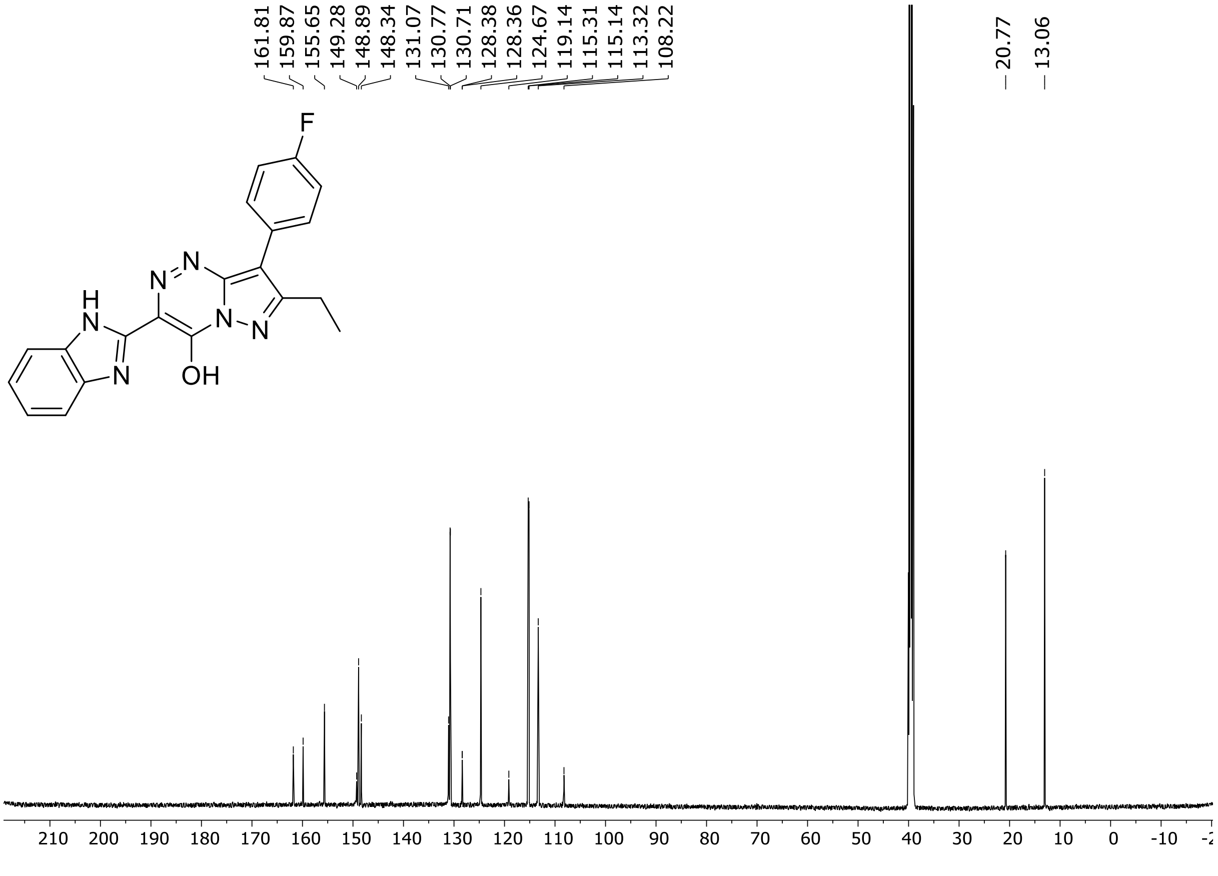

$^{19}$F (471 MHz) NMR spectrum of **18** DMSO-$d_6$

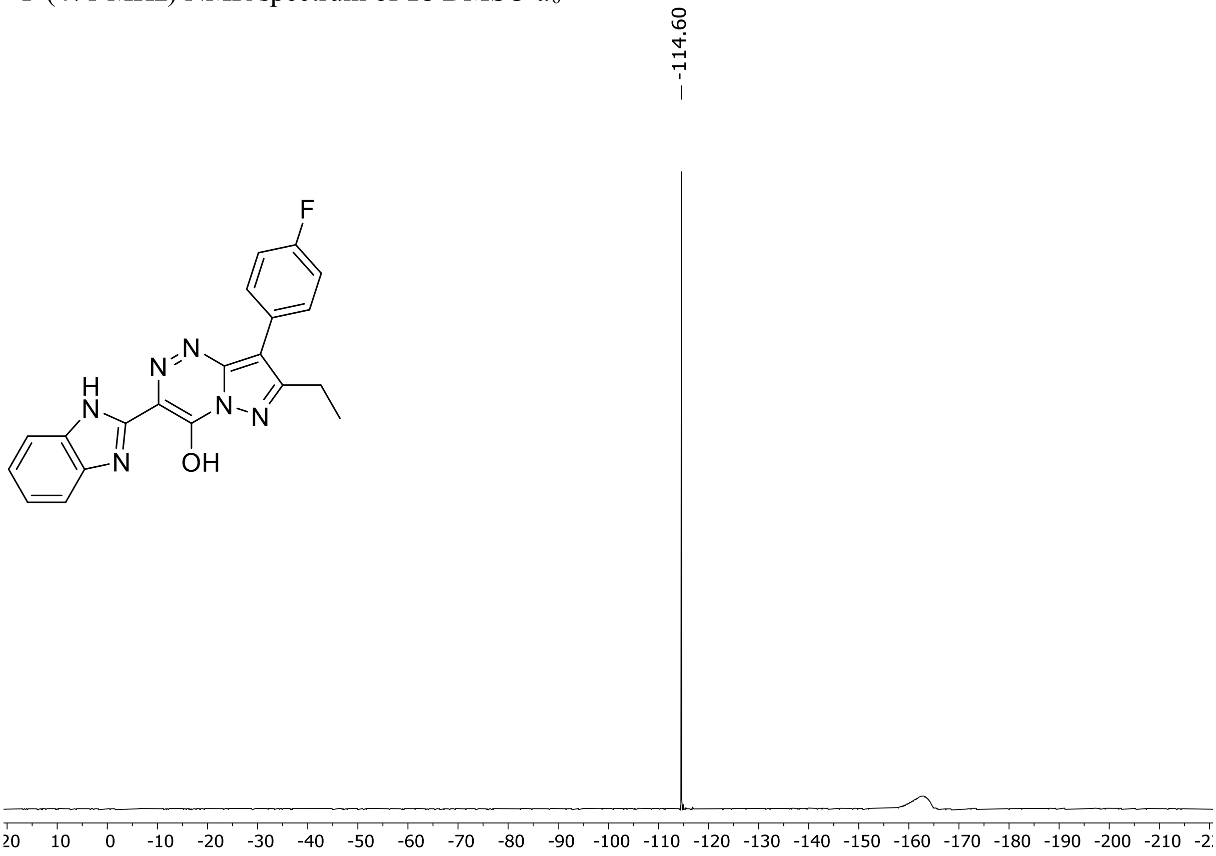

HRMS spectrum of **18**

**$C_{20}H_{15}FN_6O$** **mono m/z = 374,1291**

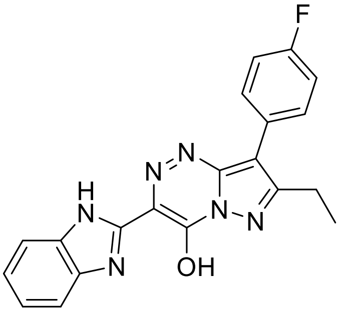

**ESI -** (MMI)
nitrogen flow 5 L/min, gas temperature 300°C, vaporizer 200°C,
nebulizer 45 psi, Vcap -2000 V, skimmer **60** V, fragmentor **25**V,
dissolved in DMSO

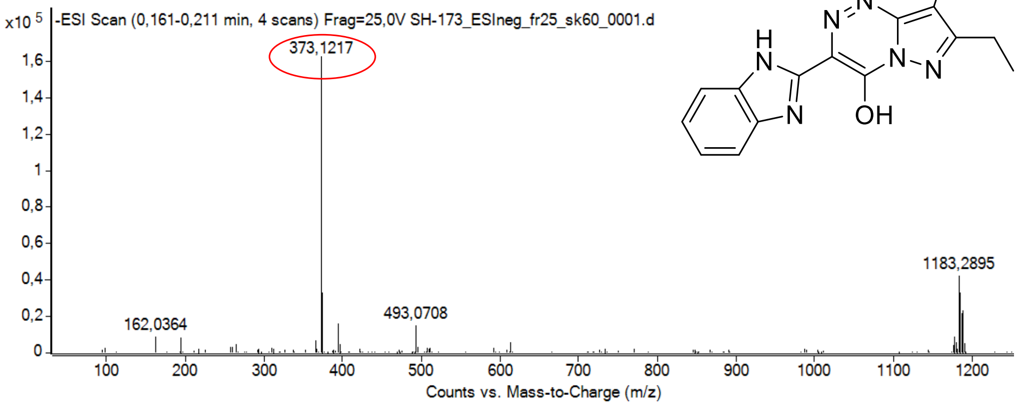

calculated mass:
[M-H]$^-$ = 373,1219 observed: [M-H]$^-$ = 373,1217 max. mass error = 0,5 ppm

$^{1}H$ (500 MHz) and $^{13}C$ NMR (126 MHz) spectra of **19** in DMSO-$d_6$

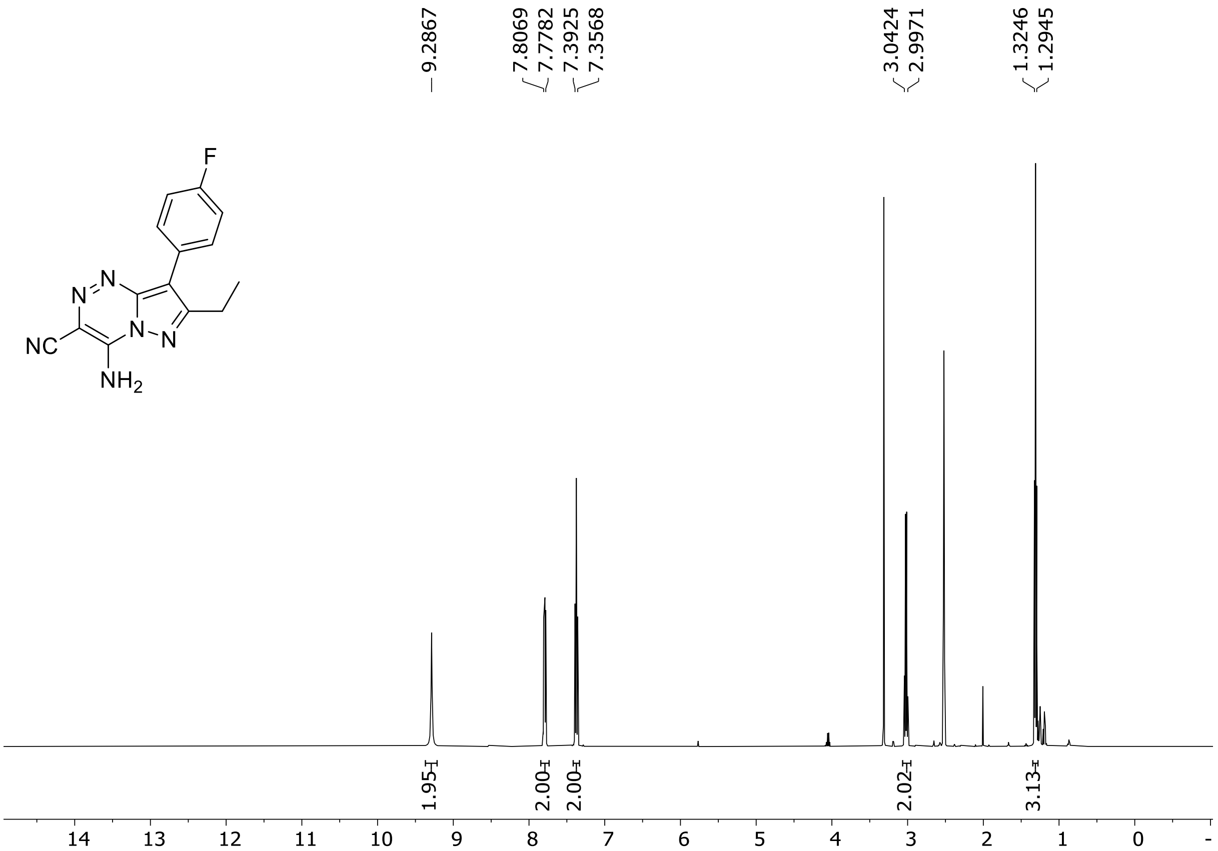

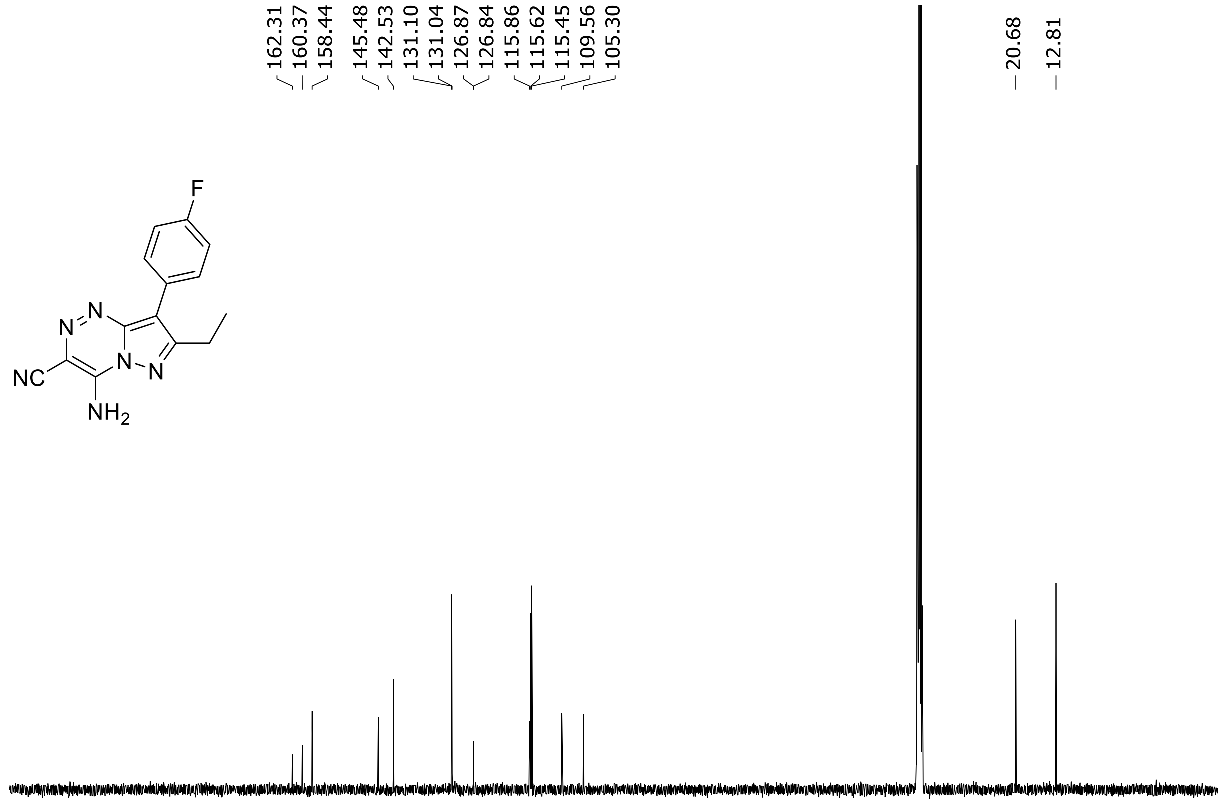

HRMS spectrum of **19**

**$C_{14}H_{11}FN_6$** **mono m/z = 282,1029**

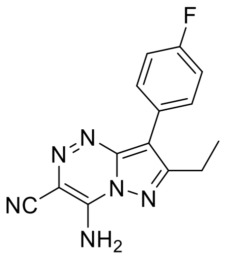

**ESI -** (MMI)
nitrogen flow 5 L/min, gas temperature 300°C, vaporizer 200°C,
nebulizer 30 psi, skimmer **55** V, fragmentor **10** V, dissolved in DMSO

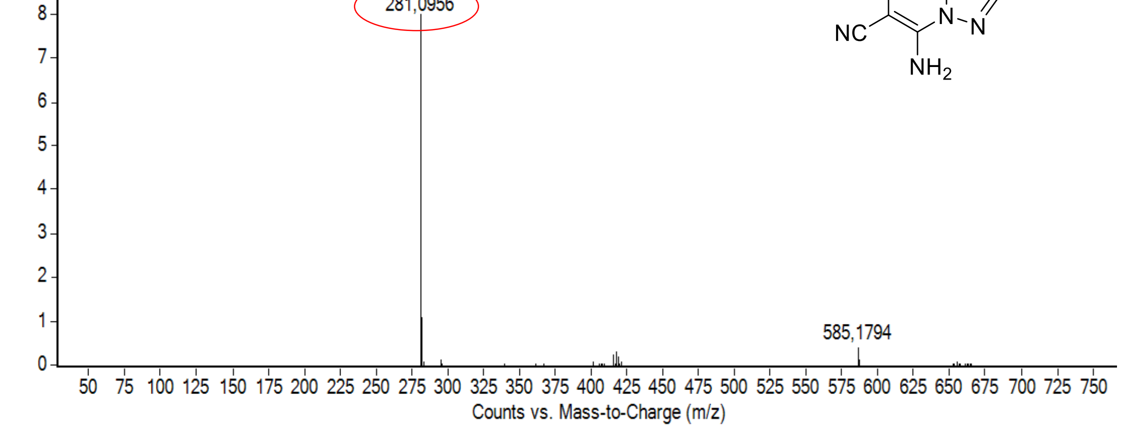

calculated mass: $[M-H]^-$ = 281,0956 observed: $[M-H]^-$ = 281,0956 max. mass error ≤ 0,1 ppm

[1]H (500 MHz) and [13]C NMR (126 MHz) spectra of **20** in DMSO-$d_6$

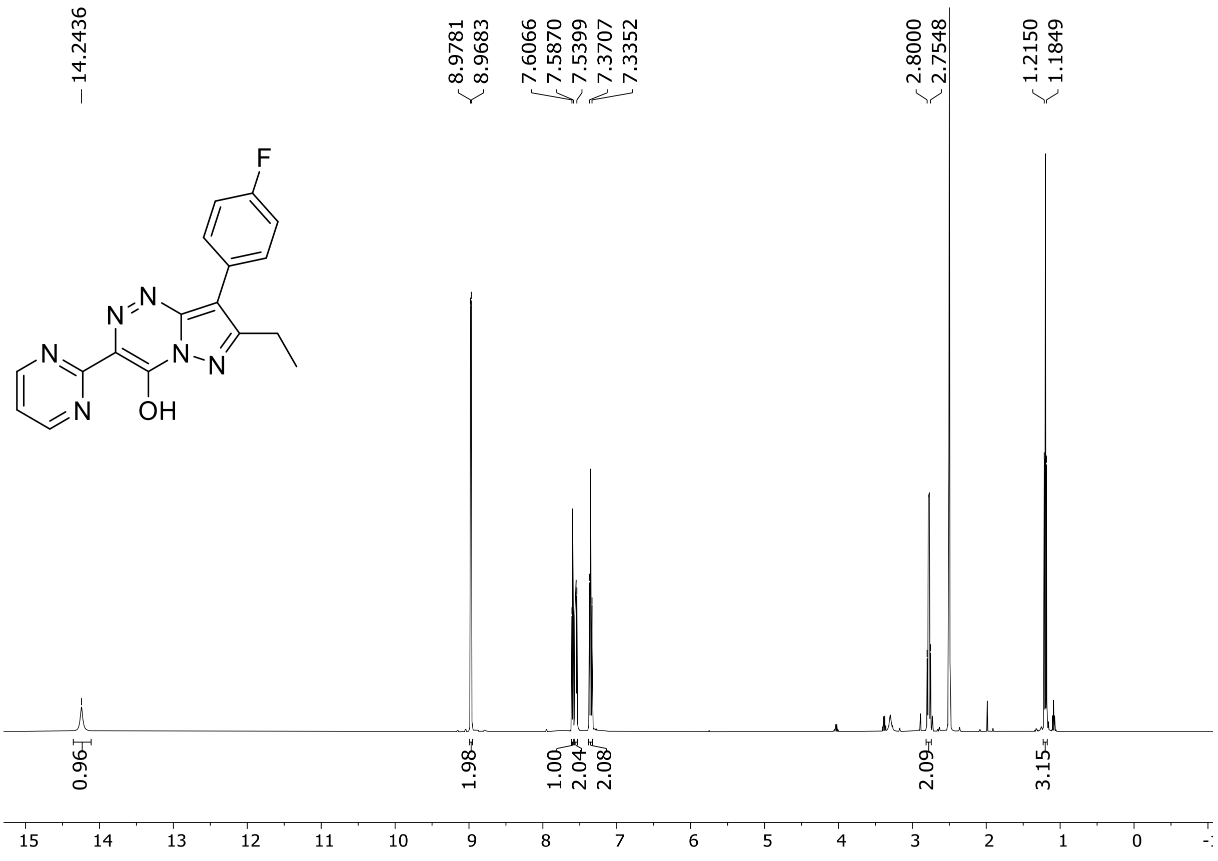

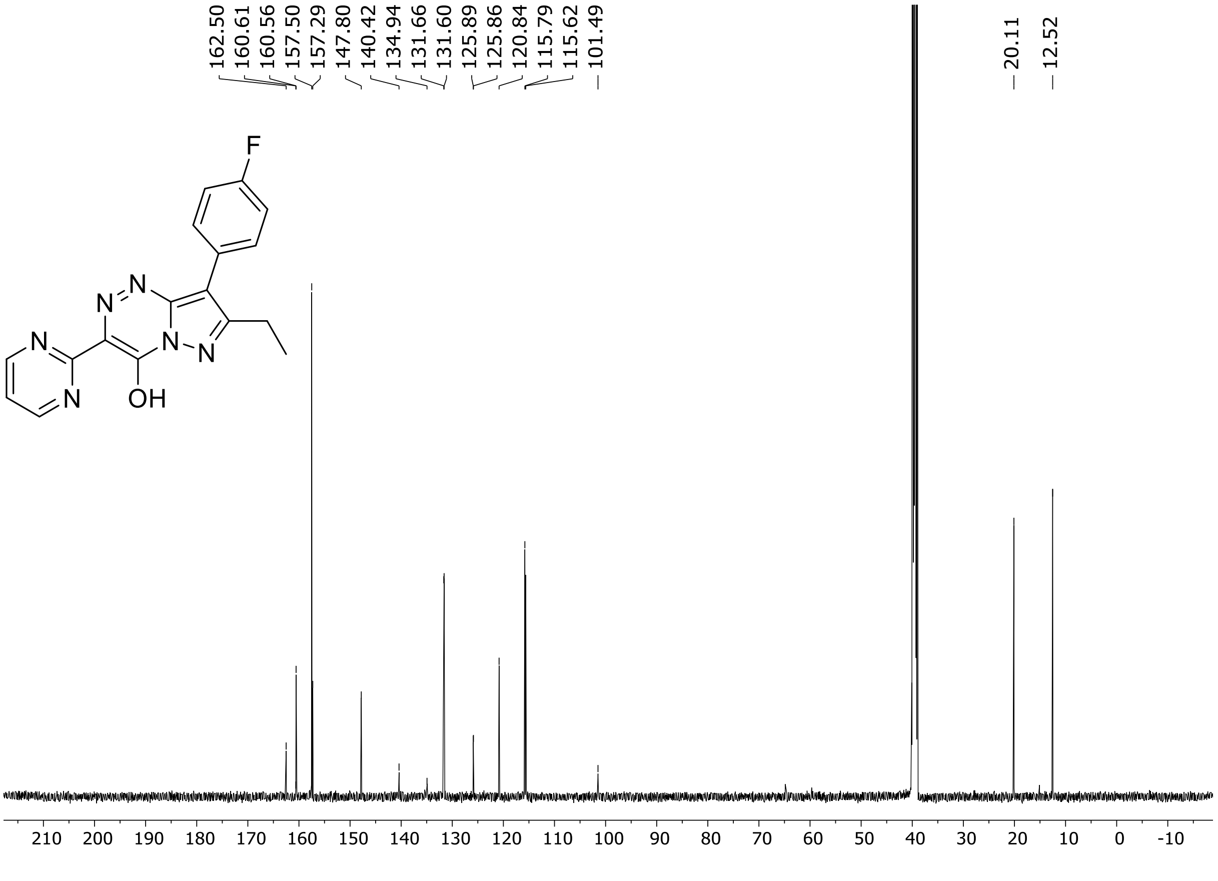

HRMS spectrum of **20**

**$C_{17}H_{13}FN_6O$** **exact mass: 336.1135**

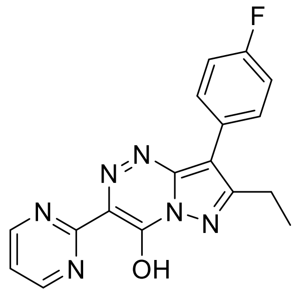

**APCI +** (MMI)
nitrogen flow 3 L/min, gas temperature 325°C, nebulizer 45 psig, skimmer 65 V, vaporizer 200°C, fragmentor 15 V, dissolved in methanol

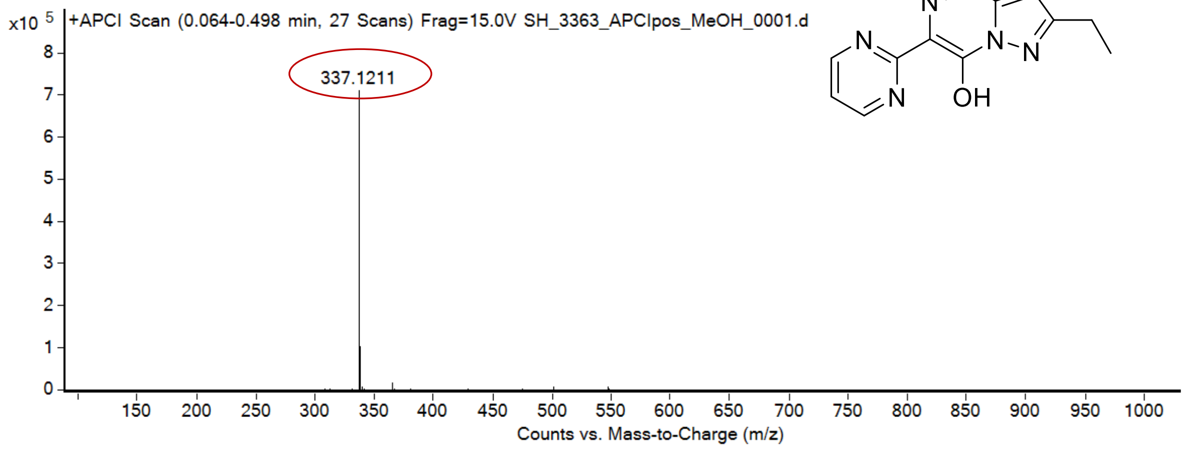

expected mass: $[M+H]^+$ = 337.1208 observed mass: $[M+H]^+$ = 337.1211 mass accuracy = 0.8 ppm

$^{1}H$ (500 MHz) and $^{13}C$ NMR (126 MHz) spectra of **21** in DMSO-$d_6$

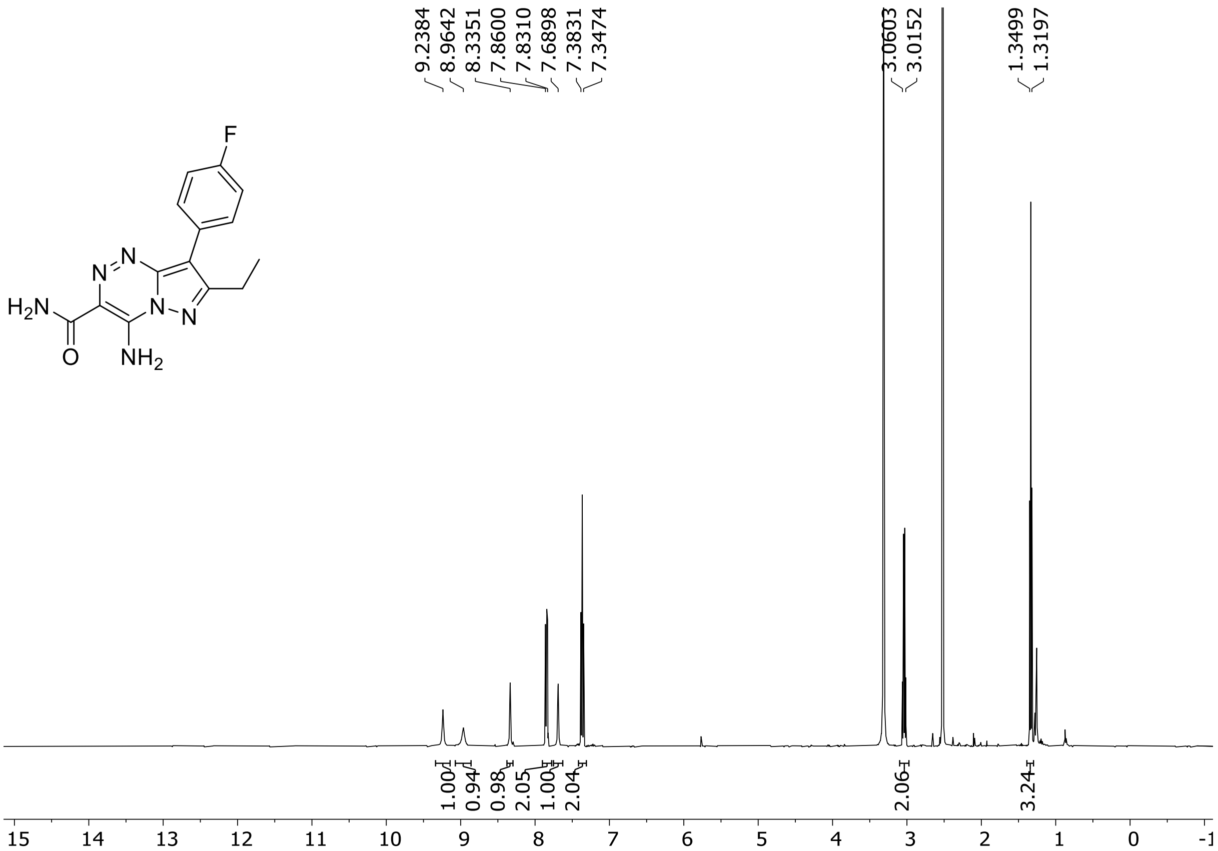

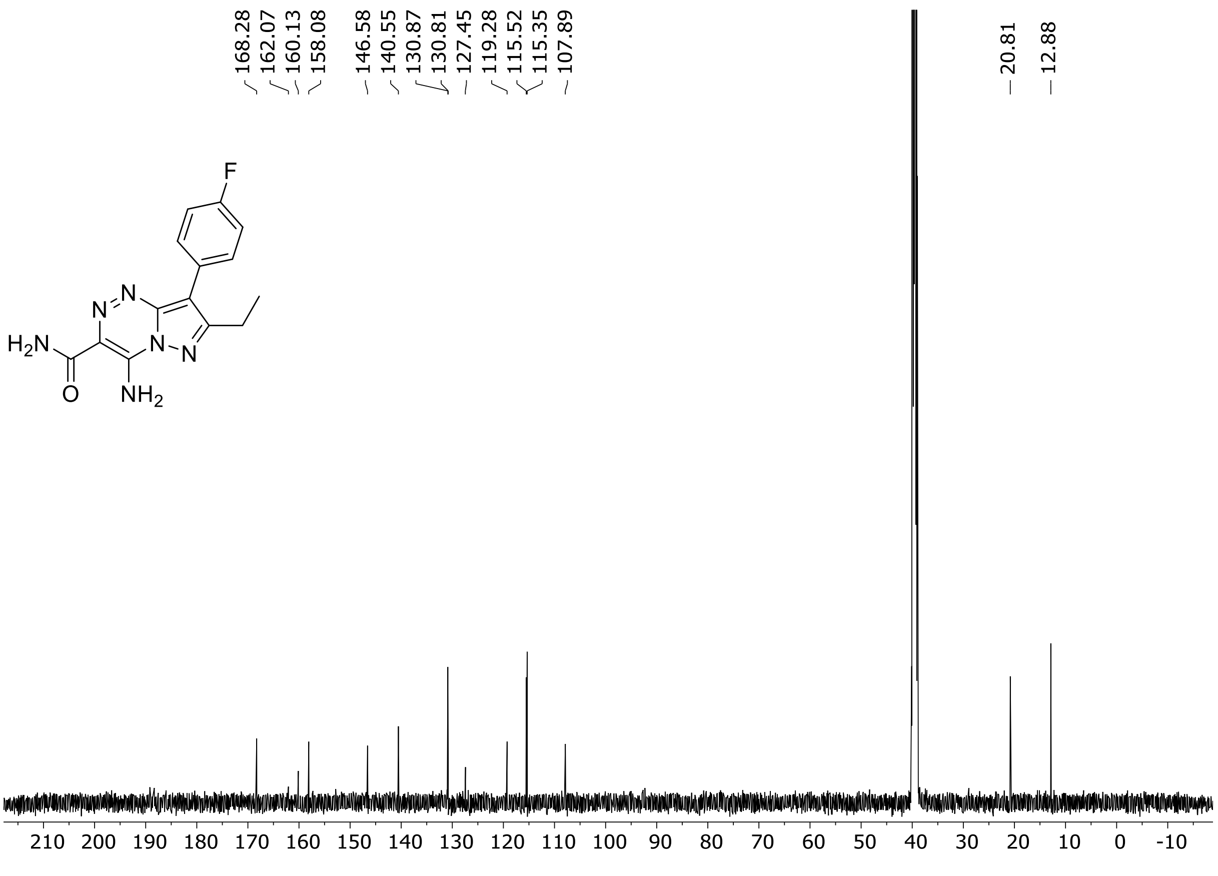

HRMS spectrum of **21**

$C_{14}H_{13}FN_6O$ **mono m/z = 300.1135**

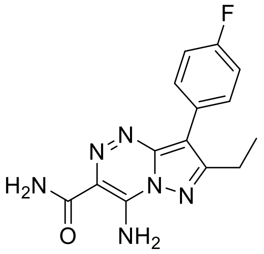

**APCI +** (MMI)
nitrogen flow 5 L/min, gas temperature 300°C, nebulizer 45 psi, vaporizer 200°C,
skimmer **65** V, fragmentor **25** V, dissolved in methanol

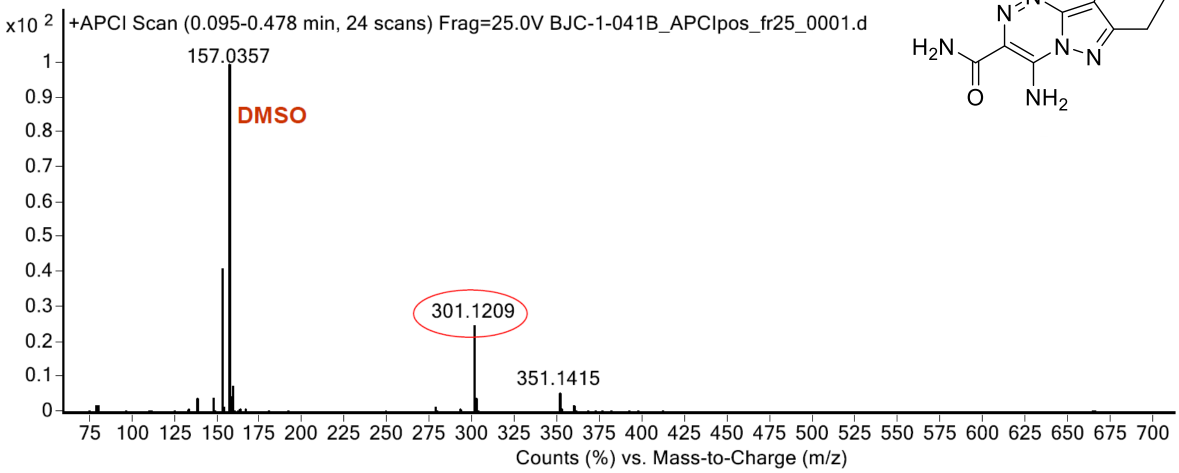

calculated mass: $[M+H]^+$ = 301.1208 observed: $[M+H]^+$ = 301.1209 max. mass error = 0.3 ppm

$^{1}H$ (500 MHz) and $^{13}C$ NMR (126 MHz) spectra of **22** in DMSO-$d_6$

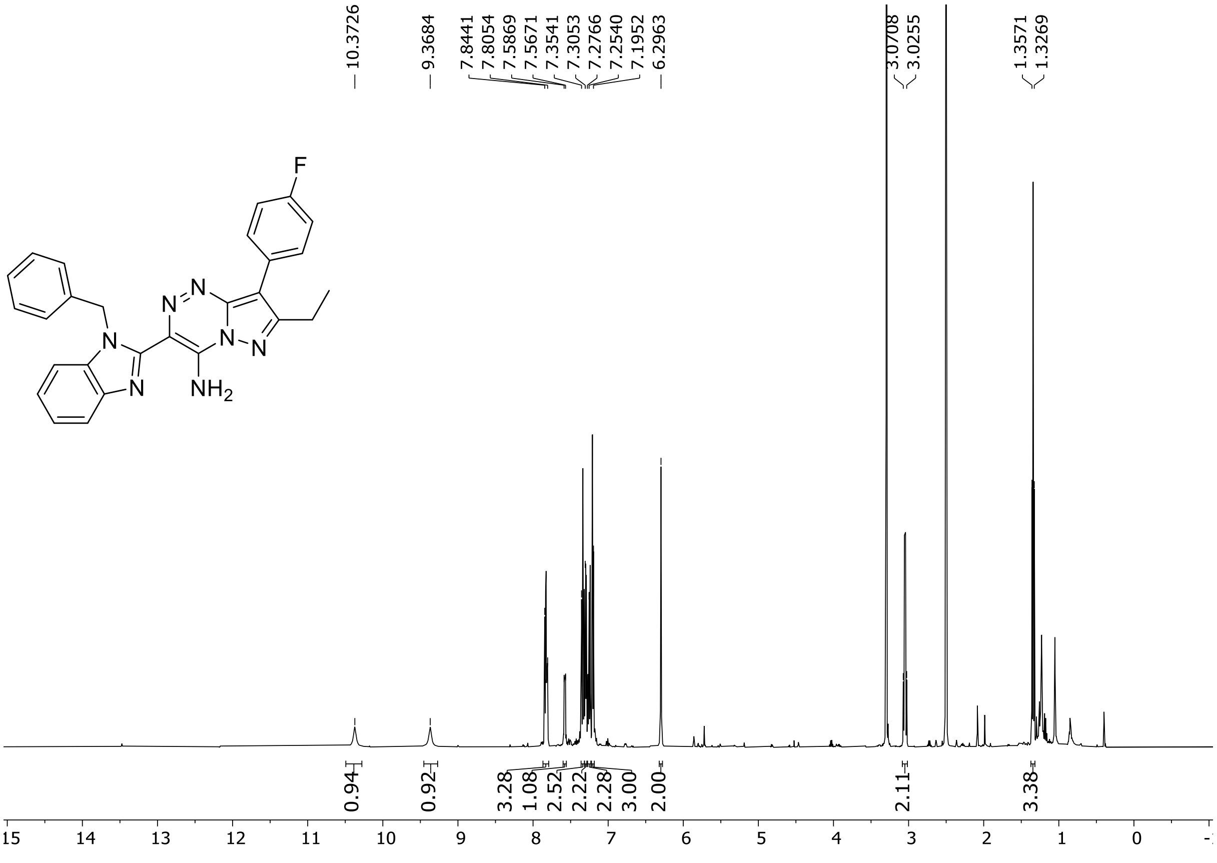

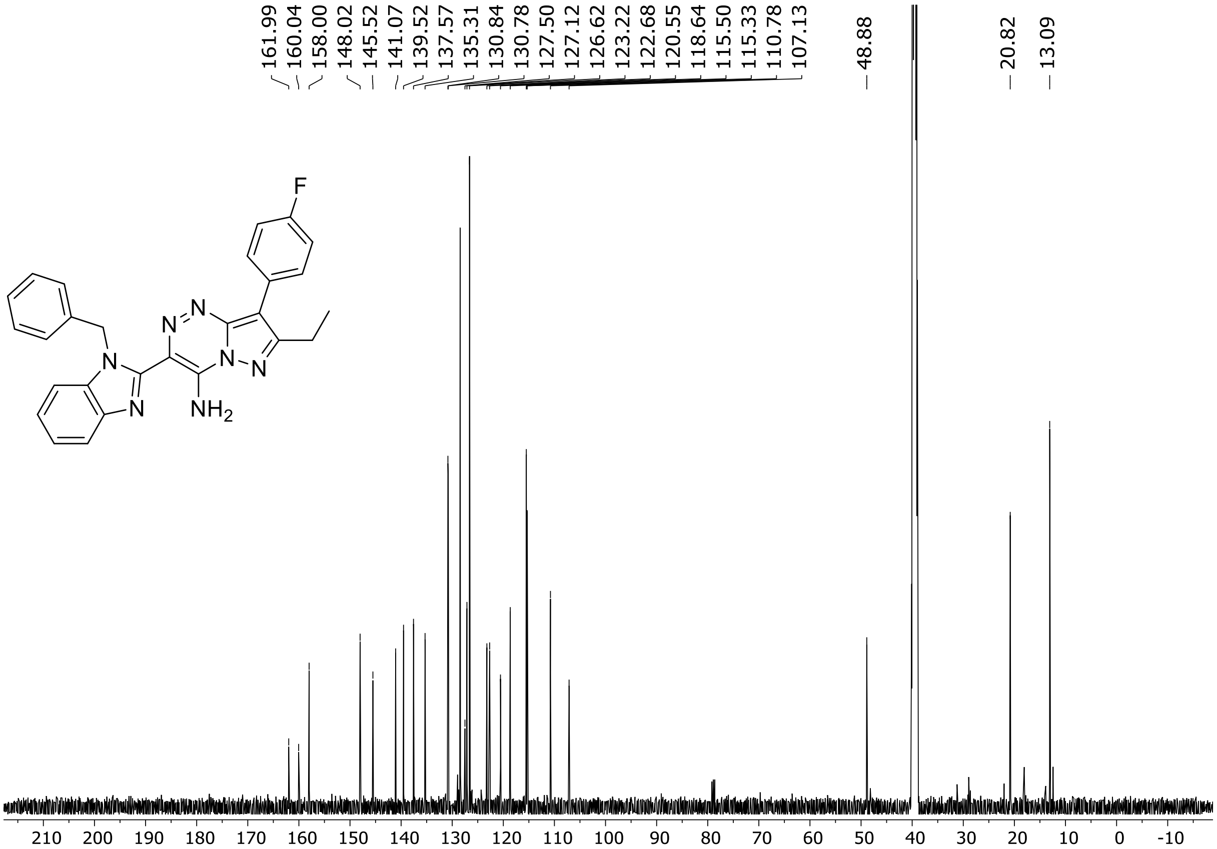

HRMS spectrum of **22**

$C_{27}H_{22}FN_7$ **mono m/z = 463.1921**

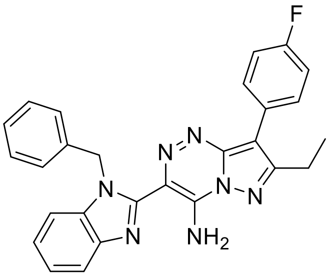

**APCI +** (MMI)
nitrogen flow 5 L/min, gas temperature 300°C, nebulizer 45 psi, vaporizer 200°C
skimmer **65** V, fragmentor **30** V, dissolved in methanol

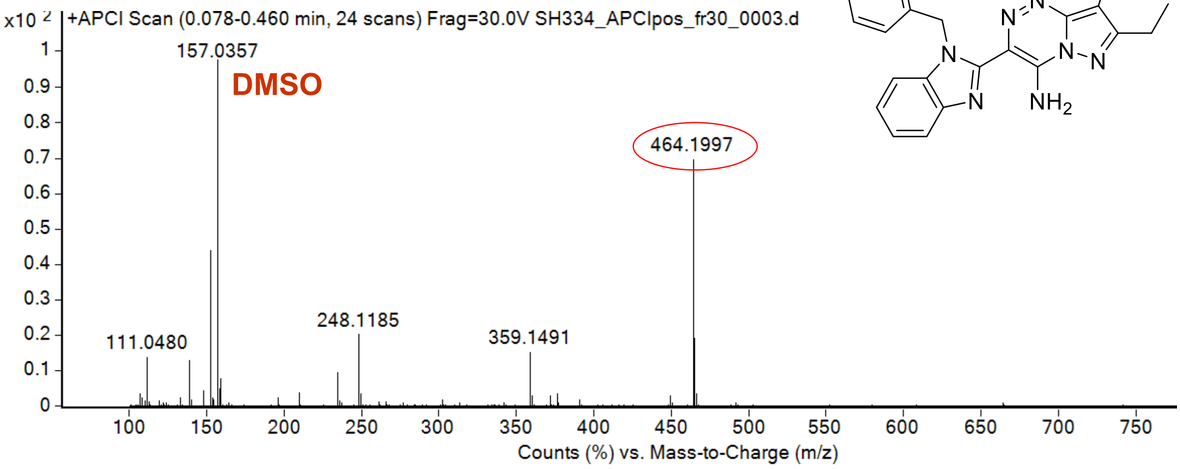

calculated mass: $[M+H]^+$ = 464.1993 observed: $[M+H]^+$ = 464.1997 max. mass error = 0.8 ppm

[1]H (500 MHz) and [13]C NMR (126 MHz) spectra of **23** in DMSO-$d_6$

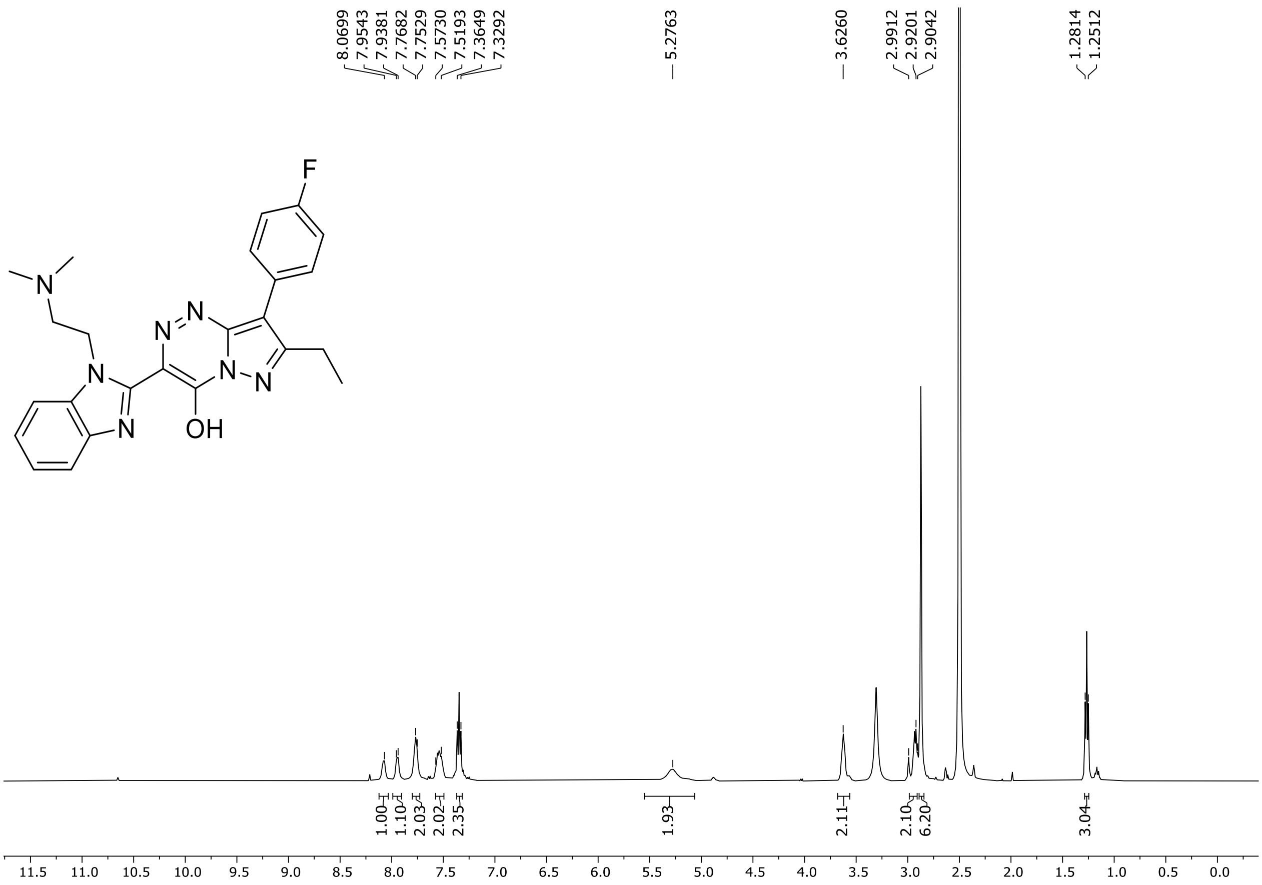

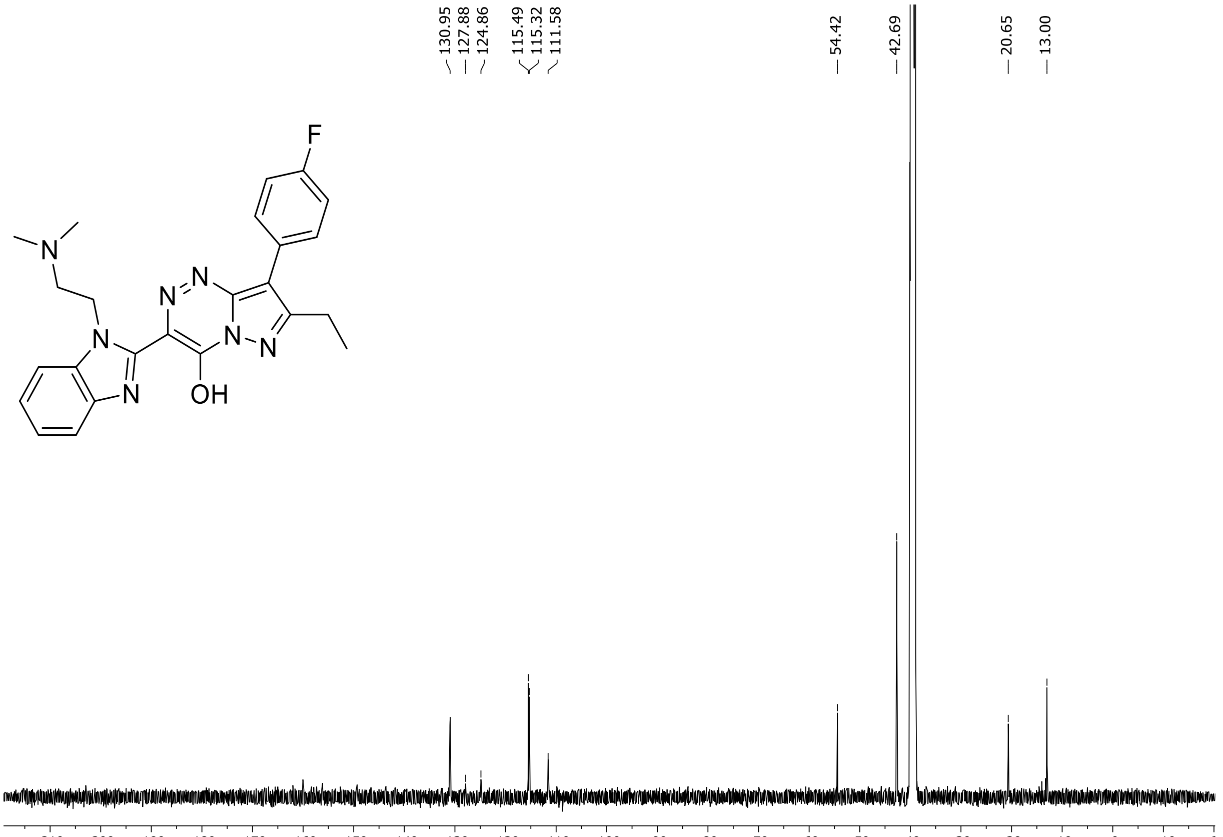

$^{19}F$ (471 MHz) NMR spectrum of **23** in DMSO-$d_6$

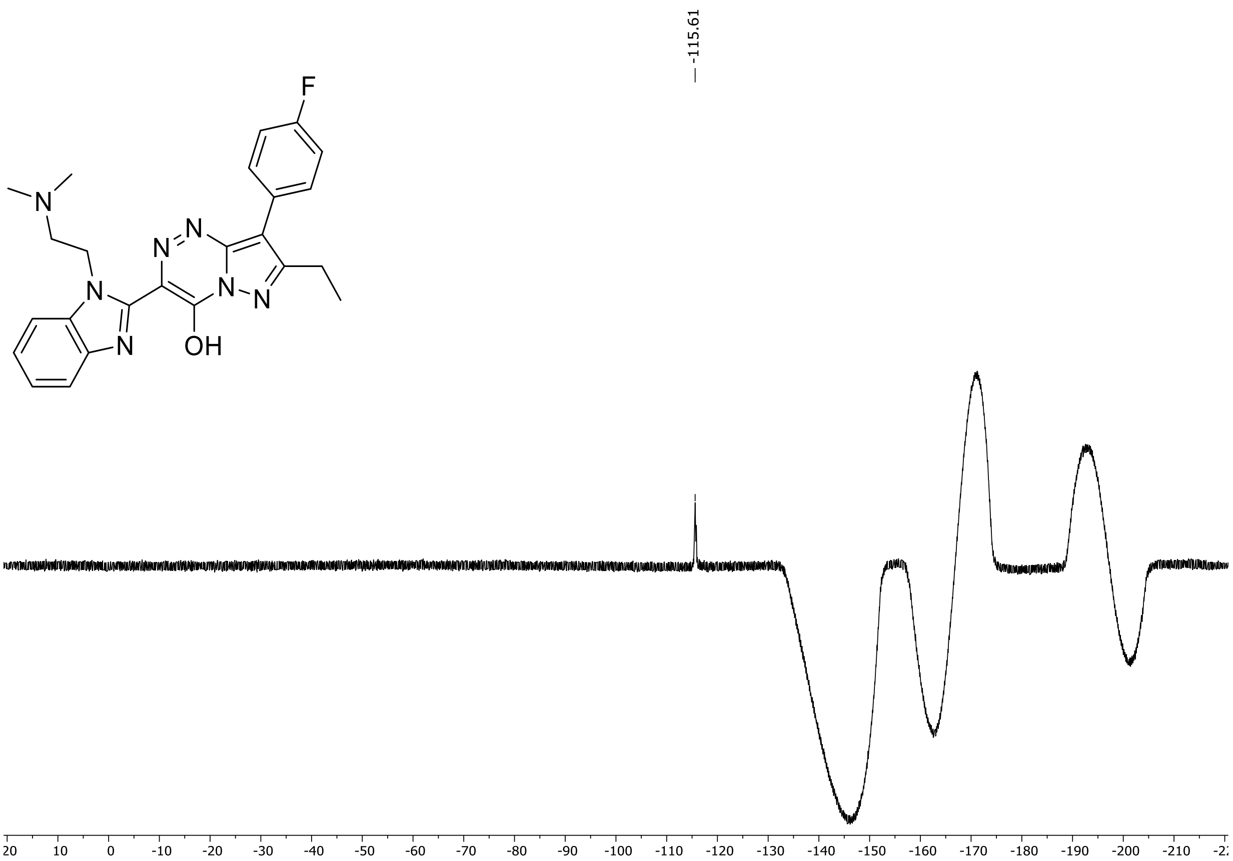

HRMS spectrum of **23**

**$C_{24}H_{24}FN_7O$** **exact mass: 445.2026**

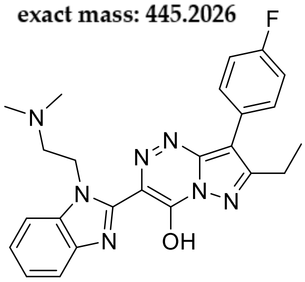

**APCI +** (MMI)
nitrogen flow 5 L/min, gas temperature 325°C, nebulizer 45 psig, skimmer 65 V, vaporizer 200°C, fragmentor 15 V, dissolved in methanol

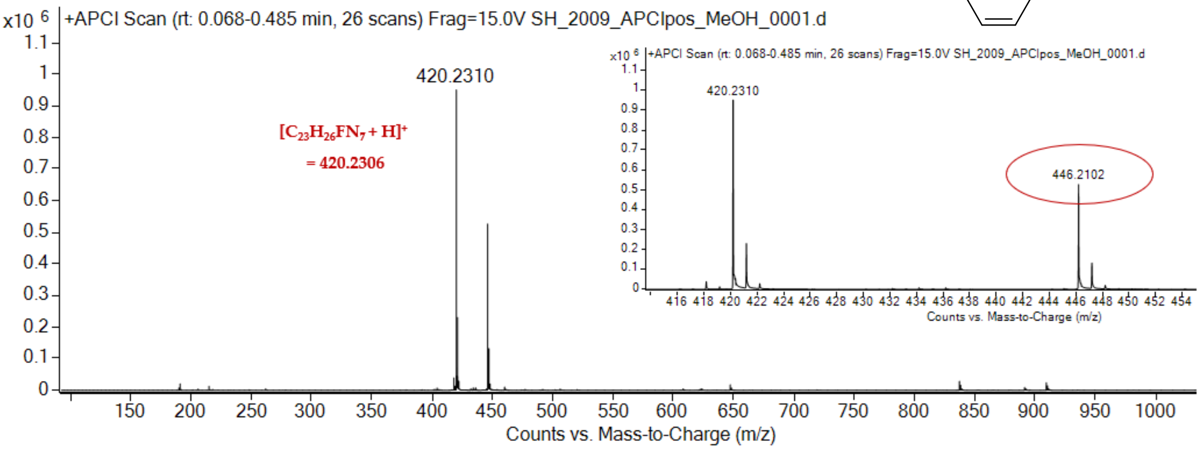

expected mass: $[M+H]^+ = 446.2099$ observed mass : $[M+H]^+ = 446.2102$ mass accuracy = 0.7 ppm

[1]H (500 MHz) and [13]C NMR (126 MHz) spectra of **24** in DMSO-$d_6$

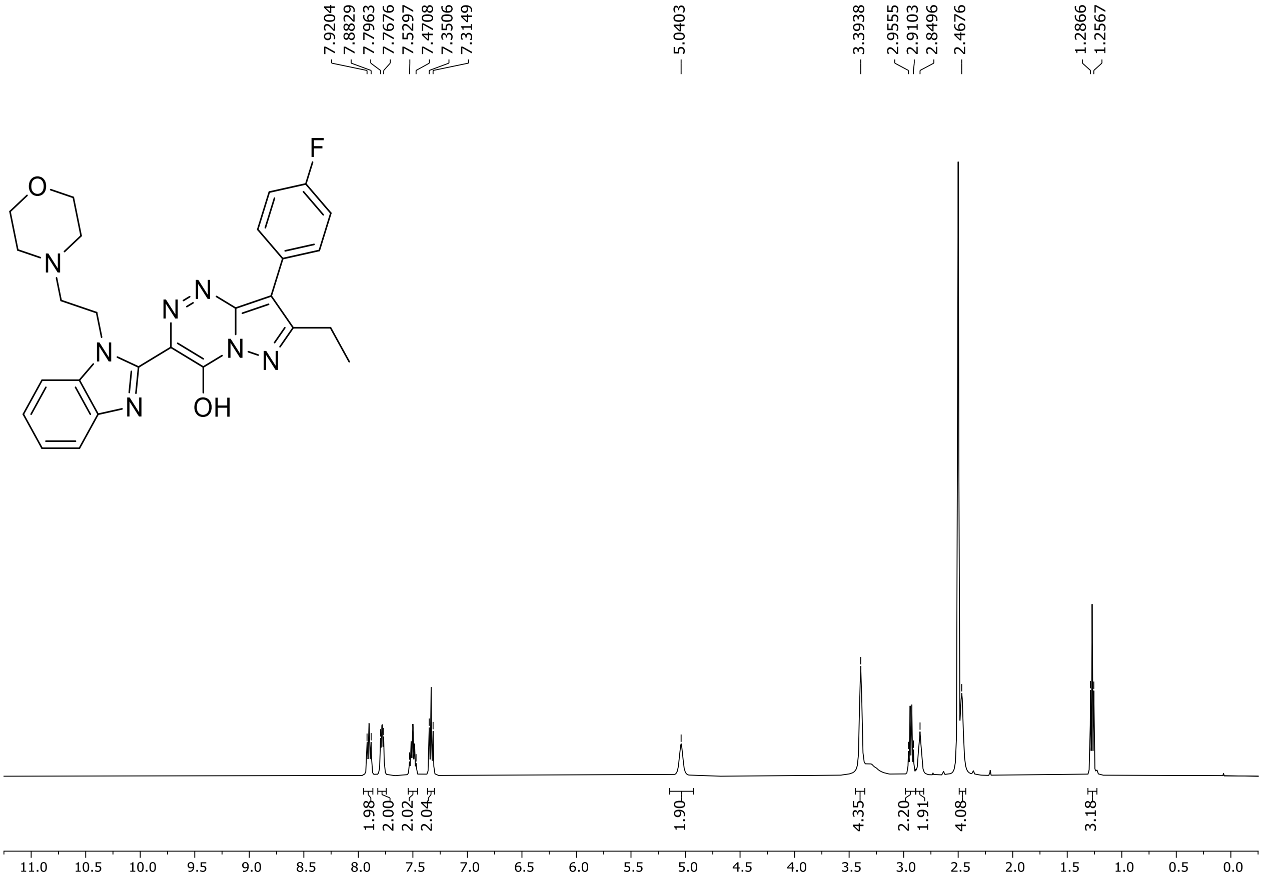

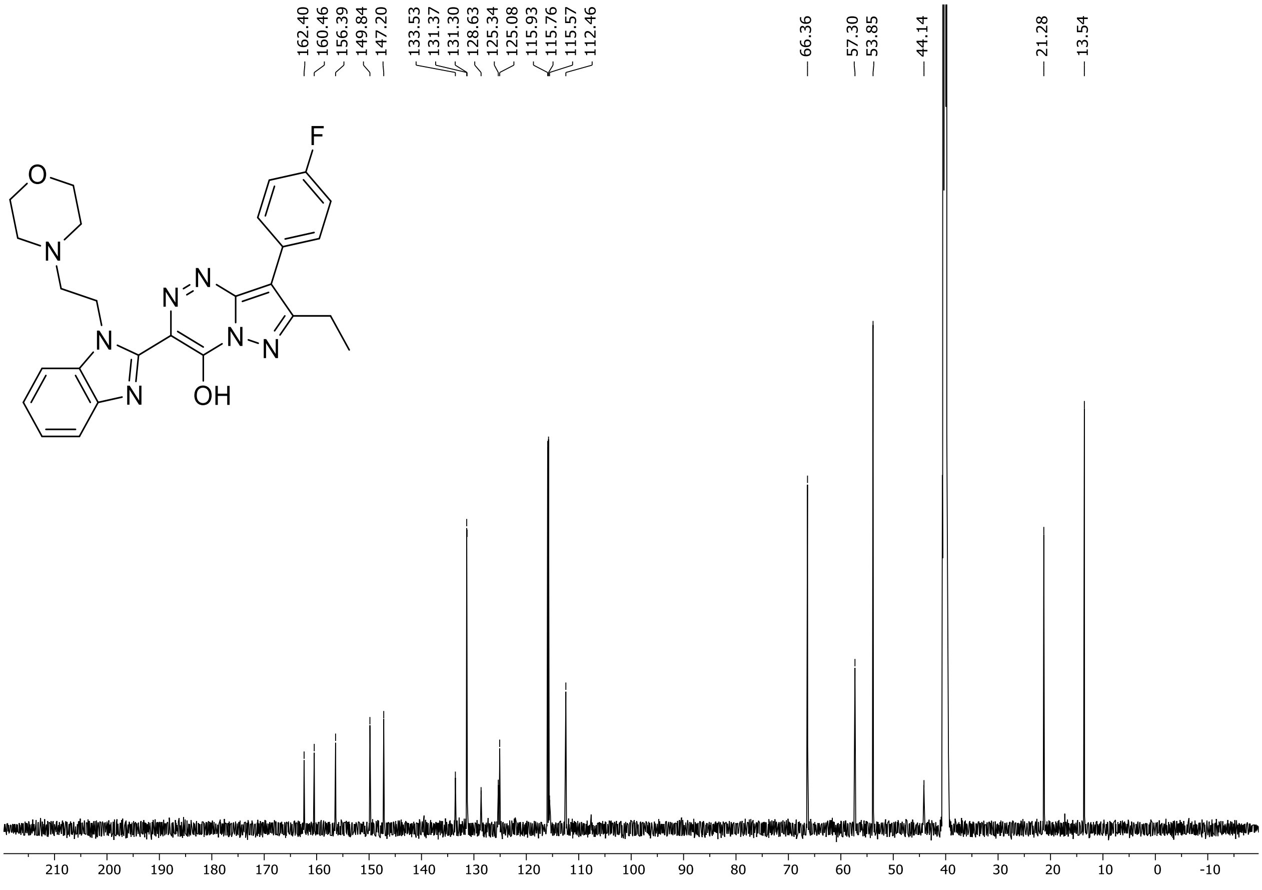

$^{19}F$ (471 MHz) NMR spectrum of **24** in DMSO-$d_6$

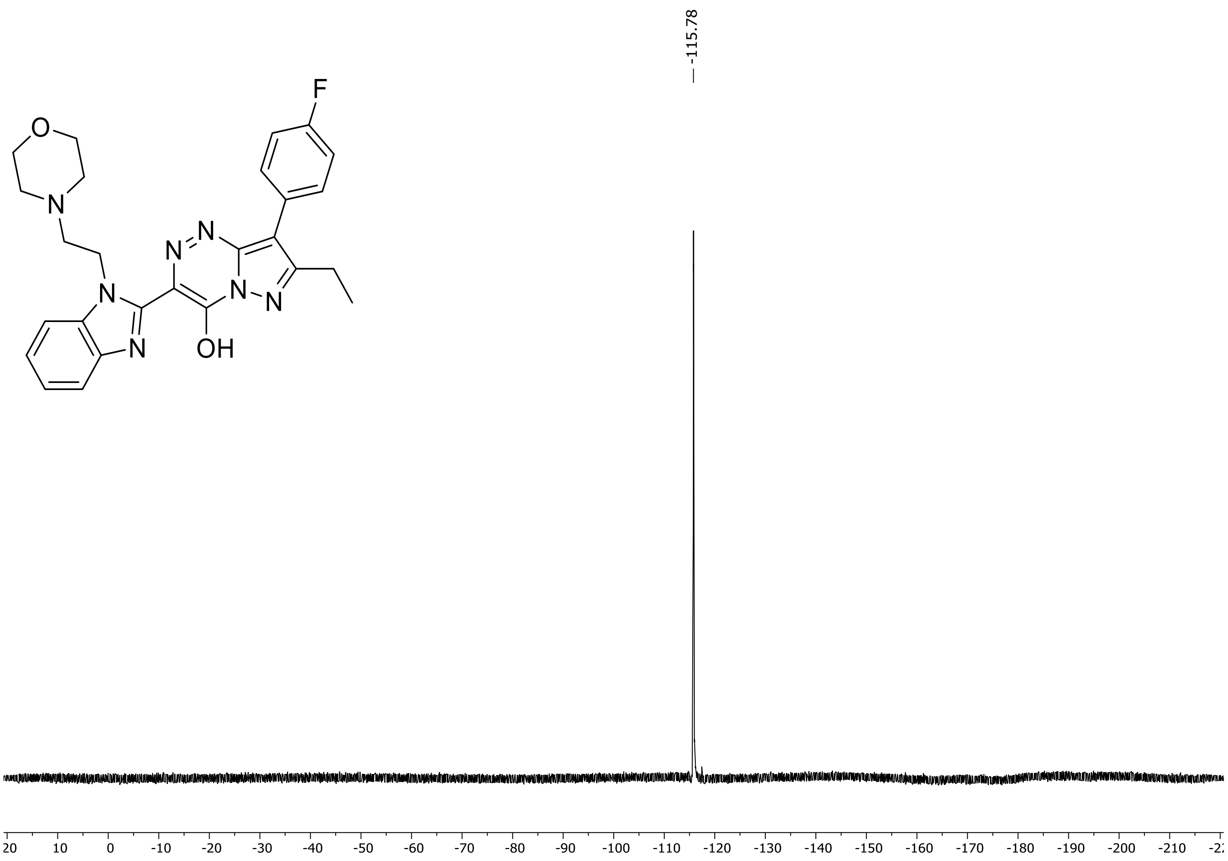

HRMS spectrum of **24**

**$C_{26}H_{26}FN_7O_2$** **exact mass: 487.2132**

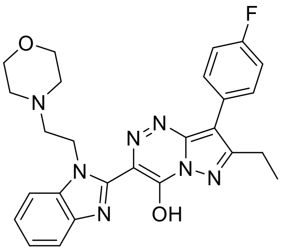

**APCI +** (MMI)
nitrogen flow 5 L/min, gas temperature 325°C, nebulizer 45 psig, skimmer 65 V, vaporizer 200°C, fragmentor 20 V, dissolved in methanol

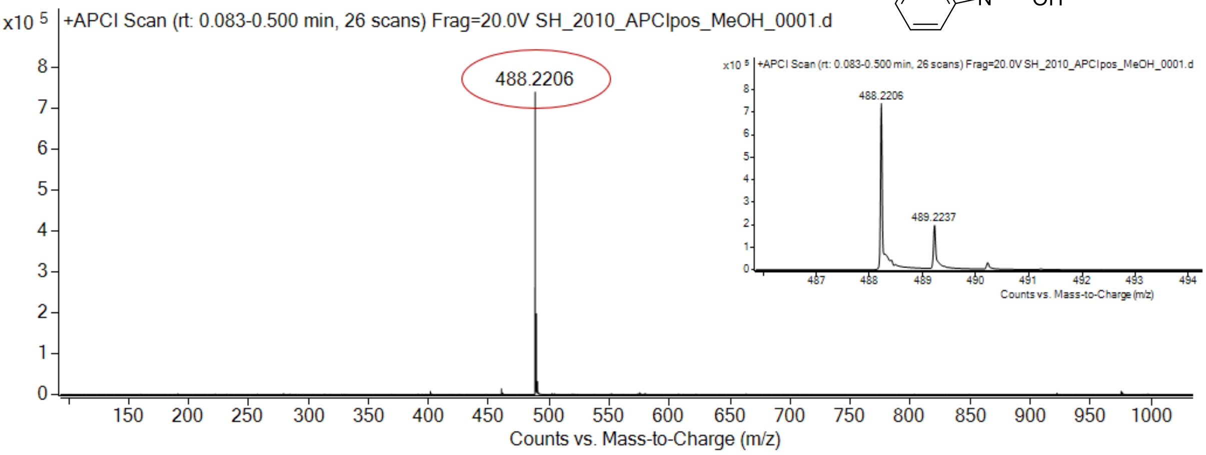

expected mass: $[M+H]^+$ = 488.2205 observed mass : $[M+H]^+$ = 488.2206 mass accuracy = 0.2 ppm

[1]H (500 MHz) and [13]C NMR (126 MHz) spectra of **25** in DMSO-$d_6$

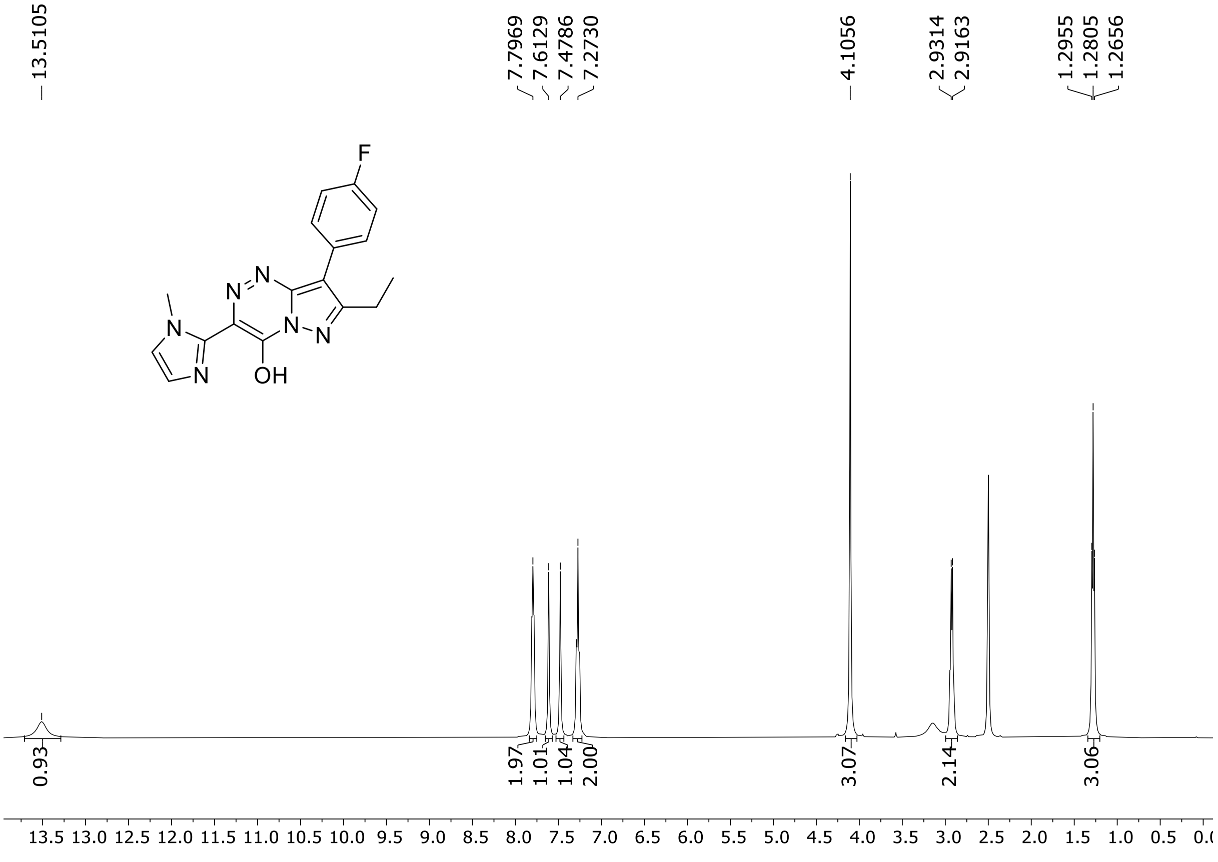

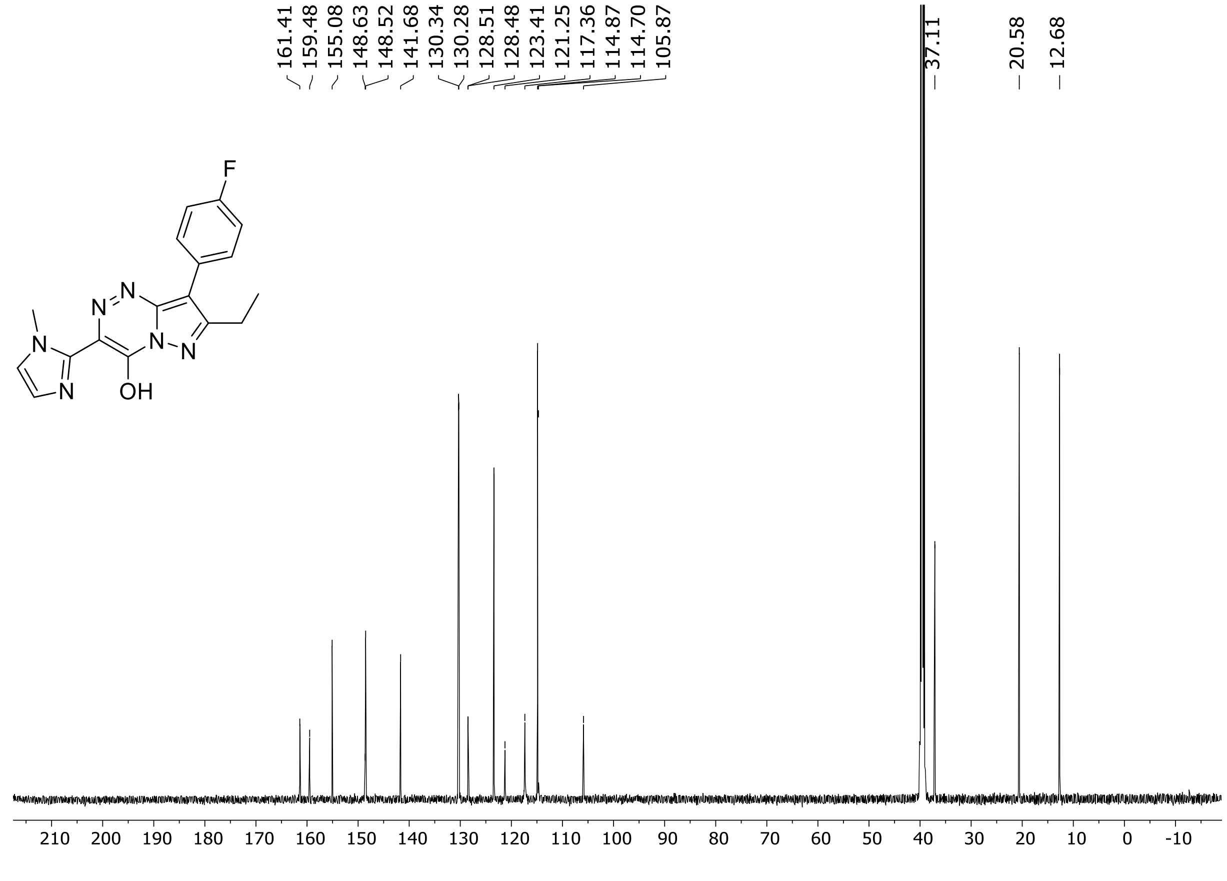

HRMS spectrum of **25**

**$C_{17}H_{15}FN_6O$** **exact mass: 338.1291**

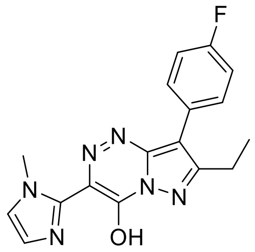

**APCI +** (MMI)
nitrogen flow 3 L/min, gas temperature 325°C, nebulizer 45 psig, skimmer 65 V, vaporizer 200°C, fragmentor 15 V, dissolved in methanol

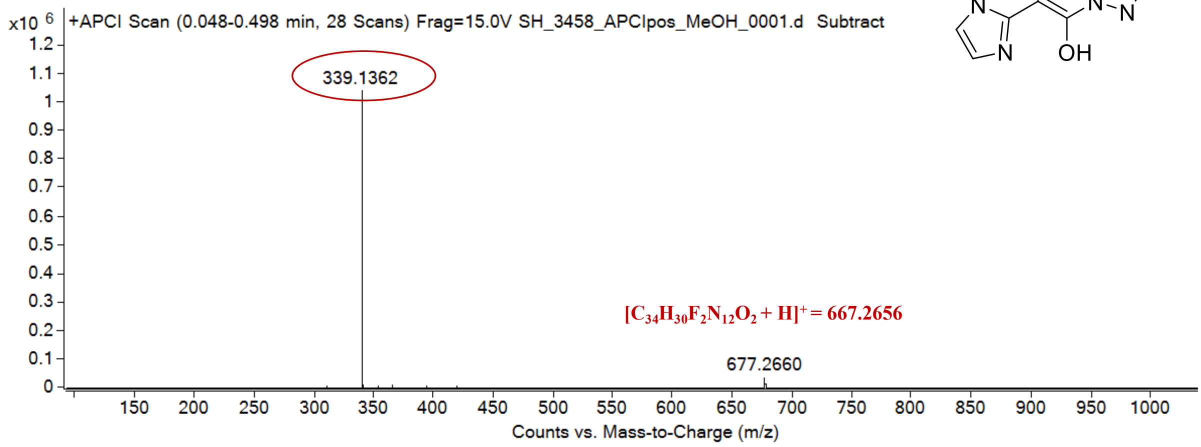

expected mass: $[M+H]^+$ = 339.1364 observed mass: $[M+H]^+$ = 339.1362 mass accuracy = - 0.6 ppm

$^{1}H$ (500 MHz) and $^{13}C$ NMR (126 MHz) spectra of **26** in MeOD-$d_4$

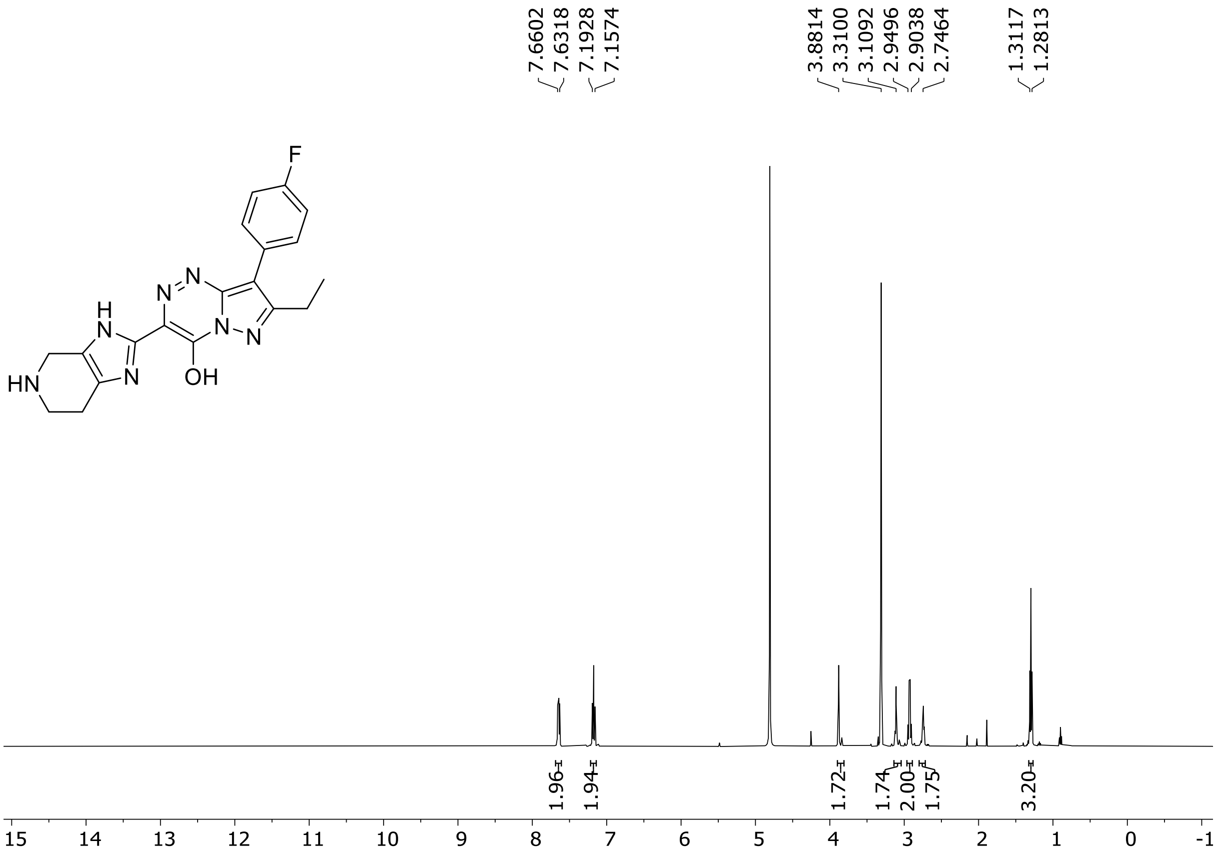

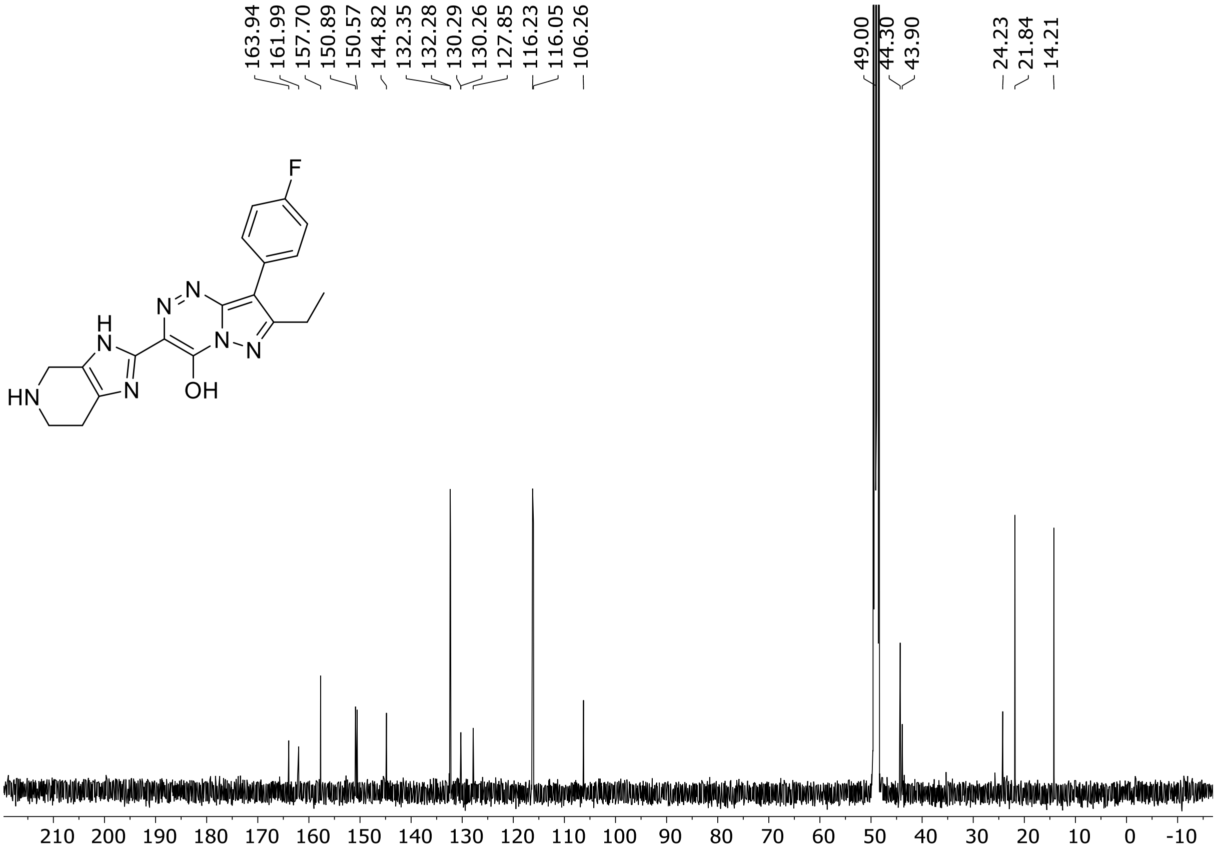

$^{19}F$ (471 MHz) NMR spectrum of **26** in MeOD-$d_4$

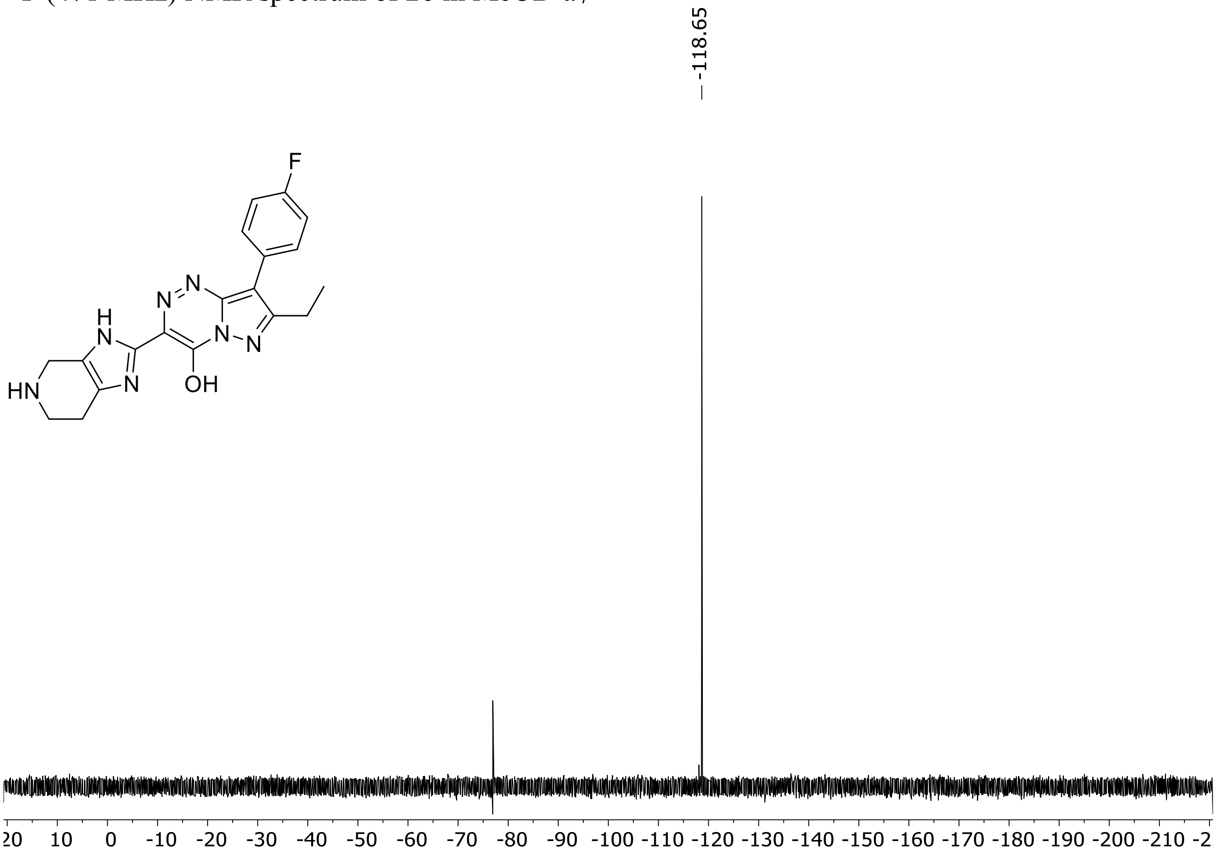

HRMS spectrum of **26**

**$C_{19}H_{18}FN_7O$** **exact mass: 379.1557**

**ESI -** (MMI)
nitrogen flow 3 L/min, gas temperature 325°C, nebulizer 45 psig, skimmer -65 V, Vcap 2500V, fragmentor -110 V, dissolved in methanol

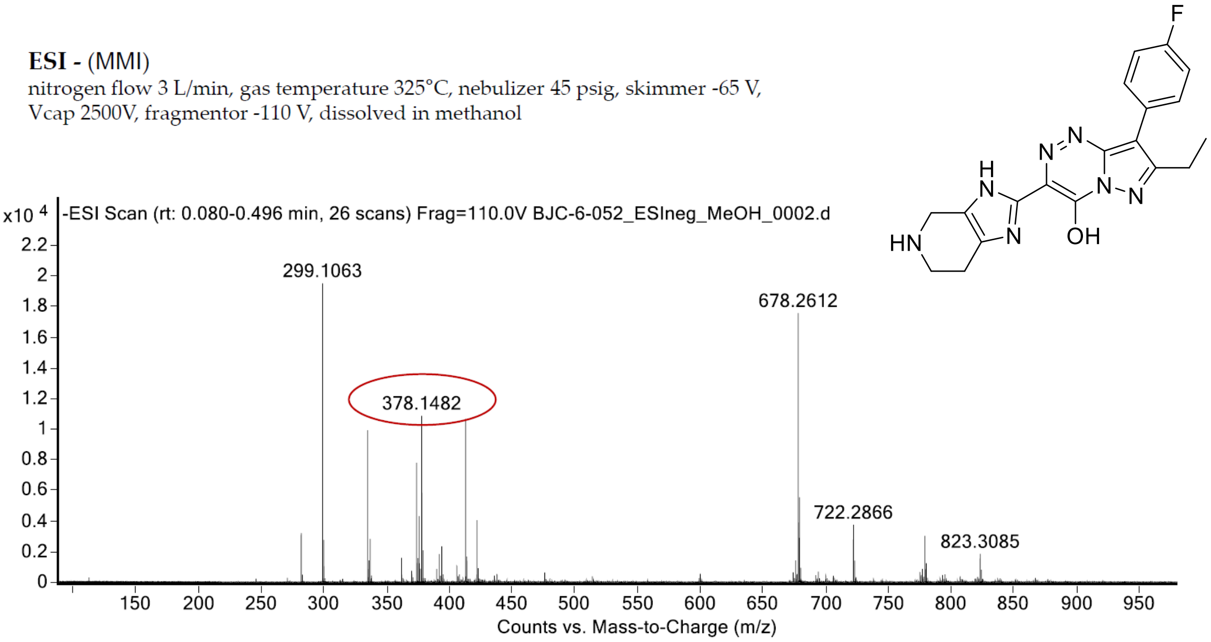

expected mass: [M-H]$^-$ = 378.1484 observed mass: [M-H]$^-$ = 378.1482 mass accuracy = - 0.5 ppm

$^{1}H$ (500 MHz) and $^{13}C$ NMR (126 MHz) spectra of **27** in DMSO-$d_6$

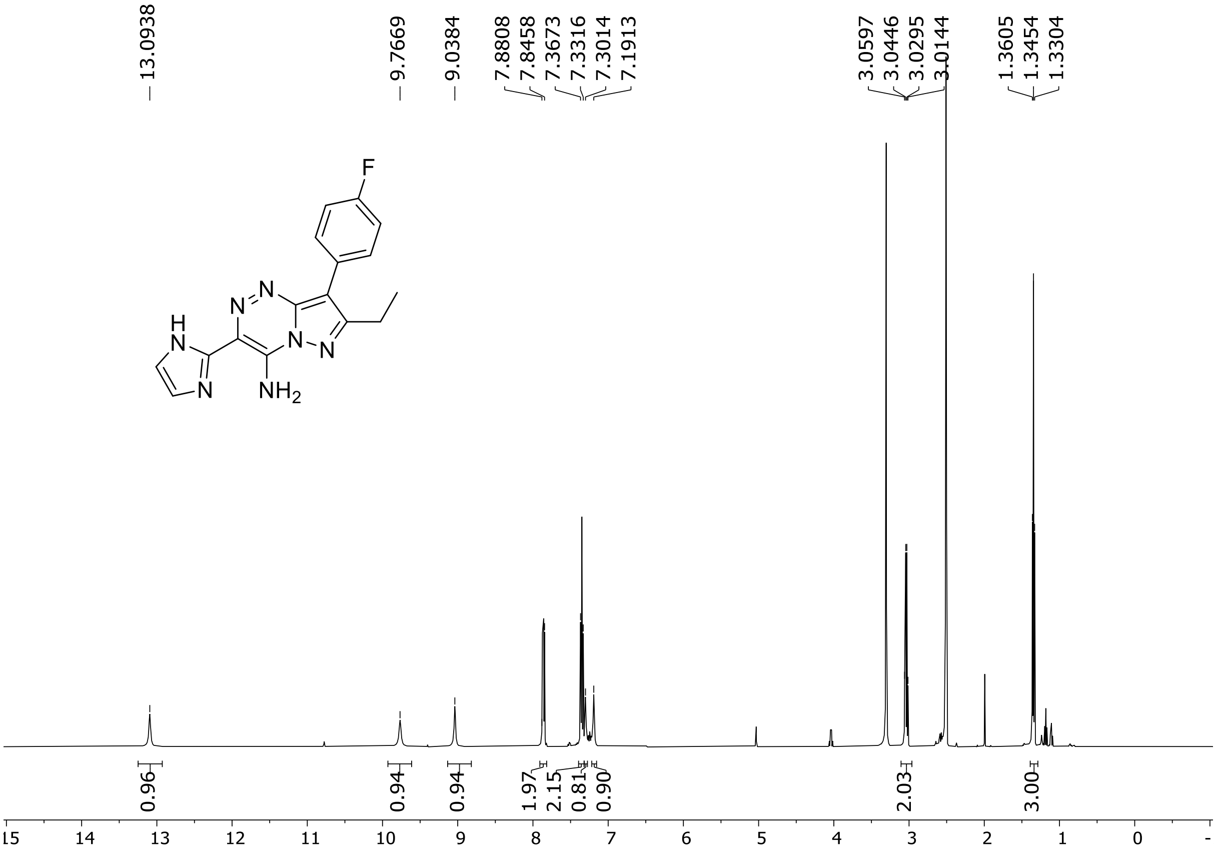

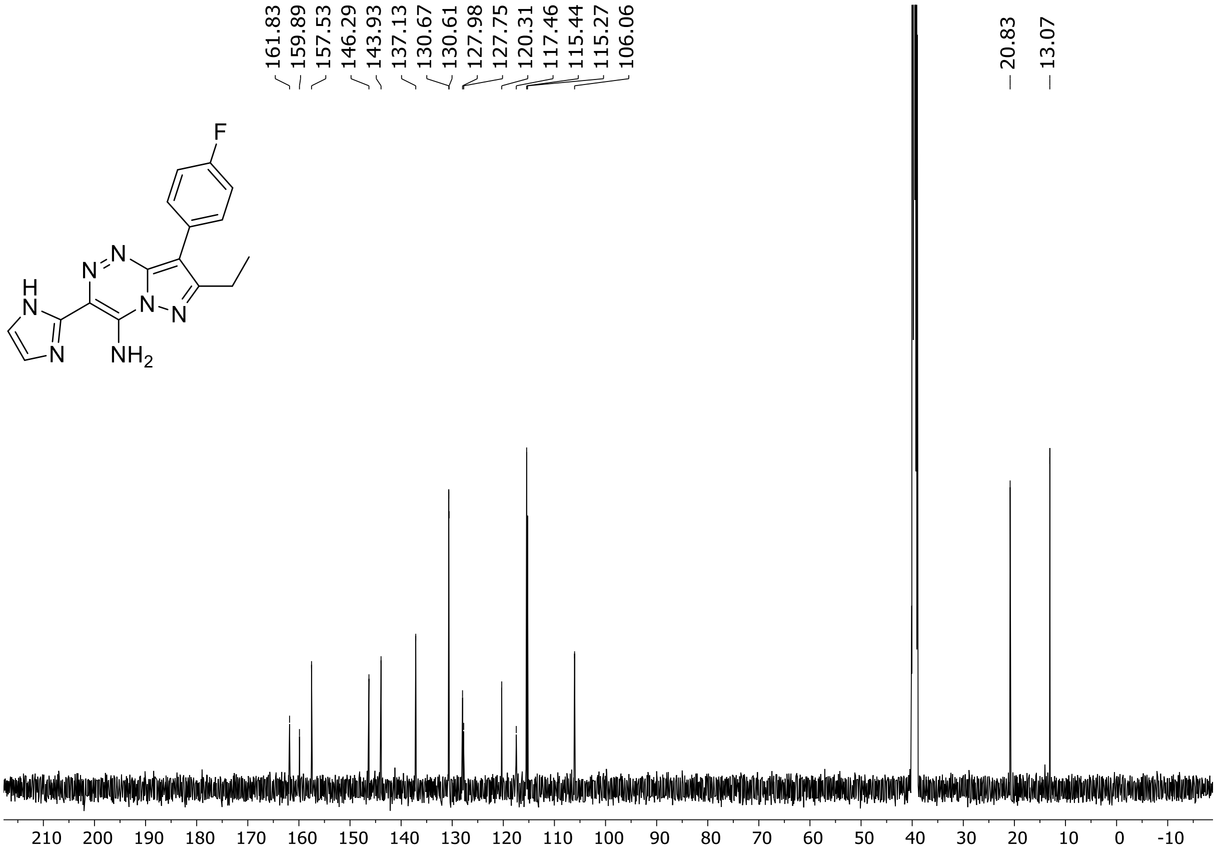

$^{19}$F (471 MHz) NMR spectrum of **27** DMSO-$d_6$

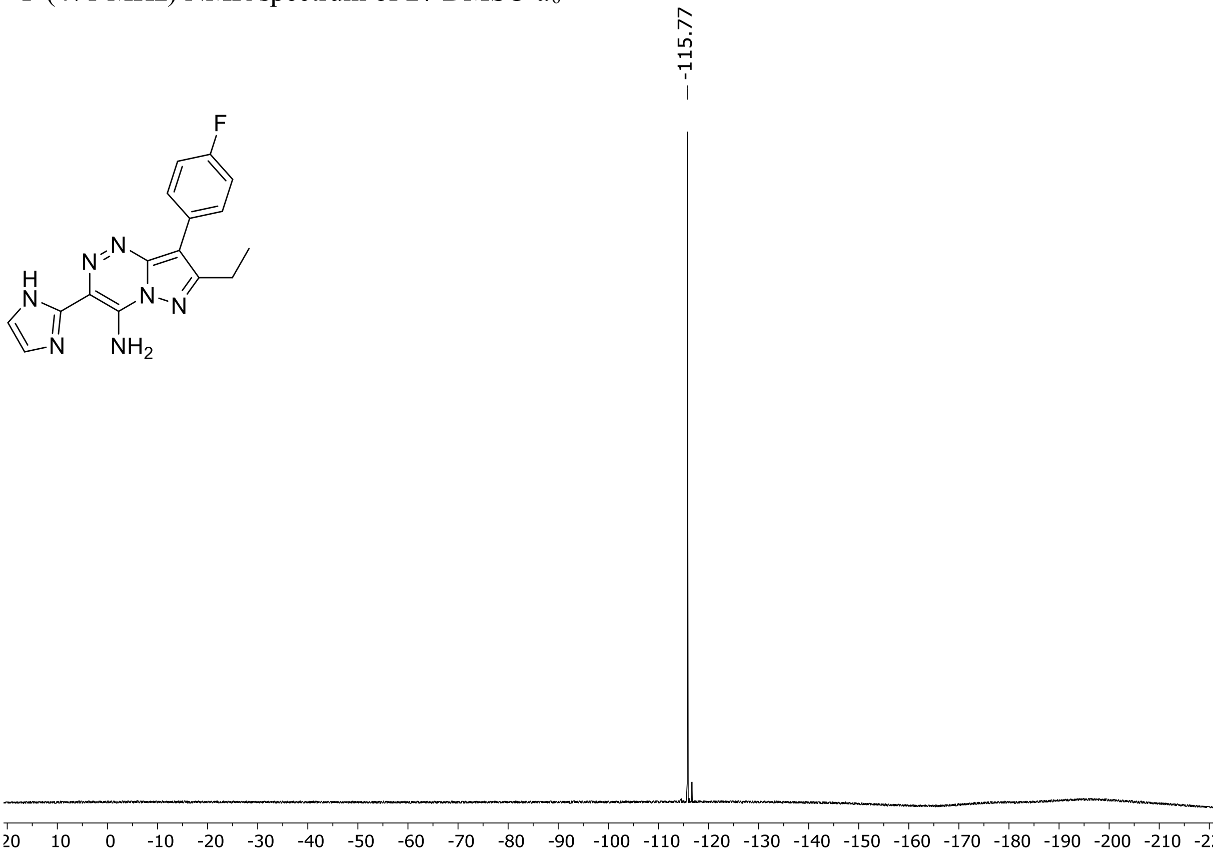

HRMS spectrum of **27**

**$C_{16}H_{14}FN_7$** **exact mass: 323.1295**

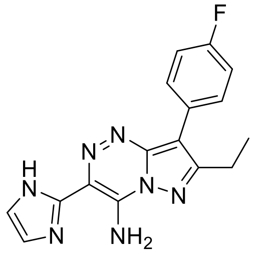

**APCI +** (MMI)
nitrogen flow 5 L/min, gas temperature 325°C, nebulizer 45 psig, skimmer 65 V, vaporizer 200°C, fragmentor 5 V, dissolved in methanol

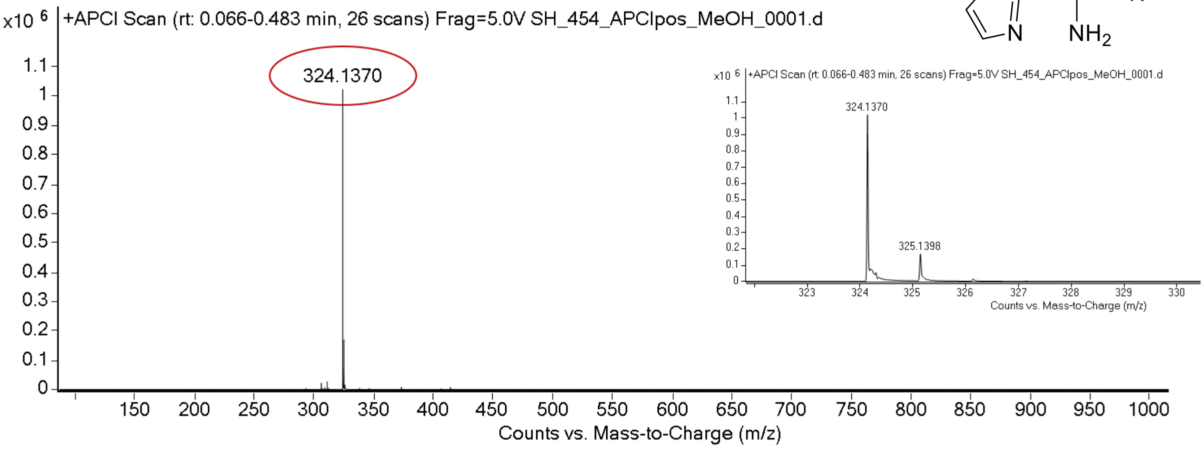

expected mass: [M+H]$^+$ = 324.1367 observed mass : [M+H]$^+$ = 324.1370 mass accuracy = 0.9 ppm

$^{1}H$ (500 MHz) and $^{13}C$ NMR (126 MHz) spectra of **28** in DMSO-$d_6$

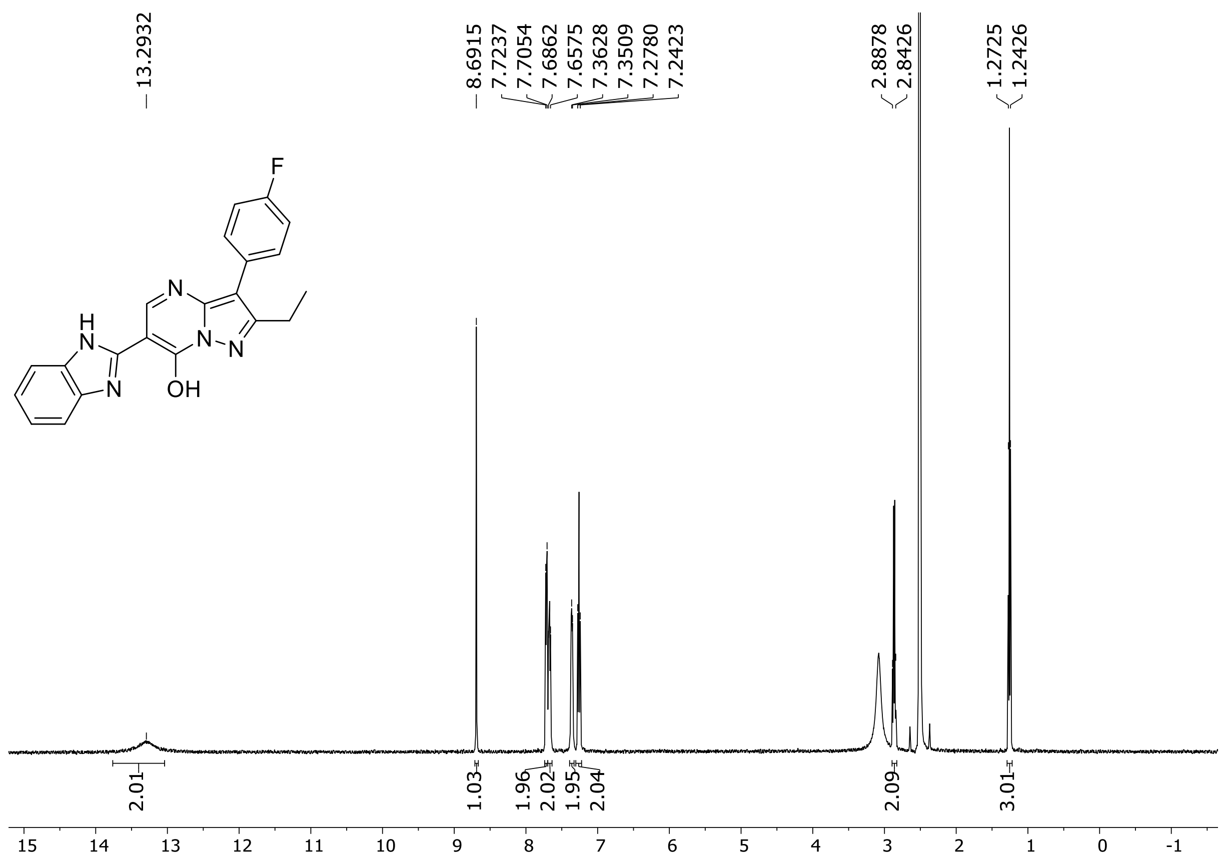

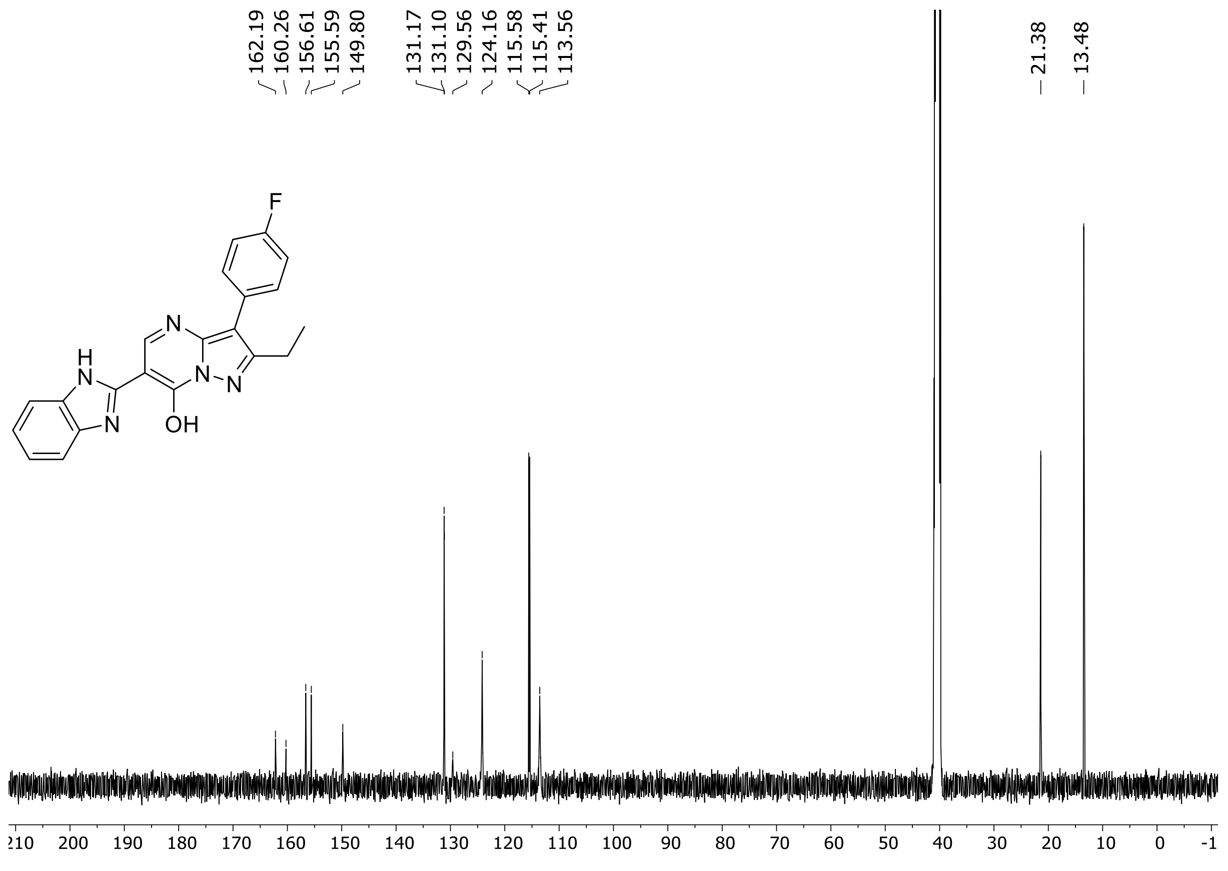

HRMS spectrum of **28**

**$C_{21}H_{16}FN_5O$** **exact mass: 373.1339**

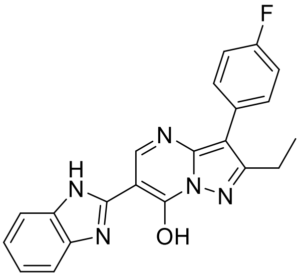

**APCI +** (MMI)
nitrogen flow 5 L/min, gas temperature 325°C, nebulizer 45 psig, skimmer 65 V, vaporizer 200°C, fragmentor 20 V, dissolved in methanol

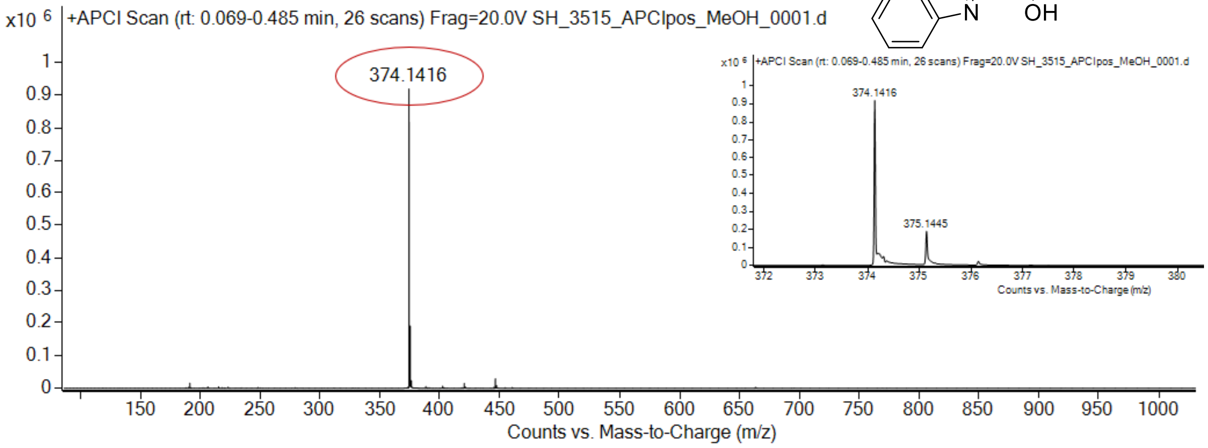

expected mass: [M+H]$^+$ = 374.1412 observed mass : [M+H]$^+$ = 374.1416 mass accuracy = 1.1 ppm

#### $^1$H NMR, $^{13}$C NMR and HRMS spectra of compound 56-117

$^1$H (500 MHz) and $^{13}$C NMR (126 MHz) spectra of **56** in DMSO-$d_6$

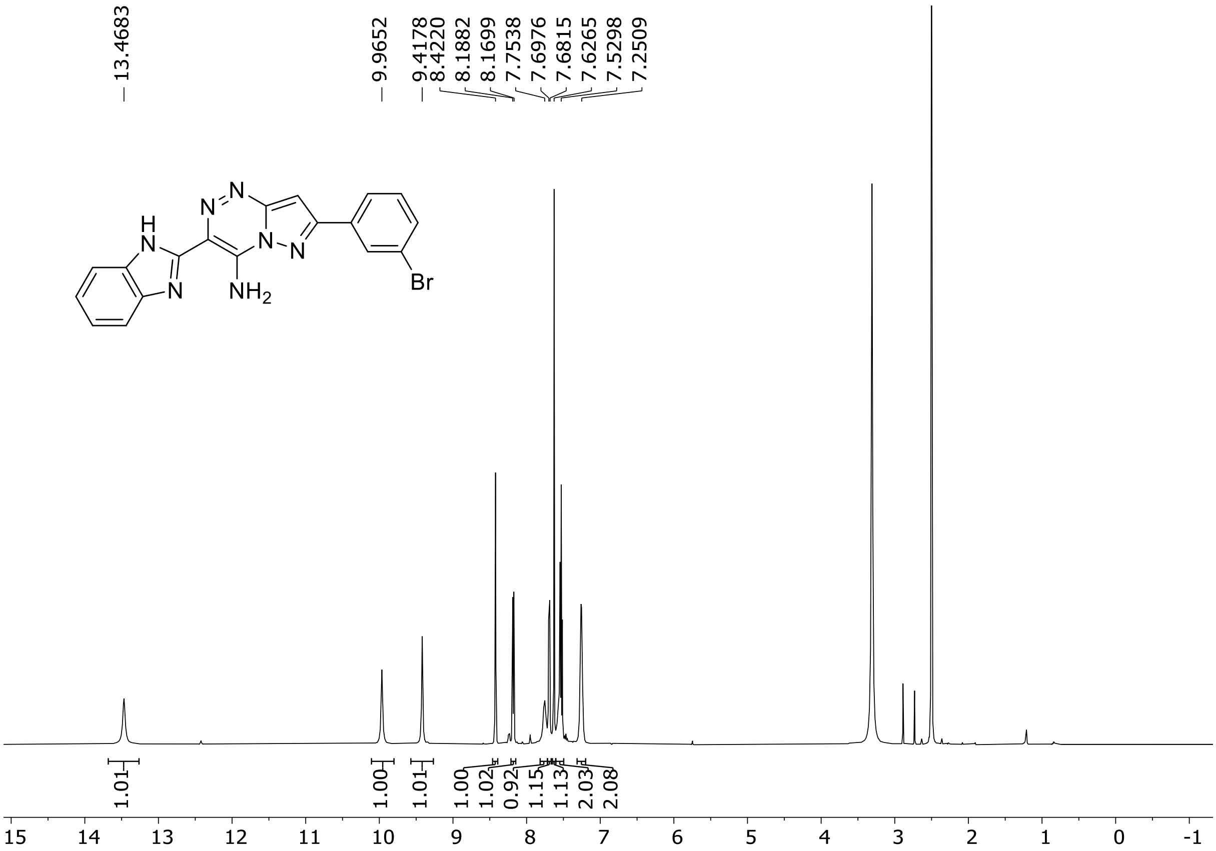

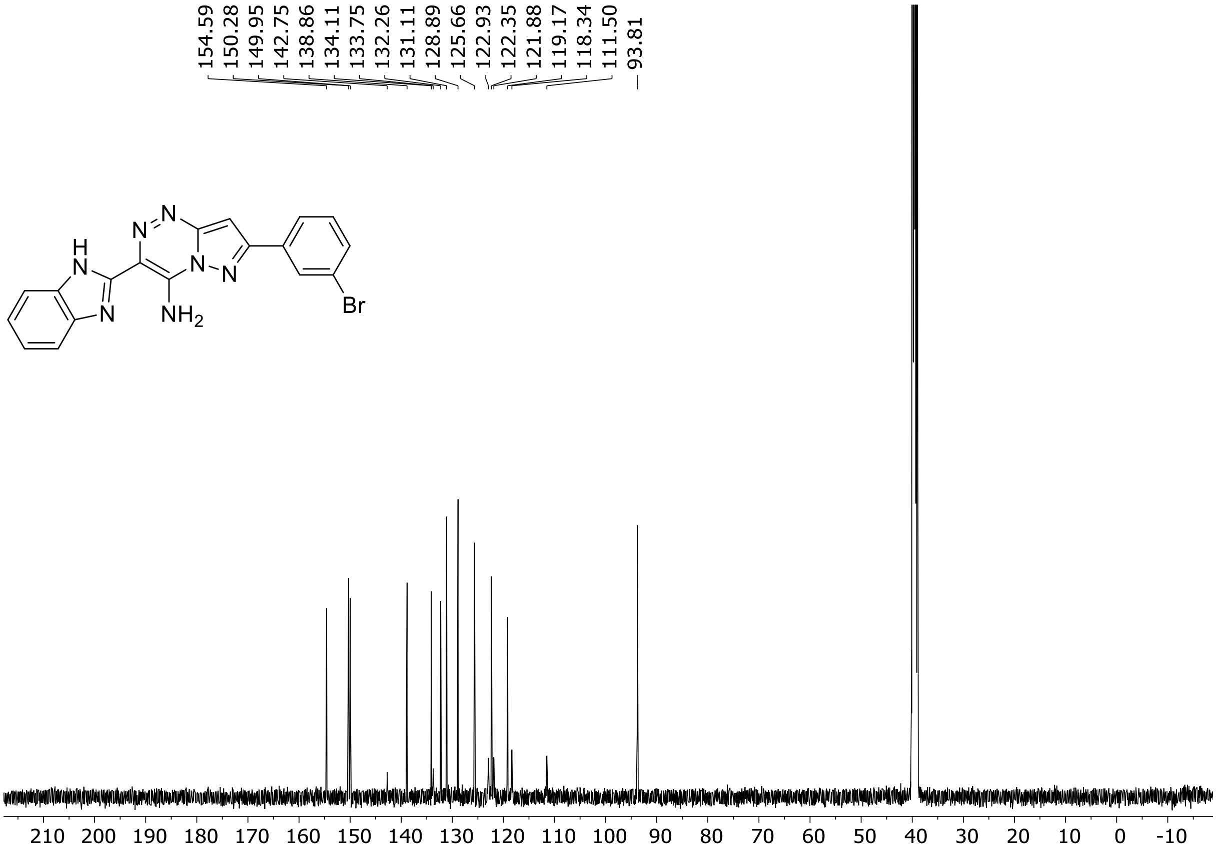

HRMS spectrum of **56**

**$C_{18}H_{12}BrN_7$ mono m/z = 405.0338**

**APCI +** (MMI)
nitrogen flow 5 L/min, gas temperature 300°C, nebulizer 45 psi,
skimmer 60 V, fragmentor 37 V, dissolved in methanol

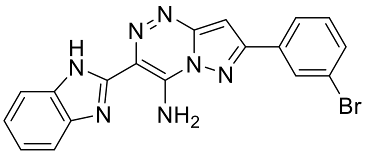

ZOOM – range of interest

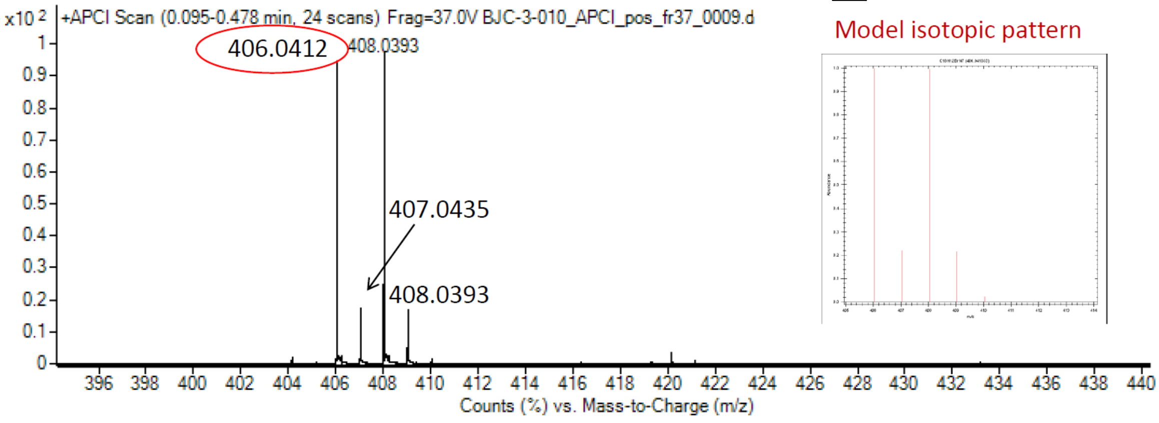

calculated mass: [M+H]+= 406. 0410 observed: $[M+H]^+$ = 406.0412 max. mass error = 0.5 ppm

$^{1}H$ (500 MHz) and $^{13}C$ NMR (126 MHz) spectra of **57** in DMSO-$d_6$

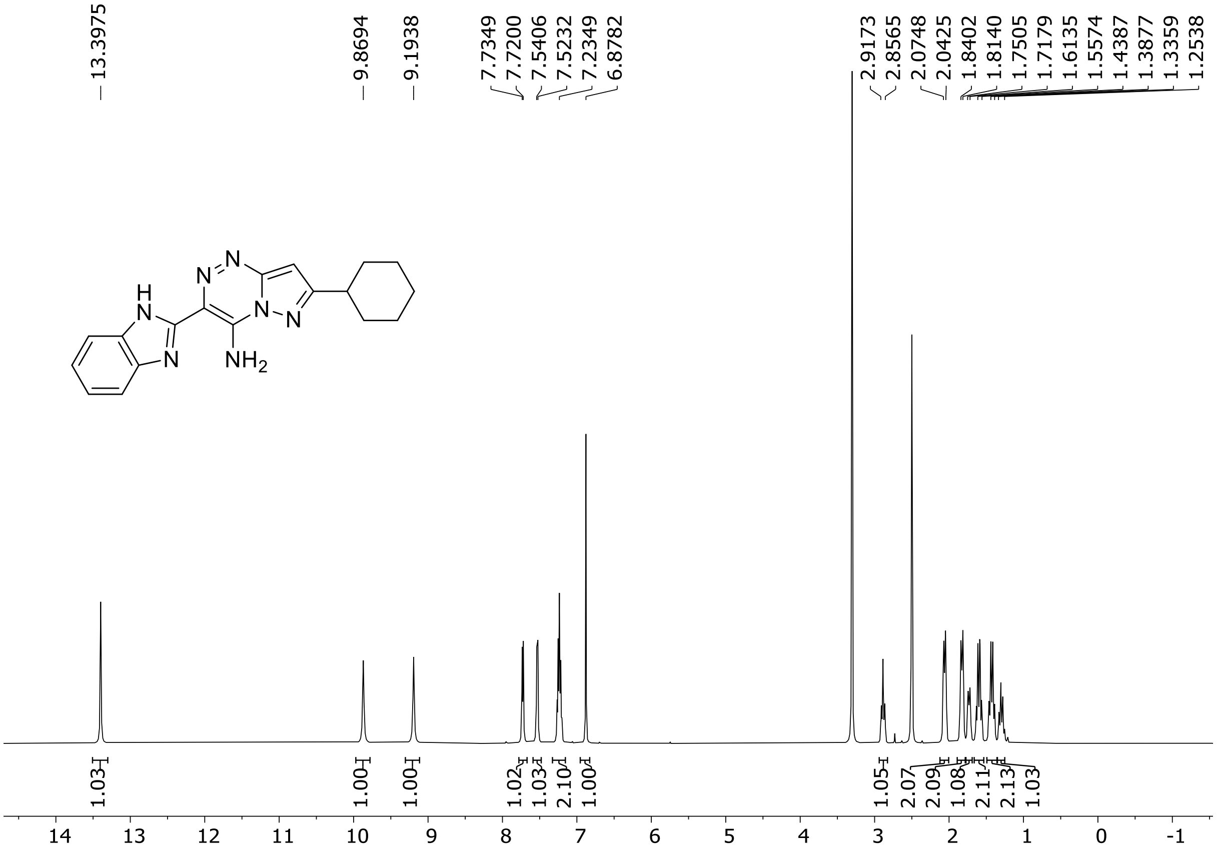

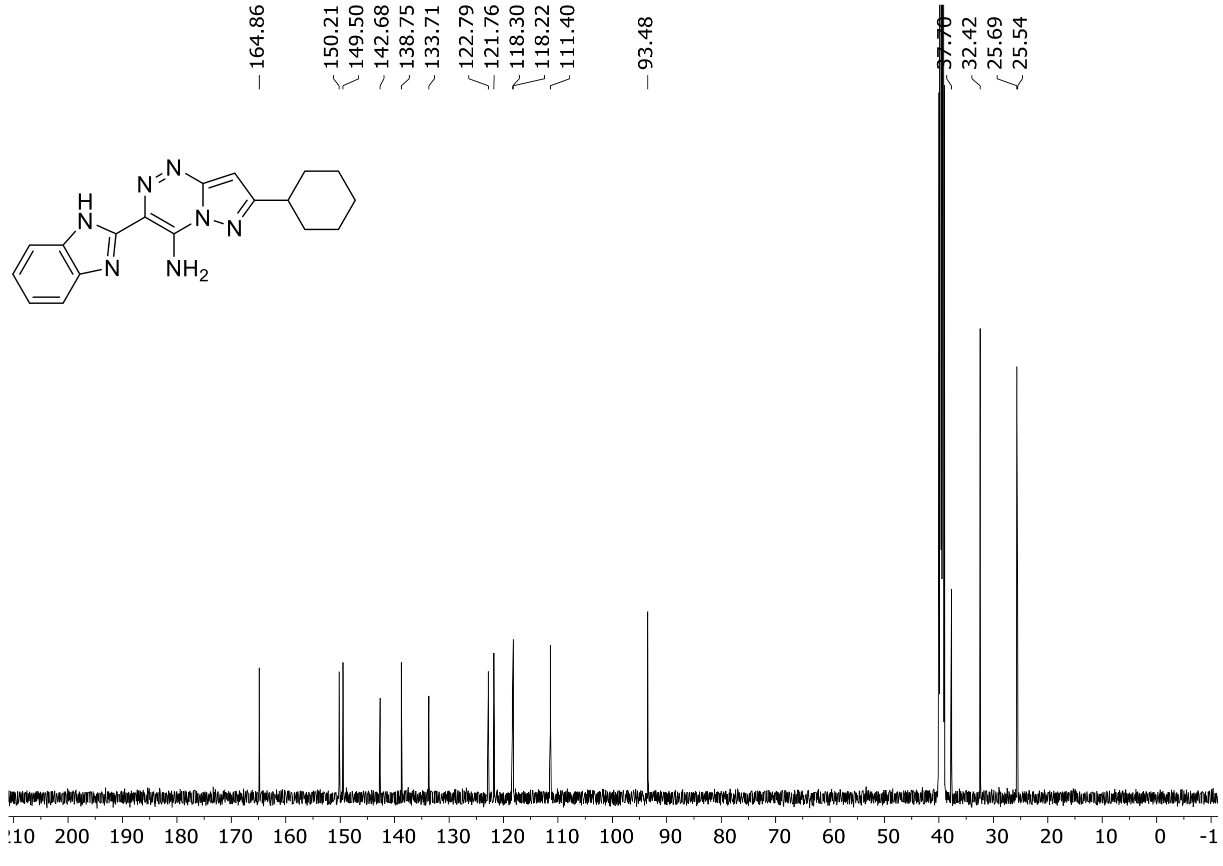

HRMS spectrum of **57**

**$C_{18}H_{19}N_7$ mono m/z = 333.1702**

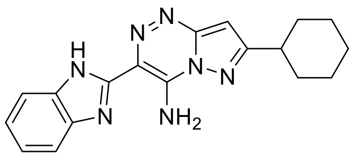

**APCI +** (MMI)
nitrogen flow 5 L/min, gas temperature 300°C, nebulizer 45 psi,
skimmer 60 V, fragmentor 17 V, dissolved in methanol

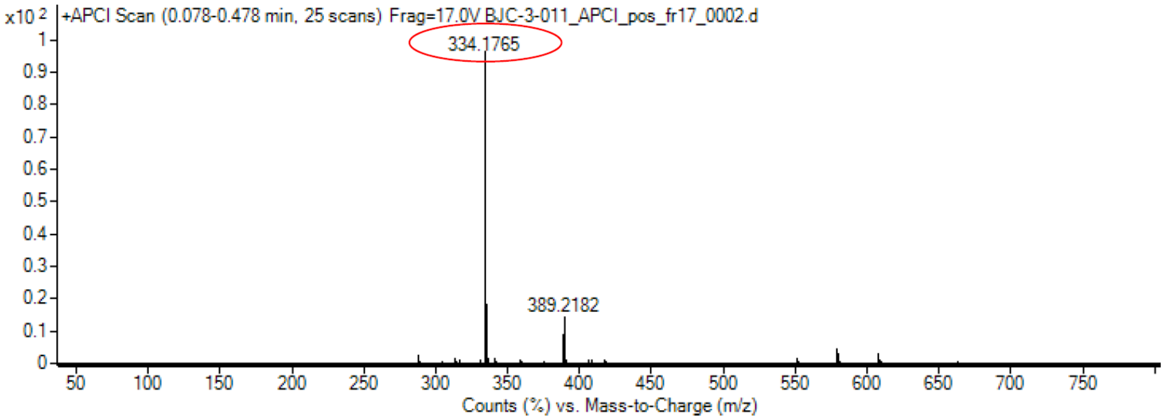

calculated mass: $[M+H]^+$ = 334.1775 observed: $[M+H]^+$ = 334.1765 max. mass error = 3.0 ppm

$^{1}H$ (500 MHz) and $^{13}C$ NMR (126 MHz) spectra of **58** in DMSO-$d_6$

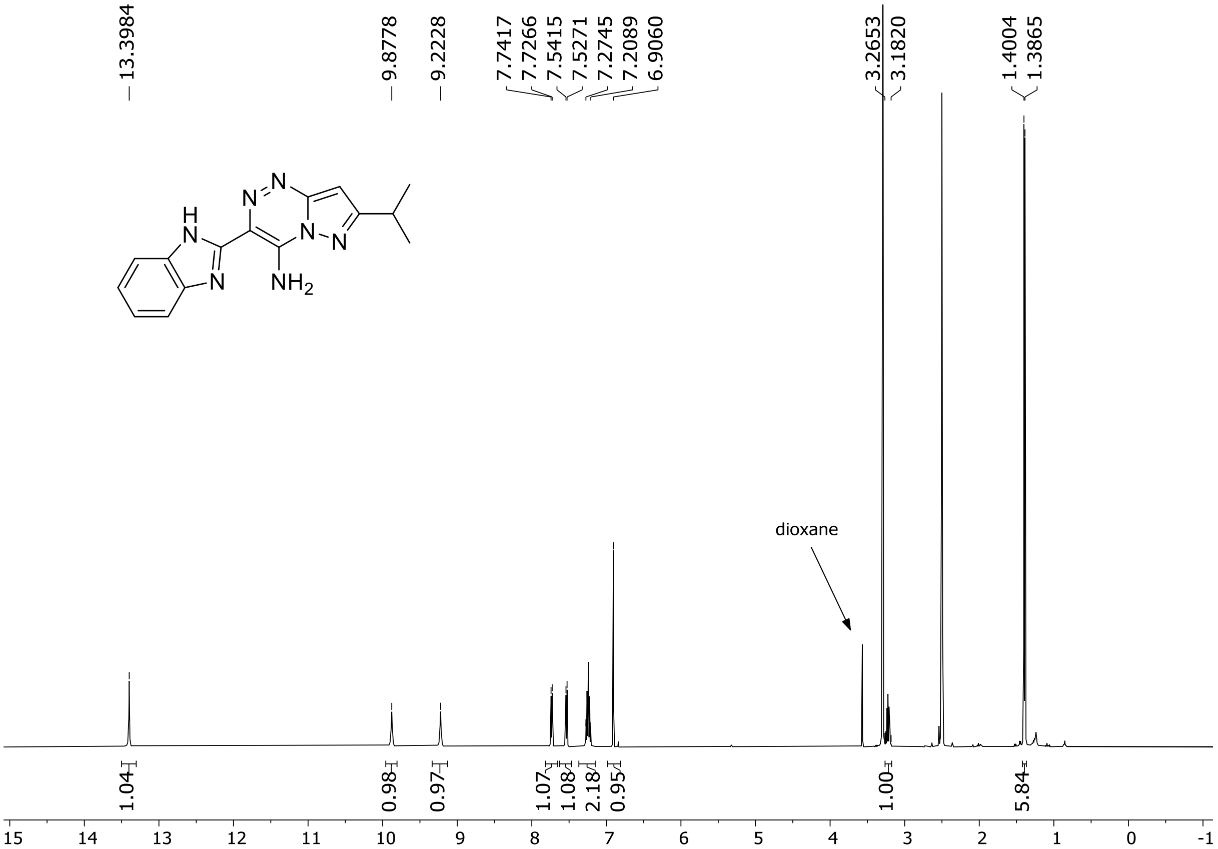

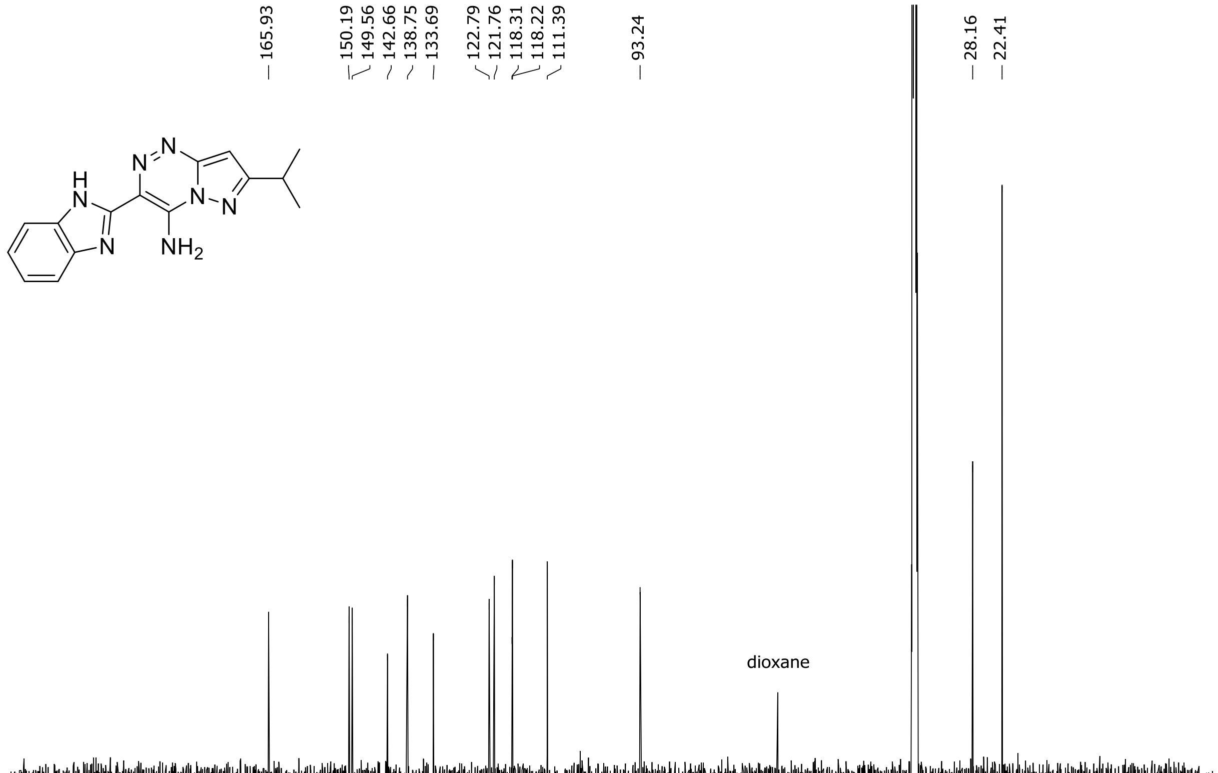

HRMS spectrum of **58**

$C_{15}H_{15}N_7$ **mono m/z = 293.1389**

**APCI +** (MMI)
nitrogen flow 5 L/min, gas temperature 300°C, nebulizer 45 psi, vaporizer 200°C
skimmer **65** V, fragmentor **20** V, dissolved in methanol

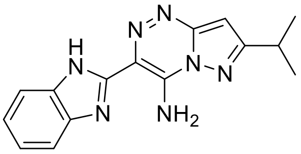

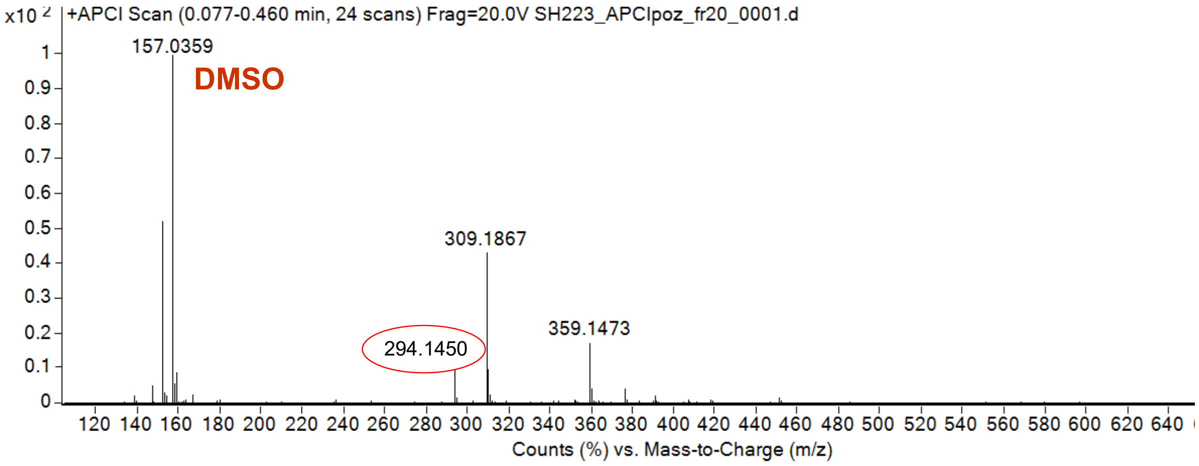

calculated mass: $[M+H]^+$ = 294.1462 observed: $[M+H]^+$ = 294.1450 max. mass error = 4 ppm

$^{1}H$ (500 MHz) and $^{13}C$ NMR (126 MHz) spectra of **59** in DMSO-$d_6$

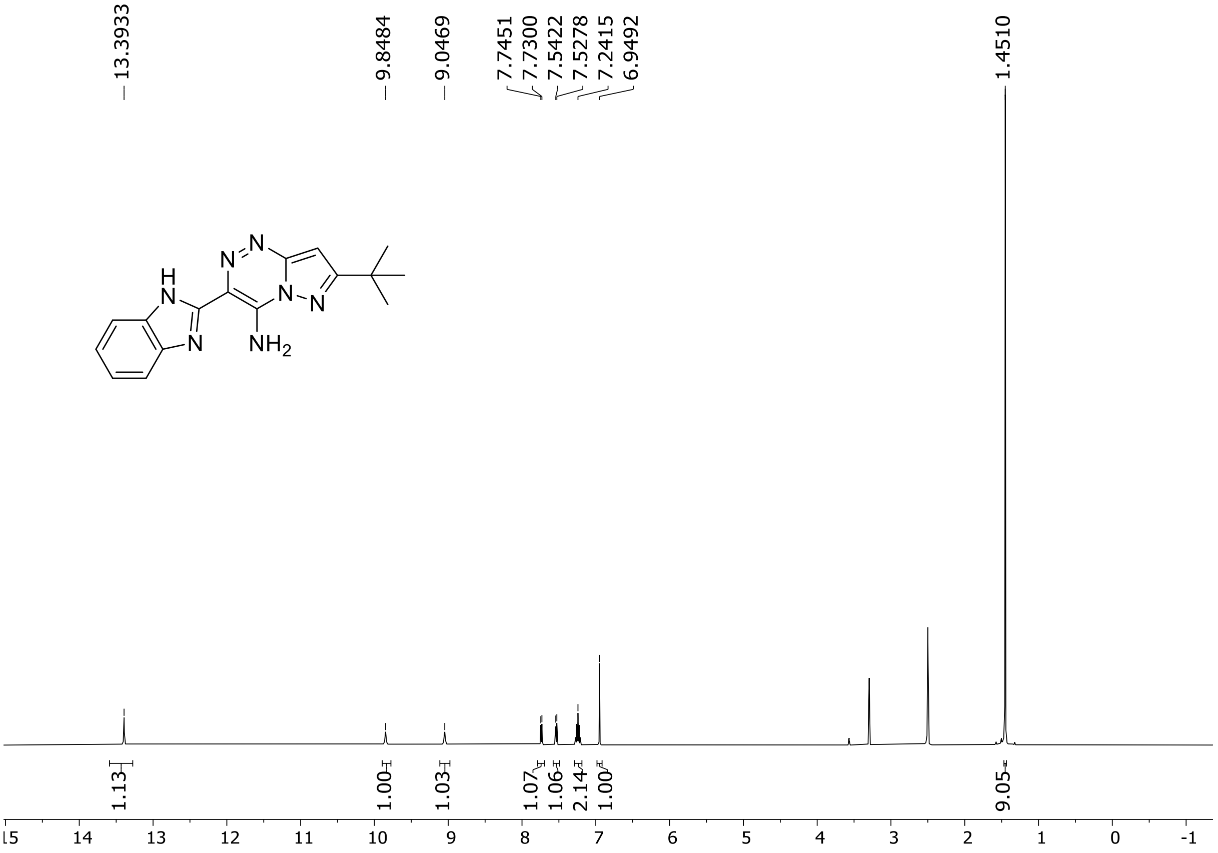

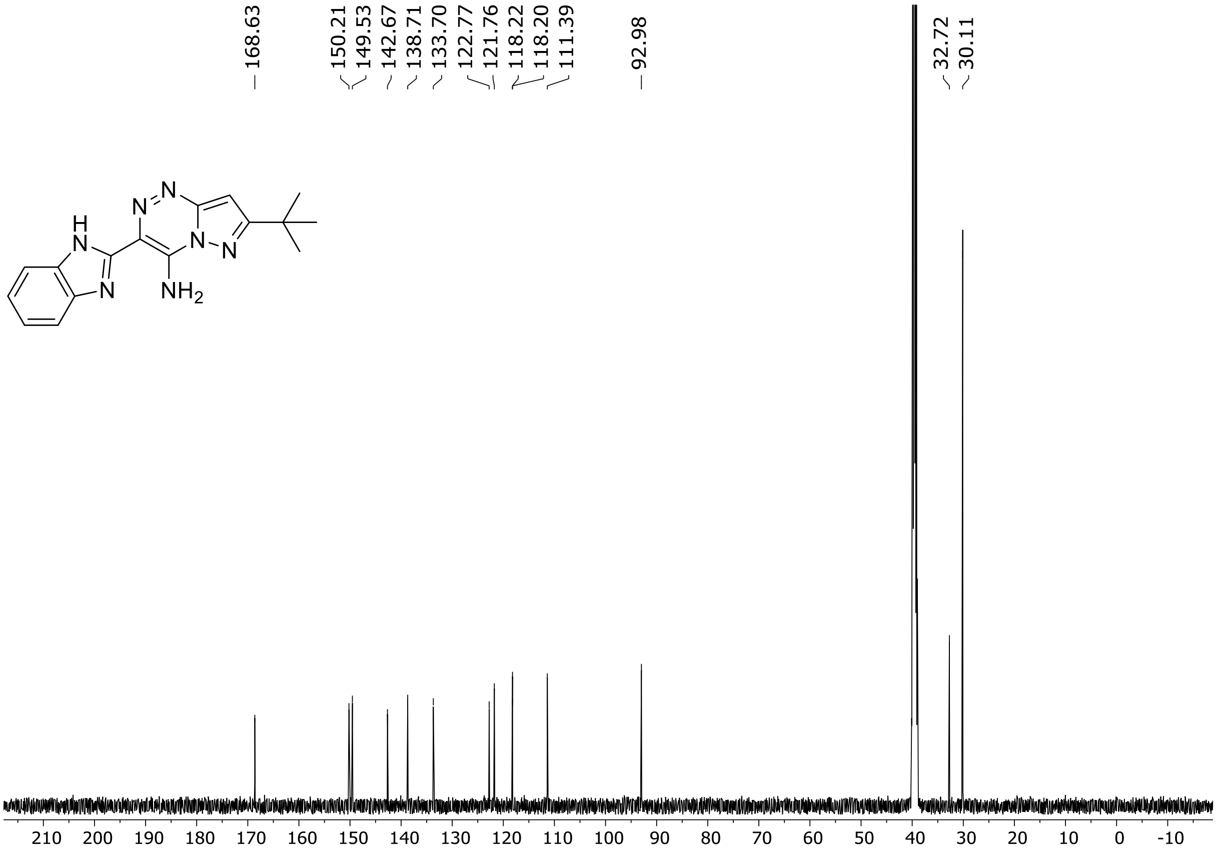

HRMS spectrum of **59**

$C_{16}H_{17}N_7$ **mono m/z = 307.1545**

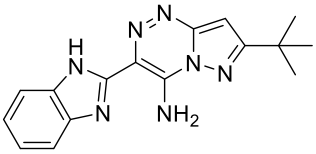

**ESI -** (MMI)
nitrogen flow 5 L/min, gas temperature 300°C, vaporizer 250°C, nebulizer 45 psi, skimmer **40** V, fragmentor **40** V, dissolved in DMSO

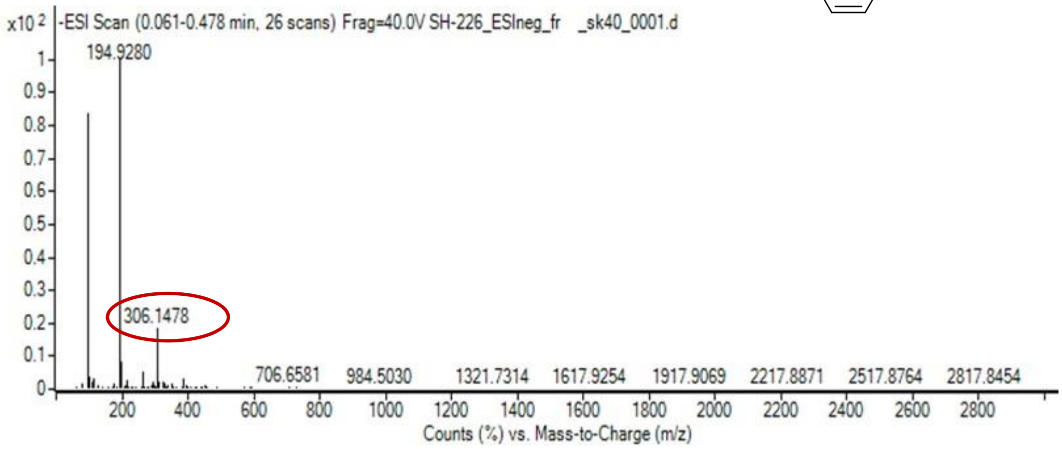

calculated mass: [M-H] $^-$ = 306.1473 observed: [M-H] $^-$ =306.1478 m ax. mass error = 1.6 ppm

[1]H (500 MHz) and [13]C NMR (126 MHz) spectra of **60** in DMSO-$d_6$

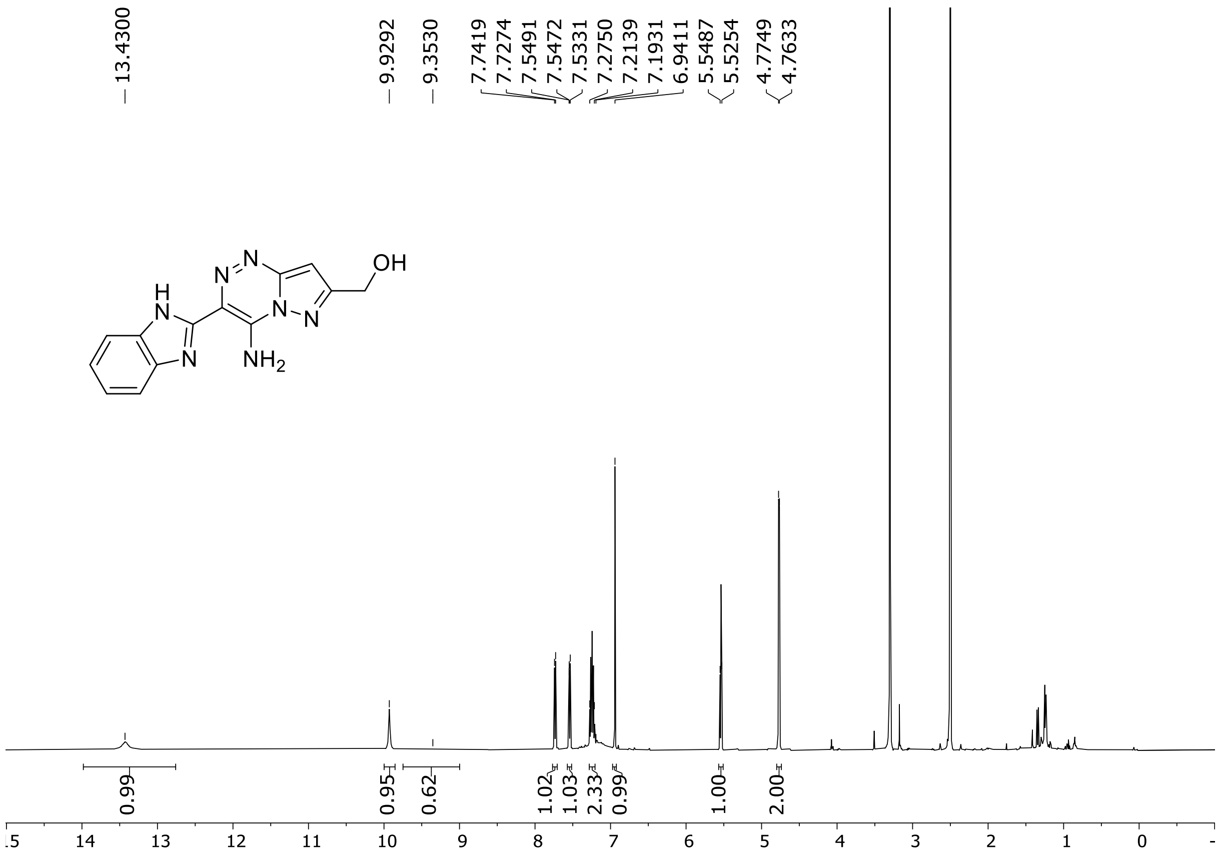

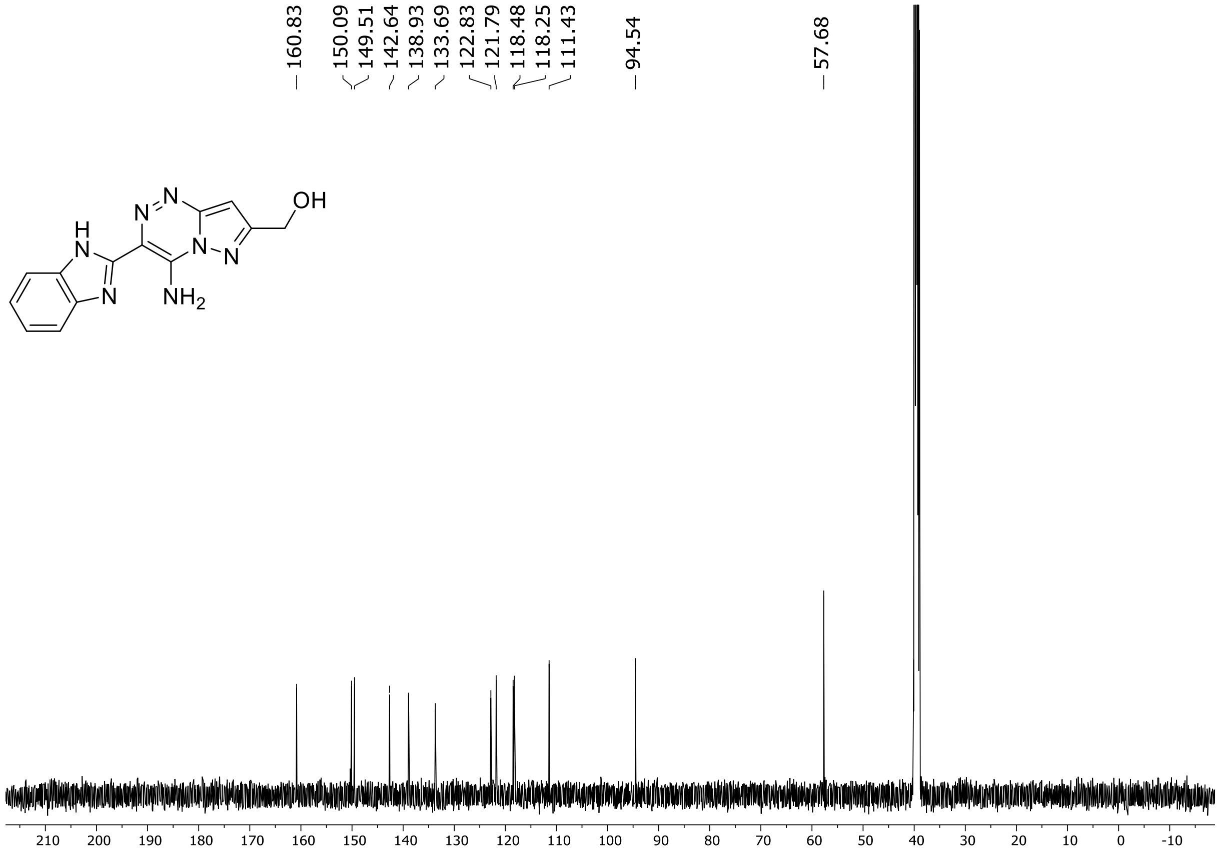

HRMS spectrum of **60**

$C_{13}H_{11}N_7O$ **mono m/z = 281.1025**

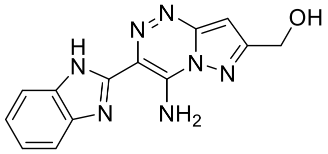

**APCI +** (MMI)
nitrogen flow 5 L/min, gas temperature 300°C, nebulizer 45 psi, vaporizer 200°C
skimmer **65** V, fragmentor **20** V, dissolved in methanol

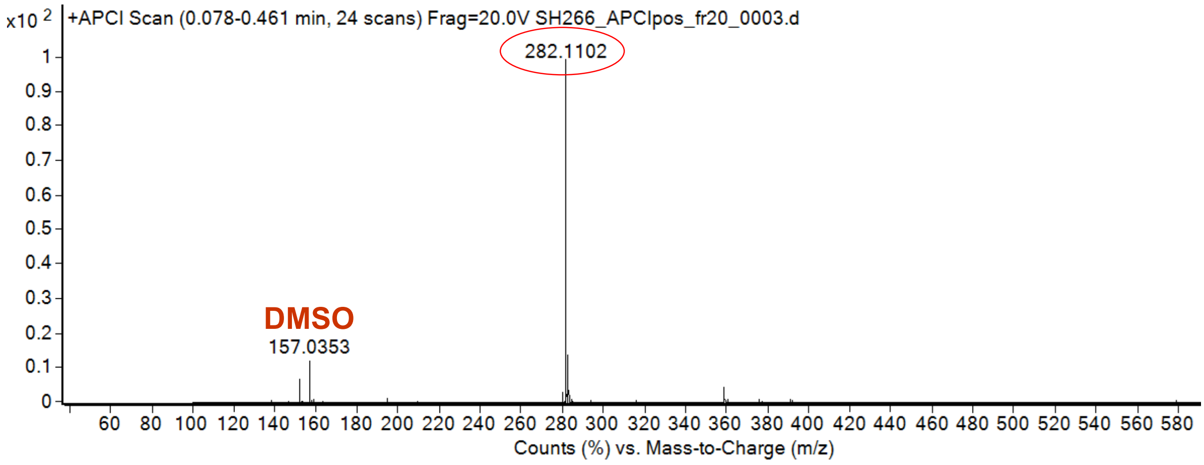

calculated mass: $[M+H]^+$ = 282.1098 observed: $[M+H]^+$ = 282.1102 max. mass error = 1.4 ppm

$^{1}H$ (500 MHz) and $^{13}C$ NMR (126 MHz) spectra of **61** in DMSO-$d_6$

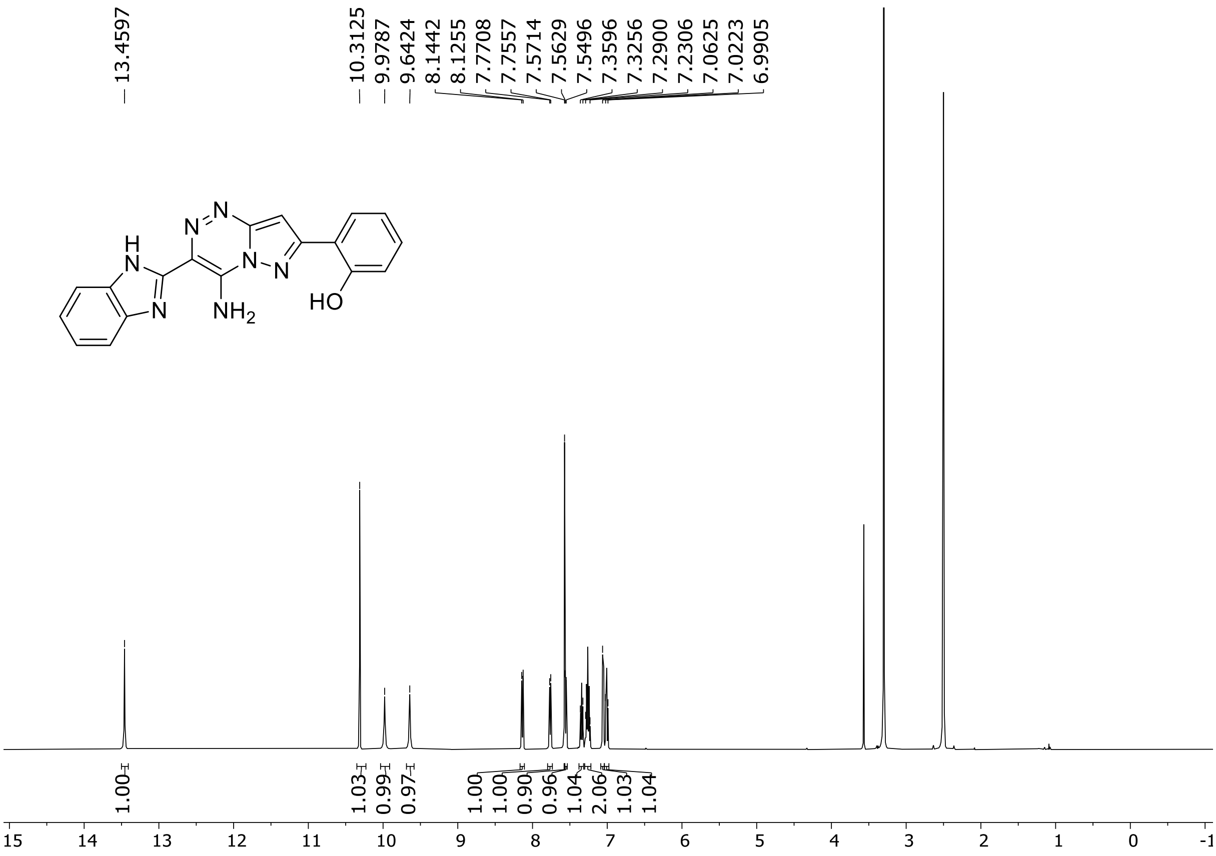

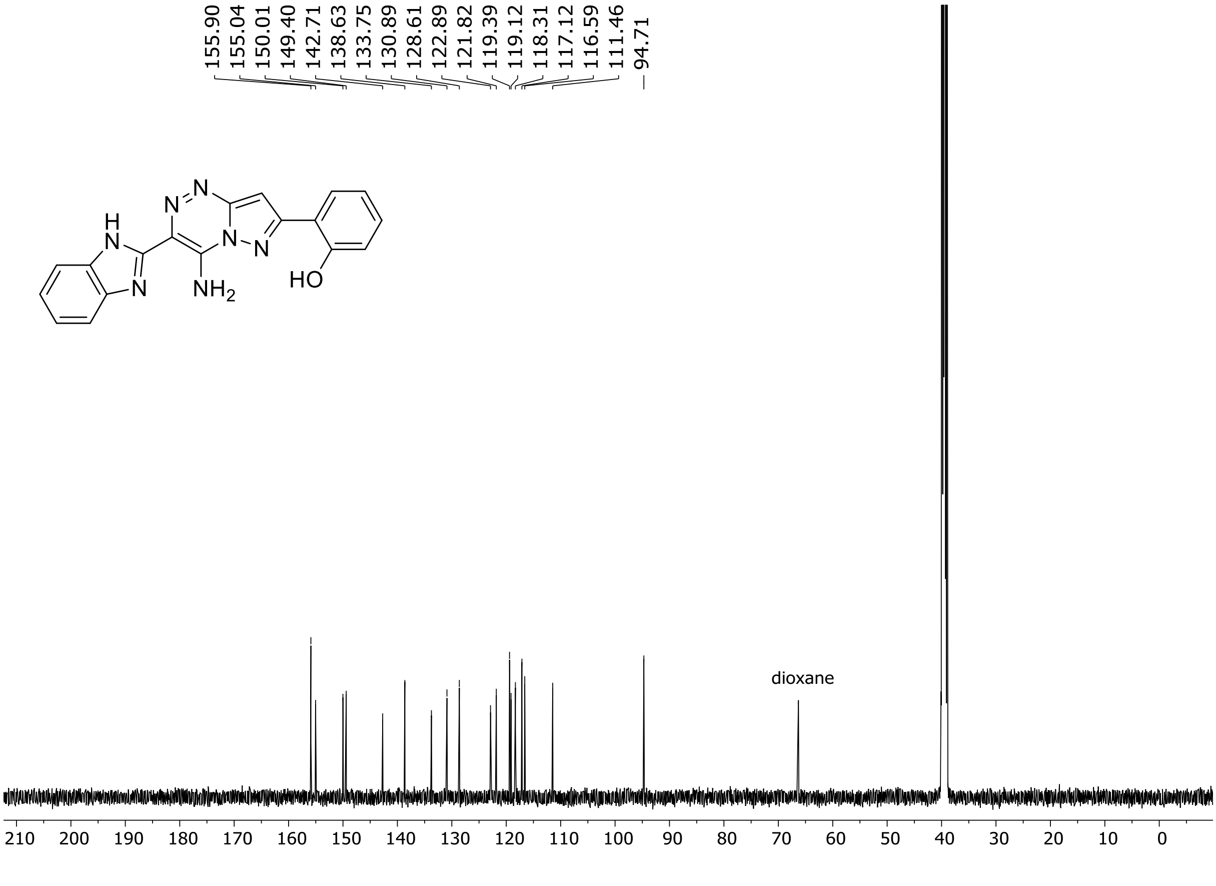

HRMS spectrum of **61**

$C_{18}H_{13}N_7O$ **mono m/z = 343.1182**

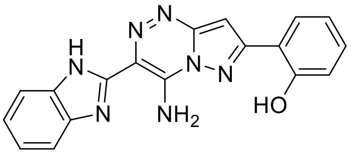

**APCI +** (MMI)
nitrogen flow 5 L/min, gas temperature 300°C, nebulizer 45 psi, vaporizer 200°C
skimmer **65** V, fragmentor 25 V, dissolved in methanol

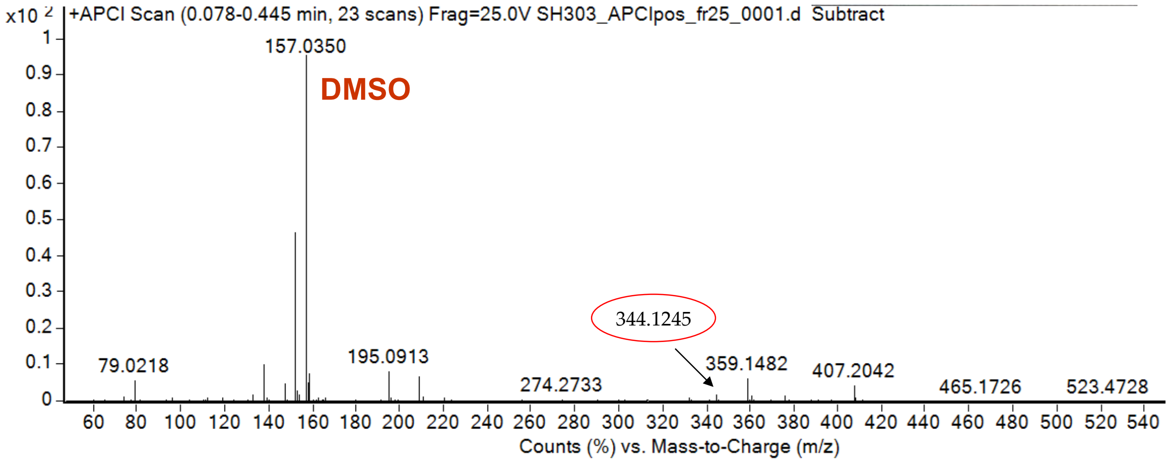

calculated mass: $[M+H]^+$ = 344.1254 observed: $[M+H]^+$ = 344.1245 max. mass error = 2.6 ppm

$^{1}H$ (500 MHz) and $^{13}C$ NMR (126 MHz) spectra of **62** in DMSO-$d_6$

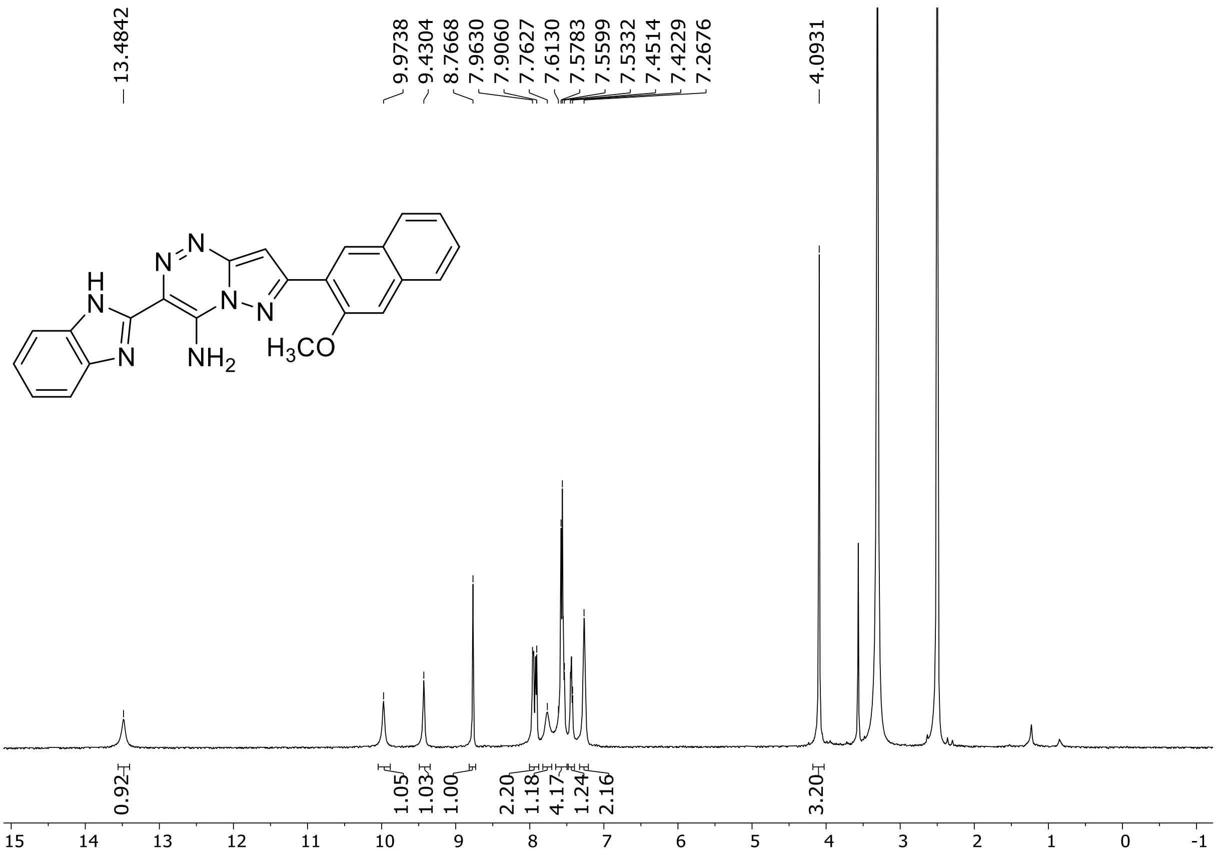

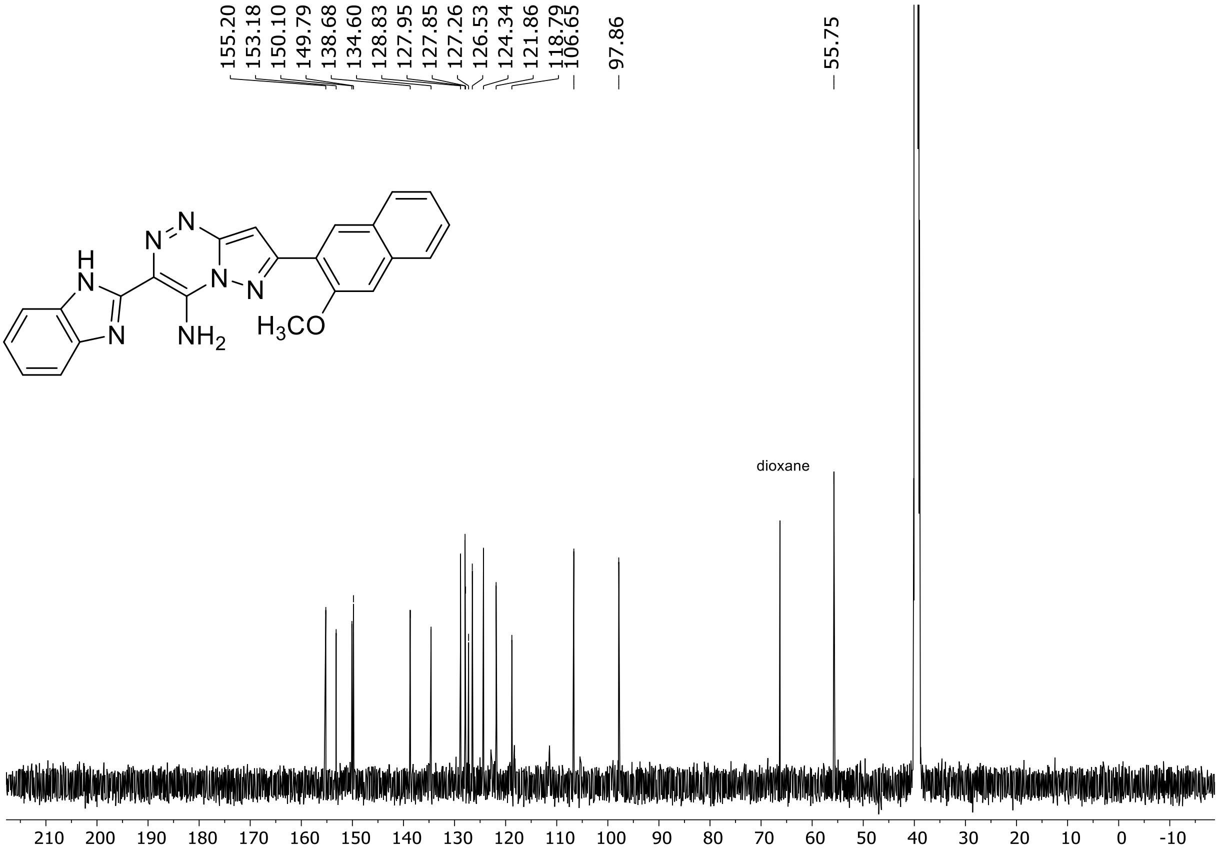

HRMS spectrum of **62**

**$C_{23}H_{17}N_7O$** **exact mass: 407.1495**

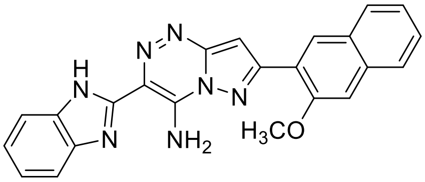

**APCI +** (MMI)
nitrogen flow 5 L/min, gas temperature 325°C, nebulizer 45 psig, skimmer 65 V, vaporizer 200°C, fragmentor 5 V, dissolved in methanol

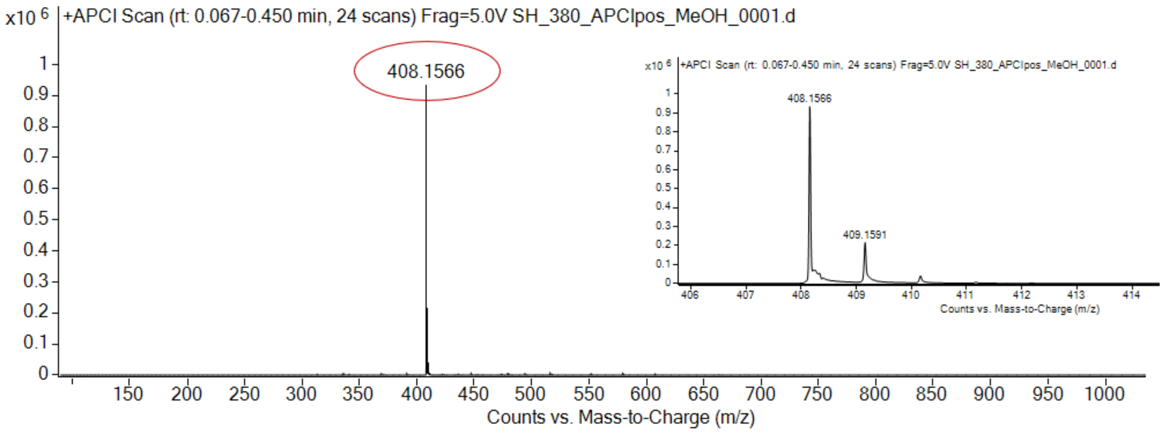

expected mass: $[M+H]^+$ = 408.1567 observed mass : $[M+H]^+$ = 408.1566 mass accuracy = - 0.2 ppm

$^{1}H$ (500 MHz) and $^{13}C$ NMR (126 MHz) spectra of **63** in DMSO-$d_6$

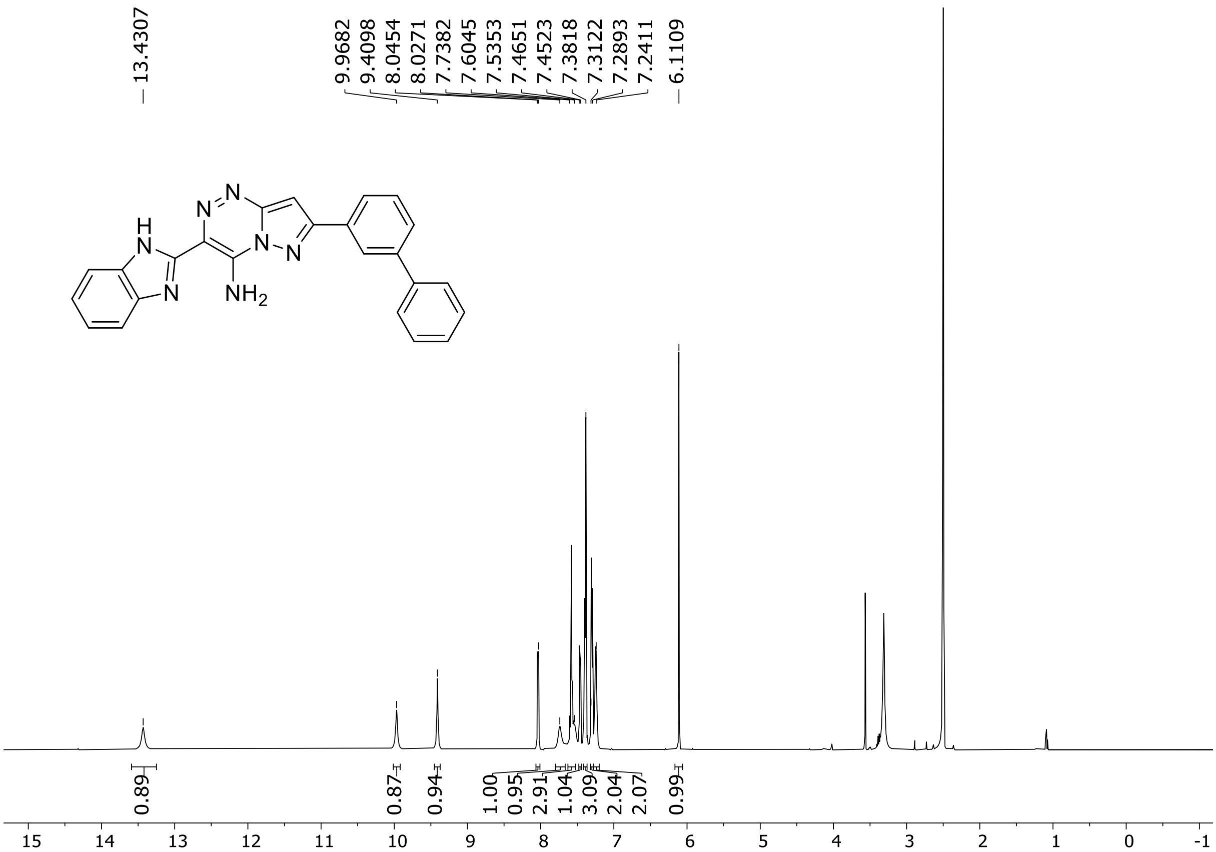

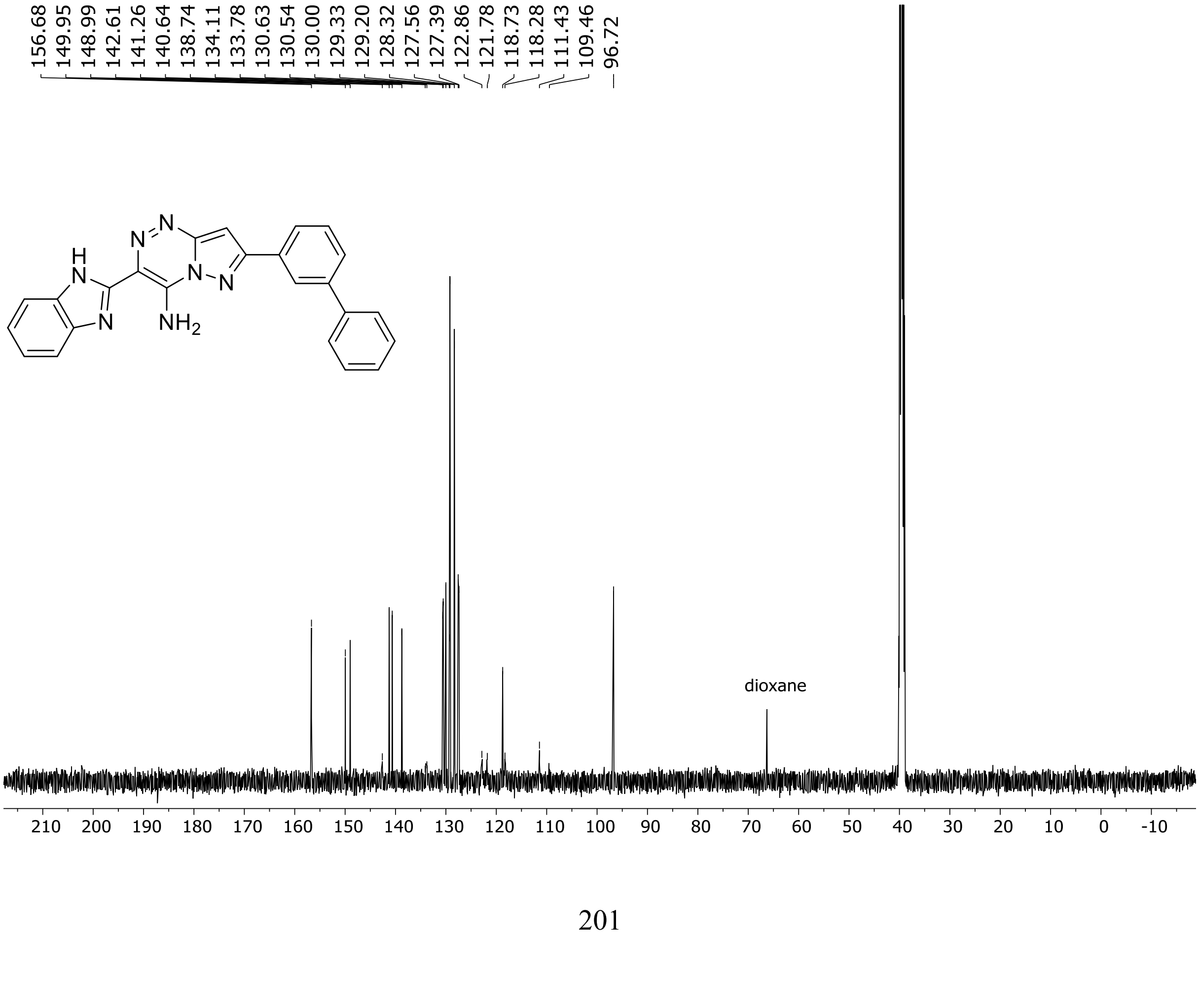

HRMS spectrum of **63**

$C_{24}H_{17}N_7$ **mono m/z = 403.1545**

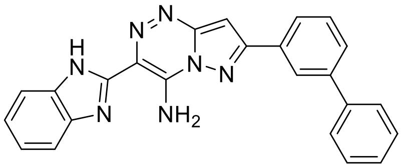

**APCI +** (MMI)
nitrogen flow 5 L/min, gas temperature 300°C, nebulizer 45 psi, vaporizer 200°C
skimmer **65** V, fragmentor **23** V, dissolved in methanol

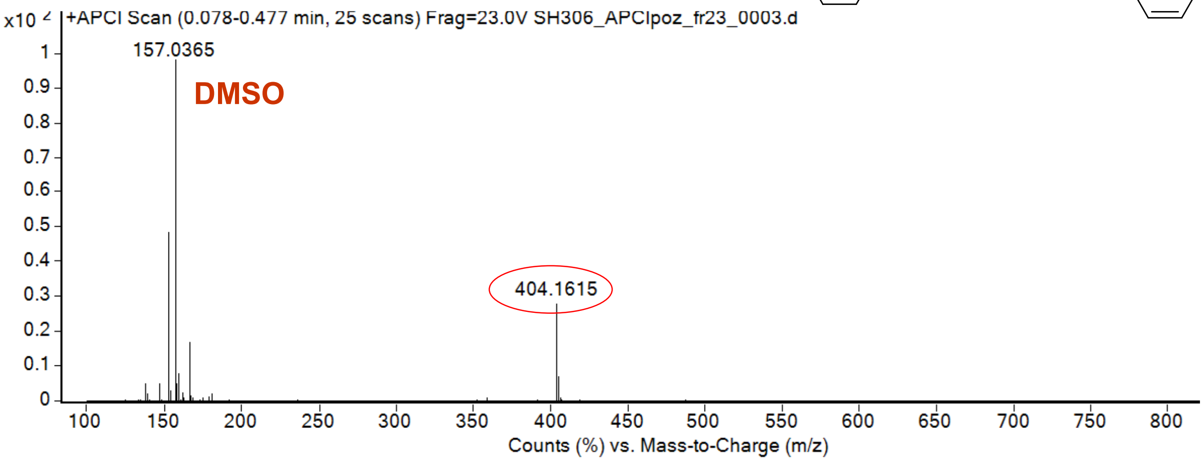

calculated mass: $[M+H]^+$ = 404.1618 observed: $[M+H]^+$ = 404.1615 max. mass error = 0.7 ppm

[1]H (500 MHz) and [13]C NMR (126 MHz) spectra of **64** in DMSO-$d_6$

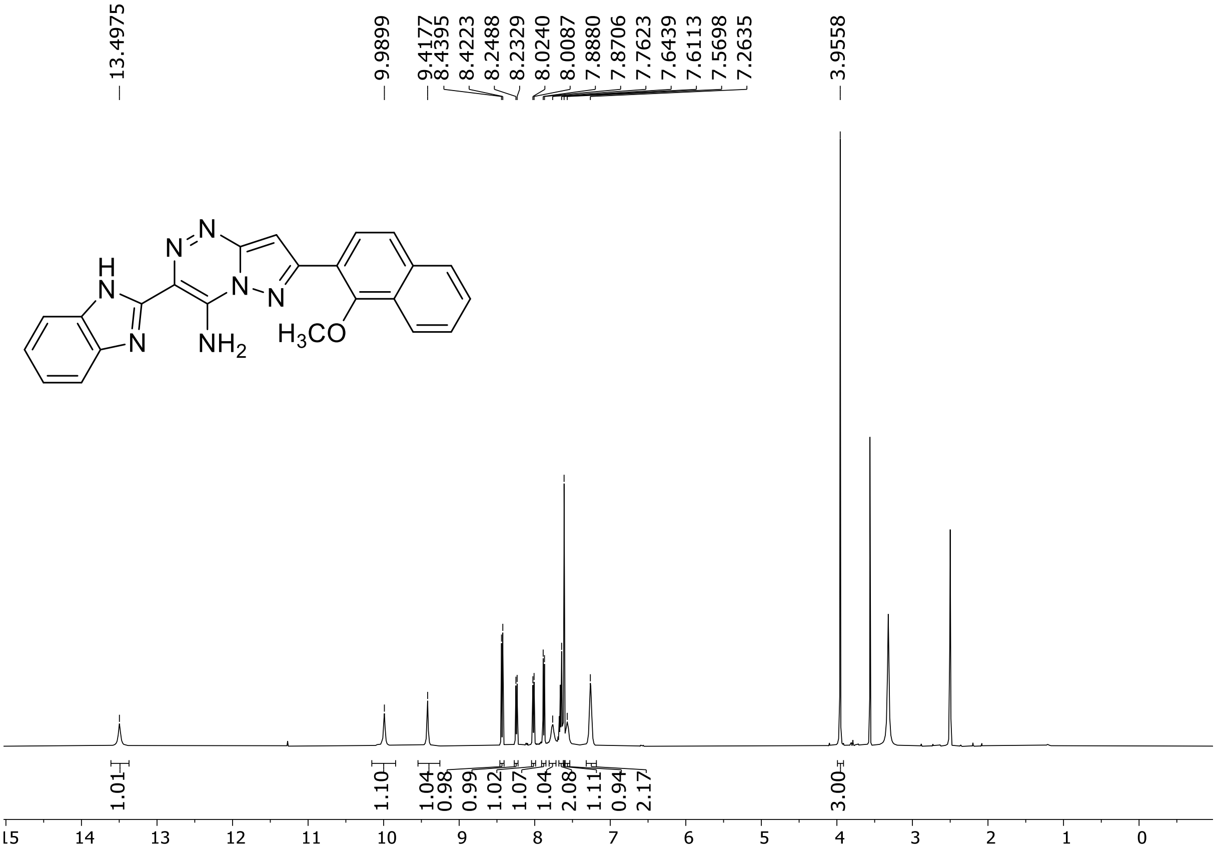

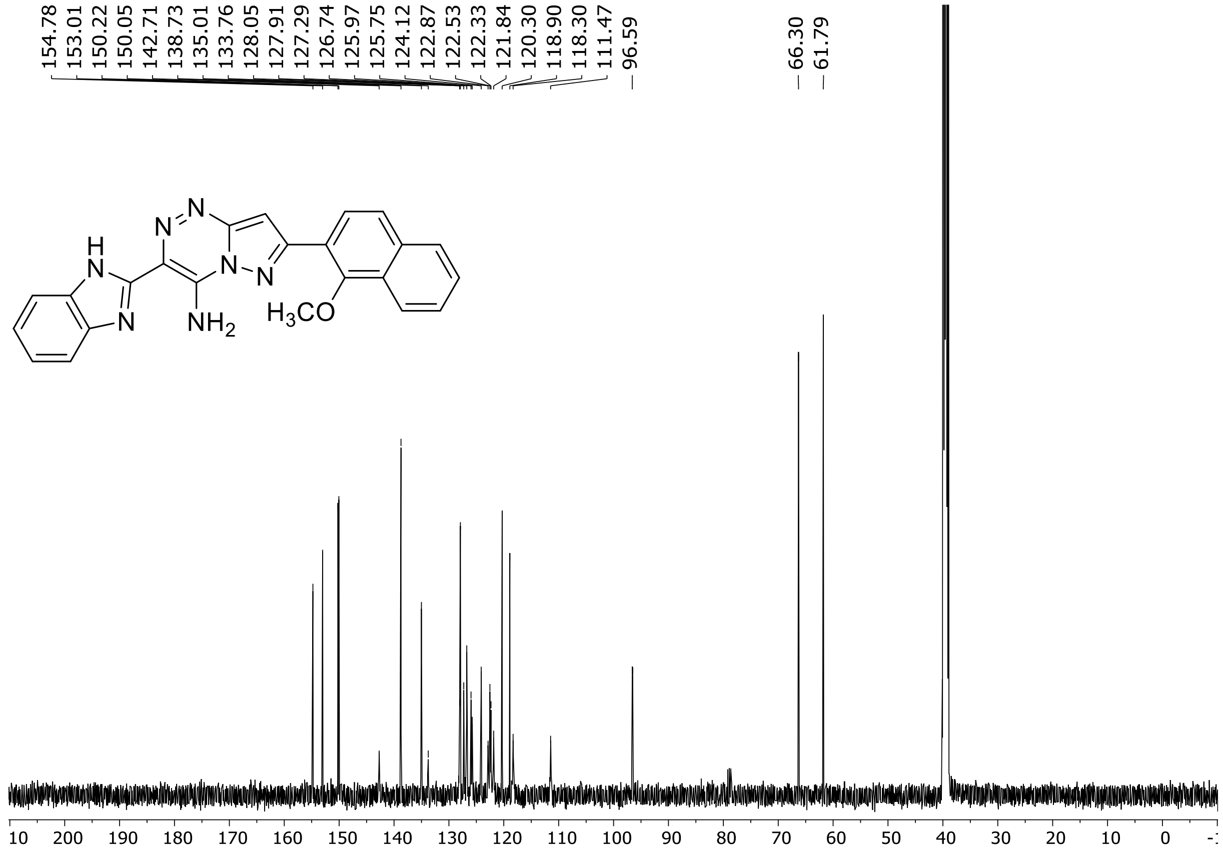

HRMS spectrum of **64**

**$C_{23}H_{17}N_7O$** **exact mass: 407.1495**

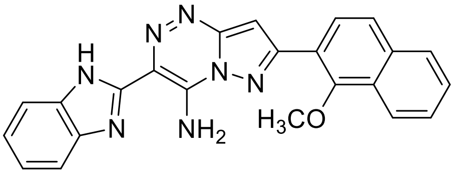

**APCI +** (MMI)
nitrogen flow 5 L/min, gas temperature 325°C, nebulizer 45 psig, skimmer 65 V, vaporizer 200°C, fragmentor 10 V, dissolved in methanol

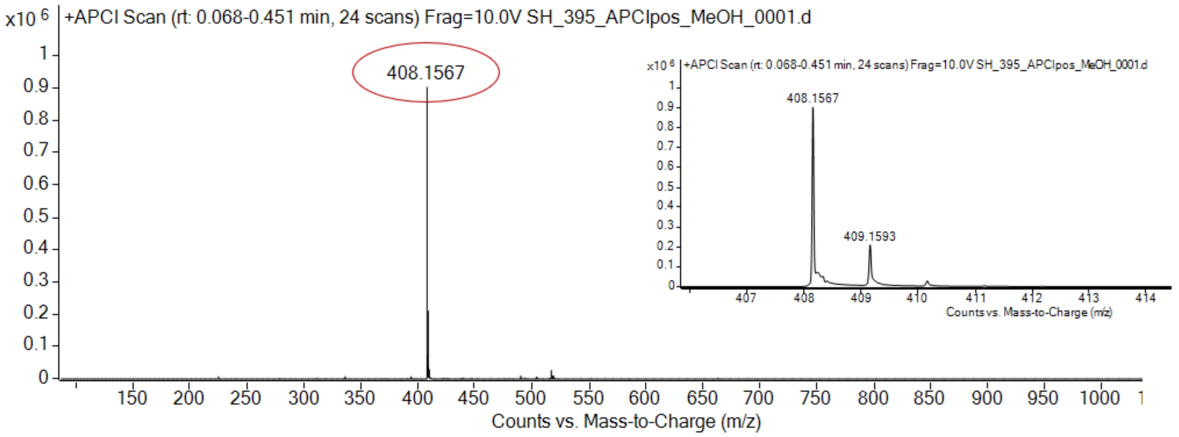

expected mass: $[M+H]^+ = 408.1567$ observed mass : $[M+H]^+ = 408.1567$ mass accuracy < 0.1 ppm

[1]H (500 MHz) and [13]C NMR (126 MHz) spectra of **65** in DMSO-$d_6$

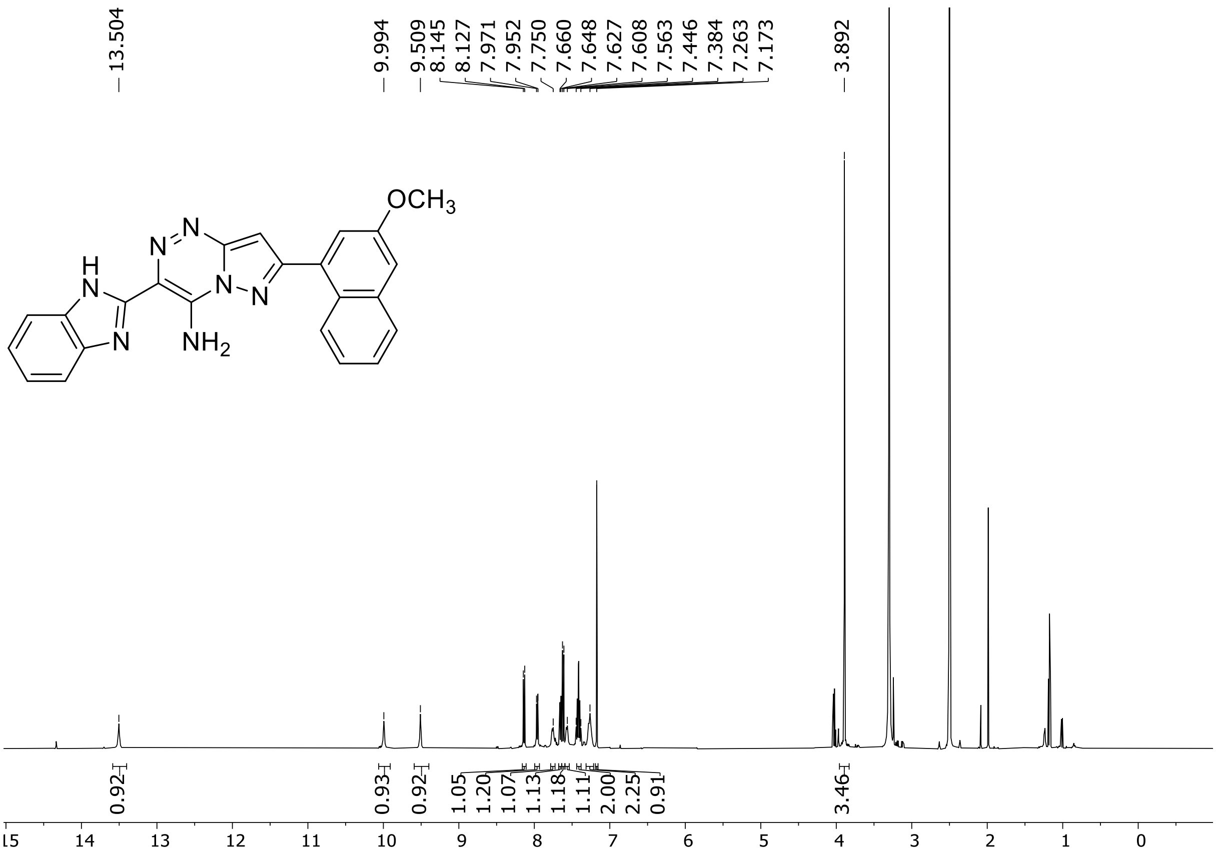

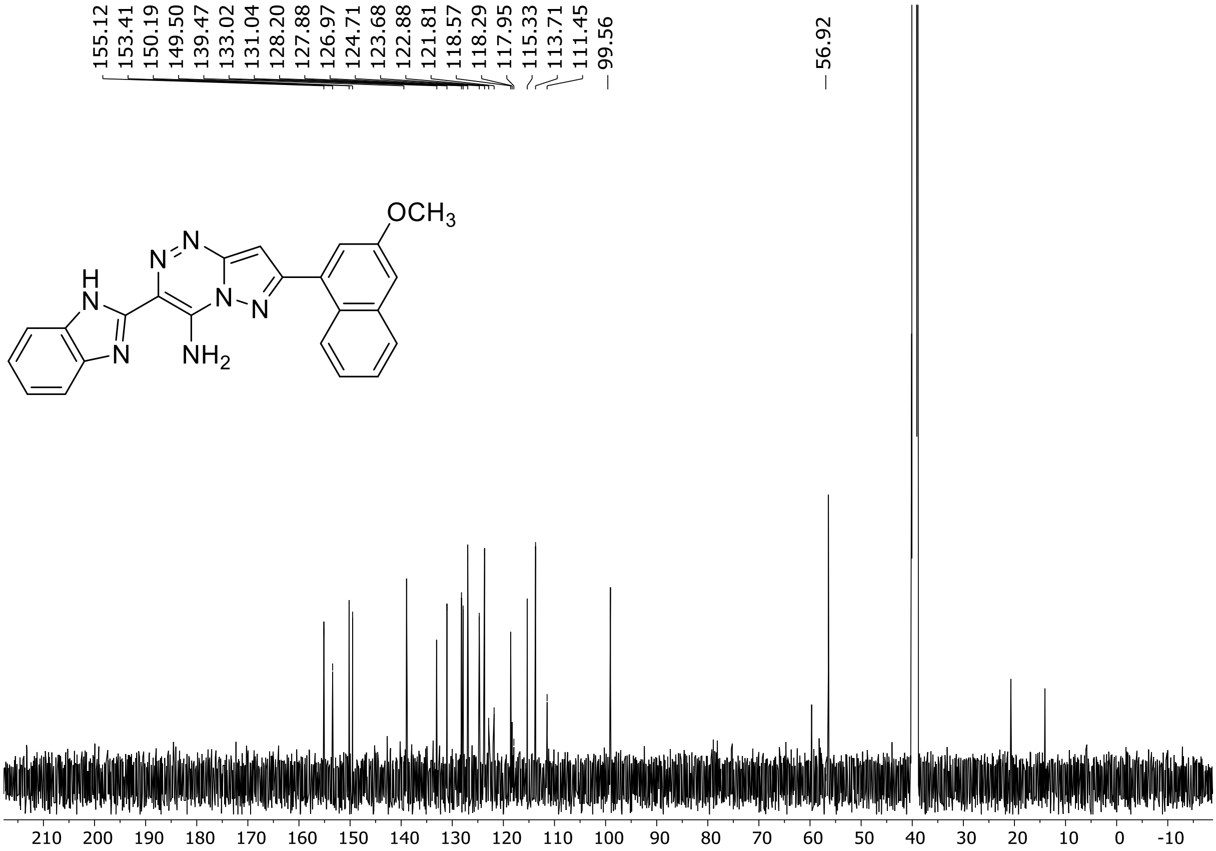

HRMS spectrum of **65**

**$C_{23}H_{17}N_7O$** **exact mass: 407.1607**

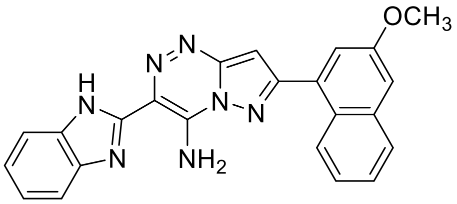

**APCI +** (MMI)
nitrogen flow 5 L/min, gas temperature 325°C, nebulizer 45 psig, skimmer 65 V, vaporizer 200°C, fragmentor 10 V, dissolved in methanol

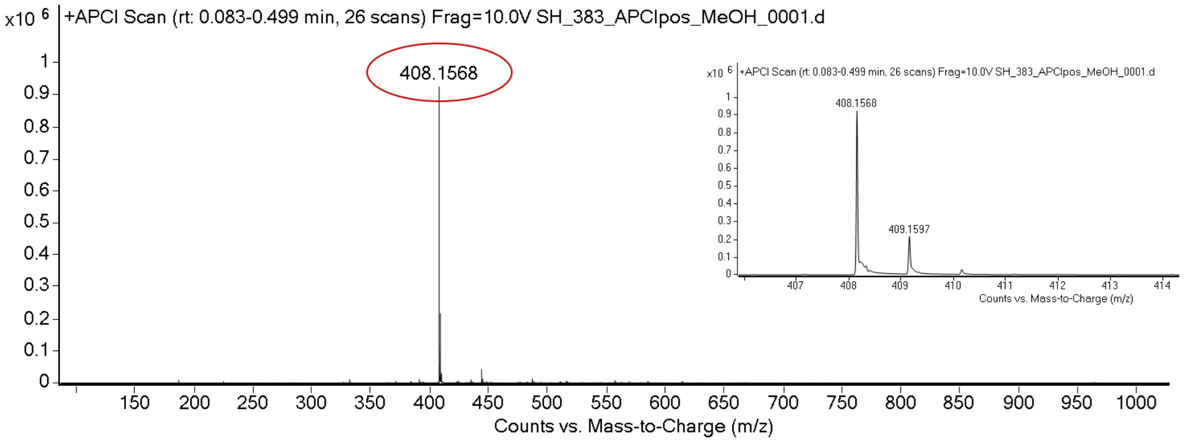

expected mass: $[M+H]^+$ = 408.1567 observed mass : $[M+H]^+$ = 408.1568 mass accuracy = 0.2 ppm

[1]H (500 MHz) and [13]C NMR (126 MHz) spectra of **66** in DMSO-$d_6$

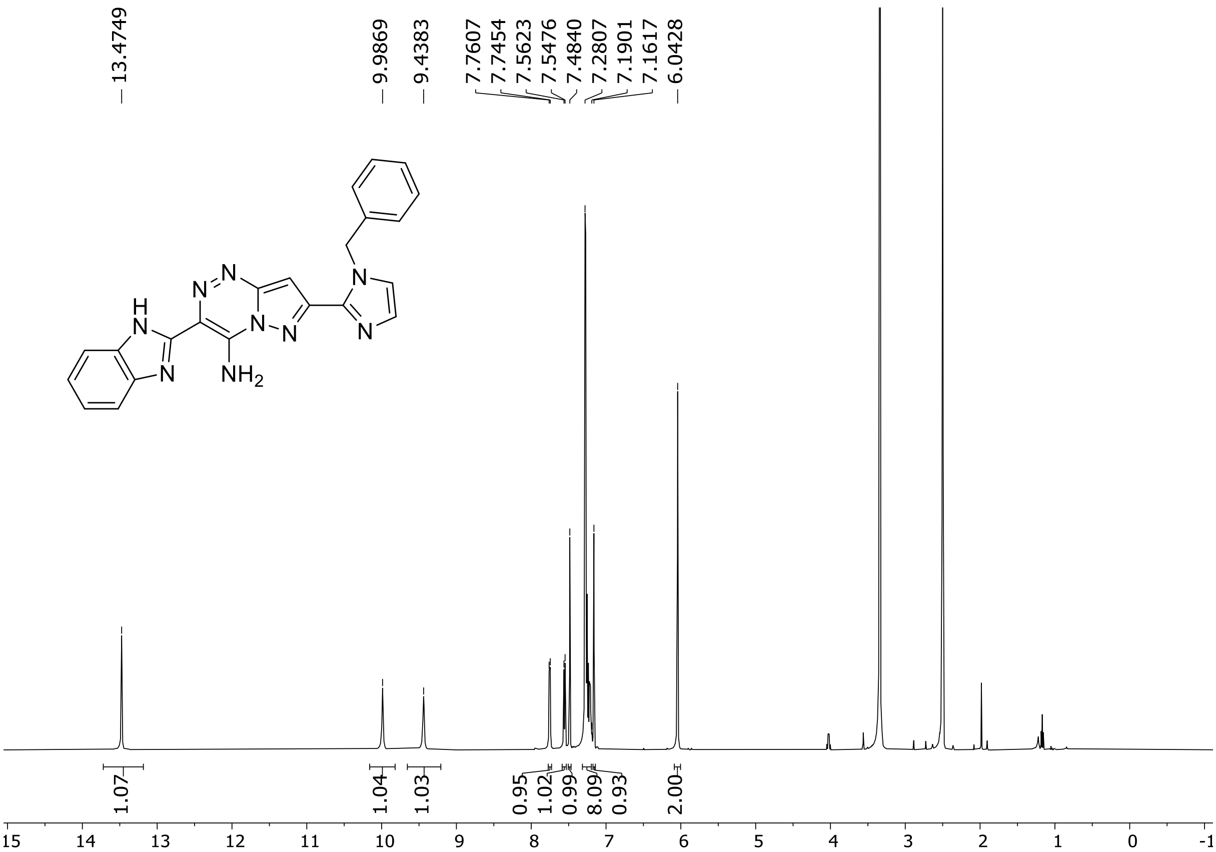

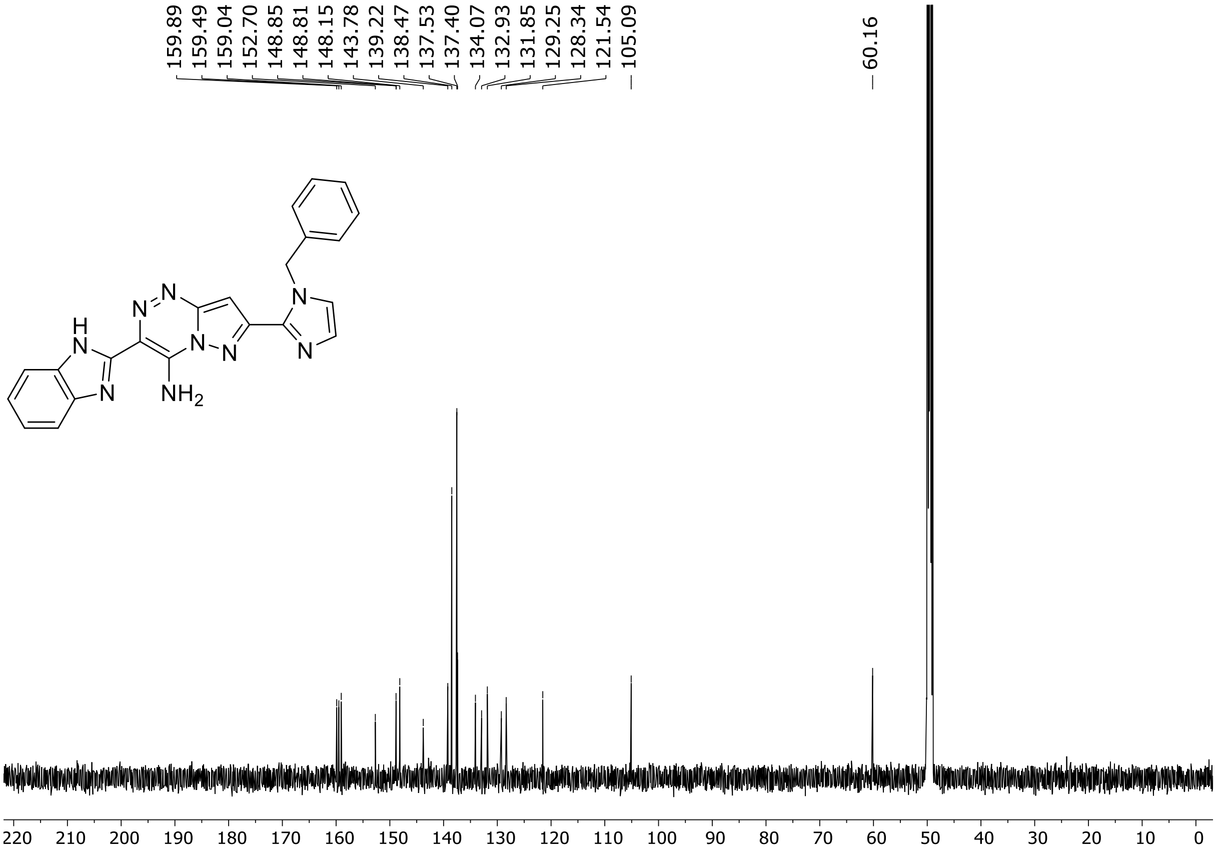

$^{1}H$ (500 MHz) and $^{13}C$ NMR (126 MHz) spectra of **67** in DMSO-$d_6$

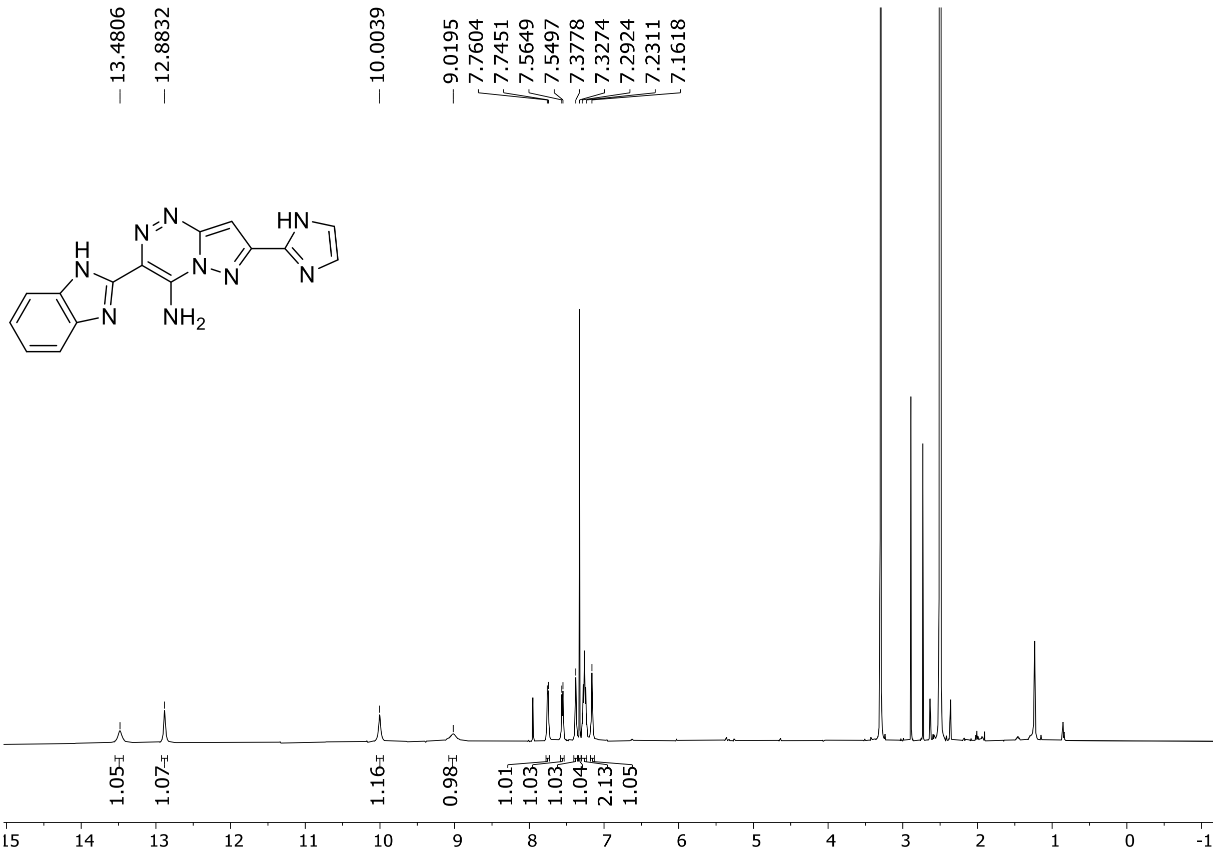

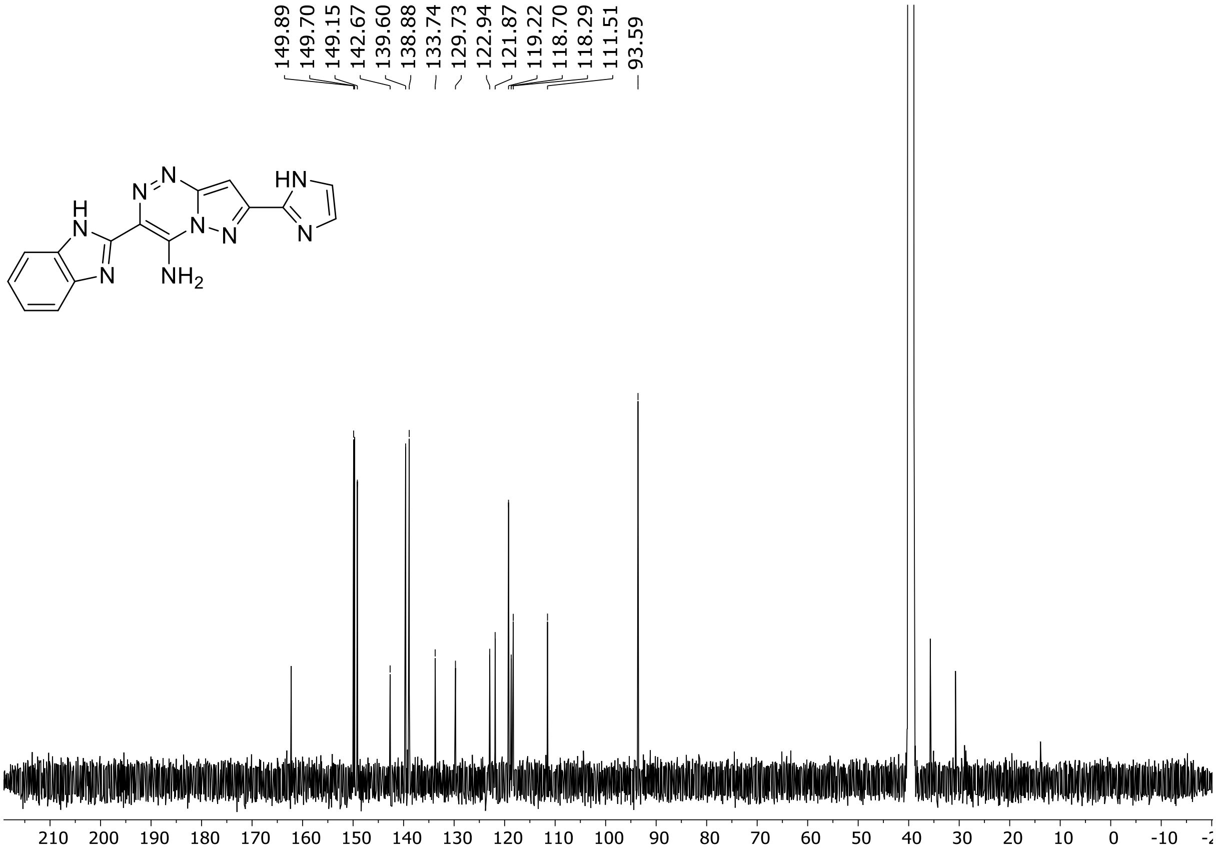

HRMS spectrum of **67**

**$C_{15}H_{11}N_9$** **exact mass: 317.1137**

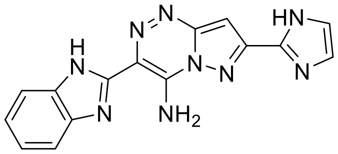

**APCI +** (MMI)
nitrogen flow 5 L/min, gas temperature 325°C, nebulizer 45 psig, skimmer 65 V, vaporizer 200°C, fragmentor 20 V, dissolved in methanol

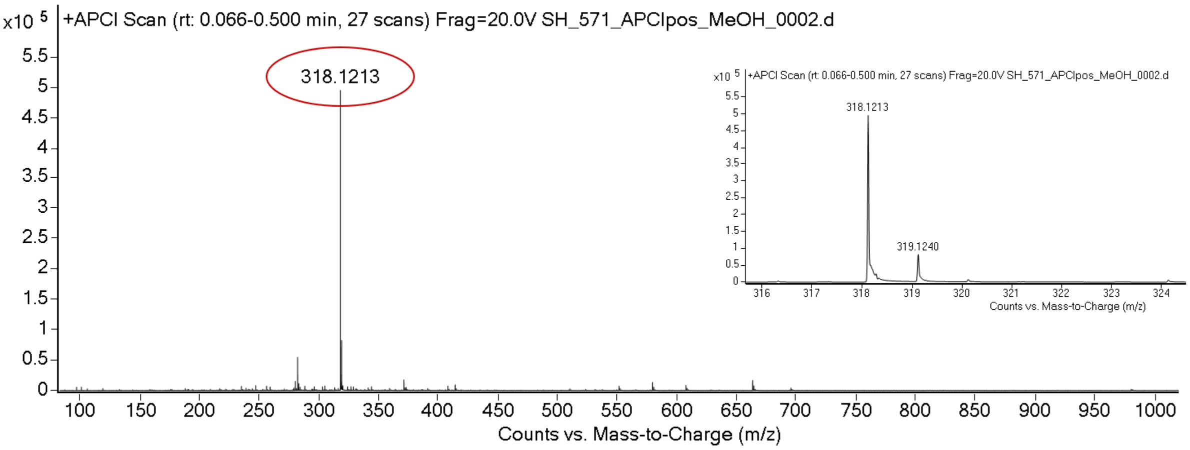

expected mass: [M+H]$^+$ = 318.1210 observed mass : [M+H]$^+$ = 318.1213 mass accuracy = 0.9 ppm

[1]H (500 MHz) and [13]C NMR (126 MHz) spectra of **68** in DMSO-$d_6$

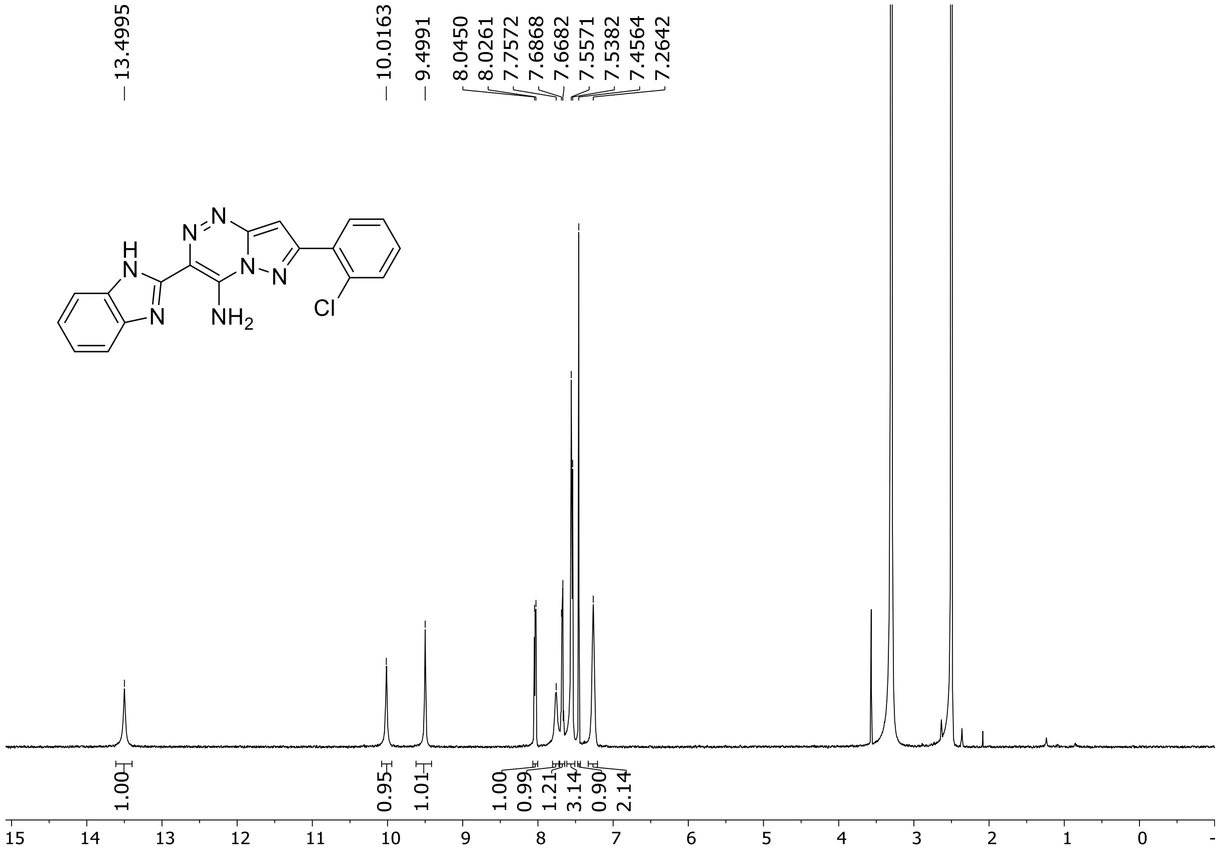

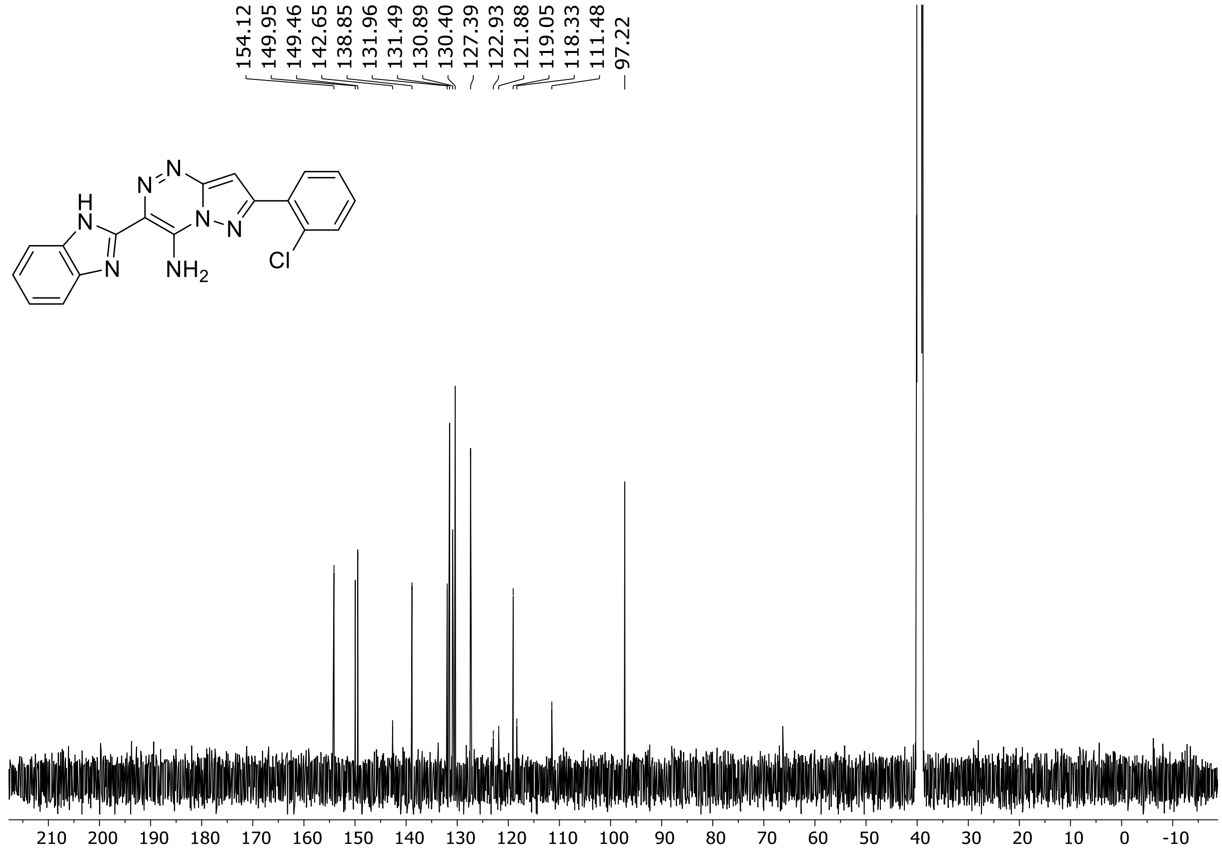

HRMS spectrum of **68**

$C_{18}H_{12}ClN_7$ **mono m/z = 361.0843**

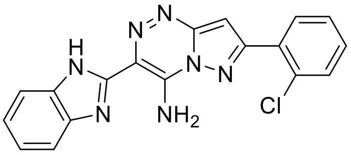

**APCI +** (MMI)
nitrogen flow 5 L/min, gas temperature 300°C, nebulizer 45 psi, vaporizer 200°C
skimmer **65** V, fragmentor 22 V, dissolved in methanol

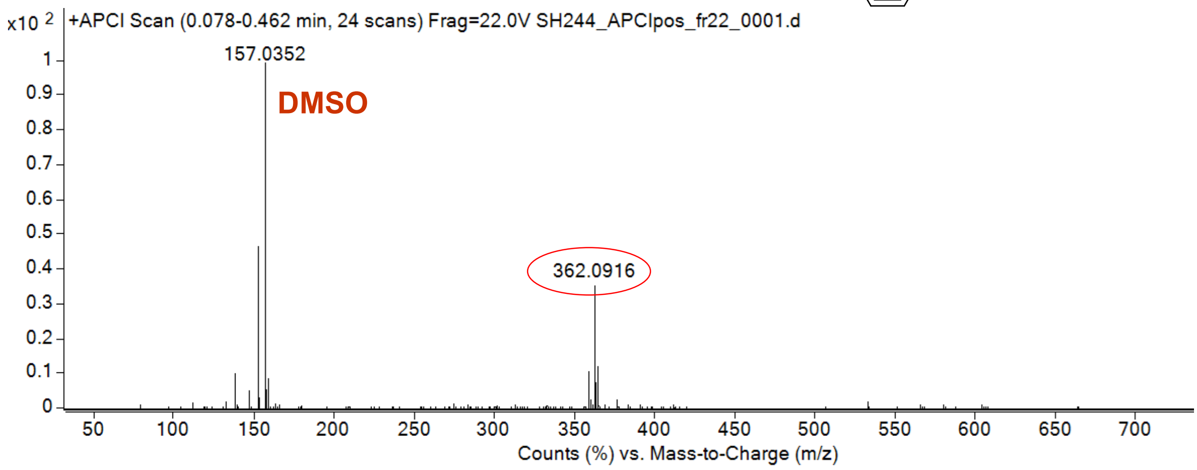

calculated mass: $[M+H]^+$ = 362.0915 observed: $[M+H]^+$ = 362.0916 max. mass error = 0.2 ppm

[1]H (500 MHz) and [13]C NMR (126 MHz) spectra of **69** in DMSO-$d_6$

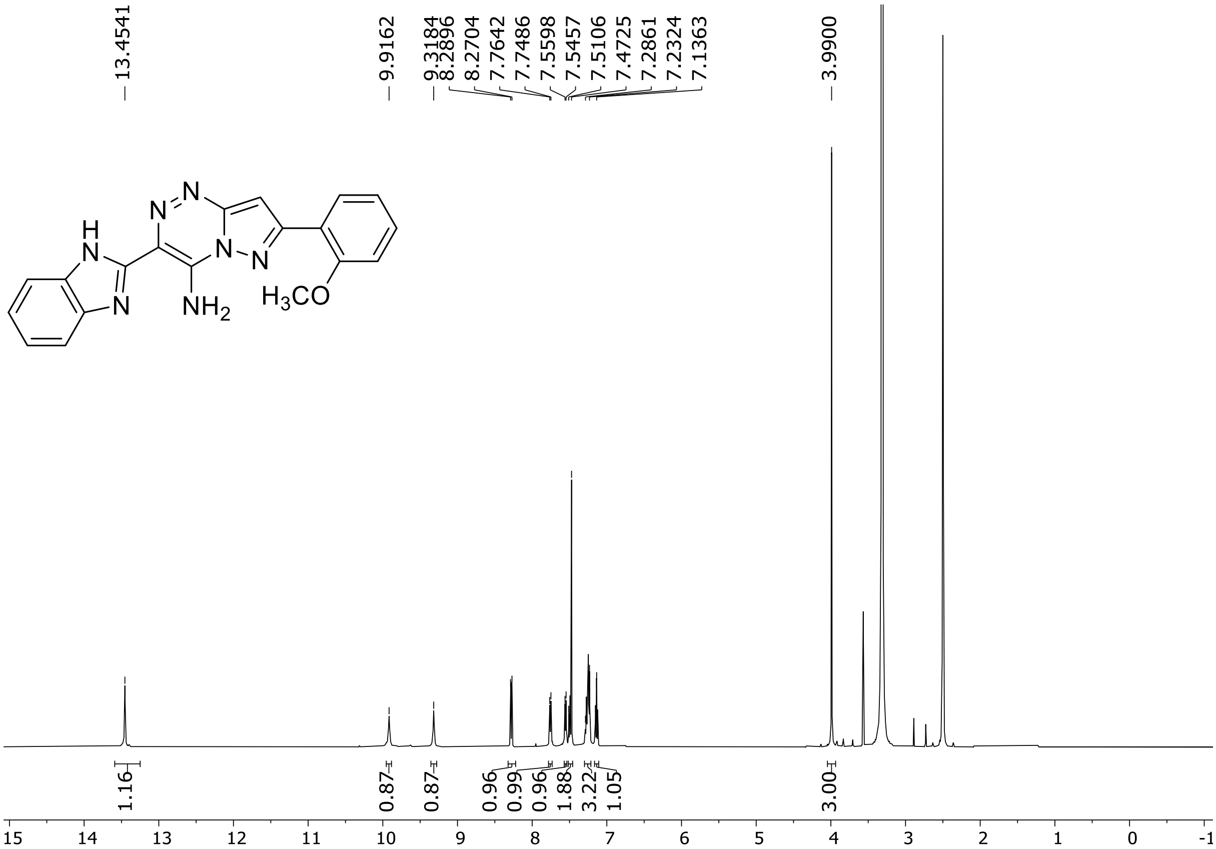

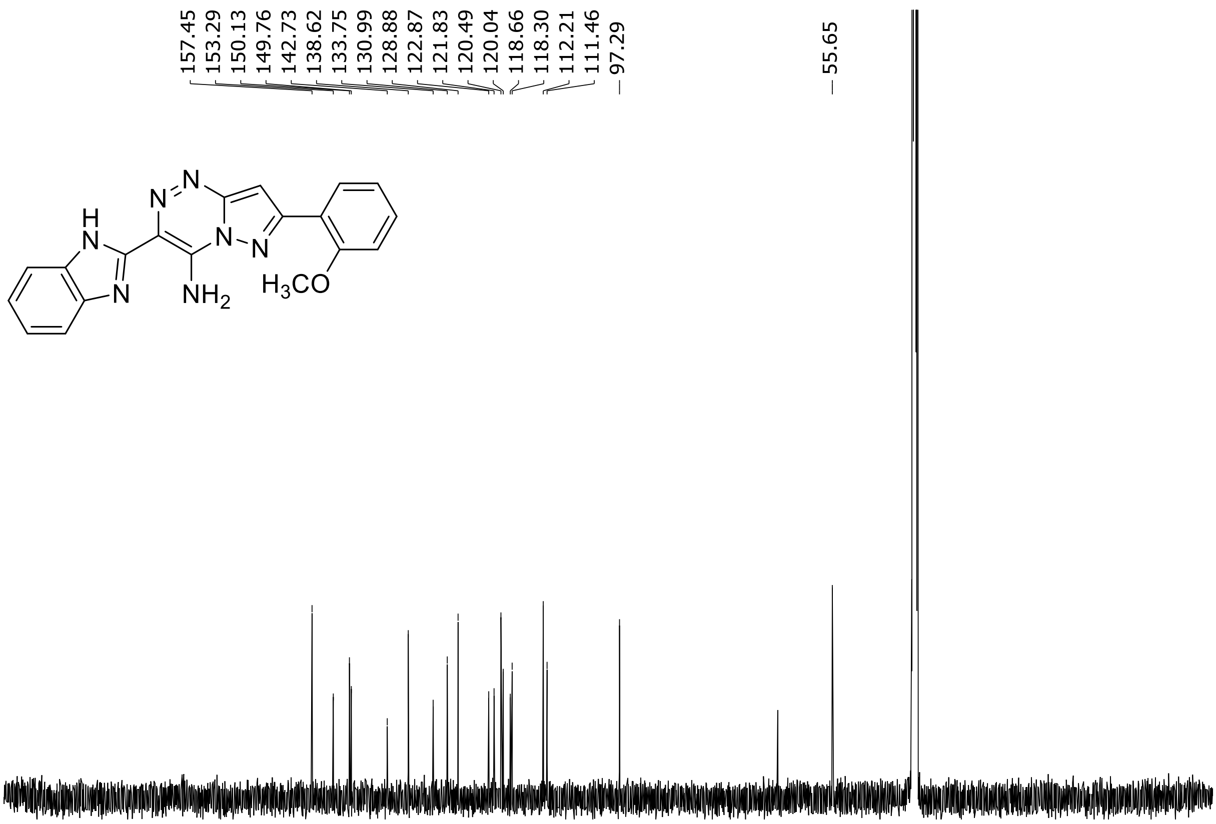

HRMS spectrum of **69**

**$C_{19}H_{15}N_7O$** **mono m/z = 357.1338**

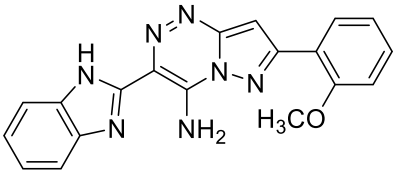

**APCI +** (MMI)
nitrogen flow 5 L/min, gas temperature 300°C, nebulizer 45 psi, vaporizer 200°C
skimmer **65** V, fragmentor **28** V, dissolved in methanol

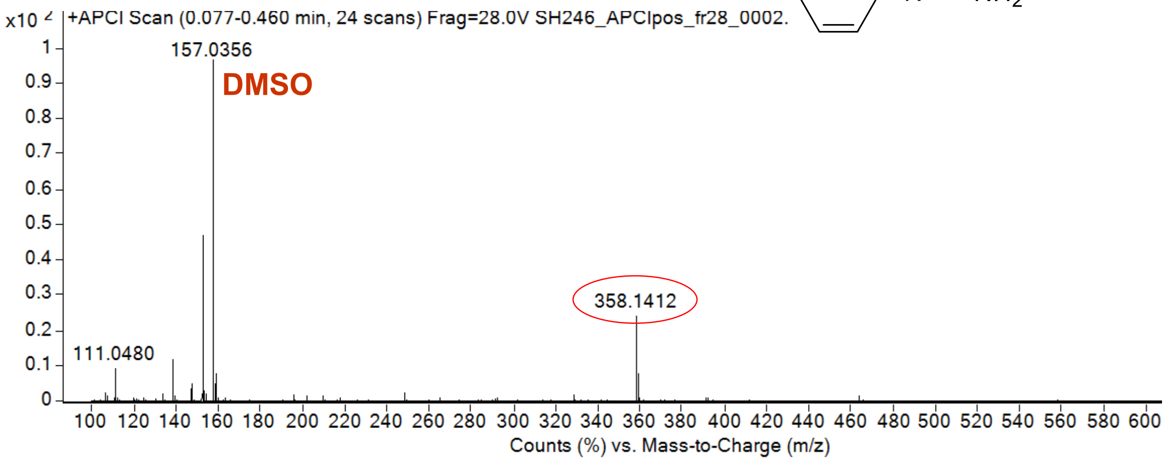

calculated mass: $[M+H]^+$ = 358.1411 observed: $[M+H]^+$ = 358.1412 max. mass error = 0.2 ppm

[1]H (500 MHz) and [13]C NMR (126 MHz) spectra of **70** in DMSO-$d_6$

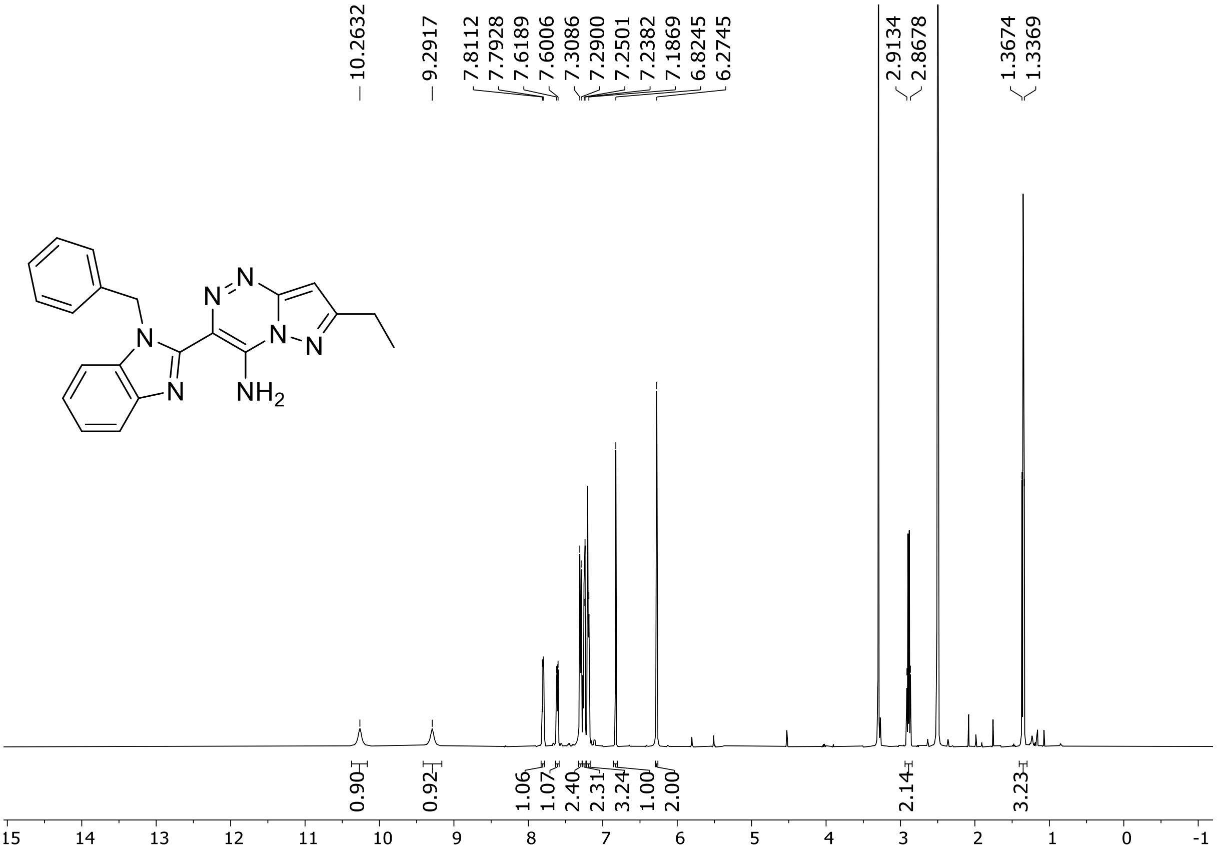

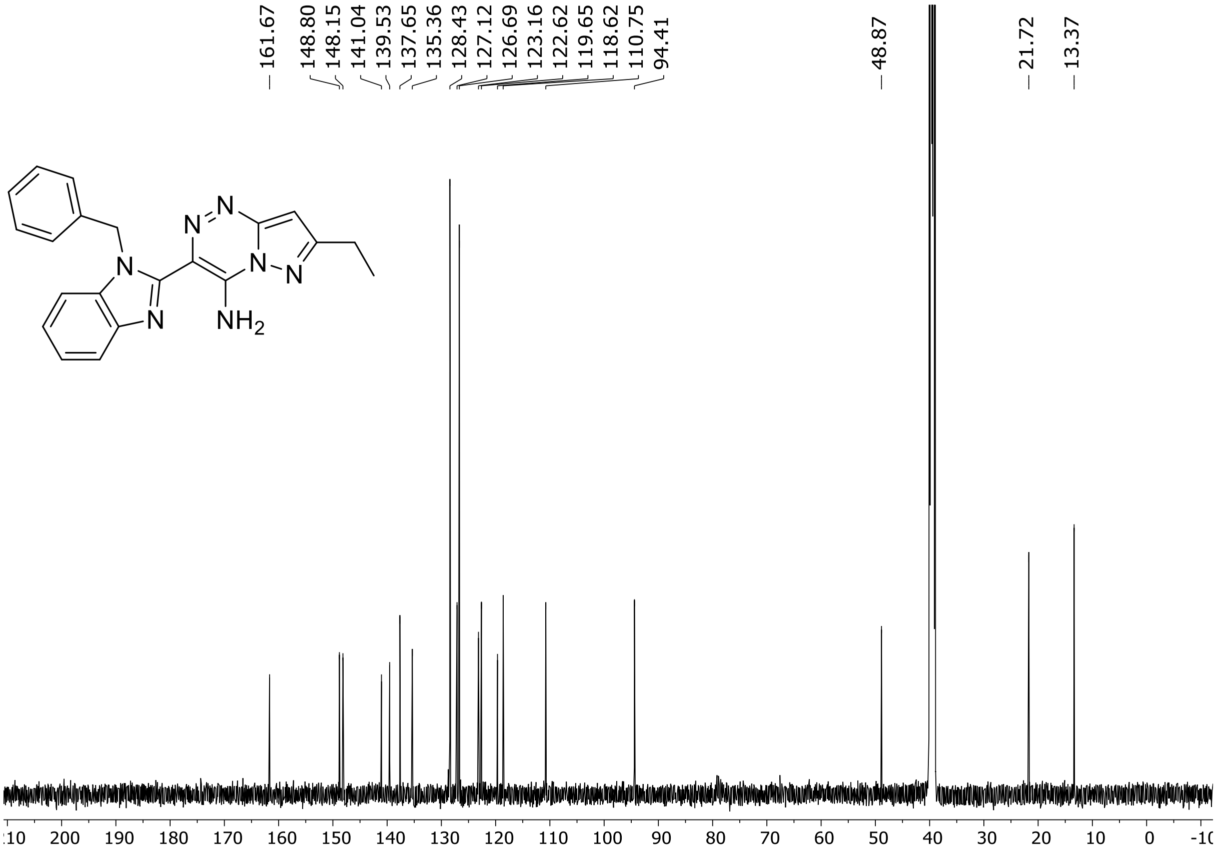

HRMS spectrum of **70**

$C_{21}H_{19}N_7$ **mono m/z = 369.1702**

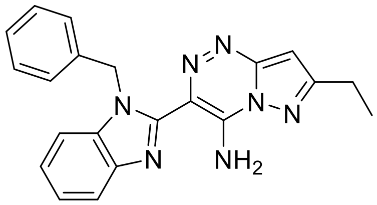

**APCI +** (MMI)
nitrogen flow 5 L/min, gas temperature 300°C, nebulizer 45 psi, vaporizer 200°C
skimmer **65** V, fragmentor **10** V, dissolved in methanol

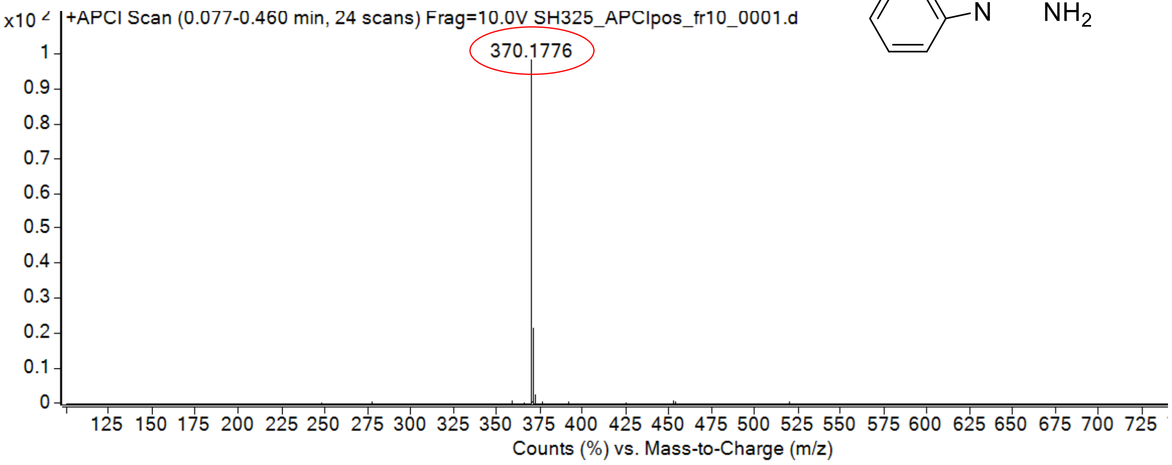

calculated mass: $[M+H]^+$ = 370.1775 observed: $[M+H]^+$ = 370.1776 max. mass error = 0.2 ppm

[1]H (500 MHz) and [13]C NMR (126 MHz) spectra of **71** in DMSO-$d_6$

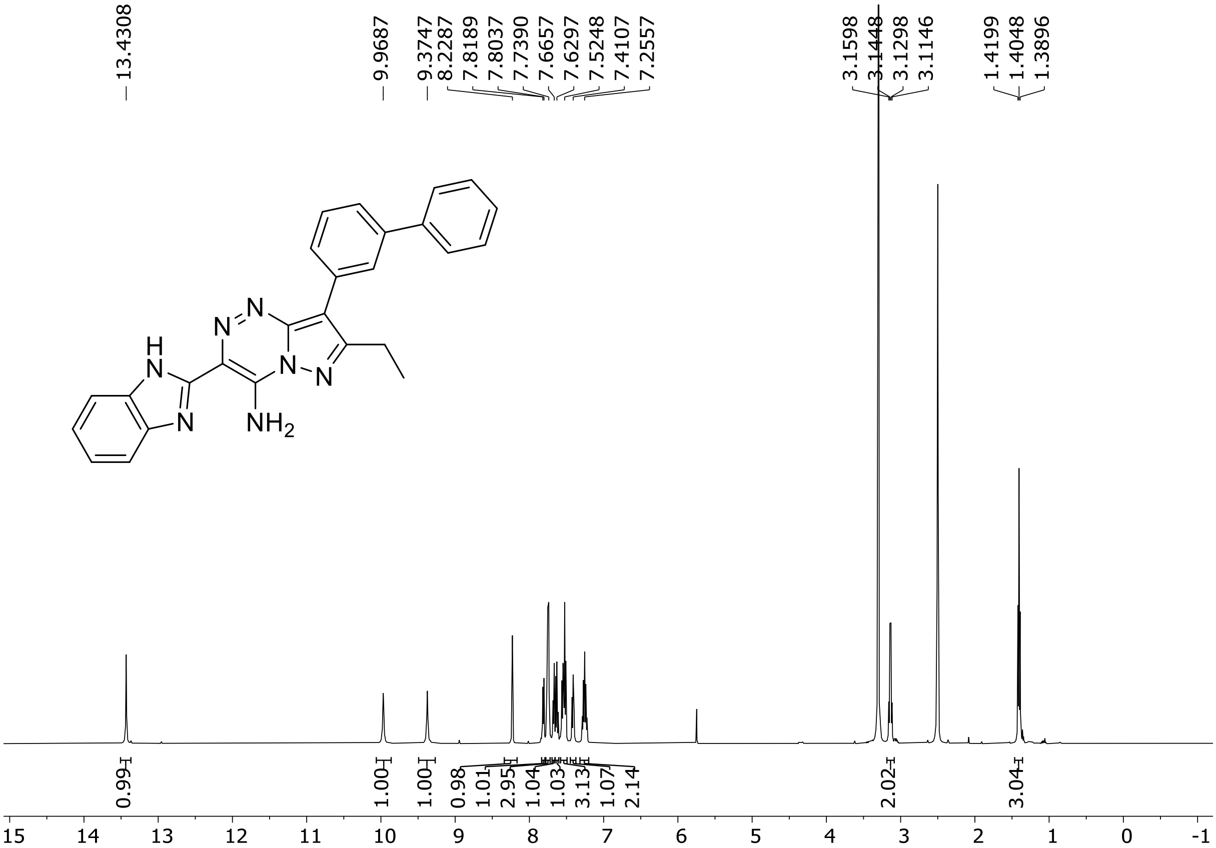

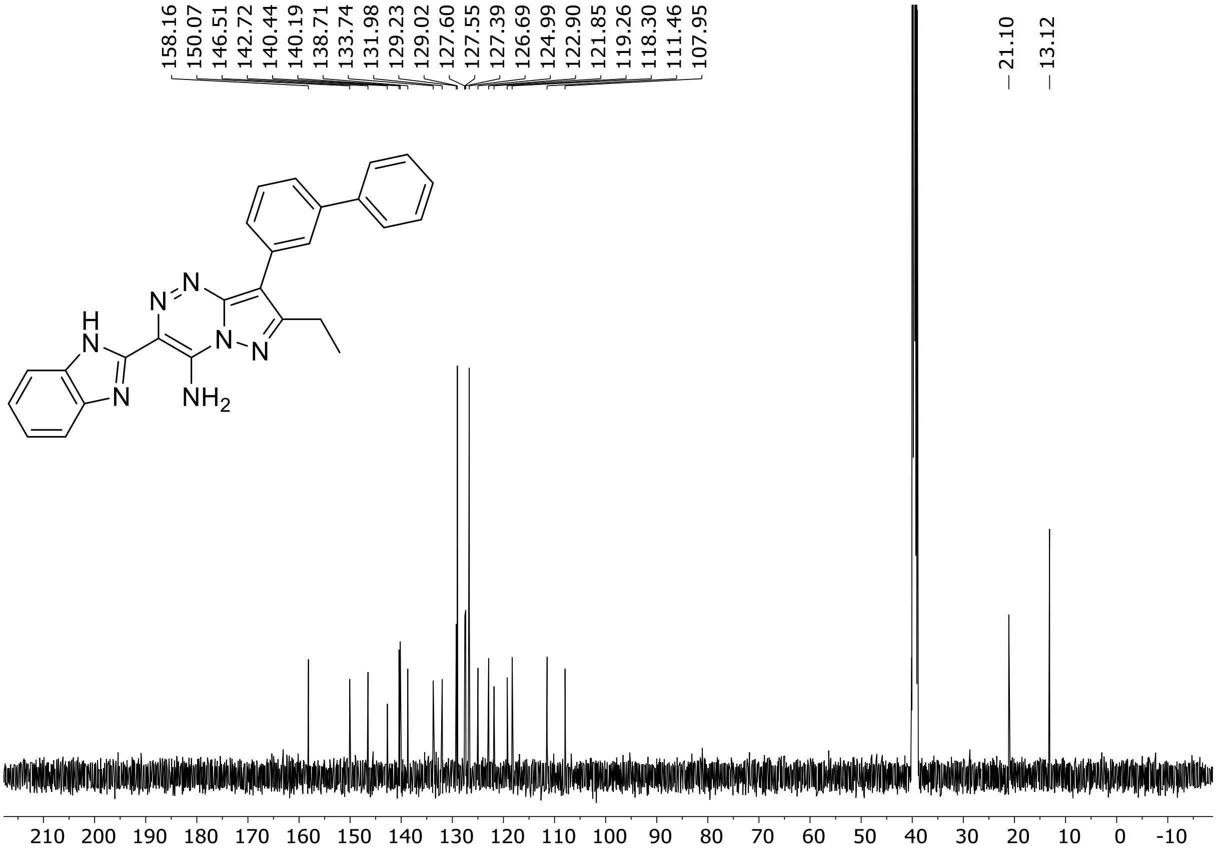

HRMS spectrum of **71**

**$C_{26}H_{21}N_7$** **mono m/z = 431.49**

**ESI +** (MMI)

nitrogen flow 5 L/min, gas temperature 300°C, vaporizer 200°C, nebulizer 45 psi,
skimmer **65** V, fragmentor 35 V, dissolved in MeOH

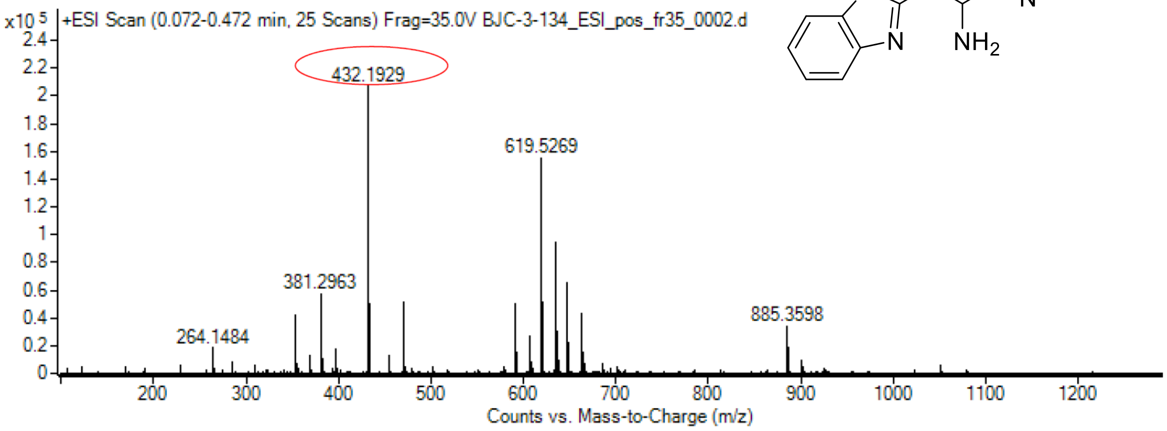

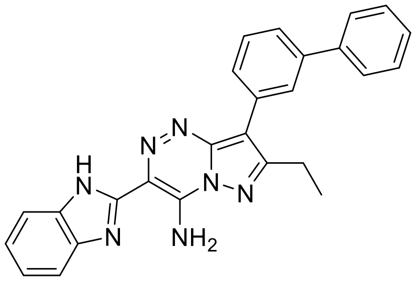

calculated mass: $[M+H]^+$ = 432.1931 observed: $[M+H]^+$ = 432.1929 max. mass error = 0.5 ppm

$^{1}H$ (500 MHz) and $^{13}C$ NMR (126 MHz) spectra of **72** in DMSO-$d_6$

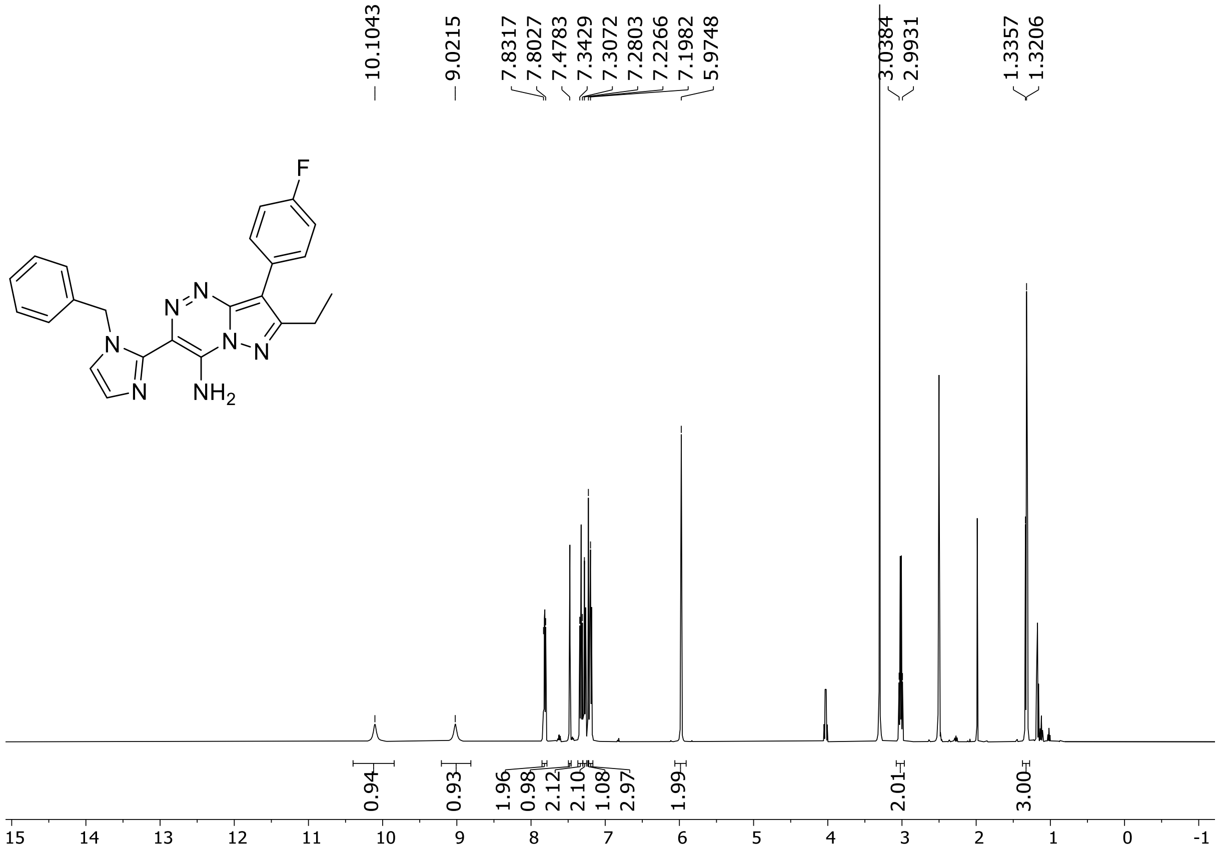

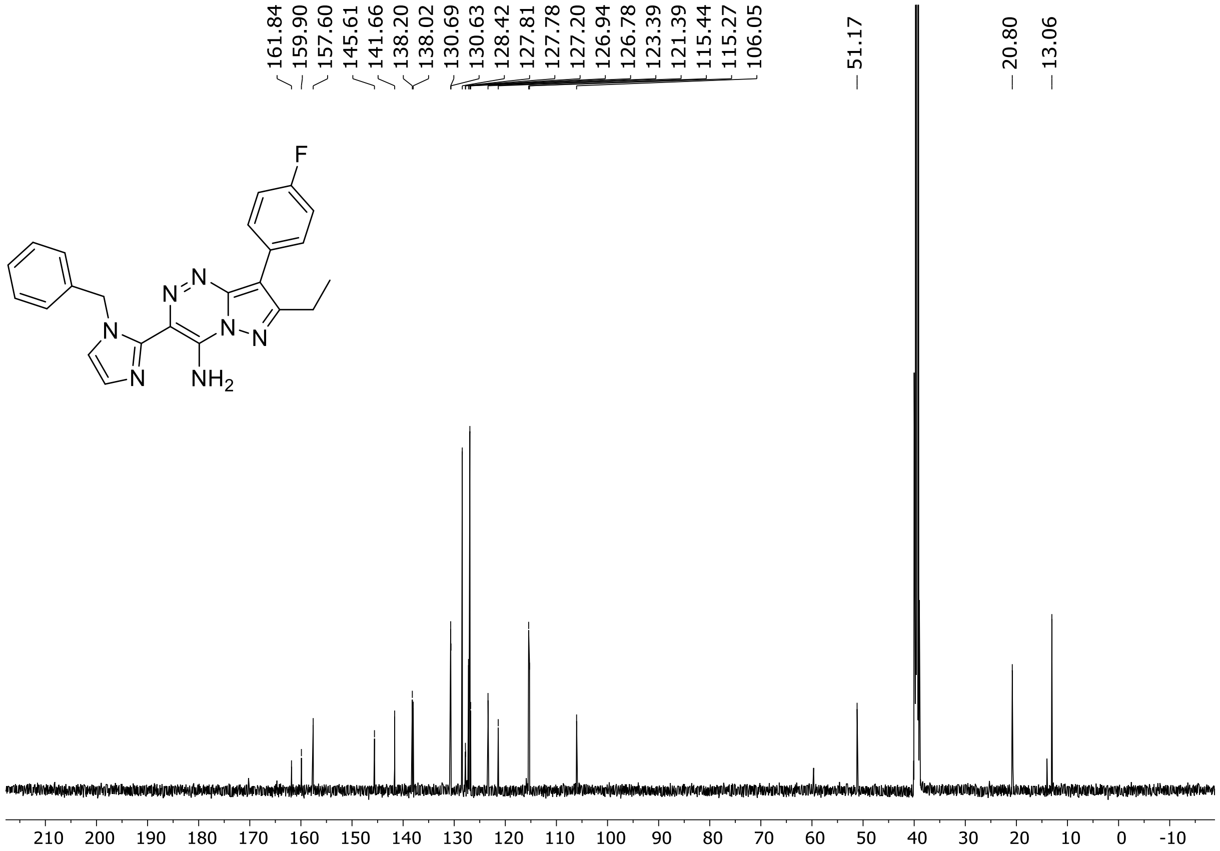

$^{19}F$ (471 MHz) NMR spectrum of **72** DMSO-$d_6$

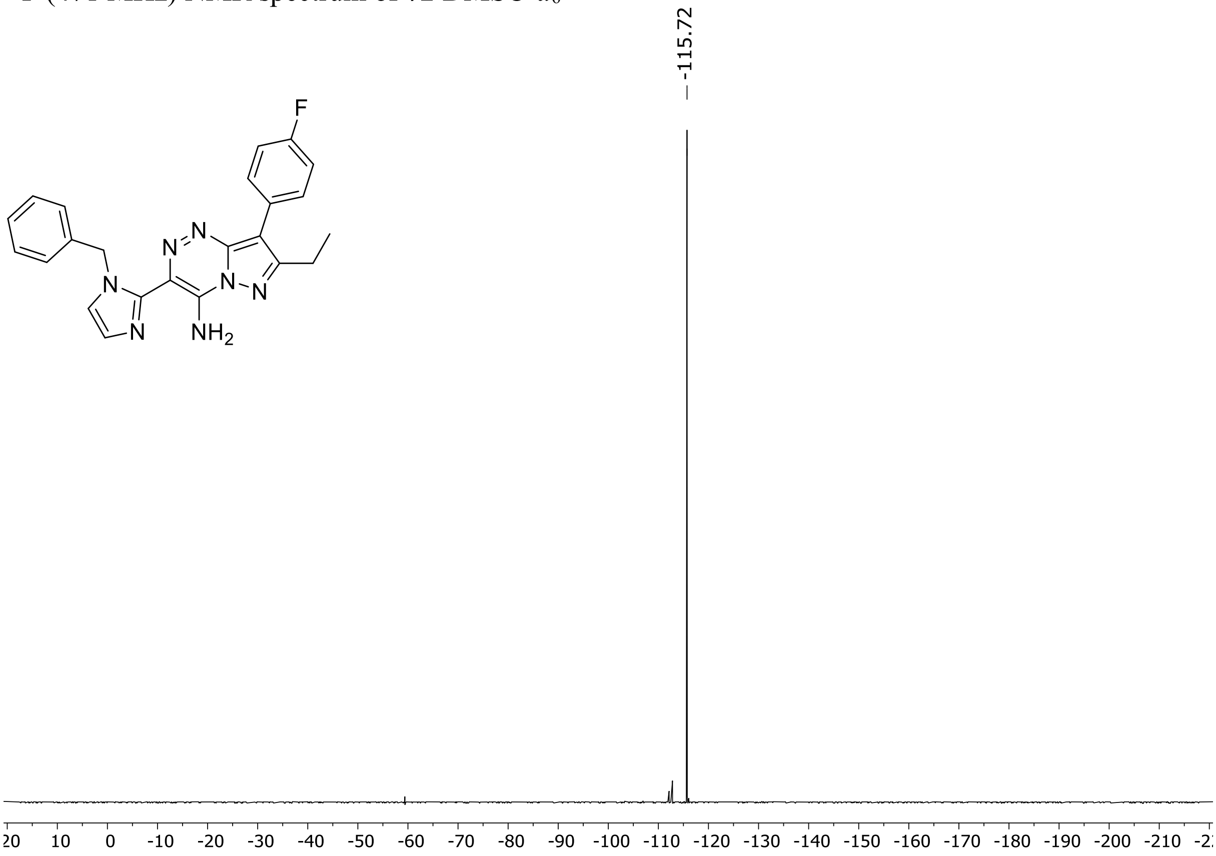

HRMS spectrum of **72**

**$C_{23}H_{20}FN_7$** **exact mass: 413.1764**

**APCI +** (MMI)
nitrogen flow 5 L/min, gas temperature 325°C, nebulizer 45 psig, skimmer 65 V, vaporizer 200°C, fragmentor 10 V, dissolved in methanol

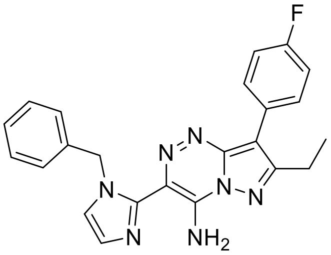

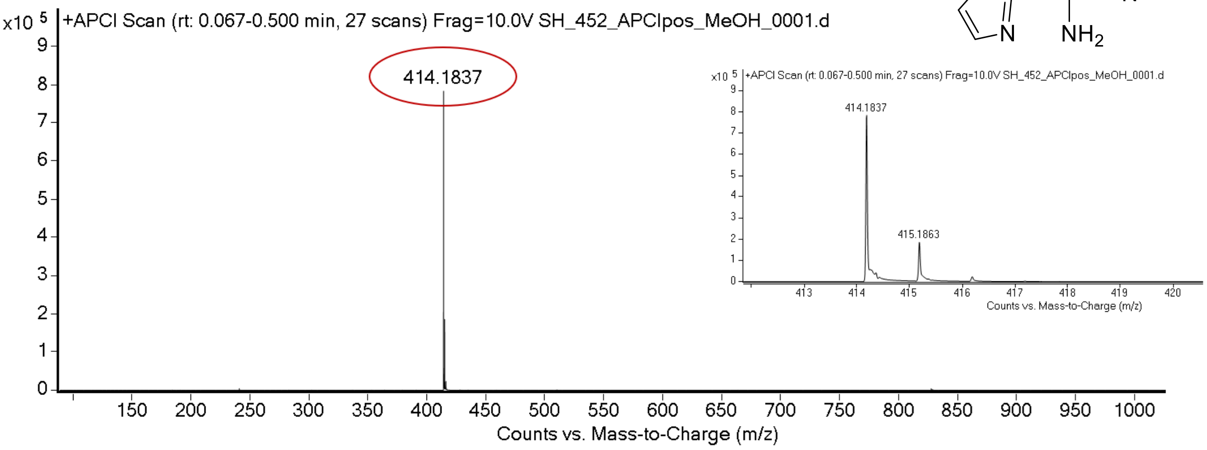

expected mass: [M+H]+ = 414.1837 observed mass : [M+H]+ = 414.1837 mass accuracy < 0.1 ppm

[1]H (500 MHz) and [13]C NMR (126 MHz) spectra of **73** in DMSO-$d_6$

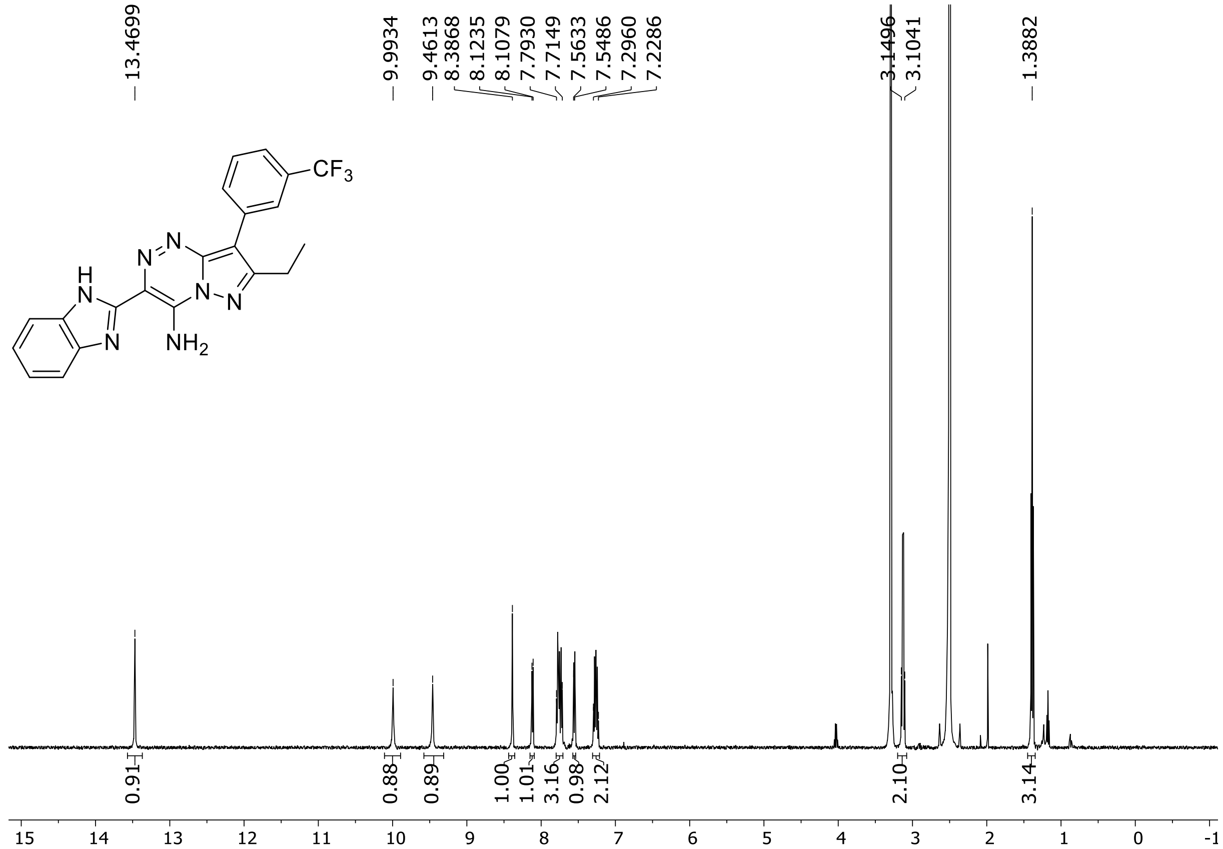

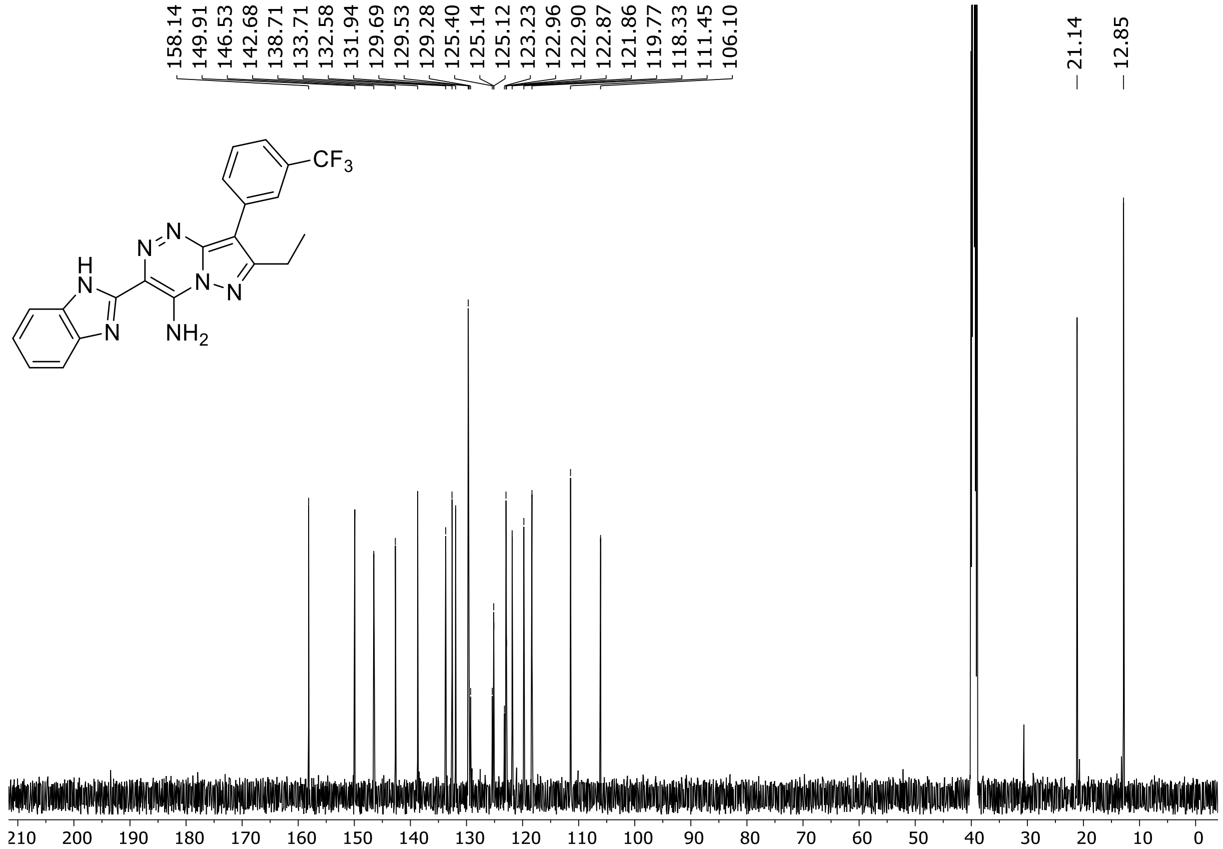

$^{19}F$ (471 MHz) NMR spectrum of **73** DMSO-$d_6$

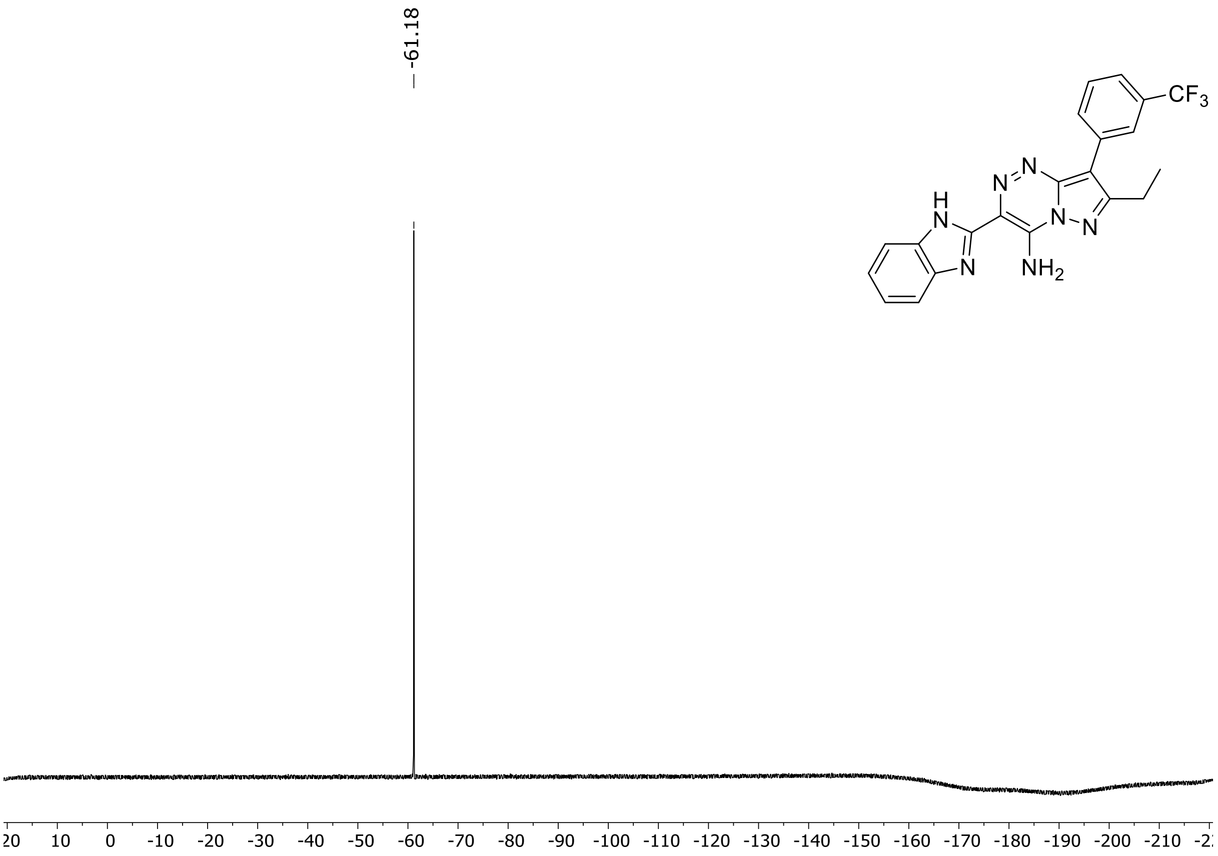

HRMS spectrum of **73**

**$C_{21}H_{16}F_3N_7$** **mono *m/z* 423.1419**

**APCI +** (MMI)
nitrogen flow 5 L/min, gas temperature 325°C, nebulizer 45 psi, skimmer 65 V, vaporizer 250°C, fragmentor 35 V, dissolved in methanol

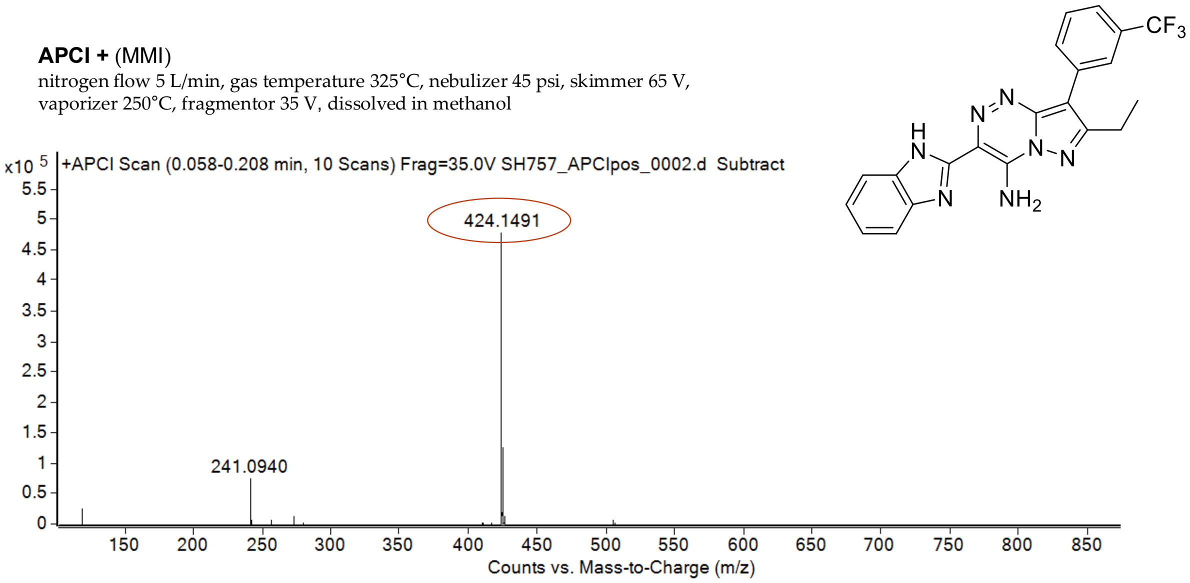

calculated mass: $[M+H]^+$ = 424.1492 observed: $[M+H]^+$ = 424.1491 mass accuracy = - 0.2 ppm

$^{1}$H (500 MHz) and $^{13}$C NMR (126 MHz) spectra of **74** in DMSO-$d_6$

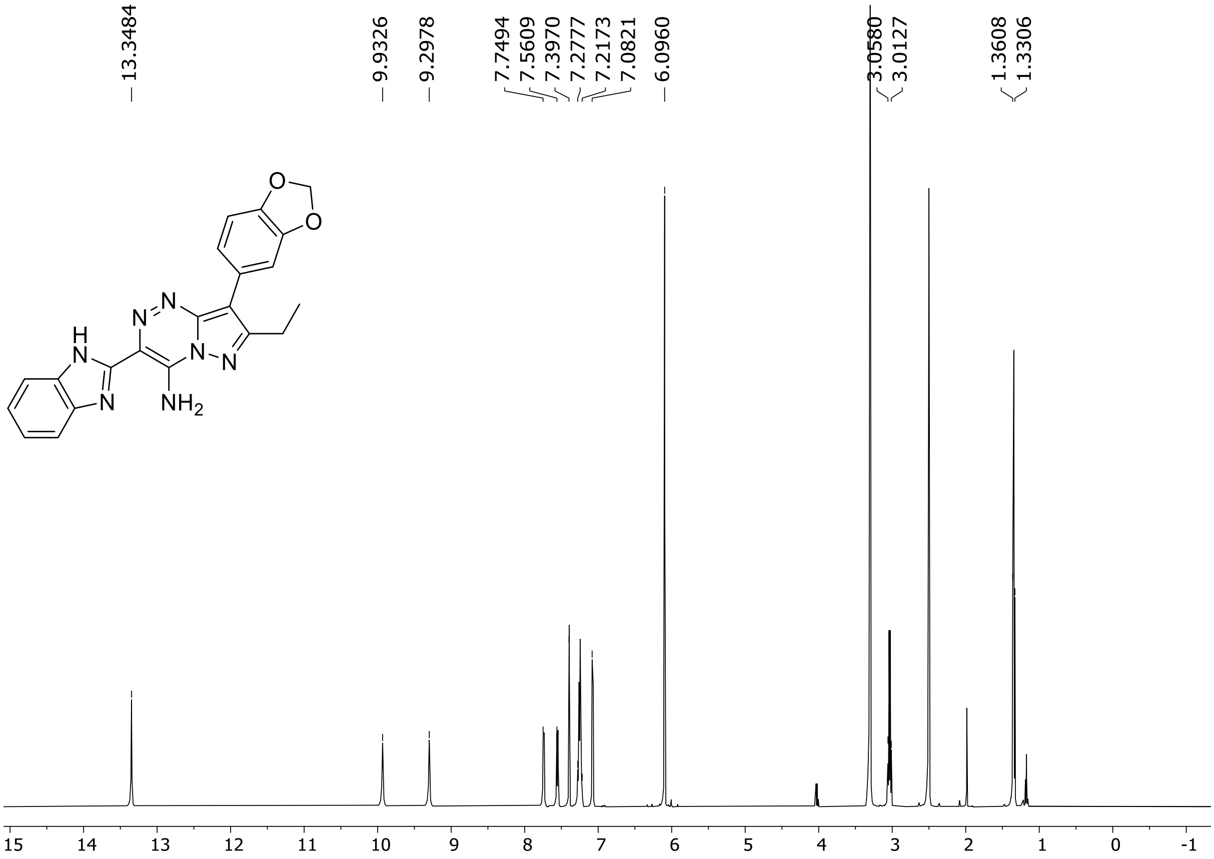

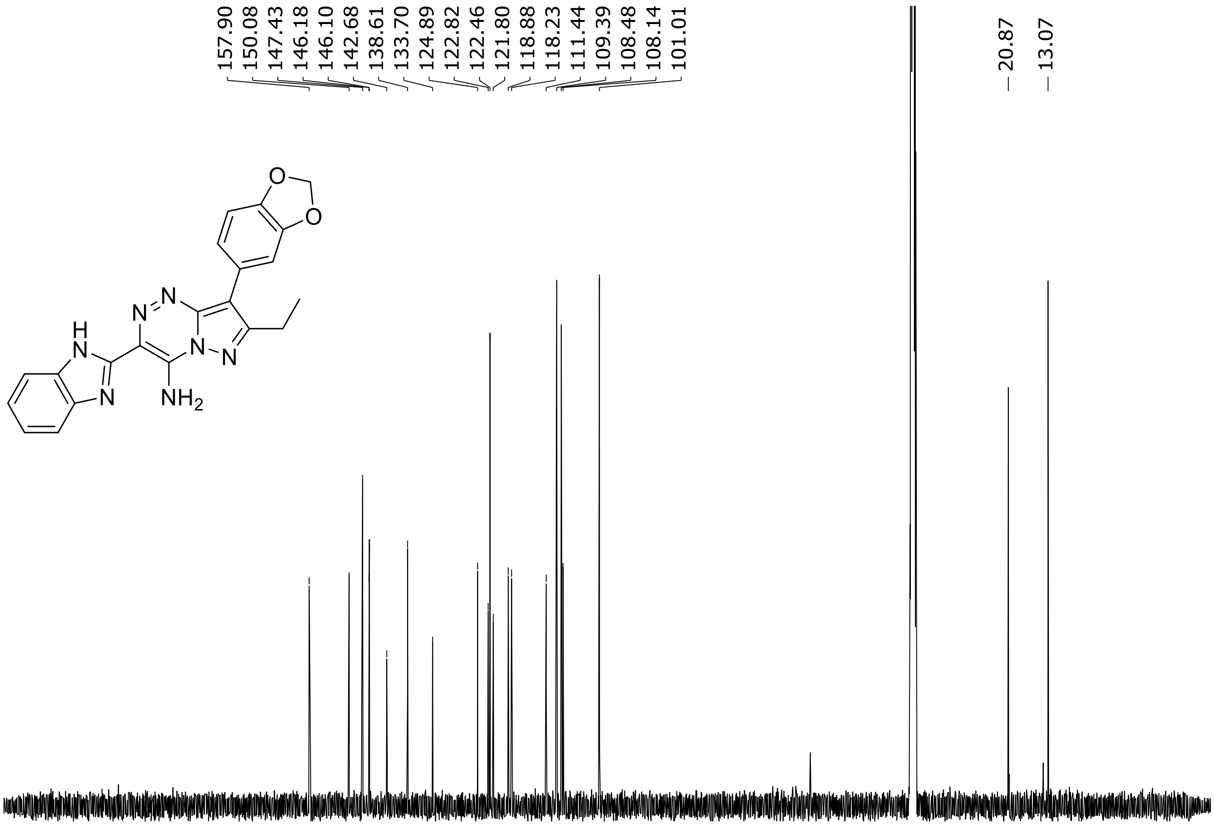

HRMS spectrum of **74**

$C_{21}H_{17}N_7O_2$ **mono *m*/*z* 399.1444**

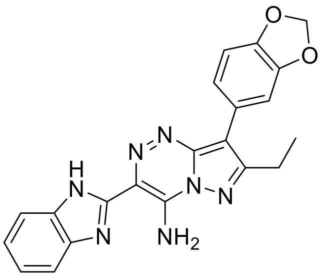

**APCI +** (MMI)
nitrogen flow 5 L/min, gas temperature 325°C, nebulizer 45 psi, skimmer 65 V,
vaporizer 250°C, fragmentor 35 V, dissolved in methanol

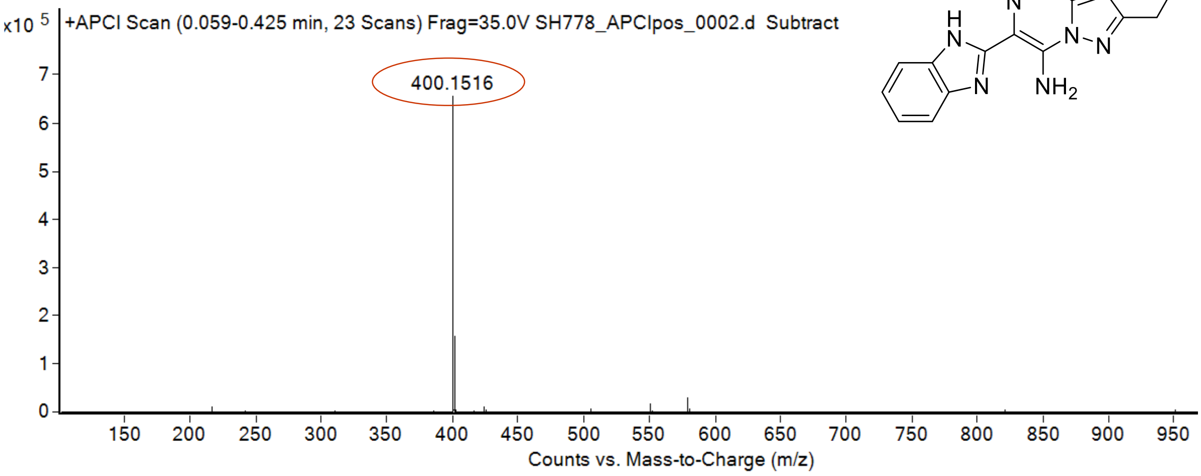

calculated mass: $[M+H]^+$ = 400.1516 observed: $[M+H]^+$ = 400.1516 mass accuracy < 0.1 ppm

[1]H (500 MHz) and [13]C NMR (126 MHz) spectra of **75** in DMSO-$d_6$

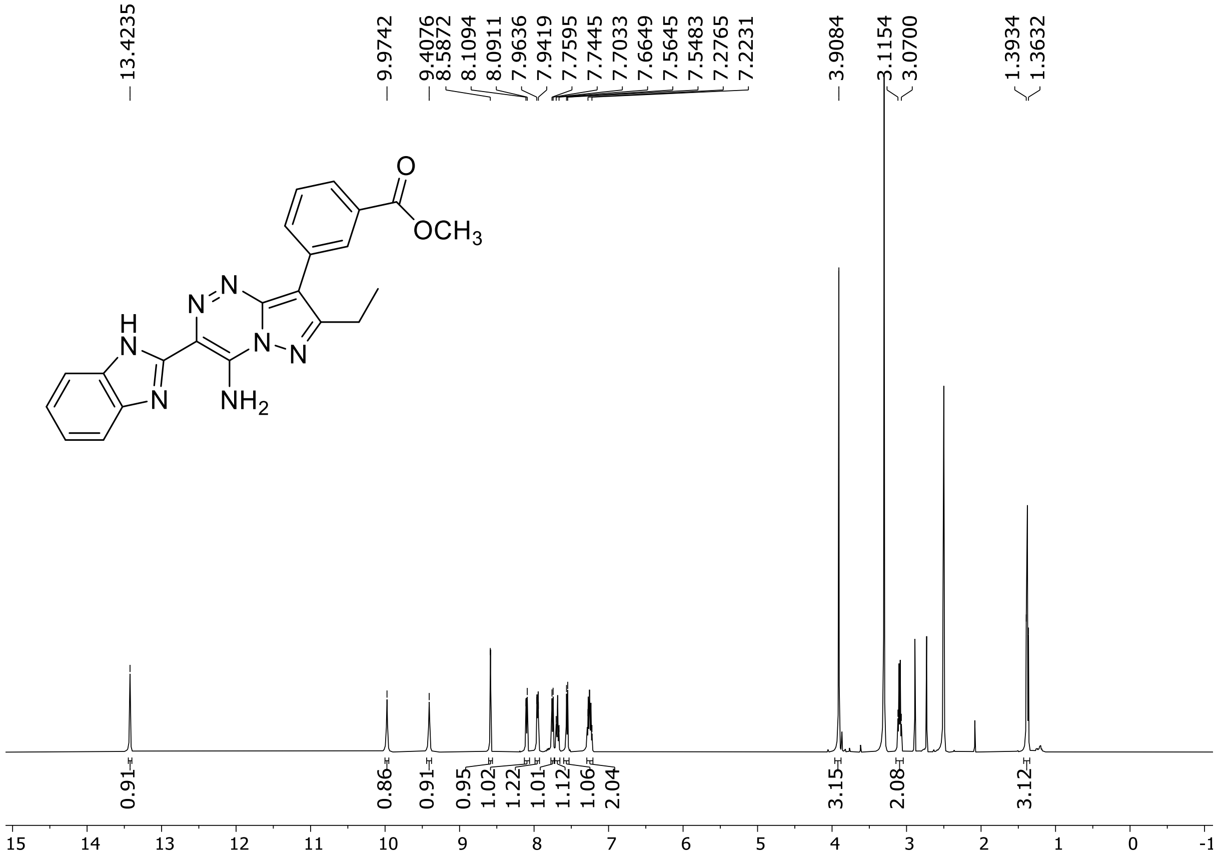

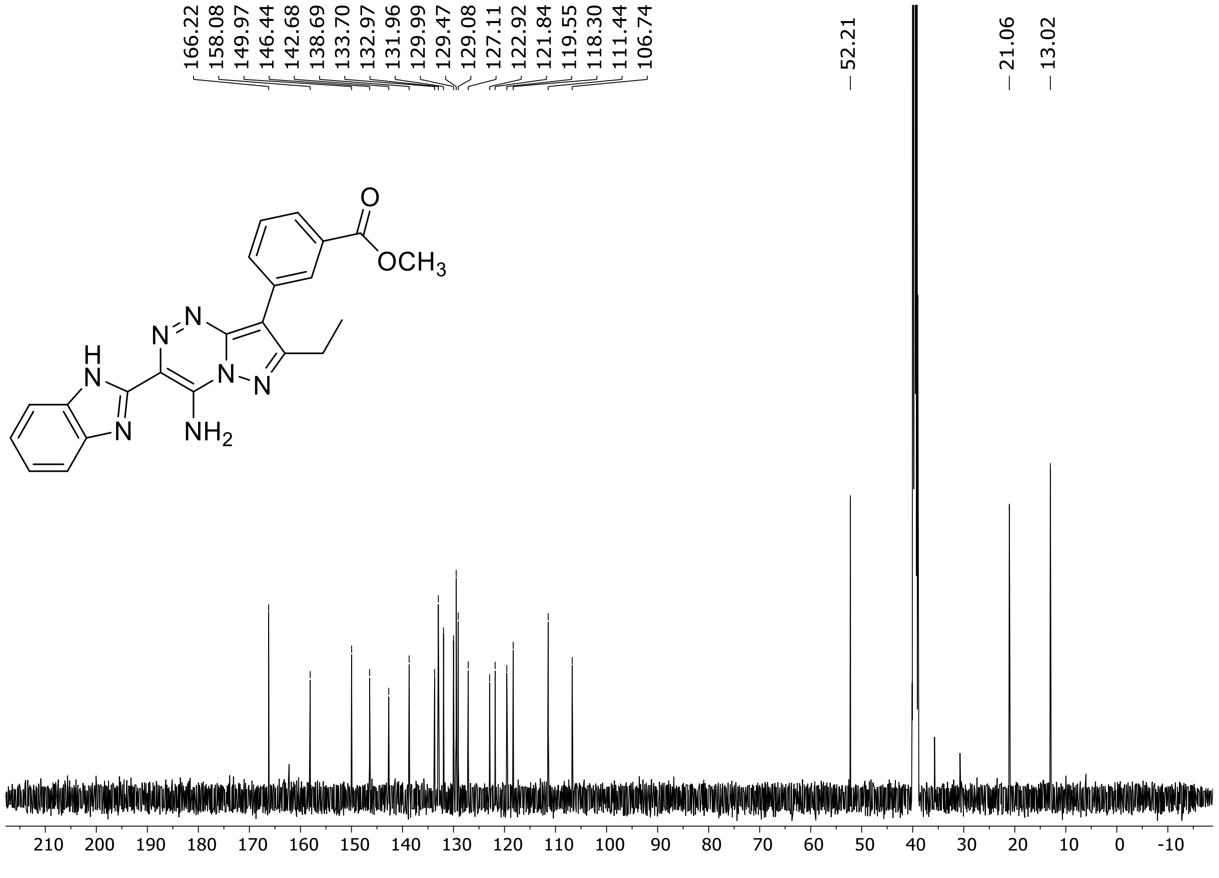

HRMS spectrum of **75**

$C_{22}H_{19}N_7O_2$ mono *m/z* **413.4410**

**APCI +** (MMI)
nitrogen flow 5 L/min, gas temperature 325°C, nebulizer 45 psi, skimmer 65 V, vaporizer 250°C, fragmentor 35 V, dissolved in methanol

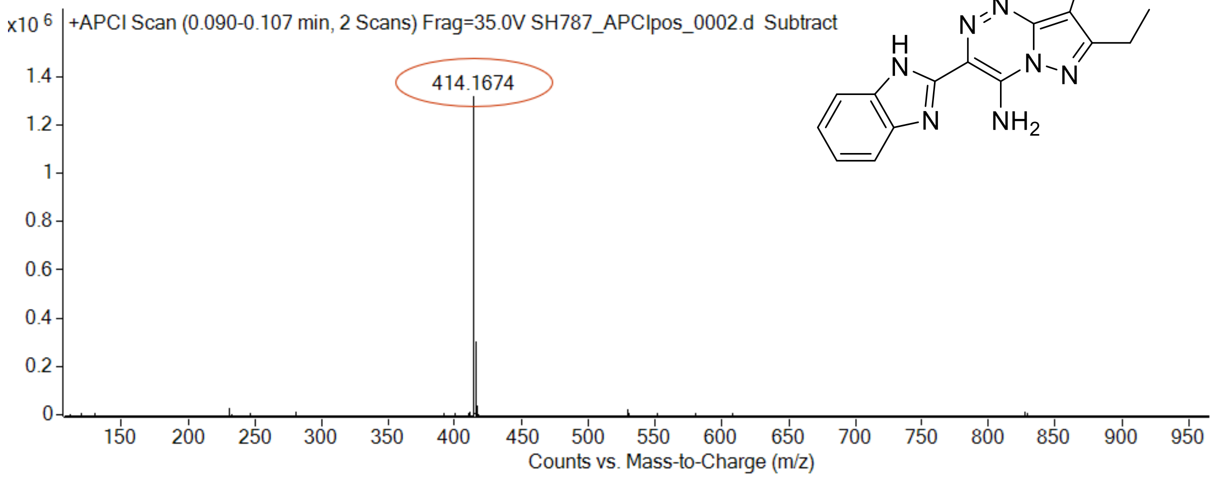

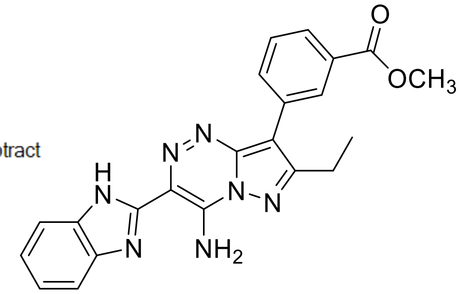

calculated mass: $[M+H]^+$ = 414.1672 observed: $[M+H]^+$ = 414.1674 mass accuracy = 0.2 ppm

[1]H (500 MHz) and [13]C NMR (126 MHz) spectra of **76** in DMSO-$d_6$

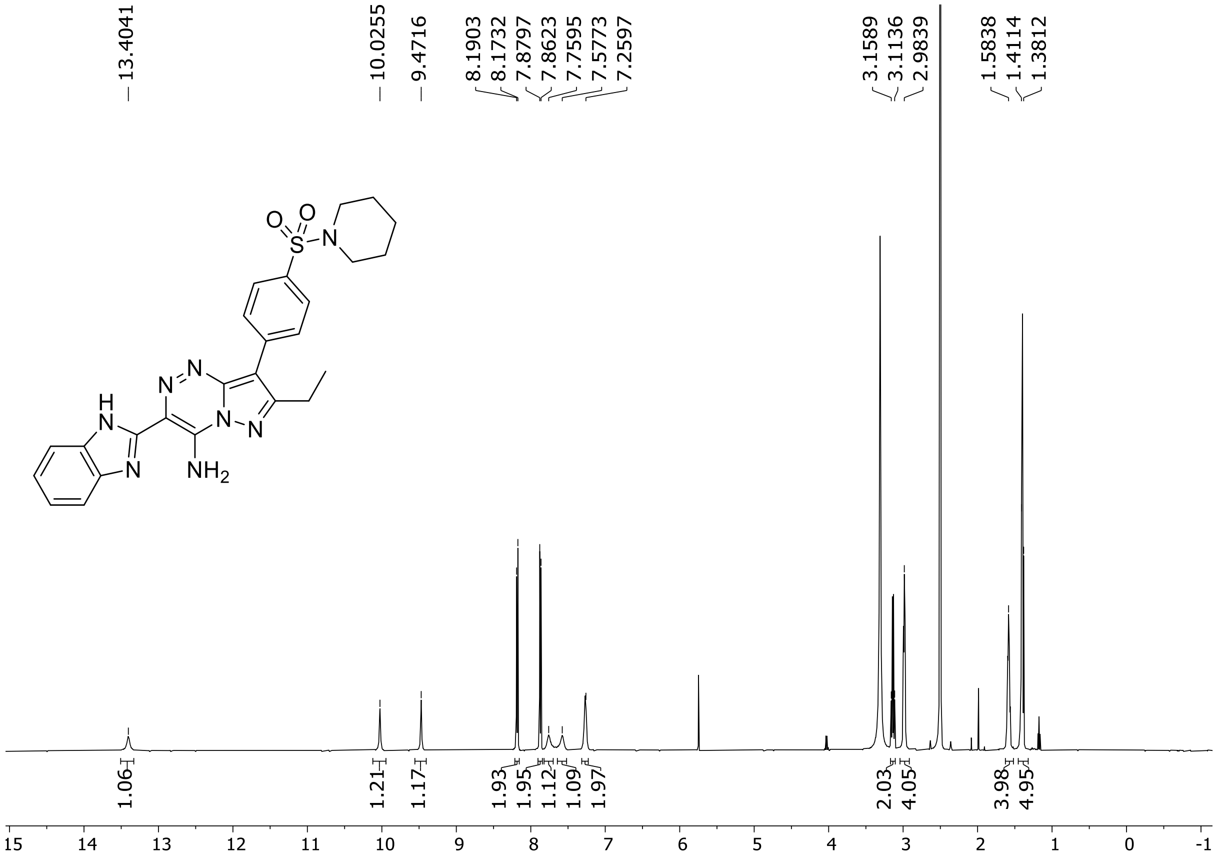

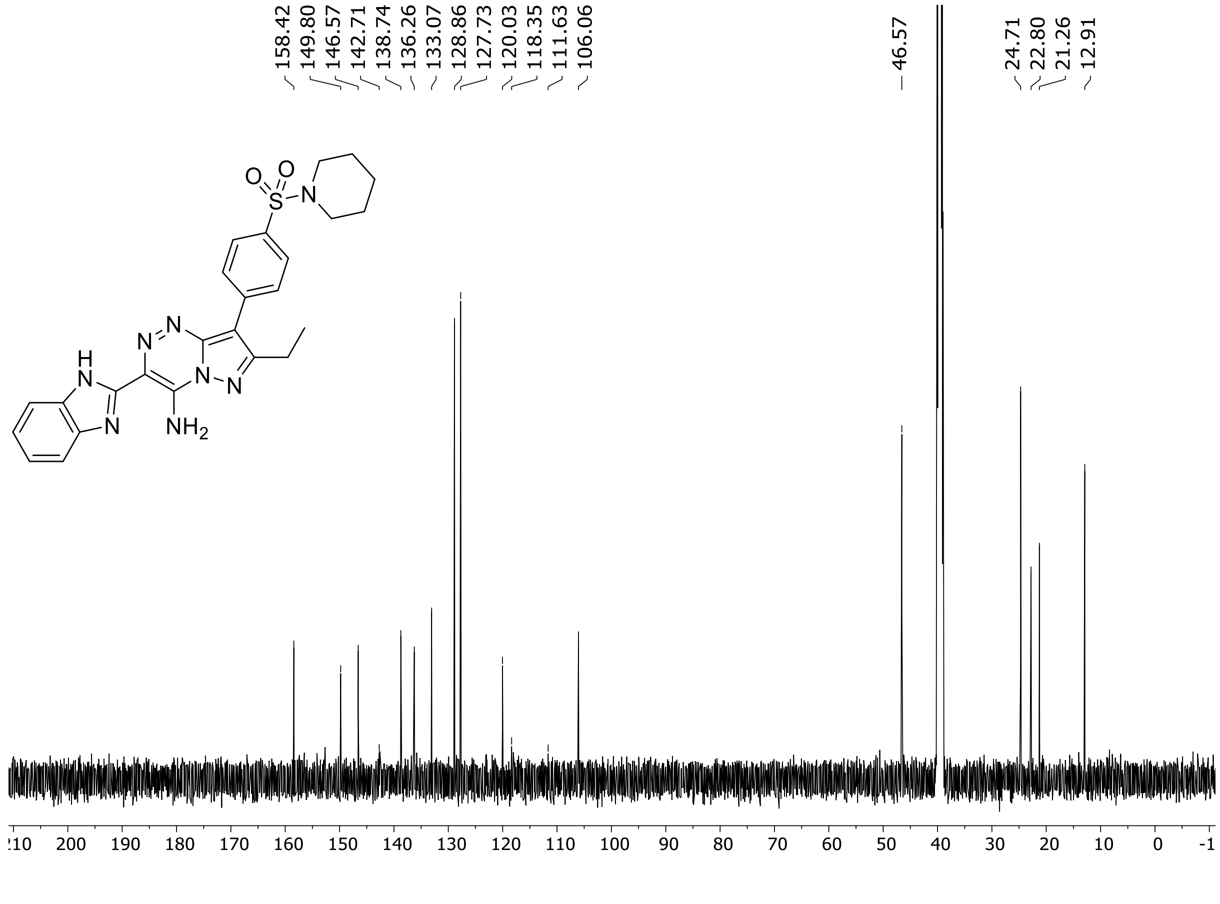

HRMS spectrum of **76**

**$C_{25}H_{26}N_8O_2S$** **exact mass: 502.1899**

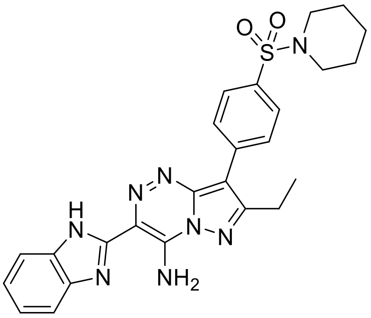

**APCI +** (MMI)
nitrogen flow 5 L/min, gas temperature 325°C, nebulizer 45 psig, skimmer 65 V,
vaporizer 200°C, fragmentor 20 V, dissolved in methanol

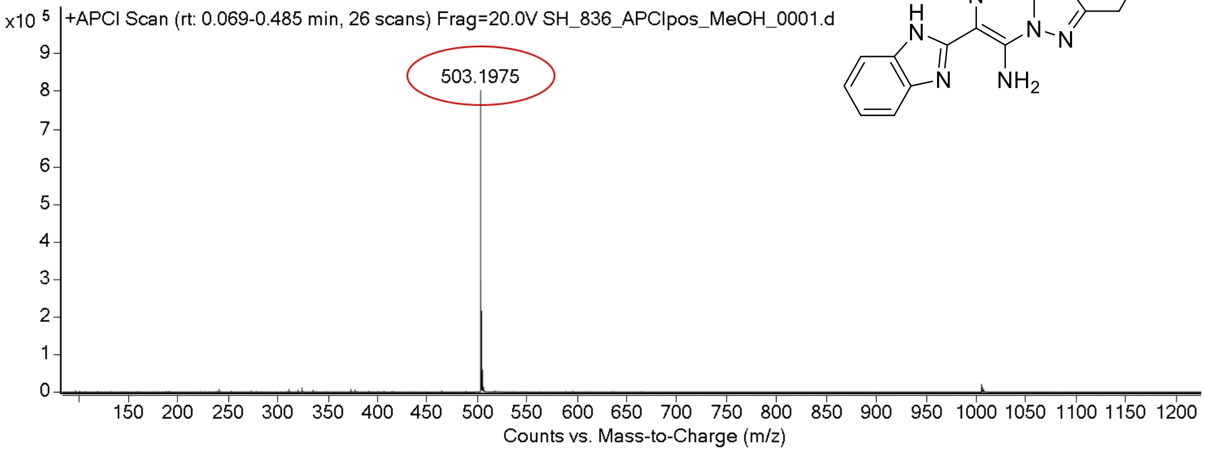

expected mass: $[M+H]^+$ = 503.1972 observed mass : $[M+H]^+$ = 503.1975 mass accuracy = 0.4 ppm

$^{1}H$ (500 MHz) and $^{13}C$ NMR (126 MHz) spectra of **77** in DMSO-$d_6$

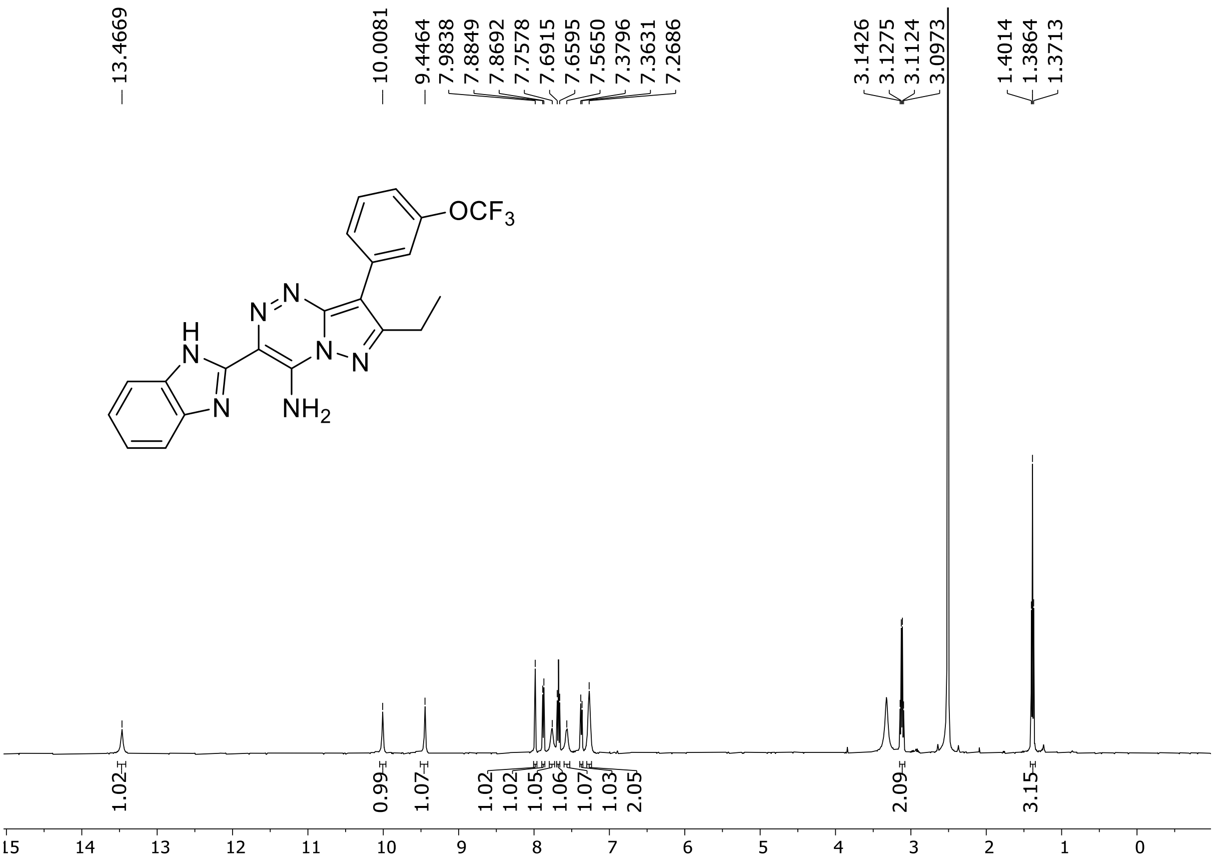

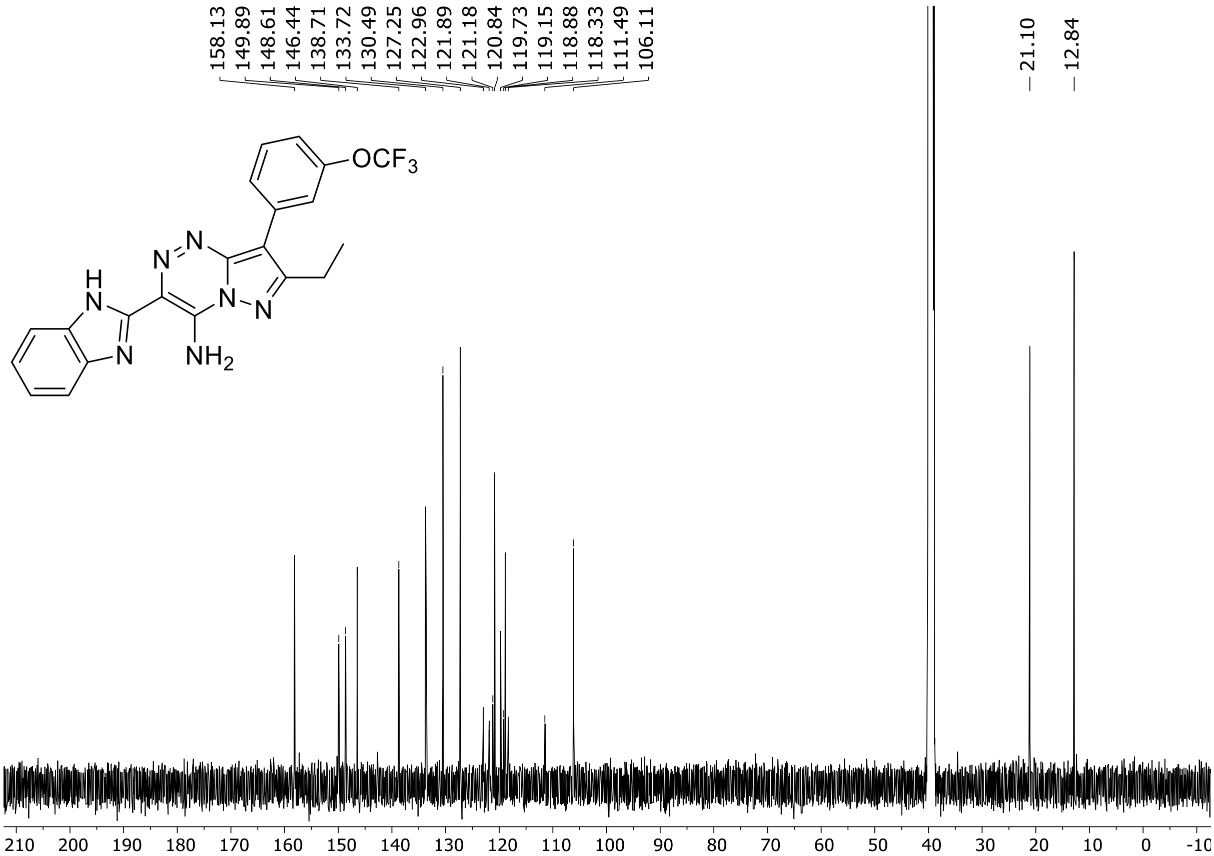

$^{19}F$ (471 MHz) NMR spectrum of **77** DMSO-$d_6$

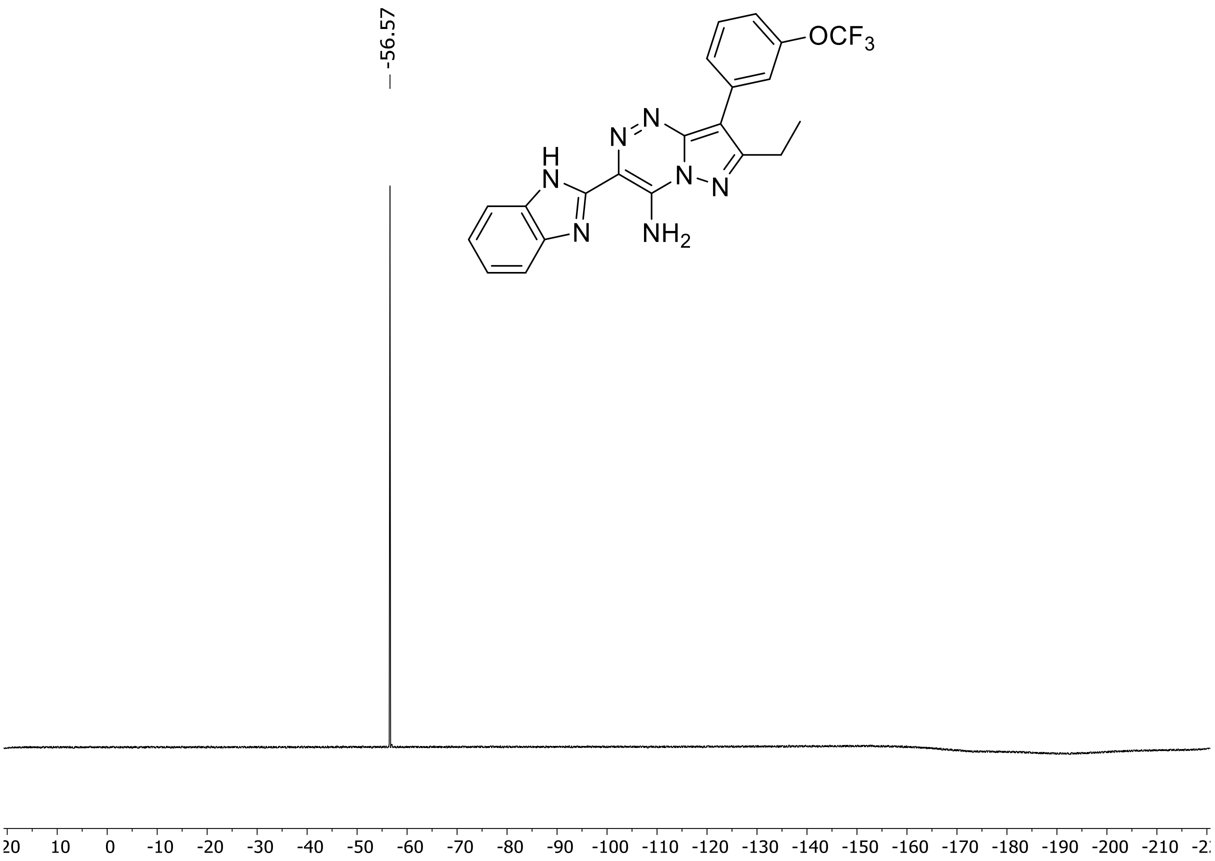

HRMS spectrum of **77**

**$C_{21}H_{16}F_3N_7O$** **mono *m/z* 439.1368**

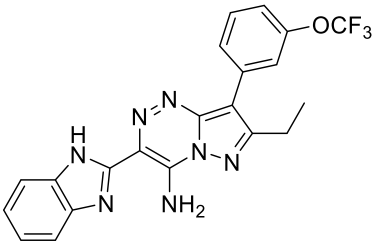

**APCI +** (MMI)
nitrogen flow 5 L/min, gas temperature 325°C, nebulizer 45 psi, skimmer 65 V, vaporizer 250°C, fragmentor 35 V, dissolved in methanol

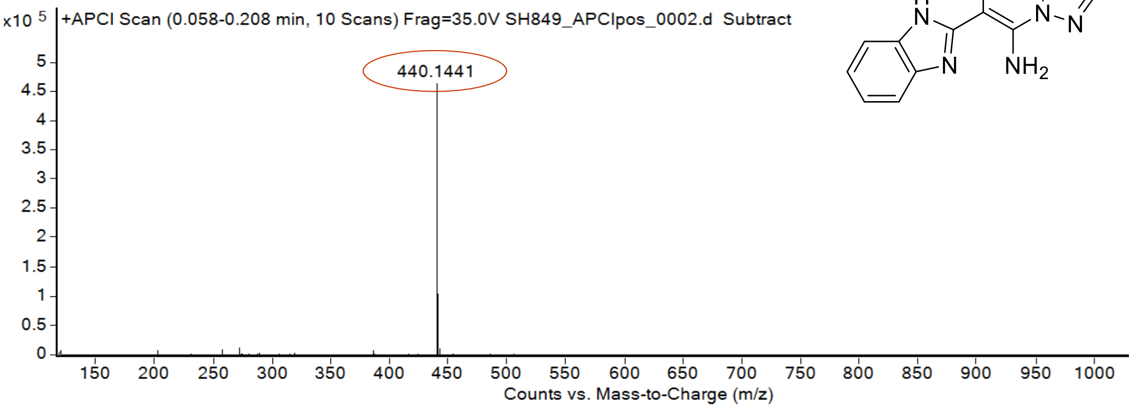

calculated mass: $[M+H]^+ = 440.1441$ observed: $[M+H]^+ = 440.1441$ mass accuracy < 0.1 ppm

$^{1}H$ (500 MHz) and $^{13}C$ NMR (126 MHz) spectra of **78** in DMSO-$d_6$

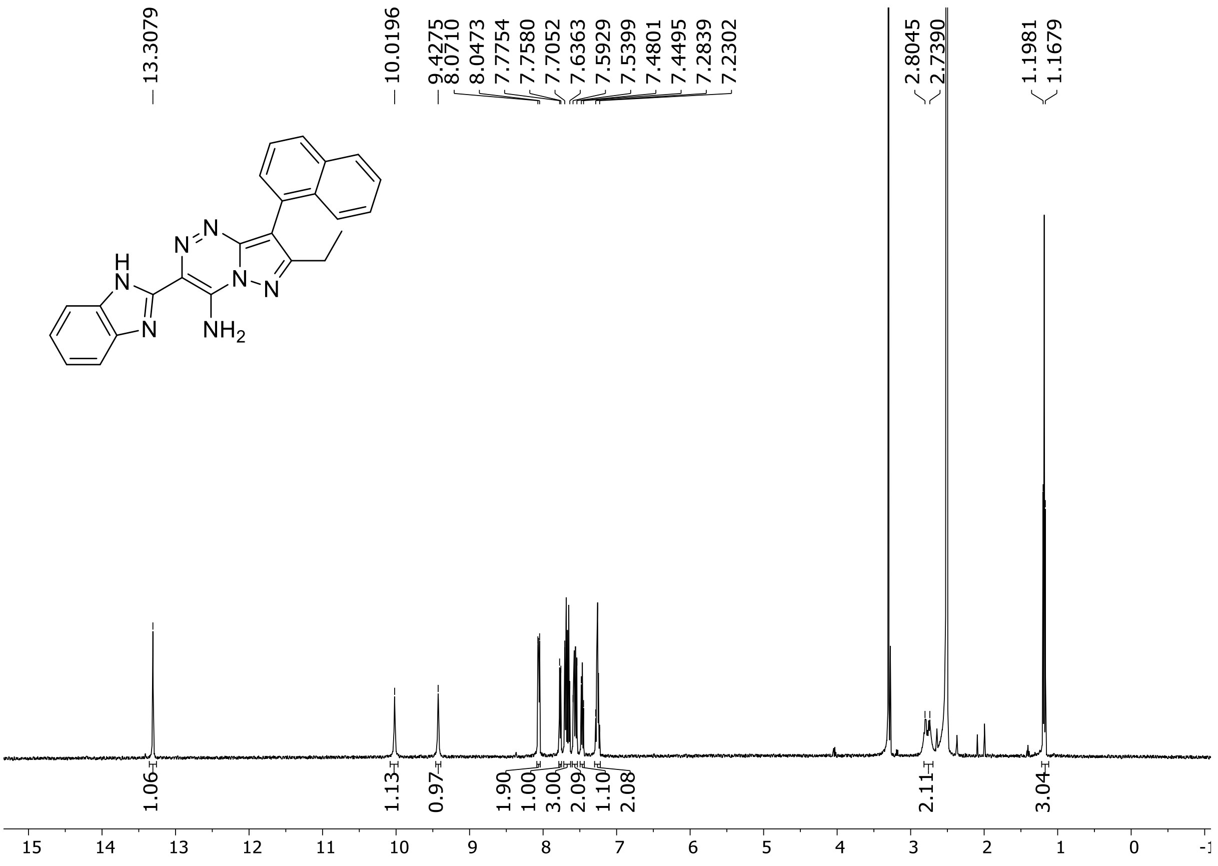

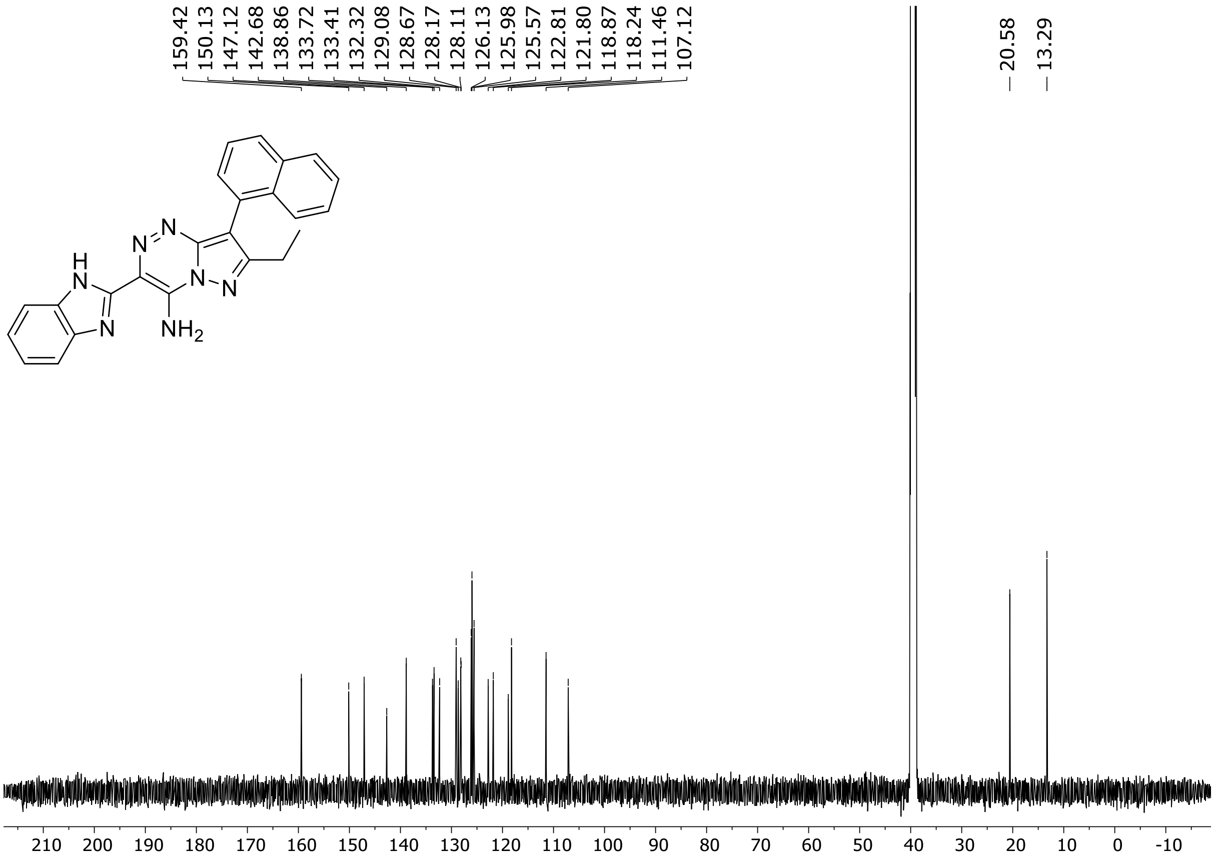

HRMS spectrum of **78**

**$C_{24}H_{19}N_7$** **exact mass: 405.1702**

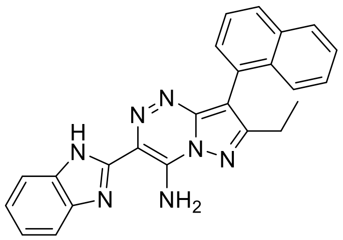

**APCI +** (MMI)
nitrogen flow 5 L/min, gas temperature 325°C, nebulizer 45 psig, skimmer 65 V,
vaporizer 200°C, fragmentor 10 V, dissolved in methanol

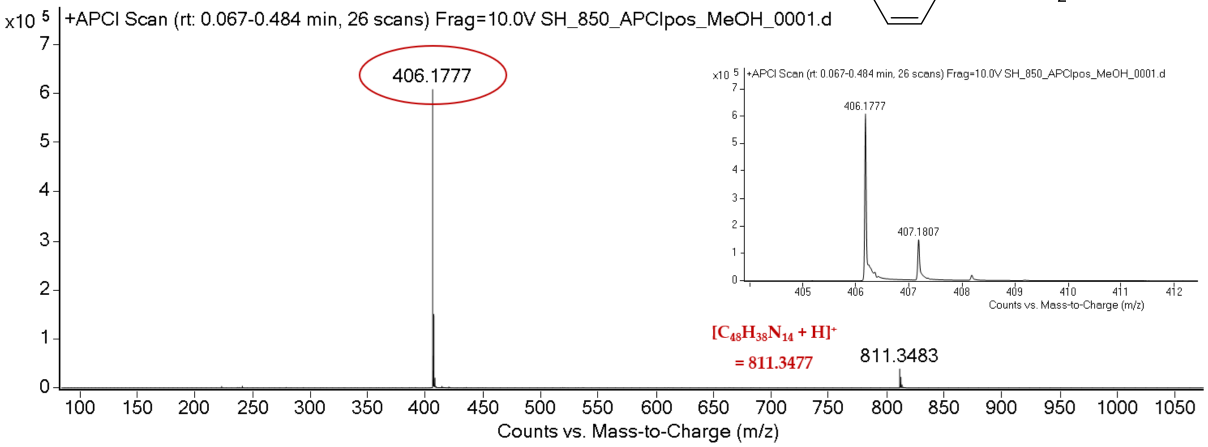

expected mass: $[M+H]^+ = 406.1775$ observed mass : $[M+H]^+ = 406.1777$ mass accuracy = 0.5 ppm

$^{1}H$ (500 MHz) and $^{13}C$ NMR (126 MHz) spectra of **79** in DMSO-$d_6$

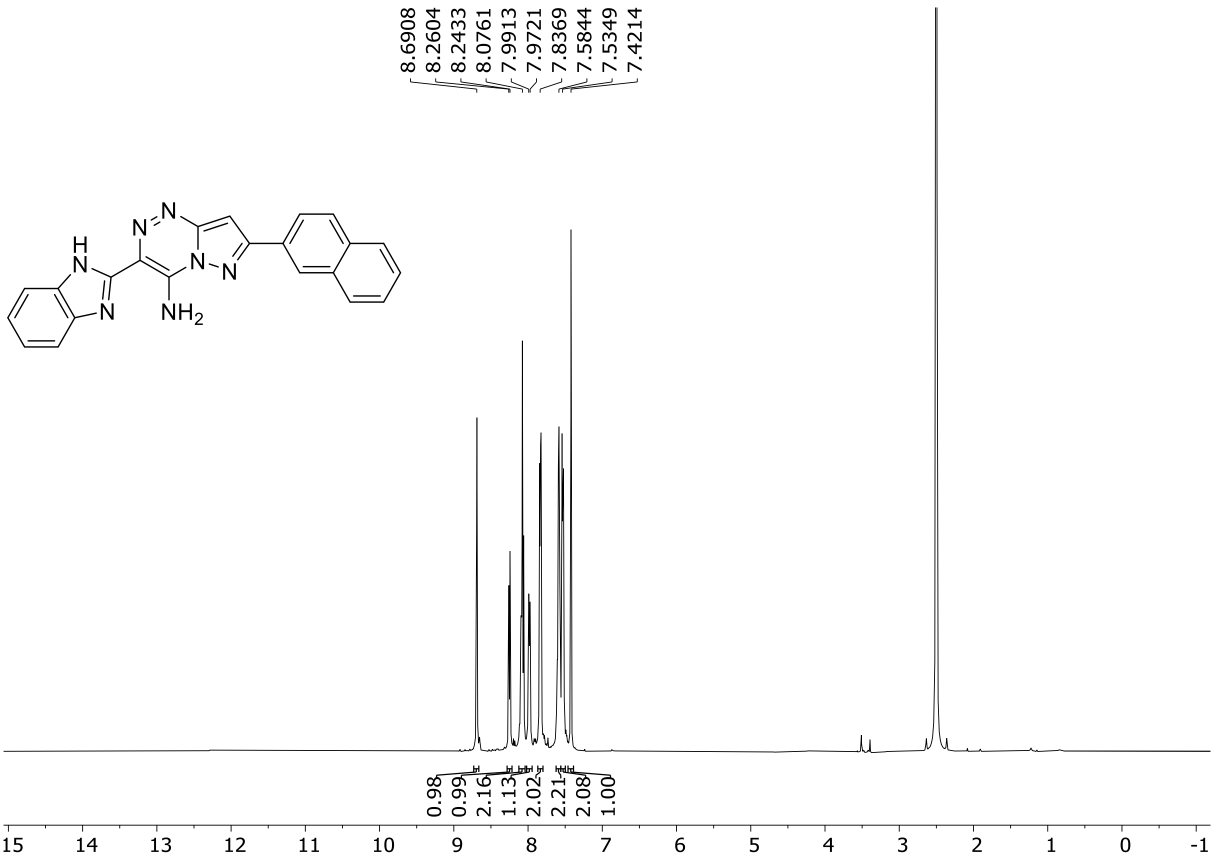

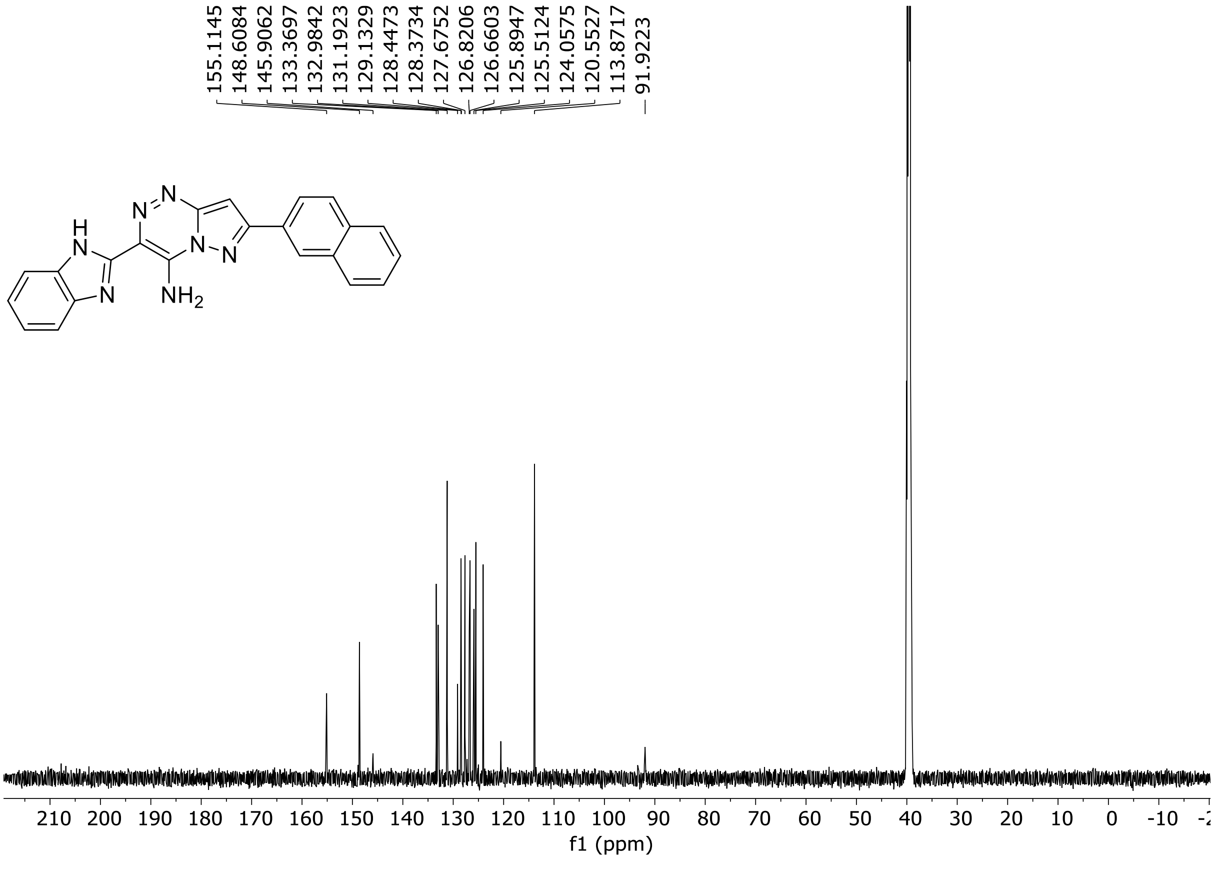

HRMS spectrum of **79**

**$C_{22}H_{14}N_6O$** **mono m/z = 378.1229**

**ESI +** (MMI)
nitrogen flow 5 L/min, gas temperature 300°C, nebulizer 45 psi,
skimmer 60 V, fragmentor 20 V, dissolved in methanol

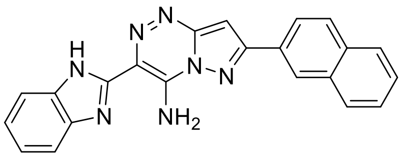

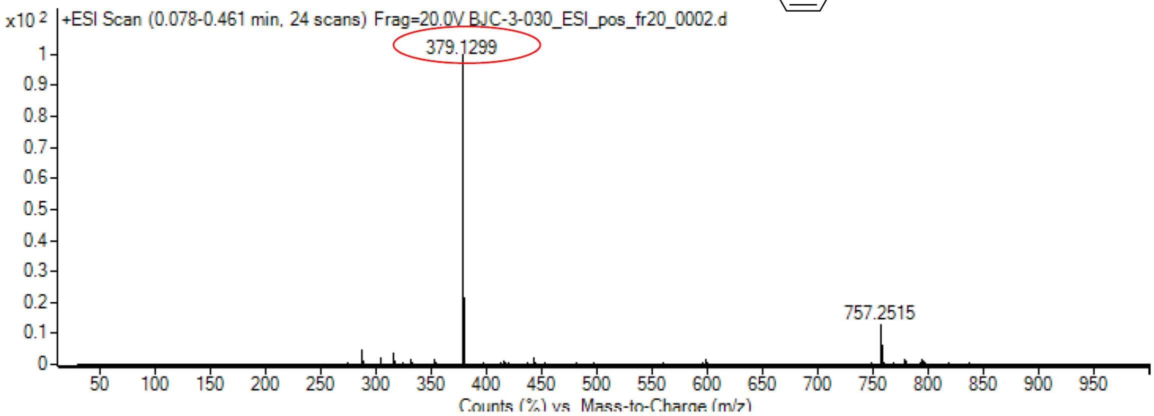

calculated mass: $[M+H]^+$ = 379.1302 observed: $[M+H]^+$ = 379.1299 max. mass error = 0.8ppm

$^{1}$H (500 MHz) and $^{13}$C NMR (126 MHz) spectra of **80** in DMSO-$d_6$

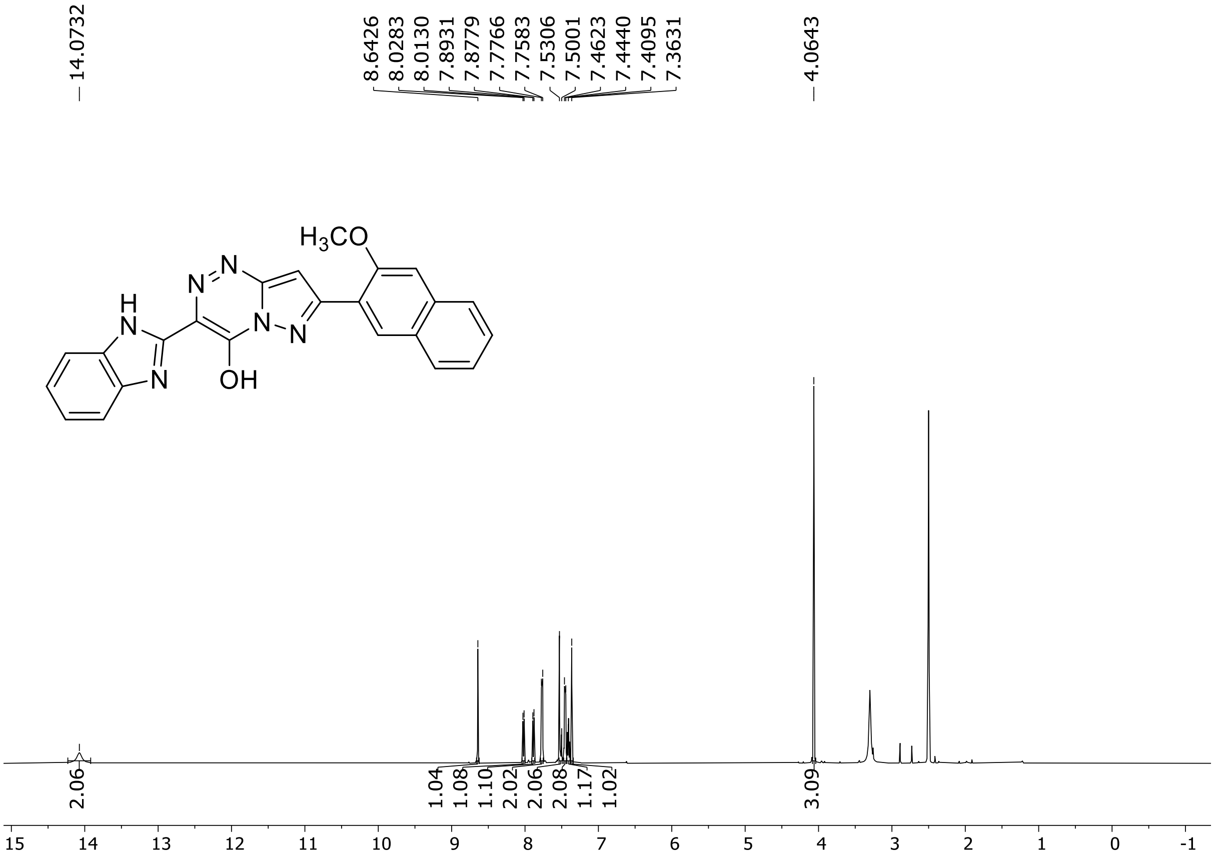

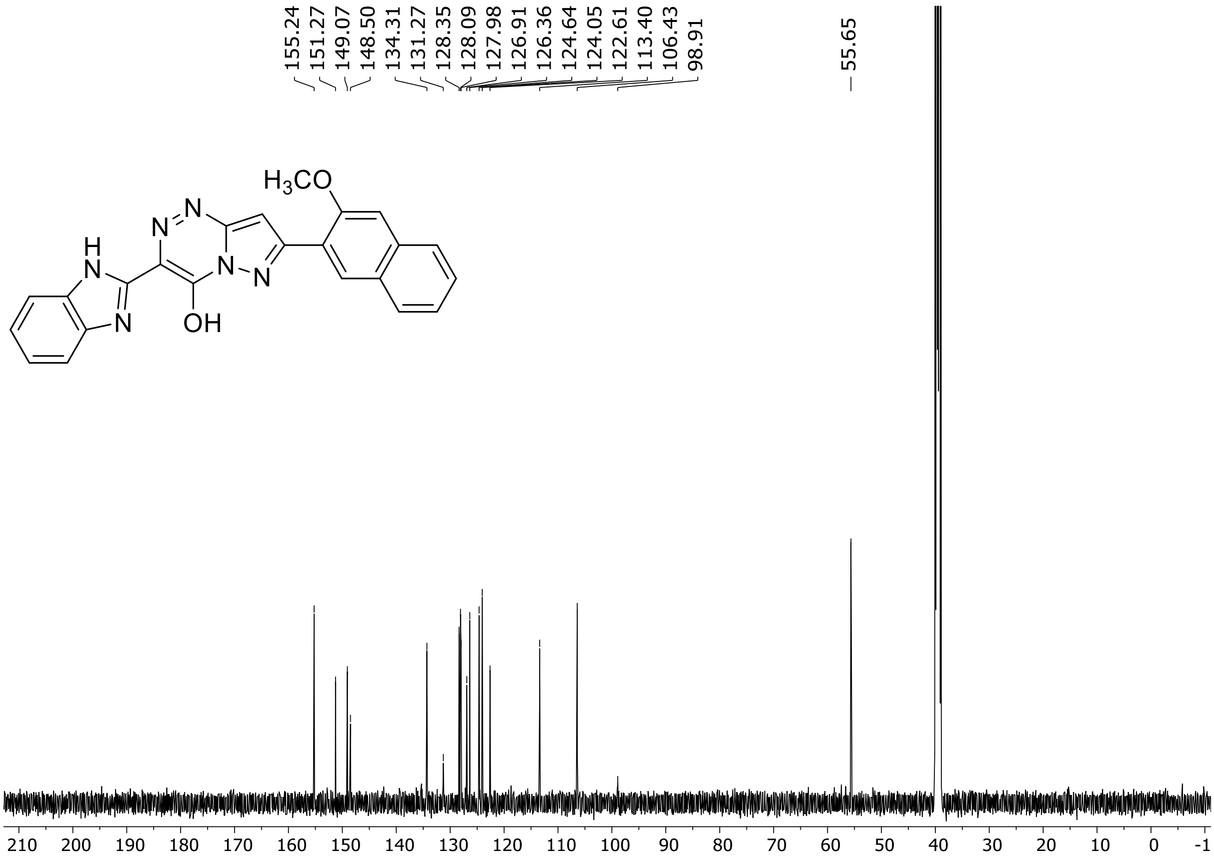

HRMS spectrum of **80**

**$C_{23}H_{17}N_7O$** **exact mass: 407.1495**

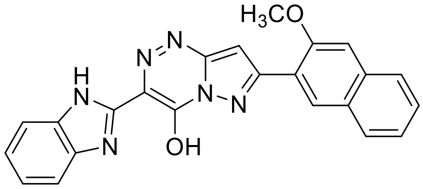

**APCI +** (MMI)
nitrogen flow 5 L/min, gas temperature 325°C, nebulizer 45 psig, skimmer 65 V, vaporizer 200°C, fragmentor 5 V, dissolved in methanol

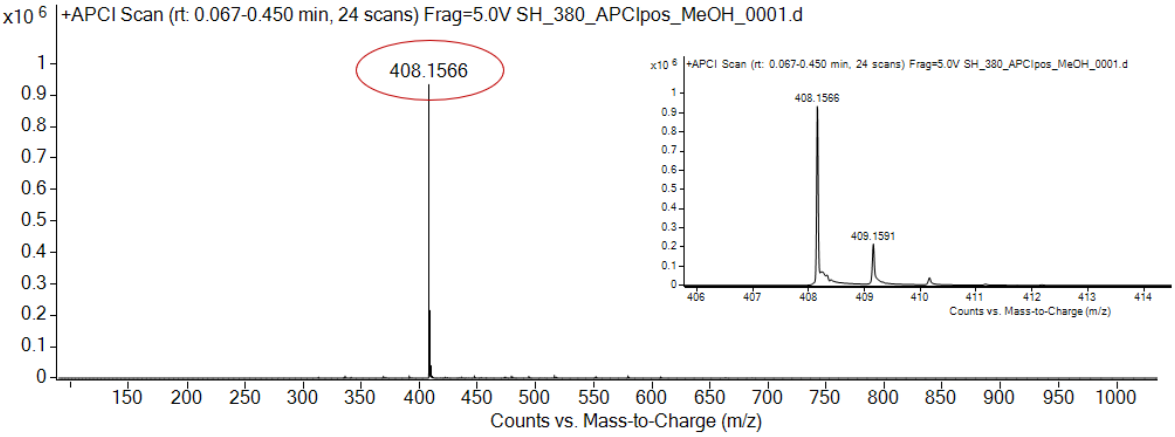

expected mass: [M+H]$^+$ = 408.1567 observed mass : [M+H]$^+$ = 408.1566 mass accuracy = - 0.2 ppm

[1]H (300 MHz) NMR spectrum of **81** in DMSO-$d_6$

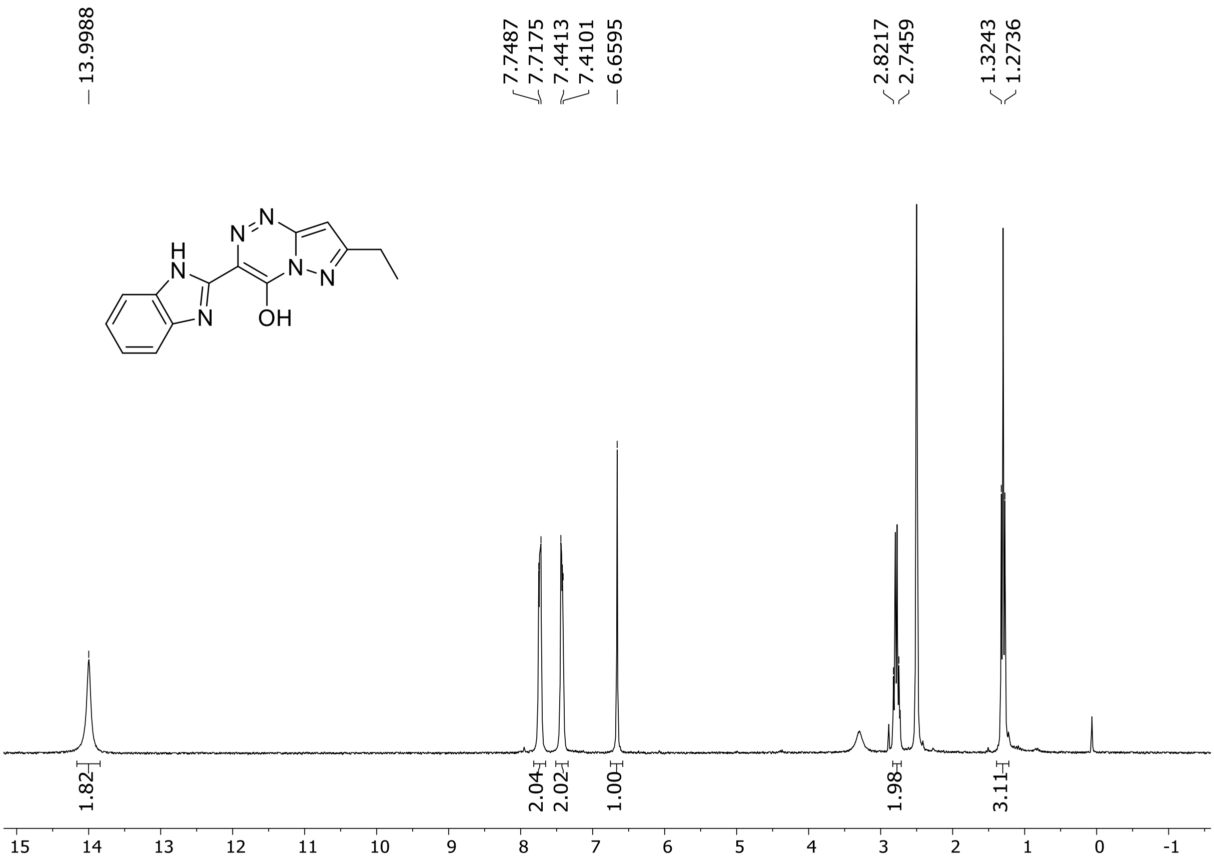

HRMS spectrum of **81**

**$C_{14}H_{12}N_6O$** **exact mass: 280.1073**

**APCI +** (MMI)
nitrogen flow 3 L/min, gas temperature 325°C, nebulizer 45 psig, skimmer 65 V,
vaporizer 200°C, fragmentor 20 V, dissolved in methanol

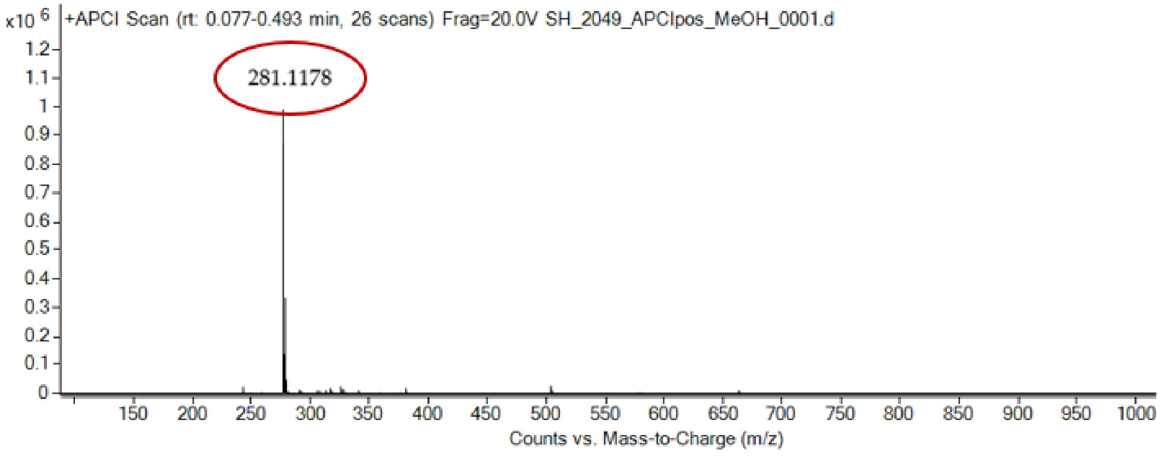

expected mass: $[M+H]^+ = 281.1175$ observed mass: $[M+H]^+ = 281.1178$ mass accuracy < 0.1 ppm

[1]H (500 MHz) and [13]C NMR (126 MHz) spectra of **82** in DMSO-$d_6$

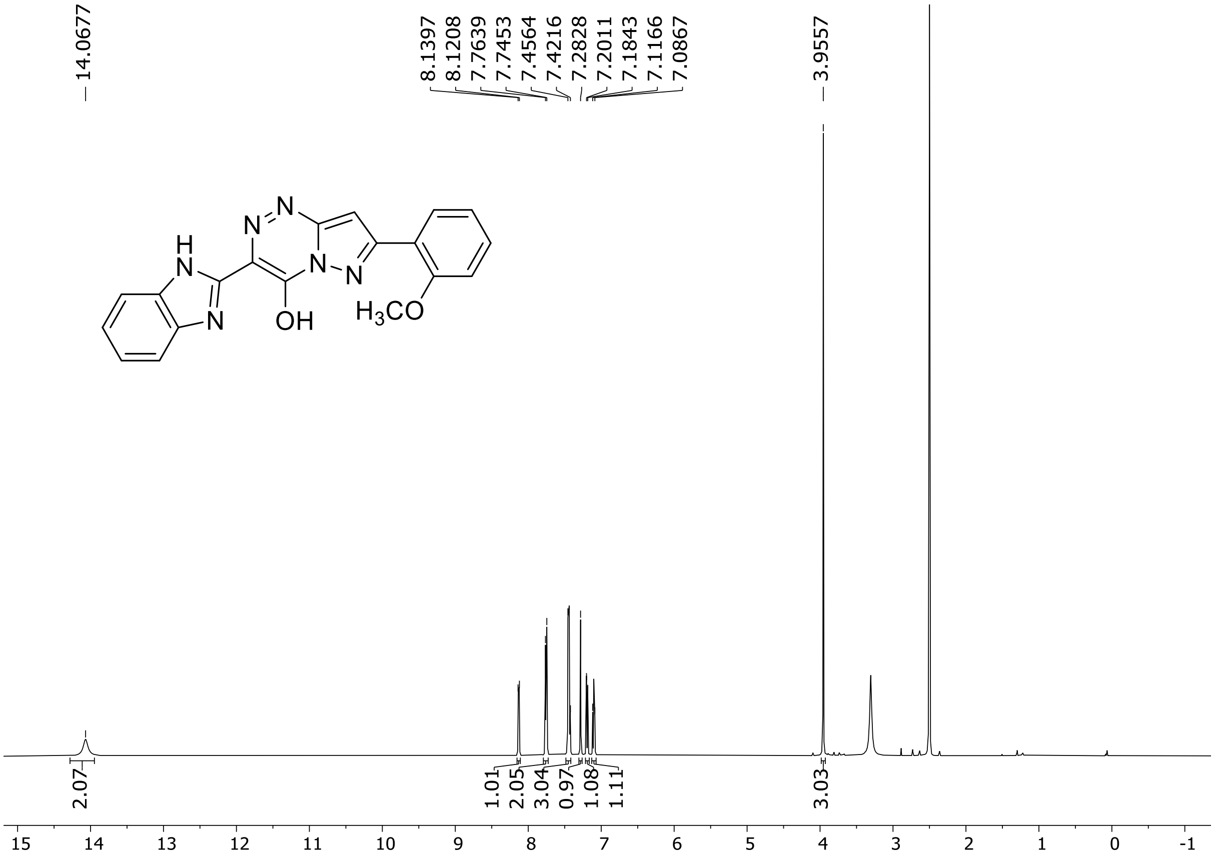

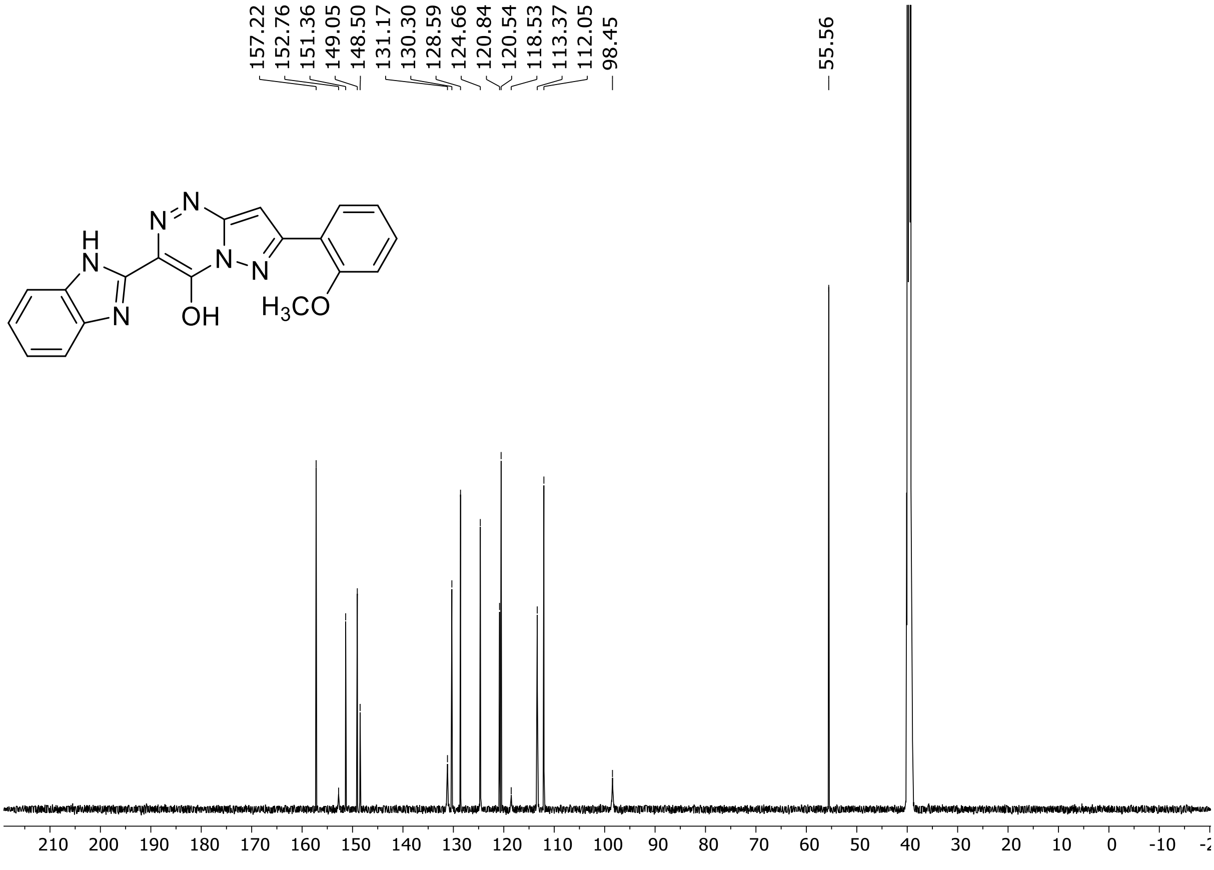

HRMS spectrum of **82**

**$C_{19}H_{14}N_6O_2$** **exact mass: 358.1178**

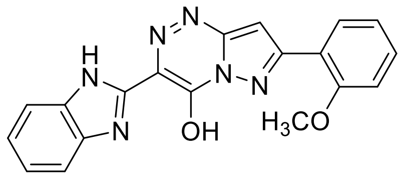

**APCI +** (MMI)
nitrogen flow 5 L/min, gas temperature 325°C, nebulizer 45 psig, skimmer 65 V, vaporizer 200°C, fragmentor 20 V, dissolved in methanol

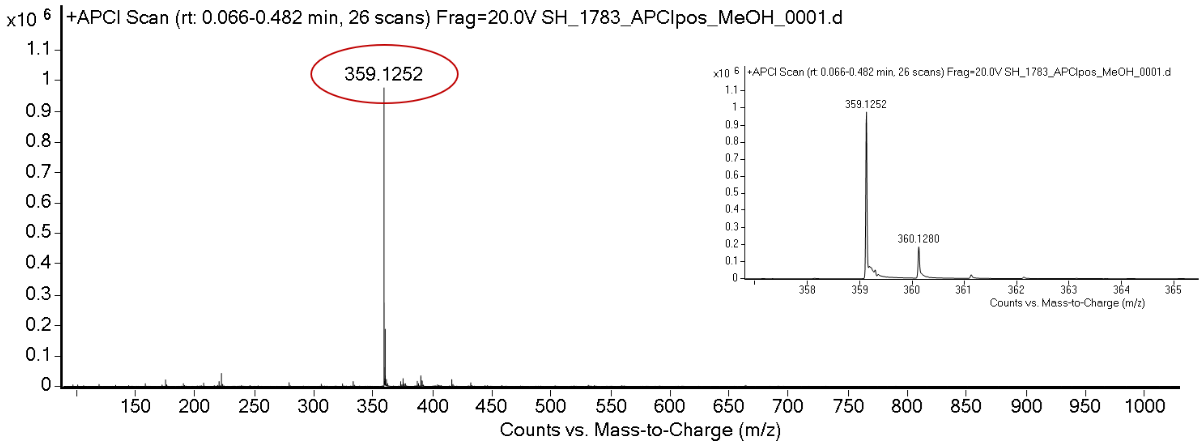

expected mass: [M+H]$^+$ = 359.1251 observed mass : [M+H]$^+$ = 359.1252 mass accuracy = 0.3 ppm

[1]H (500 MHz) and [13]C NMR (126 MHz) spectra of **83** in DMSO-$d_6$

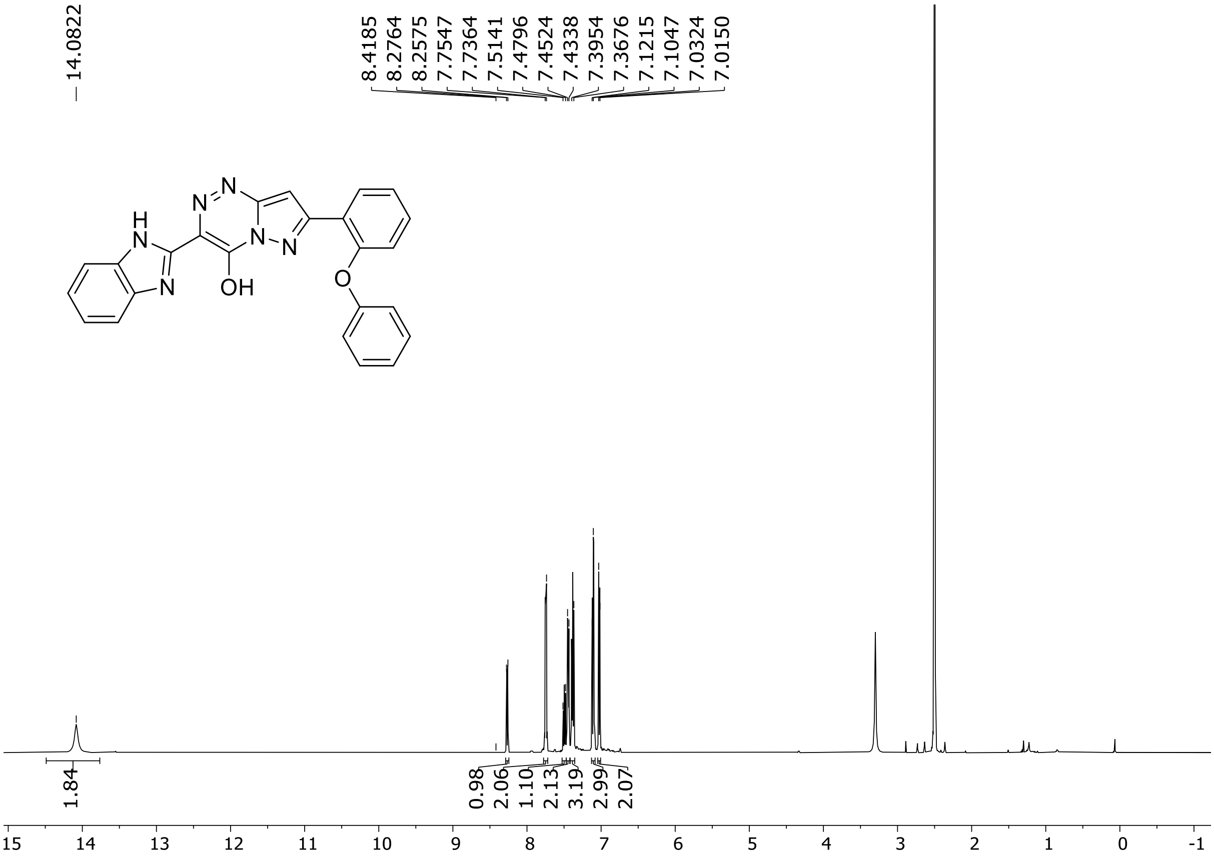

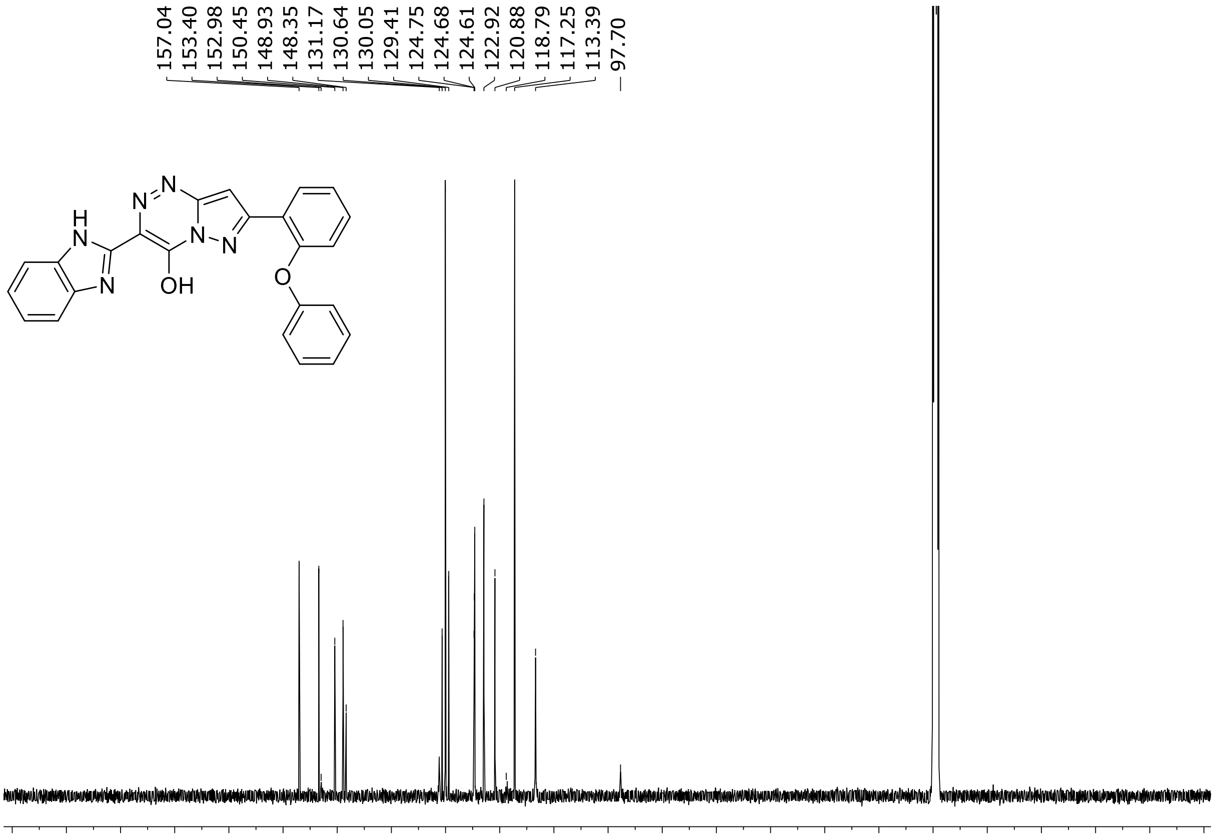

HRMS spectrum of **83**

**$C_{24}H_{16}N_6O_2$** **exact mass: 420.1335**

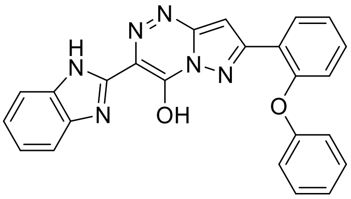

**APCI +** (MMI)
nitrogen flow 5 L/min, gas temperature 325°C, nebulizer 45 psig, skimmer 65 V, vaporizer 200°C, fragmentor 10 V, dissolved in methanol

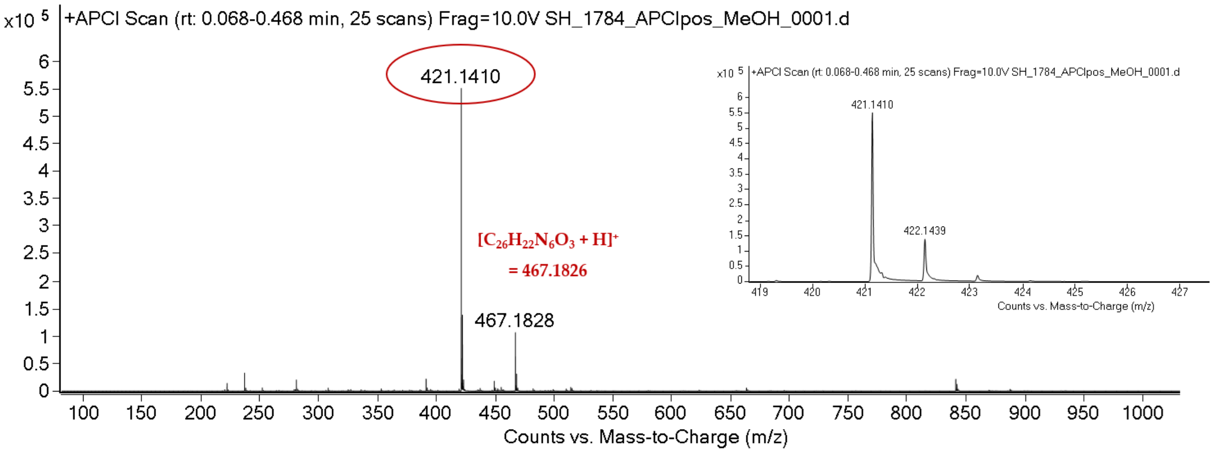

expected mass: $[M+H]^+$ = 421.1408 observed mass : $[M+H]^+$ = 421.1410 mass accuracy = 0.5 ppm

[1]H (500 MHz) and [13]C NMR (126 MHz) spectra of **84** in DMSO-$d_6$

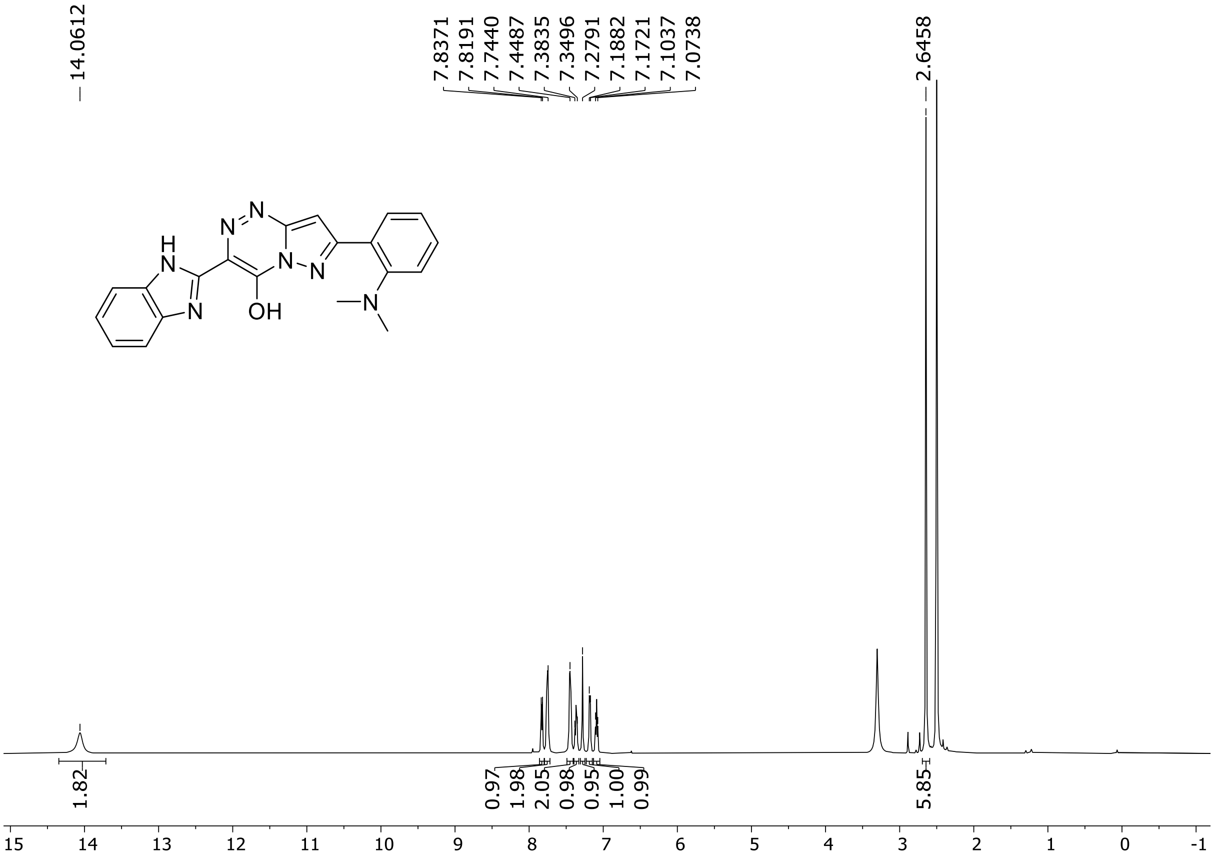

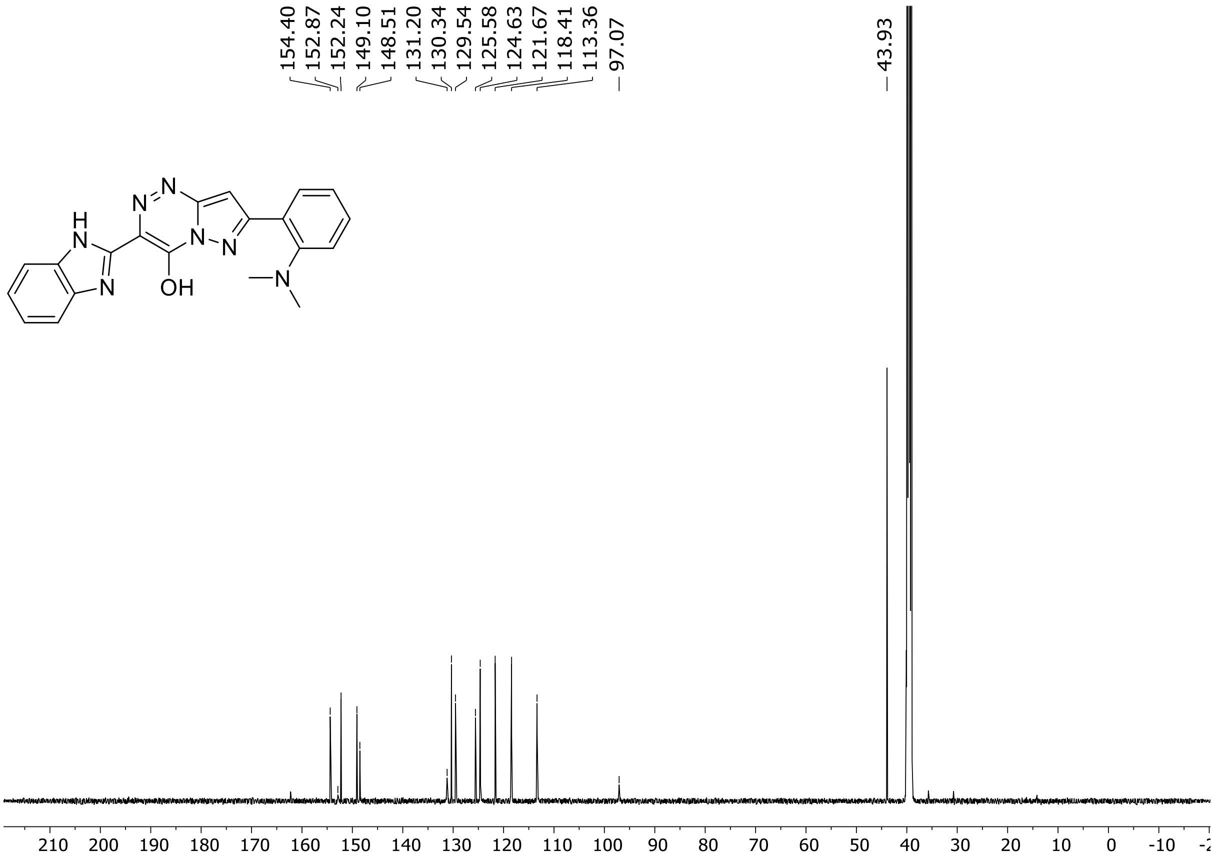

HRMS spectrum of **84**

**$C_{20}H_{17}N_7O$** **exact mass: 371.1495**

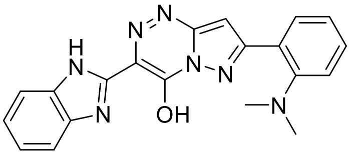

**APCI +** (MMI)
nitrogen flow 5 L/min, gas temperature 325°C, nebulizer 45 psig, skimmer 65 V, vaporizer 200°C, fragmentor 10 V, dissolved in methanol

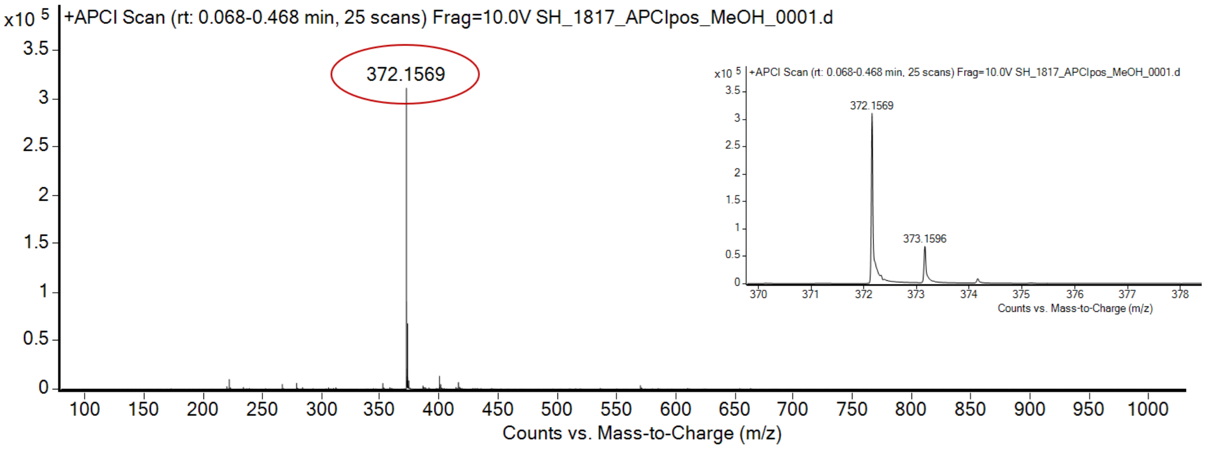

expected mass: $[M+H]^+$ = 372.1567 observed mass : $[M+H]^+$ = 372.1569 mass accuracy = 0.5 ppm

[1]H (500 MHz) and [13]C NMR (126 MHz) spectra of **85** in DMSO-$d_6$

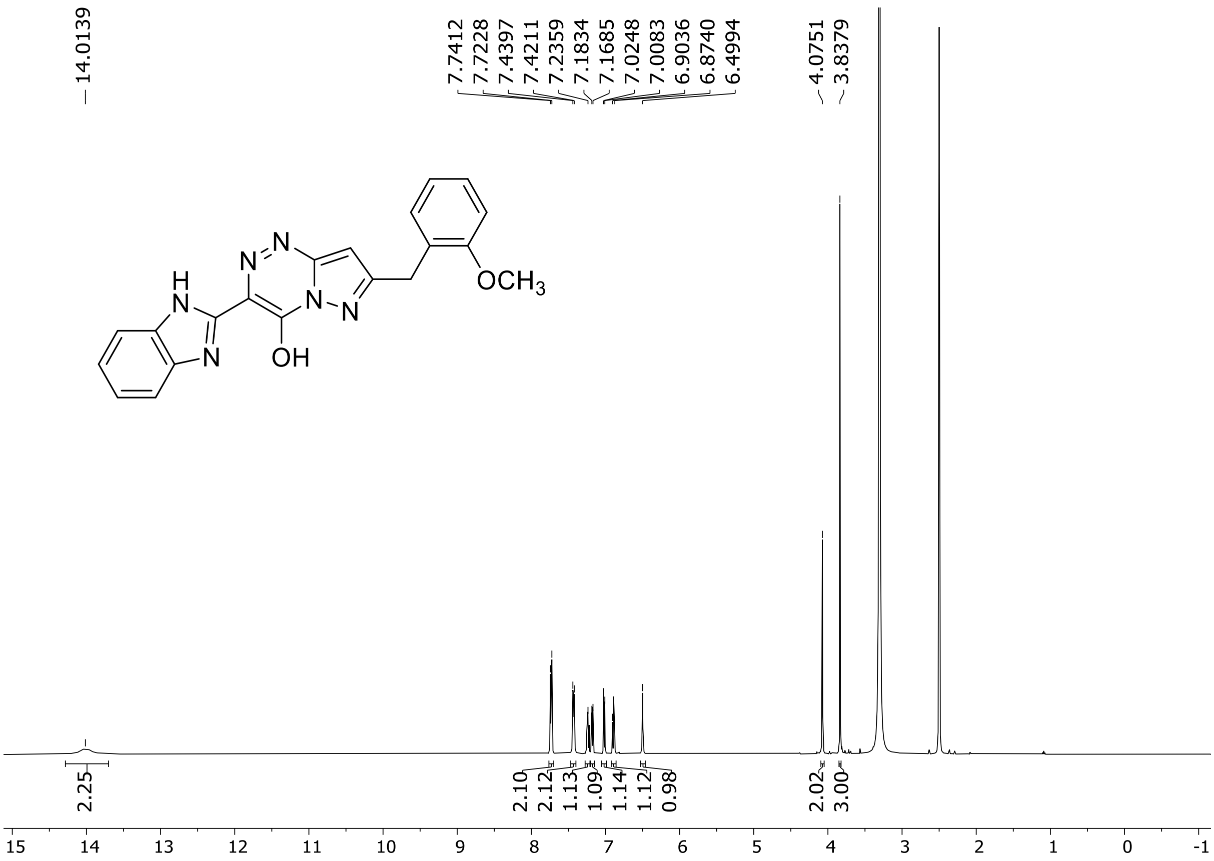

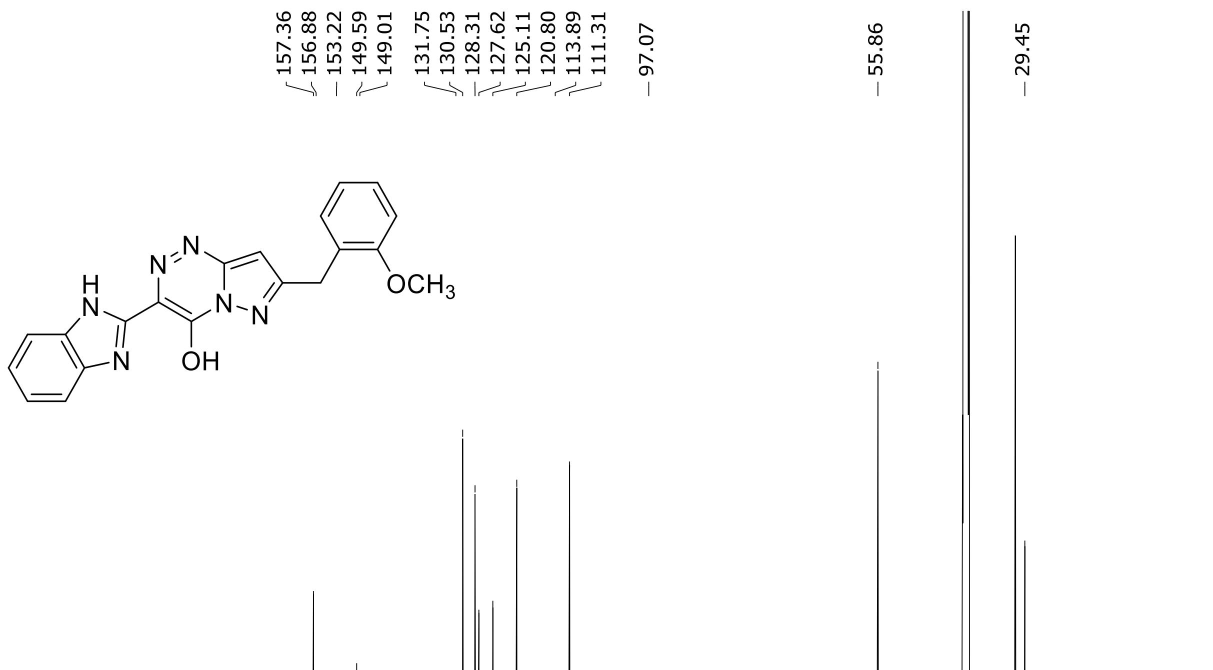

HRMS spectrum of **85**

**$C_{20}H_{16}N_6O_2$** **exact mass: 372.1335**

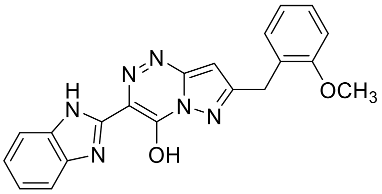

**APCI +** (MMI)
nitrogen flow 3 L/min, gas temperature 325°C, nebulizer 45 psig, skimmer 65 V, vaporizer 200°C, fragmentor 20 V, dissolved in methanol

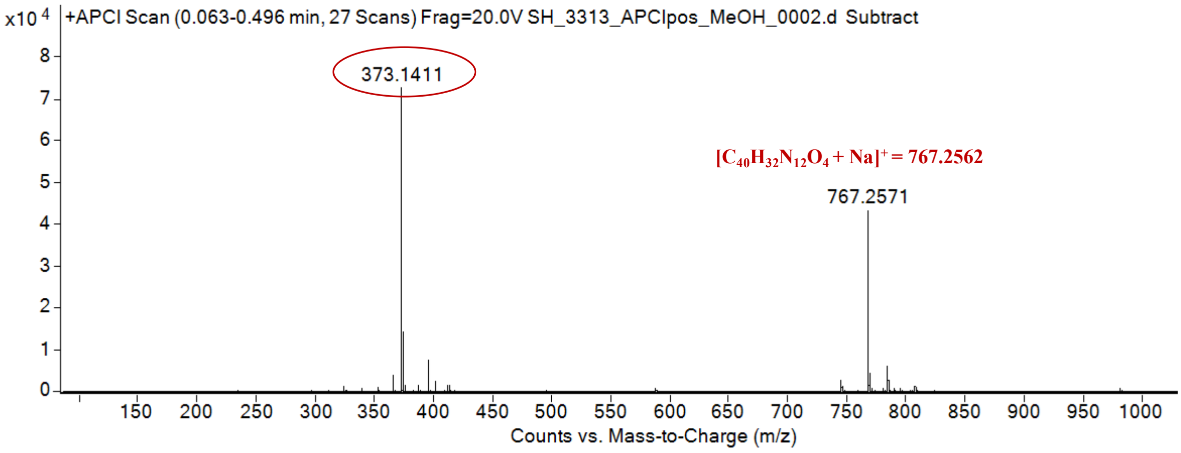

expected mass: $[M+H]^+$ = 373.1408 observed mass: $[M+H]^+$ = 373.1411 mass accuracy = 0.8 ppm

$^{1}H$ (500 MHz) and $^{13}C$ NMR (126 MHz) spectra of **86** in DMSO-$d_6$

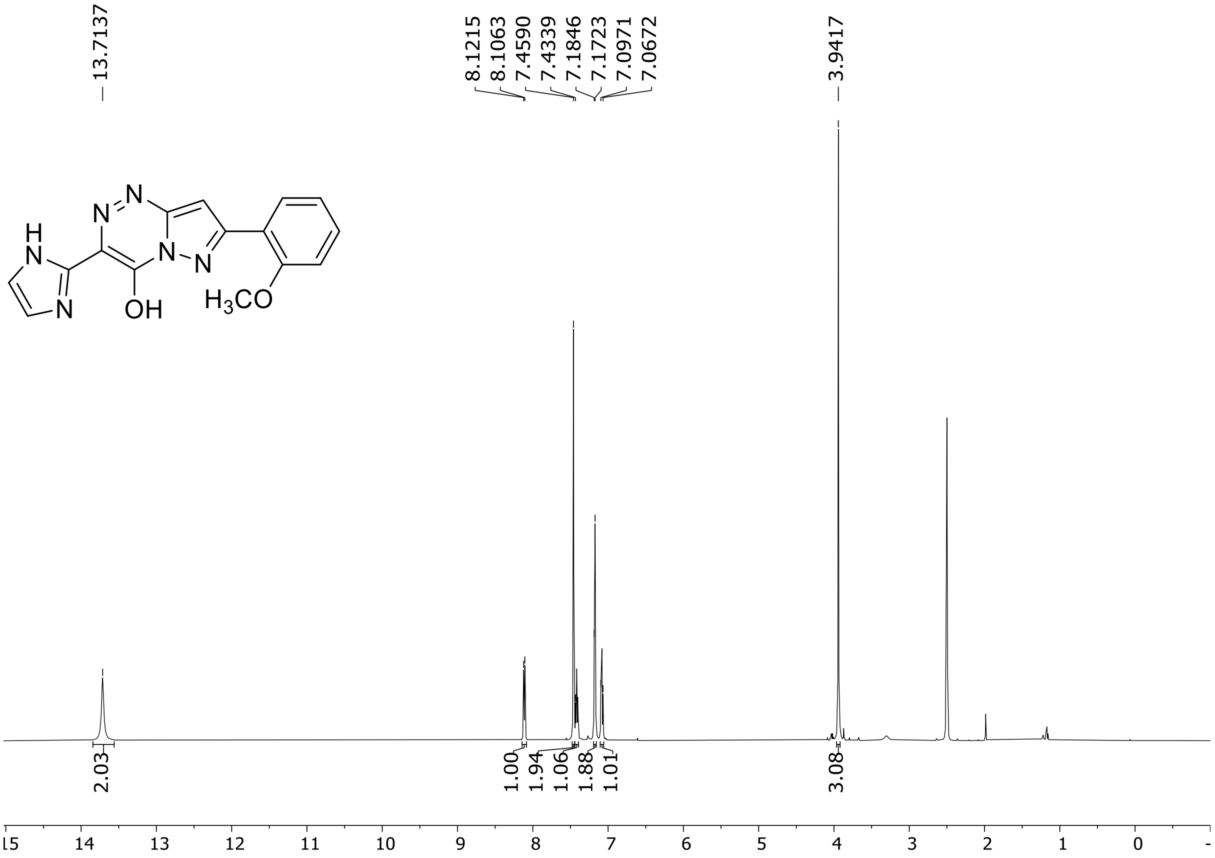

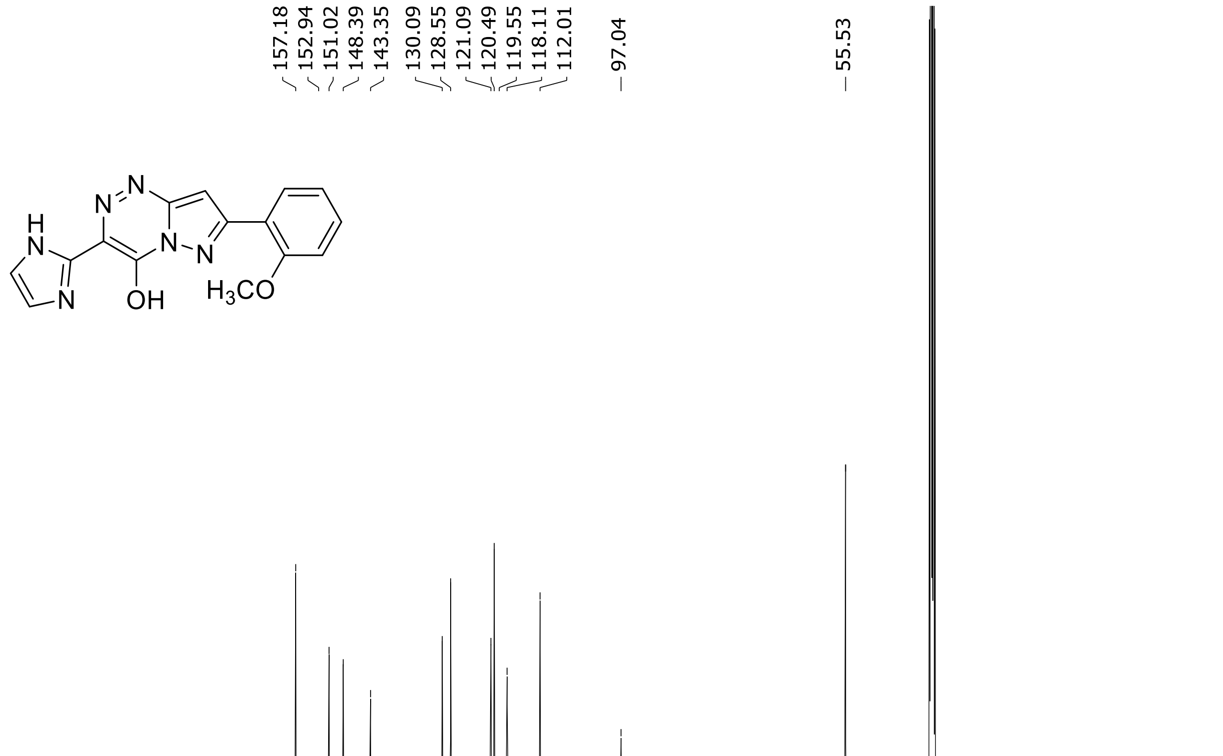

HRMS spectrum of **86**

**$C_{15}H_{12}N_6O_2$** **exact mass: 308.1022**

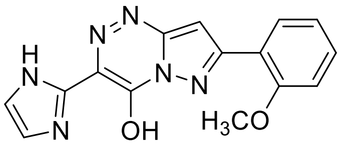

**APCI +** (MMI)
nitrogen flow 3 L/min, gas temperature 325°C, nebulizer 45 psig, skimmer 65 V, vaporizer 200°C, fragmentor 15 V, dissolved in methanol

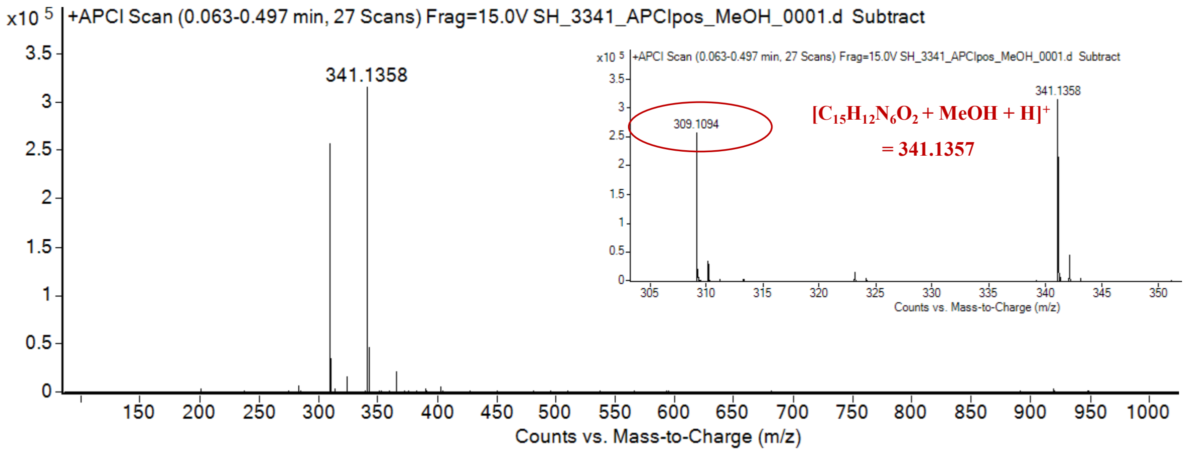

expected mass: $[M+H]^+$ = 309.1095 observed mass: $[M+H]^+$ = 309.1094 mass accuracy = - 0.3 ppm

[1]H (500 MHz) and [13]C NMR (126 MHz) spectra of **87** in DMSO-$d_6$

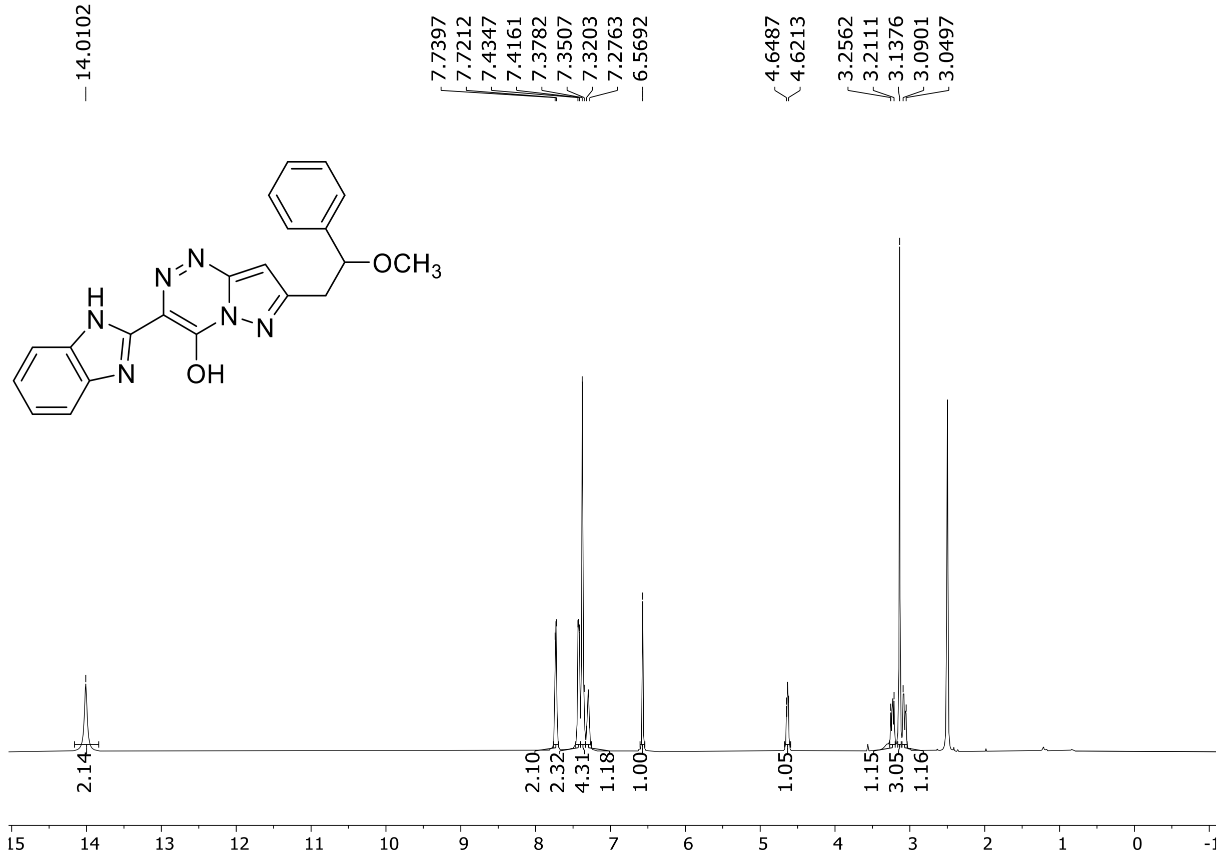

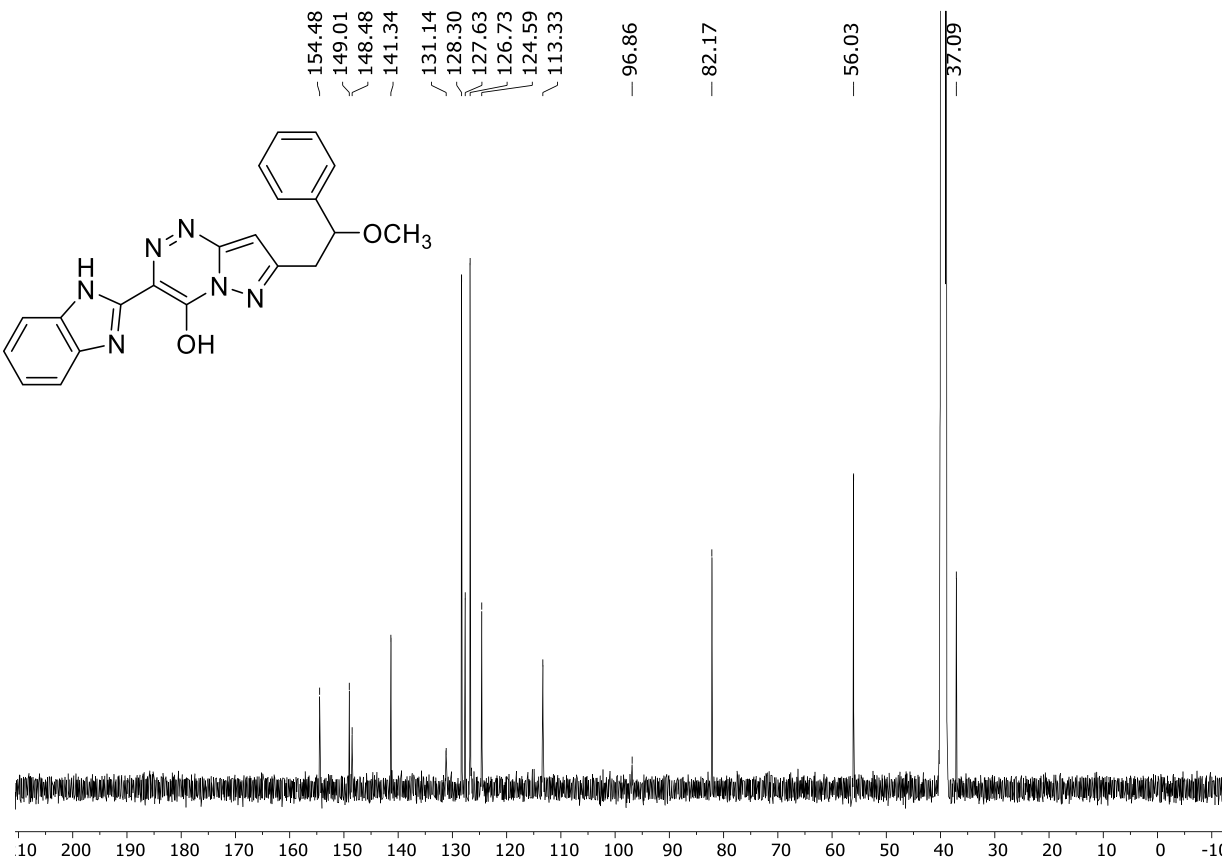

HRMS spectrum of **87**

**$C_{21}H_{18}N_6O_2$** **exact mass: 386.1491**

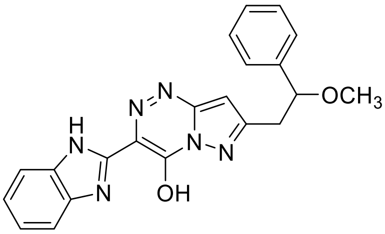

**APCI +** (MMI)
nitrogen flow 3 L/min, gas temperature 325°C, nebulizer 45 psig, skimmer 65 V, vaporizer 200°C, fragmentor 15 V, dissolved in methanol

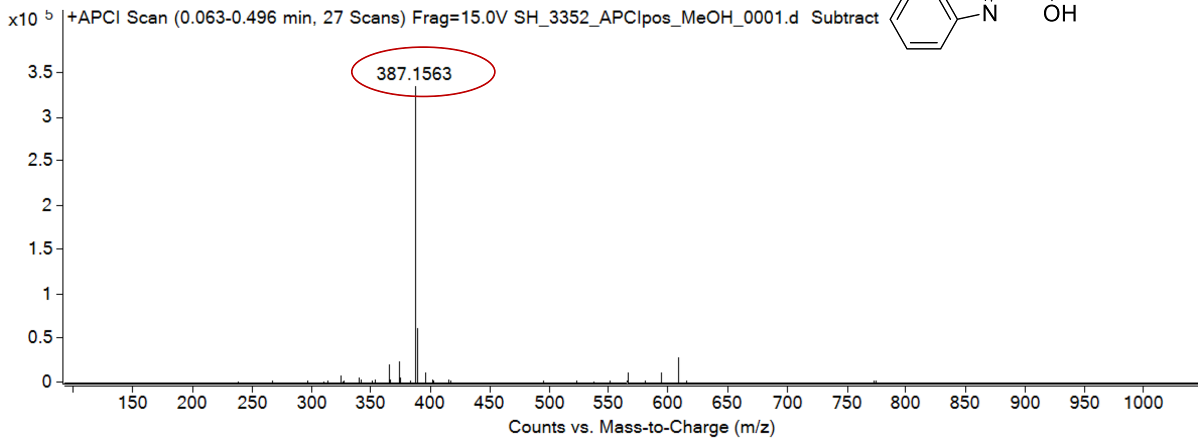

expected mass: $[M+H]^+ = 387.1564$ observed mass: $[M+H]^+ = 387.1563$ mass accuracy = - 0.3 ppm

[1]H (500 MHz) and [13]C NMR (126 MHz) spectra of **88** in DMSO-$d_6$

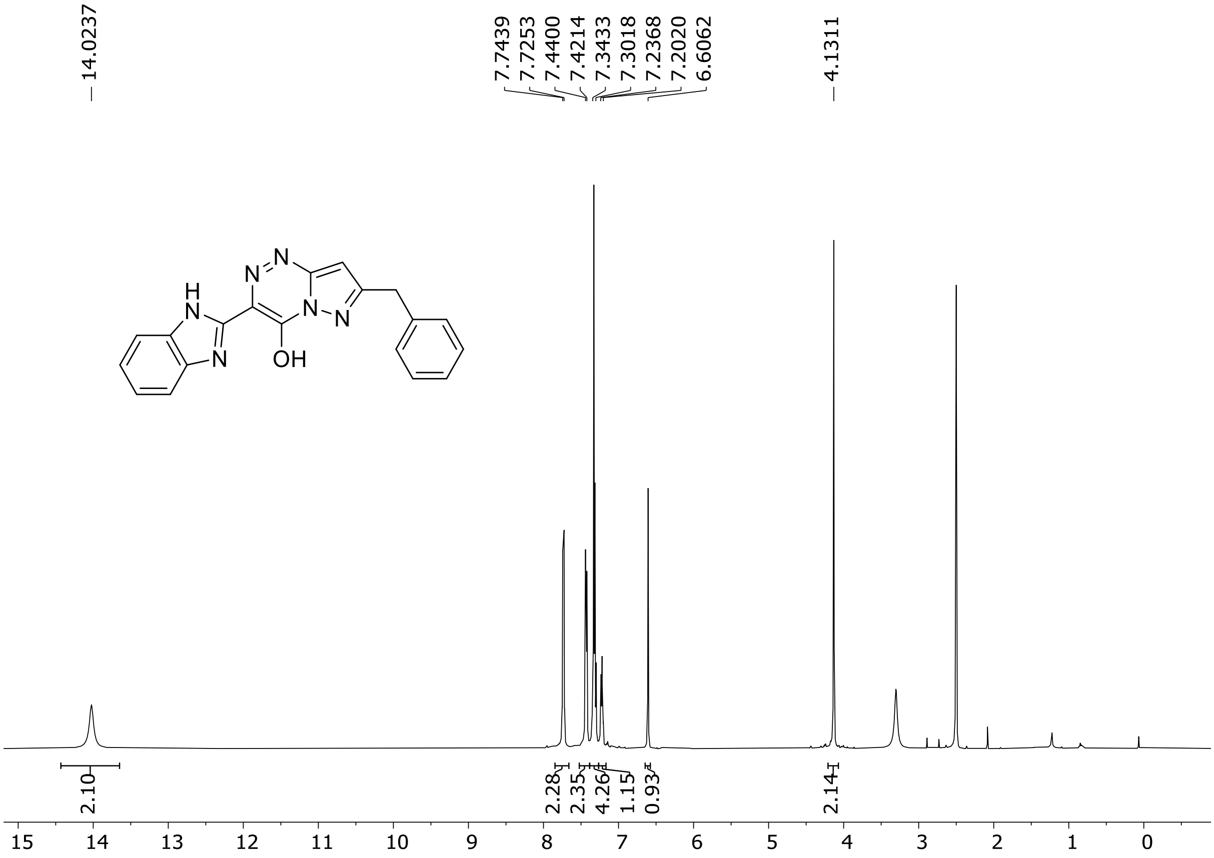

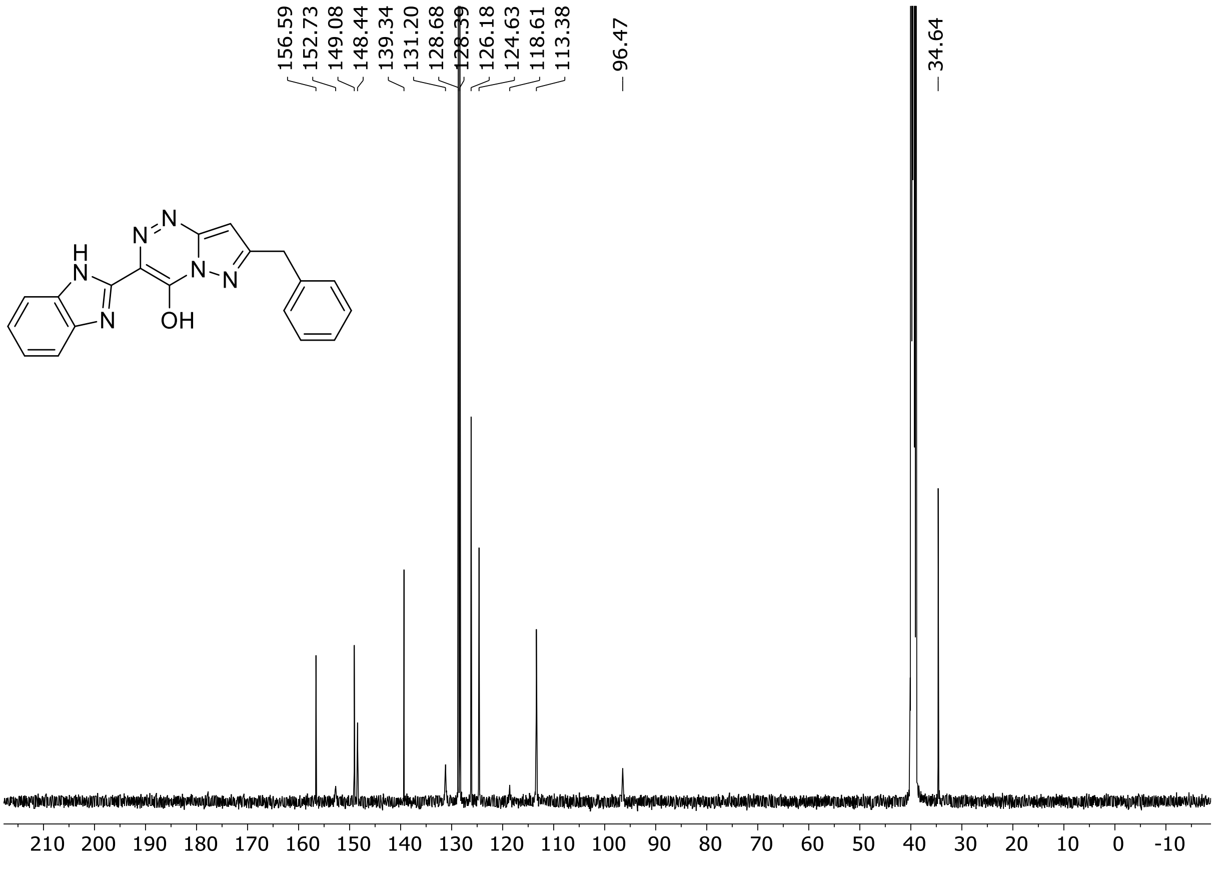

HRMS spectrum of **88**

$C_{19}H_{14}N_6O$ **exact mass: 342.1229**

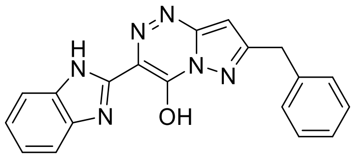

**APCI +** (MMI)
nitrogen flow 5 L/min, gas temperature 325°C, nebulizer 45 psig, skimmer 65 V, vaporizer 200°C, fragmentor 20 V, dissolved in methanol

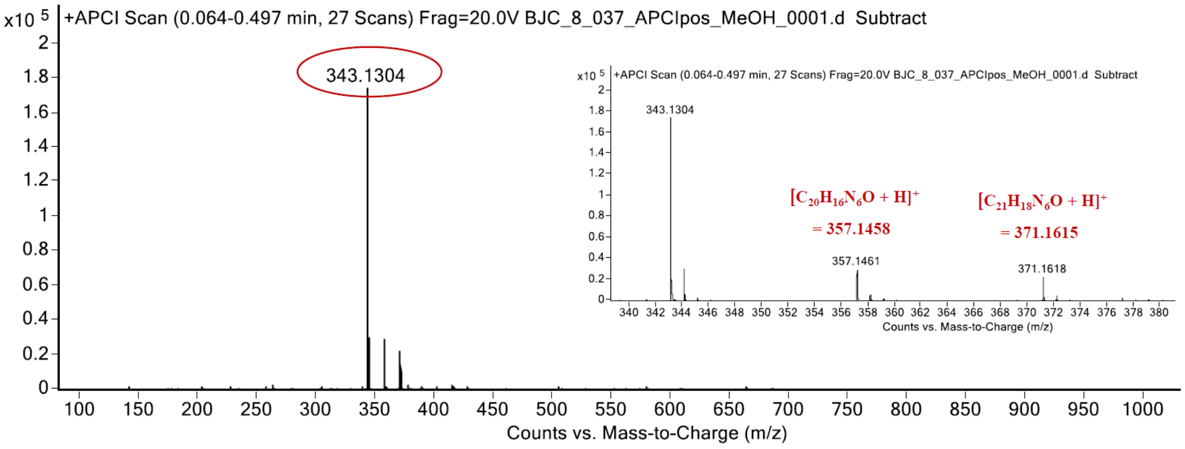

expected mass: $[M+H]^+$ = 343.1302 observed mass: $[M+H]^+$ = 343.1304 mass accuracy = 0.6 ppm

$^{1}$H (500 MHz) and $^{13}$C NMR (126 MHz) spectra of **89** in DMSO-$d_6$

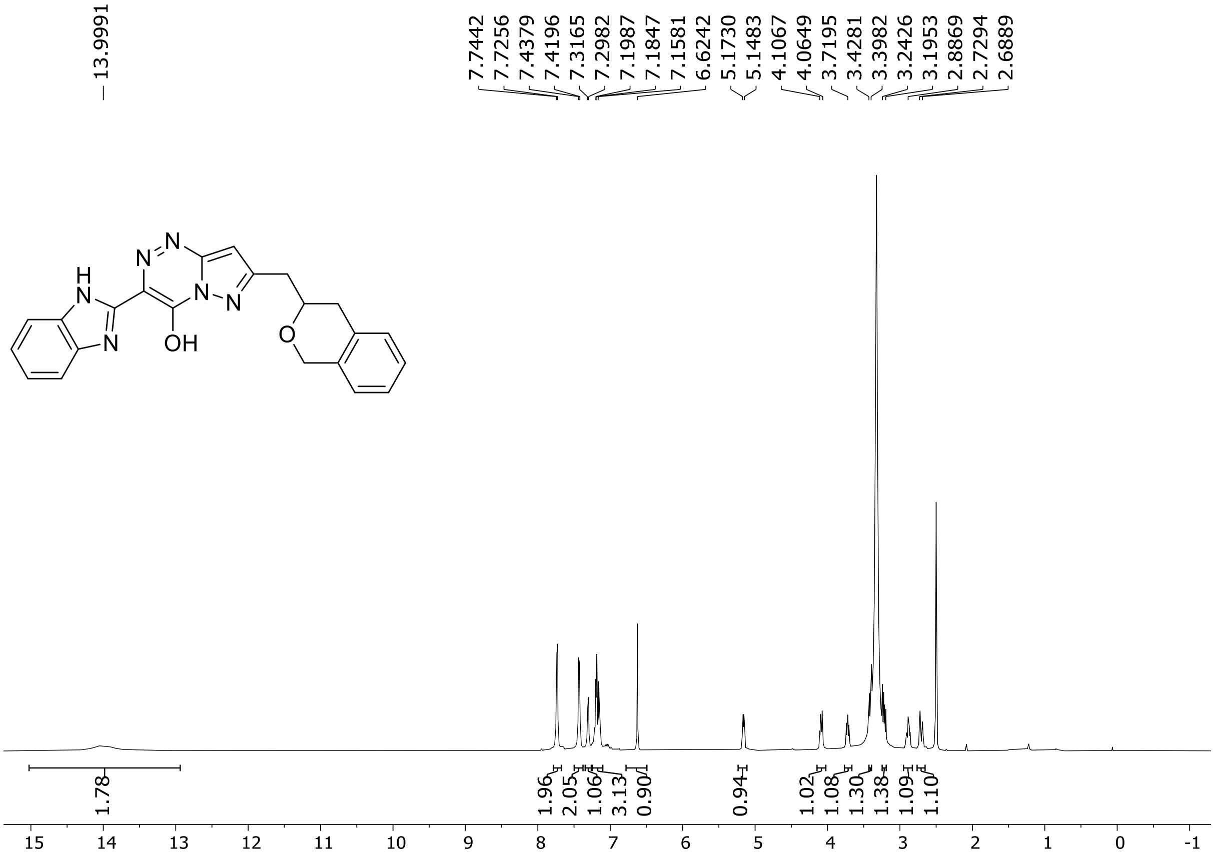

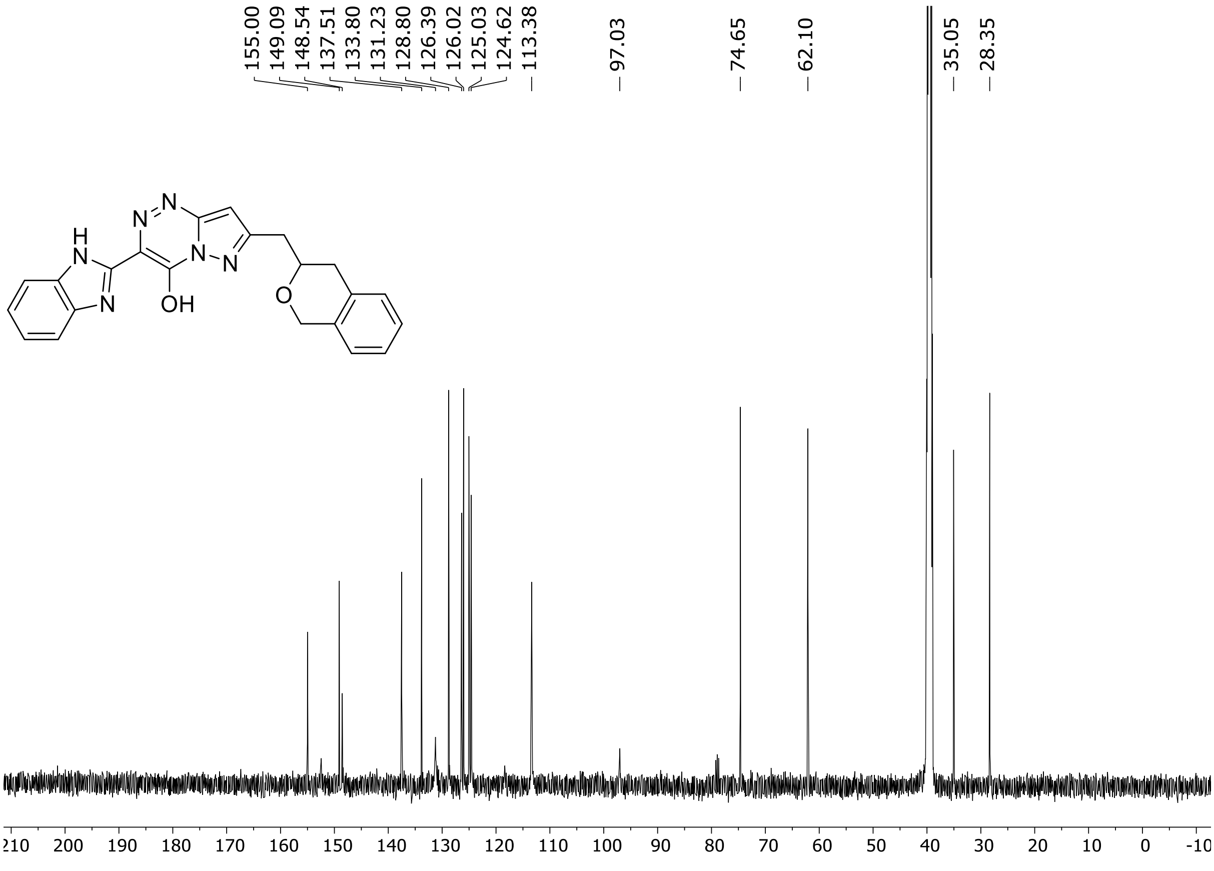

HRMS spectrum of **89**

**$C_{22}H_{18}N_6O_2$** **exact mass: 398.1491**

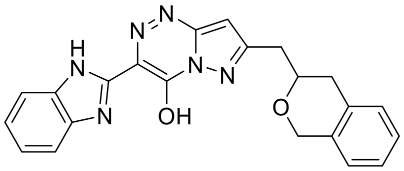

**APCI +** (MMI)
nitrogen flow 3 L/min, gas temperature 325°C, nebulizer 45 psig, skimmer 65 V, vaporizer 200°C, fragmentor 20 V, dissolved in methanol

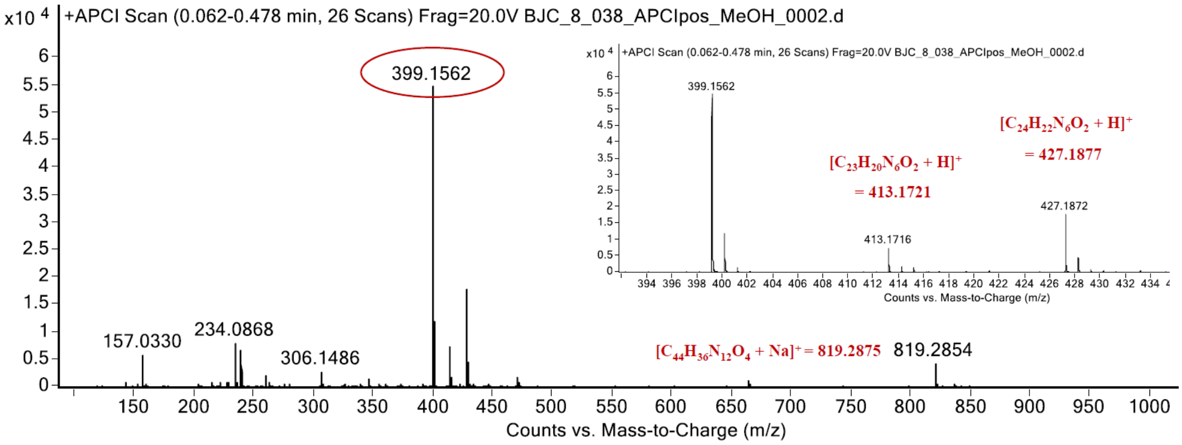

expected mass: $[M+H]^+$ = 399.1564 observed mass: $[M+H]^+$ = 399.1562 mass accuracy = - 0.5 ppm

$^{1}H$ (500 MHz) and $^{13}C$ NMR (126 MHz) spectra of **90** in DMSO-$d_6$

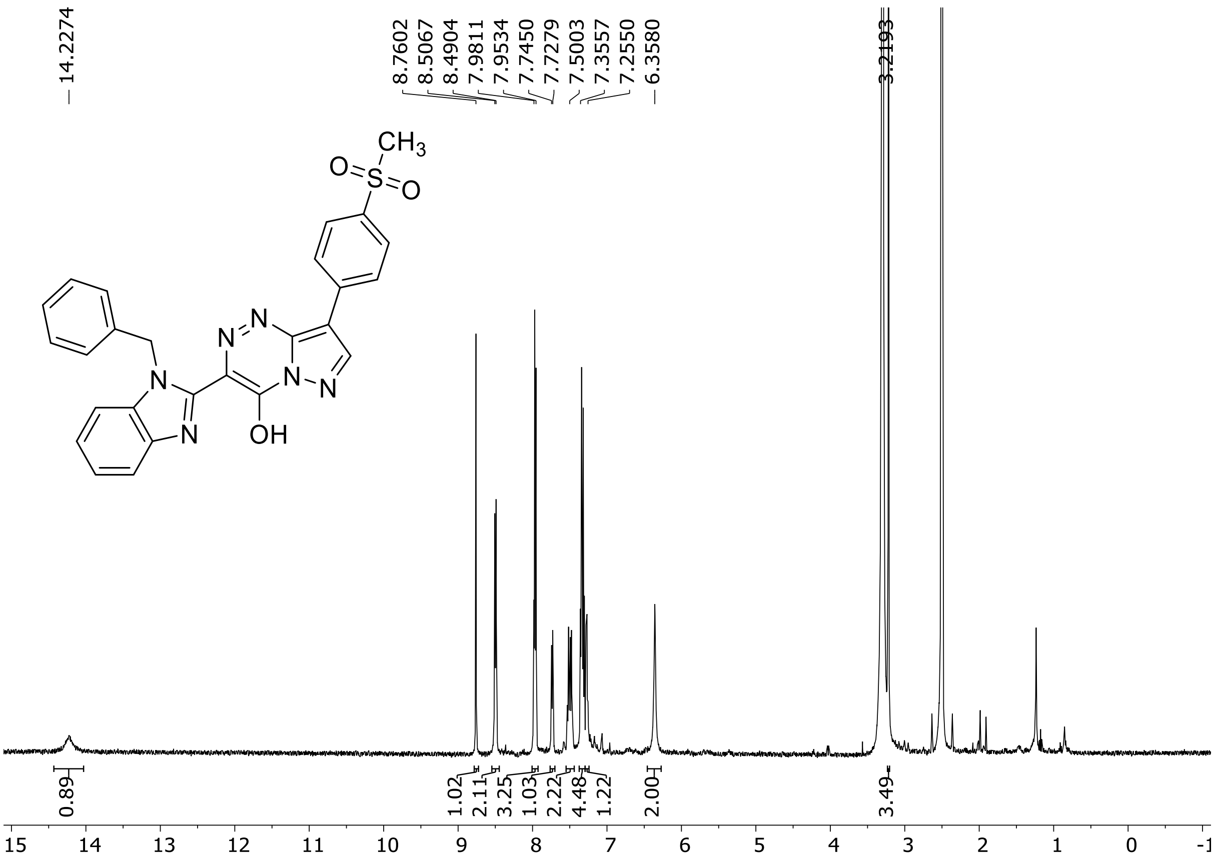

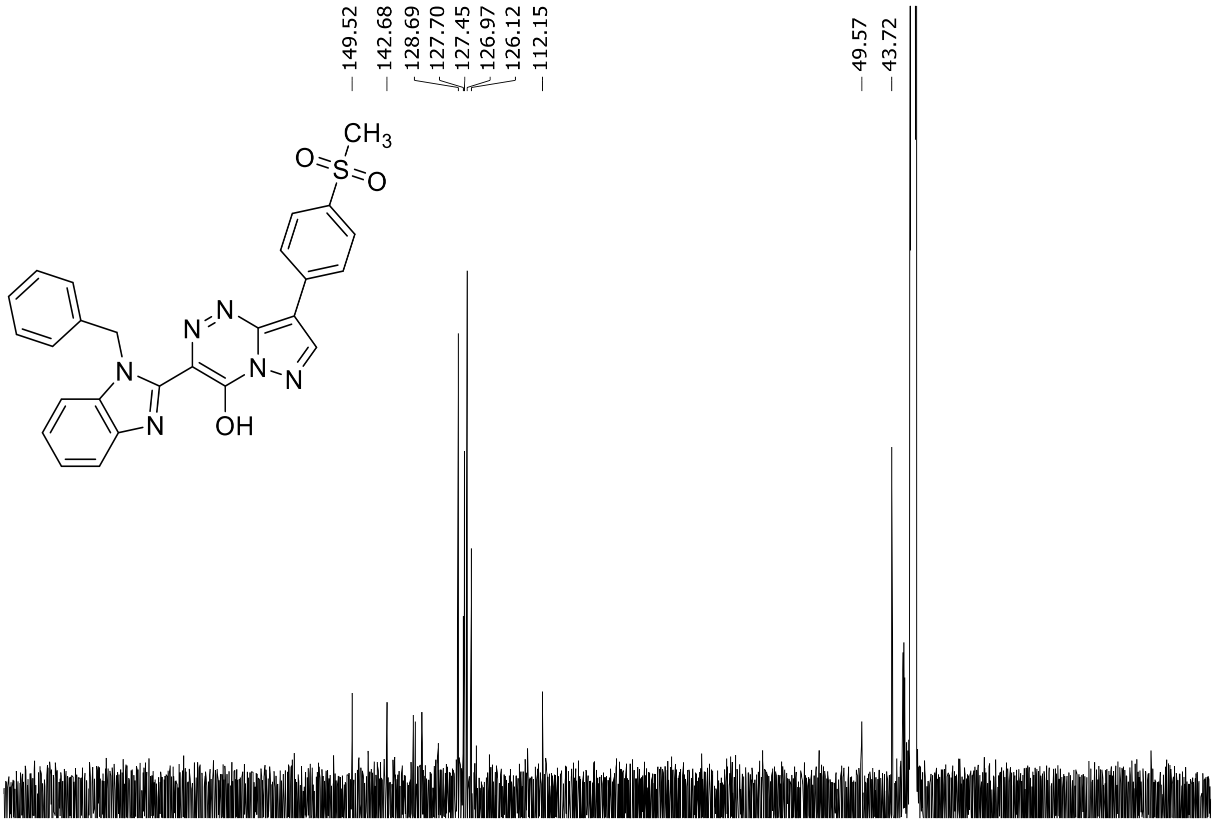

HRMS spectrum of **90**

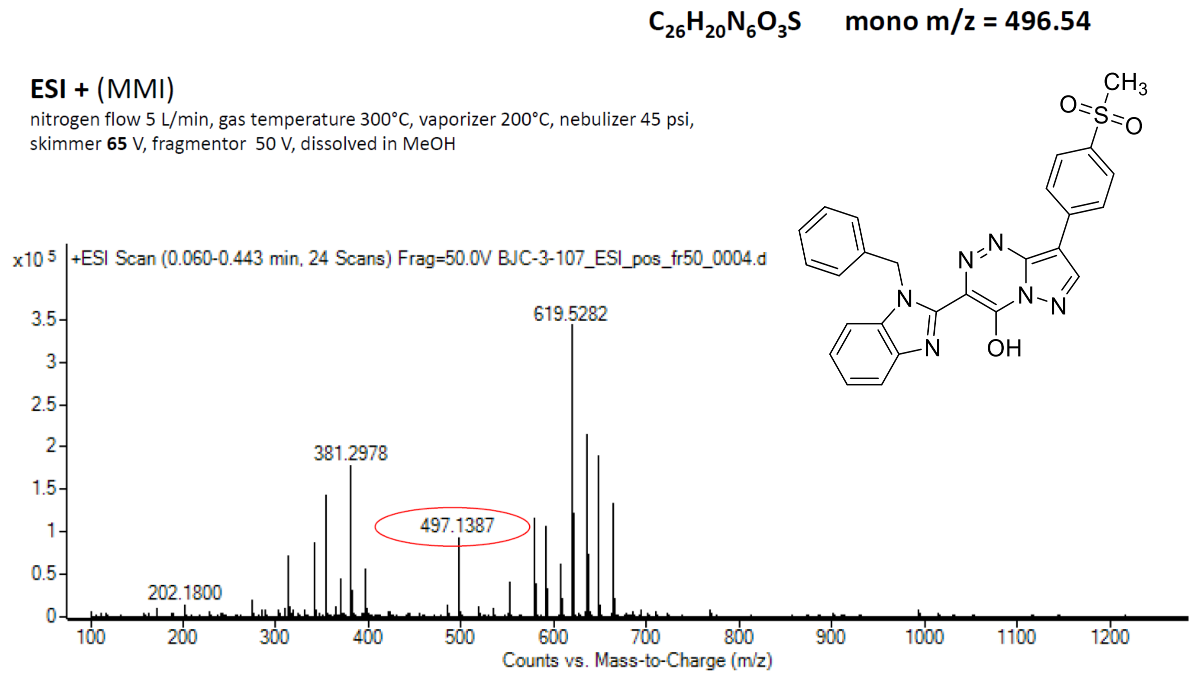

calculated mass: $[M+H]^+$ = 497.1390  observed: $[M+H]^+$ = 497.1387  max. mass error = 0.6 ppm

$^{1}H$ (500 MHz) and $^{13}C$ NMR (126 MHz) spectra of **91** in DMSO-$d_6$

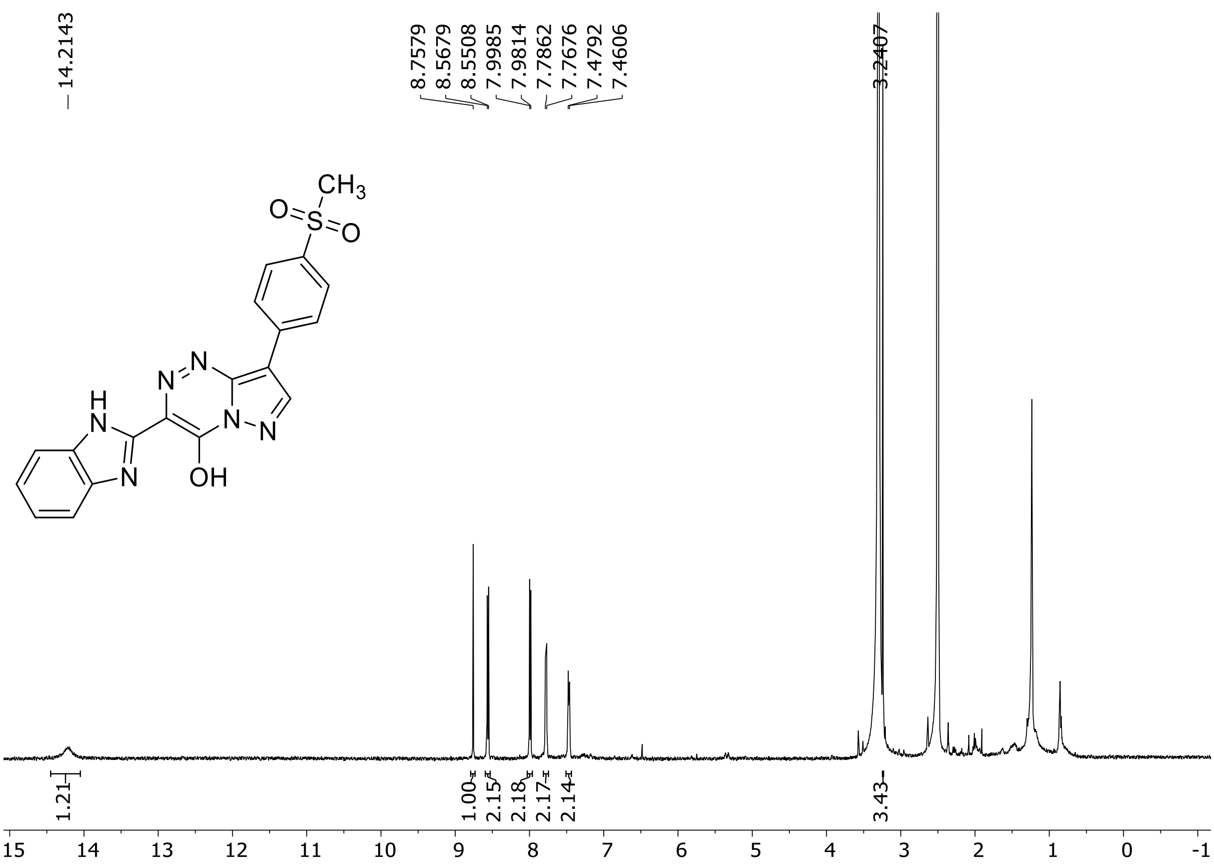

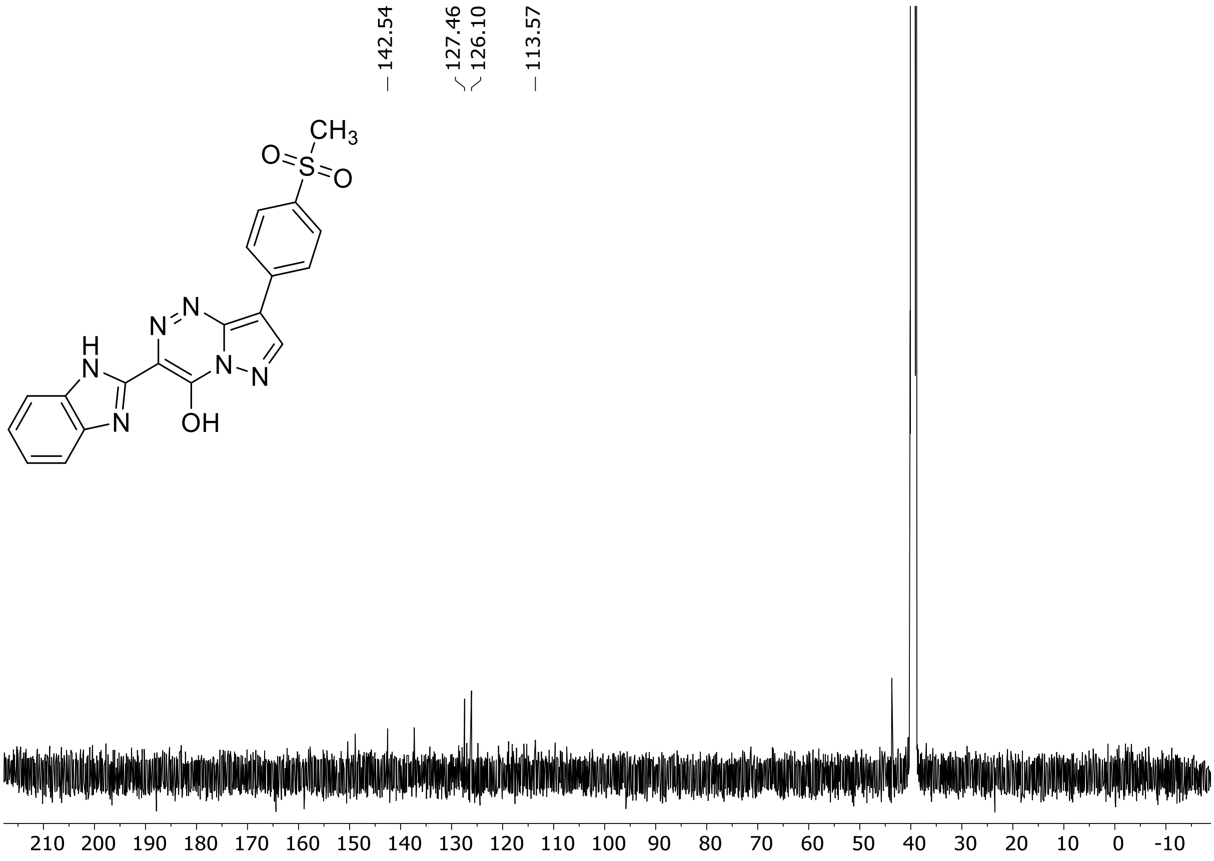

HRMS spectrum of **91**

**$C_{19}H_{14}N_6O_3S$ mono m/z = 406.42**

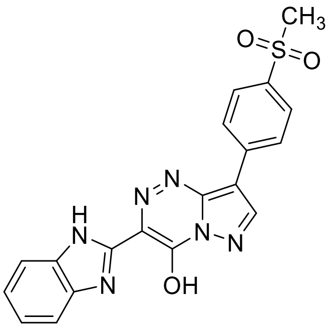

**APCI -** (MMI)

nitrogen flow 5 L/min, gas temperature 300°C, vaporizer 200°C, nebulizer 45 psi,
skimmer **65** V, fragmentor 80 V, dissolved in MeOH

**ZOOM – range of interest**

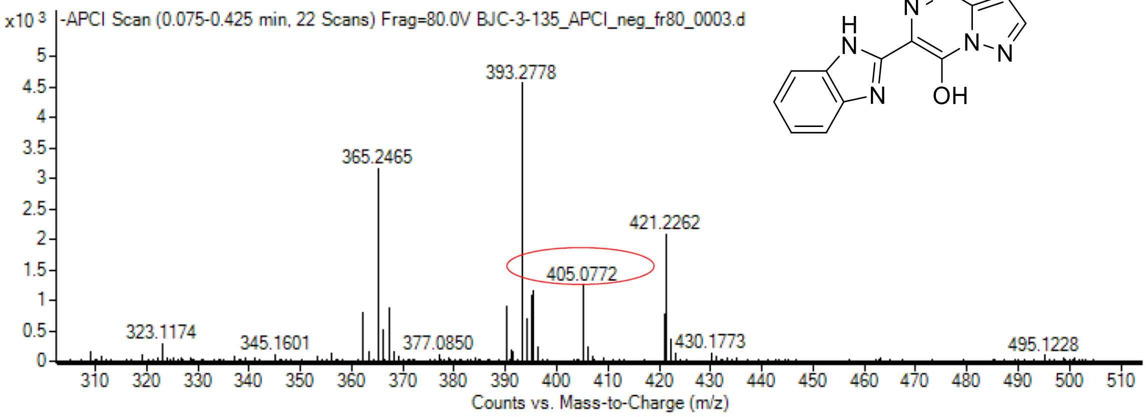

calculated mass: $[M+H]^+$ = 407.0775 observed: $[M+H]^+$ = 405.0772 max. mass error = 0.7 ppm

$^{1}H$ (500 MHz) and $^{13}C$ NMR (126 MHz) spectra of **92** in DMSO-$d_6$

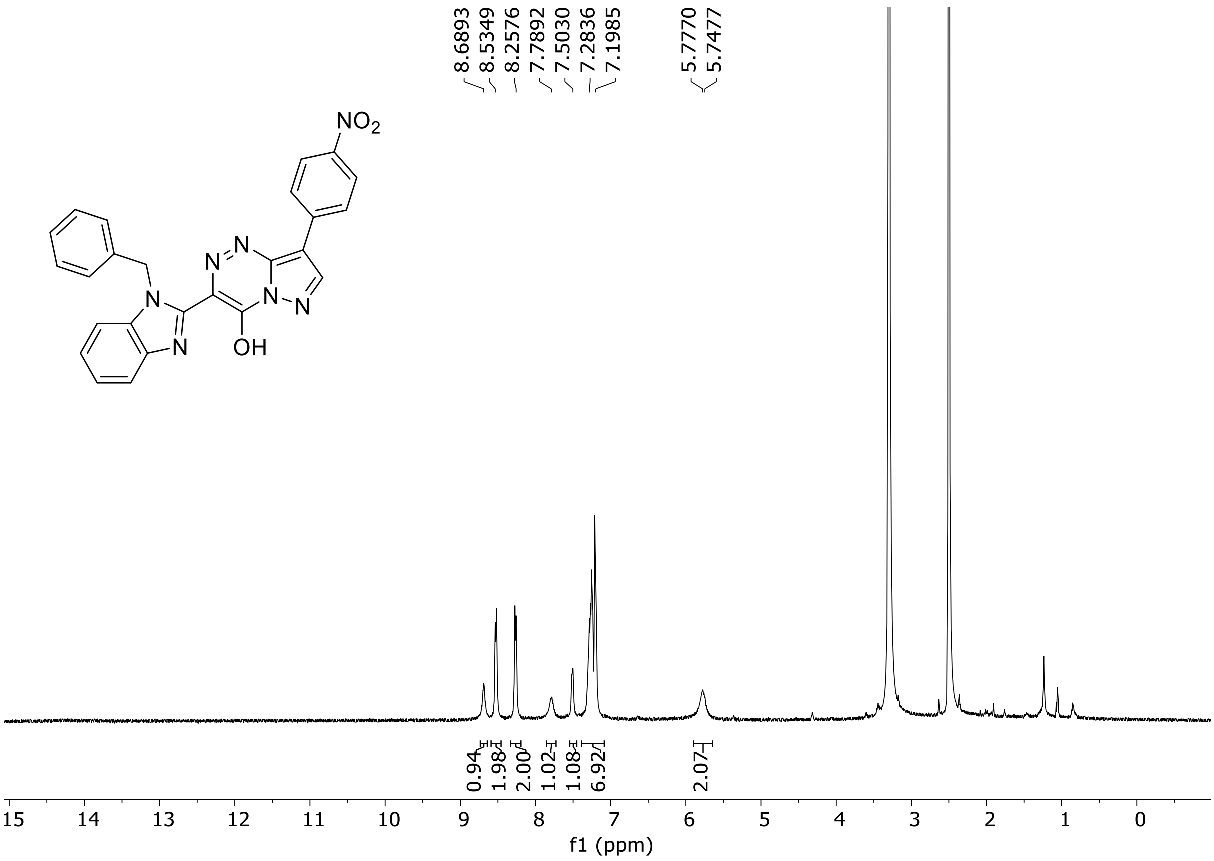

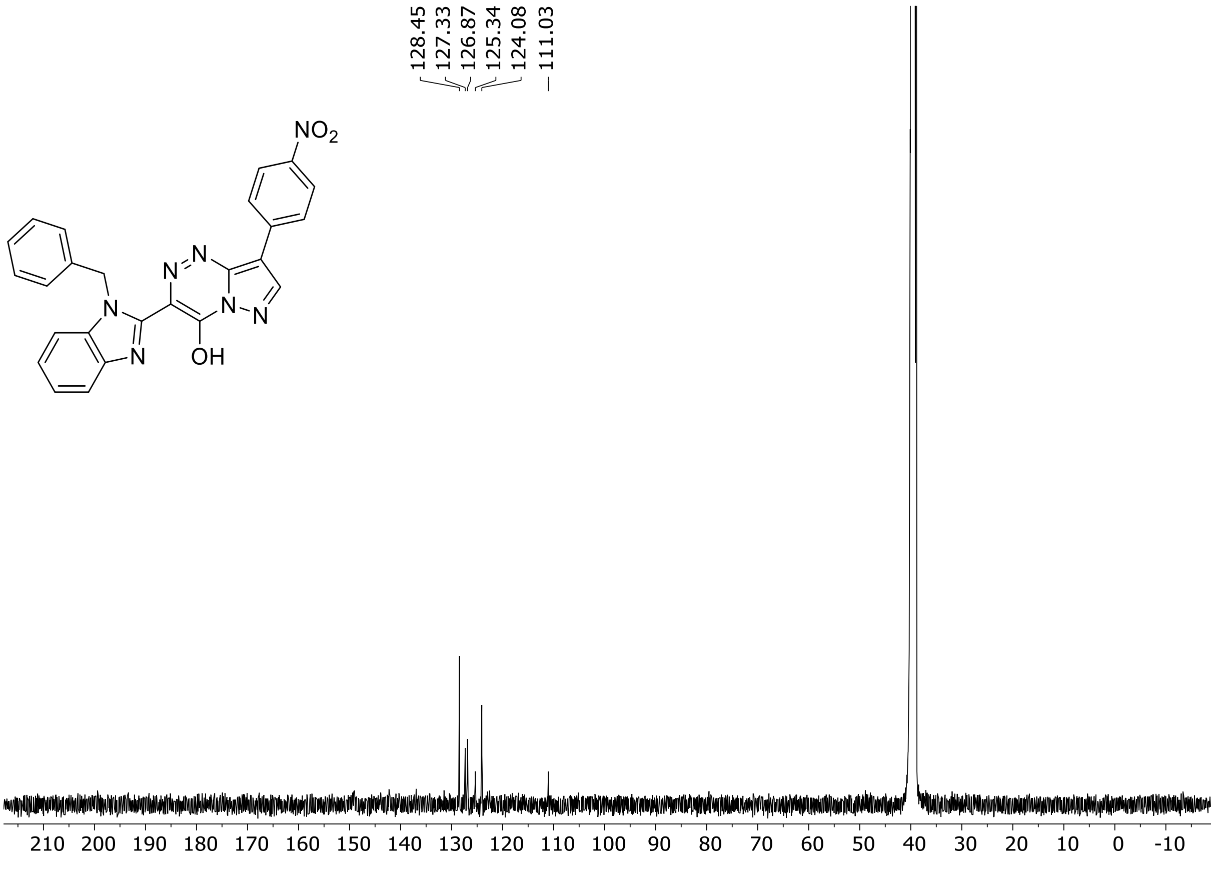

HRMS spectrum of **92**

**$C_{25}H_{17}N_7O_3$ mono m/z = 463.45**

**ESI +** (MMI)

nitrogen flow 5 L/min, gas temperature 300°C, vaporizer 200°C, nebulizer 45 psi, skimmer **65** V, fragmentor 50 V, dissolved in MeOH

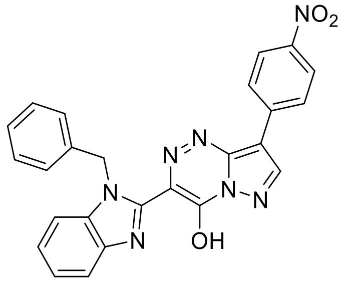

**ZOOM – range of interest**

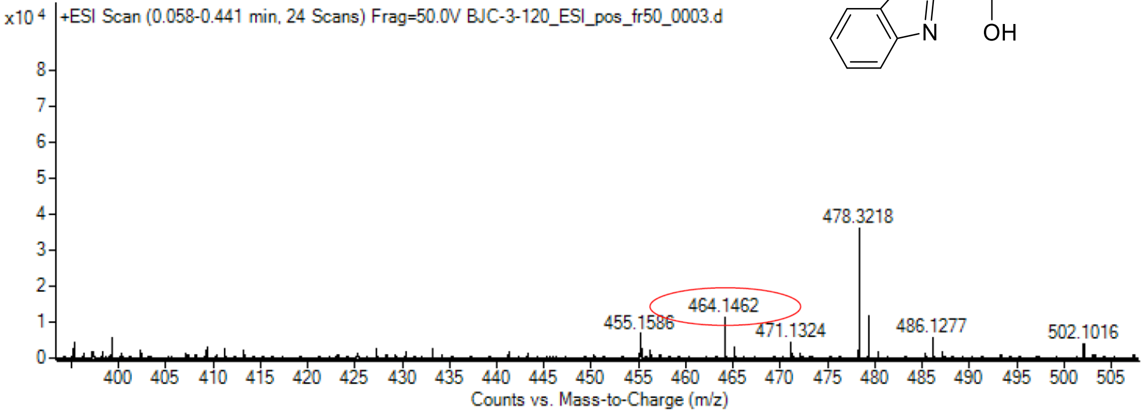

calculated mass: $[M+H]^+$ = 464.1466 observed: $[M+H]^+$ = 464.1462 max. mass error = 0.9 ppm

$^{1}H$ (500 MHz) and $^{19}F$ (282 MHz) NMR spectra of **93** in DMSO-$d_6$

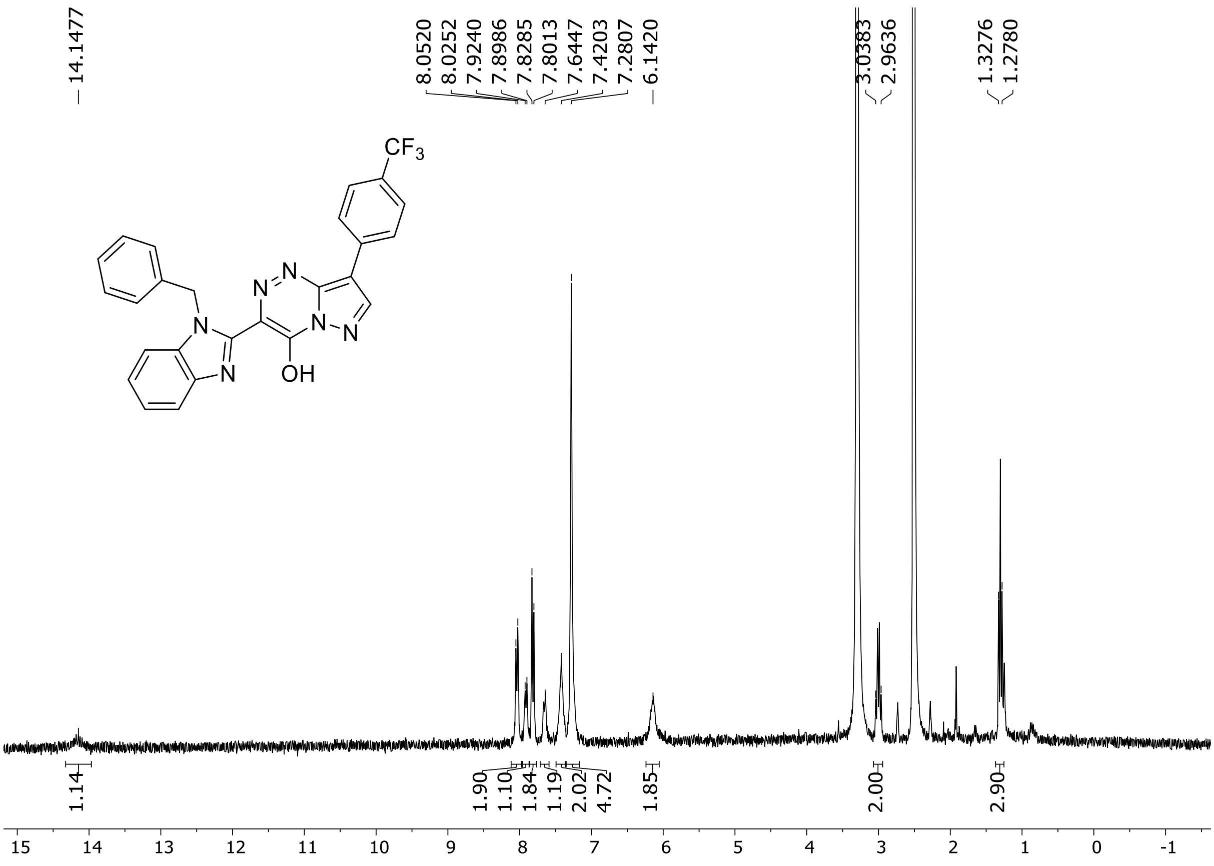

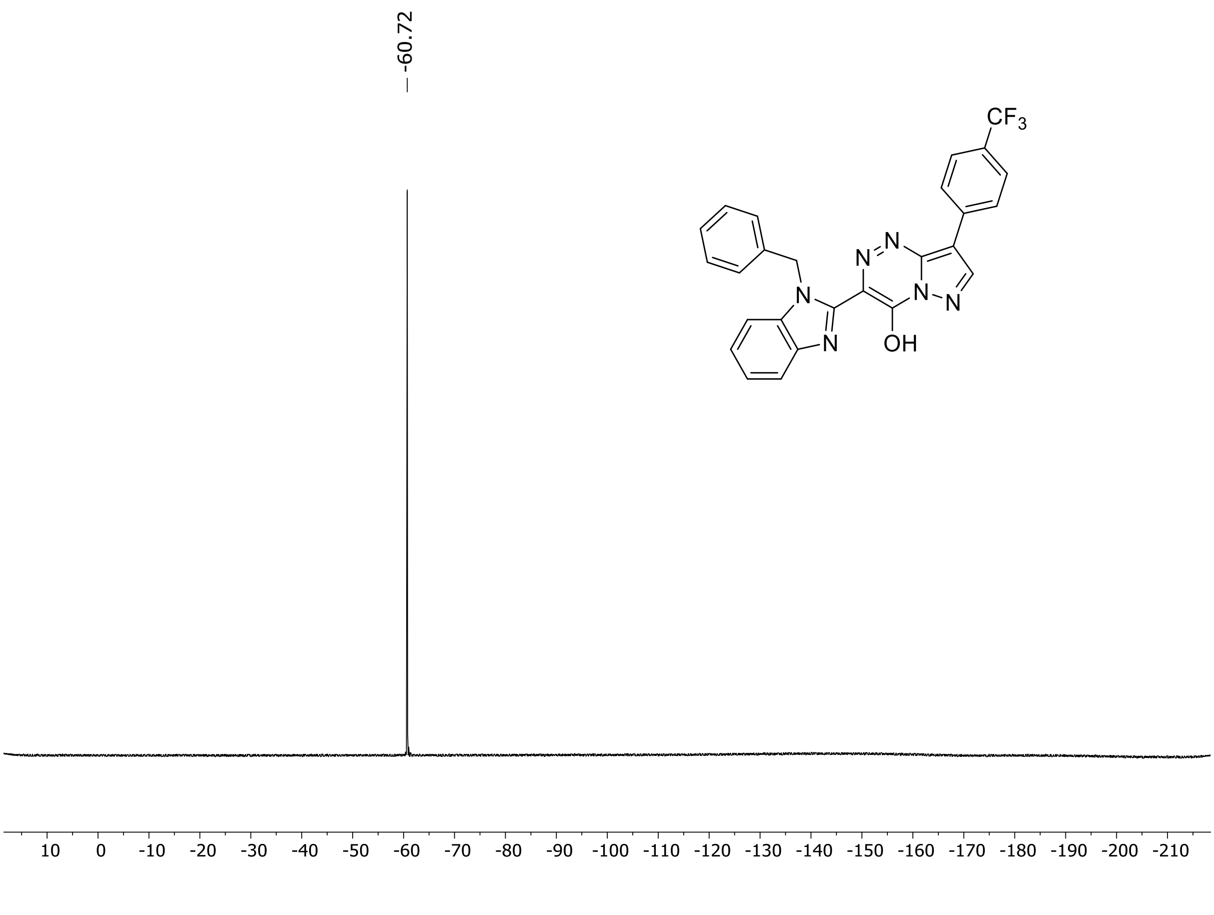

HRMS spectrum of **93**

**$C_{28}H_{21}F_3N_6O$** **mono *m*/*z* 514.1729**

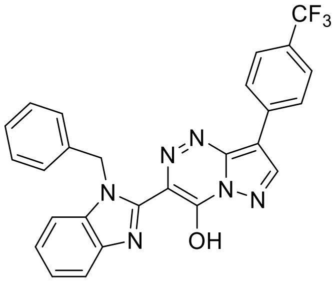

**APCI +** (MMI)
nitrogen flow 5 L/min, gas temperature 325°C, nebulizer 45 psi, skimmer 65 V, vaporizer 250°C, fragmentor 35 V, dissolved in methanol

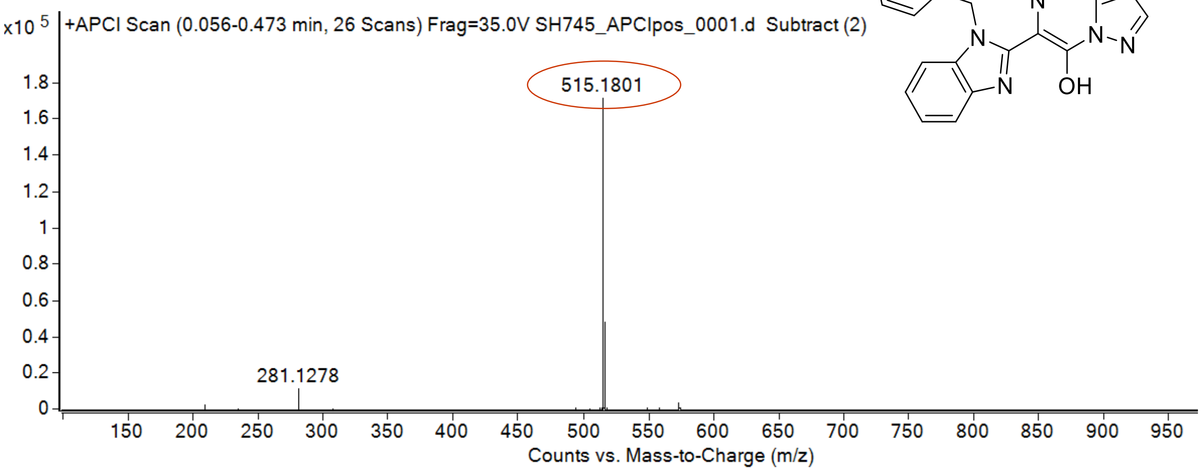

calculated mass: $[M+H]^+ = 515.1802$ observed: $[M+H]^+ = 515.1801$ mass accuracy = - 0.1 ppm

[1]H (500 MHz) and [13]C NMR (126 MHz) spectra of **94** in DMSO-$d_6$

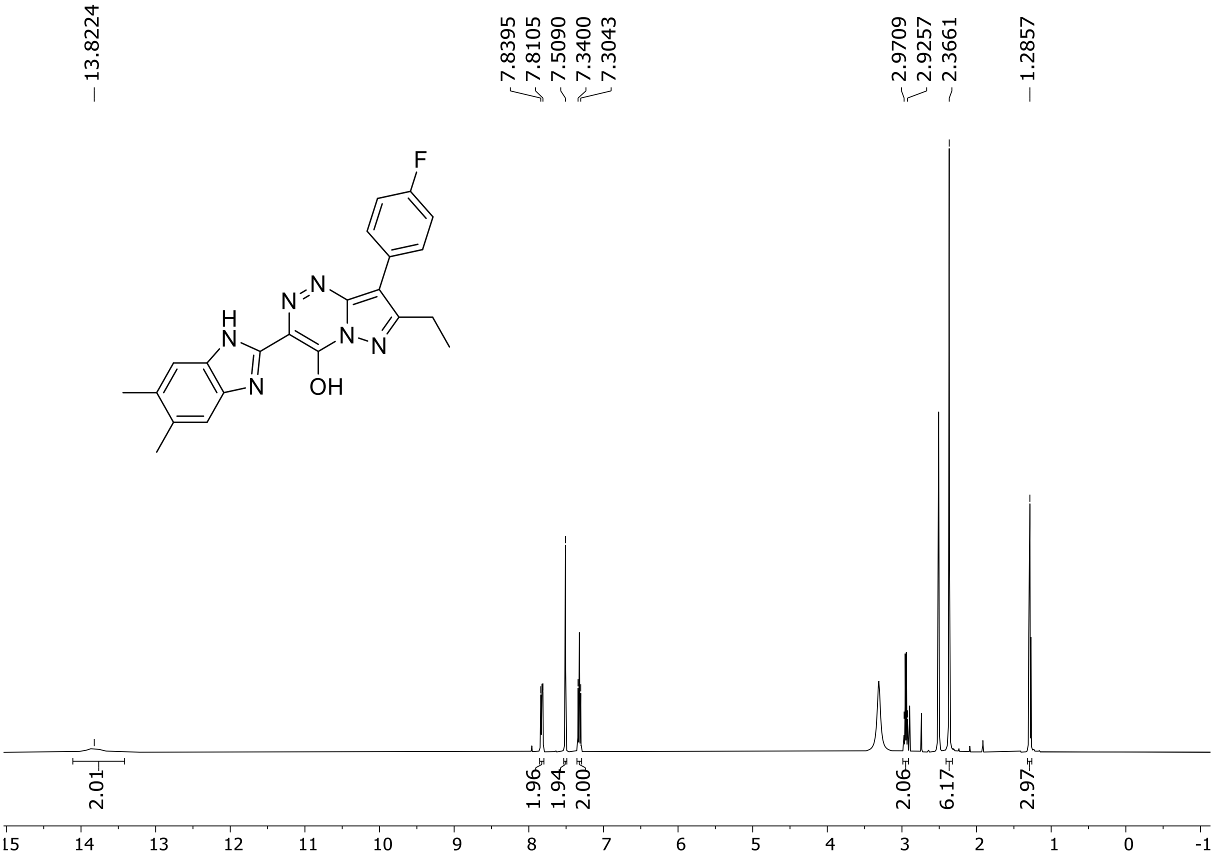

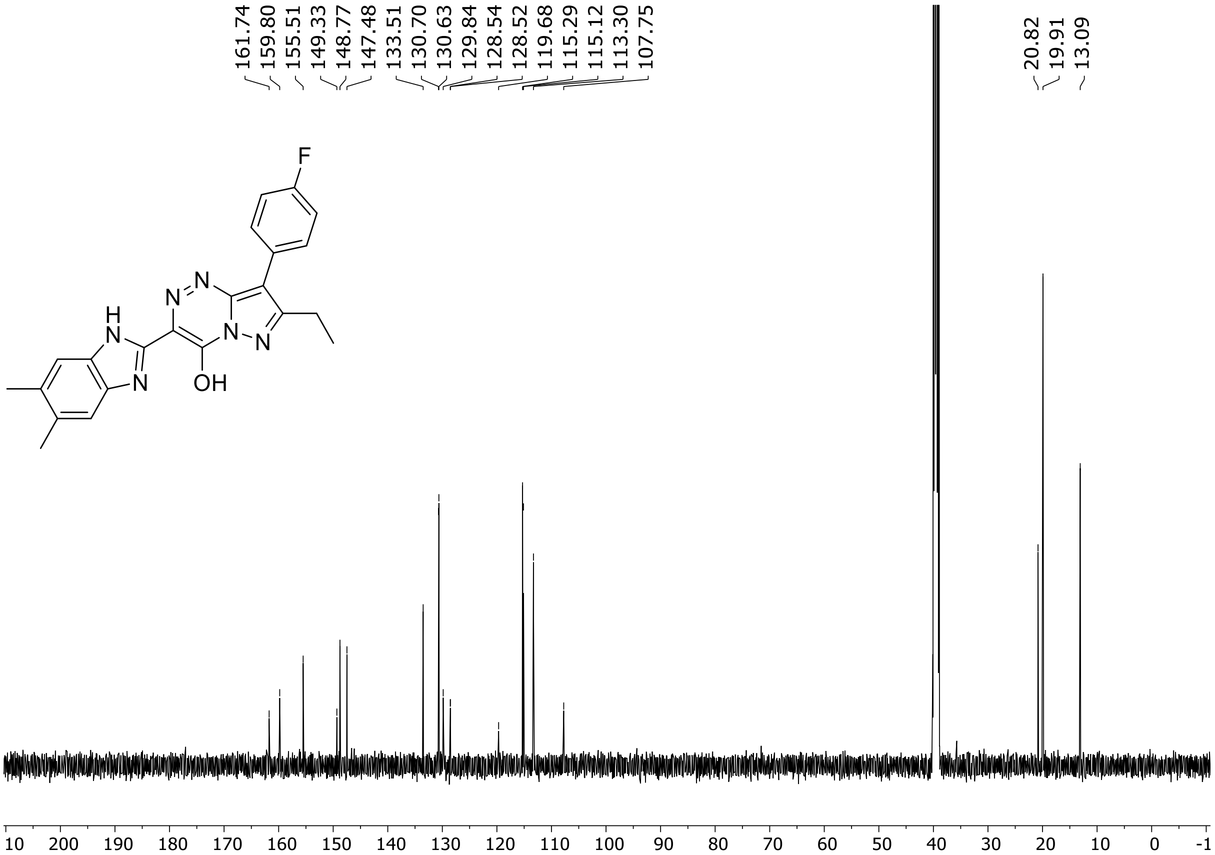

$^{19}$F (471 MHz) NMR spectrum of **94** DMSO-$d_6$

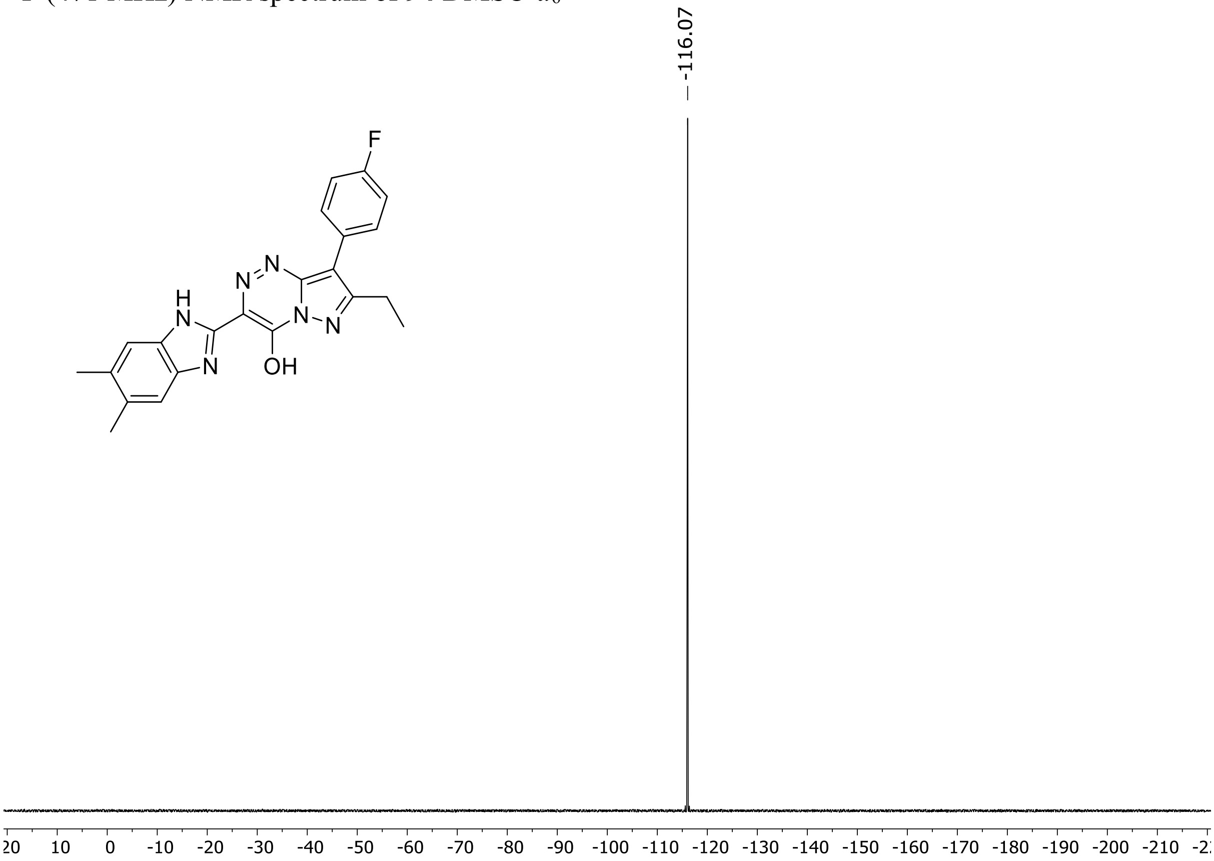

HRMS spectrum of **94**

**$C_{22}H_{19}FN_6O$** **exact mass: 402.1604**

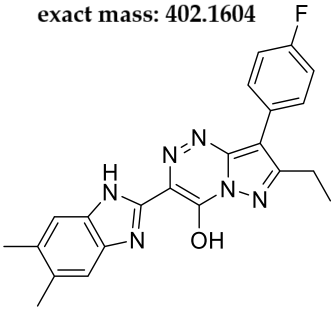

**APCI +** (MMI)
nitrogen flow 5 L/min, gas temperature 325°C, nebulizer 45 psig, skimmer 65 V, vaporizer 200°C, fragmentor 10 V, dissolved in methanol

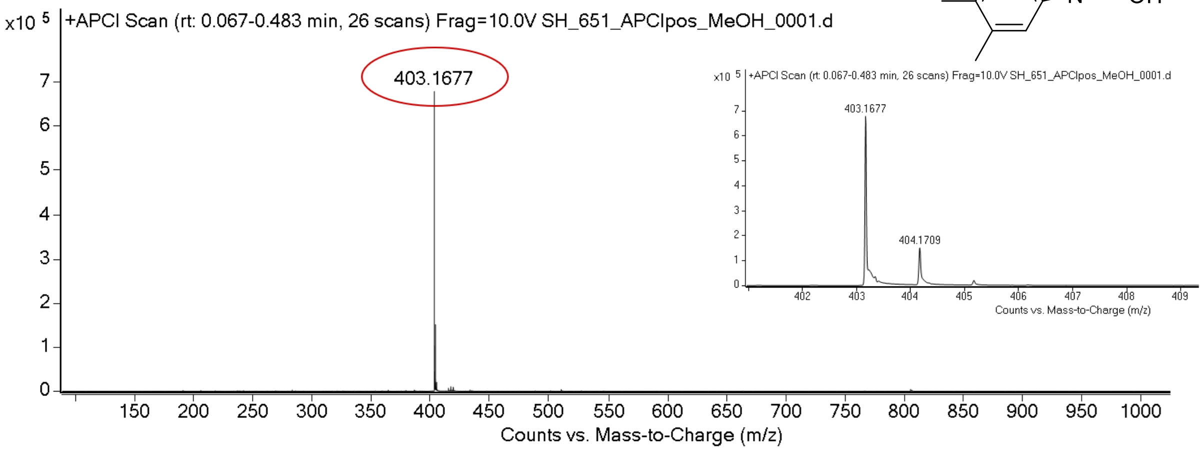

expected mass: [M+H]$^+$ = 403.1677 observed mass : [M+H]$^+$ = 403.1677 mass accuracy < 0.1 ppm

$^{1}$H (500 MHz) and $^{13}$C NMR (126 MHz) spectra of **95** in DMSO-$d_6$

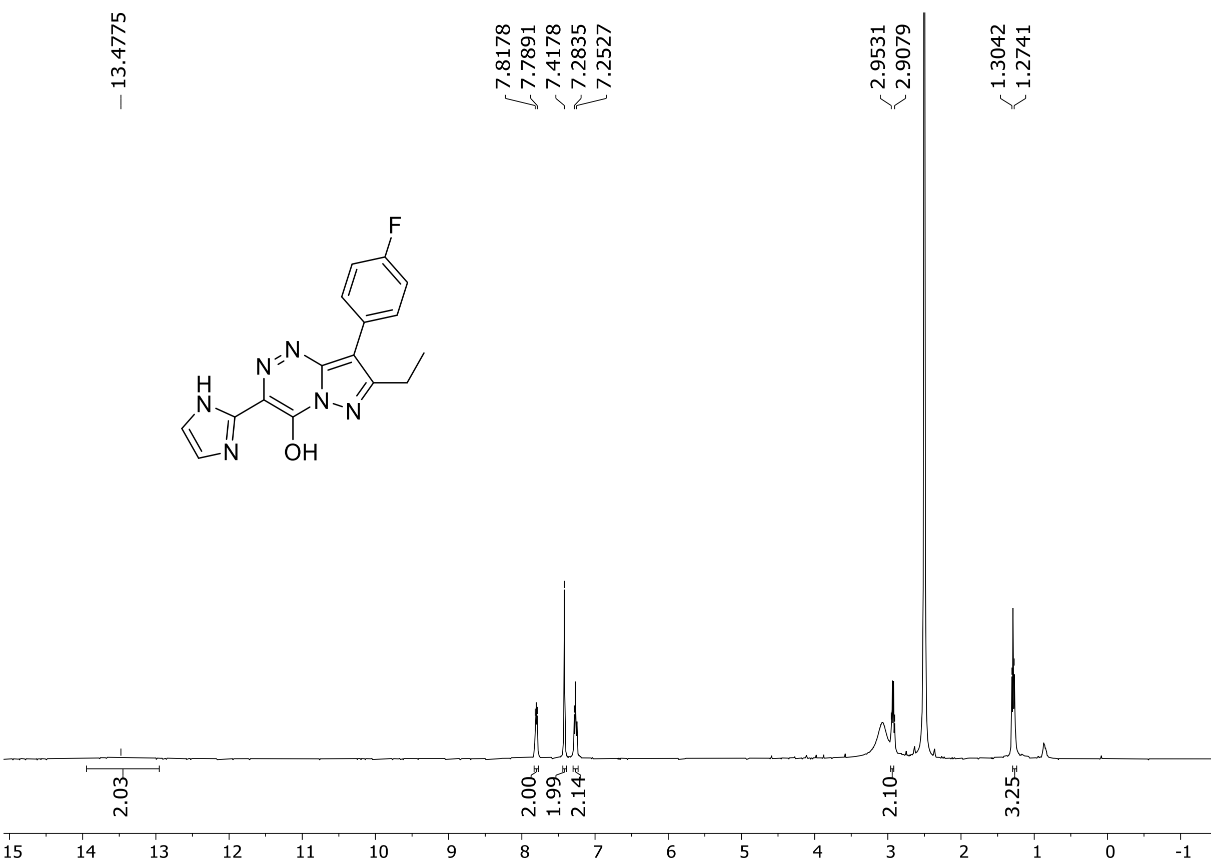

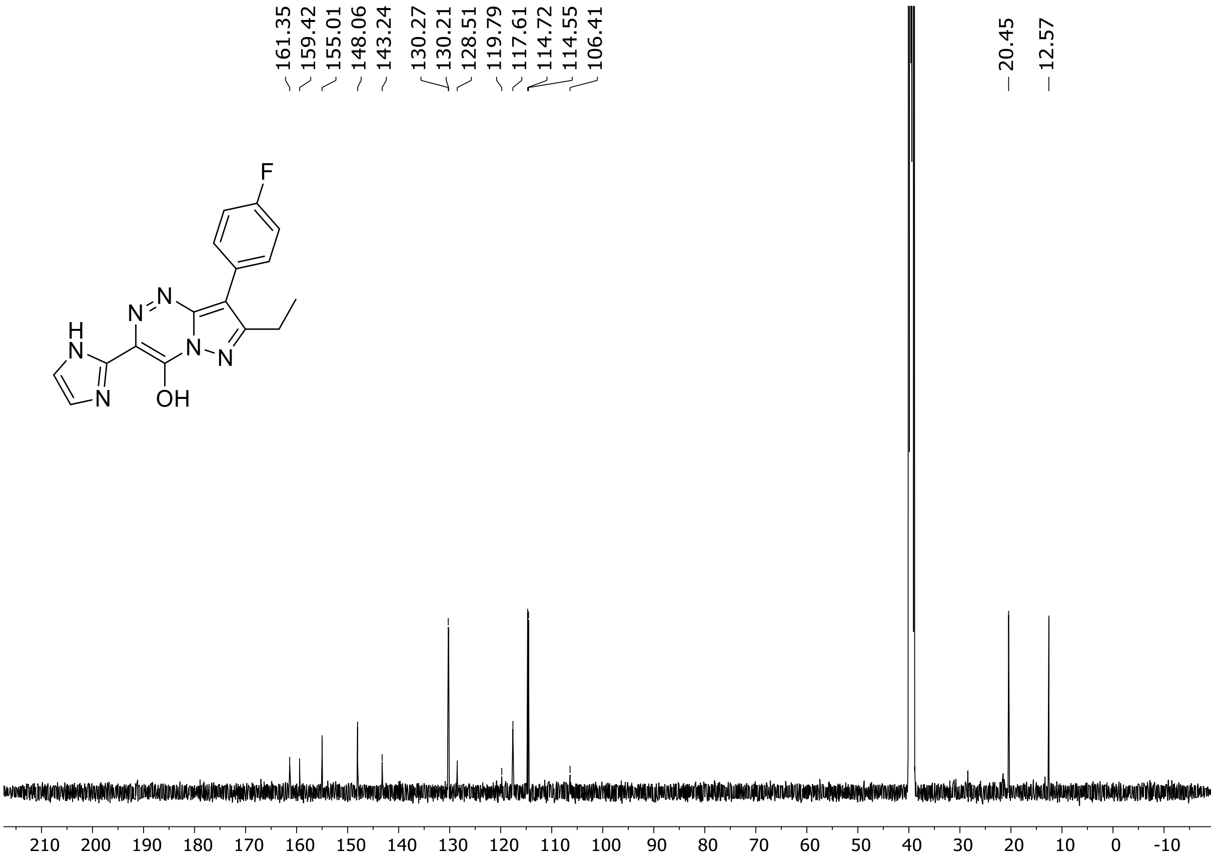

HRMS spectrum of **95**

**$C_{16}H_{13}FN_6O$** **exact mass: 324.1135**

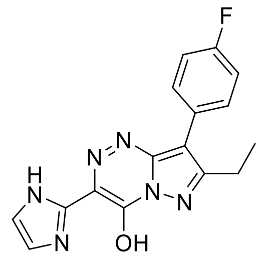

**APCI +** (MMI)
nitrogen flow 5 L/min, gas temperature 325°C, nebulizer 45 psig, skimmer 65 V, vaporizer 200°C, fragmentor 20 V, dissolved in methanol

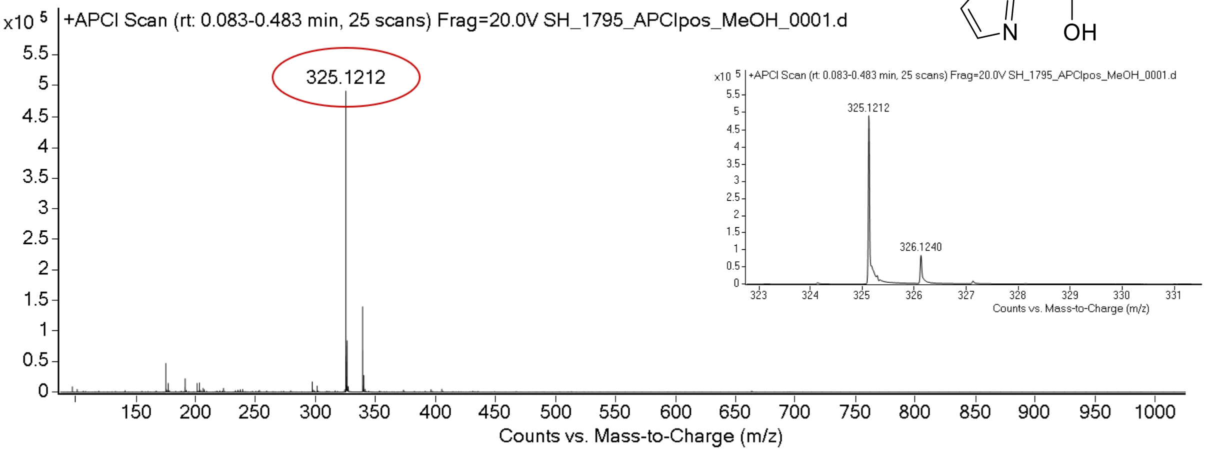

expected mass: $[M+H]^+$ = 325.1208 observed mass : $[M+H]^+$ = 325.1212 mass accuracy = 1.2 ppm

$^{1}H$ (500 MHz) and $^{13}C$ NMR (126 MHz) spectra of **96** in DMSO-$d_6$

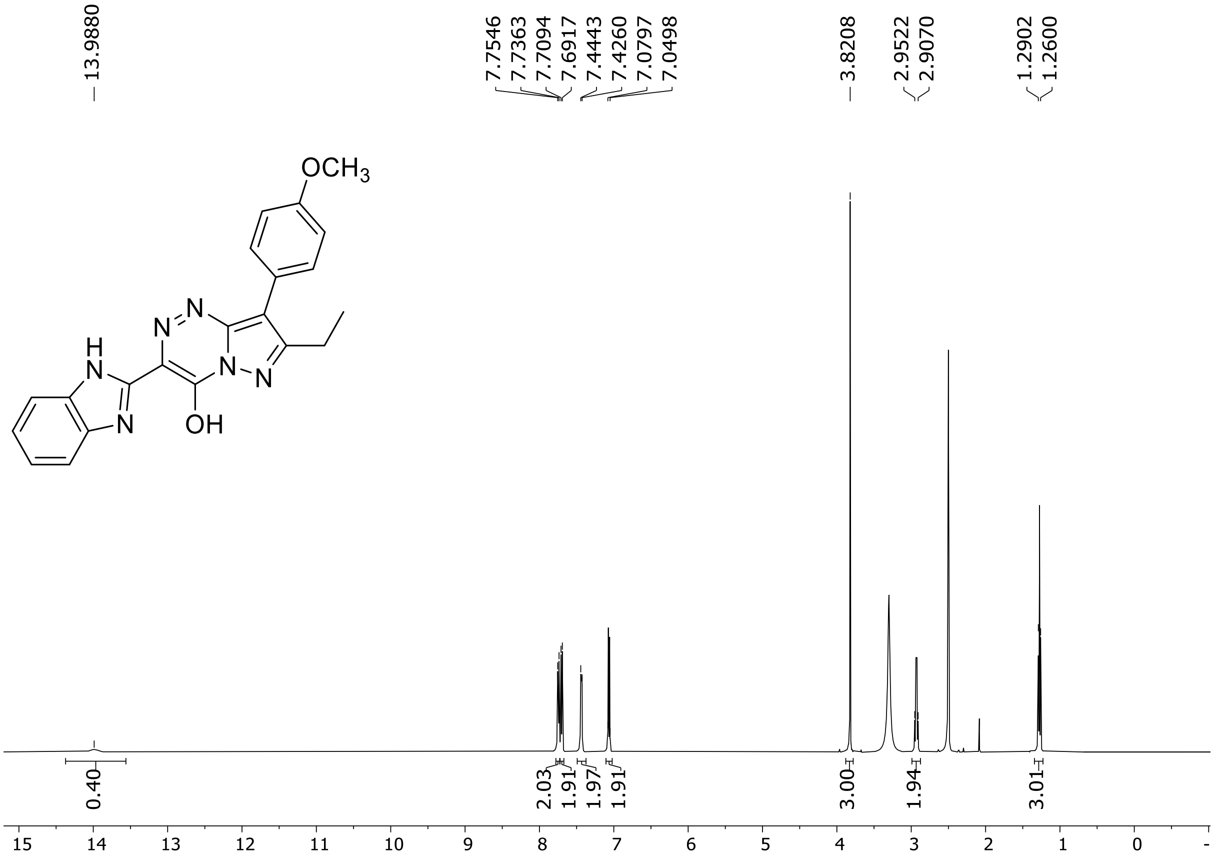

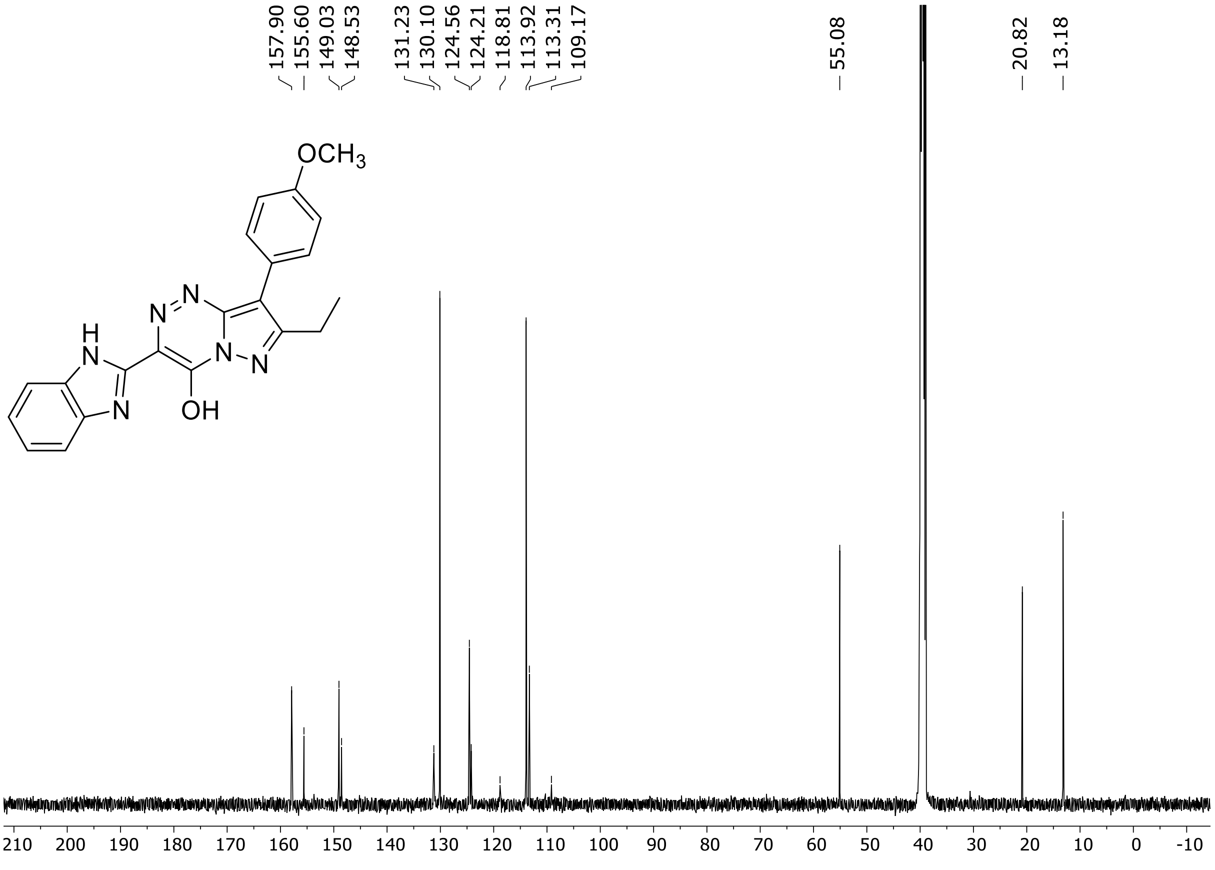

HRMS spectrum of **96**

**$C_{21}H_{18}N_6O_2$** **exact mass: 386.1491**

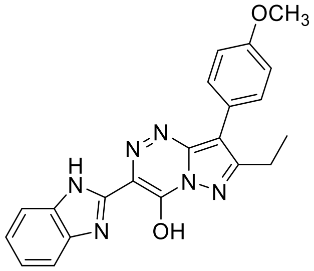

**APCI +** (MMI)
nitrogen flow 5 L/min, gas temperature 325°C, nebulizer 45 psig, skimmer 65 V, vaporizer 200°C, fragmentor 20 V, dissolved in methanol

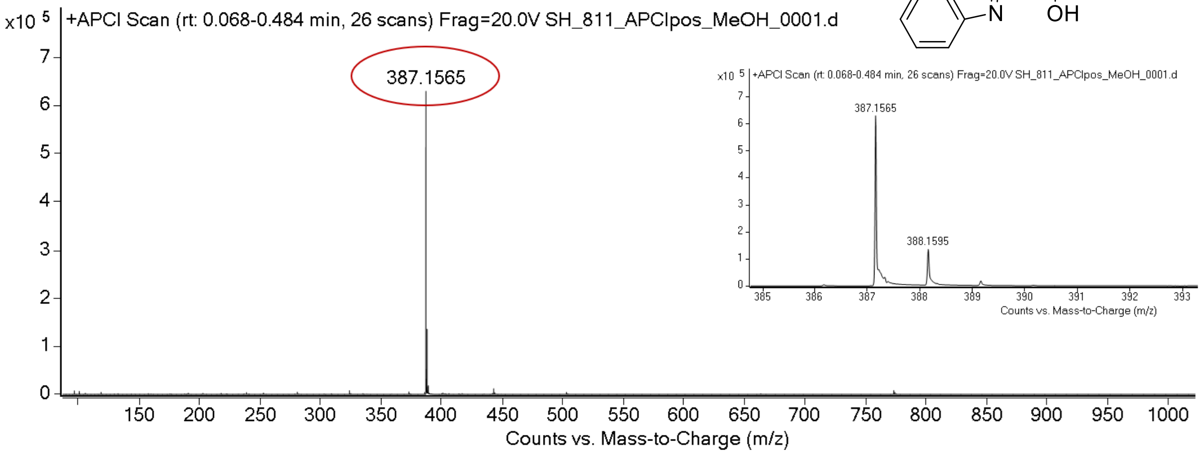

expected mass: $[M+H]^+$ = 387.1564 observed mass : $[M+H]^+$ = 387.1565 mass accuracy = 0.3 ppm

$^{1}H$ (500 MHz) and $^{13}C$ NMR (126 MHz) spectra of **97** in DMSO-$d_6$

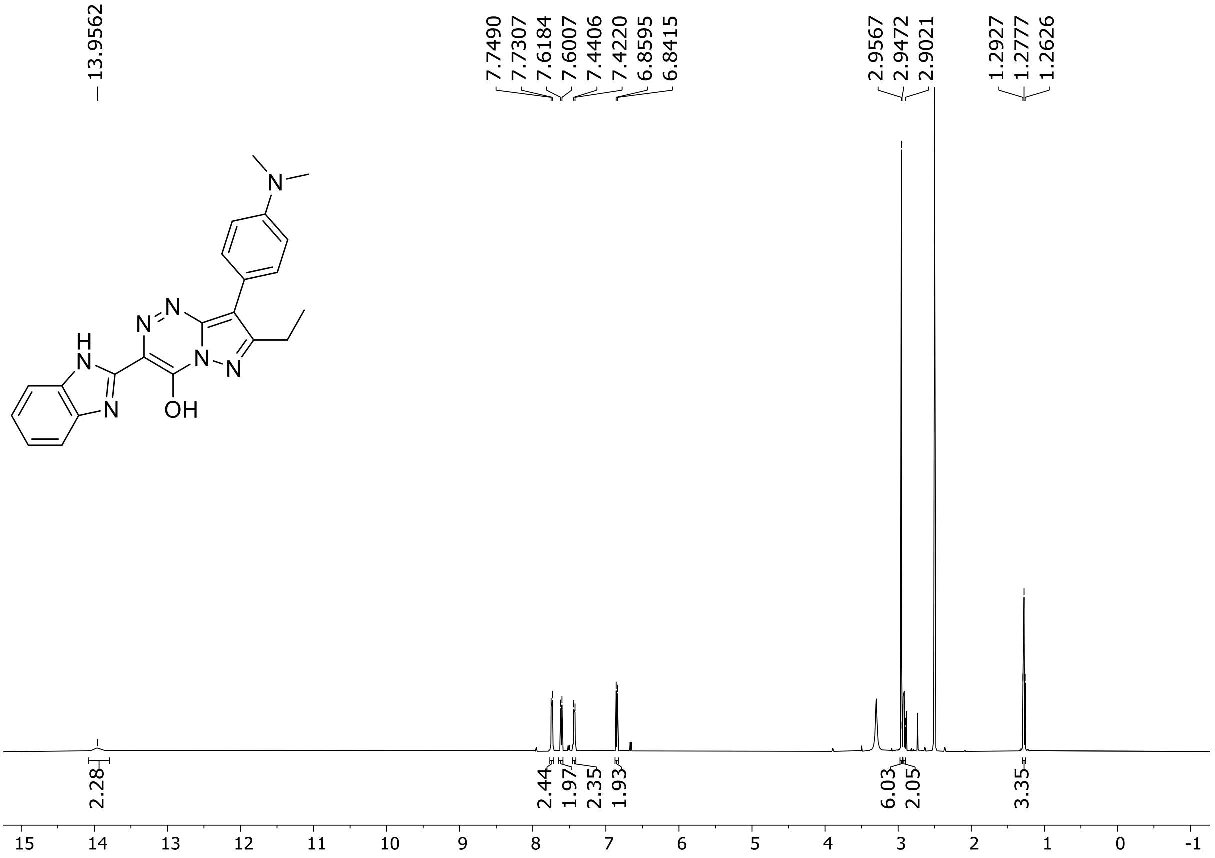

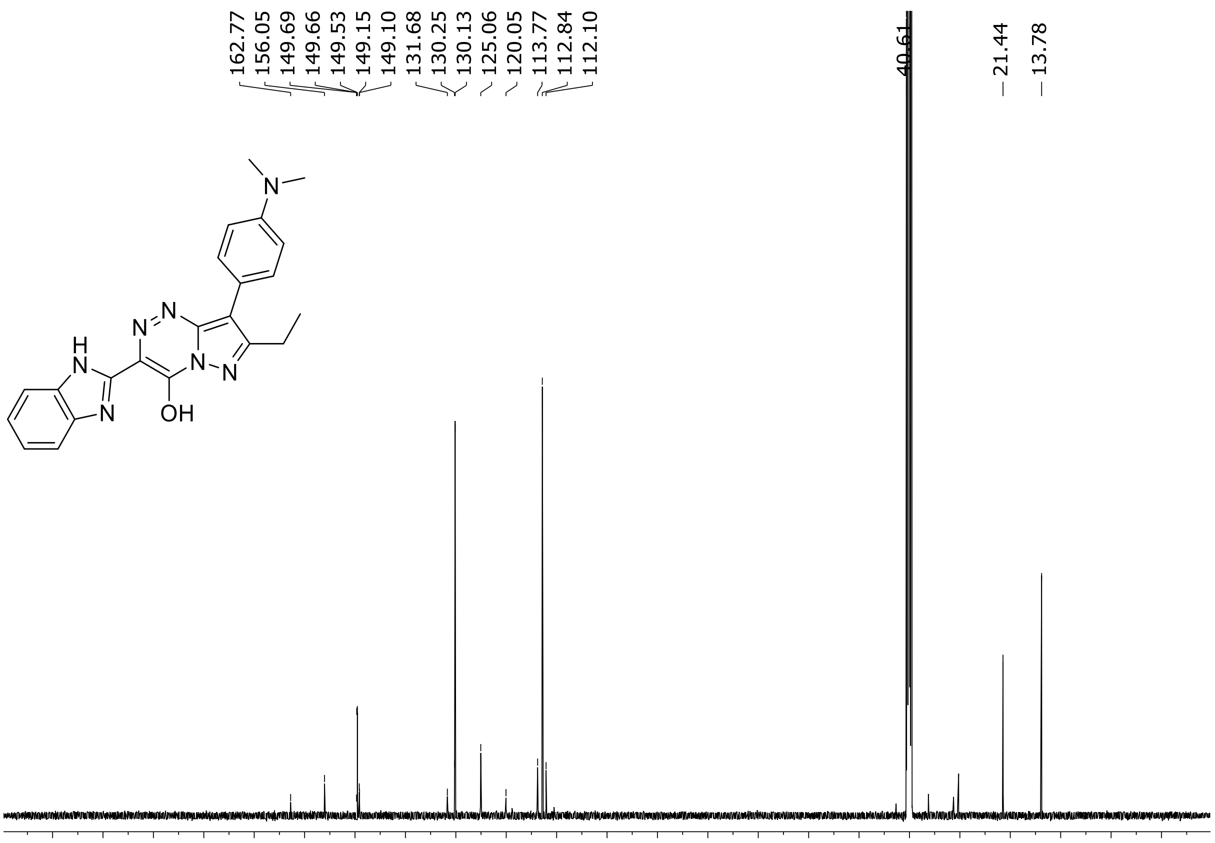

HRMS spectrum of **97**

**$C_{22}H_{21}N_7O$** **exact mass: 399.1808**

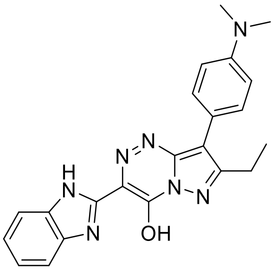

**APCI +** (MMI)
nitrogen flow 5 L/min, gas temperature 325°C, nebulizer 45 psig, skimmer 65 V, vaporizer 200°C, fragmentor 10 V, dissolved in methanol

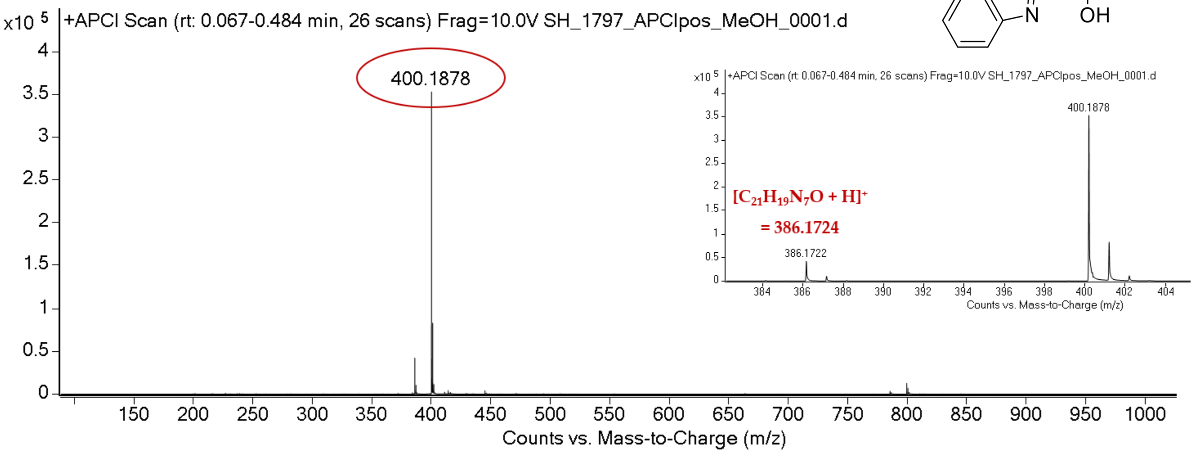

expected mass: $[M+H]^+$ = 400.1880 observed mass : $[M+H]^+$ = 400.1878 mass accuracy = - 0.5 ppm

$^{1}H$ (500 MHz) and $^{13}C$ NMR (126 MHz) spectra of **98** in DMSO-$d_6$

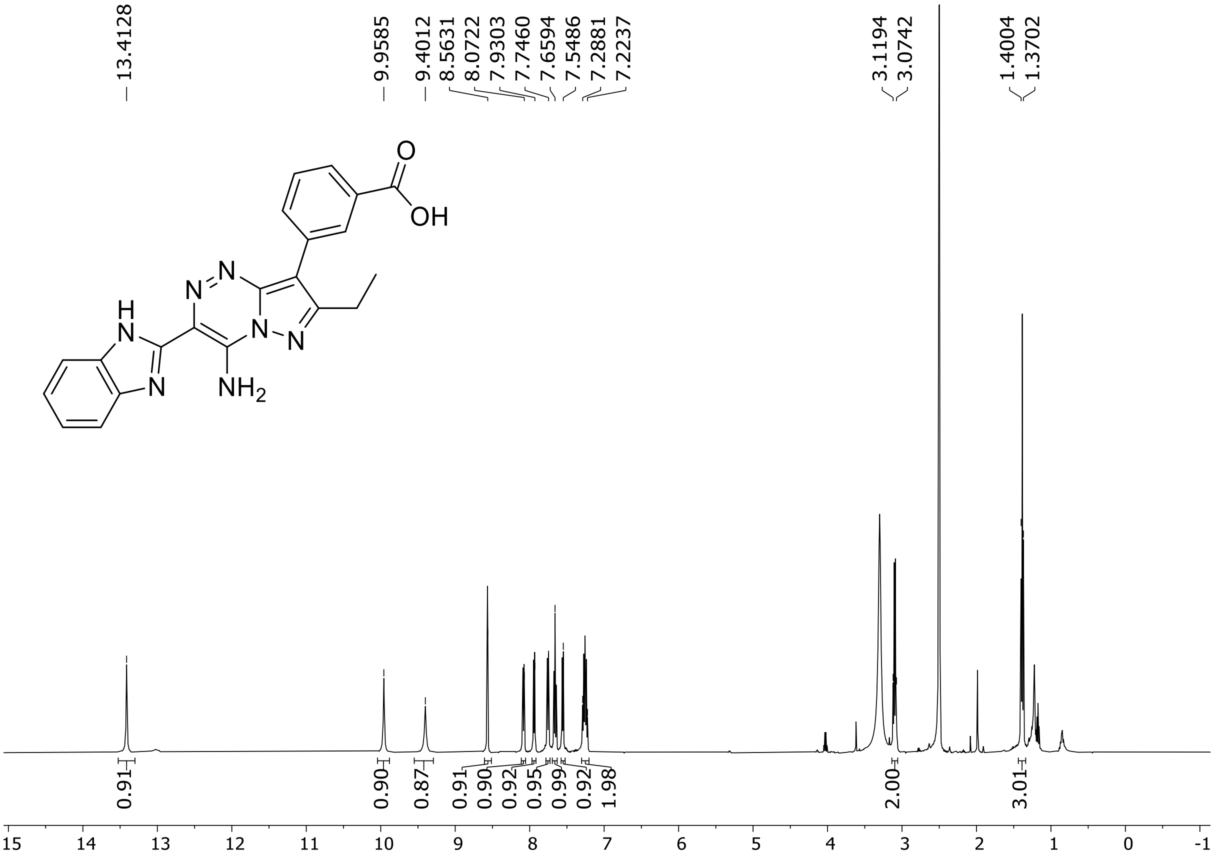

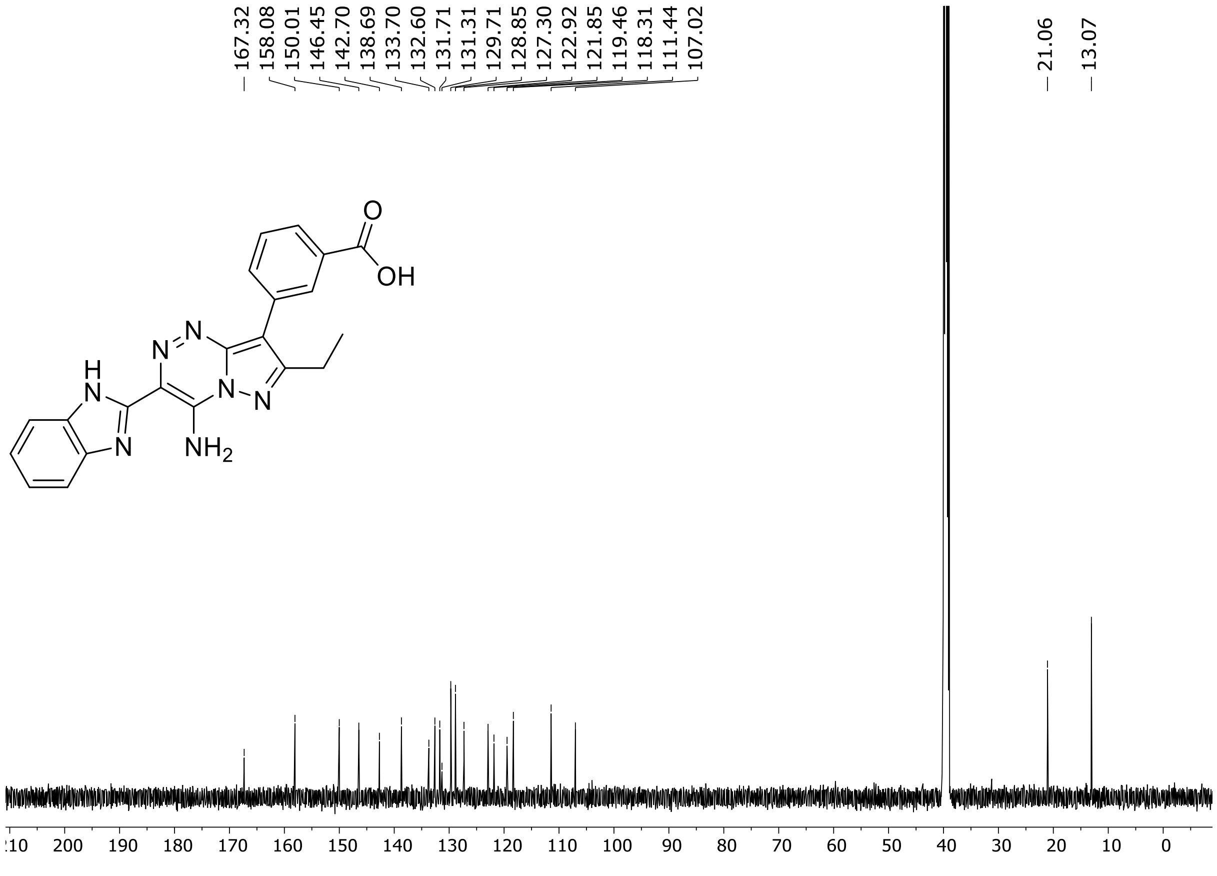

HRMS spectrum of **98**

$C_{21}H_{17}N_7O_2$ **exact mass: 399.1444**

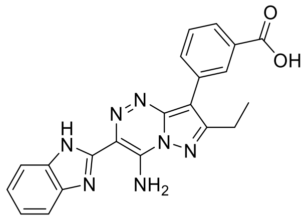

**APCI +** (MMI)
nitrogen flow 5 L/min, gas temperature 325°C, nebulizer 45 psig, skimmer 65 V, vaporizer 200°C, fragmentor 10 V, dissolved in methanol

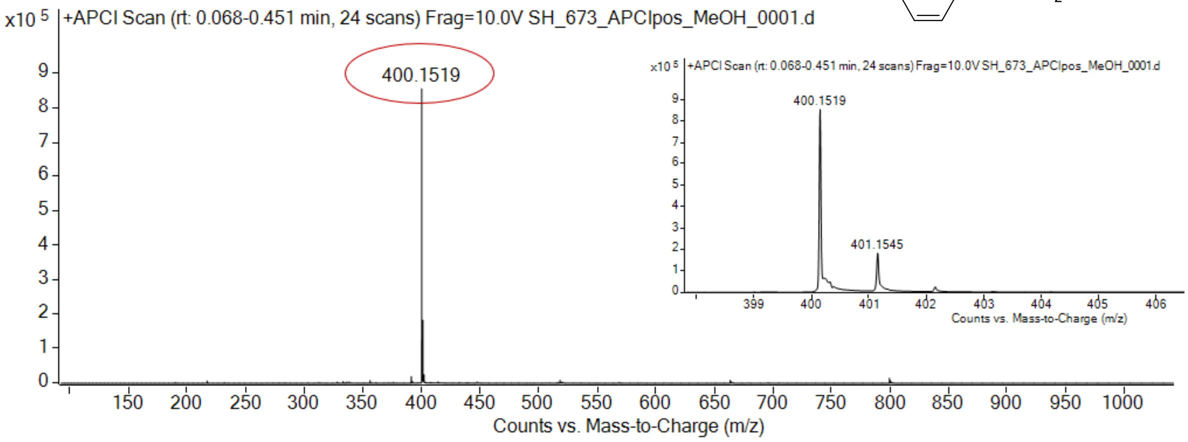

expected mass: [M+H]$^+$ = 400.1516 observed mass : [M+H]$^+$ = 400.1519 mass accuracy = 0.7 ppm

[1]H (500 MHz) and [13]C NMR (126 MHz) spectra of **99** in DMSO-$d_6$

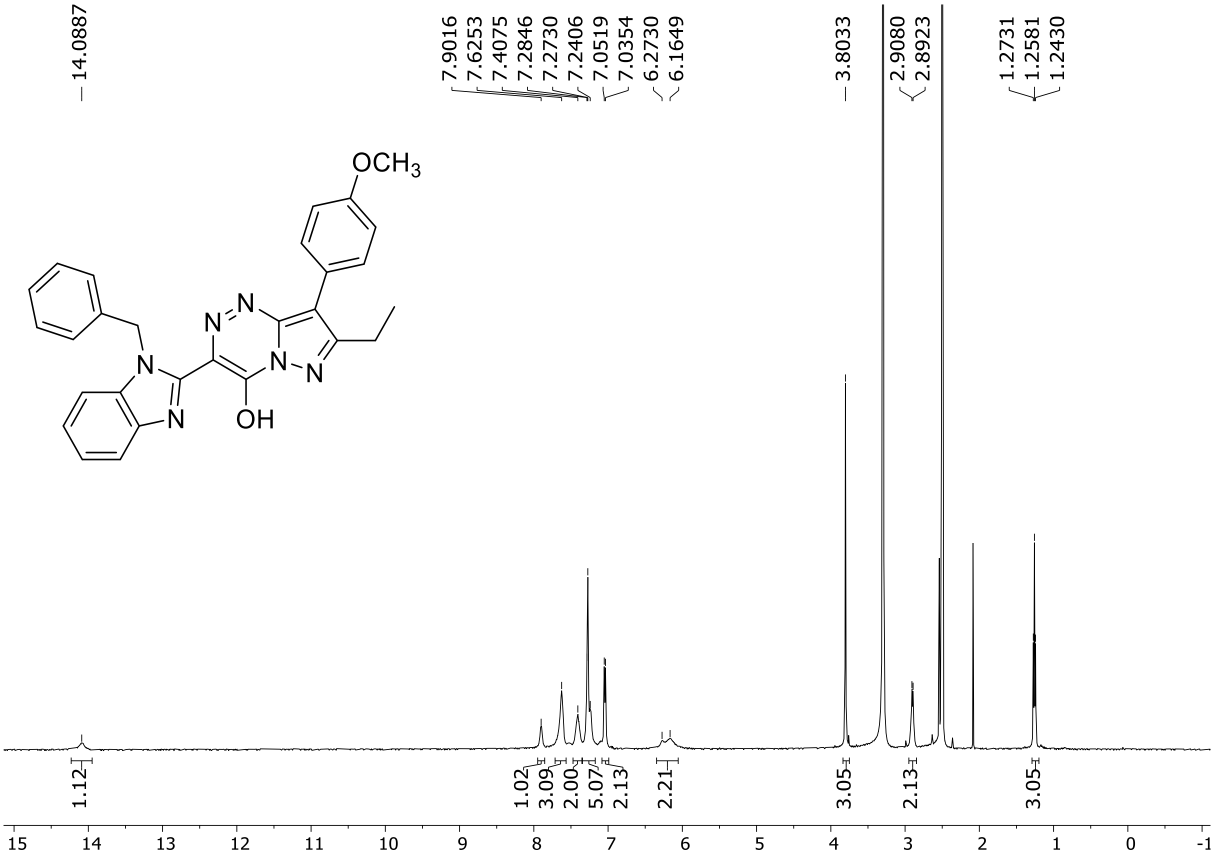

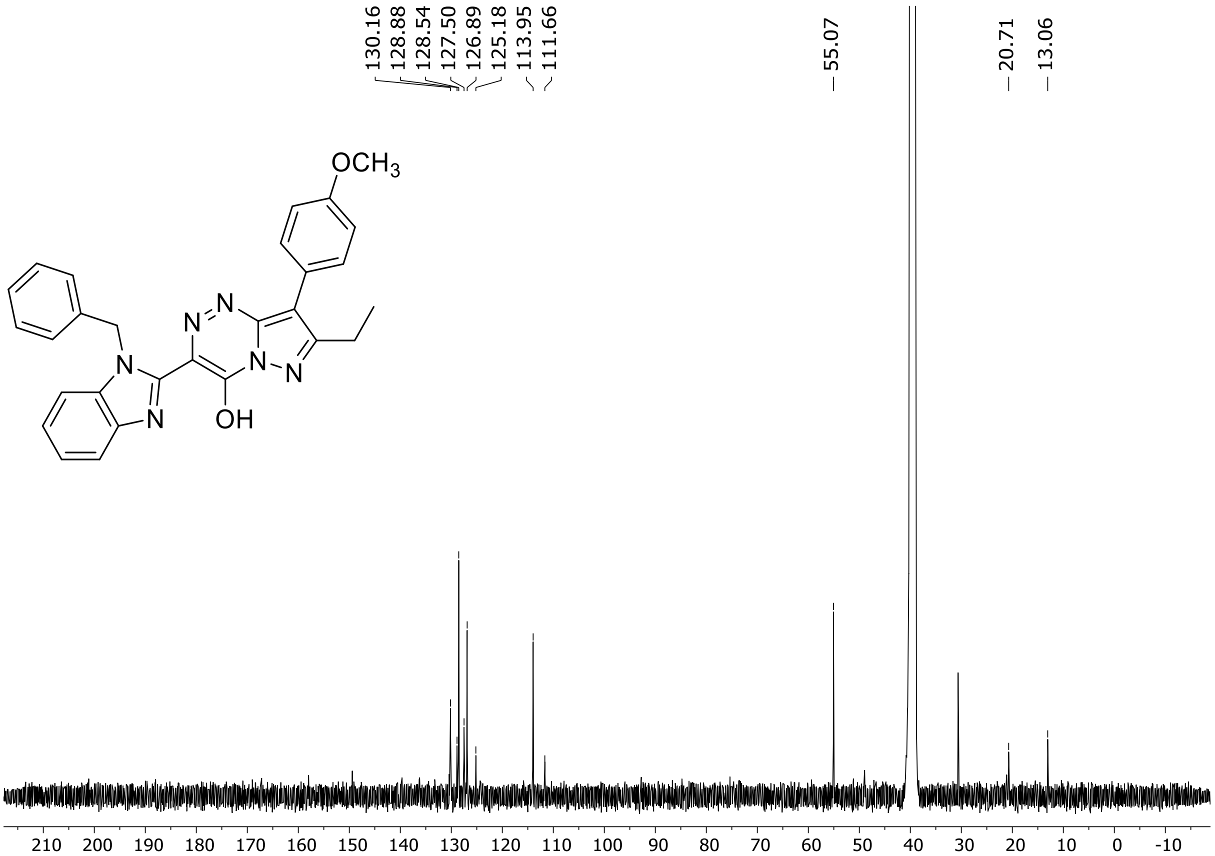

HRMS spectrum of **99**

**$C_{28}H_{24}N_6O_2$** **mono *m*/*z* 476.1961**

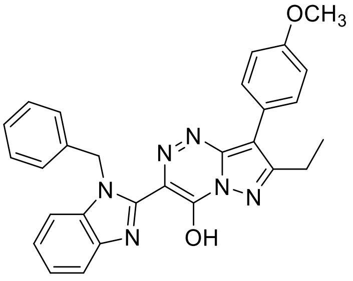

**APCI +** (MMI)
nitrogen flow 5 L/min, gas temperature 325°C, nebulizer 45 psi, skimmer 65 V, vaporizer 250°C, fragmentor 45 V, dissolved in methanol

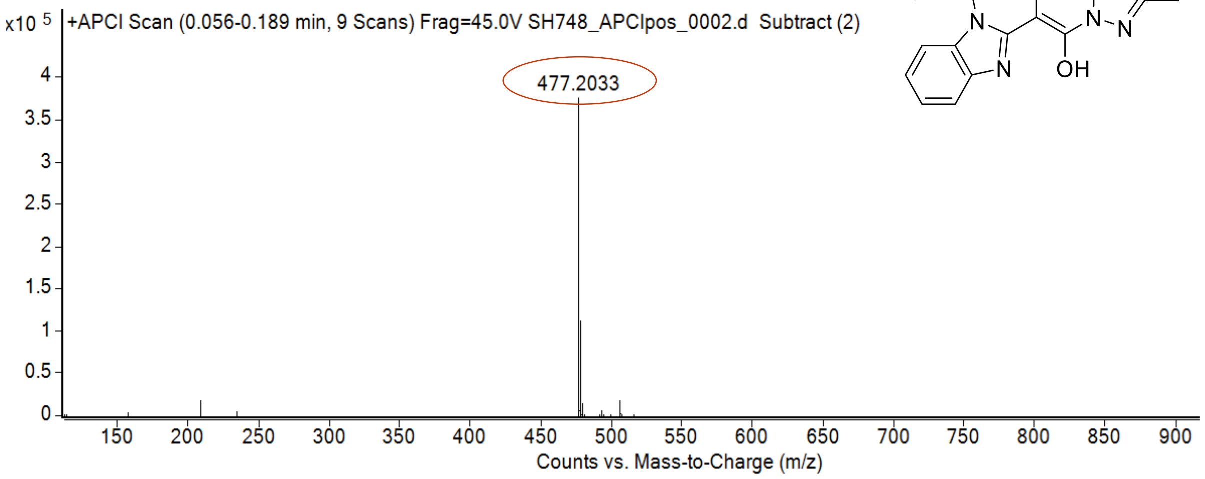

calculated mass: $[M+H]^+ = 477.2034$ observed: $[M+H]^+ = 477.2033$ mass accuracy = - 0.2 ppm

[1]H (500 MHz) and [13]C NMR (126 MHz) spectra of **100** in DMSO-$d_6$

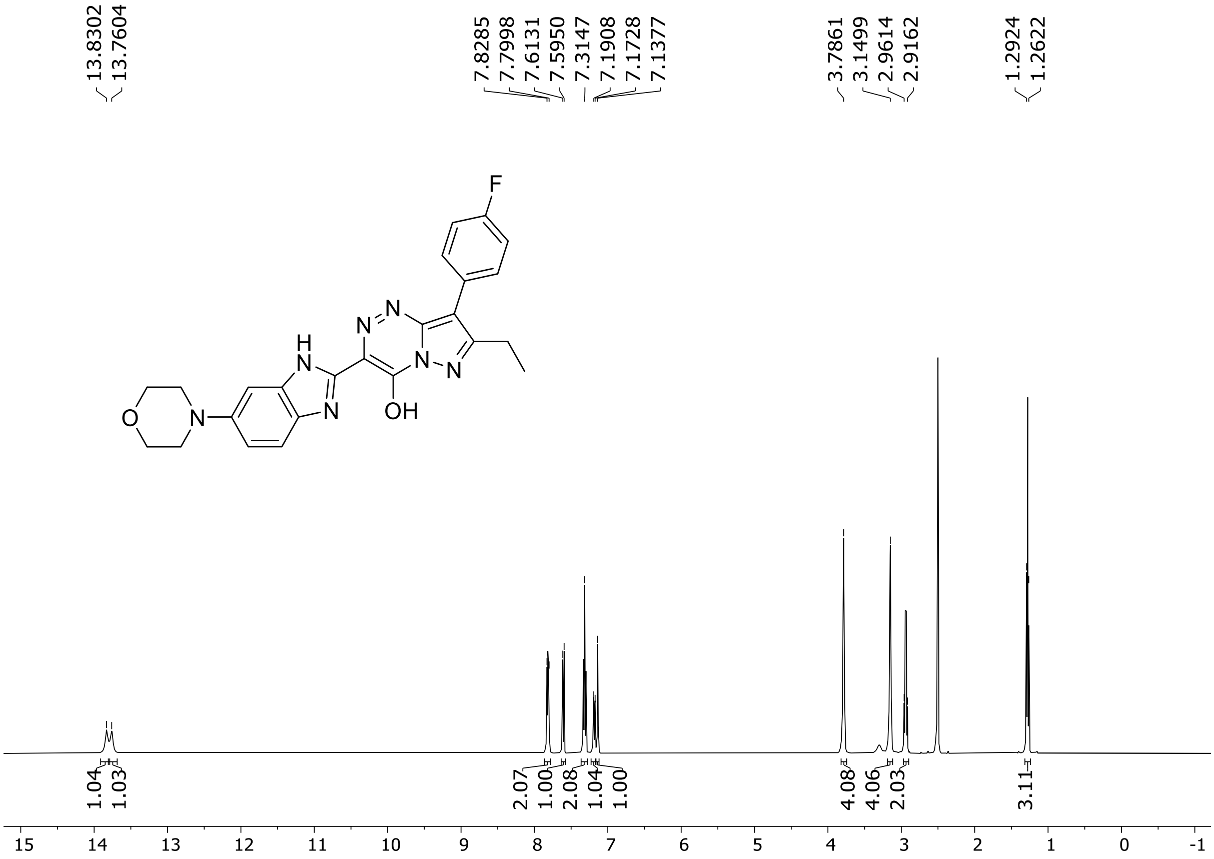

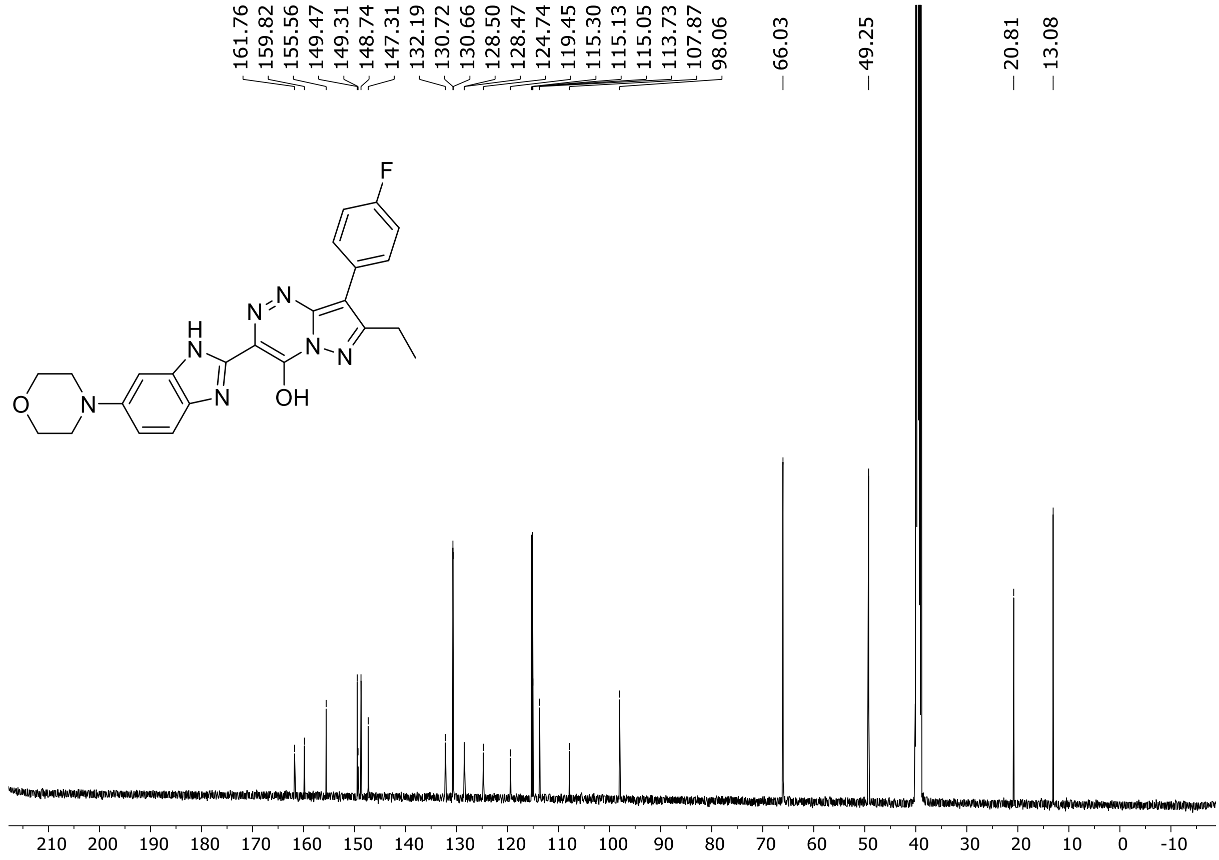

$^{19}F$ (471 MHz) NMR spectrum of **100** in DMSO-$d_6$

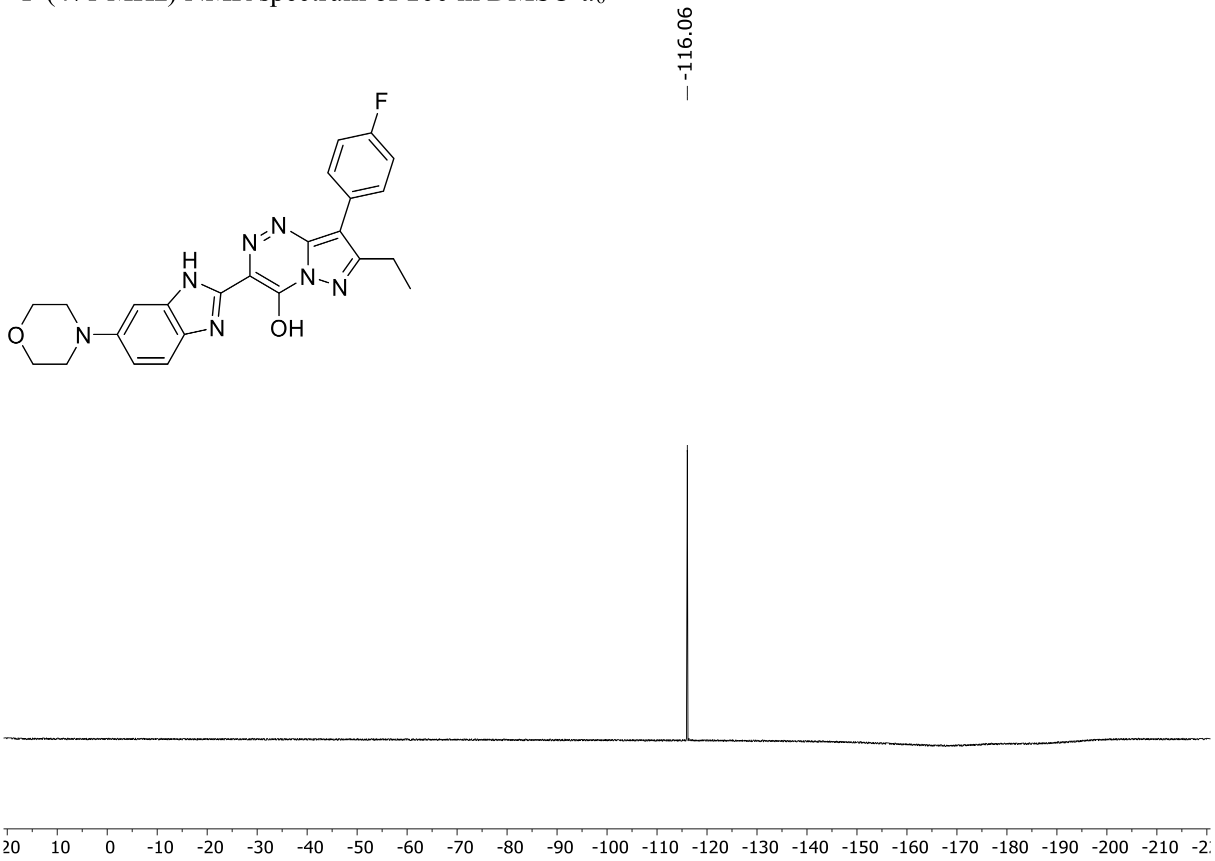

HRMS spectrum of **100**

**$C_{24}H_{22}FN_7O_2$** **exact mass: 459.1819**

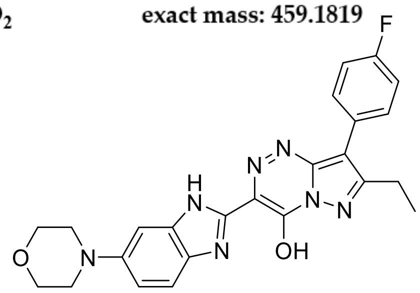

**APCI +** (MMI)
nitrogen flow 5 L/min, gas temperature 325°C, nebulizer 45 psig, skimmer 65 V,
vaporizer 200°C, fragmentor 20 V, dissolved in methanol

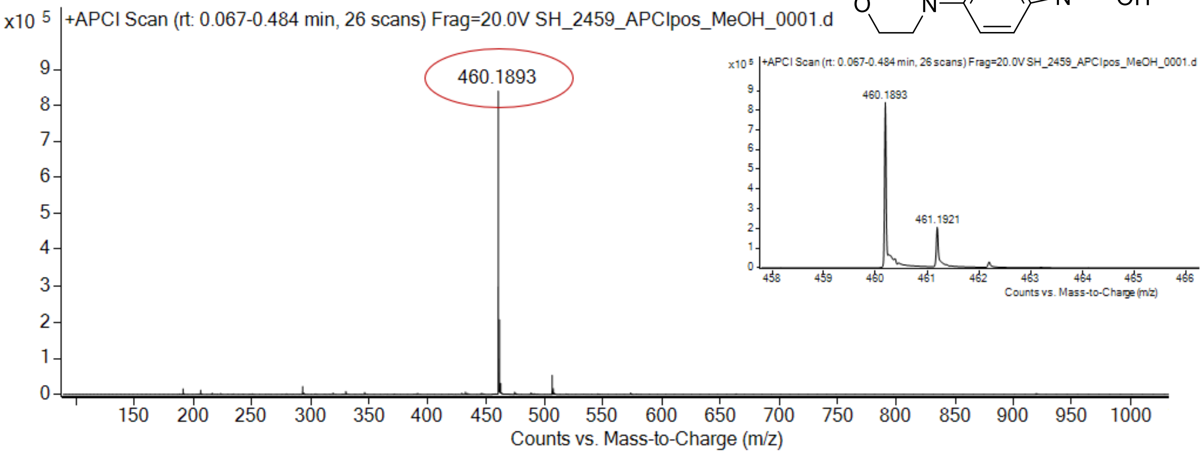

expected mass: [M+H]$^+$ = 460.1892 observed mass : [M+H]$^+$ = 460.1893 mass accuracy = 0.2 ppm

$^{1}H$ (500 MHz) and $^{13}C$ NMR (126 MHz) spectra of **101** in DMSO-$d_6$

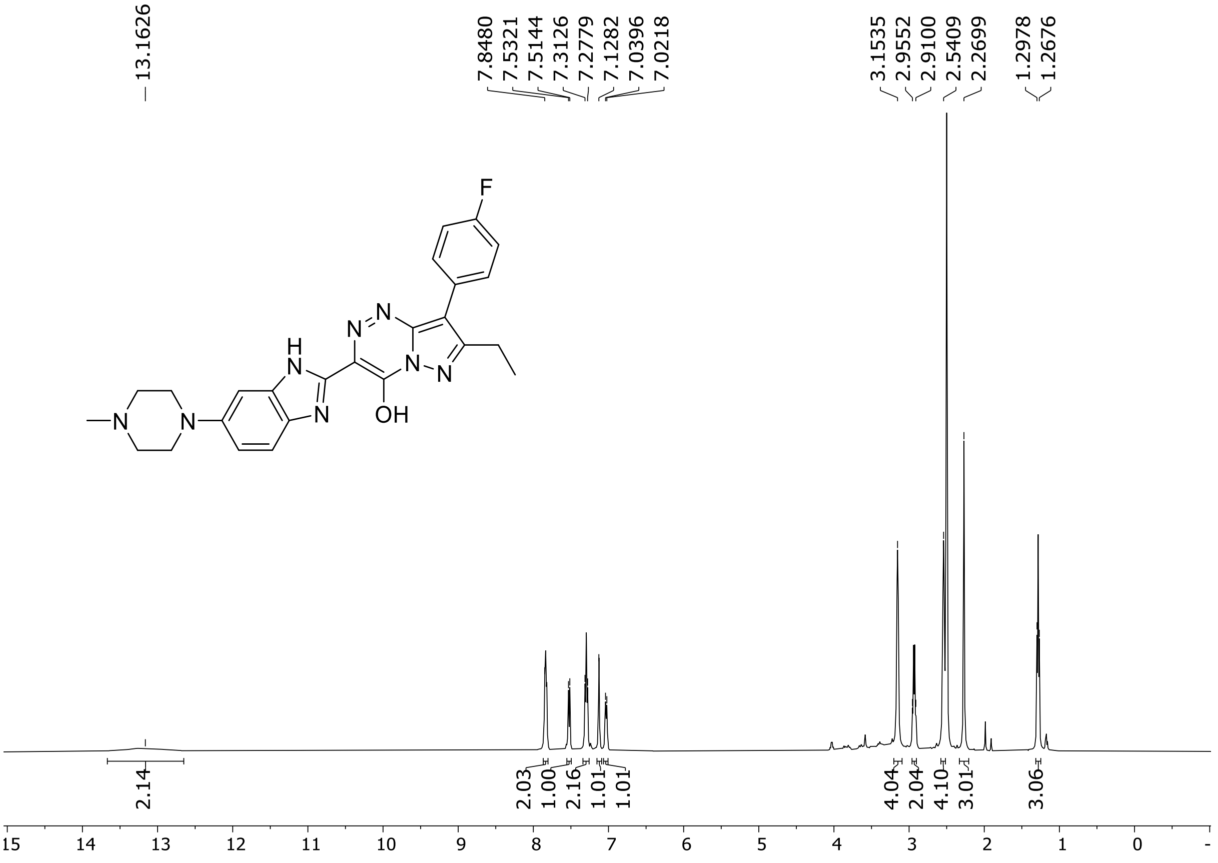

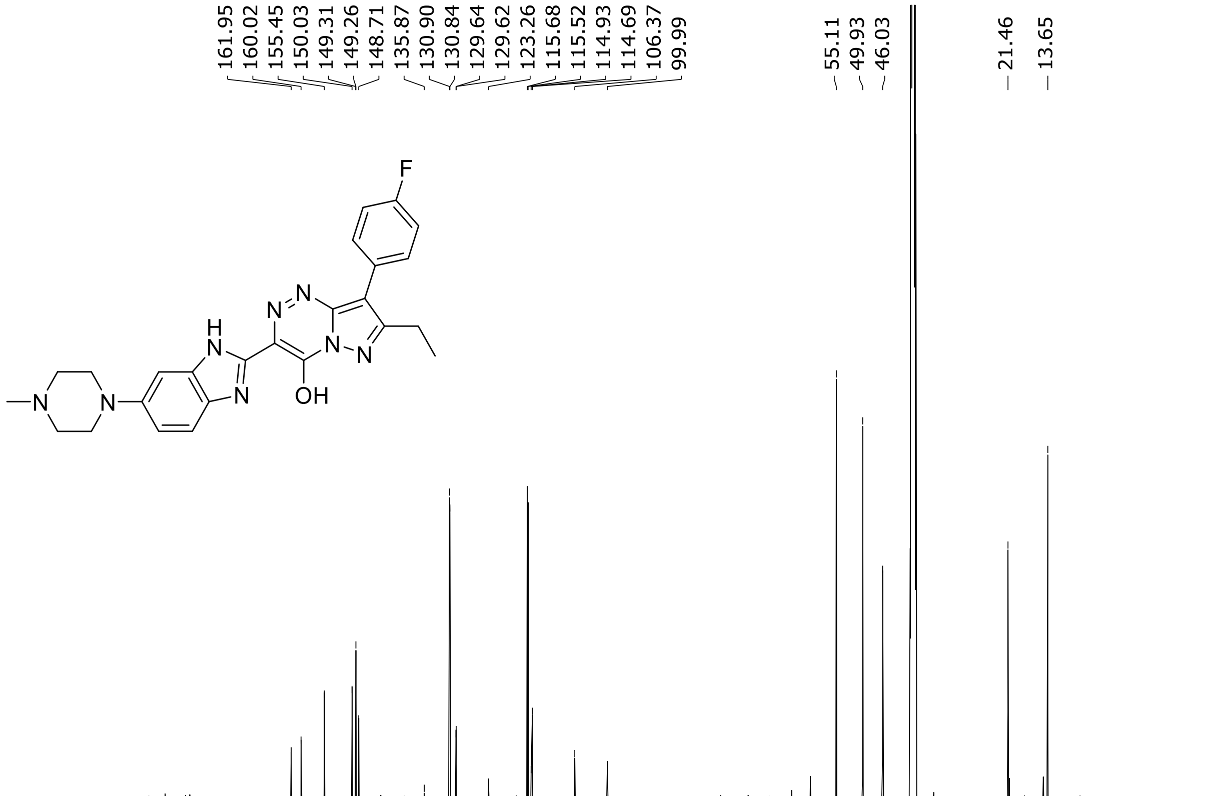

$^{19}$F (471 MHz) NMR spectrum of **101** in DMSO-$d_6$

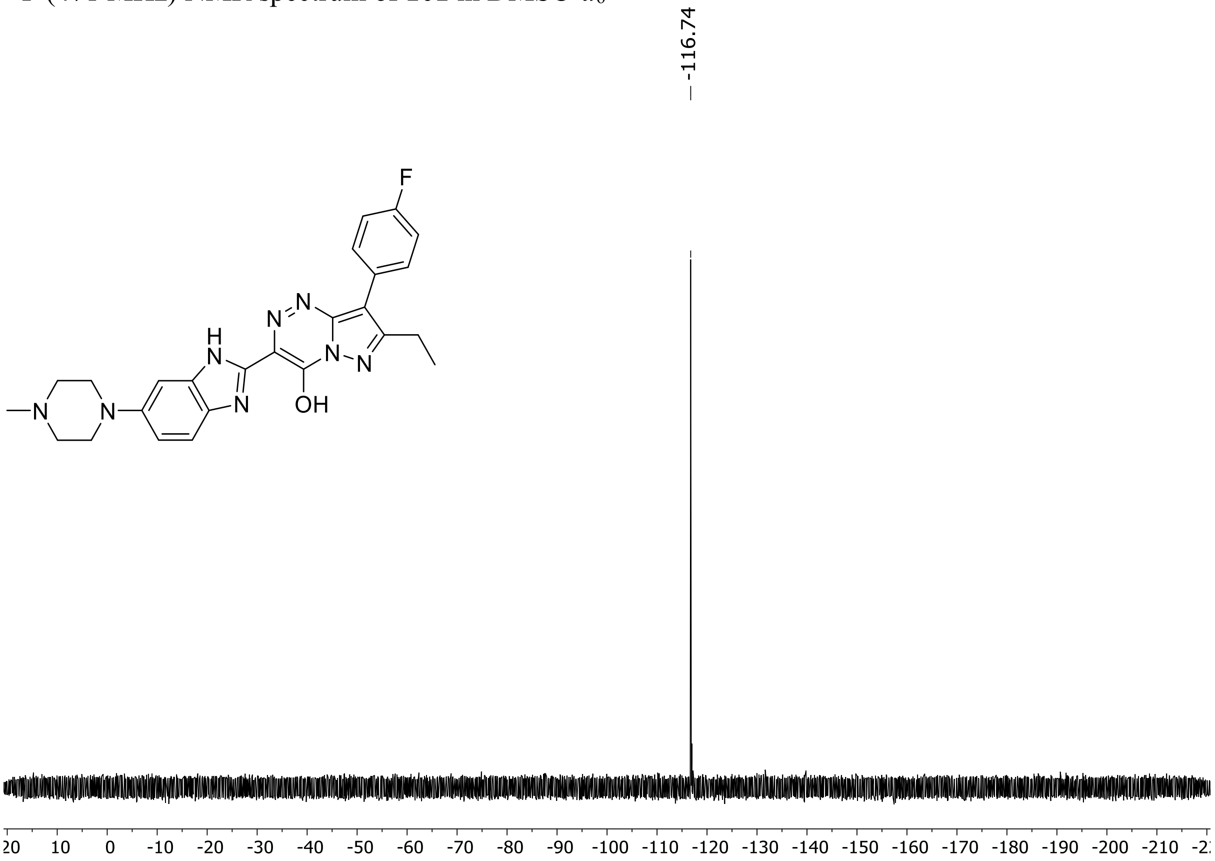

HRMS spectrum of **101**

**$C_{25}H_{25}FN_8O$** **exact mass: 472.2135**

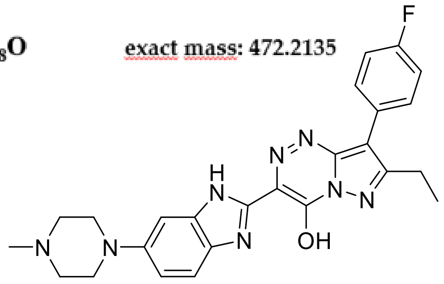

**APCI +** (MMI)
nitrogen flow 5 L/min, gas temperature 325°C, nebulizer 45 psig, skimmer 65 V, vaporizer 200°C, fragmentor 10 V, dissolved in methanol

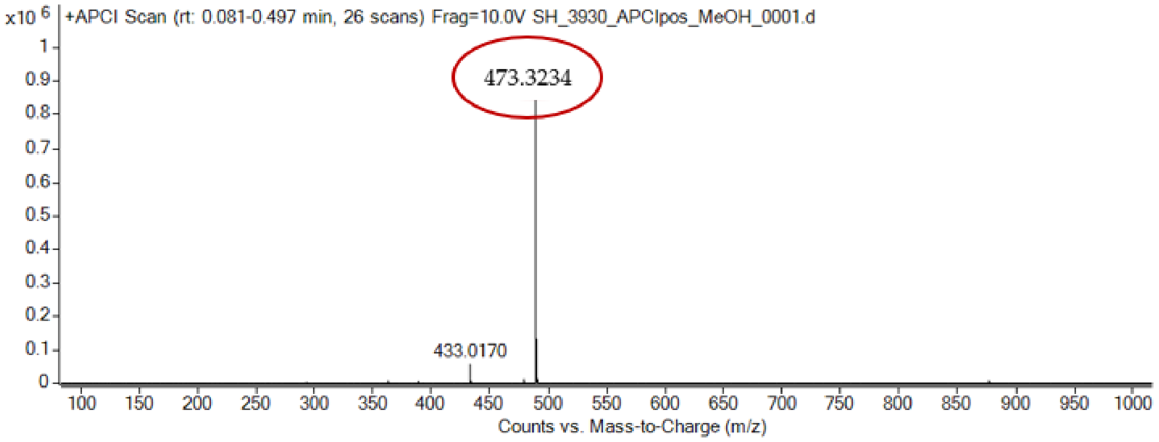

expected mass: $[M+H]^+ = 473.3223$ observed mass: $[M+H]^+ = 473.3234$ mass accuracy = 0.8 ppm

$^{1}H$ (500 MHz) and $^{13}C$ NMR (126 MHz) spectra of **102** in DMSO-$d_6$

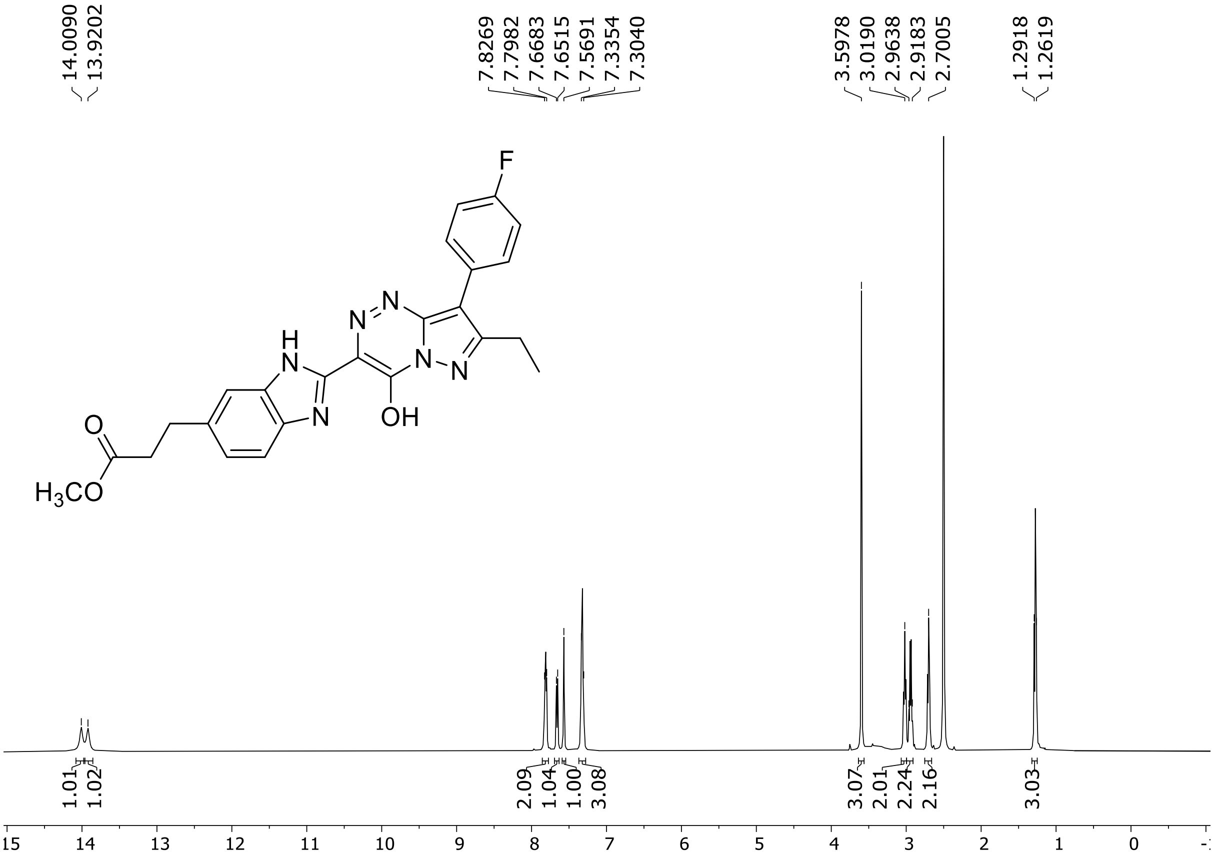

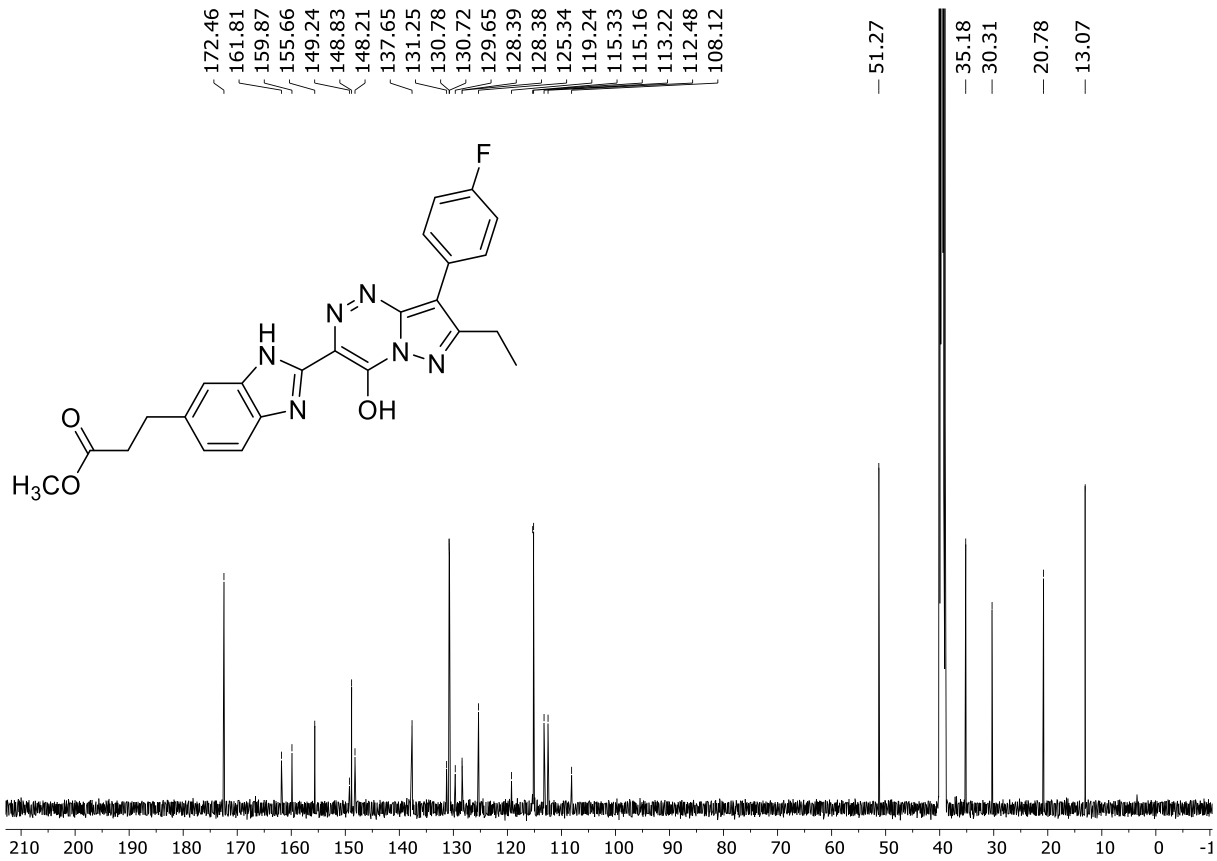

$^{19}$F (471 MHz) NMR spectrum of **102** in DMSO-$d_6$

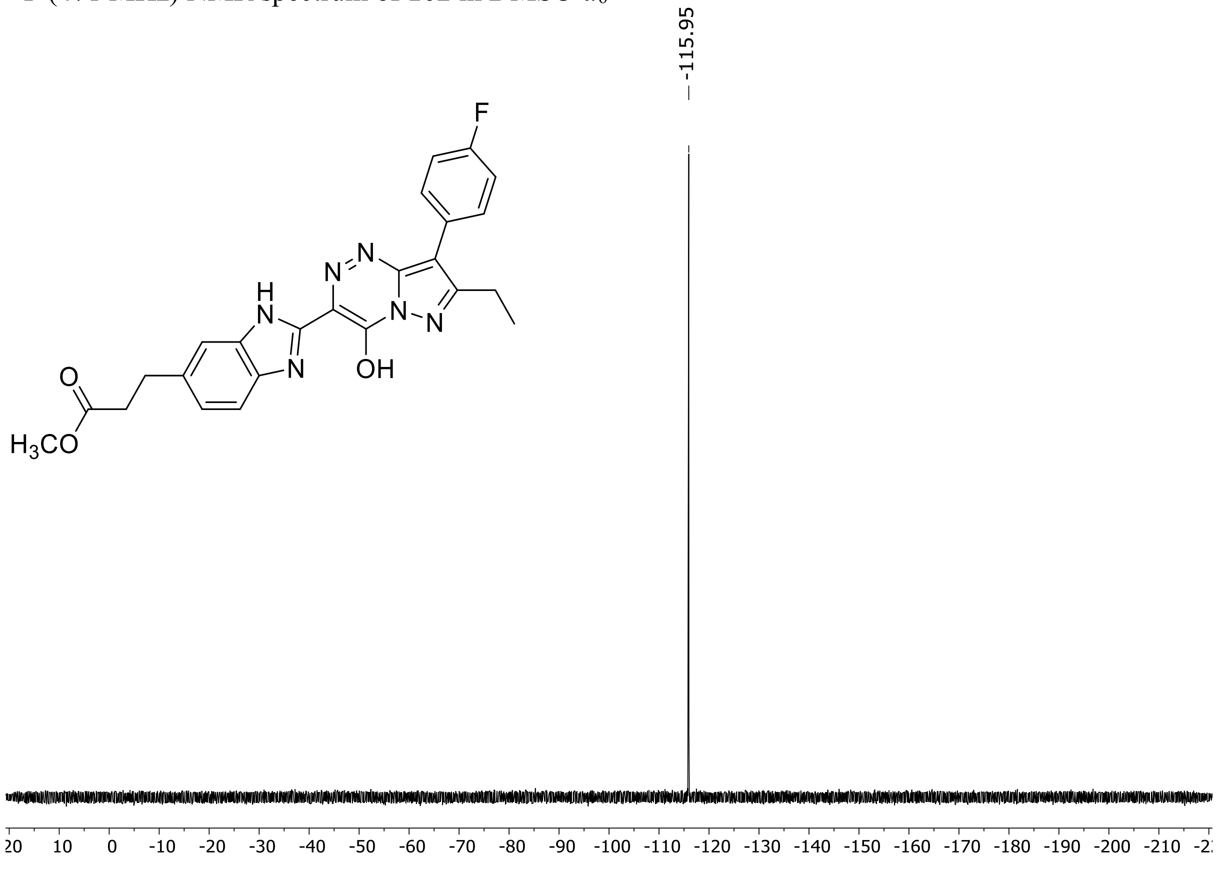

HRMS spectrum of **102**

$C_{24}H_{21}FN_6O_3$ **exact mass: 460.1659**

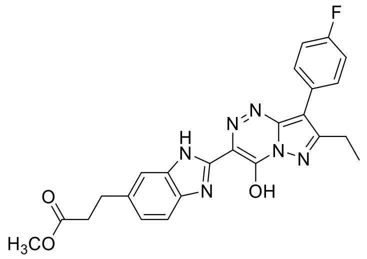

**APCI +** (MMI)
nitrogen flow 5 L/min, gas temperature 325°C, nebulizer 45 psig, skimmer 65 V, vaporizer 200°C, fragmentor 20 V, dissolved in methanol

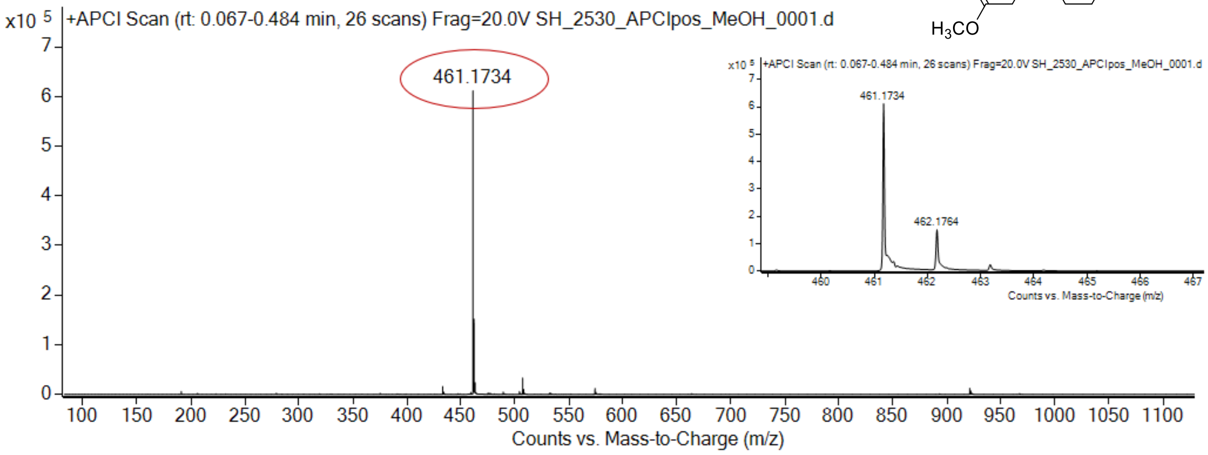

expected mass: $[M+H]^+$ = 461.1732 observed mass : $[M+H]^+$ = 461.1734 mass accuracy = 0.4 ppm

$^1$H (500 MHz) and $^{13}$C NMR (126 MHz) spectra of **103** in DMSO-$d_6$

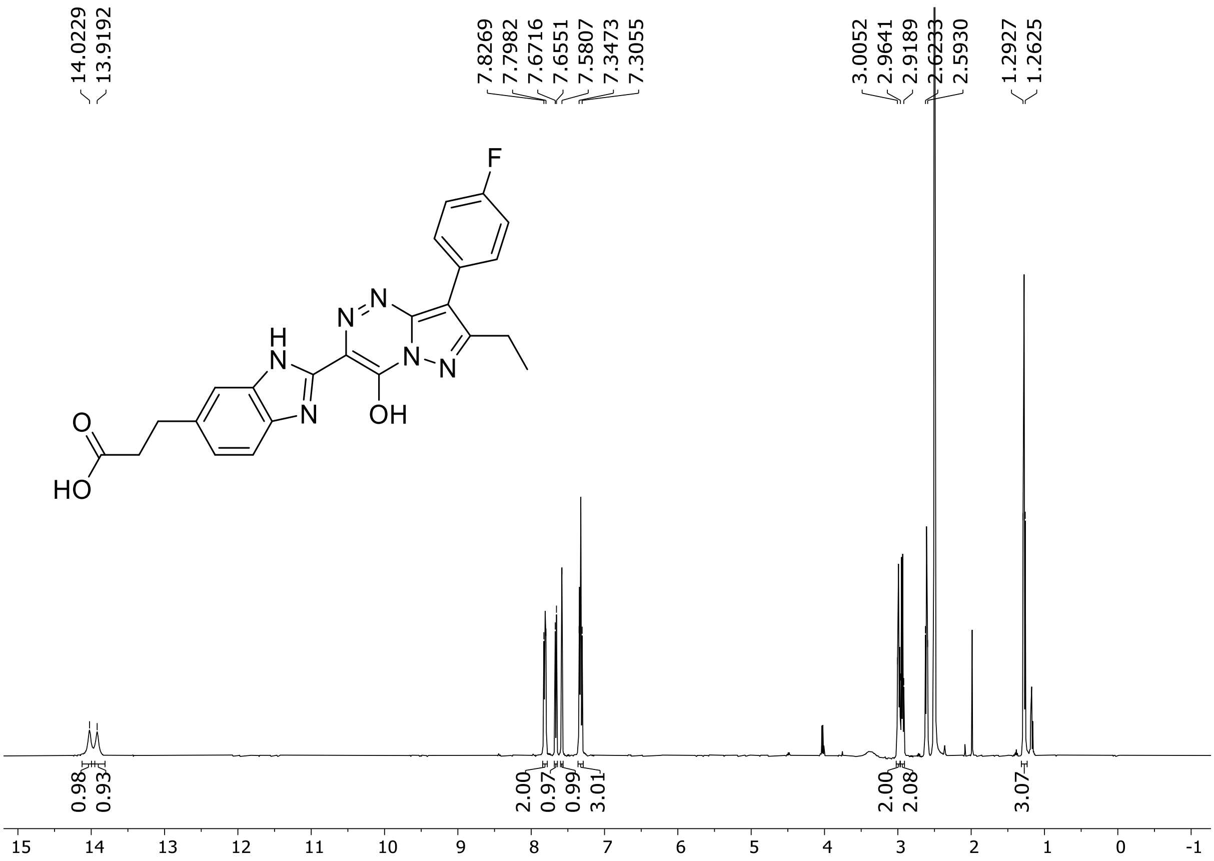

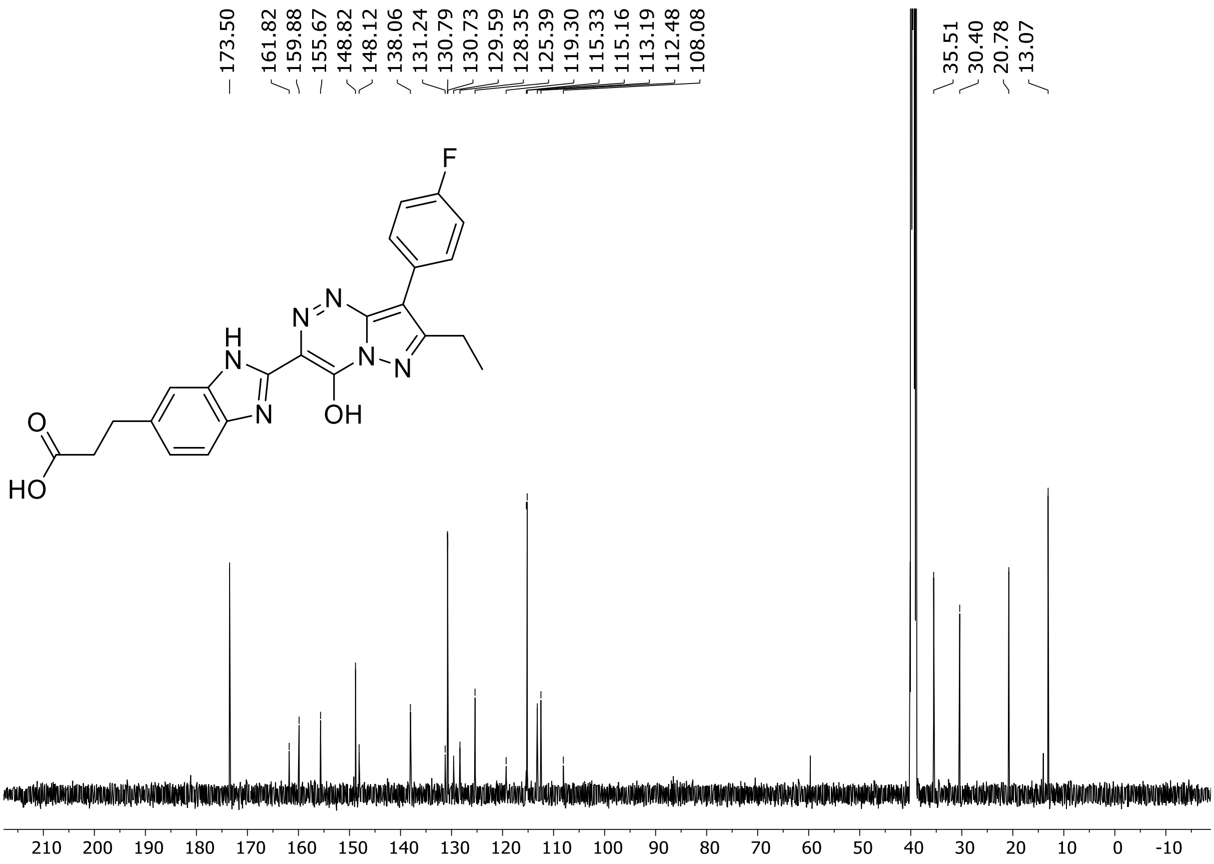

[19]F (471 MHz) NMR spectrum of **103** in DMSO-$d_6$

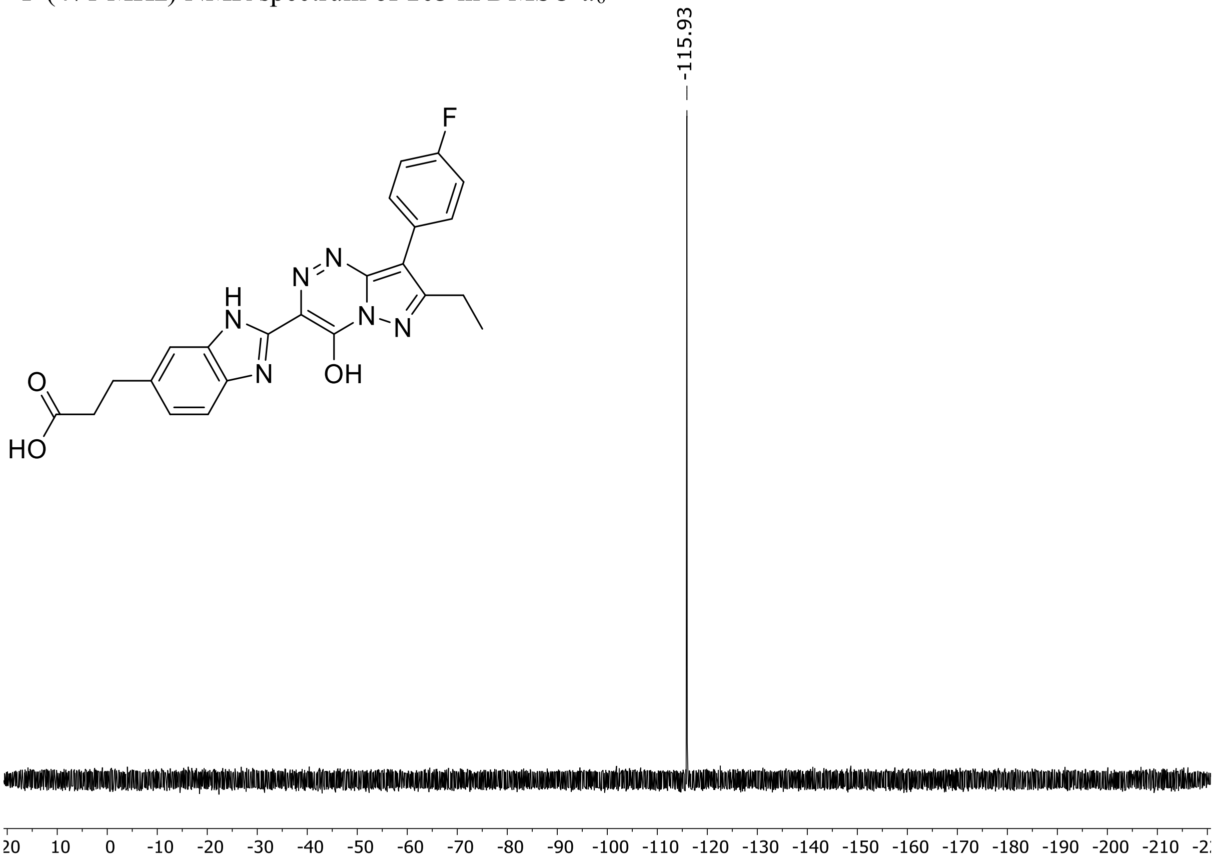

HRMS spectrum of **103**

**$C_{23}H_{19}FN_6O_3$** **exact mass: 446.1503**

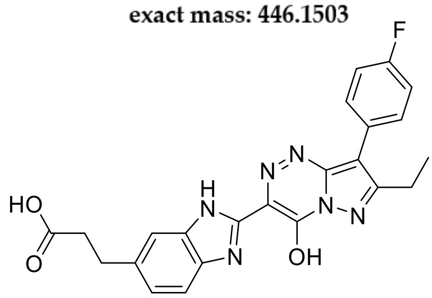

**APCI +** (MMI)
nitrogen flow 5 L/min, gas temperature 325°C, nebulizer 45 psig, skimmer 65 V, vaporizer 200°C, fragmentor 22 V, dissolved in methanol

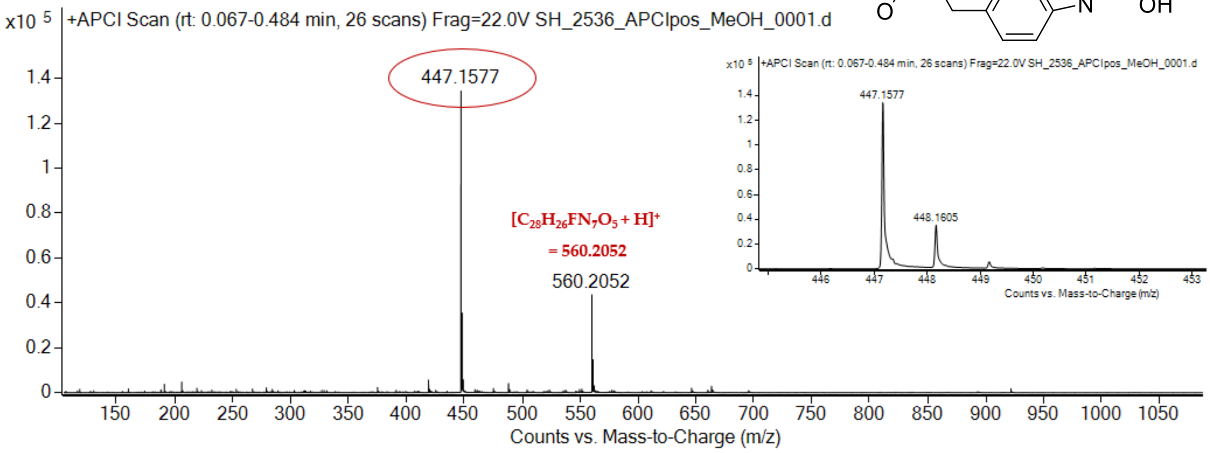

expected mass: $[M+H]^+$ = 447.1575 observed mass : $[M+H]^+$ = 447.1577 mass accuracy = 0.4 ppm

$^1$H (500 MHz) and $^{13}$C NMR (126 MHz) spectra of **104** in DMSO-$d_6$

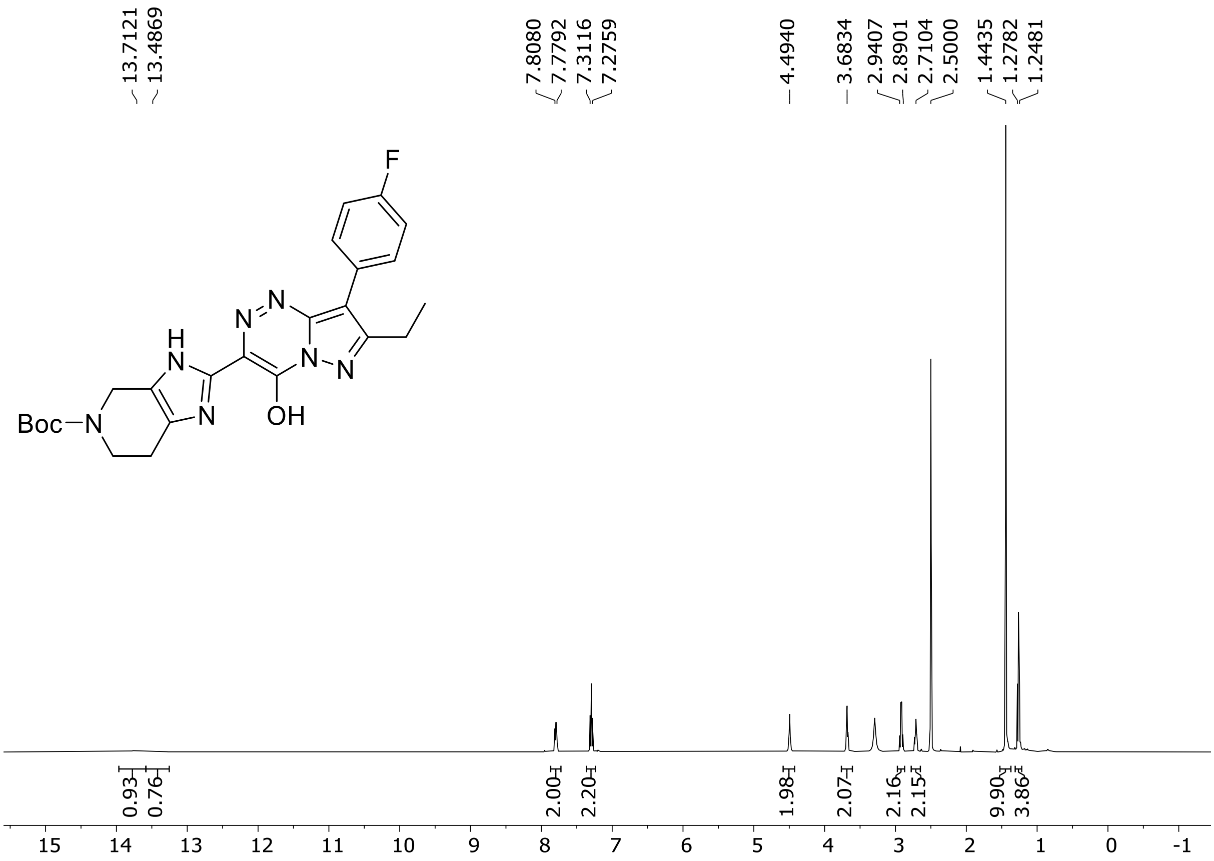

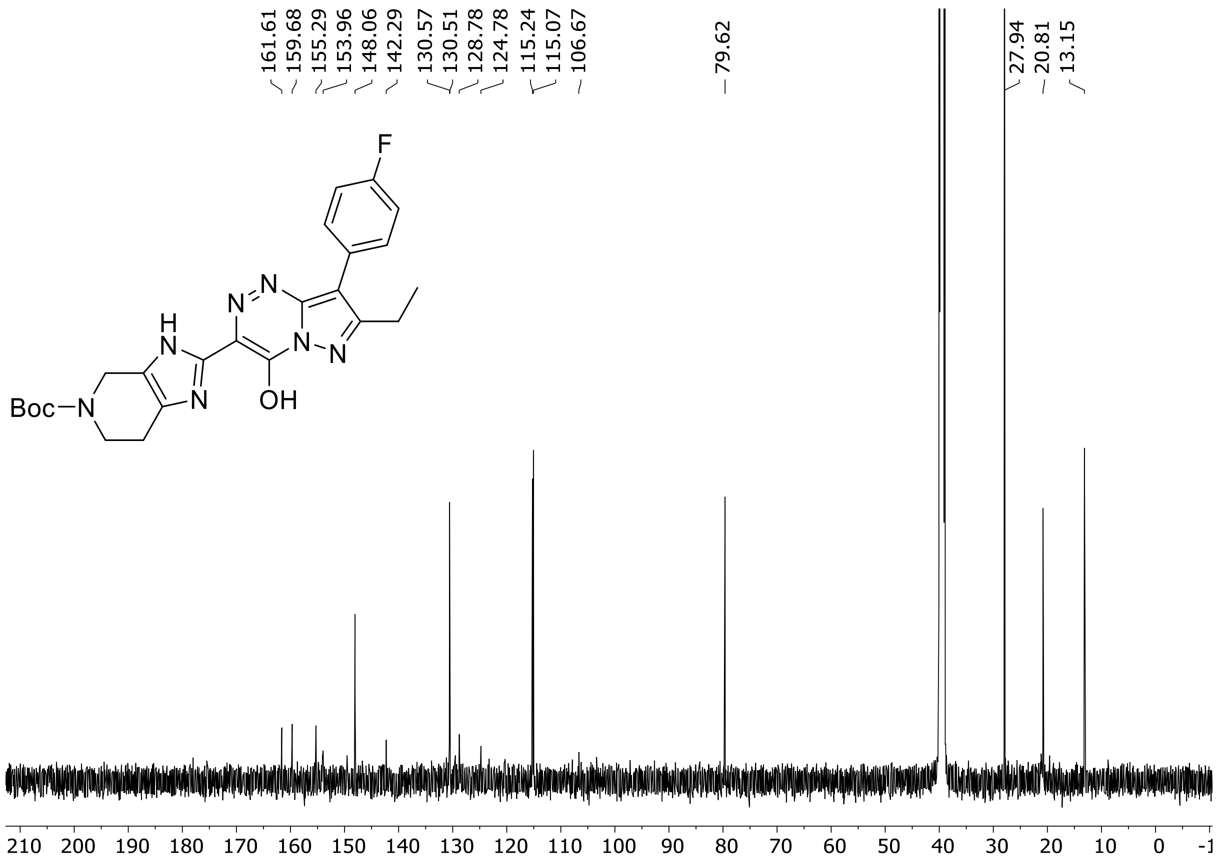

[19]F (471 MHz) NMR spectrum of **104** in DMSO-$d_6$

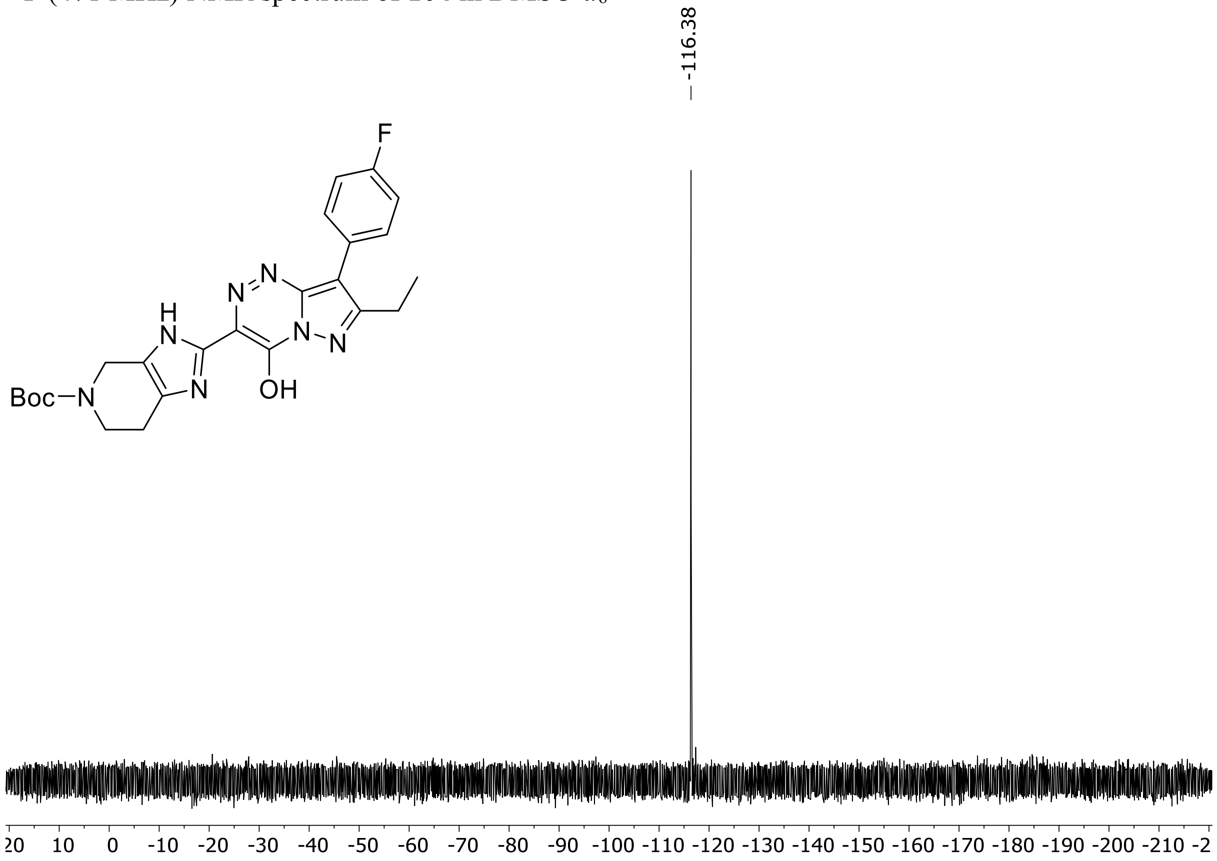

HRMS spectrum of **104**

$C_{24}H_{26}FN_7O_3$ **exact mass: 479.2081**

**APCI +** (MMI)
nitrogen flow 5 L/min, gas temperature 325°C, nebulizer 45 psig, skimmer 65 V,
vaporizer 200°C, fragmentor 30 V, dissolved in methanol

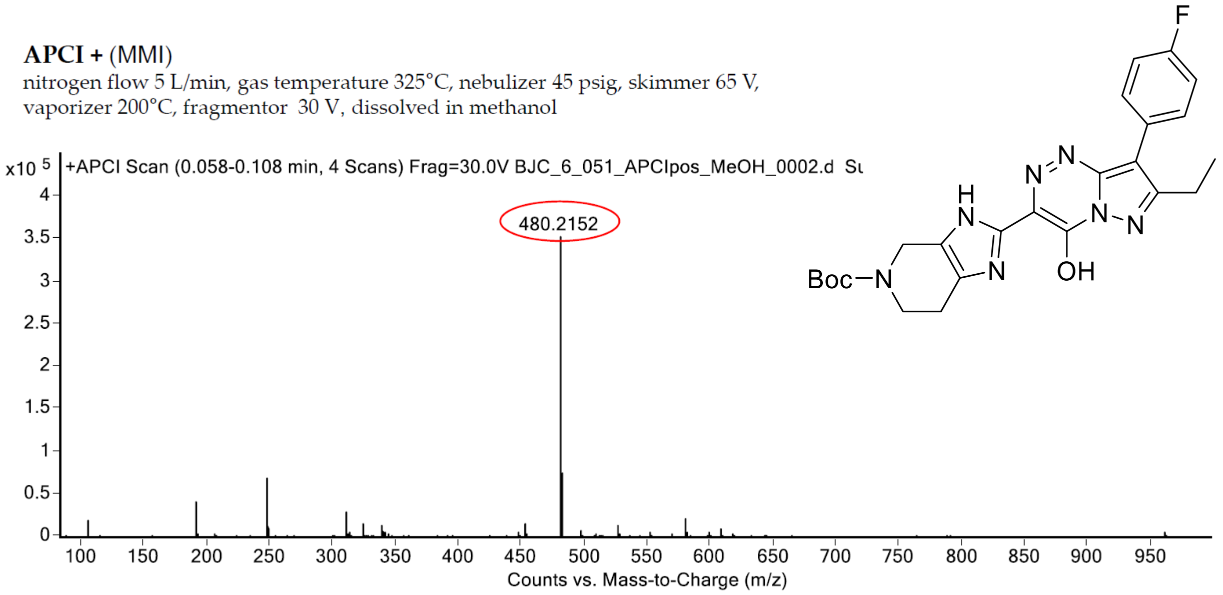

expected mass: [M+H]$^+$ = 480.2154 observed mass: [M+H]$^+$ = 480.2152 mass accuracy = - 0.4 ppm

$^1$H (500 MHz) and $^{13}$C NMR (126 MHz) spectra of **105** in DMSO-$d_6$

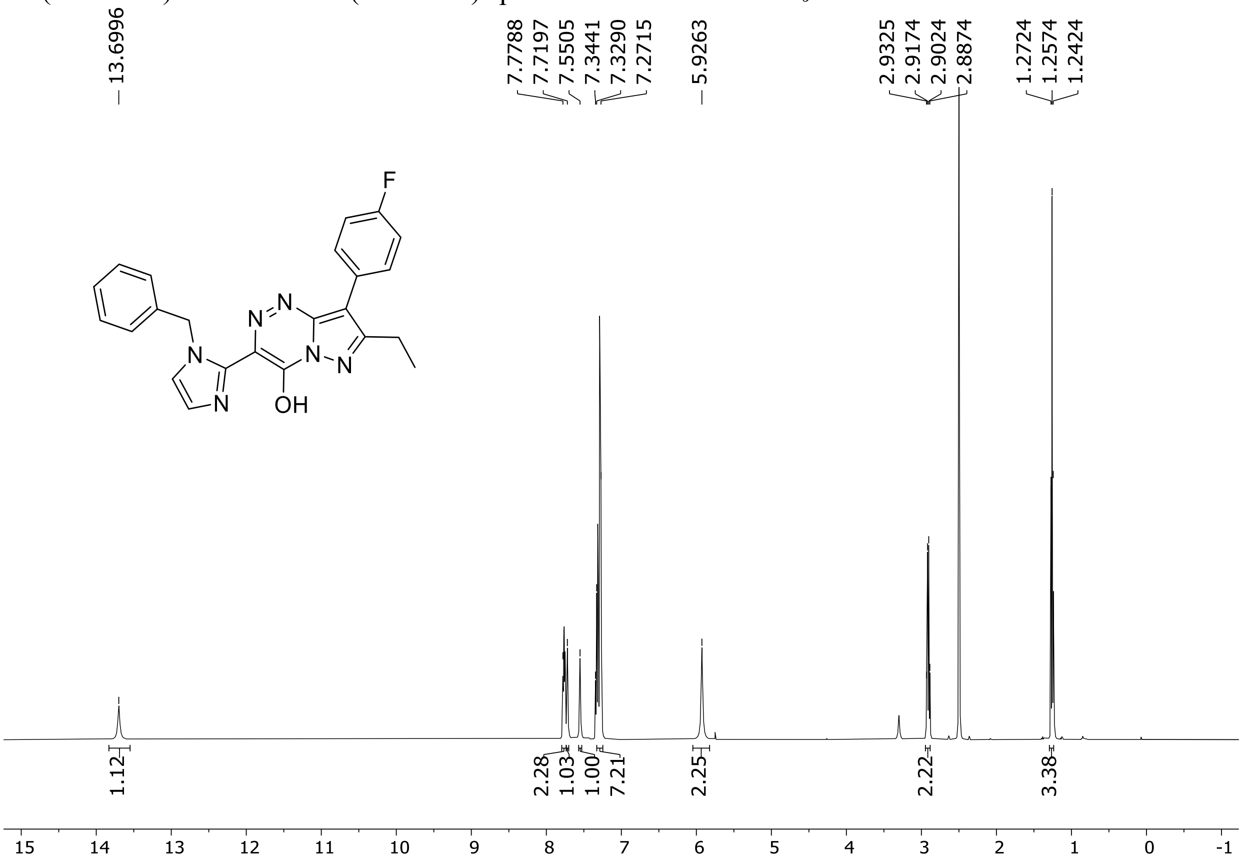

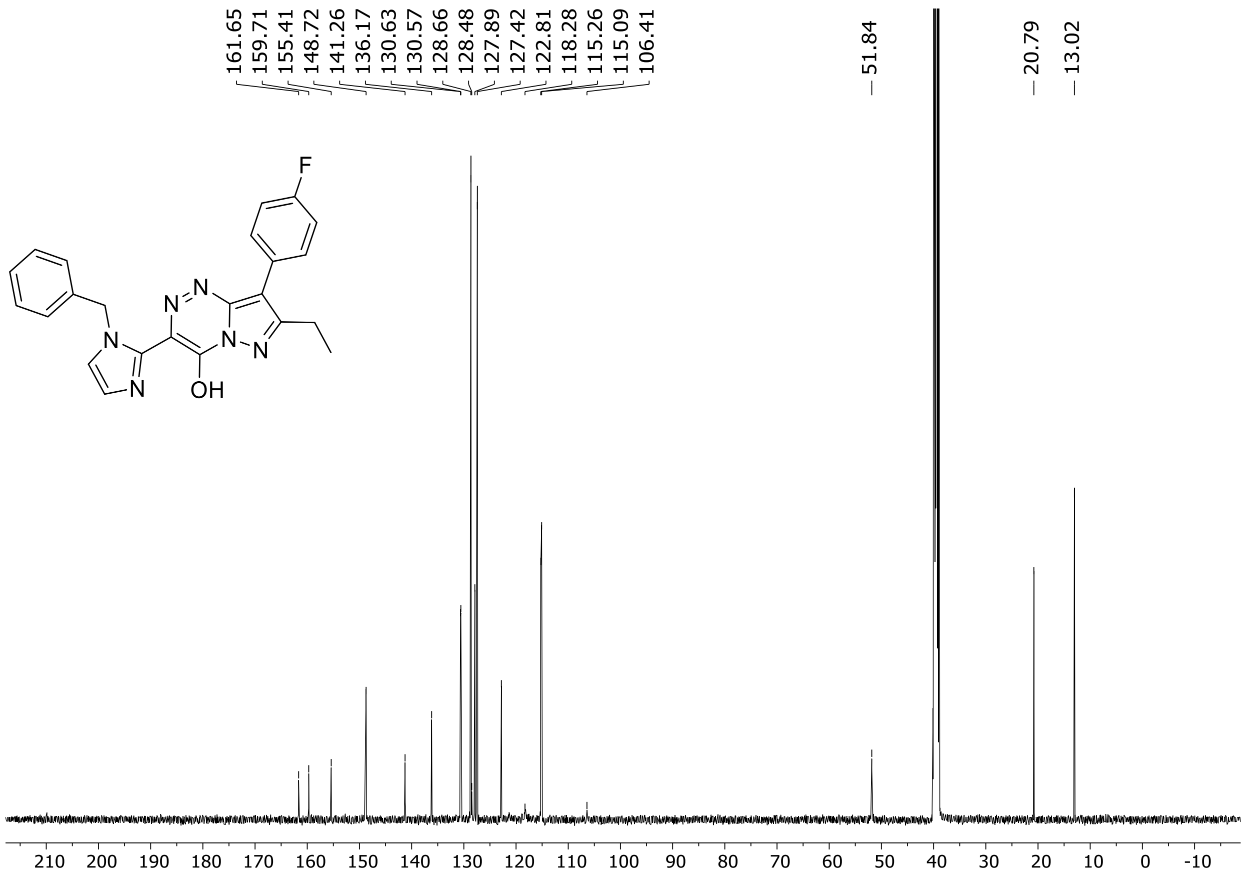

$^{19}$F (471 MHz) NMR spectrum of **105** in DMSO-$d_6$

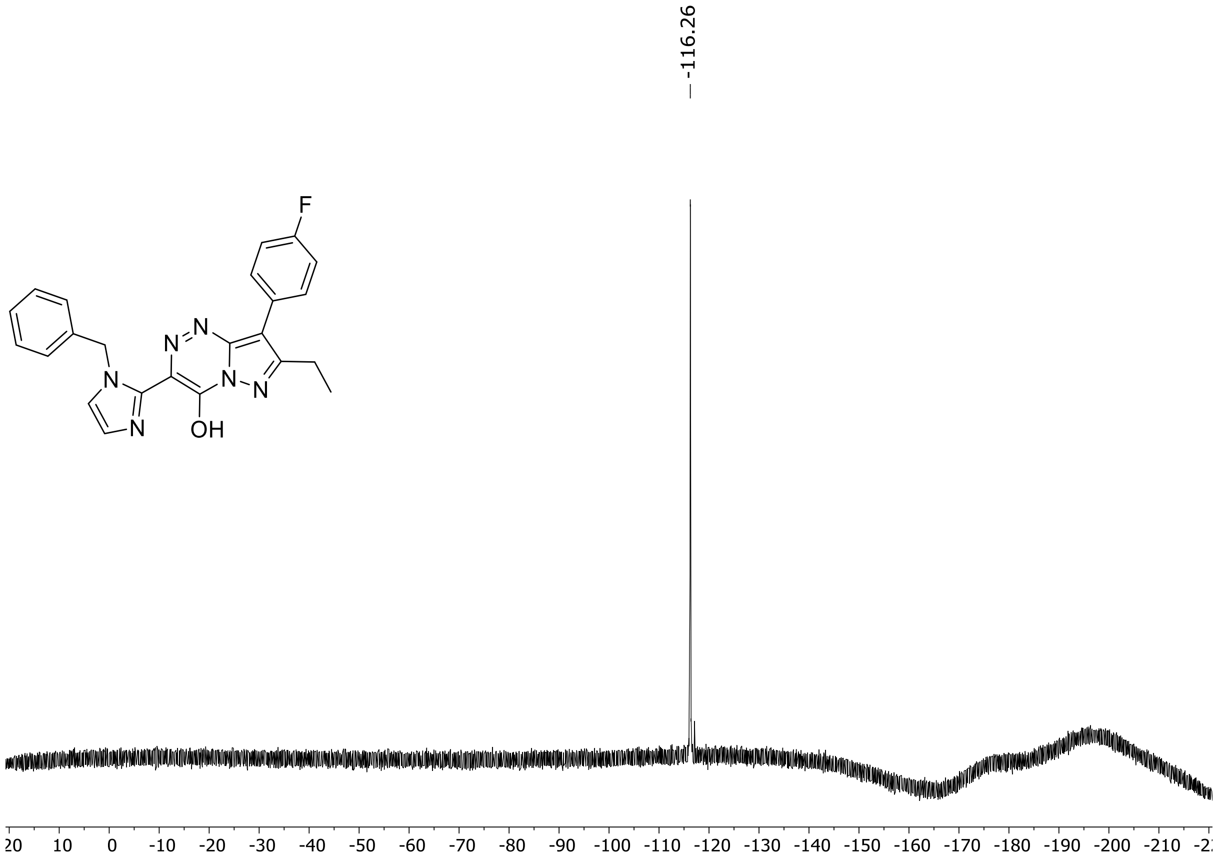

HRMS spectra of **105**

$C_{23}H_{19}FN_6O$ **exact mass: 414.1604**

**APCI +** (MMI)
nitrogen flow 3 L/min, gas temperature 325°C, nebulizer 45 psig, skimmer 65 V, vaporizer 200°C, fragmentor 15 V, dissolved in methanol

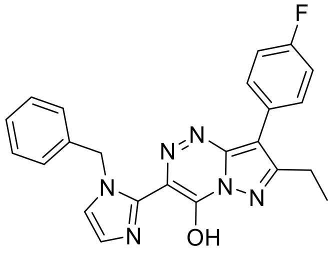

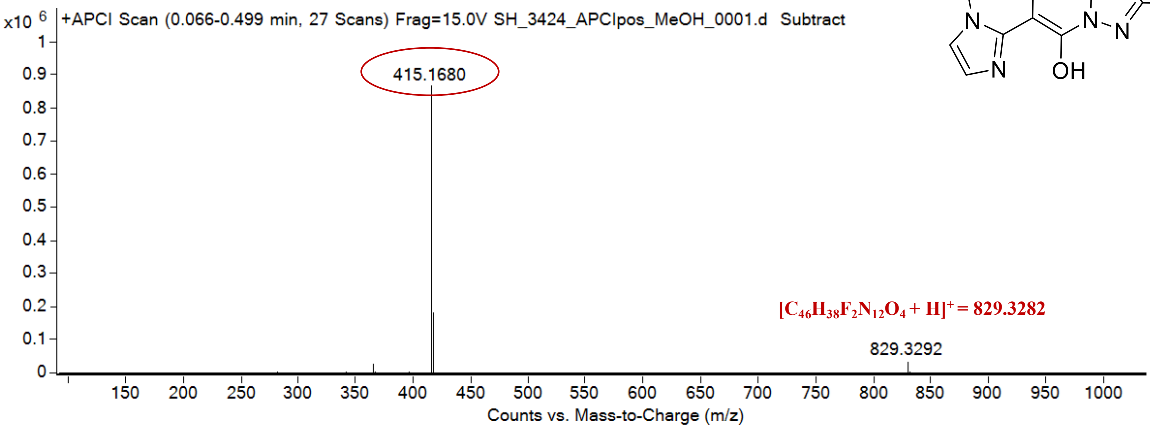

expected mass: $[M+H]^+ = 415.1677$ observed mass: $[M+H]^+ = 415.1680$ mass accuracy = 0.7 ppm

$^{1}H$ (500 MHz) and $^{13}C$ NMR (126 MHz) spectra of **106** in DMSO-$d_6$

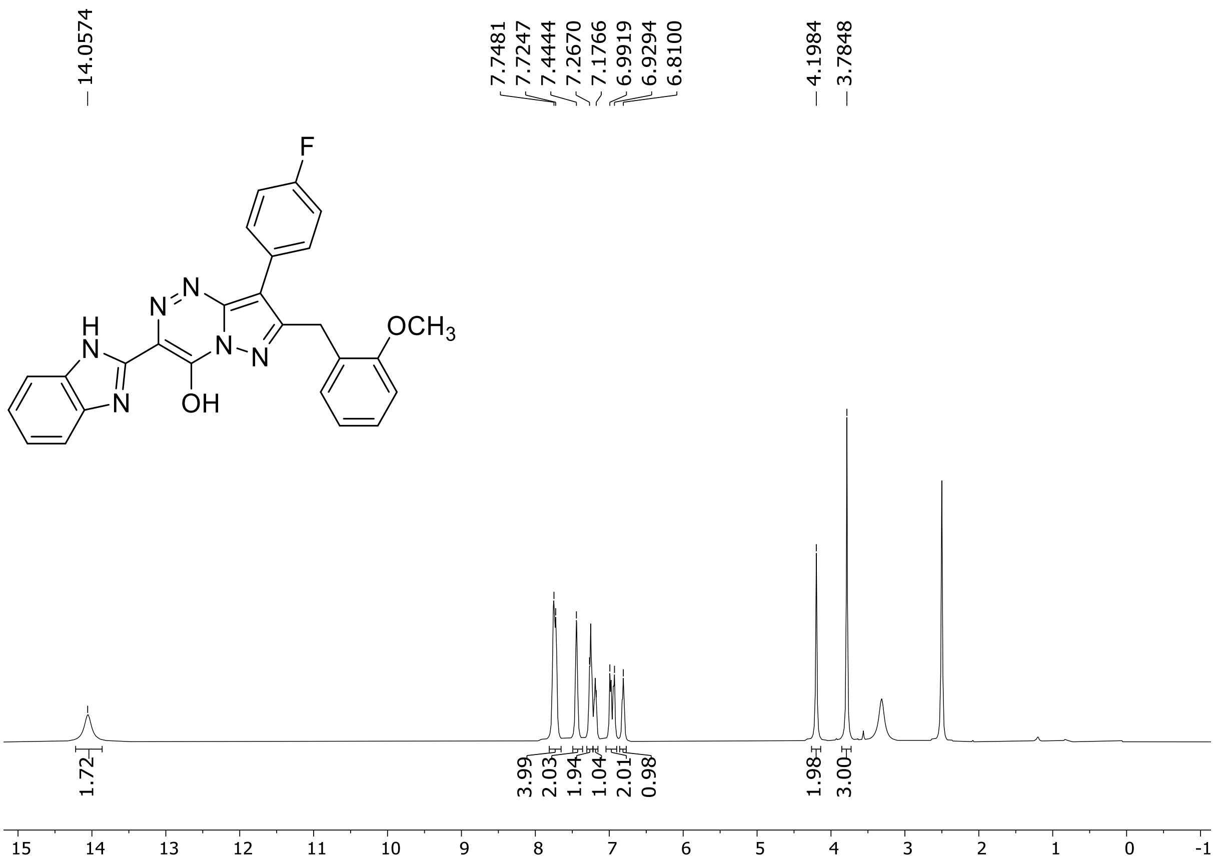

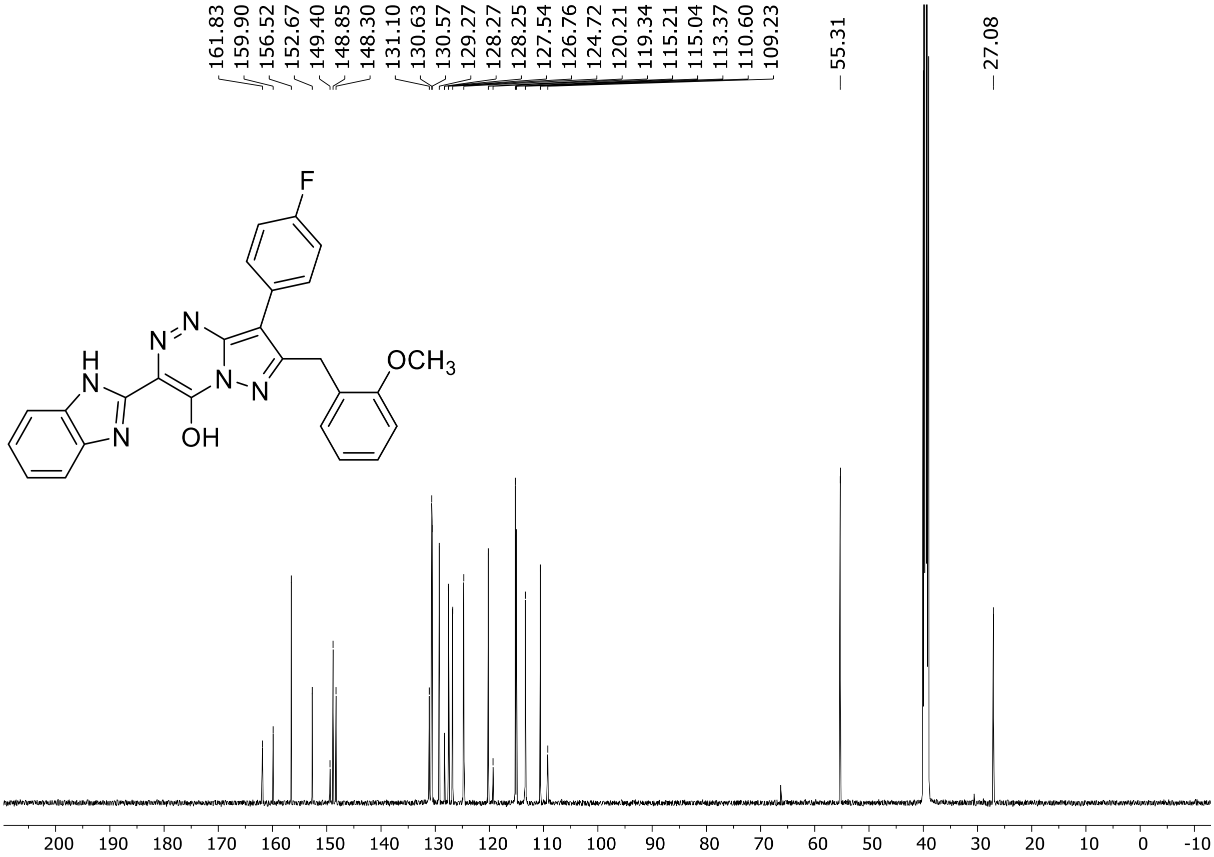

$^{19}$F (471 MHz) NMR spectrum of **106** in DMSO-$d_6$

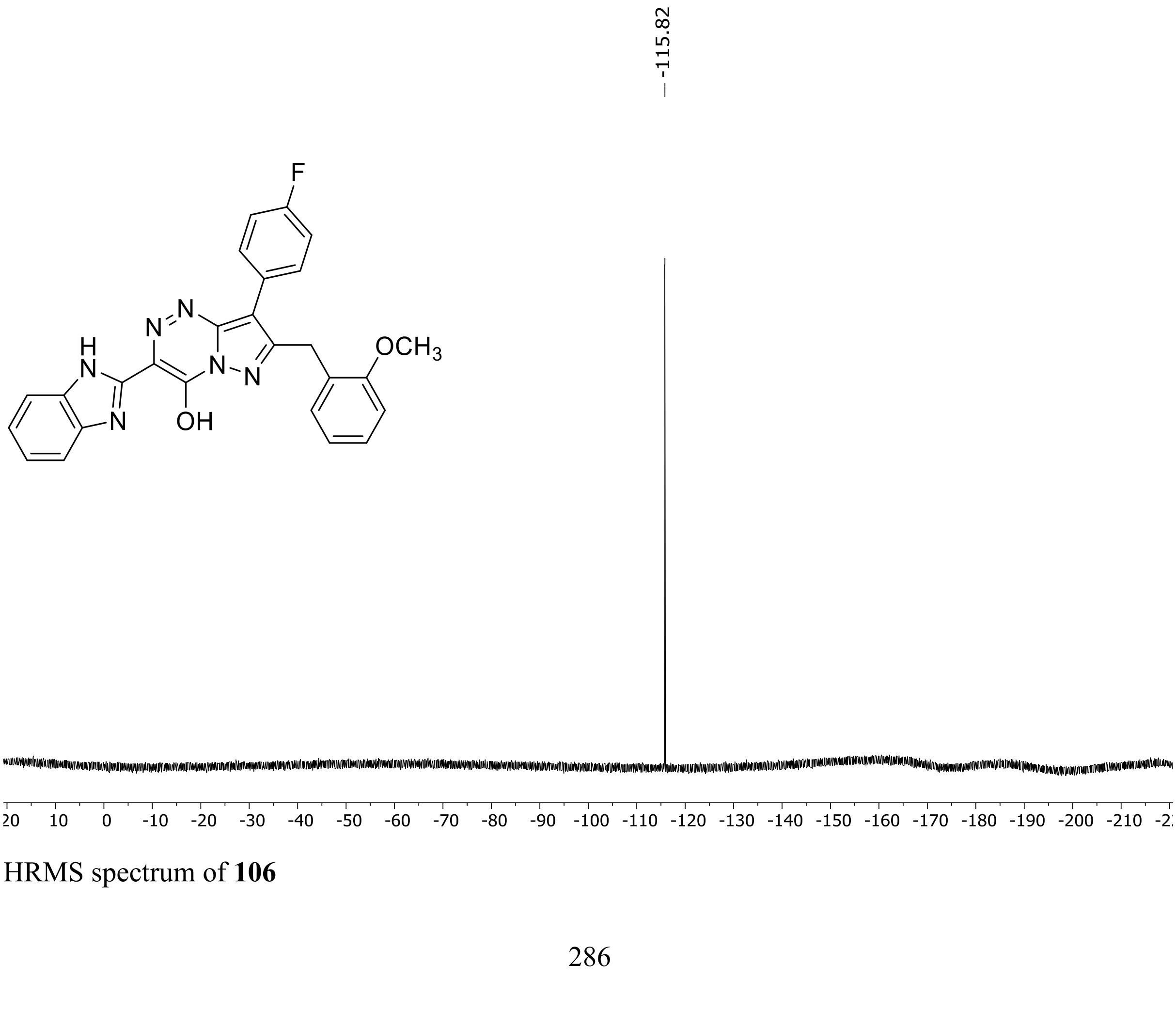

HRMS spectrum of **106**

**$C_{26}H_{19}FN_6O_2$** **exact mass: 466.1554**

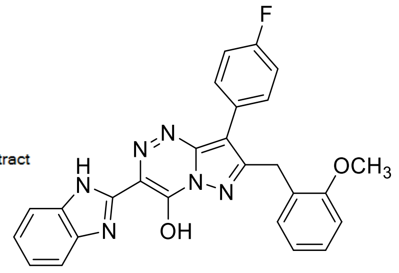

**APCI +** (MMI)
nitrogen flow 3 L/min, gas temperature 325°C, nebulizer 45 psig, skimmer 65 V, vaporizer 200°C, fragmentor 20 V, dissolved in methanol

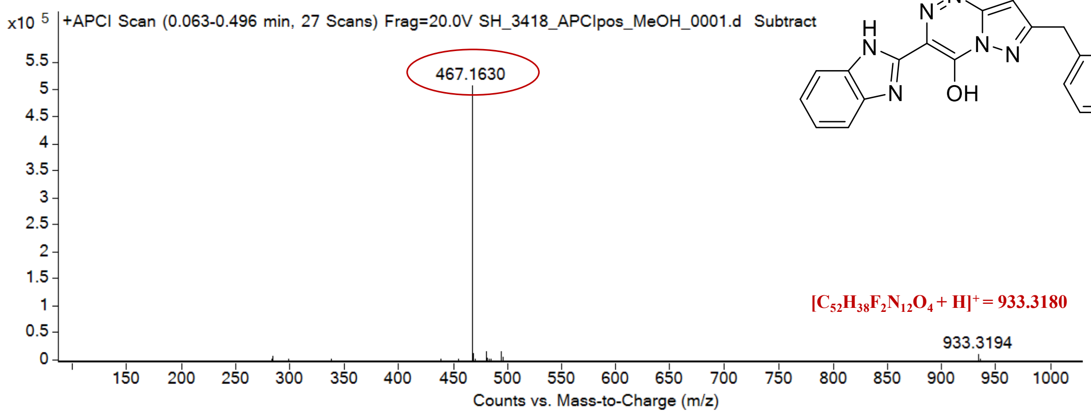

expected mass: $[M+H]^+ = 467.1626$ observed mass: $[M+H]^+ = 467.1630$ mass accuracy = 0.9 ppm

[1]H (500 MHz) and [13]C NMR (126 MHz) spectra of **107** in DMSO-$d_6$

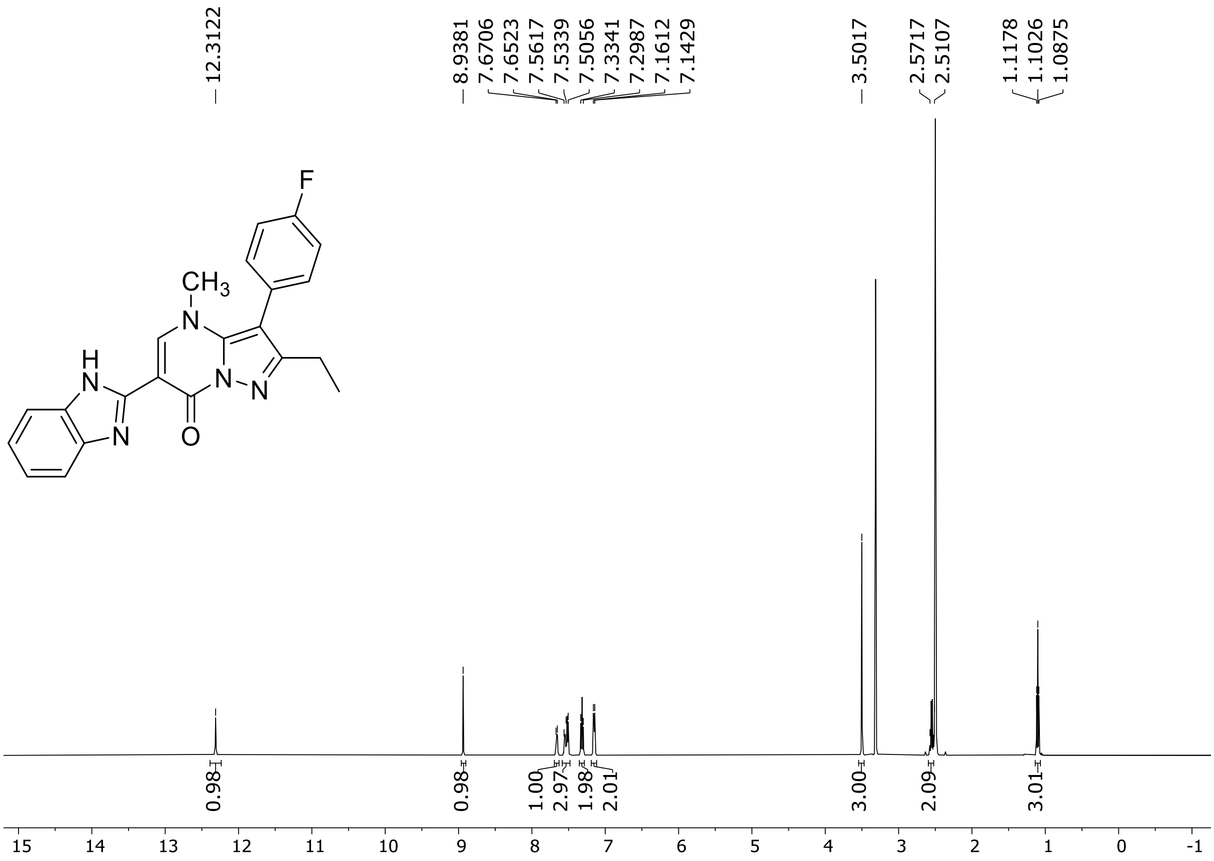

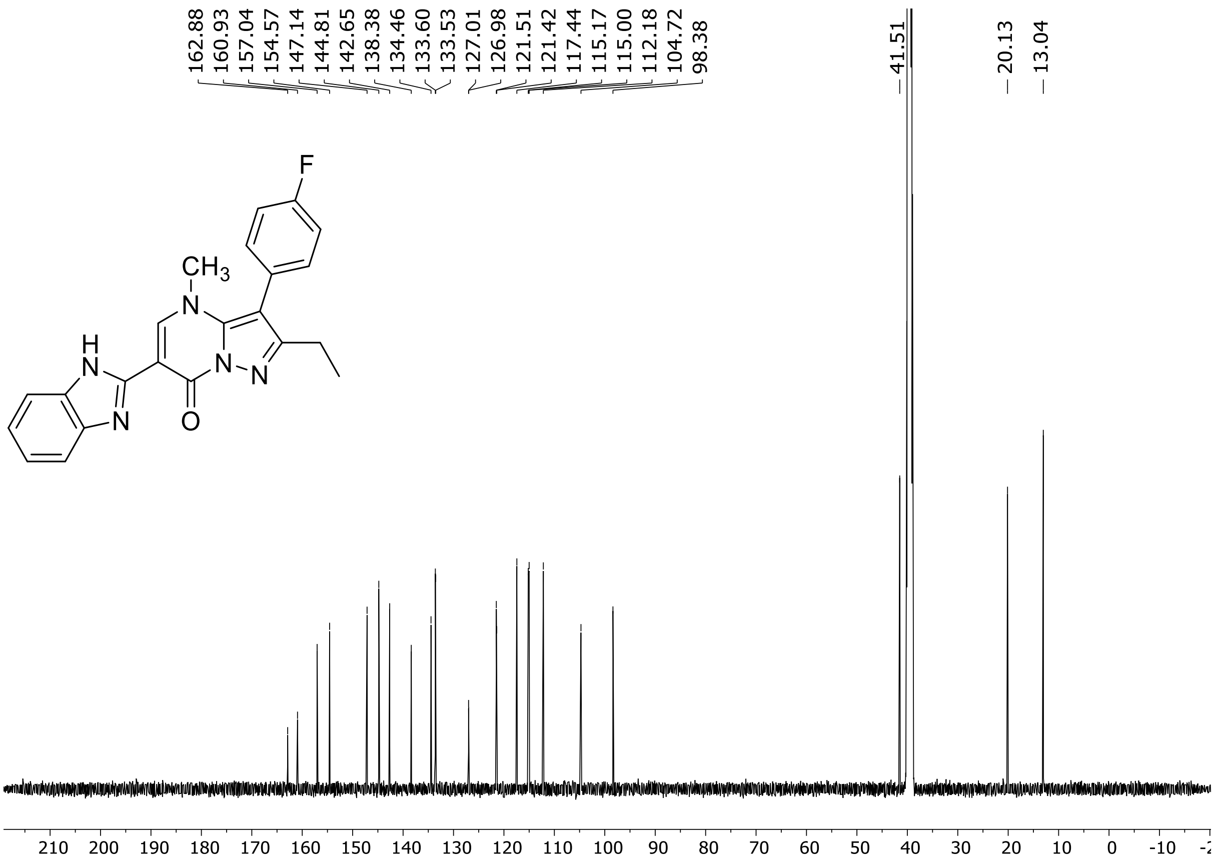

[19]F (471 MHz) NMR spectrum of **107** in DMSO-$d_6$

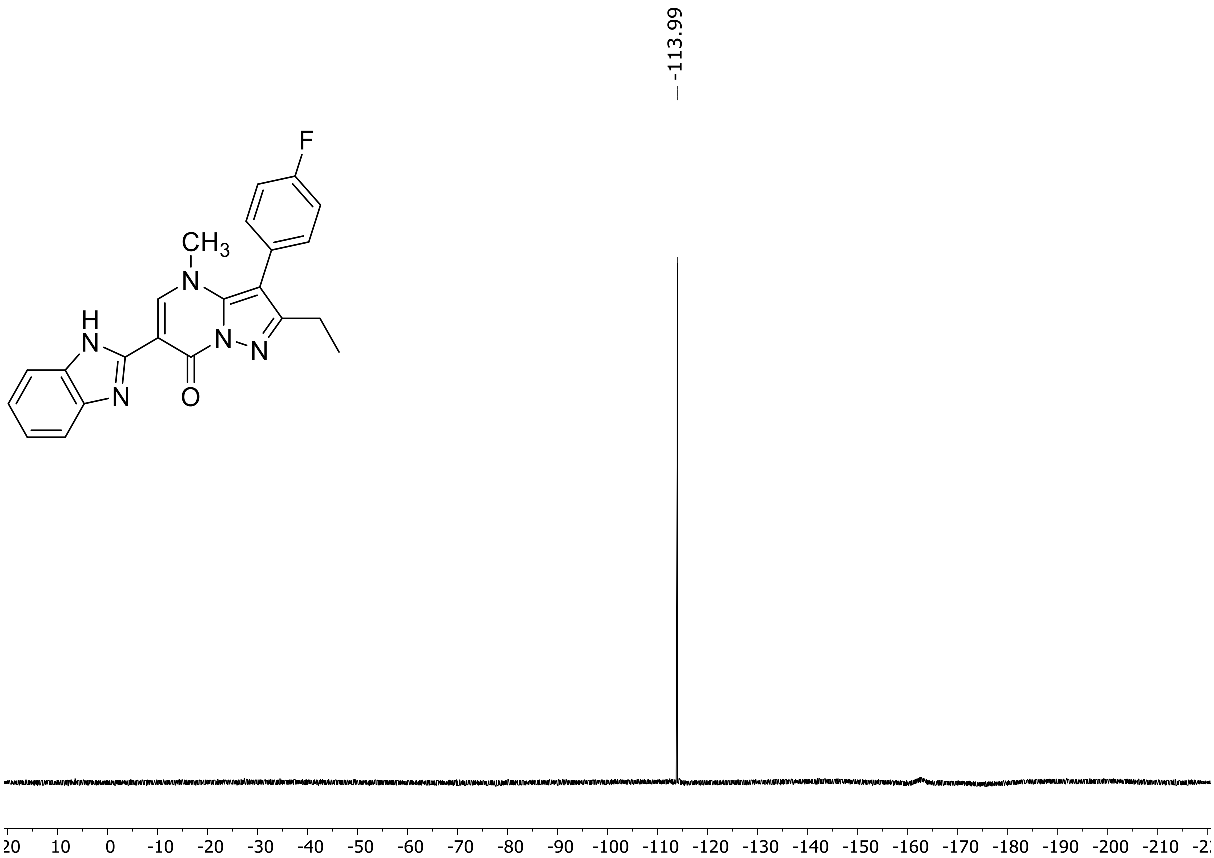

HRMS spectrum of **107**

$C_{22}H_{18}FN_5O$ **exact mass: 387.1495**

**APCI +** (MMI)
nitrogen flow 5 L/min, gas temperature 325°C, nebulizer 45 psig,
skimmer 65 V, vaporizer 200°C, fragmentor 15 V, dissolved in methanol

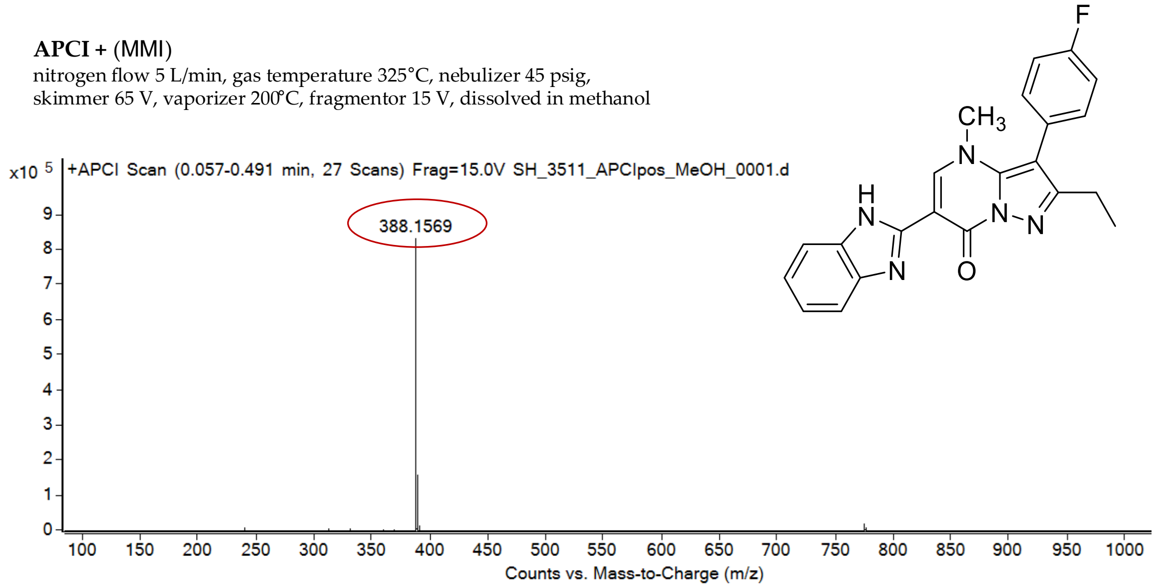

expected mass: [M+H]$^+$ = 388.1568 observed mass: [M+H]$^+$ = 388.1569 mass accuracy = 0.3 ppm

$^1$H (500 MHz) and $^{13}$C NMR (126 MHz) spectra of **108** in DMSO-$d_6$

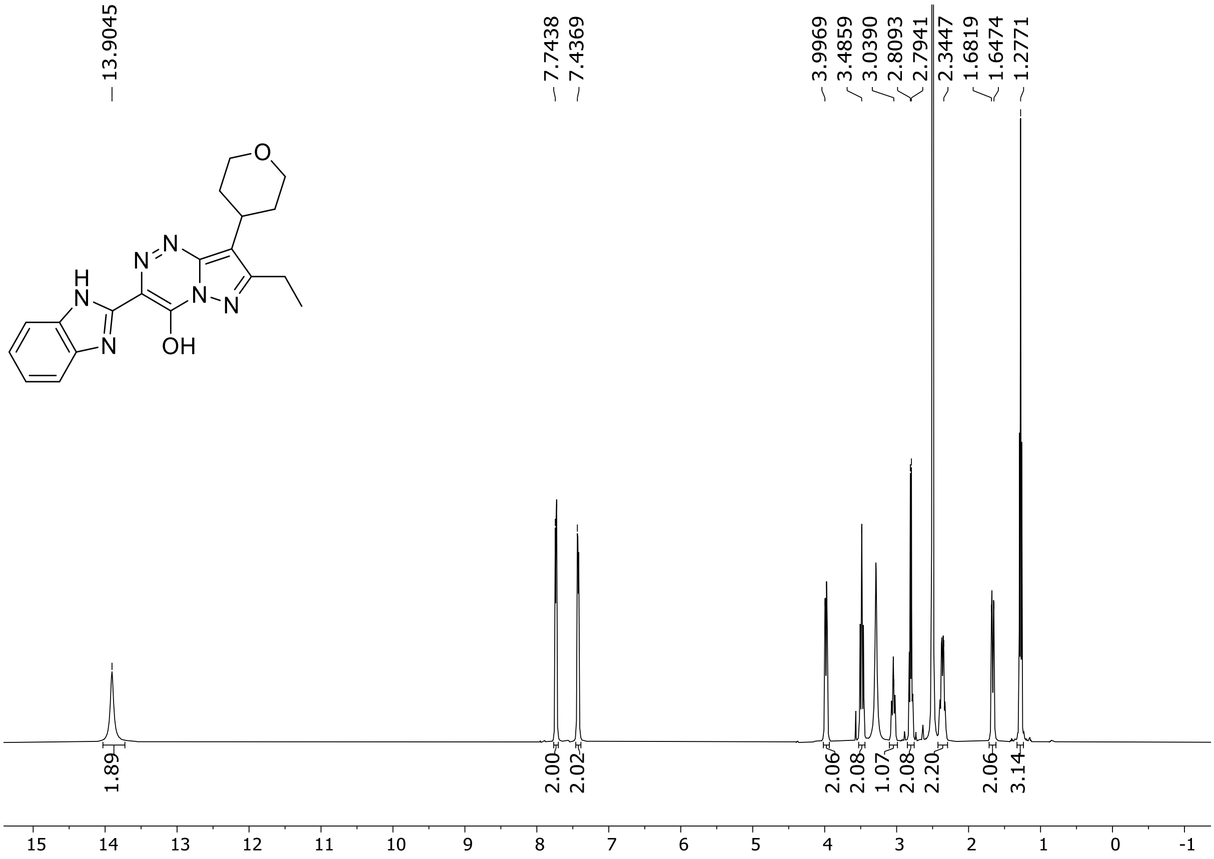

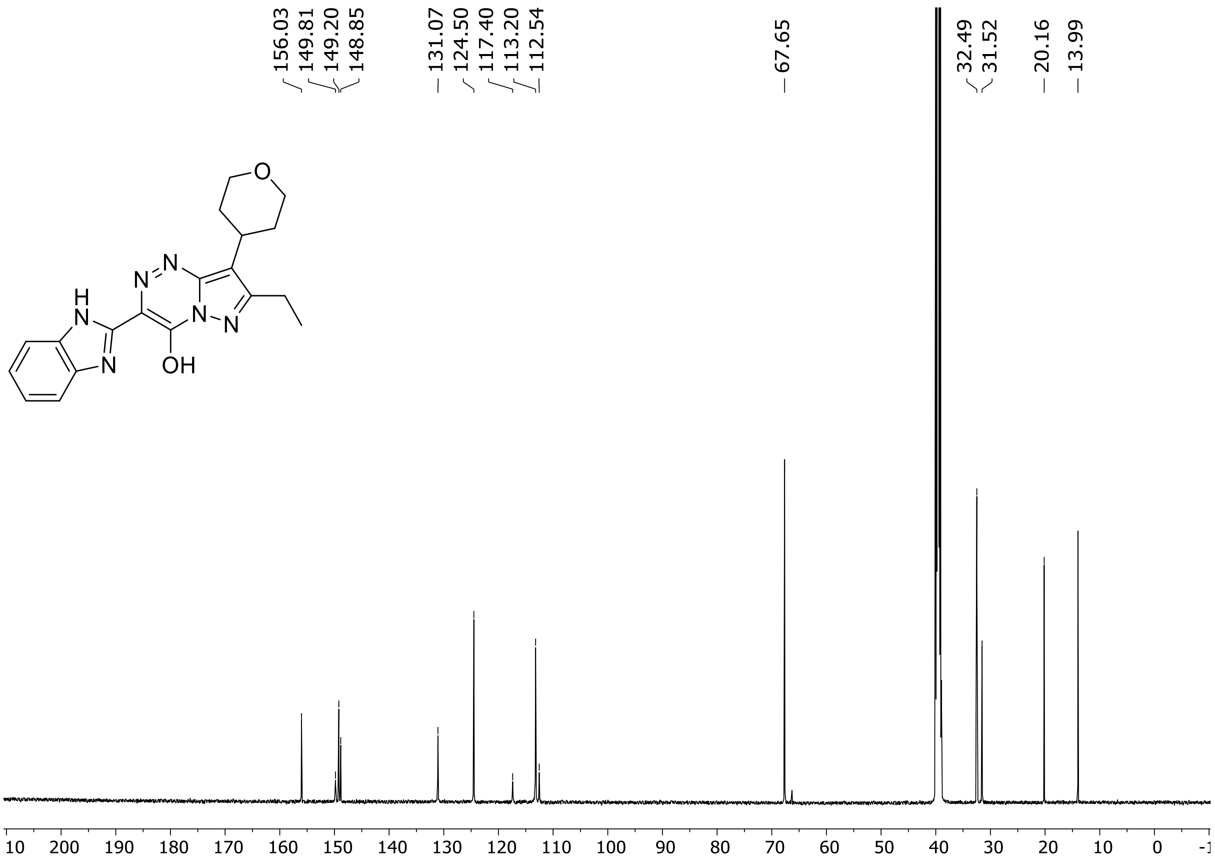

HRMS spectrum of **108**

**$C_{19}H_{20}N_6O_2$** **exact mass: 364.1648**

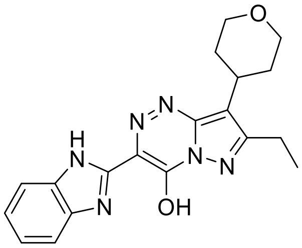

**APCI +** (MMI)
nitrogen flow 5 L/min, gas temperature 325°C, nebulizer 45 psig, skimmer 65 V, vaporizer 200°C, fragmentor 20 V, dissolved in methanol

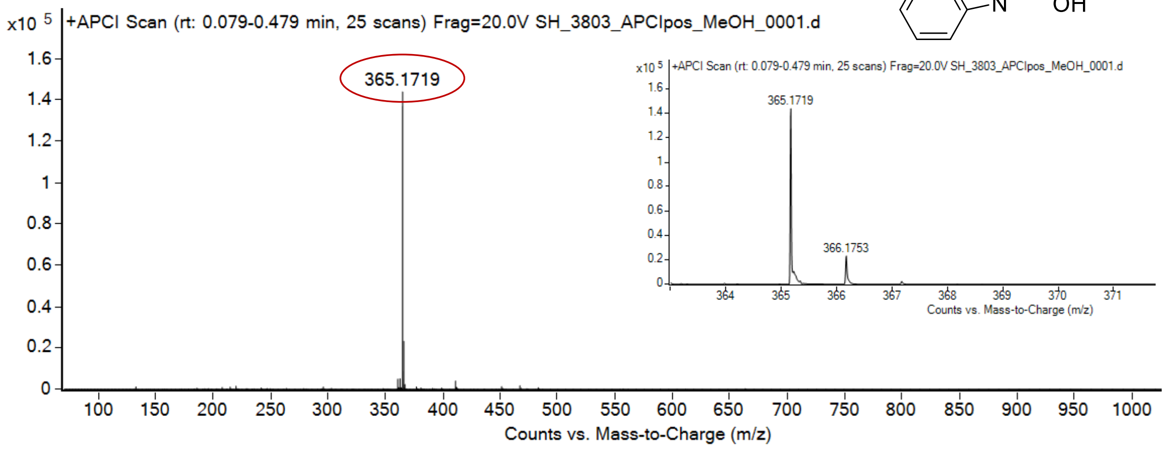

expected mass: $[M+H]^+$ = 365.1721 observed mass : $[M+H]^+$ = 365.1719 mass accuracy = - 0.6 ppm

$^1$H (500 MHz) and $^{13}$C NMR (126 MHz) spectra of **109** in DMSO-$d_6$

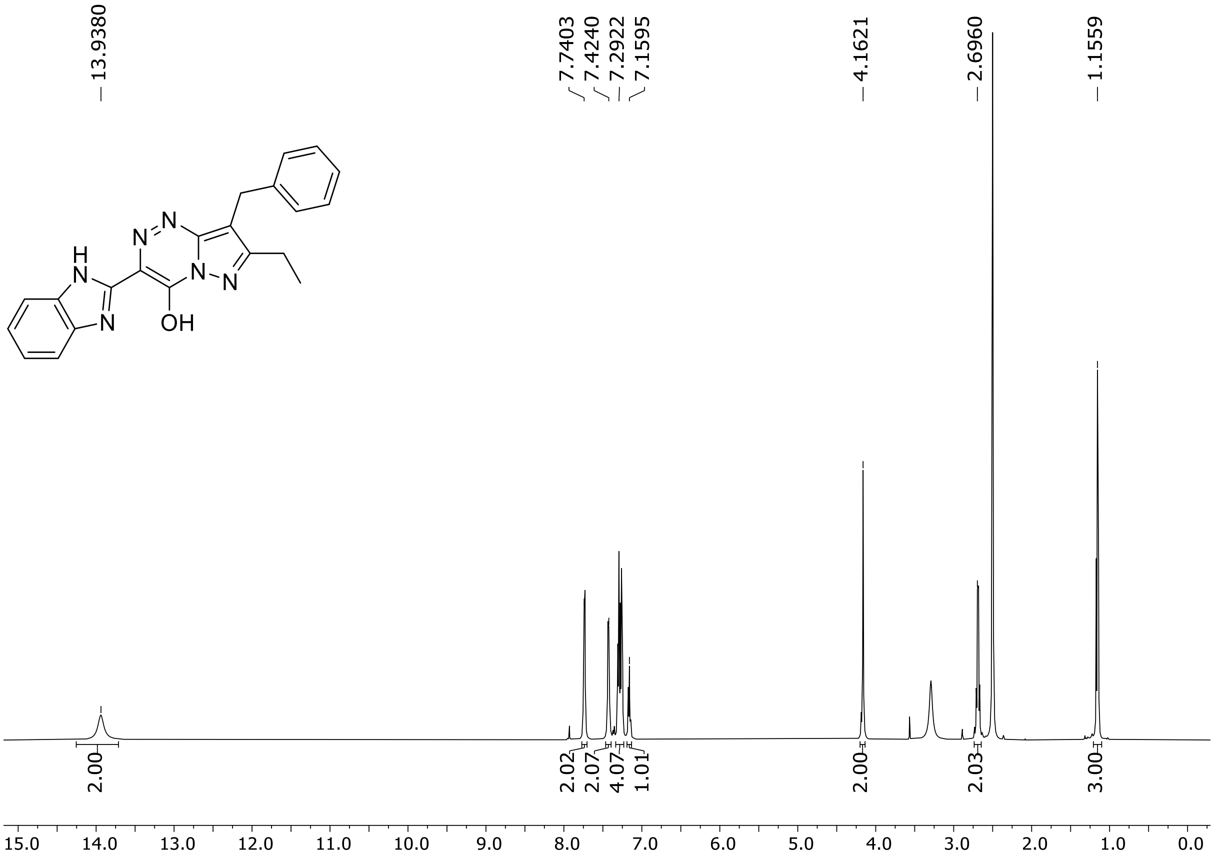

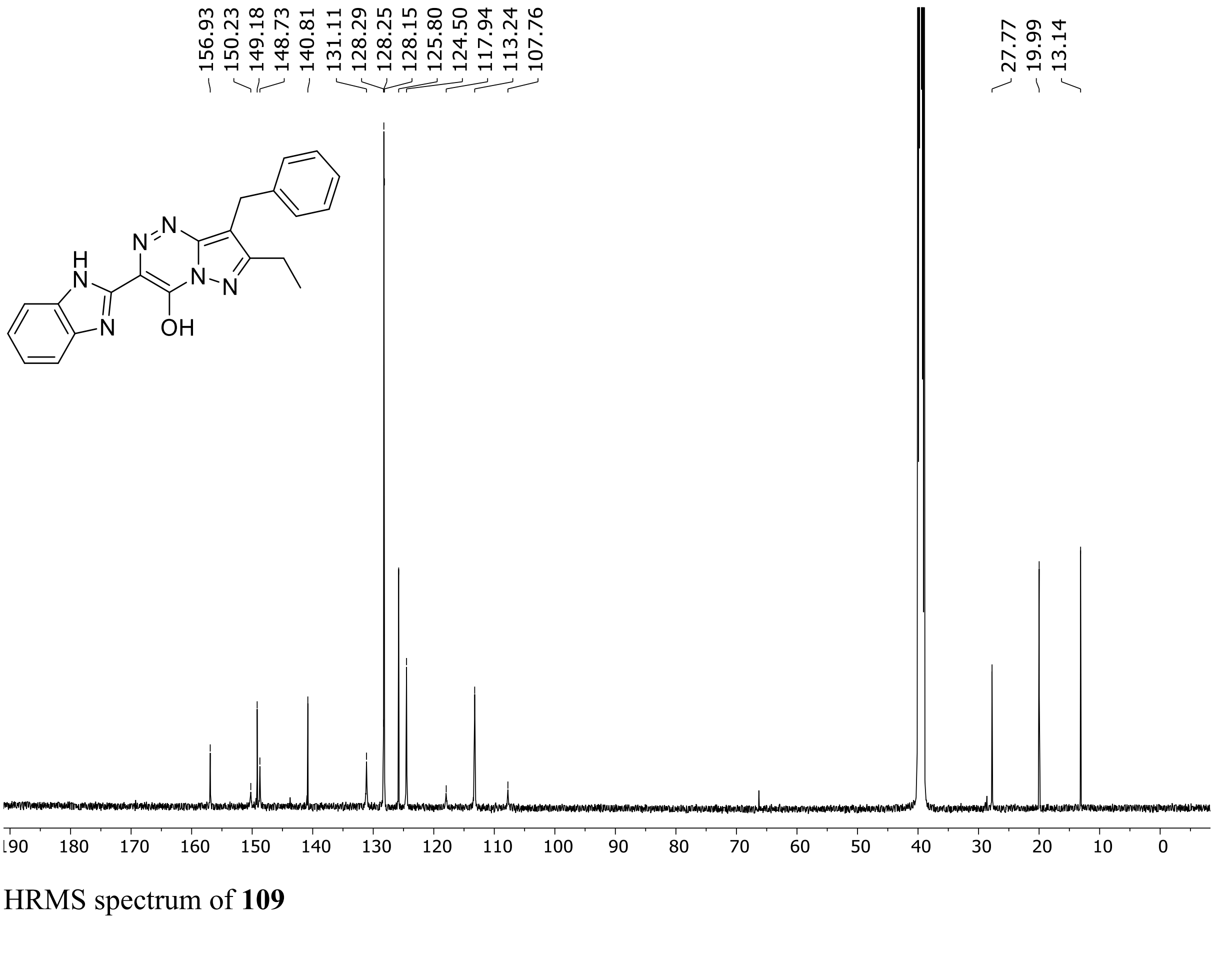

HRMS spectrum of **109**

**$C_{21}H_{18}N_6O$** **exact mass: 370.1542**

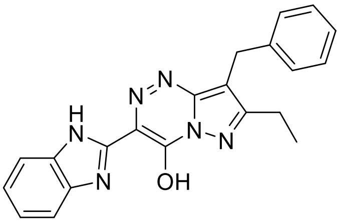

**APCI +** (MMI)
nitrogen flow 5 L/min, gas temperature 325°C, nebulizer 45 psig, skimmer 65 V, vaporizer 200°C, fragmentor 20 V, dissolved in methanol

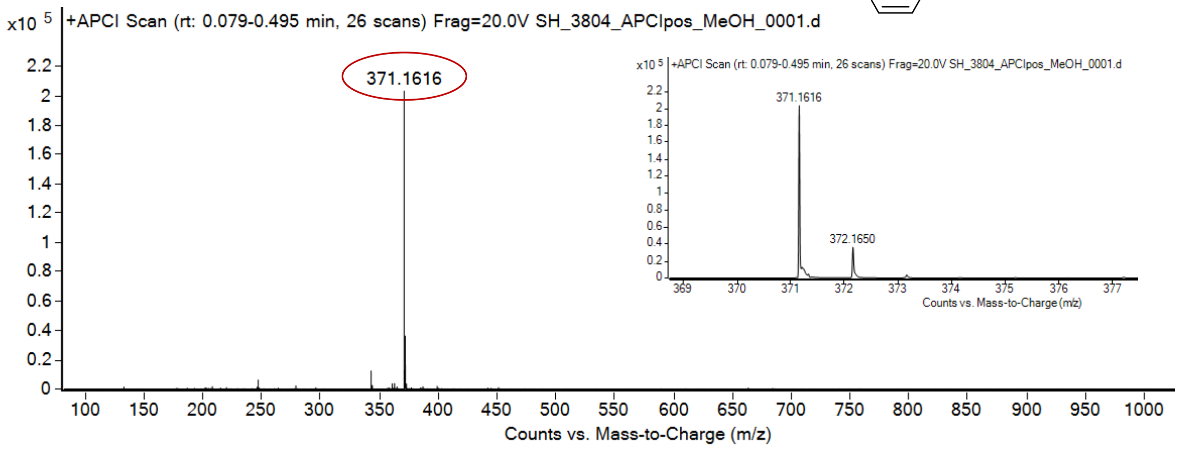

expected mass: $[M+H]^+$ = 371.1615 observed mass : $[M+H]^+$ = 371.1616 mass accuracy = 0.3 ppm

$^{1}$H (500 MHz) and $^{13}$C NMR (126 MHz) spectra of **110** in DMSO-$d_6$

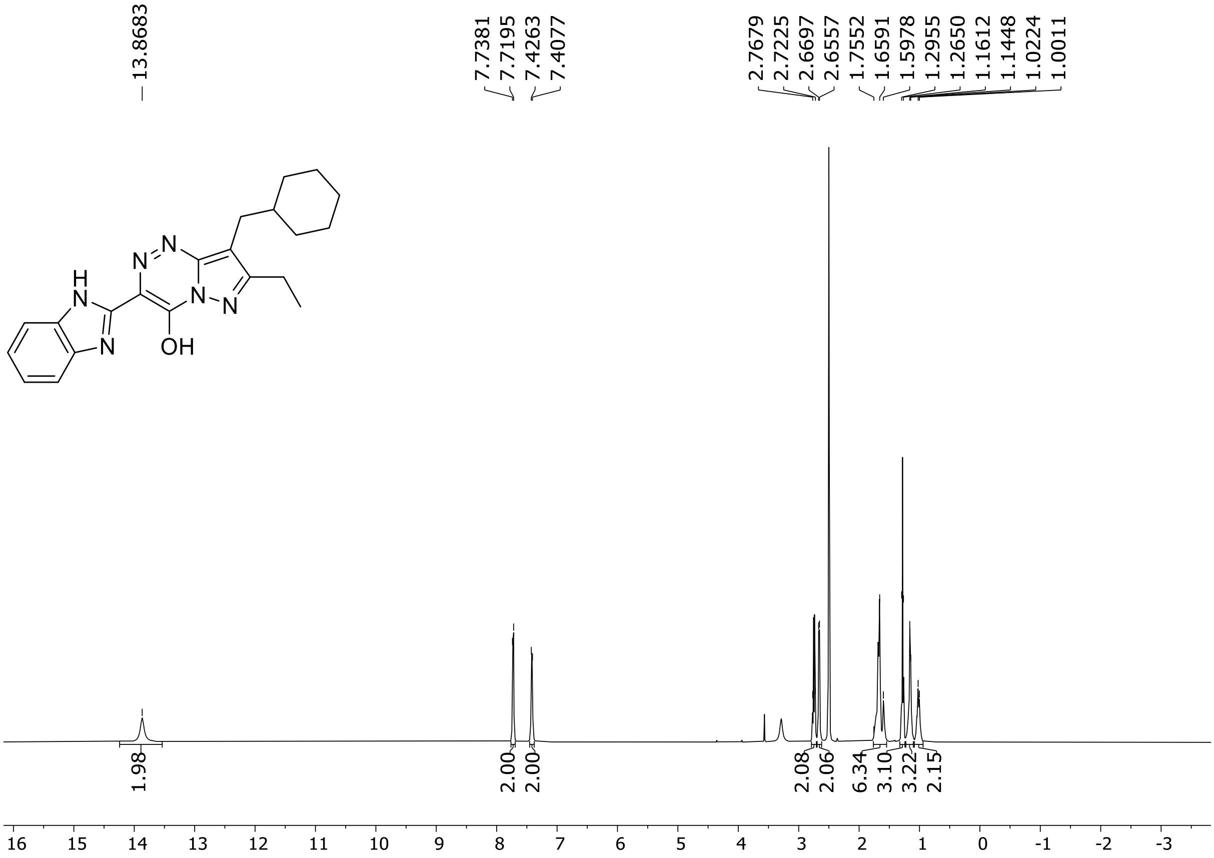

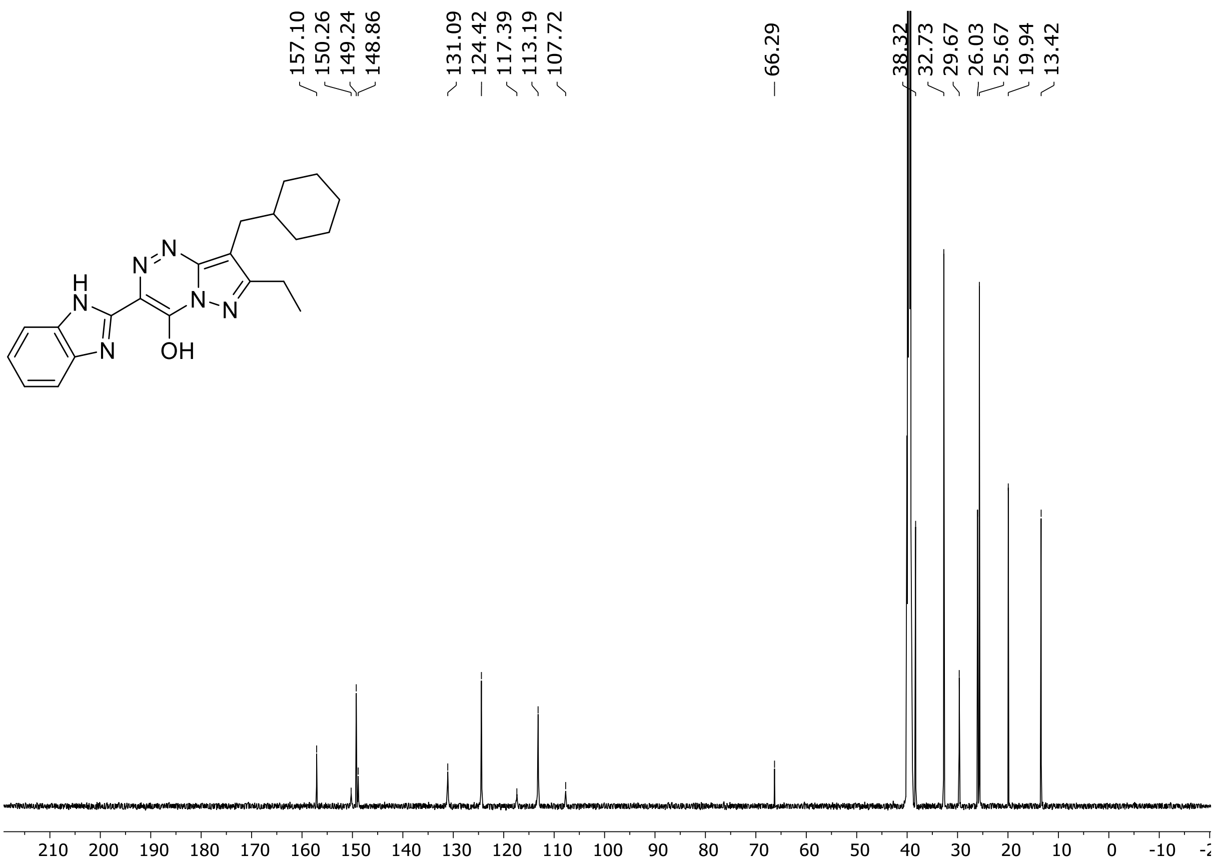

HRMS spectrum of **110**

**$C_{21}H_{24}N_6O$** **exact mass: 376.2012**

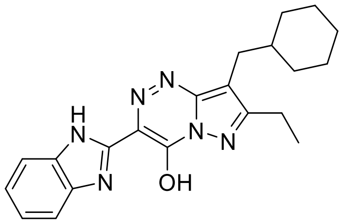

**APCI +** (MMI)
nitrogen flow 5 L/min, gas temperature 325°C, nebulizer 45 psig, skimmer 65 V, vaporizer 200°C, fragmentor 20 V, dissolved in methanol

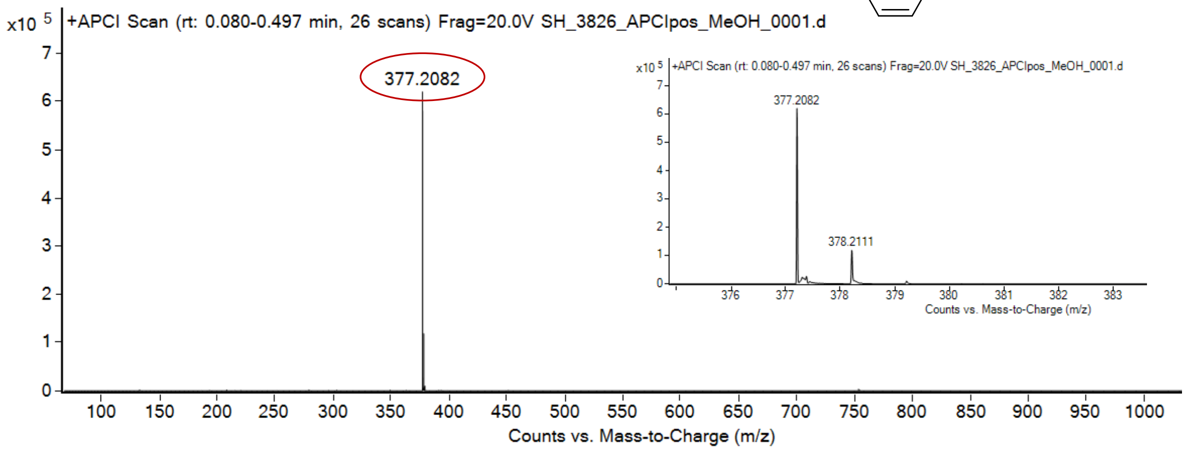

expected mass: $[M+H]^+$ = 377.2084 observed mass : $[M+H]^+$ = 377.2082 mass accuracy = - 0.5 ppm

[1]H (500 MHz) and [13]C NMR (126 MHz) spectra of **111** in DMSO-$d_6$

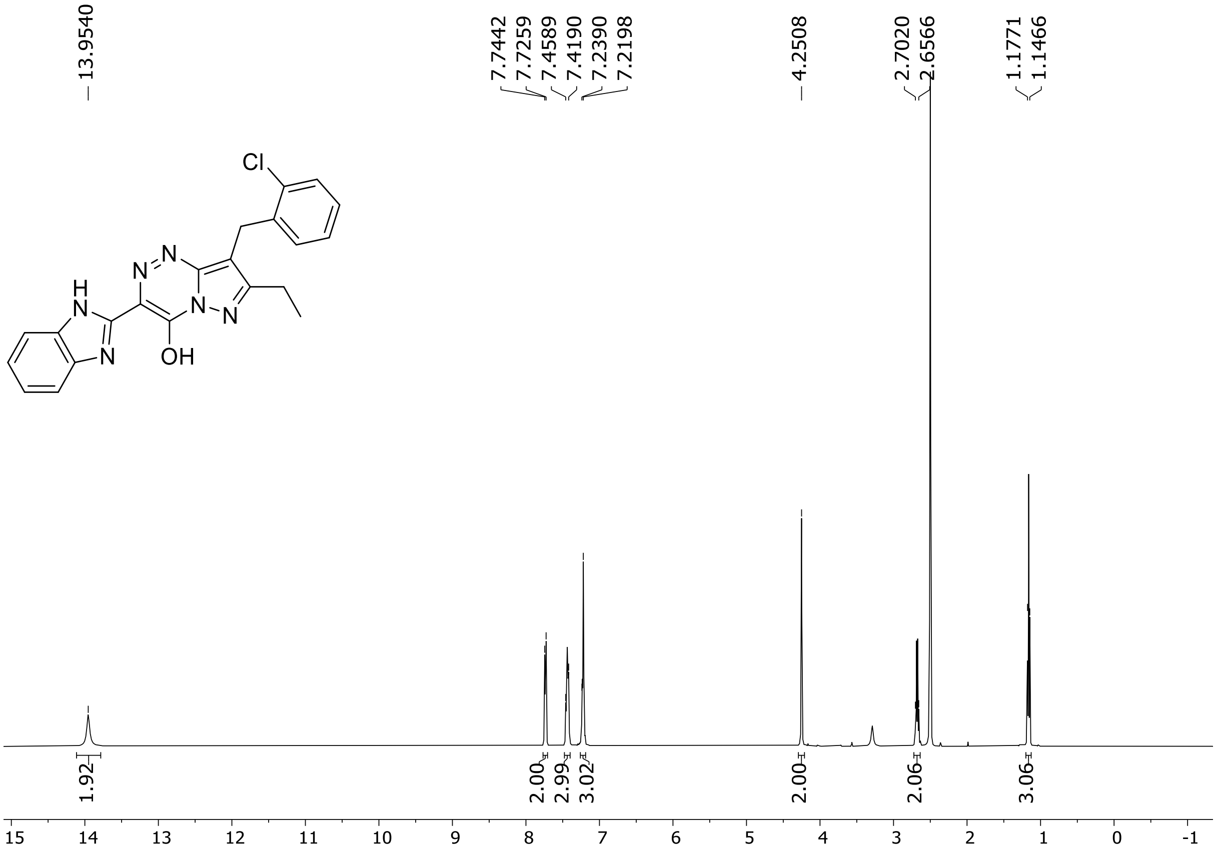

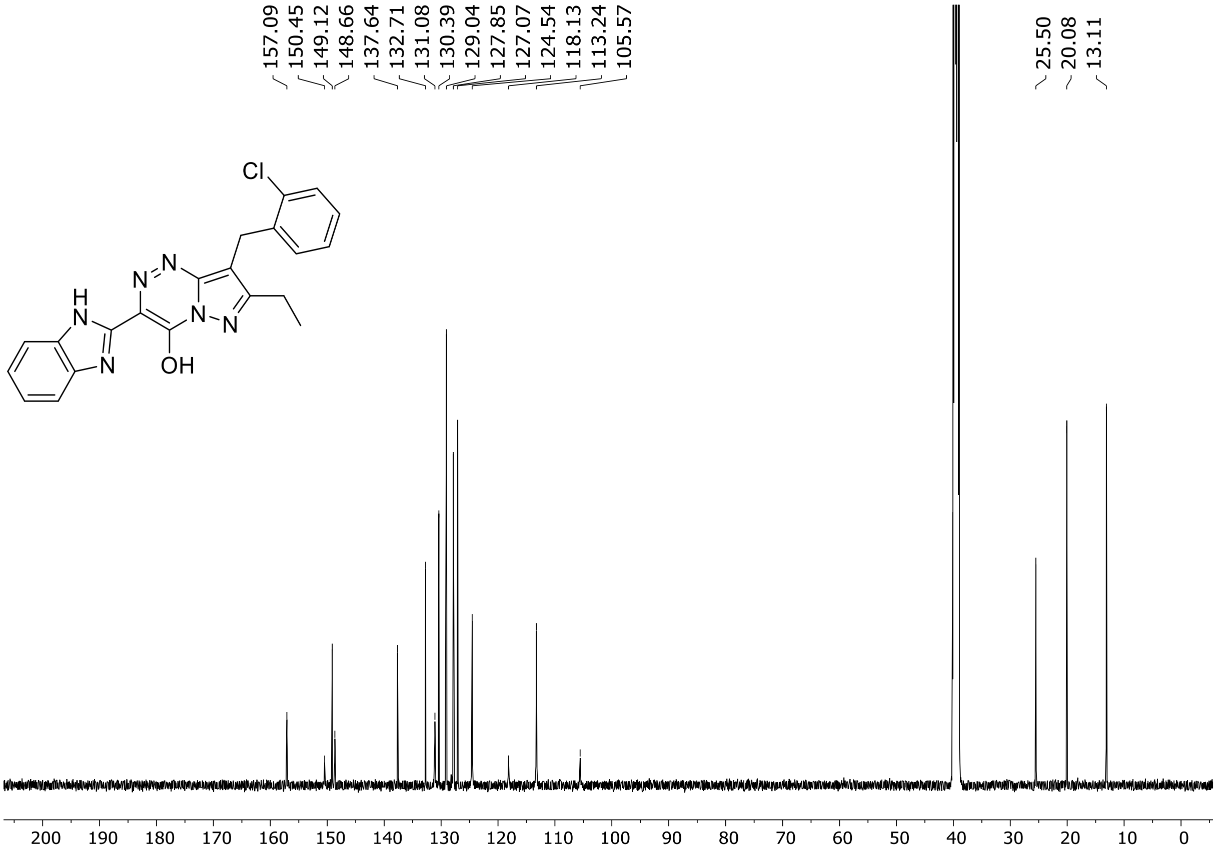

HRMS spectrum of **111**

**$C_{21}H_{17}ClN_6O$** **exact mass: 404.1152**

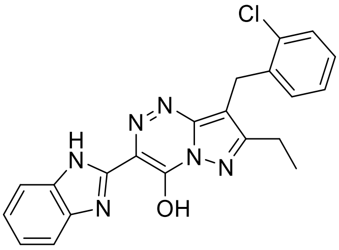

**APCI +** (MMI)
nitrogen flow 5 L/min, gas temperature 325°C, nebulizer 45 psig, skimmer 65 V, vaporizer 200°C, fragmentor 25 V, dissolved in methanol

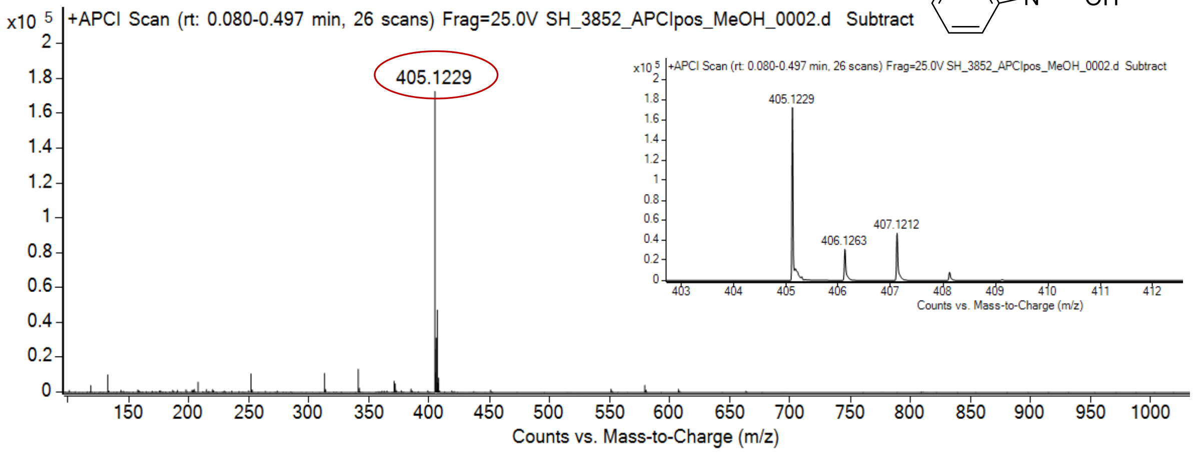

expected mass: [M+H]$^+$ = 405.1225 observed mass : [M+H]$^+$ = 405.1229 mass accuracy = 1.0 ppm

[1]H (500 MHz) and [13]C NMR (126 MHz) spectra of **112** in DMSO-$d_6$

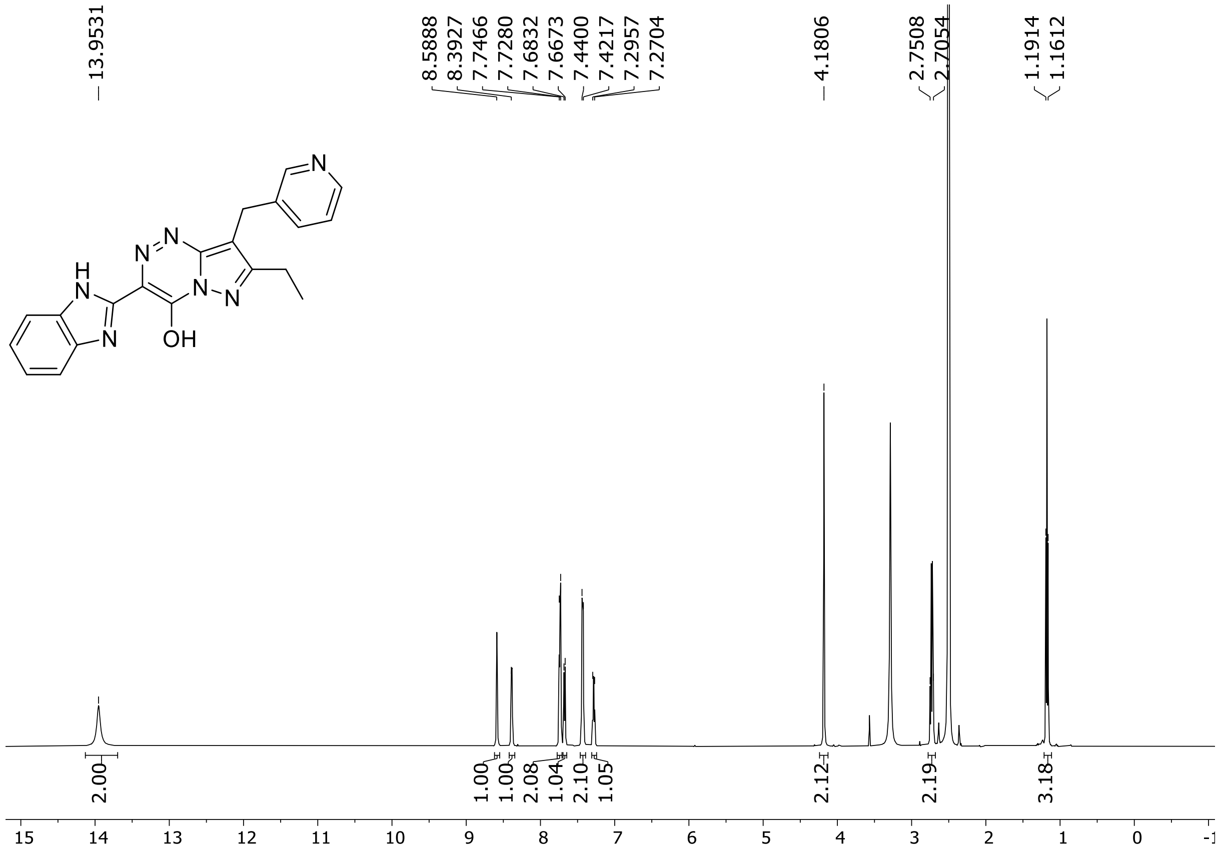

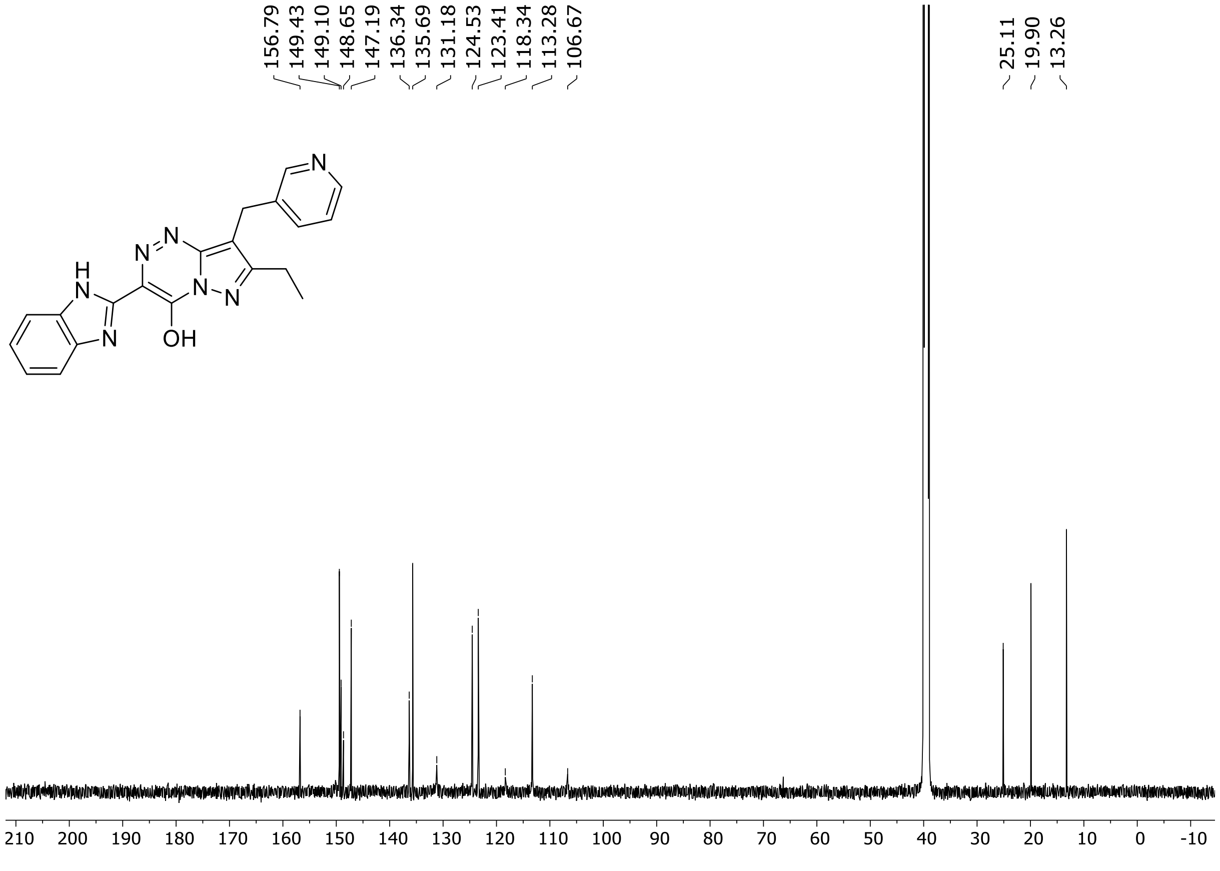

HRMS spectrum of **112**

**$C_{20}H_{17}N_7O$** **exact mass: 371.1495**

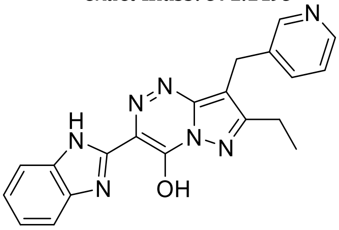

**APCI +** (MMI)
nitrogen flow 5 L/min, gas temperature 325°C, nebulizer 45 psig, skimmer 65 V, vaporizer 200°C, fragmentor 20 V, dissolved in methanol

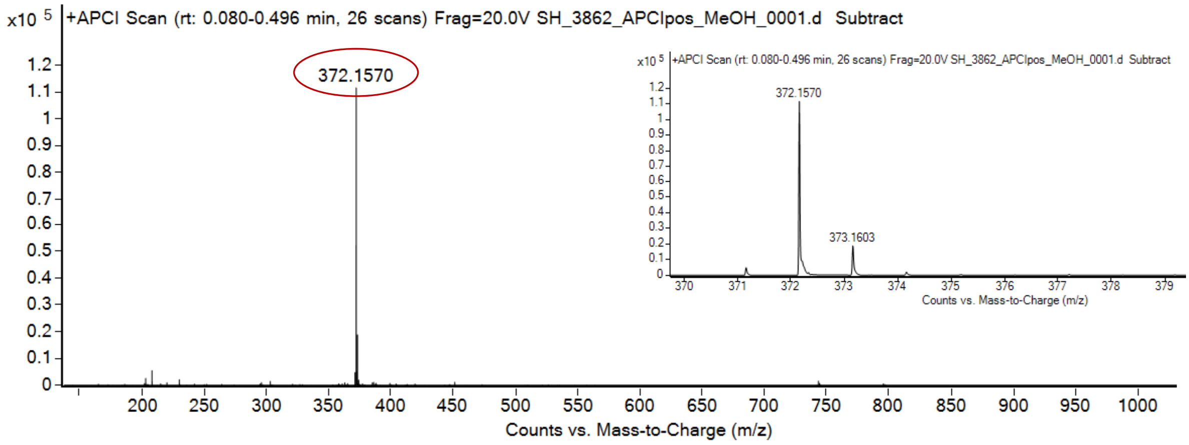

expected mass: $[M+H]^+$ = 372.1567 observed mass : $[M+H]^+$ = 372.1570 mass accuracy = 0.8 ppm

$^{1}H$ (500 MHz) and $^{13}C$ NMR (126 MHz) spectra of **113** in DMSO-$d_6$

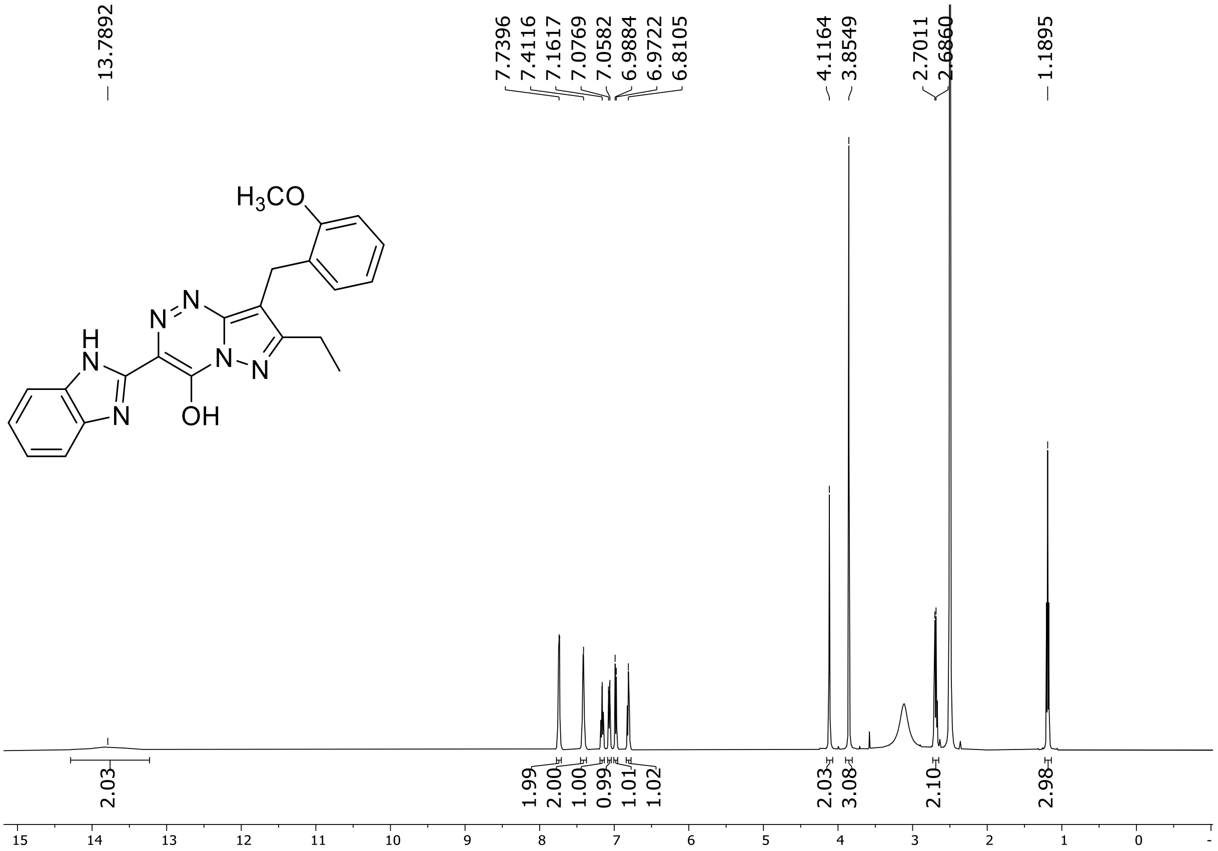

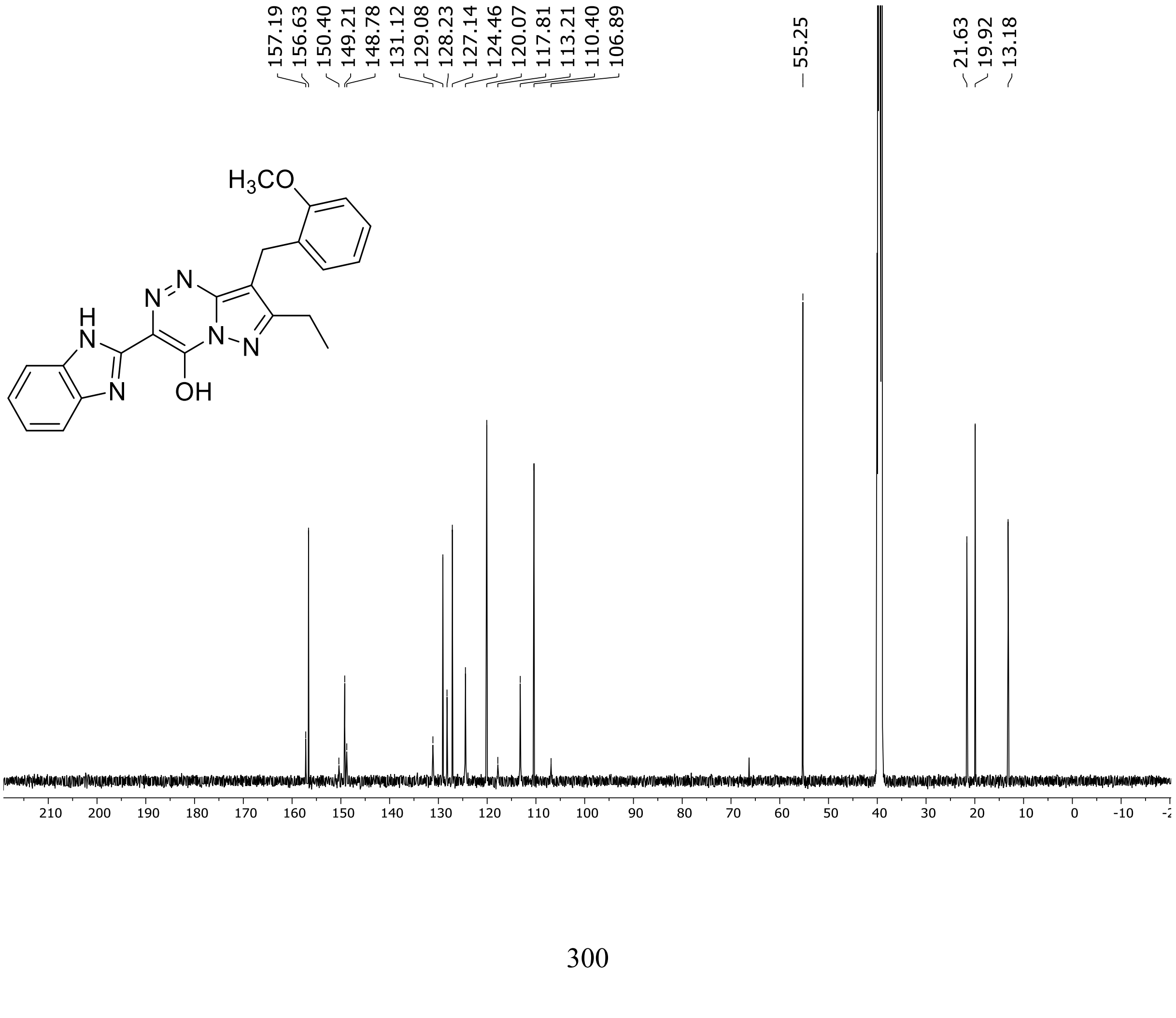

HRMS spectrum of **113**

$C_{22}H_{20}N_6O_2$ **exact mass: 400.1648**

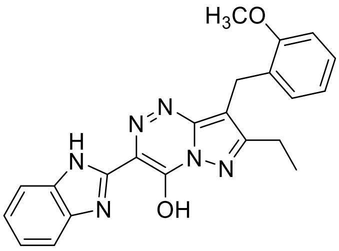

**APCI +** (MMI)
nitrogen flow 5 L/min, gas temperature 325°C, nebulizer 45 psig, skimmer 65 V, vaporizer 200°C, fragmentor 20 V, dissolved in methanol

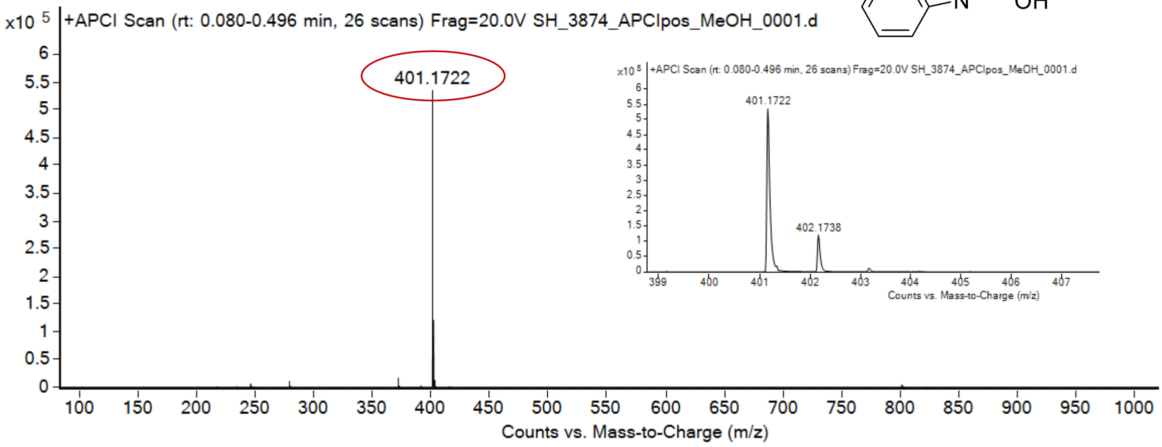

expected mass: $[M+H]^+$ = 401.1721 observed mass: $[M+H]^+$ = 401.1722 mass accuracy = 0.2 ppm

$^{1}H$ (500 MHz) and $^{13}C$ NMR (126 MHz) spectra of **114** in DMSO-$d_6$

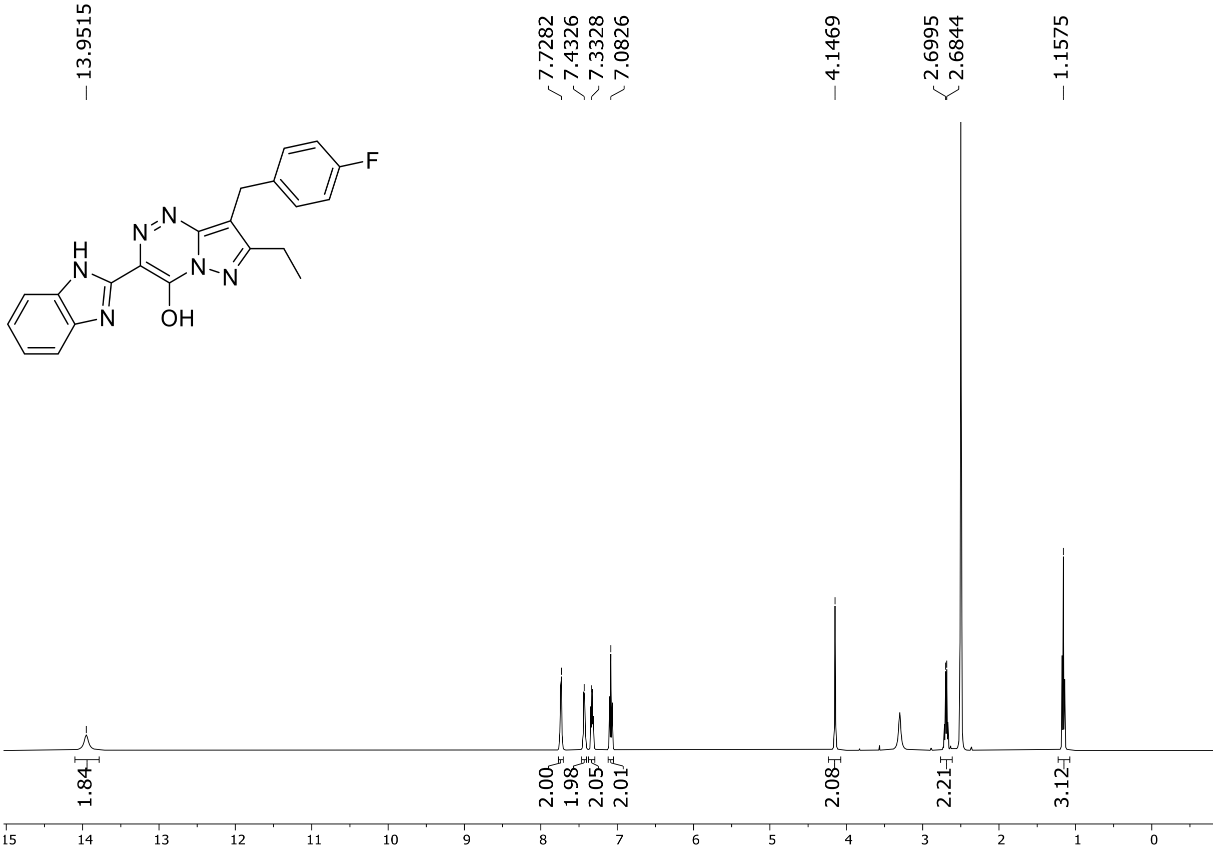

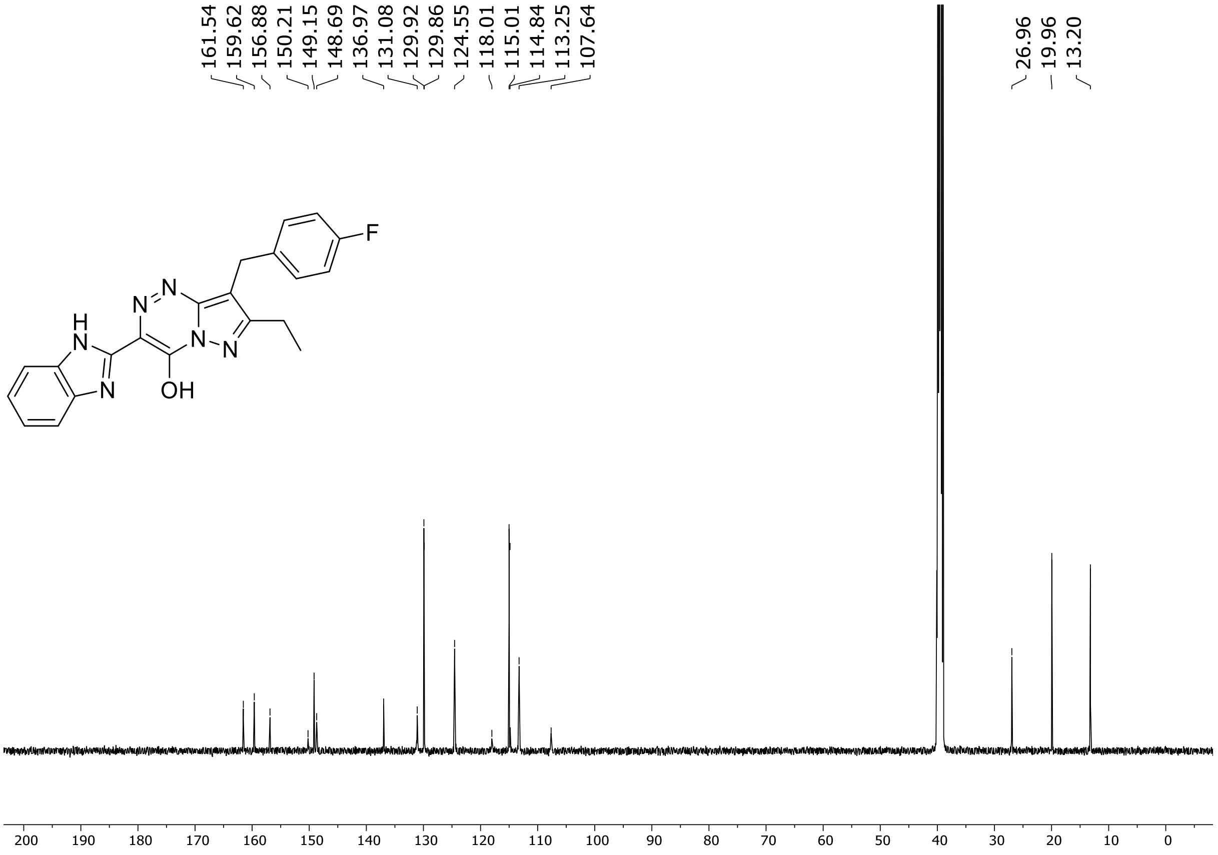

$^{19}$F (471 MHz) NMR spectrum of **114** in DMSO-$d_6$

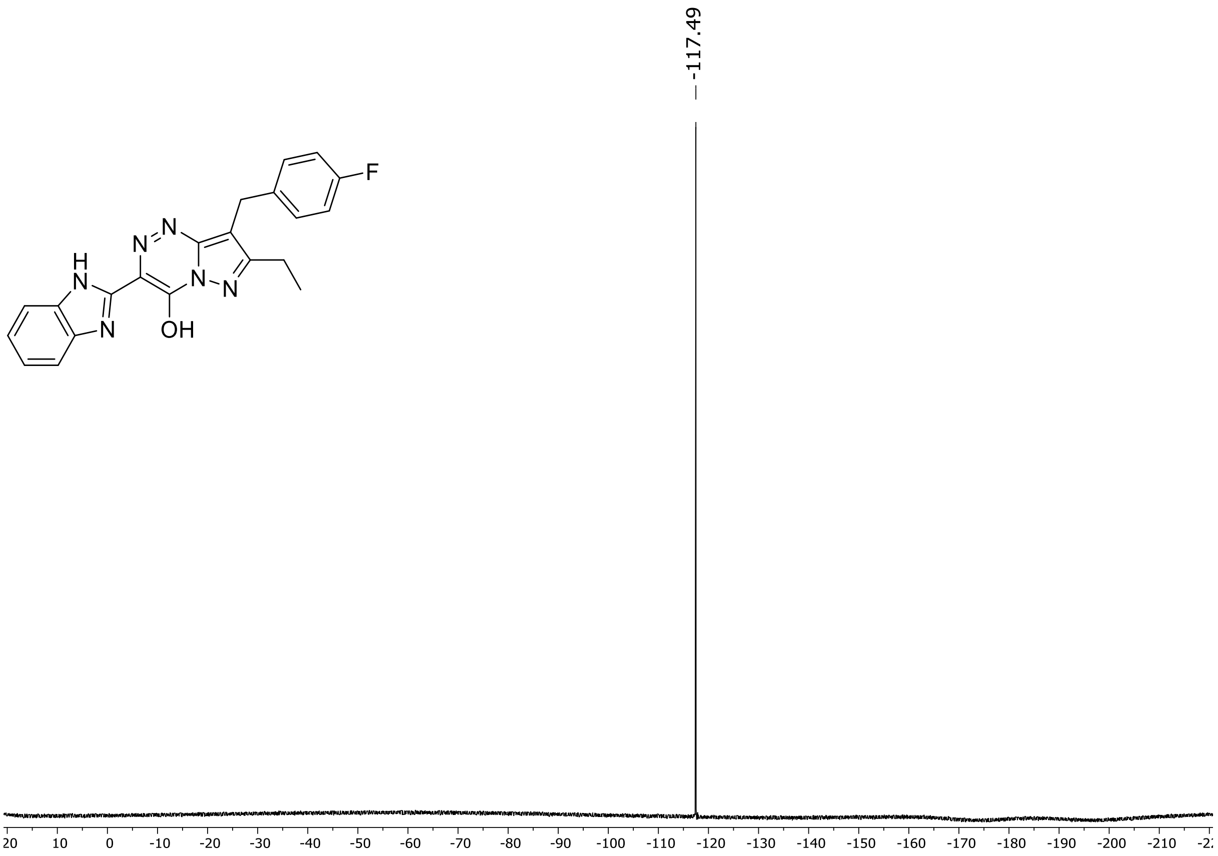

HRMS spectrum of **114**

**$C_{21}H_{17}FN_6O$** **exact mass: 388.1448**

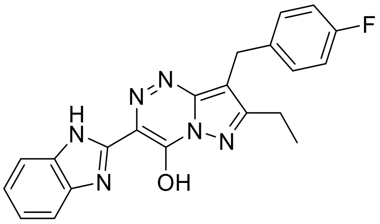

**APCI +** (MMI)
nitrogen flow 5 L/min, gas temperature 325°C, nebulizer 45 psig, skimmer 65 V, vaporizer 200°C, fragmentor 15 V, dissolved in methanol

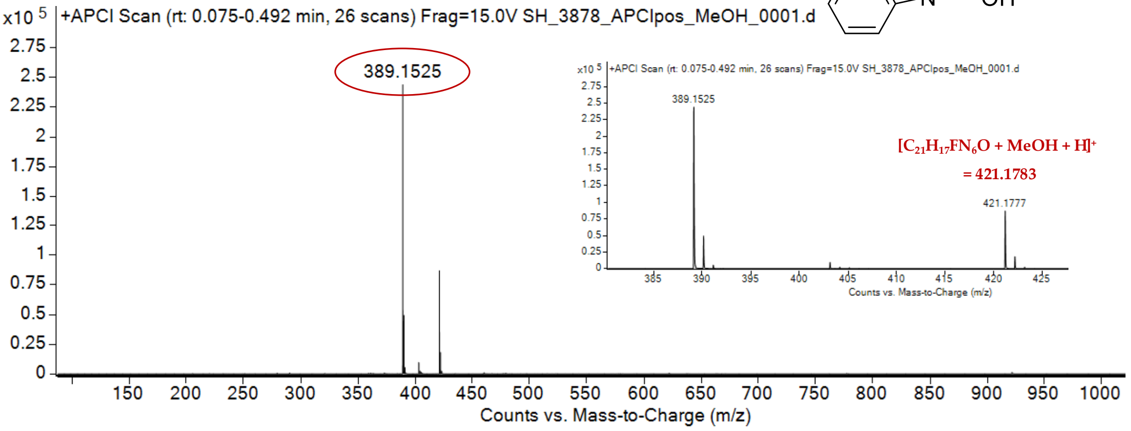

expected mass: $[M+H]^+ = 389.1521$ observed mass: $[M+H]^+ = 389.1525$ mass accuracy = 1.0 ppm

$^{1}$H (500 MHz) and $^{13}$C NMR (126 MHz) spectra of **115** in DMSO-$d_6$

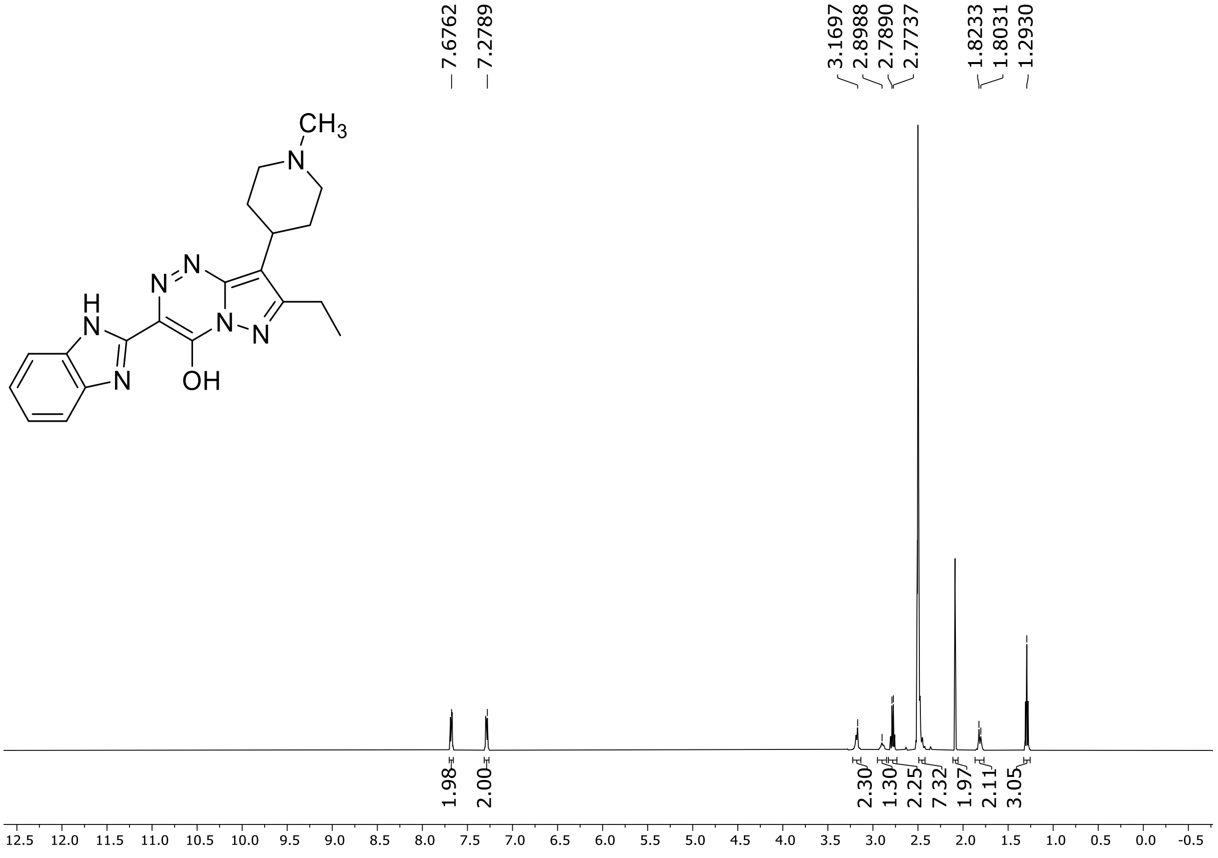

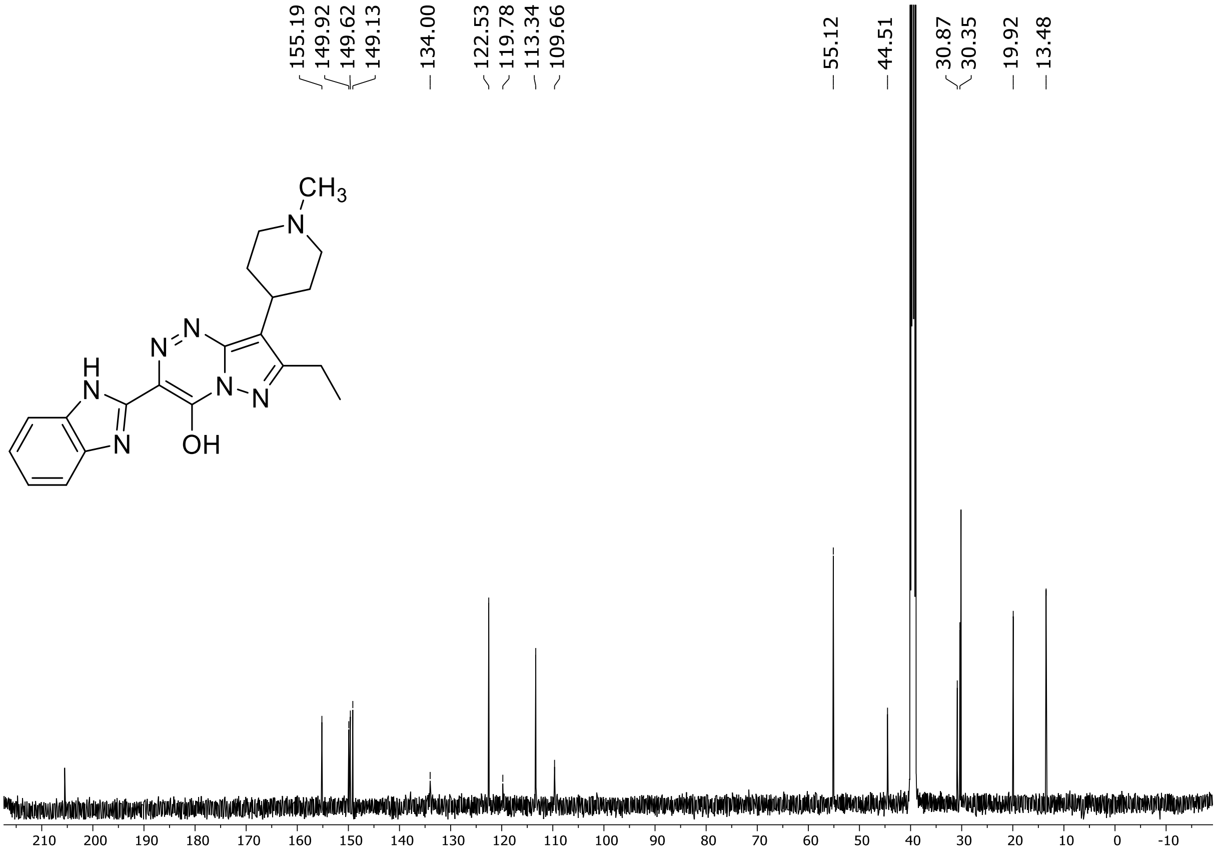

HRMS spectrum of **115**

**$C_{20}H_{23}N_7O$** **exact mass: 377.1964**

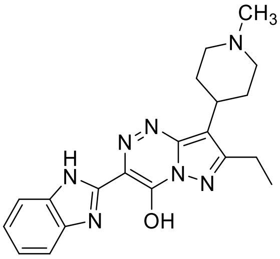

**APCI +** (MMI)
nitrogen flow 5 L/min, gas temperature 325°C, nebulizer 45 psig, skimmer 65 V, vaporizer 200°C, fragmentor 5 V, dissolved in methanol

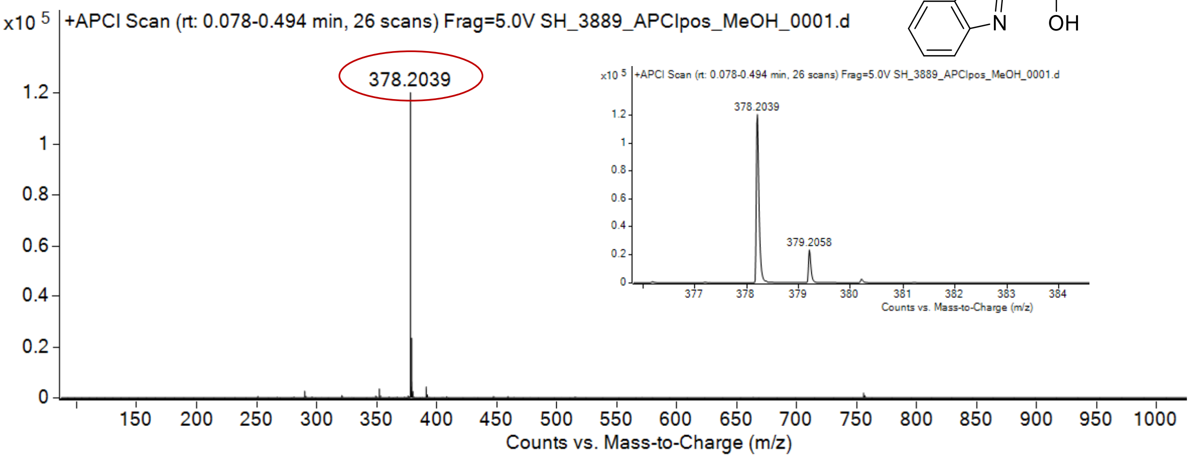

expected mass: [M+H]$^+$ = 378.2037 observed mass: [M+H]$^+$ = 378.2039 mass accuracy = 0.5 ppm

[1]H (500 MHz) and [13]C NMR (126 MHz) spectra of **116** in DMSO-$d_6$

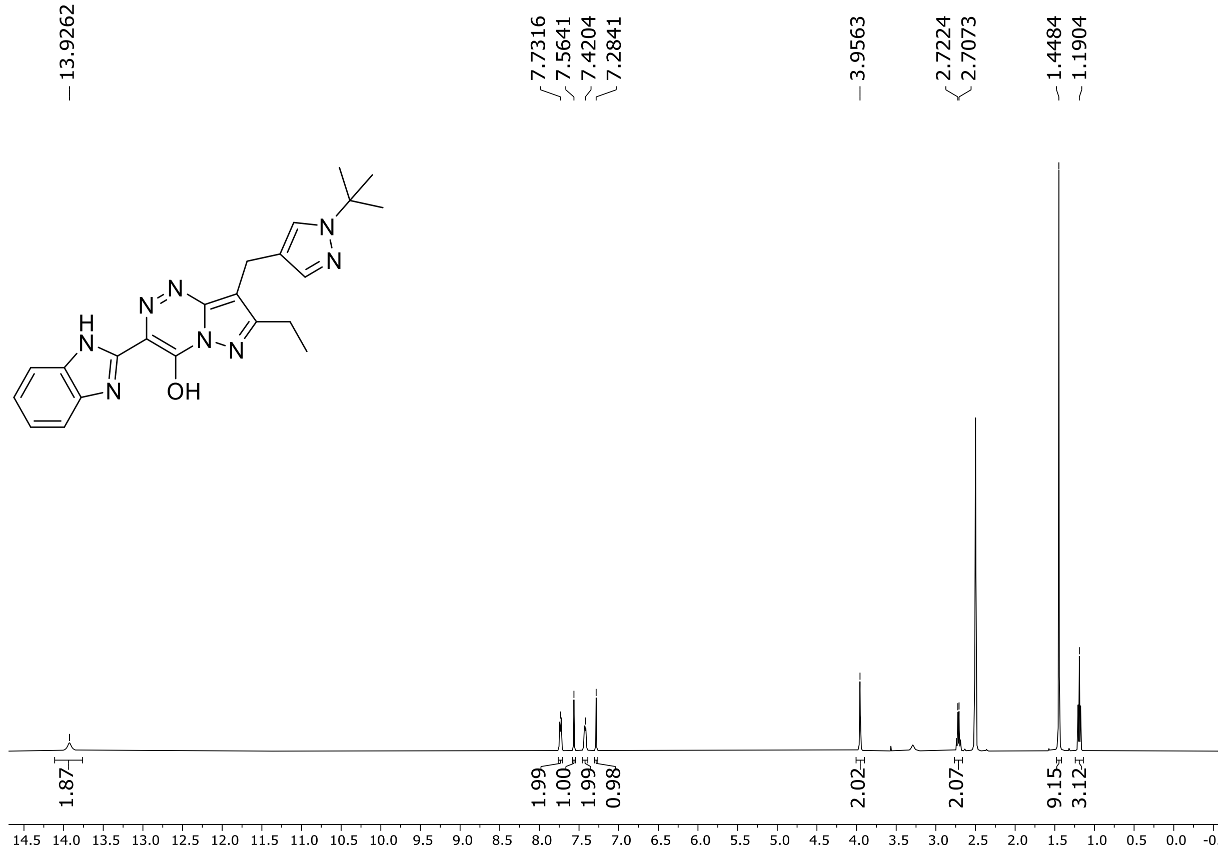

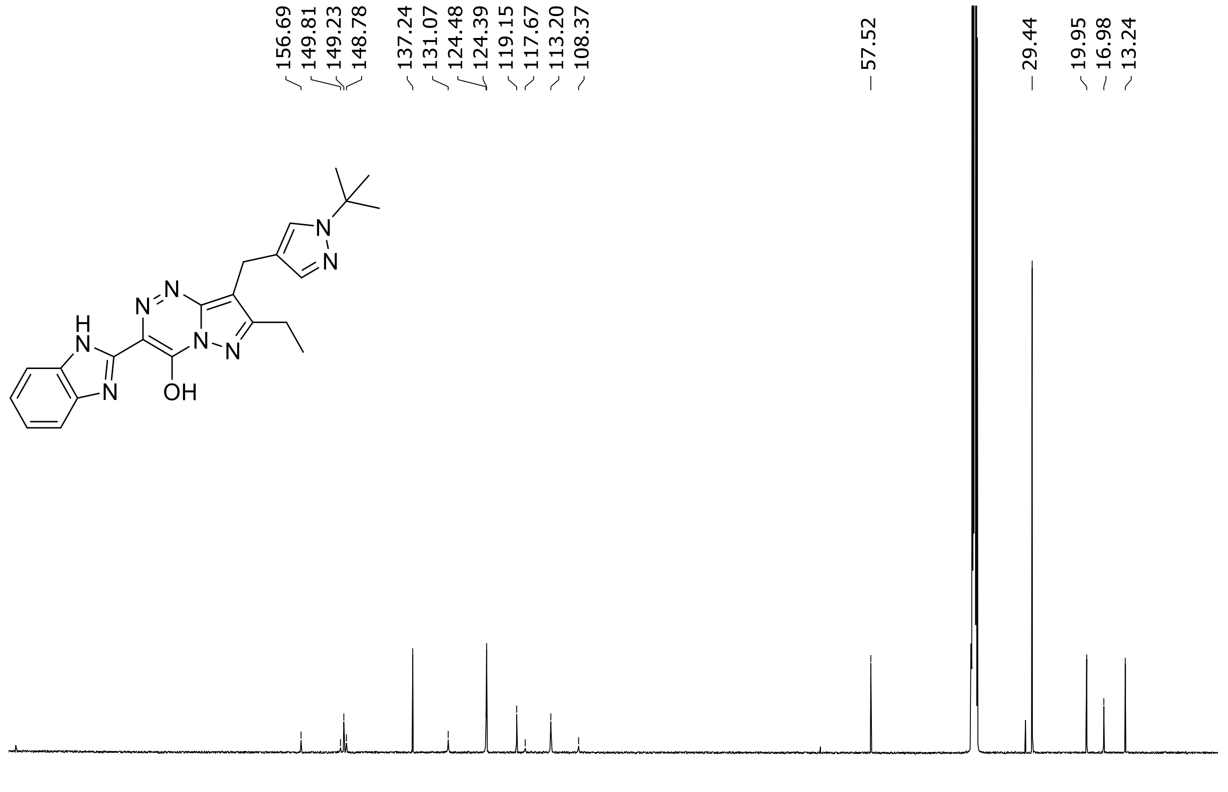

HRMS spectrum of **116**

**$C_{22}H_{24}N_8O$** **exact mass: 416.2073**

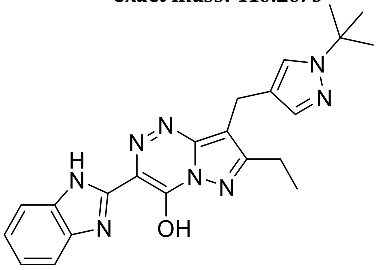

**APCI +** (MMI)
nitrogen flow 5 L/min, gas temperature 325°C, nebulizer 45 psig, skimmer 65 V, vaporizer 200°C, fragmentor 10 V, dissolved in methanol

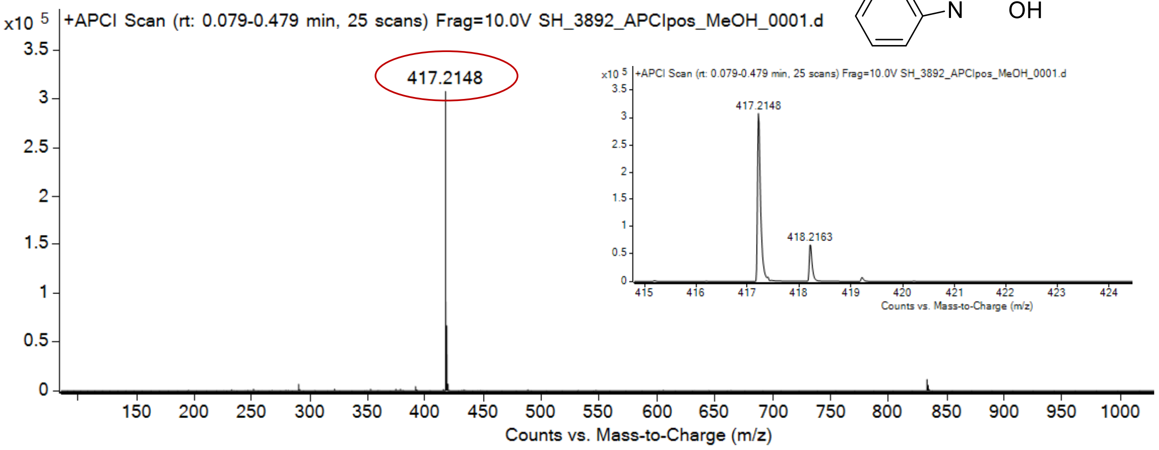

expected mass: [M+H]$^+$ = 417.2146 observed mass: [M+H]$^+$ = 417.2148 mass accuracy = 0.5 ppm

[1]H (500 MHz) and [13]C NMR (126 MHz) spectra of **117** in chloroform-*d*

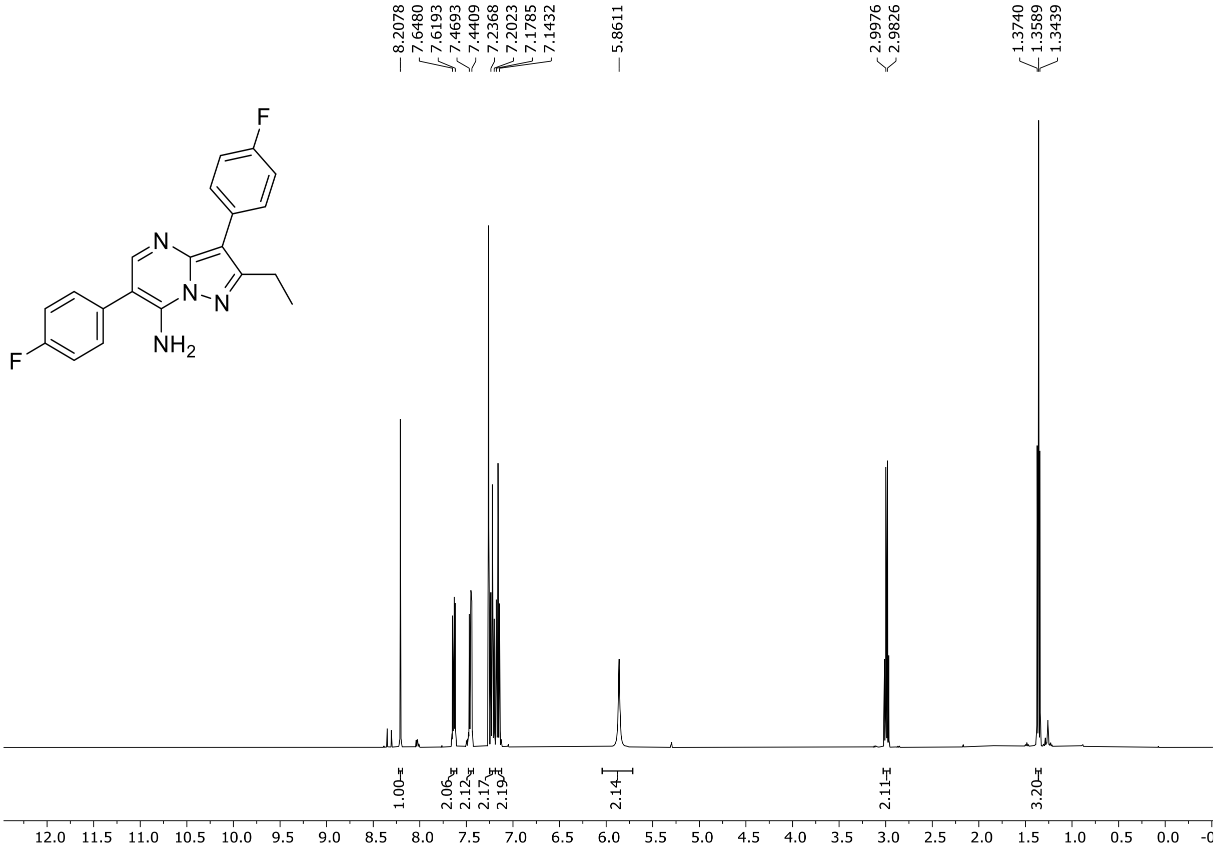

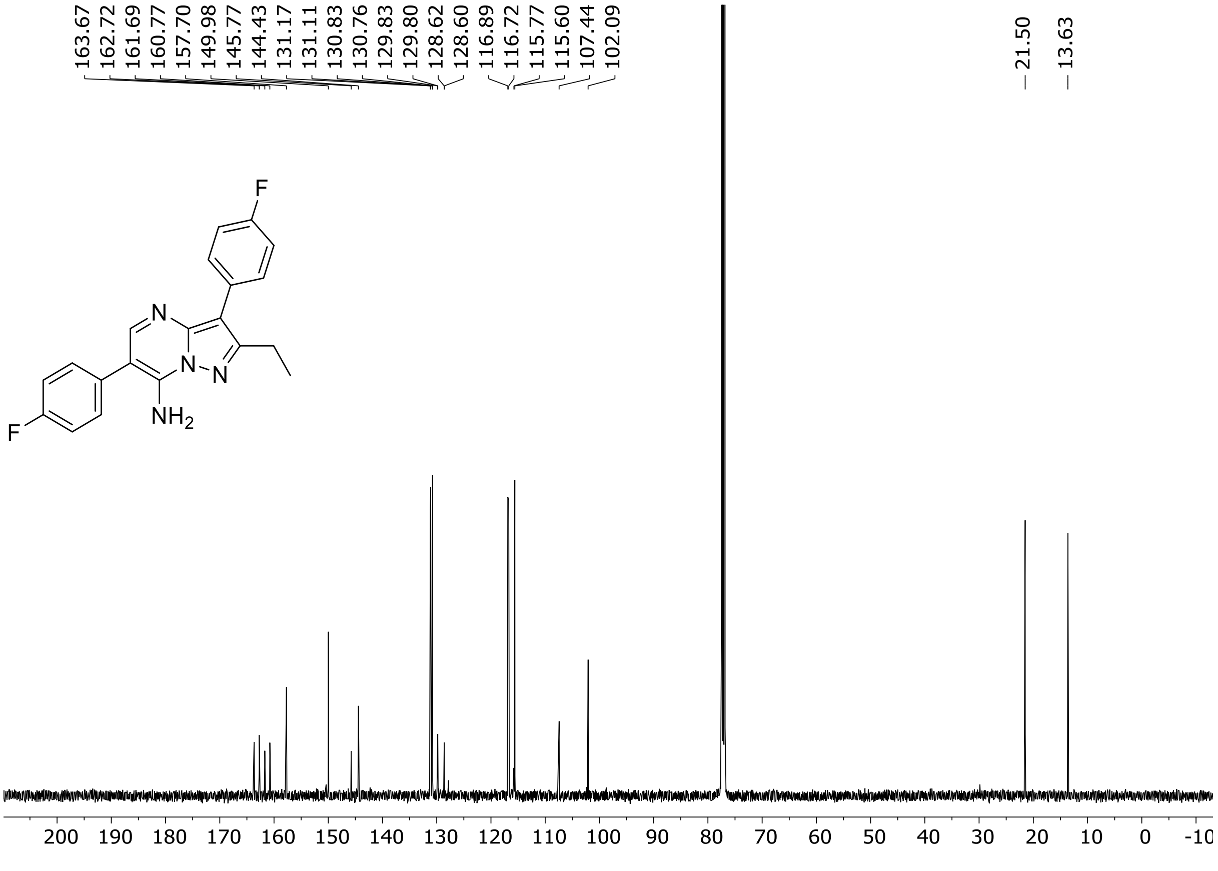

$^{19}$F (282 MHz) NMR spectrum of **117** chloroform-*d*

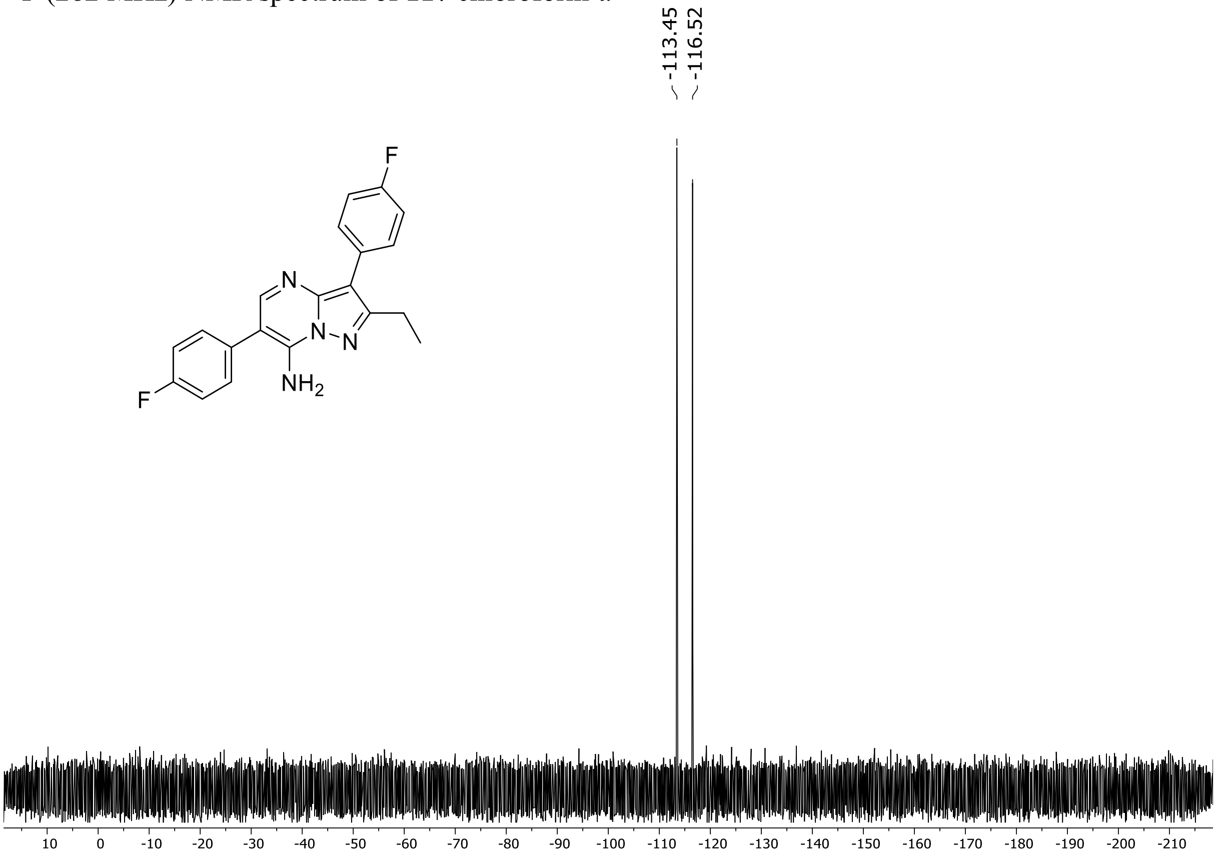

HRMS spectrum of **117**

**$C_{20}H_{16}N_4F_2$** **mono m/z = 350.1343**

**APCI +** (MMI)
nitrogen flow 5 L/min, gas temperature 300°C, nebulizer 45 psi, vaporizer 200°C, skimmer **65** V, fragmentor **15** V, dissolved in methanol

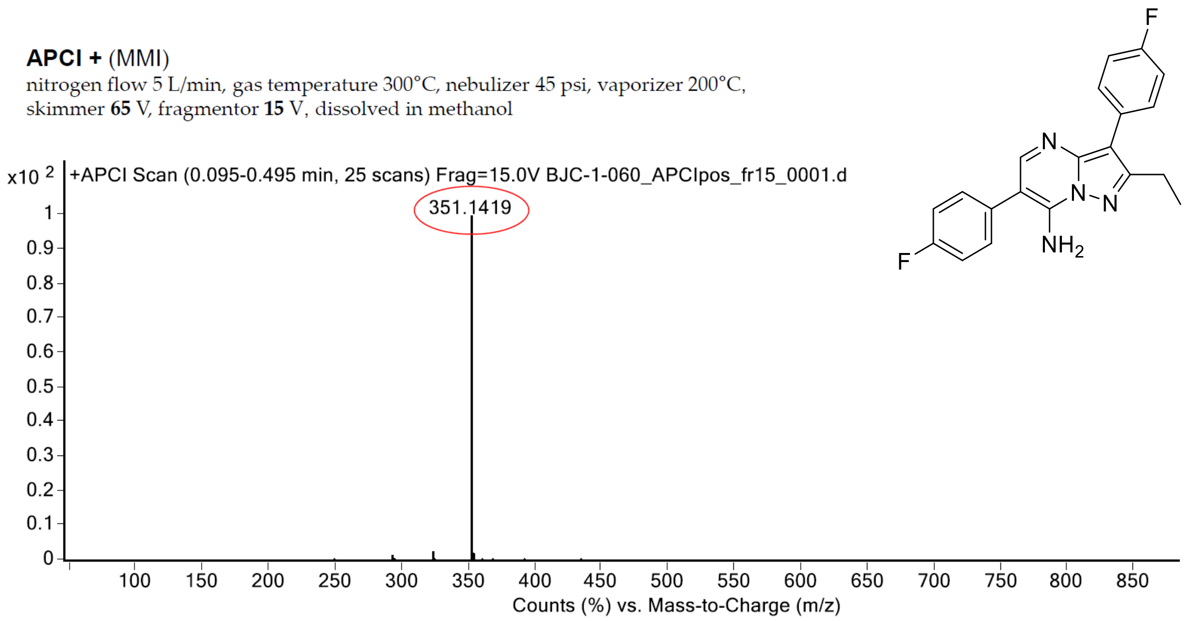

calculated mass: [M+H]$^+$ = 351.1416 observed: [M+H]$^+$ = 351.1419 max. mass error = 0.8 ppm

### [1]H NMR, [13]C NMR, HRMS and IR spectra of compound S109-S188

[1]H (500 MHz) and [13]C NMR (126 MHz) spectra of **S109** in chloroform-*d*

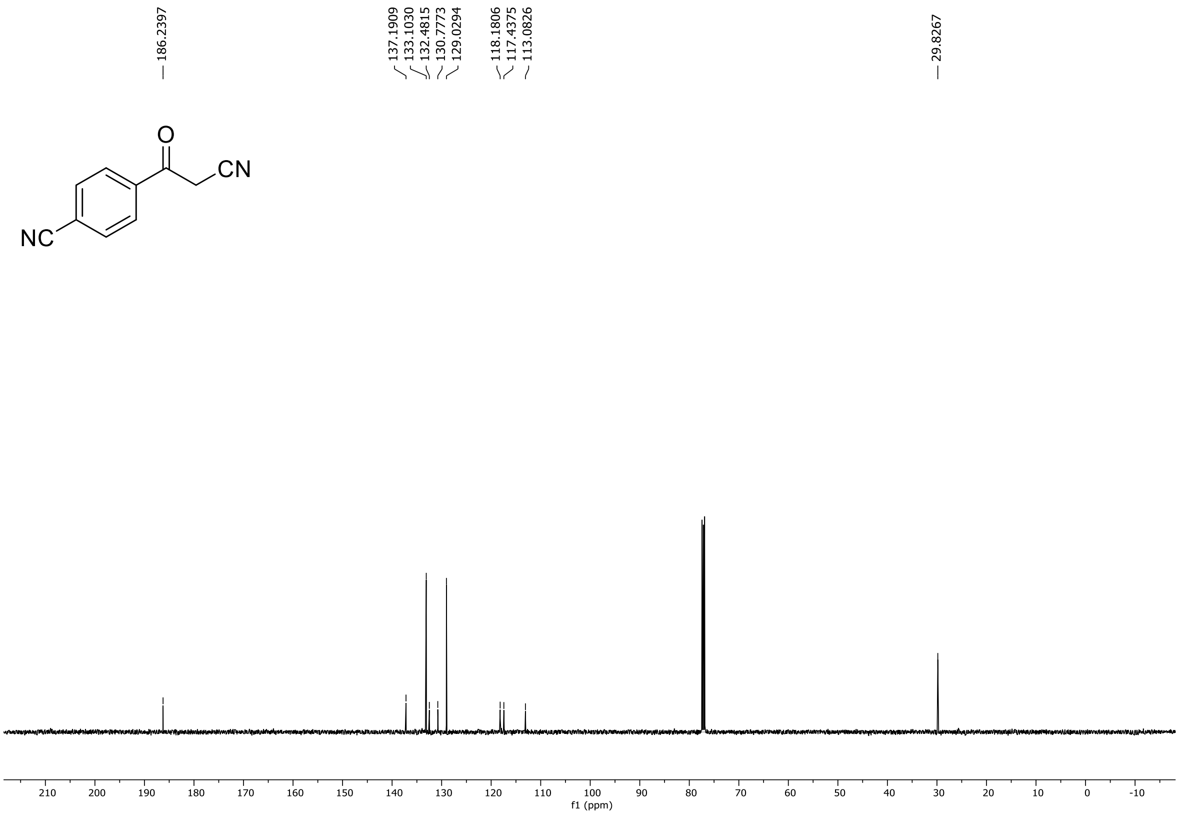

HRMS spectrum of **S109**

**NAR-A-107** $C_{10}H_6N_2O$ ***m/z* 170.0480**

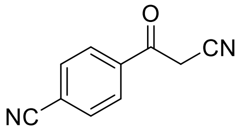

**APCI-** (MMI)
nitrogen flow 5 L/min, gas temperature 325°C, nebulizer 45 psi, skimmer 65 V, vaporizer 200°C, fragmentor 50 V, dissolved in MeOH

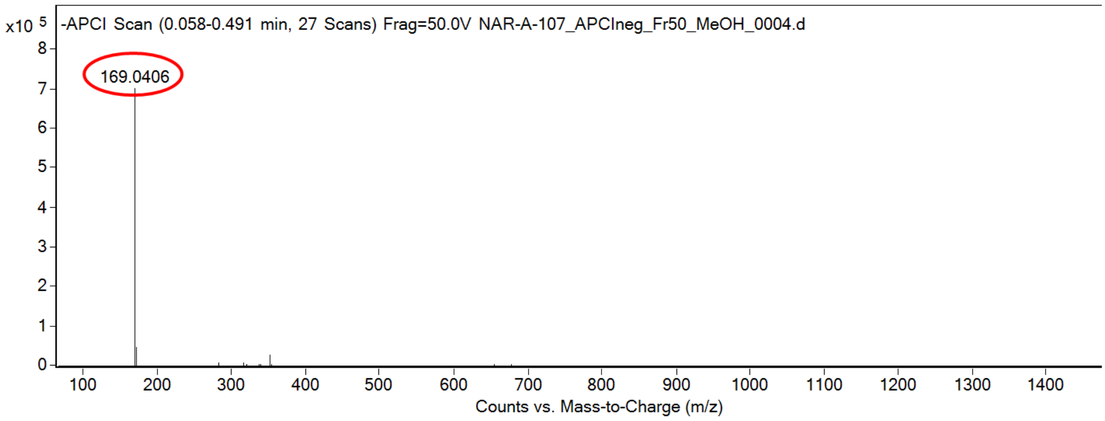

calculated mass: $[M-H]^-$ = 169.0407 observed: $[M-H]^-$ = 169.0406 mass accuracy = -0.6 ppm

FT-IR spectrum (neat) of **S109**

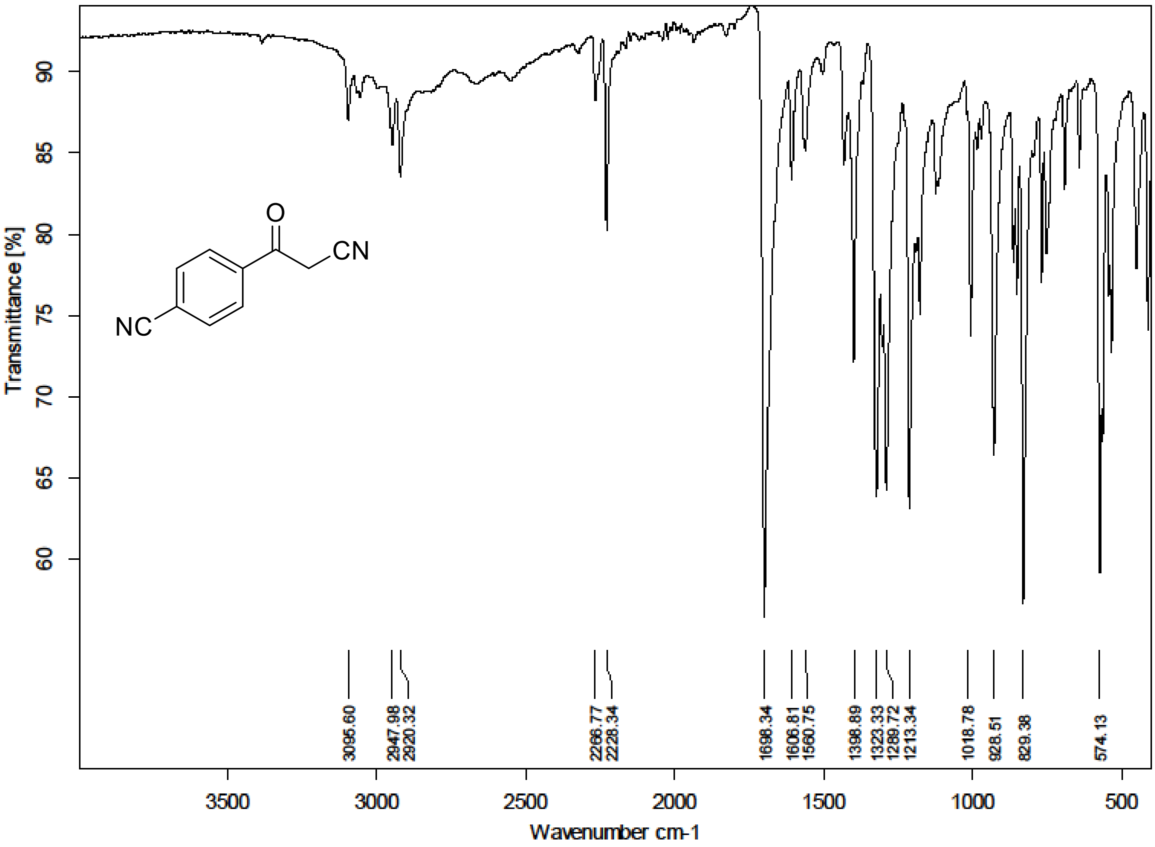

$^{1}H$ (500 MHz) and $^{13}C$ NMR (126 MHz) spectra of **S110** in chloroform-*d*

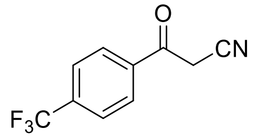

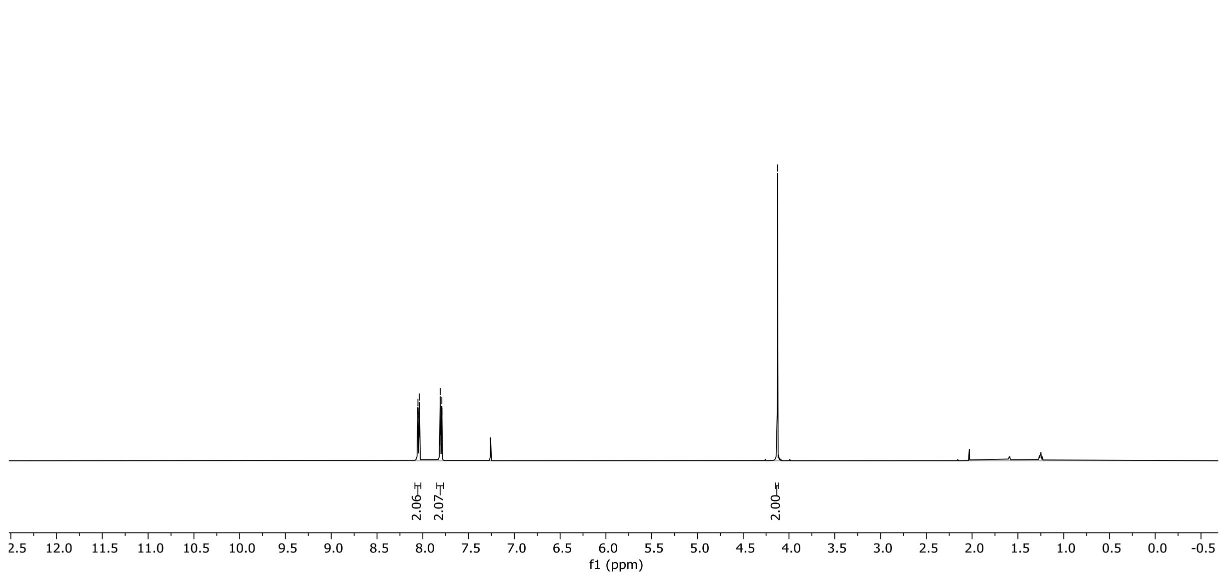

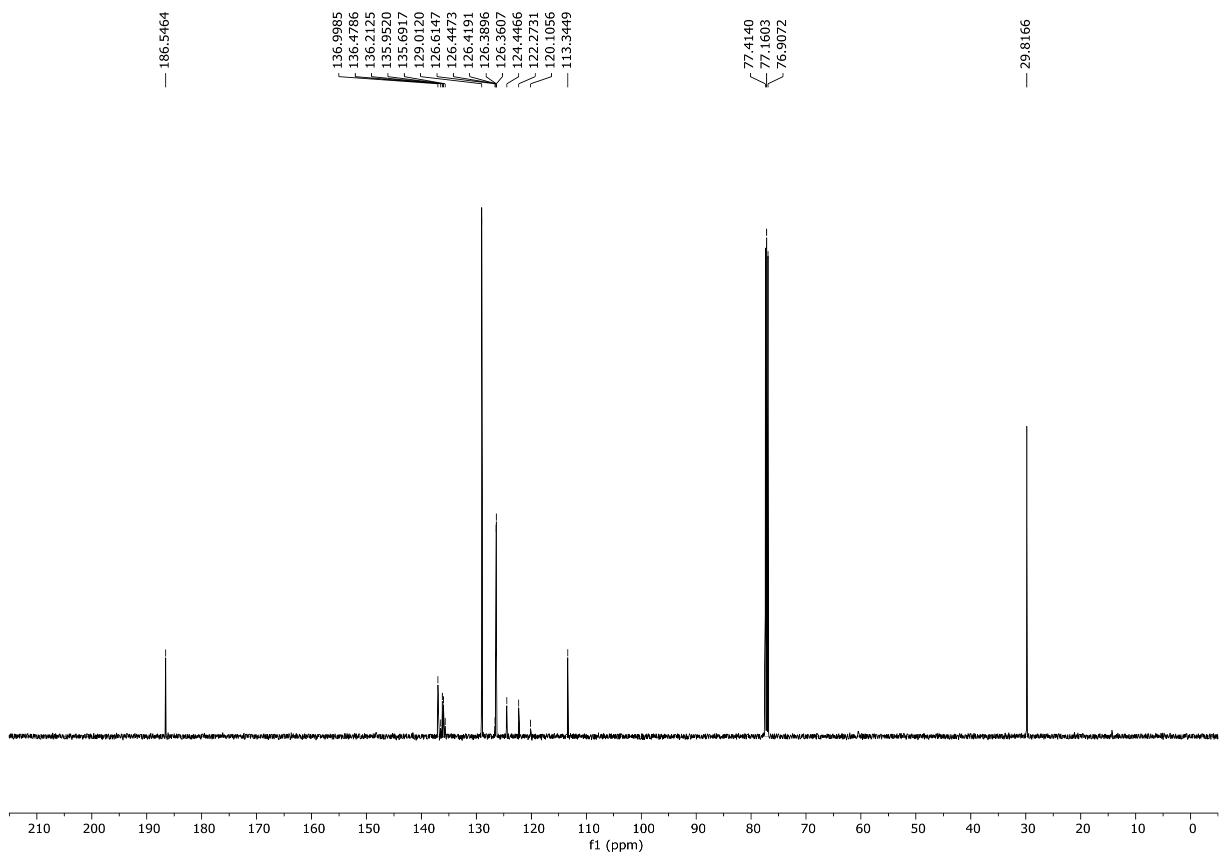

$^{19}$F NMR (471 MHz) spectrum of **S110** in chloroform-*d*

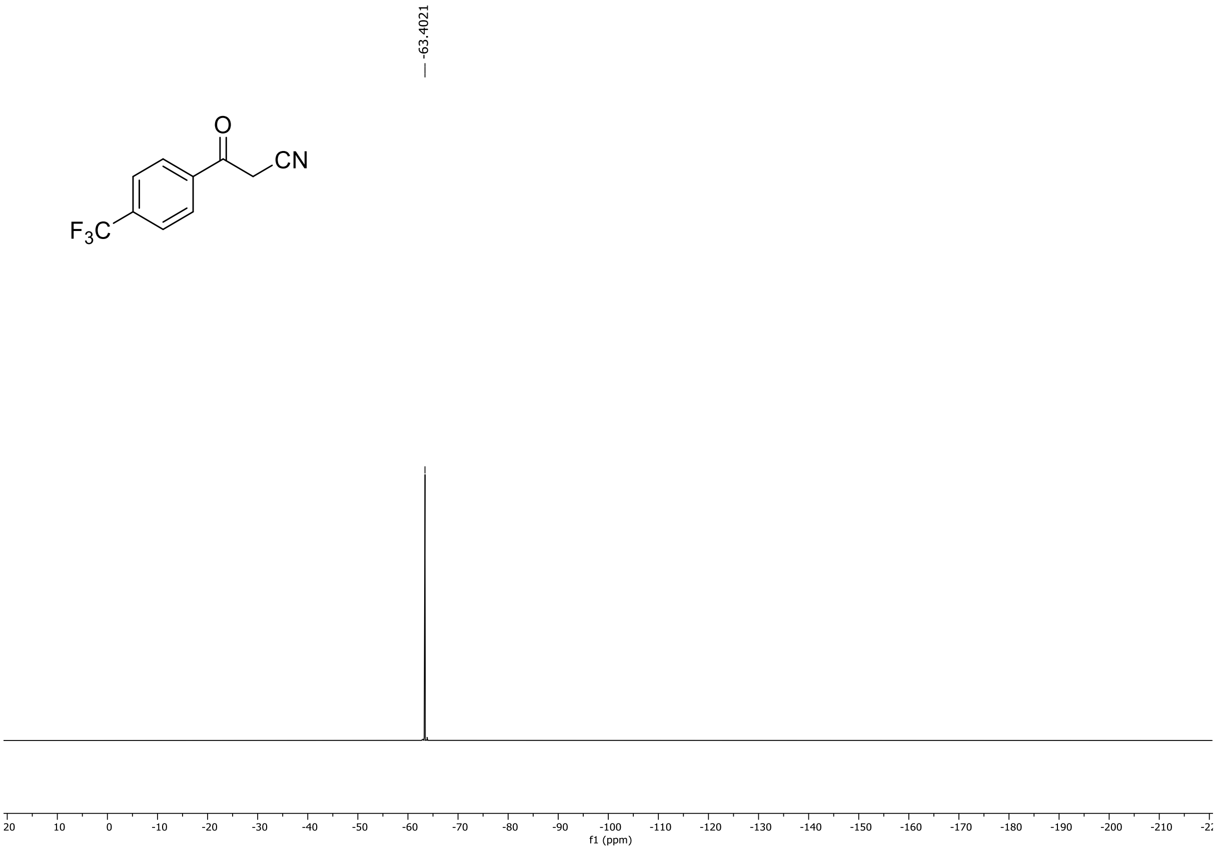

HRMS spectrum of **S110**

**NAR-A-110** **$C_{10}H_6F_3NO$** ***m/z* 213.0401**

**APCI-** (MMI)
nitrogen flow 5 L/min, gas temperature 325°C, nebulizer 45 psi, skimmer 65 V, vaporizer 200°C, fragmentor 45 V, dissolved in MeOH

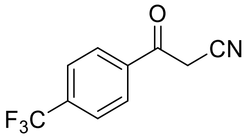

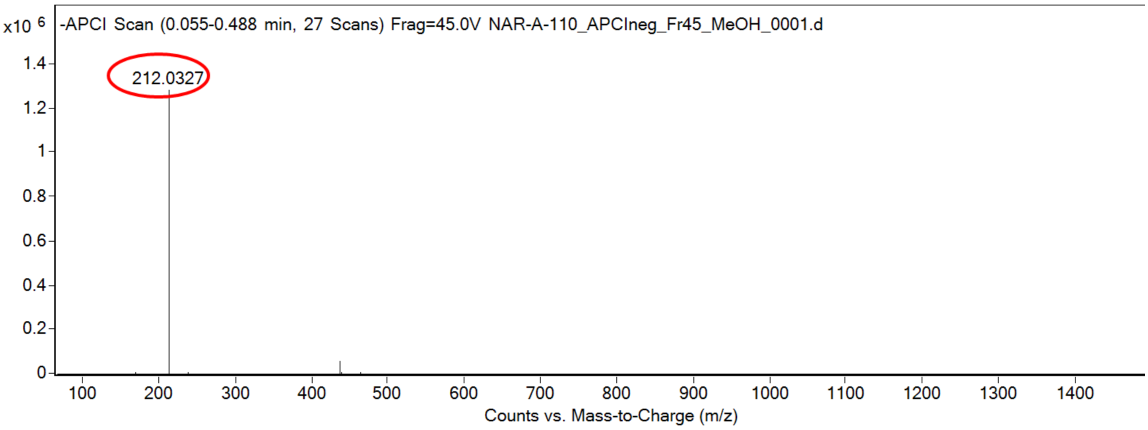

calculated mass: [M-H]$^-$ = 212.0329 observed: [M-H]$^-$ = 212.0327 mass accuracy = -0.9 ppm

FT-IR spectrum (neat) of **S110**

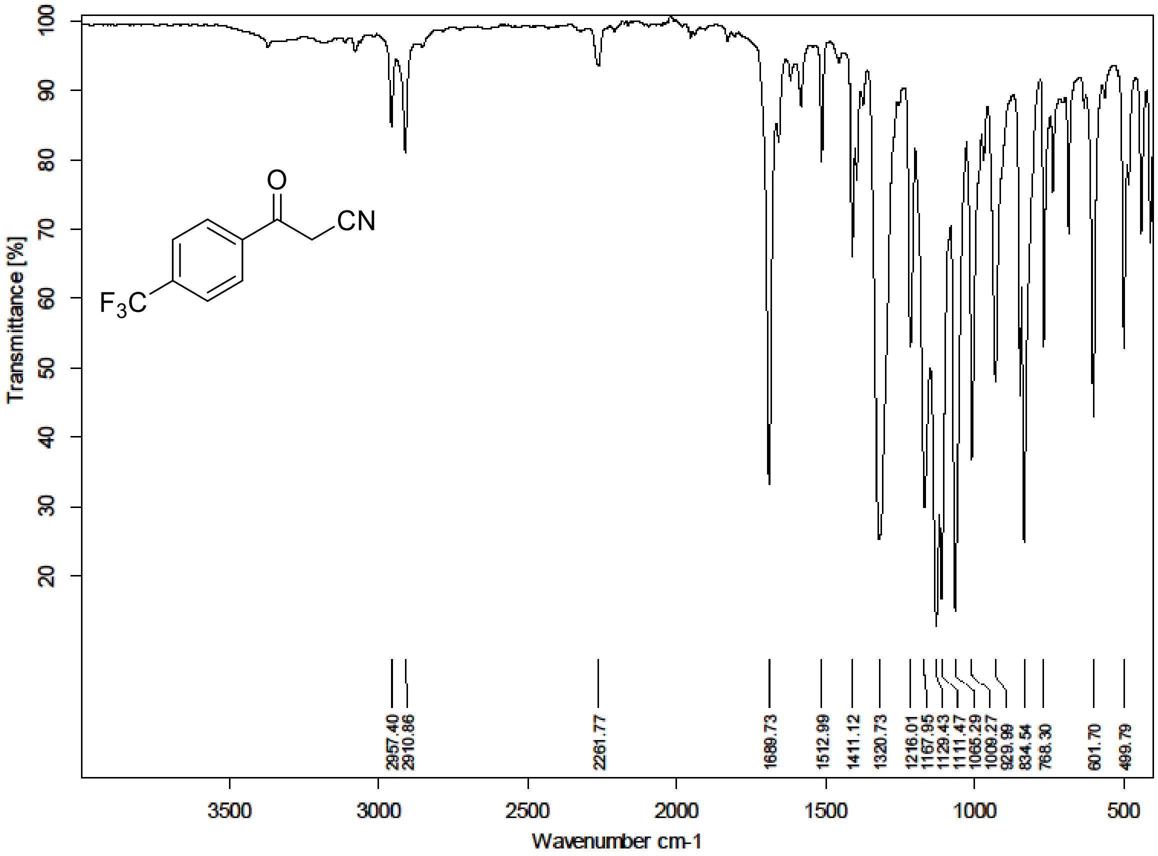

$^{1}H$ (500 MHz) and $^{13}C$ NMR (126 MHz) spectra of **S111** in chloroform-*d*

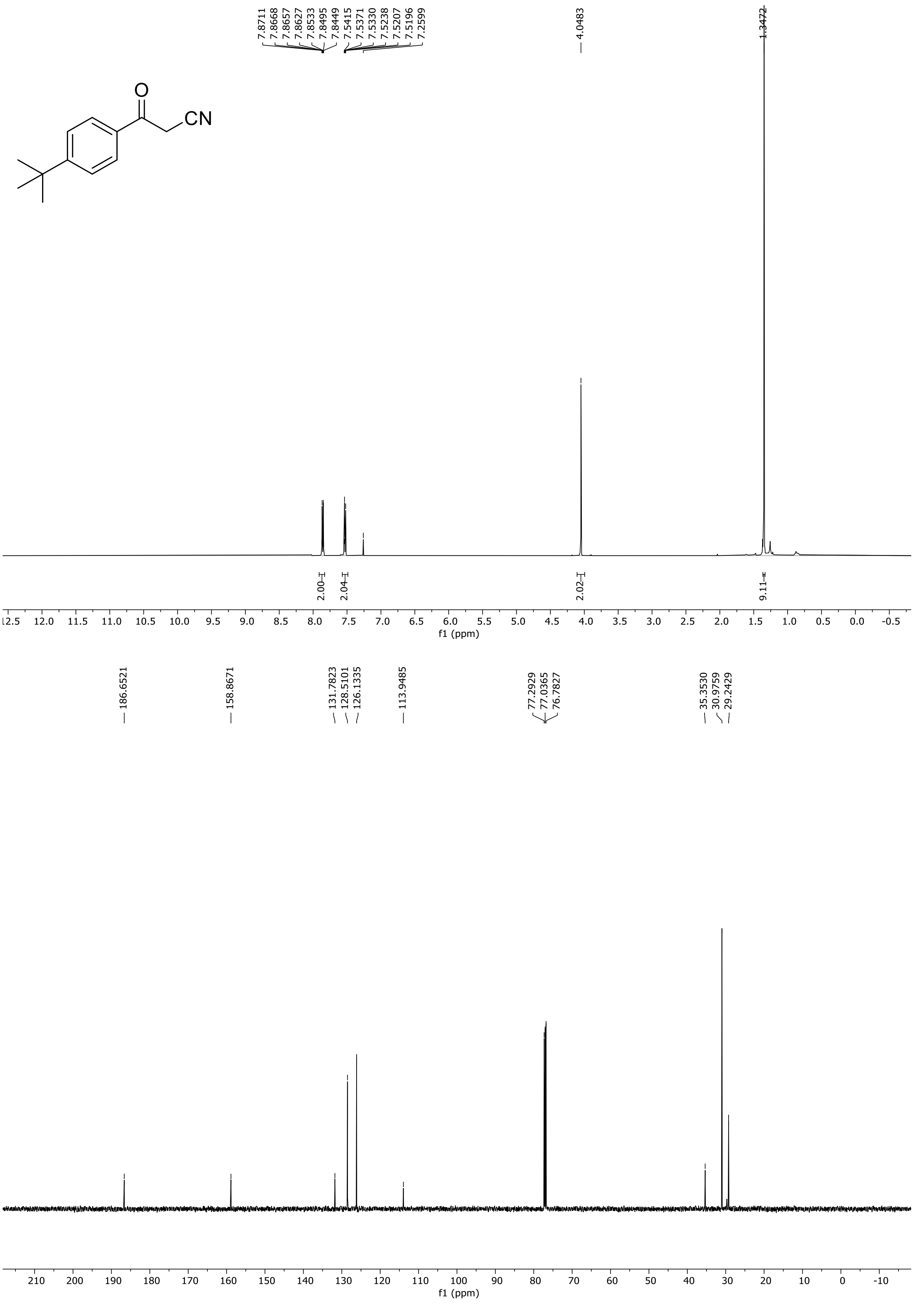

HRMS spectrum of **S111**

**NAR-A-108** $C_{13}H_{15}NO$ *m/z* **201.1154**

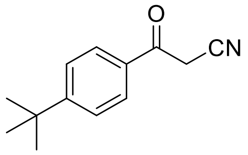

**APCI-** (MMI)
nitrogen flow 5 L/min, gas temperature 325°C, nebulizer 45 psi, skimmer 65 V, vaporizer 200°C, fragmentor 60 V, dissolved in MeOH

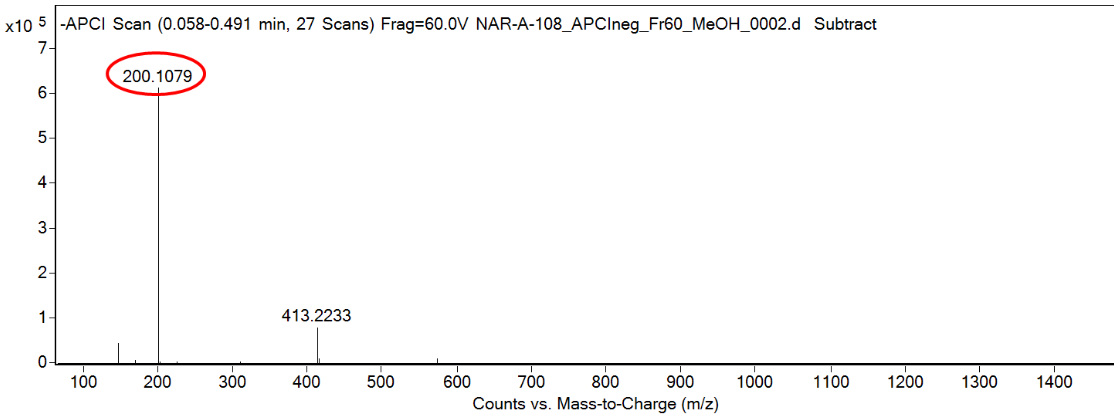

calculated mass: $[M-H]^{-}$ = 200.1081 observed: $[M-H]^{-}$ = 200.1079 mass accuracy = -1.0 ppm

FT-IR spectrum (neat) of **S111**

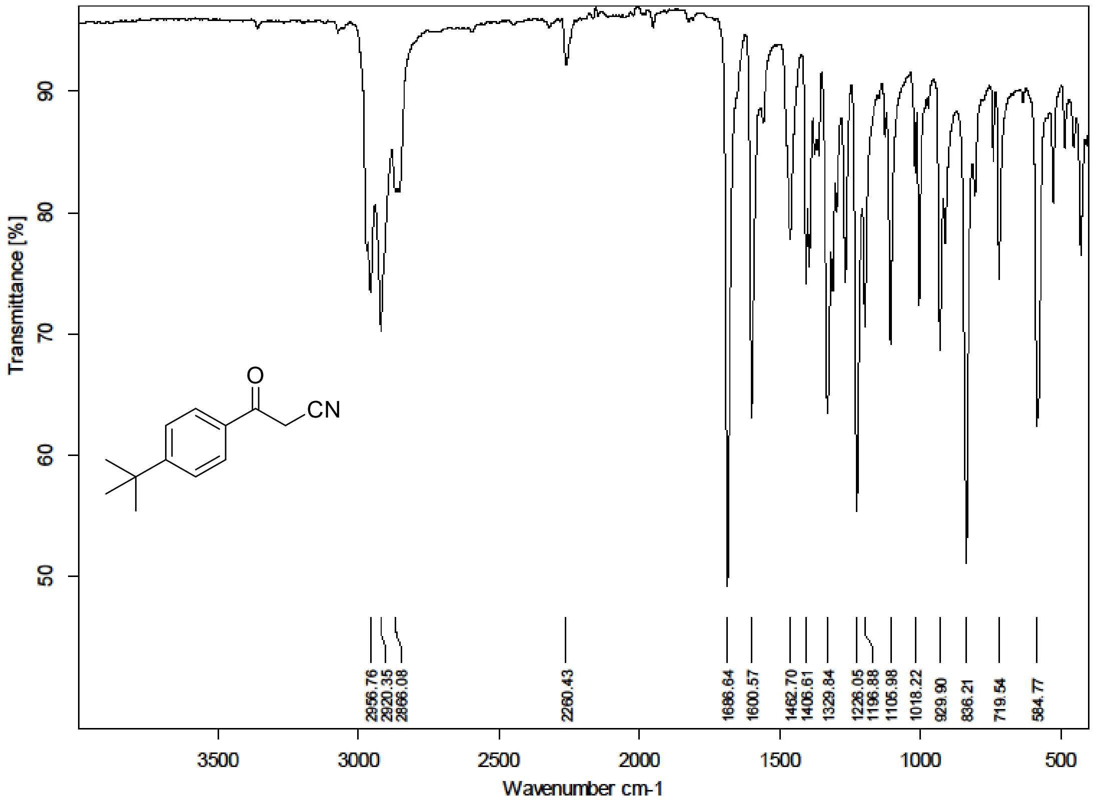

[1]H (500 MHz) and [13]C NMR (126 MHz) spectra of **S112** in chloroform-*d*

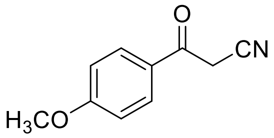

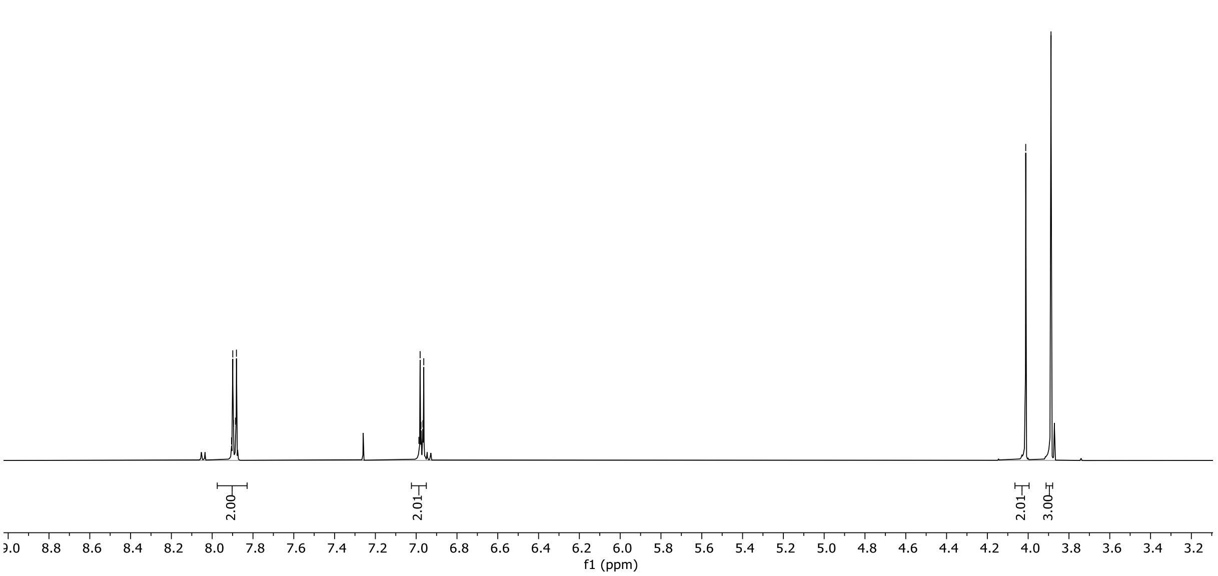

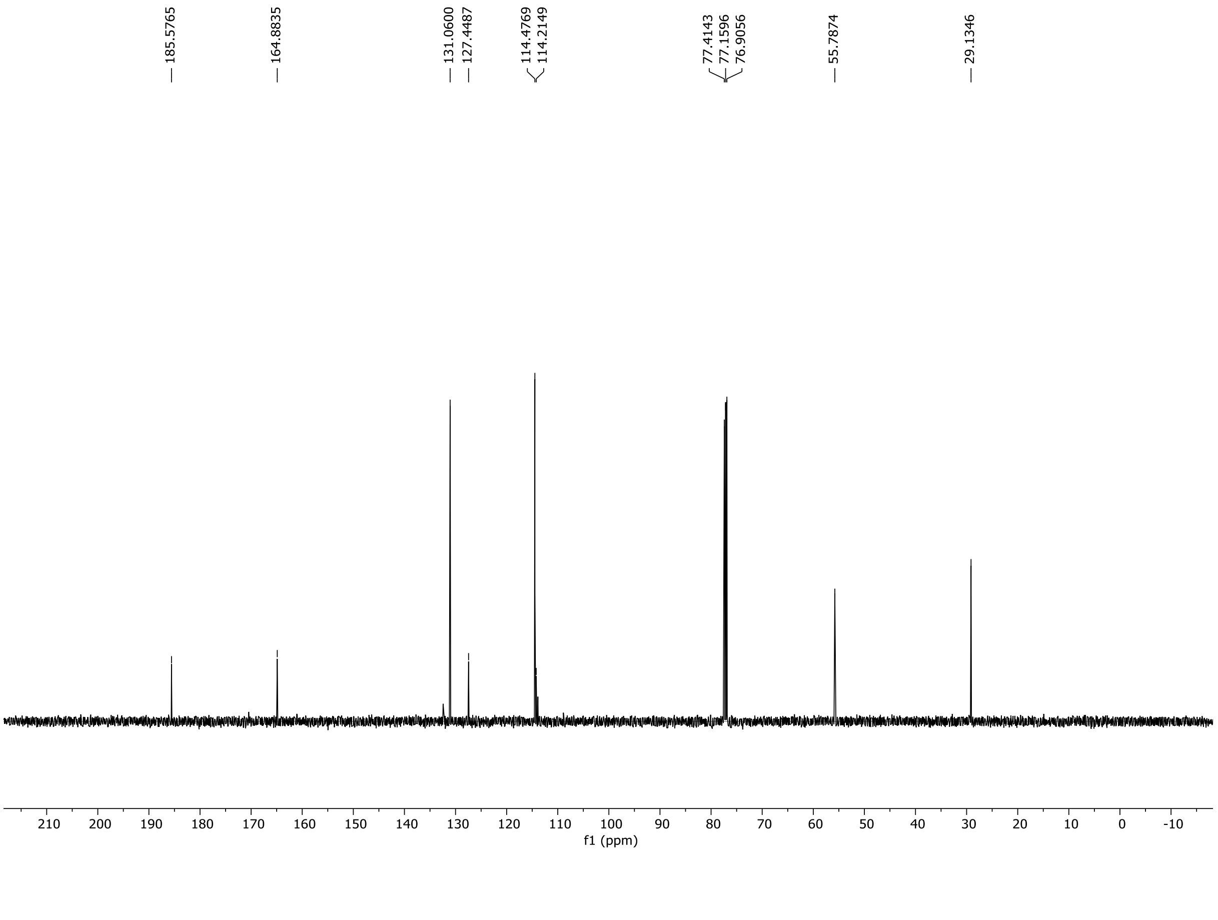

HRMS spectrum of **S112**

**NAR-A-112** $C_{10}H_9NO_2$ *m/z* **175.0633**

**APCI-** (MMI)
nitrogen flow 5 L/min, gas temperature 325°C, nebulizer 45 psi, skimmer 65 V, vaporizer 200°C, fragmentor 150 V, dissolved in MeOH

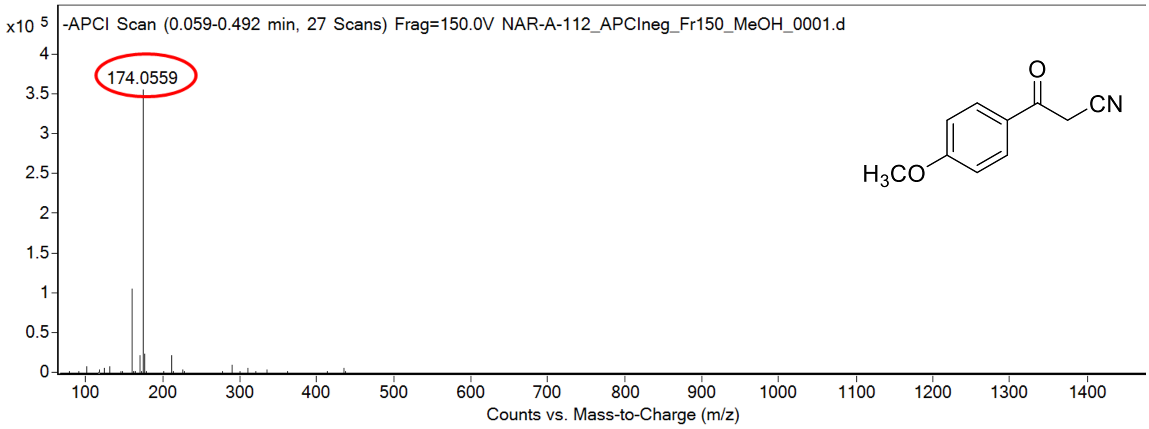

calculated mass: [M-H]$^-$ = 174.0561 observed: [M-H]$^-$ = 174.0559 mass accuracy = -1.1 ppm

FT-IR spectrum (neat) of **S112**

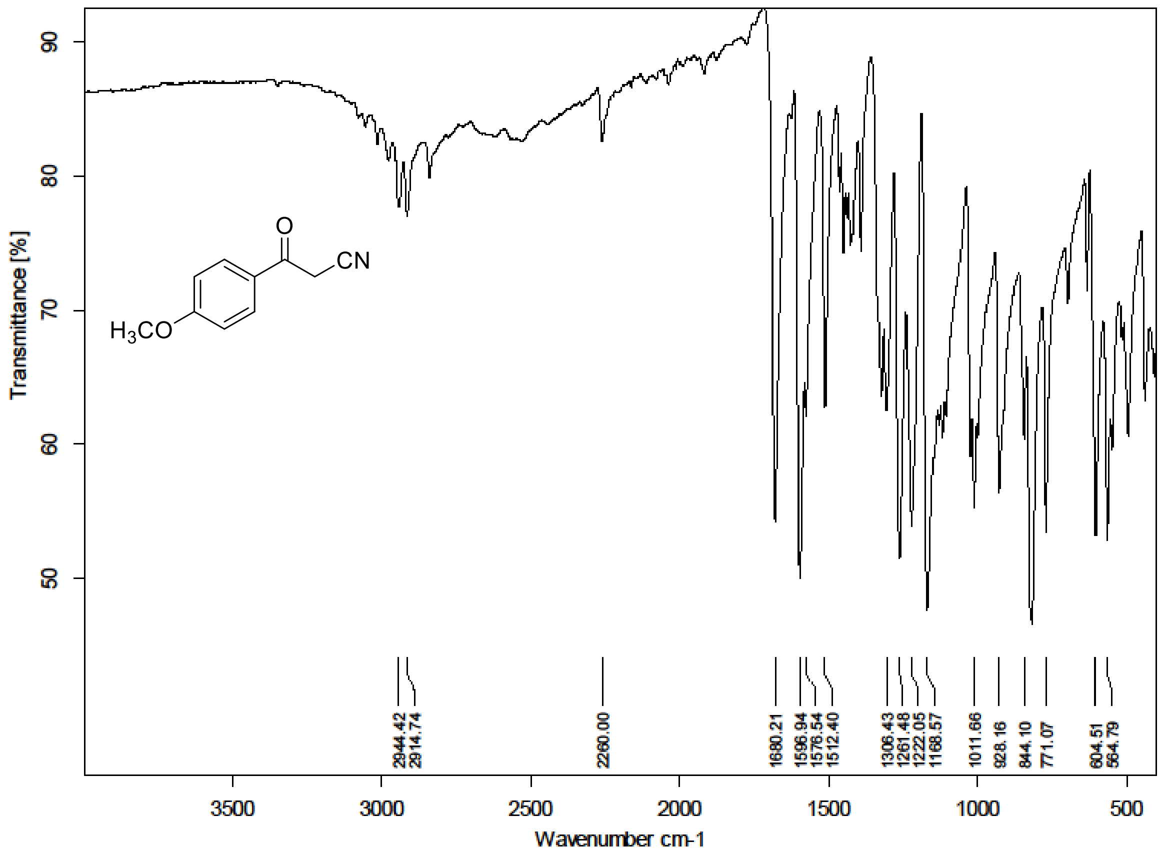

[1]H (300 MHz) and [13]C NMR (75 MHz) spectra of **S113** in chloroform-*d*

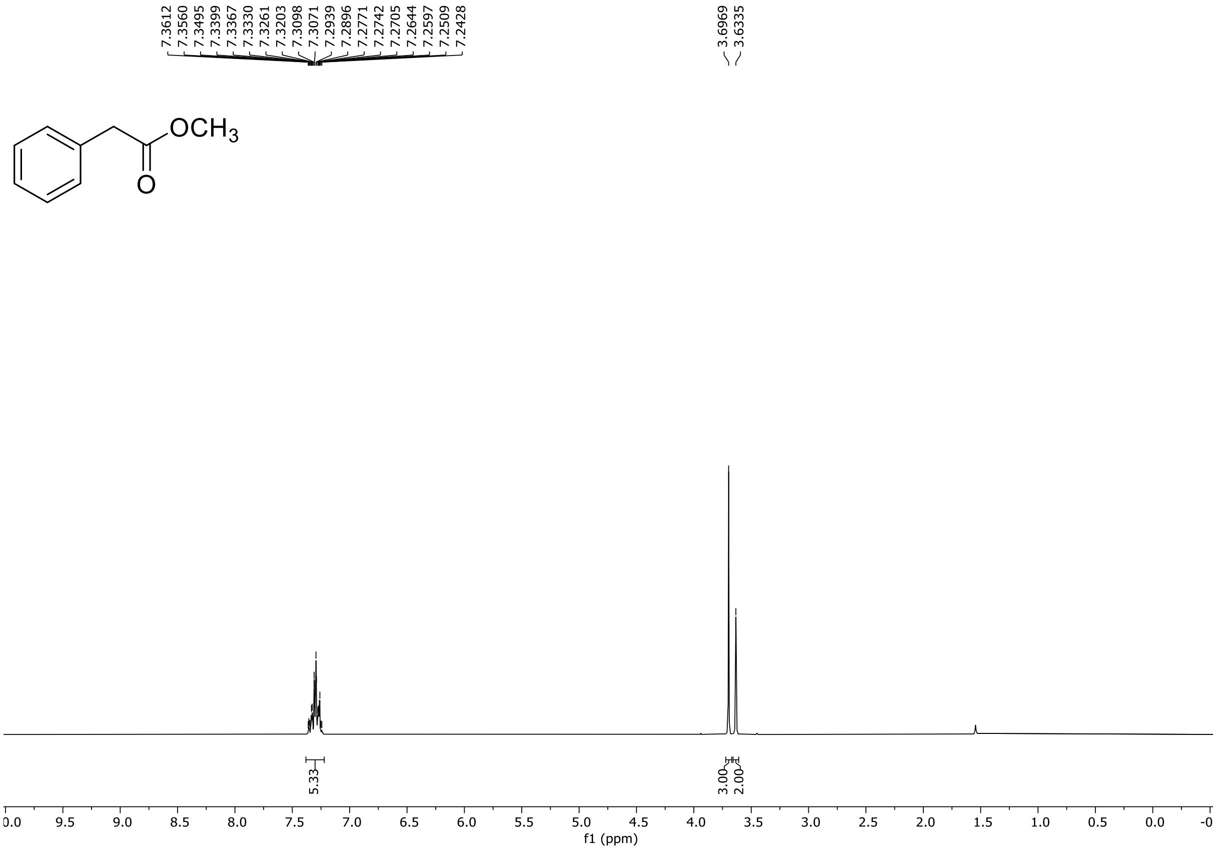

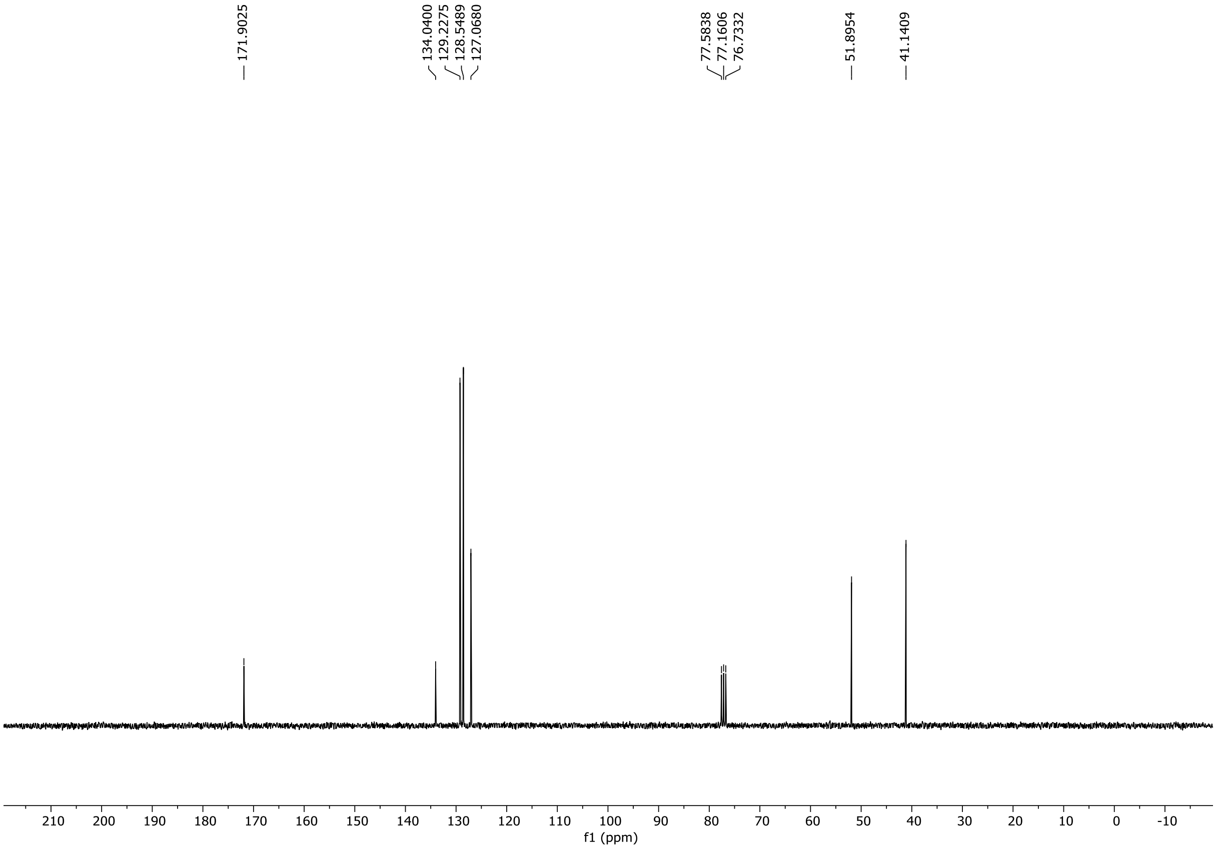

HRMS spectrum of **S113**

**NAR-A-182** $C_9H_{10}O_2$ mono *m/z* **150.0681**

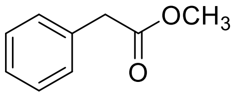

**APCI +** (MMI)
nitrogen flow 5 L/min, gas temperature 325°C, nebulizer 45 psi, skimmer **65** V, vaporizer 200°C, fragmentor 50 V, dissolved in methanol

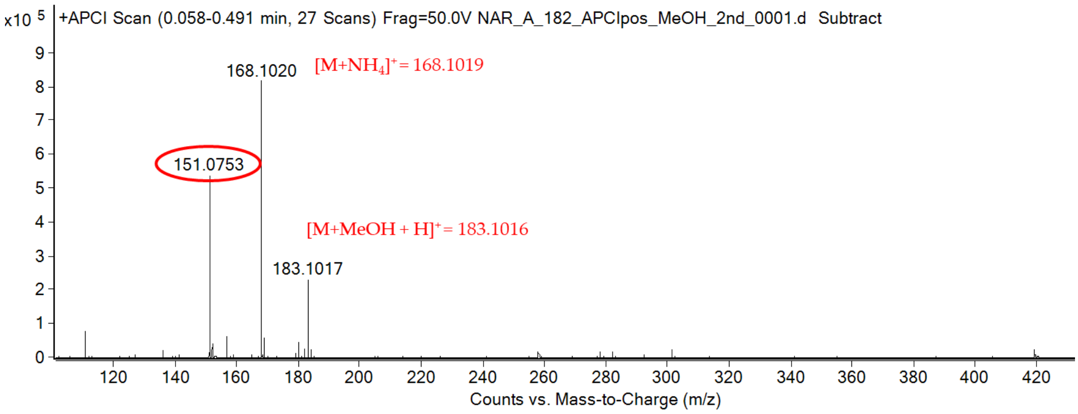

calculated mass: $[M+H]^+ = 151.0754$ observed: $[M+H]^+ = 151.0753$ mass accuracy = - 0.7 ppm

FT-IR spectrum (neat) of **S113**

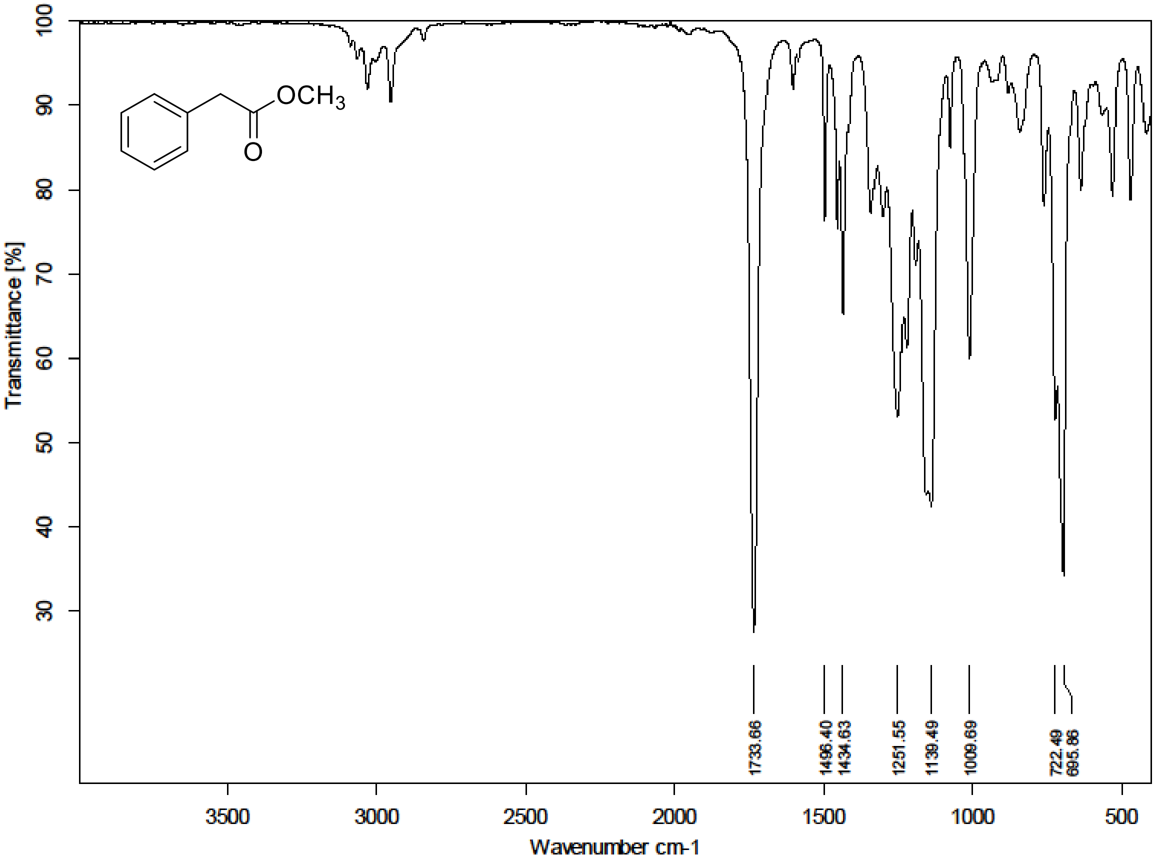

$^{1}H$ (300 MHz) and $^{13}C$ NMR (75 MHz) spectra of **S114** in chloroform-*d*

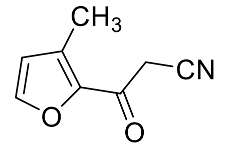

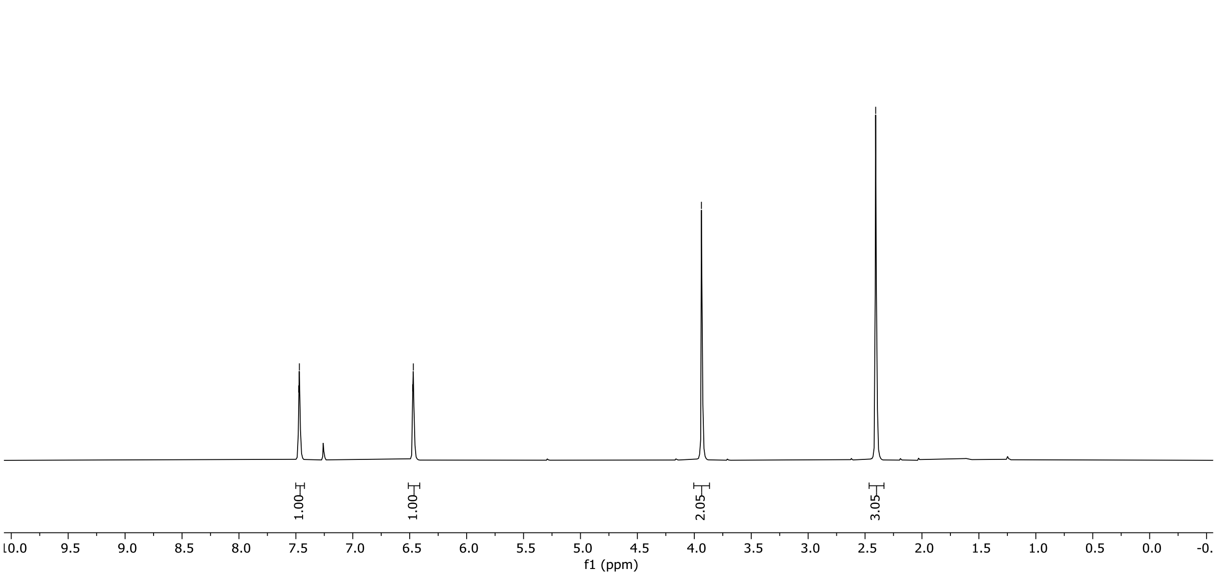

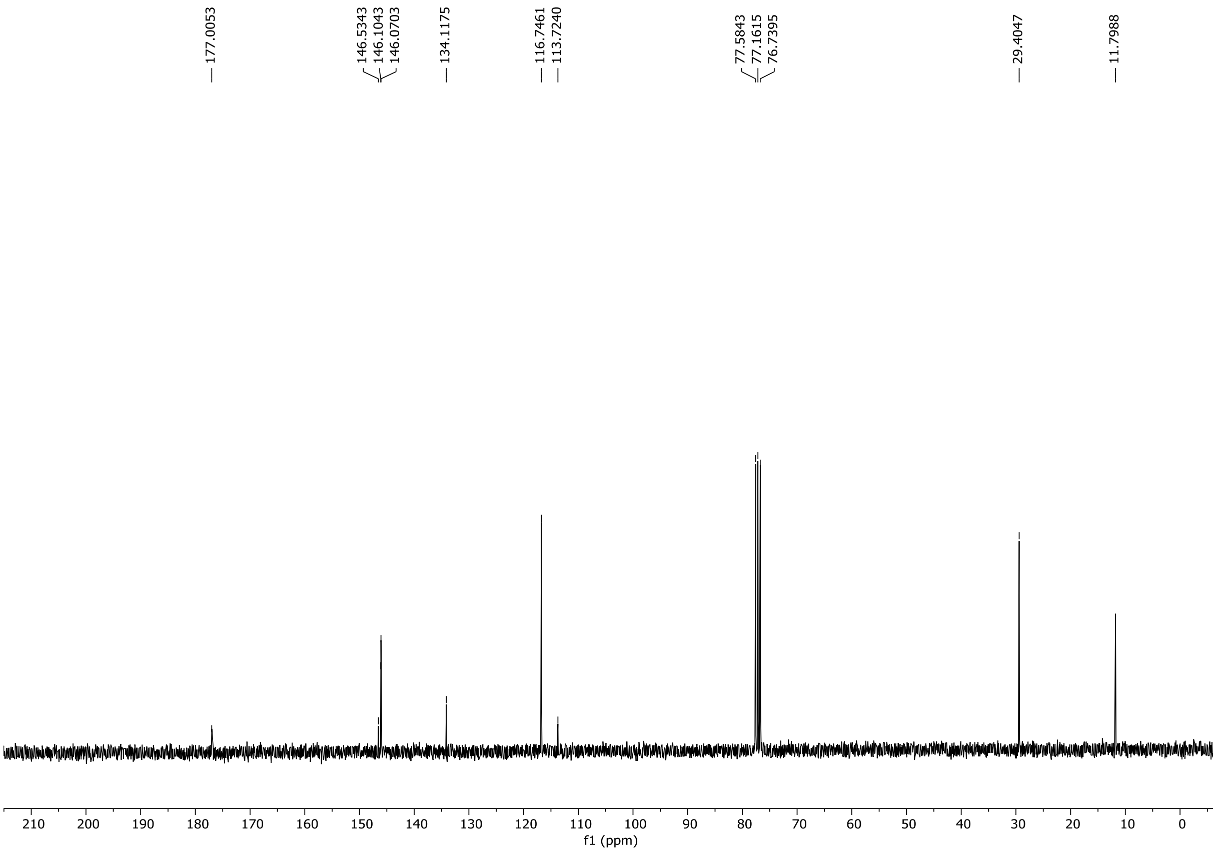

HRMS spectrum of **S114**

**NAR-A-174** $C_8H_7NO_2$ mono *m/z* 149.0477

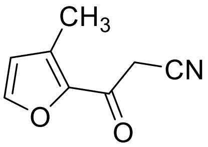

**APCI +** (MMI)
nitrogen flow 5 L/min, gas temperature 325°C, nebulizer 45 psi, skimmer **65** V, vaporizer 200°C, fragmentor 50 V, dissolved in methanol

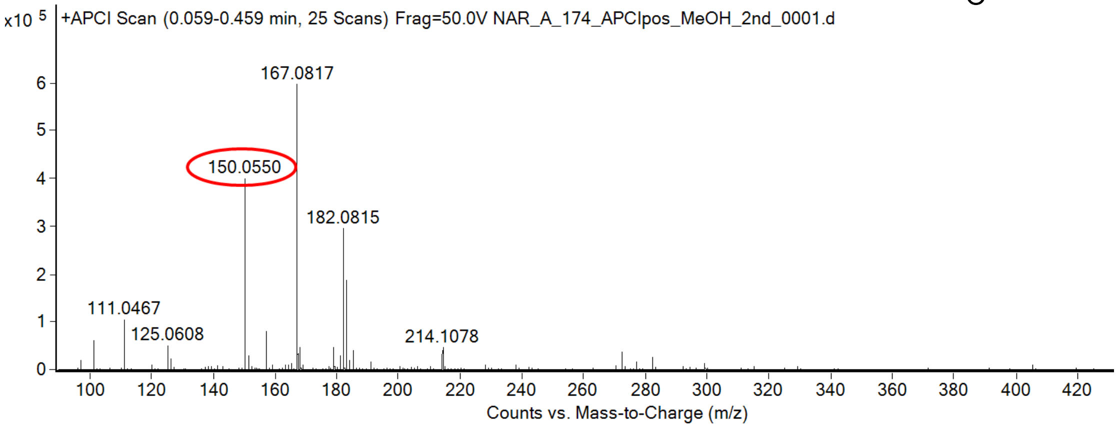

calculated mass: $[M+H]^+$ = 150.0550 observed: $[M+H]^+$ = 150.0550 mass accuracy = < 0.1 ppm

FT-IR spectrum (neat) of **S114**

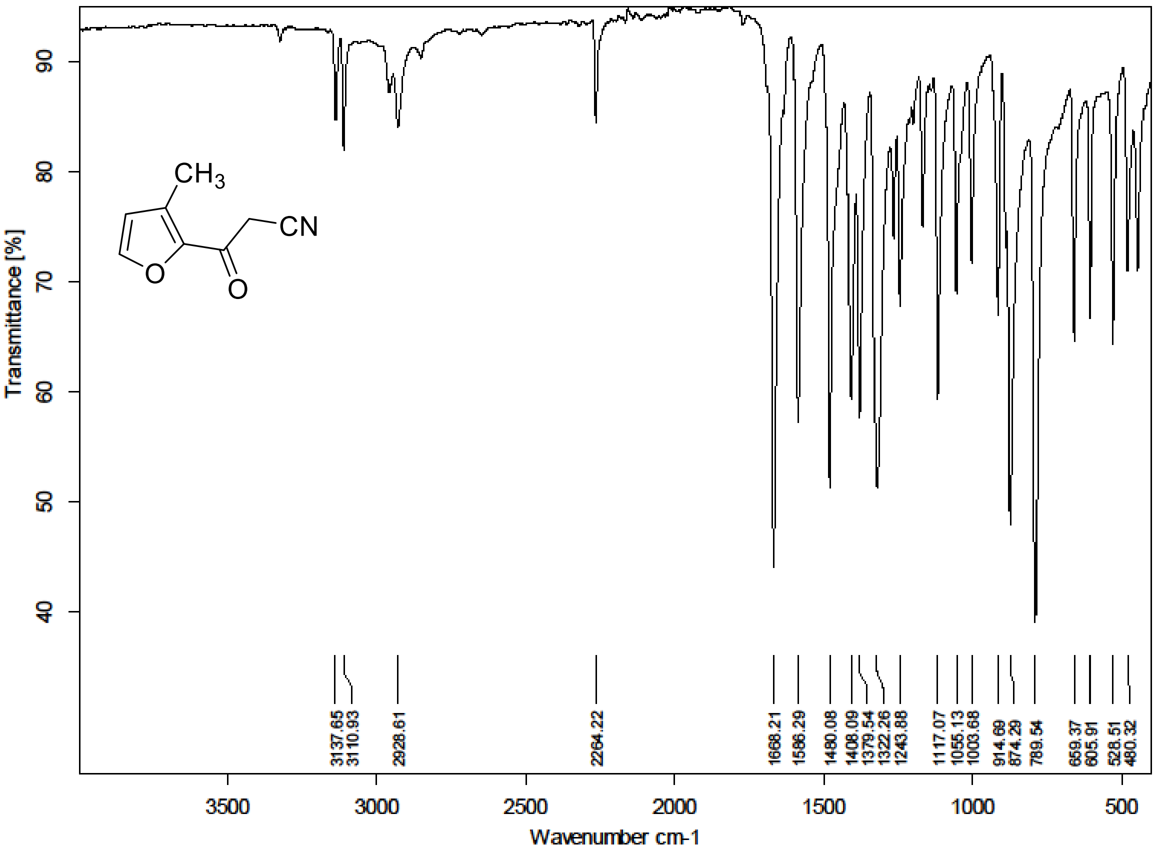

[1]H (300 MHz) and [13]C NMR (75 MHz) spectra of **S115** in chloroform-*d*

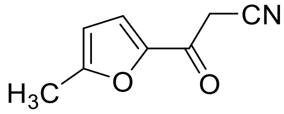

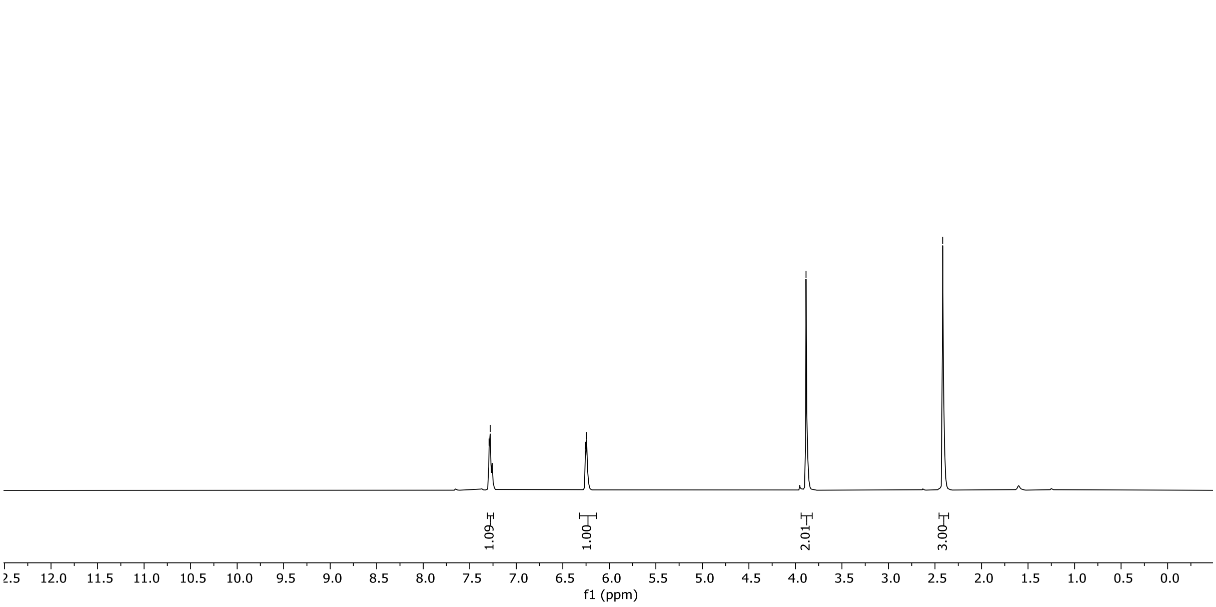

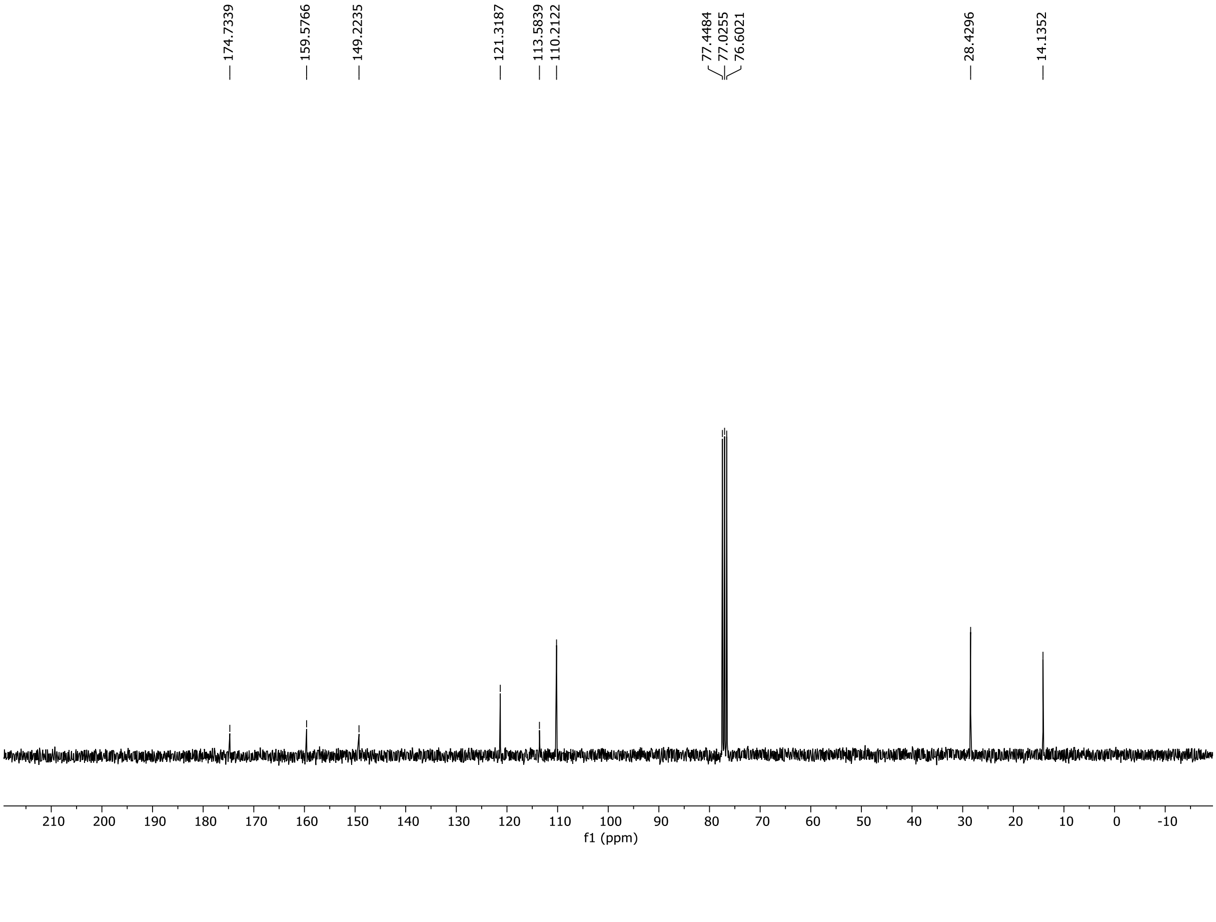

HRMS spectrum of **S115**

**NAR-A-173** $C_8H_7NO_2$ mono *m/z* 149.0477

**APCI +** (MMI)
nitrogen flow 5 L/min, gas temperature 325°C, nebulizer 45 psi, skimmer **65** V, vaporizer 200°C, fragmentor 45 V, dissolved in methanol

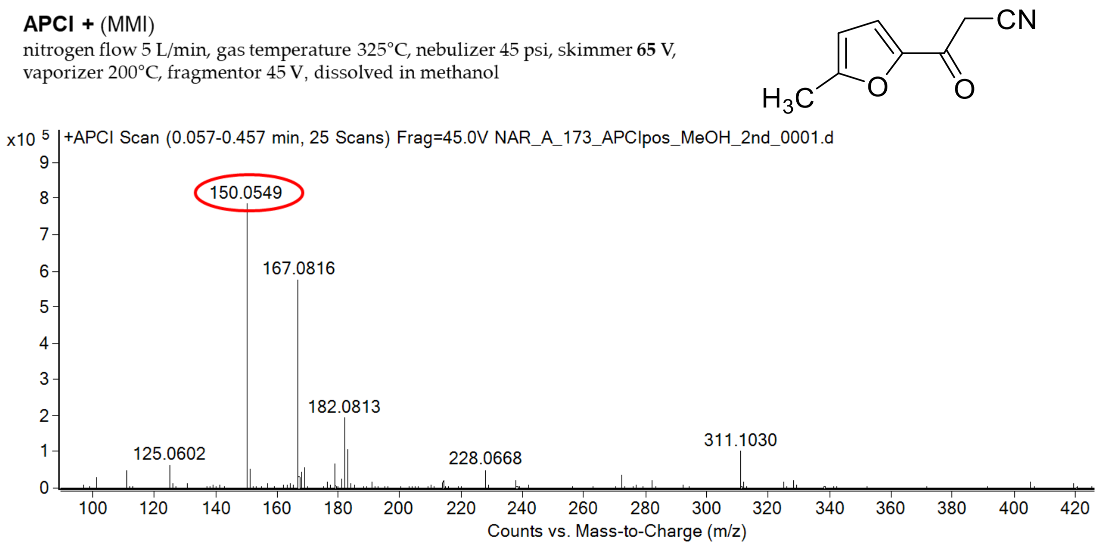

calculated mass: $[M+H]^+$ = 150.0550 observed: $[M+H]^+$ = 150.0549 mass accuracy = - 0.7 ppm

FT-IR spectrum (neat) of **S115**

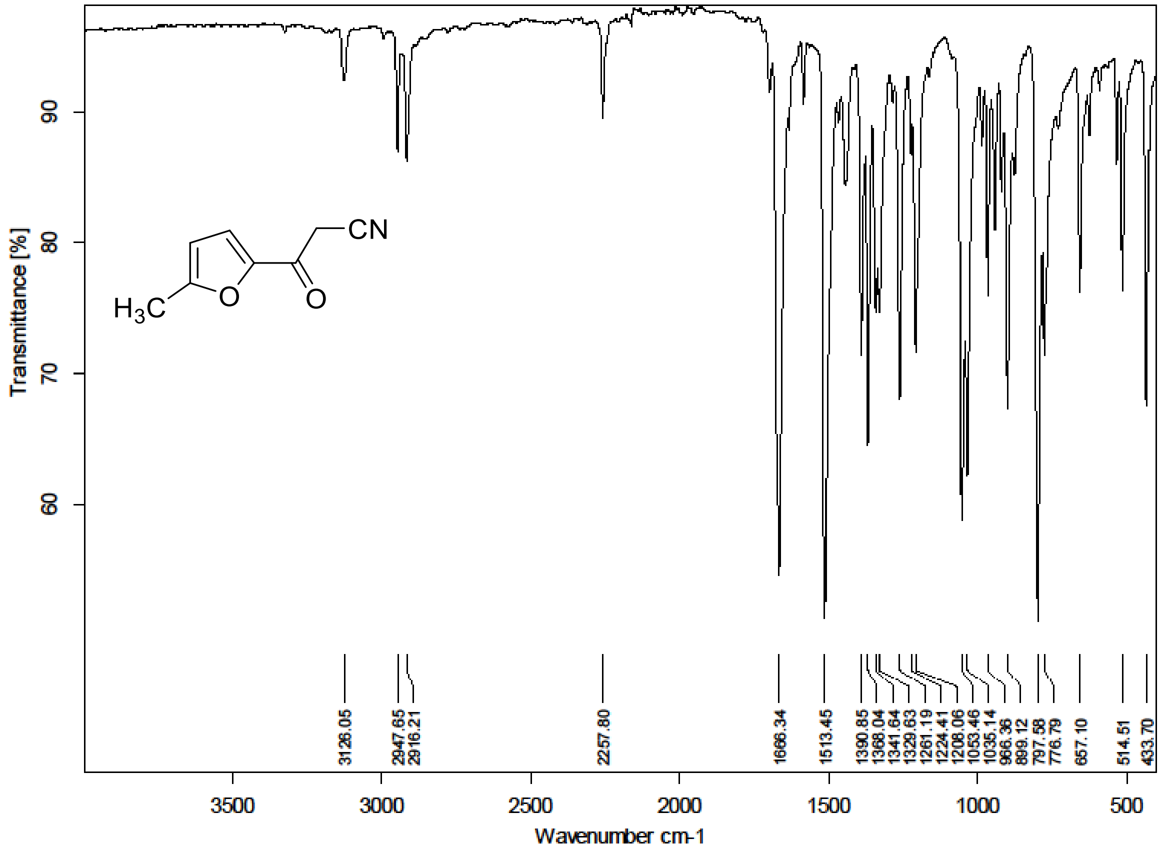

$^{1}H$ (500 MHz) and $^{13}C$ NMR (126 MHz) spectra of **S116** in chloroform-*d*

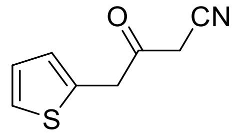

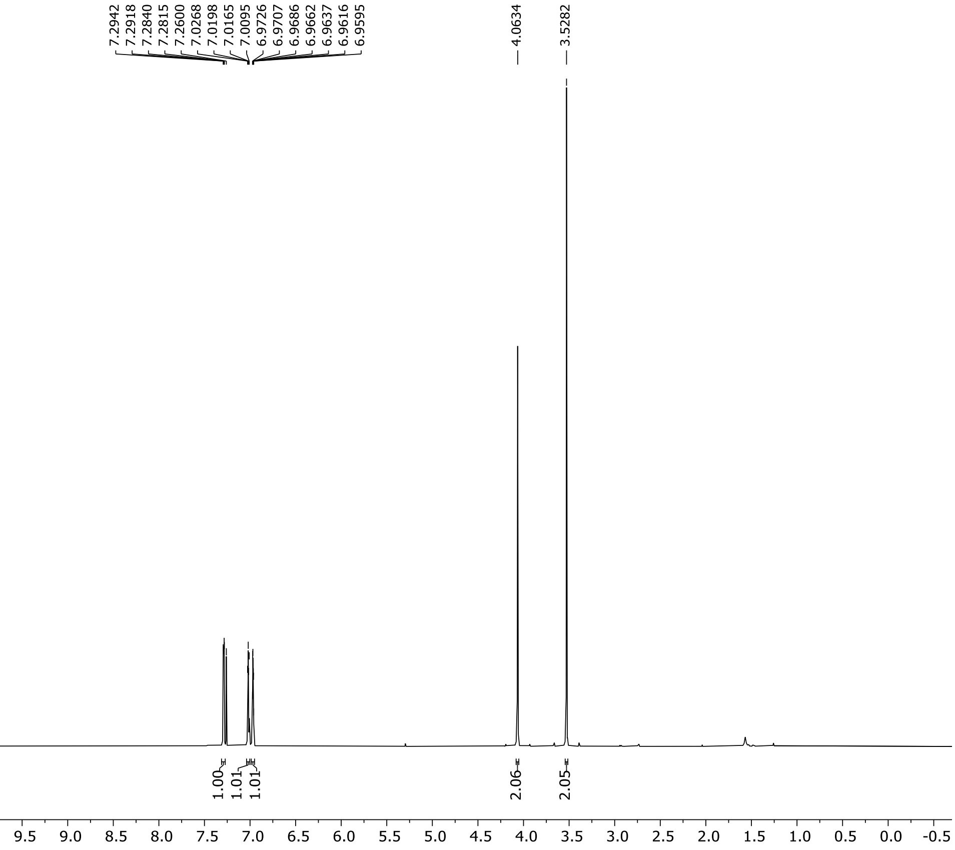

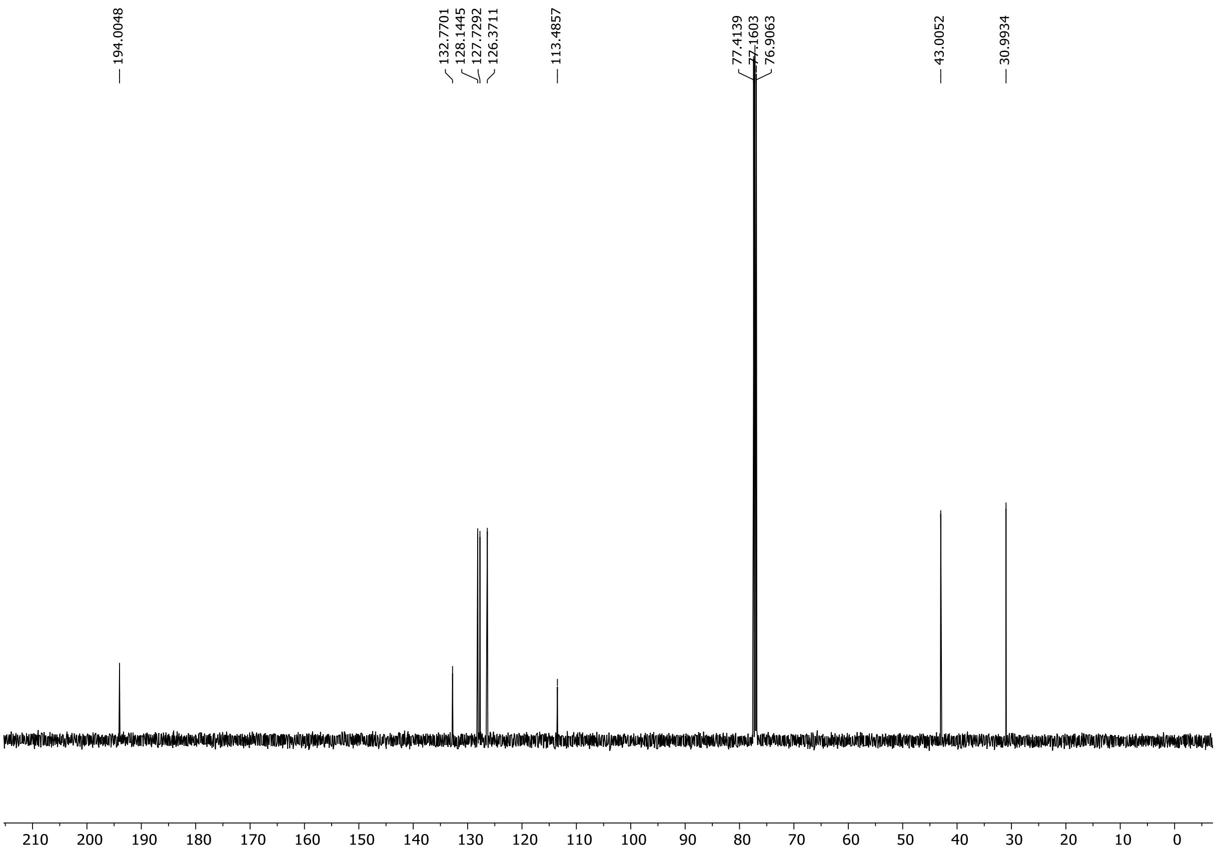

HRMS spectrum of **S116**

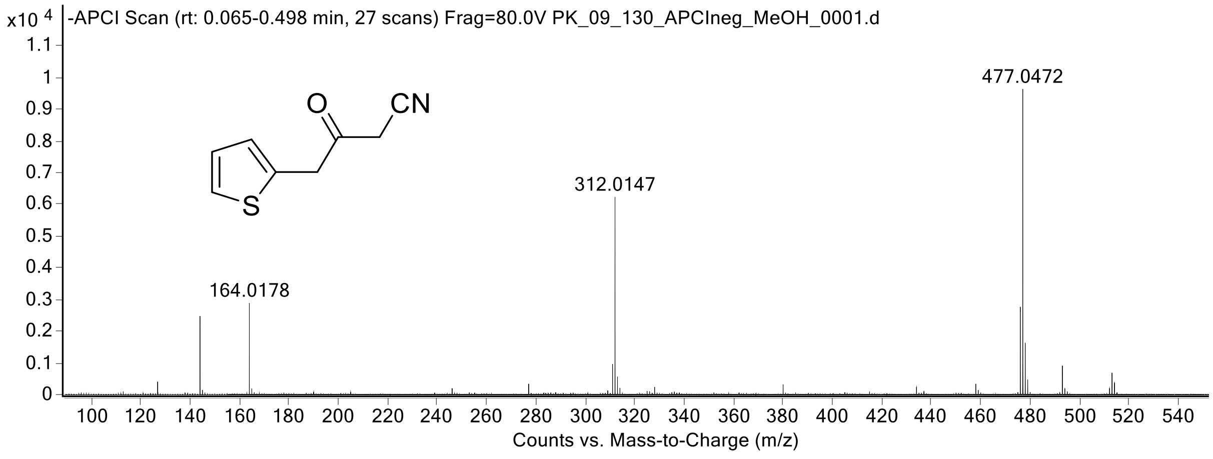

$^{1}H$ (300 MHz) spectrum of **S117** in DMSO-$d_6$

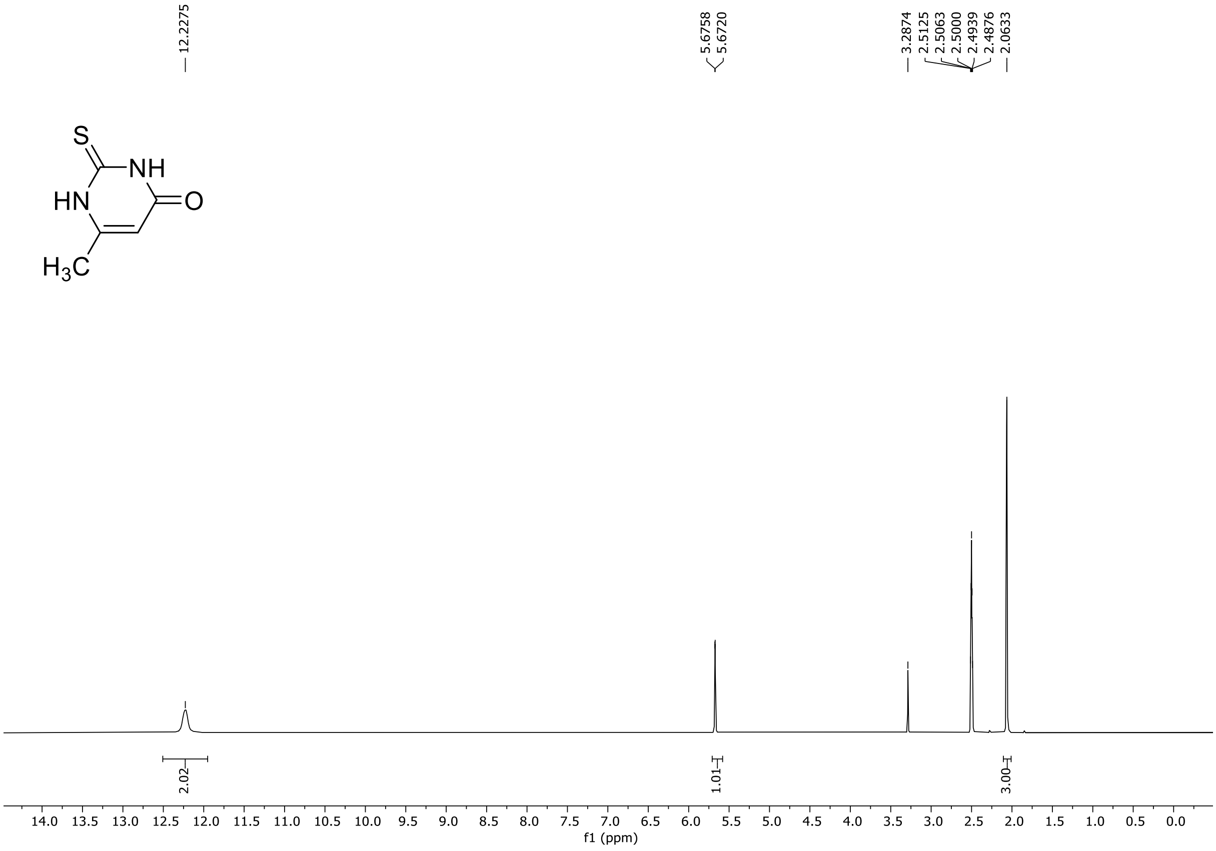

HRMS spectrum of **S117**

**NAR-A-127**

**$C_5H_6N_2OS$**

**mono *m/z* 142.0201**

**APCI -** (MMI)
nitrogen flow 5 L/min, gas temperature 325°C, nebulizer 45 psi, skimmer 65 V,
vaporizer 200°C, fragmentor 38 V, dissolved in methanol

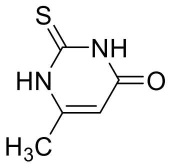

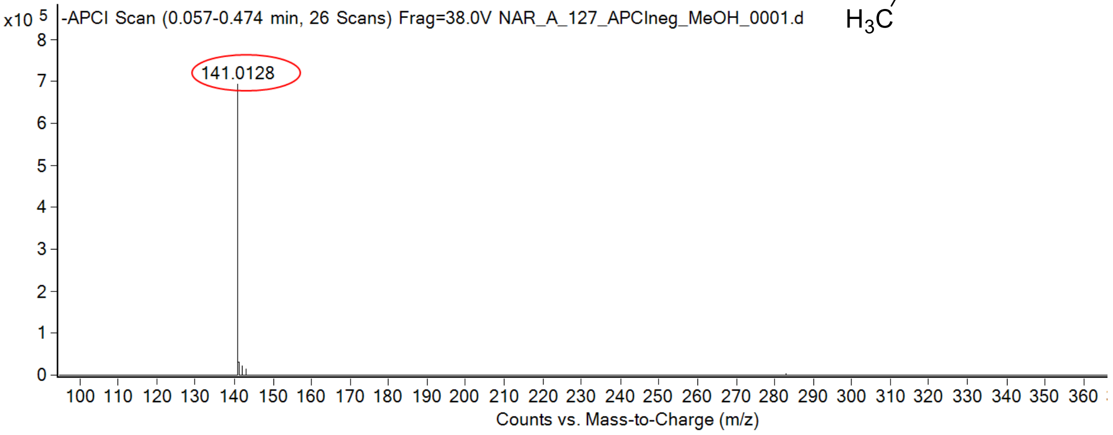

calculated mass: [M-H]$^-$ = 141.0128 observed: [M-H]$^-$ = 141.0128 mass accuracy = < 0.1 ppm

FT-IR spectrum (neat) of **S117**

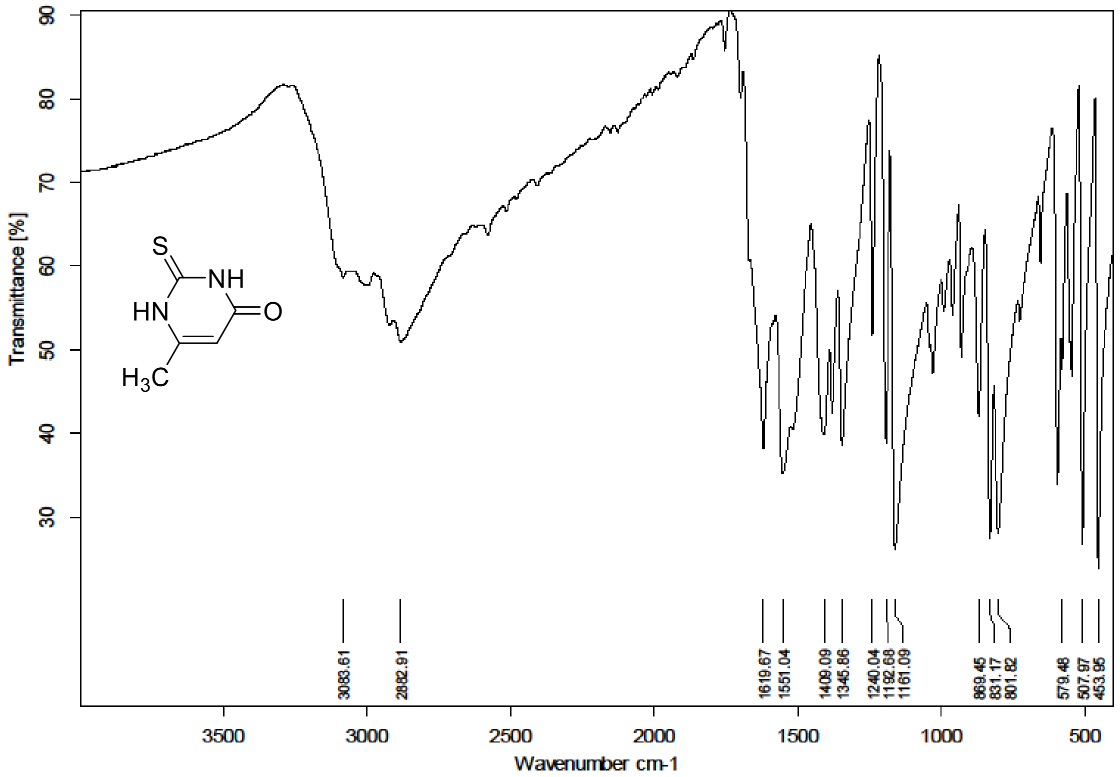

$^{1}H$ (300 MHz) spectrum of **S118** in DMSO-$d_6$

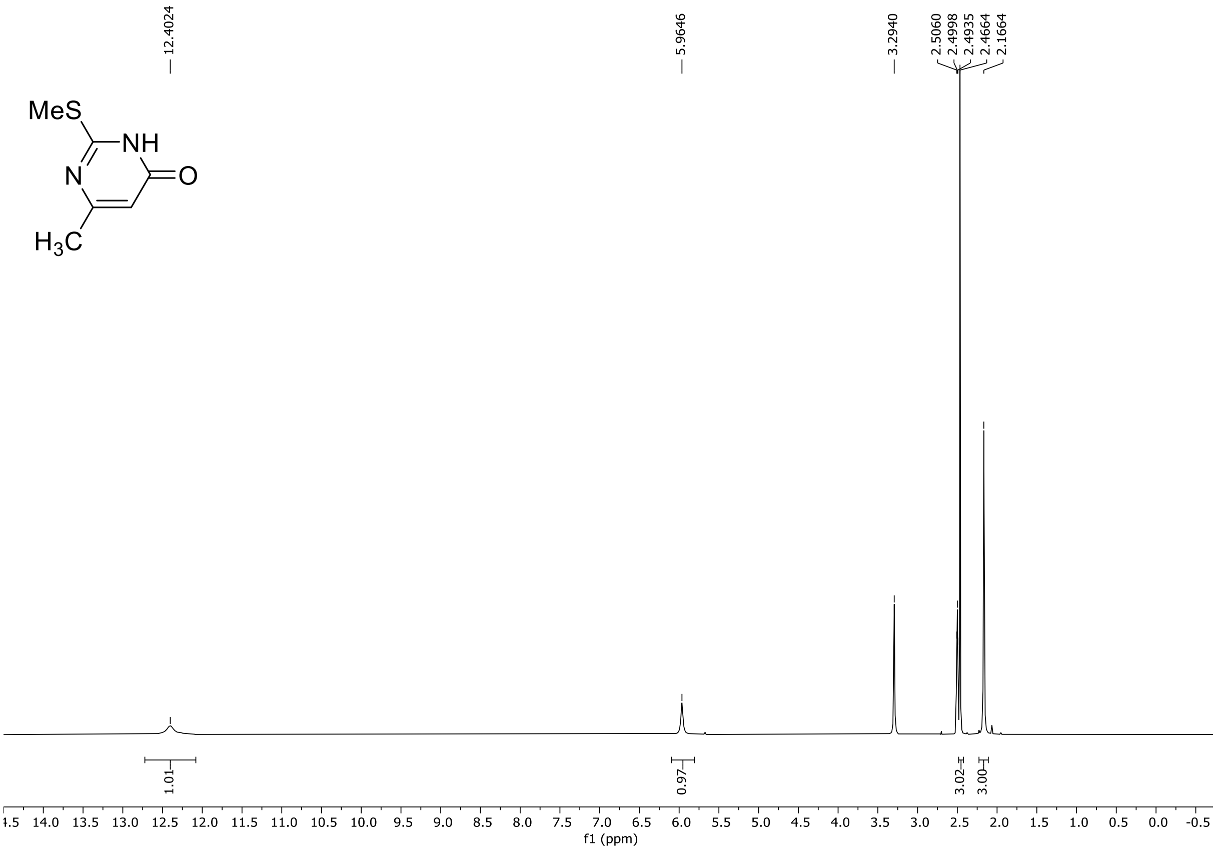

HRMS spectrum of **S118**

**NAR-A-130** **$C_6H_8N_2OS$** ***m/z* 156.0357**

**APCI+** (MMI)
nitrogen flow 5 L/min, gas temperature 325°C, nebulizer 45 psi, skimmer 65 V, vaporizer 200°C, fragmentor 30 V, dissolved in MeOH

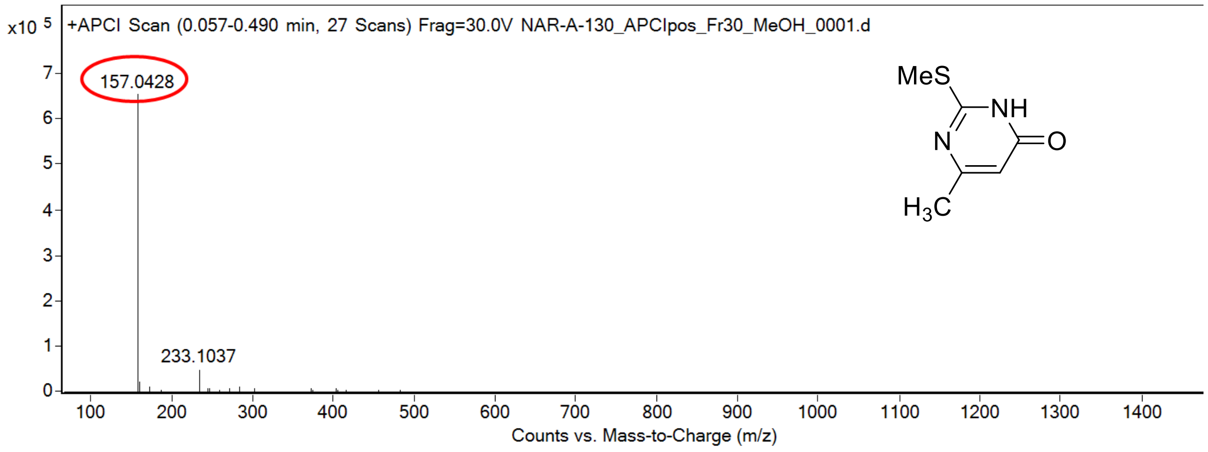

calculated mass: $[M+H]^+$ = 157.0430 observed: $[M+H]^+$ = 157.0428 mass accuracy = -1.3 ppm

FT-IR spectrum (neat) of **S118**

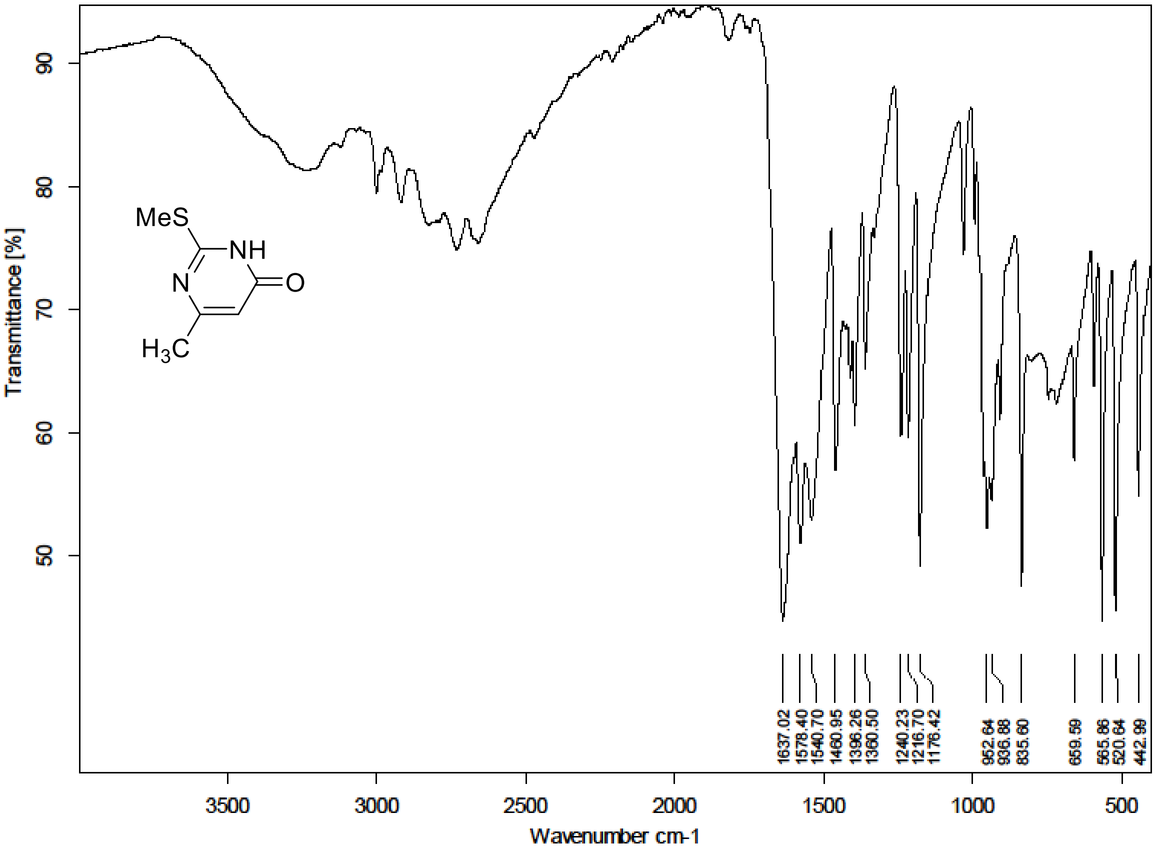

$^{1}H$ (300 MHz) spectrum of **S119** in DMSO-$d_6$

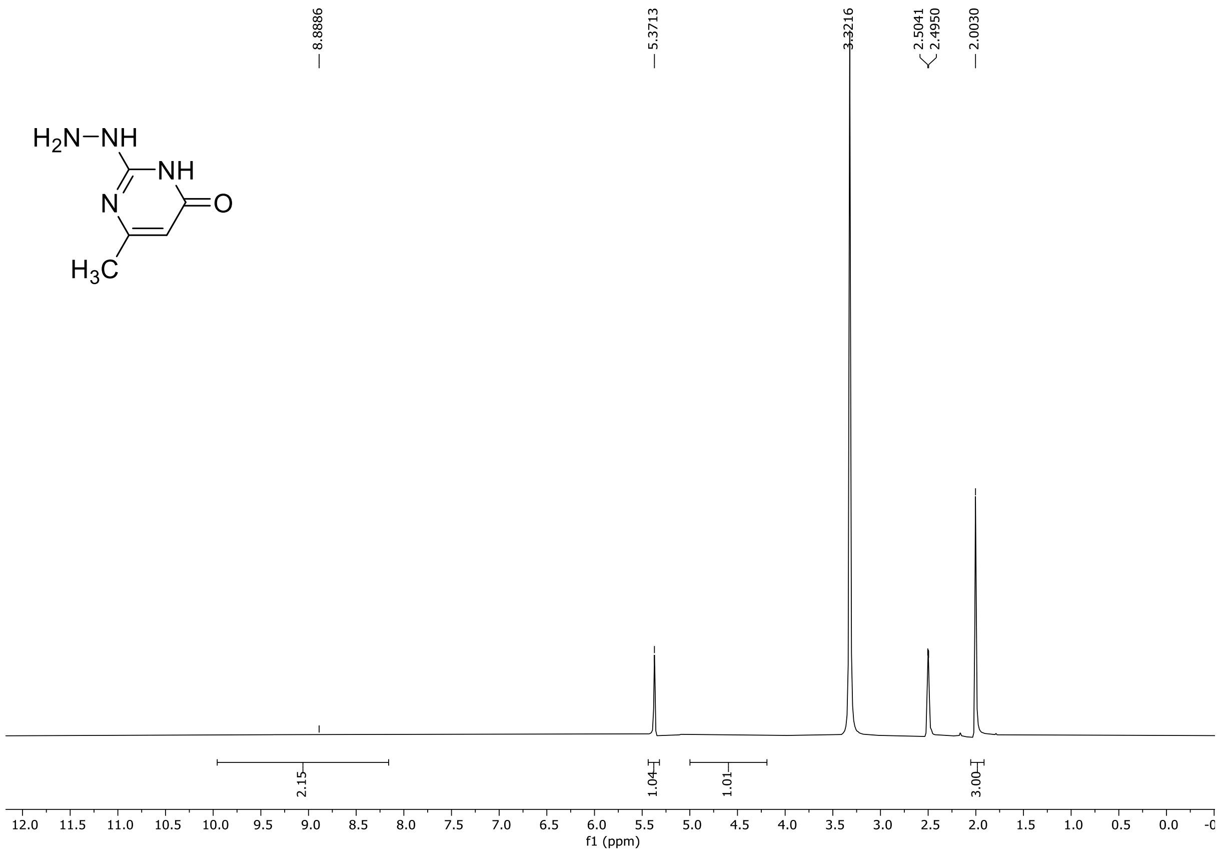

HRMS spectrum of **S119**

**NAR-A-133** **$C_5H_8N_4O$** **mono *m/z* 140.0698**

**APCI +** (MMI)
nitrogen flow 5 L/min, gas temperature 325°C, nebulizer 45 psi, skimmer 65 V, vaporizer 200°C, fragmentor 32 V, dissolved in methanol

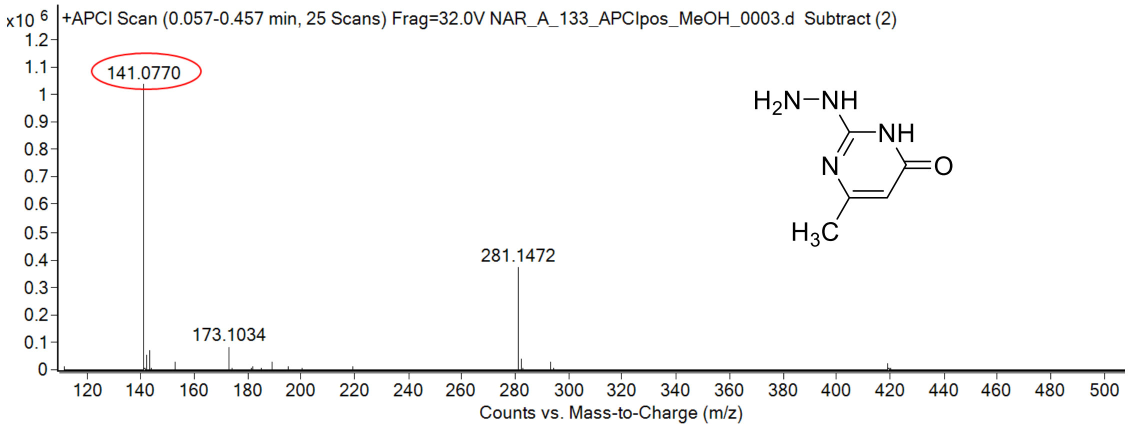

calculated mass: $[M+H]^+ = 141.0771$ observed: $[M+H]^+ = 141.0770$ mass accuracy = - 0.7 ppm

FT-IR spectrum (neat) of **S119**

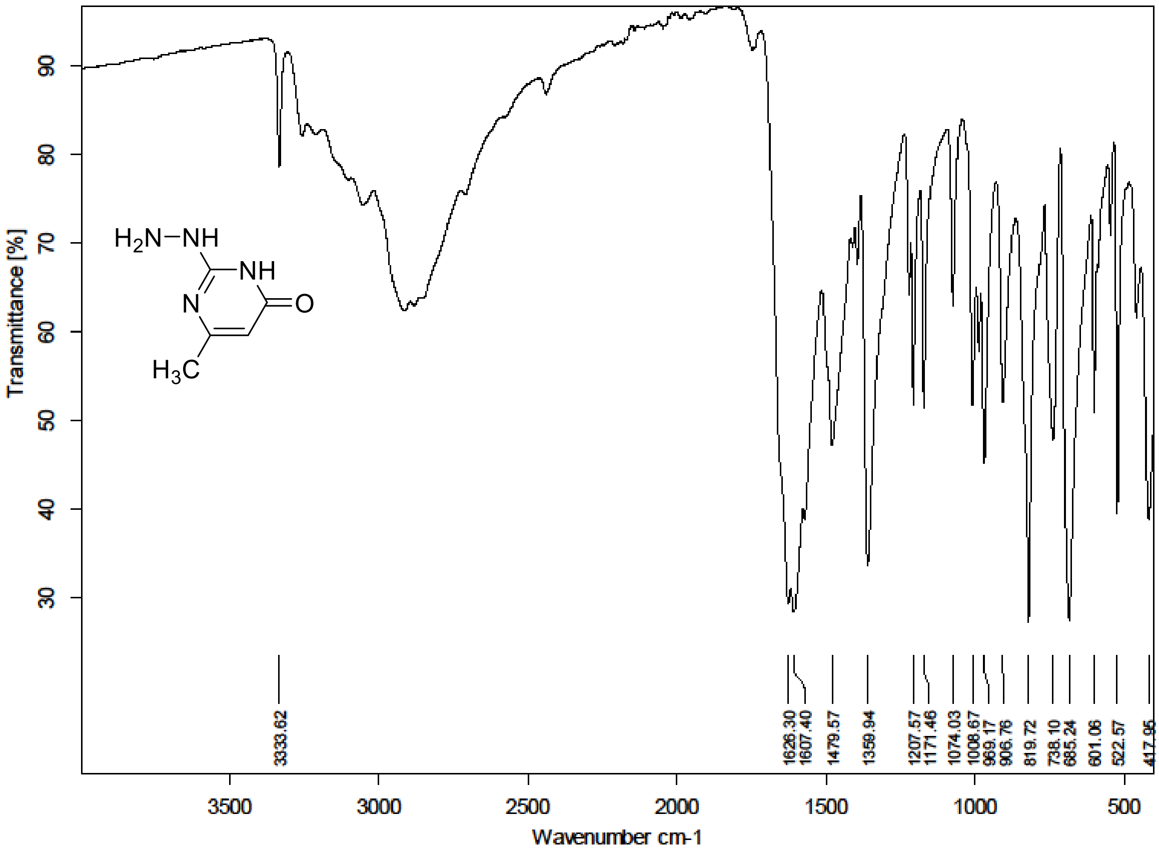

$^1$H (500 MHz) and $^{13}$C NMR (126 MHz) spectra of **S120** in DMSO-$d_6$

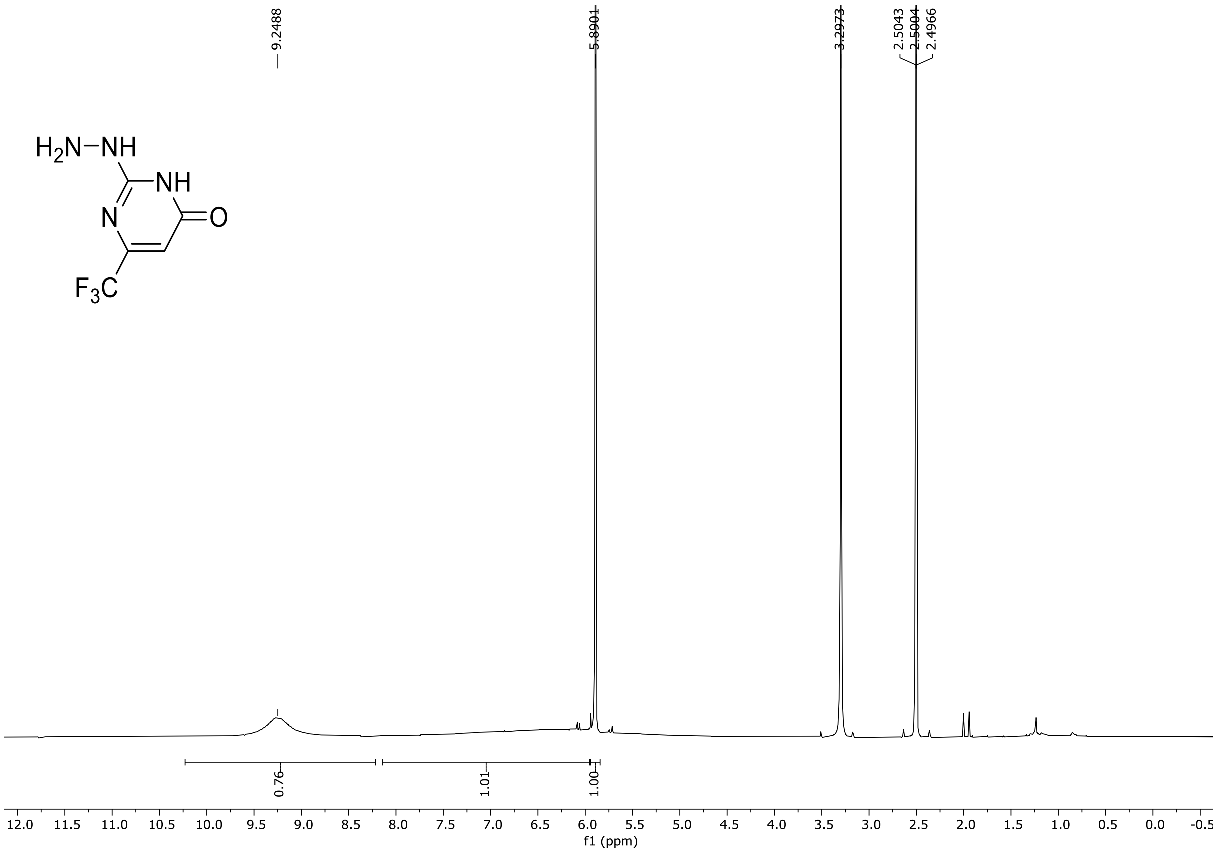

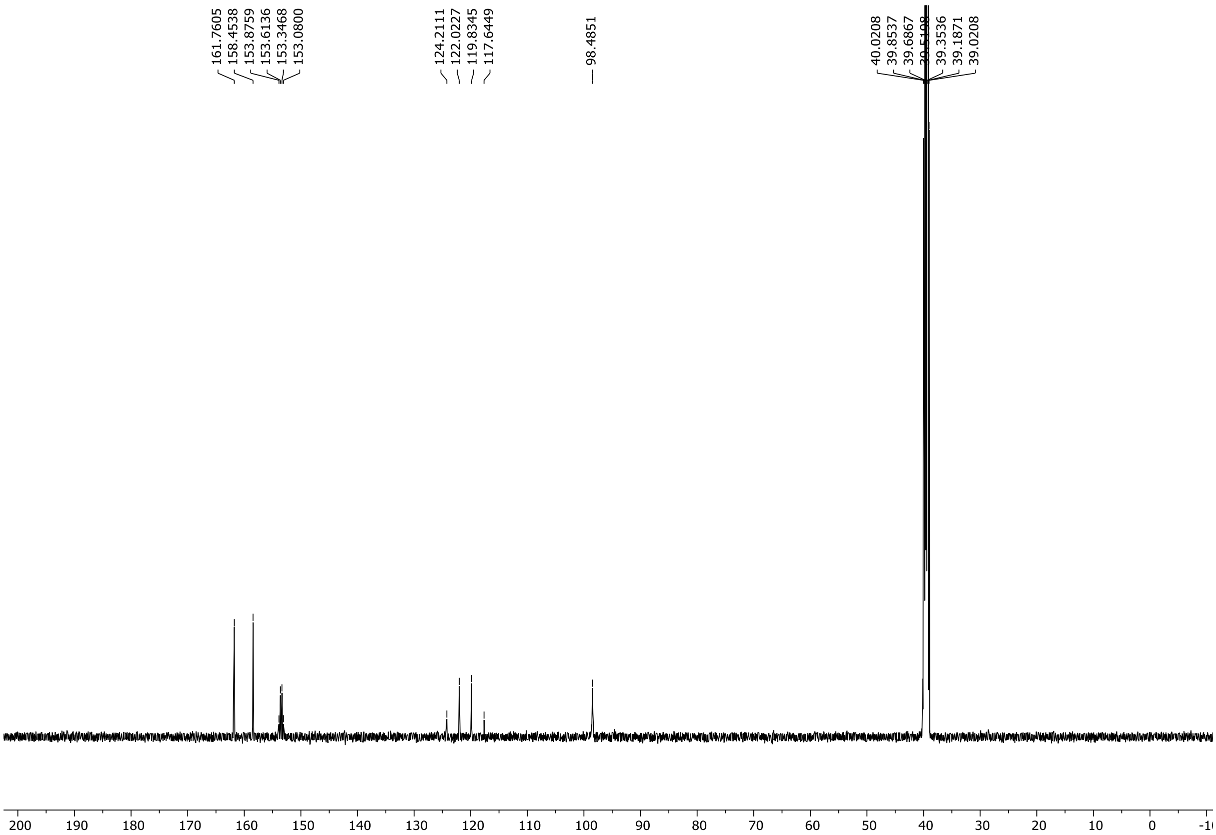

$^{19}$F NMR (471 MHz) spectrum of **S120** in DMSO-$d_6$

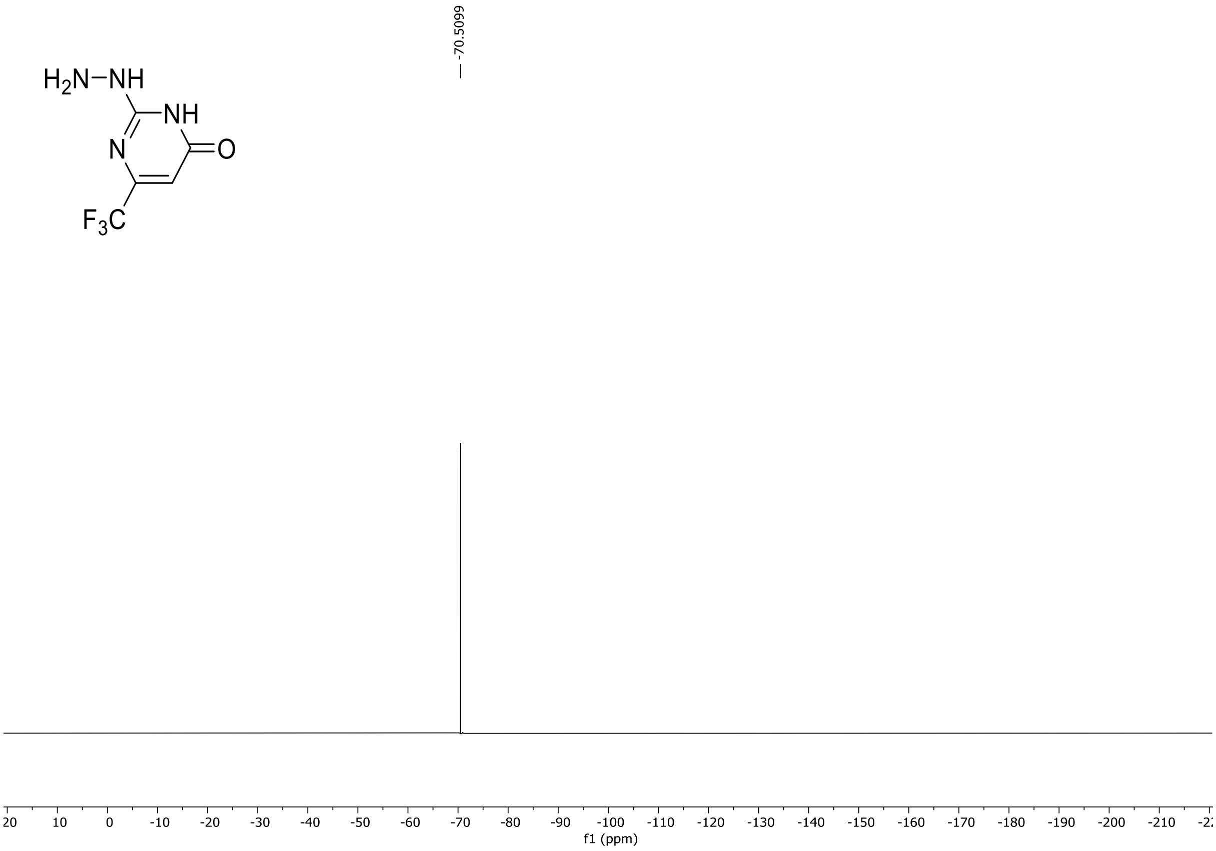

HRMS spectrum of **S120**

**NAR-A-89** **$C_5H_5F_3N_4O$** ***m/z* 194.0415**

**APCI+** (MMI)
nitrogen flow 5 L/min, gas temperature 325°C, nebulizer 45 psi, skimmer 65 V,
vaporizer 200°C, fragmentor 30 V, dissolved in MeOH

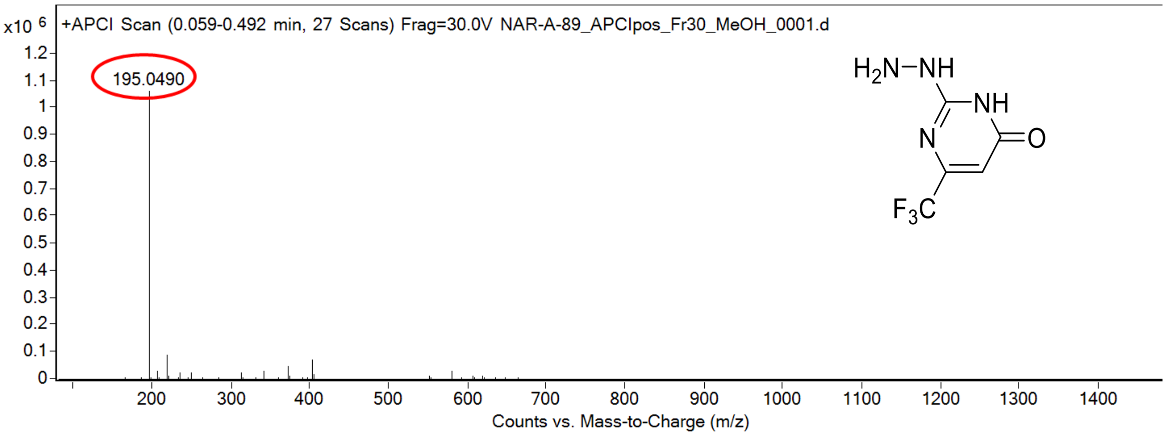

calculated mass: $[M+H]^+$ = 195.0488 observed: $[M+H]^+$ = 195.0490 mass accuracy = +1.0 ppm

FT-IR spectrum (neat) of **S120**

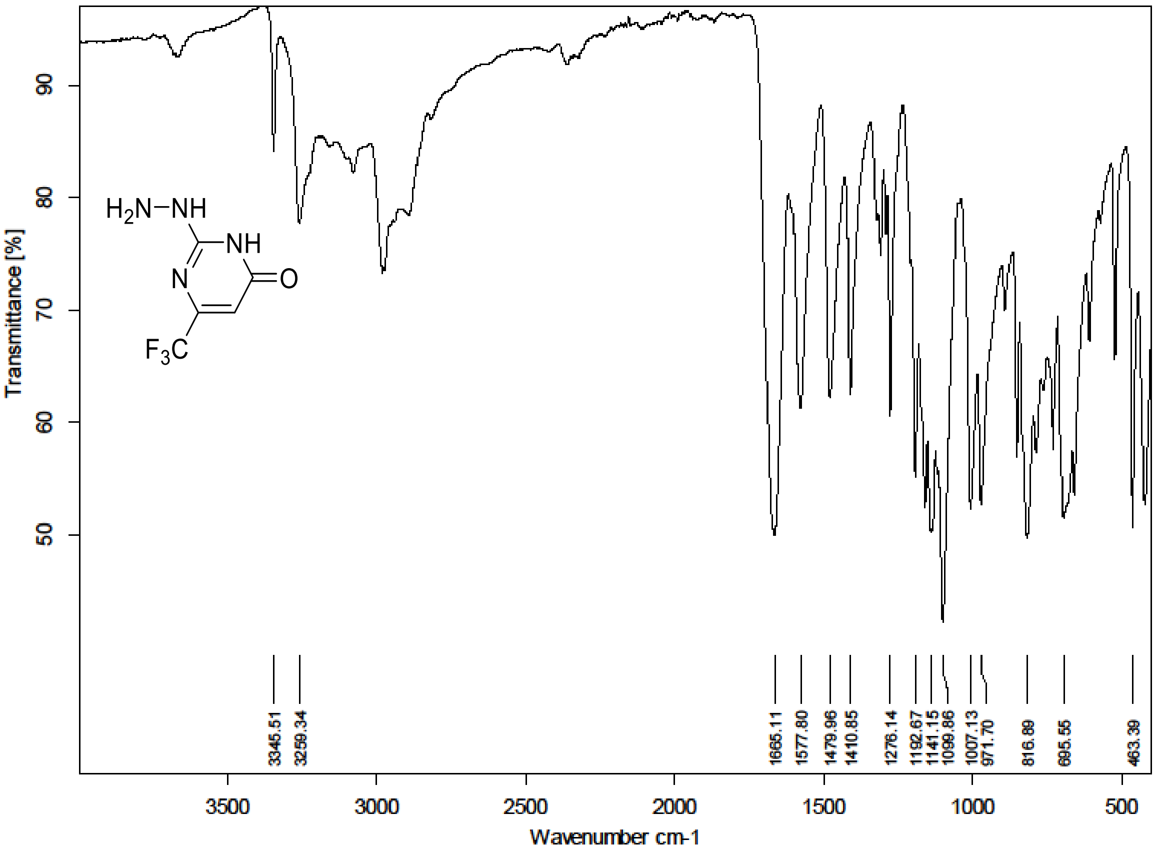

$^{1}$H (300 MHz) and $^{13}$C NMR (75 MHz) spectra of **S121** in DMSO-$d_6$

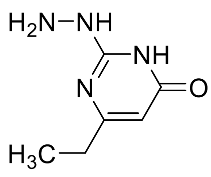

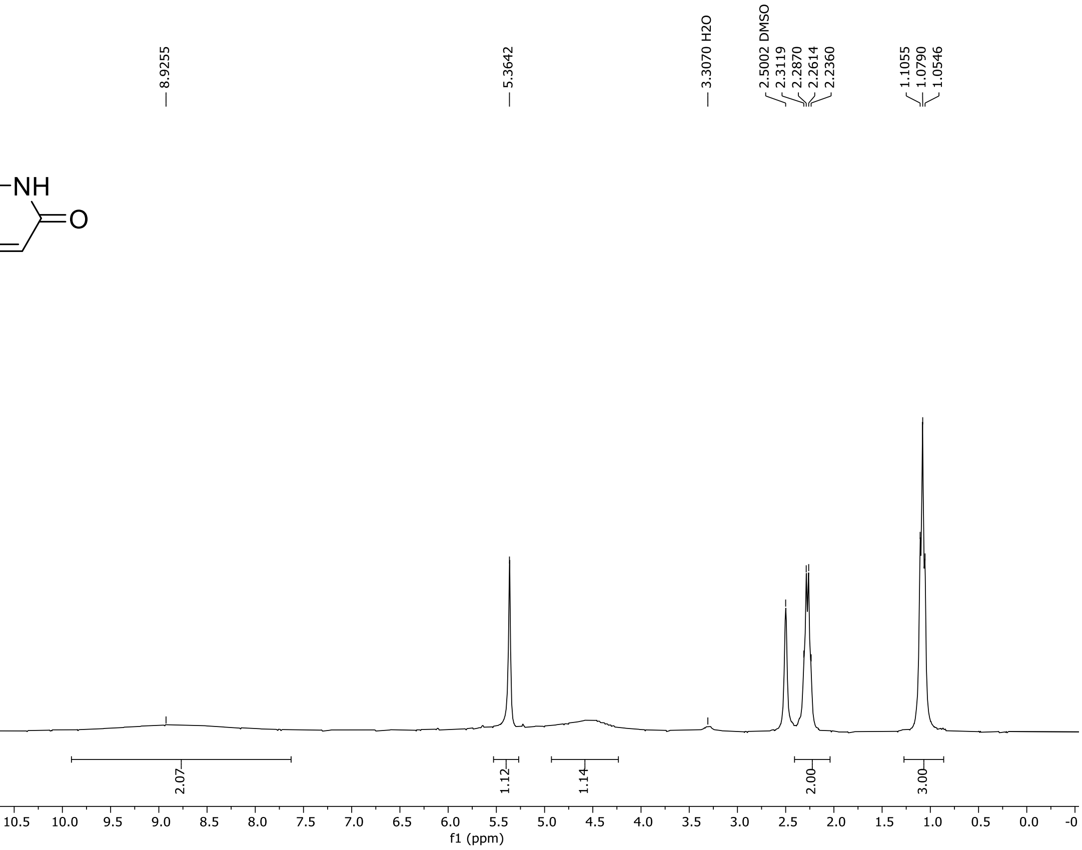

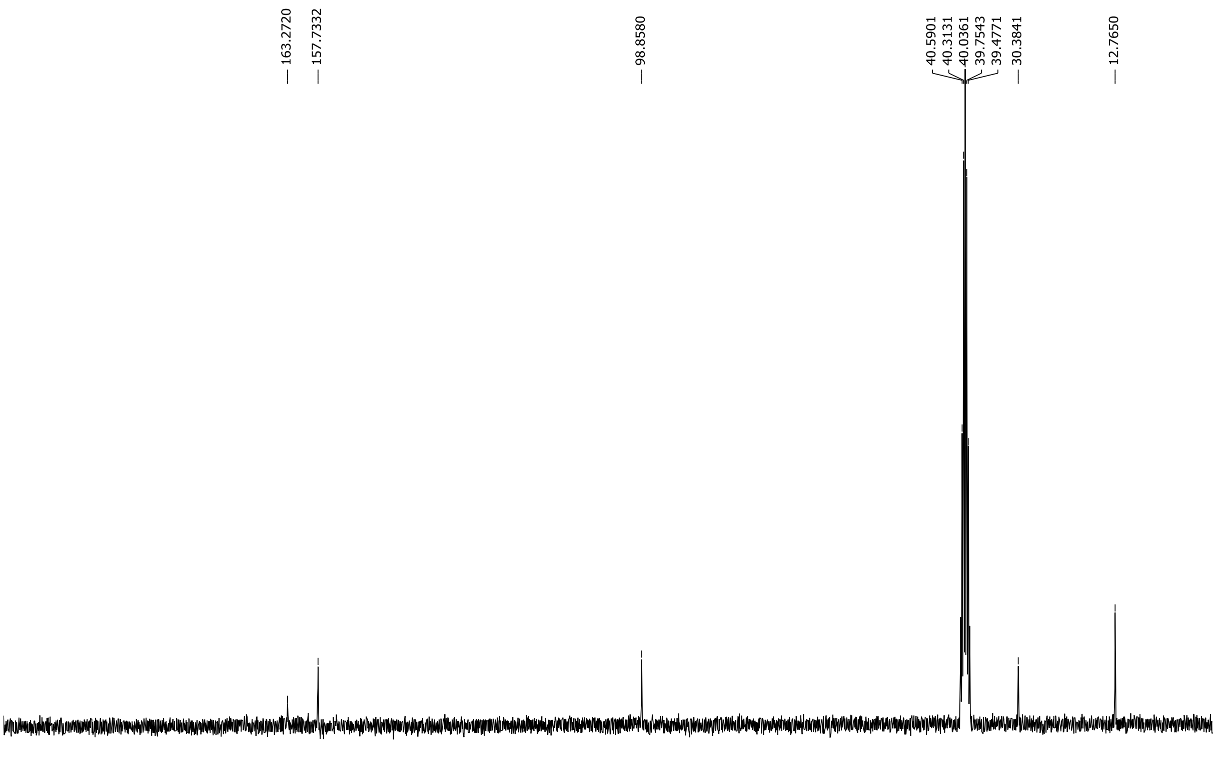

HRMS spectrum of **S121**

**NAR-A-164** $C_6H_{10}N_4O$ mono *m/z* 154.0855

**APCI +** (MMI)
nitrogen flow 5 L/min, gas temperature 325°C, nebulizer 45 psi, skimmer **65** V, vaporizer 200°C, fragmentor 20 V, dissolved in methanol

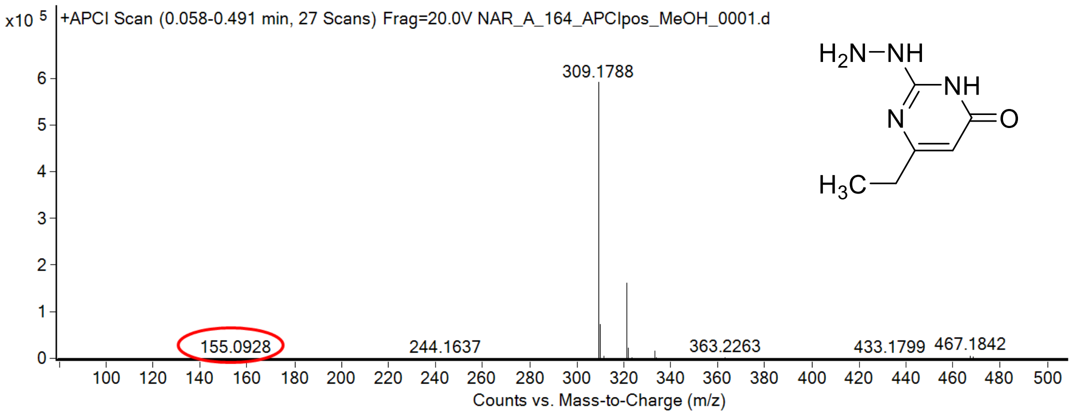

calculated mass: $[M+H]^+$ = 155.0927 observed: $[M+H]^+$ = 155.0928 mass accuracy = 0.6 ppm

FT-IR spectrum (neat) of **S121**

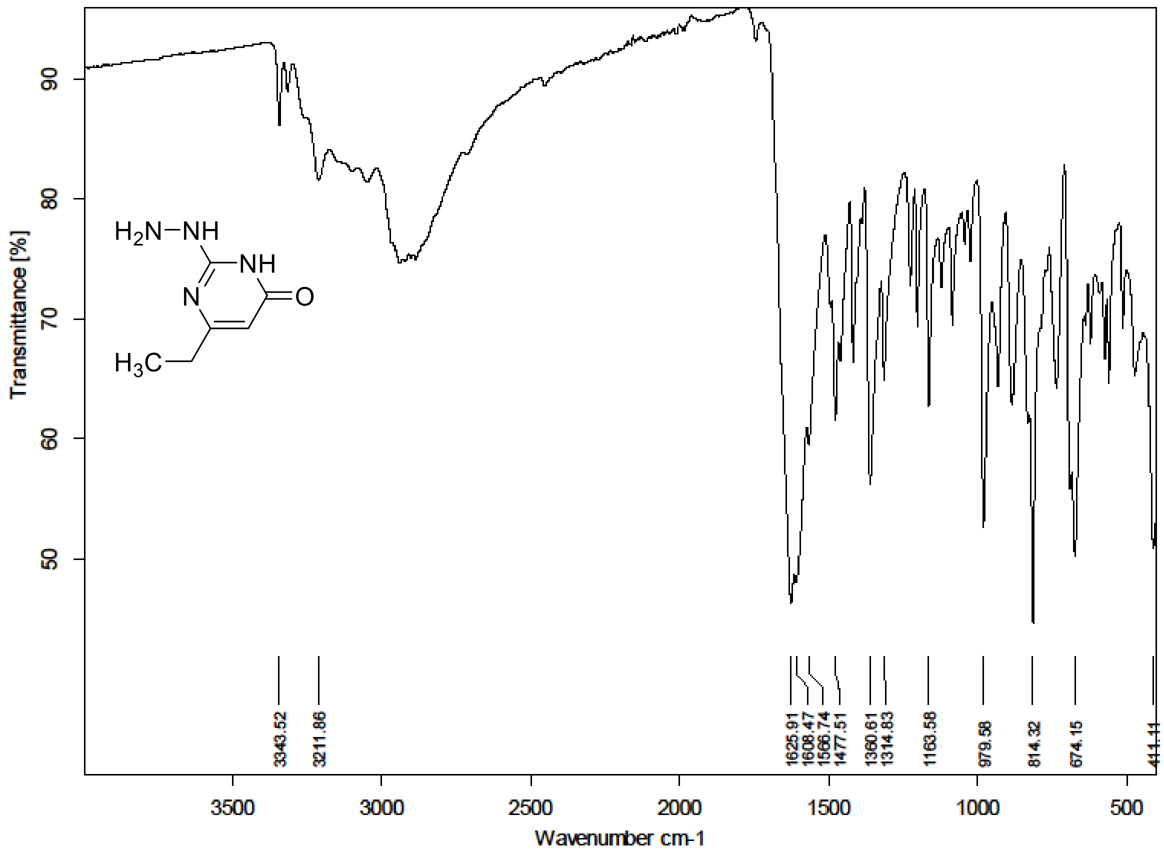

[1]H (300 MHz) and [13]C NMR (75 MHz) spectra of **S122** in Chloroform-*d*

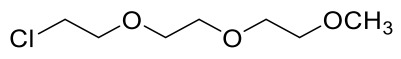

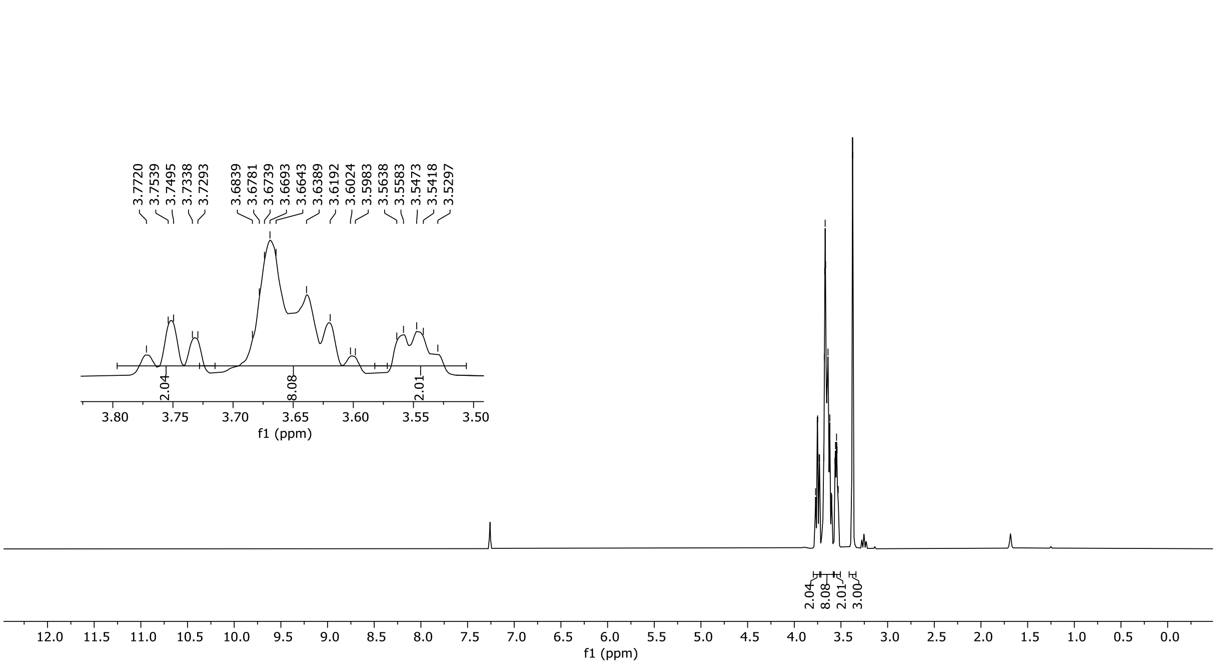

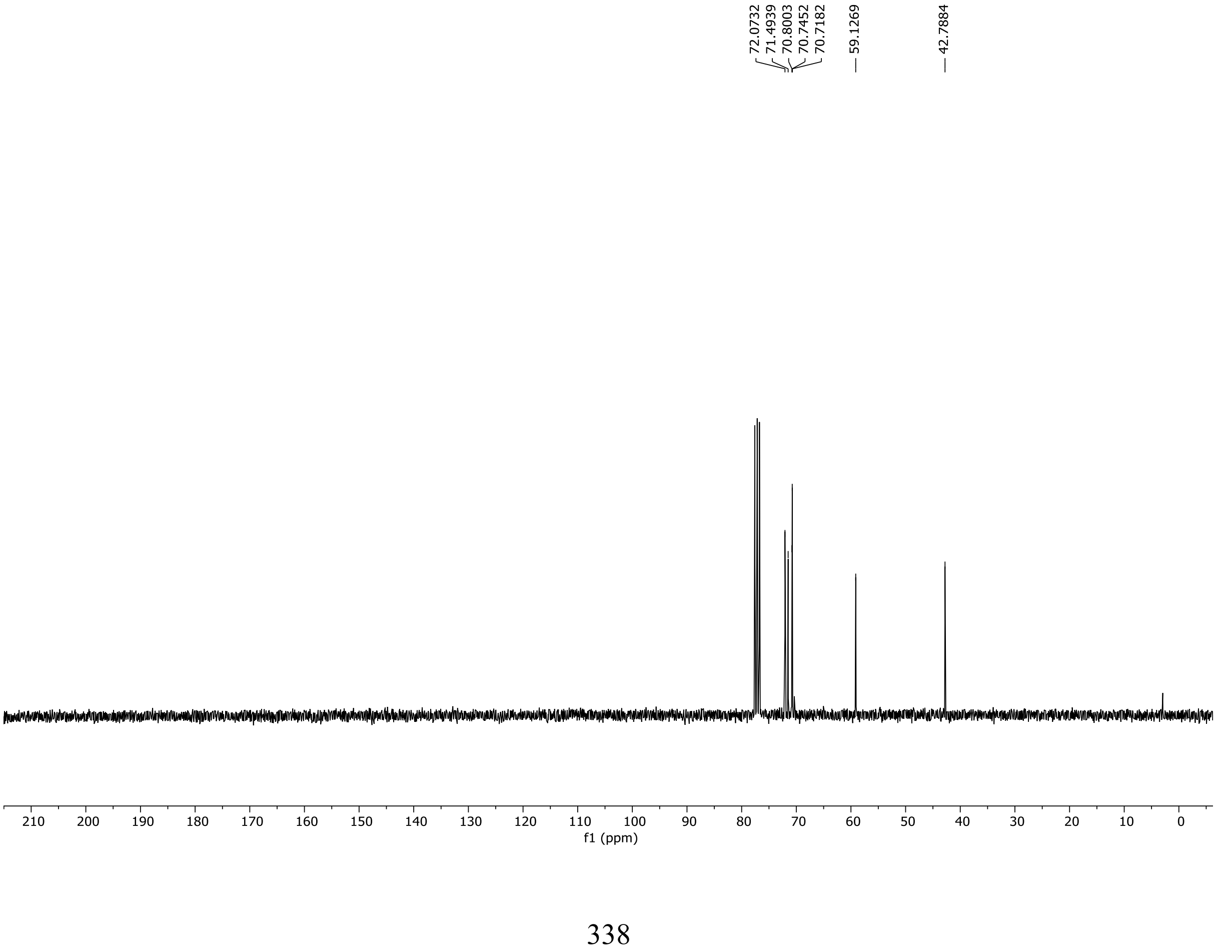

HRMS spectrum of **S122**

**NAR-A-185**

$C_7H_{15}ClO_3$ mono *m/z* 182.0710

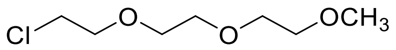

**APCI +** (MMI)
nitrogen flow 5 L/min, gas temperature 325°C, nebulizer 45 psi, skimmer **65** V, vaporizer 200°C, fragmentor 35 V, dissolved in methanol

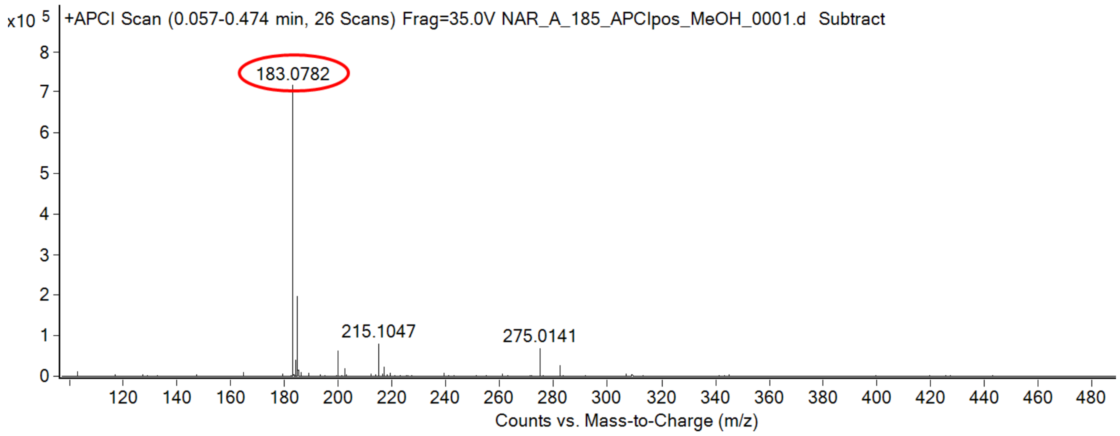

calculated mass: $[M+H]^+$ = 183.0782 observed: $[M+H]^+$ = 183.0782 mass accuracy = < 0.1 ppm

FT-IR spectrum (neat) of **S122**

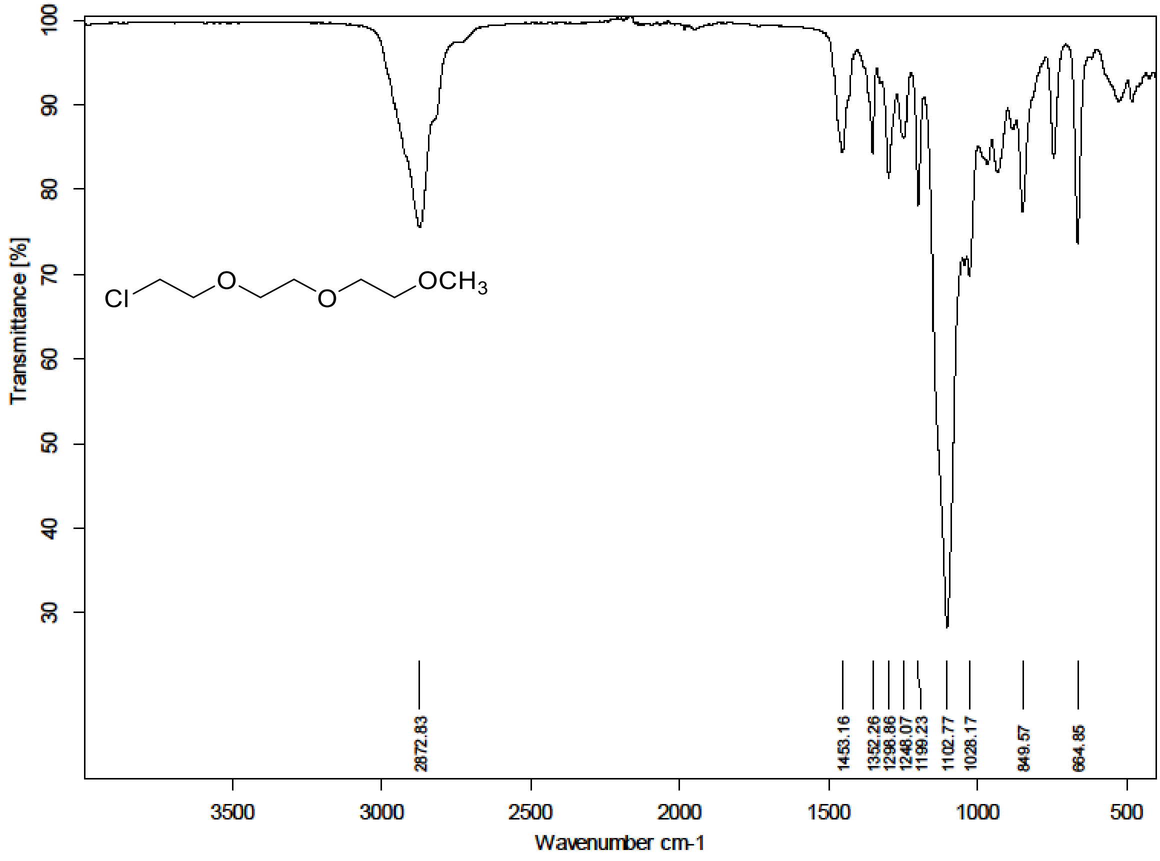

[1]H (300 MHz) and [13]C NMR (75 MHz) spectra of **S123** in Chloroform-*d*

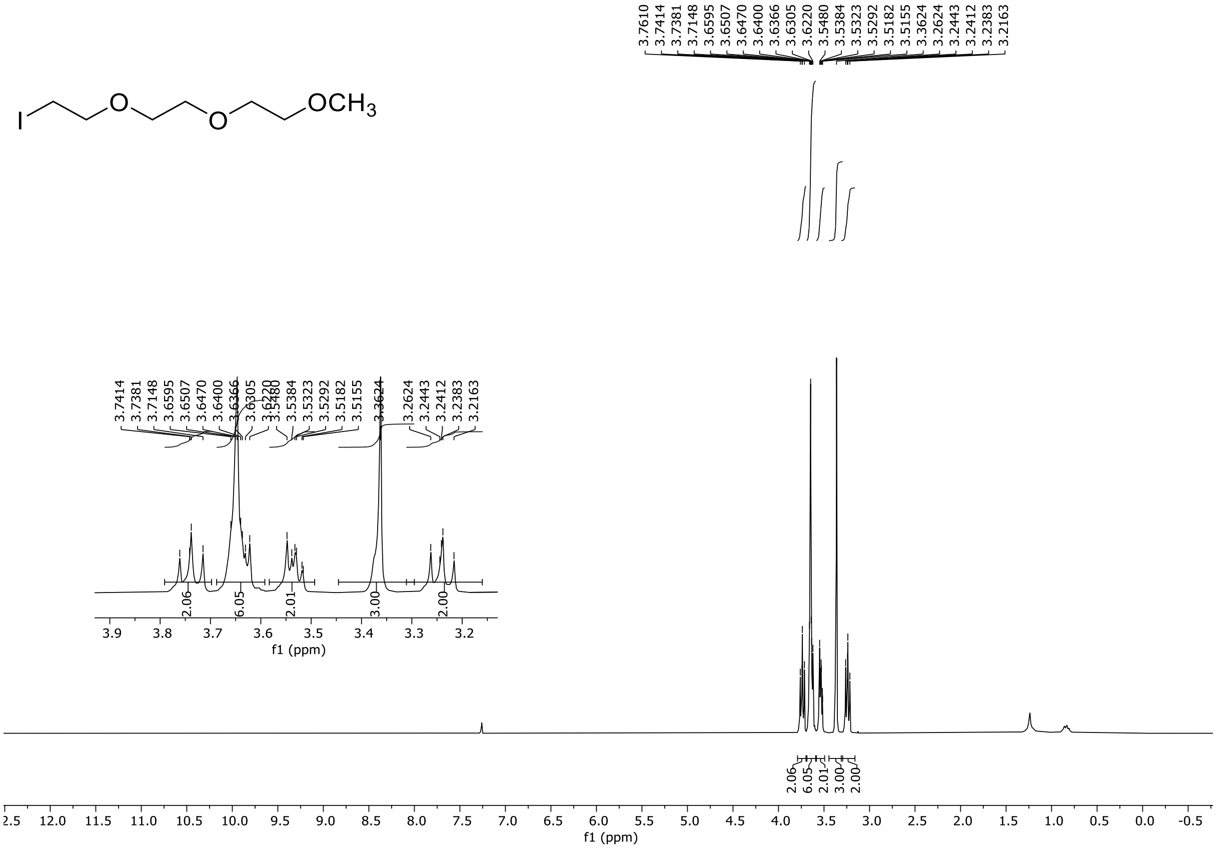

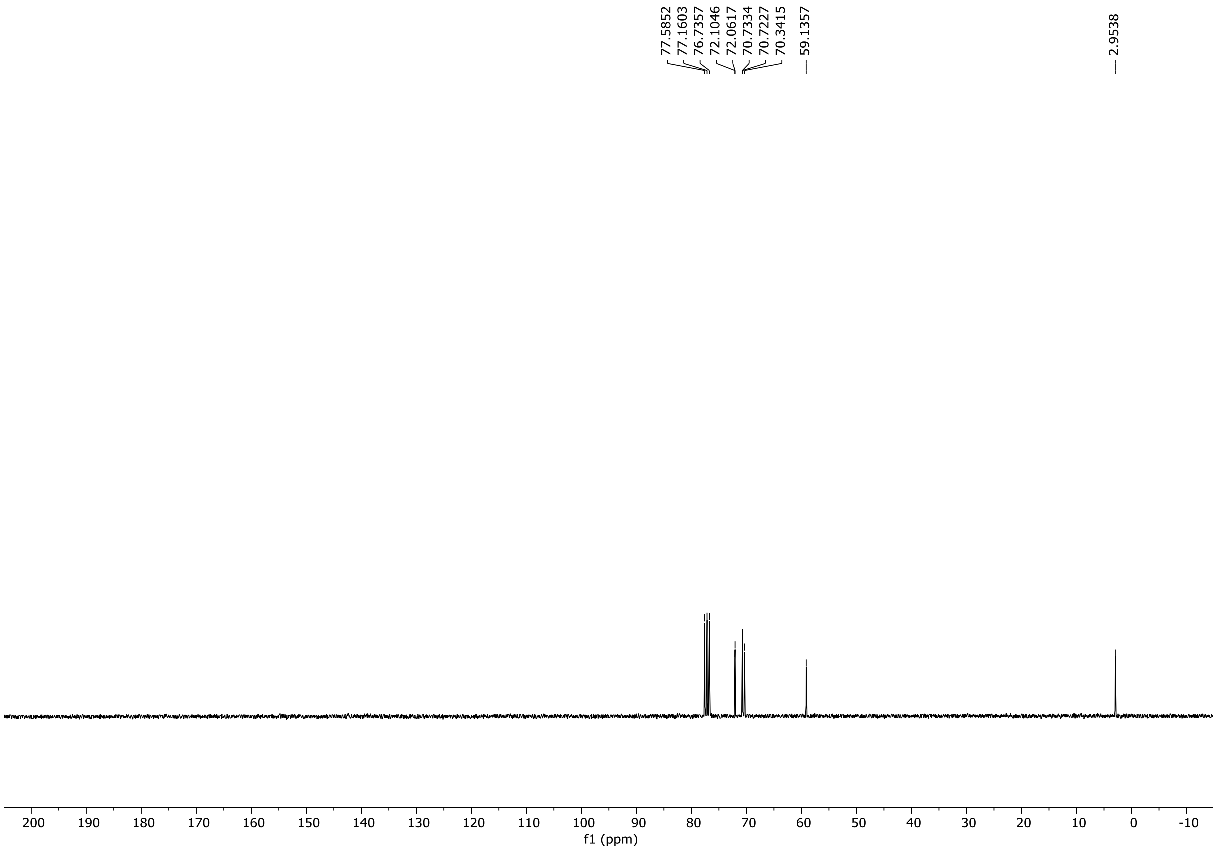

HRMS spectrum of **S123**

**NAR-A-212** $C_7H_{15}IO_3$ mono *m/z* **274.0066**

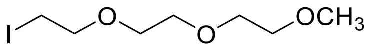

**APCI +** (MMI)
nitrogen flow 5 L/min, gas temperature 325°C, nebulizer 45 psig, skimmer 65 V, vaporizer 200°C, fragmentor 30 V, dissolved in methanol

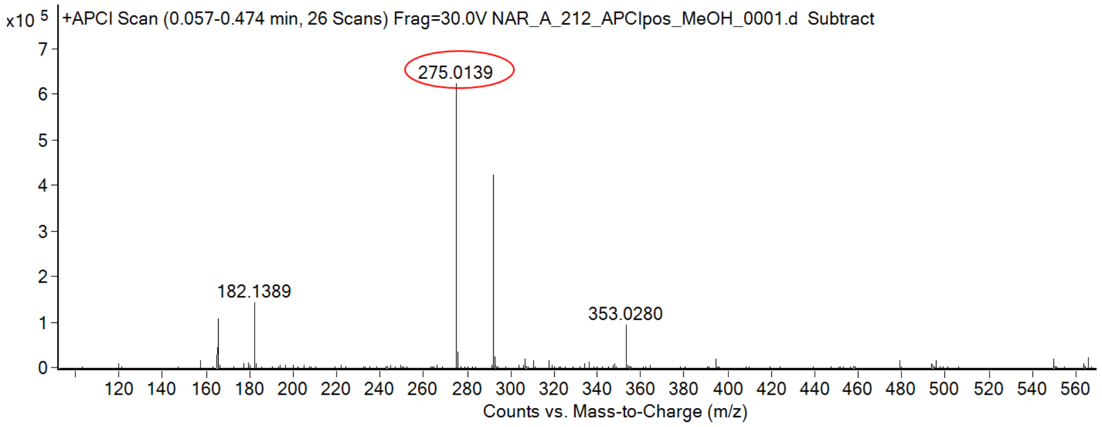

calculated mass: $[M+H]^+$ = 275.0139 observed: $[M+H]^+$ = 275.0139 mass accuracy = < 0.1 ppm

FT-IR spectrum (neat) of **S123**

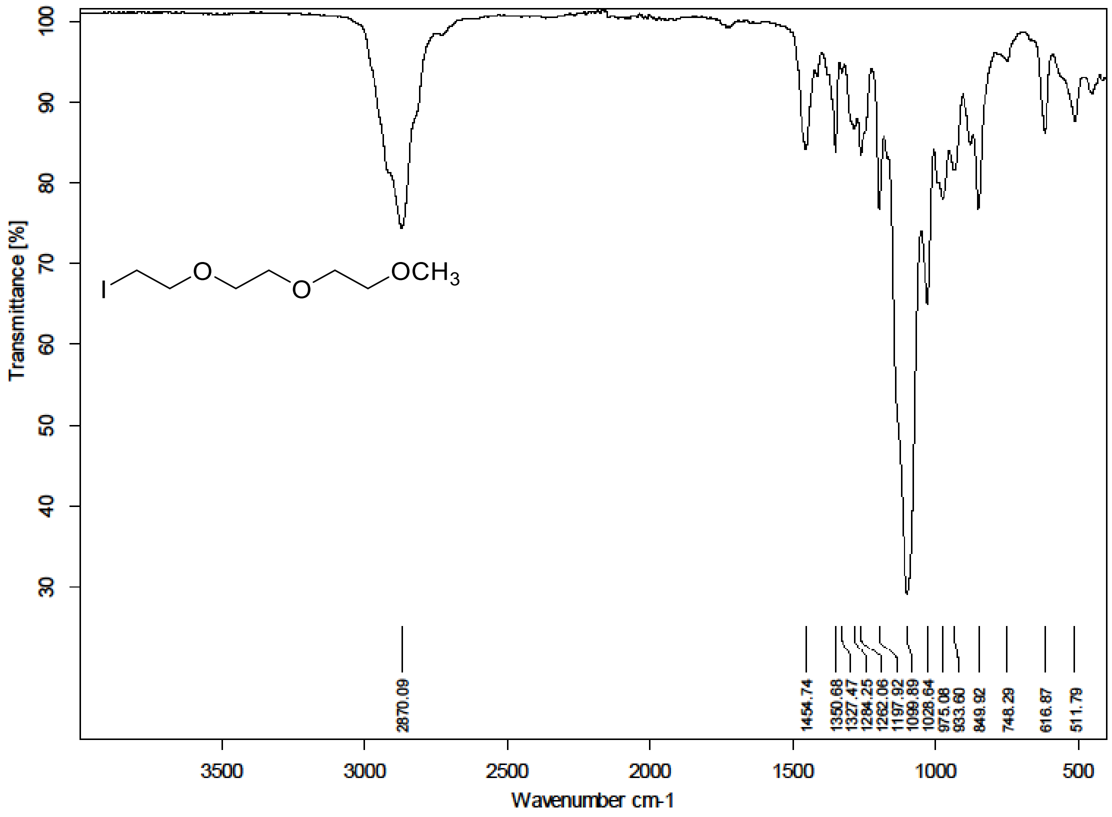

$^{1}H$ (300 MHz) and $^{13}C$ NMR (75 MHz) spectra of **S124** in Chloroform-*d*

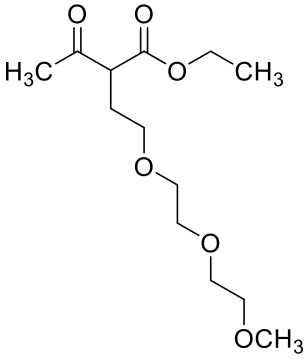

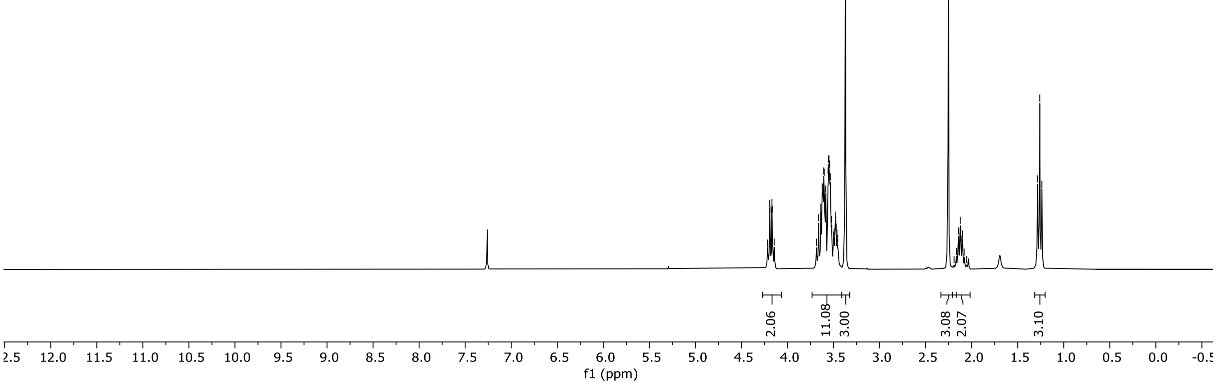

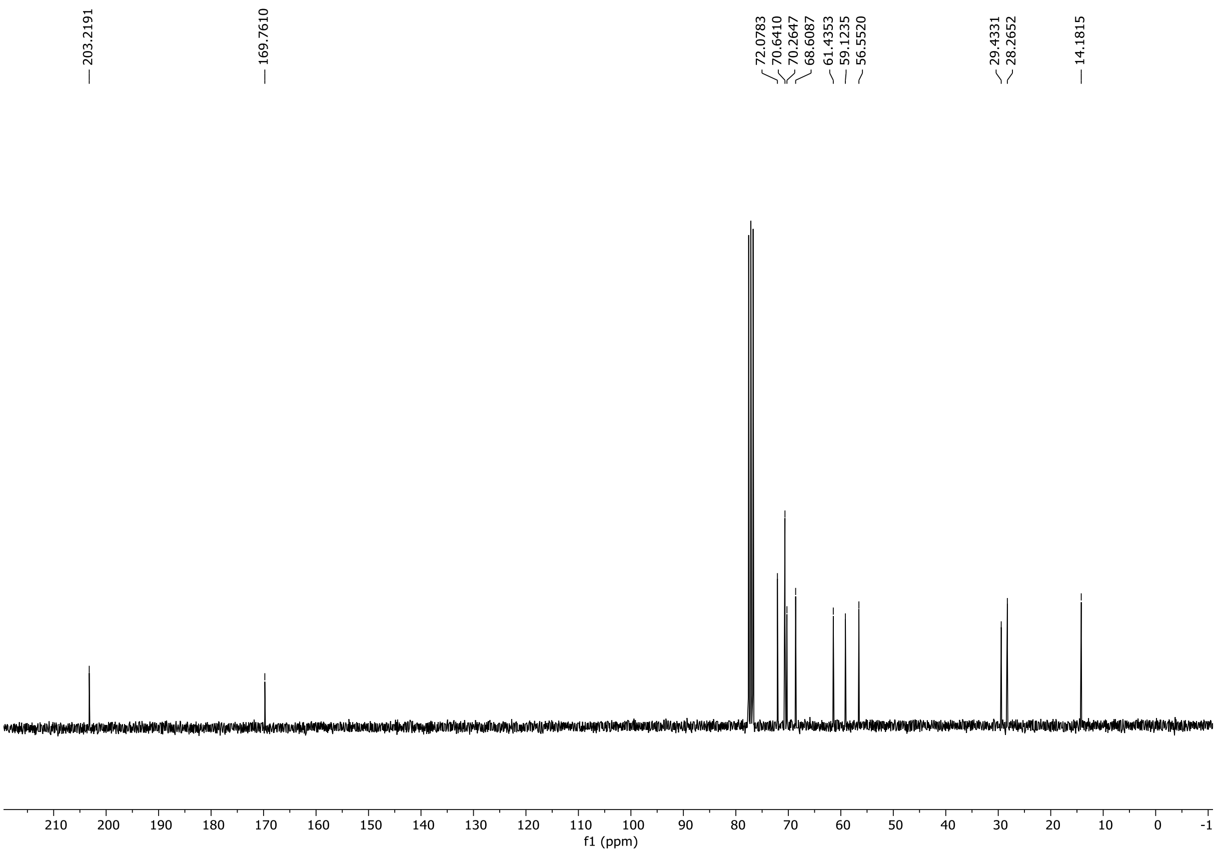

HRMS spectrum of **S124**

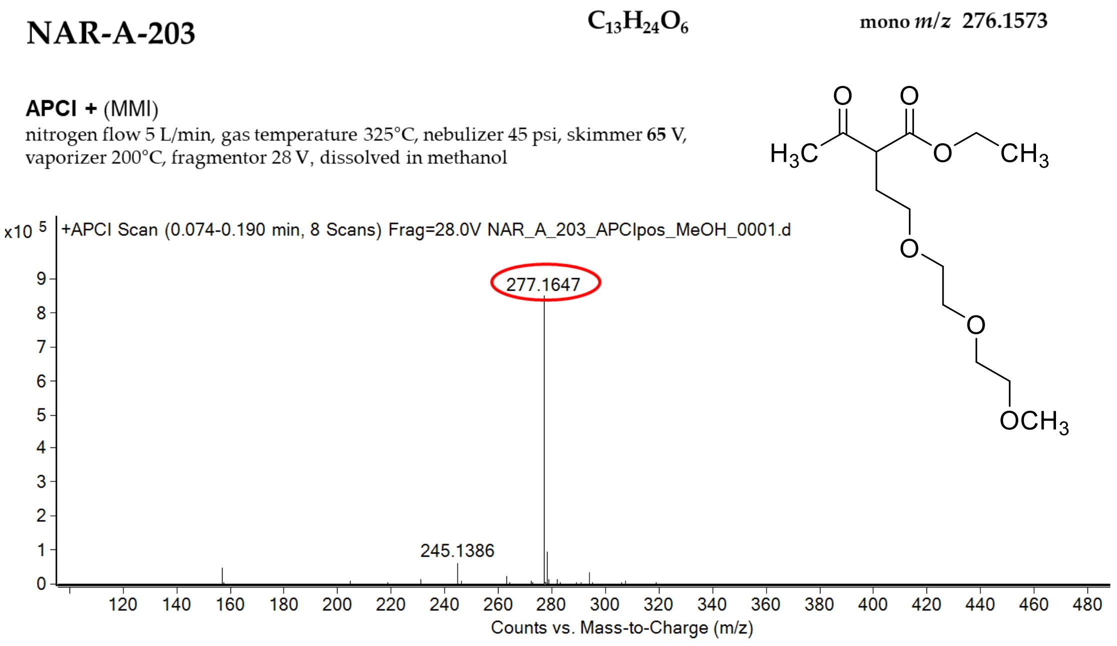

calculated mass: $[M+H]^+$ = 277.1646 observed: $[M+H]^+$ = 277.1647 mass accuracy = 0.4 ppm

*Miroslava Bittová*

FT-IR spectrum (neat) of **S124**

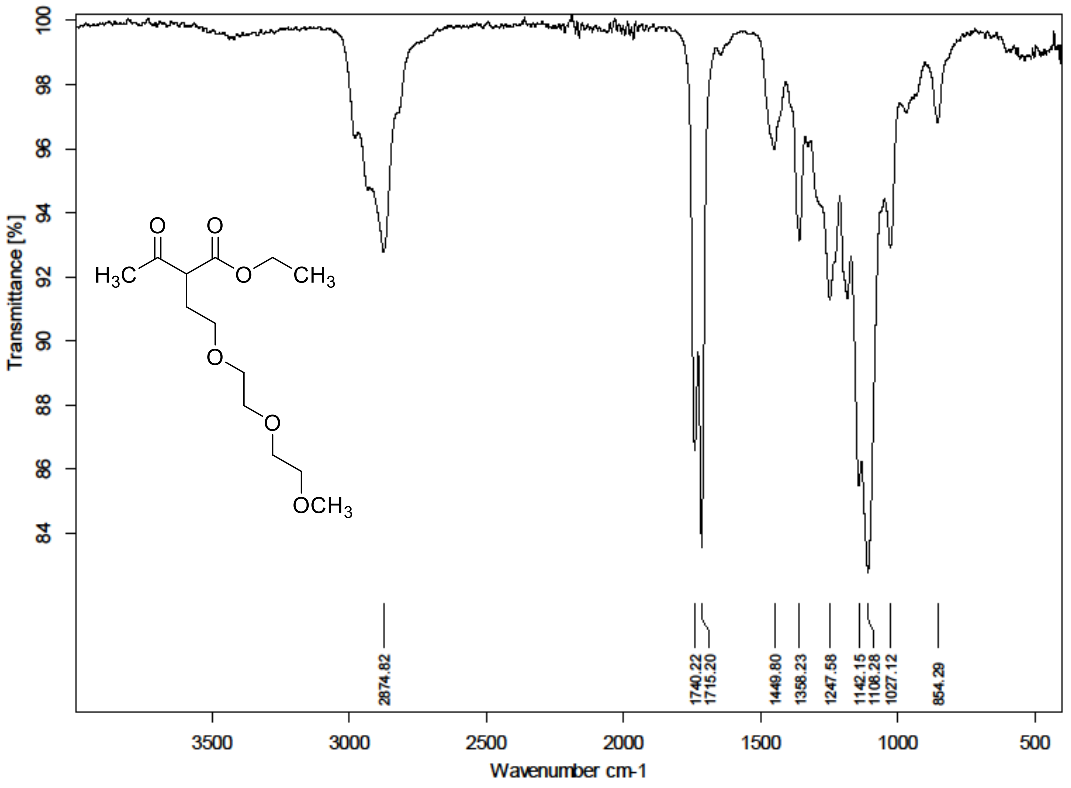

[1]H (300 MHz) and [13]C NMR (75 MHz) spectra of **S125** in Chloroform-*d*

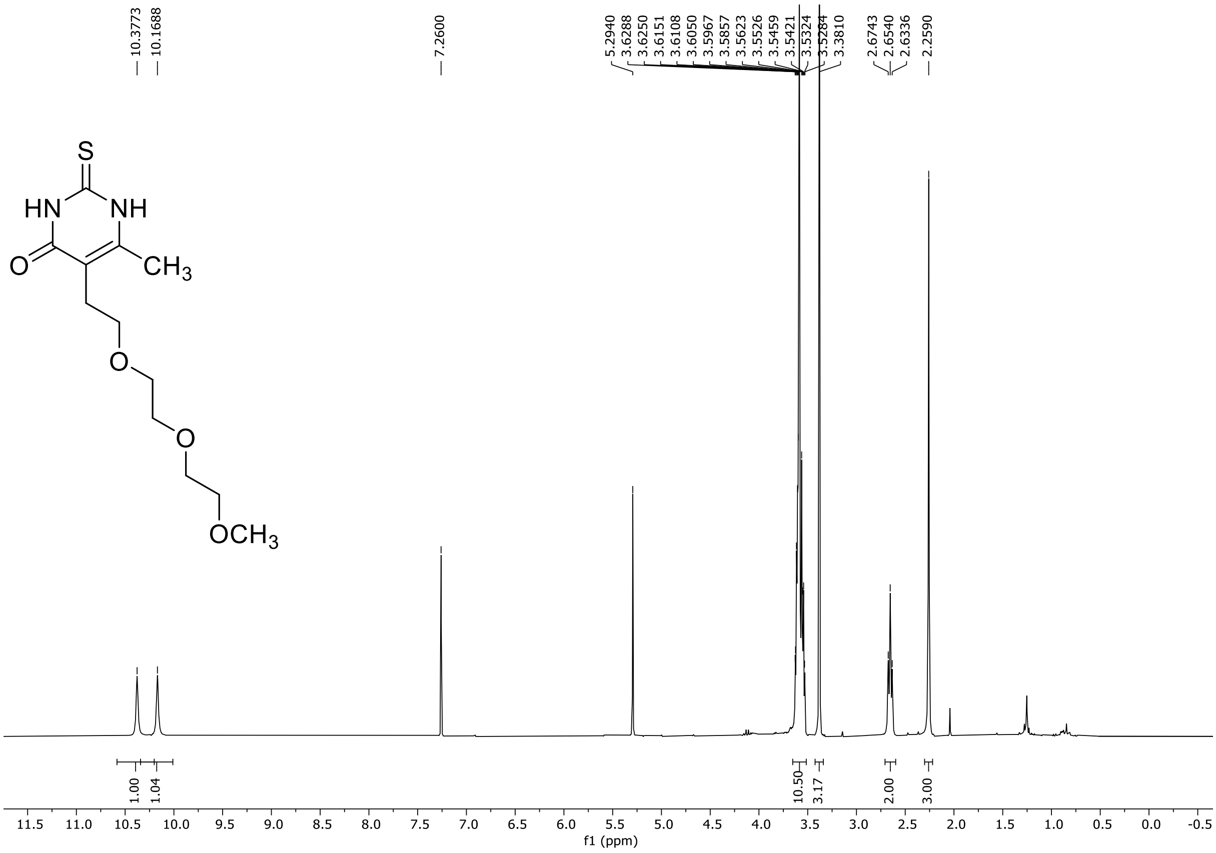

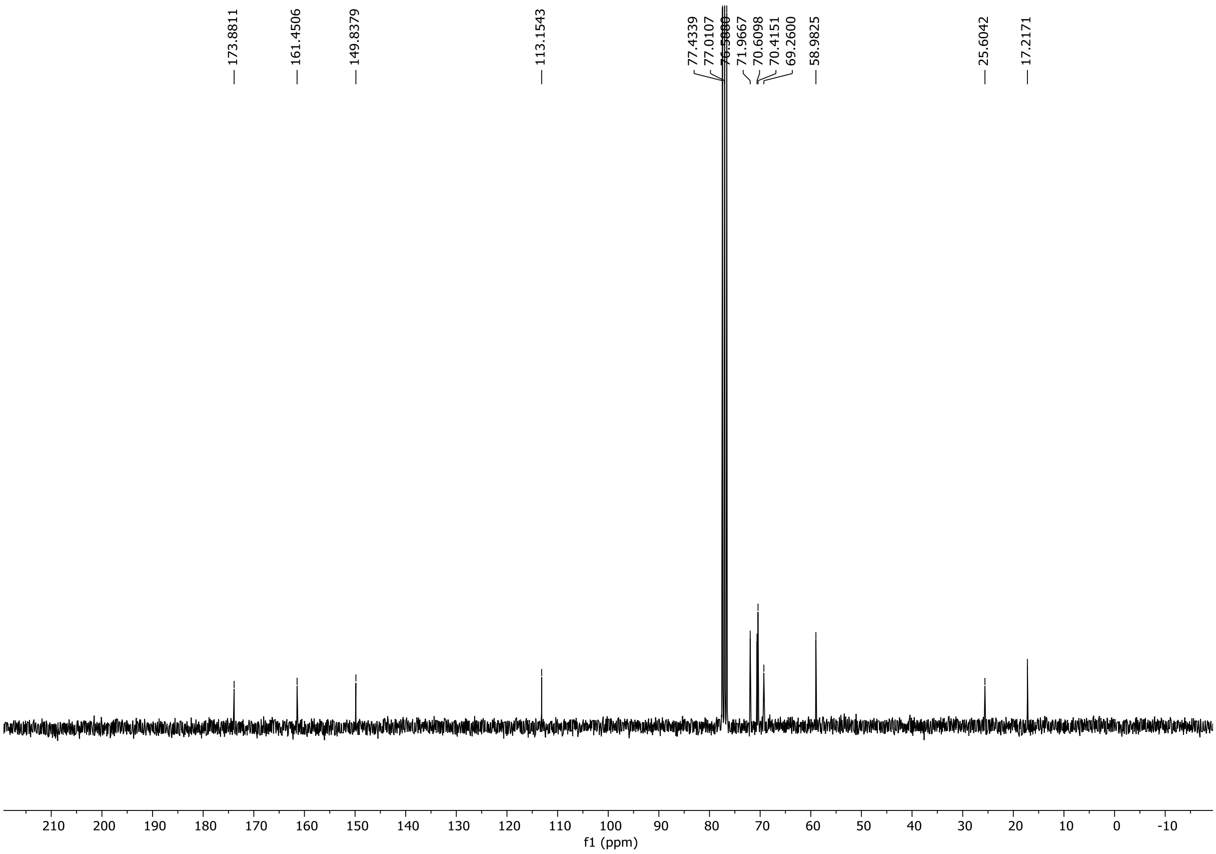

HRMS spectrum of **S125**

**NAR-A-213** $C_{12}H_{20}N_2O_4S$ mono *m/z* 288.1144

**APCI +** (MMI)
nitrogen flow 5 L/min, gas temperature 325°C, nebulizer 45 psig, skimmer 65 V, vaporizer 200°C, fragmentor 22 V, dissolved in methanol

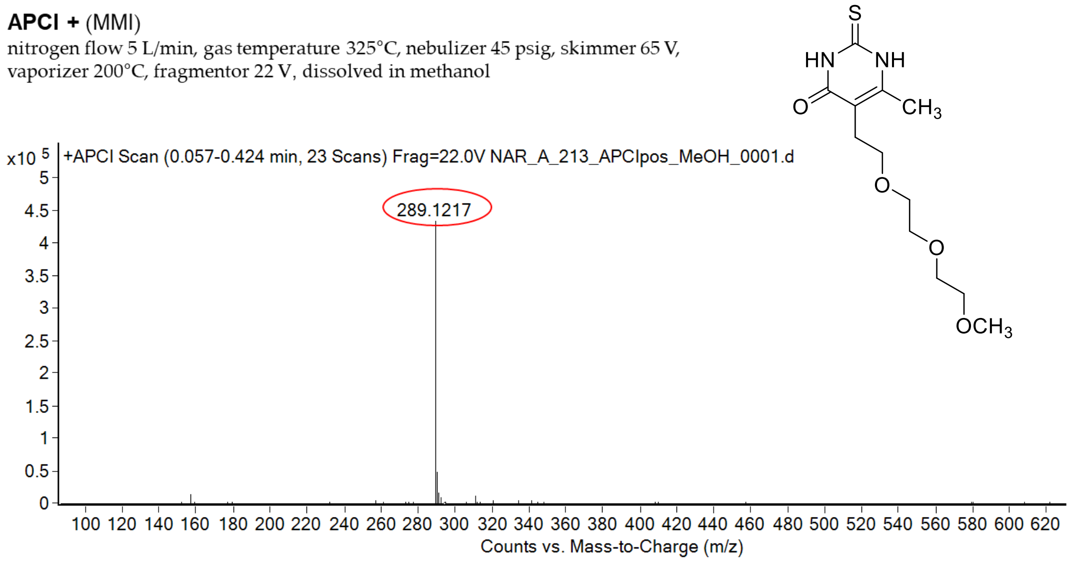

calculated mass: $[M+H]^+$ = 289.1217 observed: $[M+H]^+$ = 289.1217 mass accuracy = < 0.1 ppm

FT-IR spectrum (neat) of **S125**

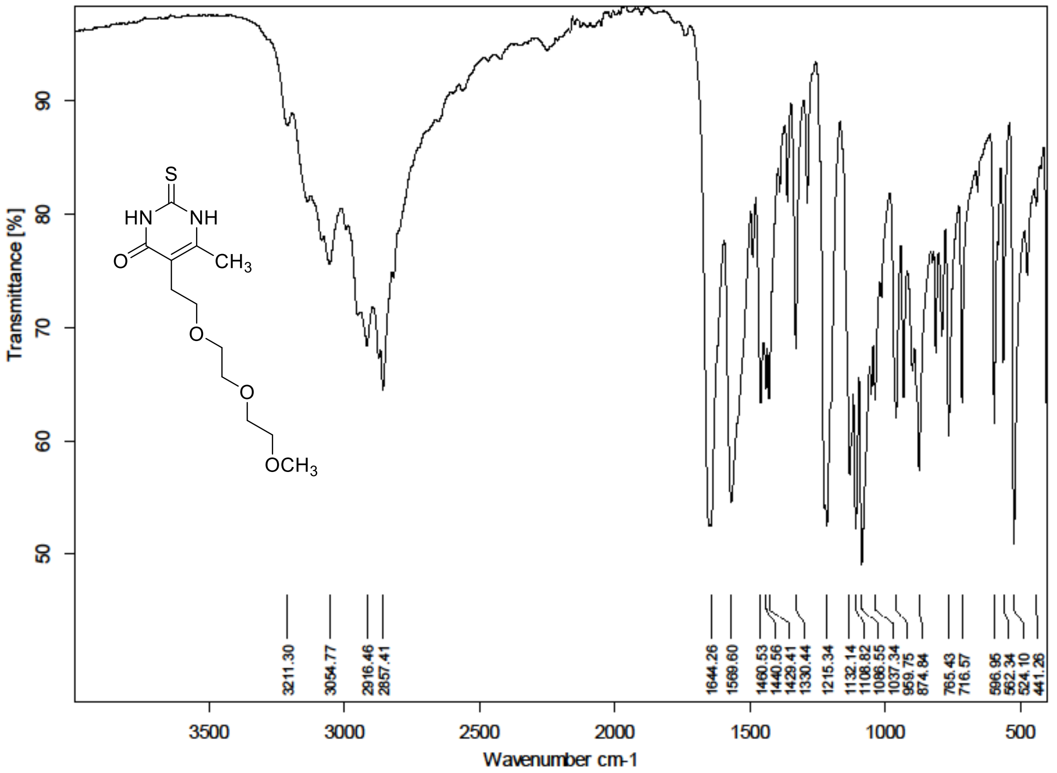

$^{1}H$ (300 MHz) spectrum of **S126** in DMSO-$d_6$

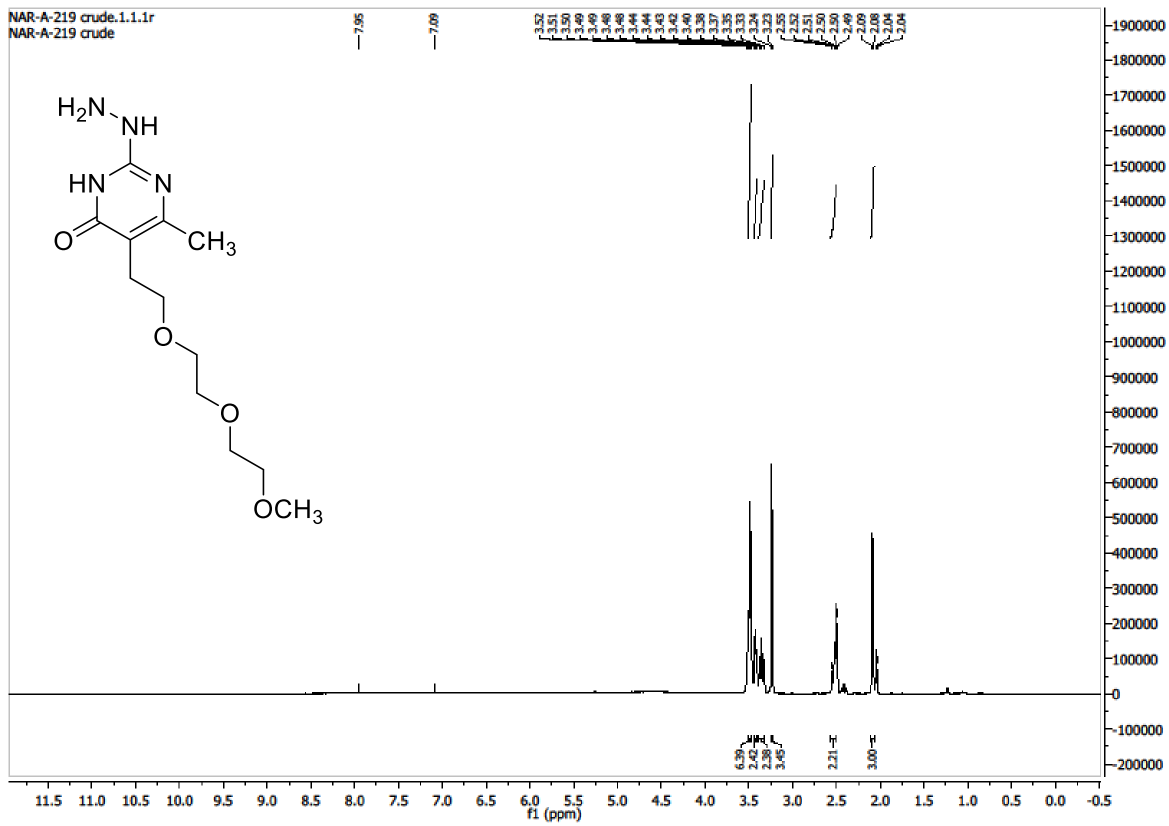

HRMS spectrum of **S126**

**NAR-A-219** $C_{12}H_{22}N_4O_4$ **mono *m/z* 286.1641**

**APCI +** (MMI)
nitrogen flow 5 L/min, gas temperature 325°C, nebulizer 45 psig, skimmer 65 V, vaporizer 200°C, fragmentor 20 V, dissolved in methanol

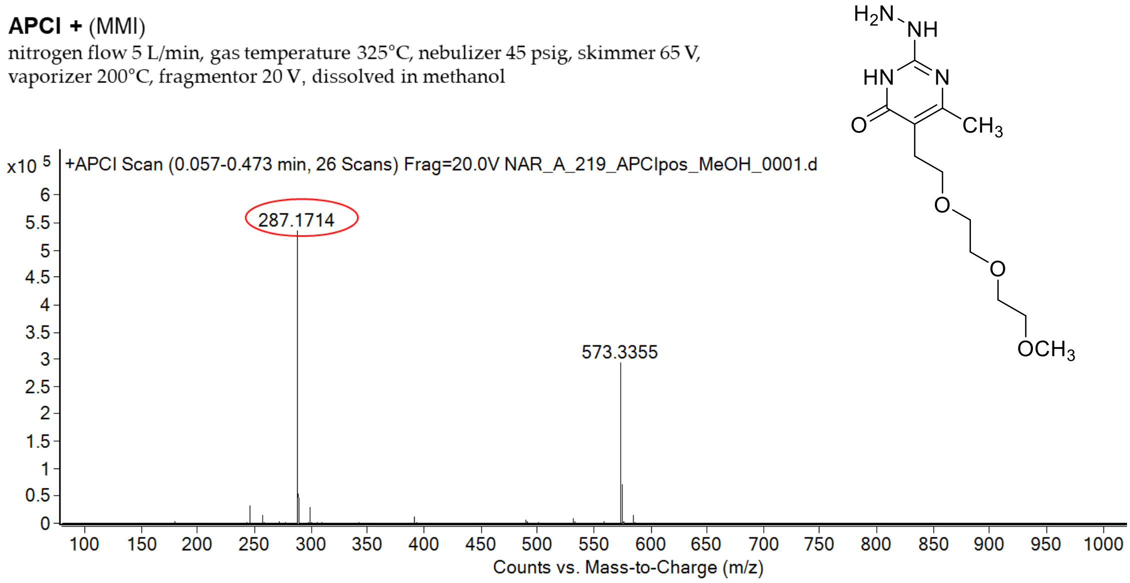

calculated mass: $[M+H]^+ = 287.1714$ observed: $[M+H]^+ = 287.1714$ mass accuracy = < 0.1 ppm

$^{1}H$ (500 MHz) and $^{13}C$ NMR (126 MHz) spectra of **S127** in Chloroform-*d*

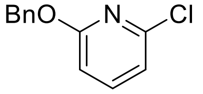

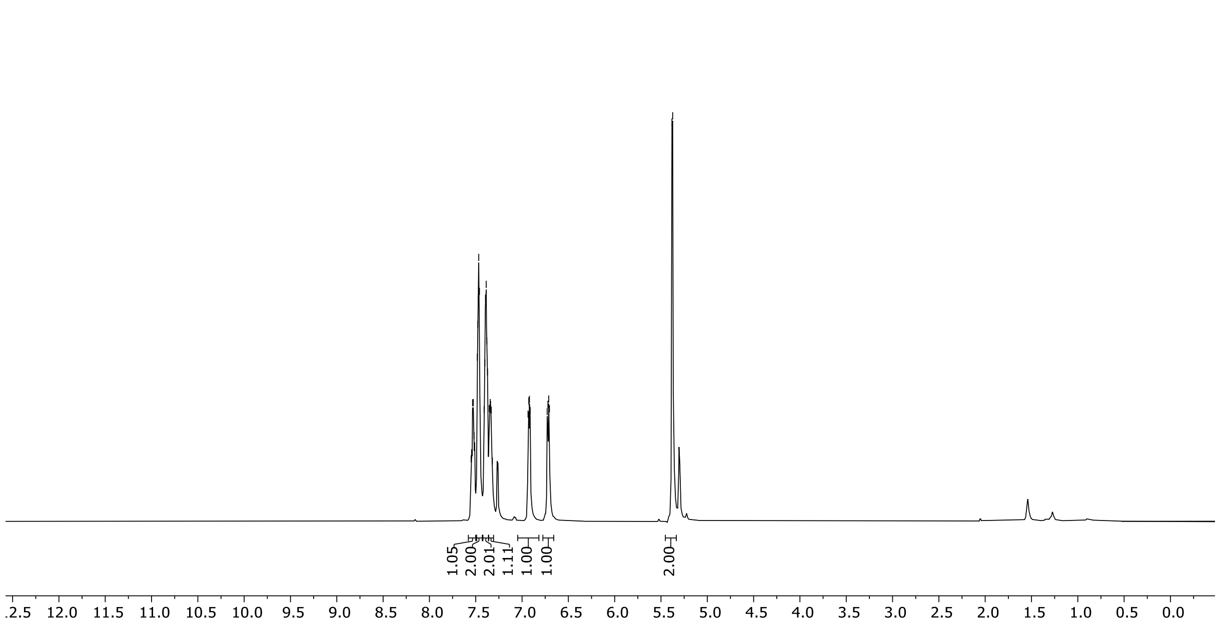

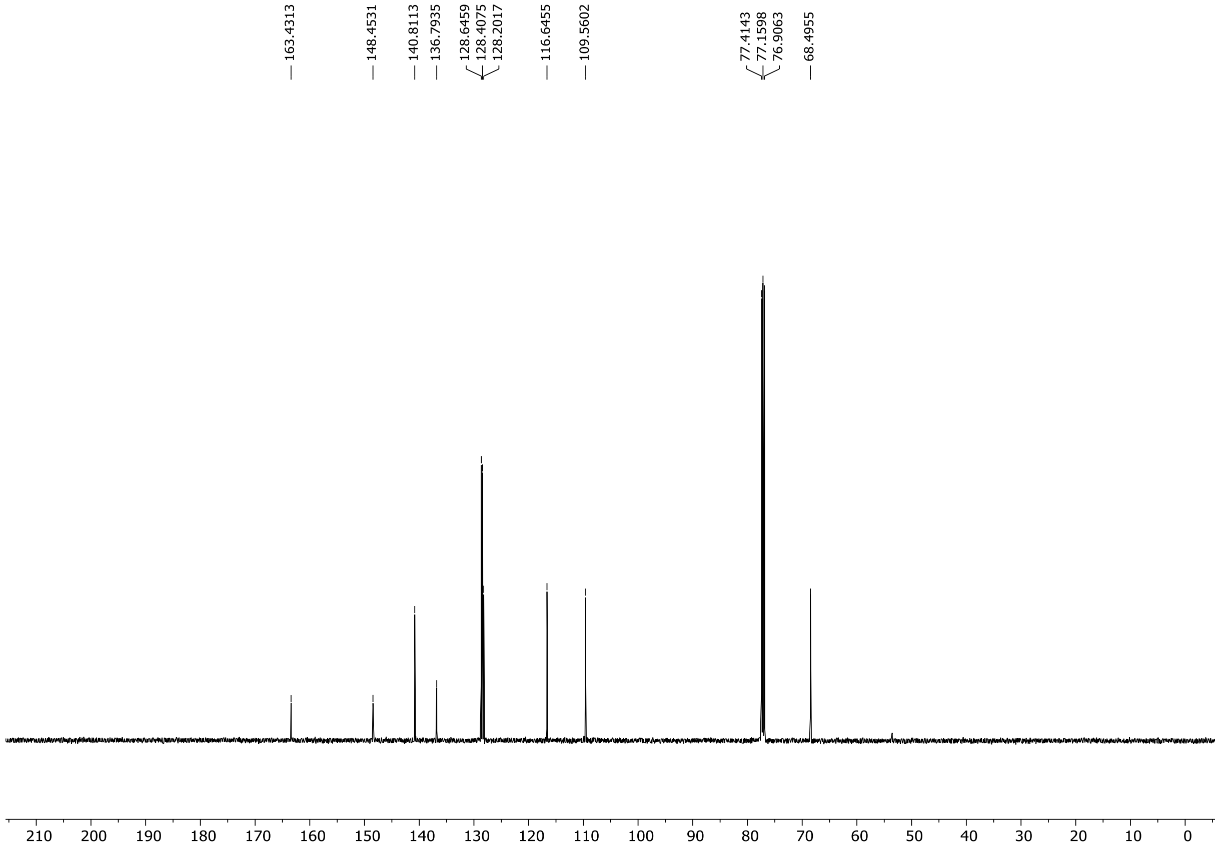

$^{1}$H (300 MHz) spectrum of **S128** in Chloroform-*d*

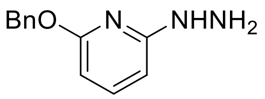

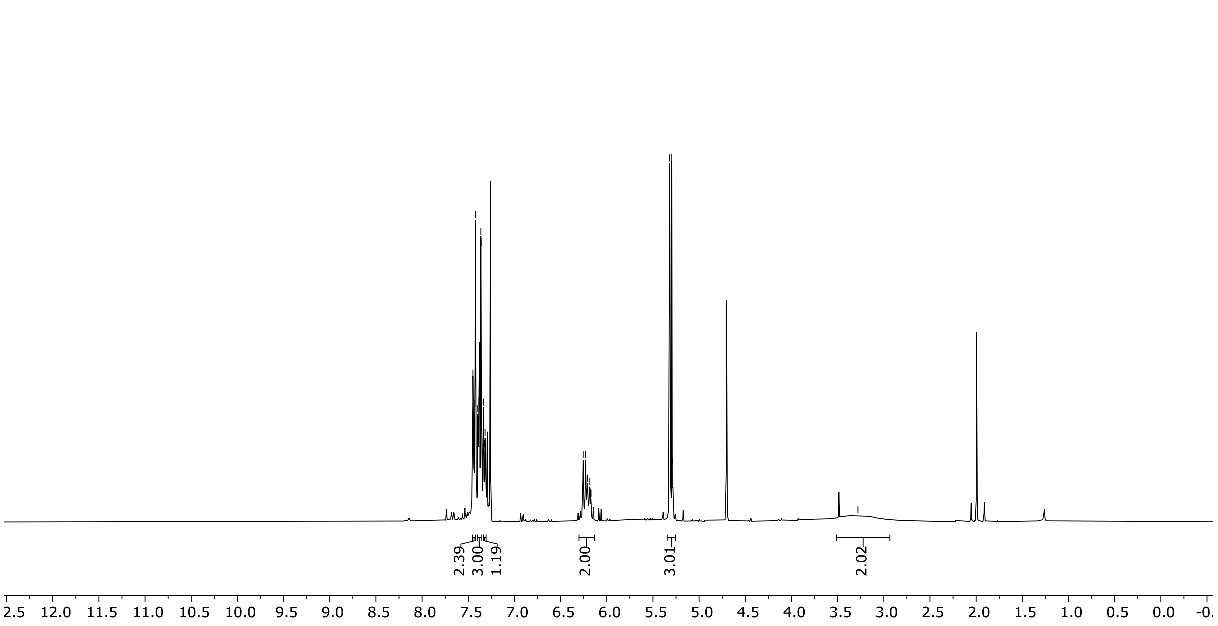

[1]H (300 MHz) NMR spectrum of **S129** in DMSO-$d_6$

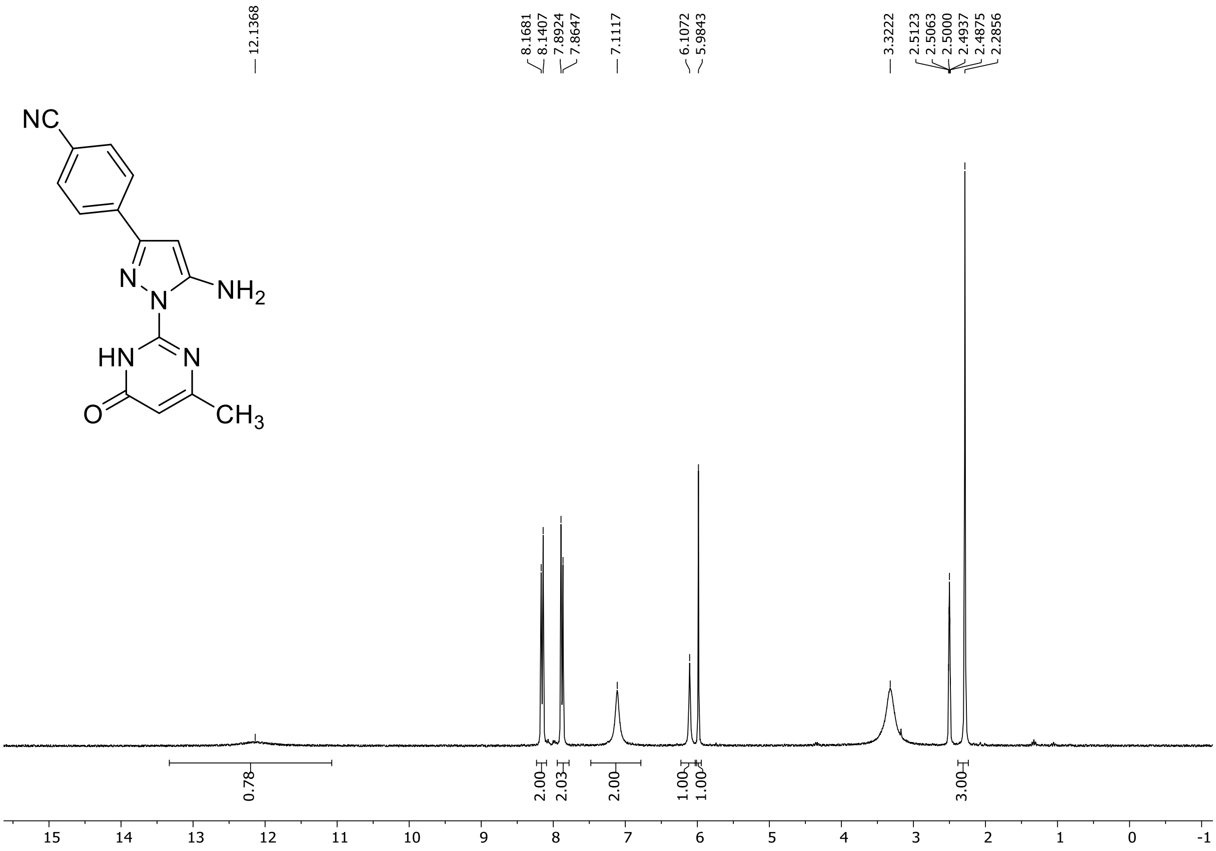

HRMS spectrum of **S129**

**NAR-A-114** **$C_{15}H_{12}N_6O$** ***m/z* 292.1073**

**APCI-** (MMI)
nitrogen flow 5 L/min, gas temperature 325°C, nebulizer 45 psi, skimmer 65 V, vaporizer 200°C, fragmentor 60 V, dissolved in DMSO, MeOH

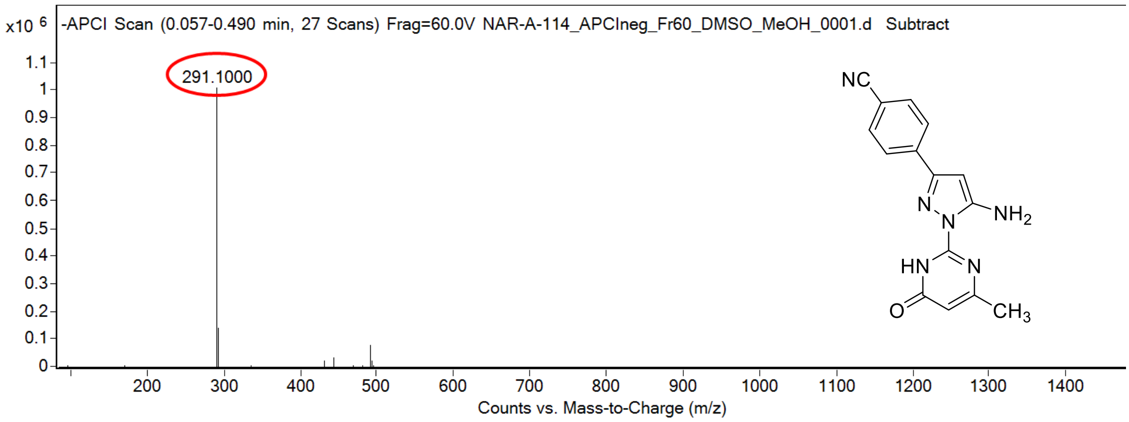

calculated mass: $[M-H]^-$ = 291.1000 observed: $[M-H]^-$ = 291.1000 mass accuracy < 0.1 ppm

FT-IR spectrum (neat) of **S129**

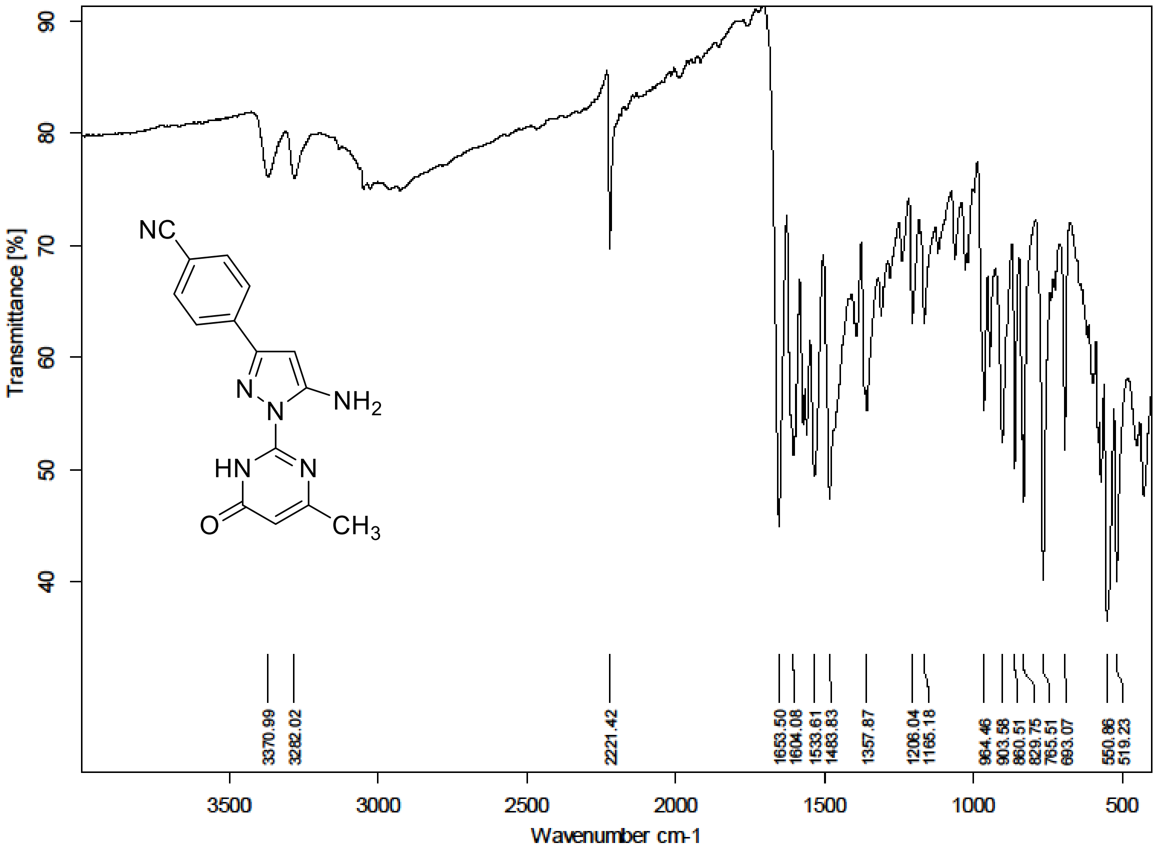

[1]H (300 MHz) and [13]C NMR (75 MHz) spectra of **S130** in Chloroform-*d*

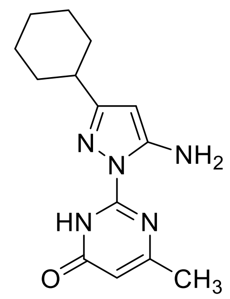

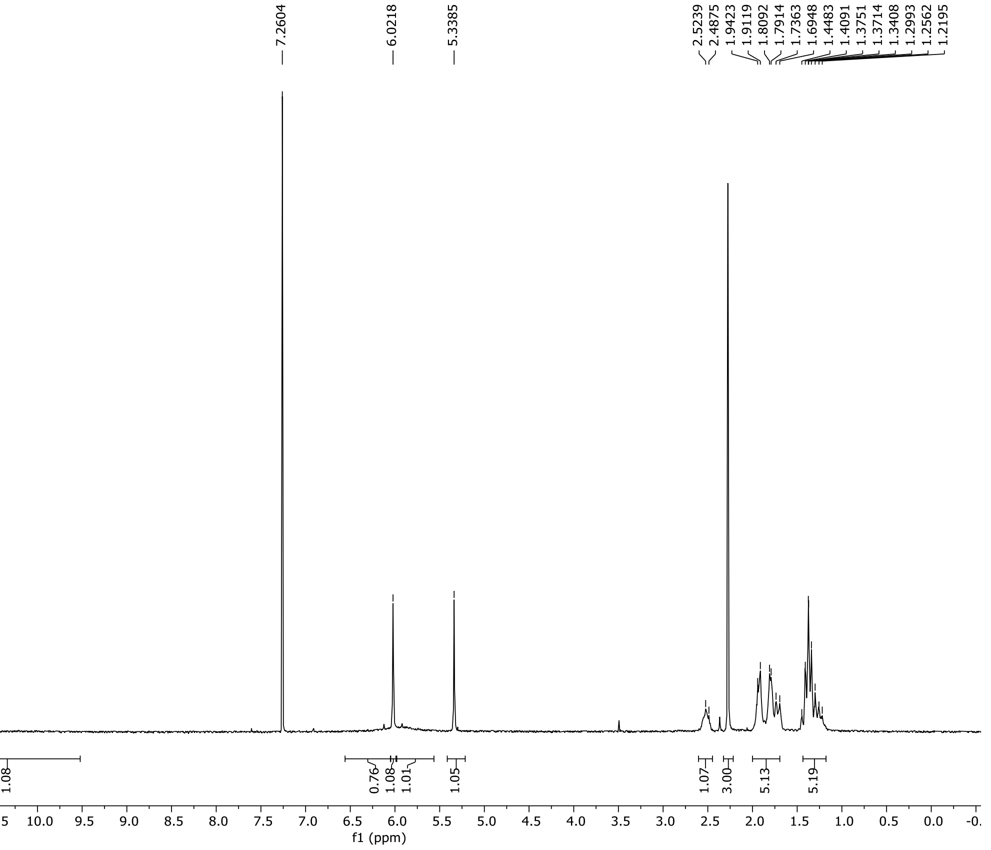

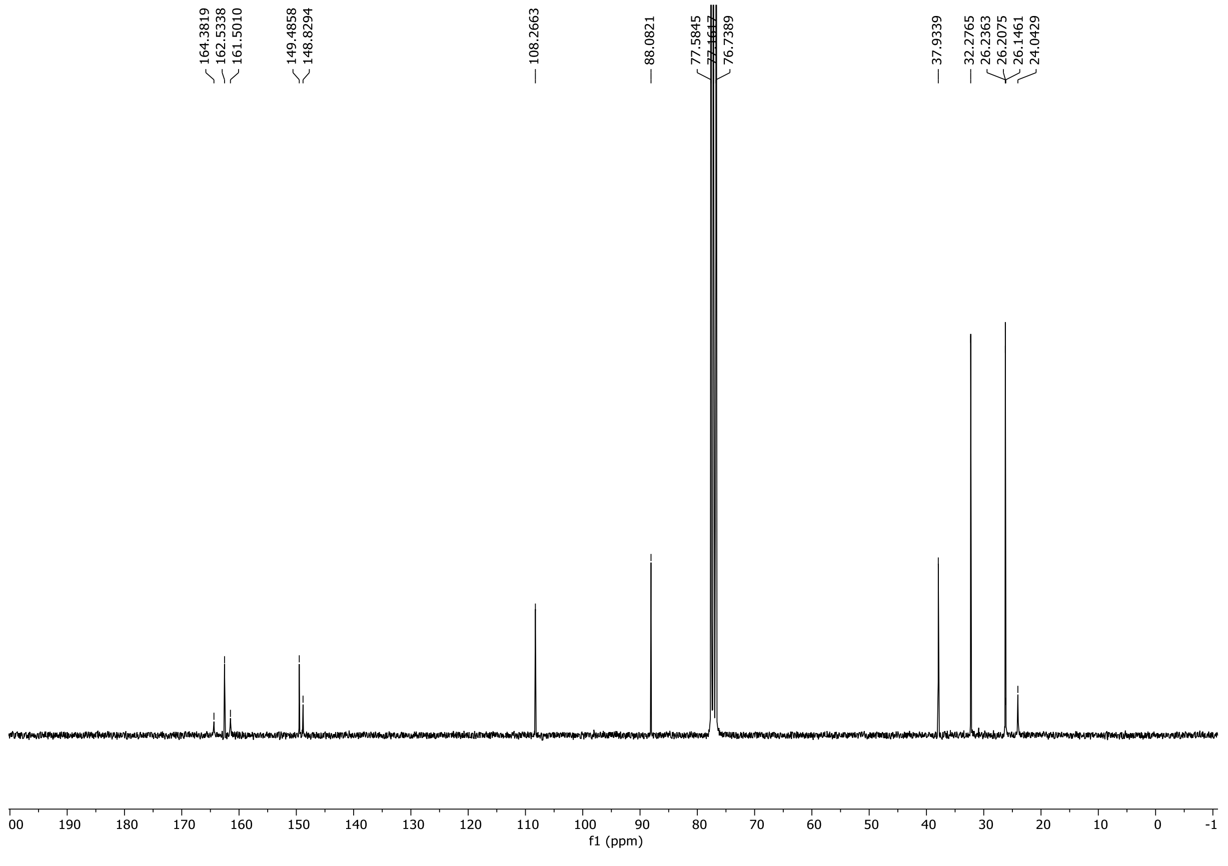

HRMS spectrum of **S130**

**NAR-A-116** $C_{14}H_{19}N_5O$ *m/z* **273.1590**

**APCI+** (MMI)
nitrogen flow 5 L/min, gas temperature 325°C, nebulizer 45 psi, skimmer 65 V, vaporizer 200°C, fragmentor 20 V, dissolved in DMSO, MeOH

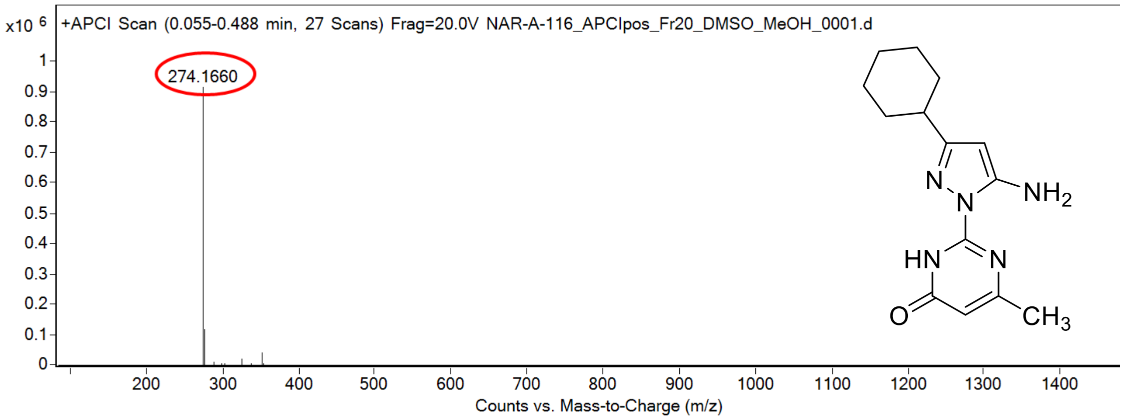

calculated mass: $[M+H]^+ = 274.1662$ observed: $[M+H]^+ = 274.1660$ mass accuracy = -0.7 ppm

FT-IR spectrum (neat) of **S130**

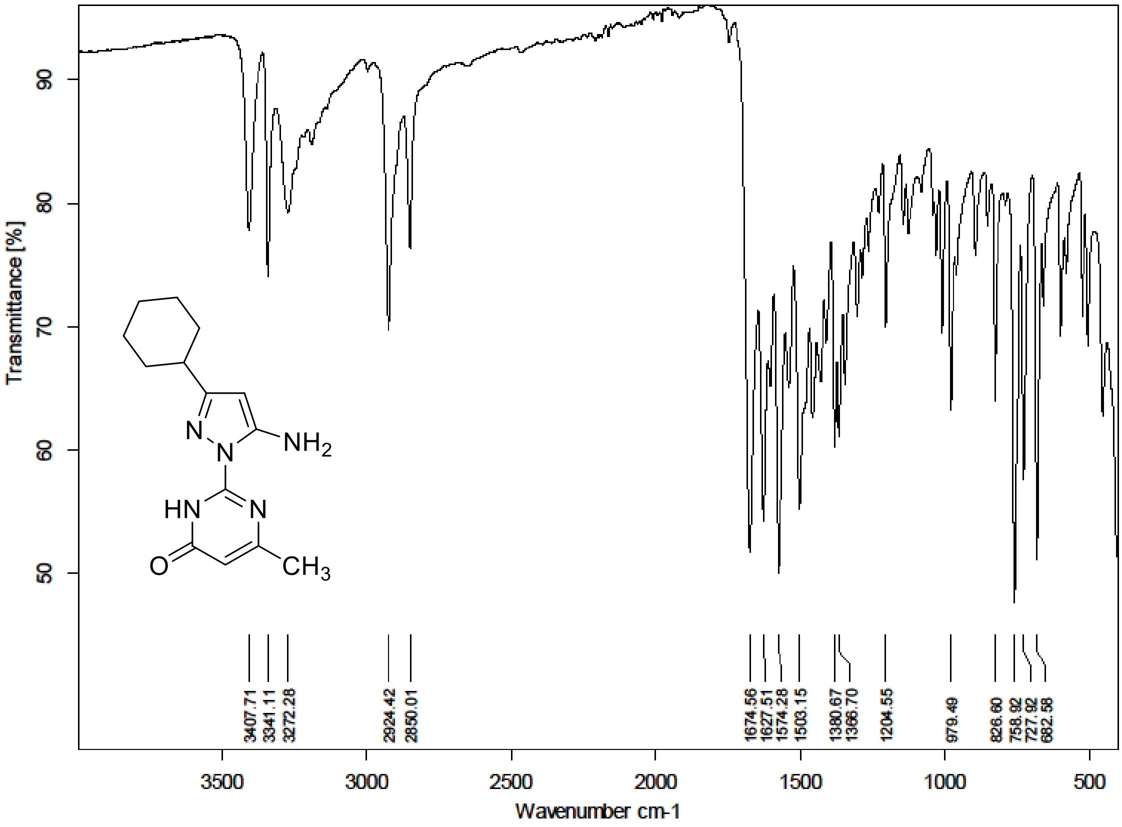

$^{1}H$ (500 MHz) and $^{13}C$ NMR (126 MHz) spectra of **S131** in DMSO-$d_6$

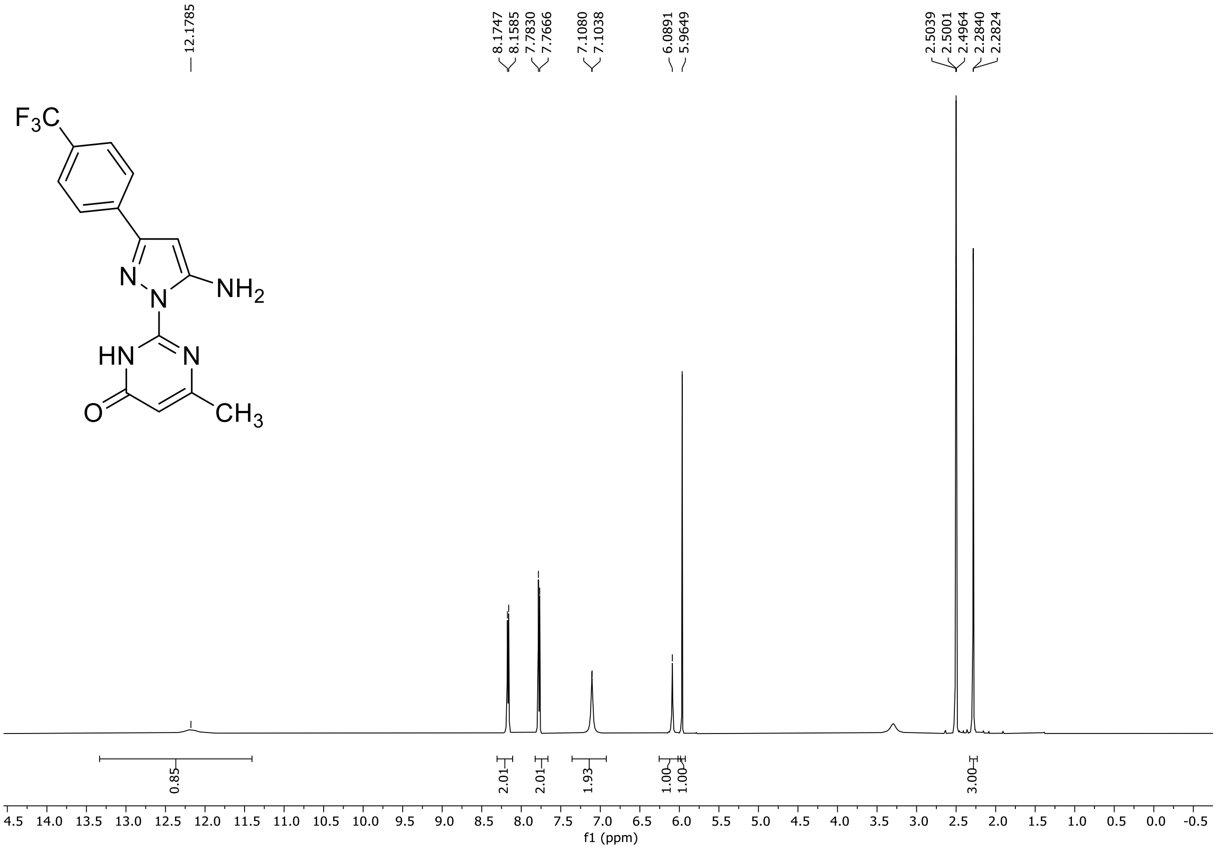

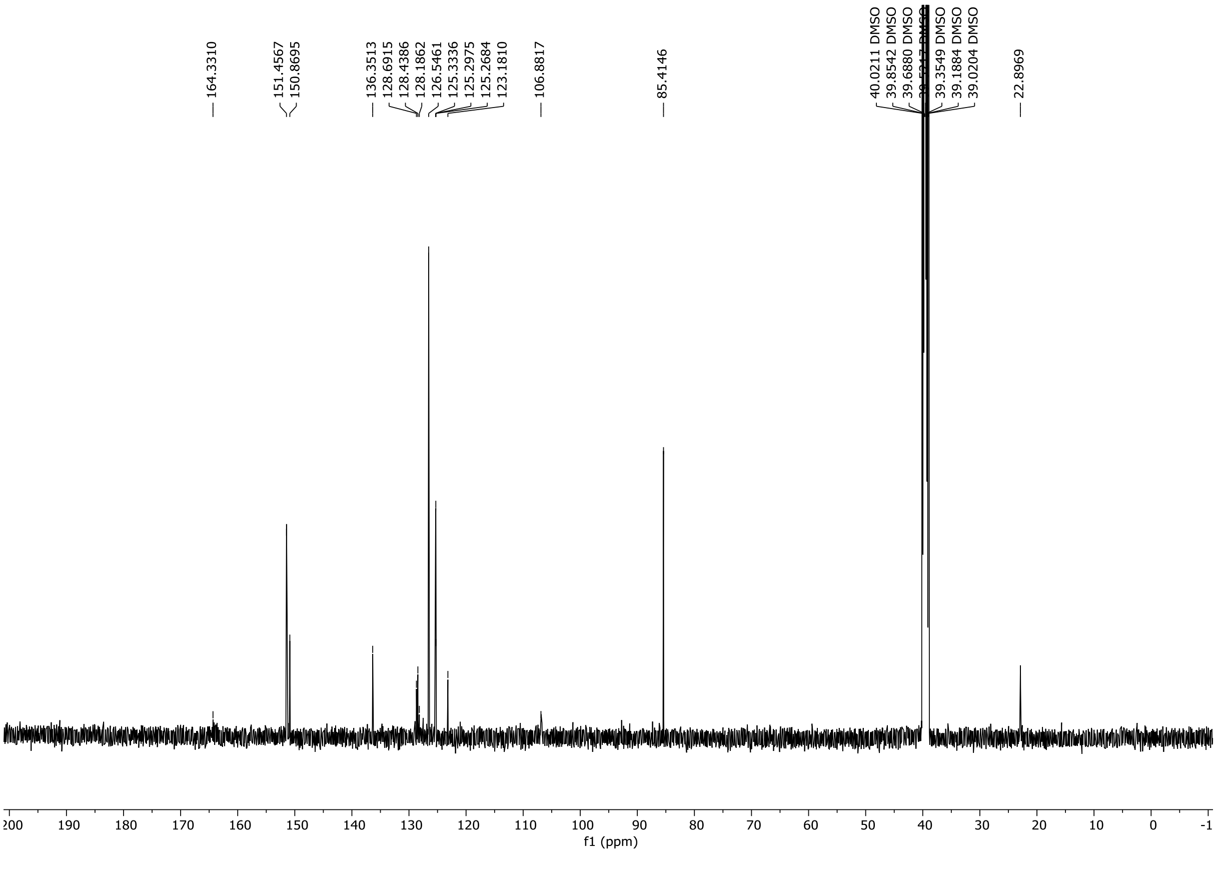

$^{19}$F NMR (282 MHz) spectrum of **S131** in DMSO-$d_6$

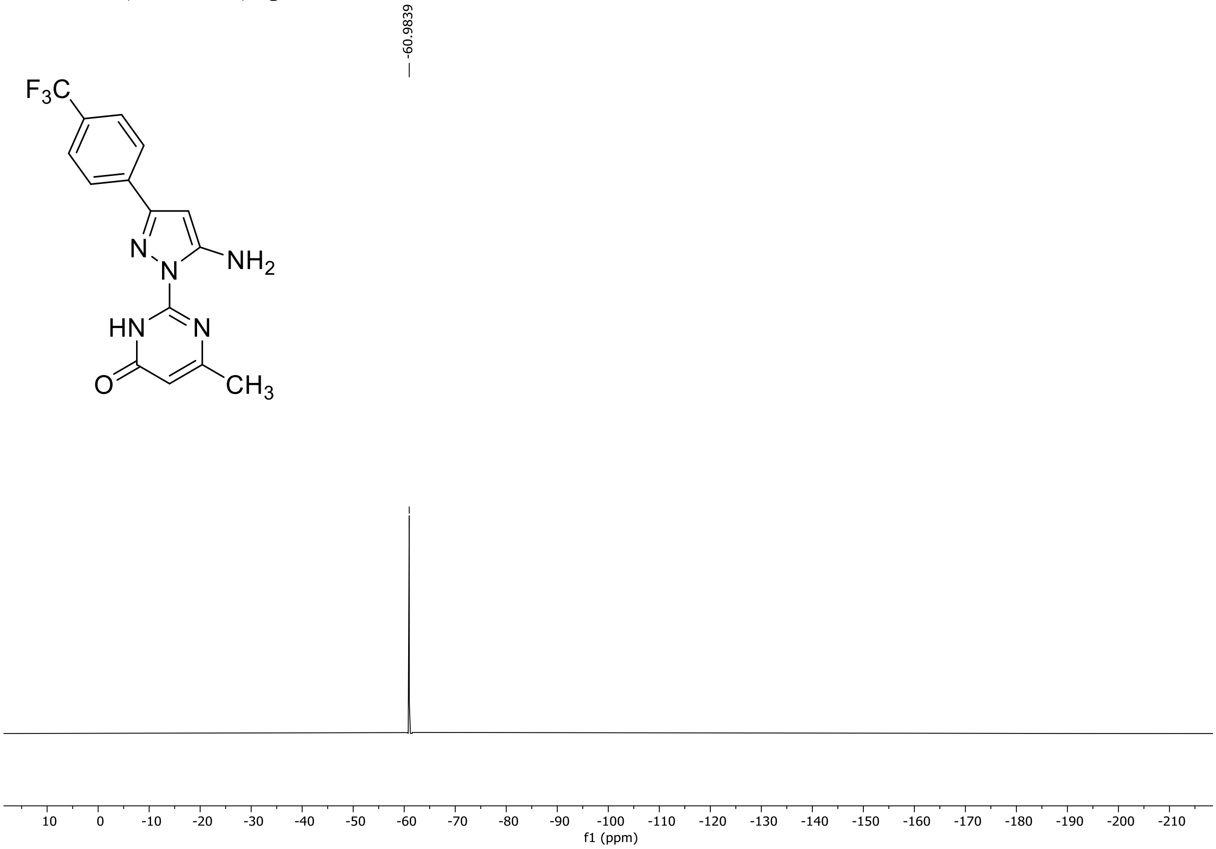

HRMS spectrum of **S131**

**NAR-A-117** **$C_{15}H_{12}F_3N_5O$** ***m/z* 335.0994**

**APCI+** (MMI)
nitrogen flow 5 L/min, gas temperature 325°C, nebulizer 45 psi, skimmer 65 V,
vaporizer 200°C, fragmentor 30 V, dissolved in DMSO, MeOH

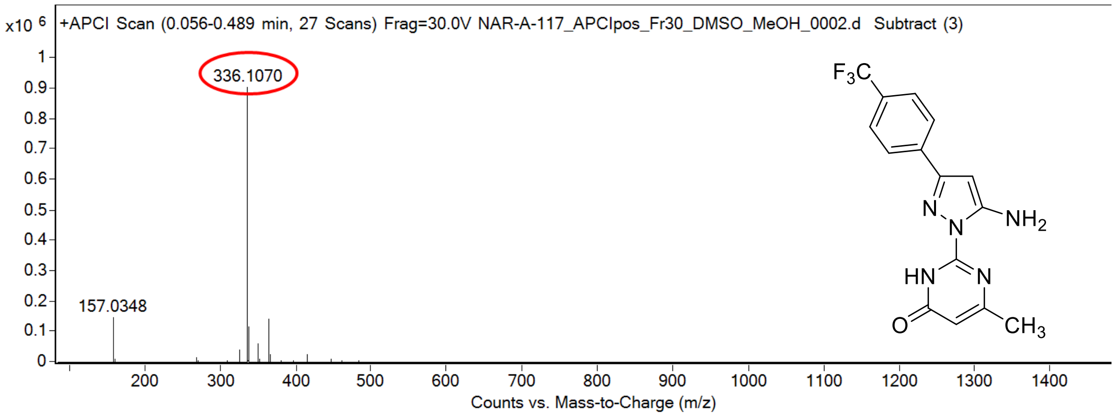

calculated mass: $[M+H]^+$ = 336.1067 observed: $[M+H]^+$ = 336.1070 mass accuracy = +0.9 ppm

FT-IR spectrum (neat) of **S131**

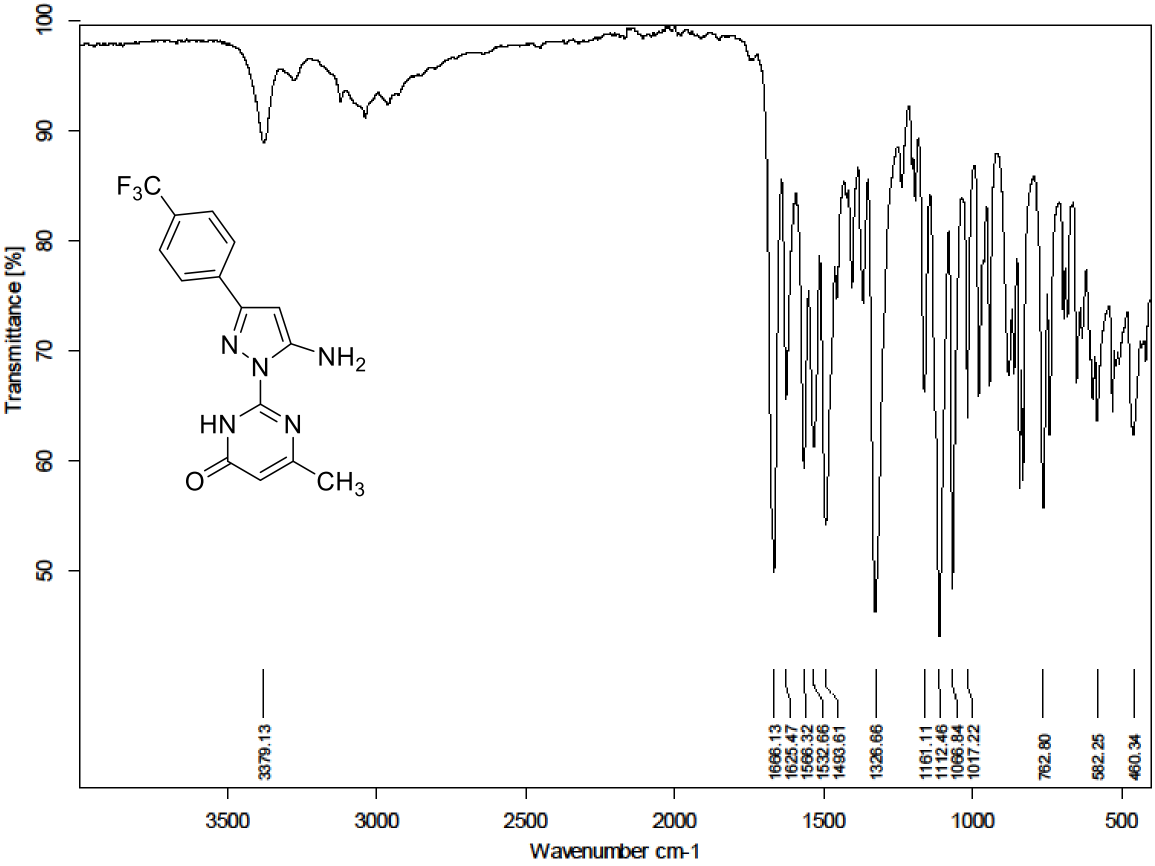

$^{1}H$ (500 MHz) and $^{13}C$ NMR (126 MHz) spectra of **S132** in DMSO-$d_6$

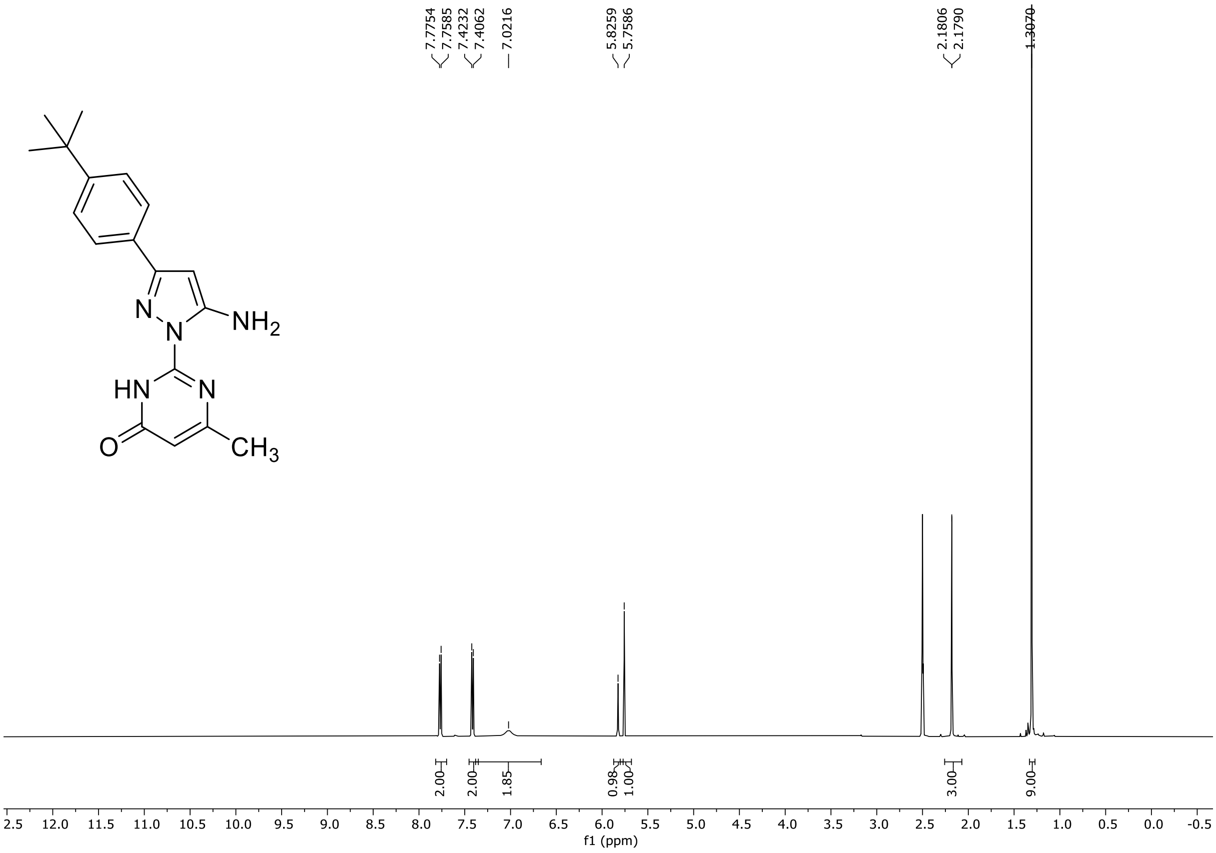

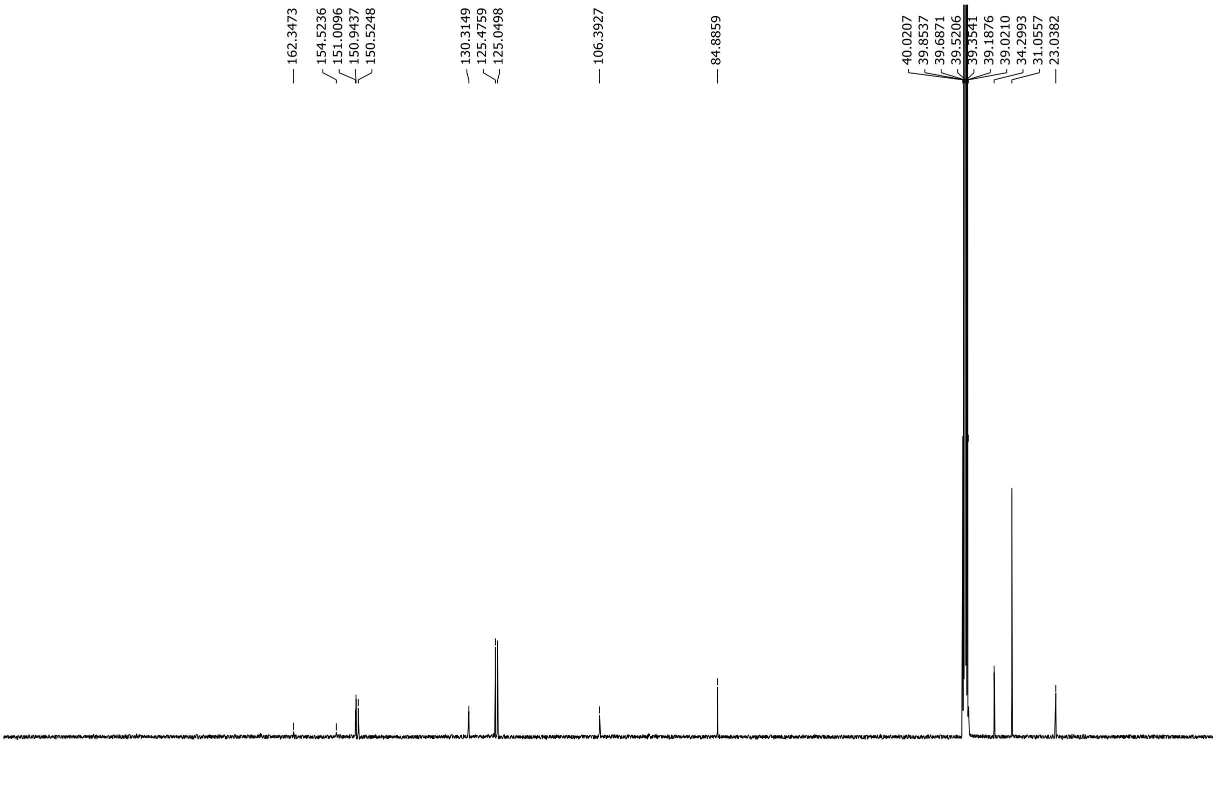

HRMS spectrum of **S132**

**NAR-A-115** $C_{28}H_{21}N_5O$ *m/z* **323.1746**

**APCI+** (MMI)
nitrogen flow 5 L/min, gas temperature 325°C, nebulizer 45 psi, skimmer 65 V, vaporizer 200°C, fragmentor 15 V, dissolved in DMSO, MeOH

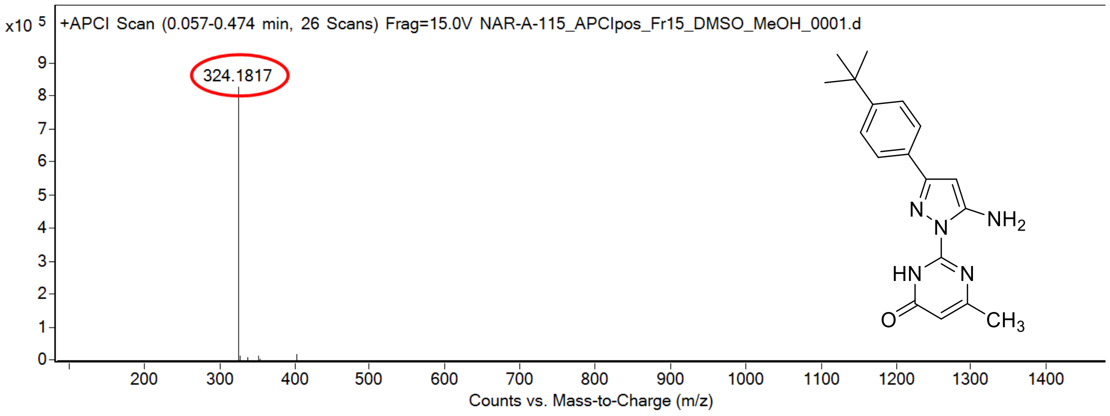

calculated mass: $[M+H]^+$ = 324.1819 observed: $[M+H]^+$ = 324.1817 mass accuracy = -0.6 ppm

FT-IR spectrum (neat) of **S132**

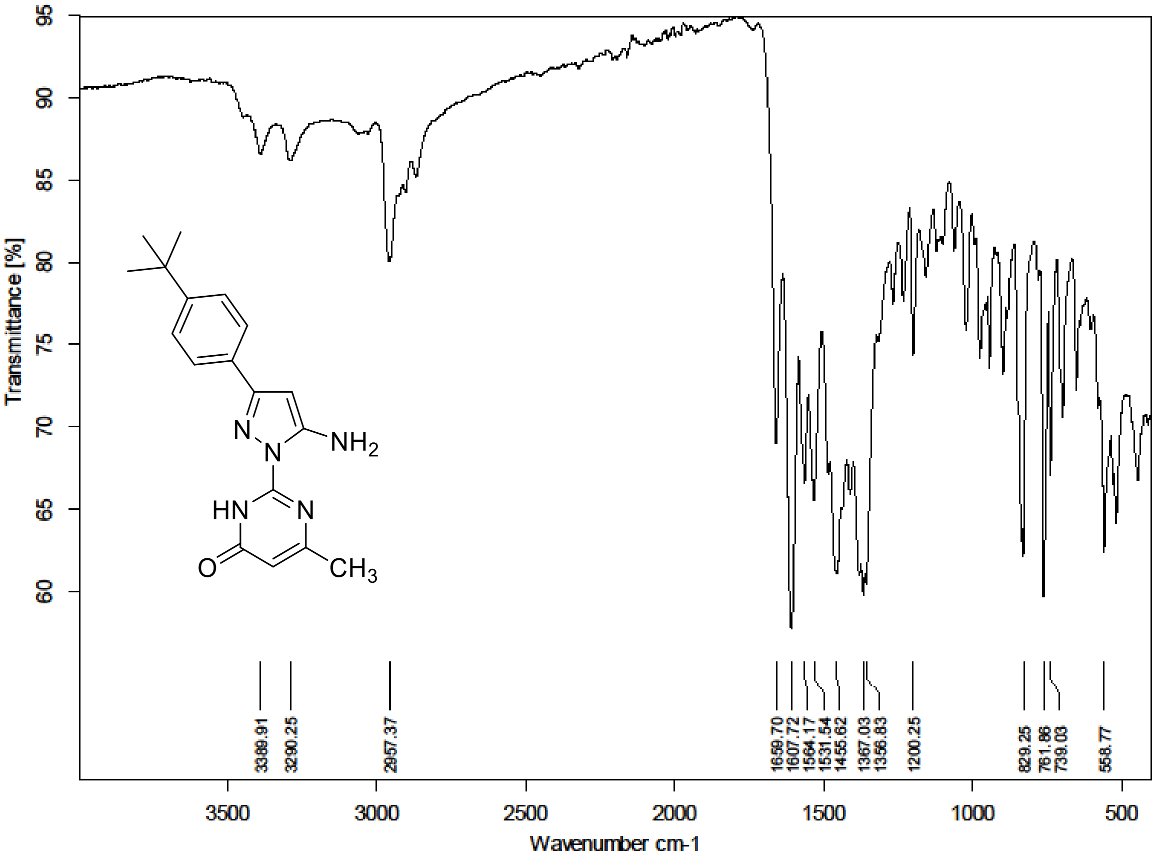

[1]H (500 MHz) and [13]C NMR (126 MHz) spectra of **S133** in DMSO-$d_6$

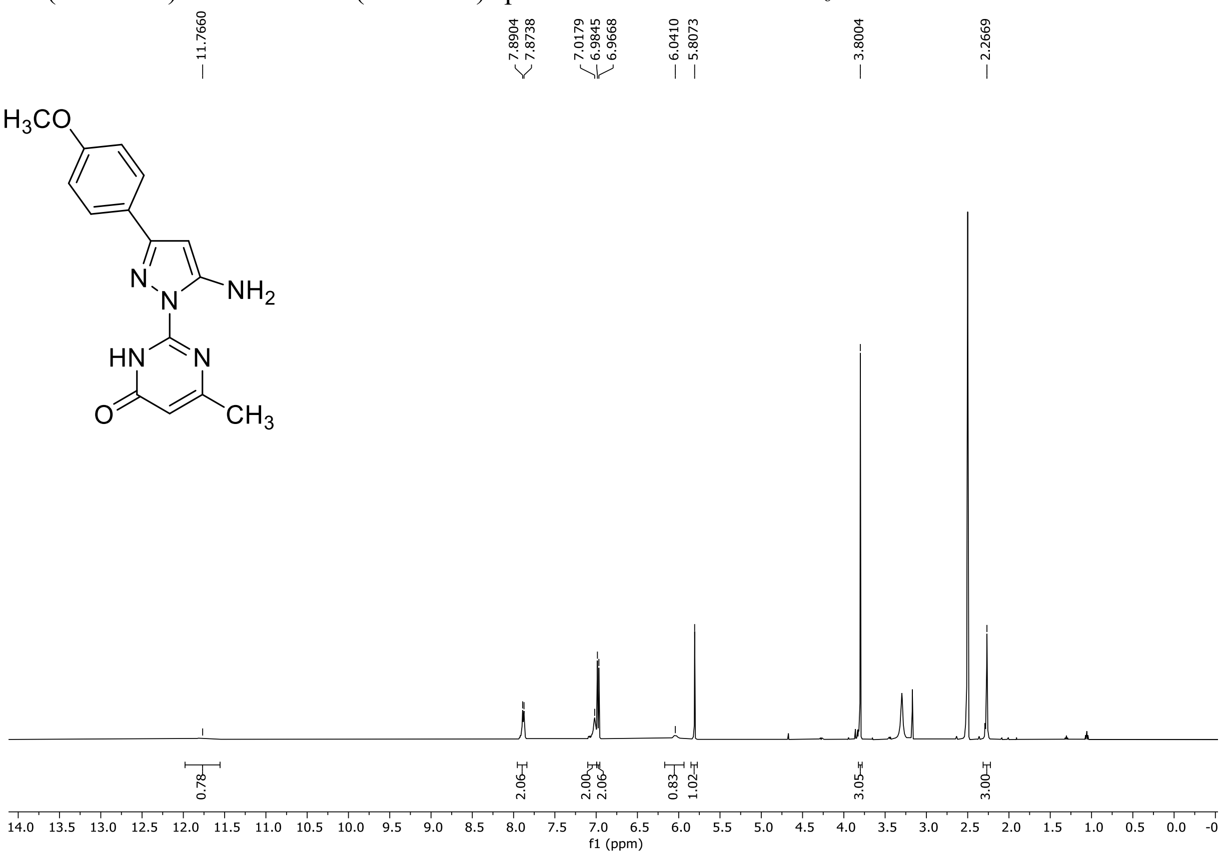

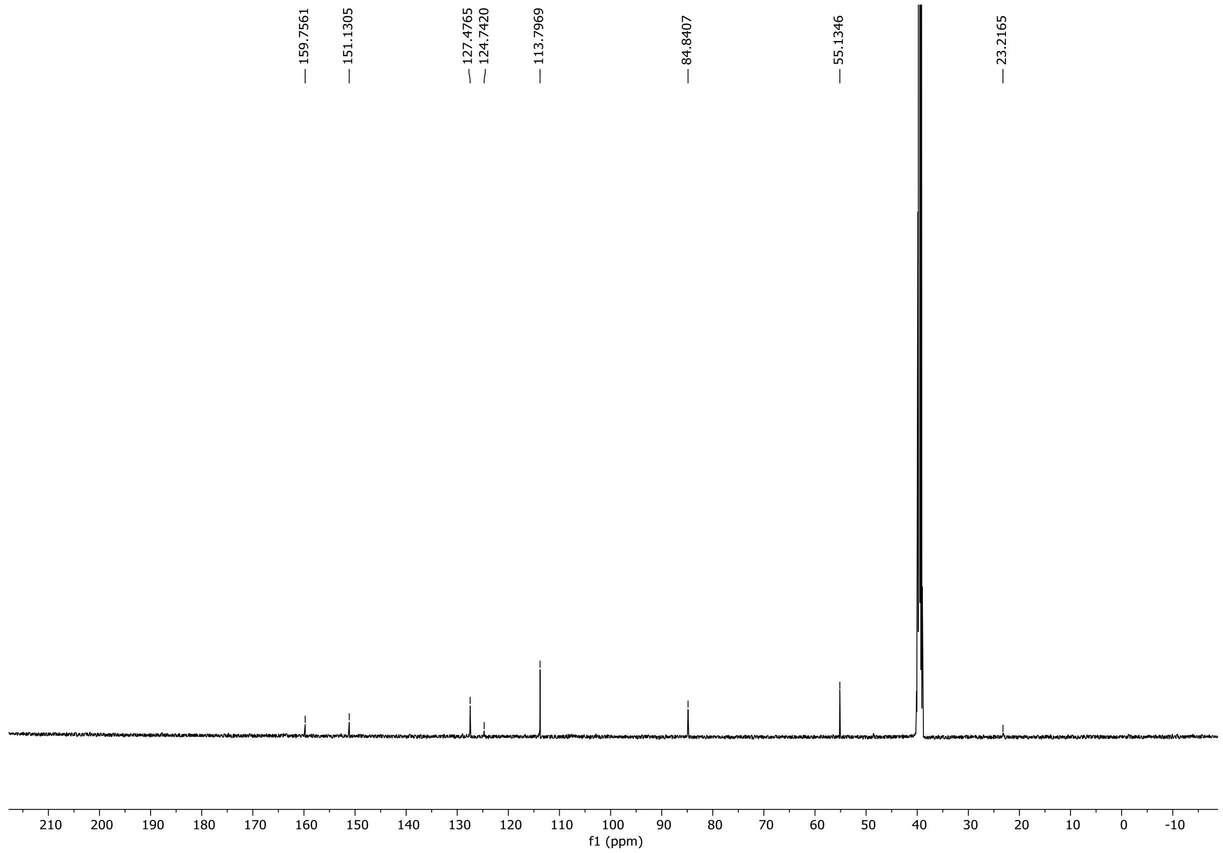

HRMS spectrum of **S133**

**NAR-A-119** $C_{15}H_{15}N_5O_2$ *m/z* **297.1226**

**APCI+** (MMI)
nitrogen flow 5 L/min, gas temperature 325°C, nebulizer 45 psi, skimmer 65 V, vaporizer 200°C, fragmentor 25 V, dissolved in DMSO, MeOH

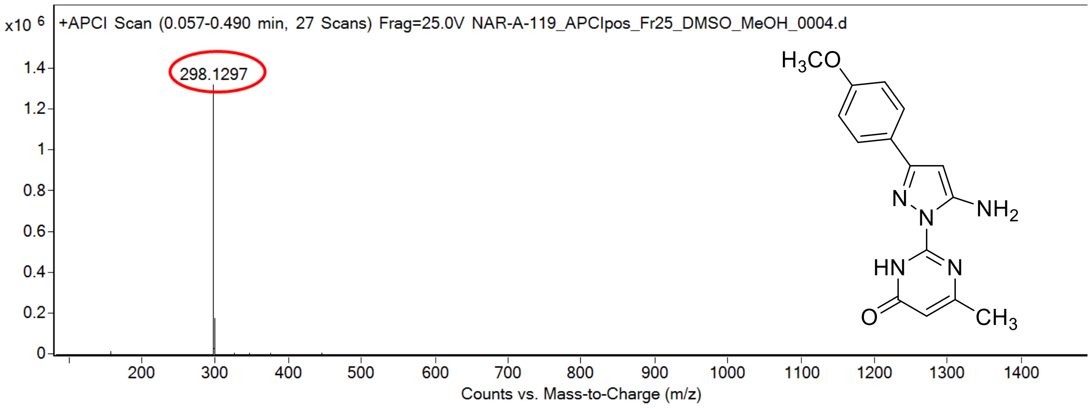

calculated mass: $[M+H]^+$ = 298.1299 observed: $[M+H]^+$ = 298.1297 mass accuracy = -0.7 ppm

FT-IR spectrum (neat) of **S133**

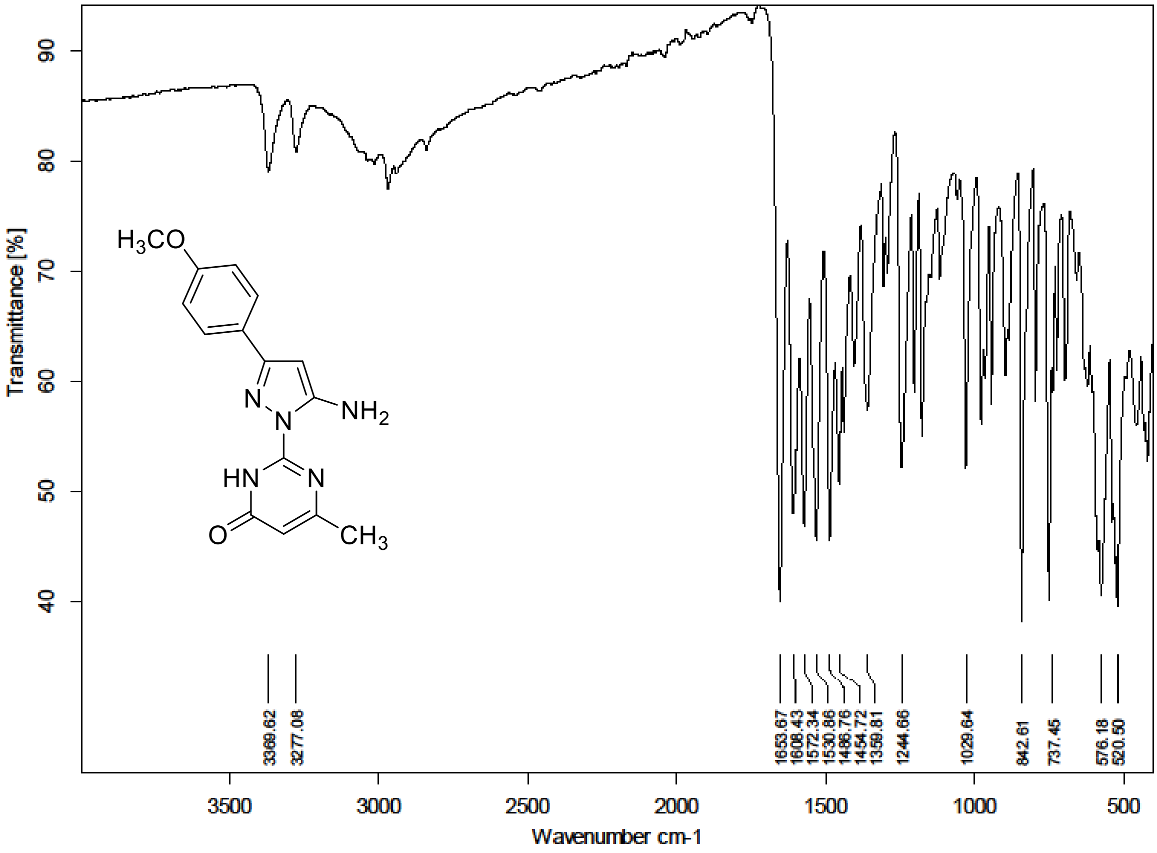

[1]H (300 MHz) and [13]C NMR (75 MHz) spectra of **S134** in DMSO-$d_6$

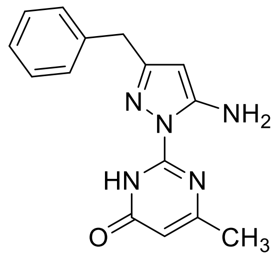

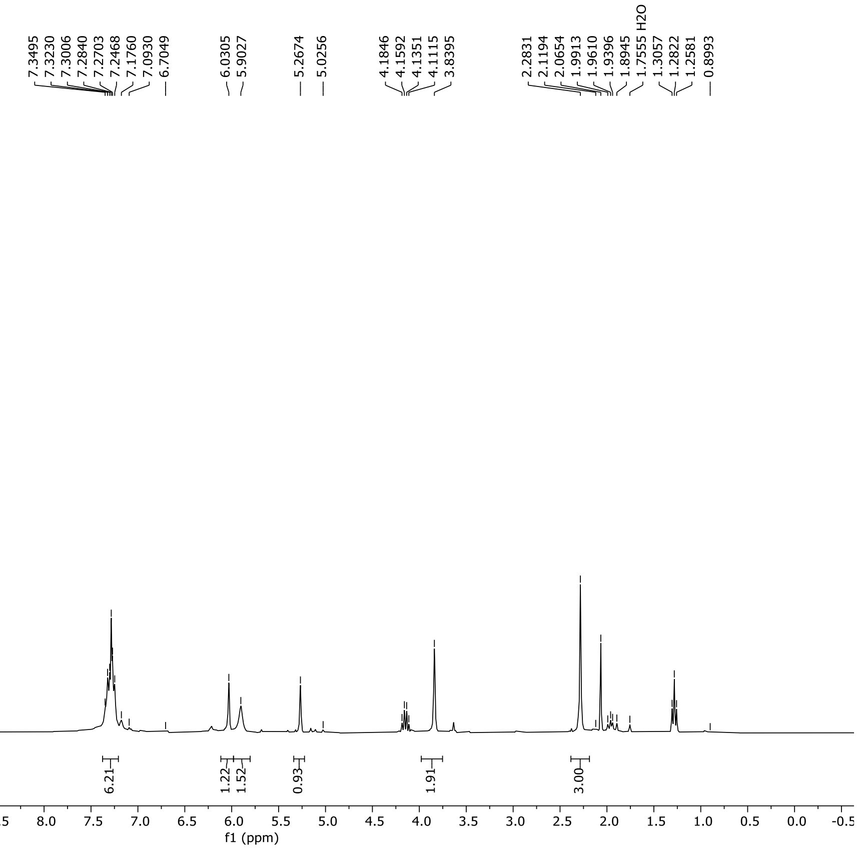

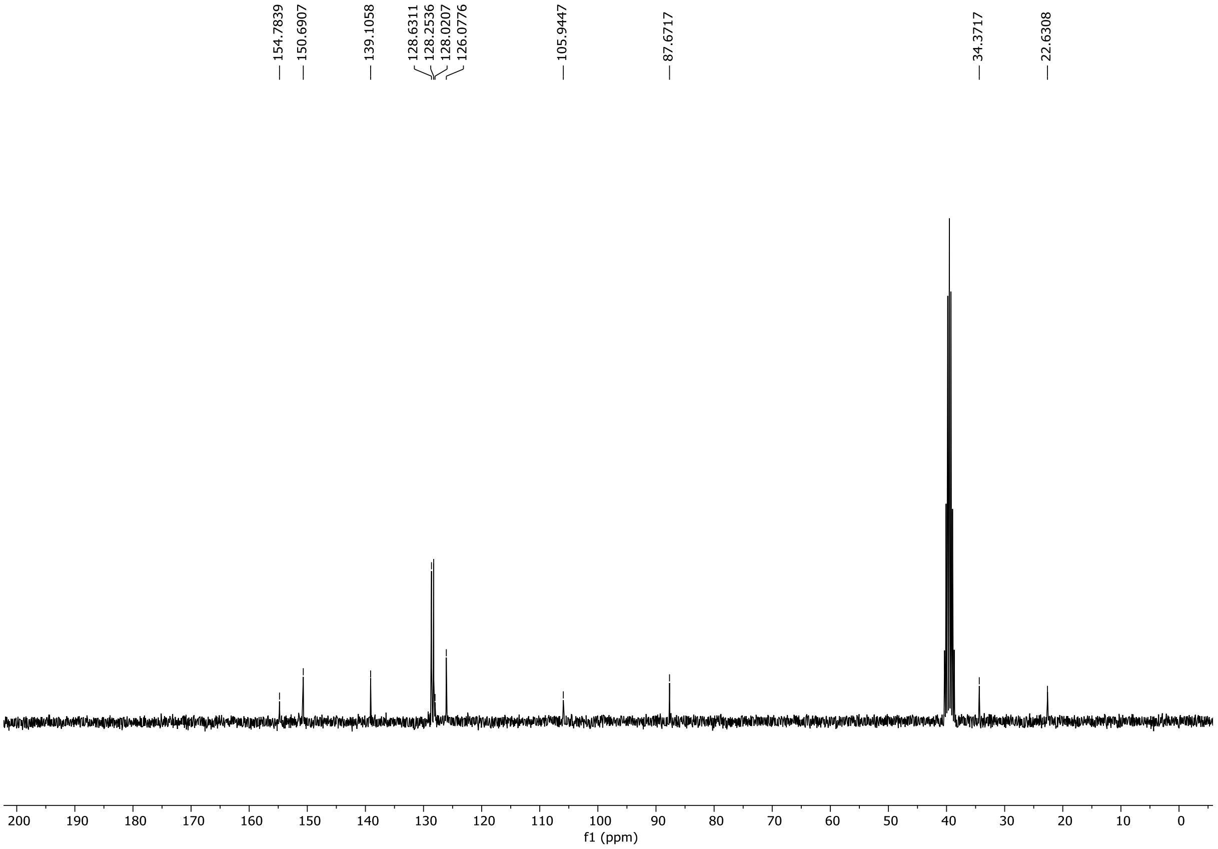

HRMS spectrum of **S134**

**NAR-A-184** $C_{15}H_{15}N_5O$ mono *m/z* 281.1277

**APCI +** (MMI)
nitrogen flow 5 L/min, gas temperature 325°C, nebulizer 45 psi, skimmer **65** V, vaporizer 200°C, fragmentor 25 V, dissolved in methanol

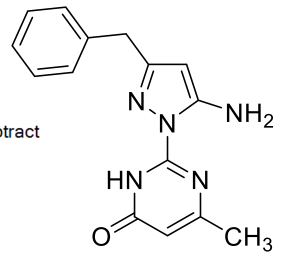

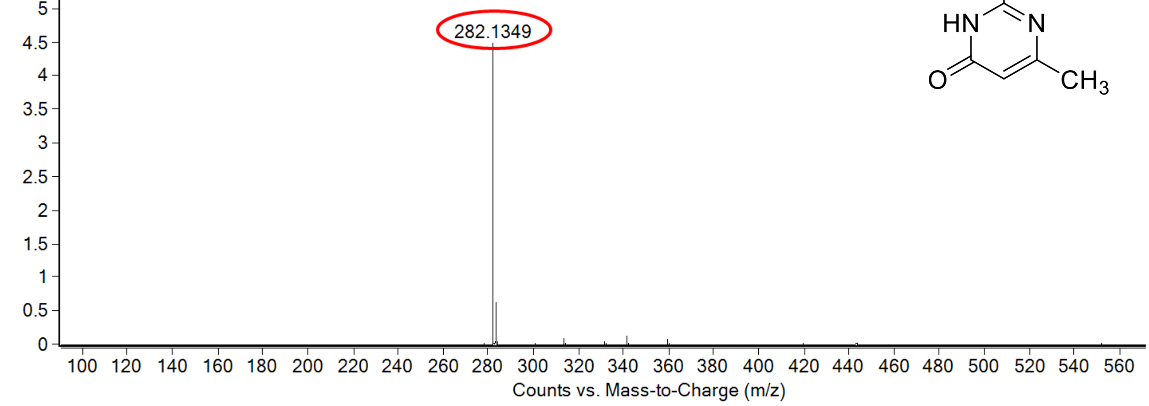

calculated mass: $[M+H]^+$ = 282..1349 observed: $[M+H]^+$ = 282.1349 mass accuracy = < 0.1 ppm

FT-IR spectrum (neat) of **S134**

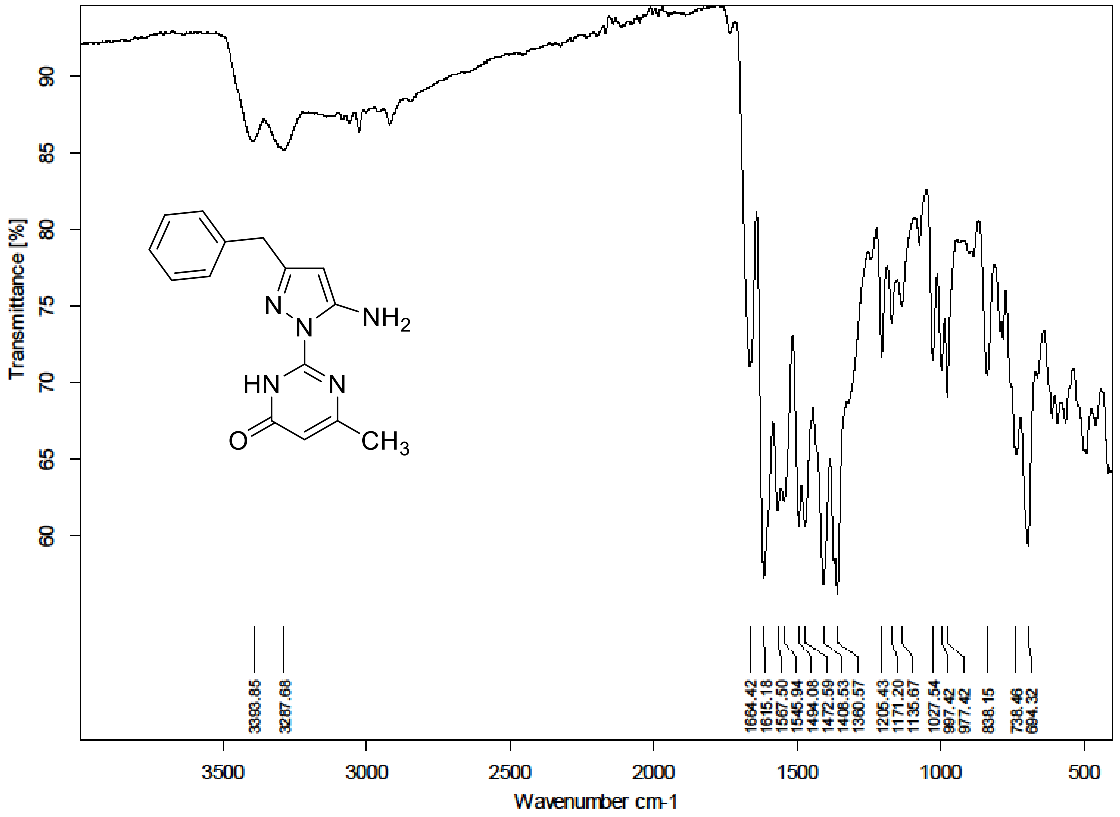

$^{1}H$ (300 MHz) and $^{13}C$ NMR (75 MHz) spectra of **S135** in DMSO-$d_6$

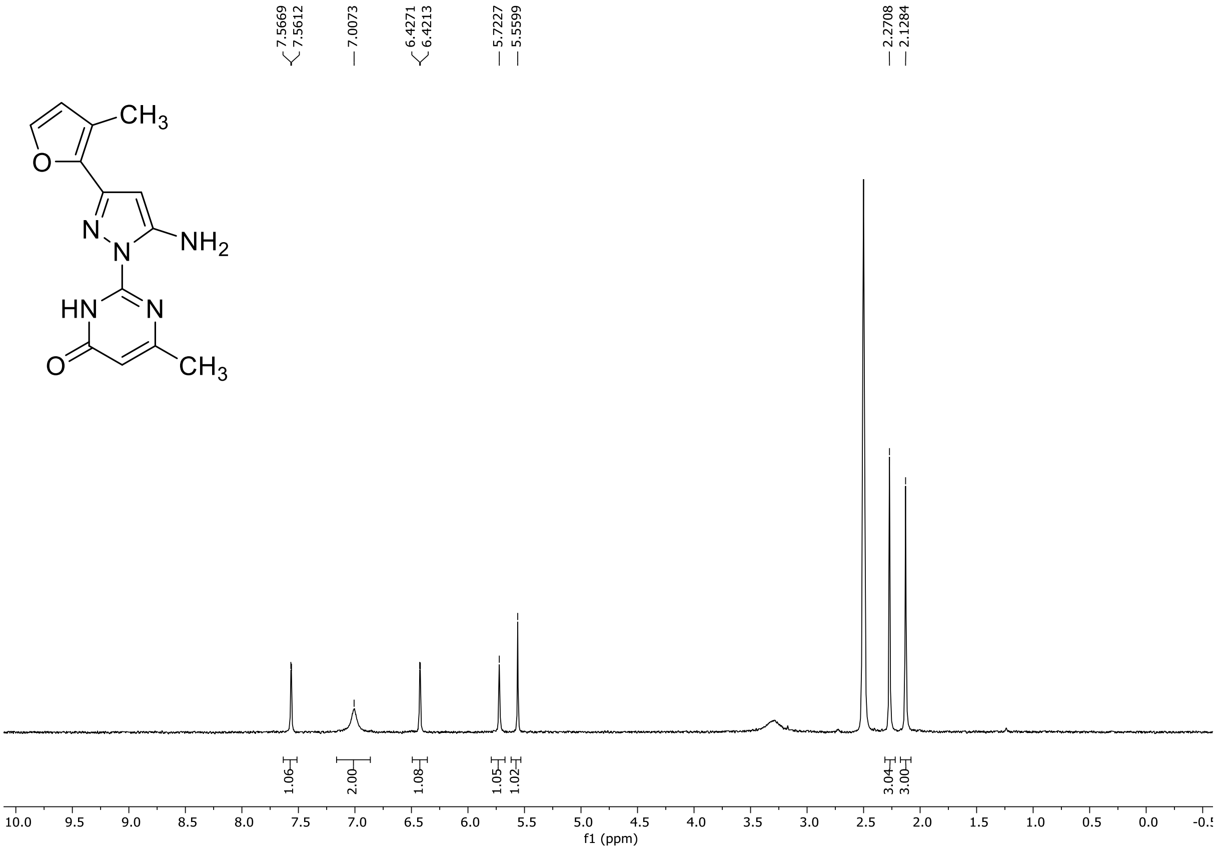

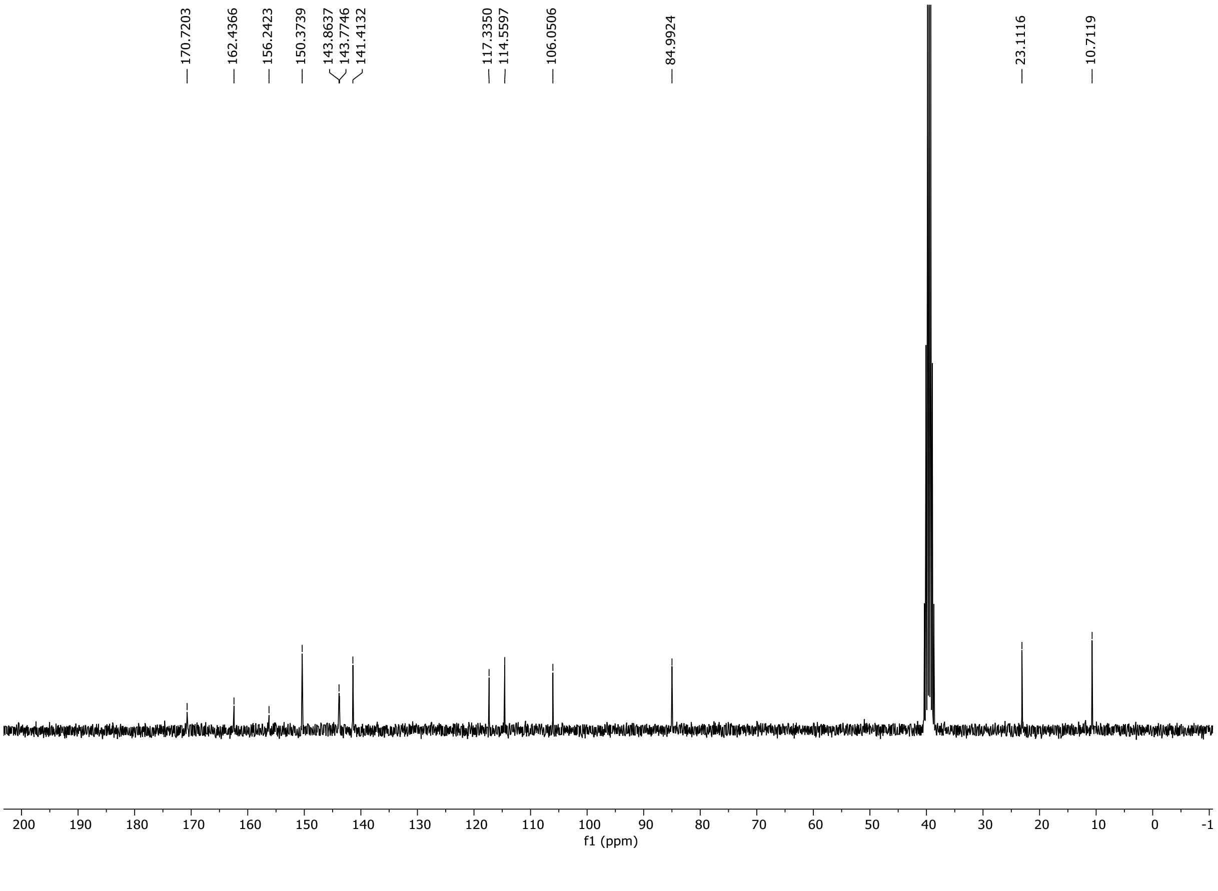

HRMS spectrum of **S135**

**NAR-A-178** $C_{13}H_{13}N_5O_2$ **mono *m/z* 271.1069**

**APCI +** (MMI)
nitrogen flow 5 L/min, gas temperature 325°C, nebulizer 45 psi, skimmer **65** V, vaporizer 200°C, fragmentor 23 V, dissolved in methanol

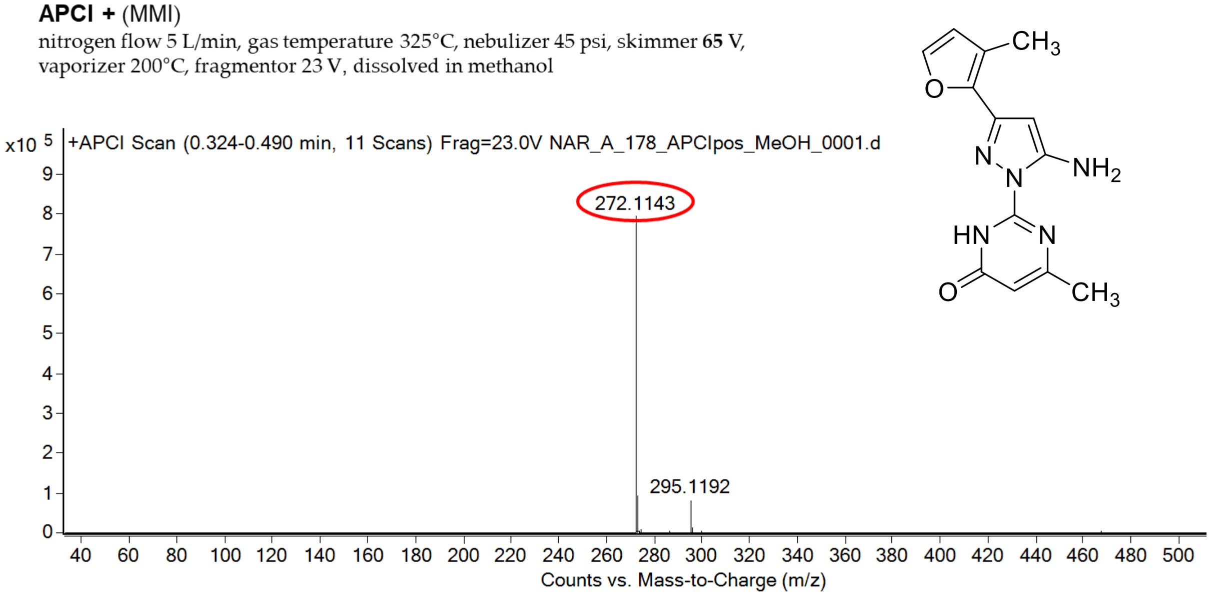

calculated mass: [M+H]$^+$ = 272..1142 observed: [M+H]$^+$ = 272.1143 mass accuracy = 0.4 ppm

FT-IR spectrum (neat) of **S135**

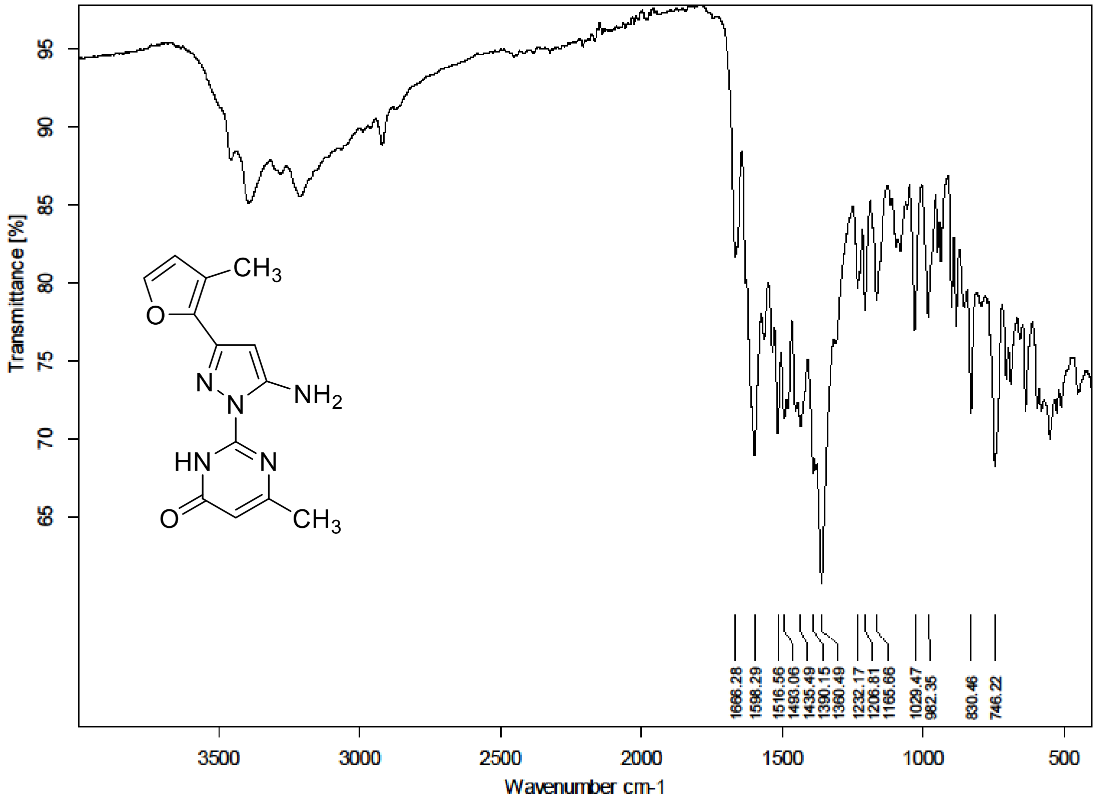

[1]H (300 MHz) and [13]C NMR (75 MHz) spectra of **S136** in DMSO-$d_6$

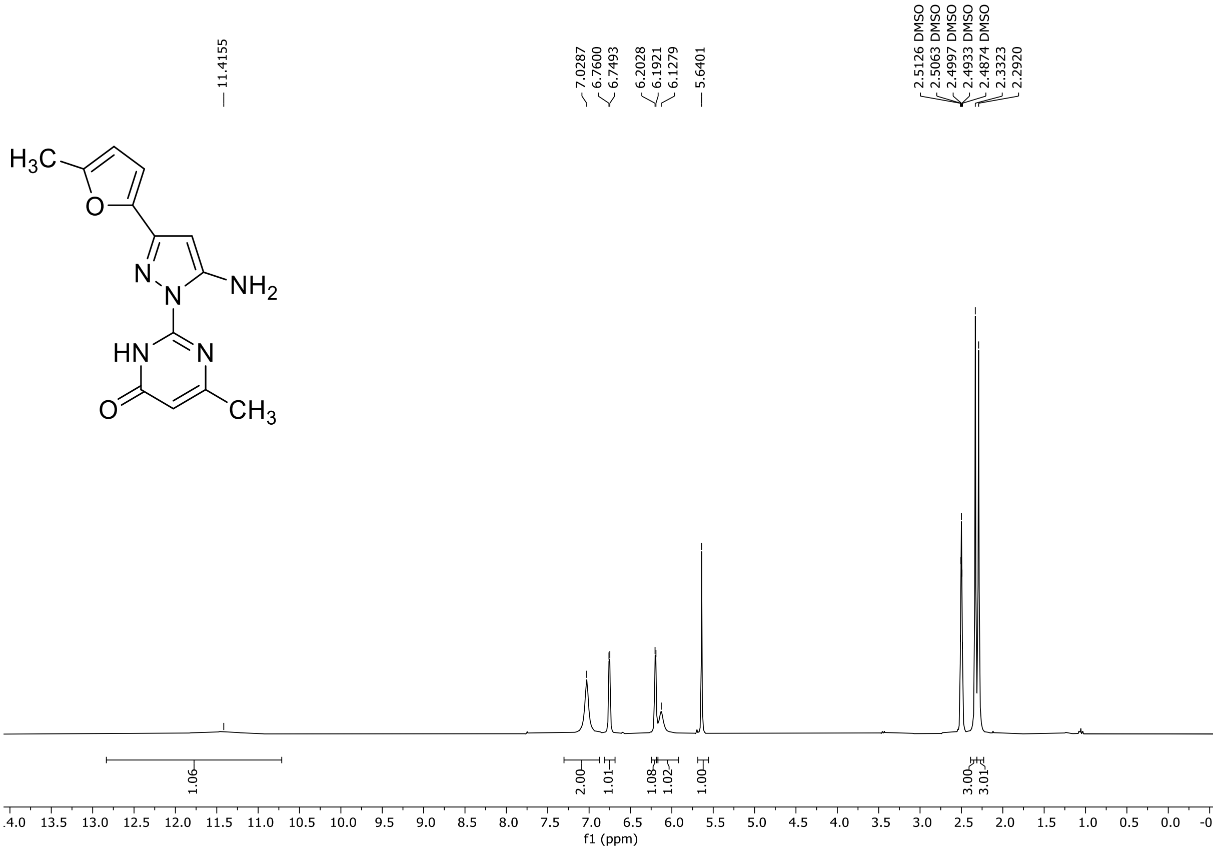

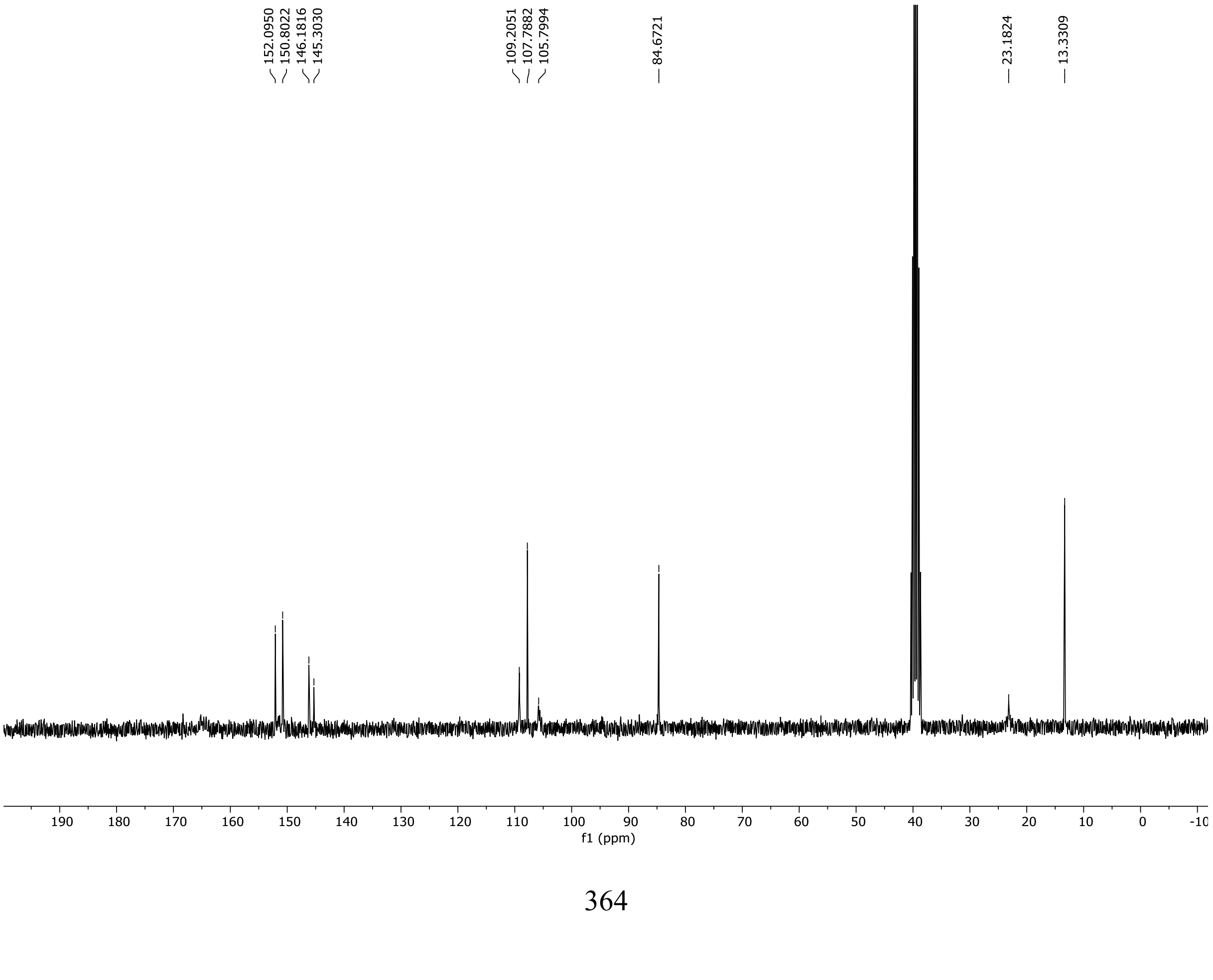

HRMS spectrum of **S136**

**NAR-A-177** $C_{13}H_{13}N_5O_2$ mono *m/z* 271.1069

**APCI +** (MMI)
nitrogen flow 5 L/min, gas temperature 325°C, nebulizer 45 psi, skimmer **65** V, vaporizer 200°C, fragmentor 20 V, dissolved in methanol

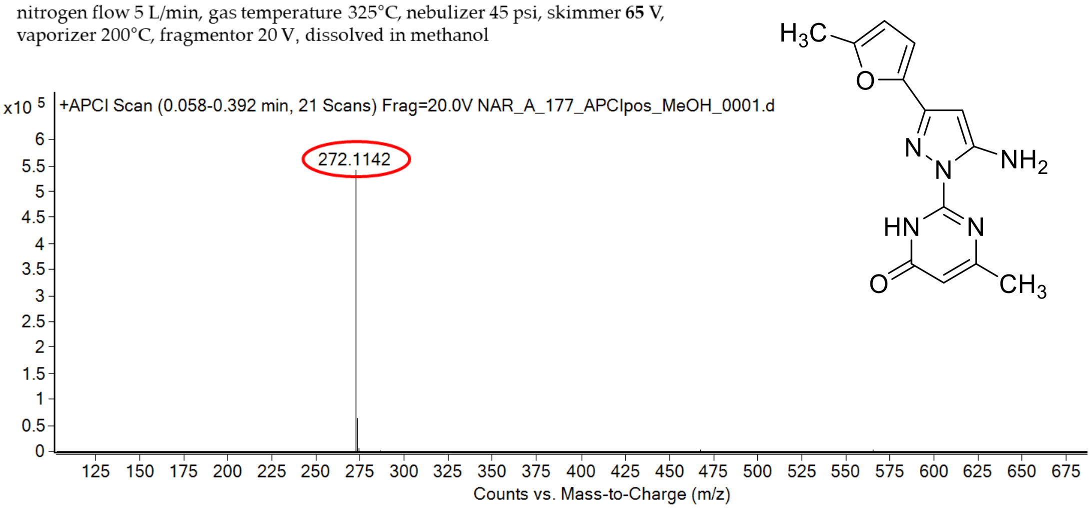

calculated mass: $[M+H]^+$ = 272..1142 observed: $[M+H]^+$ = 272.1142 mass accuracy = < 0.1 ppm

FT-IR spectrum (neat) of **S136**

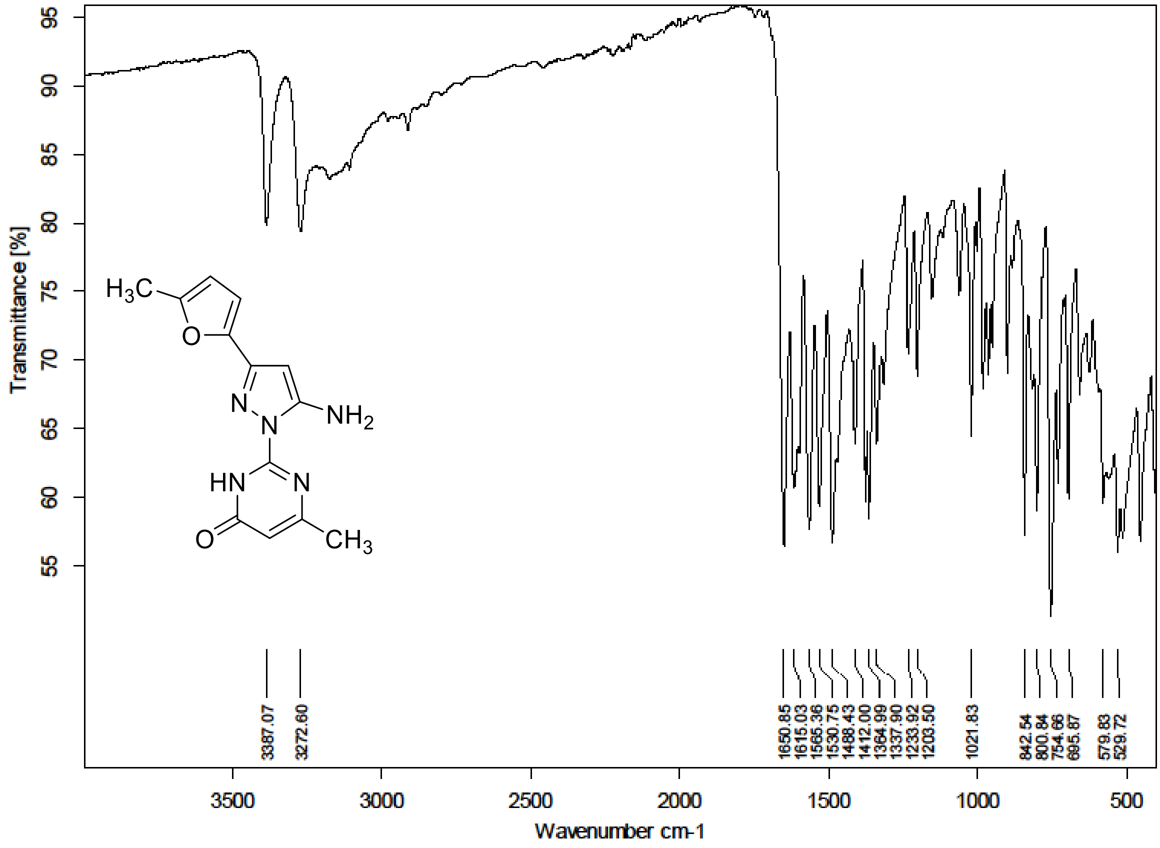

$^{1}H$ (500 MHz) and $^{13}C$ NMR (126 MHz) spectra of **S137** in DMSO-$d_6$

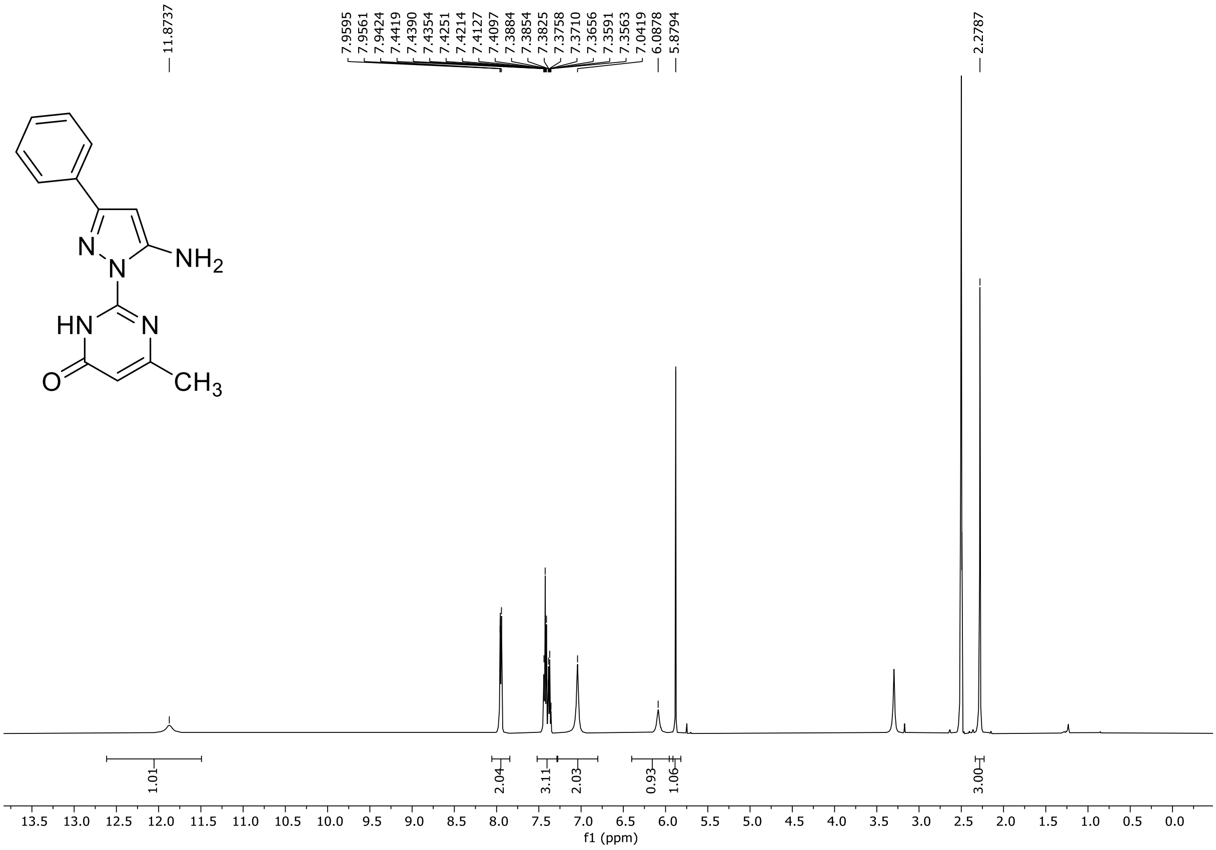

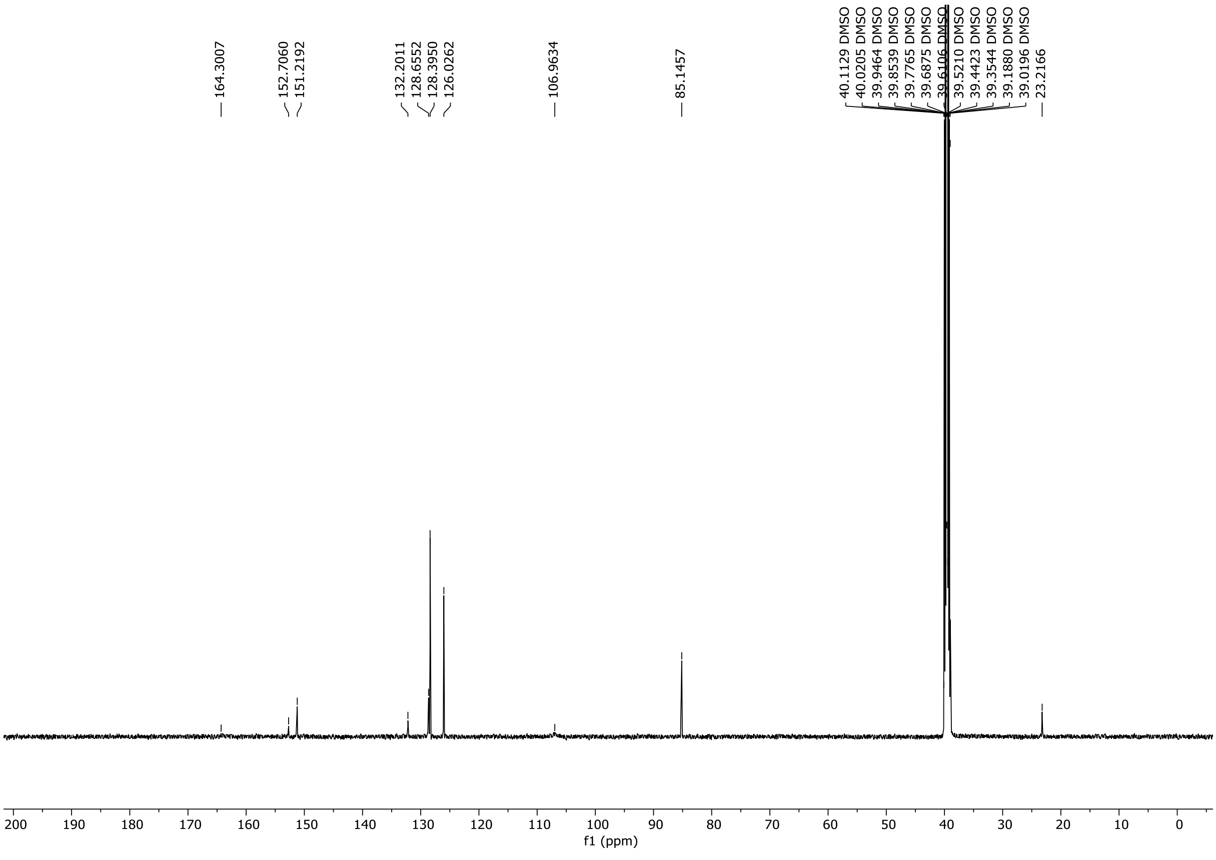

HRMS spectrum of **S137**

**NAR-A-1** $C_{14}H_{13}N_5O$ 267.1120 *mono m/z*

**APCI +** (MMI)
nitrogen flow 5 L/min, gas temperature 325°C, nebulizer 45 psi, skimmer **65** V, vaporizer 200°C, fragmentor 35 V, dissolved in methanol

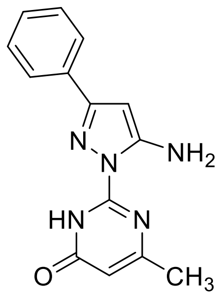

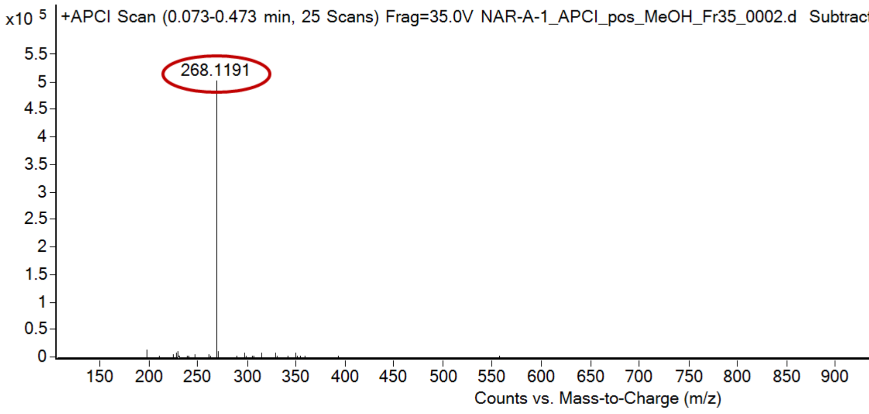

calculated mass: $[M+H]^+$ = 268.1193 observed: $[M+H]^+$ = 268.1193 mass accuracy = 0.7 ppm

FT-IR spectrum (neat) of **S137**

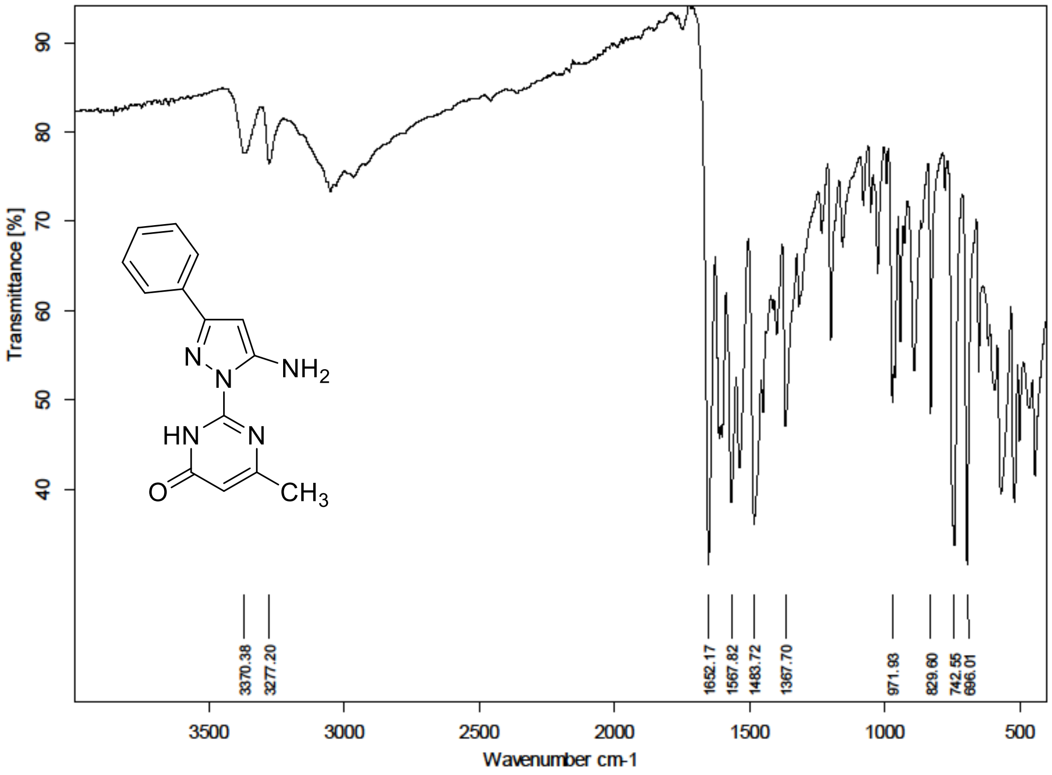

$^{1}H$ (500 MHz) and $^{13}C$ NMR (126 MHz) spectra of **S138** in DMSO-$d_6$

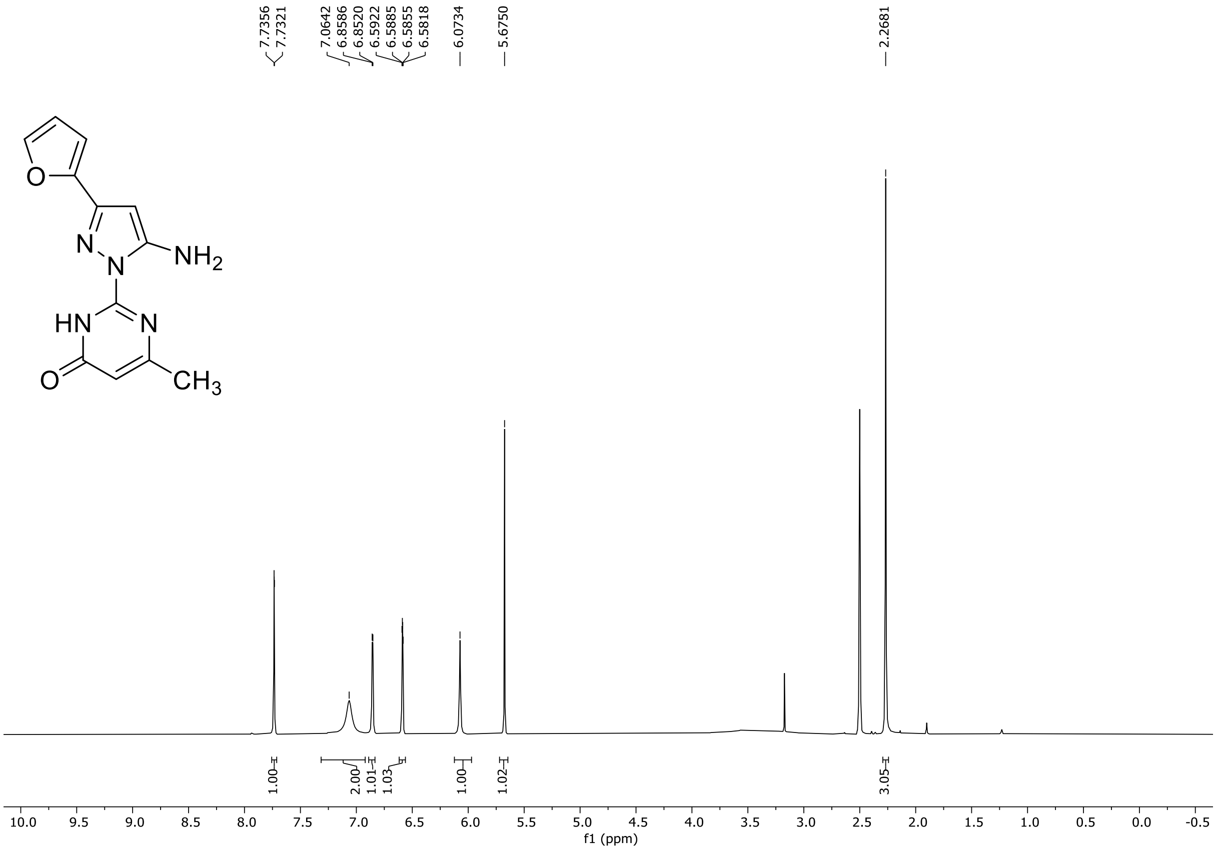

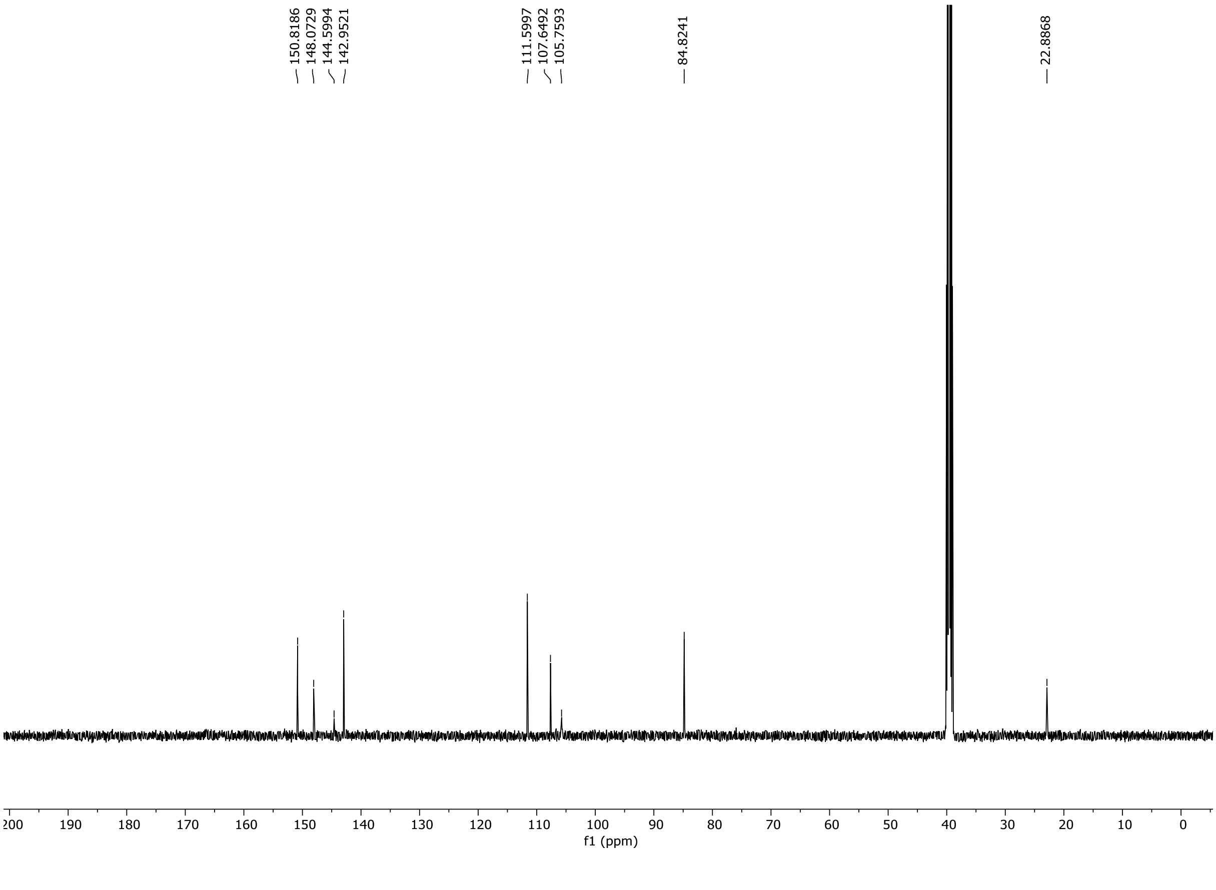

HRMS spectrum of **S138**

NAR-A-33 **$C_{12}H_{11}N_5O_2$** mono *m/z 257.0913*

**APCI +** (MMI)
nitrogen flow 5 L/min, gas temperature 325°C, nebulizer 45 psi, skimmer **65** V,
vaporizer 200°C, fragmentor 20 V, dissolved in methanol

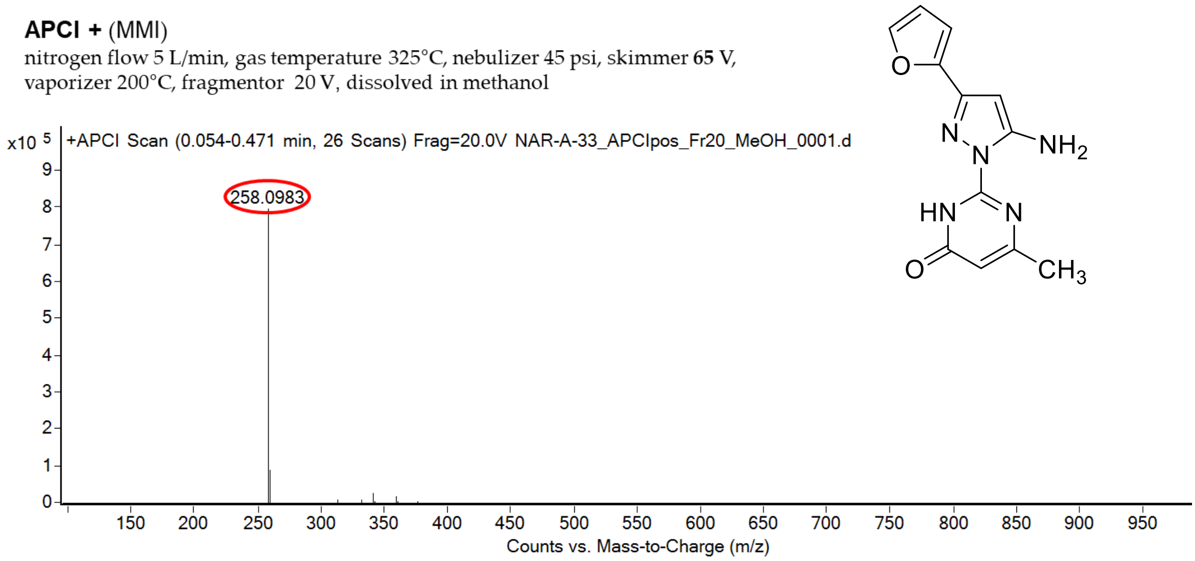

calculated mass: $[M+H]^+ = 258.0986$ observed: $[M+H]^+ = 258.0983$ mass accuracy = -1.2 ppm

FT-IR spectrum (neat) of **S138**

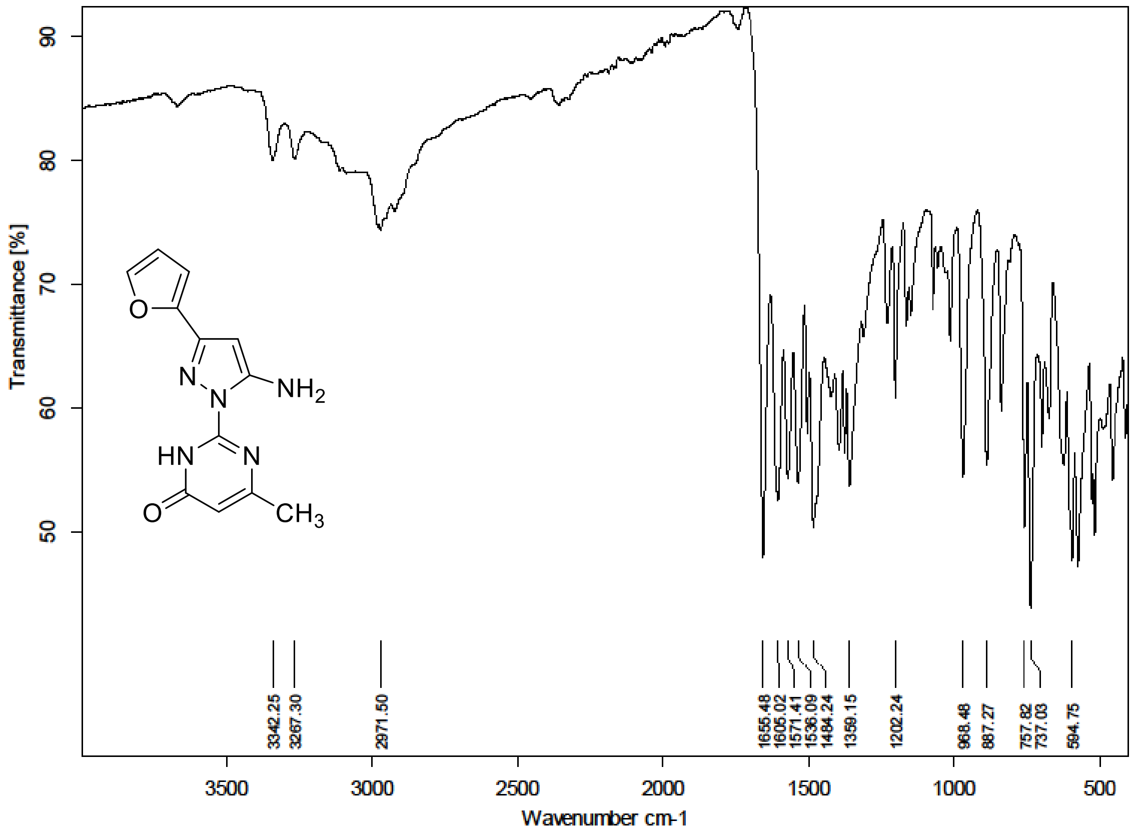

$^{1}H$ (500 MHz) and $^{13}C$ NMR (126 MHz) spectra of **S140** in DMSO-$d_6$

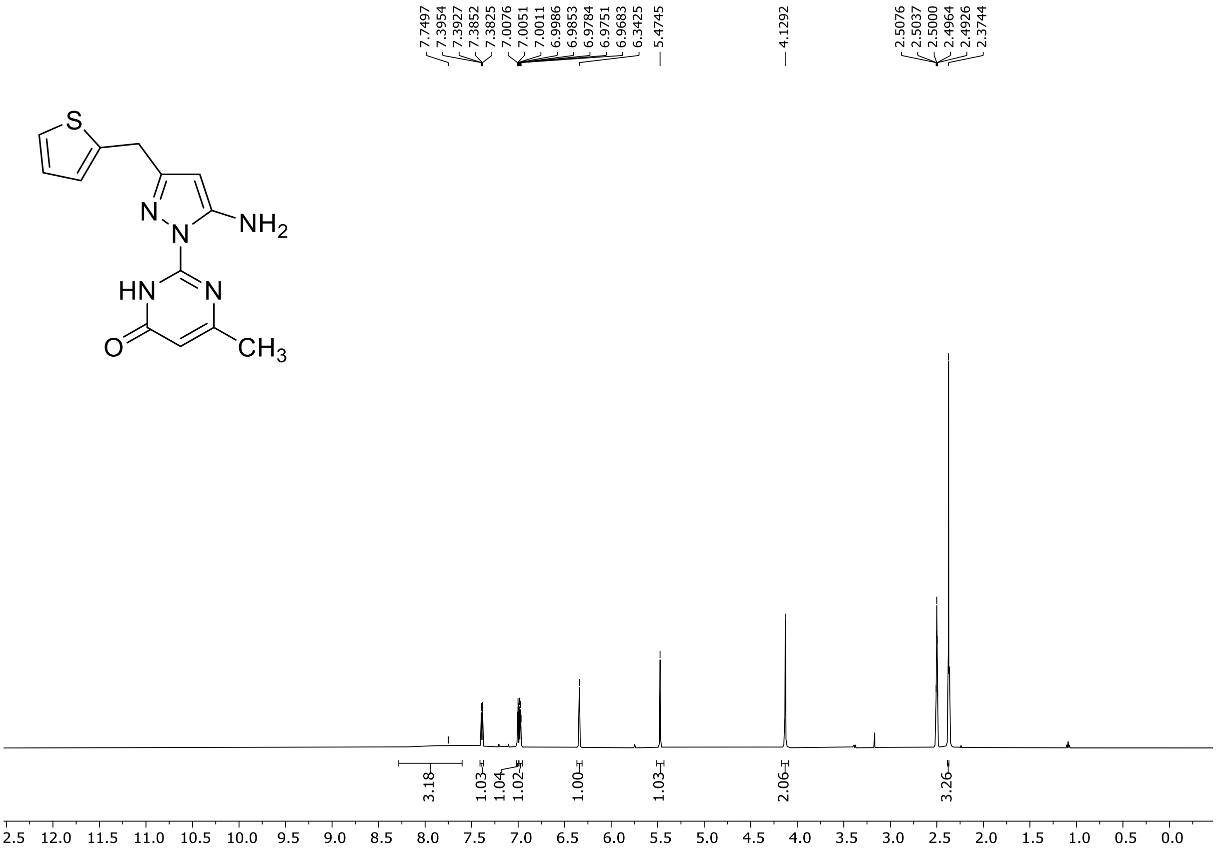

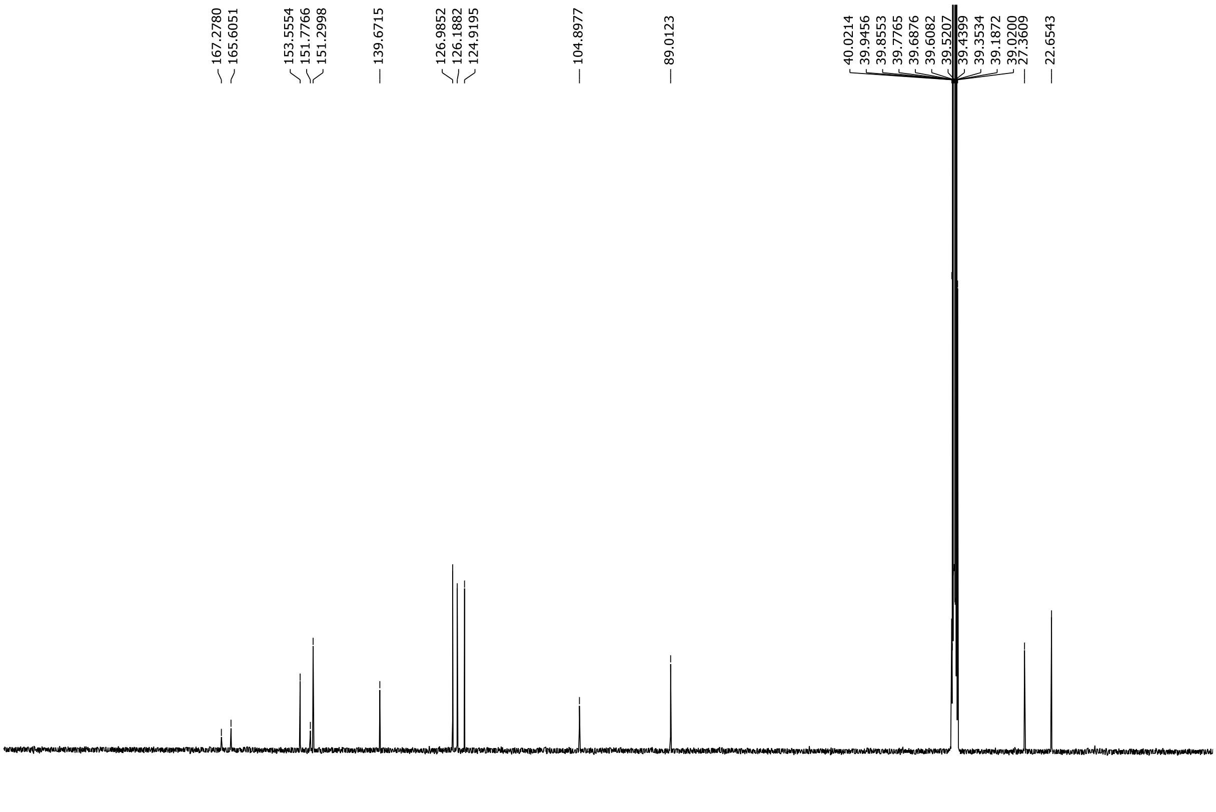

HRMS spectrum of **S140**

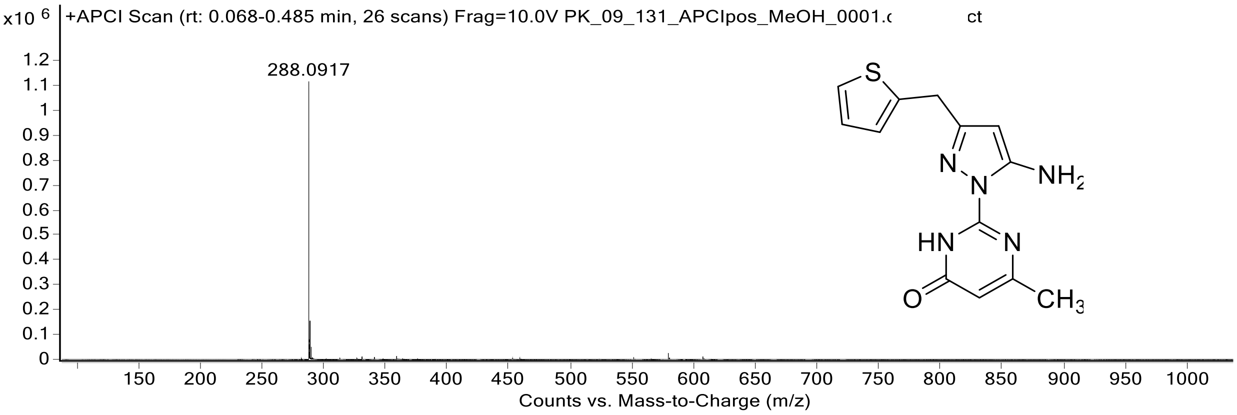

[1]H (500 MHz) and [13]C NMR (126 MHz) spectra of **S141** in DMSO-$d_6$

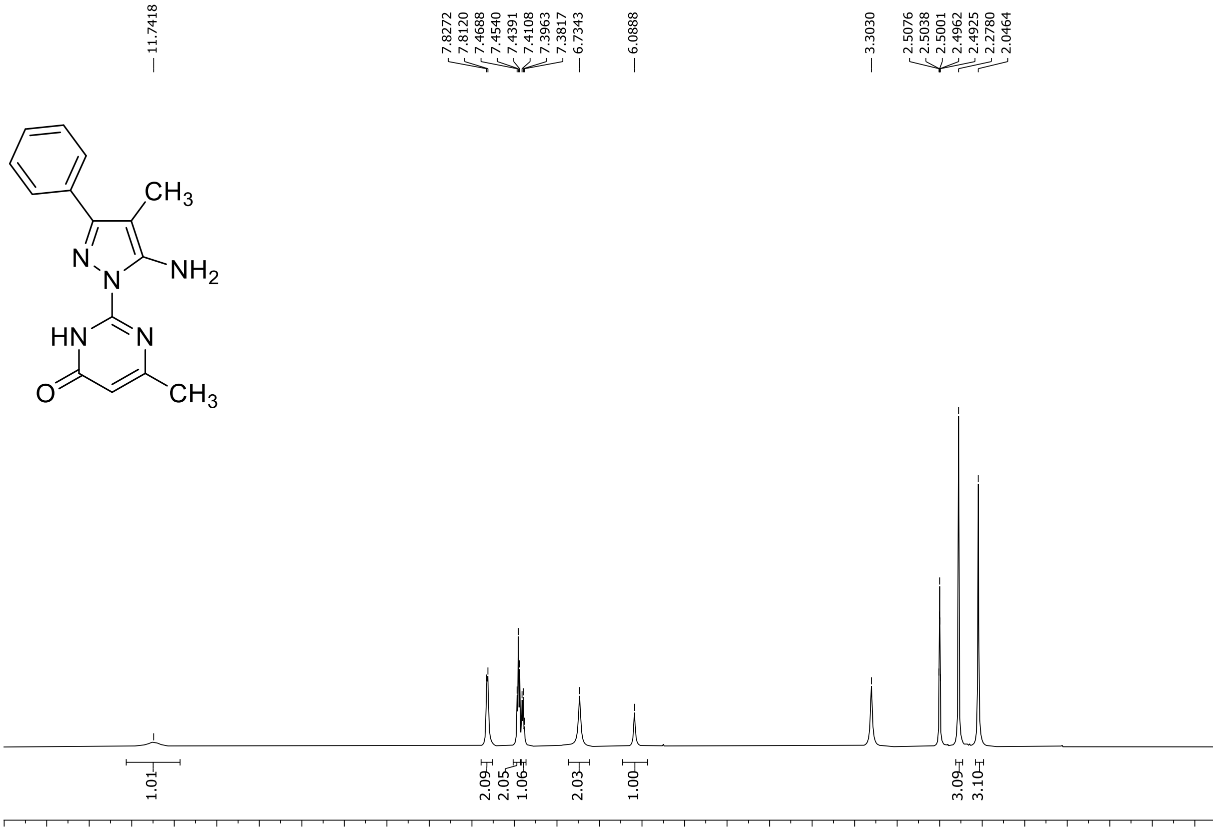

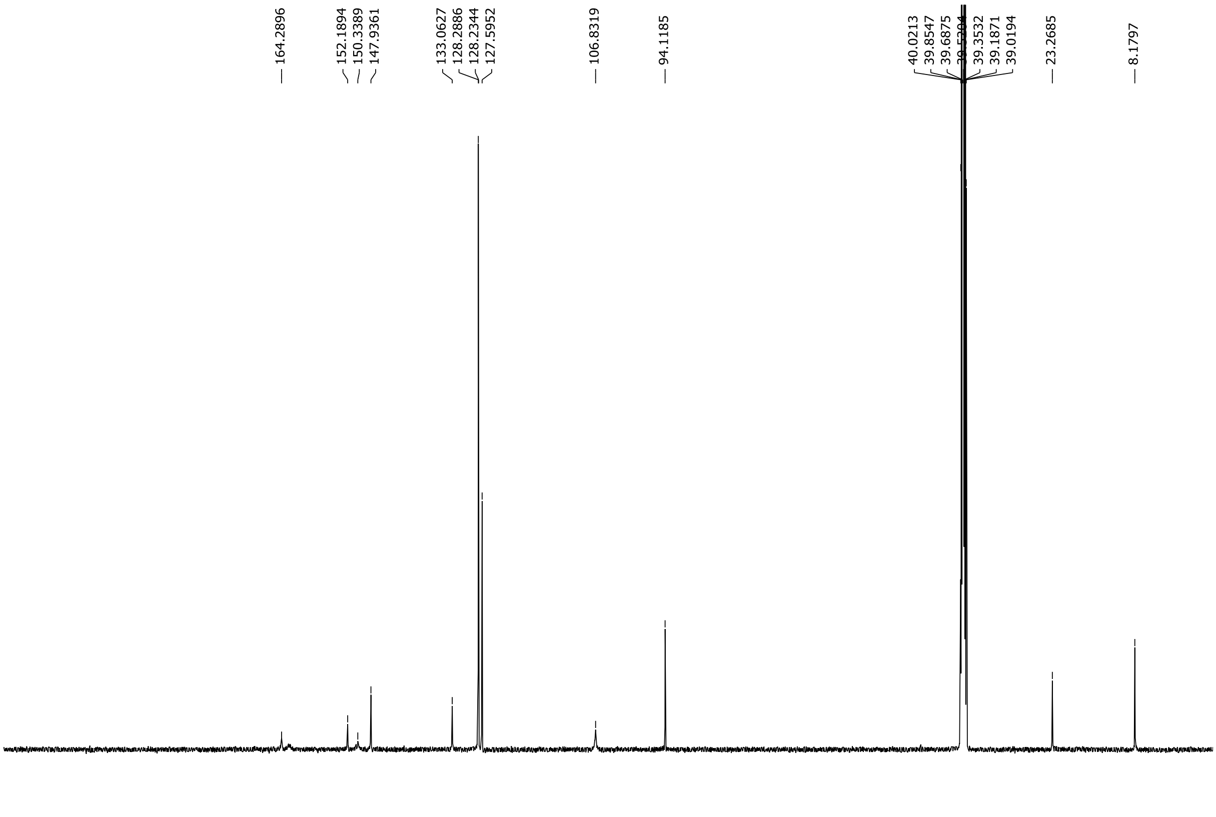

HRMS spectrum of **S141**

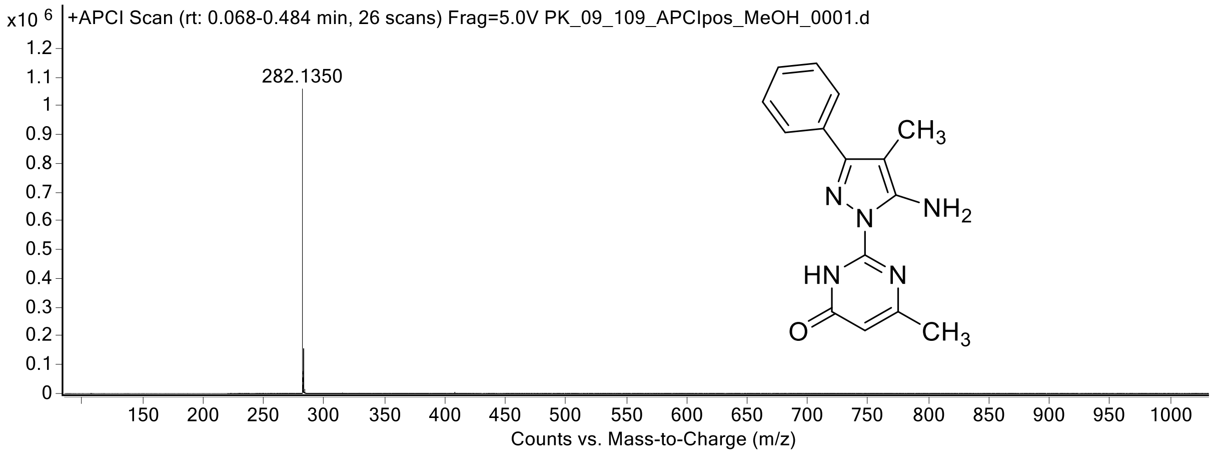

[1]H (500 MHz) and [13]C NMR (126 MHz) spectra of **S142** in DMSO-$d_6$

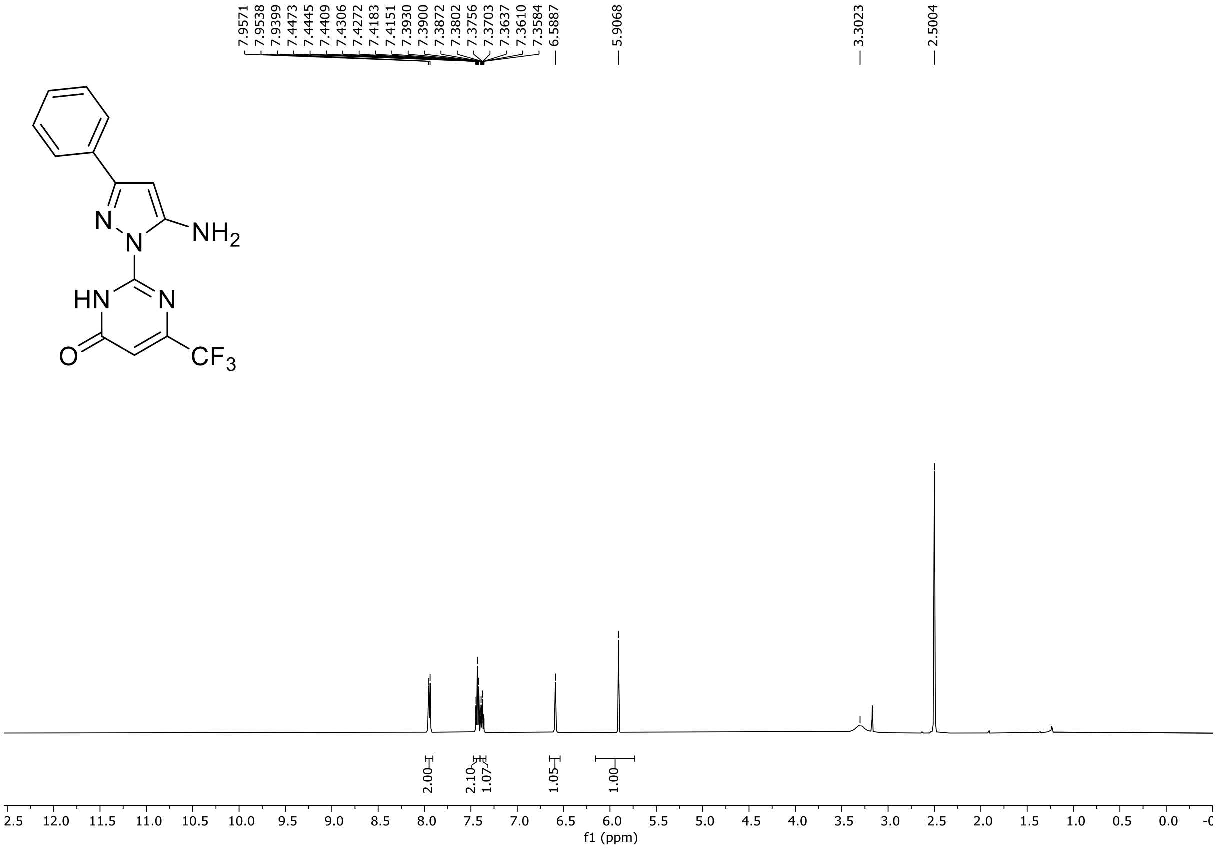

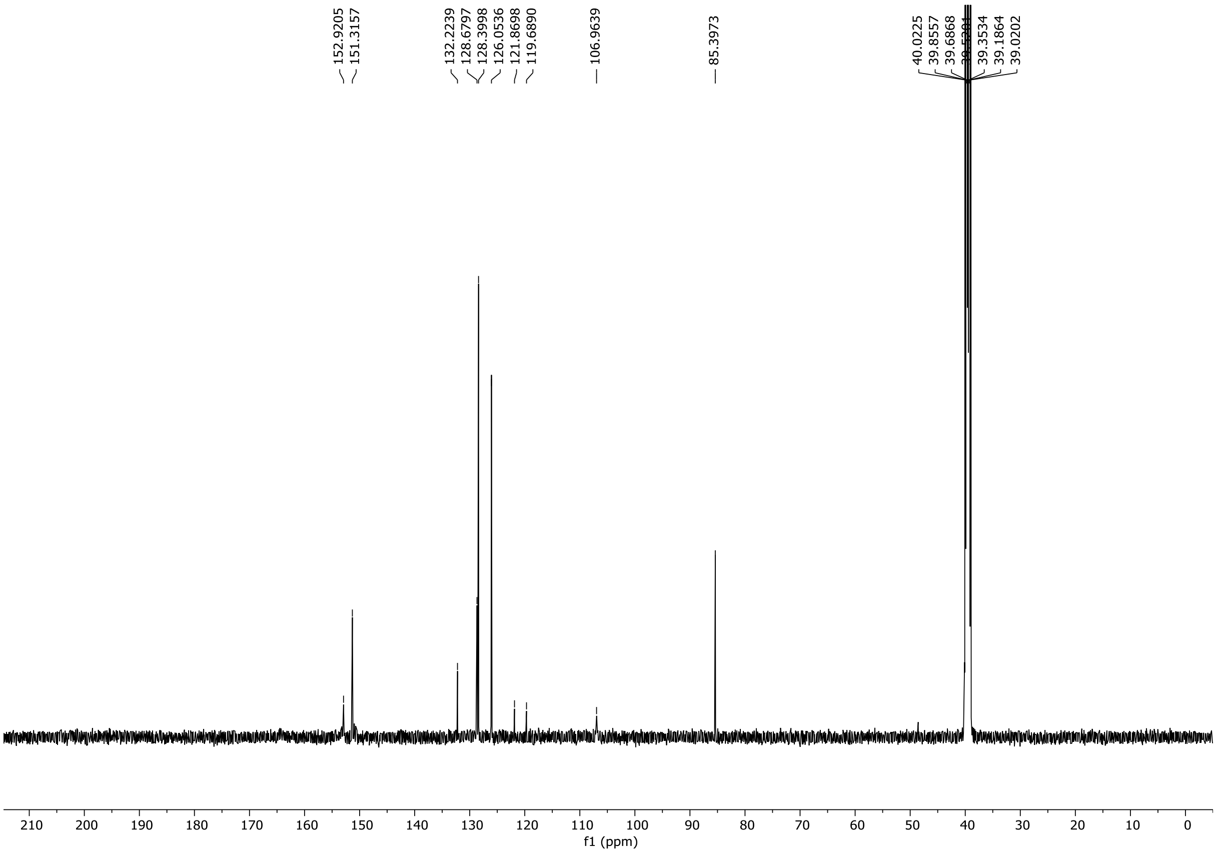

$^{19}F$ NMR (471 MHz) spectrum of **S142** in DMSO-$d_6$

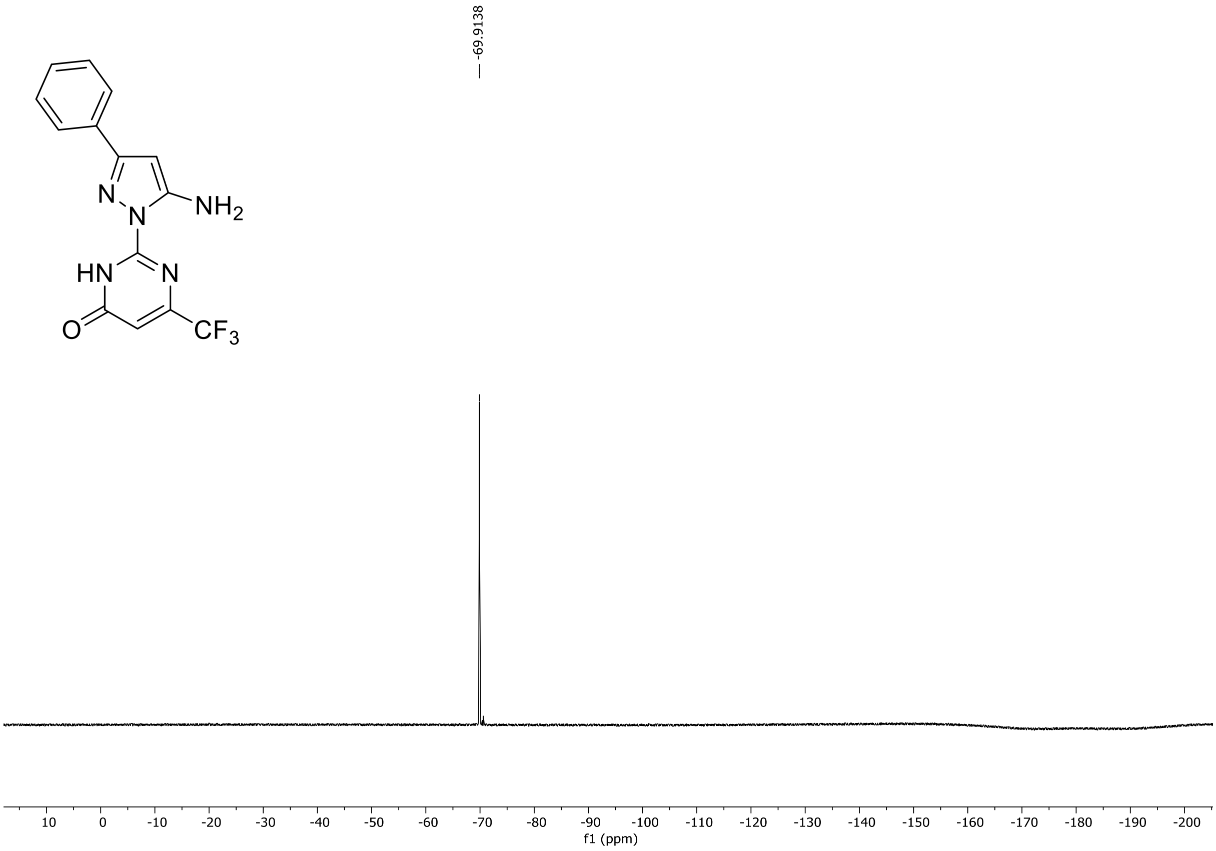

HRMS spectrum of **S142**

**NAR-A-90** **$C_{14}H_{10}F_3N_5O$** ***m/z* 321.0837**

**APCI+** (MMI)
nitrogen flow 5 L/min, gas temperature 325°C, nebulizer 45 psi, skimmer 65 V, vaporizer 200°C, fragmentor 30 V, dissolved in MeOH

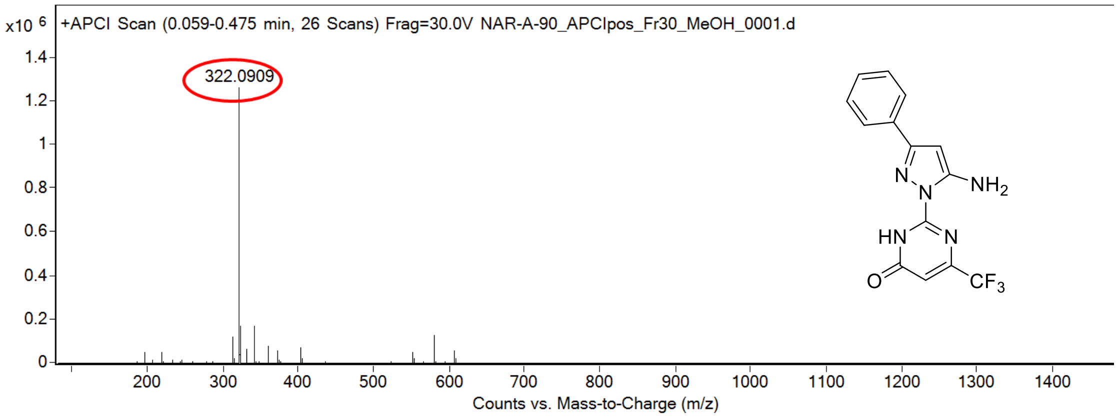

calculated mass: $[M+H]^+$ = 322.0910 observed: $[M+H]^+$ = 322.0909 mass accuracy = -0.3 ppm

FT-IR spectrum (neat) of **S142**

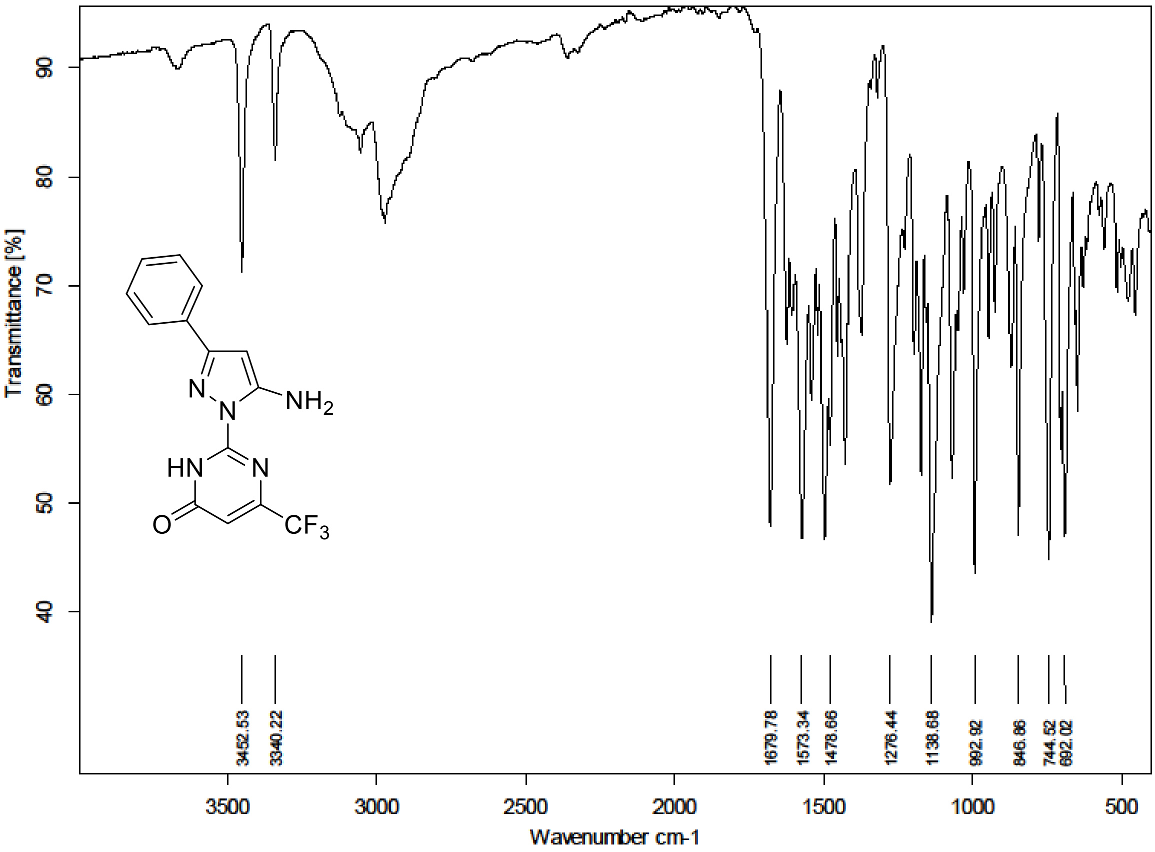

[1]H (300 MHz) and [13]C NMR (75 MHz) spectra of **S143** in DMSO-$d_6$

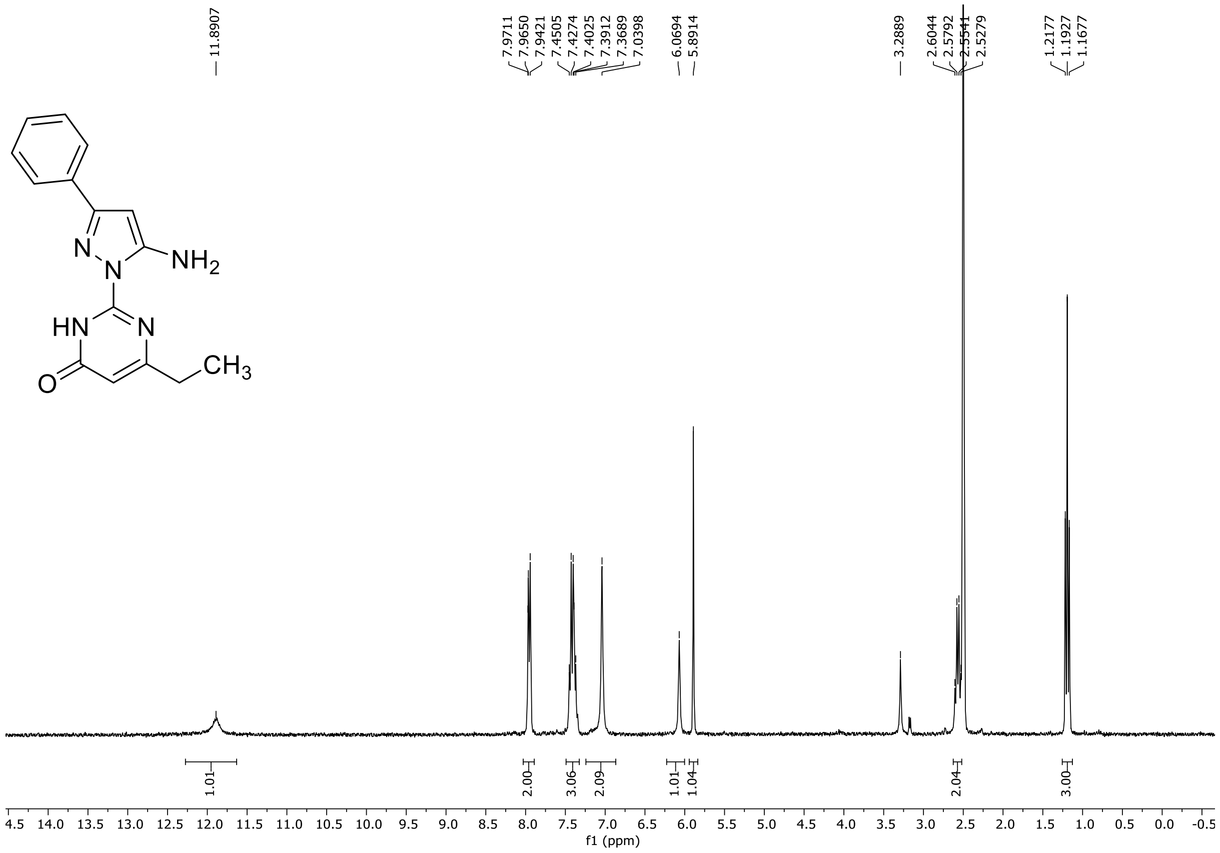

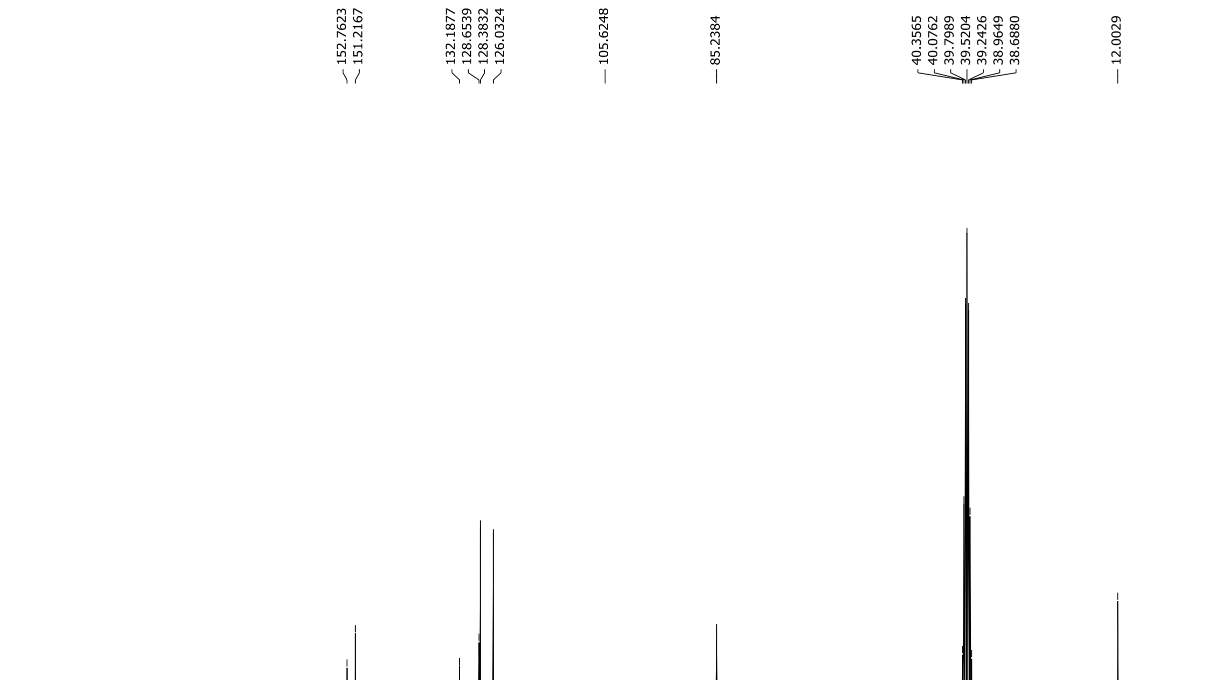

HRMS spectrum of **S143**

**NAR-A-167** $C_{15}H_{15}N_5O$ mono *m/z* 281.1277

**APCI +** (MMI)
nitrogen flow 5 L/min, gas temperature 325°C, nebulizer 45 psi, skimmer **65** V, vaporizer 200°C, fragmentor 20 V, dissolved in methanol

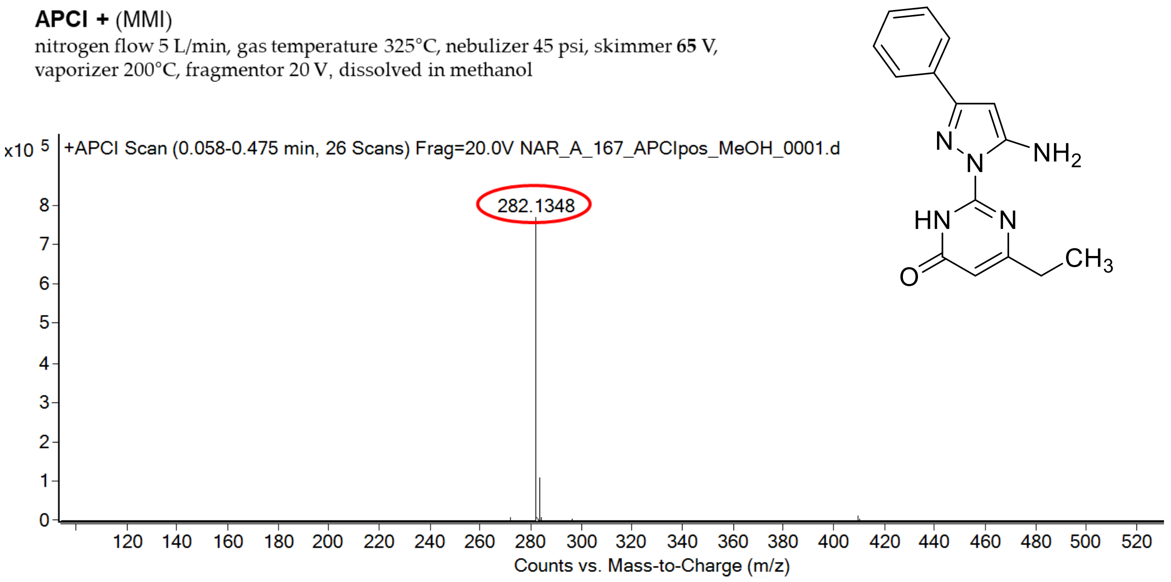

calculated mass: $[M+H]^+$ = 282..1349 observed: $[M+H]^+$ = 282.1348 mass accuracy = - 0.4 ppm

FT-IR spectrum (neat) of **S143**

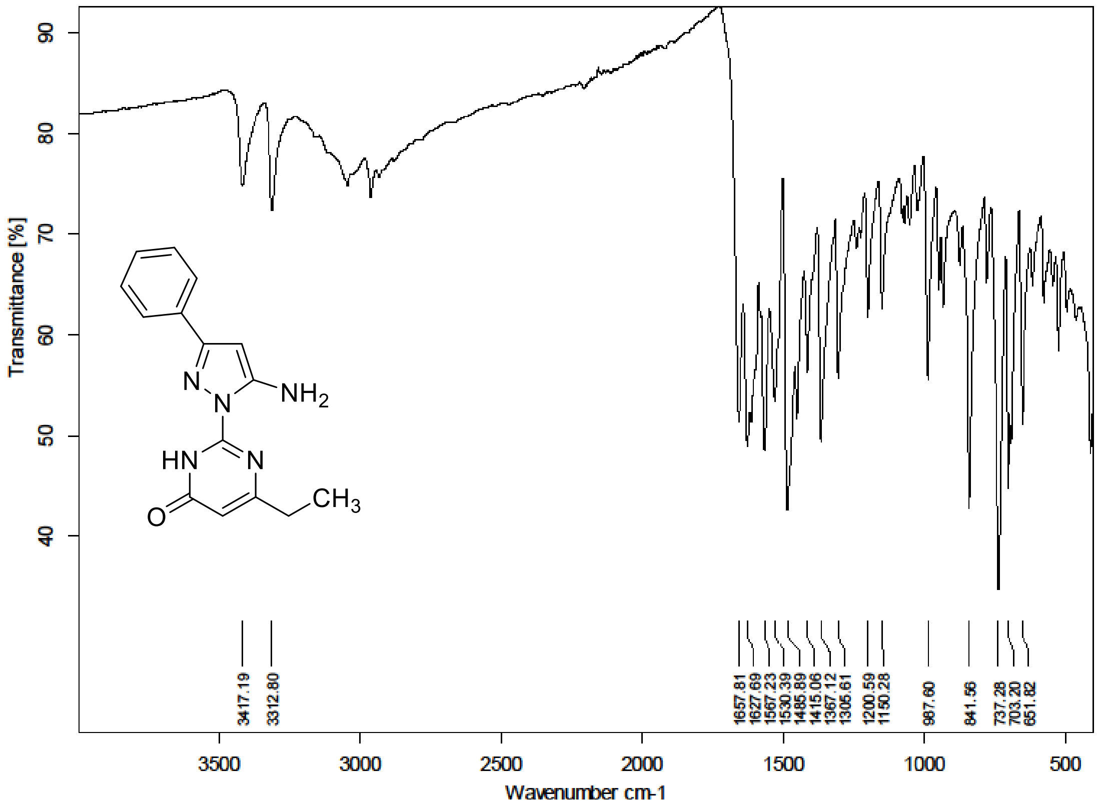

$^1H$ (500 MHz) and $^{13}C$ NMR (126 MHz) spectra of **S144** in DMSO-$d_6$

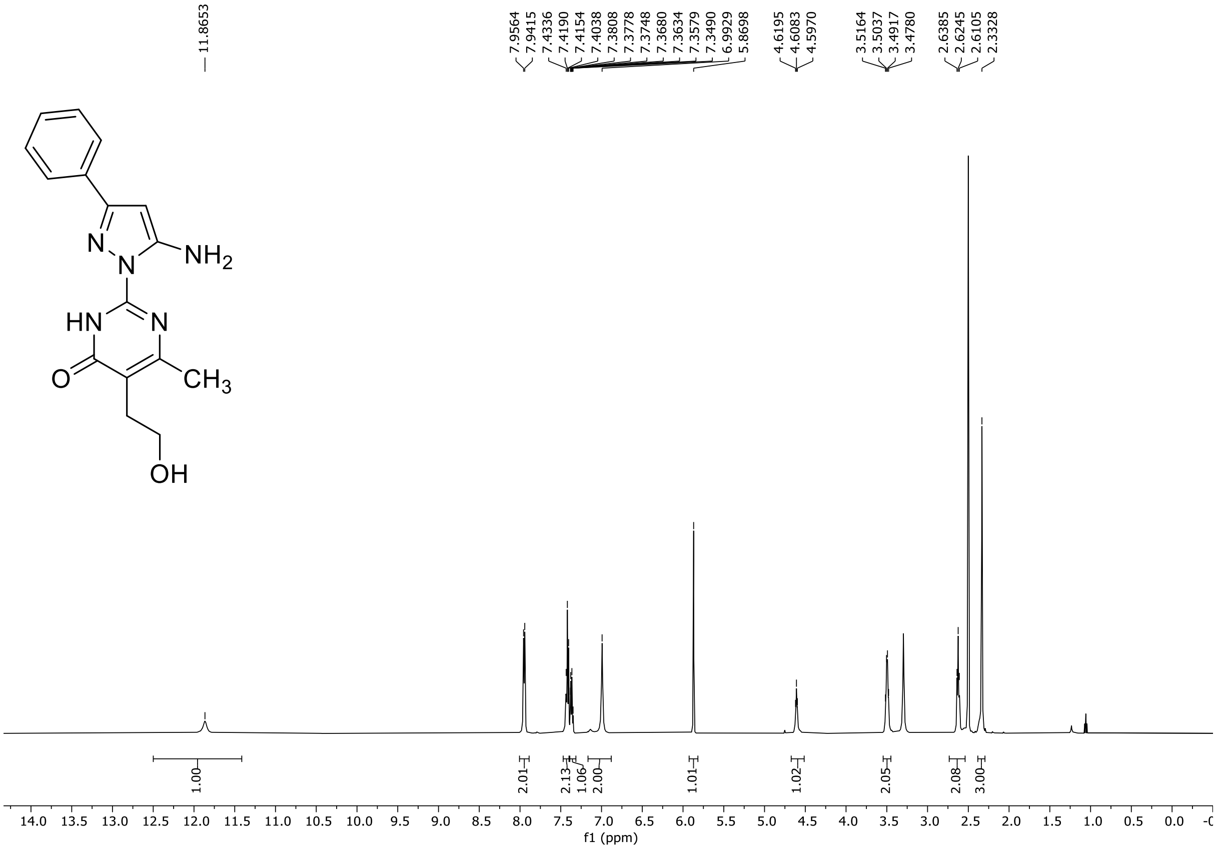

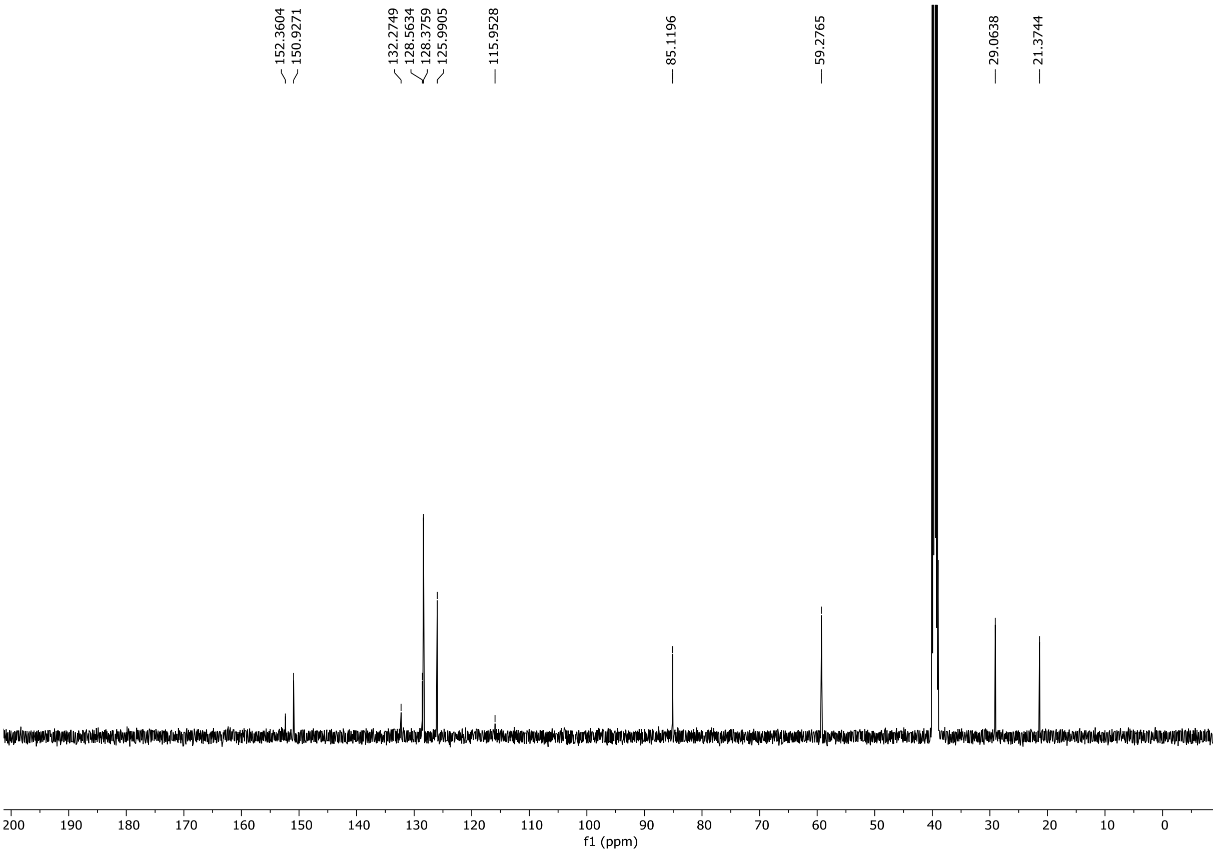

HRMS spectrum of **S144**

NAR-A-15 $C_{16}H_{17}N_5O_2$ mono *m/z 311.1382*

**APCI +** (MMI)
nitrogen flow 5 L/min, gas temperature 325°C, nebulizer 45 psi, skimmer **65** V, vaporizer 200°C, fragmentor 30 V, dissolved in methanol

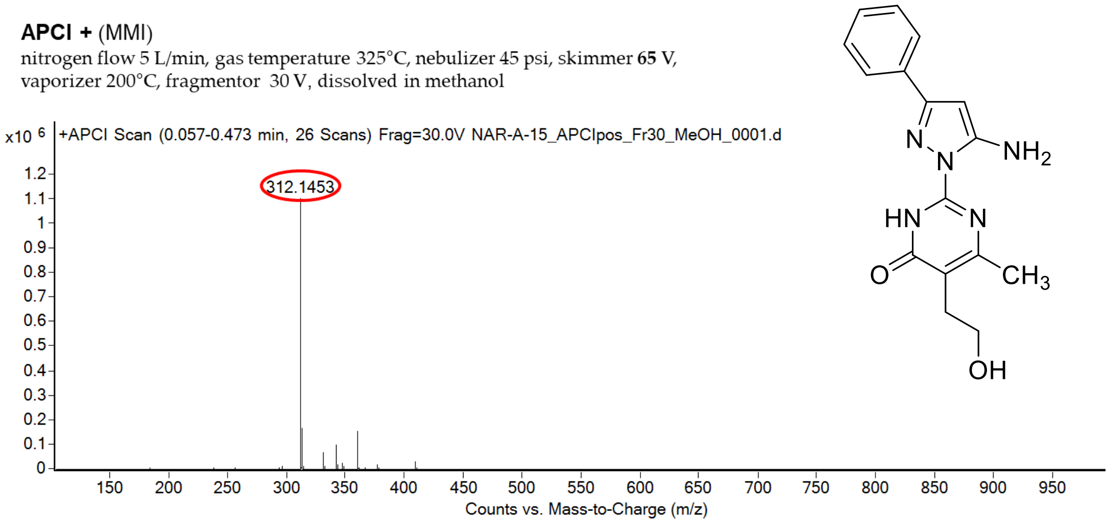

calculated mass: $[M+H]^+$ = 312.1455 observed: $[M+H]^+$ = 312.1453 mass accuracy = -0.6 ppm

FT-IR spectrum (neat) of **S144**

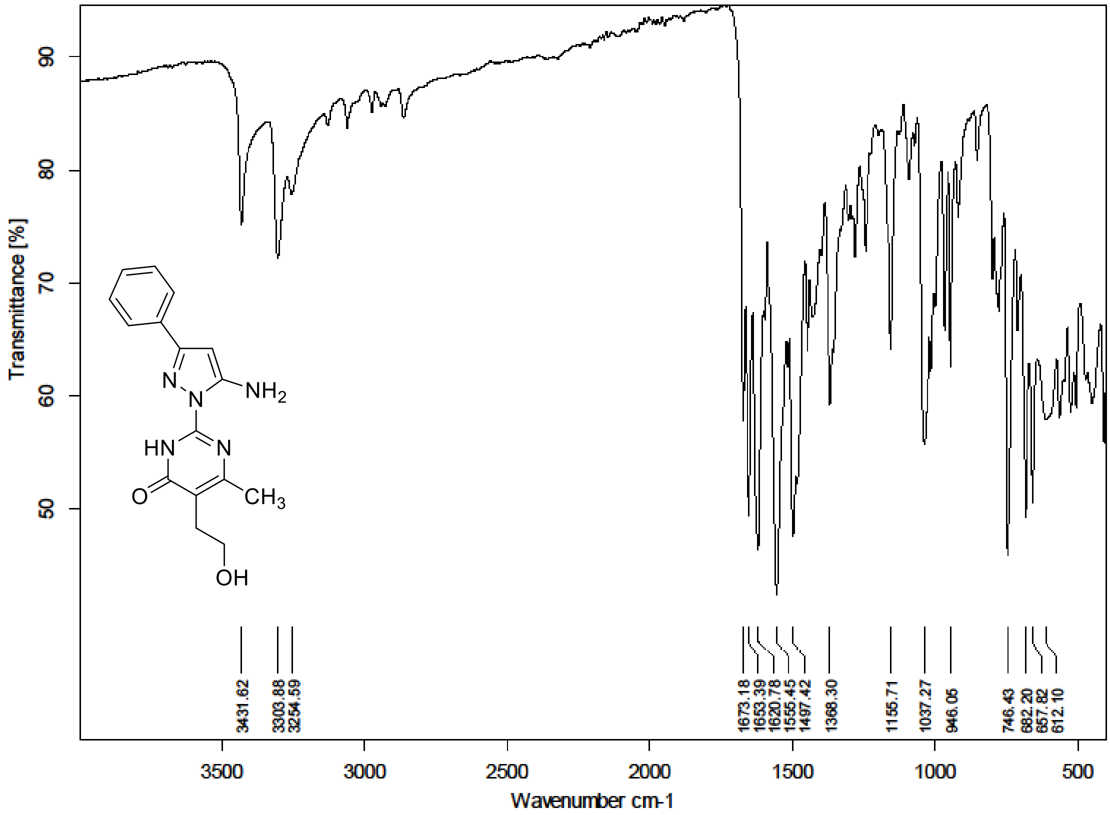

[1]H (500 MHz) and [13]C NMR (126 MHz) spectra of **S145** in Chloroform-*d*

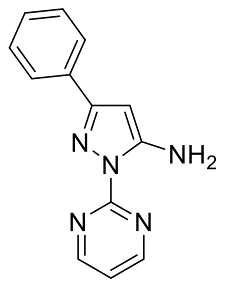

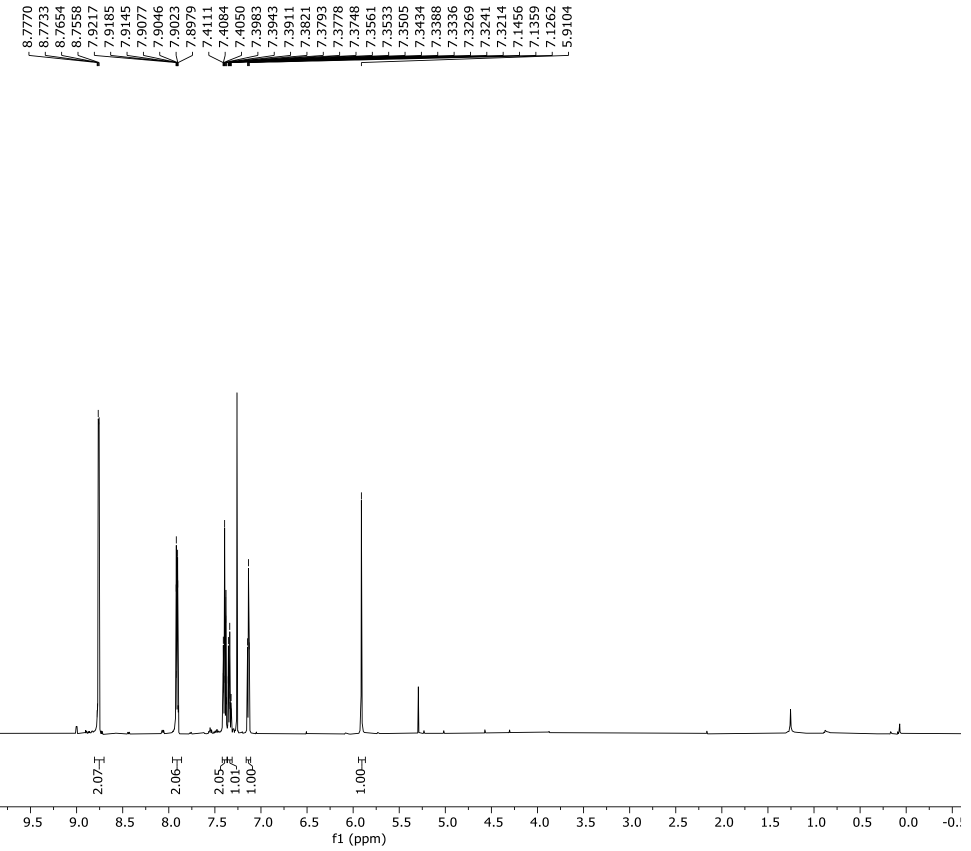

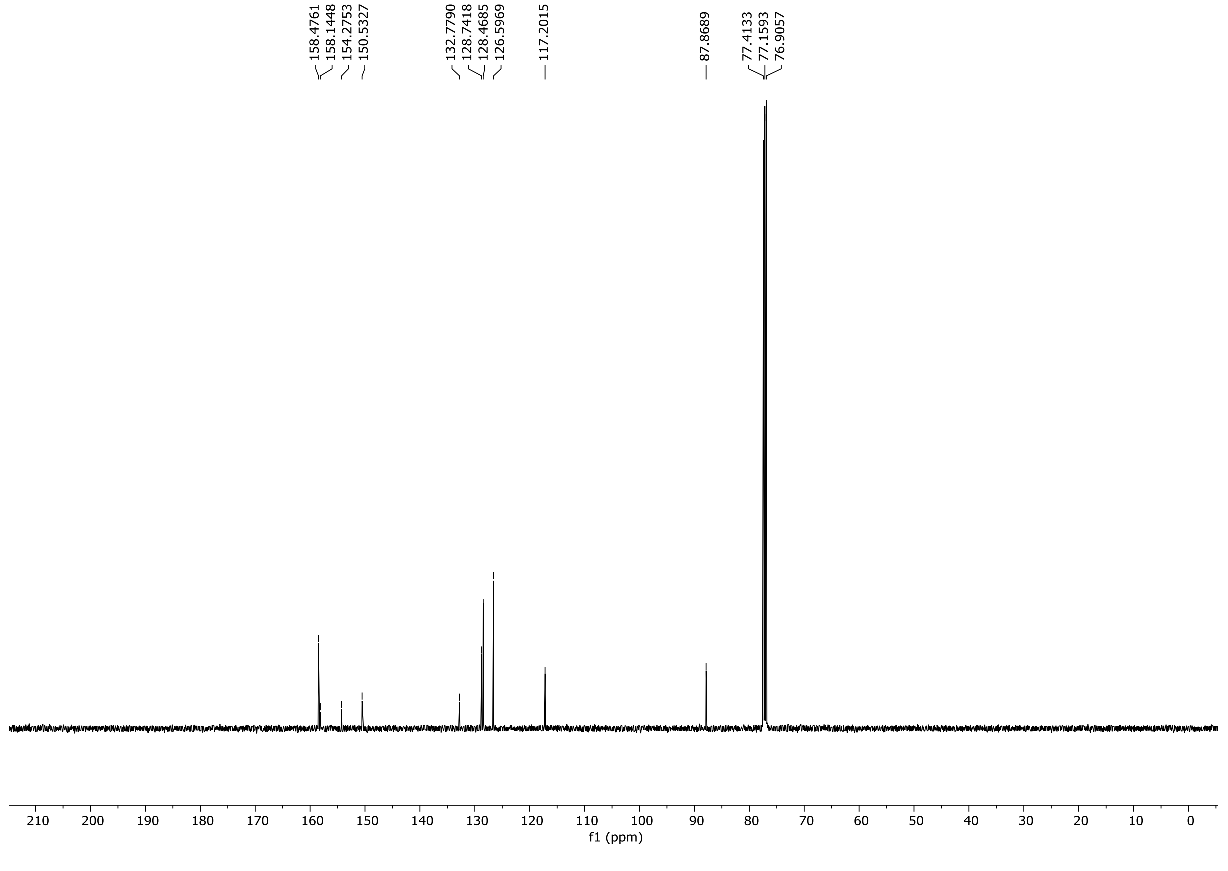

HRMS spectrum of **S145**

**NAR-A-9** $C_{13}H_{11}N_5$ *mono m/z* **237.1014**

**APCI +** (MMI)
nitrogen flow 5 L/min, gas temperature 325°C, nebulizer 45 psi, skimmer **65** V, vaporizer 200°C, fragmentor 25 V, dissolved in methanol

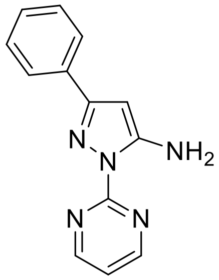

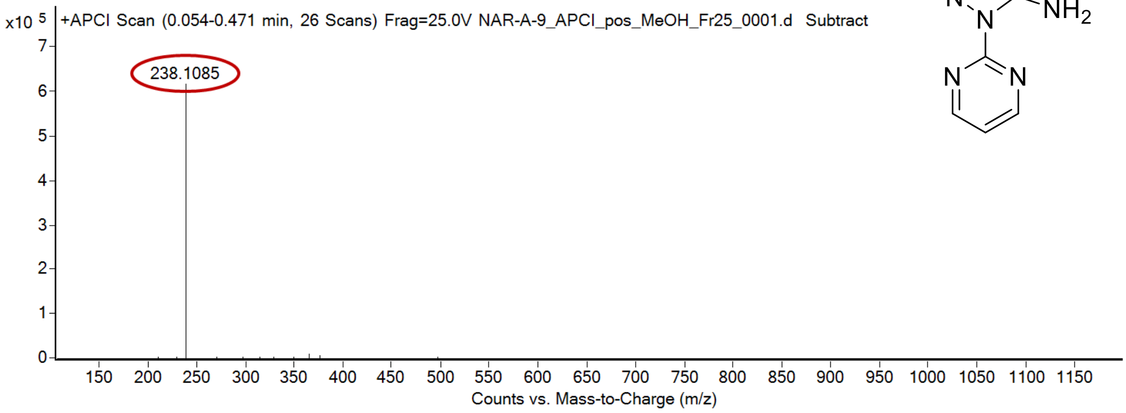

calculated mass: $[M+H]^+$ = 238.1087 observed: $[M+H]^+$ = 238.1085 mass accuracy = 0.8 ppm

FT-IR spectrum (neat) of **S145**

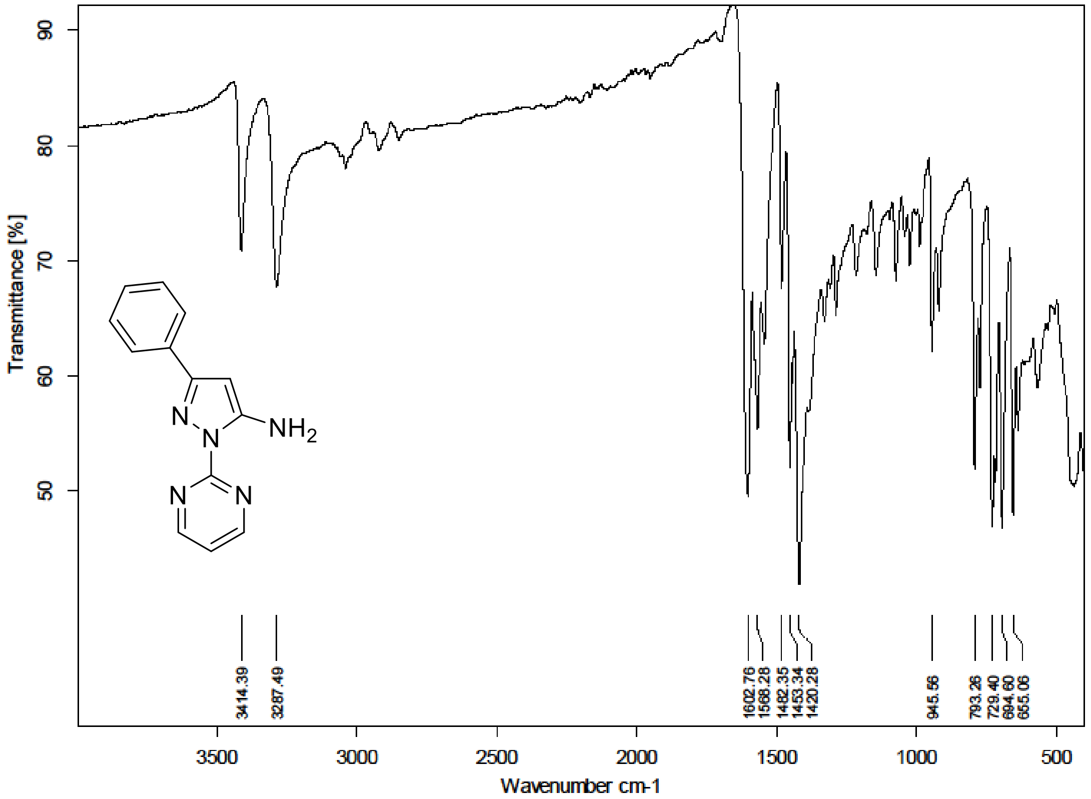

$^{1}H$ (500 MHz) and $^{13}C$ NMR (126 MHz) spectra of **S146** in Chloroform-*d*

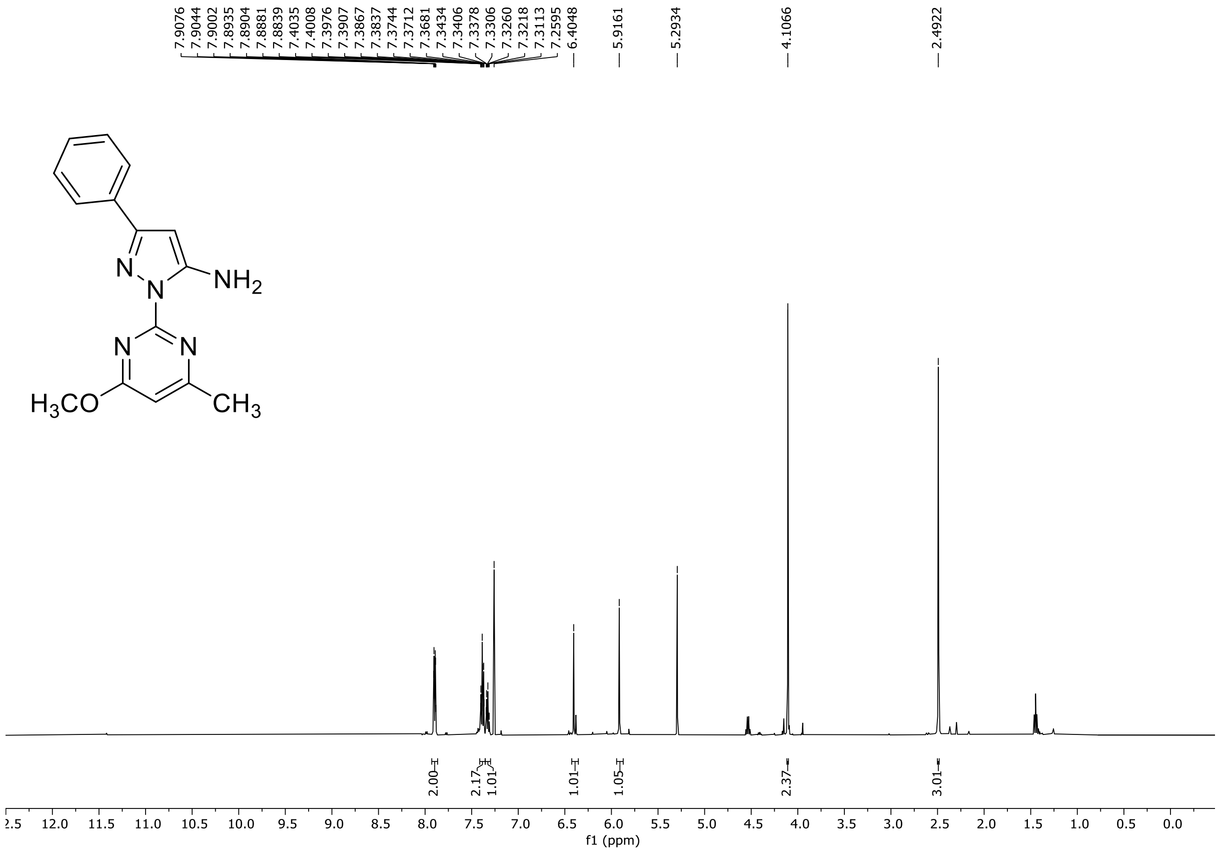

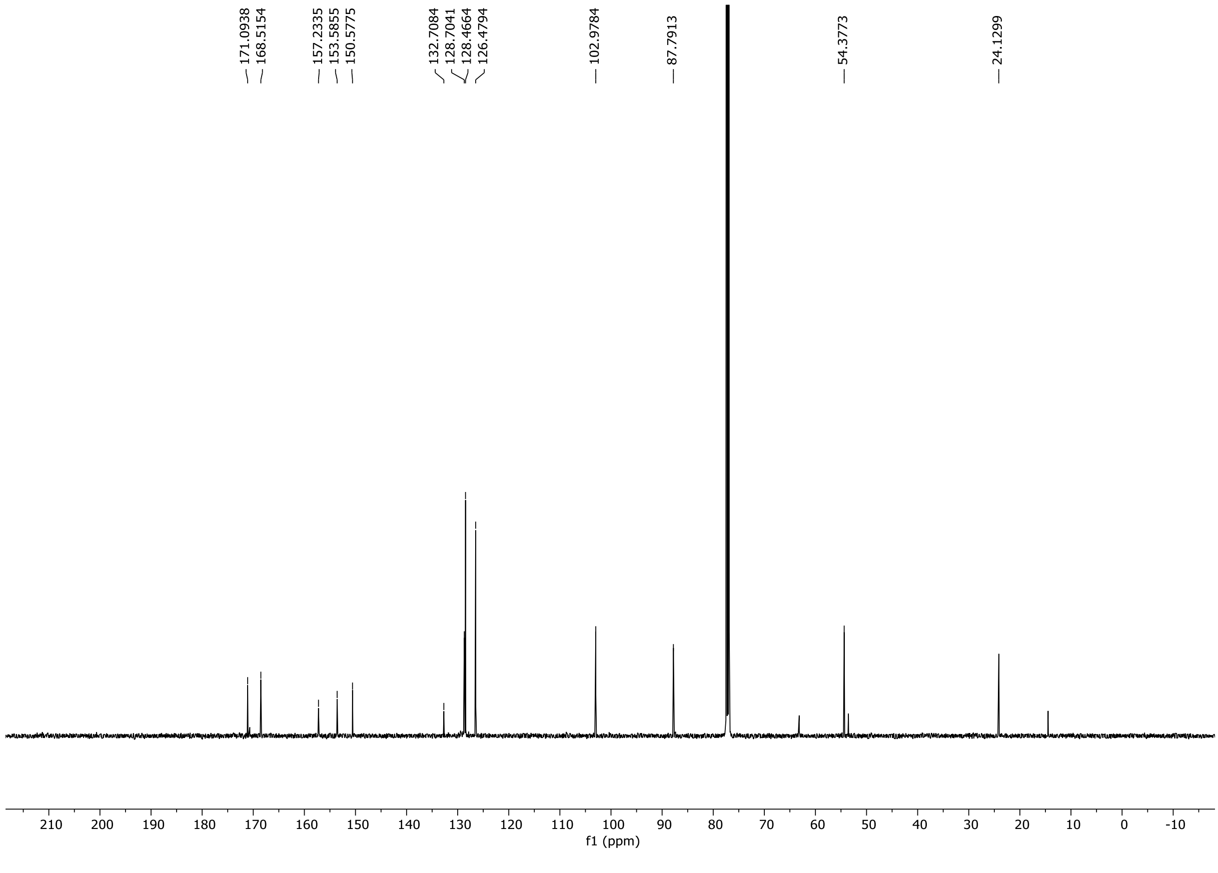

HRMS spectrum of **S146**

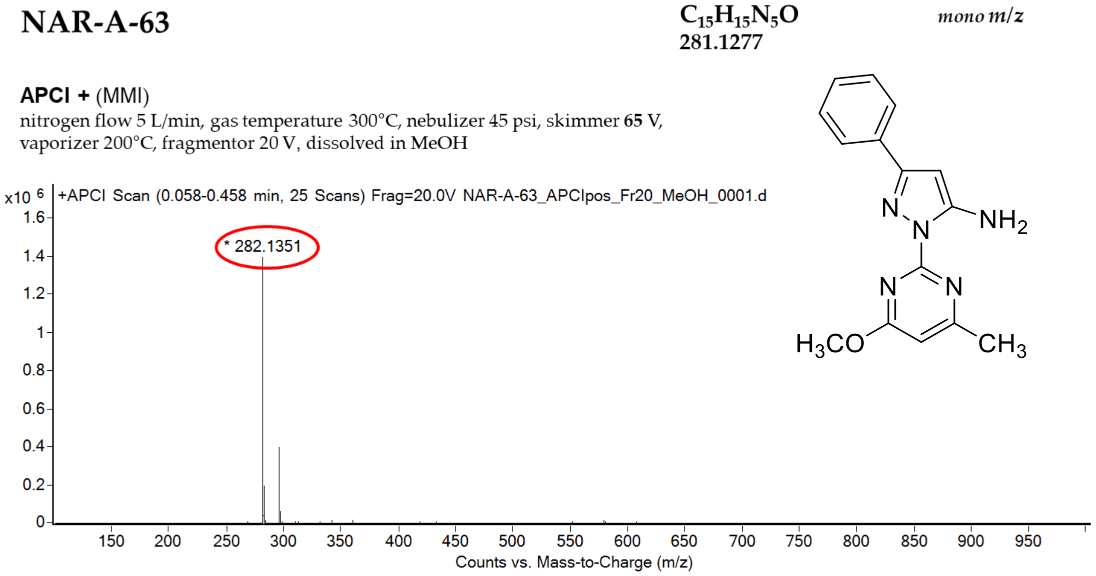

calculated mass: [M+H]$^+$ = 282.1349 observed: [M+H]$^+$ = 282.1351 mass accuracy = 0.4 ppm

FT-IR spectrum (neat) of **S146**

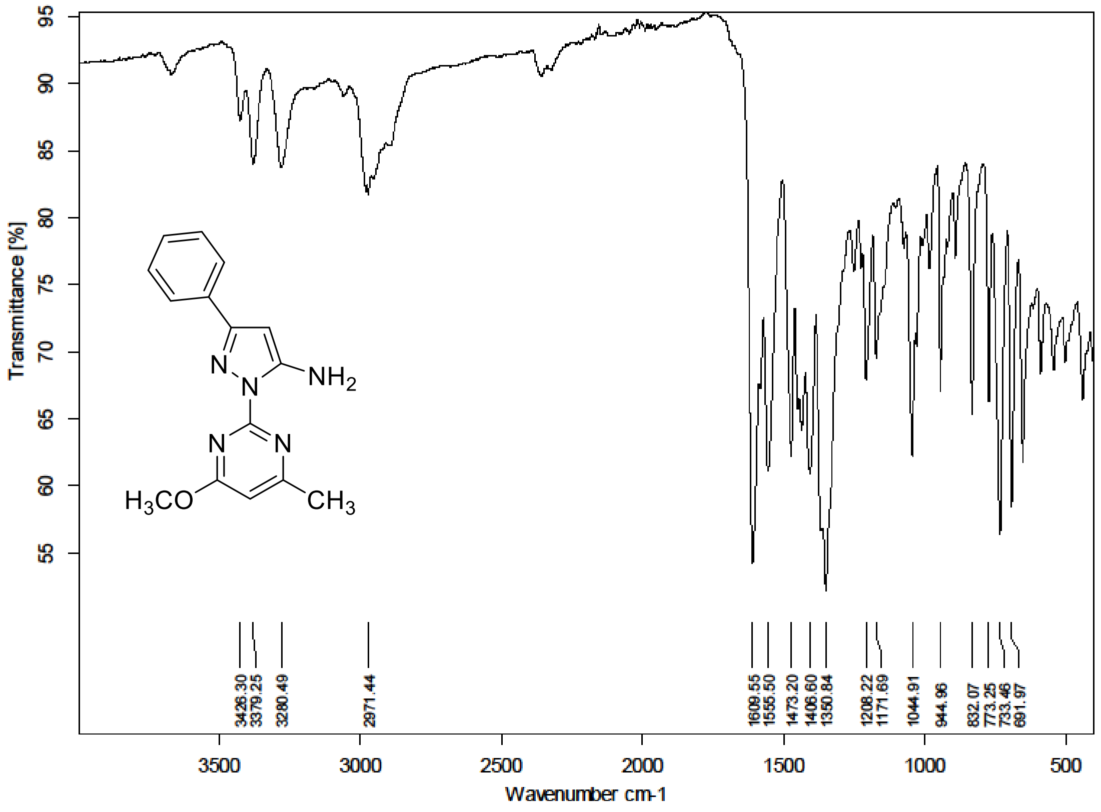

[1]H (500 MHz) and [13]C NMR (126 MHz) spectra of **S147** in DMSO-$d_6$

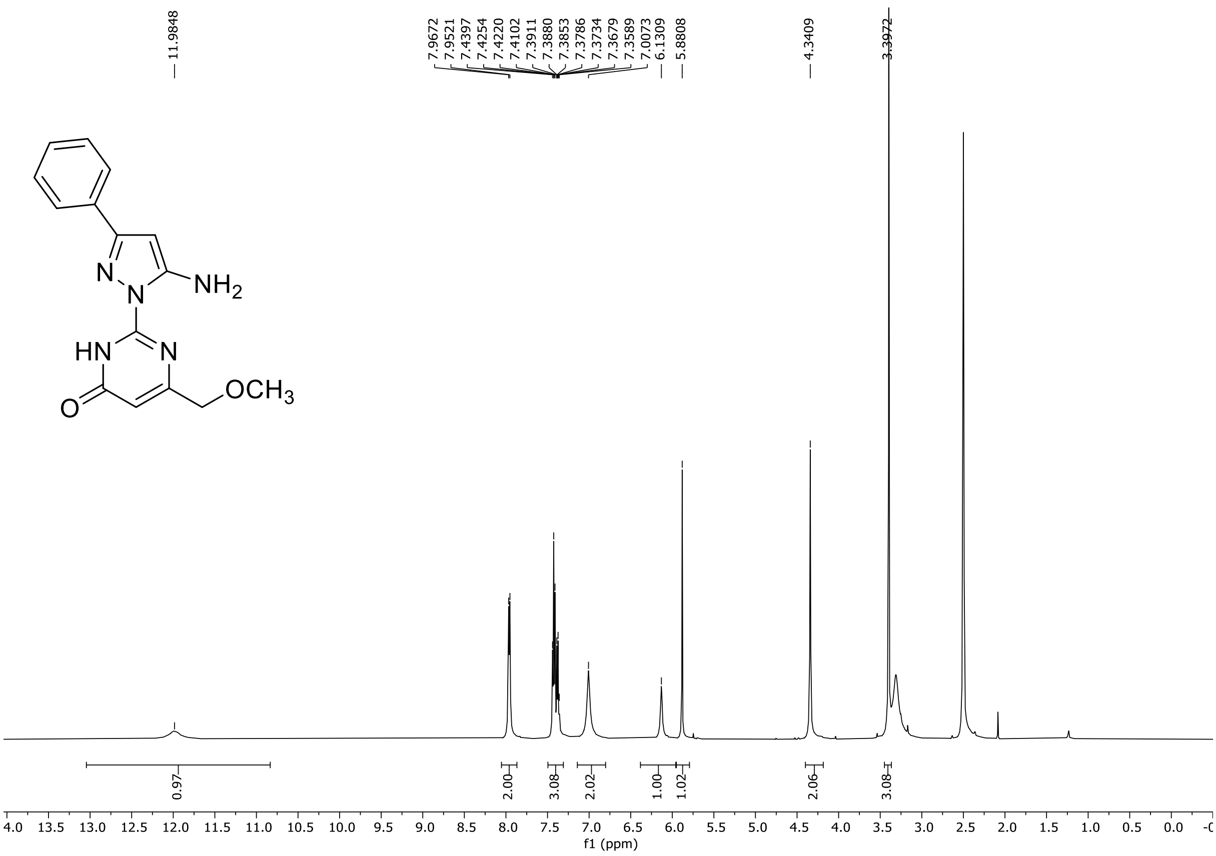

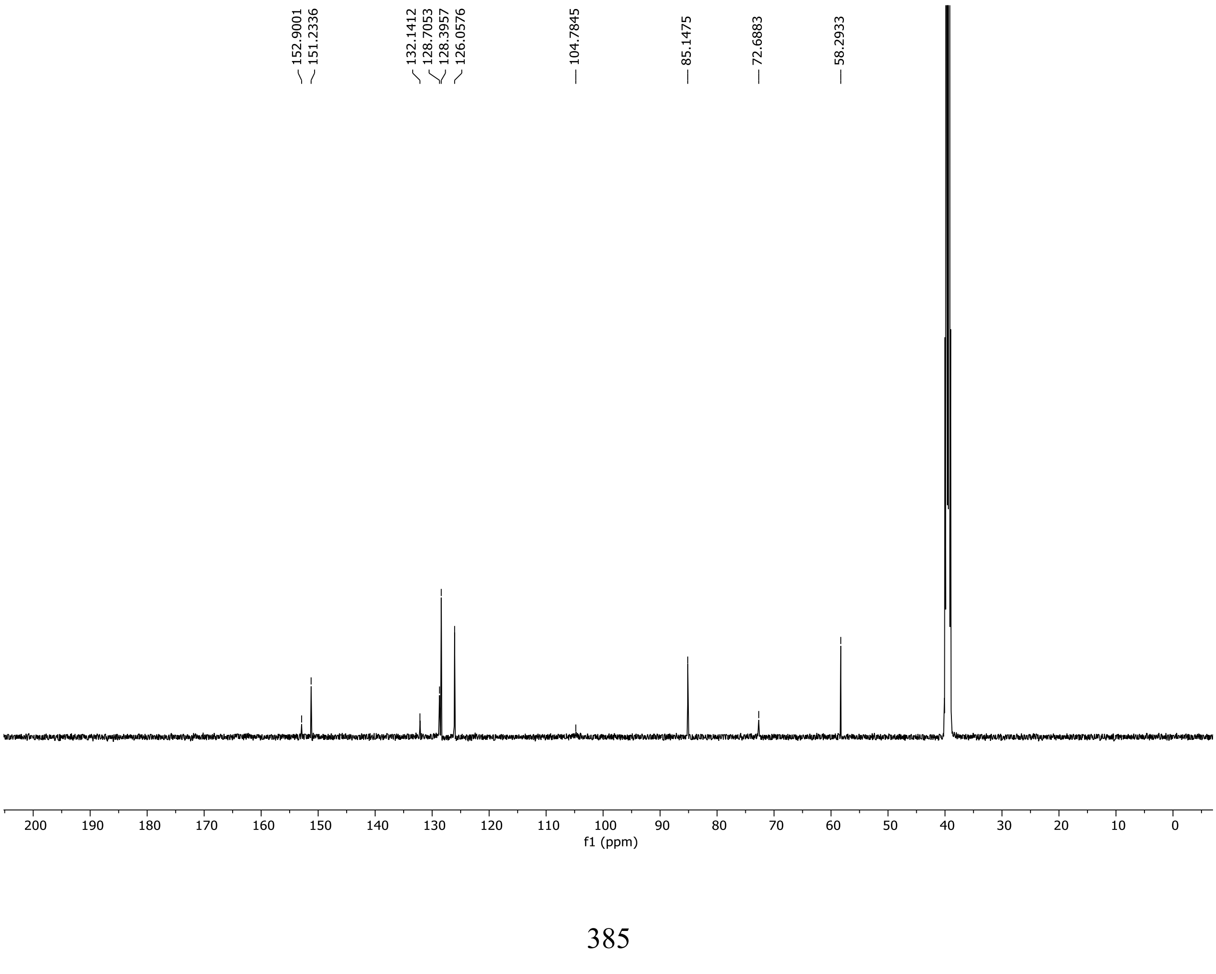

HRMS spectrum of **S147**

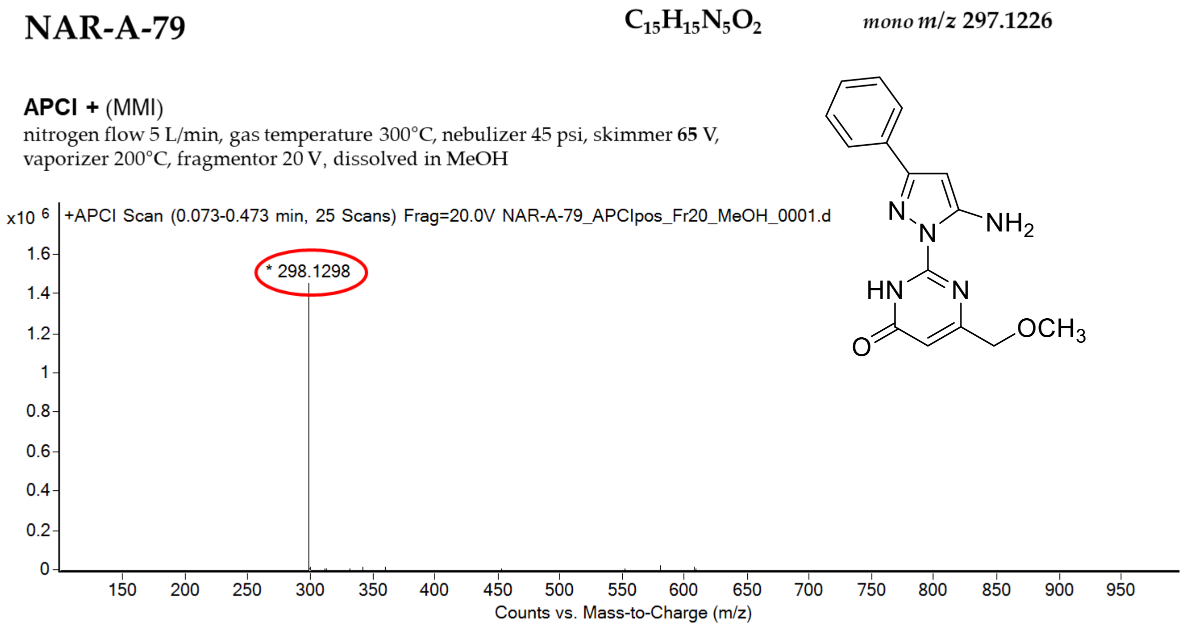

calculated mass: $[M+H]^+$ = 298.1299 observed: $[M+H]^+$ = 298.1298 mass accuracy = -0.3 ppm

FT-IR spectrum (neat) of **S147**

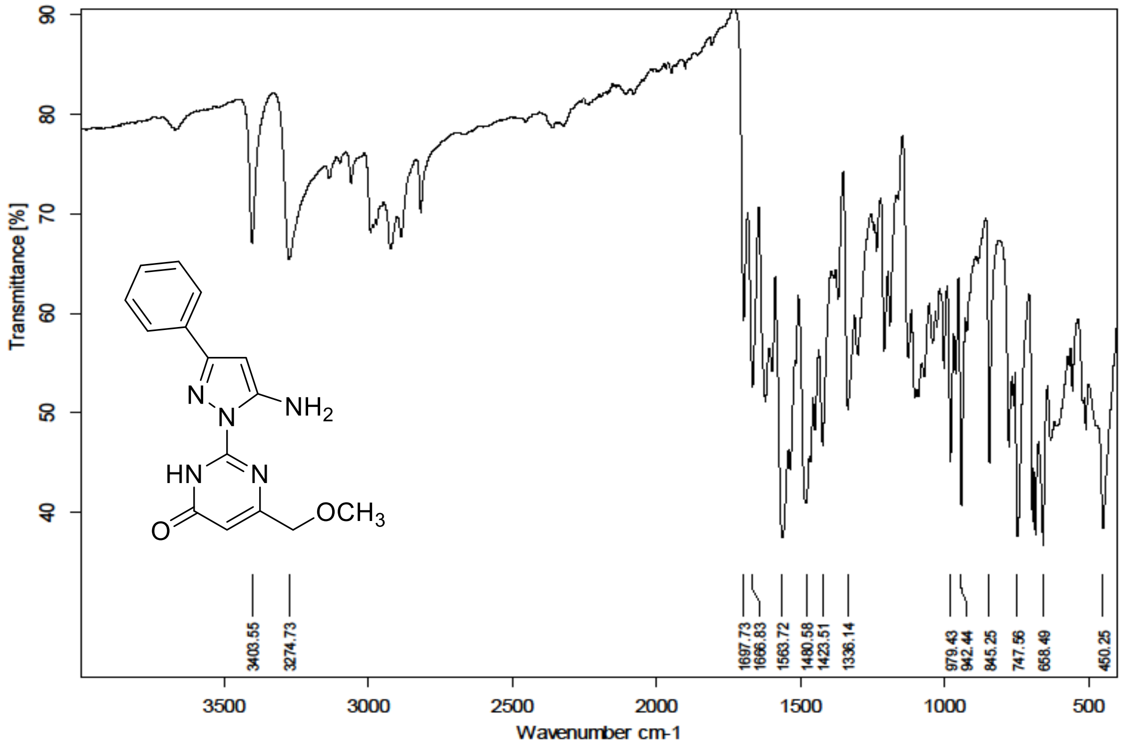

$^{1}H$ (300 MHz) and $^{13}C$ NMR (75 MHz) spectra of **S148** in Chloroform-*d*

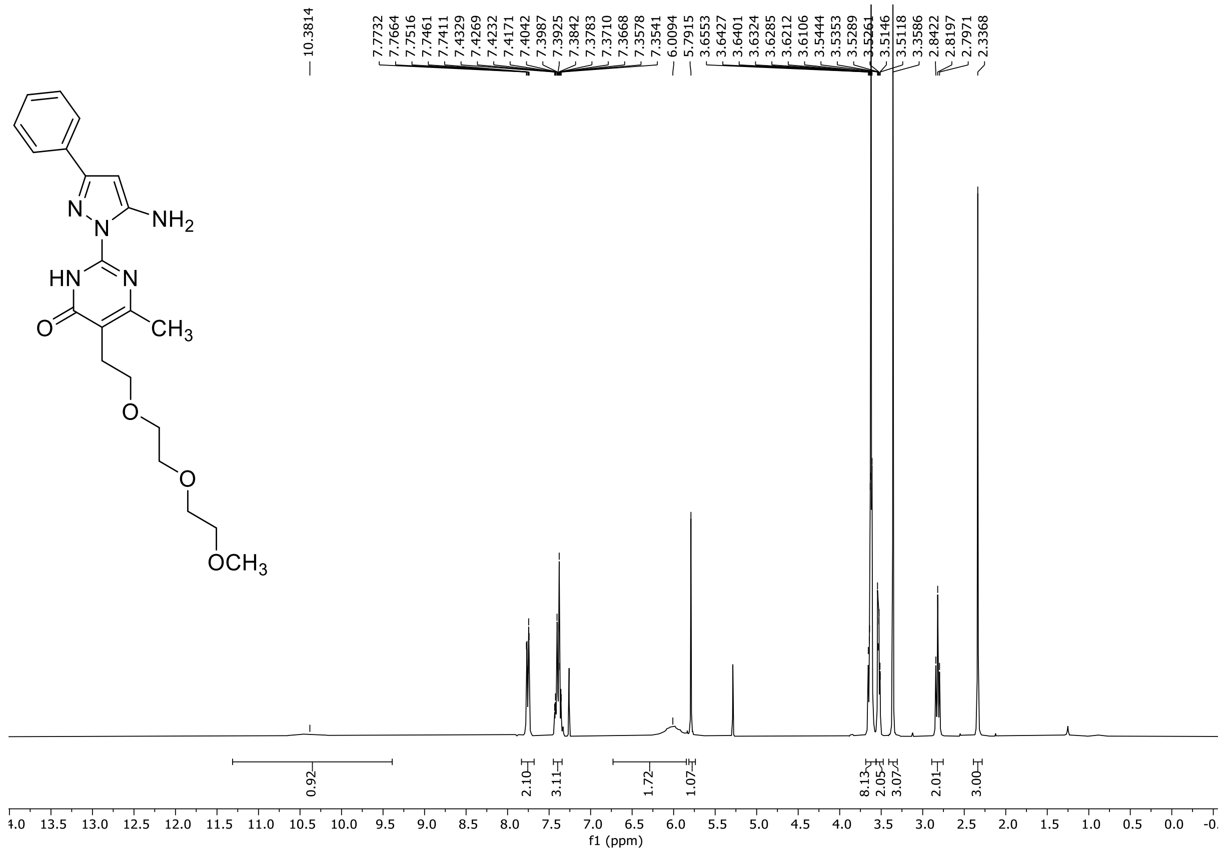

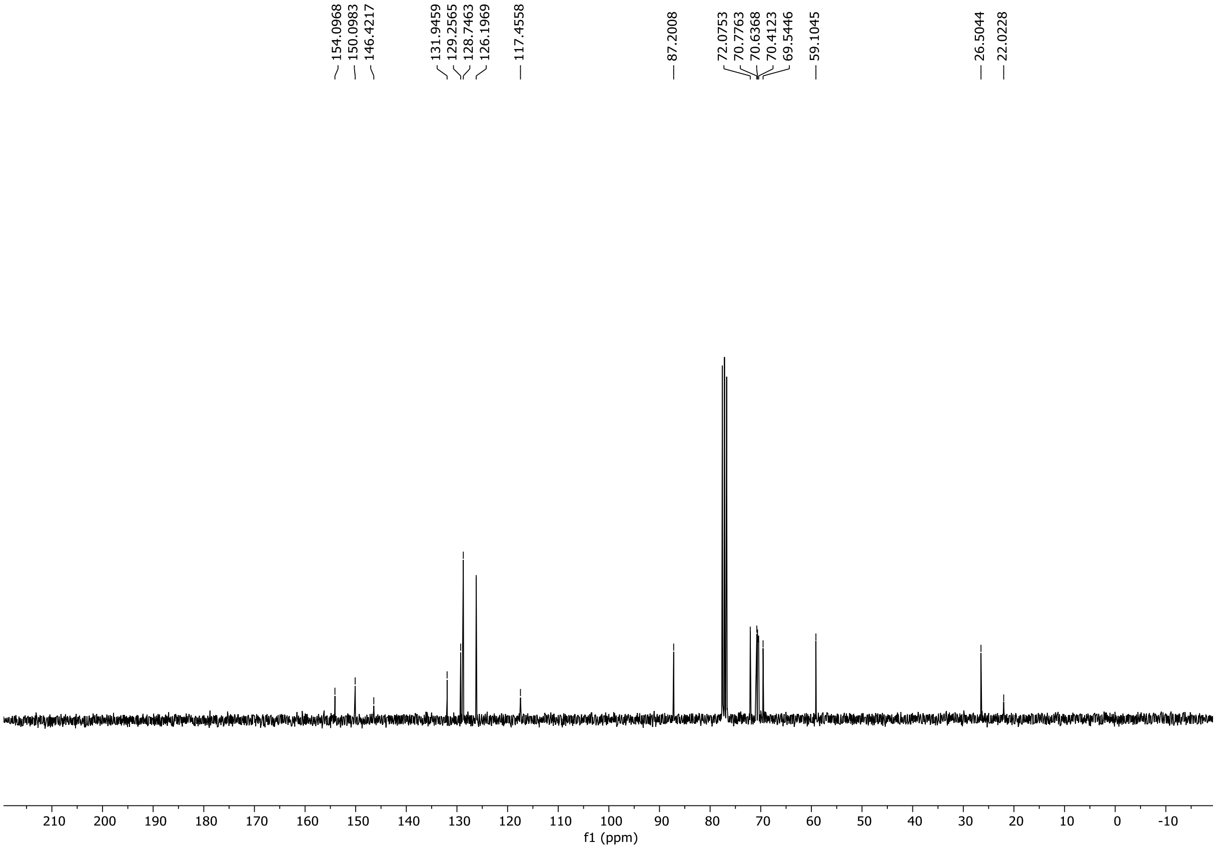

HRMS spectrum of **S148**

**NAR-A-220** $C_{21}H_{27}N_5O_4$ mono *m/z* 413.2063

**APCI +** (MMI)
nitrogen flow 5 L/min, gas temperature 325°C, nebulizer 45 psig, skimmer 65 V, vaporizer 200°C, fragmentor 10 V, dissolved in methanol

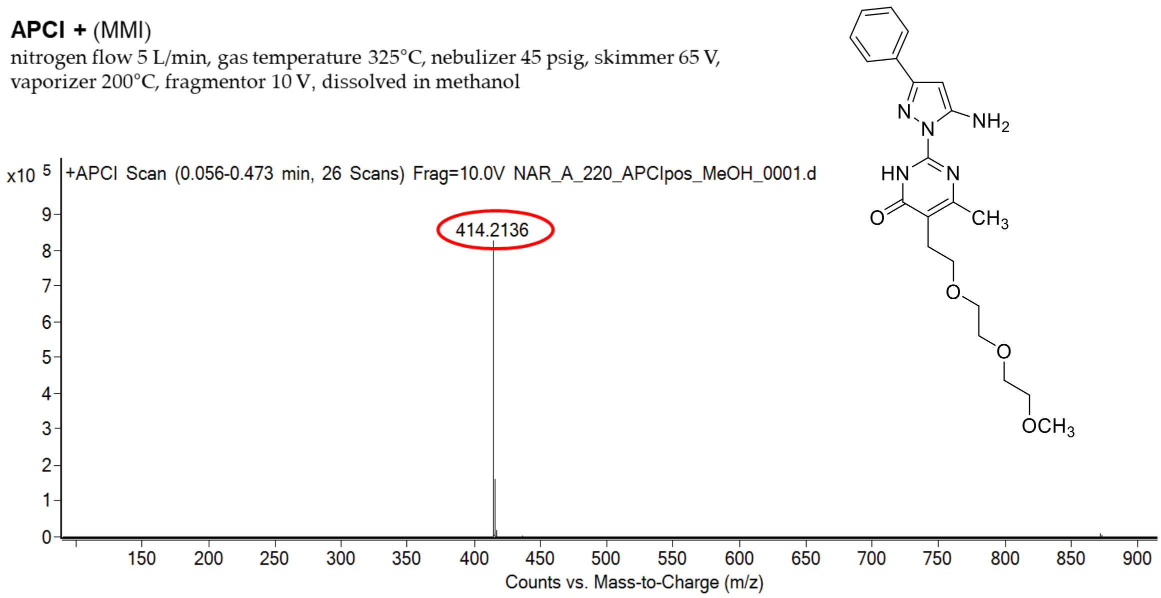

calculated mass: $[M+H]^+ = 414.2136$ observed: $[M+H]^+ = 414.2136$ mass accuracy = < 0.1 ppm

FT-IR spectrum (neat) of **S148**

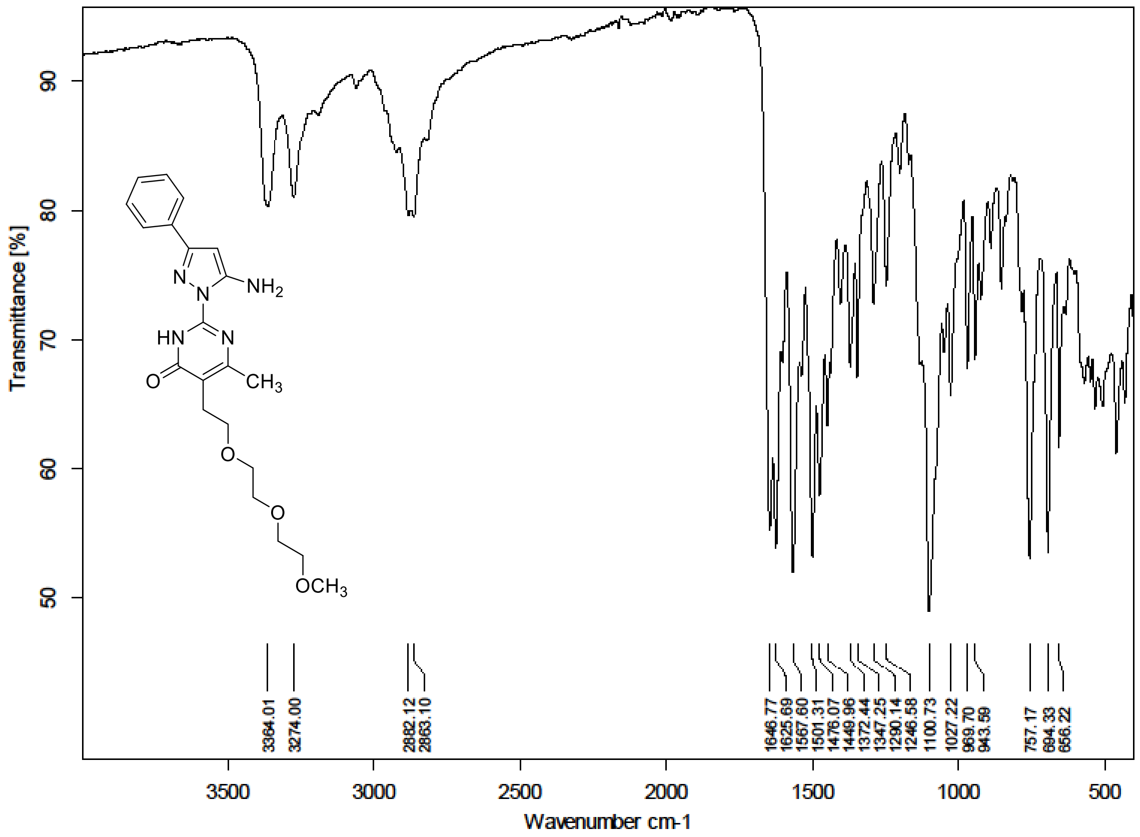

$^{1}H$ (300 MHz) and $^{13}C$ NMR (75 MHz) spectra of **S149** in DMSO-$d_6$

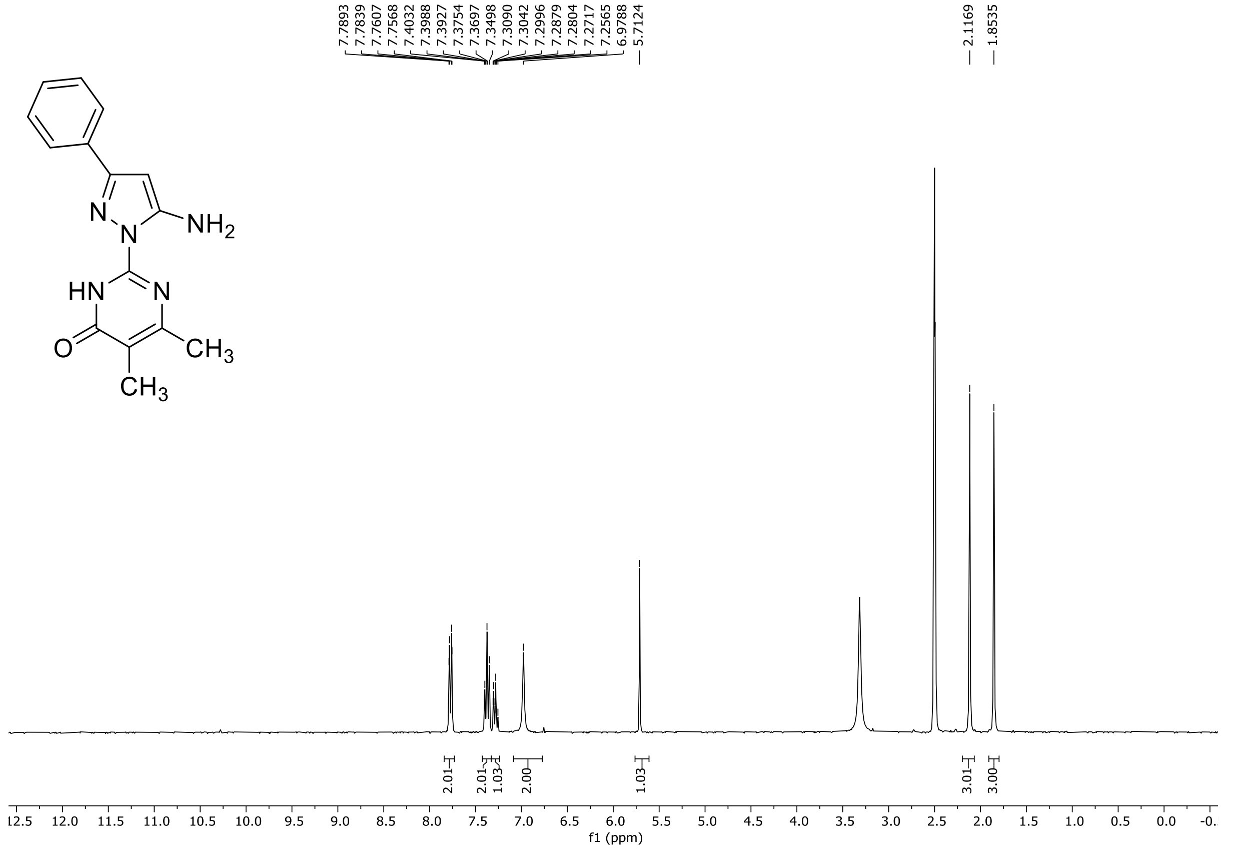

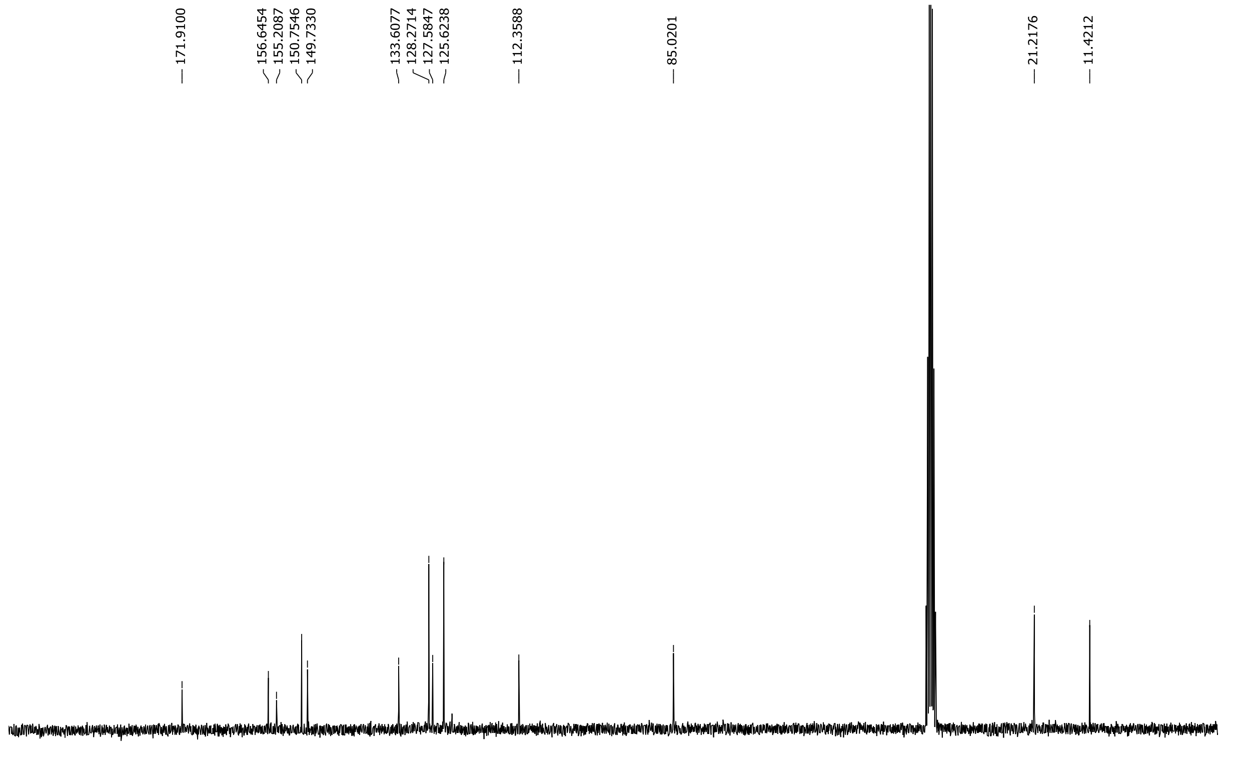

HRMS spectrum of **S149**

**NAR-A-222** $C_{15}H_{15}N_5O$ mono *m/z* 281.1277

**APCI +** (MMI)
nitrogen flow 5 L/min, gas temperature 325°C, nebulizer 45 psig, skimmer 65 V, vaporizer 200°C, fragmentor 24 V, dissolved in methanol

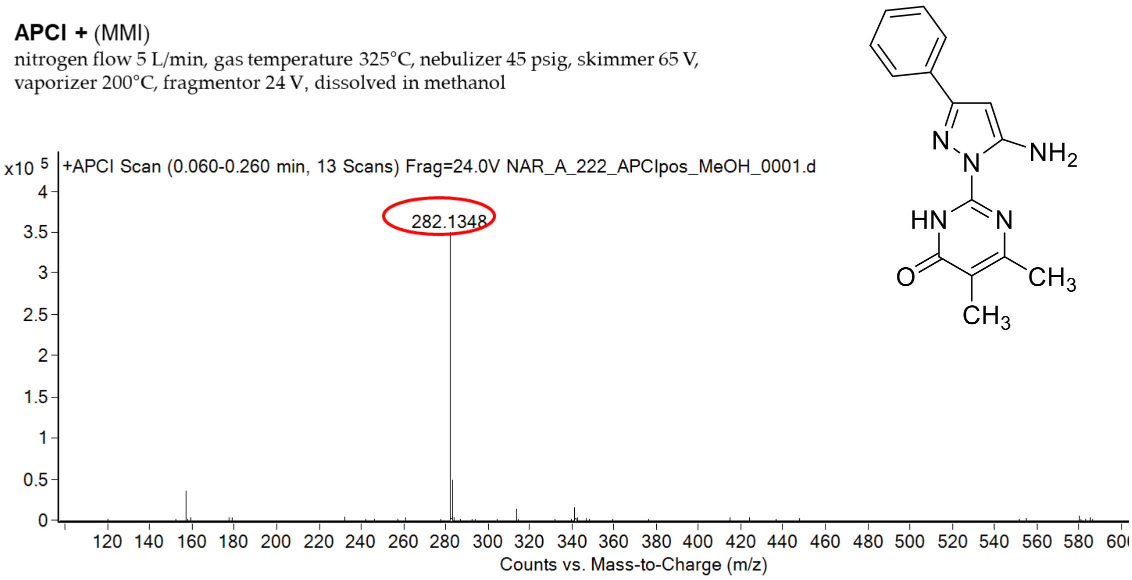

calculated mass: $[M+H]^+$ = 282.1349 observed: $[M+H]^+$ = 282.1348 mass accuracy = - 0.4 ppm

FT-IR spectrum (neat) of **S149**

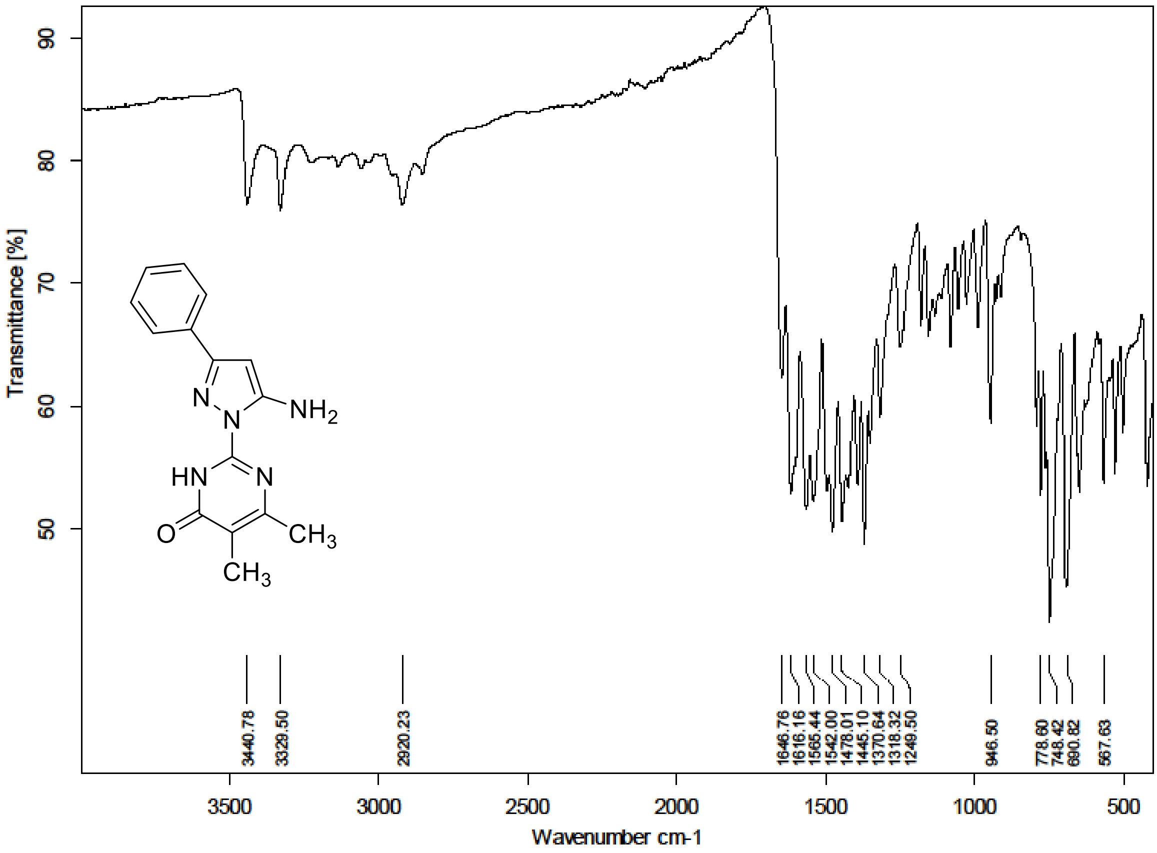

[1]H (300 MHz) and [13]C NMR (75 MHz) spectra of **S150** in Chloroform-*d*

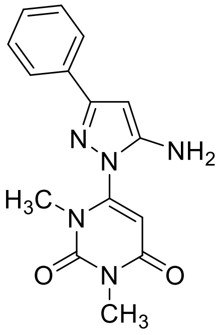

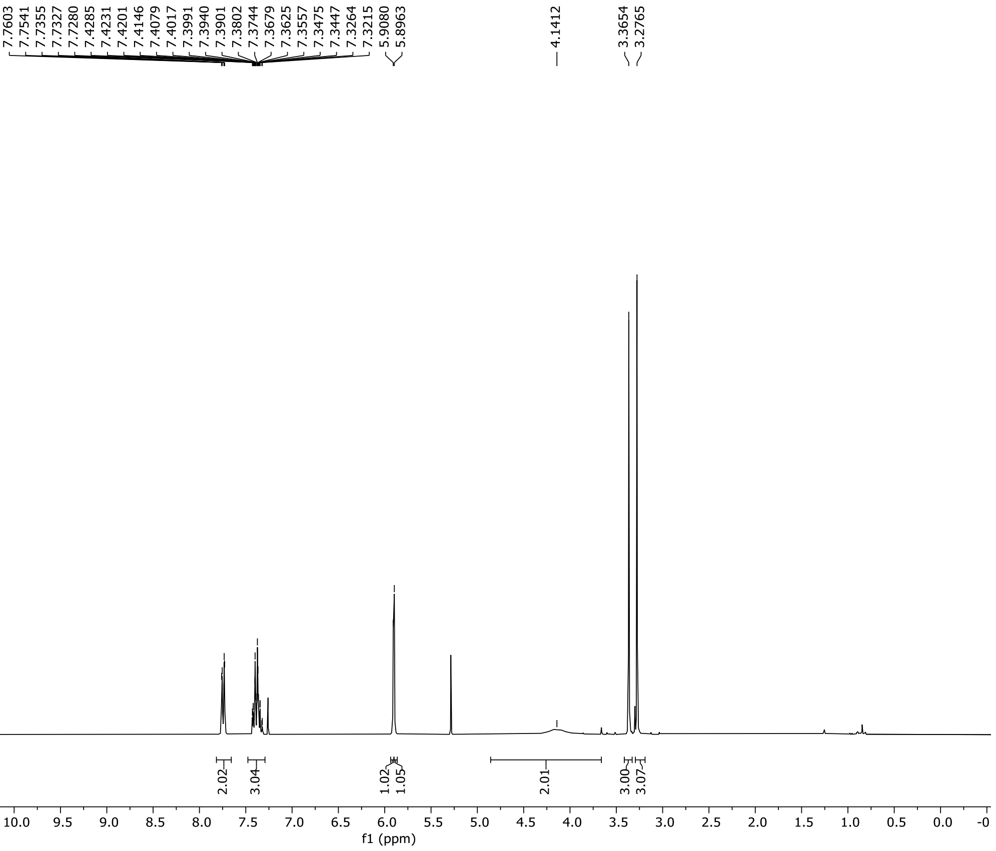

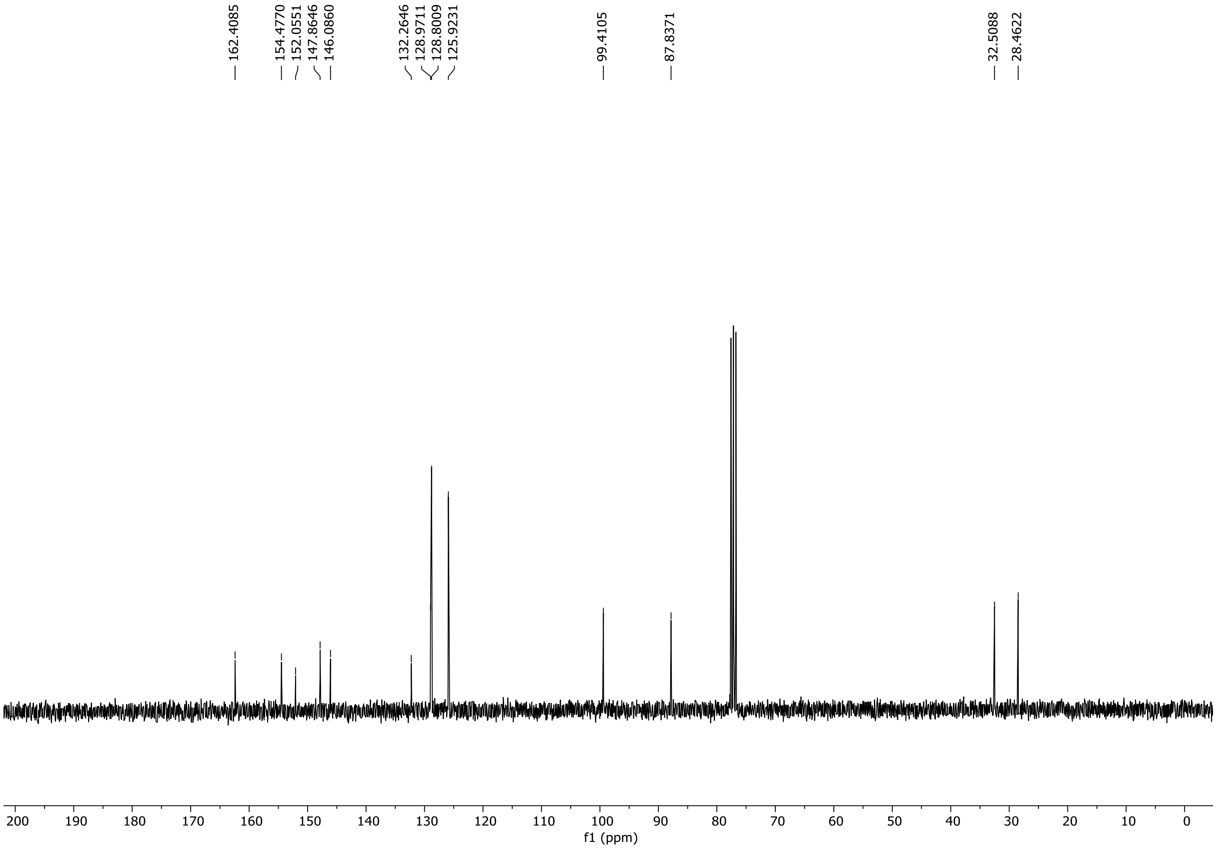

HRMS spectrum of **S150**

**NAR-A-223** $C_{15}H_{15}N_5O_2$ mono *m/z* 297.1226

**APCI +** (MMI)
nitrogen flow 5 L/min, gas temperature 325°C, nebulizer 45 psig, skimmer 65 V, vaporizer 200°C, fragmentor 20 V, dissolved in methanol

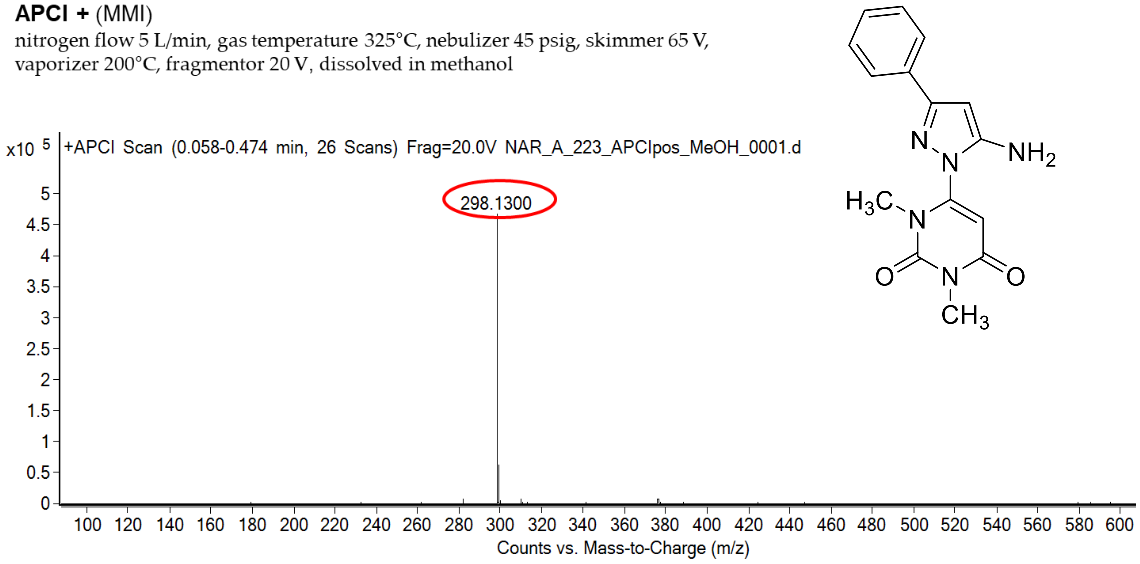

calculated mass: $[M+H]^+$ = 298.1299 observed: $[M+H]^+$ = 298.1300 mass accuracy = 0.3 ppm

FT-IR spectrum (neat) of **S150**

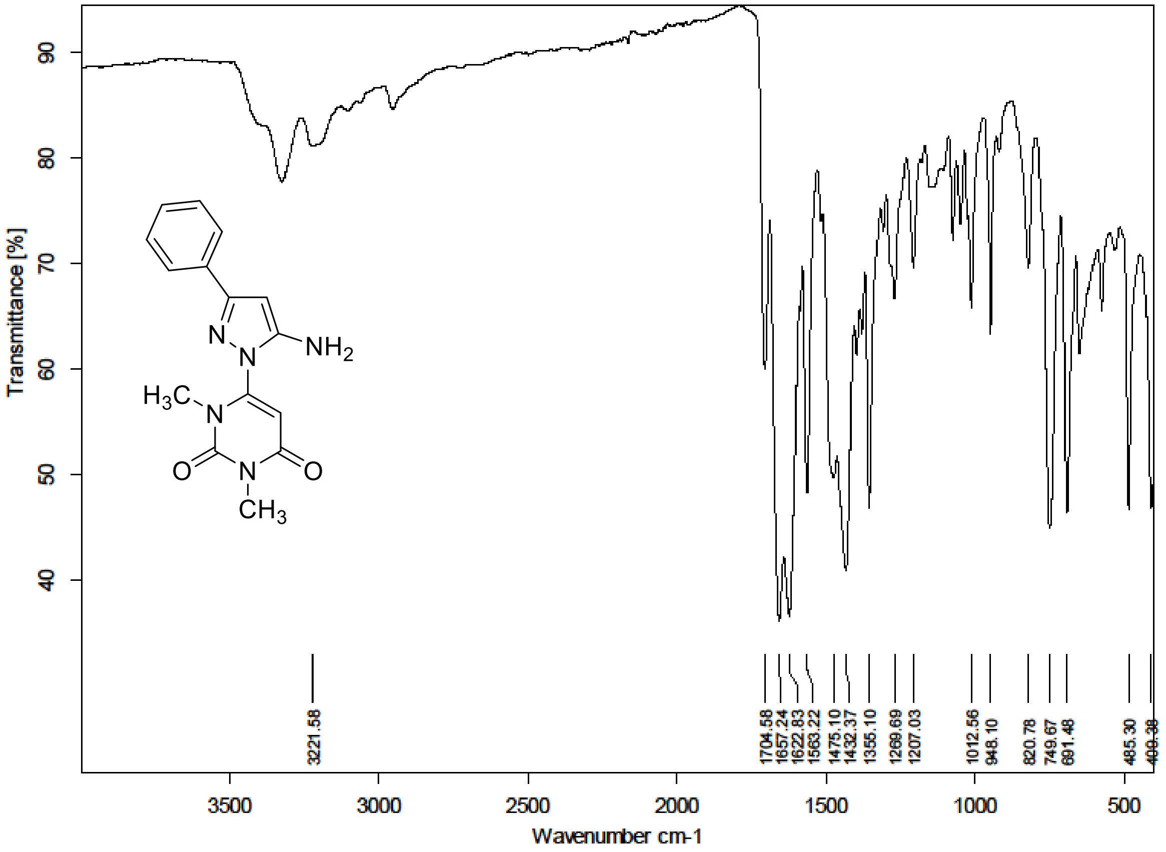

$^{1}H$ (500 MHz) and $^{13}C$ NMR (126 MHz) spectra of **S151** in Chloroform-*d*

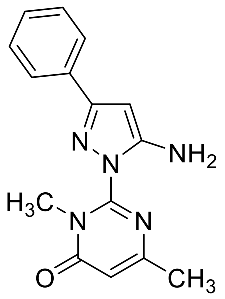

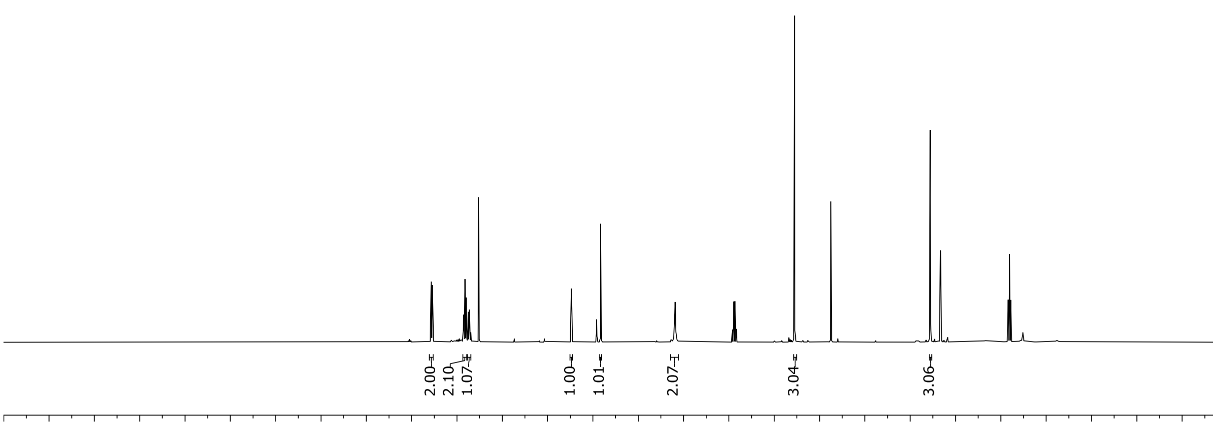

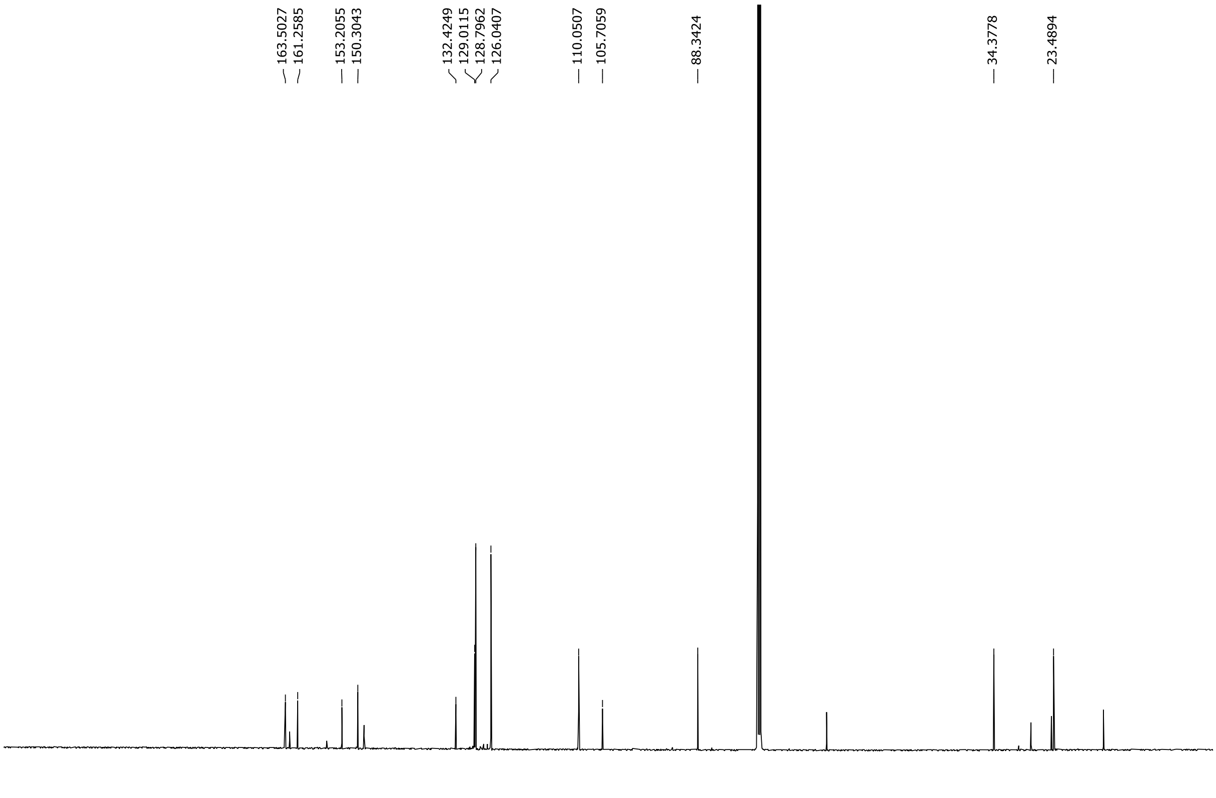

HRMS spectrum of **S151**

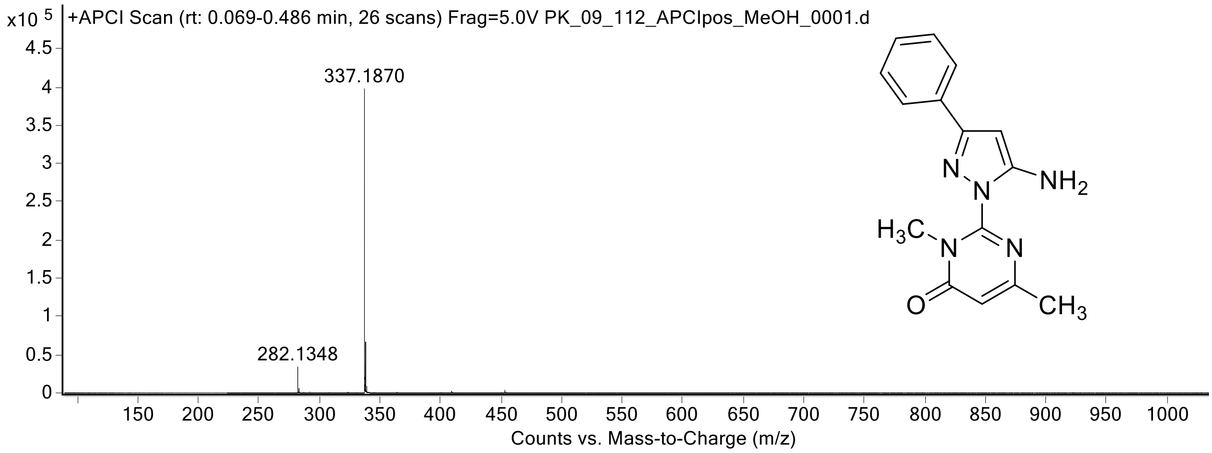

$^1H$ (500 MHz) and $^{13}C$ NMR (126 MHz) spectra of **S152** in Chloroform-*d*

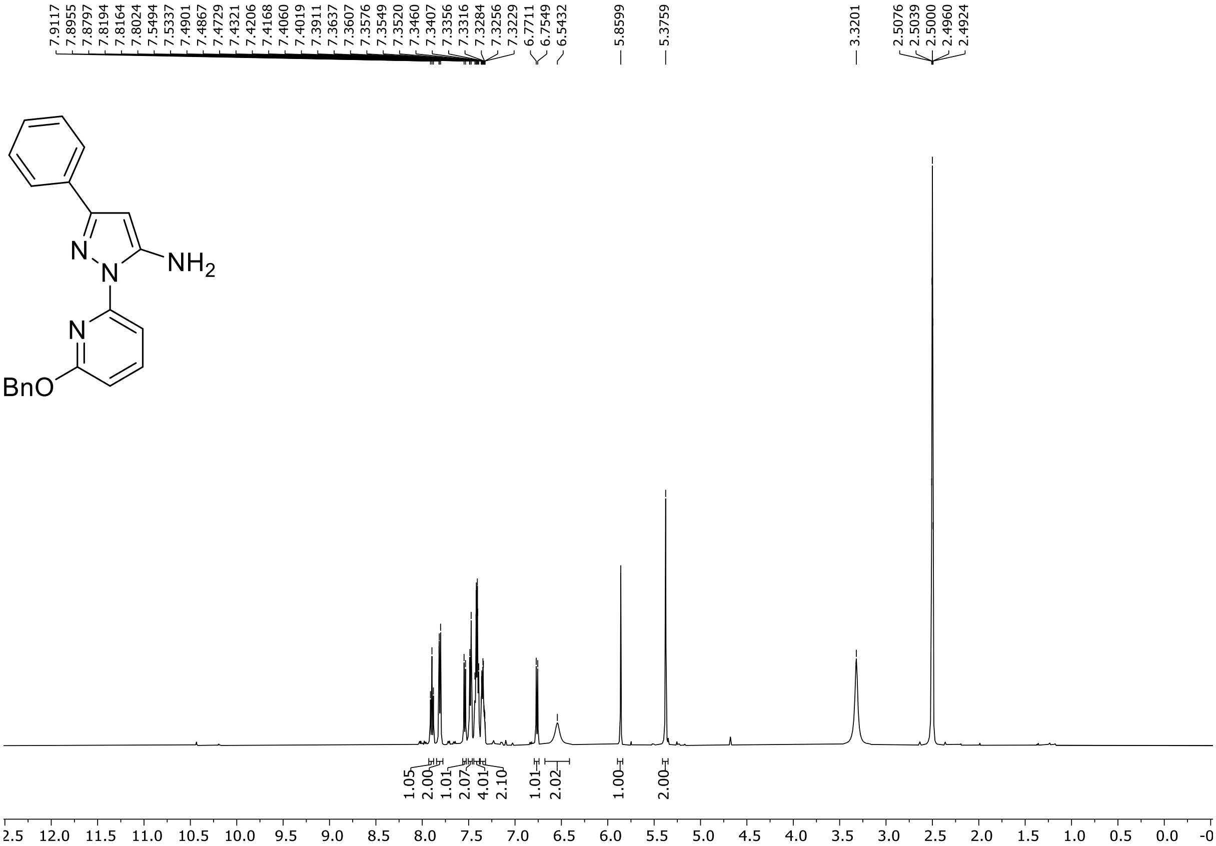

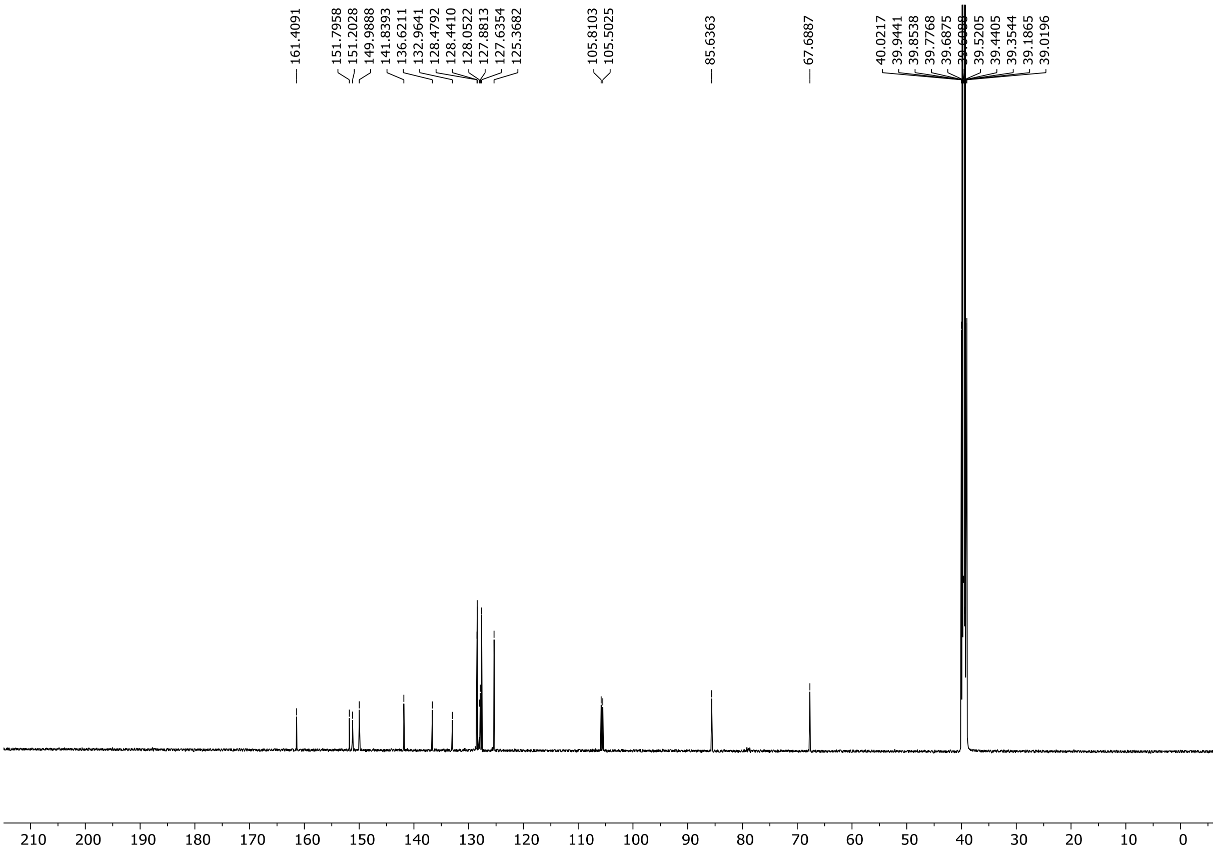

HRMS spectrum of **S152**

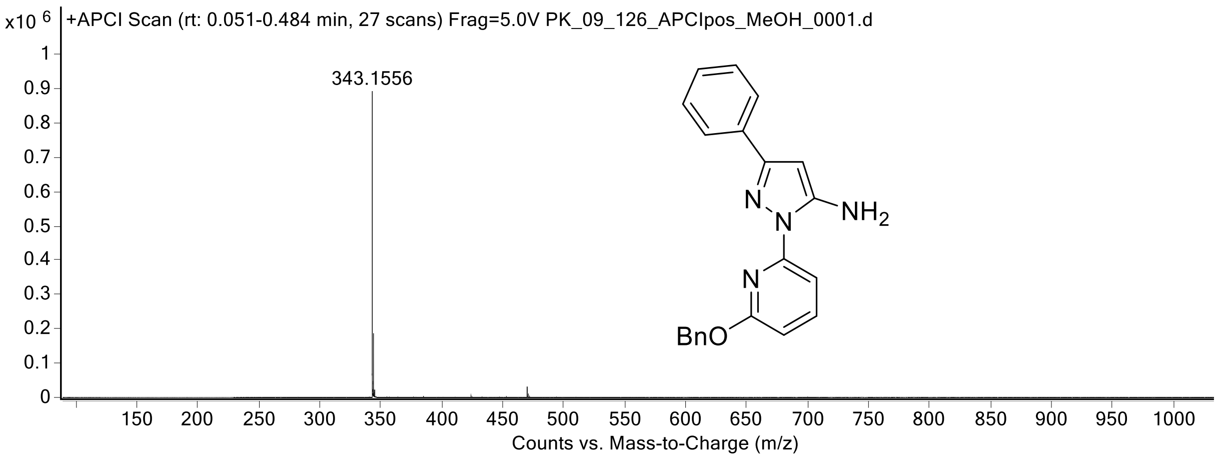

[1]H (300 MHz) and [13]C NMR (126 MHz) spectra of **S153** in DMSO-$d_6$

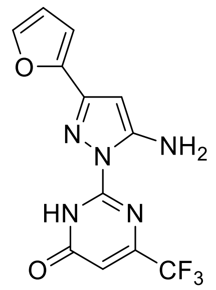

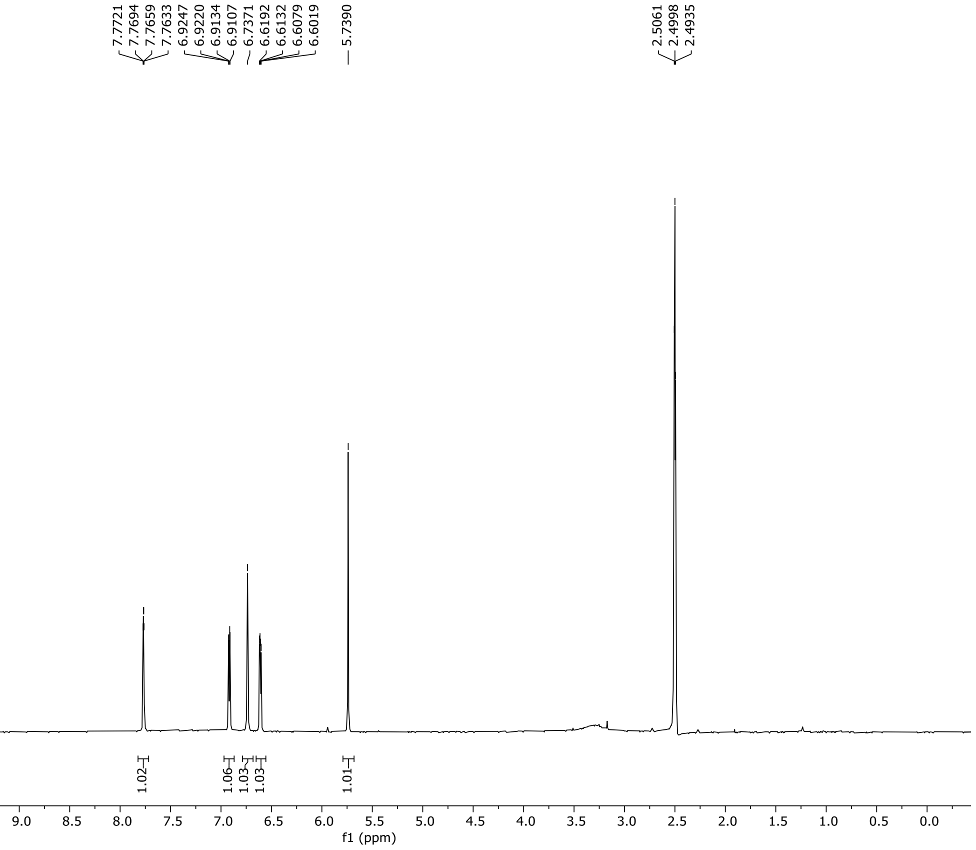

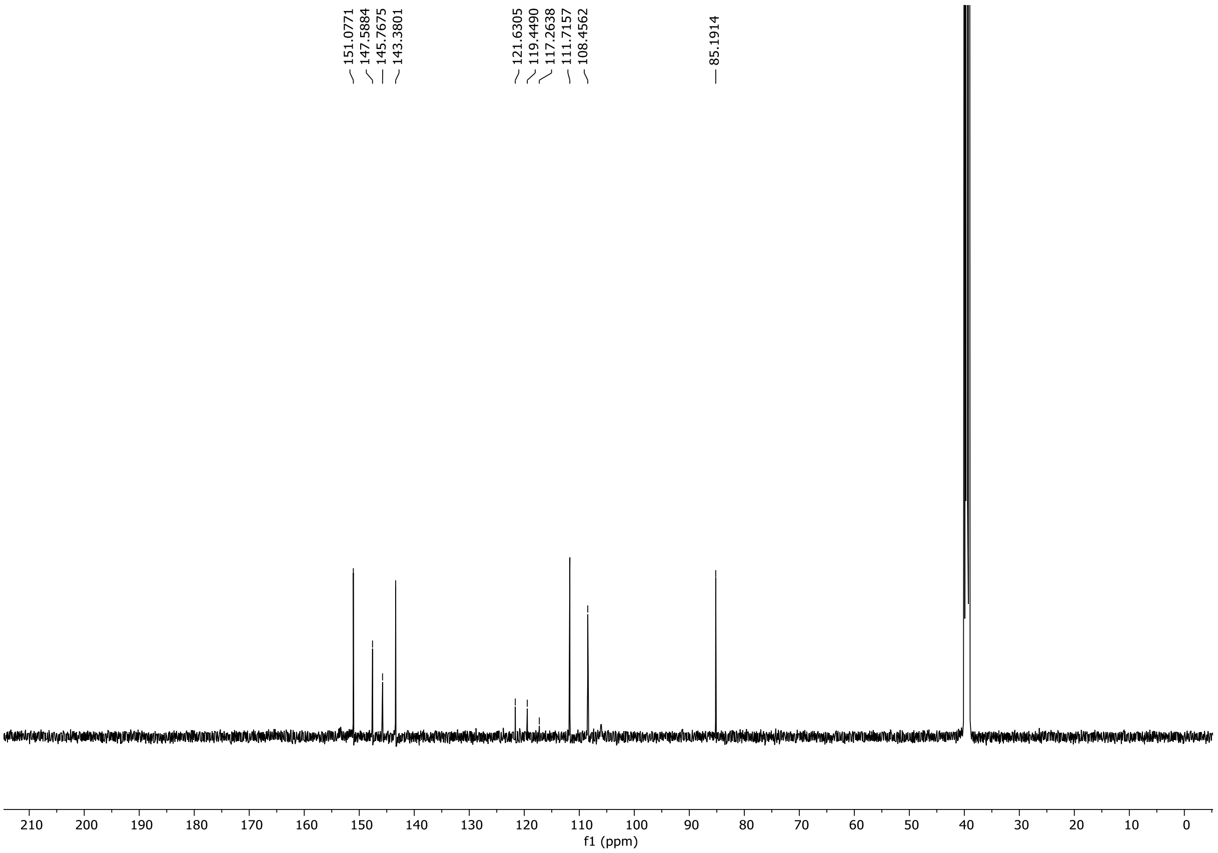

$^{19}F$ NMR (282 MHz) spectrum of **S153** in DMSO-$d_6$

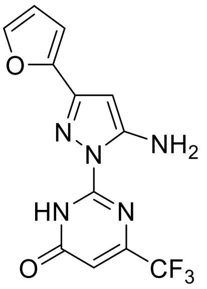

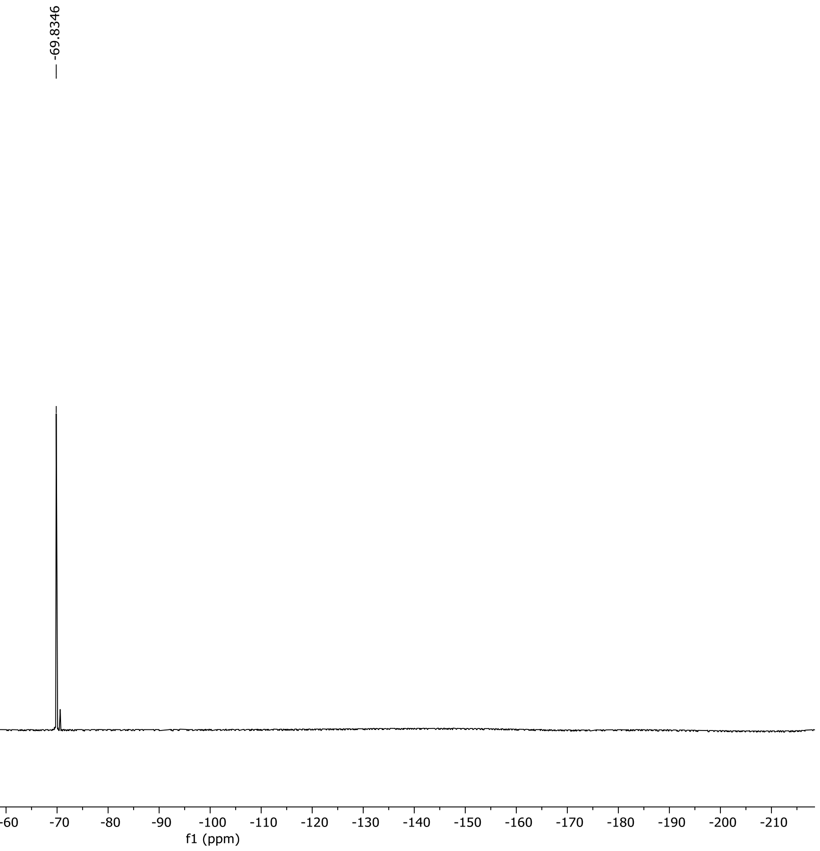

HRMS spectrum of **S153**

**NAR-A-102** **$C_{12}H_8F_3N_5O_2$** ***m/z* 311.0630**

**APCI+** (MMI)
nitrogen flow 5 L/min, gas temperature 325°C, nebulizer 45 psi, skimmer 65 V, vaporizer 200°C, fragmentor 22 V, dissolved in MeOH

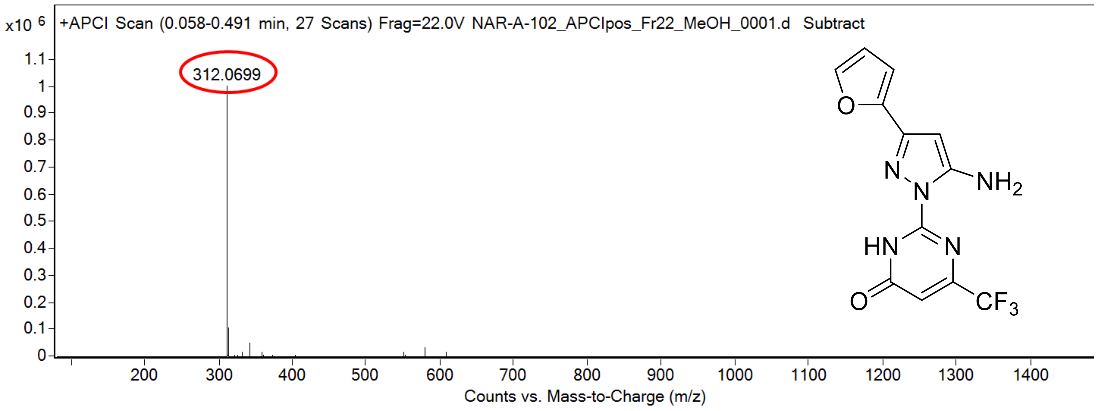

calculated mass: $[M+H]^+ = 312.0703$ observed: $[M+H]^+ = 312.0699$ mass accuracy = -1.3 ppm

FT-IR spectrum (neat) of **S153**

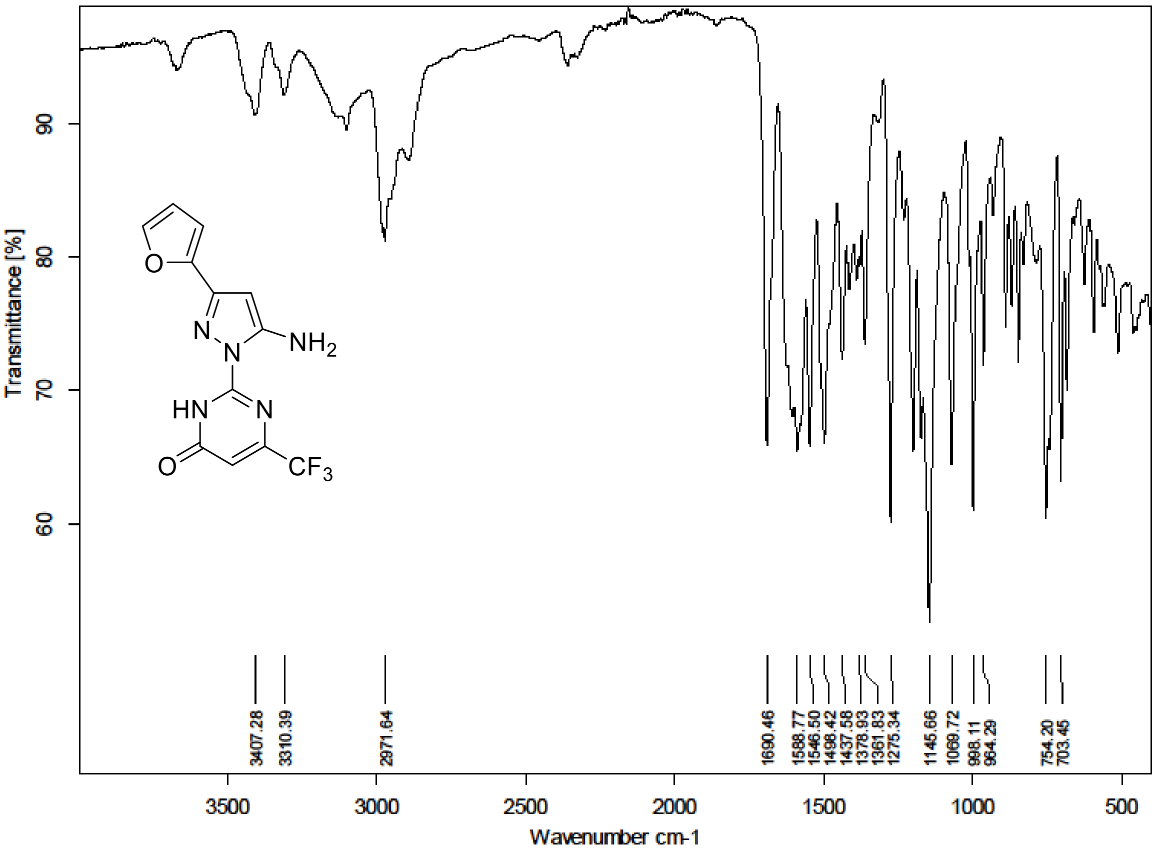

[1]H (300 MHz) and [13]C NMR (126 MHz) spectra of **S154** in DMSO-$d_6$

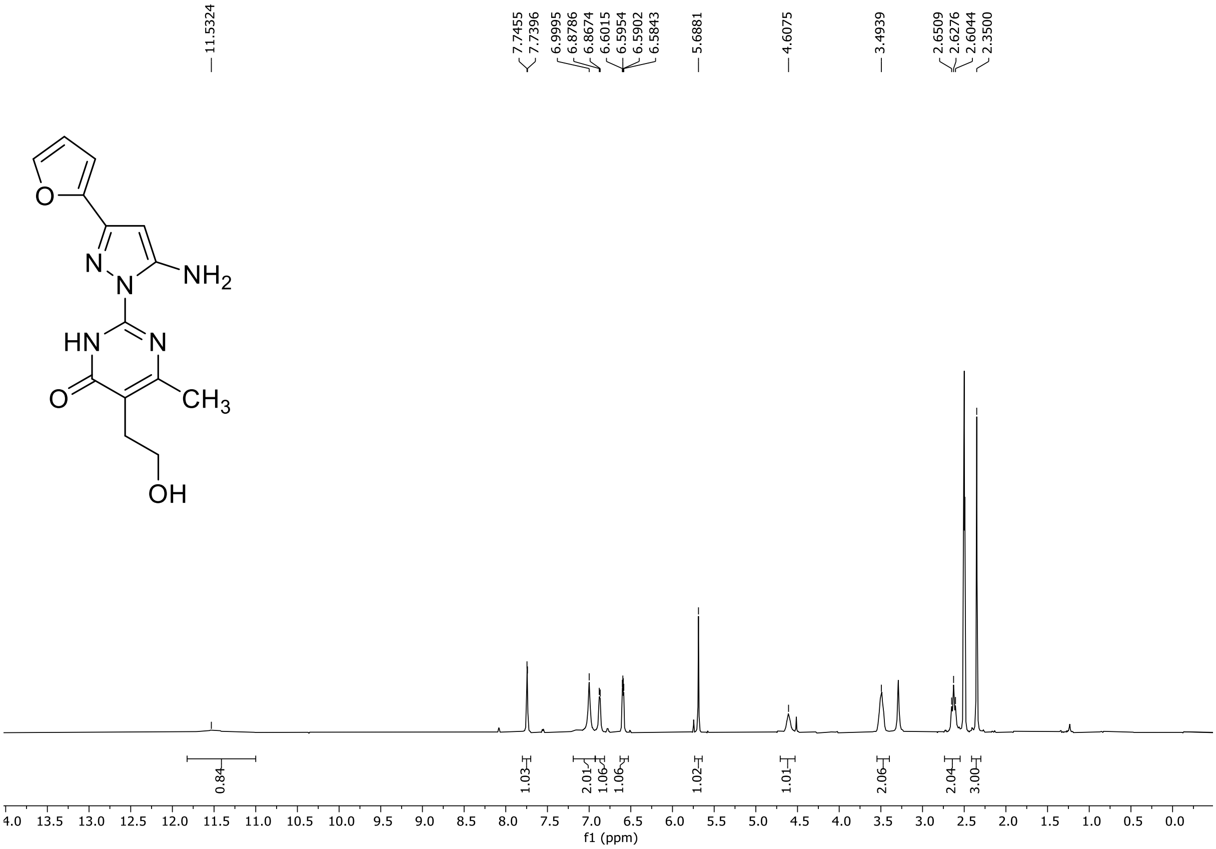

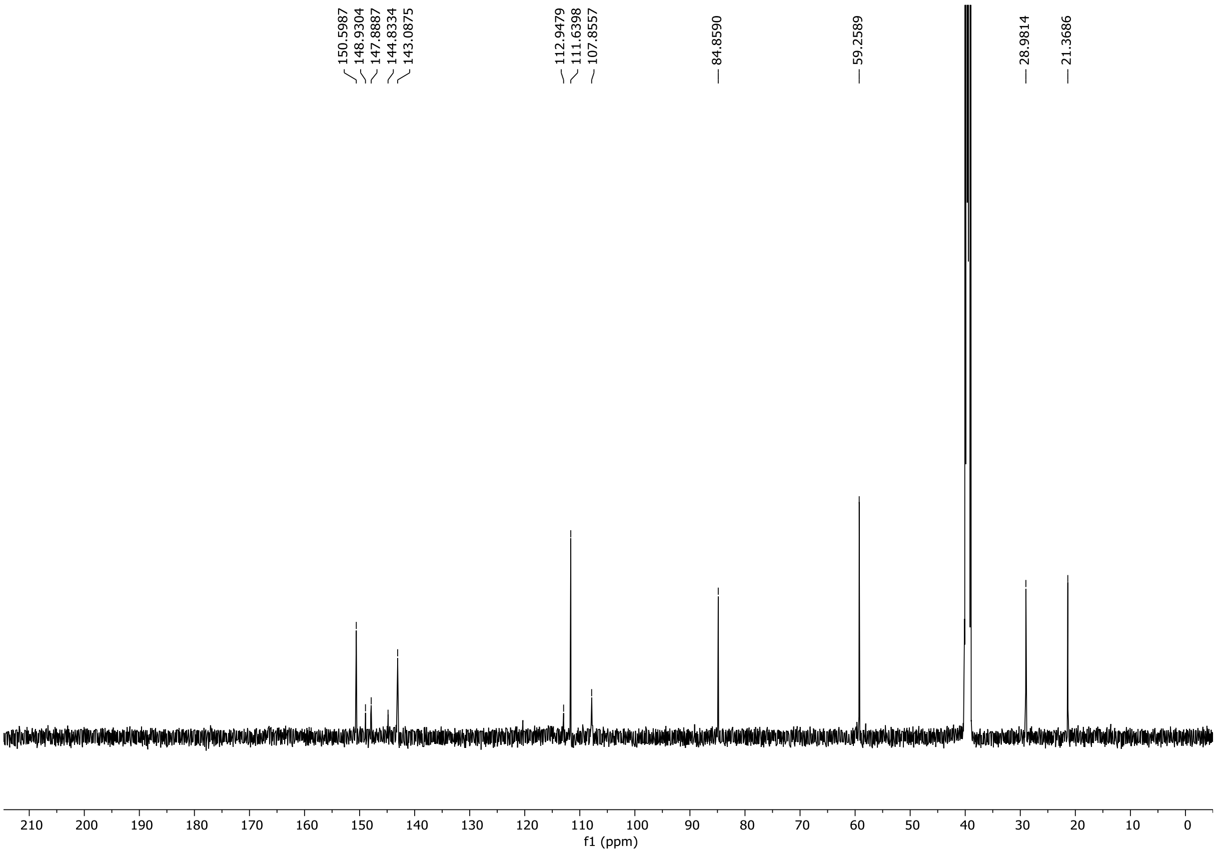

HRMS spectrum of **S154**

**NAR-A-101** $C_{14}H_{15}N_5O_3$ *m*/*z* **301.1175**

**APCI+** (MMI)
nitrogen flow 5 L/min, gas temperature 325°C, nebulizer 45 psi, skimmer 65 V, vaporizer 200°C, fragmentor 20 V, dissolved in MeOH

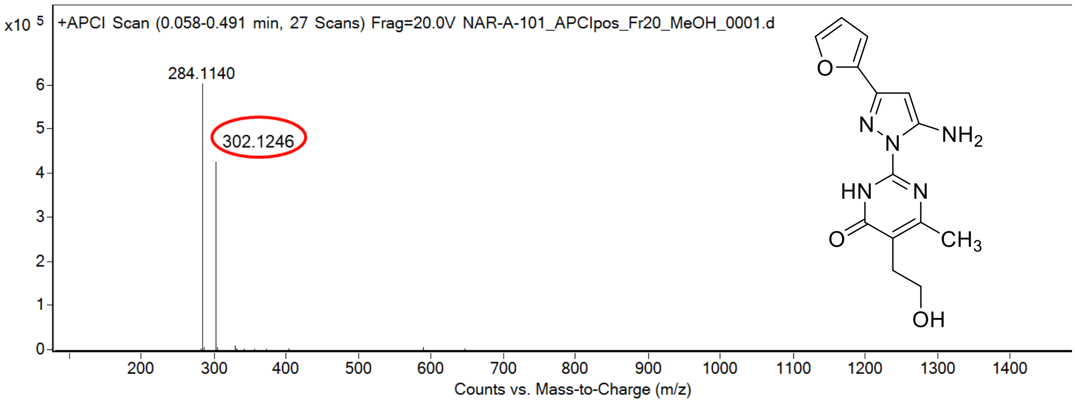

calculated mass: $[M+H]^+ = 302.1248$ observed: $[M+H]^+ = 302.1246$ mass accuracy = -0.7 ppm

FT-IR spectrum (neat) of **S154**

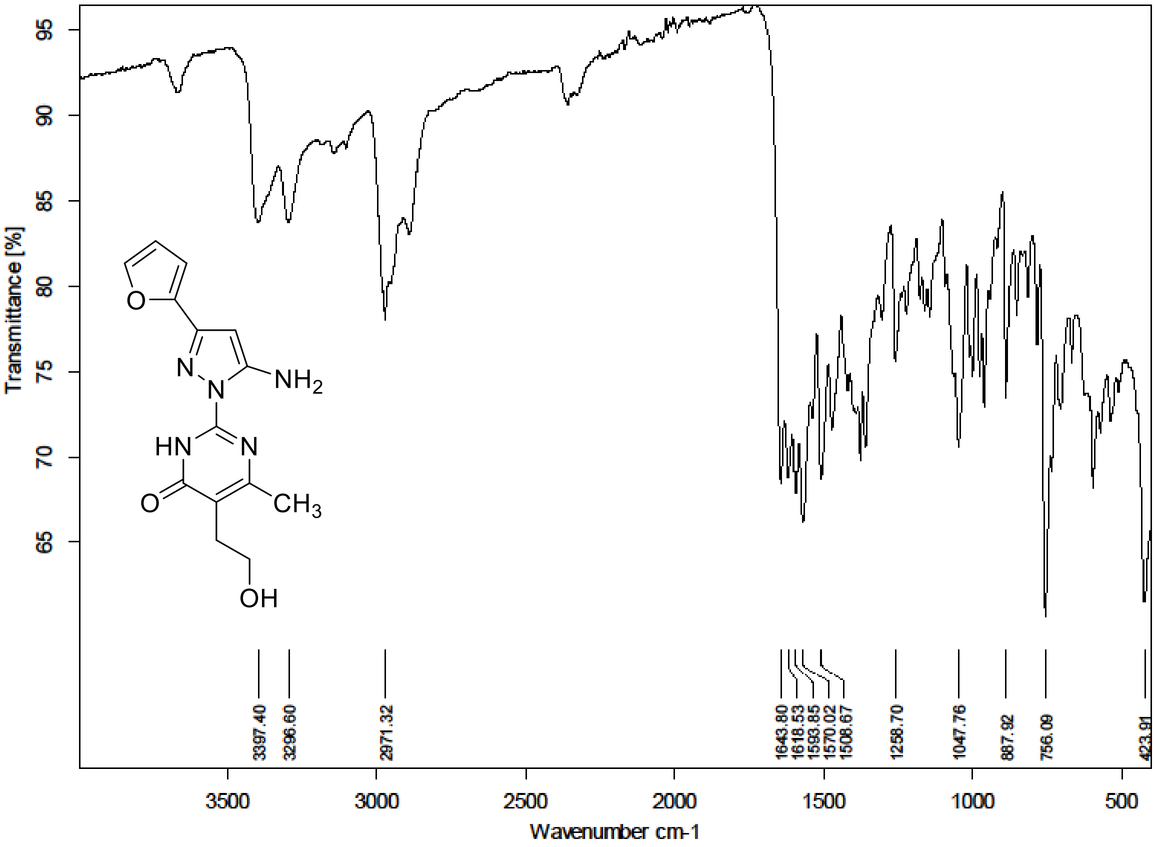

$^{1}H$ (500 MHz) and $^{13}C$ NMR (126 MHz) spectra of **S155** in Chloroform-*d*

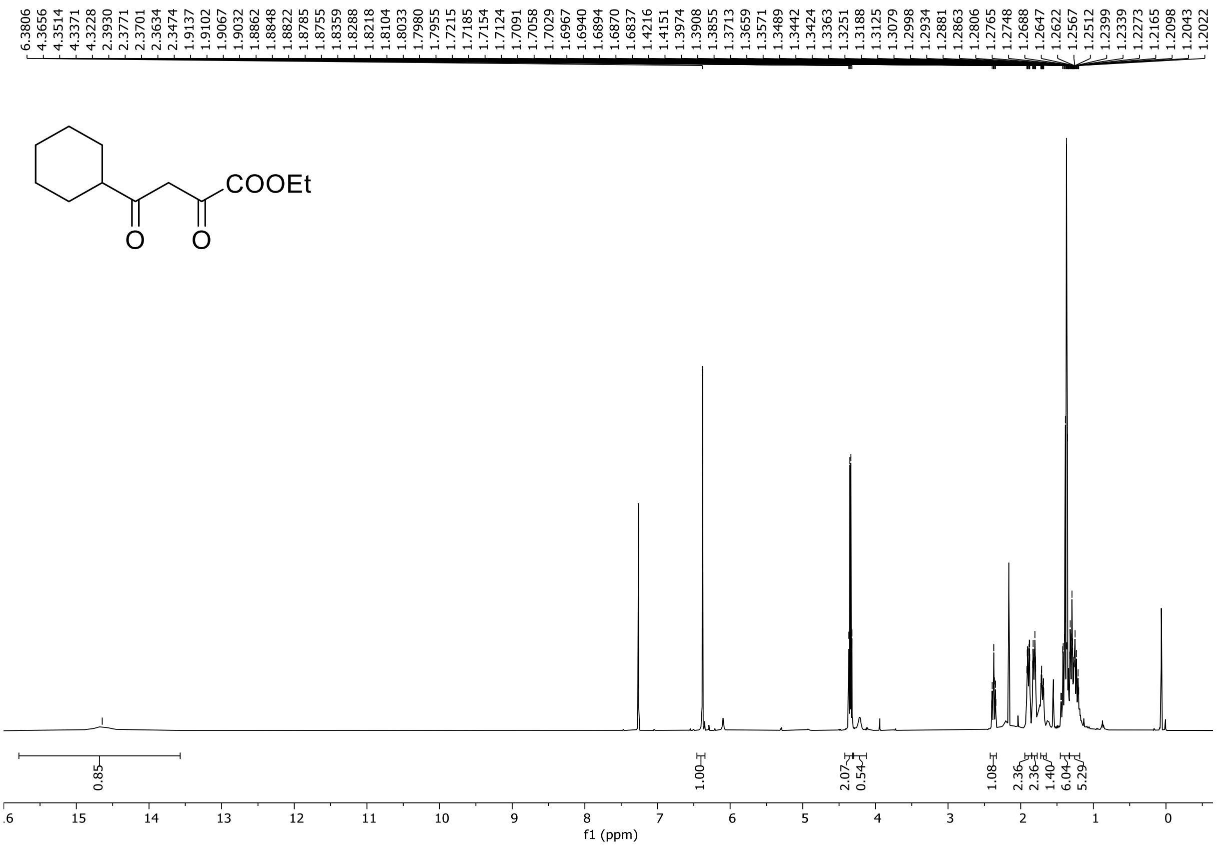

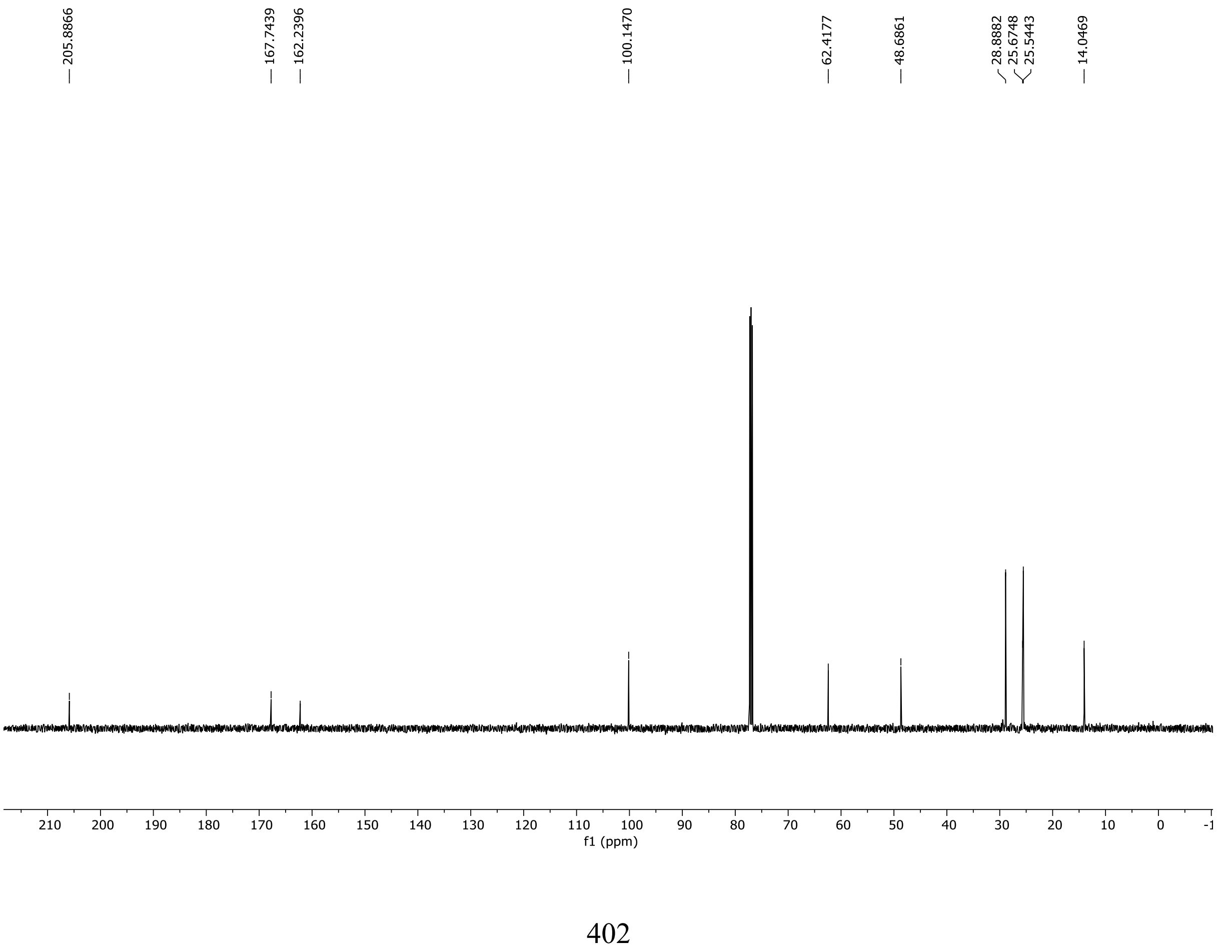

HRMS spectrum of **S155**

**NAR-A-5**

$C_{12}H_{18}O_4$ *mono m/z* **226.1205**

**APCI** + (MMI)
nitrogen flow 5 L/min, gas temperature 325°C, nebulizer 45 psi, skimmer **65** V, vaporizer 200°C, fragmentor 30 V, dissolved in methanol

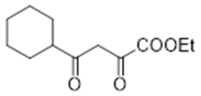

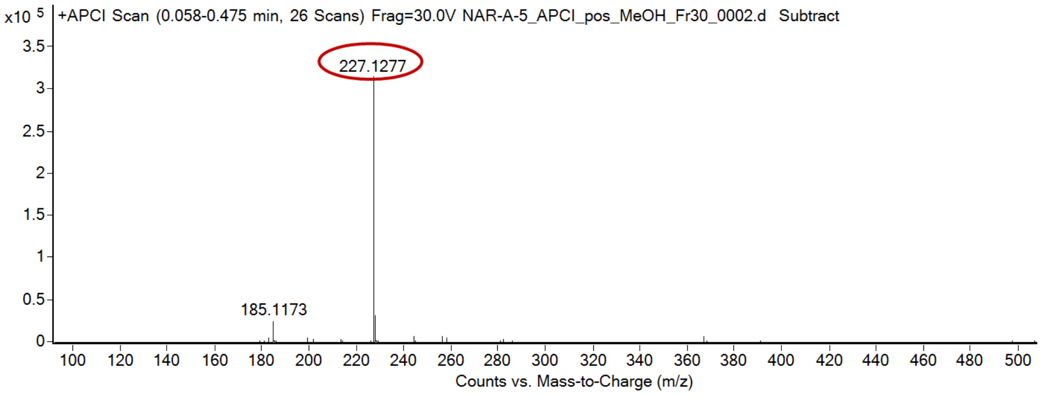

calculated mass: $[M+H]^+$ = 227.1278    observed: $[M+H]^+$ = 227.1277    mass accuracy = - 0.4 ppm

FT-IR spectrum (neat) of **S155**

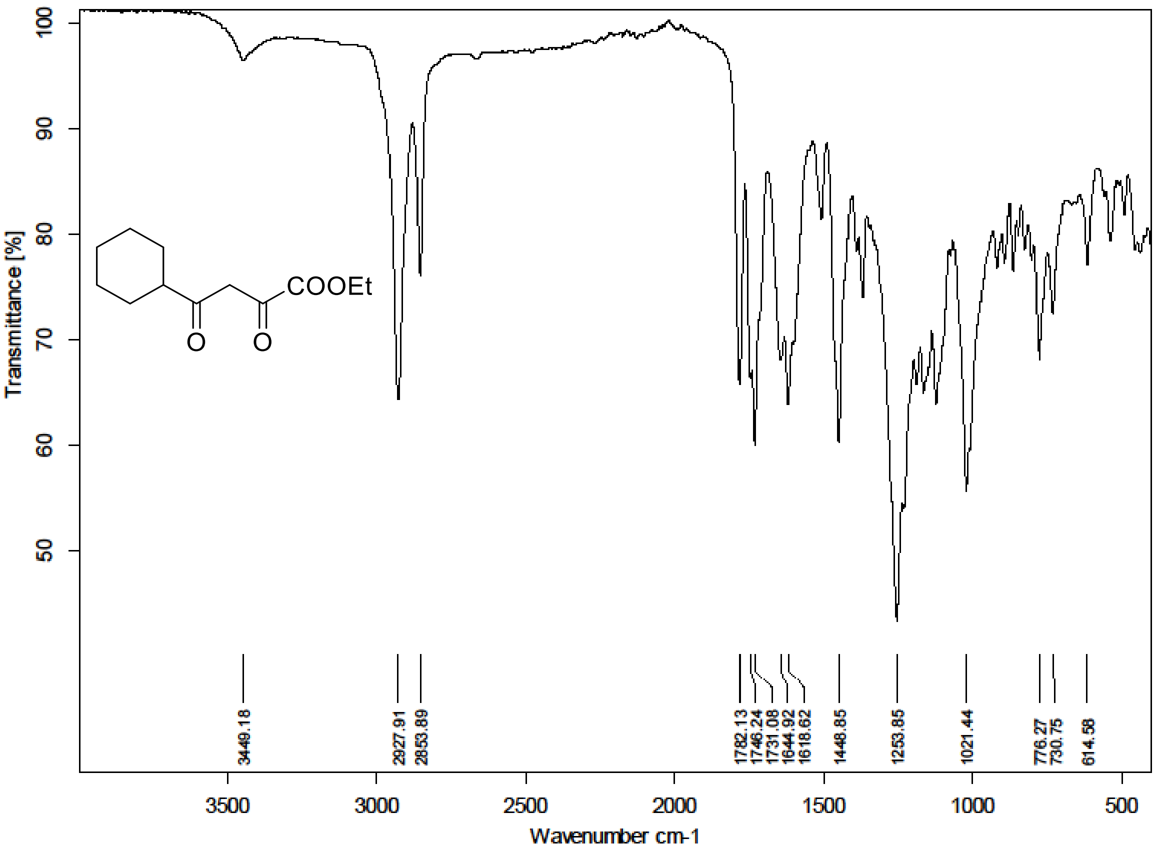

$^{1}$H (500 MHz) and $^{13}$C NMR (126 MHz) spectra of **S156** in Chloroform-*d*

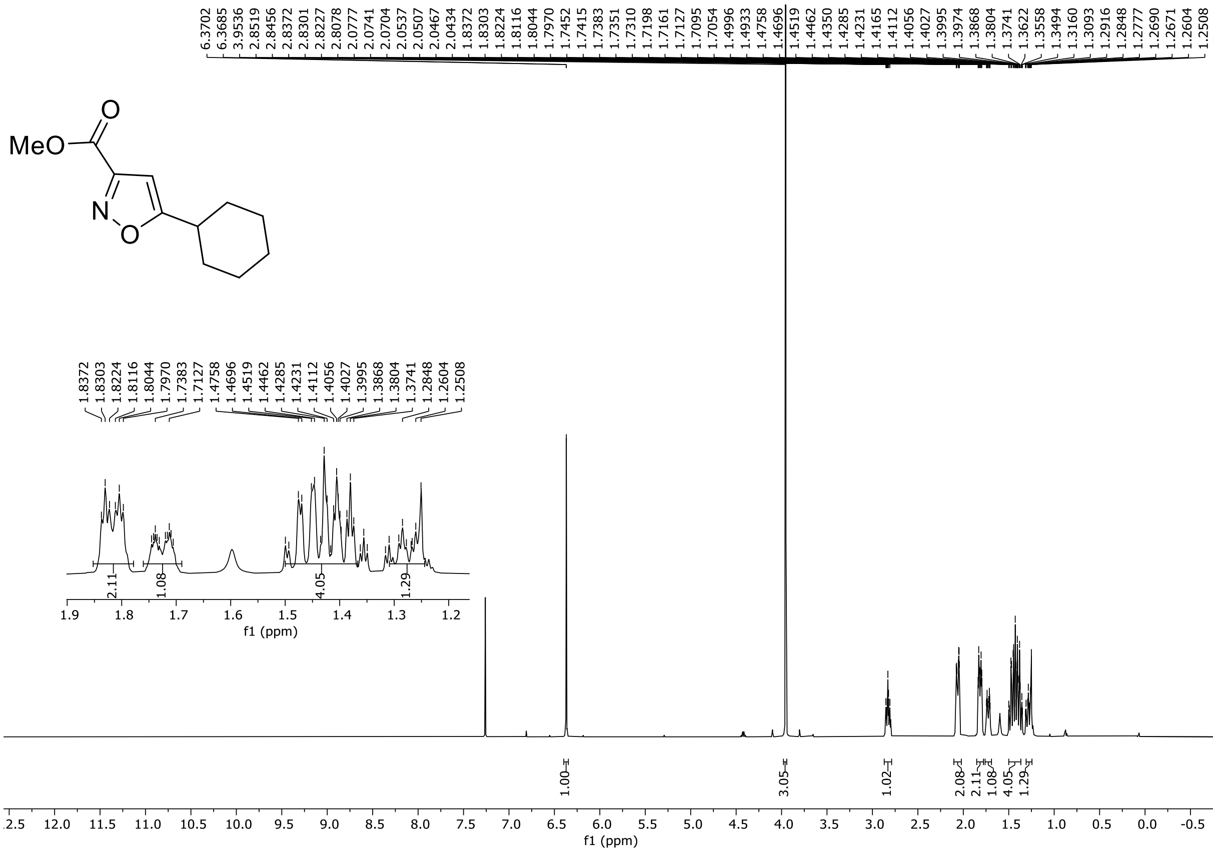

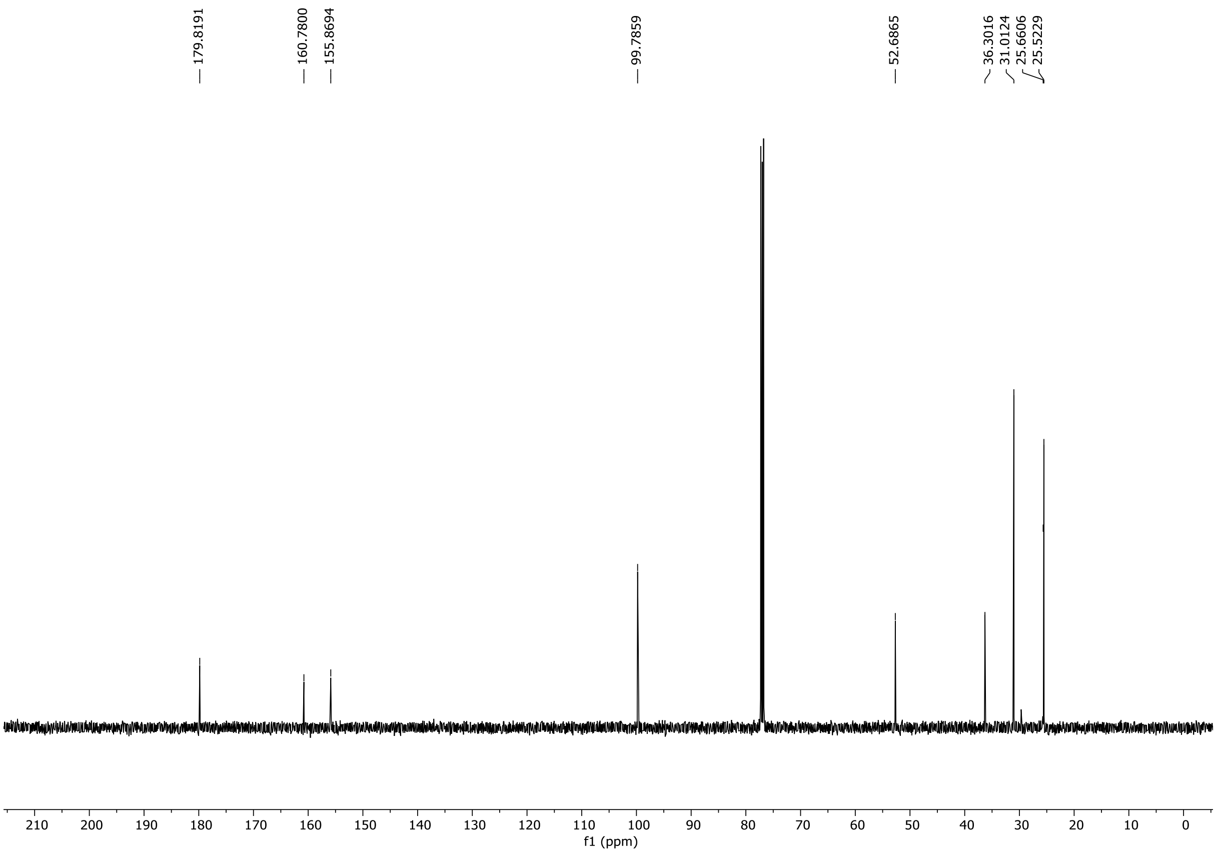

HRMS spectrum of **S156**

**NAR-A-6** $C_{11}H_{15}NO_3$ *mono m/z* **209.1052**

**APCI +** (MMI)
nitrogen flow 5 L/min, gas temperature 325°C, nebulizer 45 psi, skimmer **65** V, vaporizer 200°C, fragmentor 22 V, dissolved in methanol

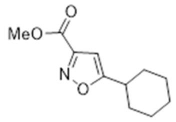

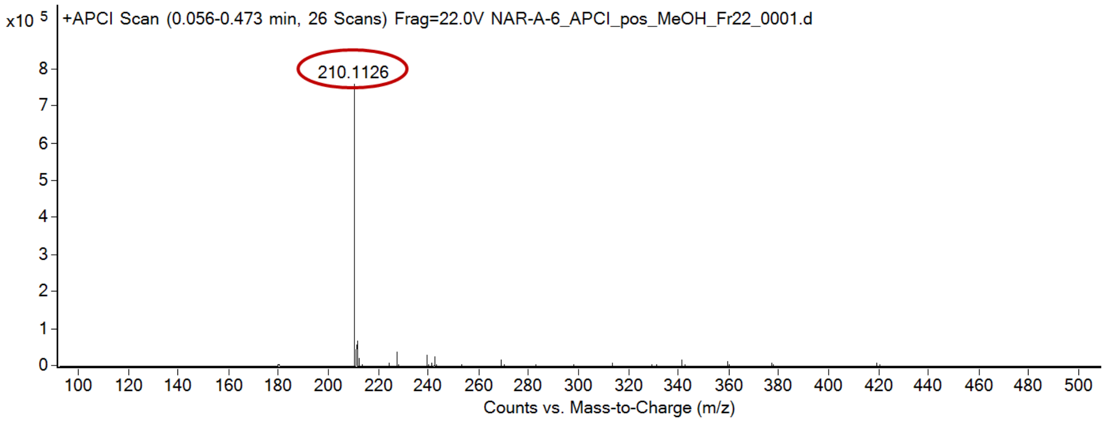

calculated mass: $[M+H]^+ = 210.1125$ observed: $[M+H]^+ = 210.1126$ mass accuracy = 0.5 ppm

FT-IR spectrum (neat) of **S156**

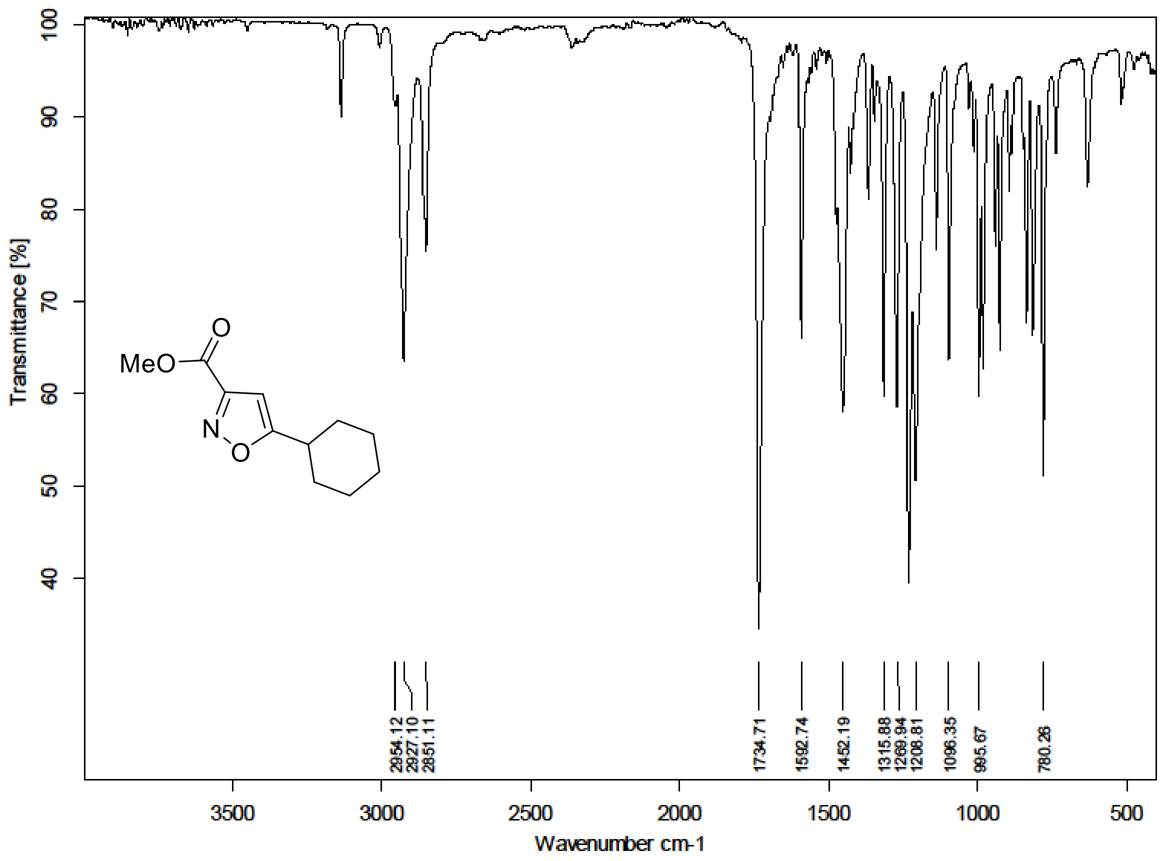

$^{1}H$ (500 MHz) and $^{13}C$ NMR (126 MHz) spectra of **S157** in Chloroform-*d*

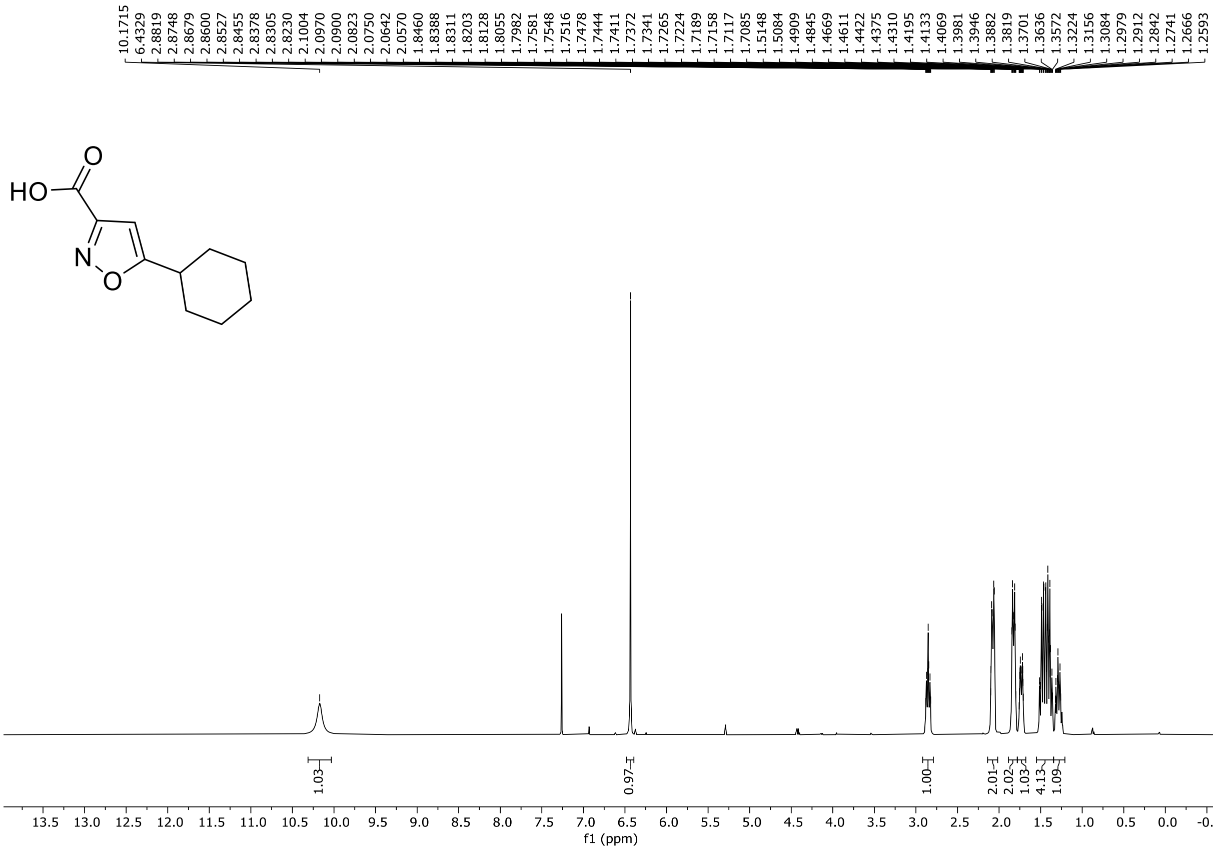

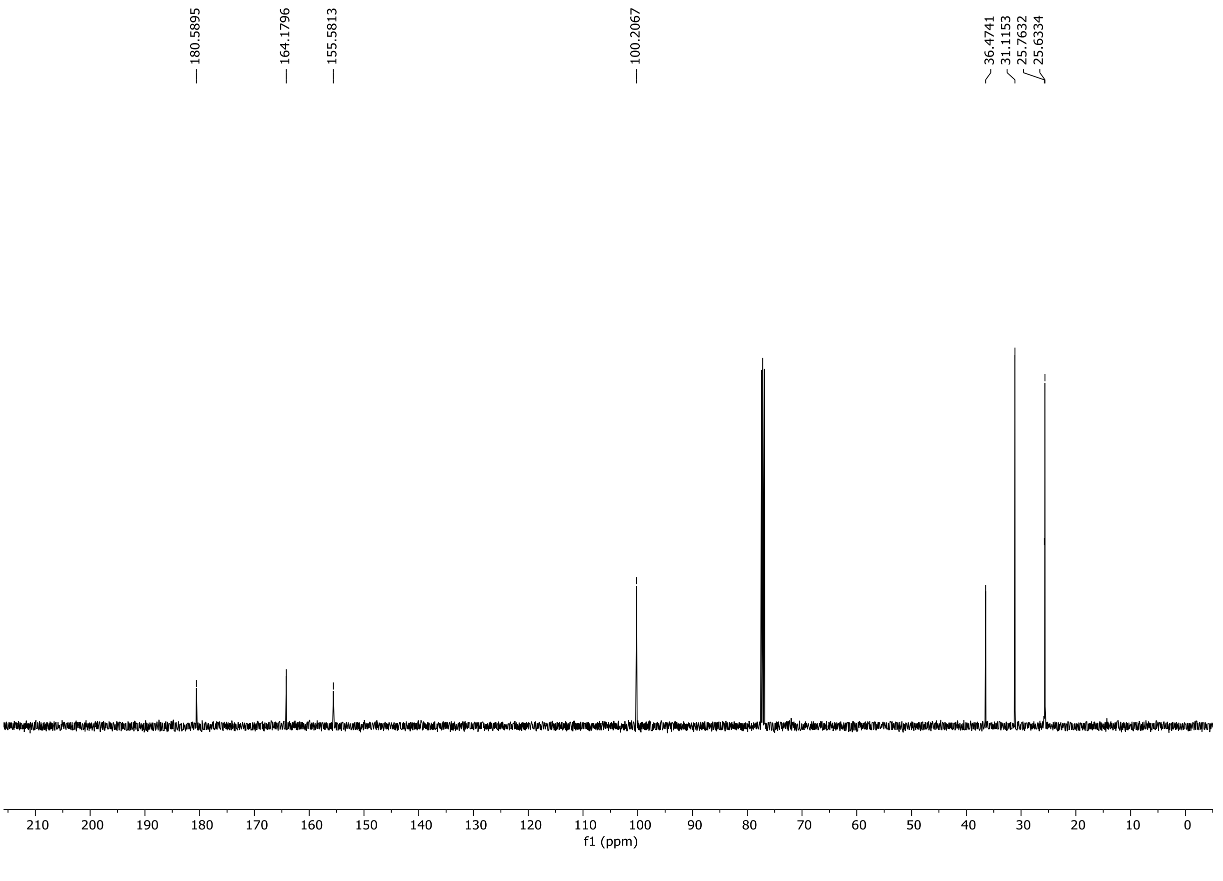

HRMS spectrum of **S157**

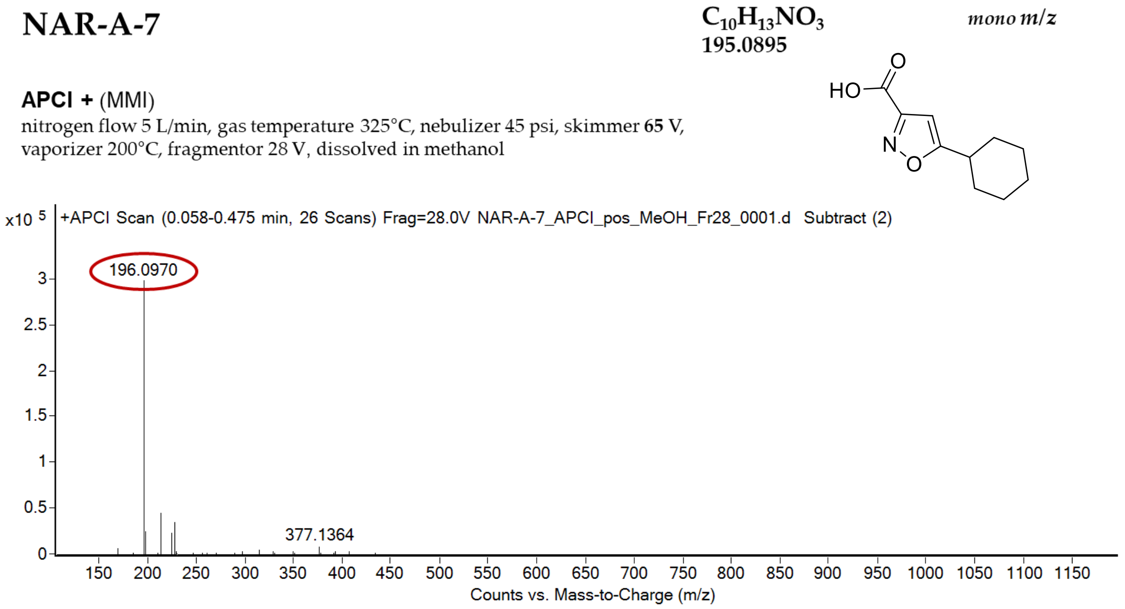

calculated mass: $[M+H]^+$ = 196.0968 observed: $[M+H]^+$ = 196.0970 mass accuracy = 1.0 ppm

FT-IR spectrum (neat) of **S157**

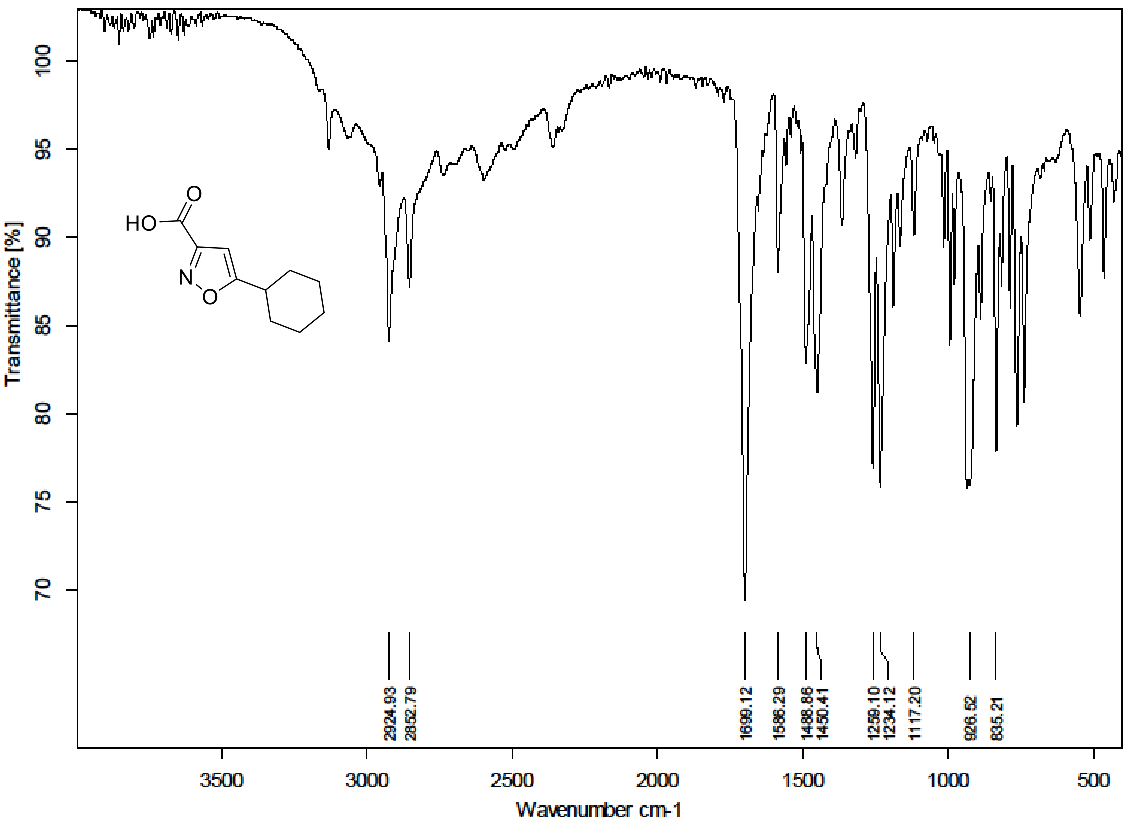

[1]H (500 MHz) and [13]C NMR (126 MHz) spectra of **S159** in Chloroform-*d*

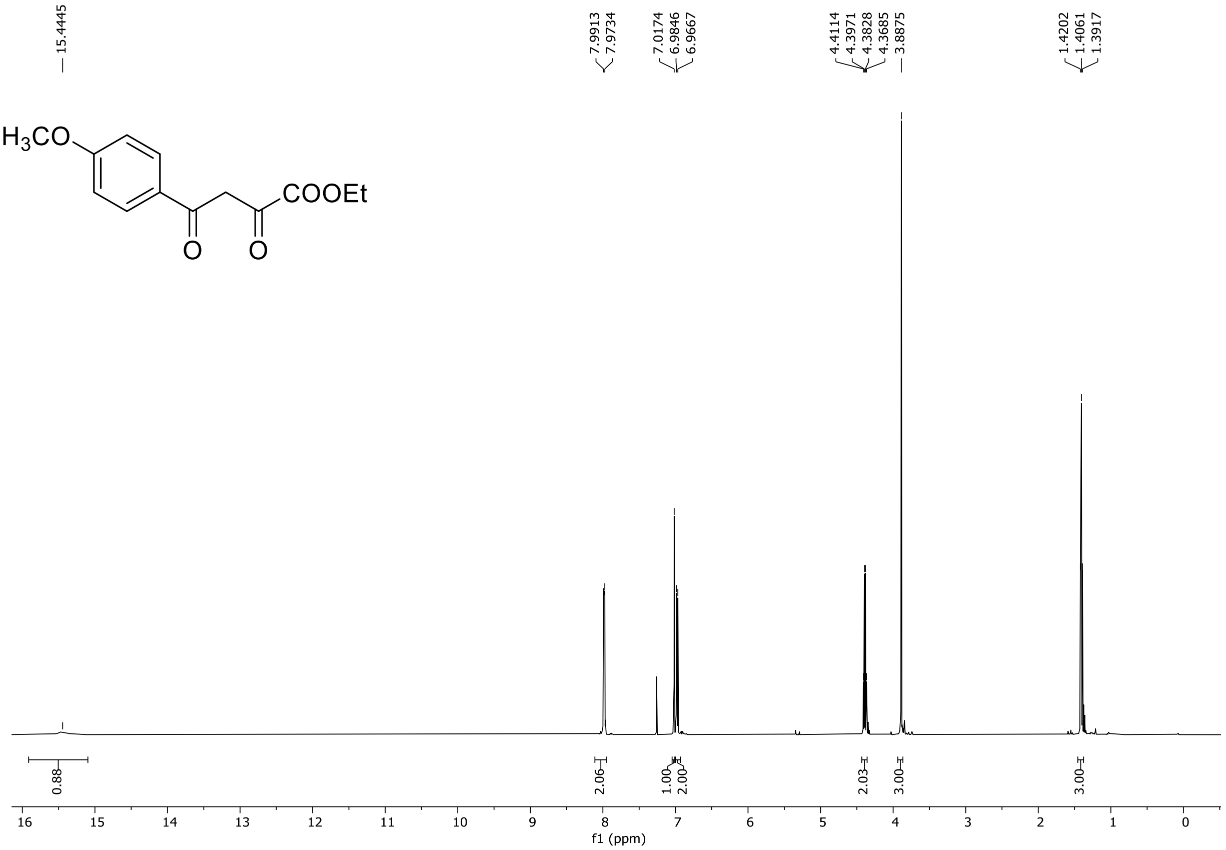

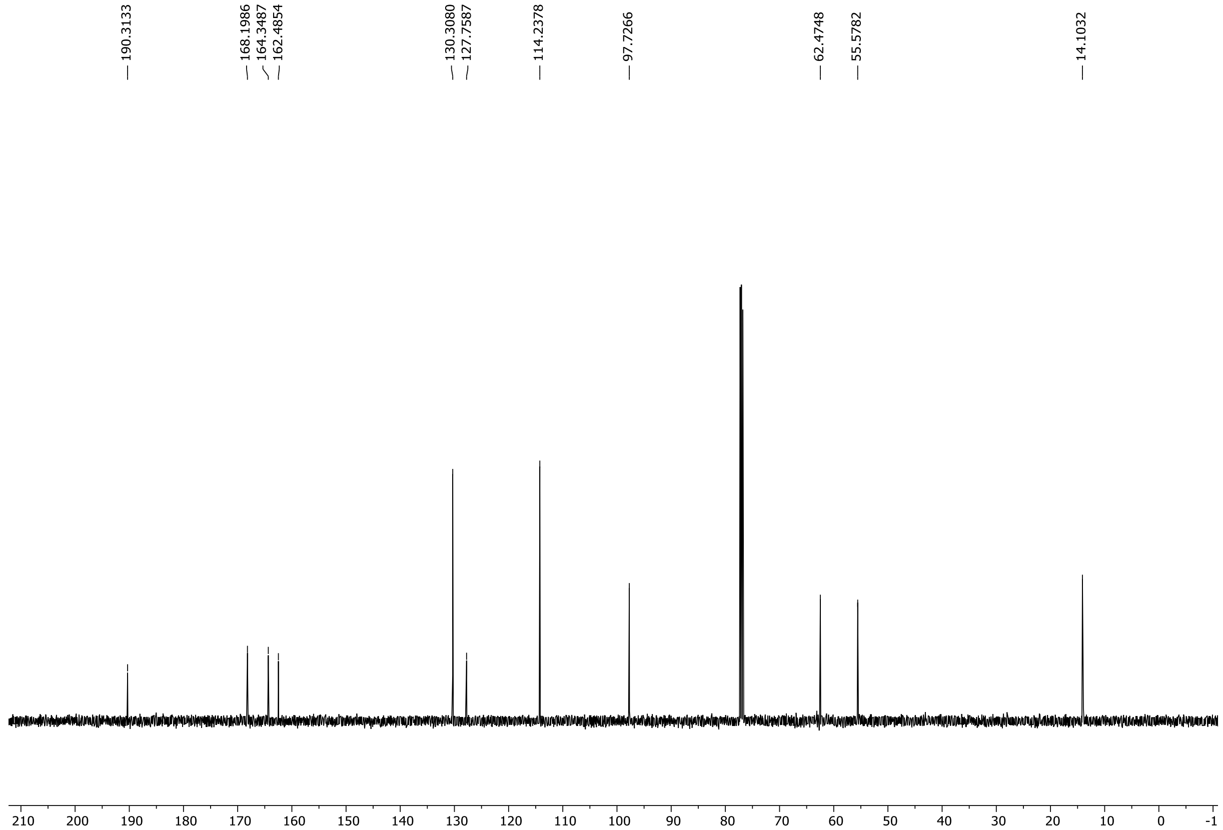

HRMS spectrum of **S159**

NAR-A-19 **$C_{13}H_{14}O_5$** **mono** ***m/z 250.0841***

**APCI +** (MMI)
nitrogen flow 5 L/min, gas temperature 325°C, nebulizer 45 psi, skimmer **65** V, vaporizer 200°C, fragmentor 20 V, dissolved in methanol

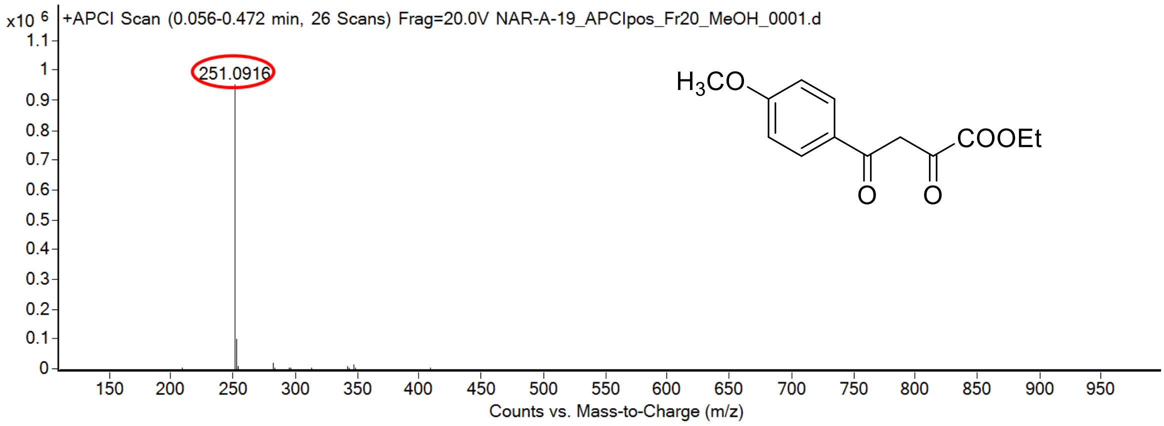

calculated mass: $[M+H]^+ = 251.0914$ observed: $[M+H]^+ = 251.0916$ mass accuracy = 0.8 ppm

FT-IR spectrum (neat) of **S159**

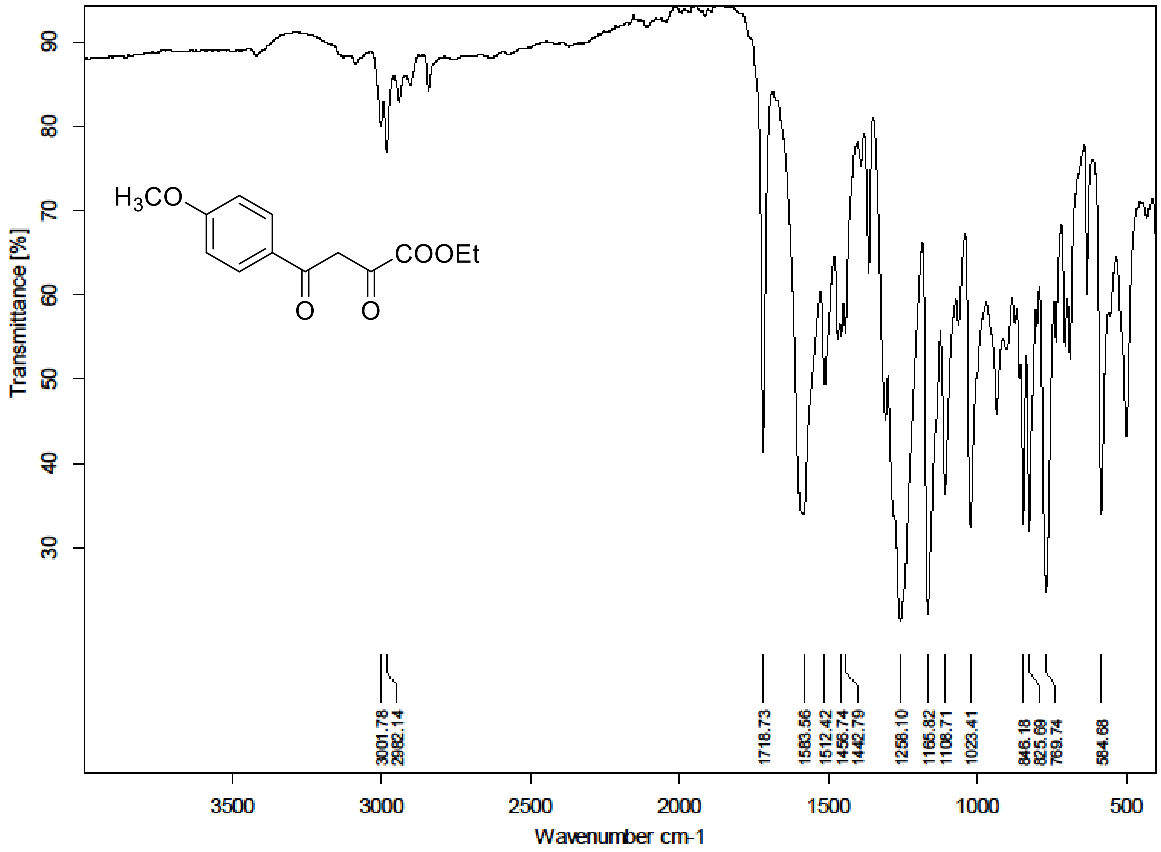

$^{1}H$ (500 MHz) and $^{13}C$ NMR (126 MHz) spectra of **S160** in Chloroform-*d*

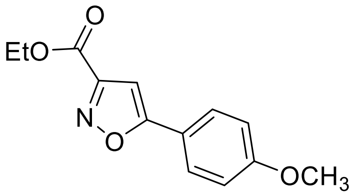

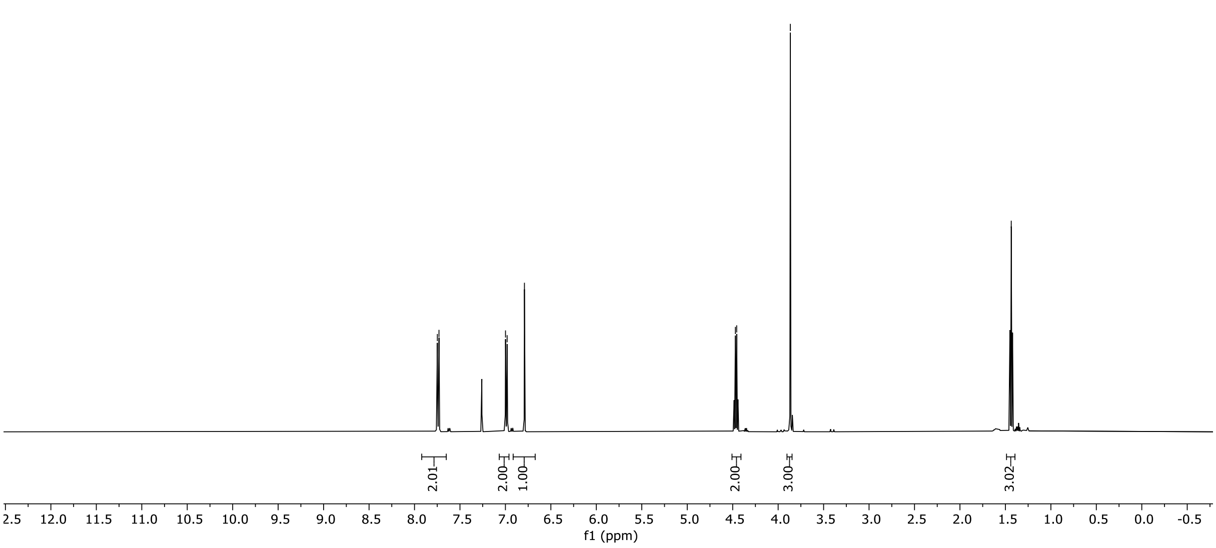

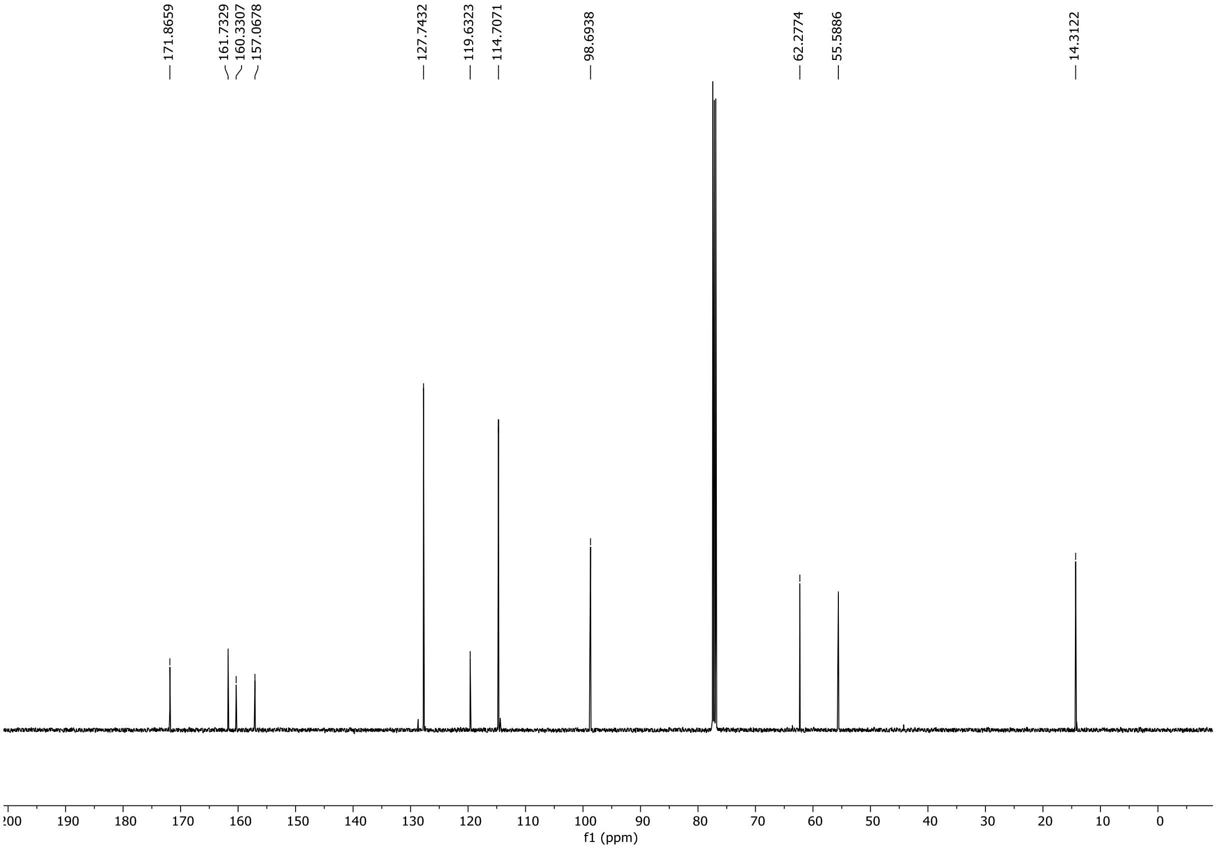

HRMS spectrum of **S160**

NAR-A-25 $C_{13}H_{13}NO_4$ mono *m/z 247.0845*

**APCI +** (MMI)
nitrogen flow 5 L/min, gas temperature 325°C, nebulizer 45 psi, skimmer **65** V, vaporizer 200°C, fragmentor 20 V, dissolved in methanol

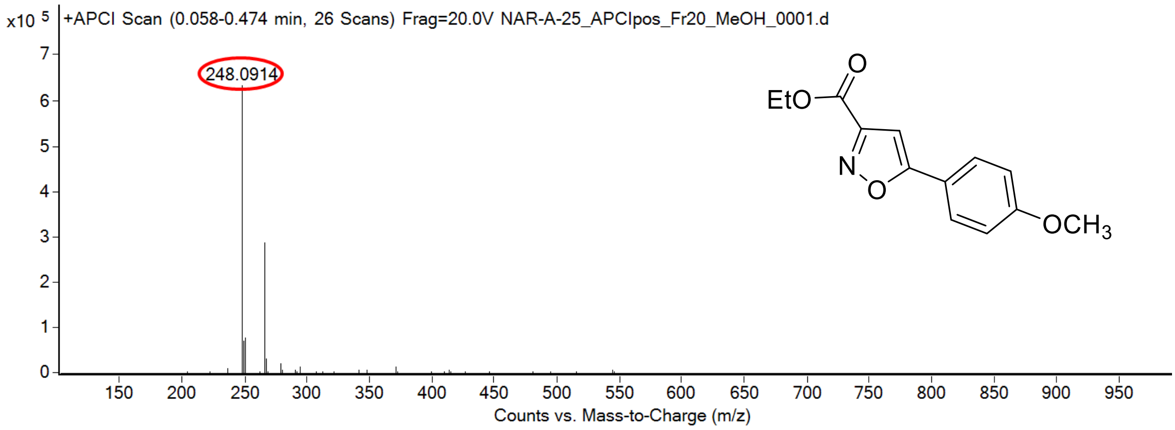

calculated mass: $[M+H]^+$ = 248.0917 observed: $[M+H]^+$ = 248.0914 mass accuracy = -1.2 ppm

FT-IR spectrum (neat) of **S160**

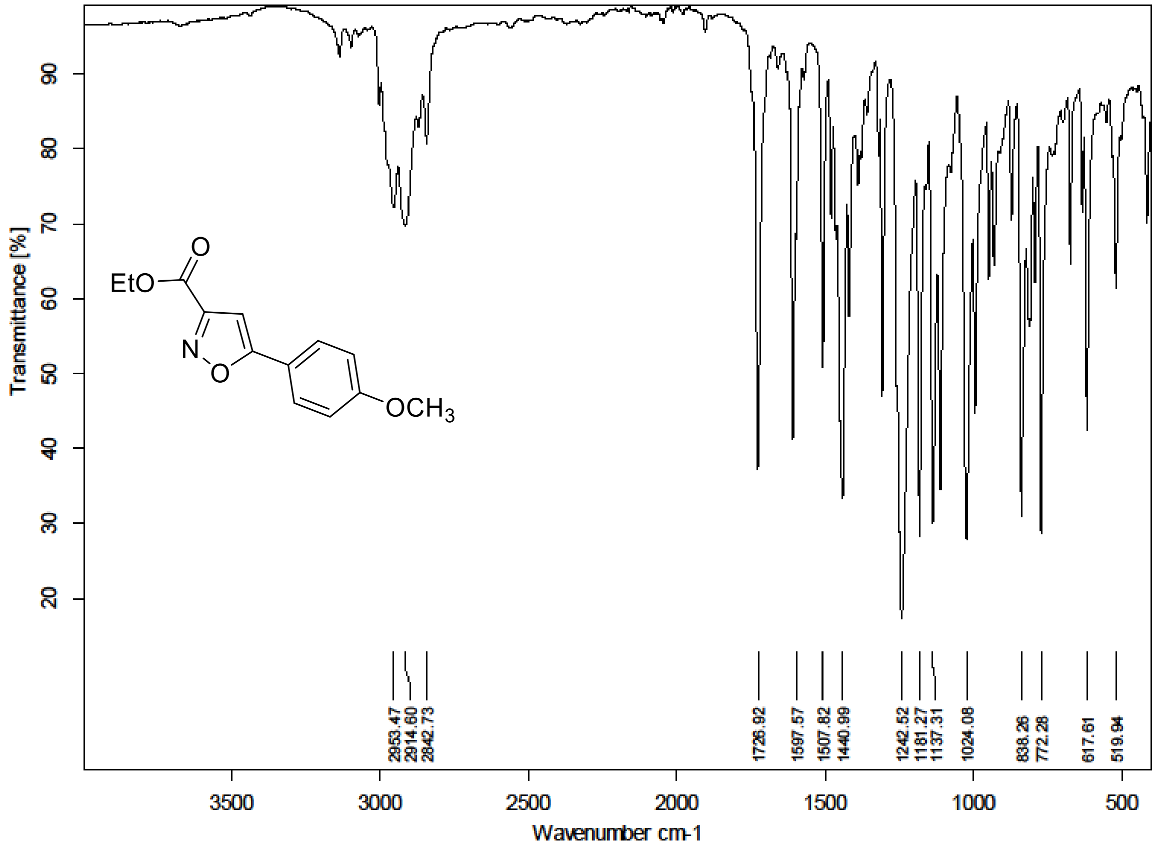

[1]H (500 MHz) and [13]C NMR (126 MHz) spectra of **S161** in Chloroform-*d*

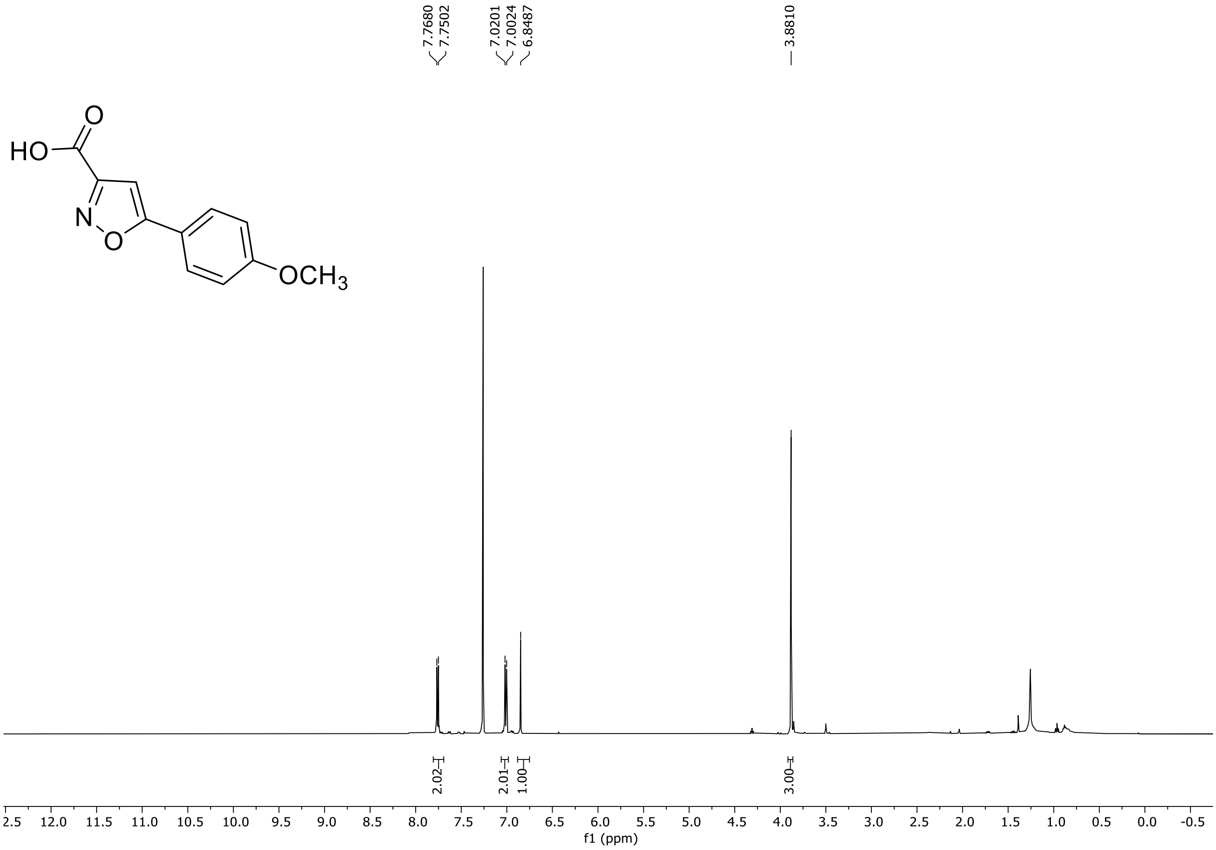

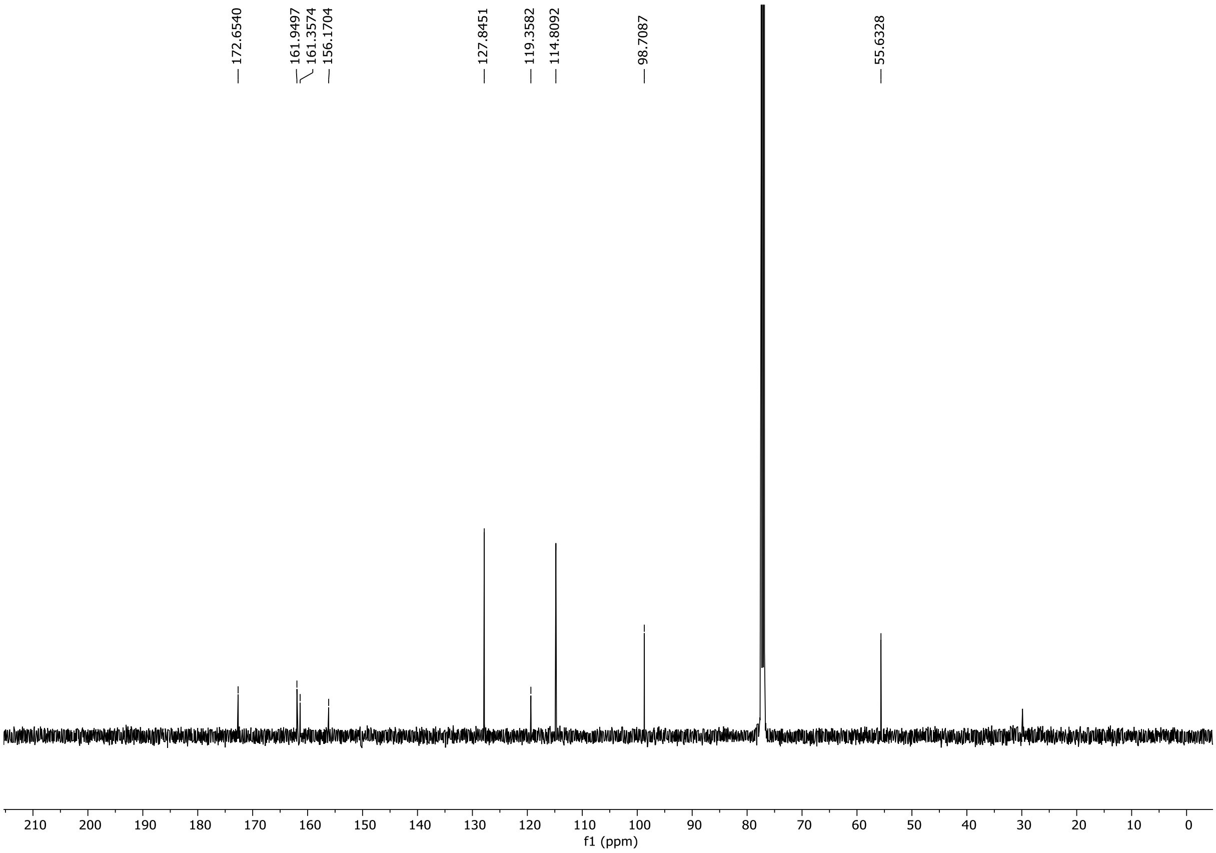

HRMS spectrum of **S161**

NAR-A-30 **$C_{11}H_9NO_4$** mono *m/z 219.0532*

**APCI +** (MMI)
nitrogen flow 5 L/min, gas temperature 325°C, nebulizer 45 psi, skimmer **65** V, vaporizer 200°C, fragmentor 25 V, dissolved in methanol

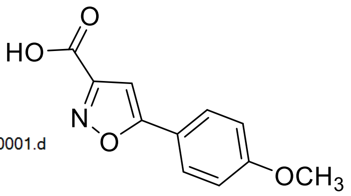

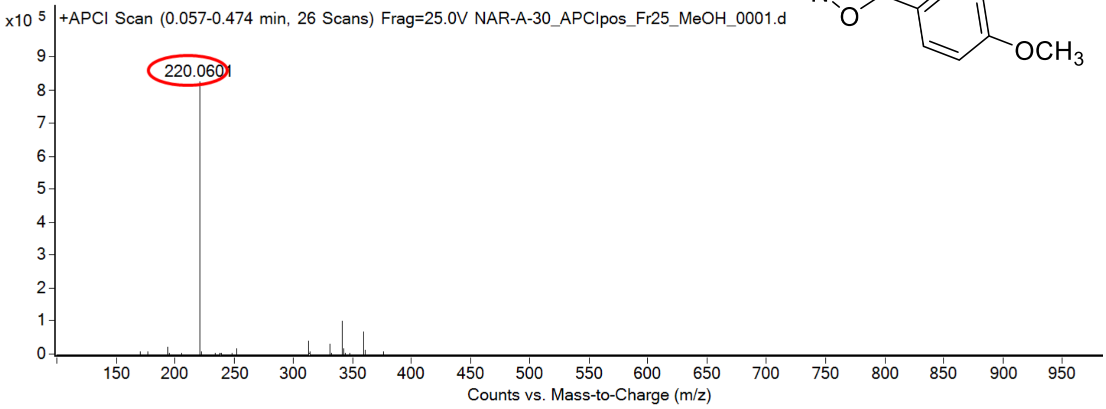

calculated mass: $[M+H]^+ = 220.0604$ observed: $[M+H]^+ = 220.0601$ mass accuracy = -1.3 ppm

FT-IR spectrum (neat) of **S161**

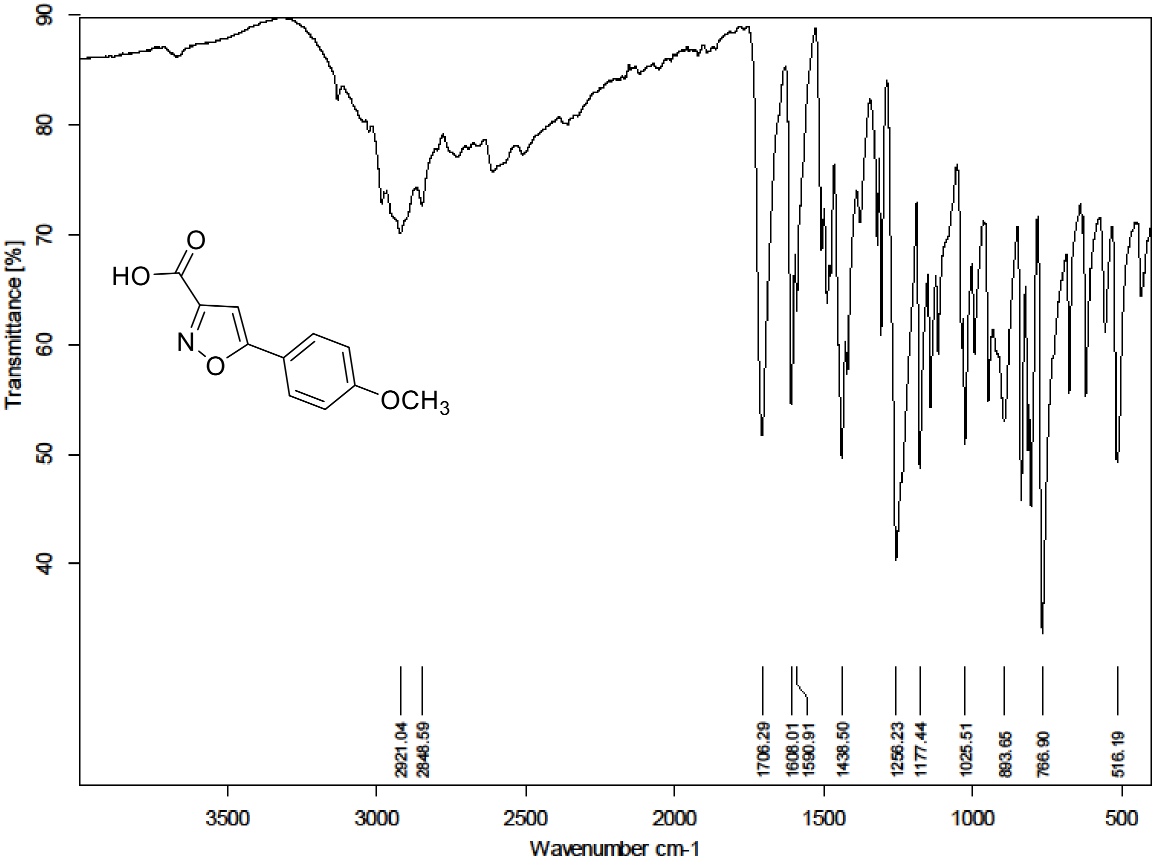

$^{1}H$ (500 MHz) and $^{13}C$ NMR (126 MHz) spectra of **S163** in Chloroform-*d*

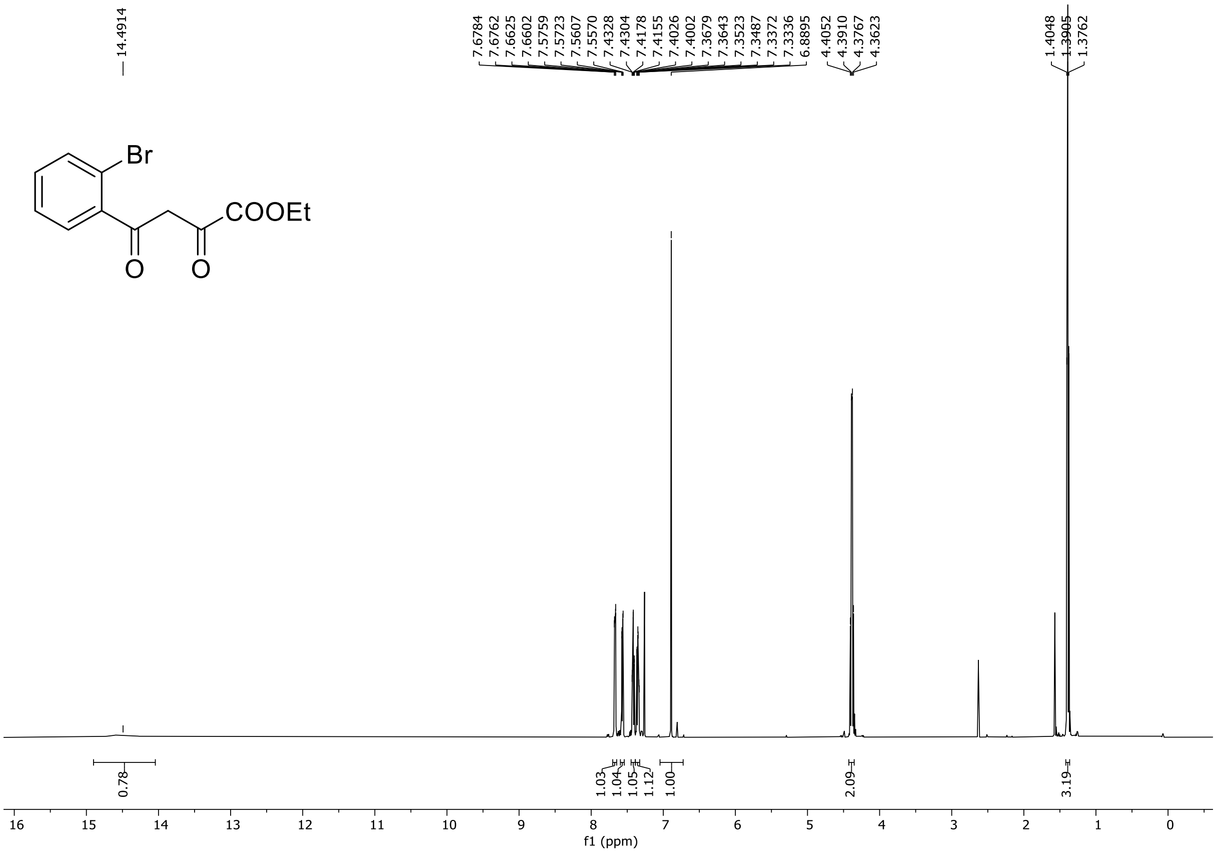

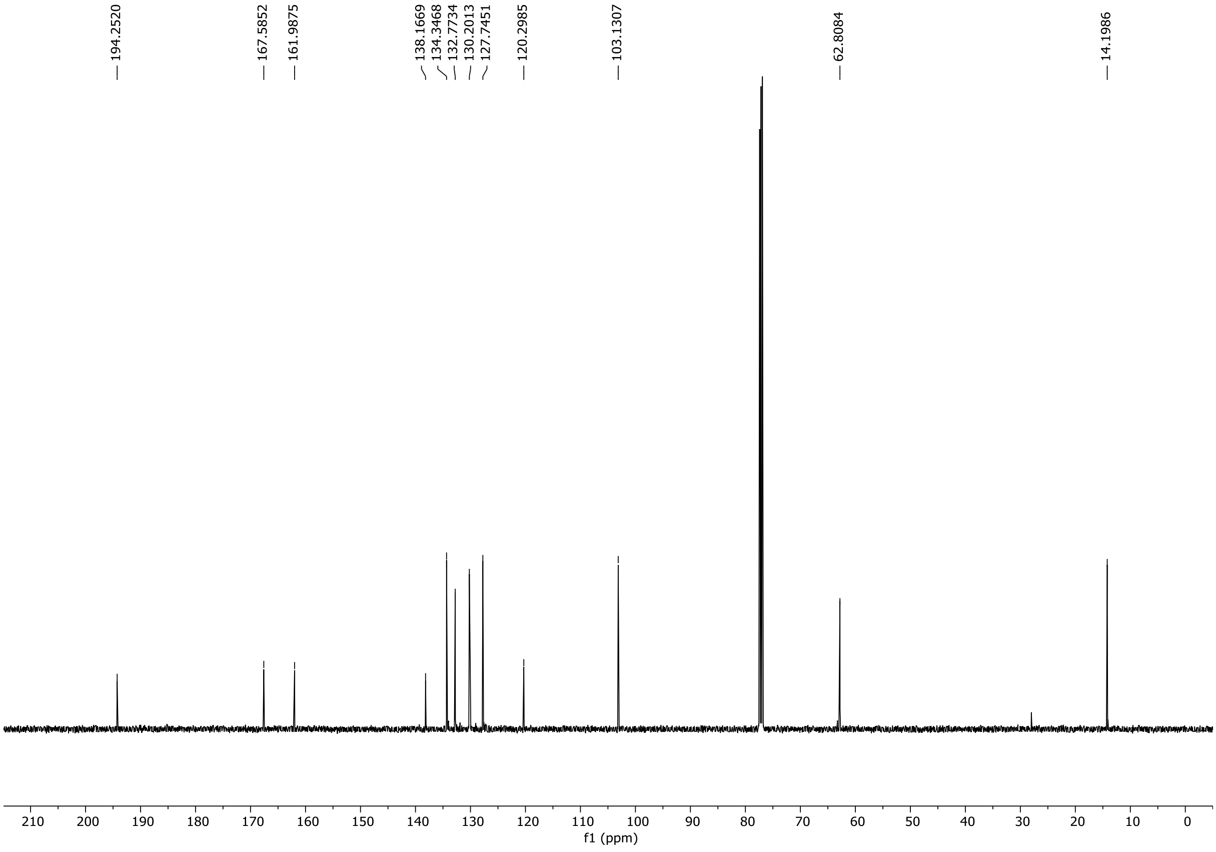

HRMS spectrum of **S163**

NAR-A-21 $C_{12}H_{11}BrO_4$ **mono *m/z* 297.9841**

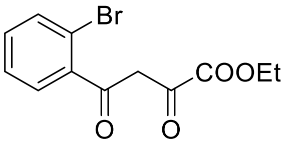

**APCI +** (MMI)
nitrogen flow 5 L/min, gas temperature 325°C, nebulizer 45 psi, skimmer **65** V, vaporizer 200°C, fragmentor 28 V, dissolved in methanol

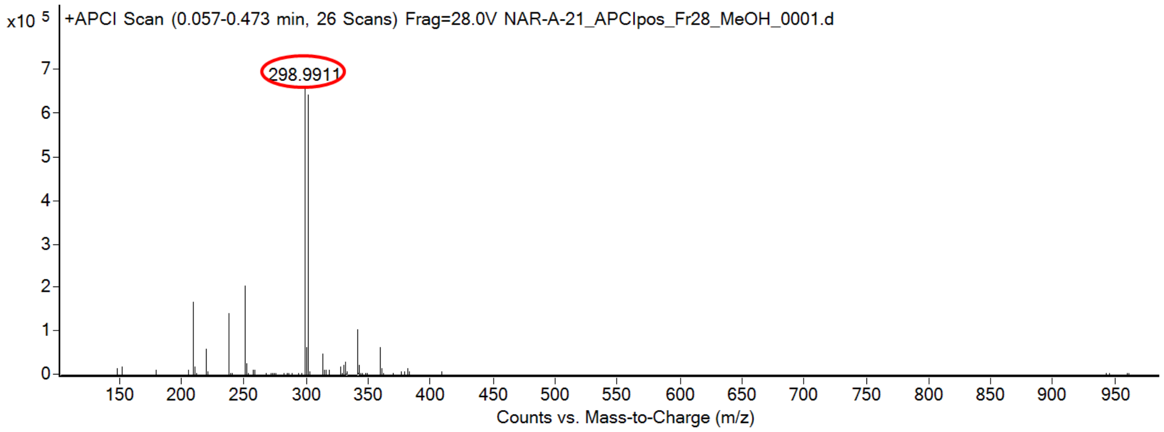

calculated mass: $[M+H]^+$ = 298.9913 observed: $[M+H]^+$ =298.9911 mass accuracy = -0.7 ppm

FT-IR spectrum (neat) of **S163**

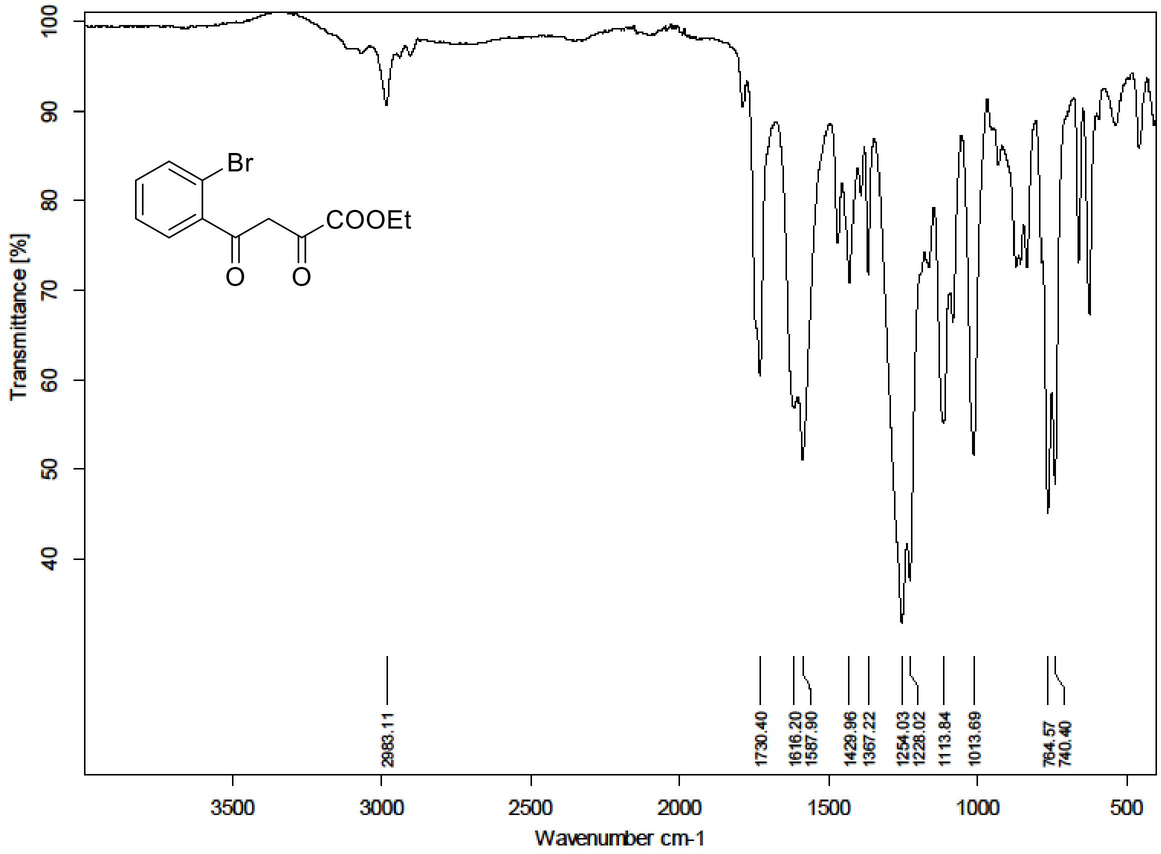

[1]H (500 MHz) and [13]C NMR (126 MHz) spectra of **S164** in Chloroform-*d*

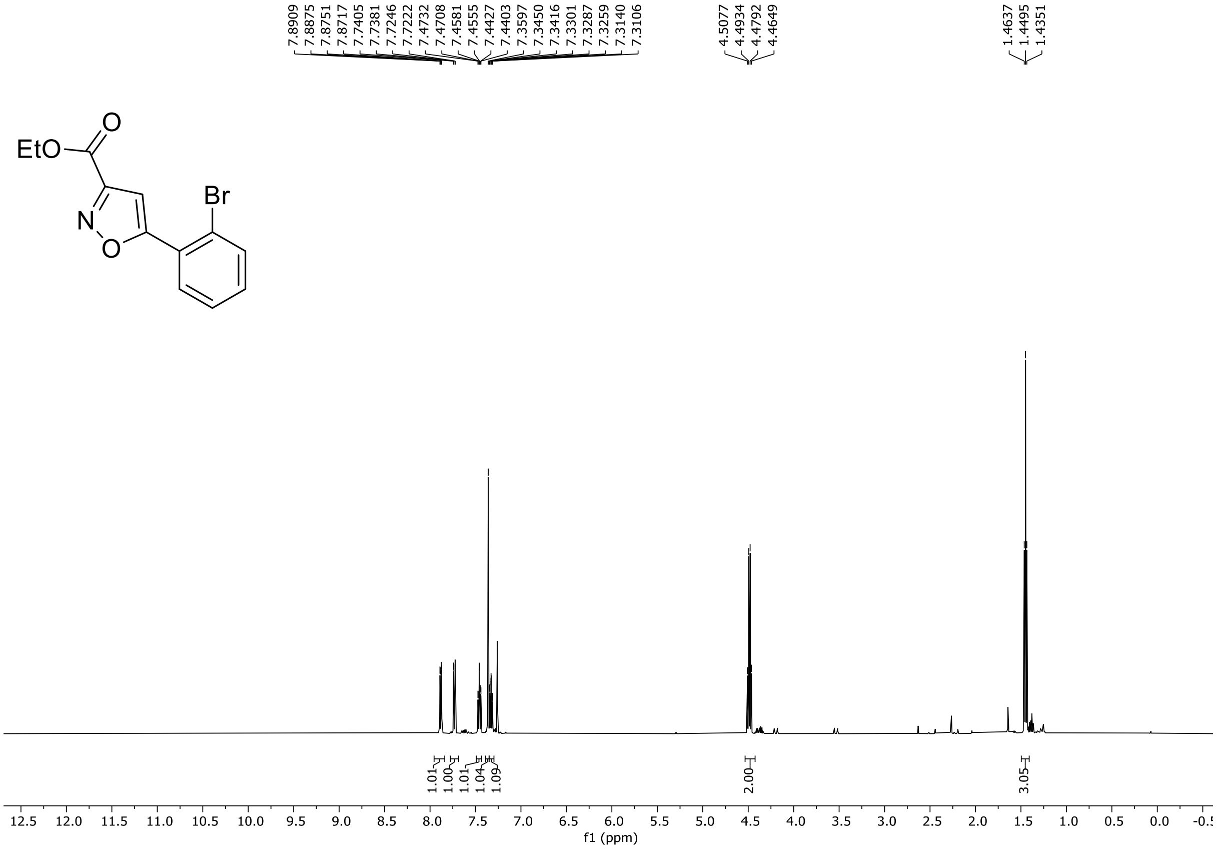

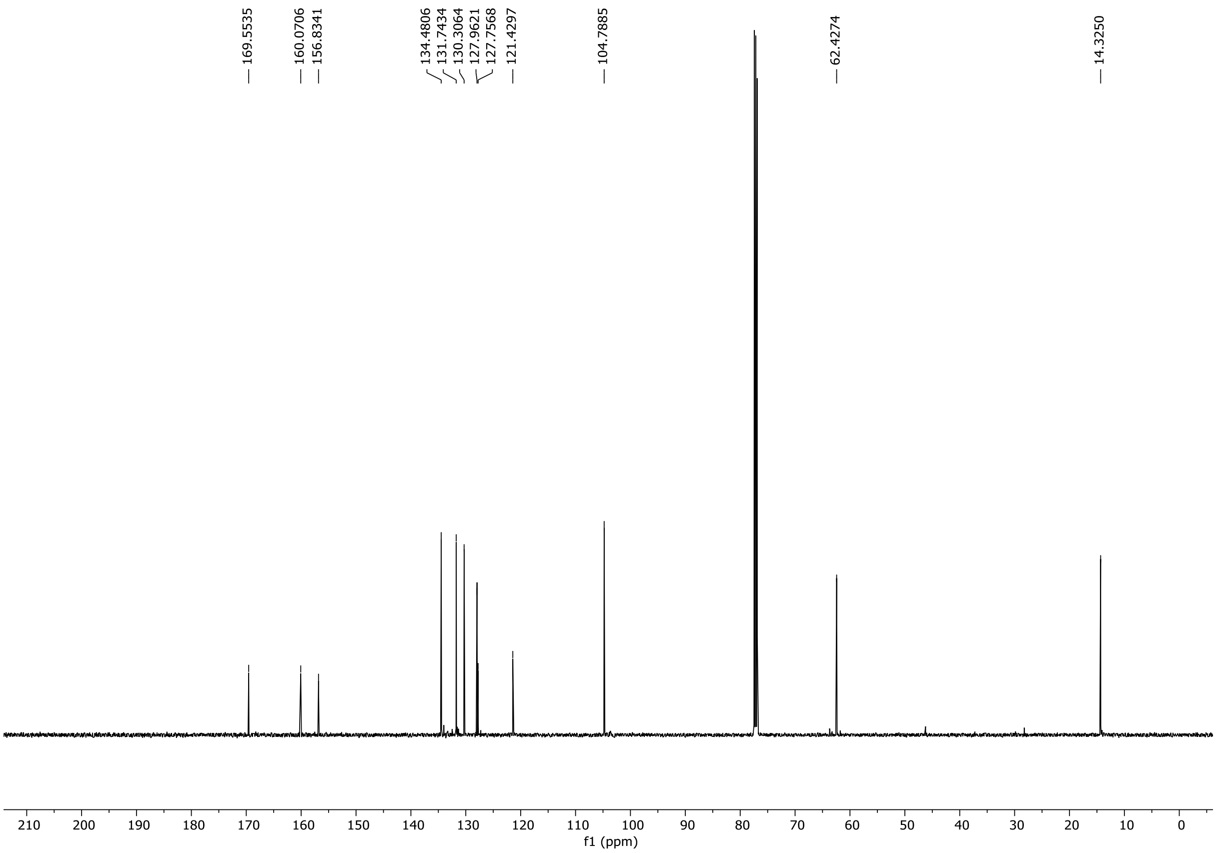

HRMS spectrum of **S164**

NAR-A-26 $C_{12}H_{10}BrNO_3$ mono *m/z 294.9844*

**APCI +** (MMI)

nitrogen flow 5 L/min, gas temperature 325°C, nebulizer 45 psi, skimmer **65** V, vaporizer 200°C, fragmentor 25 V, dissolved in methanol

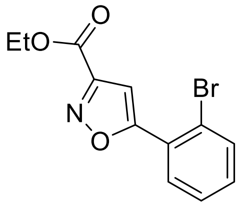

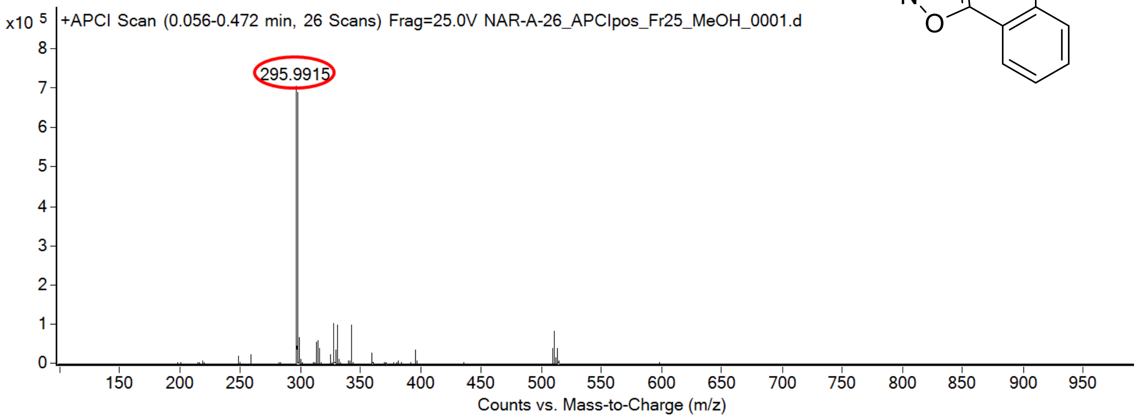

calculated mass: $[M+H]^+$ = 295.9917 observed: $[M+H]^+$ = 295.9915 mass accuracy = -0.7 ppm

FT-IR spectrum (neat) of **S164**

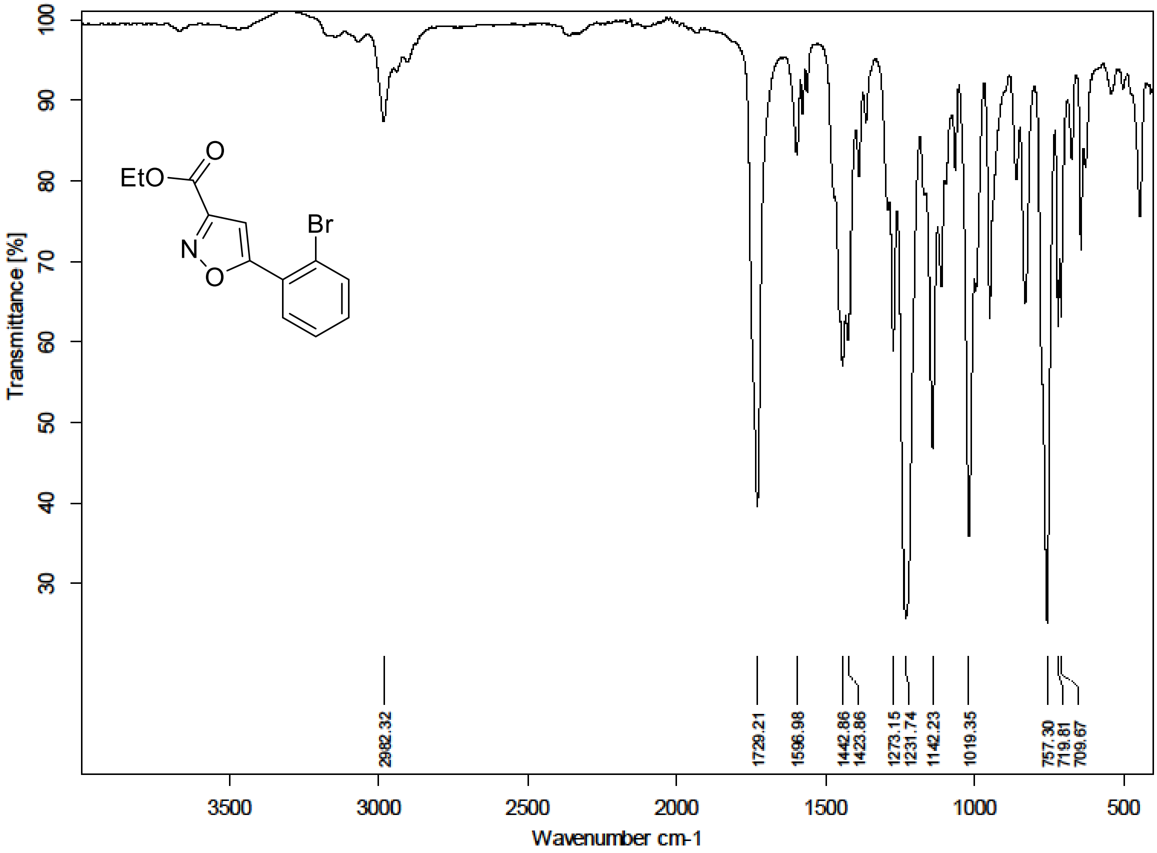

$^{1}H$ (500 MHz) and $^{13}C$ NMR (126 MHz) spectra of **S165** in Chloroform-*d*

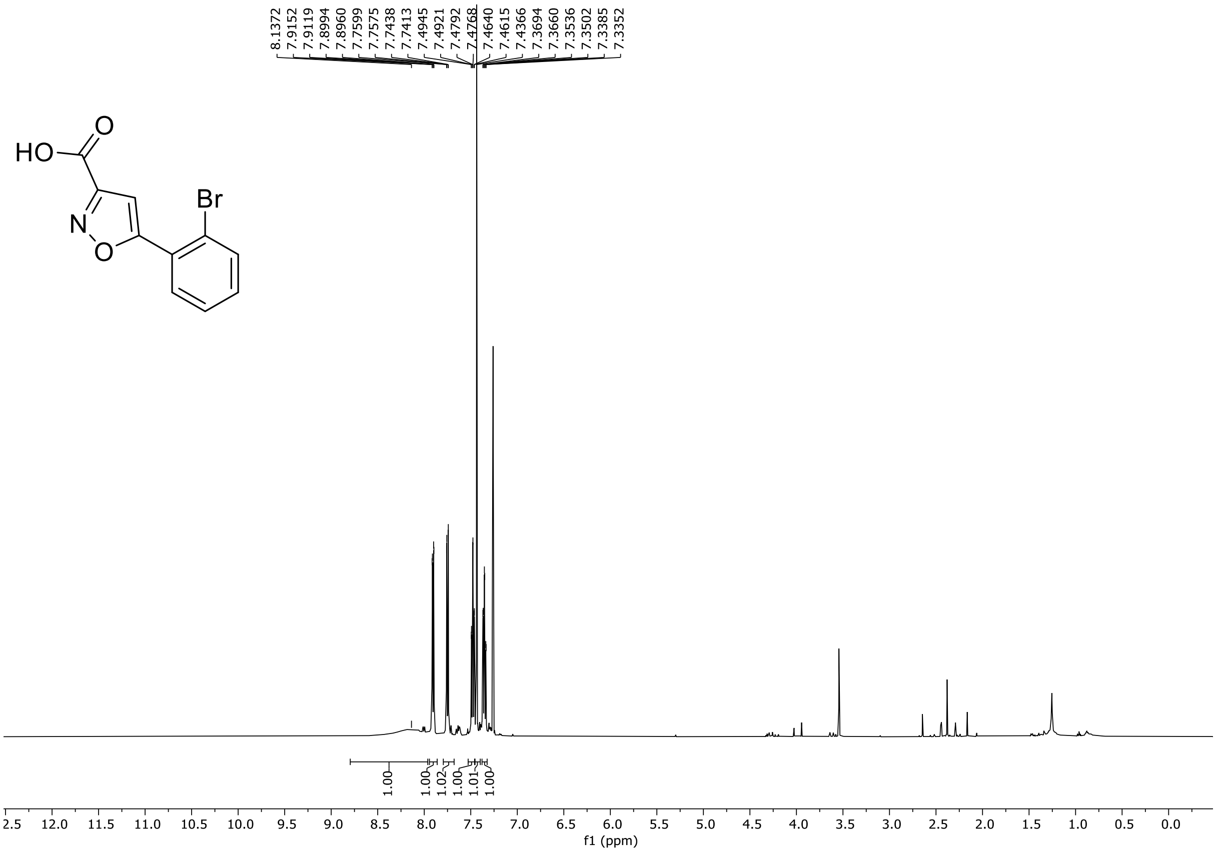

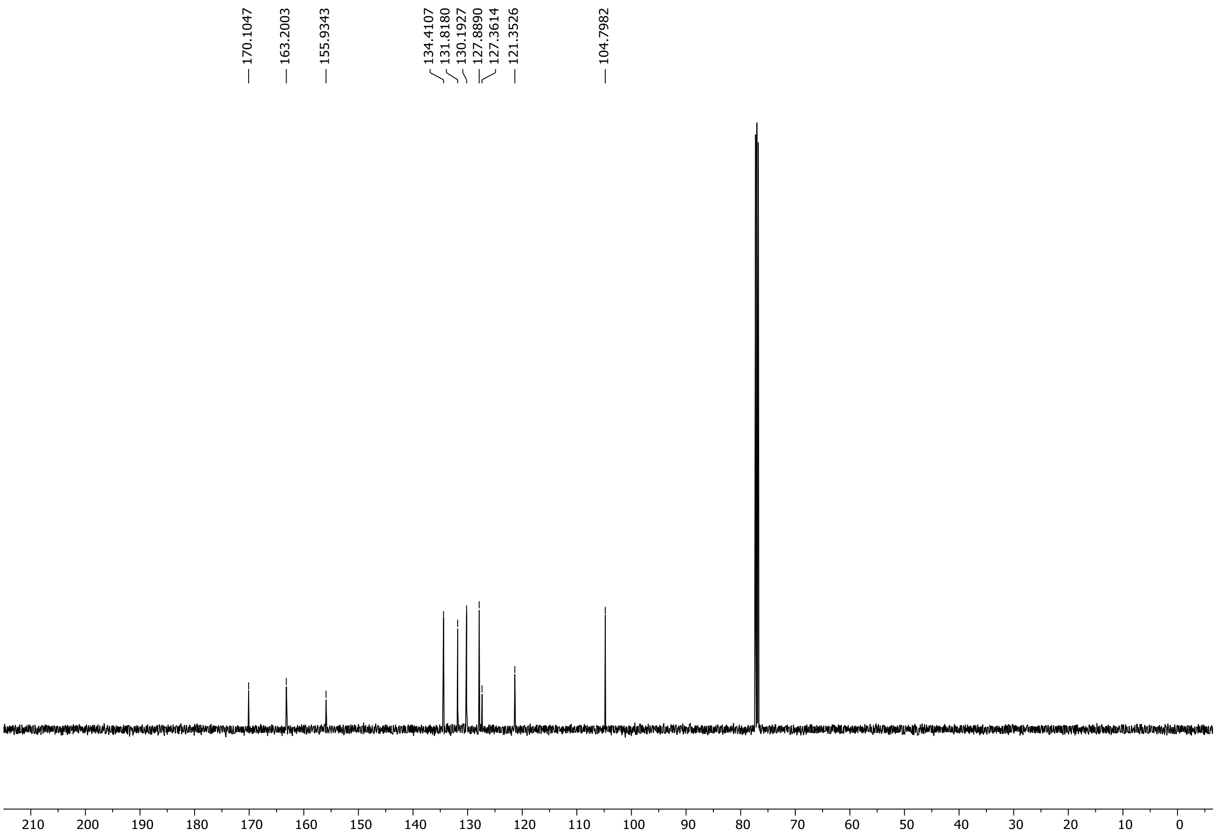

HRMS spectrum of **S165**

NAR-A-31 $C_{10}H_6BrNO_3$ mono *m/z 266.9531*

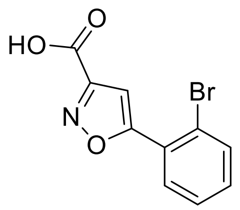

**APCI +** (MMI)
nitrogen flow 5 L/min, gas temperature 325°C, nebulizer 45 psi, skimmer **65** V, vaporizer 200°C, fragmentor 35 V, dissolved in methanol

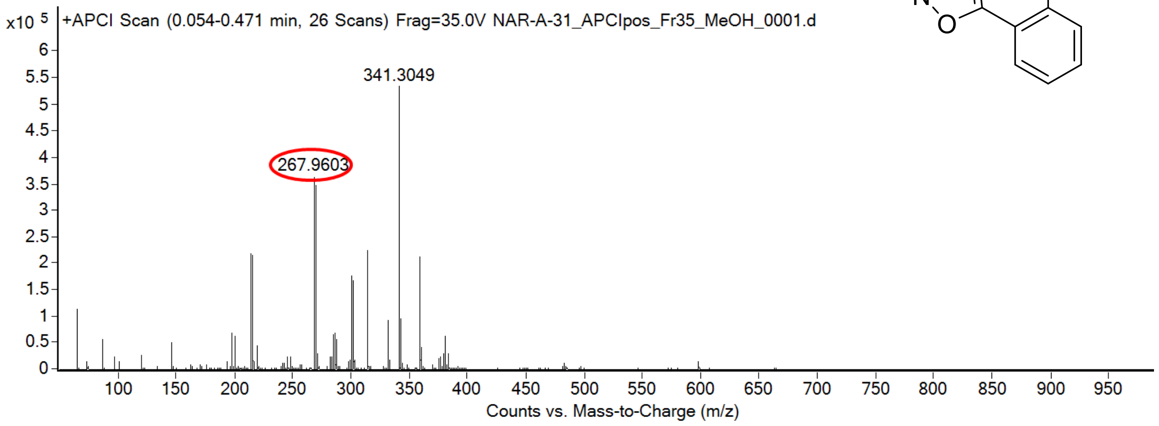

calculated mass: $[M+H]^+ = 267.9604$ observed: $[M+H]^+ = 267.9603$ mass accuracy = -0.3 ppm

FT-IR spectrum (neat) of **S165**

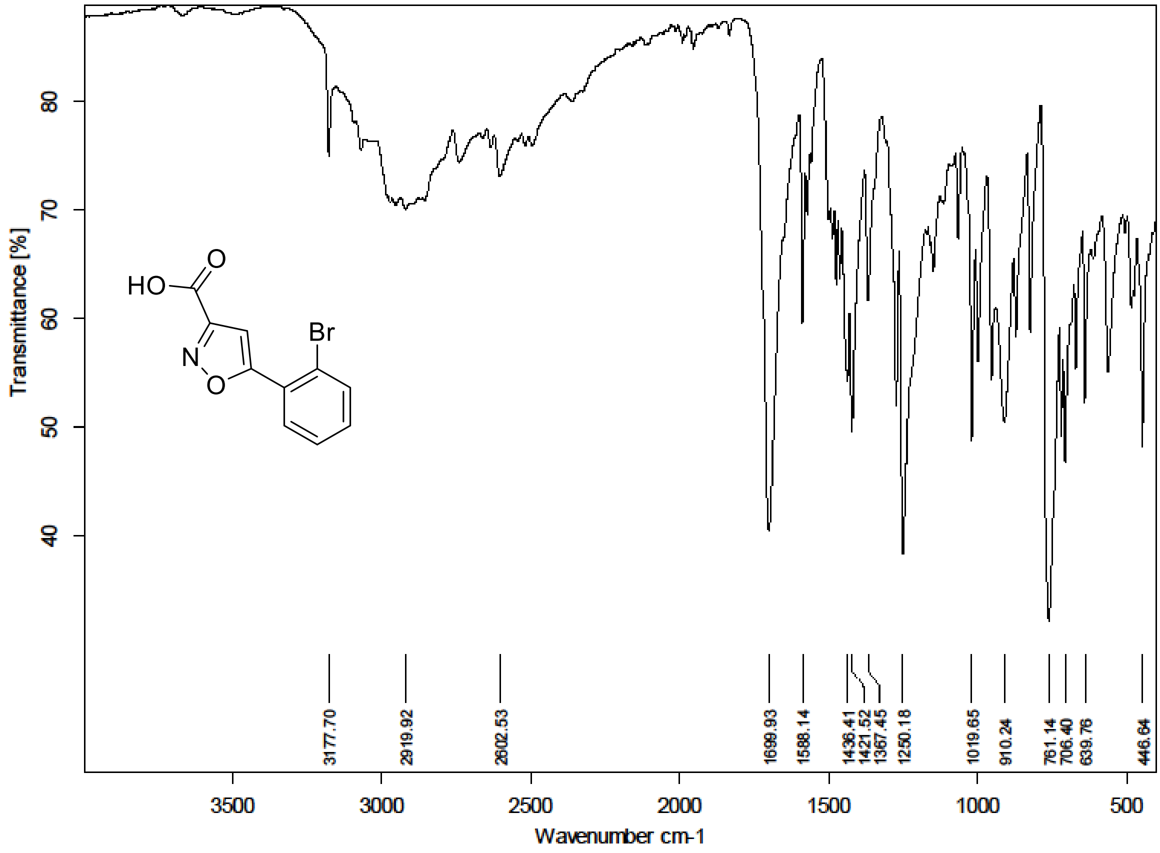

$^{1}H$ (500 MHz) and $^{13}C$ NMR (126 MHz) spectra of **S167** in Chloroform-*d*

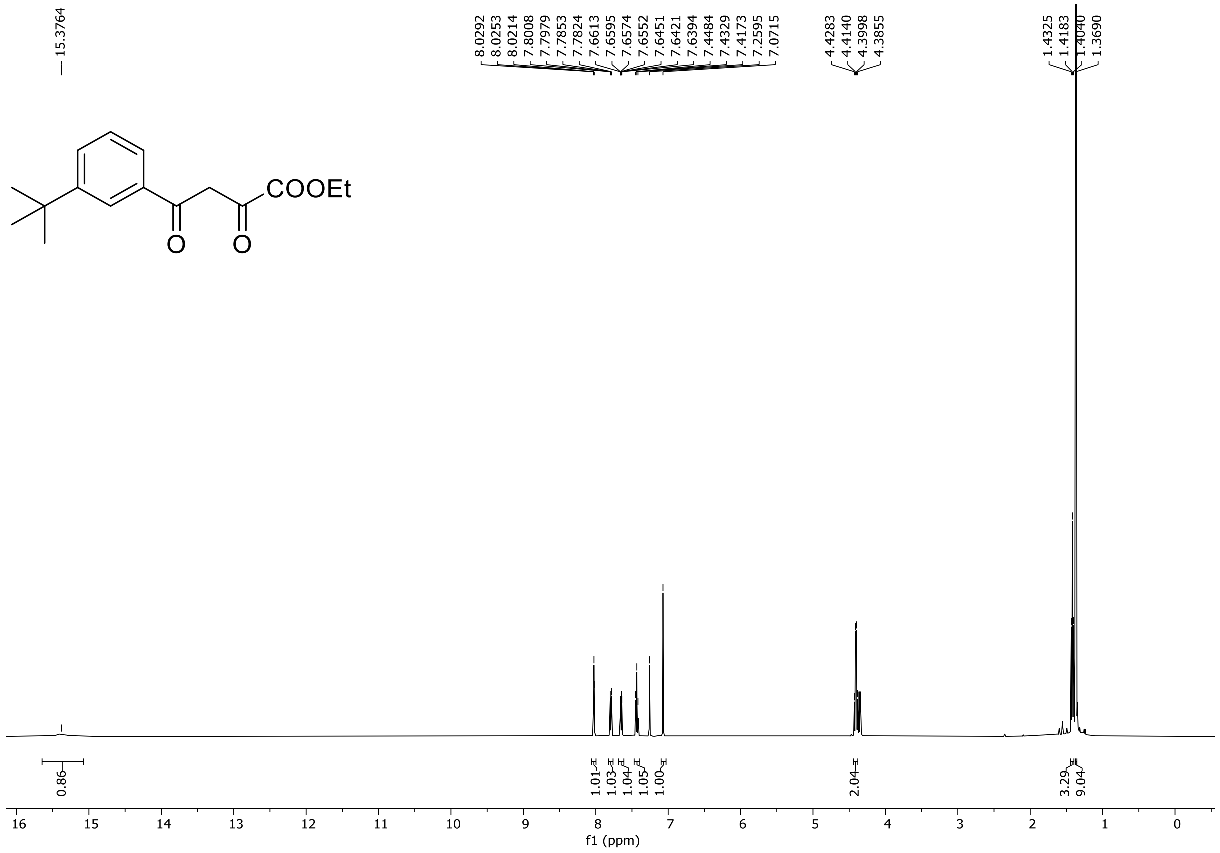

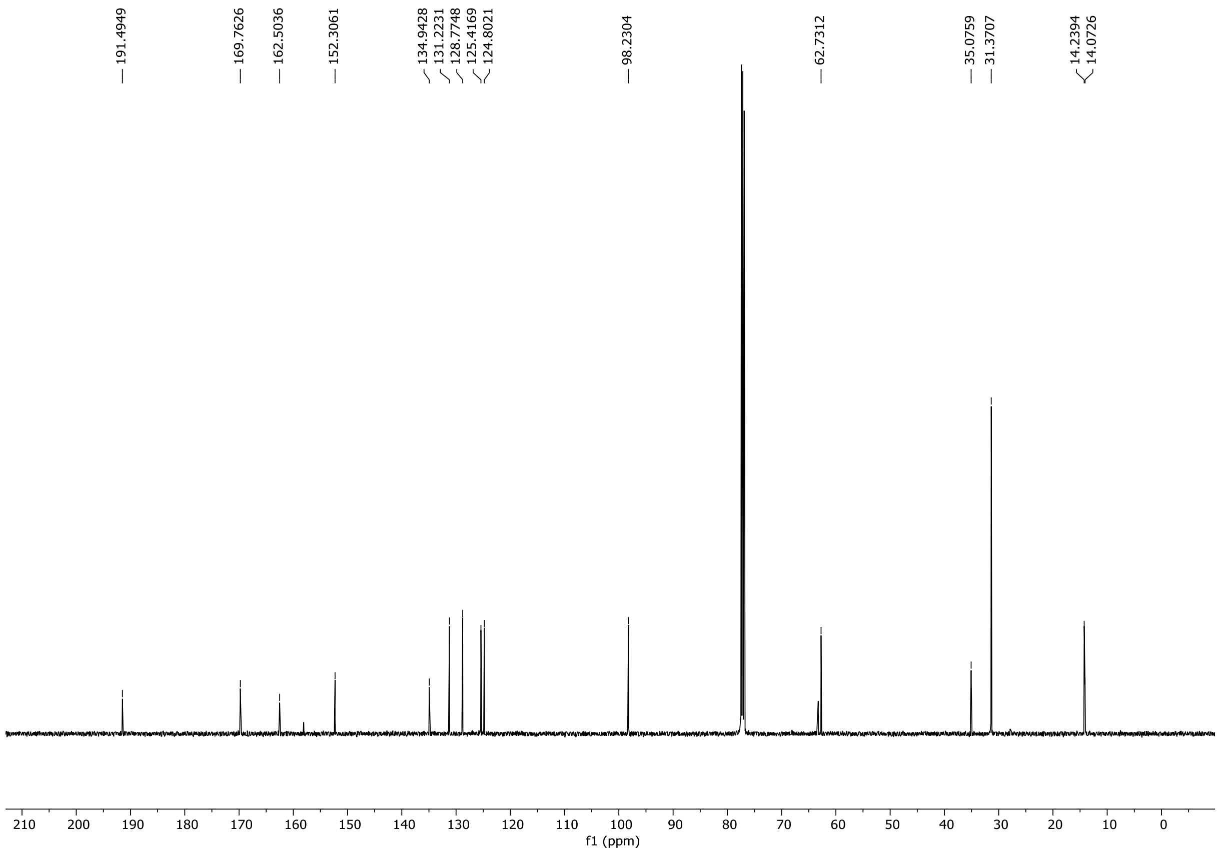

HRMS spectrum of **S167**

**NAR-A-48**

$C_{16}H_{20}O_4$ *mono m/z* **276.1362**

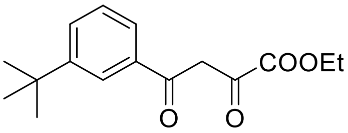

**APCI +** (MMI)
nitrogen flow 5 L/min, gas temperature 300°C, nebulizer 45 psi, skimmer **65** V, vaporizer 200°C, fragmentor 15 V, dissolved in MeOH

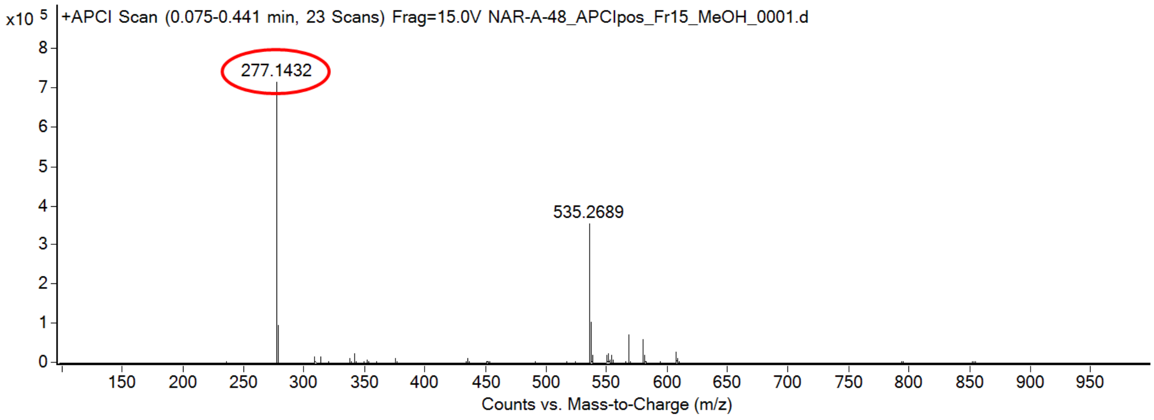

calculated mass: $[M+H]^+$ = 277.1434 observed: $[M+H]^+$ = 277.1432 mass accuracy = -0.7 ppm

FT-IR spectrum (neat) of **S167**

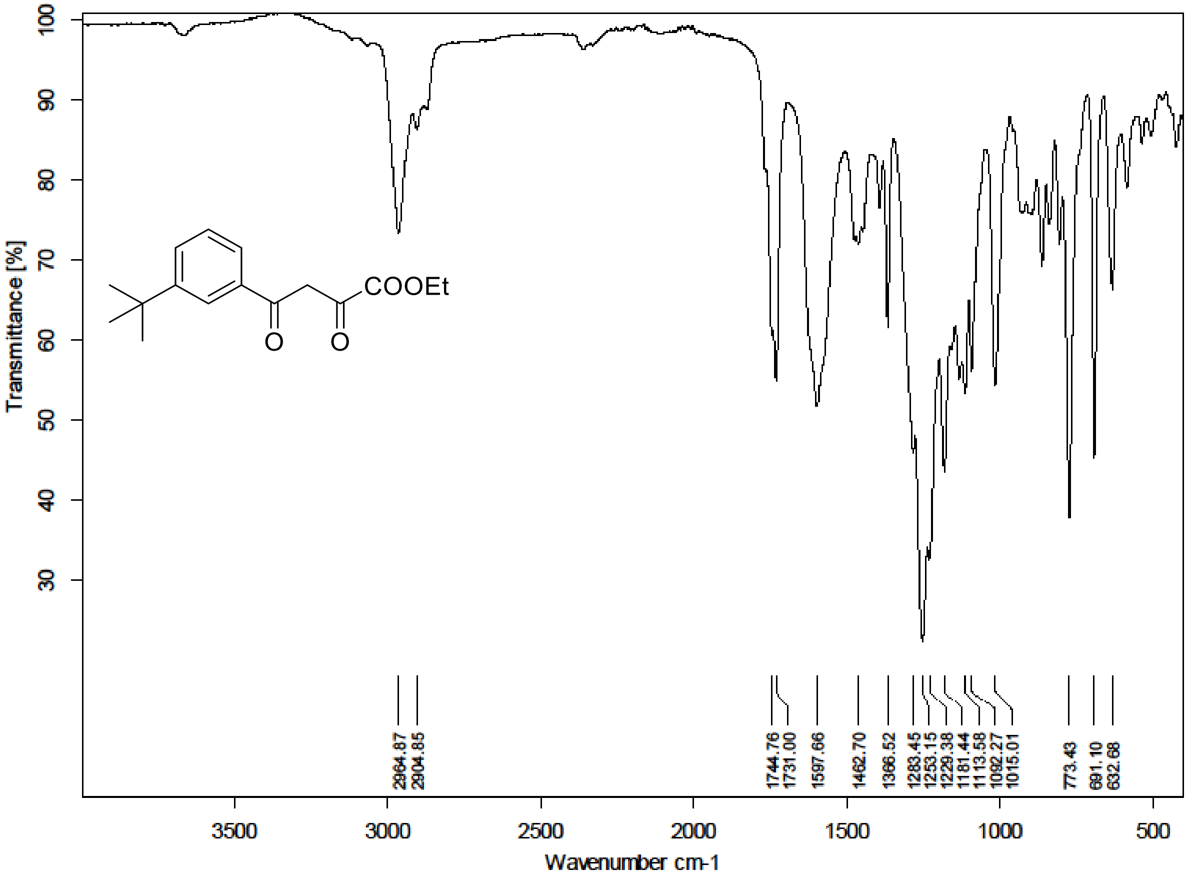

[1]H (300 MHz) and [13]C NMR (126 MHz) spectra of **S168** in Chloroform-*d*

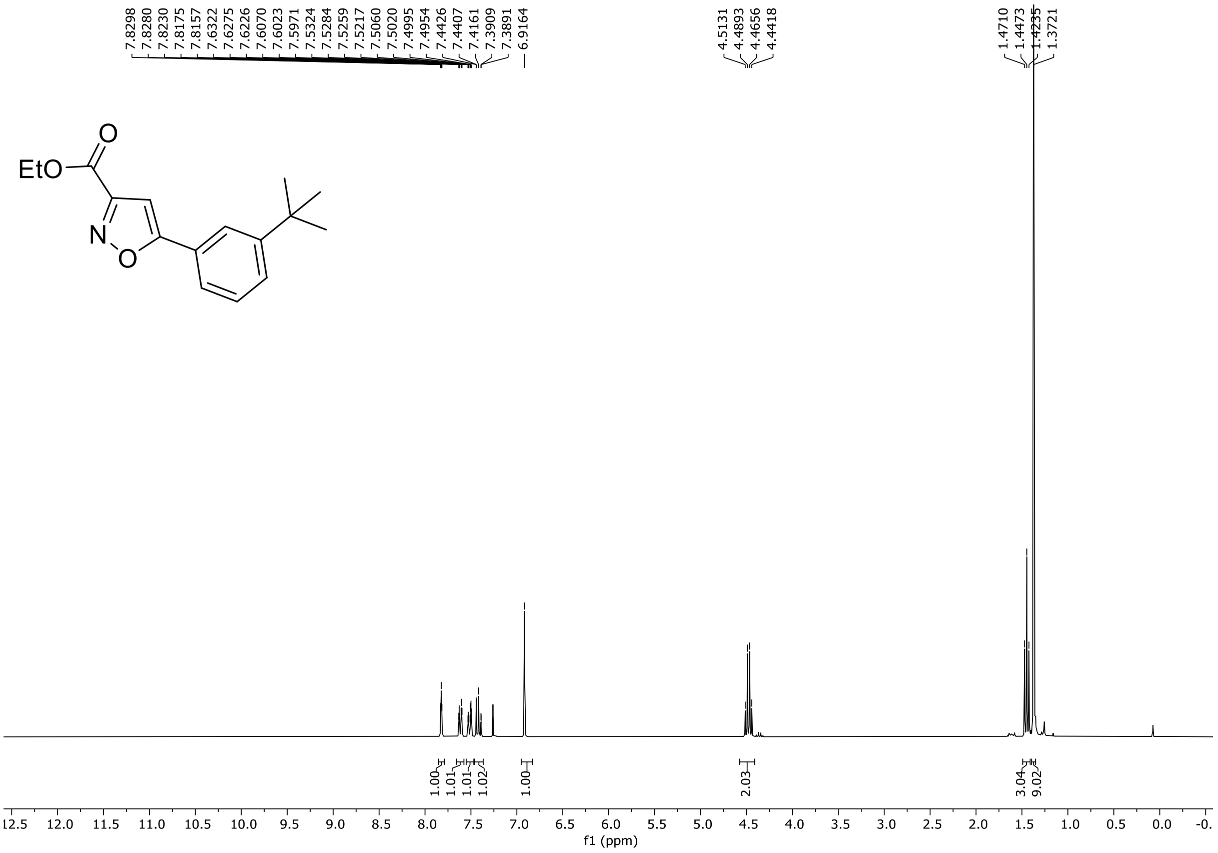

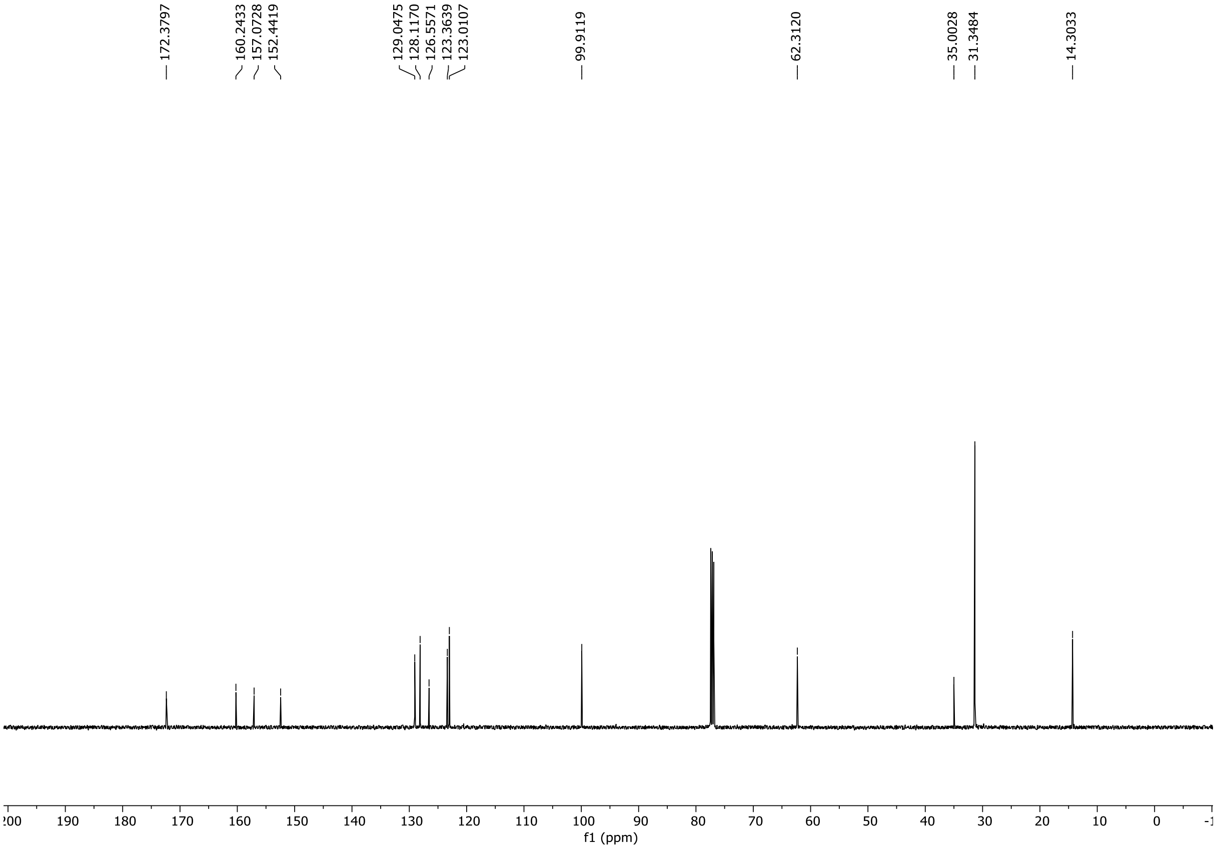

HRMS spectrum of **S168**

**NAR-A-51** $C_{16}H_{19}NO_3$ 273.1365 *mono m/z*

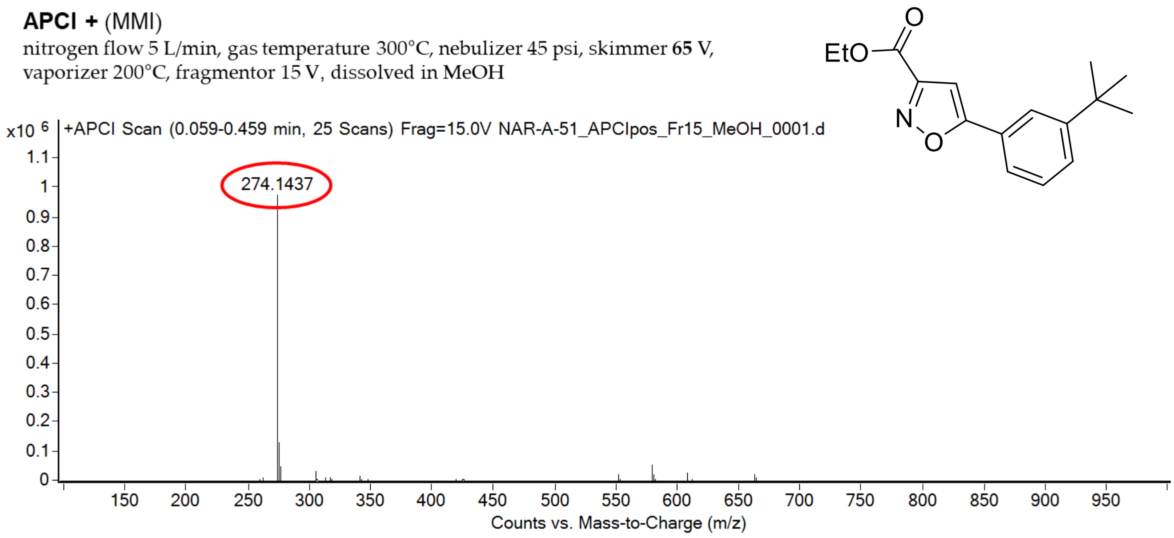

calculated mass: $[M+H]^+$ = 274.1438 observed: $[M+H]^+$ = 274.1437 mass accuracy = -0.4 ppm

FT-IR spectrum (neat) of **S168**

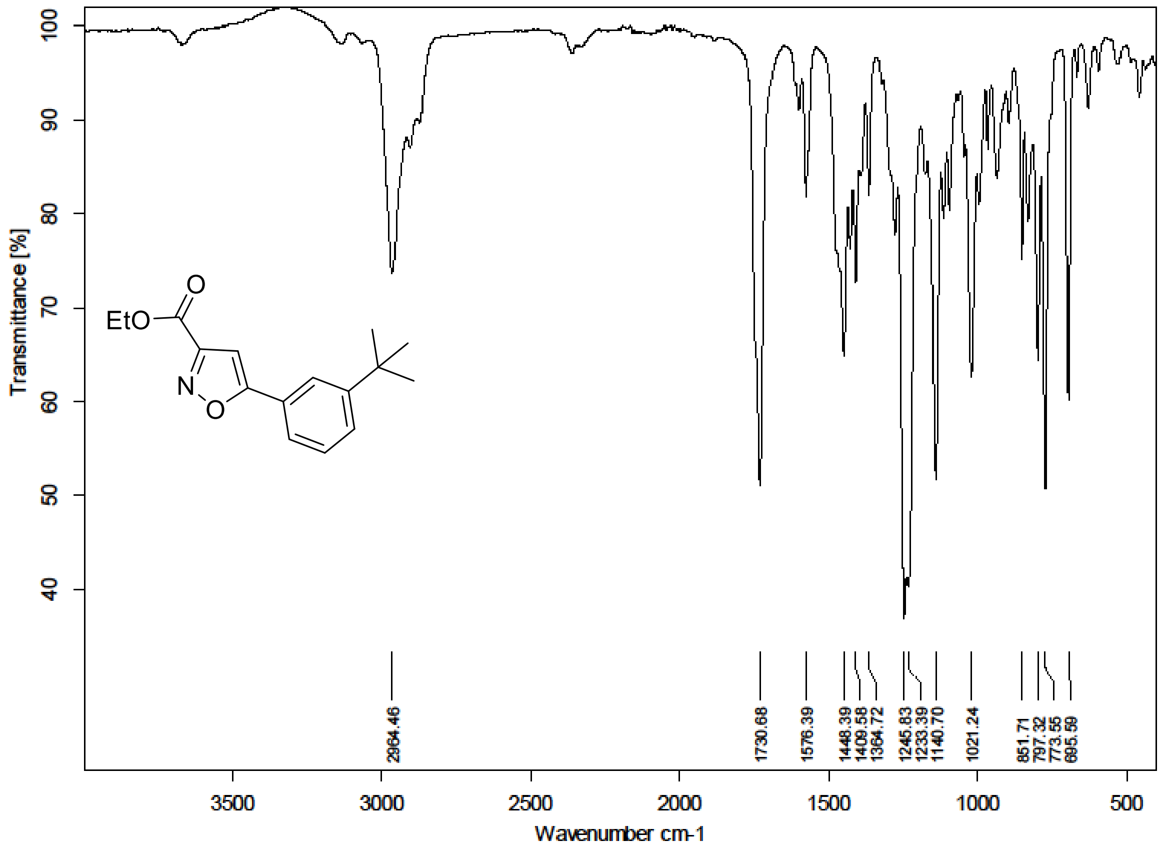

$^{1}H$ (300 MHz) and $^{13}C$ NMR (126 MHz) spectra of **S169** in DMSO-$d_6$

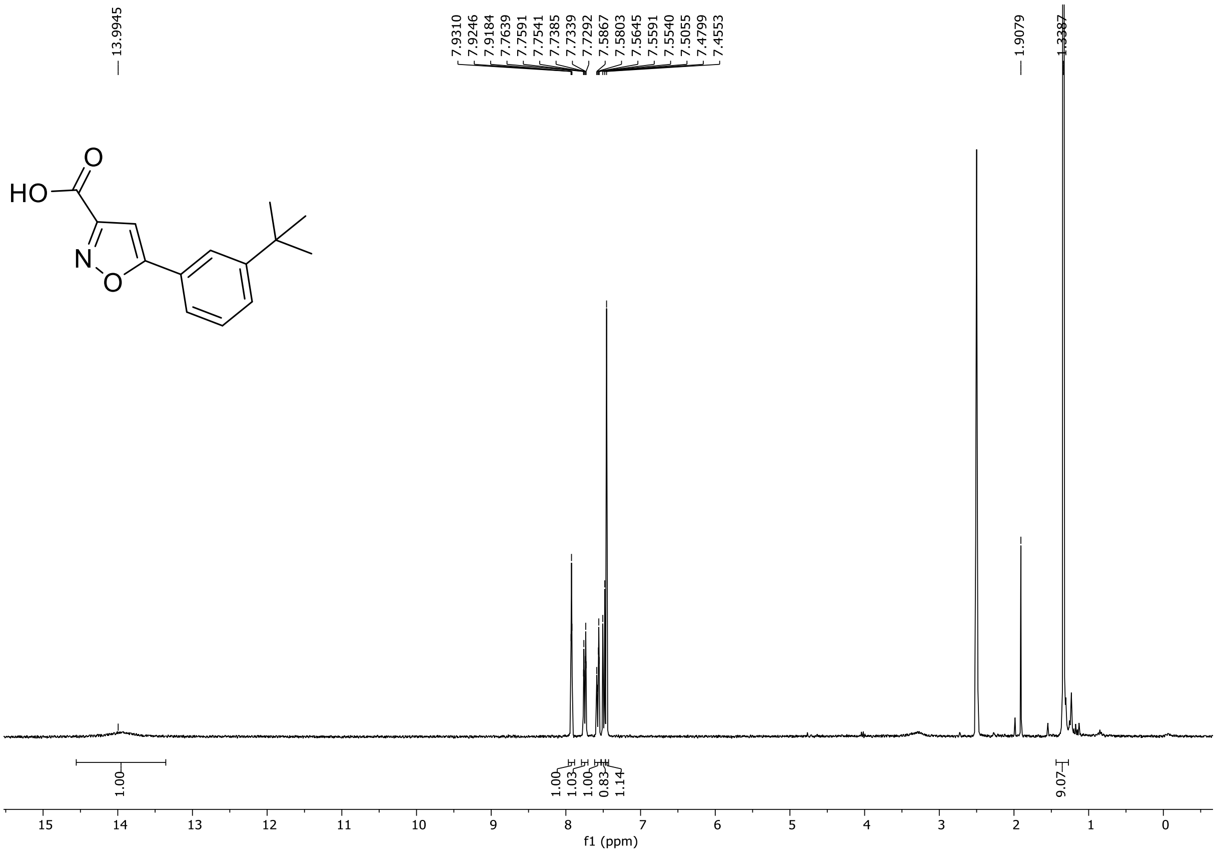

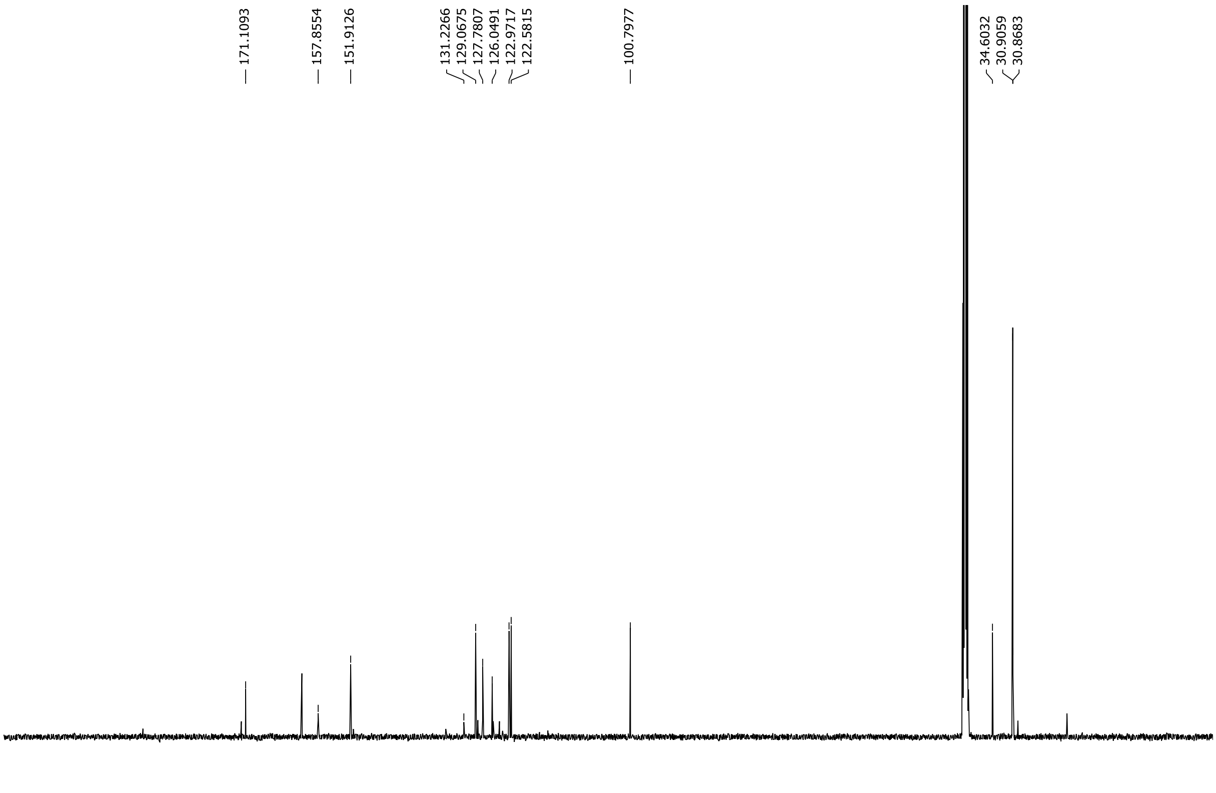

HRMS spectrum of **S169**

**NAR-A-81** $C_{14}H_{15}NO_3$ *mono m/z* **245.1052**

**APCI +** (MMI)
nitrogen flow 5 L/min, gas temperature 300°C, nebulizer 45 psi, skimmer **65** V, vaporizer 200°C, fragmentor 20 V, dissolved in MeOH

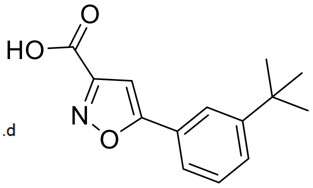

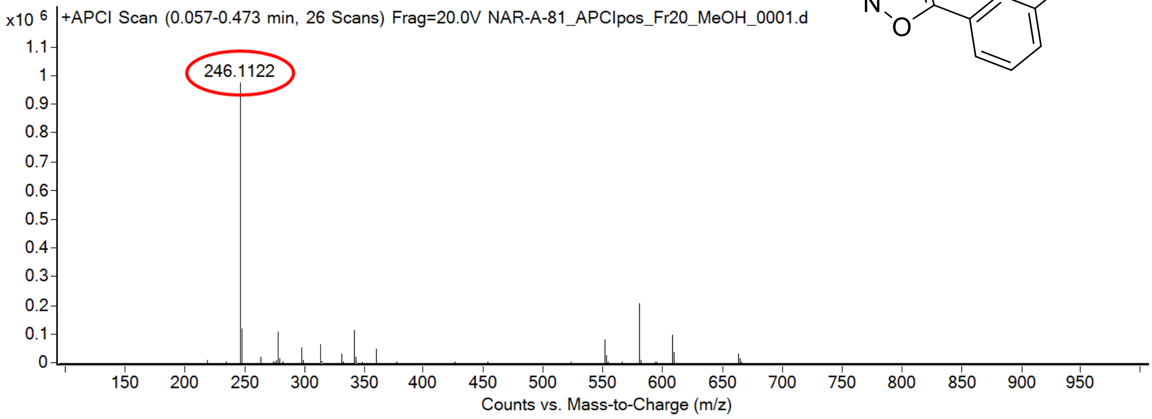

calculated mass: $[M+H]^+$ = 246.1125 observed: $[M+H]^+$ = 246.1122 mass accuracy = -1.2 ppm

FT-IR spectrum (neat) of **S169**

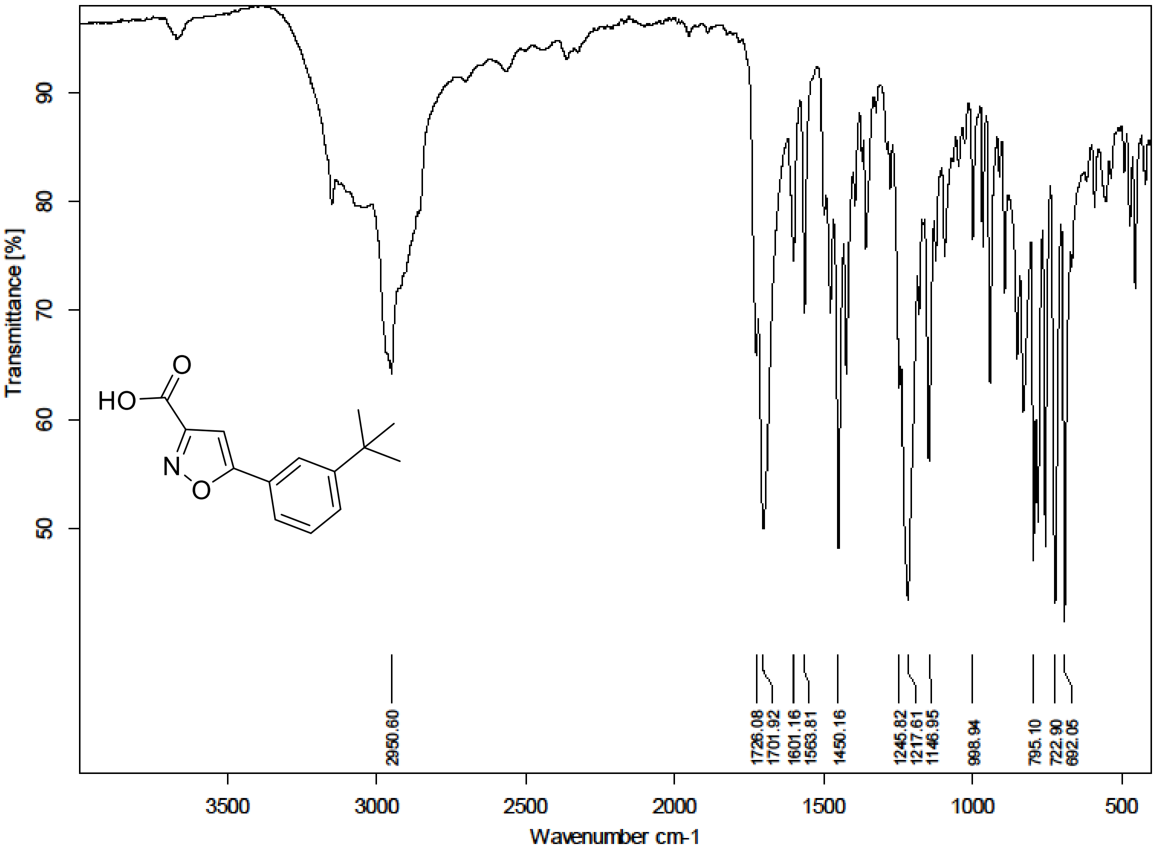

$^{1}H$ (500 MHz) and $^{13}C$ NMR (126 MHz) spectra of **S171** in Chloroform-*d*

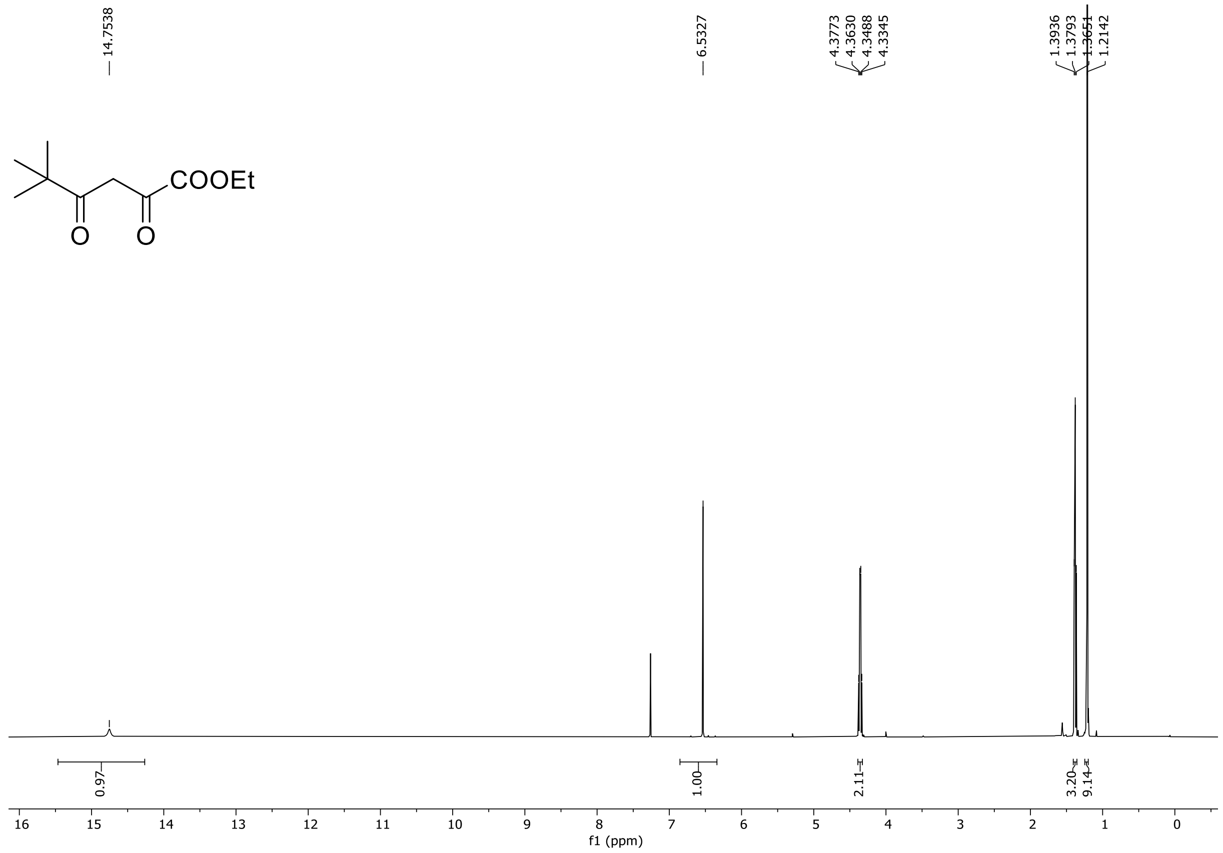

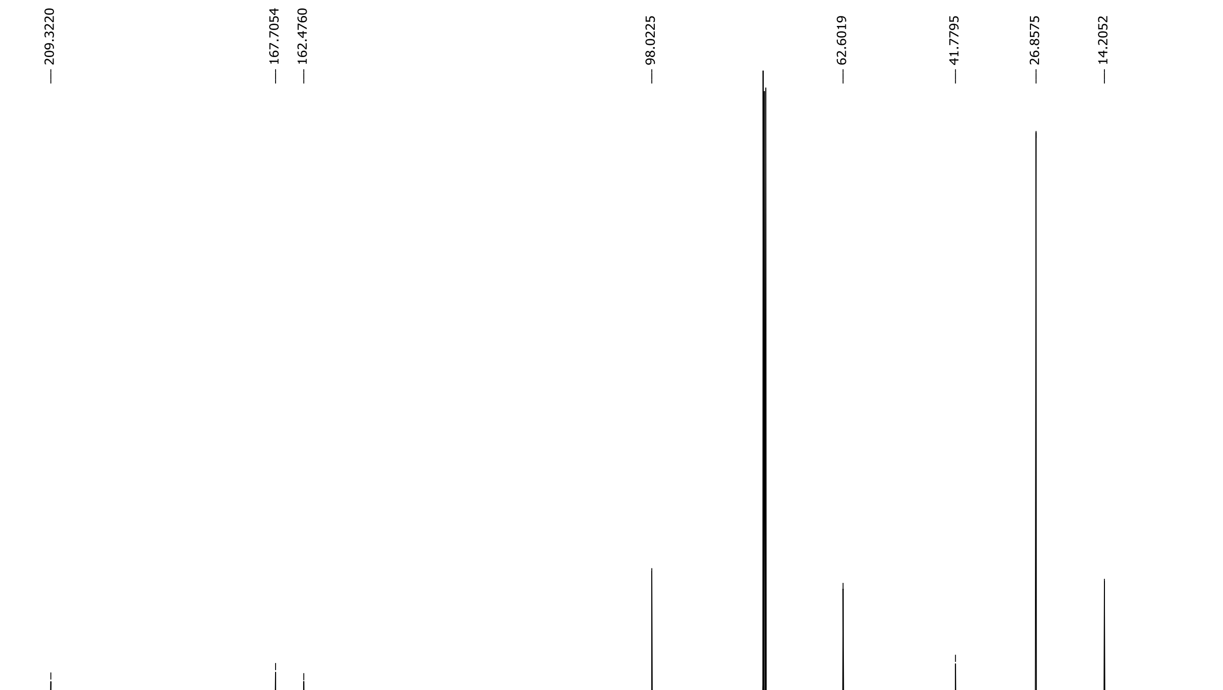

HRMS spectrum of **S171**

NAR-A-18 $C_{10}H_{16}O_4$ mono *m/z 200.1049*

**APCI +** (MMI)
nitrogen flow 5 L/min, gas temperature 325°C, nebulizer 45 psi, skimmer **65** V, vaporizer 200°C, fragmentor 30 V, dissolved in methanol

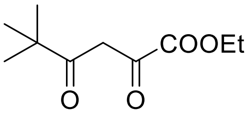

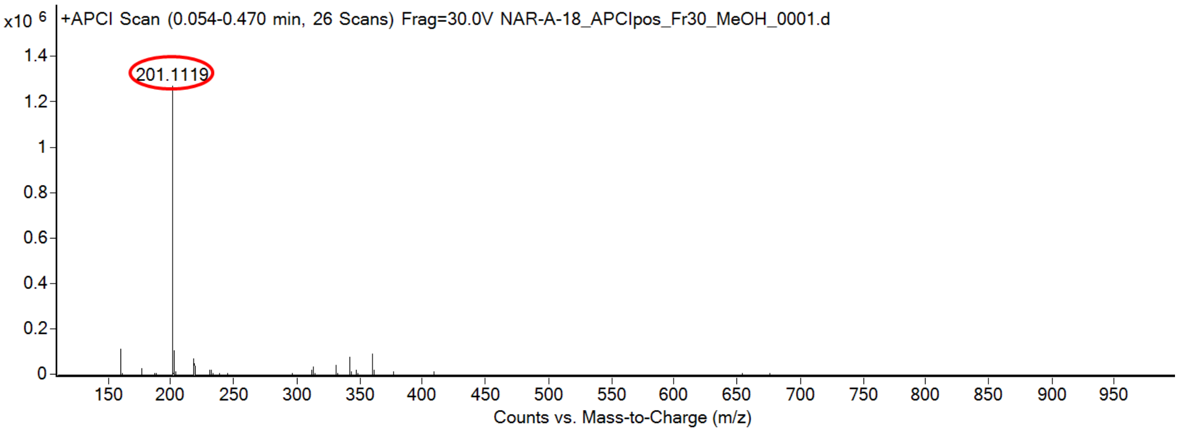

calculated mass: $[M+H]^+$ = 201.1121 observed: $[M+H]^+$ = 201.1119 mass accuracy = -1.0 ppm

FT-IR spectrum (neat) of **S171**

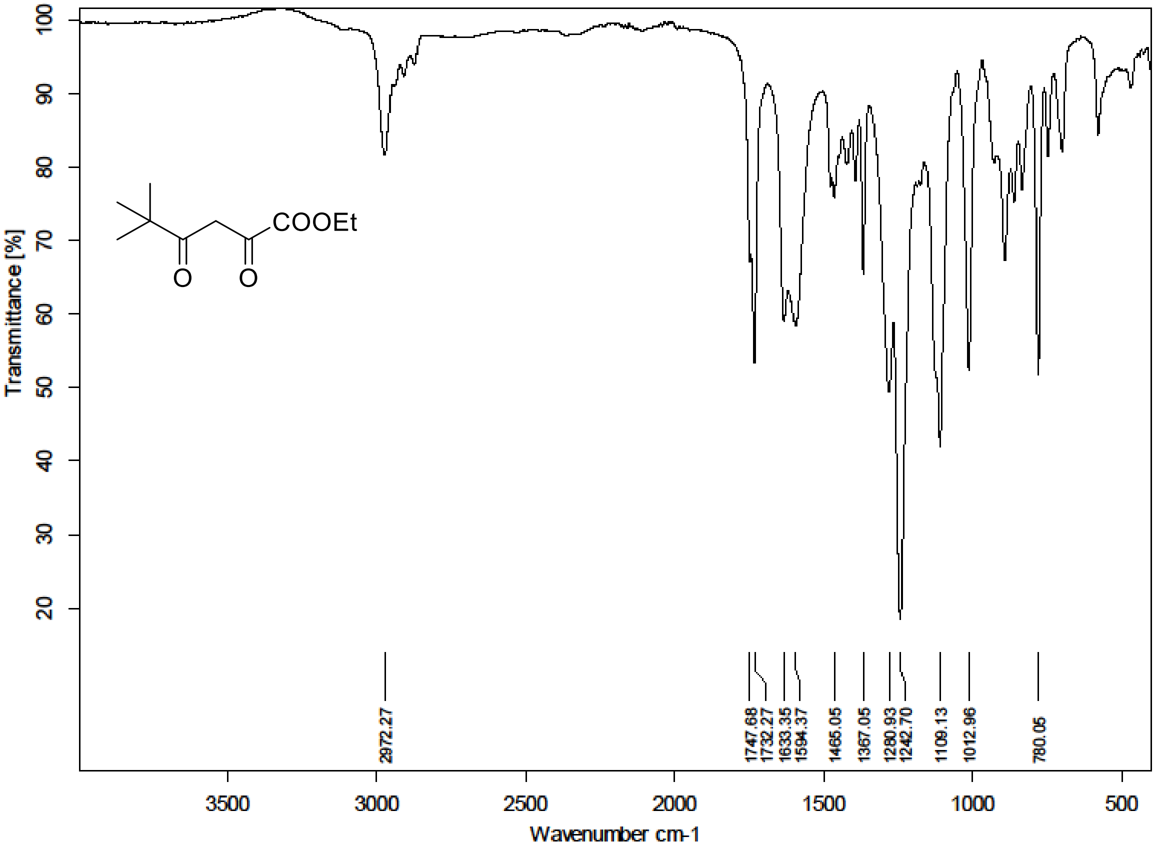

[1]H (500 MHz) and [13]C NMR (126 MHz) spectra of **S172** in Chloroform-*d*

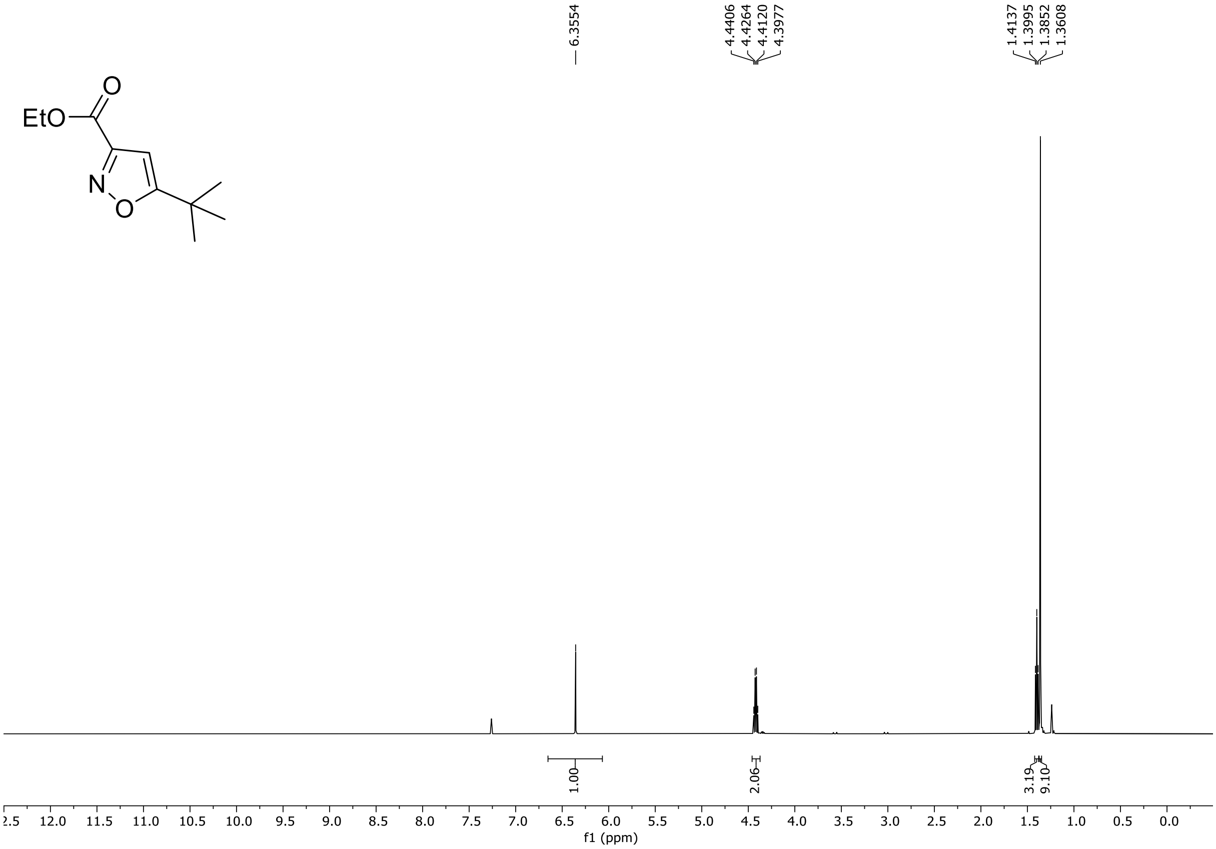

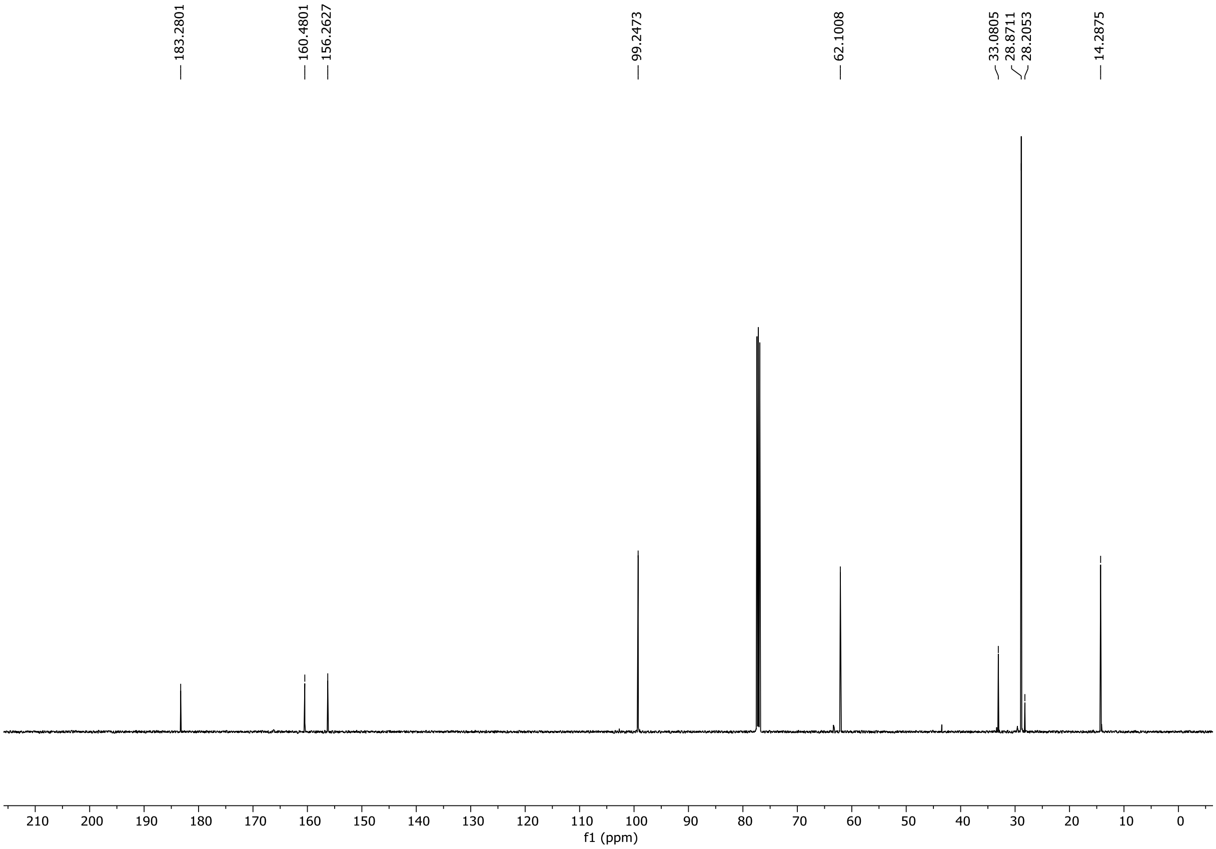

HRMS spectrum of **S172**

NAR-A-24 $C_{10}H_{15}NO_3$ **mono *m/z 197.1052***

**APCI +** (MMI)
nitrogen flow 5 L/min, gas temperature 325°C, nebulizer 45 psi, skimmer **65** V, vaporizer 200°C, fragmentor 23 V, dissolved in methanol

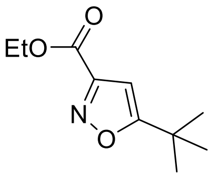

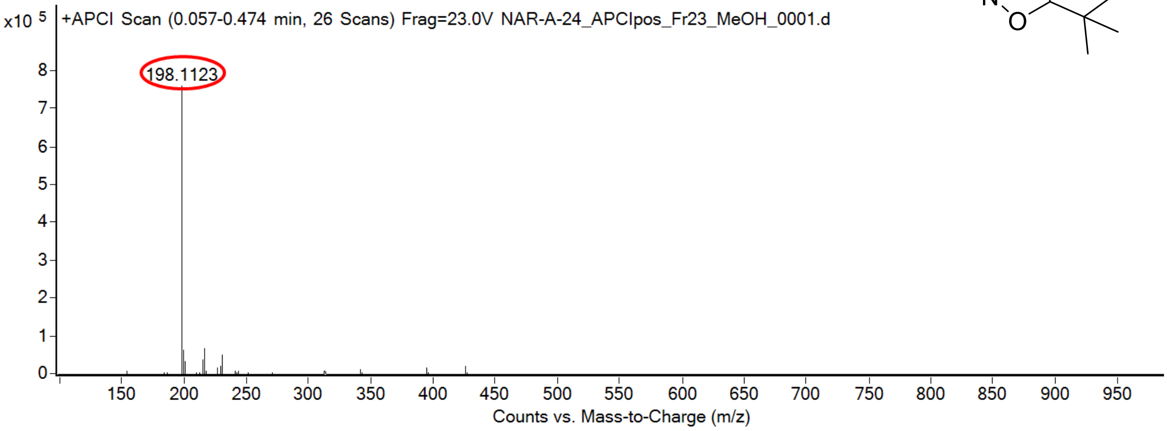

calculated mass: $[M+H]^+$ = 198.1125 observed: $[M+H]^+$ = 198.1123 mass accuracy = -1.0 ppm

FT-IR spectrum (neat) of **S172**

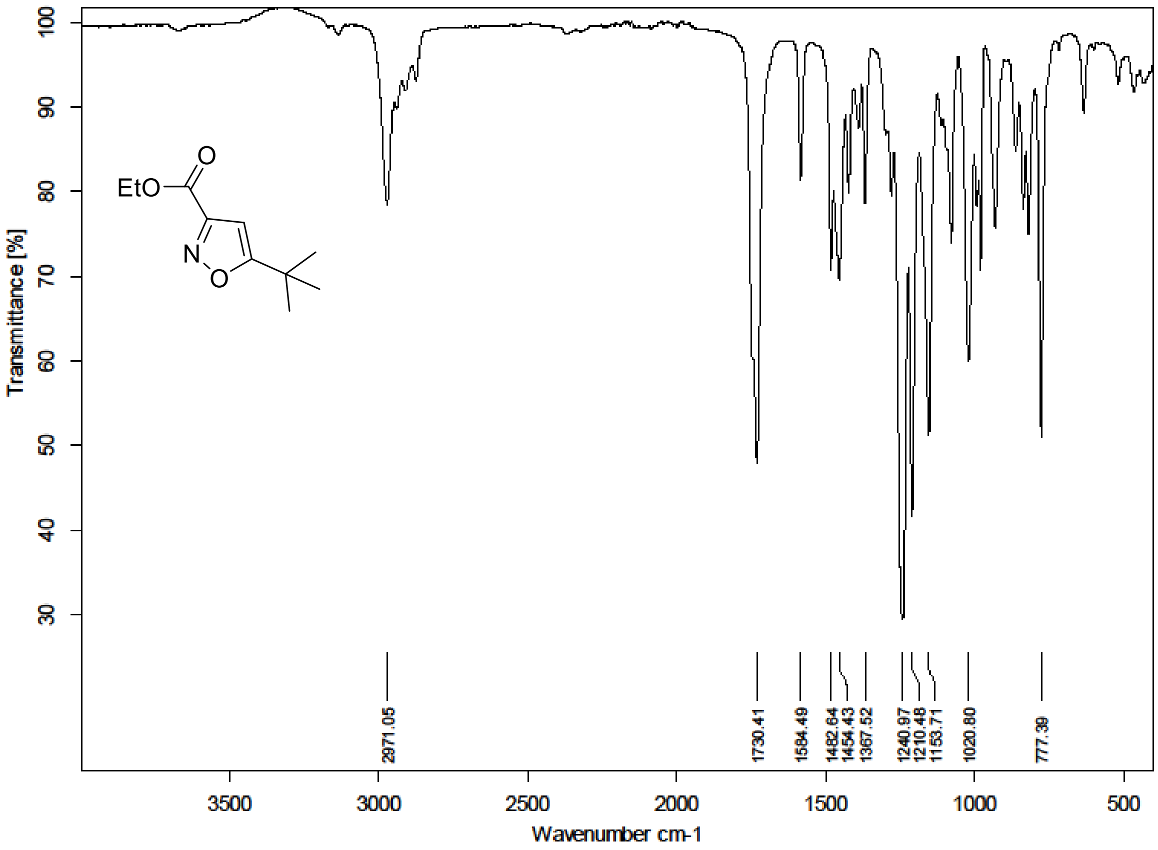

$^{1}H$ (500 MHz) and $^{13}C$ NMR (126 MHz) spectra of **S173** in Chloroform-*d*

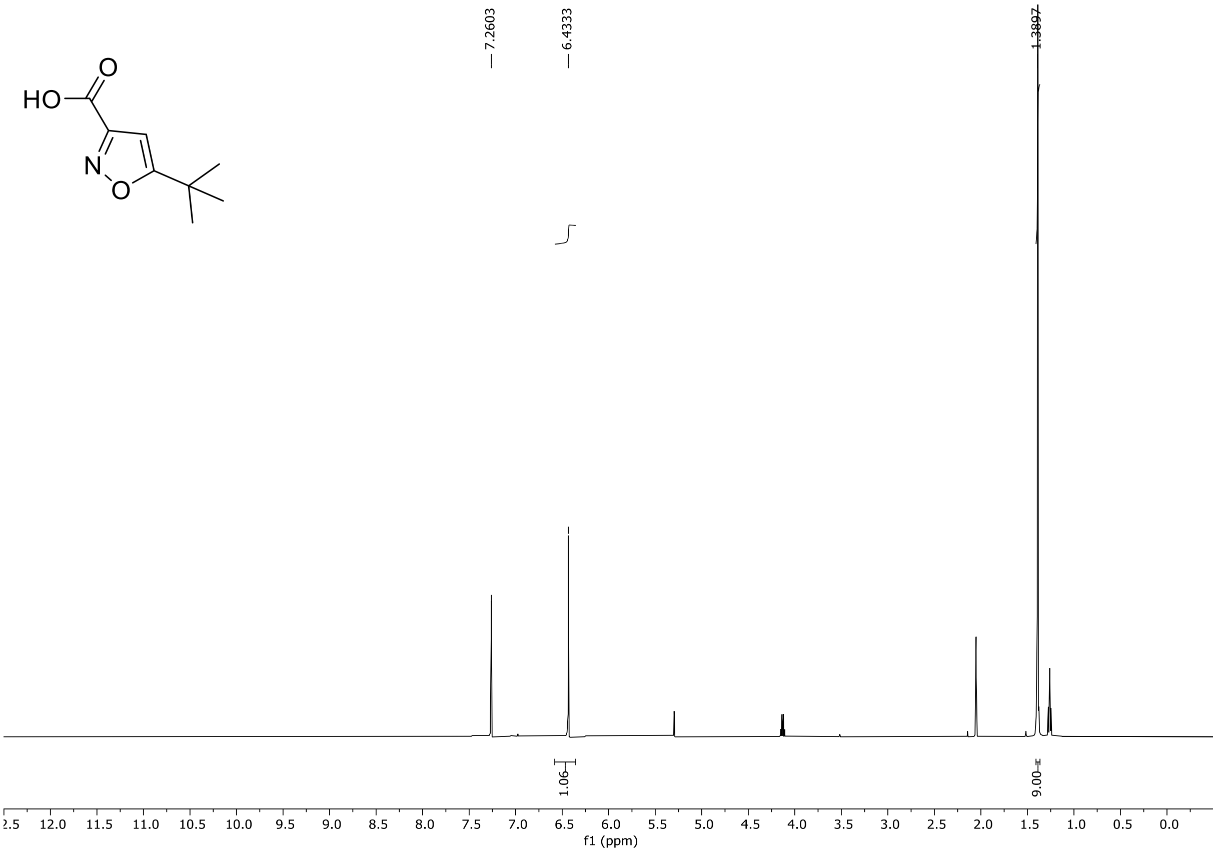

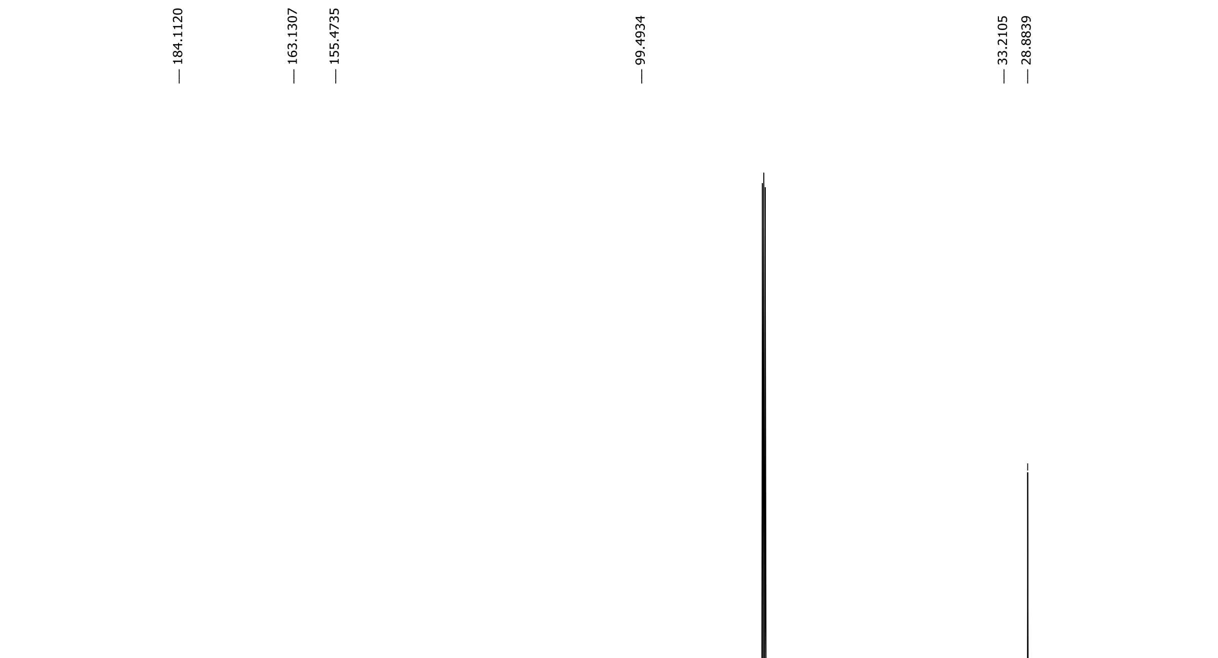

HRMS spectrum of **S173**

NAR-A-29 $C_8H_{11}NO_3$ mono *m/z 169.0739*

**APCI +** (MMI)
nitrogen flow 5 L/min, gas temperature 325°C, nebulizer 45 psi, skimmer **65** V, vaporizer 200°C, fragmentor 30 V, dissolved in methanol

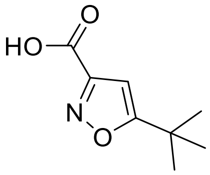

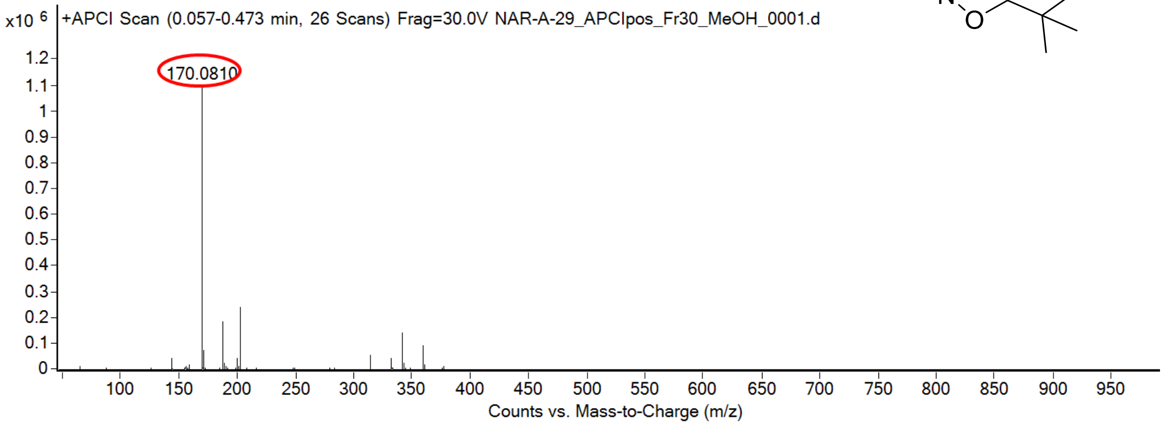

calculated mass: $[M+H]^+$ = 170.0812 observed: $[M+H]^+$ = 170.0810 mass accuracy = -1.2 ppm

FT-IR spectrum (neat) of **S173**

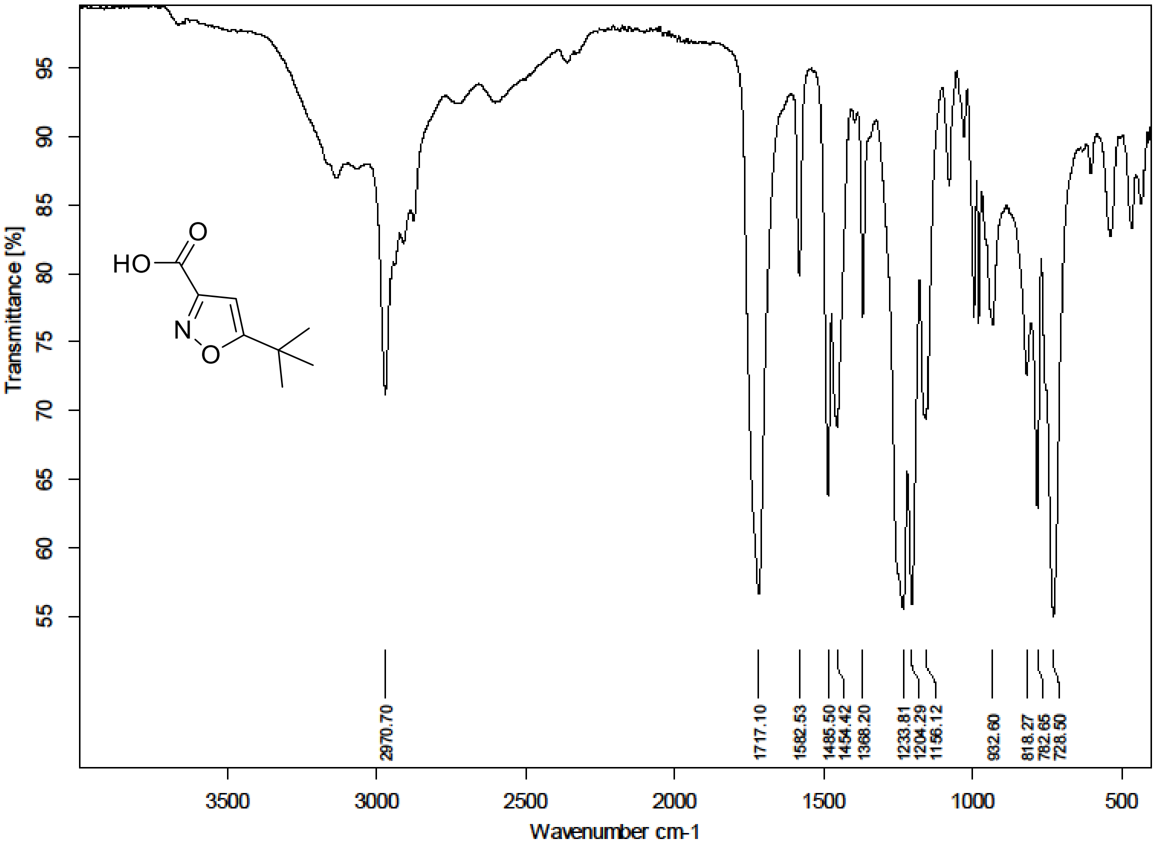

$^{1}H$ (500 MHz) and $^{13}C$ NMR (126 MHz) spectra of **S174** in Chloroform-*d*

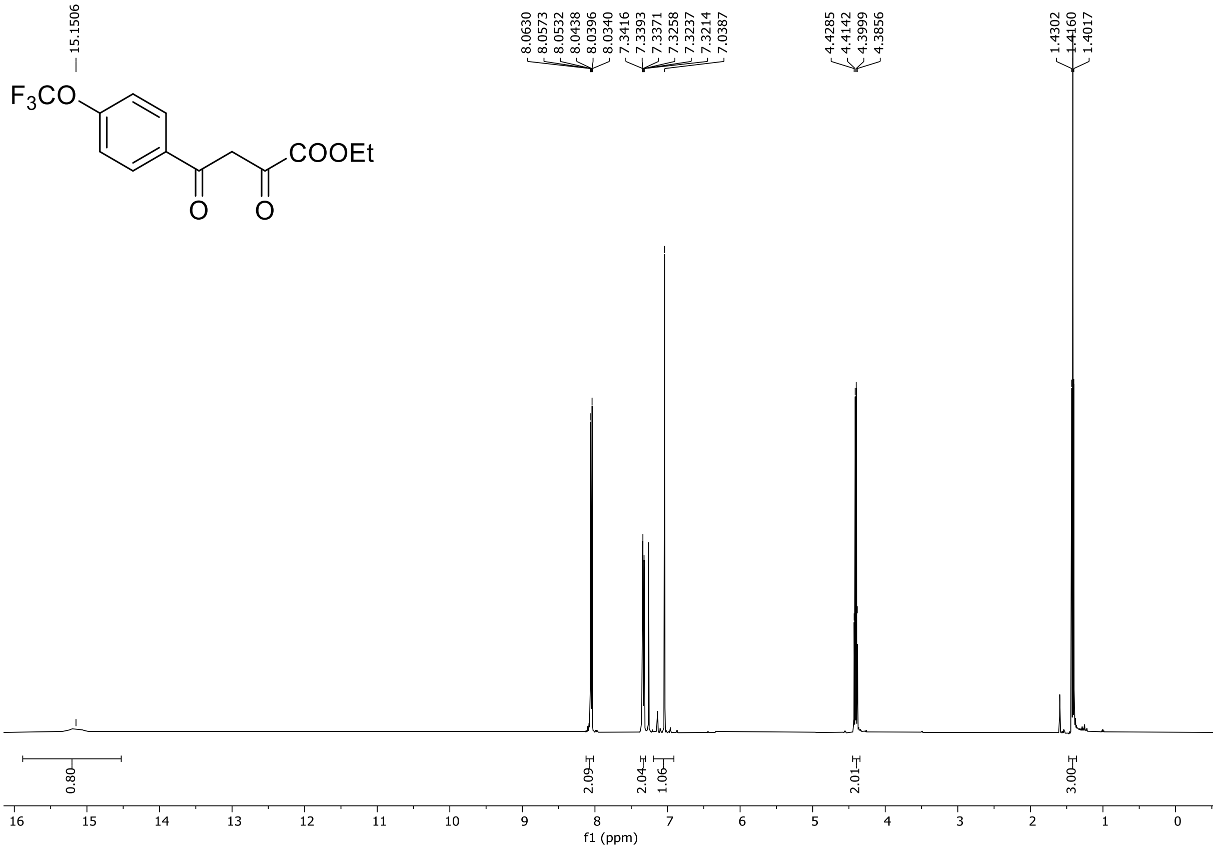

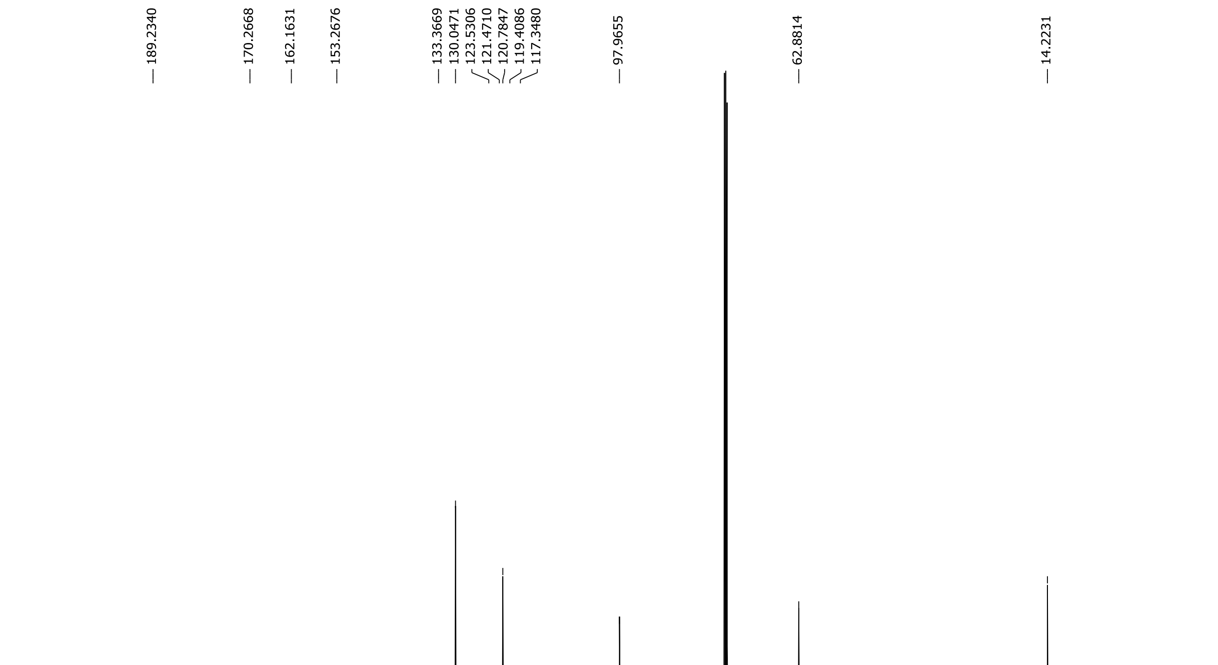

$^{19}$F NMR (282 MHz) spectrum of **S174** in Chloroform-*d*

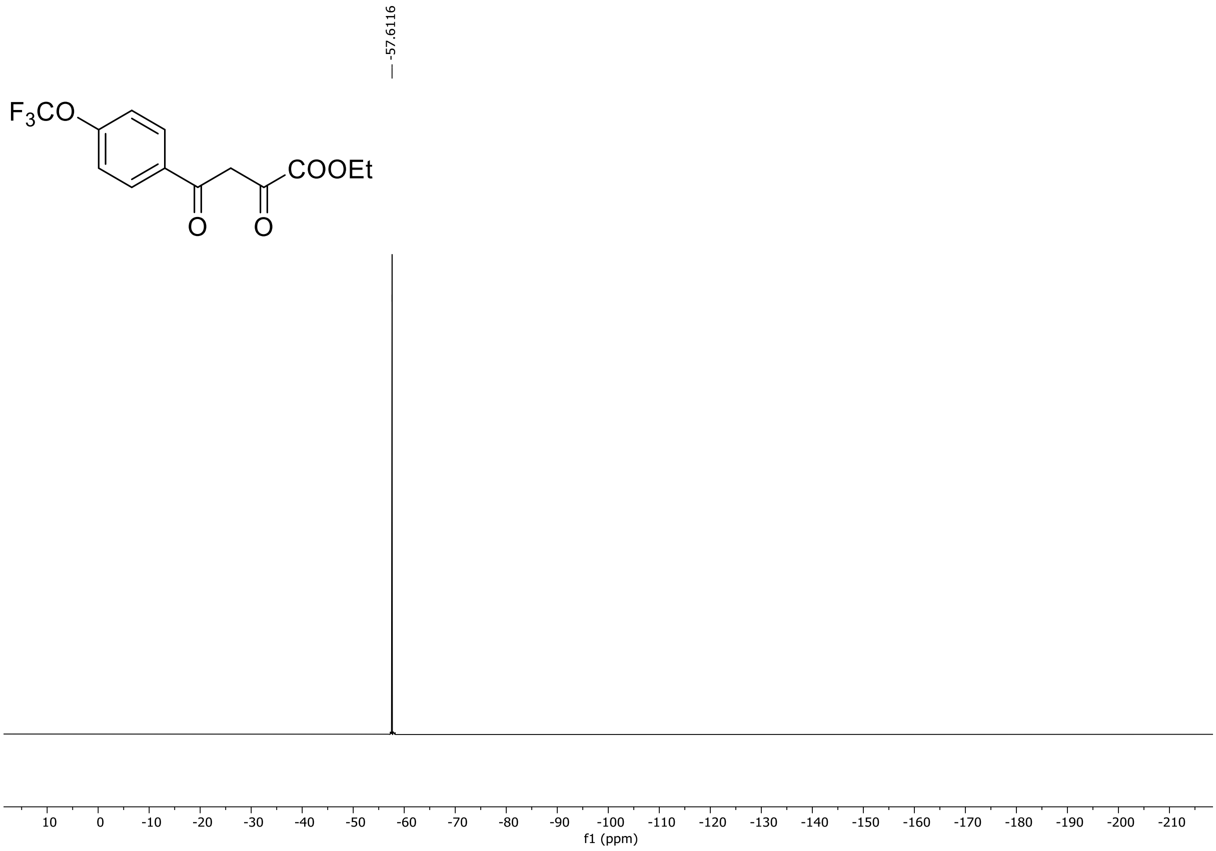

HRMS spectrum of **S174**

**NAR-A-49** **$C_{13}H_{11}F_3O_5$ 304.0559** *mono m/z*

**APCI +** (MMI)
nitrogen flow 5 L/min, gas temperature 300°C, nebulizer 45 psi, skimmer **65** V, vaporizer 200°C, fragmentor 20 V, dissolved in MeOH

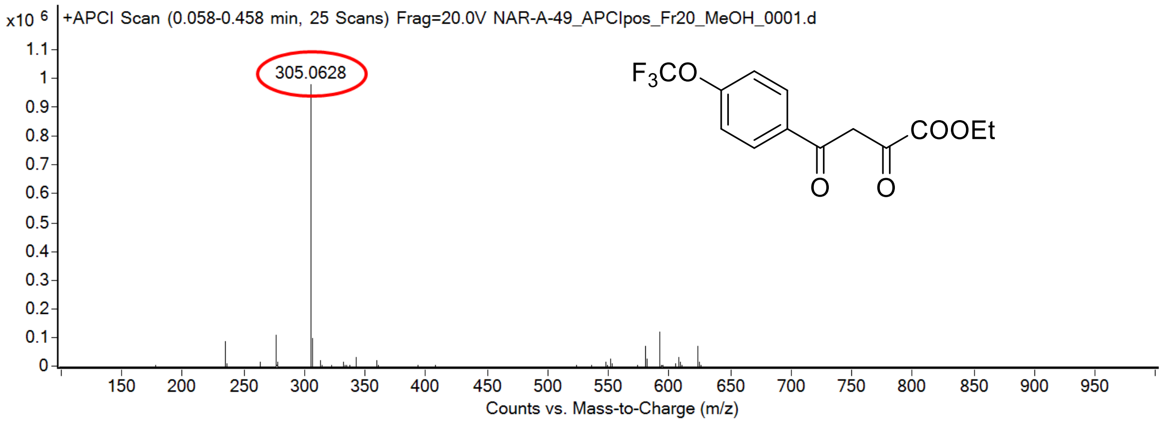

calculated mass: [M+H]$^+$ = 305.0631 observed: [M+H]$^+$ = 305.0628 mass accuracy = -1.0 ppm

FT-IR spectrum (neat) of **S174**

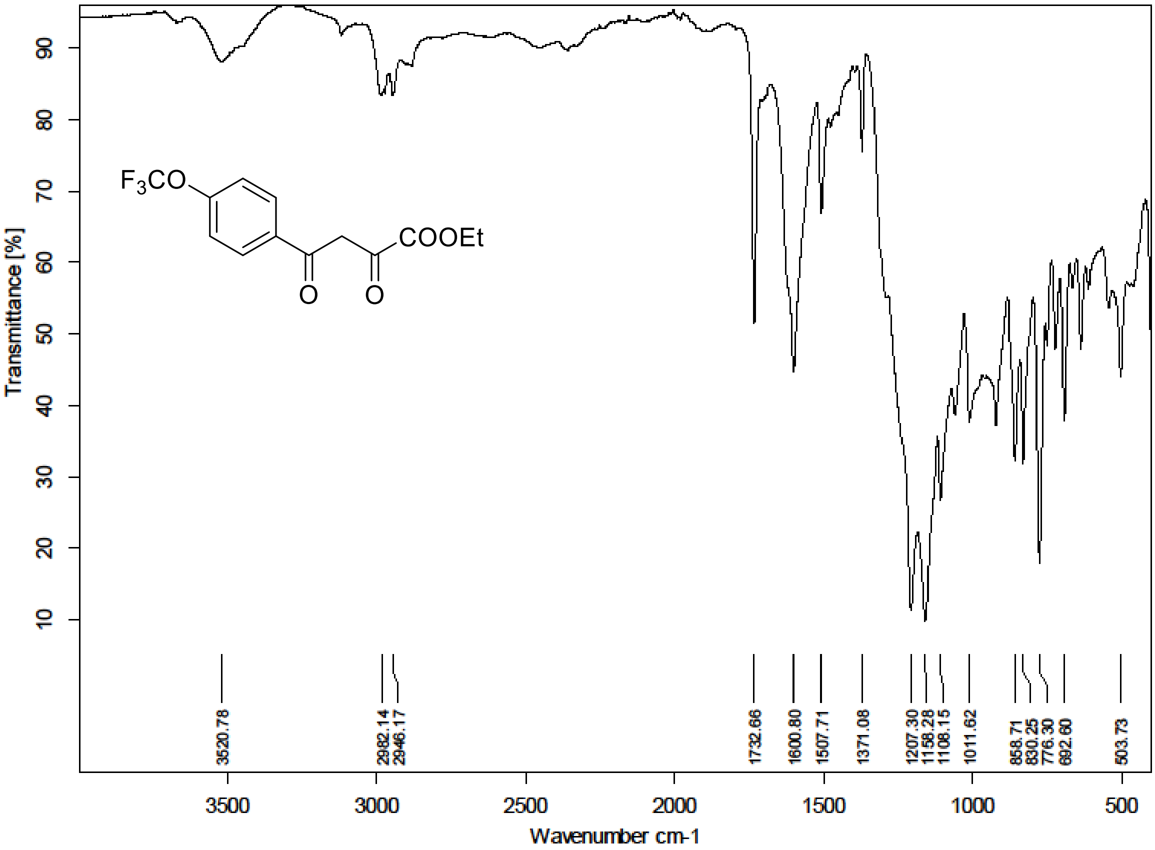

$^{1}H$ (500 MHz) and $^{13}C$ NMR (126 MHz) spectra of **S175** in Chloroform-*d*

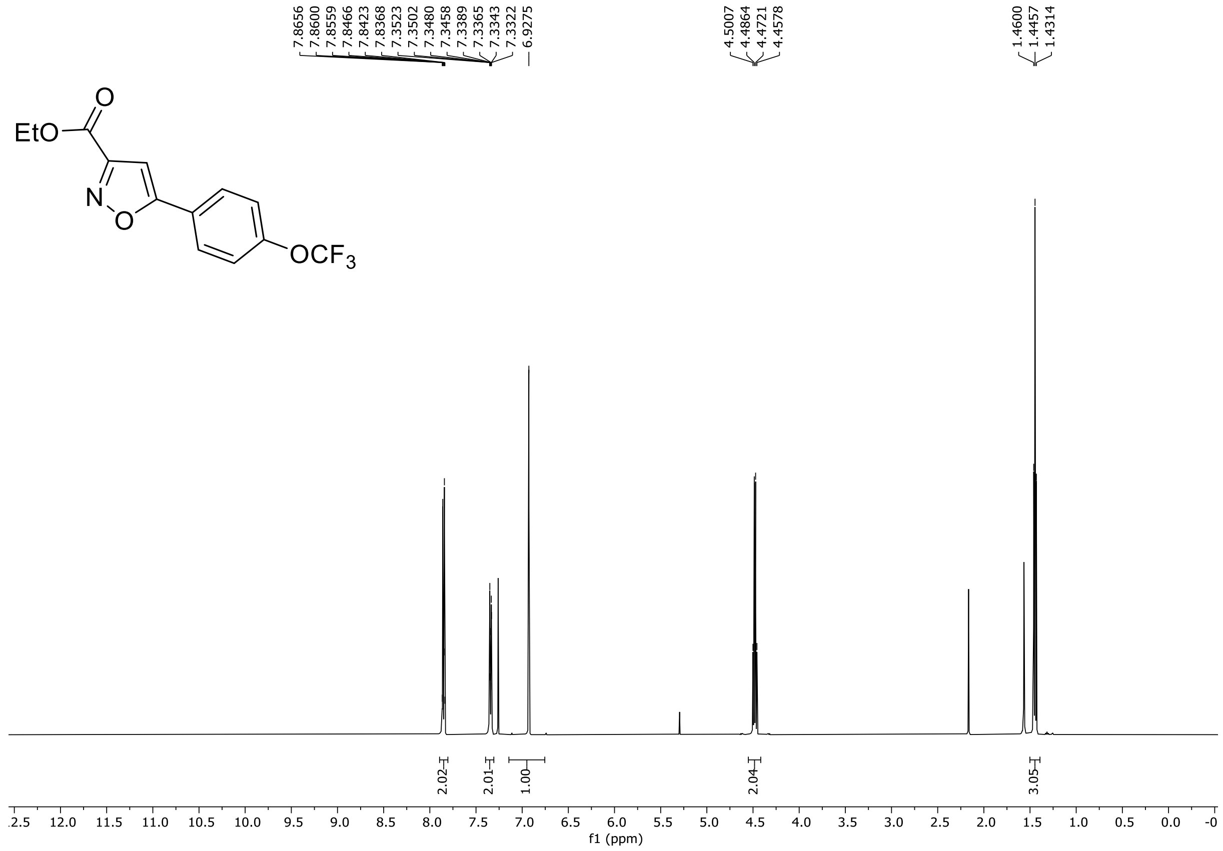

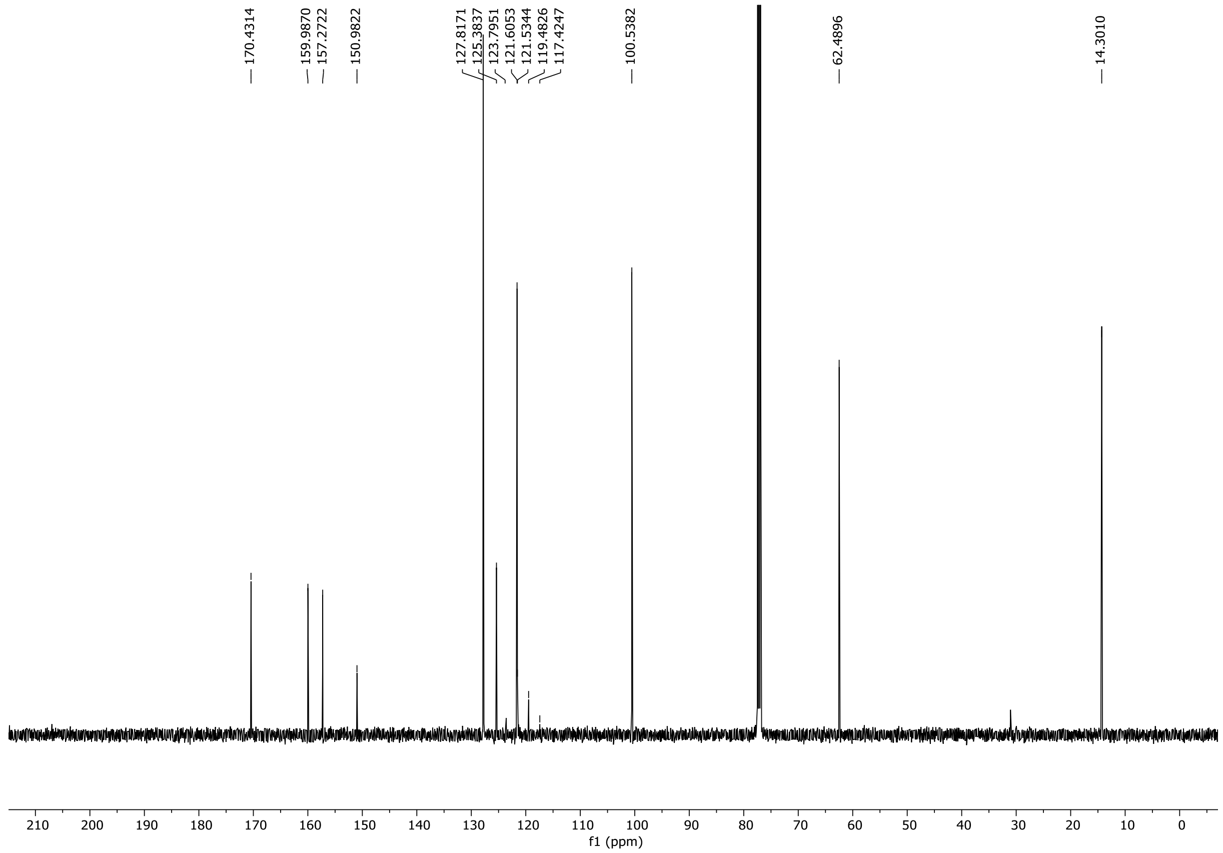

[19]F NMR (471 MHz) spectrum of **S175** in Chloroform-*d*

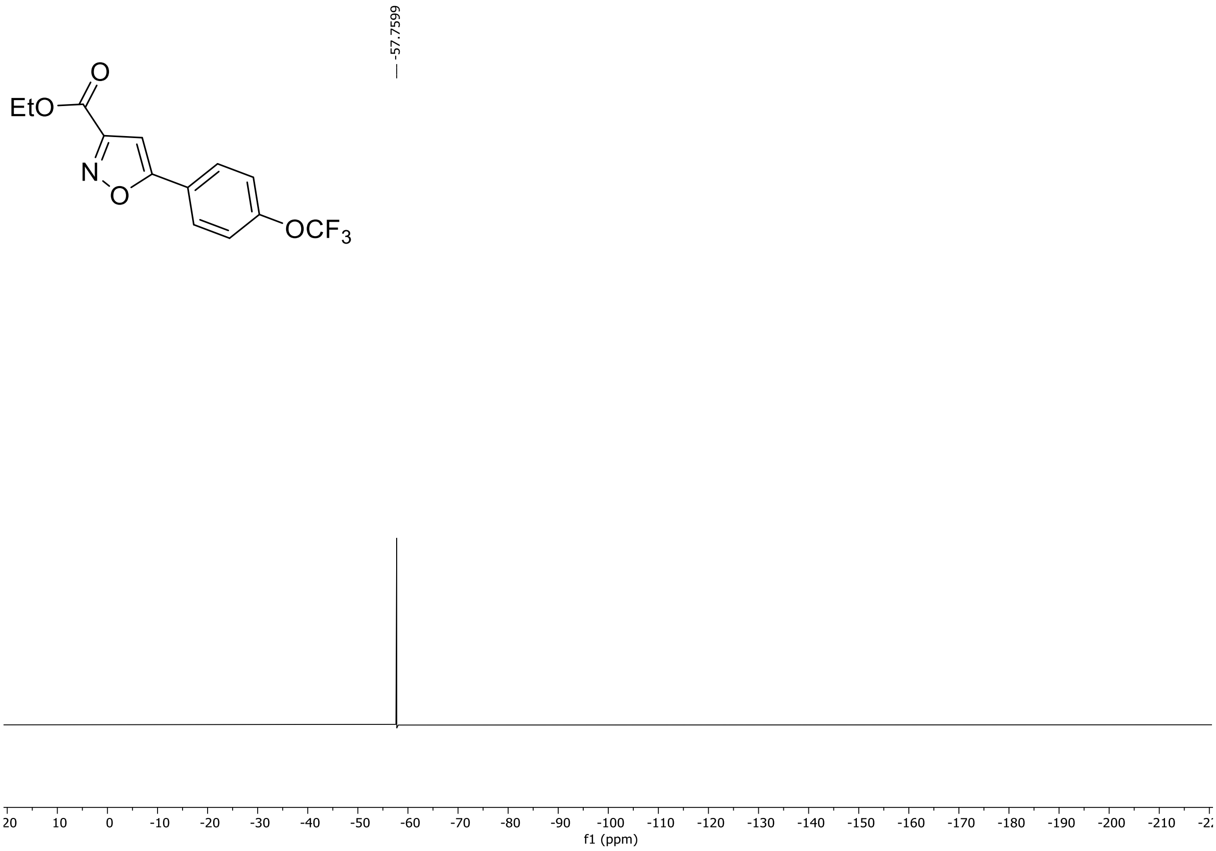

HRMS spectrum of **S175**

**NAR-A-52** $C_{13}H_{10}F_3NO_4$ 301.0562 *mono m/z*

**APCI +** (MMI)
nitrogen flow 5 L/min, gas temperature 300°C, nebulizer 45 psi, skimmer **65** V, vaporizer 200°C, fragmentor 25 V, dissolved in MeOH

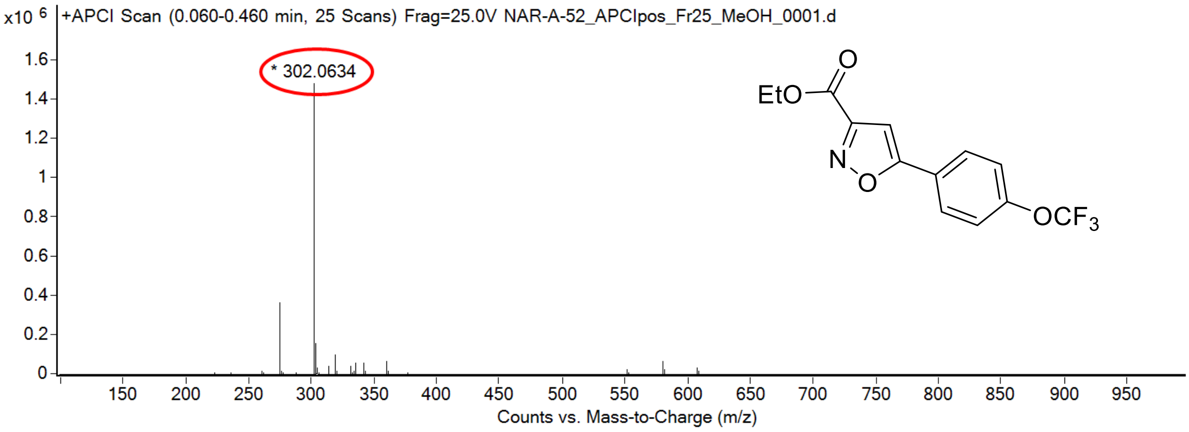

calculated mass: $[M+H]^+$ = 302.0635 observed: $[M+H]^+$ = 302.0634 mass accuracy = -0.3 ppm

FT-IR spectrum (neat) of **S175**

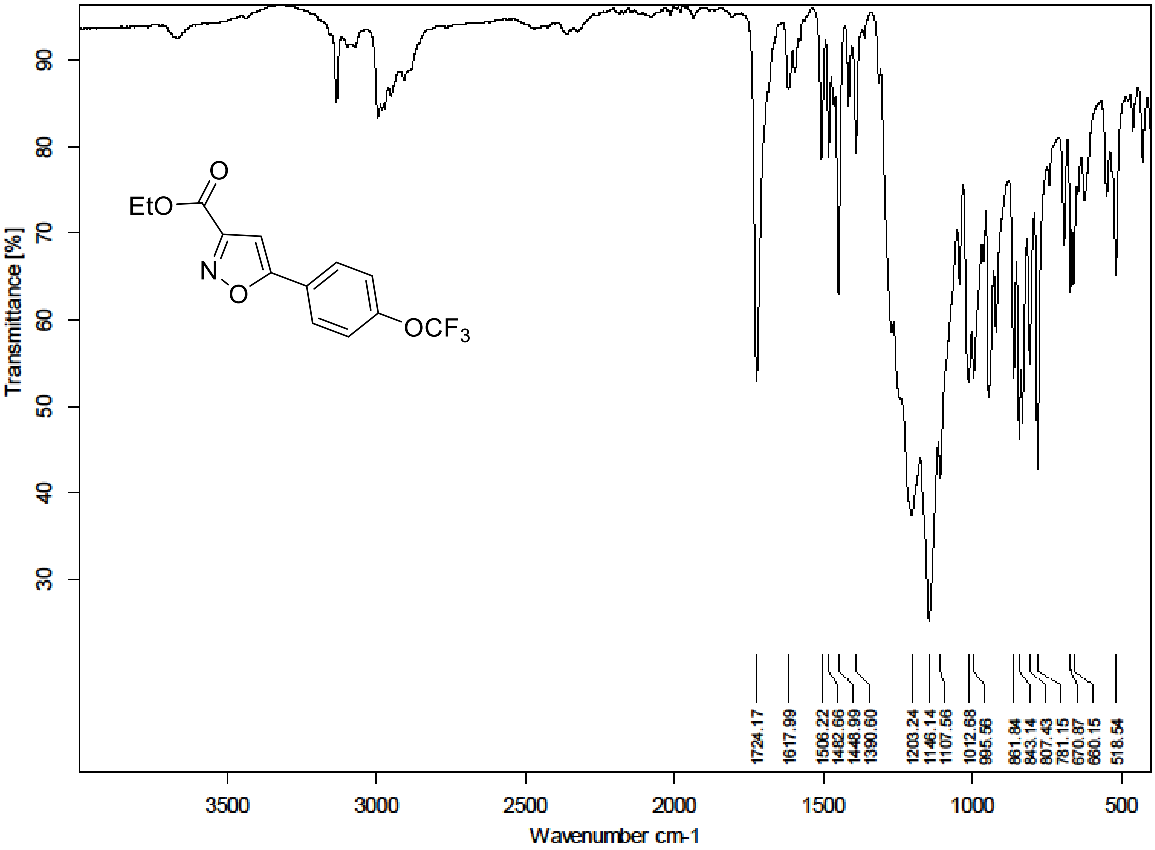

[1]H (500 MHz) and [13]C NMR (126 MHz) spectra of **S176** in DMSO-$d_6$

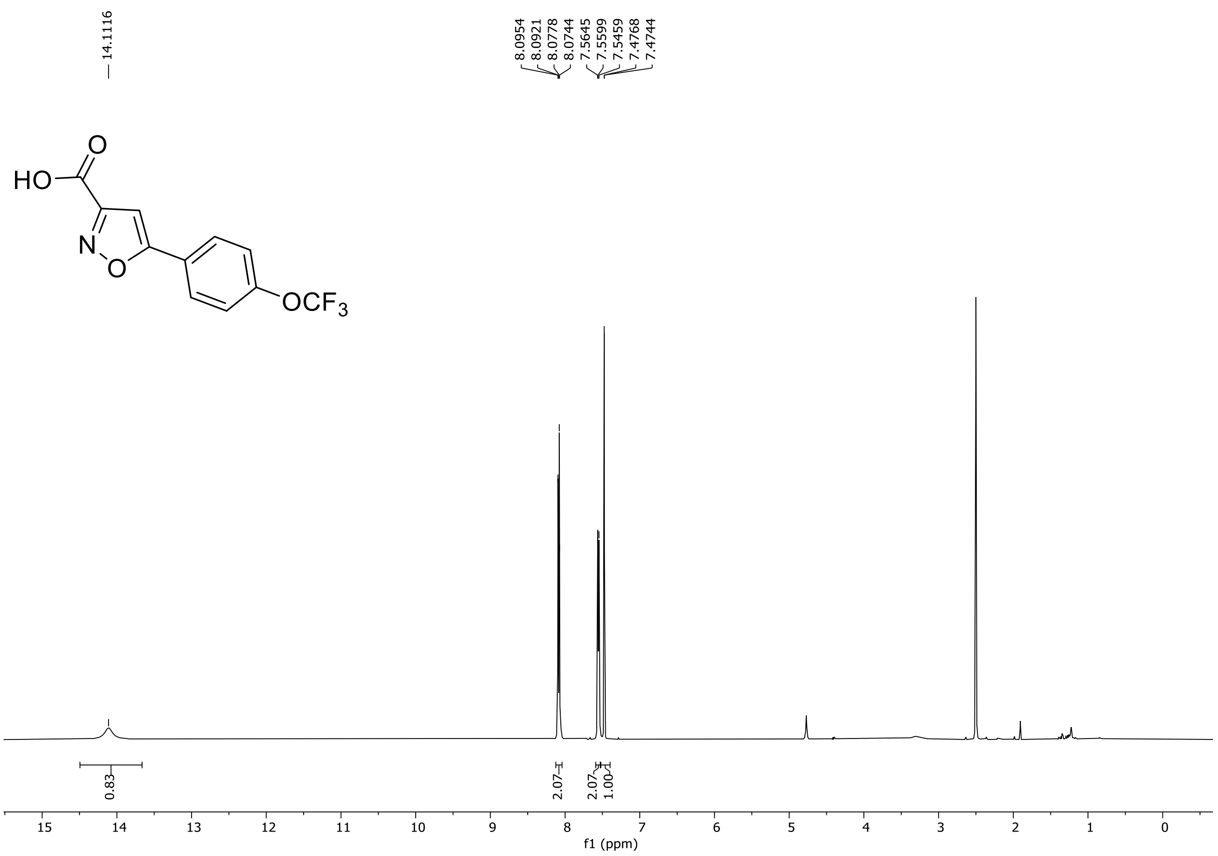

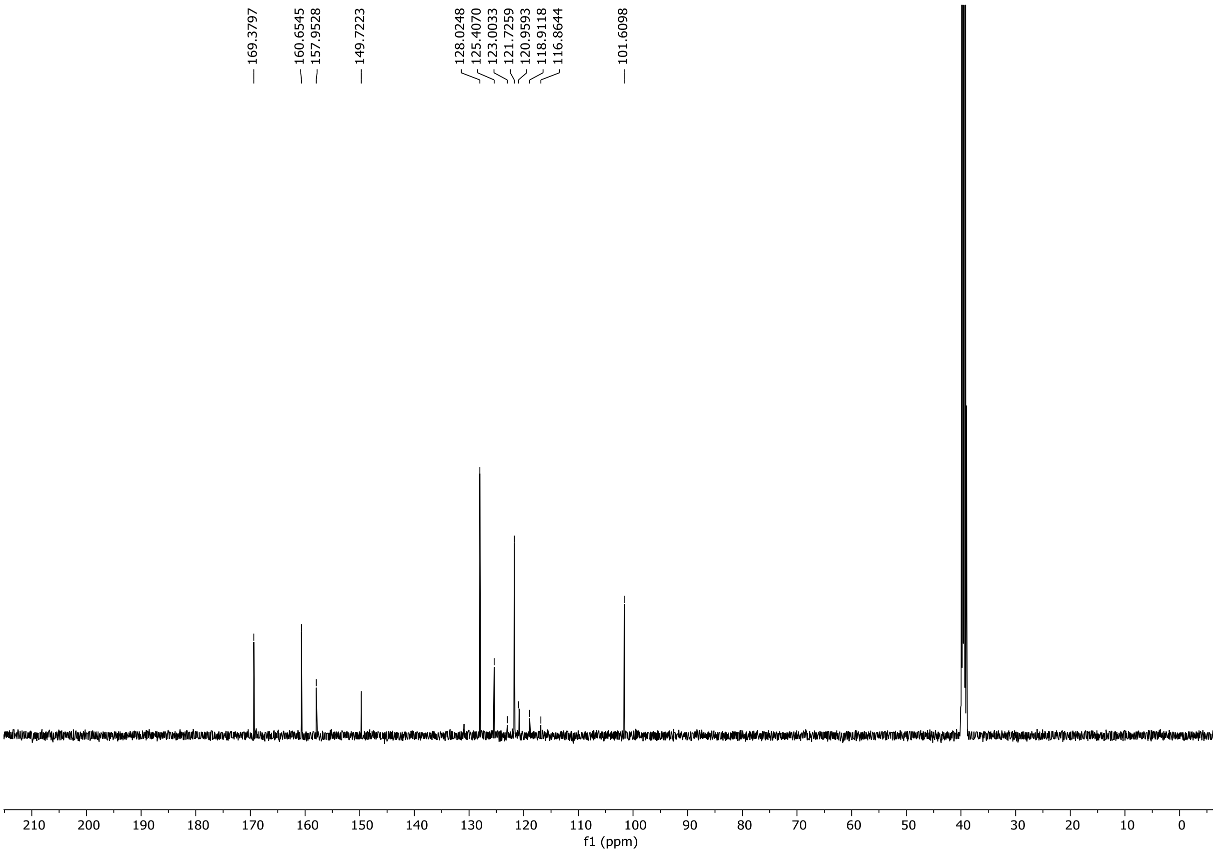

$^{19}F$ NMR (471 MHz) spectrum of **S176** in DMSO-$d_6$

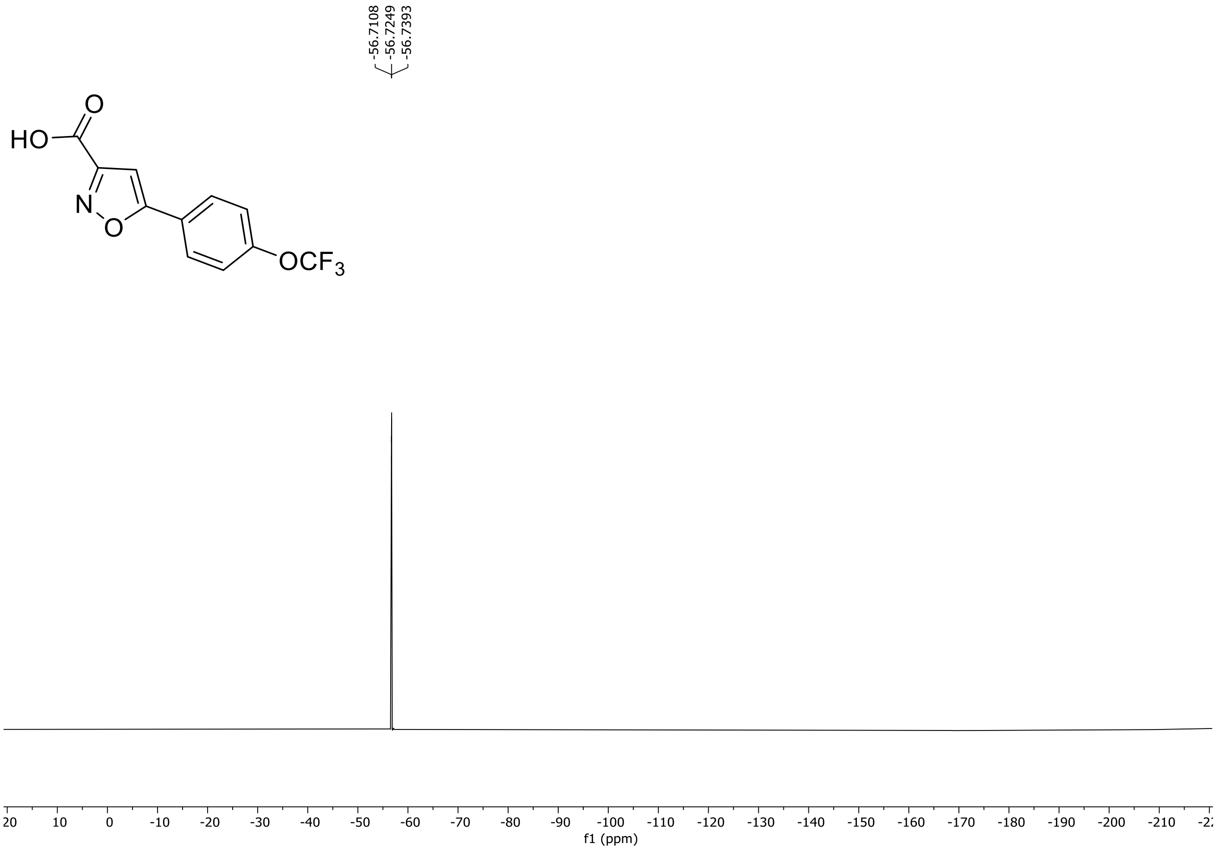

HRMS spectrum of **S176**

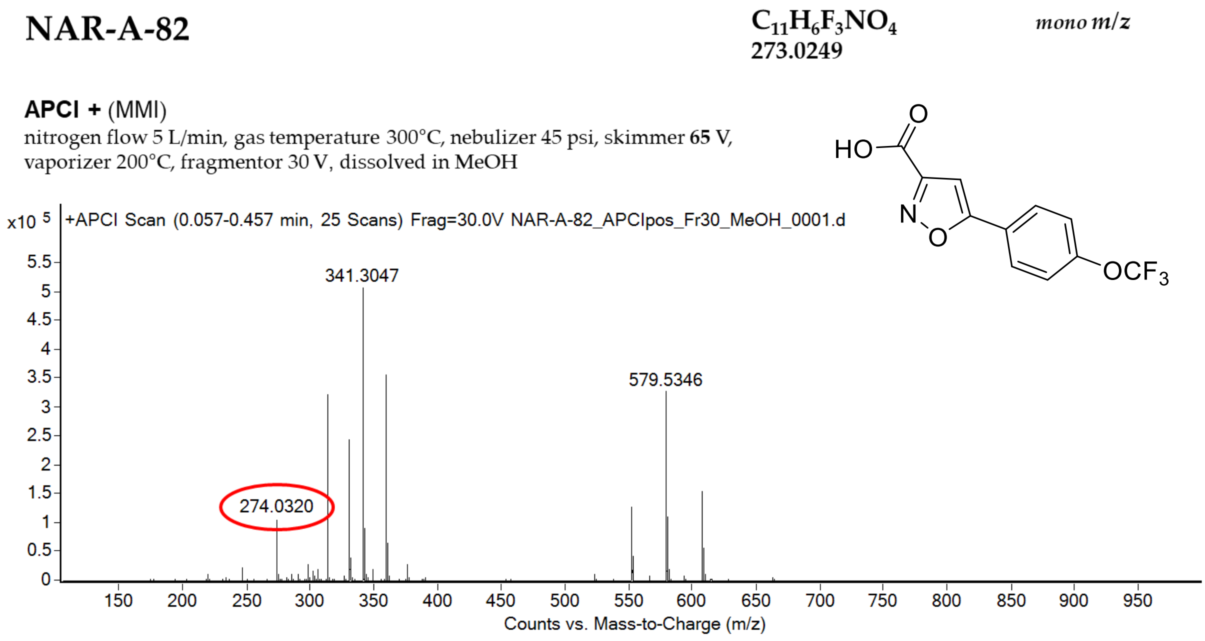

calculated mass: $[M+H]^+$ = 274.0322 observed: $[M+H]^+$ = 274.0320 mass accuracy = -0.7 ppm

FT-IR spectrum (neat) of **S176**

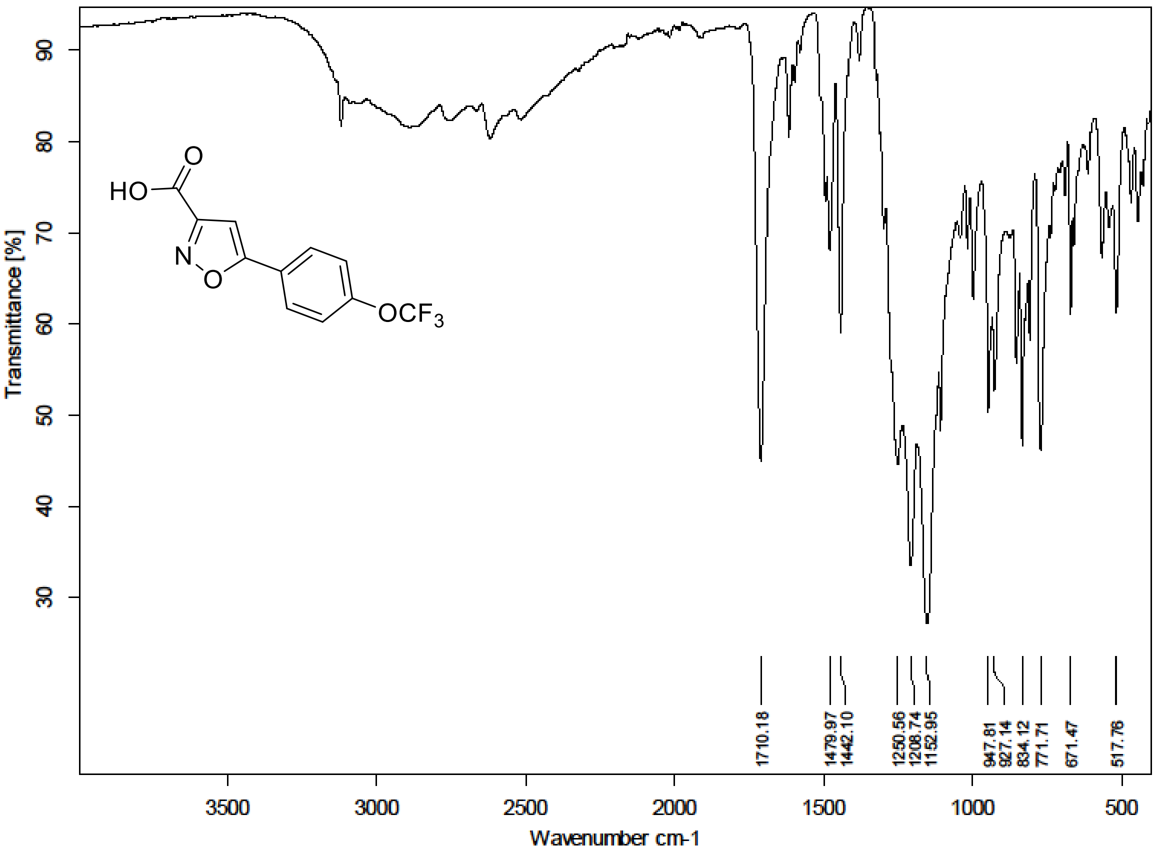

[1]H (300 MHz) and [13]C NMR (75 MHz) spectra of **S185** in DMSO-$d_6$

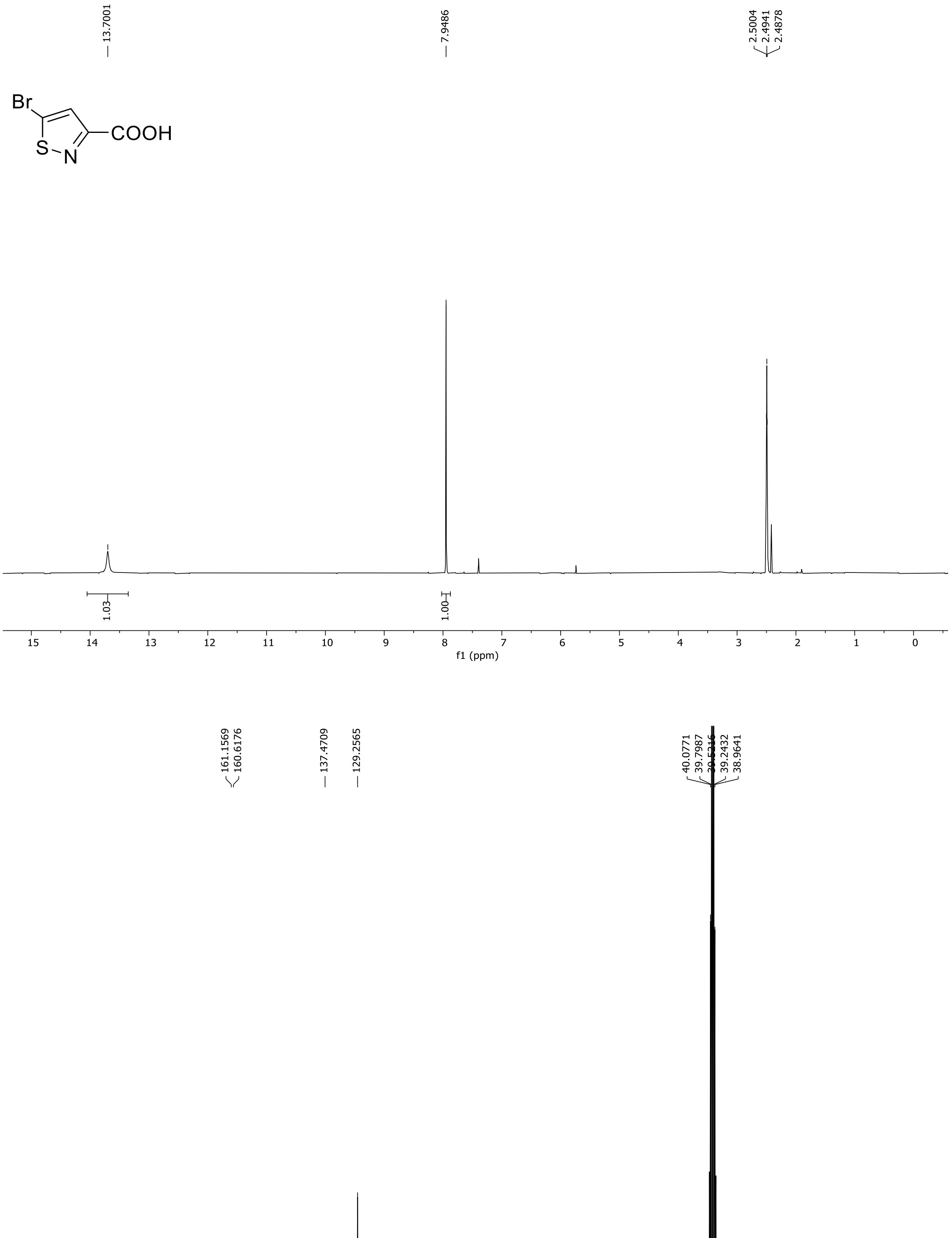

HRMS spectrum of **S185**

**NAR-A-136** $C_4H_2BrNO_2S$ $m/z_{100\%}$ **208.8969**

**APCI-** (MMI)
nitrogen flow 5 L/min, gas temperature 325°C, nebulizer 45 psi, skimmer 65 V,
vaporizer 200°C, fragmentor 90 V, dissolved in MeOH

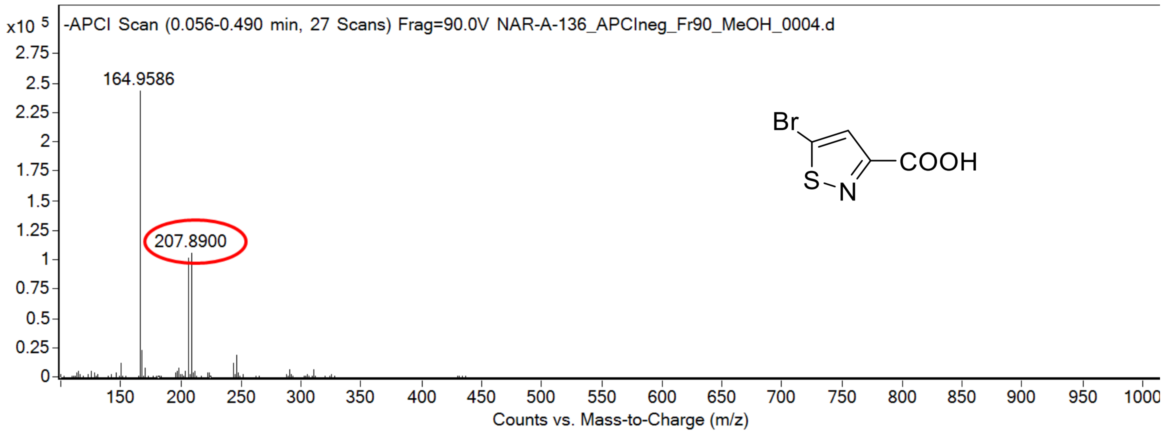

calculated mass: [M-H]- = 207.8896 observed: [M-H]- = 207.8900 mass accuracy = -1.9 ppm

FT-IR spectrum (neat) of **S185**

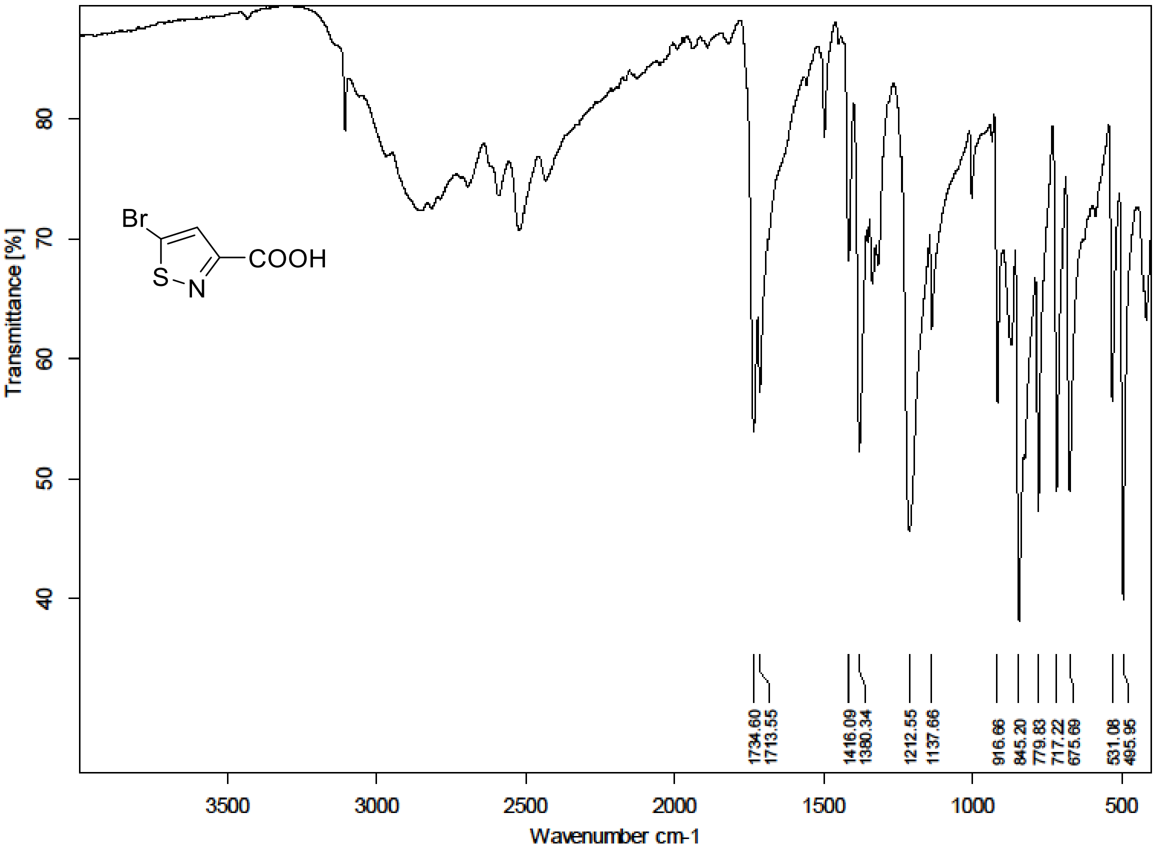

$^{1}H$ (300 MHz) and $^{13}C$ NMR (75 MHz) spectra of **S186** in Chloroform-*d*

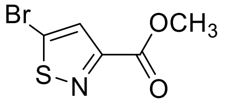

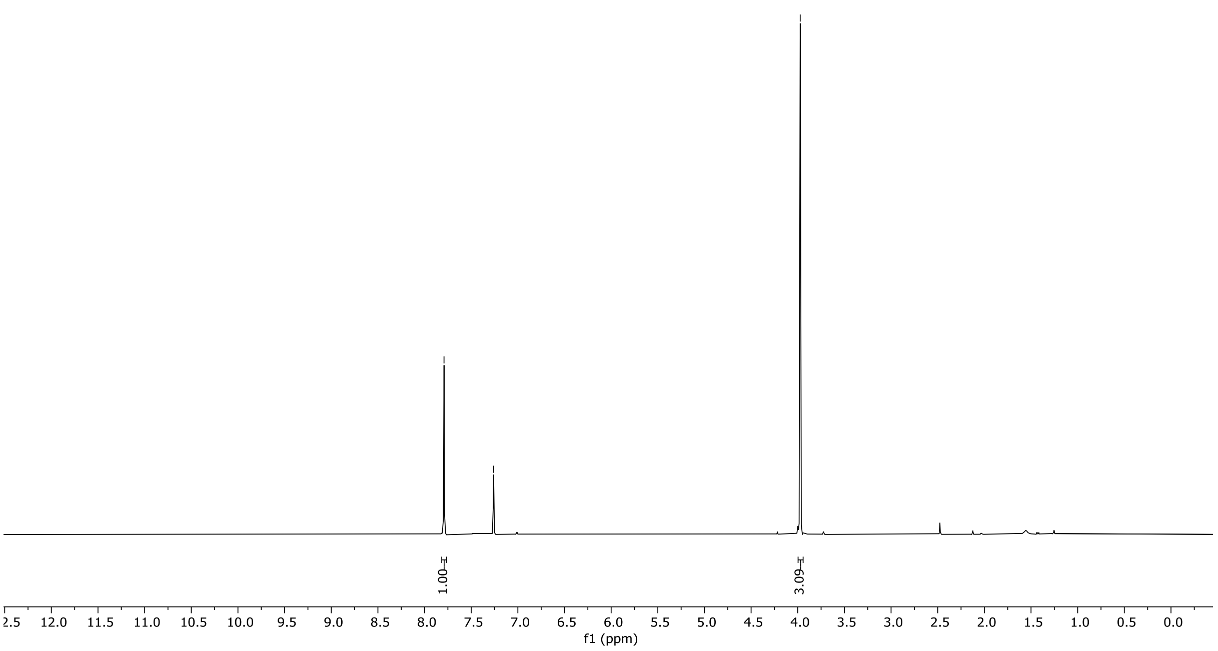

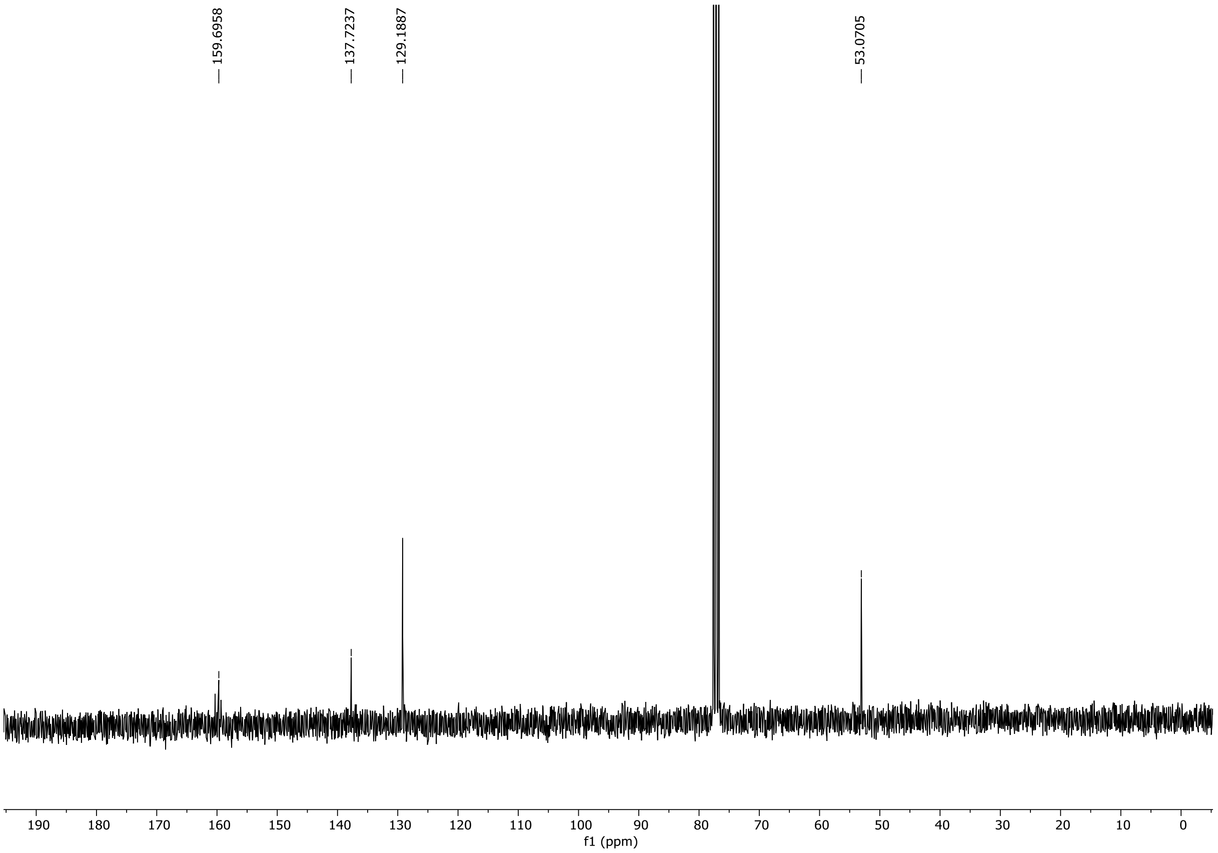

HRMS spectrum of **S186**

**NAR-A-144** $C_5H_4BrNO_2S$ $m/z_{100\%}$ **222.9125**

**APCI+** (MMI)
nitrogen flow 5 L/min, gas temperature 325°C, nebulizer 45 psi, skimmer 65 V, vaporizer 200°C, fragmentor 35 V, dissolved in MeOH

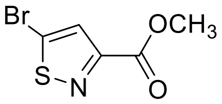

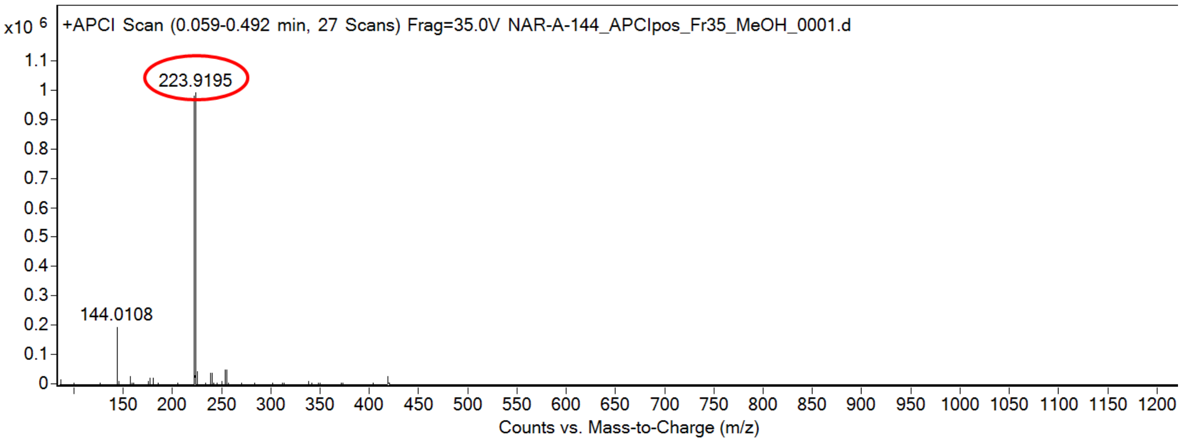

calculated mass: $[M+H]^+ = 223.9198$ observed: $[M+H]^+ = 223.9195$ mass accuracy = -1.3 ppm

FT-IR spectrum (neat) of **S186**

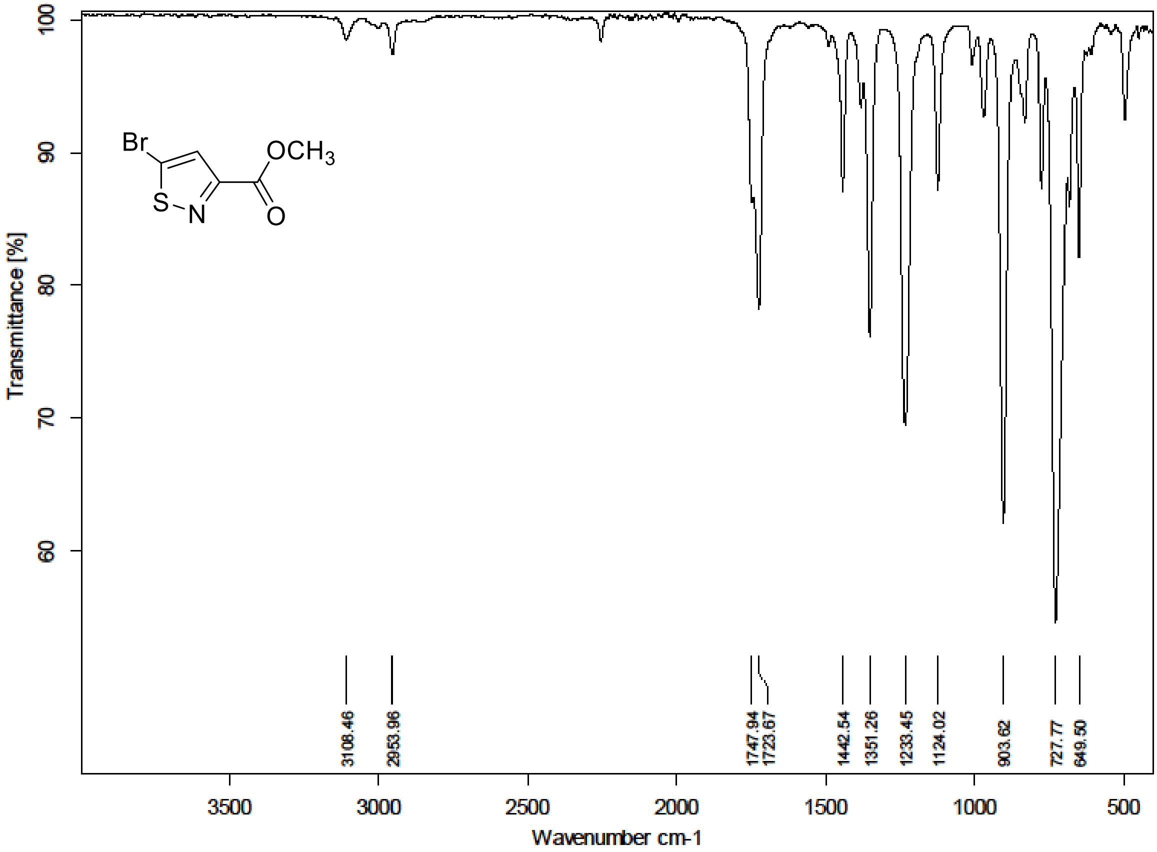

[1]H (300 MHz) and [13]C NMR (75 MHz) spectra of **S187** in Chloroform-*d*

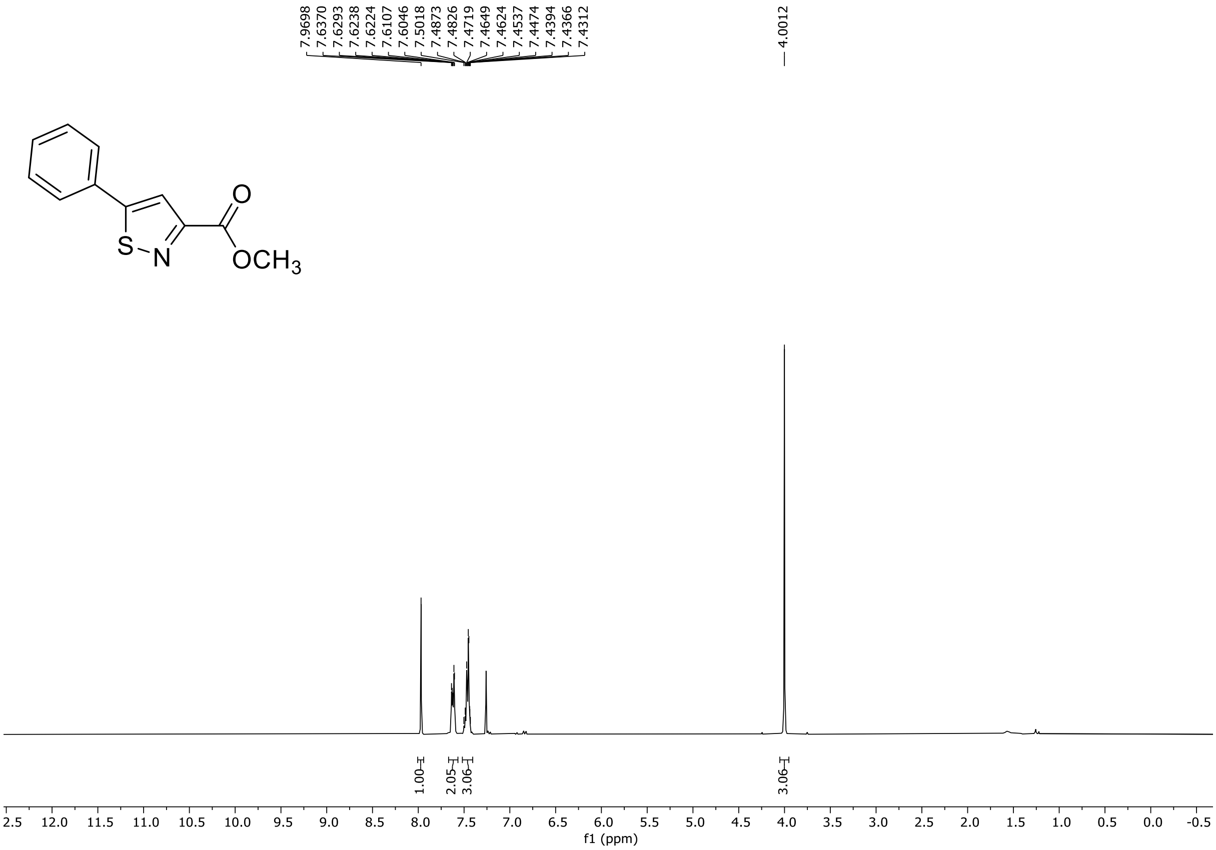

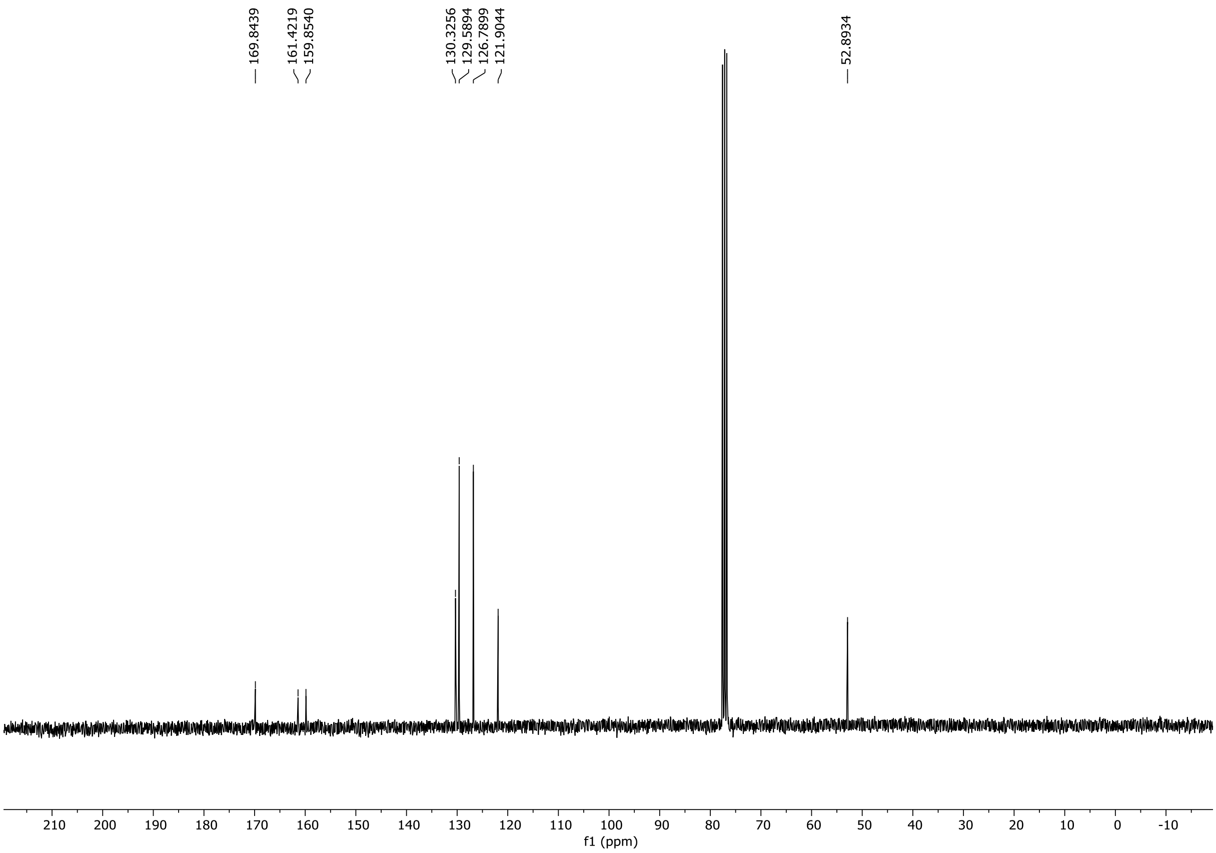

HRMS spectrum of **S187**

**NAR-A-145** $C_{11}H_9NO_2S$ *m/z* **219.0354**

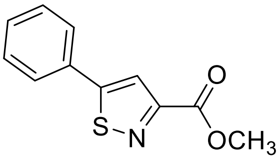

**APCI+** (MMI)
nitrogen flow 5 L/min, gas temperature 325°C, nebulizer 45 psi, skimmer 65 V, vaporizer 200°C, fragmentor 25 V, dissolved in MeOH

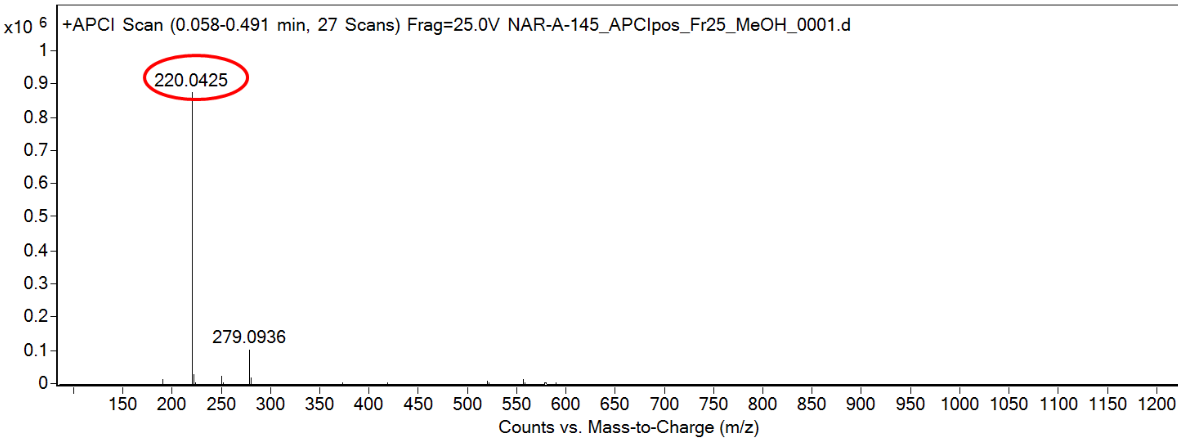

calculated mass: $[M+H]^+ = 220.0427$ observed: $[M+H]^+ = 220.0425$ mass accuracy = -0.9 ppm

FT-IR spectrum (neat) of **S187**

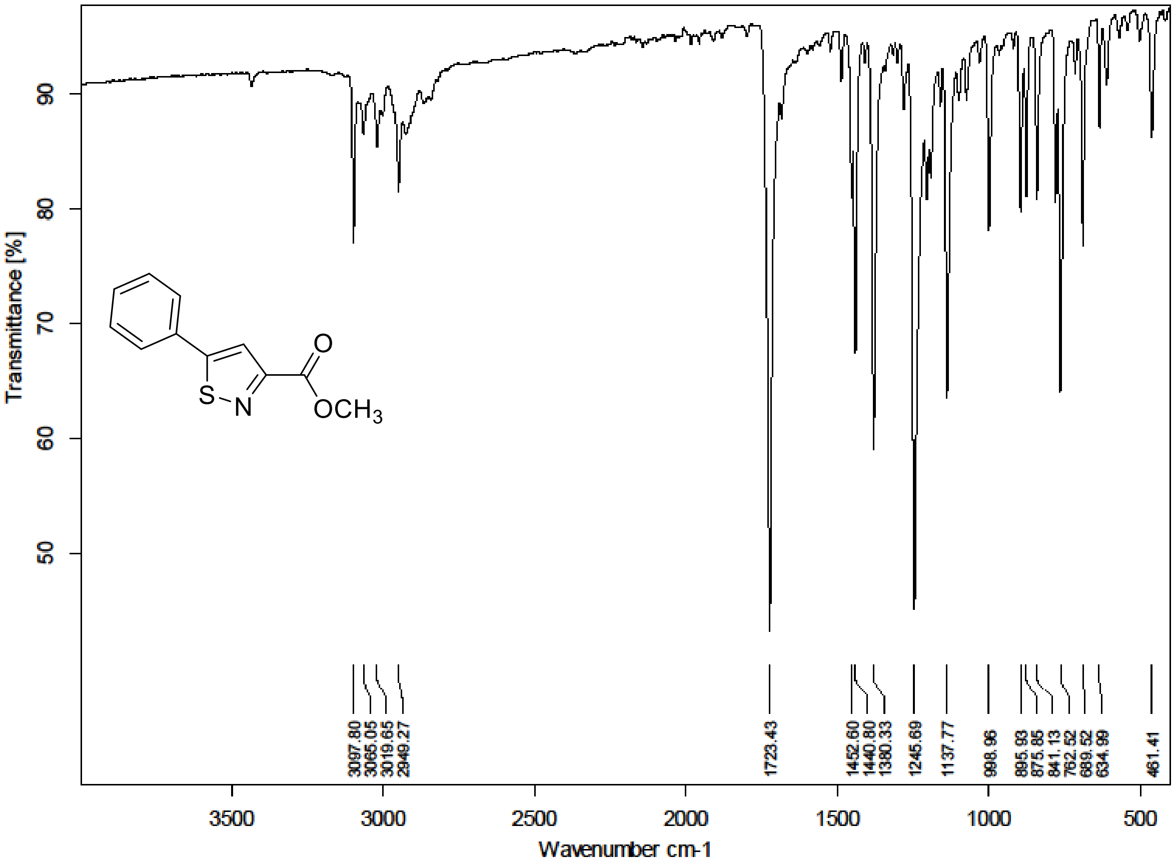

$^{1}H$ (300 MHz) and $^{13}C$ NMR (75 MHz) spectra of **S188** in Chloroform-*d*

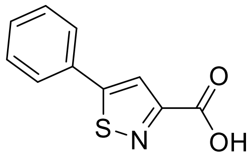

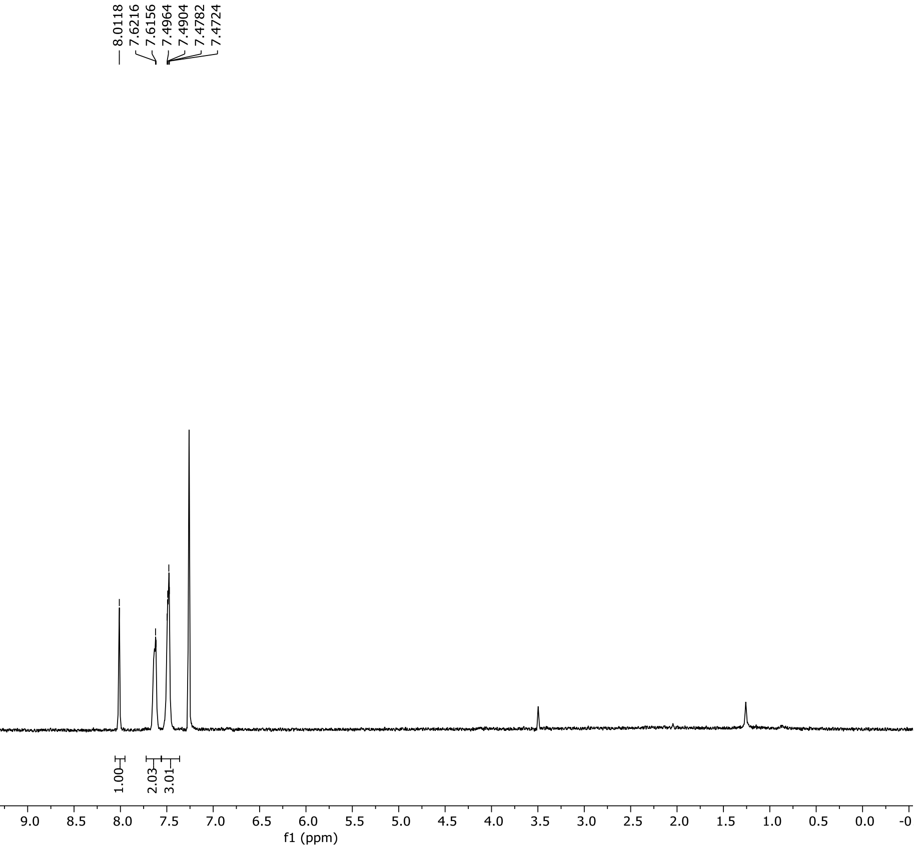

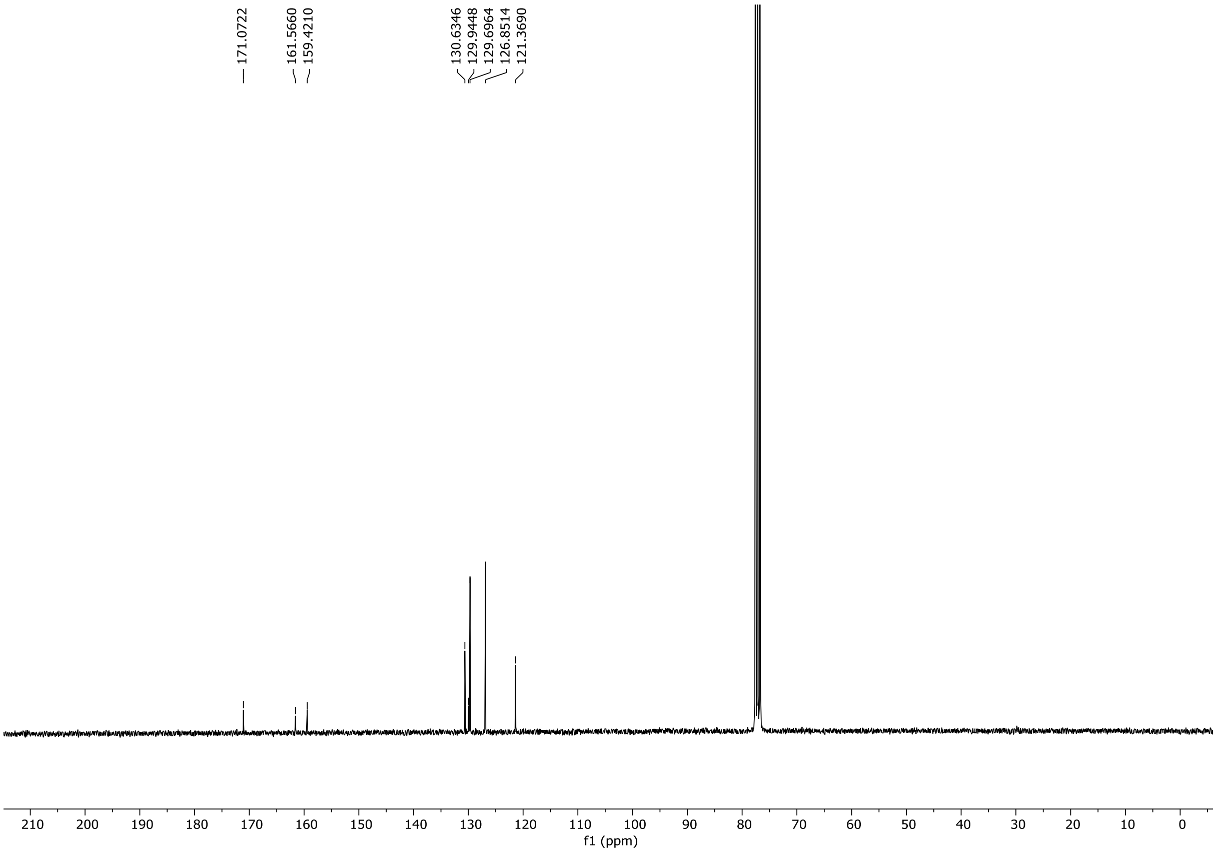

HRMS spectrum of **S188**

**NAR-A-195** $C_{10}H_7NO_2S$ mono *m/z* 205.0197

**APCI +** (MMI)
nitrogen flow 5 L/min, gas temperature 325°C, nebulizer 45 psi, skimmer **65** V, vaporizer 200°C, fragmentor 38 V, dissolved in methanol

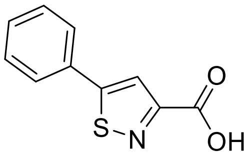

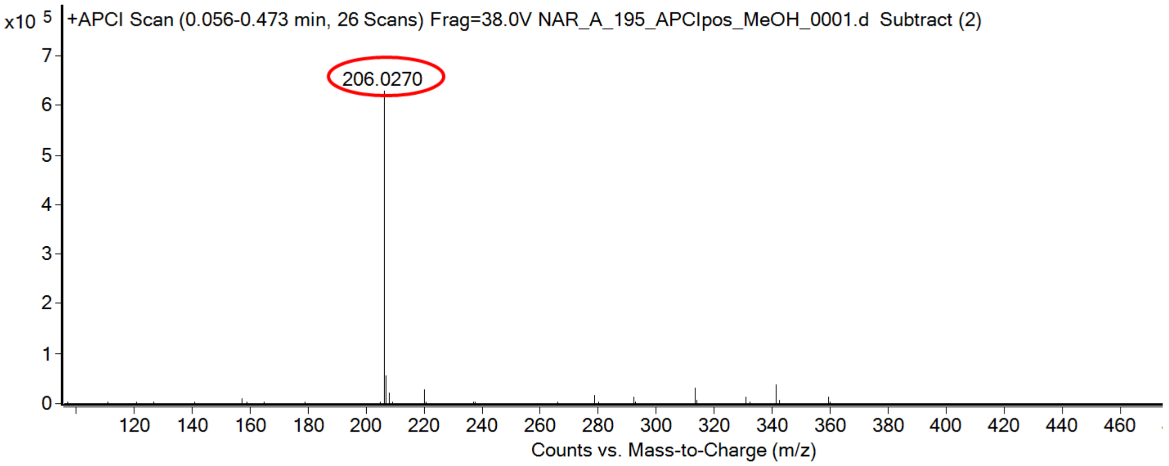

calculated mass: $[M+H]^+ = 206.0270$ observed: $[M+H]^+ = 206.0270$ mass accuracy = < 0.1 ppm

FT-IR spectrum (neat) of **S188**

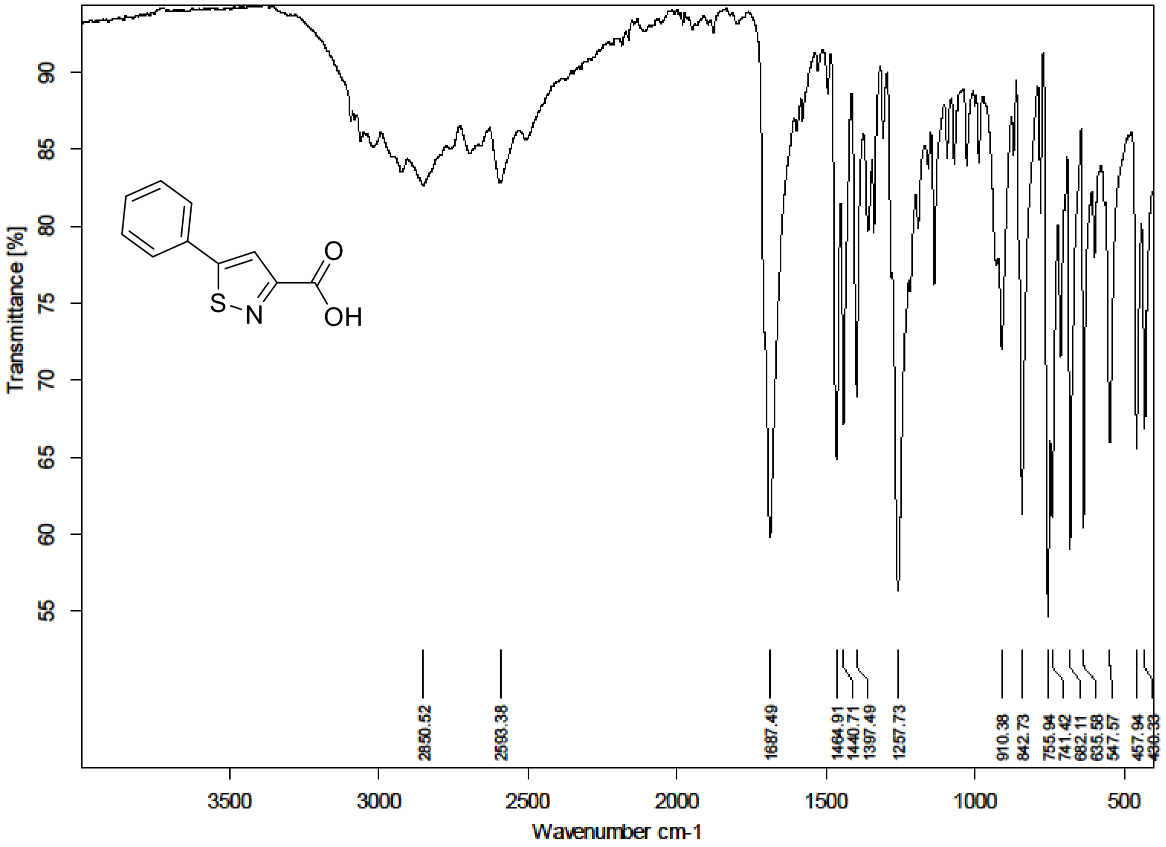

**[1]H NMR, [13]C NMR, HRMS and IR spectra of compound 2, S190 and 29-55**

[1]H (500 MHz) and [13]C NMR (126 MHz) spectra of **2** in DMSO-$d_6$

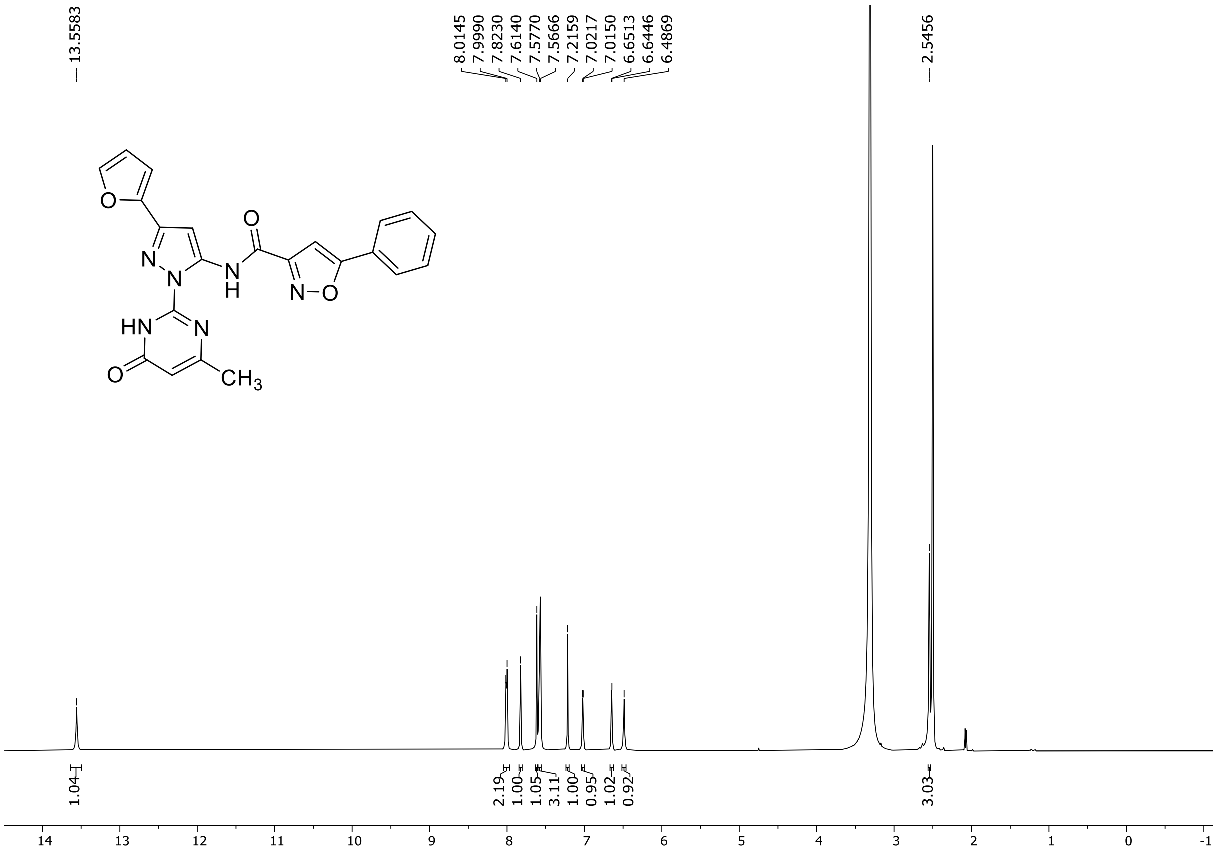

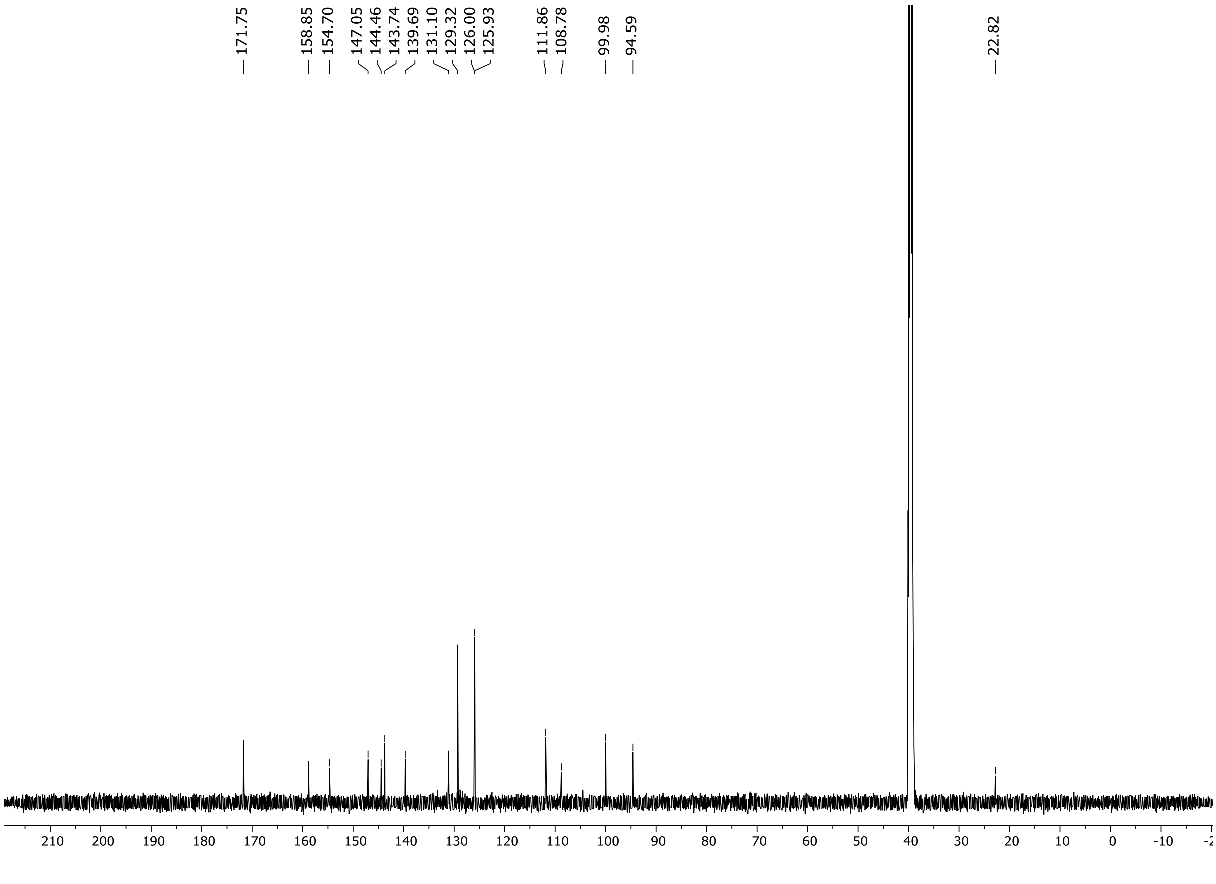

HRMS spectrum of **2**

**$C_{22}H_{16}N_6O_4$** **exact mass: 428.1233**

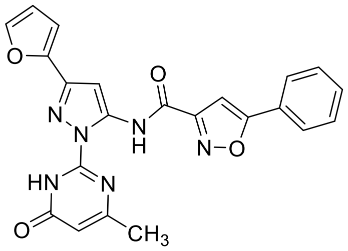

**APCI +** (MMI)
nitrogen flow 5 L/min, gas temperature 325°C, nebulizer 45 psig, skimmer 65 V, vaporizer 200°C, fragmentor 20 V, dissolved in methanol

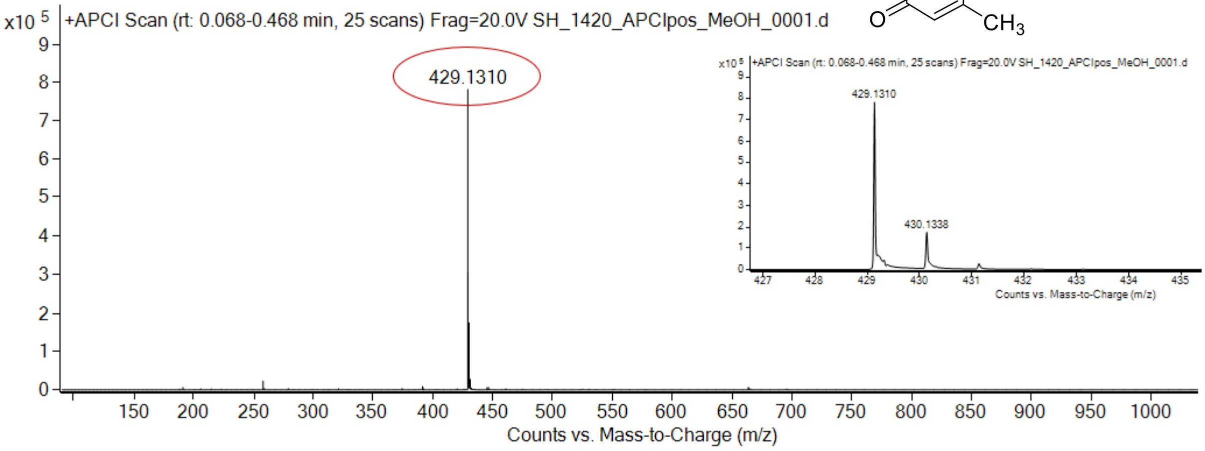

expected mass: $[M+H]^+$ = 429.1306 observed mass : $[M+H]^+$ = 429.1310 mass accuracy = 0.9 ppm

$^{1}H$ (500 MHz) and $^{13}C$ NMR (126 MHz) spectra of **29** in DMSO-$d_6$

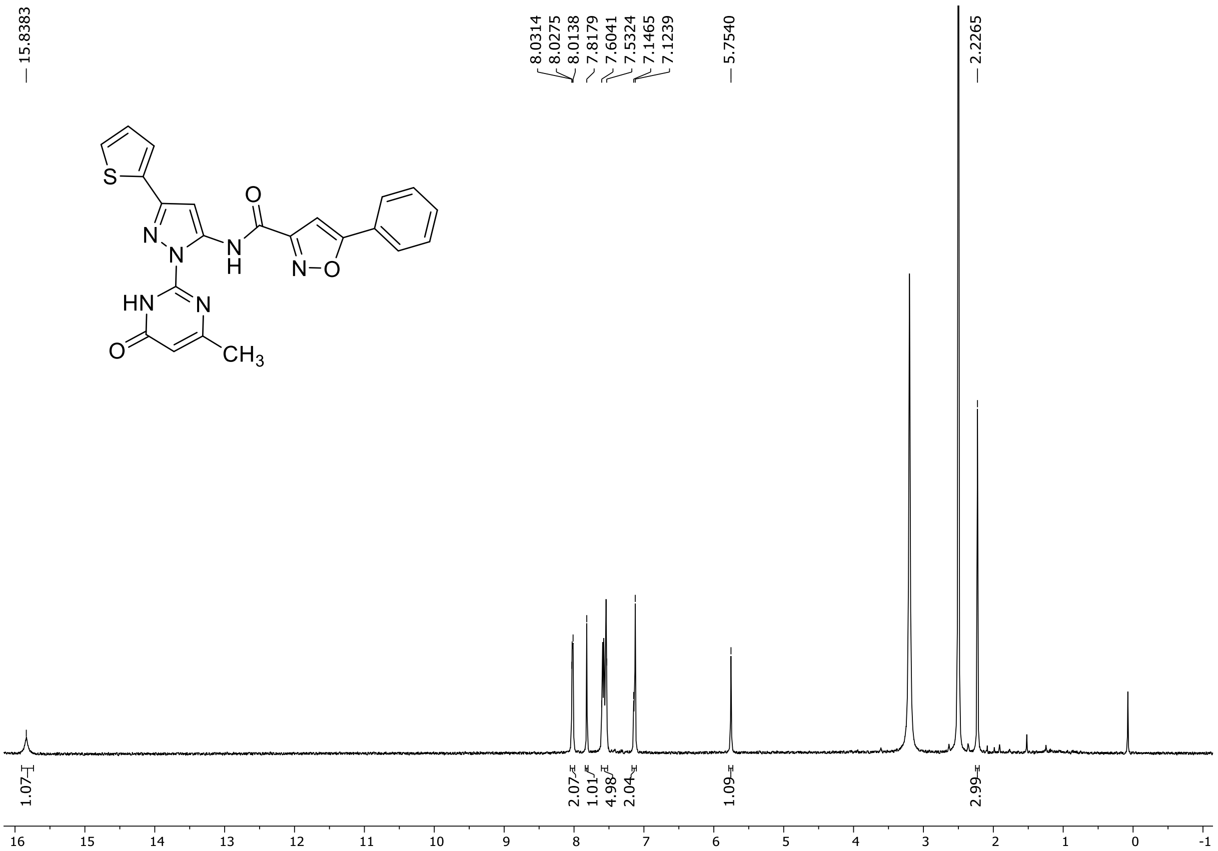

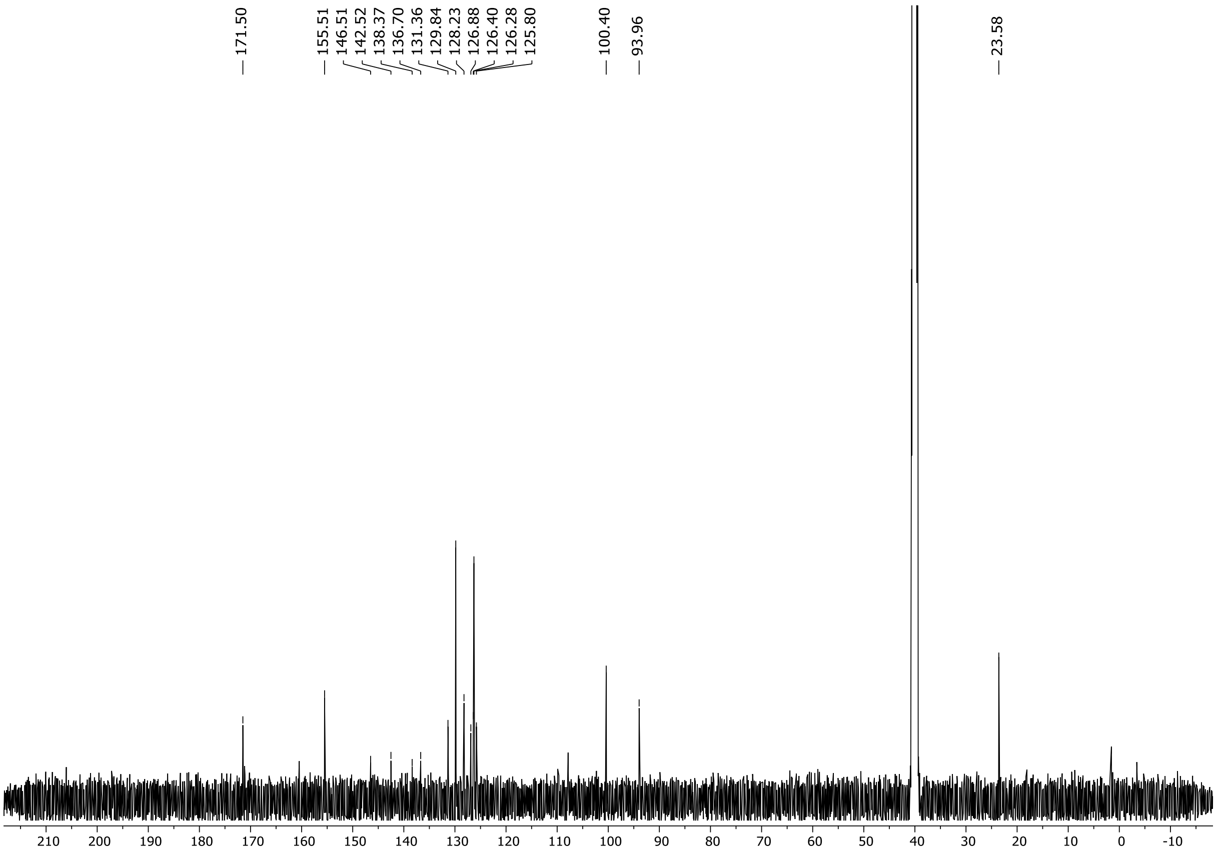

HRMS spectrum of **29**

**$C_{22}H_{16}N_6O_3S$** **exact mass: 444.1005**

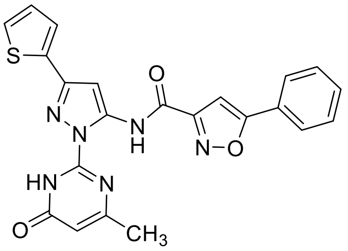

**APCI +** (MMI)
nitrogen flow 5 L/min, gas temperature 325°C, nebulizer 45 psig, skimmer 65 V, vaporizer 200°C, fragmentor 20 V, dissolved in methanol

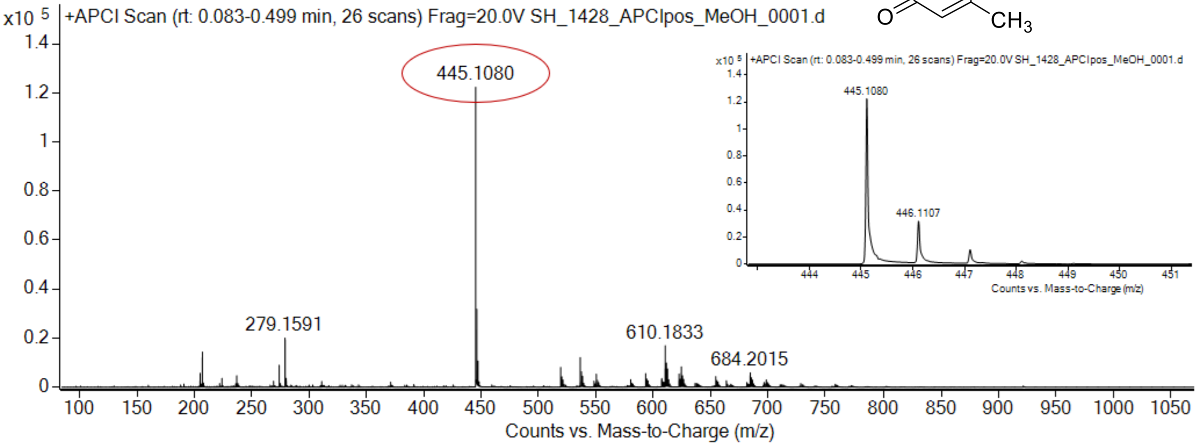

expected mass: $[M+H]^+$ = 445.1077 observed mass : $[M+H]^+$ = 445.1080 mass accuracy = 0.7 ppm

$^{1}H$ (500 MHz) and $^{13}C$ NMR (126 MHz) spectra of **30** in DMSO-$d_6$

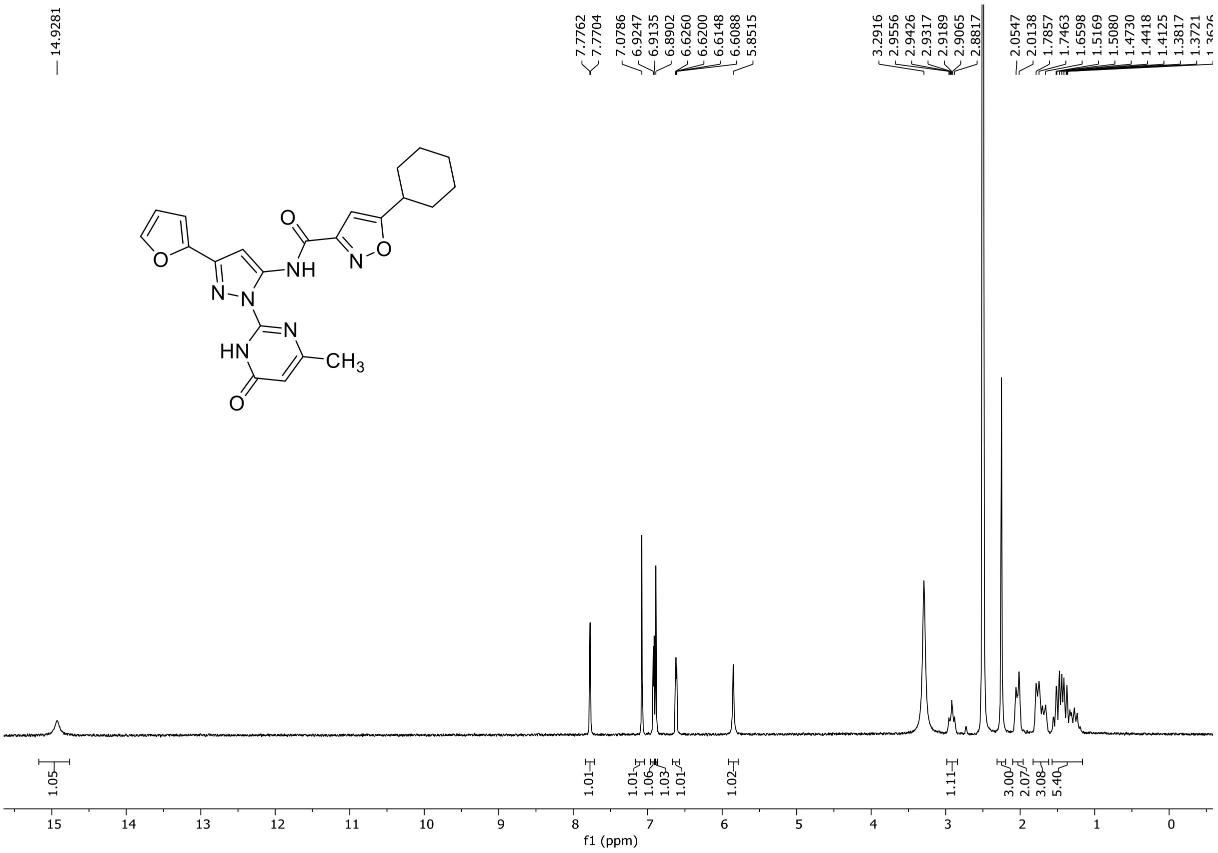

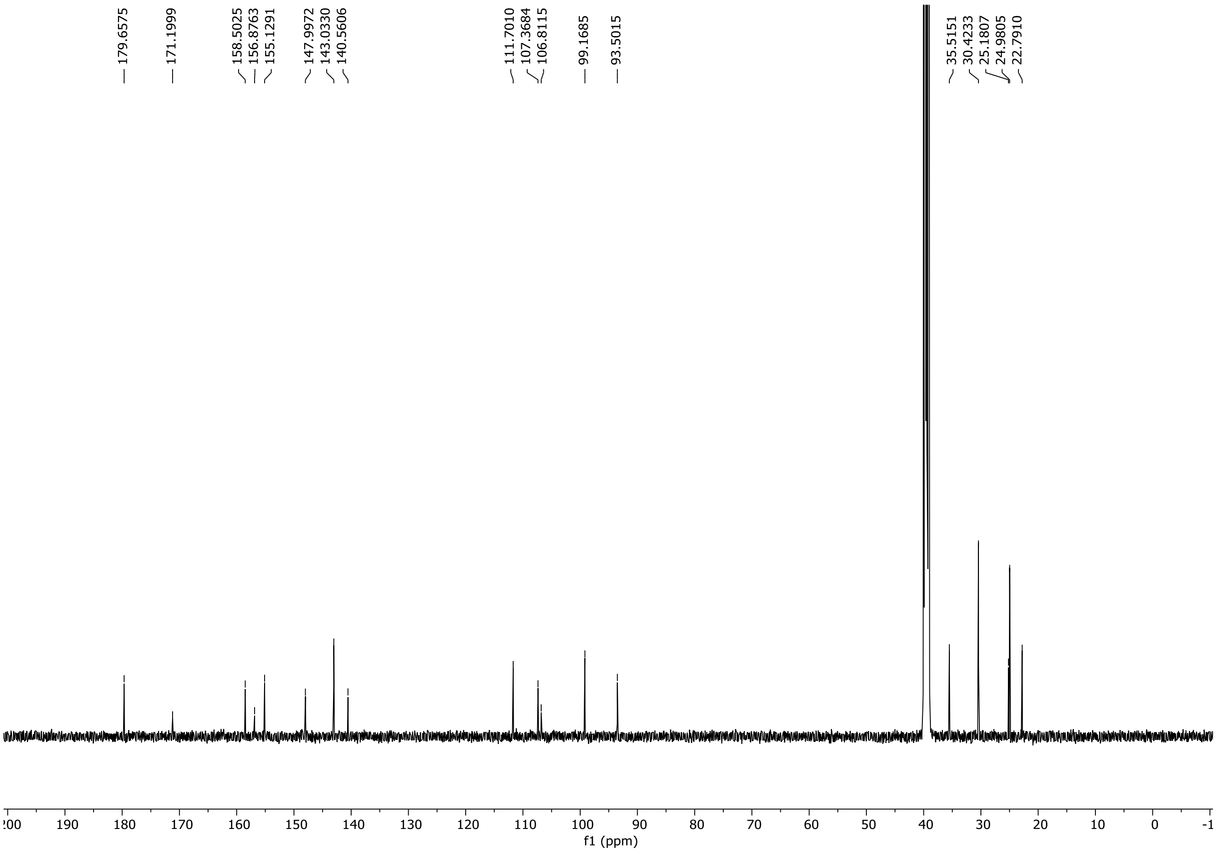

HRMS spectrum of **30**

NAR-A-12 $C_{22}H_{22}N_6O_4$ mono *m/z 434.1703*

**APCI +** (MMI)
nitrogen flow 5 L/min, gas temperature 325°C, nebulizer 45 psi, skimmer **65** V, vaporizer 200°C, fragmentor 30 V, dissolved in methanol

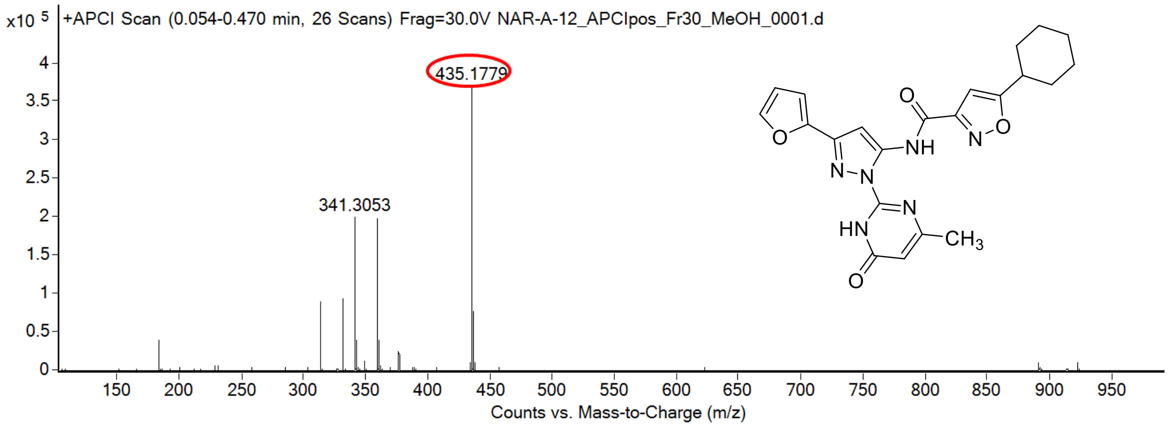

calculated mass: $[M+H]^+ = 435.1775$ observed: $[M+H]^+ = 435.1779$ mass accuracy = 0.9 ppm

FT-IR spectrum (neat) of **30**

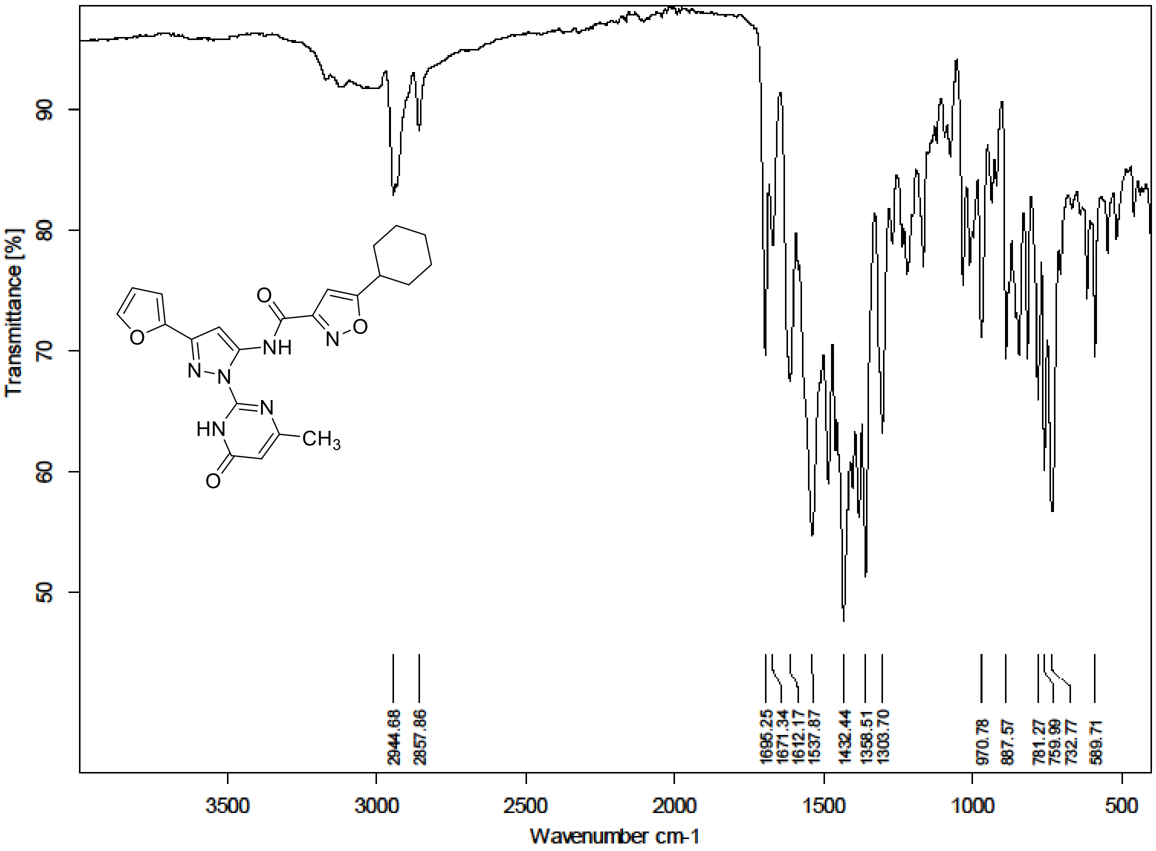

$^{1}H$ NMR (300 MHz) spectrum of **31** in DMSO-$d_6$

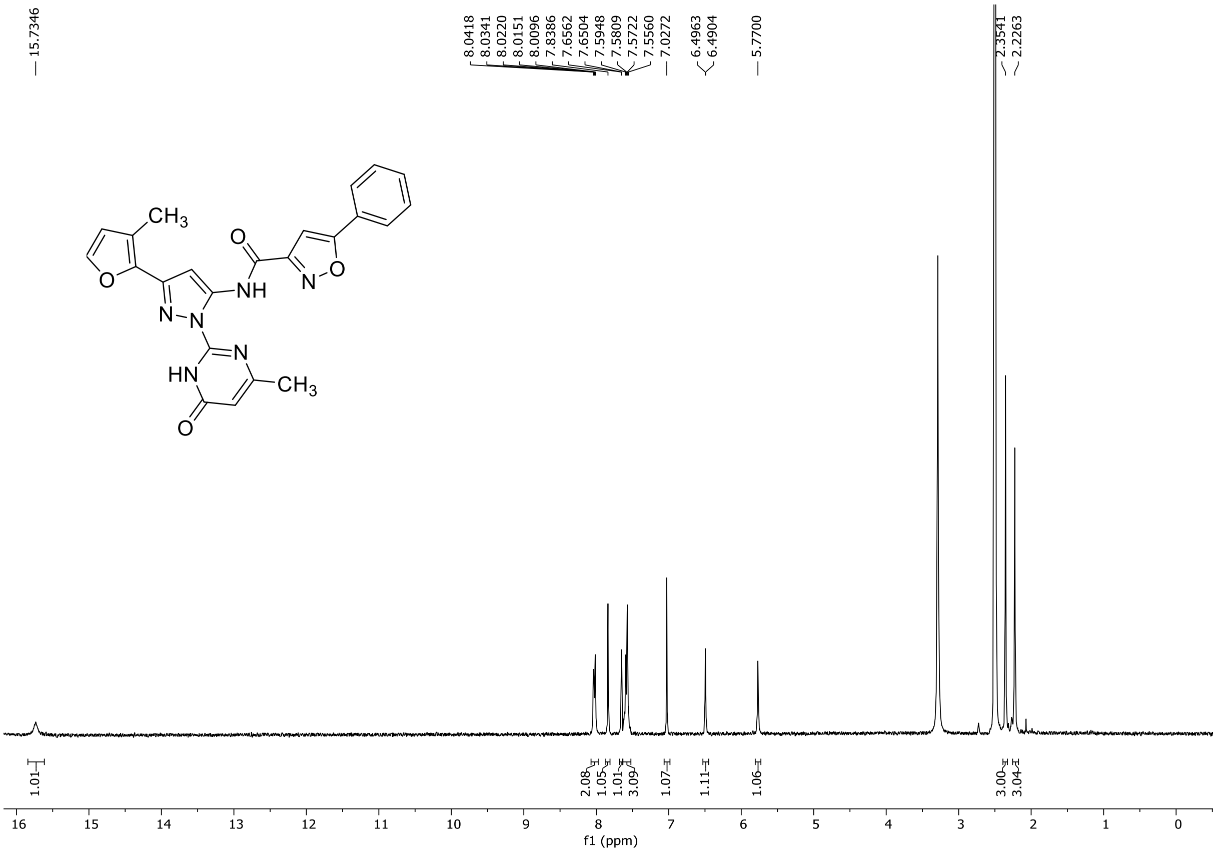

HRMS spectrum of **31**

**NAR-A-181** $C_{23}H_{18}N_6O_4$ **mono** ***m/z*** **442.1390**

**APCI +** (MMI)
nitrogen flow 5 L/min, gas temperature 325°C, nebulizer 45 psi, skimmer **65** V, vaporizer 200°C, fragmentor 25 V, dissolved in methanol

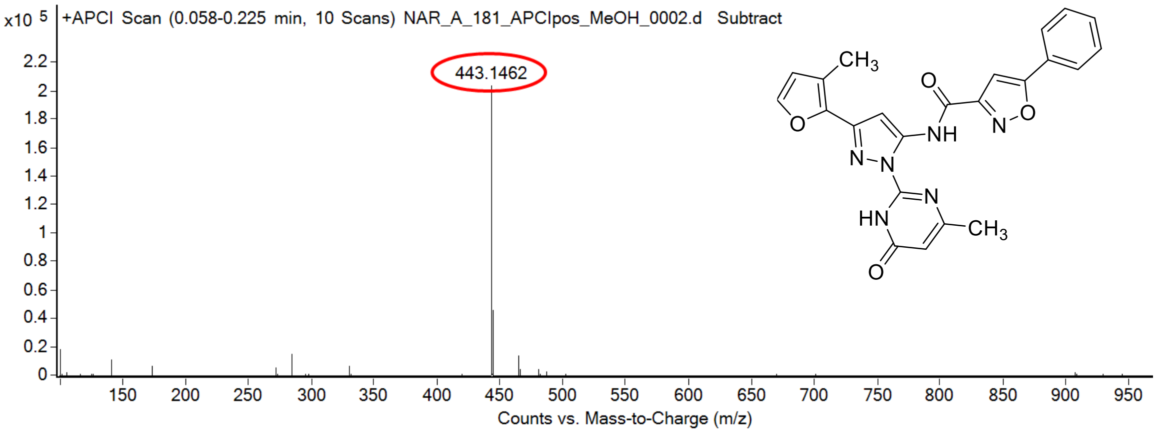

calculated mass: $[M+H]^+$ = 443.1462 observed: $[M+H]^+$ = 443.1462 mass accuracy = < 0.1 ppm

FT-IR spectrum (neat) of **31**

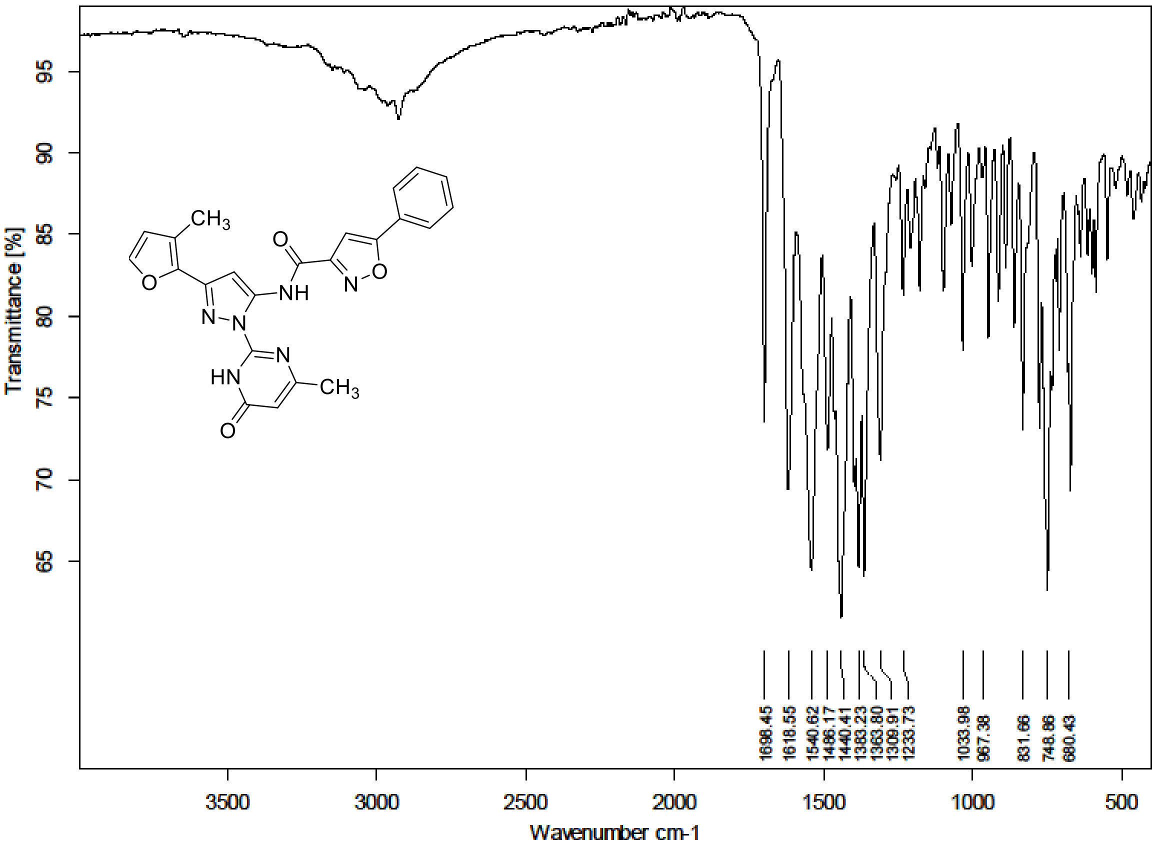

<sup></sup>

[1]H (500 MHz) and [13]C NMR (126 MHz) spectra of **32** in DMSO-$d_6$

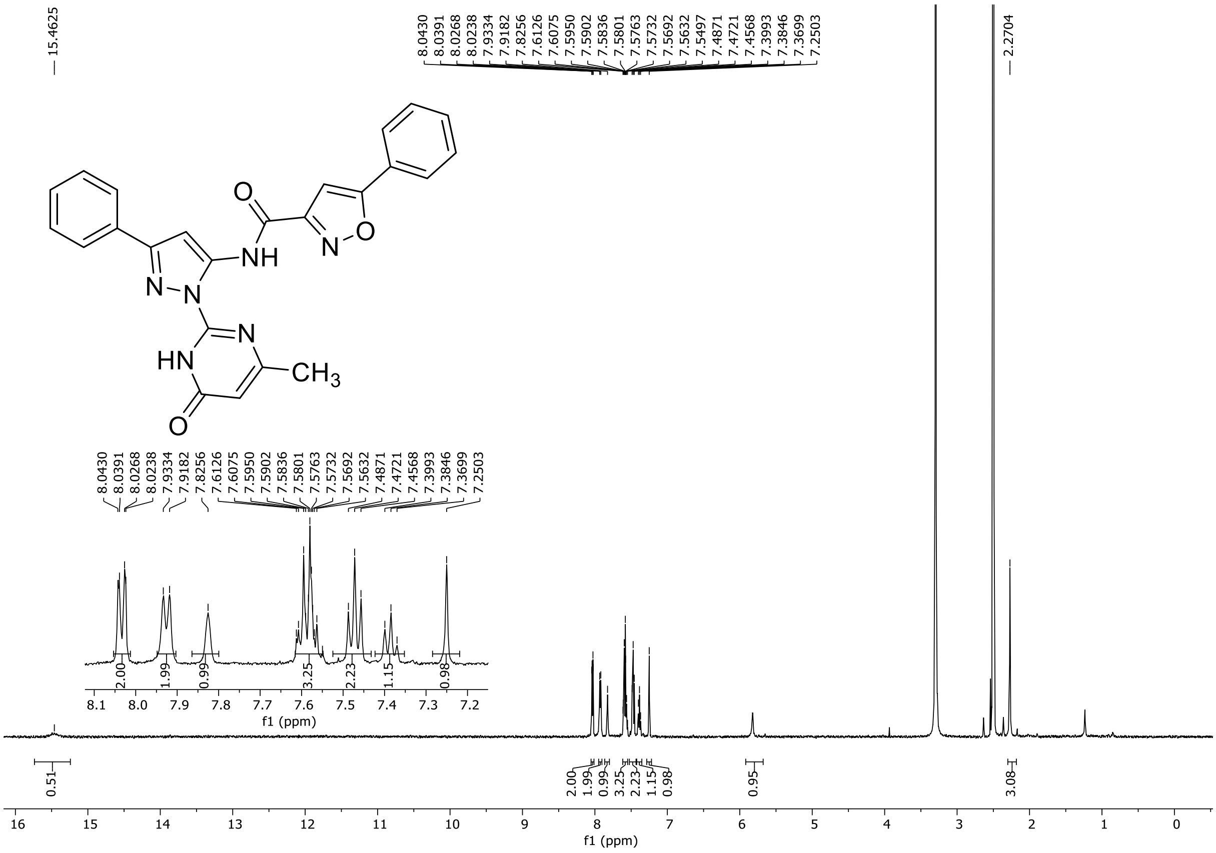

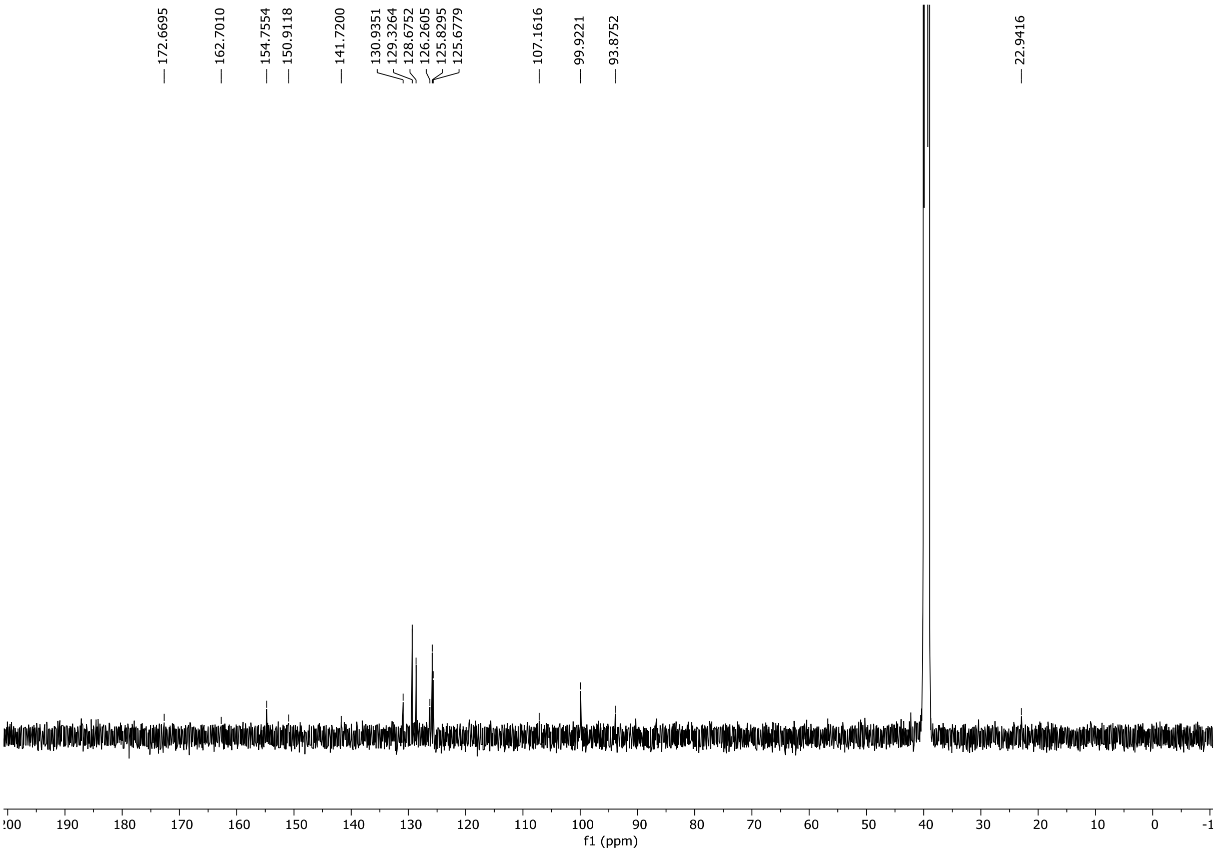

HRMS spectrum of **32**

**NAR-A-8** $C_{24}H_{18}N_6O_3$ *mono m/z* **438.1440**

**ESI -** (MMI)
nitrogen flow 5 L/min, gas temperature 325°C, nebulizer 45 psi, skimmer **65** V, vaporizer 200°C, fragmentor 90 V, dissolved in methanol

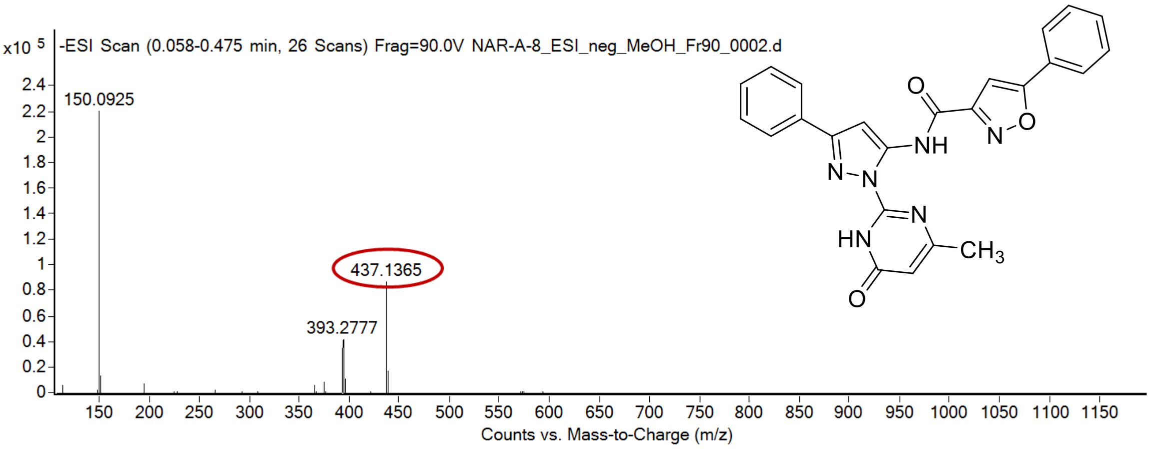

calculated mass: [M-H]$^-$ = 437.1368 observed: [M-H]$^-$ = 437.1365 mass accuracy = 0.7 ppm

FT-IR spectrum (neat) of **32**

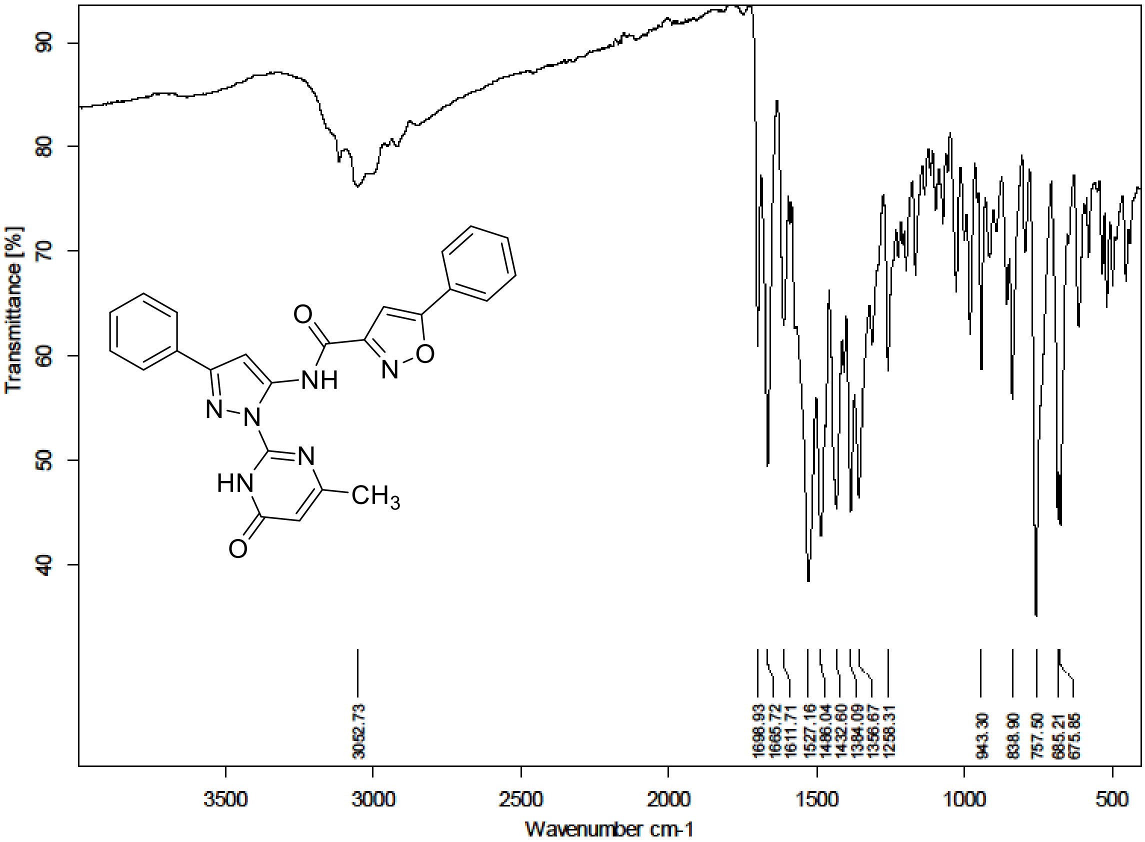

$^{1}H$ (300 MHz) and $^{13}C$ NMR (176 MHz) spectra of **33** in DMSO-$d_6$

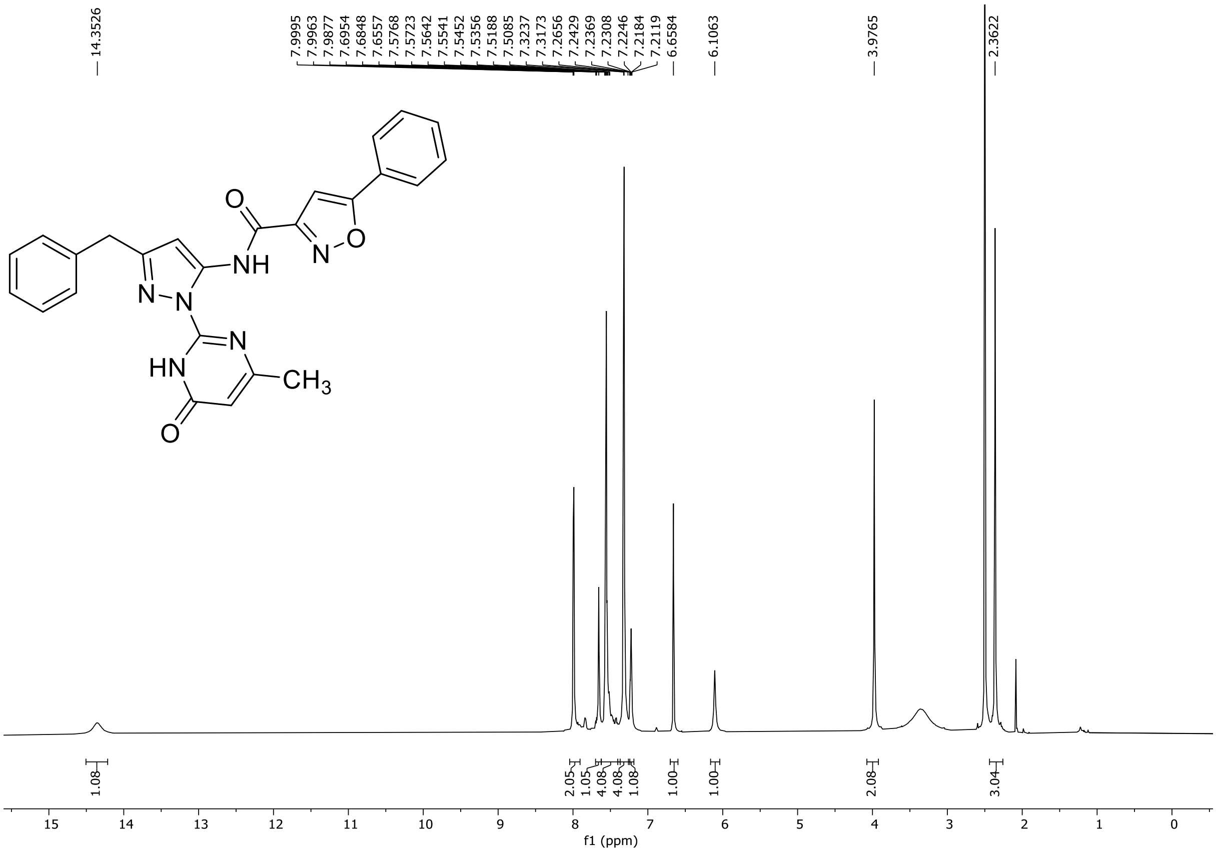

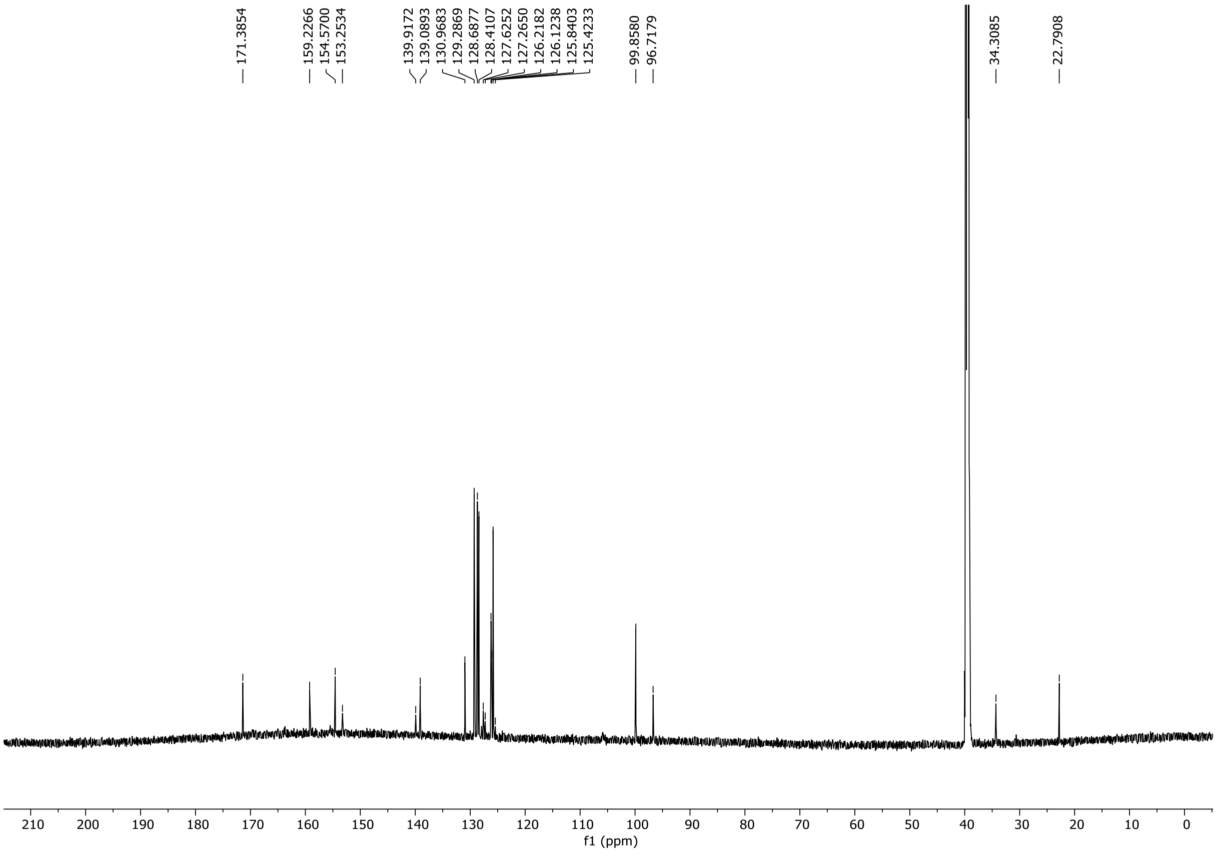

HRMS spectrum of **33**

**NAR-A-189** $C_{25}H_{20}N_6O_3$ mono *m/z* 452.1597

**APCI +** (MMI)
nitrogen flow 5 L/min, gas temperature 325°C, nebulizer 45 psi, skimmer **65** V,
vaporizer 200°C, fragmentor 35 V, dissolved in methanol

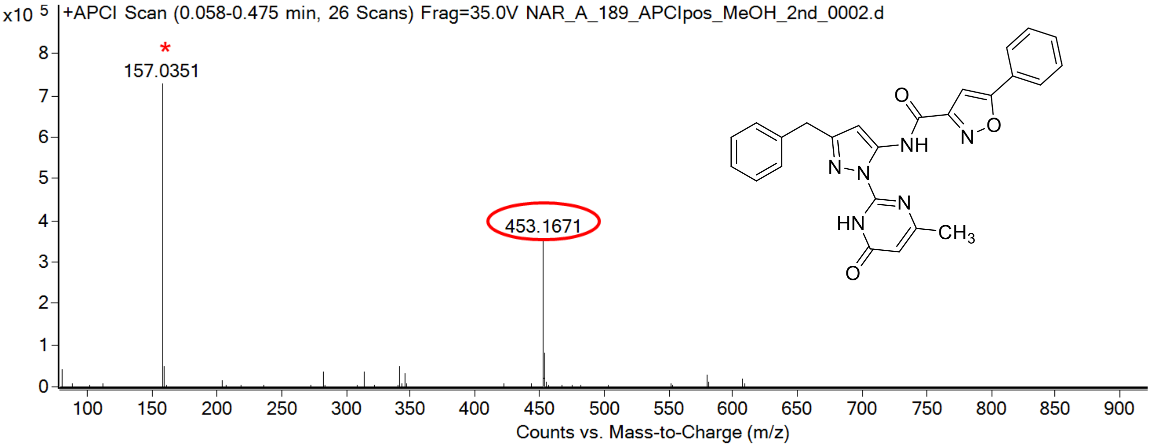

calculated mass: $[M+H]^+$ = 453.1670 observed: $[M+H]^+$ = 453.1671 mass accuracy = 0.2 ppm

* DMSO

FT-IR spectrum (neat) of **33**

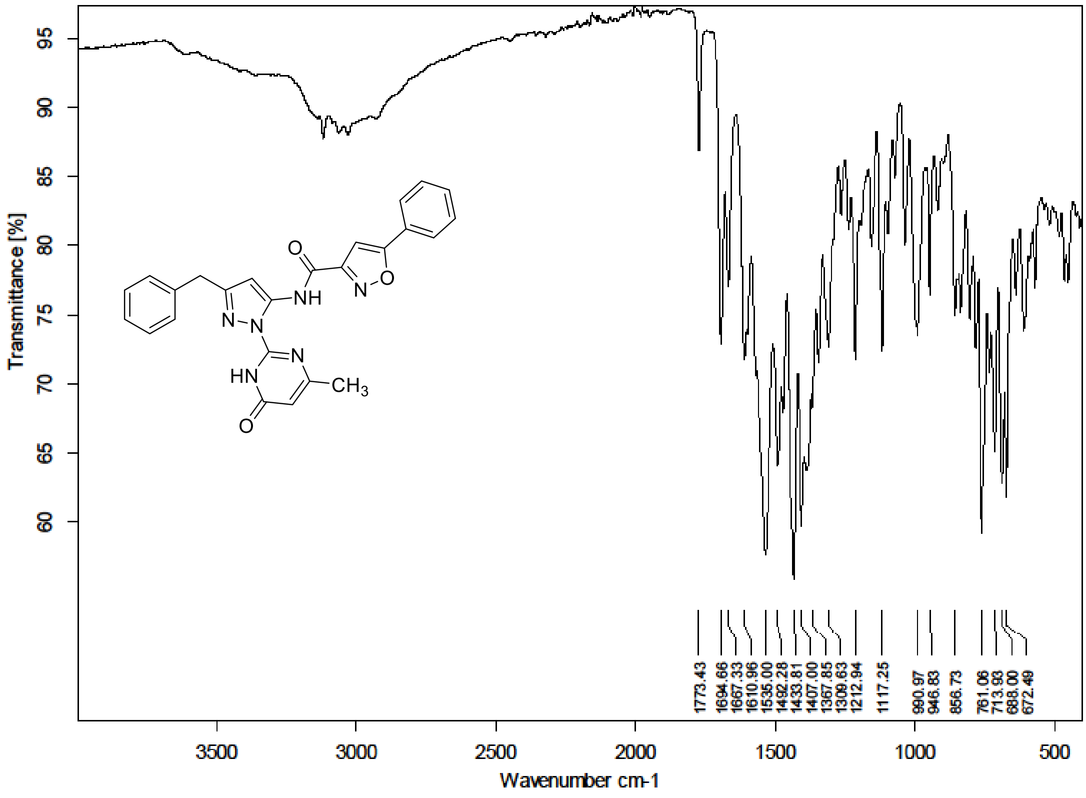

$^{1}H$ NMR (300 MHz) spectrum of **34** in DMSO-$d_6$

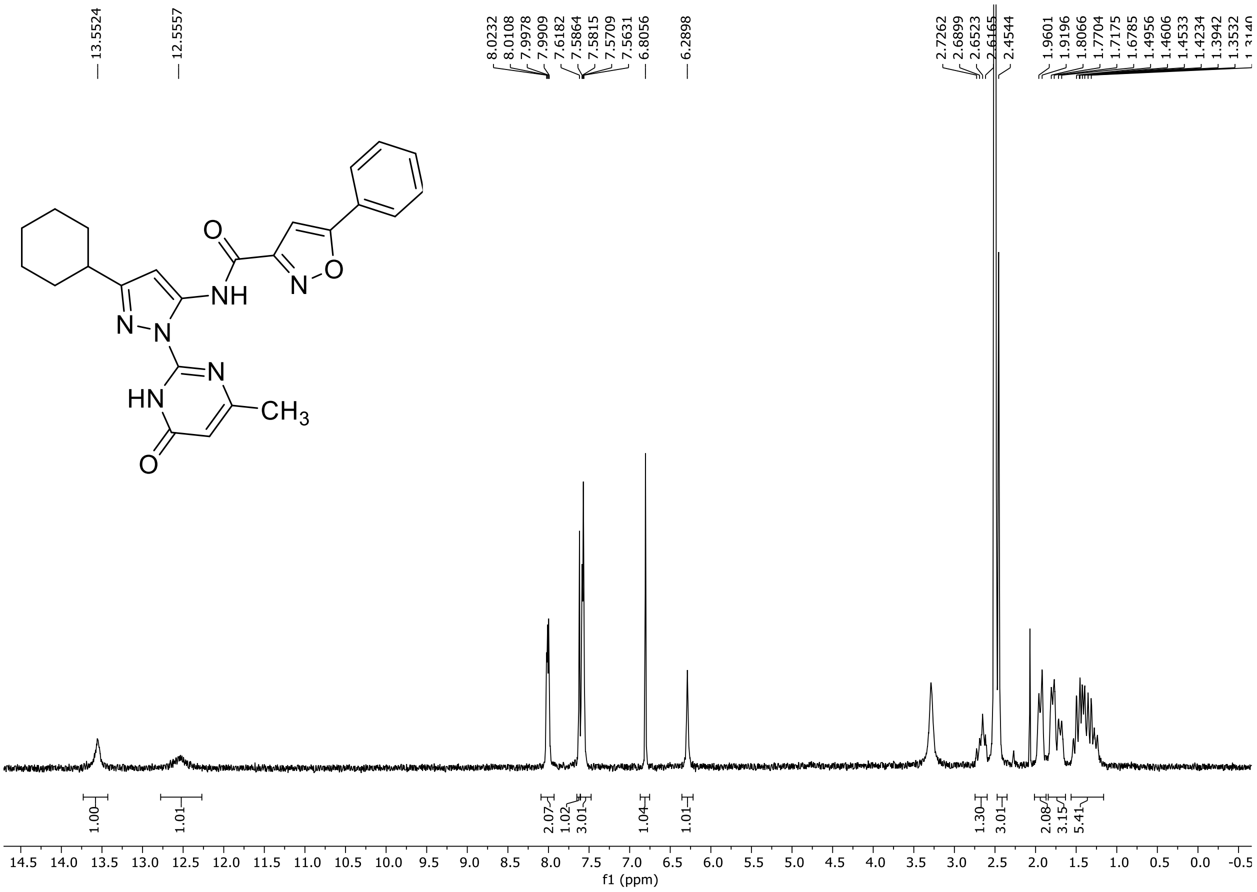

HRMS spectrum of **34**

**NAR-A-122** **$C_{24}H_{24}N_6O_3$** ***m/z* 444.1910**

**ESI-** (MMI)
nitrogen flow 5 L/min, gas temperature 325°C, nebulizer 45 psi, skimmer 65 V, fragmentor 100 V, dissolved in DMSO, MeOH

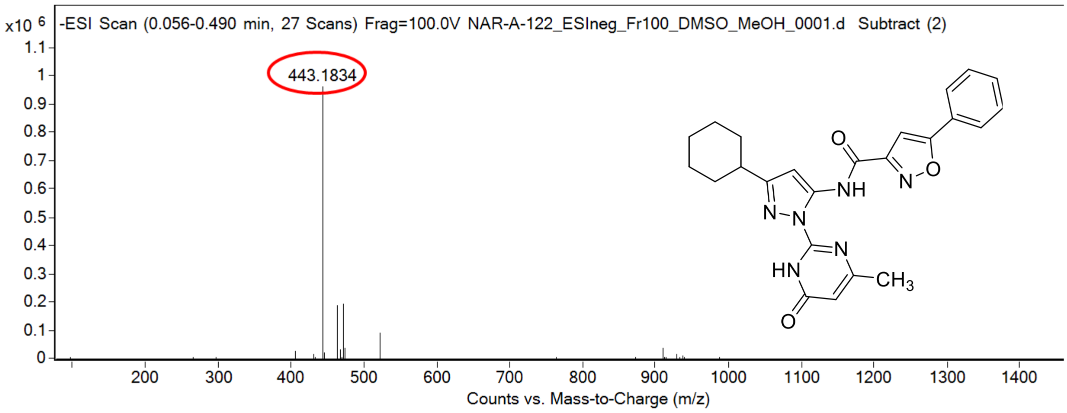

calculated mass: $[M-H]^-$ = 443.1837 observed: $[M-H]^-$ = 443.1834 mass accuracy = -0.7 ppm

FT-IR spectrum (neat) of **34**

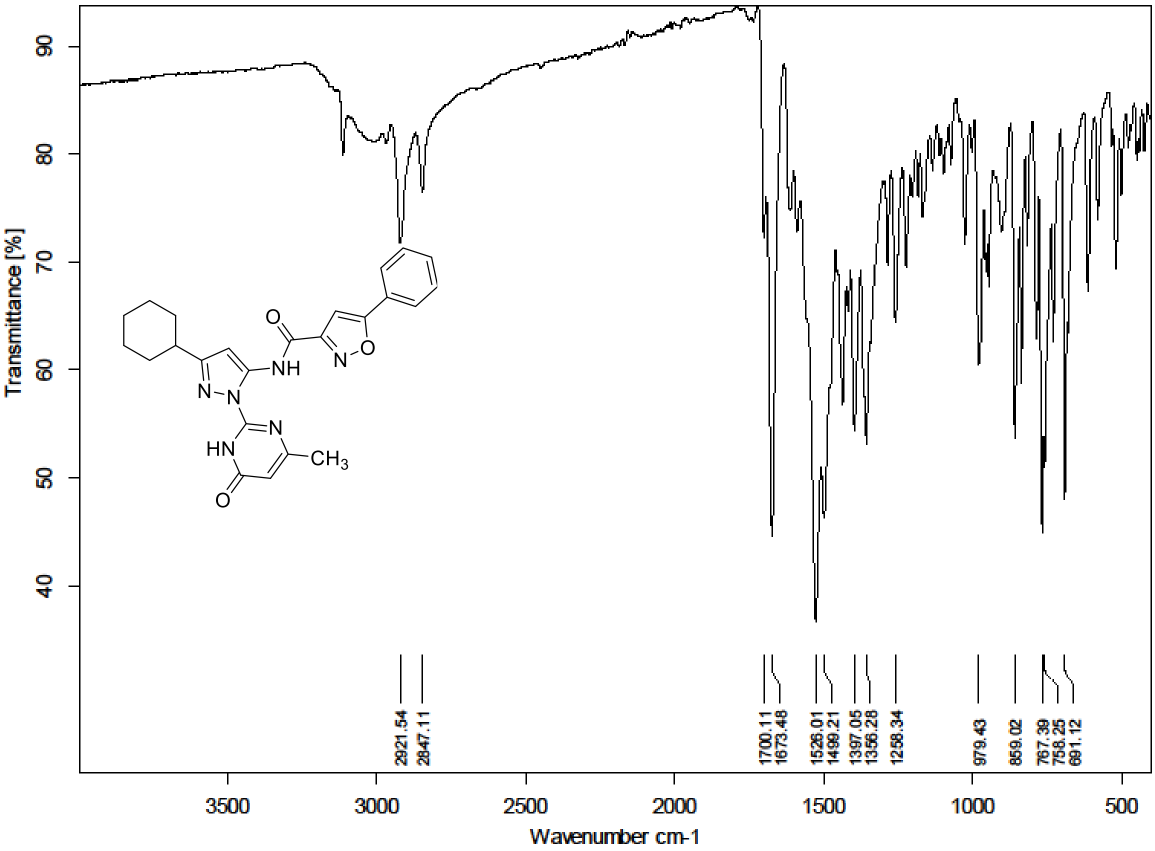

[1]H (500 MHz) and [13]C NMR (126 MHz) spectra of **35** in DMSO-$d_6$

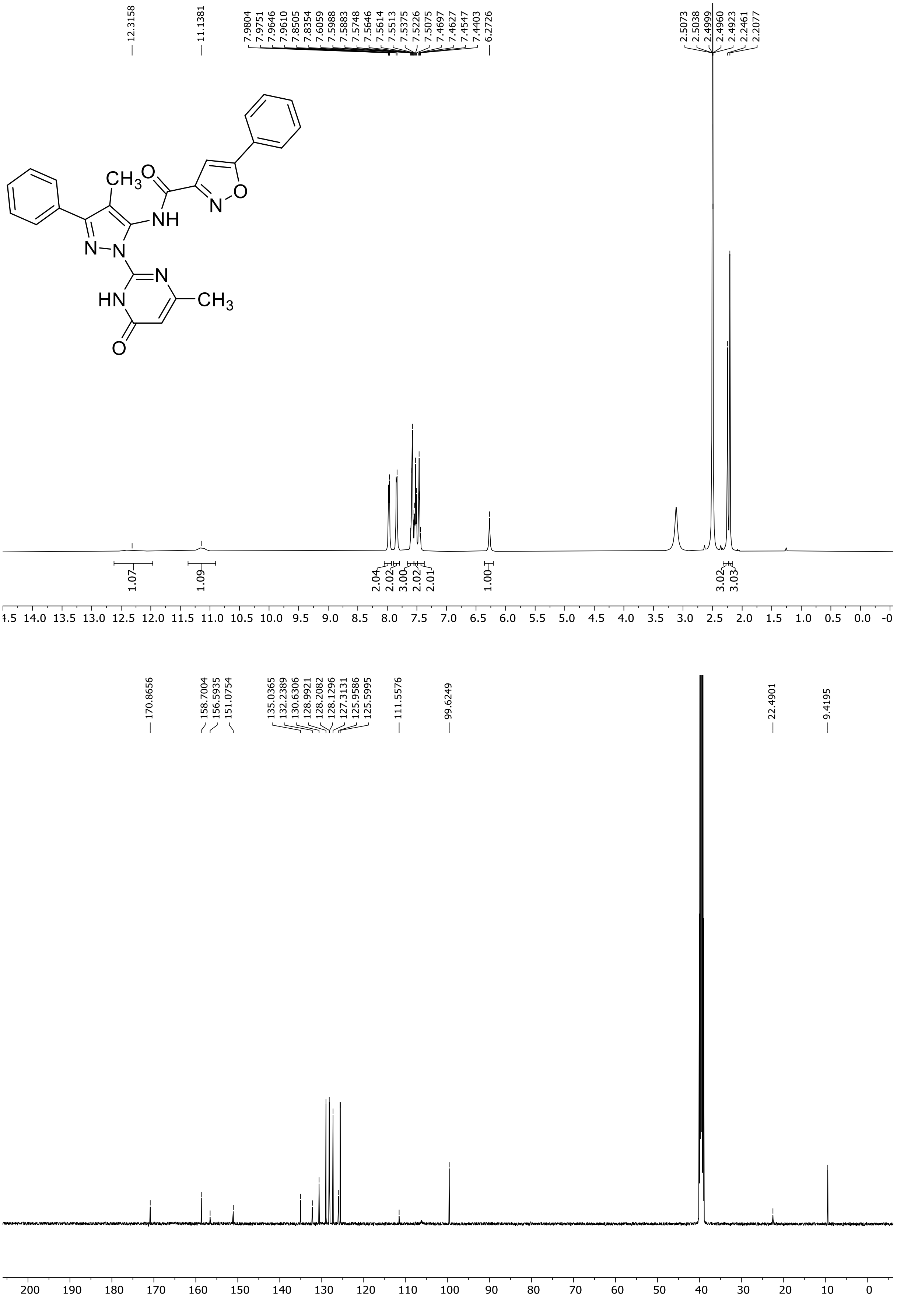

HRMS spectrum of **35**

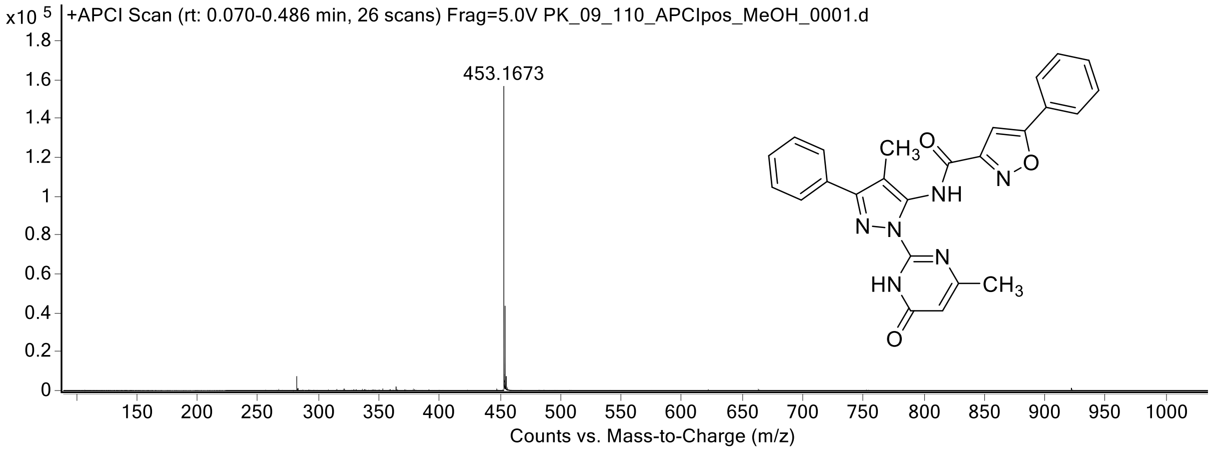

[1]H (500 MHz) and [13]C NMR (126 MHz) spectra of **36** in DMSO-$d_6$

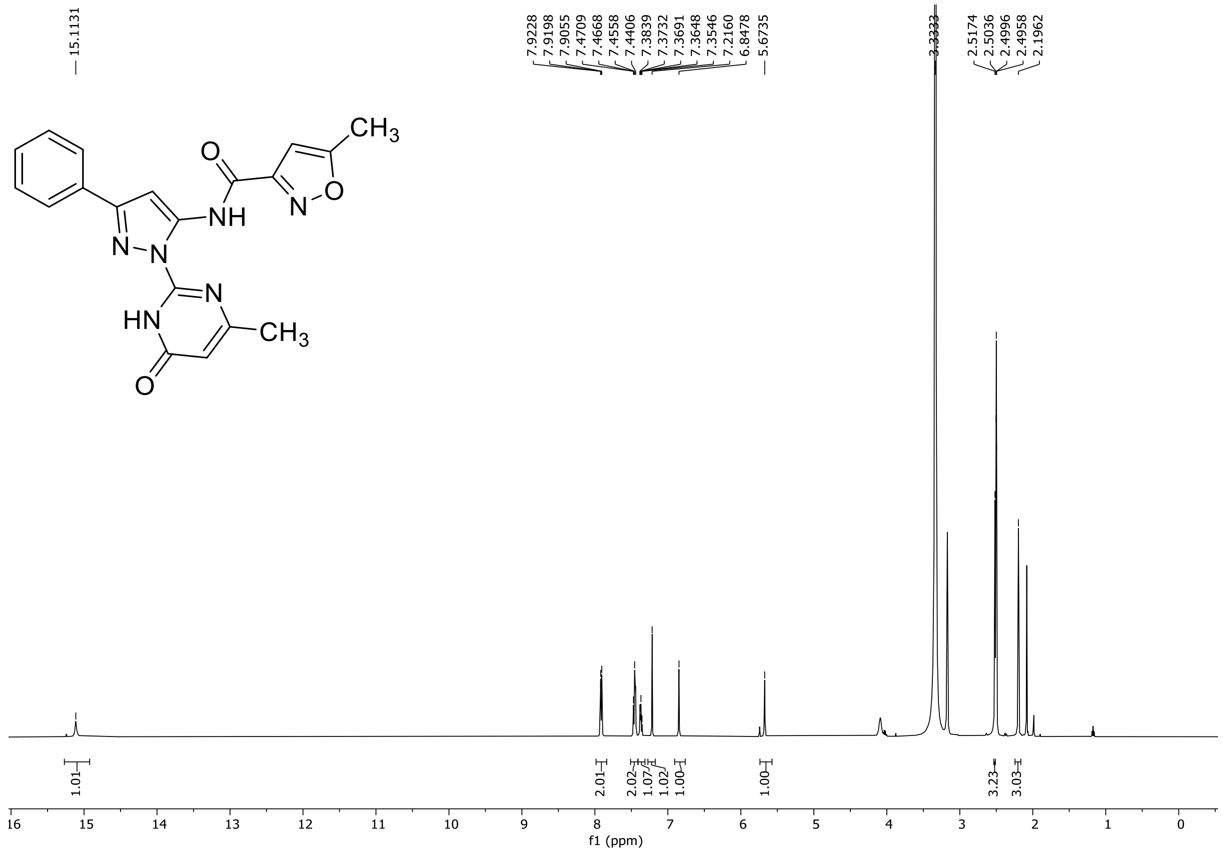

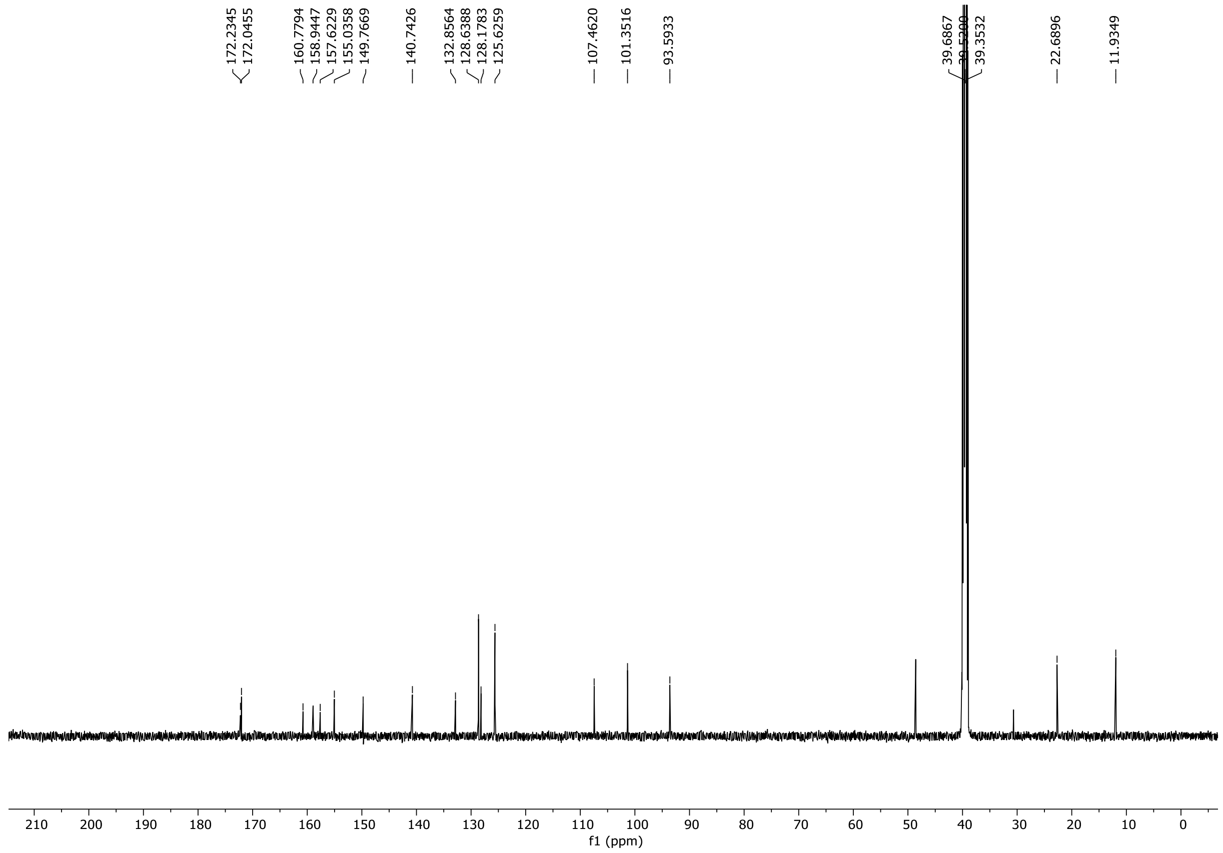

$^{1}H$ NMR (500 MHz) spectrum of **36** in Methanol-$d_4$

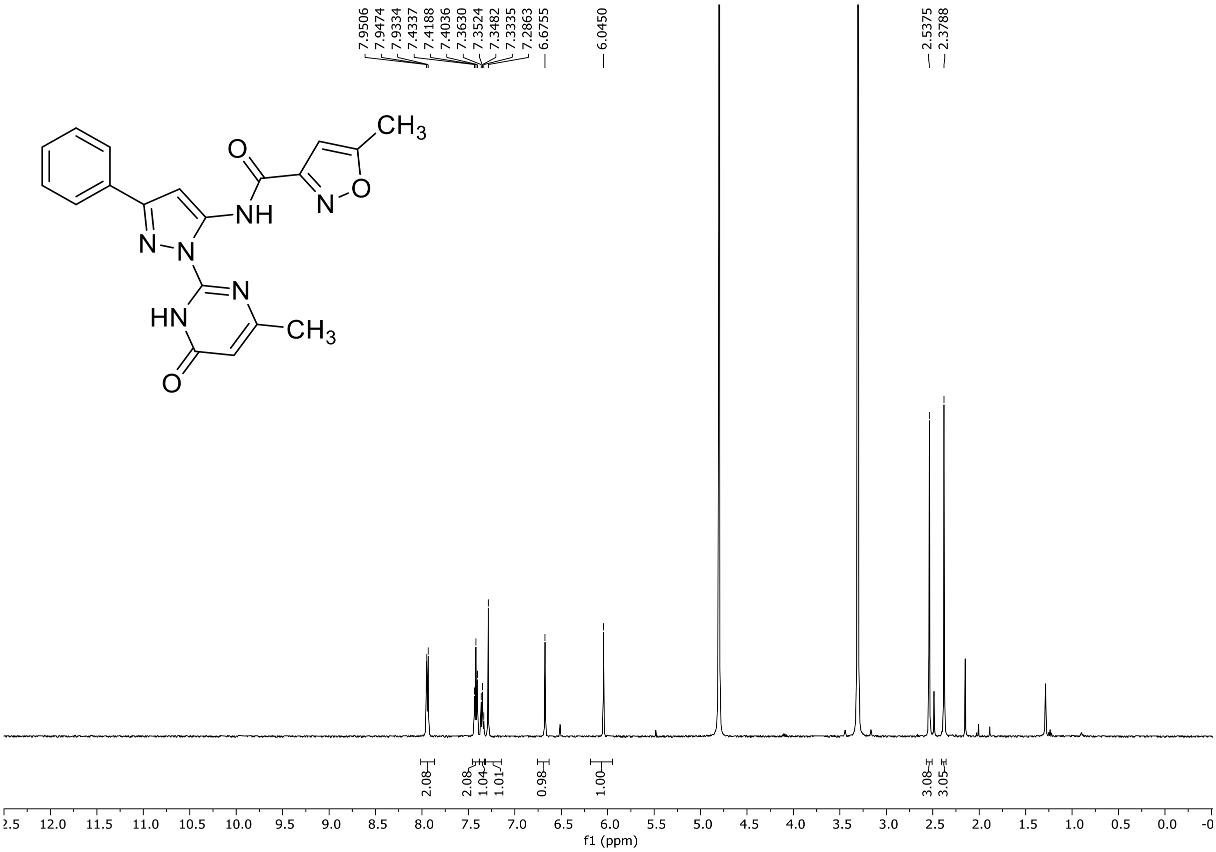

HRMS spectrum of **36**

**NAR-A-3**

$C_{19}H_{16}N_6O_3$ *mono m/z*
**376.1284**

**APCI +** (MMI)
nitrogen flow 5 L/min, gas temperature 325°C, nebulizer 45 psi, skimmer **65** V, vaporizer 200°C, fragmentor 38 V, dissolved in methanol

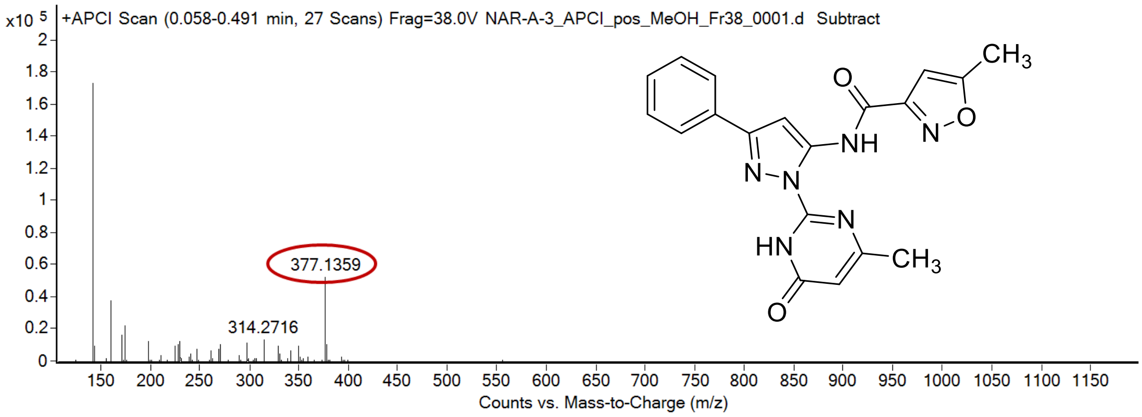

calculated mass: $[M+H]^+ = 377.1357$ observed: $[M+H]^+ = 377.1359$ mass accuracy = 0.5 ppm

FT-IR spectrum (neat) of **36**

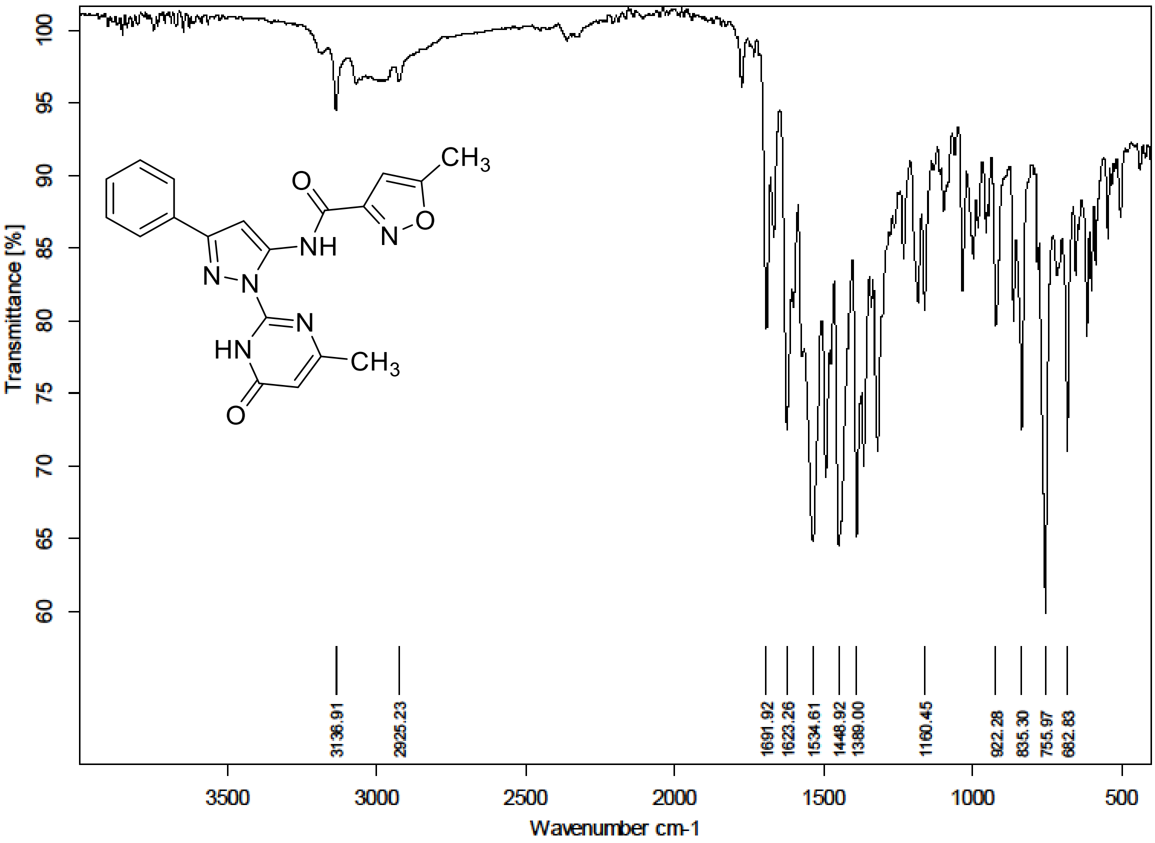

$^{1}$H NMR (300 MHz) spectrum of **37** in Trifluoroacetic Acid-*d*

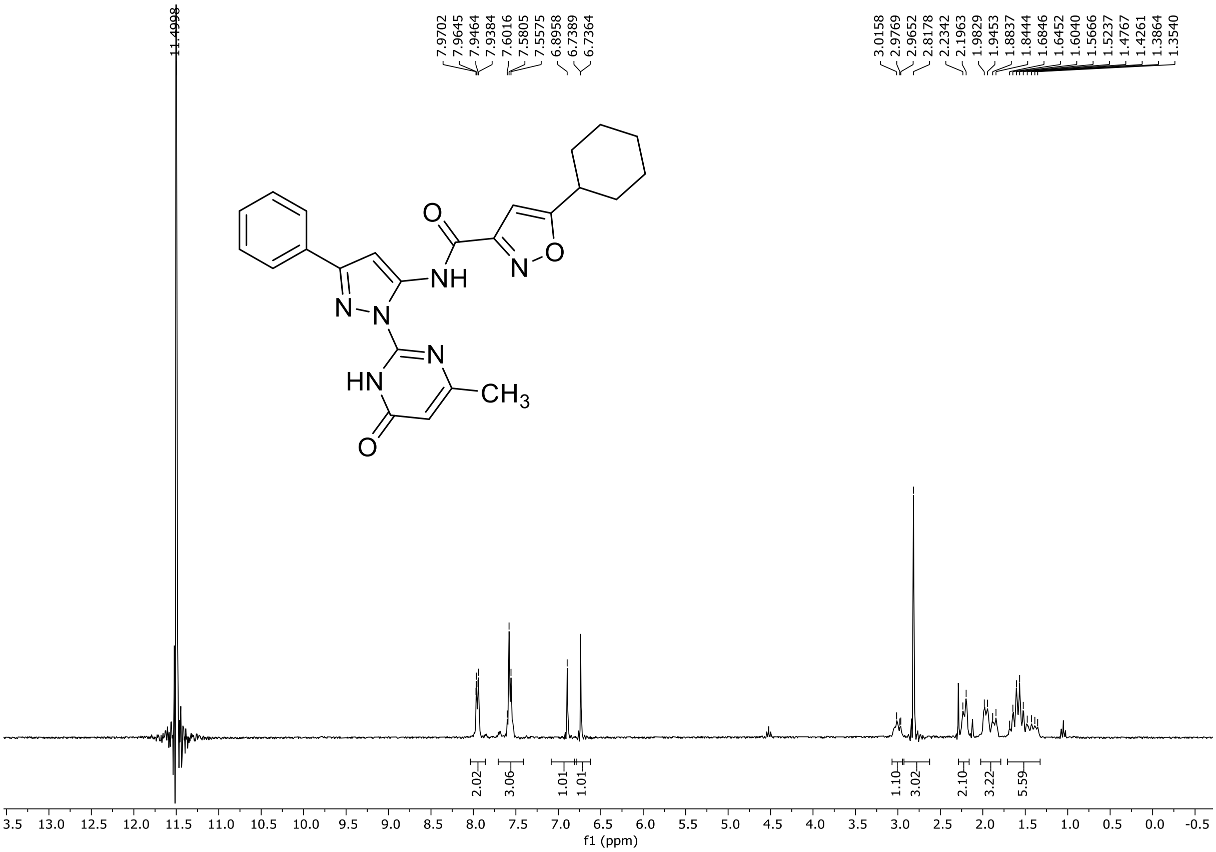

HRMS spectrum of **37**

**NAR-A-11** $C_{24}H_{24}N_6O_3$ ***m/z* 444.1910**

**ESI-** (MMI)
nitrogen flow 5 L/min, gas temperature 325°C, nebulizer 45 psi, skimmer 65 V, fragmentor 150 V, dissolved in DMSO, MeOH

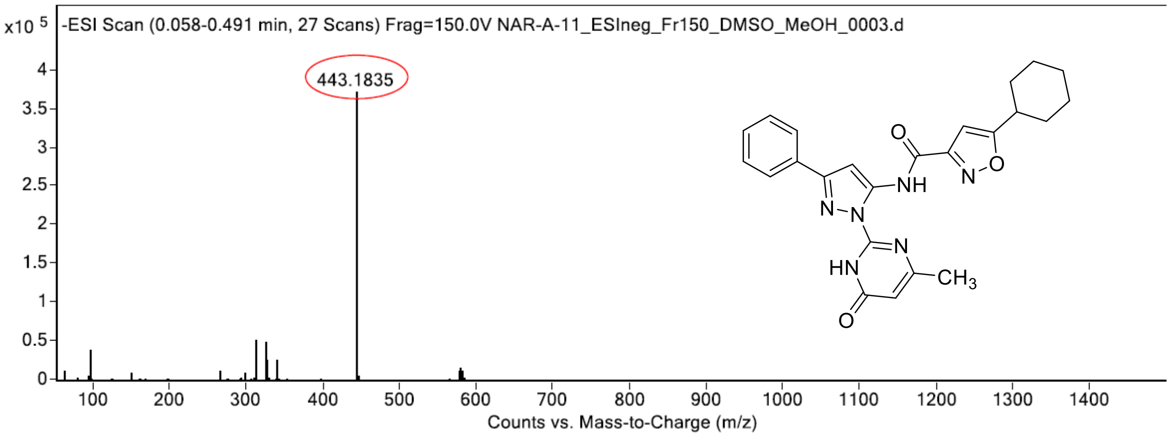

calculated mass: $[M-H]^-$ = 443.1837 observed: $[M-H]^-$ = 443.1835 mass accuracy = -0.5 ppm

FT-IR spectrum (neat) of **37**

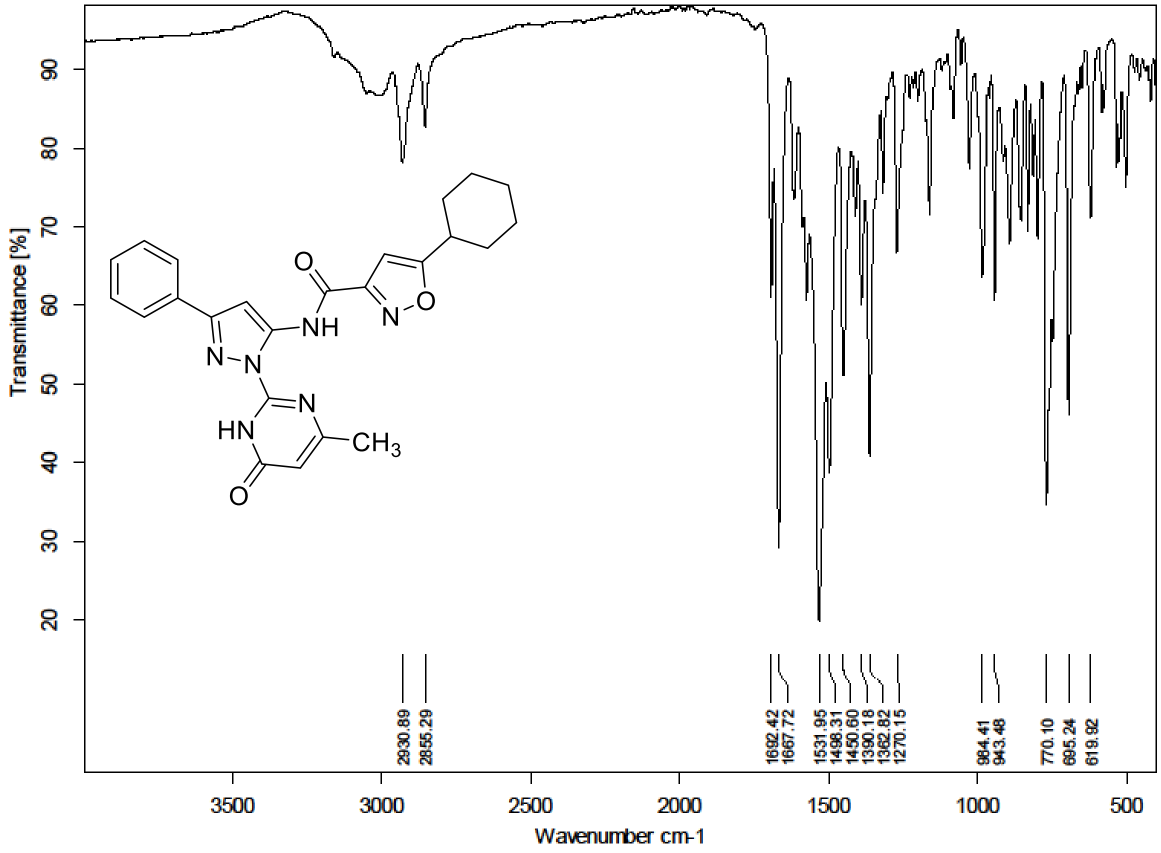

$^1$H NMR (300 MHz) spectrum of **38** in DMSO-$d_6$

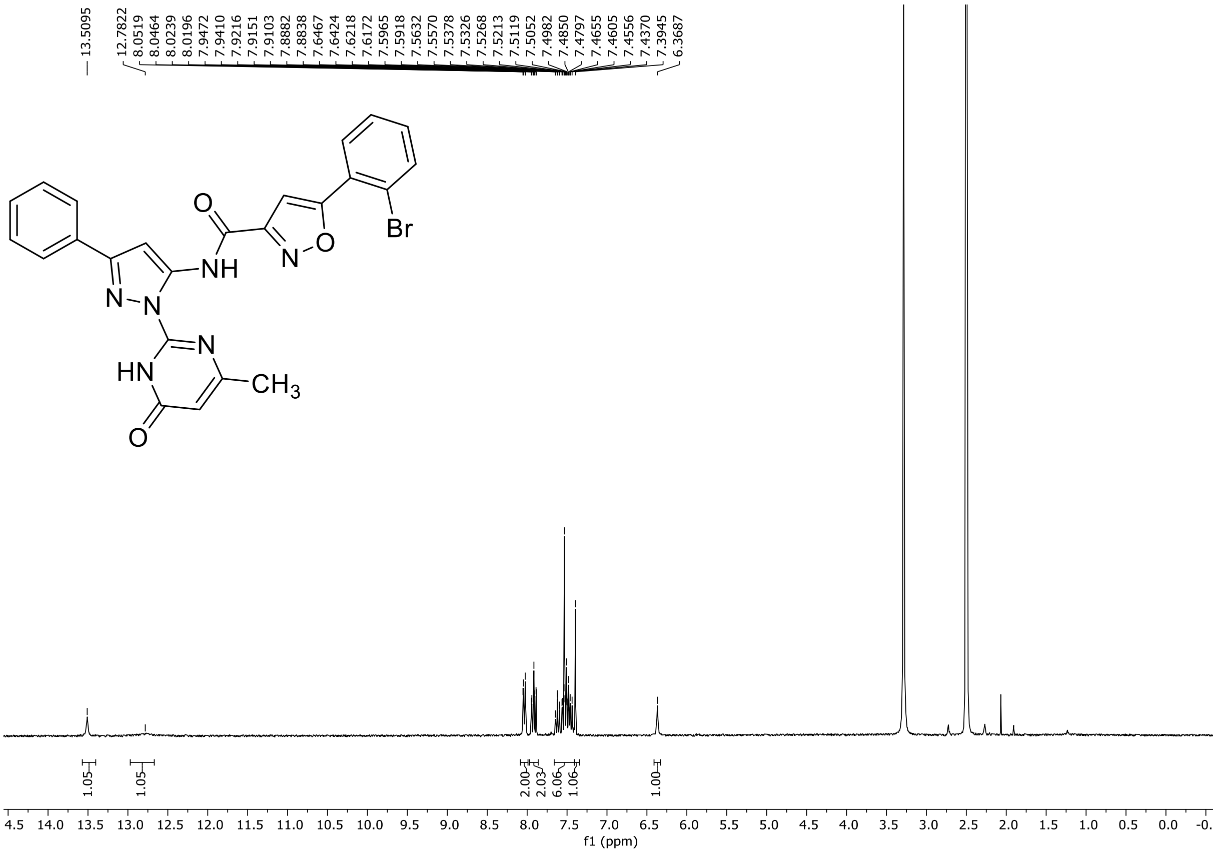

HRMS spectrum of **38**

**NAR-A-95** **$C_{24}H_{17}BrN_6O_3$** ***m/z*$_{100\%}$ 516.0546**

**Mixed Scan (APCI+ESI, both in negative mode)** (MMI)
nitrogen flow 5 L/min, gas temperature 325°C, nebulizer 45 psi, skimmer 65 V,
vaporizer 200°C, fragmentor 150 V, dissolved in DMSO, MeOH

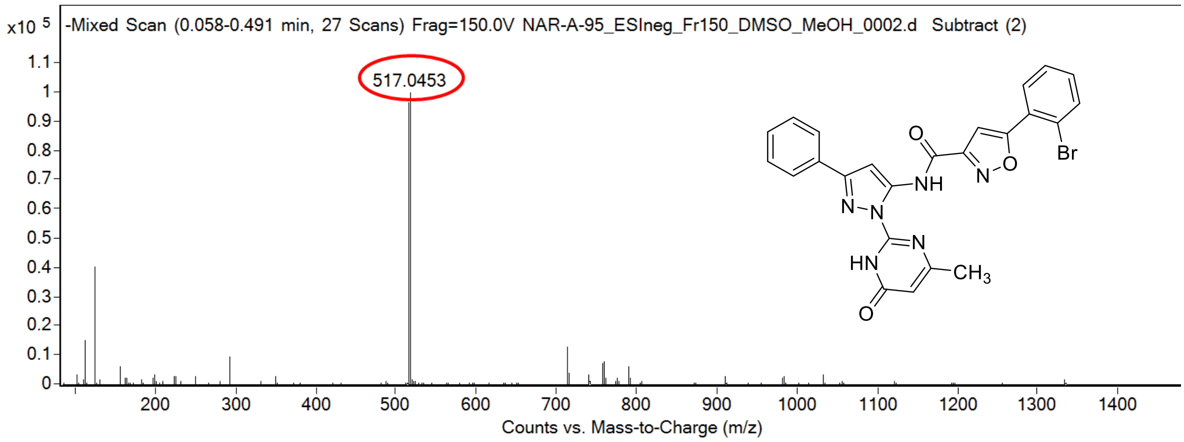

calculated mass: [M-H]$^-$ = 517.0456 observed: [M-H]$^-$ = 517.0453 mass accuracy = -0.6 ppm

FT-IR spectrum (neat) of **38**

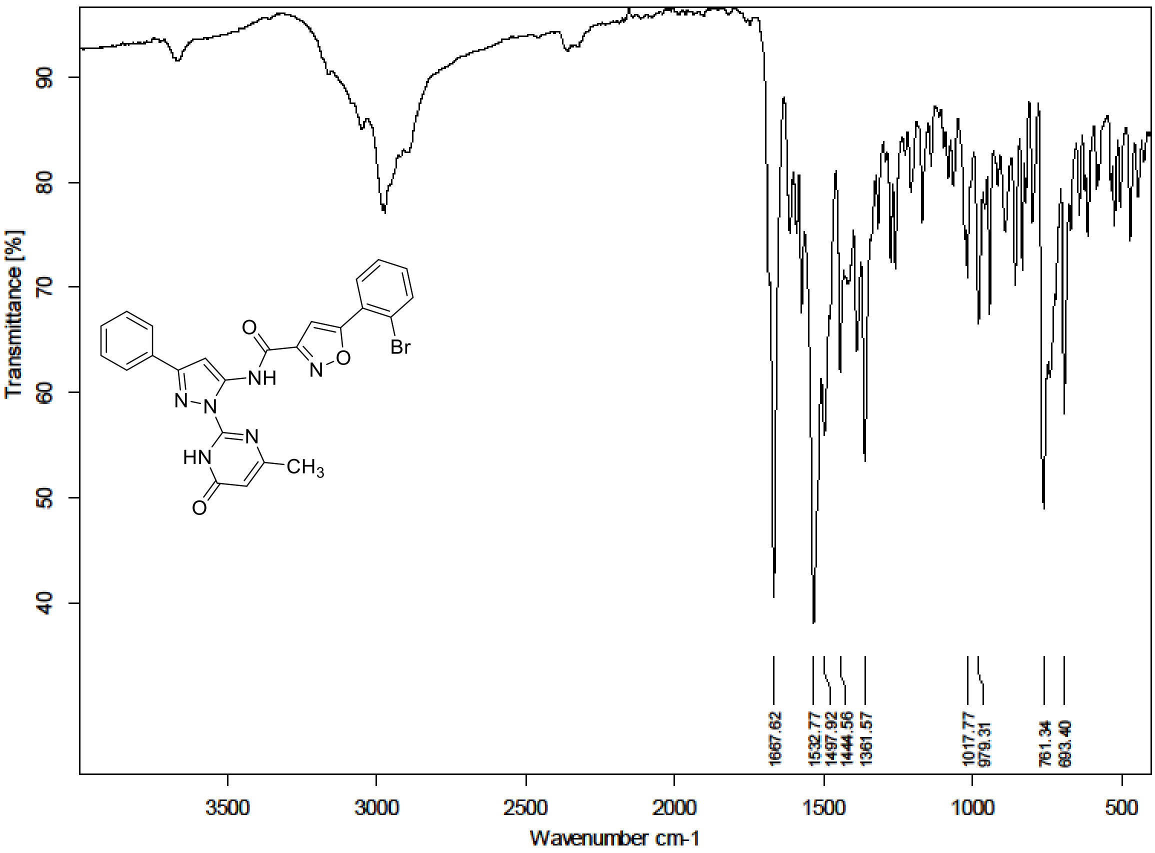

$^{1}H$ NMR (300 MHz) spectrum of **39** in DMSO-$d_6$

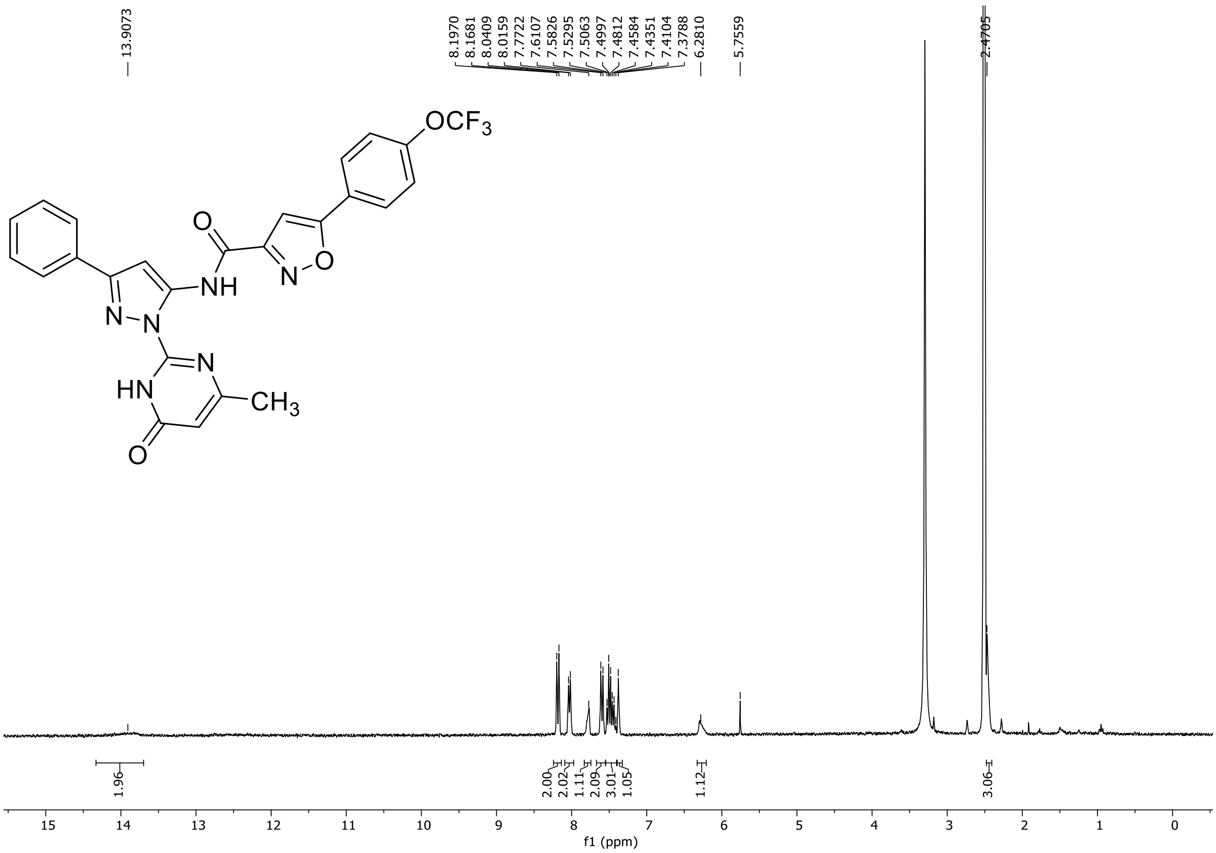

HRMS spectrum of **39**

**NAR-A-87** $C_{25}H_{17}F_3N_6O_4$ *m/z* **522.1263**

**ESI-** (MMI)
nitrogen flow 5 L/min, gas temperature 325°C, nebulizer 45 psi, skimmer 65 V, fragmentor 60 V, dissolved in DMSO, MeOH

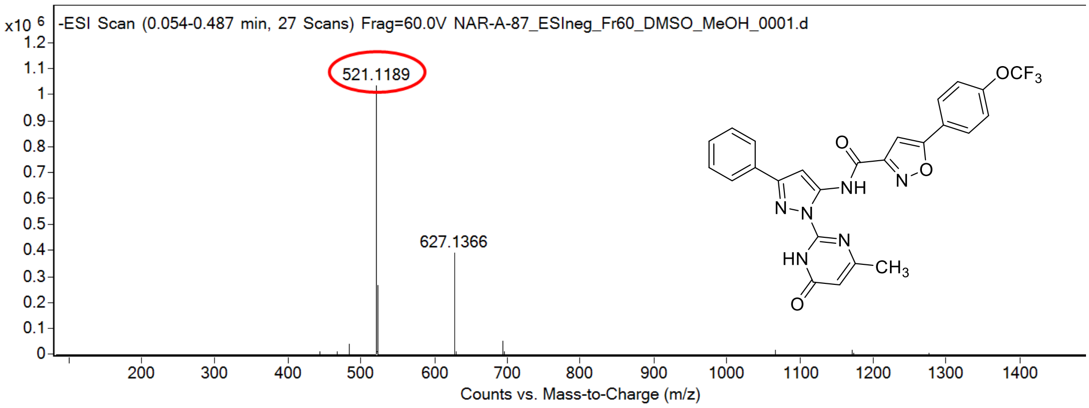

calculated mass: [M-H]⁻ = 521.1191 observed: [M-H]⁻ = 521.1189 mass accuracy = -0.4 ppm

FT-IR spectrum (neat) of **39**

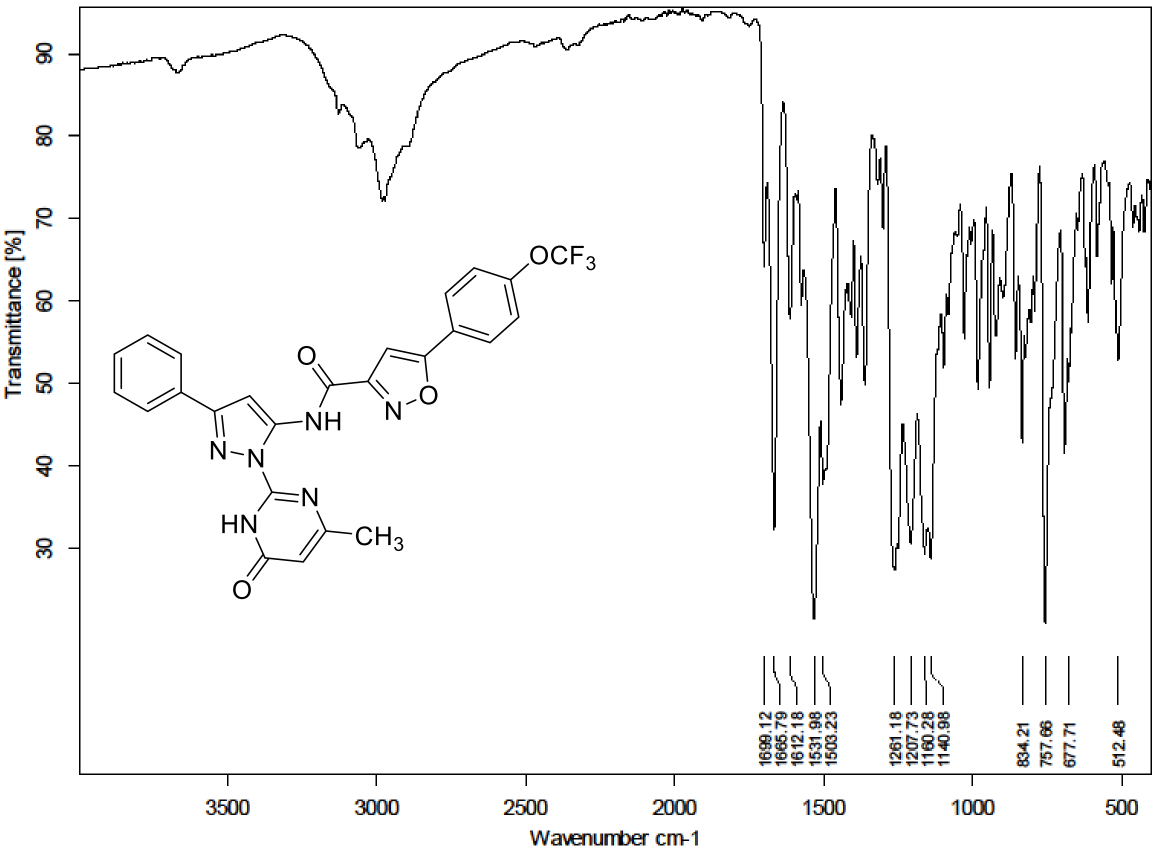

$^{1}H$ NMR (300 MHz) spectrum of **40** in DMSO-$d_6$

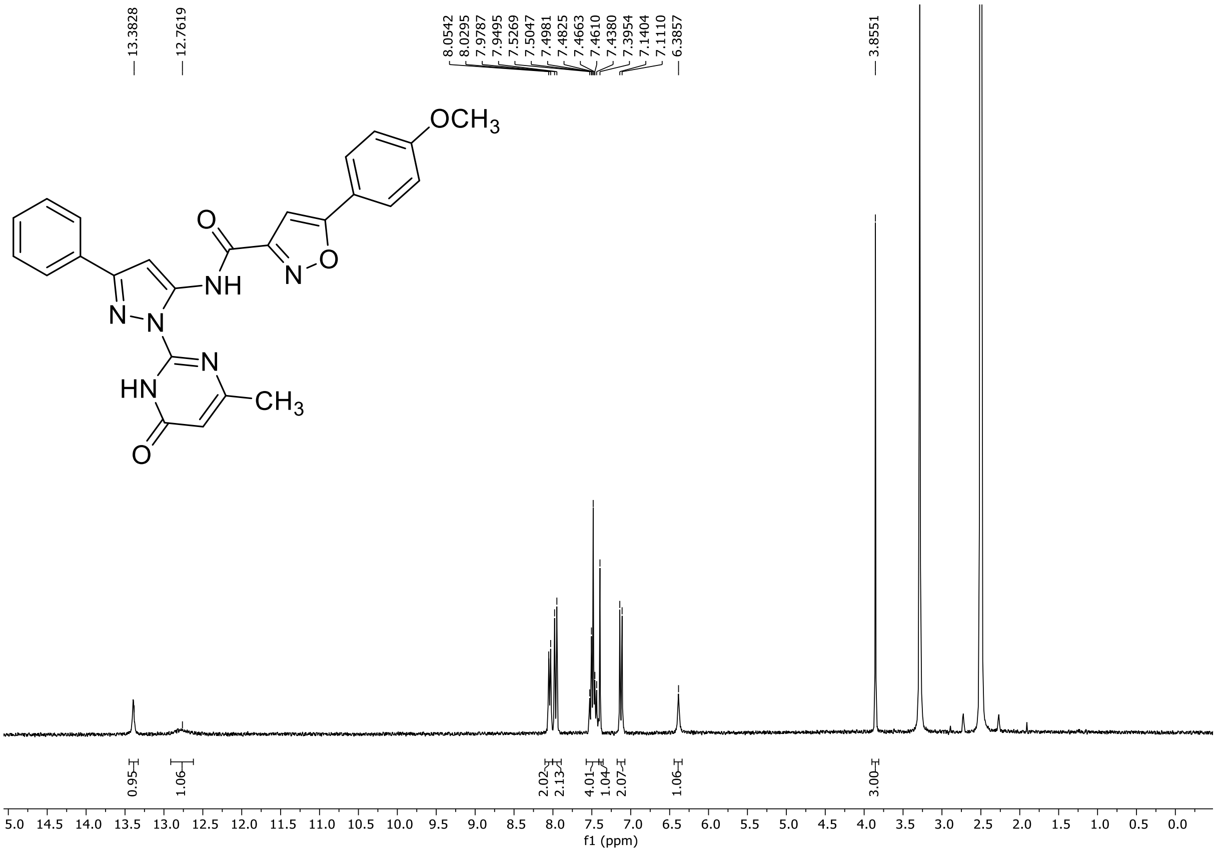

HRMS spectrum of **40**

### NAR-A-131-1

$C_{25}H_{20}N_6O_4$ *m/z* **468.1546**

**ESI-** (MMI)
nitrogen flow 5 L/min, gas temperature 325°C, nebulizer 45 psi, skimmer 65 V, fragmentor 120 V, dissolved in DMSO, MeOH

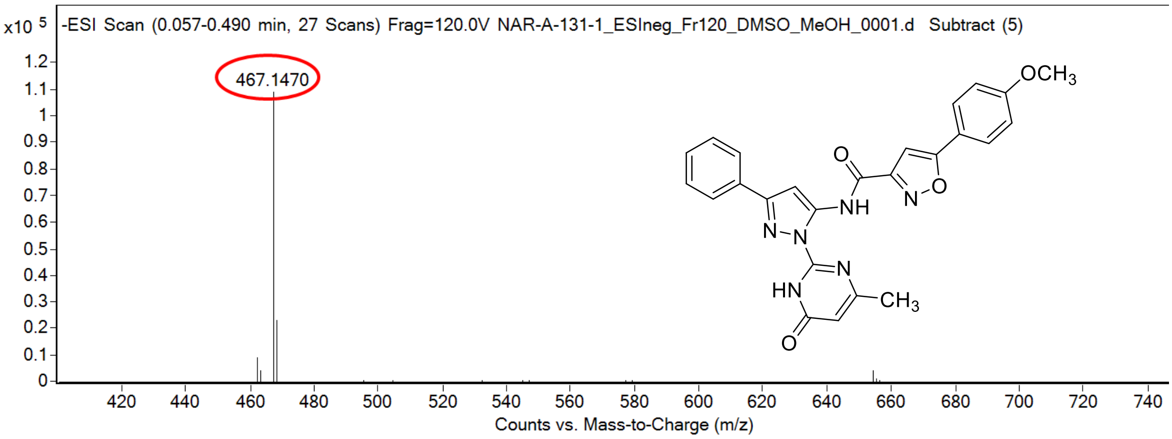

calculated mass: [M-H]$^-$ = 467.1473 observed: [M-H]$^-$ = 467.1470 mass accuracy = -0.6 ppm

FT-IR spectrum (neat) of **40**

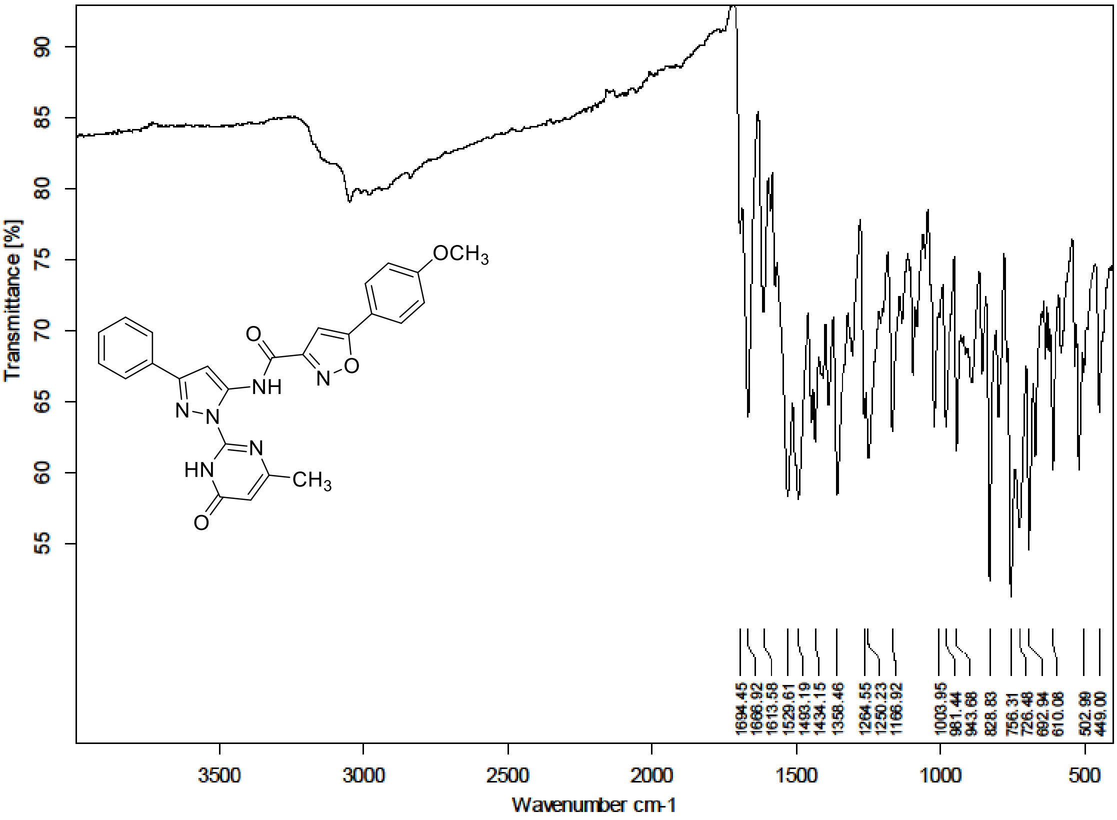

[1]H NMR (300 MHz) spectrum of **41** in Trifluoroacetic Acid-*d*

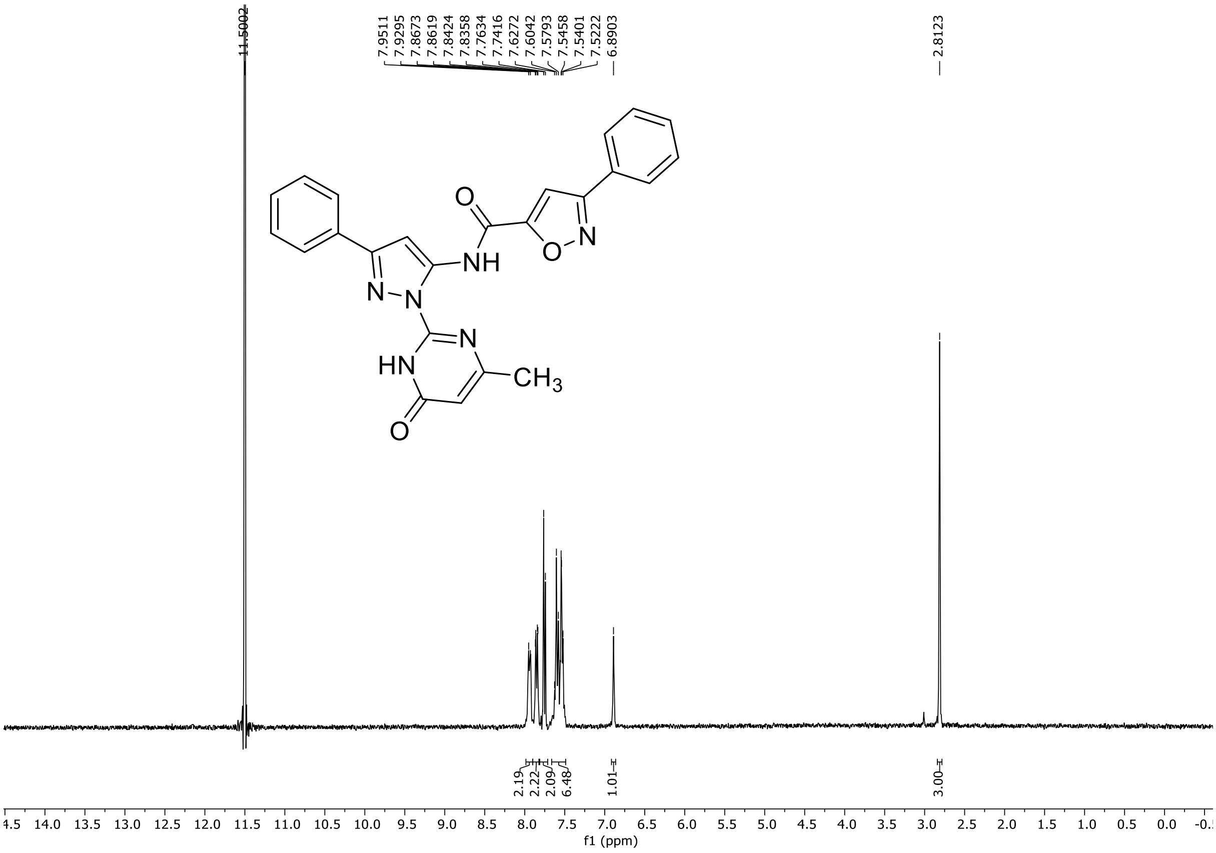

HRMS spectrum of **41**

**NAR-A-153** $C_{24}H_{18}N_6O_3$ **mono *m/z* 438.1440**

**APCI +** (MMI)
nitrogen flow 5 L/min, gas temperature 325°C, nebulizer 45 psi, skimmer 65 V, vaporizer 200°C, fragmentor 33 V, dissolved in methanol / $CH_2Cl_2$

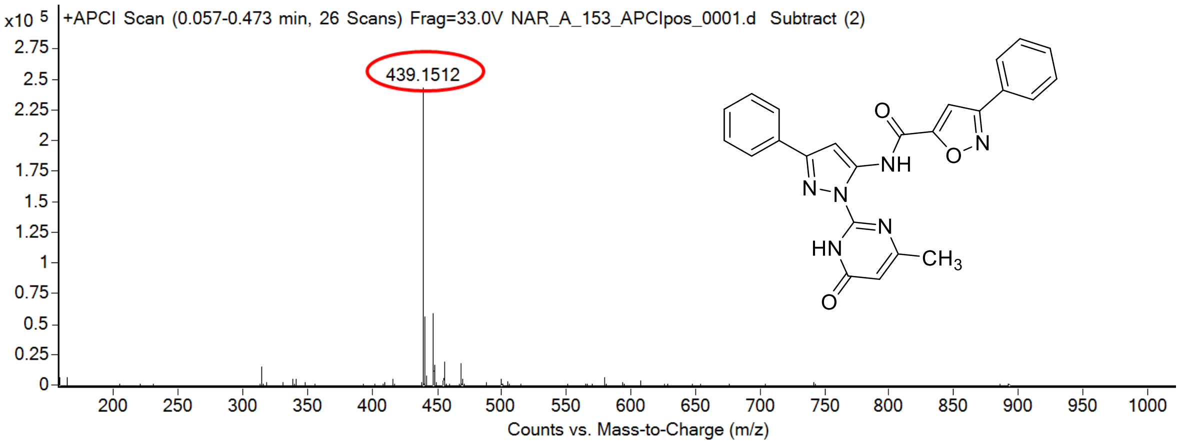

calculated mass: $[M+H]^+$ = 439.1513 observed: $[M+H]^+$ = 439.1512 mass accuracy = - 0.2 ppm

FT-IR spectrum (neat) of **41**

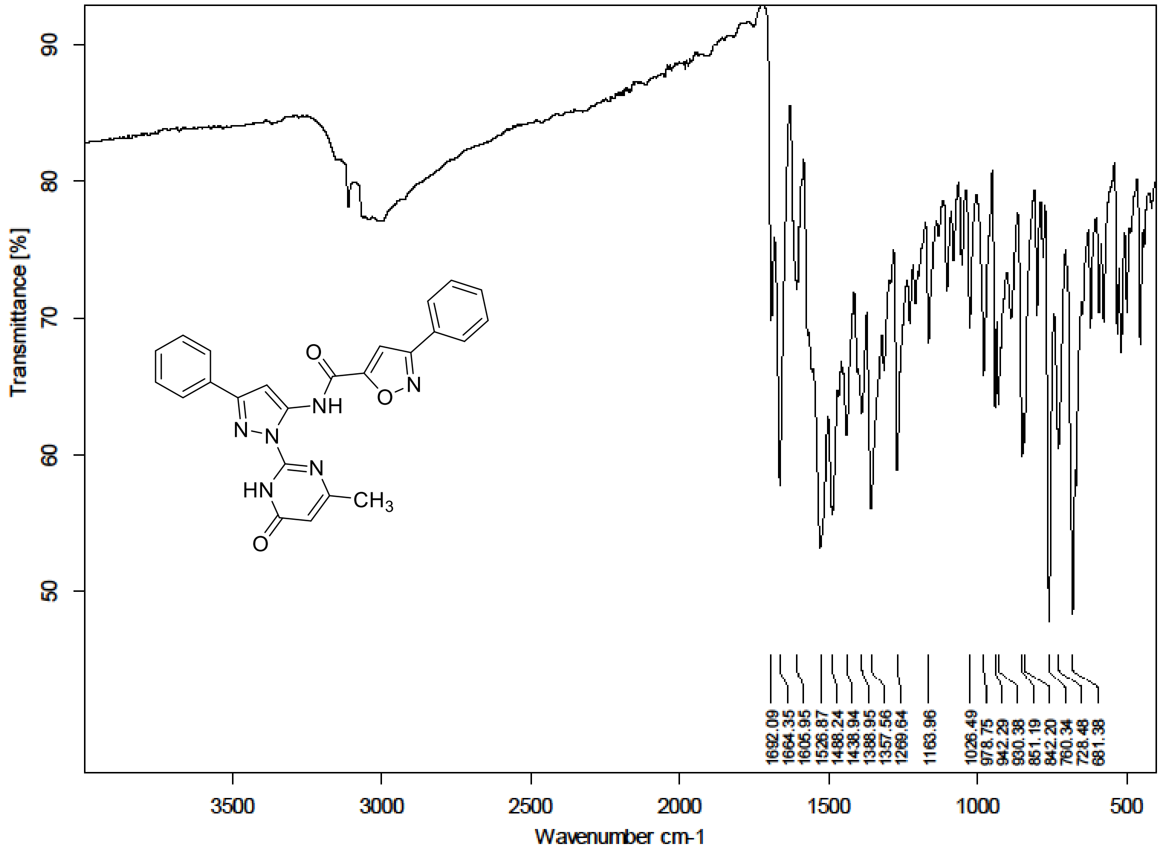

[1]H (700 MHz) and [13]C NMR (176 MHz) spectra of **42** in DMSO-$d_6$

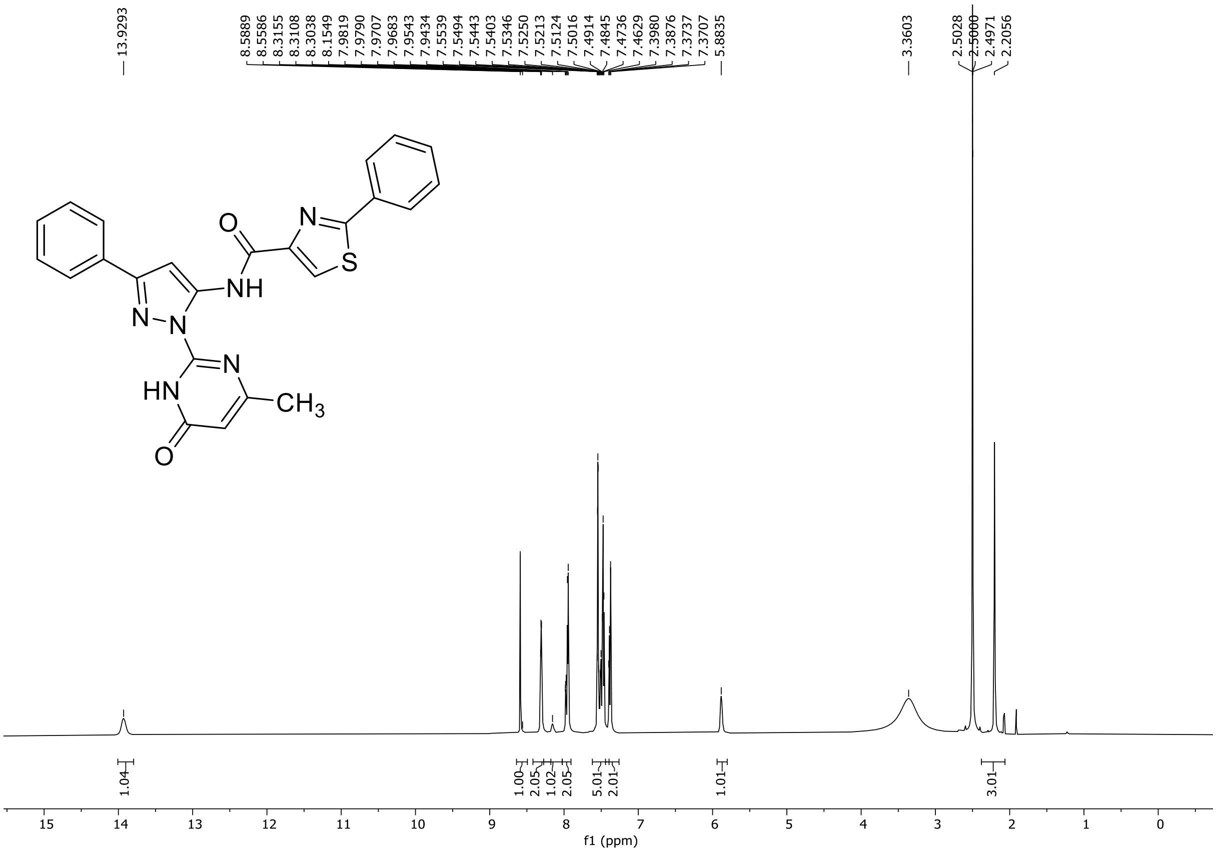

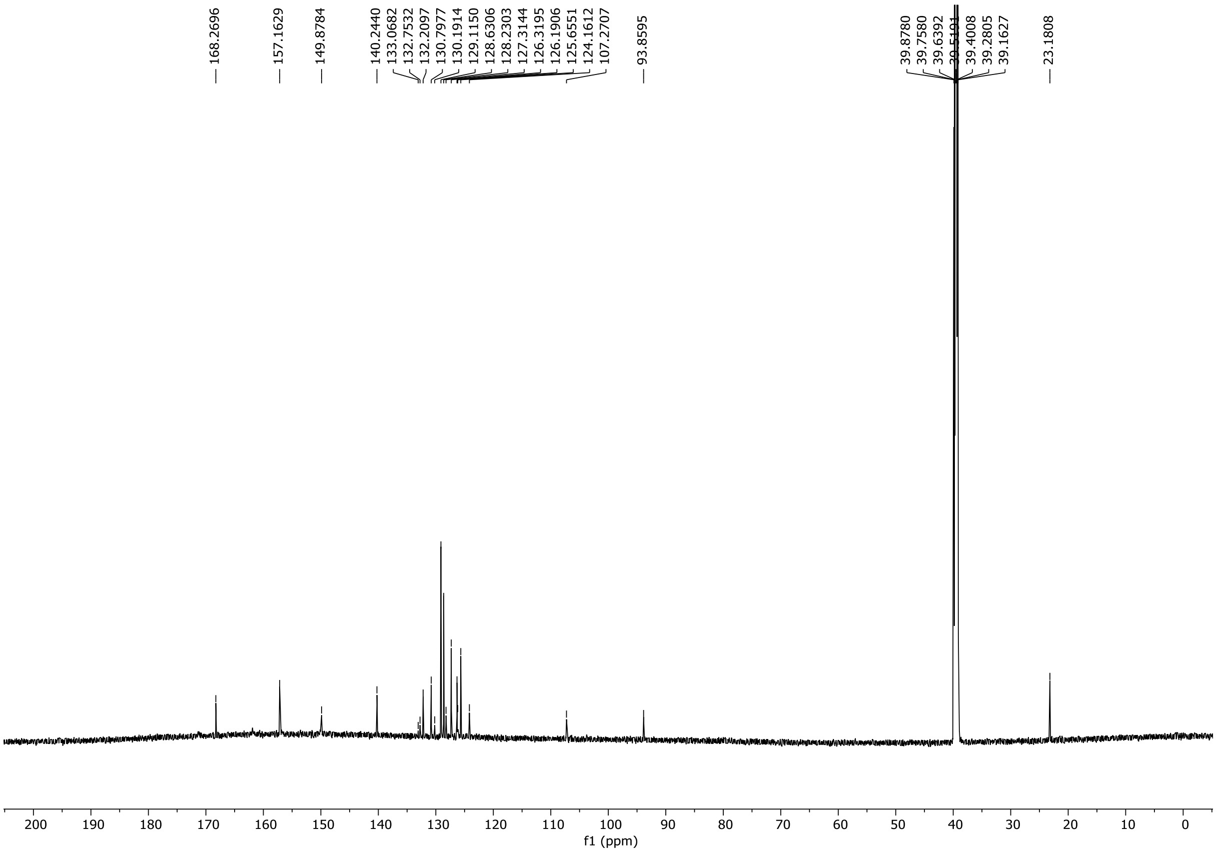

HRMS spectrum of **42**

**NAR-A-154** $C_{24}H_{18}N_6O_2S$ **mono** ***m/z*** **454.1212**

**APCI +** (MMI)
nitrogen flow 5 L/min, gas temperature 325°C, nebulizer 45 psi, skimmer **65** V, vaporizer 200°C, fragmentor 30 V, dissolved in methanol

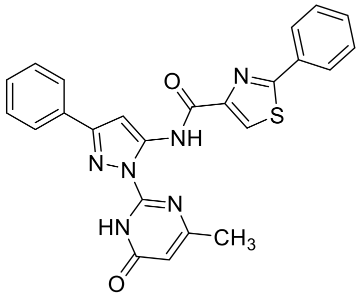

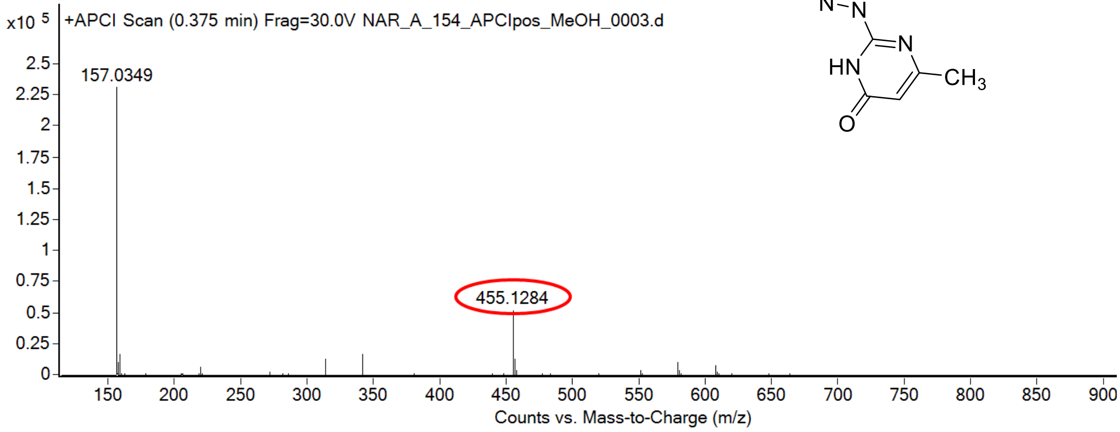

calculated mass: $[M+H]^+ = 455.1285$ observed: $[M+H]^+ = 455.1284$ mass accuracy = - 0.2 ppm

FT-IR spectrum (neat) of **42**

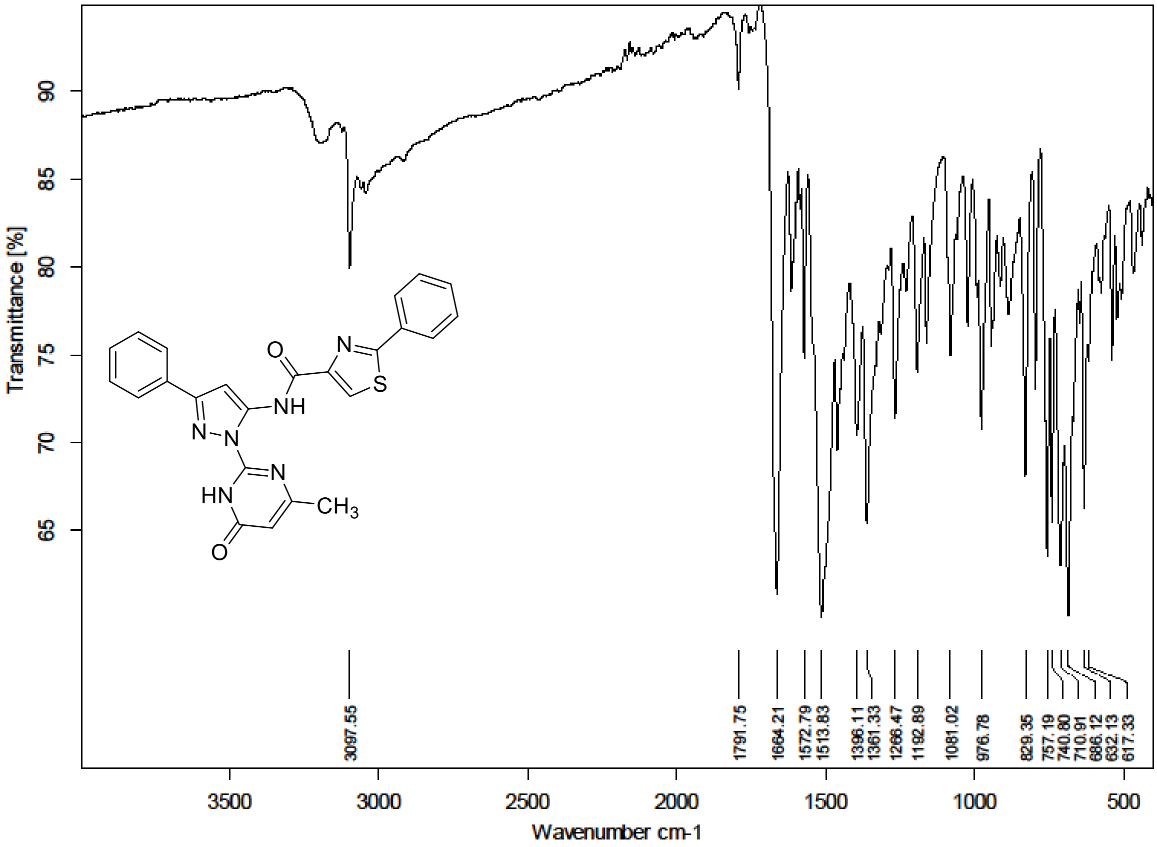

[1]H NMR (300 MHz) spectrum of **43** in DMSO-$d_6$

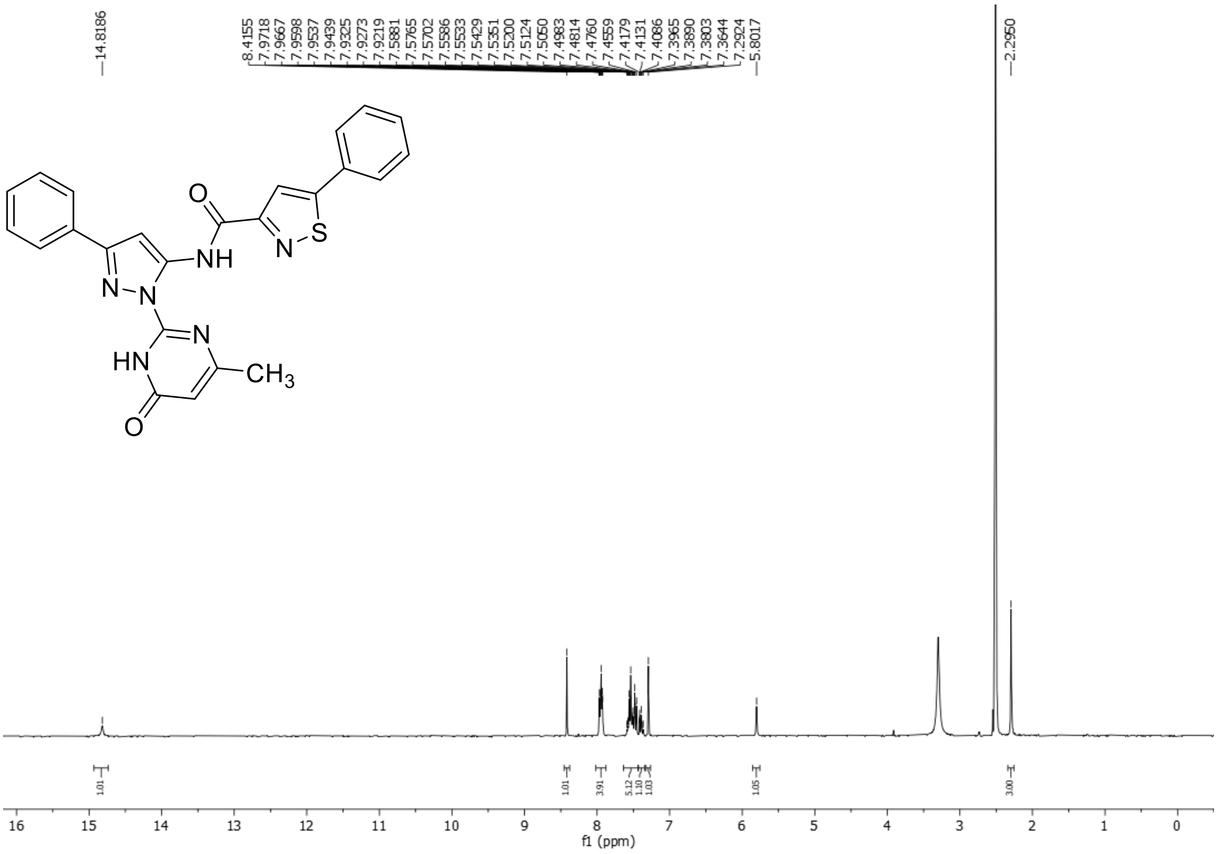

HRMS spectrum of **43**

**NAR-A-155** $C_{24}H_{18}N_6O_2S$ mono *m/z* 454.1212

**APCI +** (MMI)
nitrogen flow 5 L/min, gas temperature 325°C, nebulizer 45 psi, skimmer 65 V, vaporizer 200°C, fragmentor 22 V, dissolved in methanol / $CH_2Cl_2$

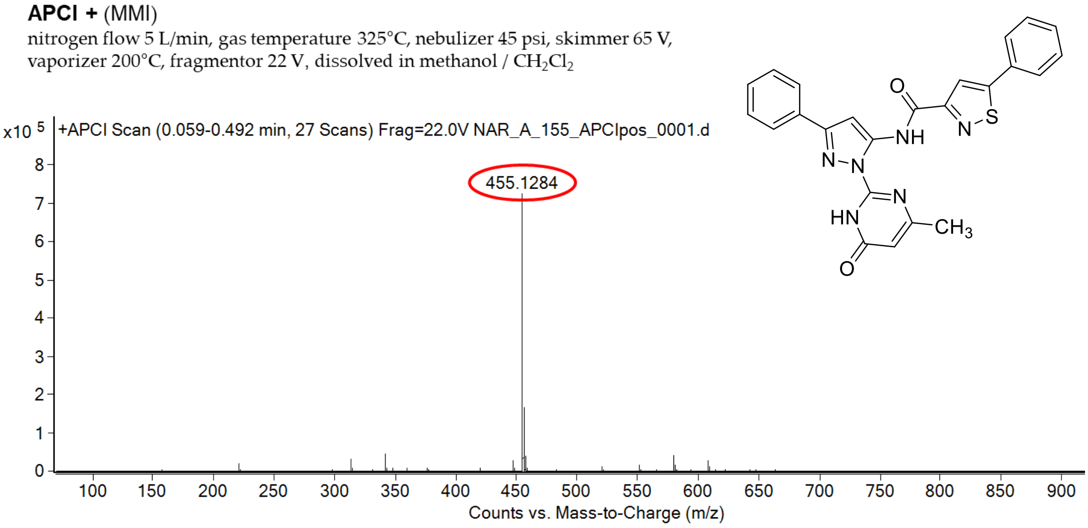

calculated mass: $[M+H]^+$ = 455.1285 observed: $[M+H]^+$ = 455.1284 mass accuracy = - 0.2 ppm

FT-IR spectrum (neat) of **43**

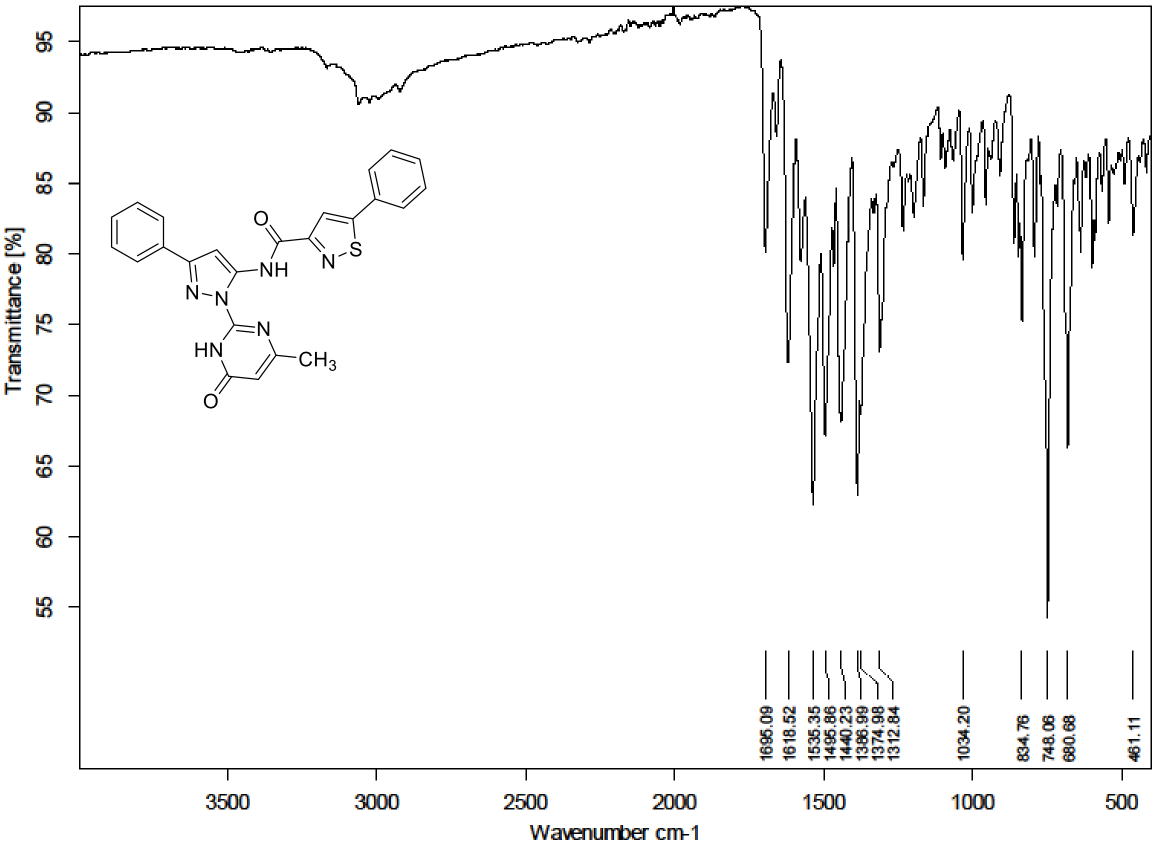

[1]H (300 MHz) and [13]C NMR (176 MHz) spectra of **44** in DMSO-*d*

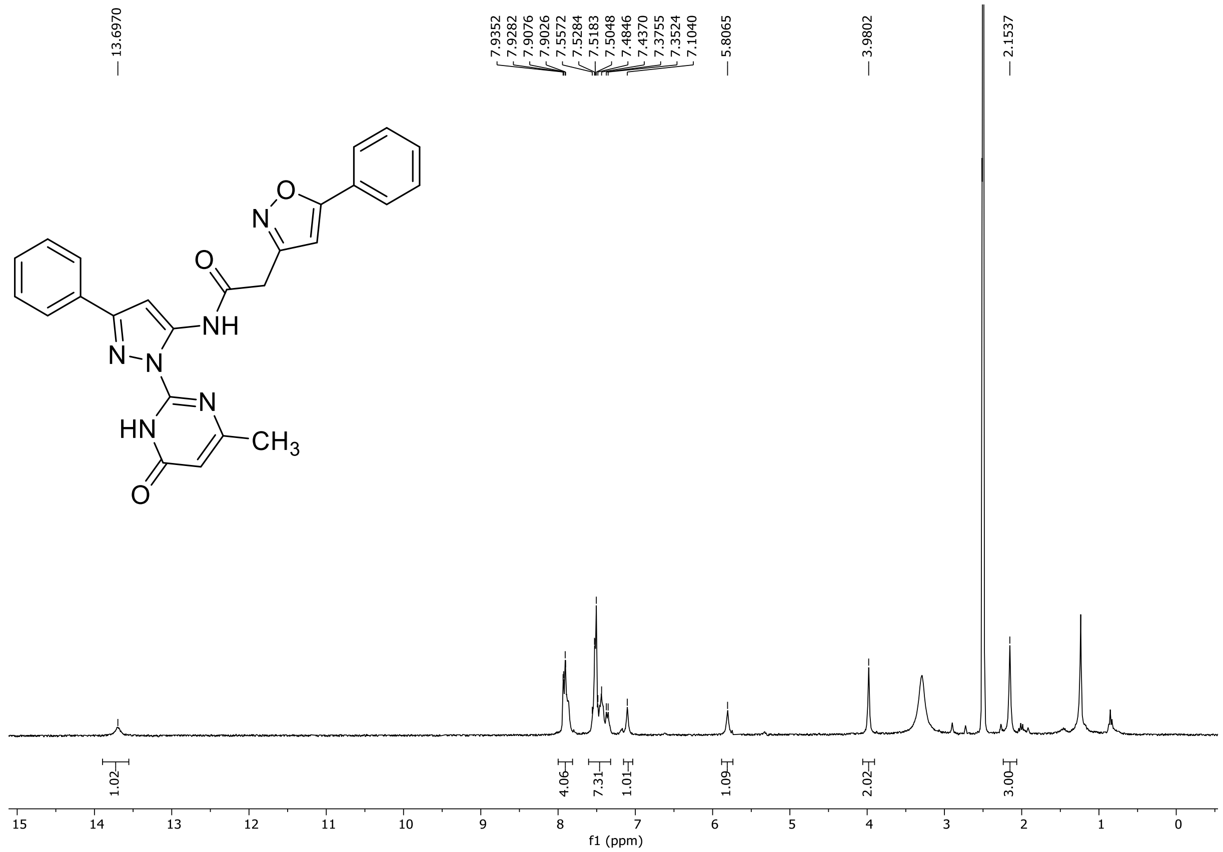

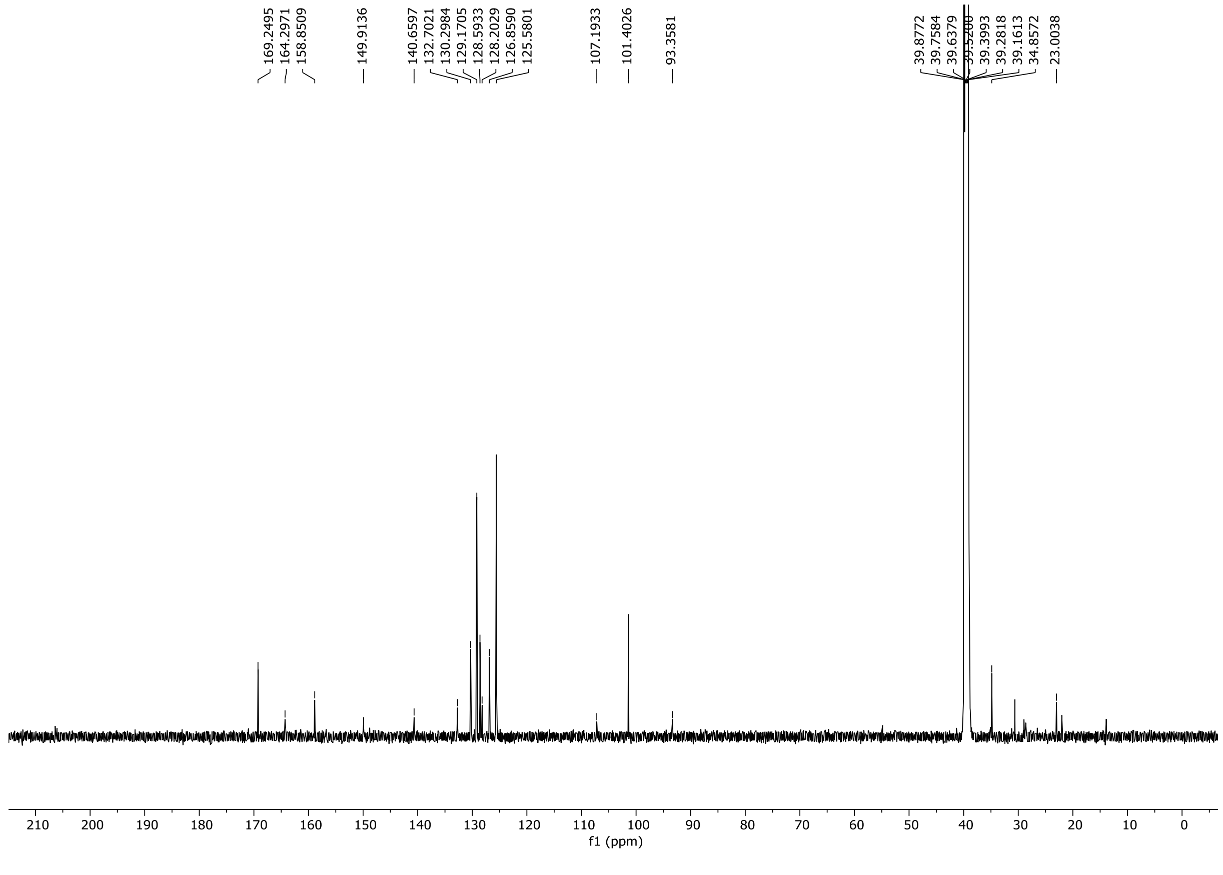

HRMS spectrum of **44**

**NAR-A-159** $C_{25}H_{20}N_6O_3$ **mono *m/z* 452.1597**

**APCI +** (MMI)
nitrogen flow 5 L/min, gas temperature 325°C, nebulizer 45 psi, skimmer **65** V, vaporizer 200°C, fragmentor 30 V, dissolved in methanol

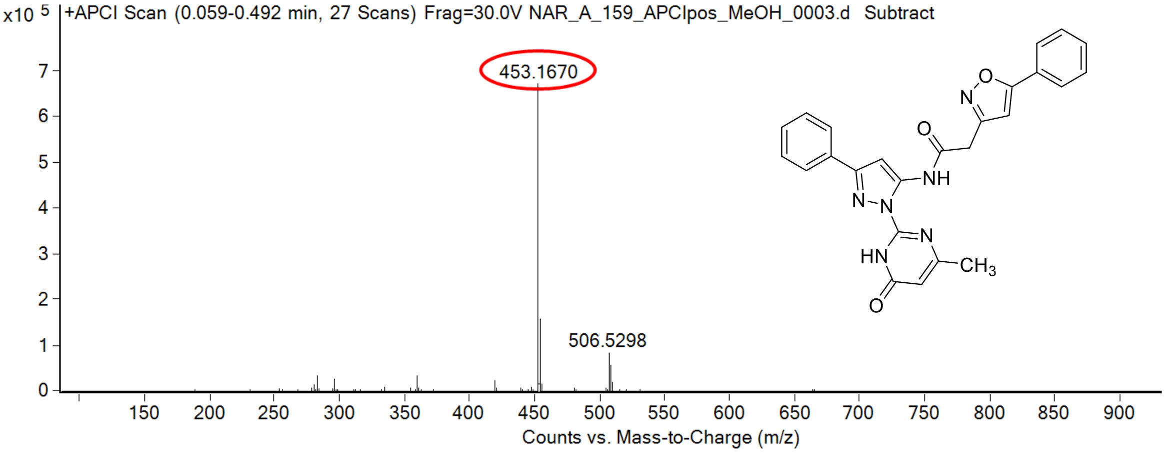

calculated mass: $[M+H]^+$ = 453.1670 observed: $[M+H]^+$ = 453.1670 mass accuracy = < 0.1 ppm

FT-IR spectrum (neat) of **44**

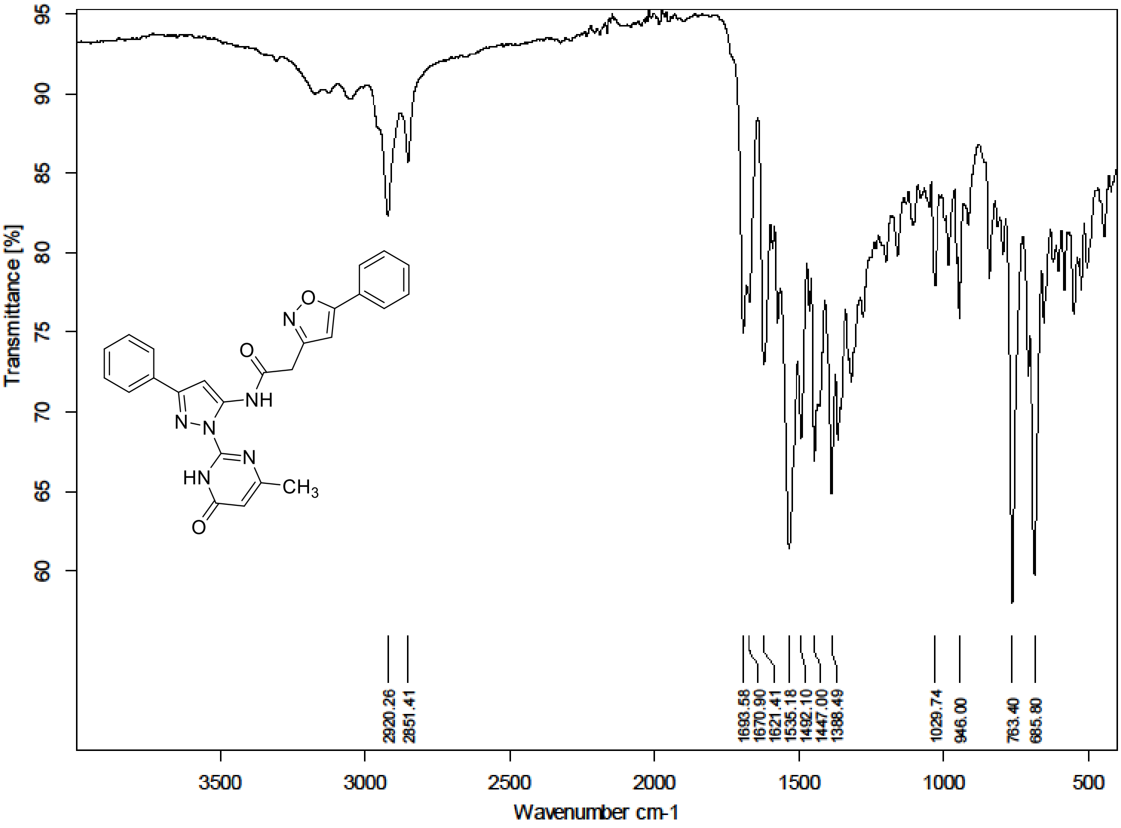

[1]H NMR (300 MHz) spectrum of **S190** in Chloroform-*d*

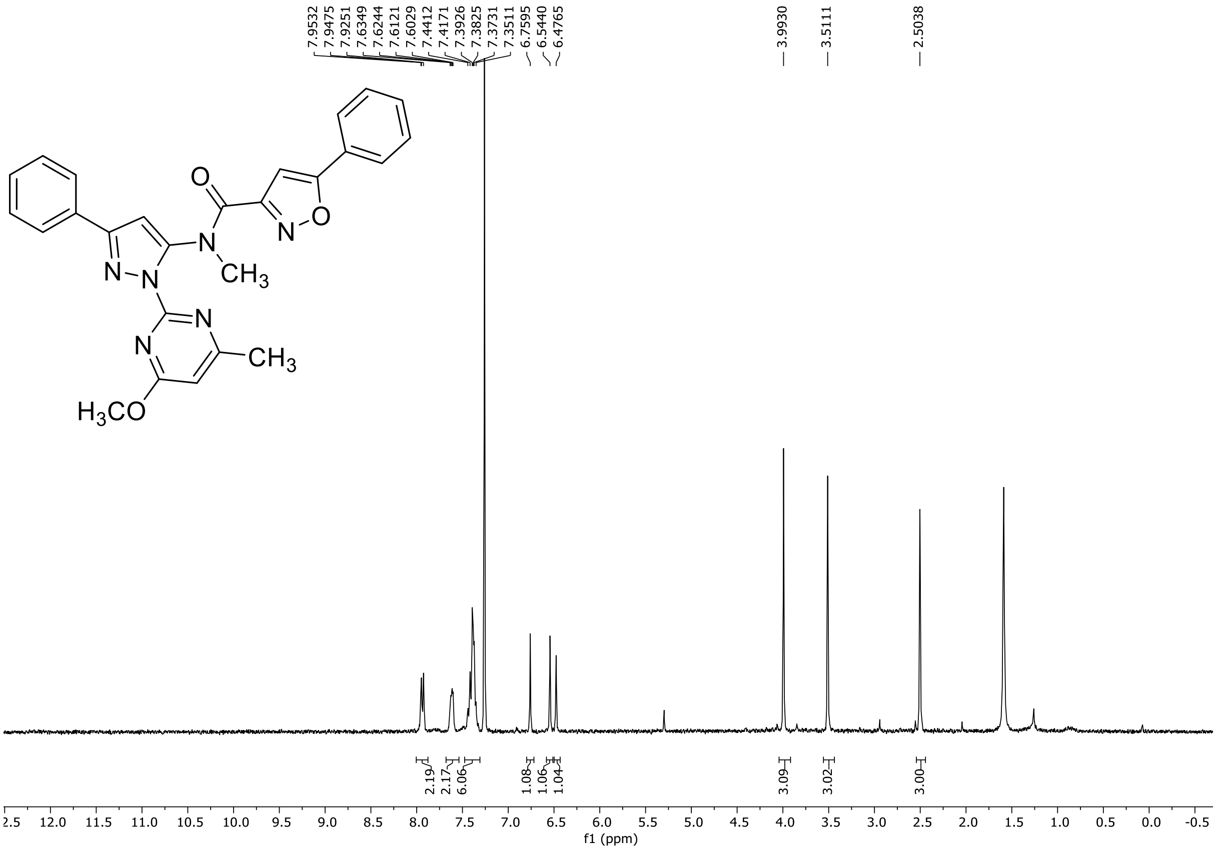

HRMS spectrum of **S190**

**NAR-A-165** **$C_{26}H_{22}N_6O_3$** **mono *m/z* 466.1753**

**APCI +** (MMI)
nitrogen flow 5 L/min, gas temperature 325°C, nebulizer 45 psi, skimmer **65** V, vaporizer 200°C, fragmentor 15 V, dissolved in methanol

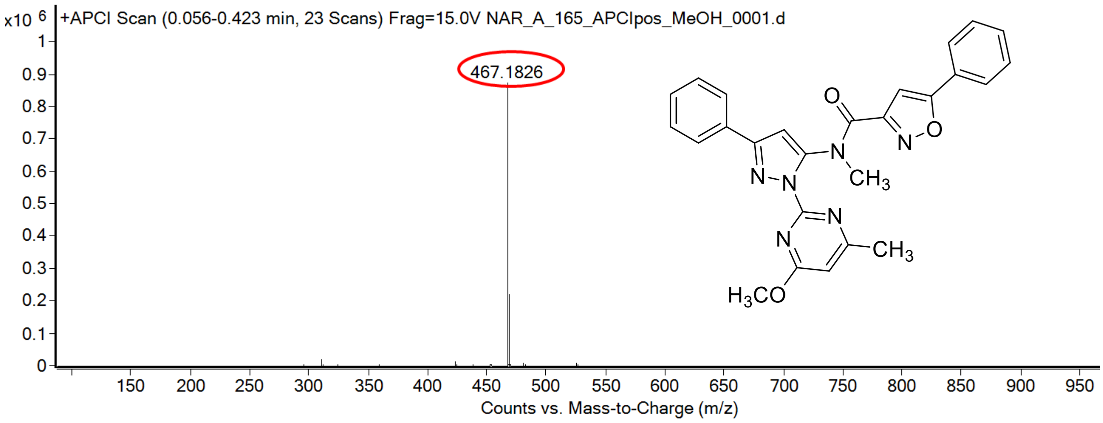

calculated mass: $[M+H]^+ = 467.1826$ observed: $[M+H]^+ = 467.1826$ mass accuracy = < 0.1 ppm

$^{1}H$ (300 MHz) and $^{13}C$ NMR (75 MHz) spectra of **45** in Chloroform-*d*

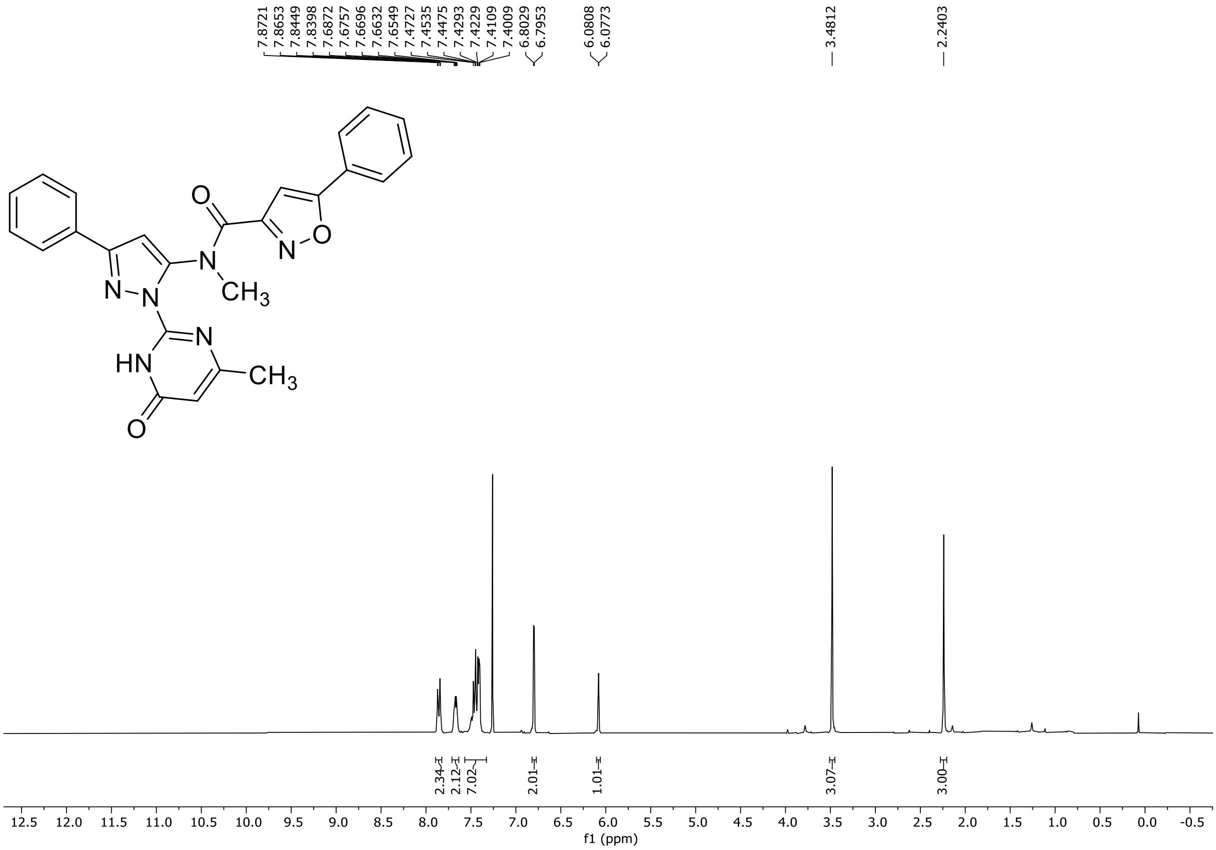

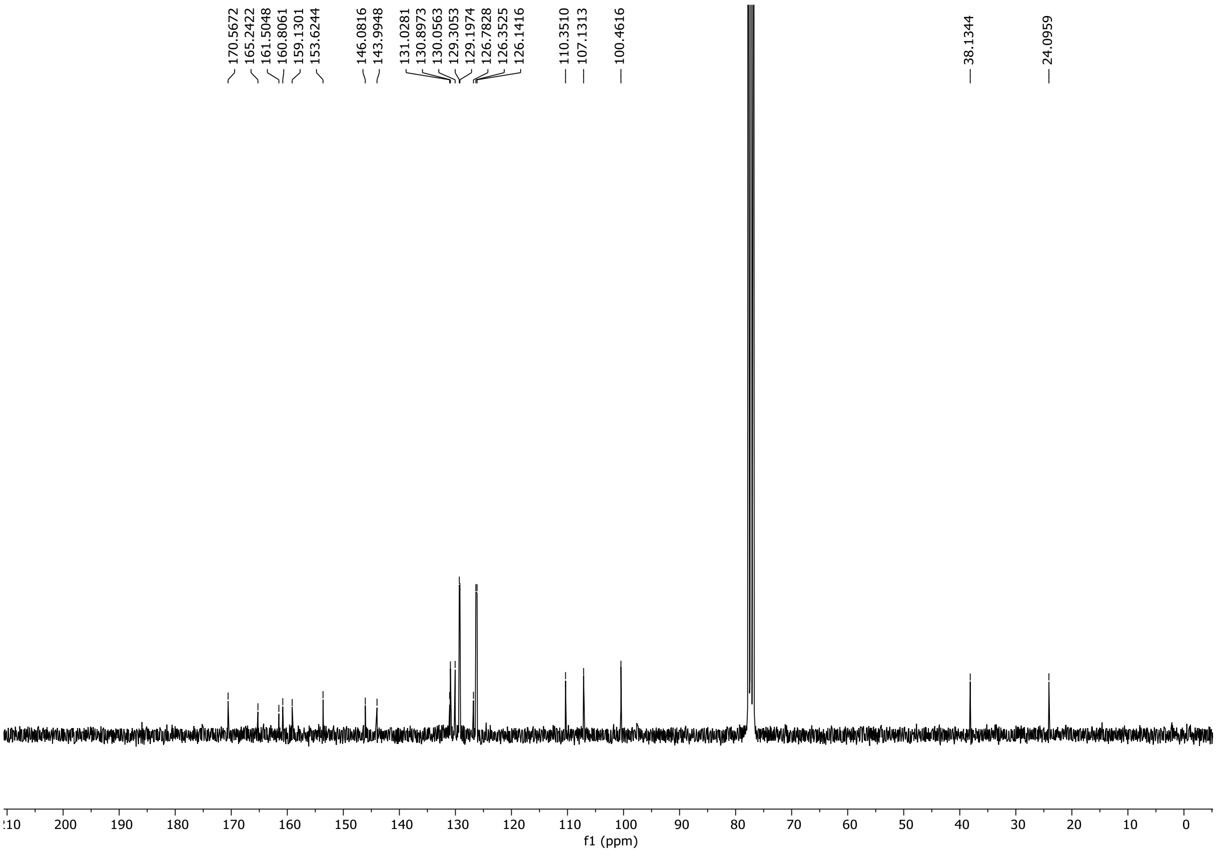

HRMS spectrum of **45**

**NAR-A-176** $C_{25}H_{20}N_6O_3$ mono *m/z* 452.1597

**APCI +** (MMI)
nitrogen flow 5 L/min, gas temperature 325°C, nebulizer 45 psi, skimmer **65** V, vaporizer 200°C, fragmentor 15 V, dissolved in methanol

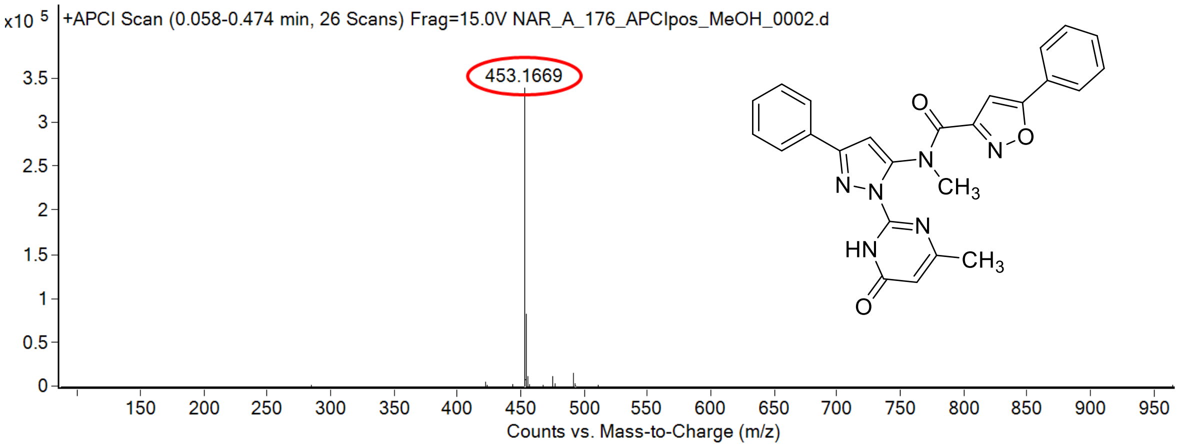

calculated mass: $[M+H]^+$ = 453.1670 observed: $[M+H]^+$ = 453.1669 mass accuracy = - 0.2 ppm

FT-IR spectrum (neat) of **45**

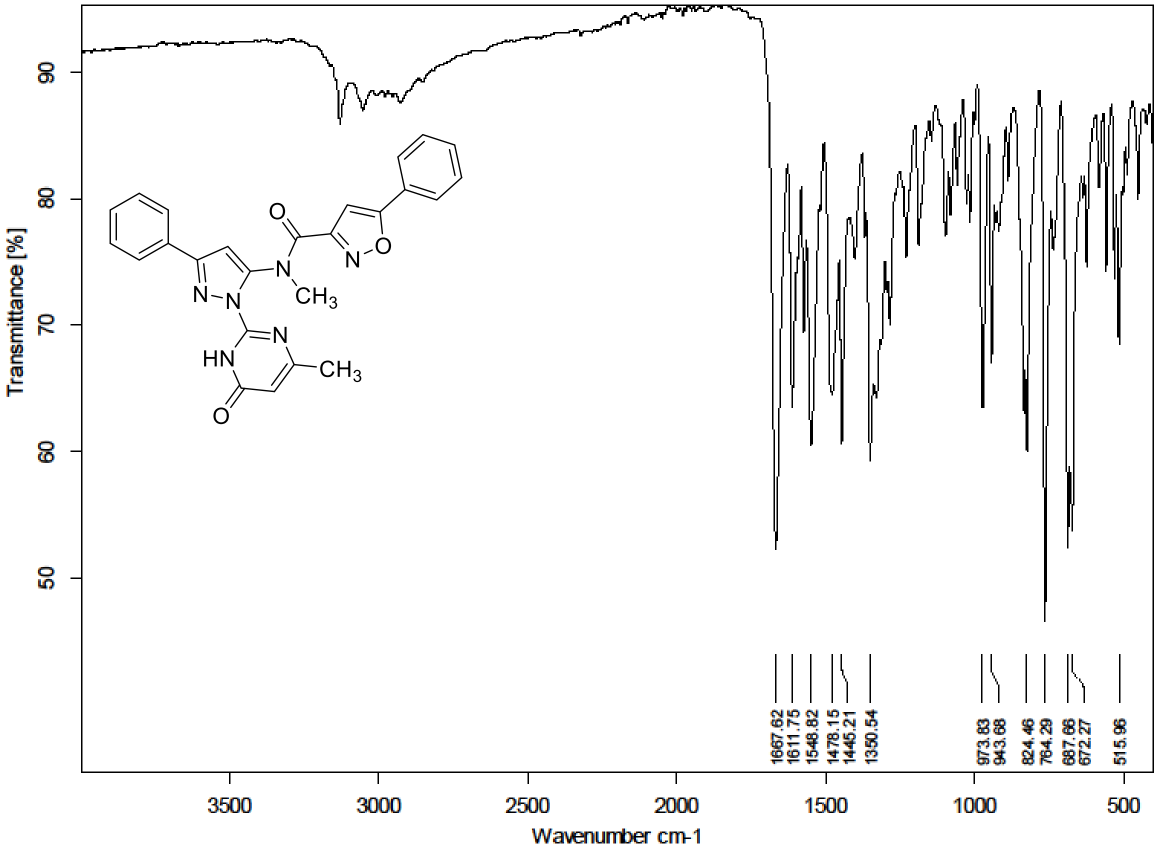

$^{1}H$ (300 MHz) and $^{13}C$ NMR (126 MHz) spectra of **46** Chloroform-*d*

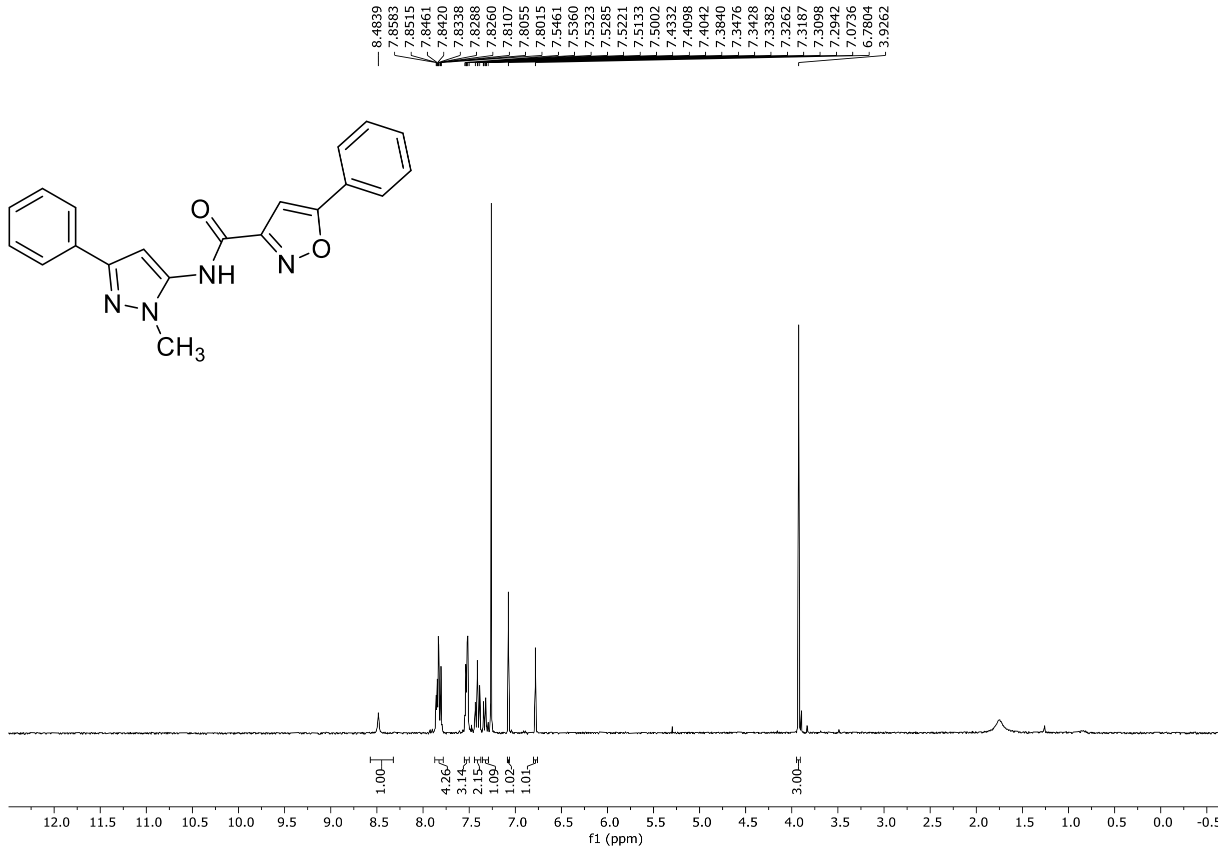

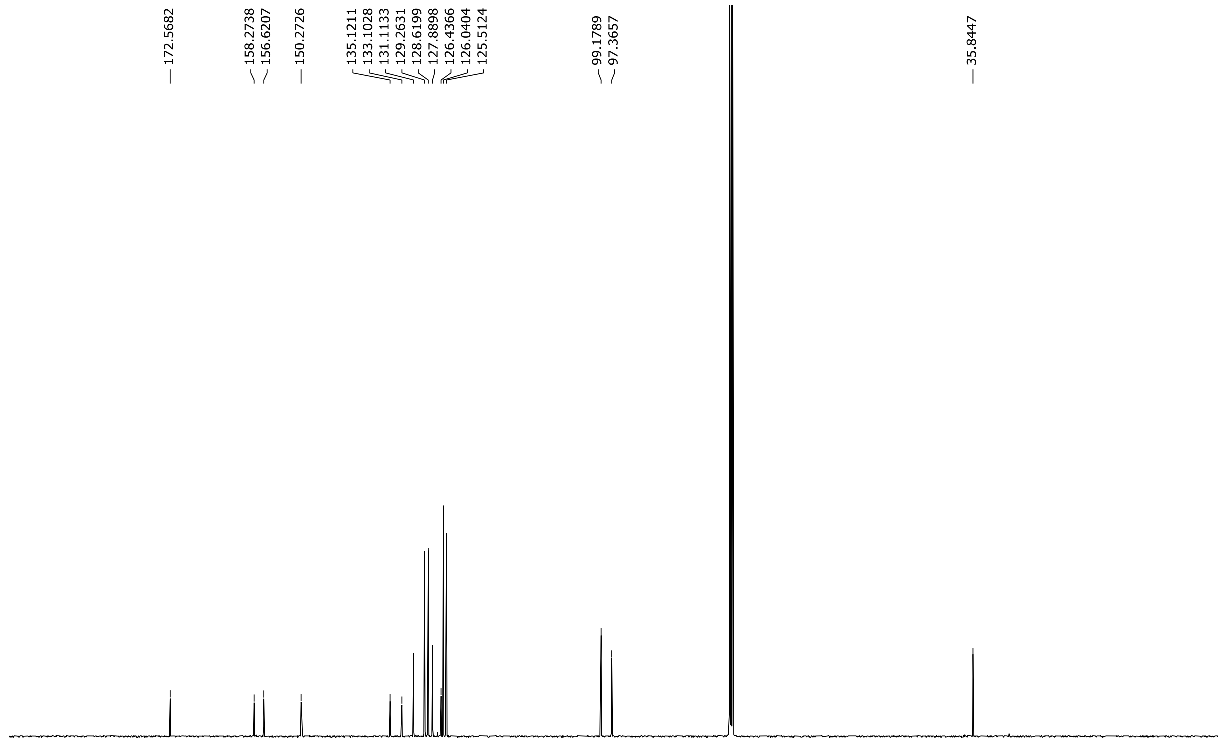

HRMS spectrum of **46**

**NAR-A-88** $C_{20}H_{16}N_4O_2$ *m/z* **344.1273**

**APCI+** (MMI)
nitrogen flow 5 L/min, gas temperature 325°C, nebulizer 45 psi, skimmer 65 V, vaporizer 200°C, fragmentor 20 V, dissolved in MeOH

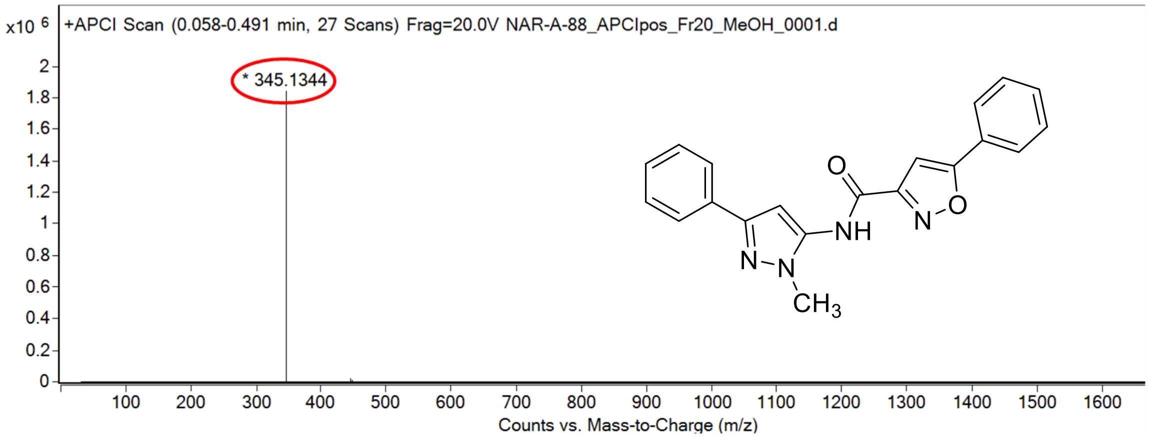

calculated mass: $[M+H]^+ = 345.1346$ observed: $[M+H]^+ = 345.1344$ mass accuracy = -0.6 ppm

FT-IR spectrum (neat) of **46**

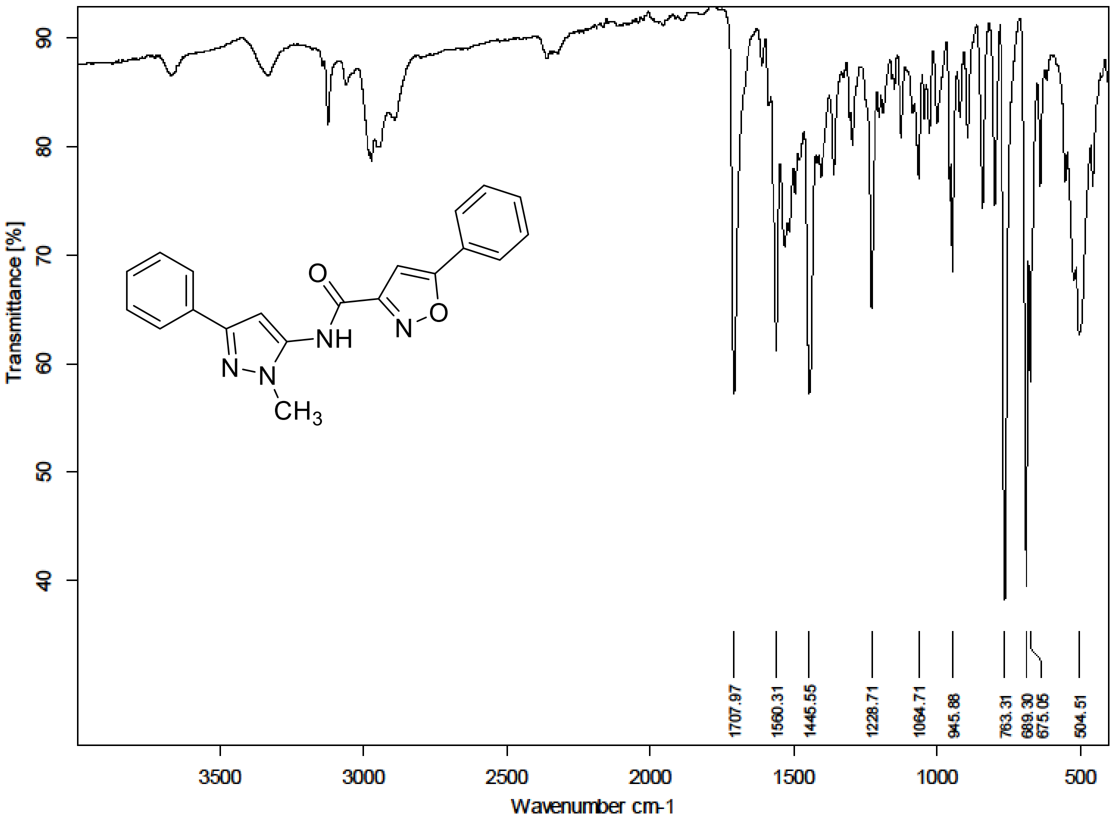

$^{1}H$ (500 MHz) and $^{13}C$ NMR (126 MHz) spectra of **47** in Chloroform-*d*

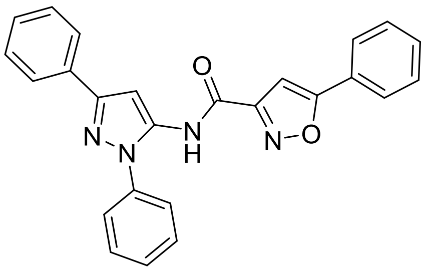

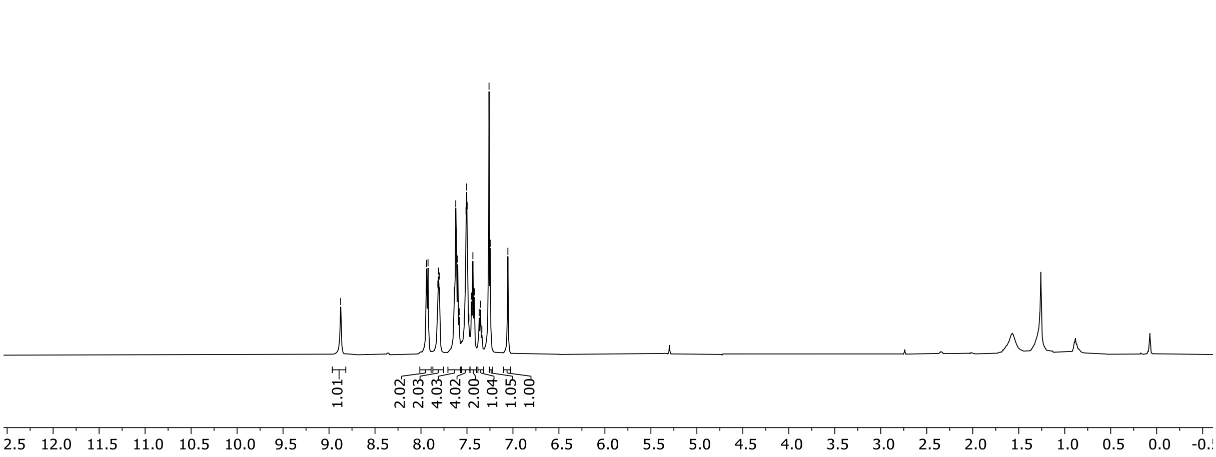

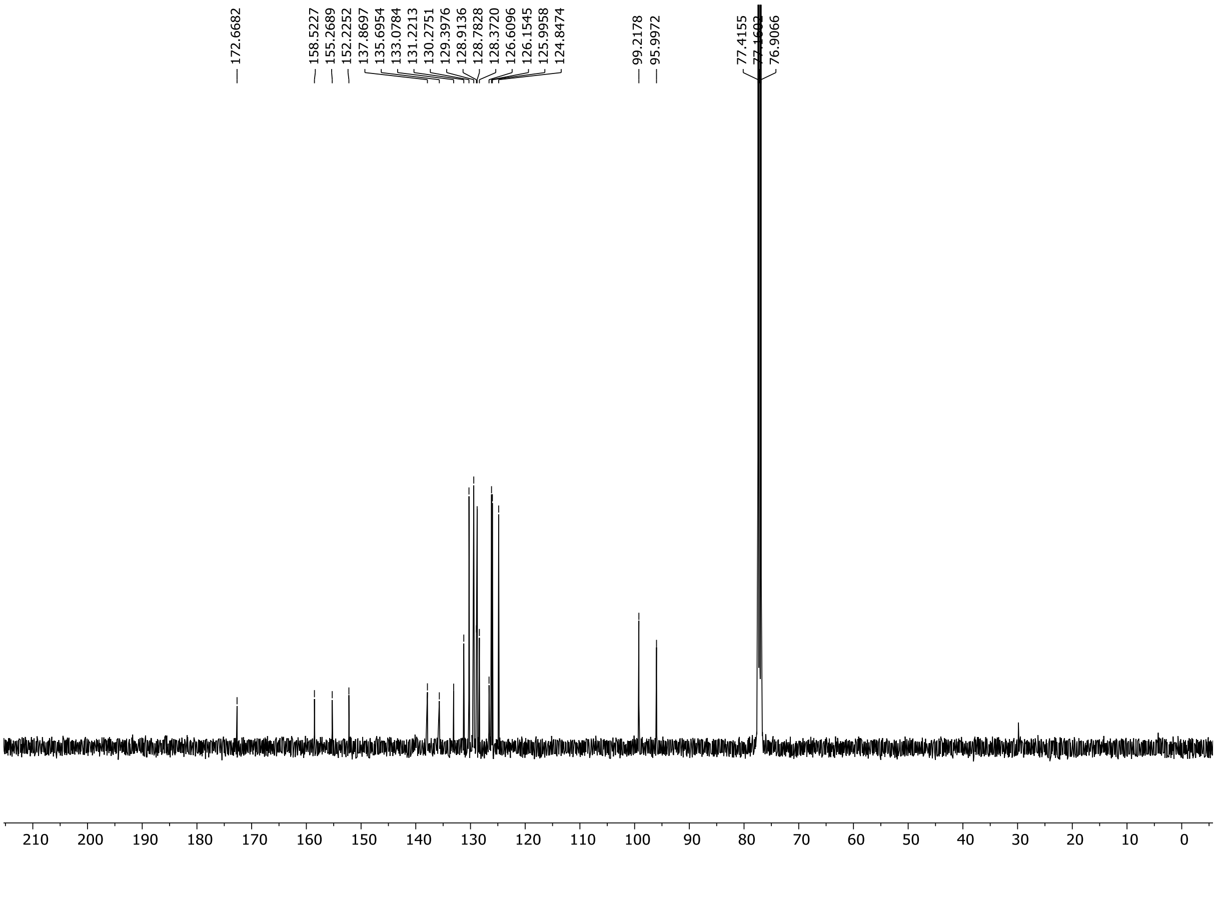

HRMS spectrum of **47**

$C_{25}H_{18}N_4O_2$ **exact mass: 406.1430**

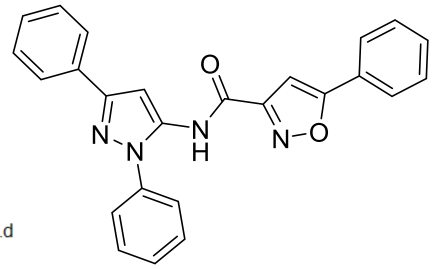

**APCI +** (MMI)
nitrogen flow 5 L/min, gas temperature 325°C, nebulizer 45 psig, skimmer 65 V, vaporizer 200°C, fragmentor 5 V, dissolved in methanol

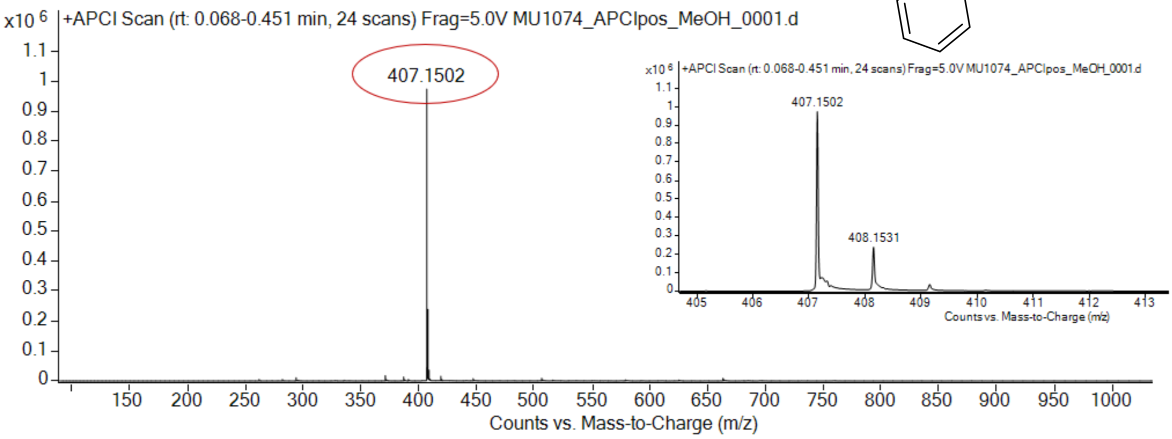

expected mass: [M+H]$^+$ = 407.1503 observed mass : [M+H]$^+$ = 407.1502 mass accuracy = - 0.2 ppm

[1]H (500 MHz) and [13]C NMR (126 MHz) spectra of **48** in Chloroform-*d*

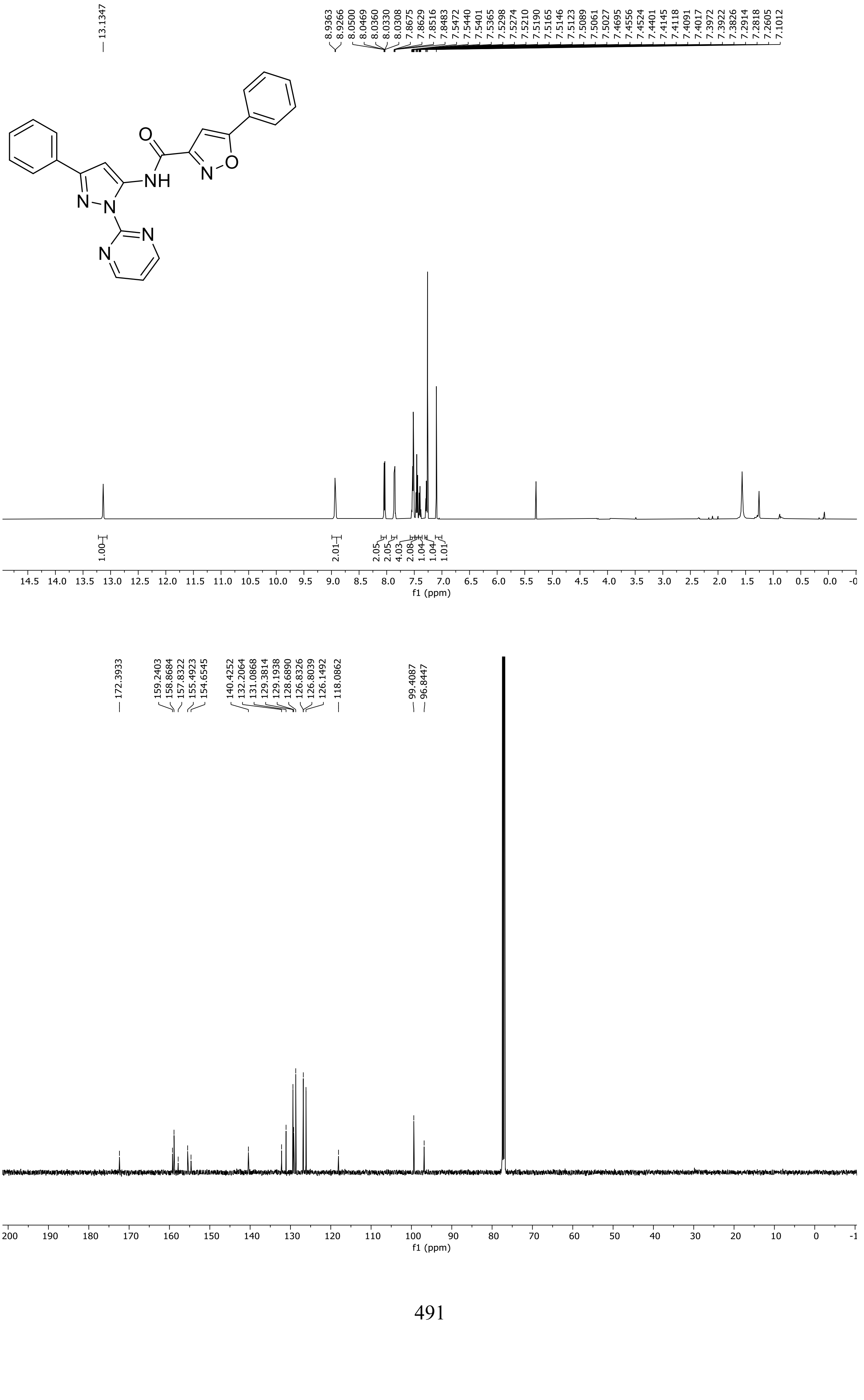

HRMS spectrum of **48**

NAR-A-14 $C_{23}H_{16}N_6O_2$ mono *m/z 408.1335*

**APCI +** (MMI)
nitrogen flow 5 L/min, gas temperature 325°C, nebulizer 45 psi, skimmer **65** V, vaporizer 200°C, fragmentor 20 V, dissolved in methanol

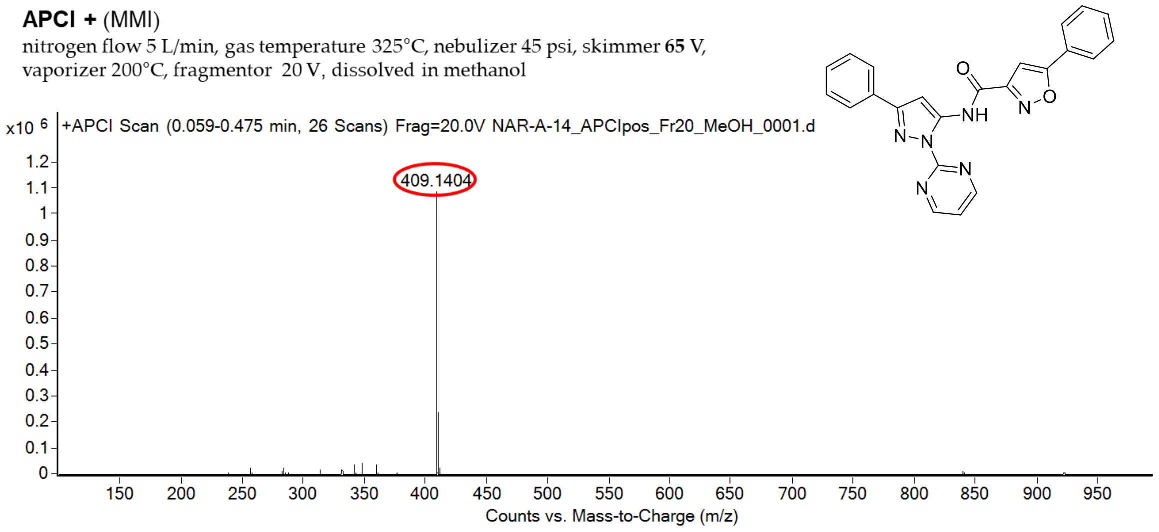

calculated mass: $[M+H]^+ = 409.1408$ observed: $[M+H]^+ = 409.1404$ mass accuracy = -0.9 ppm

FT-IR spectrum (neat) of **48**

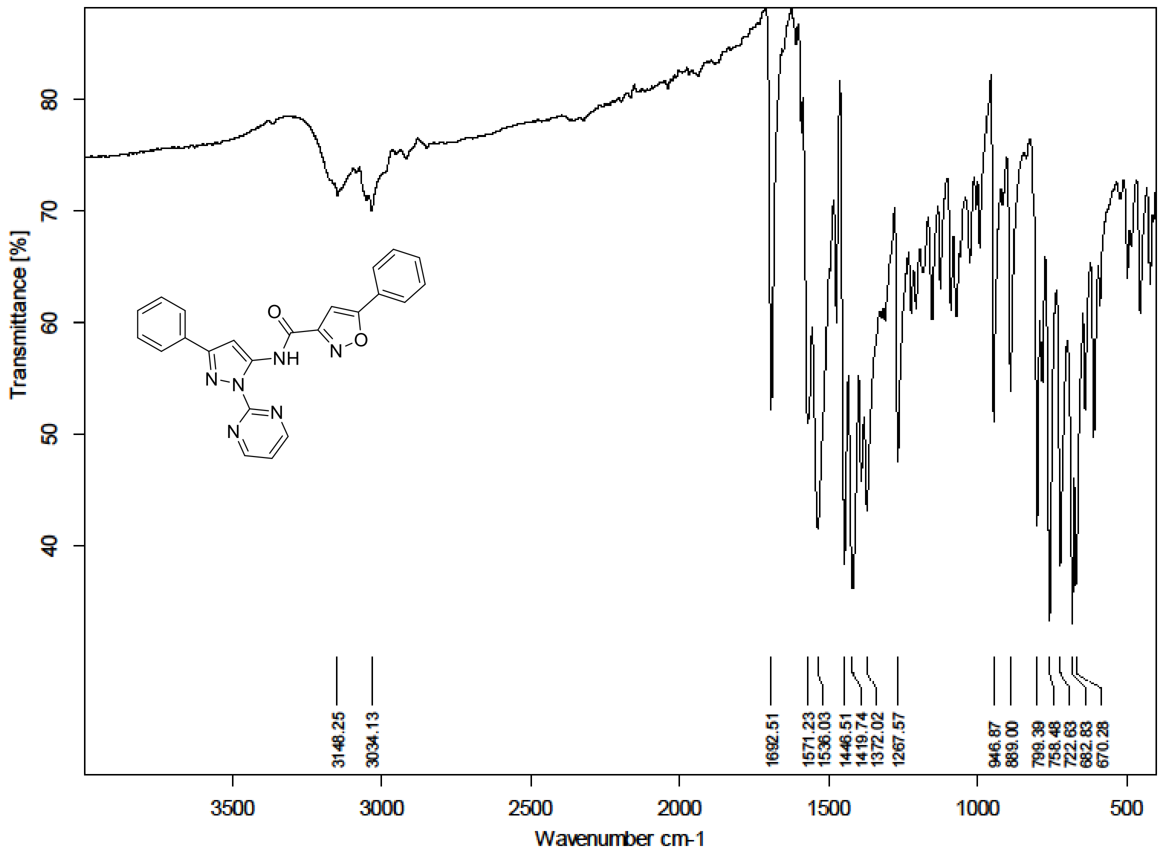

[1]H (500 MHz) and [13]C NMR (126 MHz) spectra of **49** in Chloroform-*d*

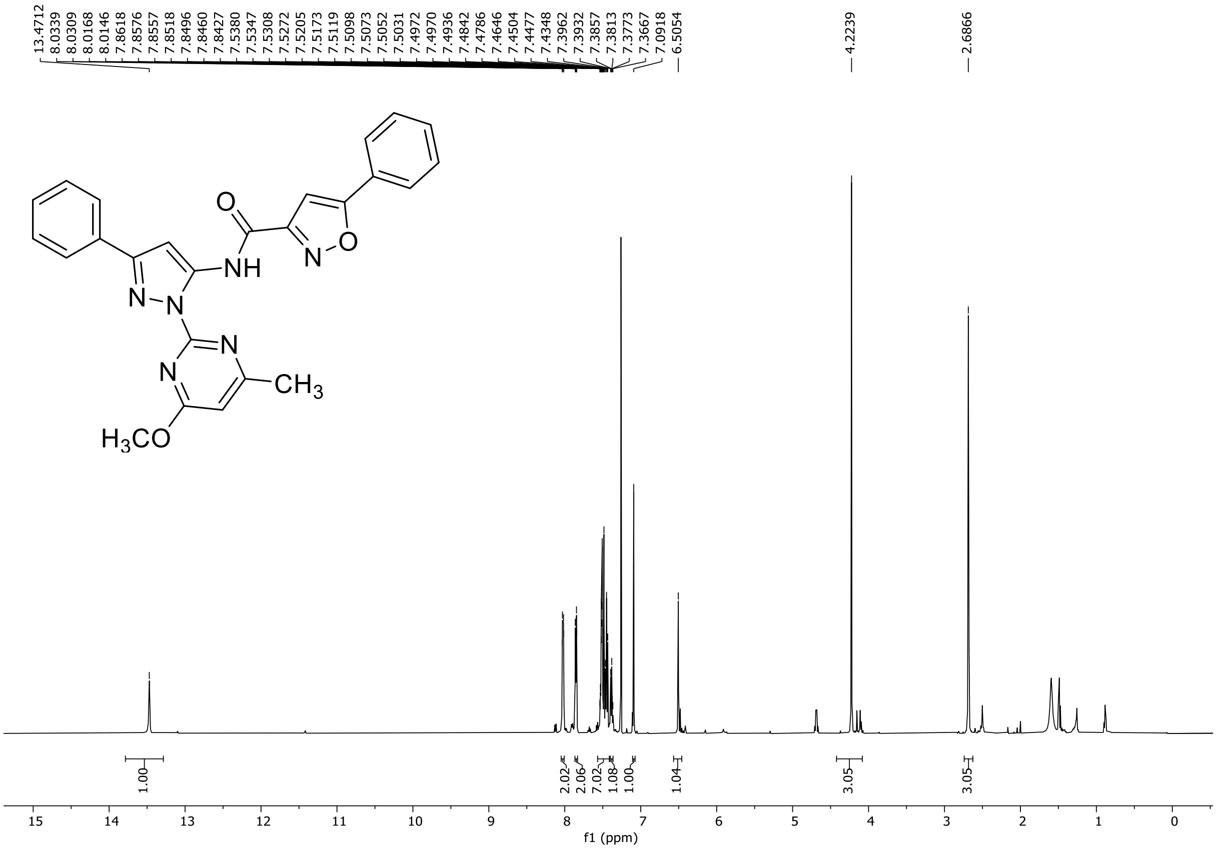

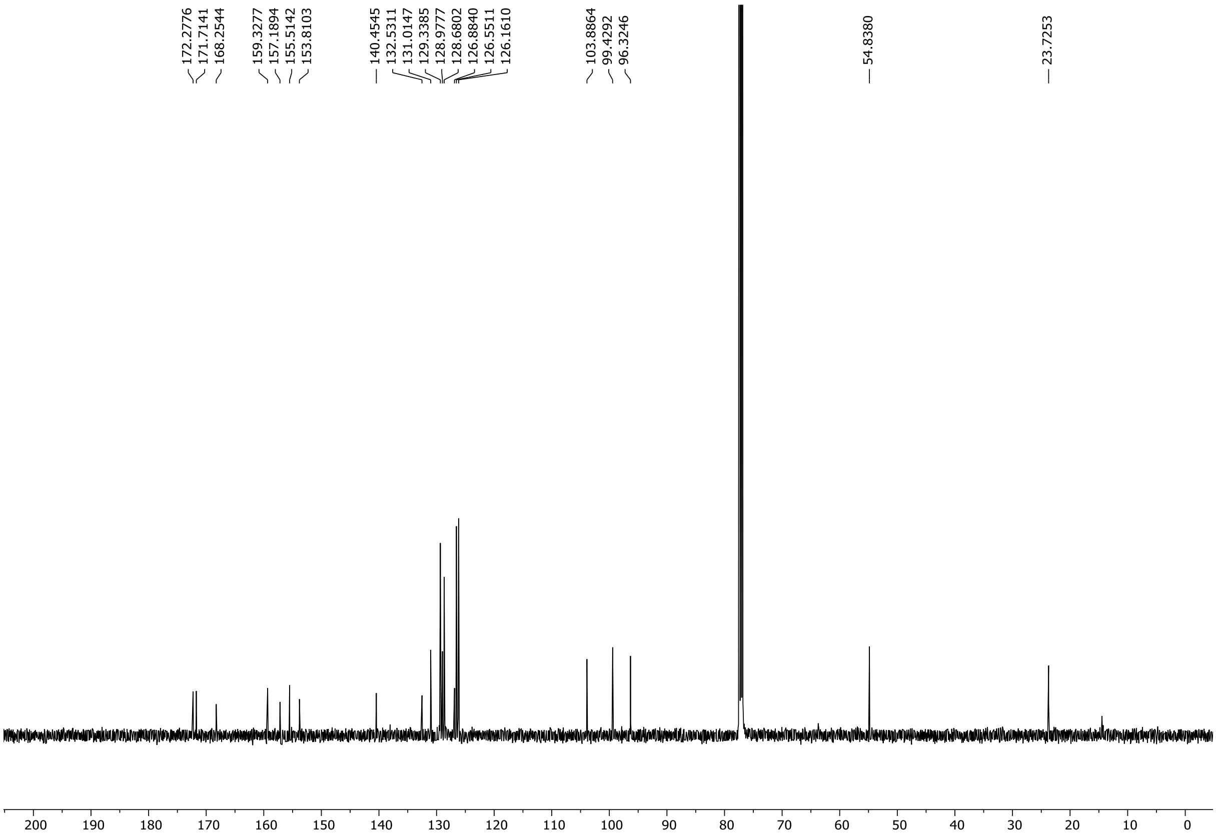

HRMS spectrum of **49**

**NAR-A-68** $C_{25}H_{20}N_6O_3$ *mono m/z* 452.1597

**APCI +** (MMI)
nitrogen flow 5 L/min, gas temperature 300°C, nebulizer 45 psi, skimmer **65** V, vaporizer 200°C, fragmentor 15 V, dissolved in MeOH

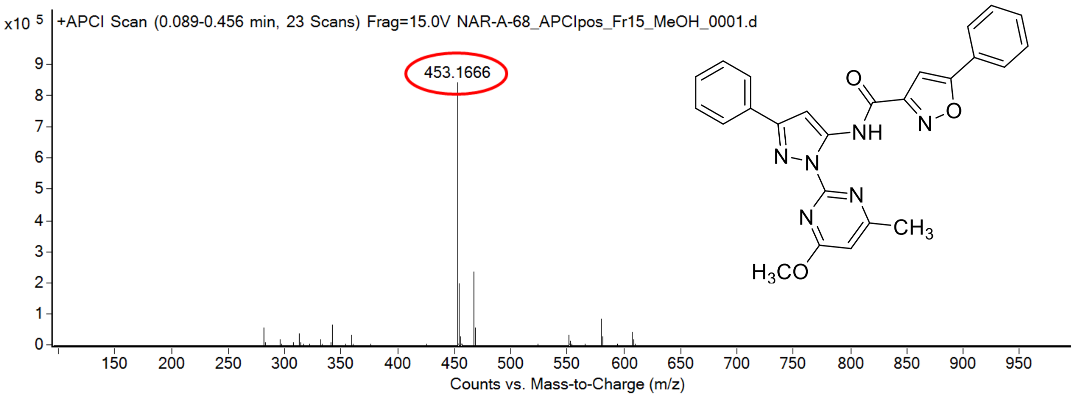

calculated mass: $[M+H]^+$ = 453.1670 observed: $[M+H]^+$ = 453.1666 mass accuracy = -0.9 ppm

FT-IR spectrum (neat) of **49**

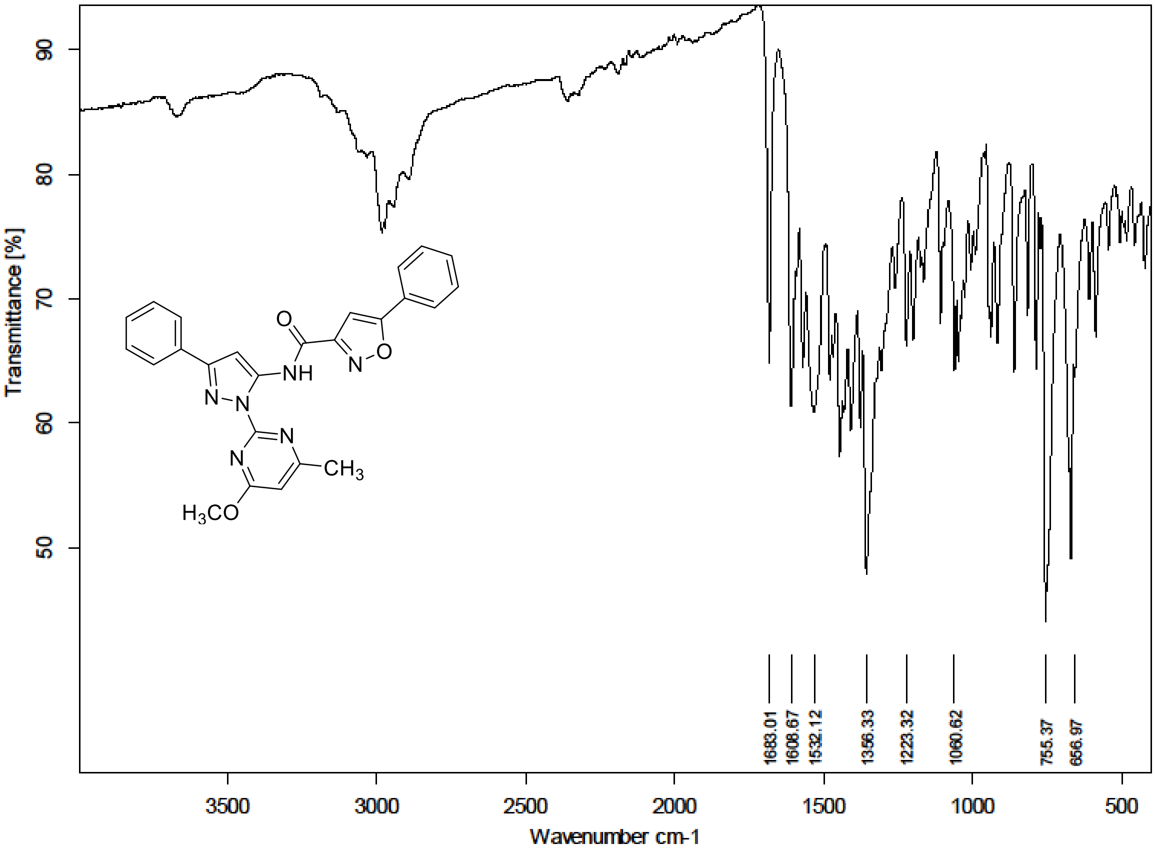

$^{1}$H NMR (300 MHz) spectrum of **50** DMSO-$d_6$

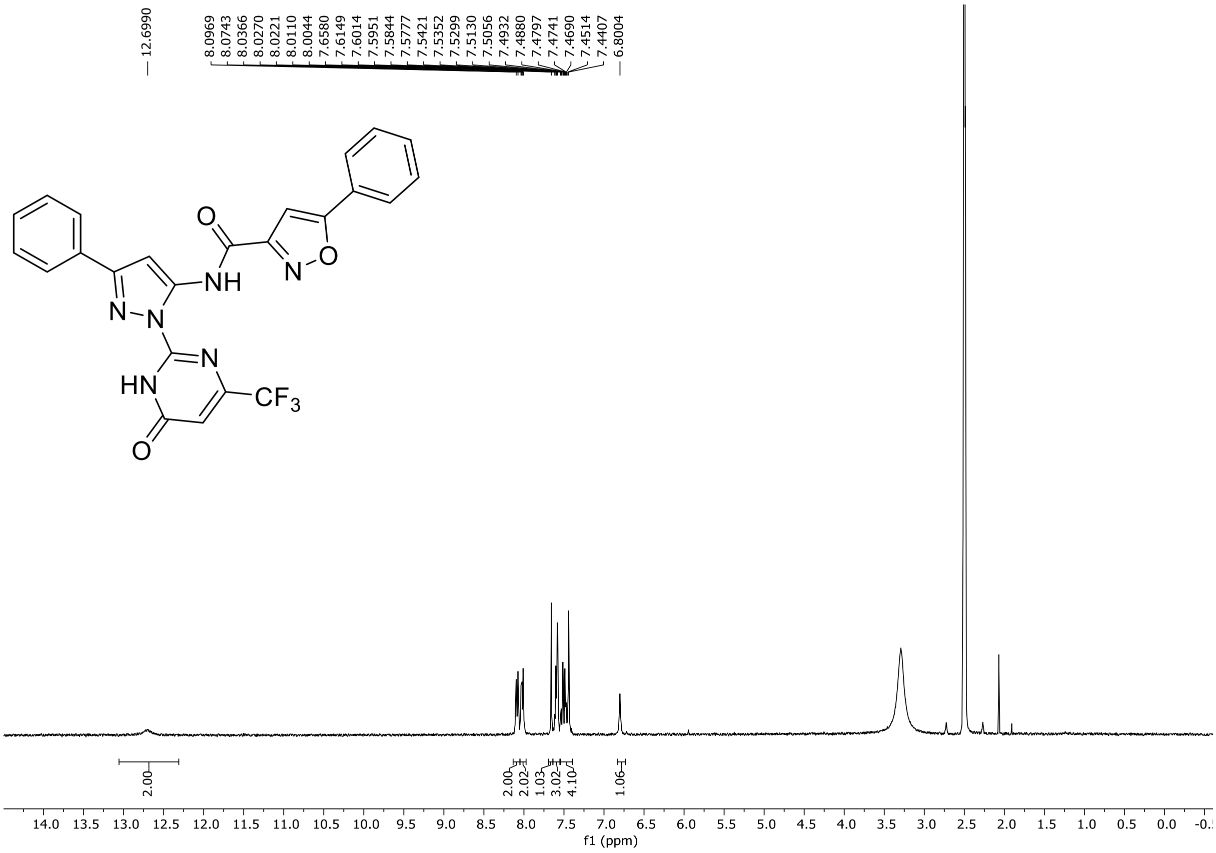

HRMS spectrum of **50**

**NAR-A-96** $C_{24}H_{15}F_3N_6O_3$ ***m/z* 492.1158**

**ESI-** (MMI)
nitrogen flow 5 L/min, gas temperature 325°C, nebulizer 45 psi, skimmer 65 V, fragmentor 60 V, dissolved in DMSO, MeOH

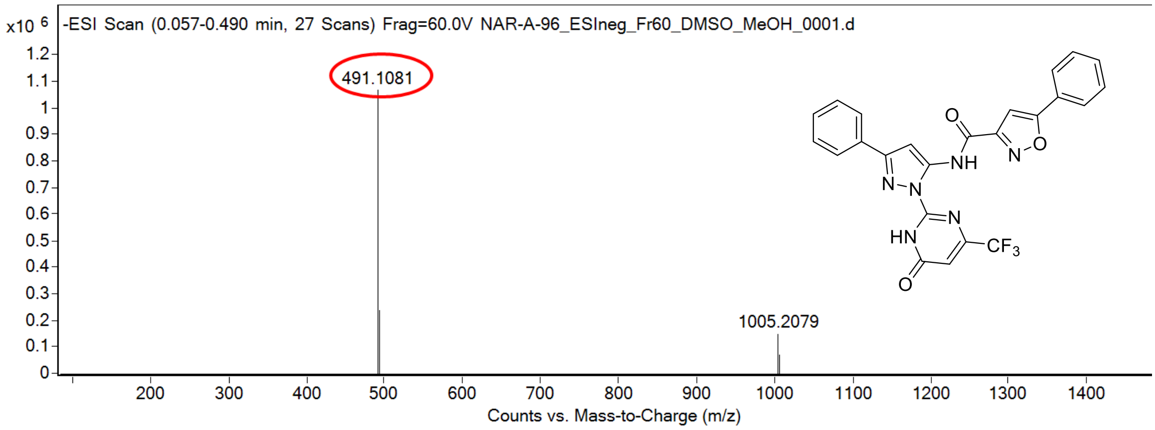

calculated mass: $[M-H]^-$ = 491.1085 observed: $[M-H]^-$ = 491.1081 mass accuracy = -0.8 ppm

FT-IR spectrum (neat) of **50**

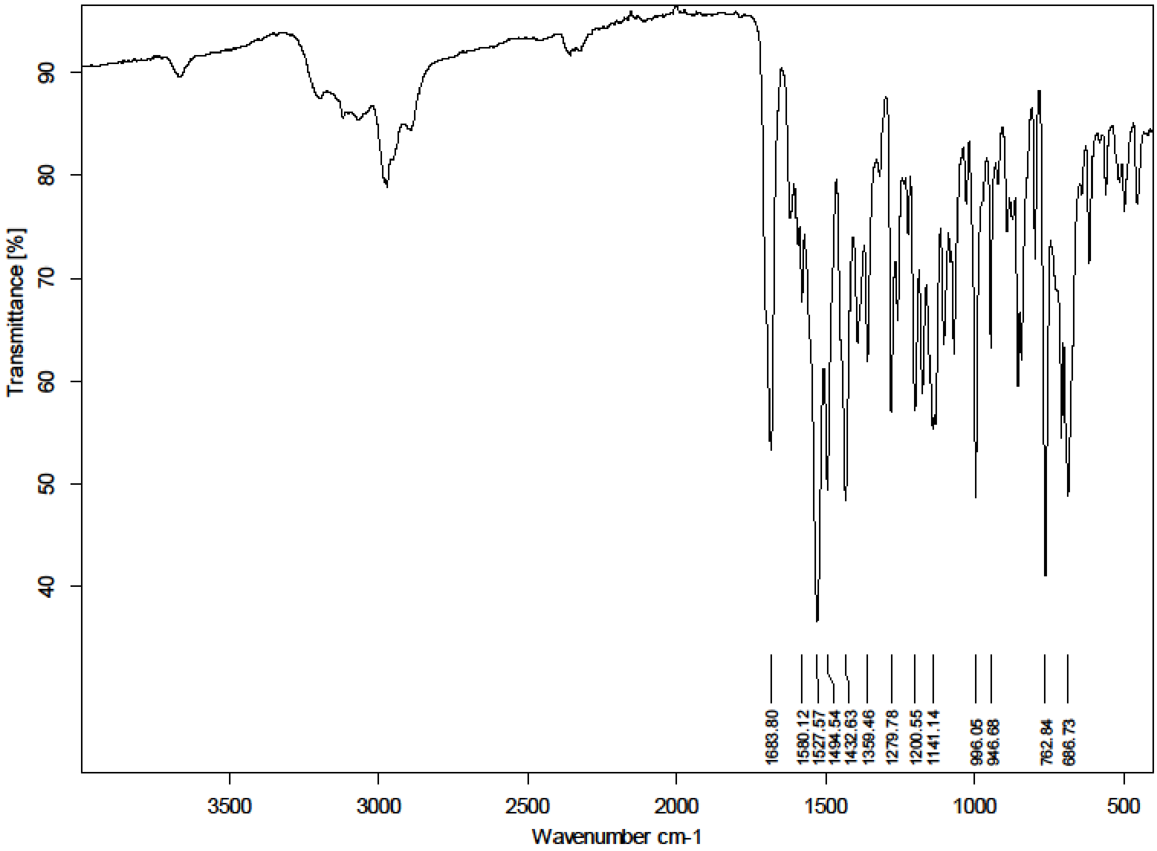

$^{1}H$ NMR (300 MHz) spectrum of **51** DMSO-$d_6$

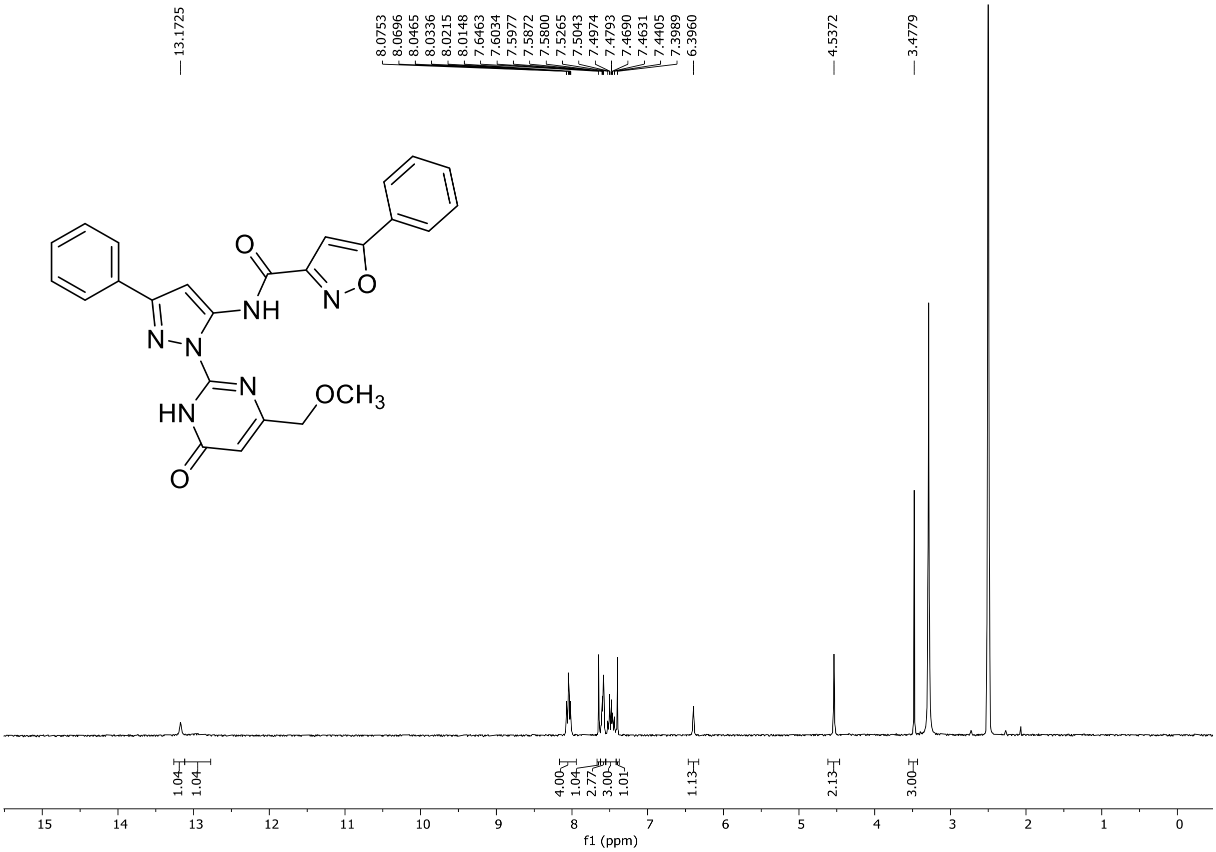

HRMS spectrum of **51**

**NAR-A-80** **$C_{25}H_{20}N_6O_4$** ***m/z* 468.1546**

**ESI-** (MMI)
nitrogen flow 5 L/min, gas temperature 325°C, nebulizer 45 psi, skimmer 65 V, fragmentor 150 V, dissolved in MeOH

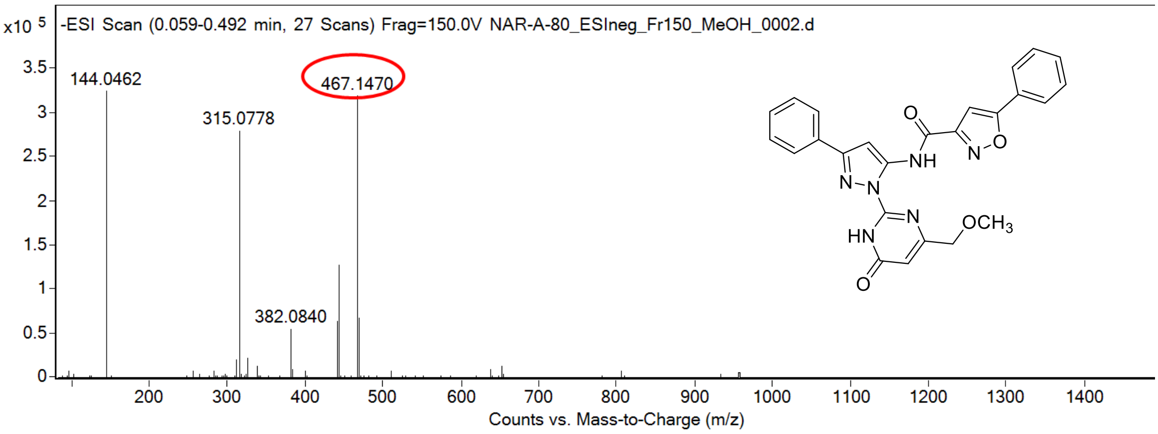

calculated mass: $[M\text{-}H]^-$ = 467.1473 observed: $[M\text{-}H]^-$ = 467.1470 mass accuracy = -0.6 ppm

FT-IR spectrum (neat) of **51**

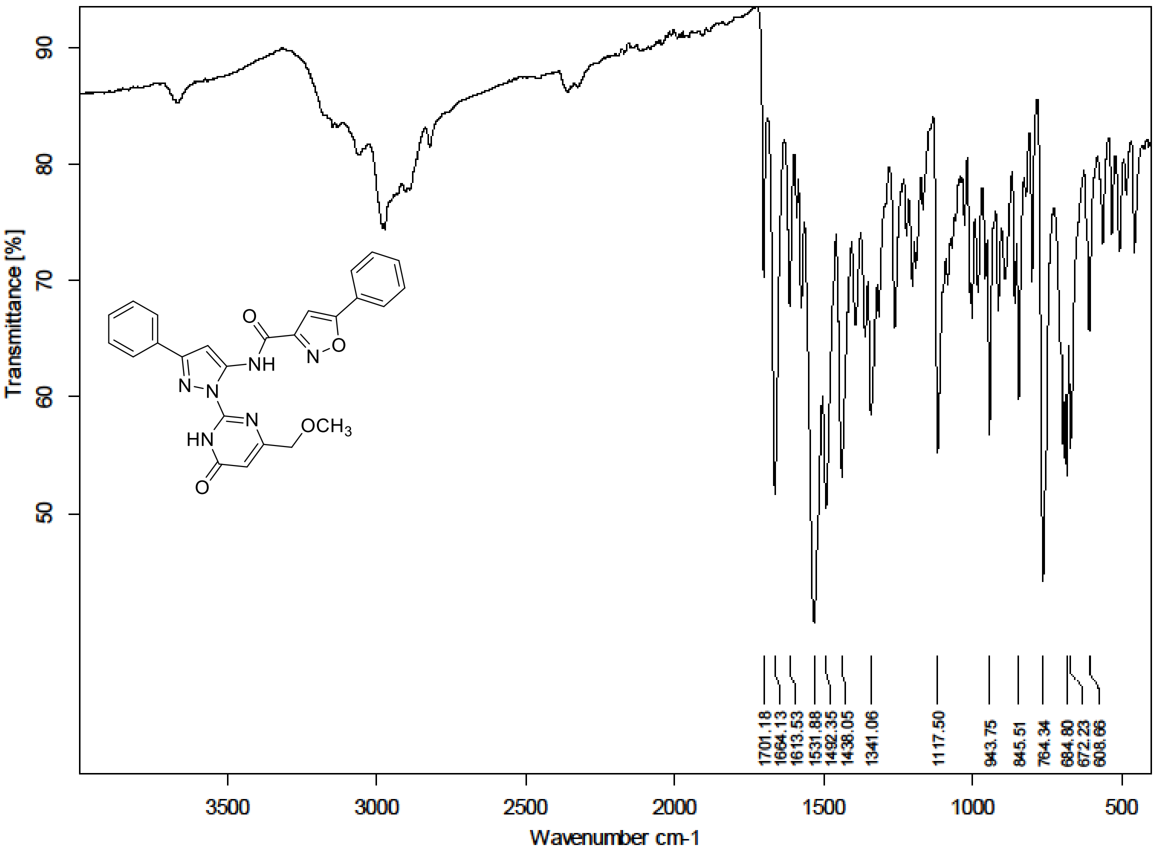

[1]H NMR (300 MHz) spectrum of **52** in DMSO-$d_6$ and [13]C NMR (75 MHz) spectrum of **52** in Trifluoroacetic Acid-*d*

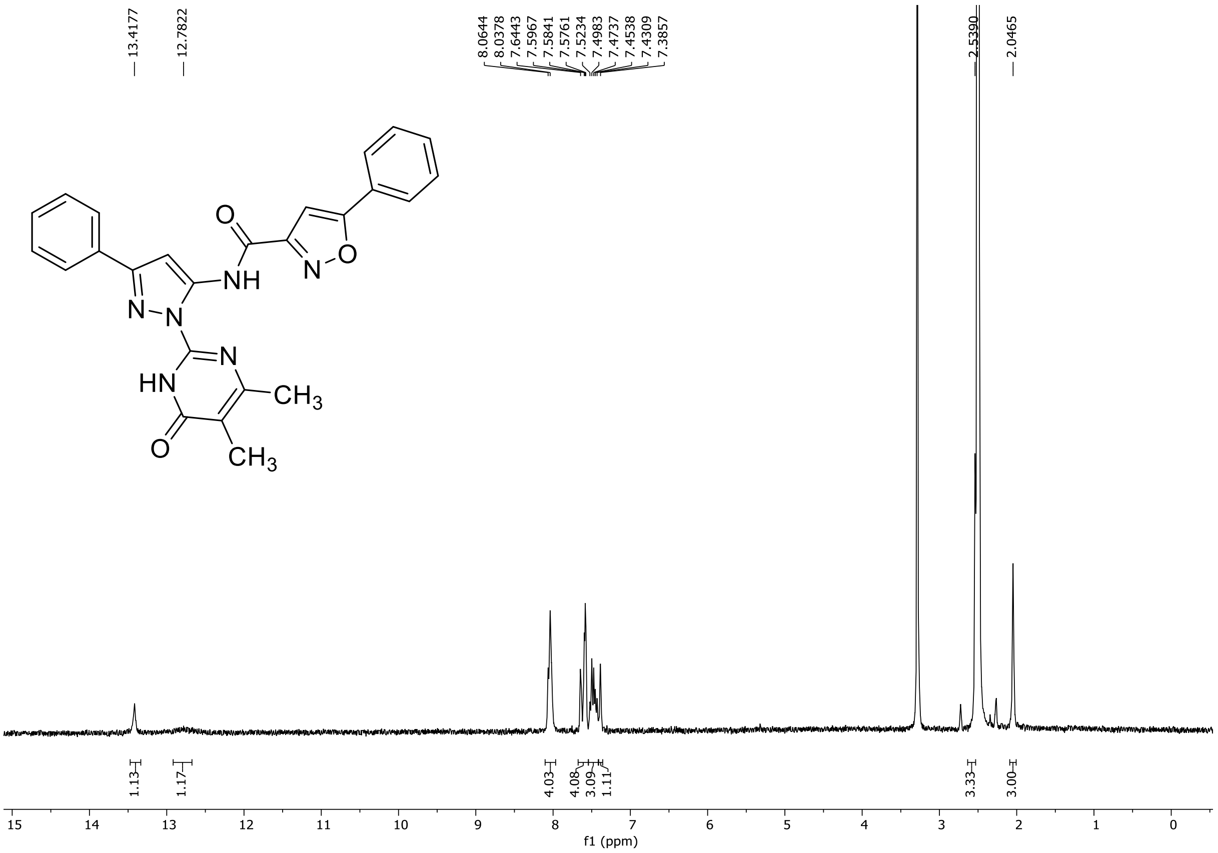

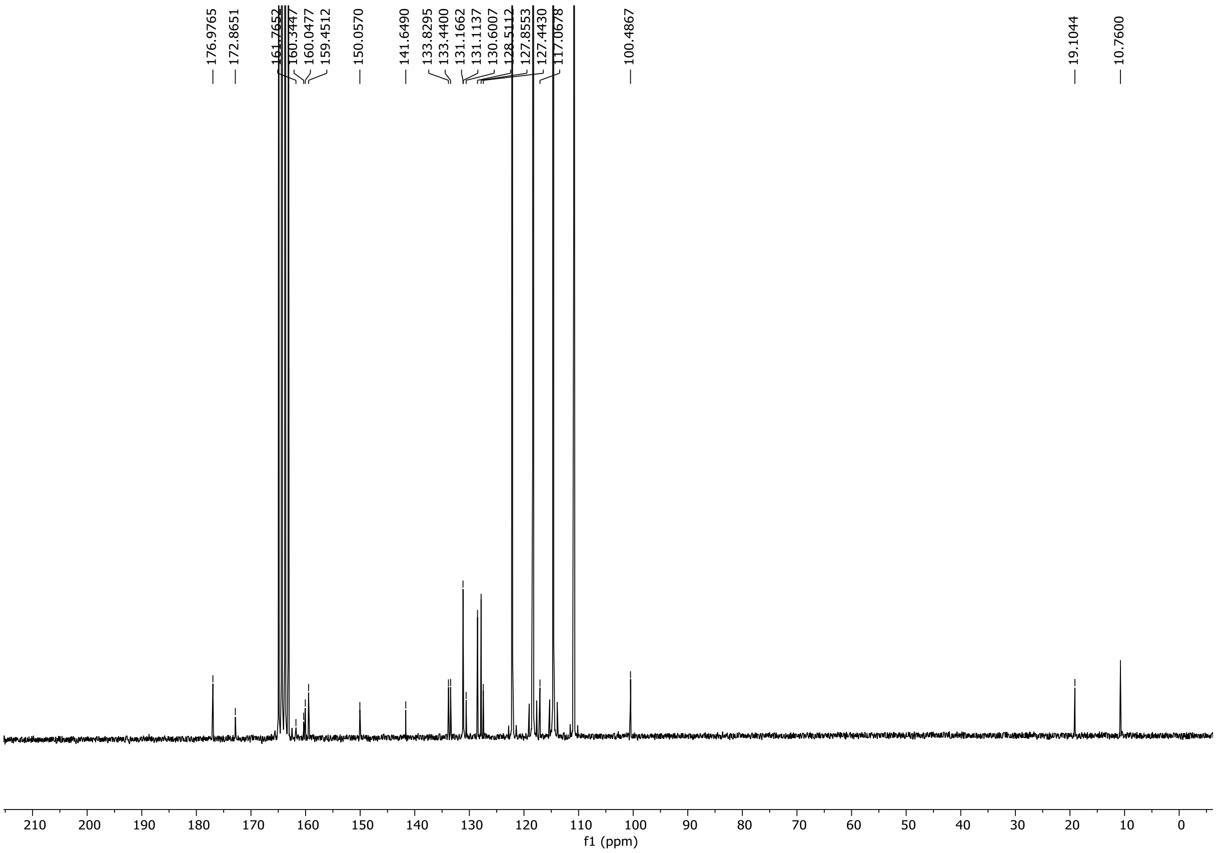

HRMS spectrum of **52**

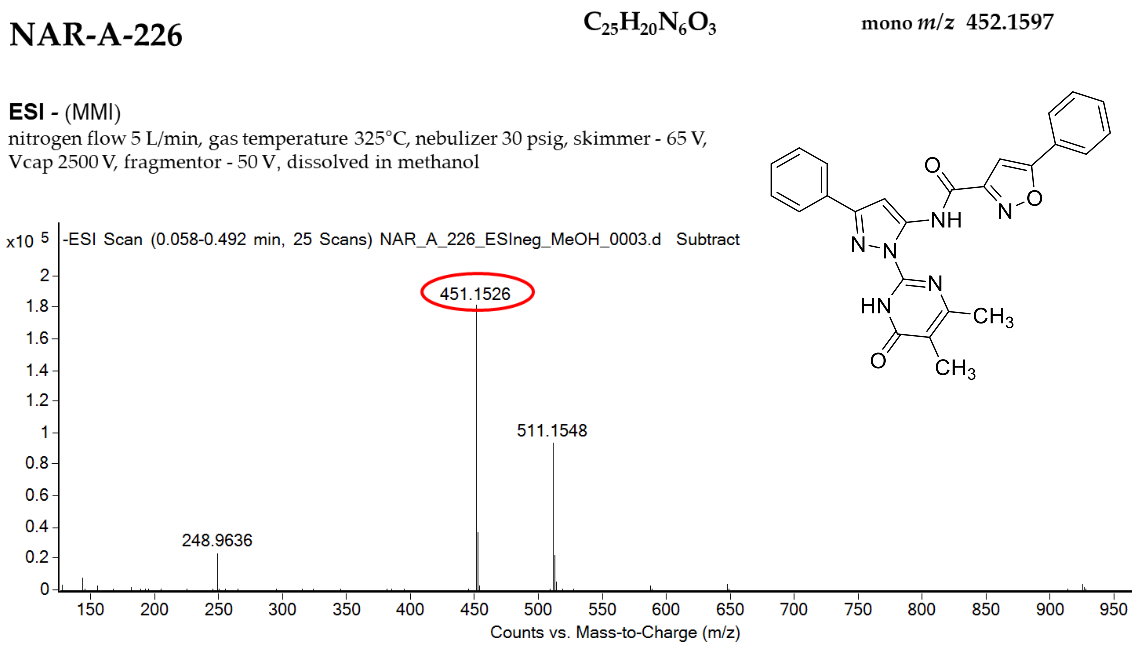

calculated mass: $[M\text{-}H]^-$ = 451.1524 observed: $[M\text{-}H]^-$ = 451.1526 mass accuracy = 0.4 ppm

FT-IR spectrum (neat) of **52**

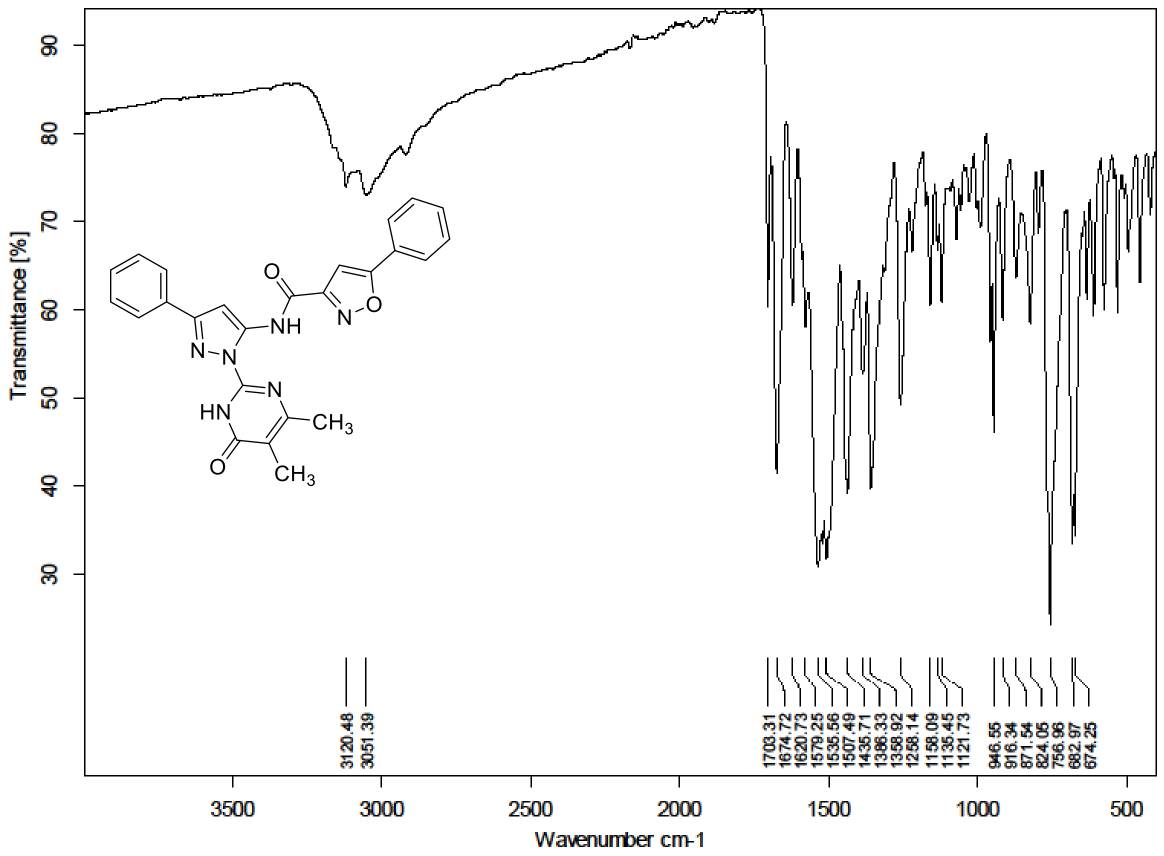

$^{1}H$ (500 MHz) and $^{13}C$ NMR (126 MHz) spectra of **53** in DMSO-$d_6$

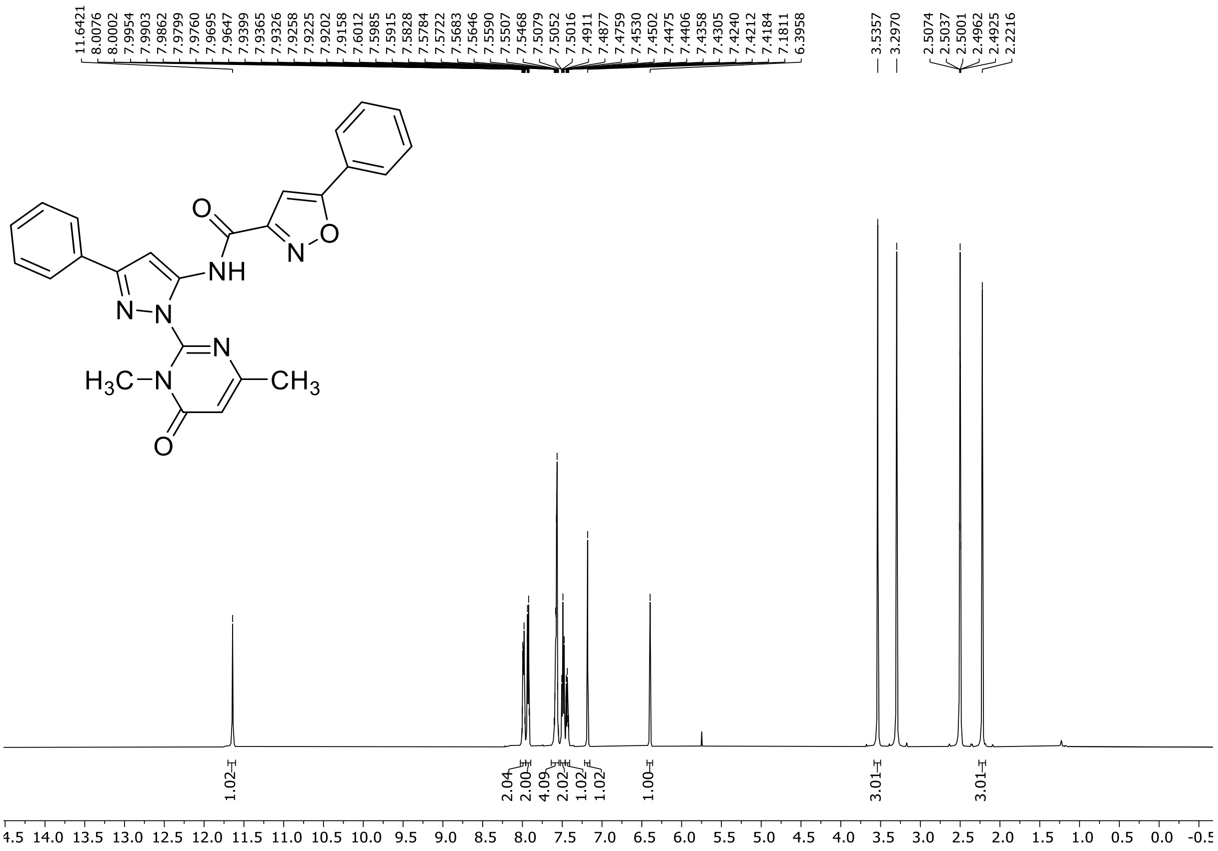

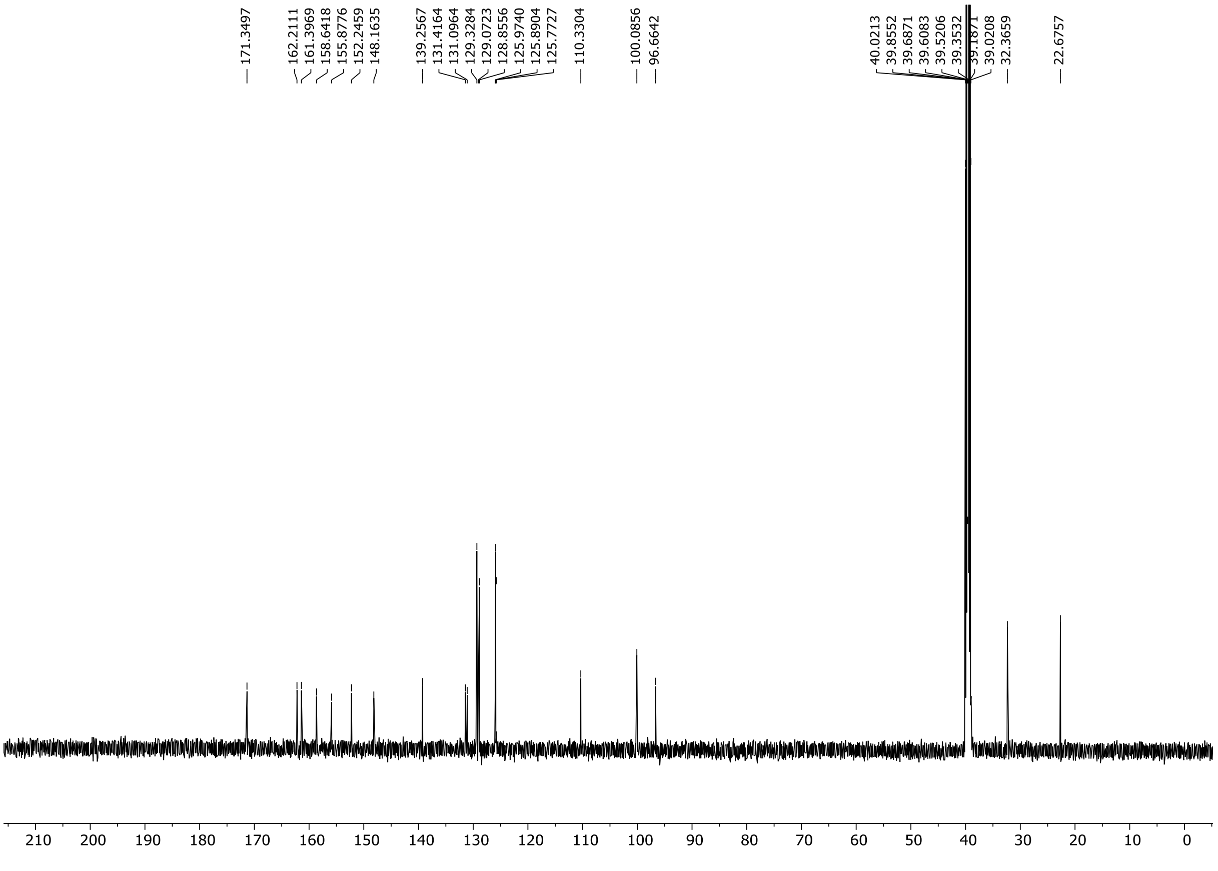

HRMS spectrum of **53**

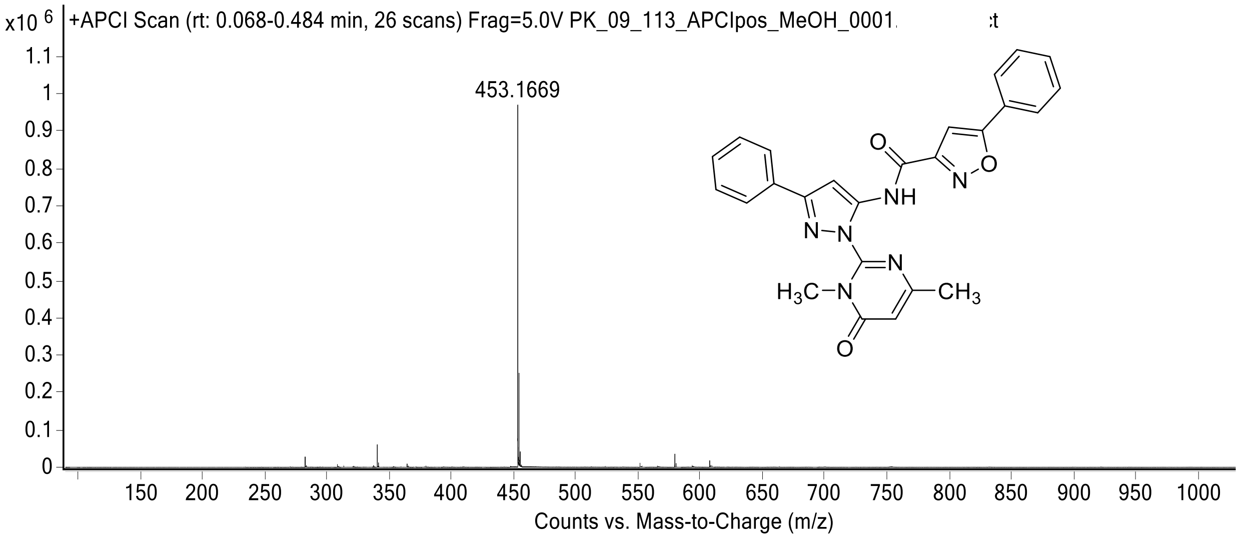

[1]H (500 MHz) and [13]C NMR (126 MHz) spectra of **54** in DMSO-$d_6$

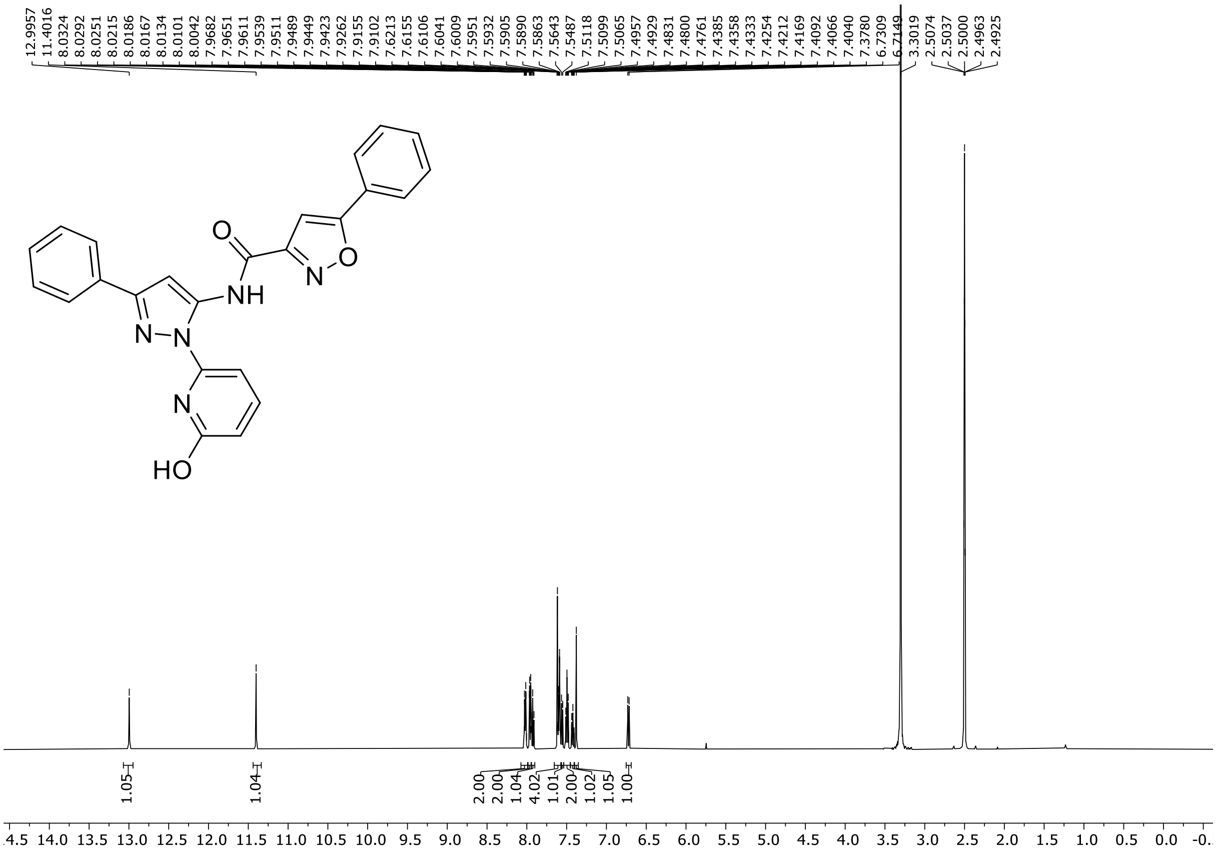

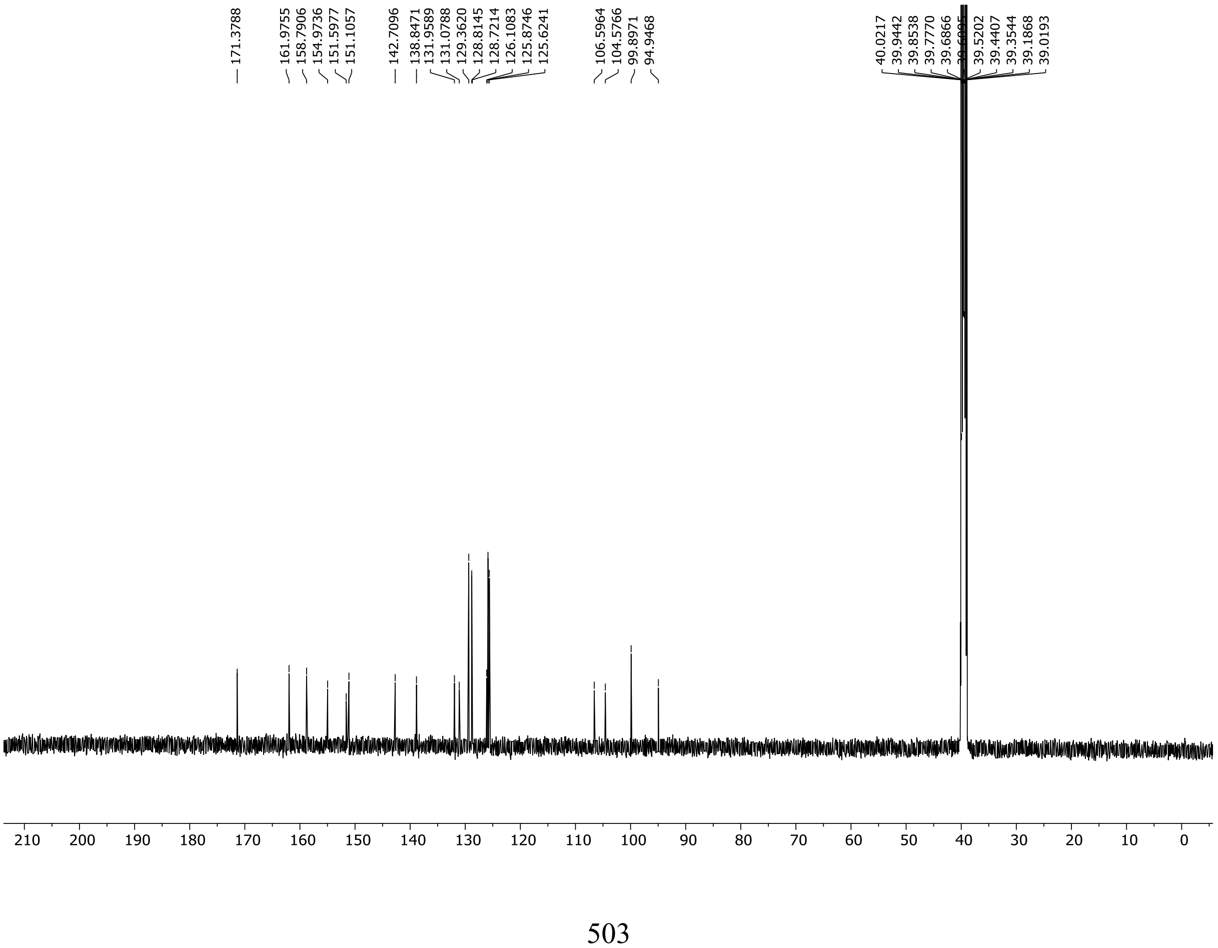

HRMS spectrum of **54**

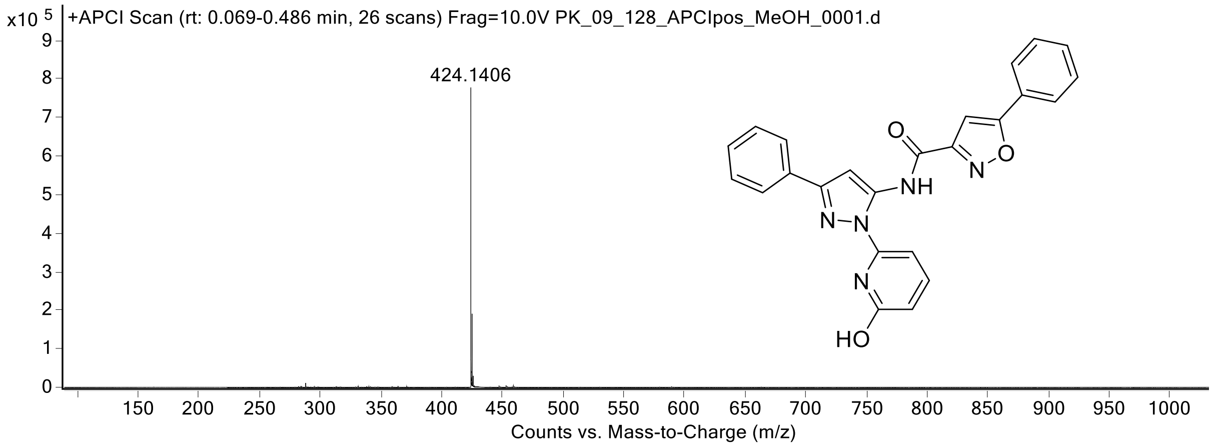

[1]H NMR (300 MHz) spectrum of **55** DMSO-$d_6$

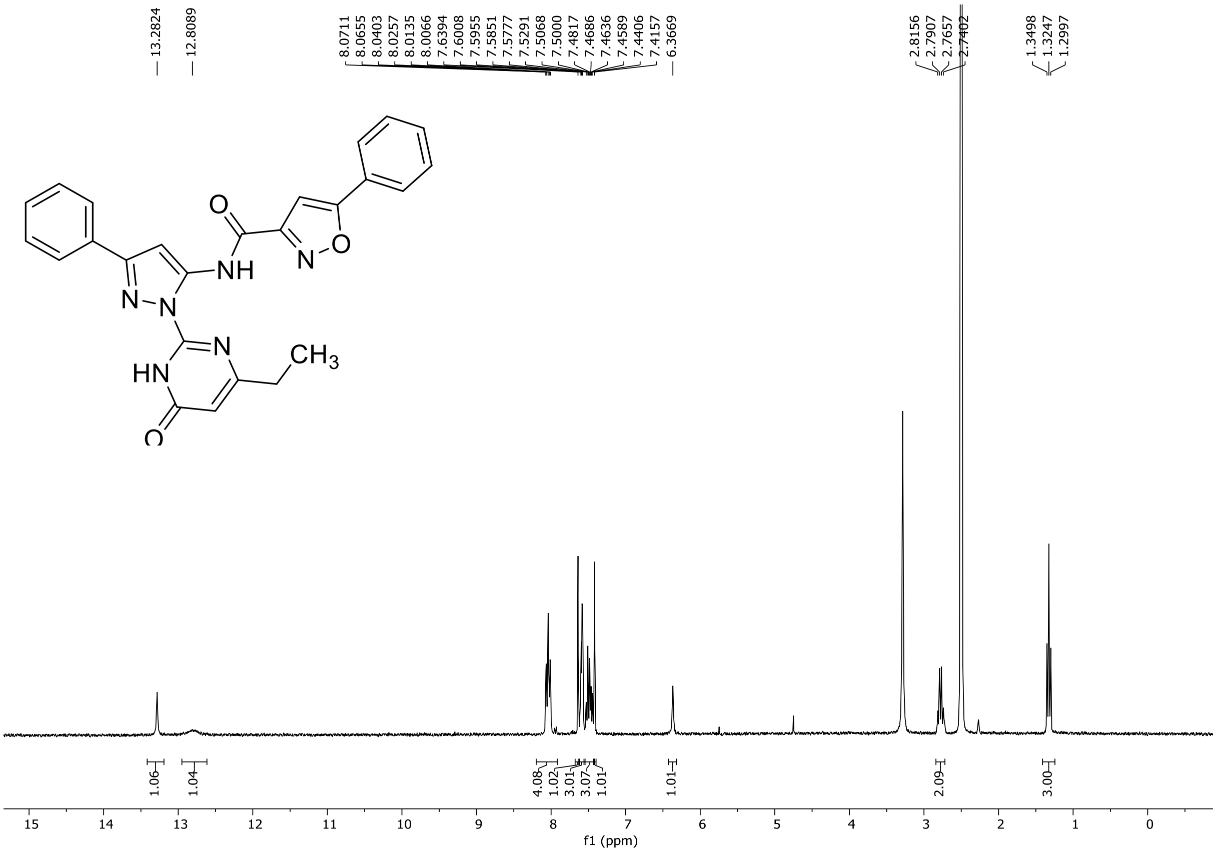

HRMS spectrum of **55**

**NAR-A-170** **$C_{25}H_{20}N_6O_3$** **mono *m/z* 452.1597**

**APCI +** (MMI)
nitrogen flow 5 L/min, gas temperature 325°C, nebulizer 45 psi, skimmer **65** V, vaporizer 200°C, fragmentor 25 V, dissolved in methanol

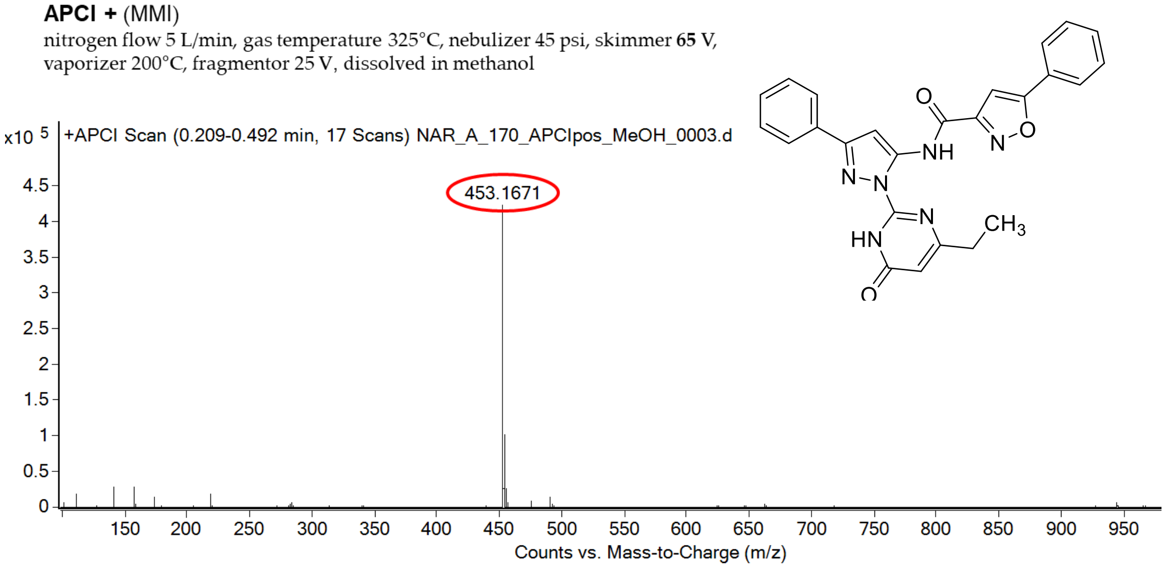

calculated mass: [M+H]$^+$ = 453.1670 observed: [M+H]$^+$ = 453.1671 mass accuracy = 0.2 ppm

FT-IR spectrum (neat) of **55**

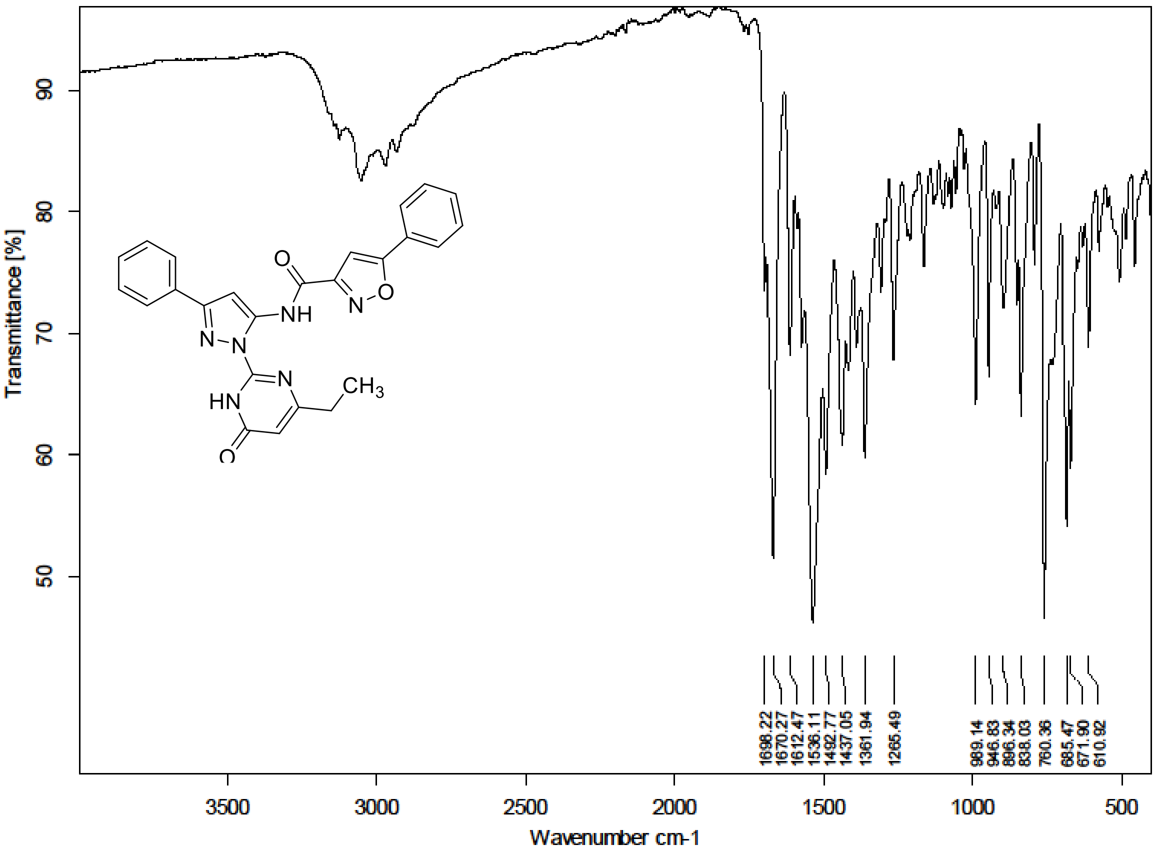

**$^{1}$H NMR, $^{13}$C NMR, HRMS and IR spectra of compound 118-141**

$^{1}$H (500 MHz) and $^{13}$C NMR (126 MHz) spectra of **118** in DMSO-$d_6$

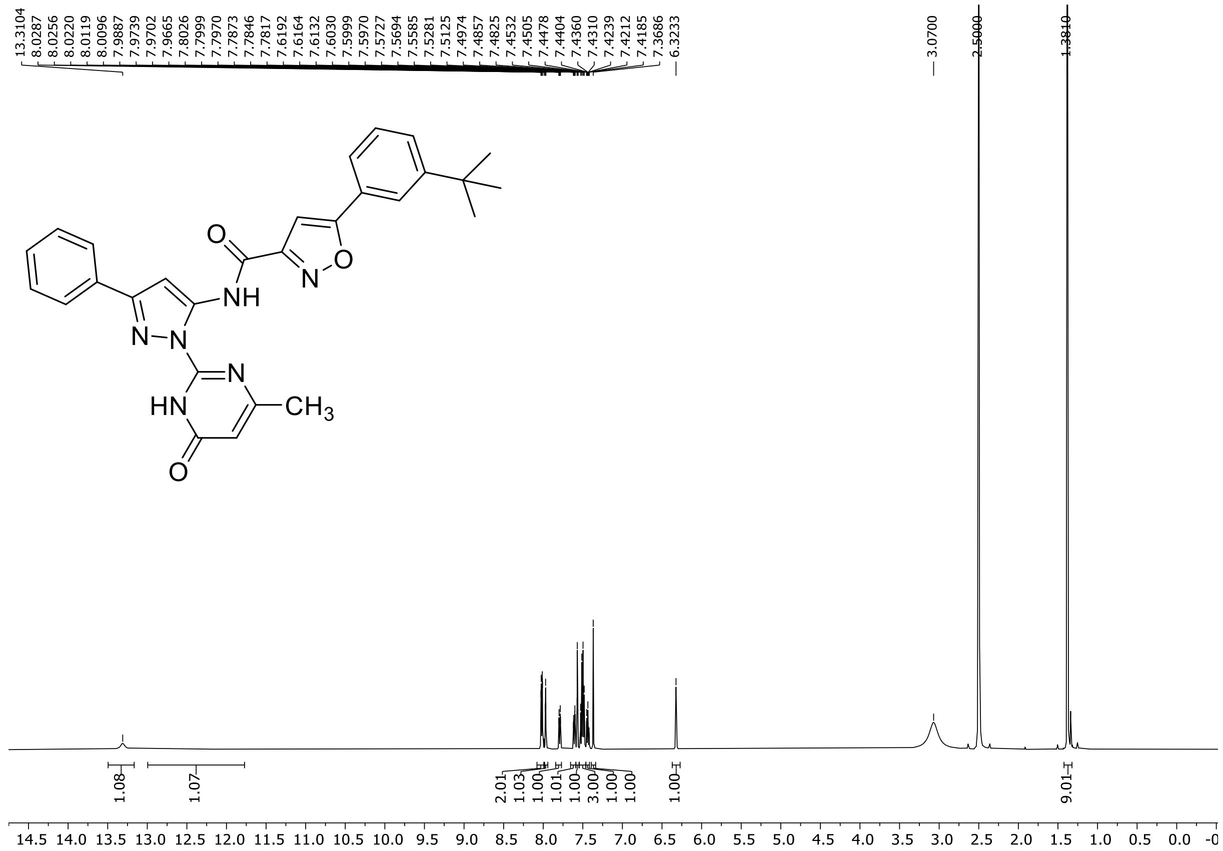

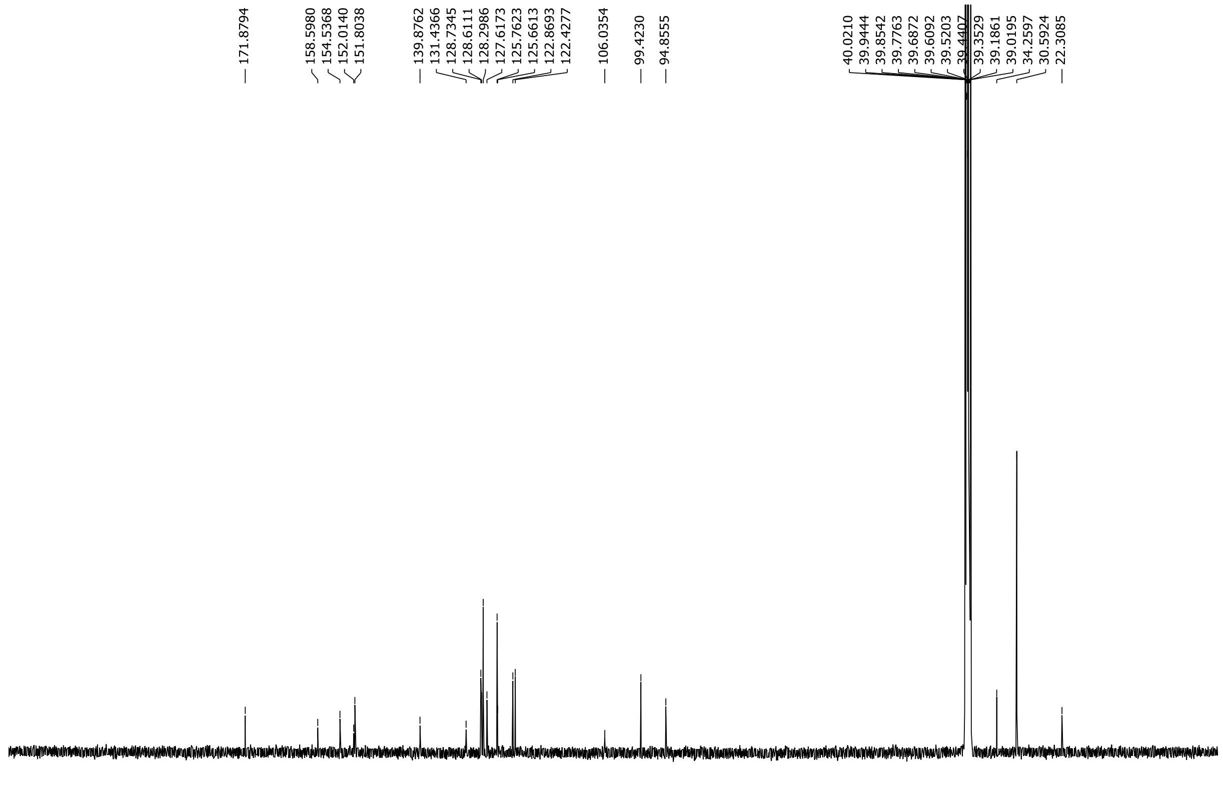

HRMS spectrum of **118**

**NAR-A-98** $C_{28}H_{26}N_6O_3$ *m/z* **494.2066**

**APCI+** (MMI)
nitrogen flow 5 L/min, gas temperature 325°C, nebulizer 45 psi, skimmer 65 V,
vaporizer 200°C, fragmentor 28 V, dissolved in DMSO, MeOH

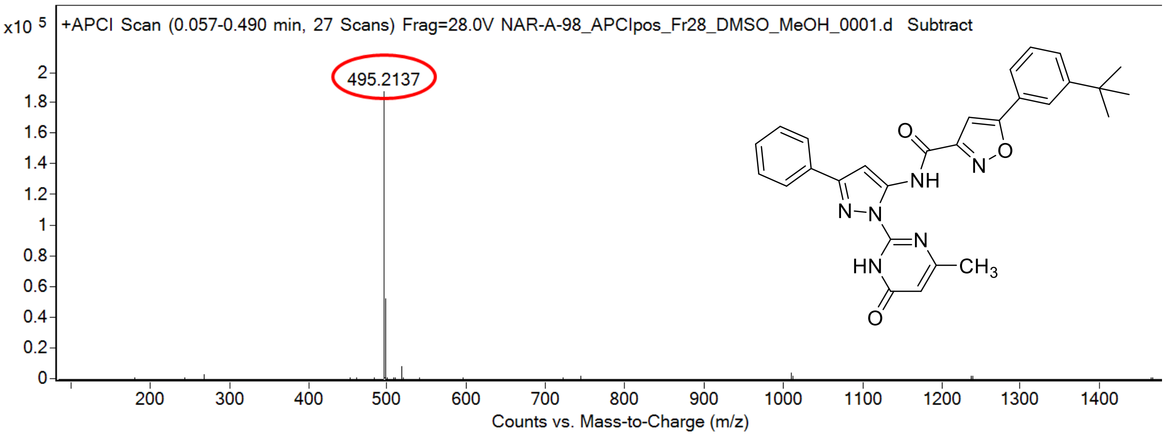

calculated mass: $[M+H]^+$ = 495.2139 observed: $[M+H]^+$ = 495.2137 mass accuracy = -0.4 ppm

FT-IR spectrum (neat) of **118**

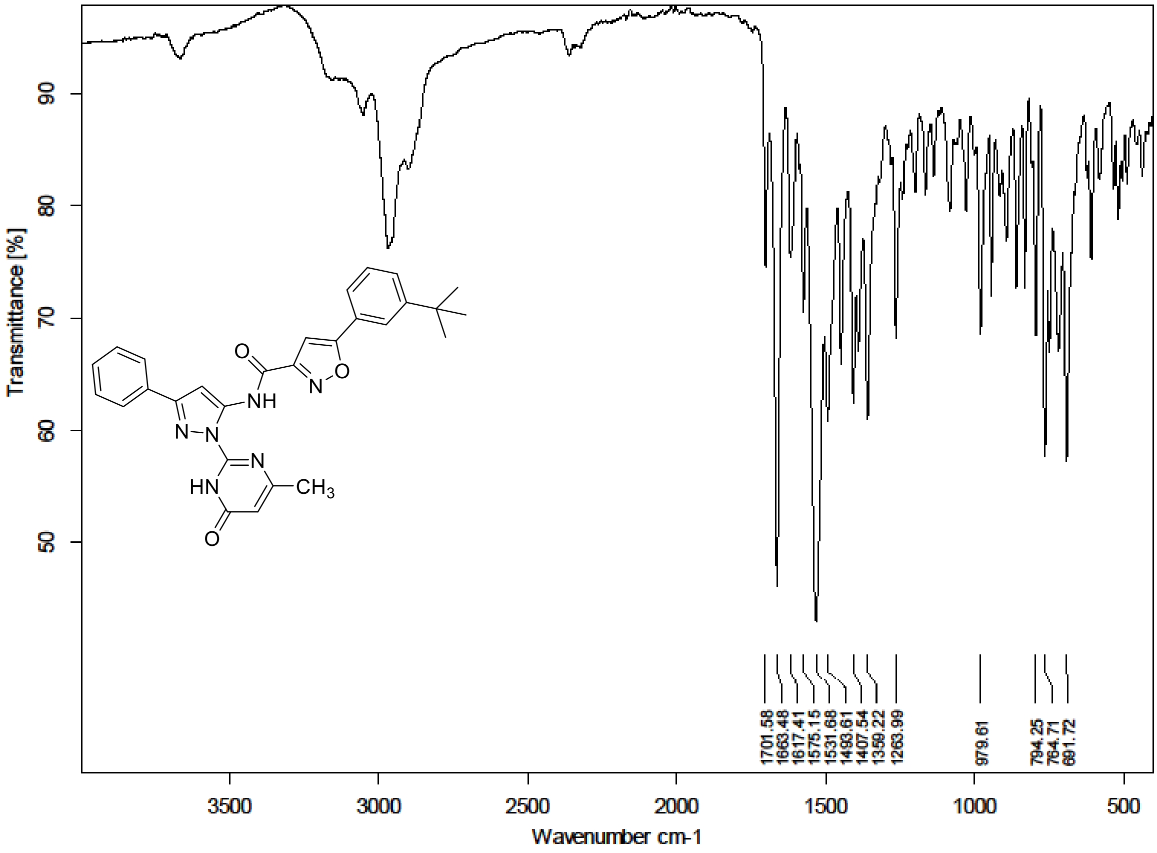

$^1$H NMR (300 MHz) spectrum of **119** in DMSO-$d_6$

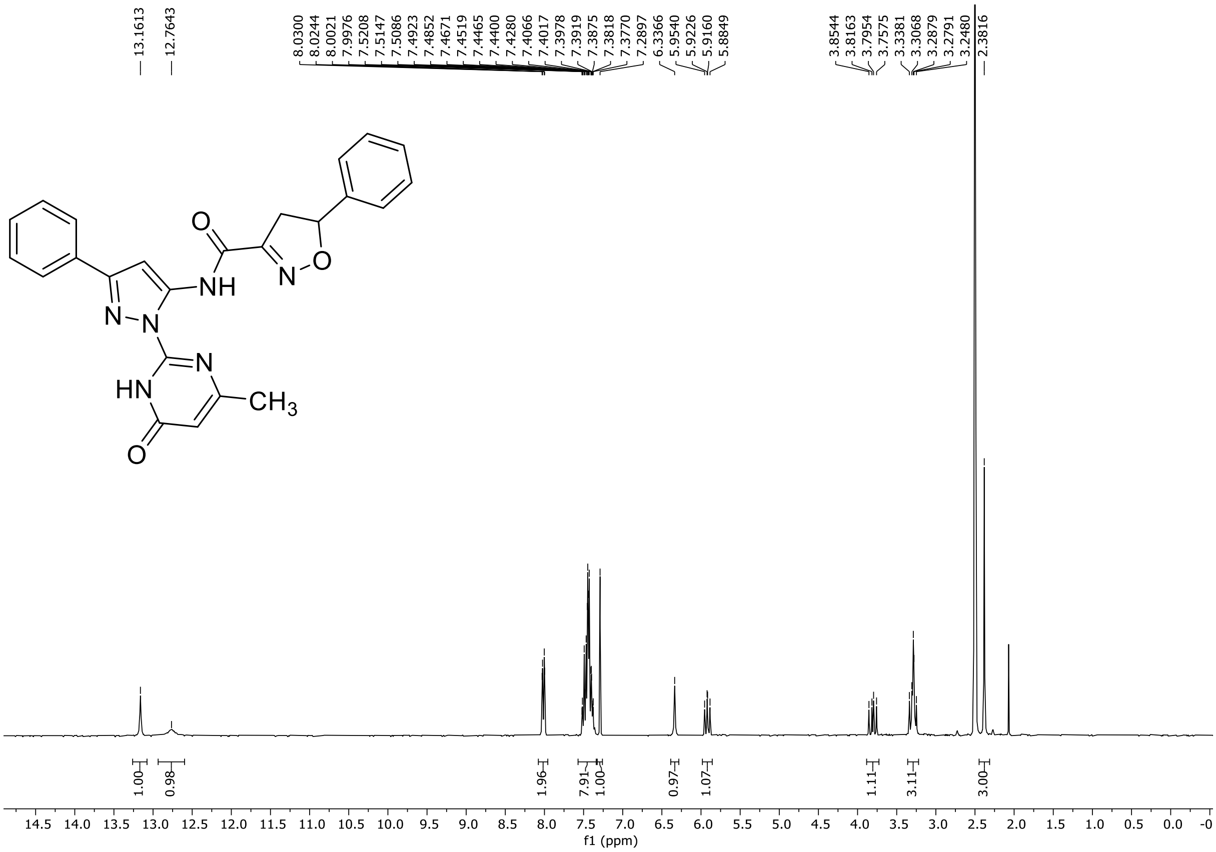

HRMS spectrum of **119**

**NAR-A-225** $C_{24}H_{20}N_6O_3$ mono *m/z* 440.1597

**ESI -** (MMI)
nitrogen flow 5 L/min, gas temperature 325°C, nebulizer 30 psig, skimmer - 65 V, Vcap 2500 V, fragmentor - 90 V, dissolved in methanol

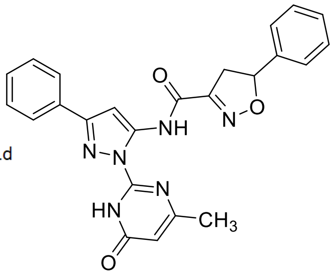

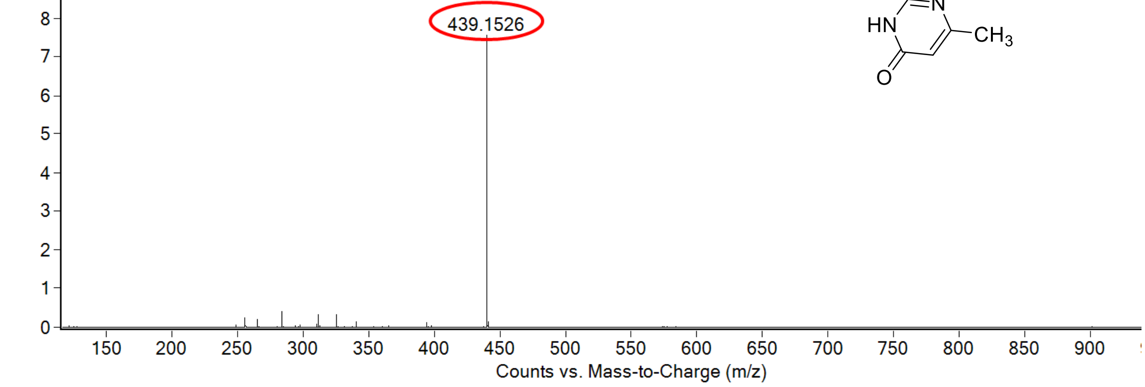

calculated mass: [M-H]$^-$ = 439.1524 observed: [M-H]$^-$ = 439.1526 mass accuracy = 0.5 ppm

FT-IR spectrum (neat) of **119**

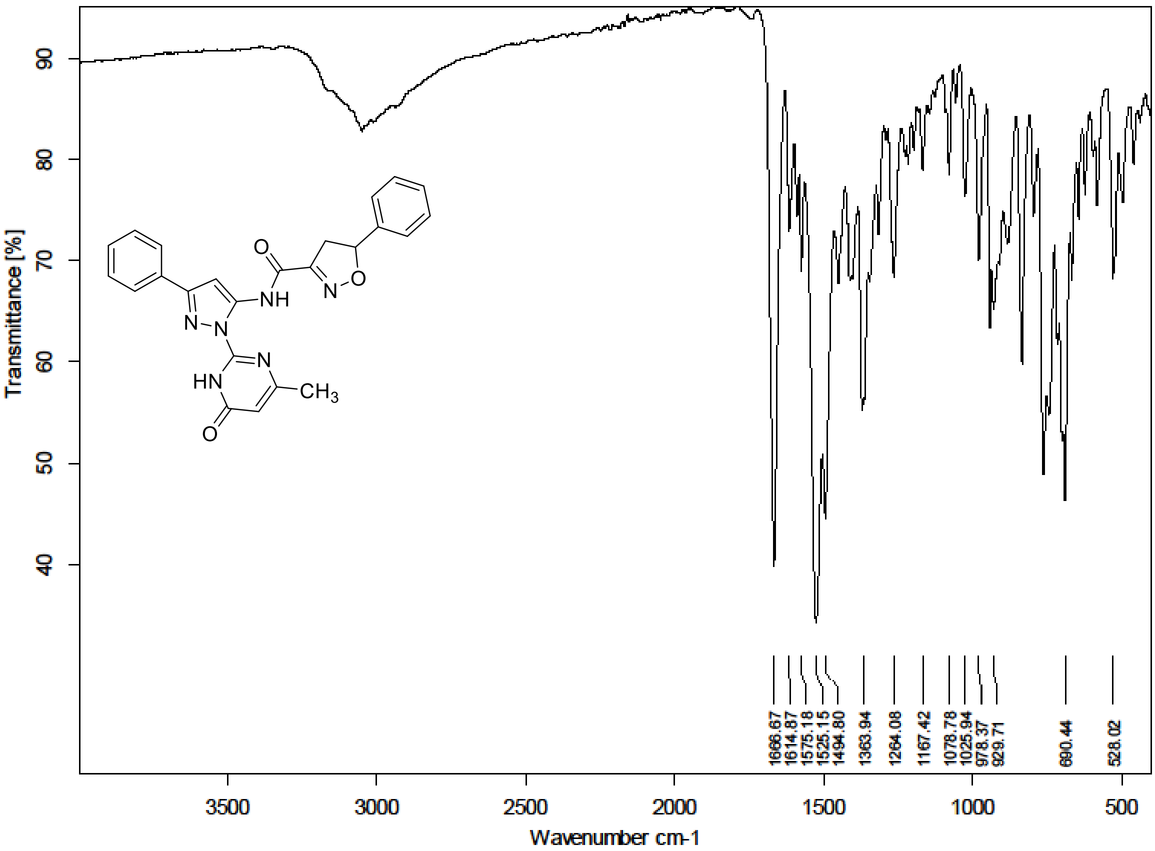

$^{1}H$ NMR (300 MHz) spectrum of **120** in DMSO-$d_6$

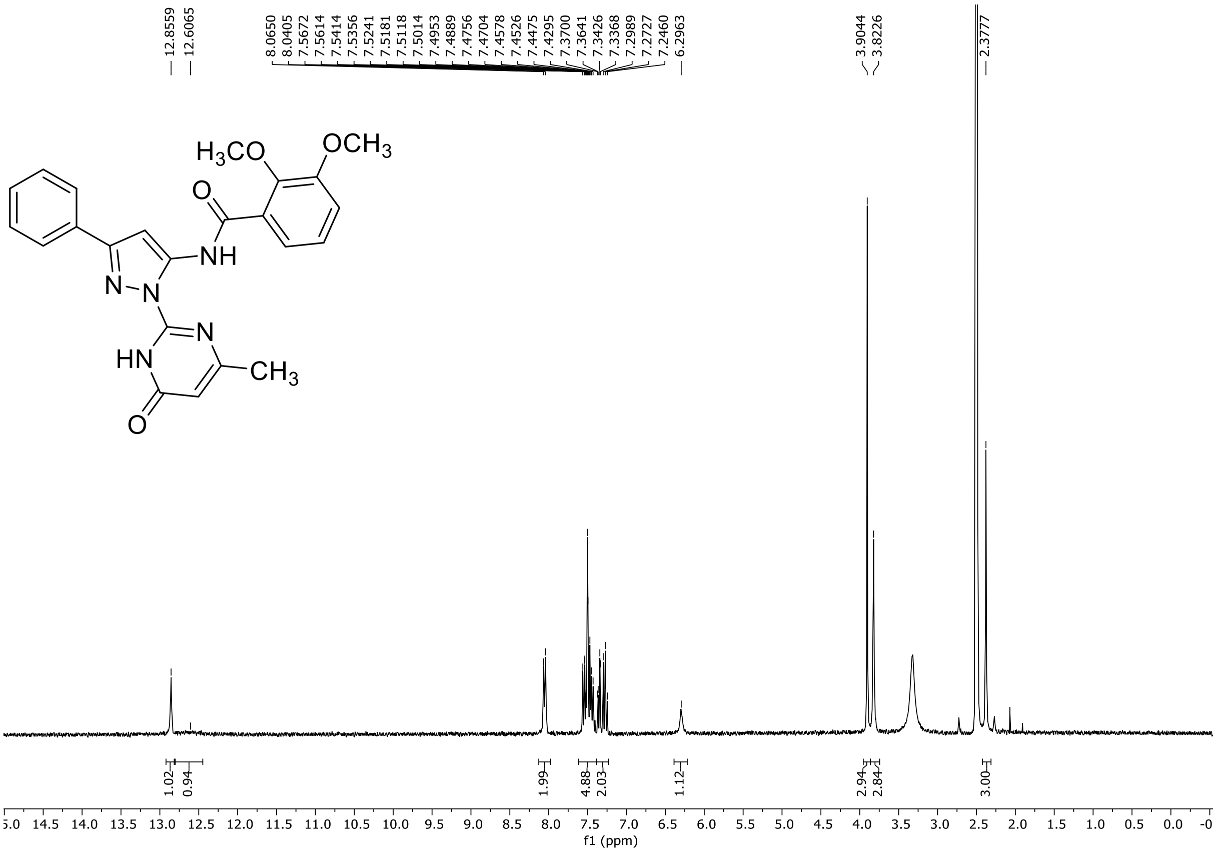

HRMS spectrum of **120**

**NAR-A-62** $C_{23}H_{21}N_5O_4$ **431.1594** *mono m/z*

**APCI +** (MMI)
nitrogen flow 5 L/min, gas temperature 300°C, nebulizer 45 psi, skimmer **65** V, vaporizer 200°C, fragmentor 30 V, dissolved in MeOH

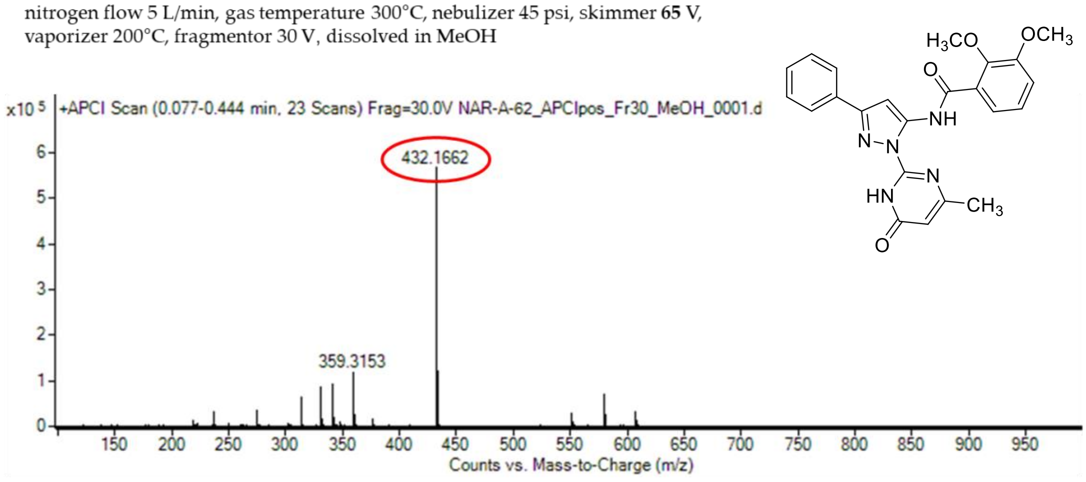

calculated mass: $[M+H]^+$ = 432.1666 observed: $[M+H]^+$ = 432.1662 mass accuracy = -0.9 ppm

FT-IR spectrum (neat) of **120**

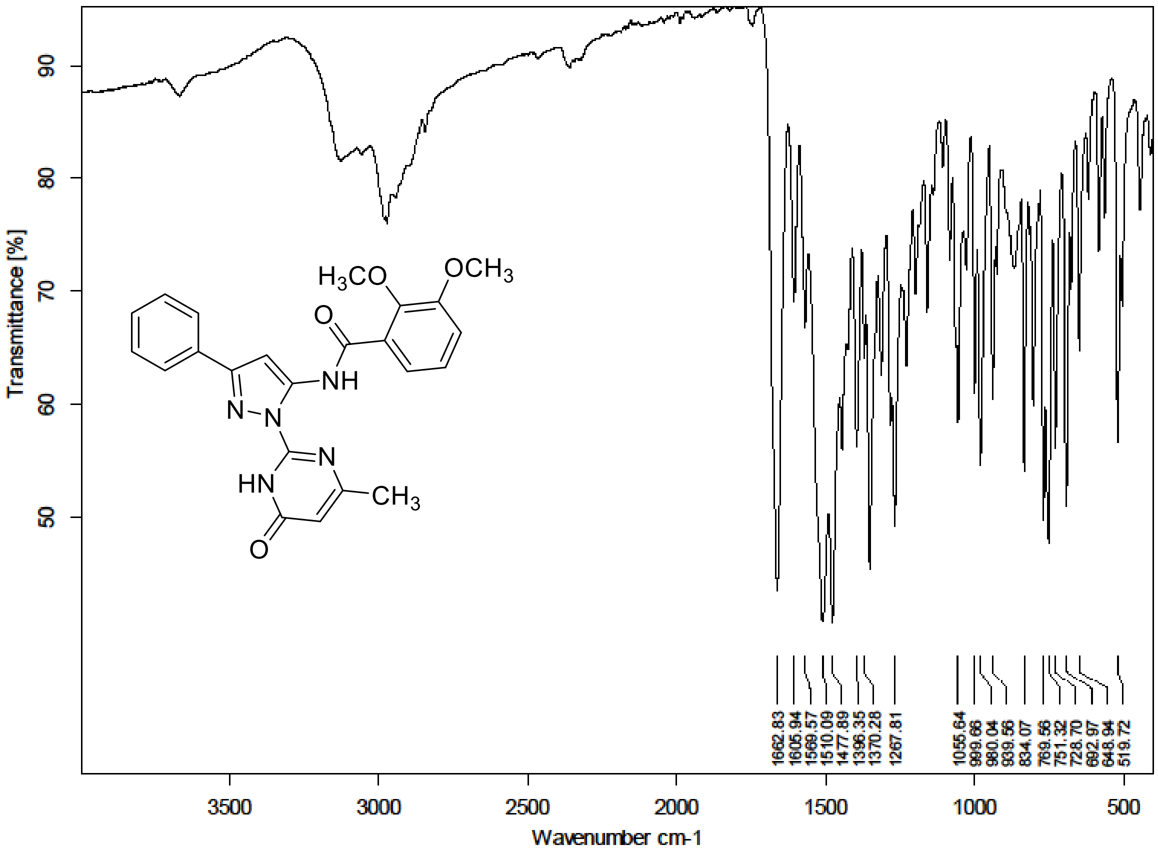

$^{1}H$ NMR (300 MHz) spectrum of **121** in DMSO-$d_6$

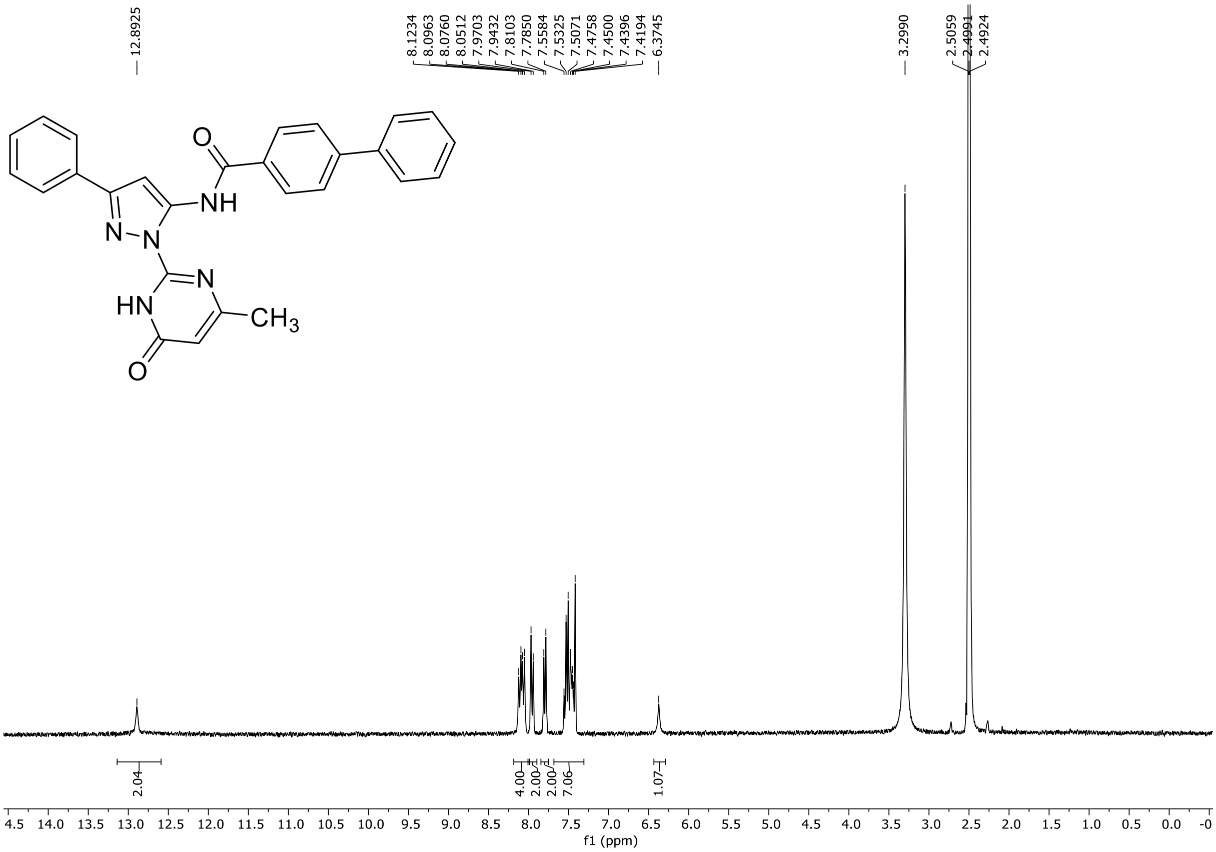

HRMS spectrum of **121**

### NAR-A-75 RC

$C_{27}H_{21}N_5O_2$ *mono m/z*
**447.1695**

**APCI +** (MMI)
nitrogen flow 5 L/min, gas temperature 300°C, nebulizer 45 psi, skimmer **65** V,
vaporizer 200°C, fragmentor 35 V, dissolved in MeOH

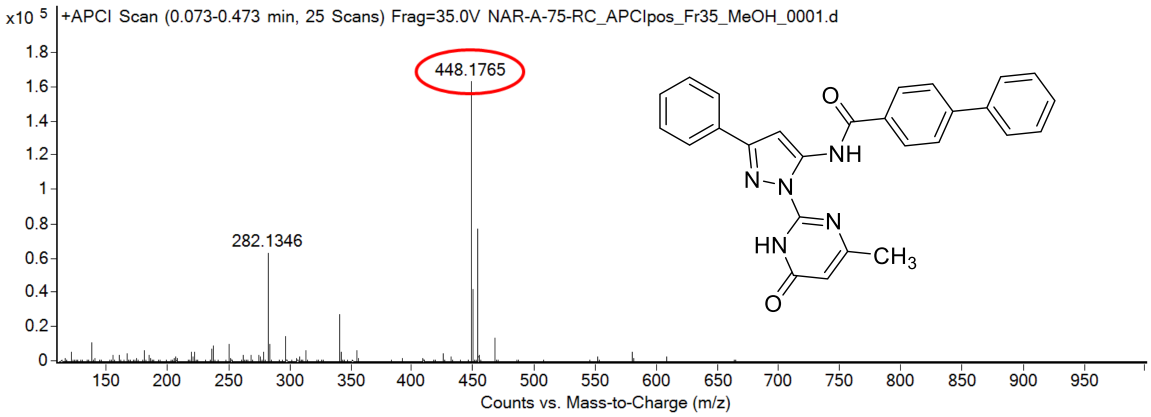

calculated mass: $[M+H]^+$ = 448.1768 observed: $[M+H]^+$ = 448.1765 mass accuracy = -0.7 ppm

FT-IR spectrum (neat) of **121**

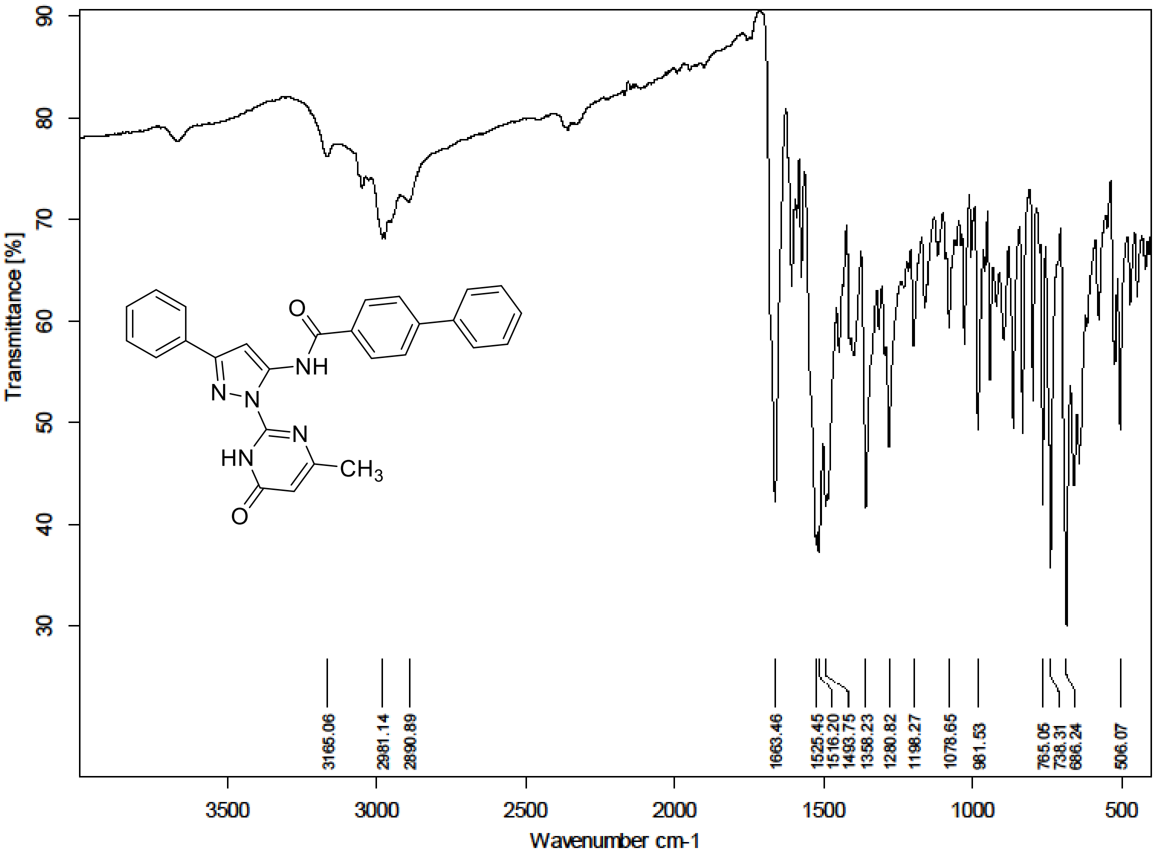

$^{1}H$ NMR (300 MHz) spectrum of **122** in DMSO-$d_6$

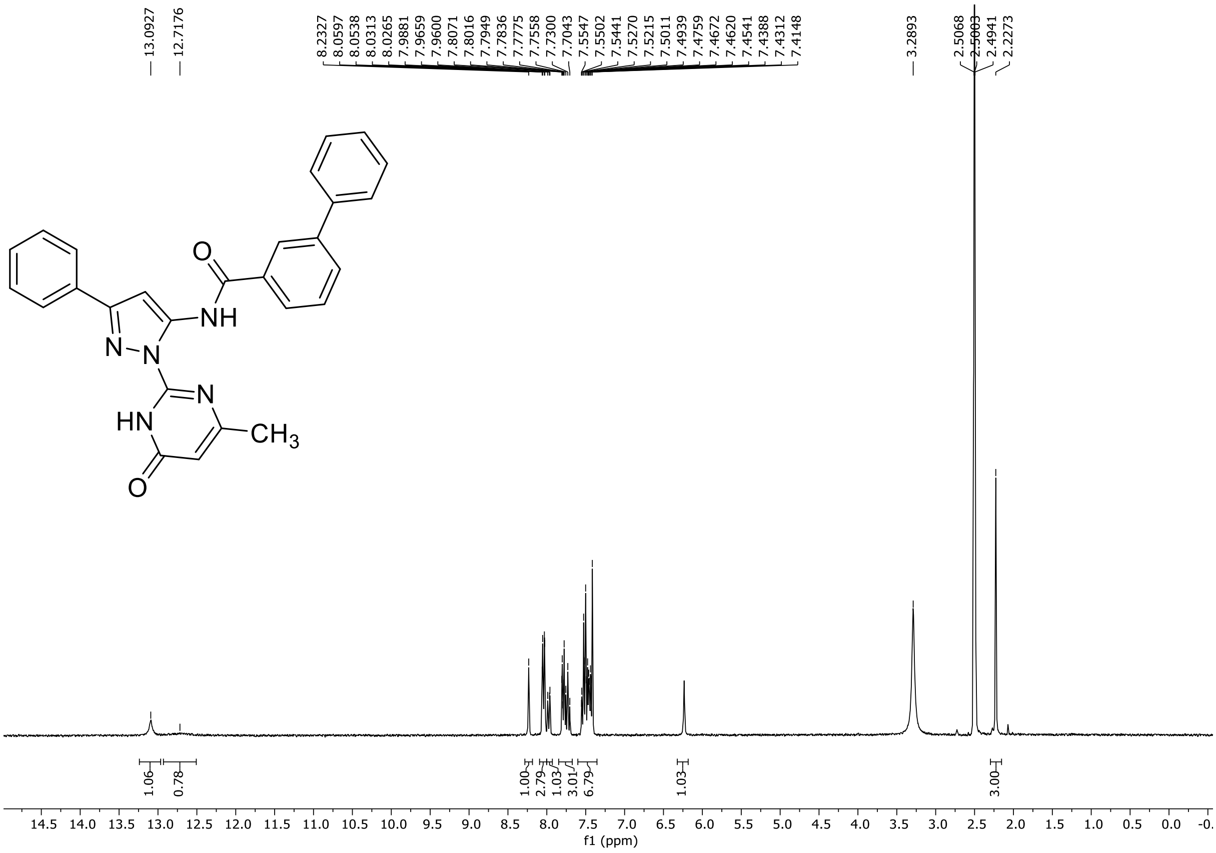

HRMS spectrum of **122**

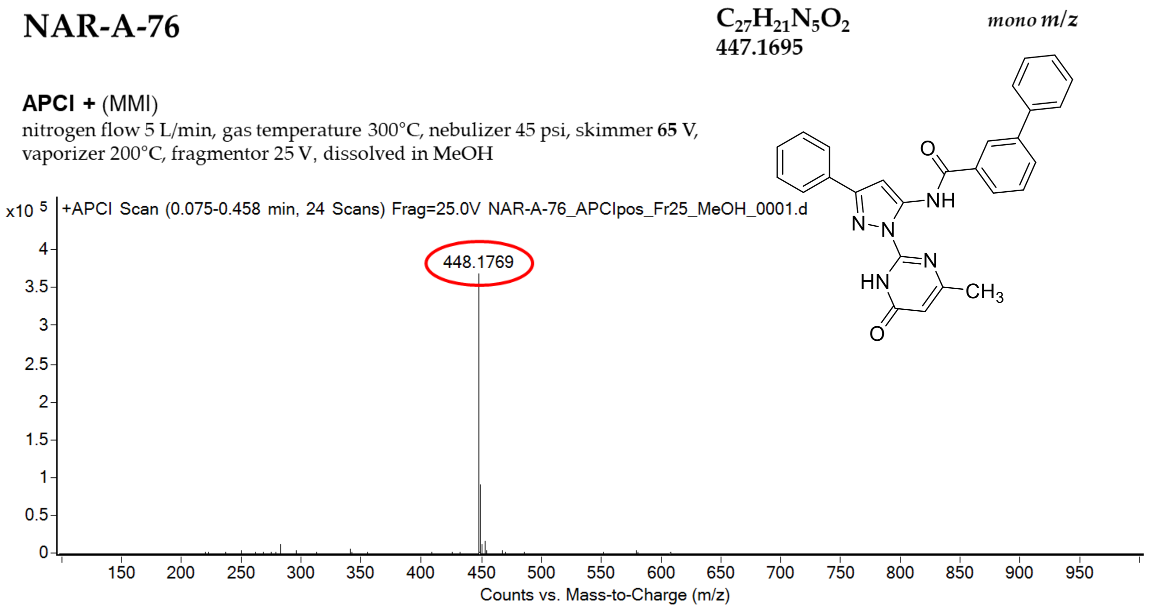

calculated mass: $[M+H]^+$ = 448.1768 observed: $[M+H]^+$ = 448.1769 mass accuracy = 0.2 ppm

FT-IR spectrum (neat) of **122**

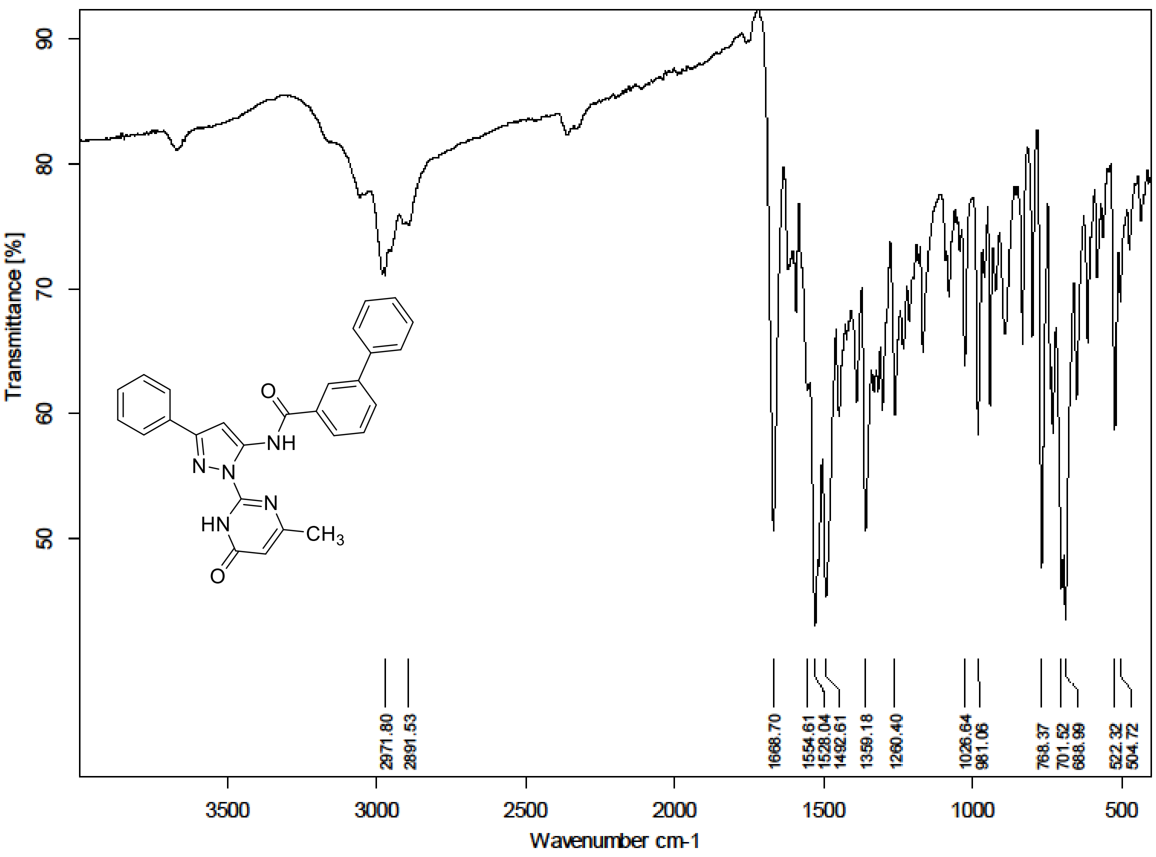

[1]H (500 MHz) and [13]C NMR (126 MHz) spectra of **123** in Methanol-$d_4$

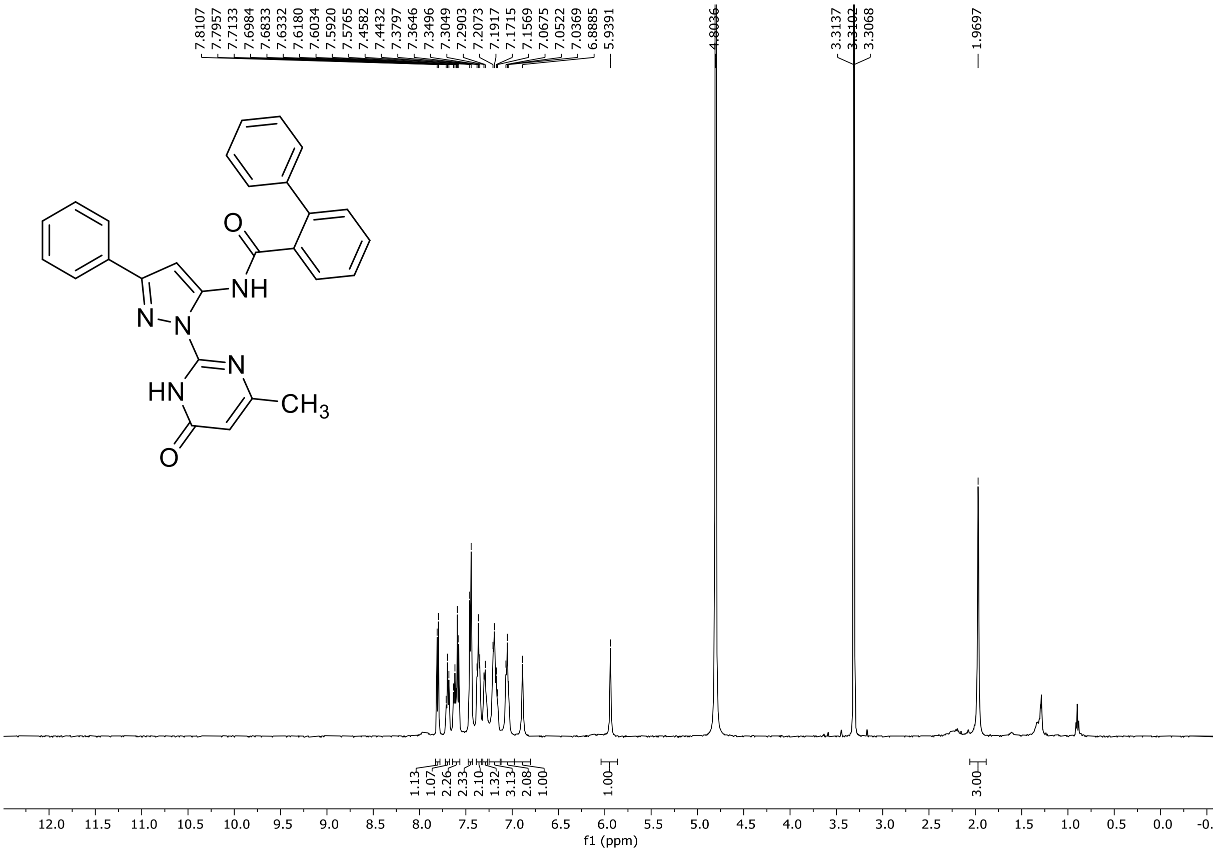

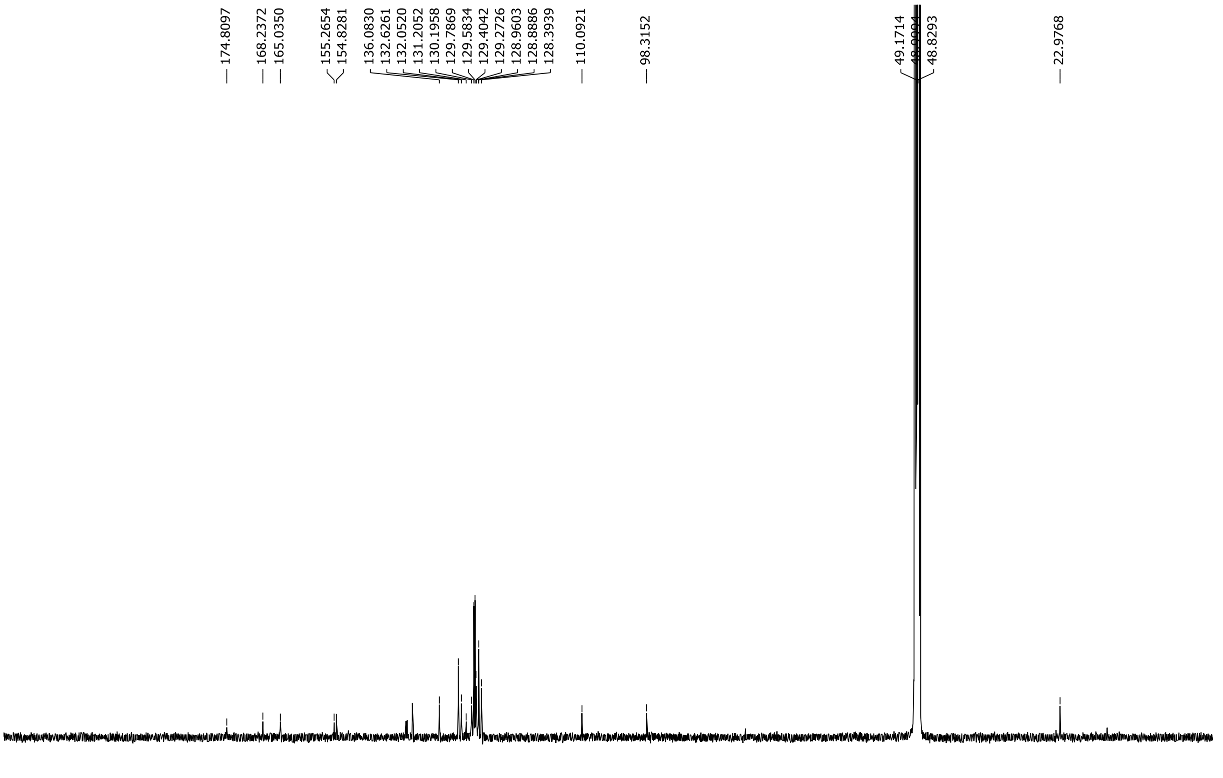

HRMS spectrum of **123**

### NAR-A-83-1

$C_{27}H_{21}N_5O_2$ *mono m/z* 447.1695

**APCI +** (MMI)
nitrogen flow 5 L/min, gas temperature 300°C, nebulizer 45 psi, skimmer **65** V, vaporizer 200°C, fragmentor 30 V, dissolved in MeOH

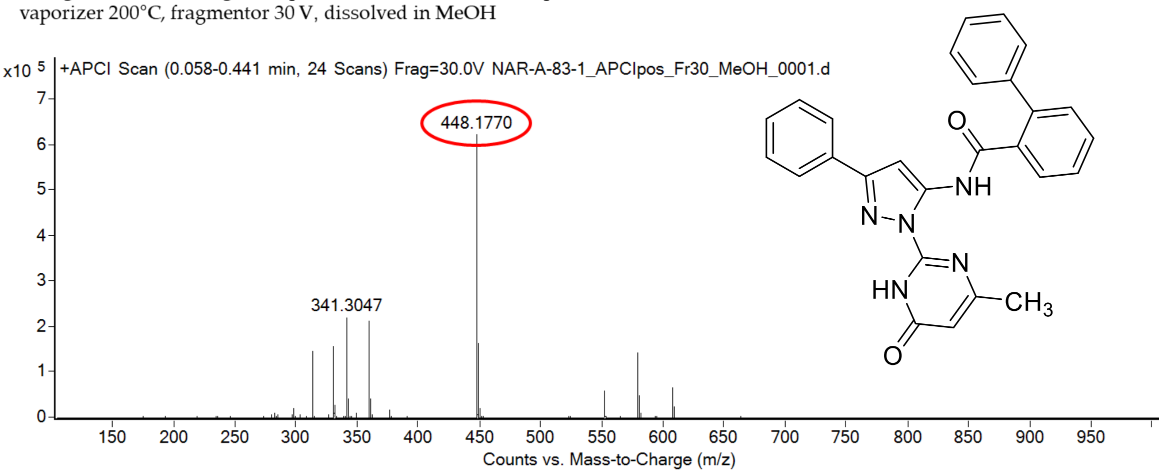

calculated mass: $[M+H]^+$ = 448.1768 observed: $[M+H]^+$ = 448.1770 mass accuracy = 0.4 ppm

FT-IR spectrum (neat) of **123**

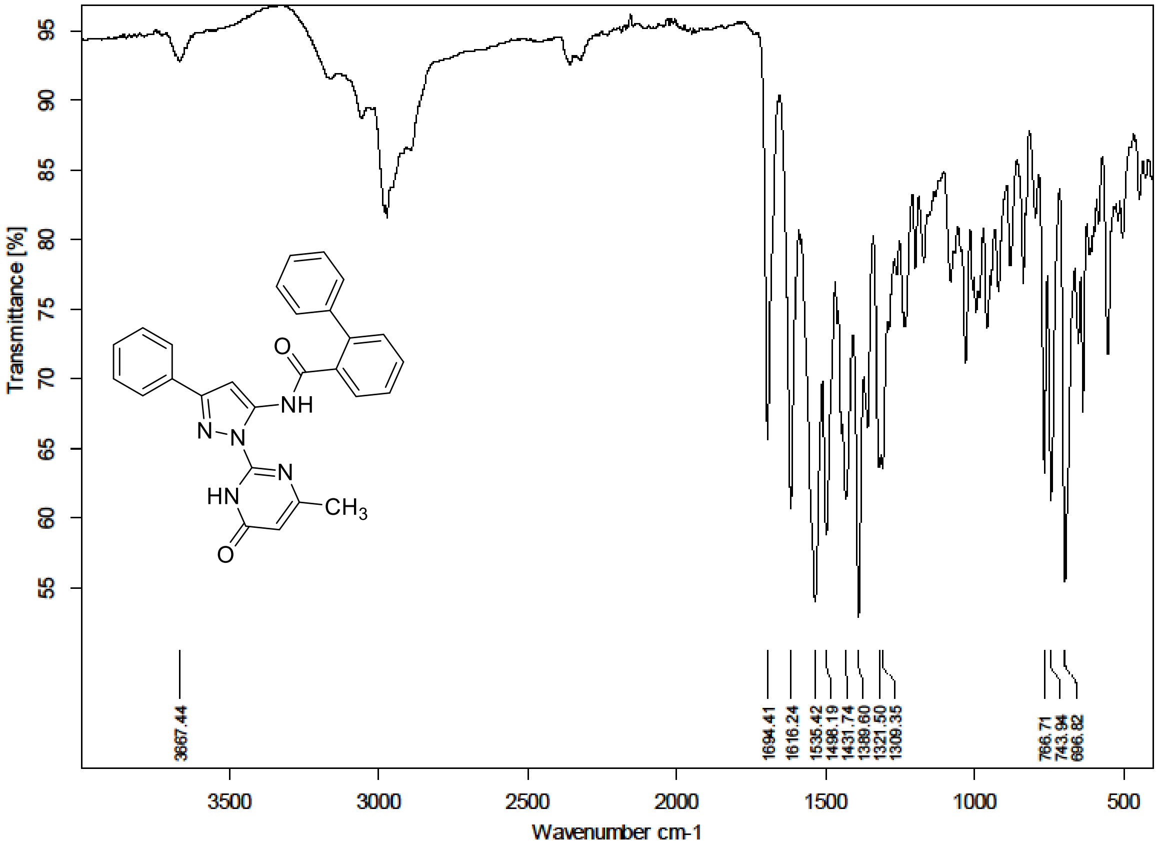

$^{1}H$ (500 MHz) and $^{13}C$ NMR (126 MHz) spectra of **124** in DMSO-$d_6$

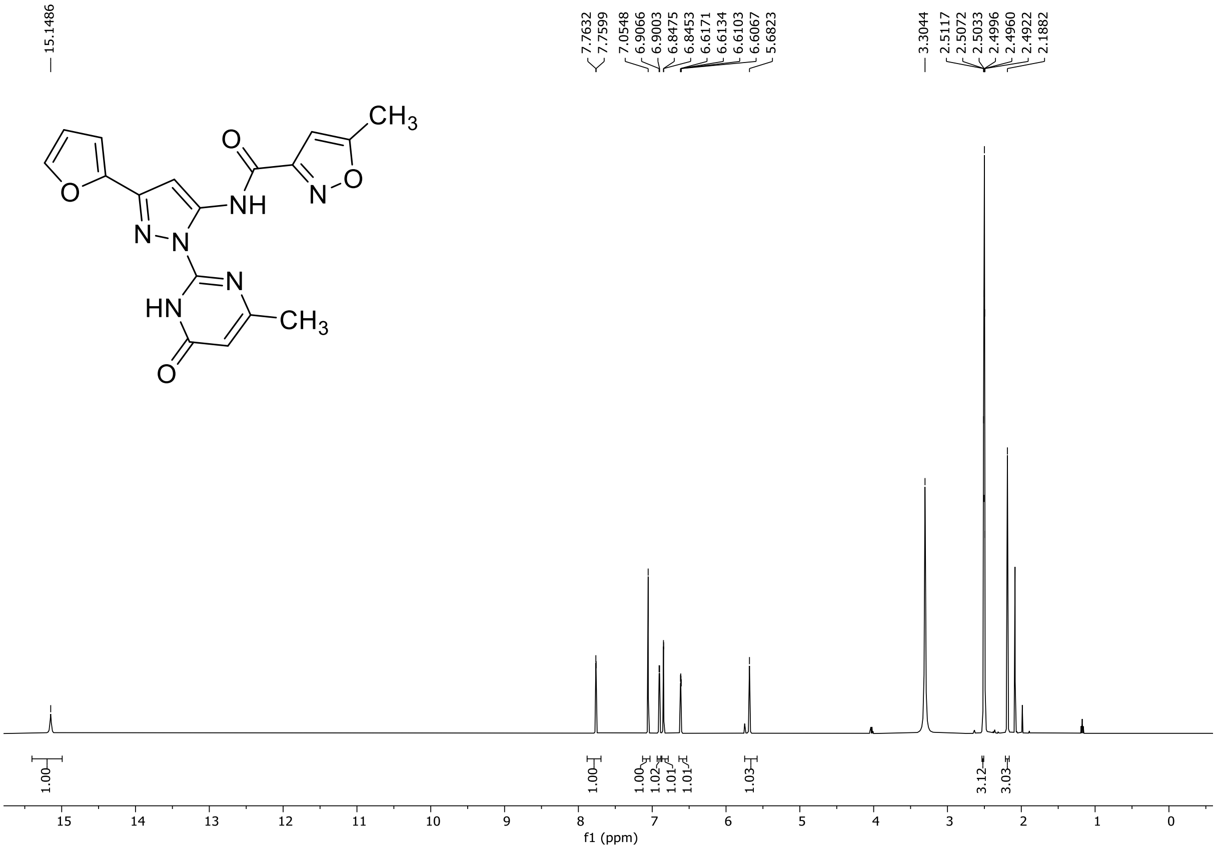

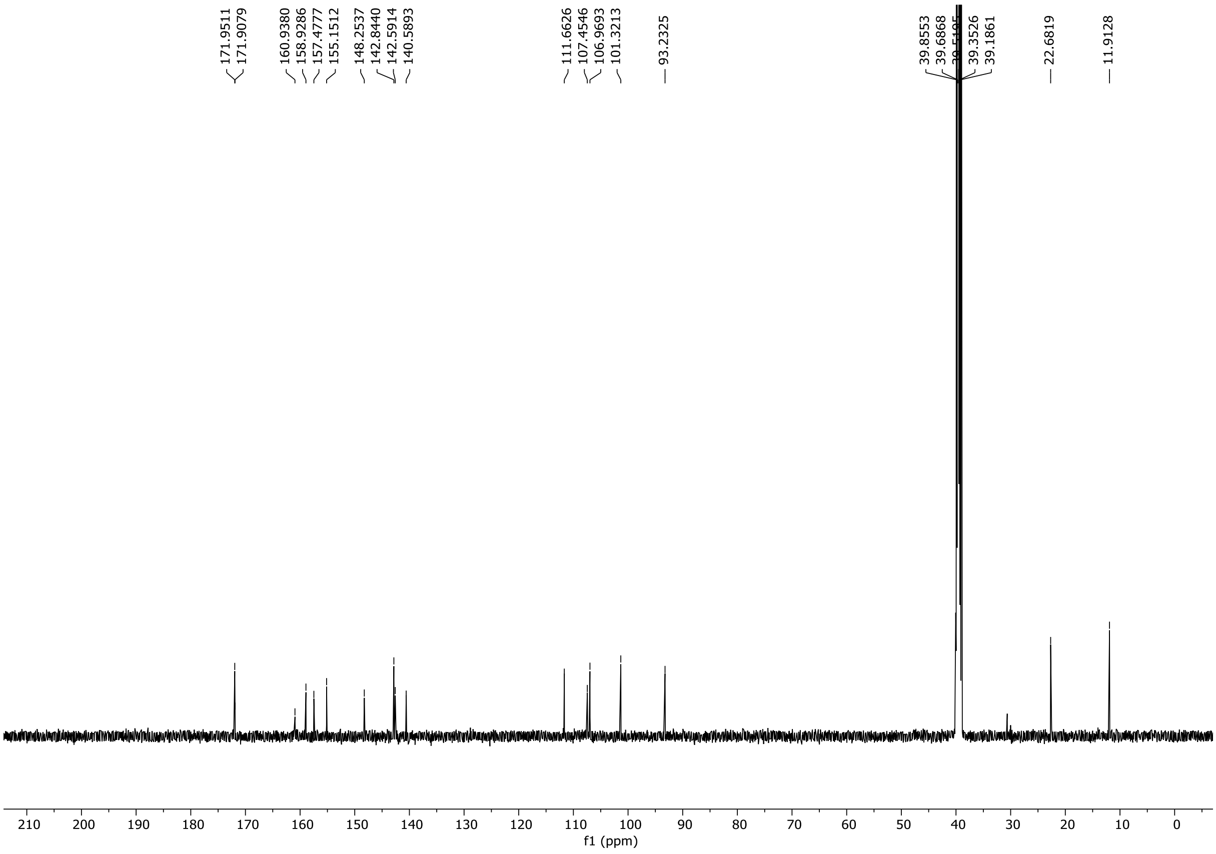

HRMS spectrum of **124**

**NAR-A-4** $C_{17}H_{14}N_6O_4$ *mono m/z* **366.1077**

**ESI -** (MMI)
nitrogen flow 5 L/min, gas temperature 325°C, nebulizer 45 psi, skimmer **65** V, vaporizer 200°C, fragmentor 35 V, dissolved in methanol

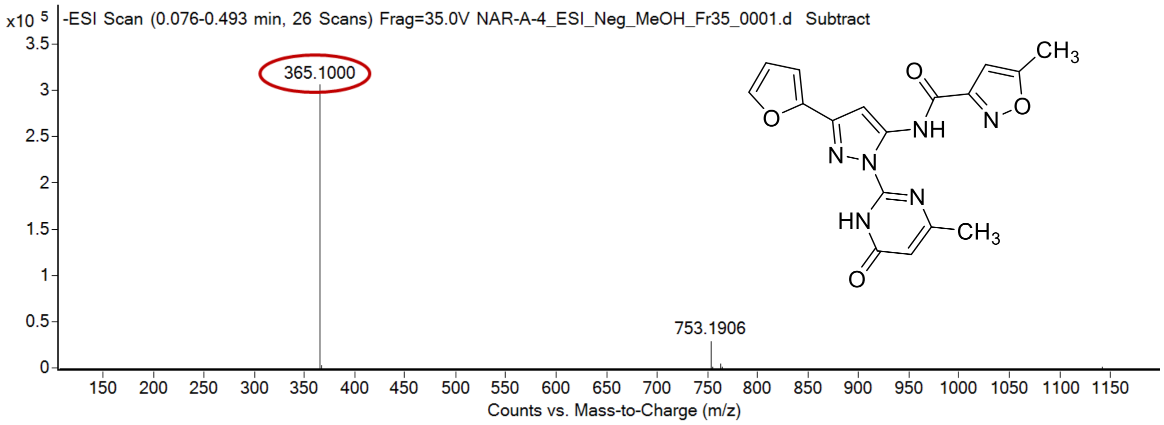

calculated mass: [M-H]$^-$ = 365.1004 observed: [M-H]$^-$ = 365.1000 mass accuracy = 1.1 ppm

FT-IR spectrum (neat) of **124**

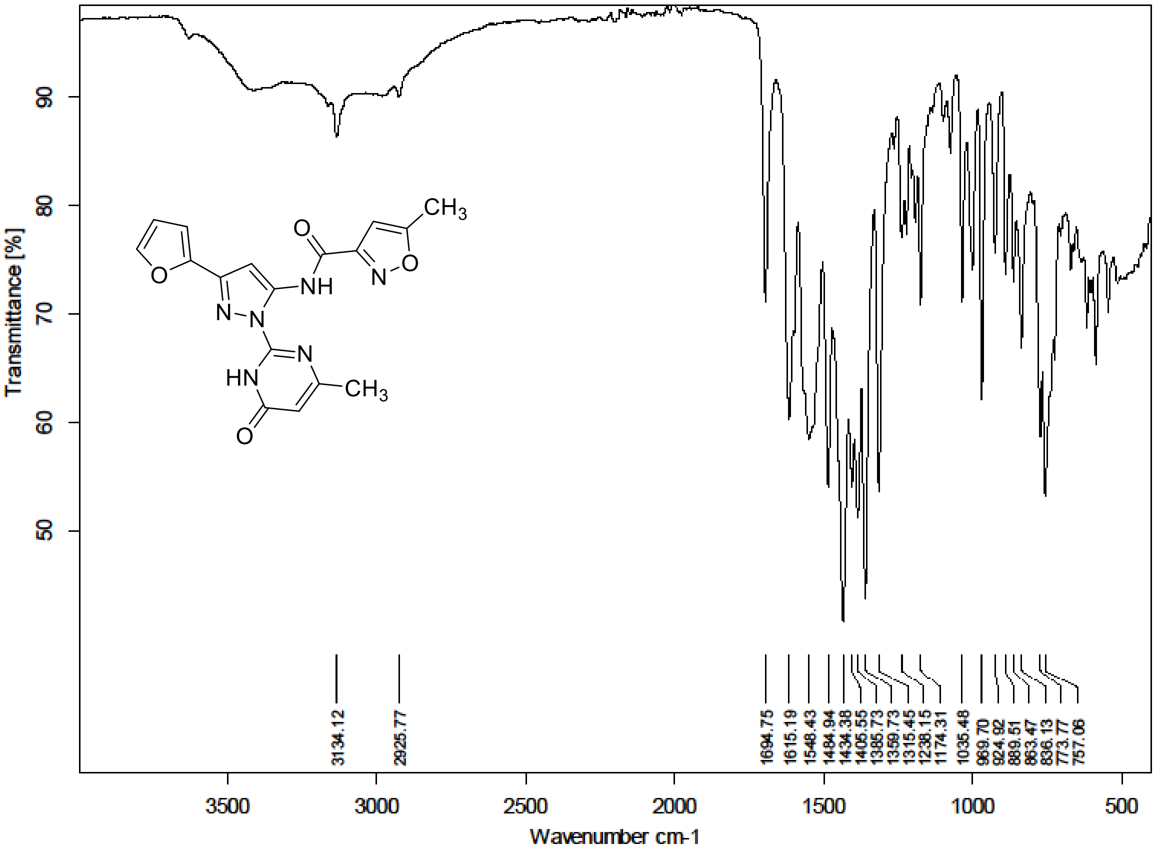

$^1$H (300 MHz) and $^{13}$C NMR (176 MHz) spectra of **125** in DMSO-$d_6$

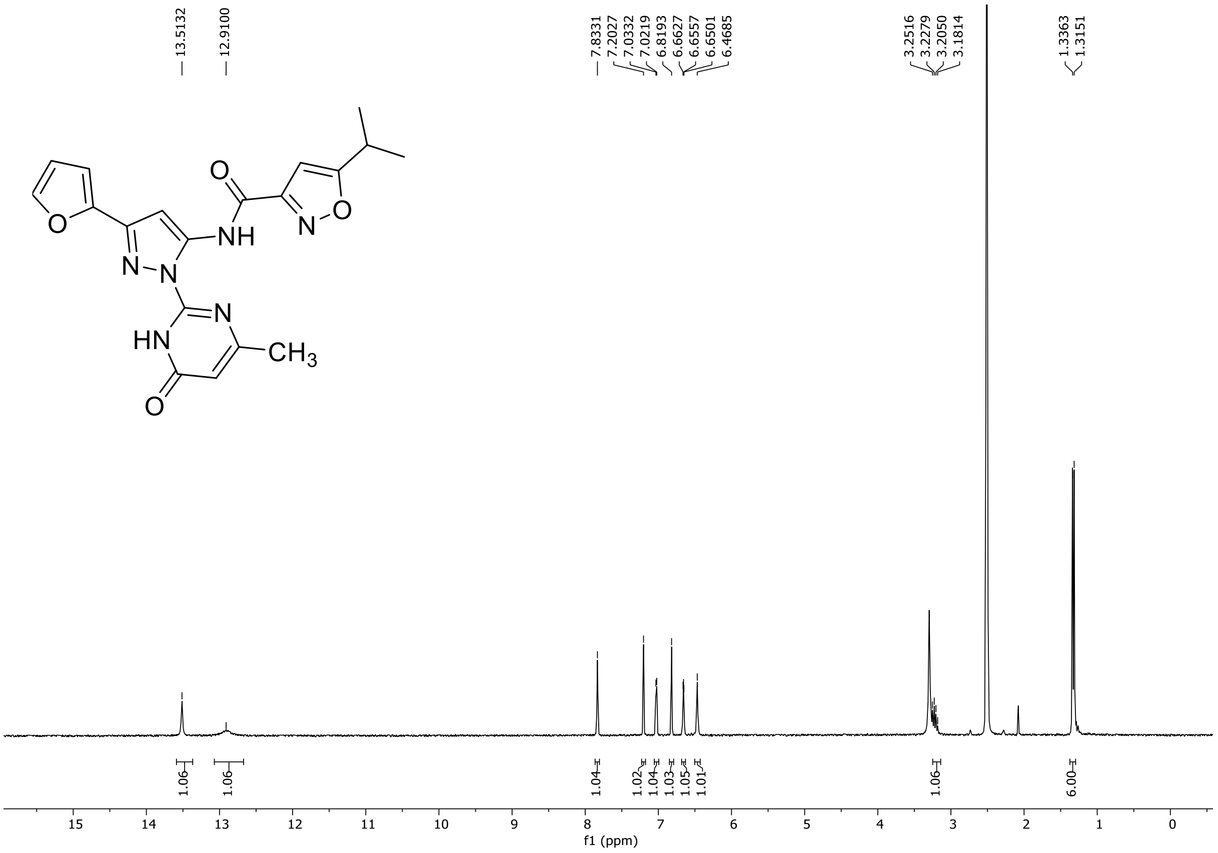

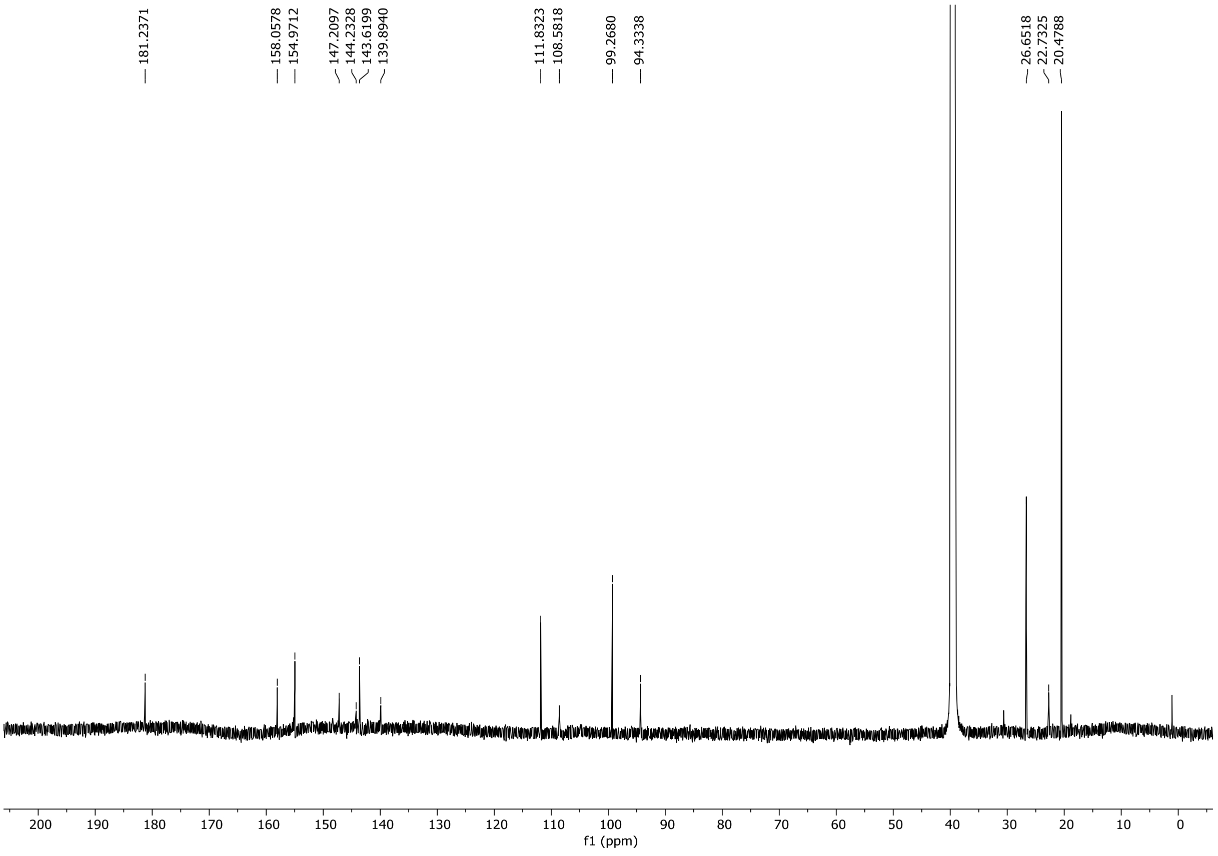

### HRMS spectrum of **125**

NAR-A-34 **$C_{19}H_{18}N_6O_4$** mono *m/z 394.1390*

**APCI +** (MMI)
nitrogen flow 5 L/min, gas temperature 325°C, nebulizer 45 psi, skimmer **65** V, vaporizer 200°C, fragmentor 25 V, dissolved in methanol

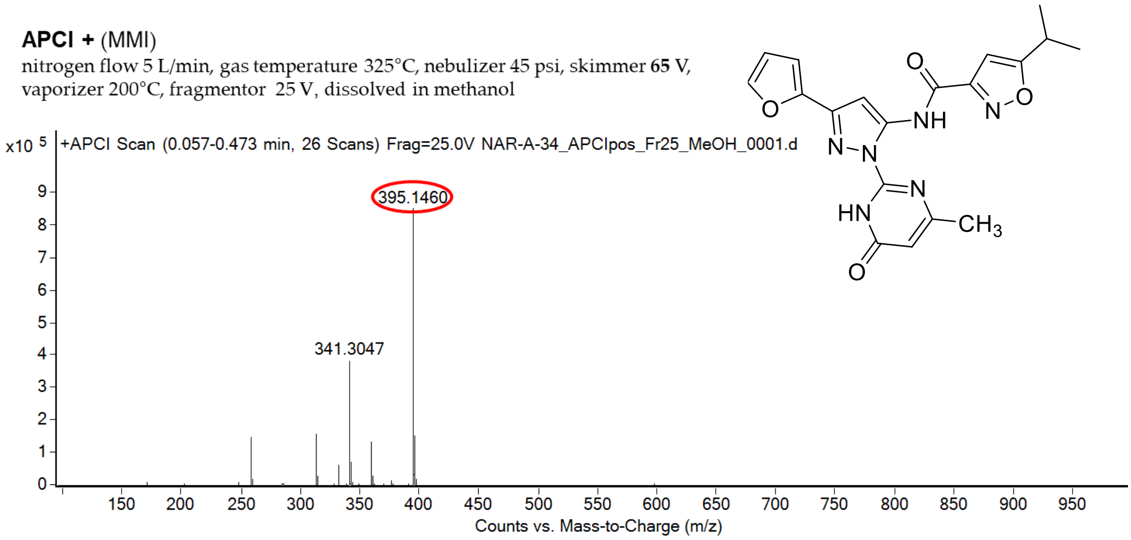

calculated mass: $[M+H]^+$ = 395.1462 observed: $[M+H]^+$ = 395.1460 mass accuracy = -0.5 ppm

### FT-IR spectrum (neat) of **125**

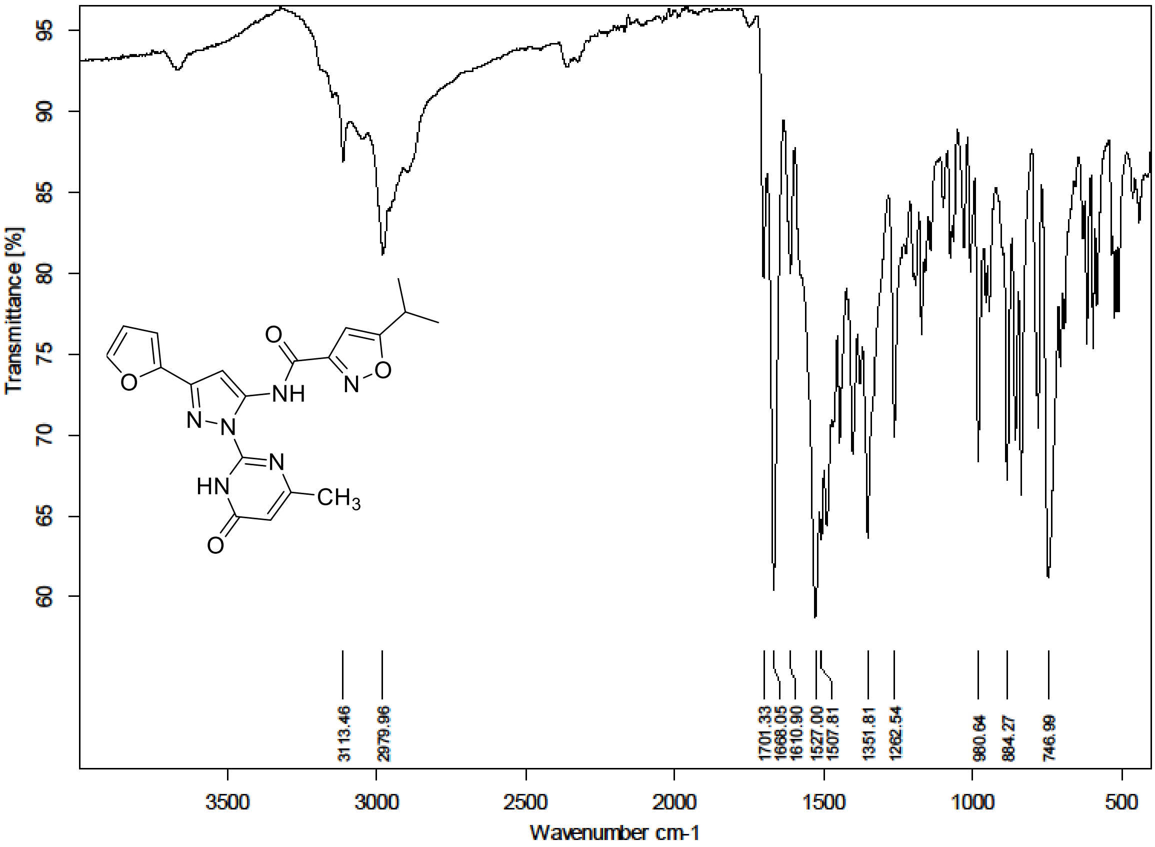

$^{1}H$ (300 MHz) and $^{13}C$ NMR (126 MHz) spectra of **126** in DMSO-$d_6$

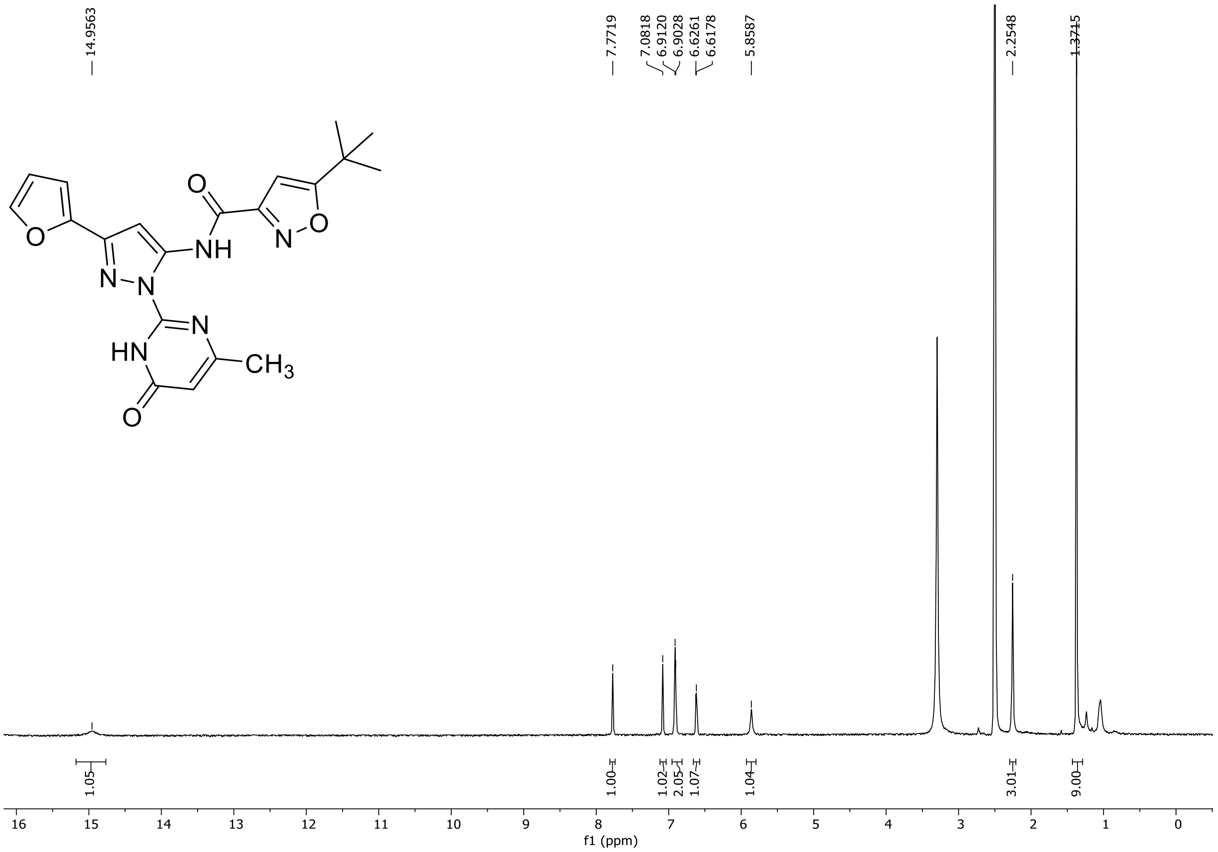

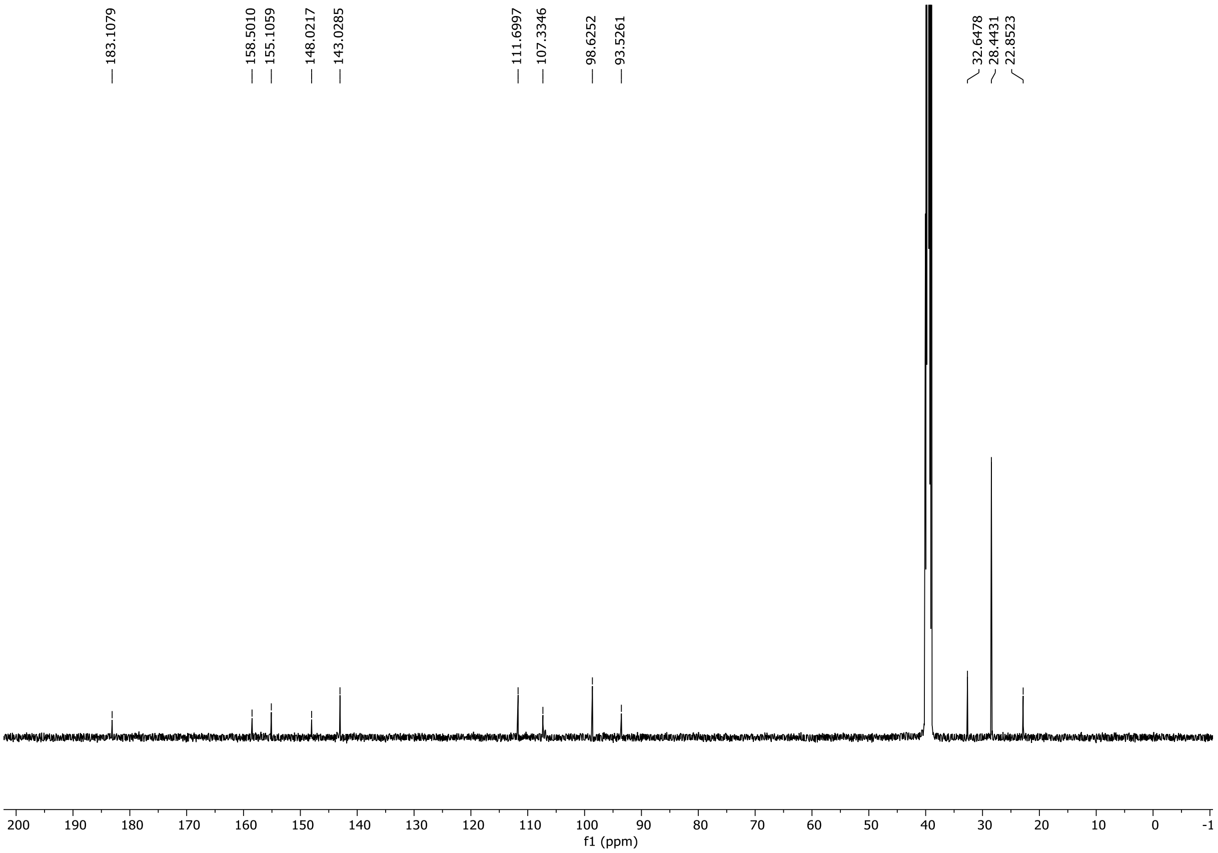

HRMS spectrum of **126**

**NAR-A-35**

$C_{20}H_{20}N_6O_4$
**408.1546**

*mono m/z*

**APCI +** (MMI)
nitrogen flow 5 L/min, gas temperature 300°C, nebulizer 45 psi, skimmer **65** V, vaporizer 200°C, fragmentor 15 V, dissolved in MeOH

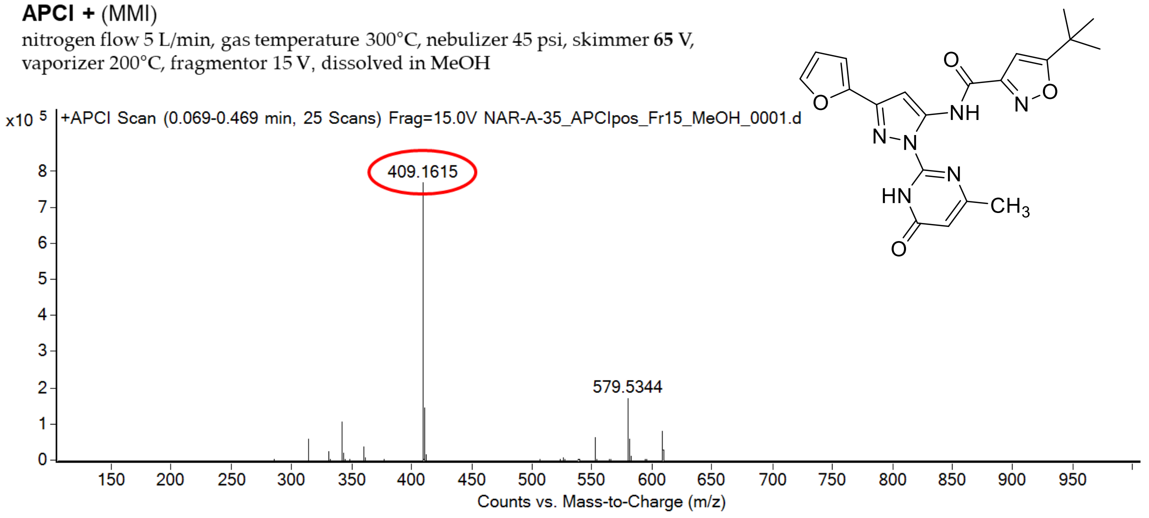

calculated mass: [M+H]$^+$ = 409.1619 observed: [M+H]$^+$ = 409.1615 mass accuracy = -1.0 ppm

FT-IR spectrum (neat) of **126**

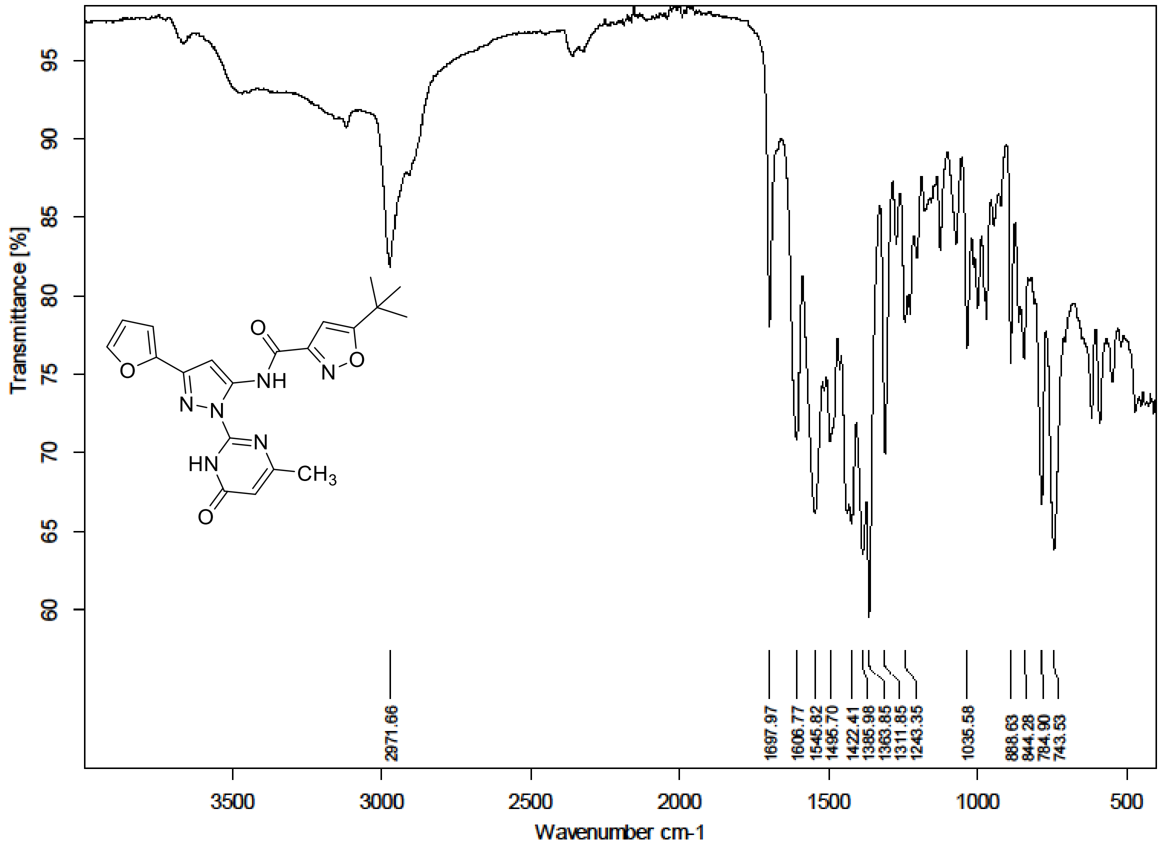

[1]H (300 MHz) and [13]C NMR (75 MHz) spectra of **127** in DMSO-$d_6$

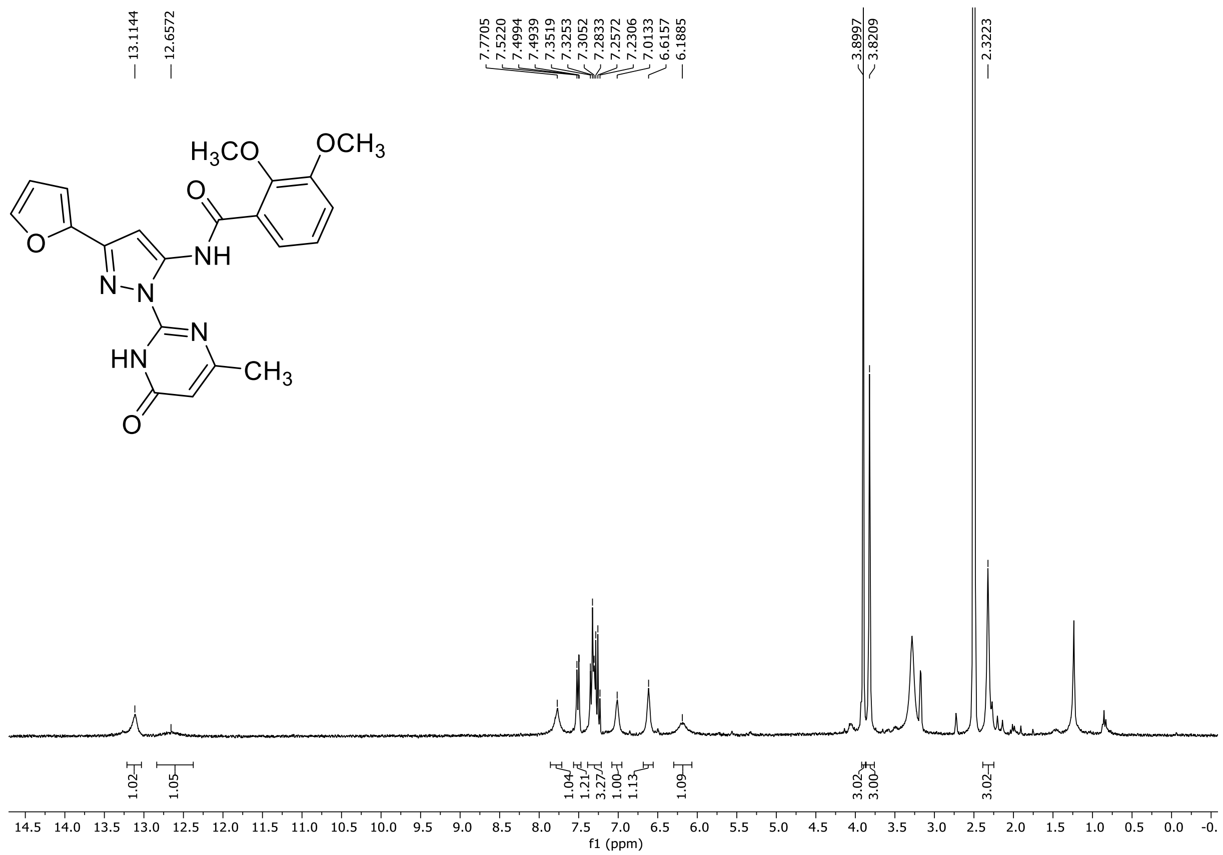

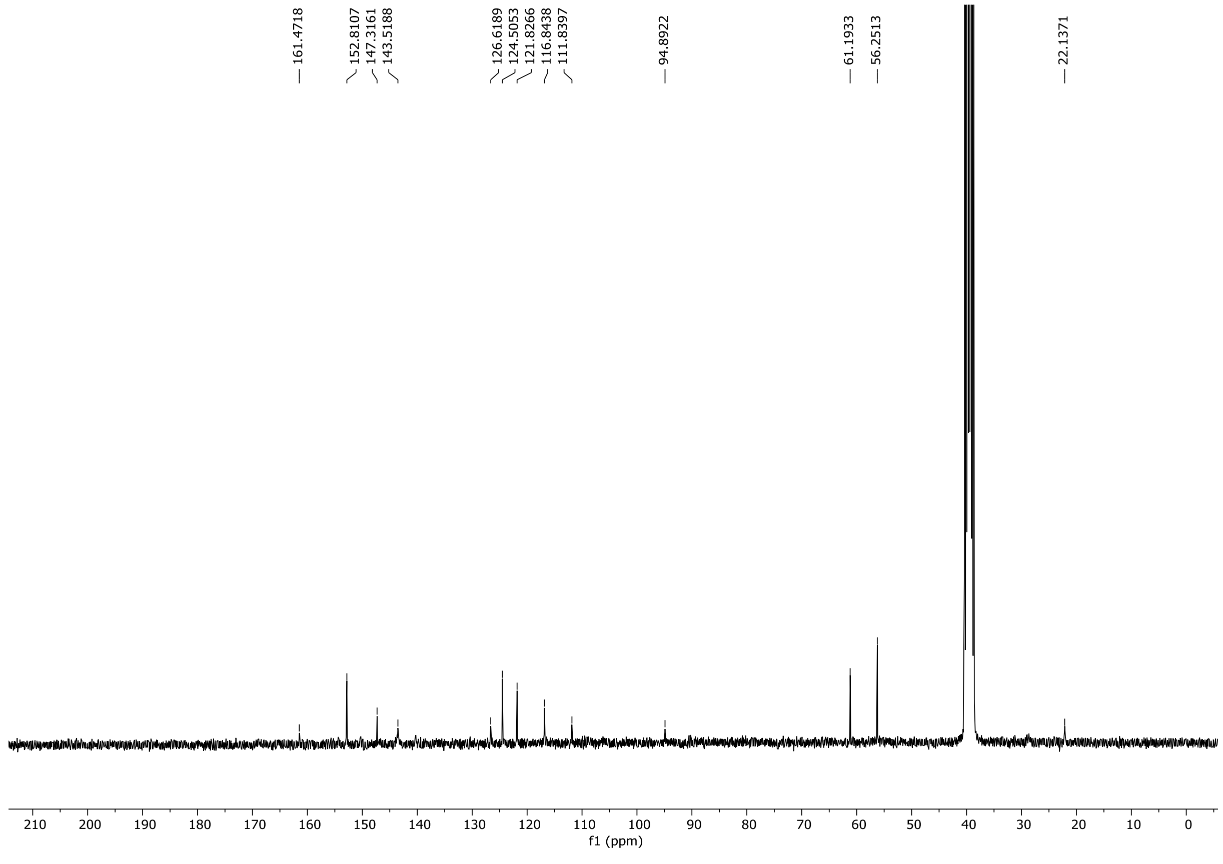

HRMS spectrum of **127**

**NAR-A-169** $C_{21}H_{19}N_5O_5$ mono *m/z* 421.1386

**APCI +** (MMI)
nitrogen flow 5 L/min, gas temperature 325°C, nebulizer 45 psi, skimmer **65** V, vaporizer 200°C, fragmentor 25 V, dissolved in methanol

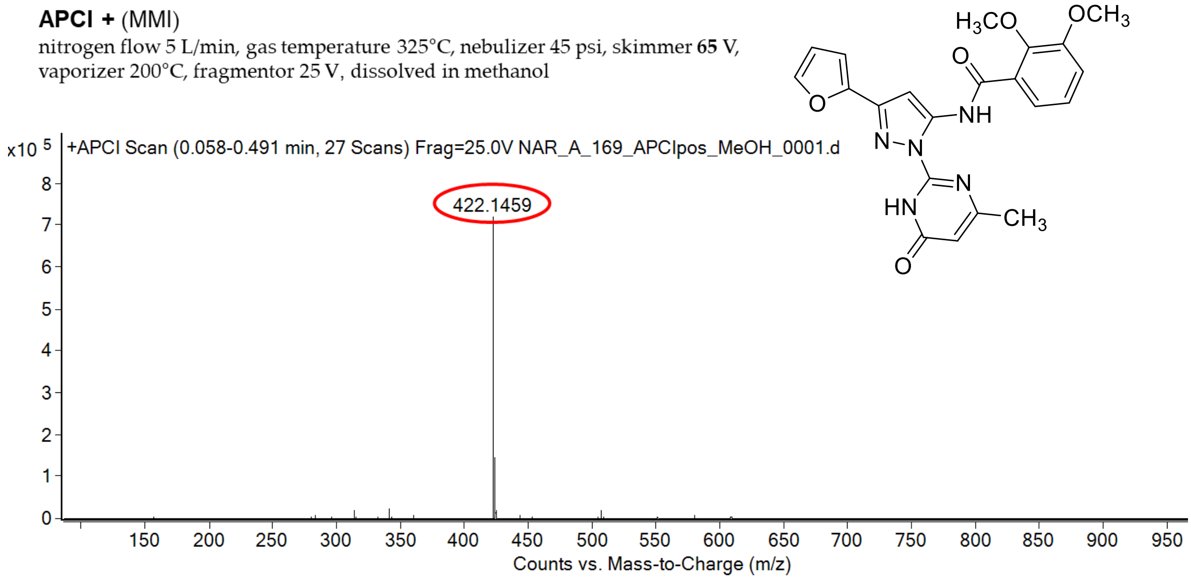

calculated mass: $[M+H]^+$ = 422.1459 observed: $[M+H]^+$ = 422.1459 mass accuracy = < 0.1 ppm

FT-IR spectrum (neat) of **127**

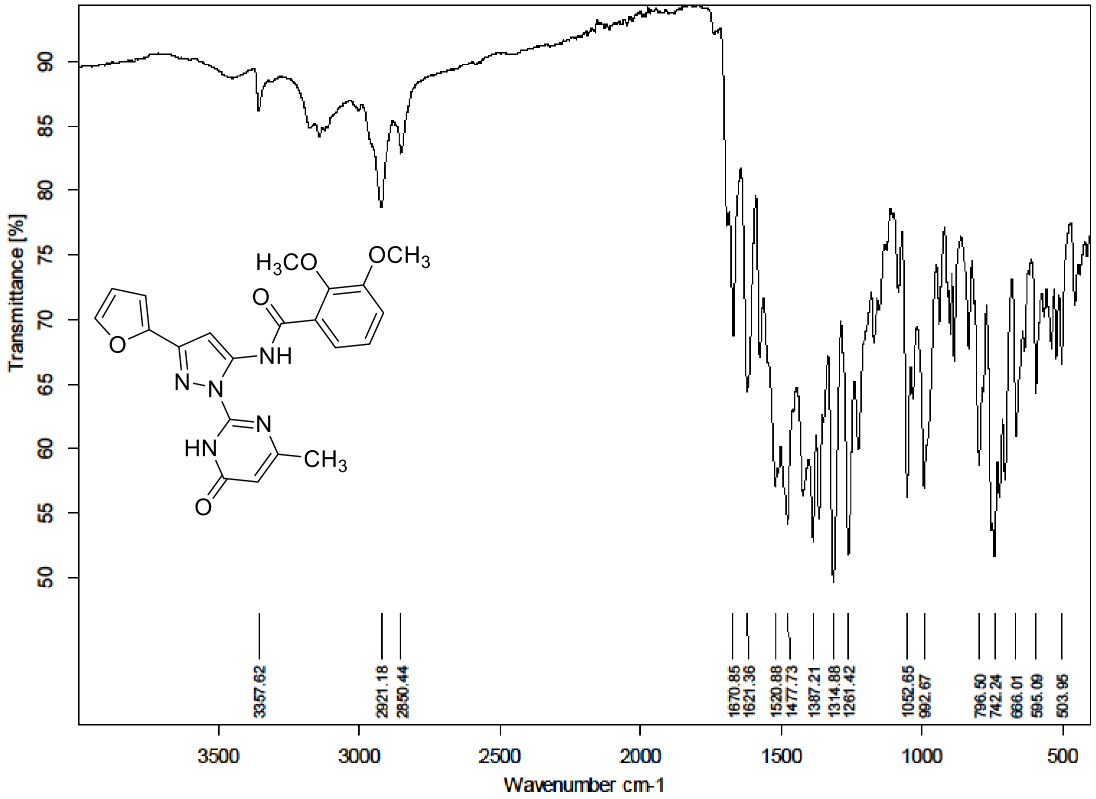

[1]H NMR (300 MHz) spectrum of **128** in DMSO-$d_6$

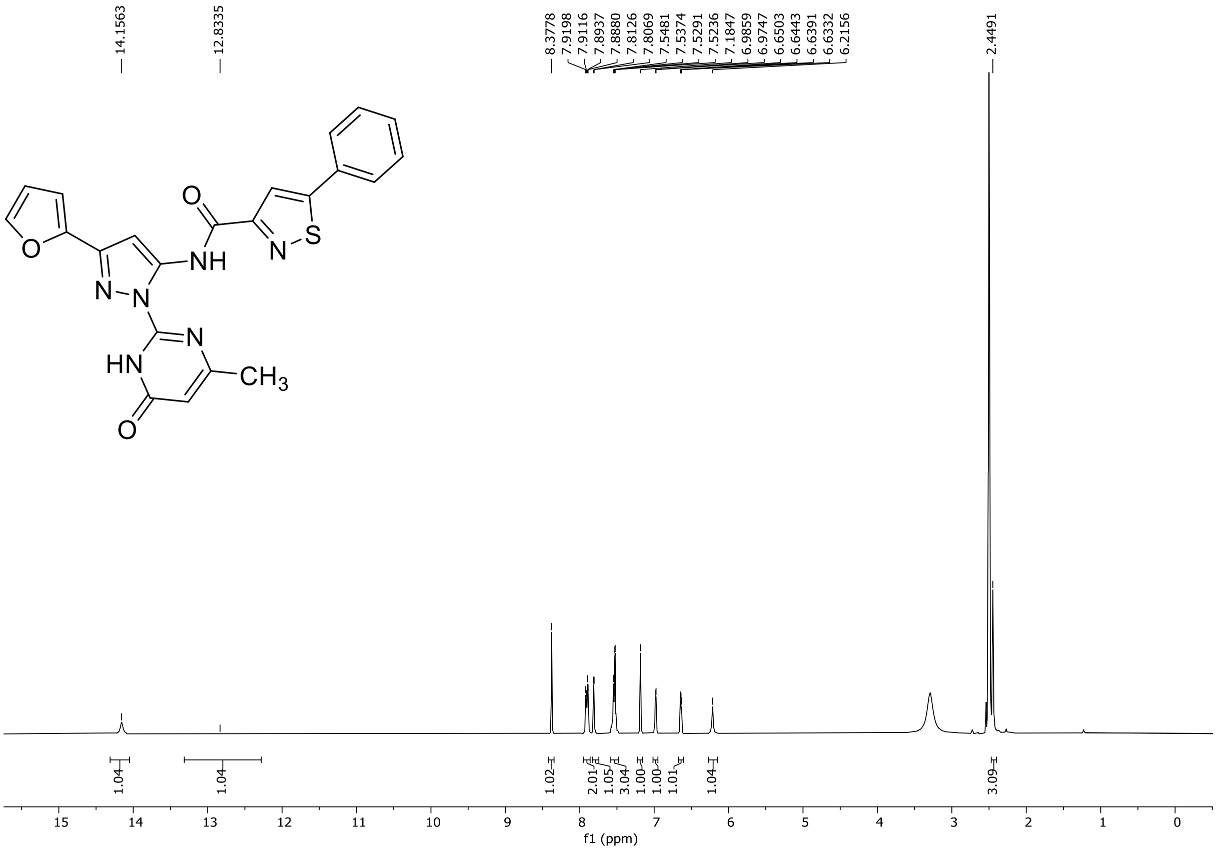

HRMS spectrum of **128**

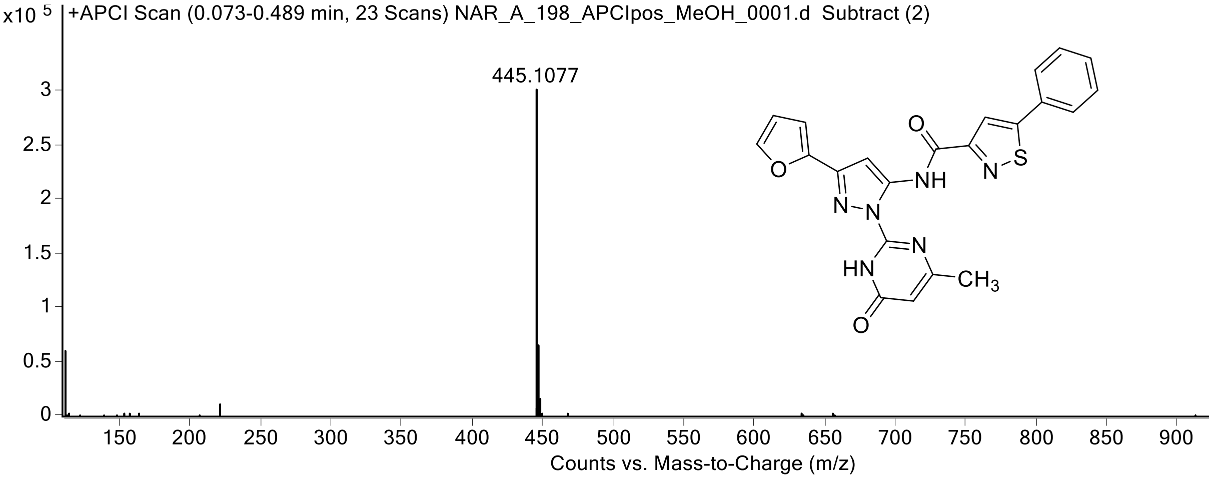

FT-IR spectrum (neat) of **128**

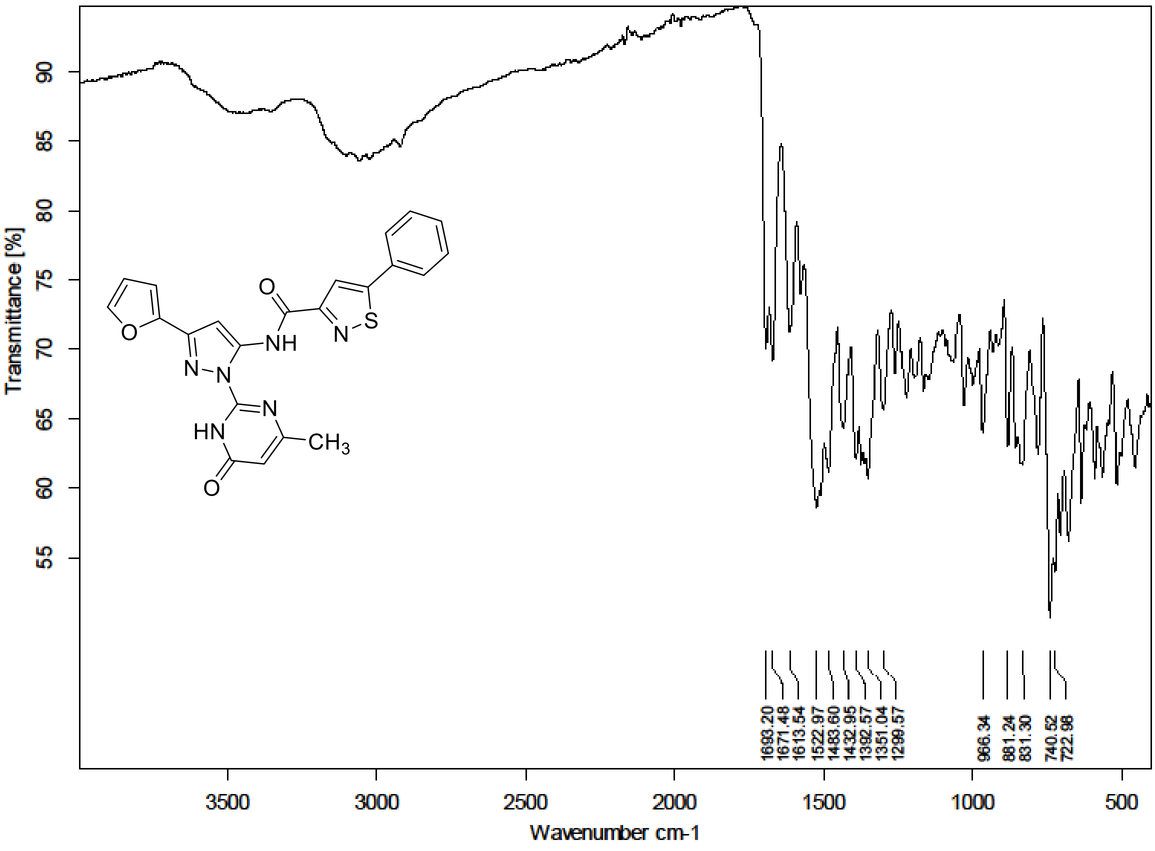

$^1H$ NMR (300 MHz) spectrum of **129** in DMSO-$d_6$

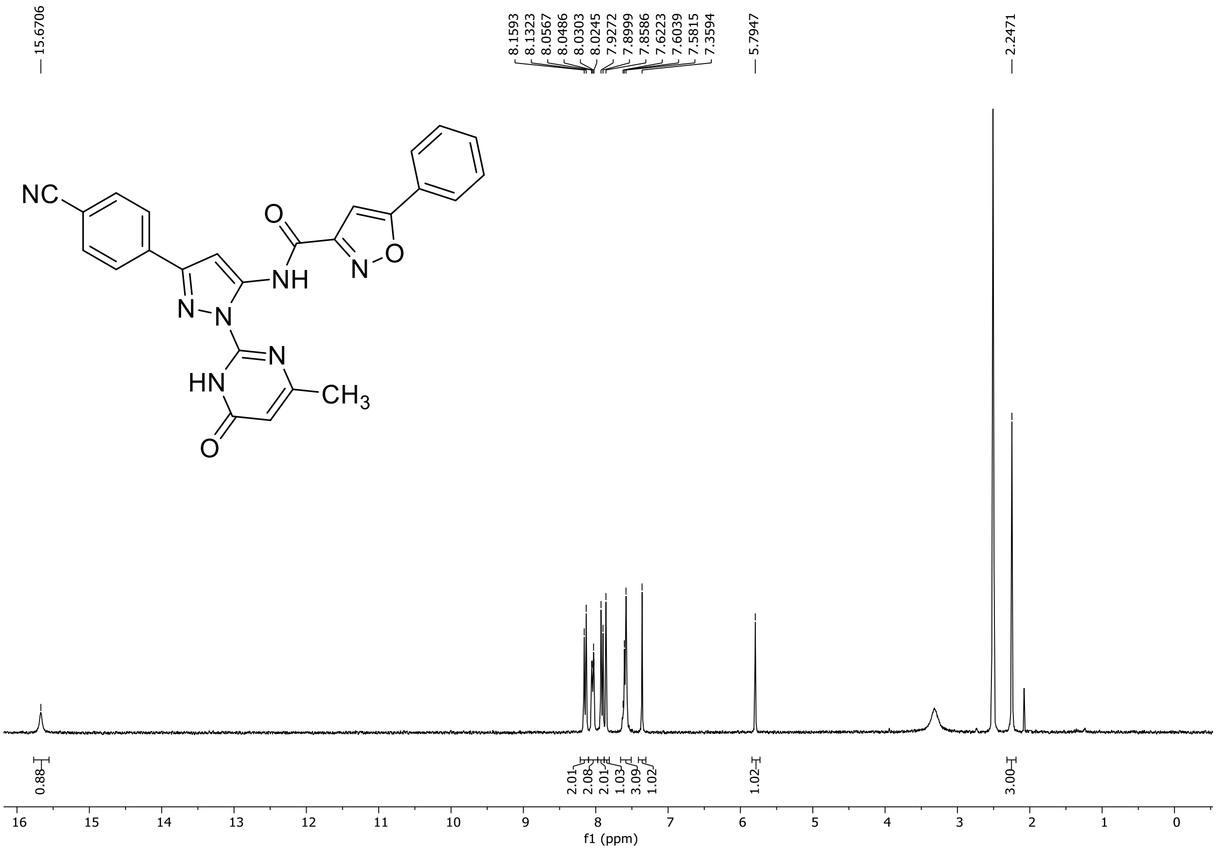

HRMS spectrum of **129**

**NAR-A-120** $C_{25}H_{17}N_7O_3$ ***m/z* 463.1393**

**ESI-** (MMI)
nitrogen flow 5 L/min, gas temperature 325°C, nebulizer 45 psi, skimmer 65 V, fragmentor 100 V, dissolved in DMSO, MeOH

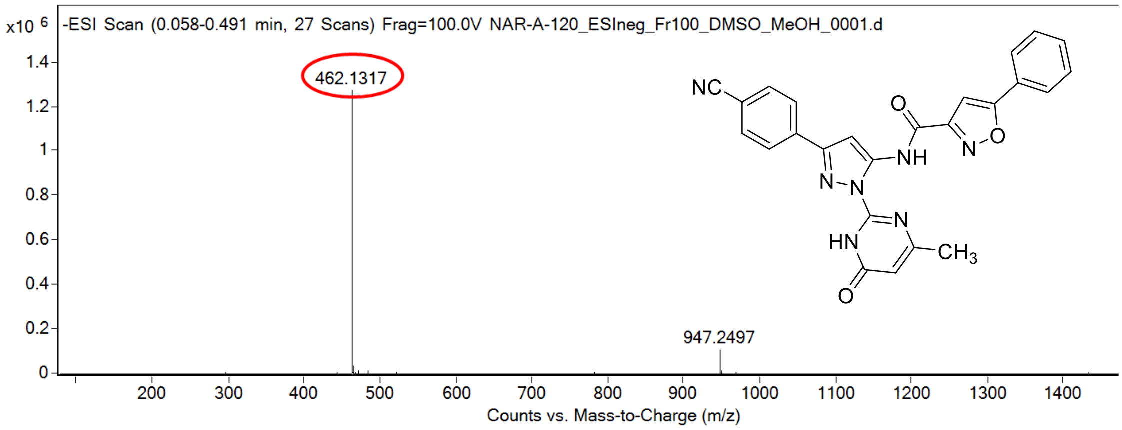

calculated mass: [M-H]$^-$ = 462.1320 observed: [M-H]$^-$ = 462.1317 mass accuracy = -0.6 ppm

FT-IR spectrum (neat) of **129**

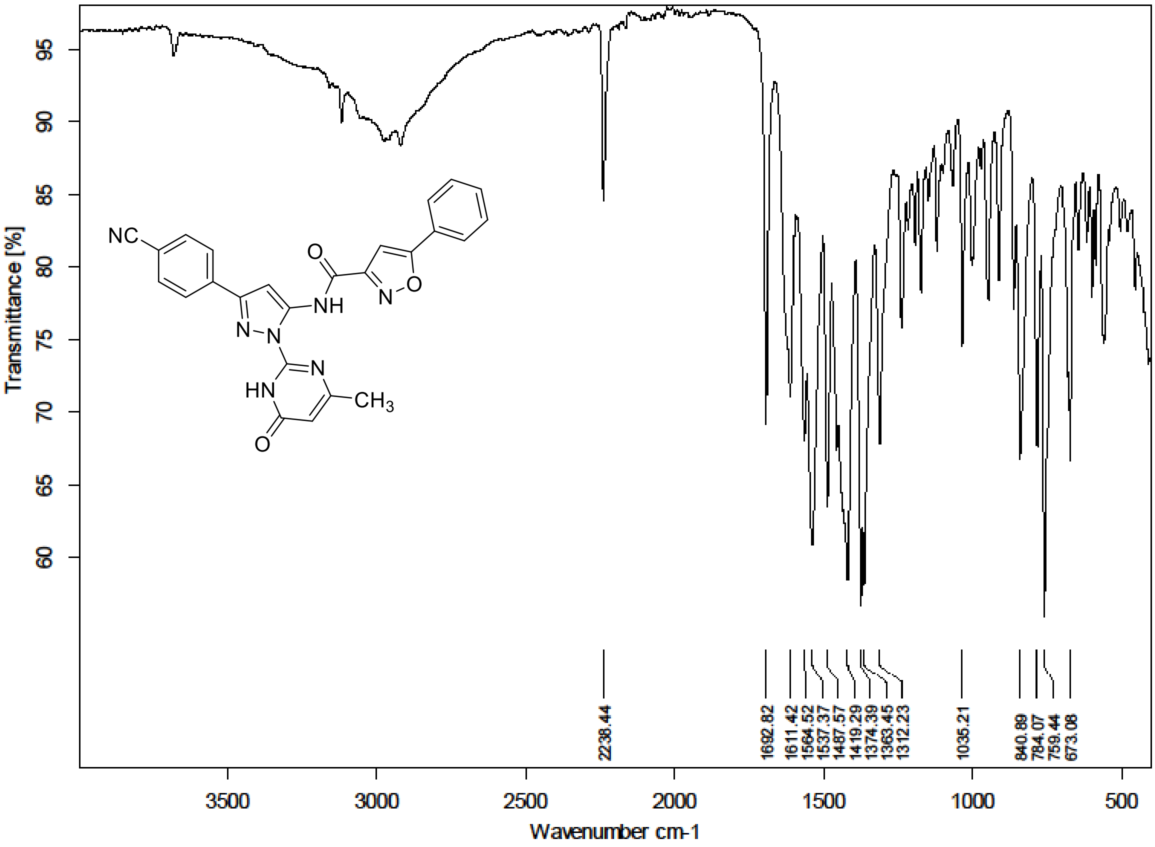

$^{1}H$ NMR (300 MHz) spectrum of **130** in Trifluoroacetic Acid-*d*

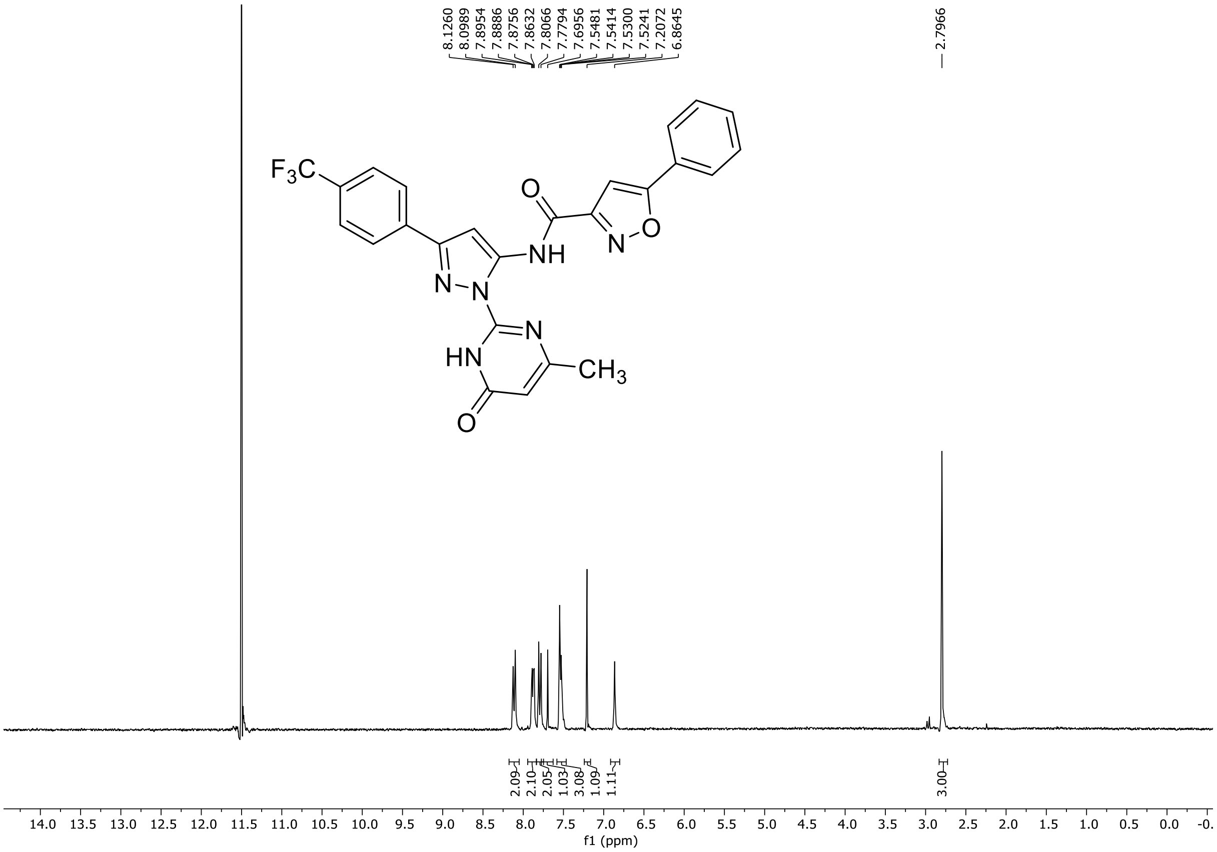

HRMS spectrum of **130**

**NAR-A-123** **$C_{25}H_{17}F_3N_6O_3$** ***m/z* 506.1314**

**ESI-** (MMI)
nitrogen flow 5 L/min, gas temperature 325°C, nebulizer 45 psi, skimmer 65 V, fragmentor 150 V, dissolved in DMSO, MeOH

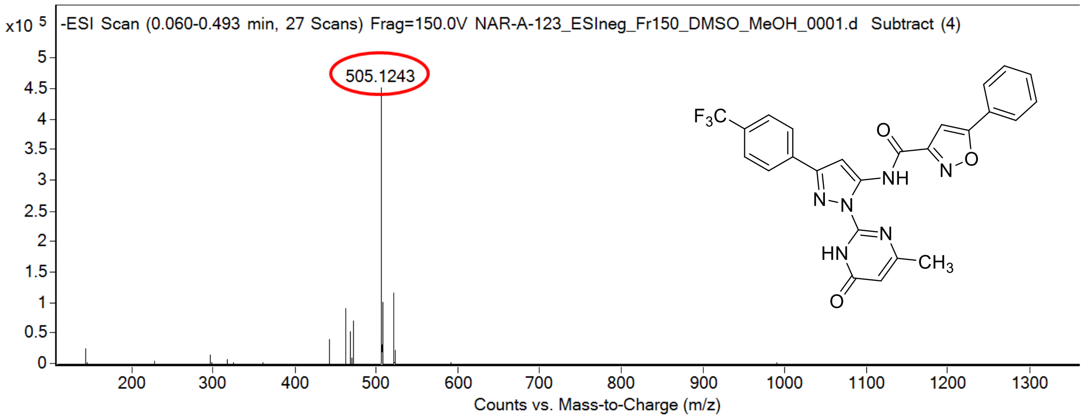

calculated mass: $[M\text{-}H]^-$ = 505.1241 observed: $[M\text{-}H]^-$ = 505.1243 mass accuracy = +0.4 ppm

FT-IR spectrum (neat) of **130**

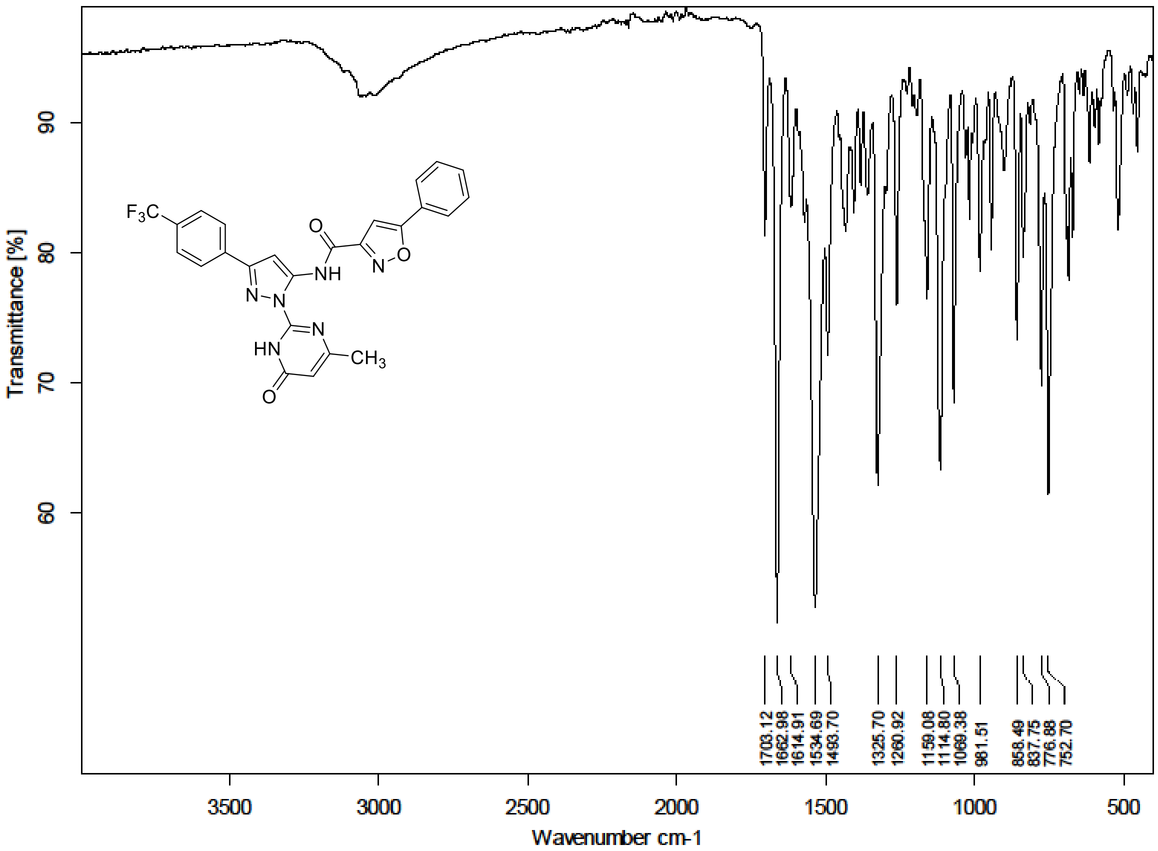

$^1$H NMR (300 MHz) spectrum of **131** in DMSO-$d_6$

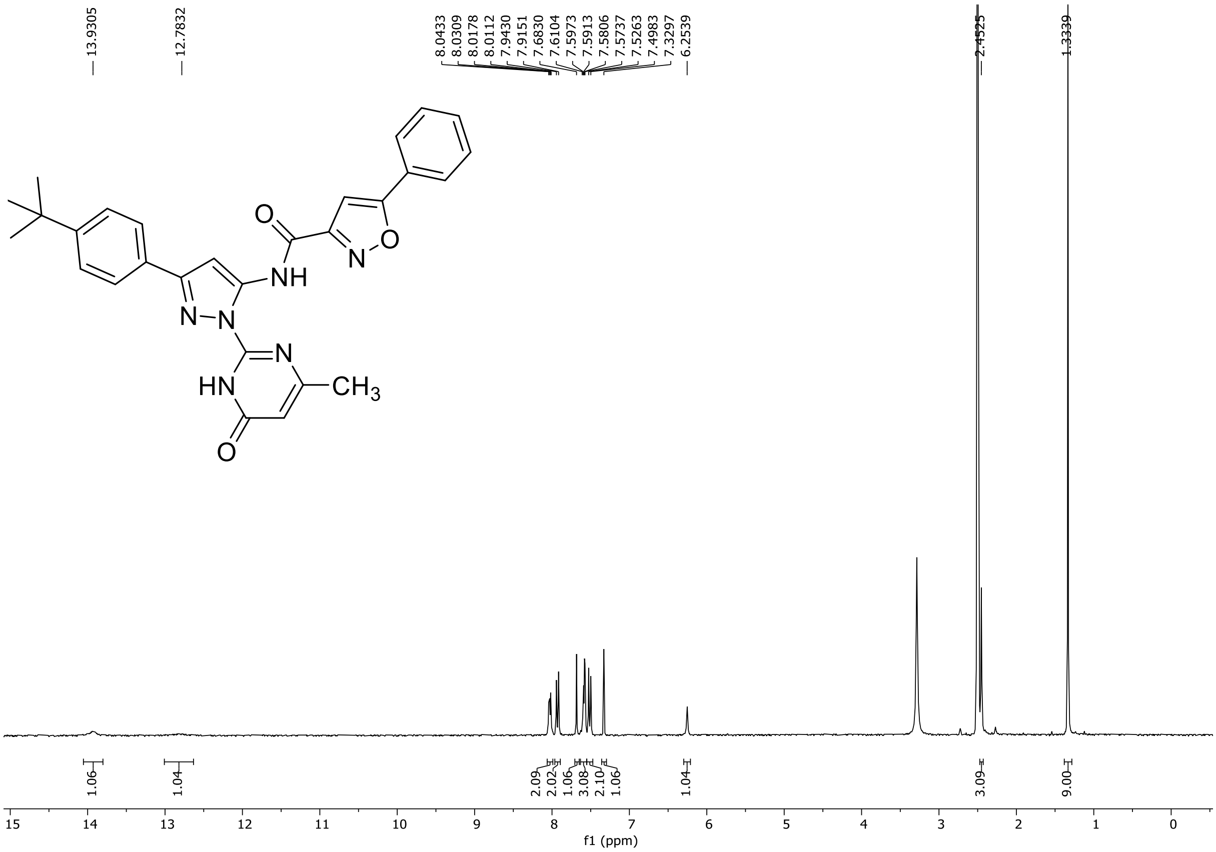

HRMS spectrum of **131**

**NAR-A-151** $C_{28}H_{26}N_6O_3$ mono *m/z* **494.2066**

**APCI +** (MMI)
nitrogen flow 5 L/min, gas temperature 325°C, nebulizer 45 psi, skimmer **65** V,
vaporizer 200°C, fragmentor 60 V, dissolved in methanol

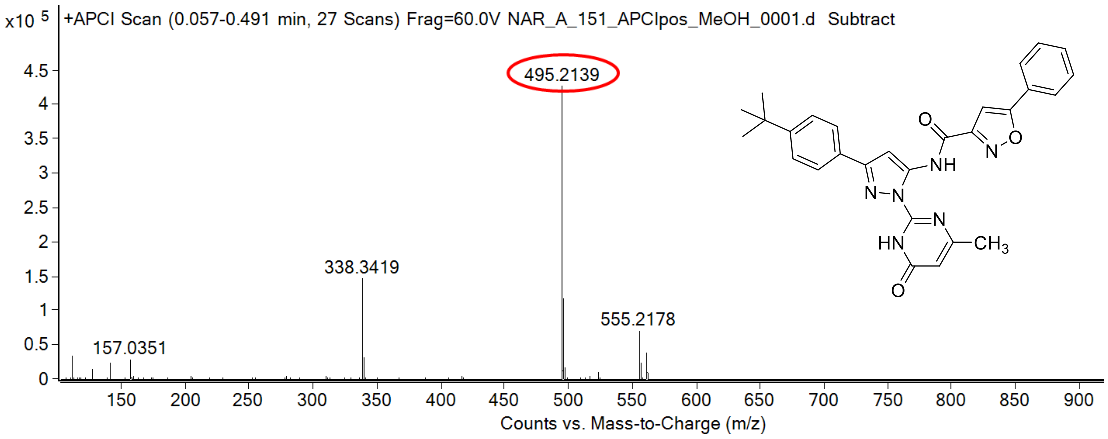

calculated mass: [M+H]$^+$ = 495.2139 observed: [M+H]$^+$ = 495.2139 mass accuracy = < 0.1 ppm

FT-IR spectrum (neat) of **131**

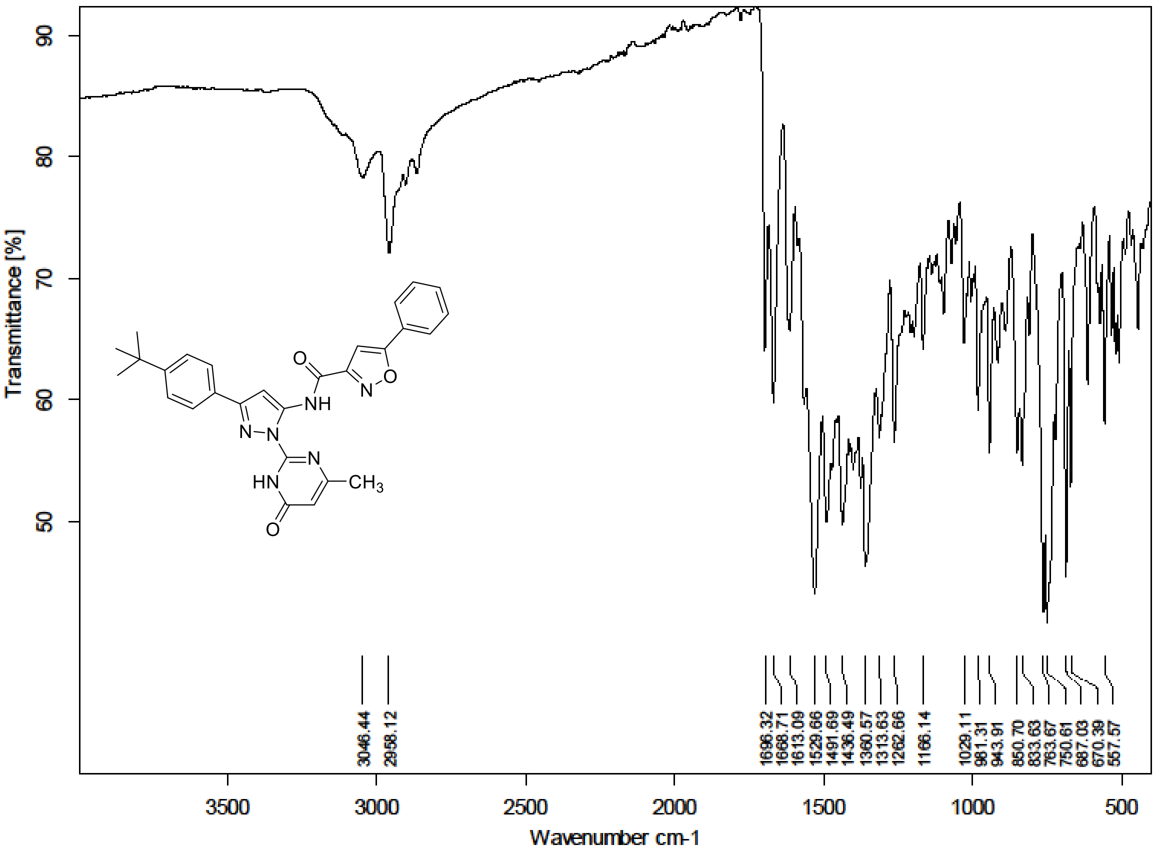

$^1$H NMR (300 MHz) spectrum of **132** in DMSO-$d_6$

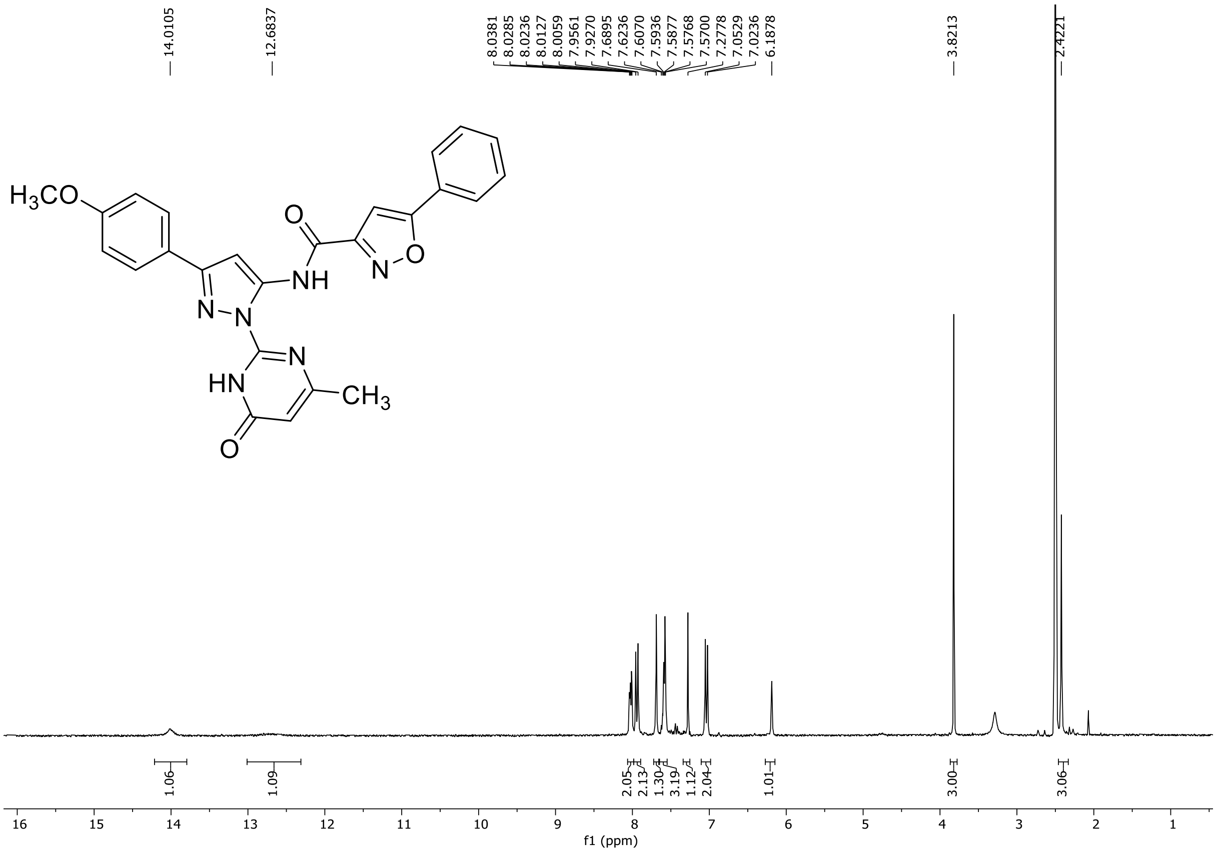

HRMS spectrum of **132**

**NAR-A-152** **$C_{25}H_{20}N_6O_4$** **mono *m/z* 468.1546**

**APCI +** (MMI)
nitrogen flow 5 L/min, gas temperature 325°C, nebulizer 45 psi, skimmer 65 V, vaporizer 200°C, fragmentor 25 V, dissolved in methanol/ $CH_2Cl_2$

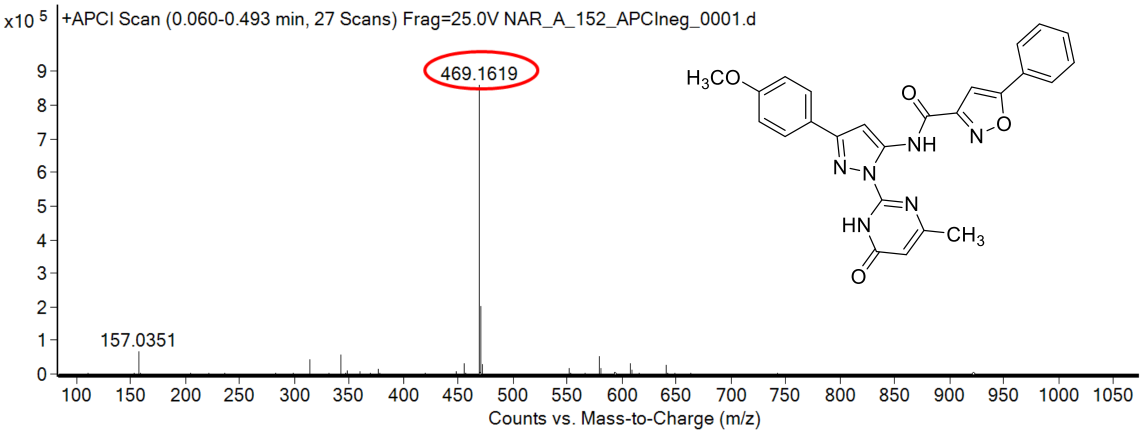

calculated mass: [M+H]$^+$ = 469.1619 observed: [M+H]$^+$ = 469.1619 mass accuracy = < 0.1 ppm

FT-IR spectrum (neat) of **132**

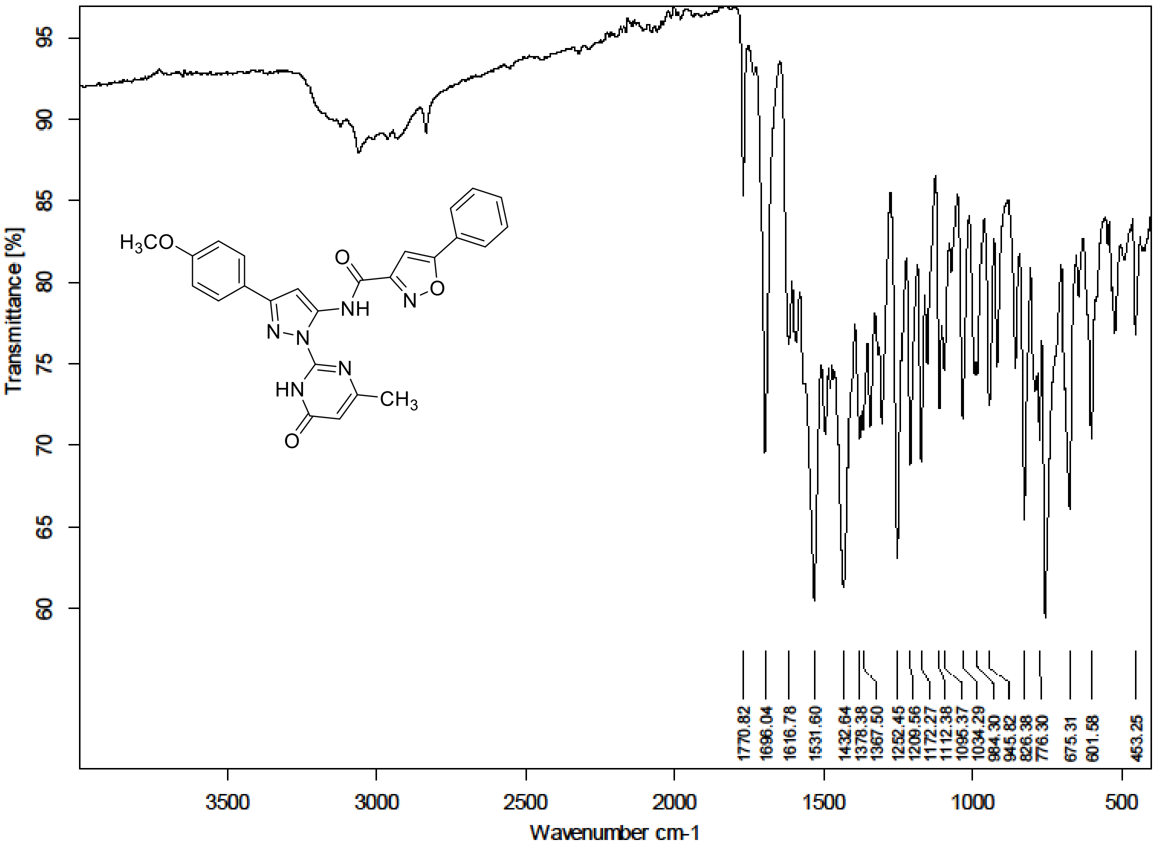

$^1H$ NMR (300 MHz) spectrum of **133** in DMSO-$d_6$

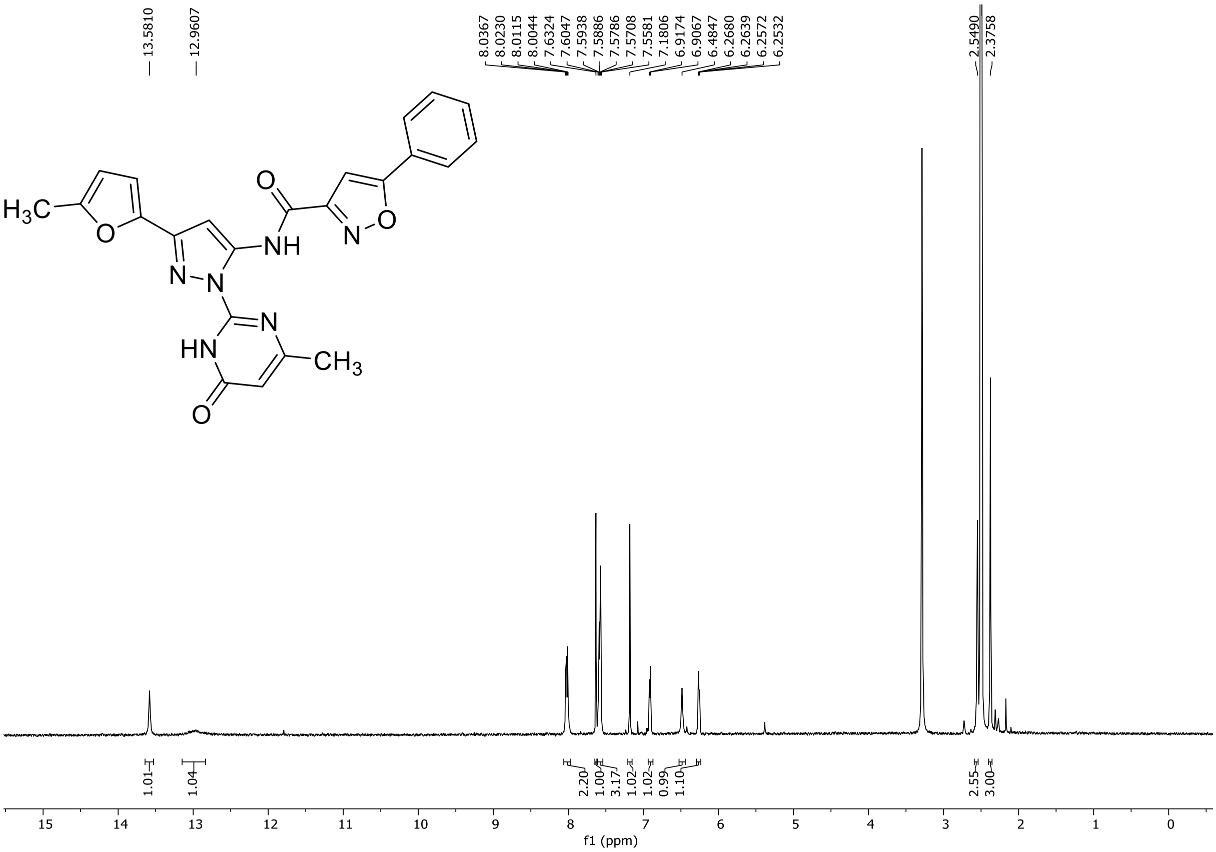

HRMS spectrum of **133**

**NAR-A-180** $C_{23}H_{18}N_6O_4$ mono *m/z* **442.1390**

**APCI +** (MMI)
nitrogen flow 5 L/min, gas temperature 325°C, nebulizer 45 psi, skimmer **65** V,
vaporizer 200°C, fragmentor 38 V, dissolved in methanol

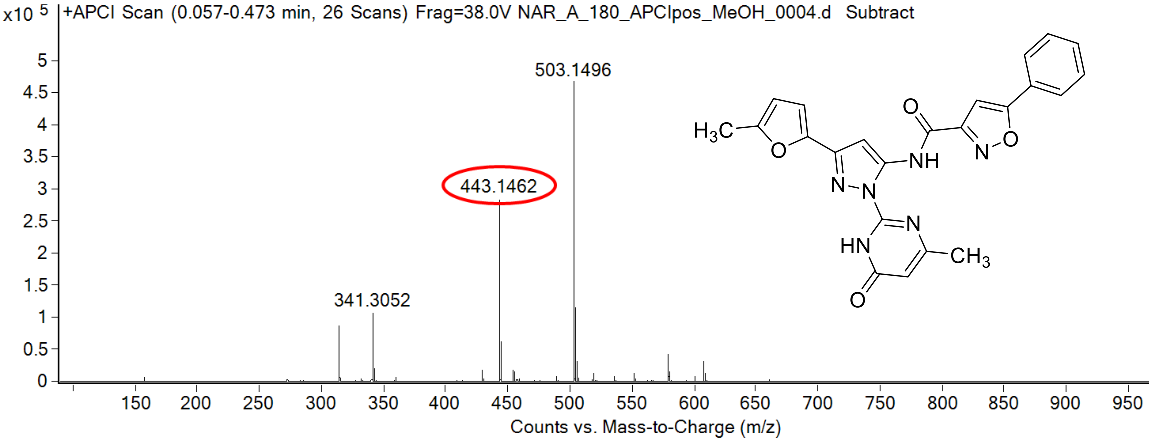

calculated mass: [M+H]$^+$ = 443.1462 observed: [M+H]$^+$ = 443.1462 mass accuracy = < 0.1 ppm

FT-IR spectrum (neat) of **133**

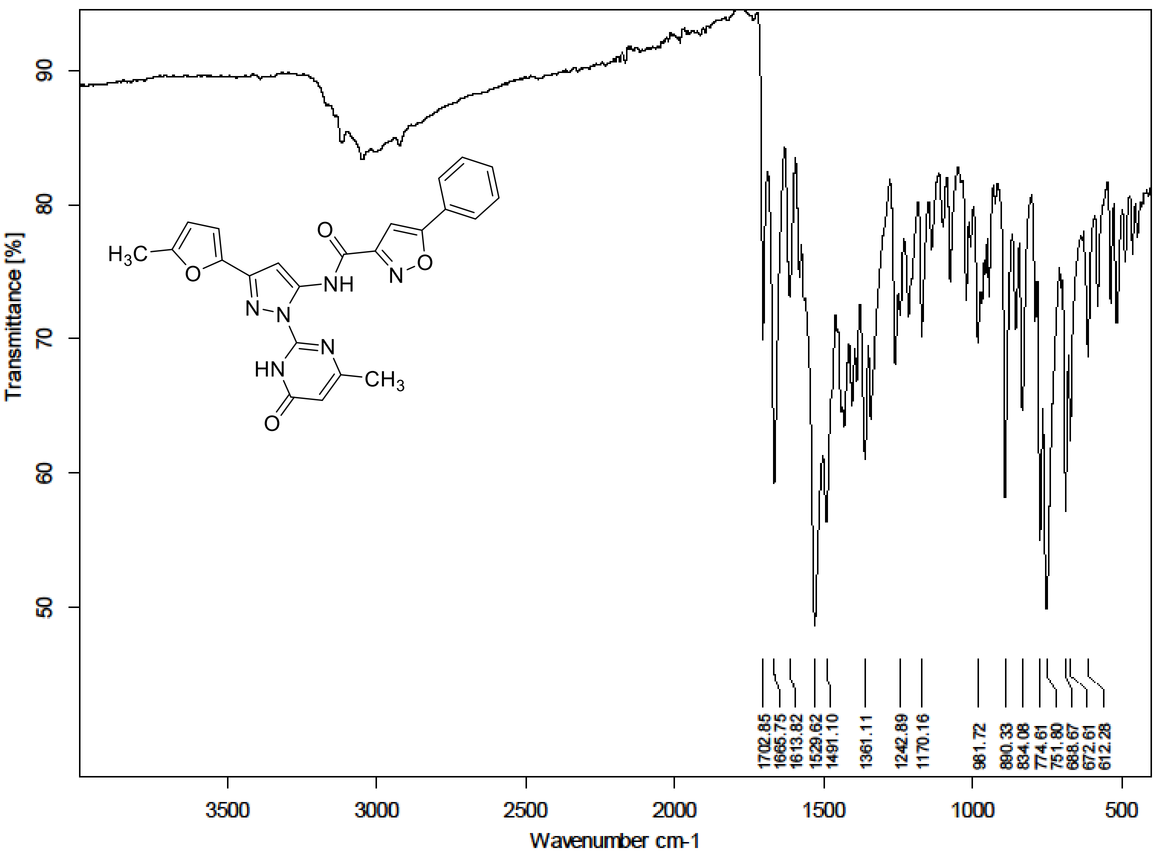

[1]H (500 MHz) and [13]C NMR (126 MHz) spectra of **134** in DMSO-$d_6$

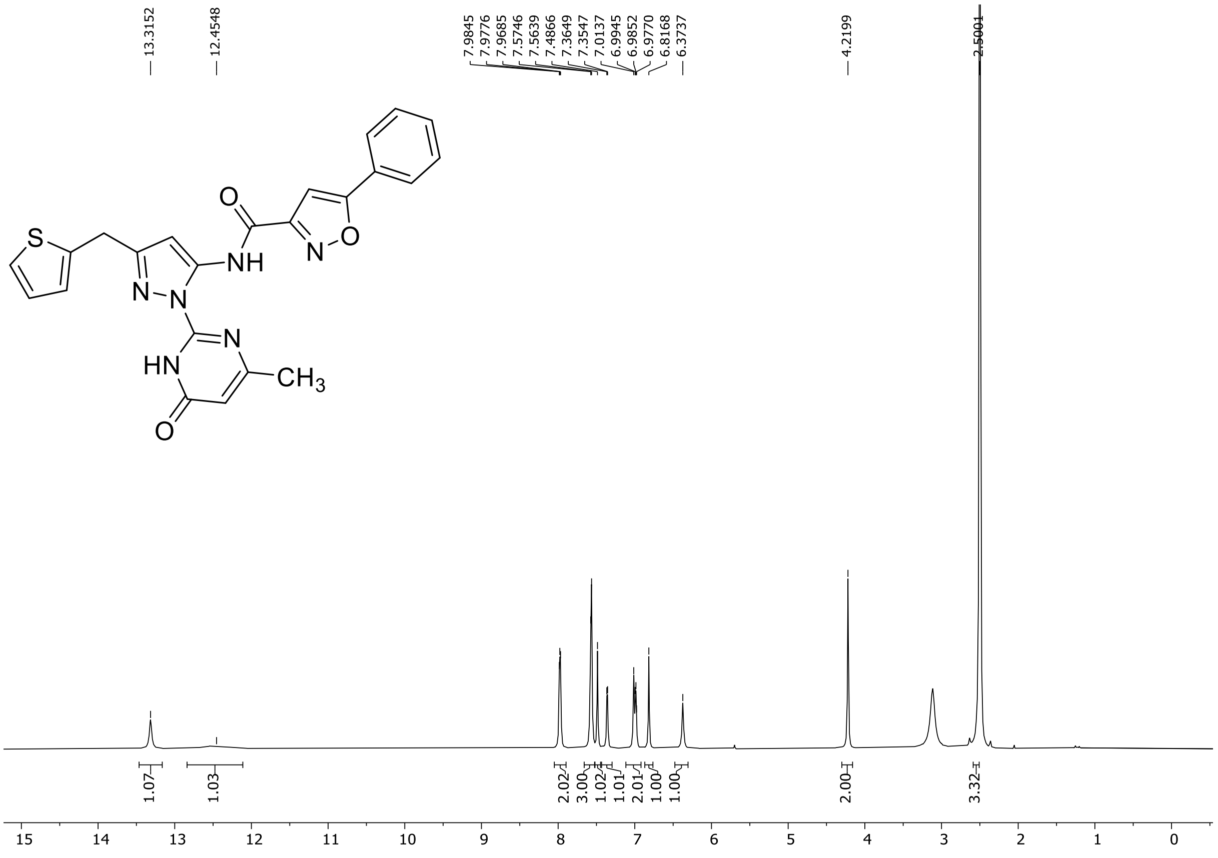

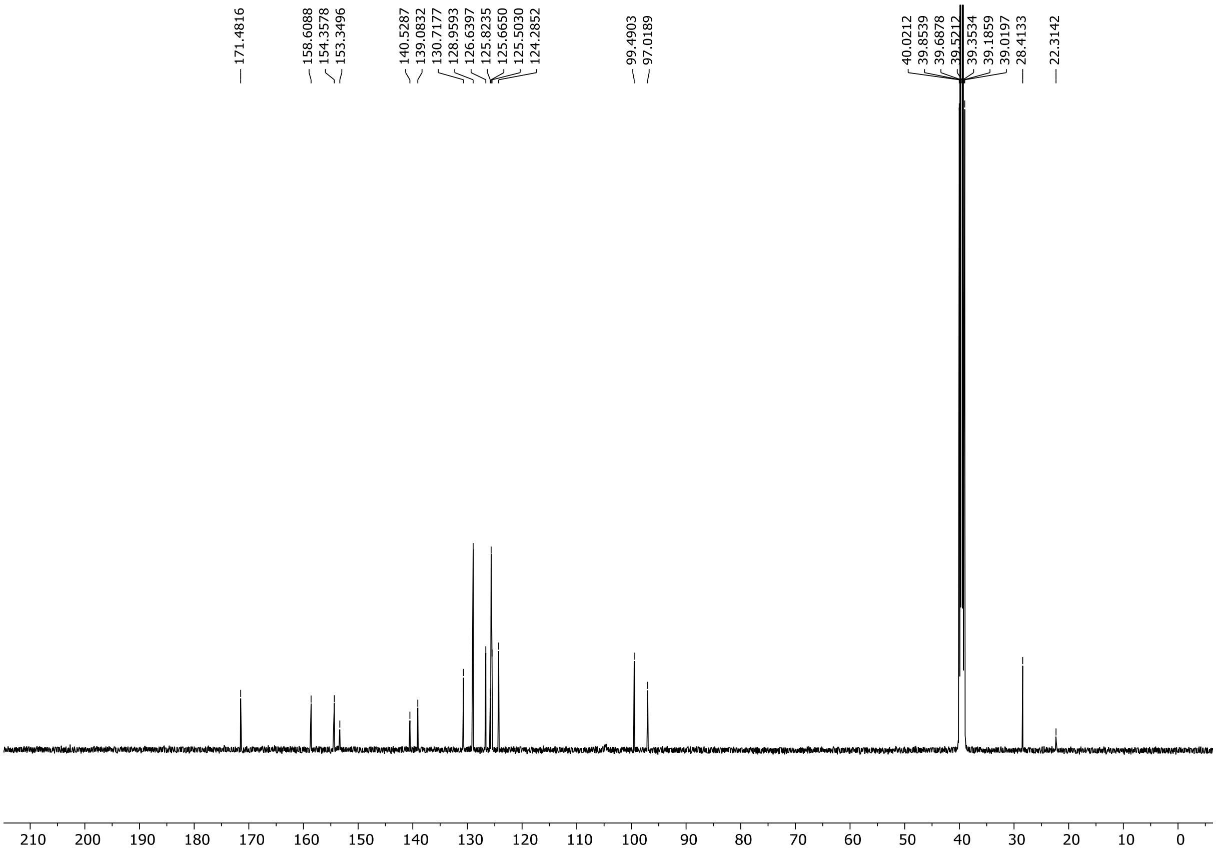

HRMS spectrum of **134**

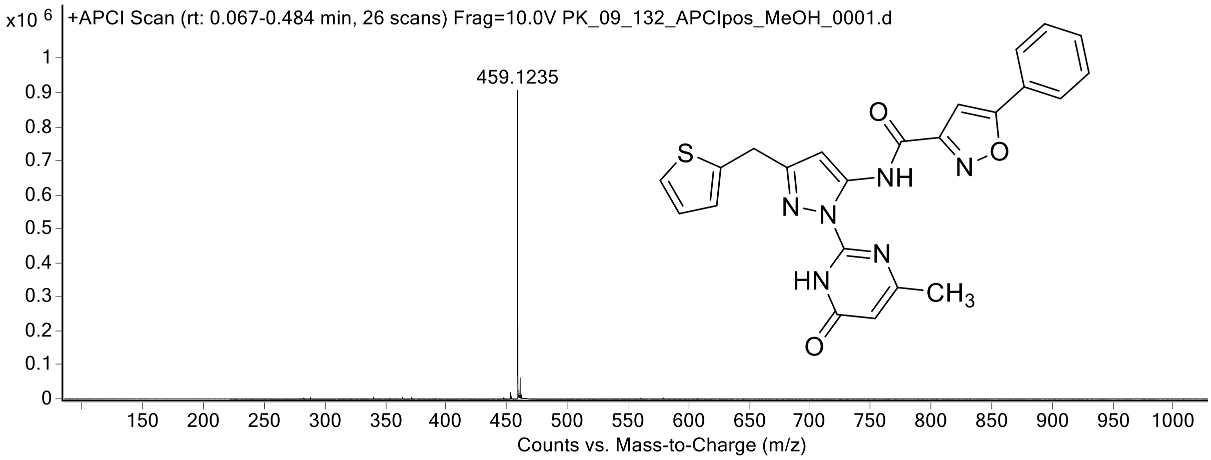

[1]H (300 MHz) and [13]C NMR (75 MHz) spectra of **135** DMSO-$d_6$

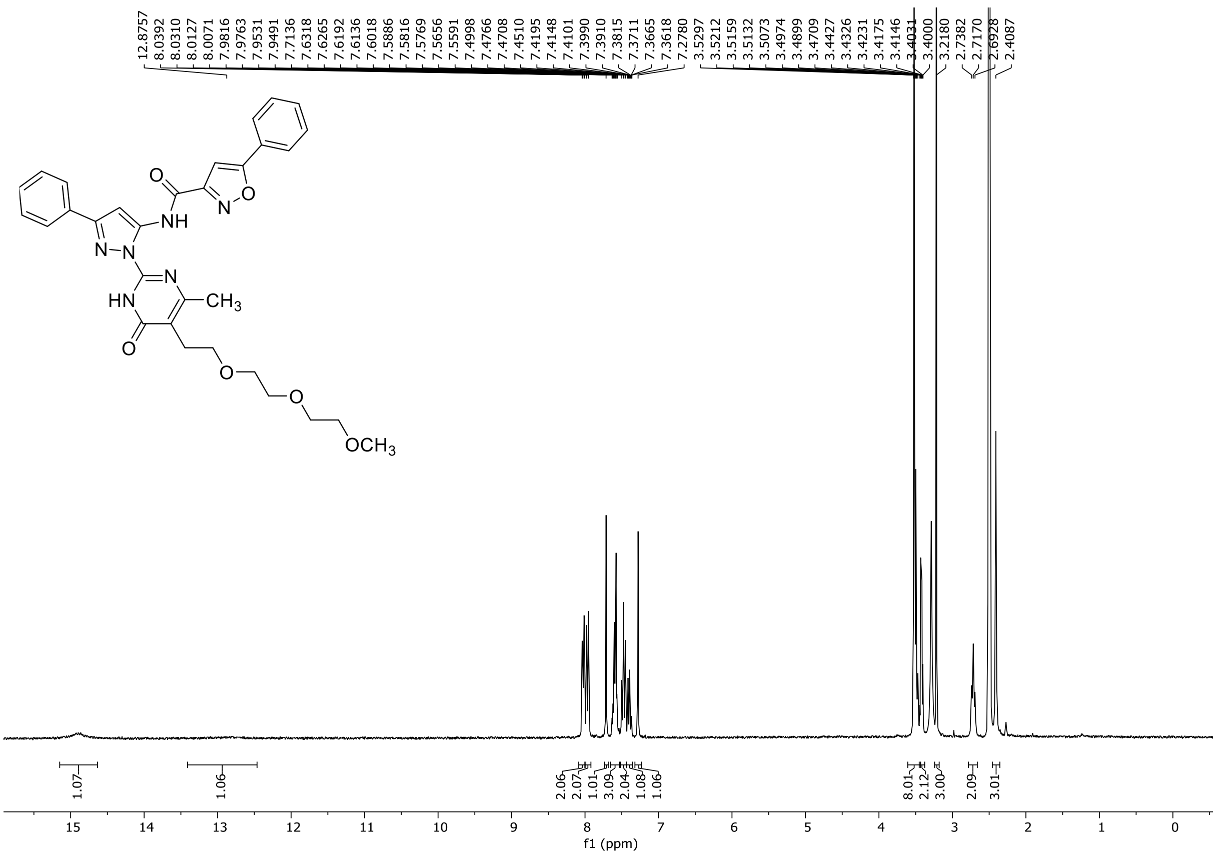

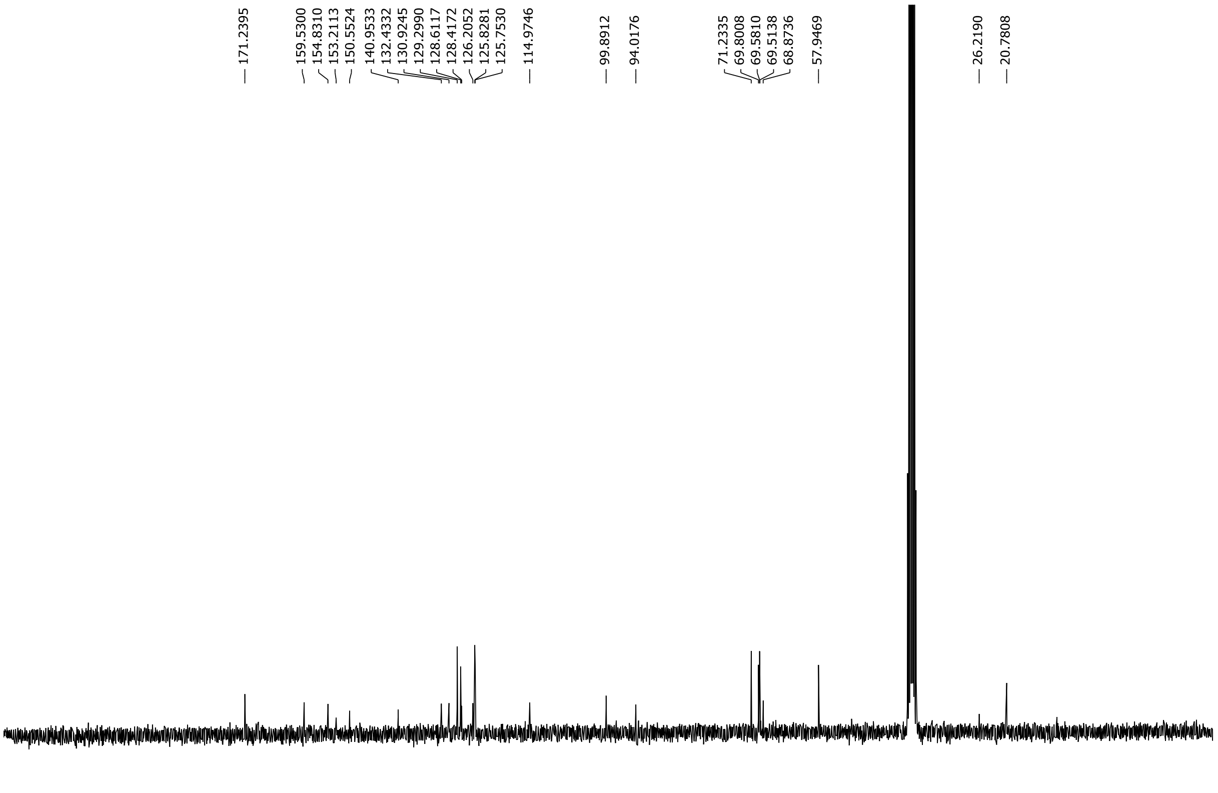

HRMS spectrum of **135**

**NAR-A-221** $C_{31}H_{32}N_6O_6$ mono *m/z* 584.2383

**APCI +** (MMI)
nitrogen flow 5 L/min, gas temperature 325°C, nebulizer 45 psig, skimmer 65 V, vaporizer 200°C, fragmentor 20 V, dissolved in methanol

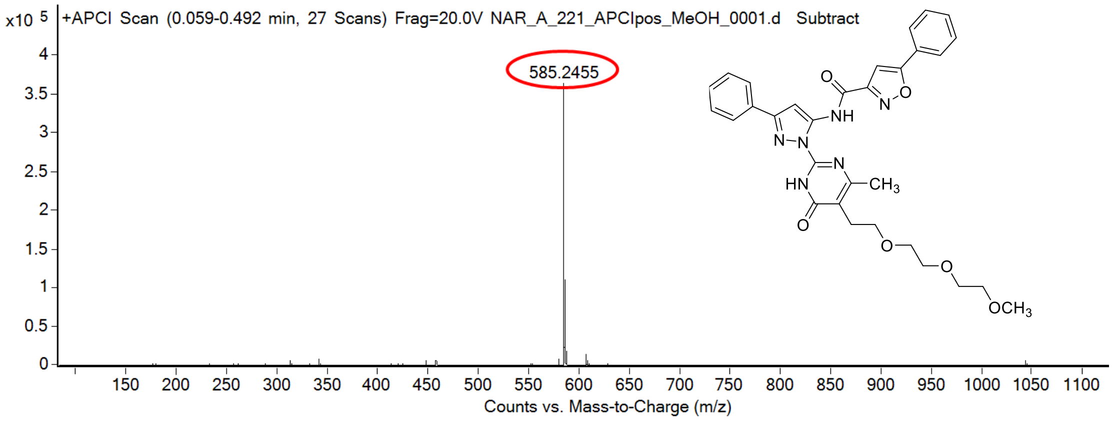

calculated mass: $[M+H]^+$ = 585.2456 observed: $[M+H]^+$ = 585.2455 mass accuracy = - 0.2 ppm

FT-IR spectrum (neat) of **135**

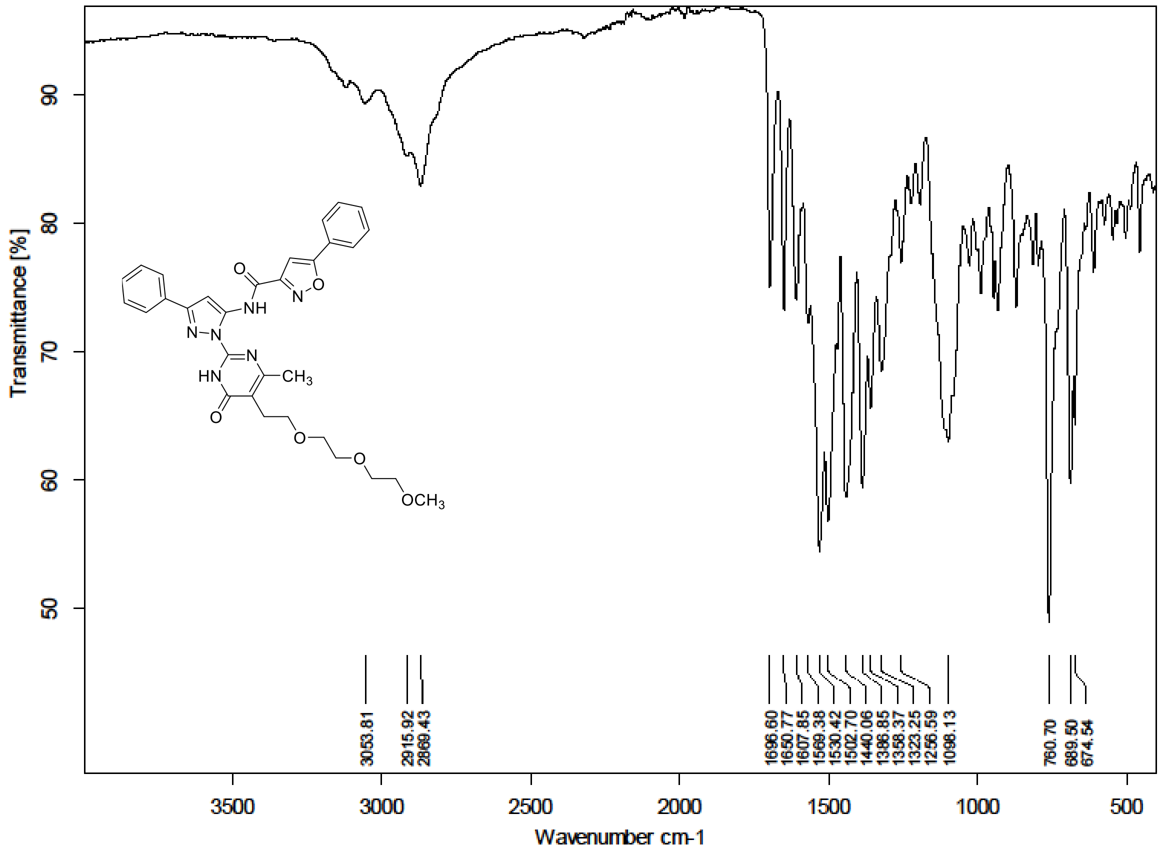

$^{1}H$ NMR (300 MHz) spectrum of **136** in Chloroform-*d*

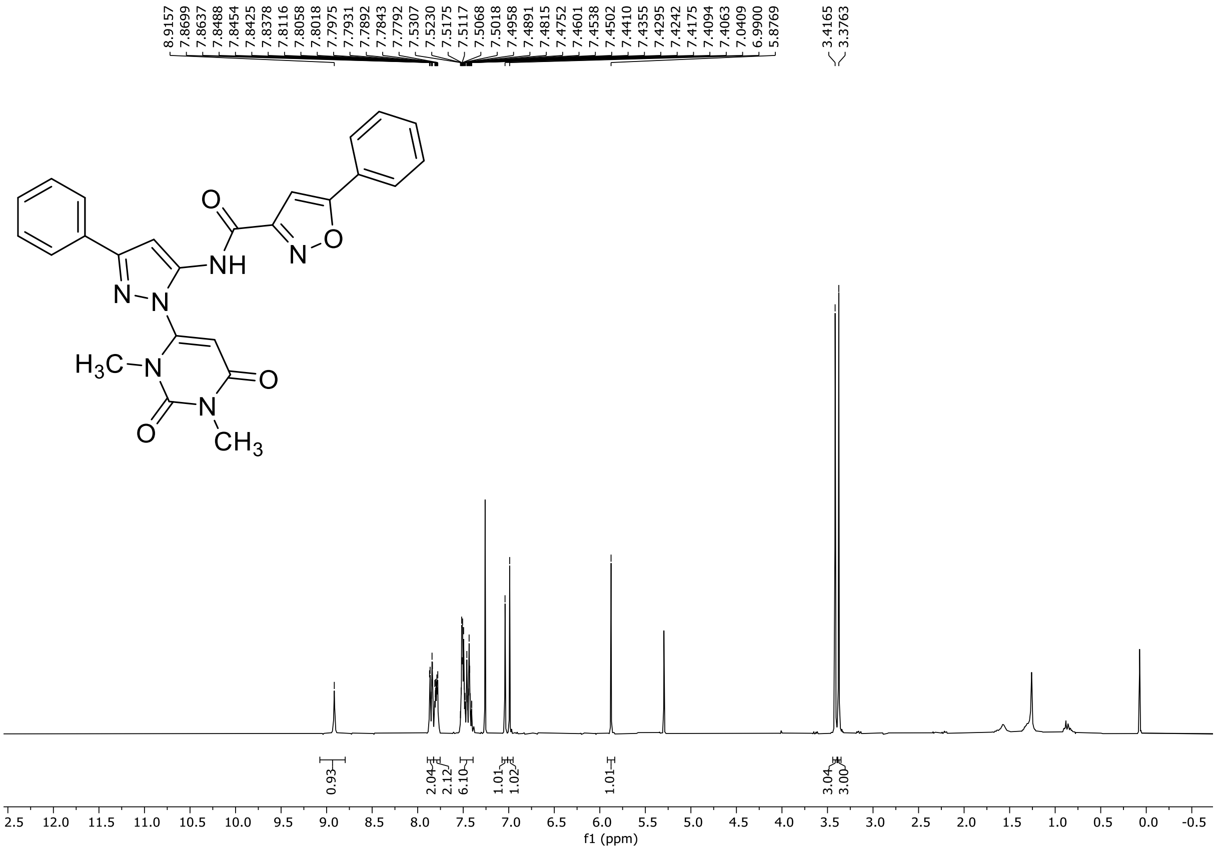

HRMS spectrum of **136**

**NAR-A-227** **$C_{25}H_{20}N_{6}O_{4}$** **mono *m/z* 468.1546**

**APCI +** (MMI)
nitrogen flow 5 L/min, gas temperature 325°C, nebulizer 45 psig, skimmer 65 V, vaporizer 200°C, fragmentor 25 V, dissolved in methanol

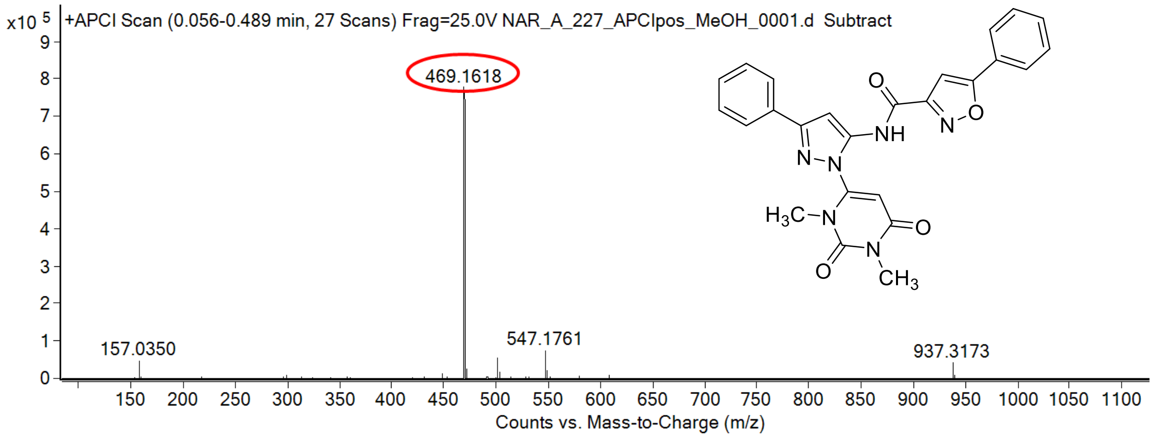

calculated mass: $[M+H]^+ = 469.1619$ observed: $[M+H]^+ = 469.1618$ mass accuracy = - 0.2 ppm

FT-IR spectrum (neat) of **136**

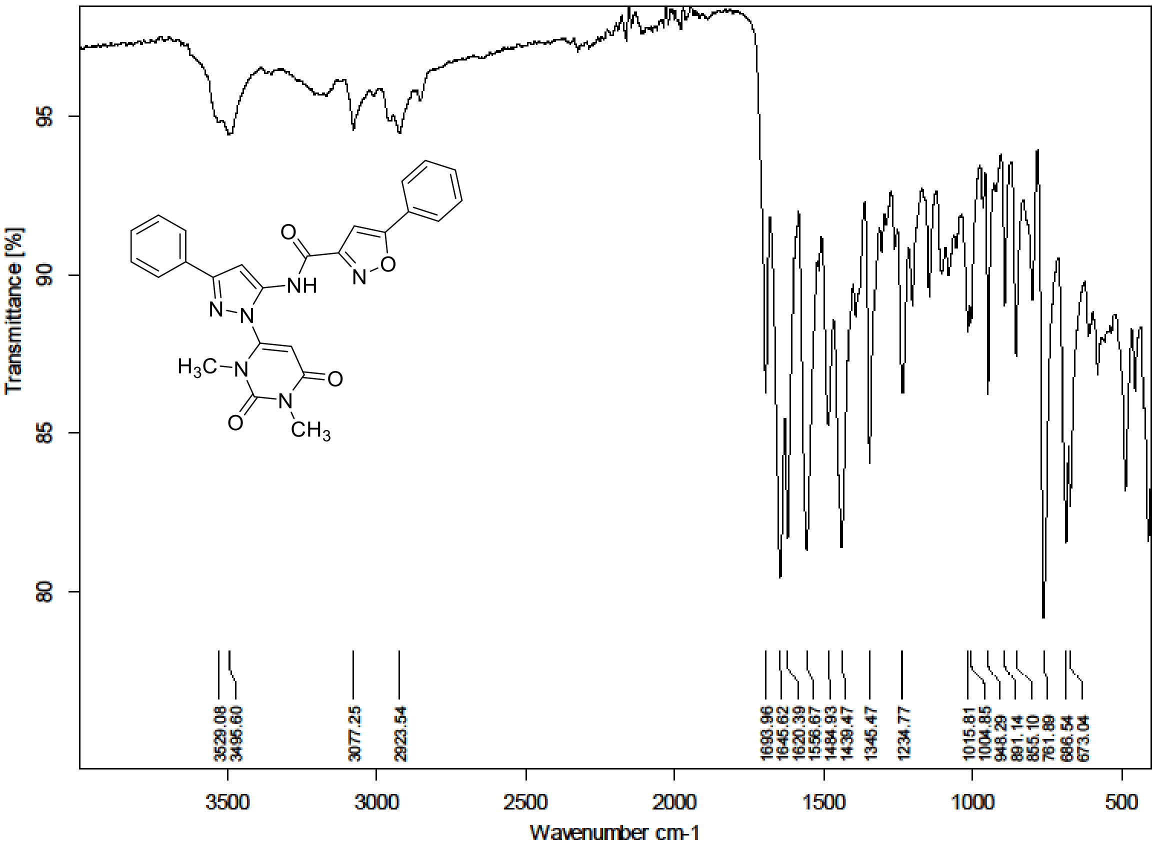

[1]H (500 MHz) and [13]C NMR (126 MHz) spectra of **137** in Chloroform-*d*

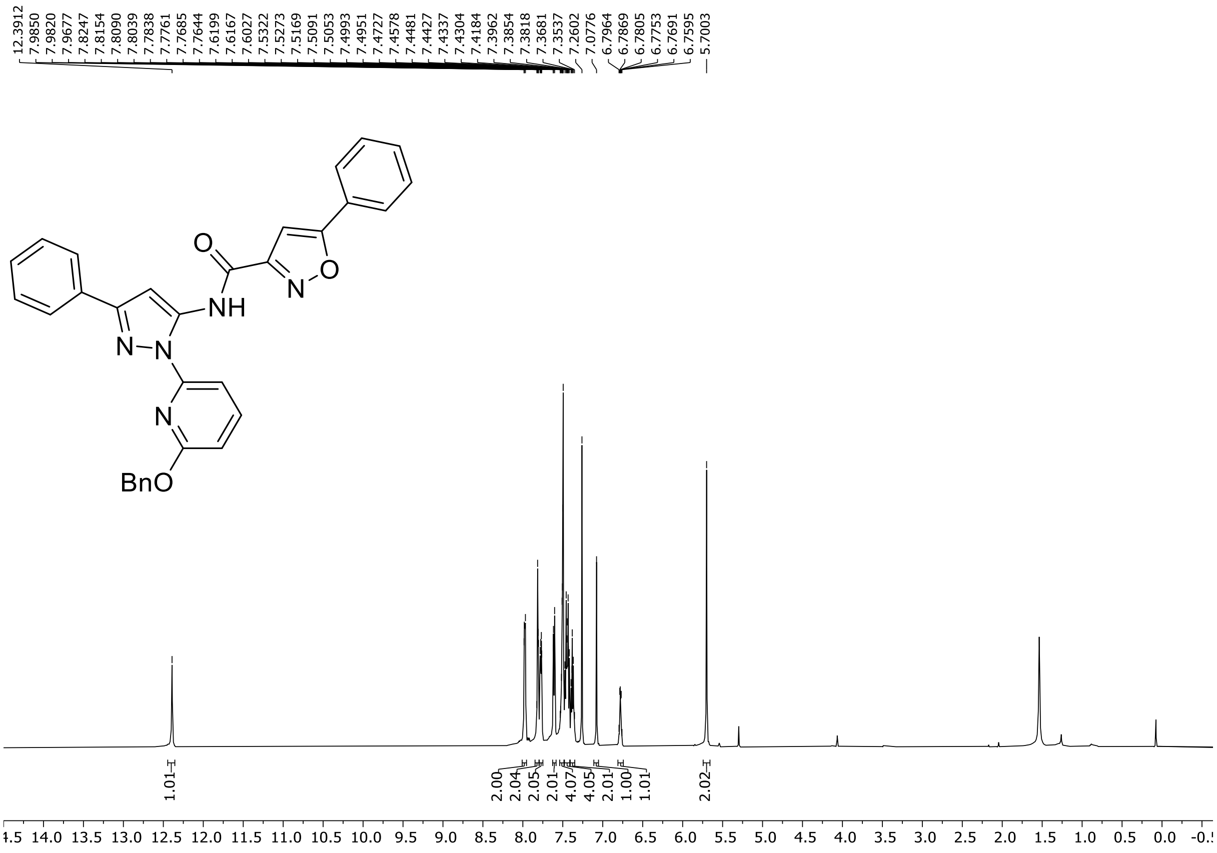

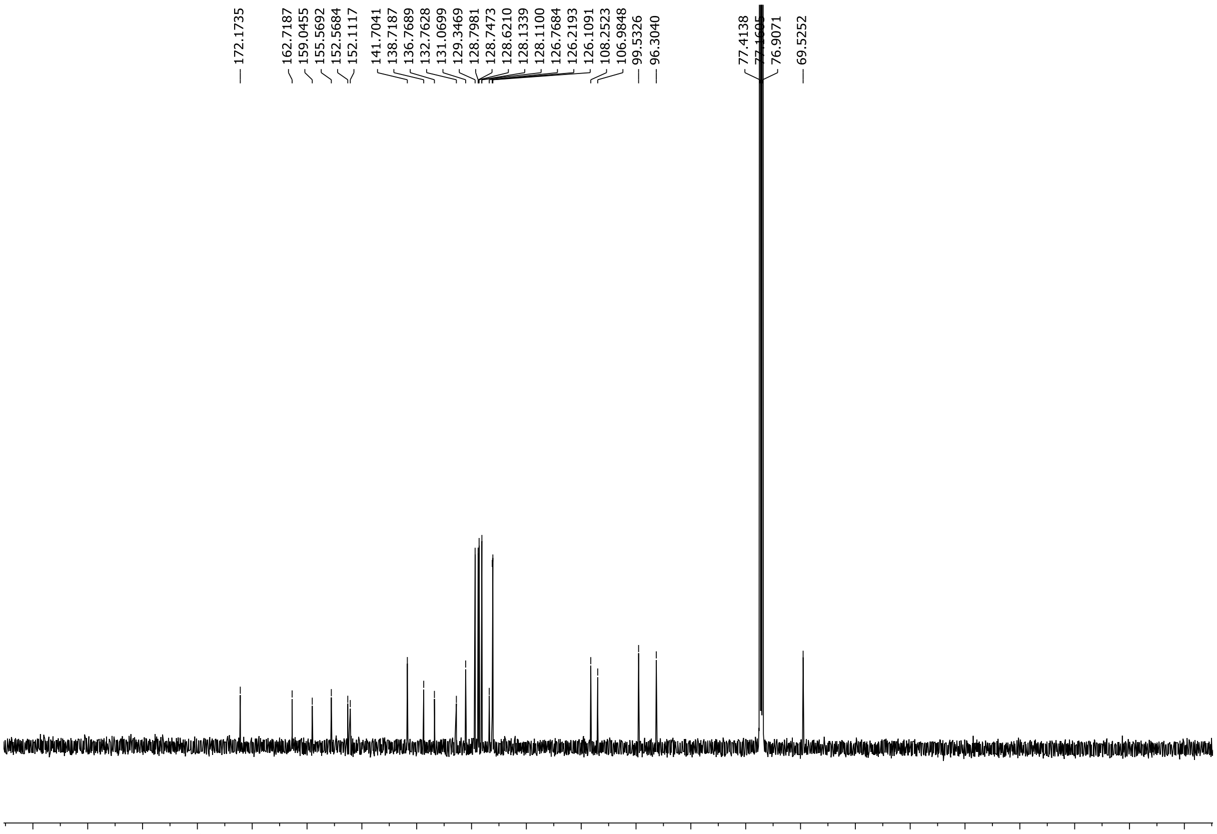

HRMS spectrum of **137**

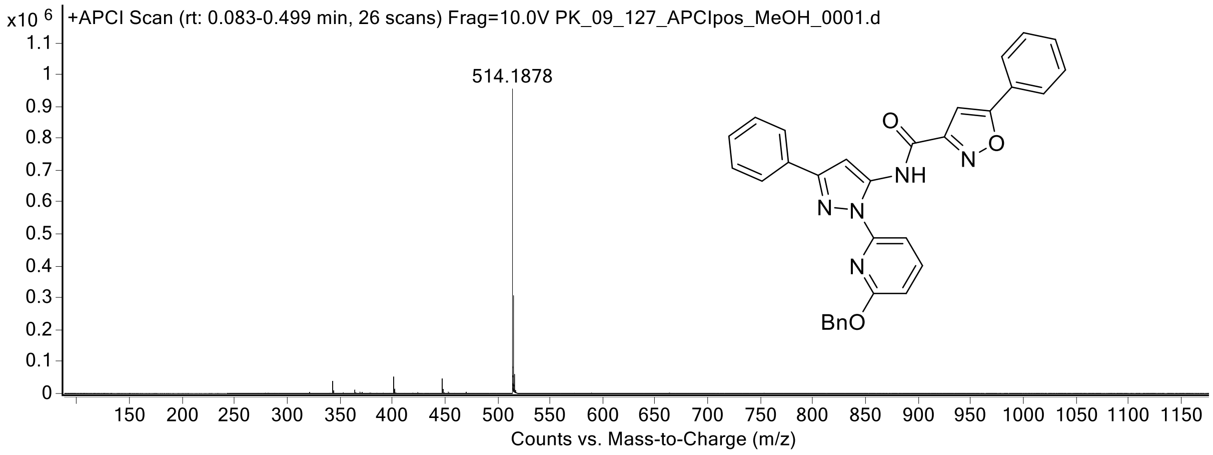

$^{1}H$ (300 MHz) and $^{13}C$ NMR (126 MHz) spectra of **138** in DMSO-$d_6$

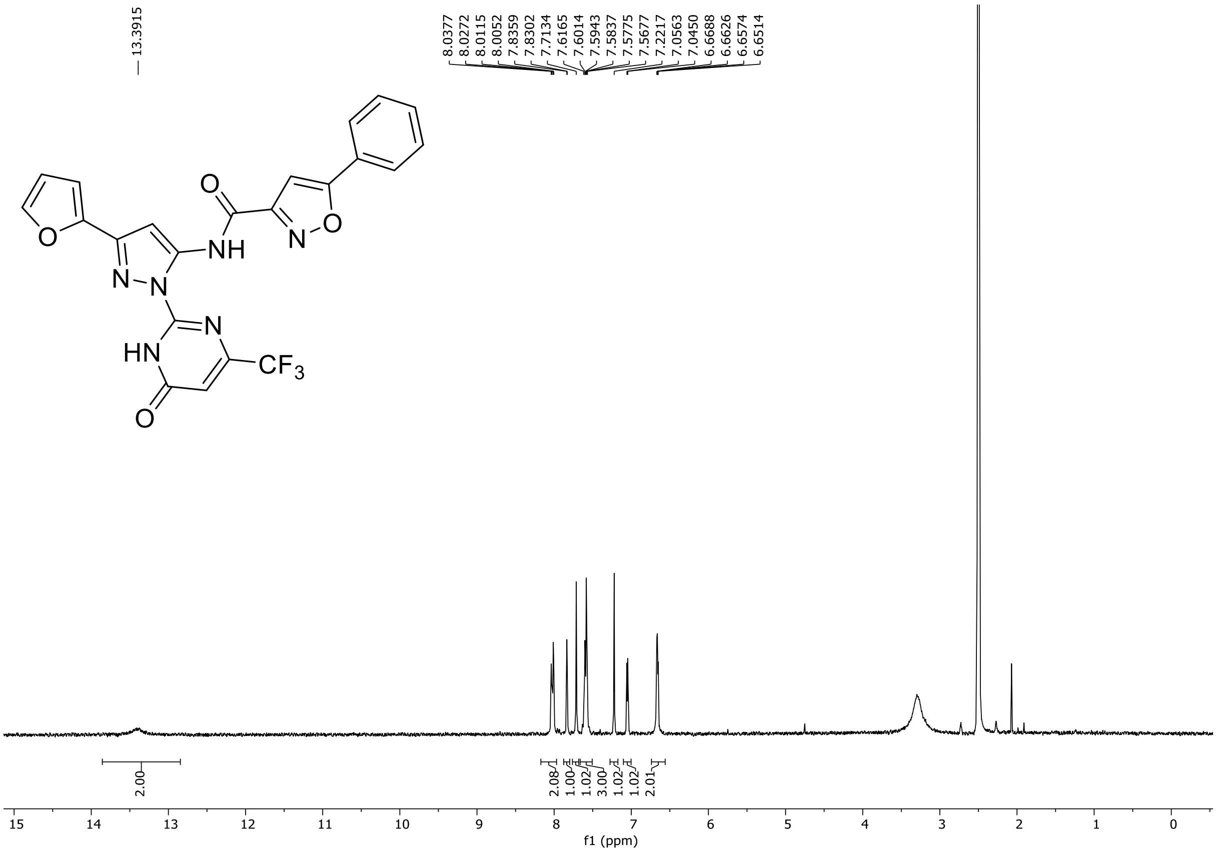

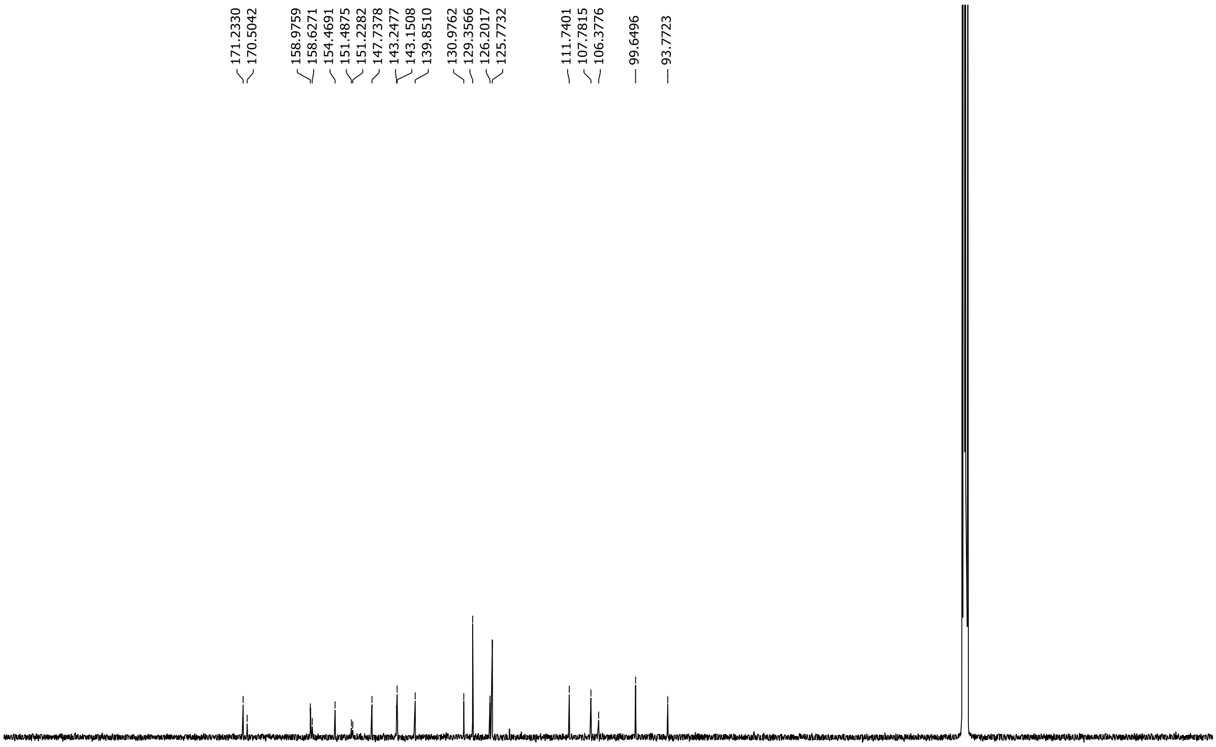

[19]F NMR (471 MHz) spectrum of **138** in DMSO-$d_6$

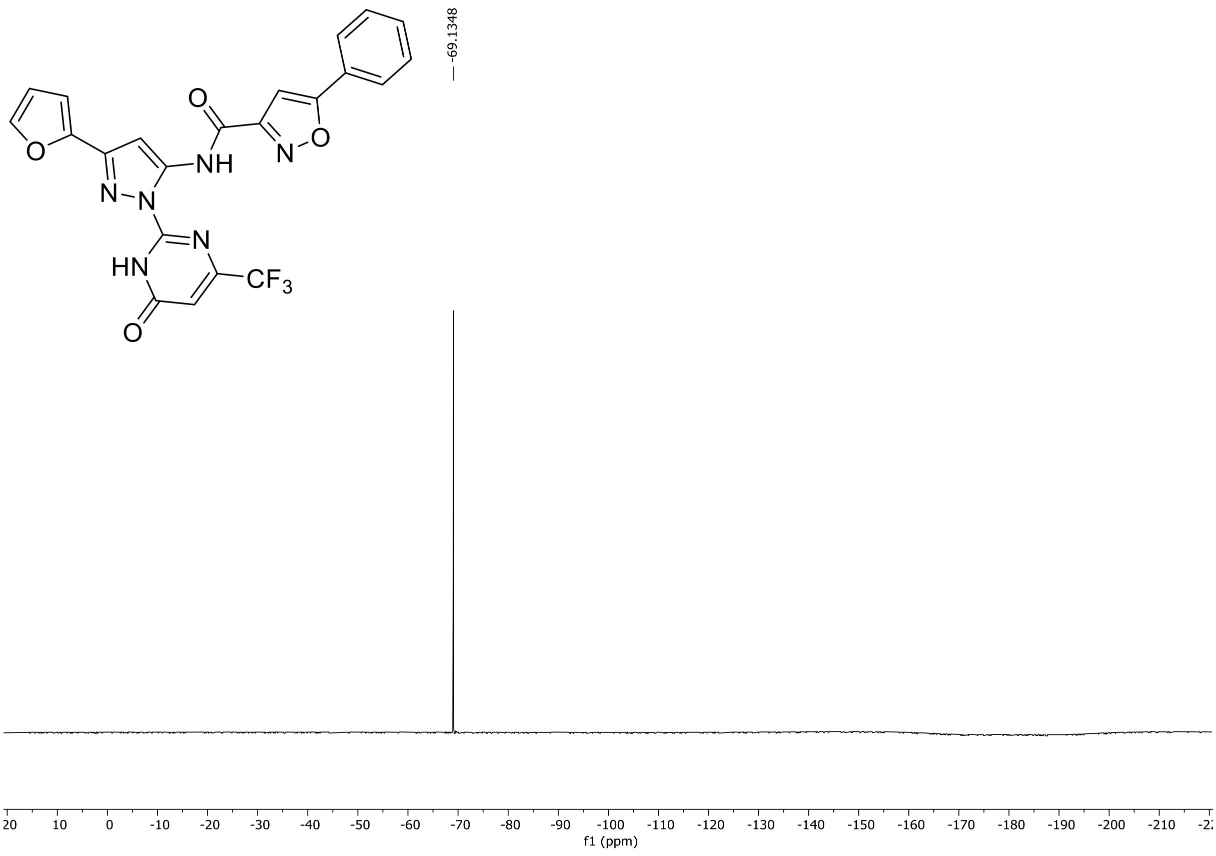

HRMS spectrum of **138**

**NAR-A-104** $C_{22}H_{13}F_3N_6O_4$ *m/z* **482.0950**

**ESI-** (MMI)
nitrogen flow 5 L/min, gas temperature 325°C, nebulizer 45 psi, skimmer 65 V, fragmentor 30 V, dissolved in DMSO, MeOH

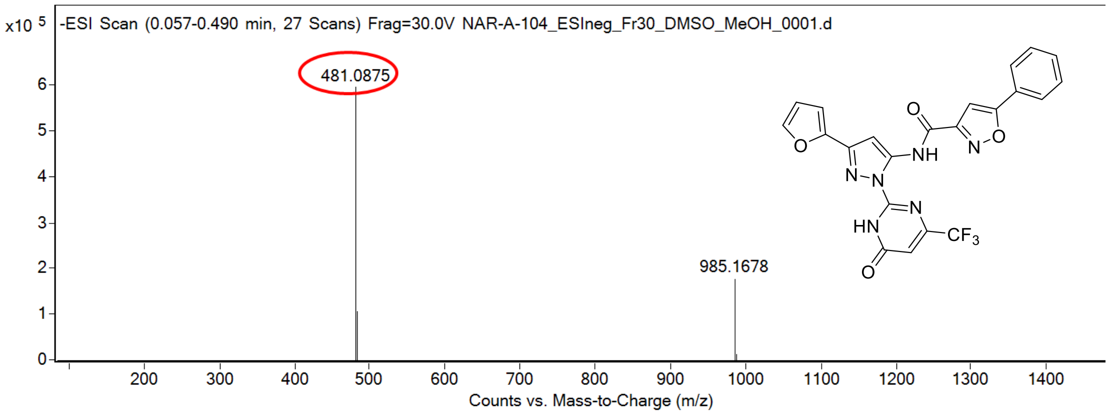

calculated mass: $[M-H]^-$ = 481.0878 observed: $[M-H]^-$ = 481.0875 mass accuracy = -0.6 ppm

FT-IR spectrum (neat) of **138**

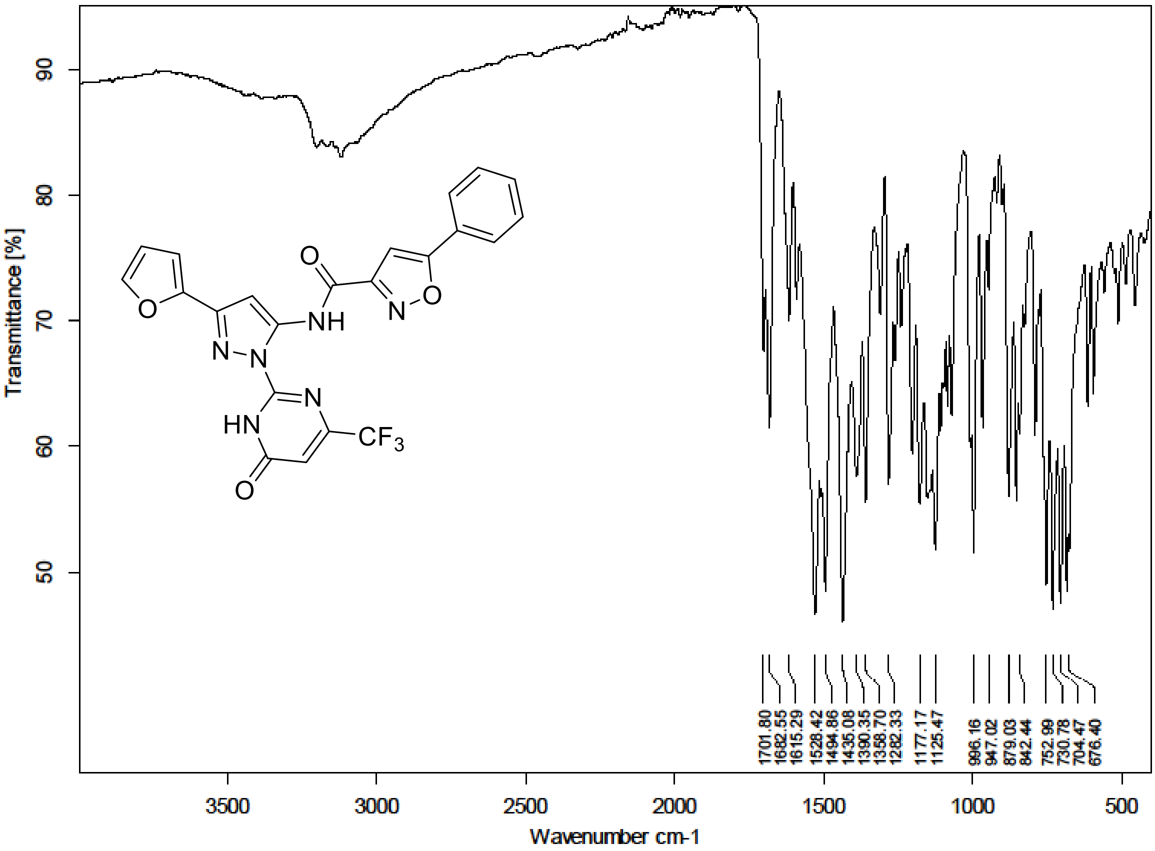

[1]H (500 MHz) and [13]C NMR (126 MHz) spectra of **139** in Chloroform-*d*

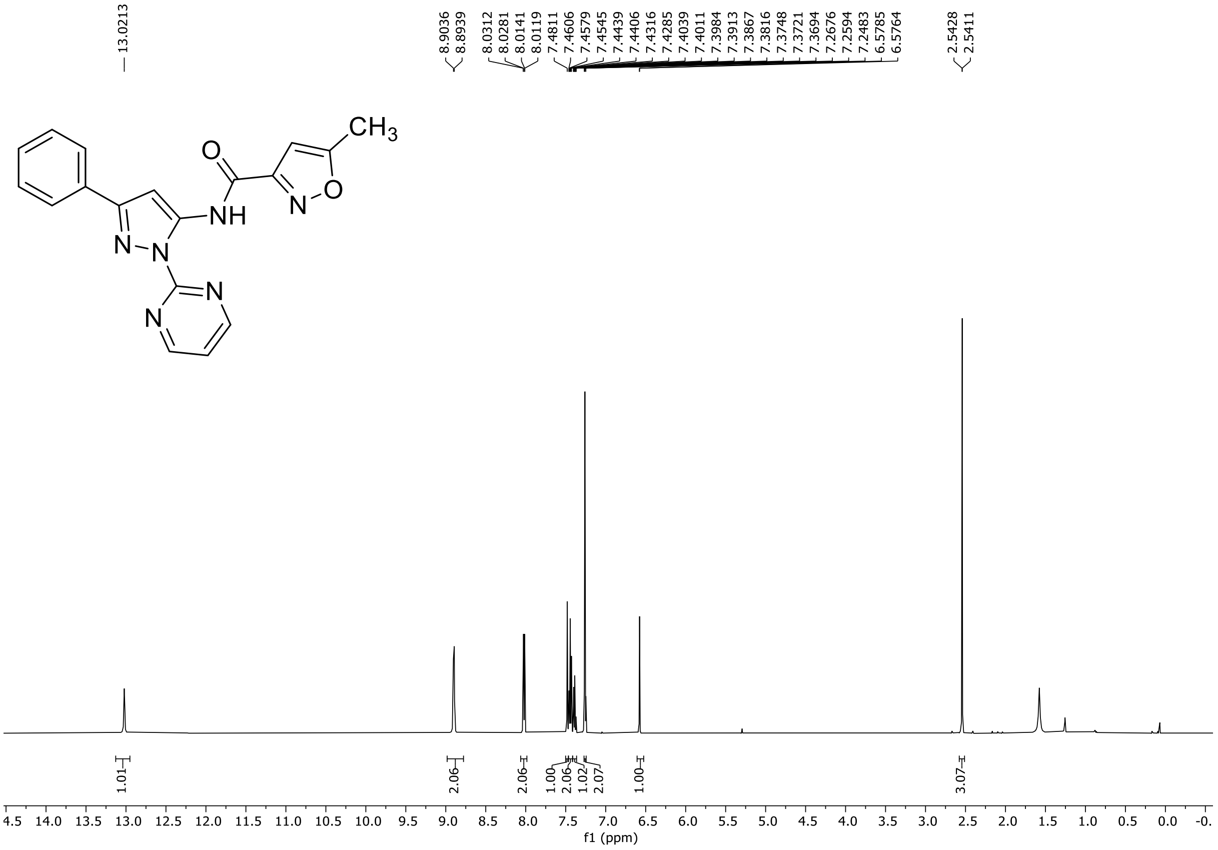

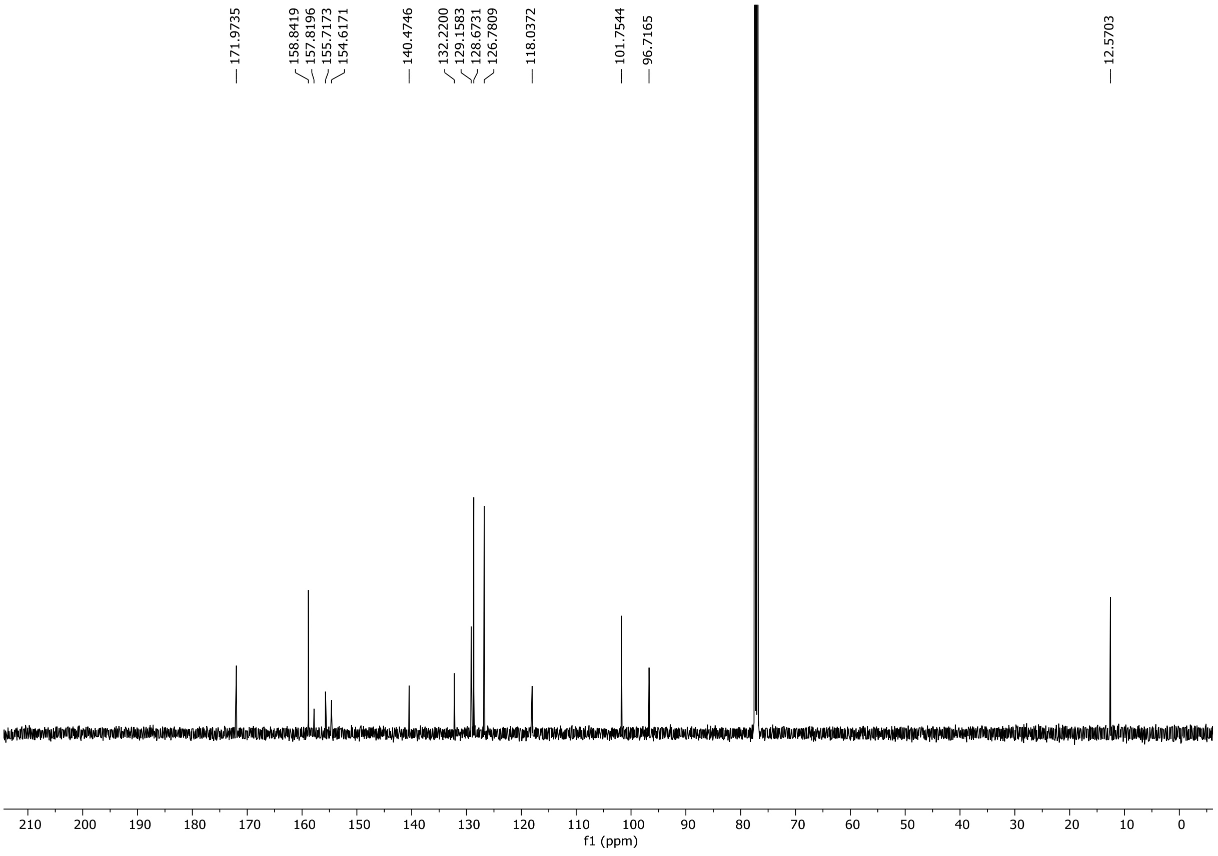

HRMS spectrum of **139**

NAR-A-13 $C_{18}H_{14}N_6O_2$ mono *m/z 346.1178*

**APCI +** (MMI)
nitrogen flow 5 L/min, gas temperature 325°C, nebulizer 45 psi, skimmer **65** V,
vaporizer 200°C, fragmentor 25 V, dissolved in methanol

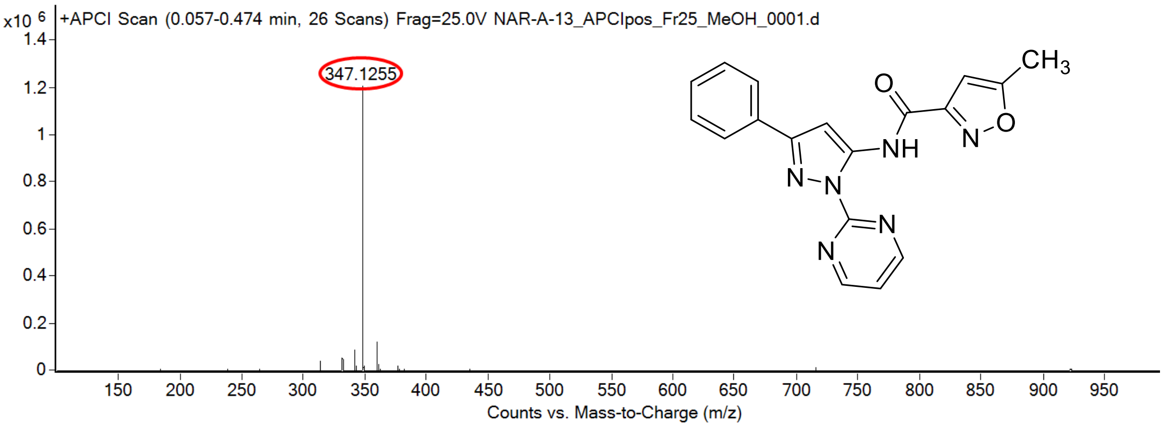

calculated mass: $[M+H]^+ = 347.1251$ observed: $[M+H]^+ = 347.1255$ mass accuracy = 1.1 ppm

FT-IR spectrum (neat) of **139**

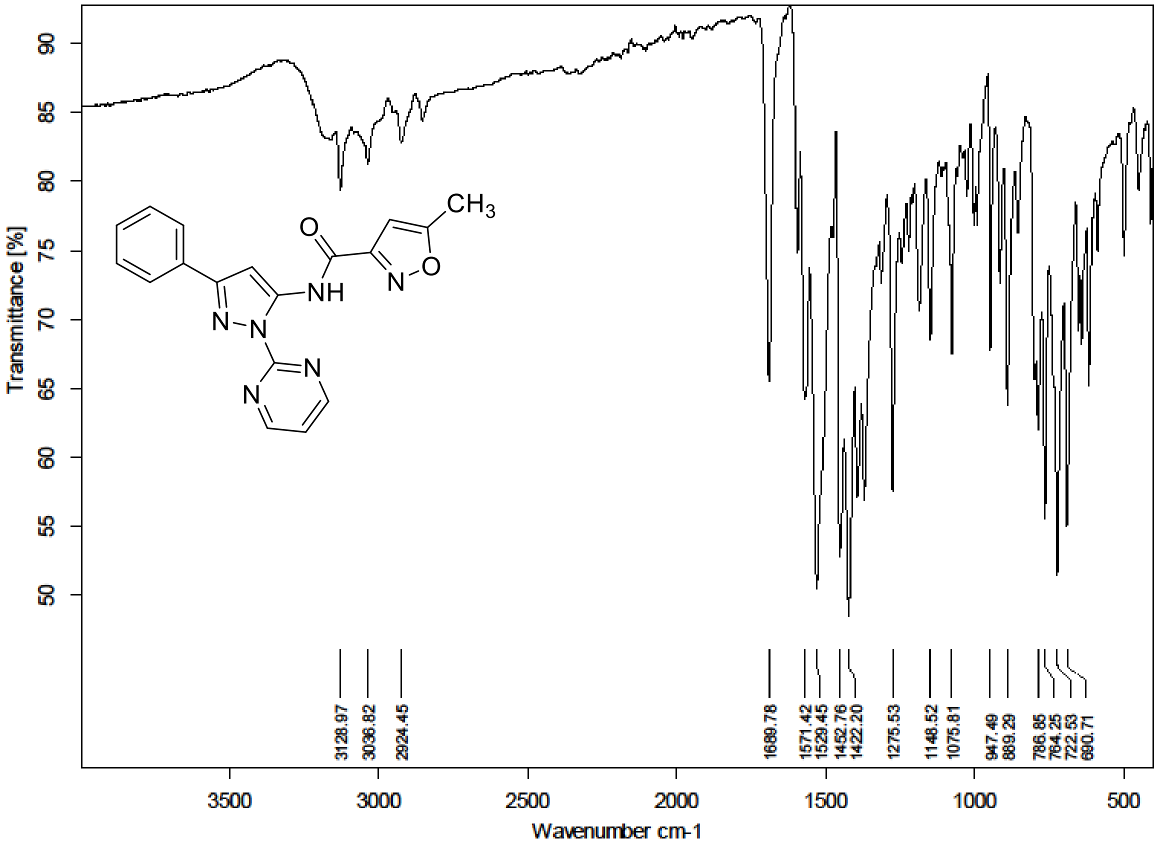

[1]H (500 MHz) and [13]C NMR (126 MHz) spectra of **140** in Chloroform-*d*

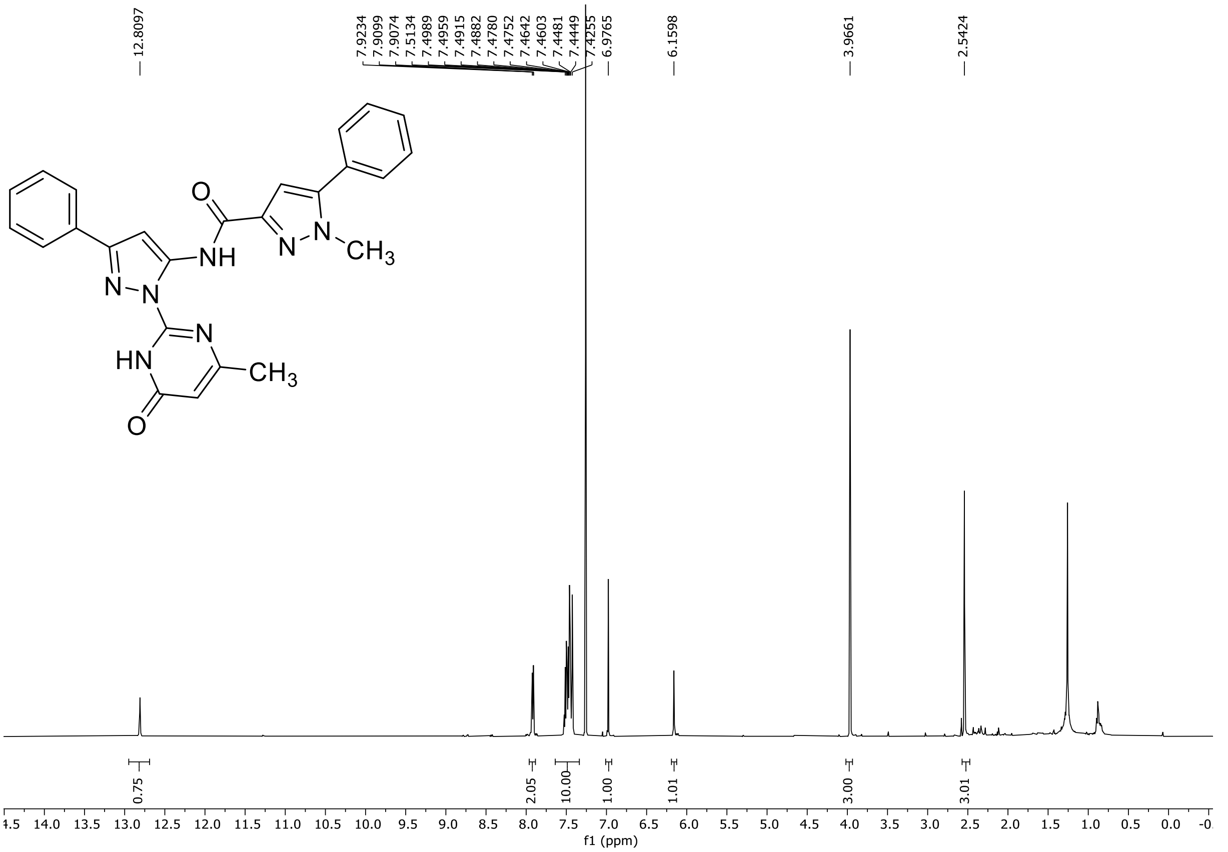

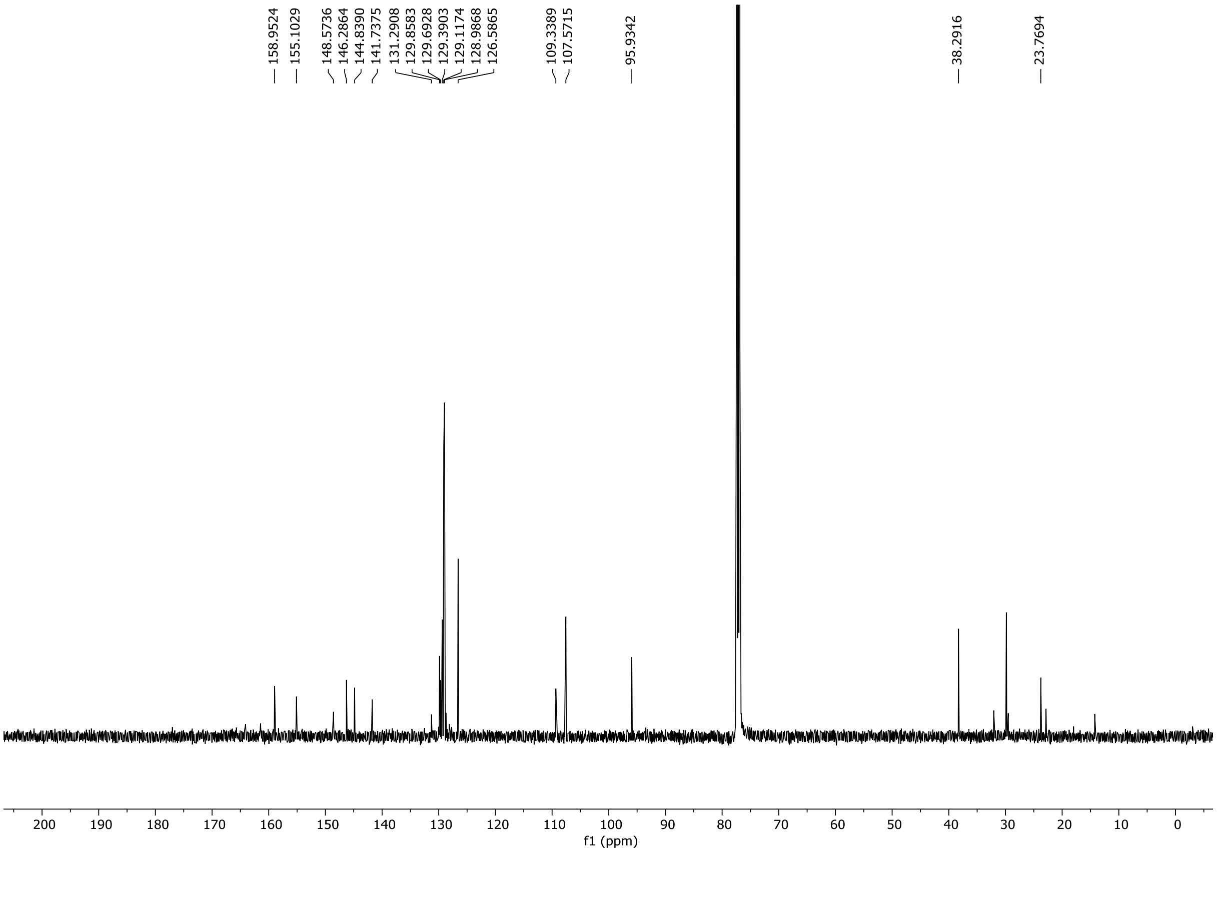

HRMS spectrum of **140**

**NAR-A-43** $C_{25}H_{21}N_7O_2$ *mono m/z*

**451.1757**

**APCI +** (MMI)
nitrogen flow 5 L/min, gas temperature 300°C, nebulizer 45 psi, skimmer **65** V, vaporizer 200°C, fragmentor 30 V, dissolved in MeOH

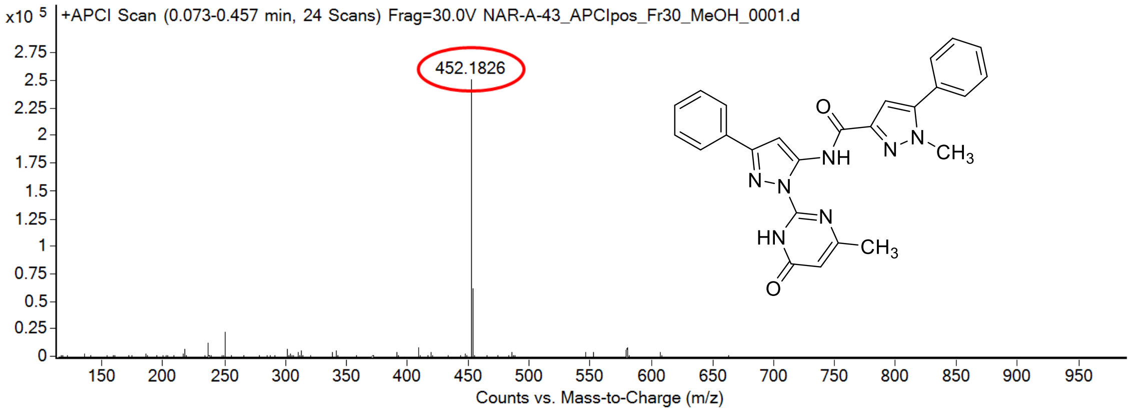

calculated mass: $[M+H]^+$ = 452.1829 observed: $[M+H]^+$ = 452.1826 mass accuracy = -0.7 ppm

FT-IR spectrum (neat) of **140**

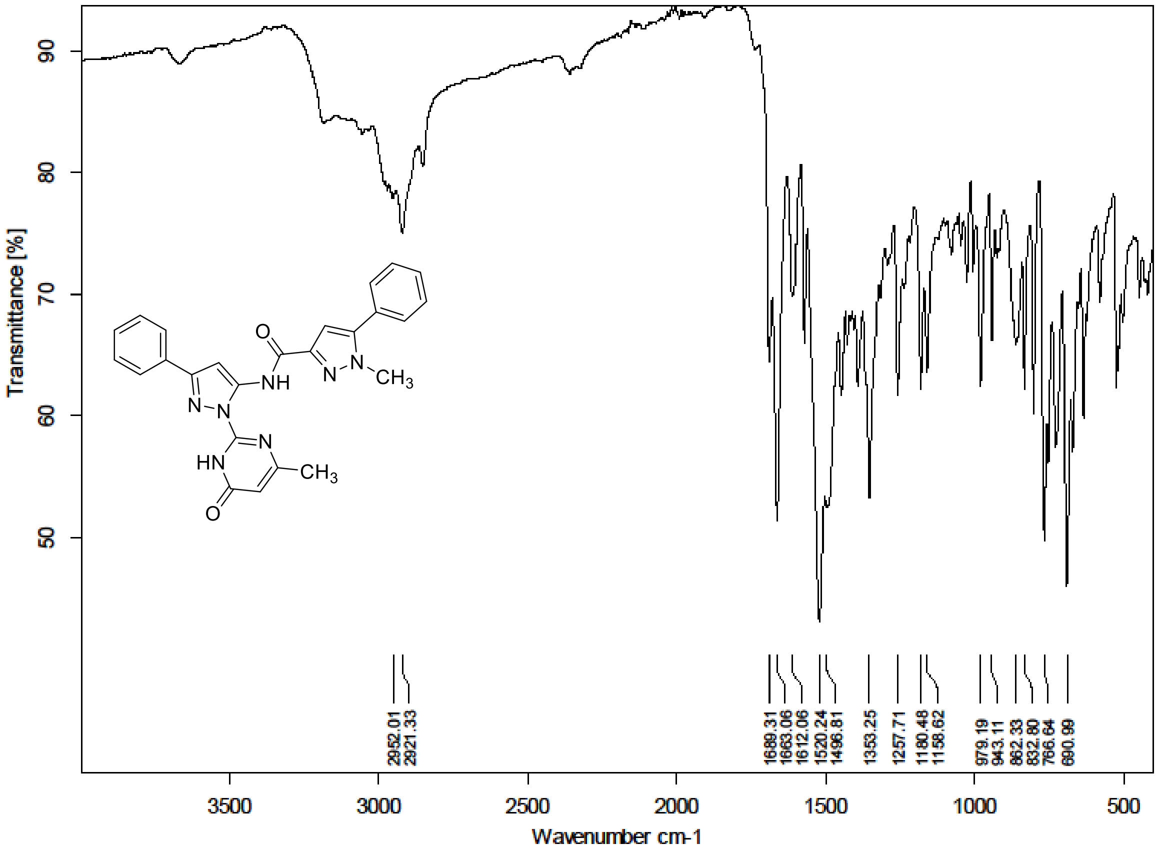

$^{1}H$ (300 MHz) and $^{13}C$ NMR (126 MHz) spectra of **141** in Chloroform-*d*

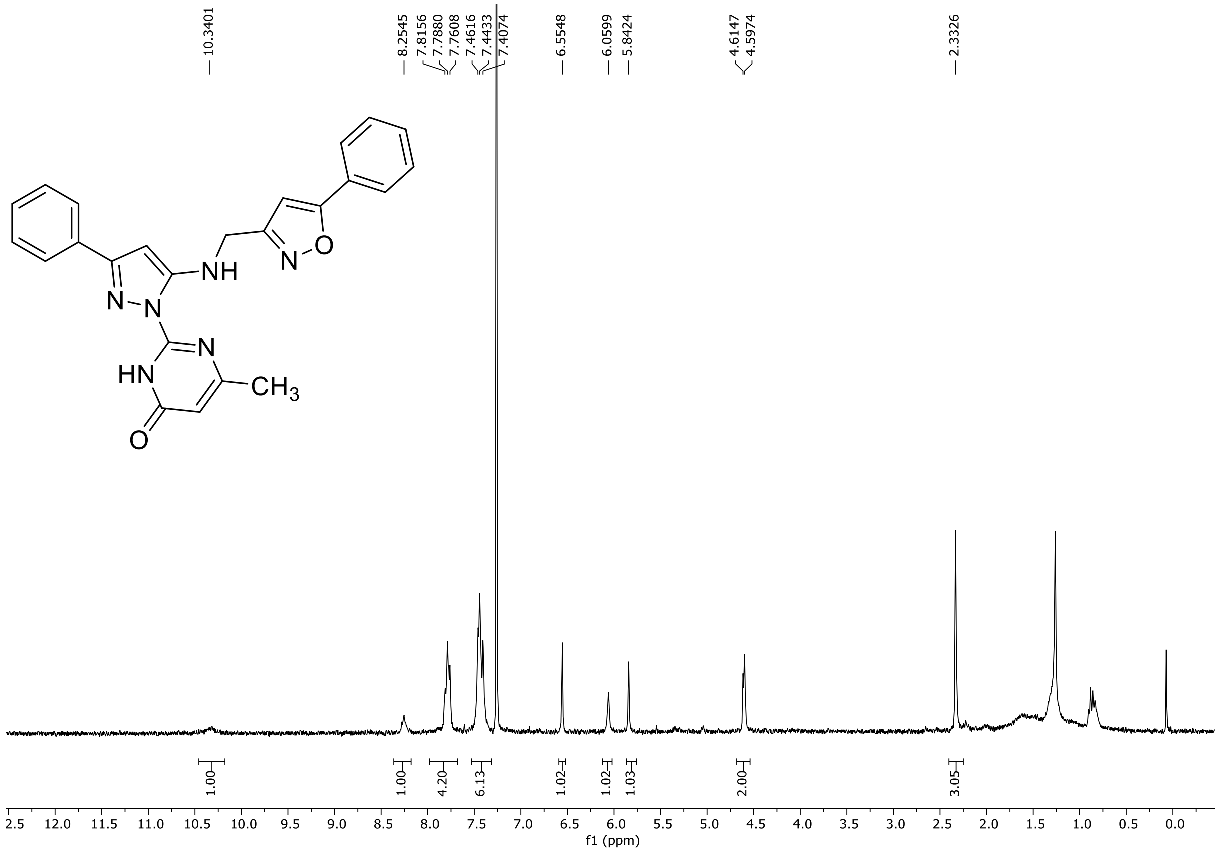

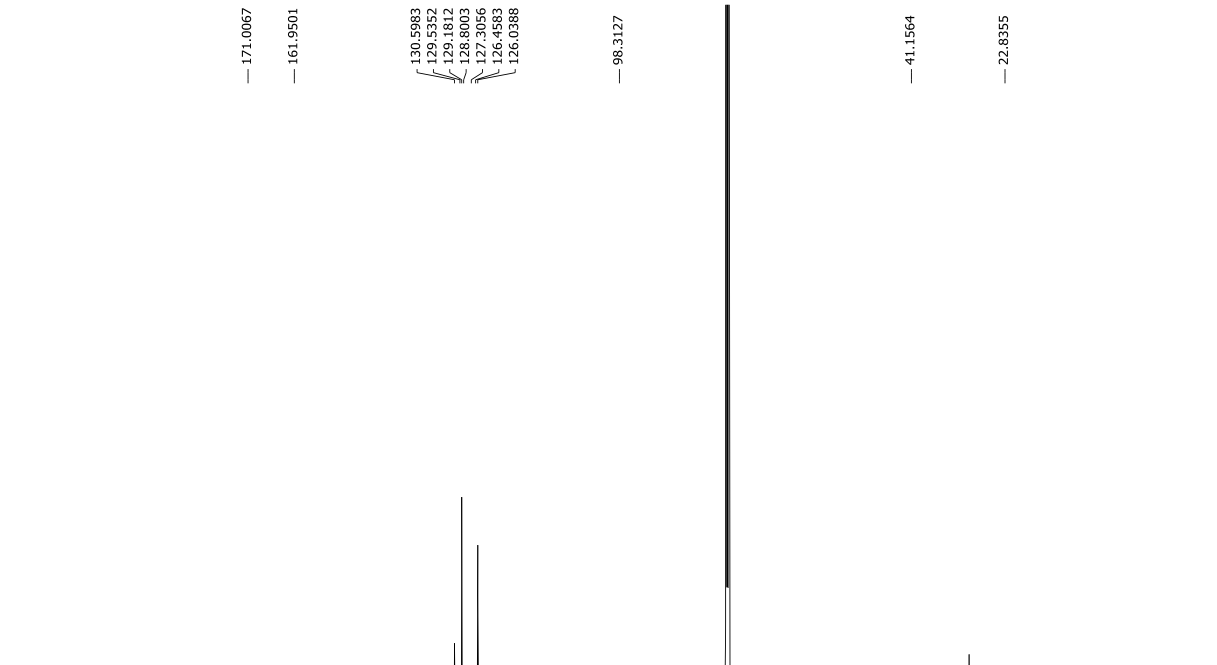

HRMS spectrum of **141**

**NAR-A-67-1** $C_{24}H_{20}N_6O_2$ *mono m/z* **424.1648**

**APCI +** (MMI)
nitrogen flow 5 L/min, gas temperature 300°C, nebulizer 45 psi, skimmer **65** V,
vaporizer 200°C, fragmentor 35 V, dissolved in MeOH

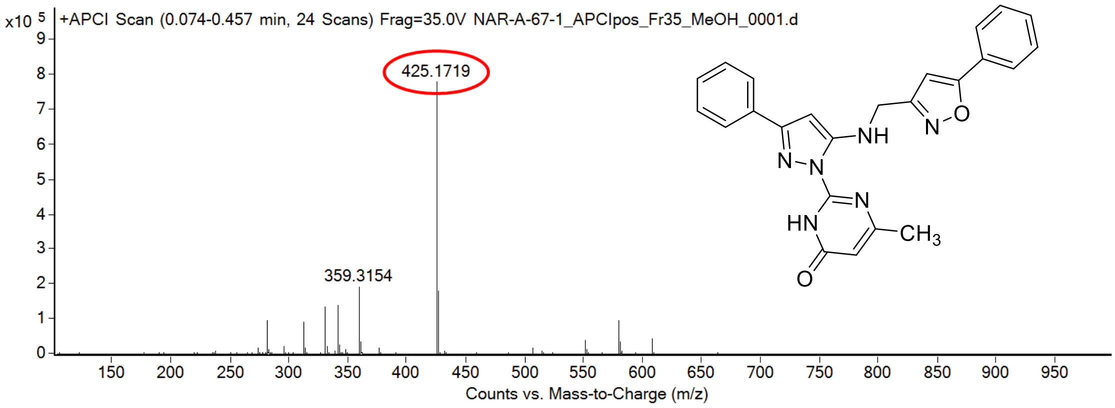

calculated mass: $[M+H]^+$ = 425.1721 observed: $[M+H]^+$ = 425.1719 mass accuracy = -0.5 ppm

FT-IR spectrum (neat) of **141**

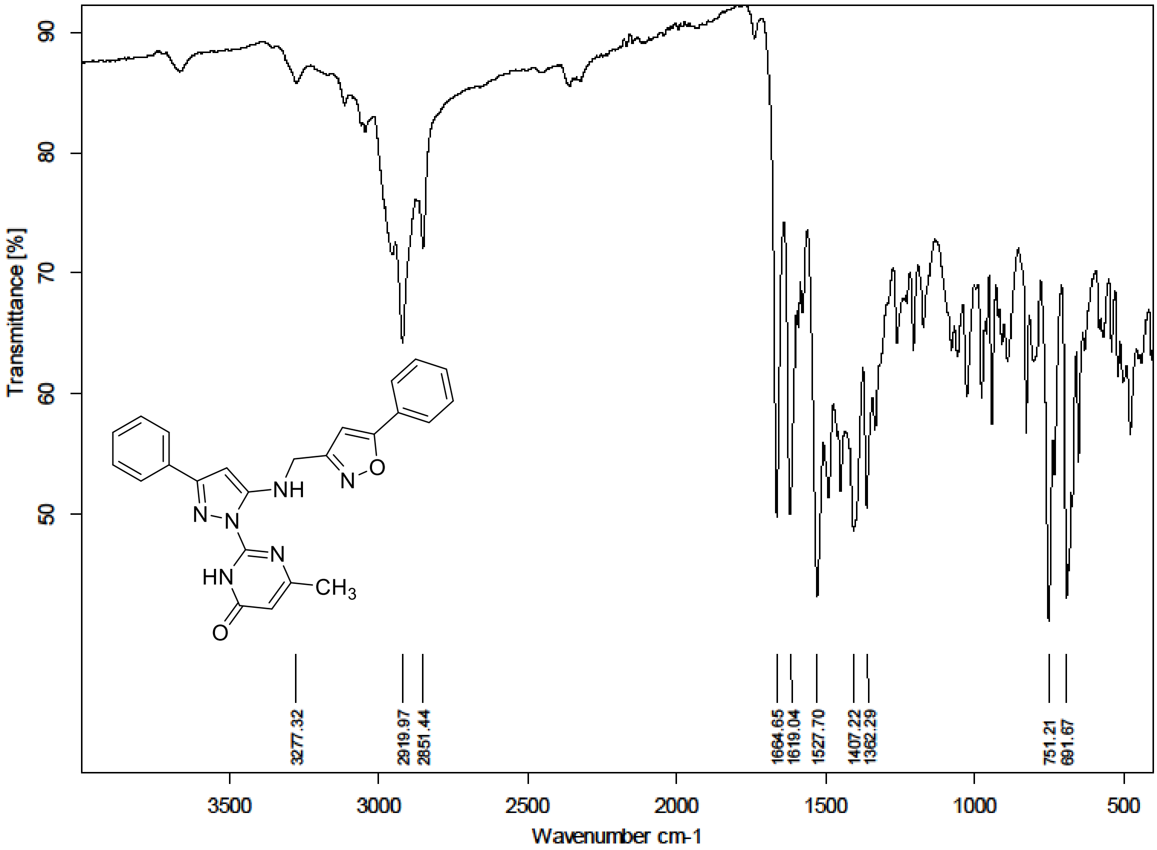

**[1]H NMR, [13]C NMR, HRMS and IR spectra of compound S191-S192**

[1]H (500 MHz) and [13]C NMR (126 MHz) spectra of **S191** in Chloroform-*d*

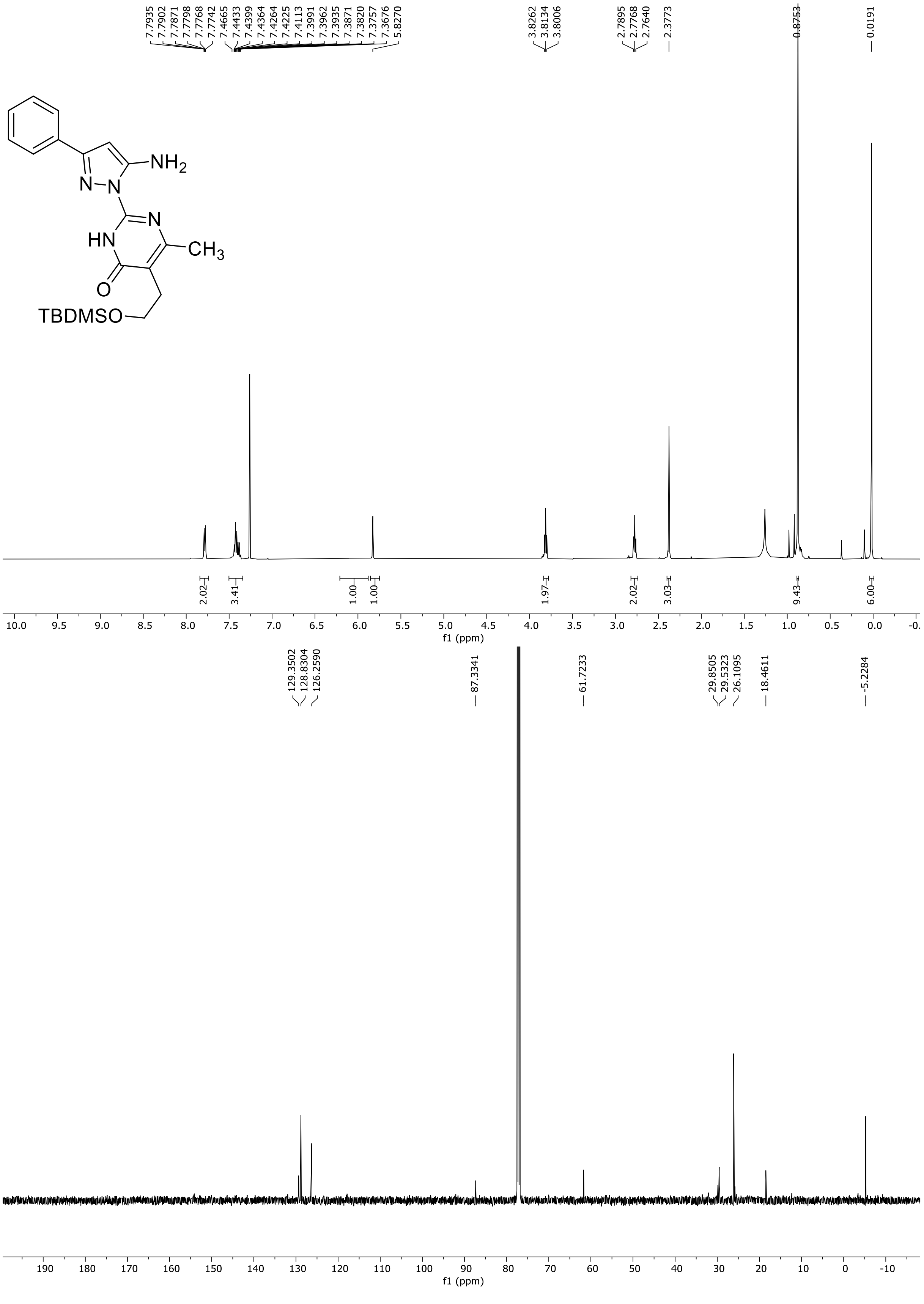

HRMS spectrum of **S191**

NAR-A-22 $C_{22}H_{31}N_5O_2Si$ **mono** ***m/z 425.2247***

**APCI +** (MMI)
nitrogen flow 5 L/min, gas temperature 325°C, nebulizer 45 psi, skimmer **65** V, vaporizer 200°C, fragmentor 25 V, dissolved in methanol

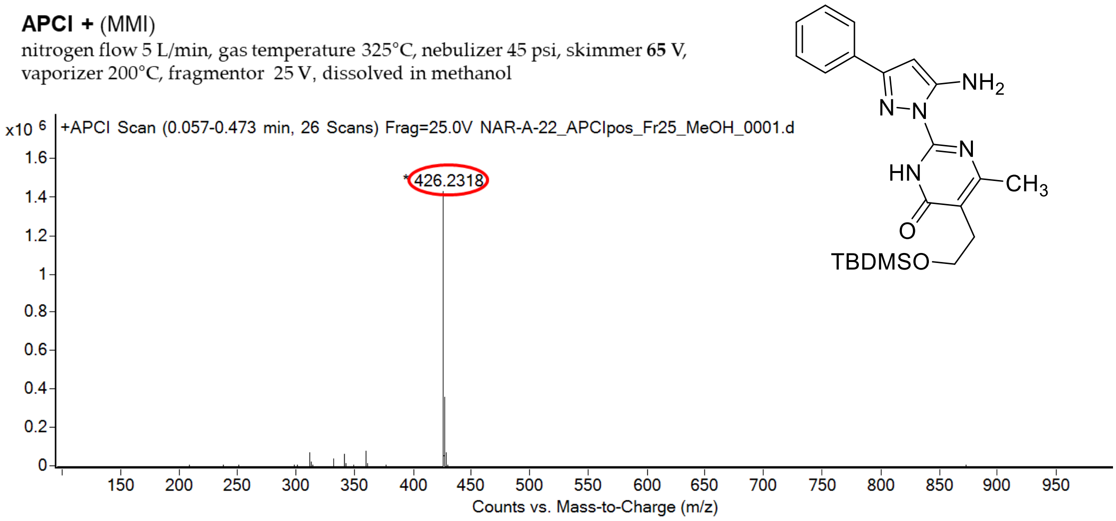

calculated mass: $[M+H]^+ = 426.2320$ observed: $[M+H]^+ = 426.2318$ mass accuracy = -0.5 ppm

FT-IR spectrum (neat) of **S191**

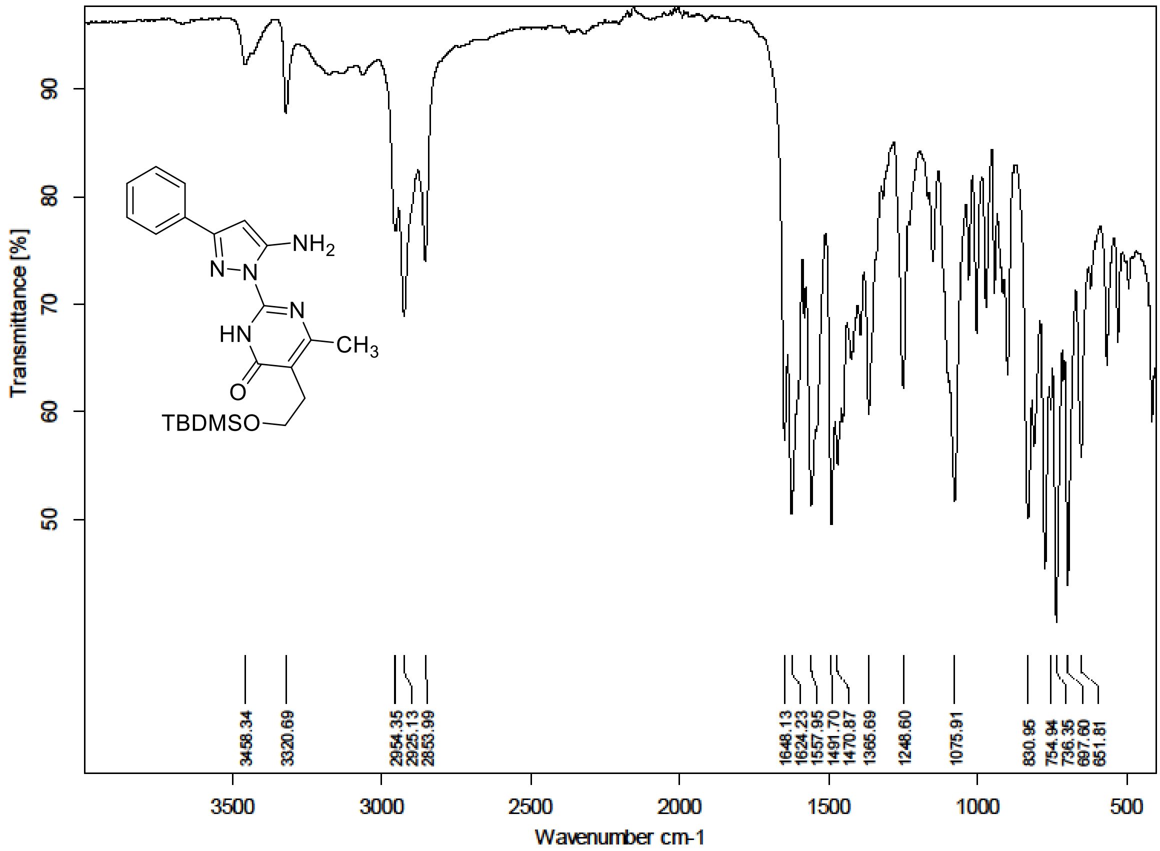

HRMS spectrum of **S192**

NAR-A-28 $C_{32}H_{36}N_6O_4Si$ mono *m/z 596.2567*

**APCI +** (MMI)
nitrogen flow 5 L/min, gas temperature 325°C, nebulizer 45 psi, skimmer **65** V, vaporizer 200°C, fragmentor 30 V, dissolved in methanol

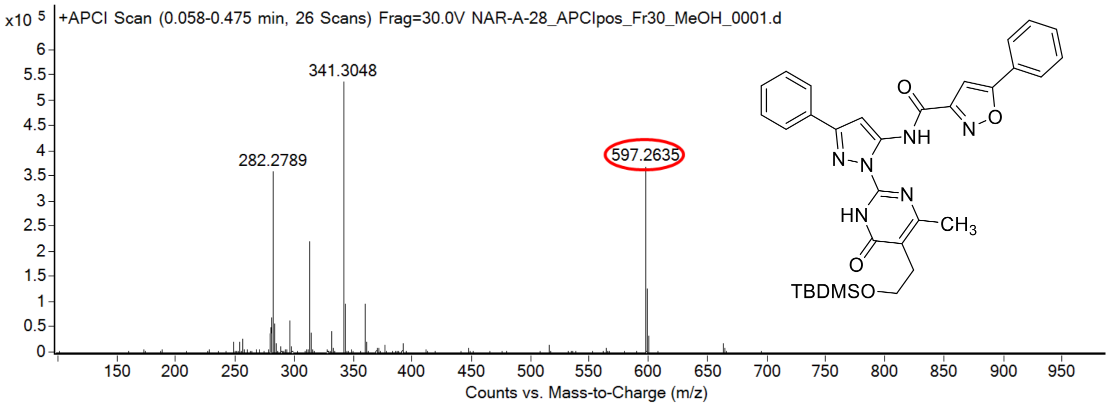

calculated mass: $[M+H]^+$ = 597.2640 observed: $[M+H]^+$ = 597.2635 mass accuracy = -0.8 ppm

**[1]H and [13]C NMR HRMS and IR spectra of compound 142**

[1]H (500 MHz) and [13]C NMR (126 MHz) spectra of **142** in DMSO-$d_6$

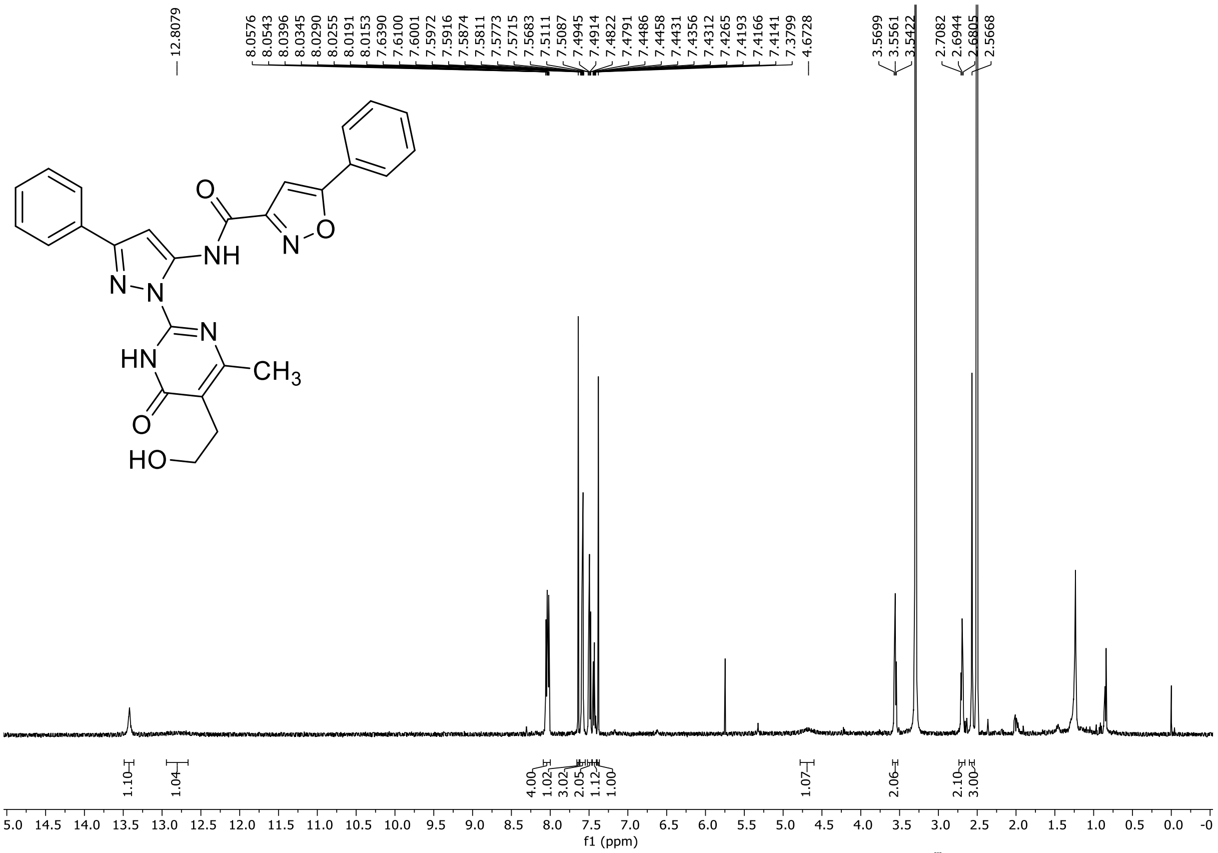

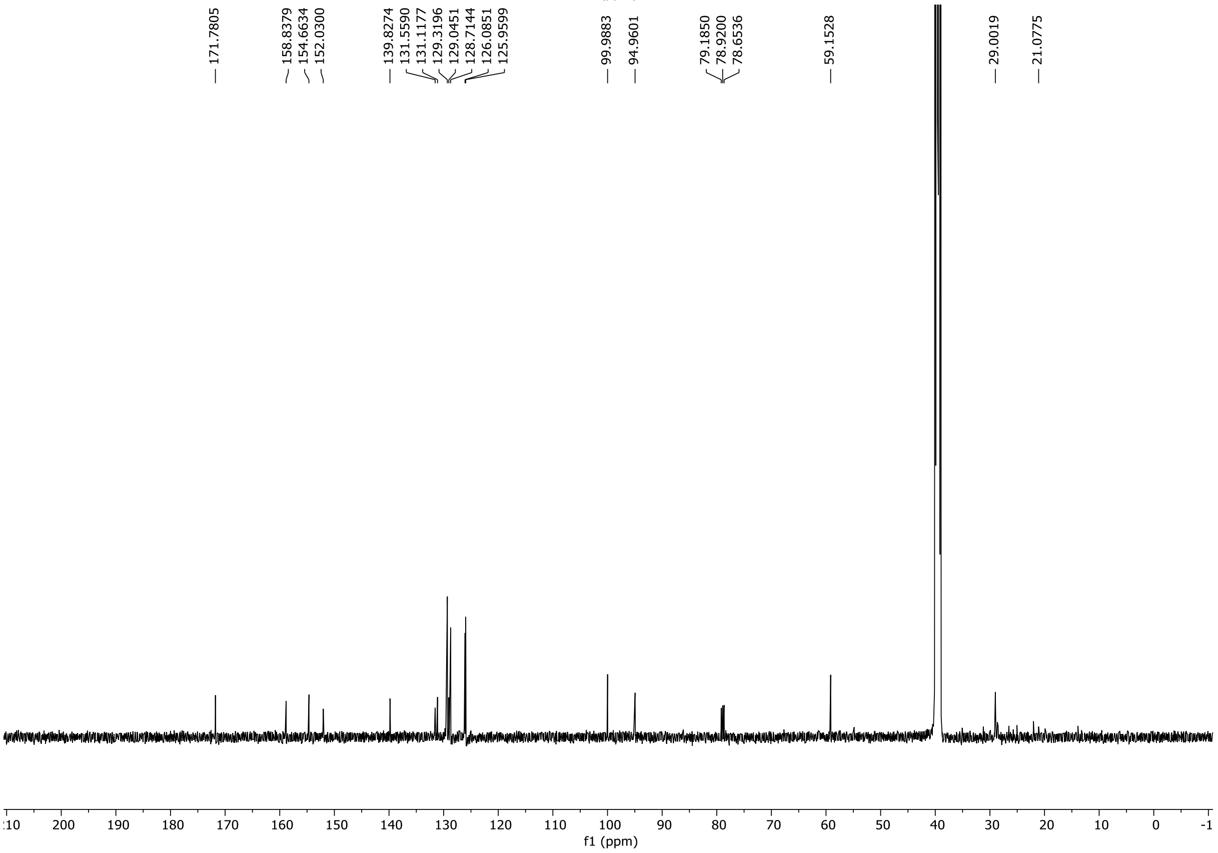

HRMS spectrum of **142**

NAR-A-32 $C_{26}H_{22}N_6O_4$ **mono *m/z* 482.1703**

**APCI +** (MMI)
nitrogen flow 5 L/min, gas temperature 325°C, nebulizer 45 psi, skimmer **65** V,
vaporizer 200°C, fragmentor 25 V, dissolved in methanol

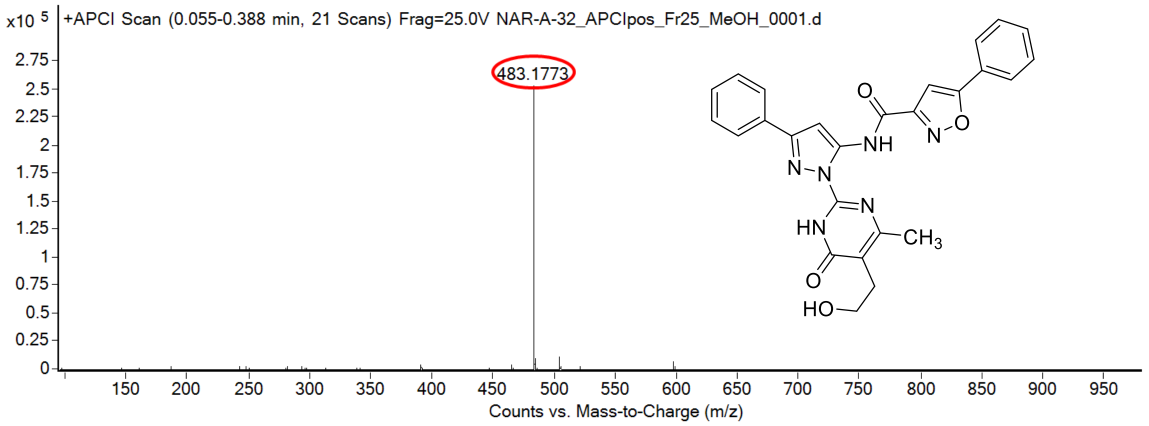

calculated mass: $[M+H]^+ = 483.1775$ observed: $[M+H]^+ = 483.1773$ mass accuracy = -0.4 ppm

FT-IR spectrum (neat) of **142**

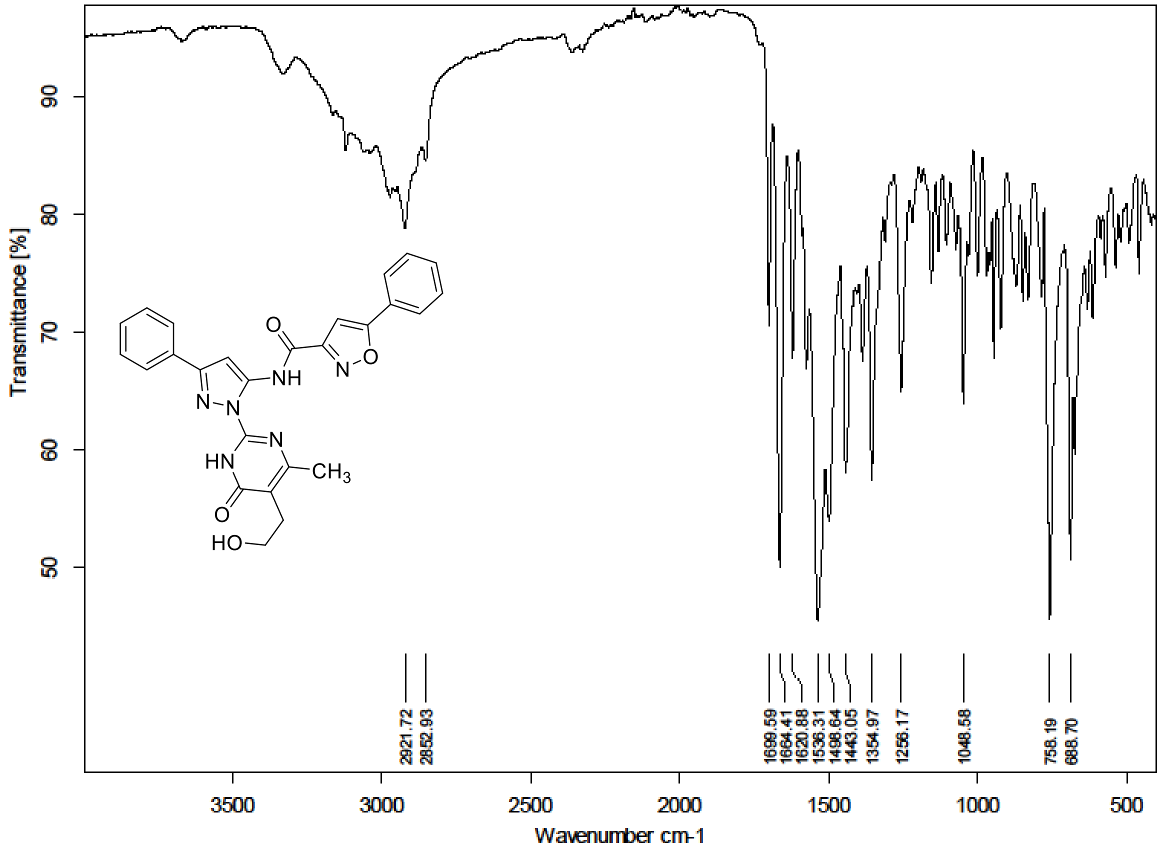

### $^{1}$H and $^{13}$C NMR HRMS and IR spectra of compound S193-S194

$^{1}$H (300 MHz) and $^{13}$C NMR (126 MHz) spectra of **S193** in Chloroform-*d*

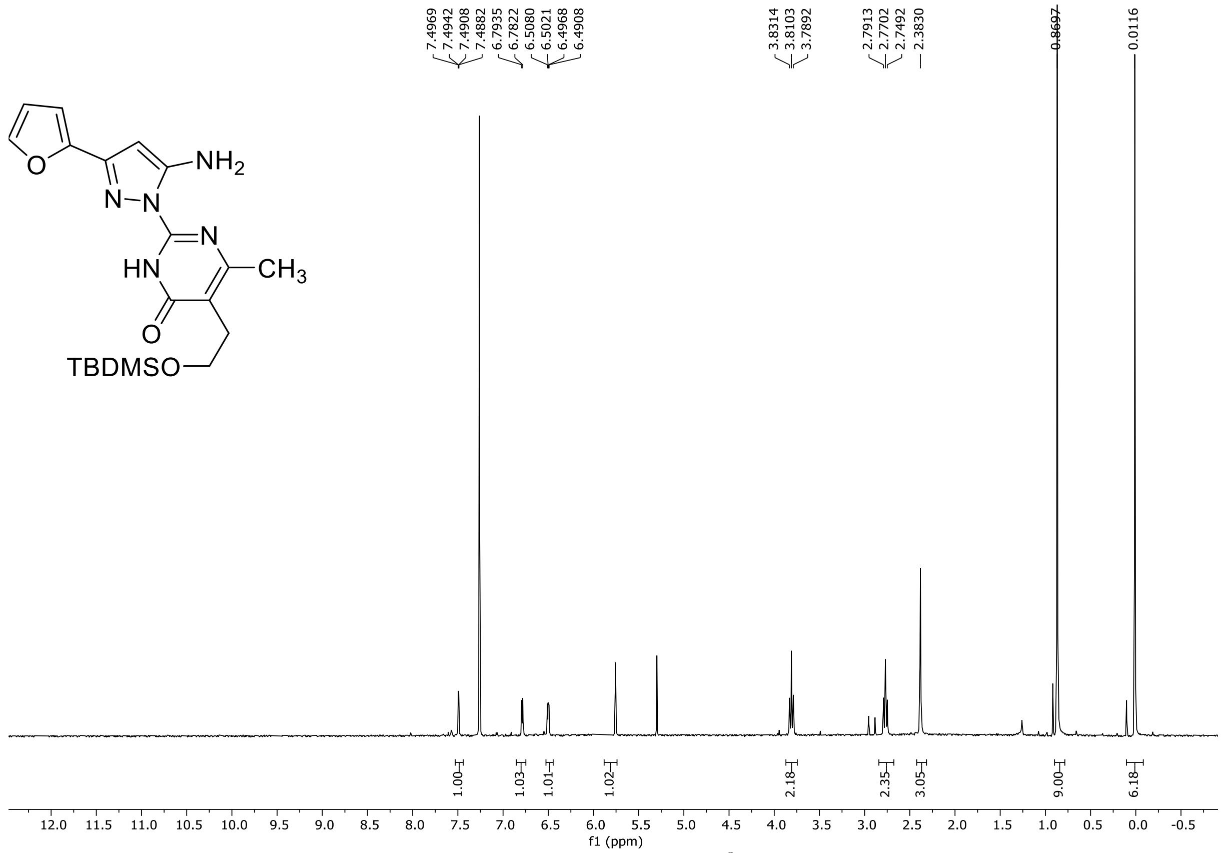

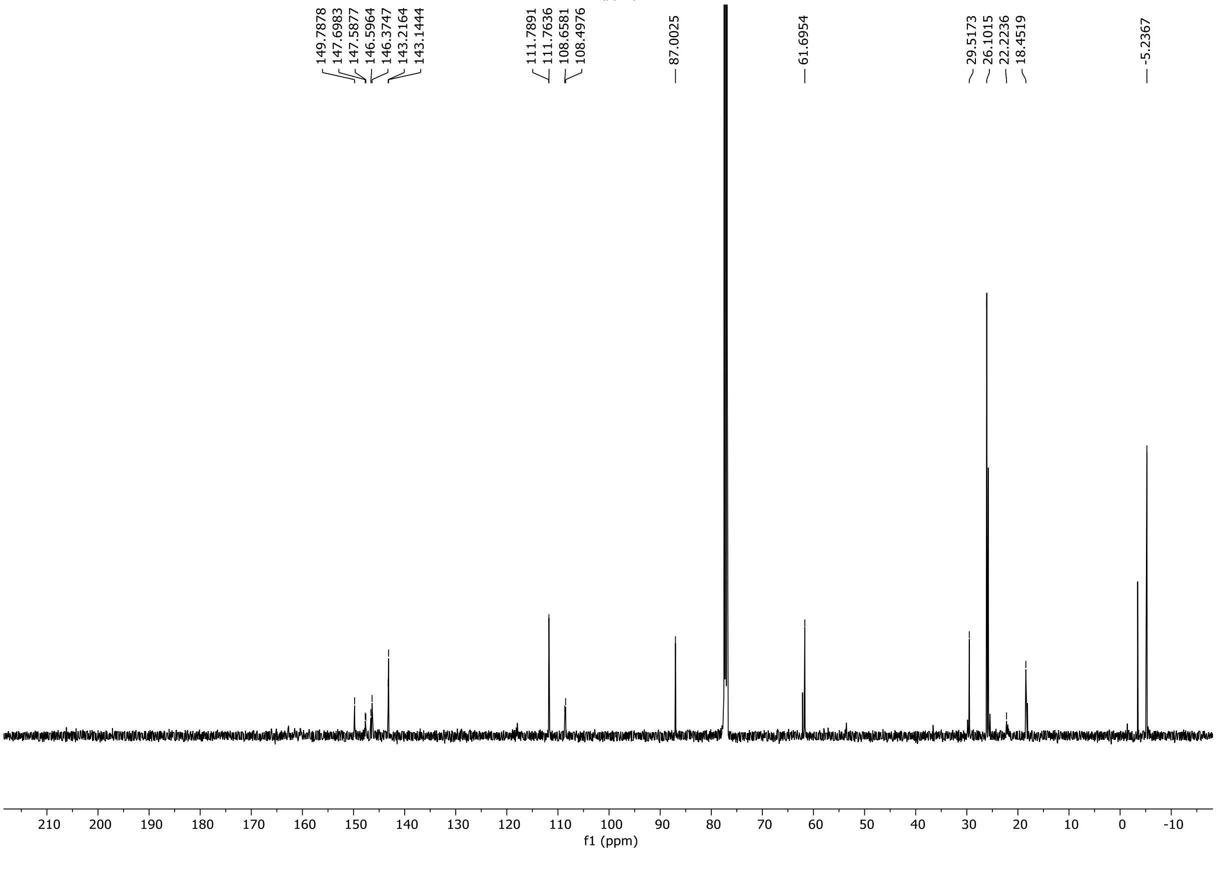

HRMS spectrum of **S193**

**NAR-A-103** $C_{20}H_{29}N_5O_3Si$ *m/z* **415.2040**

**APCI+** (MMI)
nitrogen flow 5 L/min, gas temperature 325°C, nebulizer 45 psi, skimmer 65 V, vaporizer 200°C, fragmentor 10 V, dissolved in MeOH

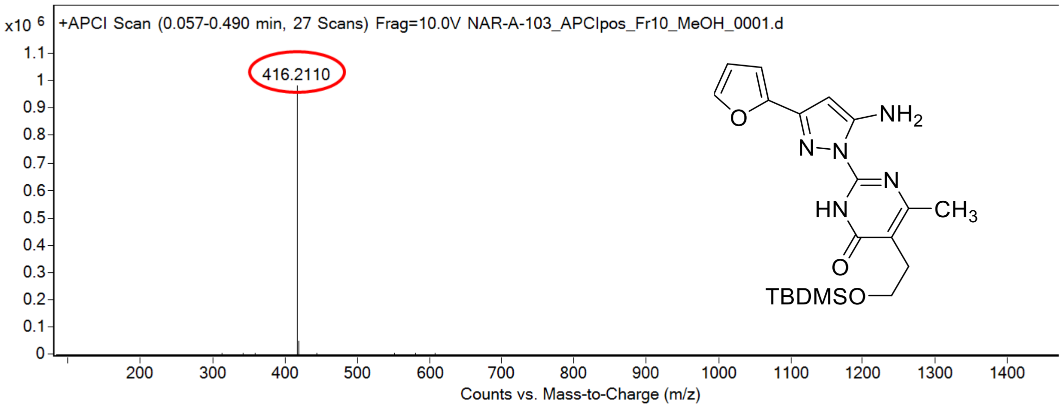

calculated mass: $[M+H]^+ = 416.2112$ observed: $[M+H]^+ = 416.2110$ mass accuracy = -0.5 ppm

FT-IR spectrum (neat) of **S193**

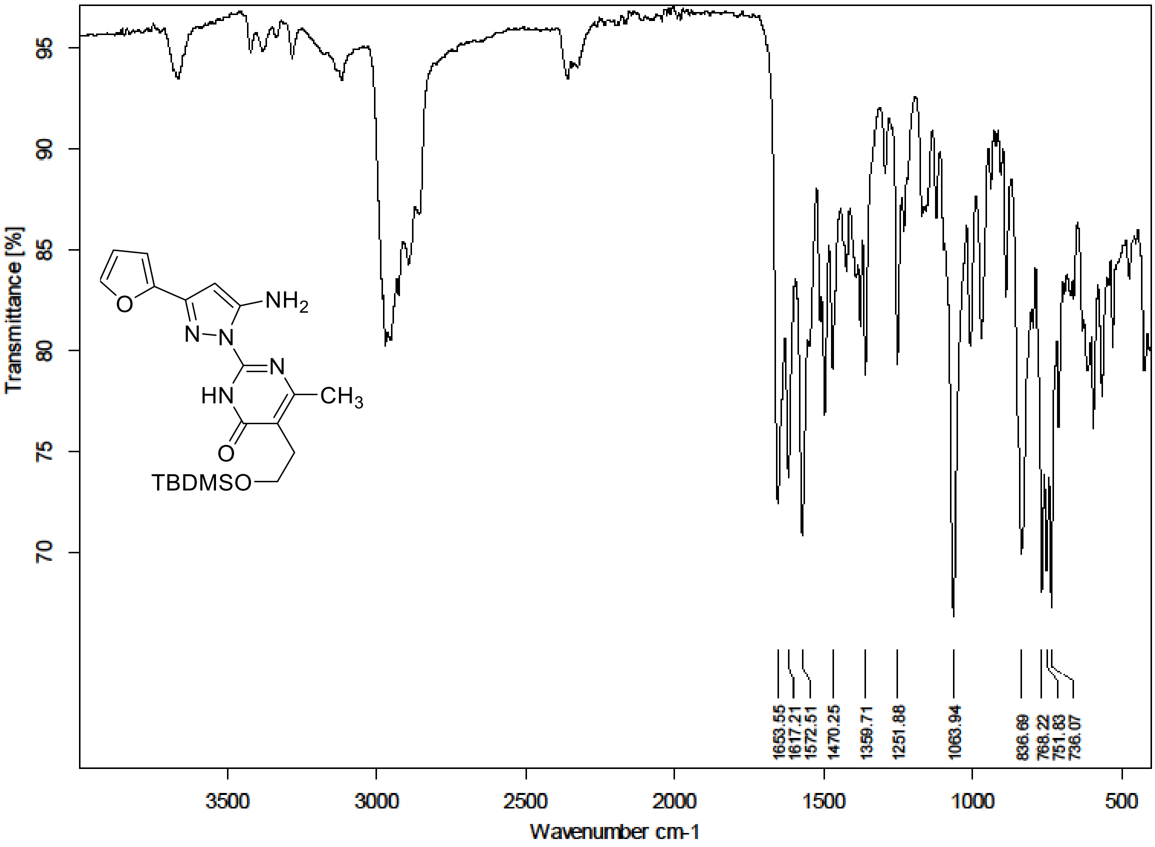

$^{1}H$ (500 MHz) and $^{13}C$ NMR (126 MHz) spectra of **S194** in Chloroform-*d*

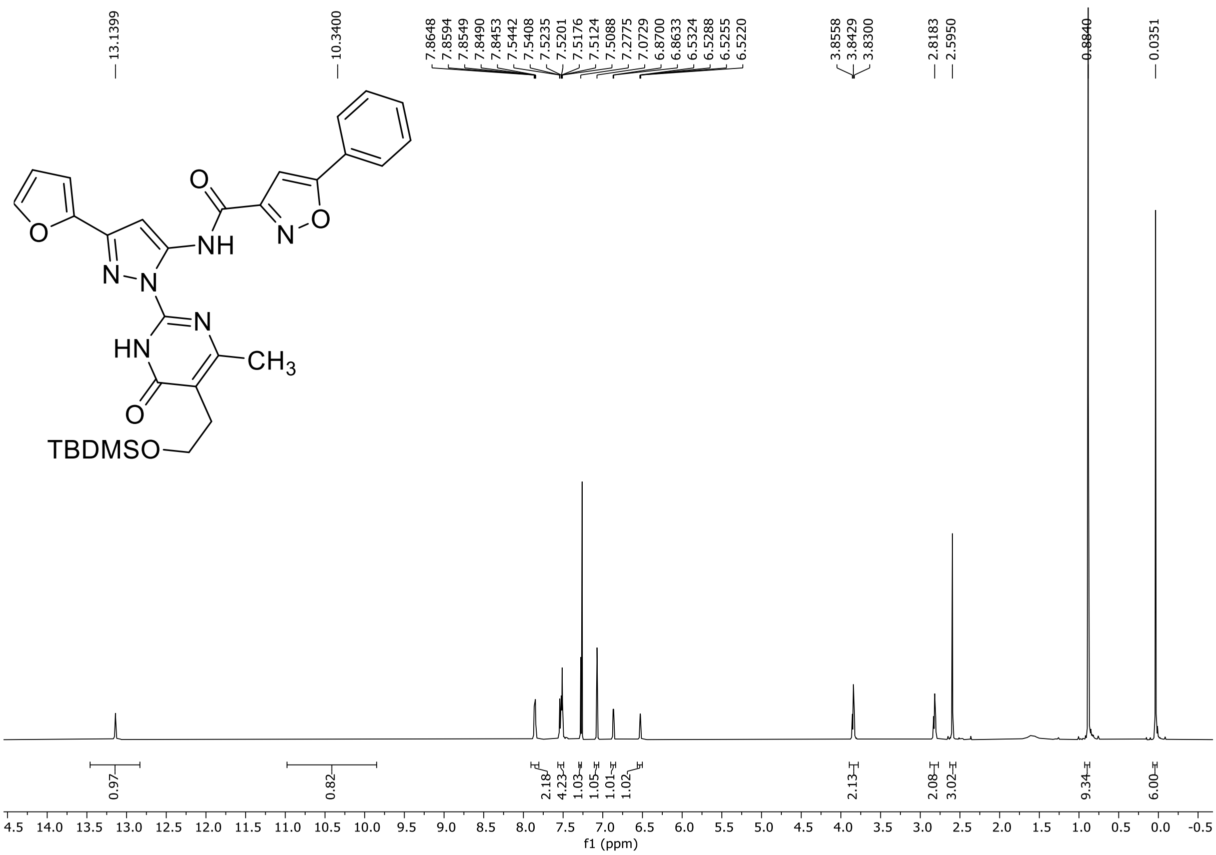

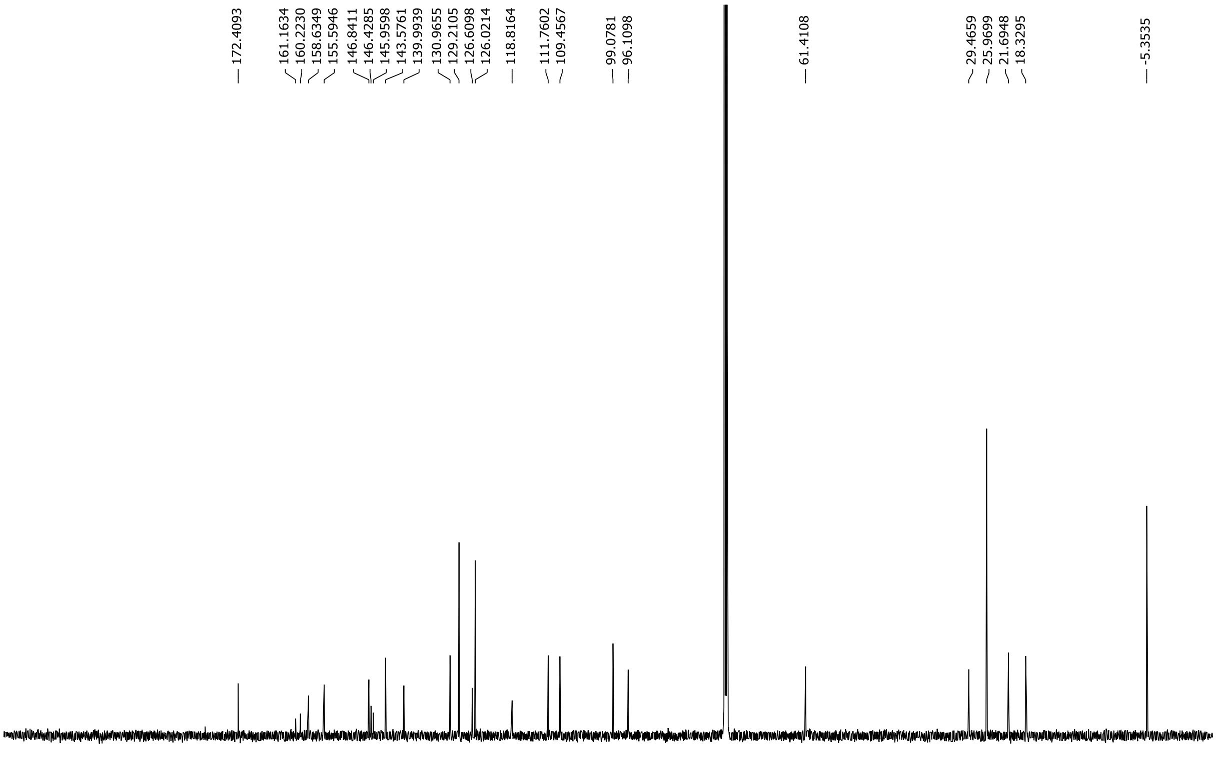

HRMS spectrum of **S194**

**NAR-A-105** $C_{30}H_{34}N_6O_5Si$ *m/z* **586.2360**

**APCI+** (MMI)
nitrogen flow 5 L/min, gas temperature 325°C, nebulizer 45 psi, skimmer 65 V,
vaporizer 200°C, fragmentor 25 V, dissolved in MeOH

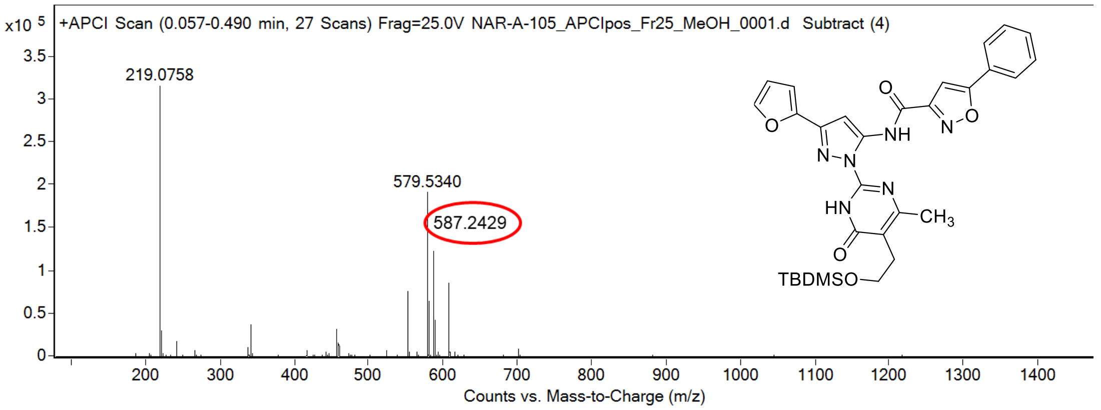

calculated mass: $[M+H]^+$ = 587.2433 observed: $[M+H]^+$ = 587.2429 mass accuracy = -0.7 ppm

FT-IR spectrum (neat) of **S194**

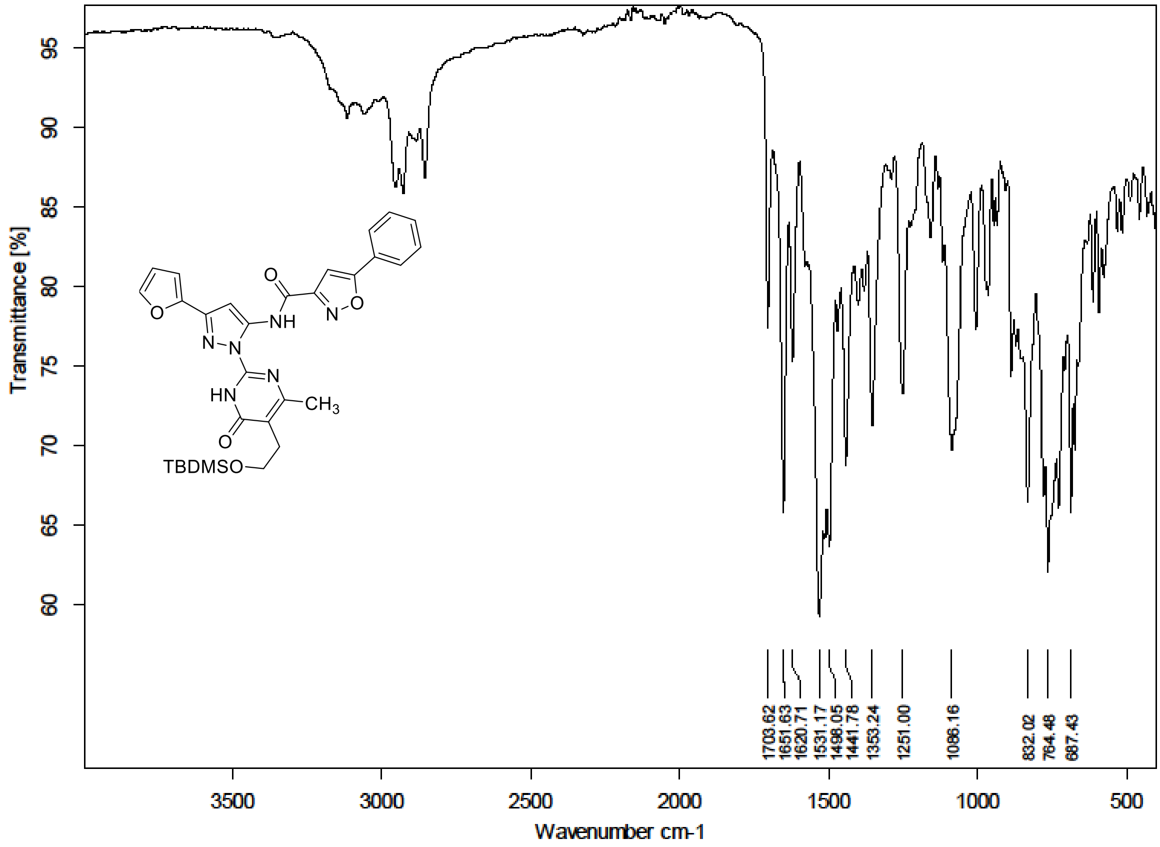

### [1]H and [13]C NMR, HRMS and IR spectra of compound 143

[1]H (300 MHz) and [13]C NMR (75 MHz) spectra of **143** in DMSO-$d_6$

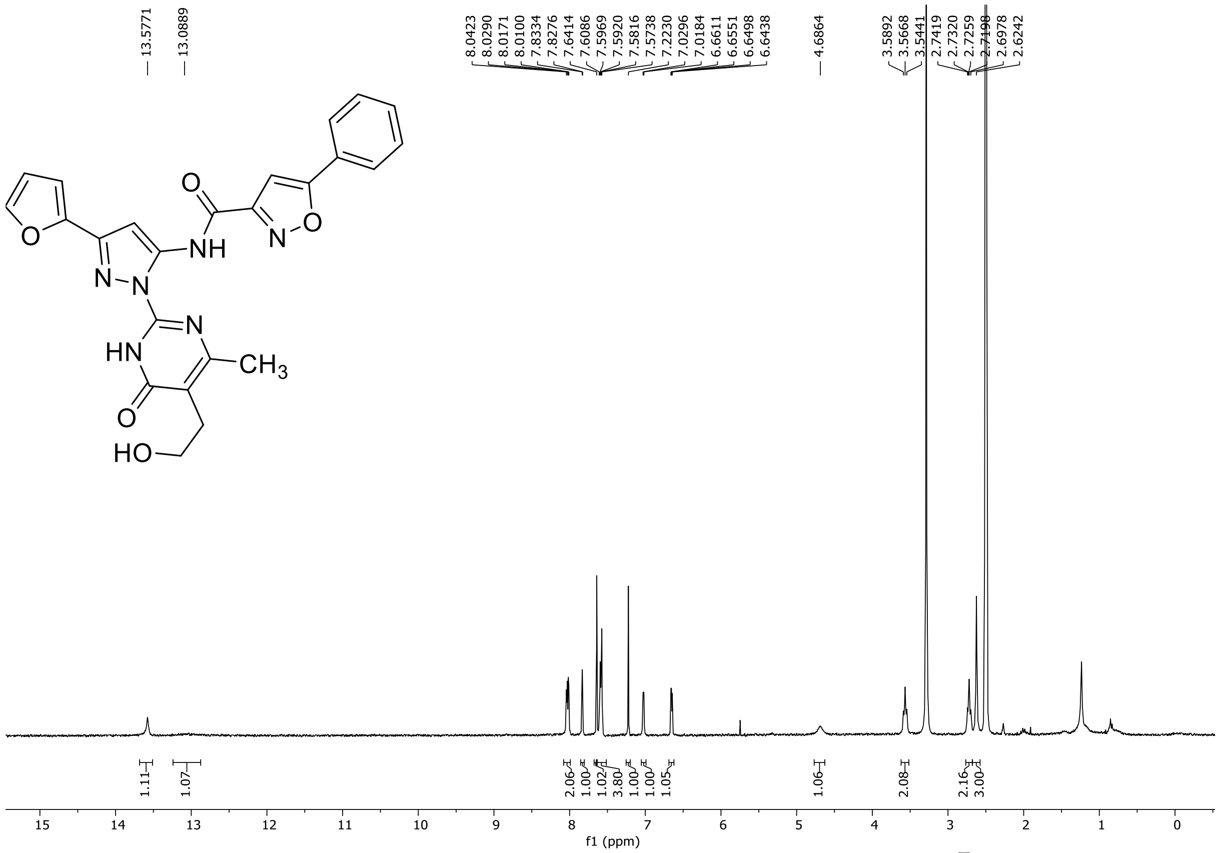

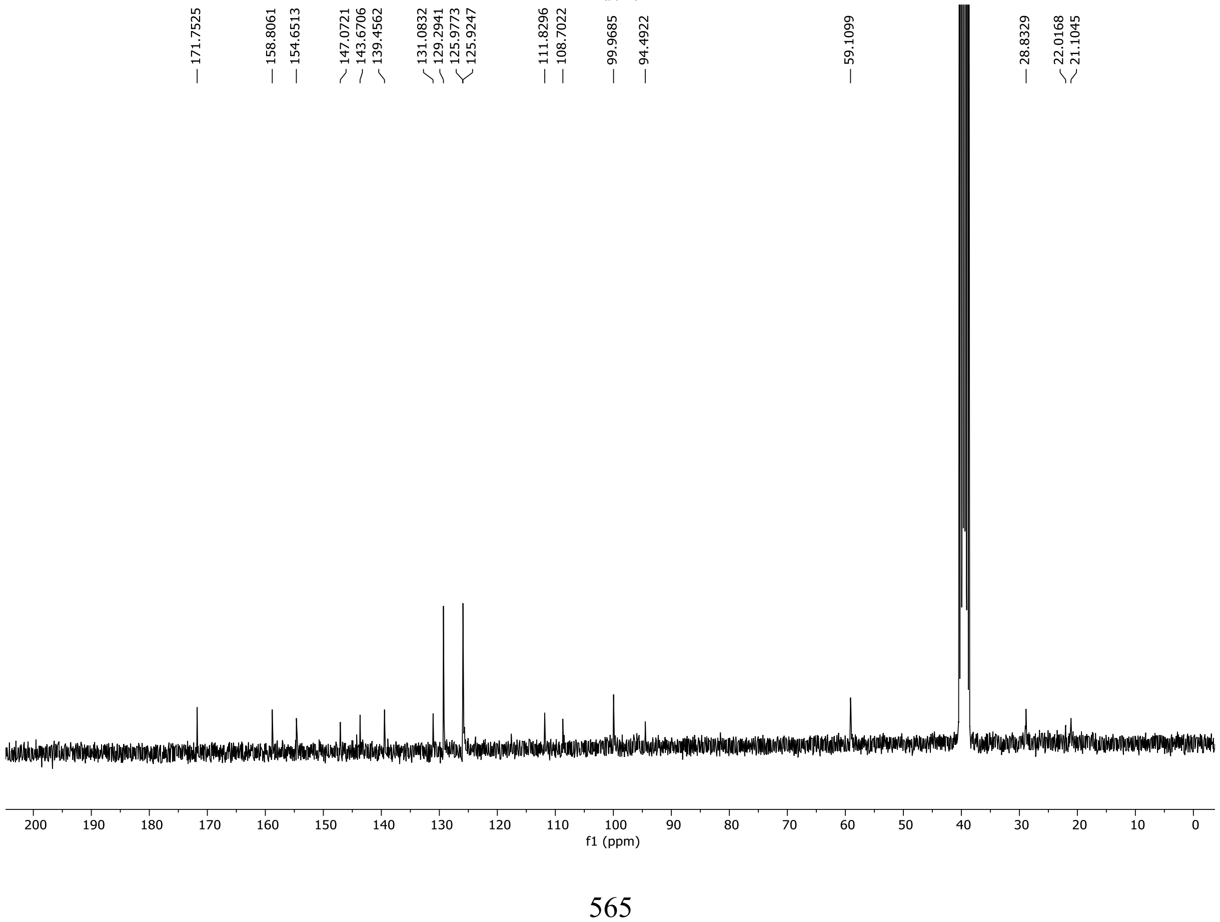

HRMS spectrum of **143**

**NAR-A-126** $C_{24}H_{20}N_6O_5$ *m/z* **472.1495**

**ESI-** (MMI)
nitrogen flow 5 L/min, gas temperature 325°C, nebulizer 45 psi, skimmer 65 V, fragmentor 120 V, dissolved in DMSO, MeOH

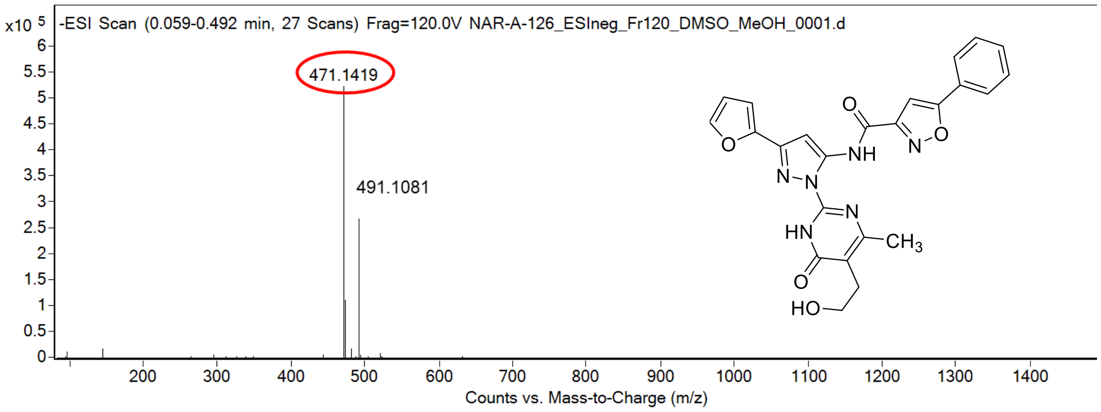

calculated mass: $[M-H]^- = 471.1422$ observed: $[M-H]^- = 471.1419$ mass accuracy = -0.6 ppm

FT-IR spectrum (neat) of **143**

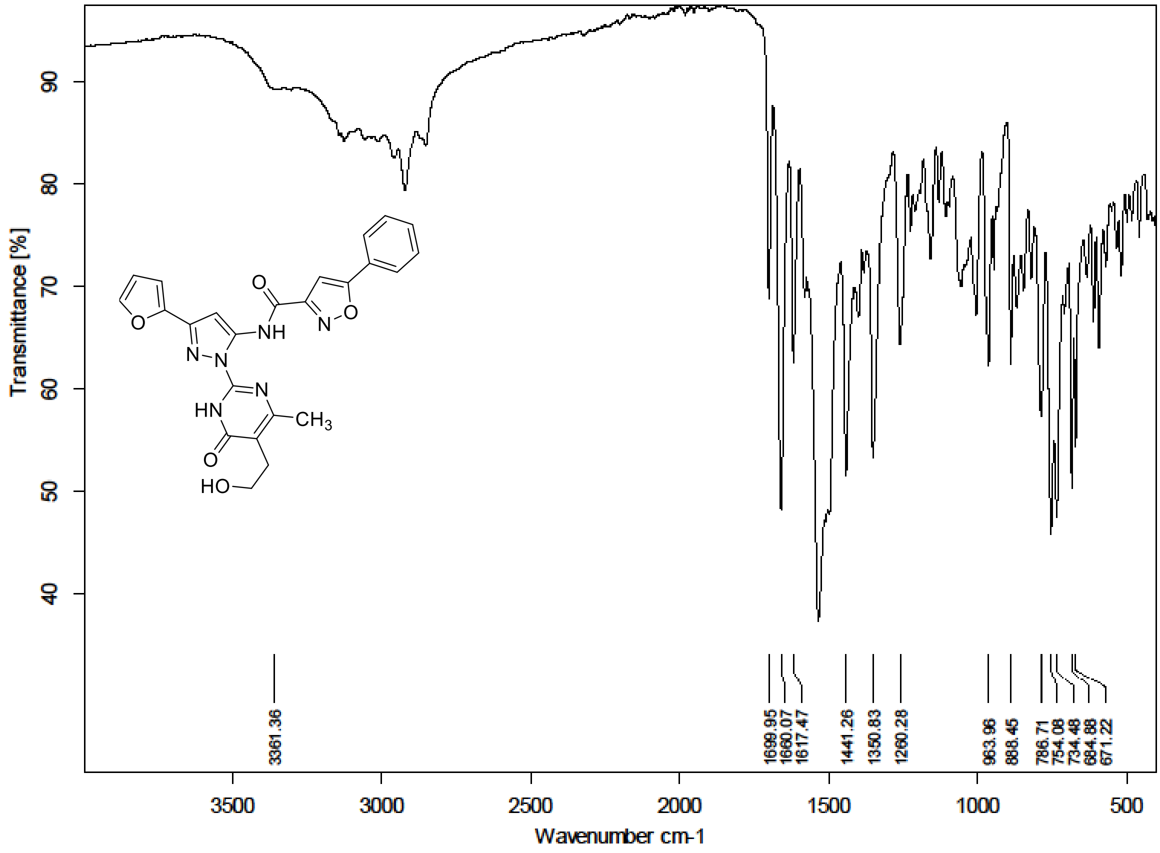

**HPLC analytical chromatogram of compound 18**

| | |
|---|---|
| Instrument | : Ultimate 3000 LC Analytical Systems, Thermo Scientific |
| Column | : Agilent ZORBAX Eclipse Plus C18 (particle size 5 µm, 4.6 × 250 mm) |
| Sample preparation | : 1 mg of compound **18** dissolved in methanol (1 mL) + 10 µL TFA |
| Injection volume | : 10 µL |
| Flow rate | : 1 mL/min |
| Mobile phase | : water:methanol = 10:90 + 0.1% Trifluoroacetic acid (Isocratic) |
| Retention times | : 3.68 min. |

### Chromatogram and Results

**Injection Details**

| | | | |
|---|---|---|---|
| *Compound Name:* | **MU262=18** | *Run Time (min):* | **9.58** |
| *Vial Number:* | **129** | *Injection Volume:* | **10.00** |
| *Mobile phase:* | **Water:MeOH + 0.1% TFA = 10:90** | | |
| *Mobile phase:* | **10:90 isocretic** | *Channel:* | **UV_VIS_3** |
| *Column:* | **Agilent ZORBAX Eclipse Plus C18, 5um, 4.60 X 25(** | *Wavelength:* | **337** |
| *Instrument Method:* | **PK analytical method_Isocratic** | *Flow rate:* | **1 mL/min** |
| *Processing Method:* | **PK processing method** | *Dilution Factor:* | **1.0000** |
| *Injection Date/Time:* | **25/Jul/25 03:34** | *Sample Weight:* | **1 mg** |

**Chromatogram**

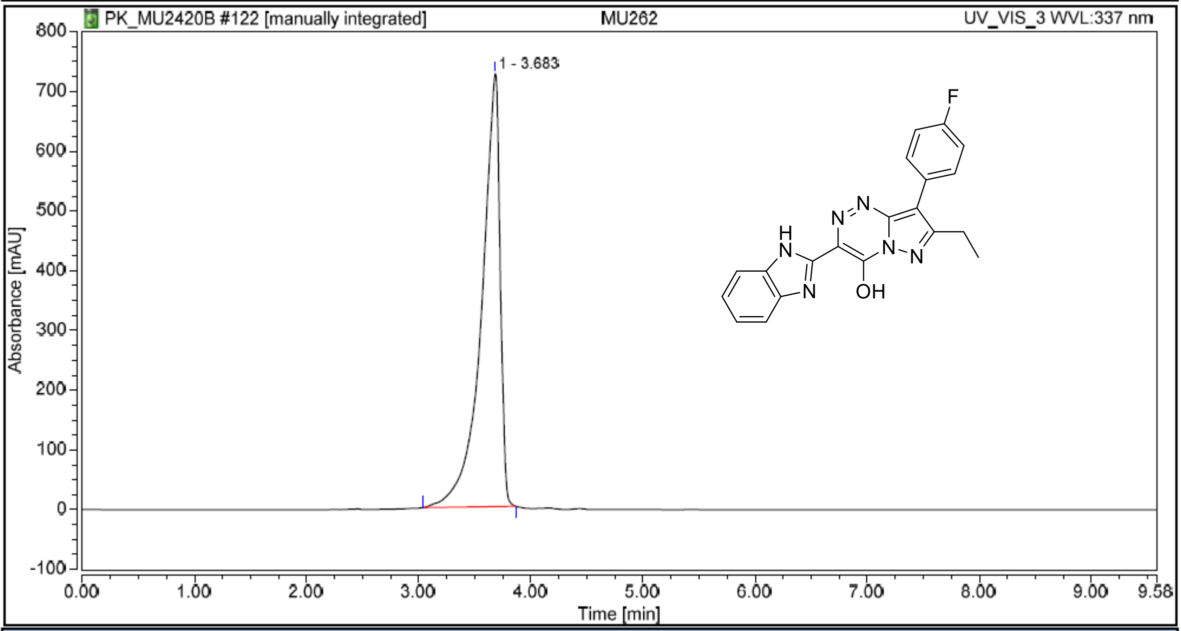

**Integration Results**

| No. | Peak Name | Retention Time min | Area mAU*min | Height mAU | Relative Area % | Relative Height % | Amount n.a. |
|---|---|---|---|---|---|---|---|
| 1 | | 3.683 | 138.375 | 725.147 | 100.00 | 100.00 | n.a. |
| **Total:** | | | **138.375** | **725.147** | **100.00** | **100.00** | |

**HPLC analytical chromatogram of compound 32**

| | |
|---|---|
| Instrument | : Ultimate 3000 LC Analytical Systems, Thermo Scientific |
| Column | : Agilent ZORBAX Eclipse Plus C18 (particle size 5 µm, 4.6 × 250 mm) |
| Sample preparation | : 1 mg of compound **32** dissolved in methanol (1 mL) + 10 µL TFA |
| Injection volume | : 10 µL |
| Flow rate | : 1 mL/min |
| Mobile phase | : water:methanol = 10:90 + 0.1% Trifluoroacetic acid (Isocratic) |
| Retention times | : 2.99 min. |

### Chromatogram and Results

**Injection Details**

| | | | |
|---|---|---|---|
| *Compound Name:* | MU876=32 | *Run Time (min):* | 6.20 |
| *Vial Number:* | 136 | *Injection Volume:* | 10.00 |
| *Mobile phase:* | Water:MeOH + 0.1% TFA = 10:90 | | |
| *Mobile phase:* | 10:90 isocretic | *Channel:* | UV_VIS_1 |
| *Column:* | Agilent ZORBAX Eclipse Plus C18, 5um, 4.60 X 25( | *Wavelength:* | 270 |
| *Instrument Method:* | PK analytical method_Isocratic | *Flow rate:* | 1 mL/min |
| *Processing Method:* | PK processing method | *Dilution Factor:* | 1.0000 |
| *Injection Date/Time:* | 25/Jul/25 04:53 | *Sample Weight:* | 1 mg |

**Chromatogram**

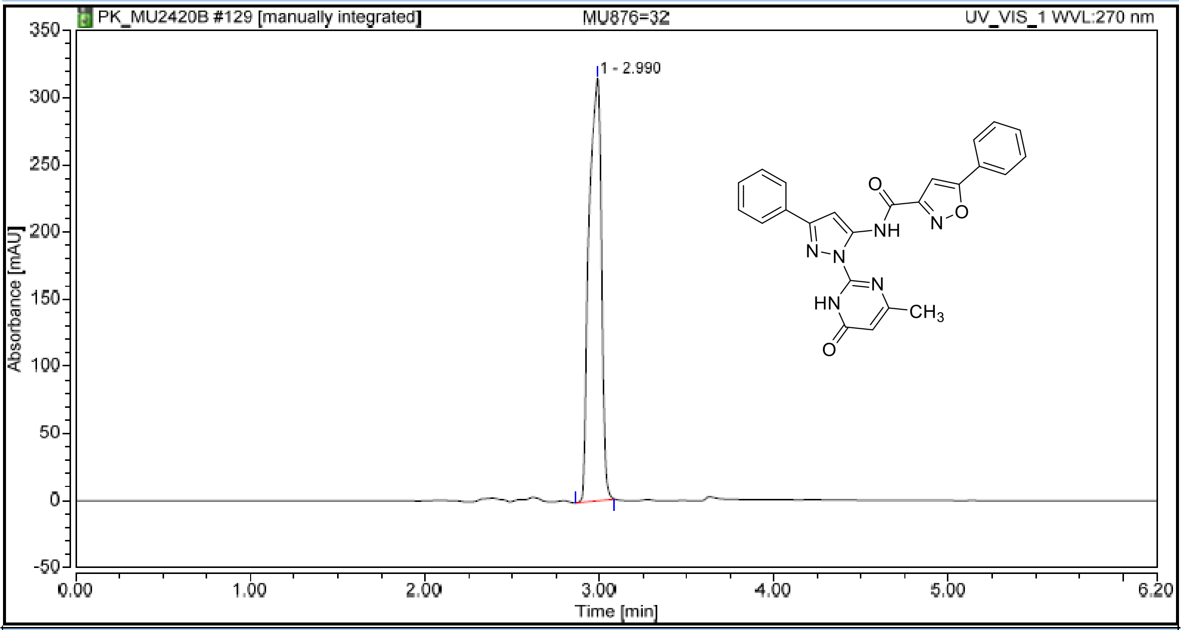

**Integration Results**

| No. | Peak Name | Retention Time min | Area mAU*min | Height mAU | Relative Area % | Relative Height % | Amount n.a. |
|---|---|---|---|---|---|---|---|
| 1 | | 2.990 | 26.190 | 314.973 | 100.00 | 100.00 | n.a. |
| Total: | | | 26.190 | 314.973 | 100.00 | 100.00 | |
