## Supporting information_Experimental procedures for "Discovery of two structurally distinct classes of inhibitors targeting the nuclease MUS81 and enhancing efficacy of chemotherapy in cancer cells"

**Experimental procedures for all intermediates and final compounds**

Republic

2 Department of Chemistry, Faculty of Science, Masaryk University, 62500 Brno, Czech

Republic

3 International Clinical Research Center, St. Anne’s University Hospital, Brno 656 91, Czech Republic

4 NCBR, Faculty of Science, Masaryk University, 62500 Brno, Czech Republic

5 GIMM - Gulbenkian Institute for Molecular Medicine, 1649-035 Lisbon, Portugal

6 Loschmidt Laboratories, Department of Experimental Biology and RECETOX, Faculty of

Science, Masaryk University, 62500 Brno, Czech Republic

7 Faculty of Medicine, University of Lisbon, 1649-028 Lisbon, Portugal

### General information

All commercially available reagents were used as supplied without further purification. The reaction solvents were purchased anhydrous and were stored under nitrogen. Unless noted otherwise, the reactions were carried out in oven-dried glassware under the atmosphere of nitrogen. Analytical thin-layer chromatography (TLC) was performed using aluminum plates pre-coated with silica gel (silica gel 60 F_254_, Merck). TLC plates were visualized by exposure to ultraviolet light (λ = 254 nm) and/or by submersion in aqueous ceric ammonium molybdate (CAM). All solutions were concentrated by rotary evaporation at 40 °C, unless noted otherwise. As indicated, either manual column chromatography was carried out using silica gel (pore size 60 Å, 230-400 mesh particle size, 40-63 μm particle size) or column chromatography was carried out using the Biotage Selekt purification system. Purification by preparative thin layer chromatography was performed using plates from Merck (PLC Silica gel 60 F_254_, 1 mm). Reverse phase column chromatography was carried out using C_18_-reversed phase silica gel (pore size 90 Å, 230-400 mesh particle size, 40-63 µm particle size). Nuclear magnetic resonance spectra were recorded using Bruker Avance 500 MHz and 300 MHz instruments at 30 °C. NMR spectra were obtained in indicated deuterated solvents; chemical shifts are quoted in parts per million (*δ*) referenced to the appropriate deuterated solvent employed. Multiplicities are indicated by s (singlet), d (doublet), t (triplet), q (quartet), p (pentet), quin (quintet), sept (septet), m (multiplet) or (br) broad, or combinations thereof. Homonuclear and heteronuclear two dimensional NMR experiments were used where appropriate to facilitate assignment of chemical shifts. Coupling constant values are given in Hz. IR spectra (4000-400 cm^–1^) were collected on Alpha Bruker FT-IR Spectrometer (Platinum ATR); solid samples were measured neat and oily samples as films. High-resolution mass spectra were obtained on Agilent 6224 Accurate-Mass TOF LC-MS with dual electrospray/chemical ionization mode or on MALDI-TOF Ultraflextreme (Bruker Daltonics) with positive ions detection. Melting points were determined with Stuart SMP40 automatic melting point apparatus. Microwave reactions were carried out using a CEM Discover SP microwave reactor equipped with a single-mode cavity and a pressure- and temperature-controlled system. Reactions were performed in sealed microwave tubes (10 mL or appropriate size) under magnetic stirring. The purity of the synthesized target compounds were determined by HPLC analysis with UV detection (Ultimate 3000 LC Analytical Systems, Thermo Scientific) using Agilent ZORBAX Eclipse Plus C18 column and ^1^H-NMR. All final compounds reported herein were >95% pure (unless stated otherwise).

### General procedure A1: Formation of ketonitrile by deprotonation of phenylacetonitrile with NaH

A solution of the appropriate phenylacetonitrile (1 eq.; unless stated otherwise) in anhydrous THF (3 mL per 1 mmol of phenylacetonitrile) was added under N_2_ to NaH (60% suspension in mineral oil, 2 eq.; unless stated otherwise). The mixture was stirred for 20 min, then the appropriate ester (1.1 eq.; unless stated otherwise) was added and the reaction mixture was stirred at 25 °C for additional 2 h. The mixture was cooled to 0 °C and saturated aqueous solution of NH_4_Cl (6 mL per 1 mmol of phenylacetonitrile) was added. The mixture was extracted with EtOAc (3 × 9 mL per 1 mmol of phenylacetonitrile). The organic extracts were combined, dried over MgSO_4_, filtered, and the solvent was evaporated. The residue obtained after the workup was purified using column chromatography or preparative TLC (unless stated otherwise).

### General procedure A2: Formation of ketonitrile by deprotonation of CH_3_CN with NaH

A solution of the appropriate ester (1 eq.; unless stated otherwise) and CH_3_CN (3 eq.; unless stated otherwise) in anhydrous THF (1 mL per 1 mmol of ester) was added under nitrogen to NaH (60% suspension in mineral oil, 3 eq.; unless stated otherwise) and the mixture was refluxed for 4 h. The reaction mixture was cooled to 0 °C, quenched with saturated aqueous solution of NH_4_Cl (2 mL per 1 mmol of ester), and extracted with EtOAc (3 ×2 mL per 1 mmol of ester). The organic extracts were combined, dried over MgSO_4_, filtered, and the solvent was evaporated. The residue obtained after the workup was purified using column chromatography or preparative TLC (unless stated otherwise).

### General procedure A3: Formation of ketonitrile by deprotonation of CH_3_CN with *n*-BuLi

*n-*BuLi solution in hexane (2.7 M, 1.05 eq.; unless stated otherwise) was added under nitrogen to a solution of CH_3_CN (1 eq.; unless stated otherwise) in anhydrous THF (3 mL per 1 mmol of ester) at -78 °C. The reaction mixture was stirred at -78 °C for 30 min, then a solution of the appropriate ester (1 eq.; unless stated otherwise) in anhydrous THF (6 mL per 1 mmol of ester) was added dropwise and the mixture was stirred at -78 °C for 2 h. The reaction mixture was quenched with saturated aqueous solution of NH_4_Cl (6 mL per 1 mmol of ester) and extracted with dichloromethane (3 × 50 mL per 1 mmol of ester). The organic extracts were combined, dried over MgSO_4_, filtered, and the solvent was evaporated. The residue obtained after the workup was purified using column chromatography or preparative TLC (unless stated otherwise).

### General procedure A4: Formation of ketonitrile by deprotonation of alkylacetonitrile with LDA

*n-*BuLi solution in hexane (2.5 M, 1.2 eq.; unless stated otherwise) was added under nitrogen to a solution of diisopropylamine (1.3 eq.; unless stated otherwise) in anhydrous THF (1.5 mL per 1 mmol of alkylacetonitrile) at -78 °C. The reaction mixture was stirred at -78 °C for 30 min and then added to a solution of alkylacetonitrile (1 eq.; unless stated otherwise) in anhydrous THF (1.5 mL per 1 mmol of alkylacetonitrile) at -78 °C. The reaction mixture was stirred at -78 °C for 30 min and the appropriate acylchloride (1.6 eq.; unless stated otherwise) was added at -78 °C. The mixture was allowed to warm to 25 °C and stirred for 18 h. The reaction mixture was quenched with saturated aqueous solution of NH_4_Cl (15 mL per 1 mmol of alkylacetonitrile) and extracted with EtOAc (3 × 15 mL per 1 mmol of alkylacetonitrile). The organic extracts were combined, dried over MgSO_4_, filtered, and the solvent was evaporated. The residue obtained after the workup was purified using column chromatography or preparative TLC (unless stated otherwise).

### General procedure A5: Formation of ketonitrile by deprotonation of CH_3_CN with NaH

NaH (60% suspension in mineral oil, 2 eq) was added to a solution of acetonitrile (1.2 eq) in anhydrous THF (10 mL per 0.62 mmol of ester) under nitrogen and stirred at room temperature for 15 min. Then a solution of appropriate ester (1.0 eq) in THF (5.0 mL per 0.62 mmol of ester) was added and the reaction mixture was refluxed for 2 to 48 h. The reaction mixture was cooled to 0°C, quenched with 1 N aqueous HCl solution (20 mL per 6.20 mmol of ester) and extracted with EtOAc (3 × 15 mL per 6.2 mmol of ester). The organic extracts were washed with brine (10 mL per 6.20 mmol of ester), dried over MgSO_4_, filtered, and the solvent was evaporated *in vacuo*. The crude material was purified by column chromatography on silica gel.

### General procedure B1: Formation of aminopyrazole using hydrazine hydrate and methanesulfonic acid

A mixture of appropriate ketonitrile (1 eq; unless stated otherwise), N_2_H_4_.H_2_O (64% in H_2_O, 1.3 eq.; unless stated otherwise) and CH_3_SO_3_H (0.1 eq.; unless stated otherwise) in absolute EtOH (5 mL per 1 mmol of the substrate) was refluxed for 4 h (unless stated otherwise). The solvent was evaporated *in vacuo* and the residue was quenched with saturated aqueous solution of NaHCO_3_ (20 mL per 1 mmol of the substrate) and extracted with EtOAc (20 mL per 1 mmol of the substrate) (unless stated otherwise). The organic extracts were washed with brine (2 × 20 mL per 1 mmol of the substrate), dried over MgSO_4_, filtered, and the solvent was removed *in vacuo*. The crude material was purified by flash column chromatography on silica gel.

### General procedure B2: Formation of aminopyrazole using arylhydrazine and methanesulfonic acid

A mixture of appropriate ketonitrile (1 to 1.4 eq) and arylhydrazine (1.0 eq; unless stated otherwise) and CH_3_SO_3_H (0.1 eq.; unless stated otherwise) in absolute EtOH (2 mL per 0.29 mmol) was refluxed for 4 (unless stated otherwise). The reaction mixture was cooled to 25 °C, concentrated *in vacuo* (unless mentioned otherwise), and the residue was purified by flash column chromatography on silica gel (unless mentioned otherwise).

### General procedure C: Diazotization of aminopyrazoles and cyclization to pyrazolotriazines

Diazotization step:

Appropriate aminopyrazole (1 eq.; unless stated otherwise) was dissolved in EtOH (5 mL per 1 mmol of aminopyrazole) and H_2_O (1.67 mL per 1 mmol of aminopyrazole), and 35% aqueous HCl (4 eq.; unless stated otherwise) was added. The solution was cooled to -10 °C and a pre-cooled solution (0 °C) of NaNO_2_ (2 eq.; unless stated otherwise) in H_2_O (1 mL per 1 mmol of aminopyrazole) was added. The reaction mixture turned yellow and was stirred for 20 min at -5 °C, then a pre-cooled solution (- 5 °C) of KOAc (8 eq.; unless stated otherwise) and 2-(cyanomethyl)-benzimidazole (or alkyl benzimidazole-2-acetate) (1.05 eq.; unless stated otherwise) in EtOH (5 mL per 1 mmol of aminopyrazole) and H_2_O (1.67 mL per 1 mmol of aminopyrazole) was added. The resulting mixture was allowed to warm up to 25 °C and stirred from 1-16 h. Cold H_2_O (5 mL per 1 mmol of aminopyrazole) was added and the precipitate was filtered and washed with water (2.5 mL per 1 mmol of aminopyrazole). The product was dried under vacuum to yield a solid, which was used directly in the next step without additional purification.

Cyclization step:

The dried solid was dissolved in anhydrous DMF (5 mL per 1 mmol of aminopyrazole), KOAc (0.05 eq.; unless stated otherwise) was added and the mixture was refluxed under N_2_ for 3 h, unless stated otherwise. The solution was poured into water (5 mL per 1 mmol of aminopyrazole), the precipitate was collected by filtration, dissolved in dioxane (2.5 mL per 1 mmol of aminopyrazole) at 50 °C, and the solution was poured into water (10 mL per 1 mmol of aminopyrazole). The precipitate was collected by filtration, washed with water (5 mL per 1 mmol of aminopyrazole), then with Et_2_O (2.5 mL per 1 mmol of aminopyrazole) and dried under vacuum to yield the product.

### General procedure D: Suzuki reaction

Degassed dioxane (4 mL per 0.1 mmol of arylhalide) and H_2_O (1 mL per 0.1 mmol of arylhalide) were added to a mixture of arylhalide (1 eq.; unless stated otherwise), appropriate boronic acid or ester (1.2 eq.; unless stated otherwise), Pd(dppf)Cl_2_ (0.05 eq.; unless stated otherwise) and K_3_PO_4_ (4 eq.: unless stated otherwise). The mixture was stirred under N_2_ at 120 °C for 4 h (unless stated otherwise). The reaction mixture was quenched with saturated aqueous solution of NH_4_Cl (10 mL per 0.1 mmol of arylhalide) and extracted with EtOAc (3 × 10 mL per 0.1 mmol of arylhalide). The organic extracts were combined, dried over MgSO_4_, filtered, and the solvent was evaporated. The residue obtained after the workup was purified using column chromatography or preparative TLC (unless stated otherwise).

### General procedure E: Preparation of diketo ester using *t*-BuOK and diethyl oxalate

*t-*BuOK (1.0 M in THF, 1.2 eq) was added dropwise to a solution of appropriate ketone (1.0 eq) and diethyl oxalate (1.1 eq) in anhydrous toluene (10 mL per 7.92 mmol of ketone) at room temperature under nitrogen atmosphere. The reaction mixture was stirred at room temperature for 16 h. The reaction mixture was acidified (pH~3-4) by the careful addition of aqueous 1 N HCl (20 mL per 7.92 mmol of ketone) and extracted with EtOAc (3 × 20 mL per 7.92 mmol of ketone). The organic extracts were washed with water (3 × 15 mL per 7.92 mmol of ketone) followed by brine (3 × 15 mL per 7.92 mmol of ketone), dried over MgSO_4_, filtered, and the solvent was evaporated *in vacuo*. The crude material was purified by column chromatography on silica gel (unless mentioned otherwise).

### General procedure F: Preparation of isoxazole esters

NH_2_OH•HCl (1.2 to 1.5 eq) was added to a solution of appropriate diketoester (1.0 eq) in EtOH/methanol (16 mL/ 6.93 mmol of diketo ester) and the mixture was stirred at 50 °C for 16 h. The solvent was evaporated *in vacuo* and the residue was purified by column chromatography on silica gel.

### General procedure G: Hydrolysis of isoxazole and isothiazole esters to corresponding carboxylic acids

NaOH (2.0 M in H_2_O, 1.5 eq) was added to a solution of appropriate isoxazole ester (1.0 eq) in EtOH (15 mL per 6.76 mmol of isoxazole ester) at room temperature and the mixture was stirred for 1 h. The reaction mixture was acidified (pH~3-4) by the careful addition of aqueous 1 N HCl and extracted with EtOAc (3 × 20 mL). The combined organic extracts were washed with water (3 × 20 mL), brine (20 mL), dried over MgSO_4_, filtered, and the solvent was evaporated *in* vacuo to provide corresponding isoxazole acid.

### **General procedure H: Preparation of acid chlorides**

SOCl_2_ (2.0 eq to 10 volumes) and appropriate acid (1.0 eq) were mixed and under nitrogen atmosphere and the mixture was refluxed for 20-240 minutes. The excess of SOCl_2_ was evaporated *in vacuo*. The obtained residue was treated with toluene (2 × 2 mL/0.53 mmol of acid), the solvents were evaporated and the resulting product was dried *in vacuo*.

### General procedure I: Preparation of amides using acid chlorides and trimethylamine

Et_3_N (1.0 to 2 eq.; unless stated otherwise) was added to a mixture of appropriate amine (1.0 eq; unless stated otherwise) in acetonitrile (0.5 mL to 1.0 mL per 0.074 mmol of amine reacted). The reaction mixture was heated to reflux, a solution of appropriate acid chloride (1.0 to 2.0 eq; unless stated otherwise) in acetonitrile (0.4 to 0.5 mL 0.074 mmol of acid chloride) was added dropwise, and the resulting mixture was refluxed for additional 2-5 h. The reaction mixture was cooled to room temperature, diluted with saturated aqueous NaHCO_3_ solution (3 mL per 0.074 mmol of amine reacted) and filtered. The obtained solid was washed with water (3 mL per 0.074 mmol of amine reacted) and then with a mixture of EtOAc:hexane (1:4, 2 mL per 0.074 mmol of amine reacted) to obtained pure compound (unless stated otherwise).

### Experimental procedures for compound 1 and its analogs

2-(4-Fluorophenyl)-3-oxopentanenitrile **(S1)**

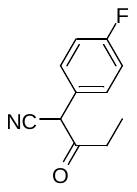

The compound was prepared according to General procedure A1 using 4-fluorophenylacetonitrile (1 mL, 8.33 mmol; 1 eq.), NaH (60% suspension in mineral oil; 667 mg, 16.66 mmol; 2 eq.), methyl propionate (0.8 mL, 8.33 mmol, 1 eq.) and THF (24 mL). Reaction time: 15 min for the formation of the sodium salt, then additional 1 h 30 min for the alkylation step. The residue obtained after the workup was purified by column chromatography on silica gel (hexane:EtOAc, gradient 1:0 to 1:1). The product was obtained as an orange oil (2.92 g, 92%).

^1^H NMR (500 MHz, Chloroform-*d*) *δ* (ppm) 7.40 – 7.34 (m, 2H), 7.15 – 7.09 (m, 2H), 4.67 (s, 1H), 2.74 – 2.52 (m, 2H), 1.05 (t, *J* = 7.2 Hz, 3H).

^13^C NMR (126 MHz, Chloroform-*d*) *δ* (ppm) 199.35, 163.24 (d, *J* = 249.8 Hz), 129.96 (d, *J* = 8.2 Hz), 125.92 (d, *J* = 3.5 Hz), 116.81 (d, *J* = 21.9 Hz), 116.36, 49.80, 33.40, 7.71.

HRMS (APCI): calcd. for C_11_H_9_FNO [M-H]^-^ = 190.0674, found [M-H]^-^ = 190.0672.

3-(3-Bromophenyl)-3-oxopropanenitrile **(S2)**

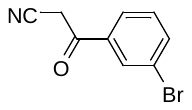

The compound was prepared according to General procedure A3 using *n-*BuLi solution in hexane (2.36 M, 2.27 mL, 5.35 mmol, 2 eq.), CH_3_CN (0.42 mL, 8.02 mmol, 3 eq.) and THF (5 mL) (reaction time: 30 min for the formation of the lithium salt) and then using methyl 3-bromobenzoate (575 mg, 2.67 mmol; 1 eq.) and THF (5 mL) (reaction time: 2 h for the alkylation step). The residue obtained after the workup was purified by column chromatography on silica gel (hexane:EtOAc, gradient 5:1 to 1:3). The product was obtained as a beige solid (590 mg, 99%).

^1^H NMR (500 MHz, Chloroform-*d*) *δ* (ppm) 8.05 (dd, *J* = 1.9 Hz, 1H), 7.84 (ddd, *J* = 7.8, 1.8, 1.0 Hz, 1H), 7.79 (ddd, *J* = 8.0, 2.0, 1.0 Hz, 1H), 7.42 (dd, *J* = 7.9 Hz, 1H), 4.06 (s, 2H).

^13^C NMR (126 MHz, Chloroform-*d*) *δ* (ppm) 186.03, 137.76, 136.07, 131.61, 130.86, 127.11, 123.68, 113.37, 29.58.

3-Cyclohexyl-3-oxopropanenitrile **(S3)**

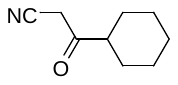

The compound was prepared according to General procedure A3 using *n-*BuLi (2.36 M, 2.57 mL, 6.08 mmol, 2 eq.), CH_3_CN (0.48 mL, 9.11 mmol, 3 eq.) in THF (5 mL) and methyl cyclohexanecarboxylate (432 mg, 3.04 mmol; 1 eq.) in THF (5 mL). Reaction time: 30 min (deprotonation step) and 1 h (alkylation step). The residue obtained after workup was purified by column chromatography on silica gel (hexane:EtOAc, gradient 1:0 to 1:1). The product was obtained as a yellow oil (460 mg, 99%).

^1^H NMR (500 MHz, Chloroform-*d*) *δ* (ppm) 3.48 (s, 2H), 2.55 (tt, *J* = 11.1, 3.4 Hz, 1H), 1.94 – 1.86 (m, 2H), 1.84 – 1.77 (m, 2H), 1.72 – 1.65 (m, 1H), 1.44 – 1.15 (m, 5H).

^13^C NMR (126 MHz, Chloroform-*d*) *δ* (ppm) 200.49, 114.05, 50.13, 30.38, 28.29, 25.65, 25.38.

HRMS (APCI): calcd. for C_9_H_17_N_2_O [M+NH_4_]^+^ = 169.1335, found [M+NH_4_]^+^ = 169.1340.

HRMS (APCI): calcd. for C_9_H_12_NO [M-H]^−^ = 150.0924, found [M-H]^−^ = 150.0923.

3-Oxo-3-(2-phenoxyphenyl)propanenitrile **(S4)**

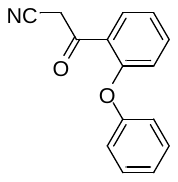

The compound was prepared according to General procedure A3 using *n-*BuLi (2.36 M, 1.92 mL, 4.52 mmol, 2 eq.), CH_3_CN (0.35 mL, 6.78 mmol, 3 eq.) in THF (5 mL) and methyl 2-phenoxybenzoate (516 mg, 2.26 mmol; 1 eq.) in THF (5 mL). Reaction time: 30 min (deprotonation step) and 1 h (alkylation step). The residue obtained after workup was purified by column chromatography on silica gel (hexane:EtOAc, gradient 1:0 to 1:1). The product was obtained as a yellow solid (528 mg, 99%).

^1^H NMR (500 MHz, Chloroform-*d*) *δ* (ppm) 7.96 (dd, *J* = 7.9, 1.8 Hz, 1H), 7.49 (ddd, *J* = 8.4, 7.3, 1.8 Hz, 1H), 7.46 – 7.41 (m, 2H), 7.29 – 7.22 (m, 1H), 7.19 (ddd, *J* = 8.1, 7.3, 1.1 Hz, 1H), 7.12 – 7.07 (m, 2H), 6.85 (dd, *J* = 8.4, 1.0 Hz, 1H), 4.16 (s, 2H).

^13^C NMR (126 MHz, Chloroform-*d*) *δ* (ppm) 187.80, 157.69, 154.89, 135.63, 131.58, 130.57, 126.26, 125.37, 123.51, 120.09, 117.96, 114.40, 34.15.

HRMS (APCI): calcd. for C_15_H_12_NO_2_ [M+H]^+^ = 238.0863, found [M+H]^+^ = 238.0865.

3-(Naphthalen-2-yl)-3-oxopropanenitrile **(S5)**

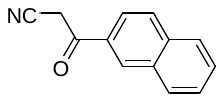

The compound was prepared according to General procedure A3 using *n-*BuLi (2.36 M, 2.37 mL, 5.61 mmol, 2 eq.), CH_3_CN (0.44 mL, 8.41 mmol, 3 eq.) in THF (5 mL) and methyl 2-naphthoate (522 mg, 2.80 mmol; 1 eq.) in THF (10 mL). Reaction time: 30 min (deprotonation step) and 1 h (alkylation step). The residue obtained after workup was purified by column chromatography on silica gel (hexane:EtOAc, gradient 1:0 to 1:1). The product was obtained as an off-white solid (540 mg, 99%).

^1^H NMR (300 MHz, Chloroform-*d*) *δ* (ppm)) δ 8.18 (t, *J* = 1.8 Hz, 2H), 7.98 (t, *J* = 1.3 Hz, 1H), 7.96 (t, *J* = 1.3 Hz, 1H), 7.70 (dd, *J* = 2.0, 1.1 Hz, 1H), 7.67 (dd, *J* = 2.1, 1.1 Hz, 1H), 7.32 (t, *J* = 7.9 Hz, 2H).

^13^C NMR (126 MHz, Chloroform-*d*) *δ* (ppm) 187.09, 136.34, 132.46, 131.83, 130.87, 129.89, 129.68, 129.38, 128.10, 127.58, 123.56, 113.97, 29.56.

HRMS (APCI): calcd. for C_13_H_8_NO [M-H]^-^ = 194.0611, found [M-H]^-^ = 194.0605.

2-([1,1'-Biphenyl]-3-yl)-3-oxopentanenitrile **(S6)**

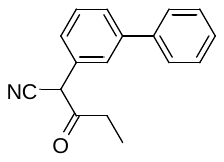

This compound was synthesized according to the procedure reported in *J. Org. Chem.* 2018, 83, 24, 15380–15405.

4-Methyl-3-oxopentanenitrile **(S7)**

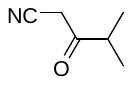

The compound was prepared according to General procedure A2 using sodium hydride (60% suspension in mineral oil, 1.40 g, 39.42 mmol, 2 eq.), CH_3_CN (2.7 mL, 59.13 mmol, 3 eq.) in THF (25 mL) and methyl isobutyrate (2.25 mL, 19.71 mmol; 1 eq.). Reaction time: 6 h. The residue obtained after workup was purified by column chromatography on silica gel (hexane:EtOAc, 4:1). The product was obtained as yellow oil (1.89 g, 99%).

^1^H NMR (500 MHz, Chloroform-*d*) *δ* (ppm) 3.51 (s, 2H), 2.81 (hept, *J* = 6.9 Hz, 1H), 1.18 (d, *J* = 6.9 Hz, 6H).

^13^C NMR (126 MHz, Chloroform-*d*) *δ* (ppm) 201.23, 113.98, 40.68, 30.18, 17.98.

HRMS (APCI): calcd. for C_6_H_8_NO [M-H]^-^ = 110.0611, found [M-H]^-^ = 110.1618.

4,4-Dimethyl-3-oxopentanenitrile **(S8)**

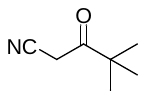

The compound was prepared according to General procedure A2 using sodium hydride (60% suspension in mineral oil, 1.40 g, 33.82 mmol, 2 eq.), CH_3_CN (2.7 mL, 50.73 mmol, 3 eq.) in THF (25 mL) and methyl pivalate (2.25 mL, 16.91 mmol; 1 eq.). Reaction time: 6 h. The residue obtained after workup was purified by column chromatography on silica gel (hexane:EtOAc, 4:1). The product was obtained as a white solid (2.40 g, 93%).

^1^H NMR (500 MHz, Chloroform-*d*) *δ* (ppm) 3.62 (s, 2H), 1.19 (s, 9H).

^13^C NMR (126 MHz, Chloroform-*d*) *δ* 202.87, 114.24, 44.79, 27.58, 26.26.

HRMS (APCI): calcd. for C_7_H_10_NO [M-H]^-^ = 124.0768, found [M-H]^-^ = 124.0774.

2-(4-Fluorophenyl)-4-methyl-3-oxopentanenitrile **(S9)**

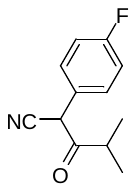

The compound was prepared according to General procedure A1 using 4-fluorophenylacetonitrile (0.2 mL, 1.31 mmol; 1 eq.), NaH (60% suspension in mineral oil; 105 mg, 2.62 mmol; 2 eq.), methyl isobutyrate (0.18 mL, 1.57 mmol, 1.2 eq.) and THF (4 mL). Reaction time: 15 min for the formation of the sodium salt, then additional 1.5 h for the alkylation step. The residue obtained after the workup was purified by column chromatography on silica gel (hexane:EtOAc, gradient 1:0 to 4:1). The product was obtained as a white solid (266 mg, 83%).

^1^H NMR (500 MHz, Chloroform-*d*) *δ* (ppm) 7.42 – 7.33 (m, 2H), 7.16 – 7.09 (m, 2H), 4.77 (s, 1H), 2.94 (hept, *J* = 6.8 Hz, 1H), 1.11 (dd, *J* = 6.9, 4.5 Hz, 6H).

^13^C NMR (126 MHz, Chloroform-*d*) *δ* (ppm) 202.47, 163.23 (d, *J* = 249.6 Hz), 130.16 (d, *J* = 8.5 Hz), 125.83 (d, *J* = 3.7 Hz), 116.79 (d, *J* = 22.1 Hz), 116.33, 48.40, 38.92, 18.97, 18.70.

HRMS (APCI): calcd. for C_12_H_11_FNO [M-H]^-^ = 204.0830, found [M-H]^-^ = 204.0838.

3-(3-Methoxynaphthalen-2-yl)-3-oxopropanenitrile **(S10)**

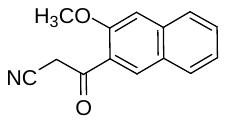

The compound was prepared according to General procedure A2 using sodium hydride (60% suspension in mineral oil, 136 mg, 3.40 mmol, 2 eq.), CH_3_CN (0.25 mL, 5.10 mmol, 3 eq.) in THF (5 mL) and methyl 3-methoxy-2-naphthoate (365 mg, 1.70 mmol; 1 eq.). Reaction time: 6 h. The residue obtained after workup was purified by column chromatography on silica gel (hexane:EtOAc, 5:2). The product was obtained as yellow oil (100 mg, 26%).

^1^H NMR (500 MHz, Chloroform-*d*) *δ* (ppm) 8.37 (s, 1H), 7.88 (d, *J* = 8.2 Hz, 1H), 7.76 (d, *J* = 8.3 Hz, 1H), 7.57 (ddd, *J* = 8.2, 6.9, 1.2 Hz, 1H), 7.41 (ddd, *J* = 8.1, 6.9, 1.1 Hz, 1H), 7.23 (s, 1H), 4.16 (s, 2H), 4.06 (s, 3H).

^13^C NMR (126 MHz, Chloroform-*d*) *δ* (ppm) 188.79, 155.26, 137.19, 133.51, 129.84, 129.64, 128.08, 126.65, 126.22, 125.20, 114.61, 107.01, 55.95, 34.14.

3-([1,1'-Biphenyl]-3-yl)-3-oxopropanenitrile **(S11)**

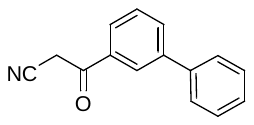

The compound was prepared according to General procedure A2 using sodium hydride (60% suspension in mineral oil, 188 mg, 4.72 mmol, 2 eq.), CH_3_CN (0.25 mL, 4.72 mmol, 2 eq.) in THF (5 mL) and methyl[1,1'-biphenyl]-3-carboxylate (500 mg, 2.36 mmol; 1 eq.). Reaction time: 16 h. The residue obtained after workup was purified by column chromatography on silica gel (hexane:EtOAc, 4:1). The product was obtained as yellow oil (0.44 g, 76%).

^1^H NMR (500 MHz, Chloroform-*d*) *δ* (ppm) ^1^H NMR (500 MHz, CDCl_3_) δ 7.65 – 7.58 (m, 2H), 7.54 – 7.44 (m, 5H), 7.39 – 7.34 (m, 2H), 3.20 (s, 2H).

^13^C NMR (126 MHz, Chloroform-*d*) *δ* (ppm) 193.68, 140.99, 139.69, 137.54, 132.43, 130.62, 129.54, 129.02, 128.10, 113.75, 32.25.

HRMS (APCI): calcd. for C_15_H_10_NO [M-H]^-^ = 220.0768, found [M- H]^-^ = 220.0765.

3-(1-Methoxynaphthalen-2-yl)-3-oxopropanenitrile **(S12)**

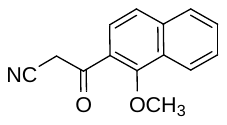

The compound was prepared according to General procedure A3 using *n-*BuLi (2.7 M, 1.03 mL, 7.76 mmol, 1.2 eq.), CH_3_CN (0.25 mL, 4.6 mmol, 2 eq.) in THF (4 mL) and methyl 1-methoxy-2-naphthoate (500 mg, 2.30 mmol; 1 eq.) in THF (4 mL). Reaction time: 30 min (deprotonation step) and 1 h (alkylation step). The residue obtained after workup was purified by column chromatography on silica gel (hexane:EtOAc, 5:1). The product was obtained as a yellow oil (260 mg, 50%).

^1^H NMR (500 MHz, Chloroform-*d*) *δ* (ppm) 8.20 (d, *J* = 8.1 Hz, 1H), 7.89 (d, *J* = 7.5 Hz, 1H), 7.78 (d, *J* = 8.7 Hz, 1H), 7.67 (d, *J* = 8.5 Hz, 1H), 7.66 – 7.57 (m, 2H), 4.27 (s, 2H), 4.06 (s, 3H).

^13^C NMR (126 MHz, Chloroform-*d*) *δ* (ppm) 188.79, 158.84, 137.94, 129.45, 128.54, 127.64, 127.27, 125.23, 125.11, 125.07, 123.64, 114.47, 64.53, 33.18.

3-(3-Methoxynaphthalen-1-yl)-3-oxopropanenitrile **(S13)**

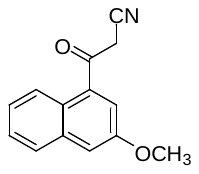

The compound was prepared according to General procedure A3 using *n-*BuLi (2.7 M, 1 mL, 2.87 mmol, 1.05 eq.), CH_3_CN (0.28 mL, 5.46 mmol, 2 eq.) in THF (4 mL) and methyl 3-methoxy-1-naphthoate (590 mg, 2.73 mmol; 1 eq.) in THF (4 mL). Reaction time: 30 min (deprotonation step) and 1 h (alkylation step). The residue obtained after workup was purified by column chromatography on silica gel (hexane:EtOAc, 5:2). The product was obtained as a yellow oil (210 mg, 36%).

^1^H NMR (500 MHz, Chloroform-*d*) *δ* (ppm) 8.00 (d, *J* = 9.0 Hz, 1H), 7.97 – 7.93 (m, 1H), 7.81 (d, *J* = 8.2 Hz, 1H), 7.54 (ddd, *J* = 8.5, 6.9, 1.4 Hz, 1H), 7.41 (ddd, *J* = 8.0, 6.8, 1.1 Hz, 1H), 7.30 (d, *J* = 9.1 Hz, 1H), 4.05 (s, 3H), 4.03 (s, 2H).

^13^C NMR (126 MHz, Chloroform-*d*) *δ* (ppm) 192.52, 156.41, 134.48, 130.95, 129.11, 128.97, 128.54, 124.85, 123.67, 120.33, 114.65, 112.25, 56.64, 34.39.

Methyl 2-((tert-butyldiphenylsilyl)oxy)acetate **(S14)**

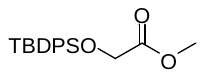

Under nitrogen atmosphere, methyl 2-hydroxyacetate (2.3 g, 25.70 mmol; 1eq.) was dissolved in dry dichloromethane (30 mL). To the solution was added Et_3_N (7.2 mL, 51.4 mmol; 2 eq.), DMAP (1.7 g, 12.50 mmol; 0.5 eq.) and TBDPSCl (7.3 mL, 28.27 mmol; 1.1 eq.), and the reaction mixture was stirred for 16 h at room temperature. The mixture was poured into water (50 mL) and extracted with dichloromethane (3 ×25 mL). The combined organic extracts were washed with hydrochloric acid (10%, 2 × 75 mL), brine (75 mL) and dried over MgSO_4_. The solvent was evaporated, and the product was dried under *in vacuo*. The obtained colorless oil (8.2 g, 98%) was used in the next step without further purification.

^1^H NMR (500 MHz, Chloroform-*d*) *δ* (ppm) 7.71 – 7.68 (m, 4H), 7.46 – 7.41 (m, 2H), 7.42 – 7.37 (m, 4H), 4.25 (s, 2H), 3.69 (s, 3H), 1.10 (s, 9H).

^13^C NMR (126 MHz, Chloroform-*d*) *δ* (ppm) 171.80, 135.74, 130.04, 127.93, 62.30, 51.77, 26.83, 19.41.

4-((*tert*-Butyldiphenylsilyl)oxy)-3-oxobutanenitrile **(S15)**

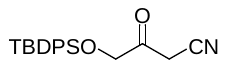

The compound was prepared according to General procedure A3 using *n-*BuLi (2.7 M, 0.6 mL, 1.6 mmol, 1.1 eq.), CH_3_CN (0.12 mL, 2.28 mmol, 1.5 eq.) in THF (5 mL) and methyl 2-((*tert*-butyldiphenylsilyl)oxy)acetate (500 mg, 1.52 mmol; 1 eq.) in THF (5 mL). Reaction time: 30 min (deprotonation step) and 1 h (alkylation step). The residue obtained after workup was purified by column chromatography on silica gel (hexane:EtOAc, 5:1). The product was obtained as a colorless oil (370 mg, 71%).

^1^H NMR (500 MHz, Chloroform-*d*) *δ* (ppm) 7.65 – 7.60 (m, 4H), 7.50 – 7.46 (m, 2H), 7.45 – 7.40 (m, 4H), 4.27 (s, 2H), 3.73 (s, 2H), 1.12 (s, 9H).

^13^C NMR (126 MHz, Chloroform-*d*) *δ* (ppm) 198.24, 135.57, 131.78, 130.56, 128.28, 113.45, 69.15, 29.50, 26.88, 19.28.

HRMS (APCI): calcd. for C_20_H_22_NO_2_Si [M-H]^-^ = 336.1425, found [M-H]^-^ = 336.1429.

3-(1-Benzyl-1*H*-imidazol-2-yl)-3-oxopropanenitrile **(S16)**

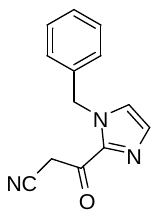

The compound was prepared according to General procedure A2 using sodium hydride (60%, 105 mg, 2.62 mmol, 2 eq.), CH_3_CN (0.2 mL, 3.9 mmol, 3 eq.) in THF (5 mL) and methyl 1-benzyl-1*H*-imidazole-2-carboxylate (280 mg, 1.30 mmol; 1 eq.). Reaction time: 2 h. The residue obtained after workup was purified by column chromatography on silica gel (hexane:EtOAc, 0:1 to 1:0). The product was obtained as yellow oil (220 mg, 76%).

^1^H NMR (500 MHz, Chloroform-*d*) *δ* (ppm) 7.42 – 7.28 (m, 3H), 7.23 – 7.16 (m, 4H), 5.60 (s, 2H), 4.30 (d, *J* = 0.8 Hz, 2H).

3-(2-chlorophenyl)-3-oxopropanenitrile **(S17)**

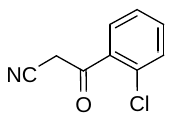

The compound was prepared according to General procedure A2 using sodium hydride (60%, 1.2 g, 30 mmol, 2 eq.), CH_3_CN (2.4 mL, 45 mmol, 3 eq.) in THF (25 mL) and methyl 2-chlorobenzoate (2.55 g, 15 mmol; 1 eq.). Reaction time: 4 h. The residue obtained after workup was purified by column chromatography on silica gel (hexane:EtOAc, gradient 4:1 to 2:1). The product was obtained as a yellow solid (2.26 g, 84%).

^1^H NMR (500 MHz, Chloroform-*d*) *δ* (ppm) 7.64 (dd, *J* = 7.7, 1.4 Hz, 1H), 7.54 – 7.45 (m, 2H), 7.44 – 7.30 (m, 1H), 4.14 (s, 2H).

^13^C NMR (126 MHz, Chloroform-*d*) *δ* (ppm) 189.50, 135.86, 133.88, 131.85, 131.14, 130.57, 127.63, 113.45, 33.05.

HRMS (APCI): calcd. for C_9_H_5_ClON [M-H]^-^ = 178.0065, found [M-H]^-^ = 178.0067.

3-(Naphthalen-1-yl)-3-oxopropanenitrile **(S18)**

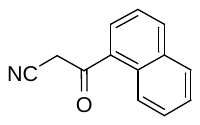

The compound was prepared according to General procedure A3 using *n-*BuLi (2.7 M, 1 mL, 2.82 mmol, 1.05 eq.), CH_3_CN (0.18 mL, 3.50 mmol, 1.3 eq.) in THF (4 mL) and methyl 1-naphthoate (500 mg, 2.69 mmol; 1 eq.) in THF (4 mL). Reaction time: 30 min (deprotonation step) and 1 h (alkylation step). The residue obtained after workup was purified by column chromatography on silica gel (hexane:EtOAc, 2:1). The product was obtained as a yellow solid (310 mg, 59%).

^1^H NMR (500 MHz, Chloroform-*d*) *δ* (ppm) 8.81 (d, *J* = 8.7 Hz, 1H), 8.11 (d, *J* = 8.2 Hz, 1H), 7.91 (t, *J* = 8.2 Hz, 1H), 7.71 – 7.65 (m, 1H), 7.63 – 7.57 (m, 1H), 7.56 – 7.52 (m, 1H), 4.19 (s, 1H).

^13^C NMR (126 MHz, Chloroform-*d*) *δ* (ppm) 189.60, 135.34, 134.26, 131.70, 130.51, 129.63, 129.32, 128.85, 127.29, 125.77, 124.31, 114.16, 31.90.

HRMS (APCI): calcd. for C_13_H_8_NO [M-H]^-^ = 194.0611, found [M-H]^-^ = 194.0608.

3-(2-(Dimethylamino)phenyl)-3-oxopropanenitrile **(S19)**

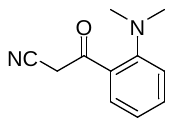

The compound was prepared according to General procedure A2 using sodium hydride (60%, 222 mg, 5.58 mmol, 2 eq.), CH_3_CN (0.44 mL, 8.37 mmol, 3 eq.) in THF (5 mL) and methyl 2-(dimethylamino)benzoate (500 mg, 2.79 mmol; 1 eq.). Reaction time: 4 h. The residue obtained after the workup was purified by column chromatography on silica gel (hexane:EtOAc, gradient 9:1 to 1:1). The product was obtained as a yellow oil (340 mg, 33%).

^1^H NMR (300 MHz, Chloroform-*d*) *δ* (ppm) 7.59 – 7.39 (m, 2H), 7.23 – 7.05 (m, 2H), 4.18 (s, 2H), 2.85 (s, 6H).

4-(2-Methoxyphenyl)-3-oxobutanenitrile **(S20)**

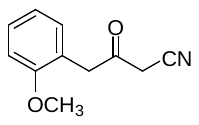

The compound was prepared according to General procedure A3 using *n-*BuLi solution in hexanes (2.5 M, 4.8 mL, 12.10 mmol, 1.3 eq.), CH_3_CN (0.73 mL, 13.98 mmol, 1.5 eq.) and THF (15 mL) (reaction time: 30 min for the formation of the lithium salt) and then using methyl 2-(2-methoxyphenyl)acetate (1.68 g, 9.32 mmol; 1 eq.) and THF (5 mL) (reaction time: 2 h for the alkylation step). The residue obtained after the workup (1.30 g, 74%) was used in the next step without further purification.

5-Methoxy-3-oxo-5-phenylpentanenitrile **(S21)**

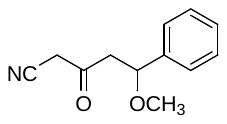

The compound was prepared according to General procedure A3 using *n-*BuLi solution in hexanes (2.5 M, 1.2 mL, 3.12 mmol, 1.3 eq.), CH_3_CN (0.19 mL, 3.60 mmol, 1.5 eq.) and THF (5 mL) (reaction time: 30 min for the formation of the lithium salt) and then using ethyl 3-methoxy-3-phenylpropanoate (500 mg, 2.40 mmol; 1 eq.) and THF (3 mL) (reaction time: 2 h for the alkylation step). The residue obtained after the workup was purified by column chromatography on silica gel (cyclohexane:EtOAc, gradient 9:1 to 1:1). The product was obtained as a colorless oil (290 mg, 60%).

^1^H NMR (500 MHz, Chloroform-*d*) *δ* (ppm) 7.41 – 7.36 (m, 2H), 7.35 – 7.29 (m, 3H), 4.61 (dd, *J* = 9.5, 3.8 Hz, 1H), 3.60 (d, *J* = 19.4 Hz, 1H), 3.49 (d, *J* = 19.5 Hz, 1H), 3.20 (s, 3H), 3.05 (dd, *J* = 15.1, 9.5 Hz, 1H), 2.70 (dd, *J* = 15.1, 3.9 Hz, 1H).

^13^C NMR (126 MHz, Chloroform-*d*) *δ* (ppm) 195.99, 139.88, 129.00, 128.58, 126.52, 113.68, 79.96, 56.97, 50.49, 33.55.

3-Oxo-4-phenylbutanenitrile **(S22)**

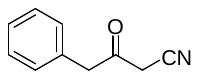

The compound was prepared according to General procedure A3 using *n-*BuLi (2.5 M, 5.68 mL, 14.23 mmol, 1.67 eq.), CH_3_CN (1.11 mL, 21.31 mmol, 2.5 eq.) in THF (12 mL) and methyl 2-phenylacetate (1.28 g, 8.52 mmol; 1 eq.) in THF (30 mL). Reaction time: 30 min (deprotonation step) and 2 h (alkylation step). The residue obtained after workup was loaded on ISOLUTE^®^ HM-N and purified by column chromatography on silica gel using Biotage Selekt purification system (cyclohexane:EtOAc, gradient 1:0 to 1:1). The product was obtained as a pale-yellow oil (1.22 g, 90%).

^1^H NMR (500 MHz, Chloroform-*d*) *δ* (ppm) 7.40 – 7.35 (m, 2H), 7.35 – 7.30 (m, 1H), 7.24 – 7.20 (m, 2H), 3.86 (s, 2H), 3.46 (s, 2H).

^13^C NMR (126 MHz, Chloroform-*d*) *δ* (ppm) 195.33, 132.07, 129.55, 129.38, 128.09, 113.70, 49.30, 31.25.

Methyl 2-(isochroman-3-yl)acetate **(S23)**

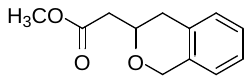

To a solution of 2-(isochroman-1-yl)acetic acid (600 mg, 3.12 mmol, 1 eq.) in MeOH (30 mL) was added conc. H_2_SO_4_ (34 µL, 0.62 mmol, 96%, 0.2 eq.) and the mixture was refluxed for 16 h. The mixture was poured into H_2_O (30 mL) and the resulting solution was extracted with dichloromethane (4 × 60 mL). The organic extracts were combined, dried over MgSO_4_, filtered, and the solvent was evaporated *in vacuo*. The residue was loaded on ISOLUTE^®^ HM-N and purified by column chromatography using Biotage Selekt purification system (cyclohexane:EtOAc, gradient 95:5 to 0:1). The product was obtained as a colorless oil (567 mg, 88%).

^1^H NMR (500 MHz, Chloroform-*d*) *δ* (ppm) 7.21 – 7.16 (m, 2H), 7.14 – 7.10 (m, 1H), 7.07 – 7.02 (m, 1H), 5.25 (dd, *J* = 9.7, 3.5 Hz, 1H), 4.13 (ddd, *J* = 11.4, 5.3, 4.3 Hz, 1H), 3.82 (ddd, *J* = 11.4, 9.0, 3.9 Hz, 1H), 3.75 (s, 3H), 3.03 – 2.94 (m, 1H), 2.89 (dd, *J* = 15.2, 3.5 Hz, 1H), 2.77 (dd, *J* = 15.2, 9.7 Hz, 1H), 2.75 – 2.69 (m, 1H).

^13^C NMR (126 MHz, Chloroform-*d*) *δ* (ppm) 171.87, 136.82, 134.09, 129.24, 126.87, 126.43, 124.62, 73.07, 63.26, 52.00, 41.72, 28.95.

HRMS (APCI): calcd. for C_12_H_15_O_3_ [M+H]^+^ = 207.1016, found [M+H]^+^ = 207.1013.

4-(Isochroman-3-yl)-3-oxobutanenitrile **(S24)**

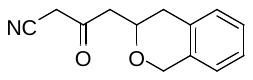

The compound was prepared according to General procedure A3 using *n-*BuLi (2.5 M, 1.36 mL, 3.40 mmol, 1.3 eq.), CH_3_CN (0.274 mL, 5.24 mmol, 2 eq.) in THF (6 mL) and methyl 2-(isochroman-3-yl)acetate (540 mg, 2.62 mmol; 1 eq.) in THF (18 mL). Reaction time: 30 min (deprotonation step) and 2 h (alkylation step). The residue obtained after the workup was loaded on ISOLUTE^®^ HM-N and purified by column chromatography on silica gel using Biotage Selekt purification system (cyclohexane:EtOAc, gradient 1:0 to 1:1). The product was obtained as a white solid (516 mg, 92%).

^1^H NMR (500 MHz, Chloroform-*d*) *δ* (ppm) 7.23 – 7.18 (m, 2H), 7.14 (dd, *J* = 5.3, 3.8 Hz, 1H), 7.06 – 6.99 (m, 1H), 5.20 (t, *J* = 6.4 Hz, 1H), 4.15 (ddd, *J* = 11.3, 5.6, 3.0 Hz, 1H), 3.78 (ddd, *J* = 11.4, 10.3, 3.6 Hz, 1H), 3.67 (d, *J* = 19.5 Hz, 1H), 3.56 (d, *J* = 19.5 Hz, 1H), 3.11 – 2.97 (m, 3H), 2.70 (dt, *J* = 16.4, 3.4 Hz, 1H).

^13^C NMR (126 MHz, Chloroform-*d*) *δ* (ppm) 196.42, 135.80, 133.86, 129.44, 127.31, 126.73, 124.43, 113.83, 73.22, 64.04, 48.51, 33.50, 28.80.

HRMS (APCI): calcd. for C_13_H_14_NO_2_ [M+H]^+^ = 216.1019, found [M+H]^+^ = 216.1020.

2-(4-Fluorophenyl)-4-(2-methoxyphenyl)-3-oxobutanenitrile **(S25)**

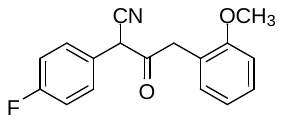

The compound was prepared according to General procedure A1 using 4-fluorophenylacetonitrile (0.82 mL, 6.84 mmol; 1.1 eq.), NaH (60% suspension in mineral oil; 497 mg, 12.43 mmol; 2 eq.), methyl 2-(2-methoxyphenyl)acetate (1 mL, 6.22 mmol, 1 eq.) and THF (10 mL). Reaction time: 15 min for the formation of the sodium salt, then an additional 1.5 h for the alkylation step. The residue obtained after the workup (1.5 g, 85%) was used in the next step without further purification.

2-Cyclohexyl-3-oxopentanenitrile **(S26)**

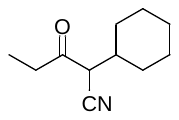

The compound was prepared according to General procedure A4 using *n-*BuLi (2.5 M, 1.15 mL, 2.88 mmol, 1.2 eq.), diisopropylamine (0.437 mL, 3.12 mmol, 1.3 eq.) and THF (3 mL) (reaction time: 30 min for the formation of LDA), then deprotonation of cyclohexylacetonitrile (295 mg, 2.40 mmol; 1 eq.) in THF (3 mL) (reaction time: 30 min) and final addition of propionyl chloride (0.346 mL, 3.84 mmol, 1.6 eq.) (reaction time: 16 h). The residue obtained after workup was loaded on ISOLUTE^®^ HM-N and purified by column chromatography on silica gel using Biotage Selekt purification system (hexane:EtOAc, gradient 1:0 to 4:1). The product was obtained as a colorless oil (319 mg, 74%) containing ca. 10 % impurities (by ^1^H NMR), was used as such in the next step.

^1^H NMR (500 MHz, Chloroform-*d*) *δ* (ppm) 2.61 (q, *J* = 7.6 Hz, 1H), 2.51 (q, *J* = 7.6 Hz, 1H), 2.27 – 2.20 (m, 1H), 1.87 – 1.59 (m, 5H), 1.44 – 1.30 (m, 1H), 1.31 – 1.13 (m, 5H), 1.12 – 1.02 (m, 2H).

HRMS (APCI): calcd. for C_11_H_18_NO [M+H]^+^ = 180.1383, found [M+H]^+^ = 180.1380.

3-Oxo-2-(tetrahydro-2*H*-pyran-4-yl)pentanenitrile **(S27)**

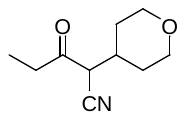

The compound was prepared according to General procedure A4 using *n-*BuLi (2.5 M, 1.15 mL, 2.88 mmol, 1.2 eq.), diisopropylamine (437 µL, 3.12 mmol, 1.3 eq.) and THF (3 mL) (reaction time: 30 min for the formation of LDA), then deprotonation of 2-(tetrahydro-2*H*-pyran-4-yl)acetonitrile (300 mg, 2.40 mmol; 1 eq.) in THF (3 mL) (reaction time: 30 min) and final addition of propionyl chloride (0.346 mL, 3.84 mmol, 1.6 eq.) (reaction time: 16 h). The residue obtained after workup was loaded on ISOLUTE^®^ HM-N and purified by column chromatography on silica gel using Biotage Selekt purification system (hexane:EtOAc, gradient 1:0 to 1:1). The product was obtained as a colorless oil (471 mg, quantitative) containing ca. 20 % impurities (by ^1^H NMR), was used as such in the next step.

^1^H NMR (500 MHz, Chloroform-*d*) *δ* (ppm) 3.99 (dd, *J* = 11.9, 4.7 Hz, 0H), 3.36 (td, *J* = 12.0, 2.1 Hz, 1H), 2.63 (q, *J* = 7.6 Hz, 1H), 2.52 (q, *J* = 7.6 Hz, 1H), 2.31 (q, *J* = 7.4 Hz, 1H), 1.78 – 1.70 (m, 1H), 1.55 – 1.48 (m, 1H), 1.38 (t, *J* = 7.0 Hz, 2H), 1.24 (t, *J* = 7.6 Hz, 1H), 1.19 (qd, *J* = 5.2, 4.6, 2.7 Hz, 2H), 1.10 (dt, *J* = 14.5, 7.5 Hz, 3H).

2-Benzyl-3-oxopentanenitrile **(S28)**

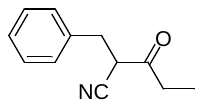

The compound was prepared according to General procedure A4 using *n-*BuLi (2.5 M, 2.3 mL, 5.75 mmol, 1.2 eq.), diisopropylamine (0.874 mL, 6.23 mmol, 1.3 eq.) and THF (3 mL) (reaction time: 30 min for the formation of LDA), then deprotonation of 3-phenylpropanenitrile (629 mg, 4.79 mmol; 1 eq.) in THF (3 mL) (reaction time: 30 min) and final addition of propionyl chloride (0.67 mL, 7.67 mmol, 1.6 eq.) Reaction time: 16 h. The residue obtained after workup was loaded on ISOLUTE^®^ HM-N and purified by column chromatography on silica gel using Biotage Selekt purification system (hexane:EtOAc, gradient 1:0 to 1:1). The product was obtained as a colorless oil (315 mg, 35%) containing ca. 10 % impurities (by ^1^H NMR), was used as such in the next step.

^1^H NMR (500 MHz, Chloroform-*d*) *δ* 7.39 – 7.16 (m, 5H), 3.43 (s, 1H), 2.96 (t, *J* = 7.4 Hz, 1H), 2.71 (q, *J* = 7.5 Hz, 1H), 2.62 (t, *J* = 7.4 Hz, 1H), 2.52 (q, *J* = 7.5 Hz, 1H), 1.18 (dt, *J* = 46.4, 7.6 Hz, 3H).

2-(Cyclohexylmethyl)-3-oxopentanenitrile **(S29)**

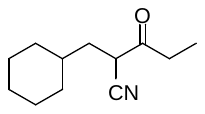

The compound was prepared according to General procedure A4 using *n-*BuLi (2.5 M, 2.6 mL, 6.57 mmol, 1.2 eq.), diisopropylamine (1.7 mL, 7.11 mmol, 1.3 eq.) and THF (3 mL) (reaction time: 30 min for the formation of LDA), then deprotonation of 3-cyclohexylpropanenitrile (750 mg, 5.47 mmol; 1 eq.) in THF (3 mL) (reaction time: 30 min) and final addition of propionyl chloride (57 µL, 6.56 mmol, 1.20 eq.) (reaction time: 16 h). The crude product obtained after the workup (1.10 g, quantitative) was used in the next step without further purification.

2-(2-Chlorobenzyl)-3-oxopentanenitrile **(S30)**

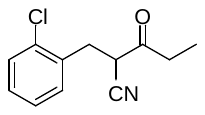

The compound was prepared according to General procedure A4 using *n-*BuLi (2.5 M, 1.4 mL, 3.62 mmol, 1.2 eq.), diisopropylamine (0.56 mL, 3.93 mmol, 1.3 eq.) and THF (3 mL) (reaction time: 30 min for the formation of LDA), then deprotonation of 3-(2-chlorophenyl)propanenitrile (500 mg, 3.01 mmol; 1 eq.) in THF (3 mL) (reaction time: 30 min) and final addition of propionyl chloride (0.32 mL, 3.62 mmol, 1.2 eq.) (reaction time: 16 h). The crude product obtained after the workup (670 mg, quantitative) was used in the next step without further purification.

3-Oxo-2-(pyridin-3-ylmethyl)pentanenitrile **(S31)**

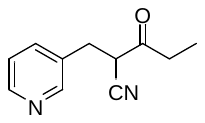

The compound was prepared according to General procedure A4 using *n-*BuLi (2.5 M, 1.5 mL, 3.64 mmol, 1.2 eq.), diisopropylamine (0.56 mL, 3.94 mmol, 1.3 eq.) and THF (3 mL) (reaction time: 30 min for the formation of LDA), then deprotonation of 3-(pyridin-3-yl)propanenitrile (400 mg, 3.03 mmol; 1 eq.) in THF (3 mL) (reaction time: 30 min) and final addition of propionyl chloride (0.32 mL, 3.64 mmol, 1.2 eq.) (reaction time: 16 h). The crude product obtained after the workup (560 mg, quantitative) was used in the next step without further purification.

2-(2-Methoxybenzyl)-3-oxopentanenitrile **(S32)**

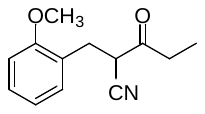

The compound was prepared according to General procedure A4 using *n-*BuLi (2.5 M, 1.5 mL, 3.72 mmol, 1.2 eq.), diisopropylamine (0.57 mL, 4.03 mmol, 1.3 eq.) and THF (3 mL) (reaction time: 30 min for the formation of LDA), then deprotonation of 3-(2-methoxyphenyl)propanenitrile (500 mg, 3.01 mmol; 1 eq.) in THF (3 mL) (reaction time: 30 min) and final addition of propionyl chloride (0.33 mL, 3.44 mmol, 1.2 eq.) (reaction time: 16 h). The crude product obtained after the workup (670 mg, quantitative) was used in the next step without further purification.

2-(4-Fluorobenzyl)-3-oxopentanenitrile **(S33)**

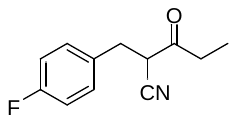

The compound was prepared according to General procedure A4 using *n-*BuLi (2.5 M, 1.9 mL, 4.83 mmol, 1.2 eq.), diisopropylamine (0.74 mL, 5.23 mmol, 1.3 eq.) and THF (3 mL) (reaction time: 30 min for the formation of LDA), then deprotonation of 3-(4-fluorophenyl)propanenitrile (600 mg, 4.02 mmol; 1 eq.) in THF (3 mL) (reaction time: 30 min) and final addition of propionyl chloride (0.42 mL, 4.83 mmol, 1.2 eq.) (reaction time: 16 h). The crude product obtained after the workup (820 mg, quantitative) was used in the next step without further purification.

2-(1-Methylpiperidin-4-yl)-3-oxopentanenitrile **(S34)**

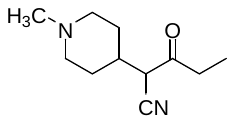

The compound was prepared according to General procedure A4 using *n-*BuLi (2.5 M, 1.7 mL, 4.34 mmol, 1.2 eq.), diisopropylamine (0.67 mL, 4.7 mmol, 1.3 eq.) and THF (3 mL) (reaction time: 30 min for the formation of LDA), then deprotonation of 2-(1-methylpiperidin-4-yl)acetonitrile (500 mg, 3.62 mmol; 1 eq.) in THF (3 mL) (reaction time: 30 min) and final addition of propionyl chloride (0.38 mL, 4.34 mmol, 1.2 eq.) (reaction time: 16 h). The crude product obtained after the workup (700 mg, quantitative) was used in the next step without further purification.

3-(1-(*tert*-Butyl)-1*H*-pyrazol-4-yl)acrylonitrile **(S35)**

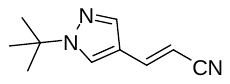

Diethyl cyanomethylphosphonate (0.45 mL, 2.76 mmol, 1.05 eq.) was slowly added at 0 °C to a stirred mixture sodium hydride in mineral oil (60%, 116 mg, 2.89 mmol, 1.1 eq.) in THF (8 mL). After 30 min, 1-*tert*-butyl-1*H*-pyrazole-4-carbaldehyde (400 mg, 2.63 mmol, 1 eq.) was added and the mixture was stirred at 23 °C for additional 16 h. The mixture was poured into water (30 mL) and extracted with EtOAc (3 × 20 mL). The combined organic extracts were washed with brine (30 mL), dried over MgSO_4_, and the solvent was evaporated. The residue was purified by column chromatography (hexane:EtOAc, gradient 9:1 to 3:7). The product was obtained as a colorless solid (350 mg, 76%) – as a mixture of *E* and *Z* isomers (5:1).

The NMR spectra of *E*-isomer:

^1^H NMR (500 MHz, Chloroform-*d*) *δ* (ppm) 7.68 (s, 1H), 7.66 (s, 1H), 7.24 (d, *J* = 16.6 Hz, 1H), 5.54 (d, *J* = 16.5 Hz, 1H), 1.58 (s, 9H).

^13^C NMR (126 MHz, Chloroform-*d*) *δ* (ppm) 141.49, 137.60, 126.17, 118.84, 117.26, 92.78, 59.41, 29.74.

The NMR spectra of *Z*-isomer:

^1^H NMR (500 MHz, Chloroform-*d*) *δ* (ppm) 8.17 (s, 1H), 7.86 (s, 1H), 7.01 (d, *J* = 11.6 Hz, 1H), 5.12 (d, *J* = 11.6 Hz, 1H), 1.60 (s, 9H).

^13^C NMR (126 MHz, Chloroform-*d*) *δ* (ppm) 140.19, 137.60, 126.84, 118.72, 116.85, 90.64, 59.41, 29.74.

HRMS (APCI): calcd. for C_10_H_14_N_3_ [M+H]^+^ =176.1182, found [M+H]^+^ = 176.1184.

3-(1-(*tert*-Butyl)-1*H*-pyrazol-4-yl)propanenitrile **(S36)**

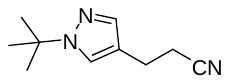

A misxture of ethyl 3-(1-(*tert*-butyl)-1*H*-pyrazol-4-yl)acrylonitrile (350 mg, 2.0 mmol, 1eq.) and Pd/C (50 mg) in Methanol (5 mL) was stirred in a pressure vessel at 23 °C under hydrogen atmosphere (15 bar) for 3 h. The mixture was filtered through a micro HPLC filter, and the filtrate was concentrated under vacuum. The crude product was dried under vacuum and used in the next step without further purification. The product was obtained as a colorless oil (350 mg, 99%).

^1^H NMR (126 MHz, DMSO-*d*_6_) *δ* (ppm) 7.44 (s, 1H), 7.40 (s, 1H), 2.83 (t, *J* = 7.2 Hz, 2H), 2.55 (t, *J* = 7.2 Hz, 2H), 1.57 (s, 9H).

^13^C NMR (126 MHz, DMSO-*d*_6_) *δ* (ppm) 137.85, 124.42, 119.54, 116.99, 58.54, 29.94, 21.02, 19.62.

HRMS (APCI): calcd. for C_10_H_16_N_3_ [M+H]^+^ = 178.1339, found [M+H]^+^ = 178.1342.

2-((1-(*tert*-Butyl)-1*H*-pyrazol-4-yl)methyl)-3-oxopentanenitrile **(S37)**

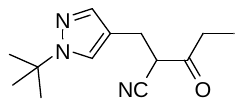

The compound was prepared according to General procedure A4 using *n-*BuLi (2.5 M, 0.92 mL, 2.3 mmol, 1.2 eq.), diisopropylamine (0.35 mL, 2.49 mmol, 1.3 eq.) and THF (3 mL) (reaction time: 30 min for the formation of LDA), then deprotonation of 3-(1-(*tert*-butyl)-1*H*-pyrazol-4-yl)propanenitrile (0.34 g, 1.92 mmol; 1 eq.) in THF (3 mL) (reaction time: 30 min) and final addition of propionyl chloride (0.2 mL, 2.3 mmol, 1.2 eq.) (reaction time: 16 h). The crude product obtained after the workup (450 mg, quantitative) was used in the next step without further purification.

3-Ethyl-4-(4-fluorophenyl)-1*H*-pyrazol-5-amine **(S38)**

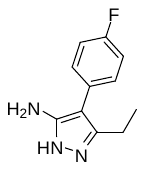

The compound was prepared according to General procedure B1 using 2-(4-fluorophenyl)-3-oxopentanenitrile (1.89 g, 9.88 mmol; 1 eq.), N_2_H_4_.H_2_O (64% in H_2_O, 0.48 mL, 9.88 mmol; 1 eq.), CH_3_SO_3_H (64 µL, 0.99 mmol, 0.1 eq.) and EtOH (20 mL). Reaction time: 45 min at reflux. The residue was sonicated in EtOH (10 mL) and the precipitate was collected by filtration to afford the product as a white crystalline solid (1.1 g, 54%). The filtrate was concentrated in a vacuum and the residue was purified by column chromatography on silica gel (hexane:EtOAc, gradient 2:1 to 0:1) to afford additional product as a white solid (727 mg, 36%).

^1^H NMR (500 MHz, DMSO-*d*_6_) *δ* (ppm) 11.42 (s, 1H), 7.36 – 7.28 (m, 2H), 7.22 – 7.13 (m, 2H), 4.38 (s, 2H), 2.53 (q, *J* = 7.6 Hz, 2H), 1.09 (t, *J* = 7.6 Hz, 3H).

^13^C NMR (126 MHz, DMSO-*d*_6_) *δ* (ppm) 160.12 (d, *J* = 242.2 Hz), 150.84, 143.06, 130.42 (d, *J* = 3.1 Hz), 129.92 (d, *J* = 7.9 Hz), 115.10, 102.76, 18.37, 13.24.

^19^F NMR (471 MHz, DMSO-*d*_6_) *δ* (ppm) -117.79.

HRMS (APCI): calcd. for C_11_H_11_FN_3_ [M-H]^-^ = 204.0942, found [M-H]^-^ = 204.0948

Mp = 163-166 °C.

4-(4-Fluorophenyl)-1*H*-pyrazol-5-amine **(S39)**

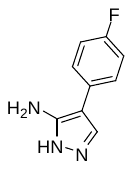

The compound was prepared according to General procedure B1 using 2-(4-fluorophenyl)-3-oxopropanenitrile (382 mg, 2.34 mmol; 1 eq.), N_2_H_4_.H_2_O (64% in H_2_O, 0.114 mL, 2.34 mmol; 1 eq.), CH_3_SO_3_H (15 µL, 0.23 mmol, 0.1 eq.) and EtOH (10 mL). Reaction time: 1 h at reflux. The residue was sonicated in EtOH (5 mL) and the precipitate was collected by filtration and dried under vacuum to afford the product as a yellow solid (222 mg, 54 %). The filtrate was concentrated in a vacuum and the residue was purified by column chromatography on silica gel (hexane:EtOAc, gradient 2:1 to 0:1) to afford additional product as a yellow solid (136 mg, 33%).

^1^H NMR (500 MHz, DMSO-*d*_6_) *δ* (ppm) 7.63 (s, 1H), 7.55 – 7.48 (m, 2H), 7.17 – 7.10 (m, 2H), 4.71 (s, 2H).

^13^C NMR (126 MHz, DMSO-*d*_6_) *δ* (ppm) 159.76 (d, *J* = 241.3 Hz), 130.48 (d, *J* = 3.3 Hz), 127.18 (d, *J* = 7.5 Hz), 115.14 (d, *J* = 21.0 Hz).

HRMS (APCI): calcd. for C_9_H_7_FN_3_ [M-H]^-^ = 176.0629, found [M-H]^-^ = 176.0627.

3-(3-Bromophenyl)-1*H*-pyrazol-5-amine **(S40)**

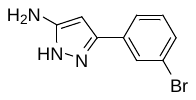

The compound was prepared according to General procedure B1 using 3-(3-bromophenyl)-3-oxopropanenitrile (575 mg, 2.57 mmol; 1 eq.), N_2_H_4_.H_2_O (64% in H_2_O, 0.39 mL, 7.7 mmol; 3 eq.), CH_3_SO_3_H (17 µL, 0.26 mmol, 0.1 eq.) and EtOH (13 mL). Reaction time: 2 h at reflux. The residue obtained after the workup was purified by column chromatography on silica gel (hexane:EtOAc:MeOH, gradient 1:1:0 to 0:8:1). The product was obtained as an off-white solid (504 mg, 82%).

^1^H NMR (500 MHz, DMSO-*d*_6_) *δ* (ppm) 11.66 (s, 1H), 7.84 (t, *J* = 1.8 Hz, 1H), 7.65 (dt, *J* = 7.8, 1.3 Hz, 1H), 7.43 (d, *J* = 7.8 Hz, 1H), 7.32 (t, *J* = 7.9 Hz, 1H), 5.78 (s, 1H), 4.89 (s, 2H).

^13^C NMR (126 MHz, DMSO-*d*_6_) *δ* (ppm) 130.69, 129.64, 127.01, 123.67, 121.97.

HRMS (APCI): calcd. For C_9_H_9_BrN_3_ [M+H]^+^ = 237.9974, found [M+H]^+^ = 237.9979.

3-Cyclohexyl-1*H*-pyrazol-5-amine **(S41)**

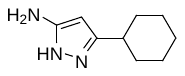

The compound was prepared according to General procedure B1 using 3-cyclohexyl-3-oxopropanenitrile (355 mg, 2.35 mmol; 1 eq.), N_2_H_4_.H_2_O (64% in H_2_O, 0.23 mL, 4.70 mmol; 2 eq.), CH_3_SO_3_H (15 µL, 0.24 mmol, 0.1 eq.) and EtOH (12 mL). Reaction time: 2 h at reflux. The residue obtained after the workup was purified by column chromatography on silica gel (hexane:EtOAc:MeOH, gradient 1:1:0 to 0:9:1). The product was obtained as a pink wax (386 mg, 99%).

^1^H NMR (500 MHz, Chloroform-*d*) *δ* (ppm) 5.42 (s, 1H), 2.61 – 2.44 (m, 1H), 1.98 – 1.88 (m, 2H), 1.82 – 1.73 (m, 2H), 1.73 – 1.64 (m, 1H), 1.43 – 1.28 (m, 4H), 1.28 – 1.17 (m, 1H).

^13^C NMR (126 MHz, Chloroform-*d*) *δ* (ppm) 154.56, 151.23, 90.07, 35.78, 32.71, 26.15, 26.03.

HRMS (APCI): calcd. for C_9_H_16_N_3_ [M+H]^+^ = 166.1339, found [M+H]^+^ = 166.1335.

3-(2-phenoxyphenyl)-1H-pyrazol-5-amine **(S42)**

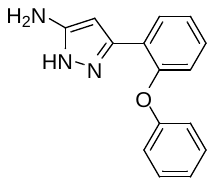

The compound was prepared according to General procedure B1 using 3-oxo-3-(2-phenoxyphenyl)propanenitrile (461 mg, 1.94 mmol; 1 eq.), N_2_H_4_.H_2_O (64% in H_2_O, 0.189 mL, 3.89 mmol; 2 eq.), CH_3_SO_3_H (13 µL, 0.19 mmol, 0.1 eq.) and EtOH (10 mL). Reaction time: 2 h at reflux. The residue obtained after the workup was purified by column chromatography on silica gel (hexane:EtOAc:MeOH, gradient 1:1:0 to 0:9:1). The product was obtained as an off-white solid (392 mg, 80%).

^1^H NMR (500 MHz, DMSO-*d*_6_) *δ* (ppm) 11.69 (s, 1H), 7.86 (s, 1H), 7.40 – 7.32 (m, 2H), 7.29 (td, *J* = 7.7, 1.7 Hz, 1H), 7.21 (td, *J* = 7.5, 1.3 Hz, 1H), 7.08 (t, *J* = 7.4 Hz, 1H), 6.98 – 6.88 (m, 3H), 5.78 (s, 1H), 4.66 (s, 2H).

^13^C NMR (126 MHz, DMSO-*d*_6_) *δ* (ppm) 157.16, 152.10, 129.91, 128.66, 127.75, 124.23, 122.73, 120.62, 117.44.

HRMS (APCI): calcd. for C_15_H_14_N_3_O [M+H]^+^ = 252.1131, found [M+H]^+^ = 252.1138.

3-(Naphthalen-2-yl)-1*H*-pyrazol-5-amine **(S43)**

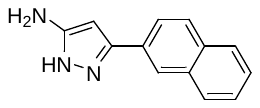

The compound was prepared according to General procedure B1 using 3-(naphthalen-2-yl)-3-oxopropanenitrile (472 mg, 2.42 mmol; 1 eq.), N_2_H_4_.H_2_O (64% in H_2_O, 0.24 mL, 4.83 mmol; 2 eq.), CH_3_SO_3_H (16 µL, 0.24 mmol, 0.1 eq.) and EtOH (12 mL). Reaction time: 2 h at reflux. The residue obtained after the workup was purified by column chromatography on silica gel (hexane:EtOAc:MeOH, gradient 1:1:0 to 0:9:1). The product was obtained as a beige solid (390 mg, 77%).

^1^H NMR (500 MHz, DMSO-*d*_6_) *δ* (ppm) 12.35 – 11.35 (m, 1H), 8.15 (d, *J* = 1.4 Hz, 1H), 7.95 – 7.81 (m, 4H), 7.58 – 7.43 (m, 2H), 5.90 (s, 1H), 4.83 (s, 2H).

^13^C NMR (126 MHz, DMSO-*d*_6_) *δ* (ppm) 133.13, 132.22, 127.97, 127.81, 127.53, 126.31, 125.68, 123.46, 122.79.

HRMS (APCI): calcd. for C_13_H_12_N_3_ [M+H]^+^ = 210.1026, found [M+H]^+^ = 210.1031.

3-Isopropyl-1*H*-pyrazol-5-amine **(S44)**

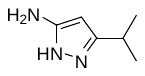

The compound was prepared according to General procedure B1 using 4-methyl-3-oxopentanenitrile (2.46 g, 22 mmol; 1 eq.), N_2_H_4_.H_2_O (64% in H_2_O, 3.4 mL, 44 mmol; 2 eq.), CH_3_SO_3_H (150 µL, 2.2 mmol, 0.1 eq.) and EtOH (50 mL). Reaction time: 2 h at reflux. The residue obtained after workup was purified by column chromatography on silica gel (hexane:EtOAc:MeOH, gradient 1:1:0 to 0:9:1). The product was obtained as a dark red wax (2.42 g, 98%).

^1^H NMR (500 MHz, Chloroform-*d*) *δ* (ppm) 5.43 (s, 1H), 2.87 (hept, *J* = 6.9 Hz, 1H), 1.23 (d, *J* = 6.9 Hz, 6H).

^13^C NMR (126 MHz, Chloroform-*d*) *δ* (ppm) 154.53, 152.19, 89.94, 26.27, 22.31.

3-(*tert*-butyl)-1*H*-pyrazol-5-amine **(S45)**

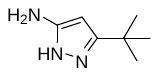

The compound was prepared according to General procedure B1 using 4,4-dimethyl-3-oxopentanenitrile (2.6 g, 20.8 mmol; 1 eq.), N_2_H_4_.H_2_O (64% in H_2_O, 3.2 mL, 41.6 mmol; 2 eq.), CH_3_SO_3_H (130 µL, 2.0 mmol, 0.1 eq.) and EtOH (25 mL). Reaction time: 2 h at reflux. The residue obtained after the workup was purified by column chromatography on silica gel (hexane:EtOAc:MeOH, gradient 1:1:0 to 0:9:1). The product was obtained as a dark red wax (1.63 g, 56%).

^1^H NMR (500 MHz, DMSO-*d*_6_) *δ* (ppm) 5.18 (s, 1H), 1.18 (s, 9H).

^13^C NMR (126 MHz, DMSO-*d*_6_) *δ* (ppm) 154.04, 153.31, 87.37, 30.64, 30.05.

HRMS (APCI): calcd. for C_7_H_14_N_3_ [M+H]^+^ = 140.1182, found [M+H]^+^ = 140.1191.

4-(4-Fluorophenyl)-3-isopropyl-1*H*-pyrazol-5-amine **(S46)**

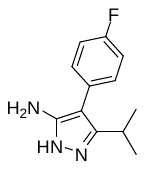

The compound was prepared according to General procedure B1 using 2-(4-fluorophenyl)-4-methyl-3-oxopentanenitrile (205 mg, 0.73 mmol; 1 eq.), N_2_H_4_.H_2_O (64% in H_2_O, 0.17 mL, 2.19 mmol; 3 eq.), CH_3_SO_3_H (6 µL, 0.07 mmol, 0.1 eq.) and EtOH (2 mL). Reaction time: 45 min at reflux. The residue obtained after the workup was purified by column chromatography on silica gel (hexane:EtOAc:MeOH, gradient 1:1:0 to 0:9:1). The product was obtained as a white solid (80 mg, 50%).

^1^H NMR (500 MHz, chloroform-*d*) *δ* (ppm) 7.30 – 7.26 (m, 2H), 7.14 – 7.09 (m, 2H), 4.97 (s, 2H), 3.04 (hept, *J* = 7.0 Hz, 1H), 1.24 (d, *J* = 7.0 Hz, 6H).

^13^C NMR (126 MHz, chloroform-*d*) *δ* (ppm) 162.01 (d, *J* = 246.9 Hz), 151.59, 148.37, 131.25 (d, *J* = 8.1 Hz), 128.34 (d, *J* = 2.8 Hz), 116.06 (d, *J* = 21.5 Hz), 104.47, 25.30, 22.07.

3-(((*tert*-Butyldiphenylsilyl)oxy)methyl)-1*H*-pyrazol-5-amine **(S47)**

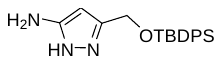

The compound was prepared according to General procedure B1 using 4-((*tert*-butyldiphenylsilyl)oxy)-3-oxobutanenitrile (350 mg, 1.0 mmol; 1 eq.), N_2_H_4_.H_2_O (64% in H_2_O, 0.24 mL, 3.0 mmol; 3 eq.), CH_3_SO_3_H (7 µL, 0.1 mmol, 0.1 eq.) and EtOH (5 mL). Reaction time: 45 min at reflux. The residue obtained after the workup was purified by column chromatography on silica gel (hexane:EtOAc:MeOH, gradient 1:1:0 to 0:9:1). The product was obtained as a colorless oil (360 mg, 98%).

^1^H NMR (500 MHz, chloroform-*d*) *δ* (ppm) 11.27 (s, 1H), 7.69 – 7.61 (m, 4H), 7.50 – 7.40 (m, 6H), 5.34 (s, 1H), 4.65 (s, 2H), 4.54 (s, 2H), 1.00 (s, 9H).

^13^C NMR (126 MHz, chloroform-*d*) *δ* (ppm) 135.00, 133.00, 129.80, 127.82, 59.68, 26.57, 18.77.

HRMS (APCI): calcd. for C_20_H_26_N_3_OSi [M+H]^+^ = 352.1840, found [M+H]^+^ = 352.1843

3-(3-Methoxynaphthalen-2-yl)-1*H*-pyrazol-5-amine **(S48)**

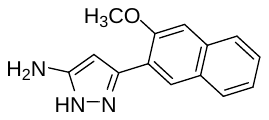

The compound was prepared according to General procedure B1 using 3-(3-methoxynaphthalen-2-yl)-3-oxopropanenitrile (100 mg, 0.44 mmol; 1 eq.), N_2_H_4_.H_2_O (64% in H_2_O, 70 µL, 0.88 mmol; 2 eq.), CH_3_SO_3_H (7 µL, 0.1 mmol, 0.4 eq.) and EtOH (5 mL). Reaction time: 45 min at reflux. The residue obtained after the workup was purified by column chromatography on silica gel (hexane:EtOAc:MeOH, gradient 1:1:0 to 0:9:1). The product was obtained as a colorless wax (93 mg, 88%).

^1^H NMR (500 MHz, chloroform-*d*) *δ* (ppm) 11.71 (s, 1H), 8.16 (s, 1H), 7.81 (t, *J* = 8.6 Hz, 2H), 7.47 – 7.41 (m, 1H), 7.41 (s, 1H), 7.38 – 7.32 (m, 1H), 5.99 (s, 1H), 4.63 (s, 2H), 3.97 (s, *J* = 21.0 Hz, 3H).

^13^C NMR (126 MHz, chloroform-*d*) *δ* (ppm) 135.00, 133.00, 129.80, 127.82, 59.68, 26.57, 18.77.

3-([1,1'-Biphenyl]-3-yl)-1H-pyrazol-5-amine **(S49)**

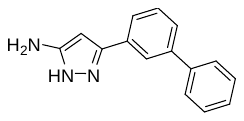

The compound was prepared according to General procedure B1 using 3-([1,1'-biphenyl]-3-yl)-3-oxopropanenitrile (424 mg, 1.92 mmol; 1 eq.), N_2_H_4_.H_2_O (64% in H_2_O, 300 µL, 3.84 mmol; 2 eq.), CH_3_SO_3_H (14 µL, 0.2 mmol, 0.1 eq.) and EtOH (5 mL). Reaction time: 45 min at reflux. The residue obtained after the workup was purified by column chromatography on silica gel (hexane:EtOAc:MeOH, gradient 1:1:0 to 0:9:1). The product was obtained as a black wax (350 mg, 77%).

^1^H NMR (500 MHz, DMSO-*d*_6_) *δ* (ppm) 11.42 (s, 1H), 7.58 (s, 1H), 7.44 – 7.25 (m, 6H), 7.21 (t, *J* = 7.4 Hz, 3H), 4.75 (s, 1H), 4.49 (s, 1H).

^13^C NMR (126 MHz, DMSO-*d*_6_) *δ* (ppm) 147.03, 141.41, 139.91, 137.80, 130.40, 128.92, 128.78, 127.92, 127.49, 127.24, 126.85, 91.00.

HRMS (APCI): calcd. for C_15_H_14_N_3_ [M+H]^+^ = 236.1182, found [M+H]^+^ = 236.1184

3-(1-Methoxynaphthalen-2-yl)-1*H*-pyrazol-5-amine **(S50)**

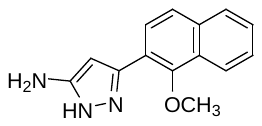

The compound was prepared according to General procedure B1 using 3-(1-methoxynaphthalen-2-yl)-3-oxopropanenitrile (260 mg, 1.16 mmol; 1 eq.), N_2_H_4_.H_2_O (64% in H_2_O, 110 µL, 2.32 mmol; 2 eq.), CH_3_SO_3_H (7 µL, 0.1 mmol, 0.1 eq.) and EtOH (5 mL). Reaction time: 45 min at reflux. The residue obtained after the workup was purified by column chromatography on silica gel (hexane:EtOAc:MeOH, gradient 1:1:0 to 0:9:1). The product was obtained as a yellow wax (186 mg, 67%).

^1^H NMR (500 MHz, DMSO-*d*_6_) *δ* (ppm) 11.42 (s, 1H), 7.58 (s, 1H), 7.44 – 7.25 (m, 4H), 7.24 – 7.16 (m, 2H), 4.75 (s, 1H), 4.49 (s, 1H), 3.31 (s, 3H).

^13^C NMR (126 MHz, DMSO-*d*_6_) *δ* (ppm) 147.03, 141.41, 139.91, 137.80, 130.39, 128.91, 128.78, 127.91, 127.48, 127.24, 126.85, 91.00, 59.69.

3-(3-Methoxynaphthalen-1-yl)-1*H*-pyrazol-5-amine **(S51)**

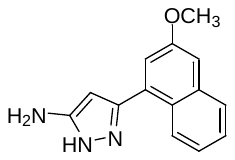

The compound was prepared according to General procedure B1 using 3-(3-methoxynaphthalen-1-yl)-3-oxopropanenitrile (220 mg, 0.97 mmol; 1 eq.), N_2_H_4_.H_2_O (64% in H_2_O, 62 µL, 1.94 mmol; 2 eq.), CH_3_SO_3_H (7 µL, 0.1 mmol, 0.1 eq.) and EtOH (5 mL). Reaction time: 45 min at reflux. The residue obtained after the workup was purified by column chromatography on silica gel (hexane:EtOAc:MeOH, gradient 1:1:0 to 0:9:1). The product was obtained as a yellow wax (200 mg, 86%).

^1^H NMR (500 MHz, DMSO-*d*_6_) *δ* (ppm) 11.50 (s, 1H), 7.98 (d, *J* = 9.0 Hz, 1H), 7.88 (d, *J* = 8.0 Hz, 1H), 7.70 (d, *J* = 8.3 Hz, 1H), 7.49 (d, *J* = 9.1 Hz, 1H), 7.44 – 7.39 (m, 1H), 7.35 (t, *J* = 7.4 Hz, 1H), 5.51 (s, 1H), 4.61 (s, *J* = 99.3 Hz, 1H), 3.84 (s, 3H).

^13^C NMR (126 MHz, DMSO-*d*_6_) *δ* (ppm) 155.28, 133.27, 129.87, 128.32, 127.75, 126.50, 124.67, 123.69, 123.46, 113.99, 56.86.

3-(1-benzyl-1*H*-imidazol-2-yl)-1*H*-pyrazol-5-amine **(S52)**

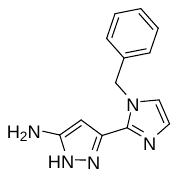

The compound was prepared according to General procedure B1 using 3-(1-benzyl-1*H*-imidazol-2-yl)-3-oxopropanenitrile (200 mg, 0.88 mmol; 1 eq.), N_2_H_4_.H_2_O (64% in H_2_O, 60 µL, 1.7 mmol; 2 eq.), CH_3_SO_3_H (7 µL, 0.1 mmol, 0.1 eq.) and EtOH (5 mL). Reaction time: 45 min at reflux. The residue obtained after the workup was purified by column chromatography on silica gel (hexane:EtOAc:MeOH, gradient 1:1:0 to 0:9:1). The product was obtained as a yellow wax (84 mg, 40%).

^1^H NMR (500 MHz, DMSO-*d*_6_) *δ* (ppm) 11.67 (s, 1H), 7.76 (s, 1H), 7.32 – 7.27 (m, 2H), 7.19 – 7.13 (m, 3H), 6.95 – 6.92 (m, 1H), 5.66 (s, 2H), 5.19 (s, 1H).

^13^C NMR (126 MHz, DMSO-*d*_6_) *δ* (ppm) 137.84, 129.13, 129.00, 128.92, 128.17, 127.92, 127.70, 127.48, 119.58, 113.33, 49.36.

3-(2-chlorophenyl)-1*H*-pyrazol-5-amine **(S53)**

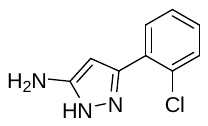

The compound was prepared according to General procedure B1 using 3-(2-chlorophenyl)-3-oxopropanenitrile (735 mg, 4.2 mmol; 1 eq.), N_2_H_4_.H_2_O (64% in H_2_O, 0.54 mL, 8.4 mmol; 2 eq.), CH_3_SO_3_H (10 µL, 0.13 mmol, 0.03 eq.) and EtOH (10 mL). Reaction time: 45 min at reflux. The residue obtained after the workup was purified by column chromatography on silica gel (hexane:EtOAc:MeOH, gradient 1:1:0 to 0:9:1). The product was obtained as a yellow wax (760 mg, 93%).

^1^H NMR (500 MHz, DMSO-*d*_6_) *δ* (ppm) 11.74 (s, 1H), 7.67 (d, *J* = 6.0 Hz, 1H), 7.48 (d, *J* = 7.7 Hz, 1H), 7.38 – 7.33 (m, 1H), 7.33 – 7.29 (m, 1H), 5.84 (s, 1H), 4.82 (s, 2H).

^13^C NMR (126 MHz, DMSO-*d*_6_) *δ* (ppm) 159.06, 143.81, 130.64, 130.23, 129.89, 128.82, 127.15, 92.73.

HRMS (APCI): calcd. for C_9_H_9_ClN_3_ [M+H]^+^ = 194.0480, found [M+H]^+^ = 194.0476.

3-(Naphthalen-1-yl)-1*H*-pyrazol-5-amine **(S54)**

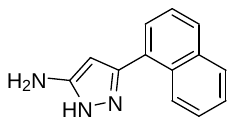

The compound was prepared according to General procedure B1 using 3-(naphthalen-1-yl)-3-oxopropanenitrile (280 mg, 1.43 mmol; 1 eq.), N_2_H_4_.H_2_O (64% in H_2_O, 0.23 mL, 2.9 mmol; 2 eq.), CH_3_SO_3_H (10 µL, 0.14 mmol, 0.1 eq.) and EtOH (5 mL). Reaction time: 45 min at reflux. The residue obtained after the workup was purified by column chromatography on silica gel (hexane:EtOAc:MeOH, gradient 1:1:0 to 0:9:1). The product was obtained as a yellow wax (185 mg, 62%).

^1^H NMR (500 MHz, DMSO-*d*_6_) *δ* (ppm) 11.75 (s, 1H), 8.45 (s, 1H), 7.95 (dd, *J* = 6.0, 3.4 Hz, 1H), 7.90 (d, *J* = 8.0 Hz, 1H), 7.60 – 7.56 (m, 1H), 7.56 – 7.51 (m, 3H), 5.71 (s, 1H), 4.83 (s, 2H).

^13^C NMR (126 MHz, DMSO-*d*_6_) *δ* (ppm) 133.46, 130.54, 128.19, 127.78, 126.25, 126.18, 125.82, 125.41.

HRMS (APCI): calcd. for C_13_H_12_N_3_ [M+H]^+^ = 210.1026, found [M+H]^+^ = 210.1028.

3-(2-(Dimethylamino)phenyl)-1*H*-pyrazol-5-amine **(S55)**

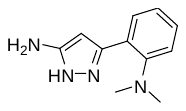

The compound was prepared according to General procedure B1 using 3-(2-(dimethylamino)phenyl)-3-oxopropanenitrile (280 mg, 1.5 mmol; 1 eq.), N_2_H_4_.H_2_O (64% in H_2_O, 0.23 mL, 3.0 mmol; 2 eq.), CH_3_SO_3_H (10 µL, 0.14 mmol, 0.1 eq.) and EtOH (5 mL). Reaction time: 45 min at reflux. The residue obtained after the workup was purified by column chromatography on silica gel (hexane:EtOAc:MeOH, gradient 1:1:0 to 0:9:1). The product was obtained as a yellow wax (215 mg, 70%).

^1^H NMR (500 MHz, DMSO-*d*_6_) *δ* (ppm) 11.75 (s, 1H), 7.98 – 7.94 (m, 1H), 7.90 (d, *J* = 8.1 Hz, 1H), 7.60 – 7.56 (m, 1H), 7.55 – 7.51 (m, 1H), 5.71 (s, 1H), 4.83 (s, 2H), 3.32 (s, 6H).

^13^C NMR (126 MHz, DMSO-*d*_6_) *δ* (ppm) 133.45, 130.53, 128.18, 127.77, 126.25, 125.82, 125.40, 59.70.

3-(2-Methoxybenzyl)-1*H*-pyrazol-5-amine **(S56)**

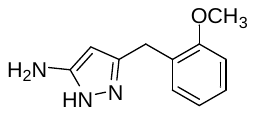

The compound was prepared according to General procedure B1 using 4-(2-methoxyphenyl)-3-oxobutanenitrile (1.3 g, 6.87 mmol; 1 eq.), N_2_H_4_.H_2_O (64% in H_2_O, 1.7 mL, 27.48 mmol; 4 eq.), CH_3_SO_3_H (45 µL, 0.7 mmol, 0.1 eq.) and EtOH (10 mL). Reaction time: 45 min at reflux. The residue obtained after the workup was purified by column chromatography on silica gel (dichloromethane:MeOH, gradient 19:1 to 9:1). The product was obtained as a colorless wax (300 mg, 22%).

^1^H NMR (500 MHz, Chloroform-*d*) *δ* (ppm) 7.23 – 7.17 (m, 1H), 7.11 (dd, *J* = 7.4, 1.7 Hz, 1H), 6.89 – 6.83 (m, 2H), 5.77 (s, 2H), 5.42 (s, 1H), 3.84 (s, 2H), 3.80 (s, 3H).

^13^C NMR (126 MHz, Chloroform-*d*) *δ* (ppm) 157.14, 154.29, 144.27, 130.17, 128.08, 126.86, 120.78, 110.63, 92.03, 55.46, 27.15.

3-(2-Methoxy-2-phenylethyl)-1*H*-pyrazol-5-amine **(S57)**

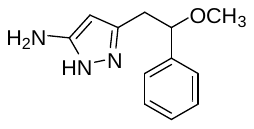

The compound was prepared according to General procedure B1 using 5-methoxy-3-oxo-5-phenylpentanenitrile (350 mg, 1.72 mmol; 1 eq.), N_2_H_4_.H_2_O (64% in H_2_O, 0.42 mL, 6.89 mmol; 4 eq.), CH_3_SO_3_H (11 µL, 0.17 mmol, 0.1 eq.) and EtOH (5 mL). Reaction time: 45 min at reflux. The residue obtained after the workup was purified by column chromatography on silica gel (dichloromethane:MeOH, gradient 19:1 to 9:1). The product was obtained as a colorless wax (350 mg, 94%).

^1^H NMR (500 MHz, Chloroform-*d*) *δ* (ppm7.40 – 7.27 (m, 5H), 5.42 (s, 1H), 4.35 (dd, *J* = 8.6, 4.0 Hz, 1H), 3.25 (s, 3H), 3.02 – 2.81 (m, 2H).

^13^C NMR (126 MHz, Chloroform-*d*) *δ* (ppm) 154.37, 142.31, 140.96, 128.68, 128.15, 126.62, 92.72, 83.27, 56.80, 34.98.

3-Benzyl-1*H*-pyrazol-5-amine **(S58)**

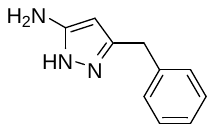

The compound was prepared according to General procedure B1 using 3-oxo-4-phenylbutanenitrile (619 mg, 3.89 mmol; 1 eq.), N_2_H_4_.H_2_O (64% in H_2_O, 0.403 mL, 7.78 mmol; 2 eq.), CH_3_SO_3_H (25 µL, 0.23 mmol, 0.1 eq.) and EtOH (19 mL). Reaction time: 2 h at reflux. The residue obtained after the workup was loaded on ISOLUTE^®^ HM-N and purified by column chromatography on silica gel using Biotage Selekt purification system (hexane:EtOAc:MeOH, gradient 1:1:0 to 0:9:1). The product was obtained as a pale-yellow wax (203 mg, 30%).

^1^H NMR (500 MHz, Chloroform-*d*) *δ* (ppm) 7.32 – 7.27 (m, 2H), 7.25 – 7.18 (m, 3H), 5.45 (s, 1H), 3.88 (s, 2H).

^13^C NMR (126 MHz, Chloroform-*d*) *δ* (ppm) 154.55, 144.46, 137.92, 128.84, 126.90, 92.65, 32.73.

HRMS (APCI): calcd. for C_10_H_12_N_3_ [M+H]^+^ = 174.1026, found [M+H]^+^ = 174.1023.

3-(Isochroman-3-ylmethyl)-1*H*-pyrazol-5-amine **(S59)**

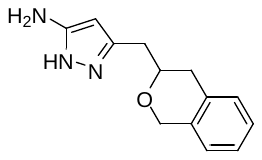

The compound was prepared according to General procedure B1 using 4-(isochroman-3-yl)-3-oxobutanenitrile (494 mg, 2.30 mmol; 1 eq.), N_2_H_4_.H_2_O (64% in H_2_O, 0.238 mL, 4.59 mmol; 2 eq.), CH_3_SO_3_H (15 µL, 0.23 mmol, 0.1 eq.) and EtOH (11 mL). Reaction time: 2 h at reflux. The residue obtained after the workup was loaded on ISOLUTE^®^ HM-N and purified by column chromatography on silica gel using Biotage Selekt purification system (hexane:EtOAc:MeOH, gradient 1:1:0 to 0:9:1). The product was obtained as a pale-yellow wax (523 mg, 99%).

^1^H NMR (500 MHz, Chloroform-*d*) *δ* (ppm) 7.21 – 7.13 (m, 2H), 7.13 – 7.05 (m, 2H), 5.44 (s, 1H), 5.21 (bs, 3H), 4.99 (ddd, *J* = 8.3, 3.2, 1.4 Hz, 1H), 4.18 (ddd, *J* = 11.3, 5.5, 3.2 Hz, 1H), 3.79 (ddd, *J* = 11.3, 10.1, 3.6 Hz, 1H), 3.18 (dd, *J* = 15.6, 3.2 Hz, 1H), 3.07 – 2.92 (m, 2H), 2.67 (dt, *J* = 16.3, 3.5 Hz, 1H).

^13^C NMR (126 MHz, Chloroform-*d*) *δ* (ppm) 154.30, 142.17, 136.64, 134.15, 129.18, 126.83, 126.40, 124.73, 92.94, 75.37, 63.72, 32.50, 29.05.

HRMS (APCI): calcd. for C_13_H_16_N_3_O [M+H]^+^ = 230.1288, found [M+H]^+^ = 230.1290.

4-(4-Fluorophenyl)-3-(2-methoxybenzyl)-1*H*-pyrazol-5-amine **(S60)**

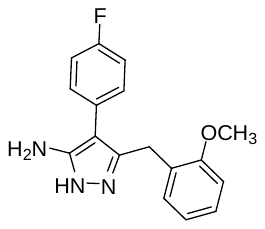

The compound was prepared according to General procedure B1 using 2-(4-fluorophenyl)-4-(2-methoxyphenyl)-3-oxobutanenitrile (580 mg, 2.05 mmol; 1 eq.), N_2_H_4_.H_2_O (64% in H_2_O, 0.23 mL, 4.1 mmol; 2 eq.), CH_3_SO_3_H (15 µL, 0.23 mmol, 0.1 eq.) and EtOH (11 mL). Reaction time: 2 h at reflux. The residue obtained after the workup was loaded on ISOLUTE^®^ HM-N and purified by column chromatography on silica gel using Biotage Selekt purification system (hexane:EtOAc:MeOH, gradient 1:1:0 to 0:4:1). The product was obtained as an off-white foam (0.2 g, 33%).

^1^H NMR (500 MHz, Chloroform-*d*) *δ* (ppm) 7.17 – 7.07 (m, 2H), 7.01 (td, *J* = 7.8, 1.8 Hz, 1H), 6.93 – 6.87 (m, 2H), 6.85 (dd, *J* = 7.7, 1.8 Hz, 1H), 6.73 – 6.63 (m, 2H), 5.90 (s, 3H), 3.71 (s, 2H), 3.61 (s, 3H).

^13^C NMR (126 MHz, Chloroform-*d*) *δ* (ppm) 161.46 (d, *J* = 245.3 Hz), 157.18, 152.00, 140.15, 130.70 (d, *J* = 7.5 Hz), 130.06, 129.06, 128.16, 126.36, 120.77, 115.62 (d, *J* = 21.6 Hz), 110.59, 105.58, 55.39, 25.88.

^19^F NMR (471 MHz, Chloroform-*d*) *δ* (ppm) -116.08.

4-Cyclohexyl-3-ethyl-1*H*-pyrazol-5-amine **(S61)**

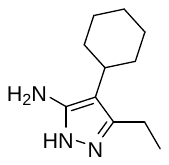

The compound was prepared according to General procedure B1 using 2-cyclohexyl-3-oxopentanenitrile (304 mg, 1.70 mmol; 1 eq.), N_2_H_4_.H_2_O (64% in H_2_O, 0.134 mL, 2.21 mmol; 1.3 eq.), CH_3_SO_3_H (33 µL, 0.51 mmol, 0.3 eq.) and EtOH (7 mL). Reaction time: 4 h at reflux. The residue obtained after the workup was loaded on ISOLUTE^®^ HM-N and purified twice by column chromatography on silica gel using Biotage Selekt purification system (cyclohexane:EtOAc:MeOH, gradient 1:2:0 to 0:9:1). The product was obtained as a pale pink solid (146 mg, 45%).

^1^H NMR (500 MHz, Chloroform-*d*) *δ* (ppm) 5.62 (s, 2H), 2.59 (q, *J* = 7.5 Hz, 2H), 2.38 – 2.29 (m, 1H), 1.82 (dt, *J* = 12.8, 3.0 Hz, 2H), 1.77 – 1.70 (m, 3H), 1.53 (qd, *J* = 12.7, 3.3 Hz, 2H), 1.33 (qt, *J* = 12.5, 3.2 Hz, 2H), 1.27 – 1.18 (m, 4H).

^13^C NMR (126 MHz, Chloroform-*d*) *δ* (ppm) 152.30, 142.96, 108.52, 34.50, 32.79, 27.28, 26.28, 18.82, 13.80.

HRMS (APCI): calcd. for C_11_H_20_N_3_ [M+H]^+^ = 194.1652, found [M+H]^+^ = 194.1654.

3-Ethyl-4-(tetrahydro-2*H*-pyran-4-yl)-1*H*-pyrazol-5-amine **(S62)**

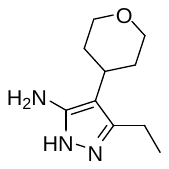

The compound was prepared according to General procedure B1 using 3-oxo-2-(tetrahydro-2*H*-pyran-4-yl)pentanenitrile (454 g, 2.51 mmol; 1 eq.), N_2_H_4_.H_2_O (64% in H_2_O, 0.214 mL, 3.5 mmol; 1.4 eq.), CH_3_SO_3_H (49 µL, 0.75 mmol, 0.3 eq.) and EtOH (10 mL). Reaction time: 4 h at reflux. The residue obtained after the workup was loaded on ISOLUTE^®^ HM-N and purified twice by column chromatography on silica gel using Biotage Selekt purification system (EtOAc:MeOH, gradient 1:0 to 9:1). The product was obtained as a white solid (160 mg, 33%).

^1^H NMR (500 MHz, Chloroform-*d*) *δ* (ppm) 5.25 (s, 2H), 4.06 (dd, *J* = 11.7, 4.7 Hz, 2H), 3.47 (td, *J* = 11.9, 2.0 Hz, 2H), 2.62 (q, *J* = 7.5 Hz, 3H), 2.02 – 1.88 (m, 2H), 1.64 – 1.58 (m, 2H), 1.23 (t, *J* = 7.6 Hz, 3H).

^13^C NMR (126 MHz, Chloroform-*d*) *δ* (ppm) 152.28, 143.29, 106.72, 68.80, 32.28, 31.62, 18.77, 13.82.

HRMS (APCI): calcd. for C_10_H_18_N_3_O [M+H]^+^ = 196.1444, found [M+H]^+^ = 196.1442

4-Benzyl-3-ethyl-1*H*-pyrazol-5-amine **(S63)**

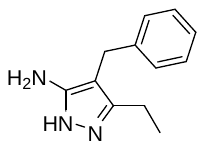

The compound was prepared according to General procedure B1 using 2-benzyl-3-oxopentanenitrile (302 mg, 1.61 mmol; 1 eq.), N_2_H_4_.H_2_O (64% in H_2_O, 0.147 mL, 2.42 mmol; 1.5 eq.), CH_3_SO_3_H (21 µL, 0.33 mmol, 0.2 eq.) and EtOH (7 mL). Reaction time: 3 h at reflux. The residue obtained after the workup was loaded on ISOLUTE^®^ HM-N and purified by column chromatography on silica gel using Biotage Selekt purification system (cyclohexane:EtOAc:MeOH, gradient 1:1:0 to 0:9:1). The product was obtained as a yellow solid (131 mg, 40%).

^1^H NMR (500 MHz, Chloroform-*d*) *δ* (ppm) 7.30 – 7.24 (m, 2H), 7.21 – 7.16 (m, 3H), 5.29 (bs, 2H), 3.70 (s, 2H), 2.55 (q, *J* = 7.6 Hz, 2H), 1.18 (t, *J* = 7.6 Hz, 3H).

^13^C NMR (126 MHz, Chloroform-*d*) *δ* (ppm) 153.46, 144.06, 140.32, 128.68, 128.24, 126.26, 101.87, 28.44, 18.42, 13.38.

HRMS (APCI): calcd. for C_12_H_16_N_3_ [M+H]^+^ = 202.1339, found [M+H]^+^ = 202.1340.

4-(Cyclohexylmethyl)-3-ethyl-1*H*-pyrazol-5-amine **(S64)**

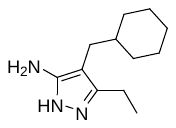

The compound was prepared according to General procedure B1 using 2-(cyclohexylmethyl)-3-oxopentanenitrile (1.1 g, 5.47 mmol; 1 eq.), N_2_H_4_.H_2_O (60 % in H_2_O, 1.7 mL, 7.1 mmol; 1.3 eq.), CH_3_SO_3_H (35 µL, 0.55 mmol, 0.1 eq.) and EtOH (10 mL). Reaction time: 4 h at reflux. The residue obtained after workup was loaded on ISOLUTE^®^ HM-N and purified by column chromatography on silica gel using Biotage Selekt purification system (dichloromethane:MeOH, gradient 19:1 to 9:1). The product was obtained as a yellow oil (330 mg, 29%).

^1^H NMR (500 MHz, Chloroform-*d*) *δ* (ppm) 2.55 – 2.46 (m, 2H), 2.14 (dd, *J* = 7.2, 2.1 Hz, 2H), 1.73 – 1.59 (m, 5H), 1.40 – 1.32 (m, 1H), 1.22 – 1.10 (m, 6H), 0.95 – 0.84 (m, 2H).

^13^C NMR (126 MHz, Chloroform-*d*) *δ* (ppm) 153.24, 143.78, 102.31, 39.05, 33.49, 30.45, 26.63, 26.41, 18.40, 13.22.

HRMS (APCI): calcd. for C_12_H_22_N_3_ [M+H]^+^ = 208.1808, found [M+H]^+^ = 208.1808.

4-(2-Chlorobenzyl)-3-ethyl-1H-pyrazol-5-amine **(S65)**

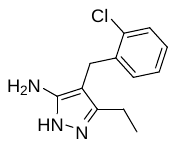

The compound was prepared according to General procedure B1 using 2-(2-chlorobenzyl)-3-oxopentanenitrile (670 mg, 3.01 mmol; 1 eq.), N_2_H_4_.H_2_O (80 % in H_2_O, 0.37 mL, 6.03 mmol; 2 eq.), CH_3_SO_3_H (20 µL, 0.30 mmol, 0.1 eq.) and EtOH (10 mL). Reaction time: 4 h at reflux. The residue obtained after the workup was loaded on ISOLUTE^®^ HM-N and purified by column chromatography on silica gel using Biotage Selekt purification system (dichloromethane:MeOH, gradient 19:1 to 9:1). The product was obtained as a colorless wax (180 mg, 25%).

^1^H NMR (500 MHz, Chloroform-*d*) *δ* (ppm) 7.38 – 7.31 (m, 1H), 7.16 – 7.09 (m, 2H), 7.08 – 7.03 (m, 1H), 3.75 (s, 2H), 2.50 (q, *J* = 7.6 Hz, 2H), 1.12 (t, *J* = 7.6 Hz, 3H).

^13^C NMR (126 MHz, Chloroform-*d*) *δ* (ppm) 152.81, 144.97, 137.72, 133.97, 129.52, 129.33, 127.47, 126.87, 99.65, 25.97, 18.47, 13.31.

HRMS (APCI): calcd. for C_12_H_15_ClN_3_ [M+H]^+^ = 236.0949, found [M+H]^+^ = 236.0952.

3-Ethyl-4-(pyridin-3-ylmethyl)-1*H*-pyrazol-5-amine **(S66)**

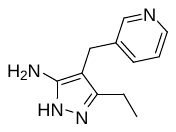

The compound was prepared according to General procedure B1 using 3-oxo-2-(pyridin-3-ylmethyl)pentanenitrile (560 mg, 3.0 mmol; 1 eq.), N_2_H_4_.H_2_O (80 % in H_2_O, 0.37 mL, 6.05 mmol; 2 eq.), CH_3_SO_3_H (20 µL, 0.30 mmol, 0.1 eq.) and EtOH (10 mL). Reaction time: 4 h at reflux. The residue obtained after the workup was loaded on ISOLUTE^®^ HM-N and purified by column chromatography on silica gel using Biotage Selekt purification system (dichloromethane:MeOH, gradient 19:1 to 9:1). The product was obtained as colorless wax (200 mg, 33%).

^1^H NMR (500 MHz, Chloroform-*d*) *δ* (ppm) 7.38 – 7.31 (m, 1H), 7.16 – 7.09 (m, 2H), 7.08 – 7.04 (m, 1H), 3.75 (s, 2H), 2.50 (q, *J* = 7.6 Hz, 2H), 1.12 (t, *J* = 7.6 Hz, 3H).

^13^C NMR (126 MHz, Chloroform-*d*) *δ* (ppm) 152.81, 144.97, 137.72, 133.97, 129.52, 129.33, 127.47, 126.87, 99.65, 25.97, 18.47, 13.31.

HRMS (APCI): calcd. for C_11_H_15_N_4_ [M+H]^+^ = 203.1291, found [M+H]^+^ = 203.1292.

3-Ethyl-4-(2-methoxybenzyl)-1*H*-pyrazol-5-amine **(S67)**

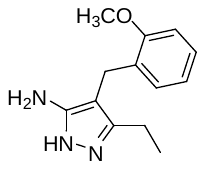

The compound was prepared according to General procedure B1 using 2-(2-methoxybenzyl)-3-oxopentanenitrile (670 mg, 3.01 mmol; 1 eq.), N_2_H_4_.H_2_O (80 % in H_2_O, 0.37 mL, 6.0 mmol; 2 eq.), CH_3_SO_3_H (20 µL, 0.30 mmol, 0.1 eq.) and EtOH (10 mL). Reaction time: 4 h at reflux. The residue obtained after the workup was loaded on ISOLUTE^®^ HM-N and purified by column chromatography on silica gel using Biotage Selekt purification system (dichloromethane:MeOH, gradient 19:1 to 9:1). The product was obtained as colorless wax (200 mg, 28%).

^1^H NMR (500 MHz, Chloroform-*d*) *δ* (ppm) 7.17 (td, *J* = 7.8, 1.8 Hz, 1H), 7.09 – 7.02 (m, 1H), 6.89 – 6.83 (m, 2H), 3.85 (s, 3H), 3.63 (s, 2H), 2.57 (q, *J* = 7.6 Hz, 2H), 1.17 (t, *J* = 7.6 Hz, 3H).

^13^C NMR (126 MHz, Chloroform-*d*) *δ* (ppm) 157.29, 153.76, 144.00, 129.30, 128.85, 127.40, 120.75, 110.40, 101.71, 55.44, 22.71, 18.41, 13.22.

3-Ethyl-4-(4-fluorobenzyl)-1*H*-pyrazol-5-amine **(S68)**

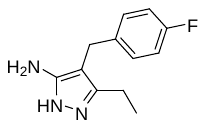

The compound was prepared according to General procedure B1 using 2-(4-fluorobenzyl)-3-oxopentanenitrile (820 mg, 4.02 mmol; 1 eq.), N_2_H_4_.H_2_O (80 % in H_2_O, 0.49 mL, 8.1 mmol; 2 eq.), CH_3_SO_3_H (25 µL, 0.40 mmol, 0.1 eq.) and EtOH (10 mL). Reaction time: 4 h at reflux. The residue obtained after the workup was loaded on ISOLUTE^®^ HM-N and purified by column chromatography on silica gel using Biotage Selekt purification system (dichloromethane:MeOH, gradient 19:1 to 9:1). The product was obtained as colorless wax (200 mg, 23%).

^1^H NMR (500 MHz, Chloroform-*d*) *δ* (ppm) 7.16 – 7.08 (m, 2H), 6.99 – 6.91 (m, 2H), 3.66 (s, 2H), 2.53 (q, *J* = 7.6 Hz, 2H), 1.17 (t, *J* = 7.6 Hz, 3H).

^13^C NMR (126 MHz, Chloroform-*d*) *δ* (ppm) 161.56 (d, *J* = 243.7 Hz), 153.50, 143.98, 135.95, 129.56 (d, *J* = 8.2 Hz), 115.40 (d, *J* = 21.1 Hz), 101.81, 27.65, 18.42, 13.39.

^19^F NMR (471 MHz, Chloroform-*d*) *δ* (ppm) -117.30.

HRMS (APCI): calcd. for C_12_H_15_FN_3_ [M+H]^+^ = 220.1245, found [M+H]^+^ = 220.1245.

3-Ethyl-4-(1-methylpiperidin-4-yl)-1*H*-pyrazol-5-amine **(S69)**

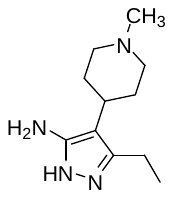

The compound was prepared according to General procedure B1 using 2-(1-methylpiperidin-4-yl)-3-oxopentanenitrile (700 mg, 3.62 mmol; 1 eq.), N_2_H_4_.H_2_O (80 % in H_2_O, 0.44 mL, 7.2 mmol; 2 eq.), CH_3_SO_3_H (25 µL, 0.4 mmol, 0.1 eq.) and EtOH (10 mL). Reaction time: 4 h at reflux. The residue obtained after the workup was loaded on ISOLUTE^®^ HM-N and purified by column chromatography on silica gel using Biotage Selekt purification system (dichloromethane:MeOH, gradient 19:1 to 4:1). The product was obtained as a pale yellow foam (160 mg, 23%).

^1^H NMR (500 MHz, Chloroform-*d*) *δ* (ppm) 4.00 (s, 2H), 3.13 – 3.02 (m, 2H), 2.59 (q, *J* = 7.6 Hz, 2H), 2.39 (s, 3H), 2.31 – 2.24 (m, 1H), 2.20 – 2.04 (m, 4H), 1.79 – 1.66 (m, 2H), 1.20 (t, *J* = 7.6 Hz, 3H).

^13^C NMR (126 MHz, Chloroform-*d*) *δ* (ppm) 152.88, 142.89, 106.59, 56.71, 46.42, 31.79, 31.36, 18.78, 13.89.

4-((1-(*tert*-Butyl)-1*H*-pyrazol-4-yl)methyl)-3-ethyl-1*H*-pyrazol-5-amine **(S70)**

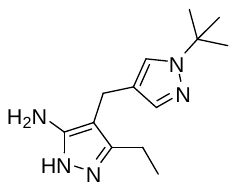

The compound was prepared according to General procedure B1 using 2-((1-(*tert*-butyl)-1*H*-pyrazol-4-yl)methyl)-3-oxopentanenitrile (450 mg, 1.92 mmol; 1 eq.), N_2_H_4_.H_2_O (80 % in H_2_O, 0.23 mL, 3.8 mmol; 2 eq.), CH_3_SO_3_H (25 µL, 0.4 mmol, 0.1 eq.) and EtOH (6 mL). Reaction time: 4 h at reflux. The residue obtained after the workup was loaded on ISOLUTE^®^ HM-N and purified by column chromatography on silica gel using Biotage Selekt purification system (dichloromethane:MeOH, gradient 19:1 to 4:1). The product was obtained as a pale yellow foam (200 mg, 42%).

^1^H NMR (500 MHz, Chloroform-*d*) *δ* (ppm) 7.34 (s, 1H), 7.24 (s, 1H), 3.52 (s, 2H), 2.56 (q, *J* = 7.6 Hz, 2H), 1.55 (s, 9H), 1.19 (t, *J* = 7.6 Hz, 3H).

^13^C NMR (126 MHz, Chloroform-*d*) *δ* (ppm) 153.21, 143.33, 137.89, 124.13, 119.16, 102.02, 58.13, 29.83, 18.24, 17.46, 13.32.

4-([1,1'-Biphenyl]-3-yl)-3-ethyl-1*H*-pyrazol-5-amine **(S71)**

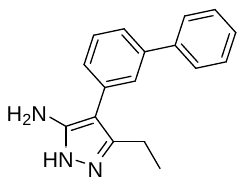

This compound was synthesized according to the procedure reported in *J. Org. Chem.* **2018**, *83*, 24, 15380–15405.

3-Ethyl-4-(4-(trifluoromethyl)phenyl)-1*H*-pyrazol-5-amine **(S72)**

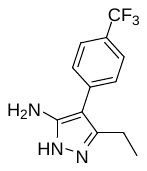

This compound was synthesized according to the procedure reported in *J. Org. Chem.* **2018**, *83*, 24, 15380–15405.

3-Ethyl-4-(4-methoxyphenyl)-1*H*-pyrazol-5-amine **(S73)**

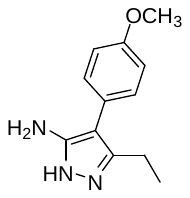

This compound was synthesized according to the procedure reported in *J. Org. Chem.* **2018**, *83*, 24, 15380–15405.

3-Ethyl-4-(3-(trifluoromethyl)phenyl)-1*H*-pyrazol-5-amine **(S74)**

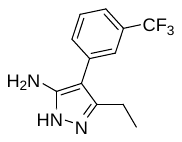

This compound was synthesized according to the procedure reported in *J. Org. Chem.* **2018**, *83*, 24, 15380–15405.

4-(Benzo[*d*][1,3]dioxol-5-yl)-3-ethyl-1*H*-pyrazol-5-amine **(S75)**

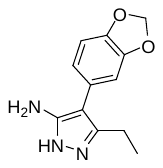

This compound was synthesized according to the procedure reported in *J. Org. Chem.* **2018**, *83*, 24, 15380–15405.

Methyl 3-(5-amino-3-ethyl-1*H*-pyrazol-4-yl)benzoate **(S76)**

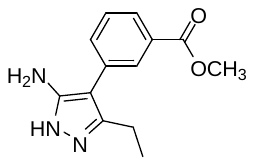

This compound was synthesized according to the procedure reported in *J. Org. Chem.* **2018**, *83*, 24, 15380–15405.

3-Ethyl-4-(4-(methylsulfonyl)phenyl)-1*H*-pyrazol-5-amine **(S77)**

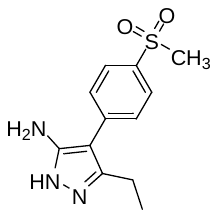

This compound was synthesized according to the procedure reported in *J. Org. Chem.* **2018**, *83*, 24, 15380–15405.

3-Ethyl-4-(4-(piperidin-1-ylsulfonyl)phenyl)-1*H*-pyrazol-5-amine **(S78)**

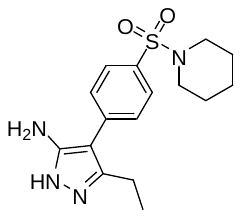

This compound was synthesized according to the procedure reported in *J. Org. Chem.* **2018**, *83*, 24, 15380–15405.

3-Ethyl-4-(3-(trifluoromethoxy)phenyl)-1*H*-pyrazol-5-amine **(S79)**

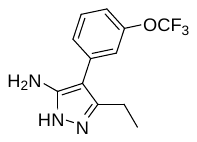

This compound was synthesized according to the procedure reported in *J. Org. Chem.* **2018**, *83*, 24, 15380–15405.

3-Ethyl-4-(naphthalen-1-yl)-1*H*-pyrazol-5-amine **(S80)**

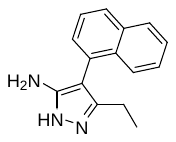

This compound was synthesized according to the procedure reported in *J. Org. Chem.* **2018**, *83*, 24, 15380–15405.

3-Ethyl-4-(4-morpholinophenyl)-1*H*-pyrazol-5-amine **(S81)**

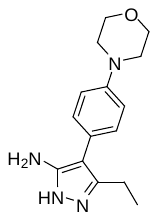

This compound was synthesized according to the procedure reported in *J. Org. Chem.* **2018**, 83, 24, 15380–15405.

*N*-(3-ethyl-4-(4-fluorophenyl)-1*H*-pyrazol-5-yl)formamide **(S82)**

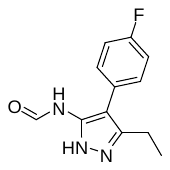

3-Ethyl-4-(4-fluorophenyl)-1*H*-pyrazol-5-amine (400 mg, 1.95 mmol; 1 eq.) was mixed with formic acid (4 mL). The mixture was stirred in the microwave reactor at 110 °C for 3 h, then poured into water (50 mL) and extracted with EtOAc (3 × 25 mL). The combined organic extracts were washed with brine (50 mL), dried over MgSO_4_, filtered, and the solvent was evaporated. The residue was purified by column chromatography on silica gel (hexane:EtOAc, gradient 1:0 to 0:1). The product was obtained as a white solid (250 mg, 55%).

^1^H NMR (500 MHz, DMSO-*d*_6_) *δ* (ppm) 12.53 (s, 1H), 9.69 (d, *J* = 10.9 Hz, 1H), 8.69 – 7.85 (m, 1H), 7.37 – 7.16 (m, 4H), 2.61 (q, *J* = 7.6 Hz, 2H), 1.13 (t, *J* = 7.6 Hz, 3H).

^13^C NMR (126 MHz, DMSO-*d*_6_) *δ* (ppm) 163.35, 160.81 (d, *J* = 242.9 Hz), 143.10 (d, *J* = 88.9 Hz), 130.97 (d, *J* = 8.1 Hz), 130.42 (d, *J* = 8.1 Hz), 128.24 (d, *J* = 3.2 Hz), 115.31 (d, *J* = 21.2 Hz), 115.03, 17.72, 13.27 (d, *J* = 8.1 Hz).

^19^F NMR (471 MHz, DMSO-*d*_6_) *δ* (ppm) -116.23.

3-Ethyl-4-(4-fluorophenyl)-*N*-methyl-1*H*-pyrazol-5-amine **(S83)**

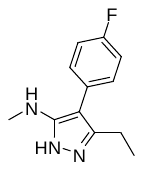

*N*-(3-Ethyl-4-(4-fluorophenyl)-1*H*-pyrazol-5-yl)formamide (500 mg, 2.14 mmol, 1 eq.) was dissolved in THF (6 mL) and the solution was cooled to 0 °C. Solution of LiAlH_4_ in THF (2 M, 1.1 mL, 1 eq.) was added and the mixture was stirred under reflux for 16 h. The mixture was cooled to 23 °C, quenched with aqueous solution of sodium hydroxide (1 M, 50 mL) and extracted with EtOAc (3 × 30 mL). The combined organic extracts were washed with brine (50 mL), dried over MgSO_4_, filtered, and the solvent was evaporated. The residue was purified by column chromatography on silica gel (dichloromethane:MeOH, gradient 9:1 to 5:1). The product was obtained as a white solid (300 mg, 64%).

^1^H NMR (500 MHz, DMSO-*d*_6_) *δ* (ppm) 11.45 (s, 1H), 7.33 – 7.26 (m, 2H), 7.21 – 7.14 (m, 2H), 4.60 (s, 1H), 2.66 (s, 1H), 2.52 (q, *J* = 7.5 Hz, 2H), 1.09 (t, q, *J* = 7.5 Hz, 3H).

^13^C NMR (126 MHz, DMSO-*d*_6_) *δ* (ppm) 160.13 (d, *J* = 241.7 Hz), 154.39, 141.79^*^, 130.22 (d, *J* = 3.1 Hz), 130.05 (d, *J* = 7.9 Hz), 115.17 (d, *J* = 21.1 Hz), 102.35, 30.58, 17.97, 13.35.

* - detected by HMBC

^19^F NMR (471 MHz, DMSO-*d*_6_) *δ* (ppm) -117.66.

2-((2-Methyl-1,3-dioxolan-2-yl)methyl)-1*H*-benzo[*d*]imidazole **(S84)**

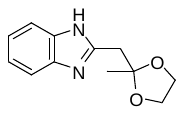

Potassium ethoxide (980 mg, 11.5 mmol, 1 eq.) was added to a solution of benzene-1,2-diamine (2 g, 18.5 mmol, 1.65 eq.) and ethyl 2-(2-methyl-1,3-dioxolan-2-yl)acetate (2 g, 11.2 mmol, 1eq.) in anhydrous EtOH (10 mL). The reaction mixture was refluxed for 72 h. The solvent was evaporated and the residue was purified by column flash chromatography on silica gel (hexane:EtOAc:MeOH, gradient 1:1:0 to 2:4:1) to afford the product as a brown solid (840 mg, 34%).

^1^H NMR (500 MHz, DMSO-*d*_6_) *δ* (ppm) 11.97 (s, 1H), 7.53 (d, *J* = 7.5 Hz, 1H), 7.44 (d, *J* = 7.5 Hz, 1H), 7.16 – 7.07 (m, 2H), 3.91 – 3.89 (m, 4H), 3.10 (s, 2H), 1.35 (s, 3H).

^13^C NMR (126 MHz, DMSO-*d*_6_) *δ* (ppm) 150.32, 143.14, 134.44, 121.43, 120.71, 118.16, 110.92, 107.96, 64.15, 38.42, 24.07.

1-(1*H*-benzo[*d*]imidazol-2-yl)propan-2-one **(S85)**

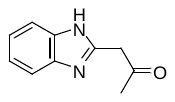

Water (0.5 mL) and hydrochloric acid (35%, 0.5 mL) were added to a solution of 2-((2-methyl-1,3-dioxolan-2-yl)methyl)-1*H*-benzo[*d*]imidazole (230 mg, 1.06 mmol, 1.0 eq.) in THF (5 mL). The reaction mixture was refluxed for 3 h. The solvent was evaporated and the crude product (183 mg) was used in the next step without further purification.

Ethyl 2-(6-morpholino-1*H*-benzo[*d*]imidazol-2-yl)acetate **(S86)**

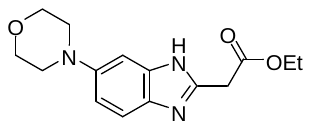

Pd/C (10%, 10 mg) was added to a solution of 5-morpholino-2-nitroaniline (350 mg, 1.57 mmol, 1 eq.) in EtOH (5 mL), and the mixture was bubbled with H_2_ for 5 min. The mixture was then stirred at reflux under an atmosphere of H_2_ for 1 h. The mixture was filtered through an micro HPLC filter and the solvent was evaporated. The crude diamine was used in the next step without further purification.

Ethyl 3-ethoxy-3-iminopropanoate hydrochloride (306 mg, 1.57 mmol, 1 eq.) was dissolved in EtOH (5 mL), the solution was cooled to 0 °C and a solution of the diamine in EtOH (5 mL) was slowly added. The mixture was stirred at 0 °C for 45 min, then under reflux for additional 2 h. The mixture was poured into water (20 mL) neutralised with saturated solution of sodium bicarbonate (10 mL) and extracted with EtOAc (3×10 mL). The combined organic extracts were washed with brine (30 mL), dried over MgSO_4_, and the solvent was evaporated. The crude product was purified by column flash chromatography on silica gel (dichloromethane:MeOH, 10:1) to afford the products as a pale brown foam (320 mg, 70%).

^1^H NMR (500 MHz, DMSO-*d*_6_) *δ* (ppm) 12.06 (s, 1H), 7.36 (d, *J* = 8.7 Hz, 1H), 6.95 (s, 1H), 6.90 (dd, *J* = 8.7, 2.2 Hz, 1H), 4.16 – 4.08 (m, 2H), 3.89 (s, 2H), 3.79 – 3.73 (m, 4H), 3.08 – 3.03 (m, 4H), 1.20 (t, *J* = 7.1 Hz, 3H).

^13^C NMR (126 MHz, DMSO-*d*_6_) *δ* (ppm) 168.79, 147.43, 146.83, 112.89, 66.25, 60.68, 50.41, 35.09, 14.00.

Methyl-3-(4-amino-3-nitrophenyl)acrylate **(S87)**

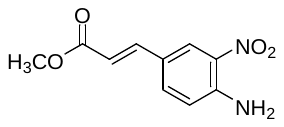

A mixture of 4-bromo-2-nitroaniline (2 g, 9.20 mmol, 1 eq.), triethylamine (3.8 mL, 27.60 mmol, 3 eq.), methyl acrylate (1.65 mL, 18.40 mmol, 2 eq.) and Pd(dppf)Cl_2_ (0.32 g, 0.46 mmol, 0.05 eq.) in degassed DMF (12 mL) and water (0.2 mL) was stirred at 100 °C for 16 h. The mixture was poured into brine solution (100 mL) and the solid was collected by filtration. The solid was suspended in dichloromethane:MeOH (10:1 mL), the mixture was filtered, and the solid was dried under *vacuum*. The product was obtained as a pale brown solid (1.18 g, 57%).

^1^H NMR (500 MHz, DMSO-*d*_6_) *δ* (ppm) 8.23 (d, *J* = 2.1 Hz, 1H), 7.85 – 7.80 (m, 3H), 7.58 (d, *J* = 16.0 Hz, 1H), 7.04 (d, *J* = 8.8 Hz, 1H), 6.44 (d, *J* = 16.0 Hz, 1H), 3.70 (s, 3H).

^13^C NMR (126 MHz, DMSO-*d*_6_) *δ* (ppm) 166.77, 147.29, 143.18, 133.52, 129.90, 127.40, 121.64, 119.87, 114.89, 51.23.

Methyl 3-(2-(2-ethoxy-2-oxoethyl)-1*H*-benzo[*d*]imidazol-6-yl)propanoate **(S88)**

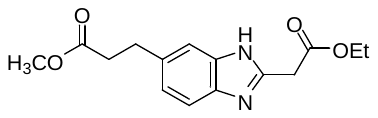

Pd/C (10%, 10 mg) was added to a solution of methyl 3-(4-amino-3-nitrophenyl)acrylate (400 mg, 1.78 mmol, 1 eq.) in EtOH (10 mL), and the mixture was bubbled with H_2_ for 5 min. The mixture was then stirred for 1 h at reflux under an atmosphere of H_2_. The mixture was filtered through an HPLC filter and the solvent was evaporated. The crude diamine was used in the next step without further purification.

Ethyl 3-ethoxy-3-iminopropanoate hydrochloride (350 mg, 1.78 mmol, 1 eq.) was dissolved in EtOH (5 mL) and the solution was cooled to 0 °C. A solution of the diamine in EtOH (5 mL) was slowly added and the mixture was stirred at 0 °C for 45 min, then under reflux for additional 2 h. The mixture was poured into water (20 mL), neutralized with saturated solution of sodium bicarbonate (10 mL) and extracted with EtOAc (3 × 10 mL). The combined organic extracts were washed with brine (30 mL), dried over MgSO_4_, and the solvent was evaporated. The residue was purified by column flash chromatography on silica gel (dichloromethane:MeOH, 20:1 to 10:1) to afford the products as a white solid (430 mg, 87%).

^1^H NMR (500 MHz, DMSO-*d*_6_) *δ* (ppm) 7.50 (d, *J* = 8.3 Hz, 1H), 7.40 (d, *J* = 1.5 Hz, 1H), 7.10 (dd, *J* = 8.3, 1.7 Hz, 1H), 4.24 (q, *J* = 7.1 Hz, 2H), 4.10 (s, 2H), 3.66 (s, 3H), 3.06 (t, *J* = 7.8 Hz, 2H), 2.72 – 2.63 (m, 2H), 1.30 (t, *J* = 7.2 Hz, 3H).

Ethyl 2-(1-benzyl-1*H*-imidazol-2-yl)acetate **(S89)**

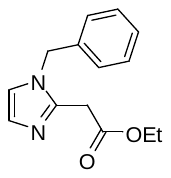

1-Benzyl-2-methylimidazole (3.32 g, 17.32 mmol; 1eq.) was dissolved in dry THF (30 mL). To the solution was added TEA (7.2 mL, 51.97 mmol; 3 eq.) and ethyl chloroformate (4.6 mL, 48.5 mmol; 2.8 eq.). The reaction mixture was stirred at 23 °C for 24 h, then poured into water (50 mL) and extracted with EtOAc (3 × 25 mL). The combined organic extracts were washed with brine (50 mL), dried over MgSO_4_, and the solvent was evaporated. The residue was purified by column flash chromatography (EtOAc:dichloromethane:MeOH, 1:1:0.5). The product was obtained as a yellow wax (1.0 g, 24%).

^1^H NMR (500 MHz, Chloroform-*d*) *δ* (ppm) 7.37 – 7.26 (m, 3H), 7.12 – 7.06 (m, 2H), 7.02 (d, *J* = 1.4 Hz, 1H), 6.86 (d, *J* = 1.3 Hz, 1H), 5.13 (s, 2H), 4.11 (q, *J* = 7.1 Hz, 2H), 3.75 (s, 2H), 1.22 (t, *J* = 7.2 Hz, 3H).

^13^C NMR (126 MHz, Chloroform-*d*) *δ* (ppm) 169.01, 141.49, 136.06, 129.10, 128.27, 128.02, 127.13, 121.11, 61.55, 50.17, 34.00, 14.21.

*tert*-Butyl 2-(2-ethoxy-2-oxoethyl)-3,4,6,7-tetrahydro-5*H*-imidazo[4,5-*c*]pyridine-5-carboxylate **(S90)**

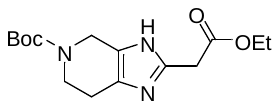

To a cold (0 °C) solution of 2-(4,5,6,7-tetrahydro-3*H*-imidazo[4,5-*c*]pyridine-2-yl)acetate dihydrochloride (262 mg, 0.93 mmol) in dichloromethane (5 mL) was added triethylamine (0.388 mL, 2.79 mmol, 3 eq.) followed by Boc_2_O (203 mg, 0.93 mmol, 1 eq.). The mixture was stirred at 0 °C for 30 min, then at 25 °C for additional 4 h. Saturated aqueous solution of NH_4_Cl (10 mL) was added and the mixture was extracted with dichloromethane (3 × 30 mL). The combined organic extracts were washed with brine (20 mL), dried over MgSO_4_, filtered, and the solvent was evaporated *in vacuo*. The residue was purified by column chromatography on silica gel (hexane:EtOAc:MeOH, gradient 2:1:0 to 0:1:0 to 0:20:1). The product was obtained as a yellow wax (177 mg, 62%).

^1^H NMR (500 MHz, Chloroform-*d*) *δ* (ppm) 10.24 (s, 1H), 4.39 (s, 2H), 4.13 (q, *J* = 7.1 Hz, 2H), 3.75 (s, 2H), 3.65 (t, *J* = 5.9 Hz, 2H), 2.59 (t, *J* = 5.6 Hz, 2H), 1.42 (s, 9H), 1.22 (t, *J* = 7.2 Hz, 3H).

^13^C NMR (126 MHz, Chloroform-*d*) *δ* (ppm) 170.17, 155.17, 139.77, 80.02, 77.36, 61.50, 43.03, 42.16, 40.99, 34.29, 28.46, 22.31, 14.11.

HRMS (APCI): calcd. for C_15_H_24_N_3_O_4_ [M+H]^+^ = 310.1761, found [M+H]^+^ = 310.1758.

*N*-(2-morpholinoethyl)-2-nitroaniline **(S91)**

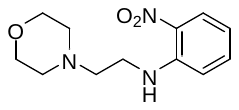

Sodium hydride (60%, 580 mg, 14.50 mmol, 2 eq.) was added to a solution of 2-nitroaniline (1 g, 7.25 mmol, 1 eq.) in anhydrou THF (50 mL) at 0 °C and the mixture was stirred for 30 min at that temperature. TBAI (130 mg, 0.36 mmol, 0.05 eq.) and 4-(2-chloroethyl)morpholine hydrochloride (1.34 g, 7.2 mmol, 1eq) were added and the mixture was stirred at 23 °C for 16 h. The mixture was poured into water (50 mL) and extracted with EtOAc (3 × 50 mL). The combined organic extracts were washed with brine (50 mL), dried over MgSO_4_, filtered, and the solvent was evaporated. The crude product (1.76 g, 97%) was used in the next step without further purification.

^1^H NMR (300 MHz, Chloroform-*d*) *δ* (ppm) 8.46 (s, 1H), 8.17 (dd, *J* = 8.6, 1.6 Hz, 1H), 7.49 – 7.36 (m, 1H), 6.82 (d, *J* = 1.2 Hz, 1H), 6.70 – 6.58 (m, 1H), 3.82 – 3.68 (m, 4H), 3.59 (t, *J* = 6.9 Hz, 2H), 2.72 (t, *J* = 6.7 Hz, 2H), 2.54 – 2.49 (m, 4H).

Ethyl 2-(1-(2-morpholinoethyl)-1*H*-benzo[*d*]imidazol-2-yl)acetate **(S92)**

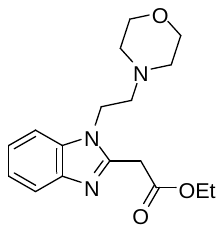

*N*-(2-morpholinoethyl)-2-nitroaniline (1.76 g, 7.0 mmol, 1 eq.) was dissolved in EtOH (20 mL), Pd/C (10%, 20 mg) was added, and the mixture was then stirred in a high-pressure apparatus at 60 °C under hydrogen (25 bar) for 16 h. The mixture was filtered through an HPLC filter and the solvent was evaporated. The crude diamine was used in the next step without further purification.

Ethyl 3-ethoxy-3-iminopropanoate hydrochloride (1.37 g, 7.0 mmol, 1 eq.) was dissolved in EtOH (5 mL), the solution was cooled to 0 °C and a solution of the diamine in EtOH (5 mL) was slowly added. The mixture was stirred for 45 min at 0 °C, then under reflux for additional 2 h. The mixture was poured into water (20 mL) neutralised with saturated solution of sodium bicarbonate (10 mL) and extracted with EtOAc (3 × 20 mL). The combined organic extracts were washed with brine (30 mL), dried over MgSO_4_, and the solvent was evaporated. The residue was purified by column flash chromatography on silica gel (dichloromethane:MeOH, 10:1) to afford the product as a light brown foam (340 mg, 15%).

^1^H NMR (300 MHz, Chloroform-*d*) *δ* (ppm) 7.77 – 7.69 (m, 1H), 7.37 – 7.31 (m, 1H), 7.30 – 7.22 (m, 1H), 4.28 (t, *J* = 6.6 Hz, 2H), 4.20 (q, *J* = 7.1 Hz, 2H), 4.13 (s, 2H), 3.73 – 3.63 (m, 4H), 2.73 (t, *J* = 6.6 Hz, 2H), 2.55 – 2.44 (m, 4H), 1.26 (t, *J* = 7.1 Hz, 3H).

^13^C NMR (75 MHz, Chloroform-*d*) *δ* (ppm) 168.60, 148.07, 142.66, 135.18, 122.86, 122.40, 119.94, 109.55, 66.96, 61.83, 57.66, 54.20, 42.19, 34.81, 14.26.

*N^1^*,*N^1^*-dimethyl-*N^2^*-(2-nitrophenyl)ethane-1,2-diamine **(S93)**

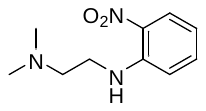

2-Nitroaniline (1 g, 7.25 mmol, 1 eq.) was dissolved in anhydrous THF (30 mL). The solution was cooled to 0 °C, sodium hydride (60%, 0.58 g, 14.5 mmol, 2 eq.) was added and the mixture was stirred at 0 °C for 30 min. Tetrabutylammonium iodide (130 mg, 0.36 mmol, 0.05 eq.) and *N^1^*,*N^1^*-dimethylethane-1,2-diamine hydrochloride (1.04 g, 7.2 mmol, 1eq) were added and the mixture was stirred at 23 °C for 16 h. The mixture was poured into water (50 mL) and extracted with EtOAc (3 × 30 mL). The combined organic extracts were washed with brine (50 mL), dried over MgSO_4_, filtered, and the solvent was evaporated. The crude product (690 mg, 46%) was used in the next step without further purification.

^1^H NMR (300 MHz, Chloroform-*d*) *δ* (ppm) 8.28 (s, 1H), 8.16 (dd, *J* = 8.6, 1.6 Hz, 1H), 7.49 – 7.38 (m, 1H), 6.84 (dd, *J* = 8.7, 1.2 Hz, 1H), 6.69 – 6.58 (m, 1H), 3.44 – 3.31 (m, 2H), 2.66 (t, *J* = 6.3 Hz, 2H), 2.33 (s, 6H).

^13^C NMR (75 MHz, Chloroform-*d*) *δ* (ppm) 145.46, 136.27, 132.24, 127.05, 115.28, 114.03, 57.54, 45.35, 40.78.

Ethyl 2-(1-(2-(dimethylamino)ethyl)-1*H*-benzo[*d*]imidazol-2-yl)acetate **(S94)**

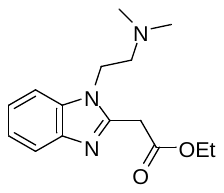

*N^1^*,*N^1^*-dimethyl-*N^2^*-(2-nitrophenyl)ethane-1,2-diamine (680 mg, 3.20 mmol, 1 eq.) was dissolved in EtOH (20 mL), Pd/C (10%, 20 mg) was added, and the mixture was stirred in a high-pressure apparatus at 60 °C under hydrogen (25 bar) for 16 h. The mixture was filtered through and HPLC filter and the solvent was evaporated. The crude diamine was used in the next step without further purification.

The diamine was dissolved in diethyl malonate (8 mL) and stirred for 3 h at 150 °C. The crude mixture was purified by column flash chromatography on silica gel (dichloromethane:MeOH, 10:1) to afford the product as a light brown foam (320 mg, 36%).

^1^H NMR (300 MHz, Chloroform-*d*) *δ* (ppm) 7.76 – 7.68 (m, 1H), 7.39 – 7.31 (m, 1H), 7.28 – 7.22 (m, 2H), 4.27 (t, *J* = 7.0 Hz, 2H), 4.20 (q, *J* = 7.2 Hz, 2H), 4.08 (s, 2H), 2.68 (t, *J* = 7.0 Hz, 2H), 2.30 (s, 6H), 1.26 (t, *J* = 7.1 Hz, 3H).

^13^C NMR (75 MHz, Chloroform-*d*) *δ* (ppm) 168.62, 148.06, 142.82, 135.27, 122.77, 122.24, 119.93, 109.54, 61.75, 58.34, 45.89, 42.76, 34.76, 14.24.

(*E*)-2-(2-(piperidin-1-yl)vinyl)-1*H*-benzo[*d*]imidazole **(S95)**

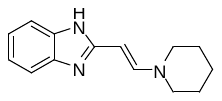

A mixture of 2-ethynyl-1*H*-benzimidazole (60 mg, 0.42 mmol, 1eq.) and piperidine (41 µL, 0.42 mmol, 1 eq.) in THF (5 mL) was stirred at 23 °C for 48 h. The solvent was evaporated and the crude product (95 mg, quantitative) was used in the next step without further purification.

^1^H NMR (300 MHz, chloroform-*d*) *δ* (ppm) 7.58 (d, *J* = 13.6 Hz, 1H), 7.47 – 7.35 (m, 2H), 7.15 – 7.03 (m, 2H), 5.16 (d, *J* = 13.5 Hz, 1H), 3.22 – 3.05 (m, 4H), 1.67 – 1.50 (m, 6H).

*N*1-benzylbenzene-1,2-diamine **(S96)**

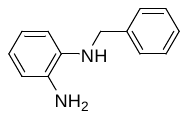

Benzyl bromide (2.7 g, 16.0 mmol) was added to a mixture *o*-diaminobenzene (8 g, 74.0 mmol) and K_2_CO_3_ (6.0 g, 44.0 mmol) in anhydrous MeOH (40 mL). The reaction mixture was stirred under N_2_ at 25 °C for 20 hrs, the solvent was evaporated and the residue was purified by column flash chromatography on silica gel (hexane:EtOAc, 2:1) to afford the product a dark red wax (2.58 g, 81%).

^1^H NMR (500 MHz, Chloroform-*d*) *δ* (ppm) δ 7.42 (d, *J* = 7.1 Hz, 2H), 7.37 (t, *J* = 7.6 Hz, 2H), 7.34 – 7.28 (m, 1H), 6.86 – 6.79 (m, 1H), 6.78 – 6.67 (m, 3H), 4.33 (s, 2H).

^13^C NMR (126 MHz, Chloroform-*d*) *δ* (ppm) 139.56, 137.82, 134.36, 128.74, 127.93, 127.40, 120.89, 119.03, 116.71, 112.24, 48.82.

1-Benzyl-2-methyl-1*H*-benzo[d]imidazole **(S97)**

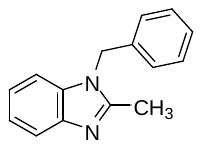

*N*1-benzylbenzene-1,2-diamine (387 mg, 1.95 mmol) was dissolved in acetic acid (2 mL) and EtOH (2 mL). The reaction mixture was refluxed for 18 h, then poured into saturated aqueous solution of sodium bicarbonate (25 mL) and water (25 mL) and extracted with EtOAc (3 × 25 mL). The combined organic layers were washed with brine (25 mL), dried over MgSO_4_, and the solvent was evaporated under reduced pressure. The residue was purified by flash column chromatography (hexane:EtOAc, gradient 1:0 to 1:1). The product was obtained as a red solid (256 mg, 60%).

^1^H NMR (500 MHz, Chloroform-*d*) *δ* (ppm) 2.59 (s, 3H), 5.34 (s, 2H), 7.09-7.06 (m, 2H), 7.85 – 7.69 (m, 3H), 7.36 – 7.19 (m, 3H), 7.75 (s, 1H).

^13^C NMR (126 MHz, Chloroform-*d*) *δ* (ppm) 14.07, 47.27, 109.49, 119.28, 122.23, 122.48, 126.39, 128.08, 129.17, 135.5, 135.98, 142.67, 152.02.

Methyl 2-(1-benzyl-1*H*-benzo[*d*]imidazol-2-yl)acetate **(S98)**

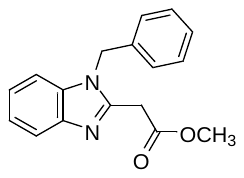

1-Benzyl-2-methyl-1*H*-benzo[*d*]imidazole (610 mg, 2.74 mmol; 1eq.) was dissolved in dry THF (5 mL). To solution was added DIPEA (1.5 mL, 8.22 mmol; 3 eq.) and methyl chloroformate (0.45 mL, 5.76 mmol; 2.1 eq.). The reaction mixture was stirred at room temperature for 18 h, then poured into water (50 mL) and extracted with EtOAc (3 × 25 mL). The combined organic extracts were washed with brine (25 mL), dried over MgSO_4_, and the solvent was evaporated *in vacuo*. The residue was purified by column flash chromatography (100% EtOAc). The product was obtained as a red solid (657 mg, 86%).

^1^H NMR (500 MHz, Chloroform-*d*) *δ* (ppm) 7.70 (d, *J* = 7.7 Hz, 1H), 7.32 – 7.09 (m, 6H), 6.97 (d, *J* = 6.4 Hz, 2H), 5.32 (s, 2H), 3.88 (s, 2H), 3.55 (s, 3H).

^13^C NMR (126 MHz, Chloroform-*d*) *δ* (ppm) 168.69, 147.93, 142.73, 135.82, 135.68, 129.13, 128.13, 126.44, 123.15, 122.50, 120.00, 110.00, 52.65, 47.53, 34.68.

HRMS (APCI): calcd. for C_18_H_19_N_2_O_2_ [M+H]^+^ = 281.1285, found [M+H]^+^ = 281.1291.

2-(1-Benzyl-1*H*-benzo[*d*]imidazol-2-yl)acetonitrile **(S99)**

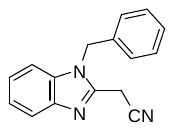

Ethyl 2-cyanoacetate (2.20 g, 20.0 mmol; 1 eq.) followed by methanesulfonic acid (0.1 mL) were added to a solution of *N*1-benzylbenzene-1,2-diamine (2.58 g, 13.0 mmol; 1.54 eq.) in ethylene glycol (15 mL). The solution was refluxed for 4 hrs under N_2_, poured into a mixture of water (100 mL) with saturated aqueous NaHCO_3_ (25 mL), and extracted with EtOAc (3 × 50 mL). The organic extracts were washed with water (100 mL), brine (25 mL), then dried over MgSO_4_, filtered, and the solvent was evaporated. The residue was purified by flash chromatography on silica gel (hexane:EtOAc, 10:1) to yield the product as a white crystalline solid (2.43 g, 75%).

MP = 135.0 – 136.0 ºC.

^1^H NMR (500 MHz, Chloroform-*d*) *δ* (ppm) 7.86 – 7.78 (m, 1H), 7.38 – 7.28 (m, 6H), 7.12 – 7.04 (m, 2H), 5.45 (s, 2H), 3.92 (s, 2H).

13C NMR (126 MHz, Chloroform-*d*) *δ* (ppm) 143.42, 142.19, 136.04, 134.83, 129.50, 128.70, 126.49, 124.07, 123.17, 120.36, 114.21, 109.95, 47.61, 18.43.

HRMS (APCI): calcd. for C_16_H_12_N_3_ [M-H]^-^ = 246.1037, found [M-H]^-^ = 246.1037.

1-Benzyl-5,6-dimethyl-1*H*-benzo[*d*]imidazole **(S100)**

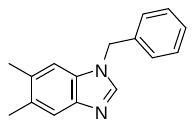

Sodium hydride (60 %, 910 mg, 22.50 mmol, 1.1 eq.) was slowly added to a stirred solution of 5,6-dimethyl-1*H*-benzo[*d*]imidazole (3 g, 20.50 mmol, 1eq.) in DMF (20 mL) at 0 °C. After 30 min was added benzyl bromide (2.7 mL, 22.50 mmol, 1.1 eq.) and the mixture was stirred at 23 °C for 16 h. The the mixture was poured into water (50 mL), the precipitate was collected by filtration, washed with water (10 mL) and diethyl ether (2 ×25 mL). The product obtained as a white solid (4.52 g, 93%), was used in the next step without further purification.

^1^H NMR (500 MHz, Chloroform-*d*) *δ* (ppm) δ 8.23 (s, 1H), 7.42 (s, *J* = 22.5 Hz, 1H), 7.33 (t, *J* = 7.3 Hz, 2H), 7.30 – 7.22 (m, 4H), 5.43 (s, 2H), 2.28 (s, *J* = 2.0 Hz, 3H), 2.27 (s, 3H).

^13^C NMR (126 MHz, Chloroform-*d*) *δ* (ppm) 143.29, 142.18, 137.16, 132.22, 130.94, 129.84, 128.60, 127.55, 127.11, 119.48, 110.51, 47.43, 20.03, 19.76.

1-Benzyl-2,5,6-trimethyl-1*H*-benzo[*d*]imidazole **(S101)**

A suspension of 1-benzyl-5,6-dimethyl-1*H*-benzo[*d*]imidazole (237 mg, 1.10 mmol, 1eq.) in dioxane/THF (2+2 mL) was cooled to -40 °C and a solution of *n*-BuLi (2.7 M, 0.47 mL, 1.26 mmol, 1.2 eq.) was added. The mixture was stirred at -40 °C for 40 min, MeI was added (80 µL, 1.26 mmol, 1.2 eq), and the mixture was stirred at 23 °C for 16 h. The reaction mixture was poured into water (25 mL) and saturated solution of NH_4_Cl (25 mL) and extracted with EtOAc (3 × 25 mL). The combined organic layers were washed with brine (50 mL) and dried over MgSO₄. The solvent was removed under reduced pressure and the residue was purified by column chromatography on silica gel (eluent: EtOAc). The product was obtained as a white solid (156 mg, 59%).

^1^H NMR (500 MHz, CDCl_3_) δ 7.40 (s, 1H), 7.23 – 7.16 (m, 3H), 7.00 – 6.92 (m, 2H), 6.90 (s, 1H), 5.18 (s, 2H), 2.43 (s, 3H), 2.28 (s, 3H), 2.24 (s, 3H).

^13^C NMR (126 MHz, CDCl_3_) δ 151.07, 141.40, 136.32, 134.19, 131.30, 130.78, 129.08, 127.88, 126.26, 119.46, 109.71, 47.08, 20.59, 20.30, 14.03.

Methyl 2-(1-benzyl-5,6-dimethyl-1*H*-benzo[*d*]imidazol-2-yl)acetate **(S102)**

Methyl chloroformate (0.18 mL, 2.38 mmol; 2.5 eq.) was added to a stirred solution of 1-benzyl-2,5,6-trimethyl-1*H*-benzo[*d*]imidazole (250 mg, 0.95 mmol; 1eq.) and DIPEA (0.5 mL, 2.85 mmol; 3 eq.) was in dry THF (5 mL) and the reaction mixture was stirred at 23 °C for 18 h. The mixture was poured into water (50 mL) and extracted with EtOAc (3 × 25 mL). The combined organic extracts were washed with brine (50 mL), dried over MgSO_4_, and the solvent was evaporated. The residue was purified by column flash chromatography (100% EtOAc). The product was obtained as a yellow solid (219 mg, 72%).

^1^H NMR (500 MHz, Chloroform-*d*) *δ* (ppm) ^1^H NMR (500 MHz, CDCl_3_) δ 7.45 (s, 1H), 7.22 – 7.17 (m, 3H), 6.98 – 6.93 (m, 2H), 6.92 (s, 1H), 5.26 (s, 2H), 3.83 (s, 2H), 3.51 (s, 3H), 2.28 (s, 3H), 2.24 (s, 3H).

^13^C NMR (126 MHz, Chloroform-*d*) *δ* (ppm) 168.79, 146.97, 141.25, 134.36, 132.24, 131.29, 129.04, 127.96, 126.32, 119.98, 110.10, 52.53, 47.33, 34.59, 20.62, 20.29.

Methyl 2-(5,6-dimethyl-1*H*-benzo[*d*]imidazol-2-yl)acetate **(S103)**

A mixture of methyl 2-(1-benzyl-5,6-dimethyl-1*H*-benzo[*d*]imidazol-2-yl)acetate (160 mg, 0.52 mmol, 1eq.) and Pd(OH)_2_/C (10 %, 35 mg, 0.03 mmol, 5%) in degassed EtOH (5 mL) was refluxed under a hydrogen atmosphere (1 bar) for 3 h. The mixture was filtered through an HPLC filter and the solvent was evaporated. The crude product was dried under vacuum and used in the next step without further purification. The product was obtained as a white solid (96 mg, 85%).

^1^H NMR (126 MHz, DMSO-*d*_6_) *δ* (ppm) 12.06 (s, 1H), 7.26 (s, 2H), 3.92 (s, 2H), 3.66 (s, 2H), 2.29 (s, 3H).

^13^C NMR (126 MHz, DMSO-*d*_6_) *δ* (ppm) 169.23, 146.58, 129.72, 51.96, 34.88, 19.85.

Ethyl-2-(1*H*-benzo[*d*]imidazol-2-yl)-3-(dimethylamino)acrylate **(S104)**

Ethyl 2-(1*H*-benzo[*d*]imidazol-2-yl)acetate (533 mg, 2.61 mmol, 1 eq.) and *t*-butoxybis(dimethylamino)methane (0.65 mL, 3.13 mmol, 1.2 eq.) were dissolved in toluene (10 mL) and the mixture was stirred at 23 °C for 5 h. The resulting precipitate was collected by filtration, washed with toluene (5 mL), diethyl ether (10 mL) and dried under *vacuum*. The product obtained as a yellow solid (500 mg, 74%), was used in the next step without further purification.

^1^H NMR (500 MHz, DMSO-*d*_6_) *δ* (ppm) 12.09 (s, 1H), 7.69 (s, 1H), 7.54 (d, *J* = 7.6 Hz, 1H), 7.40 (d, *J* = 7.5 Hz, 1H), 7.15 – 7.06 (m, 2H), 4.05 (q, *J* = 7.1 Hz, 2H), 2.73 (s, 6H), 1.13 (t, *J* = 7.1 Hz, 3H).

^13^C NMR (126 MHz, DMSO-*d*_6_) *δ* (ppm) 167.72, 152.41, 148.61, 142.91, 134.32, 121.36, 120.54, 118.21, 110.65, 87.81, 58.76, 14.54.

Ethyl 4-hydroxypyrazolo[5,1-*c*][1,2,4]triazine-3-carboxylate **(S105)**

The compound was prepared according to General procedure C using:

Diazotization step:

3-aminopyrazole (303 mg, 3.65 mmol; 1 eq.), 35% aqueous HCl (1.29 mL, 14.59 mmol; 4 eq.) in EtOH (2 mL) and H_2_O (2 mL), NaNO_2_ (302 mg, 4.38 mmol, 1.2 eq.) in EtOH (2 mL) and H_2_O (2 mL) (reaction time: 15 min), diethylmalonate (0.66 mL, 4.38 mmol, 1.2 eq.) in EtOH (3 mL) and H_2_O (3 mL) and KOAc (2.15 g, 21.88 mmol, 6 eq.). Reaction time: 6 h. EtOAc (30 mL) was added followed by water (20 mL). The organic phase was separated, washed with brine (20 mL), dried over MgSO_4_, filtered, and the solvent was evaporated. The residue was purified by column chromatography on silica gel (hexane:EtOAc, gradient 2:1; 1:1; 1:2; 0:1) to give the product (diethyl 2-(2-(1*H*-pyrazol-5-yl)hydrazono)malonate) as a yellow solid (565 mg, 61%).

Cyclization step:

the yellow solid (25 mg, 0.983 mmol) was dissolved in glacial acetic acid (10 mL) and the mixture was refluxed under N_2_ for 3 h. The solvent was evaporated, to the residue was added dichloromethane (5 mL) and the precipitate was collected by filtration and washed with cold dichloromethane (5 mL). The solid was dried *in vacuo* to afford the product as a yellow solid (190 mg, 93%).

^1^H NMR (500 MHz, DMSO-*d_6_*) *δ* (ppm) 14.58 (s, 1H), 8.11 (d, *J* = 2.1 Hz, 1H), 6.48 (d, *J* = 2.1 Hz, 1H), 4.32 (q, *J* = 7.1 Hz, 2H), 1.31 (t, *J* = 7.1 Hz, 3H).

^13^C NMR (125 MHz, DMSO-*d_6_*) *δ* (ppm) 161.48, 147.12, 144.93, 142.39, 127.15, 89.98, 61.04, 14.06.

HRMS (APCI): calcd. for C_8_H_9_N_4_O_3_ [M+H]^+^ = 209.0669, found [M+H]^+-^= 209.0670.

MP > 250 °C (dec.).

3-(1-Benzyl-1*H*-benzo[*d*]imidazol-2-yl)pyrazolo[5,1-*c*][1,2,4]triazin-4-ol **(S106)**

A mixture of ethyl 4-hydroxypyrazolo[5,1-*c*][1,2,4]triazine-3-carboxylate (532 mg, 2.55 mmol, 1.05 eq.) and *N*-benzylbenzene-1,2-diamine (250 mg, 2.43 mmol, 1 eq.) in EtOH (2 mL) and AcOH (2 mL) was refluxed for 4 h. After cooling to 25 °C, a precipitate formed. EtOH (4 mL) was added and the solid was collected by filtration and washed on filter with EtOH (10 mL). The solid was dried under vacuum to afford the product as a pale yellow solid (45 mg, 54%).

^1^H NMR (500 MHz, DMSO-*d_6_*) *δ* (ppm) 8.10 (d, *J* = 2.1 Hz, 1H), 7.86 (dd, *J* = 6.9, 2.0 Hz, 1H), 7.61 (d, *J* = 7.6 Hz, 1H), 7.43 – 7.34 (m, 2H), 7.29 – 7.21 (m, 5H), 6.63 (d, *J* = 2.2 Hz, 1H), 5.99 (s, 2H).

^13^C NMR (126 MHz, DMSO-*d_6_*) *δ* (ppm) 149.51, 146.64, 144.49, 136.38, 133.57, 128.51, 127.49, 126.93, 124.09, 123.89, 116.92, 111.64, 93.57, 48.64.

HRMS (APCI): calcd. for C_19_H_15_N_6_O [M+H]^+^ = 343.1302, found [M+H]^+-^= 343.1299.

3-(1-Benzyl-1*H*-benzo[*d*]imidazol-2-yl)-8-bromopyrazolo[5,1-*c*][1,2,4]triazin-4-ol **(S107)**

A solution of *N*-bromosuccinimide (55 mg, 0.31 mmol, 1.05 eq.) in DMF (2 mL) was added dropwise to a stirred mixture of 3-(1-benzyl-1*H*-benzo[*d*]imidazol-2-yl)pyrazolo[5,1-*c*][1,2,4]triazin-4-ol (100 mg, 0.29 mmol, 1 eq.) in DMF (20 mL) and acetonitrile (20 mL) at 0 °C. The reaction mixture was stirred at 0 °C for 10 min, then at 25 °C for 30 min. The solvent was evaporated, and the solid residue was triturated with cold EtOH (10 mL), collected by filtration, and then washed with EtOH (10 mL) and dried under vacuum. The product was obtained as a yellow solid (114 mg, 93%).

^1^H NMR (500 MHz, DMSO-d_6_) *δ* (ppm) 14.18 (s, 1H), 8.21 (s, 1H), 7.95 (d, *J* = 7.5 Hz, 1H), 7.75 (d, *J* = 7.8 Hz, 1H), 7.49 (p, *J* = 7.3 Hz, 2H), 7.31 (d, *J* = 4.4 Hz, 4H), 7.29 – 7.23 (m, 1H), 6.29 (s, 2H).

^13^C NMR (126 MHz, DMSO-d_6_) *δ* (ppm) 149.12, 146.25, 144.14, 135.86, 132.66, 128.66, 127.66, 126.95, 125.16, 112.14, 49.58.

HRMS (APCI): calcd. for C_19_H_14_BrN_6_O [M+H]^+^ = 421.0407.1245, found [M+H]^+-^= 421.0407.

3-(1*H*-benzo[*d*]imidazol-2-yl)-7-(((*tert*-butyldiphenylsilyl)oxy)methyl)pyrazolo[5,1-*c*][1,2,4]triazin-4-amine **(S108)**

The compound was prepared according to General procedure C using:

Diazotization step:

3-(((*tert*-butyldiphenylsilyl)oxy)methyl)-1*H*-pyrazol-5-amine (163 mg, 0.5 mmol; 1 eq.), 35% aqueous HCl (0.2 mL, 2.0 mmol; 4 eq.) in EtOH (4 mL) and H_2_O (4 mL), NaNO_2_ (68 mg, 1.0 mmol, 2 eq.) in H_2_O (1 mL) (reaction time: 30 min), 2-(cyanomethyl)-benzimidazole (94 mg, 0.6 mmol, 1.2 eq.) in EtOH (1 mL) and KOAc (300 mg, 3.0 mmol, 6 eq.) in H_2_O (1 mL). Reaction time: 16 h. The yellow solid (210 mg, 0.39 mmol) obtained after the filtration was used in the next step without further purification.

Cyclization step:

the dried, filtered yellow solid (210 mg, 0.39 mmol), DMF (3 mL), and KOAc (4 mg, 0.04 mmol, 0.1 eq.). Reaction time: 2 h. The product was obtained as a yellow solid (114 mg, 44%).

^1^H NMR (500 MHz, DMSO-*d*_6_) *δ* (ppm) 13.45 (s, 1H), 9.94 (s, 1H), 9.34 (s, 1H), 7.74 (d, *J* = 7.7 Hz, 1H), 7.72 – 7.66 (m, 4H), 7.54 (d, *J* = 7.7 Hz, 1H), 7.49 – 7.43 (m, 6H), 7.29 – 7.19 (m, 2H), 6.97 (s, 1H), 5.02 (s, 2H), 1.07 (s, 9H).

^13^C NMR (126 MHz, DMSO-*d*_6_) *δ* (ppm) 158.76, 150.00, 149.51, 138.95, 135.05, 134.42, 132.52, 129.99, 127.96, 127.44, 121.81, 118.77, 118.29, 111.45, 94.51, 60.35, 26.61, 18.82.

HRMS (APCI): calcd. for C_29_H_30_N_7_OSi [M+H]^+^ = 520.2276, found [M+H]^+^ = 520.2273.

### Preparation of target compound 1 and its analogs

3-(1*H*-benzo[*d*]imidazol-2-yl)-7-ethyl-8-(4-fluorophenyl)pyrazolo[5,1-*c*][1,2,4]triazin-4-amine **(1)**

The compound was prepared according to General procedure C using:

Diazotization step:

3-ethyl-4-(4-fluorophenyl)-1*H*-pyrazol-5-amine (200 mg, 0.98 mmol; 1 eq.), 35% aqueous HCl (0.344 mL, 3.90 mmol; 4 eq.) in EtOH (3 mL) and H_2_O (3 mL), NaNO_2_ (135 mg, 1.95 mmol, 2 eq.) in EtOH (2 mL) and H_2_O (2 mL) (reaction time: 15 min), 2-(cyanomethyl)-benzimidazole (184 mg, 1.17 mmol, 1.2 eq.) in EtOH (3 mL) and H_2_O (3 mL) and KOAc (765 mg, 7.8 mmol, 8 eq.). Reaction time: 1 h. The yellow solid (213 mg, 0.57 mmol) obtained after the filtration was used in the next step without further purification.

Cyclization step:

the dried yellow solid (213 mg, 0.57 mmol), DMF (2 mL), and KOAc (5 mg, 0.05 mmol, 0.05 eq.). Reaction time: 1 h. The product was obtained as a yellow solid (69 mg, 19% (2 steps)).

^1^H NMR (500 MHz, DMSO-*d*_6_) *δ* (ppm) 13.35 (s, 1H), 9.95 (s, 1H), 9.34 (s, 1H), 7.89 – 7.84 (m, 2H), 7.74 (d, *J* = 7.7 Hz, 1H), 7.56 (d, *J* = 7.6 Hz, 1H), 7.39 – 7.34 (m, 2H), 7.30 – 7.20 (m, 1H), 3.05 (q, *J* = 7.6 Hz, 2H), 1.35 (t, *J* = 7.5 Hz, 3H).

^13^C NMR (126 MHz, DMSO-*d*_6_) *δ* (ppm) 161.11 (d, *J* = 244.3 Hz), 150.08, 146.36, 142.74, 138.73, 133.76, 130.91 (d, *J* = 8.1 Hz), 127.73, 122.97, 121.93, 119.24, 118.34, 115.52 (d, *J* = 21.7 Hz), 111.54, 107.28, 20.89, 13.13.

HRMS (APCI): calcd. for C_20_H_17_FN_7_ [M+H]^+^ = 374.1524, found [M+H]^+^ = 374.1529.

Mp > 260 °C (decomp.).

3-(1*H*-benzo[*d*]imidazol-2-yl)pyrazolo[5,1-*c*][1,2,4]triazin-4-amine **(3)**

The compound was prepared according to General procedure C using:

Diazotization step:

3-aminopyrazole (250 mg, 3 mmol; 1 eq.), 35% aqueous HCl (1.06 mL, 12.03 mmol; 4 eq.) in EtOH (3 mL) and H_2_O (3 mL), NaNO_2_ (415 mg, 6.02 mmol, 2 eq.) in EtOH (2 mL) and H_2_O (2 mL) (reaction time: 20 min), 2-(cyanomethyl)-benzimidazole (567 mg, 3.61 mmol, 1.2 eq.) in EtOH (3 mL) and H_2_O (3 mL) and KOAc (1.77 g, 18 mmol, 6 eq.). Reaction time: 2 h. The yellow solid (768 mg, 3.06 mmol) obtained after the filtration was used in the next step without further purification.

Cyclization step:

the dried yellow solid (99 mg, 0.39 mmol), DMF (1 mL), and KOAc (2 mg, 0.02 mmol, 0.05 eq.). Reaction time: 1 h. The product was obtained as an orange solid (76 mg, 75% (2 steps)).

^1^H NMR (500 MHz, DMSO-*d*_6_) *δ* (ppm) 13.35 (s, 1H), 9.95 (s, 1H), 9.34 (s, 1H), 7.89 – 7.84 (m, 2H), 7.74 (d, *J* = 7.7 Hz, 1H), 7.56 (d, *J* = 7.6 Hz, 1H), 7.39 – 7.34 (m, 2H), 7.30 – 7.20 (m, 1H), 3.05 (q, *J* = 7.6 Hz, 2H), 1.35 (t, *J* = 7.5 Hz, 3H).

^13^C NMR (126 MHz, DMSO-*d*_6_) *δ* (ppm) 161.11 (d, *J* = 244.3 Hz), 150.08, 146.36, 142.74, 138.73, 133.76, 130.91 (d, *J* = 8.1 Hz), 127.73, 122.97, 121.93, 119.24, 118.34, 115.52 (d, *J* = 21.7 Hz), 111.54, 107.28, 20.89, 13.13.

HRMS (APCI): calcd. for C_12_H_8_N_7_ [M-H]^-^ = 250.0846, found [M-H]^-^ = 250.0844.

3-(1*H*-benzo[*d*]imidazol-2-yl)-8-(4-fluorophenyl)pyrazolo[5,1-*c*][1,2,4]triazin-4-amine **(4)**

The compound was prepared according to General procedure C using:

Diazotization step:

4-(4-fluorophenyl)-1*H*-pyrazol-5-amine (222 mg, 1.25 mmol; 1 eq.), 35% aqueous HCl (0.44 mL, 5.01 mmol; 4 eq.) in EtOH (3 mL) and H_2_O (3 mL), NaNO_2_ (173 mg, 2.50 mmol, 2 eq.) in EtOH (2 mL) and H_2_O (2 mL) (reaction time: 20 min), 2-(cyanomethyl)-benzimidazole (236 mg, 1.50 mmol, 1.2 eq.) in EtOH (3 mL) and H_2_O (3 mL) and KOAc (738 mg, 7.52 mmol, 6 eq.) Reaction time: 2 h. The yellow solid (247 mg, 0.72 mmol) obtained after the filtration was used in the next step without further purification.

Cyclization step:

the dried yellow solid (238 mg, 0.69 mmol), DMF (3 mL), and KOAc (3 mg, 0.03 mmol, 0.05 eq.). Reaction time: 2 h. The product was obtained as a yellow solid (235 mg, 57% (2 steps)).

^1^H NMR (500 MHz, DMSO-*d*_6_) *δ* (ppm) 13.45 (s, 1H), 10.04 (s, 1H), 9.59 (s, 1H), 8.95 (s, 1H), 8.40 – 8.32 (m, 2H), 7.75 (s, 1H), 7.59 (s, 1H), 7.40 – 7.32 (m, 2H), 7.26 (s, 2H).

^13^C NMR (126 MHz, DMSO-*d*_6_) *δ* (ppm) 160.92 (d, *J* = 245.0 Hz), 149.92, 145.05, 143.63, 139.39, 127.92 (d, *J* = 7.4 Hz), 119.29, 115.66 (d, *J* = 21.7 Hz), 108.94.

^19^F NMR (282 MHz, DMSO-*d*_6_) *δ* (ppm) -115.55.

HRMS (APCI): calcd. for C_18_H_11_FN_7_ [M-H]^-^ = 344.1065, found [M-H]^-^ = 344.1069.

3-(1*H*-benzo[*d*]imidazol-2-yl)-7-ethylpyrazolo[5,1-*c*][1,2,4]triazin-4-amine **(5)**

The compound was prepared according to General procedure C using:

Diazotization step:

3-ethyl-1*H*-pyrazol-5-amine (850 mg, 7.65 mmol; 1 eq.), 35% aqueous HCl (3.0 mL, 30.60 mmol; 4 eq.) in EtOH (20 mL) and H_2_O (20 mL), NaNO_2_ (1.10 g, 15.30 mmol, 2 eq.) in H_2_O (3 mL) (reaction time: 30 min), 2-(cyanomethyl)-benzimidazole (1.50 g, 9.18 mmol, 1.2 eq.) in EtOH (3 mL) and KOAc (4.50 g, 45.90 mmol, 6 eq.) in H_2_O (3 mL). Reaction time: 16 h. The yellow solid (1.90 g, 6.50 mmol) obtained after the filtration was used in the next step without further purification.

Cyclization step:

the dried yellow solid (1.9 g, 6.50 mmol), DMF (10 mL), and KOAc (32 mg, 0.30 mmol, 0.05 eq.). Reaction time: 2 h. The product was obtained as a yellow solid (1.25 g, 59%).

^1^H NMR (500 MHz, DMSO-*d*_6_) *δ* (ppm) 13.40 (s, 1H), 9.89 (s, 1H), 9.29 (s, 1H), 7.73 (d, *J* = 7.7 Hz, 1H), 7.56 – 7.51 (m, 1H), 7.29 – 7.19 (m, 2H), 6.88 (s, 1H), 2.90 (q, *J* = 7.6 Hz, 2H), 1.36 (t, *J* = 7.6 Hz, 3H).

^13^C NMR (126 MHz, DMSO-*d*_6_) *δ* (ppm) 161.59, 150.17, 149.62, 142.65, 138.70, 133.69, 122.78, 121.75, 118.34, 118.22, 111.39, 94.53, 21.75, 13.37.

HRMS (APCI): calcd. for C_14_H_12_N_7_ [M-H]^-^ = 278.1160, found [M-H]^-^ = 278.1159.

3-(1*H*-benzo[*d*]imidazol-2-yl)-7-ethyl-8-(4-(trifluoromethyl)phenyl)pyrazolo[5,1-*c*][1,2,4]triazin-4-amine **(6)**

The compound was prepared according to General procedure C using:

Diazotization step:

3-ethyl-4-(4-(trifluoromethyl)phenyl)*-*1*H-*pyrazol-5-amine (40 mg, 0.16 mmol; 1 eq.), 35% aqueous HCl (0.06 mL, 0.64 mmol; 4 eq.) in EtOH (3 mL) and H_2_O (3 mL), NaNO_2_ (24 mg, 0.32 mmol, 2 eq.) in H_2_O (1 mL) (reaction time: 30 min), 2-(cyanomethyl)-benzimidazole (30 mg, 0.19 mmol, 1.2 eq.) in EtOH (1 mL) and KOAc (95 mg, 0.96 mmol, 6 eq.) in H_2_O (1 mL). Reaction time: 16 h. The yellow solid (66 mg, 0.15 mmol) obtained after the filtration was used in the next step without further purification.

Cyclization step:

the dried yellow solid (66 mg, 0.15 mmol), DMF (3 mL), and KOAc (5 mg, 0.05 mmol, 0.5 eq.). Reaction time: 2 h. After the reaction completion, the reaction mixture was poured into water (20 mL) and extracted with EtOAc (3 × 20 mL). The combined organic extracts were washed with brine (50 mL) and dried over MgSO_4_. The solvent was evaporated and the residue was purified by preparative TLC (dichloromethane:MeOH, 9:1). The product was obtained as a yellow solid (20 mg, 30%).

^1^H NMR (500 MHz, DMSO-*d*_6_) *δ* (ppm) 13.40 (s, 1H), 10.00 (s, 1H), 9.43 (s, 1H), 8.11 (d, *J* = 8.0 Hz, 2H), 7.87 (d, *J* = 8.1 Hz, 2H), 7.75 (d, *J* = 7.7 Hz, 1H), 7.56 (d, *J* = 7.7 Hz, 1H), 7.30 – 7.20 (m, 2H), 3.10 (q, *J* = 7.5 Hz, 2H), 1.37 (t, *J* = 7.5 Hz, 3H).

^13^C NMR (126 MHz, DMSO-*d*_6_) *δ* (ppm) 158.33, 146.56, 142.66, 138.71, 135.82, 133.70, 129.11, 126.64 (q, *J* = 31.8 Hz), 125.36 (q, *J* = 3.8 Hz), 124.40 (q, *J* = 272.0 Hz), 122.96, 121.87, 119.85, 118.32, 111.48, 106.31, 21.10, 12.96.

^19^F NMR (471 MHz, DMSO-*d*_6_) *δ* (ppm) -60.81.

HRMS (APCI): calcd. for C_21_H_17_F_3_N_7_ [M+H]^+^ = 424.1492, found [M+H]^+^ = 424.1490.

3-(1*H*-benzo[*d*]imidazol-2-yl)-7-ethyl-8-(4-methoxyphenyl)pyrazolo[5,1-*c*][1,2,4]triazin-4-amine **(7)**

The compound was prepared according to General procedure C using:

Diazotization step:

3-ethyl-4-(4-methoxyphenyl)-1*H*-pyrazol-5-amine (55 mg, 0.25 mmol; 1 eq.), 35% aqueous HCl (0.1 mL, 1.0 mmol; 4 eq.) in EtOH (3 mL) and H_2_O (3 mL), NaNO_2_ (35 mg, 0.50 mmol, 2 eq.) in H_2_O (1 mL) (reaction time: 30 min), 2-(cyanomethyl)-benzimidazole (47 mg, 0.30 mmol, 1.2 eq.) in EtOH (1 mL) and KOAc (147 mg, 1.50 mmol, 6 eq.) in H_2_O (1 mL). Reaction time: 16 h. The yellow solid (100 mg, 0.24 mmol) obtained after the filtration was used in the next step without further purification.

Cyclization step:

the dried yellow solid (100 mg, 0.24 mmol), DMF (3 mL), and KOAc (2 mg, 0.02 mmol, 0.1 eq.). Reaction time: 2 h. The product was obtained as a yellow solid (45 mg, 46%).

^1^H NMR (500 MHz, DMSO-*d*_6_) *δ* (ppm) ^1^H NMR (500 MHz, DMSO) δ 13.34 (s, 1H), 9.92 (s, 1H), 9.27 (s, 1H), 7.78 – 7.71 (m, 3H), 7.55 (dd, *J* = 8.0, 1.3 Hz, 1H), 7.31 – 7.19 (m, 2H), 7.13 – 7.06 (m, 2H), 3.83 (s, 3H), 3.03 (q, *J* = 7.7 Hz, 2H), 1.34 (t, *J* = 7.5 Hz, 3H).

^13^C NMR (126 MHz, DMSO-*d*_6_) *δ* (ppm) 158.14, 157.86, 150.15, 146.16, 142.71, 138.62, 133.71, 130.15, 123.46, 122.79, 121.78, 118.75, 118.21, 114.06, 111.43, 108.19, 55.11, 20.86, 13.17.

HRMS (APCI): calcd. for C_21_H_20_N_7_O [M+H]^+^ = 386.1724, found [M+H]^+^ = 386.1726.

3-(1*H*-benzo[*d*]imidazol-2-yl)-7-ethyl-8-(4-(methylsulfonyl)phenyl)pyrazolo[5,1-*c*][1,2,4]triazin-4-amine **(8)**

The compound was prepared according to General procedure C using:

Diazotization step:

3-ethyl-4-(4-(methylsulfonyl)phenyl)-1*H*-pyrazol-5-amine (53 mg, 0.20 mmol; 1 eq.), 35% aqueous HCl (75 µL, 0.80 mmol; 4 eq.) in EtOH (3 mL) and H_2_O (3 mL), NaNO_2_ (30 mg, 0.40 mmol, 2 eq.) in H_2_O (1 mL) (reaction time: 30 min), 2-(cyanomethyl)-benzimidazole (37 mg, 0.22 mmol, 1.2 eq.) in EtOH (1 mL) and KOAc (125 mg, 1.20 mmol, 6 eq.) in H_2_O (1 mL). Reaction time: 16 h. The yellow solid (72 mg, 0.16 mmol) obtained after the filtration was used in the next step without further purification.

Cyclization step:

the dried yellow solid (72 mg, 0.16 mmol), DMF (3 mL), and KOAc (5 mg, 0.06 mmol, 0.4 eq.). Reaction time: 2 h. The product was obtained as a yellow solid (56 mg, 65%).

^1^H NMR (500 MHz, DMSO-*d*_6_) *δ* (ppm) 13.41 (s, 1H), 10.03 (s, 1H), 9.48 (s, 1H), 8.24 – 8.19 (m, 2H), 7.91 – 7.86 (m, 2H), 7.77 (d, *J* = 7.7 Hz, 1H), 7.57 (d, *J* = 7.7 Hz, 1H), 7.32 – 7.22 (m, 2H), 3.67 (t, *J* = 4.8 Hz, 4H), 3.15 (q, *J* = 7.5 Hz, 2H), 2.97 (t, *J* = 4.7 Hz, 4H), 1.40 (t, *J* = 7.5 Hz, 3H).

^13^C NMR (126 MHz, DMSO-*d*_6_) *δ* (ppm) 158.99, 150.33, 147.17, 143.17, 139.27, 138.80, 137.39, 134.23, 129.55, 127.76, 123.54, 122.43, 120.59, 118.88, 112.03, 106.62, 44.15, 21.71, 13.46.

HRMS (ESI): calcd. for C_21_H_18_N_7_O_2_S [M-H]^-^= 432.1248, found [M-H]^-^ = 432.1249.

3-(1*H*-benzo[*d*]imidazol-2-yl)-7-ethyl-8-(4-morpholinophenyl)pyrazolo[5,1-*c*][1,2,4]triazin-4-amine **(9)**

The compound was prepared according to General procedure C using:

Diazotization step:

3-ethyl-4-(4-morpholinophenyl)-1*H*-pyrazol-5-amine (143 mg, 0.53 mmol; 1 eq.), 35% aqueous HCl (0.185 mL, 2.1 mmol; 4 eq.) in EtOH (2 mL) and H_2_O (2 mL), NaNO_2_ (44 mg, 0.63 mmol, 1.2 eq.) in EtOH (1 mL) and H_2_O (1 mL) (reaction time: 15 min), 2-(cyanomethyl)-benzimidazole (99 mg, 0.63 mmol, 1.2 eq.) in EtOH (3 mL) and H_2_O (2 mL) and NaOAc (345 mg, 4.20 mmol, 8 eq.). Reaction time: 16 h. The mixture was poured into brine (20 mL) and extracted with EtOAc (2 × 50 mL). The combined organic extracts were dried over MgSO_4_, filtered, and the solvent was evaporated *in vacuo*. The residue was obtained as a yellow solid (296 mg, quant.) and used in the next step without further purification.

Cyclization step:

the dried yellow solid (296 mg), DMF (2 mL), and NaOAc (4 mg, 0.05 mmol, 0.1 eq.). Reaction time: 2 h. The mixture was poured into brine (20 mL) and extracted with EtOAc (3 × 50 mL). The combined organic extracts were dried over MgSO_4_, filtered, and the solvent was evaporated *in vacuo*. The residue was dissolved in boiling MeOH (25 mL) and the solution was allowed to cool to 25 °C. The resulting precipitate was collected by filtration and dried under vacuum. The product was obtained as an orange solid (47 mg, 20%).

^1^H NMR (500 MHz, DMSO-*d*_6_) *δ* (ppm) 13.34 (s, 1H), 9.90 (s, 1H), 9.25 (s, 1H), 7.76 – 7.72 (m, 1H), 7.70 (d, *J* = 8.8 Hz, 2H), 7.55 (d, *J* = 7.4 Hz, 1H), 7.25 (s, 2H), 7.09 (d, *J* = 8.9 Hz, 2H), 3.82 – 3.73 (m, 4H), 3.24 – 3.15 (m, 4H), 3.04 (q, *J* = 7.5 Hz, 2H), 1.35 (t, *J* = 7.6 Hz, 3H).

^13^C NMR (126 MHz, DMSO-*d*_6_) *δ* (ppm) 157.80, 150.20, 149.69, 146.13, 138.62, 133.72, 129.61, 122.78, 121.75, 118.59, 118.22, 114.94, 111.42, 108.52, 66.08, 48.19, 20.92, 13.20.

HRMS (APCI): calcd. for C_24_H_23_N_8_O [M-H]^-^ = 439.2000, found [M-H]^-^ = 439.2002.

3-(1*H*-benzo[d]imidazol-2-yl)-8-cyclohexyl-7-ethylpyrazolo[5,1-*c*][1,2,4]triazin-4-amine **(10)**

The compound was prepared according to General procedure C using:

Diazotization step:

4-cyclohexyl-3-ethyl-1*H*-pyrazol-5-amine (117 mg, 0.61 mmol; 1 eq.), 35% aqueous HCl (0.214 mL, 2.42 mmol; 4 eq.) in EtOH (1 mL) and H_2_O (3 mL), NaNO_2_ (84 mg, 1.21 mmol, 2 eq.) in EtOH (1 mL) and H_2_O (1 mL) (reaction time: 20 min), 2-(cyanomethyl)-benzimidazole (0.1 g, 0.70 mmol, 1.05 eq.) in EtOH (2 mL) and H_2_O (1 mL) and KOAc (475 mg, 4.84 mmol, 8 eq.). Reaction time: 16 h. The yellow solid (215 mg, 0.60 mmol) obtained after the filtration was used in the next step without further purification.

Cyclization step:

the dried yellow solid (215 mg, 0.60 mmol), DMF (3 mL), and KOAc (3 mg, 0.03 mmol, 0.05 eq.). Reaction time: 2 h. The product was obtained as a yellow solid (149 mg, 68% (2 steps)).

^1^H NMR (500 MHz, DMSO-*d*_6_) *δ* (ppm) 13.24 (s, 1H), 9.76 (s, 1H), 9.08 (s, 1H), 7.72 (d, *J* = 7.0 Hz, 1H), 7.55 (d, *J* = 7.0 Hz, 1H), 7.35 – 7.13 (m, 2H), 2.89 (q, *J* = 7.5 Hz, 3H), 2.20 – 2.07 (m, 2H), 1.90 – 1.72 (m, 5H), 1.50 – 1.36 (m, 3H), 1.33 (t, *J* = 7.6 Hz, 3H).

^13^C NMR (126 MHz, DMSO-*d*_6_) *δ* (ppm) 158.17, 150.43, 146.74, 142.77, 138.54, 133.68, 122.66, 121.70, 118.14, 117.32, 113.14, 111.35, 34.11, 32.65, 26.54, 25.71, 20.28, 14.03.

HRMS (APCI): calcd. for C_20_H_24_N_7_ [M+H]^+^ = 362.2088, found [M+H]^+^ = 362.2091.

3-(1*H*-benzo[*d*]imidazol-2-yl)-8-(4-fluorophenyl)-7-isopropylpyrazolo[5,1-*c*][1,2,4]triazin-4-amine **(11)**

The compound was prepared according to General procedure C using:

Diazotization step:

4-(4-fluorophenyl)-3-isopropyl-1*H*-pyrazol-5-amine (60 mg, 0.27 mmol; 1 eq.), 35% aqueous HCl (0.1 mL, 1.1 mmol; 4 eq.) in EtOH (2 mL) and H_2_O (2 mL), NaNO_2_ (38 mg, 0.54 mmol, 2 eq.) in H_2_O (1 mL) (reaction time: 30 min), 2-(cyanomethyl)-benzimidazole (50 mg, 0.32 mmol, 1.2 eq.) in EtOH (1 mL) and KOAc (160 mg, 1.62 mmol, 6 eq.) in H_2_O (1 mL). Reaction time: 16 h. The yellow solid (50 mg, 0.13 mmol) obtained after the filtration was used in the next step without further purification.

Cyclization step:

the dried yellow solid (40 mg, 0.1 mmol), DMF (1 mL), and KOAc (1 mg, 0.01 mmol, 0.1 eq.). Reaction time: 2 h. The product was obtained as a yellow solid (22 mg, 44%).

^1^H NMR (500 MHz, DMSO-*d*_6_) *δ* (ppm) 13.36 (s, 1H), 9.95 (s, 1H), 9.21 (s, 1H), 7.83 – 7.72 (m, 3H), 7.57 (d, *J* = 7.0 Hz, 1H), 7.42 – 7.35 (m, 2H), 7.31 – 7.22 (m, 2H), 3.46 (hept, *J* = 6.9 Hz, 1H), 1.41 (d, *J* = 6.9 Hz, 6H).

^13^C NMR (126 MHz, DMSO-*d*_6_) *δ* (ppm) 161.92, 161.18 (d, *J* = 244.1 Hz), 150.57, 146.34, 138.68, 133.72, 131.40 (d, *J* = 7.9 Hz), 127.64 (d, *J* = 3.6 Hz), 122.87, 121.84, 119.06, 118.26, 115.46 (d, *J* = 21.1 Hz), 111.46, 107.03, 26.32, 22.35.

HRMS (ESI): calcd. for C_21_H_17_FN_7_ [M-H]^-^ = 386.1535, found [M-H]^-^ = 386.1541.

3-(1*H*-benzo[*d*]imidazol-2-yl)-7-(naphthalen-1-yl)pyrazolo[5,1-*c*][1,2,4]triazin-4-amine **(12)**

The compound was prepared according to General procedure C using:

Diazotization step:

3-(naphthalen-1-yl)-1*H*-pyrazol-5-amine (120 mg, 0.57 mmol; 1 eq.), 35% aqueous HCl (0.65 mL, 2.3 mmol; 4 eq.) in EtOH (3 mL) and H_2_O (3 mL), NaNO_2_ (80 mg, 1.14 mmol, 2 eq.) in H_2_O (1 mL) (reaction time: 30 min), 2-(cyanomethyl)-benzimidazole (107 mg, 0.70 mmol, 1.2 eq.) in EtOH (1 mL) and KOAc (337 mg, 3.40 mmol, 6 eq.) in H_2_O (1 mL). Reaction time: 16 h. The yellow solid (198 mg, 0.5 mmol) obtained after the filtration was used in the next step without further purification.

Cyclization step:

the dried yellow solid (198 mg, 0.5 mmol), DMF (3 mL), and KOAc (5 mg, 0.05 mmol, 0.1 eq.). Reaction time: 2 h. The product was obtained as a yellow solid (144 mg, 67%).

^1^H NMR (500 MHz, DMSO-*d*_6_) *δ* (ppm) 13.51 (s, 1H), 10.03 (s, 1H), 9.52 (s, 1H), 8.71 – 8.64 (m, 1H), 8.10 (d, *J* = 8.2 Hz, 1H), 8.08 – 8.04 (m, 1H), 7.97 (dd, *J* = 7.1, 1.1 Hz, 1H), 7.76 (s, 1H), 7.71 – 7.66 (m, 1H), 7.65 – 7.61 (m, 2H), 7.58 (s, 1H), 7.44 (s, 1H), 7.27 (s, 2H).

^13^C NMR (126 MHz, DMSO-*d*_6_) *δ* (ppm) 157.34, 150.61, 150.07, 143.22, 139.50, 133.96, 131.11, 130.30, 130.18, 128.94, 128.91, 127.53, 126.72, 126.36, 125.90, 123.42, 122.36, 119.50, 118.85, 112.01, 97.78.

HRMS (APCI): calcd. for C_22_H_16_N_7_ [M+H]^+^ = 378.1462, found [M+H]^+^ = 378.1459.

3-(1*H*-benzo[*d*]imidazol-2-yl)-7-(2-phenoxyphenyl)pyrazolo[5,1-*c*][1,2,4]triazin-4-amine **(13)**

The compound was prepared according to General procedure C using:

Diazotization step:

3-(2-phenoxyphenyl)-1*H*-pyrazol-5-amine (213 mg, 0.85 mmol; 1 eq.), 35% aqueous HCl (0.30 mL, 3.39 mmol; 4 eq.) in EtOH (2 mL) and H_2_O (2 mL), NaNO_2_ (117 mg, 1.69 mmol, 2 eq.) in EtOH (1 mL) and H_2_O (2 mL) (reaction time: 20 min), 2-(cyanomethyl)-benzimidazole (160 mg, 1.02 mmol, 1.2 eq.) in EtOH (3 mL) and H_2_O (3 mL) and KOAc (666 mg, 6.78 mmol, 8 eq.). Reaction time: 2 h. The yellow solid (355 mg, 0.85 mmol) obtained after the filtration was used in the next step without further purification.

Cyclization step:

the dried yellow solid (345 mg, 0.82 mmol), DMF (6 mL), and KOAc (4 mg, 0.04 mmol, 0.05 eq.). Reaction time: 2 h. The product was obtained as a yellow solid (248 mg, 70%).

^1^H NMR (500 MHz, DMSO-*d*_6_) *δ* (ppm) 13.46 (s, 1H), 9.96 (s, 1H), 9.37 (s, 1H), 8.39 (d, *J* = 7.8 Hz, 1H), 7.75 (d, *J* = 7.5 Hz, 1H), 7.58 – 7.49 (m, 2H), 7.40 (t, *J* = 7.8 Hz, 3H), 7.29 (s, 1H), 7.25 (t, *J* = 8.1 Hz, 2H), 7.13 (d, *J* = 7.7 Hz, 2H), 7.06 (d, *J* = 7.9 Hz, 2H).

^13^C NMR (126 MHz, DMSO-*d*_6_) *δ* (ppm) 156.77, 153.82, 152.36, 149.99, 149.84, 142.69, 138.70, 133.74, 131.24, 130.12, 129.64, 124.41, 123.79, 123.20, 122.90, 121.83, 120.67, 118.93, 118.30, 117.53, 111.47, 96.63.

HRMS (APCI): calcd. for C_24_H_18_N_7_O [M+H]^+^ = 420.1567, found [M+H]^+^ = 420.1565.

3-(1*H*-benzo[*d*]imidazol-2-yl)-7-(naphthalen-2-yl)pyrazolo[5,1-*c*][1,2,4]triazin-4-amine **(14)**

The compound was prepared according to General procedure C using:

Diazotization step:

3-(naphthalen-2-yl)-1*H*-pyrazol-5-amine (206 mg, 0.98 mmol; 1 eq.), 35% aqueous HCl (0.35 mL, 3.94 mmol; 4 eq.) in EtOH (2 mL) and H_2_O (2 mL), NaNO_2_ (136 mg, 1.97 mmol, 2 eq.) in EtOH (1 mL) and H_2_O (2 mL) (reaction time: 20 min), 2-(cyanomethyl)-benzimidazole (186 mg, 1.18 mmol, 1.2 eq.) in EtOH (3 mL) and H_2_O (3 mL) and KOAc (773 mg, 7.87 mmol, 8 eq.). Reaction time: 2 h. The yellow solid (367 mg, 0.97 mmol) obtained after the filtration was used in the next step without further purification.

Cyclization step:

the dried yellow solid (367 mg, 0.97 mmol), DMF (6 mL), and KOAc (5 mg, 0.05 mmol, 0.05 eq.). Reaction time: 2 h. The product was obtained as an orange solid (195 mg, 52%).

^1^H NMR (500 MHz, DMSO-*d*_6_) *δ* (ppm) 13.47 (s, 1H), 9.99 (s, 1H), 9.40 (s, 1H), 8.75 (s, 1H), 8.33 (dd, *J* = 8.5, 1.8 Hz, 1H), 8.10 (d, *J* = 8.6 Hz, 1H), 8.05 (d, *J* = 7.1 Hz, 1H), 8.00 (d, *J* = 6.4 Hz, 1H), 7.77 (d, *J* = 7.6 Hz, 1H), 7.68 (s, 1H), 7.64 – 7.51 (m, 3H), 7.31 – 7.23 (m, 2H).

^13^C NMR (126 MHz, DMSO-*d*_6_) *δ* (ppm) 156.25, 150.36, 150.05, 142.72, 138.86, 133.76, 133.45, 132.95, 129.23, 128.48, 128.28, 127.76, 126.91, 126.81, 125.88, 124.21, 122.91, 121.85, 119.04, 118.31, 111.49, 93.72.

HRMS (APCI): calcd. for C_22_H_16_N_7_ [M+H]^+^ = 378.1462, found [M+H]^+^ = 378.1449.

3-(1*H*-benzo[*d*]imidazol-2-yl)-7-phenylpyrazolo[5,1-*c*][1,2,4]triazin-4-amine **(15)**

The compound was prepared according to General procedure C using:

Diazotization step:

3-phenyl-1*H*-pyrazol-5-amine (2 g, 12.50 mmol; 1 eq.), 35% aqueous HCl (4.3 mL, 50 mmol; 4 eq.) in EtOH (20 mL) and H_2_O (20 mL), NaNO_2_ (1.73 g, 25 mmol, 2 eq.) in H_2_O (3 mL) (reaction time: 30 min), 2-(cyanomethyl)-benzimidazole (2.30 g, 15 mmol, 1.2 eq.) in EtOH (10 mL) and KOAc (7.4 g, 75 mmol, 6 eq.) in H_2_O (8 mL). Reaction time: 16 h. The yellow solid (3.1 g, 9 mmol) obtained after the filtration was used in the next step without further purification.

Cyclization step:

the dried yellow solid (3.10 g, 9 mmol), DMF (10 mL), and KOAc (10 mg, 0.10 mmol, 0.01 eq.). Reaction time: 2 h. The product was obtained as a yellow solid (2.50 g, 60%).

^1^H NMR (500 MHz, DMSO-*d*_6_) *δ* (ppm) 13.46 (s, 1H), 9.95 (s, 1H), 9.34 (s, 1H), 8.19 (d, *J* = 7.4 Hz, 2H), 7.75 (s, 1H), 7.60 – 7.48 (m, 5H), 7.26 (s, 2H).

^13^C NMR (126 MHz, DMSO-*d*_6_) *δ* (ppm) 149.89, 149.70, 149.15, 142.67, 139.60, 138.88, 133.74, 129.73, 122.94, 121.87, 119.22, 118.70, 118.29, 111.51, 93.59.

HRMS (APCI): calcd. for C_18_H_14_N_7_ [M+H]^+^ = 328.1305, found [M+H]^+^ = 328.1309.

3-(1*H*-benzo[*d*]imidazol-2-yl)-7-ethyl-8-(4-fluorophenyl)pyrazolo[5,1-*c*][1,2,4]triazine **(16)**

35% aqueous HCl (0.2 mL, 2.16 mmol; 4 eq.) was added to a solution of 3-ethyl-4-(4-fluorophenyl)-1*H*-pyrazol-5-amine (110 mg, 0.54 mmol; 1 eq.) in a mixture of EtOH (2 mL) and H_2_O (2 mL). The solution was cooled to -10 °C and a pre-cooled solution (0 °C) of NaNO_2_ (68 mg, 1.08 mmol, 2 eq.) in H_2_O (0.5 mL) was added. The reaction mixture turned yellow and was stirred for 30 min at -5 °C, then a pre-cooled solution (-5 °C) of KOAc (300 mg, 3 mmol, 6 eq.) in H_2_O (0.5 mL) and (*E*)-2-(2-(piperidin-1-yl)vinyl)-1*H*-benzo[*d*]imidazole (95 mg, 0.42 mmol, 0.85 eq.) in EtOH (0.5 mL) were added. The resulting mixture was allowed to warm up to 25 °C and stirred from 16 h. The mixture was poured into water (25 mL), the solid part was collected by filtration. The compound was purified by column chromatography on silica gel (hexane:EtOAc, 1:1) and then (dichloromethane:MeOH, gradient 9:1 to 5:1). The product was obtained as a red solid (90 mg, 50%).

^1^H NMR (300 MHz, DMSO-*d*_6_) *δ* (ppm) 13.54 (s, 1H), 9.71 (s, 1H), 7.95 – 7.82 (m, 2H), 7.70 – 7.64 (m, 2H), 7.49 – 7.35 (m, 2H), 7.33 – 7.21 (m, 2H), 3.09 (q, *J* = 7.5 Hz, 2H), 1.35 (t, *J* = 7.5 Hz, 3H).

^13^C NMR (126 MHz, DMSO-*d*_6_) *δ* (ppm) 162.62 (d, *J* = 248.2 Hz), 160.92, 148.11, 146.82, 135.63, 131.35 (d, *J* = 8.0 Hz), 126.19 (d, *J* = 3.6 Hz), 124.39, 123.37, 121.13, 120.09, 116.10 (d, *J* = 22.0 Hz), 111.70, 21.47, 13.45.

^19^F NMR (282 MHz, DMSO) *δ* (ppm) -114.36.

HRMS (APCI): calcd. for C_20_H_16_FN_6_ [M+H]^+^= 359.1415, found [M+H]^+^ = 359.1418.

3-(1*H*-benzo[*d*]imidazol-2-yl)-7-ethyl-8-(4-fluorophenyl)-4-methylpyrazolo[5,1-*c*][1,2,4]triazine **(17)**

The compound was prepared according to General procedure C using:

Diazotization step:

3-ethyl-4-(4-fluorophenyl)-1*H*-pyrazol-5-amine (178 mg, 0.87 mmol; 1 eq.), 35% aqueous HCl (0.3 mL, 3.48 mmol; 4 eq.) in EtOH (5 mL) and H_2_O (5 mL), NaNO_2_ (120 mg, 1.74 mmol, 2 eq.) in EtOH (1 mL) (reaction time: 15 min), 1-(1*H*-benzo[*d*]imidazol-2-yl)propan-2-one (183 mg, 1.06 mmol, 1 eq.) in EtOH (2 mL) and KOAc (0.68 g, 6.96 mmol, 8 eq.). Reaction time: 16 h. The yellow solid (255 mg, 0.65 mmol) obtained after the filtration was used in the next step without further purification.

Cyclization step:

the dried yellow solid (255 mg), was dissolved in EtOH (5 mL) and CH_3_SO_3_H (20 µL) was added and the mixture was stirred under reflux for 4 h. The solvent was evaporated and the residue was purified by column flash chromatography on silica gel (hexane:EtOAc, gradient 1:0 to 3:2). The product was obtained as a yellow solid (151 mg, 47%).

^1^H NMR (500 MHz, DMSO-*d*_6_) *δ* (ppm) (s, 1H), 7.92 – 7.86 (m, 2H), 7.79 – 7.74 (m, 1H), 7.63 – 7.58 (m, 1H), 7.46 – 7.38 (m, 2H), 7.32 – 7.21 (m, 2H), 3.47 (s, 3H), 3.11 (q, *J* = 7.6 Hz, 2H), 1.36 (t, *J* = 7.5 Hz, 3H).

^13^C NMR (126 MHz, DMSO-*d*_6_) *δ* (ppm) 161.44 (d, *J* = 244.9 Hz), 158.49, 148.39, 146.35, 143.74, 135.81, 134.20, 134.11, 131.15 (d, *J* = 8.2 Hz), 126.82 (d, *J* = 3.2 Hz), 123.17, 121.90, 119.15, 115.69 (d, *J* = 21.5 Hz), 111.80, 109.44, 20.82, 13.53, 12.89.

^19^F NMR (282 MHz, DMSO-*d*_6_) *δ* (ppm) -114.59.

HRMS (APCI): calcd. for C_21_H_18_FN_6_ [M+H]^+^= 373.1571, found [M+H]^+^ = 373.1570.

3-(1*H*-benzo[*d*]imidazol-2-yl)-7-ethyl-8-(4-fluorophenyl)pyrazolo[5,1-*c*][1,2,4]triazin-4-ol **(18)**

The compound was prepared according to General procedure C using:

Diazotization step:

3-ethyl-4-(4-fluorophenyl)-1*H*-pyrazol-5-amine (205 mg, 1.0 mmol; 1 eq.), 35% aqueous HCl (0.34 mL, 4.0 mmol; 4 eq.) in EtOH (5 mL) and H_2_O (5 mL), NaNO_2_ (136 mg, 2.0 mmol, 2 eq.) in H_2_O (1 mL) (reaction time: 30 min), methyl 2-(1*H*-benzo[*d*]imidazol-2-yl)acetate (209 mg, 1.10 mmol, 1.1 eq.) in EtOH (5 mL) and KOAc (560 mg, 6.0 mmol, 6 eq.) in H_2_O (5 mL). Reaction time: 16 h. The yellow solid (290 mg, 0.74 mmol) obtained after the filtration was used in the next step without further purification.

Cyclization step:

the dried yellow solid (290 mg, 0.74 mmol), DMF (5 mL), and KOAc (8 mg, 0.08 mmol, 0.1 eq.). Reaction time: 2 h. The compound was purified by reversed phase column chromatography using Biotage Selekt purification system (water:MeOH:7 M NH_3_ in methanol, gradient 80:20:2 to 30:70:2). The product was obtained as an yellow solid (210 mg, 56%).

^1^H NMR (500 MHz, DMSO-*d*_6_) *δ* (ppm) 14.03 (s, 2H), 7.84 – 7.79 (m, 2H), 7.78 – 7.73 (m, 2H), 7.47 – 7.42 (m, 2H), 7.35 – 7.29 (m, 2H), 2.94 (q, *J* = 7.5 Hz, 2H), 1.28 (t, *J* = 7.5 Hz, 3H).

^13^C NMR (126 MHz, DMSO-*d*_6_) *δ* (ppm) 160.84 (d, *J* = 243.7 Hz), 155.65, 149.28, 148.89, 148.34, 131.07, 130.74 (d, *J* = 7.9 Hz), 128.37 (d, *J* = 3.2 Hz), 124.67, 119.14, 115.22 (d, *J* = 21.2 Hz), 113.32, 108.22, 20.77, 13.06.

^19^F NMR (471 MHz, DMSO-*d*_6_) *δ* (ppm) -114.60.

HRMS (APCI): calcd. for C_20_H_14_FN_6_O [M-H]^-^ = 373.1219, found [M-H]^-^ = 373.1217.

4-Amino-7-ethyl-8-(4-fluorophenyl)pyrazolo[5,1-*c*][1,2,4]triazine-3-carbonitrile **(19)**

Diazotization step (only):

The compound was prepared according to General procedure C using 3-ethyl-4-(4-fluorophenyl)-1*H*-pyrazol-5-amine (207 mg, 1.01 mmol; 1 eq.), 35% aqueous HCl (0.344 mL, 3.90 mmol; 4 eq.) in EtOH (2 mL) and H_2_O (2 mL), NaNO_2_ (139 mg, 2.02 mmol, 2 eq.) in EtOH (2 mL) and H_2_O (2 mL) (reaction time: 20 min), malononitrile (138 mg, 1.21 mmol, 1.2 eq.) in EtOH (2 mL) and H_2_O (2 mL) and KOAc (765 mg, 7.80 mmol, 8 eq.). Reaction time: 16 h. The mixture was poured into H_2_O (20 mL) and extracted with EtOAc (2 × 50 mL). The combined organic extracts were washed with brine (30 mL), dried over MgSO_4_, filtered, and the solvent was evaporated *in vacuo*. The residue was purified using column chromatography (hexane:EtOAc, gradient 10:1 to 1:2). The product was obtained as a yellow solid (188 mg, 66%).

^1^H NMR (500 MHz, DMSO-*d*_6_) *δ* (ppm) 9.29 (s, 2H), 7.85 – 7.74 (m, 2H), 7.37 (t, *J* = 8.9 Hz, 2H), 3.02 (q, *J* = 7.6 Hz, 2H), 1.31 (t, *J* = 7.5 Hz, 3H).

^13^C NMR (126 MHz, DMSO-*d*_6_) *δ* (ppm) 161.34 (d, *J* = 244.4 Hz), 158.44, 145.48, 142.53, 131.07 (d, *J* = 8.2 Hz), 126.86 (d, *J* = 3.6 Hz), 115.86, 115.54 (d, *J* = 21.7 Hz), 109.56, 105.30, 20.68, 12.81.

HRMS (APCI): calcd. for C_14_H_10_FN_6_ [M-H]^-^ = 281.0956, found [M-H]^-^ = 281.0956.

7-Ethyl-8-(4-fluorophenyl)-3-(pyrimidin-2-yl)pyrazolo[5,1-*c*][1,2,4]triazin-4-ol **(20)**

The compound was prepared according to General procedure C using:

Diazotization step:

3-ethyl-4-(4-fluorophenyl)-1*H*-pyrazol-5-amine (200 mg, 0.97 mmol; 1 eq.), 35% aqueous HCl (0.34 mL, 3.9 mmol; 4 eq.) in EtOH (3 mL) and H_2_O (3 mL), NaNO_2_ (134 mg, 1.95 mmol, 2 eq.) in H_2_O (1 mL) (reaction time: 30 min), ethyl 2-(pyrimidin-2-yl)acetate (0.19 g, 1.17 mmol, 1 eq.) in EtOH (2 mL) and KOAc (574 mg, 5.85 mmol, 6 eq.). Reaction time: 2 h. The mixture was poured into water (20 mL) and extracted with EtOAc (2 × 50 mL). The yellow solid (194 mg, 0.55 mmol) obtained after the filtration was used in the next step without further purification.

Cyclization step:

the dried yellow solid (194 mg, 0.55 mmol), DMF (3 mL), and KOAc (5 mg, 0.05 mmol, 0.1 eq.). Reaction time: 2 h. After the filtration, the solid part was suspended in hot dioxane (10 mL) and the solution was poured into water (30 mL). The precipitate was collected by filtration, washed with water (10 mL), diethyl ether (10 mL) and dried under *vacuum*. The product was obtained as a yellow solid (50 mg, 15%).

^1^H NMR (500 MHz, DMSO-*d*_6_) *δ* (ppm) 14.24 (s, 1H), 8.97 (d, *J* = 4.9 Hz, 2H), 7.60 (t, *J* = 4.9 Hz, 1H), 7.57 – 7.51 (m, 2H), 7.39 – 7.31 (m, 2H), 2.78 (q, *J* = 7.5 Hz, 2H), 1.20 (t, *J* = 7.5 Hz, 3H).

^13^C NMR (126 MHz, DMSO-*d*_6_) *δ* (ppm) 161.56 (d, *J* = 237.5 Hz), 160.56, 157.50, 157.29, 147.80, 140.42, 134.94, 131.63 (d, *J* = 8.2 Hz), 125.87 (d, *J* = 3.0 Hz), 120.84, 115.71 (d, *J* = 21.6 Hz), 101.49, 20.11, 12.52.

HRMS (APCI): calcd. for C_17_H_14_FN_6_O [M+H]^+^ = 337.1208, found [M+H]^+^ = 337.1211.

4-Amino-7-ethyl-8-(4-fluorophenyl)pyrazolo[5,1-*c*][1,2,4]triazine-3-carboxamide **(21)**

To a solution of 4-amino-7-ethyl-8-(4-fluorophenyl)pyrazolo[5,1-*c*][1,2,4]triazine-3-carbonitrile (75 mg, 0.27 mmol, 1 eq.) in EtOH (1 mL) was added KOH (60 mg, 1.06 mmol, 4 eq.) and the mixture was refluxed for 24 h. The solvent was removed *in vacuo*, H_2_O (10 mL) was added to the residue and the mixture was extracted with EtOAc (3 × 10 mL). The combined organic extracts were washed with brine (10 mL), dried over MgSO_4_, filtered and concentrated *in vacuo*. The residue was purified by column chromatography (hexane:EtOAc, gradient 10:1 to 1:2). The isolated product was further purified by reverse phase column chromatography (H_2_O:CH_3_CN:HCOOH, gradient 2:1:0.1% to 0:1:0.1%). The product was obtained as an orange solid (50 mg, 63%).

^1^H NMR (500 MHz, DMSO-*d*_6_) *δ* (ppm) 9.24 (s, 1H), 8.96 (s, 1H), 8.34 (s, 1H), 7.91 – 7.77 (m, 2H), 7.69 (s, 1H), 7.45 – 7.27 (m, 2H), 3.04 (q, *J* = 7.5 Hz, 2H), 1.33 (t, *J* = 7.5 Hz, 3H).

^13^C NMR (126 MHz, DMSO-*d*_6_) *δ* (ppm) 168.28, 161.10 (d, *J* = 244.0 Hz), 158.08, 146.58, 140.55, 130.84 (d, *J* = 8.2 Hz), 127.45, 119.28, 115.44 (d, *J* = 21.7 Hz), 107.89, 20.81, 12.88.

HRMS (APCI): calcd. for C_14_H_14_FN_6_O [M+H]^+^ = 301.1208, found [M+H]^+^ = 301.1209.

3-(1-Benzyl-1*H*-benzo[*d*]imidazol-2-yl)-7-ethyl-8-(4-fluorophenyl)pyrazolo[5,1-*c*][1,2,4]triazin-4-amine **(22)**

The compound was prepared according to General procedure C using:

Diazotization step:

3-ethyl-4-(4-fluorophenyl)-1*H*-pyrazol-5-amine (205 mg, 1.0 mmol; 1 eq.), 35% aqueous HCl (0.4 mL, 4.0 mmol; 4 eq.) in EtOH (2 mL) and H_2_O (2 mL), NaNO_2_ (140 mg, 2.0 mmol, 2 eq.) in H_2_O (1 mL) (reaction time: 20 min), 2-(1-benzyl-1*H*-benzo[*d*]imidazol-2-yl)acetonitrile (247 mg, 1.0 mmol, 1 eq.) in EtOH (1 mL) and KOAc (0.58 g, 6.0 mmol, 6 eq.) in H_2_O (1 mL). Reaction time: 16 h. The yellow solid (190 mg, 0.51 mmol) obtained after the filtration was used in the next step without further purification.

Cyclization step:

the dried yellow solid (19 mg, 0.51 mmol), DMF (3 mL), and KOAc (5 mg, 0.049 mmol, 0.1 eq.). Reaction time: 2 h. The product was poured into water (20 mL) and extracted with EtOAc (3 × 20 mL). The combined organic layers were washed with brine (30 mL), dried over MgSO_4_, the solvent was evaporated, and the residue was purified by column flash chromatography on silica gel (hexane:EtOAc, 1:1). The product was obtained as a yellow solid (60 mg, 13% (2 steps)).

^1^H NMR (500 MHz, DMSO-*d*_6_) *δ* (ppm) 10.37 (s, 1H), 9.37 (s, 1H), 7.86 – 7.78 (m, 3H), 7.61 – 7.55 (m, 1H), 7.36 – 7.31 (m, 2H), 7.32 – 7.28 (m, 2H), 7.27 – 7.23 (m, 2H), 7.20 (d, *J* = 7.3 Hz, 3H), 6.30 (s, 2H), 3.05 (q, *J* = 7.5 Hz, 2H), 1.34 (t, *J* = 7.5 Hz, 3H).

^13^C NMR (126 MHz, DMSO-*d*_6_) *δ* (ppm) 161.01 (d, *J* = 244.3 Hz), 158.00, 148.02, 145.52, 141.07, 139.52, 137.57, 135.31, 130.81 (d, *J* = 8.1 Hz), 127.51 (d, *J* = 3.0 Hz), 127.12, 126.62, 122.95 (d, *J* = 67.8 Hz), 120.55, 118.64, 115.41 (d, *J* = 21.6 Hz), 110.78, 107.13, 48.88, 20.82, 13.09.

HRMS (APCI): calcd. for C_27_H_23_N_7_F [M+H]^+^ = 464.1993, found [M+H]^+^ = 464.1997.

3-(1-(2-(Dimethylamino)ethyl)-1*H*-benzo[*d*]imidazol-2-yl)-7-ethyl-8-(4-fluorophenyl)pyrazolo[5,1-*c*][1,2,4]triazin-4-ol **(23)**

The compound was prepared according to General procedure C using:

Diazotization step:

3-ethyl-4-(4-fluorophenyl)-1*H*-pyrazol-5-amine (110 mg, 0.49 mmol; 1 eq.), 35% aqueous HCl (190 µL, 2 mmol; 4 eq.) in EtOH (3 mL) and H_2_O (3 mL), NaNO_2_ (68 mg, 1 mmol, 2 eq.) in EtOH (1 mL) (reaction time: 15 min), ethyl 2-(1-(2-(dimethylamino)ethyl)-1*H*-benzo[*d*]imidazol-2-yl)acetate (145 mg, 0.53 mmol, 1.2 eq.) in EtOH (2 mL) and KOAc (300 mg, 3.0 mmol, 6 eq.). Reaction time: 16 h. The yellow solid (185 mg, 0.40 mmol) obtained after the filtration was used in the next step without further purification.

Cyclization step:

the dried yellow solid (185 mg, 0.4 mmol), DMF (3 mL), and KOAc (4 mg, 0.04 mmol, 0.1 eq.). Reaction time: 2 h. After the filtration, the solid part was suspended in hot dioxane (5 mL) and the mixture was poured into water (20 mL). The precipitate was collected by filtration, washed with water (10 mL) and diethyl ether (10 mL), and dried under *vacuum*. The product was obtained as a yellow solid (105 mg, 48%).

^1^H NMR (500 MHz, DMSO-*d*_6_) *δ* (ppm) 8.09 – 8.05 (m, 1H), 7.97 – 7.92 (m, 1H), 7.79 – 7.73 (m, 2H), 7.59 – 7.50 (m, 2H), 7.42 – 7.27 (m, 2H), 5.28 (s, 2H), 3.63 (t, *J* = 7.0 Hz, 2H), 3.03 – 2.89 (m, 2H), 2.87 (s, 6H), 1.27 (t, *J* = 7.5 Hz, 3H).

^13^C NMR (126 MHz, DMSO-*d*_6_) *δ* (ppm) 130.95, 127.88, 124.86, 115.49, 115.32, 111.58, 54.42, 42.69, 20.65, 13.00.

^19^F NMR (471 MHz, DMSO-*d*_6_) *δ* (ppm) -115.61.

HRMS (APCI): calcd. for C_24_H_25_FN_7_O [M+H]^+^= 446.2099, found [M+H]^+^ = 446.2102.

7-Ethyl-8-(4-fluorophenyl)-3-(1-(2-morpholinoethyl)-1*H*-benzo[*d*]imidazol-2-yl)pyrazolo[5,1-*c*][1,2,4]triazin-4-ol **(24)**

The compound was prepared according to General procedure C using:

Diazotization step:

3-Ethyl-4-(4-fluorophenyl)-1*H*-pyrazol-5-amine (100 mg, 0.49 mmol; 1 eq.), 35% aqueous HCl (190 µL, 2 mmol; 4 eq.) in EtOH (3 mL) and H_2_O (3 mL), NaNO_2_ (68 mg, 1 mmol, 2 eq.) in EtOH (1 mL) (reaction time: 15 min), ethyl 2-(1-(2-morpholinoethyl)-1*H*-benzo[*d*]imidazol-2-yl)acetate (170 mg, 0.53 mmol, 1.2 eq.) in EtOH (2 mL) and KOAc (300 mg, 3.0 mmol, 6 eq.). Reaction time: 16 h. The yellow solid (177 mg, 0.35 mmol) obtained after the filtration was used in the next step without further purification.

Cyclization step:

the dried yellow solid (177 mg, 0.35 mmol), DMF (3 mL), and KOAc (3 mg, 0.03 mmol, 0.1 eq.). Reaction time: 2 h. After the filtration, the solid part was suspended in hot dioxane (5 mL) and the mixture was poured into water (20 mL). The solid precipitate was collected by filtration, washed with water (10 mL), diethyl ether (10 mL) and dried under *vacuum*. The product was obtained as a yellow solid (153 mg, 65%).

^1^H NMR (500 MHz, DMSO-*d*_6_) *δ* (ppm) 7.94 – 7.86 (m, 2H), 7.78 (dd, *J* = 8.5, 5.5 Hz, 2H), 7.55 – 7.45 (m, 2H), 7.37 – 7.30 (m, 2H), 5.04 (s, 2H), 3.39 (t, *J* = 4.5 Hz, 4H), 2.93 (q, *J* = 7.5 Hz, 2H), 2.90 – 2.79 (m, 2H), 2.47 (s, 4H), 1.27 (t, *J* = 7.5 Hz, 3H).

^13^C NMR (126 MHz, DMSO-*d*_6_) *δ* (ppm) 161.43 (d, *J* = 243.6 Hz), 156.39, 149.84, 147.20, 133.53, 131.33 (d, *J* = 8.0 Hz), 128.63, 125.34, 125.08, 115.85 (d, *J* = 21.3 Hz), 115.57, 112.46, 66.36, 57.30, 53.85, 44.14, 21.28, 13.54.

^19^F NMR (282 MHz, DMSO-*d*_6_) *δ* (ppm) -115.78.

HRMS (APCI): calcd. for C_26_H_27_FN_7_O_2_ [M+H]^+^= 488.2205, found [M+H]^+^ = 488.2206.

7-Ethyl-8-(4-fluorophenyl)-3-(1-methyl-1*H*-imidazol-2-yl)pyrazolo[5,1-*c*][1,2,4]triazin-4-ol **(25)**

The compound was prepared according to General procedure C using:

Diazotization step:

3-ethyl-4-(4-fluorophenyl)-1*H*-pyrazol-5-amine (200 mg, 0.97 mmol; 1 eq.), 35% aqueous HCl (336 µL, 3.9 mmol; 4 eq.) in EtOH (10 mL) and H_2_O (10 mL), NaNO_2_ (135 mg, 1.95 mmol, 2 eq.) in EtOH (1 mL) (reaction time: 15 min), ethyl 2-(1-methyl-1*H*-imidazol-2-yl)acetate hydrochloride (238 mg, 1.17 mmol, 1.2 eq.) in EtOH (2 mL) and KOAc (570 mg, 4.14 mmol, 6 eq.). Reaction time: 16 h. The yellow solid (214 mg, 0.6 mmol) obtained after the filtration was used in the next step without further purification.

Cyclization step:

the dried yellow solid (214 mg, 0.6 mmol), DMF (3 mL), and KOAc (7 mg, 0.07 mmol, 0.1 eq.). Reaction time: 2 h. After the filtration, the solid part was suspended in hot dioxane (5 mL) and the mixture was poured into water (30 mL). The solid precipitate was collected by filtration, washed with water (10 mL), diethyl ether (10 mL) and dried under *vacuum*. The product was obtained as a yellow solid (125 mg, 38%).

^1^H NMR (500 MHz, DMSO-*d*_6_) *δ* (ppm) 13.51 (s, 1H), 7.83 – 7.76 (m, 2H), 7.61 (s, 1H), 7.48 (s, 1H), 7.31 – 7.24 (m, 2H), 4.11 (s, 3H), 2.92 (q, *J* = 7.5 Hz, 2H), 1.28 (t, *J* = 7.4 Hz, 3H).

^13^C NMR (126 MHz, DMSO-*d*_6_) *δ* (ppm) 160.45 (d, *J* = 242.9 Hz), 155.08, 148.63, 148.52, 141.68, 130.31 (d, *J* = 7.6 Hz), 128.49 (d, *J* = 3.9 Hz), 123.41, 121.25, 117.36, 114.78 (d, *J* = 21.1 Hz), 105.87, 37.11, 20.58, 12.68.

HRMS (APCI): calcd. for C_17_H_16_FN_6_O [M+H]^+^ = 339.1364, found [M+H]^+^ = 339.1362.

7-Ethyl-8-(4-fluorophenyl)-3-(4,5,6,7-tetrahydro-3*H*-imidazo[4,5-*c*]pyridin-2-yl)pyrazolo[5,1-*c*][1,2,4]triazin-4-ol **(26)**

To a cold (0 °C) solution of *tert*-butyl 2-(7-ethyl-8-(4-fluorophenyl)-4-hydroxypyrazolo[5,1-*c*][1,2,4]triazin-3-yl)-3,4,6,7-tetrahydro-5*H*-imidazo[4,5-*c*]pyridine-5-carboxylate (121 mg, 0.25 mmol, 1 eq.) in dichloromethane (2 mL) was added TFA (1 mL) dropwise. The mixture was stirred at 0 °C for 15 min, then at 25 °C for 2 h. The solvent was removed *in vacuo*, the residue was quenched with saturated aqueous solution of NaHCO_3_ (10 mL), and the mixture was extracted with EtOAc (6 × 20 mL). The combined organic extracts were dried over MgSO_4_, filtered, and the solvent was evaporated *in vacuo*. The residue was purified by reverse phase column chromatography using Biotage Selekt purification system (H_2_O:MeOH: 7 M NH_3_ in MeOH, gradient 4:1:0.1% to 0:1:0.1%). The product was obtained as an orange solid (41 mg, 43%).

^1^H NMR (500 MHz, Methanol-*d*_4_) *δ* (ppm) 7.69 – 7.61 (m, 2H), 7.22 – 7.14 (m, 2H), 3.92 – 3.80 (m, 2H), 3.13 – 3.04 (m, 2H), 2.93 (q, *J* = 7.6 Hz, 2H), 2.82 – 2.71 (m, 2H), 1.30 (t, *J* = 7.6 Hz, 3H).

^13^C NMR (126 MHz, Methanol-*d*_4_) *δ* (ppm) 162.96 (d, *J* = 244.3 Hz), 157.70, 150.89, 150.57, 144.82, 132.32 (d, *J* = 8.1 Hz), 130.28 (d, *J* = 3.4 Hz), 127.85, 116.14 (d, *J* = 21.7 Hz), 106.26, 44.30, 43.90, 24.23, 21.84, 14.21.

^19^F NMR (471 MHz, Methanol-*d*_4_) *δ* (ppm) -118.65.

HRMS (APCI): calcd. for C_19_H_17_FN_7_O [M-H]^-^ = 378.1484, found [M-H]^-^ = 378.1482.

7-Ethyl-8-(4-fluorophenyl)-3-(1*H*-imidazol-2-yl)pyrazolo[5,1-*c*][1,2,4]triazin-4-amine **(27)**

A mixture of 3-(1-benzyl-1*H*-imidazol-2-yl)-7-ethyl-8-(4-fluorophenyl)pyrazolo[5,1-*c*][1,2,4]triazin-4-amine (50 mg, 0.12 mmol, 1eq.) and Pd(OH)_2_/C (10%, 17 mg, 0.01mmol, 10%) in degassed EtOH (5 mL) was refluxed under hydrogen atmosphere (1 bar) for 3 h. The solvent was evaporated and the residue was purified by column chromatography on silica gel (EtOAc:MeOH, 9:1). The product was obtained as a yellow solid (22 mg, 57%).

^1^H NMR (126 MHz, DMSO-*d*_6_) *δ* (ppm) 13.09 (s, 1H), 9.77 (s, 1H), 9.04 (s, 1H), 7.91 – 7.79 (m, 2H), 7.38 – 7.32 (m, 2H), 7.30 (s, 1H), 7.19 (s, 1H), 3.04 (q, *J* = 7.6 Hz, 2H), 1.35 (t, *J* = 7.5 Hz, 3H).

^13^C NMR (126 MHz, DMSO-*d*_6_) *δ* (ppm) 160.86 (d, *J* = 243.8 Hz), 157.53, 146.29, 143.93, 137.13, 130.64 (d, *J* = 7.8 Hz), 127.97 (d, *J* = 2.69 Hz), 127.75, 120.31, 117.46, 115.35 (d, *J* = 21.4 Hz), 106.06, 20.83, 13.07.

^19^F NMR (471 MHz, DMSO-*d*_6_) *δ* (ppm) -115.77.

HRMS (APCI): calcd. for C_16_H_15_FN_7_ [M+H]^+^ = 324.1367, found [M+H]^+^ = 324.1370.

6-(1*H*-benzo[*d*]imidazol-2-yl)-2-ethyl-3-(4-fluorophenyl)pyrazolo[1,5-*a*]pyrimidin-7-ol **(28)**

A mixture of 3-ethyl-4-(4-fluorophenyl)-1*H*-pyrazol-5-amine (75 mg, 0.37 mmol; 1 eq.) and ethyl-2-(1*H*-benzo[*d*]imidazol-2-yl)-3-(dimethylamino)acrylate (100 mg, 0.38 mmol, 1.05 eq.) in EtOH (3 mL) was stirred in the microwave reactor at 130 °C for 1 h. The precipitate was collected by filtration, washed with EtOH (5 mL), diethyl ether (5 mL) and dried under *vacuum*. The product was obtained as a white solid (15 mg, 11%)

^1^H NMR (500 MHz, DMSO-*d*_6_) *δ* (ppm) 13.29 (s, 2H), 8.69 (s, 1H), 7.75 – 7.70 (m, 2H), 7.69 – 7.61 (m, 2H), 7.39 – 7.33 (m, 2H), 7.31 – 7.22 (m, 2H), 2.87 (q, *J* = 7.6 Hz, 2H), 1.26 (t, *J* = 7.5 Hz, 3H).

^13^C NMR (126 MHz, DMSO-*d*_6_) *δ* (ppm) 161.23 (d, *J* = 243.6 Hz), 156.61, 155.59, 149.80, 131.13 (d, *J* = 7.9 Hz), 129.56, 124.16, 115.50 (d, *J* = 21.1 Hz), 113.56, 21.38, 13.48.

HRMS (APCI): calcd. for C_21_H_17_FN_5_O [M+H]^+^= 374.1412, found [M+H]^+^ = 374.1416.

3-(1*H*-benzo[*d*]imidazol-2-yl)-7-(3-bromophenyl)pyrazolo[5,1-*c*][1,2,4]triazin-4-amine **(56)**

The compound was prepared according to General procedure C using:

Diazotization step:

3-(3-bromophenyl)-1*H*-pyrazol-5-amine (298 mg, 1.25 mmol; 1 eq.), 35% aqueous HCl (0.44 mL, 5.01 mmol; 4 eq.) in EtOH (2 mL) and H_2_O (2 mL), NaNO_2_ (173 mg, 2.5 mmol, 2 eq.) in EtOH (1 mL) and H_2_O (2 mL) (reaction time: 20 min), 2-(cyanomethyl)-benzimidazole (236 mg, 1.5 mmol, 1.2 eq.) in EtOH (3 mL) and H_2_O (3 mL) and KOAc (982 g, 10.01 mmol, 8 eq.). Reaction time: 2 h. The yellow solid (471 mg, 1.16 mmol) obtained after the filtration was used in the next step without further purification.

Cyclization step:

the dried yellow solid (465 mg, 1.14 mmol), DMF (6 mL), and KOAc (6 mg, 0.06 mmol, 0.05 eq.). Reaction time: 2 h. The product was obtained as a brown solid (336 mg, 66%).

^1^H NMR (500 MHz, DMSO-*d*_6_) *δ* (ppm) 13.47 (s, 1H), 9.97 (s, 1H), 9.42 (s, 1H), 8.42 (t, *J* = 1.9 Hz, 1H), 8.18 (d, *J* = 7.8 Hz, 1H), 7.76 (s, 1H), 7.69 (dd, *J* = 8.2, 2.0 Hz, 1H), 7.63 (s, 1H), 7.59 – 7.49 (m, 2H), 7.26 (d, *J* = 7.2 Hz, 2H).

^13^C NMR (126 MHz, DMSO-*d*_6_) *δ* (ppm) 154.59, 150.28, 149.95, 142.75, 138.86, 134.11, 133.75, 132.26, 131.11, 128.89, 125.66, 122.93, 122.35, 121.88, 119.17, 118.34, 111.50, 93.81.

HRMS (APCI): calcd. for C_18_H_13_BrN_7_ [M+H]^+^ = 406.0410, found [M+H]^+^ = 406.0412.

3-(1*H*-benzo[*d*]imidazol-2-yl)-7-cyclohexylpyrazolo[5,1-*c*][1,2,4]triazin-4-amine **(57)**

The compound was prepared according to General procedure C using:

Diazotization step:

3-cyclohexyl-1*H*-pyrazol-5-amine (316 mg, 1.91 mmol; 1 eq.), 35% aqueous HCl (0.675 mL, 7.65 mmol; 4 eq.) in EtOH (2 mL) and H_2_O (2 mL), NaNO_2_ (264 mg, 3.82 mmol, 2 eq.) in EtOH (2 mL) and H_2_O (3 mL) (reaction time: 20 min), 2-(cyanomethyl)-benzimidazole (361 mg, 2.29 mmol, 1.2 eq.) in EtOH (4 mL) and H_2_O (4 mL) and KOAc (1.501 g, 15.30 mmol, 8 eq.). Reaction time: 2 h. The yellow solid (606 mg, 1.82 mmol) obtained after the filtration was used in the next step without further purification.

Cyclization step:

the dried yellow solid (600 mg, 1.80 mmol), DMF (10 mL), and KOAc (9 mg, 0.09 mmol, 0.05 eq.). Reaction time: 2 h. The product was obtained as a brown solid (497 mg, 78%).

^1^H NMR (500 MHz, DMSO-*d*_6_) *δ* (ppm) 13.40 (s, 1H), 9.87 (s, 1H), 9.19 (s, 1H), 7.73 (d, *J* = 7.6 Hz, 1H), 7.53 (d, *J* = 7.6 Hz, 1H), 7.30 – 7.16 (m, 2H), 6.88 (s, 1H), 2.89 (tt, *J* = 11.6, 3.6 Hz, 1H), 2.06 (d, *J* = 10.6 Hz, 2H), 1.83 (dt, *J* = 13.1, 3.5 Hz, 2H), 1.76 – 1.70 (m, 1H), 1.60 (qd, *J* = 12.5, 3.4 Hz, 2H), 1.43 (qt, *J* = 12.6, 3.4 Hz, 2H), 1.35 – 1.24 (m, 1H).

^13^C NMR (126 MHz, DMSO-*d*_6_) *δ* (ppm) 164.86, 150.21, 149.50, 142.68, 138.75, 133.71, 122.79, 121.76, 118.30, 118.22, 111.40, 93.48, 37.70, 32.42, 25.69, 25.54.

HRMS (APCI): calcd. for C_18_H_20_N_7_ [M+H]^+^ = 334.1775, found [M+H]^+^ = 334.1765.

3-(1*H*-benzo[*d*]imidazol-2-yl)-7-isopropylpyrazolo[5,1-*c*][1,2,4]triazin-4-amine **(58)**

The compound was prepared according to General procedure C using:

Diazotization step:

3-isopropyl-1*H*-pyrazol-5-amine (200 mg, 1.60 mmol; 1 eq.), 35% aqueous HCl (0.6 mL, 6.4 mmol; 4 eq.) in EtOH (6 mL) and H_2_O (6 mL), NaNO_2_ (220 mg, 3.20 mmol, 2 eq.) in H_2_O (2 mL) (reaction time: 30 min), 2-(cyanomethyl)-benzimidazole (300 mg, 1.92 mmol, 1.2 eq.) in EtOH (6 mL) and KOAc (942 mg, 9.6 mmol, 6 eq.). Reaction time: 16 h. The yellow solid (280 mg, 1.34 mmol) obtained after the filtration was used in the next step without further purification.

Cyclization step:

the dried yellow solid (280 mg, 1.34 mmol), DMF (5 mL), and KOAc (7 mg, 0.07 mmol, 0.05 eq.). Reaction time: 2 h. The product was obtained as an orange solid (22 mg, 22%).

^1^H NMR (500 MHz, DMSO-*d*_6_) *δ* (ppm) 13.40 (s, 1H), 9.88 (s, 1H), 9.22 (s, 1H), 7.73 (d, *J* = 7.6 Hz, 1H), 7.53 (d, *J* = 7.2 Hz, 1H), 7.29 – 7.19 (m, 2H), 6.91 (s, 1H), 3.22 (hept, *J* = 6.9 Hz, 1H), 1.39 (d, *J* = 6.9 Hz, 6H).

^13^C NMR (126 MHz, DMSO-*d*_6_) *δ* (ppm) 165.93, 150.19, 149.56, 142.66, 138.75, 133.69, 122.79, 121.76, 118.31, 118.22, 111.39, 93.24, 28.16, 22.41.

HRMS (APCI): calcd. for C_15_H_16_N_7_ [M+H]^+^ = 294.1462, found [M+H]^+^ = 294.1450.

3-(1*H*-benzo[*d*]imidazol-2-yl)-7-(*tert*-butyl)pyrazolo[5,1-*c*][1,2,4]triazin-4-amine **(59)**

The compound was prepared according to General procedure C using:

Diazotization step:

3-(*tert*-butyl)-1*H*-pyrazol-5-amine (222 mg, 1.60 mmol; 1 eq.), 35% aqueous HCl (0.6 mL, 6.4 mmol; 4 eq.) in EtOH (6 mL) and H_2_O (6 mL), NaNO_2_ (220 mg, 3.2 mmol, 2 eq.) in H_2_O (2 mL) (reaction time: 30 min), 2-(cyanomethyl)-benzimidazole (300 mg, 1.92 mmol, 1.2 eq.) in EtOH (6 mL) and KOAc (942 mg, 9.60 mmol, 6 eq.). Reaction time: 16 h. The yellow solid (425 mg, 1.38 mmol) obtained after the filtration was used in the next step without further purification.

Cyclization step:

the dried yellow solid (210 mg, 0.7 mmol), DMF (5 mL), and KOAc (7 mg, 0.07 mmol, 0.05 eq.). Reaction time: 2 h. The product was obtained as a yellow solid (106 mg, 22%).

^1^H NMR (500 MHz, DMSO-*d*_6_) *δ* (ppm) 13.39 (s, 1H), 9.85 (s, 1H), 9.05 (s, 1H), 7.74 (d, *J* = 7.6 Hz, 1H), 7.53 (d, *J* = 7.2 Hz, 1H), 7.29 – 7.19 (m, 2H), 6.95 (s, 1H), 1.45 (s, 9H).

^13^C NMR (126 MHz, DMSO-*d*_6_) *δ* (ppm) 168.63, 150.21, 149.53, 142.67, 138.71, 133.70, 122.77, 121.76, 118.22, 118.20, 111.39, 92.98, 32.72, 30.11.

HRMS (ESI): calcd. for C_16_H_16_N_7_ [M-H]^-^ = 306.1473, found [M-H]^-^ = 306.1478.

(4-Amino-3-(1*H*-benzo[*d*]imidazol-2-yl)pyrazolo[5,1-*c*][1,2,4]triazin-7-yl)methanol **(60)**

To solution of 3-(1*H*-benzo[*d*]imidazol-2-yl)-7-(((*tert*-butyldiphenylsilyl)oxy)methyl)pyrazolo[5,1-*c*][1,2,4]triazin-4-amine (130 mg, 0.25 mmol; 1 eq.) in THF (5 mL) was added TBAF (1M in THF, 0.5 mL, 0.5 mmol) and the mixture was stirred at 23 °C for 2 h. The solvent was evaporated and the residue was purified by column chromatography on silica gel (dichloromethane:MeOH:7M NH_3_ in MeOH, gradient 5:3:0 to 10:0:1). The product was obtained as an orange solid (52 mg, 57%).

^1^H NMR (500 MHz, DMSO-*d*_6_) *δ* (ppm) 13.43 (s, 1H), 9.93 (s, 1H), 9.42 (s, 1H), 7.73 (d, *J* = 7.2 Hz, 1H), 7.57 – 7.51 (m, 2H), 7.30 – 7.18 (m, 2H), 6.94 (s, 1H), 5.54 (t, *J* = 5.8 Hz, 1H), 4.77 (d, *J* = 5.8 Hz, 2H).

^13^C NMR (126 MHz, DMSO-*d*_6_) *δ* (ppm) 160.83, 150.09, 149.51, 142.64, 138.93, 133.69, 122.83, 121.79, 118.48, 118.25, 111.43, 94.54, 57.68.

HRMS (APCI): calcd. for C_13_H_12_N_7_O [M+H]^+^ = 282.1098, found [M+H]^+^ = 282.1102

2-(4-Amino-3-(1*H*-benzo[d]imidazol-2-yl)pyrazolo[5,1-*c*][1,2,4]triazin-7-yl)phenol **(61)**

The compound was prepared according to General procedure C using:

Diazotization step:

2-(3-Amino-1*H*-pyrazol-5-yl)phenol (117 mg, 0.67 mmol; 1 eq.), 35% aqueous HCl (0.2 mL, 2.0 mmol; 4 eq.) in EtOH (3 mL) and H_2_O (3 mL), NaNO_2_ (100 mg, 1.34 mmol, 2 eq.) in H_2_O (1 mL) (reaction time: 30 min), 2-(cyanomethyl)-benzimidazole (130 mg, 0.80 mmol, 1.2 eq.) in EtOH (1 mL) and KOAc (400 mg, 4.02 mmol, 6 eq.) in H_2_O (1 mL). Reaction time: 16 h. The yellow solid (126 mg, 0.36 mmol) obtained after the filtration was used in the next step without further purification.

Cyclization step:

the dried yellow solid (126 mg, 0.36 mmol), DMF (3 mL), and KOAc (4 mg, 0.04 mmol, 0.1 eq.). Reaction time: 2 h. The product was obtained as a yellow solid (33 mg, 14%).

^1^H NMR (500 MHz, DMSO-*d*_6_) *δ* (ppm) 13.46 (s, 1H), 10.31 (s, 1H), 9.98 (s, 1H), 9.64 (s, 1H), 8.13 (dd, *J* = 7.8, 1.6 Hz, 1H), 7.76 (d, *J* = 7.5 Hz, 1H), 7.57 (s, 1H), 7.56 (d, *J* = 6.6 Hz, 1H), 7.37 – 7.32 (m, 1H), 7.30 – 7.22 (m, 2H), 7.08 – 7.04 (m, 1H), 7.03 – 6.98 (m, 1H).

^13^C NMR (126 MHz, DMSO-*d*_6_) *δ* (ppm) 155.90, 155.04, 150.01, 149.40, 142.71, 138.63, 133.75, 130.89, 128.61, 122.89, 121.82, 119.39, 119.12, 118.31, 117.12, 116.59, 111.46, 94.71.

HRMS (APCI): calcd. for C_18_H_14_N_7_O [M+H]^+^ = 344.1254, found [M+H]^+^ = 344.1256.

3-(1*H*-benzo[*d*]imidazol-2-yl)-7-(3-methoxynaphthalen-2-yl)pyrazolo[5,1-*c*][1,2,4]triazin-4-amine **(62)**

The compound was prepared according to General procedure C using:

Diazotization step:

3-(3-methoxynaphthalen-2-yl)-1*H*-pyrazol-5-amine (90 mg, 0.38 mmol; 1 eq.), 35% aqueous HCl (0.15 mL, 1.52 mmol; 4 eq.) in EtOH (3 mL) and H_2_O (3 mL), NaNO_2_ (60 mg, 0.76 mmol, 2 eq.) in H_2_O (1 mL) (reaction time: 30 min), 2-(cyanomethyl)-benzimidazole (73 mg, 0.46 mmol, 1.2 eq.) in EtOH (1 mL) and KOAc (223 mg, 2.28 mmol, 6 eq.) in H_2_O (1 mL). Reaction time: 16 h. The yellow solid (103 mg, 0.25 mmol) obtained after the filtration was used in the next step without further purification.

Cyclization step:

the dried yellow solid (103 mg, 0.25 mmol), DMF (3 mL), and KOAc (2 mg, 0.02 mmol, 0.1 eq.). Reaction time: 2 h. The product was obtained as a yellow solid (40 mg, 25%).

^1^H NMR (500 MHz, DMSO-*d*_6_) *δ* (ppm) 13.48 (s, 1H), 9.97 (s, 1H), 9.43 (s, 1H), 8.77 (s, 1H), 7.93 (dd, *J* = 20.4, 8.1 Hz, 2H), 7.76 (s, 1H), 7.66 – 7.49 (m, 4H), 7.44 (t, *J* = 7.1 Hz, 1H), 7.27 (s, 2H), 4.09 (s, 3H).

^13^C NMR (126 MHz, DMSO-*d*_6_) *δ* (ppm) 155.20, 153.18, 150.10, 149.79, 138.68, 134.60, 128.83, 127.95, 127.85, 127.26, 126.53, 124.34, 121.86, 118.79, 106.65, 97.86, 55.75.

HRMS (APCI): calcd. for C_23_H_18_N_7_O [M+H]^+^= 408.1567, found [M+H]^+^ = 408.1566.

7-([1,1'-Biphenyl]-3-yl)-3-(1*H*-benzo[*d*]imidazol-2-yl)pyrazolo[5,1-*c*][1,2,4]triazin-4-amine **(63)**

The compound was prepared according to General procedure C using:

Diazotization step:

3-([1,1'-Biphenyl]-3-yl)-1*H*-pyrazol-5-amine (320 mg, 1.50 mmol; 1 eq.), 35% aqueous HCl (0.6 mL, 6.0 mmol; 4 eq.) in EtOH (3 mL) and H_2_O (3 mL), NaNO_2_ (228 mg, 3.0 mmol, 2 eq.) in H_2_O (1 mL) (reaction time: 30 min), 2-(cyanomethyl)-benzimidazole (130 mg, 1.8 mmol, 1.2 eq.) in EtOH (1 mL) and KOAc (880 mg, 9.0 mmol, 6 eq.) in H_2_O (1 mL). Reaction time: 16 h. The yellow solid (600 mg, 1.5 mmol) obtained after the filtration was used in the next step without further purification.

Cyclization step:

the dried yellow solid (600 mg, 1.50 mmol), DMF (3 mL), and KOAc (15 mg, 0.15 mmol, 0.1 eq.). Reaction time: 2 h. The product was obtained as a yellow solid (327 mg, 54%).

^1^H NMR (500 MHz, DMSO-*d*_6_) *δ* (ppm) 13.43 (s, 1H), 9.97 (s, 1H), 9.41 (s, 1H), 8.07 – 8.00 (m, 1H), 7.74 (s, 1H), 7.62 – 7.51 (m, 3H), 7.50 – 7.43 (m, 1H), 7.43 – 7.35 (m, 3H), 7.34 – 7.26 (m, 2H), 7.24 (d, *J* = 6.5 Hz, 2H), 6.11 (s, 1H).

^13^C NMR (126 MHz, DMSO-*d*_6_) *δ* (ppm) 156.68, 149.95, 148.99, 142.61, 141.26, 140.64, 138.74, 134.11, 133.78, 130.63, 130.54, 130.00, 129.33, 129.20, 128.32, 127.56, 127.39, 122.86, 121.78, 118.73, 118.28, 111.43, 109.46, 96.72.

HRMS (APCI): calcd. for C_24_H_18_N_7_ [M+H]^+^ = 404.1618, found [M+H]^+^ = 404.1615.

3-(1*H*-benzo[*d*]imidazol-2-yl)-7-(1-methoxynaphthalen-2-yl)pyrazolo[5,1*-c*][1,2,4]triazin-4-amine **(64)**

The compound was prepared according to General procedure C using:

Diazotization step:

3-(1-methoxynaphthalen-2-yl)-1*H*-pyrazol-5-amine (180 mg, 0.75 mmol; 1 eq.), 35% aqueous HCl (0.3 mL, 3.0 mmol; 4 eq.) in EtOH (3 mL) and H_2_O (3 mL), NaNO_2_ (103 mg, 1.50 mmol, 2 eq.) in H_2_O (1 mL) (reaction time: 30 min), 2-(cyanomethyl)-benzimidazole (138 mg, 0.90 mmol, 1.2 eq.) in EtOH (1 mL) and KOAc (450 mg, 4.5 mmol, 6 eq.) in H_2_O (1 mL). Reaction time: 16 h. The yellow solid (200 mg, 0.47 mmol) obtained after the filtration was used in the next step without further purification.

Cyclization step:

the dried yellow solid (200 mg, 0.47 mmol), DMF (3 mL), and KOAc (5 mg, 0.05 mmol, 0.1 eq.). Reaction time: 2 h. The product was obtained as a yellow solid (150 mg, 49%).

^1^H NMR (500 MHz, DMSO-*d*_6_) *δ* (ppm) 13.50 (s, 1H), 9.99 (s, *J* = 46.1 Hz, 1H), 9.42 (s, 1H), 8.43 (d, *J* = 8.6 Hz, 1H), 8.24 (d, *J* = 8.0 Hz, 1H), 8.02 (d, *J* = 7.6 Hz, 1H), 7.89 (t, *J* = 9.7 Hz, 1H), 7.76 (s, 1H), 7.70 – 7.62 (m, 2H), 7.61 (s, 1H), 7.57 (s, 1H), 7.26 (s, 2H), 3.96 (s, 3H).

^13^C NMR (126 MHz, DMSO-*d*_6_) *δ* (ppm) 154.78, 153.01, 150.22, 150.05, 142.71, 138.73, 135.01, 133.76, 128.05, 127.91, 127.29, 126.74, 125.97, 125.75, 124.12, 122.87, 122.53, 122.33, 121.84, 120.30, 118.90, 118.30, 111.47, 96.59, 66.30, 61.79.

HRMS (APCI): calcd. for C_23_H_18_N_7_O [M+H]^+^= 408.1567, found [M+H]^+^ = 408.1567.

3-(1*H*-benzo[*d*]imidazol-2-yl)-7-(3-methoxynaphthalen-1-yl)pyrazolo[5,1-*c*][1,2,4]triazin-4-amine **(65)**

The compound was prepared according to General procedure C using:

Diazotization step:

3-(3-methoxynaphthalen-1-yl)-1*H*-pyrazol-5-amine (180 mg, 0.75 mmol; 1 eq.), 35% aqueous HCl (0.3 mL, 3.0 mmol; 4 eq.) in EtOH (3 mL) and H_2_O (3 mL), NaNO_2_ (103 mg, 1.50 mmol, 2 eq.) in H_2_O (1 mL) (reaction time: 30 min), 2-(cyanomethyl)-benzimidazole (138 mg, 0.90 mmol, 1.2 eq.) in EtOH (1 mL) and KOAc (450 mg, 4.50 mmol, 6 eq.) in H_2_O (1 mL). Reaction time: 16 h. The yellow solid (300 mg, 0.71 mmol) obtained after the filtration was used in the next step without further purification.

Cyclization step:

the dried yellow solid (300 mg, 0.71 mmol), DMF (3 mL), and KOAc (7 mg, 0.07 mmol, 0.1 eq.). Reaction time: 2 h. The product was obtained as a yellow solid (30 mg, 10%).

^1^H NMR (500 MHz, DMSO-*d*_6_) *δ* (ppm) 13.50 (s, 1H), 9.99 (s, 1H), 9.51 (s, 1H), 8.14 (d, *J* = 9.1 Hz, 1H), 7.99 – 7.93 (m, 1H), 7.76 (d, *J* = 6.5 Hz, 1H), 7.68 – 7.63 (m, 1H), 7.62 (d, *J* = 9.2 Hz, 1H), 7.57 (d, *J* = 6.4 Hz, 1H), 7.46 – 7.37 (m, 1H), 7.31 – 7.22 (m, 1H), 7.17 (s, 1H), 3.89 (s, 1H).

^13^C NMR (126 MHz, DMSO-*d*_6_) *δ* (ppm) 155.12, 153.41, 150.19, 149.50, 138.95, 133.02, 131.04, 128.20, 127.88, 126.97, 124.71, 123.68, 122.88, 121.81, 118.57, 118.29, 117.95, 115.33, 113.71, 111.45, 99.05, 56.40.

HRMS (APCI): calcd. for C_23_H_18_N_7_O [M+H]^+^ = 408.1567, found [M+H]^+^ = 408.1568.

3-(1*H*-benzo[*d*]imidazol-2-yl)-7-(1-benzyl-1*H*-imidazol-2-yl)pyrazolo[5,1-*c*][1,2,4]triazin-4-amine **(66)**

The compound was prepared according to General procedure C using:

Diazotization step:

3-(1-benzyl-1*H*-imidazol-2-yl)-1*H*-pyrazol-5-amine (70 mg, 0.30 mmol; 1 eq.), 35% aqueous HCl (0.1 mL, 1.2 mmol; 4 eq.) in EtOH (4 mL) and H_2_O (4 mL), NaNO_2_ (45 mg, 0.60 mmol, 2 eq.) in H_2_O (1 mL) (reaction time: 30 min), 2-(cyanomethyl)-benzimidazole (60 mg, 0.36 mmol, 1.2 eq.) in EtOH (1 mL) and KOAc (180 mg, 1.8 mmol, 6 eq.) in H_2_O (1 mL). Reaction time: 16 h. The yellow solid (106 mg, 0.25 mmol) obtained after the filtration was used in the next step without further purification.

Cyclization step:

the dried yellow solid (106 mg, 0.25 mmol), DMF (3 mL), and KOAc (10 mg, 0.10 mmol, 0.1 eq.). Reaction time: 2 h. The product was obtained as a yellow solid (80 mg, 66%).

^1^H NMR (500 MHz, DMSO-*d*_6_) *δ* (ppm) 13.47 (s, 1H), 9.99 (s, 1H), 9.44 (s, 1H), 7.75 (d, *J* = 7.7 Hz, 1H), 7.55 (d, *J* = 7.4 Hz, 1H), 7.48 (s, 1H), 7.30 – 7.18 (m, 8H), 7.16 (s, 1H), 6.04 (s, 2H).

^13^C NMR (126 MHz, DMSO-*d*_6_) *δ* (ppm) 159.89, 159.49, 159.04, 152.70, 148.85, 148.81, 148.15, 143.78, 139.22, 138.47, 137.53, 137.40, 134.07, 132.93, 131.85, 129.25, 128.34, 121.54, 105.09, 60.16.

3-(1*H*-benzo[*d*]imidazol-2-yl)-7-(1*H*-imidazol-2-yl)pyrazolo[5,1-*c*][1,2,4]triazin-4-amine **(67)**

A mixture of 3-(1*H*-benzo[*d*]imidazol-2-yl)-7-(1-benzyl-1*H*-imidazol-2-yl)pyrazolo[5,1-*c*][1,2,4]triazin-4-amine (40 mg, 0.10 mmol, 1eq.) and Pd(OH)_2_/C (10%, 14 mg, 0.01 mmol) in degassed EtOH (5 mL) was refluxed under a hydrogen atmosphere (1 bar) for 3 h. The solvent was evaporated, and the residue was purified by column chromatography on silica gel (EtOAc:MeOH, 9:1). The product was obtained as a yellow solid (22 mg, 57%).

^1^H NMR (126 MHz, DMSO-*d*_6_) *δ* (ppm) 13.48 (s, 1H), 12.88 (s, 1H), 10.00 (s, 1H), 9.02 (s, 1H), 7.95 (s, 1H), 7.75 (d, *J* = 7.7 Hz, 1H), 7.56 (d, *J* = 7.6 Hz, 1H), 7.38 (s, 1H), 7.33 (s, 1H), 7.26 (p, *J* = 6.9 Hz, 2H), 7.16 (s, 1H).

^13^C NMR (126 MHz, DMSO-*d*_6_) *δ* (ppm) 150.42, 150.23, 149.68, 143.19, 140.13, 139.41, 134.27, 130.26, 123.47, 122.40, 119.75, 119.22, 118.82, 112.04, 94.12.

HRMS (APCI): calcd. for C_15_H_12_N_9_ [M+H]^+^ = 318.1210, found [M+H]^+^ = 318.1213.

3-(1*H*-benzo[*d*]imidazol-2-yl)-7-(2-chlorophenyl)pyrazolo[5,1-*c*][1,2,4]triazin-4-amine **(68)**

The compound was prepared according to General procedure C using:

Diazotization step:

3-(2-chlorophenyl)-1*H*-pyrazol-5-amine (350 mg, 1.81 mmol; 1 eq.), 35% aqueous HCl (0.6 mL, 7.24 mmol; 4 eq.) in EtOH (10 mL) and H_2_O (10 mL), NaNO_2_ (250 mg, 3.62 mmol, 2 eq.) in H_2_O (1 mL) (reaction time: 30 min), 2-(cyanomethyl)-benzimidazole (350 mg, 3.60 mmol, 1.2 eq.) in EtOH (1 mL) and KOAc (1.1 g, 10.90 mmol, 6 eq.) in H_2_O (1 mL). Reaction time: 16 h. The yellow solid (540 mg, 1.42 mmol) obtained after the filtration was used in the next step without further purification.

Cyclization step:

the dried yellow solid (540 mg, 1.42 mmol), DMF (5 mL), and KOAc (13 mg, 0.14 mmol, 0.1 eq.). Reaction time: 2 h. The product was obtained as a yellow solid (391 mg, 60%).

^1^H NMR (500 MHz, DMSO-*d*_6_) *δ* (ppm) 13.50 (s, 1H), 10.01 (s, 1H), 9.50 (s, 1H), 8.07 – 7.99 (m, 1H), 7.76 (s, 1H), 7.72 – 7.64 (m, 1H), 7.59 – 7.51 (m, 3H), 7.46 (s, 1H), 7.31 – 7.22 (m, 2H).

^13^C NMR (126 MHz, DMSO-*d*_6_) *δ* (ppm) 159.89, 159.49, 159.04, 152.70, 148.85, 148.81, 148.15, 143.78, 139.22, 138.47, 137.53, 137.40, 134.07, 132.93, 131.85, 129.25, 128.34, 121.54, 105.09, 60.16.

HRMS (APCI): calcd. for C_18_H_13_ClN_7_ [M+H]^+^ = 362.0915, found [M+H]^+^ = 362.0916.

3-(1*H*-benzo[*d*]imidazol-2-yl)-7-(2-methoxyphenyl)pyrazolo[5,1-*c*][1,2,4]triazin-4-amine **(69)**

The compound was prepared according to General procedure C using:

Diazotization step:

3-(2-methoxyphenyl)-1*H*-pyrazol-5-amine (350 mg, 1.85 mmol; 1 eq.), 35% aqueous HCl (0.65 mL, 7.4 mmol; 4 eq.) in EtOH (10 mL) and H_2_O (10 mL), NaNO_2_ (250 mg, 3.70 mmol, 2 eq.) in H_2_O (1 mL) (reaction time: 30 min), 2-(cyanomethyl)-benzimidazole (350 mg, 3.70 mmol, 1.2 eq.) in EtOH (1 mL) and KOAc (1.1 g, 10.90 mmol, 6 eq.) in H_2_O (1 mL). Reaction time: 16 h. The yellow solid (550 mg, 1.5 mmol) obtained after filtration was used in the next step without further purification.

Cyclization step:

the dried, filtered yellow solid (550 mg, 1.50 mmol), DMF (5 mL), and KOAc (13 mg, 0.15 mmol, 0.1 eq.). Reaction time: 2 h. The product was obtained as a yellow solid (336 mg, 51%).

^1^H NMR (500 MHz, DMSO-*d*_6_) *δ* (ppm) 13.45 (s, 1H), 9.92 (s, 1H), 9.32 (s, 1H), 8.28 (dd, *J* = 7.7, 1.8 Hz, 1H), 7.76 (d, *J* = 7.6 Hz, 1H), 7.55 (d, *J* = 7.4 Hz, 1H), 7.52 – 7.46 (m, 2H), 7.30 – 7.21 (m, 3H), 7.14 (td, *J* = 7.6, 0.9 Hz, 1H), 3.99 (s, 3H).

^13^C NMR (126 MHz, DMSO-*d*_6_) *δ* (ppm) 157.45, 153.29, 150.13, 149.76, 142.73, 138.62, 133.75, 130.99, 128.88, 122.87, 121.83, 120.49, 120.04, 118.66, 118.30, 112.21, 111.46, 97.29, 55.65.

HRMS (APCI): calcd. for C_19_H_16_N_7_O [M+H]^+^ = 358.1411, found [M+H]^+^ = 358.1412.

3-(1-benzyl-1*H*-benzo[*d*]imidazol-2-yl)-7-ethylpyrazolo[5,1-*c*][1,2,4]triazin-4-amine **(70)**

The compound was prepared according to General procedure C using:

Diazotization step:

3-ethyl-1*H*-pyrazol-5-amine (222 mg, 2.0 mmol; 1 eq.), 35% aqueous HCl (0.8 mL, 8.0 mmol; 4 eq.) in EtOH (5 mL) and H_2_O (5 mL), NaNO_2_ (140 mg, 2.0 mmol, 2 eq.) in H_2_O (1 mL) (reaction time: 20 min), 2-(1-benzyl-1*H*-benzo[*d*]imidazol-2-yl)acetonitrile (590 mg, 2.40 mmol, 1.2 eq.) in EtOH (2 mL) and KOAc (1.17 g, 12 mmol, 6 eq.) in H_2_O (2 mL). Reaction time: 16 h. The yellow solid (660 mg, 1.70 mmol) obtained after the filtration was used in the next step without further purification.

Cyclization step:

the dried yellow solid (660 mg, 1.70 mmol), DMF (5 mL), and KOAc (20 mg, 0.20 mmol, 0.1 eq.). Reaction time: 2 h. The reaction mixture was poured into water (20 mL) and extracted with EtOAc (3 × 20 mL). The combined organic layers were washed with brine (30 mL), dried over MgSO_4_, and the solvent was evaporated. The product was purified by column flash chromatography on silica gel (EtOAc:MeOH, 10:1). The product was obtained as a yellow solid (300 mg, 40%).

^1^H NMR (500 MHz, DMSO-*d*_6_) *δ* (ppm) 10.26 (s, 1H), 9.29 (s, 1H), 7.84 – 7.76 (m, 1H), 7.67 – 7.56 (m, 1H), 7.33 – 7.28 (m, 2H), 7.27 – 7.23 (m, 2H), 7.22 – 7.17 (m, 3H), 6.82 (s, 1H), 6.27 (s, 2H), 2.89 (q, *J* = 7.6 Hz, 2H), 1.35 (t, *J* = 7.6 Hz, 3H).

^13^C NMR (126 MHz, DMSO-*d*_6_) *δ* (ppm) 161.67, 148.80, 148.15, 141.04, 139.53, 137.65, 135.36, 128.43, 127.12, 126.69, 123.16, 122.62, 119.65, 118.62, 110.75, 94.41, 48.87, 21.72, 13.37.

HRMS (APCI): calcd. for C_21_H_20_N_7_ [M+H]^+^ = 370.1775, found [M+H]^+^ = 370.1776.

8-([1,1'-Biphenyl]-3-yl)-3-(1*H*-benzo[*d*]imidazol-2-yl)-7-ethylpyrazolo[5,1-*c*][1,2,4]triazin-4-amine **(71)**

The compound was prepared according to General procedure C using:

Diazotization step:

4-([1,1'-biphenyl]-3-yl)-3-ethyl-1*H*-pyrazol-5-amine (138 mg, 0.52 mmol; 1 eq.), 35% aqueous HCl (0.093 mL, 1.05 mmol; 2 eq.) in EtOH (1 mL) and H_2_O (1 mL), NaNO_2_ (38 mg, 0.55 mmol, 1.05 eq.) in H_2_O (1 mL) (reaction time: 30 min), 2-(cyanomethyl)-benzimidazole (87 mg, 0.55 mmol, 1.05 eq.) in EtOH (2 mL) and H_2_O (0.5 mL) and NaOAc (172 mg, 2.1 mmol, 4 eq.). Reaction time: 16 h. The reaction mixture was poured into brine (20 mL) and extracted with EtOAc (2 × 50 mL). The combined organic extracts were dried over MgSO_4_, filtered, and the solvent was evaporated *in vacuo*. The obtained yellow solid (126 mg, 56 %) was used in the next step without further purification.

Cyclization step:

the dried yellow solid (126 mg), DMF (2 mL) and KOAc (1.40 mg, 0.05 eq.). Reaction time: 2 h. The reaction mixture was poured into brine (20 mL) and extracted with EtOAc (3 × 50 mL). The combined organic extracts were dried over MgSO_4_, filtered and concentrated *in vacuo*. The residue was purified by column chromatography on silica gel (dichloromethane:MeOH, gradient 1:0 to 10:1). The obtained solid was triturated by EtOH (2 mL), collected by filtration and dried in a vacuum. The product was obtained as a yellow solid (98 mg, 43%).

^1^H NMR (500 MHz, DMSO-*d*_6_) *δ* (ppm) 13.43 (s, 1H), 9.97 (s, 1H), 9.37 (s, 1H), 8.23 (t, *J* = 1.8 Hz, 1H), 7.81 (d, *J* = 7.5 Hz, 1H), 7.75 (dt, *J* = 8.0, 2.3 Hz, 3H), 7.69 – 7.65 (m, 1H), 7.63 (t, *J* = 7.6 Hz, 1H), 7.57 – 7.50 (m, 3H), 7.41 (t, *J* = 7.4 Hz, 1H), 7.30 – 7.22 (m, 2H), 3.14 (q, *J* = 7.5 Hz, 2H), 1.40 (t, *J* = 7.6 Hz, 3H).

^13^C NMR (126 MHz, DMSO-*d*_6_) *δ* (ppm) 158.16, 150.07, 146.51, 142.72, 140.44, 140.19, 138.71, 133.74, 131.98, 129.23, 129.02, 127.60, 127.55, 127.39, 126.69, 124.99, 122.90, 121.85, 119.26, 118.30, 111.46, 107.95, 21.10, 13.12.

HRMS (APCI): calcd. for C_26_H_22_N_7_ [M+H]^+^ = 432.1931, found [M+H]^+^ = 432.1929.

3-(1-Benzyl-1*H*-imidazol-2-yl)-7-ethyl-8-(4-fluorophenyl)pyrazolo[5,1-*c*][1,2,4]triazin-4-amine **(72)**

The compound was prepared according to General procedure C using:

Diazotization step:

3-Ethyl-4-(4-fluorophenyl)-1*H*-pyrazol-5-amine (150 mg, 0.73 mmol; 1 eq.), 35% aqueous HCl (0.26 mL, 2.9 mmol; 4 eq.) in EtOH (3 mL) and H_2_O (3 mL), NaNO_2_ (100 mg, 1.5 mmol, 2 eq.) in H_2_O (1 mL) (reaction time: 30 min), 2-(1-benzyl-1*H*-imidazol-2-yl)acetonitrile (170 mg, 0.87 mmol, 1.2 eq.) in EtOH (1 mL) and KOAc (430 mg, 4.4 mmol, 6 eq.) in H_2_O (1 mL). Reaction time: 16 h. The yellow solid (150 mg, 0.35 mmol) obtained after the filtration was used in the next step without further purification.

Cyclization step:

the dried yellow solid (150 mg, 0.35 mmol), DMF (3 mL), and KOAc (4 mg, 0.04 mmol, 0.1 eq.). Reaction time: 2 h. The product was obtained as a yellow solid (110 mg, 36%).

^1^H NMR (500 MHz, DMSO-*d*_6_) *δ* (ppm) 10.10 (s, 1H), 9.02 (s, 1H), 7.89 – 7.75 (m, 2H), 7.48 (d, *J* = 1.1 Hz, 1H), 7.36 – 7.30 (m, 2H), 7.28 (t, *J* = 7.3 Hz, 2H), 7.23 (d, *J* = 1.1 Hz, 1H), 7.20 (t, *J* = 6.5 Hz, 3H), 5.97 (s, 2H), 3.02 (q, *J* = 7.5 Hz, 2H), 1.32 (t, *J* = 7.5 Hz, 3H).

^13^C NMR (126 MHz, DMSO-*d*_6_) *δ* (ppm) 160.87 (d, *J* = 244.1 Hz), 157.60, 145.61, 141.66, 138.20, 138.02, 130.66 (d, *J* = 8.1 Hz), 128.42, 127.80 (d, *J* = 3.4 Hz), 127.20, 126.94, 126.78, 123.39, 121.39, 115.35 (d, *J* = 21.0 Hz), 106.05, 51.17, 20.80, 13.06.

^19^F NMR (471 MHz, DMSO-*d*_6_) *δ* (ppm) -115.72.

HRMS (APCI): calcd. for C_23_H_21_FN_7_ [M+H]^+^ = 414.1837, found [M+H]^+^ = 414.1837

3-(1*H*-benzo[*d*]imidazol-2-yl)-7-ethyl-8-(3-(trifluoromethyl)phenyl)pyrazolo[5,1-*c*][1,2,4]triazin-4-amine **(73)**

The compound was prepared according to General procedure C using:

Diazotization step:

3-ethyl-4-(3-(trifluoromethyl)phenyl)-1*H*-pyrazol-5-amine (40 mg, 0.16 mmol; 1 eq.), 35% aqueous HCl (0.65 mL, 0.64 mmol; 4 eq.) in EtOH (3 mL) and H_2_O (3 mL), NaNO_2_ (22 mg, 0.32 mmol, 2 eq.) in H_2_O (1 mL) (reaction time: 30 min), 2-(cyanomethyl)-benzimidazole (30 mg, 0.19 mmol, 1.2 eq.) in EtOH (1 mL) and KOAc (94 mg, 0.96 mmol, 6 eq.) in H_2_O (1 mL). Reaction time: 16 h. The yellow solid (66 mg, 0.15 mmol) obtained after the filtration was used in the next step without further purification.

Cyclization step:

the dried yellow solid (66 mg, 0.15 mmol), DMF (3 mL), and KOAc (5 mg, 0.06 mmol, 0.4 eq.). Reaction time: 2 h. The reaction mixture was poured into water (20 mL) and extracted with EtOAc (3 × 20 mL). The combined organic layers were washed with brine (30 mL), dried over MgSO_4_, the solvent was evaporated, and the residue was purified by column flash chromatography on silica gel (EtOAc to dichloromethane:MeOH, gradient 1:0:0 to 0:10:1). The product was obtained as a yellow solid (30 mg, 45% (2 steps)).

^1^H NMR (500 MHz, DMSO-*d*_6_) *δ* (ppm) 13.47 (s, 1H), 9.99 (s, 1H), 9.46 (s, 1H), 8.39 (s, 1H), 8.12 (d, *J* = 7.7 Hz, 1H), 7.81 – 7.70 (m, 3H), 7.58 – 7.53 (m, 1H), 7.31 – 7.21 (m, 2H), 3.13 (q, *J* = 7.5 Hz, 2H), 1.39 (t, *J* = 7.5 Hz, 3H).

^13^C NMR (126 MHz, DMSO-*d*_6_) *δ* (ppm) 158.14, 149.91, 146.53, 142.68, 138.71, 133.71, 132.58, 131.94, 129.69, 129.40 (q, *J* = 31.4 Hz), 125.13 (q, *J* = 3.9 Hz), 124.31 (q, *J* = 272.5 Hz), 122.96, 122.89 (q, *J* = 4.2 Hz), 121.86, 119.77, 118.33, 111.45, 106.10, 21.14, 12.85.

^19^F NMR (471 MHz, DMSO-*d*_6_) *δ* (ppm) -61.18.

HRMS (APCI): calcd. for C_21_H_17_F_3_N_7_ [M+H]^+^ = 424.1492, found [M+H]^+^ = 424.1491.

8-(Benzo[*d*][1,3]dioxol-5-yl)-3-(1*H*-benzo[*d*]imidazol-2-yl)-7-ethylpyrazolo[5,1-*c*][1,2,4]triazin-4-amine **(74)**

The compound was prepared according to General procedure C using:

Diazotization step:

4-(benzo[*d*][1,3]dioxol-5-yl)-3-ethyl-1*H*-pyrazol-5-amine (45 mg, 0.19 mmol; 1 eq.), 35% aqueous HCl (0.065 mL, 0.76 mmol; 4 eq.) in EtOH (3 mL) and H_2_O (3 mL), NaNO_2_ (26 mg, 0.38 mmol, 2 eq.) in H_2_O (1 mL) (reaction time: 30 min), 2-(cyanomethyl)-benzimidazole (36 mg, 0.23 mmol, 1.2 eq.) in EtOH (1 mL) and KOAc (112 mg, 1.14 mmol, 6 eq.) in H_2_O (1 mL). Reaction time: 16 h. The yellow solid (71 mg, 0.17 mmol) obtained after the filtration was used in the next step without further purification.

Cyclization step:

the dried yellow solid (71 mg, 0.17 mmol), DMF (3 mL), and KOAc (5 mg, 0.06 mmol, 0.4 eq.). Reaction time: 2 h. The reaction mixture was poured into water (20 mL) and extracted with EtOAc (3 × 20 mL). The combined organic layers were washed with brine (30 mL), dried over MgSO_4_, the solvent was evaporated, and the residue was purified by column flash chromatography on silica gel (hexane:EtOAc, gradient10:1 to 0:1). The product was obtained as a yellow solid (35 mg, 46% (2 steps)).

^1^H NMR (500 MHz, DMSO-*d*_6_) *δ* (ppm) 13.35 (s, 1H), 9.93 (s, 1H), 9.30 (s, 1H), 7.74 (d, *J* = 7.7 Hz, 1H), 7.58 – 7.53 (m, 1H), 7.40 (d, *J* = 1.7 Hz, 1H), 7.31 – 7.20 (m, 3H), 7.07 (d, *J* = 8.0 Hz, 1H), 6.10 (s, 2H), 3.04 (q, *J* = 7.5 Hz, 2H), 1.35 (t, *J* = 7.5 Hz, 3H).

^13^C NMR (126 MHz, DMSO-*d*_6_) *δ* (ppm) 157.90, 150.08, 147.43, 146.18, 146.10, 142.68, 138.61, 133.70, 124.89, 122.82, 122.46, 121.80, 118.88, 118.23, 111.44, 109.39, 108.48, 108.14, 101.01, 20.87, 13.07.

HRMS (APCI): calcd. for C_21_H_18_N_7_O_2_ [M+H]^+^ = 400.1516, found [M+H]^+^ = 400.1516.

Methyl 3-(4-amino-3-(1*H*-benzo[*d*]imidazol-2-yl)-7-ethylpyrazolo[5,1-*c*][1,2,4]triazin-8-yl)benzoate **(75)**

The compound was prepared according to General procedure C using:

Diazotization step:

methyl 3-(5-amino-3-ethyl-1*H*-pyrazol-4-yl)benzoate (85 mg, 0.35 mmol; 1 eq.), 35% aqueous HCl (0.12 mL, 1.40 mmol; 4 eq.) in EtOH (3 mL) and H_2_O (3 mL), NaNO_2_ (48 mg, 0.70 mmol, 2 eq.) in H_2_O (1 mL) (reaction time: 30 min), 2-(cyanomethyl)-benzimidazole (64 mg, 0.42 mmol, 1.2 eq.) in EtOH (1 mL) and KOAc (206 mg, 2.10 mmol, 6 eq.) in H_2_O (1 mL). Reaction time: 16 h. The yellow solid (150 mg, 0.35 mmol) obtained after the filtration was used in the next step without further purification.

Cyclization step:

the dried yellow solid (150 mg, 0.35 mmol), DMF (3 mL), and KOAc (4 mg, 0.04 mmol, 0.1 eq.). Reaction time: 2 h. Reaction time: 2 h. The product was obtained as a yellow solid (90 mg, 62%).

^1^H NMR (500 MHz, DMSO-*d*_6_) *δ* (ppm) ^1^H NMR (500 MHz, DMSO) δ 13.42 (s, 1H), 9.97 (s, 1H), 9.41 (s, 1H), 8.59 (s, 1H), 8.10 (d, *J* = 7.9 Hz, 1H), 7.98 – 7.92 (m, 1H), 7.75 (d, *J* = 7.7 Hz, 1H), 7.68 (t, *J* = 7.9 Hz, 1H), 7.56 (d, *J* = 8.1 Hz, 1H), 7.31 – 7.20 (m, 2H), 3.91 (s, 3H), 3.09 (q, *J* = 7.6 Hz, 2H), 1.38 (t, *J* = 7.6 Hz, 3H).

^13^C NMR (126 MHz, DMSO-*d*_6_) *δ* (ppm) 166.22, 158.08, 149.97, 146.44, 142.68, 138.69, 133.70, 132.97, 131.96, 129.99, 129.47, 129.08, 127.11, 122.92, 121.84, 119.55, 118.30, 111.44, 106.74, 52.21, 21.06, 13.02.

HRMS (APCI): calcd. for C_22_H_20_N_7_O_2_ [M+H]^+^= 414.1672, found [M+H]^+^ = 414.1674.

3-(1*H*-benzo[*d*]imidazol-2-yl)-7-ethyl-8-(4-(piperidin-1-ylsulfonyl)phenyl)pyrazolo[5,1-*c*][1,2,4]triazin-4-amine **(76)**

The compound was prepared according to General procedure C using:

Diazotization step:

3-ethyl-4-(4-(piperidin-1-ylsulfonyl)phenyl)-1*H*-pyrazol-5-amine (57 mg, 0.2 mmol; 1 eq.), 35% aqueous HCl (75 µL, 0.80 mmol; 4 eq.) in EtOH (3 mL) and H_2_O (3 mL), NaNO_2_ (30 mg, 0.4 mmol, 2 eq.) in H_2_O (1 mL) (reaction time: 30 min), 2-(cyanomethyl)-benzimidazole (37 mg, 0.22 mmol, 1.2 eq.) in EtOH (1 mL) and KOAc (125 mg, 1.20 mmol, 6 eq.) in H_2_O (1 mL). Reaction time: 16 h. The yellow solid (88 mg, 0.17 mmol) obtained after the filtration was used in the next step without further purification.

Cyclization step:

the dried yellow solid (88 mg, 0.17 mmol), DMF (3 mL), and KOAc (5 mg, 0.06 mmol, 0.4 eq.). Reaction time: 2 h. The residue obtained after the workup was purified by column chromatography on silica gel (dichloromethane:MeOH, 20:1). The product was obtained as a yellow solid (30 mg, 30%).

^1^H NMR (500 MHz, DMSO-*d*_6_) *δ* (ppm) 13.40 (s, 1H), 10.03 (s, 1H), 9.47 (s, 1H), 8.21 – 8.15 (m, 2H), 7.90 – 7.84 (m, 2H), 7.76 (s, 1H), 7.58 (s, 1H), 7.29 – 7.24 (m, 2H), 3.14 (q, *J* = 7.5 Hz, 2H), 3.01 – 2.95 (m, 4H), 1.62 – 1.54 (m, 4H), 1.45 – 1.36 (m, 5H).

^13^C NMR (126 MHz, DMSO-*d*_6_) *δ* (ppm) 158.42, 149.80, 146.57, 142.71, 138.74, 136.26, 133.07, 128.86, 127.73, 120.03, 118.35, 111.63, 106.06, 46.57, 24.71, 22.80, 21.26, 12.91.

HRMS (APCI): calcd. for C_25_H_27_N_8_O_2_S [M+H]^+^ = 503.1972, found [M+H]^+^ = 503.1975

3-(1*H*-benzo[*d*]imidazol-2-yl)-7-ethyl-8-(3-(trifluoromethoxy)phenyl)pyrazolo[5,1-*c*][1,2,4]triazin-4-amine **(77)**

The compound was prepared according to General procedure C using:

Diazotization step:

3-ethyl-4-(3-(trifluoromethoxy)phenyl)-1*H*-pyrazol-5-amine (55 mg, 0.20 mmol; 1 eq.), 35% aqueous HCl (75 µL, 0.80 mmol; 4 eq.) in EtOH (3 mL) and H_2_O (3 mL), NaNO_2_ (30 mg, 0.4 mmol, 2 eq.) in H_2_O (1 mL) (reaction time: 30 min), 2-(cyanomethyl)-benzimidazole (37 mg, 0.22 mmol, 1.2 eq.) in EtOH (1 mL) and KOAc (125 mg, 1.20 mmol, 6 eq.) in H_2_O (1 mL). Reaction time: 16 h. The yellow solid (46 mg, 0.10 mmol) obtained after the filtration was used in the next step without further purification.

Cyclization step:

the dried yellow solid (46 mg, 0.10 mmol), DMF (3 mL), and KOAc (5 mg, 0.06 mmol, 0.6 eq.). Reaction time: 2 h. The product was obtained as a yellow solid (18 mg, 20%).

^1^H NMR (500 MHz, DMSO-*d*_6_) *δ* (ppm) 13.47 (s, 1H), 10.01 (s, 1H), 9.45 (s, 1H), 7.98 (s, 1H), 7.88 (d, *J* = 7.8 Hz, 1H), 7.79 – 7.74 (m, 1H), 7.68 (t, *J* = 8.0 Hz, 1H), 7.58 – 7.54 (m, 1H), 7.37 (d, *J* = 8.3 Hz, 1H), 7.29 – 7.25 (m, 2H), 3.12 (q, *J* = 7.6 Hz, 2H), 1.39 (t, *J* = 7.5 Hz, 3H).

^13^C NMR (126 MHz, DMSO-*d*_6_) *δ* (ppm) 158.13, 149.89, 148.61, 146.44, 138.71, 133.72, 130.49, 127.25, 122.96, 121.89, 120.84, 120.16 (q, *J* = 256.3 Hz), 119.73, 118.88, 118.32, 111.49, 106.11, 21.10, 12.84.

^19^F NMR (471 MHz, DMSO-*d*_6_) *δ* (ppm) -56.57.

HRMS (APCI): calcd. for C_21_H_17_F_3_N_7_O [M+H]^+^ = 440.1441, found [M+H]^+^ = 440.1441.

3-(1*H*-benzo[*d*]imidazol-2-yl)-7-ethyl-8-(naphthalen-1-yl)pyrazolo[5,1-*c*][1,2,4]triazin-4-amine **(78)**

The compound was prepared according to General procedure C using:

Diazotization step:

3-ethyl-4-(naphthalen-1-yl)-1*H*-pyrazol-5-amine (48 mg, 0.200 mmol; 1 eq.), 35% aqueous HCl (75 µL, 0.8 mmol; 4 eq.) in EtOH (3 mL) and H_2_O (3 mL), NaNO_2_ (30 mg, 0.40 mmol, 2 eq.) in H_2_O (1 mL) (reaction time: 30 min), 2-(cyanomethyl)-benzimidazole (37 mg, 0.22 mmol, 1.2 eq.) in EtOH (1 mL) and KOAc (125 mg, 1.20 mmol, 6 eq.) in H_2_O (1 mL). Reaction time: 16 h. The yellow solid (76 mg, 0.18 mmol) obtained after the filtration was used in the next step without further purification.

Cyclization step:

the dried yellow solid (76 mg, 0.18 mmol), DMF (3 mL), and KOAc (5 mg, 0.08 mmol, 0.3 eq.). Reaction time: 2 h. The product was obtained as a yellow solid (34 mg, 42%).

^1^H NMR (500 MHz, DMSO-*d*_6_) *δ* (ppm) 13.31 (s, 1H), 10.02 (s, 1H), 9.43 (s, 1H), 8.06 (dd, *J* = 8.1, 4.1 Hz, 2H), 7.82 – 7.74 (m, 1H), 7.72 – 7.63 (m, 3H), 7.61 – 7.52 (m, 2H), 7.50 – 7.43 (m, 1H), 7.31 – 7.21 (m, 2H), 2.88 – 2.70 (m, 2H), 1.18 (t, *J* = 7.5 Hz, 3H).

^13^C NMR (126 MHz, DMSO-*d*_6_) *δ* (ppm) 159.42, 150.13, 147.12, 142.68, 138.86, 133.72, 133.41, 132.32, 129.08, 128.67, 128.17, 128.11, 126.13, 125.98, 125.57, 122.81, 121.80, 118.87, 118.24, 111.46, 107.12, 20.58, 13.29.

HRMS (APCI): calcd. for C_24_H_20_N_7_ [M+H]^+^ = 406.1775, found [M+H]^+^ = 406.1777.

3-(1*H*-benzo[*d*]imidazol-2-yl)-7-(naphthalen-2-yl)pyrazolo[5,1-*c*][1,2,4]triazin-4-ol **(79)**

To 3-(1*H*-benzo[*d*]imidazol-2-yl)-7-(naphthalen-2-yl)pyrazolo[5,1-*c*][1,2,4]triazin-4-amine (73 mg, 0.19 mmol; 1 eq.) was added 23 % aqueous HCl (5 mL) and the mixture was refluxed for 1 h. The reaction mixture was poured into water (10 mL) and neutralized by addition of saturated aqueous solution of NaHCO_3_ (until neutral pH, 10 mL). The precipitate was collected by filtration, washed with water (6 mL), then with Et_2_O (5 mL) and dried under vacuum. The product was obtained as an orange solid (44 mg, 60%).

^1^H NMR (500 MHz, DMSO-*d*_6_) δ 8.69 (s, 1H), 8.25 (dd, *J* = 8.5, 1.7 Hz, 1H), 8.11 – 8.04 (m, 2H), 8.02 – 7.94 (m, 1H), 7.86 – 7.80 (m, 2H), 7.62 – 7.55 (m, 2H), 7.55 – 7.50 (m, 2H), 7.42 (s, 1H).

^13^C NMR (126 MHz, DMSO-*d*_6_) *δ* (ppm) 155.11, 148.61, 145.91, 133.37, 132.98, 131.19, 129.13, 128.45, 128.37, 127.68, 126.82, 126.66, 125.89, 125.51, 124.06, 120.55, 113.87, 91.92.

HRMS (APCI): calcd. for C_22_H_15_N_6_O [M+H]^+^ = 379.1302, found [M+H]^+^ = 379.1299.

3-(1*H*-benzo[*d*]imidazol-2-yl)-7-(3-methoxynaphthalen-2-yl)pyrazolo[5,1-*c*][1,2,4]triazin-4-ol **(80)**

The compound was prepared according to General procedure C using:

Diazotization step:

3-(3-methoxynaphthalen-2-yl)-1*H*-pyrazol-5-amine (240 mg, 1.0 mmol; 1 eq.), 35% aqueous HCl (0.4 mL, 4.0 mmol; 4 eq.) in EtOH (3 mL) and H_2_O (3 mL), NaNO_2_ (136 mg, 2.0 mmol, 2 eq.) in H_2_O (1 mL) (reaction time: 30 min), 2-(cyanomethyl)-benzimidazole (172 mg, 1.10 mmol, 1.2 eq.) in EtOH (1 mL) and KOAc (590 mg, 6.0 mmol, 6 eq.) in H_2_O (1 mL). Reaction time: 16 h. The yellow solid (340 mg, 0.80 mmol) obtained after the filtration was used in the next step without further purification.

Cyclization step:

the dried yellow solid (340 mg, 0.80 mmol), DMF (3 mL), and KOAc (8 mg, 0.08 mmol, 0.1 eq.). Reaction time: 2 h. The product was obtained as a yellow solid (224 mg, 55%).

^1^H NMR (500 MHz, DMSO-*d*_6_) *δ* (ppm) 14.07 (s, 2H), 8.64 (s, 1H), 8.02 (d, *J* = 8.1 Hz, 1H), 7.89 (d, *J* = 8.2 Hz, 1H), 7.80 – 7.72 (m, 2H), 7.54 – 7.50 (m, 2H), 7.47 – 7.44 (m, 2H), 7.43 – 7.38 (m, 1H), 7.36 (s, 1H), 4.06 (s, 3H).

^13^C NMR (126 MHz, DMSO-*d*_6_) *δ* (ppm) 155.24, 151.27, 149.07, 148.50, 134.31, 131.27, 128.35, 128.09, 127.98, 126.91, 126.36, 124.64, 124.05, 122.61, 113.40, 106.43, 98.91, 55.65.

HRMS (APCI): calcd. for C_23_H_18_N_7_O [M+H]^+^= 408.1567, found [M+H]^+^ = 408.1566.

3-(1*H*-benzo[*d*]imidazol-2-yl)-7-ethylpyrazolo[5,1-*c*][1,2,4]triazin-4-ol **(81)**

The compound was prepared according to General procedure C using:

Diazotization step:

3-ethyl-1*H*-pyrazol-5-amine (56 mg, 0.50 mmol; 1 eq.), 35% aqueous HCl (0.2 mL, 2.0 mmol; 4 eq.) in EtOH (2 mL) and H_2_O (2 mL), NaNO_2_ (68 mg, 1 mmol, 2 eq.) in H_2_O (0.5 mL) (reaction time: 30 min), ethyl 2-(1*H*-benzo[*d*]imidazol-2-yl)acetate (100 mg, 0.50 mmol, 1 eq.) in EtOH (0.5 mL) and KOAc (300 mg, 3 mmol, 6 eq.) in H_2_O (0.5 mL). Reaction time: 16 h. The yellow solid (120 mg, 0.40 mmol) obtained after the filtration was used in the next step without further purification.

Cyclization step:

the dried yellow solid (119 mg, 0.40 mmol), DMF (3 mL), and KOAc (4 mg, 0.04 mmol, 0.1 eq.). Reaction time: 2 h. The product was obtained as a yellow solid (91 mg, 65%).

^1^H NMR (300 MHz, DMSO-*d*_6_) *δ* (ppm) 14.00 (s, 2H), 7.73 (dd, *J* = 6.1, 3.2 Hz, 2H), 7.43 (dd, *J* = 6.1, 3.2 Hz, 2H), 6.66 (s, 1H), 2.78 (q, *J* = 7.6 Hz, 2H), 1.30 (t, *J* = 7.6 Hz, 3H).

HRMS (APCI): calcd. for C_14_H_13_N_6_O [M+H]^+^= 281.1175, found [M+H]^+^ = 281.1178.

3-(1*H*-benzo[*d*]imidazol-2-yl)-7-(2-methoxyphenyl)pyrazolo[5,1-*c*][1,2,4]triazin-4-ol **(82)**

The compound was prepared according to General procedure C using:

Diazotization step:

3-(2-methoxyphenyl)-1*H*-pyrazol-5-amine (95 mg, 0.50 mmol; 1 eq.), 35% aqueous HCl (0.2 mL, 2.0 mmol; 4 eq.) in EtOH (2 mL) and H_2_O (2 mL), NaNO_2_ (68 mg, 1 mmol, 2 eq.) in H_2_O (0.50 mL) (reaction time: 30 min), ethyl 2-(1*H*-benzo[*d*]imidazol-2-yl)acetate (100 mg, 0.50 mmol, 1 eq.) in EtOH (0.5 mL) and KOAc (300 mg, 3 mmol, 6 eq.) in H_2_O (0.5 mL). Reaction time: 16 h. The yellow solid (170 mg, 0.45 mmol) obtained after the filtration was used in the next step without further purification.

Cyclization step:

the dried yellow solid (170 mg, 0.45 mmol), DMF (3 mL), and KOAc (4 mg, 0.04 mmol, 0.1 eq.). Reaction time: 2 h. The product was obtained as a yellow solid (125 mg, 70%).

^1^H NMR (500 MHz, DMSO-*d*_6_) *δ* (ppm14.07 (s, 2H), 8.13 (dd, *J* = 7.7, 1.8 Hz, 1H), 7.79 – 7.72 (m, 2H), 7.48 – 7.40 (m, 3H), 7.28 (s, 1H), 7.19 (dd, *J* = 8.5, 1.1 Hz, 1H), 7.10 (td, *J* = 7.4, 1.1 Hz, 1H), 3.96 (s, 3H).

^13^C NMR (126 MHz, DMSO-*d*_6_) *δ* (ppm) 157.22, 152.76, 151.36, 149.05, 148.50, 131.17, 130.30, 128.59, 124.66, 120.84, 120.54, 118.53, 113.37, 112.05, 98.45, 55.56.

HRMS (APCI): calcd. for C_19_H_15_N_6_O_2_ [M+H]^+^ = 359.1251, found [M+H]^+^ = 359.1252.

3-(1*H*-benzo[*d*]imidazol-2-yl)-7-(2-phenoxyphenyl)pyrazolo[5,1-*c*][1,2,4]triazin-4-ol **(83)**

The compound was prepared according to General procedure C using:

Diazotization step:

3-(2-phenoxyphenyl)-1*H*-pyrazol-5-amine (125 mg, 0.5 mmol; 1 eq.), 35% aqueous HCl (0.2 mL, 2.0 mmol; 4 eq.) in EtOH (2 mL) and H_2_O (2 mL), NaNO_2_ (68 mg, 1 mmol, 2 eq.) in H_2_O (0.5 mL) (reaction time: 30 min), ethyl 2-(1*H*-benzo[*d*]imidazol-2-yl)acetate (100 mg, 0.5 mmol, 1 eq.) in EtOH (0.5 mL) and KOAc (300 mg, 3 mmol, 6 eq.) in H_2_O (0.5 mL). Reaction time: 16 h. The yellow solid (175 mg, 0.4 mmol) obtained after the filtration was used in the next step without further purification.

Cyclization step:

the dried yellow solid (170 mg, 0.45 mmol), DMF (3 mL), and KOAc (4 mg, 0.04 mmol, 0.1 eq.). Reaction time: 2 h. The product was obtained as a yellow solid (137 mg, 65%).

^1^H NMR (500 MHz, DMSO-*d*_6_) *δ* (ppm) 14.08 (s, 2H), 8.27 (dd, *J* = 7.8, 1.7 Hz, 1H), 7.80 – 7.71 (m, 2H), 7.55 – 7.47 (m, 1H), 7.46 – 7.43 (m, 2H), 7.41 – 7.33 (m, 3H), 7.14 – 7.07 (m, 3H), 7.06 – 6.98 (m, 2H).

^13^C NMR (126 MHz, DMSO-*d*_6_) *δ* (ppm) 157.04, 153.40, 152.98, 150.45, 148.93, 148.35, 131.17, 130.64, 130.05, 129.41, 124.75, 124.68, 124.61, 122.92, 120.88, 118.79, 117.25, 113.39, 97.70.

HRMS (APCI): calcd. for C_24_H_17_N_6_O_2_ [M+H]^+^ = 421.1408, found [M+H]^+^ = 421.1410.

3-(1*H*-benzo[*d*]imidazol-2-yl)-7-(2-(dimethylamino)phenyl)pyrazolo[5,1-*c*][1,2,4]triazin-4-ol **(84)**

The compound was prepared according to General procedure C using:

Diazotization step:

3-(2-(dimethylamino)phenyl)-1*H*-pyrazol-5-amine (101 mg, 0.50 mmol; 1 eq.), 35% aqueous HCl (0.2 mL, 2.0 mmol; 4 eq.) in EtOH (2 mL) and H_2_O (2 mL), NaNO_2_ (68 mg, 1 mmol, 2 eq.) in H_2_O (0.5 mL) (reaction time: 30 min), ethyl 2-(1*H*-benzo[*d*]imidazol-2-yl)acetate (100 mg, 0.50 mmol, 1 eq.) in EtOH (0.5 mL) and KOAc (300 mg, 3 mmol, 6 eq.) in H_2_O (0.5 mL). Reaction time: 16 h. The yellow solid (175 mg, 0.45 mmol) obtained after the filtration was used in the next step without further purification.

Cyclization step:

the dried yellow solid (175 mg, 0.45 mmol), DMF (3 mL), and KOAc (4 mg, 0.04 mmol, 0.1 eq.). Reaction time: 2 h. The product was obtained as a yellow solid (102 mg, 55%).

^1^H NMR (500 MHz, DMSO-*d*_6_) *δ* (ppm) 14.06 (s, 2H), 7.86 – 7.80 (m, 1H), 7.78 – 7.73 (m, 2H), 7.48 – 7.40 (m, 2H), 7.39 – 7.34 (m, 1H), 7.28 (s, 1H), 7.18 (d, *J* = 8.1 Hz, 1H), 7.09 (t, *J* = 7.4 Hz, 1H), 2.65 (s, 6H).

^13^C NMR (126 MHz, DMSO-*d*_6_) *δ* (ppm) 154.40, 152.87, 152.24, 149.10, 148.51, 131.20, 130.34, 129.54, 125.58, 124.63, 121.67, 118.41, 113.36, 97.07, 43.93.

HRMS (APCI): calcd. for C_14_H_12_N_7_ [M+H]^+^= 372.1567, found [M+H]^+^ = 372.1569.

3-(1*H*-benzo[*d*]imidazol-2-yl)-7-(2-methoxybenzyl)pyrazolo[5,1-*c*][1,2,4]triazin-4-ol **(85)**

The compound was prepared according to General procedure C using:

Diazotization step:

3-(2-methoxybenzyl)-1*H*-pyrazol-5-amine (300 mg, 1.48 mmol; 1 eq.), 35% aqueous HCl (510 µL, 5.9 mmol; 4 eq.) in EtOH (5 mL) and H_2_O (5 mL), NaNO_2_ (203 mg, 2.98 mmol, 2 eq.) in EtOH (1 mL) (reaction time: 15 min), ethyl 2-(1*H*-1,3-benzodiazol-2-yl)acetate (362 mg, 1.77 mmol, 1.2 eq.) in EtOH (2 mL) and KOAc (870 mg, 8.86 mmol, 6 eq.). Reaction time: 16 h. The yellow solid (526 mg, 1.35 mmol) obtained after the filtration was used in the next step without further purification.

Cyclization step:

the dried yellow solid (526 mg, 1.35 mmol), DMF (5 mL), and KOAc (10 mg, 0.10 mmol, 0.1 eq.). Reaction time: 2 h. After the filtration, the solid part was suspended in hot dioxane (10 mL) and the mixture was poured into water (30 mL). The solid was collected by filtration, washed with water (10 mL), diethyl ether (10 mL) and dried under *vacuum*. The product was obtained as a yellow solid (320 mg, 58%).

^1^H NMR (500 MHz, DMSO-*d*_6_) *δ* (ppm) 14.01 (s, 2H), 7.77 – 7.70 (m, 2H), 7.47 – 7.40 (m, 2H), 7.27 – 7.21 (m, 1H), 7.18 (dd, *J* = 7.5, 1.8 Hz, 1H), 7.02 (dd, *J* = 8.2, 1.1 Hz, 1H), 6.89 (d, *J* = 1.1 Hz, 1H), 6.50 (s, 1H), 4.08 (s, 2H), 3.84 (s, 3H).

^13^C NMR (126 MHz, DMSO-*d*_6_) *δ* (ppm) 157.36, 156.88, 153.22, 149.59, 149.01, 131.75, 130.53, 128.31, 127.62, 125.11, 120.80, 113.89, 111.31, 97.07, 55.86, 29.45.

HRMS (APCI): calcd. for C_20_H_17_N_6_O_2_ [M+H]^+^ = 373.1408., found [M+H]^+^ = 373.1411.

3-(1*H*-imidazol-2-yl)-7-(2-methoxyphenyl)pyrazolo[5,1-*c*][1,2,4]triazin-4-ol **(86)**

The compound was prepared according to General procedure C using:

Diazotization step:

3-(2-Methoxyphenyl)-1*H*-pyrazol-5-amine (230 mg, 1.22 mmol; 1 eq.), 35% aqueous HCl (420 µL, 4.86 mmol; 4 eq.) in EtOH (5 mL) and H_2_O (5 mL), NaNO_2_ (168 mg, 2.43 mmol, 2 eq.) in EtOH (1 mL) (reaction time: 15 min), ethyl 2-(1*H*-imidazol-2-yl)acetate (224 mg, 1.46 mmol, 1.2 eq.) in EtOH (2 mL) and KOAc (720 mg, 7.29 mmol, 6 eq.). Reaction time: 16 h. The yellow solid (397 mg, 1.22 mmol) obtained after the filtration was used in the next step without further purification.

Cyclization step:

the dried yellow solid (397 mg, 1.22 mmol), DMF (5 mL), and KOAc (10 mg, 0.10 mmol, 0.1 eq.). Reaction time: 2 h. After the filtration, the solid part was suspended in hot dioxane (10 mL), and the mixture was poured into water (30 mL). The solid was collected by filtration, washed with water (10 mL), diethyl ether (10 mL) and dried under *vacuum*. The product was obtained as a yellow solid (300 mg, 80%).

^1^H NMR (500 MHz, DMSO-*d*_6_) *δ* (ppm) 8.14 – 8.09 (m, 2H), 7.46 (s, 1H), 7.44 – 7.37 (m, 2H), 7.20 – 7.15 (m, 2H), 7.08 (t, *J* = 7.5 Hz, 1H), 3.94 (s, 3H).

^13^C NMR (126 MHz, DMSO-*d*_6_) *δ* (ppm) 157.18, 152.94, 151.02, 148.39, 143.35, 130.09, 128.55, 121.09, 120.49, 119.55, 118.11, 112.01, 97.04, 55.53.

HRMS (APCI): calcd. for C_15_H_13_N_6_O_2_ [M+H]^+^ = 309.1095., found [M+H]^+^ = 309.1094.

3-(1*H*-benzo[*d*]imidazol-2-yl)-7-(2-methoxy-2-phenylethyl)pyrazolo[5,1-*c*][1,2,4]triazin-4-ol **(87)**

The compound was prepared according to General procedure C using:

Diazotization step:

3-(2-methoxy-2-phenylethyl)-1*H*-pyrazol-5-amine (150 mg, 0.69 mmol; 1 eq.), 35% aqueous HCl (240 µL, 2.76 mmol; 4 eq.) in EtOH (5 mL) and H_2_O (5 mL), NaNO_2_ (95 mg, 1.38 mmol, 2 eq.) in EtOH (1 mL) (reaction time: 15 min), ethyl 2-(1*H*-1,3-benzodiazol-2-yl)acetate (169 mg, 0.83 mmol, 1.2 eq.) in EtOH (2 mL) and KOAc (0.41 g, 4.14 mmol, 6 eq.). Reaction time: 16 h. The yellow solid (222 mg, 0.55 mmol) obtained after the filtration was used in the next step without further purification.

Cyclization step:

the dried yellow solid (222 mg, 0.55 mmol), DMF (5 mL), and KOAc (5 mg, 0.05 mmol, 0.1 eq.). Reaction time: 2 h. After the filtration, the solid part was suspended in hot dioxane (10 mL), and the mixture was poured into water (30 mL). The solid was collected by filtration, washed with water (10 mL), diethyl ether (10 mL) and dried under *vacuum*. The product was obtained as a yellow solid (150 mg, 56%).

^1^H NMR (500 MHz, DMSO-*d*_6_) *δ* (ppm14.01 (s, 2H), 7.73 (dd, *J* = 6.0, 3.2 Hz, 2H), 7.45 – 7.40 (m, 2H), 7.40 – 7.33 (m, 4H), 7.34 – 7.26 (m, 1H), 6.57 (s, 1H), 4.63 (dd, *J* = 8.0, 5.7 Hz, 1H), 3.23 (dd, *J* = 14.5, 8.1 Hz, 1H), 3.14 (s, 3H), 3.07 (dd, *J* = 14.5, 5.7 Hz, 1H).

^13^C NMR (126 MHz, DMSO-*d*_6_) *δ* (ppm) 154.48, 149.01, 148.48, 141.34, 131.14, 128.30, 127.63, 126.73, 124.59, 113.33, 96.86, 82.17, 56.03, 37.09.

HRMS (APCI): calcd. for C_21_H_19_N_6_O_2_ [M-H]^-^ = 387.1564, found [M-H]^-^ = 387.1563.

3-(1*H*-benzo[*d*]imidazol-2-yl)-7-benzylpyrazolo[5,1-*c*][1,2,4]triazin-4-ol **(88)**

The compound was prepared according to General procedure C using:

Diazotization step:

3-benzyl-1*H*-pyrazol-5-amine (165 mg, 0.95 mmol; 1 eq.), 35% aqueous HCl (0.336 mL, 3.81 mmol; 4 eq.) in EtOH (2 mL) and H_2_O (2 mL), NaNO_2_ (0.131 g, 1.91 mmol, 2 eq.) in EtOH (1 mL) and H_2_O (2 mL) (reaction time: 30 min), ethyl 2-(1*H*-benzo[*d*]imidazol-2-yl)acetate (214 mg, 1.05 mmol, 1.1 eq.) in EtOH (3 mL) and H_2_O (3 mL) and KOAc (748 mg, 7.62 mmol, 8 eq.). Reaction time: 2 h. The yellow solid (384 mg, quant.) obtained after the filtration was used in the next step without further purification.

Cyclization step:

the dried yellow solid (384 mg), DMF (4 mL), and KOAc (9 mg, 0.1 mmol, 0.1 eq.). Reaction time: 2 h. The product was obtained as a yellow solid (272 mg, 83%).

^1^H NMR (500 MHz, DMSO-*d*_6_) *δ* (ppm) 14.02 (s, 2H), 7.78 – 7.70 (m, 2H), 7.47 – 7.40 (m, 2H), 7.36 – 7.28 (m, 4H), 7.25 – 7.18 (m, 1H), 6.61 (s, 1H), 4.13 (s, 2H).

^13^C NMR (126 MHz, DMSO-*d*_6_) *δ* (ppm) 156.59, 152.73, 149.08, 148.44, 139.34, 131.20, 128.68, 128.39, 126.18, 124.63, 118.61, 113.38, 96.47, 34.64.

HRMS (APCI): calcd. for C_19_H_15_N_6_O [M+H]^+^ = 343.1302, found [M+H]^+^ = 343.1304.

3-(1*H*-benzo[*d*]imidazol-2-yl)-7-(isochroman-3-ylmethyl)pyrazolo[5,1-*c*][1,2,4]triazin-4-ol **(89)**

The compound was prepared according to General procedure C using:

Diazotization step:

3-(isochroman-3-ylmethyl)-1*H*-pyrazol-5-amine (202 mg, 0.88 mmol; 1 eq.), 35% aqueous HCl (0.311 mL, 3.52 mmol; 4 eq.) in EtOH (2 mL) and H_2_O (2 mL), NaNO_2_ (122 mg, 1.76 mmol, 2 eq.) in EtOH (1 mL) and H_2_O (2 mL) (reaction time: 30 min), ethyl 2-(1*H*-benzo[*d*]imidazol-2-yl)acetate (198 mg, 0.97 mmol, 1.1 eq.) in EtOH (3 mL) and H_2_O (3 mL) and KOAc (692 mg, 7.05 mmol, 8 eq.). Reaction time: 2 h. The yellow solid (215 mg, quant.) obtained after the filtration was used in the next step without further purification.

Cyclization step:

the dried yellow solid (215 mg), DMF (5 mL), and KOAc (9 mg, 0.09 mmol, 0.1 eq.). Reaction time: 2 h. The product was obtained as a yellow solid (245 mg, 70% (2 steps)).

^1^H NMR (500 MHz, DMSO-*d*_6_) *δ* (ppm) 14.00 (s, 2H), 7.78 – 7.70 (m, 2H), 7.47 – 7.40 (m, 2H), 7.33 – 7.28 (m, 1H), 7.24 – 7.11 (m, 3H), 6.62 (s, 1H), 5.16 (dd, *J* = 8.8, 3.6 Hz, 1H), 4.09 (dt, *J* = 10.3, 4.8 Hz, 1H), 3.72 (ddd, *J* = 11.8, 8.7, 4.1 Hz, 1H), 3.41 (dd, *J* = 15.0, 3.7 Hz, 1H), 3.22 (dd, *J* = 15.0, 8.7 Hz, 1H), 2.93 – 2.84 (m, 1H), 2.75 – 2.65 (m, 1H).

^13^C NMR (126 MHz, DMSO-*d*_6_) *δ* (ppm) 155.00, 149.09, 148.54, 137.51, 133.80, 131.23, 128.80, 126.39, 126.02, 125.03, 124.62, 113.38, 97.03, 74.65, 62.10, 35.05, 28.35 ^19^F NMR (471 MHz, Methanol-*d*_4_) *δ* (ppm) -116.85.

HRMS (APCI): calcd. for C_22_H_19_N_6_O_2_ [M+H]^+^ = 399.1564, found [M+H]^+^ = 399.1562.

3-(1-Benzyl-1*H*-benzo[*d*]imidazol-2-yl)-8-(4-(methylsulfonyl)phenyl)pyrazolo[5,1-*c*][1,2,4]triazin-4-ol **(90)**

The compound was prepared according to General procedure D using 3-(1-benzyl-1*H*-benzo[*d*]imidazol-2-yl)-8-bromopyrazolo[5,1-*c*][1,2,4]triazin-4-ol (38 mg, 0.09 mmol, 1 eq.), (4-(methylsulfonyl)phenyl)boronic acid (22 mg, 0.11 mmol, 1.2 eq.), K_3_PO_4_ (77 mg, 0.36 mmol, 4 eq.), Pd(dppf)Cl_2_ (3 mg, 0.005 mmol, 0.05 eq.), dioxane (4 mL) and H_2_O (1 mL). Reaction time: 4 h at 120 °C. The residue obtained after the workup was purified using column chromatography on silica gel (EtOAc:MeOH, gradient 1:0 to 9:1). The product was obtained as a yellow solid (44 mg, 98%).

^1^H NMR (500 MHz, DMSO-*d*_6_) *δ* (ppm)) 14.23 (s, 1H), 8.76 (s, 1H), 8.50 (d, *J* = 8.2 Hz, 2H), 8.01 – 7.93 (m, 3H), 7.74 (d, *J* = 8.0 Hz, 1H), 7.50 (dt, *J* = 18.9, 7.3 Hz, 2H), 7.37 – 7.29 (m, 4H), 7.28 – 7.24 (m, 1H), 6.36 (s, 2H), 3.22 (s, 3H).

^13^C NMR (126 MHz, DMSO-*d*_6_) *δ* (ppm) 142.68, 128.69, 127.70, 127.45, 126.97, 126.12, 124.84,* 112.15, 49.57, 43.72.

*assigned by HSQC

HRMS (ESI): calcd. for C_26_H_21_N_6_O_3_S [M+H]^+^ = 497.1390, found [M-H]^-^= 497.1387.

3-(1*H*-benzo[*d*]imidazol-2-yl)-8-(4-(methylsulfonyl)phenyl)pyrazolo[5,1-*c*][1,2,4]triazin-4-ol **(91)**

To a solution of 3-(1-benzyl-1*H*-benzo[*d*]imidazol-2-yl)-8-(4-(methylsulfonyl)phenyl)pyrazolo[5,1-*c*][1,2,4]triazin-4-ol (25 mg, 0.05 mmol, 1 eq.) in MeOH (5 mL) was added 10% Pd/C (3 mg) and the mixture was bubbled with H_2_ for 5 min. The mixture was then stirred at reflux under an atmosphere of H_2_ for 16 h. The mixture was filtered through a Silia*MetS*^®^ Thiol (Si-Thiol, 500 mg) metal scavenger cartridge, and the solvent was removed *in vacuo*. The residue was triturated with dichloromethane (1 mL), the solid was collected by filtration and the trituration was repeated with EtOH (1 mL). The product was obtained as a yellow solid (10 mg, 50%).

^1^H NMR (500 MHz, DMSO-*d*_6_) *δ* (ppm) 14.21 (s, 1H), 8.76 (s, 1H), 8.59 – 8.53 (m, 2H), 7.99 (d, *J* = 8.6 Hz, 2H), 7.81 – 7.75 (m, 2H), 7.51 – 7.41 (m, 2H), 3.24 (s, 3H).

^13^C NMR (126 MHz, DMSO-*d*_6_) *δ* (ppm) 142.54, 127.46, 126.10, 124.82,* 113.57.*

*assigned by HSQC

HRMS (APCI): calcd. for C_19_H_15_N_6_O_3_S [M+H]^+^ = 407.0775, found [M+H]^+^ = 407.0772.

3-(1-Benzyl-1*H*-benzo[*d*]imidazol-2-yl)-8-(4-nitrophenyl)pyrazolo[5,1-*c*][1,2,4]triazin-4-ol **(92)**

The compound was prepared according to General procedure D using 3-(1-benzyl-1*H*-benzo[*d*]imidazol-2-yl)-8-bromopyrazolo[5,1-*c*][1,2,4]triazin-4-ol (26 mg, 0.06 mmol, 1 eq.), 4-nitrophenylboronic acid (13 mg, 0.007 mmol, 1.2 eq.), K_3_PO_4_ (52 mg, 0.25 mmol, 4 eq.), Pd(dppf)Cl_2_ (2 mg, 0.003 mmol, 0.05 eq.), dioxane (4 mL) and H_2_O (1 mL). Reaction time: 4 h at 120 °C. The residue obtained after the workup was purified using column chromatography on silica gel (EtOAc:MeOH, gradient 1:0 to 9:1). The product was obtained as an orange solid (18 mg, 63%).

^1^H NMR (500 MHz, DMSO-*d*_6_) *δ* (ppm) 8.69 (s, 1H), 8.53 (d, *J* = 8.5 Hz, 2H), 8.27 (d, *J* = 8.4 Hz, 2H), 7.79 (s, 1H), 7.51 (d, *J* = 5.9 Hz, 1H), 7.34 – 7.15 (m, 7H), 5.78 (s, 2H).

^13^C NMR (126 MHz, DMSO-*d*_6_) *δ* (ppm) 128.45, 127.33, 126.87, 125.34, 124.08, 111.03.

HRMS (ESI): calcd. for C_25_H_18_N_7_O_3_ [M+H]^+^ = 464.1466, found [M+H]^+^ = 464.1462.

3-(1-Benzyl-1*H*-benzo[*d*]imidazol-2-yl)-7-ethyl-8-(4-(trifluoromethyl)phenyl)pyrazolo[5,1-*c*][1,2,4]triazin-4-amine **(93)**

The compound was prepared according to General procedure C using:

Diazotization step:

3-ethyl-4-(4-(trifluoromethyl)phenyl)-1H-pyrazol-5-amine (40 mg, 0.16 mmol; 1 eq.), 35% aqueous HCl (0.06 mL, 0.64 mmol; 4 eq.) in EtOH (3 mL) and H_2_O (3 mL), NaNO_2_ (24 mg, 0.32 mmol, 2 eq.) in H_2_O (1 mL) (reaction time: 30 min), ethyl 2-(1-benzyl-1*H*-benzo[*d*]imidazol-2-yl)acetate (40 mg, 0.19 mmol, 1.2 eq.) in EtOH (1 mL) and KOAc (95 mg, 0.96 mmol, 6 eq.) in H_2_O (1 mL). Reaction time: 16 h. The yellow solid (75 mg, 0.14 mmol) obtained after the filtration was used in the next step without further purification.

Cyclization step:

the dried yellow solid (75 mg, 0.14 mmol), DMF (3 mL), and KOAc (5 mg, 0.05 mmol, 0.5 eq.). Reaction time: 2 h. The product was obtained as a yellow solid (32 mg, 40%).

^1^H NMR (500 MHz, DMSO-*d*_6_) *δ* (ppm) 14.15 (s, 1H), 8.04 (d, *J* = 8.1 Hz, 2H), 7.91 (d, *J* = 7.6 Hz, 1H), 7.81 (d, *J* = 8.1 Hz, 2H), 7.64 (s, 1H), 7.46 – 7.36 (m, 2H), 7.32 – 7.21 (m, 5H), 6.14 (s, 2H), 3.00 (q, *J* = 7.5 Hz, 2H), 1.30 (t, *J* = 7.4 Hz, 3H).

^19^F NMR (471 MHz, DMSO-*d*_6_) *δ* (ppm) -60.72.

HRMS (APCI): calcd. for C_28_H_22_F_3_N_6_O [M+H]^+^ = 515.1802, found [M+H]^+^ = 515.1801

3-(5,6-Dimethyl-1*H*-benzo[*d*]imidazol-2-yl)-7-ethyl-8-(4-fluorophenyl)pyrazolo[5,1-*c*][1,2,4]triazin-4-ol **(94)**

The compound was prepared according to General procedure C using:

Diazotization step:

3-ethyl-4-(4-fluorophenyl)-1*H*-pyrazol-5-amine (93 mg, 0.44 mmol; 1.1 eq.), 35% aqueous HCl (0.16 mL, 1.76 mmol; 4 eq.) in EtOH (3 mL) and H_2_O (3 mL), NaNO_2_ (60 mg, 0.88 mmol, 2 eq.) in H_2_O (1 mL) (reaction time: 30 min), methyl 2-(5,6-dimethyl-1*H*-benzo[*d*]imidazol-2-yl)acetate (90 mg, 0.4 mmol, 1 eq.) in EtOH (1 mL) and KOAc (240 mg, 2.40 mmol, 6 eq.) in H_2_O (1 mL). Reaction time: 16 h. The yellow solid (147 mg, 0.35 mmol) obtained after the filtration was used in the next step without further purification.

Cyclization step:

the dried yellow solid (147 mg, 0.35 mmol), DMF (3 mL), and KOAc (4 mg, 0.04 mmol, 0.1 eq.). Reaction time: 2 h. The product was obtained as a yellow solid (91 mg, 57%).

^1^H NMR (500 MHz, DMSO-*d*_6_) *δ* (ppm) 13.82 (s, 2H), 7.86 – 7.79 (m, 2H), 7.51 (s, 2H), 7.36 – 7.28 (m, 2H), 2.95 (q, *J* = 7.5 Hz, 2H), 2.37 (s, 6H), 1.29 (t, *J* = 7.5 Hz, 3H).

^13^C NMR (126 MHz, DMSO-*d*_6_) *δ* (ppm) 160.77 (d, *J* = 243.5 Hz), 155.51, 149.33, 148.77, 147.48, 133.51, 130.67 (d, *J* = 8.0 Hz), 129.84, 128.53 (d, *J* = 3.2 Hz), 119.68, 115.21 (d, *J* = 21.2 Hz), 113.30, 107.75, 20.82, 19.91, 13.09.

^19^F NMR (471 MHz, DMSO-*d*_6_) *δ* (ppm) -116.07.

HRMS (APCI): calcd. for C_22_H_20_FN_6_O [M+H]^+^ = 403.1677, found [M+H]^+^ = 403.1677

7-Ethyl-8-(4-fluorophenyl)-3-(1*H*-imidazol-2-yl)pyrazolo[5,1-*c*][1,2,4]triazin-4-ol **(95)**

The compound was prepared according to General procedure C using:

Diazotization step:

3-ethyl-4-(4-fluorophenyl)-1*H*-pyrazol-5-amine (117 mg, 0.50 mmol; 1 eq.), 35% aqueous HCl (0.2 mL, 2.0 mmol; 4 eq.) in EtOH (2 mL) and H_2_O (2 mL), NaNO_2_ (68 mg, 1 mmol, 2 eq.) in H_2_O (0.5 mL) (reaction time: 30 min), ethyl 2-(1*H*-imidazol-2-yl)acetate hydrobromide (100 mg, 0.50 mmol, 1 eq.) in EtOH (0.50 mL) and KOAc (300 mg, 3 mmol, 6 eq.) in H_2_O (0.5 mL). Reaction time: 16 h. The yellow solid (102 mg, 0.30 mmol) obtained after the filtration was used in the next step without further purification.

Cyclization step:

the dried yellow solid (102 mg, 0.30 mmol), DMF (3 mL), and KOAc (3 mg, 0.03 mmol, 0.1 eq.). Reaction time: 2 h. The product was obtained as a yellow solid (85 mg, 52%).

^1^H NMR (500 MHz, DMSO-*d*_6_) *δ* (ppm) 13.48 (s, 2H), 7.84 – 7.77 (m, 2H), 7.42 (s, 2H), 7.33 – 7.23 (m, 2H), 2.93 (q, *J* = 7.5 Hz, 2H), 1.29 (t, *J* = 7.5 Hz, 3H).

^13^C NMR (126 MHz, DMSO-*d*_6_) *δ* (ppm) 160.39 (d, *J* = 243.5 Hz), 155.01, 148.06, 143.24, 130.24 (d, *J* = 7.9 Hz), 128.51, 119.79, 117.61, 114.64 (d, *J* = 21.3 Hz), 106.41, 20.45, 12.57.

HRMS (APCI): calcd. for C_16_H_14_FN_6_O [M+H]^+^ = 325.1208, found [M+H]^+^ = 325.1212.

3-(1*H*-benzo[*d*]imidazol-2-yl)-7-ethyl-8-(4-methoxyphenyl)pyrazolo[5,1-*c*][1,2,4]triazin-4-ol **(96)**

The compound was prepared according to General procedure C using:

Diazotization step:

3-ethyl-4-(4-methoxyphenyl)-1*H*-pyrazol-5-amine (100 mg, 0.50 mmol; 1 eq.), 35% aqueous HCl (0.2 mL, 2.0 mmol; 4 eq.) in EtOH (2 mL) and H_2_O (2 mL), NaNO_2_ (68 mg, 1 mmol, 2 eq.) in H_2_O (0.5 mL) (reaction time: 30 min), ethyl 2-(1*H*-benzo[*d*]imidazol-2-yl)acetate (100 mg, 0.5 mmol, 1 eq.) in EtOH (0.5 mL) and KOAc (300 mg, 3 mmol, 6 eq.) in H_2_O (0.5 mL). Reaction time: 16 h. The yellow solid (182 mg, 0.45 mmol) obtained after the filtration was used in the next step without further purification.

Cyclization step:

the dried yellow solid (182 mg, 0.45 mmol), DMF (3 mL), and KOAc (4 mg, 0.04 mmol, 0.1 eq.). Reaction time: 2 h. The product was obtained as a yellow solid (80 mg, 41%).

^1^H NMR (500 MHz, DMSO-*d*_6_) *δ* (ppm) 13.99 (s, 1H), 7.78 – 7.72 (m, 2H), 7.72 – 7.68 (m, 2H), 7.47 – 7.40 (m, 2H), 7.10 – 7.03 (m, 2H), 3.82 (s, 3H), 2.93 (q, *J* = 7.5 Hz, 2H), 1.28 (t, *J* = 7.6 Hz, 3H).

^13^C NMR (126 MHz, DMSO-*d*_6_) *δ* (ppm) 157.90, 155.60, 149.03, 148.53, 131.23, 130.10, 124.56, 124.21, 118.81, 113.92, 113.31, 109.17, 55.08, 20.82, 13.18.

HRMS (APCI): calcd. for C_21_H_19_N_6_O_2_ [M+H]^+^ = 387.1564, found [M+H]^+^= 387.1565.

3-(1*H*-benzo[*d*]imidazol-2-yl)-8-(4-(dimethylamino)phenyl)-7-ethylpyrazolo[5,1-*c*][1,2,4]triazin-4-ol **(97)**

The compound was prepared according to General procedure C using:

Diazotization step:

4-(4-(dimethylamino)phenyl)-3-ethyl-1*H*-pyrazol-5-amine (115 mg, 0.50 mmol; 1 eq.), 35% aqueous HCl (0.2 mL, 2.0 mmol; 4 eq.) in EtOH (2 mL) and H_2_O (2 mL), NaNO_2_ (68 mg, 1 mmol, 2 eq.) in H_2_O (0.5 mL) (reaction time: 30 min), ethyl 2-(1*H*-benzo[*d*]imidazol-2-yl)acetate (100 mg, 0.50 mmol, 1 eq.) in EtOH (0.5 mL) and KOAc (300 mg, 3 mmol, 6 eq.) in H_2_O (0.5 mL). Reaction time: 16 h. The yellow solid (179 mg, 0.43 mmol) obtained after the filtration was used in the next step without further purification.

Cyclization step:

the dried yellow solid (179 mg, 0.43 mmol), DMF (3 mL), and KOAc (4 mg, 0.04 mmol, 0.1 eq.). Reaction time: 2 h. The product was obtained as a yellow solid (120 mg, 60%).

^1^H NMR (500 MHz, DMSO-*d*_6_) *δ* (ppm) 13.96 (s, 2H), 7.79 – 7.70 (m, 2H), 7.64 – 7.58 (m, 2H), 7.48 – 7.39 (m, 2H), 6.88 – 6.82 (m, 2H), 2.96 (s, 6H), 2.92 (q, *J* = 7.6 Hz, 2H), 1.28 (t, *J* = 7.5 Hz, 3H).

^13^C NMR (126 MHz, DMSO-*d*_6_) *δ* (ppm) 162.77, 156.05, 149.69, 149.66, 149.53, 149.15, 149.10, 131.68, 130.25, 130.13, 125.06, 120.05, 113.77, 112.84, 112.10, 40.61, 21.44, 13.78.

HRMS (APCI): calcd. for C_22_H_22_N_7_O [M+H]^+^ = 400.1880, found [M+H]^+^ = 400.1878.

3-(4-amino-3-(1*H*-benzo[*d*]imidazol-2-yl)-7-ethylpyrazolo[5,1-*c*][1,2,4]triazin-8-yl)benzoic acid **(98)**

Methyl 3-(4-amino-3-(1*H*-benzo[*d*]imidazol-2-yl)-7-ethylpyrazolo[5,1-*c*][1,2,4]triazin-8-yl)benzoate (45 mg, 0.11 mmol, 1 eq.) was dissolved in methanol (3 mL) and an aqueous solution of sodium hydroxide (3 M, 1.50 mL) was added. The mixture was stirred at 23 °C for 16 h, diluted with water (25 mL), neutralized with aqueous solution of potassium hydrogen sulfate (3 M, 1.50 mL). The mixture was extracted with EtOAc (3 × 25 mL), the combined organic extracts were washed with brine (50 mL), dried over MgSO_4_, filtered, and the solvent was evaporated. To the residue was added EtOAc (1 mL) and the solid was collected by filtration and dried under vacuum. The product was obtained as a yellow solid (10 mg, 23%).

1H NMR (500 MHz, DMSO-*d*_6_) *δ* (ppm) 13.41 (s, 1H), 9.96 (s, 1H), 9.40 (s, 1H), 8.59 – 8.55 (m, 1H), 8.08 (dt, *J* = 7.8, 1.5 Hz, 1H), 7.98 – 7.91 (m, 1H), 7.78 – 7.73 (m, 1H), 7.66 (t, *J* = 7.7 Hz, 1H), 7.62 – 7.53 (m, 1H), 7.31 – 7.21 (m, 2H), 3.10 (q, *J* = 7.5 Hz, 2H), 1.38 (t, *J* = 7.5 Hz, 3H).

^13^C NMR (126 MHz, DMSO-*d*_6_) *δ* (ppm) 167.32, 158.08, 150.01, 146.45, 142.70, 138.69, 133.70, 132.60, 131.71, 131.31, 129.71, 128.85, 127.30, 122.92, 121.85, 119.46, 118.31, 111.44, 107.02, 21.06, 13.07.

HRMS (APCI): calcd. for C_21_H_18_N_7_O_2_ [M+H]^+^= 400.1516, found [M+H]^+^ = 400.1519.

3-(1-Benzyl-1*H*-benzo[*d*]imidazol-2-yl)-7-ethyl-8-(4-methoxyphenyl)pyrazolo[5,1-*c*][1,2,4]triazin-4-ol **(99)**

The compound was prepared according to General procedure C using:

Diazotization step:

3-ethyl-4-(4-methoxyphenyl)-1*H*-pyrazol-5-amine (55 mg, 0.25 mmol; 1 eq.), 35% aqueous HCl (0.1 mL, 1.0 mmol; 4 eq.) in EtOH (3 mL) and H_2_O (3 mL), NaNO_2_ (35 mg, 0.5 mmol, 2 eq.) in H_2_O (1 mL) (reaction time: 30 min), methyl 2-(1-benzyl-1*H*-benzo[*d*]imidazol-2-yl)acetate (63 mg, 0.30 mmol, 1.2 eq.) in EtOH (1 mL) and KOAc (147 mg, 1.50 mmol, 6 eq.) in H_2_O (1 mL). Reaction time: 16 h. The yellow solid (120 mg, 0.24 mmol) obtained after the filtration was used in the next step without further purification.

Cyclization step:

the dried yellow solid (120 mg, 0.24 mmol), DMF (3 mL), and KOAc (2 mg, 0.02 mmol, 0.1 eq.). Reaction time: 2 h. The product was obtained as a yellow solid (90 mg, 76%).

1H NMR (500 MHz, DMSO-*d*6) *δ* (ppm) 14.09 (s, 1H), 7.91 – 7.88 (m, 1H), 7.64 – 7.61 (m, 3H), 7.43 – 7.39 (m, 2H), 7.30 – 7.23 (m, 5H), 7.04 (d, *J* = 8.3 Hz, 2H), 6.33 – 6.08 (m, 2H), 3.80 (s, 3H), 2.90 (d, *J* = 7.9 Hz, 2H), 1.26 (t, *J* = 7.5 Hz, 3H).

^13^C NMR (126 MHz, DMSO-*d*_6_) *δ* (ppm) 130.16, 128.88, 128.54, 127.50, 126.89, 125.18, 113.95, 111.66, 55.07, 20.71, 13.06.

HRMS (APCI): calcd. for C_28_H_25_N_6_O_2_ [M+H]^+^ = 477.2034, found [M+H]^+^ = 477.2033.

7-Ethyl-8-(4-fluorophenyl)-3-(6-morpholino-1*H*-benzo[*d*]imidazol-2-yl)pyrazolo[5,1-*c*][1,2,4]triazin-4-ol **(100)**

The compound was prepared according to General procedure C using:

Diazotization step:

3-ethyl-4-(4-fluorophenyl)-1*H*-pyrazol-5-amine (70 mg, 0.35 mmol; 1 eq.), 35% aqueous HCl (125 µL, 1.4 mmol; 4 eq.) in EtOH (3 mL) and H_2_O (3 mL), NaNO_2_ (36 mg, 0.52 mmol, 1.5 eq.) in EtOH (1 mL) (reaction time: 15 min), ethyl 2-(6-morpholino-1*H*-benzo[*d*]imidazol-2-yl)acetate (100 mg, 0.35 mmol, 1 eq.) in EtOH (2 mL) and KOAc (210 mg, 2.10 mmol, 6 eq.). Reaction time: 16 h. The yellow solid (143 mg, 0.30 mmol) obtained after the filtration was used in the next step without further purification.

Cyclization step:

the dried yellow solid (143 mg, 0.30 mmol), DMF (3 mL), and KOAc (3 mg, 0.03 mmol, 0.1 eq.). Reaction time: 2 h. After the filtration, the solid part was suspended in dichloromethane:MeOH (3:0.6 mL), the solid was collected by filtration and dried under vacuum. The product was obtained as a yellow solid (80 mg, 50%).

^1^H NMR (500 MHz, DMSO-*d*_6_) *δ* (ppm) 13.83 (s, 1H), 13.76 (s, 1H), 7.85 – 7.78 (m, 2H), 7.60 (d, *J* = 8.8 Hz, 1H), 7.36 – 7.27 (m, 2H), 7.18 (dd, *J* = 9.0, 2.2 Hz, 1H), 7.14 (d, *J* = 2.2 Hz, 1H), 3.81 – 3.76 (m, 4H), 3.18 – 3.12 (m, 4H), 2.94 (q, *J* = 7.5 Hz, 2H), 1.28 (t, *J* = 7.5 Hz, 3H).

^13^C NMR (126 MHz, DMSO-*d*_6_) *δ* (ppm) 160.79 (d, *J* = 243.7 Hz), 155.56, 149.47, 149.31, 148.74, 147.31, 132.19, 130.69 (d, *J* = 8.0 Hz), 128.48 (d, *J* = 3.2 Hz), 124.74, 119.45, 115.22 (d, *J* = 21.4 Hz), 115.05, 113.73, 98.06, 66.03, 49.25, 20.81, 13.08.

^19^F NMR (282 MHz, DMSO-*d*_6_) *δ* (ppm) -116.06.

HRMS (APCI): calcd. for C_24_H_23_FN_7_O_2_ [M+H]^+^= 460.1892, found [M+H]^+^ = 460.1893.

7-Ethyl-8-(4-fluorophenyl)-3-(6-(4-methylpiperazin-1-yl)-1*H*-benzo[*d*]imidazol-2-yl)pyrazolo[5,1-*c*][1,2,4]triazin-4-ol **(101)**

The compound was prepared according to General procedure C using:

Diazotization step:

3-ethyl-4-(4-fluorophenyl)-1*H*-pyrazol-5-amine (70 mg, 0.35 mmol; 1 eq.), 35% aqueous HCl (125 µL, 1.4 mmol; 4 eq.) in EtOH (3 mL) and H_2_O (3 mL), NaNO_2_ (36 mg, 0.52 mmol, 1.5 eq.) in EtOH (1 mL) (reaction time: 15 min), ethyl 2-(5-(4-methylpiperazin-1-yl)-1*H*-benzo[*d*]imidazol-2-yl)acetate (100 mg, 0.35 mmol, 1 eq.) in EtOH (2 mL) and KOAc (210 mg, 2.1 mmol, 6 eq.). Reaction time: 16 h. The yellow solid (123 mg, 0.25 mmol) obtained after the filtration was used in the next step without further purification.

Cyclization step:

the dried yellow solid (123 mg, 0.25 mmol), DMF (3 mL), and KOAc (3 mg, 0.03 mmol, 0.1 eq.). Reaction time: 2 h. After the reaction was completed, the solvent was evaporated, and the residue was suspended in EtOAc:MeOH (10:1 mL), filtered and the obtained solid was washed with water (3 mL), EtOAc (2 mL) and dried under vacuum. The product was obtained as a yellow solid (35 mg, 21%).

^1^H NMR (500 MHz, DMSO-*d*_6_) *δ* (ppm) 13.16 (s, 2H), 7.87 – 7.80 (m, 2H), 7.52 (d, *J* = 8.8 Hz, 1H), 7.30 (t, *J* = 8.7 Hz, 2H), 7.15 – 7.11 (m, 1H), 7.03 (d, *J* = 8.9 Hz, 1H), 3.18 – 3.13 (m, 4H), 2.93 (q, *J* = 7.5 Hz, 2H), 2.57 – 2.51 (m, 4H), 2.27 (s, 3H), 1.28 (t, *J* = 7.5 Hz, 3H).

^13^C NMR (126 MHz, DMSO-*d*_6_) *δ* (ppm) 160.99 (d, *J* = 242.9 Hz), 155.45, 150.03, 149.28 (d, *J* = 5.2 Hz), 148.71, 135.87, 130.87 (d, *J* = 7.7 Hz), 129.63 (d, *J* = 3.4 Hz), 123.26, 115.60 (d, *J* = 21.0 Hz), 114.93, 114.69, 106.37, 99.99, 55.11, 49.93, 46.03, 21.46, 13.65.

^19^F NMR (282 MHz, DMSO-*d*_6_) *δ* (ppm) -116.74.

HRMS (APCI): calcd. for C_25_H_26_FN_8_O [M+H]^+^= 473.3223, found [M+H]^+^ = 473.3234.

Methyl 3-(2-(7-ethyl-8-(4-fluorophenyl)-4-hydroxypyrazolo[5,1-*c*][1,2,4]triazin-3-yl)-1*H*-benzo[*d*]imidazol-6-yl)propanoate **(102)**

The compound was prepared according to General procedure C using:

Diazotization step:

3-ethyl-4-(4-fluorophenyl)-1*H*-pyrazol-5-amine (300 mg, 0.69 mmol; 1 eq.), 35% aqueous HCl (250 µL, 2.8 mmol; 4 eq.) in EtOH (5 mL) and H_2_O (5 mL), NaNO_2_ (72 mg, 1.04 mmol, 1.5 eq.) in EtOH (1 mL) (reaction time: 15 min), methyl 3-(2-(2-ethoxy-2-oxoethyl)-1*H*-benzo[*d*]imidazol-6-yl)propanoate (200 mg, 0.69 mmol, 1 eq.) in EtOH (2 mL) and KOAc (420 mg, 4.2 mmol, 6 eq.). Reaction time: 16 h. The yellow solid (310 mg, 0.65 mmol) obtained after the filtration was used in the next step without further purification.

Cyclization step:

the dried yellow solid (310 mg, 0.65 mmol), DMF (5 mL), and KOAc (6 mg, 0.06 mmol, 0.1 eq.). Reaction time: 2 h. After filtration, the residue was suspended in EtOAc:MeOH (20:0.5 mL), the solid was collected by filtration, washed with EtOAc (2 × 5 mL), and dried under vacuum. The product was obtained as a yellow solid (200 mg, 63%).

^1^H NMR (500 MHz, DMSO-*d*_6_) *δ* (ppm) 14.01 (s, 1H), 13.92 (s, 1H), 7.84 – 7.78 (m, 2H), 7.66 (d, *J* = 8.3 Hz, 1H), 7.57 (s, 1H), 7.36 – 7.29 (m, 3H), 3.60 (s, 3H), 3.02 (t, *J* = 7.5 Hz, 2H), 2.94 (q, *J* = 7.5 Hz, 2H), 2.70 (t, *J* = 7.5 Hz, 2H), 1.28 (t, *J* = 7.5 Hz, 3H).

^13^C NMR (126 MHz, DMSO-*d*_6_) *δ* (ppm) 172.46, 160.84 (d, *J* = 243.9 Hz), 155.66, 149.24, 148.83, 148.21, 137.65, 131.25, 130.75 (d, *J* = 7.9 Hz), 129.65, 128.79 – 128.13 (m), 125.34, 119.24, 115.24 (d, *J* = 21.3 Hz), 113.22, 112.48, 108.12, 51.27, 35.18, 30.31, 20.78, 13.07.

^19^F NMR (282 MHz, DMSO-*d*_6_) *δ* (ppm) -115.95.

HRMS (APCI): calcd. for C_24_H_22_FN_6_O_3_ [M+H]^+^= 461.1732, found [M+H]^+^ = 461.1734.

3-(2-(7-Ethyl-8-(4-fluorophenyl)-4-hydroxypyrazolo[5,1-*c*][1,2,4]triazin-3-yl)-1*H*-benzo[*d*]imidazol-6-yl)propanoic acid **(103)**

Methyl 3-(2-(7-ethyl-8-(4-fluorophenyl)-4-hydroxypyrazolo[5,1-*c*][1,2,4]triazin-3-yl)-1*H*-benzo[*d*]imidazol-6-yl)propanoate (90 mg, 0.19 mmol, 1 eq.) was dissolved in THF:H_2_O (8:2 mL, 4:1), LiOH (70 mg, 2.92 mmol, 15 eq.) was added. The mixture was stirred at 23 °C for 16 h, then neutralized with aqueous hydrochloric acid (2 M, 1.50 mL). The THF was evaporated, and the resulting solid was collected by filtration and washed on the filter with water (5 mL) and then with mixture of EtOAc:MeOH (1:0.2 mL) and dried under vacuum. The product was obtained as a yellow solid (30 mg, 35%).

^1^H NMR (500 MHz, DMSO-*d*_6_) *δ* (ppm) 14.02 (s, 1H), 13.92 (s, 1H), 7.85 – 7.77 (m, 2H), 7.66 (d, *J* = 8.3 Hz, 1H), 7.58 (s, 1H), 7.37 – 7.28 (m, 3H), 2.99 (t, *J* = 7.5 Hz, 2H), 2.94 (q, *J* = 7.5 Hz, 2H), 2.61 (t, *J* = 7.5 Hz, 2H), 1.28 (t, *J* = 7.5 Hz, 3H).

^13^C NMR (126 MHz, DMSO-*d*_6_) *δ* (ppm) 173.50, 160.85 (d, *J* = 243.7 Hz), 155.67, 148.82, 148.12, 138.06, 131.24, 130.76 (d, *J* = 7.9 Hz), 129.59, 128.35, 125.39, 119.30, 115.25 (d, *J* = 21.3 Hz), 113.19, 112.48, 108.08, 35.51, 30.40, 20.78, 13.07.

^19^F NMR (282 MHz, DMSO-*d*_6_) *δ* (ppm) -115.93.

HRMS (APCI): calcd. for C_23_H_20_FN_6_O_3_ [M+H]^+^= 447.1575, found [M+H]^+^ = 447.1577.

*tert*-Butyl 2-(7-ethyl-8-(4-fluorophenyl)-4-hydroxypyrazolo[5,1-*c*][1,2,4]triazin-3-yl)-3,4,6,7-tetrahydro-5*H*-imidazo[4,5-*c*]pyridine-5-carboxylate **(104)**

The compound was prepared according to General procedure C using:

Diazotization step:

3-ethyl-4-(4-fluorophenyl)-1*H*-pyrazol-5-amine (80 mg, 0.39 mmol; 1 eq.), 35% aqueous HCl (0.14 mL, 1.55 mmol; 4 eq.) in EtOH (1 mL) and H_2_O (3 mL), NaNO_2_ (0.054 g, 0.78 mmol, 2 eq.) in EtOH (1 mL) and H_2_O (1 mL) (reaction time: 30 min), *tert*-butyl 2-(2-ethoxy-2-oxoethyl)-3,4,6,7-tetrahydro-5*H*-imidazo[4,5-*c*]pyridine-5-carboxylate (120 mg, 0.39 mmol, 1 eq.) in EtOH (2 mL) and H_2_O (1 mL) and KOAc (305 mg, 3.10 mmol, 8 eq.) Reaction time: 2 h. The reaction mixture was poured into water (20 mL) and extracted with EtOAc (2 × 50 mL). The combined organic extracts were washed with brine (35 mL), dried over MgSO_4_, filtered, and the solvent was removed *in vacuo*. The resulting yellow solid (210 mg, quant.) was used in the next step without further purification.

Cyclization step:

the dried yellow solid (210 mg), DMF (2 mL), and KOAc (2 mg, 0.02 mmol, 0.1 eq.). Reaction time: 1.5 h. The product was obtained as an orange solid (144 mg, 77% (2 steps)).

^1^H NMR (500 MHz, DMSO-*d*_6_) *δ* (ppm) 13.71 (s, 1H), 13.49 (s, 1H), 7.79 (dd, *J* = 8.7, 5.7 Hz, 2H), 7.29 (dd, *J* = 8.9 Hz, 1H), 4.49 (s, 2H), 3.68 (t, *J* = 5.6 Hz, 2H), 2.92 (q, *J* = 7.6 Hz, 2H), 2.72 (q, *J* = 5.9, 5.4 Hz, 2H), 1.44 (s, 9H), 1.26 (t, *J* = 7.5 Hz, 3H).

^13^C NMR (126 MHz, DMSO-*d*_6_) *δ* (ppm) 160.64 (d, *J* = 243.4 Hz), 155.29, 148.06, 142.29, 130.54 (d, *J* = 7.5 Hz), 128.78, 124.78, 115.15 (d, *J* = 21.6 Hz), 79.62, 27.94, 20.81, 13.15.

^19^F NMR (471 MHz, DMSO-*d*_6_) *δ* (ppm) -116.38.

HRMS (APCI): calcd. for C_24_H_27_FN_7_O_3_ [M+H]^+^ = 480.2154, found [M+H]^+^ = 480.2152.

3-(1-Benzyl-1*H*-imidazol-2-yl)-7-ethyl-8-(4-fluorophenyl)pyrazolo[5,1-*c*][1,2,4]triazin-4-ol **(105)**

The compound was prepared according to General procedure C using:

Diazotization step:

3-ethyl-4-(4-fluorophenyl)-1*H*-pyrazol-5-amine (200 mg, 0.97 mmol; 1 eq.), 35% aqueous HCl (336 µL, 3.9 mmol; 4 eq.) in EtOH (10 mL) and H_2_O (10 mL), NaNO_2_ (135 mg, 1.95 mmol, 2 eq.) in EtOH (1 mL) (reaction time: 15 min), ethyl 2-(1-benzyl-1*H*-imidazol-2-yl)acetate (286 mg, 1.17 mmol, 1.2 eq.) in EtOH (2 mL) and KOAc (570 mg, 4.14 mmol, 6 eq.). Reaction time: 16 h. The yellow solid (302 mg, 0.7 mmol) obtained after the filtration was used in the next step without further purification.

Cyclization step:

the dried yellow solid (302 mg, 0.7 mmol), DMF (5 mL), and KOAc (7 mg, 0.07 mmol, 0.1 eq.). Reaction time: 2 h. After the filtration, the solid was suspended in hot dioxane (10 mL) and the mixture was poured into water (30 mL). The solid was collected by filtration, washed with water (10 mL), diethyl ether (10 mL) and dried under vacuum. The product was obtained as a yellow solid (200 mg, 50%).

^1^H NMR (500 MHz, DMSO-*d*_6_) *δ* (ppm) 13.70 (s, 1H), 7.79 – 7.74 (m, 2H), 7.74 – 7.70 (m, 1H), 7.55 (s, 1H), 7.37 – 7.24 (m, 7H), 5.93 (s, 2H), 2.91 (q, *J* = 7.5 Hz, 2H), 1.26 (t, *J* = 7.5 Hz, 2H).

^13^C NMR (126 MHz, DMSO-*d*_6_) *δ* (ppm) 160.68 (d, *J* = 243.1 Hz), 155.41, 148.72, 141.26, 136.17, 130.60 (d, *J* = 8.0 Hz), 128.66, 128.48, 127.89, 127.42, 122.81, 118.28, 115.18 (d, *J* = 21.2 Hz), 106.41, 51.84, 20.79, 13.02.

^19^F NMR (471 MHz, DMSO-*d*_6_) *δ* (ppm) -116.26.

HRMS (APCI): calcd. for C_23_H_20_FN_6_O [M+H]^+^ = 415.1677, found [M+H]^+^ = 415.1680.

3-(1*H*-benzo[*d*]imidazol-2-yl)-8-(4-fluorophenyl)-7-(2-methoxybenzyl)pyrazolo[5,1-*c*][1,2,4]triazin-4-ol **(106)**

The compound was prepared according to General procedure C using:

Diazotization step:

4-(4-fluorophenyl)-3-(2-methoxybenzyl)-1*H*-pyrazol-5-amine (190 mg, 0.64 mmol; 1 eq.), 35% aqueous HCl (220 µL, 2.56 mmol; 4 eq.) in EtOH (5 mL) and H_2_O (5 mL), NaNO_2_ (88 mg, 1.28 mmol, 2 eq.) in EtOH (1 mL) (reaction time: 15 min), ethyl 2-(1*H*-1,3-benzodiazol-2-yl)acetate (157 mg, 0.77 mmol, 1.2 eq.) in EtOH (2 mL) and KOAc (380 mg, 3.8 mmol, 6 eq.). Reaction time: 16 h. The yellow solid (242 mg, 0.5 mmol) obtained after the filtration was used in the next step without further purification.

Cyclization step:

the dried yellow solid (242 mg, 0.50 mmol), DMF (5 mL), and KOAc (5 mg, 0.05 mmol, 0.1 eq.). Reaction time: 2 h. After the filtration, the solid part was suspended in hot dioxane (6 mL), and the mixture was poured into water (30 mL). The solid part was collected by filtration, washed with water (10 mL), diethyl ether (10 mL) and dried under vacuum. The product was obtained as a yellow solid (200 mg, 67%).

^1^H NMR (500 MHz, DMSO-*d*_6_) *δ* (ppm) 14.06 (s, 2H), 7.78 – 7.69 (m, 4H), 7.47 – 7.42 (m, 2H), 7.29 – 7.22 (m, 2H), 7.21 – 7.12 (m, 1H), 6.96 (dd, *J* = 24.0, 7.9 Hz, 2H), 6.84 – 6.78 (m, 1H), 4.20 (s, 2H), 3.78 (s, 3H).

^13^C NMR (126 MHz, DMSO-*d*_6_) *δ* (ppm) 160.86 (d, *J* = 243.6 Hz), 156.52, 152.67, 149.40, 148.85, 148.30, 131.10, 130.60 (d, *J* = 8.0 Hz), 129.27, 128.26 (d, *J* = 2.4 Hz), 127.54, 126.76, 124.72, 120.21, 119.34, 115.21, 115.04, 113.37, 110.60, 109.23, 55.31, 27.08.

^19^F NMR (471 MHz, DMSO-*d*_6_) *δ* (ppm) -115.82.

HRMS (APCI): calcd. for C_26_H_20_FN_6_O_2_ [M+H]^+^ = 467.1626, found [M+H]^+^ = 467.1630.

6-(1*H*-benzo[*d*]imidazol-2-yl)-2-ethyl-3-(4-fluorophenyl)-4-methylpyrazolo[1,5-*a*]pyrimidin-7(4*H*)-one **(107)**

A mixture of 3-ethyl-4-(4-fluorophenyl)-*N*-methyl-1*H*-pyrazol-5-amine (75 mg, 0.34 mmol; 1 eq.) and ethyl-2-(1*H*-benzo[*d*]imidazol-2-yl)-3-(dimethylamino)acrylate (93 mg, 0.36 mmol, 1.05 eq.) in EtOH (3 mL) was stirred in the microwave reactor at 130 °C for 1 h. The precipitate was collected by filtration, washed with EtOH (5 mL), diethyl ether (5 mL) and dried under vacuum. The product was obtained as a white solid (20 mg, 15%)

^1^H NMR (500 MHz, DMSO-*d*_6_) *δ* (ppm) 12.31 (s, 1H), 8.94 (s, 1H), 7.69 – 7.63 (m, 1H), 7.59 – 7.45 (m, 3H), 7.36 – 7.27 (m, 2H), 7.19 – 7.12 (m, 2H), 3.50 (s, 3H), 2.55 (q, *J* = 7.6 Hz, 2H), 1.10 (t, *J* = 7.6 Hz, 3H).

^13^C NMR (126 MHz, DMSO-*d*_6_) *δ* (ppm) 161.91 (d, *J* = 245.1 Hz), 157.04, 154.57, 147.14, 144.81, 142.65, 138.38, 134.46, 133.57 (d, *J* = 8.3 Hz), 127.00 (d, *J* = 3.3 Hz), 121.47 (d, *J* = 10.5 Hz), 117.44, 115.09 (d, *J* = 21.4 Hz), 112.18, 104.72, 98.38, 41.51, 20.13, 13.04.

^19^F NMR (471 MHz, DMSO-*d*_6_) *δ* (ppm) -113.99.

HRMS (APCI): calcd. for C_22_H_19_FN_5_O [M+H]^+^ = 388.1568, found [M+H]^+^ = 388.1569.

3-(1*H*-benzo[*d*]imidazol-2-yl)-7-ethyl-8-(tetrahydro-2*H*-pyran-4-yl)pyrazolo[5,1-*c*][1,2,4]triazin-4-ol **(108)**

The compound was prepared according to General procedure C using:

Diazotization step:

3-ethyl-4-(tetrahydro-2*H*-pyran-4-yl)-1*H*-pyrazol-5-amine (114 mg, 0.58 mmol; 1 eq.), 35% aqueous HCl (200 µL, 2.33 mmol; 4 eq.) in EtOH (5 mL) and H_2_O (5 mL), NaNO_2_ (81 mg, 1.17 mmol, 2 eq.) in EtOH (1 mL) (reaction time: 15 min), ethyl 2-(1*H*-1,3-benzodiazol-2-yl)acetate (131 mg, 0.64 mmol, 1.1 eq.) in EtOH (2 mL) and KOAc (344 mg, 3.5 mmol, 6 eq.). Reaction time: 16 h. The yellow solid (190 mg, 0.5 mmol) obtained after the filtration was used in the next step without further purification.

Cyclization step:

the dried yellow solid (190 mg, 0.5 mmol), DMF (5 mL), and KOAc (5 mg, 0.05 mmol, 0.1 eq.). Reaction time: 2 h. After the filtration, the solid was suspended in hot dioxane (5 mL) and the mixture was poured into water (20 mL). The solid was collected by filtration, washed with water (10 mL), diethyl ether (10 mL) and dried under vacuum. The product was obtained as a yellow solid (130 mg, 61%).

^1^H NMR (500 MHz, DMSO-*d*_6_) *δ* (ppm) 13.90 (s, 2H), 7.73 (dd, *J* = 6.0, 3.2 Hz, 2H), 7.43 (dt, *J* = 6.0, 3.6 Hz, 2H), 3.98 (dd, *J* = 11.2, 4.2 Hz, 2H), 3.53 – 3.44 (m, 2H), 3.11 – 2.98 (m, 1H), 2.80 (q, *J* = 7.5 Hz, 2H), 2.42 – 2.30 (m, 2H), 1.70 – 1.63 (m, 2H), 1.28 (t, *J* = 7.5 Hz, 3H).

^13^C NMR (126 MHz, DMSO-*d*_6_) *δ* (ppm) 156.03, 149.81, 149.20, 148.85, 131.07, 124.50, 117.40, 113.20, 112.54, 67.65, 32.49, 31.52, 20.16, 13.99.

HRMS (APCI): calcd. for C_19_H_21_N_6_O_2_ [M+H]^+^ = 365.1721, found [M+H]^+^ = 365.1719.

3-(1*H*-benzo[*d*]imidazol-2-yl)-8-benzyl-7-ethylpyrazolo[5,1-*c*][1,2,4]triazin-4-ol **(109)**

The compound was prepared according to General procedure C using:

Diazotization step:

4-benzyl-3-ethyl-1*H*-pyrazol-5-amine (121 mg, 0.60 mmol; 1 eq.), 35% aqueous HCl (207 µL, 2.4 mmol; 4 eq.) in EtOH (5 mL) and H_2_O (5 mL), NaNO_2_ (83 mg, 1.20 mmol, 2 eq.) in EtOH (1 mL) (reaction time: 15 min), ethyl 2-(1*H*-1,3-benzodiazol-2-yl)acetate (135 mg, 0.60 mmol, 1.1 eq.) in EtOH (2 mL) and KOAc (354 mg, 3.60 mmol, 6 eq.) Reaction time: 16 h. The yellow solid (213 mg, 0.55 mmol) obtained after the filtration was used in the next step without further purification.

Cyclization step:

the dried yellow solid (213 mg, 0.55 mmol), DMF (5 mL), and KOAc (6 mg, 0.06 mmol, 0.1 eq.). Reaction time: 2 h. After the filtration, the solid was suspended in hot dioxane (5 mL) and the mixture was poured into water (20 mL). The solid was collected by filtration, washed with water (10 mL), diethyl ether (10 mL) and dried under vacuum. The product was obtained as a yellow solid (165 mg, 74%).

^1^H NMR (500 MHz, DMSO-*d*_6_) *δ* (ppm) 13.94 (s, 1H), 7.77 – 7.70 (m, 2H), 7.46 – 7.40 (m, 2H), 7.33 – 7.23 (m, 4H), 7.20 – 7.12 (m, 1H), 4.16 (s, 2H), 2.69 (q, *J* = 7.6 Hz, 2H), 1.16 (t, *J* = 7.6 Hz, 3H).

^13^C NMR (126 MHz, DMSO-*d*_6_) *δ* (ppm) 156.93, 150.23, 149.18, 148.73, 140.81, 131.11, 128.29, 128.25, 128.15, 125.80, 124.50, 117.94, 113.24, 107.76, 27.77, 19.99, 13.14.

HRMS (APCI): calcd. for C_21_H_18_N_6_O [M+H]^+^ = 371.1615, found [M+H]^+^ = 371.1616.

3-(1*H*-benzo[*d*]imidazol-2-yl)-8-(cyclohexylmethyl)-7-ethylpyrazolo[5,1-*c*][1,2,4]triazin-4-ol **(110)**

The compound was prepared according to General procedure C using:

Diazotization step:

4-(cyclohexylmethyl)-3-ethyl-1*H*-pyrazol-5-amine (160 mg, 0.77 mmol; 1 eq.), 35% aqueous HCl (270 µL, 3.1 mmol; 4 eq.) in EtOH (5 mL) and H_2_O (5 mL), NaNO_2_ (107 mg, 1.54 mmol, 2 eq.) in EtOH (1 mL) (reaction time: 15 min), ethyl 2-(1*H*-1,3-benzodiazol-2-yl)acetate (173 mg, 0.85 mmol, 1.1 eq.) in EtOH (2 mL) and KOAc (454 mg, 4.63 mmol, 6 eq.). Reaction time: 16 h. The yellow solid (264 mg, 0.67 mmol) obtained after the filtration was used in the next step without further purification.

Cyclization step:

the dried yellow solid (264 mg, 0.67 mmol), DMF (5 mL), and KOAc (7 mg, 0.07 mmol, 0.1 eq.). Reaction time: 2 h. After the filtration, the solid was suspended in hot dioxane (5 mL) and the mixture was poured into water (20 mL). The solid was collected by filtration, washed with water (10 mL), diethyl ether (10 mL) and dried under vacuum. The product was obtained as a yellow solid (220 mg, 76%).

^1^H NMR (500 MHz, DMSO-*d*_6_) *δ* (ppm) 13.87 (s, 2H), 7.76 – 7.69 (m, 2H), 7.45 – 7.38 (m, 2H), 2.75 (q, *J* = 7.6 Hz, 2H), 2.66 (d, *J* = 6.9 Hz, 2H), 1.77 – 1.54 (m, 6H), 1.28 (t, *J* = 7.6 Hz, 3H), 1.22 – 1.10 (m, 3H), 1.06 – 0.97 (m, 2H).

^13^C NMR (126 MHz, DMSO-*d*_6_) *δ* (ppm) 157.10, 150.26, 149.24, 148.86, 131.09, 124.42, 117.39, 113.19, 107.72, 66.29, 38.32, 32.73, 29.67, 26.03, 25.67, 19.94, 13.42.

HRMS (APCI): calcd. for C_21_H_24_N_6_O [M+H]^+^ = 377.2084, found [M+H]^+^ = 377.2082.

3-(1*H*-benzo[*d*]imidazol-2-yl)-8-(2-chlorobenzyl)-7-ethylpyrazolo[5,1-*c*][1,2,4]triazin-4-ol **(111)**

The compound was prepared according to General procedure C using:

Diazotization step:

4-(2-chlorobenzyl)-3-ethyl-1*H*-pyrazol-5-amine (130 mg, 0.55 mmol; 1 eq.), 35% aqueous HCl (190 *µ*L, 2.2 mmol; 4 eq.) in EtOH (5 mL) and H_2_O (5 mL), NaNO_2_ (76 mg, 1.1 mmol, 2 eq.) in EtOH (1 mL) (reaction time: 15 min), ethyl 2-(1*H*-1,3-benzodiazol-2-yl)acetate (124 mg, 0.6 mmol, 1.1 eq.) in EtOH (2 mL) and KOAc (325 mg, 4.63 mmol, 6 eq.). Reaction time: 16 h. The yellow solid (190 mg, 0.45 mmol) obtained after the filtration was used in the next step without further purification.

Cyclization step:

the dried yellow solid (190 mg, 0.45 mmol), DMF (5 mL), and KOAc (5 mg, 0.05 mmol, 0.1 eq.). Reaction time: 2 h. After the filtration, the solid was suspended in hot dioxane (5 mL) and the mixture was poured into water (20 mL). The solid was collected by filtration, washed with water (10 mL), diethyl ether (10 mL) and dried under vacuum. The product was obtained as a yellow solid (150 mg, 67%).

^1^H NMR (500 MHz, DMSO-*d*_6_) *δ* (ppm) 13.95 (s, 1H), 7.77 – 7.70 (m, 2H), 7.48 – 7.39 (m, 3H), 7.28 – 7.18 (m, 3H), 4.25 (s, 2H), 2.68 (q, *J* = 7.5 Hz, 2H), 1.16 (t, *J* = 7.6 Hz, 3H).

^13^C NMR (126 MHz, DMSO-*d*_6_) *δ* (ppm) 157.09, 150.45, 149.12, 148.66, 137.64, 132.71, 131.08, 130.39, 129.04, 127.85, 127.07, 124.54, 118.13, 113.24, 105.57, 25.50, 20.08, 13.11.

HRMS (APCI): calcd. for C_21_H_18_ClN_6_O [M+H]^+^ = 405.1225, found [M+H]^+^ = 405.1229.

3-(1*H*-benzo[*d*]imidazol-2-yl)-7-ethyl-8-(pyridin-3-ylmethyl)pyrazolo[5,1-*c*][1,2,4]triazin-4-ol **(112)**

The compound was prepared according to General procedure C using:

Diazotization step:

3-ethyl-4-(pyridin-3-ylmethyl)-1*H*-pyrazol-5-amine (100 mg, 0.50 mmol; 1 eq.), 35% aqueous HCl (170 µL, 2.0 mmol; 4 eq.) in EtOH (5 mL) and H_2_O (5 mL), NaNO_2_ (68 mg, 1.0 mmol, 2 eq.) in EtOH (1 mL) (reaction time: 15 min), ethyl 2-(1*H*-1,3-benzodiazol-2-yl)acetate (111 mg, 0.54 mmol, 1.1 eq.) in EtOH (2 mL) and KOAc (290 mg, 2.97 mmol, 6 eq.). Reaction time: 16 h. The yellow solid (175 mg, 0.45 mmol) obtained after the filtration was used in the next step without further purification.

Cyclization step:

the dried yellow solid (175 mg, 0.45 mmol), DMF (5 mL), and KOAc (5 mg, 0.05 mmol, 0.1 eq.). Reaction time: 2 h. After the filtration, the solid was suspended in hot dioxane (5 mL) and the mixture was poured into water (20 mL). The solid was collected by filtration, washed with water (10 mL), diethyl ether (10 mL) and dried under vacuum. The product was obtained as a yellow solid (130 mg, 71%).

^1^H NMR (500 MHz, DMSO-*d*_6_) *δ* (ppm) 13.95 (s, 1H), 8.61 – 8.57 (m, 1H), 8.41 – 8.36 (m, 1H), 7.77 – 7.70 (m, 2H), 7.69 – 7.65 (m, 1H), 7.47 – 7.40 (m, 2H), 7.28 (dd, *J* = 7.9, 4.7 Hz, 1H), 4.18 (s, 2H), 2.73 (q, *J* = 7.6 Hz, 2H), 1.18 (t, *J* = 7.6 Hz, 3H).

^13^C NMR (126 MHz, DMSO-*d*_6_) *δ* (ppm) 156.79, 149.43, 149.10, 148.65, 147.19, 136.34, 135.69, 131.18, 124.53, 123.41, 118.34, 113.28, 106.67, 25.11, 19.90, 13.26.

HRMS (APCI): calcd. for C_20_H_18_N_7_O [M+H]^+^ = 372.1567, found [M+H]^+^ = 372.1570.

3-(1*H*-benzo[*d*]imidazol-2-yl)-7-ethyl-8-(2-methoxybenzyl)pyrazolo[5,1-*c*][1,2,4]triazin-4-ol **(113)**

The compound was prepared according to General procedure C using:

Diazotization step:

3-ethyl-4-(2-methoxybenzyl)-1H-pyrazol-5-amine (110 mg, 0.48 mmol; 1 eq.), 35% aqueous HCl (160 µL, 1.9 mmol; 4 eq.) in EtOH (5 mL) and H_2_O (5 mL), NaNO_2_ (66 mg, 0.95 mmol, 2 eq.) in EtOH (1 mL) (reaction time: 15 min), ethyl 2-(1*H*-1,3-benzodiazol-2-yl)acetate (107 mg, 0.52 mmol, 1.1 eq.) in EtOH (2 mL) and KOAc (280 mg, 2.85 mmol, 6 eq.). Reaction time: 16 h. The yellow solid (167 mg, 0.4 mmol) obtained after the filtration was used in the next step without further purification.

Cyclization step:

the dried yellow solid (167 mg, 0.4 mmol), DMF (5 mL), and KOAc (4 mg, 0.04 mmol, 0.1 eq.). Reaction time: 2 h. After the filtration, the solid was suspended in hot dioxane (5 mL) and the mixture was poured into water (20 mL). The solid was collected by filtration, washed with water (10 mL), diethyl ether (10 mL) and dried under vacuum. The product was obtained as a yellow solid (100 mg, 53%).

^1^H NMR (500 MHz, DMSO-*d*_6_) *δ* (ppm) 13.79 (s, 2H), 7.77 – 7.71 (m, 2H), 7.44 – 7.39 (m, 2H), 7.20 – 7.13 (m, 1H), 7.07 (dd, *J* = 7.5, 1.8 Hz, 1H), 6.98 (d, *J* = 8.1 Hz, 1H), 6.81 (t, *J* = 7.4 Hz, 1H), 4.12 (s, 2H), 3.85 (s, 3H), 2.69 (q, *J* = 7.6 Hz, 2H), 1.19 (t, *J* = 7.6 Hz, 3H).

^13^C NMR (126 MHz, DMSO-*d*_6_) *δ* (ppm) 157.19, 156.63, 150.40, 149.21, 148.78, 131.12, 129.08, 128.23, 127.14, 124.46, 120.07, 117.81, 113.21, 110.40, 106.89, 55.25, 21.63, 19.92, 13.18.

HRMS (APCI): calcd. for C_22_H_21_N_6_O_2_ [M+H]^+^ = 401.1721, found [M+H]^+^ = 401.1722.

3-(1*H*-benzo[*d*]imidazol-2-yl)-7-ethyl-8-(4-fluorobenzyl)pyrazolo[5,1-*c*][1,2,4]triazin-4-ol **(114)**

The compound was prepared according to General procedure C using:

Diazotization step:

3-ethyl-4-(4-fluorobenzyl)-1*H*-pyrazol-5-amine (100 mg, 0.46 mmol; 1 eq.), 35% aqueous HCl (160 µL, 1.8 mmol; 4 eq.) in EtOH (5 mL) and H_2_O (5 mL), NaNO_2_ (63 mg, 0.91 mmol, 2 eq.) in EtOH (1 mL) (reaction time: 15 min), ethyl 2-(1*H*-1,3-benzodiazol-2-yl)acetate (103 mg, 0.46 mmol, 1.1 eq.) in EtOH (2 mL) and KOAc (270 mg, 2.74 mmol, 6 eq.). Reaction time: 16 h. The yellow solid (170 mg, 0.42 mmol) obtained after the filtration was used in the next step without further purification.

Cyclization step:

the dried yellow solid (170 mg, 0.42 mmol), DMF (5 mL), and KOAc (4 mg, 0.04 mmol, 0.1 eq.). Reaction time: 2 h. After the filtration, the solid was suspended in hot dioxane (5 mL) and the mixture was poured into water (20 mL). The solid part was collected by filtration, washed with water (10 mL), diethyl ether (10 mL) and dried under vacuum. The product was obtained as a yellow solid (150 mg, 85%).

^1^H NMR (500 MHz, DMSO-*d*_6_) *δ* (ppm) 13.95 (s, 1H), 7.77 – 7.70 (m, 2H), 7.47 – 7.39 (m, 2H), 7.37 – 7.29 (m, 2H), 7.12 – 7.04 (m, 2H), 4.15 (s, 2H), 2.69 (q, *J* = 7.6 Hz, 2H), 1.16 (t, *J* = 7.6 Hz, 3H).

^13^C NMR (126 MHz, DMSO-*d*_6_) *δ* (ppm) 160.58 (d, *J* = 241.5 Hz), 156.88, 150.21, 149.15, 148.69, 136.97, 131.08, 129.89 (d, *J* = 8.0 Hz), 124.55, 118.01, 114.92 (d, *J* = 21.0 Hz), 113.25, 107.64, 26.96, 19.96, 13.20.

^19^F NMR (471 MHz, DMSO-*d*_6_) *δ* (ppm) -117.49.

HRMS (APCI): calcd. for C_21_H_18_FN_6_O [M+H]^+^ 389.1521, found [M+H]^+^ = 389.1525.

3-(1*H*-benzo[*d*]imidazol-2-yl)-7-ethyl-8-(1-methylpiperidin-4-yl)pyrazolo[5,1-*c*][1,2,4]triazin-4-ol **(115)**

The compound was prepared according to General procedure C using:

Diazotization step:

3-ethyl-4-(1-methylpiperidin-4-yl)-1*H*-pyrazol-5-amine (110 mg, 0.53 mmol; 1 eq.), 35% aqueous HCl (180 µL, 2.11 mmol; 4 eq.) in EtOH (5 mL) and H_2_O (5 mL), NaNO_2_ (69 mg, 1.06 mmol, 2 eq.) in EtOH (1 mL) (reaction time: 15 min), ethyl 2-(1*H*-1,3-benzodiazol-2-yl)acetate (119 mg, 0.58 mmol, 1.1 eq.) in EtOH (2 mL) and KOAc (310 mg, 3.17 mmol, 6 eq.). Reaction time: 16 h. The yellow solid (160 mg, 0.40 mmol) obtained after the filtration was used in the next step without further purification.

Cyclization step:

the dried yellow solid (160 mg, 0.40 mmol), DMF (5 mL), and KOAc (4 mg, 0.04 mmol, 0.1 eq.). Reaction time: 2 h. After the filtration, the solid was suspended in hot dioxane (5 mL) and the mixture was poured into water (20 mL). The solid was collected by filtration, washed with water (10 mL), diethyl ether (10 mL) and dried under vacuum. The product was obtained as a yellow solid (120 mg, 60%).

^1^H NMR (500 MHz, DMSO-*d*_6_) *δ* (ppm) 7.71 – 7.64 (m, 2H), 7.32 – 7.25 (m, 2H), 3.23 – 3.13 (m, 2H), 2.95 – 2.86 (m, 1H), 2.78 (q, *J* = 7.6 Hz, 2H), 2.51 – 2.41 (m, 7H), 1.87 – 1.79 (m, 2H), 1.29 (t, *J* = 7.6 Hz, 3H).

^13^C NMR (126 MHz, DMSO-*d*_6_) *δ* (ppm) 155.19, 149.92, 149.62, 149.13, 134.00, 122.53, 119.78, 113.34, 109.66, 55.12, 44.51, 30.87, 30.35, 19.92, 13.48.

HRMS (APCI): calcd. for C_20_H_24_N_7_O [M+H]^+^ 378.2037, found [M+H]^+^ = 378.2039.

3-(1*H*-Benzo[*d*]imidazol-2-yl)-8-((1-(*tert*-butyl)-1*H*-pyrazol-4-yl)methyl)-7-ethylpyrazolo[5,1-*c*][1,2,4]triazin-4-ol **(116)**

The compound was prepared according to General procedure C using:

Diazotization step:

4-((1-(*tert*-butyl)-1*H*-pyrazol-4-yl)methyl)-3-ethyl-1*H*-pyrazol-5-amine (100 mg, 0.40 mmol; 1 eq.), 35% aqueous HCl (140 µL, 1.62 mmol; 4 eq.) in EtOH (5 mL) and H_2_O (5 mL), NaNO_2_ (56 mg, 0.80 mmol, 2 eq.) in EtOH (1 mL) (reaction time: 15 min), ethyl 2-(1*H*-1,3-benzodiazol-2-yl)acetate (91 mg, 0.45 mmol, 1.1 eq.) in EtOH (2 mL) and KOAc (230 mg, 2.43 mmol, 6 eq.). Reaction time: 16 h. The yellow solid (130 mg, 0.30 mmol) obtained after the filtration was used in the next step without further purification.

Cyclization step:

the dried yellow solid (130 mg, 0.30 mmol), DMF (5 mL), and KOAc (3 mg, 0.03 mmol, 0.1 eq.). Reaction time: 2 h. After the filtration, the solid was suspended in hot dioxane (5 mL) and the mixture was poured into water (20 mL). The solid was collected by filtration, washed with water (10 mL), diethyl ether (10 mL) and dried under vacuum. The product was obtained as a yellow solid (80 mg, 48%).

^1^H NMR (500 MHz, DMSO-*d*_6_) *δ* (ppm) 13.93 (s, 1H), 7.76 – 7.71 (m, 2H), 7.56 (s, 1H), 7.46 – 7.39 (m, 2H), 7.28 (s, 1H), 3.96 (s, 2H), 2.71 (q, *J* = 7.6 Hz, 2H), 1.45 (s, 9H), 1.19 (t, *J* = 7.6 Hz, 3H).

^13^C NMR (126 MHz, DMSO-*d*_6_) *δ* (ppm) 156.69, 149.81, 149.23, 148.78, 137.24, 131.07, 124.48, 124.39, 119.15, 117.67, 113.20, 108.37, 57.52, 29.44, 19.95, 16.98, 13.24.

HRMS (APCI): calcd. for C_22_H_25_N_8_O [M+H]^+^ 417.2146, found [M+H]^+^ = 417.2148.

2-Ethyl-3,6-bis(4-fluorophenyl)pyrazolo[1,5-*a*]pyrimidin-7-amine **(117)**

A mixture of 3-ethyl-4-(4-fluorophenyl)-1*H*-pyrazol-5-amine (41 mg, 0.20 mmol, 1eq.) and 2-(4-fluorophenyl)-3-oxopropanenitrile (33 mg, 0.20 mmol, 1 eq.) in DMF (2 mL) was refluxed for 1 h. The solvent was removed *in vacuo,* and the residue was purified twice by column chromatography on silica gel (hexane:EtOAc, gradient 3:1 to 0:1). The product was obtained as a white solid (26 mg, 37%).

^1^H NMR (500 MHz, Chloroform-*d*) *δ* (ppm) 8.21 (s, 1H), 7.63 (dd, *J* = 8.8, 5.4 Hz, 2H), 7.46 (dd, *J* = 8.8, 5.3 Hz, 2H), 7.22 (t, *J* = 8.6 Hz, 2H), 7.16 (t, *J* = 8.7 Hz, 2H), 5.86 (s, 2H), 2.99 (d, *J* = 7.5 Hz, 2H), 1.36 (t, *J* = 7.5 Hz, 3H).

^13^C NMR (126 MHz, Chloroform-*d*) *δ* (ppm) 162.68 (d, *J* = 249.0 Hz), 161.74 (d, *J* = 245.4 Hz), 157.70, 149.98, 145.77, 144.43, 131.14 (d, *J* = 8.2 Hz), 130.80 (d, *J* = 8.2 Hz), 129.82 (d, *J* = 3.1 Hz), 128.61 (d, *J* = 3.0 Hz), 116.81 (d, *J* = 21.2 Hz), 115.68 (d, *J* = 21.2 Hz), 107.44, 102.09, 21.50, 13.63.

^19^F NMR (282 MHz, Chloroform-*d*) *δ* (ppm) -113.45, -116.52.

HRMS (APCI): calcd. for C_20_H_17_N_4_F_2_ [M+H]^+^ = 351.1416, found [M+H]^+^ = 351.1419.

### Experimental procedures for compound 2 and its analogs

4-(2-Cyanoacetyl)benzonitrile (**S109**)

The compound was prepared according to General procedure A5 using NaH (60% suspension in mineral oil, 496 mg, 12.41 mmol), acetonitrile (0.388 mL, 7.45 mmol) in THF (10 mL) and methyl 4-cyanobenzoate (1 g, 6.21 mmol) in anhydrous THF (5 mL). The reaction mixture was refluxed for 2 h. The residue was purified by column chromatography (hexane:EtOAc, 8:2 to 7:3). The product was obtained as a white solid (732 mg, 69%).

^1^H NMR (500 MHz, Chloroform-*d*) *δ* (ppm) 8.07 – 7.99 (m, 2H), 7.88 – 7.81 (m, 2H), 4.11 (s, 2H).

^13^C NMR (126 MHz, Chloroform-*d*) *δ* (ppm) 186.2, 137.2, 133.1, 129.0, 118.2, 117.4, 113.1, 29.8.

HRMS (APCI): calcd. for C_10_H_5_N_2_O [M-H]^−^ = 169.0407, found [M-H]^−^ = 169.0406.

mp = 124−130 °C.

3-Oxo-3-(4-(trifluoromethyl)phenyl)propanenitrile **(S110)**

The compound was prepared according to General procedure A5 using NaH (60% suspension in mineral oil, 391 mg, 9.796 mmol), acetonitrile (0.306 mL, 5.878 mmol), THF (10 mL) and methyl 4-(trifluoromethyl)benzoate (1 g, 4.90 mmol) in anhydrous THF (5 mL). The reaction mixture was refluxed for 2 h. The residue was purified by column chromatography (hexane:EtOAc, 8:2 to 7:3). The product was obtained as a light brown solid (850 mg, 81%).

^1^H NMR (500 MHz, Chloroform-*d*) *δ* (ppm) 8.05 (dp, *J* = 7.9, 0.9 Hz, 2H), 7.85 – 7.77 (m, 2H), 4.13 (s, 2H).

^13^C NMR (126 MHz, Chloroform-*d*) *δ* (ppm) 186.55, 137.00, 136.08 (q, *J* = 33.1 Hz), 129.01, 126.40 (q, *J* = 3.6 Hz), 123.35 (*J* = 273.0 Hz), 113.34, 29.82.

^19^F NMR (471 MHz, Chloroform-*d*) *δ* (ppm) -63.40.

HRMS (APCI): calcd. for C_10_H_5_F_3_NO [M-H]^−^ = 212.0329, found [M-H]^−^ = 212.0327.

mp = 43−45 °C.

3-(4-(*tert*-Butyl)phenyl)-3-oxopropanenitrile **(S111)**

The compound was prepared according to General procedure A5 using NaH (60% suspension in mineral oil, 420 mg, 10.4 mmol), acetonitrile (0.33 mL, 6.2 mmol), THF (10 mL) and methyl 4-(trifluoromethyl)benzoate (1 g, 5.20 mmol) in anhydrous THF (5 mL). The reaction mixture was refluxed for 2 h. The residue was purified by column chromatography (hexane:EtOAc, 8:2 to 7:3). The product was obtained as a yellow solid (740 mg, 70%).

^1^H NMR (500 MHz, Chloroform-*d*) *δ* (ppm) 7.91 – 7.83 (m, 2H), 7.57 – 7.49 (m, 2H), 4.05 (s, 2H), 1.35 (s, 9H).

^13^C NMR (126 MHz, Chloroform-*d*) *δ* (ppm) 186.7, 158.9, 131.8, 128.5, 126.1, 113.9, 35.4, 31.0, 29.2.

HRMS (APCI): calcd. for C_13_H_14_NO [M-H]^−^ = 200.1081, found [M-H]^−^ = 200.1079.

mp = 71−75 °C.

3-(4-Methoxyphenyl)-3-oxopropanenitrile **(S112)**

The compound was prepared according to General procedure A5 using NaH (60% suspension in mineral oil, 481 mg, 12.03 mmol), acetonitrile (0.377 mL, 7.72 mmol), anhydrous THF (10 mL) and methyl 4-methoxybenzoate (1 g, 6.02 mmol) in anhydrous THF (5 mL). The reaction mixture was refluxed for 48 h. The residue was purified by column chromatography (hexane:EtOAc, 8:2 to 7:3). The product was obtained as a yellow solid (781 mg, 74%).

^1^H NMR (500 MHz, Chloroform-*d*) *δ* (ppm) 7.93 – 7.86 (m, 2H), 7.01 – 6.95 (m, 2H), 4.01 (s, 2H), 3.89 (s, 3H).

^13^C NMR (126 MHz, Chloroform-*d*) *δ* (ppm) 185.58, 164.88, 131.06, 127.45, 114.48, 114.21, 55.79, 29.13.

HRMS (APCI): calcd. for C_10_H_8_NO_2_ [M-H]^-^ = 174.0561, found [M-H]^-^ = 174.0559.

mp = 123−126 °C.

Methyl 2-phenylacetate **(S113)**

Conc. H_2_SO_4_ (96%, 30 µL) was added to a solution of 2-phenylacetic acid (1.50 g, 11.01 mmol) in methanol (15 mL) and the mixture was refluxed for 4 h. The reaction mixture was cooled to room temperature and the solvent was evaporated *in vacuo*. Water (15 mL) was added to the residue and the mixture was extracted with EtOAc (3 × 15 mL). The combined organic extracts were dried over MgSO_4_, ﬁltered, and the solvent was evaporated *in vacuo*. The product was obtained as a clear oil (1.25 g, 76%).

^1^H NMR (300 MHz, Chloroform-*d*) *δ* (ppm) 7.37 – 7.23 (m, 5H), 3.70 (s, 3H), 3.63 (s, 2H).

^13^C NMR (75 MHz, Chloroform-*d*) *δ* (ppm) 171.9, 134.0, 129.2, 128.5, 127.1, 51.9, 41.1.

HRMS (APCI): calcd. for C_9_H_11_O_2_ [M+H]^+^ = 151.0754, found [M+H]^+^ = 151.0753.

3-(3-Methylfuran-2-yl)-3-oxopropanenitrile **(S114)**

The compound was prepared according to General procedure A5 using NaH (60% suspension in mineral oil, 171 mg, 4.28 mmol), acetonitrile (0.134 mL, 2.57 mmol), anhydrous THF (3 mL) and methyl 3-methylfuran-2-carboxylate (300 mg, 2.14 mmol) in anhydrous THF (2 mL). The reaction mixture was refluxed for 48 h. The residue was purified by column chromatography (hexane:EtOAc, 7:3 to 8:4). The product was obtained as an off-white solid (171 mg, 54%).

^1^H NMR (300 MHz, Chloroform-*d*) *δ* (ppm) 7.47 (d, *J* = 1.8 Hz, 1H), 6.47 (d, *J* = 1.7 Hz, 1H), 3.94 (s, 2H), 2.41 (s, 3H).

13C NMR (75 MHz, Chloroform-d) *δ* (ppm) 177.01, 146.53, 146.09 (d, *J* = 2.6 Hz), 134.12, 116.75, 113.72, 29.40, 11.80.

HRMS (APCI): calcd. for C_8_H_8_NO_2_ [M+H]^+^ = 150.0550, found [M+H]^+^ = 150.0550.

mp = 92−95 °C.

3-(5-Methylfuran-2-yl)-3-oxopropanenitrile **(S115)**

The compound was prepared according to General procedure A5 using NaH (60% suspension in mineral oil, 171 mg, 4.28 mmol), acetonitrile (0.134 mL, 2.57 mmol), anhydrous THF (3 mL) and methyl 3-methylfuran-2-carboxylate (300 mg, 2.14 mmol) in anhydrous THF (2 mL). The reaction mixture was refluxed for 48 h. The residue was purified by column chromatography (hexane:EtOAc, 7:3 to 8:4). The product was obtained as a pale yellow solid (191 mg, 60%).

^1^H NMR (300 MHz, Chloroform-*d*) *δ* (ppm) 7.29 (d, *J* = 3.6 Hz, 1H), 6.30 – 6.19 (m, 1H), 3.89 (s, 2H), 2.42 (s, 3H).

^13^C NMR (75 MHz, Chloroform-*d*) *δ* (ppm) 174.7, 159.6, 149.2, 121.3, 113.6, 110.2, 28.4, 14.1.

HRMS (APCI): calcd. for C_8_H_8_NO_2_ [M+H]^+^ = 150.0550, found [M+H]^+^ = 150.0549.

mp = 97−100 °C.

3-Oxo-4-(thiophen-2-yl)butanenitrile **(S116)**

*n*-BuLi (2.7 M in heptane; 2.37 mL, 6.40 mmol) was added at -78 ºC dropwise to a solution of acetonitrile (334 µL, 6.40 mmol) in THF (5 mL) and the reaction mixture was stirred at -78 ºC for 1 h. Then, a solution of the methyl 2-(thiophen-2-yl)acetate (500 mg, 3.20 mmol; CAS: 19432-68-9) in THF (3 mL) was added dropwise and the reaction mixture was stirred at -78 ºC for 1 h and then at room temperature for additional 1 h. Aqueous saturated solution of NH_4_Cl (15 mL) was added and the mixture was extracted with EtOAc (2 × 50 mL). The combined organic extracts were dried over MgSO_4_, filtered, and the solvent was evaporated *in vacuo*. The residue was purified by column chromatography (hexane:EtOAc, 7:3). The product was obtained as an orange oil (360 mg, 68%).

^1^H NMR (500 MHz, Chloroform-*d*) *δ* (ppm) 7.29 (dd, *J* = 5.1, 1.2 Hz, 1H), 7.02 (dd, *J* = 5.2, 3.5 Hz, 1H), 6.99 – 6.95 (m, 1H), 4.06 (s, 2H), 3.53 (s, 2H).

^13^C NMR (126 MHz, Chloroform-*d*) *δ* (ppm) 194.00, 132.77, 128.14, 127.73, 126.37, 113.49, 43.01, 30.99 ppm.

HRMS (APCI): calcd. for C_8_H_6_NOS [M-H]^-^ = 164.0176, found [M-H]^-^ = 164.0178.

**Preparation of hydazine derivatives**

**Scheme S1:** synthesis of 2-hydrazinyl-6-methylpyrimidin-4(3*H*)-one **(S119)**

6-Methyl-2-thioxo-2,3-dihydropyrimidin-4(1*H*)-one **(S117)**

Thiourea (10 g, 130 mmol) was added to a solution of sodium hydroxide (10.9 g, 272 mmol) in H_2_O (205 mL) at 15 °C. The reaction mixture was stirred at 15 °C for 10 min,  ethyl acetoacetate (20.8 g, 160 mmol) was added dropwise at 15 °C. The mixture was allowed to warm to room temperature, and stirred for 3 h. The pH was adjusted to 4-5 by the careful addition of hydrochloric acid. The precipitate was collected by filtration, washed with cold water (20 mL) and dried *in vacuo*. The product was obtained as an off-white solid (7.18 g, 39%).

^1^H NMR (300 MHz, DMSO-*d*_6_) *δ* (ppm) 12.23 (s, 2H), 5.67 (d, *J* = 1.1 Hz, 1H), 2.06 (s, 3H).

HRMS (APCI): calcd. for C_5_H_5_N_2_OS [M-H]^-^ = 141.0128, found [M-H]^-^ = 141.0128.

mp >265 °C (dec.)

6-Methyl-2-(methylthio)pyrimidin-4(3*H*)-one **(S118)**

6-methyl-2-thioxo-2,3-dihydropyrimidin-4(1*H*)-one (7.18 g, 50.5 mmol) was added to a solution of NaOH (2.08 g, 52.01 mmol) in water (68 mL) and the reaction mixture was stirred at 25 °C for 20 min. lodomethane (3.92 mL, 63.12 mmol) was added dropwise and the mixture was stirred at room temperature for additional 4 h. The precipitate was collected by filtration, washed with ice cold water (2 × 20 mL) and dried *in vacuo*. The product was obtained as a white solid (7.8 g, 99%).

^1^H NMR (300 MHz, DMSO-*d*_6_) *δ* (ppm) 12.40 (s, 1H), 5.96 (s, 1H), 2.47 (s, 3H), 2.17 (s, 3H).

HRMS (APCI): calcd. for C_6_H_9_N_2_OS [M+H]^+^ = 157.0430, found [M+H]^+^ = 157.0428.

mp = 225−226 °C.

2-Hydrazinyl-6-methylpyrimidin-4(3*H*)-one **(S119)**

To a stirred solution of 6-methyl-2-(methylthio)pyrimidin-4(3*H*)-one (7.8 g, 49.93 mmol) in EtOH (20 mL) was added hydrazine hydrate (64% aq. solution, 10.18 g, 203 mmol) and the reaction mixture was stirred at 80 °C for 6 h. Then mixture was cooled to the room temperature and the resulting precipitate was collected by vacuum filtration, washed with water (2 mL) and dried *in vacuo*. The product was obtained as an off-white solid (4.15 g, 59%).

^1^H NMR (300 MHz, DMSO-*d*_6_) *δ* (ppm) 8.89 (s, 2H), 5.37 (s, 1H), 4.70 (s, 1H), 2.00 (s, 3H).

HRMS (APCI): calcd. for C_5_H_9_N_4_O [M+H]^+^ = 141.0771, found [M+H]^+^ = 141.0770.

mp = 231−233 °C.

2-Hydrazinyl-6-(trifluoromethyl)pyrimidin-4(3*H*)-one **(S120)**

To a stirred solution of 2-(methylthio)-6-(trifluoromethyl)-3,4-dihydropyrimidin-4-ol (500 mg, 2.38 mmol) in *i*-PrOH (20 mL) was added hydrazine hydrate (64% aq. solution, 930 mg, 11.89 mmol) and the mixture was stirred at 80 °C for 16 h. The solvent was evaporated *in vacuo* and the residue was purified by column chromatography on silica gel (23% aqueous NH_3_:methanol:dichloromethane; 2:8:90). The product was obtained as a white solid (171 mg, 37%).

^1^H NMR (500 MHz, DMSO-*d*_6_) *δ* (ppm) 9.25 (br s, 1H), 6.16 (br s, 1H), 5.89 (s, 1H).

^13^C NMR (126 MHz, DMSO-*d*_6_) *δ* (ppm) 161.8, 158.5, 153.5 (q, *J* = 33.3 Hz), 120.9 (q, *J* = 275.3 Hz), 98.5.

^19^F NMR (471 MHz, DMSO-*d*_6_) *δ* (ppm) -70.5.

HRMS (APCI): calcd. for C_5_H_6_F_3_N_4_O [M+H]^+^ = 195.0488, found [M+H]^+^ = 195.0490.

mp = 221−224 °C.

6-Ethyl-2-hydrazinylpyrimidin-4(3*H*)-one **(S121)**

To a stirred solution of 6-ethyl-2-(methylthio)-3,4-dihydropyrimidin-4-ol (400 mg, 2.35 mmol) in EtOH (1.2 mL) was added hydrazine hydrate (64% aq. solution, 735 mg, 9.40 mol) and the reaction mixture was stirred at 80 °C for 6 h. The solvent was evaporated *in vacuo* and the residue was mixed with EtOAc:hexane (1:4, 3 mL). The solid was collected by filtration, washed with a mixture of H_2_O:MeOH (5:95, 2 mL) and dried *in vacuo*. The product was obtained as a white solid (205 mg, 57%).

^1^H NMR (300 MHz, DMSO-*d*_6_) *δ* (ppm) 8.50 (s, 2H), 5.36 (s, 1H), 4.41 (s, 1H), 2.27 (q, *J* = 7.5 Hz, 2H), 1.08 (t, *J* = 7.6 Hz, 3H).

^13^C NMR (75 MHz, DMSO-*d*_6_) *δ* (ppm) 163.3, 157.7, 98.9, 30.4, 12.8.

HRMS (APCI): calcd. for C_6_H_11_N_4_O [M+H]^+^ = 155.0927, found [M+H]^+^ = 155.0928.

mp >180 °C (dec.)

**Scheme S2:** synthesis of 2-hydrazinyl-5-(2-(2-(2-methoxyethoxy)ethoxy)ethyl)-6-methylpyrimidin-4(3*H*)-one **(S126)**

1-Chloro-2-(2-(2-methoxyethoxy)ethoxy)ethane **(S122)**

Iodomethane (2.21 mL, 35.58 mmol) was added to a solution of 2-(2-(2-chloroethoxy)ethoxy)ethan-1-ol (3 g, 17.79 mmol) in DMF (30 mL), the mixture was cooled to 0 °C and sodium hydride (60% suspension in mineral oil, 1.067 g, 26.68 mmol) was added portion wise. The reaction mixture was stirred at 65 °C for 4 h, then poured into ice cold water (100 mL) and extracted with EtOAc (4 × 30 mL). The combined organic extracts were dried over MgSO_4_, ﬁltered, and the solvent was evaporated *in vacuo*. The resulting yellow oil was puriﬁed by ﬂash chromatography (hexane:EtOAc, 7:3). The product was obtained as a colorless oil (1.95 g, 60%).

^1^H NMR (300 MHz, Chloroform-*d*) *δ* (ppm) 3.80 – 3.72 (m, 2H), 3.72 – 3.58 (m, 8H), 3.55 (m, 2H), 3.37 (s, 3H).

^13^C NMR (75 MHz, Chloroform-*d*) *δ* (ppm) 72.1, 71.5, 70.8, 70.7, 70.7, 59.1, 42.8.

HRMS (APCI): calcd. for C_7_H_16_ClO_3_ [M+H]^+^ = 183.0782, found [M+H]^+^ = 183.0782.

1-Iodo-2-(2-(2-methoxyethoxy)ethoxy)ethane **(S123)**

NaI (5.82 g, 38.87 mmol) was added to a solution of 1-chloro-2-(2-(2-methoxyethoxy)ethoxy)ethane (3.35 g, 19.43 mmol) in acetone (40 mL) and the reaction mixture was refluxed for 48 h. The solvent was evaporated *in vacuo*, the residue was mixed with EtOAc (50 mL), and the mixture was washed with 5% aqueous Na_2_S_2_O_3_ solution (2× 20 mL) followed by water (2× 20 mL). The organic layer was dried over MgSO_4_, ﬁltered, and the solvent was evaporated *in vacuo*. The product was obtained as a light yellow oil (3.05 g, 63%).

^1^H NMR (300 MHz, Chloroform-*d*) *δ* (ppm) 3.79 – 3.68 (m, 2H), 3.69 – 3.59 (m, 6H), 3.58 – 3.49 (m, 2H), 3.36 (s, 3H), 3.30 – 3.19 (m, 2H).

^13^C NMR (75 MHz, Chloroform-*d*) *δ* (ppm) 72.1, 72.1, 70.7, 70.7, 70.3, 59.1, 3.0.

HRMS (APCI): calcd. for C_7_H_16_IO_3_ [M+H]^+^ = 275.0139, found [M+H]^+^ = 275.0139.

Ethyl 2-acetyl-4-(2-(2-methoxyethoxy)ethoxy)butanoate **(S124)**

KHMDS (1M solution in THF, 2.01 mL, 2.01 mmol) was added dropwise at -78 °C to a stirred solution of ethylacetoacetate (237 mg, 1.82 mmol) in THF (5 mL) under nitrogen atmosphere over the period of 10 min. The reaction mixture was allowed to warm to room temperature and stirred for 30 min. A solution of 1-iodo-2-(2-(2-methoxyethoxy)ethoxy)ethane (500 mg, 1.82 mmol) in THF (4 mL) was added and the reaction mixture was stirred for 16 h. The solvents were evaporated *in vacuo* and the residue was puriﬁed by column chromatography on silica gel (hexane:EtOAc, 6:4). The product was obtained as a light brown oil (360 mg, 71%).

^1^H NMR (300 MHz, Chloroform-*d*) *δ* (ppm) 4.27 – 4.06 (m, 2H), 3.73 – 3.41 (m, 11H), 3.37 (s, 3H), 2.25 (s, 3H), 2.21 – 2.04 (m, 2H), 1.26 (t, *J* = 7.1 Hz, 3H).

^13^C NMR (75 MHz, Chloroform-*d*) *δ* (ppm) 203.2, 169.8, 72.1, 70.6, 70.3, 68.6, 61.4, 59.1, 56.6, 29.4, 28.3, 14.2.

HRMS (APCI): calcd. for C_13_H_25_O_6_ [M+H]^+^ = 277.1646, found [M+H]^+^ = 277.1647.

5-(2-(2-(2-Methoxyethoxy)ethoxy)ethyl)-6-methyl-2-thioxo-2,3-dihydropyrimidin-4(1*H*)-one **(S125)**

NaOEt (2.6 M solution in EtOH, 4.17 mL, 10.86 mmol) was added to a solution of ethyl 2-acetyl-4-(2-(2-methoxyethoxy)ethoxy)butanoate (750 mg, 2.71 mmol) in EtOH (8 mL) under nitrogen atmosphere and the mixture was refluxed for 16 h. The solvent was evaporated under *in vacuo* and the residue was puriﬁed by column chromatography (dichloromethane:MeOH, 96:4). The product was obtained as a light brown gum (360 mg, 46%).

^1^H NMR (300 MHz, Chloroform-*d*) *δ* (ppm) 10.38 (s, 1H), 10.17 (s, 1H), 3.65 – 3.52 (m, 10H), 3.38 (s, 3H), 2.65 (t, *J* = 6.1 Hz, 2H), 2.26 (s, 3H).

^13^C NMR (75 MHz, Chloroform-*d*) *δ* (ppm) 173.9, 161.5, 149.8, 113.2, 72.0, 70.6, 70.4, 69.3, 59.0, 25.6, 17.2.

HRMS (APCI): calcd. for C_12_H_21_N_2_O_4_S [M+H]^+^ = 289.1217, found [M+H]^+^ = 289.1217.

2-Hydrazinyl-5-(2-(2-(2-methoxyethoxy)ethoxy)ethyl)-6-methylpyrimidin-4(3*H*)-one **(S126)**

Hydrazine hydrate (64% aq. solution, 1.38 g, 17.69 mmol) was added to a solution of 5-(2-(2-(2-methoxyethoxy)ethoxy)ethyl)-6-methyl-2-thioxo-2,3-dihydropyrimidin-4(1*H*)-one (340 mg, 1.18 mmol) in EtOH (5 mL) and the mixture was stirred at 80 °C for 16 h. The solvent was evaporated *in vacuo*, and the residue was co-evaporated with toluene (2 × 3 mL). The product was obtained as an off-white gum (340 mg), was used into the next step without further purification.

^1^H NMR (300 MHz, DMSO-*d*_6_) *δ* (ppm) 3.51 – 3.46 (m, 6H), 3.45 – 3.39 (m, 2H), 3.35 (t, *J* = 6.7 Hz, 2H), 3.24 (s, 3H), 2.56 – 2.49 (m, 2H), 2.08 (d, *J* = 1.4 Hz, 3H).

HRMS (APCI): calcd. for C_12_H_23_N_4_O_4_ [M+H]^+^ = 287.1714, found [M+H]^+^ = 287.1714.

2-(Benzyloxy)-6-chloropyridine **(S127)**

Benzyl bromide (458 µL, 3.86 mmol) and K_2_CO_3_ (1.07 g, 7.72 mmol) were added to a solution of 6-chloropyridin-2-ol (500 mg, 3.86 mmol) in DMF (5 mL) and the reaction mixture was stirred at 50 °C for 16 h. Water (35 mL) was added and the mixture was extracted with EtOAc (2 × 50 mL). The organic extracts were separated, dried over MgSO_4_, filtered, and the solvent was evaporated *in vacuo*. The residue was purified by column chromatography (hexane:EtOAc, 90:10). The product was obtained as a colorless oil (580 mg, 68%).

^1^H NMR (500 MHz, Chloroform-*d*): *δ* (ppm) 7.53 (td, *J* = 7.7, 3.7 Hz, 1H), 7.50 – 7.43 (m, 2H), 7.43 – 7.37 (m, 2H), 7.36 – 7.30 (m, 1H), 6.92 (dd, *J* = 7.5, 4.0 Hz, 1H), 6.72 (dd, *J* = 8.3, 3.9 Hz, 1H), 5.38 (d, *J* = 3.9 Hz, 2H).

^13^C NMR (126 MHz, Chloroform-*d*): *δ* (ppm) 163.43, 148.45, 140.81, 136.79, 128.65, 128.41, 128.20, 116.65, 109.56, 68.50.

2-(Benzyloxy)-6-hydrazinylpyridine **(S128)**

Hydrazine hydrate (64% aq. Solution, 794 µL, 25.49 mmol) was added to a solution of 2-(benzyloxy)-6-chloropyridine (560 mg, 2.55 mmol) in EtOH (6 mL) and the reaction mixture was stirred in the microwave reactor at 140 °C for 40 min. The solvents were evaporated *in vacuo* and the residue was purified by column chromatography (dichloromethane:MeOH, 90:10). The product was obtained as a pale yellow wax (150 mg, 27%).

^1^H NMR (300 MHz, Chloroform-*d*) *δ* (ppm) 7.46 – 7.42 (m, 2H), 7.40 – 7.35 (m, 3H), 7.34 – 7.30 (m, 1H), 6.22 (dd, *J* = 13.9, 7.9 Hz, 2H), 5.35 – 5.26 (m, 3H), 3.28 (br s, 2H).

**Preparation of aminopyrazole derivatives**

4-(5-Amino-1-(4-methyl-6-oxo-1,6-dihydropyrimidin-2-yl)-1*H*-pyrazol-3-yl)benzonitrile **(S129)**

The compound was prepared according to General procedure B2 using methanesulfonic acid (2.7 mg, 0.029 mmol), 2-hydrazinyl-6-methylpyrimidin-4(3*H*)-one (40 mg, 0.29 mmol) and 4-(2-cyanoacetyl)benzonitrile (68 mg, 0.399 mmol) in EtOH (2 mL). The reaction mixture was refluxed for 4 h. The reaction mixture was cooled to room temperature, NH_3_ (7 M solution in MeOH, 2 mL) was added, followed by saturated aqueous NaHCO_3_ solution (4 mL), and the mixture was stirred for 10 min. The precipitate was collected by filtration, washed with H_2_O (2 × 3 mL), then with a mixture of EtOAc:hexanes (1:4, 2 mL) and dried *in vacuo*. The product was obtained as a white solid (64 mg, 65%).

^1^H NMR (300 MHz, DMSO-*d*_6_) *δ* (ppm) 12.14 (s, 1H), 8.15 (d, *J* = 8.4 Hz, 2H), 7.88 (d, *J* = 8.3 Hz, 2H), 7.12 (br s, 2H), 6.11 (s, 1H), 5.98 (s, 1H), 2.29 (s, 3H).

HRMS (APCI): calcd. for C_15_H_11_N_6_O [M-H]^-^ = 291.1000, found [M-H]^-^ = 291.1000.

mp >301 °C (dec.)

2-(5-Amino-3-cyclohexyl-1*H*-pyrazol-1-yl)-6-methylpyrimidin-4(3*H*)-one **(S130)**

The compound was prepared according to General procedure B2 using methanesulfonic acid (2.7 mg, 0.029 mmol), 2-hydrazinyl-6-methylpyrimidin-4(3*H*)-one (40 mg, 0.29 mmol) and 3-cyclohexyl-3-oxopropanenitrile (43 mg, 0.29 mmol) in EtOH (2 mL). The reaction mixture was refluxed for 4 h. The crude product was purified by column chromatography on silica gel (7 NH_3_ in MeOH:MeOH: dichloromethane, 3:7:90). The product was obtained as a white solid (67 mg, 86%).

^1^H NMR (300 MHz, Chloroform-*d*) *δ* (ppm) 10.2 (s, 1H), 6.25 (s, 1H), 6.02 (s, 1H), 5.92 (s, 1H), 5.34 (s, 1H), 2.52 (t, *J* = 10.8 Hz, 1H), 2.28 (d, *J* = 0.9 Hz, 3H), 2.00 – 1.66 (m, 5H), 1.47 – 1.18 (m, 5H).

^13^C NMR (75 MHz, Chloroform-*d*) *δ* (ppm) 164.4, 162.5, 161.5, 149.5, 148.8, 108.3, 88.1, 37.9, 32.3, 26.2, 26.2, 24.0.

HRMS (APCI): calcd. for C_14_H_20_N_5_O [M+H]^+^ = 274.1662, found [M+H]^+^ = 274.1660.

mp = 221−224 °C.

2-(5-Amino-3-(4-(trifluoromethyl)phenyl)-1*H*-pyrazol-1-yl)-6-methylpyrimidin-4(3*H*)-one **(S131)**

The compound was prepared according to General procedure B2 using methanesulfonic acid (2.7 mg, 0.029 mmol), 2-hydrazinyl-6-methylpyrimidin-4(3*H*)-one (40 mg, 0.29 mmol) and 3-oxo-3-(4-(trifluoromethyl)phenyl)propanenitrile (61 mg, 0.29 mmol) in EtOH (2 mL). The reaction mixture was refluxed for 4 h. The reaction mixture was cooled to room temperature, NH_3_ (7 M solution in MeOH, 2 mL) was added, followed by saturated aqueous NaHCO_3_ solution (4 mL), and the mixture was stirred at room temperature for 10 min. The precipitated was collected by filtration, washed with H_2_O (2 × 3 mL), then with a mixture of EtOAc:hexane (1:4, 2 mL), and dried *in vacuo*. The product was obtained as a white solid (69 mg, 72%).

^1^H NMR (500 MHz, DMSO-*d*_6_) *δ* (ppm) 12.18 (s, 1H), 8.17 (d, *J* = 8.1 Hz, 2H), 7.77 (d, *J* = 8.2 Hz, 2H), 7.11 (s, 2H), 6.09 (s, 1H), 5.96 (s, 1H), 2.28 (s, 3H).

^13^C NMR (126 MHz, DMSO-*d*_6_) *δ* (ppm) 151.5, 150.9, 136.4, 128.7, 128.4, 128.2, 126.5, 125.3, 125.3, 125.3, 123.2, 85.4, 22.9.

^19^F NMR (282 MHz, DMSO-*d*_6_) *δ* (ppm) -61.0.

HRMS (APCI): calcd. for C_15_H_13_F_3_N_5_O [M+H]^+^ = 336.1067, found [M+H]^+^ = 336.1070.

mp = 284−288 °C.

2-(5-Amino-3-(4-(tert-butyl)phenyl)-1*H*-pyrazol-1-yl)-6-methylpyrimidin-4(3*H*)-one **(S132)**

The compound was prepared according to General procedure B2 using methanesulfonic acid (2.7 mg, 0.029 mmol), 2-hydrazinyl-6-methylpyrimidin-4(3*H*)-one (40 mg, 0.29 mmol) and 3-(4-(*tert*-butyl)phenyl)-3-oxopropanenitrile (57 mg, 0.29 mmol) in EtOH (2 mL). The reaction mixture was refluxed for 4 h. The reaction mixture was cooled to room temperature, NH_3_ (7 M solution in MeOH, 2 mL) was added, followed by saturated aqueous NaHCO_3_ solution (4 mL), and the mixture was stirred for 10 min. The precipitate was collected by filtration, washed with H_2_O (2 × 3 mL), then with a mixture of EtOAc:hexane (1:4, 2 mL), and dried *in vacuo*. The product was obtained as a white solid (64 mg, 69%).

^1^H NMR (500 MHz, DMSO-*d*_6_) *δ* (ppm) 7.77 (d, *J* = 8.4 Hz, 2H), 7.41 (d, *J* = 8.5 Hz, 2H), 7.02 (s, 2H), 5.83 (s, 1H), 5.76 (s, 1H), 2.18 (d, *J* = 0.8 Hz, 3H), 1.31 (s, 9H).

^13^C NMR (126 MHz, DMSO-*d*_6_) *δ* (ppm) 162.3, 154.5, 151.0, 150.9, 150.5, 130.3, 125.5, 125.0, 106.4, 84.9, 34.3, 31.1, 23.0.

HRMS (APCI): calcd. for C_28_H_22_N_5_O [M+H]^+^ = 324.1819, found [M+H]^+^ = 324.1817.

mp = 231−250 °C.

2-(5-Amino-3-(4-methoxyphenyl)-1*H*-pyrazol-1-yl)-6-methylpyrimidin-4(3*H*)-one **(S133)**

The compound was prepared according to General procedure B2 using methanesulfonic acid (2.7 mg, 0.029 mmol), 2-hydrazinyl-6-methylpyrimidin-4(3*H*)-one (40 mg, 0.29 mmol) and 3-(4-methoxyphenyl)-3-oxopropanenitrile (60 mg, 0.34 mmol) in EtOH (2 mL). The reaction mixture was refluxed for 16 h. The reaction mixture was cooled to room temperature, NH_3_ (7 M solution in MeOH, 2 mL) was added, followed by saturated aqueous NaHCO_3_ solution (4 mL), and the mixture was stirred for 10 min. The precipitate was collected by filtration and washed with H_2_O (2 × 3 mL), then with a mixture of EtOAc:hexane (1:4, 2 mL), and dried *in vacuo*. The product was obtained as a white solid (65 mg, 77%).

^1^H NMR (500 MHz, DMSO-*d*_6_) *δ* (ppm) 11.77 (s, 1H), 7.88 (d, *J* = 8.3 Hz, 2H), 7.02 (s, 2H), 6.98 (d, *J* = 8.9 Hz, 2H), 6.04 (s, 1H), 5.81 (s, 1H), 3.80 (s, 3H), 2.27 (s, 3H).

^13^C NMR (126 MHz, DMSO-*d*_6_) *δ* (ppm) 159.8, 153.2, 151.1, 127.5, 124.7, 113.8, 84.8, 55.1, 23.2.

HRMS (APCI): calcd. for C_15_H_16_N_5_O_2_ [M+H]^+^ = 298.1299, found [M+H]^+^ = 298.1297.

mp = 289−293 °C.

2-(5-Amino-3-benzyl-1*H*-pyrazol-1-yl)-6-methylpyrimidin-4(3*H*)-one **(S134)**

The compound was prepared according to General procedure B2 using methanesulfonic acid (6 mg, 0.063 mmol), 2-hydrazinyl-6-methylpyrimidin-4(3*H*)-one (88 mg, 0.63 mmol) and 3-oxo-4-phenylbutanenitrile (100 mg, 0.63 mmol) in EtOH (3 mL). The reaction mixture was refluxed for 4 h. The crude product was purified by column chromatography on silica gel (7M NH_3_ in MeOH:MeOH: dichloromethane, 2:8:90). The product was obtained as a brown solid (98 mg, 56%).

^1^H NMR (300 MHz, Chloroform-*d*) *δ* (ppm) 7.38 – 7.18 (m, 6H), 6.03 (s, 1H), 5.90 (s, 2H), 5.27 (s, 1H), 3.84 (s, 2H), 2.28 (s, 3H).

^13^C NMR (75 MHz, DMSO-*d*_6_) *δ* (ppm) 154.8, 150.7, 139.1, 128.6, 128.3, 128.0, 126.1, 105.9, 87.7, 34.4, 22.6.

HRMS (APCI): calcd. for C_15_H_16_N_5_O [M+H]^+^ = 282.1349, found [M+H]^+^ = 282.1349.

2-(5-Amino-3-(3-methylfuran-2-yl)-1*H*-pyrazol-1-yl)-6-methylpyrimidin-4(3*H*)-one **(S135)**

The compound was prepared according to General procedure B2 using methanesulfonic acid (8 mg, 0.081 mmol), 2-hydrazinyl-6-methylpyrimidin-4(3*H*)-one (113 mg, 0.81 mmol) and 3-(3-methylfuran-2-yl)-3-oxopropanenitrile (120 mg, 0.81 mmol) in EtOH (3 mL). The reaction mixture was refluxed for 16 h. The crude product was purified by column chromatography on silica gel (7M NH_3_ in MeOH:MeOH:dichloromethane, 2:8:90). The product was obtained as a pale brown solid (53 mg, 24%).

^1^H NMR (300 MHz, DMSO-*d*_6_) *δ* (ppm) 7.56 (d, *J* = 1.7 Hz, 1H), 7.01 (s, 2H), 6.42 (d, *J* = 1.7 Hz, 1H), 5.72 (s, 1H), 5.56 (s, 1H), 2.27 (s, 3H), 2.13 (s, 3H).

^13^C NMR (75 MHz, DMSO-*d*_6_) *δ* (ppm) 170.7, 162.4, 156.2, 150.4, 143.9, 143.8, 141.4, 117.3, 114.6, 106.1, 85.0, 23.1, 10.7.

HRMS (APCI): calcd. for C_13_H_14_N_5_O_2_ [M+H]^+^ = 272.1142, found [M+H]^+^ = 272.1143.

mp >220 °C (dec.)

2-(5-Amino-3-(5-methylfuran-2-yl)-1*H*-pyrazol-1-yl)-6-methylpyrimidin-4(3*H*)-one **(S136)**

The compound was prepared according to General procedure B2 using methanesulfonic acid (8 mg, 0.081 mmol), 2-hydrazinyl-6-methylpyrimidin-4(3*H*)-one (113 mg, 0.81 mmol) and 3-(5-methylfuran-2-yl)-3-oxopropanenitrile (120 mg, 0.81 mmol) in EtOH (3 mL). The reaction mixture was refluxed for 16 h. The crude product was purified by column chromatography on silica gel (7M NH_3_ in MeOH:MeOH:dichloromethane, 2:8:90). The product was obtained as a light yellow solid (130 mg, 60%).

^1^H NMR (300 MHz, DMSO-*d*_6_) *δ* (ppm) 11.42 (s, 1H), 7.03 (s, 2H), 6.75 (d, *J* = 3.2 Hz, 1H), 6.20 (d, *J* = 3.2 Hz, 1H), 6.13 (s, 1H), 5.64 (s, 1H), 2.33 (s, 3H), 2.29 (s, 3H).

^13^C NMR (75 MHz, DMSO-*d*_6_) *δ* (ppm) 152.1, 150.8, 146.2, 145.3, 109.2, 105.8, 107.8, 84.7, 13.3.

HRMS (APCI): calcd. for C_13_H_14_N_5_O_2_ [M+H]^+^ = 272.1142, found [M+H]^+^ = 272.1142.

mp = 240−243 °C.

2-(5-Amino-3-phenyl-1*H*-pyrazol-1-yl)-6-methylpyrimidin-4(3*H*)-one **(S137)**

The compound was prepared according to General procedure B2 using methanesulfonic acid (21 mg, 0.21 mmol), 2-hydrazinyl-6-methylpyrimidin-4(3*H*)-one (300 mg, 2.14 mmol) and 3-oxo-3-phenylpropanenitrile (311 mg, 2.14 mmol) in EtOH (6 mL). The reaction mixture was refluxed for 4 h. The crude product was purified by column chromatography on silica gel (7M NH_3_ in MeOH:MeOH: dichloromethane, 3:7:90). The product was obtained as a yellow solid (540 mg, 94%).

^1^H NMR (500 MHz, DMSO-*d*_6_) *δ* (ppm) 11.87 (s, 1H), 8.06 – 7.84 (m, 2H), 7.52 – 7.29 (m, 3H), 7.04 (s, 2H), 6.09 (s, 1H), 5.88 (s, 1H), 2.28 (s, 3H).

^13^C NMR (126 MHz, DMSO-*d*_6_) *δ* (ppm) 164.3, 153.2, 151.7, 132.7, 129.2, 128.9, 126.6, 107.5, 85.7, 23.7.

HRMS (APCI): calcd. for C_14_H_14_N_5_O [M+H]^+^ = 268.1193, found [M+H]^+^ = 268.1193.

mp = 239−241 °C.

2-(5-Amino-3-(furan-2-yl)-1*H*-pyrazol-1-yl)-6-methylpyrimidin-4(3*H*)-one **(S138)**

The compound was prepared according to General procedure B2 using methanesulfonic acid (14 mg, 0.14 mmol), 2-hydrazinyl-6-methylpyrimidin-4(3*H*)-one (193 mg, 1.43 mmol) and 3-(furan-2-yl)-3-oxopropanenitrile (200 mg, 1.43 mmol) in EtOH (6 mL). The reaction mixture was refluxed for 4 h. The crude product was purified by column chromatography on silica gel (7M solution NH_3_ in MeOH:MeOH:dichloromethane, 3:7:90). The product was obtained as a light brown solid (165 mg, 45%).

^1^H NMR (500 MHz, DMSO-*d*_6_) *δ* (ppm) 7.73 (d, *J* = 1.8 Hz, 1H), 7.06 (s, 2H), 6.86 (d, *J* = 3.3 Hz, 1H), 6.59 (dd, *J* = 3.3, 1.8 Hz, 1H), 6.07 (s, 1H), 5.67 (s, 1H), 2.27 (s, 3H).

^13^C NMR (126 MHz, DMSO-*d*_6_) *δ* (ppm) 150.8, 148.1, 144.6, 143.0, 111.6, 107.6, 105.8, 84.8, 22.9.

HRMS (APCI): calcd. for C_12_H_12_N_5_O_2_ [M+H]^+^ = 258.0986, found [M+H]^+^ = 258.0983.

mp >165 °C (dec).

2-(5-amino-3-(thiophen-2-yl)-1*H*-pyrazol-1-yl)-6-methylpyrimidin-4(3*H*)-one **(S139)**

The compound was prepared by General Procedure B2 using methanesulfonic acid (6 mg, 0.06 mmol), 2-hydrazinyl-6-methylpyrimidin-4(3*H*)-one (90 mg, 0.6 mmol, 1 eq.) and 3-oxo-3-(thiophen-2-yl)propanenitrile (100 mg, 0.66 mmol, 1.1 eq.) in EtOH (3 mL). The reaction mixture was refluxed for 4 h. The reaction mixture was cooled to room temperature, and NH_3_ (7 M solution in MeOH, 2 mL) was added, followed by saturated aqueous solution of NaHCO_3_ (4 mL). The mixture was stirred at room temperature for 10 min, the precipitate was collected by filtration, washed with H_2_O (2 × 3 mL), then with a mixture of EtOAc:hexane (1:4, 2 mL), and dried under vacuum. The product was obtained as a white solid (75 mg, 42%) and used directly in the next step.

2-(5-Amino-3-(thiophen-2-ylmethyl)-1*H*-pyrazol-1-yl)-6-methylpyrimidin-4(3*H*)-one **(S140)**

The compound was prepared according to General procedure B2 using methanesulfonic acid (4.63 µL, 0.71 mmol), 2-hydrazinyl-6-methylpyrimidin-4(3*H*)-one (100 mg, 0.71 mmol; CAS: 37893-08-6) and 3-oxo-4-(thiophen-2-yl)butanenitrile (123 mg, 0.75 mmol) in EtOH (5 mL). The reaction mixture was refluxed for 4 h. The solvent was evaporated *in vacuo* and the residue was dissolved in dichloromethane (2 mL). Hexane (7 mL) was added and the resulting precipitate was collected by filtration, washed with hexane (5 mL), diethyl ether (3 mL) and dried *in vacuo*. The product was obtained as an off-white solid (130 mg, 63%).

^1^H NMR (500 MHz, DMSO-*d*_6_) *δ* (ppm) 7.75 (br s, 3H), 7.39 (dd, *J* = 5.1, 1.3 Hz, 1H), 7.03 – 6.99 (m, 1H), 6.98 (dd, *J* = 5.1, 3.4 Hz, 1H), 6.34 (s, 1H), 5.47 (s, 1H), 4.13 (s, 2H), 2.37 (s, 3H).

^13^C NMR (126 MHz, DMSO-*d*_6_) *δ* (ppm) 167.28, 165.61, 153.56, 151.78, 151.30, 139.67, 126.99, 126.19, 124.92, 104.90, 89.01, 27.36, 22.65.

HRMS (APCI): calcd. for C_13_H_14_N_5_OS [M+H]^+^ = 288.0914, found [M+H]^+^ = 288.0917.

2-(5-Amino-4-methyl-3-phenyl-1*H*-pyrazol-1-yl)-6-methylpyrimidin-4(3*H*)-one **(S141)**

The compound was prepared according to General procedure B2 using methanesulfonic acid (9.28 µL, 0.14 mmol), 2-hydrazinyl-6-methylpyrimidin-4(3*H*)-one (200 mg, 1.43 mmol; CAS: 37893-08-6) and 2-methyl-3-oxo-3-phenylpropanenitrile (227 mg, 1.43 mmol; CAS: 7391-29-9) in EtOH (5 mL). The reaction mixture was refluxed for 6 h. The reaction mixture was cooled to room temperature, NH_3_ (7 M solution in MeOH, 2 mL) was added and the mixture was stirred for 2 h. The mixture was concentrated *in vacuo* to the volume of ca. 3 mL, the precipitate was collected by filtration, washed with diethyl ether (10 mL) and dried *in vacuo*. The product was obtained as a pale yellow solid (370 mg, 92%).

^1^H NMR (500 MHz, DMSO-*d*_6_) *δ* (ppm) 11.74 (br s, 1H), 7.93 – 7.74 (m, 2H), 7.51 – 7.43 (m, 2H), 7.42 – 7.36 (m, 1H), 6.73 (br s, 2H), 6.09 (s, 1H), 2.28 (s, 3H), 2.05 (s, 3H).

^13^C NMR (126 MHz, DMSO-*d*_6_) *δ* (ppm) 164.29, 163.03, 152.19, 150.34, 147.94, 133.06, 128.29, 128.23, 127.60, 106.83, 94.12, 23.27, 8.18.

HRMS (APCI): calcd. for C_15_H_16_N_5_O [M+H]^+^ = 282.1349, found [M+H]^+^ = 282.1350.

2-(5-Amino-3-phenyl-1*H*-pyrazol-1-yl)-6-(trifluoromethyl)pyrimidin-4(3*H*)-one **(S142)**

The compound was prepared according to General procedure B2 using methanesulfonic acid (3 mg, 0.03 mmol), 2-hydrazinyl-6-(trifluoromethyl)pyrimidin-4(3*H*)-one (50 mg, 0.26 mmol) and 3-oxo-3-phenylpropanenitrile (37 mg, 0.26 mmol) in EtOH (2 mL). The reaction mixture was refluxed for 4 h. The crude product was purified by column chromatography on silica gel (7M NH_3_ in MeOH:MeOH:dichloromethane, 3:7:90). The product was obtained as a light brown solid (72 mg, 87%).

^1^H NMR (500 MHz, DMSO-*d*_6_) *δ* (ppm) 7.91 – 7.99 (m, 2H), 7.40 – 7.47 (m, 2H), 7.34 – 7.40 (m, 1H), 6.59 (s, 1H), 5.91 (s, 1H).

^13^C NMR (126 MHz, DMSO-*d*_6_) *δ* (ppm) 152.9, 151.3, 132.2, 128.7, 128.4, 126.1, 121.9, 120.8 (d, *J* = 273.7 Hz), 106.96, 85.4.

^19^F NMR (471 MHz, DMSO-*d*_6_) *δ* (ppm) -69.9.

HRMS (APCI): calcd. for C_14_H_11_F_3_N_5_O [M+H]^+^ = 322.0910, found [M+H]^+^ = 322.0909.

mp >295 °C (dec.)

2-(5-Amino-3-phenyl-1*H*-pyrazol-1-yl)-6-ethylpyrimidin-4(3*H*)-one **(S143)**

The compound was prepared according to General procedure B2 using methanesulfonic acid (6 mg, 0.07 mmol), 6-ethyl-2-hydrazinylpyrimidin-4(3*H*)-one (100 mg, 0.65 mmol) and 3-oxo-3-phenylpropanenitrile (94 mg, 0.65 mmol) in EtOH (2 mL). The reaction mixture was refluxed for 4 h. The solvent was evaporated *in vacuo* and saturated aqueous NaHCO_3_ solution (4 mL) was added to the residue. The solid was collected by filtration, washed with H_2_O (2 × 3 mL) and recrystallized from MeOH (4 mL). The product was obtained as a light brown solid (79 mg, 43%).

^1^H NMR (300 MHz, DMSO-*d*_6_) *δ* (ppm) 11.89 (s, 1H), 8.03 – 7.89 (m, 2H), 7.49 – 7.33 (m, 3H), 7.04 (s, 2H), 6.07 (s, 1H), 5.89 (s, 1H), 2.57 (q, *J* = 7.7 Hz, 2H), 1.19 (t, *J* = 7.5 Hz, 3H).

^13^C NMR (75 MHz, DMSO-*d*_6_) *δ* (ppm) 152.8, 151.2, 132.2, 128.7, 128.4, 126.0, 105.6, 85.2, 12.0.

HRMS (APCI): calcd. for C_15_H_16_N_5_O [M+H]^+^ = 282.1349, found [M+H]^+^ = 282.1348.

mp = 211−213 °C.

2-(5-Amino-3-phenyl-1*H*-pyrazol-1-yl)-5-(2-hydroxyethyl)-6-methylpyrimidin-4(3*H*)-one **(S144)**

The compound was prepared according to General procedure B2 using methanesulfonic acid (5 mg, 0.05 mmol), 2-hydrazinyl-5-(2-hydroxyethyl)-6-methylpyrimidin-4(3*H*)-one (100 mg, 0.54 mmol) and 3-oxo-3-phenylpropanenitrile (79 mg, 0.54 mmol) in EtOH (3 mL). The reaction mixture was refluxed for 4 h. The solvent evaporated *in vacuo* and water (5 mL) was added to the residue. The solid was collected by filtration, washed with H_2_O (3 mL), then with a mixture of EtOAc:hexane (1:4, 2 mL), and dried *in vacuo*. The product was obtained as an off-white solid (132 mg, 78%).

^1^H NMR (500 MHz, DMSO-*d*_6_) *δ* (ppm) 11.87 (s, 1H), 7.95 (d, *J* = 7.5 Hz, 2H), 7.42 (t, *J* = 7.4 Hz, 2H), 7.39 – 7.32 (m, 1H), 6.99 (s, 2H), 5.87 (s, 1H), 4.61 (t, *J* = 5.6 Hz, 1H), 3.50 (q, *J* = 6.6 Hz, 2H), 2.62 (t, *J* = 7.0 Hz, 2H), 2.33 (s, 3H).

^13^C NMR (126 MHz, DMSO-*d*_6_) *δ* (ppm) 160.3, 150.9. 132.3, 128.6, 128.4, 126.0, 116.0, 85.1, 59.3, 29.1, 21.4.

HRMS (APCI): calcd. for C_16_H_18_N_5_O_2_ [M+H]^+^ = 312.1455, found [M+H]^+^ = 312.1453.

mp = 221−223 °C.

3-Phenyl-1-(pyrimidin-2-yl)-1*H*-pyrazol-5-amine **(S145)**

The compound was prepared according to General procedure B2 using methanesulfonic acid (7 mg, 0.069 mmol), 2-hydrazinylpyrimidine (76 mg, 0.69 mmol) and 3-oxo-3-phenylpropanenitrile (100 mg, 0.69 mmol) in EtOH (3 mL). The reaction mixture was refluxed for 6 h. The crude product was purified by column chromatography on silica gel (7M NH_3_ in MeOH:MeOH:dichloromethane, 2:8:90). The product was obtained as a light brown solid (81 mg, 50%).

^1^H NMR (500 MHz, Chloroform-*d*) *δ* (ppm) 8.76 (d, *J* = 4.8 Hz, 2H), 7.96 – 7.86 (m, 2H), 7.42 – 7.37 (m, 2H), 7.36 – 7.31 (m, 1H), 7.14 (t, *J* = 4.8 Hz, 1H), 5.91 (s, 1H).

^13^C NMR (126 MHz, Chloroform-*d*) *δ* (ppm) 158.5, 158.1, 154.3, 150.5, 132.8, 128.7, 128.5, 126.6, 117.2, 87.9.

HRMS (APCI): calcd. for C_13_H_12_N_5_ [M+H]^+^ = 238.1087, found [M+H]^+^ = 238.1085.

mp = 202−205 °C.

1-(4-Methoxy-6-methylpyrimidin-2-yl)-3-phenyl-1*H*-pyrazol-5-amine **(S146)**

The compound was prepared according to General procedure B2 using methanesulfonic acid (2.4 mg, 0.026 mmol), 2-hydrazinyl-4-methoxy-6-methylpyrimidine (40 mg, 0.26 mmol) and 3-oxo-3-phenylpropanenitrile (38 mg, 0.26 mmol) in EtOH (2 mL). The reaction mixture was refluxed for 4 h. The crude product was purified by column chromatography on silica gel (7M NH_3_ in MeOH:MeOH:dichloromethane, 2:8:90). The product was obtained as an off-white solid (63 mg, 87%).

^1^H NMR (500 MHz, Chloroform-*d*) *δ* (ppm) 7.93 – 7.87 (m, 2H), 7.38 (td, *J* = 7.4, 1.6 Hz, 2H), 7.35 – 7.30 (m, 1H), 6.40 (s, 1H), 5.92 (s, 1H), 4.11 (s, 3H), 2.49 (s, 3H).

^13^C NMR (126 MHz, Chloroform-*d*) *δ* (ppm) 171.1, 168.5, 157.2, 153.6, 150.6, 132.7, 128.7, 128.5, 126.5, 103.0, 87.8, 54.4, 24.1.

HRMS (APCI): calcd. for C_15_H_16_N_5_O [M+H]^+^ = 282.1349, found [M+H]^+^ = 282.1351.

mp = 177−180 °C.

2-(5-Amino-3-phenyl-1*H*-pyrazol-1-yl)-6-(methoxymethyl)pyrimidin-4(3*H*)-one **(S147)**

The compound was prepared according to General procedure B2 using methanesulfonic acid (2.3 mg, 0.024 mmol), 2-hydrazinyl-6-(methoxymethyl)pyrimidin-4(3*H*)-one (40 mg, 0.24 mmol) and 3-oxo-3-phenylpropanenitrile (34 mg, 0.24 mmol) in EtOH (2 mL). The reaction mixture was refluxed for 4 h. The crude product was purified by column chromatography on silica gel (7M NH_3_ in MeOH:MeOH:dichloromethane, 2:8:90). The product was obtained as a white solid (59 mg, 86%).

^1^H NMR (500 MHz, DMSO-*d*_6_) *δ* (ppm) 11.98 (s, 1H), 7.96 (d, *J* = 7.5 Hz, 2H), 7.50 – 7.31 (m, 3H), 7.01 (s, 2H), 6.13 (s, 1H), 5.88 (s, 1H), 4.34 (s, 2H), 3.40 (s, 3H).

^13^C NMR (126 MHz, DMSO-*d*_6_) *δ* (ppm) 152.9, 151.2, 132.1, 128.7, 128.4, 126.1, 104.8, 85.1, 72.7, 58.3.

HRMS (APCI): calcd. for C_15_H_16_N_5_O_2_ [M+H]^+^ = 298.1299, found [M+H]^+^ = 298.1298.

mp = 218−221 °C.

2-(5-Amino-3-phenyl-1*H*-pyrazol-1-yl)-5-(2-(2-(2-methoxyethoxy)ethoxy)ethyl)-6-methylpyrimidin-4(3*H*)-one **(S148)**

The compound was prepared according to General procedure B2 using methanesulfonic acid (11 mg, 0.12 mmol), 2-hydrazinyl-5-(2-(2-(2-ethoxyethoxy)ethoxy)ethyl)-6-methylpyrimidin-4(3*H*)-one (330 mg (crude), 1.15 mmol) and 3-oxo-3-phenylpropanenitrile (167 mg, 1.15 mmol) in EtOH (5 mL). The reaction mixture was refluxed for 4 h. The crude product was purified by column chromatography on silica gel (7M NH_3_ in MeOH:MeOH:dichloromethane, 2:8:90). The product was obtained as a light brown solid (205 mg, 43%).

^1^H NMR (300 MHz, Chloroform-*d*) *δ* (ppm) 10.38 (s, 1H), 7.83 – 7.68 (m, 2H), 7.45 – 7.34 (m, 3H), 6.01 (s, 2H), 5.79 (s, 1H), 3.69 – 3.56 (m, 8H), 3.56 – 3.48 (m, 2H), 3.36 (s, 3H), 2.82 (t, *J* = 6.8 Hz, 2H), 2.34 (s, 3H).

^13^C NMR (75 MHz, Chloroform-*d*) *δ* (ppm) 154.1, 150.1, 146.4, 131.9, 129.3, 128.7, 126.2, 117.5, 87.2, 72.1, 70.8, 70.6, 70.4, 69.5, 59.1, 26.5, 22.0.

HRMS (APCI): calcd. for C_21_H_28_N_5_O_4_ [M+H]^+^ = 414.2136, found [M+H]^+^ = 414.2136.

mp = 143−146 °C.

2-(5-Amino-3-phenyl-1*H*-pyrazol-1-yl)-5,6-dimethylpyrimidin-4(3*H*)-one **(S149)**

The compound was prepared according to General procedure B2 using methanesulfonic acid (6.2 mg, 0.065 mmol), 2-hydrazinyl-5,6-dimethylpyrimidin-4(3*H*)-one (100 mg, 0.65 mmol) and 3-oxo-3-phenylpropanenitrile (94 mg, 0.65 mmol) in EtOH (3 mL). The reaction mixture was refluxed for 4 h. The crude product was purified by column chromatography on silica gel (7M NH_3_ in MeOH:MeOH:dichloromethane, 2:8:90). The product was obtained as a white solid (102 mg, 56%).

^1^H NMR (300 MHz, DMSO-*d*_6_) *δ* (ppm) 7.82 – 7.73 (m, 2H), 7.43 – 7.33 (m, 2H), 7.33 – 7.24 (m, 1H), 6.98 (s, 2H), 5.71 (s, 1H), 2.12 (s, 3H), 1.85 (s, 3H).

^13^C NMR (75 MHz, DMSO-*d*_6_) *δ* (ppm) 171.9, 156.6, 155.2, 150.8, 149.7, 133.6, 128.3, 127.6, 125.6, 112.4, 85.0, 21.2, 11.4.

HRMS (APCI): calcd. for C_15_H_16_N_5_O [M+H]^+^ = 282.1349, found [M+H]^+^ = 282.1348.

mp = 276−281 °C.

6-(5-Amino-3-phenyl-1*H*-pyrazol-1-yl)-1,3-dimethylpyrimidine-2,4(1*H*,3*H*)-dione **(S150)**

The compound was prepared according to General procedure B2 using methanesulfonic acid (6 mg, 0.0 mmol), 6-hydrazinyl-1,3-dimethylpyrimidine-2,4(1*H*,3*H*)-dione (100 mg, 0.59 mmol) and 3-oxo-3-phenylpropanenitrile (85 mg, 0.59 mmol) in EtOH (3 mL). The reaction mixture was refluxed for 4 h. The crude product was purified by column chromatography on silica gel (7M NH_3_ in MeOH:MeOH:dichloromethane, 2:8:90). The product was obtained as a white solid (141 mg, 81%).

^1^H NMR (300 MHz, Chloroform-*d*) *δ* (ppm) 7.82 – 7.66 (m, 2H), 7.48 – 7.29 (m, 3H), 5.91 (s, 1H), 5.90 (s, 1H), 4.14 (s, 2H), 3.37 (s, 3H), 3.28 (s, 3H).

^13^C NMR (75 MHz, Chloroform-*d*) *δ* (ppm) 162.4, 154.5, 152.1, 147.9, 146.1, 132.3, 129.0, 128.8, 125.9, 99.4, 87.8, 32.5, 28.5.

HRMS (APCI): calcd. for C_15_H_16_N_5_O_2_ [M+H]^+^ = 298.1299, found [M+H]^+^ = 298.1300.

mp = 158−160 °C.

2-(5-Amino-3-phenyl-1*H*-pyrazol-1-yl)-3,6-dimethylpyrimidin-4(3*H*)-one **(S151)**

The compound was prepared according to General procedure B2 using methanesulfonic acid (9.3 µL, 0.097 mmol), 2-hydrazinyl-3,6-dimethylpyrimidin-4(3*H*)-one (150 mg, 0.97 mmol; CAS:3493-94-5) and 3-oxo-3-phenylpropanenitrile (141 mg, 0.97 mmol) in EtOH (5 mL). The reaction mixture was refluxed for 3 h and then quenched with addition of 7M NH_3_ in MeOH (2 mL). The crude product was purified by column chromatography on silica gel (EtOAc:dichloromethane, 20:80). The product was obtained as a pale yellow solid (80 mg, 29%) containing traces of residual EtOAc, which was used directly in the next step.

^1^H NMR (500 MHz, Chloroform-*d*) *δ* (ppm) 7.81 – 7.75 (m, 2H), 7.44 – 7.39 (m, 2H), 7.39 – 7.34 (m, 1H), 6.24 (d, *J* = 0.9 Hz, 1H), 5.91 (s, 1H), 5.09 (br s, 2H), 3.78 (s, 3H), 2.28 (s, 3H).

^13^C NMR (126 MHz, Chloroform-*d*) *δ* (ppm) 163.50, 161.26, 153.21, 150.30, 132.42, 129.01, 128.80, 126.04, 110.05, 105.71, 88.34, 34.38, 23.49.

HRMS (APCI): calcd. for C_15_H_16_N_5_O [M+H]^+^ = 282.1349, found [M+H]^+^ = 282.1348.

1-(6-(Benzyloxy)pyridin-2-yl)-3-phenyl-1*H*-pyrazol-5-amine **(S152)**

The compound was prepared according to General procedure B2 using methanesulfonic acid (4.5 µL, 0.07 mmol), 2-(benzyloxy)-6-hydrazinylpyridine (150 mg, 0.70 mmol) and 3-oxo-3-phenylpropanenitrile (101 mg, 0.70 mmol) in EtOH (5 mL). The reaction mixture was refluxed for 3 h. The crude product was purified by column chromatography on silica gel (EtOAc:hexane; 10:90) followed by trituration with hexanes (5 mL) and drying *in vacuo*. The product was obtained as an off-white solid (50 mg, 21%).

^1^H NMR (500 MHz, DMSO-*d*_6_) *δ* (ppm) 7.90 (t, *J* = 8.0 Hz, 1H), 7.84 – 7.78 (m, 2H), 7.54 (d, *J* = 7.9 Hz, 1H), 7.51 – 7.46 (m, 2H), 7.45 – 7.38 (m, 4H), 7.37 – 7.32 (m, 2H), 6.76 (d, *J* = 8.1 Hz, 1H), 6.54 (br s, 2H), 5.86 (br s, 1H), 5.38 (s, 2H).

^13^C NMR (126 MHz, DMSO-*d*_6_) *δ* (ppm) 161.41, 151.80, 151.20, 149.99, 141.84, 136.62, 132.96, 128.48, 128.44, 128.05, 127.88, 127.64, 125.37, 105.81, 105.50, 85.64, 67.69.

HRMS (APCI): calcd. for C_21_H_19_N_4_O [M+H]^+^ = 343.1553, found [M+H]^+^ = 343.1556.

2-(5-Amino-3-(furan-2-yl)-1*H*-pyrazol-1-yl)-6-(trifluoromethyl)pyrimidin-4(3*H*)-one **(S153)**

The compound was prepared according to General procedure B2 using methanesulfonic acid (3 mg, 0.026 mmol), 2-hydrazinyl-6-(trifluoromethyl)pyrimidin-4(3*H*)-one (50 mg, 0.26 mmol) and 3-(furan-2-yl)-3-oxopropanenitrile (35 mg, 0.26 mmol) in EtOH (3 mL). The reaction mixture was refluxed for 4 h. The crude product was purified by column chromatography on silica gel (7M NH_3_ in MeOH:MeOH:dichloromethane, 5:5:90). The product was obtained as a white solid (25 mg, 31%).

^1^H NMR (300 MHz, DMSO-*d*_6_) *δ* (ppm) 7.77 (dd, *J* = 1.8, 0.8 Hz, 1H), 6.92 (dd, *J* = 3.4, 0.8 Hz, 1H), 6.74 (s, 1H), 6.61 (dd, *J* = 3.4, 1.8 Hz, 1H), 5.74 (s, 1H).

^13^C NMR (126 MHz, DMSO-*d*_6_) *δ* (ppm) 151.1, 147.6, 145.8, 143.4, 121.6, 119.4, 117.2, 111.7, 108.5, 85.2.

^19^F NMR (282 MHz, DMSO-*d*_6_) *δ* (ppm) -69.83.

HRMS (APCI): calcd. for C_12_H_9_F_3_N_5_O_2_ [M+H]^+^ = 312.0703, found [M+H]^+^ = 312.0699.

mp >238 °C (dec.)

2-(5-Amino-3-(furan-2-yl)-1*H*-pyrazol-1-yl)-5-(2-hydroxyethyl)-6-methylpyrimidin-4(3*H*)-one **(S154)**

The compound was prepared according to General procedure B2 using methanesulfonic acid (3 mg, 0.027 mmol), 2-hydrazinyl-5-(2-hydroxyethyl)-6-methylpyrimidin-4(3*H*)-one (50 mg, 0.27 mmol) and 3-(furan-2-yl)-3-oxopropanenitrile (37 mg, 0.27 mmol) in EtOH (3 mL).The reaction mixture was refluxed for 4 h. The crude product was purified by column chromatography on silica gel (7M NH_3_ in MeOH:MeOH:dichloromethane, 7.5:7.5:85). The product was obtained as a light brown solid (55 mg, 67%).

^1^H NMR (300 MHz, DMSO-*d*_6_) *δ* (ppm) 11.53 (s, 1H), 7.74 (d, *J* = 1.8 Hz, 1H), 7.00 (s, 2H), 6.87 (d, *J* = 3.4 Hz, 1H), 6.59 (dd, *J* = 3.4, 1.8 Hz, 1H), 5.69 (s, 1H), 4.61 (s, 1H), 3.49 (s, 2H), 2.63 (t, *J* = 7.0 Hz, 2H), 2.35 (s, 3H).

^13^C NMR (126 MHz, DMSO-*d*_6_) *δ* (ppm) 150.6, 148.9, 147.9, 143.1, 112.9, 111.6, 84.9, 59.3, 29.0, 21.4.

HRMS (APCI): calcd. for C_14_H_16_N_5_O_3_ [M+H]^+^ = 302.1248, found [M+H]^+^ = 302.1246.

mp >182 °C (dec.)

**Preparation of acid chlorides**

**Scheme S3:** synthesis of 5-cyclohexylisoxazole-3-carbonyl chloride **(S158)**

Ethyl 4-cyclohexyl-2,4-dioxobutanoate **(S155)**

The compound was prepared according to General procedure E using 1-cyclohexylethan-1-one (1 g, 7.92 mmol), diethyl oxalate (1.27 g, 8.72 mmol), toluene (10 mL) and *t-*BuOK (1 M in THF, 9.5 mL, 9.5 mmol). The product was obtained as a light brown oil (1.57 g, 88%).

^1^H NMR (500 MHz, Chloroform-*d*) *δ* (ppm) 14.64 (s, 1H), 6.38 (s, 1H), 4.34 (q, *J* = 7.1 Hz, 2H), 2.37 (tt, *J* = 11.5, 3.4 Hz, 1H), 1.94 – 1.85 (m, 2H), 1.82 (dt, *J* = 13.2, 3.7 Hz, 2H), 1.70 (dtd, *J* = 9.6, 3.0, 1.6 Hz, 1H), 1.46 – 1.33 (m, 6H), 1.32 – 1.19 (m, 5H). (keto-enol tautomer)

^13^C NMR (126 MHz, Chloroform-*d*) *δ* (ppm) 205.9, 167.7, 162.2, 100.1, 62.4, 48.7, 28.9, 25.7, 25.5, 14.0.

HRMS (APCI): calcd. for C_12_H_19_O_4_ [M+H]^+^ = 227.1278, found [M+H]^+^ = 227.1277.

Ethyl 3-cyclohexyl-3-oxopropanoate **(S156)**

The compound was prepared according to General procedure F using ethyl 4-cyclohexyl-2,4-dioxobutanoate (1.57 g, 6.93 mmol), MeOH (16 mL) and HONH_2_•HCl (723 mg, 10.4 mmol). The crude product was purified by column chromatography on silica gel (hexane:EtOAc, 9:1). The product was obtained as a white solid (1.51 g, 98%).

^1^H NMR (500 MHz, Chloroform-*d*) *δ* (ppm) 6.37 (d, *J* = 0.8 Hz, 1H), 3.95 (s, 3H), 2.87 – 2.79 (m, 1H), 2.10 – 2.02 (m, 2H), 1.82 (dt, *J* = 12.7, 3.6 Hz, 2H), 1.76 – 1.69 (m, 1H), 1.51 – 1.34 (m, 4H), 1.32 – 1.23 (m, 1H).

^13^C NMR (126 MHz, Chloroform-*d*) *δ* (ppm) 179.8, 160.8, 155.9, 99.8, 52.7, 36.3, 31.0, 25.7, 25.5.

HRMS (APCI): calcd. for C_11_H_16_NO_3_ [M+H]^+^ = 210.1125, found [M+H]^+^ = 210.1126.

mp = 73−74 °C.

5-Cyclohexylisoxazole-3-carboxylic acid **(S157)**

The compound was prepared according to General procedure G using ethyl 3-cyclohexyl-3-oxopropanoate (1.51 g, 6.76 mmol), EtOH (15 mL) and NaOH (2 M in H_2_O, 5.07 mL, 10.14 mmol). The product was obtained as light brown solid (840 mg, 64%).

^1^H NMR (500 MHz, Chloroform-*d*) *δ* (ppm) 10.17 (s, 1H), 6.43 (s, 1H), 2.85 (tt, *J* = 11.3, 3.7 Hz, 1H), 2.08 (dt, *J* = 12.6, 3.6 Hz, 2H), 1.82 (dq, *J* = 11.0, 3.6 Hz, 2H), 1.73 (dddd, *J* = 12.8, 5.2, 3.5, 1.6 Hz, 1H), 1.55 – 1.34 (m, 4H), 1.29 (tt, *J* = 12.5, 3.5 Hz, 1H).

^13^C NMR (126 MHz, Chloroform-*d*) *δ* (ppm) 180.6, 164.2, 155.6, 100.2, 36.5, 31.1, 25.8, 25.6.

HRMS (APCI): calcd. for C_10_H_14_NO_3_ [M+H]^+^ = 196.0968, found [M+H]^+^ = 196.0970.

mp = 130−131 °C.

5-Cyclohexylisoxazole-3-carbonyl chloride **(S158)**

SOCl_2_ (47 µL, 0.63 mmol) was added to a solution of 5-cyclohexylisoxazole-3-carboxylic acid (100 mg, 0.53 mmol) in toluene (2 mL) and the reaction mixture was stirred at 80 °C for 3 h. The excess SOCl_2_ evaporated *in vacuo* and the residue was mixed with toluene (2 × 2 mL) and the solvent was evaporated. After drying *in vacuo*, the product was obtained as a clear gum (109 mg, quant.) and used directly in the next step.

Ethyl 4-(4-methoxyphenyl)-2,4-dioxobutanoate **(S159)**

The compound was prepared according to General procedure E using 1-(4-methoxyphenyl)ethan-1-one (1 g, 6.66 mmol), diethyl oxalate (1.07 g, 7.32 mmol), toluene (10 mL) and *t-*BuOK (1 M in THF, 7.98 mL, 7.98 mmol). The product was obtained as light brown solid (1.47 g, 89%).

^1^H NMR (500 MHz, Chloroform-*d*) *δ* (ppm) 15.44 (s, 1H), 7.98 (d, *J* = 9.0 Hz, 2H), 7.02 (s, 1H), 6.98 (d, *J* = 9.0 Hz, 2H), 4.39 (q, *J* = 7.1 Hz, 2H), 3.89 (s, 3H), 1.41 (t, *J* = 7.1 Hz, 3H).

^13^C NMR (126 MHz, Chloroform-*d*) *δ* (ppm) 190.3, 168.2, 164.3, 162.5, 130.3, 127.8, 114.2, 97.7, 62.5, 55.6, 14.1.

HRMS (APCI): calcd. for C_13_H_15_O_5_ [M+H]^+^ = 251.0914, found [M+H]^+^ = 251.0916.

Ethyl 5-(4-methoxyphenyl)isoxazole-3-carboxylate **(S160)**

The compound was prepared according to General procedure F using ethyl 4-(4-methoxyphenyl)-2,4-dioxobutanoate (1.47 g, 5.87 mmol), EtOH (15 mL) and HONH_2_•HCl (612 mg, 8.81 mmol). The crude product was purified by column chromatography on silica gel (hexane:EtOAc, 85:15). The product was obtained as a light brown solid (1.38 g, 95%).

^1^H NMR (500 MHz, Chloroform-*d*) *δ* (ppm) 7.74 (d, *J* = 8.9 Hz, 2H), 6.99 (d, *J* = 8.9 Hz, 2H), 6.79 (s, 1H), 4.46 (d, *J* = 7.1 Hz, 2H), 3.87 (s, 3H), 1.44 (s, 3H).

^13^C NMR (126 MHz, Chloroform-*d*) *δ* (ppm) 171.9, 161.7, 160.3, 157.1, 127.7, 119.6, 114.7, 98.7, 62.3, 55.6, 14.3.

HRMS (APCI): calcd. for C_13_H_14_NO_4_ [M+H]^+^ = 248.0917, found [M+H]^+^ = 248.0914.

mp = 87−89 °C.

5-(4-Methoxyphenyl)isoxazole-3-carboxylic acid **(S161)**

The compound was prepared according to General procedure G using ethyl 5-(4-methoxyphenyl)isoxazole-3-carboxylate (1.38 g, 5.58 mmol), EtOH (14 mL) and NaOH (2 M in H_2_O, 4.19 mL, 8.37 mmol). The product was obtained as a yellow solid (580 mg, 47%).

^1^H NMR (500 MHz, Chloroform-*d*) *δ* (ppm) 7.76 (d, *J* = 8.9 Hz, 2H), 7.01 (d, *J* = 8.9 Hz, 2H), 6.85 (s, 1H), 3.88 (s, 3H).

^13^C NMR (126 MHz, Chloroform-*d*) *δ* (ppm) 172.7, 161.9, 161.4, 156.2, 127.8, 119.4, 114.8, 98.7, 55.6.

HRMS (APCI): calcd. for C_11_H_10_NO_4_ [M+H]^+^ = 220.0604, found [M+H]^+^ = 220.0601.

mp > 148 °C (dec).

5-(4-Methoxyphenyl)isoxazole-3-carbonyl chloride **(S162)**

The compound was prepared according to General procedure H using 5-(4-methoxyphenyl)isoxazole-3-carboxylic acid (200 mg, 0.91 mmol) and SOCl_2_ (331 µL, 4.56 mmol). The reaction mixture was refluxed for 4 h. The product was obtained as a light brown solid (216 mg, quant.) and used directly in the next step.

Ethyl 4-(2-bromophenyl)-2,4-dioxobutanoate **(S163)**

The compound was prepared according to General procedure E using 1-(2-bromophenyl)ethan-1-one (1 g, 5.02 mmol), diethyl oxalate (807 mg, 5.53 mmol), toluene (10 mL) and *t-*BuOK (1 M in THF, 6.03 mL, 6.03 mmol). The product was obtained as a dark red oil (1.31 g, 87%).

^1^H NMR (500 MHz, Chloroform-*d*) *δ* (ppm) 14.49 (s, 1H), 7.67 (dd, *J* = 8.0, 1.1 Hz, 1H), 7.57 (dd, *J* = 7.6, 1.8 Hz, 1H), 7.42 (td, *J* = 7.6, 1.2 Hz, 1H), 7.35 (td, *J* = 7.7, 1.8 Hz, 1H), 6.89 (s, 1H), 4.38 (q, *J* = 7.2 Hz, 2H), 1.39 (t, *J* = 7.2 Hz, 3H).

^13^C NMR (126 MHz, Chloroform-*d*) *δ* (ppm) 194.3, 167.6, 162.0, 138.2, 134.3, 132.8, 130.2, 127.7, 120.3, 103.1, 62.8, 14.2.

HRMS (APCI): calcd. for C_12_H_12_BrO_4_ [M+H]^+^ = 298.9913, found [M+H]^+^ = 298.9911.

Ethyl 5-(2-bromophenyl)isoxazole-3-carboxylate **(S164)**

The compound was prepared according to General procedure F using ethyl 4-(2-bromophenyl)-2,4-dioxobutanoate (1.31 g, 4.38 mmol), EtOH (14 mL) and HONH_2_•HCl (456 mg, 6.57 mmol). The crude product was purified by column chromatography on silica gel (hexane:EtOAc, 90:10). The product was obtained as a light brown oil (1.21 g, 93%).

^1^H NMR (500 MHz, Chloroform-*d*) *δ* (ppm) 7.88 (dd, *J* = 7.9, 1.7 Hz, 1H), 7.73 (dd, *J* = 7.9, 1.2 Hz, 1H), 7.46 (td, *J* = 7.6, 1.2 Hz, 1H), 7.36 (s, 1H), 7.35 – 7.30 (m, 1H), 4.49 (q, *J* = 7.2 Hz, 2H), 1.45 (t, *J* = 7.1 Hz, 3H).

^13^C NMR (126 MHz, Chloroform-*d*) *δ* (ppm) 169.6, 160.1, 156.8, 134.5, 131.7, 130.3, 128.0, 127.8, 121.4, 104.8, 62.4, 14.3.

HRMS (APCI): calcd. for C_12_H_11_BrNO_3_ [M+H]^+^ = 295.9917, found [M+H]^+^ = 295.9915.

5-(2-Bromophenyl)isoxazole-3-carboxylic acid **(S165)**

The compound was prepared according to General procedure G using ethyl 5-(2-bromophenyl)isoxazole-3-carboxylate (1.21 g, 4.09 mmol), EtOH (13 mL) and NaOH (2 M in H_2_O, 3.06 mL, 6.13 mmol). The product was obtained as a light brown solid (930 mg, 85%).

^1^H NMR (500 MHz, Chloroform-*d*) *δ* (ppm) 8.14 (s, 1H), 7.91 (dd, *J* = 7.9, 1.7 Hz, 1H), 7.75 (dd, *J* = 8.1, 1.2 Hz, 1H), 7.48 (td, *J* = 7.6, 1.2 Hz, 1H), 7.44 (s, 1H), 7.35 (td, *J* = 7.7, 1.7 Hz, 1H).

^13^C NMR (126 MHz, Chloroform-*d*) *δ* (ppm) 170.1, 163.2, 155.9, 134.4, 131.8, 130.2, 127.9, 127.4, 121.4, 104.8.

HRMS (APCI): calcd. for C_10_H_7_BrNO_3_ [M+H]^+^ = 267.9604, found [M+H]^+^ = 267.9603.

mp = 138−141 °C.

5-(2-Bromophenyl)isoxazole-3-carbonyl chloride **(S166)**

The compound was prepared according to General procedure H using 5-(2-bromophenyl)isoxazole-3-carboxylic acid (200 mg, 0.75 mmol) and SOCl_2_ (271 µL, 3.73 mmol). The reaction mixture was refluxed for 4 h. The product was obtained as a light brown solid (213 mg, quant.) and used directly in the next step.

Ethyl 4-(3-(*tert*-butyl)phenyl)-2,4-dioxobutanoate **(S167)**

The compound was prepared according to General procedure E using 1-(3-(*tert*-butyl)phenyl)ethan-1-one (150 mg, 0.85 mmol), diethyl oxalate (137 mg, 0.94 mmol), toluene (4 mL) and *t-*BuOK (1 M in THF, 1.02 mL, 1.02 mmol). The product was obtained as a light brown gum (259 mg, crude) and used as such in the next step without further purification.

^1^H NMR (500 MHz, Chloroform-*d*) *δ* (ppm) 15.38 (s, 1H), 8.02 (d, *J* = 1.9 Hz, 1H), 7.79 (dd, *J* = 7.8, 1.4 Hz, 1H), 7.69 – 7.61 (m, 1H), 7.43 (t, *J* = 7.8 Hz, 1H), 7.07 (s, 1H), 4.41 (q, *J* = 7.1 Hz, 2H), 1.42 (t, *J* = 7.1 Hz, 3H), 1.37 (s, 9H).

^13^C NMR (126 MHz, Chloroform-*d*) *δ* (ppm) 191.5, 169.8, 162.5, 152.3, 134.9, 131.2, 128.8, 125.4, 124.8, 98.2, 62.7, 35.1, 31.4, 14.2, 14.1.

HRMS (APCI): calcd. for C_16_H_21_O_4_ [M+H]^+^ = 277.1434, found [M+H]^+^ = 277.1432.

Ethyl 5-(3-(*tert*-butyl)phenyl)isoxazole-3-carboxylate **(S168)**

The compound was prepared according to General procedure F using ethyl 4-(3-(*tert*-butyl)phenyl)-2,4-dioxobutanoate (259 mg, 0.94 mmol), EtOH (5 mL) and HONH_2_•HCl (78 mg, 1.12 mmol). The crude product was purified by column chromatography on silica gel (hexane:EtOAc, 9:1). The product was obtained as a colorless gum (175 mg, 75% over the 2 steps).

^1^H NMR (300 MHz, Chloroform-*d*) *δ* (ppm) 7.85 – 7.79 (m, 1H), 7.61 (dt, *J* = 7.6, 1.5 Hz, 1H), 7.51 (ddd, *J* = 7.9, 2.0, 1.2 Hz, 1H), 7.46 – 7.37 (m, 1H), 6.92 (s, 1H), 4.48 (q, *J* = 7.2 Hz, 2H), 1.45 (t, *J* = 7.1 Hz, 3H), 1.37 (s, 9H).

^13^C NMR (126 MHz, Chloroform-*d*) *δ* (ppm) 172.4, 160.2, 157.1, 152.4, 129.0, 128.1, 126.6, 123.4, 123.0, 99.9, 62.3, 35.0, 31.3, 14.3.

HRMS (APCI): calcd. for C_16_H_20_NO_3_ [M+H]^+^ = 274.1438, found [M+H]^+^ = 274.1437.

5-(3-(*ter*t-butyl)phenyl)isoxazole-3-carboxylic acid **(S169)**

The compound was prepared according to General procedure G using ethyl 5-(3-(*tert*-butyl)phenyl)isoxazole-3-carboxylate (175 mg, 0.64 mmol), EtOH (2 mL) and NaOH (2 M in H_2_O, 0.48 mL, 0.96 mmol). The product was obtained as off-white solid (134 mg, 85%).

^1^H NMR (300 MHz, DMSO-*d*_6_) *δ* (ppm) 13.99 (s, 1H), 7.92 (t, *J* = 1.9 Hz, 1H), 7.75 (dt, *J* = 7.5, 1.4 Hz, 1H), 7.62 – 7.53 (m, 1H), 7.49 (d, *J* = 7.7 Hz, 1H), 7.46 (s, 1H), 1.34 (s, 9H).

^13^C NMR (126 MHz, DMSO-*d*_6_) *δ* (ppm) 171.1, 157.9, 151.9, 131.2, 129.1, 127.8, 126.0, 123.0, 122.6, 100.8, 34.6, 30.9.

HRMS (APCI): calcd. for C_14_H_16_NO_3_ [M+H]^+^ = 246.1125, found [M+H]^+^ = 246.1122.

mp = 127−129 °C.

5-(3-(*tert*-butyl)phenyl)isoxazole-3-carbonyl chloride **(S170)**

The compound was prepared according to General procedure H using 5-(3-(*tert*-butyl)phenyl)isoxazole-3-carboxylic acid (110 mg, 0.45 mmol) and SOCl_2_ (163 µL, 2.24 mmol). The reaction mixture was refluxed for 4 h. The product was obtained as an off-white solid (118 mg, quant.) and used directly in the next step.

Ethyl 5,5-dimethyl-2,4-dioxohexanoate **(S171)**

The compound was prepared according to General procedure E using 3,3-dimethylbutan-2-one (1 g, 9.98 mmol), diethyl oxalate (1.60 g, 10.98 mmol), toluene (10 mL) and *t-*BuOK (1 M in THF, 11.98 mL, 11.98 mmol). The product was obtained as a light brown oil (1.63 g, 82%).

^1^H NMR (500 MHz, Chloroform-*d*) *δ* (ppm) 14.75 (s, 1H), 6.53 (s, 1H), 4.36 (q, *J* = 7.2 Hz, 2H), 1.38 (t, *J* = 7.1 Hz, 3H), 1.21 (s, 9H).

^13^C NMR (126 MHz, Chloroform-*d*) *δ* (ppm) 209.3, 167.7, 162.5, 98.0, 62.6, 41.8, 26.9, 14.2.

HRMS (APCI): calcd. for C_10_H_17_O_4_ [M+H]^+^ = 201.1121, found [M+H]^+^ = 201.1119.

Ethyl 5-(*tert*-butyl)isoxazole-3-carboxylate **(S172)**

The compound was prepared according to General procedure F using ethyl 5,5-dimethyl-2,4-dioxohexanoate (1.63 g, 7.40 mmol), EtOH (17 mL) and HONH_2_•HCl (771 mg, 11.1 mmol). The crude product was purified by column chromatography on silica gel (hexane:EtOAc, 9:1). The product was obtained as a light brown oil (1.40 g, 96%).

^1^H NMR (500 MHz, Chloroform-*d*) *δ* (ppm) 6.36 (s, 1H), 4.42 (q, *J* = 7.1 Hz, 2H), 1.40 (t, *J* = 7.1 Hz, 3H), 1.36 (s, 9H).

^13^C NMR (126 MHz, Chloroform-*d*) *δ* (ppm) 183.3, 160.5, 156.3, 99.2, 62.1, 33.1, 28.9, 14.3.

HRMS (APCI): calcd. for C_10_H_16_NO_3_ [M+H]^+^ = 198.1125, found [M+H]^+^ = 198.1123.

5-(*tert*-Butyl)isoxazole-3-carboxylic acid **(S173)**

The compound was prepared according to General procedure G using ethyl 5-(*tert*-butyl)isoxazole-3-carboxylate (1.40 g, 7.1 mmol), EtOH (14 mL) and NaOH (2 M in H_2_O, 5.32 mL, 10.65 mmol). The product was obtained as a light brown gum (1.20 g, quant.).

^1^H NMR (500 MHz, Chloroform-*d*) *δ* (ppm) 6.43 (s, 1H), 1.39 (s, 9H).

^13^C NMR (126 MHz, Chloroform-*d*) *δ* (ppm) 184.1, 163.1, 155.5, 99.5, 33.2, 28.9.

HRMS (APCI): calcd. for C_8_H_12_NO_3_ [M+H]^+^ = 170.0812, found [M+H]^+^ = 170.0810.

Ethyl 2,4-dioxo-4-(4-(trifluoromethoxy)phenyl)butanoate **(S174)**

The compound was prepared according to General procedure E using 1-(4-(trifluoromethoxy)phenyl)ethan-1-one (700 mg, 3.43 mmol), diethyl oxalate (550 mg, 3.77 mmol), toluene (7 mL) and *t-*BuOK (1 M in THF, 4.11 mL, 4.11 mmol). The product was obtained as a yellow gum (975 mg, 93%).

^1^H NMR (500 MHz, Chloroform-*d*) *δ* (ppm) 15.15 (s, 1H), 8.07 – 8.02 (m, 2H), 7.37 – 7.30 (m, 2H), 7.04 (s, 1H), 4.41 (q, *J* = 7.1 Hz, 2H), 1.42 (t, *J* = 7.1 Hz, 3H).

^13^C NMR (126 MHz, Chloroform-*d*) *δ* (ppm) 189.2, 170.3, 162.2, 153.3, 133.4, 130.0, 120.8, 120.4 (q, *J* = 259.0 Hz), 98.0, 62.9, 14.2.

^19^F NMR (282 MHz, Chloroform-*d*) *δ* (ppm) -57.6.

HRMS (APCI): calcd. for C_13_H_12_F_3_O_5_ [M+H]^+^ = 305.0631, found [M+H]^+^ = 305.0628.

Ethyl 5-(4-(trifluoromethoxy)phenyl)isoxazole-3-carboxylate **(S175)**

The compound was prepared according to General procedure F using ethyl 2,4-dioxo-4-(4-(trifluoromethoxy)phenyl)butanoate (975 mg, 3.20 mmol), EtOH (10 mL) and HONH_2_•HCl (334 mg, 4.81 mmol). The crude product was purified by column chromatography on silica gel (hexane:EtOAc, 8:2). The product was obtained as a white solid (809 mg, 84%).

^1^H NMR (500 MHz, Chloroform-*d*) *δ* (ppm) 7.80 – 7.89 (m, 2H), 7.34 (dt, *J* = 7.9, 1.1 Hz, 2H), 6.93 (s, 1H), 4.48 (q, *J* = 7.2 Hz, 2H), 1.45 (t, *J* = 7.1 Hz, 3H).

^13^C NMR (126 MHz, Chloroform-*d*) *δ* (ppm) 170.4, 160.0, 157.3, 151.0, 127.8, 125.4, 121.6, 120.5 (q, *J* = 259.2 Hz), 100.5, 62.5, 14.3.

^19^F NMR (471 MHz, Chloroform-*d*) *δ* (ppm) -57.8.

HRMS (APCI): calcd. for C_13_H_11_F_3_NO_4_ [M+H]^+^ = 302.0635, found [M+H]^+^ = 302.0634.

mp = 132−134 °C.

5-(4-(Trifluoromethoxy)phenyl)isoxazole-3-carboxylic acid **(S176)**

The compound was prepared according to General procedure G using ethyl 5-(4-(trifluoromethoxy)phenyl)isoxazole-3-carboxylate (809 mg, 2.69 mmol), EtOH (9 mL) and NaOH (2 M in H_2_O, 2.01 mL, 4.03 mmol). The product was obtained as a white solid (705 mg, 96%).

^1^H NMR (500 MHz, DMSO-*d*_6_) *δ* (ppm) 14.11 (s, 1H), 8.12 – 8.04 (m, 2H), 7.59 – 7.52 (m, 2H), 7.48 (d, *J* = 1.2 Hz, 1H).

^13^C NMR (126 MHz, DMSO-*d*_6_) *δ* (ppm) 169.4, 160.7, 158.0, 149.7, 128.0, 125.4, 121.7, 119.9 (q, *J* = 257.8 Hz), 101.6.

^19^F NMR (471 MHz, DMSO-*d*_6_) *δ* (ppm) -56.80 – -56.54 (m).

HRMS (APCI): calcd. for C_11_H_7_F_3_NO_4_ [M+H]^+^ = 274.0322, found [M+H]^+^ = 274.0320.

mp = 183−184 °C.

5-Phenylisoxazole-3-carbonyl chloride **(S177)**

The compound was prepared according to General procedure H using 5-phenylisoxazole-3-carboxylic acid (2 g, 10.57 mmol) and SOCl_2_ (3.83 mL, 52.86 mmol). The reaction mixture was refluxed for 4 h. The product was obtained as light brown solid (2.19 g, quant.) and used directly in the next step.

3-Phenylisoxazole-5-carbonyl chloride **(S178)**

The compound was prepared according to General procedure H using 3-phenylisoxazole-5-carboxylic acid (500 mg, 2.64 mmol) and SOCl_2_ (959 µL, 13.22 mmol). The reaction mixture was refluxed for 4 h. The product was obtained as a light brown solid (549 mg, quant.) and used directly in the next step.

2-Phenylthiazole-4-carbonyl chloride **(S179)**

The compound was prepared according to General procedure H using 2-phenylthiazole-4-carboxylic acid (100 mg, 0.49 mmol) and SOCl_2_ (177 µL, 2.44 mmol). The reaction mixture was refluxed for 4 h. The product was obtained as a white solid (109 mg, quant.) and used directly in the next step.

5-Phenyl-4,5-dihydroisoxazole-3-carbonyl chloride **(S180)**

The compound was prepared according to General procedure H using 5-phenyl-4,5-dihydroisoxazole-3-carboxylic acid (60 mg, 0.31 mmol) and SOCl_2_ (113 µL, 1.57 mmol). The reaction mixture was refluxed for 4 h. The product was obtained as a colorless oil (66 mg, quant.) and used directly in the next step.

2-(5-phenylisoxazol-3-yl)acetyl chloride **(S181)**

DMF (5 µL) was added to a solution of 2-(5-phenylisoxazol-3-yl)acetic acid (50 mg, 0.25 mmol) in dichloromethane (2 mL) under nitrogen atmosphere. The mixture was cooled to 0 °C, oxalyl chloride (42 µL, 0.49 mmol) was added dropwise, and the reaction mixture was stirred at room temperature for 2 h. The solvent was removed by purging nitrogen and the residue was dried *in vacuo*. The product was obtained as a brown gum (55 mg) and used directly in the next step.

[1,1'-Biphenyl]-4-carbonyl chloride **(S182)**

DMF (10 µL) was added to a solution of [1,1'-biphenyl]-4-carboxylic acid (500 mg, 2.52 mmol) in SOCl_2_ (2.5 mL) under nitrogen atmosphere and the mixture was refluxed for 16 h. The excess of SOCl_2_ was removed *in vacuo*, the residue mixed with toluene (5 mL), and the solvent was evaporated *in vacuo*. The addition and evaporation of toluene was repeated twice, and the residue was dried *in vacuo*. The product was obtained as a white solid (546 mg, quant.) and used directly in the next step.

[1,1'-Biphenyl]-3-carbonyl chloride **(S183)**

The compound was prepared according to General procedure H using [1,1'-biphenyl]-3-carboxylic acid (500 mg, 2.52 mmol) and SOCl_2_ (6.6 mL). The reaction mixture was refluxed for 2 h. The product was obtained as a brown oil (546 mg, quant.) and used directly in the next step.

2,3-Dimethoxybenzoyl chloride **(S184)**

The compound was prepared according to General procedure H using2,3-dimethoxybenzoic acid (1 g, 5.49 mmol) and SOCl_2_ (800 µL, 10.97 mmol). The reaction mixture was refluxed for 20 min. The product was obtained as a white solid (1.10 g, quant.) and used directly in the next step.

**Scheme S4:** synthesis of 5-phenylisothiazole-3-carbonyl chloride **(S189)**

5-Bromoisothiazole-3-carboxylic acid **(S185)**

CrO_3_ (2.60 g, 26.93 mmol) was added portionwise over the period of 4 h to a mixture of conc. H_2_SO_4_ (96%, 23 mL) and 5-bromo-3-methylisothiazole (1.0 g, 5.62 mmol) at room temperature. The reaction mixture was stirred at room temperature for 16 h, then poured into ice-cold water (40 mL), and extracted with EtOAc (3 × 30 mL). The combined organic extracts were washed with water (3 × 20 mL), brine (20 mL), dried over MgSO_4_, filtered, and the solvent was evaporated *in vacuo*. The product was obtained as a white solid (198 mg, 17%).

^1^H NMR (300 MHz, DMSO-*d*_6_) *δ* (ppm) 13.70 (s, 1H), 7.95 (s, 1H).

^13^C NMR (75 MHz, DMSO-*d*_6_) *δ* (ppm) 161.2, 160.6, 137.5, 129.3.

HRMS (APCI): calcd. for C_4_H_1_BrNO_2_S [M-H]^-^ = 207.8896, found [M-H]^-^ = 207.8900.

mp = 179−181 °C.

Methyl 5-bromoisothiazole-3-carboxylate **(S186)**

Conc. H_2_SO_4_ (96%, 10 µL) was added to a solution of 5-bromoisothiazole-3-carboxylic acid (190 mg, 0.91 mmol) in MeOH (2 mL) and the mixture was refluxed for 2 h. The solvent was evaporated *in vacuo*, the residue was mixed with H_2_O (4 mL), and the mixture was extracted with EtOAc (3 × 15 mL). The combined organic extracts were washed with water (3 × 10 mL), saturated aqueous NaHCO_3_ solution (10 mL), dried over MgSO_4_, filtered, and the solvent was evaporated *in vacuo*. The product was obtained as a colorless oil (172 mg, 85%).

^1^H NMR (300 MHz, Chloroform-*d*) *δ* (ppm) 7.79 (s, 1H), 3.97 (s, 3H).

^13^C NMR (75 MHz, Chloroform-*d*) *δ* (ppm) 160.3, 159.7, 137.7, 129.2, 53.1.

HRMS (APCI): calcd. for C_5_H_5_BrNO_2_S [M+H]^+^ = 223.9198, found [M+H]^+^ = 223.9195.

Methyl 5-phenylisothiazole-3-carboxylate **(S187)**

Phenyl boronic acid (98 mg, 0.80 mmol) and K_3_PO_4_ (2 M in H_2_O, 0.75 mL, 1.46 mmol) were added to a solution of methyl 5-bromoisothiazole-3-carboxylate (162 mg, 0.73 mmol) in DME (3.0 mL). The reaction mixture was deoxygenated by argon purgin for 20 min, then Pd(PPh_3_)_4_ (42 mg, 0.04 mmol) was added, and the reaction mixture was stirred at 80 °C for 2.5 h. The reaction mixture was diluted with water (10 mL), and extracted with EtOAc (3 × 5 mL). The combined organic extracts were washed with water (3 × 10 mL), brine (15 mL), dried over MgSO_4_, filtered, and the solvent was evaporated *in vacuo*. The residue was purified by column chromatography on silica gel (hexane:EtOAc, 9:1). The product methyl 5-phenylisothiazole-3-carboxylate was obtained as a white solid (66 mg, 41%).

The aqueous layer from the workup above was acidified by using aqueous 1 M HCl solution and extracted with EtOAc (3 × 5 mL). The combined organic extracts were washed with water (3 × 5 mL), brine (5 mL), dried over MgSO_4_, filtered, and the solvent was evaporated *in vacuo*. The side product 5-phenylisothiazole-3-carboxylic acid was obtained as an off-white solid (72 mg, 48%).

^1^H NMR (300 MHz, Chloroform-*d*) *δ* (ppm) 4.00 (s, 3H), 7.41 – 7.52 (m, 3H), 7.57 – 7.67 (m, 2H), 7.97 (s, 1H).

^13^C NMR (75 MHz, Chloroform-*d*) *δ* (ppm) 169.8, 161.4, 159.9, 130.3, 129.6, 126.8, 121.9, 52.9.

HRMS (APCI): calcd. for C_11_H_10_NO_2_S [M+H]^+^ = 220.0427, found [M+H]^+^ = 220.0425.

mp = 76−78 °C.

5-Phenylisothiazole-3-carboxylic acid **(S188)**

The compound was prepared according to General procedure G using methyl 5-phenylisothiazole-3-carboxylate (50 mg, 0.23 mmol), EtOH (1 mL) and NaOH (2 M in H_2_O, 0.17 mL, 0.34 mmol). The product was obtained as a white solid (42 mg, 90%).

^1^H NMR (300 MHz, Chloroform-*d*) *δ* (ppm) 8.01 (s, 1H), 7.62 (s, 2H), 7.56 – 7.41 (m, 3H).

^13^C NMR (75 MHz, Chloroform-*d*) *δ* (ppm) 171.1, 161.6, 159.4, 130.6, 129.9, 129.7, 126.9, 121.4.

HRMS (APCI): calcd. for C_10_H_8_NO_2_S [M+H]^+^ = 206.0270, found [M+H]^+^ = 206.0270.

mp = 148−150 °C.

5-Phenylisothiazole-3-carbonyl chloride **(S189)**

The compound was prepared according to General procedure H using 5-phenylisothiazole-3-carboxylic acid (40 mg, 0.19 mmol) and SOCl_2_ (71 µL, 0.97 mmol). The product was obtained as an off-white solid (44 mg, quant.) and used directly in the next step.

### Preparation of target compound 2 and its analogs

*N*-(3-(furan-2-yl)-1-(4-methyl-6-oxo-1,6-dihydropyrimidin-2-yl)-1*H*-pyrazol-5-yl)-5-phenylisoxazole-3-carboxamide **(2)**

The compound was prepared by General procedure I using Et_3_N (28 µL, 0.2 mmol), 2-(5-amino-3-(furan-2-yl)-1*H*-pyrazol-1-yl)-6-methylpyrimidin-4(3*H*)-one (50 mg, 0.2 mmol), acetonitrile (2 + 2 mL) and 5-phenylisoxazole-3-carbonyl chloride (40 mg, 0.2 mmol). The obtained solid was additionally washed with EtOAc (3 mL). The product was obtained as a white solid (30 mg, 35%).

^1^H NMR (500 MHz, DMSO-*d*_6_) *δ* (ppm) 13.56 (s, 1H), 8.04 – 7.99 (m, 2H), 7.82 (s, 1H), 7.61 (s, 1H), 7.60 – 7.53 (m, 3H), 7.22 (s, 1H), 7.02 (d, *J* = 3.3 Hz, 1H), 6.67 – 6.62 (m, 1H), 6.49 (s, 1H), 2.55 (s, 3H).

^13^C NMR (126 MHz, DMSO-*d*_6_) *δ* (ppm) 171.75, 158.85, 154.70, 147.05, 144.46, 143.74, 139.69, 131.10, 129.32, 126.00, 125.93, 111.86, 108.78, 99.98, 94.59, 22.82.

HRMS (APCI): calcd. for C_22_H_17_N_6_O_4_ [M+H]^+^= 429.1306, found [M+H]^+^ = 429.1310.

*N*-(1-(4-methyl-6-oxo-1,6-dihydropyrimidin-2-yl)-3-(thiophen-2-yl)-1*H*-pyrazol-5-yl)-5-phenylisoxazole-3-carboxamide **(29)**

The compound was prepared by General procedure I using Et_3_N (31 µL, 0.22 mmol), 2-(5-amino-3-(thiophen-2-yl)-1*H*-pyrazol-1-yl)-6-methylpyrimidin-4(3*H*)-one (60 mg, 0.22 mmol), acetonitrile (2 + 2 mL) and 5-phenylisoxazole-3-carbonyl chloride (46 mg, 0.22 mmol). The obtained solid was additionally washed with EtOAc (3 mL). The product was obtained as a white solid (33 mg, 34 %).

^1^H NMR (500 MHz, DMSO-*d*_6_) *δ* (ppm) 15.84 (s, 1H), 8.05 – 7.99 (m, 2H), 7.82 (s, 1H), 7.63 – 7.51 (m, 5H), 7.16 – 7.11 (m, 2H), 5.75 (s, 1H), 2.23 (s, 3H).

^13^C NMR (126 MHz, DMSO-*d*_6_) *δ* (ppm) 171.50, 155.51, 146.51, 142.52, 138.37, 136.70, 131.36, 129.84, 128.23, 126.88, 126.40, 126.28, 125.80, 100.40, 93.96, 23.58.

HRMS (APCI): calcd. for C_22_H_17_N_6_O_3_S [M+H]^+^= 445.1077, found [M+H]^+^ = 445.1080.

5-Cyclohexyl-*N*-(3-(furan-2-yl)-1-(4-methyl-6-oxo-1,6-dihydropyrimidin-2-yl)-1*H*-pyrazol-5-yl)isoxazole-3-carboxamide **(30)**

The compound was prepared according to General procedure I using Et_3_N (16.1 µL, 0.12 mmol), 2-(5-amino-3-(furan-2-yl)-1*H*-pyrazol-1-yl)-6-methylpyrimidin-4(3*H*)-one (30 mg, 0.12 mmol), acetonitrile (1.0 + 0.5 mL) and 5-cyclohexylisoxazole-3-carbonyl chloride (25 mg, 0.116 mmol). The product was obtained as a white solid (31 mg, 61%).

^1^H NMR (300 MHz, DMSO-*d*_6_) *δ* (ppm) 14.93 (s, 1H), 7.77 (d, *J* = 1.7 Hz, 1H), 7.08 (s, 1H), 6.92 (d, *J* = 3.4 Hz, 1H), 6.89 (s, 1H), 6.62 (dd, *J* = 3.4, 1.8 Hz, 1H), 5.85 (s, 1H), 2.92 (td, *J* = 11.2, 9.2, 5.7 Hz, 1H), 2.25 (s, 3H), 2.03 (d, *J* = 12.3 Hz, 2H), 1.72 (dd, *J* = 25.9, 11.9 Hz, 3H), 1.57 – 1.16 (m, 5H).

^13^C NMR (126 MHz, DMSO-*d*_6_) *δ* (ppm) 179.7, 171.2, 158.5, 156.9, 155.1, 148.0, 143.0, 140.6, 111.7, 107.4, 106.8, 99.2, 93.5, 35.5, 30.4, 25.2, 25.0, 22.8.

HRMS (APCI): calcd. for C_22_H_23_N_6_O_4_ [M+H]^+^= 435.1775, found [M+H]^+^ = 435.1779.

mp >285 °C (dec.)

*N*-(1-(4-Methyl-6-oxo-1,6-dihydropyrimidin-2-yl)-3-(3-methylfuran-2-yl)-1*H*-pyrazol-5-yl)-5-phenylisoxazole-3-carboxamide **(31)**

The compound was prepared according to General procedure I using Et_3_N (22 µL, 0.16 mmol), 2-(5-amino-3-(3-methylfuran-2-yl)-1*H*-pyrazol-1-yl)-6-methylpyrimidin-4(3*H*)-one (30 mg, 0.11 mmol), acetonitrile (1.0 + 0.5 mL) and 5-phenylisoxazole-3-carbonyl chloride (32 mg, 0.16 mmol). The product was obtained as an off-white solid (30 mg, 61%).

^1^H NMR (300 MHz, DMSO-*d*_6_) *δ* (ppm) 15.73 (s, 1H), 8.07 – 7.97 (m, 2H), 7.84 (s, 1H), 7.65 (d, *J* = 1.7 Hz, 1H), 7.58 (d, *J* = 6.8 Hz, 3H), 7.03 (s, 1H), 6.49 (d, *J* = 1.8 Hz, 1H), 5.77 (s, 1H), 2.35 (s, 3H), 2.23 (s, 3H).

HRMS (APCI): calcd. for C_23_H_19_N_6_O_4_ [M+H]^+^= 443.1462, found [M+H]^+^ = 443.1462.

mp >310 °C (dec.)

*N*-(1-(4-Methyl-6-oxo-1,6-dihydropyrimidin-2-yl)-3-phenyl-1*H*-pyrazol-5-yl)-5-phenylisoxazole-3-carboxamide **(32)**

The compound was prepared according to General procedure I using Et_3_N (16 µL, 0.11 mmol), 2-(5-amino-3-phenyl-1*H*-pyrazol-1-yl)-6-methylpyrimidin-4(3*H*)-one (30 mg, 0.11 mmol), acetonitrile (1.0 + 0.5 mL) and 5-phenylisoxazole-3-carbonyl chloride (11 mg, 0.11 mmol). The obtained solid was additionally washed with water (3 mL) and then with EtOAc (3 mL). The compound was purified by reversed phase column chromatography using Biotage Selekt purification system (water:MeOH:0.7 M NH_3_ in methanol, gradient 90:10:10 to 0:70:30). The product was obtained as a white solid (16 mg, 58%).

^1^H NMR (500 MHz, DMSO-*d*_6_) *δ* (ppm) 15.46 (s, 1H), 8.05 – 8.01 (m, 2H), 7.93 (d, *J* = 7.6 Hz, 2H), 7.83 (s, 1H), 7.62 – 7.55 (m, 3H), 7.47 (t, *J* = 7.6 Hz, 2H), 7.38 (t, *J* = 7.3 Hz, 1H), 7.25 (s, 1H), 5.82 (s, 1H), 2.27 (s, 3H).

^13^C NMR (126 MHz, DMSO-*d*_6_) *δ* (ppm) 172.7, 162.7, 150.9, 154.8, 141.7, 130.9, 129.3, 128.7, 126.3, 125.8, 125.7, 107.2, 99.9, 93.9, 22.9.

HRMS (APCI): calcd. for C_24_H_17_N_6_O_3_ [M-H]^-^= 437.1368, found [M-H]^-^= 437.1365.

mp >320 °C (dec.)

*N*-(3-Benzyl-1-(4-methyl-6-oxo-1,6-dihydropyrimidin-2-yl)-1*H*-pyrazol-5-yl)-5-phenylisoxazole-3-carboxamide **(33)**

The compound was prepared according to General procedure I using Et_3_N (21 *µ*L, 0.15 mmol), 2-(5-amino-3-benzyl-1*H*-pyrazol-1-yl)-6-methylpyrimidin-4(3*H*)-one (30 mg, 0.11 mmol), acetonitrile (1.0 + 0.4 mL) and 5-phenylisoxazole-3-carbonyl chloride (31 mg, 0.15 mmol). The product was obtained as a light brown solid (36 mg, 75%).

^1^H NMR (700 MHz, DMSO-*d*_6_) *δ* (ppm) 14.35 (s, 1H), 8.02 – 7.97 (m, 2H), 7.70 – 7.62 (m, 1H), 7.61 – 7.37 (m, 4H), 7.32 (d, *J* = 4.5 Hz, 4H), 7.23 (h, *J* = 4.4 Hz, 1H), 6.66 (s, 1H), 6.11 (s, 1H), 3.98 (s, 2H), 2.36 (s, 3H).

^13^C NMR (176 MHz, DMSO-*d*_6_) *δ* (ppm) 171.39, 159.23, 154.57, 153.25, 139.92, 139.09, 130.97, 129.29, 128.69, 128.41, 127.63, 127.26, 126.22, 126.12, 125.84, 125.42, 105.81, 99.86, 96.72, 34.31, 22.79.

HRMS (APCI): calcd. for C_25_H_21_N_6_O_3_ [M+H]^+^= 453.1670, found [M+H]^+^= 453.1671.

mp >240 °C (dec.)

*N*-(3-Cyclohexyl-1-(4-methyl-6-oxo-1,6-dihydropyrimidin-2-yl)-1*H*-pyrazol-5-yl)-5-phenylisoxazole-3-carboxamide **(34)**

The compound was prepared according to General procedure I using Et_3_N (10 µL, 0.07 mmol), 2-(5-amino-3-cyclohexyl-1*H*-pyrazol-1-yl)-6-methylpyrimidin-4(3*H*)-one (15 mg, 0.05 mmol) acetonitrile (0.8 + 0.3 mL) and 5-phenylisoxazole-3-carbonyl chloride (14 mg, 0.07 mmol). The product was obtained as an off white solid (21 mg, 87%).

^1^H NMR (300 MHz, DMSO-*d*_6_) *δ* (ppm) 13.55 (s, 1H), 12.56 (s, 1H), 8.01 (dd, *J* = 6.8, 2.9 Hz, 2H), 7.62 (s, 1H), 7.61 – 7.54 (m, 3H), 6.81 (s, 1H), 6.29 (s, 1H), 2.67 (q, *J* = 10.8 Hz, 1H), 2.45 (s, 3H), 1.94 (d, *J* = 12.2 Hz, 2H), 1.74 (dd, *J* = 27.2, 11.3 Hz, 3H), 1.56 – 1.16 (m, 5H).

HRMS (ESI): calcd. for C_24_H_23_N_6_O_3_ [M-H]^-^= 443.1837, found [M-H]^-^= 443.1834.

mp >276 °C (dec.)

*N*-(4-Methyl-1-(4-methyl-6-oxo-1,6-dihydropyrimidin-2-yl)-3-phenyl-1*H*-pyrazol-5-yl)-5-phenylisoxazole-3-carboxamide **(35)**

The compound was prepared according to General procedure I using Et_3_N (50 µL, 0.34 mmol), 2-(5-amino-4-methyl-3-phenyl-1*H*-pyrazol-1-yl)-6-methylpyrimidin-4(3*H*)-one (50 mg, 0.18 mmol), acetonitrile (5 mL + 2 mL) and 5-phenylisoxazole-3-carbonyl chloride (48 mg, 0.23 mmol). The reaction mixture was refluxed for 5 h. Then reaction mixture was cooled to room temperature, 7M NH_3_ in methanol (2 mL) was added and the mixture was stirred for 5 min. The solid was collected by filtration, washed with methanol (5 mL), water (10 mL), diethyl ether (15 mL), and dried *in vacuo.* The product was obtained as a white solid (40 mg, 50%).

^1^H NMR (500 MHz, DMSO-*d*_6_) *δ* (ppm) 12.32 (br s, 1H), 11.14 (br s, 1H), 8.05 – 7.93 (m, 2H), 7.90 – 7.79 (m, 2H), 7.62 – 7.55 (m, 3H), 7.55 – 7.49 (m, 2H), 7.49 – 7.42 (m, 2H), 6.27 (s, 1H), 2.25 (s, 3H), 2.21 (s, 3H).

^13^C NMR (126 MHz, DMSO-*d*_6_) *δ* (ppm) 170.87, 158.70, 156.59, 151.08, 135.04, 132.24, 130.63, 128.99, 128.21, 128.13, 127.31, 125.96, 125.60, 111.56, 106.23, 99.62, 22.49, 9.42.

HRMS (APCI): calcd. for C_25_H_21_N_6_O_3_ [M+H]^+^= 453.1670, found [M+H]^+^= 453.1671.

5-Methyl-*N*-(1-(4-methyl-6-oxo-1,6-dihydropyrimidin-2-yl)-3-phenyl-1*H*-pyrazol-5-yl)isoxazole-3-carboxamide **(36)**

The compound was prepared according to General procedure I using Et_3_N (10 µL, 0.07 mmol), 2-(5-amino-3-phenyl-1*H*-pyrazol-1-yl)-6-methylpyrimidin-4(3*H*)-one (20 mg, 0.07 mmol), acetonitrile (1 + 0.4 mL) and 5-methylisoxazole-3-carbonyl chloride (11 mg, 0.07 mmol). The obtained solid was additionally washed with EtOAc (3 mL). The product was obtained as a white solid (16 mg, 58%).

^1^H NMR (500 MHz, DMSO-*d*_6_) *δ* (ppm) 15.11 (s, 1H), 7.94 – 7.89 (m, 2H), 7.46 (t, *J* = 7.6 Hz, 2H), 7.37 (dd, *J* = 8.4, 6.3 Hz, 1H), 7.22 (s, 1H), 6.85 (s, 1H), 5.67 (s, 1H), 2.52 (s, 3H), 2.20 (s, 3H).

^13^C NMR (126 MHz, DMSO-*d*_6_) *δ* (ppm) 172.2, 172.0, 160.8, 158.9, 157.6, 155.0, 149.8, 140.7, 132.9, 128.6, 128.2, 125.6, 107.5, 101.4, 93.6, 22.7, 11.9.

^1^H NMR (500 MHz, Methanol-*d*_4_) *δ* (ppm) 8.01 – 7.86 (m, 2H), 7.42 (t, *J* = 7.5 Hz, 2H), 7.35 (t, *J* = 7.4 Hz, 1H), 7.29 (s, 1H), 6.68 (s, 1H), 6.04 (s, 1H), 2.54 (s, 3H), 2.38 (s, 3H).

HRMS (APCI): calcd. for C_19_H_17_N_6_O_3_ [M+H]^+^= 377.1357, found [M+H]^+^= 377.1359.

mp >285 °C (dec.)

5-Cyclohexyl-*N*-(1-(4-methyl-6-oxo-1,6-dihydropyrimidin-2-yl)-3-phenyl-1*H*-pyrazol-5-yl)isoxazole-3-carboxamide **(37)**

The compound was prepared according to General procedure I using Et_3_N (31 µL, 0.22 mmol), 2-(5-amino-3-phenyl-1*H*-pyrazol-1-yl)-6-methylpyrimidin-4(3*H*)-one (30 mg, 0.11 mmol), acetonitrile (1 + 0.5 mL) and 5-cyclohexylisoxazole-3-carbonyl chloride (48 mg, 0.22 mmol). The product was obtained as a white solid (30 mg, 60%).

^1^H NMR (300 MHz, trifluoroacetic acid-*d*) *δ* (ppm) 7.95 (dd, *J* = 7.5, 2.0 Hz, 2H), 7.57 (d, *J* = 6.9 Hz, 3H), 6.90 (s, 1H), 6.74 (d, *J* = 0.8 Hz, 1H), 2.99 (d, *J* = 15.2 Hz, 1H), 2.82 (s, 3H), 2.22 (d, *J* = 11.4 Hz, 2H), 1.91 (dd, *J* = 30.0, 11.5 Hz, 3H), 1.71 – 1.32 (m, 5H).

HRMS (APCI): calcd. for C_24_H_23_N_6_O_3_ [M-H]^-^= 443.1837, found [M-H]^-^= 443.1835.

mp >316 °C (dec.)

5-(2-Bromophenyl)-*N*-(1-(4-methyl-6-oxo-1,6-dihydropyrimidin-2-yl)-3-phenyl-1*H*-pyrazol-5-yl)isoxazole-3-carboxamide **(38)**

The compound was prepared according to General procedure I using Et_3_N (13 µL, 0.09 mmol), 2-(5-amino-3-phenyl-1*H*-pyrazol-1-yl)-6-methylpyrimidin-4(3*H*)-one (20 mg, 0.08 mmol), acetonitrile (1 + 0.4 mL) and 5-(2-bromophenyl)isoxazole-3-carbonyl chloride (26 mg, 0.09 mmol). The product was obtained as a light brown solid (28 mg, 72%).

^1^H NMR (300 MHz, DMSO-*d*_6_) *δ* (ppm) 13.51 (s, 1H), 12.78 (s, 1H), 8.08 – 7.99 (m, 2H), 7.97 – 7.86 (m, 2H), 7.65 – 7.42 (m, 6H), 7.39 (s, 1H), 6.37 (s, 1H), 2.50 (s, 3H, overlapped with DMSO).

HRMS (APCI): calcd. for C_24_H_16_BrN_6_O_3_ [M-H]^-^= 517.0456, found [M-H]^-^= 517.0453.

mp >295 °C (dec.)

*N*-(1-(4-Methyl-6-oxo-1,6-dihydropyrimidin-2-yl)-3-phenyl-1*H*-pyrazol-5-yl)-5-(4-(trifluoromethoxy)phenyl)isoxazole-3-carboxamide **(39)**

Et_3_N (62.6 µL, 0.449 mmol) and T_3_P/propylphosphonic anhydride (50% in EtOAc, 0.143 mL, 0.22 mmol) were added to a solution of 5-(4-(trifluoromethoxy)phenyl)isoxazole-3-carboxylic acid (20 mg, 0.75 mmol) in THF (1 mL) and the mixture was stirred at room temperature for 1 h. Then, a solution of 2-(5-amino-3-phenyl-1*H*-pyrazol-1-yl)-6-methylpyrimidin-4(3*H*)-one (20 mg, 0.075 mmol) in THF (0.4 mL) was added and the reaction mixture was refluxed for 16 h. The solvent was evaporated *in vacuo*, the residue was mixed with saturated aqueous NaHCO_3_ solution (3 mL) and the mxiture was extracted with EtOAc (3 × 3 mL). The combined organic extracts were dried over MgSO_4_, filtered, and the solvent evaporated *in vacuo*. The residue was purified by preparative TLC (dichloromethane:MeOH, 93:7). The product was obtained as a white solid (14 mg, 44%).

^1^H NMR (300 MHz, DMSO-*d*_6_) *δ* (ppm) 13.91 (s, 2H), 8.18 (d, *J* = 8.7 Hz, 2H), 8.03 (d, *J* = 7.5 Hz, 2H), 7.77 (s, 1H), 7.60 (d, *J* = 8.4 Hz, 2H), 7.55 – 7.40 (m, 3H), 7.38 (s, 1H), 6.28 (s, 1H), 2.47 (s, 3H).

HRMS (APCI): calcd. for C_25_H_16_F_3_N_6_O_4_ [M-H]^-^= 521.1191, found [M-H]^-^= 521.1189.

mp >297 °C (dec.)

5-(4-Methoxyphenyl)-*N*-(1-(4-methyl-6-oxo-1,6-dihydropyrimidin-2-yl)-3-phenyl-1*H*-pyrazol-5-yl)isoxazole-3-carboxamide **(40)**

The compound was prepared according to General procedure I using Et_3_N (13 µL, 0.09 mmol), 2-(5-amino-3-phenyl-1*H*-pyrazol-1-yl)-6-methylpyrimidin-4(3*H*)-one (20 mg, 0.08 mmol), acetonitrile (1 + 0.5 mL) and 5-(4-methoxyphenyl)isoxazole-3-carbonyl chloride (18 mg, 0.09 mmol). The crude product (15 mg) was mixed with DMSO (1.8 mL) and heated at 135 °C until a clear solution formed. The mixture was cooled to room temperature and aqueous saturated NaHCO_3_ solution (0.2 mL) was added. The resulting precipitate was collected by filtration, washed with water (3 mL) and dried *in vacuo*. The product was obtained as a white solid (30 mg, 60%).

^1^H NMR (300 MHz, DMSO-*d*_6_) *δ* (ppm) 13.38 (s, 1H), 12.76 (s, 1H), 8.04 (d, *J* = 7.4 Hz, 2H), 7.96 (d, *J* = 8.8 Hz, 2H), 7.57 – 7.41 (m, 4H), 7.40 (s, 1H), 7.13 (d, *J* = 8.8 Hz, 2H), 6.39 (s, 1H), 3.86 (s, 3H).

HRMS (APCI): calcd. for C_25_H_19_N_6_O_4_ [M-H]^-^= 467.1473, found [M-H]^-^= 467.1470.

mp >304 °C (dec.)

*N*-(1-(4-Methyl-6-oxo-1,6-dihydropyrimidin-2-yl)-3-phenyl-1*H*-pyrazol-5-yl)-3-phenylisoxazole-5-carboxamide **(41)**

The compound was prepared according to General procedure I using Et_3_N (13 µL, 0.09 mmol), 2-(5-amino-3-phenyl-1*H*-pyrazol-1-yl)-6-methylpyrimidin-4(3*H*)-one (20 mg, 0.08 mmol), acetonitrile (1 + 0.5 mL) and 3-phenylisoxazole-5-carbonyl chloride (19 mg, 0.090 mmol). The product was obtained as a white solid (13 mg, 40%).

^1^H NMR (300 MHz, Trifluoroacetic Acid-*d*) *δ* (ppm) 7.94 (d, *J* = 7.6 Hz, 2H), 7.85 (d, *J* = 7.0 Hz, 2H), 7.76 (d, *J* = 5.3 Hz, 2H), 7.56 (dd, *J* = 17.1, 7.1 Hz, 6H), 6.90 (s, 1H), 2.82 (s, 3H).

HRMS (APCI): calcd. for C_24_H_19_N_6_O_3_ [M+H]^+^= 439.1513, found [M+H]^+^= 439.1512.

mp > 300°C (dec).

*N*-(1-(4-Methyl-6-oxo-1,6-dihydropyrimidin-2-yl)-3-phenyl-1*H*-pyrazol-5-yl)-2-phenylthiazole-4-carboxamide **(42)**

The compound was prepared according to General procedure I using Et_3_N (13 µL, 0.09 mmol), 2-(5-amino-3-phenyl-1*H*-pyrazol-1-yl)-6-methylpyrimidin-4(3*H*)-one (20 mg, 0.08 mmol), acetonitrile (1 + 0.5 mL) and 2-phenylthiazole-4-carbonyl chloride (20 mg, 0.09 mmol). The product was obtained as an off-white solid (33 mg, 97%).

^1^H NMR (700 MHz, DMSO-*d*_6_) *δ* (ppm) 13.93 (br s, 1H), 8.62 – 8.52 (m, 1H), 8.37 – 8.24 (m, 2H), 8.21 – 8.09 (m, 1H), 7.96 (dd, *J* = 18.2, 6.7 Hz, 2H), 7.62 – 7.41 (m, 5H), 7.42 – 7.29 (m, 2H), 5.88 (s, 1H), 2.21 (s, 3H).

^13^C NMR (176 MHz, DMSO-*d*_6_) *δ* (ppm) 168.27, 157.16, 149.88, 140.24, 133.07, 132.75, 132.21, 130.80, 130.19, 129.12, 128.63, 128.23, 127.31, 126.32, 126.19, 125.66, 124.16, 107.27, 93.86, 23.18.

HRMS (APCI): calcd. for C_24_H_19_N_6_O_2_S [M+H]^+^= 455.1285, found [M+H]^+^= 455.1284.

mp >308 °C (dec.)

*N*-(1-(4-Methyl-6-oxo-1,6-dihydropyrimidin-2-yl)-3-phenyl-1*H*-pyrazol-5-yl)-5-phenylisothiazole-3-carboxamide **(43)**

The compound was prepared according to General procedure I using Et_3_N (15 µL, 0.11 mmol), 2-(5-amino-3-phenyl-1*H*-pyrazol-1-yl)-6-methylpyrimidin-4(3*H*)-one (20 mg, 0.08 mmol), acetonitrile (1 + 0.5 mL) and 5-phenylisothiazole-3-carbonyl chloride (23 mg, 0.11 mmol). The product was obtained as an off-white solid (19 mg, 50%).

^1^H NMR (300 MHz, DMSO-*d*_6_) *δ* (ppm) 14.81 (s, 1H), 8.41 (s, 1H), 8.01 – 7.87 (m, 4H), 7.63 – 7.42 (m, 5H), 7.42 – 7.34 (m, 1H), 7.28 (s, 1H), 5.79 (s, 1H), 2.29 (s, 3H).

HRMS (APCI): calcd. for C_24_H_19_N_6_O_2_S [M+H]^+^= 455.1285, found [M+H]^+^= 455.1284.

mp >308 °C (dec.)

*N*-(1-(4-methyl-6-oxo-1,6-dihydropyrimidin-2-yl)-3-phenyl-1*H*-pyrazol-5-yl)-2-(5-phenylisoxazol-3-yl)acetamide **(44)**

The compound was prepared according to General procedure I using Et_3_N (15 µL, 0.11 mmol), 2-(5-amino-3-phenyl-1*H*-pyrazol-1-yl)-6-methylpyrimidin-4(3*H*)-one (20 mg, 0.08 mmol), acetonitrile (1 mL+ 0.5 mL), and 2-(5-phenylisoxazol-3-yl)acetyl chloride (23 mg, 0.11 mmol). The reaction mixture was refluxed for 2 h, and then cooled to room temperature and saturated aqueous NaHCO_3_ solution (3 mL) was added. The resulting solid was collected by filtration, washed with water (3 mL), and purified by preparative TLC (dichloromethane:MeOH, 94:6). The product was obtained as a light brown solid (8 mg, 24%).

^1^H NMR (300 MHz, DMSO-*d*_6_) *δ* (ppm) 13.70 (s, 1H), 8.00 – 7.79 (m, 4H), 7.61 – 7.32 (m, 7H), 7.10 (s, 1H), 5.81 (s, 1H), 3.98 (s, 2H), 2.15 (s, 3H).

^13^C NMR (176 MHz, DMSO-*d*_6_) δ 169.25, 164.30, 158.85, 149.91, 140.66, 132.70, 130.30, 129.17, 128.59, 128.20, 126.86, 125.58, 107.19, 101.40, 93.36, 34.86, 23.00.

HRMS (APCI): calcd. for C_25_H_21_N_6_O_3_ [M+H]^+^= 453.1670, found [M+H]^+^= 453.1670.

mp = 185−188 °C.

*N*-(1-(4-methoxy-6-methylpyrimidin-2-yl)-3-phenyl-1*H*-pyrazol-5-yl)-*N*-methyl-5-phenylisoxazole-3-carboxamide **(S190)**

NaH (60%suspension in mineral oil, 17 mg, 0.43 mmol) was added to a cooled (0 °C ) solution of *N*-(1-(4-methoxy-6-methylpyrimidin-2-yl)-3-phenyl-1*H*-pyrazol-5-yl)-5-phenylisoxazole-3-carboxamide (130 mg, 0.29 mmol) in DMF (2 mL) and the mixture was stirred at room temperature for 20 min. Then, MeI (27 µL, 0.43 mmol) was added and thhe reaction mixture was stirred at room temperature for 2 h. The reaction mixture was poured into ice-cold water (10 mL) and extracted with EtOAc (3 × 20 mL). The combined organic extracts were washed with water (3 × 10 mL), brine (10 mL), dried over MgSO_4_, filtered, and the solvent evaporated *in vacuo*. The residue was purified by preparative TLC (hexane:EtOAc, 6:4). The product was obtained as a white solid (66 mg, 49%).

^1^H NMR (300 MHz, Chloroform-*d*) *δ* (ppm) 8.00 – 7.88 (m, 2H), 7.68 – 7.54 (m, 2H), 7.48 – 7.31 (m, 6H), 6.76 (s, 1H), 6.54 (s, 1H), 6.48 (s, 1H), 3.99 (s, 3H), 3.51 (s, 3H), 2.50 (s, 3H).

HRMS (APCI): calcd. for C_26_H_23_N_6_O_3_ [M+H]^+^= 467.1826, found [M+H]^+^= 467.1826.

*N*-Methyl-*N*-(1-(4-methyl-6-oxo-1,6-dihydropyrimidin-2-yl)-3-phenyl-1*H*-pyrazol-5-yl)-5-phenylisoxazole-3-carboxamide **(45)**

A mixture of *N*-(1-(4-methoxy-6-methylpyrimidin-2-yl)-3-phenyl-1*H*-pyrazol-5-yl)-N-methyl-5-phenylisoxazole-3-carboxamide (21 mg, 0.045 mmol) and pyridine•HCl (16 mg, 0.14 mmol) was heated at 150 °C for 1.5 h. The reaction mixture was cooled to room temperature, water (0.2 mL) was added and the pH was neutralized by adding saturated aqueous solution of NaHCO_3_. The solid was collected by filtration, washed with water (3 mL) and dried *in vacuo*. The resulting solid (18 mg) was mixed with MeOH (0.5 mL) and heated to reflux. The mixture was cooled to room temperature, the precipitate was collected by filtration, washed with MeOH (0.1 mL), and dried *in vacuo*. The product was obtained as an off-white solid (11 mg, 54%).

^1^H NMR (300 MHz, Chloroform-*d*) *δ* (ppm) 7.86 (dd, *J* = 7.9, 1.8 Hz, 2H), 7.67 (dd, *J* = 6.7, 3.0 Hz, 2H), 7.53 – 7.36 (m, 7H), 6.80 (d, *J* = 2.3 Hz, 2H), 6.08 (d, *J* = 1.0 Hz, 1H), 3.48 (s, 3H), 2.24 (d, *J* = 0.9 Hz, 3H).

^13^C NMR (75 MHz, Chloroform-*d*) *δ* (ppm) 170.6, 165.2, 161.5, 160.8, 159.1, 153.6, 146.1, 144.0, 131.0, 130.9, 130.1, 129.3, 129.2, 126.8, 126.4, 126.1, 110.4, 107.1, 100.5, 38.1, 24.1.

HRMS (APCI): calcd. for C_25_H_21_N_6_O_3_ [M+H]^+^= 453.1670, found [M+H]^+^= 453.1669.

mp >257 °C (dec.)

*N*-(1-Methyl-3-phenyl-1*H*-pyrazol-5-yl)-5-phenylisoxazole-3-carboxamide **(46)**

Et_3_N (145 µL, 1.04 mmol) and T_3_P/propylphosphonic anhydride (50% in EtOAc, 0.33 mL, 0.52 mmol) were added to a solution of 5-phenylisoxazole-3-carboxylic acid (33 mg, 0.17 mmol) in THF (5 mL) and the mixture was stirred at room temperature for 1 h. Then, 1-methyl-3-phenyl-1*H*-pyrazol-5-amine (30 mg, 0.17 mmol) was added and the reaction mixture was refluxed for 16 h. The reaction mixture was cooled to room temperature, quenched with saturated aqueous solution of NaHCO_3_ (5 mL), and extracted with EtOAc (3 × 15 mL). The combined organic extracts were dried over MgSO_4_, filtered, and the solvent evaporated *in vacuo*. The residue was purified by preparative TLC (hexanes:EtOAc, 6:4, run two times). The product was obtained as a white solid (25 mg, 42%).

^1^H NMR (300 MHz, Chloroform-*d*) *δ* (ppm) 8.48 (s, 1H), 7.87 – 7.78 (m, 4H), 7.53 (dt, *J* = 4.6, 2.8 Hz, 3H), 7.41 (t, *J* = 7.4 Hz, 2H), 7.36 – 7.29 (m, 1H), 7.07 (s, 1H), 6.78 (s, 1H), 3.93 (s, 3H).

^13^C NMR (126 MHz, Chloroform-*d*) *δ* (ppm) 172.6, 158.3, 156.6, 150.3, 135.1, 133.1, 131.1, 129.3, 128.6, 127.9, 126.4, 126.0, 125.5, 99.2, 97.4, 35.8.

HRMS (APCI): calcd. for C_20_H_17_N_4_O_2_ [M+H]^+^= 345.1346, found [M+H]^+^= 345.1344.

mp = 168−170 °C.

*N*-(1,3-Diphenyl-1*H*-pyrazol-5-yl)-5-phenylisoxazole-3-carboxamide **(47)**

The compound was prepared according to General procedure I using Et_3_N (83 µL, 0.6 mmol), 1,3-diphenyl-1*H*-pyrazol-5-amine (70 mg, 0.3 mmol, CAS: 5356-71-8), acetonitrile (5 mL) and 5-phenylisoxazole-3-carbonyl chloride (80 mg, 0.39 mmol). The reaction mixture was refluxed for 3 h. The crude product was purified by column chromatography on silica gel (hexane:EtOAc, 7:3). The product was obtained as a white solid (60 mg, 50%).

^1^H NMR (500 MHz, Chloroform-*d*) *δ* (ppm) 8.87 (br s, 1H), 7.93 (d, J = 7.7 Hz, 2H), 7.87 – 7.75 (m, 2H), 7.71 – 7.57 (m, 4H), 7.57 – 7.47 (m, 4H), 7.44 (t, J = 7.5 Hz, 2H), 7.38 – 7.33 (m, 1H), 7.25 (s, 1H), 7.05 (s, 1H).

^13^C NMR (126 MHz, Chloroform-*d*) *δ* (ppm) 172.67, 158.52, 155.27, 152.23, 137.87, 135.70, 133.08, 131.22, 130.28, 129.40, 128.91, 128.78, 128.37, 126.61, 126.15, 126.00, 124.85, 99.22, 96.00.

HRMS (APCI): calcd. for C_25_H_19_N_4_O_2_ [M+H]^+^= 407.1503, found [M+H]^+^ = 407.1502.

5-Phenyl-*N*-(3-phenyl-1-(pyrimidin-2-yl)-1*H*-pyrazol-5-yl)isoxazole-3-carboxamide **(48)**

The compound was prepared according to General procedure I using Et_3_N (23 µL, 0.17 mmol), 3-phenyl-1-(pyrimidin-2-yl)-1*H*-pyrazol-5-amine (20 mg, 0.084 mmol), acetonitrile (1 + 0.4 mL) and 5-phenylisoxazole-3-carbonyl chloride (35 mg, 0.17 mmol). The reaction mixture was cooled to room temperature and quenched with aqueous saturated solution of NaHCO_3_ (3 mL). The resulting precipitate was collected by filtration, washed with water (3 mL), dried *in vacuo*, and purified by reversed phase column chromatography (100% CH_3_CN). The product was obtained as a light brown solid (16 mg, 45%).

^1^H NMR (500 MHz, Chloroform-*d*) *δ* (ppm) 13.13 (s, 1H), 8.93 (d, *J* = 4.9 Hz, 2H), 8.10 – 8.01 (m, 2H), 7.86 (dd, *J* = 7.6, 2.0 Hz, 2H), 7.58 – 7.49 (m, 4H), 7.46 (t, *J* = 7.3 Hz, 2H), 7.42 – 7.36 (m, 1H), 7.29 (d, *J* = 4.8 Hz, 1H), 7.10 (s, 1H).

^13^C NMR (126 MHz, Chloroform-*d*) *δ* (ppm) 172.4, 159.2, 158.9, 157.8, 155.5, 154.7, 140.4, 132.2, 131.1, 129.4, 129.2, 128.7, 126.8, 126.8, 126.1, 118.1, 99.4, 96.8.

HRMS (APCI): calcd. for C_23_H_17_N_6_O_2_ [M+H]^+^= 409.1408, found [M+H]^+^= 409.1404.

mp >226 °C (dec.)

*N*-(1-(4-Methoxy-6-methylpyrimidin-2-yl)-3-phenyl-1*H*-pyrazol-5-yl)-5-phenylisoxazole-3-carboxamide **(49)**

The compound was prepared according to General procedure I using Et_3_N (13 µL, 0.09 mmol), 1-(4-methoxy-6-methylpyrimidin-2-yl)-3-phenyl-1*H*-pyrazol-5-amine (20 mg, 0.07 mmol), acetonitrile (1 + 0.4 mL) and 5-phenylisoxazole-3-carbonyl chloride (19 mg, 0.09 mmol). The reaction mixture was cooled to room temperature and quenched with aqeuous saturated solution of NaHCO_3_ (3 mL). The resulting precipitate was collected by filtration, washed with water (3 mL), and dried *in vacuo*. The product was obtained as an off-white solid (15 mg, 47%).

^1^H NMR (500 MHz, Chloroform-*d*) *δ* (ppm) 13.47 (s, 1H), 8.05 – 8.01 (m, 2H), 7.87 – 7.83 (m, 2H), 7.54 – 7.43 (m, 7H), 7.41 – 7.36 (m, 1H), 7.09 (s, 1H), 6.51 (s, 1H), 4.22 (s, 3H), 2.69 (s, 3H).

^13^C NMR (126 MHz, Chloroform-*d*) *δ* (ppm) 172.3, 171.7, 168.3, 159.3, 157.2, 155.5, 153.8, 140.5, 132.5, 131.0, 129.3, 129.0, 128.7, 126.9, 126.6, 126.2, 103.9, 99.4, 96.3, 54.8, 23.7.

HRMS (APCI): calcd. for C_25_H_21_N_6_O_3_ [M+H]^+^= 453.1670, found [M+H]^+^= 453.1666.

mp >206 °C (dec.)

*N*-(1-(6-Oxo-4-(trifluoromethyl)-1,6-dihydropyrimidin-2-yl)-3-phenyl-1*H*-pyrazol-5-yl)-5-phenylisoxazole-3-carboxamide **(50)**

The compound was prepared according to General procedure I using Et_3_N (10 µL, 0.08 mmol), 2-(5-amino-3-phenyl-1*H*-pyrazol-1-yl)-6-(trifluoromethyl)pyrimidin-4(3*H*)-one (20 mg, 0.06 mmol), acetonitrile (1 + 0.3 mL) and 5-phenylisoxazole-3-carbonyl chloride (16 mg, 0.08 mmol). The product was obtained as an off-white solid (21 mg, 69%).

^1^H NMR (300 MHz, DMSO-*d*_6_) *δ* (ppm) 12.70 (s, 2H), 8.09 (d, *J* = 6.8 Hz, 2H), 8.05 – 7.97 (m, 2H), 7.66 (s, 1H), 7.59 (dd, *J* = 5.2, 2.0 Hz, 3H), 7.55 – 7.39 (m, 4H), 6.80 (s, 1H).

HRMS (ESI): calcd. for C_24_H_14_F_3_N_6_O_3_ [M-H]^-^= 491.1085, found [M-H]^-^= 491.1081.

mp >250 °C (dec.)

*N*-(1-(4-(Methoxymethyl)-6-oxo-1,6-dihydropyrimidin-2-yl)-3-phenyl-1*H*-pyrazol-5-yl)-5-phenylisoxazole-3-carboxamide **(51)**

The compound was prepared according to General procedure I using Et_3_N (11.2 µL, 0.08 mmol), 2-(5-amino-3-phenyl-1*H*-pyrazol-1-yl)-6-(methoxymethyl)pyrimidin-4(3*H*)-one (20 mg, 0.07 mmol), acetonitrile (1 + 0.4 mL) and 5-phenylisoxazole-3-carbonyl chloride (17mg, 0.08 mmol). The reaction mixture was cooled to room temperature and quenched with aqueous saturated solution of NaHCO_3_ (3 mL). The resulting precipitate was collected by filtration, washed with water (3 mL), and dried *in vacuo*. The product was obtained as an off-white solid (16 mg, 49%).

^1^H NMR (300 MHz, DMSO-*d*_6_) *δ* (ppm) 13.17 (s, 1H), 8.13 – 7.94 (m, 4H), 7.65 (s, 1H), 7.59 (dd, *J* = 5.1, 1.9 Hz, 3H), 7.55 – 7.42 (m, 3H), 7.40 (s, 1H), 6.40 (s, 1H), 4.54 (s, 2H), 3.48 (s, 3H).

HRMS (ESI): calcd. for C_25_H_19_N_6_O_4_ [M-H]^-^= 467.1473, found [M-H]^-^= 467.1470.

mp >309 °C (dec.)

*N*-(1-(4,5-Dimethyl-6-oxo-1,6-dihydropyrimidin-2-yl)-3-phenyl-1*H*-pyrazol-5-yl)-5-phenylisoxazole-3-carboxamide **(52)**

The compound was prepared according to General procedure I using Et_3_N (21 µL, 0.15 mmol), 2-(5-amino-3-phenyl-1*H*-pyrazol-1-yl)-5,6-dimethylpyrimidin-4(3*H*)-one (30 mg, 0.11 mmol), acetonitrile (1+ 0.4 mL) and 5-phenylisoxazole-3-carbonyl chloride (31 mg, 0.15 mmol). The crude product (43 mg) was mixed with DMSO (1 mL) and heated to 135 °C until a clear solution obtained (10 min). Then cooled to room temperature, solid precipitated were filtered, washed with water (3 mL) and dried *in vacuo*. The product was obtained as a white solid (22 mg, 46%).

^1^H NMR (300 MHz, DMSO-*d*_6_) *δ* (ppm) 13.42 (s, 1H), 12.78 (s, 1H), 8.10 – 7.97 (m, 4H), 7.68 – 7.54 (m, 4H), 7.54 – 7.42 (m, 3H), 7.39 (s, 1H), 2.54 (s, 3H), 2.05 (s, 3H).

^13^C NMR (75 MHz, Trifluoroacetic Acid-*d*) *δ* (ppm) 177.0, 172.9, 161.8, 160.3, 160.0, 159.5, 150.1, 141.6, 133.8, 133.4, 131.2, 131.1, 130.6, 128.5, 127.9, 127.4, 117.1, 100.5, 19.1, 10.8.

HRMS (ESI): calcd. for C_25_H_19_N_6_O_3_ [M-H]^-^= 451.1524, found [M-H]^-^= 451.1526.

mp >335 °C (dec.)

*N*-(1-(1,4-Dimethyl-6-oxo-1,6-dihydropyrimidin-2-yl)-3-phenyl-1*H*-pyrazol-5-yl)-5-phenylisoxazole-3-carboxamide **(53)**

The compound was prepared according to General procedure I using Et_3_N (79 µL, 0.57 mmol), 2-(5-amino-3-phenyl-1*H*-pyrazol-1-yl)-3,6-dimethylpyrimidin-4(3*H*)-one (80 mg, 0.28 mmol), acetonitrile (5 mL) and 5-phenylisoxazole-3-carbonyl chloride (77 mg, 0.37 mmol). The reaction mixture was refluxed for 3 h and then cooled to room temperature. The precipitate was collected by filtration, washed with diethyl ether (20 mL), water (20 mL), again with diethyl ether (10 mL), and dried *in vacuo*. The product was obtained as a white solid (50 mg, 39%).

^1^H NMR (500 MHz, DMSO-*d*_6_) *δ* (ppm) 11.64 (s, 1H), 8.02 – 7.95 (m, 2H), 7.93 (dt, *J* = 6.2, 1.4 Hz, 2H), 7.64 – 7.54 (m, 4H), 7.52 – 7.47 (m, 2H), 7.46 – 7.41 (m, 1H), 7.18 (s, 1H), 6.40 (s, 1H), 3.54 (s, 3H), 2.22 (s, 3H).

^13^C NMR (126 MHz, DMSO-*d*_6_) *δ* (ppm) 71.35, 162.21, 161.40, 158.64, 155.88, 152.25, 148.16, 139.26, 131.42, 131.10, 129.33, 129.07, 128.86, 125.97, 125.89, 125.77, 110.33, 100.09, 96.66, 32.37, 22.68 ppm.

HRMS (APCI): calcd. for C_25_H_21_N_6_O_3_ [M+H]^+^= 453.1670, found [M+H]^+^= 453.1669.

*N*-(1-(6-Hydroxypyridin-2-yl)-3-phenyl-1*H*-pyrazol-5-yl)-5-phenylisoxazole-3-carboxamide **(54)**

Palladium on activated charcoal (10% Pd basis, 11 mg, 0.01 mmol) was added to solution of *N*-(1-(6-(benzyloxy)pyridin-2-yl)-3-phenyl-1*H*-pyrazol-5-yl)-5-phenylisoxazole-3-carboxamide (50 mg, 0.1 mmol) in dichloromethane:THF (1:1, 10 mL) at 23 °C and the mixture was purged with H_2_ (1 bar) for 5 min. The reaction mixture was stirred at 50 °C under H_2_ atmosphere for 4 h, filtered through Celite^®^, the filter plug was washed with dichloromethane (20 mL) and methanol (10 mL), the filtrate eas collected and the solvents were evaporated *in vacuo*. The residue was purified by column chromatography on silica gel (dichloromethane:MeOH, 95:5). The product was obtained as a white solid (29 mg, 71%).

^1^H NMR (500 MHz, DMSO-*d*_6_) *δ* (ppm) 13.00 (s, 1H), 11.40 (s, 1H), 8.07 – 7.99 (m, 2H), 7.99 – 7.94 (m, 2H), 7.93 – 7.90 (m, 1H), 7.66 – 7.56 (m, 4H), 7.57 – 7.53 (m, 1H), 7.52 – 7.46 (m, 2H), 7.45 – 7.40 (m, 1H), 7.38 (br s, 1H), 6.72 (d, *J* = 8.0 Hz, 1H).

^13^C NMR (126 MHz, DMSO-*d*_6_) *δ* (ppm) 171.38, 161.98, 158.79, 154.97, 151.60, 151.11, 142.71, 138.85, 131.96, 131.08, 129.36, 128.81, 128.72, 126.11, 125.87, 125.62, 106.60, 104.58, 99.90, 94.95.

HRMS (APCI): calcd. for C_24_H_18_N_5_O_3_ [M+H]^+^= 424.1404, found [M+H]^+^= 424.1406.

*N*-(1-(4-Ethyl-6-oxo-1,6-dihydropyrimidin-2-yl)-3-phenyl-1*H*-pyrazol-5-yl)-5-phenylisoxazole-3-carboxamide **(55)**

The compound was prepared according to General procedure I using Et_3_N (21 µL, 0.15 mmol), 2-(5-amino-3-phenyl-1*H*-pyrazol-1-yl)-6-ethylpyrimidin-4(3*H*)-one (30 mg, 0.11 mmol), acetonitrile (1 + 0.4 mL) and 5-phenylisoxazole-3-carbonyl chloride (31 mg, 0.15 mmol). The product was obtained as a white solid (36 mg, 75%).

^1^H NMR (300 MHz, DMSO-*d*_6_) *δ* (ppm) 13.28 (s, 1H), 12.81 (s, 1H), 8.10 – 7.97 (m, 4H), 7.64 (s, 1H), 7.59 (dd, *J* = 5.1, 1.9 Hz, 3H), 7.55 – 7.42 (m, 3H), 7.42 (s, 1H), 6.37 (s, 1H), 2.78 (q, *J* = 7.6 Hz, 2H), 1.32 (t, *J* = 7.5 Hz, 3H).

HRMS (APCI): calcd. for C_25_H_21_N_6_O_3_ [M+H]^+^= 453.1670, found [M+H]^+^= 453.1671.

mp >290 °C (dec.)

5-(3-(t*ert*-Butyl)phenyl)-*N*-(1-(4-methyl-6-oxo-1,6-dihydropyrimidin-2-yl)-3-phenyl-1*H*-pyrazol-5-yl)isoxazole-3-carboxamide **(118)**

The compound was prepared according to General procedure I using Et_3_N (10 µL, 0.07 mmol), 2-(5-amino-3-phenyl-1*H*-pyrazol-1-yl)-6-methylpyrimidin-4(3*H*)-one (15 mg, 0.06 mmol), acetonitrile (0.8 + 0.4 mL) and 5-(3-(*ter*t-butyl)phenyl)isoxazole-3-carbonyl chloride (18 mg, 0.07 mmol). The product was obtained as a white solid (21 mg, 74%).

^1^H NMR (500 MHz, DMSO-*d*_6_) *δ* (ppm) 13.31 (s, 1H), 12.38 (br s, 1H), 8.05 – 7.99 (m, 2H), 7.97 (t, *J* = 1.9 Hz, 1H), 7.79 (dt, *J* = 7.6, 1.4 Hz, 1H), 7.61 (dt, *J* = 8.1, 1.4 Hz, 1H), 7.58 – 7.55 (m, 1H), 7.55 – 7.46 (m, 3H), 7.46 – 7.41 (m, 1H), 7.37 (s, 1H), 6.32 (s, 1H), 2.50 (s, 3H, overlapped with DMSO), 1.38 (s, 9H).

^13^C NMR (126 MHz, DMSO-*d*_6_) *δ* (ppm) 171.88, 158.60, 154.54, 152.01, 151.80, 139.88, 131.44, 128.73, 128.61, 128.30, 127.62, 125.76, 125.66, 122.87, 122.43, 106.04, 99.42, 94.86, 34.26, 30.59, 22.31.

HRMS (APCI): calcd. for C_28_H_27_N_6_O_3_ [M+H]^+^= 495.2139, found [M+H]^+^= 495.2137.

mp >309 °C (dec.)

*N*-(1-(4-Methyl-6-oxo-1,6-dihydropyrimidin-2-yl)-3-phenyl-1*H*-pyrazol-5-yl)-5-phenyl-4,5-dihydroisoxazole-3-carboxamide **(119)**

The compound was prepared according to General procedure I using Et_3_N (19 µL, 0.14 mmol), 2-(5-amino-3-phenyl-1*H*-pyrazol-1-yl)-6-methylpyrimidin-4(3*H*)-one (30 mg, 0.11 mmol), acetonitrile (1 + 0.5 mL) and 5-phenyl-4,5-dihydroisoxazole-3-carbonyl chloride (28 mg, 0.14 mmol). The product was obtained as a white solid (35 mg, 71%).

^1^H NMR (300 MHz, DMSO-*d*_6_) *δ* (ppm) 13.16 (s, 1H), 12.76 (s, 1H), 8.08 – 7.96 (m, 2H), 7.57 – 7.34 (m, 8H), 7.29 (s, 1H), 6.34 (s, 1H), 5.92 (dd, *J* = 11.4, 9.4 Hz, 1H), 3.81 (dd, *J* = 17.7, 11.4 Hz, 1H), 3.36 – 3.22 (m, 1H), 2.38 (s, 3H).

HRMS (ESI): calcd. for C_24_H_19_N_6_O_3_ [M-H]^-^= 439.1524, found [M-H]^-^= 439.1526.

mp >294 °C (dec.)

2,3-Dimethoxy-*N*-(1-(4-methyl-6-oxo-1,6-dihydropyrimidin-2-yl)-3-phenyl-1*H*-pyrazol-5-yl)benzamide **(120)**

The compound was prepared according to General procedure I using Et_3_N (13 µL, 0.09 mmol), 2-(5-amino-3-phenyl-1*H*-pyrazol-1-yl)-6-methylpyrimidin-4(3*H*)-one (20 mg, 0.08 mmol), acetonitrile (1 + 0.5 mL) and 2,3-dimethoxybenzoyl chloride (18 mg, 0.09 mmol). The product was obtained as a white solid (13 mg, 40%).

^1^H NMR (300 MHz, DMSO-*d*_6_) *δ* (ppm) 12.86 (s, 1H), 12.61 (s, 1H), 8.05 (d, *J* = 7.4 Hz, 2H), 7.61 – 7.39 (m, 5H), 7.39 – 7.23 (m, 2H), 6.30 (s, 1H), 3.90 (s, 3H), 3.82 (s, 3H), 2.38 (s, 3H).

HRMS (APCI): calcd. for C_23_H_22_N_5_O_4_ [M+H]^+^= 432.1666, found [M+H]^+^= 432.1662.

mp = 269−272 °C

*N*-(1-(4-Methyl-6-oxo-1,6-dihydropyrimidin-2-yl)-3-phenyl-1*H*-pyrazol-5-yl)-[1,1'-biphenyl]-4-carboxamide **(121)**

The compound was prepared according to General procedure I using Et_3_N (13 µL, 0.09 mmol), 2-(5-amino-3-phenyl-1*H*-pyrazol-1-yl)-6-methylpyrimidin-4(3*H*)-one (20 mg, 0.08 mmol), acetonitrile (1 + 0.5 mL) and [1,1'-biphenyl]-4-carbonyl chloride (19 mg, 0.09 mmol). The crude product (24 mg) was combined with DMSO (2 mL) and heated at 135 °C until a clear solution formed. The solution was cooled to room temperature, the precipitate was collected by filtration, washed with water (1 mL) and dried *in vacuo*. The product was obtained as a white solid (12 mg, 36%).

^1^H NMR (300 MHz, DMSO-*d*_6_) *δ* (ppm) 12.89 (s, 2H), 8.09 (dd, *J* = 13.9, 7.8 Hz, 4H), 7.96 (d, *J* = 8.1 Hz, 2H), 7.80 (d, *J* = 7.6 Hz, 2H), 7.60 – 7.35 (m, 7H), 6.37 (s, 1H).

HRMS (APCI): calcd. for C_27_H_22_N_5_O_2_ [M+H]^+^= 448.1768, found [M+H]^+^= 448.1765.

mp >232 °C (dec.)

*N*-(1-(4-Methyl-6-oxo-1,6-dihydropyrimidin-2-yl)-3-phenyl-1*H*-pyrazol-5-yl)-[1,1'-biphenyl]-3-carboxamide **(122)**

The compound was prepared according to General procedure I using Et_3_N (13 µL, 0.09 mmol), 2-(5-amino-3-phenyl-1*H*-pyrazol-1-yl)-6-methylpyrimidin-4(3*H*)-one (20 mg, 0.08 mmol), acetonitrile (1 + 0.5 mL) and [1,1'-biphenyl]-3-carbonyl chloride (19 mg, 0.09 mmol). The product was obtained as a white solid (16 mg, 48%).

^1^H NMR (300 MHz, DMSO-*d*_6_) *δ* (ppm) 13.09 (s, 1H), 12.72 (s, 1H), 8.23 (s, 1H), 8.09 – 8.00 (m, 3H), 8.00 – 7.93 (m, 1H), 7.85 – 7.67 (m, 3H), 7.60 – 7.36 (m, 7H), 6.23 (s, 1H), 2.23 (s, 3H),

HRMS (APCI): calcd. for C_27_H_22_N_5_O_2_ [M+H]^+^= 448.1768, found [M+H]^+^= 448.1769.

mp >246 °C (dec.)

*N*-(1-(4-Methyl-6-oxo-1,6-dihydropyrimidin-2-yl)-3-phenyl-1*H*-pyrazol-5-yl)-[1,1'-biphenyl]-2-carboxamide **(123)**

Et_3_N (62 µL, 0.45 mmol) and T_3_P/propylphosphonic anhydride (50% in EtOAc, 0.143 mL, 0.22 mmol) were added to a solution of [1,1'-biphenyl]-2-carboxylic acid (15 mg, 0.75 mmol) in THF (1.0 mL), and the mixture was stirred at room temperature for 1 h. Then, 2-(5-amino-3-phenyl-1*H*-pyrazol-1-yl)-6-methylpyrimidin-4(3*H*)-one (20 mg, 0.08 mmol) was added and the reaction mixture was refluxed for 48 h. The solvent was evaporated *in vacuo*, saturated aqueous solution of NaHCO_3_ (3 mL) was added to the residue and the mixture was extracted with EtOAc (3 × 10 mL). The combined organic extracts were dried over MgSO_4_, filtered, and the solvent evaporated *in vacuo*. The residue was purified by preparative TLC (dichloromethane:MeOH, 94:6). The product was obtained as a white solid (14 mg, 42%).

^1^H NMR (500 MHz, Methanol-*d*_4_) *δ* (ppm) 7.80 (d, *J* = 7.5 Hz, 1H), 7.70 (t, *J* = 7.5 Hz, 1H), 7.64 – 7.56 (m, 2H), 7.45 (d, *J* = 7.5 Hz, 2H), 7.36 (t, *J* = 7.5 Hz, 2H), 7.30 (d, *J* = 7.3 Hz, 1H), 7.23 – 7.14 (m, 3H), 7.05 (t, *J* = 7.6 Hz, 2H), 6.89 (s, 1H), 5.94 (s, 1H), 1.97 (s, 3H).

^13^C NMR (126 MHz, Methanol-*d*_4_) *δ* (ppm) 174.8, 168.2, 165.0, 155.3, 154.8, 136.1, 132.6, 132.1, 131.2, 130.2, 129.8, 129.6, 129.4, 129.3, 129.0, 128.9, 128.4, 110.1, 98.3, 23.0.

HRMS (APCI): calcd. for C_27_H_22_N_5_O_2_ [M+H]^+^= 448.1768, found [M+H]^+^= 448.1770.

mp >238 °C (dec.)

*N*-(3-(Furan-2-yl)-1-(4-methyl-6-oxo-1,6-dihydropyrimidin-2-yl)-1*H*-pyrazol-5-yl)-5-methylisoxazole-3-carboxamide **(124)**

The compound was prepared according to General procedure I using Et_3_N (16 µL, 0.12 mmol), 2-(5-amino-3-(furan-2-yl)-1*H*-pyrazol-1-yl)-6-methylpyrimidin-4(3*H*)-one (30 mg, 0.12 mmol), acetonitrile (1 + 0.5 mL) and 5-methylisoxazole-3-carbonyl chloride (17 mg, 0.12 mmol). The obtained solid was washed with water (3 mL), then with EtOAc (2 mL), and dried *in vacuo*. The product was obtained as a white solid (32 mg, 75%).

^1^H NMR (500 MHz, DMSO-*d*_6_) *δ* (ppm) 15.15 (s, 1H), 7.76 (d, *J* = 1.6 Hz, 1H), 7.05 (s, 1H), 6.90 (d, *J* = 3.2 Hz, 1H), 6.85 (d, *J* = 1.1 Hz, 1H), 6.61 (dd, *J* = 3.3, 1.8 Hz, 1H), 5.68 (s, 1H), 2.51 (s, 3H), 2.19 (s, 3H).

^13^C NMR (126 MHz, DMSO-*d*_6_) *δ* (ppm) 172.0, 171.9, 160.9, 158.9, 157.5, 155.2, 148.3, 142.8, 142.6, 140.6, 111.7, 107.5, 107.0, 101.3, 93.2, 22.7, 11.9.

HRMS (ESI): calcd. for C_17_H_13_N_6_O_4_ [M-H]^-^= 365.1004, found [M-H]^-^= 365.1000.

mp >290 °C (dec.)

*N*-(3-(Furan-2-yl)-1-(4-methyl-6-oxo-1,6-dihydropyrimidin-2-yl)-1*H*-pyrazol-5-yl)-5-isopropylisoxazole-3-carboxamide **(125)**

The compound was prepared according to General procedure I using Et_3_N (14 µL, 0.1 mmol), 2-(5-amino-3-(furan-2-yl)-1*H*-pyrazol-1-yl)-6-methylpyrimidin-4(3*H*)-one (25 mg, 0.1 mmol), acetonitrile (1 + 0.5 mL) and 5-isopropylisoxazole-3-carbonyl chloride (17 mg, 0.1 mmol). The product was obtained as a light brown solid (26 mg, 68%).

^1^H NMR (300 MHz, DMSO-*d*_6_) *δ* (ppm) 13.50 (s, 1H), 12.90 (s, 1H), 7.82 (s, 1H), 7.19 (s, 1H), 7.02 (d, *J* = 3.4 Hz, 1H), 6.81 (s, 1H), 6.67 – 6.62 (m, 1H), 6.46 (s, 1H), 3.24 – 3.13 (m, 1H), 1.31 (d, *J* = 6.4 Hz, 6H),

^13^C NMR (176 MHz, DMSO-*d*_6_) *δ* (ppm) 181.24, 158.06, 154.97, 147.21, 144.23, 143.62, 139.89, 111.83, 108.58, 99.27, 94.33, 26.65, 22.73, 20.48.

HRMS (APCI): calcd. for C_19_H_19_N_6_O_4_ [M+H]^+^= 395.1462, found [M+H]^+^= 395.1460.

mp >250°C (dec.)

5-(t*ert*-Butyl)-N-(3-(furan-2-yl)-1-(4-methyl-6-oxo-1,6-dihydropyrimidin-2-yl)-1*H*-pyrazol-5-yl)isoxazole-3-carboxamide **(126)**

HOBt•H_2_O (10 mg, 0.08 mmol), EDCI•HCl (22 mg, 0.12 mmol) and DIPEA (27 µL, 0.16 mmol) were added to a solution of 5-(*tert*-butyl)isoxazole-3-carboxylic acid (12 mg, 0.78 mmol) in DMF (1 mL) and the mixture was stirred at room temperature for 15 min. Then, 2-(5-amino-3-(furan-2-yl)-1*H*-pyrazol-1-yl)-6-methylpyrimidin-4(3*H*)-one (20 mg, 0.08 mmol) was added and the reaction mixture was stirred at room temperature for 48 h. H_2_O (30 mL) was dded and the mixture was stirred for 15 min. The precipitate was collected by filtration, washed with water (3 mL), and dried *in vacuo*. The product was obtained as a light brown solid (6 mg, 18%).

^1^H NMR (300 MHz, DMSO-*d*_6_) *δ* (ppm) 14.96 (s, 1H), 7.77 (s, 1H), 7.08 (s, 1H), 6.91 (d, *J* = 2.7 Hz, 2H), 6.62 (d, *J* = 2.5 Hz, 1H), 5.86 (s, 1H), 2.25 (s, 3H), 1.37 (s, 9H).

^13^C NMR (126 MHz, DMSO-*d*_6_) *δ* (ppm) 183.1, 158.5, 155.1, 148.0, 143.0, 111.7, 107.3, 98.6, 93.5, 32.6, 28.4, 22.9.

HRMS (APCI): calcd. for C_20_H_21_N_6_O_4_ [M+H]^+^= 409.1619, found [M+H]^+^= 409.1615.

mp >197 °C (dec.)

*N*-(3-(furan-2-yl)-1-(4-methyl-6-oxo-1,6-dihydropyrimidin-2-yl)-1*H*-pyrazol-5-yl)-2,3-dimethoxybenzamide **(127)**

The compound was prepared according to General procedure I using Et_3_N (20 µL, 0.14 mmol), 2-(5-amino-3-(furan-2-yl)-1*H*-pyrazol-1-yl)-6-methylpyrimidin-4(3*H*)-one (30 mg, 0.12 mmol), acetonitrile (1 + 0.5 mL) and 2,3-dimethoxybenzoyl chloride (28 mg, 0.14 mmol). The reaction mixture was cooled to room temperature, quenched with saturated aqueous solution of NaHCO_3_ (3 mL) and extracted with EtOAc (3 × 5 mL). The combined organic extracts were washed with water (3 × 5 mL), brine (5 mL), dried over MgSO_4_, filtered, and the solvent evaporated *in vacuo*. The residue was purified by preparative TLC (dichloromethane:MeOH, 94:6). The product was obtained as an off-white solid (12 mg, 24%).

^1^H NMR (300 MHz, DMSO-*d*_6_) *δ* (ppm) 13.11 (s, 1H), 12.66 (s, 1H), 7.77 (s, 1H), 7.56 – 7.47 (m, 1H), 7.29 (dt, *J* = 22.4, 8.0 Hz, 3H), 7.01 (s, 1H), 6.62 (s, 1H), 6.19 (s, 1H), 3.90 (s, 3H), 3.82 (s, 3H), 2.32 (s, 3H).

^13^C NMR (75 MHz, DMSO-*d*_6_) *δ* (ppm) 161.5, 152.8, 147.3, 143.5, 126.6, 124.5, 121.8, 116.8, 111.8, 94.9, 61.2, 56.3, 22.1.

HRMS (APCI): calcd. for C_21_H_20_N_5_O_5_ [M+H]^+^= 422.1459, found [M+H]^+^= 422.1459.

mp >210 °C (dec.)

*N*-(3-(furan-2-yl)-1-(4-methyl-6-oxo-1,6-dihydropyrimidin-2-yl)-1*H*-pyrazol-5-yl)-5-phenylisothiazole-3-carboxamide **(128)**

The compound was prepared according to General procedure I using Et_3_N (20 µL, 0.14 mmol), 2-(5-amino-3-(furan-2-yl)-1*H*-pyrazol-1-yl)-6-methylpyrimidin-4(3*H*)-one (30 mg, 0.12 mmol), acetonitrile (1 + 0.5 mL) and 5-phenylisothiazole-3-carbonyl chloride (31 mg, 0.14 mmol). The crude product (30 mg) was mixed with DMSO (1.5 mL) and heated at 135 °C until a clear solution formed. The solution was cooled to room temperature and H_2_O (0.2 mL) was added. The resulting precipitate was collceted by filtration, washed with water (2 mL), and dried *in vacuo*. The product was obtained as a light brown solid (19 mg, 37%).

^1^H NMR (300 MHz, DMSO-*d*_6_) *δ* (ppm) 14.16 (s, 1H), 12.83 (s, 1H), 8.38 (s, 1H), 7.90 (dd, *J* = 7.5, 2.1 Hz, 2H), 7.81 (d, *J* = 1.7 Hz, 1H), 7.53 (dd, *J* = 4.9, 2.4 Hz, 3H), 7.18 (s, 1H), 6.98 (d, *J* = 3.4 Hz, 1H), 6.64 (dd, *J* = 3.4, 1.8 Hz, 1H), 6.22 (s, 1H), 2.45 (s, 3H).

HRMS (APCI): calcd. for C_22_H_17_N_6_O_3_S [M+H]^+^= 445.1077, found [M+H]^+^= 445.1077.

mp >289 °C (dec.)

*N*-(3-(4-cyanophenyl)-1-(4-methyl-6-oxo-1,6-dihydropyrimidin-2-yl)-1*H*-pyrazol-5-yl)-5-phenylisoxazole-3-carboxamide **(129)**

The compound was prepared according to General procedure I using Et_3_N (9 µL, 0.06 mmol), 4-(5-amino-1-(4-methyl-6-oxo-1,6-dihydropyrimidin-2-yl)-1*H*-pyrazol-3-yl)benzonitrile (15 mg, 0.05 mmol), acetonitrile (0.8+ 0.3 mL) and 5-phenylisoxazole-3-carbonyl chloride (13 mg, 0.06 mmol). The product was obtained as a white solid (21 mg, 87%).

^1^H NMR (300 MHz, DMSO-*d*_6_) *δ* (ppm) 15.67 (s, 1H), 8.15 (d, *J* = 8.1 Hz, 2H), 8.10 – 7.97 (m, 2H), 7.91 (d, *J* = 8.2 Hz, 2H), 7.86 (s, 1H), 7.59 (d, *J* = 6.7 Hz, 3H), 7.36 (s, 1H), 5.79 (s, 1H), 2.25 (s, 3H).

HRMS (ESI): calcd. for C_25_H_16_N_7_O_3_ [M-H]^-^= 462.1320, found [M-H]^-^= 462.1317.

mp >330 °C (dec.)

*N*-(1-(4-methyl-6-oxo-1,6-dihydropyrimidin-2-yl)-3-(4-(trifluoromethyl)phenyl)-1*H*-pyrazol-5-yl)-5-phenylisoxazole-3-carboxamide **(130)**

The compound was prepared according to General procedure I using Et_3_N (8 µL, 0.54 mmol), 2-(5-amino-3-(4-(trifluoromethyl)phenyl)-1*H*-pyrazol-1-yl)-6-methylpyrimidin-4(3*H*)-one (15 mg, 0.05 mmol), acetonitrile (0.8 + 0.3 mL) and 5-phenylisoxazole-3-carbonyl chloride (11 mg, 0.05 mmol). The crude product (20 mg) was mixed with DMSO (1.5 mL) and heated at 135 °C until a clear solution formed. The solution was cooled to room temperature, the resulting precipitated was collected by filtration, washed with H_2_O (1 mL), and dried *in vacuo*. The product was obtained as a white solid (12 mg, 53%).

^1^H NMR (300 MHz, Trifluoroacetic Acid-*d*) *δ* (ppm) 8.11 (d, *J* = 8.1 Hz, 2H), 7.88 (dd, *J* = 6.8, 2.8 Hz, 2H), 7.79 (d, *J* = 8.2 Hz, 2H), 7.70 (s, 1H), 7.58 – 7.47 (m, 3H), 7.21 (s, 1H), 6.86 (s, 1H), 2.80 (s, 3H).

HRMS (ESI): calcd. for C_25_H_16_F_3_N_6_O_3_ [M-H]^-^= 505.1241, found [M-H]^-^= 505.1243.

mp >350 °C (dec.)

*N*-(3-(4-(t*ert*-butyl)phenyl)-1-(4-methyl-6-oxo-1,6-dihydropyrimidin-2-yl)-1*H*-pyrazol-5-yl)-5-phenylisoxazole-3-carboxamide **(131)**

The compound was prepared according to General procedure I using Et_3_N (12 µL, 0.09 mmol), 2-(5-amino-3-(4-(*tert*-butyl)phenyl)-1*H*-pyrazol-1-yl)-6-methylpyrimidin-4(3*H*)-one (20 mg, 0.06 mmol), acetonitrile (0.8 + 0.3 mL) and 5-phenylisoxazole-3-carbonyl chloride (18 mg, 0.09 mmol). The product was obtained as a white solid (11 mg, 36%).

^1^H NMR (300 MHz, DMSO-*d*_6_) *δ* (ppm) 13.93 (s, 1H), 12.78 (s, 1H), 8.03 (dd, *J* = 6.8, 2.8 Hz, 2H), 7.93 (d, *J* = 8.4 Hz, 2H), 7.68 (s, 1H), 7.59 (dd, *J* = 5.1, 1.9 Hz, 3H), 7.51 (d, *J* = 8.4 Hz, 2H), 7.33 (s, 1H), 6.25 (s, 1H), 2.45 (s, 3H), 1.33 (s, 9H).

HRMS (APCI): calcd. for C_28_H_27_N_6_O_3_ [M+H]^+^= 495.2139, found [M+H]^+^= 495.2139.

mp >309 °C (dec.)

*N*-(3-(4-methoxyphenyl)-1-(4-methyl-6-oxo-1,6-dihydropyrimidin-2-yl)-1*H*-pyrazol-5-yl)-5-phenylisoxazole-3-carboxamide **(132)**

The compound was prepared according to General procedure I using Et_3_N (13 µL, 0.09 mmol), 2-(5-amino-3-(4-methoxyphenyl)-1*H*-pyrazol-1-yl)-6-methylpyrimidin-4(3*H*)-one (20 mg, 0.07 mmol), acetonitrile (0.8 + 0.3 mL) and 5-phenylisoxazole-3-carbonyl chloride (20 mg, 0.09 mmol). The product was obtained as an off-white solid (24 mg, 76%).

^1^H NMR (300 MHz, DMSO-*d*_6_) *δ* (ppm) 14.01 (s, 1H), 12.68 (s, 1H), 8.06 – 7.98 (m, 2H), 7.94 (d, *J* = 8.7 Hz, 2H), 7.69 (s, 1H), 7.58 (dd, *J* = 5.2, 1.9 Hz, 3H), 7.28 (s, 1H), 7.04 (d, *J* = 8.8 Hz, 2H), 6.19 (s, 1H), 3.82 (s, 3H), 2.42 (s, 3H).

HRMS (APCI): calcd. for C_25_H_21_N_6_O_4_ [M+H]^+^= 469.1619, found [M+H]^+^= 469.1619.

mp >270 °C (dec.)

*N*-(1-(4-methyl-6-oxo-1,6-dihydropyrimidin-2-yl)-3-(5-methylfuran-2-yl)-1*H*-pyrazol-5-yl)-5-phenylisoxazole-3-carboxamide **(133)**

The compound was prepared according to General procedure I using Et_3_N (22 µL, 0.16 mmol), 2-(5-amino-3-(5-methylfuran-2-yl)-1*H*-pyrazol-1-yl)-6-methylpyrimidin-4(3*H*)-one (30 mg, 0.11 mmol), acetonitrile (1 + 0.4 mL) and 5-phenylisoxazole-3-carbonyl chloride (32 mg, 0.16 mmol). The product was obtained as a light yellow solid (29 mg, 59%).

^1^H NMR (300 MHz, DMSO-*d*_6_) *δ* (ppm) 13.58 (s, 1H), 12.96 (s, 1H), 8.02 (dd, *J* = 6.6, 3.1 Hz, 2H), 7.63 (s, 1H), 7.58 (dd, *J* = 5.0, 1.9 Hz, 3H), 7.18 (s, 1H), 6.91 (d, *J* = 3.2 Hz, 1H), 6.48 (s, 1H), 6.26 (dd, *J* = 3.2, 1.2 Hz, 1H), 2.55 (s, 3H), 2.38 (s, 3H).

HRMS (APCI): calcd. for C_23_H_19_N_6_O_4_ [M+H]^+^= 443.1462, found [M+H]^+^= 443.1462.

mp >300 °C (dec.)

*N*-(1-(4-methyl-6-oxo-1,6-dihydropyrimidin-2-yl)-3-(thiophen-2-ylmethyl)-1*H*-pyrazol-5-yl)-5-phenylisoxazole-3-carboxamide **(134)**

The compound was prepared according to General procedure I using Et_3_N (78 µL, 0.56 mmol), 2-(5-amino-3-(thiophen-2-ylmethyl)-1*H*-pyrazol-1-yl)-6-methylpyrimidin-4(3*H*)-one (80 mg, 0.28 mmol), acetonitrile (5 mL) and 5-phenylisoxazole-3-carbonyl chloride (75 mg, 0.36 mmol). The reaction mixture was refluxed for 3 h, then cooled to room temperature. The resulting precipitate was collected by filtration, washed with water (5 mL), diethyl ether (15 mL), and dried *in vacuo*. The product was obtained as an off-white solid (50 mg, 39%).

^1^H NMR (500 MHz, DMSO-*d*_6_) *δ* (ppm) 13.32 (br s, 1H), 12.45 (br s, 1H), 8.10 – 7.89 (m, 2H), 7.68 – 7.52 (m, 3H), 7.49 (s, 1H), 7.36 (d, J = 5.1 Hz, 1H), 7.11 – 6.94 (m, 2H), 6.82 (s, 1H), 6.37 (s, 1H), 4.22 (s, 2H), 2.55 (s, 3H; mearged with DMSO signal).

^13^C NMR (126 MHz, DMSO-*d*_6_) *δ* (ppm) 171.48, 158.61, 154.36, 153.35, 140.53, 139.08, 130.72, 128.96, 126.64, 125.82, 125.66, 125.50, 124.29, 104.75, 99.49, 97.02, 28.41, 22.31.

HRMS (APCI): calcd. for C_23_H_19_N_6_O_3_S [M+H]^+^= 459.1234, found [M+H]^+^= 459.1235.

*N*-(1-(5-(2-(2-(2-methoxyethoxy)ethoxy)ethyl)-4-methyl-6-oxo-1,6-dihydropyrimidin-2-yl)-3-phenyl-1*H*-pyrazol-5-yl)-5-phenylisoxazole-3-carboxamide **(135)**

The compound was prepared according to General procedure I using using Et_3_N (14 µL, 0.10 mmol), 2-(5-amino-3-phenyl-1*H*-pyrazol-1-yl)-5-(2-(2-(2-methoxyethoxy)ethoxy)ethyl)-6-methylpyrimidin-4(3*H*)-one (30 mg, 0.07 mmol), acetonitrile (1 + 0.4 mL) and 5-phenylisoxazole-3-carbonyl chloride (21 mg, 0.10 mmol). The product was obtained as a white solid (25 mg, 59%).

^1^H NMR (300 MHz, DMSO-*d*_6_) *δ* (ppm) 14.87 (s, 1H), 12.88 (s, 1H), 8.02 (dd, *J* = 7.6, 2.1 Hz, 2H), 8.00 – 7.92 (m, 2H), 7.71 (s, 1H), 7.65 – 7.53 (m, 3H), 7.48 (t, *J* = 7.3 Hz, 2H), 7.43 – 7.35 (m, 1H), 7.28 (s, 1H), 3.60 – 3.45 (m, 8H), 3.45 – 3.38 (m, 2H), 3.22 (s, 3H), 2.72 (t, *J* = 6.8 Hz, 2H), 2.41 (s, 3H).

^13^C NMR (75 MHz, DMSO-*d*_6_) *δ* (ppm) 171.2, 159.5, 154.8, 153.2, 150.6, 141.0, 132.4, 130.9, 129.3, 128.6, 128.4, 126.2, 125.8, 125.8, 115.0, 99.9, 94.0, 71.2, 69.8, 69.6, 69.5, 68.9, 57.9, 20.8.

HRMS (APCI): calcd. for C_31_H_33_N_6_O_6_ [M+H]^+^= 585.2456, found [M+H]^+^= 585.2455.

mp >195 °C (dec.)

*N*-(1-(1,3-dimethyl-2,6-dioxo-1,2,3,6-tetrahydropyrimidin-4-yl)-3-phenyl-1*H*-pyrazol-5-yl)-5-phenylisoxazole-3-carboxamide **(136)**

The compound was prepared according to General procedure I using Et_3_N (20 µL, 0.14 mmol), 6-(5-amino-3-phenyl-1*H*-pyrazol-1-yl)-1,3-dimethylpyrimidine-2,4(1*H*,3*H*)-dione (30 mg, 0.10 mmol), acetonitrile (1 mL + 0.4 mL) and 5-phenylisoxazole-3-carbonyl chloride (29 mg, 0.14 mmol). The reaction mixture was cooled to room temperature, quenched with saturated aqueous solution of NaHCO_3_ (3 mL) and extracted with EtOAc (3 × 15 mL). The combined organic extracts were washed with water (3 × 10 mL), brine (10 mL), dried over MgSO_4_, filtered, and the solvent evaporated *in vacuo*. The residue was purified by preparative TLC (hexane:EtOAc, 7:3). The product was obtained as a white solid (10 mg, 21%).

^1^H NMR (300 MHz, Chloroform-*d*) *δ* (ppm) 8.92 (s, 1H), 7.89 – 7.83 (m, 2H), 7.83 – 7.75 (m, 2H), 7.56 – 7.37 (m, 6H), 7.04 (s, 1H), 6.99 (s, 1H), 5.88 (s, 1H), 3.42 (s, 3H), 3.38 (s, 3H).

HRMS (APCI): calcd. for C_25_H_21_N_6_O_4_ [M+H]^+^= 469.1619, found [M+H]^+^= 469.1618.

*N*-(1-(6-(benzyloxy)pyridin-2-yl)-3-phenyl-1*H*-pyrazol-5-yl)-5-phenylisoxazole-3-carboxamide **(137)**

The compound was prepared according to General procedure I using Et_3_N (41 µL, 0.29 mmol), 1-(6-(benzyloxy)pyridin-2-yl)-3-phenyl-1*H*-pyrazol-5-amine (50 mg, 0.15 mmol), acetonitrile (4 mL) and 5-phenylisoxazole-3-carbonyl chloride (39 mg, 0.19 mmol). The reaction mixture was refluxed for 3 h. The crude product was purified by column chromatography on silica gel (hexane:EtOAc, 8:2). The product was obtained as a white solid (50 mg, 67%).

^1^H NMR (500 MHz, Chloroform-*d*) *δ* (ppm) 12.39 (s, 1H), 8.00 – 7.95 (m, 2H), 7.85 – 7.80 (m, 2H), 7.79 – 7.75 (m, 2H), 7.64 – 7.58 (m, 2H), 7.54 – 7.48 (m, 4H), 7.48 – 7.41 (m, 4H), 7.40 – 7.35 (m, 2H), 7.08 (s, 1H), 6.81 – 6.74 (m, 1H), 5.70 (s, 2H).

^13^C NMR (126 MHz, Chloroform-*d*) *δ* (ppm) 172.17, 162.72, 159.05, 155.57, 152.57, 152.11, 141.70, 138.72, 136.77, 132.76, 131.07, 129.35, 128.80, 128.75, 128.62, 128.13, 128.11, 126.77, 126.22, 126.11, 108.25, 106.98, 99.53, 96.30, 69.53.

HRMS (APCI): calcd. for C_31_H_24_N_5_O_3_ [M+H]^+^= 514.1874, found [M+H]^+^= 514.1878.

*N*-(3-(furan-2-yl)-1-(6-oxo-4-(trifluoromethyl)-1,6-dihydropyrimidin-2-yl)-1*H*-pyrazol-5-yl)-5-phenylisoxazole-3-carboxamide **(138)**

The compound was prepared according to General procedure I using Et_3_N (8 µL, 0.06 mmol), 2-(5-amino-3-(furan-2-yl)-1*H*-pyrazol-1-yl)-6-(trifluoromethyl)pyrimidin-4(3*H*)-one (15 mg, 0.06 mmol), acetonitrile (0.8+ 0.3 mL) and 5-phenylisoxazole-3-carbonyl chloride (12 mg, 0.06 mmol). The product was obtained as an off-white solid (23 mg, 65%).

^1^H NMR (300 MHz, DMSO-*d*_6_) *δ* (ppm) 13.39 (s, 2H), 8.02 (dd, *J* = 7.2, 2.5 Hz, 2H), 7.83 (d, *J* = 1.7 Hz, 1H), 7.71 (s, 1H), 7.59 (dd, *J* = 5.2, 2.0 Hz, 3H), 7.22 (s, 1H), 7.05 (d, *J* = 3.4 Hz, 1H), 6.66 (dd, *J* = 3.4, 1.8 Hz, 2H).

^13^C NMR (126 MHz, DMSO-*d*_6_) *δ* (ppm) 171.2, 170.5, 159.0, 158.6, 154.5, 151.23, 151.49, 147.7, 143.2, 143.2, 139.9, 131.0, 129.4, 126.2, 125.8, 111.7, 107.8, 106.4, 99.6, 93.8.

^19^F NMR (471 MHz, DMSO-*d*_6_) *δ* (ppm) -69.13.

HRMS (ESI): calcd. for C_22_H_12_F_3_N_6_O_4_ [M-H]^-^= 481.0878, found [M-H]^-^= 481.0875.

mp >280 °C (dec.)

5-Methyl-*N*-(3-phenyl-1-(pyrimidin-2-yl)-1*H*-pyrazol-5-yl)isoxazole-3-carboxamide **(139)**

The compound was prepared according to General procedure I using Et_3_N (12 µL, 0.08 mmol), 3-phenyl-1-(pyrimidin-2-yl)-1*H*-pyrazol-5-amine (20 mg, 0.08 mmol), acetonitrile (0.8+ 0.3 mL) and 5-methylisoxazole-3-carbonyl chloride (12 mg, 0.08 mmol). The reaction mixture was cooled to room temperature, quenched with saturated aqueous solution of NaHCO_3_ (1 mL) and the solvents were evaporated *in vacuo*. The residue was purified by column chromatography on silica gel. (dichloromethane:MeOH, 98:2). The product was obtained as a light brown solid (16 mg, 55%).

^1^H NMR (500 MHz, Chloroform-*d*) *δ* (ppm) 13.02 (s, 1H), 8.90 (d, *J* = 4.8 Hz, 2H), 8.09 – 7.95 (m, 2H), 7.48 (s, 1H), 7.46 – 7.41 (m, 2H), 7.41 – 7.36 (m, 1H), 7.26 (t, 2H, merged with CHCl_3_ solvent residual peak), 6.58 (d, *J* = 1.1 Hz, 1H), 2.54 (d, *J* = 0.9 Hz, 3H).

^13^C NMR (126 MHz, Chloroform-*d*) *δ* (ppm) 172.0, 158.8, 157.8, 155.7, 154.6, 140.5, 132.2, 129.2, 128.7, 126.8, 118.0, 101.8, 96.7, 12.6.

HRMS (APCI): calcd. for C_18_H_15_N_6_O_2_ [M+H]^+^= 347.1251, found [M+H]^+^= 347.1255.

mp >218 °C (dec.)

1-methyl-*N*-(1-(4-methyl-6-oxo-1,6-dihydropyrimidin-2-yl)-3-phenyl-1*H*-pyrazol-5-yl)-5-phenyl-1*H*-pyrazole-3-carboxamide **(140)**

HATU (43 mg, 0.11 mmol) and DIPEA (39 µL, 0.22 mmol) were added to a solution of 1-methyl-5-phenyl-1*H*-pyrazole-3-carboxylic acid (25 mg, 0.11 mmol) in DMF (2 mL) and the mixture was stirred at room temperature for 20 min. Then, 2-(5-amino-3-phenyl-1*H*-pyrazol-1-yl)-6-methylpyrimidin-4(3*H*)-one (30 mg, 0.11 mmol) was added and the reaction mixture was stirred at room temperature for 16 h. The solvents were removed *in vacuo* and the residue was purified by column chromatography on silica gel (dichloromethane:MeOH, 98.5:1.5). The product was obtained as a white solid (4 mg, 8%).

^1^H NMR (500 MHz, Chloroform-*d*) *δ* (ppm) 12.81 (s, 1H), 7.96 – 7.85 (m, 2H), 7.63 – 7.34 (m, 10H), 6.98 (s, 1H), 6.16 (s, 1H), 3.97 (s, 3H), 2.54 (s, 3H).

^13^C NMR (126 MHz, Chloroform-*d*) *δ* (ppm) 159.0, 155.1, 148.6, 146.3, 144.8, 141.7, 131.3, 129.9, 129.7, 129.4, 129.1, 129.0, 126.6, 109.3, 107.6, 95.9, 38.3, 23.8.

HRMS (APCI): calcd. for C_25_H_22_N_7_O_2_ [M+H]^+^= 452.1829, found [M+H]^+^= 452.1826.

mp >215 °C (dec.)

6-Methyl-2-(3-phenyl-5-(((5-phenylisoxazol-3-yl)methyl)amino)-1*H*-pyrazol-1-yl)pyrimidin-4(3*H*)-one **(141)**

ZnCl_2_ (0.5 M in THF, 1.49 mL, 0.75 mmol) and 5-phenylisoxazole-3-carbaldehyde (84 mg, 0.49 mmol) were added to a solution of 2-(5-amino-3-phenyl-1*H*-pyrazol-1-yl)-6-methylpyrimidin-4(3*H*)-one (100 mg, 0.37 mmol) in THF (3 mL) and the reaction mixture was stirred at room temperature for 16 h. Then, NaBH_3_CN (31 mg, 0.49 mmol) was added and reaction mixture was refluxed for 16 h. The solvent was evaporated *in vacuo* and the residue was purified by column chromatography on silica gel (dichloromethane:MeOH, 9:1). The product was obtained as an off-white solid (6 mg, 4%).

^1^H NMR (300 MHz, Chloroform-*d*) *δ* (ppm) 10.34 (s, 1H), 8.25 (s, 1H), 8.03 – 7.69 (m, 4H), 7.53 – 7.35 (m, 6H), 6.55 (s, 1H), 6.06 (s, 1H), 5.84 (s, 1H), 4.61 (d, *J* = 5.2 Hz, 2H), 2.33 (s, 3H).

^13^C NMR (126 MHz, Chloroform-*d*) *δ* (ppm) 171.0, 162.0, 130.6, 129.5, 129.2, 128.8, 127.3, 126.5, 126.0, 98.3, 41.2, 22.8.

HRMS (APCI): calcd. for C_24_H_21_N_6_O_2_ [M+H]^+^= 425.1721, found [M+H]^+^= 425.1719.

mp >216 °C (dec.)

**Scheme S5:** synthesis of **142**

2-(5-Amino-3-phenyl-1*H*-pyrazol-1-yl)-5-(2-((tert-butyldimethylsilyl)oxy)ethyl)-6-methylpyrimidin-4(3H)-one **(S191)**

NaH (60% dispersion in mineral oil, 17 mg, 0.43 mmol) was added to a cooled (0 °C) solution of 2-(5-amino-3-phenyl-1*H*-pyrazol-1-yl)-5-(2-hydroxyethyl)-6-methylpyrimidin-4(3*H*)-one (30 mg, 0.1 mmol) in DMF (1 mL) and the reaction mixture was stirred at 0 °C for 15 min. Then, a solution of TBDMSCl (15 mg, 0.1 mmol) in DMF (0.5 mL) was added and the reaction mixture was stirred at 0 °C for 30 min. The reaction mixture was quenched with ice-cold water (10 mL) and extracted with EtOAc (3 × 10 mL). The combined organic extracts were washed with water (3 × 6 mL) brine (5 mL), dried over MgSO_4_, filtered, and the solvent was evaporated *in vacuo*. The product was obtained as a light brown solid (30 mg, 73%) and used as such in the next step.

^1^H NMR (500 MHz, Chloroform-*d*) *δ* (ppm) 7.78 (dt, *J* = 6.5, 1.5 Hz, 2H), 7.50 – 7.34 (m, 3H), 6.02 (s, 1H), 5.83 (s, 1H), 3.81 (t, *J* = 6.4 Hz, 2H), 2.78 (t, *J* = 6.4 Hz, 2H), 2.38 (s, 3H), 0.88 (s, 9H), 0.02 (s, 6H).

^13^C NMR (126 MHz, Chloroform-*d*) *δ* (ppm) 129.4, 128.8, 126.3, 87.3, 61.7, 29.9, 29.5, 26.1, 18.5, -5.2.

HRMS (APCI) m/z: [M+H]^+^ calcd for C_22_H_32_N_5_O_2_Si 426.2320; found 426.2318.

*N*-(1-(5-(2-((*tert*-butyldimethylsilyl)oxy)ethyl)-4-methyl-6-oxo-1,6-dihydropyrimidin-2-yl)-3-phenyl-1*H*-pyrazol-5-yl)-5-phenylisoxazole-3-carboxamide **(S192)**

The compound was prepared according to General procedure I using Et_3_N (7 µL, 0.05 mmol), 2-(5-amino-3-phenyl-1*H*-pyrazol-1-yl)-5-(2-((*tert*-butyldimethylsilyl)oxy)ethyl)-6-methylpyrimidin-4(3*H*)-one (20 mg, 0.05 mmol), acetonitrile (1 + 0.4 mL) and 5-phenylisoxazole-3-carbonyl chloride (10 mg, 0.05 mmol). The product was obtained as a white solid (17 mg) and used directly in the next step without further purification.

HRMS (APCI): calcd. for C_32_H_37_N_6_O_4_Si [M+H]^+^= 597.2640, found [M+H]^+^= 597.2635.

*N*-(1-(5-(2-hydroxyethyl)-4-methyl-6-oxo-1,6-dihydropyrimidin-2-yl)-3-phenyl-1H-pyrazol-5-yl)-5-phenylisoxazole-3-carboxamide **(142)**

TBAF (1 M in THF, 0.14 mL, 0.14 mmol) was added to a solution of *N*-(1-(5-(2-((*tert*-butyldimethylsilyl)oxy)ethyl)-4-methyl-6-oxo-1,6-dihydropyrimidin-2-yl)-3-phenyl-1*H*-pyrazol-5-yl)-5-phenylisoxazole-3-carboxamide (15 mg, 0.03 mmol) in THF (0.5 mL) and the reaction mixture was stirred at room temperature for 2 h. The solvent was evaporated *in vacuo* and the residue was purified by column chromatography on silica gel (dichloromethane:MeOH, 97:3). The product was obtained as a white solid (9 mg, 40%, over the 2 steps).

^1^H NMR (500 MHz, DMSO-*d*_6_) *δ* (ppm) 13.42 (s, 1H), 12.81 (s, 1H), 8.08 – 8.00 (m, 4H), 7.64 (s, 1H), 7.62 – 7.56 (m, 3H), 7.52 – 7.46 (m, 2H), 7.46 – 7.41 (m, 1H), 7.38 (s, 1H), 4.67 (s, 2H), 3.56 (t, *J* = 6.9 Hz, 2H), 2.69 (t, *J* = 6.9 Hz, 2H), 2.57 (s, 3H).

^13^C NMR (126 MHz, DMSO-*d*_6_) *δ* (ppm) 171.8, 158.8, 154.7, 152.0, 139.8, 131.6, 131.1, 129.3, 129.0, 128.7, 126.1, 126.0, 100.0, 95.0, 79.2, 78.9, 78.7, 59.2, 29.0.

HRMS (APCI): calcd. for C_26_H_23_N_6_O_4_ [M+H]^+^= 483.1775, found [M+H]^+^= 483.1773.

mp >255 °C (dec.)

**Scheme S6: synthesis of 143**

2-(5-Amino-3-(furan-2-yl)-1*H*-pyrazol-1-yl)-5-(2-((*tert*-butyldimethylsilyl)oxy)ethyl)-6-methylpyrimidin-4(3*H*)-one **(S193)**

NaH (60% dispersion in mineral oil, 4 mg, 0.17 mmol) was added to a cooled (0 °C) solution of 2-(5-amino-3-(furan-2-yl)-1*H*-pyrazol-1-yl)-5-(2-hydroxyethyl)-6-methylpyrimidin-4(3*H*)-one (47 mg, 0.16 mmol) in DMF (2 mL) and the reaction mixture was stirred at 0 °C for 15 min. Then a solution of TBDMSCl (26 mg, 0.17 mmol) in DMF (0.4 mL) was added and the reaction mixture was stirred at 0 °C for 30 min. The reaction mixture was quenched with ice-cold water (5 mL) and extracted with EtOAc (3 × 10 mL). The combined organic extracts were washed with water (2 × 8 mL), brine (5 mL), dried over MgSO_4_, filtered, and the solvent was evaporated *in vacuo*. The product was obtained as a light brown solid (31 mg, 48%).

^1^H NMR (300 MHz, Chloroform-*d*) *δ* (ppm) 7.49 (dd, *J* = 1.8, 0.8 Hz, 1H), 6.79 (d, *J* = 3.4 Hz, 1H), 6.50 (dd, *J* = 3.4, 1.8 Hz, 1H), 5.76 (s, 1H), 3.81 (t, *J* = 6.3 Hz, 2H), 2.77 (t, *J* = 6.3 Hz, 2H), 2.38 (s, 3H), 0.87 (s, 9H), 0.01 (s, 6H).

^13^C NMR (126 MHz, Chloroform-*d*) *δ* (ppm) 149.8, 147.7, 147.6, 146.6, 146.4, 143.2, 143.1, 111.8, 111.8, 108.7, 108.5, 87.0, 61.7, 29.5, 26.1, 22.2, 18.5, -5.2.

HRMS (APCI): calcd. for C_20_H_30_N_5_O_3_Si [M+H]^+^= 416.2112, found [M+H]^+^= 416.2110.

mp = 169−171 °C.

*N*-(1-(5-(2-((*tert*-butyldimethylsilyl)oxy)ethyl)-4-methyl-6-oxo-1,6-dihydropyrimidin-2-yl)-3-(furan-2-yl)-1*H*-pyrazol-5-yl)-5-phenylisoxazole-3-carboxamide **(S194)**

The compound was prepared according to General procedure I using Et_3_N (8 µL, 0.06 mmol), 2-(5-amino-3-(furan-2-yl)-1*H*-pyrazol-1-yl)-5-(2-((*tert*-butyldimethylsilyl)oxy)ethyl)-6-methylpyrimidin-4(3*H*)-one (20 mg, 0.05 mmol), acetonitrile (1 + 0.4 mL) and 5-phenylisoxazole-3-carbonyl chloride (12 mg, 0.06 mmol). The product was obtained as an off-white solid (18 mg, 64%).

^1^H NMR (500 MHz, Chloroform-*d*) *δ* (ppm) 13.14 (s, 1H), 10.34 (s, 1H), 7.91 – 7.81 (m, 2H), 7.60 – 7.47 (m, 4H), 7.28 (s, 1H), 7.07 (s, 1H), 6.87 (d, *J* = 3.3 Hz, 1H), 6.53 (dd, *J* = 3.4, 1.8 Hz, 1H), 3.84 (t, *J* = 6.4 Hz, 2H), 2.82 (s, 2H), 2.60 (s, 3H), 0.88 (s, 9H), 0.04 (s, 6H).

^13^C NMR (126 MHz, Chloroform-*d*) *δ* (ppm) 172.4, 161.2, 160.2, 158.6, 155.6, 146.8, 146.4, 146.0, 143.6, 140.0, 131.0, 129.2, 126.6, 126.0, 118.8, 111.8, 109.5, 99.1, 96.1, 61.4, 29.5, 26.0, 21.7, 18.3, -5.4.

HRMS (APCI): calcd. for C_30_H_35_N_6_O_5_Si [M+H]^+^= 587.2433, found [M+H]^+^= 587.2429.

mp >220 °C (dec.)

*N*-(3-(furan-2-yl)-1-(5-(2-hydroxyethyl)-4-methyl-6-oxo-1,6-dihydropyrimidin-2-yl)-1*H*-pyrazol-5-yl)-5-phenylisoxazole-3-carboxamide **(143)**

TBAF (1 M in THF, 128 µL, 0.128 mmol) was added to a solution of *N*-(1-(5-(2-((*tert*-butyldimethylsilyl)oxy)ethyl)-4-methyl-6-oxo-1,6-dihydropyrimidin-2-yl)-3-(furan-2-yl)-1*H*-pyrazol-5-yl)-5-phenylisoxazole-3-carboxamide (15 mg, 0.03 mmol) in THF (0.5 mL) and the reaction mixture was stirred at room temperature for 2 h. The solvent was evaporated *in vacuo* and the residue was purified by column chromatography on silica gel (dichloromethane:MeOH, 97:3). The product was obtained as a white solid (11 mg, 92%).

^1^H NMR (300 MHz, DMSO-*d*_6_) *δ* (ppm) 13.58 (s, 1H), 13.09 (s, 1H), 8.02 (dd, *J* = 6.6, 3.0 Hz, 2H), 7.83 (d, *J* = 1.7 Hz, 1H), 7.64 (s, 1H), 7.59 (dd, *J* = 5.0, 1.9 Hz, 3H), 7.22 (s, 1H), 7.02 (d, *J* = 3.4 Hz, 1H), 6.65 (dd, *J* = 3.4, 1.8 Hz, 1H), 4.69 (s, 1H), 3.57 (t, *J* = 6.8 Hz, 2H), 2.77 – 2.68 (m, 2H), 2.62 (s, 3H).

^13^C NMR (75 MHz, DMSO-*d*_6_) *δ* (ppm) 171.8, 158.8, 154.7, 147.1, 143.7, 139.5, 131.1, 129.3, 126.0, 125.9, 111.8, 108.7, 100.0, 94.5, 59.1.

HRMS (ESI): calcd. for C_24_H_19_N_6_O_5_ [M-H]^-^= 471.1422, found [M-H]^-^= 471.1419.

mp >265 °C (dec.)
